## Supplementary material for "Intergeneric chromosomal transfer in yeast results in improved phenotypes and widespread transcriptional responses": TableS6

Table S6.Classification of the transferred genes based on cis and trans effects.

**YPD**

| | Normalized read counts | | | | | | | | | | | | | | | | log2 fold change adjusted with ashr method | | | |
|---|---|---|---|---|---|---|---|---|---|---|---|---|---|---|---|---|---|---|---|---|
| W303_gene | KS.1.sc | KS.2.sc | KS.3.sc | KS.1.km | KS.2.km | KS.3.km | Km.V.1 | Km.V.2 | Km.V.3 | SC.1 | SC.2 | SC.3 | locus_tag | gene | ambiguous | Category (Fig. 5) | log2FC(KSsc/Psc) | log2(KSsc/KSkm) | log2FC(KSsc/Pkm) | Category (Fig. S14) |
| W303.asm02.HP0_G0000040 | 2040.744 | 5917.805 | 3294.239 | 3731.149 | 3481.11 | 2931.641 | 2651.037 | 2312.19 | 2573.827 | 1300.71 | 733.5658 | 1056.313 | YAL067C | SEO1 | n | km-like | 1.724676057 | 0.122205691 | 0.542048744 | km-like |
| W303.asm02.HP0_G0000100 | 28219.63 | 43918.05 | 48465.71 | 8487.015 | 21941.31 | 11284.66 | 12048.63 | 10369.36 | 21262.58 | 2686.868 | 2768.687 | 2793.268 | YAL063C | FLO9 | n | transgressive | 3.741861075 | 1.423442241 | 1.358543826 | transgressive |
| W303.asm02.HP0_G0000110 | 16139.9 | 13645.73 | 13285.06 | 7175.052 | 4928.653 | 3871.294 | 8658.424 | 8986.936 | 6755.47 | 1279.823 | 1081.486 | 1287.415 | YAL062W | GDH3 | n | transgressive | 3.494378659 | 1.385523078 | 0.792526285 | transgressive |
| W303.asm02.HP0_G0000120 | 2377.961 | 3726.799 | 2745.055 | 8360.708 | 9631.038 | 8629.331 | 2819.372 | 2954.529 | 8076.783 | 4160.373 | 5320.448 | 4494.344 | YAL061W | BDH2 | n | NA | -0.587522588 | -1.479503447 | -0.59888942 | NA |
| W303.asm02.HP0_G0000130 | 17676.76 | 18852.16 | 19021.66 | 38169.16 | 30243.15 | 29233.12 | 7249.679 | 6837.658 | 22141.88 | 14250.84 | 14012.15 | 13782.72 | YAL060W | BDH1 | n | NA | 0.35443191 | -0.742490308 | 0.56924048 | NA |
| W303.asm02.HP0_G0000140 | 1896.361 | 1287.387 | 1599.12 | 1945.535 | 1638.782 | 2045.877 | 4782.964 | 5113.116 | 2279.783 | 890.5591 | 967.2589 | 1118.573 | YAL059W | ECM1 | n | km-like | 0.636258707 | -0.198008278 | -1.28757357 | intermediate |
| W303.asm02.HP0_G0000150 | 6556.35 | 7537.84 | 7870.196 | 3499.926 | 4064.292 | 3567.548 | 4932.49 | 4298.555 | 6873.276 | 1191.526 | 1110.828 | 1353.896 | YAL058W | CNE1 | n | transgressive | 2.531241441 | 0.960692805 | 0.438326417 | transgressive |
| W303.asm02.HP0_G0000170 | 875.0199 | 964.8205 | 831.1273 | 2397.796 | 2474.74 | 2337.866 | 1630.684 | 1594.212 | 2176.113 | 1275.076 | 1098.253 | 1112.242 | YAL055W | PEX22 | n | transgressive | -0.374731842 | -1.41349954 | -1.001874305 | transgressive |
| W303.asm02.HP0_G0000180 | 1664.766 | 2145.645 | 1961.495 | 680.428 | 828.3848 | 616.3108 | 831.3291 | 815.7247 | 1022.557 | 1119.37 | 1372.816 | 1261.033 | YAL054C | ACS1 | n | transgressive | 0.604330581 | 1.417362219 | 1.092932726 | transgressive |
| W303.asm02.HP0_G0000190 | 18260.1 | 20578.76 | 21218.39 | 3760.689 | 4021.69 | 3120.747 | 4909.92 | 4219.426 | 4033.679 | 7969.459 | 8320.732 | 8602.463 | YAL053W | FLC2 | n | transgressive | 1.249314119 | 2.43596379 | 2.164094714 | transgressive |
| W303.asm02.HP0_G0000200 | 1219.02 | 3602.236 | 1606.904 | 4078.494 | 3854.119 | 3721.381 | 6392.958 | 6325.649 | 9293.484 | 3964.792 | 4561.731 | 4185.153 | YAL051W | OAF1 | n | transgressive | -0.897398149 | -0.800646438 | -1.669335683 | transgressive |
| W303.asm02.HP0_G0000210 | 1985.51 | 2224.847 | 2229.601 | 1981.186 | 5562.959 | 3567.548 | 4234.7 | 4937.404 | 4995.919 | 3352.414 | 3186.819 | 3308.234 | YAL049C | AIM2 | n | NA | -0.568320514 | -0.747576385 | -1.084556921 | NA |
| W303.asm02.HP0_G0000220 | 3852.801 | 4166.728 | 3720.614 | 1939.424 | 1815.819 | 1748.01 | 2609.659 | 2564.704 | 2235.488 | 1495.342 | 1598.125 | 1537.511 | YAL048C | GEM1 | n | transgressive | 1.330489844 | 1.086108369 | 0.659836845 | transgressive |
| W303.asm02.HP0_G0000230 | 2067.876 | 2305.489 | 2203.655 | 190.4791 | 438.3339 | 273.3716 | 1470.813 | 1363.808 | 1454.198 | 974.1083 | 1101.397 | 1168.171 | YAL047C | SPC72 | n | NA | 0.970890579 | 2.791492827 | 0.59553214 | NA |
| W303.asm02.HP0_G0000240 | 317.8367 | 696.9748 | 500.752 | 105.9349 | 224.3739 | 130.3169 | 1010.008 | 1165.986 | 614.4764 | 705.4215 | 551.2223 | 656.3704 | YAL046C | BOL3 | n | NA | -0.298677557 | 1.604840881 | -0.818777745 | NA |
| W303.asm02.HP0_G0000250 | 926.3776 | 980.6608 | 851.8839 | 264.8373 | 422.2396 | 396.8297 | 1520.655 | 1633.777 | 1217.643 | 1288.367 | 1235.534 | 1346.509 | YAL044W- | BOL1 | n | NA | -0.471540154 | 1.319205057 | -0.649769458 | NA |
| W303.asm02.HP0_G0000260 | 191.8648 | 400.3285 | 345.0778 | 4041.824 | 6700.923 | 5132.331 | 5832.47 | 6646.818 | 6707.406 | 1514.33 | 1033.28 | 1359.172 | YAL044C | GCV3 | n | transgressive | -1.952808966 | -4.017641436 | -4.304166029 | transgressive |
| W303.asm02.HP0_G0000270 | 5489.466 | 4145.848 | 4894.224 | 3336.949 | 3355.195 | 2968.874 | 4483.911 | 3902.911 | 3139.296 | 1979.548 | 1972.244 | 2302.573 | YAL043C | PTA1 | n | transgressive | 1.189198685 | 0.578641886 | 0.328365667 | transgressive |
| W303.asm02.HP0_G0000280 | 12992.54 | 9949.891 | 11775.89 | 2065.731 | 2448.232 | 2523.053 | 5849.397 | 6175.537 | 5010.998 | 8047.312 | 7671.002 | 8834.619 | YAL042W | ERV46 | n | transgressive | 0.487156962 | 2.272605997 | 1.011494803 | transgressive |
| W303.asm02.HP0_G0000290 | 3897.375 | 4630.418 | 4345.905 | 4121.275 | 3984.768 | 4232.85 | 6960.03 | 6407.105 | 6569.808 | 2426.726 | 2691.138 | 2529.453 | YAL041W | CDC24 | n | km-like | 0.744881042 | 0.042826555 | -0.626964009 | intermediate |
| W303.asm02.HP0_G0000300 | 5349.928 | 5310.833 | 5785.891 | 3434.735 | 3700.75 | 4002.591 | 8217.368 | 8904.316 | 8324.647 | 890.5591 | 972.4986 | 1084.805 | YAL040C | CLN3 | n | transgressive | 2.459628604 | 0.557946032 | -0.624837343 | intermediate |
| W303.asm02.HP0_G0000310 | 4051.449 | 1937.561 | 2994.134 | 2881.633 | 2299.596 | 2156.598 | 4082.353 | 4175.207 | 3412.606 | 11863.99 | 12770.33 | 12270.54 | YAL039C | CYC3 | n | km-like | -1.954289113 | 0.254225931 | -0.364206533 | km-like |
| W303.asm02.HP0_G0000320 | 52155.25 | 49779.7 | 55300.67 | 111052.4 | 104514.7 | 107042.1 | 78163.74 | 82534.82 | 114060.8 | 92958.99 | 101208 | 112732.1 | YAL038W | CDC19 | n | NA | -0.946928786 | -1.023638159 | -0.794764049 | NA |
| W303.asm02.HP0_G0000340 | 3064.023 | 2332.849 | 3029.593 | 3554.931 | 2545.745 | 2706.28 | 7042.787 | 7196.065 | 4372.018 | 441.4818 | 447.4751 | 495.9712 | YAL036C | RBG1 | n | km-like | 2.52612943 | -0.050213477 | -1.11100489 | intermediate |
| W303.asm02.HP0_G0000350 | 10644.62 | 12786.75 | 13634.47 | 9332.457 | 9551.513 | 8698.899 | 26207.56 | 25894.9 | 21600.92 | 15476.55 | 16174.08 | 17736.78 | YAL035W | FUN12 | n | intermediate | -0.405763002 | 0.420234975 | -0.979985519 | transgressive |
| W303.asm02.HP0_G0000360 | 157.9493 | 223.2047 | 148.7554 | 422.721 | 541.527 | 347.8384 | 931.0134 | 1028.674 | 832.182 | 705.4215 | 740.9014 | 745.0121 | YAL034W- | MTW1 | n | NA | -1.979737326 | -1.26989033 | -2.335699475 | NA |
| W303.asm02.HP0_G0000380 | 729.6677 | 949.7001 | 868.3161 | 355.4931 | 480.9365 | 460.5184 | 982.7363 | 1040.311 | 661.5988 | 712.0675 | 726.2301 | 794.6092 | YAL033W | POP5 | n | NA | 0.182874976 | 0.950060623 | -0.072335318 | NA |
| W303.asm02.HP0_G0000390 | 1906.051 | 2095.964 | 2184.628 | 1459.661 | 1316.895 | 1515.791 | 1598.71 | 1410.354 | 1292.097 | 657.0009 | 836.265 | 660.5915 | YAL032C | PRP45 | n | transgressive | 1.498214464 | 0.520846814 | 0.517865116 | transgressive |
| W303.asm02.HP0_G0000400 | 3351.82 | 3299.83 | 3173.159 | 3217.773 | 3246.322 | 2826.799 | 5985.758 | 5665.854 | 6888.356 | 1794.41 | 1974.34 | 1799.215 | YAL031C | GIP4 | n | km-like | 0.809868106 | 0.058947235 | -0.909438728 | intermediate |
| W303.asm02.HP0_G0000440 | 874.0509 | 974.1807 | 890.8024 | 369.7535 | 817.0241 | 403.6885 | 836.0312 | 902.9991 | 850.0885 | 1730.799 | 1644.235 | 1641.981 | YAL027W | SAW1 | n | intermediate | -0.823748489 | 0.751787484 | 0.078276445 | km-like |
| W303.asm02.HP0_G0000450 | 1792.676 | 3192.548 | 2668.083 | 5736.782 | 7188.487 | 6584.433 | 9563.106 | 9119.593 | 9054.102 | 3471.092 | 3552.554 | 3708.176 | YAL026C | DRS2 | n | transgressive | -0.468066203 | -1.318366432 | -1.814764796 | transgressive |
| W303.asm02.HP0_G0000460 | 6532.125 | 5432.515 | 7420.471 | 3153.601 | 1003.529 | 2325.128 | 5105.527 | 4250.845 | 708.7212 | 1678.58 | 1852.778 | 2199.158 | YAL025C | MAK16 | n | NA | 1.309075908 | 1.2532109 | 0.747185187 | NA |
| W303.asm02.HP0_G0000470 | 2426.412 | 3305.59 | 2455.328 | 5013.573 | 5763.665 | 5980.86 | 9040.234 | 8485.399 | 8461.302 | 741.4996 | 831.0252 | 852.6484 | YAL024C | LTE1 | n | intermediate | 1.726278721 | -1.019182926 | -1.644169457 | intermediate |
| W303.asm02.HP0_G0000480 | 5824.745 | 8231.935 | 7259.607 | 10688.22 | 12820.56 | 11268.98 | 15193.39 | 16471.59 | 17183.66 | 6267.143 | 5662.08 | 7226.406 | YAL023C | PMT2 | n | sc-like | 0.149622251 | -0.693740295 | -1.175573643 | sc-like |
| W303.asm02.HP0_G0000490 | 5812.147 | 5271.952 | 4745.469 | 1227.419 | 1152.165 | 973.9474 | 1103.11 | 1195.077 | 1449.486 | 1519.077 | 1345.569 | 1704.242 | YAL022C | FUN26 | n | NA | 1.757846946 | 2.207866063 | 2.047082594 | NA |
| W303.asm02.HP0_G0000500 | 21211.72 | 25443.9 | 28539.41 | 5939.485 | 7623.98 | 6022.993 | 14288.7 | 13178.43 | 14115.05 | 10890.83 | 11141.82 | 12385.56 | YAL021C | CCR4 | n | NA | 1.104656537 | 1.914765902 | 0.841954355 | NA |
| W303.asm02.HP0_G0000510 | 927.3466 | 916.5794 | 1075.017 | 4094.791 | 2844.91 | 4325.933 | 1623.161 | 1760.616 | 1670.019 | 1211.464 | 934.7724 | 1214.602 | YAL020C | ATS1 | n | sc-like | -0.196215531 | -1.912183258 | -0.775326704 | sc-like |
| W303.asm02.HP0_G0000520 | 6580.576 | 6013.568 | 8096.789 | 2210.372 | 1829.074 | 2116.425 | 4553.502 | 3928.512 | 2570.057 | 1407.045 | 1461.892 | 1780.22 | YAL019W | FUN30 | n | transgressive | 2.085119512 | 1.703920729 | 0.879584527 | transgressive |
| W303.asm02.HP0_G0000540 | 2803.358 | 4431.694 | 2741.596 | 6294.978 | 7110.855 | 6304.203 | 7951.23 | 6936.569 | 7138.104 | 8566.647 | 8850.995 | 8541.258 | YAL017W | PSK1 | n | NA | -1.337968295 | -0.959450348 | -1.115922181 | NA |
| W303.asm02.HP0_G0000550 | 2765.567 | 4344.572 | 3137.7 | 8554.243 | 9827.01 | 8502.933 | 14189.96 | 14481.73 | 13858.7 | 8382.458 | 8861.474 | 9256.723 | YAL016W | TPD3 | n | transgressive | -1.336280001 | -1.365402496 | -2.015971806 | transgressive |
| W303.asm02.HP0_G0000570 | 2372.147 | 2481.173 | 2826.352 | 750.7117 | 831.225 | 845.5902 | 727.8832 | 608.5935 | 665.3686 | 1946.318 | 1911.463 | 2129.51 | YAL014C | SYN8 | n | transgressive | 0.353719814 | 1.647029867 | 1.921296246 | transgressive |
| W303.asm02.HP0_G0000580 | 5554.39 | 5405.155 | 6552.154 | 1868.121 | 1627.421 | 1596.137 | 2527.842 | 2414.592 | 2514.452 | 4141.384 | 4295.552 | 4325.502 | YAL013W | DEP1 | n | transgressive | 0.45068475 | 1.769833138 | 1.222363093 | transgressive |
| W303.asm02.HP0_G0000590 | 15024.57 | 22877.77 | 25751.11 | 11326.89 | 17349.69 | 14475.95 | 18886.41 | 18839.63 | 16064.98 | 16862.71 | 14089.7 | 18421.64 | YAL012W | CYS3 | n | NA | 0.34729272 | 0.537206766 | 0.235106833 | NA |
| W303.asm02.HP0_G0000600 | 1851.786 | 1578.273 | 1681.281 | 4004.136 | 3682.762 | 3555.79 | 6101.429 | 5643.745 | 5061.891 | 394.9601 | 448.5231 | 448.4846 | YAL011W | SWC3 | n | intermediate | 1.961426233 | -1.127848602 | -1.703244521 | intermediate |
| W303.asm02.HP0_G0000610 | 1612.44 | 2939.102 | 2323.005 | 1078.702 | 1519.494 | 1392.333 | 1822.529 | 1690.796 | 1825.523 | 1895.049 | 2109.526 | 1988.106 | YAL010C | MDM10 | n | NA | 0.189184736 | 0.763093668 | 0.354859644 | NA |
| W303.asm02.HP0_G0000620 | 873.0818 | 1253.547 | 925.3967 | 1324.186 | 1789.311 | 1675.503 | 1930.677 | 2015.457 | 2000.818 | 962.7152 | 788.0592 | 867.422 | YAL009W | SPO7 | n | sc-like | 0.214590261 | -0.636470057 | -0.943003875 | sc-like |
| W303.asm02.HP0_G0000630 | 2269.431 | 3196.868 | 2751.109 | 1251.865 | 3153.543 | 2393.716 | 1637.267 | 2224.915 | 3109.137 | 3591.668 | 3672.021 | 3425.367 | YAL008W | FUN14 | n | NA | -0.344935 | 0.230422953 | 0.223645251 | NA |
| W303.asm02.HP0_G0000660 | 5847.032 | 4261.77 | 6009.889 | 7948.173 | 8336.865 | 6393.367 | 13952.03 | 16942.87 | 14302.6 | 10761.71 | 11973.89 | 13633.93 | YAL003W | EFB1 | n | NA | -1.139525109 | -0.477870747 | -1.455926832 | NA |
| W303.asm02.HP0_G0000670 | 5837.342 | 7506.879 | 5927.728 | 1455.586 | 1624.581 | 1534.408 | 3429.703 | 3217.516 | 3630.311 | 2512.174 | 2455.349 | 2648.697 | YAL002W | VPS8 | n | transgressive | 1.322934361 | 2.044685075 | 0.898395695 | transgressive |
| W303.asm02.HP0_G0000680 | 4407.077 | 5853.004 | 5354.328 | 1699.033 | 2485.154 | 2094.869 | 4375.763 | 4000.659 | 4814.969 | 4125.244 | 4643.471 | 4448.967 | YAL001C | TFC3 | n | NA | 0.233477506 | 1.292458293 | 0.239109664 | NA |
| W303.asm02.HP0_G0000700 | 3401.24 | 3336.551 | 3308.942 | 3286.019 | 3399.691 | 3173.658 | 6926.175 | 6318.667 | 6117.433 | 3357.161 | 3845.98 | 4148.219 | YAR002W | NUP60 | n | conserved | -0.173804527 | 0.018249546 | -0.94010598 | sc-like |
| W303.asm02.HP0_G0000710 | 3030.108 | 3882.322 | 3438.67 | 2603.554 | 4292.453 | 3392.159 | 6461.609 | 7019.189 | 6714.945 | 4621.793 | 4550.204 | 4996.646 | YAR002C-. | ERP1 | n | km-like | -0.437760752 | 0.006733307 | -0.944881027 | transgressive |
| W303.asm02.HP0_G0000720 | 2327.572 | 2878.621 | 2342.897 | 1159.172 | 2123.505 | 1482.477 | 3335.661 | 3311.773 | 3843.305 | 2978.341 | 2646.077 | 3209.039 | YAR003W | SWD1 | n | NA | -0.216018854 | 0.644604958 | -0.461593894 | NA |
| W303.asm02.HP0_G0000730 | 1066.885 | 2693.577 | 2013.386 | 2512.898 | 5038.473 | 3456.827 | 8286.019 | 9120.757 | 10715.64 | 4090.116 | 4627.752 | 4649.466 | YAR007C | RFA1 | n | transgressive | -1.115459024 | -0.874395662 | -2.160853116 | transgressive |
| W303.asm02.HP0_G0000740 | 267.4479 | 692.6547 | 475.6712 | 1474.94 | 2130.132 | 1987.088 | 3089.272 | 3660.87 | 2368.373 | 400.6566 | 472.6259 | 501.2475 | YAR008W | SEN34 | n | sc-like | 0.059060217 | -1.864245145 | -2.542488338 | sc-like |
| W303.asm02.HP0_G0000750 | 2697.736 | 3460.393 | 2907.648 | 1145.93 | 1376.539 | 1235.561 | 4705.849 | 4509.177 | 5069.43 | 2317.542 | 2354.746 | 2831.257 | YAR014C | BUD14 | n | sc-like | 0.267445783 | 1.255195698 | -0.647512984 | sc-like |
| W303.asm02.HP0_G0000760 | 408.924 | 1334.188 | 888.2078 | 4880.136 | 16591.36 | 12901.37 | 7678.509 | 6607.254 | 10347.14 | 4107.205 | 3052.682 | 4131.335 | YAR015W | ADE1 | n | transgressive | -1.832428499 | -3.496050591 | -2.963961205 | transgressive |
| W303.asm02.HP0_G0000770 | 949.634 | 1288.107 | 1059.449 | 538.8419 | 653.2406 | 464.4377 | 1578.961 | 1506.938 | 1341.104 | 2151.393 | 2222.704 | 2522.066 | YAR018C | KIN3 | n | intermediate | -1.037702658 | 0.975900548 | -0.416563152 | transgressive |
| W303.asm02.HP0_G0000780 | 4528.204 | 7152.632 | 6245.995 | 1343.54 | 1295.12 | 924.9561 | 2370.793 | 2038.73 | 2451.308 | 1748.838 | 2093.806 | 1707.407 | YAR019C | CDC15 | n | NA | 1.638686768 | 2.280382994 | 1.350894423 | NA |
| W303.asm02.HP0_G0000810 | 2285.905 | 3335.111 | 3008.837 | 979.8978 | 1671.917 | 986.6852 | 909.3838 | 1018.201 | 885.9015 | 3930.613 | 3729.658 | 3694.458 | YAR027W | UIP3 | n | sc-like | -0.375907036 | 1.209114647 | 1.569526439 | sc-like |
| W303.asm02.HP0_G0000840 | 3658.029 | 2913.902 | 2271.979 | 2261.303 | 3234.014 | 1025.878 | 1322.227 | 786.6333 | 937.7362 | 598.1367 | 587.9006 | 621.5469 | YAR035W | YAT1 | n | km-like | 2.109328479 | 0.381411127 | 1.432026702 | transgressive |
| W303.asm02.HP0_G0000850 | 16264.9 | 15440.73 | 16728.06 | 10879.72 | 11962.82 | 9688.523 | 14734.46 | 13476.33 | 20680.15 | 5033.842 | 5575.1 | 5585.48 | YAR042W | SWH1 | n | transgressive | 1.542675328 | 0.562376981 | -0.013353135 | km-like |
| W303.asm02.HP0_G0004930 | 14422.81 | 11209.92 | 13596.41 | 1137.782 | 1097.255 | 1051.354 | 1236.649 | 1039.147 | 1226.125 | 35.12866 | 44.01395 | 27.43671 | YCL073C | GEX1 | y | transgressive | 8.505556007 | 3.555536232 | 3.45987369 | transgressive |
| W303.asm02.HP0_G0004980 | 3253.95 | 2609.335 | 3495.751 | 5335.452 | 4601.086 | 5681.033 | 2067.978 | 2022.439 | 5239.071 | 5203.789 | 3153.285 | 4401.481 | YCL064C | CHA1 | n | NA | -0.402257999 | -0.681901624 | 0.0040627 | NA |
| W303.asm02.HP0_G0004990 | 3016.541 | 2575.495 | 2709.596 | 4897.452 | 4049.145 | 4518.959 | 2801.504 | 2497.212 | 2773.626 | 7798.563 | 6933.244 | 6911.939 | YCL063W | VAC17 | n | transgressive | -1.367485037 | -0.692003939 | 0.039406349 | km-like |
| W303.asm02.HP0_G0005000 | 11102 | 8308.976 | 9649.206 | 3848.289 | 4273.519 | 4006.51 | 3547.255 | 3593.378 | 3703.822 | 8667.285 | 7833.434 | 6952.039 | YCL061C | MRC1 | n | transgressive | 0.302218921 | 1.244706215 | 1.403918612 | transgressive |
| W303.asm02.HP0_G0005010 | 17449.04 | 11724.73 | 15332.18 | 9521.918 | 9224.893 | 9317.169 | 6951.566 | 6769.002 | 3059.188 | 15597.13 | 17321.58 | 17135.28 | YCL059C | KRR1 | n | sc-like | -0.156422444 | 0.627225062 | 1.34286645 | sc-like |
| W303.asm02.HP0_G0005030 | 1109.521 | 676.0943 | 934.0453 | 12856.83 | 10128.07 | 11257.23 | 5541.881 | 6893.514 | 4413.486 | 5199.042 | 7031.752 | 5869.345 | YCL057C-/ | MIC10 | n | NA | -2.645116778 | -3.597818618 | -2.557043918 | NA |
| W303.asm02.HP0_G0005040 | 2084.35 | 2746.858 | 2601.489 | 9657.392 | 8933.302 | 9038.898 | 5193.926 | 5735.674 | 6699.866 | 12314.97 | 5654.744 | 5860.903 | YCL057W | PRD1 | n | NA | -1.581019214 | -1.810684247 | -1.186399039 | NA |
| W303.asm02.HP0_G0005060 | 5317.95 | 3024.784 | 4584.605 | 365.6791 | 314.3129 | 329.2217 | 135.4201 | 123.3478 | 283.677 | 5282.591 | 6588.468 | 6069.844 | YCL055W | KAR4 | n | sc-like | -0.434778231 | 3.587176318 | 4.519051082 | sc-like |
| W303.asm02.HP0_G0005070 | 16786.23 | 13804.85 | 15048.51 | 13094.16 | 11548.16 | 12018.55 | 7075.702 | 5829.93 | 1309.061 | 32709.53 | 29027.2 | 26951.29 | YCL054W | SPB1 | n | km-like | -0.826403181 | 0.258520722 | 1.520847372 | intermediate |
| W303.asm02.HP0_G0005080 | 5050.502 | 3584.236 | 4360.608 | 3844.215 | 2973.665 | 3433.312 | 1704.977 | 1583.739 | 1562.58 | 11106.35 | 13303.74 | 11738.69 | YCL052C | PBN1 | n | intermediate | -1.445717673 | 0.325544524 | 1.398183065 | intermediate |
| W303.asm02.HP0_G0005100 | 21572.2 | 14275.74 | 20050.84 | 14904.23 | 12825.29 | 14050.71 | 12415.39 | 13699.75 | 19740.52 | 24197.95 | 25965.08 | 27222.49 | YCL050C | APA1 | n | km-like | -0.448851767 | 0.396001515 | 0.277927531 | km-like |
| W303.asm02.HP0_G0005130 | 20927.8 | 13836.53 | 18136.91 | 173.1628 | 147.6892 | 170.4898 | 183.3814 | 137.3117 | 106.4967 | 300.0178 | 685.36 | 355.6219 | YCL048W | SPS22 | n | transgressive | 5.271247427 | 6.696728028 | 6.900418078 | transgressive |
| W303.asm02.HP0_G0005150 | 8938.187 | 9155.714 | 9247.913 | 8543.039 | 8635.083 | 8091.406 | 6359.103 | 6295.393 | 7753.523 | 20803.76 | 14617.87 | 15199.94 | YCL045C | EMC1 | n | km-like | -0.866921424 | 0.089465123 | 0.414754272 | intermediate |
| W303.asm02.HP0_G0005160 | 1184.135 | 3139.987 | 2297.059 | 6251.178 | 5125.572 | 5844.664 | 3482.366 | 3410.684 | 3583.189 | 2972.644 | 1352.905 | 1252.591 | YCL044C | MGR1 | n | sc-like | 0.219057107 | -1.282700185 | -0.613481065 | conserved |

| | | | | | | | | | | | | | | | | | | | |
|---|---|---|---|---|---|---|---|---|---|---|---|---|---|---|---|---|---|---|---|
| W303.asm02.HP0_G0005170 | 14771.65 | 17757.02 | 18136.91 | 32634.06 | 33064.39 | 33048.56 | 20640.28 | 22606.4 | 25677.95 | 34491.6 | 25283.92 | 23061.61 | YCL043C | PDI1 | n | NA | -0.689643039 | -0.945706394 | -0.435947831 | NA |
| W303.asm02.HP0_G0005180 | 1814.964 | 2939.822 | 2250.357 | 703.8859 | 683.5358 | 670.2012 | 463.6258 | 445.6813 | 514.5768 | 9661.332 | 8597.391 | 5917.886 | YCL040W | GLK1 | n | intermediate | -1.726503519 | 1.721394101 | 2.238622192 | intermediate |
| W303.asm02.HP0_G0005190 | 2731.651 | 3101.106 | 3047.755 | 4341.294 | 4165.592 | 3771.352 | 3828.44 | 3790.036 | 9357.57 | 3207.152 | 2701.618 | 2526.287 | YCL039W | GID7 | n | conserved | 0.067825283 | -0.420562647 | -0.885867499 | sc-like |
| W303.asm02.HP0_G0005200 | 4728.79 | 3492.794 | 4114.123 | 3421.494 | 3003.013 | 3002.188 | 2293.678 | 2529.794 | 3039.396 | 3620.151 | 4671.766 | 3247.029 | YCL038C | ATG22 | n | sc-like | 0.092795308 | 0.370028759 | 0.636327264 | sc-like |
| W303.asm02.HP0_G0005210 | 941.8819 | 1607.074 | 1320.636 | 7596.755 | 12656.77 | 7994.403 | 17599.91 | 15455.71 | 8791.159 | 1322.547 | 732.5178 | 565.6182 | YCL037C | SRO9 | n | sc-like | 0.495806986 | -2.701002684 | -3.262318541 | sc-like |
| W303.asm02.HP0_G0005230 | 5849.939 | 3302.71 | 4912.386 | 6690.196 | 4833.034 | 5841.725 | 3609.323 | 4625.543 | 4454.954 | 7350.435 | 11335.69 | 8713.265 | YCL035C | GRX1 | n | km-like | -0.911249238 | -0.266067557 | 0.142720263 | km-like |
| W303.asm02.HP0_G0005240 | 3455.505 | 3710.959 | 3349.59 | 4784.387 | 4698.599 | 4631.639 | 3035.668 | 3012.712 | 6800.708 | 6667.8 | 5755.347 | 5876.731 | YCL034W | LSB5 | n | NA | -0.746699975 | -0.38336404 | -0.276155458 | NA |
| W303.asm02.HP0_G0005250 | 1667.674 | 930.2597 | 1222.907 | 2716.619 | 1924.693 | 2331.987 | 1215.96 | 1390.572 | 1211.989 | 3624.898 | 6794.915 | 4309.673 | YCL033C | MXR2 | n | transgressive | -1.847849891 | -0.830371878 | 0.000465928 | km-like |
| W303.asm02.HP0_G0005260 | 3714.232 | 3085.265 | 3239.753 | 3775.968 | 3731.992 | 3330.43 | 3489.889 | 3115.114 | 2950.806 | 5643.372 | 6355.823 | 5058.906 | YCL032W | STE50 | n | km-like | -0.752572137 | -0.085941251 | 0.070084803 | km-like |
| W303.asm02.HP0_G0005270 | 17690.32 | 10887.35 | 15038.13 | 11246.42 | 7977.109 | 10407.72 | 5336.87 | 5575.089 | 3349.462 | 15119.57 | 17226.22 | 17163.77 | YCL031C | RRP7 | n | sc-like | -0.171520505 | 0.528313107 | 1.554888188 | sc-like |
| W303.asm02.HP0_G0005280 | 5326.671 | 5346.113 | 4844.062 | 10684.15 | 12379.38 | 10616.42 | 9637.399 | 9853.862 | 11168.01 | 13392.57 | 8830.036 | 8824.067 | YCL030C | HIS4 | n | NA | -0.97011966 | -1.09444053 | -0.961317533 | NA |
| W303.asm02.HP0_G0005290 | 2892.508 | 3002.464 | 2880.838 | 4950.419 | 3851.279 | 4511.121 | 2726.27 | 2627.541 | 2698.23 | 6901.358 | 6276.179 | 5566.486 | YCL029C | BIK1 | n | transgressive | -1.080513549 | -0.595294152 | 0.122965706 | km-like |
| W303.asm02.HP0_G0005310 | 1000.023 | 820.0974 | 908.0996 | 660.0559 | 857.7333 | 663.3425 | 1207.496 | 1182.277 | 1827.408 | 1369.068 | 972.4986 | 1225.154 | YCL027W | FUS1 | n | NA | -0.369348114 | 0.290496516 | -0.611490827 | NA |
| W303.asm02.HP0_G0005320 | 3865.398 | 2121.885 | 2958.675 | 3067.019 | 2440.658 | 2715.099 | 1575.199 | 1868.836 | 1906.573 | 1183.931 | 1934.518 | 1258.923 | YCL026C-[illegible] | HBN1 | n | NA | 0.977291737 | 0.098125291 | 0.713545977 | NA |
| W303.asm02.HP0_G0005340 | 7583.506 | 8186.574 | 7705.873 | 10506.91 | 10116.71 | 10394 | 13030.43 | 14480.57 | 15387.36 | 18960.93 | 17690.46 | 11147.74 | YCL025C | AGP1 | n | NA | -0.993115795 | -0.381514795 | -0.849911111 | NA |
| W303.asm02.HP0_G0005370 | 2929.33 | 2391.891 | 2339.437 | 3259.535 | 2861.951 | 2762.131 | 1612.816 | 1653.559 | 2732.158 | 4276.203 | 3400.601 | 4009.98 | YCL018W | LEU2 | n | km-like | -0.582921773 | -0.180917611 | 0.342494495 | km-like |
| W303.asm02.HP0_G0005380 | 5276.283 | 4726.9 | 5121.681 | 7605.922 | 16595.15 | 11133.77 | 17615.9 | 17820.27 | 13139.62 | 8677.729 | 7480.275 | 6432.852 | YCL017C | NFS1 | n | transgressive | -0.540380418 | -1.173625776 | -1.614278906 | transgressive |
| W303.asm02.HP0_G0005390 | 3282.051 | 2318.449 | 2707.002 | 2974.326 | 2154.747 | 2535.791 | 1561.093 | 1689.632 | 1414.615 | 2226.398 | 2455.349 | 2186.494 | YCL016C | DCC1 | n | conserved | 0.26557352 | 0.091554233 | 0.817146552 | sc-like |
| W303.asm02.HP0_G0005400 | 9754.097 | 7793.445 | 8597.541 | 5838.643 | 9110.339 | 7448.64 | 6268.823 | 6298.884 | 5524.633 | 4908.519 | 2821.084 | 2886.13 | YCL014W | BUD3 | n | km-like | 1.236751709 | 0.18794971 | 0.511203568 | transgressive |
| W303.asm02.HP0_G0005430 | 1071.73 | 1918.841 | 1396.744 | 1772.372 | 1604.7 | 1501.094 | 1310.942 | 1310.28 | 1003.708 | 1777.32 | 903.3338 | 959.2295 | YCL010C | SGF29 | n | NA | 0.24854002 | -0.124732241 | 0.262650793 | NA |
| W303.asm02.HP0_G0005440 | 3346.006 | 9412.04 | 7573.55 | 11167.98 | 12687.07 | 10797.69 | 11432.66 | 11758.77 | 13271.56 | 9778.111 | 2638.741 | 2544.227 | YCL009C | ILV6 | n | conserved | 0.36169216 | -0.650635882 | -0.737144372 | conserved |
| W303.asm02.HP0_G0005450 | 676.3719 | 1199.545 | 732.5336 | 633.5722 | 778.2083 | 622.1898 | 515.3488 | 395.6439 | 736.9947 | 601.9344 | 307.0497 | 311.3011 | YCL008C | STP22 | n | NA | 1.001785397 | 0.308775391 | 0.620691402 | NA |
| W303.asm02.HP0_G0005470 | 4530.142 | 3155.107 | 3980.935 | 1857.935 | 1355.711 | 1508.933 | 1030.698 | 1096.166 | 1899.976 | 1804.854 | 2027.785 | 1826.651 | YCL005W | LDB16 | n | NA | 0.990576302 | 1.258661386 | 1.479569671 | NA |
| W303.asm02.HP0_G0005480 | 3988.463 | 3312.79 | 3489.697 | 6761.499 | 5731.476 | 5929.909 | 3177.671 | 3029.004 | 2578.539 | 1655.794 | 1801.428 | 1572.334 | YCL004W | PGS1 | n | intermediate | 1.08497983 | -0.763870738 | 0.293347902 | transgressive |
| W303.asm02.HP0_G0005500 | 4065.984 | 2268.768 | 3759.532 | 8883.253 | 6451.934 | 8402.991 | 3465.439 | 3558.468 | 3836.708 | 5211.385 | 6052.965 | 6188.032 | YCL001W | RER1 | n | transgressive | -0.755230891 | -1.198609034 | -0.102699894 | km-like |
| W303.asm02.HP0_G0005530 | 7517.613 | 5168.99 | 6700.91 | 16645.02 | 13870.47 | 16240.62 | 9863.099 | 10170.38 | 9726.068 | 7338.093 | 8000.059 | 7604.189 | YCR002C | CDC10 | n | sc-like | -0.236520477 | -1.251208344 | -0.608749351 | sc-like |
| W303.asm02.HP0_G0005540 | 5511.753 | 2813.1 | 4460.931 | 7891.131 | 6562.701 | 7110.6 | 4939.073 | 5017.696 | 3394.699 | 3872.698 | 5720.765 | 4613.588 | YCR003W | MRPL32 | n | NA | -0.141907898 | -0.721863968 | -0.060018416 | NA |
| W303.asm02.HP0_G0005550 | 4784.992 | 2967.183 | 3891.855 | 10816.56 | 9219.213 | 9913.883 | 6140.927 | 7087.845 | 9806.176 | 7154.854 | 10280.4 | 7752.98 | YCR004C | YCP4 | n | NA | -1.058583922 | -1.316029149 | -0.949122892 | NA |
| W303.asm02.HP0_G0005580 | 1519.414 | 905.0592 | 1382.906 | 715.0606 | 633.3593 | 661.3828 | 389.3329 | 475.9364 | 477.8214 | 592.4401 | 704.2231 | 767.1725 | YCR007C | DFP3 | n | transgressive | 0.850133981 | 0.897801443 | 1.46510749 | transgressive |
| W303.asm02.HP0_G0005590 | 15140.85 | 11515.2 | 13577.39 | 786.3629 | 3623.118 | 1734.293 | 8037.749 | 7567.272 | 6561.326 | 9267.321 | 7019.176 | 7873.279 | YCR008W | SAT4 | n | NA | 0.636802817 | 2.511649688 | 0.779952659 | NA |
| W303.asm02.HP0_G0005600 | 6963.336 | 4163.128 | 5932.917 | 5224.424 | 7844.567 | 5282.244 | 10163.09 | 12268.45 | 12422.41 | 9753.426 | 11468.78 | 11002.12 | YCR009C | RVS161 | n | NA | -0.873972866 | -0.084590276 | -0.996922107 | NA |
| W303.asm02.HP0_G0005610 | 1340.147 | 1333.468 | 1279.988 | 997.2141 | 1119.976 | 959.25 | 796.5337 | 790.1242 | 790.7142 | 270.5856 | 402.4132 | 241.6541 | YCR010C | ADY2 | n | transgressive | 2.065343018 | 0.34427592 | 0.720149678 | transgressive |
| W303.asm02.HP0_G0005620 | 10301.59 | 9425 | 9814.394 | 5132.75 | 6264.482 | 5406.682 | 4120.91 | 4189.171 | 6595.254 | 6890.914 | 4634.04 | 5307.947 | YCR011C | ADP1 | n | NA | 0.77459116 | 0.789007298 | 0.956756895 | NA |
| W303.asm02.HP0_G0005630 | 24926.92 | 17377.57 | 21579.04 | 47393.64 | 48123 | 47547.05 | 48914.88 | 57679.07 | 79179.81 | 73417.01 | 75082.55 | 79259.37 | YCR012W | PGK1 | n | NA | -1.788850192 | -1.139897632 | -1.509354356 | NA |
| W303.asm02.HP0_G0005640 | 6531.156 | 4141.528 | 5636.271 | 888.2234 | 1605.646 | 950.4316 | 1942.903 | 1886.291 | 1293.039 | 2232.094 | 2511.939 | 2324.733 | YCR014C | POL4 | n | NA | 1.140867652 | 2.164687345 | 1.605584141 | NA |
| W303.asm02.HP0_G0005650 | 3420.62 | 1893.64 | 3071.971 | 282.1535 | 537.7401 | 368.4147 | 631.0202 | 552.7379 | 376.0369 | 2020.373 | 2363.13 | 2236.092 | YCR015C | CTO1 | n | sc-like | 0.314475428 | 2.715665589 | 2.320995605 | sc-like |
| W303.asm02.HP0_G0005660 | 13143.71 | 7098.631 | 11852 | 3929.777 | 3871.16 | 3946.741 | 3005.574 | 2561.213 | 416.5622 | 7490 | 8750.392 | 8554.976 | YCR016W | RBP95 | n | sc-like | 0.300483545 | 1.263140113 | 2.131103926 | sc-like |
| W303.asm02.HP0_G0005670 | 6457.511 | 6341.174 | 6048.808 | 6003.657 | 8447.631 | 6911.695 | 9031.77 | 8824.024 | 11553.48 | 10606.01 | 5738.58 | 6721.993 | YCR017C | CWH43 | n | NA | -0.274051373 | -0.14947933 | -0.618670013 | NA |
| W303.asm02.HP0_G0005800 | 5527.257 | 4328.732 | 5025.682 | 871.9257 | 2933.902 | 1602.996 | 3373.278 | 3394.392 | 3227.886 | 4538.243 | 3419.464 | 3569.938 | YCR028C | FEN2 | n | NA | 0.33222234 | 1.373335983 | 0.538245492 | NA |
| W303.asm02.HP0_G0005810 | 3591.167 | 1813.718 | 2925.81 | 6477.308 | 6678.201 | 6507.027 | 5733.726 | 6794.603 | 5752.705 | 6270.941 | 7899.455 | 7946.092 | YCR028C-[illegible] | RIM1 | n | NA | -1.351114748 | -1.200811878 | -1.0981122 | NA |
| W303.asm02.HP0_G0005840 | 8659.11 | 5608.199 | 7355.606 | 4477.787 | 5024.272 | 4518.959 | 4397.392 | 4333.465 | 6365.297 | 4859.149 | 5001.871 | 5037.801 | YCR032W | BPH1 | n | NA | 0.515940127 | 0.608332601 | 0.505267082 | NA |
| W303.asm02.HP0_G0005850 | 7523.427 | 8576.102 | 7700.684 | 3489.74 | 10906.28 | 6283.627 | 9824.542 | 9117.266 | 7878.869 | 6338.35 | 3473.958 | 3472.854 | YCR033W | SNT1 | n | km-like | 0.748435654 | 0.165735012 | -0.158962424 | km-like |
| W303.asm02.HP0_G0005860 | 11965.39 | 7125.271 | 9861.961 | 21997.79 | 17901.63 | 20964.37 | 8223.951 | 8911.298 | 8661.101 | 14394.21 | 15818.82 | 16222.48 | YCR034W | ELO2 | n | transgressive | -0.658673678 | -1.049212713 | 0.162747139 | km-like |
| W303.asm02.HP0_G0005870 | 8927.528 | 6248.292 | 7976.573 | 5514.726 | 4443.929 | 5299.881 | 2638.812 | 2624.05 | 1237.435 | 8035.919 | 8565.952 | 8512.766 | YCR035C | RRP43 | n | sc-like | -0.108933097 | 0.566400454 | 1.756822787 | sc-like |
| W303.asm02.HP0_G0005880 | 10184.34 | 6719.182 | 8620.892 | 4892.359 | 3631.639 | 4826.625 | 2027.54 | 2236.552 | 2170.459 | 5671.855 | 8180.306 | 5963.263 | YCR036W | RBK1 | n | sc-like | 0.348909942 | 0.910199698 | 1.937810522 | sc-like |
| W303.asm02.HP0_G0005890 | 4583.437 | 3976.644 | 3986.125 | 6415.173 | 16881.06 | 10248 | 25651.77 | 26690.84 | 19331.5 | 7860.276 | 5645.313 | 6557.373 | YCR037C | PHO87 | n | transgressive | -0.621432159 | -1.344594334 | -2.393261674 | transgressive |
| W303.asm02.HP0_G0005900 | 1218.051 | 1036.102 | 1150.259 | 8708.053 | 9467.255 | 9415.152 | 4565.727 | 4214.772 | 3881.945 | 828.8466 | 710.5108 | 640.5416 | YCR038C | BUD5 | n | intermediate | 0.630819096 | -2.999518922 | -1.87792524 | intermediate |
| W303.asm02.HP0_G0005920 | 6918.762 | 6454.217 | 6640.37 | 5226.461 | 9855.412 | 7079.246 | 7347.482 | 6892.35 | 6733.794 | 6838.696 | 5101.426 | 5234.079 | YCR042C | TAF2 | n | NA | 0.209059758 | -0.119815627 | -0.065507707 | NA |
| W303.asm02.HP0_G0005930 | 2012.643 | 1026.022 | 1744.416 | 3701.61 | 2712.368 | 3277.519 | 1073.957 | 1287.006 | 902.8656 | 5159.166 | 7589.262 | 6089.893 | YCR043C | | n | transgressive | -1.886527271 | -0.979700129 | 0.529165461 | intermediate |
| W303.asm02.HP0_G0005940 | 1199.64 | 897.859 | 997.1798 | 6574.075 | 6873.227 | 6512.906 | 5283.266 | 5706.582 | 4505.846 | 3900.231 | 4206.476 | 4186.208 | YCR044C | PER1 | n | transgressive | -1.963881431 | -2.666791662 | -2.300731675 | transgressive |
| W303.asm02.HP0_G0005950 | 3417.713 | 4397.853 | 3965.368 | 56.02326 | 53.96335 | 44.09219 | 66.76965 | 73.3105 | 73.51098 | 1109.876 | 553.3182 | 518.1316 | YCR045C | RRT12 | n | transgressive | 2.316379709 | 6.209510164 | 5.748061146 | transgressive |
| W303.asm02.HP0_G0005960 | 1978.727 | 1399.71 | 1588.742 | 6273.587 | 5107.584 | 5962.244 | 3459.796 | 3450.248 | 2172.344 | 3848.962 | 5668.367 | 3846.415 | YCR046C | IMG1 | n | transgressive | -1.369705102 | -1.751121543 | -0.842881172 | transgressive |
| W303.asm02.HP0_G0005970 | 3007.82 | 3708.799 | 3146.349 | 6332.666 | 4509.253 | 5601.667 | 2642.573 | 2470.447 | 565.4691 | 5094.605 | 2822.132 | 3161.553 | YCR047C | BUD23 | n | conserved | -0.142129299 | -0.644918634 | 0.717212482 | conserved |
| W303.asm02.HP0_G0005990 | 7225.939 | 5486.516 | 6168.158 | 5722.522 | 4478.011 | 5283.224 | 3652.582 | 3445.593 | 1455.14 | 7677.986 | 7388.055 | 7592.581 | YCR051W | | n | conserved | -0.241786894 | 0.243962026 | 1.085825108 | sc-like |
| W303.asm02.HP0_G0006010 | 23857.13 | 19559.93 | 21052.34 | 24393.55 | 20546.78 | 22804.48 | 16199.63 | 16400.61 | 15291.23 | 22485.19 | 21364.58 | 22320.82 | YCR053W | THR4 | n | conserved | -0.037115066 | -0.05161051 | 0.425408531 | sc-like |
| W303.asm02.HP0_G0006020 | 6526.311 | 3857.842 | 5132.925 | 3843.196 | 2597.815 | 3474.464 | 1729.428 | 1375.445 | 1037.636 | 4189.805 | 3954.967 | 4393.039 | YCR054C | CTR86 | n | sc-like | 0.290073868 | 0.620954184 | 1.843596344 | sc-like |
| W303.asm02.HP0_G0006030 | 9201.759 | 14483.83 | 12163.34 | 8901.587 | 10059.9 | 8936.017 | 7302.342 | 6510.67 | 1819.868 | 13918.55 | 5967.034 | 6170.093 | YCR057C | PWP2 | n | NA | 0.383492818 | 0.293924285 | 1.063033142 | NA |
| W303.asm02.HP0_G0006040 | 7736.61 | 5066.747 | 6164.699 | 2251.117 | 1723.04 | 1963.572 | 1587.425 | 1665.196 | 1490.953 | 4996.815 | 6607.332 | 5564.375 | YCR059C | YIH1 | n | sc-like | 0.138629351 | 1.643808501 | 1.961326687 | sc-like |
| W303.asm02.HP0_G0006050 | 2723.899 | 1865.56 | 2199.331 | 1538.093 | 1171.099 | 1377.636 | 1054.208 | 991.4372 | 799.1963 | 2524.517 | 3585.041 | 2897.738 | YCR060W | TAH1 | n | sc-like | -0.391389696 | 0.713470623 | 1.223507565 | sc-like |
| W303.asm02.HP0_G0006060 | 502.9184 | 1278.027 | 735.9931 | 6443.694 | 6664.001 | 6111.177 | 4801.772 | 4885.039 | 5164.617 | 1101.331 | 931.6285 | 807.2723 | YCR061W | TVS1 | n | sc-like | -0.160467107 | -2.840223886 | -2.461959352 | sc-like |
| W303.asm02.HP0_G0006070 | 1256.811 | 959.7804 | 1129.503 | 282.1535 | 213.96 | 222.4206 | 322.5632 | 330.4791 | 211.1084 | 606.6815 | 607.8116 | 531.85 | YCR063W | BUD31 | n | transgressive | 0.904257653 | 2.170446601 | 1.907579887 | transgressive |
| W303.asm02.HP0_G0006080 | 2687.077 | 3125.586 | 3105.7 | 2467.061 | 2940.529 | 2894.407 | 2525.962 | 3047.622 | 2691.633 | 2744.783 | 1415.782 | 1462.587 | YCR065W | HCM1 | n | km-like | 0.629652256 | 0.082942699 | 0.10605264 | km-like |
| W303.asm02.HP0_G0006090 | 2841.15 | 2495.573 | 2667.218 | 3723.001 | 3060.763 | 3542.072 | 2735.675 | 2423.901 | 2308.056 | 1765.927 | 1566.687 | 1583.942 | YCR066W | RAD18 | n | intermediate | 0.691949362 | -0.362555904 | 0.098701646 | km-like |
| W303.asm02.HP0_G0006100 | 7050.548 | 7894.247 | 7579.604 | 7701.671 | 11981.76 | 8762.587 | 10552.43 | 10131.98 | 11626.99 | 8616.017 | 5228.228 | 5040.967 | YCR067C | SED4 | n | conserved | 0.239177244 | -0.300124346 | -0.502016004 | sc-like |
| W303.asm02.HP0_G0006110 | 1598.874 | 4930.665 | 3294.239 | 3259.535 | 3625.959 | 3178.557 | 2532.544 | 2679.906 | 2997.928 | 1278.873 | 426.5161 | 480.1424 | YCR068W | ATG15 | n | km-like | 1.925191621 | -0.027650368 | 0.233574433 | km-like |
| W303.asm02.HP0_G0006120 | 1403.133 | 3120.546 | 2265.06 | 8837.415 | 7629.661 | 7424.144 | 4324.04 | 4499.868 | 3968.65 | 4026.504 | 3554.65 | 3439.086 | YCR069W | CPR4 | n | NA | -0.661413951 | -1.75700785 | -0.881858419 | NA |
| W303.asm02.HP0_G0006130 | 2694.829 | 1623.634 | 2011.657 | 5590.103 | 3973.407 | 4911.87 | 2264.525 | 2385.5 | 1335.449 | 3148.288 | 4537.628 | 3738.779 | YCR071C | IMG2 | n | transgressive | -0.801108357 | -1.146015606 | 0.077322916 | km-like |
| W303.asm02.HP0_G0006140 | 4383.82 | 4634.738 | 4390.878 | 5012.554 | 5673.726 | 4751.178 | 4975.749 | 4192.662 | 919.8297 | 5561.722 | 3747.473 | 3965.659 | YCR072C | RSA4 | n | NA | 0.012311775 | -0.163609978 | 0.36937027 | NA |
| W303.asm02.HP0_G0006170 | 937.0368 | 588.2525 | 887.343 | 2295.935 | 1828.127 | 2194.811 | 1032.578 | 1098.494 | 1160.154 | 1087.09 | 1457.7 | 1281.083 | YCR075C | ERS1 | n | NA | -0.641614699 | -1.358840431 | -0.438469024 | NA |
| W303.asm02.HP0_G0006180 | 534.8959 | 265.6856 | 356.321 | 1310.944 | 770.6345 | 1069.97 | 393.0945 | 477.1001 | 362.8426 | 456.6726 | 689.5518 | 565.6182 | YCR075W-[illegible] | EGO2 | n | NA | -0.524238937 | -1.381159501 | -0.088528028 | NA |
| W303.asm02.HP0_G0006190 | 2275.245 | 1570.353 | 1935.549 | 1174.451 | 1260.092 | 1050.374 | 911.2646 | 950.7091 | 1462.68 | 1468.758 | 1505.906 | 1576.555 | YCR076C | FUB1 | n | sc-like | 0.329725159 | 0.710535425 | 0.776429401 | sc-like |
| W303.asm02.HP0_G0006200 | 2309.161 | 4641.938 | 3154.132 | 3407.233 | 16996.56 | 7518.208 | 22063.14 | 21291.46 | 21410.54 | 3316.336 | 2402.952 | 2216.042 | YCR077C | PAT1 | n | sc-like | 0.290358237 | -1.307514047 | -2.418436439 | sc-like |
| W303.asm02.HP0_G0006210 | 4128.001 | 4175.369 | 3971.422 | 1220.289 | 1599.019 | 1270.835 | 1053.268 | 1008.892 | 998.0529 | 1482.05 | 1133.883 | 987.7214 | YCR079W | PTC6 | n | NA | 1.729810233 | 1.558995883 | 1.970598033 | NA |
| W303.asm02.HP0_G0006220 | 12131.09 | 8470.98 | 10452.66 | 7123.103 | 5792.066 | 6608.929 | 3617.786 | 3429.302 | 3566.225 | 4012.263 | 4074.434 | 3698.679 | YCR081W | SRB8 | n | NA | 1.370732321 | 0.659411229 | 1.526138293 | NA |
| W303.asm02.HP0_G0006230 | 3907.065 | 2286.768 | 3204.294 | 4431.95 | 3621.225 | 4278.902 | 1637.267 | 1598.867 | 1665.306 | 4419.566 | 5967.034 | 5002.978 | YCR082W | AHC2 | n | km-like | -0.68608584 | -0.37025503 | 0.916493866 | intermediate |
| W303.asm02.HP0_G0006240 | 1921.555 | 966.2605 | 1539.445 | 3759.67 | 2814.615 | 3144.263 | 2170.484 | 2553.067 | 1915.998 | 2102.973 | 3312.573 | 2530.509 | YCR083W | TRX3 | n | NA | -0.794017563 | -1.089782763 | -0.561058177 | NA |
| W303.asm02.HP0_G0006250 | 2873.127 | 12063.86 | 5452.057 | 12572.64 | 24127.3 | 19032.15 | 16257 | 16953.34 | 15946.23 | 4571.473 | 2005.778 | 1879.414 | YCR084C | TUP1 | n | intermediate | 1.071027125 | -1.282814594 | -1.119580376 | intermediate |
| W303.asm02.HP0_G0006260 | 1780.079 | 1174.345 | 1475.446 | 2109.531 | 2210.604 | 1873.428 | 1950.426 | 2157.423 | 1385.399 | 1080.444 | 1360.241 | 1127.015 | YCR086W | CSM1 | n | sc-like | 0.297845489 | -0.463837846 | -0.302274344 | conserved |
| W303.asm02.HP0_G0006270 | 5395.471 | 3256.629 | 4558.66 | 4694.75 | 3961.099 | 4163.282 | 2877.678 | 2998.748 | 1127.168 | 4703.443 | 5085.707 | 4670.572 | YCR087C-A | | n | conserved | -0.118559882 | 0.03482777 | 0.864578339 | sc-like |
| W303.asm02.HP0_G0006280 | 1680.271 | 8077.851 | 4438.445 | 7266.727 | 12704.11 | 9424.95 | 14045.14 | 13553.13 | 18787.71 | 4222.085 | 1396.919 | 1020.434 | YCR088W | ABP1 | n | conserved | 0.869129679 | -0.871679831 | -1.449048417 | sc-like |

| | | | | | | | | | | | | | | | | | | | | |
|---|---|---|---|---|---|---|---|---|---|---|---|---|---|---|---|---|---|---|---|---|
| W303.asm02.HP0_G0006300 | 6328.632 | 3533.115 | 5560.164 | 5460.74 | 3815.304 | 5069.622 | 2286.155 | 2574.013 | 2317.481 | 3139.743 | 4063.954 | 4047.969 | YCR090C | | n | conserved | 0.432339138 | 0.083271306 | 1.070169126 | sc-like |
| W303.asm02.HP0_G0006320 | 4029.161 | 2983.743 | 3647.966 | 4510.382 | 3783.115 | 4431.755 | 1941.962 | 2149.278 | 2051.71 | 2900.488 | 1741.695 | 1818.209 | YCR092C | MSH3 | n | NA | 0.693647008 | -0.222912545 | 0.77461591 | NA |
| W303.asm02.HP0_G0006330 | 13005.14 | 13347.64 | 13577.39 | 15215.92 | 22065.33 | 18071.92 | 19615.23 | 18921.09 | 19060.08 | 13128.63 | 7874.304 | 8869.443 | YCR093W | CDC39 | n | sc-like | 0.400738181 | -0.451128282 | -0.514724602 | sc-like |
| W303.asm02.HP0_G0006340 | 6723.021 | 4327.292 | 5801.459 | 5113.396 | 4870.903 | 4764.896 | 4445.354 | 4700.017 | 4246.673 | 4176.513 | 4371.004 | 4387.762 | YCR094W | CDC50 | n | NA | 0.371500379 | 0.163801849 | 0.326218273 | NA |
| W303.asm02.HP0_G0006350 | 3158.986 | 1657.475 | 2643.867 | 2671.8 | 2947.156 | 2631.814 | 3225.632 | 3097.659 | 2701.057 | 2988.785 | 3720.226 | 3766.216 | YCR095C | OCA4 | n | NA | -0.467526451 | -0.11852047 | -0.267104002 | NA |
| W303.asm02.HP0_G0006450 | 4854.761 | 3515.835 | 4085.583 | 5242.759 | 6096.912 | 5605.587 | 5096.123 | 5144.535 | 3696.283 | 6069.663 | 4691.677 | 4953.381 | YCR106W | RDS1 | n | NA | -0.32358558 | -0.428622541 | -0.158690342 | NA |

**NaCl**

| | Normalized read counts | | | | | | | | | | | | | | | | log2 fold change adjusted with ashr method | |
|---|---|---|---|---|---|---|---|---|---|---|---|---|---|---|---|---|---|---|
| W303_gene | KS.1_NaCl.sc | KS.2_NaCl.sc | KS.3_NaCl.sc | KS.1_NaCl.km | KS.2_NaCl.km | KS.3_NaCl.km | Km.V.1_NaCl | Km.V.2_NaCl | Km.V.3_NaCl | SC.1_NaCl | SC.2_NaCl | SC.3_NaCl | locus_tag | gene | ambiguous | Category | log2FC(KSsc/Psc) | log2(KSsc/KSkm) |
| W303.asm02.HP0_G0000040 | 1218.074547 | 2178.317417 | 1257.587841 | 425.3047173 | 582.3405146 | 692.5540811 | 294.4088771 | 224.3043998 | 356.6121468 | 173.1105445 | 283.049605 | 210.632487 | YAL067C | SEO1 | n | NA | 2.697049826 | 1.384503218 |
| W303.asm02.HP0_G0000100 | 4854.305948 | 6601.697353 | 6584.957438 | 665.5134941 | 1931.122539 | 1501.231279 | 3336.031877 | 3848.799613 | 4032.863836 | 506.4129362 | 820.5000695 | 509.2625463 | YAL063C | FLO9 | n | transgressi | 3.16442673 | 2.033146949 |
| W303.asm02.HP0_G0000110 | 3392.43657 | 2982.725918 | 2810.388333 | 484.5770129 | 486.4142069 | 536.6771052 | 521.9887453 | 671.5937617 | 538.4144177 | 227.3690734 | 250.9630101 | 180.6758666 | YAL062W | GDH3 | n | transgressi | 3.781068646 | 2.583455742 |
| W303.asm02.HP0_G0000120 | 456.103246 | 336.3259681 | 402.6447789 | 270.364857 | 194.7594733 | 264.6770129 | 559.9187233 | 294.234595 | 508.6967388 | 305.742504 | 261.2765584 | 248.0782624 | YAL061W | BDH2 | n | transgressi | 0.529477608 | 0.678115213 |
| W303.asm02.HP0_G0000130 | 1733.55218 | 1718.03626 | 1533.841882 | 1190.645235 | 428.2770507 | 1447.877414 | 2554.854949 | 818.0513404 | 2397.517448 | 1377.994384 | 1183.766162 | 1517.490051 | YAL060W | BDH1 | n | NA | 0.256593621 | 0.600242301 |
| W303.asm02.HP0_G0000140 | 1829.810656 | 1341.836594 | 1286.477152 | 3212.142474 | 2730.508436 | 2222.031523 | 3815.575171 | 3880.466116 | 3998.775911 | 993.8784496 | 2385.86752 | 1185.158793 | YAL059W | ECM1 | n | sc-like | -0.03147475 | -0.818431773 |
| W303.asm02.HP0_G0000150 | 1206.379592 | 1188.409542 | 1127.585939 | 2068.291156 | 1792.562316 | 1993.969907 | 856.1337898 | 774.5098981 | 848.7019474 | 299.7137786 | 444.6285293 | 321.0975246 | YAL058W | CNE1 | n | intermedia | 1.70911446 | -0.721561798 |
| W303.asm02.HP0_G0000170 | 289.6750399 | 230.5739884 | 239.2396108 | 649.9155215 | 466.0662022 | 471.8155447 | 488.5742408 | 460.4837384 | 492.0898006 | 574.4514089 | 569.5370594 | 634.7058941 | YAL055W | PEX22 | n | NA | -1.208772459 | -1.045509911 |
| W303.asm02.HP0_G0000180 | 380.5358443 | 342.3937046 | 362.0191846 | 173.6574274 | 106.5847864 | 216.5539196 | 93.01875565 | 85.76344698 | 84.7827898 | 377.2259627 | 499.6341205 | 362.2878776 | YAL054C | ACS1 | n | sc-like | -0.18351776 | 1.089197999 |
| W303.asm02.HP0_G0000190 | 2383.971602 | 3777.599405 | 3384.563399 | 1648.185763 | 2020.266178 | 1608.985161 | 1535.261016 | 1709.991189 | 1509.483278 | 1194.548882 | 1632.97849 | 1329.325029 | YAL053W | FLC2 | n | NA | 1.17246863 | 0.831933258 |
| W303.asm02.HP0_G0000200 | 646.8209741 | 798.3407645 | 694.2462667 | 676.9520072 | 796.47904 | 781.4771882 | 608.6858379 | 862.9122204 | 631.9377013 | 318.6612013 | 552.3478121 | 307.9915032 | YAL051W | OAF1 | n | km-like | 0.832175278 | -0.064754558 |
| W303.asm02.HP0_G0000210 | 471.3966488 | 365.7978313 | 355.6996477 | 1907.112106 | 1722.797729 | 1401.84663 | 2025.641446 | 1882.837521 | 1998.076881 | 1255.697383 | 1606.621644 | 1321.835874 | YAL049C | AIM2 | n | transgressi | -1.781661516 | -2.051214633 |
| W303.asm02.HP0_G0000220 | 332.8564123 | 644.0468925 | 664.4541643 | 677.9918721 | 763.5346515 | 683.1386933 | 615.9105956 | 812.7735898 | 730.7052812 | 180.8617629 | 252.1089599 | 163.8252676 | YAL048C | GEM1 | n | km-like | 1.402153561 | -0.338049775 |
| W303.asm02.HP0_G0000230 | 671.1104961 | 919.6954952 | 690.6351028 | 470.0189052 | 528.0791689 | 472.8616989 | 408.1988112 | 530.4139336 | 476.3569118 | 400.4796179 | 454.9420776 | 375.393899 | YAL047C | SPC72 | n | NA | 0.876770766 | 0.619629555 |
| W303.asm02.HP0_G0000240 | 408.423814 | 403.9378895 | 314.1712624 | 566.7263348 | 457.3456288 | 363.0155078 | 337.7574234 | 407.7062326 | 382.8336282 | 431.4844915 | 740.2835822 | 452.1577387 | YAL046C | BOL3 | n | km-like | -0.505340397 | -0.270268632 |
| W303.asm02.HP0_G0000250 | 703.4965254 | 489.7530206 | 472.1596847 | 932.7587565 | 670.5152016 | 541.9078762 | 550.8877762 | 661.0382605 | 648.5446395 | 755.3131718 | 1001.560141 | 873.4227126 | YAL044W- | BOL1 | n | km-like | -0.636258841 | -0.337119061 |
| W303.asm02.HP0_G0000260 | 199.7138474 | 142.1583989 | 149.8633034 | 2515.433035 | 1984.414932 | 1774.277525 | 3321.582362 | 2324.849132 | 3289.047815 | 945.6486461 | 1388.891179 | 1174.861205 | YAL044C | GCV3 | n | transgressi | -2.770846854 | -3.619480252 |
| W303.asm02.HP0_G0000270 | 1074.136639 | 787.0721109 | 1139.322222 | 958.7553774 | 1221.849233 | 1247.015808 | 1275.169738 | 1452.700848 | 1394.10876 | 440.0969564 | 474.4232245 | 313.6083695 | YAL043C | PTA1 | n | km-like | 1.258432639 | -0.170738783 |
| W303.asm02.HP0_G0000280 | 2863.464758 | 2252.863895 | 2643.372001 | 2838.830998 | 2449.512181 | 2529.600858 | 2135.819001 | 2187.627617 | 2194.737992 | 2192.733564 | 2338.883578 | 2009.901998 | YAL042W | ERV46 | n | NA | 0.244439338 | -0.008878366 |
| W303.asm02.HP0_G0000290 | 994.9707892 | 1205.745932 | 1166.405951 | 1276.954017 | 1311.961825 | 1573.415919 | 1495.524848 | 1563.53361 | 1543.571204 | 532.2503309 | 838.8352666 | 550.4528993 | YAL041W | CDC24 | n | km-like | 0.794260424 | -0.286466303 |
| W303.asm02.HP0_G0000300 | 1194.684637 | 1719.769899 | 1569.05073 | 2273.144528 | 1180.184271 | 2102.769944 | 1152.348856 | 2192.905367 | 1361.768933 | 249.7614821 | 735.699783 | 182.5481554 | YAL040C | CLN3 | n | km-like | 1.636750727 | -0.235987449 |
| W303.asm02.HP0_G0000310 | 958.9863122 | 702.1237994 | 707.7881315 | 1666.90333 | 1599.740748 | 1479.26204 | 1455.788681 | 1765.40757 | 1560.178142 | 1265.171094 | 1402.642577 | 1186.094938 | YAL039C | CYC3 | n | transgressi | -0.691432728 | -0.98731722 |
| W303.asm02.HP0_G0000320 | 5895.156945 | 6983.964755 | 4687.290789 | 18733.16499 | 17839.38638 | 17445.66746 | 38080.79485 | 33620.59065 | 37319.28635 | 17393.73411 | 20941.08697 | 15154.30533 | YAL038W | CDC19 | n | transgressi | -1.580135441 | -1.597396033 |
| W303.asm02.HP0_G0000340 | 2971.418189 | 2662.002701 | 2637.052464 | 2889.784375 | 2540.593726 | 2444.862368 | 2943.185676 | 3137.622722 | 2978.760285 | 924.9787304 | 1992.806733 | 1101.841943 | YAL036C | RBG1 | n | km-like | 1.003855133 | 0.060241321 |
| W303.asm02.HP0_G0000350 | 5353.590566 | 6209.028117 | 6262.661056 | 5750.452535 | 6616.977329 | 6637.848406 | 8281.378537 | 8774.260345 | 8556.943424 | 2315.030565 | 3271.68673 | 2316.957357 | YAL035W | FUN12 | n | km-like | 1.157296697 | -0.079236169 |
| W303.asm02.HP0_G0000360 | 218.6056978 | 227.1067104 | 238.3368198 | 335.8763415 | 330.4128378 | 242.7077747 | 260.0912779 | 302.1512209 | 234.2452337 | 354.8335539 | 498.4881707 | 399.7336531 | YAL034W- | MTW1 | n | transgressi | -0.851825624 | -0.393892121 |
| W303.asm02.HP0_G0000380 | 642.3229145 | 575.5681516 | 471.2568937 | 354.5939086 | 292.6236862 | 279.3231717 | 315.1800555 | 394.5118561 | 294.5546409 | 453.8769003 | 687.5698906 | 520.4962789 | YAL033W | POP5 | n | sc-like | 0.024867447 | 0.840738666 |
| W303.asm02.HP0_G0000390 | 510.0799615 | 529.6267178 | 542.5773814 | 297.4013427 | 295.530544 | 349.4155032 | 219.4520157 | 296.8734703 | 223.7566411 | 190.3354743 | 234.9197126 | 178.8035778 | YAL032C | PRP45 | n | NA | 1.373153899 | 0.734473055 |
| W303.asm02.HP0_G0000400 | 406.6245901 | 611.9745708 | 651.8150905 | 724.7857896 | 718.9628317 | 759.50795 | 617.7167851 | 684.7881382 | 661.6553802 | 283.3500952 | 435.4609307 | 265.8650058 | YAL031C | GIP4 | n | km-like | 0.74213161 | -0.374472984 |
| W303.asm02.HP0_G0000440 | 191.61734 | 206.3030423 | 188.6833157 | 248.5276954 | 245.1450087 | 202.953915 | 186.0375113 | 196.5962092 | 203.6535054 | 624.4037053 | 830.8136178 | 661.8540813 | YAL027W | SAW1 | n | km-like | -1.825569653 | -0.231496289 |
| W303.asm02.HP0_G0000450 | 980.5769984 | 1251.687366 | 1352.380894 | 1751.132381 | 2450.481134 | 2345.477719 | 2140.334475 | 2327.488007 | 2174.634856 | 699.33215 | 923.6355531 | 750.787798 | YAL026C | DRS2 | n | intermedia | 0.583421969 | -0.849322772 |
| W303.asm02.HP0_G0000460 | 5161.073614 | 4547.335125 | 4479.648863 | 3271.414769 | 2512.494101 | 2211.569981 | 4573.271637 | 4540.184939 | 4767.939365 | 2336.561728 | 4796.945937 | 2972.258427 | YAL025C | MAK16 | n | NA | 0.470906924 | 0.791509988 |
| W303.asm02.HP0_G0000470 | 735.8825547 | 1133.799913 | 904.596566 | 1584.754008 | 1741.207828 | 1590.154386 | 1484.687711 | 1661.171996 | 1567.170537 | 150.7181358 | 390.7688878 | 171.3144227 | YAL024C | LTE1 | n | intermedia | 1.86482414 | -0.769835897 |
| W303.asm02.HP0_G0000480 | 3844.041756 | 5840.629827 | 5529.594777 | 8381.310566 | 9823.241493 | 9703.080216 | 6883.387918 | 7252.948739 | 7083.29617 | 2207.374754 | 2962.280279 | 2176.535699 | YAL023C | PMT2 | n | intermedia | 1.028900561 | -0.854451347 |
| W303.asm02.HP0_G0000490 | 2131.180651 | 2861.371187 | 2370.729124 | 583.3641721 | 423.4322877 | 455.0770775 | 225.7736788 | 270.4847174 | 242.9857275 | 701.9158894 | 757.4728295 | 678.7046802 | YAL022C | FUN26 | n | transgressi | 1.758831174 | 2.307287423 |
| W303.asm02.HP0_G0000500 | 1489.757348 | 2260.66527 | 2484.480788 | 1980.94251 | 2708.222527 | 2821.47788 | 3629.53766 | 3492.551448 | 3707.717467 | 750.1456929 | 1187.204011 | 614.1107176 | YAL021C | CCR4 | n | km-like | 1.235718735 | -0.237420412 |
| W303.asm02.HP0_G0000510 | 702.5969135 | 667.4510192 | 561.5359921 | 294.2817482 | 218.9832884 | 240.6154663 | 123.723976 | 146.4575787 | 100.5156786 | 327.2736662 | 456.0880274 | 303.3107812 | YAL020C | ATS1 | n | transgressi | 0.810751702 | 1.324975515 |
| W303.asm02.HP0_G0000520 | 716.0910924 | 1145.068567 | 1048.140332 | 1058.582401 | 1420.484517 | 1163.323472 | 1918.173175 | 1963.323217 | 1879.206166 | 329.8574057 | 825.0838688 | 356.6710113 | YAL019W | FUN30 | n | km-like | 0.887178999 | -0.282192897 |
| W303.asm02.HP0_G0000540 | 626.1298999 | 1197.944556 | 961.472398 | 700.8688984 | 1265.4521 | 935.2618558 | 1410.633945 | 1401.24278 | 1457.040315 | 356.5560469 | 561.5154107 | 340.7565567 | YAL017W | PSK1 | n | km-like | 1.089747106 | -0.049944603 |
| W303.asm02.HP0_G0000550 | 641.4233026 | 968.2373875 | 877.5128365 | 3063.441802 | 3156.847582 | 3494.155032 | 2839.329784 | 3078.248028 | 2886.111051 | 2758.572508 | 3237.308235 | 2888.941577 | YAL016W | TPD3 | n | NA | -1.812333511 | -1.944382524 |
| W303.asm02.HP0_G0000570 | 1015.661863 | 1002.910168 | 1051.751496 | 200.6939131 | 150.1876535 | 227.0154616 | 106.5651764 | 143.8187034 | 102.2637774 | 873.303941 | 1060.003581 | 989.5046166 | YAL014C | SYN8 | n | sc-like | 0.070512949 | 2.381814918 |
| W303.asm02.HP0_G0000580 | 1384.502753 | 1318.432468 | 1596.13446 | 739.3438973 | 569.7441308 | 565.9694228 | 493.0897144 | 511.9418066 | 488.5936031 | 707.0833684 | 630.2723997 | 569.175787 | YAL013W | DEP1 | n | transgressi | 1.157357383 | 1.181939833 |
| W303.asm02.HP0_G0000590 | 5638.767547 | 5350.009987 | 4899.44667 | 6815.274125 | 6027.854146 | 5508.001869 | 10299.79522 | 9691.269508 | 9931.823098 | 2153.977472 | 2444.310961 | 1808.630955 | YAL012W | CYS3 | n | km-like | 1.296638337 | -0.193524356 |
| W303.asm02.HP0_G0000600 | 413.8214855 | 461.1479769 | 541.6745904 | 738.3040325 | 1007.710708 | 1001.16957 | 853.4245057 | 994.8559849 | 948.3435766 | 195.5029533 | 296.8010028 | 188.1650217 | YAL011W | SWC3 | n | intermedia | 1.034303434 | -0.928600995 |
| W303.asm02.HP0_G0000610 | 718.7899282 | 1125.131718 | 1002.097992 | 416.9857986 | 460.2524866 | 424.7386057 | 399.167864 | 468.4003643 | 468.4904673 | 554.6427396 | 521.4071671 | 512.0709794 | YAL010C | MDM10 | n | NA | 0.826269365 | 1.107594425 |
| W303.asm02.HP0_G0000620 | 556.8597816 | 488.0193815 | 416.1866436 | 1014.908078 | 624.9744292 | 753.2310248 | 333.2419498 | 385.2757926 | 371.4709862 | 642.4898816 | 550.0559125 | 683.3854022 | YAL009W | SPO7 | n | sc-like | -0.352269129 | -0.690623103 |
| W303.asm02.HP0_G0000630 | 625.2302879 | 730.7288431 | 644.5927626 | 822.5330841 | 505.793259 | 579.5694274 | 417.2297583 | 281.0402186 | 442.268986 | 960.2898365 | 893.8408578 | 1161.755184 | YAL008W | FUN14 | n | km-like | -0.573460603 | 0.058712214 |
| W303.asm02.HP0_G0000660 | 7315.644175 | 4041.979354 | 5341.814253 | 10497.4355 | 7829.137036 | 7980.064246 | 8271.444495 | 9048.703375 | 7874.310859 | 12878.21877 | 16360.72555 | 13142.53104 | YAL003W | EFB1 | n | transgressi | -1.305249402 | -0.630750104 |
| W303.asm02.HP0_G0000670 | 2683.542373 | 2794.626085 | 2683.997596 | 562.5668754 | 655.9809125 | 750.0925622 | 436.1947474 | 564.7193124 | 429.1582453 | 382.3934416 | 519.1152674 | 386.6276316 | YAL002W | VPS8 | n | NA | 2.631343512 | 2.025861991 |
| W303.asm02.HP0_G0000680 | 765.5697483 | 1084.391201 | 869.3877176 | 606.2411985 | 770.3173197 | 818.0925853 | 625.8446375 | 712.4963287 | 736.8236268 | 639.0448956 | 707.0510375 | 689.0022685 | YAL001C | TFC3 | n | NA | 0.411311188 | 0.293688834 |
| W303.asm02.HP0_G0000700 | 2123.084143 | 2295.33805 | 1988.848538 | 1424.614823 | 1196.656465 | 1373.600466 | 1257.107843 | 1311.52102 | 1351.280341 | 1287.563503 | 1337.323437 | 1158.010606 | YAR002W | NUP60 | n | NA | 0.755342581 | 0.675992367 |
| W303.asm02.HP0_G0000710 | 3496.791553 | 2395.022293 | 2288.575145 | 3890.134346 | 2840.969033 | 2582.954723 | 2717.411997 | 2856.582503 | 2798.706113 | 2825.749734 | 3911.126728 | 3260.590898 | YAR002C-. | ERP1 | n | NA | -0.280541169 | -0.165049804 |
| W303.asm02.HP0_G0000720 | 1220.773382 | 1127.732177 | 1129.391521 | 798.6161929 | 782.9137036 | 813.9079685 | 601.4610802 | 694.0242017 | 714.9723923 | 1874.072362 | 2233.456195 | 2067.006805 | YAR003W | SWD1 | n | intermedia | -0.821657486 | 0.532992607 |
| W303.asm02.HP0_G0000730 | 1281.946993 | 1286.360146 | 1168.211533 | 2028.776292 | 2239.249466 | 2137.293033 | 1890.177239 | 1859.087643 | 2037.409103 | 1567.468612 | 1762.47082 | 1696.293628 | YAR007C | RFA1 | n | transgressi | -0.424577699 | -0.773102633 |
| W303.asm02.HP0_G0000740 | 605.4388256 | 489.7530206 | 434.2424633 | 477.297959 | 321.6922643 | 372.4308956 | 343.1759917 | 459.1643007 | 397.6924676 | 674.3560017 | 1174.598563 | 776.0636964 | YAR008W | SEN34 | n | km-like | -0.748909237 | 0.353988189 |
| W303.asm02.HP0_G0000750 | 917.6041636 | 900.6254661 | 1042.723587 | 813.1743006 | 716.0559739 | 758.4617958 | 773.0490761 | 851.0372815 | 752.5565156 | 942.2036602 | 1183.766162 | 1014.780515 | YAR014C | BUD14 | n | sc-like | -0.131740502 | 0.316307942 |
| W303.asm02.HP0_G0000760 | 229.4010409 | 183.7657351 | 211.2530903 | 1080.419563 | 944.7287883 | 829.6002815 | 3195.149102 | 1005.411486 | 2861.637668 | 997.3234355 | 1257.10695 | 1146.776874 | YAR015W | ADE1 | n | transgressi | -2.308233196 | -2.082626779 |
| W303.asm02.HP0_G0000770 | 1089.430041 | 1025.447475 | 1069.807316 | 637.4371435 | 491.2589699 | 520.9847922 | 325.1140974 | 455.2059878 | 360.1083443 | 535.6953168 | 610.7912528 | 548.5806105 | YAR018C | KIN3 | n | transgressi | 0.898089176 | 0.934634914 |
| W303.asm02.HP0_G0000780 | 3297.077705 | 3290.446842 | 2854.625092 | 277.6439108 | 319.7543591 | 359.8770452 | 183.3282271 | 296.8734703 | 229.8749868 | 138.6606849 | 203.9790676 | 104.8481713 | YAR019C | CDC15 | n | transgressi | 4.365098044 | 3.240660892 |
| W303.asm02.HP0_G0000810 | 1153.302488 | 870.2867834 | 877.5128365 | 251.6472899 | 413.7427617 | 224.9231532 | 380.202875 | 339.095475 | 357.4861962 | 1152.347804 | 1359.096484 | 1197.32867 | YAR027W | UIP3 | n | sc-like | -0.343016494 | 1.660713705 |
| W303.asm02.HP0_G0000840 | 307.6672784 | 487.152562 | 449.5899101 | 268.2851273 | 93.98840253 | 233.2923869 | 53.28258819 | 76.52738346 | 81.28659228 | 92.15337444 | 214.2926159 | 109.5288932 | YAR035W | YAT1 | n | transgressi | 1.441914572 | 0.955886226 |
| W303.asm02.HP0_G0000850 | 1254.059024 | 1593.214251 | 1332.519492 | 2645.416139 | 2903.950952 | 2927.139455 | 2065.377613 | 2103.183607 | 2122.191893 | 291.9625601 | 709.3429372 | 282.7156047 | YAR042W | SWH1 | n | intermedia | 1.619556673 | -0.960930063 |
| W303.asm02.HP0_G0004930 | 2216.643783 | 3294.78094 | 3138.101461 | 204.8533724 | 268.3998711 | 293.9693305 | 316.986245 | 208.471148 | 346.9976036 | 28.42113417 | 12.60544799 | 17.78674334 | YCL073C | GEX1 | y | transgressi | 7.11308831 | 3.422309578 |
| W303.asm02.HP0_G0004980 | 586.5469752 | 746.3315941 | 594.9392585 | 996.1905114 | 1167.587887 | 949.9080146 | 415.4235689 | 407.7062326 | 488.5936031 | 645.0736211 | 313.9902501 | 700.2360011 | YCL064C | CHA1 | n | NA | 0.20497287 | -0.654386981 |
| W303.asm02.HP0_G0004990 | 666.6124365 | 765.4016233 | 700.5658036 | 482.4972832 | 411.8048564 | 476.0001615 | 485.8649567 | 583.1914394 | 529.6739239 | 1251.39115 | 1097.819925 | 1166.435906 | YCL063W | VAC17 | n | intermedia | -0.715789265 | 0.631905166 |
| W303.asm02.HP0_G0005000 | 1996.238862 | 1688.564397 | 1709.886124 | 765.3405182 | 742.2176942 | 692.5540811 | 646.6158159 | 774.5098981 | 750.8084169 | 887.9451313 | 974.0573451 | 897.7624667 | YCL061C | MRC1 | n | transgressi | 0.960157738 | 1.284099526 |
| W303.asm02.HP0_G0005010 | 4205.68575 | 3458.609827 | 3940.682645 | 3557.377599 | 2802.210929 | 3706.524335 | 4140.689268 | 4558.657066 | 4346.647564 | 4225.27528 | 3382.843862 | 3700.57876 | YCL059C | KRR1 | n | NA | 0.036482214 | 0.189316508 |
| W303.asm02.HP0_G0005030 | 218.6056978 | 176.8311791 | 183.2665698 | 3828.78232 | 3379.706681 | 3287.0165 | 3504.007494 | 4128.520394 | 3490.953221 | 2632.83052 | 2620.787233 | 2510.739245 | YCL057C-/ | MIC10 | n | transgressi | -3.73092215 | -4.166929699 |
| W303.asm02.HP0_G0005040 | 367.0416654 | 660.5164631 | 430.6312994 | 1410.056716 | 2020.266178 | 1559.815914 | 1375.413251 | 1186.174444 | 1389.738513 | 3872.164219 | 4584.945221 | 3613.517332 | YCL057W | PRD1 | n | transgressi | -2.986318951 | -1.730358458 |
| W303.asm02.HP0_G0005060 | 736.7821667 | 728.995204 | 764.6639635 | 64.47161974 | 45.54077236 | 64.86156047 | 55.08877762 | 51.45806819 | 48.07271587 | 1656.177 | 1286.901645 | 1872.288773 | YCL055W | KAR4 | n | intermedia | -1.092048951 | 3.634624326 |
| W303.asm02.HP0_G0005070 | 4184.095064 | 3945.762389 | 3778.180268 | 4200.014066 | 4194.59582 | 4929.478596 | 6853.585792 | 6335.939575 | 6882.264813 | 5038.291967 | 4720.167299 | 4648.893024 | YCL054W | SPB1 | n | km-like | -0.273381516 | -0.151502849 |
| W303.asm02.HP0_G0005080 | 641.4233026 | 654.4487266 | 626.5369429 | 553.2080919 | 486.4142069 | 469.7232363 | 425.3576108 | 562.0804371 | 490.3417018 | 3196.085725 | 3137.610601 | 3216.592112 | YCL052C | PBN1 | n | intermedia | -2.298217081 | 0.344093098 |
| W303.asm02.HP0_G0005100 | 4916.379171 | 4496.192775 | 4761.31965 | 2958.415454 | 3457.222889 | 3117.539519 | 4505.539533 | 3664.078342 | 4828.248772 | 8054.377175 | 7598.793241 | 8075.181478 | YCL050C | APA1 | n | intermedia | -0.736677187 | 0.565430298 |
| W303.asm02.HP0_G0005130 | 3893.520412 | 3422.203407 | 3693.317916 | 17.67770219 | 17.44114686 | 27.20000923 | 31.60831503 | 25.06931527 | 20.97718511 | 106.7945648 | 89.38408578 | 121.6987702 | YCL048W | SPS22 | n | transgressi | 5.09366003 | 7.441861726 |
| W303.asm02.HP0_G0005150 | 2134.779098 | 2125.441427 | 2026.765759 | 1191.6851 | 1329.402972 | 1115.200378 | 1308.584242 | 1329.993147 | 1304.081674 | 5623.93958 | 5665.575899 | 5404.361543 | YCL045C | EMC1 | n | intermedia | -1.403621558 | 0.785341114 |
| W303.asm02.HP0_G0005160 | 590.1454229 | 583.3695271 | 612.9950782 | 1062.741861 | 889.4984899 | 910.154155 | 827.234759 | 947.3562297 | 838.2133548 | 911.1987865 | 805.6027219 | 815.3817607 | YCL044C | MGR1 | n | NA | -0.500319264 | -0.673265891 |
| W303.asm02.HP0_G0005170 | 5452.547878 | 5727.076472 | 5010.489961 | 8153.580167 | 8622.709218 | 8220.679713 | 8509.8615 | 8162.041277 | 8327.068437 | 8937.154828 | 10414.39194 | 8852.181319 | YCL043C | PDI1 | n | NA | -0.796178147 | -0.622416623 |
| W303.asm02.HP0_G0005180 | 947.2913571 | 913.6277587 | 863.9709717 | 127.9033746 | 115.3053598 | 128.6769667 | 133.6580178 | 170.2074563 | 114.5004687 | 8775.240488 | 9055.29546 | 9814.537748 | YCL040W | GLK1 | n | intermedia | -3.327371571 | 2.853136363 |

| | | | | | | | | | | | | | | | | | |
|---|---|---|---|---|---|---|---|---|---|---|---|---|---|---|---|---|---|
| W303.asm02.HP0_G0005190 | 660.315153 | 720.327009 | 661.7457913 | 229.8101284 | 390.4878992 | 230.1539242 | 166.1694276 | 209.7905857 | 186.1725178 | 1127.371656 | 1008.43584 | 1139.287718 | YCL039W | GID7 | n | intermedia | -0.66424191 | 1.231185487 |
| W303.asm02.HP0_G0005200 | 825.8437472 | 733.3293016 | 729.4551151 | 323.3979635 | 419.5564773 | 334.7693444 | 288.0872141 | 296.8734703 | 330.3906654 | 1150.625311 | 1227.312255 | 1095.288932 | YCL038C | ATG22 | n | intermedia | -0.59523818 | 1.073017249 |
| W303.asm02.HP0_G0005210 | 410.2230378 | 417.8070016 | 393.616869 | 3940.047858 | 3737.250191 | 4258.893753 | 5285.813367 | 5276.431146 | 5393.75872 | 108.5170578 | 91.67598542 | 89.86986111 | YCL037C | SRO9 | n | intermedia | 2.050034466 | -3.279044908 |
| W303.asm02.HP0_G0005230 | 902.3107609 | 1120.79762 | 1052.654287 | 1770.889813 | 1704.387629 | 1985.600674 | 2099.695212 | 2348.59901 | 2095.096362 | 3492.354517 | 3516.919991 | 3520.839038 | YCL035C | GRX1 | n | transgressi | -1.763453774 | -0.820471984 |
| W303.asm02.HP0_G0005240 | 806.9518968 | 901.4922856 | 791.747693 | 422.1851228 | 961.2009826 | 608.8617451 | 773.9521708 | 416.9422961 | 748.1862687 | 2785.271149 | 2698.711821 | 3170.721037 | YCL034W | LSB5 | n | km-like | -1.705463125 | 0.28858401 |
| W303.asm02.HP0_G0005250 | 186.2196685 | 324.190495 | 221.1837911 | 376.4310701 | 348.8229372 | 394.4001338 | 372.9781173 | 435.4144231 | 370.5969369 | 1843.067489 | 1977.909385 | 1875.097206 | YCL033C | MXR2 | n | transgressi | -2.918205907 | -0.595052406 |
| W303.asm02.HP0_G0005260 | 532.5702597 | 509.6898692 | 488.4099224 | 593.7628205 | 485.4452543 | 525.169409 | 518.3763664 | 577.9136889 | 519.1853313 | 1719.047994 | 1573.3891 | 1693.485195 | YCL032W | STE50 | n | km-like | -1.693830732 | -0.056467121 |
| W303.asm02.HP0_G0005270 | 3686.609669 | 3229.769477 | 3399.008055 | 2274.184393 | 1933.060444 | 2143.569958 | 2183.683021 | 2600.6116 | 2267.28409 | 3357.138818 | 2896.961139 | 3558.284813 | YCL031C | RRP7 | n | sc-like | 0.070991599 | 0.691828384 |
| W303.asm02.HP0_G0005280 | 1066.040131 | 1110.395786 | 944.3193693 | 2170.19791 | 2944.646962 | 2351.754644 | 2389.588616 | 2133.530673 | 2476.181892 | 4694.654617 | 4978.006008 | 4164.906376 | YCL030C | HIS4 | n | transgressi | -2.125215643 | -1.243100167 |
| W303.asm02.HP0_G0005290 | 696.29963 | 746.3315941 | 726.7467422 | 686.3107907 | 583.3094673 | 659.0771467 | 666.4838997 | 721.7323923 | 642.4262938 | 1259.142369 | 1129.90652 | 1246.944323 | YCL029C | BIK1 | n | km-like | -0.74138023 | 0.159233922 |
| W303.asm02.HP0_G0005310 | 106.1542072 | 108.3524382 | 93.89026234 | 247.4878306 | 382.7362783 | 302.3385641 | 416.3266636 | 460.4837384 | 477.2309611 | 432.345738 | 214.2926159 | 558.8781988 | YCL027W | FUS1 | n | NA | -1.870418201 | -1.519679648 |
| W303.asm02.HP0_G0005320 | 445.3079029 | 452.4797818 | 517.2992339 | 537.6101194 | 530.9860267 | 485.4155493 | 598.7517961 | 486.8724913 | 537.5403683 | 626.9874448 | 632.5642994 | 641.2589048 | YCL026C-I | HBN1 | n | km-like | -0.422330233 | -0.121137305 |
| W303.asm02.HP0_G0005340 | 1967.45128 | 1624.419753 | 1703.566587 | 3034.325587 | 2018.328273 | 2321.416172 | 2608.137537 | 4434.629928 | 2804.824458 | 4295.897493 | 5184.276975 | 3826.022108 | YCL025C | AGP1 | n | NA | -1.28553873 | -0.449743475 |
| W303.asm02.HP0_G0005370 | 526.2729762 | 501.0216741 | 502.8545781 | 270.364857 | 326.5370273 | 208.184686 | 153.5261016 | 199.2350845 | 191.4168141 | 1036.079528 | 704.7591379 | 1103.714232 | YCL018W | LEU2 | n | intermedia | -0.870638849 | 0.89577268 |
| W303.asm02.HP0_G0005380 | 953.5886406 | 1065.321172 | 1130.294312 | 3639.526921 | 3326.414288 | 3257.724182 | 3167.153166 | 3376.440936 | 3323.13574 | 2345.174193 | 2524.527448 | 2213.981474 | YCL017C | NFS1 | n | transgressi | -1.16207702 | -1.690903064 |
| W303.asm02.HP0_G0005390 | 453.4044103 | 403.07107 | 398.130824 | 424.2648525 | 333.3196956 | 348.369349 | 307.9552978 | 381.3174796 | 384.5817269 | 567.561437 | 434.3149809 | 572.9203646 | YCL016C | DCC1 | n | km-like | -0.324519283 | 0.163587213 |
| W303.asm02.HP0_G0005400 | 1658.88439 | 2087.301369 | 1869.680128 | 1742.813463 | 1533.851971 | 1567.138993 | 1540.679584 | 1847.212704 | 1728.869672 | 647.6573605 | 799.8729728 | 626.2805946 | YCL014W | BUD3 | n | km-like | 1.422270847 | 0.198455556 |
| W303.asm02.HP0_G0005430 | 502.8830661 | 501.0216741 | 538.9662175 | 373.3114756 | 282.9341602 | 348.369349 | 316.0831503 | 428.8172349 | 335.6349617 | 603.7337896 | 567.2451598 | 634.7058941 | YCL010C | SGF29 | n | sc-like | -0.224434061 | 0.608756258 |
| W303.asm02.HP0_G0005440 | 2912.943413 | 3249.706326 | 3099.281448 | 2431.203983 | 3350.638103 | 2565.170101 | 2099.695212 | 1628.186055 | 2333.711843 | 2414.935158 | 2568.073541 | 1979.945378 | YCL009C | ILV6 | n | NA | 0.403129772 | 0.131599274 |
| W303.asm02.HP0_G0005450 | 400.3273067 | 442.0779478 | 430.6312994 | 48.87364722 | 58.1371562 | 40.80001384 | 50.57330404 | 44.86087996 | 45.45056773 | 327.2736662 | 380.4553395 | 307.0553588 | YCL008C | STP22 | n | transgressi | 0.32608111 | 3.066091749 |
| W303.asm02.HP0_G0005470 | 842.0367619 | 722.9274675 | 719.5244143 | 106.0662131 | 111.4295494 | 94.1538781 | 82.18161907 | 100.2772611 | 88.27898732 | 682.9684666 | 661.2130448 | 668.407092 | YCL005W | LDB16 | n | sc-like | 0.180786896 | 2.851538264 |
| W303.asm02.HP0_G0005480 | 782.6623748 | 760.2007062 | 739.3858159 | 937.9580807 | 837.1750493 | 822.2772021 | 797.4326334 | 923.6063521 | 906.3892064 | 446.9869284 | 394.2067373 | 519.5601345 | YCL004W | PGS1 | n | km-like | 0.735948014 | -0.172673867 |
| W303.asm02.HP0_G0005500 | 618.0333925 | 597.2386392 | 574.1750659 | 1666.90333 | 1406.91918 | 1465.662036 | 1664.40356 | 1696.796812 | 1632.724241 | 1548.521189 | 1340.761287 | 1403.280435 | YCL001W | RER1 | n | transgressi | -1.255518369 | -1.334958296 |
| W303.asm02.HP0_G0005530 | 1455.572095 | 1611.417461 | 1481.480005 | 3198.624231 | 2831.279507 | 2817.293264 | 2680.385114 | 3036.026023 | 2816.1871 | 2726.706388 | 2549.738344 | 2783.157261 | YCR002C | CDC10 | n | NA | -0.820948611 | -0.953828689 |
| W303.asm02.HP0_G0005540 | 610.8364971 | 718.59337 | 727.6495331 | 1911.271566 | 1659.81581 | 1884.123716 | 2002.160983 | 2016.100723 | 2055.76414 | 1085.170578 | 1144.803868 | 1231.029868 | YCR003W | MRPL32 | n | transgressi | -0.744822981 | -1.398037312 |
| W303.asm02.HP0_G0005550 | 756.573629 | 819.1444326 | 691.5378938 | 2145.241154 | 2648.147465 | 2337.108485 | 3032.592053 | 2306.377005 | 2920.198976 | 2408.045186 | 2652.873828 | 2380.615175 | YCR004C | YCP4 | n | NA | -1.694879521 | -1.636591817 |
| W303.asm02.HP0_G0005580 | 316.6633976 | 241.842642 | 303.3377706 | 133.1026988 | 97.86421294 | 130.7692751 | 107.4682711 | 164.9297057 | 129.3593081 | 347.0823355 | 265.8603577 | 389.4360648 | YCR007C | DFP3 | n | sc-like | -0.213785255 | 1.216390712 |
| W303.asm02.HP0_G0005590 | 3079.37162 | 2414.959142 | 2417.674255 | 479.3776887 | 434.0907663 | 493.7847829 | 568.0465757 | 668.9548864 | 588.2352323 | 1520.961302 | 1207.831108 | 1557.744259 | YCR008W | SAT4 | n | transgressi | 0.870850765 | 2.467741421 |
| W303.asm02.HP0_G0005600 | 940.0944617 | 813.9435156 | 980.4310087 | 1865.517513 | 2165.609069 | 1924.92373 | 1802.577051 | 1795.754636 | 1875.709968 | 2782.687409 | 2503.900352 | 2728.860887 | YCR009C | RVS161 | n | transgressi | -1.542440878 | -1.11580027 |
| W303.asm02.HP0_G0005610 | 417.4199332 | 380.5337629 | 384.5889592 | 276.604046 | 279.0583498 | 260.4923961 | 205.0025003 | 205.8322727 | 203.6535054 | 82.67966305 | 74.48673815 | 104.8481713 | YCR010C | ADY2 | n | transgressi | 2.143522442 | 0.527961385 |
| W303.asm02.HP0_G0005620 | 2296.709245 | 2085.56773 | 2139.614632 | 722.70606 | 684.080538 | 647.5694505 | 661.9684261 | 699.3019523 | 706.2318985 | 1906.799729 | 1808.308812 | 2014.58272 | YCR011C | ADP1 | n | transgressi | 0.185300957 | 1.66013638 |
| W303.asm02.HP0_G0005630 | 5355.38979 | 7005.635243 | 5910.572573 | 17572.67584 | 20788.87811 | 17972.92918 | 26834.55636 | 23682.5863 | 26640.15103 | 31325.25734 | 29211.4068 | 31447.89838 | YCR012W | PGK1 | n | transgressi | -2.313505833 | -1.611567617 |
| W303.asm02.HP0_G0005640 | 934.6967902 | 726.3947455 | 718.6216233 | 285.9628295 | 280.996255 | 319.0770313 | 259.1881832 | 287.6374068 | 304.169184 | 459.0443792 | 324.3037984 | 498.964958 | YCR014C | POL4 | n | transgressi | 0.870385093 | 1.398656658 |
| W303.asm02.HP0_G0005650 | 407.5242021 | 397.0033335 | 355.6996477 | 73.83040325 | 93.98840253 | 63.81540627 | 85.79399793 | 77.8468211 | 75.16824663 | 416.8433012 | 413.6878842 | 481.1782147 | YCR015C | CTO1 | n | sc-like | -0.175232799 | 2.292609846 |
| W303.asm02.HP0_G0005660 | 2602.577299 | 2484.304702 | 2693.025505 | 1515.083064 | 1547.417308 | 1866.339095 | 2336.306028 | 2115.058546 | 2359.933324 | 1426.224188 | 1103.549674 | 1358.345505 | YCR016W | RBP95 | n | transgressi | 0.988781221 | 0.649935793 |
| W303.asm02.HP0_G0005670 | 1339.522156 | 1451.055852 | 1223.281783 | 1500.524956 | 1271.265816 | 1280.492742 | 1444.951544 | 1737.69938 | 1602.132512 | 2698.285253 | 2641.41433 | 2677.372945 | YCR017C | CWH43 | n | km-like | -0.990084843 | -0.011052328 |
| W303.asm02.HP0_G0005800 | 1026.457207 | 786.2052914 | 811.6090947 | 667.5932237 | 591.0610881 | 628.7386749 | 689.9643623 | 773.1904604 | 694.8692566 | 847.4665462 | 724.2402848 | 792.9142954 | YCR028C | FEN2 | n | NA | 0.147218651 | 0.46833841 |
| W303.asm02.HP0_G0005810 | 531.6706477 | 501.0216741 | 541.6745904 | 2406.247227 | 2246.032135 | 2550.523942 | 2440.16192 | 2641.514167 | 2470.937595 | 2277.13572 | 1943.530891 | 2235.512795 | YCR028C- | RIM1 | n | transgressi | -2.026021473 | -2.184289638 |
| W303.asm02.HP0_G0005840 | 1226.171054 | 898.0250076 | 1017.445439 | 486.6567425 | 640.4776709 | 402.7693674 | 319.6955291 | 366.8036655 | 355.7380974 | 554.6427396 | 556.9316114 | 647.8119155 | YCR032W | BPH1 | n | transgressi | 0.817279882 | 1.012714637 |
| W303.asm02.HP0_G0005850 | 1697.567703 | 1650.424338 | 1597.940042 | 1944.54724 | 1626.871421 | 1819.262156 | 1684.271644 | 1962.003779 | 1772.572141 | 822.490398 | 794.1432237 | 851.8913917 | YCR033W | SNT1 | n | km-like | 0.995530461 | -0.11083972 |
| W303.asm02.HP0_G0005860 | 2902.14807 | 2468.701951 | 2571.148723 | 5991.701176 | 5153.858898 | 5569.724967 | 5529.64894 | 6045.663293 | 5867.493484 | 4268.337605 | 3866.434685 | 4043.207605 | YCR034W | ELO2 | n | transgressi | -0.613106155 | -1.066774965 |
| W303.asm02.HP0_G0005870 | 2059.211697 | 2199.987905 | 2242.532804 | 1466.209417 | 1178.246366 | 1569.231302 | 1733.038758 | 1740.338255 | 1780.438586 | 1470.147759 | 1252.523151 | 1550.255104 | YCR035C | RRP43 | n | transgressi | 0.598183436 | 0.617788106 |
| W303.asm02.HP0_G0005880 | 2155.470173 | 1798.650474 | 2001.487612 | 819.4134896 | 783.8826562 | 800.3079639 | 851.6183163 | 823.329091 | 870.5531819 | 1534.741245 | 1657.043436 | 1587.70088 | YCR036W | RBK1 | n | transgressi | 0.316199055 | 1.302594155 |
| W303.asm02.HP0_G0005890 | 669.3112723 | 521.8253423 | 537.1606355 | 3044.724235 | 3039.604317 | 3466.955023 | 4939.928091 | 3464.843258 | 5055.50161 | 1222.108769 | 1106.987524 | 1256.305767 | YCR037C | PHO87 | n | transgressi | -1.03382381 | -2.436869427 |
| W303.asm02.HP0_G0005900 | 215.0072501 | 225.3730714 | 206.7391353 | 1753.212111 | 1418.546611 | 1497.046662 | 1332.064705 | 1616.311116 | 1426.448587 | 147.2731498 | 121.4706807 | 180.6758666 | YCR038C | BUD5 | n | intermedia | 0.510532845 | -2.8246304 |
| W303.asm02.HP0_G0005920 | 1227.070666 | 1222.215503 | 1093.279882 | 1637.787114 | 1461.180526 | 1424.862022 | 1295.940916 | 1558.25586 | 1453.544118 | 1005.074654 | 960.3059472 | 1039.120269 | YCR042C | TAF2 | n | intermedia | 0.235370893 | -0.348660829 |
| W303.asm02.HP0_G0005930 | 360.744382 | 263.5131296 | 338.546619 | 942.11754 | 775.1620827 | 803.4464265 | 740.5376663 | 956.5922932 | 708.8540467 | 1199.716361 | 1204.393258 | 1122.437119 | YCR043C | | n | transgressi | -1.845196124 | -1.367487472 |
| W303.asm02.HP0_G0005940 | 351.7482627 | 303.3868269 | 328.6159182 | 1970.543861 | 1539.665687 | 1562.954376 | 1435.920597 | 1954.087153 | 1500.742784 | 890.5288708 | 725.3862346 | 964.2287181 | YCR044C | PER1 | n | transgressi | -1.365932821 | -2.335681205 |
| W303.asm02.HP0_G0005950 | 1316.132246 | 1288.960605 | 1254.879468 | 8.318918676 | 10.65847864 | 13.60000461 | 5.41856829 | 9.236063521 | 9.614543173 | 316.0774619 | 285.3415046 | 310.7999363 | YCR045C | RRT12 | n | transgressi | 2.064693452 | 6.85015872 |
| W303.asm02.HP0_G0005960 | 417.4199332 | 492.3534791 | 520.9103978 | 1559.797252 | 1367.192123 | 1536.800521 | 1530.745542 | 1735.060504 | 1557.555994 | 1236.74996 | 1299.507093 | 1295.623831 | YCR046C | IMG1 | n | transgressi | -1.41017648 | -1.630797043 |
| W303.asm02.HP0_G0005970 | 1358.414007 | 1299.362439 | 1279.254824 | 1979.902645 | 1599.740748 | 2083.939169 | 2699.350103 | 2689.013922 | 2790.839668 | 1721.631734 | 1288.047595 | 1474.427409 | YCR047C | BUD23 | n | sc-like | -0.186411524 | -0.516167932 |
| W303.asm02.HP0_G0005990 | 2336.292169 | 1950.343887 | 1936.486661 | 1497.405362 | 1255.762574 | 1438.462027 | 1867.599871 | 1840.615516 | 1874.835919 | 2821.443502 | 2218.558847 | 2505.122378 | YCR051W | | n | intermedia | -0.275565855 | 0.562663281 |
| W303.asm02.HP0_G0006010 | 5649.56289 | 5821.559798 | 5107.088597 | 5309.549845 | 5417.414006 | 5295.632566 | 5836.701143 | 5509.971609 | 5976.749656 | 5857.337379 | 5486.807727 | 5613.121742 | YCR053W | THR4 | n | NA | -0.032659701 | 0.04105575 |
| W303.asm02.HP0_G0006020 | 825.8437472 | 773.2029988 | 770.9835004 | 509.5337689 | 499.0105908 | 621.4155955 | 648.4220054 | 624.0940065 | 674.7661209 | 557.2264791 | 410.2500347 | 527.985434 | YCR054C | CTR86 | n | transgressi | 0.652511614 | 0.53174018 |
| W303.asm02.HP0_G0006030 | 3654.22364 | 3293.914121 | 3231.991723 | 4352.874197 | 3950.419764 | 4980.740162 | 6141.044062 | 5631.359872 | 6149.811433 | 2125.556338 | 2106.255765 | 2036.114041 | YCR057C | PWP2 | n | intermedia | 0.694508068 | -0.379911836 |
| W303.asm02.HP0_G0006040 | 2342.589453 | 1947.743429 | 2111.628112 | 475.2182294 | 423.4322877 | 459.2616943 | 503.926851 | 563.3998748 | 499.956245 | 1745.746635 | 1539.010605 | 1644.805687 | YCR059C | YIH1 | n | transgressi | 0.373437806 | 2.225516087 |
| W303.asm02.HP0_G0006050 | 691.8015704 | 619.7759464 | 647.3011356 | 128.9432395 | 148.2497483 | 168.4308264 | 182.4251324 | 163.6102681 | 180.0541722 | 1129.955395 | 1060.003581 | 1089.672066 | YCR060W | TAH1 | n | intermedia | -0.739234073 | 2.122126241 |
| W303.asm02.HP0_G0006060 | 460.6013057 | 591.1709027 | 706.8853405 | 1233.279694 | 866.2436274 | 1140.308079 | 1317.615189 | 1562.214173 | 1123.153452 | 456.4606397 | 443.4825795 | 452.1577387 | YCR061W | TVS1 | n | sc-like | 0.369013965 | -0.856383889 |
| W303.asm02.HP0_G0006070 | 392.2307993 | 311.1882024 | 354.7968567 | 49.91351205 | 71.70249265 | 72.18463988 | 130.9487337 | 73.88850816 | 120.6188144 | 151.5793823 | 138.6599279 | 138.5493692 | YCR063W | BUD31 | n | NA | 1.267169115 | 2.395984133 |
| W303.asm02.HP0_G0006080 | 846.5348215 | 763.6679843 | 683.4127749 | 795.4965984 | 921.4739258 | 889.231071 | 957.2803979 | 865.5510956 | 1003.408688 | 656.2698254 | 495.0503213 | 626.2805946 | YCR065W | HCM1 | n | km-like | 0.360253414 | -0.167996007 |
| W303.asm02.HP0_G0006090 | 436.3117837 | 395.2696945 | 393.616869 | 695.6695743 | 620.1296662 | 687.3233101 | 713.4448249 | 720.4129566 | 726.3350343 | 292.8238066 | 248.6711104 | 295.8216261 | YCR066W | RAD18 | n | intermedia | 0.539509058 | -0.702868329 |
| W303.asm02.HP0_G0006100 | 1465.467826 | 1441.520837 | 1229.60132 | 1891.514134 | 1975.694358 | 1950.031431 | 1761.034694 | 1930.337276 | 1804.037919 | 1511.48759 | 1524.113258 | 1424.811756 | YCR067C | SED4 | n | sc-like | -0.107771745 | -0.48836295 |
| W303.asm02.HP0_G0006110 | 706.1953612 | 780.1375549 | 638.2732257 | 333.7966119 | 262.5861555 | 301.2924099 | 223.9674893 | 325.9010985 | 275.3255545 | 499.5229643 | 459.5258769 | 525.1770008 | YCR068W | ATG15 | n | transgressi | 0.508277301 | 1.224358556 |
| W303.asm02.HP0_G0006120 | 515.4776331 | 505.3557716 | 389.1029141 | 1839.520892 | 1818.724037 | 1712.554427 | 1669.822128 | 1625.54718 | 1647.58308 | 1135.98412 | 1103.549674 | 1140.223863 | YCR069W | CPR4 | n | transgressi | -1.251612662 | -1.918386377 |
| W303.asm02.HP0_G0006130 | 392.2307993 | 389.2019579 | 431.5340904 | 1158.409426 | 891.4363951 | 1149.723467 | 1164.992182 | 1308.882145 | 1156.367329 | 858.6627506 | 898.4246571 | 924.9106539 | YCR071C | IMG2 | n | transgressi | -1.132733627 | -1.385879103 |
| W303.asm02.HP0_G0006140 | 1543.734063 | 1414.649433 | 1440.854411 | 2400.008038 | 2271.224902 | 2820.431726 | 3445.306338 | 3162.692037 | 3600.209394 | 738.088242 | 657.7751954 | 751.7239424 | YCR072C | RSA4 | n | intermedia | 1.024350936 | -0.760516384 |
| W303.asm02.HP0_G0006170 | 122.3472218 | 94.48332609 | 88.47351644 | 422.1851228 | 332.350743 | 355.6924284 | 347.6914653 | 393.1924184 | 387.2038751 | 362.5847723 | 287.6334042 | 364.1601664 | YCR075C | ERS1 | n | NA | -1.704838128 | -1.835135811 |
| W303.asm02.HP0_G0006180 | 54.87632743 | 74.54647746 | 70.41769676 | 127.9033746 | 150.1876535 | 153.7846676 | 129.1425442 | 149.096454 | 133.729555 | 101.6270858 | 116.8868814 | 139.4855136 | YCR075W- | EGO2 | n | NA | -0.818077795 | -1.084241666 |
| W303.asm02.HP0_G0006190 | 372.439337 | 387.4683189 | 308.7545165 | 120.6243208 | 124.0259332 | 97.29234071 | 78.56924021 | 71.24963287 | 83.03469104 | 540.0015493 | 438.8987802 | 561.6866319 | YCR076C | FUB1 | n | intermedia | -0.520917725 | 1.618086382 |
| W303.asm02.HP0_G0006200 | 1371.908186 | 1553.340554 | 1792.942894 | 3358.763415 | 3124.872146 | 3512.985807 | 3634.053133 | 4149.631396 | 4005.768386 | 1384.884356 | 1442.75082 | 1246.008178 | YCR077C | PAT1 | n | sc-like | 0.210116356 | -1.072499246 |
| W303.asm02.HP0_G0006210 | 1211.777263 | 1327.100663 | 1177.239443 | 226.6905339 | 205.4179519 | 172.6154432 | 166.1694276 | 191.3184586 | 185.2984684 | 525.360359 | 516.8233678 | 533.6023003 | YCR079W | PTC6 | n | transgressi | 1.22805636 | 2.603606408 |
| W303.asm02.HP0_G0006220 | 1770.436269 | 1627.020212 | 1577.175849 | 621.839171 | 457.3456288 | 511.5694044 | 440.7102209 | 514.5806819 | 487.7195537 | 447.8481748 | 437.7528304 | 483.0505035 | YCR081W | SRB8 | n | NA | 1.843228909 | 1.631350924 |
| W303.asm02.HP0_G0006230 | 565.8559009 | 576.4349711 | 564.244365 | 760.141194 | 630.7881448 | 604.6771283 | 557.2094392 | 511.9418066 | 566.3839978 | 1089.47681 | 1074.900929 | 1125.245553 | YCR082W | AHC2 | n | transgressi | -0.940316896 | -0.216912762 |
| W303.asm02.HP0_G0006240 | 263.5862941 | 240.109003 | 228.406119 | 998.2702411 | 778.0689405 | 831.6925899 | 782.9831179 | 907.7731003 | 846.9538486 | 733.7820096 | 696.7374892 | 666.5348032 | YCR083W | TRX3 | n | transgressi | -1.50278269 | -1.817342145 |
| W303.asm02.HP0_G0006250 | 2559.395927 | 2196.520627 | 2623.5106 | 5222.201199 | 4519.194942 | 4148.001407 | 4130.755226 | 4359.421982 | 4199.807268 | 1347.850757 | 1659.335336 | 1041.928702 | YCR084C | TUP1 | n | intermedia | 0.850348855 | -0.89359563 |
| W303.asm02.HP0_G0006260 | 398.5280828 | 307.7209244 | 269.0317132 | 692.5499798 | 648.2292917 | 726.0310156 | 646.6158159 | 687.4270135 | 634.5598494 | 425.4557661 | 355.2444435 | 392.244498 | YCR086W | CSM1 | n | sc-like | -0.2641683 | -1.067145978 |
| W303.asm02.HP0_G0006270 | 1703.864986 | 1432.852642 | 1745.997663 | 1942.467511 | 1161.774171 | 1779.508296 | 2227.031567 | 3286.719176 | 2303.994164 | 1181.630184 | 836.5433669 | 1062.523879 | YCR087C-A | | n | km-like | 0.640574406 | -0.000141335 |
| W303.asm02.HP0_G0006280 | 1543.734063 | 2115.906413 | 1854.332681 | 2377.131012 | 2799.304071 | 2569.354718 | 2638.842757 | 2339.362946 | 2748.885298 | 1312.539651 | 1678.816483 | 1039.120269 | YCR088W | ABP1 | n | intermedia | 0.442490931 | -0.473871688 |
| W303.asm02.HP0_G0006300 | 1292.742336 | 1051.45206 | 1194.392472 | 932.7587565 | 755.7830307 | 990.7080285 | 851.6183163 | 952.6339803 | 872.3012806 | 863.8302296 | 792.9972739 | 891.209456 | YCR090C | | n | NA | 0.467400767 | 0.394758072 |
| W303.asm02.HP0_G0006320 | 678.3073915 | 604.1731953 | 612.0922872 | 775.7391665 | 672.4531068 | 723.9387072 | 547.2753973 | 663.6771358 | 659.9072814 | 478.8530485 | 434.3149809 | 418.4565408 | YCR092C | MSH3 | n | km-like | 0.50246927 | -0.184237235 |
| W303.asm02.HP0_G0006330 | 2483.828525 | 2484.304702 | 2494.411489 | 3896.373535 | 3351.607055 | 3665.724321 | 3696.366669 | 4201.089464 | 3944.584849 | 1163.544008 | 1290.339495 | 1116.820253 | YCR093W | CDC39 | n | intermedia | 1.058524365 | -0.544269766 |

| | | | | | | | | | | | | | | | | | |
|---|---|---|---|---|---|---|---|---|---|---|---|---|---|---|---|---|---|
| W303.asm02.HP0_G0006340 | 925.7006709 | 912.7609392 | 860.3598078 | 940.0378104 | 856.5541014 | 1003.261879 | 930.1875565 | 1046.314053 | 988.5498481 | 813.0166866 | 766.640428 | 808.82875 | YCR094W | CDC50 | n | conserved | 0.174301528 | -0.04359029 |
| W303.asm02.HP0_G0006350 | 409.3234259 | 283.4499783 | 324.1019633 | 402.4276909 | 421.4943825 | 406.9539842 | 415.4235689 | 412.9839831 | 428.2841959 | 538.2790563 | 420.5635831 | 671.2155251 | YCR095C | OCA4 | n | km-like | -0.667114047 | -0.254471185 |
| W303.asm02.HP0_G0006450 | 727.7860474 | 579.0354296 | 702.3713856 | 783.0182204 | 571.682036 | 691.5079269 | 764.9212236 | 741.5239569 | 695.743306 | 906.0313076 | 843.4190658 | 917.4214988 | YCR106W | RDS1 | n | km-like | -0.403903326 | -0.021675624 |

**TM**

| W303_gene | KS.1_TM.sc | KS.2_TM.sc | KS.3_TM.sc | KS.1_TM.km | KS.2_TM.km | KS.3_TM.km | Km.V.1_TM | Km.V.2_TM | Km.V.3_TM | SC.1_TM | SC.2_TM | SC.3_TM | locus_tag | gene | ambiguous | Category | log2FC(KSsc/Psc) | log2(KSsc/KSkm) |
|---|---|---|---|---|---|---|---|---|---|---|---|---|---|---|---|---|---|---|
| | Normalized read counts | | | | | | | | | | | | | | | | log2 fold change adjusted with ashr method | |
| W303.asm02.HP0_G0000040 | 1054.193968 | 1278.940416 | 1573.285446 | 691.0526547 | 600.8635871 | 1028.036511 | 820.1716397 | 703.4956843 | 356.3941189 | 253.6268541 | 249.2413544 | 275.4761803 | YAL067C | SEO1 | n | transgressi | 2.16839965 | 0.667184386 |
| W303.asm02.HP0_G0000100 | 5833.206623 | 5327.391488 | 8464.866518 | 577.5660017 | 642.0656616 | 813.9973635 | 665.4057759 | 684.0313768 | 822.8362801 | 710.1551914 | 981.4623669 | 792.1131688 | YAL063C | FLO9 | n | NA | 2.918541723 | 3.215306047 |
| W303.asm02.HP0_G0000110 | 2171.726339 | 2116.298252 | 3158.958967 | 762.9950865 | 534.482467 | 1073.438754 | 1433.988794 | 1285.571165 | 1402.355329 | 213.2344282 | 338.6820318 | 301.2127092 | YAL062W | GDH3 | n | transgressi | 3.04319267 | 1.594486095 |
| W303.asm02.HP0_G0000120 | 590.0015623 | 440.6664329 | 825.2363396 | 513.7297594 | 353.6511398 | 688.6006913 | 305.1598105 | 321.6245092 | 291.7787546 | 243.2939082 | 283.825083 | 255.45888 | YAL061W | BDH2 | n | NA | 1.181687495 | 0.203881287 |
| W303.asm02.HP0_G0000130 | 1386.503671 | 1424.607698 | 2073.573066 | 1535.109636 | 820.6079846 | 1878.788071 | 501.0216946 | 446.7522 | 448.26909 | 1035.173308 | 1409.585076 | 1496.531498 | YAL060W | BDH1 | n | conserved | 0.284325904 | 0.156713748 |
| W303.asm02.HP0_G0000140 | 1055.929267 | 1162.589819 | 1647.613893 | 1576.653857 | 1967.399059 | 2174.983659 | 2520.847374 | 2641.584585 | 2634.085712 | 1662.664932 | 1860.36609 | 2080.846026 | YAL059W | ECM1 | n | transgressi | -0.517350367 | -0.533554503 |
| W303.asm02.HP0_G0000150 | 870.2523044 | 953.7084331 | 1355.064752 | 1819.839542 | 1484.419186 | 2450.640137 | 1031.772425 | 1001.948399 | 1064.134281 | 353.1988783 | 435.2779635 | 338.3876955 | YAL058W | CNE1 | n | intermedia | 1.440564282 | -0.824370377 |
| W303.asm02.HP0_G0000170 | 474.6041979 | 415.0143326 | 762.3430389 | 602.8978439 | 534.482467 | 795.620265 | 681.9033095 | 716.4718793 | 702.6920871 | 507.2537081 | 667.8237248 | 631.9747665 | YAL055W | PEX22 | n | NA | -0.12308785 | -0.183684008 |
| W303.asm02.HP0_G0000180 | 431.2217301 | 417.7627719 | 608.9215023 | 363.7652537 | 262.0909742 | 421.59226 | 360.2459654 | 375.3830726 | 306.9279806 | 459.3464135 | 553.3396577 | 555.7183844 | YAL054C | ACS1 | n | sc-like | -0.100897014 | 0.429977977 |
| W303.asm02.HP0_G0000190 | 5225.852073 | 5389.689445 | 7730.158414 | 2923.294588 | 3003.173433 | 3922.970029 | 3536.006515 | 3343.226526 | 3768.893048 | 1558.396114 | 1891.372192 | 1797.744207 | YAL053W | FLC2 | n | transgressi | 1.756944579 | 0.872266339 |
| W303.asm02.HP0_G0000200 | 780.0167714 | 713.6780686 | 1002.481096 | 868.3755499 | 747.3598521 | 1192.349392 | 1099.974331 | 1153.955371 | 1129.759261 | 532.6163936 | 691.6745721 | 628.1619474 | YAL051W | OAF1 | n | km-like | 0.414505325 | -0.138054487 |
| W303.asm02.HP0_G0000210 | 337.5155996 | 393.0268183 | 565.0867776 | 1656.702479 | 1794.579247 | 2222.547914 | 2176.34031 | 2176.29495 | 2244.374295 | 1384.614752 | 1849.633209 | 1610.916071 | YAL049C | AIM2 | n | transgressi | -1.845256066 | -2.085194277 |
| W303.asm02.HP0_G0000220 | 598.6780559 | 521.2873184 | 852.8712747 | 735.6366969 | 796.5734412 | 1048.575621 | 913.7306647 | 864.7713748 | 904.6151006 | 219.8099402 | 289.7877948 | 259.2716991 | YAL048C | GEM1 | n | km-like | 1.309360129 | -0.341116806 |
| W303.asm02.HP0_G0000230 | 631.6487314 | 692.6067007 | 872.8827795 | 563.3801701 | 587.1295622 | 770.7571317 | 603.3245537 | 578.3679934 | 641.105568 | 335.3510626 | 443.62576 | 366.030634 | YAL047C | SPC72 | n | km-like | 0.910041977 | 0.159341666 |
| W303.asm02.HP0_G0000240 | 394.7804571 | 408.6013076 | 476.4643993 | 518.7961279 | 505.8699152 | 696.1677319 | 573.5955177 | 564.4649166 | 544.1825215 | 647.2181573 | 871.7484693 | 751.1253635 | YAL046C | BOL3 | n | transgressi | -0.800245808 | -0.390202186 |
| W303.asm02.HP0_G0000250 | 511.0454709 | 495.6352184 | 730.8963885 | 546.1545174 | 580.2625498 | 732.9219289 | 664.5313925 | 763.7423503 | 776.393987 | 886.7546305 | 1203.275247 | 1044.712434 | YAL044W- | BOL1 | n | km-like | -0.820320321 | -0.077967285 |
| W303.asm02.HP0_G0000260 | 203.0299494 | 185.9777252 | 317.3252899 | 1902.927985 | 2187.143457 | 2765.212823 | 2590.798047 | 2514.603151 | 2792.595277 | 1373.342447 | 1642.130837 | 1698.610911 | YAL044C | GCV3 | n | transgressi | -2.671197922 | -3.22415817 |
| W303.asm02.HP0_G0000270 | 1134.885358 | 973.8636546 | 1741.000915 | 1203.76914 | 1141.068564 | 1543.676275 | 1402.510991 | 1387.527061 | 1522.499522 | 512.8898605 | 835.9721983 | 733.0144727 | YAL043C | PTA1 | n | km-like | 0.846391868 | -0.011324738 |
| W303.asm02.HP0_G0000280 | 2105.784988 | 1867.106423 | 2968.373208 | 3481.608389 | 3332.79003 | 4711.023254 | 3611.203488 | 3578.651959 | 3707.306528 | 2422.606136 | 3027.865066 | 2986.390563 | YAL042W | ERV46 | n | NA | -0.27110861 | -0.703854026 |
| W303.asm02.HP0_G0000290 | 1665.886764 | 1649.063573 | 2318.475767 | 1844.158111 | 1665.250513 | 2477.665282 | 2200.823046 | 2281.958333 | 2187.835852 | 743.9721053 | 1084.02101 | 879.8080082 | YAL041W | CDC24 | n | km-like | 1.019417198 | -0.069827474 |
| W303.asm02.HP0_G0000300 | 1411.665503 | 1323.831591 | 2027.832483 | 2048.839396 | 2056.670221 | 3275.447558 | 2306.623439 | 2338.497512 | 2246.393526 | 728.0030071 | 1070.903044 | 1010.397063 | YAL040C | CLN3 | n | intermedia | 0.72780217 | -0.591664705 |
| W303.asm02.HP0_G0000310 | 583.0603675 | 518.5388791 | 899.5647859 | 1104.468319 | 1125.045535 | 1478.815927 | 1394.641541 | 1352.305933 | 1466.970693 | 1211.772747 | 1574.155923 | 1802.510231 | YAL039C | CYC3 | n | NA | -1.148565368 | -0.855861046 |
| W303.asm02.HP0_G0000320 | 9366.274802 | 9582.891653 | 12280.39343 | 16124.22418 | 13822.15151 | 20802.87552 | 21171.44553 | 19528.26162 | 21736.00279 | 21592.09951 | 31044.26286 | 28190.07804 | YAL038W | CDC19 | n | NA | -1.329177094 | -0.67385232 |
| W303.asm02.HP0_G0000340 | 2320.094379 | 2152.944109 | 3281.886782 | 1872.529774 | 2087.571777 | 2669.003307 | 2799.775683 | 2562.800484 | 2814.806809 | 1485.126134 | 1706.528125 | 1695.751296 | YAL036C | RBG1 | n | km-like | 0.643617045 | 0.189080214 |
| W303.asm02.HP0_G0000350 | 5047.116306 | 4926.119351 | 7365.186684 | 4823.182751 | 4920.214402 | 6775.744322 | 7441.877212 | 6933.000947 | 7746.776414 | 2577.600324 | 3636.061673 | 3361.00004 | YAL035W | FUN12 | n | km-like | 0.825050956 | 0.055173346 |
| W303.asm02.HP0_G0000360 | 222.9858846 | 321.5673968 | 434.5355322 | 254.3316955 | 304.4375508 | 419.4302484 | 363.743499 | 368.8949702 | 382.6441106 | 416.1359124 | 558.1098271 | 427.9889444 | YAL034W- | MTW1 | n | NA | -0.490194068 | 0.000552325 |
| W303.asm02.HP0_G0000380 | 342.7214958 | 453.4924826 | 615.5920039 | 345.5263273 | 368.5296667 | 492.9386424 | 473.0414254 | 498.65702 | 439.1825544 | 610.5831672 | 748.9166056 | 827.3817455 | YAL033W | POP5 | n | km-like | -0.603575827 | 0.182834522 |
| W303.asm02.HP0_G0000390 | 493.6924838 | 535.9456613 | 782.3545436 | 327.287401 | 337.6281108 | 442.1313701 | 442.438006 | 407.8235851 | 359.4229641 | 208.5376356 | 373.2657604 | 264.9909277 | YAL032C | PRP45 | n | transgressi | 1.046878819 | 0.667396575 |
| W303.asm02.HP0_G0000400 | 614.2957443 | 607.4050828 | 958.6463714 | 851.1498973 | 765.6718853 | 1173.972293 | 998.5458555 | 955.6048096 | 907.6439458 | 373.8647701 | 566.4576237 | 441.3338113 | YAL031C | GIP4 | n | km-like | 0.628746528 | -0.304694546 |
| W303.asm02.HP0_G0000440 | 193.4858065 | 209.7975324 | 330.6662931 | 273.5838955 | 304.4375508 | 408.6201904 | 406.5882862 | 396.7011237 | 376.5864202 | 760.8805622 | 850.2827067 | 811.1772643 | YAL027W | SAW1 | n | km-like | -1.660674376 | -0.378709139 |
| W303.asm02.HP0_G0000450 | 1101.047033 | 1059.981419 | 1904.904668 | 2585.87445 | 2522.482564 | 3533.807943 | 2825.132801 | 2634.169611 | 2864.277947 | 906.4811636 | 1312.989145 | 1169.58226 | YAL026C | DRS2 | n | sc-like | 0.250282079 | -1.0479605 |
| W303.asm02.HP0_G0000460 | 3332.641178 | 3991.649993 | 5823.347888 | 1191.609856 | 1467.251655 | 1746.905364 | 2392.313013 | 2407.086024 | 2598.749184 | 3017.220205 | 3613.403368 | 3167.499471 | YAL025C | MAK16 | n | NA | 0.407548939 | 1.536899243 |
| W303.asm02.HP0_G0000470 | 736.6343036 | 1024.251708 | 1314.088813 | 1457.087562 | 1339.067423 | 1969.592558 | 1534.542887 | 1457.969317 | 1635.57641 | 294.019279 | 418.5823703 | 342.2005146 | YAL024C | LTE1 | n | intermedia | 1.483436141 | -0.593642909 |
| W303.asm02.HP0_G0000480 | 6676.561797 | 7539.885115 | 10715.68434 | 12326.4744 | 11838.72942 | 15792.41366 | 11467.53844 | 11274.46838 | 12426.34225 | 4233.689746 | 6488.623011 | 5295.052531 | YAL023C | PMT2 | n | intermedia | 0.613507028 | -0.648642512 |
| W303.asm02.HP0_G0000490 | 1542.680556 | 1907.416866 | 2503.343954 | 587.6987386 | 580.2625498 | 809.6733403 | 458.1769074 | 409.6773287 | 435.1440942 | 742.0933878 | 988.6176211 | 870.2759605 | YAL022C | FUN26 | n | transgressi | 1.151737342 | 1.549155404 |
| W303.asm02.HP0_G0000500 | 2097.976144 | 1899.171548 | 3627.799936 | 3294.152757 | 3000.884429 | 4413.74666 | 3910.242615 | 3660.216676 | 3964.758371 | 1147.896354 | 1651.671176 | 1626.167348 | YAL021C | CCR4 | n | km-like | 0.746181392 | -0.437358696 |
| W303.asm02.HP0_G0000510 | 287.191937 | 323.3996896 | 485.0407585 | 248.2520534 | 241.4899369 | 349.1648717 | 289.4209091 | 260.4509714 | 269.5672231 | 414.257195 | 485.3647428 | 503.2921217 | YAL020C | ATS1 | n | conserved | -0.340977232 | 0.332550038 |
| W303.asm02.HP0_G0000520 | 1153.973644 | 1136.021573 | 1889.657808 | 1052.791361 | 960.2372373 | 1392.335464 | 1358.791821 | 1444.993112 | 1572.980275 | 731.7604419 | 1041.089485 | 1006.584243 | YAL019W | FUN30 | n | km-like | 0.562093145 | 0.246341282 |
| W303.asm02.HP0_G0000540 | 975.2378766 | 1201.984116 | 1617.120171 | 1062.924098 | 1205.16068 | 1586.916507 | 1396.390308 | 1423.675061 | 1462.932233 | 557.0397202 | 881.2888082 | 710.1375581 | YAL017W | PSK1 | n | km-like | 0.785537959 | -0.017877409 |
| W303.asm02.HP0_G0000550 | 903.22298 | 1012.341805 | 1493.239427 | 3707.568422 | 3457.540755 | 5214.771954 | 4319.454051 | 4299.758207 | 4192.931376 | 2870.680245 | 3408.286081 | 2831.018185 | YAL016W | TPD3 | n | transgressi | -1.368053345 | -1.816348167 |
| W303.asm02.HP0_G0000570 | 1126.208865 | 1087.465812 | 1867.740445 | 377.9510853 | 352.5066378 | 501.5866888 | 477.4133425 | 448.6059436 | 431.1056339 | 918.692827 | 1101.909146 | 989.4265575 | YAL014C | SYN8 | n | sc-like | 0.420510576 | 1.680225982 |
| W303.asm02.HP0_G0000580 | 1301.474035 | 1326.58003 | 2165.05423 | 617.0836755 | 597.4300809 | 892.9107865 | 696.8835787 | 697.0075818 | 683.5094008 | 489.4058925 | 717.9105041 | 550.9523605 | YAL013W | DEP1 | n | NA | 1.386175819 | 1.14301332 |
| W303.asm02.HP0_G0000590 | 4790.292097 | 5697.514645 | 7919.791245 | 4724.895203 | 5357.414193 | 6374.691172 | 6863.909778 | 6780.993974 | 7163.218905 | 2501.512268 | 4281.227093 | 3443.928856 | YAL012W | CYS3 | n | km-like | 0.812448463 | 0.128911922 |
| W303.asm02.HP0_G0000600 | 404.3246001 | 491.0544862 | 757.5783949 | 1072.043561 | 1010.595328 | 1528.542194 | 1433.114411 | 1357.867164 | 1418.50917 | 226.3854512 | 329.1416929 | 288.8210471 | YAL011W | SWC3 | n | intermedia | 0.922783671 | -1.085270334 |
| W303.asm02.HP0_G0000610 | 860.7081615 | 1037.993905 | 1458.933991 | 505.6235699 | 560.8060146 | 717.7878478 | 533.3738808 | 585.7829677 | 699.6632419 | 422.7114235 | 577.190505 | 448.0062447 | YAL010C | MDM10 | n | NA | 1.163805043 | 0.876608866 |
| W303.asm02.HP0_G0000620 | 336.6479503 | 411.3497468 | 546.0282016 | 791.3667497 | 702.7242714 | 1075.600766 | 647.0437243 | 630.2728133 | 643.1247981 | 454.6496199 | 579.5755897 | 555.7183844 | YAL009W | SPO7 | n | NA | -0.284582748 | -0.957264919 |
| W303.asm02.HP0_G0000630 | 730.5607581 | 818.1187615 | 1240.713296 | 1465.193752 | 1357.379456 | 2065.802074 | 918.9769651 | 830.4771187 | 859.1824226 | 929.0257729 | 1075.673214 | 1141.939322 | YAL008W | FUN14 | n | NA | -0.166528275 | -0.776800299 |
| W303.asm02.HP0_G0000660 | 3993.789987 | 3572.054929 | 6121.614602 | 6785.893883 | 6223.80226 | 8711.8257 | 7798.625644 | 7219.404328 | 8678.651122 | 14120.44026 | 18501.10226 | 14087.41338 | YAL003W | EFB1 | n | transgressi | -1.709870823 | -0.631866666 |
| W303.asm02.HP0_G0000670 | 2049.38778 | 2286.701488 | 3460.084468 | 1136.893077 | 945.3587103 | 1493.950008 | 1128.828984 | 1140.979166 | 1104.518884 | 442.4379566 | 628.4698267 | 580.5017086 | YAL002W | VPS8 | n | transgressi | 2.164618311 | 1.085384516 |
| W303.asm02.HP0_G0000680 | 918.8406684 | 936.3016509 | 1389.370188 | 930.1852449 | 757.6603708 | 1237.751635 | 943.4597006 | 953.7510661 | 995.4804567 | 575.8268946 | 722.6806736 | 691.0734626 | YAL001C | TFC3 | n | km-like | 0.676988607 | 0.118884799 |
| W303.asm02.HP0_G0000700 | 1451.577373 | 1638.985962 | 2328.005055 | 964.6365502 | 965.9597476 | 1404.226527 | 1178.668838 | 1231.812601 | 1253.941914 | 1012.628699 | 1361.883382 | 896.0124894 | YAR002W | NUP60 | n | NA | 0.69732582 | 0.663957365 |
| W303.asm02.HP0_G0000710 | 2943.934266 | 3156.12445 | 4204.321859 | 3004.356483 | 2897.879243 | 3898.106896 | 3306.043678 | 3315.420372 | 3547.787348 | 4276.900247 | 5688.427084 | 5327.461493 | YAR002C-. | ERP1 | n | km-like | -0.551224166 | 0.057993679 |
| W303.asm02.HP0_G0000720 | 1786.490025 | 1612.417716 | 2468.085588 | 903.840129 | 862.9545613 | 1231.2656 | 1186.538289 | 1100.196808 | 1173.172709 | 2273.248099 | 2753.580322 | 2462.127936 | YAR003W | SWD1 | n | sc-like | -0.339647972 | 0.940441027 |
| W303.asm02.HP0_G0000730 | 1769.137038 | 1563.861955 | 2465.226802 | 2456.175418 | 2226.056527 | 3561.914094 | 2721.081176 | 2626.754637 | 2671.441469 | 1587.516235 | 1942.651514 | 1667.155153 | YAR007C | RFA1 | n | sc-like | 0.151609548 | -0.467245074 |
| W303.asm02.HP0_G0000740 | 465.060055 | 469.0669719 | 646.0857254 | 383.0174538 | 365.0961605 | 601.0392219 | 618.1890717 | 569.0992756 | 570.4325132 | 898.0269352 | 1116.219654 | 1088.559854 | YAR008W | SEN34 | n | km-like | -0.935015717 | 0.18623828 |
| W303.asm02.HP0_G0000750 | 1126.208865 | 975.6959474 | 1700.024977 | 1150.065635 | 1023.184851 | 1573.944437 | 1447.104545 | 1406.991369 | 1369.038032 | 924.3289793 | 1161.536264 | 1045.665639 | YAR014C | BUD14 | n | NA | 0.267804536 | 0.016073112 |
| W303.asm02.HP0_G0000760 | 275.9124953 | 258.3532931 | 431.6767458 | 1349.680551 | 1360.812962 | 2018.237819 | 1591.623774 | 1761.05639 | 1635.57641 | 947.8129473 | 1113.834569 | 923.6554279 | YAR015W | ADE1 | n | transgressi | -1.566385722 | -2.235939727 |
| W303.asm02.HP0_G0000770 | 602.1486533 | 590.914447 | 931.0114362 | 411.389117 | 420.0322599 | 563.2040191 | 470.4182752 | 458.8015332 | 481.5863873 | 452.7709025 | 614.1593183 | 500.4325074 | YAR018C | KIN3 | n | NA | 0.420855706 | 0.568918662 |
| W303.asm02.HP0_G0000780 | 3054.125734 | 3098.407225 | 4231.956795 | 445.8404224 | 422.3212641 | 621.578332 | 474.7901923 | 497.7301482 | 482.5960023 | 195.3866135 | 307.6759303 | 265.9441325 | YAR019C | CDC15 | n | transgressi | 3.715778223 | 2.745344985 |
| W303.asm02.HP0_G0000810 | 968.2966817 | 975.6959474 | 1455.122275 | 250.2786007 | 320.4605798 | 375.1090108 | 296.4159764 | 364.2606112 | 290.7691395 | 1382.736034 | 1611.124736 | 1599.477614 | YAR027W | UIP3 | n | intermedia | -0.417552615 | 1.800522203 |
| W303.asm02.HP0_G0000840 | 400.8540026 | 354.5486682 | 548.886988 | 187.4556321 | 141.9182568 | 255.1173676 | 132.9062785 | 128.835178 | 117.1153478 | 185.0536676 | 238.5084731 | 224.9563271 | YAR035W | YAT1 | n | conserved | 0.960017847 | 1.111616211 |
| W303.asm02.HP0_G0000850 | 1435.959685 | 1397.123305 | 2212.70067 | 4049.041654 | 3521.632871 | 5478.537368 | 3740.612303 | 3628.703036 | 3880.96032 | 557.0397202 | 942.1084688 | 793.0663736 | YAR042W | SWH1 | n | intermedia | 1.087760074 | -1.325476238 |
| W303.asm02.HP0_G0004930 | 5132.145943 | 4363.605444 | 3594.447428 | 274.5971692 | 258.657468 | 190.2570199 | 201.1081846 | 213.1805104 | 233.2210806 | 16.90845694 | 48.89423699 | 61.95831044 | YCL073C | GEX1 | y | transgressi | 6.607988893 | 4.073876481 |
| W303.asm02.HP0_G0004980 | 810.3844988 | 700.8520186 | 549.8399168 | 1121.693972 | 931.6246855 | 669.142587 | 239.5810547 | 302.1602017 | 191.8268629 | 650.9755921 | 506.8305054 | 582.4081181 | YCL064C | CHA1 | n | conserved | 0.232034974 | -0.348963685 |
| W303.asm02.HP0_G0004990 | 628.178134 | 705.4327507 | 507.9110497 | 641.402244 | 670.6782134 | 461.5894744 | 675.0239934 | 744.2780428 | 538.1248311 | 1901.262047 | 1862.751175 | 1887.345456 | YCL063W | VAC17 | n | km-like | -1.569304284 | 0.043231594 |
| W303.asm02.HP0_G0005000 | 2140.490962 | 1991.702338 | 1485.615997 | 1104.468319 | 1104.444498 | 793.4582534 | 1017.782291 | 1083.513116 | 1072.211202 | 2178.372869 | 1872.291514 | 1935.005695 | YCL061C | MRC1 | n | sc-like | -0.088568263 | 0.879685566 |
| W303.asm02.HP0_G0005010 | 4810.248032 | 4784.116655 | 3461.037397 | 3034.754693 | 3346.524055 | 2240.925013 | 2723.704326 | 2832.520173 | 3003.604826 | 4248.719485 | 3222.249472 | 3871.9178 | YCL059C | KRR1 | n | sc-like | 0.195703179 | 0.569086562 |
| W303.asm02.HP0_G0005030 | 262.0301056 | 279.424661 | 177.2447565 | 4106.798254 | 4771.429132 | 3200.858158 | 3450.31694 | 3593.481908 | 3803.21996 | 2904.497159 | 3316.460319 | 2515.507404 | YCL057C-/ | MIC10 | n | NA | -3.558379878 | -4.032187049 |
| W303.asm02.HP0_G0005040 | 1422.077295 | 1267.946658 | 956.7405138 | 3284.02002 | 2662.111816 | 2257.1401 | 2129.99799 | 2262.494025 | 2173.701241 | 5702.846782 | 4864.380309 | 4840.373852 | YCL057W | PRD1 | n | transgressi | -2.027353704 | -1.141159738 |
| W303.asm02.HP0_G0005060 | 1069.811656 | 1130.524694 | 742.3315341 | 84.10171604 | 92.70466772 | 67.02235928 | 90.06149135 | 76.00348631 | 49.47113832 | 2092.891225 | 1623.05016 | 1824.433941 | YCL055W | KAR4 | n | intermedia | -0.87499206 | 3.529623684 |
| W303.asm02.HP0_G0005070 | 4287.05547 | 4182.208451 | 3095.112738 | 2820.953945 | 2899.023745 | 2065.802074 | 2946.672096 | 3021.602017 | 3085.383647 | 7479.174119 | 5342.589798 | 6683.871889 | YCL054W | SPB1 | n | intermedia | -0.729453105 | 0.540542974 |
| W303.asm02.HP0_G0005080 | 1193.017865 | 1198.31953 | 829.0480548 | 1212.888604 | 1304.732361 | 868.0476532 | 1064.998995 | 1115.026757 | 970.24008 | 3778.100767 | 4438.642685 | 3694.621712 | YCL052C | PBN1 | n | km-like | -1.832962182 | -0.057189606 |
| W303.asm02.HP0_G0005100 | 7914.697429 | 7484.916329 | 5428.835366 | 4501.974992 | 5117.068758 | 3257.07046 | 3909.368231 | 3994.81739 | 3938.508379 | 8983.087428 | 9423.469773 | 8384.389209 | YCL050C | APA1 | n | sc-like | -0.350857682 | 0.668176947 |
| W303.asm02.HP0_G0005130 | 5996.324702 | 5315.481584 | 4164.29885 | 36.47785274 | 42.34657661 | 24.86313328 | 23.6083521 | 23.17179461 | 18.17307122 | 108.9656114 | 127.6020331 | 98.18009192 | YCL048W | SPS22 | n | transgressi | 5.513152151 | 7.191333457 |
| W303.asm02.HP0_G0005150 | 2417.271107 | 2161.189427 | 1590.438165 | 2140.034027 | 2018.901653 | 1499.355037 | 1628.976295 | 1727.689006 | 1530.576443 | 6080.468987 | 5668.153864 | 5513.336424 | YCL045C | EMC1 | n | km-like | -1.44210137 | 0.101177668 |
| W303.asm02.HP0_G0005160 | 1729.225167 | 1832.292859 | 1330.288603 | 1285.844309 | 1296.720846 | 819.4023925 | 872.6346443 | 1087.220603 | 941.9708581 | 2923.284333 | 1822.204735 | 2445.923455 | YCL044C | MGR1 | n | intermedia | -0.531717296 | 0.479181681 |
| W303.asm02.HP0_G0005170 | 11489.41278 | 11949.29788 | 8744.074656 | 33885.89865 | 35616.90444 | 25129.06071 | 24633.12946 | 25711.4233 | 27780.5682 | 23579.78256 | 21963.05275 | 22187.74757 | YCL043C | PDI1 | n | NA | -1.044584815 | -1.52718507 |
| W303.asm02.HP0_G0005180 | 1401.253711 | 1519.886927 | 992.9518081 | 230.013127 | 206.0103727 | 144.8547765 | 186.2436666 | 187.2281004 | 195.8653231 | 9475.311396 | 11549.77281 | 8591.234646 | YCL040W | GLK1 | n | intermedia | -2.85834661 | 2.694827974 |
| W303.asm02.HP0_G0005190 | 1090.635241 | 1226.720069 | 789.977974 | 940.3179817 | 912.1681503 | 646.4414653 | 615.5659215 | 633.9803005 | 562.3555927 | 1545.245092 | 1093.561349 | 1264.902738 | YCL039W | GID7 | n | conserved | -0.316208784 | 0.268113245 |
| W303.asm02.HP0_G0005200 | 1553.092348 | 1577.604152 | 1094.91519 | 762.9950865 | 789.7064287 | 553.4749669 | 595.455103 | 681.2507615 | 546.2017516 | 1363.009501 | 1527.64677 | 1149.56496 | YCL038C | ATG22 | n | sc-like | 0.062806366 | 0.974284926 |

| | | | | | | | | | | | | | | | | | | |
|---|---|---|---|---|---|---|---|---|---|---|---|---|---|---|---|---|---|---|
| W303.asm02.HP0_G0005210 | 603.883952 | 652.2962578 | 473.6056129 | 3174.586462 | 3422.061191 | 2363.078667 | 3516.77008 | 3591.628164 | 3602.306561 | 340.0478562 | 275.4772865 | 249.7396513 | YCL037C | SRO9 | n | intermedia | 0.965133862 | -2.329348213 |
| W303.asm02.HP0_G0005230 | 1549.62175 | 1506.14473 | 1068.233183 | 2649.710692 | 2845.232148 | 1875.545054 | 2379.197262 | 2562.800469 | 2436.201158 | 4404.653032 | 4980.056919 | 3974.863916 | YCL035C | GRX1 | n | transgressi | -1.645287862 | -0.811030674 |
| W303.asm02.HP0_G0005240 | 1634.651387 | 1671.967234 | 1140.655772 | 1421.622983 | 1697.296571 | 929.6649835 | 905.861214 | 942.6286046 | 800.6247487 | 3294.331027 | 3368.932183 | 2936.823915 | YCL034W | LSB5 | n | km-like | -1.06614794 | 0.107796087 |
| W303.asm02.HP0_G0005250 | 494.5601331 | 604.6566435 | 324.9487203 | 692.0659284 | 753.0823625 | 422.6732658 | 556.1078495 | 643.2490183 | 586.5863543 | 2067.52854 | 2437.556595 | 1913.081985 | YCL033C | MXR2 | n | km-like | -2.088414708 | -0.334672998 |
| W303.asm02.HP0_G0005260 | 944.8701491 | 1027.916294 | 714.6965989 | 1300.030141 | 1469.540659 | 899.3968213 | 1246.870744 | 1334.695369 | 1078.268892 | 1868.384492 | 1743.496939 | 1765.335245 | YCL032W | STE50 | n | transgressi | -0.965097845 | -0.407455676 |
| W303.asm02.HP0_G0005270 | 3494.023958 | 3479.524139 | 2571.954827 | 2404.49846 | 2575.129659 | 1781.49755 | 2052.177866 | 2368.157409 | 2127.258948 | 4589.7067 | 4246.643364 | 4174.083714 | YCL031C | RRP7 | n | intermedia | -0.433106891 | 0.466825819 |
| W303.asm02.HP0_G0005280 | 2759.124953 | 2747.523142 | 2004.962192 | 2868.577809 | 3146.236192 | 2074.45012 | 2355.58891 | 2550.75115 | 2476.585761 | 8093.514721 | 6979.950466 | 7064.200595 | YCL030C | HIS4 | n | km-like | -1.515696635 | -0.08624554 |
| W303.asm02.HP0_G0005290 | 1260.694515 | 1277.108123 | 931.964365 | 1074.070108 | 1118.178523 | 755.6230506 | 828.9154738 | 878.6744515 | 875.3362637 | 2390.667939 | 1826.974904 | 2092.284483 | YCL029C | BIK1 | n | km-like | -0.834115123 | 0.197005485 |
| W303.asm02.HP0_G0005310 | 417.3393404 | 417.7627719 | 287.7844972 | 182.3892637 | 201.4323644 | 147.0167881 | 149.5195633 | 150.1532291 | 122.1634232 | 495.9814035 | 292.1728796 | 407.0184393 | YCL027W | FUS1 | n | sc-like | -0.087108022 | 1.038596315 |
| W303.asm02.HP0_G0005320 | 1114.061774 | 884.9974509 | 683.2499486 | 1180.463846 | 1312.743875 | 813.9973635 | 1099.099948 | 1199.372089 | 1127.740031 | 512.8898605 | 685.7118602 | 503.2921217 | YCL026C-I | HBN1 | n | km-like | 0.629055197 | -0.253463953 |
| W303.asm02.HP0_G0005340 | 2411.197561 | 2492.834435 | 1696.213262 | 2233.255207 | 2794.874056 | 1629.075733 | 2665.995021 | 2791.737814 | 2607.83572 | 7306.332115 | 7807.574868 | 5850.770915 | YCL025C | AGP1 | n | km-like | -1.610161895 | -0.009542404 |
| W303.asm02.HP0_G0005370 | 841.6198757 | 923.4756009 | 560.3221336 | 591.7518333 | 718.7473004 | 415.1062252 | 479.1621093 | 440.2640975 | 419.9986907 | 1650.453269 | 1589.658973 | 1335.439891 | YCL018W | LEU2 | n | km-like | -0.934484892 | 0.380756718 |
| W303.asm02.HP0_G0005380 | 1766.53409 | 1506.14473 | 1251.195513 | 4033.842549 | 4533.372702 | 2923.039669 | 3436.326806 | 3208.830117 | 3474.085448 | 2802.107058 | 2627.170831 | 2620.359929 | YCL017C | NFS1 | n | NA | -0.805346649 | -1.314608304 |
| W303.asm02.HP0_G0005390 | 684.5753422 | 660.5415757 | 530.7813408 | 677.8800967 | 771.3943956 | 511.3157409 | 668.0289261 | 695.1538382 | 636.0574926 | 638.7639288 | 583.1532168 | 593.8465754 | YCL016C | DCC1 | n | NA | 0.045502472 | -0.05068883 |
| W303.asm02.HP0_G0005400 | 1866.313766 | 1875.351741 | 1349.347179 | 2362.954239 | 2342.795739 | 1680.964011 | 1809.099278 | 1750.860801 | 1983.893608 | 1221.166334 | 1190.157281 | 1160.050212 | YCL014W | BUD3 | n | km-like | 0.495583124 | -0.287837072 |
| W303.asm02.HP0_G0005430 | 731.4284074 | 752.1562186 | 557.4633472 | 437.7342329 | 521.8929442 | 313.4916805 | 466.0463581 | 436.5556104 | 454.3267804 | 829.4537487 | 647.5505045 | 699.6523056 | YCL010C | SGF29 | n | sc-like | -0.089552108 | 0.650266516 |
| W303.asm02.HP0_G0005440 | 3219.846761 | 3998.063018 | 2497.626381 | 3245.51562 | 4235.802163 | 2712.243539 | 2405.428764 | 2422.842844 | 2486.681912 | 3764.949745 | 3085.1071 | 3270.445586 | YCL009C | ILV6 | n | conserved | -0.056755023 | -0.054791455 |
| W303.asm02.HP0_G0005450 | 639.4575756 | 635.8056221 | 450.7353217 | 166.1768847 | 154.5077795 | 137.2877359 | 146.8964131 | 126.9814345 | 138.3172643 | 430.2262932 | 288.5952525 | 318.3703952 | YCL008C | STP22 | n | transgressi | 0.70387493 | 1.864261431 |
| W303.asm02.HP0_G0005470 | 1550.4894 | 1768.162609 | 1154.949704 | 240.1458639 | 254.0794597 | 183.7709851 | 176.6254491 | 181.6668697 | 155.4807204 | 607.7650911 | 527.1037256 | 539.5139032 | YCL005W | LDB16 | n | transgressi | 1.372842537 | 2.671368977 |
| W303.asm02.HP0_G0005480 | 1190.414917 | 1138.770012 | 799.507262 | 1622.251173 | 1539.355285 | 1090.734847 | 1206.649107 | 1140.979166 | 1153.990022 | 464.0432071 | 456.743726 | 455.6318829 | YCL004W | PGS1 | n | intermedia | 1.145707356 | -0.403626102 |
| W303.asm02.HP0_G0005500 | 724.4872126 | 818.1187615 | 524.1108392 | 3602.187958 | 3660.117622 | 2504.690427 | 2692.226523 | 2740.759866 | 2922.835621 | 1807.326175 | 1617.087448 | 1552.77058 | YCL001W | RER1 | n | transgressi | -1.226304798 | -2.195759837 |
| W303.asm02.HP0_G0005530 | 2107.520287 | 2006.360681 | 1468.463279 | 5264.970079 | 5427.228819 | 3725.145969 | 4171.683255 | 4180.191747 | 4290.864038 | 2810.561287 | 2785.778966 | 2703.288745 | YCR002C | CDC10 | n | transgressi | -0.554184606 | -1.33919518 |
| W303.asm02.HP0_G0005540 | 868.5170057 | 887.7458902 | 568.8984928 | 1891.781974 | 2099.016798 | 1382.606412 | 1804.727361 | 1753.641416 | 1705.239849 | 1347.979761 | 1462.05694 | 1228.680956 | YCR003W | MRPL32 | n | NA | -0.766758614 | -1.17477113 |
| W303.asm02.HP0_G0005550 | 963.9584349 | 1061.813712 | 769.9664693 | 3605.227779 | 4106.473429 | 2542.525629 | 2868.851972 | 2722.222431 | 2993.508676 | 2894.164213 | 3885.303027 | 2621.313134 | YCR004C | YCP4 | n | NA | -1.691122687 | -1.829806606 |
| W303.asm02.HP0_G0005580 | 367.8833271 | 363.7101325 | 233.4675557 | 181.37599 | 193.4208499 | 137.2877359 | 161.7609311 | 125.1276909 | 152.4518752 | 180.356874 | 170.5335583 | 180.1557027 | YCR007C | DFP3 | n | NA | 0.822427814 | 0.877615684 |
| W303.asm02.HP0_G0005590 | 3062.802228 | 2928.920135 | 2125.98415 | 757.928718 | 748.5043542 | 580.5001118 | 911.9818978 | 797.1097345 | 795.5766733 | 2381.274352 | 2277.755918 | 2164.728046 | YCR008W | SAT4 | n | sc-like | 0.243063537 | 1.924695438 |
| W303.asm02.HP0_G0005600 | 1578.254179 | 1505.228584 | 1074.903685 | 3304.285494 | 3616.626543 | 2249.573059 | 2646.758586 | 2596.167868 | 2783.508742 | 3308.421408 | 3560.931504 | 2858.661123 | YCR009C | RVS161 | n | NA | -1.183624043 | -1.108590485 |
| W303.asm02.HP0_G0005610 | 378.2951194 | 409.517454 | 268.7259212 | 571.4863596 | 571.1065332 | 353.4888949 | 327.0193958 | 338.3082013 | 388.701801 | 74.20933878 | 93.01830452 | 51.4730579 | YCR010C | ADY2 | n | km-like | 2.179911618 | -0.446425357 |
| W303.asm02.HP0_G0005620 | 4433.688211 | 4414.909644 | 3183.735116 | 1387.171678 | 1363.101966 | 968.5811921 | 1080.737896 | 1048.291988 | 1016.682373 | 2821.833591 | 2826.325407 | 2709.961178 | YCR011C | ADP1 | n | transgressi | 0.509682001 | 1.661281048 |
| W303.asm02.HP0_G0005630 | 11288.98578 | 11944.71715 | 7629.147962 | 23916.29884 | 27539.00882 | 18078.74091 | 20743.87205 | 18911.89189 | 21700.66626 | 48331.8848 | 48920.47292 | 45721.42028 | YCR012W | PGK1 | n | transgressi | -2.15444235 | -1.141077538 |
| W303.asm02.HP0_G0005640 | 1396.915464 | 1414.530087 | 933.8702226 | 688.0128336 | 804.5849557 | 530.7738453 | 631.3048229 | 588.563583 | 617.8844214 | 587.0991992 | 500.8677936 | 526.1690363 | YCR014C | POL4 | n | NA | 1.171761806 | 0.860107282 |
| W303.asm02.HP0_G0005650 | 684.5753422 | 586.3337149 | 442.1589625 | 178.336169 | 222.0334017 | 149.1787997 | 200.2338012 | 193.7162029 | 161.5384108 | 510.0717843 | 549.7620306 | 462.3043163 | YCR015C | CTO1 | n | sc-like | 0.164637031 | 1.597455451 |
| W303.asm02.HP0_G0005660 | 2301.006093 | 2543.222488 | 1630.461174 | 1028.472793 | 1134.201552 | 815.0783693 | 1234.629377 | 1139.125423 | 1280.191906 | 1734.056195 | 1322.529483 | 1444.105236 | YCR016W | RBP95 | n | NA | 0.505665834 | 1.090074878 |
| W303.asm02.HP0_G0005670 | 1619.033699 | 1702.200066 | 1189.255141 | 2178.538428 | 2187.143457 | 1500.436043 | 1722.53532 | 1649.831776 | 1791.05713 | 3011.584052 | 2854.946423 | 2840.550232 | YCR017C | CWH43 | n | km-like | -0.919456327 | -0.339672091 |
| W303.asm02.HP0_G0005800 | 1204.297307 | 1212.061726 | 860.4947051 | 1681.021047 | 1738.498645 | 1243.156664 | 1469.838514 | 1519.142854 | 1474.037999 | 951.5703825 | 815.6989781 | 888.3868512 | YCR028C | FEN2 | n | sc-like | 0.29324469 | -0.477077605 |
| W303.asm02.HP0_G0005810 | 1075.885202 | 1135.105426 | 755.6725373 | 3178.639557 | 3591.447498 | 2297.137314 | 2881.09334 | 2893.693711 | 3041.970199 | 2931.738561 | 2947.964728 | 2622.266339 | YCR028C-. | RIM1 | n | NA | -1.47164066 | -1.57493305 |
| W303.asm02.HP0_G0005840 | 2733.095473 | 2465.350042 | 1882.034377 | 1167.291288 | 1257.807776 | 898.3158155 | 897.9917633 | 908.3343486 | 918.7497116 | 1180.77391 | 1266.479992 | 1117.155997 | YCR032W | BPH1 | n | transgressi | 0.961682797 | 1.065721508 |
| W303.asm02.HP0_G0005850 | 2308.814937 | 2188.67382 | 1773.400494 | 2583.847902 | 2505.315033 | 1889.598129 | 2202.571813 | 2209.662334 | 2074.758964 | 1545.245092 | 1385.734229 | 1385.006539 | YCR033W | SNT1 | n | km-like | 0.524366263 | -0.128454121 |
| W303.asm02.HP0_G0005860 | 2734.830771 | 2682.476746 | 1704.789621 | 6118.146523 | 6945.983067 | 4440.771805 | 4954.256408 | 4824.367637 | 5073.315715 | 4477.923012 | 4163.165398 | 3939.595339 | YCR034W | ELO2 | n | NA | -0.790302168 | -1.262111344 |
| W303.asm02.HP0_G0005870 | 2978.64024 | 3023.283217 | 2286.076188 | 1427.702625 | 1444.361613 | 989.1203023 | 1357.917438 | 1435.724394 | 1424.56686 | 2895.103571 | 2194.277953 | 2440.204226 | YCR035C | RRP43 | n | sc-like | 0.13365237 | 1.074041284 |
| W303.asm02.HP0_G0005880 | 3582.524192 | 3839.569686 | 2631.036413 | 1214.915151 | 1141.068564 | 755.6230506 | 875.2577946 | 868.4788619 | 872.3074185 | 1554.63868 | 1880.639311 | 1584.226338 | YCR036W | RBK1 | n | transgressi | 0.967082482 | 1.652282067 |
| W303.asm02.HP0_G0005890 | 905.825928 | 877.6682795 | 620.3566479 | 3549.497726 | 3725.35424 | 2649.545203 | 4232.01571 | 4021.696672 | 3926.392998 | 1765.055033 | 1552.69016 | 1689.078863 | YCR037C | PHO87 | n | transgressi | -1.026463882 | -2.007701423 |
| W303.asm02.HP0_G0005900 | 311.4861189 | 276.6762217 | 223.9382677 | 2581.821355 | 2464.112958 | 1800.955654 | 1517.055218 | 1616.464392 | 1718.364845 | 231.0822448 | 224.1979647 | 222.0967128 | YCR038C | BUD5 | n | sc-like | 0.253243266 | -3.03032042 |
| W303.asm02.HP0_G0005920 | 1812.519505 | 1791.982416 | 1362.688182 | 2298.104723 | 2428.633394 | 1707.989156 | 1789.862843 | 1956.626337 | 1877.884026 | 1768.812467 | 1824.589819 | 1554.67699 | YCR042C | TAF2 | n | NA | -0.049720352 | -0.33630418 |
| W303.asm02.HP0_G0005930 | 227.3241314 | 287.6699789 | 182.0094005 | 1455.061015 | 1634.348957 | 976.1482327 | 996.7970867 | 1053.853219 | 1043.94198 | 1291.618238 | 1540.764736 | 1291.592471 | YCR043C | | n | NA | -2.493463151 | -2.4872248 |
| W303.asm02.HP0_G0005940 | 286.3242876 | 295.9152967 | 182.9623293 | 2081.264154 | 2442.367419 | 1519.894148 | 1652.584647 | 1694.321622 | 1676.970627 | 965.6607629 | 1033.934231 | 975.1284858 | YCR044C | PER1 | n | transgressi | -1.895351746 | -2.924748985 |
| W303.asm02.HP0_G0005950 | 1724.019271 | 1837.789738 | 1236.901581 | 20.26547374 | 24.03454348 | 11.89106374 | 7.8694507 | 8.341846059 | 10.09615068 | 401.1061729 | 362.5328792 | 387.9543438 | YCR045C | RRT12 | n | transgressi | 1.996483052 | 6.378717415 |
| W303.asm02.HP0_G0005960 | 707.1342254 | 831.860958 | 526.9696256 | 1834.025374 | 2007.456632 | 1234.508618 | 1538.914804 | 1606.268802 | 1620.432184 | 1823.295273 | 2110.799987 | 1705.283344 | YCR046C | IMG1 | n | NA | -1.397051346 | -1.259738409 |
| W303.asm02.HP0_G0005970 | 1295.400489 | 1671.051087 | 1027.257245 | 1379.065488 | 1494.719704 | 1043.170592 | 1440.983862 | 1352.305933 | 1522.499522 | 1728.420043 | 1093.561349 | 1558.489809 | YCR047C | BUD23 | n | NA | -0.128030152 | 0.021905418 |
| W303.asm02.HP0_G0005990 | 2330.506171 | 2846.466956 | 1795.317857 | 1202.755867 | 1299.00985 | 932.9080009 | 1244.247594 | 1164.150961 | 1299.374592 | 2482.725094 | 2460.2149 | 2419.233721 | YCR051W | | n | sc-like | -0.075833636 | 0.995043092 |
| W303.asm02.HP0_G0006010 | 7623.167245 | 7549.962726 | 5762.360445 | 7504.304928 | 8669.603185 | 5504.481507 | 6067.34649 | 6223.01716 | 6620.045999 | 7875.583498 | 6810.60945 | 7090.890328 | YCR053W | THR4 | n | NA | -0.055041625 | -0.040296392 |
| W303.asm02.HP0_G0006020 | 977.8408246 | 888.6620366 | 724.2258869 | 864.3224552 | 759.9493749 | 589.1481582 | 707.3761796 | 705.3494279 | 719.8555432 | 883.9365544 | 633.2399961 | 765.4234351 | YCR054C | CTR86 | n | NA | 0.175535461 | 0.189448163 |
| W303.asm02.HP0_G0006030 | 4432.820562 | 4592.642051 | 3299.99243 | 2812.847756 | 2919.624782 | 2136.067451 | 2735.07131 | 2586.89915 | 2924.854851 | 3068.884934 | 1772.117955 | 2810.04768 | YCR057C | PWP2 | n | NA | 0.659446141 | 0.612474121 |
| W303.asm02.HP0_G0006040 | 2066.740767 | 2641.250156 | 1627.602388 | 624.1765913 | 738.2038356 | 392.4051035 | 561.35415 | 554.269327 | 516.9229146 | 1665.483008 | 1649.286092 | 1513.689184 | YCR059C | YIH1 | n | sc-like | 0.37455232 | 1.80176844 |
| W303.asm02.HP0_G0006050 | 980.4437727 | 1215.726312 | 761.3901101 | 287.7697272 | 231.1894183 | 178.3659561 | 233.4603708 | 200.2043054 | 224.134545 | 1095.292266 | 1167.498976 | 967.5028476 | YCR060W | TAH1 | n | sc-like | -0.12168679 | 2.032590182 |
| W303.asm02.HP0_G0006060 | 519.7219645 | 567.0946399 | 415.4769562 | 1504.711426 | 1144.502071 | 894.7990647 | 1280.971697 | 1219.763268 | 1745.624452 | 605.8863736 | 565.2650813 | 540.467108 | YCR061W | TVS1 | n | sc-like | -0.180244994 | -1.221785752 |
| W303.asm02.HP0_G0006070 | 311.4861189 | 289.5022717 | 191.5386885 | 52.69023174 | 37.76856833 | 51.88827815 | 69.95067289 | 58.39292241 | 58.55767392 | 231.0822448 | 188.4216938 | 178.2492931 | YCR063W | BUD31 | n | sc-like | 0.385623829 | 2.390186436 |
| W303.asm02.HP0_G0006080 | 1025.561539 | 1282.605001 | 718.5083141 | 837.9773393 | 923.613171 | 637.7934189 | 1130.577751 | 1099.269936 | 1085.336198 | 793.7581174 | 732.2210125 | 721.5760154 | YCR065W | HCM1 | n | conserved | 0.412293636 | 0.288943255 |
| W303.asm02.HP0_G0006090 | 699.3253812 | 620.2311328 | 531.7342696 | 868.3755499 | 832.0530054 | 599.9582161 | 685.5165943 | 679.3970179 | 684.5190159 | 450.89285 | 393.5389807 | 394.6267773 | YCR066W | RAD18 | n | km-like | 0.560511967 | -0.273550775 |
| W303.asm02.HP0_G0006100 | 2580.389186 | 2524.89956 | 1817.235219 | 6067.482839 | 5516.49998 | 3827.841519 | 3858.653993 | 3902.130212 | 4281.777502 | 2127.647498 | 1982.005412 | 2070.360773 | YCR067C | SED4 | n | sc-like | 0.157981271 | -1.124893468 |
| W303.asm02.HP0_G0006110 | 1278.915151 | 1238.629973 | 926.2467922 | 1135.879803 | 1106.733502 | 779.4051781 | 807.0558885 | 805.4515806 | 765.2882213 | 563.6152313 | 451.9735566 | 473.7427737 | YCR068W | ATG15 | n | km-like | 1.169715406 | 0.155317897 |
| W303.asm02.HP0_G0006120 | 1064.60576 | 1130.524694 | 816.6599804 | 2986.117556 | 3288.154449 | 2191.198746 | 2504.23409 | 2396.890434 | 2448.316539 | 1637.302247 | 1650.478634 | 1454.590488 | YCR069W | CPR4 | n | transgressi | -0.634161708 | -1.459584404 |
| W303.asm02.HP0_G0006130 | 479.8100941 | 462.6539469 | 336.3838659 | 1404.39733 | 1390.570016 | 927.5029719 | 1092.979264 | 1157.662859 | 1204.470776 | 1298.193749 | 1567.000668 | 1241.072618 | YCR071C | IMG2 | n | NA | -1.62862866 | -1.502886619 |
| W303.asm02.HP0_G0006140 | 1560.033543 | 1591.346348 | 1133.98527 | 1551.322015 | 1719.04211 | 1259.371751 | 1843.200231 | 1741.592083 | 1923.316704 | 1169.501605 | 670.2088095 | 959.8772094 | YCR072C | RSA4 | n | km-like | 0.58812878 | -0.062793407 |
| W303.asm02.HP0_G0006170 | 222.1182352 | 271.1793431 | 141.986391 | 825.8180551 | 924.7576731 | 615.0922972 | 687.2653612 | 734.0824532 | 747.1151501 | 401.1061729 | 460.3213531 | 372.7030674 | YCR075C | ERS1 | n | transgressi | -0.91301861 | -1.842689911 |
| W303.asm02.HP0_G0006180 | 131.0150528 | 120.0151823 | 69.5638023 | 290.8095482 | 342.2061191 | 193.5000373 | 264.9381736 | 305.8676888 | 260.4806875 | 169.0845694 | 228.9681342 | 134.4018734 | YCR075W- | EGO2 | n | transgressi | -0.677749599 | -1.299012831 |
| W303.asm02.HP0_G0006190 | 488.4865876 | 685.2775293 | 458.3587521 | 355.6590642 | 349.0731315 | 267.0084313 | 285.9233754 | 266.0122021 | 221.1056998 | 533.5557523 | 547.3769458 | 518.5433981 | YCR076C | FUB1 | n | sc-like | 0.028261838 | 0.718844217 |
| W303.asm02.HP0_G0006200 | 1979.108182 | 1836.873591 | 1398.899476 | 4901.204825 | 5299.044587 | 3660.285621 | 4034.405059 | 4005.01298 | 4601.825478 | 2687.505294 | 1817.434565 | 2112.301784 | YCR077C | PAT1 | n | sc-like | -0.330967704 | -1.374567763 |
| W303.asm02.HP0_G0006210 | 1584.327725 | 1780.072513 | 1234.995723 | 387.0705485 | 393.7087123 | 259.4413908 | 340.1351469 | 312.3557913 | 321.0575915 | 661.308538 | 628.4698267 | 540.467108 | YCR079W | PTC6 | n | transgressi | 1.285905204 | 2.099196065 |
| W303.asm02.HP0_G0006220 | 2390.373977 | 2313.269734 | 1677.154686 | 1351.707099 | 1171.97012 | 936.1510183 | 930.3439495 | 931.5061432 | 920.7689417 | 958.1458932 | 797.8108426 | 839.7734076 | YCR081W | SRB8 | n | NA | 1.258896238 | 0.858562292 |
| W303.asm02.HP0_G0006230 | 1146.1648 | 1134.18928 | 740.4256765 | 1211.87533 | 1420.32707 | 865.8856416 | 1085.109813 | 1121.514859 | 1107.547729 | 1590.334311 | 1727.993888 | 1393.585382 | YCR082W | AHC2 | n | km-like | -0.61523857 | -0.172458937 |
| W303.asm02.HP0_G0006240 | 356.6038855 | 355.4648146 | 234.4204845 | 1195.662951 | 1406.593045 | 898.3158155 | 1097.351181 | 1069.610039 | 1172.163094 | 788.1219651 | 884.8664353 | 689.167053 | YCR083W | TRX3 | n | transgressi | -1.269561718 | -1.840321865 |
| W303.asm02.HP0_G0006250 | 4194.216989 | 4269.242361 | 2962.655635 | 6604.517893 | 7376.315845 | 4767.235555 | 5330.241274 | 5232.191222 | 5444.85406 | 3419.265736 | 2269.408122 | 2732.838093 | YCR084C | TUP1 | n | intermedia | 0.422532022 | -0.683424555 |
| W303.asm02.HP0_G0006260 | 348.7950413 | 391.1945254 | 245.85563 | 875.4684658 | 1122.756581 | 692.9247145 | 906.7355974 | 842.5264519 | 886.4420294 | 447.1347501 | 397.1166078 | 390.8139581 | YCR086W | CSM1 | n | sc-like | -0.311314729 | -1.40793416 |
| W303.asm02.HP0_G0006270 | 1416.871399 | 1586.765616 | 1062.51561 | 966.6630976 | 1187.993149 | 669.142587 | 1188.287056 | 1125.222346 | 1291.297672 | 1073.687016 | 857.4379609 | 1043.75923 | YCR087C-A | | n | NA | 0.431448964 | 0.484179823 |
| W303.asm02.HP0_G0006280 | 2904.890045 | 2866.622178 | 2075.478923 | 4014.590349 | 4478.436602 | 2866.827368 | 3285.058476 | 3146.729708 | 3173.220158 | 2175.554793 | 1758.999989 | 1883.532637 | YCR088W | ABP1 | n | intermedia | 0.416431369 | -0.500348311 |
| W303.asm02.HP0_G0006300 | 1605.151309 | 1577.604152 | 1116.832552 | 1320.295614 | 1418.038066 | 888.5867633 | 1094.728031 | 1143.759782 | 1122.691955 | 1111.261364 | 1075.673214 | 1016.116291 | YCR090C | | n | NA | 0.409510129 | 0.205736324 |
| W303.asm02.HP0_G0006320 | 1047.252773 | 1103.956448 | 691.8263078 | 1278.751393 | 1290.998336 | 902.6398387 | 1000.294622 | 1105.758039 | 1114.615035 | 833.2111836 | 628.4698267 | 773.0490733 | YCR092C | MSH3 | n | conserved | 0.33268752 | -0.243130864 |
| W303.asm02.HP0_G0006330 | 4091.834365 | 3996.230725 | 3082.724663 | 6581.212598 | 7031.820722 | 4986.679731 | 5844.37872 | 5629.819218 | 5768.940497 | 3180.668622 | 3149.504388 | 2904.414952 | YCR093W | CDC39 | n | sc-like | 0.267072119 | -0.715689363 |
| W303.asm02.HP0_G0006340 | 1534.871171 | 1496.067119 | 1087.291759 | 1860.37049 | 1873.5489 | 1231.2656 | 1602.744793 | 1521.92347 | 1492.21107 | 1089.656114 | 1136.492874 | 1063.77653 | YCR094W | CDC50 | n | conserved | 0.313011815 | -0.229286839 |
| W303.asm02.HP0_G0006350 | 678.5017967 | 617.4826935 | 448.8294641 | 758.9419917 | 867.5325695 | 551.3129554 | 640.048657 | 573.7336345 | 610.8171159 | 823.8175964 | 728.6433854 | 783.5343258 | YCR095C | OCA4 | n | NA | -0.404890103 | -0.274414588 |
| W303.asm02.HP0_G0006450 | 944.0024997 | 927.1401867 | 639.4152239 | 1712.432531 | 1872.405388 | 1228.022583 | 1606.242326 | 1609.976289 | 1767.835983 | 1811.08361 | 1504.988465 | 1583.273133 | YCR106W | RDS1 | n | NA | -0.93091268 | -0.910142187 |

**YPD_gw**

| | Normalized read counts (genome-wide) | | | | | | | | | | | | | |
|---|---|---|---|---|---|---|---|---|---|---|---|---|---|---|
| W303_gene | SC.RI.1 | SC.RI.2 | SC.RI.3 | Km.V.1 | Km.V.2 | Km.V.3 | KS.1.1 | KS.1.2 | KS.1.3 | padj.sc.v.km | log2FC.sc.v.km | locus_tag | gene | ambiguous |
| W303.asm02.HP0_G0000040 | 1450.691 | 778.961 | 1078.988 | 1958.405 | 1724.403 | 2074.644 | 4521.875 | 3948.362 | 3578.684 | 0.000989678 | -0.799089962 | YAL067C | SEO1 | n |
| W303.asm02.HP0_G0000100 | 2996.683 | 2940.021 | 2853.228 | 8900.703 | 7733.345 | 17138.79 | 10285.63 | 24886.39 | 13775.3 | 5.40E-07 | -1.941946417 | YAL063C | FLO9 | n |
| W303.asm02.HP0_G0000110 | 1427.395 | 1148.411 | 1315.05 | 6396.251 | 6702.348 | 5445.276 | 8695.628 | 5590.202 | 4725.729 | 1.72E-20 | -2.252716089 | YAL062W | GDH3 | n |
| W303.asm02.HP0_G0000120 | 4640.094 | 5649.693 | 4590.819 | 2082.759 | 2203.452 | 6510.326 | 10132.56 | 10923.77 | 10533.91 | 0.294674605 | 0.462843037 | YAL061W | BDH2 | n |
| W303.asm02.HP0_G0000130 | 15894.07 | 14879.27 | 14078.58 | 5355.567 | 5099.443 | 17847.56 | 46258.18 | 34302.54 | 35685.18 | 0.161603944 | 0.664229828 | YAL060W | BDH1 | n |
| W303.asm02.HP0_G0000140 | 993.247 | 1027.116 | 1142.585 | 3533.326 | 3813.3 | 1837.629 | 2357.844 | 1858.748 | 2497.424 | 4.90E-08 | -1.537845626 | YAL059W | ECM1 | n |
| W303.asm02.HP0_G0000150 | 1328.918 | 1179.569 | 1382.959 | 3643.786 | 3205.81 | 5540.234 | 4241.65 | 4609.823 | 4354.943 | 1.71E-14 | -1.670796651 | YAL058W | CNE1 | n |
| W303.asm02.HP0_G0000170 | 1422.101 | 1166.216 | 1136.117 | 1204.638 | 1188.944 | 1754.066 | 2905.95 | 2806.913 | 2853.857 | 0.464267741 | -0.155290737 | YAL055W | PEX22 | n |
| W303.asm02.HP0_G0000180 | 1248.442 | 1457.77 | 1288.103 | 614.1291 | 608.3576 | 824.2361 | 824.628 | 939.5749 | 752.337 | 4.89E-08 | 0.964385337 | YAL054C | ACS1 | n |
| W303.asm02.HP0_G0000190 | 8888.396 | 8835.643 | 8787.123 | 3627.113 | 3146.797 | 3251.365 | 4557.675 | 4561.502 | 3809.529 | 2.76E-35 | 1.402769463 | YAL053W | FLC2 | n |
| W303.asm02.HP0_G0000200 | 4421.961 | 4844.024 | 4274.992 | 4722.68 | 4717.592 | 7491.053 | 4942.83 | 4371.44 | 4542.728 | 0.142813352 | -0.322394302 | YAL051W | OAF1 | n |
| W303.asm02.HP0_G0000210 | 3738.971 | 3384.029 | 3379.248 | 3128.307 | 3682.256 | 4026.982 | 2401.05 | 6309.648 | 4354.943 | 0.90337307 | -0.045313166 | YAL049C | AIM2 | n |
| W303.asm02.HP0_G0000220 | 1667.765 | 1697.022 | 1570.515 | 1927.837 | 1912.725 | 1801.924 | 2350.437 | 2059.548 | 2133.814 | 0.053574234 | -0.193223177 | YAL048C | GEM1 | n |
| W303.asm02.HP0_G0000230 | 1086.43 | 1169.554 | 1193.247 | 1086.536 | 1017.111 | 1172.163 | 230.8465 | 497.1694 | 333.7075 | 0.801954069 | 0.074340392 | YAL047C | SPC72 | n |
| W303.asm02.HP0_G0000240 | 786.7617 | 585.3335 | 670.4601 | 746.1251 | 869.5782 | 495.3013 | 128.3852 | 254.4906 | 159.0792 | 0.897190289 | -0.047246501 | YAL046C | BOL3 | n |
| W303.asm02.HP0_G0000250 | 1436.926 | 1311.993 | 1375.413 | 1123.356 | 1218.451 | 981.4867 | 320.963 | 478.9148 | 484.4142 | 0.115393122 | 0.311762282 | YAL044W- | BOL1 | n |
| W303.asm02.HP0_G0000260 | 1688.943 | 1097.222 | 1388.348 | 4308.629 | 4957.116 | 5406.533 | 4898.389 | 7600.356 | 6265.09 | 1.66E-15 | -1.813277164 | YAL044C | GCV3 | n |
| W303.asm02.HP0_G0000270 | 2207.804 | 2094.292 | 2352 | 3312.407 | 2910.744 | 2530.443 | 4044.134 | 3805.547 | 3624.136 | 0.004064098 | -0.395722798 | YAL043C | PTA1 | n |
| W303.asm02.HP0_G0000280 | 8975.225 | 8145.706 | 9024.264 | 4321.134 | 4605.64 | 4039.137 | 2503.511 | 2776.847 | 3079.917 | 1.17E-17 | 1.011920145 | YAL042W | ERV46 | n |
| W303.asm02.HP0_G0000290 | 2706.545 | 2857.674 | 2583.751 | 5141.594 | 4778.341 | 5295.622 | 4994.678 | 4519.624 | 5167.084 | 1.09E-19 | -0.901183819 | YAL041W | CDC24 | n |
| W303.asm02.HP0_G0000300 | 993.247 | 1032.68 | 1108.092 | 6070.43 | 6640.731 | 6710.118 | 4162.643 | 4197.484 | 4886.004 | 9.74E-123 | -2.631518619 | YAL040C | CLN3 | n |
| W303.asm02.HP0_G0000310 | 13232 | 13560.6 | 12533.94 | 3015.763 | 3113.819 | 2750.746 | 3492.324 | 2608.26 | 2632.582 | 3.33E-49 | 2.146863442 | YAL039C | CYC3 | n |
| W303.asm02.HP0_G0000320 | 103677.8 | 107471 | 115152.1 | 57742.03 | 61553.47 | 91939.17 | 134587.2 | 118543.2 | 130667.4 | 0.0016842 | 0.627356095 | YAL038W | CDC19 | n |
| W303.asm02.HP0_G0000340 | 492.3879 | 475.1662 | 506.6178 | 5202.729 | 5366.738 | 3524.084 | 4308.311 | 2887.448 | 3303.585 | 3.39E-49 | -3.256999232 | YAL036C | RBG1 | n |
| W303.asm02.HP0_G0000350 | 17261.11 | 17174.98 | 18117.51 | 19360.35 | 19312.1 | 17411.51 | 11310.24 | 10833.57 | 10618.84 | 0.287452671 | -0.093789078 | YAL035W | FUN12 | n |
| W303.asm02.HP0_G0000360 | 786.7617 | 786.7506 | 761.0046 | 687.769 | 767.1728 | 670.7839 | 512.3063 | 614.2136 | 424.6099 | 0.445435395 | 0.135579789 | YAL034W- | MTW1 | n |
| W303.asm02.HP0_G0000380 | 794.174 | 771.1714 | 811.6663 | 725.9784 | 775.8512 | 533.2846 | 430.8311 | 545.4904 | 562.1596 | 0.273395311 | 0.224379847 | YAL033W | POP5 | n |
| W303.asm02.HP0_G0000390 | 732.7579 | 888.0155 | 674.7717 | 1181.017 | 1051.825 | 1041.5 | 1769 | 1493.656 | 1850.342 | 0.001557763 | -0.51284077 | YAL032C | PRP45 | n |
| W303.asm02.HP0_G0000400 | 2001.319 | 2096.518 | 1837.837 | 4421.868 | 4225.525 | 5552.389 | 3899.7 | 3682.06 | 3450.703 | 1.47E-18 | -1.258469747 | YAL031C | GIP4 | n |
| W303.asm02.HP0_G0000440 | 1930.372 | 1745.985 | 1677.228 | 617.6026 | 673.4458 | 685.2175 | 448.1137 | 926.6893 | 492.7867 | 2.51E-07 | 1.437830286 | YAL027W | SAW1 | n |
| W303.asm02.HP0_G0000450 | 3871.334 | 3772.397 | 3787.776 | 7064.568 | 6801.282 | 7298.098 | 6952.552 | 8153.363 | 8037.687 | 1.14E-20 | -0.888623876 | YAL026C | DRS2 | n |
| W303.asm02.HP0_G0000460 | 1872.133 | 1967.433 | 2246.365 | 3771.614 | 3170.229 | 571.2678 | 3821.929 | 1138.228 | 2838.308 | 0.656228957 | -0.303933956 | YAL025C | MAK16 | n |
| W303.asm02.HP0_G0000470 | 826.9999 | 882.4515 | 870.9514 | 6678.306 | 6328.308 | 6820.269 | 6076.077 | 6537.294 | 7300.899 | 3.03E-171 | -2.941949891 | YAL024C | LTE1 | n |
| W303.asm02.HP0_G0000480 | 6989.79 | 6012.466 | 7381.529 | 11223.83 | 12284.31 | 13850.96 | 12953.33 | 14541.4 | 13756.17 | 6.60E-11 | -0.873996062 | YAL023C | PMT2 | n |
| W303.asm02.HP0_G0000490 | 1694.238 | 1428.837 | 1740.825 | 814.902 | 891.2743 | 1168.364 | 1487.54 | 1306.815 | 1188.908 | 5.93E-05 | 0.758891769 | YAL022C | FUN26 | n |
| W303.asm02.HP0_G0000500 | 12146.63 | 11831.3 | 12651.43 | 10555.52 | 9828.316 | 11377.5 | 7198.213 | 8647.311 | 7352.33 | 0.091101949 | 0.205695808 | YAL021C | CCR4 | n |
| W303.asm02.HP0_G0000510 | 1351.155 | 992.6188 | 1240.675 | 1199.08 | 1313.046 | 1346.126 | 4962.582 | 3226.769 | 5280.712 | 0.674216182 | -0.106022 | YAL020C | ATS1 | n |
| W303.asm02.HP0_G0000520 | 1569.288 | 1552.358 | 1818.434 | 3363.816 | 2929.836 | 2071.605 | 2678.807 | 2074.581 | 2583.542 | 0.000449425 | -0.759909417 | YAL019W | FUN30 | n |
| W303.asm02.HP0_G0000540 | 9554.443 | 9398.72 | 8724.605 | 5873.825 | 5173.209 | 5753.7 | 7629.044 | 8065.311 | 7695.607 | 1.72E-15 | 0.720030155 | YAL017W | PSK1 | n |
| W303.asm02.HP0_G0000550 | 9349.017 | 9409.848 | 9455.428 | 10482.57 | 10800.3 | 11170.87 | 10367.11 | 11146.04 | 10379.62 | 0.004008878 | -0.201934775 | YAL016W | TPD3 | n |
| W303.asm02.HP0_G0000570 | 2170.742 | 2029.75 | 2175.223 | 537.7103 | 453.8816 | 536.3232 | 909.8067 | 942.7963 | 1032.221 | 1.89E-71 | 2.059613541 | YAL014C | SYN8 | n |
| W303.asm02.HP0_G0000580 | 4618.916 | 4561.373 | 4418.354 | 1867.397 | 1800.773 | 2026.785 | 2264.024 | 1845.862 | 1948.421 | 8.31E-29 | 1.255581293 | YAL013W | DEP1 | n |
| W303.asm02.HP0_G0000590 | 18807.1 | 14961.61 | 18817.08 | 13951.98 | 14050.37 | 12949.24 | 13727.34 | 19678.46 | 17670.95 | 0.046312199 | 0.360765159 | YAL012W | CYS3 | n |
| W303.asm02.HP0_G0000600 | 440.5019 | 476.279 | 458.1118 | 4507.318 | 4209.036 | 4080.159 | 4852.714 | 4177.082 | 4340.59 | 6.61E-195 | -3.218674689 | YAL011W | SWC3 | n |
| W303.asm02.HP0_G0000610 | 2113.562 | 2240.069 | 2030.783 | 1346.36 | 1260.975 | 1471.47 | 1307.307 | 1723.449 | 1699.636 | 1.38E-05 | 0.646205821 | YAL010C | MDM10 | n |
| W303.asm02.HP0_G0000620 | 1073.723 | 836.8266 | 886.0421 | 1426.252 | 1503.103 | 1612.768 | 1604.815 | 2029.482 | 2045.304 | 1.62E-05 | -0.699455646 | YAL009W | SPO7 | n |
| W303.asm02.HP0_G0000630 | 4005.814 | 3899.256 | 3498.896 | 1209.501 | 1659.315 | 2506.134 | 1517.167 | 3576.828 | 2922.034 | 0.002827769 | 1.08525429 | YAL008W | FUN14 | n |
| W303.asm02.HP0_G0000660 | 12002.62 | 12714.87 | 13926.6 | 10306.81 | 12635.79 | 11528.67 | 9632.594 | 9455.882 | 7804.45 | 0.27702364 | 0.164905591 | YAL003W | EFB1 | n |
| W303.asm02.HP0_G0000670 | 2801.846 | 2607.294 | 2705.554 | 2533.63 | 2399.585 | 2926.228 | 1764.062 | 1842.641 | 1873.068 | 0.711501617 | 0.045995669 | YAL002W | VPS8 | n |
| W303.asm02.HP0_G0000680 | 4600.915 | 4930.823 | 4544.469 | 3232.514 | 2983.643 | 3881.127 | 2059.101 | 2818.725 | 2557.228 | 0.004817769 | 0.479205775 | YAL001C | TFC3 | n |
| W303.asm02.HP0_G0000700 | 3744.266 | 4083.981 | 4237.265 | 5116.584 | 4712.385 | 4930.983 | 3982.41 | 3856.016 | 3874.117 | 0.000852465 | -0.290984505 | YAR002W | NUP60 | n |

| | | | | | | | | | | | | | | |
|---|---|---|---|---|---|---|---|---|---|---|---|---|---|---|
| W303.asm02.HP0_G0000710 | 5154.719 | 4831.784 | 5103.905 | 4773.395 | 5234.826 | 5412.61 | 3155.313 | 4868.609 | 4140.844 | 0.877497891 | -0.031209945 | YAR002C- | ERP1 | n |
| W303.asm02.HP0_G0000720 | 3321.765 | 2809.823 | 3277.925 | 2464.158 | 2469.88 | 3097.912 | 1404.83 | 2408.533 | 1809.676 | 0.34393391 | 0.22837464 | YAR003W | SWD1 | n |
| W303.asm02.HP0_G0000730 | 4561.736 | 4914.131 | 4749.272 | 6121.144 | 6802.15 | 8637.387 | 3045.445 | 5714.763 | 4219.785 | 0.018924435 | -0.599958215 | YAR007C | RFA1 | n |
| W303.asm02.HP0_G0000740 | 446.8553 | 501.8734 | 512.0073 | 2282.143 | 2730.233 | 1909.037 | 1787.517 | 2416.05 | 2425.659 | 1.18E-30 | -2.244298296 | YAR008W | SEN34 | n |
| W303.asm02.HP0_G0000750 | 2584.772 | 2500.465 | 2892.033 | 3476.36 | 3362.89 | 4086.236 | 1388.782 | 1561.305 | 1508.262 | 0.000422997 | -0.453769806 | YAR014C | BUD14 | n |
| W303.asm02.HP0_G0000760 | 4580.796 | 3241.59 | 4220.018 | 5672.357 | 4927.61 | 8340.358 | 5914.361 | 18818.34 | 15748.84 | 0.125991187 | -0.653338266 | YAR015W | ADE1 | n |
| W303.asm02.HP0_G0000770 | 2399.464 | 2360.252 | 2576.205 | 1166.428 | 1123.856 | 1081.003 | 653.0363 | 740.922 | 566.944 | 2.46E-19 | 1.12159721 | YAR018C | KIN3 | n |
| W303.asm02.HP0_G0000780 | 1950.491 | 2223.377 | 1744.059 | 1751.379 | 1520.46 | 1975.888 | 1628.27 | 1468.958 | 1129.104 | 0.410771596 | 0.173198831 | YAR019C | CDC15 | n |
| W303.asm02.HP0_G0000810 | 4383.841 | 3960.46 | 3773.763 | 671.7905 | 759.3622 | 714.0847 | 1187.563 | 1896.331 | 1204.457 | 1.63E-33 | 2.498194875 | YAR027W | UIP3 | n |
| W303.asm02.HP0_G0000840 | 667.1062 | 624.2816 | 634.8891 | 976.7709 | 586.6615 | 755.8663 | 2740.53 | 3668.101 | 1252.3 | 0.511443257 | -0.268149805 | YAR035W | YAT1 | n |
| W303.asm02.HP0_G0000850 | 5614.281 | 5920.103 | 5705.378 | 10884.81 | 10050.48 | 16669.32 | 13185.41 | 13568.54 | 11826.88 | 6.08E-08 | -1.125190814 | YAR042W | SWH1 | n |
| W303.asm02.HP0_G0000910 | 4589.267 | 5091.066 | 4438.834 | 853.8061 | 831.3931 | 744.4713 | 669.0844 | 642.1324 | 600.4343 | 7.62E-117 | 2.538733576 | YBL107C | MIX23 | n |
| W303.asm02.HP0_G0000920 | 1465.516 | 1295.301 | 1504.763 | 3445.792 | 3439.26 | 3572.703 | 5550.191 | 4827.804 | 5273.536 | 1.42E-35 | -1.293613277 | YBL106C | SRO77 | n |
| W303.asm02.HP0_G0000930 | 3399.065 | 3660.004 | 3310.262 | 9038.257 | 8482.293 | 11066.03 | 12286.71 | 11262.01 | 11828.08 | 5.98E-29 | -1.463079003 | YBL105C | PKC1 | n |
| W303.asm02.HP0_G0000940 | 1182.79 | 1379.874 | 1159.831 | 3210.283 | 3132.911 | 3704.884 | 3265.181 | 3583.27 | 3423.193 | 4.47E-31 | -1.432806566 | YBL104C | SEA4 | n |
| W303.asm02.HP0_G0000950 | 2056.381 | 2422.569 | 2014.614 | 3289.481 | 2917.687 | 4093.073 | 3730.578 | 2627.588 | 2847.877 | 0.001128423 | -0.665706152 | YBL103C | RTG3 | n |
| W303.asm02.HP0_G0000960 | 2250.16 | 2016.396 | 1977.965 | 2031.35 | 2298.915 | 2025.266 | 1750.483 | 2233.504 | 1776.185 | 0.879665215 | -0.025183382 | YBL102W | SFT2 | n |
| W303.asm02.HP0_G0000970 | 2873.851 | 3135.874 | 2708.788 | 2542.661 | 2824.827 | 4724.354 | 3818.225 | 4347.816 | 3228.232 | 0.45423023 | -0.21105078 | YBL101C | ECM21 | n |
| W303.asm02.HP0_G0000980 | 12483.36 | 11357.25 | 12035.94 | 39916.31 | 44005.69 | 43185.41 | 12043.52 | 23763.19 | 19235.43 | 1.30E-14 | -1.824929402 | YBL099W | ATP1 | n |
| W303.asm02.HP0_G0001000 | 392.8514 | 420.6389 | 404.2163 | 3329.08 | 3123.365 | 3710.962 | 4007.1 | 3691.724 | 3814.313 | 3.19E-167 | -3.061504912 | YBL097W | BRN1 | n |
| W303.asm02.HP0_G0001010 | 388.6158 | 412.8493 | 376.1906 | 1622.162 | 1801.641 | 2892.043 | 836.9728 | 2147.6 | 1271.438 | 1.56E-11 | -2.423086232 | YBL095W | MRX3 | n |
| W303.asm02.HP0_G0001020 | 2476.764 | 2641.79 | 2369.246 | 2921.976 | 2905.537 | 5028.22 | 9625.187 | 5734.092 | 7046.133 | 0.066627944 | -0.535872835 | YBL093C | ROX3 | n |
| W303.asm02.HP0_G0001030 | 12896.33 | 12434.44 | 12822.82 | 10779.91 | 13078.39 | 10721.15 | 10312.79 | 8531.34 | 8061.608 | 0.359757796 | 0.14196395 | YBL092W | RPL32 | n |
| W303.asm02.HP0_G0001050 | 13072.1 | 12567.98 | 13996.66 | 10030.31 | 10409.77 | 10907.26 | 6268.654 | 7613.241 | 6739.935 | 0.00259346 | 0.338514229 | YBL091C | MAP2 | n |
| W303.asm02.HP0_G0001060 | 4612.563 | 4504.62 | 4287.927 | 1735.401 | 1826.808 | 1267.121 | 607.3608 | 473.5458 | 608.8069 | 4.30E-16 | 1.473014126 | YBL090W | MRP21 | n |
| W303.asm02.HP0_G0001080 | 3396.947 | 3138.1 | 3163.666 | 2369.677 | 2328.421 | 1967.531 | 1555.436 | 1770.696 | 1499.89 | 1.80E-05 | 0.541097067 | YBL088C | TEL1 | n |
| W303.asm02.HP0_G0001100 | 5776.292 | 5970.179 | 5621.301 | 1645.783 | 1559.513 | 1883.209 | 3007.176 | 2448.264 | 3255.742 | 5.99E-35 | 1.770961129 | YBL086C | | n |
| W303.asm02.HP0_G0001120 | 900.0639 | 973.7012 | 865.5618 | 3189.442 | 3046.995 | 3187.553 | 4726.797 | 5113.435 | 4321.453 | 3.60E-64 | -1.782768421 | YBL084C | CDC27 | n |
| W303.asm02.HP0_G0001130 | 1450.691 | 1181.795 | 1649.203 | 1683.992 | 1604.641 | 1365.117 | 1233.239 | 1969.349 | 1834.793 | 0.63487677 | -0.12016882 | YBL082C | ALG3 | n |
| W303.asm02.HP0_G0001140 | 770.8783 | 756.7049 | 987.3657 | 1542.27 | 1633.28 | 1267.121 | 982.6406 | 554.0808 | 952.0831 | 0.001567998 | -0.820810772 | YBL081W | | n |
| W303.asm02.HP0_G0001150 | 1760.949 | 1727.068 | 1617.943 | 1749.295 | 1517.857 | 1385.628 | 1519.636 | 1390.571 | 1705.616 | 0.379094701 | 0.133841311 | YBL080C | PET112 | n |
| W303.asm02.HP0_G0001160 | 3532.486 | 3585.446 | 3678.907 | 12210.33 | 11639.5 | 11847.73 | 9888.129 | 10205.39 | 10736.05 | 1.98E-140 | -1.725276353 | YBL079W | NUP170 | n |
| W303.asm02.HP0_G0001170 | 3345.061 | 3397.383 | 3089.29 | 266.771 | 255.1457 | 644.9553 | 548.1061 | 441.3318 | 261.9425 | 9.20E-15 | 3.074609553 | YBL078C | ATG8 | n |
| W303.asm02.HP0_G0001180 | 20885.72 | 19549.69 | 21086.08 | 19580.57 | 20778.75 | 21118.68 | 16237.02 | 16282.03 | 15149.6 | 0.991251526 | 0.001055946 | YBL076C | ILS1 | n |
| W303.asm02.HP0_G0001190 | 2453.468 | 2468.193 | 2509.375 | 3094.96 | 3440.996 | 8904.789 | 9586.918 | 7294.323 | 11295.82 | 0.008664297 | -1.055115053 | YBL075C | SSA3 | n |
| W303.asm02.HP0_G0001200 | 456.3853 | 400.6085 | 355.7103 | 490.4696 | 477.3134 | 330.4541 | 466.6308 | 424.151 | 413.8452 | 0.670341253 | -0.09811743 | YBL074C | AAR2 | n |
| W303.asm02.HP0_G0001210 | 10489.45 | 11215.93 | 11460.34 | 30121.5 | 35015.71 | 33082.63 | 27710.22 | 24514.85 | 22683.74 | 7.10E-41 | -1.566308891 | YER102W | RPS8B | y |
| W303.asm02.HP0_G0001220 | 492.3879 | 440.6693 | 477.5142 | 767.6613 | 814.0363 | 415.5365 | 1927.012 | 1176.885 | 1443.674 | 0.107950249 | -0.501497901 | YBL071W- | KTI11 | n |
| W303.asm02.HP0_G0001260 | 1844.602 | 1997.478 | 1780.708 | 8578.355 | 7956.38 | 7923.302 | 10783.12 | 10894.77 | 12011.08 | 9.11E-111 | -2.121109316 | YBL066C | SEF1 | n |
| W303.asm02.HP0_G0001270 | 5839.826 | 4872.957 | 5012.282 | 4867.876 | 5236.562 | 5582.016 | 894.993 | 4337.078 | 2705.543 | 0.994422139 | 0.003554325 | YBL064C | PRX1 | n |
| W303.asm02.HP0_G0001290 | 2514.884 | 2477.096 | 2456.557 | 6437.934 | 7313.309 | 7924.062 | 1814.675 | 4524.993 | 2865.818 | 3.61E-07 | -1.541012383 | YBL061C | SKT5 | n |
| W303.asm02.HP0_G0001300 | 1152.082 | 1186.246 | 1241.752 | 1652.73 | 1614.187 | 1758.624 | 892.524 | 1745.999 | 998.7304 | 0.067777008 | -0.489310289 | YBL060W | YEL1 | n |
| W303.asm02.HP0_G0001310 | 2111.444 | 2163.286 | 1931.615 | 1827.103 | 2094.972 | 1602.133 | 1124.605 | 954.6081 | 941.3184 | 0.290575857 | 0.168230965 | YBL059C-/ | CMC2 | n |
| W303.asm02.HP0_G0001330 | 9917.645 | 9131.648 | 9628.971 | 4188.444 | 4471.125 | 6358.393 | 4176.222 | 5953.147 | 4761.612 | 1.10E-05 | 0.933269796 | YBL058W | SHP1 | n |
| W303.asm02.HP0_G0001340 | 1639.175 | 1653.623 | 1613.631 | 2520.43 | 2707.669 | 1909.797 | 4121.906 | 3705.684 | 4406.375 | 0.000990516 | -0.540703203 | YBL057C | PTH2 | n |
| W303.asm02.HP0_G0001350 | 10525.45 | 10517.09 | 10332.85 | 8849.294 | 9164.417 | 12690.96 | 9507.912 | 13726.38 | 11165.45 | 0.893767654 | 0.031170901 | YBL056W | PTC3 | n |
| W303.asm02.HP0_G0001360 | 1818.129 | 1758.226 | 1861.551 | 3186.663 | 3321.233 | 3317.455 | 2143.045 | 2453.633 | 2643.346 | 1.66E-15 | -0.853359247 | YBL055C | | n |
| W303.asm02.HP0_G0001380 | 892.6516 | 903.5947 | 909.7562 | 1474.188 | 1405.905 | 1825.474 | 559.2163 | 655.018 | 724.8271 | 4.78E-07 | -0.798347343 | YBL052C | SAS3 | n |
| W303.asm02.HP0_G0001390 | 7238.632 | 7938.725 | 7799.758 | 14605.71 | 14026.94 | 17805.78 | 19118.28 | 18738.88 | 18589.54 | 3.89E-17 | -1.015156004 | YBL051C | PIN4 | n |
| W303.asm02.HP0_G0001410 | 2192.979 | 2186.655 | 2029.705 | 111.1546 | 106.7446 | 134.4606 | 1941.826 | 746.291 | 1016.672 | 2.66E-36 | 4.184733949 | YBL049W | MOH1 | n |
| W303.asm02.HP0_G0001420 | 22225.22 | 22268.27 | 23221.42 | 14566.81 | 13987.02 | 22477.72 | 19344.19 | 26067.57 | 22390.7 | 0.074231822 | 0.408076046 | YBL047C | EDE1 | n |
| W303.asm02.HP0_G0001430 | 2656.777 | 2735.266 | 2617.166 | 2075.812 | 2254.655 | 2038.18 | 1467.789 | 1450.704 | 1502.282 | 4.90E-05 | 0.331052461 | YBL046W | PSY4 | n |
| W303.asm02.HP0_G0001440 | 19320.67 | 20123.9 | 18604.73 | 24271.3 | 24380.3 | 24367.76 | 6036.573 | 15747.28 | 12814.85 | 0.323215072 | -0.33100017 | YBL045C | COR1 | n |
| W303.asm02.HP0_G0001460 | 119.6556 | 114.6185 | 103.4794 | 722.5048 | 898.217 | 1053.655 | 351.8248 | 1194.066 | 1236.751 | 2.93E-12 | -2.985081535 | YBL043W | ECM13 | n |

| | | | | | | | | | | | | | | |
|---|---|---|---|---|---|---|---|---|---|---|---|---|---|---|
| W303.asm02.HP0_G0001470 | 849.2368 | 911.3843 | 866.6398 | 4605.273 | 4427.733 | 2508.413 | 12765.69 | 4660.292 | 6122.756 | 3.07E-08 | -2.135194561 | YBL042C | FUI1 | n |
| W303.asm02.HP0_G0001480 | 9170.063 | 9672.47 | 9018.874 | 3751.467 | 4044.146 | 4763.857 | 2443.022 | 4165.27 | 3038.054 | 6.79E-08 | 1.149501368 | YBL041W | PRE7 | n |
| W303.asm02.HP0_G0001500 | 283.7849 | 326.0508 | 225.2832 | 201.4677 | 251.6743 | 332.7331 | 109.8681 | 190.0626 | 212.903 | 0.80727298 | 0.087836577 | YBL039W- | MIN6 | n |
| W303.asm02.HP0_G0001510 | 953.0088 | 974.814 | 1279.479 | 13577.53 | 13480.2 | 7960.526 | 14653.2 | 8189.872 | 14043.23 | 3.44E-31 | -3.448662673 | YBL039C | URA7 | n |
| W303.asm02.HP0_G0001520 | 3498.601 | 3182.612 | 3038.629 | 2049.413 | 2121.007 | 1517.05 | 478.9756 | 526.162 | 438.9629 | 5.38E-06 | 0.773261141 | YBL038W | MRPL16 | n |
| W303.asm02.HP0_G0001530 | 2287.221 | 2275.679 | 2170.911 | 5153.404 | 4990.094 | 7023.859 | 5234.166 | 6172.202 | 5551.027 | 2.32E-16 | -1.350207525 | YBL037W | APL3 | n |
| W303.asm02.HP0_G0001540 | 3672.261 | 3425.203 | 3526.922 | 2277.974 | 2400.452 | 2094.395 | 2709.668 | 1817.943 | 2313.227 | 0.000113156 | 0.649634745 | YBL036C | | n |
| W303.asm02.HP0_G0001550 | 894.7694 | 824.5858 | 927.0027 | 2695.499 | 2907.272 | 2598.813 | 464.1619 | 667.9036 | 535.8458 | 1.33E-25 | -1.631731244 | YBL035C | POL12 | n |
| W303.asm02.HP0_G0001560 | 1509.99 | 1656.961 | 1631.956 | 2291.869 | 2332.761 | 3014.349 | 1044.364 | 1578.486 | 1138.672 | 0.001097988 | -0.670716334 | YBL034C | STU1 | n |
| W303.asm02.HP0_G0001570 | 3934.868 | 4119.591 | 3719.868 | 4427.426 | 4614.319 | 4784.368 | 9878.254 | 6679.036 | 7625.038 | 0.187142681 | -0.231746785 | YBL033C | RIB1 | n |
| W303.asm02.HP0_G0001580 | 2388.875 | 1801.625 | 2223.729 | 6601.887 | 6750.079 | 7343.678 | 5824.244 | 6313.944 | 6866.72 | 9.50E-33 | -1.689854751 | YBL032W | HEK2 | n |
| W303.asm02.HP0_G0001590 | 545.3328 | 585.3335 | 555.1237 | 1725.675 | 1740.024 | 2071.605 | 1903.558 | 1535.534 | 1704.42 | 1.69E-35 | -1.71592144 | YBL031W | SHE1 | n |
| W303.asm02.HP0_G0001600 | 12509.83 | 11609.86 | 12118.94 | 27457.96 | 30754.6 | 22787.66 | 4965.051 | 9008.108 | 7108.329 | 6.04E-07 | -1.160387955 | YBL030C | PET9 | n |
| W303.asm02.HP0_G0001610 | 1836.13 | 1701.473 | 1449.789 | 361.2524 | 510.2914 | 470.9921 | 2085.025 | 1214.468 | 1502.282 | 6.36E-15 | 1.894057011 | YBL029C-A | | n |
| W303.asm02.HP0_G0001620 | 1386.098 | 1145.073 | 1101.624 | 491.859 | 663.8995 | 467.1937 | 908.5722 | 507.9074 | 546.6105 | 2.07E-05 | 1.16303747 | YBL029W | | n |
| W303.asm02.HP0_G0001630 | 845.0012 | 852.4059 | 924.8469 | 2450.959 | 2177.417 | 673.8225 | 3108.403 | 1125.342 | 2303.658 | 0.043732316 | -1.015781021 | YBL028C | | n |
| W303.asm02.HP0_G0001640 | 1062.075 | 1030.454 | 962.5738 | 45.15655 | 52.07055 | 38.7429 | 102.4613 | 67.6494 | 78.94157 | 2.12E-111 | 4.494423623 | NA | NA | NA |
| W303.asm02.HP0_G0001650 | 1142.552 | 1270.819 | 1150.13 | 1151.145 | 1254.9 | 1109.11 | 1003.627 | 899.8443 | 907.828 | 0.875565603 | 0.020000414 | YBL026W | LSM2 | n |
| W303.asm02.HP0_G0001660 | 628.9858 | 624.2816 | 618.7204 | 273.7182 | 303.7449 | 251.449 | 474.0377 | 515.424 | 398.2961 | 2.18E-15 | 1.176527563 | YBL025W | RRN10 | n |
| W303.asm02.HP0_G0001670 | 4184.768 | 4217.517 | 4658.728 | 10078.25 | 9934.193 | 5982.359 | 7231.543 | 4938.406 | 6915.76 | 6.30E-05 | -0.992947743 | YBL024W | NCL1 | n |
| W303.asm02.HP0_G0001680 | 2242.747 | 2213.362 | 2143.963 | 4753.943 | 4741.892 | 4038.377 | 2466.477 | 1831.903 | 2047.696 | 1.14E-12 | -1.036039499 | YBL023C | MCM2 | n |
| W303.asm02.HP0_G0001690 | 19158.65 | 20355.36 | 19276.27 | 8672.836 | 8658.465 | 10865.48 | 4171.285 | 7023.725 | 5078.574 | 6.41E-07 | 1.06004008 | YBL022C | PIM1 | n |
| W303.asm02.HP0_G0001700 | 982.658 | 890.2411 | 939.9376 | 694.7161 | 654.3533 | 765.7419 | 727.1046 | 495.0218 | 574.1205 | 0.020252345 | 0.411333857 | YBL021C | HAP3 | n |
| W303.asm02.HP0_G0001710 | 897.9461 | 964.7988 | 1052.04 | 2343.278 | 2154.853 | 2962.692 | 1698.635 | 3117.241 | 2414.894 | 3.64E-08 | -1.356009816 | YBL020W | RFT1 | n |
| W303.asm02.HP0_G0001720 | 2104.032 | 2067.585 | 1948.862 | 1739.569 | 1783.416 | 1801.924 | 1181.391 | 1636.471 | 1525.008 | 0.191092501 | 0.200924487 | YBL019W | APN2 | n |
| W303.asm02.HP0_G0001730 | 141.8924 | 122.4082 | 137.9725 | 359.1682 | 458.2208 | 328.1751 | 292.5701 | 370.461 | 339.6879 | 7.48E-14 | -1.508049876 | YBL018C | POP8 | n |
| W303.asm02.HP0_G0001740 | 6139.495 | 5688.641 | 5720.469 | 7323.003 | 7508.573 | 10675.57 | 10834.97 | 16668.6 | 13412.89 | 0.016348308 | -0.539546254 | YBL017C | PEP1 | n |
| W303.asm02.HP0_G0001750 | 4860.345 | 4661.525 | 4999.347 | 160.4794 | 170.0971 | 362.36 | 407.3761 | 699.0438 | 454.512 | 5.12E-38 | 4.389083469 | YBL016W | FUS3 | n |
| W303.asm02.HP0_G0001760 | 4834.931 | 4347.715 | 4758.973 | 12322.18 | 13164.3 | 19910.05 | 15406.22 | 28058.39 | 20693.45 | 2.33E-10 | -1.703178346 | YBL015W | ACH1 | n |
| W303.asm02.HP0_G0001770 | 832.2944 | 915.8355 | 814.9001 | 885.7631 | 678.6528 | 584.1821 | 1593.705 | 839.7115 | 1405.399 | 0.391606552 | 0.254138231 | YBL014C | RRN6 | n |
| W303.asm02.HP0_G0001780 | 250.959 | 262.6211 | 214.5041 | 1742.348 | 1848.505 | 1670.502 | 1124.605 | 1340.102 | 1450.85 | 2.44E-80 | -2.853376444 | YBL013W | FMT1 | n |
| W303.asm02.HP0_G0001790 | 8081.515 | 7982.124 | 8330.09 | 7117.367 | 6967.908 | 8714.113 | 7226.605 | 7124.663 | 8385.747 | 0.504220299 | 0.097491983 | YBL011W | SCT1 | n |
| W303.asm02.HP0_G0001800 | 690.402 | 745.5769 | 810.5884 | 2275.89 | 2406.527 | 2555.512 | 1603.581 | 1980.087 | 2045.304 | 7.95E-34 | -1.687862876 | YBL010C | LAA2 | n |
| W303.asm02.HP0_G0001810 | 1245.265 | 1290.85 | 1522.009 | 5176.33 | 5278.218 | 5908.671 | 6688.375 | 4556.133 | 5058.241 | 3.44E-29 | -2.011571088 | YBL009W | ALK2 | n |
| W303.asm02.HP0_G0001830 | 1567.17 | 1620.239 | 1678.306 | 2843.473 | 2674.691 | 2992.319 | 1920.84 | 2238.873 | 1930.48 | 4.89E-14 | -0.806750595 | YBL008W | HIR1 | n |
| W303.asm02.HP0_G0001840 | 12337.23 | 11225.94 | 11834.38 | 15306.68 | 14752.45 | 19047.07 | 13543.4 | 17417.03 | 15259.64 | 0.002678007 | -0.472261262 | YBL007C | SLA1 | n |
| W303.asm02.HP0_G0001850 | 2854.791 | 2750.845 | 2771.307 | 974.6867 | 1095.217 | 862.979 | 1148.06 | 653.9442 | 743.9644 | 2.07E-11 | 1.514293618 | YBL006C | LDB7 | n |
| W303.asm02.HP0_G0001870 | 2487.353 | 2447.05 | 2631.179 | 9617.65 | 9059.408 | 3632.716 | 8794.386 | 4924.446 | 9019.672 | 3.91E-05 | -1.560147286 | YBL004W | UTP20 | n |
| W303.asm02.HP0_G0001880 | 4453.728 | 3880.338 | 4236.187 | 6130.175 | 7476.463 | 5234.849 | 10454.75 | 11557.31 | 9303.144 | 0.000733323 | -0.583790163 | YBL003C | HTA2 | y |
| W303.asm02.HP0_G0001920 | 1453.868 | 1276.383 | 1467.036 | 2735.097 | 2810.942 | 2177.958 | 2312.168 | 2215.249 | 2050.089 | 8.08E-10 | -0.879719103 | YBR002C | RER2 | n |
| W303.asm02.HP0_G0001930 | 5833.473 | 5587.376 | 5375.538 | 4724.764 | 4402.565 | 3998.875 | 1702.338 | 1882.371 | 1656.577 | 0.00126505 | 0.355647958 | YBR003W | COQ1 | n |
| W303.asm02.HP0_G0001940 | 2146.388 | 2500.465 | 2023.237 | 1069.168 | 1005.829 | 1084.041 | 1035.723 | 1254.198 | 1041.789 | 4.47E-15 | 1.077934166 | YBR004C | GPI18 | n |
| W303.asm02.HP0_G0001950 | 2783.845 | 3372.901 | 2453.323 | 1563.111 | 1470.993 | 1611.249 | 4452.744 | 3604.746 | 3277.271 | 5.48E-07 | 0.890096355 | YBR005W | RCR1 | n |
| W303.asm02.HP0_G0001960 | 3257.172 | 3130.31 | 2853.228 | 1045.548 | 1072.653 | 869.816 | 356.7627 | 584.1472 | 305.0015 | 4.46E-11 | 1.628875896 | YBR006W | UGA2 | n |
| W303.asm02.HP0_G0001970 | 2426.996 | 2566.12 | 2583.751 | 2913.639 | 2582.699 | 3766.417 | 2486.229 | 2159.412 | 1913.735 | 0.1290506 | -0.289950163 | YBR007C | DSF2 | n |
| W303.asm02.HP0_G0001990 | 5535.922 | 5261.325 | 5324.876 | 6599.803 | 7909.517 | 5968.685 | 8952.399 | 8722.477 | 6416.992 | 0.062268592 | -0.344963496 | YNL030W | HHF2 | y |
| W303.asm02.HP0_G0002000 | 2191.92 | 1826.107 | 1926.225 | 5709.872 | 7326.327 | 4776.771 | 5966.208 | 4467.008 | 4476.944 | 2.81E-14 | -1.583245847 | YNL031C | HHT2 | y |
| W303.asm02.HP0_G0002010 | 22894.45 | 22958.21 | 24066.5 | 19630.59 | 22899.76 | 23628.61 | 27744.78 | 28216.24 | 25335.46 | 0.504293427 | 0.079782688 | YBR011C | IPP1 | n |
| W303.asm02.HP0_G0002020 | 2260.749 | 2168.85 | 2120.249 | 2013.287 | 2233.827 | 2407.377 | 3105.934 | 2476.183 | 2639.758 | 0.881966092 | -0.022724645 | YBR014C | GRX7 | n |
| W303.asm02.HP0_G0002030 | 6796.012 | 6032.496 | 7002.104 | 9562.768 | 9486.387 | 8698.92 | 6163.724 | 7168.688 | 6863.132 | 1.62E-05 | -0.484624049 | YBR015C | MNN2 | n |
| W303.asm02.HP0_G0002040 | 5385.559 | 5244.633 | 4885.089 | 3372.152 | 3910.498 | 3719.318 | 3870.073 | 3493.071 | 3136.133 | 5.82E-05 | 0.496091527 | YBR016W | CPP1 | n |
| W303.asm02.HP0_G0002050 | 6044.194 | 6069.219 | 6277.749 | 8435.938 | 8242.768 | 6441.956 | 5836.589 | 5361.483 | 6153.854 | 0.019946539 | -0.330162849 | YBR017C | KAP104 | n |
| W303.asm02.HP0_G0002060 | 227.6632 | 233.6883 | 195.1017 | 1818.072 | 1691.425 | 1965.252 | 21152.7 | 8066.385 | 18179.29 | 7.44E-21 | -3.060102383 | YBR018C | GAL7 | n |
| W303.asm02.HP0_G0002070 | 400.2637 | 447.3462 | 399.9047 | 2561.418 | 2352.721 | 4131.816 | 22378.53 | 9604.067 | 13893.72 | 1.71E-17 | -2.858293614 | YBR019C | GAL10 | n |

| | | | | | | | | | | | | | | |
|---|---|---|---|---|---|---|---|---|---|---|---|---|---|---|
| W303.asm02.HP0_G0002090 | 751.8181 | 679.9216 | 749.1475 | 732.9255 | 667.3709 | 899.4429 | 1947.999 | 1282.117 | 1596.773 | 0.735169866 | -0.076740298 | YBR021W | FUR4 | n |
| W303.asm02.HP0_G0002110 | 2282.986 | 2343.56 | 2458.713 | 7549.48 | 6662.427 | 6832.424 | 6127.924 | 6520.113 | 6595.209 | 4.60E-68 | -1.570680235 | YBR023C | CHS3 | n |
| W303.asm02.HP0_G0002130 | 20747 | 21133.21 | 21193.87 | 15604.02 | 16880.4 | 16383.69 | 14634.68 | 14641.26 | 13984.62 | 1.70E-08 | 0.368218667 | YBR025C | OLA1 | n |
| W303.asm02.HP0_G0002140 | 10223.67 | 10182.13 | 9904.916 | 3207.504 | 3196.264 | 6000.591 | 3187.41 | 18529.49 | 7469.547 | 0.018924435 | 1.288975744 | YBR026C | ETR1 | n |
| W303.asm02.HP0_G0002150 | 1904.959 | 1655.848 | 1837.837 | 1731.233 | 1632.412 | 1182.798 | 1265.335 | 1065.21 | 1381.477 | 0.222732578 | 0.247889608 | YBR028C | YPK3 | n |
| W303.asm02.HP0_G0002160 | 4209.122 | 4898.552 | 4730.948 | 5327.778 | 5449.183 | 5263.716 | 4858.886 | 5893.014 | 5937.363 | 0.096816895 | -0.213053493 | YBR029C | CDS1 | n |
| W303.asm02.HP0_G0002170 | 894.7694 | 931.4147 | 1119.949 | 2480.831 | 2224.28 | 1216.223 | 2355.375 | 1547.346 | 2651.719 | 0.00147505 | -1.007109692 | YBR030W | RKM3 | n |
| W303.asm02.HP0_G0002200 | 1736.594 | 1817.205 | 1935.927 | 4105.078 | 3796.811 | 2022.987 | 5550.191 | 4134.13 | 5235.261 | 0.002120536 | -0.854317518 | YBR034C | HMT1 | n |
| W303.asm02.HP0_G0002230 | 3299.528 | 3004.564 | 2924.37 | 3213.757 | 3532.987 | 3514.968 | 6051.387 | 4013.864 | 4927.867 | 0.413029798 | -0.153038643 | YBR037C | SCO1 | n |
| W303.asm02.HP0_G0002240 | 4557.5 | 4033.905 | 4583.274 | 10857.02 | 10231 | 9074.194 | 1262.866 | 4021.381 | 3418.409 | 0.001389253 | -1.194970834 | YBR038W | CHS2 | n |
| W303.asm02.HP0_G0002250 | 6237.972 | 6951.67 | 6636.693 | 21056.85 | 22136.06 | 22736.76 | 6039.042 | 13308.68 | 10559.03 | 6.88E-11 | -1.733511876 | YBR039W | ATP3 | n |
| W303.asm02.HP0_G0002270 | 4046.052 | 3943.768 | 3925.749 | 3584.735 | 3342.929 | 5964.887 | 13917.45 | 11932.06 | 15138.84 | 0.67402414 | -0.113726112 | YBR041W | FAT1 | n |
| W303.asm02.HP0_G0002280 | 1602.114 | 1473.349 | 1595.307 | 1658.982 | 1740.892 | 2622.362 | 3624.413 | 4049.3 | 3809.529 | 0.077734047 | -0.366638456 | YBR042C | CST26 | n |
| W303.asm02.HP0_G0002290 | 1538.58 | 1636.931 | 1540.334 | 1595.763 | 1412.848 | 1369.675 | 2677.572 | 1800.762 | 1363.536 | 0.698777536 | 0.107059002 | YBR043C | QDR3 | n |
| W303.asm02.HP0_G0002300 | 4510.908 | 4438.965 | 4687.831 | 2548.914 | 2222.545 | 1769.259 | 697.4773 | 862.2613 | 937.7301 | 3.53E-09 | 1.059994086 | YBR044C | TCM62 | n |
| W303.asm02.HP0_G0002460 | 6196.675 | 6006.902 | 5602.977 | 1909.08 | 2249.448 | 3848.461 | 2850.398 | 5044.712 | 4232.942 | 0.000208196 | 1.153081817 | YBR053C | | n |
| W303.asm02.HP0_G0002480 | 2275.573 | 2122.112 | 2362.779 | 3176.242 | 2970.625 | 3211.862 | 3145.437 | 2666.245 | 2745.014 | 1.62E-05 | -0.469256475 | YBR055C | PRP6 | n |
| W303.asm02.HP0_G0002490 | 7839.027 | 7016.213 | 6679.809 | 1129.608 | 1272.257 | 2409.656 | 636.9881 | 2115.386 | 802.5726 | 2.91E-06 | 2.162116741 | YBR056W | MRX18 | n |
| W303.asm02.HP0_G0002510 | 2378.286 | 2350.237 | 2434.999 | 638.4441 | 657.8246 | 1312.7 | 887.5862 | 1532.312 | 961.6518 | 1.27E-05 | 1.457123922 | YBR057C | MUM2 | n |
| W303.asm02.HP0_G0002520 | 3837.449 | 4057.274 | 4263.135 | 6707.484 | 6565.229 | 9275.505 | 7703.112 | 9508.498 | 8755.337 | 4.03E-07 | -0.891150786 | YBR058C | UBP14 | n |
| W303.asm02.HP0_G0002530 | 2350.755 | 1774.918 | 1911.135 | 1346.36 | 1644.562 | 1603.652 | 3360.236 | 2521.282 | 2822.759 | 0.033811959 | 0.394164363 | YBR058C-A | TSC3 | n |
| W303.asm02.HP0_G0002540 | 5857.828 | 5852.222 | 5820.715 | 7211.154 | 6892.405 | 10002.5 | 5124.298 | 7747.466 | 5854.833 | 0.037118408 | -0.459523895 | YBR059C | AKL1 | n |
| W303.asm02.HP0_G0002550 | 2461.939 | 2817.613 | 2562.192 | 2209.892 | 2019.47 | 1941.703 | 3812.053 | 2688.795 | 4218.589 | 0.080545183 | 0.345560178 | YBR060C | ORC2 | n |
| W303.asm02.HP0_G0002580 | 1416.806 | 1429.95 | 1543.567 | 1668.013 | 1738.289 | 1318.018 | 750.5596 | 984.6745 | 850.416 | 0.563044653 | -0.105652686 | YBR061C | TRM7 | n |
| W303.asm02.HP0_G0002590 | 3242.348 | 3272.749 | 2959.941 | 740.5674 | 801.0186 | 891.8463 | 877.7104 | 1448.556 | 1135.084 | 3.35E-23 | 1.961228502 | YBR062C | | n |
| W303.asm02.HP0_G0002610 | 1058.899 | 1117.253 | 977.6645 | 511.3111 | 386.1899 | 279.5566 | 1744.31 | 951.3867 | 1430.517 | 1.05E-06 | 1.421439422 | YBR065C | ECM2 | n |
| W303.asm02.HP0_G0002660 | 1050.428 | 888.0155 | 1001.379 | 734.315 | 906.0276 | 1172.163 | 1270.273 | 1119.973 | 1495.105 | 0.790255297 | 0.064122238 | YBR070C | ALG14 | n |
| W303.asm02.HP0_G0002670 | 4208.063 | 4808.415 | 4451.769 | 287.6125 | 330.648 | 328.9348 | 76.53733 | 143.8892 | 83.7259 | 5.40E-71 | 3.830237237 | YBR071W | | n |
| W303.asm02.HP0_G0002680 | 9393.49 | 8748.844 | 8525.191 | 157.0058 | 204.8108 | 87.36143 | 824.628 | 238.3836 | 1076.476 | 4.21E-34 | 5.892358746 | YBR072W | HSP26 | n |
| W303.asm02.HP0_G0002700 | 6113.022 | 6270.636 | 6633.459 | 621.7709 | 570.1725 | 689.0158 | 150.6057 | 191.1364 | 129.1771 | 4.45E-110 | 3.337327569 | YBR073W | RDH54 | n |
| W303.asm02.HP0_G0002710 | 2565.712 | 2382.508 | 2523.388 | 6258.698 | 6312.686 | 8204.378 | 7566.086 | 9851.041 | 8748.161 | 8.20E-20 | -1.475401492 | YBR074W | PFF1 | n |
| W303.asm02.HP0_G0002730 | 1629.645 | 1551.245 | 1555.424 | 612.7396 | 749.8159 | 777.8966 | 1296.197 | 1097.424 | 1208.045 | 1.98E-17 | 1.146656705 | YBR077C | SLM4 | n |
| W303.asm02.HP0_G0002740 | 22297.23 | 23314.3 | 23139.5 | 18637.84 | 19693.95 | 22216.39 | 27729.97 | 24835.92 | 25177.58 | 0.09520322 | 0.183302043 | YBR078W | ECM33 | n |
| W303.asm02.HP0_G0002750 | 29539.04 | 30081.25 | 33097.23 | 31669.33 | 31185.05 | 29699.09 | 23806.81 | 23577.43 | 24353.47 | 0.978596391 | 0.002553174 | YBR079C | RPG1 | n |
| W303.asm02.HP0_G0002760 | 7531.947 | 7192.035 | 7353.503 | 6256.614 | 6725.779 | 8496.089 | 5932.878 | 10138.82 | 7414.527 | 0.877831791 | 0.039689761 | YBR080C | SEC18 | n |
| W303.asm02.HP0_G0002770 | 4420.902 | 4657.074 | 4377.393 | 6537.279 | 6108.743 | 6553.627 | 3014.583 | 4058.964 | 3868.137 | 0.000233132 | -0.512948404 | YBR081C | SPT7 | n |
| W303.asm02.HP0_G0002790 | 1533.285 | 1668.089 | 1630.878 | 1624.941 | 1540.42 | 3315.936 | 757.9665 | 1678.349 | 924.5732 | 0.30106787 | -0.423626628 | YBR083W | TEC1 | n |
| W303.asm02.HP0_G0002800 | 7110.505 | 6656.778 | 7787.901 | 7668.971 | 7245.617 | 6159.361 | 5315.641 | 5059.745 | 5891.911 | 0.830966744 | 0.032572785 | YBR084W | MIS1 | n |
| W303.asm02.HP0_G0002830 | 5297.67 | 5244.633 | 4766.519 | 955.9294 | 1201.094 | 1404.62 | 1673.946 | 1871.633 | 1721.165 | 7.98E-39 | 2.104053731 | YBR085C-A | | n |
| W303.asm02.HP0_G0002840 | 13682.03 | 14005.72 | 13878.09 | 8222.66 | 7758.512 | 11228.6 | 14384.08 | 12463.6 | 14226.23 | 0.000318557 | 0.611245727 | YBR086C | IST2 | n |
| W303.asm02.HP0_G0002850 | 2307.34 | 2300.16 | 2314.273 | 2760.802 | 2902.933 | 2866.215 | 2544.249 | 3495.219 | 3038.054 | 0.033408071 | -0.30133762 | YBR087W | RFC5 | n |
| W303.asm02.HP0_G0002860 | 1310.917 | 1342.038 | 1528.477 | 5952.328 | 7158.833 | 6691.886 | 856.7243 | 1770.696 | 1420.948 | 1.72E-18 | -2.243626443 | YBR088C | POL30 | n |
| W303.asm02.HP0_G0002880 | 909.594 | 823.473 | 805.1989 | 1476.967 | 1544.76 | 1049.857 | 1227.066 | 978.2317 | 922.181 | 0.000808118 | -0.681576208 | YBR091C | TIM12 | n |
| W303.asm02.HP0_G0002920 | 1441.161 | 1737.083 | 1493.983 | 2340.499 | 2371.814 | 2197.71 | 1955.405 | 2266.792 | 1910.147 | 9.81E-06 | -0.564689318 | YBR095C | RXT2 | n |
| W303.asm02.HP0_G0002930 | 1660.353 | 1541.23 | 1637.346 | 1557.554 | 1730.478 | 2184.036 | 1702.338 | 2621.146 | 2260.599 | 0.432213713 | -0.177339908 | YBR096W | | n |
| W303.asm02.HP0_G0002940 | 2170.742 | 2148.819 | 2042.64 | 2321.047 | 2168.738 | 3309.859 | 2125.763 | 2875.636 | 2436.424 | 0.1753436 | -0.293954056 | YBR097W | VPS15 | n |
| W303.asm02.HP0_G0002950 | 1350.096 | 1396.566 | 1402.361 | 1024.706 | 1063.107 | 1330.932 | 791.2973 | 715.1508 | 801.3765 | 0.054709948 | 0.279263159 | YBR098W | MMS4 | n |
| W303.asm02.HP0_G0002960 | 10730.88 | 13352.5 | 9964.201 | 2182.103 | 2201.716 | 1982.725 | 1528.278 | 1129.638 | 1491.517 | 3.30E-44 | 2.418979145 | YBR101C | FES1 | n |
| W303.asm02.HP0_G0002970 | 2353.932 | 2212.249 | 2362.779 | 5126.31 | 4843.429 | 6817.23 | 4395.959 | 4114.801 | 4262.845 | 1.10E-16 | -1.276655979 | YBR102C | EXO84 | n |
| W303.asm02.HP0_G0002980 | 2184.508 | 2187.768 | 2321.818 | 3768.835 | 3917.441 | 4200.186 | 4312.015 | 3815.211 | 4150.413 | 1.48E-18 | -0.82830915 | YBR103W | SIF2 | n |
| W303.asm02.HP0_G0002990 | 410.8527 | 344.9684 | 464.5793 | 1451.262 | 1498.764 | 2982.443 | 1609.753 | 1442.113 | 1254.692 | 7.43E-14 | -2.28121216 | YBR104W | YMC2 | n |
| W303.asm02.HP0_G0003000 | 1454.927 | 1541.23 | 1319.362 | 2697.583 | 2742.382 | 4768.415 | 11636.14 | 8998.443 | 17068.12 | 5.84E-05 | -1.242176233 | YBR105C | VID24 | n |
| W303.asm02.HP0_G0003010 | 7879.265 | 7833.009 | 8432.491 | 6112.807 | 6953.154 | 8464.183 | 5440.323 | 6507.228 | 6243.56 | 0.322011282 | 0.165380582 | YBR106W | SND3 | n |
| W303.asm02.HP0_G0003030 | 5170.602 | 4629.254 | 4822.57 | 4619.168 | 4495.424 | 6105.425 | 4761.363 | 5816.774 | 5510.36 | 0.752849828 | -0.057805376 | YBR108W | AIM3 | n |

| | | | | | | | | | | | | | | |
|---|---|---|---|---|---|---|---|---|---|---|---|---|---|---|
| W303.asm02.HP0_G0003040 | 6907.196 | 7293.3 | 7286.673 | 3603.493 | 4601.301 | 4083.197 | 2947.922 | 2803.692 | 2584.738 | 7.86E-11 | 0.806453947 | YBR109C | CMD1 | n |
| W303.asm02.HP0_G0003050 | 2473.587 | 2743.055 | 2599.919 | 4555.948 | 5148.91 | 5664.819 | 3054.086 | 3897.894 | 3622.939 | 8.16E-12 | -0.97546476 | YBR110W | ALG1 | n |
| W303.asm02.HP0_G0003060 | 5097.538 | 4802.851 | 4725.558 | 2601.712 | 2742.382 | 3284.79 | 2484.994 | 3625.149 | 3244.977 | 1.80E-05 | 0.761300171 | YBR111C | YSA1 | n |
| W303.asm02.HP0_G0003080 | 6258.091 | 6229.462 | 6096.66 | 10115.76 | 9808.356 | 9791.317 | 6369.881 | 6448.169 | 6383.502 | 2.76E-37 | -0.677196537 | YBR112C | CYC8 | n |
| W303.asm02.HP0_G0003090 | 3190.462 | 3060.204 | 2896.345 | 935.0879 | 791.4724 | 408.6996 | 2612.145 | 926.6893 | 1824.029 | 1.93E-07 | 2.098839028 | YBR114W | RAD16 | n |
| W303.asm02.HP0_G0003100 | 3491.189 | 3355.096 | 3466.559 | 5955.107 | 5031.751 | 5909.431 | 3140.5 | 3833.466 | 2529.718 | 5.29E-05 | -0.712319618 | YBR115C | LYS2 | n |
| W303.asm02.HP0_G0003120 | 25766.18 | 28015.89 | 29307.3 | 99386.79 | 110541.4 | 118050.4 | 115724.4 | 102517.8 | 95087.51 | 1.14E-57 | -1.980865555 | YPR080W | TEF1 | y |
| W303.asm02.HP0_G0003130 | 3120.574 | 3042.399 | 2898.5 | 728.7572 | 794.0759 | 575.8258 | 374.0453 | 513.2764 | 381.5509 | 2.06E-31 | 2.110630646 | YBR119W | MUD1 | n |
| W303.asm02.HP0_G0003140 | 2920.443 | 2856.561 | 2485.661 | 1907.691 | 1868.465 | 1295.988 | 1641.849 | 1205.877 | 1495.105 | 0.00066553 | 0.704041233 | YBR120C | CBP6 | n |
| W303.asm02.HP0_G0003150 | 23822.04 | 24370.35 | 24700.31 | 18435.68 | 19360.7 | 20096.93 | 13418.72 | 15145.95 | 12628.26 | 0.0010747 | 0.332388514 | YBR121C | GRS1 | n |
| W303.asm02.HP0_G0003160 | 4622.093 | 4427.837 | 4430.211 | 2125.831 | 1992.566 | 1519.329 | 651.8018 | 574.483 | 623.1599 | 1.37E-16 | 1.257624616 | YBR122C | MRPL36 | n |
| W303.asm02.HP0_G0003170 | 2379.345 | 2474.87 | 2325.052 | 2157.094 | 2088.029 | 2332.17 | 2071.446 | 2108.943 | 1846.754 | 0.237169597 | 0.126170071 | YBR123C | TFC1 | n |
| W303.asm02.HP0_G0003180 | 3070.806 | 3039.061 | 3184.147 | 2057.749 | 2107.989 | 2516.009 | 5498.343 | 4145.942 | 4809.455 | 0.001647739 | 0.476069544 | YBR125C | PTC4 | n |
| W303.asm02.HP0_G0003190 | 29732.82 | 27186.85 | 26927.27 | 7245.195 | 7889.556 | 13433.91 | 28654.59 | 23897.42 | 26372.46 | 2.07E-10 | 1.553322201 | YBR126C | TPS1 | n |
| W303.asm02.HP0_G0003210 | 34892.83 | 34125.17 | 36549.78 | 12237.42 | 12298.2 | 15603.51 | 9469.643 | 10401.9 | 9494.517 | 3.17E-27 | 1.395079429 | YBR127C | VMA2 | n |
| W303.asm02.HP0_G0003220 | 1169.024 | 1122.817 | 1019.703 | 357.0841 | 436.5248 | 676.8612 | 582.6713 | 1093.128 | 493.9828 | 0.001005056 | 1.17129858 | YBR128C | ATG14 | n |
| W303.asm02.HP0_G0003230 | 1004.895 | 896.9179 | 838.6141 | 883.6789 | 880.8602 | 945.0228 | 2380.064 | 1728.818 | 1965.167 | 0.931295759 | 0.016379476 | YBR129C | OPY1 | n |
| W303.asm02.HP0_G0003240 | 2812.435 | 2790.906 | 2895.267 | 1777.084 | 1845.901 | 1697.85 | 1872.696 | 1618.216 | 1699.636 | 4.06E-13 | 0.67577701 | YBR130C | SHE3 | n |
| W303.asm02.HP0_G0003250 | 414.0294 | 413.9621 | 400.9826 | 1804.178 | 1812.923 | 2112.627 | 1251.756 | 1606.405 | 1449.654 | 5.84E-56 | -2.221061192 | YBR131W | CCZ1 | n |
| W303.asm02.HP0_G0003260 | 1356.449 | 1781.595 | 1163.065 | 1713.17 | 1468.39 | 1577.823 | 4603.35 | 3018.452 | 1674.518 | 0.709334488 | -0.146153752 | YBR132C | AGP2 | n |
| W303.asm02.HP0_G0003270 | 2512.767 | 2366.929 | 2843.527 | 3440.234 | 3442.731 | 2815.317 | 5108.25 | 3541.392 | 4518.807 | 0.082025333 | -0.328486979 | YBR133C | HSL7 | n |
| W303.asm02.HP0_G0003280 | 1589.407 | 1425.499 | 1627.644 | 1825.019 | 1950.91 | 1968.291 | 1171.515 | 1094.202 | 1229.575 | 0.003665562 | -0.306875709 | YBR135W | CKS1 | n |
| W303.asm02.HP0_G0003290 | 1596.819 | 1458.883 | 1460.568 | 1448.483 | 1478.804 | 1427.41 | 1390.017 | 1080.243 | 992.75 | 0.757081278 | 0.052651771 | YBR136W | MEC1 | n |
| W303.asm02.HP0_G0003310 | 1892.252 | 2037.539 | 1855.083 | 1419.305 | 1398.094 | 1908.278 | 2168.969 | 1782.508 | 1684.087 | 0.118051584 | 0.291655291 | YBR138C | | n |
| W303.asm02.HP0_G0003320 | 7605.011 | 7844.137 | 7236.011 | 1248.405 | 1305.235 | 1702.408 | 543.1682 | 1256.346 | 804.9648 | 1.11E-16 | 2.414156354 | YBR139W | ATG42 | n |
| W303.asm02.HP0_G0003330 | 7445.117 | 7585.967 | 7405.243 | 7930.185 | 7870.464 | 10365.62 | 7906.8 | 11681.87 | 10024.38 | 0.271829928 | -0.221884088 | YBR140C | IRA1 | n |
| W303.asm02.HP0_G0003340 | 519.9193 | 564.1903 | 590.6948 | 742.6516 | 617.9039 | 206.6288 | 793.7662 | 1097.424 | 704.4937 | 0.842269794 | 0.095747185 | YBR141C | BMT2 | n |
| W303.asm02.HP0_G0003350 | 2956.445 | 3253.831 | 3362.002 | 4473.972 | 3592 | 1104.552 | 6064.966 | 2378.467 | 5013.985 | 0.914105155 | 0.061846387 | YBR142W | MAK5 | n |
| W303.asm02.HP0_G0003360 | 10243.79 | 10203.28 | 10349.02 | 10270.68 | 10589.41 | 7250.999 | 7061.186 | 4644.185 | 6116.775 | 0.574437714 | 0.131620764 | YBR143C | SUP45 | n |
| W303.asm02.HP0_G0003380 | 12384.88 | 12141.78 | 12145.89 | 6354.569 | 6516.629 | 4711.44 | 2336.858 | 1941.43 | 1877.852 | 1.21E-10 | 1.060579359 | YBR146W | MRPS9 | n |
| W303.asm02.HP0_G0003420 | 15501.22 | 15084.02 | 15212.55 | 644.6966 | 605.7541 | 806.7638 | 1233.239 | 1439.966 | 1450.85 | 1.73E-223 | 4.476101018 | YBR149W | ARA1 | n |
| W303.asm02.HP0_G0003440 | 9259.01 | 9414.3 | 8997.316 | 1799.315 | 1818.998 | 1679.618 | 1452.975 | 917.0251 | 961.6518 | 6.34E-37 | 2.384875075 | YBR151W | APD1 | n |
| W303.asm02.HP0_G0003460 | 735.9346 | 807.8938 | 706.0311 | 5729.324 | 6894.141 | 12014.1 | 32932.04 | 31639.51 | 31349.37 | 1.41E-38 | -3.45298827 | YBR153W | RIB7 | n |
| W303.asm02.HP0_G0003470 | 4310.777 | 4250.901 | 4303.017 | 6035.694 | 6159.078 | 4160.683 | 4973.692 | 3685.281 | 4449.434 | 0.089893493 | -0.346328231 | YBR154C | RPB5 | n |
| W303.asm02.HP0_G0003480 | 1000.659 | 935.866 | 1204.026 | 3093.571 | 3363.758 | 1805.723 | 3528.124 | 2081.024 | 3349.036 | 2.95E-06 | -1.39558782 | YBR155W | CNS1 | n |
| W303.asm02.HP0_G0003490 | 1443.279 | 1555.696 | 1663.215 | 1538.796 | 1385.944 | 1938.664 | 1523.34 | 1177.959 | 1208.045 | 0.764801283 | -0.061142761 | YBR156C | SLI15 | n |
| W303.asm02.HP0_G0003500 | 588.7477 | 590.8975 | 529.2539 | 315.4011 | 328.0445 | 602.414 | 399.9693 | 836.4901 | 410.2569 | 0.249094028 | 0.455831501 | YBR157C | ICS2 | n |
| W303.asm02.HP0_G0003510 | 2096.619 | 2114.323 | 2360.623 | 2346.056 | 2678.162 | 3094.114 | 2610.911 | 2416.05 | 3017.721 | 0.059389929 | -0.304857779 | YBR158W | AMN1 | n |
| W303.asm02.HP0_G0003520 | 5046.711 | 5033.201 | 4954.075 | 4382.269 | 4758.381 | 5145.968 | 4509.53 | 6588.836 | 5225.692 | 0.689810936 | 0.073591842 | YBR159W | IFA38 | n |
| W303.asm02.HP0_G0003530 | 4102.174 | 4458.995 | 4331.043 | 3320.048 | 3468.767 | 4147.009 | 2829.412 | 4253.322 | 3927.941 | 0.230852915 | 0.237430746 | YBR160W | CDC28 | n |
| W303.asm02.HP0_G0003550 | 14492.09 | 14587.71 | 15836.66 | 10678.48 | 11594.38 | 11337.99 | 5508.219 | 9366.757 | 7093.976 | 0.042893426 | 0.418328529 | YBR162C | TOS1 | n |
| W303.asm02.HP0_G0003560 | 1221.969 | 928.0763 | 925.9248 | 1554.775 | 1900.575 | 2323.814 | 1383.844 | 1326.143 | 1382.673 | 3.63E-06 | -0.909585848 | YBR162W- | YSY6 | n |
| W303.asm02.HP0_G0003570 | 1069.488 | 1056.049 | 1047.729 | 1130.998 | 1089.142 | 1409.178 | 1417.175 | 1487.213 | 1175.751 | 0.24657362 | -0.193881966 | YBR163W | EXO5 | n |
| W303.asm02.HP0_G0003580 | 2672.66 | 2674.062 | 2682.918 | 4294.735 | 4400.829 | 4783.608 | 4079.934 | 5087.664 | 4408.767 | 9.70E-11 | -0.747316669 | YBR164C | ARL1 | n |
| W303.asm02.HP0_G0003590 | 825.941 | 992.6188 | 779.329 | 759.3247 | 646.5427 | 758.1453 | 1265.335 | 970.7151 | 1106.378 | 0.137483626 | 0.262941735 | YBR165W | UBS1 | n |
| W303.asm02.HP0_G0003600 | 5312.495 | 5028.749 | 5240.799 | 2680.91 | 2550.589 | 3021.946 | 2560.297 | 2795.101 | 2884.955 | 2.56E-18 | 0.916693751 | YBR166C | TYR1 | n |
| W303.asm02.HP0_G0003610 | 992.1881 | 1068.289 | 997.0669 | 488.3854 | 512.8949 | 278.0372 | 935.7306 | 447.7746 | 825.2982 | 0.000146201 | 1.25717685 | YBR167C | POP7 | n |
| W303.asm02.HP0_G0003620 | 787.8206 | 809.0066 | 722.1998 | 323.043 | 307.2163 | 326.6558 | 925.8548 | 396.2322 | 647.0816 | 1.19E-05 | 1.276916813 | YBR168W | PEX32 | n |
| W303.asm02.HP0_G0003640 | 3700.851 | 3958.234 | 3658.427 | 6004.432 | 5915.215 | 8388.217 | 6387.164 | 9682.454 | 7509.017 | 7.94E-05 | -0.843504753 | YBR170C | NPL4 | n |
| W303.asm02.HP0_G0003650 | 3094.102 | 3197.078 | 3063.421 | 2283.532 | 2345.778 | 2366.355 | 3847.853 | 3293.344 | 3554.763 | 4.38E-06 | 0.419260317 | YBR171W | SEC66 | n |
| W303.asm02.HP0_G0003670 | 7455.706 | 7168.666 | 7106.662 | 3433.287 | 3719.573 | 4270.075 | 3104.7 | 4411.17 | 3654.038 | 1.33E-08 | 0.927854098 | YBR173C | UMP1 | n |
| W303.asm02.HP0_G0003680 | 443.6786 | 340.5172 | 445.1769 | 1112.935 | 1214.98 | 1052.135 | 583.9058 | 1272.453 | 781.0431 | 5.88E-07 | -1.458901939 | YBR175W | SWD3 | n |
| W303.asm02.HP0_G0003690 | 1211.38 | 1133.945 | 1177.078 | 944.8139 | 1038.807 | 1017.951 | 365.404 | 668.9774 | 541.8262 | 0.325834234 | 0.23098159 | YBR176W | ECM31 | n |
| W303.asm02.HP0_G0003700 | 15715.12 | 16332.59 | 15984.33 | 6462.944 | 6514.894 | 9264.87 | 8026.544 | 12276.75 | 9622.498 | 2.14E-07 | 1.110653571 | YBR177C | EHT1 | n |

| | | | | | | | | | | | | | | |
|---|---|---|---|---|---|---|---|---|---|---|---|---|---|---|
| W303.asm02.HP0_G0003710 | 5798.529 | 6063.655 | 5732.326 | 4406.584 | 3984.265 | 3961.651 | 6113.111 | 5434.501 | 6900.21 | 3.08E-05 | 0.510216477 | YBR179C | FZO1 | n |
| W303.asm02.HP0_G0003720 | 265.7836 | 283.7644 | 223.1274 | 3572.23 | 3165.022 | 2468.91 | 2243.038 | 2260.349 | 3783.215 | 8.34E-43 | -3.574800534 | YBR180W | DTR1 | n |
| W303.asm02.HP0_G0003730 | 10556.16 | 11239.29 | 12242.9 | 35470.82 | 40478.78 | 37464.38 | 32081.49 | 28536.23 | 26277.97 | 1.08E-46 | -1.736354513 | YPL090C | RPS6A | y |
| W303.asm02.HP0_G0003780 | 3660.613 | 3594.348 | 3229.419 | 3757.025 | 3656.221 | 2772.016 | 1473.961 | 1772.844 | 1657.773 | 0.819447937 | 0.041769191 | YBR185C | MBA1 | n |
| W303.asm02.HP0_G0003790 | 538.9794 | 543.0471 | 580.9936 | 91.70253 | 95.46268 | 88.88076 | 58.02024 | 248.0478 | 142.334 | 1.71E-09 | 2.590998429 | YBR186W | PCH2 | n |
| W303.asm02.HP0_G0003800 | 2919.384 | 2537.187 | 2824.125 | 2386.35 | 2839.581 | 2124.022 | 2046.756 | 2412.828 | 2088.363 | 0.294575044 | 0.172296577 | YBR187W | GDT1 | n |
| W303.asm02.HP0_G0003820 | 13397.19 | 12471.16 | 13852.22 | 17511.01 | 20199.9 | 18304.88 | 15833.35 | 10395.46 | 8926.377 | 0.030904574 | -0.495934432 | YBR189W | RPS9B | y |
| W303.asm02.HP0_G0003840 | 3483.777 | 3215.996 | 3184.147 | 1711.781 | 1561.249 | 940.4648 | 3119.513 | 2044.515 | 2717.504 | 2.27E-06 | 1.230056228 | YBR192W | RIM2 | n |
| W303.asm02.HP0_G0003850 | 2096.619 | 2366.929 | 2266.845 | 1168.513 | 1172.455 | 885.0093 | 1229.535 | 1566.674 | 1334.83 | 1.59E-10 | 1.061014817 | YBR193C | MED8 | n |
| W303.asm02.HP0_G0003860 | 1074.782 | 1053.823 | 994.9111 | 815.5967 | 830.5253 | 856.9017 | 791.2973 | 847.2281 | 782.2391 | 0.000612406 | 0.319616717 | YBR194W | AIM4 | n |
| W303.asm02.HP0_G0003870 | 424.6184 | 334.9532 | 408.5279 | 1237.984 | 1248.825 | 1152.411 | 569.0921 | 586.2948 | 480.8259 | 1.23E-29 | -1.638895142 | YBR195C | MSI1 | n |
| W303.asm02.HP0_G0003880 | 44516.1 | 43634.05 | 46148.57 | 28421.53 | 31131.25 | 49461.76 | 91302.86 | 81578.73 | 82388.68 | 0.199058037 | 0.300925004 | YBR196C | PGI1 | n |
| W303.asm02.HP0_G0003900 | 978.4224 | 791.2018 | 865.5618 | 856.585 | 899.9527 | 1366.637 | 1143.122 | 1310.036 | 1081.26 | 0.301506032 | -0.245058494 | YBR197C | | n |
| W303.asm02.HP0_G0003910 | 5354.851 | 5085.502 | 5284.993 | 6377.494 | 6072.294 | 7551.826 | 6018.056 | 5955.294 | 6400.247 | 0.002895773 | -0.347065526 | YBR198C | TAF5 | n |
| W303.asm02.HP0_G0003930 | 3416.007 | 3428.541 | 3322.119 | 3756.33 | 3462.692 | 3229.334 | 3139.265 | 2549.201 | 2856.249 | 0.76544726 | -0.039517717 | YBR200W | BEM1 | n |
| W303.asm02.HP0_G0003950 | 1621.174 | 1604.66 | 1469.192 | 689.1584 | 747.2124 | 796.8882 | 1792.455 | 1640.766 | 1461.615 | 1.10E-17 | 1.072286525 | YBR201W | DER1 | n |
| W303.asm02.HP0_G0003960 | 1444.338 | 1447.755 | 1553.269 | 6219.794 | 6132.175 | 5702.802 | 2529.435 | 2359.138 | 2499.816 | 4.05E-128 | -2.021990608 | YBR202W | MCM7 | n |
| W303.asm02.HP0_G0003970 | 3584.372 | 4119.591 | 3533.389 | 1228.953 | 1205.433 | 2515.25 | 7614.23 | 4033.193 | 5484.047 | 0.000672967 | 1.182855331 | YBR203W | COS111 | n |
| W303.asm02.HP0_G0003980 | 4309.718 | 4413.37 | 4235.109 | 227.8669 | 237.7888 | 181.5598 | 474.0377 | 234.0884 | 264.3346 | 7.10E-59 | 4.323786374 | YBR204C | LDH1 | n |
| W303.asm02.HP0_G0003990 | 5621.693 | 5367.041 | 5563.094 | 6747.083 | 7425.261 | 7616.397 | 5076.153 | 5632.081 | 5431.419 | 1.30E-05 | -0.396486703 | YBR205W | KTR3 | n |
| W303.asm02.HP0_G0004000 | 2987.153 | 3510.888 | 2847.839 | 2576.007 | 2492.444 | 2374.712 | 3009.645 | 2712.419 | 3026.093 | 0.007738108 | 0.328293064 | YBR207W | FTH1 | n |
| W303.asm02.HP0_G0004010 | 4833.873 | 5103.307 | 5255.89 | 14532.77 | 15980.45 | 23087.73 | 6360.005 | 11516.5 | 5340.517 | 4.67E-09 | -1.818845581 | YBR208C | DUR12 | n |
| W303.asm02.HP0_G0004030 | 696.7553 | 899.1435 | 717.8882 | 323.7377 | 303.7449 | 305.3852 | 419.7209 | 354.354 | 324.1389 | 4.37E-15 | 1.309851016 | YBR211C | AME1 | n |
| W303.asm02.HP0_G0004040 | 9076.88 | 9637.973 | 9457.584 | 1248.405 | 1284.407 | 986.8043 | 661.6776 | 613.1398 | 391.1196 | 1.24E-45 | 3.000870274 | YBR212W | NGR1 | n |
| W303.asm02.HP0_G0004050 | 1256.913 | 998.1828 | 1244.986 | 1723.591 | 1895.368 | 1751.027 | 220.9707 | 1203.73 | 718.8467 | 0.226950099 | -0.617495488 | YBR213W | MET8 | n |
| W303.asm02.HP0_G0004060 | 16814.25 | 15827.37 | 15577.96 | 5521.604 | 4929.345 | 6201.142 | 6626.652 | 5302.424 | 4882.416 | 7.83E-23 | 1.533858143 | YBR214W | SDS24 | n |
| W303.asm02.HP0_G0004070 | 2529.709 | 2812.049 | 2626.867 | 2819.853 | 2650.391 | 2848.742 | 2061.57 | 2958.319 | 2553.64 | 0.720874706 | -0.062173081 | YBR215W | HPC2 | n |
| W303.asm02.HP0_G0004090 | 768.7605 | 902.4819 | 841.8478 | 546.0469 | 584.058 | 876.653 | 454.2861 | 746.291 | 501.1593 | 0.254786935 | 0.324518783 | YBR217W | ATG12 | n |
| W303.asm02.HP0_G0004110 | 1609.526 | 1547.907 | 1789.331 | 1548.522 | 1576.002 | 2124.022 | 2589.925 | 1952.168 | 2313.227 | 0.681028982 | -0.085450377 | YBR220C | | n |
| W303.asm02.HP0_G0004120 | 13732.86 | 13296.86 | 13935.22 | 20012.69 | 20423.81 | 23877.78 | 4758.894 | 12560.24 | 11200.13 | 0.054119348 | -0.650748016 | YBR221C | PDB1 | n |
| W303.asm02.HP0_G0004140 | 1716.475 | 1813.866 | 1688.007 | 792.6711 | 691.6705 | 749.0293 | 759.201 | 449.9222 | 840.8473 | 1.63E-07 | 1.224150802 | YBR223C | TDP1 | n |
| W303.asm02.HP0_G0004150 | 1831.895 | 1936.274 | 1691.241 | 2448.874 | 2096.708 | 2443.081 | 10080.71 | 6610.312 | 7641.783 | 0.065057568 | -0.356422081 | YBR225W | | n |
| W303.asm02.HP0_G0004160 | 2517.002 | 2652.918 | 2724.957 | 2343.278 | 2356.192 | 2337.488 | 1660.366 | 1243.46 | 1211.633 | 0.296026973 | 0.165959028 | YBR227C | MCX1 | n |
| W303.asm02.HP0_G0004170 | 1239.97 | 1224.082 | 1225.584 | 900.3521 | 851.3535 | 666.9855 | 1188.798 | 749.5124 | 852.8081 | 0.004878515 | 0.609225144 | YBR228W | SLX1 | n |
| W303.asm02.HP0_G0004180 | 2541.357 | 2380.282 | 2382.181 | 5178.414 | 5090.764 | 5580.496 | 3384.925 | 4814.919 | 4590.572 | 9.00E-13 | -1.117725523 | YBR229C | ROT2 | n |
| W303.asm02.HP0_G0004190 | 2712.898 | 2926.668 | 2484.583 | 1020.538 | 1157.702 | 1931.827 | 1228.301 | 3970.912 | 2881.367 | 0.0200636 | 0.983050453 | YBR230C | OM14 | n |
| W303.asm02.HP0_G0004210 | 1584.112 | 1416.596 | 1427.153 | 796.1447 | 763.7014 | 800.6865 | 1331.996 | 1221.984 | 1181.731 | 3.97E-19 | 0.90729778 | YBR231C | SWC5 | n |
| W303.asm02.HP0_G0004220 | 1049.369 | 1068.289 | 1127.494 | 792.6711 | 819.2433 | 1014.912 | 469.0998 | 726.9626 | 570.5322 | 0.140870082 | 0.304946852 | YBR233W | PBP2 | n |
| W303.asm02.HP0_G0004230 | 360.0256 | 292.6668 | 331.9963 | 145.1957 | 164.0222 | 121.5463 | 86.41312 | 56.9114 | 72.96114 | 3.10E-08 | 1.194520536 | YBR233W- | DAD3 | n |
| W303.asm02.HP0_G0004240 | 11035.84 | 10539.34 | 11255.54 | 7086.105 | 7712.516 | 9514.799 | 8832.655 | 12860.9 | 10749.21 | 0.021923051 | 0.433304275 | YBR234C | ARC40 | n |
| W303.asm02.HP0_G0004250 | 1924.019 | 1764.903 | 1908.979 | 1581.174 | 1566.456 | 2255.444 | 4229.305 | 3088.249 | 3631.312 | 0.821616967 | 0.05106814 | YBR235W | VHC1 | n |
| W303.asm02.HP0_G0004260 | 2406.877 | 2499.352 | 2376.792 | 3173.463 | 3313.423 | 3144.252 | 3162.72 | 2918.588 | 2961.505 | 1.36E-07 | -0.403063911 | YBR236C | ABD1 | n |
| W303.asm02.HP0_G0004270 | 2101.914 | 2142.143 | 1961.796 | 2964.354 | 2880.369 | 3066.006 | 2073.915 | 2730.673 | 2595.503 | 0.000103212 | -0.521974666 | YBR237W | PRP5 | n |
| W303.asm02.HP0_G0004280 | 1488.812 | 1314.218 | 1475.659 | 17422.79 | 17038.35 | 26683.98 | 29038.51 | 27622.43 | 34513.01 | 6.37E-77 | -3.836963241 | YBR238C | | n |
| W303.asm02.HP0_G0004290 | 1219.851 | 1110.576 | 1305.349 | 2152.231 | 2186.963 | 2892.043 | 2641.772 | 3067.846 | 2626.601 | 1.85E-09 | -0.991948007 | YBR239C | ERT1 | n |
| W303.asm02.HP0_G0004300 | 185.3073 | 156.905 | 191.868 | 156.3111 | 151.8724 | 189.9162 | 103.6957 | 835.4163 | 212.903 | 0.890607117 | 0.100613858 | YBR240C | THI2 | n |
| W303.asm02.HP0_G0004310 | 2833.613 | 2836.531 | 2801.488 | 550.9099 | 533.7231 | 730.0377 | 1373.969 | 795.6857 | 723.631 | 4.58E-17 | 2.222819957 | YBR241C | VVS1 | n |
| W303.asm02.HP0_G0004320 | 1652.941 | 1880.634 | 1836.759 | 3322.133 | 3416.696 | 2528.164 | 1039.426 | 1930.692 | 1941.245 | 0.003302728 | -0.787085379 | YBR242W | | n |
| W303.asm02.HP0_G0004330 | 1609.526 | 1426.611 | 1585.606 | 1579.09 | 1659.315 | 2551.713 | 1001.158 | 2503.028 | 2108.697 | 0.383702981 | -0.325152103 | YBR243C | ALG7 | n |
| W303.asm02.HP0_G0004350 | 5047.77 | 5279.13 | 5594.354 | 10470.76 | 10622.39 | 12336.19 | 11633.67 | 12590.3 | 13135.4 | 3.72E-22 | -1.070174897 | YBR245C | ISW1 | n |
| W303.asm02.HP0_G0004360 | 3258.231 | 3248.267 | 3321.041 | 1221.311 | 1135.138 | 642.6763 | 892.524 | 568.0402 | 892.2789 | 1.01E-09 | 1.712291183 | YBR246W | RRT2 | n |
| W303.asm02.HP0_G0004370 | 2996.683 | 3360.66 | 3739.27 | 4806.741 | 4057.164 | 1239.773 | 7482.141 | 2854.16 | 6638.268 | 0.998298462 | -0.001014422 | YBR247C | ENP1 | n |
| W303.asm02.HP0_G0004380 | 2017.202 | 1625.803 | 2201.093 | 6406.672 | 6324.836 | 8131.45 | 3592.317 | 4208.222 | 3181.584 | 7.86E-23 | -1.835839611 | YBR248C | HIS7 | n |
| W303.asm02.HP0_G0004390 | 18113.52 | 18108.62 | 18000.02 | 12030.4 | 12572.43 | 11105.54 | 22409.39 | 16100.56 | 16826.51 | 0.000100544 | 0.602634013 | YBR249C | ARO4 | n |

| | | | | | | | | | | | | | | |
|---|---|---|---|---|---|---|---|---|---|---|---|---|---|---|
| W303.asm02.HP0_G0004400 | 225.5454 | 291.554 | 242.5298 | 1388.043 | 1085.671 | 2093.636 | 530.8234 | 7883.839 | 1392.242 | 0.004954712 | -2.588024445 | YBR250W | SPO23 | n |
| W303.asm02.HP0_G0004410 | 4853.992 | 4857.378 | 4484.106 | 6033.61 | 6127.836 | 4759.299 | 1385.079 | 1489.361 | 1331.242 | 0.07270295 | -0.253331828 | YBR251W | MRPS5 | n |
| W303.asm02.HP0_G0004420 | 1107.608 | 905.8203 | 950.7167 | 740.5674 | 852.2214 | 832.5924 | 812.2833 | 689.3796 | 588.4735 | 0.097722336 | 0.289899083 | YBR253W | SRB6 | n |
| W303.asm02.HP0_G0004430 | 776.1728 | 682.1472 | 780.4069 | 659.2856 | 757.6265 | 582.6628 | 323.4319 | 522.9406 | 403.0804 | 0.478810004 | 0.163441504 | YBR254C | TRS20 | n |
| W303.asm02.HP0_G0004450 | 711.5799 | 711.0801 | 673.6938 | 1020.538 | 1004.094 | 976.169 | 732.0425 | 968.5675 | 1010.691 | 0.000697096 | -0.517506235 | YBR255C-/ | RCF3 | n |
| W303.asm02.HP0_G0004460 | 3841.685 | 3596.574 | 3430.988 | 1979.246 | 2085.426 | 2611.727 | 2975.08 | 3523.138 | 2898.112 | 6.21E-06 | 0.703126991 | YBR256C | RIB5 | n |
| W303.asm02.HP0_G0004470 | 694.6376 | 760.0433 | 723.2777 | 1136.556 | 1128.195 | 857.6614 | 1199.908 | 1177.959 | 1220.006 | 0.00045451 | -0.519694991 | YBR257W | POP4 | n |
| W303.asm02.HP0_G0004480 | 621.5735 | 586.4463 | 579.9157 | 661.3698 | 744.6089 | 526.4476 | 714.7599 | 581.9996 | 712.8663 | 0.566583509 | -0.111517737 | YBR258C | SHG1 | n |
| W303.asm02.HP0_G0004500 | 2526.532 | 2678.513 | 2741.125 | 7100.694 | 6512.29 | 8571.296 | 5116.891 | 6837.958 | 5357.262 | 5.69E-19 | -1.481253041 | YBR260C | RGD1 | n |
| W303.asm02.HP0_G0004510 | 2339.107 | 2360.252 | 2645.191 | 1567.28 | 1697.5 | 862.2193 | 1671.477 | 875.1469 | 1538.164 | 0.011536198 | 0.831664446 | YBR261C | TAE1 | n |
| W303.asm02.HP0_G0004520 | 2083.913 | 1947.402 | 1835.681 | 876.0371 | 1070.05 | 692.8141 | 114.806 | 328.5828 | 285.8642 | 0.001356203 | 1.152841863 | YBR262C | MIC12 | n |
| W303.asm02.HP0_G0004530 | 9235.715 | 8100.081 | 9990.071 | 7119.451 | 7758.512 | 6983.597 | 5877.326 | 7306.135 | 7194.447 | 0.026105635 | 0.321929724 | YBR263W | SHM1 | n |
| W303.asm02.HP0_G0004540 | 831.2355 | 797.8786 | 755.615 | 409.8825 | 409.6217 | 391.9869 | 532.0579 | 429.52 | 423.4139 | 1.70E-13 | 0.977131191 | YBR264C | YPT10 | n |
| W303.asm02.HP0_G0004550 | 2126.269 | 1929.598 | 2096.535 | 1972.299 | 2232.959 | 1753.306 | 1538.153 | 2057.401 | 1630.263 | 0.801671716 | 0.046410277 | YBR265W | TSC10 | n |
| W303.asm02.HP0_G0004560 | 1447.515 | 1711.489 | 1668.605 | 4872.739 | 4031.128 | 1289.911 | 8222.825 | 3034.559 | 6782.994 | 0.032231455 | -1.0783218 | YBR267W | REI1 | n |
| W303.asm02.HP0_G0004570 | 3068.688 | 2647.354 | 2603.153 | 1086.536 | 1119.517 | 787.7722 | 548.1061 | 491.8004 | 471.2572 | 2.70E-17 | 1.474651401 | YBR268W | MRPL37 | n |
| W303.asm02.HP0_G0004580 | 2438.644 | 2336.883 | 2256.066 | 729.4519 | 800.1508 | 437.5668 | 277.7564 | 405.8964 | 366.0018 | 3.27E-13 | 1.83800787 | YBR269C | SDH8 | n |
| W303.asm02.HP0_G0004600 | 634.2803 | 528.5807 | 696.33 | 1743.043 | 1641.09 | 1515.531 | 2365.25 | 1748.146 | 2286.913 | 1.48E-15 | -1.397864704 | YBR271W | EFM2 | n |
| W303.asm02.HP0_G0004610 | 2782.786 | 2764.199 | 2600.997 | 1700.665 | 1579.473 | 1811.8 | 1328.293 | 1811.5 | 1651.792 | 3.21E-06 | 0.678074501 | YBR272C | HSM3 | n |
| W303.asm02.HP0_G0004620 | 5956.305 | 5847.771 | 5715.08 | 2469.021 | 2292.84 | 3808.958 | 4488.544 | 4585.126 | 4220.982 | 5.77E-07 | 1.031356016 | YBR273C | UBX7 | n |
| W303.asm02.HP0_G0004630 | 1778.95 | 1708.15 | 1788.253 | 1155.313 | 1032.733 | 1254.206 | 2133.169 | 1690.161 | 2066.834 | 9.84E-06 | 0.615579583 | YBR274W | CHK1 | n |
| W303.asm02.HP0_G0004640 | 3232.818 | 3330.615 | 3332.898 | 2516.262 | 2382.228 | 1809.521 | 1639.38 | 1364.8 | 1841.97 | 0.001799511 | 0.560992716 | YBR275C | RIF1 | n |
| W303.asm02.HP0_G0004650 | 2811.376 | 2678.513 | 2809.034 | 3878.6 | 3512.159 | 4836.785 | 8891.91 | 6116.364 | 8018.549 | 0.003902125 | -0.559185777 | YBR276C | PPS1 | n |
| W303.asm02.HP0_G0004670 | 4691.98 | 5140.03 | 4657.65 | 4609.442 | 4188.208 | 4069.523 | 4091.044 | 3484.481 | 3985.353 | 0.148660363 | 0.171198006 | YBR279W | PAF1 | n |
| W303.asm02.HP0_G0004680 | 3597.079 | 4050.597 | 3196.004 | 904.5204 | 853.0892 | 2380.789 | 2470.181 | 2379.541 | 1868.284 | 0.000378823 | 1.389639377 | YBR280C | SAF1 | n |
| W303.asm02.HP0_G0004690 | 6456.105 | 4712.714 | 5571.718 | 3183.884 | 3042.656 | 4165.241 | 5555.129 | 7193.386 | 6881.073 | 0.00062442 | 0.687889101 | YBR281C | DUG2 | n |
| W303.asm02.HP0_G0004700 | 4514.085 | 4227.532 | 4125.162 | 2348.835 | 2147.91 | 1822.435 | 1452.975 | 1510.837 | 1747.479 | 1.75E-13 | 1.025762598 | YBR282W | MRPL27 | n |
| W303.asm02.HP0_G0004710 | 9336.31 | 9425.428 | 9832.696 | 12088.76 | 12433.58 | 14274.86 | 8615.388 | 12899.56 | 11145.11 | 0.012236185 | -0.440217612 | YBR283C | SSH1 | n |
| W303.asm02.HP0_G0004740 | 26521.18 | 26741.73 | 26255.74 | 5891.888 | 6064.484 | 7916.465 | 3908.342 | 8818.045 | 6800.935 | 1.95E-12 | 2.00049302 | YBR286W | APE3 | n |
| W303.asm02.HP0_G0004750 | 4267.362 | 4908.567 | 4093.903 | 2202.25 | 2244.241 | 2550.954 | 6897.001 | 6861.582 | 7323.624 | 2.83E-15 | 0.923188292 | YBR287W | | n |
| W303.asm02.HP0_G0004760 | 3480.6 | 3481.955 | 3457.936 | 3875.127 | 4052.825 | 4647.628 | 3426.897 | 3903.263 | 3750.92 | 0.016751927 | -0.271183317 | YBR288C | APM3 | n |
| W303.asm02.HP0_G0004770 | 3463.658 | 3397.383 | 3696.154 | 5575.792 | 4887.689 | 5360.953 | 5964.974 | 5068.336 | 5537.87 | 7.23E-08 | -0.584046695 | YBR289W | SNF5 | n |
| W303.asm02.HP0_G0004780 | 4046.052 | 4102.899 | 3656.271 | 3073.424 | 3448.806 | 3220.978 | 3047.914 | 4501.369 | 3442.331 | 0.121459794 | 0.277028138 | YBR290W | BSD2 | n |
| W303.asm02.HP0_G0004790 | 1053.604 | 1219.63 | 1221.272 | 1121.967 | 944.2127 | 1266.361 | 699.9462 | 604.5494 | 543.0223 | 0.721096752 | 0.068015414 | YBR291C | CTP1 | n |
| W303.asm02.HP0_G0004800 | 1408.335 | 1249.676 | 1215.883 | 830.8805 | 708.1595 | 995.1606 | 560.4508 | 1840.493 | 816.9256 | 0.146004206 | 0.612177018 | YBR293W | VBA2 | n |
| W303.asm02.HP0_G0004810 | 912.7707 | 827.9242 | 931.3144 | 857.2797 | 815.772 | 1054.414 | 1369.031 | 1459.294 | 1064.515 | 0.885207293 | -0.029746816 | YBR294W | SUL1 | n |
| W303.asm02.HP0_G0004830 | 394.9692 | 417.3005 | 371.879 | 989.9705 | 886.0672 | 981.4867 | 973.9993 | 912.7299 | 610.003 | 2.24E-10 | -1.271192829 | YBR296C | PHO89 | n |
| W303.asm02.HP0_G0004980 | 411.9116 | 422.8645 | 488.2933 | 1527.681 | 1508.31 | 4222.976 | 3786.129 | 2795.101 | 3216.271 | 4.16E-10 | -2.455904425 | YCL064C | CHA1 | n |
| W303.asm02.HP0_G0004990 | 587.6888 | 590.8975 | 675.8497 | 2069.559 | 1862.39 | 2235.693 | 2370.188 | 1563.453 | 1773.793 | 3.25E-20 | -1.733823966 | YCL063W | VAC17 | n |
| W303.asm02.HP0_G0005000 | 2128.386 | 2223.377 | 2195.703 | 2620.469 | 2679.898 | 2985.482 | 729.5736 | 1226.28 | 938.9262 | 0.099406916 | -0.339721171 | YCL061C | MRC1 | n |
| W303.asm02.HP0_G0005010 | 3960.281 | 3924.85 | 4342.9 | 5135.342 | 5048.24 | 2465.871 | 6994.525 | 3357.772 | 5882.343 | 0.903633622 | -0.04887087 | YCL059C | KRR1 | n |
| W303.asm02.HP0_G0005030 | 2715.016 | 2511.593 | 2394.038 | 4093.962 | 5141.099 | 3557.509 | 1866.523 | 2914.293 | 2526.13 | 0.000598094 | -0.74721974 | YCL057C-/ | MIC10 | n |
| W303.asm02.HP0_G0005040 | 8885.219 | 7767.354 | 8958.511 | 3836.917 | 4277.596 | 5400.456 | 1702.338 | 9163.809 | 3268.898 | 0.079444631 | 0.922215254 | YCL057W | PRD1 | n |
| W303.asm02.HP0_G0005060 | 978.4224 | 1001.521 | 963.6517 | 100.0391 | 91.99131 | 228.659 | 129.6197 | 422.0034 | 227.256 | 2.10E-09 | 2.806333461 | YCL055W | KAR4 | n |
| W303.asm02.HP0_G0005070 | 2260.749 | 2671.836 | 2873.708 | 5227.044 | 4347.891 | 1055.174 | 8052.468 | 2823.02 | 6805.72 | 0.454109894 | -0.445452738 | YCL054W | SPB1 | n |
| W303.asm02.HP0_G0005080 | 7711.959 | 7581.516 | 7359.97 | 1259.52 | 1181.134 | 1259.524 | 1182.625 | 1301.446 | 1208.045 | 1.79E-237 | 2.613580294 | YCL052C | PBN1 | n |
| W303.asm02.HP0_G0005100 | 10509.57 | 10607.22 | 10254.16 | 9171.642 | 10217.11 | 15911.94 | 26200.46 | 31080.07 | 29375.83 | 0.480981055 | -0.170267173 | YCL050C | APA1 | n |
| W303.asm02.HP0_G0005130 | 691.4609 | 598.6871 | 545.4225 | 135.4696 | 102.4054 | 85.8421 | 214.7983 | 432.7414 | 305.0015 | 5.84E-17 | 2.501927435 | YCL048W | SPS22 | n |
| W303.asm02.HP0_G0005150 | 7266.163 | 6723.546 | 7359.97 | 4697.671 | 4695.028 | 6249.761 | 2351.671 | 5011.424 | 3841.823 | 0.121804491 | 0.448742723 | YCL045C | EMC1 | n |
| W303.asm02.HP0_G0005160 | 1737.653 | 1780.482 | 1606.086 | 2572.534 | 2543.646 | 2888.245 | 2434.381 | 2129.345 | 2105.108 | 1.66E-08 | -0.643557107 | YCL044C | MGR1 | n |
| W303.asm02.HP0_G0005170 | 21444.82 | 21506 | 23047.87 | 15247.63 | 16859.58 | 20697.82 | 16602.43 | 27474.24 | 19355.04 | 0.163836391 | 0.321766091 | YCL043C | PDI1 | n |
| W303.asm02.HP0_G0005180 | 42779.51 | 39452.15 | 39667.09 | 342.4951 | 332.3837 | 414.7769 | 7704.347 | 4432.646 | 7530.547 | 1.33E-197 | 6.80547806 | YCL040W | GLK1 | n |
| W303.asm02.HP0_G0005190 | 4952.469 | 4450.093 | 5024.139 | 2828.189 | 2826.563 | 7542.71 | 8216.653 | 8934.015 | 7516.194 | 0.757081278 | 0.128466715 | YCL039W | GID7 | n |
| W303.asm02.HP0_G0005200 | 2179.214 | 1789.385 | 1976.887 | 1694.413 | 1886.69 | 2449.918 | 3788.598 | 5091.959 | 3822.686 | 0.932395718 | -0.020530269 | YCL038C | ATG22 | n |

| | | | | | | | | | | | | | | |
|---|---|---|---|---|---|---|---|---|---|---|---|---|---|---|
| W303.asm02.HP0_G0005210 | 343.0832 | 318.2612 | 459.1897 | 13001.61 | 11526.68 | 7086.152 | 9593.09 | 5650.335 | 8300.825 | 3.33E-60 | -4.818254102 | YCL037C | SRO9 | n |
| W303.asm02.HP0_G0005230 | 6182.91 | 5942.359 | 5331.344 | 2666.321 | 3449.674 | 3590.935 | 3591.082 | 5544.029 | 4323.845 | 3.90E-05 | 0.846786178 | YCL035C | GRX1 | n |
| W303.asm02.HP0_G0005240 | 4965.176 | 4751.662 | 4637.169 | 2242.544 | 2246.844 | 5481.74 | 7527.817 | 9656.683 | 7343.958 | 0.155022258 | 0.525602237 | YCL034W | LSB5 | n |
| W303.asm02.HP0_G0005250 | 2903.5 | 2547.202 | 2562.192 | 898.268 | 1037.072 | 976.9287 | 454.2861 | 904.1395 | 808.553 | 5.63E-09 | 1.460335776 | YCL033C | MXR2 | n |
| W303.asm02.HP0_G0005260 | 3036.921 | 3484.181 | 3202.471 | 2578.092 | 2323.214 | 2378.51 | 3647.868 | 3240.728 | 3368.173 | 8.25E-05 | 0.417278284 | YCL032W | STE50 | n |
| W303.asm02.HP0_G0005270 | 2899.265 | 3006.789 | 3004.136 | 3942.514 | 4157.833 | 2699.848 | 3175.065 | 2932.548 | 3064.368 | 0.143305604 | -0.277470617 | YCL031C | RRP7 | n |
| W303.asm02.HP0_G0005280 | 5757.232 | 4900.777 | 5011.204 | 7119.451 | 7348.89 | 9002.026 | 12391.64 | 11558.38 | 10931.01 | 3.46E-05 | -0.582882312 | YCL030C | HIS4 | n |
| W303.asm02.HP0_G0005290 | 1786.362 | 1794.949 | 2041.562 | 2013.982 | 1959.588 | 2174.92 | 1193.735 | 1581.707 | 1062.123 | 0.503524202 | -0.128958702 | YCL029C | BIK1 | n |
| W303.asm02.HP0_G0005310 | 360.0256 | 388.3677 | 351.3987 | 892.0155 | 881.728 | 1472.99 | 461.6929 | 845.0805 | 871.9455 | 4.26E-07 | -1.561866407 | YCL027W | FUS1 | n |
| W303.asm02.HP0_G0005320 | 1032.426 | 754.4793 | 794.4198 | 1163.65 | 1393.755 | 1536.802 | 539.4647 | 1225.206 | 721.2388 | 0.039187963 | -0.665338092 | YCL026C-I | HBN1 | n |
| W303.asm02.HP0_G0005340 | 3994.166 | 4617.013 | 4006.592 | 9625.987 | 10799.43 | 12403.04 | 21381.07 | 16985.37 | 14960.62 | 9.31E-15 | -1.37949977 | YCL025C | AGP1 | n |
| W303.asm02.HP0_G0005370 | 868.2969 | 1053.823 | 855.8607 | 1191.438 | 1233.204 | 2202.268 | 5555.129 | 3718.569 | 4348.963 | 0.011790872 | -0.736108286 | YCL018W | LEU2 | n |
| W303.asm02.HP0_G0005380 | 6320.566 | 6455.361 | 6392.007 | 13013.42 | 13290.14 | 10591.24 | 8404.293 | 10765.92 | 8830.691 | 1.21E-10 | -0.944712437 | YCL017C | NFS1 | n |
| W303.asm02.HP0_G0005390 | 1074.782 | 1088.32 | 1167.377 | 1153.229 | 1260.107 | 1140.257 | 265.4117 | 868.7041 | 448.5316 | 0.824664091 | -0.09350995 | YCL016C | DCC1 | n |
| W303.asm02.HP0_G0005400 | 2042.616 | 2102.082 | 2117.016 | 4630.978 | 4697.632 | 4453.154 | 1662.835 | 2810.134 | 2231.893 | 4.20E-09 | -1.138128895 | YCL014W | BUD3 | n |
| W303.asm02.HP0_G0005430 | 1626.468 | 1662.525 | 1609.32 | 968.4343 | 977.1907 | 809.0428 | 470.3342 | 548.7118 | 462.8846 | 8.95E-11 | 0.830541516 | YCL010C | SGF29 | n |
| W303.asm02.HP0_G0005440 | 10026.71 | 8312.626 | 9793.892 | 8445.664 | 8769.549 | 10697.6 | 9756.041 | 8340.204 | 7867.843 | 0.952095314 | 0.011363553 | YCL009C | ILV6 | n |
| W303.asm02.HP0_G0005450 | 2321.106 | 1950.741 | 2465.18 | 380.7044 | 295.0665 | 594.0577 | 251.8325 | 343.616 | 246.3934 | 7.39E-18 | 2.406957538 | YCL008C | STP22 | n |
| W303.asm02.HP0_G0005470 | 1007.013 | 1066.064 | 1013.236 | 761.4089 | 817.5077 | 1531.484 | 1985.033 | 2496.585 | 2521.346 | 0.973353763 | -0.011278856 | YCL005W | LDB16 | n |
| W303.asm02.HP0_G0005480 | 1622.233 | 1663.638 | 1439.01 | 2347.446 | 2258.994 | 2078.442 | 1415.941 | 1547.346 | 1697.244 | 4.81E-05 | -0.500709061 | YCL004W | PGS1 | n |
| W303.asm02.HP0_G0005500 | 2403.7 | 2507.141 | 2333.675 | 2560.029 | 2653.862 | 3092.595 | 1510.995 | 2561.013 | 2179.266 | 0.382242279 | -0.197355738 | YCL001W | RER1 | n |
| W303.asm02.HP0_G0005530 | 6280.328 | 6512.114 | 6256.19 | 7286.183 | 7584.944 | 7839.739 | 3952.783 | 5626.712 | 4499.669 | 0.105448727 | -0.253689409 | YCR002C | CDC10 | n |
| W303.asm02.HP0_G0005540 | 3679.673 | 3660.004 | 3119.472 | 3648.649 | 3742.137 | 2736.312 | 1868.992 | 1526.943 | 1647.008 | 0.810577156 | 0.046586854 | YCR003W | MRPL32 | n |
| W303.asm02.HP0_G0005550 | 5347.438 | 5378.169 | 4900.179 | 4536.496 | 5286.029 | 7904.31 | 10575.73 | 12686.95 | 12081.65 | 0.45089137 | -0.181998475 | YCR004C | YCP4 | n |
| W303.asm02.HP0_G0005580 | 831.2355 | 780.0738 | 725.4335 | 287.6125 | 354.9476 | 385.15 | 569.0921 | 2.1476 | 587.2774 | 0.399873602 | 1.185088798 | YCR007C | DFP3 | n |
| W303.asm02.HP0_G0005590 | 1955.786 | 2246.746 | 1879.875 | 5937.739 | 5643.58 | 5288.785 | 16132.09 | 9765.137 | 13752.58 | 1.98E-13 | -1.471802427 | YCR008W | SAT4 | n |
| W303.asm02.HP0_G0005600 | 10225.78 | 10428.06 | 9851.021 | 7507.797 | 9149.664 | 10013.14 | 11104.09 | 11591.67 | 9817.46 | 0.198819499 | 0.193839581 | YCR009C | RVS161 | n |
| W303.asm02.HP0_G0005610 | 302.845 | 433.9925 | 362.1778 | 588.4246 | 589.2651 | 637.3586 | 1522.105 | 1271.379 | 1286.987 | 2.56E-05 | -0.724459333 | YCR010C | ADY2 | n |
| W303.asm02.HP0_G0005620 | 6106.669 | 6400.834 | 5692.443 | 3044.246 | 3124.233 | 5316.133 | 8709.208 | 7067.751 | 8328.335 | 0.006697361 | 0.66420725 | YCR011C | ADP1 | n |
| W303.asm02.HP0_G0005630 | 78844.54 | 77566.71 | 82225.14 | 36134.97 | 43016.35 | 63823.22 | 126094 | 97324.93 | 108475.3 | 0.001356203 | 0.739057728 | YCR012W | PGK1 | n |
| W303.asm02.HP0_G0005640 | 2253.336 | 2237.844 | 2114.86 | 1435.284 | 1406.773 | 1042.26 | 2614.614 | 2440.747 | 2028.559 | 1.01E-05 | 0.766165276 | YCR014C | POL4 | n |
| W303.asm02.HP0_G0005650 | 955.1266 | 1044.92 | 975.5087 | 466.1545 | 412.2252 | 303.1062 | 475.2721 | 612.066 | 449.7277 | 1.46E-10 | 1.332228427 | YCR015C | CTO1 | n |
| W303.asm02.HP0_G0005660 | 1516.343 | 1531.215 | 1860.473 | 2220.313 | 1910.121 | 335.7718 | 2950.391 | 1319.7 | 2328.776 | 0.833431175 | 0.136095126 | YCR016W | RBP95 | n |
| W303.asm02.HP0_G0005670 | 11599.18 | 10579.4 | 11371.95 | 6672.054 | 6580.85 | 9312.729 | 25225.22 | 18161.18 | 21049.89 | 0.003863929 | 0.572197417 | YCR017C | CWH43 | n |
| W303.asm02.HP0_G0005690 | 1377.627 | 1399.904 | 1290.258 | 1290.088 | 1350.363 | 1263.322 | 1034.488 | 1274.601 | 1338.418 | 0.684066318 | 0.059502271 | YCR019W | MAK32 | n |
| W303.asm02.HP0_G0005700 | 376.9679 | 297.118 | 327.6847 | 118.1017 | 148.4011 | 94.19841 | 234.5499 | 464.9554 | 285.8642 | 2.20E-06 | 1.475393964 | YCR020C | PET18 | n |
| W303.asm02.HP0_G0005710 | 938.1843 | 714.4185 | 784.7186 | 168.816 | 222.1677 | 163.3279 | 140.7299 | 219.0552 | 226.0599 | 3.51E-19 | 2.138382368 | YCR020C-. | MAK31 | n |
| W303.asm02.HP0_G0005720 | 830.1766 | 643.1992 | 676.9276 | 738.4833 | 856.5606 | 673.8225 | 574.03 | 516.4978 | 485.6102 | 0.684393907 | -0.076662267 | YCR020W- | HTL1 | n |
| W303.asm02.HP0_G0005760 | 4003.696 | 4450.093 | 3925.749 | 2476.663 | 2239.902 | 2303.303 | 824.628 | 839.7115 | 796.5922 | 1.64E-17 | 0.818110978 | YCR024C | SLM5 | n |
| W303.asm02.HP0_G0005780 | 2854.791 | 3024.594 | 2903.89 | 1065.695 | 1015.376 | 1373.474 | 1980.095 | 1510.837 | 1586.008 | 3.11E-15 | 1.346087503 | YCR026C | NPP1 | n |
| W303.asm02.HP0_G0005790 | 784.6439 | 810.1194 | 704.9532 | 1647.867 | 1632.412 | 1432.727 | 3563.924 | 2335.515 | 2936.387 | 1.71E-08 | -1.035218309 | YCR027C | RHB1 | n |
| W303.asm02.HP0_G0005800 | 620.5146 | 659.8912 | 651.0577 | 2491.947 | 2531.497 | 2601.851 | 3458.994 | 2955.097 | 2430.443 | 1.45E-39 | -1.981158381 | YCR028C | FEN2 | n |
| W303.asm02.HP0_G0005810 | 4276.892 | 4041.695 | 3909.58 | 4235.684 | 5067.332 | 4636.993 | 2910.888 | 4506.738 | 3520.076 | 0.338360396 | -0.188944958 | YCR028C-. | RIM1 | n |
| W303.asm02.HP0_G0005840 | 4052.405 | 3844.729 | 3660.583 | 3248.493 | 3231.846 | 5130.775 | 1655.428 | 3752.931 | 2719.896 | 0.985973269 | -0.006660998 | YCR032W | BPH1 | n |
| W303.asm02.HP0_G0005850 | 5360.145 | 5760.973 | 5421.888 | 7257.7 | 6799.546 | 6350.796 | 6666.155 | 4960.956 | 5210.143 | 0.044857453 | -0.302950787 | YCR033W | SNT1 | n |
| W303.asm02.HP0_G0005860 | 4891.053 | 4913.018 | 5130.852 | 6075.293 | 6645.938 | 6981.318 | 7773.477 | 7099.965 | 7553.273 | 1.56E-05 | -0.399610509 | YCR034W | ELO2 | n |
| W303.asm02.HP0_G0005870 | 2752.078 | 2805.372 | 2834.904 | 1949.373 | 1956.985 | 997.4396 | 2592.393 | 1298.224 | 2075.206 | 0.020897812 | 0.775211573 | YCR035C | RRP43 | n |
| W303.asm02.HP0_G0005880 | 3816.271 | 3891.466 | 3658.427 | 1497.808 | 1667.993 | 1749.508 | 1462.851 | 1794.32 | 1629.067 | 1.58E-24 | 1.209606673 | YCR036W | RBK1 | n |
| W303.asm02.HP0_G0005890 | 4040.757 | 3940.43 | 3975.333 | 18949.77 | 19905.7 | 15582.24 | 16655.51 | 13556.72 | 15129.27 | 1.03E-58 | -2.186786975 | YCR037C | PHO87 | n |
| W303.asm02.HP0_G0005900 | 406.6171 | 447.3462 | 367.5674 | 3372.847 | 3143.326 | 3129.059 | 1669.008 | 2091.762 | 1535.772 | 2.53E-80 | -2.981396421 | YCR038C | BUD5 | n |
| W303.asm02.HP0_G0005920 | 3413.889 | 3577.656 | 3435.3 | 5427.817 | 5140.231 | 5427.804 | 3360.236 | 3759.374 | 3435.154 | 5.10E-14 | -0.617535297 | YCR042C | TAF2 | n |
| W303.asm02.HP0_G0005930 | 712.6388 | 620.9432 | 666.1485 | 793.3658 | 959.8338 | 727.7587 | 446.8793 | 438.1104 | 387.5313 | 0.05432147 | -0.310197365 | YCR043C | | n |
| W303.asm02.HP0_G0005940 | 1873.192 | 1595.757 | 1730.046 | 3902.915 | 4255.9 | 3631.957 | 4178.691 | 4053.595 | 3802.352 | 1.18E-25 | -1.181088432 | YCR044C | PER1 | n |
| W303.asm02.HP0_G0005950 | 624.7502 | 565.3031 | 639.2007 | 49.32485 | 54.67408 | 59.25384 | 53.08234 | 157.8486 | 125.5889 | 3.28E-23 | 3.48675019 | YCR045C | RRT12 | n |

| | | | | | | | | | | | | | |
|---|---|---|---|---|---|---|---|---|---|---|---|---|---|
| W303.asm02.HP0_G0005960 | 5811.236 | 5607.406 | 5490.874 | 2555.861 | 2573.153 | 1751.027 | 806.1109 | 1048.029 | 991.5539 | 7.09E-12 | 1.297398132 | YCR046C | IMG1 | n |
| W303.asm02.HP0_G0005970 | 1794.833 | 1762.677 | 1911.135 | 1952.152 | 1842.43 | 455.7988 | 2554.125 | 982.5269 | 1669.734 | 0.518136013 | 0.363609174 | YCR047C | BUD23 | n |
| W303.asm02.HP0_G0005990 | 4282.186 | 3910.384 | 4353.679 | 2698.277 | 2569.682 | 1172.922 | 3538 | 1929.618 | 3241.389 | 0.006518484 | 0.961943291 | YCR051W | | n |
| W303.asm02.HP0_G0006010 | 13869.46 | 13477.14 | 14680.06 | 11967.18 | 12231.37 | 12325.56 | 15144.52 | 14320.2 | 13519.34 | 0.015178324 | 0.202478533 | YCR053W | THR4 | n |
| W303.asm02.HP0_G0006020 | 670.2829 | 652.1016 | 678.0055 | 1277.583 | 1025.79 | 836.3907 | 2122.059 | 1802.91 | 2636.17 | 0.002742385 | -0.650605399 | YCR054C | CTR86 | n |
| W303.asm02.HP0_G0006030 | 2823.024 | 2837.644 | 3347.989 | 5394.471 | 4855.579 | 1466.912 | 7413.011 | 2887.448 | 6214.854 | 0.471042618 | -0.379211373 | YCR057C | PWP2 | n |
| W303.asm02.HP0_G0006040 | 2681.132 | 1979.674 | 2646.269 | 1172.681 | 1241.883 | 1201.789 | 3113.341 | 2204.511 | 3090.682 | 1.60E-07 | 1.014903673 | YCR059C | YIH1 | n |
| W303.asm02.HP0_G0006050 | 2294.633 | 1932.936 | 1833.525 | 778.7768 | 739.4018 | 644.1956 | 1256.694 | 1400.235 | 1591.988 | 2.52E-21 | 1.486924374 | YCR060W | TAH1 | n |
| W303.asm02.HP0_G0006060 | 2505.354 | 2329.093 | 2333.675 | 3547.221 | 3643.203 | 4162.962 | 3686.137 | 8651.606 | 4960.162 | 0.02962776 | -0.663448506 | YCR061W | TVS1 | n |
| W303.asm02.HP0_G0006070 | 334.612 | 273.7491 | 330.9184 | 238.2876 | 246.4673 | 170.1649 | 309.8527 | 242.6788 | 264.3346 | 0.017099665 | 0.521085343 | YCR063W | BUD31 | n |
| W303.asm02.HP0_G0006080 | 490.2701 | 467.3766 | 533.5655 | 1866.008 | 2272.88 | 2169.602 | 1499.885 | 2107.869 | 1382.673 | 1.39E-26 | -2.080600396 | YCR065W | HCM1 | n |
| W303.asm02.HP0_G0006090 | 1153.141 | 1191.81 | 1032.638 | 2020.929 | 1807.716 | 1860.419 | 2062.805 | 2347.327 | 2279.737 | 2.10E-11 | -0.752555791 | YCR066W | RAD18 | n |
| W303.asm02.HP0_G0006100 | 5138.835 | 5295.822 | 5434.823 | 7795.41 | 7556.305 | 9371.982 | 9099.301 | 8740.731 | 8334.316 | 3.78E-08 | -0.639680298 | YCR067C | SED4 | n |
| W303.asm02.HP0_G0006110 | 1154.2 | 1081.643 | 1114.559 | 1870.871 | 1998.641 | 2416.493 | 2781.268 | 2601.817 | 2361.07 | 6.86E-11 | -0.907718503 | YCR068W | ATG15 | n |
| W303.asm02.HP0_G0006120 | 5367.558 | 5207.91 | 5321.642 | 3194.305 | 3355.947 | 3198.948 | 4430.524 | 4537.879 | 4552.297 | 9.87E-31 | 0.705698536 | YCR069W | CPR4 | n |
| W303.asm02.HP0_G0006130 | 4375.369 | 4062.838 | 3755.439 | 1672.876 | 1779.077 | 1076.445 | 871.538 | 857.9661 | 997.5343 | 7.60E-12 | 1.429192956 | YCR071C | IMG2 | n |
| W303.asm02.HP0_G0006140 | 335.6709 | 349.4196 | 420.385 | 3675.743 | 3126.837 | 741.4327 | 6510.611 | 2423.566 | 5283.104 | 2.83E-07 | -2.770665762 | YCR072C | RSA4 | n |
| W303.asm02.HP0_G0006170 | 240.37 | 164.6946 | 206.9587 | 762.7983 | 819.2433 | 935.1471 | 302.4459 | 609.9184 | 486.8063 | 4.51E-13 | -2.0396974 | YCR075C | ERS1 | n |
| W303.asm02.HP0_G0006180 | 184.2484 | 111.2801 | 123.9597 | 290.3913 | 355.8154 | 292.4709 | 224.6741 | 351.1326 | 352.8449 | 3.80E-05 | -1.160586835 | YCR075W- | EGO2 | n |
| W303.asm02.HP0_G0006190 | 1605.29 | 1652.51 | 1576.983 | 673.1799 | 709.0273 | 1178.999 | 835.7383 | 1334.733 | 926.9654 | 0.000999482 | 0.916571867 | YCR076C | FUB1 | n |
| W303.asm02.HP0_G0006200 | 13159.99 | 12695.95 | 13756.29 | 16298.74 | 15878.91 | 17258.06 | 10700.41 | 11826.83 | 10958.52 | 0.000182164 | -0.319617435 | YCR077C | PAT1 | n |
| W303.asm02.HP0_G0006210 | 1357.508 | 1364.294 | 1262.233 | 778.0821 | 752.4195 | 804.4848 | 446.8793 | 542.269 | 465.2768 | 8.60E-12 | 0.77054624 | YCR079W | PTC6 | n |
| W303.asm02.HP0_G0006220 | 1450.691 | 1542.343 | 1422.841 | 2672.573 | 2557.532 | 2874.571 | 2610.911 | 2868.12 | 2619.425 | 8.41E-20 | -0.876295687 | YCR081W | SRB8 | n |
| W303.asm02.HP0_G0006230 | 2353.932 | 2322.416 | 2333.675 | 1209.501 | 1192.416 | 1342.327 | 1523.34 | 978.2317 | 1176.947 | 3.84E-07 | 0.904698338 | YCR082W | AHC2 | n |
| W303.asm02.HP0_G0006240 | 1621.174 | 1738.196 | 1506.918 | 1603.405 | 1904.046 | 1544.398 | 1202.377 | 1044.807 | 917.3967 | 0.757081278 | -0.053729356 | YCR083W | TRX3 | n |
| W303.asm02.HP0_G0006250 | 10537.1 | 10089.77 | 11623.11 | 12009.56 | 12643.6 | 12853.53 | 6500.735 | 8669.861 | 6919.348 | 0.146884349 | -0.217832876 | YCR084C | TUP1 | n |
| W303.asm02.HP0_G0006260 | 603.5723 | 492.971 | 596.0843 | 1440.841 | 1608.98 | 1116.707 | 872.7725 | 1541.977 | 1188.908 | 2.20E-07 | -1.299345697 | YCR086W | CSM1 | n |
| W303.asm02.HP0_G0006270 | 1634.94 | 1573.501 | 1690.163 | 2125.831 | 2236.43 | 908.5589 | 1402.361 | 545.4904 | 1090.829 | 0.820632381 | -0.105614033 | YCR087C-A | | n |
| W303.asm02.HP0_G0006280 | 13893.81 | 13157.76 | 13971.87 | 10375.59 | 10107.76 | 15143.91 | 12473.12 | 16243.37 | 13978.64 | 0.338704932 | 0.203457205 | YCR088W | ABP1 | n |
| W303.asm02.HP0_G0006300 | 1328.918 | 1348.715 | 1279.479 | 1688.855 | 1919.668 | 1868.015 | 1672.711 | 2089.615 | 1971.147 | 0.000194293 | -0.468569554 | YCR090C | | n |
| W303.asm02.HP0_G0006320 | 1491.988 | 1449.98 | 1426.075 | 1434.589 | 1602.905 | 1653.79 | 317.2596 | 588.4424 | 324.1389 | 0.713317967 | -0.102930447 | YCR092C | MSH3 | n |
| W303.asm02.HP0_G0006330 | 6055.842 | 5984.646 | 5760.352 | 14490.39 | 14111.12 | 15363.46 | 12864.44 | 15388.63 | 14504.91 | 1.89E-39 | -1.304423698 | YCR093W | CDC39 | n |
| W303.asm02.HP0_G0006340 | 2594.302 | 2569.458 | 2678.607 | 3283.923 | 3505.216 | 3423.049 | 3007.176 | 3668.101 | 3186.369 | 0.000424418 | -0.380817743 | YCR094W | CDC50 | n |
| W303.asm02.HP0_G0006350 | 1334.212 | 1462.221 | 1348.466 | 2382.876 | 2310.197 | 2177.199 | 3382.456 | 2687.721 | 2961.505 | 2.91E-09 | -0.729154841 | YCR095C | OCA4 | n |
| W303.asm02.HP0_G0006450 | 822.7643 | 756.7049 | 756.6929 | 3764.667 | 3836.732 | 2979.405 | 1627.036 | 3442.603 | 2576.366 | 4.20E-16 | -2.179196569 | YCR106W | RDS1 | n |
| W303.asm02.HP0_G0006500 | 160.9526 | 173.597 | 148.7516 | 748.2093 | 290.7272 | 284.1146 | 9342.492 | 4680.694 | 5419.458 | 0.000706795 | -1.453695353 | YJR159W | SOR1 | y |
| W303.asm02.HP0_G0006530 | 389.6747 | 360.5476 | 341.6975 | 136.1644 | 156.2117 | 129.143 | 71.59944 | 210.4648 | 110.0398 | 0.000169421 | 1.37376002 | YDL244W | THI13 | y |
| W303.asm02.HP0_G0006570 | 408.7349 | 405.0597 | 438.7094 | 1406.8 | 1343.42 | 1523.128 | 5121.829 | 1884.519 | 5124.025 | 3.98E-07 | -1.770564221 | YDL240W | LRG1 | n |
| W303.asm02.HP0_G0006590 | 1057.84 | 1060.5 | 1022.937 | 1126.135 | 1085.671 | 2649.71 | 2962.735 | 6367.634 | 5579.733 | 0.136113749 | -0.630099395 | YDL238C | GUD1 | n |
| W303.asm02.HP0_G0006600 | 4291.716 | 3973.814 | 4137.019 | 496.0273 | 459.0887 | 1581.622 | 15430.91 | 7502.64 | 21537.89 | 7.94E-06 | 2.28951907 | YDL237W | AIM6 | n |
| W303.asm02.HP0_G0006610 | 4733.277 | 4275.383 | 4730.948 | 6581.046 | 7131.93 | 9945.529 | 51431.85 | 23750.31 | 47774 | 0.011864826 | -0.784012181 | YDL236W | PHO13 | n |
| W303.asm02.HP0_G0006620 | 1898.605 | 1971.884 | 1875.564 | 2761.497 | 2716.347 | 3120.702 | 5139.111 | 2246.389 | 4997.24 | 0.06072327 | -0.581535169 | YDL235C | YPD1 | n |
| W303.asm02.HP0_G0006630 | 10281.91 | 12465.6 | 9654.841 | 1199.775 | 1142.081 | 1704.687 | 5341.565 | 2943.286 | 5474.478 | 3.93E-27 | 3.001269632 | YDL234C | GYP7 | n |
| W303.asm02.HP0_G0006640 | 2421.701 | 2380.282 | 2002.757 | 1629.109 | 1298.292 | 2061.73 | 5503.281 | 1528.017 | 4295.139 | 0.314657821 | 0.447695054 | YDL233W | MFG1 | n |
| W303.asm02.HP0_G0006650 | 2608.068 | 2613.97 | 2726.035 | 3096.35 | 2828.299 | 3596.252 | 31859.28 | 8769.724 | 24842.67 | 0.552034212 | -0.260506702 | YDL231C | BRE4 | n |
| W303.asm02.HP0_G0006660 | 2050.028 | 1948.515 | 1898.2 | 1398.464 | 1345.156 | 1456.277 | 6757.506 | 2651.212 | 4980.495 | 0.127338271 | 0.489546344 | YDL230W | PTP1 | n |
| W303.asm02.HP0_G0006670 | 34254.31 | 33638.87 | 38801.53 | 71589.11 | 80609.55 | 75316.19 | 126849.5 | 60802.85 | 107474.2 | 2.94E-05 | -1.092469573 | YDL229W | SSB1 | y |
| W303.asm02.HP0_G0006690 | 3697.674 | 3123.633 | 3566.805 | 4805.352 | 5068.2 | 4610.405 | 9867.143 | 4402.58 | 8973.025 | 0.110659703 | -0.479499588 | YDL226C | GCS1 | n |
| W303.asm02.HP0_G0006700 | 4343.602 | 4418.934 | 4349.367 | 6843.649 | 6694.537 | 7734.905 | 4366.331 | 5401.214 | 5330.948 | 1.83E-08 | -0.698182306 | YDL225W | SHS1 | n |
| W303.asm02.HP0_G0006710 | 1349.037 | 1309.767 | 1460.568 | 9374.5 | 8328.685 | 8270.469 | 9131.397 | 9241.122 | 10099.74 | 2.04E-148 | -2.656569529 | YDL224C | WHI4 | n |
| W303.asm02.HP0_G0006730 | 1052.545 | 1372.084 | 1040.183 | 29.17808 | 29.50665 | 37.22357 | 377.7488 | 292.0736 | 173.4322 | 5.52E-60 | 5.174515541 | YDL222C | FMP45 | n |
| W303.asm02.HP0_G0006740 | 816.4109 | 843.5034 | 798.7314 | 1829.188 | 1712.253 | 1597.575 | 1190.032 | 1609.626 | 1511.851 | 4.17E-13 | -1.063757509 | YDL220C | CDC13 | n |
| W303.asm02.HP0_G0006750 | 1149.964 | 1217.405 | 1202.948 | 990.6652 | 1326.063 | 1557.312 | 3898.466 | 3486.628 | 3932.725 | 0.562764887 | -0.117513187 | YDL219W | DTD1 | n |
| W303.asm02.HP0_G0006770 | 1284.444 | 1250.789 | 1258.999 | 950.3717 | 928.5915 | 587.9804 | 765.3733 | 361.8706 | 700.9054 | 0.053689595 | 0.621122241 | YDL217C | TIM22 | n |

| | | | | | | | | | | | | | | |
|---|---|---|---|---|---|---|---|---|---|---|---|---|---|---|
| W303.asm02.HP0_G0006780 | 1182.79 | 1259.691 | 1024.015 | 269.5499 | 237.7888 | 245.3717 | 182.702 | 238.3836 | 190.1774 | 1.02E-43 | 2.202172971 | YDL216C | RRI1 | n |
| W303.asm02.HP0_G0006790 | 11266.68 | 11017.85 | 10884.74 | 10370.03 | 10327.33 | 19138.99 | 5692.155 | 24678.07 | 8496.983 | 0.633784569 | -0.264238579 | YDL215C | GDH2 | n |
| W303.asm02.HP0_G0006810 | 2722.429 | 2697.431 | 2916.825 | 2204.334 | 2116.668 | 619.8863 | 2591.159 | 987.8959 | 2277.345 | 0.139028624 | 0.754721224 | YDL213C | NOP6 | n |
| W303.asm02.HP0_G0006820 | 4846.579 | 4604.772 | 4782.687 | 2262.69 | 2568.814 | 2260.762 | 3075.072 | 2690.943 | 2977.054 | 3.57E-23 | 1.00539056 | YDL212W | SHR3 | n |
| W303.asm02.HP0_G0006840 | 374.8501 | 367.2245 | 328.7626 | 446.0078 | 399.2076 | 455.0391 | 1538.153 | 1442.113 | 1190.104 | 0.081861119 | -0.280802714 | YDL210W | UGA4 | n |
| W303.asm02.HP0_G0006850 | 276.3726 | 260.3955 | 314.7498 | 1094.873 | 1048.354 | 881.211 | 1124.605 | 885.8849 | 1062.123 | 1.48E-28 | -1.828398223 | YDL209C | CWC2 | n |
| W303.asm02.HP0_G0006860 | 3813.094 | 3913.722 | 4313.796 | 7515.439 | 7585.811 | 4190.31 | 5851.402 | 4208.222 | 5930.186 | 0.011003923 | -0.680046926 | YDL208W | NHP2 | n |
| W303.asm02.HP0_G0006870 | 2801.846 | 2960.052 | 2978.266 | 2979.638 | 2851.731 | 3177.677 | 3392.332 | 3127.979 | 3533.233 | 0.679323537 | -0.043881762 | YDL207W | GLE1 | n |
| W303.asm02.HP0_G0006880 | 1108.667 | 1089.433 | 1011.08 | 865.6163 | 896.4813 | 1540.6 | 1417.175 | 2238.873 | 1696.048 | 0.897831493 | -0.041491952 | YDL206W | YCX1 | n |
| W303.asm02.HP0_G0006890 | 1277.032 | 1107.237 | 1121.027 | 4769.921 | 4839.958 | 3160.205 | 651.8018 | 857.9661 | 858.7885 | 1.05E-18 | -1.865086238 | YDL205C | HEM3 | n |
| W303.asm02.HP0_G0006910 | 1979.082 | 1930.71 | 2048.029 | 2575.313 | 2474.219 | 2626.161 | 1751.717 | 1908.142 | 1952.01 | 1.87E-05 | -0.365625361 | YDL203C | ACK1 | n |
| W303.asm02.HP0_G0006920 | 4104.291 | 4169.667 | 4115.461 | 4125.224 | 4305.367 | 3195.909 | 1834.427 | 1677.275 | 1800.107 | 0.554316398 | 0.091761102 | YDL202W | MRPL11 | n |
| W303.asm02.HP0_G0006930 | 1530.109 | 1274.158 | 1624.411 | 2428.728 | 2438.637 | 1296.748 | 1503.588 | 799.9809 | 1388.654 | 0.161603944 | -0.476963344 | YDL201W | TRM8 | n |
| W303.asm02.HP0_G0006940 | 762.4071 | 738.9001 | 623.0321 | 141.7221 | 149.2689 | 150.4136 | 172.8262 | 177.177 | 214.0991 | 1.59E-46 | 2.267510149 | YDL200C | MGT1 | n |
| W303.asm02.HP0_G0006950 | 986.8936 | 1036.018 | 851.549 | 399.4618 | 389.6613 | 1018.71 | 1055.474 | 1667.611 | 901.8476 | 0.119882569 | 0.668904682 | YDL199C | | n |
| W303.asm02.HP0_G0006960 | 4723.747 | 4308.767 | 4357.991 | 5101.301 | 4997.037 | 3690.451 | 1909.73 | 1604.257 | 1766.617 | 0.819047437 | -0.042257583 | YDL198C | GGC1 | n |
| W303.asm02.HP0_G0006980 | 16709.42 | 15854.08 | 17507.42 | 17786.82 | 16935.08 | 22651.68 | 12796.55 | 16249.81 | 14479.8 | 0.252618797 | -0.196421775 | YDL195W | SEC31 | n |
| W303.asm02.HP0_G0006990 | 1787.421 | 2017.509 | 1645.969 | 1068.473 | 870.446 | 909.3185 | 2570.173 | 1720.227 | 2231.893 | 9.56E-07 | 0.936027257 | YDL194W | SNF3 | n |
| W303.asm02.HP0_G0007000 | 1986.494 | 2141.03 | 1761.305 | 1702.055 | 1741.76 | 1691.773 | 983.8751 | 1126.416 | 972.4166 | 0.116486397 | 0.197443378 | YDL193W | NUS1 | n |
| W303.asm02.HP0_G0007030 | 6559.877 | 6367.449 | 6125.763 | 6551.868 | 6235.448 | 8293.259 | 3881.183 | 5435.575 | 4121.707 | 0.463220561 | -0.14591111 | YDL190C | UFD2 | n |
| W303.asm02.HP0_G0007040 | 1651.882 | 1781.595 | 1876.642 | 3797.318 | 3218.828 | 4240.448 | 4114.499 | 4571.166 | 4520.003 | 2.84E-15 | -1.084147018 | YDL189W | RBS1 | n |
| W303.asm02.HP0_G0007060 | 429.9129 | 332.7276 | 362.1778 | 188.2681 | 164.8901 | 141.2976 | 398.7348 | 395.1584 | 349.2566 | 9.50E-11 | 1.185016704 | YDL186W | | n |
| W303.asm02.HP0_G0007070 | 30703.83 | 27910.17 | 30732.3 | 8351.183 | 8847.654 | 11150.36 | 7141.427 | 8110.411 | 6212.462 | 1.47E-24 | 1.656104611 | YDL185W | VMA1 | n |
| W303.asm02.HP0_G0007080 | 1989.671 | 2343.56 | 1806.577 | 1321.35 | 1278.332 | 1516.291 | 1111.026 | 1530.165 | 1315.693 | 0.001234861 | 0.576813883 | YDL183C | MRX19 | n |
| W303.asm02.HP0_G0007090 | 1942.02 | 2439.261 | 2497.518 | 1136.556 | 949.4197 | 1028.586 | 1308.541 | 1330.438 | 1273.83 | 5.47E-16 | 1.142533878 | YDL182W | LYS20 | n |
| W303.asm02.HP0_G0007100 | 1544.933 | 1363.182 | 1191.091 | 6116.976 | 7056.427 | 7496.37 | 15841.99 | 12907.08 | 14339.85 | 1.59E-52 | -2.334009358 | YDL181W | INH1 | n |
| W303.asm02.HP0_G0007110 | 1176.436 | 1294.188 | 1047.729 | 516.8688 | 437.3926 | 510.4946 | 772.7802 | 859.0399 | 767.8861 | 6.25E-20 | 1.263037877 | YDL180W | AIT1 | n |
| W303.asm02.HP0_G0007130 | 6482.578 | 6060.316 | 6174.269 | 5090.185 | 4922.403 | 5710.399 | 2845.46 | 6818.63 | 4750.847 | 0.425580962 | 0.251481686 | YDL178W | DLD2 | n |
| W303.asm02.HP0_G0007140 | 354.7311 | 322.7124 | 339.5417 | 378.6203 | 488.5953 | 383.6306 | 460.4585 | 489.6528 | 297.825 | 0.225056764 | -0.297694686 | YDL177C | | n |
| W303.asm02.HP0_G0007150 | 1189.143 | 1015.988 | 1172.766 | 373.7573 | 402.6789 | 574.3065 | 506.134 | 460.6602 | 407.8648 | 9.27E-11 | 1.322431583 | YDL176W | GID12 | n |
| W303.asm02.HP0_G0007160 | 13276.47 | 14804.71 | 13638.8 | 5825.89 | 5880.501 | 5669.377 | 16765.38 | 9160.587 | 10306.66 | 4.99E-08 | 1.263661877 | YDL174C | DLD1 | n |
| W303.asm02.HP0_G0007170 | 2705.486 | 3649.989 | 3626.09 | 1549.217 | 1818.13 | 1268.64 | 4544.095 | 1763.179 | 2213.952 | 0.002122641 | 1.106447609 | YDL173W | PAR32 | n |
| W303.asm02.HP0_G0007180 | 8735.914 | 8236.956 | 8216.909 | 17489.48 | 18806.15 | 19784.71 | 7017.98 | 8907.17 | 5354.87 | 2.06E-09 | -1.154647916 | YDL171C | GLT1 | n |
| W303.asm02.HP0_G0007190 | 1158.435 | 1254.127 | 1153.364 | 2103.6 | 2193.906 | 2667.182 | 3288.636 | 1945.725 | 2515.365 | 7.71E-06 | -0.965811585 | YDL170W | UGA3 | n |
| W303.asm02.HP0_G0007200 | 2009.79 | 2112.097 | 1706.332 | 45.85127 | 43.39213 | 104.8337 | 1702.338 | 886.9587 | 611.1991 | 4.46E-29 | 4.907486218 | YDL169C | UGX2 | n |
| W303.asm02.HP0_G0007210 | 3004.096 | 2733.04 | 2730.346 | 11378.06 | 14042.56 | 14464.01 | 6768.616 | 13209.89 | 7988.647 | 2.44E-18 | -2.235806418 | YDL168W | SFA1 | n |
| W303.asm02.HP0_G0007220 | 1590.466 | 1643.608 | 1772.084 | 3413.835 | 2864.748 | 1347.645 | 2856.571 | 1596.74 | 2668.464 | 0.103851859 | -0.607285509 | YDL167C | NRP1 | n |
| W303.asm02.HP0_G0007230 | 1484.576 | 1419.935 | 1519.853 | 2523.904 | 2592.246 | 1205.588 | 1989.971 | 1041.586 | 1876.656 | 0.156352441 | -0.514809843 | YDL166C | FAP7 | n |
| W303.asm02.HP0_G0007240 | 1390.334 | 1210.728 | 1186.779 | 2204.334 | 2397.849 | 2257.723 | 2259.086 | 2425.714 | 2503.404 | 1.44E-15 | -0.856448827 | YDL165W | CDC36 | n |
| W303.asm02.HP0_G0007250 | 1620.115 | 1632.48 | 1814.123 | 3328.385 | 3333.383 | 3697.288 | 5976.084 | 3653.067 | 5761.538 | 5.35E-07 | -1.031771993 | YDL164C | CDC9 | n |
| W303.asm02.HP0_G0007260 | 7854.911 | 8728.814 | 8330.09 | 10037.95 | 9758.021 | 12145.52 | 11820.08 | 11794.62 | 10956.13 | 0.003911296 | -0.358525968 | YDL161W | ENT1 | n |
| W303.asm02.HP0_G0007270 | 759.2304 | 606.4767 | 626.2658 | 188.9628 | 200.4716 | 210.4271 | 165.4194 | 157.8486 | 138.7458 | 3.55E-28 | 1.732270709 | YDL160C-A | MHF2 | n |
| W303.asm02.HP0_G0007280 | 3265.644 | 3958.234 | 3171.212 | 11232.87 | 10491.35 | 11677.56 | 8612.919 | 7881.691 | 7824.784 | 4.77E-43 | -1.684130086 | YDL160C | DHH1 | n |
| W303.asm02.HP0_G0007290 | 1244.206 | 1418.822 | 1233.129 | 5152.71 | 5003.112 | 5423.246 | 5041.588 | 5713.689 | 4633.631 | 6.90E-61 | -1.999655644 | YDL159W | STE7 | n |
| W303.asm02.HP0_G0007300 | 1165.847 | 1181.795 | 1186.779 | 922.583 | 1081.332 | 785.4932 | 1765.297 | 1027.627 | 1490.321 | 0.154775204 | 0.341781207 | YDL157C | DMO2 | n |
| W303.asm02.HP0_G0007310 | 488.1523 | 501.8734 | 537.8772 | 2030.655 | 2194.774 | 2182.516 | 859.1933 | 1316.479 | 1578.831 | 1.87E-20 | -2.068274758 | YDL156W | CMR1 | n |
| W303.asm02.HP0_G0007320 | 1285.503 | 1494.492 | 1330.141 | 1936.869 | 1812.923 | 2207.585 | 1964.047 | 1584.929 | 1658.969 | 0.000258598 | -0.535731566 | YDL155W | CLB3 | n |
| W303.asm02.HP0_G0007330 | 1587.289 | 1705.925 | 1409.906 | 80.58707 | 85.04857 | 29.62692 | 77.7718 | 93.42059 | 112.4319 | 2.07E-39 | 4.59150205 | YDL154W | MSH5 | n |
| W303.asm02.HP0_G0007340 | 5148.366 | 5825.515 | 5839.039 | 6767.925 | 5511.668 | 1942.462 | 9725.179 | 4470.229 | 8592.67 | 0.624353832 | 0.241431603 | YDL153C | SAS10 | n |
| W303.asm02.HP0_G0007350 | 3400.124 | 3527.58 | 3810.412 | 3632.671 | 3107.744 | 1469.191 | 5200.835 | 2878.858 | 4722.141 | 0.304181713 | 0.387341833 | YDL150W | RPC53 | n |
| W303.asm02.HP0_G0007360 | 1968.493 | 1956.305 | 1753.76 | 1411.663 | 1409.376 | 2177.958 | 1519.636 | 2519.135 | 1675.714 | 0.521612218 | 0.183841017 | YDL149W | ATG9 | n |
| W303.asm02.HP0_G0007370 | 2362.403 | 2714.123 | 2921.136 | 4573.316 | 3718.705 | 929.0698 | 6610.603 | 2393.5 | 5768.715 | 0.734027316 | -0.205367493 | YDL148C | NOP14 | n |
| W303.asm02.HP0_G0007380 | 14461.38 | 16513.97 | 15040.08 | 7295.214 | 7085.934 | 9619.633 | 3514.545 | 6744.537 | 4354.943 | 0.000342505 | 0.93902537 | YDL147W | RPN5 | n |
| W303.asm02.HP0_G0007390 | 2474.646 | 2385.846 | 2412.363 | 499.5009 | 429.582 | 601.6544 | 434.5345 | 415.5606 | 501.1593 | 2.19E-47 | 2.247468023 | YDL146W | LDB17 | n |

| | | | | | | | | | | | | | | |
|---|---|---|---|---|---|---|---|---|---|---|---|---|---|---|
| W303.asm02.HP0_G0007400 | 16492.35 | 16083.32 | 16903.79 | 16304.99 | 16319.78 | 20507.15 | 14166.81 | 15433.73 | 14594.62 | 0.450365262 | -0.102755291 | YDL145C | COP1 | n |
| W303.asm02.HP0_G0007410 | 2433.349 | 2376.944 | 2403.74 | 3275.587 | 3191.057 | 4061.167 | 2663.993 | 4496 | 3630.116 | 0.012210482 | -0.54534804 | YDL144C | | n |
| W303.asm02.HP0_G0007420 | 9371.254 | 9350.87 | 9764.788 | 11050.85 | 11010.32 | 11740.62 | 6145.207 | 7134.327 | 6707.641 | 0.007120491 | -0.246807108 | YDL143W | CCT4 | n |
| W303.asm02.HP0_G0007430 | 1553.404 | 1628.028 | 1430.387 | 2761.497 | 2725.893 | 2111.868 | 840.6762 | 1593.519 | 1556.106 | 0.006981469 | -0.720524764 | YDL142C | CRD1 | n |
| W303.asm02.HP0_G0007440 | 1454.927 | 1389.889 | 1558.658 | 3979.334 | 3768.172 | 5455.152 | 3361.47 | 5517.184 | 4255.668 | 4.16E-12 | -1.584127404 | YDL141W | BPL1 | n |
| W303.asm02.HP0_G0007450 | 7608.187 | 7213.179 | 7600.344 | 16269.56 | 14846.18 | 17241.35 | 15376.6 | 14368.52 | 12900.97 | 2.54E-24 | -1.108852135 | YDL140C | RPO21 | n |
| W303.asm02.HP0_G0007460 | 1081.136 | 1087.207 | 1114.559 | 436.2817 | 406.1503 | 437.5668 | 372.8109 | 400.5274 | 325.3349 | 1.39E-32 | 1.358048413 | YDL139C | SCM3 | n |
| W303.asm02.HP0_G0007480 | 6877.547 | 6830.375 | 6745.562 | 11449.62 | 12415.36 | 13623.83 | 7045.138 | 8619.392 | 7391.801 | 3.89E-13 | -0.874093687 | YDL137W | ARF2 | y |
| W303.asm02.HP0_G0007500 | 3679.673 | 3782.412 | 3488.117 | 3499.98 | 4034.6 | 4564.065 | 3180.003 | 4290.905 | 3590.645 | 0.416217241 | -0.14383779 | YDL135C | RDI1 | n |
| W303.asm02.HP0_G0007510 | 6427.515 | 6436.443 | 6720.77 | 6584.52 | 7032.128 | 7751.618 | 7643.857 | 7040.906 | 6772.229 | 0.253660047 | -0.12570481 | YDL134C | PPH21 | n |
| W303.asm02.HP0_G0007520 | 926.5364 | 1030.454 | 811.6663 | 805.8707 | 692.5383 | 877.4126 | 593.7816 | 512.2026 | 362.4136 | 0.349302197 | 0.220415013 | YDL133W | SRF1 | n |
| W303.asm02.HP0_G0007530 | 8720.031 | 9407.623 | 8835.629 | 6237.856 | 5983.774 | 7545.749 | 5769.927 | 6268.844 | 5199.379 | 0.001141885 | 0.44782718 | YDL132W | CDC53 | n |
| W303.asm02.HP0_G0007540 | 4398.665 | 4788.384 | 4991.802 | 6596.33 | 6524.44 | 7458.387 | 7011.807 | 5508.594 | 3521.272 | 0.033923005 | -0.537452118 | YDL131W | LYS21 | n |
| W303.asm02.HP0_G0007560 | 2267.102 | 1921.808 | 2001.679 | 9781.603 | 11930.23 | 10692.28 | 9074.612 | 6767.087 | 6903.799 | 1.36E-48 | -2.387933197 | YDL130W | RPP1B | y |
| W303.asm02.HP0_G0007570 | 1148.905 | 1375.422 | 1216.961 | 2694.109 | 2433.43 | 2263.801 | 4034.258 | 2673.762 | 3614.567 | 1.43E-07 | -0.982445194 | YDL129W | | n |
| W303.asm02.HP0_G0007580 | 6618.117 | 7078.53 | 6987.014 | 7174.334 | 8295.707 | 8199.82 | 4341.642 | 7679.817 | 6183.756 | 0.401231672 | -0.194540353 | YDL128W | VCX1 | n |
| W303.asm02.HP0_G0007590 | 1110.785 | 1288.624 | 1255.765 | 4030.743 | 4998.773 | 3448.118 | 4498.42 | 4018.159 | 4032 | 1.46E-26 | -1.771260393 | YDL127W | PCL2 | n |
| W303.asm02.HP0_G0007600 | 40174.62 | 41575.37 | 40903.46 | 18296.74 | 19572.45 | 28615.81 | 14274.21 | 28153.96 | 18618.25 | 0.00193152 | 0.883486748 | YDL126C | CDC48 | n |
| W303.asm02.HP0_G0007620 | 24730.58 | 24306.92 | 24044.94 | 2252.27 | 2153.985 | 5237.888 | 3509.607 | 10337.47 | 6007.932 | 2.65E-11 | 2.921782003 | YDL124W | | n |
| W303.asm02.HP0_G0007630 | 1816.011 | 1911.793 | 1886.343 | 735.7044 | 792.3402 | 1064.29 | 2781.268 | 1832.976 | 2289.305 | 1.26E-07 | 1.114816998 | YDL123W | SNA4 | n |
| W303.asm02.HP0_G0007640 | 5286.022 | 5817.726 | 5915.571 | 12013.73 | 12378.04 | 12325.56 | 10574.5 | 9410.783 | 8906.044 | 1.04E-27 | -1.109299667 | YDL122W | UBP1 | n |
| W303.asm02.HP0_G0007650 | 1071.605 | 1115.027 | 1123.182 | 1670.792 | 1807.716 | 1442.603 | 1947.999 | 1502.246 | 1818.048 | 0.000151955 | -0.572066219 | YDL121C | EXP1 | n |
| W303.asm02.HP0_G0007660 | 3664.848 | 3618.83 | 3367.391 | 1096.957 | 1070.918 | 1248.129 | 767.8423 | 496.0956 | 632.7286 | 5.48E-21 | 1.640536867 | YDL120W | YFH1 | n |
| W303.asm02.HP0_G0007670 | 2156.977 | 1908.454 | 1766.695 | 614.1291 | 599.6792 | 547.7182 | 316.0251 | 246.974 | 346.8645 | 3.61E-27 | 1.72720196 | YDL119C | HEM25 | n |
| W303.asm02.HP0_G0007680 | 1966.375 | 1999.704 | 2142.885 | 5709.872 | 5389.302 | 5258.398 | 3286.167 | 3474.817 | 3124.172 | 2.21E-58 | -1.421032351 | YDL117W | CYK3 | n |
| W303.asm02.HP0_G0007690 | 2602.773 | 2514.931 | 2589.14 | 4595.547 | 4354.834 | 4086.236 | 4412.007 | 4296.274 | 4732.906 | 5.08E-18 | -0.758398574 | YDL116W | NUP84 | n |
| W303.asm02.HP0_G0007700 | 3723.088 | 4162.99 | 3505.364 | 3509.011 | 2968.889 | 3903.916 | 4041.665 | 3135.496 | 3309.565 | 0.46051015 | 0.133743878 | YDL115C | IWR1 | n |
| W303.asm02.HP0_G0007720 | 3846.979 | 4128.493 | 3691.842 | 3255.44 | 3346.401 | 6707.839 | 2189.955 | 4376.809 | 2316.815 | 0.639255532 | -0.190017122 | YDL113C | ATG20 | n |
| W303.asm02.HP0_G0007730 | 3193.639 | 3479.73 | 3806.101 | 5140.205 | 4754.041 | 2506.893 | 2396.112 | 1389.497 | 2845.485 | 0.512294055 | -0.242908387 | YDL112W | TRM3 | n |
| W303.asm02.HP0_G0007740 | 1628.586 | 1661.412 | 1768.851 | 1851.419 | 1855.447 | 1066.569 | 4040.43 | 2770.404 | 3753.313 | 0.767913303 | 0.08383191 | YDL111C | RRP42 | n |
| W303.asm02.HP0_G0007750 | 6588.468 | 6211.657 | 5760.352 | 964.9607 | 953.7589 | 1535.282 | 2603.504 | 3338.444 | 2337.149 | 2.59E-25 | 2.425807704 | YDL110C | TMA17 | n |
| W303.asm02.HP0_G0007770 | 1688.943 | 1725.955 | 1762.383 | 1001.086 | 1024.922 | 999.7186 | 1351.748 | 1185.475 | 1217.614 | 5.83E-18 | 0.775080111 | YDL108W | KIN28 | n |
| W303.asm02.HP0_G0007780 | 4456.905 | 4543.568 | 4132.707 | 1493.64 | 1502.235 | 1492.741 | 1575.188 | 1521.574 | 1508.262 | 5.31E-92 | 1.548887954 | YDL107W | MSS2 | n |
| W303.asm02.HP0_G0007790 | 5108.127 | 5782.116 | 5308.707 | 6553.257 | 6181.642 | 7317.849 | 11030.02 | 9076.831 | 10017.21 | 0.017632191 | -0.307974665 | YDL106C | PHO2 | n |
| W303.asm02.HP0_G0007800 | 514.6248 | 550.8367 | 481.8258 | 1215.059 | 1227.997 | 1178.999 | 928.3238 | 949.2391 | 918.5928 | 2.48E-38 | -1.227285393 | YDL105W | NSE4 | n |
| W303.asm02.HP0_G0007810 | 1370.215 | 1545.681 | 1299.96 | 1419.305 | 1396.359 | 1212.425 | 1909.73 | 1374.464 | 1859.911 | 0.734027316 | 0.065673356 | YDL104C | QRI7 | n |
| W303.asm02.HP0_G0007820 | 4924.938 | 4432.288 | 4744.96 | 3565.978 | 3829.789 | 3783.889 | 998.6887 | 1638.619 | 989.1617 | 0.138489611 | 0.335077009 | YDL103C | QRI1 | n |
| W303.asm02.HP0_G0007830 | 1932.49 | 1908.454 | 2051.263 | 4201.643 | 4630.808 | 4063.446 | 1540.622 | 1835.124 | 1815.656 | 2.90E-23 | -1.129888767 | YDL102W | POL3 | n |
| W303.asm02.HP0_G0007840 | 2287.221 | 2331.319 | 2273.312 | 1887.544 | 1834.619 | 1893.844 | 1493.712 | 1722.375 | 1685.283 | 0.001118035 | 0.295194741 | YDL101C | DUN1 | n |
| W303.asm02.HP0_G0007850 | 12787.26 | 13163.33 | 13245.36 | 6209.373 | 6435.052 | 8057.763 | 3986.114 | 6603.87 | 5165.888 | 1.10E-05 | 0.920918812 | YDL100C | GET3 | n |
| W303.asm02.HP0_G0007860 | 7631.483 | 7859.716 | 7560.462 | 3190.831 | 3213.621 | 3488.38 | 2033.177 | 3319.116 | 2630.189 | 3.98E-11 | 1.220408848 | YDL099W | BUG1 | n |
| W303.asm02.HP0_G0007870 | 648.046 | 645.4248 | 491.527 | 315.4011 | 340.1943 | 323.6171 | 332.0733 | 450.996 | 368.394 | 2.83E-06 | 0.866611214 | YDL098C | SNU23 | n |
| W303.asm02.HP0_G0007880 | 14437.02 | 15392.27 | 14798.63 | 6173.248 | 6599.074 | 8260.593 | 5813.134 | 7238.485 | 5816.558 | 7.46E-12 | 1.08530024 | YDL097C | RPN6 | n |
| W303.asm02.HP0_G0007890 | 5373.911 | 5003.155 | 5529.679 | 14800.23 | 14780.23 | 21421.78 | 19576.27 | 25620.87 | 22352.42 | 1.20E-18 | -1.680920157 | YDL095W | PMT1 | n |
| W303.asm02.HP0_G0007910 | 2468.293 | 2462.629 | 2404.818 | 1988.278 | 2361.399 | 1771.538 | 1730.731 | 1379.833 | 1597.969 | 0.10094382 | 0.261407239 | YDL092W | SRP14 | n |
| W303.asm02.HP0_G0007920 | 4535.263 | 4691.571 | 4439.912 | 1062.221 | 1076.993 | 1479.827 | 3873.777 | 2929.326 | 3327.507 | 5.42E-27 | 1.916917748 | YDL091C | UBX3 | n |
| W303.asm02.HP0_G0007930 | 1663.53 | 1525.651 | 1546.801 | 1001.086 | 1003.226 | 983.7657 | 525.8855 | 717.2984 | 653.062 | 1.40E-06 | 0.664515372 | YDL090C | RAM1 | n |
| W303.asm02.HP0_G0007940 | 2316.87 | 2373.605 | 2230.196 | 1181.017 | 1067.446 | 1271.679 | 1224.597 | 1514.058 | 1406.595 | 8.07E-15 | 0.974881105 | YDL089W | NUR1 | n |
| W303.asm02.HP0_G0007950 | 4642.212 | 4992.027 | 4698.61 | 6187.142 | 5844.919 | 6397.895 | 5689.686 | 5405.509 | 5397.929 | 4.93E-06 | -0.362883542 | YDL088C | ASM4 | n |
| W303.asm02.HP0_G0007960 | 796.2918 | 823.473 | 757.7708 | 1517.26 | 1391.152 | 1237.494 | 2264.024 | 1464.663 | 1725.95 | 2.93E-05 | -0.802332431 | YDL087C | LUC7 | n |
| W303.asm02.HP0_G0007970 | 8745.444 | 8156.834 | 8588.788 | 2693.414 | 3060.013 | 4366.552 | 7217.964 | 6989.364 | 7605.9 | 2.90E-12 | 1.332807428 | YDL086W | | n |
| W303.asm02.HP0_G0007980 | 511.4481 | 466.2638 | 435.4757 | 1076.115 | 1419.79 | 1017.191 | 1238.177 | 601.328 | 788.2196 | 4.33E-06 | -1.313538442 | YDL085C-A | | n |
| W303.asm02.HP0_G0008000 | 12137.1 | 11424.02 | 12607.24 | 16746.83 | 17918.34 | 18857.91 | 10104.16 | 11962.13 | 10301.87 | 3.51E-07 | -0.565409436 | YDL084W | SUB2 | n |
| W303.asm02.HP0_G0008030 | 8601.434 | 8210.249 | 8401.232 | 10142.16 | 12029.17 | 10774.32 | 11276.91 | 7310.43 | 8477.846 | 0.039321641 | -0.385891207 | YDL081C | RPP1A | n |

| | | | | | | | | | | | | | | |
|---|---|---|---|---|---|---|---|---|---|---|---|---|---|---|
| W303.asm02.HP0_G0008040 | 2097.678 | 1937.387 | 2319.663 | 2691.33 | 2423.016 | 2558.55 | 3944.142 | 3961.248 | 3435.154 | 0.028221522 | -0.272054049 | YDL080C | THI3 | n |
| W303.asm02.HP0_G0008050 | 2959.622 | 3732.336 | 2842.449 | 3395.772 | 3320.365 | 5210.54 | 12481.76 | 7904.241 | 6843.995 | 0.298842441 | -0.323014504 | YDL079C | MRK1 | n |
| W303.asm02.HP0_G0008070 | 10449.21 | 10373.53 | 10304.82 | 3879.295 | 4177.794 | 4844.381 | 8894.379 | 7317.947 | 5722.067 | 4.75E-12 | 1.270673649 | YDL078C | MDH3 | n |
| W303.asm02.HP0_G0008080 | 3115.28 | 3283.877 | 2904.968 | 1904.912 | 1842.43 | 2022.227 | 1352.983 | 2108.943 | 1660.165 | 0.000137187 | 0.689347848 | YDL077C | VAM6 | n |
| W303.asm02.HP0_G0008090 | 3795.093 | 3633.296 | 3383.56 | 2550.303 | 2557.532 | 2834.309 | 2167.735 | 2283.972 | 2370.639 | 5.91E-06 | 0.445005922 | YDL076C | RXT3 | n |
| W303.asm02.HP0_G0008110 | 3787.681 | 3981.603 | 4094.981 | 5181.888 | 4971.87 | 3898.599 | 2768.923 | 2274.308 | 2646.935 | 0.12681786 | -0.244199167 | YDL074C | BRE1 | n |
| W303.asm02.HP0_G0008120 | 1017.602 | 945.8812 | 1047.729 | 2417.612 | 1920.535 | 2062.489 | 2909.653 | 2311.891 | 2791.661 | 2.34E-13 | -1.088097474 | YDL073W | AHK1 | n |
| W303.asm02.HP0_G0008130 | 8632.142 | 8557.443 | 8060.612 | 955.2347 | 1230.601 | 1488.183 | 1006.096 | 1753.515 | 1278.614 | 1.19E-30 | 2.781006177 | YDL072C | YET3 | n |
| W303.asm02.HP0_G0008160 | 3306.941 | 2806.485 | 2722.801 | 5210.371 | 5349.381 | 6087.952 | 3331.843 | 4502.443 | 4939.828 | 6.41E-07 | -0.913791908 | YDL067C | COX9 | n |
| W303.asm02.HP0_G0008170 | 3562.135 | 2916.652 | 3432.066 | 3921.673 | 4102.292 | 4243.487 | 1165.343 | 2995.902 | 1886.225 | 0.360681342 | -0.307735739 | YDL066W | IDP1 | n |
| W303.asm02.HP0_G0008180 | 3744.266 | 4029.454 | 3733.881 | 2844.168 | 2786.642 | 2809.24 | 2141.811 | 3474.817 | 3387.311 | 0.028179541 | 0.447241871 | YDL065C | PEX19 | n |
| W303.asm02.HP0_G0008190 | 2571.006 | 2279.017 | 2325.052 | 2240.46 | 2497.651 | 2048.815 | 2752.875 | 2710.271 | 2660.092 | 0.519439265 | 0.080597849 | YDL064W | UBC9 | n |
| W303.asm02.HP0_G0008200 | 1032.426 | 1083.869 | 1308.583 | 3298.512 | 2804.867 | 1017.191 | 3670.088 | 1695.53 | 3304.781 | 0.018796553 | -1.055948273 | YDL063C | SYO1 | n |
| W303.asm02.HP0_G0008220 | 3355.65 | 3347.307 | 3839.516 | 13372.59 | 12577.64 | 6226.971 | 12328.68 | 6806.818 | 11856.78 | 1.31E-06 | -1.609821768 | YDL060W | TSR1 | n |
| W303.asm02.HP0_G0008230 | 810.0575 | 697.7265 | 657.5252 | 137.5538 | 170.965 | 111.6707 | 153.0747 | 175.0294 | 154.2949 | 5.23E-33 | 2.368438345 | YDL059C | RAD59 | n |
| W303.asm02.HP0_G0008240 | 3143.87 | 2978.969 | 3187.38 | 5686.252 | 5102.046 | 6694.924 | 2409.691 | 3411.462 | 3145.702 | 2.08E-07 | -0.909119113 | YDL058W | USO1 | n |
| W303.asm02.HP0_G0008250 | 1845.66 | 1794.949 | 1669.683 | 1928.532 | 1785.152 | 3002.954 | 7808.042 | 8170.544 | 10404.74 | 0.16900795 | -0.339021714 | YDL057W | | n |
| W303.asm02.HP0_G0008260 | 3730.5 | 3945.994 | 3557.103 | 4109.246 | 3880.992 | 4869.45 | 3303.45 | 3641.256 | 3346.644 | 0.137317971 | -0.195140648 | YDL056W | MBP1 | n |
| W303.asm02.HP0_G0008280 | 1752.477 | 1980.786 | 1789.331 | 1576.311 | 1611.584 | 1994.12 | 2330.685 | 2715.64 | 1880.245 | 0.651272492 | 0.091771851 | YDL054C | MCH1 | n |
| W303.asm02.HP0_G0008290 | 3039.039 | 3338.404 | 3196.004 | 1010.117 | 1157.702 | 764.9823 | 812.2833 | 877.2945 | 889.8867 | 5.37E-24 | 1.707173803 | YDL053C | PBP4 | n |
| W303.asm02.HP0_G0008300 | 4074.642 | 3534.257 | 4140.253 | 5198.561 | 5161.927 | 6609.842 | 7432.762 | 8048.13 | 7737.469 | 0.000222581 | -0.53044669 | YDL052C | SLC1 | n |
| W303.asm02.HP0_G0008310 | 2652.541 | 2793.131 | 2881.254 | 6789.461 | 7199.622 | 5835.744 | 3226.913 | 2985.164 | 3432.762 | 1.35E-25 | -1.251411892 | YDL051W | LHP1 | n |
| W303.asm02.HP0_G0008340 | 5099.656 | 5691.979 | 5405.719 | 4204.422 | 4121.384 | 4510.129 | 5977.319 | 4613.044 | 4771.18 | 0.015893283 | 0.335522739 | YDL047W | SIT4 | n |
| W303.asm02.HP0_G0008350 | 12857.15 | 12842.84 | 12679.46 | 2201.555 | 2301.518 | 3260.481 | 4061.416 | 4131.982 | 3796.372 | 1.04E-42 | 2.305523829 | YDL046W | NPC2 | n |
| W303.asm02.HP0_G0008360 | 2482.059 | 2544.977 | 2226.962 | 2055.665 | 2454.259 | 2025.266 | 1759.124 | 1554.862 | 1306.124 | 0.379094701 | 0.150760529 | YDL045W- | MRP10 | n |
| W303.asm02.HP0_G0008370 | 4113.821 | 4185.246 | 4013.059 | 1042.074 | 1231.469 | 1008.075 | 2156.624 | 1927.471 | 1401.811 | 1.33E-25 | 1.907911388 | YDL045C | FAD1 | n |
| W303.asm02.HP0_G0008380 | 1830.836 | 1883.973 | 1779.63 | 2456.516 | 2436.034 | 1251.927 | 4183.629 | 1743.851 | 3303.585 | 0.690332504 | -0.161300424 | YDL044C | MTF2 | n |
| W303.asm02.HP0_G0008390 | 868.2969 | 951.4452 | 783.6407 | 595.3717 | 613.5647 | 460.3568 | 797.4696 | 758.1027 | 691.3367 | 0.000108102 | 0.641352428 | YDL043C | PRP11 | n |
| W303.asm02.HP0_G0008400 | 680.8719 | 738.9001 | 633.8112 | 7048.59 | 7509.441 | 7687.806 | 8473.423 | 9823.122 | 6793.759 | 4.73E-101 | -3.437408505 | YDL042C | SIR2 | n |
| W303.asm02.HP0_G0008410 | 10148.49 | 9449.909 | 10151.76 | 11418.35 | 11150.04 | 11841.65 | 7104.393 | 8124.37 | 8145.334 | 0.029853785 | -0.209935081 | YDL040C | NAT1 | n |
| W303.asm02.HP0_G0008440 | 1927.196 | 1964.094 | 1794.72 | 3630.587 | 3206.678 | 4342.243 | 5909.423 | 2976.573 | 4551.101 | 0.000171266 | -0.975411928 | YDL035C | GPR1 | n |
| W303.asm02.HP0_G0008450 | 1606.349 | 1720.391 | 1667.527 | 594.677 | 512.8949 | 315.2608 | 420.9553 | 315.6972 | 452.1199 | 1.51E-12 | 1.811321609 | YDL033C | SLM3 | n |
| W303.asm02.HP0_G0008460 | 2892.911 | 3106.941 | 3605.609 | 5570.234 | 4925.874 | 1278.516 | 7411.776 | 2412.828 | 6208.874 | 0.622998311 | -0.293752822 | YDL031W | DBP10 | n |
| W303.asm02.HP0_G0008470 | 1364.92 | 1445.529 | 1340.92 | 1597.847 | 1293.085 | 1288.391 | 2114.652 | 1781.434 | 2001.049 | 0.951116887 | -0.010202484 | YDL030W | PRP9 | n |
| W303.asm02.HP0_G0008480 | 10076.48 | 10112.03 | 9721.672 | 5272.895 | 5910.008 | 6940.296 | 7514.238 | 7892.429 | 7329.605 | 8.81E-09 | 0.722841587 | YDL029W | ARP2 | n |
| W303.asm02.HP0_G0008490 | 2469.352 | 2621.76 | 2440.389 | 1697.192 | 1540.42 | 1987.283 | 2545.484 | 1816.869 | 2258.207 | 0.002237987 | 0.527363282 | YDL028C | MPS1 | n |
| W303.asm02.HP0_G0008510 | 2760.549 | 3181.499 | 2629.023 | 3993.228 | 3683.991 | 4773.732 | 8620.326 | 4436.941 | 5027.142 | 0.053574234 | -0.538747452 | YDL025C | RTK1 | n |
| W303.asm02.HP0_G0008520 | 666.0473 | 819.0218 | 624.11 | 4113.414 | 4059.767 | 13763.6 | 18231.93 | 37288.78 | 19396.9 | 4.86E-12 | -3.378632628 | YDL024C | DIA3 | n |
| W303.asm02.HP0_G0008530 | 6589.527 | 6957.234 | 6401.708 | 5575.792 | 5507.329 | 14362.22 | 24704.28 | 14398.58 | 18774.94 | 0.405022517 | -0.351132621 | YDL022W | GPD1 | n |
| W303.asm02.HP0_G0008550 | 6854.251 | 8380.507 | 6828.561 | 5392.387 | 4940.627 | 5659.501 | 6898.236 | 6913.124 | 6144.285 | 0.000445548 | 0.464150065 | YDL020C | RPN4 | n |
| W303.asm02.HP0_G0008570 | 1181.731 | 1237.435 | 1107.014 | 483.5224 | 529.3839 | 629.762 | 309.8527 | 730.184 | 539.434 | 0.000233132 | 1.102112664 | YDL018C | ERP3 | n |
| W303.asm02.HP0_G0008580 | 1497.283 | 1591.306 | 1392.66 | 922.583 | 857.4284 | 1161.527 | 572.7955 | 590.59 | 456.9042 | 0.000578554 | 0.607062979 | YDL017W | CDC7 | n |
| W303.asm02.HP0_G0008590 | 8134.46 | 8366.041 | 8080.014 | 3419.393 | 4016.375 | 3971.527 | 2999.77 | 5686.844 | 3998.51 | 2.04E-06 | 1.107601027 | YDL015C | TSC13 | n |
| W303.asm02.HP0_G0008600 | 10464.04 | 10343.49 | 11193.02 | 16775.31 | 17657.99 | 11081.99 | 13168.12 | 7945.046 | 11597.23 | 0.047628471 | -0.508245232 | YDL014W | NOP1 | n |
| W303.asm02.HP0_G0008610 | 2613.362 | 2944.472 | 2648.425 | 2121.663 | 1858.051 | 2194.671 | 3244.195 | 2841.275 | 2867.014 | 0.00078746 | 0.410067938 | YDL013W | SLX5 | n |
| W303.asm02.HP0_G0008630 | 1397.746 | 1160.652 | 1258.999 | 530.7631 | 649.1462 | 503.6576 | 628.3468 | 547.638 | 583.6892 | 5.30E-15 | 1.182280249 | YDL008W | APC11 | n |
| W303.asm02.HP0_G0008650 | 15643.11 | 16794.4 | 16245.18 | 8007.298 | 8464.936 | 10533.51 | 4147.83 | 10119.49 | 6310.541 | 0.006281415 | 0.850142917 | YDL007W | RPT2 | n |
| W303.asm02.HP0_G0008660 | 1219.851 | 1196.261 | 1238.519 | 2246.017 | 2214.734 | 1988.042 | 2147.983 | 1910.29 | 1916.127 | 6.95E-16 | -0.819257564 | YDL006W | PTC1 | n |
| W303.asm02.HP0_G0008670 | 2911.971 | 3418.526 | 3105.459 | 1513.092 | 1342.552 | 1589.978 | 2482.525 | 1692.309 | 2143.383 | 3.76E-10 | 1.08556867 | YDL005C | MED2 | n |
| W303.asm02.HP0_G0008680 | 4485.495 | 4341.038 | 4124.084 | 7973.257 | 8842.447 | 7679.45 | 1497.416 | 3455.488 | 2544.071 | 0.001161085 | -0.919461933 | YDL004W | ATP16 | n |
| W303.asm02.HP0_G0008690 | 1425.278 | 1508.959 | 1653.514 | 3412.446 | 3803.754 | 3698.047 | 1143.122 | 1250.977 | 1286.987 | 2.21E-31 | -1.250233173 | YDL003W | MCD1 | n |
| W303.asm02.HP0_G0008700 | 1971.669 | 2245.633 | 1888.499 | 2866.399 | 2969.757 | 2979.405 | 1687.525 | 2247.463 | 2047.696 | 0.000252012 | -0.5299098 | YDL002C | NHP10 | n |
| W303.asm02.HP0_G0008710 | 1053.604 | 1099.448 | 922.6911 | 2402.328 | 2579.228 | 2615.525 | 1341.872 | 1651.504 | 1139.868 | 3.72E-15 | -1.304503982 | YDL001W | RMD1 | n |
| W303.asm02.HP0_G0008720 | 21830.26 | 20651.37 | 20336.93 | 5888.414 | 5986.378 | 9827.021 | 19278.77 | 21775.59 | 23190.88 | 3.33E-12 | 1.53336312 | YDR001C | NTH1 | n |

| | | | | | | | | | | | | | | |
|---|---|---|---|---|---|---|---|---|---|---|---|---|---|---|
| W303.asm02.HP0_G0008730 | 11779.19 | 11202.57 | 11697.48 | 11894.24 | 13059.29 | 12352.15 | 9628.89 | 8795.495 | 8512.532 | 0.265513663 | -0.105250051 | YDR002W | YRB1 | n |
| W303.asm02.HP0_G0008750 | 2024.614 | 2100.969 | 1982.277 | 1631.193 | 1519.592 | 1838.388 | 966.5924 | 1089.907 | 1011.887 | 0.012744602 | 0.291567347 | YDR004W | RAD57 | n |
| W303.asm02.HP0_G0008770 | 2753.137 | 3224.898 | 2665.672 | 6920.067 | 5848.391 | 9036.211 | 20420.65 | 13748.93 | 17825.24 | 1.76E-09 | -1.334975298 | YDR006C | SOK1 | n |
| W303.asm02.HP0_G0008780 | 607.8079 | 436.2181 | 480.7479 | 1876.428 | 2048.976 | 2303.303 | 2845.46 | 1101.719 | 2626.601 | 2.68E-09 | -2.030219961 | YDR007W | TRP1 | n |
| W303.asm02.HP0_G0008790 | 718.9922 | 816.7962 | 742.6801 | 895.4891 | 836.6002 | 1189.635 | 16467.87 | 5665.368 | 10948.96 | 0.348697547 | -0.358841065 | YDR009W | GAL3 | n |
| W303.asm02.HP0_G0008800 | 4122.293 | 4514.635 | 4112.227 | 10804.92 | 10650.16 | 5997.552 | 8851.172 | 17349.39 | 8627.356 | 0.001043144 | -1.106542194 | YDR011W | SNQ2 | n |
| W303.asm02.HP0_G0008810 | 19824.7 | 19640.94 | 22079.91 | 51113.74 | 59928.86 | 51884.33 | 54589.63 | 42496.71 | 41609.38 | 4.30E-21 | -1.404488398 | YDR012W | RPL4B | y |
| W303.asm02.HP0_G0008820 | 1255.854 | 1058.274 | 1207.259 | 792.6711 | 825.3182 | 790.8109 | 251.8325 | 298.5164 | 249.9816 | 1.52E-05 | 0.548224637 | YDR013W | PSF1 | n |
| W303.asm02.HP0_G0008830 | 1340.566 | 1447.755 | 1201.87 | 425.861 | 505.9522 | 502.1383 | 307.3838 | 221.2028 | 214.0991 | 2.19E-16 | 1.477007976 | YDR014W | RAD61 | n |
| W303.asm02.HP0_G0008850 | 601.4545 | 539.7087 | 508.7736 | 448.7866 | 495.5381 | 549.2375 | 334.5422 | 431.6676 | 345.6684 | 0.424285078 | 0.144086019 | YDR016C | DAD1 | n |
| W303.asm02.HP0_G0008860 | 3916.866 | 4284.285 | 3841.672 | 2890.714 | 2422.148 | 2960.413 | 4983.568 | 3191.333 | 3389.703 | 0.008826014 | 0.541527743 | YDR017C | KCS1 | n |
| W303.asm02.HP0_G0008880 | 770.8783 | 555.2879 | 710.3428 | 4933.179 | 4863.389 | 8132.21 | 12724.95 | 21130.24 | 21494.83 | 4.66E-27 | -3.138012798 | YDR019C | GCV1 | n |
| W303.asm02.HP0_G0008890 | 757.1126 | 717.7569 | 759.9266 | 747.5146 | 779.3226 | 227.1397 | 1119.667 | 491.8004 | 938.9262 | 0.489926456 | 0.349624301 | YDR020C | DAS2 | n |
| W303.asm02.HP0_G0008900 | 587.6888 | 637.6352 | 755.615 | 1117.104 | 1077.86 | 273.4793 | 1986.267 | 433.8152 | 1423.34 | 0.630096951 | -0.317392578 | YDR021W | FAL1 | n |
| W303.asm02.HP0_G0008910 | 865.1202 | 770.0586 | 739.4464 | 74.33463 | 104.1411 | 145.8556 | 196.2812 | 186.8412 | 194.9617 | 3.01E-32 | 2.873897301 | YDR022C | ATG31 | n |
| W303.asm02.HP0_G0008920 | 19065.47 | 19923.6 | 19500.47 | 13654.65 | 14132.82 | 13499.24 | 8738.835 | 8096.451 | 8372.59 | 1.98E-14 | 0.502531873 | YDR023W | SES1 | n |
| W303.asm02.HP0_G0008950 | 4334.072 | 4504.62 | 4011.982 | 4554.559 | 4290.613 | 5425.525 | 2684.979 | 4334.93 | 3675.567 | 0.495215134 | -0.151246972 | YDR027C | VPS54 | n |
| W303.asm02.HP0_G0008960 | 15171.9 | 15511.34 | 15402.26 | 7328.561 | 6989.604 | 10817.62 | 10985.58 | 9798.424 | 9519.635 | 4.45E-06 | 0.87454626 | YDR028C | REG1 | n |
| W303.asm02.HP0_G0008990 | 8094.222 | 7849.701 | 7533.514 | 2396.771 | 2927.233 | 3426.087 | 6746.395 | 8560.333 | 6445.698 | 1.53E-15 | 1.423996323 | YDR032C | PST2 | n |
| W303.asm02.HP0_G0009010 | 2473.587 | 2837.644 | 2619.322 | 2710.782 | 2210.395 | 1969.051 | 4404.6 | 2515.913 | 3377.742 | 0.424904686 | 0.202760265 | YDR034C | LYS14 | n |
| W303.asm02.HP0_G0009030 | 9395.608 | 9861.646 | 9439.259 | 5414.618 | 5786.774 | 6067.441 | 9144.977 | 8803.012 | 9055.554 | 9.65E-22 | 0.732778572 | YDR035W | ARO3 | n |
| W303.asm02.HP0_G0009040 | 6818.249 | 6829.262 | 6842.574 | 5954.412 | 6034.109 | 6571.099 | 7159.944 | 7484.386 | 7346.35 | 0.061551277 | 0.142753721 | YDR036C | EHD3 | n |
| W303.asm02.HP0_G0009050 | 15181.43 | 14617.76 | 16249.5 | 24526.95 | 26023.99 | 27154.97 | 14093.98 | 15822.44 | 14324.31 | 1.99E-15 | -0.754845298 | YDR037W | KRS1 | n |
| W303.asm02.HP0_G0009090 | 33.88476 | 43.39925 | 43.11641 | 1368.591 | 1449.297 | 1094.677 | 543.1682 | 910.5823 | 252.3738 | 3.26E-32 | -5.022671087 | YDR040C | ENA1 | y |
| W303.asm02.HP0_G0009100 | 6352.333 | 6765.832 | 6123.608 | 2868.483 | 3136.383 | 2462.833 | 2144.28 | 1759.958 | 1810.872 | 1.97E-17 | 1.184326714 | YDR041W | RSM10 | n |
| W303.asm02.HP0_G0009120 | 1495.165 | 1985.238 | 1445.477 | 711.3893 | 870.446 | 579.6241 | 2343.03 | 2262.496 | 1902.97 | 2.05E-08 | 1.188795248 | YDR043C | NRG1 | n |
| W303.asm02.HP0_G0009140 | 751.8181 | 766.7201 | 781.4849 | 1233.121 | 1411.98 | 720.9217 | 1783.814 | 885.8849 | 1475.968 | 0.104302538 | -0.549266703 | YDR045C | RPC11 | n |
| W303.asm02.HP0_G0009150 | 3959.222 | 2834.305 | 3270.379 | 10224.14 | 9987.132 | 11639.58 | 32343.19 | 16939.19 | 26079.42 | 4.50E-11 | -1.662116144 | YDR046C | BAP3 | n |
| W303.asm02.HP0_G0009160 | 6652.002 | 5768.762 | 5775.443 | 5377.798 | 5681.765 | 5230.291 | 2851.633 | 3000.197 | 2277.345 | 0.294216395 | 0.159722135 | YDR047W | HEM12 | n |
| W303.asm02.HP0_G0009170 | 2163.33 | 2178.865 | 2115.938 | 2641.311 | 2560.135 | 2777.334 | 1765.297 | 2100.353 | 1753.46 | 0.005864 | -0.305143132 | YDR049W | VMS1 | n |
| W303.asm02.HP0_G0009180 | 36772.38 | 36302.92 | 38316.47 | 22236.47 | 26522.14 | 37920.94 | 67476.3 | 54329.98 | 62440.39 | 0.105053931 | 0.361882926 | YDR050C | TPI1 | n |
| W303.asm02.HP0_G0009190 | 3173.519 | 3261.621 | 3225.107 | 2594.07 | 2958.475 | 5417.928 | 11353.45 | 7846.256 | 8194.374 | 0.577100791 | -0.183499024 | YDR051C | DET1 | n |
| W303.asm02.HP0_G0009200 | 2603.832 | 2957.826 | 2987.967 | 5660.547 | 5577.624 | 6840.78 | 5419.337 | 5009.277 | 4694.631 | 2.54E-16 | -1.080446625 | YDR052C | DBF4 | n |
| W303.asm02.HP0_G0009210 | 7866.558 | 9141.663 | 8141.455 | 5633.453 | 5466.54 | 7192.505 | 12568.17 | 8824.488 | 10783.9 | 0.015635802 | 0.45925577 | YDR054C | CDC34 | n |
| W303.asm02.HP0_G0009230 | 4693.039 | 4502.394 | 4555.248 | 1599.237 | 1615.055 | 1971.33 | 4233.008 | 3590.787 | 4033.196 | 9.80E-29 | 1.406870465 | YDR056C | EMC10 | n |
| W303.asm02.HP0_G0009240 | 2584.772 | 2642.903 | 2570.816 | 2578.092 | 2541.911 | 3923.668 | 2502.277 | 3283.68 | 3200.722 | 0.351499948 | -0.213757163 | YDR057W | YOS9 | n |
| W303.asm02.HP0_G0009250 | 1922.96 | 1759.339 | 1570.515 | 1423.473 | 1569.927 | 1680.378 | 1698.635 | 1730.965 | 1535.772 | 0.225056764 | 0.168717742 | YDR058C | TGL2 | n |
| W303.asm02.HP0_G0009270 | 4521.497 | 4866.28 | 5283.916 | 9896.926 | 8854.597 | 2972.568 | 11310.24 | 4221.108 | 9048.378 | 0.255720832 | -0.566259748 | YDR060W | MAK21 | n |
| W303.asm02.HP0_G0009280 | 5930.892 | 6241.703 | 5671.963 | 1260.215 | 1223.658 | 2149.851 | 1391.251 | 1354.062 | 1017.868 | 8.11E-15 | 1.945161804 | YDR061W | | n |
| W303.asm02.HP0_G0009290 | 5359.086 | 5808.823 | 5568.484 | 6047.504 | 5902.197 | 7725.789 | 8031.482 | 7995.514 | 8290.06 | 0.096278577 | -0.233461408 | YDR062W | LCB2 | n |
| W303.asm02.HP0_G0009300 | 4166.766 | 3846.954 | 3758.673 | 1900.049 | 2424.752 | 1935.625 | 853.0209 | 874.0731 | 891.0828 | 3.19E-12 | 0.911550707 | YDR063W | AIM7 | n |
| W303.asm02.HP0_G0009320 | 1127.727 | 1330.91 | 1110.247 | 714.1682 | 611.829 | 578.8644 | 601.1884 | 425.2248 | 515.5123 | 4.24E-07 | 0.905269983 | YDR065W | RRG1 | n |
| W303.asm02.HP0_G0009340 | 2092.384 | 1946.29 | 1804.422 | 840.6065 | 810.5649 | 935.9068 | 1269.038 | 1424.933 | 1406.595 | 5.76E-24 | 1.175169432 | YDR067C | OCA6 | n |
| W303.asm02.HP0_G0009350 | 3749.56 | 3832.488 | 3747.894 | 2565.587 | 2646.052 | 2847.223 | 2947.922 | 3745.414 | 3493.762 | 5.11E-05 | 0.491508111 | YDR068W | DOS2 | n |
| W303.asm02.HP0_G0009360 | 2136.858 | 2533.849 | 2007.069 | 2778.17 | 2416.941 | 3660.824 | 3600.958 | 3526.359 | 3131.349 | 0.040635907 | -0.407428858 | YDR069C | DOA4 | n |
| W303.asm02.HP0_G0009370 | 1635.999 | 1636.931 | 1426.075 | 27.78865 | 17.35685 | 24.30927 | 85.17864 | 197.5792 | 89.70632 | 5.47E-64 | 6.072509876 | YDR070C | FMP16 | n |
| W303.asm02.HP0_G0009380 | 6079.137 | 6329.614 | 5954.376 | 4924.148 | 5585.434 | 6056.806 | 4473.73 | 5694.361 | 4655.16 | 0.32907377 | 0.148598956 | YDR071C | PAA1 | n |
| W303.asm02.HP0_G0009390 | 2304.164 | 2627.324 | 2198.937 | 5415.312 | 5465.672 | 4867.171 | 11222.59 | 6428.84 | 8554.395 | 1.06E-07 | -1.1431431 | YDR072C | IPT1 | n |
| W303.asm02.HP0_G0009400 | 1356.449 | 1297.526 | 1222.35 | 919.8042 | 944.2127 | 983.7657 | 1357.92 | 1075.948 | 1381.477 | 0.001151097 | 0.444935649 | YDR073W | SNF11 | n |
| W303.asm02.HP0_G0009410 | 29524.21 | 30664.35 | 27870.44 | 11621.91 | 11294.97 | 21156.66 | 30781.59 | 30332.7 | 34840.74 | 0.000101173 | 0.998549247 | YDR074W | TPS2 | n |
| W303.asm02.HP0_G0009420 | 707.3443 | 747.8025 | 808.4326 | 2027.182 | 1572.531 | 1229.897 | 4029.32 | 2136.862 | 3778.43 | 0.000146709 | -1.093416429 | YDR075W | PPH3 | n |
| W303.asm02.HP0_G0009430 | 901.1228 | 891.3539 | 914.0678 | 1845.166 | 1895.368 | 2158.207 | 2305.996 | 2570.677 | 2289.305 | 1.25E-24 | -1.1239265 | YDR076W | RAD55 | n |
| W303.asm02.HP0_G0009440 | 65220.75 | 68913.56 | 71475.14 | 3982.808 | 3762.097 | 6538.433 | 12345.97 | 22936.37 | 16749.96 | 6.86E-42 | 3.847479869 | YDR077W | SED1 | n |
| W303.asm02.HP0_G0009450 | 1029.25 | 787.8634 | 910.8341 | 509.2269 | 600.547 | 513.5333 | 278.9909 | 328.5828 | 344.4723 | 5.56E-06 | 0.749846496 | YDR078C | SHU2 | n |

| | | | | | | | | | | | | | |
|---|---|---|---|---|---|---|---|---|---|---|---|---|---|
| W303.asm02.HP0_G0009460 | 2166.507 | 2084.277 | 1865.862 | 2025.792 | 2222.545 | 1675.82 | 2049.225 | 1725.596 | 1940.049 | 0.786410426 | 0.046324206 | YDR079W | PET100 | n |
| W303.asm02.HP0_G0009470 | 685.1075 | 617.6048 | 575.604 | 103.5127 | 131.0442 | 69.88914 | 125.9163 | 121.3394 | 118.4123 | 7.03E-29 | 2.628608958 | YDR079C- | TFB5 | n |
| W303.asm02.HP0_G0009480 | 4777.751 | 5058.795 | 4573.573 | 3424.951 | 3359.418 | 4350.599 | 5031.712 | 5569.8 | 5920.617 | 0.012032295 | 0.371922675 | YDR080W | VPS41 | n |
| W303.asm02.HP0_G0009490 | 2021.438 | 2066.472 | 2030.783 | 3170.684 | 2902.065 | 2992.319 | 6701.954 | 4971.694 | 6348.816 | 5.45E-05 | -0.567197302 | YDR081C | PDC2 | n |
| W303.asm02.HP0_G0009500 | 991.1292 | 909.1587 | 882.8084 | 473.7964 | 444.3354 | 477.829 | 782.6559 | 687.232 | 810.9452 | 3.91E-17 | 0.994916344 | YDR082W | STN1 | n |
| W303.asm02.HP0_G0009510 | 2438.644 | 2492.675 | 2791.787 | 2878.209 | 2529.761 | 568.9888 | 4381.145 | 1794.32 | 3835.842 | 0.528524021 | 0.369771265 | YDR083W | RRP8 | n |
| W303.asm02.HP0_G0009520 | 1807.54 | 1857.265 | 1758.071 | 1805.567 | 1867.597 | 2198.469 | 2494.87 | 2583.563 | 2538.091 | 0.333566196 | -0.114730321 | YDR084C | TVP23 | n |
| W303.asm02.HP0_G0009530 | 7772.317 | 7889.762 | 7266.192 | 5458.385 | 4262.842 | 5666.338 | 7087.11 | 4174.934 | 4991.26 | 0.0118816 | 0.57529843 | YDR085C | AFR1 | n |
| W303.asm02.HP0_G0009540 | 3412.831 | 3402.947 | 3366.313 | 1692.329 | 2149.646 | 1731.276 | 1387.548 | 1422.785 | 1376.693 | 8.35E-13 | 0.870021008 | YDR086C | SSS1 | n |
| W303.asm02.HP0_G0009550 | 1773.655 | 1688.12 | 1985.51 | 2688.551 | 2970.625 | 719.4024 | 3331.843 | 807.4975 | 2106.304 | 0.719281692 | -0.227666072 | YDR087C | RRP1 | n |
| W303.asm02.HP0_G0009560 | 1123.492 | 1240.774 | 1105.936 | 1788.894 | 1891.029 | 651.0326 | 2856.571 | 752.7338 | 1847.95 | 0.563657918 | -0.319636904 | YDR088C | SLU7 | n |
| W303.asm02.HP0_G0009570 | 2647.247 | 2108.759 | 2256.066 | 5131.868 | 4825.204 | 4810.196 | 709.822 | 991.1173 | 757.1214 | 8.48E-11 | -1.07445135 | YDR089W | VTC5 | n |
| W303.asm02.HP0_G0009580 | 706.2854 | 812.345 | 748.0696 | 115.3229 | 96.33052 | 166.3666 | 164.1849 | 225.498 | 136.3536 | 2.52E-22 | 2.582552479 | YDR090C | ILT1 | n |
| W303.asm02.HP0_G0009590 | 8133.401 | 8601.955 | 9183.794 | 10811.17 | 10636.28 | 6438.158 | 9705.427 | 7940.75 | 9874.872 | 0.674498176 | -0.105495873 | YDR091C | RLI1 | n |
| W303.asm02.HP0_G0009600 | 2849.496 | 2790.906 | 2750.827 | 2987.279 | 3460.956 | 4508.61 | 2541.78 | 3995.61 | 3109.819 | 0.097230647 | -0.384851148 | YDR092W | UBC13 | n |
| W303.asm02.HP0_G0009630 | 1970.61 | 2085.39 | 2242.053 | 4861.624 | 5430.091 | 5666.338 | 1036.957 | 2463.297 | 1850.342 | 3.68E-06 | -1.341297224 | YDR097C | MSH6 | n |
| W303.asm02.HP0_G0009650 | 15993.61 | 16692.02 | 16502.8 | 14932.23 | 17017.52 | 19869.03 | 18450.43 | 21831.43 | 18877.8 | 0.634727277 | -0.075143381 | YDR099W | BMH2 | n |
| W303.asm02.HP0_G0009660 | 1857.308 | 1576.84 | 1552.191 | 1525.597 | 2012.527 | 2512.971 | 5050.229 | 6195.826 | 4902.75 | 0.224492224 | -0.279057728 | YDR100W | TVP15 | n |
| W303.asm02.HP0_G0009670 | 2068.029 | 2132.127 | 2345.532 | 6326.085 | 5583.699 | 1586.18 | 7366.101 | 2192.699 | 5701.734 | 0.057449392 | -1.04390959 | YDR101C | ARX1 | n |
| W303.asm02.HP0_G0009680 | 3166.107 | 3014.579 | 3094.68 | 558.5518 | 394.0005 | 597.0964 | 1578.891 | 1173.663 | 1432.909 | 6.53E-41 | 2.58039134 | YDR103W | STE5 | n |
| W303.asm02.HP0_G0009690 | 883.1215 | 1070.515 | 894.6654 | 1626.33 | 1590.755 | 2645.912 | 3863.901 | 3915.075 | 3258.134 | 9.32E-06 | -1.041676366 | YDR104C | SPO71 | n |
| W303.asm02.HP0_G0009700 | 3195.756 | 2957.826 | 3023.538 | 1053.19 | 1112.574 | 1247.369 | 2282.541 | 2450.411 | 1932.872 | 2.46E-27 | 1.427057042 | YDR105C | TMS1 | n |
| W303.asm02.HP0_G0009710 | 290.1382 | 208.0939 | 216.6599 | 213.2779 | 218.6963 | 154.2119 | 91.35101 | 141.7416 | 84.92199 | 0.345644365 | 0.287003973 | YDR106W | ARP10 | n |
| W303.asm02.HP0_G0009730 | 1034.544 | 950.3324 | 1090.845 | 2176.546 | 1964.795 | 1757.864 | 1229.535 | 1276.748 | 1047.77 | 2.73E-11 | -0.939634003 | YDR108W | TRS85 | n |
| W303.asm02.HP0_G0009740 | 1297.151 | 1215.179 | 1311.817 | 3847.338 | 3367.229 | 5713.438 | 18404.76 | 12846.94 | 12917.71 | 3.97E-13 | -1.757317812 | YDR109C | | n |
| W303.asm02.HP0_G0009750 | 1256.913 | 1169.554 | 1310.739 | 2280.753 | 2309.329 | 1871.814 | 2014.66 | 1768.548 | 2010.618 | 2.51E-09 | -0.789849254 | YDR110W | FOB1 | n |
| W303.asm02.HP0_G0009770 | 1640.234 | 1718.165 | 1637.346 | 1305.372 | 1143.816 | 1265.601 | 1613.456 | 1266.01 | 1245.124 | 0.002868219 | 0.427046889 | YDR113C | PDS1 | n |
| W303.asm02.HP0_G0009790 | 10017.18 | 9760.381 | 9674.244 | 4066.174 | 3965.172 | 3075.122 | 1156.701 | 1689.087 | 1290.575 | 1.65E-14 | 1.406964589 | YDR116C | MRPL1 | n |
| W303.asm02.HP0_G0009800 | 2851.614 | 2842.095 | 2900.656 | 4595.547 | 4188.208 | 5000.872 | 2814.599 | 3925.813 | 3235.408 | 9.57E-06 | -0.681658762 | YDR117C | TMA64 | n |
| W303.asm02.HP0_G0009830 | 932.8898 | 1052.71 | 841.8478 | 43.0724 | 32.11017 | 47.09921 | 59.25471 | 195.4316 | 137.5497 | 2.60E-31 | 4.528490365 | YDR119W- | COX26 | n |
| W303.asm02.HP0_G0009840 | 2405.818 | 2799.808 | 2872.631 | 5384.745 | 5013.526 | 3035.62 | 3636.758 | 1749.22 | 3076.329 | 0.023897555 | -0.733770118 | YDR120C | TRM1 | n |
| W303.asm02.HP0_G0009850 | 1670.942 | 1543.456 | 1573.749 | 1541.575 | 1563.852 | 1435.766 | 1229.535 | 1158.63 | 1272.634 | 0.448264539 | 0.076633063 | YDR121W | DPB4 | n |
| W303.asm02.HP0_G0009860 | 6271.857 | 5719.799 | 5876.766 | 9150.106 | 9004.734 | 14033.28 | 6731.582 | 7572.437 | 5418.262 | 0.000137184 | -0.849120742 | YDR122W | KIN1 | n |
| W303.asm02.HP0_G0009870 | 1254.795 | 2062.021 | 1135.039 | 1103.904 | 1263.579 | 1169.884 | 507.3684 | 504.686 | 437.7669 | 0.201143979 | 0.331719015 | YDR123C | INO2 | n |
| W303.asm02.HP0_G0009890 | 1413.63 | 1432.175 | 1254.687 | 1761.105 | 1577.738 | 1649.232 | 654.2707 | 692.601 | 715.2584 | 0.009666313 | -0.283091387 | YDR125C | ECM18 | n |
| W303.asm02.HP0_G0009900 | 612.0435 | 536.3703 | 592.8506 | 566.1937 | 553.6835 | 678.3805 | 246.8946 | 352.2064 | 253.5699 | 0.826400944 | -0.046494532 | YDR126W | SWF1 | n |
| W303.asm02.HP0_G0009910 | 13791.1 | 12174.05 | 13080.44 | 18142.51 | 18004.26 | 21319.23 | 25510.39 | 20371.06 | 22595.23 | 4.09E-05 | -0.55754525 | YDR127W | ARO1 | n |
| W303.asm02.HP0_G0009920 | 2403.7 | 2428.133 | 2235.586 | 1192.133 | 1053.561 | 1527.686 | 1156.701 | 1330.438 | 1145.849 | 7.71E-08 | 0.905039112 | YDR128W | MTC5 | n |
| W303.asm02.HP0_G0009930 | 25607.35 | 27519.58 | 25331.97 | 12180.46 | 12745.14 | 16995.22 | 16083.95 | 19787.99 | 17813.28 | 5.46E-08 | 0.904266178 | YDR129C | SAC6 | n |
| W303.asm02.HP0_G0009940 | 1451.75 | 1490.041 | 1448.711 | 506.4481 | 448.6746 | 594.0577 | 682.6636 | 599.1804 | 604.0226 | 1.59E-27 | 1.501991972 | YDR130C | FIN1 | n |
| W303.asm02.HP0_G0009950 | 1628.586 | 1623.577 | 1571.593 | 999.6965 | 883.4637 | 989.843 | 836.9728 | 1144.671 | 873.1416 | 1.12E-06 | 0.747241851 | YDR131C | | n |
| W303.asm02.HP0_G0009960 | 1781.068 | 2177.752 | 1844.304 | 2463.463 | 2287.633 | 2721.878 | 1494.947 | 2213.102 | 1210.437 | 0.145332675 | -0.364932365 | YDR132C | MRX16 | n |
| W303.asm02.HP0_G0009970 | 11744.25 | 12585.78 | 12130.8 | 30661.99 | 34356.15 | 40673.96 | 41299.3 | 36548.93 | 35254.59 | 6.11E-29 | -1.535447807 | YLR110C | CCW12 | y |
| W303.asm02.HP0_G0009980 | 5845.121 | 5778.778 | 6080.491 | 5354.177 | 5566.342 | 7046.649 | 3144.203 | 7424.253 | 4699.415 | 0.952592663 | -0.021255678 | YDR135C | YCF1 | n |
| W303.asm02.HP0_G0009990 | 2878.087 | 2986.759 | 2761.606 | 1775 | 1731.346 | 2341.286 | 1132.012 | 1735.261 | 1359.948 | 0.006842489 | 0.560851179 | YDR137W | RGP1 | n |
| W303.asm02.HP0_G0010000 | 2258.631 | 2157.722 | 2335.831 | 3358.953 | 3319.498 | 3184.514 | 1477.664 | 2376.319 | 2076.402 | 0.00364808 | -0.546655209 | YDR138W | HPR1 | n |
| W303.asm02.HP0_G0010010 | 1547.051 | 1291.962 | 1354.933 | 430.0293 | 627.4501 | 347.9264 | 269.1151 | 329.6566 | 258.3542 | 6.58E-11 | 1.57842946 | YDR139C | RUB1 | n |
| W303.asm02.HP0_G0010020 | 1134.081 | 1214.066 | 1108.092 | 2038.297 | 2350.118 | 1669.743 | 1557.905 | 1428.154 | 1703.224 | 5.54E-07 | -0.809492318 | YDR140W | MTQ2 | n |
| W303.asm02.HP0_G0010030 | 5418.385 | 5580.699 | 5196.605 | 7637.015 | 7630.071 | 9904.507 | 7284.626 | 9611.583 | 9036.417 | 0.000143166 | -0.636204159 | YDR141C | DOP1 | n |
| W303.asm02.HP0_G0010040 | 673.4596 | 792.3146 | 609.0192 | 634.9706 | 593.6043 | 762.7033 | 985.1095 | 938.5011 | 793.0039 | 0.766179734 | 0.058791452 | YDR142C | PEX7 | n |
| W303.asm02.HP0_G0010050 | 2411.112 | 2187.768 | 2546.024 | 6863.101 | 6314.422 | 6899.274 | 7662.374 | 7738.876 | 8286.472 | 1.19E-52 | -1.490537676 | YDR143C | SAN1 | n |
| W303.asm02.HP0_G0010070 | 8267.881 | 8743.28 | 8565.074 | 5823.805 | 5560.267 | 5809.155 | 4679.887 | 5596.645 | 5372.811 | 5.52E-09 | 0.572896352 | YDR145W | TAF12 | n |
| W303.asm02.HP0_G0010100 | 10100.83 | 9987.392 | 9396.143 | 8150.41 | 8103.046 | 7780.485 | 3066.431 | 4721.498 | 4464.983 | 0.112662563 | 0.294877583 | YDR148C | KGD2 | n |
| W303.asm02.HP0_G0010110 | 5956.305 | 6548.836 | 6050.31 | 10203.99 | 9809.224 | 12943.93 | 17048.07 | 19903.96 | 19868.16 | 2.14E-08 | -0.8287828 | YDR150W | NUM1 | n |

| | | | | | | | | | | | | | | |
|---|---|---|---|---|---|---|---|---|---|---|---|---|---|---|
| W303.asm02.HP0_G0010130 | 3180.932 | 3337.291 | 3495.663 | 6385.831 | 7081.595 | 5793.202 | 6074.842 | 5726.575 | 6171.795 | 6.22E-18 | -0.943562734 | YDR152W | GIR2 | n |
| W303.asm02.HP0_G0010140 | 3990.989 | 4482.364 | 4204.927 | 8676.31 | 8718.346 | 10465.14 | 14907.5 | 10216.13 | 12523 | 3.70E-11 | -1.135849243 | YDR153C | ENT5 | n |
| W303.asm02.HP0_G0010150 | 21217.15 | 20845 | 21036.49 | 10285.27 | 12733.85 | 12108.29 | 12354.61 | 14723.94 | 11934.53 | 2.02E-10 | 0.845071506 | YDR155C | CPR1 | n |
| W303.asm02.HP0_G0010160 | 579.2176 | 669.9064 | 628.4216 | 3888.326 | 3926.987 | 1949.299 | 5867.451 | 2990.533 | 5419.458 | 1.24E-12 | -2.37873357 | YDR156W | RPA14 | n |
| W303.asm02.HP0_G0010170 | 8465.895 | 8179.09 | 8132.832 | 10914.69 | 11122.27 | 12258.71 | 17799.87 | 12654.73 | 13703.54 | 0.002753531 | -0.468973885 | YDR158W | HOM2 | n |
| W303.asm02.HP0_G0010180 | 2251.219 | 2437.035 | 2115.938 | 5450.743 | 4955.381 | 5639.75 | 4442.869 | 4226.477 | 4059.51 | 2.31E-33 | -1.237897852 | YDR159W | SAC3 | n |
| W303.asm02.HP0_G0010190 | 2523.356 | 2618.422 | 2430.687 | 701.6633 | 573.6439 | 818.9185 | 1251.756 | 1241.313 | 1101.594 | 3.16E-33 | 1.853540176 | YDR160W | SSY1 | n |
| W303.asm02.HP0_G0010200 | 2618.657 | 2569.458 | 2738.97 | 5047.807 | 4904.178 | 2815.317 | 4689.763 | 3635.887 | 4857.298 | 0.006193912 | -0.687575377 | YDR161W | ACL4 | n |
| W303.asm02.HP0_G0010210 | 2455.586 | 2557.218 | 2344.455 | 2353.698 | 2422.148 | 2525.125 | 3861.432 | 3713.2 | 3847.803 | 0.903484114 | 0.011091322 | YDR162C | NBP2 | n |
| W303.asm02.HP0_G0010220 | 2226.864 | 1890.65 | 1741.903 | 1085.841 | 923.3844 | 1011.114 | 1234.473 | 681.863 | 999.9265 | 4.87E-05 | 0.955890042 | YDR163W | CWC15 | n |
| W303.asm02.HP0_G0010230 | 4939.762 | 4800.625 | 5001.503 | 2910.166 | 2838.713 | 3642.592 | 2430.678 | 3140.865 | 2997.387 | 3.33E-05 | 0.650452612 | YDR164C | SEC1 | n |
| W303.asm02.HP0_G0010240 | 2182.39 | 2514.931 | 2530.933 | 5464.637 | 5170.606 | 1999.437 | 3731.812 | 2047.736 | 3864.548 | 0.046832264 | -0.805669504 | YDR165W | TRM82 | n |
| W303.asm02.HP0_G0010250 | 3990.989 | 4201.938 | 3961.32 | 4756.721 | 4601.301 | 6185.949 | 4368.8 | 4339.226 | 4079.844 | 0.015730868 | -0.354953893 | YDR166C | SEC5 | n |
| W303.asm02.HP0_G0010260 | 2806.082 | 2283.468 | 2536.323 | 2268.248 | 2416.074 | 2190.873 | 2293.651 | 2476.183 | 1985.5 | 0.308191474 | 0.149694395 | YDR167W | TAF10 | n |
| W303.asm02.HP0_G0010270 | 9208.183 | 9837.164 | 8762.332 | 4426.731 | 4279.331 | 5467.306 | 3652.806 | 4644.185 | 4401.59 | 1.96E-10 | 0.972254534 | YDR168W | CDC37 | n |
| W303.asm02.HP0_G0010280 | 3724.147 | 3979.378 | 3631.479 | 762.7983 | 837.468 | 2079.962 | 1964.047 | 2941.138 | 1873.068 | 3.08E-05 | 1.622877846 | YDR169C | STB3 | n |
| W303.asm02.HP0_G0010300 | 9558.679 | 9785.975 | 9800.359 | 10423.52 | 11049.37 | 12203.25 | 8349.976 | 12432.46 | 10957.33 | 0.245474175 | -0.208472533 | YDR170C | SEC7 | n |
| W303.asm02.HP0_G0010310 | 35385.22 | 40489.28 | 33720.26 | 3373.542 | 3731.723 | 10001.74 | 12938.51 | 11322.15 | 12300.53 | 2.93E-12 | 2.67951018 | YDR171W | HSP42 | n |
| W303.asm02.HP0_G0010320 | 11133.26 | 11439.6 | 12093.07 | 14794.67 | 15064.88 | 12318.72 | 11342.34 | 10401.9 | 11481.21 | 0.01817502 | -0.282966699 | YDR172W | SUP35 | n |
| W303.asm02.HP0_G0010330 | 1170.083 | 889.1283 | 978.7424 | 781.5557 | 812.3006 | 774.8579 | 950.5443 | 1205.877 | 1063.319 | 0.025556278 | 0.35934406 | YDR173C | ARG82 | n |
| W303.asm02.HP0_G0010340 | 10456.62 | 10994.48 | 10670.23 | 4779.647 | 5212.262 | 4652.186 | 11732.43 | 5144.575 | 10905.9 | 9.77E-05 | 1.133227269 | YDR174W | HMO1 | n |
| W303.asm02.HP0_G0010350 | 5906.537 | 6631.183 | 5999.648 | 5095.048 | 5202.716 | 4355.157 | 6629.12 | 2806.913 | 6363.169 | 0.304181713 | 0.33925106 | YDR175C | RSM24 | n |
| W303.asm02.HP0_G0010360 | 3293.175 | 3569.867 | 3267.146 | 4370.459 | 4016.375 | 4546.593 | 10783.12 | 4082.587 | 9340.222 | 0.309417391 | -0.352461436 | YDR176W | NGG1 | n |
| W303.asm02.HP0_G0010370 | 5449.093 | 5188.993 | 5366.915 | 3761.193 | 3926.12 | 4068.764 | 2713.372 | 3347.034 | 3219.859 | 6.64E-05 | 0.445172826 | YDR177W | UBC1 | n |
| W303.asm02.HP0_G0010390 | 253.0768 | 272.6363 | 253.3089 | 370.9784 | 340.1943 | 473.2711 | 1662.835 | 597.0328 | 1330.046 | 0.105610239 | -0.604679988 | YDR179C | CSN9 | n |
| W303.asm02.HP0_G0010400 | 417.2061 | 431.7669 | 406.3721 | 1154.618 | 1107.367 | 1036.183 | 3918.218 | 1422.785 | 3869.333 | 8.05E-05 | -1.393609315 | YDR179W- | NVJ3 | n |
| W303.asm02.HP0_G0010410 | 707.3443 | 803.4426 | 852.6269 | 3330.469 | 2732.836 | 3701.086 | 8503.051 | 3922.591 | 10193.03 | 4.57E-10 | -2.046716978 | YDR180W | SCC2 | n |
| W303.asm02.HP0_G0010420 | 1611.644 | 1820.543 | 1592.073 | 1488.082 | 1423.262 | 1543.639 | 3530.593 | 1429.228 | 3688.724 | 0.636605676 | 0.173456471 | YDR181C | SAS4 | n |
| W303.asm02.HP0_G0010430 | 4668.684 | 4444.529 | 4185.525 | 3122.749 | 3075.634 | 3748.945 | 7950.007 | 4172.787 | 9547.145 | 0.169331196 | 0.418900075 | YDR182W | CDC1 | n |
| W303.asm02.HP0_G0010450 | 1036.662 | 958.122 | 930.2365 | 1595.763 | 1575.134 | 938.9455 | 3384.925 | 1742.777 | 3765.273 | 0.151233177 | -0.490596072 | YDR183W | PLP1 | n |
| W303.asm02.HP0_G0010460 | 954.0677 | 1186.246 | 1228.818 | 2401.634 | 1817.262 | 450.4811 | 9193.121 | 1387.35 | 7219.565 | 0.521508348 | -0.470875944 | YDR184C | ATC1 | n |
| W303.asm02.HP0_G0010480 | 4632.682 | 4943.064 | 4630.702 | 1559.638 | 1319.121 | 1472.23 | 5972.381 | 1745.999 | 4905.142 | 8.80E-06 | 1.707079883 | YDR186C | SND1 | n |
| W303.asm02.HP0_G0010490 | 8907.456 | 8381.62 | 9459.739 | 8431.77 | 8550.852 | 8923.78 | 10916.45 | 6735.947 | 12273.02 | 0.856586462 | 0.046170729 | YDR188W | CCT6 | n |
| W303.asm02.HP0_G0010500 | 4621.034 | 4810.64 | 4823.648 | 5148.541 | 4913.724 | 5565.303 | 10544.87 | 5110.214 | 11484.8 | 0.678693249 | -0.132603164 | YDR189W | SLY1 | n |
| W303.asm02.HP0_G0010510 | 7259.81 | 7727.293 | 7677.954 | 6166.3 | 6330.043 | 5794.722 | 9548.649 | 4256.543 | 9901.186 | 0.336114607 | 0.309333954 | YDR190C | RVB1 | n |
| W303.asm02.HP0_G0010520 | 213.8975 | 196.9658 | 203.725 | 1189.354 | 1120.385 | 1346.126 | 3483.683 | 1318.626 | 3414.821 | 6.95E-14 | -2.572493756 | YDR191W | HST4 | n |
| W303.asm02.HP0_G0010530 | 545.3328 | 625.3944 | 554.0458 | 2954.628 | 2825.695 | 2885.206 | 8621.56 | 3426.496 | 9035.221 | 1.94E-12 | -2.328933727 | YDR192C | NUP42 | n |
| W303.asm02.HP0_G0010540 | 19168.18 | 20550.1 | 19415.32 | 3802.876 | 2916.819 | 1442.603 | 4893.451 | 1175.811 | 5130.006 | 2.37E-08 | 2.856913365 | YDR194C | MSS116 | n |
| W303.asm02.HP0_G0010560 | 2494.765 | 2750.845 | 2728.191 | 1606.878 | 1291.35 | 1109.87 | 6995.759 | 1990.825 | 5494.811 | 0.015395324 | 0.992289711 | YDR195W | REF2 | n |
| W303.asm02.HP0_G0010570 | 3551.546 | 3537.596 | 3280.081 | 1299.119 | 1174.191 | 1059.732 | 4374.973 | 1500.099 | 4014.059 | 1.37E-05 | 1.553291057 | YDR196C | CAB5 | n |
| W303.asm02.HP0_G0010580 | 2784.904 | 2991.21 | 2716.334 | 2001.477 | 1744.363 | 1428.169 | 2589.925 | 904.1395 | 2508.189 | 0.059982722 | 0.714865119 | YDR197W | CBS2 | n |
| W303.asm02.HP0_G0010590 | 1563.993 | 1678.104 | 1671.839 | 2470.411 | 2262.465 | 2036.661 | 4620.633 | 2218.471 | 5324.967 | 0.146784947 | -0.462206113 | YDR198C | RKM2 | n |
| W303.asm02.HP0_G0010610 | 643.8104 | 603.1383 | 481.8258 | 826.7122 | 989.3405 | 823.4764 | 2381.299 | 946.0177 | 2219.933 | 0.082709218 | -0.610398234 | YDR201W | SPC19 | n |
| W303.asm02.HP0_G0010620 | 3139.635 | 3236.026 | 2761.606 | 596.7612 | 539.798 | 764.9823 | 3056.555 | 789.2429 | 2644.542 | 1.15E-07 | 2.264516514 | YDR202C | RAV2 | n |
| W303.asm02.HP0_G0010630 | 4431.491 | 4135.17 | 3926.827 | 2623.943 | 2991.453 | 3129.059 | 13823.63 | 5917.711 | 14733.37 | 0.123645372 | 0.514761528 | YDR204W | COQ4 | n |
| W303.asm02.HP0_G0010640 | 989.0114 | 1061.613 | 1007.846 | 3849.422 | 3597.207 | 4079.399 | 2866.447 | 3181.669 | 2680.425 | 2.22E-68 | -1.914229147 | YDR205W | MSC2 | n |
| W303.asm02.HP0_G0010650 | 3968.752 | 3646.65 | 4155.344 | 13942.95 | 13757.91 | 12099.94 | 9662.221 | 9273.336 | 9926.304 | 1.76E-67 | -1.757543241 | YDR206W | EBS1 | n |
| W303.asm02.HP0_G0010660 | 1985.435 | 2141.03 | 2063.12 | 4220.401 | 4023.318 | 7831.383 | 2977.549 | 5067.262 | 3369.37 | 9.70E-06 | -1.376938092 | YDR207C | UME6 | n |
| W303.asm02.HP0_G0010670 | 3013.626 | 3449.684 | 3070.966 | 6034.999 | 5417.941 | 6449.553 | 5648.949 | 4893.306 | 4134.864 | 3.88E-09 | -0.909055341 | YDR208W | MSS4 | n |
| W303.asm02.HP0_G0010700 | 4933.409 | 5322.529 | 5414.343 | 9342.543 | 9479.444 | 8593.326 | 7764.836 | 6672.593 | 8608.219 | 1.63E-10 | -0.806952687 | YDR211W | GCD6 | n |
| W303.asm02.HP0_G0010710 | 9000.639 | 8950.261 | 9460.817 | 10935.53 | 11562.27 | 12074.87 | 6442.715 | 9198.17 | 7647.763 | 0.031404402 | -0.334827247 | YDR212W | TCP1 | n |
| W303.asm02.HP0_G0010720 | 1551.287 | 1553.471 | 1544.645 | 4122.446 | 4114.441 | 7408.249 | 17367.8 | 15258.7 | 15964.14 | 5.97E-13 | -1.750618144 | YDR213W | UPC2 | n |
| W303.asm02.HP0_G0010730 | 6973.907 | 6365.224 | 6590.343 | 4222.485 | 4444.222 | 5901.834 | 2362.781 | 4448.753 | 4357.335 | 0.104304853 | 0.452049833 | YDR214W | AHA1 | n |
| W303.asm02.HP0_G0010750 | 4511.967 | 6160.468 | 4181.213 | 1096.262 | 818.3755 | 892.6059 | 2466.477 | 2790.806 | 1678.106 | 3.81E-22 | 2.403273262 | YDR216W | ADR1 | n |

| | | | | | | | | | | | | | | |
|---|---|---|---|---|---|---|---|---|---|---|---|---|---|---|
| W303.asm02.HP0_G0010760 | 1394.57 | 1575.727 | 1536.022 | 1938.953 | 1619.394 | 2081.481 | 1777.641 | 1621.438 | 2126.638 | 0.053261231 | -0.3240083 | YDR217C | RAD9 | n |
| W303.asm02.HP0_G0010770 | 301.7861 | 327.1636 | 299.659 | 189.6575 | 244.7316 | 127.6237 | 161.716 | 118.118 | 120.8045 | 0.009666313 | 0.726013308 | YDR218C | SPR28 | n |
| W303.asm02.HP0_G0010780 | 2514.884 | 2972.292 | 2311.039 | 2337.025 | 2070.672 | 2608.688 | 3029.397 | 1614.995 | 1797.715 | 0.602056345 | 0.152339051 | YDR219C | MFB1 | n |
| W303.asm02.HP0_G0010790 | 2448.174 | 2792.019 | 2663.516 | 5411.839 | 5198.377 | 6299.139 | 3777.488 | 4252.248 | 4444.649 | 6.74E-18 | -1.097319848 | YDR221W | GTB1 | n |
| W303.asm02.HP0_G0010800 | 645.9282 | 692.1625 | 659.681 | 321.6536 | 339.3264 | 252.2087 | 930.7927 | 749.5124 | 825.2982 | 4.01E-12 | 1.130048157 | YDR222W | | n |
| W303.asm02.HP0_G0010820 | 5641.812 | 5816.613 | 5434.823 | 7935.048 | 9823.977 | 7218.333 | 15269.2 | 10704.71 | 13307.63 | 0.002302997 | -0.564119797 | YDR224C | HTB1 | y |
| W303.asm02.HP0_G0010840 | 8663.909 | 8006.606 | 8818.383 | 11751.82 | 13369.98 | 14260.42 | 17552.97 | 13149.75 | 13931.99 | 4.30E-05 | -0.62764617 | YDR226W | ADK1 | n |
| W303.asm02.HP0_G0010850 | 2277.691 | 2484.885 | 2325.052 | 2362.73 | 1935.289 | 2143.014 | 2540.546 | 2183.035 | 2619.425 | 0.323215072 | 0.137622266 | YDR227W | SIR4 | n |
| W303.asm02.HP0_G0010860 | 3515.544 | 3532.032 | 3888.022 | 1695.802 | 1720.064 | 1537.561 | 735.746 | 812.8665 | 649.4738 | 2.69E-21 | 1.142616657 | YDR228C | PCF11 | n |
| W303.asm02.HP0_G0010870 | 4717.394 | 5015.396 | 4342.9 | 1804.873 | 1526.535 | 1854.341 | 1207.315 | 1344.398 | 1379.085 | 1.54E-30 | 1.440170841 | YDR229W | IVY1 | n |
| W303.asm02.HP0_G0010880 | 2145.329 | 1976.335 | 1920.836 | 1893.101 | 2258.126 | 1631 | 1807.269 | 1529.091 | 1642.224 | 0.717366993 | 0.063845673 | YDR231C | COX20 | n |
| W303.asm02.HP0_G0010890 | 3879.805 | 3453.023 | 4360.147 | 5149.931 | 4813.055 | 3436.723 | 3209.63 | 2476.183 | 3963.823 | 0.445435395 | -0.196550956 | YDR232W | HEM1 | n |
| W303.asm02.HP0_G0010900 | 18152.7 | 19935.84 | 19982.3 | 13856.11 | 14918.21 | 16600.95 | 14743.31 | 17204.42 | 15008.47 | 0.003637305 | 0.355914739 | YDR233C | RTN1 | n |
| W303.asm02.HP0_G0010910 | 5119.775 | 5471.644 | 6014.739 | 4576.79 | 4123.12 | 4547.352 | 4405.834 | 3647.698 | 3040.446 | 0.056830361 | 0.325962326 | YDR234W | LYS4 | n |
| W303.asm02.HP0_G0010920 | 2252.278 | 2341.334 | 2170.911 | 2383.571 | 2120.139 | 2199.989 | 1841.834 | 1985.456 | 2005.833 | 0.908788216 | 0.012617249 | YDR235W | PRP42 | n |
| W303.asm02.HP0_G0010930 | 1286.562 | 1207.389 | 1187.857 | 1304.677 | 1388.548 | 1473.749 | 934.4961 | 1173.663 | 919.7888 | 0.222161451 | -0.17843068 | YDR236C | FMN1 | n |
| W303.asm02.HP0_G0010940 | 9291.836 | 9202.867 | 8637.294 | 4137.035 | 4123.12 | 3139.694 | 1610.987 | 1520.501 | 1461.615 | 1.82E-19 | 1.251012765 | YDR237W | MRPL7 | n |
| W303.asm02.HP0_G0010950 | 13165.29 | 13158.88 | 13740.12 | 11148.11 | 11511.06 | 14045.44 | 8088.268 | 10427.67 | 9091.437 | 0.431447846 | 0.126354403 | YDR238C | SEC26 | n |
| W303.asm02.HP0_G0010960 | 2652.541 | 2686.303 | 2685.074 | 2751.076 | 2579.228 | 3600.051 | 1213.487 | 2882.079 | 1857.519 | 0.650498956 | -0.154435704 | YDR239C | | n |
| W303.asm02.HP0_G0010970 | 930.772 | 904.7075 | 988.4436 | 1667.319 | 1501.368 | 1242.052 | 940.6685 | 937.4273 | 923.3771 | 4.76E-06 | -0.643439648 | YDR240C | SNU56 | n |
| W303.asm02.HP0_G0010980 | 1986.494 | 1846.137 | 1842.148 | 567.5831 | 546.7408 | 426.9315 | 913.5101 | 1014.741 | 577.7087 | 5.10E-16 | 1.880473044 | YDR242W | AMD2 | n |
| W303.asm02.HP0_G0010990 | 776.1728 | 803.4426 | 791.186 | 1406.8 | 1353.834 | 1093.157 | 1398.658 | 1175.811 | 1608.733 | 3.62E-05 | -0.700940167 | YDR243C | PRP28 | n |
| W303.asm02.HP0_G0011000 | 3000.919 | 3418.526 | 2931.916 | 1445.704 | 1332.138 | 1786.731 | 1599.877 | 1422.785 | 1144.653 | 1.59E-08 | 1.034527163 | YDR244W | PEX5 | n |
| W303.asm02.HP0_G0011010 | 3660.613 | 3626.62 | 3855.685 | 5536.193 | 6121.761 | 5167.239 | 2150.452 | 2849.865 | 2532.111 | 2.31E-05 | -0.594420183 | YDR245W | MNN10 | n |
| W303.asm02.HP0_G0011020 | 4291.716 | 4732.744 | 4175.824 | 2201.555 | 2221.677 | 2443.081 | 2815.833 | 3017.378 | 3034.466 | 4.80E-22 | 0.942888767 | YDR246W | TRS23 | n |
| W303.asm02.HP0_G0011050 | 2059.558 | 1861.717 | 1832.447 | 294.5596 | 339.3264 | 310.7028 | 493.7892 | 398.3798 | 436.5708 | 8.41E-89 | 2.608266948 | YDR248C | | n |
| W303.asm02.HP0_G0011060 | 291.1971 | 322.7124 | 268.3996 | 1210.196 | 1222.79 | 738.394 | 919.6825 | 994.3387 | 764.2979 | 6.36E-15 | -1.845808144 | YDR249C | | n |
| W303.asm02.HP0_G0011100 | 1585.171 | 1067.177 | 1234.207 | 825.3228 | 846.1465 | 653.3116 | 639.4571 | 991.1173 | 772.6705 | 0.0019582 | 0.741594875 | YDR253C | MET32 | n |
| W303.asm02.HP0_G0011110 | 1473.987 | 1488.928 | 1449.789 | 762.7983 | 667.3709 | 577.3451 | 1111.026 | 834.3425 | 1194.888 | 3.06E-10 | 1.135944958 | YDR254W | CHL4 | n |
| W303.asm02.HP0_G0011120 | 2428.055 | 2951.149 | 2317.507 | 1570.753 | 1583.813 | 2170.362 | 3491.09 | 3303.009 | 3518.88 | 0.00326995 | 0.531371995 | YDR255C | RMD5 | n |
| W303.asm02.HP0_G0011130 | 1272.796 | 1332.023 | 1353.855 | 7985.762 | 7093.745 | 7584.492 | 3152.844 | 5407.656 | 3966.216 | 8.40E-36 | -2.517348755 | YDR256C | CTA1 | n |
| W303.asm02.HP0_G0011140 | 2771.138 | 2868.802 | 2838.137 | 2378.013 | 2124.478 | 866.7773 | 1203.611 | 599.1804 | 1034.613 | 0.105610239 | 0.6590076 | YDR257C | RKM4 | n |
| W303.asm02.HP0_G0011150 | 32233.94 | 33733.46 | 28850.26 | 2113.326 | 2158.324 | 4221.456 | 1034.488 | 1776.065 | 873.1416 | 5.89E-24 | 3.48077247 | YDR258C | HSP78 | n |
| W303.asm02.HP0_G0011170 | 1138.316 | 1254.127 | 1215.883 | 772.5243 | 884.3315 | 1367.396 | 1930.716 | 1380.907 | 1637.439 | 0.338600638 | 0.254749826 | YDR260C | SWM1 | n |
| W303.asm02.HP0_G0011190 | 3920.043 | 4160.764 | 3896.645 | 1678.434 | 1608.112 | 1845.985 | 5021.837 | 3831.318 | 4204.236 | 1.12E-19 | 1.222444735 | YDR262W | | n |
| W303.asm02.HP0_G0011210 | 3485.895 | 3321.712 | 3591.597 | 4077.984 | 4245.486 | 4421.248 | 1991.205 | 4303.79 | 2687.601 | 0.30634536 | -0.29341542 | YDR264C | AKR1 | n |
| W303.asm02.HP0_G0011220 | 1687.885 | 1836.122 | 1623.333 | 1397.074 | 1397.226 | 1858.14 | 1265.335 | 1895.257 | 1399.419 | 0.518929766 | 0.145765847 | YDR265W | PEX10 | n |
| W303.asm02.HP0_G0011230 | 2623.951 | 2721.912 | 2775.619 | 4080.068 | 4287.142 | 3887.964 | 4867.527 | 3792.661 | 4065.491 | 2.31E-06 | -0.593527923 | YDR266C | HEL2 | n |
| W303.asm02.HP0_G0011240 | 2156.977 | 2094.292 | 2385.415 | 1834.745 | 1753.042 | 1577.823 | 1677.649 | 1774.991 | 1783.362 | 0.000973346 | 0.361478542 | YDR267C | CIA1 | n |
| W303.asm02.HP0_G0011250 | 2340.166 | 2276.792 | 2174.145 | 1589.511 | 1474.464 | 1759.383 | 1323.355 | 1298.224 | 1600.361 | 0.000215236 | 0.493407385 | YDR268W | MSW1 | n |
| W303.asm02.HP0_G0011260 | 4874.111 | 5044.329 | 4314.874 | 3687.553 | 3338.59 | 3264.279 | 4431.758 | 3280.459 | 4040.373 | 0.002180844 | 0.467886566 | YDR270W | CCC2 | n |
| W303.asm02.HP0_G0011290 | 5469.212 | 5419.343 | 4930.361 | 2881.683 | 3265.691 | 3761.859 | 4274.98 | 4923.373 | 4924.279 | 1.31E-06 | 0.674882433 | YDR276C | PMP3 | n |
| W303.asm02.HP0_G0011300 | 5341.085 | 6162.694 | 5019.828 | 5757.113 | 4688.953 | 4715.998 | 30769.24 | 19376.72 | 28130.71 | 0.592017895 | 0.123993469 | YDR277C | MTH1 | n |
| W303.asm02.HP0_G0011310 | 968.8923 | 952.558 | 970.1191 | 1329.687 | 1352.966 | 979.2077 | 875.2414 | 686.1582 | 931.7497 | 0.080005084 | -0.34061293 | YDR279W | RNH202 | n |
| W303.asm02.HP0_G0011320 | 1545.992 | 1631.367 | 1592.073 | 2505.841 | 2509.801 | 1414.496 | 2473.884 | 1788.951 | 2546.464 | 0.110433693 | -0.431003452 | YDR280W | RRP45 | n |
| W303.asm02.HP0_G0011340 | 2015.084 | 2373.605 | 1965.03 | 1099.041 | 1128.195 | 1096.196 | 608.5952 | 817.1617 | 584.8852 | 1.72E-08 | 0.934922702 | YDR282C | MRX10 | n |
| W303.asm02.HP0_G0011350 | 2259.69 | 2381.395 | 2264.689 | 2923.366 | 2893.387 | 3172.359 | 2194.893 | 2466.518 | 2377.816 | 3.82E-05 | -0.380492204 | YDR283C | GCN2 | n |
| W303.asm02.HP0_G0011360 | 2664.189 | 2511.593 | 2591.296 | 1597.152 | 1576.87 | 2349.643 | 3693.543 | 5235.848 | 4339.394 | 0.02404924 | 0.491696394 | YDR284C | DPP1 | n |
| W303.asm02.HP0_G0011370 | 1253.736 | 1288.624 | 1322.596 | 806.5654 | 734.1948 | 626.7233 | 444.4103 | 644.28 | 462.8846 | 8.06E-06 | 0.83425677 | YDR285W | ZIP1 | n |
| W303.asm02.HP0_G0011380 | 582.3943 | 459.587 | 467.813 | 268.1604 | 331.5158 | 233.217 | 302.4459 | 286.7046 | 287.0602 | 5.55E-06 | 0.860031571 | YDR286C | MGP12 | n |
| W303.asm02.HP0_G0011400 | 1715.416 | 1626.916 | 1588.84 | 1808.346 | 1776.474 | 2093.636 | 3421.959 | 2238.873 | 2817.975 | 0.294216395 | -0.203595886 | YDR288W | NSE3 | n |
| W303.asm02.HP0_G0011410 | 2947.974 | 3202.642 | 2928.682 | 1694.413 | 1740.892 | 1670.502 | 2086.26 | 1812.574 | 1668.538 | 5.09E-13 | 0.830482599 | YDR289C | RTT103 | n |
| W303.asm02.HP0_G0011420 | 2106.15 | 1927.372 | 1938.082 | 2256.438 | 2254.655 | 2462.073 | 2187.486 | 2414.976 | 2093.148 | 0.036906474 | -0.223660769 | YDR291W | HRQ1 | n |
| W303.asm02.HP0_G0011430 | 5916.067 | 6535.482 | 6514.889 | 4523.991 | 4553.57 | 4687.131 | 4521.875 | 4264.06 | 4384.845 | 3.70E-09 | 0.462435383 | YDR292C | SRP101 | n |

| | | | | | | | | | | | | | |
|---|---|---|---|---|---|---|---|---|---|---|---|---|---|
| W303.asm02.HP0_G0011440 | 11159.73 | 11204.8 | 11063.67 | 17734.71 | 16803.17 | 20572.48 | 18160.33 | 19662.35 | 17633.87 | 4.78E-11 | -0.721280869 | YDR293C | SSD1 | y |
| W303.asm02.HP0_G0011460 | 10759.47 | 10137.62 | 11047.5 | 1428.336 | 1384.209 | 2220.5 | 2514.622 | 3641.256 | 2922.034 | 1.14E-31 | 2.665997969 | YDR294C | DPL1 | n |
| W303.asm02.HP0_G0011470 | 2593.243 | 3082.46 | 2676.451 | 2930.313 | 2989.717 | 3777.052 | 3260.243 | 4286.609 | 3756.901 | 0.236802482 | -0.215462364 | YDR295C | HDA2 | n |
| W303.asm02.HP0_G0011480 | 9176.416 | 9481.068 | 9278.65 | 4007.123 | 3950.419 | 3157.926 | 1613.456 | 1534.46 | 1525.008 | 1.47E-28 | 1.329582843 | YDR296W | MHR1 | n |
| W303.asm02.HP0_G0011490 | 5017.062 | 4921.92 | 5329.188 | 4815.078 | 5332.024 | 12371.14 | 50228.24 | 32603.79 | 35115.84 | 0.144727187 | -0.560571035 | YDR297W | SUR2 | n |
| W303.asm02.HP0_G0011500 | 4657.037 | 4780.595 | 4705.078 | 13485.13 | 15116.95 | 15143.91 | 5087.264 | 9932.649 | 7890.568 | 3.35E-12 | -1.629086239 | YDR298C | ATP5 | n |
| W303.asm02.HP0_G0011510 | 1246.324 | 1277.496 | 1562.97 | 2739.266 | 2065.465 | 492.2627 | 5656.356 | 1667.611 | 4636.023 | 0.561623312 | -0.374224222 | YDR299W | BFR2 | n |
| W303.asm02.HP0_G0011520 | 2132.622 | 2270.115 | 2301.338 | 3729.931 | 3775.115 | 3371.392 | 4545.33 | 3236.433 | 4230.55 | 6.96E-06 | -0.698093135 | YDR300C | PRO1 | n |
| W303.asm02.HP0_G0011530 | 1759.89 | 1915.131 | 1611.476 | 2918.502 | 2705.065 | 2842.665 | 2008.488 | 1970.423 | 2241.462 | 9.79E-10 | -0.679643313 | YDR301W | CFT1 | n |
| W303.asm02.HP0_G0011540 | 931.8309 | 804.5554 | 819.2117 | 898.268 | 991.0762 | 813.6008 | 783.8904 | 1037.291 | 824.1021 | 0.660625547 | -0.080425133 | YDR302W | GPI11 | n |
| W303.asm02.HP0_G0011560 | 4149.824 | 3717.869 | 3813.646 | 2229.344 | 2535.836 | 2997.637 | 4247.822 | 4683.915 | 4146.824 | 4.39E-05 | 0.589670899 | YDR304C | CPR5 | n |
| W303.asm02.HP0_G0011580 | 3115.28 | 3289.441 | 2888.799 | 1397.074 | 1371.191 | 1160.008 | 1703.573 | 1545.198 | 1423.34 | 9.57E-22 | 1.242324261 | YDR306C | PFU1 | n |
| W303.asm02.HP0_G0011590 | 1974.846 | 2135.466 | 2020.004 | 1150.45 | 1202.83 | 1845.985 | 2338.092 | 3350.256 | 2789.269 | 0.021272889 | 0.545775489 | YDR307W | PMT7 | n |
| W303.asm02.HP0_G0011600 | 1839.307 | 1757.113 | 1777.474 | 830.8805 | 805.3579 | 723.2007 | 708.5876 | 668.9774 | 698.5132 | 6.12E-35 | 1.187466491 | YDR308C | SRB7 | n |
| W303.asm02.HP0_G0011610 | 1322.564 | 1481.139 | 1647.047 | 4482.309 | 5108.121 | 5547.831 | 578.9679 | 1625.733 | 1184.123 | 1.35E-07 | -1.766075873 | YDR309C | GIC2 | n |
| W303.asm02.HP0_G0011620 | 5377.088 | 5750.957 | 5738.794 | 10179.68 | 9816.167 | 8580.412 | 9756.041 | 7898.872 | 9415.576 | 2.45E-09 | -0.760644969 | YDR310C | SUM1 | n |
| W303.asm02.HP0_G0011630 | 2153.8 | 2413.666 | 2224.807 | 1844.471 | 1689.689 | 2151.37 | 1734.435 | 1618.216 | 2130.226 | 0.127083296 | 0.256411693 | YDR311W | TFB1 | n |
| W303.asm02.HP0_G0011640 | 2637.717 | 2980.082 | 3164.744 | 6800.576 | 5516.007 | 1956.136 | 7568.555 | 3727.16 | 7359.507 | 0.129459266 | -0.70055996 | YDR312W | SSF2 | n |
| W303.asm02.HP0_G0011650 | 1488.812 | 1893.988 | 1478.893 | 1285.92 | 1227.997 | 1150.892 | 1683.821 | 1758.884 | 1706.812 | 0.00324896 | 0.407452676 | YDR313C | PIB1 | n |
| W303.asm02.HP0_G0011670 | 760.2893 | 809.0066 | 772.8616 | 446.0078 | 451.2781 | 419.3349 | 1145.591 | 997.5601 | 1166.182 | 2.38E-13 | 0.831105549 | YDR315C | IPK1 | n |
| W303.asm02.HP0_G0011680 | 7668.544 | 8390.522 | 7416.022 | 4719.207 | 4363.512 | 4109.026 | 2862.743 | 2724.23 | 3073.937 | 3.59E-15 | 0.831392606 | YDR316W | OMS1 | n |
| W303.asm02.HP0_G0011690 | 795.2329 | 719.9825 | 714.6544 | 327.906 | 334.9872 | 598.6157 | 167.8883 | 377.9776 | 204.5304 | 0.026394415 | 0.821727487 | YDR317W | HIM1 | n |
| W303.asm02.HP0_G0011710 | 1331.036 | 1283.06 | 1187.857 | 251.4872 | 245.5994 | 322.0978 | 797.4696 | 474.6196 | 705.6898 | 4.93E-23 | 2.214210377 | YDR319C | YFT2 | n |
| W303.asm02.HP0_G0011720 | 3077.16 | 3172.597 | 3031.083 | 5295.126 | 5003.112 | 5888.16 | 4507.061 | 4843.911 | 4372.884 | 9.99E-16 | -0.802563999 | YDR320C | SWA2 | n |
| W303.asm02.HP0_G0011730 | 762.4071 | 457.3614 | 452.7223 | 239.6771 | 321.1017 | 209.6674 | 365.404 | 391.937 | 278.6876 | 0.000134876 | 1.119455465 | YDR320C- | DAD4 | n |
| W303.asm02.HP0_G0011740 | 2263.925 | 2052.006 | 2427.454 | 4491.34 | 5326.817 | 3310.618 | 983.8751 | 1031.922 | 1300.144 | 6.82E-06 | -0.961086323 | YDR321W | ASP1 | n |
| W303.asm02.HP0_G0011750 | 9201.83 | 8968.066 | 8503.633 | 6089.187 | 6357.814 | 4483.541 | 3392.332 | 2574.972 | 3363.389 | 0.000435666 | 0.655813462 | YDR322W | MRPL35 | n |
| W303.asm02.HP0_G0011760 | 2611.244 | 2052.006 | 1969.342 | 4417.7 | 5310.328 | 3802.121 | 1280.149 | 2664.098 | 2365.855 | 0.000473369 | -1.028471353 | YDR322C- | TIM11 | n |
| W303.asm02.HP0_G0011770 | 769.8194 | 776.7354 | 784.7186 | 1893.101 | 1807.716 | 2090.597 | 1586.298 | 2108.943 | 1853.931 | 6.30E-21 | -1.312911344 | YDR323C | PEP7 | n |
| W303.asm02.HP0_G0011780 | 3658.495 | 3676.696 | 4253.433 | 6116.281 | 5493.443 | 1798.886 | 8066.047 | 3640.182 | 6684.915 | 0.674216182 | -0.210447941 | YDR324C | UTP4 | n |
| W303.asm02.HP0_G0011800 | 3033.745 | 3297.23 | 3164.744 | 8154.578 | 7824.468 | 11592.48 | 21029.25 | 16847.92 | 18332.38 | 1.10E-16 | -1.537863466 | YDR326C | YSP2 | n |
| W303.asm02.HP0_G0011810 | 9095.94 | 8632 | 8942.343 | 3570.146 | 3957.362 | 4229.053 | 5402.054 | 4545.395 | 4833.377 | 7.36E-26 | 1.181871459 | YDR328C | SKP1 | n |
| W303.asm02.HP0_G0011820 | 4416.667 | 4845.137 | 4556.326 | 2324.52 | 2217.338 | 3331.129 | 1757.89 | 1753.515 | 1283.398 | 0.000133866 | 0.811513362 | YDR329C | PEX3 | n |
| W303.asm02.HP0_G0011830 | 3828.978 | 3957.122 | 3679.985 | 4300.988 | 4255.9 | 4324.771 | 2799.785 | 4762.303 | 3521.272 | 0.438475966 | -0.167955414 | YDR330W | UBX5 | n |
| W303.asm02.HP0_G0011840 | 1223.028 | 1278.609 | 1185.701 | 2480.831 | 2545.382 | 2624.641 | 1743.076 | 2508.397 | 1812.068 | 1.79E-10 | -1.053056008 | YDR331W | GPI8 | n |
| W303.asm02.HP0_G0011850 | 1740.829 | 1836.122 | 1693.397 | 2948.375 | 2863.88 | 2325.333 | 1815.91 | 1481.844 | 1838.382 | 2.98E-05 | -0.62673007 | YDR332W | IRC3 | n |
| W303.asm02.HP0_G0011860 | 2957.504 | 2946.698 | 2985.811 | 3188.052 | 2961.947 | 2427.128 | 1610.987 | 1450.704 | 1609.93 | 0.72855565 | 0.051641941 | YDR333C | RQC1 | n |
| W303.asm02.HP0_G0011870 | 2680.073 | 2938.908 | 2472.726 | 3248.493 | 3043.524 | 2638.315 | 5230.462 | 4651.701 | 5524.713 | 0.338704932 | -0.14235946 | YDR334W | SWR1 | n |
| W303.asm02.HP0_G0011880 | 7562.655 | 7435.739 | 7377.217 | 5281.927 | 5207.923 | 6685.808 | 4086.106 | 5474.232 | 4316.668 | 0.025420715 | 0.381541566 | YDR335W | MSN5 | n |
| W303.asm02.HP0_G0011890 | 948.7732 | 837.9394 | 950.7167 | 474.4911 | 447.8067 | 197.5128 | 398.7348 | 166.439 | 343.2762 | 0.001571392 | 1.289631074 | YDR336W | MRX8 | n |
| W303.asm02.HP0_G0011900 | 7017.322 | 7518.086 | 6894.313 | 3518.737 | 3694.406 | 2640.594 | 1459.147 | 1296.077 | 1325.261 | 4.35E-13 | 1.120939949 | YDR337W | MRPS28 | n |
| W303.asm02.HP0_G0011910 | 1216.675 | 1393.227 | 1149.052 | 3404.109 | 3745.608 | 4170.559 | 6451.356 | 6194.752 | 5978.029 | 4.43E-36 | -1.590588334 | YDR338C | | n |
| W303.asm02.HP0_G0011920 | 2257.572 | 2222.264 | 2220.495 | 3200.557 | 3539.062 | 1670.502 | 3687.371 | 2089.615 | 3310.761 | 0.337696779 | -0.327843539 | YDR339C | FCF1 | n |
| W303.asm02.HP0_G0011940 | 13601.55 | 13054.27 | 14288.78 | 14320.88 | 14148.44 | 12773 | 6421.729 | 4822.435 | 6193.325 | 0.951116887 | -0.010446916 | YDR341C | | n |
| W303.asm02.HP0_G0011960 | 8671.322 | 9471.052 | 9541.661 | 11180.07 | 12004 | 14097.1 | 12744.7 | 15270.51 | 12667.73 | 0.002611814 | -0.429392235 | YDR346C | SVF1 | n |
| W303.asm02.HP0_G0011970 | 11139.61 | 11252.65 | 11215.66 | 2903.913 | 2933.308 | 2114.147 | 764.1388 | 928.8369 | 879.122 | 1.09E-38 | 2.079579159 | YDR347W | MRP1 | n |
| W303.asm02.HP0_G0011980 | 4603.033 | 4519.086 | 4729.87 | 1554.08 | 1273.125 | 1454.758 | 1665.304 | 1408.826 | 1545.341 | 5.88E-47 | 1.693161365 | YDR348C | PAL1 | n |
| W303.asm02.HP0_G0012000 | 5152.601 | 5062.133 | 5651.483 | 3529.158 | 2991.453 | 4262.478 | 2641.772 | 3801.252 | 2576.366 | 0.00945521 | 0.557114309 | YDR349C | YPS7 | n |
| W303.asm02.HP0_G0012010 | 1462.339 | 1476.687 | 1356.011 | 538.405 | 471.2385 | 281.8356 | 491.3203 | 308.1806 | 391.1196 | 2.73E-10 | 1.73351003 | YDR350C | ATP22 | n |
| W303.asm02.HP0_G0012030 | 1209.262 | 1240.774 | 1267.622 | 2199.471 | 2001.245 | 3372.911 | 6214.338 | 5186.454 | 5345.301 | 2.52E-06 | -1.026680499 | YDR352W | YPQ2 | n |
| W303.asm02.HP0_G0012040 | 14047.35 | 15098.49 | 14197.15 | 9127.181 | 10056.56 | 12482.05 | 4792.225 | 7104.26 | 5372.811 | 0.023650583 | 0.452877866 | YDR353W | TRR1 | n |
| W303.asm02.HP0_G0012050 | 1477.164 | 1358.73 | 1486.438 | 1971.604 | 2200.849 | 2425.609 | 1271.507 | 2027.334 | 1800.107 | 0.002295541 | -0.610143298 | YDR354W | TRP4 | n |
| W303.asm02.HP0_G0012060 | 1785.303 | 1908.454 | 1912.213 | 1904.912 | 1851.108 | 2398.261 | 771.5457 | 1197.287 | 764.2979 | 0.574335982 | -0.134683055 | YDR356W | SPC110 | n |
| W303.asm02.HP0_G0012070 | 1571.406 | 1556.809 | 1381.881 | 143.1115 | 173.5685 | 140.538 | 391.328 | 357.5754 | 375.5705 | 9.10E-129 | 3.306005133 | YDR357C | CNL1 | n |

| | | | | | | | | | | | | | | |
|---|---|---|---|---|---|---|---|---|---|---|---|---|---|---|
| W303.asm02.HP0_G0012090 | 2453.468 | 2341.334 | 2594.53 | 3843.17 | 3439.26 | 4453.914 | 5764.989 | 4164.196 | 4980.495 | 8.01E-05 | -0.667527639 | YDR359C | EAF1 | n |
| W303.asm02.HP0_G0012100 | 1444.338 | 1653.623 | 1708.488 | 2302.984 | 2316.272 | 1235.974 | 3346.657 | 2097.131 | 3032.074 | 0.360630057 | -0.284732638 | YDR361C | BCP1 | n |
| W303.asm02.HP0_G0012110 | 2152.741 | 2161.06 | 2208.638 | 1566.585 | 1602.037 | 1713.044 | 1664.07 | 1619.29 | 1381.477 | 0.000124467 | 0.418067851 | YDR362C | TFC6 | n |
| W303.asm02.HP0_G0012120 | 2127.328 | 2341.334 | 2109.47 | 1220.616 | 1087.407 | 1088.599 | 892.524 | 1022.258 | 1088.437 | 3.67E-15 | 0.953123465 | YDR363W | ESC2 | n |
| W303.asm02.HP0_G0012130 | 3097.279 | 3071.332 | 2792.865 | 1335.244 | 1652.372 | 1378.032 | 1007.33 | 1210.173 | 973.6126 | 6.08E-13 | 1.037997456 | YDR363W- | SEM1 | n |
| W303.asm02.HP0_G0012140 | 914.8885 | 926.9635 | 1067.131 | 1388.738 | 1383.341 | 1431.968 | 1936.888 | 2616.85 | 1968.755 | 0.000999796 | -0.531266767 | YDR364C | CDC40 | n |
| W303.asm02.HP0_G0012150 | 2711.84 | 2996.774 | 3326.431 | 5489.647 | 5050.843 | 1669.743 | 6566.162 | 2708.123 | 5492.419 | 0.378044036 | -0.434485026 | YDR365C | ESF1 | n |
| W303.asm02.HP0_G0012170 | 4444.198 | 4165.216 | 4101.448 | 440.45 | 495.5381 | 1182.798 | 18529.44 | 16726.58 | 12587.59 | 1.16E-12 | 2.584695988 | YDR368W | YPR1 | n |
| W303.asm02.HP0_G0012180 | 1854.132 | 1829.445 | 1639.501 | 2902.524 | 2617.413 | 2888.245 | 3450.352 | 3310.525 | 3038.054 | 6.53E-10 | -0.65975374 | YDR369C | XRS2 | n |
| W303.asm02.HP0_G0012190 | 875.7092 | 837.9394 | 808.4326 | 1926.448 | 1937.024 | 1842.187 | 4899.624 | 4890.085 | 5352.477 | 1.44E-40 | -1.177633275 | YDR370C | DXO1 | n |
| W303.asm02.HP0_G0012200 | 1970.61 | 1931.823 | 1942.394 | 193.1311 | 131.9121 | 151.1733 | 195.0467 | 332.878 | 407.8648 | 7.28E-40 | 3.615681225 | YDR371W | CTS2 | n |
| W303.asm02.HP0_G0012210 | 6219.971 | 5891.17 | 6510.577 | 5337.504 | 5535.967 | 6353.075 | 4587.302 | 6292.468 | 5543.851 | 0.503524202 | 0.112365549 | YDR372C | VPS74 | n |
| W303.asm02.HP0_G0012220 | 1302.445 | 1107.237 | 1157.675 | 1804.178 | 1845.033 | 2033.622 | 2330.685 | 1944.652 | 1990.284 | 2.36E-07 | -0.671600926 | YDR373W | FRQ1 | n |
| W303.asm02.HP0_G0012230 | 148.2458 | 117.9569 | 103.4794 | 44.46183 | 39.92076 | 93.43875 | 32.0963 | 225.498 | 43.05904 | 0.160341129 | 1.055472249 | YDR374C | PHO92 | n |
| W303.asm02.HP0_G0012240 | 513.5659 | 455.1358 | 449.4885 | 355.6947 | 407.0181 | 318.2995 | 304.9149 | 394.0846 | 314.5702 | 0.027963002 | 0.392792663 | YDR374W- | WIP1 | n |
| W303.asm02.HP0_G0012250 | 8379.065 | 8096.743 | 8161.936 | 2370.371 | 2343.175 | 1738.872 | 1070.288 | 981.4531 | 1003.515 | 8.84E-41 | 1.933007362 | YDR375C | BCS1 | n |
| W303.asm02.HP0_G0012260 | 3347.179 | 3238.252 | 3184.147 | 1818.072 | 1807.716 | 1932.587 | 745.6217 | 1029.774 | 724.8271 | 2.57E-07 | 0.813617902 | YDR376W | ARH1 | n |
| W303.asm02.HP0_G0012280 | 1434.808 | 1370.971 | 1363.556 | 2063.307 | 2369.21 | 1901.441 | 2040.584 | 2082.098 | 1932.872 | 2.46E-07 | -0.602878934 | YDR378C | LSM6 | n |
| W303.asm02.HP0_G0012300 | 1746.124 | 1394.34 | 1431.465 | 937.1721 | 1061.371 | 847.7857 | 1335.7 | 1385.202 | 1257.085 | 4.22E-06 | 0.68428526 | YDR379C- | SDH6 | n |
| W303.asm02.HP0_G0012310 | 1139.375 | 1153.975 | 1116.715 | 6104.471 | 6967.04 | 2576.782 | 1988.736 | 1586.002 | 1656.577 | 3.92E-11 | -2.198104982 | YDR380W | ARO10 | n |
| W303.asm02.HP0_G0012320 | 7757.492 | 6863.759 | 7107.739 | 2514.872 | 2606.999 | 3182.995 | 2170.204 | 2999.123 | 2264.188 | 9.95E-16 | 1.387599882 | YDR381W | YRA1 | n |
| W303.asm02.HP0_G0012330 | 2258.631 | 2094.292 | 1910.057 | 687.769 | 781.0583 | 631.2813 | 334.5422 | 504.686 | 283.472 | 4.03E-12 | 1.57665719 | YDR381C- | COI1 | n |
| W303.asm02.HP0_G0012340 | 10882.3 | 9338.629 | 9909.228 | 10674.31 | 13337.87 | 11382.05 | 11860.82 | 9081.126 | 8462.297 | 0.210485651 | -0.232256009 | YDR382W | RPP2B | n |
| W303.asm02.HP0_G0012350 | 1043.015 | 924.7379 | 915.1457 | 823.9333 | 884.3315 | 714.8444 | 770.3112 | 915.9513 | 1051.358 | 0.15074072 | 0.251029354 | YDR383C | NKP1 | n |
| W303.asm02.HP0_G0012360 | 788.8795 | 827.9242 | 859.0944 | 566.8884 | 467.7671 | 1516.291 | 9058.564 | 6696.216 | 8215.903 | 0.92915821 | -0.043220987 | YDR384C | ATO3 | n |
| W303.asm02.HP0_G0012370 | 27731.5 | 29410.23 | 31374.73 | 91509.4 | 102641.5 | 103303.8 | 92633.63 | 82055.5 | 75304.27 | 7.91E-49 | -1.748649239 | YOR133W | EFT1 | y |
| W303.asm02.HP0_G0012380 | 1459.162 | 1544.568 | 1478.893 | 514.7847 | 595.34 | 610.7704 | 344.418 | 379.0514 | 282.2759 | 5.15E-22 | 1.381769661 | YDR386W | MUS81 | n |
| W303.asm02.HP0_G0012390 | 2650.423 | 2452.614 | 2396.194 | 780.8609 | 730.7234 | 1004.277 | 2594.862 | 1781.434 | 2011.814 | 5.14E-16 | 1.575499614 | YDR387C | CIN10 | n |
| W303.asm02.HP0_G0012400 | 13105.99 | 13148.86 | 13159.13 | 9587.083 | 10519.12 | 13468.09 | 17409.77 | 16677.19 | 16551.41 | 0.133584625 | 0.231358547 | YDR388W | RVS167 | n |
| W303.asm02.HP0_G0012410 | 1793.774 | 1979.674 | 1952.095 | 5384.745 | 5025.676 | 5421.726 | 6324.206 | 5308.867 | 6024.677 | 2.25E-43 | -1.46752478 | YDR389W | SAC7 | n |
| W303.asm02.HP0_G0012420 | 4365.839 | 4945.289 | 4792.388 | 2792.759 | 2724.158 | 3698.807 | 4716.922 | 4567.945 | 5366.83 | 0.000155272 | 0.613810481 | YDR390C | UBA2 | n |
| W303.asm02.HP0_G0012430 | 5332.614 | 4521.312 | 4500.275 | 186.8786 | 241.2602 | 204.3498 | 383.9211 | 404.8226 | 230.8443 | 6.98E-86 | 4.50564739 | YDR391C | | n |
| W303.asm02.HP0_G0012440 | 2904.559 | 2686.303 | 2633.334 | 2481.526 | 2489.84 | 3089.556 | 5721.783 | 4872.904 | 5000.829 | 0.854871663 | 0.028913663 | YDR392W | SPT3 | n |
| W303.asm02.HP0_G0012450 | 2378.286 | 2605.068 | 2297.027 | 2223.092 | 1938.76 | 2399.781 | 799.9386 | 1211.246 | 1084.848 | 0.457262125 | 0.149805471 | YDR393W | SHE9 | n |
| W303.asm02.HP0_G0012460 | 12910.09 | 14191.56 | 13595.68 | 6153.101 | 6083.576 | 8470.26 | 4221.898 | 6754.202 | 5513.949 | 7.90E-06 | 0.974804745 | YDR394W | RPT3 | n |
| W303.asm02.HP0_G0012470 | 6835.191 | 7258.803 | 7516.267 | 9598.198 | 9292.858 | 8108.66 | 6148.91 | 7539.149 | 7208.8 | 0.013670246 | -0.321239153 | YDR395W | SXM1 | n |
| W303.asm02.HP0_G0012480 | 1580.936 | 1619.126 | 1555.424 | 3264.471 | 3441.863 | 2869.253 | 4928.017 | 5404.435 | 5527.106 | 1.62E-20 | -1.009665215 | YDR397C | NCB2 | n |
| W303.asm02.HP0_G0012490 | 2535.004 | 2868.802 | 3178.757 | 4539.97 | 3900.952 | 1204.068 | 4840.369 | 1379.833 | 3741.352 | 0.775601774 | -0.168374835 | YDR398W | UTP5 | n |
| W303.asm02.HP0_G0012500 | 3702.969 | 3477.504 | 3835.204 | 6888.111 | 6954.022 | 7911.147 | 9379.527 | 12748.15 | 11503.94 | 2.00E-11 | -0.981662125 | YDR399W | HPT1 | n |
| W303.asm02.HP0_G0012510 | 1985.435 | 1780.482 | 1864.785 | 904.5204 | 942.477 | 1136.458 | 1103.619 | 1742.777 | 1437.693 | 3.73E-06 | 0.916363506 | YDR400W | URH1 | n |
| W303.asm02.HP0_G0012520 | 354.7311 | 341.63 | 348.165 | 3128.307 | 2954.136 | 2983.203 | 1078.929 | 1830.829 | 3421.997 | 4.88E-18 | -3.117553277 | YDR402C | DIT2 | n |
| W303.asm02.HP0_G0012530 | 1239.97 | 1221.856 | 1257.921 | 5191.614 | 5272.143 | 4440.999 | 3710.826 | 2123.976 | 6603.582 | 1.90E-08 | -2.002488776 | YDR403W | DIT1 | n |
| W303.asm02.HP0_G0012540 | 1709.063 | 1743.76 | 1724.656 | 2189.745 | 2275.483 | 1756.345 | 1698.635 | 1844.788 | 1728.342 | 0.050104651 | -0.264902247 | YDR404C | RPB7 | n |
| W303.asm02.HP0_G0012550 | 6033.605 | 5551.766 | 5483.329 | 5845.342 | 6551.343 | 4548.112 | 4981.099 | 3709.979 | 3803.548 | 0.961917911 | 0.010571205 | YDR405W | MRP20 | n |
| W303.asm02.HP0_G0012570 | 2422.76 | 2405.877 | 2523.388 | 4763.669 | 4279.331 | 5741.545 | 4387.317 | 4913.708 | 4838.161 | 2.37E-13 | -1.007961436 | YDR407C | TRS120 | n |
| W303.asm02.HP0_G0012580 | 2864.321 | 2371.38 | 2650.581 | 1925.058 | 1917.064 | 2453.717 | 1591.236 | 3280.459 | 2076.402 | 0.269252225 | 0.324947401 | YDR408C | ADE8 | n |
| W303.asm02.HP0_G0012590 | 1641.293 | 1671.428 | 1595.307 | 3154.706 | 3043.524 | 3534.719 | 3852.791 | 3543.54 | 3546.39 | 1.16E-23 | -0.98787341 | YDR409W | SIZ1 | n |
| W303.asm02.HP0_G0012600 | 881.0037 | 736.6745 | 845.0816 | 1269.941 | 1297.425 | 1242.052 | 1725.793 | 1683.718 | 1669.734 | 7.83E-10 | -0.628775814 | YDR410C | STE14 | n |
| W303.asm02.HP0_G0012610 | 2921.502 | 2802.034 | 2740.048 | 1043.464 | 945.0805 | 1734.314 | 1683.821 | 1555.936 | 1336.026 | 1.55E-06 | 1.184716182 | YDR411C | DFM1 | n |
| W303.asm02.HP0_G0012620 | 1754.595 | 1807.189 | 1969.342 | 3751.467 | 3358.551 | 1902.2 | 4762.597 | 2617.924 | 4421.924 | 0.027686021 | -0.704317685 | YDR412W | RRP17 | n |
| W303.asm02.HP0_G0012630 | 714.7566 | 698.8393 | 736.2126 | 1024.012 | 987.6048 | 1012.633 | 966.5924 | 1100.645 | 1048.966 | 3.63E-07 | -0.492545046 | YDR414C | ERD1 | n |
| W303.asm02.HP0_G0012640 | 2408.995 | 2467.081 | 2318.585 | 3927.23 | 4034.6 | 6859.012 | 6572.335 | 9580.443 | 8011.373 | 5.21E-05 | -1.042647129 | YDR415C | | n |
| W303.asm02.HP0_G0012650 | 2099.796 | 2205.572 | 2108.392 | 2397.465 | 2317.14 | 2105.031 | 3620.71 | 2874.562 | 3370.566 | 0.518136013 | -0.088585157 | YDR416W | SYF1 | n |
| W303.asm02.HP0_G0012660 | 1446.456 | 1260.804 | 1387.27 | 1144.197 | 972.8515 | 633.5603 | 713.5254 | 1331.512 | 1487.929 | 0.084908309 | 0.573893414 | YDR419W | RAD30 | n |

| | | | | | | | | | | | | | | |
|---|---|---|---|---|---|---|---|---|---|---|---|---|---|---|
| W303.asm02.HP0_G0012670 | 2699.133 | 2754.183 | 2965.331 | 6892.974 | 5645.316 | 5164.96 | 10663.38 | 7169.762 | 8815.141 | 1.33E-08 | -1.072407118 | YDR420W | HKR1 | n |
| W303.asm02.HP0_G0012680 | 2628.187 | 2694.092 | 2478.115 | 1775.694 | 1690.557 | 2237.972 | 2240.569 | 1961.832 | 1709.204 | 0.0072638 | 0.451399526 | YDR421W | ARO80 | n |
| W303.asm02.HP0_G0012720 | 2965.975 | 3267.185 | 2655.971 | 2344.667 | 2177.417 | 4409.093 | 2326.982 | 3177.374 | 2170.893 | 0.98513419 | -0.006857117 | YDR425W | SNX41 | n |
| W303.asm02.HP0_G0012730 | 7456.765 | 8301.498 | 7809.459 | 4228.042 | 4476.332 | 5563.024 | 3686.137 | 5617.047 | 4504.454 | 0.000213389 | 0.724086545 | YDR427W | RPN9 | n |
| W303.asm02.HP0_G0012750 | 10100.83 | 10629.48 | 10346.86 | 10581.22 | 11718.48 | 8638.146 | 7873.469 | 8204.905 | 7824.784 | 0.9669376 | 0.006527838 | YDR429C | TIF35 | n |
| W303.asm02.HP0_G0012760 | 13093.28 | 13794.29 | 12746.29 | 4225.264 | 4097.952 | 3890.243 | 2031.943 | 2101.426 | 1928.088 | 9.95E-100 | 1.698202433 | YDR430C | CYM1 | n |
| W303.asm02.HP0_G0012770 | 18843.1 | 20136.14 | 20092.25 | 10458.95 | 11245.5 | 10100.5 | 11501.59 | 12089.91 | 11097.27 | 1.24E-26 | 0.893271576 | YDR432W | NPL3 | n |
| W303.asm02.HP0_G0012780 | 1902.841 | 1990.802 | 2029.705 | 6632.455 | 6262.352 | 7740.223 | 6367.412 | 8923.277 | 7212.388 | 5.59E-29 | -1.800652484 | YDR434W | GPI17 | n |
| W303.asm02.HP0_G0012790 | 3940.162 | 3802.442 | 3567.883 | 1514.481 | 1496.16 | 1879.41 | 2326.982 | 2688.795 | 2247.442 | 5.57E-18 | 1.209653222 | YDR435C | PPM1 | n |
| W303.asm02.HP0_G0012810 | 442.6197 | 509.663 | 440.8652 | 484.9119 | 526.7804 | 370.7163 | 618.471 | 1388.423 | 678.1798 | 0.977573304 | 0.011276354 | YDR437W | GPI19 | n |
| W303.asm02.HP0_G0012830 | 1215.616 | 1254.127 | 1254.687 | 1565.195 | 1595.095 | 1482.106 | 785.1249 | 1349.767 | 1070.495 | 0.134321291 | -0.317830554 | YDR439W | LRS4 | n |
| W303.asm02.HP0_G0012840 | 1297.151 | 1265.255 | 1269.778 | 1364.422 | 1204.565 | 1319.537 | 1066.585 | 1420.637 | 1090.829 | 0.901675359 | -0.021323115 | YDR440W | DOT1 | n |
| W303.asm02.HP0_G0012860 | 1969.552 | 2157.722 | 1914.368 | 4610.831 | 4339.213 | 5571.38 | 3952.783 | 4907.266 | 4107.354 | 4.22E-17 | -1.265266197 | YDR443C | SSN2 | n |
| W303.asm02.HP0_G0012870 | 2282.986 | 2214.475 | 2163.366 | 1230.342 | 1054.429 | 1988.042 | 1175.218 | 1374.464 | 1035.809 | 0.013921412 | 0.640368644 | YDR444W | | n |
| W303.asm02.HP0_G0012880 | 32.82586 | 43.39925 | 30.18148 | 409.1878 | 386.1899 | 404.9012 | 509.8374 | 1157.556 | 708.0819 | 1.03E-24 | -3.497116312 | YDR446W | ECM11 | n |
| W303.asm02.HP0_G0012900 | 767.7016 | 863.5339 | 798.7314 | 2518.346 | 2427.356 | 2778.094 | 5235.4 | 4007.421 | 4398.002 | 1.61E-32 | -1.668576939 | YDR448W | ADA2 | n |
| W303.asm02.HP0_G0012910 | 1566.111 | 1782.708 | 1859.395 | 2531.546 | 2051.58 | 603.1737 | 5072.45 | 2053.105 | 3473.429 | 0.991883289 | 0.006062556 | YDR449C | UTP6 | n |
| W303.asm02.HP0_G0012920 | 9873.172 | 9288.553 | 9956.656 | 17144.2 | 20219.86 | 18316.27 | 15837.06 | 13878.86 | 12926.08 | 7.30E-15 | -0.935190522 | YML026C | RPS18B | y |
| W303.asm02.HP0_G0012940 | 8813.214 | 8955.825 | 8746.163 | 7368.159 | 7833.147 | 9925.778 | 8036.42 | 9848.893 | 8495.787 | 0.651942442 | 0.077577829 | YDR452W | PPN1 | n |
| W303.asm02.HP0_G0012960 | 4571.266 | 4400.017 | 4542.313 | 3774.393 | 4404.301 | 3963.17 | 6919.222 | 4598.011 | 4816.632 | 0.439831614 | 0.154500448 | YDR454C | GUK1 | n |
| W303.asm02.HP0_G0012970 | 3201.051 | 3164.807 | 3107.615 | 1985.499 | 2132.289 | 3169.321 | 4974.927 | 4717.203 | 4370.492 | 0.06475758 | 0.37853809 | YDR456W | NHX1 | n |
| W303.asm02.HP0_G0012980 | 11422.34 | 11459.63 | 11162.84 | 11003.61 | 10795.96 | 9685.724 | 5298.359 | 7358.751 | 7117.898 | 0.501543612 | 0.112756016 | YDR457W | TOM1 | n |
| W303.asm02.HP0_G0013000 | 448.9731 | 370.5629 | 409.6059 | 2123.747 | 2361.399 | 1980.446 | 1897.385 | 2971.204 | 2445.992 | 1.41E-34 | -2.394814464 | YDR459C | PFA5 | n |
| W303.asm02.HP0_G0013010 | 1870.015 | 1848.363 | 1829.214 | 1598.542 | 1299.16 | 1252.687 | 2064.039 | 1868.412 | 2078.795 | 0.001603578 | 0.418113473 | YDR460W | TFB3 | n |
| W303.asm02.HP0_G0013030 | 4103.232 | 4226.42 | 3935.45 | 2038.297 | 2088.029 | 1378.791 | 871.538 | 836.4901 | 844.4355 | 6.96E-11 | 1.155801352 | YDR462W | MRPL28 | n |
| W303.asm02.HP0_G0013040 | 4860.345 | 5134.466 | 5171.813 | 8159.441 | 7717.723 | 8262.872 | 20100.93 | 14912.93 | 17704.44 | 5.35E-07 | -0.670564433 | YDR463W | STP1 | n |
| W303.asm02.HP0_G0013050 | 2027.791 | 2212.249 | 2328.286 | 3381.878 | 3033.977 | 2841.905 | 2920.763 | 2799.396 | 2803.622 | 8.06E-06 | -0.495338833 | YDR464W | SPP41 | n |
| W303.asm02.HP0_G0013060 | 3690.262 | 3657.778 | 4037.851 | 2810.127 | 2805.735 | 1266.361 | 6714.299 | 2322.629 | 5854.833 | 0.099493922 | 0.726319048 | YDR465C | RMT2 | n |
| W303.asm02.HP0_G0013070 | 3417.066 | 3577.656 | 3437.455 | 2125.831 | 1932.685 | 2705.166 | 4647.791 | 3223.547 | 3837.039 | 0.001227085 | 0.625050143 | YDR466W | PKH3 | n |
| W303.asm02.HP0_G0013080 | 2754.196 | 2918.878 | 2639.802 | 2067.475 | 1943.967 | 2449.918 | 1249.287 | 1840.493 | 1843.166 | 0.067115045 | 0.363423114 | YDR468C | TLG1 | n |
| W303.asm02.HP0_G0013090 | 1138.316 | 1028.228 | 1039.105 | 723.8942 | 714.2344 | 758.905 | 766.6078 | 647.5014 | 788.2196 | 6.94E-06 | 0.545051688 | YDR469W | SDC1 | n |
| W303.asm02.HP0_G0013100 | 3878.746 | 4134.057 | 3722.024 | 3156.79 | 3480.048 | 3168.561 | 2863.978 | 2563.16 | 2704.347 | 0.009096924 | 0.259321012 | YDR470C | UGO1 | n |
| W303.asm02.HP0_G0013120 | 1652.941 | 1633.592 | 1666.449 | 1431.115 | 1173.323 | 1447.161 | 1235.708 | 1748.146 | 1597.969 | 0.101301022 | 0.289474752 | YDR472W | TRS31 | n |
| W303.asm02.HP0_G0013130 | 2621.833 | 2934.457 | 2631.179 | 1342.886 | 1159.438 | 736.115 | 3365.174 | 1581.707 | 2807.21 | 2.95E-05 | 1.33805945 | YDR473C | PRP3 | n |
| W303.asm02.HP0_G0013140 | 5614.281 | 5861.125 | 5162.112 | 1893.796 | 1727.874 | 1527.686 | 2166.5 | 1699.825 | 1247.516 | 1.89E-15 | 1.691917829 | YDR475C | JIP4 | n |
| W303.asm02.HP0_G0013150 | 2461.939 | 2064.247 | 2245.287 | 1409.579 | 1740.892 | 1572.506 | 3931.797 | 2708.123 | 3378.938 | 0.004304819 | 0.52007904 | YDR476C | | n |
| W303.asm02.HP0_G0013160 | 8862.982 | 9201.755 | 9115.886 | 6310.801 | 5857.937 | 7443.194 | 8825.248 | 7633.644 | 6732.759 | 0.001896085 | 0.470799472 | YDR477W | SNF1 | n |
| W303.asm02.HP0_G0013170 | 493.4468 | 525.2422 | 501.2282 | 514.7847 | 467.7671 | 429.2105 | 1280.149 | 702.2652 | 1087.241 | 0.688177472 | 0.106263526 | YDR478W | SNM1 | n |
| W303.asm02.HP0_G0013180 | 3990.989 | 4021.664 | 3698.31 | 1722.896 | 1678.407 | 1974.368 | 2397.347 | 2093.91 | 1986.696 | 4.00E-20 | 1.123224324 | YDR479C | PEX29 | n |
| W303.asm02.HP0_G0013200 | 3021.038 | 3041.286 | 2900.656 | 4781.036 | 4873.804 | 7100.585 | 5448.964 | 9912.247 | 7672.881 | 0.000492632 | -0.902587591 | YDR481C | PHO8 | n |
| W303.asm02.HP0_G0013210 | 980.5402 | 899.1435 | 826.7571 | 681.5165 | 707.2917 | 461.1164 | 1598.643 | 1080.243 | 1383.87 | 0.019539493 | 0.549156503 | YDR482C | CWC21 | n |
| W303.asm02.HP0_G0013220 | 10819.83 | 10620.58 | 10867.49 | 5406.281 | 5319.875 | 5843.34 | 4839.135 | 5649.261 | 4990.064 | 4.84E-24 | 0.96332533 | YDR483W | KRE2 | n |
| W303.asm02.HP0_G0013230 | 2787.021 | 3092.475 | 2861.851 | 2348.141 | 2185.227 | 2842.665 | 2057.867 | 2476.183 | 2258.207 | 0.107933261 | 0.244850658 | YDR484W | VPS52 | n |
| W303.asm02.HP0_G0013240 | 3809.918 | 4274.27 | 3909.58 | 2159.178 | 2054.183 | 2050.335 | 2515.856 | 2083.172 | 2420.875 | 2.84E-17 | 0.937044715 | YDR485C | VPS72 | n |
| W303.asm02.HP0_G0013250 | 4536.322 | 4895.213 | 4349.367 | 1623.552 | 1680.143 | 2060.97 | 2298.589 | 2617.924 | 2483.071 | 1.32E-24 | 1.361077722 | YDR486C | VPS60 | n |
| W303.asm02.HP0_G0013260 | 6194.557 | 6226.124 | 6193.672 | 5102.69 | 5298.179 | 6115.3 | 8272.204 | 10186.07 | 6945.662 | 0.339490872 | 0.172539107 | YDR487C | RIB3 | n |
| W303.asm02.HP0_G0013270 | 946.6554 | 863.5339 | 920.5353 | 831.5752 | 824.4504 | 928.3102 | 539.4647 | 651.7966 | 679.3759 | 0.589652523 | 0.079491202 | YDR488C | PAC11 | n |
| W303.asm02.HP0_G0013280 | 850.2957 | 761.1561 | 882.8084 | 622.4657 | 635.2607 | 613.809 | 496.2582 | 493.948 | 492.7867 | 8.59E-05 | 0.414870615 | YDR489W | SLD5 | n |
| W303.asm02.HP0_G0013310 | 886.2982 | 1009.311 | 992.7552 | 871.174 | 948.5519 | 694.3335 | 2092.432 | 1736.334 | 1751.067 | 0.259753444 | 0.200509867 | YDR492W | IZH1 | n |
| W303.asm02.HP0_G0013320 | 3051.746 | 2617.309 | 2592.374 | 1954.236 | 2161.796 | 1532.244 | 924.6203 | 1182.254 | 1159.006 | 0.003128367 | 0.548764773 | YDR493W | MZM1 | n |
| W303.asm02.HP0_G0013330 | 12139.21 | 12941.88 | 12017.62 | 5755.723 | 5944.721 | 4535.198 | 1939.357 | 2528.799 | 2257.011 | 2.81E-14 | 1.192234527 | YDR494W | RSM28 | n |
| W303.asm02.HP0_G0013340 | 2854.791 | 3095.813 | 2843.527 | 2753.855 | 2554.061 | 3363.795 | 2559.063 | 3461.931 | 2846.681 | 0.919450817 | 0.020121448 | YDR495C | VPS3 | n |
| W303.asm02.HP0_G0013350 | 2509.59 | 2676.287 | 3087.135 | 6278.844 | 5311.196 | 1347.645 | 8480.83 | 3007.714 | 6791.367 | 0.256219173 | -0.645096667 | YDR496C | PUF6 | n |
| W303.asm02.HP0_G0013360 | 12792.56 | 13498.28 | 14753.36 | 2258.522 | 1913.593 | 1742.671 | 3991.051 | 2094.984 | 2032.147 | 2.52E-24 | 2.794721275 | YDR497C | ITR1 | n |

| | | | | | | | | | | | | | | |
|---|---|---|---|---|---|---|---|---|---|---|---|---|---|---|
| W303.asm02.HP0_G0013370 | 2457.704 | 2617.309 | 2529.855 | 1369.286 | 1287.878 | 1251.168 | 1135.715 | 1257.42 | 1198.476 | 1.33E-27 | 0.96000158 | YDR498C | SEC20 | n |
| W303.asm02.HP0_G0013380 | 1091.725 | 1169.554 | 1005.69 | 1682.602 | 1590.755 | 1700.889 | 1011.033 | 1547.346 | 1389.85 | 0.000876412 | -0.606647877 | YDR499W | LCD1 | n |
| W303.asm02.HP0_G0013400 | 774.055 | 695.5009 | 756.6929 | 4294.04 | 4762.72 | 4967.447 | 1471.492 | 2340.884 | 1629.067 | 2.08E-42 | -2.655131269 | YDR501W | PLM2 | n |
| W303.asm02.HP0_G0013420 | 624.7502 | 574.2055 | 682.3171 | 1020.538 | 1003.226 | 971.611 | 2575.111 | 1230.575 | 2068.03 | 0.013234957 | -0.670990307 | YDR503C | LPP1 | n |
| W303.asm02.HP0_G0013450 | 857.708 | 854.6315 | 862.3281 | 1586.732 | 1987.359 | 2499.297 | 2978.784 | 2277.53 | 2530.914 | 4.51E-10 | -1.238003711 | YDR506C | GMC1 | n |
| W303.asm02.HP0_G0013460 | 4623.152 | 5061.021 | 5366.915 | 7015.938 | 7740.287 | 5938.298 | 3119.513 | 3623.001 | 3328.703 | 0.001397109 | -0.45933137 | YDR507C | GIN4 | n |
| W303.asm02.HP0_G0013480 | 3687.085 | 3508.663 | 3740.348 | 3486.78 | 3952.155 | 3429.886 | 3740.453 | 3762.595 | 3436.35 | 0.938210864 | 0.009177403 | YDR510W | SMT3 | n |
| W303.asm02.HP0_G0013490 | 3571.665 | 3024.594 | 3215.406 | 1756.242 | 1893.632 | 1842.946 | 4397.193 | 3601.525 | 4146.824 | 9.22E-12 | 0.837174168 | YDR511W | SDH7 | n |
| W303.asm02.HP0_G0013500 | 3995.225 | 3775.735 | 3520.455 | 388.3463 | 439.9962 | 284.1146 | 214.7983 | 195.4316 | 206.9226 | 4.02E-79 | 3.344357485 | YDR512C | EMI1 | n |
| W303.asm02.HP0_G0013510 | 1498.342 | 1474.462 | 1408.829 | 2139.726 | 2067.201 | 1207.867 | 1944.295 | 1203.73 | 1812.068 | 0.296316865 | -0.305416911 | YDR514C | | n |
| W303.asm02.HP0_G0013520 | 2874.91 | 3369.563 | 2912.513 | 61.13502 | 58.14545 | 73.68747 | 196.2812 | 302.8116 | 144.7262 | 5.37E-93 | 5.567541917 | YDR515W | SLF1 | n |
| W303.asm02.HP0_G0013540 | 5296.611 | 4930.823 | 5214.929 | 2193.914 | 2105.386 | 2692.251 | 3176.299 | 2721.009 | 3235.408 | 1.49E-16 | 1.143120971 | YDR517W | GRH1 | n |
| W303.asm02.HP0_G0013560 | 2299.928 | 2284.581 | 2368.169 | 1757.632 | 1847.637 | 1829.272 | 1512.23 | 2077.803 | 1749.871 | 0.012035317 | 0.355501116 | YDR519W | FPR2 | n |
| W303.asm02.HP0_G0013570 | 877.827 | 933.6404 | 866.6398 | 1570.058 | 1365.116 | 1526.166 | 2362.781 | 1940.356 | 2527.326 | 1.65E-07 | -0.736648564 | YDR520C | URC2 | n |
| W303.asm02.HP0_G0013590 | 499.8002 | 517.4526 | 476.4363 | 379.315 | 360.1546 | 188.3968 | 144.4334 | 318.9186 | 293.0407 | 0.068671172 | 0.686929578 | YDR523C | SPS1 | n |
| W303.asm02.HP0_G0013620 | 894.7694 | 949.2196 | 782.5628 | 409.8825 | 558.8906 | 708.7671 | 733.277 | 1713.785 | 1003.515 | 0.066784927 | 0.646998216 | YDR525W- | SNA2 | n |
| W303.asm02.HP0_G0013630 | 1622.233 | 1600.208 | 1694.475 | 2156.399 | 1815.527 | 1518.57 | 6400.743 | 3305.156 | 5417.066 | 0.579833879 | -0.15925061 | YDR527W | RBA50 | n |
| W303.asm02.HP0_G0013640 | 87.88859 | 86.79851 | 113.1806 | 1428.336 | 1445.826 | 1326.374 | 1788.752 | 1444.261 | 1391.046 | 5.17E-107 | -3.866741185 | YDR528W | HLR1 | n |
| W303.asm02.HP0_G0013650 | 6681.651 | 6373.013 | 6191.516 | 10013.64 | 11402.58 | 9425.159 | 3419.49 | 7011.914 | 6204.089 | 0.01064702 | -0.68027912 | YDR529C | QCR7 | n |
| W303.asm02.HP0_G0013670 | 2016.143 | 1722.617 | 1740.825 | 2307.847 | 2416.074 | 2435.485 | 3966.362 | 2565.308 | 2888.544 | 0.046847646 | -0.385699067 | YDR531W | CAB1 | n |
| W303.asm02.HP0_G0013680 | 1462.339 | 1426.611 | 1327.985 | 174.3738 | 166.6258 | 100.2757 | 213.5638 | 197.5792 | 178.2166 | 6.73E-50 | 3.256618842 | YDR532C | KRE28 | n |
| W303.asm02.HP0_G0013690 | 6523.875 | 6121.52 | 5745.261 | 533.542 | 555.4192 | 436.8072 | 318.4941 | 782.8001 | 374.3744 | 2.92E-29 | 3.591455315 | YDR533C | HSP31 | n |
| W303.asm02.HP0_G0013710 | 654.3994 | 588.6719 | 656.4473 | 2988.669 | 2452.523 | 2485.623 | 8540.085 | 11548.72 | 6757.876 | 9.74E-22 | -2.061130009 | YDR536W | STL1 | n |
| W303.asm02.HP0_G0013750 | 1702.709 | 1571.276 | 1579.138 | 287.6125 | 276.8418 | 294.7499 | 344.418 | 423.0772 | 362.4136 | 3.01E-96 | 2.497335276 | YDR541C | | n |
| W303.asm02.HP0_G0013840 | 195.8963 | 189.1762 | 188.6343 | 2491.252 | 2436.902 | 1972.089 | 2468.946 | 3356.699 | 2396.953 | 7.03E-81 | -3.588175799 | YEL065W | SIT1 | n |
| W303.asm02.HP0_G0013850 | 284.8438 | 304.9076 | 291.0357 | 885.0684 | 823.5825 | 953.3791 | 1743.076 | 1888.814 | 1724.754 | 2.64E-43 | -1.596462025 | YEL064C | AVT2 | n |
| W303.asm02.HP0_G0013870 | 1517.402 | 1432.175 | 1390.504 | 1795.841 | 1668.861 | 1969.81 | 1885.04 | 1857.674 | 2133.814 | 0.006391683 | -0.324616686 | YEL062W | NPR2 | n |
| W303.asm02.HP0_G0013880 | 1619.056 | 1981.899 | 1730.046 | 2021.624 | 1935.289 | 2306.342 | 1264.1 | 1221.984 | 1074.084 | 0.112283658 | -0.232726202 | YEL061C | CIN8 | n |
| W303.asm02.HP0_G0013890 | 31256.57 | 30980.39 | 29836.55 | 3251.272 | 3450.542 | 7039.052 | 10367.11 | 6538.368 | 5437.399 | 1.98E-15 | 2.744303684 | YEL060C | PRB1 | n |
| W303.asm02.HP0_G0013900 | 940.3021 | 695.5009 | 731.901 | 864.2269 | 956.3625 | 980.727 | 2087.494 | 1264.936 | 1479.556 | 0.323287516 | -0.242369915 | YEL059C-A | SOM1 | n |
| W303.asm02.HP0_G0013910 | 10220.49 | 8838.981 | 10598.01 | 4933.179 | 4718.46 | 6398.655 | 3686.137 | 5193.97 | 4705.396 | 4.19E-06 | 0.885777327 | YEL058W | PCM1 | n |
| W303.asm02.HP0_G0013930 | 2916.207 | 3079.121 | 2733.58 | 3959.882 | 3894.877 | 4344.522 | 3034.335 | 3833.466 | 3856.176 | 0.000411304 | -0.482976972 | YEL056W | HAT2 | n |
| W303.asm02.HP0_G0013940 | 4293.834 | 4517.974 | 4682.442 | 5263.864 | 4814.79 | 3568.145 | 4772.473 | 4158.827 | 5521.125 | 0.942211286 | -0.01623981 | YEL055C | POL5 | n |
| W303.asm02.HP0_G0013950 | 10741.47 | 10997.82 | 11394.59 | 20510.8 | 23910.8 | 21014.6 | 17997.38 | 15213.6 | 14747.72 | 4.59E-16 | -0.981752246 | YEL054C | RPL12A | y |
| W303.asm02.HP0_G0013960 | 3968.752 | 4025.003 | 4377.393 | 5698.756 | 5285.161 | 5515.925 | 4544.095 | 3895.746 | 3898.039 | 9.64E-05 | -0.415547839 | YEL053C | MAK10 | n |
| W303.asm02.HP0_G0013970 | 4686.686 | 4619.239 | 4410.808 | 4288.483 | 4247.221 | 3988.999 | 5711.907 | 2952.95 | 4451.826 | 0.612704083 | 0.131161438 | YEL052W | AFG1 | n |
| W303.asm02.HP0_G0013980 | 5840.885 | 5150.045 | 5608.366 | 4729.627 | 5013.526 | 5819.031 | 4436.696 | 4372.513 | 3902.823 | 0.500820506 | 0.093124465 | YEL051W | VMA8 | n |
| W303.asm02.HP0_G0013990 | 10891.83 | 10942.18 | 10114.03 | 6198.257 | 6551.343 | 5021.383 | 2865.212 | 2332.293 | 2441.208 | 8.37E-09 | 0.846249053 | YEL050C | RML2 | n |
| W303.asm02.HP0_G0014010 | 141.8924 | 135.7618 | 167.0761 | 605.7925 | 772.3798 | 619.1267 | 309.8527 | 566.9664 | 464.0807 | 2.61E-17 | -2.166509139 | YEL048C | TCA17 | n |
| W303.asm02.HP0_G0014020 | 7135.918 | 6298.456 | 6768.198 | 4116.888 | 4511.913 | 12810.22 | 35858.97 | 29008.71 | 29566.01 | 0.853517801 | -0.08570487 | YEL047C | FRD1 | n |
| W303.asm02.HP0_G0014030 | 5102.833 | 5380.395 | 5614.834 | 14408.41 | 14121.53 | 14432.11 | 24154.93 | 18692.71 | 22146.7 | 4.55E-32 | -1.416191365 | YEL046C | GLY1 | n |
| W303.asm02.HP0_G0014050 | 2093.443 | 1807.189 | 1858.317 | 1941.037 | 2157.456 | 1976.647 | 2462.774 | 2411.755 | 2267.776 | 0.50997592 | -0.076677545 | YEL044W | IES6 | n |
| W303.asm02.HP0_G0014060 | 4939.762 | 4925.259 | 5066.178 | 7411.232 | 7037.335 | 8378.341 | 6995.759 | 6869.098 | 6726.778 | 1.04E-10 | -0.61246865 | YEL043W | GTA1 | n |
| W303.asm02.HP0_G0014070 | 4192.18 | 3796.878 | 4753.584 | 5065.175 | 5319.007 | 5961.848 | 4729.266 | 4820.288 | 5128.81 | 0.005933884 | -0.359224202 | YEL042W | GDA1 | n |
| W303.asm02.HP0_G0014090 | 3947.574 | 4621.464 | 4694.299 | 13832.49 | 16016.9 | 11233.16 | 2968.908 | 6090.593 | 5525.91 | 5.76E-09 | -1.631077112 | YEL040W | UTR2 | n |
| W303.asm02.HP0_G0014110 | 2108.267 | 1794.949 | 1915.446 | 2060.528 | 2517.611 | 2499.297 | 1496.181 | 1459.294 | 1432.909 | 0.031547583 | -0.282149212 | YEL038W | UTR4 | n |
| W303.asm02.HP0_G0014120 | 8213.877 | 8524.059 | 8850.72 | 7149.324 | 7440.014 | 8911.626 | 5008.257 | 7228.821 | 6817.681 | 0.525239472 | 0.122783534 | YEL037C | RAD23 | n |
| W303.asm02.HP0_G0014130 | 2412.171 | 2024.186 | 2643.036 | 10317.92 | 9903.819 | 8713.353 | 6842.684 | 7314.725 | 6937.289 | 1.90E-51 | -2.031044364 | YEL036C | ANP1 | n |
| W303.asm02.HP0_G0014160 | 1809.658 | 1789.385 | 2020.004 | 6153.101 | 6053.202 | 6043.132 | 2215.879 | 3605.82 | 2942.367 | 6.42E-20 | -1.699448889 | YEL032W | MCM3 | n |
| W303.asm02.HP0_G0014170 | 8259.41 | 7637.156 | 9347.637 | 11856.72 | 12200.13 | 17422.91 | 21429.22 | 24185.2 | 21568.99 | 0.000146201 | -0.716453323 | YEL031W | SPF1 | n |
| W303.asm02.HP0_G0014190 | 578.1587 | 498.535 | 543.2667 | 2170.293 | 2248.58 | 2320.016 | 1194.97 | 1188.697 | 1046.574 | 2.64E-78 | -2.056104323 | YEL029C | BUD16 | n |
| W303.asm02.HP0_G0014210 | 9163.709 | 8821.176 | 8910.005 | 3536.8 | 3943.476 | 4627.877 | 4258.932 | 5023.236 | 4326.237 | 3.16E-17 | 1.151409045 | YEL027W | VMA3 | n |
| W303.asm02.HP0_G0014220 | 2433.349 | 2055.344 | 2387.571 | 8724.94 | 10190.21 | 6095.549 | 6002.008 | 4067.554 | 4984.083 | 1.18E-15 | -1.862788426 | YEL026W | SNU13 | n |
| W303.asm02.HP0_G0014240 | 5944.657 | 5822.177 | 5526.445 | 14437.59 | 15607.28 | 13670.92 | 4281.153 | 10559.75 | 8845.044 | 9.33E-06 | -1.337937596 | YEL024W | RIP1 | n |

| | | | | | | | | | | | | | | |
|---|---|---|---|---|---|---|---|---|---|---|---|---|---|---|
| W303.asm02.HP0_G0014250 | 624.7502 | 659.8912 | 704.9532 | 797.5341 | 796.6794 | 601.6544 | 972.7648 | 1362.652 | 869.5533 | 0.555408039 | -0.142260905 | YEL023C | | n |
| W303.asm02.HP0_G0014260 | 3815.212 | 3868.098 | 4142.409 | 8749.95 | 8313.063 | 10523.63 | 6814.291 | 7663.71 | 7255.448 | 1.52E-22 | -1.222076854 | YEL022W | GEA2 | n |
| W303.asm02.HP0_G0014270 | 3477.423 | 3483.068 | 3385.716 | 12015.12 | 12664.43 | 16525.74 | 2149.218 | 2375.245 | 2472.306 | 1.97E-40 | -1.993729495 | YEL021W | URA3 | n |
| W303.asm02.HP0_G0014280 | 3846.979 | 3642.199 | 3648.726 | 2057.054 | 2411.734 | 1594.536 | 1249.287 | 1060.914 | 1236.751 | 6.14E-07 | 0.877529697 | YEL020W- | TIM9 | n |
| W303.asm02.HP0_G0014300 | 321.9052 | 372.7885 | 313.6719 | 706.5263 | 657.8246 | 670.0242 | 853.0209 | 738.7744 | 763.1018 | 2.76E-15 | -1.013508999 | YEL019C | MMS21 | n |
| W303.asm02.HP0_G0014310 | 1229.381 | 1238.548 | 1226.662 | 932.3091 | 943.3448 | 812.8411 | 917.2135 | 561.5974 | 757.1214 | 0.021539819 | 0.458677135 | YEL018W | EAF5 | n |
| W303.asm02.HP0_G0014320 | 2401.582 | 2198.896 | 2494.284 | 6755.42 | 6427.242 | 7434.078 | 2426.974 | 4172.787 | 3910 | 7.84E-13 | -1.538985233 | YEL017W | GTT3 | n |
| W303.asm02.HP0_G0014340 | 6253.856 | 6449.797 | 6309.008 | 3904.999 | 3739.533 | 3685.133 | 1161.639 | 1427.08 | 1176.947 | 4.86E-12 | 0.74679497 | YEL015W | EDC3 | n |
| W303.asm02.HP0_G0014350 | 5599.456 | 5980.195 | 5666.574 | 8082.328 | 7883.481 | 10594.28 | 10153.54 | 9092.938 | 9323.477 | 2.43E-05 | -0.623013524 | YEL013W | VAC8 | n |
| W303.asm02.HP0_G0014360 | 3213.758 | 3111.393 | 2732.502 | 2775.391 | 2843.052 | 4347.561 | 5276.138 | 5130.616 | 4252.08 | 0.559498073 | -0.137900016 | YEL012W | UBC8 | n |
| W303.asm02.HP0_G0014370 | 11673.3 | 14024.64 | 11605.86 | 1802.094 | 1746.099 | 4750.943 | 11379.37 | 9998.151 | 7855.882 | 1.62E-08 | 2.168257627 | YEL011W | GLC3 | n |
| W303.asm02.HP0_G0014380 | 20213.32 | 22858.05 | 23289.33 | 15833.28 | 17463.59 | 17767.04 | 16809.82 | 11958.91 | 13105.5 | 0.021610133 | 0.378033817 | YEL009C | GCN4 | n |
| W303.asm02.HP0_G0014420 | 2369.815 | 2264.551 | 2220.495 | 929.5302 | 943.3448 | 1014.912 | 654.2707 | 871.9255 | 724.8271 | 3.89E-21 | 1.247185185 | YEL005C | VAB2 | n |
| W303.asm02.HP0_G0014430 | 1587.289 | 1474.462 | 1348.466 | 1352.612 | 1365.116 | 1516.291 | 3205.927 | 3014.156 | 3296.408 | 0.635132582 | 0.058843963 | YEL004W | YEA4 | n |
| W303.asm02.HP0_G0014440 | 1272.796 | 1057.161 | 1145.818 | 1092.788 | 1309.574 | 796.1285 | 881.4138 | 952.4605 | 1027.436 | 0.601305422 | 0.120377305 | YEL003W | GIM4 | n |
| W303.asm02.HP0_G0014450 | 8703.088 | 8924.667 | 9365.961 | 4617.083 | 4765.323 | 5499.212 | 2625.724 | 5436.649 | 4425.512 | 0.000953216 | 0.85909261 | YEL002C | WBP1 | n |
| W303.asm02.HP0_G0014460 | 5266.962 | 5213.474 | 5067.256 | 2690.636 | 2862.145 | 3445.079 | 1655.428 | 3380.322 | 3073.937 | 0.003560212 | 0.789053158 | YEL001C | IRC22 | n |
| W303.asm02.HP0_G0014470 | 4322.425 | 4778.369 | 4857.063 | 1812.514 | 1839.826 | 1662.146 | 2507.215 | 2519.135 | 2387.384 | 2.86E-51 | 1.393151721 | YER001W | MNN1 | n |
| W303.asm02.HP0_G0014480 | 3964.517 | 4230.871 | 4244.81 | 2821.242 | 2712.008 | 701.1704 | 2517.091 | 715.1508 | 1837.186 | 0.077441672 | 0.996693885 | YER002W | NOP16 | n |
| W303.asm02.HP0_G0014490 | 6300.447 | 5980.195 | 6693.822 | 9605.145 | 10190.21 | 11267.35 | 11778.11 | 11300.67 | 11265.92 | 6.26E-13 | -0.71108249 | YER003C | PMI40 | n |
| W303.asm02.HP0_G0014500 | 7041.676 | 6648.988 | 7071.091 | 3192.915 | 3649.278 | 3085.758 | 3857.728 | 4043.931 | 4907.534 | 1.86E-15 | 1.064527184 | YER004W | FMP52 | n |
| W303.asm02.HP0_G0014510 | 2806.082 | 2558.33 | 2676.451 | 3124.833 | 3039.184 | 3906.195 | 5423.04 | 4503.517 | 5705.322 | 0.043304344 | -0.324695474 | YER005W | YND1 | n |
| W303.asm02.HP0_G0014520 | 6221.03 | 6971.701 | 7421.411 | 5982.201 | 5071.672 | 1576.304 | 6360.005 | 2541.684 | 5883.539 | 0.16633827 | 0.706755915 | YER006W | NUG1 | n |
| W303.asm02.HP0_G0014530 | 754.9948 | 781.1866 | 773.9395 | 1488.082 | 1442.354 | 1341.568 | 1346.81 | 1515.132 | 1589.596 | 2.24E-16 | -0.887078361 | YER007W | PAC2 | n |
| W303.asm02.HP0_G0014540 | 2104.032 | 2196.67 | 2233.43 | 2853.199 | 3142.458 | 2337.488 | 2663.993 | 2780.068 | 2892.132 | 0.012240738 | -0.350709195 | YER007C-/ | TMA20 | n |
| W303.asm02.HP0_G0014550 | 4661.272 | 5009.832 | 4614.533 | 6305.244 | 6248.466 | 8276.546 | 3650.337 | 3973.06 | 3225.839 | 0.0006633 | -0.544148393 | YER008C | SEC3 | n |
| W303.asm02.HP0_G0014560 | 6134.2 | 5648.58 | 6130.075 | 2936.565 | 3712.63 | 2895.842 | 3786.129 | 2573.898 | 2646.935 | 3.52E-06 | 0.908315613 | YER009W | NTF2 | n |
| W303.asm02.HP0_G0014570 | 2359.226 | 2015.283 | 2229.118 | 411.272 | 437.3926 | 647.2343 | 525.8855 | 1710.563 | 490.3946 | 3.71E-06 | 2.142240601 | YER010C | | n |
| W303.asm02.HP0_G0014590 | 8552.725 | 9151.678 | 8455.127 | 2134.168 | 2357.928 | 2623.882 | 1411.003 | 1740.63 | 1407.791 | 1.31E-43 | 1.878285784 | YER012W | PRE1 | n |
| W303.asm02.HP0_G0014600 | 1513.166 | 1748.211 | 1409.906 | 2081.37 | 2024.677 | 1849.783 | 2756.578 | 1950.021 | 2701.954 | 0.057239588 | -0.350564768 | YER013W | PRP22 | n |
| W303.asm02.HP0_G0014610 | 4537.381 | 4553.583 | 4376.315 | 3311.712 | 3420.167 | 4686.371 | 5989.663 | 6167.907 | 6167.011 | 0.15197382 | 0.238096039 | YER014W | HEM14 | n |
| W303.asm02.HP0_G0014620 | 1792.716 | 1771.58 | 1648.125 | 1642.309 | 1753.042 | 1227.618 | 2110.949 | 1376.612 | 1539.361 | 0.460344499 | 0.173239766 | YER015W | FAA2 | n |
| W303.asm02.HP0_G0014630 | 1988.612 | 1811.641 | 1969.342 | 1365.117 | 1365.116 | 1476.028 | 723.4012 | 904.1395 | 800.1804 | 0.000150973 | 0.455961065 | YER016W | BIM1 | n |
| W303.asm02.HP0_G0014640 | 8680.852 | 8312.626 | 8797.903 | 7769.705 | 6856.824 | 7616.397 | 3242.961 | 2975.5 | 2656.503 | 0.068449719 | 0.213465251 | YER017C | AFG3 | n |
| W303.asm02.HP0_G0014650 | 1382.922 | 1133.945 | 1237.441 | 603.7083 | 656.0889 | 579.6241 | 940.6685 | 434.889 | 549.0027 | 0.000312028 | 1.02945728 | YER018C | SPC25 | n |
| W303.asm02.HP0_G0014670 | 2457.704 | 2099.856 | 2073.899 | 1815.988 | 2282.426 | 1856.62 | 1256.694 | 984.6745 | 1099.201 | 0.372882775 | 0.155567634 | YER019C-/ | SBH2 | n |
| W303.asm02.HP0_G0014680 | 6683.769 | 7453.544 | 6113.906 | 4731.712 | 4848.636 | 5876.765 | 12858.27 | 5920.933 | 8325.943 | 0.188511718 | 0.3897311 | YER020W | GPA2 | n |
| W303.asm02.HP0_G0014690 | 12878.33 | 14423.02 | 13470.64 | 7684.255 | 7872.199 | 11141.24 | 6532.832 | 11547.64 | 8328.335 | 0.016365277 | 0.610855971 | YER021W | RPN3 | n |
| W303.asm02.HP0_G0014700 | 2898.206 | 3085.798 | 2928.682 | 3590.293 | 3279.577 | 3362.276 | 2583.752 | 3800.178 | 2928.014 | 0.254786935 | -0.19925246 | YER022W | SRB4 | n |
| W303.asm02.HP0_G0014710 | 5688.404 | 4888.536 | 5580.341 | 4976.252 | 5962.946 | 6348.517 | 4778.645 | 5574.095 | 4499.669 | 0.56013125 | -0.097468821 | YER023W | PRO3 | n |
| W303.asm02.HP0_G0014720 | 1050.428 | 1056.049 | 1048.807 | 1505.45 | 1345.156 | 1268.64 | 3561.455 | 2664.098 | 2704.347 | 0.016777719 | -0.384798044 | YER024W | YAT2 | n |
| W303.asm02.HP0_G0014730 | 13746.62 | 14132.58 | 14965.7 | 18644.1 | 19839.75 | 16909.37 | 14087.81 | 12596.75 | 13098.32 | 0.000361996 | -0.370560063 | YER025W | GCD11 | n |
| W303.asm02.HP0_G0014740 | 8856.629 | 9304.132 | 8982.225 | 6228.13 | 5908.272 | 7453.829 | 4982.333 | 8105.042 | 7151.388 | 0.023949312 | 0.470423041 | YER026C | CHO1 | n |
| W303.asm02.HP0_G0014750 | 6191.381 | 5967.954 | 6197.983 | 3868.179 | 3563.361 | 4335.406 | 5485.998 | 4948.07 | 4642.003 | 5.68E-08 | 0.64150798 | YER027C | GAL83 | n |
| W303.asm02.HP0_G0014770 | 2173.919 | 2093.179 | 2050.185 | 2750.381 | 2664.277 | 2463.592 | 4166.347 | 4192.115 | 4043.961 | 0.000128905 | -0.31859356 | YER029C | SMB1 | n |
| W303.asm02.HP0_G0014780 | 4101.115 | 4221.968 | 4135.941 | 2257.827 | 2181.756 | 1793.568 | 2454.133 | 2423.566 | 2730.661 | 4.66E-16 | 0.999142644 | YER030W | CHZ1 | n |
| W303.asm02.HP0_G0014800 | 1079.018 | 1103.899 | 1167.377 | 4437.152 | 3880.992 | 4561.786 | 3450.352 | 3374.953 | 3671.979 | 1.31E-77 | -1.94297158 | YER032W | FIR1 | n |
| W303.asm02.HP0_G0014810 | 3273.056 | 3655.553 | 3333.976 | 2388.434 | 2022.073 | 2283.552 | 2652.883 | 2319.408 | 2316.815 | 2.20E-07 | 0.616062021 | YER033C | ZRG8 | n |
| W303.asm02.HP0_G0014820 | 902.1817 | 867.9851 | 904.3666 | 1285.225 | 1300.028 | 1348.405 | 785.1249 | 1175.811 | 1163.79 | 0.002004349 | -0.556548656 | YER034W | | n |
| W303.asm02.HP0_G0014840 | 6290.917 | 6251.718 | 6871.677 | 11667.76 | 12075.16 | 9271.707 | 8515.395 | 7143.991 | 8281.688 | 1.40E-07 | -0.765960846 | YER036C | ARB1 | n |
| W303.asm02.HP0_G0014860 | 702.0498 | 597.5743 | 645.6682 | 793.3658 | 759.3622 | 945.0228 | 955.4822 | 1066.283 | 1016.672 | 0.011868519 | -0.360693535 | YER038C | KRE29 | n |
| W303.asm02.HP0_G0014870 | 212.8386 | 205.8683 | 248.9972 | 6036.389 | 6415.092 | 6486.017 | 4214.491 | 5958.516 | 4863.279 | 6.69E-182 | -4.825702044 | YER039C | HVG1 | n |
| W303.asm02.HP0_G0014890 | 1431.631 | 1482.251 | 1316.128 | 4874.128 | 3848.882 | 4241.208 | 7156.241 | 5528.996 | 6951.642 | 1.30E-25 | -1.615969467 | YER040W | GLN3 | n |
| W303.asm02.HP0_G0014900 | 1277.032 | 1208.502 | 1261.155 | 2223.786 | 2107.122 | 2687.693 | 2002.315 | 2156.19 | 2187.638 | 1.07E-12 | -0.905656665 | YER041W | YEN1 | n |

| | | | | | | | | | | | | | | |
|---|---|---|---|---|---|---|---|---|---|---|---|---|---|---|
| W303.asm02.HP0_G0014910 | 6637.177 | 4767.241 | 5650.405 | 1584.648 | 1585.548 | 1157.729 | 739.4494 | 1035.143 | 867.1611 | 4.26E-21 | 1.978503817 | YER042W | MXR1 | n |
| W303.asm02.HP0_G0014920 | 38343.78 | 41071.27 | 44058.5 | 23433.47 | 26540.36 | 21618.54 | 21824.25 | 21406.2 | 20411.18 | 7.70E-12 | 0.786352537 | YER043C | SAH1 | n |
| W303.asm02.HP0_G0014930 | 8286.941 | 7983.237 | 7957.133 | 1372.064 | 1657.579 | 1504.896 | 2525.732 | 2191.626 | 2042.912 | 3.84E-85 | 2.418087331 | YER044C | ERG28 | n |
| W303.asm02.HP0_G0014960 | 461.6798 | 353.8708 | 341.6975 | 641.223 | 682.1242 | 1105.312 | 1324.59 | 1357.283 | 1479.556 | 2.32E-05 | -1.069320674 | YER046W | SPO73 | n |
| W303.asm02.HP0_G0014970 | 1536.462 | 1990.802 | 1498.295 | 4703.923 | 4141.344 | 5066.963 | 6937.739 | 5925.228 | 6466.032 | 3.12E-20 | -1.469203384 | YER047C | SAP1 | n |
| W303.asm02.HP0_G0014980 | 7352.993 | 7434.626 | 7076.48 | 4810.909 | 5030.015 | 6600.726 | 6676.03 | 8265.038 | 7038.956 | 0.014246969 | 0.411202338 | YER048C | CAJ1 | n |
| W303.asm02.HP0_G0014990 | 1799.069 | 1704.812 | 1568.359 | 2371.066 | 2527.157 | 2123.263 | 2015.895 | 2198.068 | 2035.736 | 5.41E-05 | -0.468943841 | YER048W- | ISD11 | n |
| W303.asm02.HP0_G0015000 | 2074.383 | 2171.075 | 2514.764 | 4913.727 | 4811.319 | 6985.116 | 21850.17 | 17762.8 | 21671.85 | 1.42E-11 | -1.305608894 | YER049W | TPA1 | n |
| W303.asm02.HP0_G0015010 | 4122.293 | 4201.938 | 3910.658 | 2070.949 | 1992.566 | 1542.879 | 1387.548 | 1218.763 | 1304.928 | 4.42E-15 | 1.125850561 | YER050C | RSM18 | n |
| W303.asm02.HP0_G0015020 | 1165.847 | 1217.405 | 1033.716 | 1433.199 | 1401.566 | 1886.247 | 1954.171 | 2729.599 | 2118.265 | 0.020115715 | -0.466483429 | YER051W | JHD1 | n |
| W303.asm02.HP0_G0015030 | 3465.775 | 2827.628 | 3199.237 | 5634.843 | 5615.809 | 5679.253 | 6962.428 | 7344.792 | 8029.314 | 1.66E-13 | -0.834602249 | YER052C | HOM3 | n |
| W303.asm02.HP0_G0015050 | 3699.792 | 4030.567 | 3296.249 | 591.2034 | 573.6439 | 2006.274 | 8988.199 | 7938.603 | 4798.69 | 0.0001909 | 1.797859975 | YER054C | GIP2 | n |
| W303.asm02.HP0_G0015060 | 4082.055 | 3549.836 | 4189.837 | 3482.612 | 3784.661 | 3621.321 | 1860.351 | 2166.928 | 1856.323 | 0.358005265 | 0.118776415 | YER055C | HIS1 | n |
| W303.asm02.HP0_G0015070 | 1940.961 | 2276.792 | 2097.613 | 20790.08 | 19671.39 | 21738.56 | 24904.26 | 37201.8 | 31312.29 | 1.28E-86 | -3.300032799 | YER056C | FCY2 | n |
| W303.asm02.HP0_G0015100 | 1451.75 | 1209.615 | 1244.986 | 1160.176 | 1204.565 | 1274.717 | 759.201 | 644.28 | 690.1407 | 0.460104115 | 0.102333439 | YER058W | PET117 | n |
| W303.asm02.HP0_G0015110 | 3216.934 | 3527.58 | 3145.342 | 2195.998 | 2401.32 | 3667.661 | 6290.875 | 4881.494 | 5143.163 | 0.283953662 | 0.258922732 | YER059W | PCL6 | n |
| W303.asm02.HP0_G0015130 | 1299.269 | 1492.267 | 1324.752 | 2635.753 | 2466.408 | 2804.682 | 2380.064 | 2624.367 | 2032.147 | 8.34E-12 | -0.941964488 | YER061C | CEM1 | n |
| W303.asm02.HP0_G0015150 | 5961.6 | 5695.317 | 5664.418 | 2291.869 | 2202.584 | 2871.532 | 2002.315 | 1581.707 | 2242.658 | 1.29E-12 | 1.23354036 | YER063W | THO1 | n |
| W303.asm02.HP0_G0015170 | 650.1638 | 655.44 | 655.3694 | 19.45205 | 20.82822 | 15.19329 | 1708.511 | 265.2286 | 990.3578 | 9.14E-21 | 5.144751286 | YER065C | ICL1 | n |
| W303.asm02.HP0_G0015190 | 14773.75 | 15371.13 | 12563.04 | 4872.044 | 4347.891 | 10301.05 | 45700.19 | 33562.69 | 39672.92 | 0.000712012 | 1.129457348 | YER067W | RGI1 | n |
| W303.asm02.HP0_G0015200 | 3017.861 | 3526.468 | 3097.914 | 4435.068 | 4116.177 | 4580.018 | 4162.643 | 3590.787 | 3450.703 | 0.000404419 | -0.445706916 | YER068W | MOT2 | n |
| W303.asm02.HP0_G0015210 | 1436.926 | 1353.166 | 1386.192 | 4645.567 | 5230.487 | 5721.034 | 2557.828 | 4339.226 | 2780.896 | 3.88E-18 | -1.900946665 | YER069W | ARG56 | n |
| W303.asm02.HP0_G0015220 | 2640.893 | 2530.51 | 3346.911 | 16722.51 | 19704.36 | 22527.85 | 8589.464 | 10812.09 | 6446.895 | 3.28E-34 | -2.790949053 | YER070W | RNR1 | n |
| W303.asm02.HP0_G0015230 | 1542.815 | 1285.286 | 1375.413 | 854.5009 | 1096.085 | 1271.679 | 1498.65 | 1621.438 | 1410.183 | 0.039190471 | 0.383916564 | YER071C | TDA2 | n |
| W303.asm02.HP0_G0015240 | 4787.281 | 4384.437 | 4569.261 | 4257.22 | 5516.007 | 5735.468 | 2162.797 | 2303.301 | 2247.442 | 0.249535068 | -0.174444418 | YER072W | VTC1 | n |
| W303.asm02.HP0_G0015250 | 643.8104 | 783.4122 | 663.9926 | 14737.01 | 15974.38 | 9966.8 | 10967.06 | 3727.16 | 3442.331 | 1.27E-23 | -4.281867386 | YER073W | ALD5 | n |
| W303.asm02.HP0_G0015270 | 509.3303 | 460.6998 | 450.5664 | 217.4462 | 275.9739 | 179.2808 | 164.1849 | 185.7674 | 179.4126 | 3.54E-08 | 1.080705171 | YER074W- | YOS1 | n |
| W303.asm02.HP0_G0015280 | 1097.019 | 1218.518 | 1020.781 | 3337.416 | 2677.294 | 2903.438 | 5444.026 | 3936.551 | 5749.577 | 2.28E-14 | -1.418681234 | YER075C | PTP3 | n |
| W303.asm02.HP0_G0015290 | 802.6452 | 731.1105 | 718.9661 | 901.7415 | 933.7985 | 1079.483 | 1512.23 | 1354.062 | 1330.046 | 0.004293392 | -0.371687065 | YER076C | | n |
| W303.asm02.HP0_G0015300 | 2747.842 | 3007.902 | 2817.657 | 1313.014 | 1133.402 | 723.2007 | 628.3468 | 426.2986 | 572.9244 | 1.03E-08 | 1.435465027 | YER077C | MRX1 | n |
| W303.asm02.HP0_G0015310 | 3461.54 | 3554.288 | 3372.781 | 2443.317 | 2390.906 | 1841.427 | 1071.523 | 1676.202 | 1466.399 | 0.001788697 | 0.638030908 | YER078C | ICP55 | n |
| W303.asm02.HP0_G0015330 | 4096.879 | 4763.903 | 3683.219 | 400.1565 | 361.0225 | 624.4443 | 2915.825 | 2370.95 | 2477.091 | 2.40E-42 | 3.177977116 | YER079W | | n |
| W303.asm02.HP0_G0015340 | 9246.304 | 9183.95 | 8985.459 | 8490.821 | 7877.407 | 7247.96 | 1494.947 | 3367.437 | 2637.366 | 0.471930873 | 0.215221318 | YER080W | AIM9 | n |
| W303.asm02.HP0_G0015350 | 8486.014 | 6788.088 | 7448.359 | 6633.15 | 6322.233 | 8132.21 | 3267.65 | 7300.766 | 4506.846 | 0.741410092 | 0.107723205 | YER081W | SER3 | n |
| W303.asm02.HP0_G0015360 | 4136.058 | 4351.053 | 4531.534 | 4163.434 | 3770.776 | 1024.788 | 4563.847 | 2132.567 | 4220.982 | 0.294575044 | 0.53917356 | YER082C | UTP7 | n |
| W303.asm02.HP0_G0015370 | 3893.571 | 3487.52 | 3604.532 | 5647.347 | 5328.553 | 7028.417 | 6992.056 | 6185.088 | 6556.934 | 4.73E-07 | -0.712748072 | YER083C | GET2 | n |
| W303.asm02.HP0_G0015400 | 8464.836 | 7816.317 | 9080.315 | 8203.208 | 8296.574 | 8910.866 | 4792.225 | 6207.637 | 5718.479 | 0.985973269 | -0.002782986 | YER086W | ILV1 | n |
| W303.asm02.HP0_G0015410 | 6107.728 | 5794.357 | 5429.433 | 3459.686 | 3046.127 | 3664.622 | 1862.82 | 2074.581 | 2102.716 | 3.96E-11 | 0.768837777 | YER087W | AIM10 | n |
| W303.asm02.HP0_G0015430 | 7871.853 | 8647.579 | 8056.3 | 3417.309 | 2581.831 | 1674.301 | 3120.748 | 1203.73 | 2387.384 | 5.00E-06 | 1.679244511 | YER088C | DOT6 | n |
| W303.asm02.HP0_G0015450 | 6303.624 | 5781.003 | 6567.706 | 6646.349 | 6374.303 | 7797.957 | 5772.396 | 5871.538 | 5408.693 | 0.211214018 | -0.158539732 | YER090W | TRP2 | n |
| W303.asm02.HP0_G0015460 | 31471.53 | 21564.98 | 28028.9 | 6104.471 | 5311.196 | 7498.649 | 1404.83 | 8463.691 | 4891.985 | 4.17E-05 | 2.099603273 | YER091C | MET6 | n |
| W303.asm02.HP0_G0015480 | 1906.018 | 1838.348 | 1906.823 | 476.5753 | 491.1989 | 502.1383 | 822.1591 | 587.3686 | 623.1599 | 2.19E-38 | 1.942966871 | YER092W | IES5 | n |
| W303.asm02.HP0_G0015490 | 2634.54 | 2656.257 | 2630.101 | 3515.958 | 3572.908 | 4156.125 | 4829.259 | 3968.765 | 4344.178 | 3.93E-05 | -0.50556421 | YER093C | TSC11 | n |
| W303.asm02.HP0_G0015500 | 1718.593 | 1618.013 | 1527.399 | 602.3189 | 612.6968 | 448.9618 | 375.2798 | 242.6788 | 339.6879 | 5.59E-14 | 1.54768735 | YER093C-/ | AIM11 | n |
| W303.asm02.HP0_G0015510 | 6761.068 | 6918.286 | 6832.872 | 3160.264 | 3516.498 | 4097.631 | 2536.842 | 4023.528 | 2942.367 | 3.23E-06 | 0.928903928 | YER094C | PUP3 | n |
| W303.asm02.HP0_G0015520 | 2363.462 | 2257.874 | 2375.714 | 3836.223 | 4971.87 | 4497.215 | 2712.137 | 3053.887 | 2186.442 | 3.01E-08 | -0.927042161 | YER095W | RAD51 | n |
| W303.asm02.HP0_G0015540 | 2636.658 | 2927.78 | 2524.466 | 3569.452 | 3357.683 | 5075.319 | 3359.001 | 3306.23 | 2876.583 | 0.003368636 | -0.569388938 | YER098W | UBP9 | n |
| W303.asm02.HP0_G0015550 | 3510.249 | 3190.402 | 3151.809 | 4091.878 | 4091.01 | 4511.648 | 5969.912 | 4722.572 | 4689.847 | 0.009088374 | -0.365638083 | YER099C | PRS2 | n |
| W303.asm02.HP0_G0015560 | 6206.205 | 6021.368 | 6082.647 | 3256.829 | 3454.013 | 3000.675 | 3193.582 | 2724.23 | 2658.895 | 2.91E-16 | 0.915004857 | YER100W | UBC6 | n |
| W303.asm02.HP0_G0015570 | 2667.366 | 2697.431 | 2557.881 | 2676.741 | 2738.911 | 4034.579 | 3749.095 | 3669.174 | 3593.037 | 0.187351025 | -0.254395827 | YER101C | AST2 | n |
| W303.asm02.HP0_G0015630 | 2252.278 | 2158.835 | 2334.753 | 3726.457 | 3842.807 | 3589.415 | 2935.577 | 3287.975 | 2884.955 | 6.07E-15 | -0.72596429 | YER107C | GLE2 | n |
| W303.asm02.HP0_G0015650 | 4695.157 | 4111.801 | 5387.395 | 17260.22 | 17337.76 | 9476.816 | 11153.46 | 4681.768 | 9925.108 | 5.44E-06 | -1.634627492 | YER110C | KAP123 | n |
| W303.asm02.HP0_G0015660 | 1176.436 | 1397.679 | 1235.285 | 5474.363 | 5165.399 | 5938.298 | 1899.854 | 1975.792 | 1808.479 | 3.60E-80 | -2.121907484 | YER111C | SWI4 | n |
| W303.asm02.HP0_G0015670 | 4097.938 | 4644.833 | 4111.149 | 3957.103 | 4228.997 | 4096.871 | 3396.035 | 3359.92 | 3430.37 | 0.497236523 | 0.065598542 | YER112W | LSM4 | n |

| | | | | | | | | | | | | | | |
|---|---|---|---|---|---|---|---|---|---|---|---|---|---|---|
| W303.asm02.HP0_G0015680 | 953.0088 | 783.4122 | 929.1585 | 5889.109 | 5918.686 | 5256.119 | 8604.277 | 6346.158 | 7859.47 | 1.23E-65 | -2.678249381 | YER113C | TMN3 | n |
| W303.asm02.HP0_G0015690 | 2776.432 | 2777.552 | 2790.709 | 6049.588 | 5713.007 | 7009.425 | 4593.474 | 4941.627 | 4323.845 | 1.48E-24 | -1.169714688 | YER114C | BOI2 | n |
| W303.asm02.HP0_G0015700 | 1217.734 | 1179.569 | 1201.87 | 172.2896 | 122.3658 | 252.9683 | 638.2226 | 355.4278 | 468.8651 | 4.01E-19 | 2.715138881 | YER115C | SPR6 | n |
| W303.asm02.HP0_G0015710 | 1656.118 | 1658.074 | 1654.592 | 555.7729 | 486.8597 | 725.4797 | 1322.121 | 917.0251 | 1053.75 | 2.22E-13 | 1.490255922 | YER116C | SLX8 | n |
| W303.asm02.HP0_G0015730 | 1136.198 | 1146.185 | 1257.921 | 2593.375 | 2890.783 | 2941.421 | 1614.691 | 1903.847 | 1662.557 | 6.27E-27 | -1.250736778 | YER118C | SHO1 | n |
| W303.asm02.HP0_G0015740 | 2473.587 | 1984.125 | 2488.895 | 2261.301 | 2409.131 | 3790.726 | 3673.792 | 3761.521 | 3353.82 | 0.242814543 | -0.284523199 | YER119C | AVT6 | n |
| W303.asm02.HP0_G0015770 | 7292.635 | 7955.417 | 8040.132 | 5522.993 | 5731.232 | 5858.534 | 3231.851 | 3458.71 | 2712.719 | 0.000286727 | 0.44453521 | YER122C | GLO3 | n |
| W303.asm02.HP0_G0015780 | 2008.731 | 2389.185 | 2195.703 | 4779.647 | 4345.287 | 5098.869 | 5197.132 | 4116.949 | 4679.082 | 9.63E-16 | -1.109316316 | YER123W | YCK3 | n |
| W303.asm02.HP0_G0015790 | 5765.703 | 5585.15 | 5871.377 | 3390.909 | 3707.423 | 4703.084 | 4416.945 | 6035.829 | 5592.89 | 0.002810203 | 0.545331627 | YER124C | DSE1 | n |
| W303.asm02.HP0_G0015800 | 14582.09 | 13926.71 | 15169.43 | 9678.091 | 9521.968 | 12739.58 | 14231.01 | 11085.91 | 11857.98 | 0.008170354 | 0.451556297 | YER125W | RSP5 | n |
| W303.asm02.HP0_G0015810 | 3190.462 | 3238.252 | 3524.766 | 4549.001 | 4947.57 | 1667.464 | 5566.239 | 1953.242 | 3992.529 | 0.744530791 | -0.165564583 | YER126C | NSA2 | n |
| W303.asm02.HP0_G0015820 | 884.1804 | 985.942 | 1122.104 | 2408.581 | 1965.663 | 407.1802 | 4779.88 | 1344.398 | 3520.076 | 0.294216395 | -0.676232537 | YER127W | LCP5 | n |
| W303.asm02.HP0_G0015830 | 831.2355 | 839.0522 | 837.5362 | 684.2954 | 720.3093 | 475.5501 | 972.7648 | 637.8372 | 814.5334 | 0.075707573 | 0.415810528 | YER128W | VFA1 | n |
| W303.asm02.HP0_G0015840 | 4002.637 | 3595.461 | 4123.006 | 2401.634 | 2201.716 | 2495.498 | 2191.19 | 1776.065 | 2243.854 | 8.52E-08 | 0.723344808 | YER129W | SAK1 | n |
| W303.asm02.HP0_G0015850 | 1918.724 | 1534.553 | 1792.565 | 4949.852 | 4933.685 | 5215.098 | 7874.704 | 8436.846 | 8475.454 | 1.04E-42 | -1.524994966 | YER130C | COM2 | n |
| W303.asm02.HP0_G0015860 | 13137.76 | 13764.24 | 13636.64 | 16386.27 | 19732.14 | 17959.99 | 16664.15 | 14246.1 | 13861.42 | 0.000761366 | -0.415715678 | YER131W | RPS26B | y |
| W303.asm02.HP0_G0015880 | 1066.311 | 1130.606 | 935.626 | 664.8433 | 750.6838 | 919.9539 | 638.2226 | 922.3941 | 697.3172 | 0.046202853 | 0.42369855 | YER139C | RTR1 | n |
| W303.asm02.HP0_G0015890 | 860.8846 | 777.8482 | 838.6141 | 1273.415 | 1308.707 | 1362.079 | 2813.364 | 2365.581 | 2563.209 | 1.38E-09 | -0.670675645 | YER140W | EMP65 | n |
| W303.asm02.HP0_G0015900 | 16426.7 | 16738.76 | 15753.66 | 2971.996 | 3192.793 | 3390.383 | 4683.591 | 4300.569 | 4655.16 | 2.69E-157 | 2.356164845 | YER141W | COX15 | n |
| W303.asm02.HP0_G0015910 | 2446.056 | 2858.787 | 2320.741 | 523.1213 | 492.0667 | 929.0698 | 325.9009 | 802.1285 | 468.8651 | 8.44E-08 | 1.971509373 | YER142C | MAG1 | n |
| W303.asm02.HP0_G0015920 | 7693.958 | 8115.66 | 7482.852 | 2928.923 | 2905.537 | 4270.834 | 1639.38 | 4126.613 | 2728.268 | 0.000245205 | 1.204738473 | YER143W | DDI1 | n |
| W303.asm02.HP0_G0015930 | 3012.567 | 3069.106 | 3263.912 | 10626.38 | 12366.76 | 8911.626 | 16449.35 | 8328.392 | 10875.99 | 7.27E-12 | -1.77139713 | YER144C | UBP5 | n |
| W303.asm02.HP0_G0015940 | 932.8898 | 762.2689 | 849.3932 | 1317.877 | 1530.006 | 1526.166 | 1046.833 | 1045.881 | 980.7891 | 7.76E-10 | -0.780926412 | YER146W | LSM5 | n |
| W303.asm02.HP0_G0015950 | 954.0677 | 983.7164 | 1029.404 | 1463.767 | 1405.905 | 1911.316 | 4915.672 | 3338.444 | 4652.768 | 0.000742201 | -0.688303478 | YER147C | SCC4 | n |
| W303.asm02.HP0_G0015960 | 4307.6 | 4292.075 | 4341.822 | 3792.455 | 4189.076 | 3473.946 | 4453.979 | 3935.477 | 4412.355 | 0.12754056 | 0.176139164 | YER148W | SPT15 | n |
| W303.asm02.HP0_G0015970 | 1373.392 | 1403.243 | 1555.424 | 1101.82 | 1176.794 | 1137.218 | 988.8129 | 1077.021 | 936.534 | 0.001578821 | 0.343132756 | YER149C | PEA2 | n |
| W303.asm02.HP0_G0015990 | 4813.754 | 4796.174 | 5189.059 | 8385.919 | 7930.345 | 9974.396 | 12768.16 | 10809.94 | 11411.84 | 2.52E-10 | -0.82908281 | YER151C | UBP3 | n |
| W303.asm02.HP0_G0016000 | 4024.874 | 3674.47 | 3951.619 | 2737.876 | 2868.22 | 3224.776 | 561.6853 | 1915.659 | 1391.046 | 0.322068642 | 0.399828558 | YER152C | | n |
| W303.asm02.HP0_G0016010 | 343.0832 | 346.0812 | 365.4115 | 1456.125 | 1523.931 | 1258.764 | 783.8904 | 521.8668 | 496.375 | 2.12E-22 | -2.006917665 | YER153C | PET122 | n |
| W303.asm02.HP0_G0016020 | 4222.888 | 3958.234 | 4345.056 | 3347.837 | 3328.176 | 2292.668 | 728.3391 | 809.6451 | 825.2982 | 0.007440975 | 0.482024163 | YER154W | OXA1 | n |
| W303.asm02.HP0_G0016030 | 7144.39 | 7557.034 | 7041.987 | 8322.005 | 8009.319 | 8855.41 | 6611.838 | 10951.69 | 8754.141 | 0.304181713 | -0.212100458 | YER155C | BEM2 | n |
| W303.asm02.HP0_G0016040 | 5739.231 | 5263.55 | 5745.261 | 4235.684 | 4775.737 | 2106.55 | 3168.892 | 2899.26 | 3841.823 | 0.053772139 | 0.591151701 | YER156C | MYG1 | n |
| W303.asm02.HP0_G0016050 | 2204.627 | 2238.956 | 2233.43 | 3622.25 | 3561.626 | 4126.498 | 4040.43 | 4273.724 | 4502.061 | 1.92E-14 | -0.760449198 | YER157W | COG3 | n |
| W303.asm02.HP0_G0016070 | 4886.818 | 4606.998 | 4568.183 | 1994.53 | 2363.135 | 1871.814 | 4231.774 | 3122.61 | 3577.488 | 6.46E-14 | 1.174865728 | YER159C | BUR6 | n |
| W303.asm02.HP0_G0016080 | 4024.874 | 4306.541 | 4103.604 | 1910.469 | 1819.866 | 1795.847 | 1228.301 | 1266.01 | 1434.105 | 8.67E-35 | 1.169866116 | YER161C | SPT2 | n |
| W303.asm02.HP0_G0016090 | 3022.097 | 3266.072 | 2835.982 | 2381.487 | 2104.518 | 2033.622 | 3773.784 | 2818.725 | 3036.858 | 0.001832731 | 0.484738615 | YER162C | RAD4 | n |
| W303.asm02.HP0_G0016100 | 3702.969 | 3453.023 | 3256.367 | 1263.689 | 1294.821 | 1378.032 | 1881.337 | 1565.6 | 1690.067 | 8.49E-36 | 1.403360292 | YER163C | GCG1 | n |
| W303.asm02.HP0_G0016110 | 4231.359 | 3866.985 | 4159.655 | 9000.048 | 8792.98 | 9362.866 | 9681.972 | 8916.835 | 9230.183 | 1.27E-47 | -1.147508528 | YER164W | CHD1 | n |
| W303.asm02.HP0_G0016120 | 30418.98 | 28603.45 | 34172.99 | 26663.21 | 27979.24 | 24543.24 | 20718.16 | 21368.62 | 21631.19 | 0.033063381 | 0.235037913 | YER165W | PAB1 | n |
| W303.asm02.HP0_G0016130 | 5506.273 | 5302.499 | 5590.042 | 10009.47 | 9983.66 | 10500.08 | 8617.857 | 10590.89 | 9140.476 | 3.15E-17 | -0.894889111 | YER166W | DNF1 | n |
| W303.asm02.HP0_G0016140 | 2447.115 | 2980.082 | 2644.114 | 5532.719 | 4649.9 | 4714.479 | 6210.634 | 4610.897 | 4961.358 | 7.79E-08 | -0.884285422 | YER167W | BCK2 | n |
| W303.asm02.HP0_G0016150 | 1193.379 | 1191.81 | 1305.349 | 4182.191 | 3901.82 | 3596.252 | 4107.092 | 3153.75 | 4398.002 | 2.24E-26 | -1.662191826 | YER168C | CCA1 | n |
| W303.asm02.HP0_G0016160 | 2721.37 | 2701.882 | 2708.788 | 2987.974 | 2406.527 | 3042.457 | 9636.297 | 5807.11 | 7666.901 | 0.821020326 | -0.05322142 | YER169W | RPH1 | n |
| W303.asm02.HP0_G0016170 | 2366.639 | 2233.392 | 2252.832 | 1495.724 | 1562.984 | 1398.543 | 192.5778 | 486.4314 | 240.4129 | 0.057802738 | 0.62058313 | YER170W | ADK2 | n |
| W303.asm02.HP0_G0016180 | 3876.628 | 3597.687 | 4052.942 | 3704.226 | 3589.397 | 3746.666 | 4049.072 | 3569.311 | 3438.742 | 0.591177267 | 0.062256667 | YER171W | RAD3 | n |
| W303.asm02.HP0_G0016190 | 3556.841 | 3763.494 | 3509.675 | 6127.396 | 5681.765 | 4644.589 | 4844.072 | 4737.605 | 5425.438 | 1.39E-05 | -0.603425452 | YER172C | BRR2 | n |
| W303.asm02.HP0_G0016200 | 864.0613 | 938.0916 | 999.2227 | 847.5537 | 747.2124 | 882.7303 | 834.5038 | 908.4347 | 846.8277 | 0.169575129 | 0.17646274 | YER173W | RAD24 | n |
| W303.asm02.HP0_G0016210 | 1473.987 | 1372.084 | 1498.295 | 8249.059 | 8847.654 | 9312.729 | 6794.54 | 11233.02 | 8746.965 | 4.88E-42 | -2.603798265 | YER174C | GRX4 | n |
| W303.asm02.HP0_G0016220 | 1322.564 | 1349.828 | 1200.792 | 166.7319 | 148.4011 | 383.6306 | 660.4431 | 502.5384 | 714.0623 | 4.42E-12 | 2.470159569 | YER175C | TMT1 | n |
| W303.asm02.HP0_G0016250 | 43097.18 | 45539.17 | 45708.78 | 16109.77 | 17712.67 | 19748.24 | 4666.308 | 11621.74 | 8654.866 | 1.10E-05 | 1.32643122 | YER178W | PDA1 | n |
| W303.asm02.HP0_G0016260 | 896.8872 | 873.5491 | 820.2896 | 96.56554 | 98.93405 | 131.422 | 288.8667 | 198.653 | 94.49066 | 1.20E-16 | 2.986226558 | YER179W | DMC1 | n |
| W303.asm02.HP0_G0016270 | 785.7028 | 824.5858 | 759.9266 | 1748.601 | 1885.822 | 1847.504 | 1291.259 | 1533.386 | 1324.065 | 5.47E-28 | -1.209569237 | YER180C | ISC10 | n |
| W303.asm02.HP0_G0016280 | 398.1459 | 310.4716 | 307.2044 | 373.7573 | 442.5997 | 334.2524 | 319.7285 | 389.7894 | 288.2563 | 0.426371421 | -0.178626217 | YER180C-/ | SLO1 | n |
| W303.asm02.HP0_G0016290 | 9719.631 | 9037.06 | 9013.485 | 2054.97 | 2383.096 | 2648.191 | 2652.883 | 3023.821 | 2657.699 | 9.22E-53 | 1.970611815 | YER182W | FMP10 | n |

| | | | | | | | | | | | | | | |
|---|---|---|---|---|---|---|---|---|---|---|---|---|---|---|
| W303.asm02.HP0_G0016300 | 1142.552 | 1038.244 | 1102.702 | 690.5478 | 741.1375 | 840.9487 | 4774.942 | 1761.032 | 2505.797 | 0.136206905 | 0.53091151 | YER183C | FAU1 | n |
| W303.asm02.HP0_G0016320 | 132.3623 | 127.9722 | 131.505 | 9371.026 | 10823.73 | 8126.892 | 10730.04 | 15086.89 | 11759.9 | 5.34E-225 | -6.175345631 | YER185W | PUG1 | n |
| W303.asm02.HP0_G0016410 | 3406.477 | 3416.3 | 3542.013 | 3015.763 | 2797.056 | 3372.151 | 4398.428 | 4844.985 | 4338.198 | 0.129450799 | 0.174181731 | YFL049W | SWP82 | n |
| W303.asm02.HP0_G0016420 | 3472.129 | 3456.361 | 3460.092 | 3939.735 | 4018.111 | 4524.562 | 2331.92 | 2812.282 | 2786.876 | 0.028324805 | -0.264898902 | YFL048C | EMP47 | n |
| W303.asm02.HP0_G0016430 | 2773.256 | 2733.04 | 2848.917 | 3992.534 | 3770.776 | 4710.68 | 5092.201 | 5239.07 | 4979.299 | 3.50E-07 | -0.578263814 | YFL047W | RGD2 | n |
| W303.asm02.HP0_G0016440 | 1882.722 | 1639.156 | 1676.15 | 599.54 | 529.3839 | 563.6711 | 275.2875 | 343.616 | 245.1973 | 3.67E-26 | 1.618239991 | YFL046W | FMP32 | n |
| W303.asm02.HP0_G0016450 | 9940.941 | 8928.005 | 10301.59 | 7946.858 | 9193.924 | 8761.972 | 3512.076 | 4512.107 | 3854.98 | 0.243249608 | 0.171472745 | YFL045C | SEC53 | n |
| W303.asm02.HP0_G0016460 | 3972.988 | 3781.299 | 3724.18 | 500.1956 | 592.7364 | 599.3754 | 198.7502 | 400.5274 | 263.1386 | 1.57E-30 | 2.762138435 | YFL044C | OTU1 | n |
| W303.asm02.HP0_G0016470 | 5066.83 | 4894.1 | 4954.075 | 3208.199 | 3061.748 | 4125.739 | 4841.603 | 5375.442 | 5086.947 | 0.000300407 | 0.5207075 | YFL042C | LAM5 | n |
| W303.asm02.HP0_G0016500 | 1489.87 | 1465.559 | 1440.088 | 5216.623 | 4918.931 | 5836.503 | 4395.959 | 4604.454 | 4426.708 | 4.77E-83 | -1.861529556 | YFL041W | FET5 | n |
| W303.asm02.HP0_G0016510 | 432.0307 | 465.151 | 377.2685 | 53.49314 | 48.59918 | 39.50256 | 38.26867 | 96.64199 | 61.0003 | 7.82E-23 | 3.169241737 | YFL040W | | n |
| W303.asm02.HP0_G0016520 | 30813.95 | 32243.42 | 32777.09 | 26527.04 | 28789.81 | 32760.54 | 27627.51 | 31757.63 | 28462.02 | 0.341655678 | 0.121778661 | YFL039C | ACT1 | n |
| W303.asm02.HP0_G0016530 | 10835.71 | 10763.01 | 10926.78 | 4904.696 | 5373.681 | 6044.651 | 4231.774 | 5301.35 | 4285.57 | 3.51E-13 | 0.994698322 | YFL038C | YPT1 | n |
| W303.asm02.HP0_G0016540 | 8753.916 | 8478.434 | 8392.608 | 4486.477 | 3556.419 | 2174.16 | 6257.544 | 5056.524 | 6723.19 | 4.61E-07 | 1.326523373 | YFL036W | RPO41 | n |
| W303.asm02.HP0_G0016570 | 1282.326 | 1286.398 | 1173.844 | 1875.039 | 1851.108 | 2018.429 | 1776.407 | 1706.268 | 1467.595 | 9.99E-08 | -0.618257302 | YFL034W | MIL1 | n |
| W303.asm02.HP0_G0016580 | 3898.865 | 3818.022 | 3825.503 | 2612.827 | 2370.946 | 3274.154 | 4147.83 | 3502.735 | 3322.722 | 0.003385726 | 0.482978414 | YFL033C | RIM15 | n |
| W303.asm02.HP0_G0016590 | 7534.064 | 7894.213 | 7101.272 | 15674.88 | 16206.09 | 18631.53 | 30701.35 | 32481.37 | 27015.96 | 1.60E-21 | -1.164831461 | YFL031W | HAC1 | n |
| W303.asm02.HP0_G0016600 | 836.53 | 743.3513 | 708.187 | 193.8258 | 141.4583 | 243.0927 | 520.9476 | 1297.15 | 508.3358 | 4.67E-07 | 1.983380084 | YFL030W | AGX1 | n |
| W303.asm02.HP0_G0016610 | 2641.952 | 2660.708 | 2660.282 | 1352.612 | 1356.438 | 1350.684 | 492.5548 | 950.3129 | 821.7099 | 2.91E-05 | 0.971918673 | YFL029C | CAK1 | n |
| W303.asm02.HP0_G0016620 | 3024.215 | 2700.769 | 2931.916 | 2062.612 | 2186.095 | 2818.356 | 2008.488 | 2965.835 | 2301.266 | 0.162048706 | 0.292762042 | YFL028C | CAF16 | n |
| W303.asm02.HP0_G0016630 | 1449.632 | 1514.523 | 1331.219 | 902.4363 | 820.1112 | 1379.551 | 428.3622 | 642.1324 | 440.159 | 0.080079611 | 0.469378485 | YFL027C | GYP8 | n |
| W303.asm02.HP0_G0016640 | 2068.029 | 2292.371 | 2483.505 | 158.3953 | 182.2469 | 148.1346 | 183.9365 | 183.6198 | 116.0202 | 9.99E-82 | 3.808977282 | YFL026W | STE2 | n |
| W303.asm02.HP0_G0016650 | 1736.594 | 1439.965 | 1681.54 | 2002.867 | 1752.174 | 2507.653 | 1565.312 | 1670.833 | 1621.89 | 0.034287714 | -0.366490254 | YFL025C | BST1 | n |
| W303.asm02.HP0_G0016670 | 1343.742 | 1565.712 | 1358.167 | 4234.295 | 3725.648 | 4207.023 | 3779.957 | 2900.334 | 4187.491 | 2.82E-19 | -1.511614333 | YFL024C | EPL1 | y |
| W303.asm02.HP0_G0016680 | 1289.739 | 1163.99 | 1275.168 | 3072.035 | 2446.448 | 1135.699 | 6160.021 | 2216.323 | 4814.239 | 0.061480266 | -0.835526614 | YFL023W | BUD27 | n |
| W303.asm02.HP0_G0016690 | 11709.3 | 11254.87 | 13816.65 | 13890.15 | 14437.43 | 14755.73 | 9698.021 | 9756.546 | 9897.598 | 0.039399869 | -0.228153068 | YFL022C | FRS2 | n |
| W303.asm02.HP0_G0016700 | 4411.372 | 4026.115 | 4299.784 | 3565.978 | 3629.317 | 6231.529 | 3198.52 | 2169.076 | 1641.028 | 0.826400944 | -0.076069112 | YFL021W | GAT1 | n |
| W303.asm02.HP0_G0016720 | 13521.08 | 13912.24 | 13963.25 | 23184.76 | 24855.01 | 27095.72 | 6588.383 | 20801.65 | 14475.01 | 0.018231502 | -0.859981683 | YFL018C | LPD1 | n |
| W303.asm02.HP0_G0016740 | 1738.712 | 1407.694 | 1402.361 | 1813.209 | 1943.967 | 2367.875 | 5331.689 | 3969.838 | 4614.493 | 0.022396513 | -0.429149627 | YFL017C | GNA1 | n |
| W303.asm02.HP0_G0016750 | 9508.91 | 10291.19 | 8865.811 | 2980.332 | 2553.193 | 1970.57 | 806.1109 | 653.9442 | 797.7882 | 5.96E-26 | 1.93349661 | YFL016C | MDJ1 | n |
| W303.asm02.HP0_G0016770 | 7552.066 | 7663.863 | 6439.435 | 450.1761 | 399.2076 | 1121.265 | 1038.192 | 3617.632 | 3508.115 | 7.08E-12 | 3.457889574 | YFL014W | HSP12 | n |
| W303.asm02.HP0_G0016780 | 6549.289 | 6799.216 | 6420.033 | 6594.246 | 6207.678 | 8001.547 | 10031.33 | 11020.41 | 10263.6 | 0.601612994 | -0.073678685 | YFL013C | IES1 | n |
| W303.asm02.HP0_G0016810 | 9265.364 | 8406.102 | 8655.618 | 4426.037 | 5025.676 | 6056.046 | 6704.423 | 9935.871 | 6340.443 | 0.000436302 | 0.763581213 | YFL010C | WWM1 | n |
| W303.asm02.HP0_G0016820 | 2160.153 | 2063.134 | 2133.184 | 4297.514 | 3994.679 | 3811.997 | 7384.618 | 4715.055 | 5427.831 | 4.03E-07 | -0.929223667 | YFL009W | CDC4 | n |
| W303.asm02.HP0_G0016830 | 2010.849 | 2302.386 | 2123.483 | 4163.434 | 4682.01 | 4516.206 | 876.4759 | 1761.032 | 1423.34 | 2.09E-05 | -1.053695184 | YFL008W | SMC1 | n |
| W303.asm02.HP0_G0016840 | 8131.283 | 8597.503 | 8441.114 | 8397.034 | 8369.473 | 10241.8 | 6376.054 | 11328.59 | 8959.868 | 0.681028982 | -0.101711153 | YFL007W | BLM10 | n |
| W303.asm02.HP0_G0016850 | 7571.126 | 7850.814 | 7746.94 | 7309.108 | 8407.658 | 8091.188 | 7494.486 | 6135.693 | 6450.483 | 0.763355181 | -0.039184302 | YFL005W | SEC4 | n |
| W303.asm02.HP0_G0016860 | 6844.721 | 6987.28 | 7306.075 | 14860.67 | 15267.09 | 20561.08 | 4010.803 | 8032.023 | 5303.438 | 2.39E-06 | -1.26182541 | YFL004W | VTC2 | n |
| W303.asm02.HP0_G0016870 | 546.3917 | 515.227 | 438.7094 | 2634.364 | 2568.814 | 3328.85 | 3309.622 | 3545.687 | 3462.664 | 5.03E-61 | -2.507632816 | YFL003C | MSH4 | n |
| W303.asm02.HP0_G0016880 | 1901.782 | 1967.433 | 2197.859 | 1819.462 | 1660.183 | 639.6376 | 2921.998 | 1378.759 | 2553.64 | 0.198004462 | 0.55861287 | YFL002C | SPB4 | n |
| W303.asm02.HP0_G0016890 | 1925.078 | 1962.982 | 2055.575 | 3446.487 | 3095.594 | 1988.042 | 3215.802 | 2392.426 | 2984.23 | 0.027874618 | -0.521242554 | YFL001W | DEG1 | n |
| W303.asm02.HP0_G0016900 | 2289.339 | 2454.84 | 2384.337 | 2902.524 | 2133.157 | 981.4867 | 2139.342 | 1657.947 | 2362.267 | 0.53263629 | 0.244450742 | YFR001W | LOC1 | n |
| W303.asm02.HP0_G0016910 | 6213.618 | 6731.336 | 6600.044 | 9457.171 | 9188.717 | 10010.1 | 6992.056 | 7401.703 | 7120.29 | 1.07E-12 | -0.552116026 | YFR002W | NIC96 | n |
| W303.asm02.HP0_G0016920 | 3300.587 | 3256.057 | 2939.461 | 663.4539 | 769.7763 | 922.9925 | 1327.059 | 1306.815 | 1133.888 | 4.10E-38 | 2.011141977 | YFR003C | YPI1 | n |
| W303.asm02.HP0_G0016930 | 7715.136 | 8171.3 | 7564.773 | 6330.948 | 6663.295 | 7739.463 | 4566.316 | 6479.309 | 5356.066 | 0.311698988 | 0.177684353 | YFR004W | RPN11 | n |
| W303.asm02.HP0_G0016940 | 1615.879 | 1611.336 | 1628.722 | 1053.884 | 986.7369 | 997.4396 | 1043.13 | 1348.693 | 1081.26 | 1.95E-07 | 0.676377916 | YFR005C | SAD1 | n |
| W303.asm02.HP0_G0016950 | 9062.055 | 8470.644 | 8601.723 | 4479.53 | 4478.067 | 3951.775 | 3455.29 | 7586.396 | 3826.274 | 0.000619362 | 1.017537777 | YFR006W | | n |
| W303.asm02.HP0_G0016960 | 3331.295 | 3144.777 | 2987.967 | 1257.436 | 1356.438 | 1831.551 | 703.6497 | 1647.209 | 1211.633 | 0.000412596 | 1.090141358 | YFR007W | YFH7 | n |
| W303.asm02.HP0_G0016970 | 1413.63 | 1537.891 | 1301.038 | 528.679 | 513.7628 | 723.2007 | 249.3636 | 285.6308 | 220.0795 | 1.91E-11 | 1.267863435 | YFR008W | FAR7 | n |
| W303.asm02.HP0_G0016980 | 7471.589 | 7764.015 | 7778.2 | 7243.805 | 7196.15 | 6501.969 | 5399.585 | 5199.339 | 5741.205 | 0.139985577 | 0.136106116 | YFR009W | GCN20 | n |
| W303.asm02.HP0_G0016990 | 10160.13 | 10608.34 | 9552.44 | 5359.735 | 5460.465 | 7977.998 | 2752.875 | 6537.294 | 4585.787 | 0.033298068 | 0.689712739 | YFR010W | UBP6 | n |
| W303.asm02.HP0_G0017000 | 2494.765 | 2717.461 | 2336.909 | 5152.015 | 5487.368 | 3902.397 | 1035.723 | 2263.57 | 1999.853 | 0.001468256 | -0.94581212 | YFR011C | MIC19 | n |
| W303.asm02.HP0_G0017020 | 3753.796 | 3904.82 | 3969.943 | 7071.516 | 6605.149 | 7311.772 | 2425.74 | 4287.683 | 3462.664 | 5.02E-05 | -0.851941543 | YFR013W | IOC3 | n |
| W303.asm02.HP0_G0017050 | 9414.668 | 9481.068 | 9439.259 | 7819.03 | 7859.182 | 13057.12 | 18645.48 | 19355.24 | 20673.12 | 0.937778362 | -0.020252313 | YFR016C | AIP5 | n |

| | | | | | | | | | | | | | | |
|---|---|---|---|---|---|---|---|---|---|---|---|---|---|---|
| W303.asm02.HP0_G0017070 | 5668.285 | 5264.663 | 5940.363 | 1665.929 | 1627.205 | 2260.002 | 3612.068 | 3446.898 | 3640.881 | 5.60E-25 | 1.603280092 | YFR018C | | n |
| W303.asm02.HP0_G0017080 | 4395.489 | 4161.877 | 4005.514 | 4159.96 | 3837.6 | 5430.842 | 10637.45 | 8377.787 | 9260.085 | 0.619153337 | -0.09617145 | YFR019W | FAB1 | n |
| W303.asm02.HP0_G0017100 | 1124.55 | 1038.244 | 1139.351 | 3788.287 | 3479.181 | 4289.826 | 4130.547 | 4120.17 | 3828.666 | 5.08E-55 | -1.807351365 | YFR021W | ATG18 | n |
| W303.asm02.HP0_G0017120 | 497.6824 | 624.2816 | 465.6572 | 277.8865 | 255.1457 | 408.6996 | 918.448 | 883.7373 | 575.3166 | 0.007231723 | 0.752964558 | YFR023W | PES4 | n |
| W303.asm02.HP0_G0017140 | 2591.125 | 2346.898 | 2529.855 | 902.4363 | 1025.79 | 1785.972 | 1270.273 | 2046.663 | 1867.088 | 0.000798827 | 1.007651339 | YFR025C | HIS2 | n |
| W303.asm02.HP0_G0017160 | 458.5031 | 510.7758 | 459.1897 | 162.5636 | 158.8152 | 178.5212 | 133.3231 | 111.6752 | 117.2163 | 2.64E-25 | 1.513922795 | YFR027W | ECO1 | n |
| W303.asm02.HP0_G0017170 | 1799.069 | 1843.912 | 2057.73 | 3025.489 | 3038.317 | 2762.9 | 1211.018 | 1453.925 | 1163.79 | 1.27E-06 | -0.630708174 | YFR028C | CDC14 | n |
| W303.asm02.HP0_G0017180 | 884.1804 | 990.3932 | 918.3794 | 1828.493 | 1734.817 | 1418.294 | 1186.329 | 1059.841 | 1175.751 | 2.14E-09 | -0.834979736 | YFR029W | PTR3 | n |
| W303.asm02.HP0_G0017190 | 3632.023 | 2668.498 | 3151.809 | 3436.761 | 3286.52 | 4618.761 | 4202.146 | 5181.085 | 4578.611 | 0.208082547 | -0.262930252 | YFR030W | MET10 | n |
| W303.asm02.HP0_G0017200 | 2562.535 | 2853.223 | 2679.685 | 2903.219 | 2568.814 | 3120.702 | 3681.199 | 2381.688 | 3107.427 | 0.679255423 | -0.086118031 | YFR031C | SMC2 | n |
| W303.asm02.HP0_G0017220 | 29.64916 | 45.62486 | 26.94775 | 614.8238 | 629.1858 | 235.496 | 244.4257 | 337.1732 | 211.7069 | 4.34E-19 | -3.856618958 | YFR032C | RRT5 | n |
| W303.asm02.HP0_G0017250 | 5442.739 | 5185.654 | 4879.699 | 11758.77 | 11886.84 | 10727.98 | 2675.103 | 5721.206 | 5409.889 | 3.11E-05 | -1.148274974 | YFR033C | QCR6 | n |
| W303.asm02.HP0_G0017260 | 1185.967 | 1139.509 | 1315.05 | 11538.54 | 11855.6 | 11748.97 | 3425.663 | 6333.272 | 4175.53 | 1.60E-47 | -3.27100151 | YFR034C | PHO4 | n |
| W303.asm02.HP0_G0017280 | 229.781 | 227.0115 | 206.9587 | 487.6907 | 457.353 | 409.4592 | 432.0656 | 425.2248 | 447.3355 | 3.11E-14 | -1.029451577 | YFR036W | CDC26 | n |
| W303.asm02.HP0_G0017290 | 13451.19 | 12246.38 | 13383.33 | 5060.312 | 5047.372 | 5038.096 | 4245.353 | 4045.004 | 4235.335 | 7.58E-80 | 1.367570715 | YFR037C | RSC8 | n |
| W303.asm02.HP0_G0017300 | 1574.582 | 1610.224 | 1746.214 | 1351.223 | 1437.147 | 1335.49 | 674.0223 | 689.3796 | 635.1208 | 0.007781306 | 0.258234907 | YFR038W | IRC5 | n |
| W303.asm02.HP0_G0017320 | 6435.986 | 6947.219 | 6352.124 | 6925.625 | 6695.405 | 8075.995 | 8662.298 | 7933.234 | 7671.685 | 0.273085389 | -0.136774104 | YFR040W | SAP155 | n |
| W303.asm02.HP0_G0017330 | 2562.535 | 2796.47 | 2605.309 | 1489.471 | 1543.024 | 1571.746 | 1588.767 | 2423.566 | 2041.716 | 3.45E-06 | 0.790603318 | YFR041C | ERJ5 | n |
| W303.asm02.HP0_G0017340 | 892.6516 | 984.8292 | 818.1338 | 666.9275 | 611.829 | 584.9418 | 920.9169 | 799.9809 | 837.259 | 4.75E-05 | 0.53184335 | YFR042W | KEG1 | n |
| W303.asm02.HP0_G0017350 | 1751.418 | 1833.897 | 1772.084 | 663.4539 | 614.4325 | 792.3302 | 880.1793 | 707.6342 | 824.1021 | 2.72E-21 | 1.371291139 | YFR043C | IRC6 | n |
| W303.asm02.HP0_G0017360 | 9042.995 | 8267.001 | 9197.807 | 9537.063 | 10383.74 | 10598.84 | 8346.273 | 9576.148 | 6577.268 | 0.236802482 | -0.203295043 | YFR044C | DUG1 | n |
| W303.asm02.HP0_G0017370 | 2519.12 | 2664.046 | 2389.727 | 2765.665 | 2995.792 | 2569.945 | 1472.726 | 2195.921 | 1877.852 | 0.460662806 | -0.137656032 | YFR045W | MRX20 | n |
| W303.asm02.HP0_G0017380 | 1223.028 | 1290.85 | 1274.09 | 627.3287 | 552.8157 | 557.5938 | 304.9149 | 548.7118 | 306.1976 | 4.90E-06 | 1.123990106 | YFR046C | CNN1 | n |
| W303.asm02.HP0_G0017400 | 1070.547 | 1159.539 | 1119.949 | 2214.755 | 2121.007 | 2057.172 | 2097.37 | 2047.736 | 1875.46 | 3.91E-24 | -0.932574696 | YFR048W | RMD8 | n |
| W303.asm02.HP0_G0017410 | 2756.313 | 2825.403 | 2503.985 | 3404.804 | 4141.344 | 3452.676 | 1801.096 | 2669.467 | 2562.013 | 0.020252345 | -0.443808099 | YFR049W | KGD4 | n |
| W303.asm02.HP0_G0017420 | 8419.304 | 8664.271 | 7973.301 | 4483.698 | 5086.425 | 6397.136 | 4328.063 | 6909.903 | 5470.89 | 0.002829064 | 0.650103982 | YFR050C | PRE4 | n |
| W303.asm02.HP0_G0017430 | 7951.27 | 7537.004 | 8057.378 | 10463.81 | 10329.06 | 12202.49 | 8864.751 | 9998.151 | 9854.539 | 1.14E-05 | -0.486808892 | YFR051C | RET2 | n |
| W303.asm02.HP0_G0017440 | 3949.692 | 4063.951 | 3909.58 | 3785.508 | 4317.517 | 5243.965 | 2967.673 | 6635.01 | 4985.279 | 0.609581431 | -0.162728842 | YFR052W | RPN12 | n |
| W303.asm02.HP0_G0017570 | 1369.156 | 1416.596 | 1412.062 | 347.3581 | 402.6789 | 414.7769 | 483.9135 | 743.0696 | 321.7467 | 8.04E-11 | 1.84977461 | YGL256W | ADH4 | n |
| W303.asm02.HP0_G0017600 | 38900.76 | 42641.44 | 41618.11 | 47979.88 | 49084.3 | 81549.24 | 146291.2 | 100756.8 | 118947 | 0.036648969 | -0.536304525 | YGL253W | HXK2 | n |
| W303.asm02.HP0_G0017610 | 6604.351 | 6922.737 | 6860.898 | 4965.136 | 5008.319 | 4861.094 | 3425.663 | 3064.625 | 2791.661 | 7.25E-06 | 0.458761821 | YGL252C | RTG2 | n |
| W303.asm02.HP0_G0017620 | 2169.683 | 2220.039 | 2148.275 | 723.1995 | 691.6705 | 1484.385 | 2710.903 | 1994.046 | 2392.169 | 0.000146574 | 1.173051731 | YGL251C | HFM1 | n |
| W303.asm02.HP0_G0017630 | 1211.38 | 1202.938 | 1032.638 | 625.9392 | 710.763 | 970.0917 | 799.9386 | 892.3277 | 737.984 | 0.004486147 | 0.57951697 | YGL250W | RMR1 | n |
| W303.asm02.HP0_G0017640 | 672.4007 | 580.8823 | 593.9285 | 85.45009 | 72.03093 | 131.422 | 156.7781 | 412.3392 | 251.1777 | 6.87E-13 | 2.675727676 | YGL249W | ZIP2 | n |
| W303.asm02.HP0_G0017650 | 6374.57 | 6031.383 | 5705.378 | 541.8786 | 520.7055 | 635.8393 | 567.8576 | 672.1988 | 621.9638 | 4.06E-163 | 3.414217893 | YGL248W | PDE1 | n |
| W303.asm02.HP0_G0017660 | 541.0972 | 477.3918 | 475.3584 | 33.34637 | 31.24233 | 11.39497 | 45.6755 | 17.1808 | 39.47078 | 9.15E-24 | 4.297710029 | YGL247W | BRR6 | n |
| W303.asm02.HP0_G0017670 | 1578.818 | 1688.12 | 1732.202 | 3012.984 | 2758.004 | 1738.113 | 2709.668 | 1595.667 | 2204.383 | 0.033380519 | -0.58697756 | YGL246C | RAI1 | n |
| W303.asm02.HP0_G0017680 | 20158.25 | 18413.52 | 20505.08 | 23082.64 | 23848.31 | 26088.4 | 27471.96 | 22497.18 | 23698.02 | 0.012629 | -0.305676238 | YGL245W | GUS1 | n |
| W303.asm02.HP0_G0017690 | 7884.56 | 8043.328 | 7971.145 | 6545.615 | 6241.523 | 5752.94 | 4767.535 | 4878.273 | 4784.337 | 3.64E-06 | 0.366245579 | YGL244W | RTF1 | n |
| W303.asm02.HP0_G0017700 | 1019.719 | 994.8444 | 944.2493 | 1497.808 | 1514.385 | 1632.519 | 1669.008 | 2254.98 | 2222.325 | 1.62E-05 | -0.650564626 | YGL243W | TAD1 | n |
| W303.asm02.HP0_G0017710 | 1473.987 | 1283.06 | 1392.66 | 2609.354 | 2918.554 | 2898.88 | 2352.906 | 3042.075 | 2429.247 | 6.75E-13 | -1.021769501 | YGL242C | ANK1 | n |
| W303.asm02.HP0_G0017720 | 3081.395 | 3119.182 | 3137.796 | 4678.218 | 4370.455 | 4565.584 | 3905.873 | 4894.38 | 4473.355 | 1.04E-06 | -0.543956864 | YGL241W | KAP114 | n |
| W303.asm02.HP0_G0017730 | 555.9218 | 575.3183 | 551.89 | 1292.867 | 1262.711 | 1355.242 | 1176.453 | 1341.176 | 1137.476 | 3.55E-29 | -1.216564785 | YGL240W | DOC1 | n |
| W303.asm02.HP0_G0017740 | 5392.971 | 4937.5 | 5394.94 | 7777.347 | 7978.944 | 6966.884 | 6569.866 | 7303.987 | 6584.444 | 2.67E-07 | -0.530998183 | YGL238W | CSE1 | n |
| W303.asm02.HP0_G0017750 | 1915.548 | 1830.558 | 1792.565 | 840.6065 | 854.8249 | 670.0242 | 469.0998 | 634.6158 | 395.9039 | 1.10E-09 | 1.227535675 | YGL237C | HAP2 | n |
| W303.asm02.HP0_G0017760 | 4033.345 | 3697.839 | 4093.903 | 1913.943 | 1604.641 | 1087.08 | 3598.489 | 2570.677 | 4052.334 | 7.83E-08 | 1.360296938 | YGL236C | MTO1 | n |
| W303.asm02.HP0_G0017770 | 7480.06 | 5339.221 | 7752.33 | 19364.52 | 17773.41 | 24233.3 | 18318.35 | 43692.92 | 30903.23 | 8.63E-07 | -1.576901051 | YGL234W | ADE57 | n |
| W303.asm02.HP0_G0017780 | 4150.883 | 4264.255 | 3953.774 | 3699.363 | 3563.361 | 4472.146 | 4816.914 | 4989.948 | 5141.967 | 0.567826193 | 0.075826457 | YGL233W | SEC15 | n |
| W303.asm02.HP0_G0017790 | 2725.605 | 2641.79 | 2602.075 | 2126.526 | 2197.377 | 1851.303 | 1370.265 | 1510.837 | 1686.479 | 0.003697019 | 0.36812349 | YGL232W | TAN1 | n |
| W303.asm02.HP0_G0017800 | 4686.686 | 4889.649 | 4632.858 | 1030.959 | 1144.684 | 1229.137 | 1035.723 | 1247.756 | 1056.142 | 1.72E-65 | 2.061457599 | YGL231C | EMC4 | n |
| W303.asm02.HP0_G0017830 | 6186.086 | 6352.983 | 6506.266 | 4661.545 | 4374.794 | 5980.84 | 5645.245 | 5795.298 | 5284.301 | 0.021440833 | 0.342758818 | YGL228W | SHE10 | n |
| W303.asm02.HP0_G0017840 | 7078.738 | 7423.498 | 6719.692 | 2767.054 | 2639.109 | 5123.178 | 4151.533 | 4000.979 | 3231.82 | 0.000236121 | 1.01110262 | YGL227W | VID30 | n |
| W303.asm02.HP0_G0017850 | 1444.338 | 1296.414 | 1326.907 | 261.2133 | 330.648 | 187.6372 | 182.702 | 307.1068 | 244.0012 | 4.42E-20 | 2.384723918 | YGL226C-, | OST5 | n |
| W303.asm02.HP0_G0017860 | 1279.15 | 1111.689 | 1137.195 | 1286.614 | 1470.125 | 842.4681 | 1080.164 | 941.7225 | 1125.515 | 0.913848869 | -0.028537027 | YGL226W | MTC3 | n |

| | | | | | | | | | | | | | |
|---|---|---|---|---|---|---|---|---|---|---|---|---|---|
| W303.asm02.HP0_G0017890 | 1739.771 | 1631.367 | 1777.474 | 1046.937 | 1173.323 | 1865.736 | 483.9135 | 1355.136 | 720.0428 | 0.420430594 | 0.333501978 | YGL224C | SDT1 | n |
| W303.asm02.HP0_G0017900 | 2715.016 | 2975.631 | 2856.462 | 1471.409 | 1556.909 | 1719.881 | 1467.789 | 1460.368 | 1292.967 | 1.47E-14 | 0.848109227 | YGL223C | COG1 | n |
| W303.asm02.HP0_G0017920 | 4059.818 | 3845.842 | 4028.15 | 2474.579 | 2412.602 | 2493.979 | 2833.116 | 2667.319 | 3027.289 | 4.93E-17 | 0.693073375 | YGL221C | NIF3 | n |
| W303.asm02.HP0_G0017930 | 2437.585 | 1956.305 | 2174.145 | 933.6985 | 1098.689 | 749.789 | 1269.038 | 1078.095 | 1137.476 | 5.72E-12 | 1.239757782 | YGL220W | BOL2 | n |
| W303.asm02.HP0_G0017940 | 6637.177 | 6653.439 | 6823.171 | 4691.418 | 4056.296 | 3395.701 | 4099.685 | 4195.336 | 4262.845 | 1.13E-07 | 0.727916773 | YGL219C | MDM34 | n |
| W303.asm02.HP0_G0017950 | 913.8296 | 890.2411 | 975.5087 | 1376.233 | 1239.279 | 1751.787 | 992.5164 | 1174.737 | 648.2777 | 0.009804957 | -0.651962111 | YGL216W | KIP3 | n |
| W303.asm02.HP0_G0017960 | 8493.427 | 8850.109 | 9218.288 | 17066.4 | 16785.81 | 17659.16 | 16917.22 | 13979.8 | 13949.93 | 4.42E-18 | -0.955548323 | YGL215W | CLG1 | n |
| W303.asm02.HP0_G0017970 | 1272.796 | 1373.197 | 1287.025 | 2562.808 | 2698.99 | 2312.419 | 1623.332 | 2629.736 | 1812.068 | 2.85E-06 | -0.945443604 | YGL213C | SKI8 | n |
| W303.asm02.HP0_G0017980 | 1051.486 | 1020.439 | 970.1191 | 2380.097 | 2374.417 | 2395.982 | 919.6825 | 1363.726 | 1124.319 | 8.54E-15 | -1.232992976 | YGL212W | VAM7 | n |
| W303.asm02.HP0_G0017990 | 1387.157 | 1279.722 | 1513.386 | 1869.481 | 1556.042 | 1178.999 | 4357.69 | 3074.289 | 4171.942 | 0.569830338 | -0.139549011 | YGL211W | NCS6 | n |
| W303.asm02.HP0_G0018010 | 497.6824 | 439.5565 | 459.1897 | 4849.813 | 4012.904 | 3987.48 | 8119.13 | 5954.221 | 8328.335 | 6.86E-75 | -3.201912305 | YGL209W | MIG2 | n |
| W303.asm02.HP0_G0018030 | 9659.274 | 9624.619 | 9928.63 | 11045.99 | 11361.79 | 12710.71 | 7120.441 | 9327.026 | 7823.588 | 0.055306394 | -0.265640969 | YGL207W | SPT16 | n |
| W303.asm02.HP0_G0018040 | 17729.14 | 18661.68 | 17753.18 | 18050.81 | 18146.59 | 28047.58 | 25701.73 | 33282.43 | 31077.86 | 0.273294841 | -0.246787964 | YGL206C | CHC1 | n |
| W303.asm02.HP0_G0018050 | 446.8553 | 558.6263 | 495.8387 | 79.19764 | 65.08819 | 63.81183 | 225.9086 | 287.7784 | 244.0012 | 1.25E-49 | 2.846985532 | YGL205W | POX1 | n |
| W303.asm02.HP0_G0018070 | 2986.094 | 2800.921 | 3113.004 | 3110.244 | 3035.713 | 3674.498 | 5203.304 | 3701.388 | 4087.02 | 0.435653193 | -0.141999513 | YGL203C | KEX1 | n |
| W303.asm02.HP0_G0018080 | 13508.37 | 12364.34 | 13733.65 | 12891.85 | 13756.17 | 12352.15 | 7547.568 | 4586.199 | 4411.159 | 0.932995457 | 0.022262644 | YGL202W | ARO8 | n |
| W303.asm02.HP0_G0018090 | 2455.586 | 2205.572 | 2547.102 | 4623.336 | 4708.913 | 4365.793 | 2551.656 | 2767.182 | 2698.366 | 2.19E-22 | -0.926100542 | YGL201C | MCM6 | n |
| W303.asm02.HP0_G0018100 | 6930.492 | 6773.622 | 7128.22 | 3827.886 | 4527.534 | 3982.922 | 3351.594 | 4371.44 | 3869.333 | 2.35E-08 | 0.755825454 | YGL200C | EMP24 | n |
| W303.asm02.HP0_G0018110 | 3628.846 | 2932.232 | 3163.666 | 1580.479 | 1791.227 | 1817.118 | 2188.721 | 2521.282 | 2408.914 | 3.98E-12 | 0.906547941 | YGL198W | YIP4 | n |
| W303.asm02.HP0_G0018120 | 5341.085 | 5157.834 | 5218.163 | 2915.724 | 2633.034 | 3305.301 | 3246.664 | 2245.316 | 2572.777 | 1.19E-06 | 0.827836472 | YGL197W | MDS3 | n |
| W303.asm02.HP0_G0018130 | 2753.137 | 2669.61 | 2588.062 | 4753.248 | 5225.28 | 7647.544 | 969.0614 | 3348.108 | 2314.423 | 0.005276018 | -1.137688484 | YGL196W | DSD1 | n |
| W303.asm02.HP0_G0018140 | 12995.86 | 11816.84 | 12675.15 | 18395.39 | 18314.95 | 16319.87 | 11100.38 | 13495.52 | 13282.52 | 3.42E-05 | -0.500377388 | YGL195W | GCN1 | n |
| W303.asm02.HP0_G0018150 | 491.329 | 458.4742 | 410.6838 | 93.78668 | 116.2909 | 103.3144 | 55.55129 | 42.952 | 55.01988 | 1.76E-31 | 2.121535519 | YGL194C-A | | n |
| W303.asm02.HP0_G0018160 | 1044.074 | 1268.594 | 943.1714 | 2152.925 | 2138.364 | 1821.676 | 2387.471 | 1590.298 | 1880.245 | 7.10E-06 | -0.908933915 | YGL194C | HOS2 | n |
| W303.asm02.HP0_G0018180 | 2513.826 | 2671.836 | 2515.842 | 215.362 | 187.454 | 184.5985 | 119.7439 | 510.055 | 144.7262 | 5.77E-15 | 3.712513901 | YGL192W | IME4 | n |
| W303.asm02.HP0_G0018190 | 5916.067 | 6023.594 | 5649.327 | 12345.11 | 13964.45 | 13566.85 | 9494.333 | 13357 | 13203.57 | 9.56E-14 | -1.180845352 | YGL191W | COX13 | n |
| W303.asm02.HP0_G0018200 | 3302.705 | 4118.478 | 3424.521 | 5862.015 | 6112.215 | 7467.503 | 5147.753 | 5744.83 | 4424.316 | 7.87E-07 | -0.842076742 | YGL190C | CDC55 | n |
| W303.asm02.HP0_G0018220 | 2242.747 | 2305.724 | 2244.209 | 9323.091 | 10736.08 | 9869.563 | 6155.083 | 9298.034 | 9382.085 | 7.79E-32 | -2.139459179 | YGL187C | COX4 | n |
| W303.asm02.HP0_G0018230 | 2350.755 | 2172.188 | 2063.12 | 2744.823 | 2743.25 | 2423.33 | 4699.639 | 6691.921 | 5486.439 | 0.120449875 | -0.264478405 | YGL186C | TPN1 | n |
| W303.asm02.HP0_G0018240 | 1968.493 | 2058.683 | 1835.681 | 2193.914 | 2111.461 | 2991.559 | 2140.576 | 2544.906 | 2316.815 | 0.080231752 | -0.315785525 | YGL185C | | n |
| W303.asm02.HP0_G0018250 | 3139.635 | 1625.803 | 2235.586 | 182.7103 | 208.2822 | 293.9902 | 714.7599 | 866.5565 | 486.8063 | 3.38E-25 | 3.353578004 | YGL184C | STR3 | n |
| W303.asm02.HP0_G0018260 | 130.2445 | 100.1521 | 109.9468 | 16.67319 | 19.09254 | 12.9143 | 70.36497 | 51.5424 | 52.62771 | 4.51E-19 | 2.811654994 | YGL183C | MND1 | n |
| W303.asm02.HP0_G0018270 | 6187.145 | 6360.773 | 6141.932 | 3210.978 | 3448.806 | 4242.727 | 4604.585 | 4662.439 | 4354.943 | 2.18E-09 | 0.777612625 | YGL181W | GTS1 | n |
| W303.asm02.HP0_G0018280 | 1918.724 | 1972.997 | 1838.915 | 2104.99 | 1978.681 | 3495.977 | 4550.268 | 3081.806 | 2478.287 | 0.199230422 | -0.40348045 | YGL180W | ATG1 | n |
| W303.asm02.HP0_G0018310 | 2864.321 | 3507.55 | 2993.356 | 10624.99 | 8836.372 | 10614.79 | 13865.6 | 9807.015 | 14132.93 | 1.11E-19 | -1.683305667 | YGL178W | MPT5 | n |
| W303.asm02.HP0_G0018330 | 304.9628 | 323.8252 | 288.8799 | 375.1467 | 353.2119 | 399.5836 | 254.3015 | 173.9556 | 184.197 | 0.120967442 | -0.298164301 | YGL175C | SAE2 | n |
| W303.asm02.HP0_G0018340 | 1136.198 | 1129.493 | 1165.221 | 603.0136 | 586.6615 | 543.1602 | 686.367 | 678.6416 | 692.5328 | 2.69E-27 | 0.985302428 | YGL174W | BUD13 | n |
| W303.asm02.HP0_G0018350 | 17460.18 | 17646.8 | 18005.41 | 16888.55 | 17029.67 | 18914.89 | 16108.64 | 20432.27 | 19988.96 | 0.959198143 | 0.007603611 | YGL173C | XRN1 | n |
| W303.asm02.HP0_G0018360 | 2597.479 | 2329.093 | 2536.323 | 9546.789 | 9515.025 | 9748.016 | 8124.067 | 8516.307 | 8882.122 | 7.96E-129 | -1.94863747 | YGL172W | NUP49 | n |
| W303.asm02.HP0_G0018370 | 2204.627 | 2238.956 | 2330.442 | 4790.068 | 4482.407 | 1764.701 | 7511.769 | 2845.57 | 6364.365 | 0.120554586 | -0.704280716 | YGL171W | ROK1 | n |
| W303.asm02.HP0_G0018390 | 2329.577 | 2267.889 | 2427.454 | 2211.281 | 1871.936 | 1599.854 | 4083.637 | 2650.138 | 3628.92 | 0.153825624 | 0.305706986 | YGL169W | SUA5 | n |
| W303.asm02.HP0_G0018400 | 5050.947 | 4827.332 | 5082.346 | 7714.823 | 7231.732 | 9149.401 | 5945.222 | 8429.329 | 7227.938 | 4.35E-05 | -0.68763955 | YGL167C | PMR1 | n |
| W303.asm02.HP0_G0018430 | 2983.977 | 2834.305 | 2872.631 | 2117.495 | 2279.822 | 3037.899 | 1610.987 | 1704.12 | 1587.204 | 0.185530391 | 0.225155135 | YGL163C | RAD54 | n |
| W303.asm02.HP0_G0018440 | 212.8386 | 260.3955 | 228.5169 | 3246.409 | 2916.819 | 3793.005 | 4586.068 | 4212.517 | 6605.974 | 1.52E-69 | -3.826951067 | YGL162W | SUT1 | n |
| W303.asm02.HP0_G0018450 | 8276.352 | 8318.19 | 7821.316 | 1575.616 | 1723.535 | 1692.533 | 2225.755 | 2453.633 | 2682.817 | 8.26E-110 | 2.290492686 | YGL161C | YIP5 | n |
| W303.asm02.HP0_G0018460 | 3590.726 | 3539.821 | 3413.741 | 1710.391 | 1655.844 | 1725.198 | 1539.388 | 2397.795 | 2115.873 | 1.45E-09 | 1.050276096 | YGL160W | AIM14 | n |
| W303.asm02.HP0_G0018470 | 1250.559 | 1232.984 | 1140.429 | 2865.704 | 2812.678 | 3984.441 | 3972.534 | 3892.525 | 3705.469 | 8.08E-18 | -1.414926603 | YGL159W | | n |
| W303.asm02.HP0_G0018510 | 1708.004 | 1727.068 | 1734.357 | 1690.244 | 2032.487 | 2105.031 | 940.6685 | 1392.719 | 1126.711 | 0.366160255 | -0.172780323 | YGL155W | CDC43 | n |
| W303.asm02.HP0_G0018520 | 892.6516 | 773.397 | 862.3281 | 423.7768 | 457.353 | 422.3735 | 239.4878 | 385.4942 | 380.3548 | 2.55E-06 | 0.956187898 | YGL154C | LYS5 | n |
| W303.asm02.HP0_G0018530 | 2339.107 | 2381.395 | 2121.327 | 2043.16 | 2172.21 | 2515.25 | 1152.998 | 1526.943 | 1345.595 | 0.893767654 | 0.023661626 | YGL153W | PEX14 | n |
| W303.asm02.HP0_G0018540 | 3983.577 | 4092.883 | 4149.954 | 4503.845 | 4152.626 | 4887.682 | 3061.493 | 3836.687 | 3664.802 | 0.277094917 | -0.147759421 | YGL151W | NUT1 | n |
| W303.asm02.HP0_G0018550 | 8625.789 | 9064.88 | 8459.439 | 8148.326 | 7703.838 | 9082.55 | 11023.84 | 10431.97 | 11472.84 | 0.535957407 | 0.068591834 | YGL150C | INO80 | n |
| W303.asm02.HP0_G0018560 | 9019.699 | 8201.346 | 9430.636 | 8705.488 | 8777.359 | 8588.008 | 4076.23 | 3985.945 | 4006.882 | 0.729424257 | 0.031835741 | YGL148W | ARO2 | n |
| W303.asm02.HP0_G0018580 | 334.612 | 308.246 | 308.2823 | 1034.432 | 993.6797 | 1181.278 | 6805.65 | 4106.211 | 6257.913 | 1.98E-16 | -1.754558025 | YGL146C | RRT6 | n |

| | | | | | | | | | | | | | | |
|---|---|---|---|---|---|---|---|---|---|---|---|---|---|---|
| W303.asm02.HP0_G0018590 | 2511.708 | 2469.306 | 2580.517 | 2228.649 | 2199.113 | 2607.929 | 2768.923 | 2585.71 | 2642.15 | 0.35880876 | 0.103905647 | YGL145W | TIP20 | n |
| W303.asm02.HP0_G0018600 | 634.2803 | 622.056 | 693.0962 | 1310.929 | 1270.521 | 1121.265 | 1207.315 | 1077.021 | 1096.809 | 5.92E-15 | -0.92549367 | YGL144C | ROG1 | n |
| W303.asm02.HP0_G0018610 | 5310.377 | 5143.368 | 5137.32 | 1223.395 | 1065.711 | 1013.393 | 580.2024 | 429.52 | 535.8458 | 5.37E-56 | 2.238770763 | YGL143C | MRF1 | n |
| W303.asm02.HP0_G0018620 | 1471.869 | 1485.59 | 1455.179 | 1530.46 | 1576.002 | 1817.118 | 2839.288 | 2642.622 | 2694.778 | 0.152240182 | -0.158074788 | YGL142C | GPI10 | n |
| W303.asm02.HP0_G0018630 | 3977.224 | 3770.171 | 3607.765 | 2227.26 | 1897.104 | 3055.371 | 2513.387 | 2834.832 | 2920.838 | 0.00080432 | 0.66121058 | YGL141W | HUL5 | n |
| W303.asm02.HP0_G0018640 | 2522.297 | 2549.428 | 2493.206 | 2584.344 | 2640.845 | 4053.57 | 18098.61 | 12389.5 | 13931.99 | 0.229070741 | -0.294629305 | YGL140C | | n |
| W303.asm02.HP0_G0018660 | 15.88348 | 21.14323 | 12.93492 | 323.043 | 323.7053 | 223.3414 | 260.4738 | 234.0884 | 300.2172 | 7.37E-39 | -4.124963471 | YGL138C | | n |
| W303.asm02.HP0_G0018670 | 12840.21 | 12477.84 | 12902.58 | 7397.337 | 7815.79 | 8493.81 | 5219.352 | 6532.999 | 5719.675 | 5.94E-09 | 0.689064835 | YGL137W | SEC27 | n |
| W303.asm02.HP0_G0018680 | 2782.786 | 2718.574 | 2508.297 | 555.7729 | 542.4016 | 355.523 | 359.2317 | 265.2286 | 328.9232 | 4.34E-32 | 2.462118044 | YGL136C | MRM2 | n |
| W303.asm02.HP0_G0018690 | 10108.25 | 9209.544 | 10504.23 | 27361.4 | 31912.3 | 30567.38 | 27679.36 | 22090.21 | 20528.4 | 1.23E-26 | -1.590974608 | YPL220W | RPL1A | y |
| W303.asm02.HP0_G0018700 | 1595.76 | 1602.434 | 1602.852 | 2236.986 | 1930.95 | 1689.494 | 2929.405 | 1846.936 | 2377.816 | 0.176369851 | -0.287017417 | YGL134W | PCL10 | n |
| W303.asm02.HP0_G0018710 | 4721.629 | 4762.79 | 4718.013 | 4848.424 | 4613.451 | 4760.058 | 4373.738 | 4232.919 | 4673.101 | 0.980758632 | -0.002110889 | YGL133W | ITC1 | n |
| W303.asm02.HP0_G0018720 | 3151.283 | 3239.365 | 2783.164 | 3213.757 | 2961.079 | 3054.611 | 2552.89 | 2345.179 | 2773.72 | 0.948578123 | -0.008873424 | YGL131C | SNT2 | n |
| W303.asm02.HP0_G0018730 | 2956.445 | 3000.113 | 2895.267 | 3724.373 | 3612.828 | 3774.014 | 3582.441 | 3551.056 | 3529.645 | 1.08E-07 | -0.328159147 | YGL130W | CEG1 | n |
| W303.asm02.HP0_G0018740 | 9054.643 | 10105.35 | 9184.872 | 5821.027 | 5249.579 | 4456.193 | 1198.673 | 1664.39 | 1658.969 | 1.02E-06 | 0.868290712 | YGL129C | RSM23 | n |
| W303.asm02.HP0_G0018750 | 1517.402 | 1544.568 | 1433.62 | 616.2132 | 755.8908 | 680.6595 | 588.8437 | 775.2835 | 590.8657 | 5.23E-13 | 1.13165886 | YGL128C | CWC23 | n |
| W303.asm02.HP0_G0018760 | 363.2023 | 387.2549 | 313.6719 | 1017.759 | 1061.371 | 909.3185 | 1301.135 | 1020.11 | 1221.202 | 1.35E-21 | -1.489756705 | YGL127C | SOH1 | n |
| W303.asm02.HP0_G0018770 | 2669.484 | 2195.557 | 2466.258 | 746.8198 | 731.5912 | 953.3791 | 1294.962 | 1632.176 | 1318.085 | 1.28E-20 | 1.591947463 | YGL126W | SCS3 | n |
| W303.asm02.HP0_G0018780 | 4490.789 | 3657.778 | 4106.838 | 2068.865 | 2035.091 | 2402.06 | 3019.521 | 2251.758 | 1814.46 | 1.55E-05 | 0.913567053 | YGL125W | MET13 | n |
| W303.asm02.HP0_G0018790 | 2313.694 | 2442.599 | 2236.664 | 839.9118 | 765.4371 | 1220.781 | 1625.801 | 1270.305 | 1070.495 | 2.08E-08 | 1.306883076 | YGL124C | MON1 | n |
| W303.asm02.HP0_G0018800 | 22264.4 | 21145.45 | 24256.21 | 25689.21 | 30800.6 | 26693.85 | 24248.75 | 19331.62 | 18361.09 | 0.061013763 | -0.297846387 | YGL123W | RPS2 | n |
| W303.asm02.HP0_G0018810 | 9253.716 | 10095.33 | 9533.037 | 8492.21 | 8726.156 | 9030.893 | 6595.79 | 9113.34 | 7486.292 | 0.370517771 | 0.13789602 | YGL122C | NAB2 | n |
| W303.asm02.HP0_G0018830 | 3407.536 | 3593.236 | 4196.304 | 5238.854 | 4763.588 | 1558.072 | 9562.229 | 4068.628 | 7571.214 | 0.931882536 | -0.046080819 | YGL120C | PRP43 | n |
| W303.asm02.HP0_G0018850 | 1008.072 | 1024.89 | 1057.43 | 2532.24 | 2081.086 | 3003.714 | 10189.34 | 7564.921 | 8087.922 | 1.75E-12 | -1.30160445 | YGL117W | ARO5 | n |
| W303.asm02.HP0_G0018860 | 1892.252 | 1839.461 | 2196.781 | 3042.857 | 2845.656 | 3220.218 | 1580.126 | 2781.142 | 1675.714 | 0.010680919 | -0.619602985 | YGL116W | CDC20 | n |
| W303.asm02.HP0_G0018870 | 5334.732 | 5927.893 | 5574.951 | 3301.986 | 3512.159 | 3545.355 | 3159.017 | 4302.716 | 3195.937 | 4.61E-06 | 0.700756079 | YGL115W | SNF4 | n |
| W303.asm02.HP0_G0018880 | 1767.302 | 1607.998 | 1672.917 | 1590.9 | 1568.191 | 1820.916 | 1750.483 | 2871.341 | 1777.381 | 0.940199165 | 0.019618842 | YGL114W | | n |
| W303.asm02.HP0_G0018890 | 964.6567 | 990.3932 | 897.8991 | 1928.532 | 1727.007 | 1569.467 | 1232.004 | 1797.541 | 1717.577 | 1.42E-06 | -0.873105378 | YGL113W | SLD3 | n |
| W303.asm02.HP0_G0018900 | 3348.238 | 3402.947 | 3399.729 | 4131.477 | 3999.886 | 3980.643 | 3549.11 | 4371.44 | 3914.784 | 0.017102555 | -0.254877555 | YGL112C | TAF6 | n |
| W303.asm02.HP0_G0018910 | 2913.03 | 3025.707 | 3362.002 | 3424.256 | 3063.484 | 1302.065 | 4467.558 | 2307.596 | 3680.351 | 0.535724763 | 0.255755807 | YGL111W | NSA1 | n |
| W303.asm02.HP0_G0018920 | 2476.764 | 2539.413 | 2363.857 | 2150.841 | 1959.588 | 1880.93 | 1955.405 | 2014.449 | 2257.011 | 0.006087415 | 0.300498993 | YGL110C | CUE3 | n |
| W303.asm02.HP0_G0018930 | 1908.135 | 1751.549 | 1705.254 | 664.1486 | 871.3139 | 790.8109 | 1053.006 | 1345.471 | 1076.476 | 9.20E-14 | 1.206344377 | YGL108C | | n |
| W303.asm02.HP0_G0018950 | 5356.969 | 4664.863 | 5000.425 | 3900.831 | 4827.808 | 4037.617 | 1772.703 | 3631.591 | 3143.31 | 0.412412406 | 0.234822576 | YGL106W | MLC1 | n |
| W303.asm02.HP0_G0018960 | 18064.81 | 18079.68 | 18595.03 | 12105.43 | 13260.63 | 12793.51 | 10360.93 | 10382.57 | 10981.25 | 2.95E-13 | 0.520622446 | YGL105W | ARC1 | n |
| W303.asm02.HP0_G0018990 | 3519.779 | 3437.443 | 3561.415 | 3177.632 | 3642.335 | 3196.669 | 3423.194 | 4285.536 | 4051.138 | 0.611159265 | 0.070739221 | YGL100W | SEH1 | n |
| W303.asm02.HP0_G0019000 | 1531.168 | 1612.449 | 1796.876 | 5409.06 | 5062.993 | 1530.724 | 7003.166 | 2813.356 | 5645.518 | 0.00776694 | -1.280636373 | YGL099W | LSG1 | n |
| W303.asm02.HP0_G0019010 | 2262.867 | 2433.697 | 2155.82 | 1177.544 | 1273.125 | 974.6497 | 1334.465 | 1661.168 | 1176.947 | 1.70E-08 | 1.000547033 | YGL098W | USE1 | n |
| W303.asm02.HP0_G0019020 | 3178.814 | 2964.503 | 3160.433 | 3310.322 | 3151.136 | 2344.325 | 3224.444 | 2514.839 | 3625.332 | 0.717734912 | 0.079370855 | YGL097W | SRM1 | n |
| W303.asm02.HP0_G0019040 | 4906.937 | 5520.608 | 5047.853 | 3304.07 | 2975.832 | 3893.281 | 4066.354 | 4897.601 | 4655.16 | 3.70E-05 | 0.605068979 | YGL095C | VPS45 | n |
| W303.asm02.HP0_G0019050 | 2470.411 | 2612.858 | 2461.947 | 3156.79 | 2671.219 | 2521.327 | 2849.164 | 2696.312 | 3090.682 | 0.278509635 | -0.146343786 | YGL094C | PAN2 | n |
| W303.asm02.HP0_G0019060 | 3035.863 | 3054.64 | 2956.708 | 1326.908 | 1148.156 | 1139.497 | 1085.102 | 978.2317 | 907.828 | 7.43E-31 | 1.323145262 | YGL093W | SPC105 | n |
| W303.asm02.HP0_G0019070 | 4957.764 | 5155.609 | 5179.358 | 10101.87 | 9552.343 | 9386.416 | 8074.688 | 8823.414 | 8981.397 | 1.70E-30 | -0.925292744 | YGL092W | NUP145 | n |
| W303.asm02.HP0_G0019080 | 2099.796 | 2099.856 | 1780.708 | 1206.027 | 1234.072 | 1204.828 | 2603.504 | 1888.814 | 1999.853 | 8.95E-06 | 0.714380116 | YGL091C | NBP35 | n |
| W303.asm02.HP0_G0019110 | 3315.412 | 3227.124 | 3065.576 | 2195.303 | 2287.633 | 3325.052 | 4673.715 | 4752.638 | 4828.592 | 0.115190188 | 0.299280748 | YGL087C | MMS2 | n |
| W303.asm02.HP0_G0019120 | 3122.692 | 3238.252 | 3131.329 | 2161.957 | 2061.994 | 2528.164 | 3719.467 | 3924.739 | 4456.61 | 0.000124294 | 0.491272133 | YGL086W | MAD1 | n |
| W303.asm02.HP0_G0019130 | 591.9244 | 551.9495 | 582.0715 | 680.8218 | 812.3006 | 642.6763 | 685.1326 | 761.3241 | 698.5132 | 0.029448751 | -0.30625651 | YGL085W | LCL3 | n |
| W303.asm02.HP0_G0019140 | 3173.519 | 3230.462 | 2967.487 | 2605.88 | 2272.88 | 2558.55 | 3268.885 | 4256.543 | 3859.764 | 0.016682263 | 0.333289611 | YGL084C | GUP1 | n |
| W303.asm02.HP0_G0019150 | 3113.162 | 2940.021 | 2840.293 | 1837.524 | 1702.707 | 1709.245 | 1897.385 | 1803.984 | 1845.558 | 7.00E-20 | 0.760265614 | YGL083W | SCY1 | n |
| W303.asm02.HP0_G0019160 | 6600.116 | 6385.254 | 6342.423 | 4015.459 | 4030.261 | 5946.655 | 6425.432 | 8268.259 | 7731.489 | 0.020546839 | 0.466015892 | YGL082W | | n |
| W303.asm02.HP0_G0019170 | 383.3213 | 392.8189 | 352.4766 | 300.8121 | 357.5511 | 644.1956 | 1351.748 | 1300.372 | 917.3967 | 0.548276121 | -0.206798551 | YGL081W | | n |
| W303.asm02.HP0_G0019180 | 2627.128 | 2733.04 | 2376.792 | 3181.8 | 3783.793 | 3192.87 | 1199.908 | 1774.991 | 1505.87 | 0.030886442 | -0.392745266 | YGL080W | MPC1 | n |
| W303.asm02.HP0_G0019190 | 1123.492 | 1145.073 | 1094.079 | 332.769 | 328.0445 | 443.6441 | 482.679 | 578.7782 | 471.2572 | 8.11E-22 | 1.605892503 | YGL079W | KXD1 | n |
| W303.asm02.HP0_G0019200 | 5582.514 | 5726.476 | 6084.803 | 6292.739 | 5573.285 | 2165.044 | 7454.983 | 4256.543 | 7418.115 | 0.463300999 | 0.309946551 | YGL078C | DBP3 | n |
| W303.asm02.HP0_G0019210 | 3419.184 | 3463.038 | 3934.372 | 3154.011 | 2889.048 | 3271.116 | 4312.015 | 3942.993 | 3152.878 | 0.183084333 | 0.215665439 | YGL077C | HNM1 | n |

| | | | | | | | | | | | | | | |
|---|---|---|---|---|---|---|---|---|---|---|---|---|---|---|
| W303.asm02.HP0_G0019230 | 429.9129 | 559.7391 | 418.2291 | 175.0685 | 170.0971 | 237.0154 | 35.79972 | 82.68259 | 23.92169 | 0.001539 | 1.273744151 | YGL075C | MPS2 | n |
| W303.asm02.HP0_G0019240 | 7675.957 | 8003.267 | 7677.954 | 11414.88 | 10716.12 | 13967.95 | 15028.48 | 14337.38 | 12982.3 | 3.45E-06 | -0.628124865 | YGL073W | HSF1 | n |
| W303.asm02.HP0_G0019250 | 620.5146 | 770.0586 | 634.8891 | 6612.308 | 5576.756 | 5262.956 | 6283.468 | 4070.776 | 4530.767 | 8.82E-52 | -3.10726019 | YGL071W | AFT1 | n |
| W303.asm02.HP0_G0019260 | 1951.55 | 1719.278 | 1987.666 | 1367.896 | 1467.522 | 1014.152 | 1378.906 | 1128.564 | 1245.124 | 0.002278018 | 0.55595927 | YGL070C | RPB9 | n |
| W303.asm02.HP0_G0019270 | 8528.37 | 8165.736 | 8152.234 | 4052.279 | 4784.416 | 3647.909 | 1804.8 | 1707.342 | 1803.695 | 1.59E-15 | 0.993081738 | YGL068W | MNP1 | n |
| W303.asm02.HP0_G0019280 | 1668.824 | 1640.269 | 1773.162 | 2270.332 | 2355.325 | 2036.661 | 2704.731 | 3434.012 | 3026.093 | 0.003291605 | -0.390437693 | YGL067W | NPY1 | n |
| W303.asm02.HP0_G0019290 | 3600.256 | 3821.36 | 3543.091 | 3518.043 | 3471.37 | 5137.612 | 4129.312 | 4215.739 | 3644.469 | 0.4708225 | -0.145404693 | YGL066W | SGF73 | n |
| W303.asm02.HP0_G0019300 | 2303.105 | 2241.182 | 2401.584 | 4867.181 | 5115.064 | 6417.647 | 4207.084 | 6741.316 | 5426.635 | 1.20E-09 | -1.239450636 | YGL065C | ALG2 | n |
| W303.asm02.HP0_G0019310 | 2591.125 | 2566.12 | 2689.386 | 1432.505 | 1247.09 | 518.8509 | 1493.712 | 681.863 | 1137.476 | 0.001332184 | 1.294687026 | YGL064C | MRH4 | n |
| W303.asm02.HP0_G0019330 | 4537.381 | 4917.469 | 4847.362 | 18100.13 | 17384.62 | 25575.63 | 32013.59 | 23014.75 | 16601.65 | 1.48E-15 | -2.094006319 | YGL062W | PYC1 | n |
| W303.asm02.HP0_G0019340 | 985.8347 | 929.1891 | 951.7947 | 811.4285 | 886.9351 | 900.9622 | 956.7166 | 830.0473 | 870.7494 | 0.214767834 | 0.141902453 | YGL061C | DUO1 | n |
| W303.asm02.HP0_G0019350 | 1802.246 | 1790.497 | 1864.785 | 3886.937 | 3651.013 | 4952.254 | 6285.937 | 6152.874 | 6378.718 | 7.43E-18 | -1.194549323 | YGL060W | YBP2 | n |
| W303.asm02.HP0_G0019360 | 2243.806 | 2471.532 | 1974.731 | 2260.606 | 2121.875 | 2995.358 | 2828.178 | 3751.857 | 2963.897 | 0.516280589 | -0.141278188 | YGL059W | PKP2 | n |
| W303.asm02.HP0_G0019370 | 1529.05 | 1624.69 | 1483.204 | 2978.943 | 2936.779 | 3853.779 | 2560.297 | 3111.872 | 3106.231 | 9.00E-12 | -1.075190657 | YGL058W | RAD6 | n |
| W303.asm02.HP0_G0019400 | 16614.12 | 17138.25 | 18565.92 | 20161.36 | 20644.24 | 26893.65 | 52463.87 | 55745.25 | 67405.33 | 0.031757765 | -0.371826871 | YGL055W | OLE1 | n |
| W303.asm02.HP0_G0019410 | 3587.549 | 3574.318 | 3423.443 | 3767.446 | 3978.19 | 4263.238 | 2708.434 | 3494.145 | 3203.114 | 0.17524814 | -0.182003663 | YGL054C | ERV14 | n |
| W303.asm02.HP0_G0019420 | 3050.687 | 3143.664 | 2663.516 | 1340.802 | 1708.782 | 1743.43 | 2961.501 | 3966.617 | 2689.994 | 7.52E-06 | 0.886222186 | YGL053W | PRM8 | n |
| W303.asm02.HP0_G0019430 | 594.0422 | 685.4856 | 563.747 | 439.0606 | 543.2694 | 164.0876 | 186.4054 | 298.5164 | 254.766 | 0.109940459 | 0.685508318 | YGL051W | MST27 | y |
| W303.asm02.HP0_G0019460 | 15455.69 | 15319.94 | 15607.06 | 8772.181 | 9528.911 | 10952.84 | 7858.656 | 10958.13 | 8607.023 | 4.59E-05 | 0.664967534 | YGL048C | RPT6 | n |
| W303.asm02.HP0_G0019470 | 1494.106 | 1431.063 | 1382.959 | 644.6966 | 806.2257 | 735.3553 | 692.5394 | 679.7154 | 615.9834 | 4.68E-15 | 0.979759644 | YGL047W | ALG13 | n |
| W303.asm02.HP0_G0019480 | 4337.249 | 5338.108 | 4078.812 | 1783.336 | 1691.425 | 2042.738 | 4116.968 | 1788.951 | 2447.189 | 1.24E-05 | 1.317742268 | YGL045W | RIM8 | n |
| W303.asm02.HP0_G0019490 | 2943.738 | 2934.457 | 2843.527 | 1838.914 | 1938.76 | 1719.121 | 1809.738 | 1734.187 | 1725.95 | 1.14E-15 | 0.666347806 | YGL044C | RNA15 | n |
| W303.asm02.HP0_G0019500 | 2848.438 | 3044.625 | 3067.732 | 3031.741 | 3103.405 | 2658.066 | 2799.785 | 2389.205 | 2570.385 | 0.832829888 | 0.027285346 | YGL043W | DST1 | n |
| W303.asm02.HP0_G0019520 | 8404.479 | 8153.496 | 8011.028 | 5386.134 | 5796.32 | 4865.652 | 6514.314 | 5363.631 | 5144.359 | 3.87E-06 | 0.614504511 | YGL040C | HEM2 | n |
| W303.asm02.HP0_G0019540 | 1443.279 | 1592.419 | 1343.076 | 3917.504 | 3445.335 | 4126.498 | 3383.691 | 3694.946 | 4045.157 | 4.31E-26 | -1.391918317 | YGL038C | OCH1 | n |
| W303.asm02.HP0_G0019550 | 23027.87 | 25223.87 | 22172.61 | 1571.448 | 1652.372 | 2576.782 | 2167.735 | 2991.607 | 2291.698 | 3.93E-56 | 3.601774229 | YGL037C | PNC1 | n |
| W303.asm02.HP0_G0019560 | 5118.716 | 5220.151 | 4931.439 | 209.8043 | 154.476 | 181.5598 | 411.0795 | 728.0364 | 383.9431 | 7.92E-76 | 4.804720128 | YGL036W | VIR1 | n |
| W303.asm02.HP0_G0019570 | 1327.859 | 1543.456 | 1327.985 | 5370.85 | 5225.28 | 5803.838 | 7617.933 | 4531.436 | 5902.676 | 1.99E-22 | -1.965538321 | YGL035C | MIG1 | n |
| W303.asm02.HP0_G0019600 | 13323.06 | 12910.72 | 13501.9 | 20940.13 | 24843.73 | 22192.84 | 19646.64 | 16810.34 | 15729.7 | 6.62E-10 | -0.774571166 | YGL031C | RPL24A | y |
| W303.asm02.HP0_G0019610 | 9480.32 | 8235.843 | 9015.64 | 12259.66 | 14460.86 | 11988.27 | 10841.14 | 8945.827 | 8157.295 | 0.000608738 | -0.534052368 | YGL030W | RPL30 | n |
| W303.asm02.HP0_G0019620 | 972.069 | 800.1042 | 1068.209 | 1793.757 | 1997.773 | 625.204 | 1898.62 | 620.6564 | 1369.517 | 0.218653443 | -0.636824341 | YGL029W | CGR1 | n |
| W303.asm02.HP0_G0019630 | 2527.591 | 2352.462 | 2922.214 | 6625.508 | 6391.66 | 8181.588 | 8298.128 | 9203.539 | 10951.35 | 3.33E-17 | -1.442009868 | YGL028C | SCW11 | n |
| W303.asm02.HP0_G0019640 | 5422.62 | 4982.012 | 5401.408 | 3577.093 | 3165.889 | 4687.89 | 1666.539 | 3877.492 | 2825.151 | 0.144009035 | 0.467515229 | YGL027C | CWH41 | n |
| W303.asm02.HP0_G0019650 | 8077.279 | 6568.867 | 7924.795 | 11075.16 | 11370.47 | 13304.77 | 5394.647 | 7084.932 | 5496.007 | 8.25E-05 | -0.663473804 | YGL026C | TRP5 | n |
| W303.asm02.HP0_G0019660 | 2224.746 | 2227.828 | 2226.962 | 1097.651 | 1061.371 | 1012.633 | 993.7508 | 638.911 | 679.3759 | 5.12E-09 | 1.074470889 | YGL025C | PGD1 | n |
| W303.asm02.HP0_G0019670 | 3116.339 | 2940.021 | 3157.199 | 2517.651 | 2115.8 | 2737.831 | 6171.131 | 2885.3 | 4769.984 | 0.267227374 | 0.321782637 | YGL023C | PIB2 | n |
| W303.asm02.HP0_G0019680 | 6442.34 | 5449.388 | 6420.033 | 7634.236 | 7906.913 | 9615.075 | 4697.17 | 8912.539 | 7229.134 | 0.073691781 | -0.458135628 | YGL022W | STT3 | n |
| W303.asm02.HP0_G0019700 | 2772.197 | 2622.873 | 2720.645 | 4587.905 | 5283.425 | 5613.162 | 8572.181 | 8001.957 | 7714.744 | 4.25E-17 | -0.931893252 | YGL020C | GET1 | n |
| W303.asm02.HP0_G0019710 | 4971.529 | 4900.777 | 4859.219 | 4465.635 | 4799.169 | 4295.903 | 4467.558 | 3191.333 | 3799.96 | 0.452271258 | 0.119532373 | YGL019W | CKB1 | n |
| W303.asm02.HP0_G0019720 | 2251.219 | 2356.913 | 2257.144 | 1746.516 | 2072.408 | 2968.769 | 2286.244 | 3853.868 | 2931.603 | 0.957562864 | 0.016417285 | YGL018C | JAC1 | n |
| W303.asm02.HP0_G0019730 | 1557.64 | 1661.412 | 1605.008 | 1604.794 | 1487.482 | 2260.002 | 1751.717 | 2364.507 | 1761.832 | 0.518136013 | -0.150025705 | YGL017W | ATE1 | n |
| W303.asm02.HP0_G0019740 | 2074.383 | 1891.762 | 2057.73 | 3945.293 | 3811.564 | 3265.039 | 1789.986 | 1827.607 | 1284.595 | 4.41E-07 | -0.871581994 | YGL016W | KAP122 | n |
| W303.asm02.HP0_G0019760 | 2752.078 | 2930.006 | 2806.878 | 7313.972 | 6865.502 | 6022.621 | 3941.673 | 3693.872 | 3939.902 | 1.12E-32 | -1.250915859 | YGL014W | PUF4 | n |
| W303.asm02.HP0_G0019770 | 2697.015 | 2932.232 | 2682.918 | 7642.572 | 7588.415 | 9294.497 | 5356.379 | 6160.39 | 4453.022 | 1.90E-22 | -1.561021538 | YGL013C | PDR1 | n |
| W303.asm02.HP0_G0019780 | 9969.531 | 9409.848 | 9808.982 | 6540.058 | 6841.203 | 8532.553 | 8314.176 | 6756.349 | 6866.72 | 0.007437718 | 0.413557785 | YGL012W | ERG4 | n |
| W303.asm02.HP0_G0019790 | 10705.47 | 11538.64 | 11189.79 | 4228.737 | 4833.015 | 5660.261 | 3180.003 | 5124.173 | 3760.489 | 9.89E-09 | 1.183358317 | YGL011C | SCL1 | n |
| W303.asm02.HP0_G0019800 | 355.79 | 342.7428 | 302.8927 | 458.5127 | 561.4941 | 589.4997 | 1107.322 | 871.9255 | 1039.397 | 7.69E-05 | -0.683852778 | YGL010W | MPO1 | n |
| W303.asm02.HP0_G0019810 | 8508.251 | 9980.716 | 9319.611 | 2835.831 | 2614.81 | 3643.352 | 8774.635 | 6235.556 | 9285.203 | 3.13E-15 | 1.612478187 | YGL009C | LEU1 | n |
| W303.asm02.HP0_G0019820 | 41356.35 | 36365.24 | 40633.98 | 65843.11 | 71828.72 | 67480.25 | 40664.78 | 62275.03 | 56513.79 | 8.18E-06 | -0.793561302 | YGL008C | PMA1 | n |
| W303.asm02.HP0_G0019830 | 7394.29 | 7937.612 | 6704.601 | 8425.517 | 8869.351 | 7039.052 | 3249.133 | 9160.587 | 6466.032 | 0.705103301 | -0.143069117 | YGL006W | PMC1 | n |
| W303.asm02.HP0_G0019840 | 3547.311 | 3731.223 | 3414.819 | 1285.92 | 1273.993 | 1326.374 | 746.8562 | 1165.073 | 840.8473 | 1.64E-16 | 1.460217043 | YGL005C | COG7 | n |
| W303.asm02.HP0_G0019850 | 3171.402 | 3112.505 | 2841.371 | 1484.608 | 1627.205 | 1734.314 | 1215.956 | 1902.773 | 1412.576 | 1.16E-06 | 0.913112257 | YGL004C | RPN14 | n |
| W303.asm02.HP0_G0019860 | 1223.028 | 1439.965 | 1205.104 | 605.7925 | 544.1373 | 661.6679 | 840.6762 | 413.413 | 594.4539 | 3.05E-05 | 1.094132194 | YGL003C | CDH1 | n |
| W303.asm02.HP0_G0019880 | 8194.817 | 8604.18 | 8632.982 | 5879.383 | 6175.567 | 7696.162 | 10448.58 | 8537.783 | 8700.317 | 0.018300973 | 0.364706896 | YGL001C | ERG26 | n |

| | | | | | | | | | | | | | | |
|---|---|---|---|---|---|---|---|---|---|---|---|---|---|---|
| W303.asm02.HP0_G0019890 | 2419.584 | 2482.66 | 2460.869 | 3369.373 | 3831.525 | 3094.114 | 1603.581 | 1541.977 | 1275.026 | 0.000521519 | -0.483415554 | YGR001C | EFM5 | n |
| W303.asm02.HP0_G0019900 | 3933.809 | 4117.365 | 3821.191 | 2560.029 | 2337.968 | 2784.93 | 2760.282 | 2399.943 | 2529.718 | 1.66E-08 | 0.627648066 | YGR002C | SWC4 | n |
| W303.asm02.HP0_G0019910 | 1251.618 | 1365.407 | 1288.103 | 2733.013 | 2580.096 | 2920.151 | 3570.096 | 3167.71 | 3950.667 | 1.04E-18 | -1.076287656 | YGR003W | CUL3 | n |
| W303.asm02.HP0_G0019930 | 6888.136 | 7271.044 | 6796.223 | 4655.988 | 4558.777 | 5398.936 | 6190.883 | 5114.509 | 4760.416 | 0.000250681 | 0.519962006 | YGR005C | TFG2 | n |
| W303.asm02.HP0_G0019940 | 745.4647 | 717.7569 | 644.5903 | 684.9901 | 756.7587 | 692.0545 | 699.9462 | 836.4901 | 657.8464 | 0.918048052 | -0.017188096 | YGR006W | PRP18 | n |
| W303.asm02.HP0_G0019950 | 3499.66 | 3860.308 | 3791.01 | 2114.021 | 2715.479 | 3113.106 | 1464.085 | 2096.057 | 1885.029 | 0.017030839 | 0.489581817 | YGR007W | ECT1 | n |
| W303.asm02.HP0_G0019960 | 8950.871 | 8347.123 | 7419.255 | 565.4989 | 673.4458 | 1697.091 | 11538.62 | 8662.344 | 9079.476 | 1.31E-15 | 3.073547478 | YGR008C | STF2 | n |
| W303.asm02.HP0_G0019970 | 5387.677 | 5784.342 | 5256.968 | 4760.89 | 4554.438 | 5550.869 | 7305.612 | 7379.153 | 7000.682 | 0.224025526 | 0.144111386 | YGR009C | SEC9 | n |
| W303.asm02.HP0_G0019990 | 1708.004 | 1838.348 | 1793.642 | 3060.919 | 3414.96 | 3066.766 | 2982.487 | 2862.751 | 2975.858 | 1.34E-21 | -0.837339774 | YGR012W | MCY1 | n |
| W303.asm02.HP0_G0020000 | 4568.089 | 4550.245 | 4564.949 | 2428.728 | 2245.109 | 2056.412 | 1910.964 | 1878.076 | 1908.951 | 7.24E-30 | 1.023485389 | YGR013W | SNU71 | n |
| W303.asm02.HP0_G0020010 | 1118.197 | 1130.606 | 1381.881 | 6880.469 | 6860.295 | 7549.547 | 2441.788 | 4989.948 | 3754.509 | 9.90E-24 | -2.551878159 | YGR014W | MSB2 | n |
| W303.asm02.HP0_G0020020 | 1079.018 | 1111.689 | 1055.274 | 2007.035 | 2186.963 | 2375.471 | 2299.823 | 1621.438 | 1943.637 | 1.40E-10 | -1.017078984 | YGR015C | EAT1 | n |
| W303.asm02.HP0_G0020030 | 494.5057 | 487.407 | 406.3721 | 923.9725 | 871.3139 | 951.1001 | 1420.879 | 1122.121 | 977.2009 | 4.17E-08 | -0.984400636 | YGR016W | | n |
| W303.asm02.HP0_G0020050 | 7322.285 | 7837.46 | 6953.598 | 1280.362 | 1239.279 | 1623.403 | 640.6915 | 920.2465 | 485.6102 | 9.23E-24 | 2.416095597 | YGR019W | UGA1 | n |
| W303.asm02.HP0_G0020060 | 6778.011 | 6586.671 | 6617.29 | 1501.282 | 1748.703 | 1792.049 | 1671.477 | 1286.412 | 1507.066 | 3.99E-50 | 1.986881989 | YGR020C | VMA7 | n |
| W303.asm02.HP0_G0020070 | 6428.574 | 6537.708 | 6185.048 | 994.1388 | 1100.424 | 812.8411 | 335.7767 | 260.9334 | 391.1196 | 1.19E-51 | 2.719917137 | YGR021W | DPC29 | n |
| W303.asm02.HP0_G0020090 | 1676.237 | 1454.431 | 1595.307 | 433.5029 | 392.2648 | 93.43875 | 775.2491 | 216.9076 | 538.2379 | 3.27E-05 | 2.362181083 | YGR024C | THG1 | n |
| W303.asm02.HP0_G0020100 | 5959.482 | 5740.942 | 5628.847 | 2391.908 | 2558.4 | 3131.338 | 4614.46 | 3932.255 | 4406.375 | 1.27E-15 | 1.100520911 | YGR026W | | n |
| W303.asm02.HP0_G0020120 | 4366.898 | 4454.544 | 3907.424 | 3060.225 | 2794.453 | 2875.331 | 3852.791 | 3438.307 | 3264.114 | 1.22E-06 | 0.543883646 | YGR028W | MSP1 | n |
| W303.asm02.HP0_G0020140 | 352.6133 | 403.9469 | 394.5151 | 475.8806 | 579.7188 | 309.1835 | 533.2924 | 417.7082 | 392.3157 | 0.388171262 | -0.245161594 | YGR030C | POP6 | n |
| W303.asm02.HP0_G0020170 | 3127.987 | 3147.002 | 3078.511 | 4291.956 | 4157.833 | 3529.402 | 2554.125 | 3027.042 | 3065.564 | 0.006154703 | -0.356960483 | YGR033C | TIM21 | n |
| W303.asm02.HP0_G0020210 | 1036.662 | 997.07 | 825.6792 | 909.3834 | 965.9087 | 1160.008 | 1011.033 | 1734.187 | 1492.713 | 0.741405288 | -0.086097761 | YGR036C | CAX4 | n |
| W303.asm02.HP0_G0020220 | 6134.2 | 6010.24 | 5822.871 | 3600.714 | 4406.904 | 4616.482 | 3899.7 | 5597.719 | 4449.434 | 0.00415608 | 0.509276402 | YGR037C | ACB1 | n |
| W303.asm02.HP0_G0020230 | 1898.605 | 1876.183 | 1903.589 | 3216.536 | 3205.81 | 4150.807 | 4328.063 | 5640.671 | 4493.689 | 5.82E-08 | -0.896880275 | YGR038W | ORM1 | n |
| W303.asm02.HP0_G0020240 | 1137.257 | 1397.679 | 1403.439 | 1480.44 | 1534.346 | 1484.385 | 2752.875 | 1141.449 | 2050.089 | 0.561308396 | -0.192069539 | YGR040W | KSS1 | n |
| W303.asm02.HP0_G0020250 | 1631.763 | 1465.559 | 1571.593 | 3195 | 2922.894 | 3000.675 | 1651.725 | 1888.814 | 1719.969 | 6.14E-22 | -0.965768003 | YGR041W | BUD9 | n |
| W303.asm02.HP0_G0020260 | 2164.389 | 2285.694 | 2202.17 | 486.3013 | 470.3706 | 663.1872 | 1049.302 | 994.3387 | 937.7301 | 1.67E-37 | 2.037597855 | YGR042W | MTE1 | n |
| W303.asm02.HP0_G0020280 | 5314.613 | 5587.376 | 5216.007 | 1097.651 | 1113.442 | 1996.399 | 2656.586 | 2658.729 | 2163.717 | 1.21E-14 | 1.937572811 | YGR044C | RME1 | n |
| W303.asm02.HP0_G0020290 | 2138.975 | 2504.916 | 2008.147 | 2417.612 | 2185.227 | 2063.249 | 3483.683 | 2975.5 | 3167.231 | 0.984560557 | -0.003260265 | YGR046W | TAM41 | n |
| W303.asm02.HP0_G0020300 | 1479.281 | 1562.373 | 1450.867 | 1687.466 | 1647.165 | 1975.888 | 1103.619 | 1165.073 | 897.0632 | 0.113941858 | -0.241451896 | YGR047C | TFC4 | n |
| W303.asm02.HP0_G0020310 | 3117.398 | 3456.361 | 3128.095 | 1457.514 | 1605.509 | 2134.658 | 1172.749 | 1923.176 | 1386.262 | 0.000116456 | 0.900398419 | YGR048W | UFD1 | n |
| W303.asm02.HP0_G0020340 | 657.5761 | 880.2259 | 640.2786 | 716.9471 | 556.2871 | 535.5636 | 1729.497 | 708.708 | 968.8283 | 0.467864142 | 0.267731564 | YGR052W | FMP48 | n |
| W303.asm02.HP0_G0020350 | 687.2253 | 695.5009 | 544.3446 | 879.5106 | 777.5869 | 609.251 | 1141.888 | 690.4534 | 1095.613 | 0.392199427 | -0.234100441 | YGR053C | MCO32 | n |
| W303.asm02.HP0_G0020360 | 8983.697 | 8644.241 | 9652.685 | 15737.4 | 15462.35 | 15591.36 | 5741.534 | 6376.224 | 6457.659 | 7.27E-19 | -0.778351957 | YGR054W | | n |
| W303.asm02.HP0_G0020370 | 3170.343 | 2274.566 | 2856.462 | 1115.714 | 1372.927 | 1550.475 | 4818.148 | 7901.02 | 5656.283 | 1.91E-05 | 1.039469554 | YGR055W | MUP1 | n |
| W303.asm02.HP0_G0020380 | 2120.974 | 2386.959 | 2263.611 | 6849.901 | 6606.885 | 8295.538 | 6771.085 | 6721.988 | 6213.658 | 2.96E-41 | -1.68370976 | YGR056W | RSC1 | n |
| W303.asm02.HP0_G0020390 | 1307.74 | 1439.965 | 1277.324 | 1342.886 | 1455.372 | 1564.149 | 1936.888 | 1859.821 | 2087.167 | 0.344068766 | -0.116051464 | YGR057C | LST7 | n |
| W303.asm02.HP0_G0020400 | 2631.363 | 2561.669 | 2718.489 | 1071.252 | 1012.772 | 762.7033 | 1809.738 | 1456.073 | 1776.185 | 1.64E-18 | 1.474629744 | YGR058W | PEF1 | n |
| W303.asm02.HP0_G0020410 | 355.79 | 373.9013 | 308.2823 | 42.37768 | 39.92076 | 26.58826 | 72.83391 | 76.23979 | 84.92199 | 1.34E-44 | 3.25242605 | YGR059W | SPR3 | n |
| W303.asm02.HP0_G0020420 | 11223.27 | 13073.19 | 13951.39 | 11853.94 | 12497.8 | 16135.28 | 34431.92 | 23866.28 | 24486.24 | 0.726030496 | -0.082084713 | YGR060W | ERG25 | n |
| W303.asm02.HP0_G0020430 | 12905.86 | 9976.264 | 12992.05 | 12674.4 | 11458.12 | 15085.42 | 12501.51 | 32935.59 | 22582.07 | 0.727521287 | -0.128569472 | YGR061C | ADE6 | n |
| W303.asm02.HP0_G0020440 | 2628.187 | 2465.968 | 2468.414 | 1518.649 | 1490.953 | 1384.869 | 1293.728 | 1485.065 | 1608.733 | 1.09E-11 | 0.783202391 | YGR062C | COX18 | n |
| W303.asm02.HP0_G0020450 | 887.3571 | 820.1346 | 795.4977 | 801.7024 | 827.9218 | 648.7536 | 507.3684 | 446.7008 | 438.9629 | 0.392386838 | 0.13598499 | YGR063C | SPT4 | n |
| W303.asm02.HP0_G0020460 | 945.5965 | 811.2322 | 787.9523 | 1737.485 | 1728.742 | 1605.171 | 1331.996 | 2724.23 | 1343.203 | 0.000778093 | -0.994802186 | YGR065C | VHT1 | n |
| W303.asm02.HP0_G0020480 | 432.0307 | 472.9406 | 419.307 | 1217.837 | 1015.376 | 1412.217 | 1190.032 | 1436.744 | 1072.888 | 1.55E-15 | -1.461331313 | YGR067C | | n |
| W303.asm02.HP0_G0020490 | 695.6965 | 581.9951 | 685.5508 | 2000.088 | 1747.835 | 2071.605 | 1841.834 | 1972.57 | 1942.441 | 1.49E-36 | -1.567648177 | YGR068C | ART5 | n |
| W303.asm02.HP0_G0020520 | 2659.954 | 2713.01 | 2723.879 | 4367.68 | 3755.155 | 2774.295 | 2230.693 | 2553.496 | 2401.737 | 0.022317535 | -0.428577036 | YGR072W | UPF3 | n |
| W303.asm02.HP0_G0020530 | 659.6939 | 737.7873 | 637.0449 | 971.2132 | 1030.997 | 1155.45 | 1239.411 | 1136.08 | 1050.162 | 2.18E-06 | -0.634254033 | YGR075C | PRP38 | n |
| W303.asm02.HP0_G0020540 | 4091.585 | 3987.167 | 3590.519 | 1972.994 | 2022.941 | 1509.454 | 795.0007 | 846.1543 | 948.4949 | 3.89E-12 | 1.083895113 | YGR076C | MRPL25 | n |
| W303.asm02.HP0_G0020550 | 2271.338 | 2481.547 | 2212.95 | 964.9607 | 983.2656 | 1033.904 | 1096.212 | 1300.372 | 967.6322 | 8.59E-19 | 1.223935626 | YGR077C | PEX8 | n |
| W303.asm02.HP0_G0020560 | 1446.456 | 1424.386 | 1545.723 | 2598.238 | 2643.448 | 2459.034 | 1493.712 | 2034.851 | 1899.382 | 1.81E-08 | -0.802016623 | YGR078C | PAC10 | n |
| W303.asm02.HP0_G0020570 | 5822.884 | 5856.674 | 5851.974 | 2115.411 | 2186.963 | 2683.895 | 1356.686 | 1819.017 | 1413.772 | 1.20E-16 | 1.327350334 | YGR079W | | n |
| W303.asm02.HP0_G0020580 | 1867.897 | 1651.397 | 1877.719 | 1290.088 | 1346.892 | 315.2608 | 1924.544 | 1005.077 | 1796.519 | 0.085516629 | 0.870419122 | YGR081C | SLX9 | n |
| W303.asm02.HP0_G0020590 | 9414.668 | 10016.33 | 9548.128 | 6769.314 | 7266.445 | 6075.798 | 5565.005 | 5628.859 | 6687.307 | 2.17E-05 | 0.527039469 | YGR082W | TOM20 | n |

| | | | | | | | | | | | | | |
|---|---|---|---|---|---|---|---|---|---|---|---|---|---|
| W303.asm02.HP0_G0020600 | 4201.71 | 4489.041 | 4774.064 | 8558.208 | 8371.209 | 8436.076 | 7595.713 | 7072.046 | 7249.467 | 2.88E-29 | -0.913724076 | YGR083C | GCD2 | n |
| W303.asm02.HP0_G0020610 | 7628.306 | 7867.506 | 7214.453 | 4538.581 | 4163.908 | 3484.582 | 1330.762 | 1555.936 | 1446.066 | 2.91E-11 | 0.897940303 | YGR084C | MRP13 | n |
| W303.asm02.HP0_G0020620 | 6431.751 | 6680.147 | 6638.849 | 18445.41 | 21391.45 | 19716.34 | 17028.32 | 14757.23 | 13473.89 | 2.58E-39 | -1.592247588 | YGR085C | RPL11B | y |
| W303.asm02.HP0_G0020630 | 22458.18 | 24992.41 | 22967.03 | 9368.942 | 10393.28 | 14091.02 | 8395.651 | 11514.36 | 9801.911 | 1.06E-07 | 1.056644251 | YGR086C | PIL1 | n |
| W303.asm02.HP0_G0020650 | 9278.07 | 11145.82 | 9953.422 | 1187.27 | 1244.486 | 2038.18 | 1659.132 | 11092.35 | 2154.148 | 7.41E-06 | 2.764661804 | YGR088W | CTT1 | n |
| W303.asm02.HP0_G0020660 | 3584.372 | 3656.665 | 3761.906 | 7142.377 | 6884.595 | 8187.665 | 6952.552 | 7663.71 | 8007.785 | 3.09E-20 | -1.013667417 | YGR089W | NNF2 | n |
| W303.asm02.HP0_G0020670 | 2971.27 | 3252.718 | 3350.145 | 17028.19 | 17376.81 | 10389.17 | 12978.02 | 9670.642 | 12761.02 | 2.45E-22 | -2.226097351 | YGR090W | UTP22 | n |
| W303.asm02.HP0_G0020680 | 1440.102 | 1407.694 | 1474.581 | 1526.291 | 1542.156 | 1182.038 | 1467.789 | 635.6896 | 999.9265 | 0.944851079 | 0.024229145 | YGR091W | PRP31 | n |
| W303.asm02.HP0_G0020690 | 3438.244 | 3655.553 | 3435.3 | 3677.827 | 3743.873 | 4093.073 | 2493.636 | 2166.928 | 2318.011 | 0.220122493 | -0.129140956 | YGR092W | DBF2 | n |
| W303.asm02.HP0_G0020710 | 26639.77 | 25230.55 | 27442.51 | 15861.06 | 15824.24 | 16758.96 | 14706.28 | 13621.15 | 14154.46 | 2.79E-21 | 0.711228857 | YGR094W | VAS1 | n |
| W303.asm02.HP0_G0020720 | 2647.247 | 2509.367 | 2857.54 | 2656.595 | 2900.33 | 1808.002 | 2836.819 | 2100.353 | 2728.268 | 0.613802459 | 0.121978993 | YGR095C | RRP46 | n |
| W303.asm02.HP0_G0020730 | 884.1804 | 903.5947 | 911.912 | 1160.871 | 1116.045 | 745.9907 | 125.9163 | 437.0366 | 376.7666 | 0.720874706 | -0.163138136 | YGR096W | TPC1 | n |
| W303.asm02.HP0_G0020740 | 8903.22 | 9093.813 | 9921.085 | 6959.666 | 6329.175 | 12684.88 | 22215.58 | 21762.7 | 24483.85 | 0.727464594 | 0.104130298 | YGR097W | ASK10 | n |
| W303.asm02.HP0_G0020750 | 1187.025 | 1241.886 | 1098.39 | 2455.127 | 2445.58 | 2644.393 | 1454.209 | 1740.63 | 1617.106 | 1.09E-22 | -1.097087862 | YGR098C | ESP1 | n |
| W303.asm02.HP0_G0020760 | 1748.242 | 1610.224 | 1719.267 | 1387.348 | 1302.632 | 1118.986 | 822.1591 | 1194.066 | 976.0048 | 0.019539493 | 0.414756696 | YGR099W | TEL2 | n |
| W303.asm02.HP0_G0020770 | 6068.548 | 6207.206 | 5762.508 | 4608.747 | 4298.424 | 5709.639 | 7631.513 | 6793.932 | 7641.783 | 0.036352716 | 0.303357436 | YGR100W | MDR1 | n |
| W303.asm02.HP0_G0020780 | 3547.311 | 3818.022 | 3255.289 | 2275.195 | 2399.585 | 2310.9 | 1060.412 | 1304.667 | 1010.691 | 8.35E-06 | 0.604446171 | YGR101W | PCP1 | n |
| W303.asm02.HP0_G0020790 | 1406.217 | 1269.706 | 1193.247 | 607.8766 | 634.3929 | 547.7182 | 434.5345 | 452.0698 | 413.8452 | 5.05E-20 | 1.112585443 | YGR102C | GTF1 | n |
| W303.asm02.HP0_G0020800 | 7298.989 | 8002.155 | 8544.594 | 10235.25 | 9340.589 | 3693.489 | 8468.485 | 3349.182 | 7331.997 | 0.944851079 | 0.035302547 | YGR103W | NOP7 | n |
| W303.asm02.HP0_G0020810 | 1290.798 | 1221.856 | 1266.544 | 1581.869 | 1485.746 | 1897.642 | 738.2149 | 1112.457 | 941.3184 | 0.039869988 | -0.39386697 | YGR104C | SRB5 | n |
| W303.asm02.HP0_G0020820 | 1255.854 | 1112.801 | 1073.599 | 636.36 | 758.4944 | 794.6092 | 1004.861 | 946.0177 | 1010.691 | 1.27E-06 | 0.653551034 | YGR105W | VMA21 | n |
| W303.asm02.HP0_G0020830 | 4784.104 | 4673.766 | 4684.597 | 4519.128 | 4879.878 | 5894.997 | 8148.757 | 7694.85 | 8218.295 | 0.409288282 | -0.112902011 | YGR106C | VOA1 | n |
| W303.asm02.HP0_G0020860 | 378.0268 | 534.1447 | 339.5417 | 3169.99 | 3014.017 | 2895.082 | 3370.112 | 2799.396 | 2981.838 | 2.03E-52 | -2.85918095 | YGR110W | CLD1 | n |
| W303.asm02.HP0_G0020870 | 7287.341 | 7309.992 | 7010.728 | 1509.618 | 1556.042 | 1423.611 | 1124.605 | 2527.725 | 1160.202 | 1.14E-13 | 2.267035027 | YGR111W | | n |
| W303.asm02.HP0_G0020880 | 5984.895 | 5718.686 | 5541.536 | 2699.667 | 2831.77 | 3305.301 | 3895.997 | 3226.769 | 3585.861 | 2.83E-14 | 0.964628224 | YGR112W | SHY1 | n |
| W303.asm02.HP0_G0020890 | 2233.217 | 2577.248 | 2316.429 | 1448.483 | 1569.927 | 1513.252 | 1178.922 | 1147.892 | 1120.731 | 1.68E-11 | 0.653443844 | YGR113W | DAM1 | n |
| W303.asm02.HP0_G0020900 | 10213.08 | 10498.17 | 9967.435 | 15078.12 | 14605.79 | 15133.28 | 11863.29 | 13103.58 | 11538.63 | 1.97E-11 | -0.546846854 | YGR116W | SPT6 | n |
| W303.asm02.HP0_G0020910 | 1921.901 | 1767.129 | 1937.005 | 1617.299 | 1636.751 | 1550.475 | 1650.491 | 1421.711 | 1912.539 | 0.113637183 | 0.227815901 | YGR117C | | n |
| W303.asm02.HP0_G0020920 | 8218.113 | 8265.889 | 8787.123 | 259.8238 | 304.6127 | 192.9548 | 419.7209 | 359.723 | 464.0807 | 3.45E-163 | 5.061784297 | YPR132W | RPS23B | y |
| W303.asm02.HP0_G0020930 | 5438.504 | 5370.379 | 5099.593 | 8260.175 | 7893.896 | 7282.905 | 6664.92 | 7543.445 | 7086.8 | 5.04E-09 | -0.559016067 | YGR119C | NUP57 | n |
| W303.asm02.HP0_G0020940 | 1631.763 | 1666.976 | 1639.501 | 704.4422 | 676.0493 | 832.5924 | 711.0565 | 1234.87 | 857.5925 | 1.44E-07 | 1.157813459 | YGR120C | COG2 | n |
| W303.asm02.HP0_G0020950 | 1728.123 | 1741.534 | 1683.696 | 968.4343 | 994.5475 | 1344.606 | 1433.223 | 1068.431 | 1424.536 | 0.000852465 | 0.63967992 | YGR122W | | n |
| W303.asm02.HP0_G0020980 | 2515.943 | 2532.736 | 2728.191 | 3576.399 | 3578.983 | 3143.492 | 1248.052 | 2093.91 | 1888.617 | 0.056163393 | -0.405213734 | YGR123C | PPT1 | n |
| W303.asm02.HP0_G0020990 | 23158.11 | 24878.9 | 25565.87 | 25587.09 | 26308.65 | 25721.48 | 18596.1 | 15885.8 | 15675.88 | 0.501491223 | -0.076611993 | YGR124W | ASN2 | n |
| W303.asm02.HP0_G0021000 | 2769.02 | 3034.609 | 2665.672 | 8987.543 | 9121.025 | 10611 | 4235.477 | 8736.436 | 5738.813 | 7.82E-12 | -1.761730787 | YGR125W | VSB1 | n |
| W303.asm02.HP0_G0021010 | 718.9922 | 748.9153 | 602.5518 | 298.0332 | 341.93 | 333.4928 | 1352.983 | 1365.874 | 1115.947 | 2.25E-12 | 1.08964351 | YGR126W | | n |
| W303.asm02.HP0_G0021020 | 2496.883 | 2195.557 | 2100.847 | 486.996 | 533.7231 | 700.4108 | 1757.89 | 1893.109 | 1395.83 | 2.14E-24 | 1.98085342 | YGR127W | | n |
| W303.asm02.HP0_G0021030 | 3039.039 | 3246.042 | 3601.298 | 4501.761 | 3974.719 | 1502.617 | 6588.383 | 2526.651 | 5688.577 | 0.979816212 | -0.013468989 | YGR128C | UTP8 | n |
| W303.asm02.HP0_G0021040 | 658.635 | 647.6504 | 696.33 | 245.2348 | 204.8108 | 101.0354 | 382.6867 | 243.7526 | 374.3744 | 2.82E-08 | 1.861234373 | YGR129W | SYF2 | n |
| W303.asm02.HP0_G0021050 | 18903.46 | 19958.09 | 18247.94 | 10092.14 | 10663.18 | 21081.45 | 13573.03 | 19437.93 | 15047.94 | 0.159580924 | 0.448954497 | YGR130C | | n |
| W303.asm02.HP0_G0021070 | 14506.91 | 15156.35 | 14689.76 | 5575.792 | 5857.069 | 6306.736 | 2482.525 | 3965.543 | 3114.604 | 2.09E-13 | 1.322067498 | YGR132C | PHB1 | n |
| W303.asm02.HP0_G0021080 | 1267.502 | 1115.027 | 1144.741 | 1654.814 | 1867.597 | 2276.715 | 3692.309 | 3642.329 | 3326.311 | 2.94E-06 | -0.717108076 | YGR133W | PEX4 | n |
| W303.asm02.HP0_G0021090 | 1158.435 | 1224.082 | 1096.235 | 2053.581 | 2036.826 | 2212.903 | 1481.368 | 1523.722 | 1527.4 | 5.51E-22 | -0.8577597 | YGR134W | CAF130 | n |
| W303.asm02.HP0_G0021100 | 9465.496 | 9803.78 | 9192.418 | 6259.392 | 7031.26 | 7527.517 | 3782.426 | 6362.265 | 4829.789 | 0.030761782 | 0.451195944 | YGR135W | PRE9 | n |
| W303.asm02.HP0_G0021130 | 1752.477 | 1919.582 | 2015.692 | 1973.689 | 1875.408 | 2287.35 | 1938.123 | 2044.515 | 2294.09 | 0.4521849 | -0.109716268 | YGR140W | CBF2 | n |
| W303.asm02.HP0_G0021140 | 3309.058 | 3416.3 | 2856.462 | 992.7494 | 1098.689 | 1773.057 | 2339.327 | 2967.983 | 2790.465 | 7.89E-08 | 1.310004532 | YGR141W | VPS62 | n |
| W303.asm02.HP0_G0021150 | 16974.15 | 22222.64 | 15018.52 | 484.9119 | 619.6396 | 679.1402 | 465.3964 | 416.6344 | 753.5331 | 1.17E-74 | 4.926015862 | YGR142W | BTN2 | n |
| W303.asm02.HP0_G0021170 | 144.0102 | 158.0178 | 129.3492 | 896.1838 | 932.0629 | 1277.756 | 1097.447 | 2125.05 | 1032.221 | 2.05E-19 | -2.848208294 | YGR144W | THI4 | n |
| W303.asm02.HP0_G0021180 | 1513.166 | 1635.818 | 1802.266 | 4494.813 | 4036.336 | 1088.599 | 7743.85 | 3183.817 | 6279.443 | 0.062438715 | -0.95820544 | YGR145W | ENP2 | n |
| W303.asm02.HP0_G0021190 | 626.868 | 750.0281 | 604.7076 | 4905.391 | 4742.759 | 4917.309 | 3842.915 | 3052.813 | 4735.298 | 2.32E-54 | -2.877990404 | YGR146C | ECL1 | n |
| W303.asm02.HP0_G0021210 | 5136.718 | 5225.715 | 5159.956 | 1363.728 | 1440.619 | 869.816 | 490.0858 | 384.4204 | 425.806 | 4.05E-23 | 2.079010444 | YGR147C | NAT2 | n |
| W303.asm02.HP0_G0021230 | 4796.811 | 5347.011 | 4750.35 | 1853.503 | 1852.844 | 2239.491 | 4158.94 | 4352.111 | 4633.631 | 1.60E-27 | 1.324699493 | YGR149W | GPC1 | n |
| W303.asm02.HP0_G0021240 | 8285.882 | 8584.15 | 8391.53 | 2172.377 | 1486.614 | 619.8863 | 3044.211 | 1371.243 | 2270.168 | 2.94E-09 | 2.561561277 | YGR150C | CCM1 | n |
| W303.asm02.HP0_G0021250 | 1308.799 | 1246.338 | 1428.231 | 2293.258 | 2567.946 | 1823.955 | 2257.851 | 1259.567 | 1225.986 | 0.007018672 | -0.746895347 | YGR152C | RSR1 | n |

| | | | | | | | | | | | | | | |
|---|---|---|---|---|---|---|---|---|---|---|---|---|---|---|
| W303.asm02.HP0_G0021280 | 18822.98 | 15957.57 | 18654.31 | 16329.3 | 17262.26 | 14397.16 | 19277.53 | 16232.63 | 16721.26 | 0.283534731 | 0.155114423 | YGR155W | CYS4 | n |
| W303.asm02.HP0_G0021290 | 2961.74 | 2757.522 | 3006.291 | 3823.023 | 3585.057 | 3401.778 | 3403.442 | 2643.695 | 2973.466 | 0.019160915 | -0.309051922 | YGR156W | PTI1 | n |
| W303.asm02.HP0_G0021300 | 12715.26 | 13034.24 | 12867.01 | 18808.05 | 19162.83 | 28777.62 | 39745.1 | 47016.33 | 32491.63 | 0.000344517 | -0.789521271 | YGR157W | CHO2 | n |
| W303.asm02.HP0_G0021310 | 1461.28 | 1585.742 | 1794.72 | 1226.869 | 1302.632 | 508.2156 | 2098.604 | 1022.258 | 1661.361 | 0.095485523 | 0.672618316 | YGR158C | MTR3 | n |
| W303.asm02.HP0_G0021320 | 2560.417 | 2660.708 | 3244.51 | 12491 | 12919.57 | 5174.835 | 18067.75 | 11831.13 | 17403.03 | 9.99E-08 | -1.853143492 | YGR159C | NSR1 | n |
| W303.asm02.HP0_G0021330 | 4299.129 | 5037.652 | 3900.957 | 1417.221 | 1626.337 | 2461.313 | 2720.779 | 2015.522 | 1176.947 | 0.000166565 | 1.265877118 | YGR161C | RTS3 | n |
| W303.asm02.HP0_G0021360 | 19047.47 | 19455.11 | 20484.6 | 23816.26 | 23738.96 | 20130.35 | 24699.34 | 19851.34 | 24230.28 | 0.157908649 | -0.198443659 | YGR162W | TIF4631 | n |
| W303.asm02.HP0_G0021370 | 2740.43 | 2637.339 | 2567.582 | 2188.356 | 2304.122 | 2495.498 | 1819.613 | 2173.371 | 1864.695 | 0.123645372 | 0.185318559 | YGR163W | GTR2 | n |
| W303.asm02.HP0_G0021380 | 9884.819 | 9619.055 | 9613.881 | 3712.563 | 3683.124 | 2620.083 | 1994.909 | 1566.674 | 1794.126 | 5.81E-19 | 1.539654326 | YGR165W | MRPS35 | n |
| W303.asm02.HP0_G0021390 | 1645.529 | 1590.193 | 1615.787 | 1669.403 | 1571.663 | 1831.551 | 859.1933 | 1284.265 | 995.1422 | 0.737051691 | -0.064381401 | YGR166W | TRS65 | n |
| W303.asm02.HP0_G0021400 | 9444.318 | 9623.506 | 8911.083 | 4617.083 | 4944.967 | 6960.047 | 4324.359 | 5734.092 | 5223.3 | 0.000131403 | 0.759939465 | YGR167W | CLC1 | n |
| W303.asm02.HP0_G0021410 | 2065.911 | 1859.491 | 1966.108 | 730.1467 | 839.2037 | 894.8849 | 809.8143 | 981.4531 | 1083.652 | 1.17E-16 | 1.25789113 | YGR168C | PEX35 | n |
| W303.asm02.HP0_G0021420 | 2165.448 | 2088.728 | 2132.106 | 1654.814 | 1303.499 | 325.1365 | 2630.662 | 963.1985 | 2222.325 | 0.089136134 | 0.959737715 | YGR169C | PUS6 | n |
| W303.asm02.HP0_G0021430 | 2246.983 | 2005.268 | 1821.668 | 2574.618 | 2849.995 | 2253.925 | 2804.723 | 3095.765 | 2855.053 | 0.019621665 | -0.33799231 | YGR169C- | LSO2 | n |
| W303.asm02.HP0_G0021440 | 4119.116 | 4130.719 | 4097.136 | 3371.457 | 2912.479 | 3896.32 | 6635.293 | 5104.845 | 5394.34 | 0.097459347 | 0.278260288 | YGR170W | PSD2 | n |
| W303.asm02.HP0_G0021450 | 3549.428 | 3665.568 | 3516.143 | 1747.211 | 1551.702 | 1706.966 | 1275.211 | 1290.708 | 1401.811 | 5.31E-34 | 1.099556244 | YGR171C | MSM1 | n |
| W303.asm02.HP0_G0021460 | 1983.317 | 1897.326 | 1933.771 | 2912.25 | 3145.929 | 2957.374 | 3668.854 | 4371.44 | 3774.842 | 1.64E-09 | -0.632589839 | YGR172C | YIP1 | n |
| W303.asm02.HP0_G0021470 | 1657.176 | 1520.087 | 1643.813 | 2609.354 | 2648.655 | 2109.589 | 2367.719 | 2049.884 | 2127.834 | 6.09E-06 | -0.611707913 | YGR173W | RBG2 | n |
| W303.asm02.HP0_G0021480 | 4532.086 | 3875.887 | 3674.596 | 1688.855 | 1728.742 | 1383.349 | 1101.15 | 1171.516 | 1193.692 | 3.25E-21 | 1.331678319 | YGR174C | CBP4 | n |
| W303.asm02.HP0_G0021490 | 10885.48 | 12302.02 | 13958.94 | 6351.095 | 6335.25 | 8886.557 | 33285.1 | 21841.09 | 23297.33 | 0.000769832 | 0.783991535 | YGR175C | ERG1 | n |
| W303.asm02.HP0_G0021510 | 17676.2 | 16929.05 | 17575.32 | 9116.76 | 8937.042 | 11682.88 | 5678.576 | 7839.813 | 6412.208 | 2.43E-06 | 0.811257522 | YGR178C | PBP1 | n |
| W303.asm02.HP0_G0021520 | 1319.388 | 1437.739 | 1393.738 | 729.4519 | 636.9964 | 609.251 | 443.1758 | 329.6566 | 480.8259 | 2.86E-10 | 1.070620424 | YGR179C | OKP1 | n |
| W303.asm02.HP0_G0021550 | 2430.173 | 2353.575 | 2220.495 | 1445.01 | 1678.407 | 1230.657 | 514.7753 | 619.5826 | 522.6888 | 1.24E-05 | 0.686214104 | YGR181W | TIM13 | n |
| W303.asm02.HP0_G0021560 | 3284.704 | 3295.005 | 3034.317 | 4483.698 | 4973.605 | 4896.798 | 1906.026 | 3569.311 | 2880.171 | 0.012744602 | -0.578227893 | YGR183C | QCR9 | n |
| W303.asm02.HP0_G0021570 | 9408.315 | 10001.86 | 9327.156 | 9390.478 | 9728.515 | 12194.14 | 10351.06 | 10669.28 | 10184.66 | 0.37733788 | -0.123853301 | YGR184C | UBR1 | n |
| W303.asm02.HP0_G0021580 | 6987.673 | 6457.586 | 7182.115 | 11134.22 | 11728.89 | 12199.45 | 11601.58 | 11739.85 | 10622.42 | 4.04E-17 | -0.76531651 | YGR185C | TYS1 | n |
| W303.asm02.HP0_G0021590 | 9087.469 | 9281.876 | 9168.704 | 7105.557 | 6757.89 | 8265.911 | 8838.827 | 8068.533 | 8113.04 | 0.00480131 | 0.315407338 | YGR186W | TFG1 | n |
| W303.asm02.HP0_G0021600 | 1121.374 | 1175.118 | 1128.572 | 2453.043 | 2557.532 | 983.006 | 2578.814 | 1569.895 | 2523.738 | 0.031134799 | -0.80725068 | YGR187C | HGH1 | n |
| W303.asm02.HP0_G0021610 | 506.1536 | 475.1662 | 392.3593 | 1243.542 | 1209.772 | 1148.613 | 622.1744 | 791.3905 | 586.0813 | 1.74E-16 | -1.390720382 | YGR188C | BUB1 | n |
| W303.asm02.HP0_G0021620 | 11245.5 | 12683.71 | 12363.63 | 8905.566 | 10199.75 | 8875.921 | 4528.047 | 5576.243 | 5096.515 | 0.003442189 | 0.375277035 | YGR189C | CRH1 | n |
| W303.asm02.HP0_G0021640 | 102148.8 | 109037.8 | 110854.4 | 63508.87 | 74394.06 | 85927.94 | 156947.2 | 126337.9 | 132840.7 | 0.000790858 | 0.5248391 | YGR192C | TDH3 | y |
| W303.asm02.HP0_G0021650 | 7457.824 | 8342.672 | 7677.954 | 9803.834 | 9819.638 | 12614.23 | 3038.038 | 4773.041 | 5645.518 | 0.069474307 | -0.45742055 | YGR193C | PDX1 | n |
| W303.asm02.HP0_G0021660 | 7498.062 | 7273.27 | 6824.249 | 1163.65 | 1088.275 | 1771.538 | 9274.596 | 5717.985 | 8635.729 | 4.90E-22 | 2.424131622 | YGR194C | XKS1 | n |
| W303.asm02.HP0_G0021670 | 2438.644 | 2269.002 | 2392.961 | 1935.479 | 1976.945 | 966.2934 | 1648.022 | 1176.885 | 1595.576 | 0.07099177 | 0.541506193 | YGR195W | SKI6 | n |
| W303.asm02.HP0_G0021680 | 3365.18 | 3555.4 | 3378.17 | 2885.851 | 2849.127 | 3306.82 | 1119.667 | 1962.906 | 1236.751 | 0.444505289 | 0.187766564 | YGR196C | FYV8 | n |
| W303.asm02.HP0_G0021690 | 1285.503 | 1328.685 | 1116.715 | 4571.232 | 4345.287 | 4052.051 | 1857.882 | 4299.495 | 1979.52 | 1.70E-08 | -1.797436942 | YGR197C | SNG1 | n |
| W303.asm02.HP0_G0021700 | 3289.998 | 3528.693 | 3181.991 | 3479.138 | 3495.67 | 3042.457 | 2298.589 | 2750.002 | 2124.246 | 0.988438634 | -0.002389489 | YGR198W | YPP1 | n |
| W303.asm02.HP0_G0021710 | 2417.466 | 2325.755 | 2135.34 | 2701.056 | 2513.272 | 3049.294 | 3883.652 | 4221.108 | 4487.708 | 0.038238091 | -0.264797112 | YGR199W | PMT6 | n |
| W303.asm02.HP0_G0021720 | 3344.002 | 3401.834 | 3604.532 | 4850.508 | 4799.169 | 4112.065 | 2513.387 | 2155.116 | 2468.718 | 0.000598094 | -0.410968369 | YGR200C | ELP2 | n |
| W303.asm02.HP0_G0021740 | 7069.208 | 6650.101 | 6463.149 | 7375.801 | 7815.79 | 7583.732 | 7232.778 | 7926.791 | 6262.698 | 0.158854311 | -0.174324206 | YGR202C | PCT1 | n |
| W303.asm02.HP0_G0021750 | 713.6977 | 585.3335 | 673.6938 | 791.9764 | 874.7853 | 1027.826 | 1117.198 | 1856.6 | 1132.692 | 0.076734251 | -0.449708717 | YGR203W | YCH1 | n |
| W303.asm02.HP0_G0021770 | 1902.841 | 2112.097 | 1799.032 | 1772.916 | 1905.782 | 3305.301 | 3151.61 | 3382.47 | 2798.837 | 0.337972643 | -0.264568235 | YGR205W | TDA10 | n |
| W303.asm02.HP0_G0021780 | 728.5223 | 636.5224 | 601.4739 | 95.87083 | 110.216 | 135.98 | 143.1989 | 128.856 | 141.138 | 4.03E-43 | 2.524558984 | YGR206W | MVB12 | n |
| W303.asm02.HP0_G0021790 | 3155.518 | 2982.308 | 2872.631 | 3088.708 | 3115.555 | 3375.95 | 851.7864 | 1904.921 | 1194.888 | 0.77364847 | -0.088454653 | YGR207C | CIR1 | n |
| W303.asm02.HP0_G0021800 | 1123.492 | 938.0916 | 1206.181 | 4234.295 | 4848.636 | 4700.805 | 1727.028 | 2840.201 | 2204.383 | 9.39E-24 | -2.076471153 | YGR208W | SER2 | n |
| W303.asm02.HP0_G0021810 | 4828.578 | 4253.127 | 4322.42 | 3985.586 | 5241.769 | 4326.29 | 2386.236 | 4807.402 | 3017.721 | 0.959210133 | -0.01594026 | YGR209C | TRX2 | n |
| W303.asm02.HP0_G0021820 | 4781.987 | 3879.226 | 4382.783 | 4552.475 | 5196.641 | 2765.939 | 4293.497 | 2223.84 | 3726.999 | 0.866534167 | 0.059768361 | YGR210C | | n |
| W303.asm02.HP0_G0021830 | 9298.19 | 12140.66 | 8037.976 | 11993.58 | 12617.56 | 8250.717 | 10072.07 | 7767.869 | 10057.87 | 0.551463358 | -0.156832463 | YGR211W | ZPR1 | n |
| W303.asm02.HP0_G0021840 | 1197.614 | 1337.587 | 1190.013 | 1208.111 | 1301.764 | 1538.321 | 1240.645 | 1725.596 | 1401.811 | 0.53033421 | -0.119965949 | YGR212W | SLI1 | n |
| W303.asm02.HP0_G0021860 | 17290.76 | 16643.06 | 18450.59 | 2161.262 | 2249.448 | 1927.269 | 4110.795 | 3237.507 | 3097.858 | 5.08E-100 | 3.047130174 | YGR214W | RPS0A | n |
| W303.asm02.HP0_G0021870 | 2207.804 | 2036.427 | 1859.395 | 1428.336 | 1464.918 | 1077.964 | 836.9728 | 764.5455 | 690.1407 | 0.000262047 | 0.620225433 | YGR215W | RSM27 | n |
| W303.asm02.HP0_G0021880 | 1998.142 | 2128.789 | 2035.094 | 1529.765 | 1356.438 | 1466.912 | 1822.082 | 2188.404 | 1798.911 | 3.76E-05 | 0.500957959 | YGR216C | GPI1 | n |
| W303.asm02.HP0_G0021890 | 1124.55 | 1143.96 | 1040.183 | 5120.058 | 4565.719 | 5373.868 | 5272.435 | 5374.369 | 6402.639 | 2.65E-66 | -2.186484823 | YGR217W | CCH1 | n |
| W303.asm02.HP0_G0021900 | 8571.785 | 8775.552 | 8505.789 | 13697.72 | 13496.69 | 14922.85 | 10296.74 | 10146.34 | 10123.66 | 3.74E-23 | -0.704109988 | YGR218W | CRM1 | n |

| | | | | | | | | | | | | | | |
|---|---|---|---|---|---|---|---|---|---|---|---|---|---|---|
| W303.asm02.HP0_G0021910 | 5620.634 | 5553.992 | 5376.616 | 4067.563 | 4478.067 | 3332.649 | 1393.72 | 1394.866 | 1449.654 | 0.000353159 | 0.478741706 | YGR220C | MRPL9 | n |
| W303.asm02.HP0_G0021930 | 1967.434 | 1802.738 | 1698.786 | 814.2073 | 860.0319 | 810.5621 | 1081.398 | 700.1176 | 876.7298 | 1.36E-10 | 1.138299704 | YGR222W | PET54 | n |
| W303.asm02.HP0_G0021940 | 2045.792 | 2262.325 | 2016.77 | 1397.074 | 1549.967 | 2136.177 | 1190.032 | 2224.913 | 1523.811 | 0.269294293 | 0.315296286 | YGR223C | HSV2 | n |
| W303.asm02.HP0_G0021960 | 497.6824 | 438.4437 | 434.3978 | 98.64969 | 116.2909 | 130.6623 | 278.9909 | 325.3614 | 412.6491 | 9.20E-22 | 1.989038534 | YGR225W | AMA1 | n |
| W303.asm02.HP0_G0021970 | 1778.95 | 1810.528 | 1916.524 | 1360.254 | 1409.376 | 1374.993 | 1614.691 | 1900.626 | 1819.244 | 4.79E-05 | 0.40992348 | YGR227W | DIE2 | n |
| W303.asm02.HP0_G0021980 | 4972.588 | 5501.69 | 5494.108 | 12937.7 | 13180.79 | 15937.76 | 10544.87 | 14660.59 | 13853.05 | 3.91E-17 | -1.397115717 | YGR229C | SMI1 | n |
| W303.asm02.HP0_G0021990 | 523.096 | 497.4222 | 489.3712 | 2250.186 | 2374.417 | 2388.386 | 2001.081 | 2060.622 | 2257.011 | 1.28E-115 | -2.215291952 | YGR230W | BNS1 | n |
| W303.asm02.HP0_G0022000 | 14900.82 | 16103.35 | 15460.47 | 4567.064 | 4743.627 | 5341.962 | 2259.086 | 2508.397 | 2273.756 | 1.83E-60 | 1.664977362 | YGR231C | PHB2 | n |
| W303.asm02.HP0_G0022010 | 1645.529 | 1360.956 | 1333.375 | 2102.211 | 2417.809 | 2455.996 | 1835.661 | 2110.017 | 1484.341 | 0.000119502 | -0.6845764 | YGR232W | NAS6 | n |
| W303.asm02.HP0_G0022020 | 3362.003 | 3596.574 | 3471.949 | 7486.261 | 8520.478 | 15084.66 | 1667.773 | 3003.418 | 2008.226 | 6.02E-07 | -1.575707357 | YGR233C | PHO81 | n |
| W303.asm02.HP0_G0022030 | 41944.04 | 48774.08 | 47146.71 | 18439.16 | 20746.64 | 30693.49 | 45600.2 | 50768.19 | 53380.05 | 3.38E-06 | 0.980317764 | YGR234W | YHB1 | n |
| W303.asm02.HP0_G0022040 | 5010.709 | 4700.473 | 4514.288 | 3193.61 | 3795.943 | 2690.732 | 806.1109 | 2027.334 | 1353.967 | 0.091564047 | 0.555401711 | YGR235C | MIC26 | n |
| W303.asm02.HP0_G0022060 | 6025.134 | 6243.929 | 5711.846 | 13021.76 | 13826.47 | 17327.95 | 38209.41 | 23659.03 | 27789.82 | 6.14E-10 | -1.296804982 | YGR237C | | n |
| W303.asm02.HP0_G0022080 | 714.7566 | 583.1079 | 607.9413 | 1665.235 | 1553.438 | 1808.002 | 6243.965 | 4689.284 | 6945.662 | 3.02E-14 | -1.399131573 | YGR239C | PEX21 | n |
| W303.asm02.HP0_G0022090 | 44582.81 | 44036.89 | 46557.09 | 33709.02 | 34161.75 | 63859.69 | 91771.96 | 89355.19 | 103338.1 | 0.902566886 | 0.037257024 | YGR240C | PFK1 | n |
| W303.asm02.HP0_G0022130 | 10525.45 | 11667.72 | 10140.98 | 15657.51 | 17710.93 | 21661.84 | 4134.25 | 10382.57 | 7650.155 | 0.017571242 | -0.767165938 | YGR244C | LSC2 | n |
| W303.asm02.HP0_G0022140 | 2621.833 | 2681.851 | 3184.147 | 7588.384 | 7157.965 | 1959.935 | 13629.82 | 5306.719 | 12292.16 | 0.05895333 | -0.976917252 | YGR245C | SDA1 | n |
| W303.asm02.HP0_G0022150 | 2456.645 | 2529.398 | 2597.763 | 3060.919 | 2969.757 | 2727.956 | 3447.883 | 2904.629 | 3066.76 | 0.062451663 | -0.207825157 | YGR246C | BRF1 | n |
| W303.asm02.HP0_G0022160 | 1930.372 | 1918.47 | 1680.462 | 2013.982 | 2276.351 | 2228.856 | 1678.883 | 2263.57 | 1940.049 | 0.130934439 | -0.237456249 | YGR247W | CPD1 | n |
| W303.asm02.HP0_G0022180 | 2739.371 | 3974.927 | 2489.972 | 2962.27 | 2136.628 | 1720.64 | 8738.835 | 4617.34 | 6113.187 | 0.200229641 | 0.432519399 | YGR249W | MGA1 | n |
| W303.asm02.HP0_G0022190 | 8178.934 | 9200.642 | 7274.816 | 3667.406 | 3576.379 | 3963.17 | 2946.687 | 3239.654 | 2412.502 | 2.29E-13 | 1.137408616 | YGR250C | RIE1 | n |
| W303.asm02.HP0_G0022200 | 596.16 | 628.7328 | 727.5893 | 724.5889 | 551.08 | 148.8943 | 1839.365 | 729.1102 | 1760.636 | 0.433855781 | 0.454699483 | YGR251W | NOP19 | n |
| W303.asm02.HP0_G0022210 | 2871.733 | 2702.995 | 2783.164 | 3930.009 | 3693.538 | 3328.85 | 2923.232 | 2437.526 | 2516.561 | 0.001343619 | -0.39005796 | YGR252W | GCN5 | n |
| W303.asm02.HP0_G0022220 | 8532.606 | 8614.195 | 8313.921 | 5479.921 | 5915.215 | 7421.164 | 5088.498 | 6651.117 | 5650.302 | 0.009476171 | 0.43629797 | YGR253C | PUP2 | n |
| W303.asm02.HP0_G0022240 | 7257.692 | 6528.806 | 6847.963 | 3281.144 | 3368.097 | 3285.549 | 2920.763 | 2955.097 | 3126.564 | 8.41E-42 | 1.054623769 | YGR255C | COQ6 | n |
| W303.asm02.HP0_G0022260 | 3012.567 | 3030.158 | 2843.527 | 1254.657 | 1315.649 | 1306.623 | 913.5101 | 714.077 | 826.4943 | 2.34E-25 | 1.196805847 | YGR257C | MTM1 | n |
| W303.asm02.HP0_G0022270 | 4830.696 | 4849.588 | 4589.741 | 1091.399 | 985.8691 | 814.3605 | 2519.56 | 1523.722 | 2318.011 | 4.83E-27 | 2.302936909 | YGR258C | RAD2 | n |
| W303.asm02.HP0_G0022290 | 5074.243 | 4437.852 | 4616.689 | 6533.111 | 6845.542 | 5297.141 | 32776.5 | 34538.77 | 25984.93 | 0.018468749 | -0.4024815 | YGR260W | TNA1 | n |
| W303.asm02.HP0_G0022300 | 5046.711 | 5173.414 | 5108.216 | 5969.001 | 6127.836 | 8165.635 | 3352.829 | 5675.033 | 4094.197 | 0.089328862 | -0.402615245 | YGR261C | APL6 | n |
| W303.asm02.HP0_G0022310 | 2938.444 | 2551.654 | 2747.593 | 2450.264 | 2903.801 | 2270.638 | 2415.864 | 2434.304 | 2176.873 | 0.451115264 | 0.111883984 | YGR262C | BUD32 | n |
| W303.asm02.HP0_G0022320 | 1368.097 | 1283.06 | 1236.363 | 821.1545 | 840.0716 | 813.6008 | 543.1682 | 348.985 | 423.4139 | 0.000193784 | 0.651613999 | YGR263C | SAY1 | n |
| W303.asm02.HP0_G0022330 | 9239.95 | 8118.999 | 9613.881 | 14632.81 | 14670.01 | 16807.58 | 14976.63 | 14122.62 | 14244.17 | 3.55E-12 | -0.773567222 | YGR264C | MES1 | n |
| W303.asm02.HP0_G0022340 | 3392.711 | 3598.8 | 3493.507 | 10902.18 | 11815.68 | 15472.85 | 5719.314 | 10012.11 | 8357.041 | 1.15E-15 | -1.864896039 | YGR266W | | n |
| W303.asm02.HP0_G0022350 | 5227.783 | 5227.941 | 5291.461 | 3965.44 | 4582.208 | 3026.504 | 2544.249 | 2601.817 | 2296.482 | 0.010728726 | 0.444306412 | YGR267C | FOL2 | n |
| W303.asm02.HP0_G0022360 | 2006.613 | 1666.976 | 1423.919 | 1408.19 | 1540.42 | 1389.427 | 2047.991 | 1853.379 | 1990.284 | 0.152944971 | 0.233004421 | YGR268C | HUA1 | n |
| W303.asm02.HP0_G0022370 | 8285.882 | 8676.512 | 8224.454 | 15637.37 | 15443.26 | 20323.31 | 23832.74 | 11566.97 | 21951.74 | 0.000202867 | -1.029211839 | YGR270W | YTA7 | n |
| W303.asm02.HP0_G0022380 | 2626.069 | 2575.022 | 2583.751 | 5648.737 | 5531.628 | 5252.321 | 3930.562 | 5367.926 | 5217.32 | 4.06E-14 | -1.077828189 | YGR271W | SLH1 | n |
| W303.asm02.HP0_G0022390 | 858.7669 | 757.8177 | 815.978 | 2146.673 | 2157.456 | 689.0158 | 2813.364 | 968.5675 | 2523.738 | 0.038135825 | -1.037433479 | YGR271C- | EFG1 | n |
| W303.asm02.HP0_G0022410 | 2490.53 | 2638.452 | 2446.856 | 6000.263 | 5806.734 | 5987.677 | 5213.18 | 5356.114 | 4598.944 | 8.72E-40 | -1.232064939 | YGR274C | TAF1 | n |
| W303.asm02.HP0_G0022420 | 1834.013 | 1682.556 | 1518.775 | 1705.528 | 1987.359 | 1743.43 | 1239.411 | 2076.729 | 1767.813 | 0.638296281 | -0.110395291 | YGR275W | RTT102 | n |
| W303.asm02.HP0_G0022430 | 2922.56 | 3098.039 | 2980.422 | 8514.441 | 8313.931 | 6667.576 | 6680.968 | 4841.764 | 6710.033 | 1.54E-15 | -1.384260771 | YGR276C | RNH70 | n |
| W303.asm02.HP0_G0022440 | 2586.89 | 2310.176 | 2542.79 | 1679.824 | 1776.474 | 1772.298 | 2313.403 | 2745.706 | 2172.089 | 5.68E-05 | 0.509025106 | YGR277C | CAB4 | n |
| W303.asm02.HP0_G0022450 | 1209.262 | 1191.81 | 1152.286 | 1895.88 | 1740.892 | 1997.158 | 2436.85 | 2360.212 | 2272.56 | 1.35E-12 | -0.66532081 | YGR278W | CWC22 | n |
| W303.asm02.HP0_G0022460 | 16256.21 | 14844.77 | 16849.89 | 21645.97 | 24529.57 | 22005.2 | 18141.82 | 17050.87 | 16863.59 | 3.12E-07 | -0.507757065 | YGR279C | SCW4 | n |
| W303.asm02.HP0_G0022470 | 3089.866 | 2977.856 | 3040.785 | 1685.381 | 1710.518 | 281.8356 | 2257.851 | 719.446 | 2027.363 | 0.035642189 | 1.308428729 | YGR280C | PXR1 | n |
| W303.asm02.HP0_G0022480 | 5084.832 | 5254.648 | 4967.01 | 5619.559 | 6052.334 | 6224.692 | 2884.964 | 6333.272 | 3137.329 | 0.485261956 | -0.225534675 | YGR281W | YOR1 | n |
| W303.asm02.HP0_G0022490 | 25489.81 | 25356.29 | 26202.92 | 15449.79 | 16943.76 | 16639.69 | 24387.02 | 20396.83 | 20701.83 | 5.24E-10 | 0.652046667 | YGR282C | BGL2 | n |
| W303.asm02.HP0_G0022510 | 4874.111 | 4941.951 | 4504.586 | 4184.275 | 4283.671 | 5189.269 | 6025.463 | 6661.855 | 6676.543 | 0.623726767 | 0.068414381 | YGR284C | ERV29 | n |
| W303.asm02.HP0_G0022520 | 19852.23 | 19521.87 | 21161.53 | 13266.3 | 13419.45 | 11031.09 | 7451.28 | 5273.431 | 7117.898 | 5.53E-05 | 0.682584658 | YGR285C | ZUO1 | n |
| W303.asm02.HP0_G0022530 | 3457.304 | 3554.288 | 3736.037 | 1797.925 | 2088.897 | 2997.637 | 7322.894 | 6246.294 | 5826.127 | 0.002912912 | 0.642631959 | YGR286C | BIO2 | n |
| W303.asm02.HP0_G0022660 | 984.7758 | 832.3754 | 977.6645 | 314.7064 | 374.908 | 344.1281 | 1377.672 | 1457.147 | 2205.579 | 4.27E-11 | 1.435595488 | YHL047C | ARN2 | n |
| W303.asm02.HP0_G0022710 | 553.804 | 499.6478 | 535.7213 | 5218.013 | 5319.875 | 6522.48 | 7771.008 | 11125.64 | 7140.623 | 1.21E-64 | -3.424217894 | YHL040C | ARN1 | n |
| W303.asm02.HP0_G0022720 | 2487.353 | 2502.69 | 2805.8 | 5765.449 | 5548.985 | 5904.113 | 4849.01 | 4989.948 | 4913.514 | 1.08E-42 | -1.143237569 | YHL039W | EFM1 | n |
| W303.asm02.HP0_G0022750 | 1571.406 | 1216.292 | 1307.505 | 351.5264 | 334.1194 | 481.6274 | 693.7739 | 634.6158 | 491.5907 | 6.58E-17 | 1.810534675 | YHL036W | MUP3 | n |

| | | | | | | | | | | | | | |
|---|---|---|---|---|---|---|---|---|---|---|---|---|---|
| W303.asm02.HP0_G0022770 | 27179.81 | 28975.12 | 27502.88 | 4645.567 | 4620.394 | 7193.264 | 7727.802 | 7368.415 | 7890.568 | 9.34E-35 | 2.345578302 | YHL034C | SBP1 | n |
| W303.asm02.HP0_G0022790 | 2010.849 | 1853.927 | 1902.511 | 5224.96 | 3003.603 | 1266.361 | 39896.94 | 15148.1 | 10384.4 | 0.226950099 | -0.719294314 | YHL032C | GUT1 | n |
| W303.asm02.HP0_G0022800 | 3856.509 | 3474.166 | 3556.026 | 1775 | 1861.522 | 1966.012 | 1872.696 | 2319.408 | 1883.833 | 2.41E-14 | 0.958516346 | YHL031C | GOS1 | n |
| W303.asm02.HP0_G0022810 | 7000.379 | 7348.94 | 6881.378 | 4698.365 | 4586.548 | 7459.907 | 2822.005 | 8048.13 | 5768.715 | 0.387708416 | 0.34242787 | YHL030W | ECM29 | n |
| W303.asm02.HP0_G0022820 | 663.9295 | 698.8393 | 692.0183 | 4223.179 | 3932.194 | 5057.847 | 13742.15 | 9205.687 | 12818.44 | 9.96E-47 | -2.684997898 | YHL029C | OCA5 | n |
| W303.asm02.HP0_G0022830 | 850.2957 | 747.8025 | 876.3409 | 3085.929 | 2850.863 | 3463.311 | 6129.159 | 3460.857 | 4139.648 | 2.89E-17 | -1.925551096 | YHL028W | WSC4 | n |
| W303.asm02.HP0_G0022840 | 2322.165 | 2364.703 | 2235.586 | 6518.522 | 7111.969 | 7428.001 | 13000.24 | 9049.986 | 9957.402 | 6.42E-24 | -1.605031424 | YHL027W | RIM101 | n |
| W303.asm02.HP0_G0022860 | 5191.78 | 4949.741 | 4975.633 | 2271.722 | 2295.443 | 1902.2 | 2462.774 | 2056.327 | 1943.637 | 2.80E-19 | 1.224559221 | YHL025W | SNF6 | n |
| W303.asm02.HP0_G0022870 | 1657.176 | 1423.273 | 1518.775 | 1467.935 | 1227.129 | 1621.124 | 1539.388 | 1893.109 | 1658.969 | 0.601271141 | 0.091360108 | YHL024W | RIM4 | n |
| W303.asm02.HP0_G0022880 | 2090.266 | 2339.108 | 2312.117 | 3597.24 | 3238.788 | 3323.533 | 2222.052 | 1336.881 | 1684.087 | 0.00321457 | -0.591768208 | YHL023C | NPR3 | n |
| W303.asm02.HP0_G0022900 | 16335.63 | 17873.82 | 14448.31 | 76.41878 | 48.59918 | 59.25384 | 1199.908 | 917.0251 | 720.0428 | 1.95E-234 | 8.0386329 | YHL021C | AIM17 | n |
| W303.asm02.HP0_G0022910 | 1932.49 | 2035.314 | 1885.265 | 1441.536 | 1531.742 | 1299.786 | 2004.784 | 1730.965 | 1627.871 | 0.000407763 | 0.45411191 | YHL020C | OPI1 | n |
| W303.asm02.HP0_G0022920 | 4083.113 | 4290.962 | 4056.176 | 1217.143 | 1239.279 | 1474.509 | 907.3377 | 847.2281 | 767.8861 | 1.55E-42 | 1.660854083 | YHL019C | APM2 | n |
| W303.asm02.HP0_G0022930 | 366.379 | 316.0356 | 278.1008 | 394.5988 | 434.7891 | 353.244 | 380.2177 | 229.7932 | 227.256 | 0.267413237 | -0.299689109 | YHL018W | MCO14 | n |
| W303.asm02.HP0_G0022950 | 562.2752 | 442.8949 | 478.5921 | 1178.239 | 1241.015 | 2159.727 | 1359.155 | 1713.785 | 1007.103 | 6.96E-08 | -1.625696153 | YHL016C | DUR3 | n |
| W303.asm02.HP0_G0022960 | 17981.16 | 19017.78 | 19050.98 | 15880.52 | 18491.99 | 16214.28 | 18256.62 | 15326.35 | 14674.76 | 0.276689543 | 0.147986227 | YHL015W | RPS20 | n |
| W303.asm02.HP0_G0022970 | 2867.498 | 2845.433 | 2745.437 | 1178.239 | 1040.543 | 942.7438 | 330.8388 | 394.0846 | 344.4723 | 5.78E-29 | 1.419395804 | YHL014C | YLF2 | n |
| W303.asm02.HP0_G0022980 | 1050.428 | 1146.185 | 1085.456 | 2004.256 | 2009.055 | 1219.262 | 1333.231 | 906.2871 | 1254.692 | 0.00716265 | -0.672894928 | YHL013C | OTU2 | n |
| W303.asm02.HP0_G0023000 | 4516.203 | 4392.227 | 4641.481 | 7698.844 | 8327.817 | 6081.875 | 7334.005 | 5844.693 | 6591.621 | 8.95E-06 | -0.706270728 | YHL011C | PRS3 | n |
| W303.asm02.HP0_G0023010 | 1647.646 | 1824.994 | 1625.488 | 1414.442 | 1343.42 | 1555.793 | 1444.334 | 1170.442 | 1320.477 | 0.073691781 | 0.2407971 | YHL010C | ETP1 | n |
| W303.asm02.HP0_G0023020 | 1915.548 | 2006.381 | 1832.447 | 2594.07 | 2216.47 | 2935.344 | 9469.643 | 3980.576 | 6721.994 | 0.173493465 | -0.428825767 | YHL009C | YAP3 | n |
| W303.asm02.HP0_G0023030 | 992.1881 | 1102.786 | 987.3657 | 2475.968 | 2233.827 | 1521.608 | 1617.16 | 988.9697 | 1478.36 | 6.56E-05 | -1.015588848 | YHL008C | | n |
| W303.asm02.HP0_G0023040 | 2733.018 | 2834.305 | 3033.239 | 11737.23 | 11799.19 | 11678.32 | 10389.33 | 9155.218 | 9739.715 | 6.30E-123 | -2.033687287 | YHL007C | STE20 | n |
| W303.asm02.HP0_G0023050 | 348.3777 | 347.194 | 362.1778 | 113.2387 | 121.498 | 50.89753 | 75.30286 | 37.583 | 58.60813 | 3.77E-07 | 1.889669203 | YHL006C | SHU1 | n |
| W303.asm02.HP0_G0023060 | 14120.41 | 14552.1 | 15132.78 | 6419.177 | 6363.021 | 4661.302 | 2271.43 | 2306.522 | 2222.325 | 1.59E-19 | 1.328433751 | YHL004W | MRP4 | n |
| W303.asm02.HP0_G0023070 | 3851.215 | 3692.275 | 3727.413 | 3515.958 | 3552.079 | 5307.777 | 10473.27 | 8705.296 | 11128.37 | 0.540868538 | -0.134938613 | YHL003C | LAG1 | n |
| W303.asm02.HP0_G0023080 | 8016.922 | 8458.403 | 8050.911 | 4697.671 | 4166.512 | 5552.389 | 8841.296 | 7772.164 | 8132.177 | 1.54E-08 | 0.766486776 | YHL002W | HSE1 | n |
| W303.asm02.HP0_G0023090 | 3208.463 | 3124.746 | 3025.694 | 3233.209 | 3959.965 | 3548.393 | 3114.576 | 2655.507 | 2728.268 | 0.123313646 | -0.198547506 | YHL001W | RPL14B | y |
| W303.asm02.HP0_G0023100 | 3628.846 | 3464.151 | 3397.573 | 5757.807 | 6197.263 | 7465.224 | 9583.215 | 9817.753 | 8456.316 | 4.72E-11 | -0.888445113 | YHR001W | OSH7 | n |
| W303.asm02.HP0_G0023120 | 900.0639 | 974.814 | 737.2905 | 2111.937 | 1991.699 | 1590.738 | 1499.885 | 1184.401 | 967.6322 | 2.54E-07 | -1.124364767 | YHR002W | LEU5 | n |
| W303.asm02.HP0_G0023140 | 1613.762 | 1724.842 | 1527.399 | 1579.09 | 1559.513 | 1960.694 | 1735.669 | 1921.028 | 1667.342 | 0.666333178 | -0.067705586 | YHR004C | NEM1 | n |
| W303.asm02.HP0_G0023150 | 2684.308 | 2893.284 | 3011.681 | 137.5538 | 126.705 | 102.5547 | 208.626 | 333.9518 | 190.1774 | 2.00E-82 | 4.549002696 | YHR005C | GPA1 | n |
| W303.asm02.HP0_G0023160 | 6136.318 | 6351.87 | 5758.196 | 2339.109 | 2733.704 | 1778.375 | 1043.13 | 1012.593 | 1082.456 | 1.50E-16 | 1.413383639 | YHR005C- | TIM10 | n |
| W303.asm02.HP0_G0023180 | 19286.78 | 20087.18 | 22014.16 | 10080.33 | 9733.722 | 13724.1 | 49683.84 | 32496.41 | 37774.74 | 4.74E-05 | 0.872145317 | YHR007C | ERG11 | n |
| W303.asm02.HP0_G0023210 | 12564.89 | 12239.7 | 11901.21 | 5771.007 | 6954.022 | 6233.048 | 2825.709 | 5411.952 | 4796.298 | 8.18E-05 | 0.953228521 | YHR008C | SOD2 | n |
| W303.asm02.HP0_G0023220 | 8451.071 | 8569.683 | 8839.941 | 12104.04 | 11708.93 | 15846.6 | 20625.58 | 15370.37 | 18105.13 | 0.000402776 | -0.616920938 | YHR009C | TDA3 | n |
| W303.asm02.HP0_G0023240 | 4382.782 | 4257.578 | 4111.149 | 1714.559 | 1641.09 | 1671.262 | 625.8779 | 846.1543 | 653.062 | 2.30E-23 | 1.342844311 | YHR011W | DIA4 | n |
| W303.asm02.HP0_G0023260 | 3644.729 | 3471.94 | 3546.324 | 2359.951 | 2568.814 | 2342.806 | 3054.086 | 2474.035 | 2832.328 | 9.44E-07 | 0.552483848 | YHR013C | ARD1 | n |
| W303.asm02.HP0_G0023300 | 3123.751 | 3289.441 | 2900.656 | 3088.013 | 3493.066 | 3832.508 | 5099.608 | 5004.981 | 5033.123 | 0.191092543 | -0.160929517 | YHR017W | YSC83 | n |
| W303.asm02.HP0_G0023310 | 5872.652 | 6099.264 | 5749.573 | 6186.447 | 6533.118 | 7886.078 | 6240.261 | 5255.177 | 4183.903 | 0.261595107 | -0.217540364 | YHR018C | ARG4 | n |
| W303.asm02.HP0_G0023320 | 18803.92 | 17457.63 | 20382.2 | 18325.92 | 19613.24 | 18242.59 | 12502.74 | 13901.41 | 12985.89 | 0.919450817 | 0.011850314 | YHR019C | DED81 | n |
| W303.asm02.HP0_G0023330 | 14590.57 | 13372.53 | 15775.21 | 14938.48 | 16136.66 | 14595.44 | 8096.909 | 8894.285 | 8199.158 | 0.592213238 | -0.062321434 | YHR020W | | n |
| W303.asm02.HP0_G0023350 | 654.3994 | 618.7176 | 552.9679 | 3809.129 | 2832.638 | 1864.977 | 13059.49 | 7490.828 | 11368.78 | 1.46E-13 | -2.219903834 | YHR022C | | n |
| W303.asm02.HP0_G0023360 | 4189.003 | 4516.861 | 4474.405 | 5743.913 | 5689.576 | 6762.534 | 1987.502 | 4573.314 | 3722.214 | 0.120554586 | -0.465251276 | YHR023W | MYO1 | n |
| W303.asm02.HP0_G0023370 | 10470.39 | 10936.61 | 10400.75 | 2987.279 | 2903.801 | 2467.391 | 1660.366 | 1703.047 | 1668.538 | 3.23E-80 | 1.928055575 | YHR024C | MAS2 | n |
| W303.asm02.HP0_G0023380 | 9355.37 | 8531.848 | 9977.136 | 6760.977 | 6988.736 | 7168.955 | 8058.64 | 6609.239 | 7134.643 | 0.00048944 | 0.413663011 | YHR025W | THR1 | n |
| W303.asm02.HP0_G0023390 | 5297.67 | 4788.384 | 4976.711 | 2033.434 | 2242.505 | 2649.71 | 4057.713 | 3051.739 | 3134.937 | 1.56E-11 | 1.121030276 | YHR026W | VMA16 | n |
| W303.asm02.HP0_G0023400 | 18654.62 | 19922.48 | 19196.5 | 18985.9 | 18983.19 | 24610.09 | 11116.43 | 21850.75 | 16029.92 | 0.684317936 | -0.11527498 | YHR027C | RPN1 | n |
| W303.asm02.HP0_G0023410 | 7884.56 | 7624.915 | 7313.62 | 3037.994 | 3240.524 | 5099.629 | 913.5101 | 4893.306 | 2638.562 | 0.050008378 | 1.004228152 | YHR028C | DAP2 | n |
| W303.asm02.HP0_G0023430 | 13021.28 | 12530.14 | 12986.66 | 7167.386 | 6455.88 | 7157.56 | 13237.25 | 11711.94 | 11421.41 | 9.29E-20 | 0.890937842 | YHR030C | SLT2 | n |
| W303.asm02.HP0_G0023440 | 636.3981 | 569.7543 | 617.6425 | 2993.532 | 2948.061 | 2593.495 | 725.8702 | 1470.032 | 1161.398 | 3.80E-19 | -2.226404013 | YHR031C | RRM3 | n |
| W303.asm02.HP0_G0023450 | 1193.379 | 1027.116 | 1282.713 | 3971.692 | 3448.806 | 3299.223 | 8540.085 | 6236.63 | 7564.037 | 6.53E-22 | -1.613500601 | YHR032W | ERC1 | n |
| W303.asm02.HP0_G0023470 | 1178.554 | 1235.21 | 1088.689 | 1241.458 | 1314.781 | 1107.591 | 733.277 | 1160.778 | 985.5735 | 0.762981663 | -0.064902201 | YHR034C | PIH1 | n |
| W303.asm02.HP0_G0023480 | 536.8616 | 523.0166 | 558.3575 | 978.1603 | 1003.226 | 1160.768 | 809.8143 | 1189.77 | 760.7096 | 2.60E-06 | -0.957305512 | YHR035W | NEL1 | n |

| | | | | | | | | | | | | | | |
|---|---|---|---|---|---|---|---|---|---|---|---|---|---|---|
| W303.asm02.HP0_G0023490 | 634.2803 | 536.3703 | 659.681 | 2735.792 | 2442.109 | 3259.721 | 5007.023 | 4594.79 | 5491.223 | 9.23E-43 | -2.204646468 | YHR036W | BRL1 | n |
| W303.asm02.HP0_G0023500 | 4329.837 | 4669.315 | 4357.991 | 5123.532 | 4826.072 | 8508.244 | 10223.91 | 12505.47 | 7442.037 | 0.106968 | -0.466646233 | YHR037W | PUT2 | n |
| W303.asm02.HP0_G0023510 | 2973.388 | 2970.067 | 2542.79 | 1938.258 | 1888.425 | 2101.232 | 1364.093 | 1537.681 | 1525.008 | 9.25E-06 | 0.517492889 | YHR038W | RRF1 | n |
| W303.asm02.HP0_G0023520 | 9280.188 | 9108.279 | 9708.737 | 6278.844 | 6369.964 | 8237.043 | 7489.548 | 8564.628 | 9251.712 | 0.006190018 | 0.427890934 | YHR039C | MSC7 | n |
| W303.asm02.HP0_G0023530 | 3727.323 | 3122.521 | 3382.482 | 1777.779 | 2047.24 | 2100.473 | 1193.735 | 1342.25 | 1069.299 | 2.48E-08 | 0.788363812 | YHR039C- | VMA10 | n |
| W303.asm02.HP0_G0023540 | 1424.219 | 1227.42 | 1422.841 | 919.1095 | 770.6442 | 235.496 | 1527.043 | 643.2062 | 1400.615 | 0.03035819 | 1.08155043 | YHR040W | BCD1 | n |
| W303.asm02.HP0_G0023560 | 7602.893 | 6449.797 | 8122.053 | 10029.62 | 10454.9 | 13041.92 | 6643.934 | 6269.918 | 5008.005 | 0.001032164 | -0.596367268 | YHR042W | NCP1 | n |
| W303.asm02.HP0_G0023570 | 5361.204 | 4906.341 | 4765.441 | 689.8531 | 550.2122 | 2057.931 | 22120.52 | 14967.7 | 16630.36 | 1.13E-06 | 2.188375586 | YHR043C | DOG2 | n |
| W303.asm02.HP0_G0023590 | 2072.265 | 2049.78 | 2057.73 | 1868.786 | 1894.5 | 2095.155 | 3926.859 | 2525.577 | 3655.234 | 0.709906475 | 0.077035913 | YHR045W | DDE1 | n |
| W303.asm02.HP0_G0023640 | 3474.247 | 3635.522 | 3590.519 | 744.041 | 886.9351 | 638.1183 | 686.367 | 324.2876 | 351.6488 | 1.79E-14 | 2.237670947 | YHR049W | FSH1 | n |
| W303.asm02.HP0_G0023650 | 1727.064 | 1843.912 | 1766.695 | 1856.976 | 1530.874 | 1571.746 | 1662.835 | 995.4125 | 1306.124 | 0.64382639 | 0.105804325 | YHR050W | SMF2 | n |
| W303.asm02.HP0_G0023670 | 3556.841 | 3585.446 | 3437.455 | 9302.249 | 10940.02 | 10638.34 | 4992.209 | 10147.41 | 9337.83 | 2.12E-09 | -1.54538426 | YHR051W | COX6 | n |
| W303.asm02.HP0_G0023680 | 3422.361 | 3560.964 | 3686.453 | 7675.224 | 7155.362 | 2411.935 | 6971.07 | 3536.023 | 6407.424 | 0.11793457 | -0.692436979 | YHR052W | CIC1 | n |
| W303.asm02.HP0_G0023860 | 892.6516 | 923.6251 | 918.3794 | 2270.332 | 2403.924 | 2808.48 | 2797.316 | 2777.92 | 2347.914 | 5.02E-28 | -1.452210238 | YHR058C | MED6 | n |
| W303.asm02.HP0_G0023870 | 2548.769 | 2272.34 | 2234.508 | 1972.299 | 1917.064 | 1537.561 | 585.1402 | 632.4682 | 584.8852 | 0.007901771 | 0.378692158 | YHR059W | FYV4 | n |
| W303.asm02.HP0_G0023880 | 1360.685 | 1315.331 | 1263.311 | 980.9392 | 1087.407 | 1022.509 | 991.2819 | 1285.339 | 965.24 | 0.020264534 | 0.350253674 | YHR060W | VMA22 | n |
| W303.asm02.HP0_G0023900 | 2514.884 | 2295.709 | 2492.128 | 1913.943 | 1993.434 | 1310.421 | 1681.352 | 1836.198 | 1650.596 | 0.00949259 | 0.48512995 | YHR062C | RPP1 | n |
| W303.asm02.HP0_G0023910 | 3834.272 | 3872.549 | 3870.775 | 2523.904 | 2731.968 | 4036.858 | 5289.717 | 6310.722 | 6755.484 | 0.147185623 | 0.317156709 | YHR063C | PAN5 | n |
| W303.asm02.HP0_G0023920 | 15472.63 | 14565.46 | 16585.8 | 17933.4 | 18499.8 | 15583 | 11500.35 | 8877.104 | 10834.13 | 0.304735521 | -0.157876736 | YHR064C | SSZ1 | n |
| W303.asm02.HP0_G0023930 | 4154.06 | 4322.121 | 4695.377 | 2443.317 | 2067.201 | 603.9334 | 4007.1 | 1801.836 | 3979.373 | 0.005556899 | 1.364765736 | YHR065C | RRP3 | n |
| W303.asm02.HP0_G0023950 | 1388.216 | 1449.98 | 1433.62 | 1022.622 | 999.7546 | 935.1471 | 896.2275 | 1240.239 | 1388.654 | 0.003409781 | 0.530421514 | YHR067W | HTD2 | n |
| W303.asm02.HP0_G0023960 | 5983.837 | 5223.49 | 6049.232 | 7158.355 | 7730.741 | 6779.247 | 15555.6 | 10206.47 | 11731.2 | 0.084378234 | -0.328404523 | YHR068W | DYS1 | n |
| W303.asm02.HP0_G0023980 | 1376.568 | 1355.392 | 1533.866 | 4786.594 | 4753.173 | 2309.38 | 4353.987 | 1860.895 | 3648.057 | 7.10E-05 | -1.473870384 | YHR070W | TRM5 | n |
| W303.asm02.HP0_G0023990 | 2631.363 | 2603.955 | 2523.388 | 2523.209 | 2396.113 | 3913.792 | 7753.725 | 3879.639 | 4431.492 | 0.579833879 | -0.187133679 | YHR071W | PCL5 | n |
| W303.asm02.HP0_G0024000 | 6180.792 | 5961.277 | 6717.536 | 3455.518 | 3679.652 | 4792.724 | 6583.445 | 5791.003 | 6420.581 | 3.08E-05 | 0.660970274 | YHR072W | ERG7 | n |
| W303.asm02.HP0_G0024010 | 2525.473 | 1984.125 | 2210.794 | 2577.397 | 2982.775 | 1612.768 | 2198.597 | 1232.722 | 1913.735 | 0.776223805 | -0.093920267 | YHR072W- | NOP10 | n |
| W303.asm02.HP0_G0024020 | 7750.08 | 7646.058 | 7507.644 | 11356.52 | 10951.3 | 14585.56 | 9284.472 | 13221.7 | 11489.59 | 0.000149732 | -0.687787962 | YHR073W | OSH3 | n |
| W303.asm02.HP0_G0024030 | 7244.985 | 6375.239 | 7195.05 | 5937.044 | 5810.206 | 7898.233 | 4016.975 | 7109.629 | 5863.205 | 0.753910749 | 0.083440118 | YHR074W | QNS1 | n |
| W303.asm02.HP0_G0024040 | 1800.128 | 1634.705 | 1537.1 | 1385.959 | 1586.416 | 2266.839 | 3904.638 | 3830.244 | 3949.47 | 0.737105659 | -0.075484668 | YHR075C | PPE1 | n |
| W303.asm02.HP0_G0024050 | 3439.303 | 3185.95 | 3007.369 | 2607.27 | 2764.078 | 2817.596 | 1667.773 | 1783.582 | 1369.517 | 0.093422748 | 0.234334838 | YHR076W | PTC7 | n |
| W303.asm02.HP0_G0024060 | 6811.895 | 7068.514 | 7352.425 | 4267.641 | 3871.445 | 3027.263 | 2662.758 | 2385.983 | 2823.955 | 3.55E-09 | 0.927096695 | YHR077C | NMD2 | n |
| W303.asm02.HP0_G0024070 | 2723.487 | 2508.254 | 2691.542 | 1169.207 | 1139.477 | 1474.509 | 1219.659 | 1639.692 | 1454.439 | 1.14E-10 | 1.066402383 | YHR078W | | n |
| W303.asm02.HP0_G0024080 | 1786.362 | 1941.838 | 1693.397 | 3675.743 | 3160.682 | 3397.22 | 4340.407 | 3460.857 | 4814.239 | 5.64E-09 | -0.916684302 | YHR079C | IRE1 | n |
| W303.asm02.HP0_G0024090 | 26.47247 | 14.46642 | 15.09074 | 28.48336 | 39.05291 | 24.30927 | 11.11026 | 39.7306 | 16.74518 | 0.259753444 | -0.708891741 | YHR079C- | SAE3 | n |
| W303.asm02.HP0_G0024110 | 1686.826 | 1880.634 | 1824.902 | 1012.201 | 823.5825 | 455.0391 | 1060.412 | 573.4092 | 1046.574 | 0.000298854 | 1.234938236 | YHR081W | LRP1 | n |
| W303.asm02.HP0_G0024120 | 5548.629 | 6074.783 | 5195.527 | 6539.363 | 6100.065 | 9488.971 | 8389.479 | 6710.176 | 5905.068 | 0.087044914 | -0.39584269 | YHR082C | KSP1 | n |
| W303.asm02.HP0_G0024130 | 2738.312 | 2281.243 | 2735.736 | 1973.689 | 1877.143 | 2091.357 | 1378.906 | 1523.722 | 1725.95 | 0.006517866 | 0.384159869 | YHR083W | SAM35 | n |
| W303.asm02.HP0_G0024140 | 1065.252 | 875.7747 | 1042.339 | 3581.262 | 3772.511 | 4143.211 | 2305.996 | 2550.275 | 1972.343 | 1.36E-39 | -1.946051187 | YHR084W | STE12 | n |
| W303.asm02.HP0_G0024150 | 596.16 | 600.9127 | 737.2905 | 1267.857 | 802.7543 | 207.3884 | 3256.54 | 813.9403 | 2963.897 | 0.735169866 | -0.235995495 | YHR085W | IPI1 | n |
| W303.asm02.HP0_G0024160 | 1984.376 | 2139.917 | 1989.822 | 4770.616 | 4476.332 | 4813.995 | 6466.17 | 5995.025 | 6893.034 | 1.54E-37 | -1.201664159 | YHR086W | NAM8 | n |
| W303.asm02.HP0_G0024180 | 7050.148 | 7585.967 | 6487.941 | 73.63991 | 51.20271 | 219.5431 | 470.3342 | 512.2026 | 754.7292 | 1.90E-37 | 5.937731305 | YHR087W | RTC3 | n |
| W303.asm02.HP0_G0024190 | 2266.043 | 2451.501 | 2547.102 | 2520.43 | 2271.144 | 977.6884 | 4344.111 | 2452.559 | 4035.589 | 0.398758192 | 0.33250079 | YHR088W | RPF1 | n |
| W303.asm02.HP0_G0024200 | 8129.165 | 8136.804 | 8186.728 | 8542.23 | 9263.351 | 4608.126 | 7343.88 | 4094.399 | 6250.737 | 0.714879007 | 0.125629969 | YHR089C | GAR1 | n |
| W303.asm02.HP0_G0024210 | 1763.066 | 1784.933 | 1641.657 | 967.7396 | 1109.971 | 686.7368 | 2670.165 | 1158.63 | 1796.519 | 0.004602053 | 0.90879288 | YHR090C | YNG2 | n |
| W303.asm02.HP0_G0024230 | 14358.67 | 14637.79 | 13599.99 | 2723.982 | 2860.409 | 4011.029 | 5810.665 | 6684.405 | 7025.799 | 5.58E-34 | 2.150301863 | YHR092C | HXT4 | n |
| W303.asm02.HP0_G0024260 | 1034.544 | 1150.637 | 999.2227 | 32073.65 | 31148.6 | 48553.96 | 43871.94 | 57226.02 | 42934.64 | 1.18E-124 | -5.133495256 | YHR096C | HXT5 | n |
| W303.asm02.HP0_G0024280 | 5974.307 | 5409.327 | 5908.025 | 6314.275 | 6026.298 | 7307.214 | 7268.578 | 7190.164 | 7824.784 | 0.1233908 | -0.184316665 | YHR098C | SFB3 | n |
| W303.asm02.HP0_G0024290 | 6658.355 | 6133.761 | 6486.863 | 13249.63 | 12937.8 | 12996.34 | 15900.01 | 14757.23 | 17328.87 | 6.71E-29 | -1.023210914 | YHR099W | TRA1 | n |
| W303.asm02.HP0_G0024300 | 965.7156 | 905.8203 | 958.2621 | 1114.325 | 1302.632 | 660.1486 | 962.889 | 477.841 | 741.5723 | 0.737305987 | -0.120700847 | YHR100C | GEP4 | n |
| W303.asm02.HP0_G0024320 | 2374.051 | 2537.187 | 2427.454 | 4922.064 | 4456.371 | 6673.654 | 8305.535 | 5575.169 | 6406.228 | 1.29E-07 | -1.129215331 | YHR102W | KIC1 | n |
| W303.asm02.HP0_G0024330 | 3042.216 | 3486.407 | 3070.966 | 7514.744 | 7495.556 | 7866.327 | 7405.604 | 5613.826 | 6405.032 | 8.78E-21 | -1.252882942 | YHR103W | SBE22 | n |
| W303.asm02.HP0_G0024340 | 11222.21 | 9998.52 | 10185.17 | 3493.033 | 4126.591 | 9621.912 | 15922.23 | 10891.55 | 12149.82 | 0.02326094 | 0.86514491 | YHR104W | GRE3 | n |
| W303.asm02.HP0_G0024370 | 5819.707 | 5932.344 | 6594.654 | 11310.67 | 11591.77 | 11642.62 | 8095.675 | 8174.839 | 8104.667 | 1.21E-31 | -0.912937629 | YHR107C | CDC12 | n |
| W303.asm02.HP0_G0024380 | 11507.05 | 11269.34 | 11693.17 | 7852.377 | 7497.291 | 13325.28 | 11434.92 | 14166.64 | 12767 | 0.302025481 | 0.265517951 | YHR108W | GGA2 | n |

| | | | | | | | | | | | | | | |
|---|---|---|---|---|---|---|---|---|---|---|---|---|---|---|
| W303.asm02.HP0_G0024390 | 2113.562 | 2187.768 | 2117.016 | 1267.857 | 1061.371 | 1230.657 | 2099.839 | 1923.176 | 1989.088 | 6.74E-16 | 0.849509024 | YHR109W | CTM1 | n |
| W303.asm02.HP0_G0024400 | 2513.826 | 2429.245 | 2398.35 | 566.1937 | 561.4941 | 583.4224 | 518.4787 | 841.8591 | 714.0623 | 3.79E-31 | 2.101090327 | YHR110W | ERP5 | n |
| W303.asm02.HP0_G0024410 | 1970.61 | 1867.281 | 1953.173 | 2736.487 | 2390.906 | 1994.12 | 3504.669 | 2818.725 | 3301.193 | 0.071164605 | -0.298443259 | YHR111W | UBA4 | n |
| W303.asm02.HP0_G0024420 | 4037.581 | 3416.3 | 3950.541 | 2632.974 | 2803.131 | 3217.18 | 2324.513 | 3320.189 | 2859.838 | 0.025747446 | 0.398324786 | YHR112C | | n |
| W303.asm02.HP0_G0024430 | 7340.286 | 6941.655 | 7073.246 | 4851.203 | 4990.962 | 6202.662 | 3315.795 | 4499.222 | 3588.253 | 0.015178324 | 0.412478079 | YHR113W | APE4 | n |
| W303.asm02.HP0_G0024440 | 4794.693 | 4388.889 | 4327.809 | 3911.947 | 3887.067 | 5724.833 | 6951.318 | 7329.758 | 7487.488 | 0.994485197 | -0.001349653 | YHR114W | BZZ1 | n |
| W303.asm02.HP0_G0024450 | 2676.896 | 2554.992 | 2789.631 | 6874.911 | 6626.845 | 8211.215 | 12078.08 | 8786.905 | 9203.869 | 3.42E-18 | -1.436615499 | YHR115C | DMA1 | n |
| W303.asm02.HP0_G0024460 | 2933.149 | 2807.598 | 2593.452 | 716.2523 | 705.556 | 354.0037 | 162.9504 | 105.2324 | 160.2753 | 4.46E-14 | 2.230661502 | YHR116W | COX23 | n |
| W303.asm02.HP0_G0024480 | 1628.586 | 1628.028 | 1561.892 | 723.8942 | 657.8246 | 517.3316 | 645.6294 | 429.52 | 665.0229 | 2.44E-10 | 1.343179772 | YHR118C | ORC6 | n |
| W303.asm02.HP0_G0024490 | 3567.43 | 3734.561 | 3813.646 | 3200.557 | 3101.669 | 2990.04 | 4977.395 | 3187.038 | 4868.063 | 0.194733183 | 0.258469764 | YHR119W | SET1 | n |
| W303.asm02.HP0_G0024500 | 1091.725 | 1011.536 | 1004.612 | 3062.309 | 2798.792 | 2837.347 | 5652.652 | 4095.473 | 4978.103 | 2.15E-24 | -1.484859166 | YHR120W | MSH1 | n |
| W303.asm02.HP0_G0024510 | 3766.503 | 3573.205 | 3557.103 | 2874.735 | 3004.471 | 2683.135 | 2782.502 | 3067.846 | 3240.192 | 0.000644504 | 0.34798208 | YHR121W | LSM12 | n |
| W303.asm02.HP0_G0024520 | 1436.926 | 1398.791 | 1429.309 | 1220.616 | 1053.561 | 909.3185 | 1271.507 | 1494.73 | 1231.967 | 0.006657205 | 0.421650962 | YHR122W | CIA2 | n |
| W303.asm02.HP0_G0024540 | 839.7067 | 948.1068 | 848.3153 | 366.1154 | 294.1986 | 368.4373 | 235.7844 | 578.7782 | 337.2958 | 1.77E-05 | 1.357085652 | YHR124W | NDT80 | n |
| W303.asm02.HP0_G0024560 | 1115.02 | 1197.374 | 1202.948 | 2482.915 | 2417.809 | 1649.992 | 2480.056 | 2505.175 | 2526.13 | 4.52E-07 | -0.897991385 | YHR127W | | n |
| W303.asm02.HP0_G0024570 | 3143.87 | 3527.58 | 3478.416 | 5266.643 | 6029.77 | 4195.628 | 7446.342 | 5539.734 | 6956.426 | 0.001164722 | -0.610015451 | YHR128W | FUR1 | n |
| W303.asm02.HP0_G0024580 | 686.1664 | 688.8241 | 763.1604 | 1292.867 | 1208.905 | 1275.477 | 911.0411 | 1054.472 | 1054.946 | 2.15E-13 | -0.821221013 | YHR129C | ARP1 | n |
| W303.asm02.HP0_G0024590 | 1329.977 | 1177.344 | 1186.779 | 3366.594 | 3237.92 | 4678.774 | 12502.74 | 8012.695 | 10347.33 | 2.53E-13 | -1.610889107 | YHR131C | | n |
| W303.asm02.HP0_G0024600 | 5808.059 | 6295.117 | 5698.911 | 3670.88 | 3627.582 | 5082.156 | 11059.64 | 6443.873 | 7753.019 | 0.033463933 | 0.523940048 | YHR132C | ECM14 | n |
| W303.asm02.HP0_G0024610 | 2095.561 | 1901.778 | 1831.369 | 1680.518 | 1906.65 | 2294.187 | 5150.222 | 3333.075 | 3948.274 | 0.957620156 | -0.012898575 | YHR132W- | IGO2 | n |
| W303.asm02.HP0_G0024630 | 1358.567 | 1325.346 | 1187.857 | 848.2484 | 891.2743 | 670.0242 | 897.4619 | 878.3683 | 898.2593 | 2.51E-06 | 0.684574185 | YHR134W | WSS1 | n |
| W303.asm02.HP0_G0024650 | 136.5979 | 152.4538 | 120.7259 | 1560.332 | 1465.786 | 962.4951 | 501.1961 | 492.8742 | 336.0997 | 6.46E-37 | -3.283257102 | YHR136C | SPL2 | n |
| W303.asm02.HP0_G0024660 | 636.3981 | 813.4578 | 790.1081 | 4553.864 | 4238.543 | 3584.857 | 2393.643 | 2367.729 | 1807.283 | 5.72E-43 | -2.466329569 | YHR137W | ARO9 | n |
| W303.asm02.HP0_G0024670 | 4178.414 | 4130.719 | 3513.987 | 717.6418 | 804.49 | 1054.414 | 4250.291 | 2588.932 | 3398.076 | 1.36E-21 | 2.198157313 | YHR138C | | n |
| W303.asm02.HP0_G0024690 | 327.1997 | 275.9747 | 216.6599 | 1012.896 | 1020.583 | 1684.936 | 5536.612 | 4559.354 | 4283.178 | 1.28E-16 | -2.181222109 | YHR140W | | n |
| W303.asm02.HP0_G0024710 | 4984.236 | 4900.777 | 4827.96 | 1624.941 | 1455.372 | 1245.09 | 2994.832 | 2310.817 | 2338.345 | 1.72E-29 | 1.76604373 | YHR142W | CHS7 | n |
| W303.asm02.HP0_G0024720 | 7984.096 | 7914.243 | 8510.101 | 4598.326 | 4987.491 | 6249.001 | 5962.505 | 7076.342 | 7438.448 | 0.000100647 | 0.624294894 | YHR143W | DSE2 | n |
| W303.asm02.HP0_G0024730 | 1341.625 | 1402.13 | 1427.153 | 2054.276 | 2253.787 | 1385.628 | 2238.1 | 1066.283 | 1657.773 | 0.150503348 | -0.448943594 | YHR143W- | RPC10 | n |
| W303.asm02.HP0_G0024740 | 627.9269 | 655.44 | 682.3171 | 2189.051 | 2418.677 | 1392.465 | 1290.024 | 838.6377 | 993.9461 | 1.74E-10 | -1.609897807 | YHR144C | DCD1 | n |
| W303.asm02.HP0_G0024750 | 4932.35 | 5405.989 | 4463.626 | 13055.11 | 13458.5 | 13904.9 | 6271.123 | 11317.85 | 9415.576 | 4.24E-11 | -1.449234758 | YHR146W | CRP1 | n |
| W303.asm02.HP0_G0024760 | 4598.797 | 4566.937 | 4230.797 | 4990.841 | 4645.561 | 3606.128 | 2709.668 | 2094.984 | 2588.326 | 0.934396795 | 0.016664031 | YHR147C | MRPL6 | n |
| W303.asm02.HP0_G0024770 | 1471.869 | 1571.276 | 1598.541 | 1391.516 | 1369.455 | 508.9753 | 2898.543 | 1332.586 | 2265.384 | 0.244989213 | 0.505416635 | YHR148W | IMP3 | n |
| W303.asm02.HP0_G0024780 | 971.0101 | 955.8964 | 1252.532 | 6189.921 | 5861.408 | 5761.296 | 5132.939 | 4654.923 | 4700.611 | 1.88E-75 | -2.486021487 | YHR149C | SKG6 | n |
| W303.asm02.HP0_G0024790 | 1014.425 | 1010.424 | 916.2236 | 1315.792 | 1224.526 | 1612.768 | 1578.891 | 1346.545 | 1357.556 | 0.001252684 | -0.498062964 | YHR150W | PEX28 | n |
| W303.asm02.HP0_G0024800 | 533.6849 | 568.6415 | 522.7864 | 1281.057 | 1110.838 | 1233.695 | 1477.664 | 1299.298 | 1339.614 | 6.37E-24 | -1.158448869 | YHR151C | MTC6 | n |
| W303.asm02.HP0_G0024830 | 2100.855 | 2381.395 | 2549.257 | 2094.569 | 2051.58 | 3142.733 | 3265.181 | 3936.551 | 3566.723 | 0.82408182 | -0.051936199 | YHR154W | RTT107 | n |
| W303.asm02.HP0_G0024850 | 1829.777 | 2095.405 | 1716.033 | 1262.994 | 1222.79 | 1133.42 | 1318.417 | 1441.04 | 1414.968 | 7.50E-08 | 0.640146354 | YHR156C | LIN1 | n |
| W303.asm02.HP0_G0024860 | 3790.857 | 3964.911 | 3756.517 | 5876.604 | 5887.444 | 7710.596 | 3942.907 | 3900.041 | 3377.742 | 3.66E-07 | -0.758453414 | YHR158C | KEL1 | n |
| W303.asm02.HP0_G0024870 | 1813.893 | 1942.951 | 1719.267 | 2316.184 | 2133.157 | 2083 | 1023.378 | 1583.855 | 1492.713 | 0.196179104 | -0.25453602 | YHR159W | TDA11 | n |
| W303.asm02.HP0_G0024890 | 5670.403 | 6109.28 | 5307.63 | 8864.578 | 9119.289 | 14789.15 | 13675.49 | 15590.5 | 14658.01 | 2.16E-05 | -0.939597114 | YHR161C | YAP1801 | n |
| W303.asm02.HP0_G0024900 | 6192.44 | 5955.713 | 6098.816 | 4415.616 | 5211.394 | 3847.701 | 829.5659 | 1231.649 | 1118.339 | 0.023436887 | 0.437489723 | YHR162W | MPC2 | n |
| W303.asm02.HP0_G0024910 | 2726.664 | 2240.069 | 2603.153 | 3482.612 | 3732.591 | 5043.413 | 1946.764 | 3380.322 | 3920.764 | 0.015893283 | -0.695431985 | YHR163W | SOL3 | n |
| W303.asm02.HP0_G0024920 | 2130.504 | 2329.093 | 2087.912 | 1848.64 | 1989.963 | 1922.711 | 532.0579 | 1011.52 | 717.6506 | 0.460344499 | 0.184576393 | YHR164C | DNA2 | n |
| W303.asm02.HP0_G0024930 | 5936.186 | 6094.813 | 5631.003 | 5510.488 | 5303.386 | 5071.521 | 4952.706 | 4727.941 | 5273.536 | 0.092425479 | 0.152886005 | YHR165C | PRP8 | n |
| W303.asm02.HP0_G0024940 | 1559.758 | 1557.922 | 1507.996 | 780.8609 | 806.2257 | 916.1555 | 608.5952 | 555.1546 | 569.3361 | 8.30E-17 | 0.88590555 | YHR166C | CDC23 | n |
| W303.asm02.HP0_G0024950 | 325.0819 | 255.9443 | 294.2695 | 1160.176 | 1111.706 | 1056.693 | 603.6573 | 811.7927 | 721.2388 | 1.78E-32 | -1.926599792 | YHR167W | THP2 | n |
| W303.asm02.HP0_G0024960 | 1119.256 | 980.378 | 1035.872 | 1595.763 | 1536.949 | 1576.304 | 1656.663 | 1688.013 | 1690.067 | 2.30E-11 | -0.586640061 | YHR168W | MTG2 | n |
| W303.asm02.HP0_G0024970 | 562.2752 | 691.0497 | 790.1081 | 1580.479 | 1357.306 | 335.0121 | 2876.322 | 1138.228 | 2674.445 | 0.234233419 | -0.679548179 | YHR169W | DBP8 | n |
| W303.asm02.HP0_G0024980 | 2877.028 | 3140.325 | 3467.637 | 4828.277 | 4232.468 | 1577.823 | 7164.882 | 2991.607 | 5962.48 | 0.7341004 | -0.165589333 | YHR170W | NMD3 | n |
| W303.asm02.HP0_G0024990 | 887.3571 | 1047.146 | 811.6663 | 515.4794 | 542.4016 | 795.3689 | 1173.984 | 1168.294 | 827.6904 | 0.02232312 | 0.567241335 | YHR171W | ATG7 | n |
| W303.asm02.HP0_G0025000 | 825.941 | 861.3083 | 955.0284 | 1638.141 | 1707.914 | 2073.884 | 1164.108 | 1888.814 | 1841.97 | 2.58E-06 | -1.036499795 | YHR172W | SPC97 | n |
| W303.asm02.HP0_G0025020 | 88835.25 | 88892.8 | 97102.46 | 53486.2 | 63001.03 | 81802.21 | 163788.7 | 136058 | 139851 | 0.011934611 | 0.470937069 | YHR174W | ENO2 | n |
| W303.asm02.HP0_G0025030 | 1380.804 | 1221.856 | 1315.05 | 970.5184 | 1152.495 | 1119.746 | 897.4619 | 1020.11 | 1469.988 | 0.218945298 | 0.273042268 | YHR175W | CTR2 | n |
| W303.asm02.HP0_G0025050 | 1377.627 | 1237.435 | 1219.116 | 955.9294 | 986.7369 | 1591.497 | 892.524 | 1999.415 | 2042.912 | 0.760715062 | 0.11753428 | YHR176W | FMO1 | n |

| | | | | | | | | | | | | | | |
|---|---|---|---|---|---|---|---|---|---|---|---|---|---|---|
| W303.asm02.HP0_G0025060 | 958.3033 | 1111.689 | 984.132 | 468.9334 | 394.8683 | 194.4741 | 176.5297 | 237.3098 | 192.5696 | 1.51E-06 | 1.528841425 | YHR177W | ROF1 | n |
| W303.asm02.HP0_G0025070 | 914.8885 | 863.5339 | 937.7818 | 3912.641 | 4293.217 | 4747.904 | 6263.716 | 5395.845 | 5440.988 | 1.23E-75 | -2.253559847 | YHR178W | STB5 | n |
| W303.asm02.HP0_G0025080 | 12807.38 | 12336.52 | 12849.77 | 5760.586 | 6866.37 | 19299.28 | 46597.66 | 61187.27 | 52872.91 | 0.586718724 | 0.251014205 | YHR179W | OYE2 | n |
| W303.asm02.HP0_G0025100 | 1228.323 | 1231.871 | 1179.234 | 2619.08 | 2696.387 | 3129.059 | 6066.201 | 4805.255 | 5676.616 | 1.62E-19 | -1.21431418 | YHR181W | SVP26 | n |
| W303.asm02.HP0_G0025110 | 1512.107 | 1629.141 | 1481.049 | 3529.853 | 3782.926 | 3753.503 | 4140.423 | 3577.901 | 4366.904 | 3.06E-29 | -1.259484858 | YHR182W | RGD3 | n |
| W303.asm02.HP0_G0025120 | 17644.43 | 17275.13 | 19075.78 | 25386.32 | 28145 | 31896.8 | 35486.16 | 39622.14 | 33176.99 | 4.61E-07 | -0.661865306 | YHR183W | GND1 | n |
| W303.asm02.HP0_G0025130 | 244.6056 | 278.2003 | 240.374 | 403.6301 | 409.6217 | 271.9599 | 427.1277 | 299.5902 | 196.1578 | 0.122544049 | -0.507855816 | YHR184W | SSP1 | n |
| W303.asm02.HP0_G0025140 | 25.41357 | 24.48163 | 16.16865 | 56.96672 | 56.40976 | 42.54122 | 97.52337 | 66.5756 | 90.90241 | 0.000502989 | -1.238679125 | YHR185C | PFS1 | n |
| W303.asm02.HP0_G0025150 | 2502.178 | 2519.382 | 2605.309 | 4154.403 | 3755.155 | 3886.444 | 5169.973 | 3750.783 | 4728.121 | 1.37E-05 | -0.629210091 | YHR186C | KOG1 | n |
| W303.asm02.HP0_G0025160 | 1337.389 | 1249.676 | 1427.153 | 2419.002 | 2317.14 | 1397.783 | 2767.689 | 1692.309 | 2447.189 | 0.026730712 | -0.611661641 | YHR187W | IKI1 | n |
| W303.asm02.HP0_G0025170 | 4089.467 | 3737.9 | 4486.262 | 4655.988 | 4468.521 | 5408.052 | 4455.213 | 5152.092 | 4761.612 | 0.084534878 | -0.239052218 | YHR188C | GPI16 | n |
| W303.asm02.HP0_G0025180 | 524.1549 | 444.0077 | 410.6838 | 368.1996 | 340.1943 | 237.775 | 291.3356 | 355.4278 | 289.4524 | 0.01683285 | 0.543298095 | YHR189W | PTH1 | n |
| W303.asm02.HP0_G0025190 | 13061.52 | 13942.29 | 13792.94 | 6520.606 | 6788.264 | 7993.191 | 12216.35 | 10188.21 | 10798.25 | 9.56E-14 | 0.937459166 | YHR190W | ERG9 | n |
| W303.asm02.HP0_G0025200 | 732.7579 | 543.0471 | 601.4739 | 583.5616 | 723.7807 | 587.9804 | 175.2952 | 249.1216 | 215.2952 | 0.957154707 | -0.012990095 | YHR191C | CTF8 | n |
| W303.asm02.HP0_G0025210 | 1794.833 | 1839.461 | 1836.759 | 2009.814 | 1959.588 | 2207.585 | 1525.809 | 2008.006 | 2005.833 | 0.24927673 | -0.175159798 | YHR192W | LNP1 | n |
| W303.asm02.HP0_G0025220 | 9904.939 | 9873.887 | 10428.78 | 8155.967 | 10108.63 | 8254.516 | 5137.877 | 6177.571 | 5353.673 | 0.187554486 | 0.187973031 | YHR193C | EGD2 | n |
| W303.asm02.HP0_G0025230 | 2399.464 | 2210.024 | 2257.144 | 646.7807 | 665.6352 | 674.5822 | 796.2351 | 709.7818 | 605.2187 | 3.22E-46 | 1.789186668 | YHR194W | MDM31 | n |
| W303.asm02.HP0_G0025250 | 1189.143 | 1301.978 | 1445.477 | 4918.59 | 4448.561 | 1820.156 | 7153.772 | 3994.536 | 6372.737 | 6.45E-05 | -1.506846137 | YHR196W | UTP9 | n |
| W303.asm02.HP0_G0025260 | 912.7707 | 1108.35 | 1235.285 | 5655.684 | 4883.35 | 2156.688 | 7440.169 | 3169.857 | 6024.677 | 1.46E-06 | -1.963008977 | YHR197W | RIX1 | n |
| W303.asm02.HP0_G0025270 | 2863.262 | 2615.083 | 2503.985 | 1457.514 | 1497.028 | 2174.92 | 3410.849 | 3245.023 | 3055.995 | 0.000677746 | 0.637971967 | YHR198C | AIM18 | n |
| W303.asm02.HP0_G0025300 | 6145.848 | 5709.784 | 6210.918 | 5292.348 | 5795.452 | 6846.098 | 3913.28 | 6746.685 | 4761.612 | 0.967025572 | 0.010647807 | YHR200W | RPN10 | n |
| W303.asm02.HP0_G0025310 | 4523.615 | 4657.074 | 4610.222 | 1373.454 | 1227.129 | 1091.638 | 1544.326 | 1196.213 | 1590.792 | 6.43E-36 | 1.900963387 | YHR201C | PPX1 | n |
| W303.asm02.HP0_G0025320 | 3046.452 | 2562.782 | 2669.983 | 1142.113 | 1148.156 | 1686.455 | 1318.417 | 2296.858 | 1316.889 | 0.00014442 | 1.057890599 | YHR202W | SMN1 | n |
| W303.asm02.HP0_G0025340 | 1765.184 | 1722.617 | 1961.796 | 1691.634 | 1645.429 | 1463.874 | 1064.116 | 1370.169 | 1275.026 | 0.222652057 | 0.182843245 | YHR204W | MNL1 | n |
| W303.asm02.HP0_G0025350 | 4832.814 | 5266.889 | 5228.942 | 9364.079 | 8464.068 | 10652.78 | 11525.04 | 11483.22 | 10870.01 | 9.99E-15 | -0.893847308 | YHR205W | SCH9 | n |
| W303.asm02.HP0_G0025360 | 2096.619 | 2295.709 | 2306.728 | 3490.949 | 3426.242 | 2769.737 | 2401.05 | 2069.212 | 2225.913 | 9.57E-05 | -0.532102218 | YHR206W | SKN7 | n |
| W303.asm02.HP0_G0025370 | 1571.406 | 1720.391 | 1559.736 | 2579.481 | 2629.563 | 3157.166 | 4142.892 | 3422.2 | 3332.291 | 6.34E-08 | -0.786205864 | YHR207C | SET5 | n |
| W303.asm02.HP0_G0025380 | 13158.93 | 14252.76 | 14238.12 | 15067 | 16929 | 16619.18 | 22042.75 | 14807.7 | 17966.38 | 0.206088354 | -0.2230856 | YHR208W | BAT1 | n |
| W303.asm02.HP0_G0025580 | 100.5954 | 85.68571 | 112.1027 | 2363.424 | 1967.399 | 2300.264 | 21283.55 | 12060.92 | 15284.76 | 5.72E-75 | -4.473588125 | YOL156W | HXT11 | y |
| W303.asm02.HP0_G0025620 | 185.3073 | 227.0115 | 183.2447 | 939.2562 | 1037.072 | 1589.978 | 1314.714 | 1339.029 | 1013.083 | 4.16E-25 | -2.582436625 | YCL064C | CHA1 | y |
| W303.asm02.HP0_G0025630 | 896.8872 | 961.4604 | 879.5747 | 3490.949 | 3213.621 | 3280.991 | 401.2038 | 630.3206 | 1588.4 | 1.48E-05 | -1.866777853 | YIL166C | SOA1 | n |
| W303.asm02.HP0_G0025670 | 16206.44 | 20600.18 | 17393.16 | 5580.655 | 2868.22 | 20295.2 | 86292.14 | 54280.59 | 71454.08 | 0.132993993 | 0.915012125 | YIL162W | SUC2 | n |
| W303.asm02.HP0_G0025680 | 1497.283 | 1350.941 | 1446.555 | 612.7396 | 651.7497 | 421.6139 | 611.0642 | 398.3798 | 547.8066 | 3.68E-09 | 1.349188053 | YIL161W | SMU2 | n |
| W303.asm02.HP0_G0025690 | 546.3917 | 623.1688 | 558.3575 | 229.951 | 229.9783 | 183.0792 | 369.1075 | 238.3836 | 184.197 | 1.12E-07 | 1.426196788 | YIL160C | POT1 | n |
| W303.asm02.HP0_G0025700 | 1058.899 | 1073.853 | 1056.352 | 1315.098 | 1278.332 | 1661.387 | 492.5548 | 551.9332 | 360.0214 | 0.038303991 | -0.416032809 | YIL159W | BNR1 | n |
| W303.asm02.HP0_G0025720 | 8412.95 | 8579.699 | 7922.64 | 1717.338 | 1940.496 | 1820.156 | 1350.514 | 1390.571 | 1194.888 | 1.42E-102 | 2.185680928 | YIL157C | COA1 | n |
| W303.asm02.HP0_G0025740 | 2297.81 | 2266.776 | 2137.496 | 2824.716 | 2543.646 | 4260.199 | 2712.137 | 3250.392 | 2442.404 | 0.024297653 | -0.522774793 | YIL156W | UBP7 | n |
| W303.asm02.HP0_G0025750 | 2019.32 | 2141.03 | 1968.264 | 3135.254 | 3187.586 | 2966.49 | 4776.176 | 3749.709 | 2346.717 | 0.018383501 | -0.60001525 | YIL155C | GUT2 | n |
| W303.asm02.HP0_G0025760 | 8420.363 | 8180.203 | 8026.119 | 3206.81 | 3127.704 | 3373.671 | 4831.728 | 3657.363 | 3969.804 | 7.71E-26 | 1.342912738 | YIL154C | IMP21 | n |
| W303.asm02.HP0_G0025770 | 1568.229 | 1271.932 | 1541.412 | 2291.869 | 2276.351 | 2282.033 | 2712.137 | 2520.208 | 2344.325 | 1.41E-07 | -0.644532316 | YIL153W | RRD1 | n |
| W303.asm02.HP0_G0025800 | 491.329 | 617.6048 | 487.2154 | 1777.084 | 1499.632 | 1497.299 | 907.3377 | 1602.109 | 910.2202 | 1.15E-09 | -1.580836281 | YIL150C | MCM10 | n |
| W303.asm02.HP0_G0025820 | 1837.189 | 1907.342 | 1896.044 | 2980.332 | 2542.779 | 2278.234 | 5309.469 | 3918.296 | 4918.299 | 0.005540518 | -0.468014162 | YIL147C | SLN1 | n |
| W303.asm02.HP0_G0025830 | 797.3507 | 873.5491 | 823.5233 | 1294.256 | 1198.491 | 2252.406 | 2488.698 | 2988.385 | 2162.52 | 0.000641937 | -0.927868654 | YIL146C | ATG32 | n |
| W303.asm02.HP0_G0025840 | 3418.125 | 3257.17 | 3619.622 | 1927.143 | 2039.43 | 1567.948 | 506.134 | 475.6934 | 784.6313 | 5.31E-05 | 0.895477301 | YIL145C | PAN6 | n |
| W303.asm02.HP0_G0025850 | 2096.619 | 2243.408 | 2028.627 | 2231.428 | 1973.474 | 2348.123 | 3339.25 | 2250.685 | 2430.443 | 0.841428969 | -0.041313966 | YIL144W | NDC80 | n |
| W303.asm02.HP0_G0025860 | 4008.991 | 3980.491 | 3827.659 | 5602.191 | 5629.694 | 6397.136 | 10136.26 | 7292.175 | 7738.666 | 0.000253926 | -0.577079054 | YIL143C | SSL2 | n |
| W303.asm02.HP0_G0025870 | 10106.13 | 10500.39 | 10685.32 | 8688.12 | 8821.619 | 9138.006 | 4095.982 | 5420.542 | 4697.023 | 0.076743506 | 0.231773331 | YIL142W | CCT2 | n |
| W303.asm02.HP0_G0025880 | 4022.756 | 4256.465 | 4267.446 | 5463.248 | 6033.241 | 5835.744 | 2639.303 | 3480.186 | 2532.111 | 0.002205845 | -0.466117237 | YIL140W | AXL2 | n |
| W303.asm02.HP0_G0025890 | 617.3379 | 597.5743 | 584.2273 | 466.8492 | 540.6659 | 534.0442 | 894.993 | 717.2984 | 919.7888 | 0.149035466 | 0.223710659 | YIL139C | REV7 | n |
| W303.asm02.HP0_G0025900 | 4363.722 | 4686.007 | 4444.224 | 12579.92 | 13947.1 | 14682.04 | 8257.39 | 12416.35 | 10858.05 | 1.75E-21 | -1.610646469 | YIL138C | TPM2 | n |
| W303.asm02.HP0_G0025910 | 6536.582 | 6149.34 | 6505.188 | 5709.177 | 5269.54 | 4206.263 | 1002.392 | 1413.121 | 1243.928 | 0.068062953 | 0.337782215 | YIL137C | TMA108 | n |
| W303.asm02.HP0_G0025920 | 14451.85 | 14638.9 | 12798.03 | 265.3816 | 307.2163 | 1112.909 | 2103.542 | 4066.48 | 3472.233 | 3.15E-20 | 4.635237751 | YIL136W | OM45 | n |
| W303.asm02.HP0_G0025930 | 4590.326 | 4936.387 | 4491.652 | 7707.876 | 7474.728 | 8957.205 | 8431.451 | 6224.818 | 7082.015 | 2.28E-07 | -0.784130456 | YIL135C | VHS2 | n |
| W303.asm02.HP0_G0025950 | 915.9474 | 783.4122 | 826.7571 | 1224.785 | 1174.191 | 1051.376 | 1018.44 | 1157.556 | 995.1422 | 0.000608738 | -0.449696331 | YIL134W | FLX1 | n |

| | | | | | | | | | | | | | | |
|---|---|---|---|---|---|---|---|---|---|---|---|---|---|---|
| W303.asm02.HP0_G0025960 | 10864.3 | 10610.56 | 11045.35 | 24745.79 | 29116.98 | 26146.9 | 23846.32 | 20030.66 | 18944.78 | 4.48E-25 | -1.298810508 | YIL133C | RPL16A | n |
| W303.asm02.HP0_G0025970 | 490.2701 | 541.9343 | 481.8258 | 259.8238 | 275.1061 | 166.3666 | 107.3992 | 187.915 | 120.8045 | 8.90E-05 | 1.110689175 | YIL132C | CSM2 | n |
| W303.asm02.HP0_G0025990 | 1756.713 | 1915.131 | 1772.084 | 5565.371 | 5415.337 | 6841.54 | 12487.93 | 8149.068 | 11512.31 | 5.44E-19 | -1.711002602 | YIL130W | ASG1 | n |
| W303.asm02.HP0_G0026000 | 3320.706 | 3131.423 | 3032.161 | 5101.995 | 4793.962 | 5188.509 | 5787.21 | 4337.078 | 4838.161 | 7.49E-07 | -0.66948283 | YIL129C | TAO3 | n |
| W303.asm02.HP0_G0026010 | 2071.206 | 2039.765 | 2188.158 | 3459.686 | 3159.815 | 3429.886 | 2604.738 | 3046.37 | 2776.112 | 2.33E-11 | -0.674033337 | YIL128W | MET18 | n |
| W303.asm02.HP0_G0026020 | 1208.203 | 1125.042 | 1264.389 | 1483.219 | 1246.222 | 398.0643 | 2518.325 | 1107.088 | 2109.893 | 0.698777536 | 0.202023296 | YIL127C | RRT14 | n |
| W303.asm02.HP0_G0026030 | 5994.426 | 6246.154 | 6296.073 | 7327.171 | 7004.357 | 7088.431 | 7237.716 | 6470.718 | 6561.719 | 0.011693293 | -0.208649799 | YIL126W | STH1 | n |
| W303.asm02.HP0_G0026040 | 15989.37 | 15170.82 | 15320.34 | 29285.76 | 28914.78 | 30331.89 | 13176.77 | 21815.32 | 19062 | 1.17E-06 | -0.929578601 | YIL125W | KGD1 | n |
| W303.asm02.HP0_G0026050 | 5214.017 | 4761.677 | 4958.387 | 3073.424 | 3526.044 | 4405.295 | 4113.264 | 5288.465 | 3792.783 | 0.025765756 | 0.440523648 | YIL124W | AYR1 | n |
| W303.asm02.HP0_G0026080 | 2333.813 | 1885.086 | 2131.028 | 769.0508 | 833.9967 | 850.8244 | 6900.705 | 5193.97 | 6035.442 | 1.80E-19 | 1.371981645 | YIL121W | QDR2 | n |
| W303.asm02.HP0_G0026100 | 1854.132 | 1683.668 | 1971.498 | 2236.986 | 1656.711 | 3814.276 | 5989.663 | 5712.616 | 3563.135 | 0.165925289 | -0.484548132 | YIL119C | RPI1 | n |
| W303.asm02.HP0_G0026110 | 1550.228 | 1561.26 | 1608.242 | 3685.469 | 3790.736 | 3764.898 | 2324.513 | 2129.345 | 1969.951 | 1.90E-42 | -1.251955102 | YIL118W | RHO3 | n |
| W303.asm02.HP0_G0026130 | 3335.531 | 2948.924 | 3146.42 | 2899.745 | 2798.792 | 3465.59 | 2244.272 | 1924.249 | 1523.811 | 0.838340878 | 0.041375426 | YIL116W | HIS5 | n |
| W303.asm02.HP0_G0026140 | 5508.391 | 5502.803 | 5710.768 | 7821.114 | 7446.089 | 7163.637 | 4973.692 | 5472.084 | 5472.086 | 2.83E-07 | -0.423784092 | YIL115C | NUP159 | n |
| W303.asm02.HP0_G0026170 | 6716.595 | 6859.308 | 6498.72 | 3836.223 | 3389.793 | 2971.048 | 3071.369 | 2228.135 | 2692.386 | 1.41E-09 | 0.977146976 | YIL112W | HOS4 | n |
| W303.asm02.HP0_G0026190 | 1423.16 | 1311.993 | 1482.126 | 1622.162 | 1467.522 | 567.4695 | 1673.946 | 893.4015 | 1629.067 | 0.643529263 | 0.20560238 | YIL110W | HPM1 | n |
| W303.asm02.HP0_G0026200 | 13512.61 | 12415.52 | 13490.05 | 14639.06 | 14734.23 | 18879.94 | 8275.908 | 12080.25 | 10230.11 | 0.130577779 | -0.291764643 | YIL109C | SEC24 | n |
| W303.asm02.HP0_G0026210 | 4538.44 | 4404.468 | 4523.989 | 3413.835 | 3486.123 | 4408.334 | 2620.786 | 4549.69 | 3648.057 | 0.288426138 | 0.25203167 | YIL108W | | n |
| W303.asm02.HP0_G0026220 | 9298.19 | 9104.941 | 8652.385 | 3656.291 | 3657.956 | 6653.143 | 16587.61 | 11830.05 | 14855.37 | 0.000393099 | 0.953840839 | YIL107C | PFK26 | n |
| W303.asm02.HP0_G0026240 | 657.5761 | 652.1016 | 761.0046 | 996.9177 | 719.4414 | 229.4187 | 1846.772 | 779.5787 | 1801.303 | 0.877497891 | 0.089663251 | YIL104C | SHQ1 | n |
| W303.asm02.HP0_G0026250 | 1574.582 | 1505.62 | 1677.228 | 1377.622 | 1155.098 | 370.7163 | 2496.105 | 955.6819 | 1868.284 | 0.158938841 | 0.71240378 | YIL103W | DPH1 | n |
| W303.asm02.HP0_G0026260 | 340.9654 | 287.1028 | 263.0101 | 108.3757 | 113.6874 | 116.2287 | 77.7718 | 91.27299 | 95.68675 | 3.94E-14 | 1.3981686 | YIL102C-A | | n |
| W303.asm02.HP0_G0026280 | 1816.011 | 1993.027 | 1636.268 | 412.6614 | 355.8154 | 436.8072 | 862.8967 | 3022.747 | 856.3964 | 2.37E-06 | 2.17551595 | YIL101C | XBP1 | n |
| W303.asm02.HP0_G0026290 | 769.8194 | 768.9457 | 804.121 | 1178.239 | 1195.019 | 1476.028 | 978.9372 | 1096.35 | 897.0632 | 8.67E-07 | -0.716348462 | YIL099W | SGA1 | n |
| W303.asm02.HP0_G0026300 | 4174.179 | 4036.131 | 3646.57 | 1629.109 | 1622.866 | 1042.26 | 1170.28 | 731.2578 | 1080.064 | 2.21E-09 | 1.465297427 | YIL098C | FMC1 | n |
| W303.asm02.HP0_G0026310 | 5131.423 | 5523.946 | 4685.675 | 987.8863 | 932.9307 | 1735.834 | 2018.364 | 1668.685 | 1450.85 | 6.40E-15 | 2.068691409 | YIL097W | FYV10 | n |
| W303.asm02.HP0_G0026320 | 260.4891 | 300.4564 | 337.3859 | 1825.019 | 1810.319 | 811.3218 | 1317.183 | 1032.996 | 1374.301 | 4.51E-13 | -2.307418148 | YIL096C | BMT5 | n |
| W303.asm02.HP0_G0026330 | 2734.076 | 2849.884 | 2696.931 | 5559.813 | 5149.777 | 7190.985 | 5469.95 | 6519.039 | 5973.245 | 1.96E-12 | -1.112210169 | YIL095W | PRK1 | n |
| W303.asm02.HP0_G0026340 | 5183.309 | 5561.781 | 5569.562 | 1580.479 | 1453.636 | 2057.172 | 1431.989 | 1186.549 | 984.3774 | 2.83E-18 | 1.679939127 | YIL094C | LYS12 | n |
| W303.asm02.HP0_G0026350 | 8973.108 | 8943.585 | 8844.253 | 3706.311 | 3921.78 | 2839.626 | 645.6294 | 880.5159 | 710.4741 | 6.93E-15 | 1.354244131 | YIL093C | RSM25 | n |
| W303.asm02.HP0_G0026370 | 2088.148 | 2358.026 | 2483.505 | 3443.708 | 2989.717 | 935.1471 | 7371.039 | 2853.086 | 6686.111 | 0.87419794 | -0.088602112 | YIL091C | UTP25 | n |
| W303.asm02.HP0_G0026380 | 2866.439 | 2780.891 | 3049.408 | 2808.043 | 2822.224 | 2395.982 | 4740.377 | 3978.429 | 4607.317 | 0.392064473 | 0.115814133 | YIL090W | ICE2 | n |
| W303.asm02.HP0_G0026400 | 7236.514 | 6892.692 | 6714.302 | 2524.598 | 2702.462 | 3244.528 | 4324.359 | 4677.472 | 3487.782 | 2.41E-16 | 1.298920294 | YIL088C | AVT7 | n |
| W303.asm02.HP0_G0026410 | 2885.499 | 2578.361 | 2592.374 | 613.4344 | 905.1597 | 930.5892 | 832.0349 | 1319.7 | 806.1608 | 8.08E-12 | 1.718194672 | YIL087C | AIM19 | n |
| W303.asm02.HP0_G0026440 | 1909.194 | 2048.667 | 1776.396 | 2845.557 | 2822.224 | 2986.242 | 3733.047 | 2820.872 | 3498.547 | 2.31E-05 | -0.593830625 | YIL084C | SDS3 | n |
| W303.asm02.HP0_G0026450 | 2954.327 | 2573.91 | 2931.916 | 2481.526 | 2573.153 | 2726.436 | 3910.811 | 3771.185 | 3925.549 | 0.24239938 | 0.120871822 | YIL083C | CAB2 | n |
| W303.asm02.HP0_G0026460 | 24537.86 | 24368.12 | 25153.03 | 18670.5 | 19660.1 | 19493.75 | 17355.46 | 16801.75 | 16626.77 | 1.33E-09 | 0.357045364 | YIL078W | THS1 | n |
| W303.asm02.HP0_G0026470 | 1834.013 | 1918.47 | 1584.528 | 1195.606 | 1276.596 | 1381.83 | 2115.887 | 2032.703 | 2135.011 | 8.34E-05 | 0.46975465 | YIL077C | RCI37 | n |
| W303.asm02.HP0_G0026480 | 7944.917 | 8293.709 | 8017.496 | 5870.351 | 6389.057 | 7304.175 | 5493.405 | 7026.947 | 6311.737 | 0.030867501 | 0.310197082 | YIL076W | SEC28 | n |
| W303.asm02.HP0_G0026490 | 26249.04 | 26788.47 | 27069.56 | 14780.09 | 15167.28 | 19760.4 | 11250.99 | 17967.89 | 14071.93 | 0.000993092 | 0.688456047 | YIL075C | RPN2 | n |
| W303.asm02.HP0_G0026510 | 1644.47 | 1763.79 | 1662.137 | 131.9961 | 110.216 | 143.5766 | 232.0809 | 353.2802 | 327.7271 | 3.38E-83 | 3.714000797 | YIL073C | SPO22 | n |
| W303.asm02.HP0_G0026520 | 1290.798 | 1171.78 | 1141.507 | 137.5538 | 116.2909 | 140.538 | 185.171 | 353.2802 | 168.6479 | 1.02E-28 | 3.191040785 | YIL072W | HOP1 | n |
| W303.asm02.HP0_G0026530 | 2038.38 | 1958.53 | 1710.643 | 1171.291 | 1176.794 | 1804.203 | 2045.522 | 2212.028 | 2289.305 | 0.029031652 | 0.458892092 | YIL071C | PCI8 | n |
| W303.asm02.HP0_G0026540 | 12482.3 | 12629.18 | 12049.96 | 3572.925 | 3506.952 | 2386.866 | 1677.649 | 1321.848 | 1597.969 | 5.25E-26 | 1.972887018 | YIL070C | MAM33 | n |
| W303.asm02.HP0_G0026560 | 4280.069 | 4591.418 | 4165.045 | 4931.79 | 4833.883 | 5447.555 | 4067.589 | 4536.805 | 4597.748 | 0.03628841 | -0.222844571 | YIL068C | SEC6 | n |
| W303.asm02.HP0_G0026570 | 1321.506 | 1270.819 | 1301.038 | 1356.781 | 1369.455 | 1295.988 | 1943.061 | 1774.991 | 1764.224 | 0.628984694 | -0.046791222 | YIL067C | | n |
| W303.asm02.HP0_G0026590 | 2468.293 | 2267.889 | 2179.534 | 1322.045 | 1566.456 | 995.1606 | 324.6664 | 664.6822 | 480.8259 | 0.003677086 | 0.832617637 | YIL065C | FIS1 | n |
| W303.asm02.HP0_G0026600 | 929.7131 | 843.5034 | 1033.716 | 1613.131 | 1825.941 | 894.1253 | 1522.105 | 958.9033 | 1377.889 | 0.043353937 | -0.626292377 | YIL064W | EFM4 | n |
| W303.asm02.HP0_G0026610 | 3564.253 | 3298.343 | 3241.276 | 4039.08 | 4416.451 | 3917.59 | 2855.336 | 4163.122 | 3350.232 | 0.085970965 | -0.29222721 | YIL063C | YRB2 | n |
| W303.asm02.HP0_G0026620 | 6585.291 | 6081.46 | 6217.386 | 2903.913 | 3157.211 | 3730.713 | 3335.546 | 4469.155 | 3687.528 | 2.38E-09 | 0.947563847 | YIL062C | ARC15 | n |
| W303.asm02.HP0_G0026630 | 949.8321 | 1125.042 | 1111.325 | 836.4382 | 818.3755 | 723.9604 | 846.8485 | 831.1211 | 726.0232 | 0.002054923 | 0.421459469 | YIL061C | SNP1 | n |
| W303.asm02.HP0_G0026660 | 1559.758 | 1751.549 | 1602.852 | 5224.96 | 4411.244 | 3323.533 | 8843.765 | 5056.524 | 7175.31 | 2.21E-08 | -1.399072172 | YIL056W | VHR1 | n |
| W303.asm02.HP0_G0026670 | 1650.823 | 1843.912 | 1617.943 | 838.5224 | 682.9921 | 1529.205 | 4703.342 | 2402.09 | 3795.176 | 0.038340855 | 0.744778217 | YIL055C | | n |
| W303.asm02.HP0_G0026690 | 5205.546 | 5676.4 | 5940.363 | 15700.58 | 18409.54 | 34483.46 | 26394.27 | 24562.1 | 33039.44 | 8.34E-12 | -2.027699962 | YIL053W | GPP1 | n |

| | | | | | | | | | | | | | | |
|---|---|---|---|---|---|---|---|---|---|---|---|---|---|---|
| W303.asm02.HP0_G0026700 | 5873.711 | 5275.791 | 5692.443 | 20672.67 | 23797.11 | 21754.52 | 18871.39 | 14483.41 | 14997.7 | 5.17E-46 | -1.975262303 | YIL052C | RPL34B | y |
| W303.asm02.HP0_G0026710 | 6122.552 | 5718.686 | 5581.419 | 5138.121 | 6040.184 | 6610.601 | 7161.178 | 6729.504 | 7188.467 | 0.831552997 | -0.029916393 | YIL051C | MMF1 | n |
| W303.asm02.HP0_G0026730 | 787.8206 | 791.2018 | 785.7965 | 1009.423 | 1003.226 | 998.959 | 1208.549 | 1162.925 | 1212.83 | 1.37E-05 | -0.348874716 | YIL049W | DFG10 | n |
| W303.asm02.HP0_G0026740 | 1925.078 | 2276.792 | 1929.459 | 4864.402 | 4521.46 | 6570.339 | 5620.556 | 6885.205 | 6299.776 | 5.61E-14 | -1.379938426 | YIL048W | NEO1 | n |
| W303.asm02.HP0_G0026750 | 1931.431 | 1784.933 | 1946.706 | 3425.645 | 3749.08 | 3928.985 | 8361.086 | 6548.032 | 7440.841 | 5.29E-14 | -0.971262252 | YIL047C | SYG1 | n |
| W303.asm02.HP0_G0026760 | 1551.287 | 1454.431 | 1484.282 | 1451.262 | 1418.923 | 1301.305 | 2062.805 | 1328.291 | 2046.5 | 0.620950467 | 0.106165605 | YIL046W | MET30 | n |
| W303.asm02.HP0_G0026780 | 12521.48 | 12030.5 | 12927.38 | 1671.487 | 1431.94 | 1722.16 | 1770.234 | 1184.401 | 1674.518 | 1.66E-64 | 2.957122968 | YIL044C | AGE2 | y |
| W303.asm02.HP0_G0026790 | 897.9461 | 902.4819 | 805.1989 | 2173.072 | 2303.254 | 2234.174 | 2264.024 | 2931.474 | 2567.993 | 1.12E-25 | -1.364726009 | YIL042C | PKP1 | n |
| W303.asm02.HP0_G0026800 | 9764.105 | 9289.666 | 10006.24 | 9621.124 | 9953.286 | 11953.32 | 9309.162 | 12327.22 | 12112.75 | 0.496346726 | -0.117581933 | YIL041W | GVP36 | n |
| W303.asm02.HP0_G0026810 | 290.1382 | 210.3195 | 286.7241 | 207.0254 | 247.3351 | 214.9851 | 317.2596 | 170.7342 | 190.1774 | 0.449015733 | 0.234903927 | YIL040W | APQ12 | n |
| W303.asm02.HP0_G0026820 | 4869.875 | 4511.297 | 4806.401 | 2981.722 | 2878.634 | 3148.81 | 2139.342 | 2892.817 | 2792.857 | 4.70E-06 | 0.655131979 | YIL039W | TED1 | n |
| W303.asm02.HP0_G0026830 | 4121.234 | 4475.687 | 4341.822 | 10048.37 | 9062.879 | 7338.36 | 9662.221 | 9642.723 | 10549.46 | 2.80E-13 | -1.031590816 | YIL038C | NOT3 | n |
| W303.asm02.HP0_G0026840 | 415.0883 | 420.6389 | 325.5289 | 330.6849 | 298.5378 | 441.3651 | 630.8157 | 562.6712 | 572.9244 | 0.608096856 | 0.116589972 | YIL037C | PRM2 | n |
| W303.asm02.HP0_G0026850 | 5266.962 | 5434.922 | 5240.799 | 4640.704 | 4516.252 | 9064.318 | 11954.64 | 14815.22 | 15607.7 | 0.539862694 | -0.19274859 | YIL036W | CST6 | n |
| W303.asm02.HP0_G0026860 | 5374.97 | 5709.784 | 5429.433 | 7980.204 | 7564.983 | 8533.313 | 7357.46 | 6273.139 | 7115.506 | 1.56E-07 | -0.544108814 | YIL035C | CKA1 | n |
| W303.asm02.HP0_G0026870 | 9808.579 | 9712.53 | 9865.034 | 4225.264 | 4224.657 | 5601.007 | 3642.93 | 5234.775 | 4603.729 | 6.77E-09 | 1.064452471 | YIL034C | CAP2 | n |
| W303.asm02.HP0_G0026880 | 22405.24 | 22802.41 | 22352.62 | 7434.157 | 7764.587 | 14185.22 | 18485 | 20028.52 | 17777.4 | 2.70E-06 | 1.201138733 | YIL033C | BCY1 | n |
| W303.asm02.HP0_G0026900 | 1430.572 | 1631.367 | 1321.518 | 2698.972 | 2435.166 | 3160.205 | 2556.594 | 2798.323 | 2571.581 | 3.50E-10 | -0.920293342 | YIL031W | ULP2 | n |
| W303.asm02.HP0_G0026910 | 4419.843 | 4840.686 | 4715.857 | 3977.25 | 3714.366 | 4051.291 | 5763.755 | 5148.871 | 5775.891 | 0.009866859 | 0.251029178 | YIL030C | SSM4 | n |
| W303.asm02.HP0_G0026940 | 1094.901 | 1083.869 | 1126.416 | 1554.08 | 1737.421 | 1661.387 | 1643.084 | 2014.449 | 1894.598 | 5.77E-07 | -0.583207996 | YIL027C | EMC5 | n |
| W303.asm02.HP0_G0026950 | 2488.412 | 2687.415 | 2718.489 | 2526.683 | 2983.643 | 2864.695 | 803.642 | 1521.574 | 1062.123 | 0.744566404 | -0.085222838 | YIL026C | IRR1 | n |
| W303.asm02.HP0_G0026960 | 687.2253 | 679.9216 | 676.9276 | 371.6731 | 348.8727 | 660.1486 | 238.2533 | 555.1546 | 261.9425 | 0.158938841 | 0.565909829 | YIL024C | | n |
| W303.asm02.HP0_G0026970 | 3390.594 | 3305.02 | 3374.937 | 1218.532 | 1184.605 | 1520.089 | 1506.057 | 2282.899 | 2361.07 | 2.35E-11 | 1.359960122 | YIL023C | YKE4 | n |
| W303.asm02.HP0_G0026980 | 13439.54 | 12705.97 | 13609.69 | 7180.586 | 6762.229 | 5592.651 | 3931.797 | 3226.769 | 3389.703 | 1.55E-13 | 1.025038741 | YIL022W | TIM44 | n |
| W303.asm02.HP0_G0026990 | 4183.709 | 4310.993 | 4199.538 | 6606.75 | 7146.683 | 5673.935 | 4161.409 | 3998.831 | 3965.02 | 4.58E-08 | -0.613841355 | YIL021W | RPB3 | n |
| W303.asm02.HP0_G0027000 | 1067.37 | 1147.298 | 1080.066 | 1800.704 | 1836.355 | 814.3605 | 1665.304 | 942.7963 | 1288.183 | 0.228486416 | -0.434064672 | YIL020C | HIS6 | n |
| W303.asm02.HP0_G0027010 | 829.1177 | 894.6923 | 881.7305 | 2552.387 | 2618.281 | 513.5333 | 4420.648 | 1122.121 | 2822.759 | 0.070246869 | -1.125357332 | YIL019W | FAF1 | n |
| W303.asm02.HP0_G0027030 | 2623.951 | 2816.5 | 2424.22 | 2054.276 | 1883.218 | 4620.28 | 3370.112 | 4710.76 | 3215.075 | 0.757190948 | -0.121891717 | YIL017C | VID28 | n |
| W303.asm02.HP0_G0027040 | 1288.68 | 1140.621 | 1213.727 | 2173.072 | 2407.395 | 2753.784 | 3915.749 | 3833.466 | 3370.566 | 8.01E-14 | -1.00932064 | YIL016W | SNL1 | n |
| W303.asm02.HP0_G0027050 | 1570.347 | 1580.178 | 1932.693 | 370.2837 | 431.3177 | 420.8542 | 729.5736 | 722.6674 | 627.9443 | 9.17E-46 | 2.056923735 | YIL015W | BAR1 | n |
| W303.asm02.HP0_G0027070 | 2035.203 | 1781.595 | 1883.109 | 1708.307 | 1590.755 | 1978.926 | 1192.501 | 1535.534 | 1319.281 | 0.50404604 | 0.110856964 | YIL014W | MNT3 | n |
| W303.asm02.HP0_G0027110 | 3655.318 | 3633.296 | 3653.037 | 2771.917 | 3204.942 | 3220.978 | 1481.368 | 3706.757 | 3017.721 | 0.451710922 | 0.250492855 | YIL010W | DOT5 | n |
| W303.asm02.HP0_G0027120 | 767.7016 | 753.3665 | 755.615 | 240.3718 | 226.5069 | 118.5077 | 137.0265 | 95.56819 | 122.0006 | 5.21E-12 | 1.959569032 | YIL009C-A | EST3 | n |
| W303.asm02.HP0_G0027140 | 1479.281 | 1348.715 | 1421.763 | 932.3091 | 1076.125 | 608.4914 | 1365.327 | 1092.055 | 1205.653 | 0.002300297 | 0.699866269 | YIL008W | URM1 | n |
| W303.asm02.HP0_G0027150 | 2572.065 | 2538.3 | 2458.713 | 735.0097 | 682.1242 | 1004.277 | 1256.694 | 1741.703 | 1624.283 | 7.92E-17 | 1.644032575 | YIL007C | NAS2 | n |
| W303.asm02.HP0_G0027160 | 425.6773 | 436.2181 | 421.4629 | 1643.698 | 1575.134 | 1373.474 | 1304.838 | 685.0844 | 895.8672 | 1.84E-14 | -1.839329573 | YIL006W | YIA6 | n |
| W303.asm02.HP0_G0027170 | 4391.253 | 4847.363 | 4522.911 | 3577.093 | 3178.039 | 4482.021 | 5981.022 | 4278.019 | 5594.086 | 0.143435991 | 0.292280211 | YIL005W | EPS1 | n |
| W303.asm02.HP0_G0027180 | 950.891 | 735.5617 | 776.0953 | 1665.235 | 2005.584 | 1512.492 | 1985.033 | 2016.596 | 2074.01 | 2.61E-11 | -1.073190583 | YIL004C | BET1 | n |
| W303.asm02.HP0_G0027190 | 556.9807 | 513.0014 | 535.7213 | 1161.565 | 1283.539 | 1091.638 | 1385.079 | 1437.818 | 1181.731 | 3.35E-18 | -1.138672027 | YIL003W | CFD1 | n |
| W303.asm02.HP0_G0027210 | 3101.514 | 3063.542 | 3125.939 | 2601.017 | 2545.382 | 3459.513 | 2617.083 | 2775.773 | 2636.17 | 0.494378719 | 0.11043914 | YIL002C | INP51 | n |
| W303.asm02.HP0_G0027220 | 1275.973 | 1446.642 | 1371.102 | 1407.495 | 1461.447 | 1453.998 | 1020.909 | 1042.66 | 978.397 | 0.430622269 | -0.078620506 | YIL001W | | n |
| W303.asm02.HP0_G0027230 | 3514.485 | 3564.303 | 3505.364 | 7115.977 | 7038.203 | 7023.099 | 5004.554 | 4560.428 | 5449.36 | 9.32E-28 | -1.000631516 | YIR001C | SGN1 | n |
| W303.asm02.HP0_G0027240 | 3408.595 | 3517.565 | 3542.013 | 2069.559 | 1963.06 | 2288.11 | 1125.839 | 923.4679 | 1090.829 | 7.37E-10 | 0.727688112 | YIR002C | MPH1 | n |
| W303.asm02.HP0_G0027250 | 12124.39 | 12261.96 | 12112.48 | 2538.493 | 2523.686 | 4222.216 | 2423.271 | 3802.326 | 2727.072 | 1.65E-14 | 1.974942035 | YIR003W | AIM21 | n |
| W303.asm02.HP0_G0027260 | 7936.446 | 8423.906 | 8090.794 | 5811.301 | 6210.281 | 6136.571 | 6197.055 | 6342.936 | 5552.223 | 1.47E-06 | 0.429346732 | YIR004W | DJP1 | n |
| W303.asm02.HP0_G0027270 | 543.215 | 511.8886 | 483.9817 | 530.7631 | 522.4412 | 382.1113 | 1072.757 | 2151.895 | 776.2587 | 0.806851796 | 0.100750311 | YIR005W | IST3 | n |
| W303.asm02.HP0_G0027280 | 17704.79 | 16383.77 | 17621.67 | 11480.18 | 10358.57 | 17196.53 | 15172.91 | 17723.07 | 16465.3 | 0.060109158 | 0.405655246 | YIR006C | PAN1 | n |
| W303.asm02.HP0_G0027290 | 2207.804 | 2138.804 | 2123.483 | 815.5967 | 750.6838 | 1648.472 | 1291.259 | 1467.885 | 898.2593 | 0.002592864 | 1.008977566 | YIR007W | EGH1 | n |
| W303.asm02.HP0_G0027300 | 1446.456 | 1557.922 | 1385.115 | 2330.078 | 2262.465 | 2398.261 | 1573.953 | 1418.49 | 1381.477 | 1.35E-11 | -0.671619764 | YIR008C | PRI1 | n |
| W303.asm02.HP0_G0027310 | 1141.493 | 1149.524 | 1123.182 | 352.9158 | 303.7449 | 280.3162 | 227.143 | 150.332 | 190.1774 | 8.64E-27 | 1.864645358 | YIR009W | MSL1 | n |
| W303.asm02.HP0_G0027320 | 1649.764 | 1814.979 | 1676.15 | 1021.927 | 1015.376 | 1067.329 | 796.2351 | 642.1324 | 562.1596 | 1.22E-06 | 0.72750822 | YIR010W | DSN1 | n |
| W303.asm02.HP0_G0027330 | 4687.745 | 5035.426 | 4621.001 | 4437.847 | 4472.86 | 4621.8 | 5278.607 | 5206.856 | 5155.123 | 0.269016788 | 0.083974592 | YIR011C | STS1 | n |
| W303.asm02.HP0_G0027340 | 1614.821 | 1369.858 | 1676.15 | 5435.459 | 5305.989 | 2380.789 | 6280.999 | 3631.591 | 5472.086 | 1.24E-05 | -1.493324737 | YIR012W | SQT1 | n |
| W303.asm02.HP0_G0027350 | 48.70934 | 51.18886 | 38.80477 | 160.4794 | 190.9254 | 305.3852 | 939.434 | 354.354 | 385.1392 | 6.15E-07 | -2.243562078 | YIR013C | GAT4 | n |

| | | | | | | | | | | | | | | |
|---|---|---|---|---|---|---|---|---|---|---|---|---|---|---|
| W303.asm02.HP0_G0027360 | 604.6312 | 598.6871 | 490.4491 | 366.8101 | 459.9565 | 628.2426 | 722.1668 | 696.8962 | 453.316 | 0.460104115 | 0.219401596 | YIR014W | VLD1 | n |
| W303.asm02.HP0_G0027370 | 715.8155 | 640.9736 | 616.5646 | 1151.839 | 1190.68 | 1069.608 | 2147.983 | 1334.733 | 1627.871 | 7.41E-05 | -0.78988257 | YIR015W | RPR2 | n |
| W303.asm02.HP0_G0027390 | 508.2714 | 354.9836 | 349.2429 | 106.2916 | 104.1411 | 91.91942 | 387.6246 | 208.3172 | 394.7078 | 2.34E-11 | 2.003862047 | YIR017C | MET28 | n |
| W303.asm02.HP0_G0027400 | 780.4083 | 776.7354 | 612.253 | 684.2954 | 571.0404 | 409.4592 | 2077.618 | 914.8775 | 1962.774 | 0.295802046 | 0.381805832 | YIR018W | YAP5 | n |
| W303.asm02.HP0_G0027410 | 3756.973 | 3332.84 | 4006.592 | 1897.964 | 1031.865 | 1543.639 | 10126.38 | 3949.436 | 3778.43 | 0.001312868 | 1.310480619 | YIR019C | FLO11 | n |
| W303.asm02.HP0_G0027460 | 2628.187 | 2421.456 | 2477.038 | 2196.692 | 2235.562 | 2764.42 | 2804.723 | 3095.765 | 3040.446 | 0.65432168 | 0.064686164 | YIR022W | SEC11 | n |
| W303.asm02.HP0_G0027470 | 2045.792 | 2154.383 | 2336.909 | 7345.234 | 6547.004 | 7131.731 | 9738.758 | 7268.552 | 9154.829 | 7.82E-32 | -1.685376756 | YIR023W | DAL81 | n |
| W303.asm02.HP0_G0027480 | 2420.642 | 2578.361 | 2240.975 | 2654.51 | 2598.32 | 2574.503 | 2865.212 | 2558.865 | 2914.857 | 0.302057999 | -0.112644299 | YIR024C | INA22 | n |
| W303.asm02.HP0_G0027490 | 1491.988 | 1613.562 | 1461.646 | 897.5733 | 1004.962 | 837.9101 | 1682.587 | 1259.567 | 1059.731 | 0.000176516 | 0.737113523 | YIR025W | MND2 | n |
| W303.asm02.HP0_G0027500 | 1963.198 | 1846.137 | 2088.99 | 1937.563 | 1806.848 | 269.6809 | 2071.446 | 525.0882 | 1611.126 | 0.433568893 | 0.555246378 | YIR026C | YVH1 | n |
| W303.asm02.HP0_G0027510 | 486.0345 | 501.8734 | 516.319 | 911.4676 | 689.9348 | 1160.768 | 260.4738 | 471.3982 | 290.6485 | 0.002306721 | -0.877049074 | YIR027C | DAL1 | n |
| W303.asm02.HP0_G0027530 | 849.2368 | 922.5123 | 831.0687 | 1675.655 | 1777.341 | 2048.815 | 1871.461 | 2120.755 | 1845.558 | 4.01E-17 | -1.079899869 | YIR029W | DAL2 | n |
| W303.asm02.HP0_G0027540 | 618.3968 | 471.8278 | 420.385 | 863.5322 | 1029.261 | 1125.063 | 201.2191 | 510.055 | 413.8452 | 0.003597877 | -0.99823751 | YIR030C | DCG1 | n |
| W303.asm02.HP0_G0027560 | 477.5633 | 389.4805 | 429.0082 | 354.9999 | 392.2648 | 433.7685 | 478.9756 | 677.5678 | 516.7084 | 0.521626567 | 0.134623986 | YIR032C | DAL3 | n |
| W303.asm02.HP0_G0027570 | 7087.209 | 7145.298 | 6800.535 | 10003.22 | 9587.056 | 11398.77 | 16667.86 | 13141.16 | 14284.84 | 2.26E-05 | -0.559126239 | YIR033W | MGA2 | n |
| W303.asm02.HP0_G0027580 | 3268.82 | 3501.986 | 3160.433 | 2493.336 | 1967.399 | 2463.592 | 2292.417 | 1942.504 | 1228.379 | 0.035756499 | 0.520177408 | YIR034C | LYS1 | n |
| W303.asm02.HP0_G0027590 | 2788.08 | 2141.03 | 2466.258 | 1575.616 | 1872.804 | 1978.926 | 923.3859 | 1402.383 | 1053.75 | 0.034381634 | 0.446539819 | YIR035C | NRE1 | n |
| W303.asm02.HP0_G0027610 | 7652.661 | 7000.633 | 7186.427 | 3183.884 | 3993.811 | 3821.113 | 1428.285 | 2732.821 | 1464.007 | 0.000248287 | 0.989660345 | YIR037W | HYR1 | n |
| W303.asm02.HP0_G0027620 | 4606.209 | 4216.404 | 4168.279 | 2958.796 | 3197.132 | 4776.771 | 2166.5 | 9276.558 | 6904.995 | 0.619739327 | 0.248852638 | YIR038C | GTT1 | n |
| W303.asm02.HP0_G0027720 | 1018.661 | 1021.552 | 904.3666 | 91.00781 | 91.99131 | 207.3884 | 661.6776 | 471.3982 | 516.7084 | 1.85E-18 | 2.914526798 | YJL218W | | n |
| W303.asm02.HP0_G0027770 | 1443.279 | 1148.411 | 1431.465 | 1597.152 | 1693.161 | 1765.461 | 4703.342 | 9917.616 | 5322.575 | 0.288426138 | -0.32954316 | YJL212C | OPT1 | n |
| W303.asm02.HP0_G0027780 | 1379.745 | 1313.106 | 1370.024 | 1001.086 | 1076.125 | 1242.811 | 2465.243 | 1213.394 | 1527.4 | 0.304626887 | 0.291348978 | YJL210W | PEX2 | n |
| W303.asm02.HP0_G0027800 | 2878.087 | 2950.036 | 3022.46 | 3204.031 | 3648.41 | 1612.768 | 1148.06 | 923.4679 | 1184.123 | 0.847257916 | 0.064294088 | YJL208C | NUC1 | n |
| W303.asm02.HP0_G0027810 | 3125.869 | 3239.365 | 3084.979 | 1940.342 | 1597.698 | 1863.457 | 1269.038 | 1443.187 | 1528.596 | 8.61E-11 | 0.806537512 | YJL207C | LAA1 | n |
| W303.asm02.HP0_G0027820 | 903.2406 | 958.122 | 875.263 | 1280.362 | 1346.024 | 1746.469 | 5055.167 | 3061.404 | 4374.08 | 0.002924455 | -0.676210768 | YJL206C | | n |
| W303.asm02.HP0_G0027840 | 2117.797 | 2179.978 | 2009.225 | 2156.399 | 2119.271 | 2231.895 | 1868.992 | 2087.467 | 1862.303 | 0.658652473 | -0.045293242 | YJL204C | RCY1 | n |
| W303.asm02.HP0_G0027850 | 1974.846 | 2003.042 | 1920.836 | 1140.724 | 1015.376 | 986.0447 | 1298.666 | 1155.409 | 1309.712 | 1.67E-17 | 0.908108739 | YJL203W | PRP21 | n |
| W303.asm02.HP0_G0027860 | 1963.198 | 1972.997 | 2219.417 | 1977.162 | 1936.157 | 1794.328 | 932.0272 | 1020.11 | 851.612 | 0.389149584 | 0.109001173 | YJL201W | ECM25 | n |
| W303.asm02.HP0_G0027870 | 5491.449 | 6337.404 | 6575.252 | 3513.874 | 2824.827 | 2054.133 | 2761.516 | 1442.113 | 1790.538 | 0.000101173 | 1.132745898 | YJL200C | ACO2 | n |
| W303.asm02.HP0_G0027900 | 2249.101 | 2245.633 | 2332.598 | 5639.706 | 5292.972 | 3934.303 | 3035.569 | 2758.592 | 3600.214 | 2.85E-10 | -1.12272267 | YJL197W | UBP12 | n |
| W303.asm02.HP0_G0027920 | 133.4212 | 151.341 | 137.9725 | 1422.084 | 1733.949 | 1096.956 | 274.053 | 241.605 | 205.7265 | 6.84E-52 | -3.330711707 | YJL194W | CDC6 | n |
| W303.asm02.HP0_G0027930 | 541.0972 | 600.9127 | 627.3437 | 540.4892 | 552.8157 | 875.1336 | 3883.652 | 3119.389 | 3993.726 | 0.543424457 | -0.153994425 | YJL193W | | n |
| W303.asm02.HP0_G0027940 | 3180.932 | 3017.917 | 2932.993 | 2098.043 | 2217.338 | 2749.986 | 885.1172 | 2206.659 | 1545.341 | 0.261595107 | 0.370147444 | YJL192C | SOP4 | n |
| W303.asm02.HP0_G0027960 | 8154.579 | 7541.455 | 8395.842 | 8491.515 | 9685.122 | 9021.017 | 6803.181 | 5560.136 | 5460.125 | 0.202666451 | -0.174867915 | YJL190C | RPS22A | y |
| W303.asm02.HP0_G0027980 | 2342.284 | 2196.67 | 2440.389 | 3308.933 | 3541.665 | 4175.117 | 2933.108 | 2641.548 | 2064.442 | 0.000140676 | -0.659670655 | YJL187C | SWE1 | n |
| W303.asm02.HP0_G0027990 | 3303.764 | 3212.658 | 3210.016 | 4725.459 | 3940.005 | 4802.6 | 4062.651 | 6461.054 | 4439.865 | 0.023401036 | -0.469620826 | YJL186W | MNN5 | n |
| W303.asm02.HP0_G0028010 | 848.1779 | 731.1105 | 758.8487 | 808.6496 | 1092.614 | 703.4494 | 925.8548 | 757.0289 | 788.2196 | 0.482932843 | -0.155015605 | YJL184W | GON7 | n |
| W303.asm02.HP0_G0028020 | 1192.32 | 1119.478 | 1229.895 | 8832.621 | 9232.109 | 10383.1 | 5209.476 | 6157.169 | 5341.713 | 3.10E-141 | -3.005720765 | YJL183W | MNN11 | n |
| W303.asm02.HP0_G0028040 | 3067.63 | 3282.764 | 2864.007 | 2749.686 | 2667.748 | 1897.642 | 669.0844 | 803.2023 | 777.4548 | 0.07650196 | 0.33301697 | YJL180C | ATP12 | n |
| W303.asm02.HP0_G0028050 | 2026.732 | 1777.144 | 1795.798 | 1607.573 | 1766.06 | 1323.336 | 1443.099 | 1218.763 | 1286.987 | 0.114154985 | 0.253903079 | YJL179W | PFD1 | n |
| W303.asm02.HP0_G0028060 | 3538.839 | 3572.092 | 3519.377 | 3685.469 | 4506.706 | 4709.921 | 4252.76 | 4937.332 | 4126.491 | 0.050041085 | -0.279296705 | YJL178C | ATG27 | n |
| W303.asm02.HP0_G0028070 | 9285.483 | 10065.29 | 10312.37 | 14668.24 | 16680.8 | 16837.21 | 13895.23 | 12455.01 | 11610.39 | 1.83E-09 | -0.699927501 | YJL177W | RPL17B | y |
| W303.asm02.HP0_G0028080 | 3964.517 | 4569.162 | 4274.992 | 12056.8 | 10953.04 | 11824.18 | 6741.458 | 6740.242 | 6686.111 | 2.51E-59 | -1.443517312 | YJL176C | SWI3 | n |
| W303.asm02.HP0_G0028090 | 6502.697 | 6404.172 | 6153.789 | 2969.217 | 2940.25 | 3003.714 | 4139.188 | 3915.075 | 3249.761 | 1.43E-21 | 1.096568081 | YJL174W | KRE9 | n |
| W303.asm02.HP0_G0028100 | 1315.152 | 1137.283 | 1222.35 | 823.2386 | 1235.808 | 990.6027 | 370.3419 | 688.3058 | 381.5509 | 0.37733788 | 0.269363862 | YJL173C | RFA3 | n |
| W303.asm02.HP0_G0028110 | 2859.027 | 3584.333 | 2764.84 | 2646.174 | 2942.854 | 3947.977 | 2266.493 | 3184.891 | 2011.814 | 0.85575428 | -0.050634061 | YJL172W | CPS1 | n |
| W303.asm02.HP0_G0028130 | 371.6734 | 306.0204 | 306.1265 | 41.68297 | 34.7137 | 26.58826 | 11.11026 | 52.6162 | 20.33343 | 1.56E-12 | 3.254843902 | YJL170C | ASG7 | n |
| W303.asm02.HP0_G0028140 | 1039.839 | 1162.877 | 1045.573 | 4047.416 | 3850.617 | 3444.319 | 2322.044 | 2649.064 | 2771.327 | 1.44E-48 | -1.804134237 | YJL168C | SET2 | n |
| W303.asm02.HP0_G0028150 | 16084.67 | 16604.11 | 17521.43 | 11721.25 | 13041.07 | 13485.57 | 16778.96 | 15795.6 | 15071.86 | 7.25E-05 | 0.392643527 | YJL167W | ERG20 | n |
| W303.asm02.HP0_G0028160 | 3026.333 | 3116.957 | 2731.424 | 10594.42 | 12041.31 | 11334.2 | 3788.598 | 8039.54 | 7372.664 | 5.62E-13 | -1.936480466 | YJL166W | QCR8 | n |
| W303.asm02.HP0_G0028180 | 5219.312 | 5196.782 | 4798.856 | 4007.817 | 4336.609 | 4169.039 | 6619.245 | 6463.202 | 6487.561 | 0.000270782 | 0.282200389 | YJL164C | TPK1 | n |
| W303.asm02.HP0_G0028190 | 2128.386 | 1790.497 | 1710.643 | 1732.622 | 1790.359 | 2060.97 | 2041.818 | 3094.691 | 2157.736 | 0.960531662 | 0.011769603 | YJL163C | | n |
| W303.asm02.HP0_G0028200 | 1008.072 | 994.8444 | 1103.78 | 2348.835 | 2210.395 | 2585.139 | 765.3733 | 997.5601 | 760.7096 | 1.16E-15 | -1.20149151 | YJL162C | JJJ2 | n |
| W303.asm02.HP0_G0028210 | 3122.692 | 2638.452 | 2402.662 | 262.6027 | 276.8418 | 391.2273 | 232.0809 | 505.7598 | 342.0801 | 8.21E-26 | 3.132870084 | YJL161W | FMP33 | n |

| | | | | | | | | | | | | | | |
|---|---|---|---|---|---|---|---|---|---|---|---|---|---|---|
| W303.asm02.HP0_G0028220 | 105.8899 | 102.3777 | 82.99908 | 7462.641 | 8144.702 | 9093.185 | 4147.83 | 4647.406 | 3295.212 | 3.28E-239 | -6.406065891 | YJL160C | PIR5 | n |
| W303.asm02.HP0_G0028250 | 2370.874 | 2241.182 | 2546.024 | 480.0489 | 429.582 | 445.1635 | 433.3001 | 856.8923 | 598.0422 | 1.68E-23 | 2.401264281 | YJL157C | FAR1 | n |
| W303.asm02.HP0_G0028260 | 2160.153 | 2112.097 | 2084.678 | 5401.418 | 4974.473 | 6286.984 | 7654.968 | 8013.769 | 5745.989 | 5.90E-17 | -1.390273282 | YJL156C | SSY5 | n |
| W303.asm02.HP0_G0028270 | 6626.588 | 7392.34 | 6162.412 | 2831.663 | 2526.29 | 3420.77 | 4888.513 | 3368.51 | 3021.309 | 5.90E-08 | 1.200868585 | YJL155C | FBP26 | n |
| W303.asm02.HP0_G0028280 | 6297.271 | 6551.062 | 6065.4 | 3862.622 | 3915.705 | 5936.779 | 5157.629 | 5900.531 | 4987.672 | 0.022507239 | 0.46364268 | YJL154C | VPS35 | n |
| W303.asm02.HP0_G0028290 | 3507.073 | 4637.043 | 3688.609 | 5896.056 | 2845.656 | 6400.934 | 1180.156 | 2393.5 | 1272.634 | 0.37941281 | -0.355903891 | YJL153C | INO1 | n |
| W303.asm02.HP0_G0028310 | 1493.047 | 1630.254 | 1601.774 | 2511.399 | 2512.404 | 3562.067 | 3650.337 | 2870.267 | 2657.699 | 1.78E-05 | -0.861684543 | YJL149W | DAS1 | n |
| W303.asm02.HP0_G0028320 | 1357.508 | 1445.529 | 1633.034 | 4200.254 | 4767.927 | 1883.209 | 4139.188 | 1718.08 | 2974.662 | 0.00142765 | -1.290496847 | YJL148W | RPA34 | n |
| W303.asm02.HP0_G0028330 | 430.9718 | 517.4526 | 522.7864 | 2271.722 | 2349.25 | 2806.201 | 4931.72 | 4467.008 | 3798.764 | 1.49E-46 | -2.336022413 | YJL147C | SMT1 | n |
| W303.asm02.HP0_G0028340 | 1368.097 | 1642.495 | 1379.725 | 2136.252 | 2114.064 | 4209.302 | 5727.955 | 4824.583 | 5183.829 | 0.001080109 | -0.946335113 | YJL146W | IDS2 | n |
| W303.asm02.HP0_G0028350 | 2001.319 | 2026.411 | 2034.016 | 1260.215 | 1490.953 | 1506.415 | 766.6078 | 1169.368 | 1102.79 | 0.006800943 | 0.509884789 | YJL145W | SFH5 | n |
| W303.asm02.HP0_G0028370 | 7736.314 | 7245.45 | 7410.632 | 4306.545 | 4971.87 | 3682.094 | 2862.743 | 2140.083 | 2254.619 | 3.37E-06 | 0.788987323 | YJL143W | TIM17 | n |
| W303.asm02.HP0_G0028380 | 12190.04 | 12710.42 | 11561.66 | 3946.682 | 3968.644 | 5620.759 | 5240.338 | 3945.141 | 4139.648 | 7.68E-14 | 1.429565174 | YJL141C | YAK1 | n |
| W303.asm02.HP0_G0028390 | 6059.018 | 5562.894 | 5586.808 | 3074.814 | 3279.577 | 2914.833 | 3004.708 | 3086.101 | 2971.073 | 3.08E-25 | 0.892862897 | YJL140W | RPB4 | n |
| W303.asm02.HP0_G0028400 | 2072.265 | 2307.95 | 2067.432 | 2428.728 | 2462.937 | 2528.164 | 2413.395 | 2659.802 | 2608.66 | 0.033070137 | -0.2027135 | YJL139C | YUR1 | n |
| W303.asm02.HP0_G0028410 | 15068.13 | 16968 | 16794.92 | 35768.16 | 39353.19 | 38877.36 | 27037.43 | 25550 | 24370.22 | 4.48E-38 | -1.223144858 | YKR059W | TIF1 | y |
| W303.asm02.HP0_G0028420 | 1004.895 | 1060.5 | 986.2878 | 992.7494 | 863.5033 | 2392.184 | 8919.068 | 4210.37 | 5198.182 | 0.291725278 | -0.477398232 | YJL137C | GLG2 | n |
| W303.asm02.HP0_G0028450 | 1818.129 | 1684.781 | 1636.268 | 2946.291 | 2787.51 | 2992.319 | 3570.096 | 3402.872 | 3772.45 | 5.33E-17 | -0.763878614 | YJL134W | LCB3 | n |
| W303.asm02.HP0_G0028460 | 729.5812 | 926.9635 | 646.7461 | 238.2876 | 276.8418 | 338.0508 | 845.6141 | 459.5864 | 413.8452 | 6.07E-06 | 1.432889987 | YJL133C-A | DPI8 | n |
| W303.asm02.HP0_G0028490 | 2871.733 | 2849.884 | 2544.946 | 1350.528 | 1220.187 | 974.6497 | 517.2442 | 570.1878 | 638.709 | 7.20E-14 | 1.221221847 | YJL131C | AIM23 | n |
| W303.asm02.HP0_G0028500 | 13035.04 | 12353.21 | 13884.56 | 22704.02 | 23336.29 | 18396.04 | 14365.56 | 22652.88 | 17150.65 | 0.000363374 | -0.714336001 | YJL130C | URA2 | n |
| W303.asm02.HP0_G0028520 | 3563.194 | 3573.205 | 3683.219 | 8991.016 | 8278.35 | 6887.119 | 7874.704 | 7300.766 | 6794.955 | 1.13E-18 | -1.158782746 | YJL128C | PBS2 | n |
| W303.asm02.HP0_G0028530 | 1750.36 | 1599.096 | 1680.462 | 721.8101 | 775.8512 | 733.0764 | 885.1172 | 610.9922 | 726.0232 | 5.03E-14 | 1.173308535 | YJL127C-B | MCO6 | n |
| W303.asm02.HP0_G0028540 | 2322.165 | 2183.316 | 2138.574 | 1490.861 | 1633.28 | 1993.36 | 1281.383 | 1515.132 | 1233.163 | 0.018796553 | 0.376706167 | YJL127C | SPT10 | n |
| W303.asm02.HP0_G0028550 | 1585.171 | 1334.249 | 1523.087 | 840.6065 | 957.2303 | 1738.872 | 2073.915 | 2663.024 | 2735.445 | 0.291107992 | 0.328982487 | YJL126W | NIT2 | n |
| W303.asm02.HP0_G0028560 | 1731.299 | 2197.783 | 2174.145 | 2054.97 | 1735.685 | 537.0829 | 3641.696 | 1418.49 | 2722.288 | 0.343870658 | 0.495939675 | YJL125C | GCD14 | n |
| W303.asm02.HP0_G0028570 | 5943.598 | 6118.182 | 6381.228 | 3315.185 | 3576.379 | 3126.78 | 3562.689 | 2742.485 | 3314.35 | 1.48E-11 | 0.880527211 | YJL124C | LSM1 | n |
| W303.asm02.HP0_G0028580 | 11591.76 | 11823.51 | 11394.59 | 5259.001 | 5506.461 | 6457.149 | 3899.7 | 4815.993 | 4466.179 | 3.74E-15 | 1.015197658 | YJL123C | MTC1 | n |
| W303.asm02.HP0_G0028590 | 864.0613 | 801.217 | 927.0027 | 3144.98 | 3137.251 | 1008.835 | 3347.891 | 972.8627 | 2706.739 | 0.003692939 | -1.491883238 | YJL122W | ALB1 | n |
| W303.asm02.HP0_G0028600 | 4434.668 | 3829.15 | 4388.172 | 3312.407 | 4085.803 | 4637.752 | 6862.436 | 7252.445 | 6225.619 | 0.689810936 | 0.072132806 | YJL121C | RPE1 | n |
| W303.asm02.HP0_G0028620 | 1915.548 | 2024.186 | 2055.575 | 6965.224 | 7901.706 | 6724.551 | 4662.605 | 7392.039 | 5908.657 | 1.02E-23 | -1.848532361 | YJL117W | PHO86 | n |
| W303.asm02.HP0_G0028640 | 1635.999 | 1520.087 | 1530.632 | 3873.737 | 4286.274 | 4169.799 | 987.5785 | 1855.526 | 1105.182 | 7.40E-09 | -1.395461254 | YJL115W | ASF1 | n |
| W303.asm02.HP0_G0028650 | 2987.153 | 2879.93 | 2900.656 | 2730.929 | 2488.104 | 2625.401 | 1273.976 | 1415.268 | 1452.046 | 0.094996285 | 0.160296346 | YJL112W | MDV1 | n |
| W303.asm02.HP0_G0028660 | 8029.629 | 7709.488 | 8684.722 | 8659.637 | 8919.685 | 9351.471 | 4583.599 | 6648.969 | 5857.225 | 0.413335836 | -0.140955918 | YJL111W | CCT7 | n |
| W303.asm02.HP0_G0028670 | 6715.536 | 6835.939 | 7348.113 | 6689.422 | 6031.505 | 2470.429 | 12705.2 | 7554.183 | 14062.36 | 0.253220671 | 0.460224258 | YJL109C | UTP10 | n |
| W303.asm02.HP0_G0028680 | 230.8399 | 293.7796 | 239.2961 | 3301.986 | 3592 | 7262.394 | 5094.67 | 4521.772 | 4225.766 | 1.85E-42 | -4.212100694 | YJL107C | | n |
| W303.asm02.HP0_G0028690 | 419.3239 | 421.7517 | 443.0211 | 223.6986 | 234.3175 | 293.9902 | 251.8325 | 653.9442 | 397.1 | 0.024896846 | 0.771939262 | YJL106W | IME2 | n |
| W303.asm02.HP0_G0028700 | 721.11 | 893.5795 | 785.7965 | 2525.293 | 2302.386 | 2473.468 | 2781.268 | 2359.138 | 1882.637 | 5.84E-20 | -1.605039006 | YJL105W | SET4 | n |
| W303.asm02.HP0_G0028710 | 6169.144 | 6133.761 | 5752.806 | 1608.963 | 1654.976 | 1210.905 | 846.8485 | 798.9071 | 818.1217 | 1.59E-44 | 2.012691909 | YJL104W | PAM16 | n |
| W303.asm02.HP0_G0028720 | 1732.358 | 2282.356 | 1621.177 | 539.7944 | 336.7229 | 346.4071 | 2875.088 | 1079.169 | 1503.478 | 4.33E-09 | 2.203796665 | YJL103C | GSM1 | n |
| W303.asm02.HP0_G0028730 | 2087.089 | 2177.752 | 2010.302 | 1888.933 | 1674.068 | 843.2277 | 864.1312 | 406.9702 | 795.3961 | 0.182669339 | 0.510087811 | YJL102W | MEF2 | n |
| W303.asm02.HP0_G0028740 | 7327.579 | 6311.809 | 6728.315 | 13337.16 | 14149.3 | 15789.63 | 14163.11 | 24581.43 | 16899.48 | 6.41E-07 | -1.08727495 | YJL101C | GSH1 | n |
| W303.asm02.HP0_G0028750 | 3189.403 | 2894.396 | 3068.81 | 2267.553 | 2351.853 | 2727.956 | 2502.277 | 3048.518 | 3825.078 | 0.096732683 | 0.31698028 | YJL100W | LSB6 | n |
| W303.asm02.HP0_G0028760 | 3827.919 | 3997.183 | 3978.566 | 3898.747 | 4034.6 | 5054.808 | 1910.964 | 3348.108 | 2675.641 | 0.576212351 | -0.137967924 | YJL099W | CHS6 | n |
| W303.asm02.HP0_G0028780 | 2666.307 | 2464.855 | 2597.763 | 1619.383 | 1752.174 | 2266.839 | 2089.963 | 3078.584 | 2637.366 | 0.025503699 | 0.455014958 | YJL097W | PHS1 | n |
| W303.asm02.HP0_G0028790 | 4001.578 | 3812.458 | 3687.531 | 2407.191 | 2436.034 | 1533.003 | 1201.142 | 1162.925 | 1217.614 | 1.37E-05 | 0.85112352 | YJL096W | MRPL49 | n |
| W303.asm02.HP0_G0028800 | 1469.751 | 1544.568 | 1511.23 | 4199.559 | 3737.798 | 5600.248 | 4210.788 | 4275.871 | 3988.941 | 3.68E-21 | -1.580906849 | YJL095W | BCK1 | n |
| W303.asm02.HP0_G0028810 | 1285.503 | 1552.358 | 1168.455 | 2717.73 | 2727.629 | 3171.6 | 3439.242 | 4684.989 | 3869.333 | 4.48E-10 | -1.105023715 | YJL094C | KHA1 | n |
| W303.asm02.HP0_G0028820 | 1308.799 | 1332.023 | 1280.557 | 949.677 | 943.3448 | 1014.152 | 1075.226 | 990.0435 | 803.7687 | 0.001688937 | 0.431662731 | YJL093C | TOK1 | n |
| W303.asm02.HP0_G0028830 | 2231.1 | 2313.514 | 2484.583 | 2118.884 | 2121.007 | 2042.738 | 1438.161 | 1259.567 | 1142.261 | 0.195925486 | 0.161996148 | YJL092W | SRS2 | n |
| W303.asm02.HP0_G0028840 | 1777.891 | 1701.473 | 1684.774 | 1934.09 | 1815.527 | 2650.47 | 5477.357 | 5127.395 | 5785.46 | 0.081510039 | -0.309665759 | YJL091C | GWT1 | n |
| W303.asm02.HP0_G0028850 | 1247.383 | 1397.679 | 1380.803 | 1922.974 | 1695.764 | 1422.852 | 2422.036 | 1606.405 | 2389.776 | 0.136521307 | -0.324698831 | YJL090C | DPB11 | n |
| W303.asm02.HP0_G0028860 | 600.3956 | 673.2448 | 488.2933 | 654.4226 | 535.4588 | 623.6847 | 7933.959 | 2790.806 | 2542.875 | 0.932995457 | -0.041821522 | YJL089W | SIP4 | n |
| W303.asm02.HP0_G0028870 | 87.88859 | 80.1217 | 91.62236 | 2973.385 | 3131.176 | 3332.649 | 1887.509 | 2246.389 | 1615.91 | 8.99E-183 | -5.183242412 | YJL088W | ARG3 | n |

| | | | | | | | | | | | | | |
|---|---|---|---|---|---|---|---|---|---|---|---|---|---|
| W303.asm02.HP0_G0028880 | 783.585 | 844.6162 | 846.1595 | 2953.933 | 2834.374 | 2202.268 | 2141.811 | 1603.183 | 2107.501 | 1.05E-21 | -1.691291635 | YJL087C | TRL1 | n |
| W303.asm02.HP0_G0028890 | 1823.424 | 1887.311 | 1886.343 | 2061.223 | 1935.289 | 2282.033 | 1699.869 | 2524.504 | 2157.736 | 0.365961435 | -0.165852939 | YJL085W | EXO70 | n |
| W303.asm02.HP0_G0028910 | 897.9461 | 1058.274 | 895.7433 | 655.812 | 764.5693 | 1425.131 | 746.8562 | 1101.719 | 801.3765 | 0.992921844 | 0.003223725 | YJL083W | TAX4 | n |
| W303.asm02.HP0_G0028930 | 5295.552 | 5165.624 | 5566.328 | 3497.896 | 3367.229 | 3474.706 | 3536.765 | 3445.824 | 3505.723 | 3.12E-21 | 0.63221119 | YJL081C | ARP4 | n |
| W303.asm02.HP0_G0028940 | 17322.52 | 15698.29 | 18031.28 | 13697.02 | 13943.63 | 15341.43 | 4212.022 | 7581.027 | 5924.206 | 0.288697356 | 0.248237948 | YJL080C | SCP160 | n |
| W303.asm02.HP0_G0028990 | 3017.861 | 3356.209 | 3440.689 | 3057.446 | 3418.432 | 3844.663 | 1328.293 | 1898.478 | 1431.713 | 0.717152345 | -0.072466617 | YJL074C | SMC3 | n |
| W303.asm02.HP0_G0029000 | 1453.868 | 1575.727 | 1574.827 | 1233.816 | 1099.556 | 1381.07 | 1159.17 | 1286.412 | 1188.908 | 0.015967588 | 0.309378686 | YJL073W | JEM1 | n |
| W303.asm02.HP0_G0029010 | 1076.9 | 923.6251 | 972.2749 | 733.6202 | 726.3842 | 698.8914 | 953.0132 | 650.7228 | 865.9651 | 0.008148333 | 0.461619499 | YJL072C | PSF2 | n |
| W303.asm02.HP0_G0029020 | 1220.91 | 1156.201 | 1167.377 | 1349.139 | 1077.86 | 2059.451 | 1292.493 | 1992.973 | 1535.772 | 0.242814543 | -0.340139885 | YJL071W | ARG2 | n |
| W303.asm02.HP0_G0029040 | 949.8321 | 1000.408 | 1194.324 | 4072.426 | 3453.145 | 1365.117 | 5266.262 | 2056.327 | 4079.844 | 0.000756078 | -1.499442187 | YJL069C | UTP18 | n |
| W303.asm02.HP0_G0029050 | 5901.242 | 5828.854 | 5541.536 | 2339.109 | 2750.193 | 3374.43 | 2684.979 | 4129.835 | 3255.742 | 6.89E-07 | 1.029088162 | YJL068C | | n |
| W303.asm02.HP0_G0029060 | 10494.75 | 10919.92 | 10656.22 | 2301.595 | 2662.541 | 2549.434 | 1848.006 | 1734.187 | 1591.988 | 9.09E-97 | 2.094008168 | YJL066C | MPM1 | n |
| W303.asm02.HP0_G0029070 | 1945.197 | 1887.311 | 1886.343 | 987.8863 | 1107.367 | 1452.479 | 598.7194 | 659.3132 | 587.2774 | 5.24E-05 | 0.688919616 | YJL065C | DLS1 | n |
| W303.asm02.HP0_G0029080 | 7590.186 | 6785.863 | 7490.398 | 2157.788 | 2235.562 | 1450.2 | 1151.763 | 888.0325 | 1084.848 | 1.29E-21 | 1.90388935 | YJL063C | MRPL8 | n |
| W303.asm02.HP0_G0029090 | 4436.786 | 3691.162 | 3669.206 | 1667.319 | 1818.998 | 1349.924 | 1881.337 | 1391.645 | 1560.89 | 1.38E-12 | 1.286658591 | YJL062W-/ | COA3 | n |
| W303.asm02.HP0_G0029100 | 1426.337 | 1421.047 | 1573.749 | 2057.054 | 1818.13 | 2162.765 | 1156.701 | 1456.073 | 1281.006 | 0.001128423 | -0.449856332 | YJL062W | LAS21 | n |
| W303.asm02.HP0_G0029110 | 3046.452 | 3039.061 | 3178.757 | 6100.997 | 5861.408 | 6495.132 | 5565.005 | 5798.52 | 5698.146 | 1.29E-39 | -0.994541603 | YJL061W | NUP82 | n |
| W303.asm02.HP0_G0029120 | 5958.423 | 4176.344 | 5127.619 | 3264.471 | 3354.211 | 3966.969 | 2960.266 | 6249.516 | 3089.486 | 0.105610239 | 0.527883719 | YJL060W | BNA3 | n |
| W303.asm02.HP0_G0029130 | 1175.378 | 1005.972 | 1096.235 | 469.6281 | 515.4985 | 517.3316 | 509.8374 | 684.0106 | 399.4922 | 8.96E-08 | 1.125600116 | YJL059W | YHC3 | n |
| W303.asm02.HP0_G0029140 | 445.7964 | 486.2942 | 450.5664 | 673.8747 | 788.8688 | 754.347 | 953.0132 | 1030.848 | 1098.005 | 4.45E-08 | -0.680599542 | YJL058C | BIT61 | n |
| W303.asm02.HP0_G0029150 | 3520.838 | 3621.056 | 3253.133 | 1021.927 | 1306.971 | 2563.108 | 3607.13 | 4086.883 | 3392.095 | 0.00083094 | 1.087404912 | YJL057C | IKS1 | n |
| W303.asm02.HP0_G0029160 | 299.6683 | 358.322 | 286.7241 | 5233.297 | 5269.54 | 7731.107 | 14176.69 | 13766.12 | 14591.03 | 1.41E-108 | -4.27093305 | YJL056C | ZAP1 | n |
| W303.asm02.HP0_G0029180 | 7657.955 | 8164.624 | 8205.052 | 3643.091 | 3171.097 | 1620.365 | 1334.465 | 898.7705 | 1350.379 | 9.02E-07 | 1.510295484 | YJL054W | TIM54 | n |
| W303.asm02.HP0_G0029190 | 4806.341 | 4851.814 | 4782.687 | 1177.544 | 1169.852 | 1425.131 | 1260.397 | 1628.954 | 1376.693 | 4.14E-43 | 1.936458585 | YJL053W | PEP8 | n |
| W303.asm02.HP0_G0029210 | 1947.315 | 1845.025 | 1807.655 | 1065 | 951.1554 | 1422.852 | 499.9616 | 744.1434 | 478.4337 | 0.002791854 | 0.703282131 | YJL051W | IRC8 | n |
| W303.asm02.HP0_G0029220 | 1843.543 | 1770.467 | 2249.598 | 5978.032 | 5390.17 | 2173.4 | 7420.418 | 3121.536 | 6585.64 | 0.004992512 | -1.207532268 | YJL050W | MTR4 | n |
| W303.asm02.HP0_G0029230 | 1823.424 | 1921.808 | 1823.824 | 1372.759 | 1262.711 | 1472.99 | 1755.421 | 1872.707 | 1778.577 | 8.09E-06 | 0.438308571 | YJL049W | CHM7 | n |
| W303.asm02.HP0_G0029250 | 2443.938 | 2799.808 | 2427.454 | 847.5537 | 687.3313 | 615.3283 | 777.718 | 775.2835 | 655.4542 | 1.92E-29 | 1.834346126 | YJL047C | RTT101 | n |
| W303.asm02.HP0_G0029260 | 1909.194 | 1928.485 | 1894.966 | 1315.098 | 1201.962 | 947.3018 | 233.3154 | 516.4978 | 419.8256 | 0.0118971 | 0.726574399 | YJL046W | AIM22 | n |
| W303.asm02.HP0_G0029280 | 3648.965 | 3357.322 | 3518.299 | 1126.83 | 1166.38 | 894.1253 | 1394.955 | 988.9697 | 1021.456 | 2.01E-21 | 1.723484474 | YJL044C | GYP6 | n |
| W303.asm02.HP0_G0029300 | 11328.1 | 11144.71 | 10944.02 | 15254.58 | 13863.78 | 15440.94 | 12666.93 | 11668.98 | 11907.02 | 4.58E-07 | -0.415212893 | YJL042W | MHP1 | n |
| W303.asm02.HP0_G0029310 | 3238.112 | 3436.331 | 3376.015 | 10078.25 | 9482.915 | 8883.518 | 5482.295 | 7472.574 | 7394.193 | 5.69E-24 | -1.500925336 | YJL039C | NUP192 | n |
| W303.asm02.HP0_G0029340 | 5468.153 | 5794.357 | 5056.476 | 1636.751 | 1569.927 | 2698.329 | 1288.79 | 1858.748 | 1169.77 | 5.32E-08 | 1.466482935 | YJL036W | SNX4 | n |
| W303.asm02.HP0_G0029350 | 1025.014 | 950.3324 | 967.9633 | 628.7181 | 603.1505 | 483.1467 | 534.5268 | 356.5016 | 510.728 | 8.69E-05 | 0.779235514 | YJL035C | TAD2 | n |
| W303.asm02.HP0_G0029360 | 12956.68 | 13776.48 | 12919.83 | 15758.94 | 15997.81 | 21671.71 | 3402.208 | 10088.35 | 6274.658 | 0.255214261 | -0.430179581 | YJL034W | KAR2 | n |
| W303.asm02.HP0_G0029370 | 2388.875 | 2592.827 | 2964.253 | 4178.718 | 3293.462 | 826.5151 | 4444.103 | 1788.951 | 3709.057 | 0.919450817 | -0.062675091 | YJL033W | HCA4 | n |
| W303.asm02.HP0_G0029390 | 1610.585 | 1570.163 | 1541.412 | 532.1526 | 459.9565 | 436.0475 | 185.171 | 294.2212 | 264.3346 | 4.10E-21 | 1.724815086 | YJL030W | MAD2 | n |
| W303.asm02.HP0_G0029400 | 3185.167 | 3290.554 | 3035.395 | 4743.522 | 4320.988 | 5608.604 | 2797.316 | 3893.599 | 3205.506 | 0.000248422 | -0.625552623 | YJL029C | VPS53 | n |
| W303.asm02.HP0_G0029430 | 17544.89 | 17302.95 | 18053.92 | 10497.16 | 12554.21 | 18958.95 | 21531.68 | 25819.52 | 23360.72 | 0.164335563 | 0.332577679 | YJL026W | RNR2 | n |
| W303.asm02.HP0_G0029440 | 223.4276 | 208.0939 | 238.2181 | 1422.084 | 1293.953 | 900.9622 | 1688.759 | 1051.25 | 1452.046 | 7.37E-23 | -2.433042914 | YJL025W | RRN7 | n |
| W303.asm02.HP0_G0029460 | 1306.681 | 1301.978 | 1208.337 | 1204.638 | 1239.279 | 1216.983 | 518.4787 | 1646.135 | 703.2976 | 0.89256113 | 0.060249079 | YJL023C | PET130 | n |
| W303.asm02.HP0_G0029470 | 13918.16 | 13389.23 | 13814.5 | 7848.903 | 7532.005 | 13375.41 | 12364.48 | 12788.96 | 12613.91 | 0.030191423 | 0.516012648 | YJL020C | BBC1 | n |
| W303.asm02.HP0_G0029480 | 874.6503 | 875.7747 | 958.2621 | 858.6691 | 878.2566 | 1219.262 | 290.1012 | 656.0918 | 349.2566 | 0.721691129 | -0.126164601 | YJL019W | MPS3 | n |
| W303.asm02.HP0_G0029490 | 7876.089 | 7236.547 | 7275.893 | 3569.452 | 3900.084 | 4853.497 | 1036.957 | 4793.443 | 2126.638 | 0.070317677 | 0.861406959 | YJL016W | TPH3 | n |
| W303.asm02.HP0_G0029500 | 12979.98 | 13363.63 | 13658.2 | 8430.38 | 8385.962 | 8885.037 | 3398.504 | 4804.181 | 4093.001 | 1.01E-05 | 0.63821312 | YJL014W | CCT3 | n |
| W303.asm02.HP0_G0029520 | 7509.71 | 7642.72 | 8576.931 | 11612.18 | 13109.63 | 17576.36 | 3918.218 | 9097.233 | 5435.007 | 0.00915594 | -0.833919048 | YJL012C | VTC4 | n |
| W303.asm02.HP0_G0029530 | 174.7183 | 155.7922 | 160.6086 | 1614.52 | 1405.905 | 655.5906 | 1431.989 | 511.1288 | 916.2006 | 7.91E-12 | -2.903943211 | YJL011C | RPC17 | n |
| W303.asm02.HP0_G0029540 | 1633.881 | 1542.343 | 1897.122 | 4367.68 | 4075.388 | 1474.509 | 4540.392 | 2157.264 | 4252.08 | 0.026073132 | -0.967038345 | YJL010C | NOP9 | n |
| W303.asm02.HP0_G0029550 | 10167.55 | 11140.25 | 10906.29 | 10003.22 | 10334.27 | 10692.28 | 4830.493 | 7217.009 | 6120.363 | 0.766712774 | 0.05403893 | YJL008C | CCT8 | n |
| W303.asm02.HP0_G0029560 | 597.2189 | 614.2664 | 610.0971 | 1933.395 | 1929.214 | 1342.327 | 1723.324 | 1437.818 | 1690.067 | 3.76E-17 | -1.514653463 | YJL006C | CTK2 | n |
| W303.asm02.HP0_G0029570 | 9376.548 | 9245.154 | 8606.035 | 3683.385 | 3478.313 | 3167.042 | 3517.014 | 2913.219 | 3254.545 | 3.22E-34 | 1.39836073 | YJL005W | CYR1 | n |
| W303.asm02.HP0_G0029580 | 1428.454 | 1310.88 | 1297.804 | 856.585 | 886.9351 | 915.3959 | 596.2505 | 1311.11 | 968.8283 | 0.029678623 | 0.602535635 | YJL004C | SYS1 | n |
| W303.asm02.HP0_G0029590 | 1081.136 | 1037.131 | 883.8863 | 1134.471 | 1157.702 | 1199.51 | 4349.049 | 2445.042 | 2865.818 | 0.384796709 | -0.217909412 | YJL003W | COX16 | n |
| W303.asm02.HP0_G0029600 | 7277.811 | 7080.755 | 7680.11 | 5122.837 | 4949.306 | 6543.751 | 2270.196 | 4911.561 | 3988.941 | 0.165690065 | 0.407464365 | YJL002C | OST1 | n |

| | | | | | | | | | | | | | | |
|---|---|---|---|---|---|---|---|---|---|---|---|---|---|---|
| W303.asm02.HP0_G0029610 | 7566.89 | 8575.247 | 7716.759 | 295.2544 | 339.3264 | 312.9818 | 178.9986 | 211.5386 | 200.9422 | 0 | 4.655800559 | YJL001W | PRE3 | n |
| W303.asm02.HP0_G0029620 | 4555.382 | 3976.039 | 4230.797 | 5602.191 | 5962.946 | 7761.493 | 8438.858 | 11163.22 | 9366.536 | 0.00099336 | -0.598690379 | YJR001W | AVT1 | n |
| W303.asm02.HP0_G0029630 | 5431.091 | 5478.321 | 6000.726 | 5443.101 | 4389.547 | 1754.066 | 8462.313 | 3816.285 | 7604.704 | 0.226114426 | 0.545405746 | YJR002W | MPP10 | n |
| W303.asm02.HP0_G0029640 | 1790.598 | 1786.046 | 1974.731 | 1585.342 | 1622.866 | 1710.765 | 460.4585 | 608.8446 | 601.6304 | 0.236189227 | 0.174524848 | YJR003C | MRX12 | n |
| W303.asm02.HP0_G0029650 | 1381.863 | 1414.371 | 1436.854 | 1272.025 | 1177.662 | 1805.723 | 2123.294 | 1737.408 | 2216.344 | 0.974016996 | -0.007741776 | YJR004C | SAG1 | n |
| W303.asm02.HP0_G0029660 | 4750.22 | 5246.858 | 4860.297 | 4328.082 | 4098.82 | 4515.446 | 4604.585 | 4491.705 | 4700.611 | 0.022012621 | 0.198891608 | YJR005W | APL1 | n |
| W303.asm02.HP0_G0029680 | 4264.185 | 4158.539 | 4297.628 | 1790.978 | 1816.394 | 2211.384 | 554.2784 | 885.8849 | 631.5325 | 9.12E-09 | 1.128341675 | YJR006W | POL31 | n |
| W303.asm02.HP0_G0029690 | 7675.957 | 8398.312 | 8320.388 | 12738.32 | 14039.09 | 11551.46 | 7448.811 | 7390.965 | 7488.684 | 3.47E-10 | -0.651827703 | YJR007W | SUI2 | n |
| W303.asm02.HP0_G0029700 | 7960.8 | 8303.724 | 7469.917 | 426.5557 | 500.7451 | 568.2291 | 1776.407 | 1444.261 | 1441.282 | 2.45E-151 | 3.988807023 | YJR008W | MHO1 | n |
| W303.asm02.HP0_G0029720 | 4978.942 | 3687.824 | 4290.082 | 1071.252 | 1091.746 | 2181.757 | 2650.414 | 3907.558 | 3059.584 | 5.33E-07 | 1.576333561 | YJR010W | MET3 | n |
| W303.asm02.HP0_G0029730 | 468.0332 | 410.6237 | 402.0605 | 995.5282 | 1234.072 | 1201.789 | 1573.953 | 1097.424 | 1260.673 | 9.87E-15 | -1.42137276 | YJR010C-A | SPC1 | n |
| W303.asm02.HP0_G0029750 | 1768.361 | 1536.779 | 1537.1 | 1198.385 | 1163.777 | 1081.003 | 1313.479 | 1826.534 | 1596.773 | 0.002024689 | 0.49196723 | YJR013W | GPI14 | n |
| W303.asm02.HP0_G0029760 | 6494.226 | 6170.484 | 6367.215 | 4643.483 | 5168.87 | 3724.636 | 2146.749 | 2143.305 | 2395.757 | 0.000980072 | 0.491623252 | YJR014W | TMA22 | n |
| W303.asm02.HP0_G0029800 | 5980.66 | 6261.733 | 6579.563 | 8383.834 | 7903.442 | 12184.26 | 6425.432 | 4953.439 | 5557.008 | 0.003720828 | -0.597125162 | YJR016C | ILV3 | n |
| W303.asm02.HP0_G0029810 | 4570.207 | 3687.824 | 4216.784 | 1645.088 | 1917.064 | 1996.399 | 3067.666 | 3267.573 | 2890.936 | 5.44E-18 | 1.16651829 | YJR017C | ESS1 | n |
| W303.asm02.HP0_G0029820 | 1706.945 | 1976.335 | 1813.045 | 1321.35 | 1300.028 | 2658.066 | 2301.058 | 2385.983 | 1809.676 | 0.865842532 | 0.058008951 | YJR019C | TES1 | n |
| W303.asm02.HP0_G0029830 | 1102.314 | 959.2348 | 916.2236 | 67.38747 | 78.97367 | 93.43875 | 135.792 | 112.749 | 105.2554 | 5.75E-83 | 3.636324365 | YJR021C | REC107 | y |
| W303.asm02.HP0_G0029840 | 427.7951 | 377.2397 | 305.0486 | 760.7142 | 980.662 | 579.6241 | 1144.357 | 892.3277 | 1088.437 | 1.44E-05 | -1.063530694 | YJR022W | LSM8 | n |
| W303.asm02.HP0_G0029850 | 2006.613 | 1753.775 | 2037.25 | 1242.152 | 1319.988 | 1074.925 | 1371.5 | 834.3425 | 1306.124 | 0.002092765 | 0.672823369 | YJR024C | MDE1 | n |
| W303.asm02.HP0_G0029870 | 1190.202 | 1116.14 | 1156.598 | 4737.269 | 5515.139 | 4982.64 | 1480.133 | 1539.829 | 1180.535 | 3.32E-55 | -2.137169294 | YJR030C | RBH2 | n |
| W303.asm02.HP0_G0029890 | 1531.168 | 1367.633 | 1642.735 | 1202.554 | 1105.631 | 681.4192 | 1006.096 | 659.3132 | 940.1223 | 0.026042653 | 0.603233553 | YJR032W | CPR7 | n |
| W303.asm02.HP0_G0029900 | 3470.011 | 3170.371 | 3054.797 | 3149.843 | 2993.189 | 3992.797 | 2199.831 | 3256.835 | 2808.406 | 0.766435113 | -0.064157284 | YJR033C | RAV1 | n |
| W303.asm02.HP0_G0029910 | 2656.777 | 2415.892 | 2449.012 | 1500.587 | 1497.896 | 1504.136 | 1801.096 | 1347.619 | 1419.752 | 1.01E-07 | 0.740321214 | YJR034W | PET191 | n |
| W303.asm02.HP0_G0029920 | 2406.877 | 2432.584 | 2200.015 | 3407.583 | 3234.449 | 2756.823 | 3414.553 | 4987.801 | 3805.94 | 0.023373487 | -0.417040251 | YJR035W | RAD26 | n |
| W303.asm02.HP0_G0029930 | 1590.466 | 1958.53 | 1394.816 | 239.6771 | 199.6038 | 395.0256 | 562.9197 | 813.9403 | 504.7476 | 7.22E-17 | 2.566384955 | YJR036C | HUL4 | n |
| W303.asm02.HP0_G0029950 | 1227.264 | 1063.838 | 1112.403 | 1802.788 | 1737.421 | 1514.771 | 2106.011 | 2047.736 | 2207.972 | 1.35E-06 | -0.570558072 | YJR040W | GEF1 | n |
| W303.asm02.HP0_G0029960 | 1775.773 | 1821.656 | 1991.978 | 4581.653 | 3969.512 | 1608.21 | 8246.28 | 3657.363 | 7522.174 | 0.044687772 | -0.86204904 | YJR041C | URB2 | n |
| W303.asm02.HP0_G0029970 | 5459.682 | 5006.493 | 5550.159 | 5659.158 | 5270.408 | 4561.786 | 4205.85 | 4731.162 | 5181.437 | 0.752712006 | 0.048069823 | YJR042W | NUP85 | n |
| W303.asm02.HP0_G0029980 | 1329.977 | 1307.542 | 1412.062 | 1872.955 | 1823.337 | 2036.661 | 693.7739 | 566.9664 | 435.3747 | 0.006274335 | -0.501535192 | YJR043C | POL32 | n |
| W303.asm02.HP0_G0029990 | 3517.662 | 3483.068 | 3203.549 | 678.7377 | 847.0143 | 1071.887 | 1271.507 | 2460.076 | 1648.204 | 4.11E-13 | 1.974028325 | YJR044C | VPS55 | n |
| W303.asm02.HP0_G0030000 | 54921.9 | 55213.87 | 56556.87 | 20988.07 | 20913.27 | 19680.63 | 11806.5 | 9823.122 | 10617.64 | 3.92E-51 | 1.436612578 | YJR045C | SSC1 | n |
| W303.asm02.HP0_G0030010 | 1332.095 | 1344.264 | 1320.44 | 1648.561 | 1495.293 | 1176.72 | 3423.194 | 1734.187 | 3176.8 | 0.712328028 | -0.112416197 | YJR046W | TAH11 | n |
| W303.asm02.HP0_G0030020 | 9008.051 | 9232.913 | 8920.784 | 10159.53 | 11828.69 | 8550.785 | 3173.83 | 7068.825 | 7656.136 | 0.630096951 | -0.169064233 | YJR048W | CYC1 | n |
| W303.asm02.HP0_G0030030 | 5822.884 | 6009.127 | 5764.663 | 1779.863 | 1848.505 | 1985.763 | 2043.053 | 1870.559 | 1880.245 | 1.04E-83 | 1.648245346 | YJR049C | UTR1 | n |
| W303.asm02.HP0_G0030040 | 1726.005 | 1741.534 | 1612.554 | 1950.763 | 1973.474 | 1917.393 | 5274.904 | 4118.023 | 5173.065 | 0.137303605 | -0.201501992 | YJR050W | ISY1 | n |
| W303.asm02.HP0_G0030060 | 2789.139 | 2661.821 | 2609.62 | 2219.618 | 2006.452 | 2201.508 | 3242.961 | 3310.525 | 3203.114 | 8.87E-05 | 0.326199177 | YJR052W | RAD7 | n |
| W303.asm02.HP0_G0030070 | 715.8155 | 768.9457 | 761.0046 | 2135.557 | 1988.227 | 2026.785 | 2139.342 | 2178.74 | 1588.4 | 7.49E-22 | -1.453673542 | YJR053W | BFA1 | n |
| W303.asm02.HP0_G0030080 | 632.1625 | 638.748 | 704.9532 | 1133.082 | 639.5999 | 704.2091 | 3502.2 | 3137.643 | 2851.465 | 0.197879879 | -0.326786526 | YJR054W | KCH1 | n |
| W303.asm02.HP0_G0030090 | 435.2074 | 482.9558 | 450.5664 | 183.4051 | 172.7007 | 79.00512 | 1139.419 | 473.5458 | 1076.476 | 5.18E-05 | 1.653450546 | YJR055W | HIT1 | n |
| W303.asm02.HP0_G0030110 | 1243.147 | 1210.728 | 1211.571 | 1002.475 | 1183.737 | 900.9622 | 211.0949 | 397.306 | 307.3937 | 0.325834234 | 0.247936389 | YJR057W | CDC8 | n |
| W303.asm02.HP0_G0030120 | 938.1843 | 961.4604 | 835.3804 | 1116.409 | 1186.341 | 1128.102 | 1417.175 | 1200.508 | 1281.006 | 0.005866828 | -0.326823119 | YJR058C | APS2 | n |
| W303.asm02.HP0_G0030140 | 2535.004 | 2792.019 | 2593.452 | 4188.444 | 4659.446 | 4753.222 | 8317.88 | 6490.047 | 7749.43 | 2.78E-09 | -0.780035581 | YJR060W | CBF1 | n |
| W303.asm02.HP0_G0030150 | 1950.491 | 1714.827 | 1689.085 | 1706.223 | 1522.196 | 1811.8 | 3744.157 | 3293.344 | 2456.757 | 0.677212151 | 0.087138158 | YJR061W | MNN14 | n |
| W303.asm02.HP0_G0030160 | 3035.863 | 3471.94 | 2984.733 | 1385.264 | 1381.605 | 1716.082 | 1154.232 | 1579.56 | 1359.948 | 1.36E-10 | 1.082256173 | YJR062C | NTA1 | n |
| W303.asm02.HP0_G0030180 | 190.6018 | 183.6122 | 185.4005 | 1770.137 | 1855.447 | 616.8477 | 2330.685 | 722.6674 | 1772.597 | 1.77E-09 | -2.922331649 | YJR063W | RPA12 | n |
| W303.asm02.HP0_G0030190 | 7329.697 | 6966.137 | 7746.94 | 8609.617 | 8896.254 | 8929.858 | 3919.452 | 5437.723 | 4169.55 | 0.093768003 | -0.262166249 | YJR064W | CCT5 | n |
| W303.asm02.HP0_G0030200 | 12704.67 | 11804.6 | 12141.58 | 7492.514 | 8224.544 | 10161.27 | 11143.59 | 12907.08 | 11849.61 | 0.000644504 | 0.502121065 | YJR065C | ARP3 | n |
| W303.asm02.HP0_G0030220 | 888.416 | 627.62 | 695.252 | 727.3678 | 897.3492 | 493.782 | 672.7878 | 674.3464 | 674.5916 | 0.824925732 | 0.062435596 | YJR067C | YAE1 | n |
| W303.asm02.HP0_G0030230 | 3203.169 | 3059.091 | 3122.706 | 2623.943 | 2696.387 | 2602.611 | 1561.608 | 2011.227 | 1746.283 | 0.044853281 | 0.244388456 | YJR068W | RFC2 | n |
| W303.asm02.HP0_G0030240 | 1400.923 | 1033.792 | 1285.947 | 1925.058 | 2077.615 | 1354.482 | 2571.407 | 2206.659 | 2721.092 | 0.01648781 | -0.525691992 | YJR069C | HAM1 | n |
| W303.asm02.HP0_G0030250 | 6026.193 | 5904.524 | 6446.981 | 5492.426 | 5846.655 | 4684.092 | 21751.42 | 11481.07 | 14940.29 | 0.450717195 | 0.197817927 | YJR070C | LIA1 | n |
| W303.asm02.HP0_G0030260 | 4078.878 | 3846.954 | 4183.369 | 3875.821 | 3889.67 | 2548.675 | 4173.754 | 2984.09 | 3668.391 | 0.310210729 | 0.231511392 | YJR072C | NPA3 | n |
| W303.asm02.HP0_G0030270 | 10407.92 | 10324.57 | 10842.7 | 2555.166 | 2941.986 | 3155.647 | 2777.564 | 5038.269 | 3085.898 | 2.03E-15 | 1.86759495 | YJR073C | OPI3 | n |
| W303.asm02.HP0_G0030280 | 1615.879 | 1693.684 | 1690.163 | 1654.814 | 1936.157 | 1822.435 | 2235.631 | 2096.057 | 2041.716 | 0.294473886 | -0.114213288 | YJR074W | MOG1 | n |

| | | | | | | | | | | | | | | |
|---|---|---|---|---|---|---|---|---|---|---|---|---|---|---|
| W303.asm02.HP0_G0030290 | 4699.392 | 4747.211 | 5110.372 | 7295.214 | 7630.939 | 8236.284 | 4276.215 | 5832.881 | 5072.594 | 2.75E-06 | -0.670064582 | YJR075W | HOC1 | n |
| W303.asm02.HP0_G0030300 | 4738.572 | 4875.183 | 4989.646 | 8609.617 | 8919.685 | 9962.242 | 5397.116 | 6986.142 | 6040.226 | 1.98E-12 | -0.912683406 | YJR076C | CDC11 | n |
| W303.asm02.HP0_G0030310 | 4541.617 | 4314.331 | 4246.966 | 21923.85 | 24050.52 | 21225.79 | 3518.248 | 10748.74 | 7800.862 | 7.17E-12 | -2.3585718 | YJR077C | MIR1 | n |
| W303.asm02.HP0_G0030330 | 4315.012 | 4533.553 | 4173.668 | 5101.995 | 5220.941 | 6653.902 | 11496.65 | 10233.31 | 9889.225 | 0.009372174 | -0.382616032 | YJR080C | AIM24 | n |
| W303.asm02.HP0_G0030340 | 2059.558 | 1745.985 | 1791.487 | 391.8199 | 420.9036 | 402.6222 | 497.4927 | 330.7304 | 460.4925 | 4.91E-36 | 2.203684151 | YJR082C | EAF6 | n |
| W303.asm02.HP0_G0030350 | 1615.879 | 1747.098 | 1547.879 | 1294.256 | 1374.663 | 1319.537 | 533.2924 | 773.1359 | 613.5913 | 0.068778093 | 0.300215456 | YJR083C | ACF4 | n |
| W303.asm02.HP0_G0030360 | 3247.642 | 3144.777 | 2932.993 | 2438.454 | 3086.916 | 2132.379 | 720.9323 | 1881.297 | 1379.085 | 0.425159521 | 0.284315959 | YJR085C | TMH11 | n |
| W303.asm02.HP0_G0030370 | 964.6567 | 791.2018 | 732.9789 | 15.28376 | 9.546268 | 25.8286 | 19.75157 | 94.49439 | 47.84337 | 2.94E-27 | 5.613178789 | YJR086W | STE18 | n |
| W303.asm02.HP0_G0030380 | 4140.294 | 4714.939 | 4462.548 | 1727.759 | 1797.302 | 1909.797 | 1187.563 | 1664.39 | 1603.949 | 1.00E-16 | 1.293051374 | YJR088C | EMC2 | n |
| W303.asm02.HP0_G0030390 | 1428.454 | 1508.959 | 1403.439 | 2949.07 | 2488.104 | 2961.173 | 1896.151 | 1930.692 | 1594.38 | 2.42E-13 | -0.952409358 | YJR089W | BIR1 | n |
| W303.asm02.HP0_G0030400 | 3015.744 | 3535.37 | 2981.499 | 4007.123 | 4017.243 | 5958.05 | 5932.878 | 5977.844 | 4970.926 | 0.006920987 | -0.552719365 | YJR090C | GRR1 | n |
| W303.asm02.HP0_G0030410 | 6764.245 | 7251.014 | 6649.628 | 10807 | 10264.84 | 13173.34 | 13345.89 | 11515.43 | 11231.23 | 2.07E-07 | -0.728754834 | YJR091C | JSN1 | n |
| W303.asm02.HP0_G0030420 | 3526.133 | 3461.925 | 3621.778 | 7661.33 | 6863.766 | 6939.536 | 2578.814 | 2665.171 | 2780.896 | 8.28E-34 | -1.016666287 | YJR092W | BUD4 | n |
| W303.asm02.HP0_G0030430 | 2362.403 | 2553.879 | 2515.842 | 2416.917 | 2200.849 | 2146.053 | 2365.25 | 1989.751 | 2313.227 | 0.263986637 | 0.135712511 | YJR093C | FIP1 | n |
| W303.asm02.HP0_G0030440 | 830.1766 | 888.0155 | 693.0962 | 582.1721 | 487.7275 | 732.3167 | 650.5673 | 2075.655 | 868.3572 | 0.344068766 | 0.419887529 | YJR094C | IME1 | n |
| W303.asm02.HP0_G0030460 | 227.6632 | 221.4475 | 174.6214 | 6485.87 | 6902.819 | 5850.937 | 2028.239 | 2304.375 | 1889.813 | 7.33E-224 | -4.947023057 | YJR095W | SFC1 | n |
| W303.asm02.HP0_G0030470 | 6326.92 | 6536.595 | 5908.025 | 426.5557 | 570.1725 | 1328.653 | 198.7502 | 585.221 | 228.4521 | 1.06E-09 | 3.013009947 | YJR096W | | n |
| W303.asm02.HP0_G0030480 | 75.18181 | 75.67049 | 88.38863 | 535.6261 | 508.5557 | 311.4625 | 450.5827 | 388.7156 | 501.1593 | 1.93E-21 | -2.502330583 | YJR097W | JJJ3 | n |
| W303.asm02.HP0_G0030490 | 1106.549 | 1220.743 | 1088.689 | 2090.401 | 2094.972 | 1890.805 | 2567.704 | 2776.847 | 3054.799 | 6.70E-13 | -0.830957055 | YJR098C | | n |
| W303.asm02.HP0_G0030500 | 1429.513 | 1427.724 | 1263.311 | 865.6163 | 899.0848 | 1435.766 | 936.9651 | 1416.342 | 1184.123 | 0.171845343 | 0.364499359 | YJR099W | YUH1 | n |
| W303.asm02.HP0_G0030510 | 3184.108 | 3065.768 | 2918.981 | 843.3854 | 858.2962 | 942.7438 | 513.5408 | 726.9626 | 556.1792 | 1.21E-31 | 1.793794213 | YJR100C | AIM25 | n |
| W303.asm02.HP0_G0030520 | 7872.912 | 7443.528 | 7359.97 | 3471.497 | 3495.67 | 2606.409 | 1314.714 | 1144.671 | 1217.614 | 2.72E-17 | 1.244114456 | YJR101W | RSM26 | n |
| W303.asm02.HP0_G0030530 | 1587.289 | 1397.679 | 1336.609 | 552.994 | 649.1462 | 722.4411 | 388.859 | 663.6084 | 586.0813 | 2.28E-07 | 1.167265414 | YJR102C | VPS25 | n |
| W303.asm02.HP0_G0030550 | 10275.55 | 9921.737 | 10035.34 | 4281.536 | 5446.58 | 5029.739 | 6809.354 | 9456.956 | 6259.109 | 3.85E-08 | 1.034707103 | YJR104C | SOD1 | n |
| W303.asm02.HP0_G0030560 | 6155.378 | 5378.169 | 6277.749 | 9321.701 | 10073.05 | 10052.64 | 6337.785 | 6425.619 | 6025.873 | 1.16E-13 | -0.725252572 | YJR105W | ADO1 | n |
| W303.asm02.HP0_G0030570 | 1530.109 | 1644.72 | 1474.581 | 1629.109 | 1608.98 | 2290.389 | 3371.346 | 2667.319 | 2759.367 | 0.212700114 | -0.249933145 | YJR106W | ECM27 | n |
| W303.asm02.HP0_G0030580 | 1590.466 | 1484.477 | 1593.151 | 37.51467 | 35.58154 | 28.86726 | 37.03419 | 35.4354 | 43.05904 | 1.46E-202 | 5.515715933 | YJR107W | LIH1 | n |
| W303.asm02.HP0_G0030610 | 2623.951 | 2576.135 | 2584.829 | 5374.324 | 5290.368 | 4045.214 | 5642.776 | 6134.619 | 4935.044 | 2.63E-09 | -0.918015495 | YJR109C | CPA2 | n |
| W303.asm02.HP0_G0030620 | 1178.554 | 1043.808 | 1014.313 | 2255.049 | 2305.858 | 2727.956 | 1646.787 | 2260.349 | 1666.145 | 2.08E-11 | -1.171127959 | YJR110W | YMR1 | n |
| W303.asm02.HP0_G0030630 | 1614.821 | 1641.382 | 1731.124 | 960.0977 | 936.4021 | 977.6884 | 316.0251 | 610.9922 | 418.6295 | 0.000674684 | 0.795075501 | YJR111C | PXP2 | n |
| W303.asm02.HP0_G0030640 | 230.8399 | 228.1243 | 239.2961 | 769.0508 | 724.6485 | 625.9636 | 822.1591 | 693.6748 | 659.0425 | 3.02E-25 | -1.602078351 | YJR112W | NNF1 | n |
| W303.asm02.HP0_G0030650 | 957.2444 | 770.0586 | 846.1595 | 198.6888 | 219.5642 | 154.9716 | 262.9428 | 204.022 | 265.5307 | 2.21E-28 | 2.167850808 | YJR112W-A | | n |
| W303.asm02.HP0_G0030660 | 5585.691 | 5503.916 | 5525.367 | 3789.677 | 3856.692 | 2671.74 | 1541.857 | 1405.604 | 1727.146 | 9.67E-05 | 0.687351126 | YJR113C | RSM7 | n |
| W303.asm02.HP0_G0030680 | 777.2317 | 688.8241 | 762.0825 | 1195.606 | 1346.024 | 1868.775 | 2515.856 | 3310.525 | 2414.894 | 7.55E-06 | -0.984973824 | YJR116W | TDA4 | n |
| W303.asm02.HP0_G0030690 | 5150.483 | 4958.643 | 4946.53 | 5575.792 | 5333.76 | 7096.027 | 4005.865 | 6783.194 | 5458.929 | 0.269970492 | -0.258154023 | YJR117W | STE24 | n |
| W303.asm02.HP0_G0030700 | 1805.422 | 1813.866 | 1794.72 | 1249.794 | 1403.301 | 1194.952 | 1265.335 | 1755.663 | 1623.086 | 0.001630143 | 0.492844647 | YJR118C | ILM1 | n |
| W303.asm02.HP0_G0030710 | 447.9142 | 403.9469 | 472.1246 | 2726.761 | 2749.325 | 2889.764 | 9606.67 | 6264.549 | 7918.078 | 3.16E-50 | -2.659474774 | YJR119C | JHD2 | n |
| W303.asm02.HP0_G0030730 | 24512.45 | 24007.58 | 23664.44 | 42958.47 | 47469.25 | 46464.13 | 14827.26 | 28044.43 | 22632.31 | 5.05E-05 | -0.923273069 | YJR121W | ATP2 | n |
| W303.asm02.HP0_G0030740 | 3335.531 | 3290.554 | 3030.005 | 1593.679 | 1367.72 | 831.8328 | 270.3496 | 715.1508 | 309.7858 | 0.000635661 | 1.347982198 | YJR122W | IBA57 | n |
| W303.asm02.HP0_G0030750 | 26998.74 | 26886.39 | 28987.16 | 40046.22 | 43086.65 | 45468.97 | 33664.08 | 29338.36 | 28873.48 | 2.24E-09 | -0.63394107 | YJR123W | RPS5 | n |
| W303.asm02.HP0_G0030760 | 797.3507 | 717.7569 | 845.0816 | 848.2484 | 715.1022 | 333.4928 | 1960.343 | 1206.951 | 1582.42 | 0.401038223 | 0.315250777 | YJR124C | | n |
| W303.asm02.HP0_G0030770 | 7625.13 | 7671.653 | 7117.441 | 5355.567 | 5310.328 | 7829.863 | 7390.79 | 9062.871 | 7530.547 | 0.169852854 | 0.27719822 | YJR125C | ENT3 | n |
| W303.asm02.HP0_G0030780 | 3398.006 | 3298.343 | 3094.68 | 1162.26 | 1011.904 | 1638.597 | 1824.551 | 2132.567 | 1588.4 | 1.40E-10 | 1.360424991 | YJR126C | VPS70 | n |
| W303.asm02.HP0_G0030800 | 910.6529 | 903.5947 | 1044.495 | 582.8668 | 613.5647 | 225.6204 | 591.3126 | 429.52 | 543.0223 | 0.005747536 | 1.007610983 | YJR129C | EFM3 | n |
| W303.asm02.HP0_G0030810 | 3886.158 | 3863.646 | 3954.852 | 3864.011 | 3733.459 | 4026.982 | 2772.627 | 2719.935 | 2510.581 | 0.916192861 | 0.009816865 | YJR130C | STR2 | n |
| W303.asm02.HP0_G0030820 | 1408.335 | 1402.13 | 1316.128 | 2992.142 | 2839.581 | 2820.635 | 2204.769 | 2591.079 | 2730.661 | 4.78E-21 | -1.068229394 | YJR131W | MNS1 | n |
| W303.asm02.HP0_G0030830 | 2331.695 | 2127.676 | 2362.779 | 6190.616 | 5535.1 | 3249.086 | 7453.749 | 4368.218 | 6925.328 | 0.000106671 | -1.134234202 | YJR132W | NMD5 | n |
| W303.asm02.HP0_G0030840 | 2280.868 | 2351.349 | 2162.288 | 5496.594 | 6190.321 | 3868.212 | 5787.21 | 3152.677 | 4811.847 | 6.56E-06 | -1.194920984 | YJR133W | XPT1 | n |
| W303.asm02.HP0_G0030850 | 3725.206 | 3963.798 | 3854.607 | 3950.156 | 3432.317 | 4791.205 | 8257.39 | 6313.944 | 8097.491 | 0.695105856 | -0.076767661 | YJR134C | SGM1 | n |
| W303.asm02.HP0_G0030860 | 599.3367 | 594.2359 | 616.5646 | 599.54 | 593.6043 | 414.0172 | 434.5345 | 493.948 | 391.1196 | 0.408558751 | 0.171738578 | YJR135C | MCM22 | n |
| W303.asm02.HP0_G0030870 | 1558.699 | 1397.679 | 1396.972 | 1254.657 | 1416.319 | 998.959 | 655.5052 | 660.387 | 645.8855 | 0.136406595 | 0.246788739 | YJR135W-. | TIM8 | n |
| W303.asm02.HP0_G0030880 | 815.352 | 815.6834 | 848.3153 | 1369.286 | 1249.693 | 1196.472 | 1422.113 | 1405.604 | 1411.38 | 5.61E-11 | -0.622322253 | YJR136C | TTI2 | n |
| W303.asm02.HP0_G0030890 | 3988.871 | 2661.821 | 3162.588 | 4625.42 | 4785.284 | 5272.072 | 4846.541 | 4447.679 | 3994.922 | 0.001379911 | -0.581239464 | YJR137C | MET5 | n |
| W303.asm02.HP0_G0030900 | 1841.425 | 1920.695 | 1713.877 | 2257.827 | 2061.994 | 2501.576 | 2334.389 | 2006.932 | 2259.403 | 0.013170732 | -0.317182667 | YJR138W | IML1 | n |

| | | | | | | | | | | | | | | |
|---|---|---|---|---|---|---|---|---|---|---|---|---|---|---|
| W303.asm02.HP0_G0030910 | 6590.586 | 6060.316 | 6377.994 | 4586.516 | 4842.561 | 5484.019 | 3291.105 | 4164.196 | 3533.233 | 0.01047261 | 0.351644482 | YJR139C | HOM6 | n |
| W303.asm02.HP0_G0030920 | 2492.648 | 2497.126 | 2475.96 | 4287.788 | 4066.71 | 5133.054 | 5102.077 | 5771.675 | 5146.751 | 8.00E-12 | -0.85334817 | YJR140C | HIR3 | n |
| W303.asm02.HP0_G0030930 | 852.4135 | 727.7721 | 812.7442 | 994.1388 | 1002.358 | 611.53 | 1291.259 | 872.9993 | 1089.633 | 0.652452649 | -0.124018163 | YJR141W | IPA1 | n |
| W303.asm02.HP0_G0030940 | 4921.761 | 4509.071 | 4486.262 | 2270.332 | 2325.818 | 2180.997 | 1285.086 | 1625.733 | 1693.655 | 3.83E-15 | 1.038177075 | YJR142W | | n |
| W303.asm02.HP0_G0030950 | 6161.732 | 5720.912 | 6565.551 | 9321.006 | 9315.422 | 11084.27 | 6366.178 | 10508.21 | 9791.146 | 0.001217982 | -0.687985706 | YJR143C | PMT4 | n |
| W303.asm02.HP0_G0030960 | 1297.151 | 1367.633 | 1315.05 | 1941.037 | 2331.893 | 1732.035 | 1103.619 | 1670.833 | 1363.536 | 0.002621966 | -0.593278985 | YJR144W | MGM101 | n |
| W303.asm02.HP0_G0030990 | 1395.628 | 1315.331 | 1252.532 | 609.9608 | 652.6176 | 910.8379 | 780.187 | 1020.11 | 782.2391 | 2.58E-05 | 0.866814561 | YJR149W | | n |
| W303.asm02.HP0_G0031020 | 599.3367 | 638.748 | 609.0192 | 224.3933 | 227.3747 | 344.8877 | 364.1696 | 458.5126 | 199.7461 | 0.00019082 | 1.213078446 | YJR152W | DAL5 | n |
| W303.asm02.HP0_G0031030 | 455.3264 | 476.279 | 481.8258 | 368.1996 | 321.9696 | 540.1215 | 574.03 | 2032.703 | 706.8858 | 0.698096482 | 0.200058021 | YJR153W | PGU1 | n |
| W303.asm02.HP0_G0031140 | 1498.342 | 1600.208 | 1522.009 | 1057.358 | 914.706 | 1308.142 | 3803.412 | 2798.323 | 2794.053 | 0.013250049 | 0.493990181 | YKL222C | | n |
| W303.asm02.HP0_G0031150 | 387.5569 | 385.0293 | 376.1906 | 222.3092 | 215.2249 | 316.0205 | 354.2938 | 770.9883 | 510.728 | 0.057208452 | 0.608078171 | YKL221W | MCH2 | n |
| W303.asm02.HP0_G0031190 | 1113.961 | 1246.338 | 1016.469 | 3354.09 | 2121.875 | 1444.882 | 22576.04 | 11115.98 | 10004.05 | 0.009029409 | -1.035400271 | YKL217W | JEN1 | n |
| W303.asm02.HP0_G0031200 | 2340.166 | 2632.888 | 2772.385 | 6620.645 | 7000.886 | 13055.6 | 27474.43 | 23893.12 | 24927.59 | 3.11E-11 | -1.784200158 | YKL216W | URA1 | n |
| W303.asm02.HP0_G0031210 | 4499.261 | 4205.276 | 4438.834 | 5013.072 | 4599.565 | 7452.31 | 857.9588 | 6127.102 | 2739.033 | 0.533055348 | -0.376703018 | YKL215C | OXP1 | n |
| W303.asm02.HP0_G0031230 | 8836.51 | 8912.426 | 8474.53 | 4233.6 | 4427.733 | 5948.174 | 2581.283 | 4831.026 | 3329.899 | 0.000954978 | 0.843947541 | YKL213C | DOA1 | n |
| W303.asm02.HP0_G0031240 | 10357.09 | 9635.747 | 10723.05 | 10224.14 | 10608.51 | 14195.85 | 8889.441 | 12486.15 | 10129.64 | 0.358588278 | -0.189542565 | YKL212W | SAC1 | n |
| W303.asm02.HP0_G0031250 | 3049.628 | 2602.842 | 2995.512 | 6089.187 | 6100.065 | 6844.578 | 3677.495 | 4993.17 | 4266.433 | 3.82E-14 | -1.138090037 | YKL211C | TRP3 | n |
| W303.asm02.HP0_G0031260 | 18035.16 | 18598.25 | 18257.64 | 16556.48 | 16937.68 | 27706.49 | 9149.915 | 21763.78 | 13650.91 | 0.675544609 | -0.156977266 | YKL210W | UBA1 | n |
| W303.asm02.HP0_G0031270 | 1999.201 | 1948.515 | 2096.535 | 955.2347 | 866.1068 | 1372.714 | 443.1758 | 990.0435 | 436.5708 | 0.009247901 | 0.920101967 | YKL209C | STE6 | n |
| W303.asm02.HP0_G0031280 | 1385.04 | 1428.837 | 1450.867 | 1211.585 | 1218.451 | 869.816 | 1587.532 | 1104.94 | 1565.674 | 0.08517657 | 0.370127174 | YKL208W | CBT1 | n |
| W303.asm02.HP0_G0031290 | 3605.55 | 2996.774 | 3181.991 | 1954.931 | 1947.439 | 2457.515 | 2543.015 | 4719.351 | 3980.569 | 0.01321996 | 0.621467135 | YKL207W | EMC3 | n |
| W303.asm02.HP0_G0031300 | 2090.266 | 1951.854 | 1963.952 | 1460.293 | 1559.513 | 1676.58 | 1754.186 | 2027.334 | 1770.205 | 0.001641235 | 0.355068003 | YKL206C | ADD66 | n |
| W303.asm02.HP0_G0031310 | 3652.142 | 3875.887 | 4001.202 | 8319.226 | 7886.085 | 5749.142 | 3013.349 | 2707.05 | 2619.425 | 2.74E-08 | -0.929226945 | YKL205W | LOS1 | n |
| W303.asm02.HP0_G0031320 | 6043.135 | 6476.504 | 6001.804 | 8375.498 | 7591.018 | 8709.555 | 8470.954 | 7483.312 | 8158.491 | 4.77E-05 | -0.41402116 | YKL204W | EAP1 | n |
| W303.asm02.HP0_G0031330 | 2524.415 | 2509.367 | 2306.728 | 7491.124 | 7174.454 | 7731.866 | 8029.013 | 7131.105 | 7535.331 | 2.61E-72 | -1.609443801 | YKL203C | TOR2 | n |
| W303.asm02.HP0_G0031350 | 979.4813 | 1002.634 | 1046.651 | 10914.69 | 10456.63 | 12725.14 | 19126.93 | 12960.77 | 13205.97 | 8.04E-80 | -3.492836884 | YKL198C | PTK1 | n |
| W303.asm02.HP0_G0031360 | 2030.968 | 2312.401 | 1993.056 | 1990.362 | 1857.183 | 2899.64 | 883.8827 | 2205.585 | 1641.028 | 0.810369306 | -0.090661074 | YKL197C | PEX1 | n |
| W303.asm02.HP0_G0031370 | 5523.216 | 5705.333 | 5704.3 | 3606.966 | 4360.909 | 4053.57 | 2586.221 | 3085.027 | 2618.229 | 6.35E-05 | 0.494397412 | YKL196C | YKT6 | n |
| W303.asm02.HP0_G0031380 | 21436.35 | 23233.07 | 21351.24 | 8805.527 | 9328.439 | 10164.31 | 2480.056 | 6224.818 | 4552.297 | 5.10E-05 | 1.222205427 | YKL195W | MIA40 | n |
| W303.asm02.HP0_G0031390 | 2865.38 | 2627.324 | 2581.595 | 2255.049 | 2061.994 | 2069.326 | 1075.226 | 990.0435 | 1234.359 | 0.006155535 | 0.338211619 | YKL194C | MST1 | n |
| W303.asm02.HP0_G0031400 | 4471.729 | 4738.308 | 4066.955 | 2535.019 | 2546.25 | 3122.981 | 2531.904 | 3200.998 | 2511.777 | 1.37E-05 | 0.694442857 | YKL193C | SDS22 | n |
| W303.asm02.HP0_G0031410 | 7511.827 | 7066.289 | 7147.622 | 3968.219 | 4473.728 | 4041.416 | 1362.858 | 2247.463 | 1867.088 | 3.11E-05 | 0.799455435 | YKL192C | ACP1 | n |
| W303.asm02.HP0_G0031420 | 1393.511 | 1156.201 | 1723.578 | 3816.77 | 3857.56 | 1346.885 | 3193.582 | 1765.327 | 2601.483 | 0.009629181 | -1.077936196 | YKL191W | DPH2 | n |
| W303.asm02.HP0_G0031430 | 2486.294 | 2405.877 | 2522.31 | 1684.687 | 1747.835 | 1894.604 | 1650.491 | 1652.578 | 1704.42 | 1.67E-08 | 0.477124249 | YKL190W | CNB1 | n |
| W303.asm02.HP0_G0031440 | 1463.398 | 1563.486 | 1413.14 | 1731.233 | 1783.416 | 1959.935 | 2634.366 | 1728.818 | 1814.46 | 0.129377318 | -0.302201028 | YKL189W | HYM1 | n |
| W303.asm02.HP0_G0031450 | 1847.778 | 2172.188 | 1684.774 | 140.3327 | 122.3658 | 89.64042 | 309.8527 | 290.9998 | 169.844 | 2.49E-49 | 4.016474135 | YKL188C | PXA2 | n |
| W303.asm02.HP0_G0031460 | 2060.617 | 2263.438 | 2034.016 | 22339.99 | 22332.19 | 22901.61 | 70882.21 | 55131.04 | 71376.33 | 7.74E-150 | -3.409856228 | YKL187C | FAT3 | n |
| W303.asm02.HP0_G0031470 | 2672.66 | 2761.973 | 2459.791 | 1059.442 | 1187.209 | 659.3889 | 713.5254 | 661.4608 | 716.4545 | 8.79E-11 | 1.442038831 | YKL186C | MTR2 | n |
| W303.asm02.HP0_G0031480 | 5794.294 | 6586.671 | 6186.126 | 2463.463 | 2476.823 | 2889.764 | 2078.853 | 3007.714 | 3183.976 | 2.42E-11 | 1.245628846 | YKL185W | ASH1 | n |
| W303.asm02.HP0_G0031490 | 3112.103 | 3164.807 | 3396.495 | 3668.796 | 3979.058 | 3617.523 | 2803.488 | 3269.721 | 3012.936 | 0.038268749 | -0.219655097 | YKL184W | SPE1 | n |
| W303.asm02.HP0_G0031510 | 1832.954 | 1799.4 | 1850.772 | 1902.133 | 1858.051 | 1877.891 | 1204.846 | 1311.11 | 1408.987 | 0.692000143 | -0.04027137 | YKL183W | LOT5 | n |
| W303.asm02.HP0_G0031520 | 54549.17 | 52552.04 | 55891.8 | 21278.46 | 23439.56 | 27205.87 | 5935.347 | 20768.36 | 18802.45 | 0.00349935 | 1.180269136 | YKL182W | FAS1 | n |
| W303.asm02.HP0_G0031530 | 7031.087 | 6519.903 | 7873.056 | 7104.862 | 6947.079 | 4286.028 | 4058.948 | 2137.936 | 3207.898 | 0.47317173 | 0.224407014 | YKL181W | PRS1 | n |
| W303.asm02.HP0_G0031550 | 3329.178 | 3753.479 | 3178.757 | 4676.134 | 4562.248 | 6591.61 | 3778.722 | 4899.749 | 4249.688 | 0.001499446 | -0.625478488 | YKL179C | COY1 | n |
| W303.asm02.HP0_G0031560 | 435.2074 | 464.0382 | 407.45 | 196.6047 | 216.0928 | 238.5347 | 492.5548 | 506.8336 | 327.7271 | 1.93E-06 | 1.005009975 | YKL178C | STE3 | n |
| W303.asm02.HP0_G0031570 | 2612.303 | 2487.111 | 2768.073 | 2926.144 | 2735.44 | 3442.04 | 3977.472 | 3554.278 | 3835.842 | 0.112950442 | -0.210632063 | YKL176C | LST4 | n |
| W303.asm02.HP0_G0031580 | 2209.922 | 2400.313 | 2353.078 | 3148.454 | 3020.092 | 4119.661 | 5837.823 | 5330.343 | 5543.851 | 0.000192706 | -0.563245318 | YKL175W | ZRT3 | n |
| W303.asm02.HP0_G0031590 | 2341.225 | 2253.423 | 2099.769 | 1245.626 | 1326.931 | 1272.438 | 898.6964 | 1311.11 | 1145.849 | 4.93E-07 | 0.800107828 | YKL174C | TPO5 | n |
| W303.asm02.HP0_G0031600 | 4322.425 | 4216.404 | 4046.475 | 3829.275 | 3516.498 | 3551.432 | 2819.537 | 3123.684 | 3024.897 | 0.019729821 | 0.207606925 | YKL173W | SNU114 | n |
| W303.asm02.HP0_G0031610 | 4262.067 | 4950.853 | 5123.307 | 8612.396 | 8108.253 | 2480.305 | 7942.6 | 3313.747 | 6875.093 | 0.405022517 | -0.421510132 | YKL172W | EBP2 | n |
| W303.asm02.HP0_G0031620 | 1984.376 | 2523.834 | 1917.602 | 1511.008 | 1353.834 | 2212.903 | 2907.184 | 1793.246 | 2266.58 | 0.228514084 | 0.339565912 | YKL171W | NNK1 | n |
| W303.asm02.HP0_G0031630 | 7397.466 | 7295.526 | 7008.572 | 2004.951 | 2337.968 | 1638.597 | 892.524 | 856.8923 | 881.5141 | 4.10E-37 | 1.859509628 | YKL170W | MRPL38 | n |
| W303.asm02.HP0_G0031640 | 2781.727 | 2714.123 | 2386.493 | 4030.743 | 3861.031 | 4963.649 | 3408.38 | 4003.126 | 3680.351 | 1.60E-06 | -0.705734205 | YKL168C | KKQ8 | n |
| W303.asm02.HP0_G0031650 | 4066.171 | 4033.905 | 3788.854 | 1386.653 | 1510.046 | 832.5924 | 662.912 | 558.376 | 726.0232 | 1.38E-12 | 1.672803063 | YKL167C | MRP49 | n |

| | | | | | | | | | | | | | | |
|---|---|---|---|---|---|---|---|---|---|---|---|---|---|---|
| W303.asm02.HP0_G0031670 | 1951.55 | 1773.805 | 1969.342 | 4601.105 | 4722.799 | 5923.865 | 1543.091 | 4089.03 | 2689.994 | 1.17E-05 | -1.420898114 | YKL165C | MCD4 | n |
| W303.asm02.HP0_G0031680 | 21153.62 | 24454.92 | 24736.96 | 39170.87 | 35865.33 | 43568.29 | 60381.78 | 63039.57 | 66963.98 | 6.01E-10 | -0.753646584 | YKL164C | PIR1 | n |
| W303.asm02.HP0_G0031710 | 718.9922 | 898.0307 | 659.681 | 1102.515 | 1062.239 | 1138.737 | 666.6155 | 844.0067 | 919.7888 | 0.003597877 | -0.53733134 | YKL162C | | n |
| W303.asm02.HP0_G0031730 | 2669.484 | 2647.354 | 2428.532 | 4306.545 | 4555.305 | 4312.616 | 2235.631 | 2563.16 | 2422.071 | 3.95E-16 | -0.766248968 | YKL160W | ELF1 | n |
| W303.asm02.HP0_G0031740 | 1476.105 | 1539.004 | 1457.335 | 705.1369 | 745.4767 | 798.4075 | 424.6587 | 487.5052 | 449.7277 | 2.21E-21 | 0.991985325 | YKL159C | RCN1 | n |
| W303.asm02.HP0_G0031760 | 13251.06 | 12990.84 | 13137.57 | 12024.84 | 13016.77 | 20574.76 | 7875.938 | 15955.59 | 9902.382 | 0.528495371 | -0.212107917 | YKL157W | APE2 | n |
| W303.asm02.HP0_G0031770 | 5754.056 | 5276.904 | 5580.341 | 9194.568 | 11289.76 | 9359.828 | 8846.234 | 7256.74 | 6891.838 | 8.98E-09 | -0.845204836 | YKL156W | RPS27A | y |
| W303.asm02.HP0_G0031780 | 3576.96 | 3398.495 | 3405.118 | 1599.237 | 1252.297 | 569.7485 | 1388.782 | 595.959 | 1031.025 | 8.25E-05 | 1.601219829 | YKL155C | RSM22 | n |
| W303.asm02.HP0_G0031790 | 769.8194 | 672.132 | 712.4986 | 4253.052 | 4392.151 | 4995.555 | 6569.866 | 6680.109 | 7306.879 | 3.94E-118 | -2.662286412 | YKL154W | SRP102 | n |
| W303.asm02.HP0_G0031800 | 60627.25 | 62924.47 | 64931.15 | 27549.66 | 32373.13 | 48515.22 | 87971.02 | 72147.54 | 78683.21 | 0.000376411 | 0.797564555 | YKL152C | GPM1 | n |
| W303.asm02.HP0_G0031810 | 7480.06 | 6726.884 | 6615.135 | 1076.115 | 1155.966 | 2051.854 | 6835.277 | 6587.763 | 6482.777 | 5.12E-20 | 2.281074107 | YKL151C | NNR2 | n |
| W303.asm02.HP0_G0031820 | 14013.47 | 13660.75 | 13478.19 | 11515.61 | 12606.28 | 13440.75 | 2959.032 | 9853.188 | 6944.466 | 0.748860772 | 0.131681824 | YKL150W | MCR1 | n |
| W303.asm02.HP0_G0031830 | 806.8808 | 948.1068 | 828.9129 | 1324.129 | 1353.834 | 1405.38 | 1327.059 | 2192.699 | 1533.38 | 0.001685885 | -0.660275336 | YKL149C | DBR1 | n |
| W303.asm02.HP0_G0031840 | 12110.62 | 10582.74 | 11177.93 | 27143.95 | 28215.3 | 21822.89 | 12145.98 | 13263.58 | 14290.82 | 2.83E-17 | -1.188183032 | YKL148C | SDH1 | n |
| W303.asm02.HP0_G0031850 | 3244.466 | 3473.053 | 3364.158 | 2746.213 | 2872.559 | 3435.963 | 6909.346 | 4582.978 | 6066.54 | 0.43895943 | 0.154984451 | YKL146W | AVT3 | n |
| W303.asm02.HP0_G0031860 | 16712.6 | 17399.76 | 17119.37 | 6053.756 | 6822.11 | 8186.146 | 8925.24 | 9778.022 | 8421.63 | 3.82E-20 | 1.282422109 | YKL145W | RPT1 | n |
| W303.asm02.HP0_G0031870 | 722.1689 | 687.7112 | 811.6663 | 2068.865 | 2230.355 | 869.0563 | 3371.346 | 1352.988 | 2741.425 | 0.003922546 | -1.218060944 | YKL144C | RPC25 | n |
| W303.asm02.HP0_G0031880 | 1980.141 | 2206.685 | 2392.961 | 2997.005 | 2679.898 | 724.72 | 5165.035 | 1274.601 | 3560.743 | 0.953066208 | 0.039603825 | YKL143W | LTV1 | n |
| W303.asm02.HP0_G0031890 | 12256.75 | 12960.8 | 12137.27 | 1078.894 | 1281.803 | 1324.855 | 1187.563 | 1780.36 | 1564.478 | 1.92E-83 | 3.341569978 | YKL142W | MRP8 | n |
| W303.asm02.HP0_G0031900 | 6383.041 | 6283.989 | 5878.922 | 5362.514 | 5906.536 | 4573.181 | 1614.691 | 2843.422 | 2115.873 | 0.342081962 | 0.227361849 | YKL141W | SDH3 | n |
| W303.asm02.HP0_G0031920 | 1799.069 | 1878.409 | 1697.708 | 2505.146 | 2580.096 | 2367.875 | 3508.373 | 2801.544 | 2467.522 | 0.002913439 | -0.471511966 | YKL139W | CTK1 | n |
| W303.asm02.HP0_G0031930 | 1104.431 | 1002.634 | 950.7167 | 220.225 | 452.146 | 163.3279 | 322.1975 | 179.3246 | 184.197 | 2.65E-06 | 1.872740706 | YKL138C-A | HSK3 | n |
| W303.asm02.HP0_G0031940 | 5406.737 | 5113.322 | 4819.336 | 2402.328 | 2657.334 | 1899.921 | 1257.928 | 994.3387 | 1081.26 | 7.72E-12 | 1.140319861 | YKL138C | MRPL31 | n |
| W303.asm02.HP0_G0031950 | 2942.68 | 2749.732 | 2591.296 | 773.2191 | 859.1641 | 557.5938 | 939.434 | 634.6158 | 804.9648 | 3.62E-19 | 1.919680567 | YKL137W | CMC1 | n |
| W303.asm02.HP0_G0031960 | 5631.223 | 5500.577 | 5731.248 | 5638.316 | 5619.28 | 6610.601 | 4556.44 | 5044.712 | 4474.551 | 0.485261956 | -0.083547042 | YKL135C | APL2 | n |
| W303.asm02.HP0_G0031970 | 1236.794 | 1265.255 | 1185.701 | 4061.311 | 3961.701 | 4393.9 | 2091.197 | 2343.031 | 1935.264 | 1.79E-57 | -1.751582425 | YKL134C | OCT1 | n |
| W303.asm02.HP0_G0032000 | 1893.311 | 1856.153 | 1836.759 | 2368.982 | 2606.131 | 2326.093 | 2185.017 | 2065.991 | 2262.992 | 2.16E-05 | -0.385897705 | YKL130C | SHE2 | n |
| W303.asm02.HP0_G0032020 | 2012.966 | 1786.046 | 1925.148 | 2707.309 | 2815.281 | 3732.232 | 8801.793 | 5564.431 | 7619.057 | 0.001493731 | -0.693122091 | YKL128C | PMU1 | n |
| W303.asm02.HP0_G0032040 | 8477.543 | 7973.222 | 8380.751 | 9255.703 | 9916.836 | 11954.84 | 14676.65 | 13159.42 | 12734.71 | 0.016978925 | -0.325994518 | YKL126W | YPK1 | n |
| W303.asm02.HP0_G0032050 | 1679.413 | 1561.26 | 1753.76 | 1668.708 | 1515.253 | 524.9283 | 2634.366 | 1147.892 | 1910.147 | 0.363590828 | 0.429344892 | YKL125W | RRN3 | n |
| W303.asm02.HP0_G0032060 | 2526.532 | 2366.929 | 2238.819 | 1676.35 | 1582.945 | 1785.972 | 6330.378 | 4069.702 | 5402.713 | 0.007264441 | 0.499366681 | YKL124W | SSH4 | n |
| W303.asm02.HP0_G0032070 | 3055.982 | 3199.304 | 2994.434 | 2433.591 | 3056.541 | 2289.629 | 3003.473 | 3000.197 | 2969.877 | 0.077455613 | 0.250015735 | YKL122C | SRP21 | n |
| W303.asm02.HP0_G0032090 | 263.6658 | 290.4412 | 309.3602 | 2968.522 | 3507.819 | 2532.722 | 3670.088 | 1312.184 | 1429.321 | 8.67E-19 | -3.383163215 | YKL120W | OAC1 | n |
| W303.asm02.HP0_G0032100 | 1975.905 | 1831.671 | 1962.874 | 1929.227 | 1996.038 | 2048.056 | 1920.84 | 1870.559 | 1863.499 | 0.56540936 | -0.049578985 | YKL119C | VPH2 | n |
| W303.asm02.HP0_G0032110 | 14768.46 | 15377.8 | 14916.12 | 2799.706 | 3236.185 | 4497.974 | 2207.238 | 4877.199 | 3801.156 | 2.43E-12 | 2.096898921 | YKL117W | SBA1 | n |
| W303.asm02.HP0_G0032120 | 1690.002 | 1806.077 | 1833.525 | 2353.698 | 2500.254 | 2939.902 | 1702.338 | 1698.751 | 1084.848 | 0.009064385 | -0.548309235 | YKL116C | PRR1 | n |
| W303.asm02.HP0_G0032140 | 2609.126 | 2789.793 | 2693.697 | 2299.51 | 2276.351 | 2430.167 | 1448.037 | 2076.729 | 1667.342 | 0.200048987 | 0.207966197 | YKL114C | APN1 | n |
| W303.asm02.HP0_G0032150 | 1458.104 | 1554.584 | 1639.501 | 2239.765 | 2612.206 | 2149.091 | 265.4117 | 488.579 | 405.4726 | 0.010134843 | -0.589579783 | YKL113C | RAD27 | n |
| W303.asm02.HP0_G0032160 | 1257.972 | 1461.108 | 1437.932 | 2717.73 | 2725.893 | 4150.807 | 3961.424 | 2726.378 | 3011.74 | 1.87E-07 | -1.20671061 | YKL112W | ABF1 | n |
| W303.asm02.HP0_G0032180 | 6383.041 | 8781.116 | 6540.759 | 5147.152 | 4403.433 | 4444.038 | 5956.333 | 4497.074 | 5464.909 | 0.000733507 | 0.63306433 | YKL109W | HAP4 | n |
| W303.asm02.HP0_G0032190 | 688.2842 | 713.3057 | 773.9395 | 680.1271 | 689.067 | 649.5132 | 498.7271 | 296.3688 | 387.5313 | 0.626773158 | 0.107975944 | YKL108W | SLD2 | n |
| W303.asm02.HP0_G0032210 | 394.9692 | 430.6541 | 469.9688 | 1427.642 | 1398.094 | 1049.097 | 1646.787 | 1408.826 | 1971.147 | 5.57E-16 | -1.580554842 | YKL106W | AAT1 | n |
| W303.asm02.HP0_G0032220 | 26245.86 | 28477.7 | 27975 | 17797.93 | 18262.88 | 19841.68 | 10216.5 | 11063.36 | 10075.81 | 2.53E-10 | 0.564946789 | YKL104C | GFA1 | n |
| W303.asm02.HP0_G0032230 | 9243.127 | 8144.593 | 8497.166 | 4147.455 | 4577.001 | 9605.959 | 7868.531 | 16943.49 | 12079.26 | 0.214954641 | 0.497867888 | YKL103C | APE1 | n |
| W303.asm02.HP0_G0032240 | 7522.416 | 7402.355 | 8040.132 | 4819.941 | 5133.288 | 6065.922 | 2541.78 | 1776.065 | 1382.673 | 0.029628141 | 0.519640105 | YKL101W | HSL1 | n |
| W303.asm02.HP0_G0032260 | 1142.552 | 1184.021 | 1413.14 | 3318.659 | 3654.485 | 1150.892 | 4877.403 | 1994.046 | 3643.273 | 0.014869281 | -1.119236893 | YKL099C | UTP11 | n |
| W303.asm02.HP0_G0032270 | 2462.998 | 2422.569 | 2370.324 | 105.5969 | 78.97367 | 82.80344 | 402.4382 | 146.0368 | 156.687 | 2.69E-37 | 4.76092945 | YKL098W | MTC2 | n |
| W303.asm02.HP0_G0032300 | 7267.222 | 7316.669 | 7289.906 | 19217.93 | 18800.07 | 25397.11 | 42790.54 | 47007.74 | 47380.49 | 1.34E-26 | -1.535633304 | YKL096W | CWP1 | n |
| W303.asm02.HP0_G0032310 | 1278.091 | 1518.974 | 1413.14 | 562.7201 | 675.1815 | 386.6693 | 920.9169 | 555.1546 | 516.7084 | 2.48E-06 | 1.374176878 | YKL095W | YJU2 | n |
| W303.asm02.HP0_G0032320 | 5588.867 | 5153.383 | 5389.551 | 1885.46 | 2168.738 | 3003.714 | 9609.139 | 9506.351 | 6664.582 | 1.15E-07 | 1.192626758 | YKL094W | YJU3 | n |
| W303.asm02.HP0_G0032330 | 1843.543 | 1981.899 | 1790.409 | 1372.064 | 1086.539 | 1835.35 | 26823.87 | 11766.7 | 16020.35 | 0.261096551 | 0.387076903 | YKL093W | MBR1 | n |
| W303.asm02.HP0_G0032340 | 1682.59 | 1756.001 | 1584.528 | 3128.307 | 3161.55 | 3596.252 | 2088.728 | 3020.599 | 2882.563 | 8.74E-09 | -0.976850912 | YKL092C | BUD2 | n |
| W303.asm02.HP0_G0032360 | 2047.91 | 1947.402 | 1998.445 | 383.4833 | 354.0797 | 302.3465 | 382.6867 | 148.1844 | 275.0994 | 9.92E-17 | 2.526909103 | YKL090W | CUE2 | n |
| W303.asm02.HP0_G0032370 | 452.1497 | 445.1205 | 500.1503 | 984.4128 | 945.0805 | 1103.793 | 1480.133 | 512.2026 | 1062.123 | 0.001340397 | -1.118134053 | YKL089W | MIF2 | n |

| | | | | | | | | | | | | | | |
|---|---|---|---|---|---|---|---|---|---|---|---|---|---|---|
| W303.asm02.HP0_G0032380 | 4351.015 | 4756.113 | 4374.159 | 6050.283 | 5835.373 | 5120.899 | 10197.98 | 7050.57 | 9010.103 | 0.05119939 | -0.335153354 | YKL088W | CAB3 | n |
| W303.asm02.HP0_G0032390 | 7313.813 | 7269.931 | 6910.482 | 2205.029 | 2005.584 | 1540.6 | 1654.194 | 1128.564 | 1599.165 | 1.43E-21 | 1.901967529 | YKL087C | CYT2 | n |
| W303.asm02.HP0_G0032400 | 158.8348 | 142.4386 | 146.5958 | 689.8531 | 640.4678 | 867.537 | 1278.914 | 1379.833 | 1393.438 | 6.57E-43 | -2.294935033 | YKL086W | SRX1 | n |
| W303.asm02.HP0_G0032410 | 13344.24 | 13111.03 | 12717.18 | 15431.73 | 17283.95 | 22356.93 | 11238.64 | 12801.84 | 13738.22 | 0.004261058 | -0.491489353 | YKL085W | MDH1 | n |
| W303.asm02.HP0_G0032420 | 1262.207 | 1131.719 | 1115.637 | 132.6908 | 153.6081 | 142.0573 | 64.1926 | 162.1438 | 84.92199 | 8.39E-24 | 3.035084084 | YKL084W | HOT13 | n |
| W303.asm02.HP0_G0032430 | 2014.025 | 2117.661 | 2129.95 | 4681.692 | 4653.372 | 853.863 | 3860.197 | 1349.767 | 2756.974 | 0.248645586 | -0.702379493 | YKL082C | RRP14 | n |
| W303.asm02.HP0_G0032450 | 8959.342 | 8768.875 | 9208.586 | 4444.794 | 4822.601 | 6625.035 | 5830.416 | 6344.01 | 5661.067 | 1.04E-05 | 0.761244674 | YKL080W | VMA5 | n |
| W303.asm02.HP0_G0032470 | 163.0704 | 164.6946 | 227.439 | 3183.189 | 3020.96 | 964.0144 | 5845.23 | 1169.368 | 3847.803 | 9.59E-11 | -3.690486157 | YKL078W | DHR2 | n |
| W303.asm02.HP0_G0032480 | 5502.038 | 5424.907 | 5648.249 | 3695.195 | 3741.269 | 3544.595 | 2696.089 | 2567.456 | 2692.386 | 3.90E-19 | 0.594099426 | YKL077W | PSG1 | n |
| W303.asm02.HP0_G0032490 | 2454.527 | 2678.513 | 2589.14 | 1991.056 | 1982.152 | 1565.669 | 2051.694 | 1700.899 | 1794.126 | 0.001072207 | 0.479436481 | YKL075C | AAN1 | n |
| W303.asm02.HP0_G0032500 | 2494.765 | 2550.541 | 2444.7 | 2320.352 | 2025.544 | 1624.923 | 6653.81 | 3604.746 | 6277.051 | 0.2312072 | 0.326986269 | YKL074C | MUD2 | n |
| W303.asm02.HP0_G0032510 | 3548.37 | 3779.073 | 3693.998 | 3936.262 | 3874.049 | 5544.792 | 1193.735 | 2993.754 | 1885.029 | 0.43046453 | -0.277090729 | YKL073W | LHS1 | n |
| W303.asm02.HP0_G0032520 | 942.4198 | 1017.1 | 924.8469 | 1777.779 | 1632.412 | 2187.834 | 3368.877 | 2815.503 | 3026.093 | 3.80E-10 | -0.956908801 | YKL072W | STB6 | n |
| W303.asm02.HP0_G0032540 | 605.6901 | 586.4463 | 673.6938 | 948.2875 | 894.7456 | 1027.067 | 1018.44 | 872.9993 | 454.512 | 0.035229272 | -0.621321214 | YKL070W | | n |
| W303.asm02.HP0_G0032550 | 740.1702 | 793.4274 | 771.7837 | 1376.927 | 1498.764 | 1045.299 | 323.4319 | 1295.003 | 656.6503 | 0.087167089 | -0.766183689 | YKL069W | | n |
| W303.asm02.HP0_G0032560 | 362.1434 | 237.0267 | 327.6847 | 123.6595 | 140.5905 | 110.911 | 491.3203 | 565.8926 | 678.1798 | 2.01E-08 | 1.306743773 | YKL068W-A | | n |
| W303.asm02.HP0_G0032580 | 7849.616 | 7508.071 | 7065.701 | 7712.739 | 9490.726 | 6677.452 | 646.8639 | 1992.973 | 1285.791 | 0.82408182 | -0.090836496 | YKL067W | YNK1 | n |
| W303.asm02.HP0_G0032610 | 9972.708 | 10006.31 | 9460.817 | 1536.712 | 1760.852 | 2373.952 | 2093.666 | 3415.758 | 2639.758 | 4.11E-25 | 2.375995741 | YKL065C | YET1 | n |
| W303.asm02.HP0_G0032620 | 14674.22 | 14547.65 | 14600.29 | 4414.226 | 4299.292 | 5495.414 | 6977.242 | 4811.697 | 5799.813 | 5.71E-21 | 1.62483593 | YKL064W | MNR2 | n |
| W303.asm02.HP0_G0032630 | 2947.974 | 3128.085 | 2826.28 | 2507.925 | 2753.664 | 2699.088 | 2894.839 | 2696.312 | 2562.013 | 0.113941858 | 0.16146156 | YKL063C | | n |
| W303.asm02.HP0_G0032650 | 818.5287 | 796.7658 | 696.33 | 228.5616 | 208.2822 | 231.6977 | 281.4599 | 347.9112 | 281.0798 | 7.12E-33 | 1.788703886 | YKL061W | BLI1 | n |
| W303.asm02.HP0_G0032660 | 99475.06 | 106818.9 | 109496.3 | 54419.89 | 63512.19 | 84499.01 | 145702.4 | 123007 | 133840.6 | 0.000563862 | 0.641537547 | YKL060C | FBA1 | n |
| W303.asm02.HP0_G0032670 | 2408.995 | 2589.489 | 2366.013 | 2298.816 | 2245.976 | 1913.595 | 2793.613 | 2045.589 | 2325.188 | 0.255441356 | 0.189389229 | YKL059C | MPE1 | n |
| W303.asm02.HP0_G0032680 | 2486.294 | 2106.533 | 2242.053 | 1497.113 | 1872.804 | 1345.366 | 2081.322 | 1787.877 | 1748.675 | 0.001663309 | 0.536003248 | YKL058W | TOA2 | n |
| W303.asm02.HP0_G0032690 | 2500.06 | 2705.22 | 2660.282 | 6119.754 | 6098.329 | 5989.196 | 3893.528 | 4794.517 | 4369.296 | 2.11E-30 | -1.210939859 | YKL057C | NUP120 | n |
| W303.asm02.HP0_G0032700 | 15493.81 | 14394.09 | 15725.63 | 17998.71 | 21001.79 | 17308.2 | 15822.24 | 12409.91 | 13632.97 | 0.03414338 | -0.303859816 | YKL056C | TMA19 | n |
| W303.asm02.HP0_G0032710 | 104.831 | 122.4082 | 119.648 | 946.8981 | 964.173 | 1179.759 | 767.8423 | 1321.848 | 1025.044 | 2.79E-40 | -3.155840716 | YKL055C | OAR1 | n |
| W303.asm02.HP0_G0032720 | 15205.79 | 16008.76 | 15900.25 | 39778.06 | 37549.81 | 44143.35 | 28545.96 | 30675.24 | 28444.08 | 6.69E-48 | -1.366381474 | YKL054C | DEF1 | n |
| W303.asm02.HP0_G0032730 | 3607.668 | 3445.233 | 3290.86 | 1424.168 | 1759.117 | 1382.59 | 1852.944 | 1457.147 | 1444.87 | 8.51E-14 | 1.180188978 | YKL053C-/ | MDM35 | n |
| W303.asm02.HP0_G0032740 | 996.4237 | 954.7836 | 973.3529 | 537.7103 | 551.9478 | 533.2846 | 387.6246 | 302.8116 | 399.4922 | 1.14E-10 | 0.849845109 | YKL052C | ASK1 | n |
| W303.asm02.HP0_G0032750 | 2164.389 | 2358.026 | 2023.237 | 4917.896 | 5984.642 | 4231.332 | 8382.072 | 8092.156 | 11305.39 | 7.22E-10 | -1.209116891 | YKL051W | SFK1 | n |
| W303.asm02.HP0_G0032770 | 1214.557 | 1327.572 | 1260.077 | 664.1486 | 721.1771 | 670.7839 | 659.2086 | 606.697 | 654.2581 | 1.01E-19 | 0.887568671 | YKL049C | CSE4 | n |
| W303.asm02.HP0_G0032780 | 603.5723 | 658.7784 | 552.9679 | 762.7983 | 689.9348 | 774.0982 | 96.2889 | 309.2544 | 174.6283 | 0.45423023 | -0.294867272 | YKL048C | ELM1 | n |
| W303.asm02.HP0_G0032790 | 3087.749 | 2715.235 | 3134.563 | 1570.753 | 1520.46 | 1958.415 | 2036.881 | 2297.932 | 2145.775 | 8.36E-09 | 0.823621318 | YKL047W | ANR2 | n |
| W303.asm02.HP0_G0032800 | 6537.641 | 6535.482 | 6568.784 | 5154.099 | 5088.161 | 5957.29 | 6432.839 | 6644.674 | 5662.263 | 0.016131518 | 0.277947398 | YKL046C | DCW1 | n |
| W303.asm02.HP0_G0032810 | 2531.827 | 2528.285 | 2589.14 | 3956.408 | 4415.583 | 4419.729 | 798.7041 | 1715.932 | 1325.261 | 0.00505395 | -0.741797263 | YKL045W | PRI2 | n |
| W303.asm02.HP0_G0032840 | 1529.05 | 1484.477 | 1580.216 | 569.6672 | 551.08 | 726.2394 | 269.1151 | 287.7784 | 239.2169 | 4.12E-18 | 1.314209911 | YKL042W | SPC42 | n |
| W303.asm02.HP0_G0032850 | 2661.012 | 2362.477 | 2264.689 | 1734.706 | 1733.082 | 1847.504 | 1627.036 | 1388.423 | 1483.145 | 8.29E-05 | 0.455441223 | YKL041W | VPS24 | n |
| W303.asm02.HP0_G0032860 | 3299.528 | 3309.471 | 3065.576 | 2981.027 | 3062.616 | 2433.965 | 2317.106 | 3138.717 | 2194.815 | 0.325834234 | 0.190580899 | YKL040C | NFU1 | n |
| W303.asm02.HP0_G0032870 | 9817.05 | 9832.713 | 9757.243 | 5304.158 | 5300.782 | 6738.985 | 5760.051 | 8343.425 | 6311.737 | 2.99E-05 | 0.7617159 | YKL039W | PTM1 | n |
| W303.asm02.HP0_G0032880 | 2736.194 | 2723.025 | 2478.115 | 3371.457 | 2563.607 | 4112.065 | 20256.47 | 9298.034 | 13377.01 | 0.291725278 | -0.340118722 | YKL038W | RGT1 | n |
| W303.asm02.HP0_G0032900 | 18488.37 | 18490.31 | 16724.85 | 13951.98 | 14995.45 | 23477.43 | 36082.41 | 34315.42 | 35407.68 | 0.887568619 | 0.034763361 | YKL035W | UGP1 | n |
| W303.asm02.HP0_G0032910 | 2620.774 | 2717.461 | 2493.206 | 2143.894 | 2105.386 | 3134.376 | 3637.992 | 4798.812 | 3759.293 | 0.712753313 | 0.084883598 | YKL034W | TUL1 | n |
| W303.asm02.HP0_G0032920 | 4444.198 | 4058.387 | 4173.668 | 1610.352 | 1761.72 | 2364.836 | 3234.319 | 3576.828 | 3333.487 | 6.89E-12 | 1.143820676 | YKL033W-A | | n |
| W303.asm02.HP0_G0032930 | 1065.252 | 1227.42 | 1114.559 | 1035.127 | 873.9174 | 941.9841 | 606.1263 | 765.6193 | 702.1015 | 0.082451502 | 0.256443675 | YKL033W | TTI1 | n |
| W303.asm02.HP0_G0032940 | 2715.016 | 2995.661 | 2853.228 | 7002.044 | 6891.537 | 6618.198 | 7451.28 | 4868.609 | 7473.135 | 8.00E-12 | -1.260132857 | YKL032C | IXR1 | y |
| W303.asm02.HP0_G0032960 | 2362.403 | 1996.366 | 2521.232 | 5532.025 | 5098.575 | 4082.438 | 2877.557 | 8568.923 | 13422.46 | 0.018395824 | -1.096608323 | YKL029C | MAE1 | n |
| W303.asm02.HP0_G0032970 | 3191.521 | 3300.569 | 3516.143 | 6503.932 | 6016.752 | 6629.593 | 4235.477 | 4716.129 | 4187.491 | 2.83E-22 | -0.93627621 | YKL028W | TFA1 | n |
| W303.asm02.HP0_G0032980 | 2738.312 | 2832.079 | 2896.345 | 3214.452 | 2748.457 | 1325.615 | 2788.675 | 1443.187 | 2158.932 | 0.579045228 | 0.216162344 | YKL027W | TCD2 | n |
| W303.asm02.HP0_G0033000 | 4005.814 | 4006.085 | 4030.306 | 4249.579 | 3984.265 | 5158.882 | 3499.731 | 3221.4 | 3676.763 | 0.275447301 | -0.153429137 | YKL025C | PAN3 | n |
| W303.asm02.HP0_G0033010 | 1948.374 | 1671.428 | 2001.679 | 5003.346 | 5493.443 | 5442.997 | 2229.458 | 3031.337 | 2601.483 | 1.31E-23 | -1.503493106 | YKL024C | URA6 | n |
| W303.asm02.HP0_G0033040 | 1534.344 | 1610.224 | 1592.073 | 1863.229 | 1803.377 | 2174.92 | 1301.135 | 1206.951 | 887.4946 | 0.096424352 | -0.302562225 | YKL022C | CDC16 | n |
| W303.asm02.HP0_G0033050 | 3382.122 | 3392.931 | 3406.196 | 4558.727 | 4314.913 | 1998.678 | 5004.554 | 2283.972 | 4078.648 | 0.820475319 | -0.094738527 | YKL021C | MAK11 | n |
| W303.asm02.HP0_G0033070 | 3230.7 | 2626.211 | 2995.512 | 1682.602 | 1781.681 | 1994.879 | 513.5408 | 1308.962 | 967.6322 | 0.029384607 | 0.697418276 | YKL019W | RAM2 | n |

| | | | | | | | | | | | | | | |
|---|---|---|---|---|---|---|---|---|---|---|---|---|---|---|
| W303.asm02.HP0_G0033080 | 658.635 | 369.4501 | 390.2035 | 421.6927 | 536.3267 | 355.523 | 439.4724 | 342.5422 | 382.747 | 0.72677868 | 0.111511921 | YKL018C-/ | MCO12 | n |
| W303.asm02.HP0_G0033090 | 1037.721 | 1109.463 | 1041.261 | 1429.726 | 1519.592 | 1155.45 | 1738.138 | 1449.63 | 1621.89 | 0.016726795 | -0.364290036 | YKL018W | SWD2 | n |
| W303.asm02.HP0_G0033100 | 1846.719 | 1955.192 | 1861.551 | 1900.049 | 1949.174 | 2084.52 | 1343.107 | 2127.198 | 1637.439 | 0.741410092 | -0.067265922 | YKL017C | HCS1 | n |
| W303.asm02.HP0_G0033110 | 5409.913 | 5318.078 | 5138.398 | 10608.32 | 11955.4 | 10985.51 | 3924.39 | 6861.582 | 5089.339 | 1.88E-07 | -1.080288429 | YKL016C | ATP7 | n |
| W303.asm02.HP0_G0033120 | 3104.691 | 3414.075 | 3288.704 | 2296.037 | 2226.884 | 1656.069 | 374.0453 | 1115.678 | 595.65 | 0.076075827 | 0.666509984 | YKL015W | PUT3 | n |
| W303.asm02.HP0_G0033130 | 1931.431 | 2093.179 | 2263.611 | 4406.584 | 4112.706 | 1789.77 | 5230.462 | 2327.998 | 4712.572 | 0.081607244 | -0.713187037 | YKL014C | URB1 | n |
| W303.asm02.HP0_G0033140 | 7081.915 | 7283.285 | 7049.532 | 3373.542 | 3952.155 | 4256.401 | 4988.506 | 5429.132 | 4970.926 | 1.87E-14 | 0.88684598 | YKL013C | ARC19 | n |
| W303.asm02.HP0_G0033150 | 5607.928 | 6056.978 | 5447.758 | 3019.931 | 2865.616 | 3210.343 | 5139.111 | 3814.137 | 5027.142 | 6.05E-10 | 0.911702953 | YKL012W | PRP40 | n |
| W303.asm02.HP0_G0033160 | 1365.979 | 1354.279 | 1344.154 | 431.4187 | 422.6393 | 442.1248 | 277.7564 | 315.6972 | 328.9232 | 2.36E-60 | 1.648464997 | YKL011C | CCE1 | n |
| W303.asm02.HP0_G0033170 | 10092.36 | 10388 | 9922.163 | 7714.128 | 7722.063 | 9721.428 | 10615.23 | 10227.94 | 9459.831 | 0.039591678 | 0.27317455 | YKL010C | UFD4 | n |
| W303.asm02.HP0_G0033180 | 3402.242 | 3336.179 | 3865.386 | 6581.741 | 7253.428 | 2354.201 | 6900.705 | 3274.016 | 5992.382 | 0.170481616 | -0.610444866 | YKL009W | MRT4 | n |
| W303.asm02.HP0_G0033200 | 9208.183 | 9683.598 | 9070.614 | 2538.493 | 2786.642 | 3661.583 | 2735.592 | 3670.248 | 3221.055 | 1.79E-19 | 1.637634882 | YKL007W | CAP1 | n |
| W303.asm02.HP0_G0033240 | 2130.504 | 2252.31 | 2303.494 | 2354.393 | 2046.373 | 2106.55 | 6355.067 | 4628.078 | 5448.164 | 0.818582239 | 0.038956019 | YKL004W | AUR1 | n |
| W303.asm02.HP0_G0033250 | 3288.939 | 2813.162 | 2841.371 | 1980.636 | 1917.064 | 1389.427 | 1059.178 | 976.0841 | 1109.966 | 1.27E-05 | 0.758401908 | YKL003C | MRP17 | n |
| W303.asm02.HP0_G0033270 | 2330.636 | 1585.742 | 1662.137 | 1056.663 | 1061.371 | 1403.86 | 1681.352 | 1803.984 | 1728.342 | 0.001242037 | 0.66356599 | YKL001C | MET14 | n |
| W303.asm02.HP0_G0033280 | 15008.83 | 15824.04 | 15269.68 | 10569.41 | 10590.28 | 14904.62 | 6458.763 | 10522.17 | 7870.235 | 0.135589324 | 0.354267367 | YKR001C | VPS1 | n |
| W303.asm02.HP0_G0033290 | 3768.62 | 3794.653 | 3927.905 | 6585.214 | 6332.647 | 7952.929 | 3683.668 | 4548.616 | 4828.592 | 1.31E-08 | -0.860986056 | YKR002W | PAP1 | n |
| W303.asm02.HP0_G0033310 | 789.9384 | 791.2018 | 697.4079 | 724.5889 | 771.512 | 732.3167 | 864.1312 | 1078.095 | 838.4551 | 0.843226213 | 0.032337554 | YKR004C | ECM9 | n |
| W303.asm02.HP0_G0033320 | 820.6465 | 754.4793 | 726.5114 | 65.30332 | 69.4274 | 88.1211 | 98.75785 | 97.71579 | 113.628 | 2.01E-83 | 3.368937739 | YKR005C | | n |
| W303.asm02.HP0_G0033330 | 8438.364 | 8472.87 | 8028.275 | 5044.334 | 5177.548 | 4017.866 | 3310.857 | 3367.437 | 3187.565 | 9.25E-11 | 0.808561765 | YKR006C | MRPL13 | n |
| W303.asm02.HP0_G0033340 | 1843.543 | 1701.473 | 1696.631 | 1055.274 | 1053.561 | 1237.494 | 977.7027 | 638.911 | 880.3181 | 0.000480572 | 0.6474182 | YKR007W | MEH1 | n |
| W303.asm02.HP0_G0033350 | 3256.114 | 3759.043 | 3337.21 | 2931.007 | 2657.334 | 2702.127 | 2765.22 | 2687.721 | 2749.798 | 0.001267347 | 0.320126123 | YKR008W | RSC4 | n |
| W303.asm02.HP0_G0033360 | 2521.238 | 2760.86 | 2500.752 | 992.7494 | 929.4593 | 1218.502 | 3360.236 | 1918.88 | 2862.23 | 8.91E-09 | 1.309079238 | YKR009C | FOX2 | n |
| W303.asm02.HP0_G0033370 | 1029.25 | 1015.988 | 1186.779 | 7706.486 | 7604.036 | 7851.134 | 6787.133 | 7155.803 | 6747.112 | 5.51E-208 | -2.841171498 | YKR010C | TOF2 | n |
| W303.asm02.HP0_G0033380 | 4419.843 | 4786.159 | 4310.563 | 1132.387 | 1113.442 | 1731.276 | 941.903 | 1276.748 | 893.475 | 8.36E-15 | 1.764851398 | YKR011C | FPT1 | n |
| W303.asm02.HP0_G0033390 | 7066.031 | 7230.983 | 8207.208 | 5404.197 | 5856.201 | 6788.363 | 4341.642 | 3536.023 | 4055.922 | 0.032984423 | 0.318318002 | YKR013W | PRY2 | n |
| W303.asm02.HP0_G0033400 | 6384.1 | 6691.275 | 6303.619 | 2511.399 | 2710.272 | 3483.822 | 3791.067 | 3453.341 | 3304.781 | 2.31E-14 | 1.154505928 | YKR014C | YPT52 | n |
| W303.asm02.HP0_G0033420 | 7234.396 | 7758.451 | 7070.013 | 9095.918 | 9464.69 | 8343.396 | 1546.795 | 3327.706 | 2449.581 | 0.307248378 | -0.286197914 | YKR016W | MIC60 | n |
| W303.asm02.HP0_G0033430 | 1490.929 | 1648.059 | 1378.647 | 1262.299 | 1234.072 | 1708.486 | 1610.987 | 1763.179 | 1570.459 | 0.584690573 | 0.10334109 | YKR017C | HEL1 | n |
| W303.asm02.HP0_G0033440 | 14659.39 | 15232.03 | 15048.7 | 5354.872 | 5297.311 | 7139.328 | 15817.3 | 13209.89 | 15091 | 3.71E-18 | 1.336787294 | YKR018C | | n |
| W303.asm02.HP0_G0033460 | 1434.808 | 1398.791 | 1277.324 | 675.2641 | 846.1465 | 831.8328 | 504.8995 | 695.8224 | 550.1988 | 2.84E-06 | 0.805355046 | YKR020W | VPS51 | n |
| W303.asm02.HP0_G0033470 | 2929.973 | 3132.536 | 3094.68 | 3920.978 | 3870.578 | 5703.562 | 6708.127 | 5243.365 | 5031.927 | 0.006919957 | -0.559496325 | YKR021W | ALY1 | n |
| W303.asm02.HP0_G0033480 | 1949.433 | 1932.936 | 1912.213 | 640.5283 | 569.3047 | 807.5235 | 744.3873 | 705.4866 | 741.5723 | 2.80E-24 | 1.521638413 | YKR022C | NTR2 | n |
| W303.asm02.HP0_G0033490 | 2102.973 | 2263.438 | 2212.95 | 2673.268 | 2476.823 | 2330.651 | 2239.334 | 2463.297 | 2510.581 | 0.081069658 | -0.18544548 | YKR023W | RQT4 | n |
| W303.asm02.HP0_G0033500 | 510.3892 | 537.4831 | 603.6297 | 3090.792 | 2733.704 | 635.0796 | 5136.643 | 1430.302 | 4277.198 | 0.000747786 | -1.967659898 | YKR024C | DBP7 | n |
| W303.asm02.HP0_G0033510 | 900.0639 | 1021.552 | 1017.547 | 2607.964 | 2638.241 | 999.7186 | 4718.156 | 1944.652 | 3691.116 | 0.011003923 | -1.087496826 | YKR025W | RPC37 | n |
| W303.asm02.HP0_G0033520 | 3400.124 | 3326.163 | 3409.43 | 3151.927 | 3276.105 | 1877.891 | 3519.483 | 1961.832 | 2984.23 | 0.343762283 | 0.287273248 | YKR026C | GCN3 | n |
| W303.asm02.HP0_G0033540 | 4290.658 | 4466.785 | 4451.769 | 16807.27 | 15811.22 | 15844.32 | 9665.924 | 10029.29 | 10505.21 | 2.35E-151 | -1.875404227 | YKR028W | SAP190 | n |
| W303.asm02.HP0_G0033560 | 1518.461 | 1616.9 | 1447.633 | 1326.213 | 1361.645 | 1478.307 | 1783.814 | 2180.888 | 2089.559 | 0.296026973 | 0.137533159 | YKR030W | GMH1 | n |
| W303.asm02.HP0_G0033570 | 3850.156 | 4313.218 | 3658.427 | 2480.137 | 2370.078 | 3584.857 | 1020.909 | 2562.087 | 1784.558 | 0.161603944 | 0.486949962 | YKR031C | SPO14 | n |
| W303.asm02.HP0_G0033590 | 700.9909 | 732.2233 | 720.044 | 4873.434 | 4767.059 | 4352.119 | 5436.619 | 5702.951 | 6829.641 | 1.42E-103 | -2.700142739 | YKR034W | DAL80 | n |
| W303.asm02.HP0_G0033600 | 4347.838 | 4731.631 | 4384.938 | 2532.935 | 2828.299 | 3124.501 | 4823.086 | 4413.318 | 4554.689 | 4.57E-09 | 0.666140593 | YKR035W- | DID2 | n |
| W303.asm02.HP0_G0033620 | 1536.462 | 1444.416 | 1372.18 | 1212.28 | 1149.891 | 1101.514 | 1034.488 | 1303.593 | 1074.084 | 0.014055316 | 0.329623813 | YKR037C | SPC34 | n |
| W303.asm02.HP0_G0033630 | 2856.909 | 2687.415 | 3082.823 | 3495.117 | 3702.216 | 3376.709 | 2367.719 | 2867.046 | 2411.306 | 0.018702818 | -0.293452242 | YKR038C | KAE1 | n |
| W303.asm02.HP0_G0033640 | 3962.399 | 4813.979 | 4184.447 | 2636.448 | 2843.052 | 4611.924 | 2301.058 | 3356.699 | 2390.973 | 0.198108536 | 0.361006739 | YKR039W | GAP1 | n |
| W303.asm02.HP0_G0033660 | 23144.35 | 23100.64 | 24508.44 | 5240.939 | 4963.191 | 4272.354 | 3833.039 | 2637.253 | 3475.821 | 2.38E-43 | 2.289076513 | YKR042W | UTH1 | n |
| W303.asm02.HP0_G0033670 | 6234.796 | 5885.606 | 7040.909 | 3732.015 | 4171.719 | 4662.821 | 12264.49 | 8850.259 | 7257.84 | 0.006030665 | 0.608636746 | YKR043C | SHB17 | n |
| W303.asm02.HP0_G0033700 | 16002.08 | 16126.72 | 16387.47 | 3857.064 | 4705.442 | 9573.294 | 21078.63 | 21537.21 | 22814.11 | 7.62E-06 | 1.419627847 | YKR046C | PLN1 | n |
| W303.asm02.HP0_G0033710 | 13271.18 | 11323.87 | 13191.46 | 10089.36 | 10279.59 | 10098.98 | 8422.81 | 9405.414 | 8512.532 | 0.003226771 | 0.310614831 | YKR048C | NAP1 | n |
| W303.asm02.HP0_G0033720 | 3157.636 | 2999 | 2810.112 | 280.6653 | 306.3484 | 369.197 | 393.7969 | 659.3132 | 441.3551 | 5.98E-49 | 3.229346843 | YKR049C | FMP46 | n |
| W303.asm02.HP0_G0033730 | 808.9986 | 810.1194 | 831.0687 | 2086.233 | 1989.095 | 1457.037 | 2596.097 | 2000.489 | 2325.188 | 7.99E-11 | -1.175002352 | YKR050W | TRK2 | n |
| W303.asm02.HP0_G0033740 | 853.4724 | 915.8355 | 776.0953 | 996.2229 | 994.5475 | 1267.88 | 1341.872 | 1277.822 | 1088.437 | 0.033785882 | -0.35655679 | YKR051W | HFL1 | n |
| W303.asm02.HP0_G0033750 | 1493.047 | 1849.476 | 1418.53 | 5941.212 | 5918.686 | 5699.764 | 4431.758 | 4403.654 | 4845.338 | 8.01E-49 | -1.883087491 | YKR052C | MRS4 | n |
| W303.asm02.HP0_G0033770 | 2899.265 | 2630.662 | 2886.643 | 4430.9 | 4332.27 | 4345.282 | 1561.608 | 2588.932 | 2245.05 | 0.001102903 | -0.639181545 | YKR054C | DYN1 | n |

| | | | | | | | | | | | | | | |
|---|---|---|---|---|---|---|---|---|---|---|---|---|---|---|
| W303.asm02.HP0_G0033780 | 562.2752 | 694.3881 | 535.7213 | 591.2034 | 691.6705 | 384.3903 | 290.1012 | 202.9482 | 217.6873 | 0.727464594 | 0.104685456 | YKR055W | RHO4 | n |
| W303.asm02.HP0_G0033790 | 1807.54 | 2400.313 | 2538.478 | 4460.078 | 4099.688 | 1686.455 | 3319.498 | 1404.53 | 2636.17 | 0.171472289 | -0.602924791 | YKR056W | TRM2 | n |
| W303.asm02.HP0_G0033830 | 778.2906 | 628.7328 | 679.0834 | 1925.058 | 1938.76 | 650.2729 | 2255.382 | 952.4605 | 1721.165 | 0.013933935 | -1.113560091 | YKR060W | UTP30 | n |
| W303.asm02.HP0_G0033850 | 2300.987 | 2336.883 | 2138.574 | 2163.346 | 2534.1 | 1893.844 | 2746.703 | 1966.128 | 1966.363 | 0.852462053 | 0.040121889 | YKR062W | TFA2 | n |
| W303.asm02.HP0_G0033860 | 1190.202 | 1169.554 | 1220.194 | 1695.107 | 1654.108 | 925.2715 | 2331.92 | 1894.183 | 2966.289 | 0.386626694 | -0.255771518 | YKR063C | LAS1 | n |
| W303.asm02.HP0_G0033870 | 1811.776 | 1995.253 | 1889.576 | 2177.24 | 2056.787 | 2412.695 | 7316.722 | 5729.796 | 7159.761 | 0.130739766 | -0.222700739 | YKR064W | OAF3 | n |
| W303.asm02.HP0_G0033880 | 4145.588 | 4138.508 | 3941.917 | 3618.776 | 4320.988 | 3485.341 | 2588.69 | 3554.278 | 3439.939 | 0.591386521 | 0.097869855 | YKR065C | PAM17 | n |
| W303.asm02.HP0_G0033890 | 10224.73 | 9491.083 | 9628.971 | 16471.02 | 17136.42 | 20838.36 | 22609.37 | 44205.12 | 39316.49 | 0.000482866 | -0.891714354 | YKR066C | CCP1 | n |
| W303.asm02.HP0_G0033900 | 5821.825 | 5967.954 | 5404.641 | 5179.109 | 5058.654 | 6678.971 | 5969.912 | 8093.23 | 5860.813 | 0.914105155 | 0.023463993 | YKR067W | GPT2 | n |
| W303.asm02.HP0_G0033910 | 2565.712 | 2714.123 | 2688.308 | 1441.536 | 1474.464 | 1519.329 | 695.0083 | 1371.243 | 999.9265 | 0.000396607 | 0.845207002 | YKR068C | BET3 | n |
| W303.asm02.HP0_G0033920 | 3623.551 | 2285.694 | 2871.553 | 523.1213 | 478.1812 | 506.6963 | 338.2456 | 278.1142 | 328.9232 | 3.86E-39 | 2.541514235 | YKR069W | MET1 | n |
| W303.asm02.HP0_G0033930 | 2546.651 | 2097.631 | 2311.039 | 1856.976 | 1946.571 | 2164.284 | 1178.922 | 1650.43 | 1234.359 | 0.232143624 | 0.220988033 | YKR070W | | n |
| W303.asm02.HP0_G0033940 | 4778.81 | 5460.516 | 4952.997 | 4526.77 | 4692.424 | 4450.115 | 7136.489 | 6875.541 | 6455.267 | 0.119997358 | 0.152406638 | YKR071C | DRE2 | n |
| W303.asm02.HP0_G0033950 | 4056.641 | 4009.423 | 4175.824 | 7627.983 | 7563.248 | 9492.009 | 15527.2 | 13035.93 | 16440.18 | 3.01E-12 | -1.011717801 | YKR072C | SIS2 | n |
| W303.asm02.HP0_G0033970 | 3108.927 | 2957.826 | 2743.281 | 2057.749 | 2147.91 | 2243.29 | 1520.871 | 1828.681 | 1582.42 | 0.000103684 | 0.450167324 | YKR074W | AIM29 | n |
| W303.asm02.HP0_G0033980 | 3958.163 | 5044.329 | 4183.369 | 12039.43 | 10500.03 | 11880.39 | 29565.63 | 19750.4 | 26465.76 | 1.29E-13 | -1.384302528 | YKR075C | | n |
| W303.asm02.HP0_G0033990 | 4877.287 | 4964.207 | 4793.466 | 2528.072 | 2763.211 | 6624.275 | 4583.599 | 13368.81 | 7658.528 | 0.554561912 | 0.296564752 | YKR076W | ECM4 | n |
| W303.asm02.HP0_G0034010 | 2366.639 | 2474.87 | 2294.871 | 5154.099 | 5197.509 | 7149.204 | 4908.265 | 6821.851 | 5622.792 | 8.11E-12 | -1.294183901 | YKR078W | VPS501 | n |
| W303.asm02.HP0_G0034020 | 1248.442 | 1188.472 | 1441.166 | 5318.052 | 4648.165 | 2617.045 | 2426.974 | 1518.353 | 2422.071 | 1.65E-08 | -1.69808865 | YKR079C | TRZ1 | n |
| W303.asm02.HP0_G0034040 | 1913.43 | 2206.685 | 2318.585 | 3868.179 | 3784.661 | 1354.482 | 3775.019 | 1476.475 | 2852.661 | 0.304653186 | -0.48432139 | YKR081C | RPF2 | n |
| W303.asm02.HP0_G0034050 | 4945.057 | 5127.789 | 4922.816 | 9278.629 | 8635.901 | 9396.292 | 6784.664 | 7847.33 | 7644.175 | 1.14E-20 | -0.865006031 | YKR082W | NUP133 | n |
| W303.asm02.HP0_G0034060 | 584.5121 | 563.0775 | 517.3969 | 449.4813 | 453.0138 | 510.4946 | 313.5562 | 544.4166 | 495.1789 | 0.32331656 | 0.23673805 | YKR083C | DAD2 | n |
| W303.asm02.HP0_G0034080 | 26.47247 | 24.48163 | 23.71402 | 1346.36 | 1233.204 | 1517.05 | 1236.942 | 1436.744 | 1495.105 | 9.35E-142 | -5.77741869 | YKR084C | HBS1 | y |
| W303.asm02.HP0_G0034090 | 4677.156 | 4539.117 | 4486.262 | 2682.994 | 2790.114 | 1985.763 | 1224.597 | 973.9365 | 907.828 | 1.61E-06 | 0.87748614 | YKR085C | MRPL20 | n |
| W303.asm02.HP0_G0034100 | 1383.981 | 1373.197 | 1331.219 | 2180.019 | 2142.703 | 2183.276 | 1598.643 | 1688.013 | 1770.205 | 1.76E-17 | -0.670321562 | YKR086W | PRP16 | n |
| W303.asm02.HP0_G0034110 | 1385.04 | 1607.998 | 1381.881 | 1424.863 | 1533.478 | 1347.645 | 1704.807 | 1892.035 | 2093.148 | 0.880777542 | 0.023011347 | YKR087C | OMA1 | n |
| W303.asm02.HP0_G0034120 | 2476.764 | 2669.61 | 2404.818 | 741.9568 | 942.477 | 878.932 | 762.9044 | 1433.523 | 1107.574 | 4.18E-11 | 1.55894163 | YKR088C | TVP38 | n |
| W303.asm02.HP0_G0034130 | 4725.865 | 4657.074 | 4563.872 | 2821.242 | 2889.048 | 3683.614 | 4300.904 | 3632.665 | 3317.938 | 0.000405432 | 0.570122966 | YKR089C | TGL4 | n |
| W303.asm02.HP0_G0034140 | 3066.571 | 3194.853 | 3347.989 | 2055.665 | 1933.553 | 1716.082 | 1675.18 | 657.1656 | 865.9651 | 0.022934855 | 0.752131658 | YKR090W | PXL1 | n |
| W303.asm02.HP0_G0034160 | 2282.986 | 2423.681 | 2736.814 | 3336.722 | 3235.317 | 1397.023 | 3342.953 | 2410.681 | 3058.388 | 0.787628743 | -0.098416107 | YKR092C | SRP40 | y |
| W303.asm02.HP0_G0034170 | 2887.617 | 3933.753 | 2636.568 | 4117.583 | 4360.041 | 4538.236 | 2661.524 | 6228.04 | 1873.068 | 0.291107992 | -0.460682285 | YKR093W | PTR2 | n |
| W303.asm02.HP0_G0034180 | 9530.088 | 10152.09 | 9525.492 | 10314.45 | 9784.057 | 11812.78 | 10933.73 | 12515.14 | 12642.61 | 0.314404177 | -0.127762178 | YKR095W | MLP1 | n |
| W303.asm02.HP0_G0034190 | 284.8438 | 260.3955 | 230.6728 | 705.8316 | 866.1068 | 554.5552 | 901.1654 | 741.9958 | 795.3961 | 2.55E-12 | -1.45382385 | YKR095W- | PCC1 | n |
| W303.asm02.HP0_G0034200 | 4543.734 | 5158.947 | 4436.678 | 6308.023 | 6080.105 | 8841.736 | 17240.65 | 19428.26 | 17265.48 | 0.001236953 | -0.586422748 | YKR096W | ESL2 | n |
| W303.asm02.HP0_G0034210 | 852.4135 | 812.345 | 863.406 | 821.8492 | 798.4151 | 1134.179 | 8283.314 | 4419.761 | 4923.083 | 0.685794931 | -0.123730202 | YKR097W | PCK1 | n |
| W303.asm02.HP0_G0034230 | 429.9129 | 474.0534 | 455.956 | 2627.416 | 1881.483 | 1315.739 | 8142.584 | 3821.654 | 6613.15 | 2.07E-10 | -2.098768511 | YKR099W | BAS1 | n |
| W303.asm02.HP0_G0034240 | 3242.348 | 3284.99 | 3315.652 | 5641.79 | 6135.647 | 5105.706 | 7977.165 | 6010.058 | 7115.506 | 3.44E-08 | -0.778356533 | YKR100C | SKG1 | n |
| W303.asm02.HP0_G0034420 | 588.7477 | 547.4983 | 514.1631 | 329.9902 | 368.8331 | 171.6842 | 608.5952 | 636.7634 | 513.1202 | 0.001462364 | 0.923524416 | YLL057C | JLP1 | n |
| W303.asm02.HP0_G0034440 | 1313.034 | 1304.203 | 1039.105 | 894.0997 | 974.5871 | 872.8546 | 335.7767 | 441.3318 | 263.1386 | 0.05871111 | 0.415552056 | YLL055W | YCT1 | n |
| W303.asm02.HP0_G0034500 | 5997.602 | 5542.864 | 5814.247 | 11523.26 | 11508.46 | 9911.344 | 12091.66 | 14737.9 | 12822.02 | 2.74E-14 | -0.924620793 | YLL048C | YBT1 | n |
| W303.asm02.HP0_G0034520 | 28204.83 | 27552.96 | 30186.87 | 22251.06 | 25705.5 | 22803.61 | 22394.58 | 19742.89 | 17547.75 | 0.036457767 | 0.280496279 | YLL045C | RPL8B | y |
| W303.asm02.HP0_G0034530 | 3346.12 | 3258.282 | 3351.223 | 2531.546 | 2133.157 | 2417.253 | 10226.38 | 6540.515 | 7133.447 | 0.012809615 | 0.491245228 | YLL043W | FPS1 | n |
| W303.asm02.HP0_G0034540 | 681.9308 | 600.9127 | 556.2016 | 291.0861 | 361.0225 | 339.5701 | 248.1291 | 541.1952 | 227.256 | 0.01006742 | 0.891375487 | YLL042C | ATG10 | n |
| W303.asm02.HP0_G0034550 | 8599.316 | 9040.398 | 8733.228 | 12109.6 | 12548.13 | 8932.137 | 6657.513 | 6566.287 | 6377.522 | 0.024684098 | -0.34895063 | YLL041C | SDH2 | n |
| W303.asm02.HP0_G0034560 | 10079.66 | 10806.41 | 9545.972 | 4960.273 | 4688.953 | 8069.917 | 5534.143 | 9549.303 | 7979.079 | 0.005674758 | 0.780261646 | YLL040C | VPS13 | n |
| W303.asm02.HP0_G0034570 | 753.9359 | 825.6986 | 688.7846 | 1323.434 | 1482.275 | 843.9874 | 839.4417 | 1069.505 | 1321.673 | 0.010874786 | -0.68597987 | YLL038C | ENT4 | n |
| W303.asm02.HP0_G0034580 | 3921.102 | 3760.156 | 3878.321 | 1929.227 | 1791.227 | 1604.412 | 1660.366 | 1432.449 | 1590.792 | 1.71E-23 | 1.118144071 | YLL036C | PRP19 | n |
| W303.asm02.HP0_G0034590 | 468.0332 | 477.3918 | 560.5133 | 1663.15 | 1427.601 | 465.6744 | 1554.202 | 723.7412 | 1636.243 | 0.009550637 | -1.239772773 | YLL035W | GRC3 | n |
| W303.asm02.HP0_G0034600 | 835.4711 | 900.2563 | 1069.287 | 3142.896 | 2688.576 | 1356.761 | 4341.642 | 2409.607 | 4299.923 | 0.00011829 | -1.357650091 | YLL034C | RIX7 | n |
| W303.asm02.HP0_G0034610 | 409.7938 | 373.9013 | 409.6059 | 305.6751 | 281.181 | 268.9213 | 330.8388 | 263.081 | 281.0798 | 0.001362876 | 0.479148598 | YLL033W | IRC19 | n |
| W303.asm02.HP0_G0034620 | 1661.412 | 1875.07 | 1702.02 | 1091.399 | 983.2656 | 1156.21 | 222.2052 | 485.3576 | 253.5699 | 0.018061886 | 0.697130018 | YLL032C | | n |
| W303.asm02.HP0_G0034630 | 1910.253 | 1983.012 | 1919.758 | 2180.714 | 2012.527 | 2651.23 | 1823.317 | 1998.342 | 1839.578 | 0.09399421 | -0.235844253 | YLL031C | GPI13 | n |
| W303.asm02.HP0_G0034640 | 10718.17 | 10281.17 | 9926.474 | 13801.93 | 14606.66 | 18995.41 | 18227 | 17364.42 | 15184.29 | 0.00011363 | -0.616194298 | YLL029W | FRA1 | n |
| W303.asm02.HP0_G0034650 | 1038.78 | 1101.673 | 989.5215 | 11016.11 | 10565.98 | 10261.55 | 10618.94 | 12886.67 | 11258.74 | 1.08E-200 | -3.346900462 | YLL028W | TPO1 | n |

| | | | | | | | | | | | | | |
|---|---|---|---|---|---|---|---|---|---|---|---|---|---|
| W303.asm02.HP0_G0034660 | 2798.669 | 2809.823 | 2834.904 | 2554.471 | 2685.973 | 1956.136 | 2926.936 | 2538.463 | 2603.876 | 0.1484027 | 0.230625119 | YLL027W | ISA1 | n |
| W303.asm02.HP0_G0034670 | 82139.83 | 82198.19 | 74733.67 | 3527.769 | 3551.212 | 10023.01 | 11906.49 | 19706.38 | 22555.76 | 7.64E-20 | 3.805195751 | YLL026W | HSP104 | n |
| W303.asm02.HP0_G0034690 | 18396.25 | 18044.07 | 19020.8 | 39631.47 | 43037.18 | 45567.72 | 11585.53 | 22184.71 | 17233.18 | 1.57E-07 | -1.209254032 | YLL024C | SSA2 | y |
| W303.asm02.HP0_G0034700 | 5585.691 | 5124.45 | 5306.552 | 2527.377 | 2554.061 | 3710.202 | 3031.866 | 6832.589 | 3986.549 | 0.006155535 | 0.86536306 | YLL023C | POM33 | n |
| W303.asm02.HP0_G0034710 | 1531.168 | 1448.867 | 1661.06 | 3683.385 | 3716.102 | 3667.661 | 1593.705 | 2543.832 | 2286.913 | 1.68E-11 | -1.25371511 | YLL022C | HIF1 | n |
| W303.asm02.HP0_G0034720 | 3620.375 | 3587.672 | 4058.332 | 8442.191 | 8249.711 | 8209.696 | 1782.579 | 2612.555 | 2009.422 | 1.37E-12 | -1.144213819 | YLL021W | SPA2 | n |
| W303.asm02.HP0_G0034730 | 2591.125 | 3395.157 | 2640.88 | 882.2895 | 734.1948 | 1070.367 | 5237.869 | 2122.902 | 2796.445 | 6.60E-07 | 1.68282335 | YLL019C | KNS1 | n |
| W303.asm02.HP0_G0034740 | 1172.201 | 1398.791 | 1089.767 | 571.0567 | 679.5207 | 544.6795 | 304.9149 | 359.723 | 320.5506 | 2.38E-10 | 1.028598738 | YLL018C-A | COX19 | n |
| W303.asm02.HP0_G0034750 | 23568.97 | 21796.44 | 24028.77 | 17224.79 | 17814.2 | 18724.21 | 16845.62 | 17705.89 | 16171.06 | 1.80E-05 | 0.368215769 | YLL018C | DPS1 | n |
| W303.asm02.HP0_G0034780 | 6321.625 | 6065.88 | 5897.246 | 2300.9 | 2376.153 | 1735.074 | 2640.538 | 2509.47 | 1966.363 | 6.16E-18 | 1.51184446 | YLL015W | BPT1 | n |
| W303.asm02.HP0_G0034790 | 1588.348 | 1465.559 | 1490.75 | 445.313 | 596.2078 | 461.1164 | 438.2379 | 381.199 | 413.8452 | 4.06E-26 | 1.598349178 | YLL014W | EMC6 | n |
| W303.asm02.HP0_G0034810 | 1563.993 | 1823.881 | 1520.931 | 6642.876 | 6124.365 | 4859.575 | 1804.8 | 2715.64 | 2588.326 | 4.25E-19 | -1.844386108 | YLL013C | PUF3 | n |
| W303.asm02.HP0_G0034830 | 839.7067 | 865.7595 | 922.6911 | 2367.593 | 2004.716 | 569.7485 | 4509.53 | 1814.722 | 4022.432 | 0.080005084 | -0.911068758 | YLL011W | SOF1 | n |
| W303.asm02.HP0_G0034840 | 627.9269 | 734.4489 | 673.6938 | 2182.798 | 1922.271 | 2117.185 | 1011.033 | 690.4534 | 919.7888 | 4.63E-21 | -1.611928808 | YLL010C | PSR1 | n |
| W303.asm02.HP0_G0034850 | 2607.009 | 2133.24 | 2045.873 | 1113.63 | 1427.601 | 986.0447 | 760.4354 | 1005.077 | 942.5144 | 3.03E-06 | 0.944465745 | YLL009C | COX17 | n |
| W303.asm02.HP0_G0034860 | 2443.938 | 2688.528 | 3047.252 | 5309.715 | 4504.103 | 1495.02 | 5278.607 | 2158.338 | 4822.612 | 0.357347777 | -0.467332826 | YLL008W | DRS1 | n |
| W303.asm02.HP0_G0034870 | 1413.63 | 1348.715 | 1380.803 | 3406.193 | 3119.894 | 3011.311 | 2465.243 | 3116.167 | 3107.427 | 1.93E-20 | -1.202919736 | YLL007C | LMO1 | n |
| W303.asm02.HP0_G0034880 | 1544.933 | 1549.02 | 1356.011 | 1555.469 | 1373.795 | 1326.374 | 783.8904 | 1084.538 | 991.5539 | 0.721831599 | 0.064214275 | YLL006W | MMM1 | n |
| W303.asm02.HP0_G0034890 | 1309.858 | 1188.472 | 1260.077 | 1135.166 | 1259.239 | 913.8765 | 636.9881 | 708.708 | 676.9837 | 0.256456672 | 0.184482368 | YLL005C | SPO75 | n |
| W303.asm02.HP0_G0034900 | 799.4685 | 821.2474 | 886.0421 | 1738.18 | 1538.685 | 1491.222 | 2085.025 | 1398.088 | 1992.676 | 4.24E-07 | -0.927722601 | YLL004W | ORC3 | n |
| W303.asm02.HP0_G0034910 | 2198.274 | 2273.453 | 2196.781 | 2148.062 | 1910.989 | 2139.216 | 2614.614 | 2395.648 | 2463.934 | 0.288258083 | 0.105020445 | YLL003W | SFI1 | n |
| W303.asm02.HP0_G0034920 | 1236.794 | 1141.734 | 1229.895 | 1472.798 | 1537.817 | 1870.294 | 677.7257 | 1293.929 | 708.0819 | 0.123382909 | -0.435760562 | YLL002W | RTT109 | n |
| W303.asm02.HP0_G0034930 | 3957.104 | 3883.677 | 3821.191 | 3977.25 | 3533.855 | 4311.097 | 3392.332 | 2529.873 | 2656.503 | 0.912547656 | -0.01977599 | YLL001W | DNM1 | n |
| W303.asm02.HP0_G0034940 | 2929.973 | 3067.993 | 2644.114 | 1492.945 | 1359.909 | 1942.462 | 1839.365 | 1518.353 | 1629.067 | 1.09E-06 | 0.849563247 | YLR001C | | n |
| W303.asm02.HP0_G0034950 | 2526.532 | 2708.559 | 2806.878 | 4275.978 | 3382.85 | 858.421 | 5453.902 | 2180.888 | 4744.867 | 0.89256113 | -0.082849916 | YLR002C | NOC3 | n |
| W303.asm02.HP0_G0034960 | 1848.837 | 1924.034 | 2035.094 | 1549.217 | 1333.874 | 423.8929 | 3065.197 | 1188.697 | 2550.052 | 0.10453938 | 0.812506196 | YLR003C | CMS1 | n |
| W303.asm02.HP0_G0034970 | 546.3917 | 586.4463 | 541.1109 | 107.681 | 116.2909 | 142.0573 | 1046.833 | 1576.338 | 739.1801 | 2.28E-14 | 2.193371824 | YLR004C | THI73 | n |
| W303.asm02.HP0_G0034980 | 2765.843 | 2706.333 | 2837.059 | 3370.068 | 3297.802 | 2768.978 | 2647.945 | 2370.95 | 2701.954 | 0.144009035 | -0.183565875 | YLR005W | SSL1 | n |
| W303.asm02.HP0_G0034990 | 2189.803 | 2744.168 | 2159.054 | 1319.266 | 1390.284 | 1899.921 | 474.0377 | 820.3831 | 509.5319 | 0.021925136 | 0.621762359 | YLR006C | SSK1 | n |
| W303.asm02.HP0_G0035000 | 1264.325 | 1072.741 | 1211.571 | 1318.571 | 1554.306 | 1504.136 | 909.8067 | 962.1247 | 1009.495 | 0.017955048 | -0.302074779 | YLR007W | NSE1 | n |
| W303.asm02.HP0_G0035010 | 4211.24 | 3900.369 | 3945.151 | 2093.18 | 2245.976 | 1574.025 | 1270.273 | 889.1063 | 1039.397 | 1.00E-07 | 1.027951966 | YLR008C | PAM18 | n |
| W303.asm02.HP0_G0035020 | 2160.153 | 2371.38 | 2505.063 | 6057.23 | 6051.466 | 1922.711 | 5868.685 | 2602.891 | 4653.964 | 0.028305183 | -0.995701754 | YLR009W | RLP24 | n |
| W303.asm02.HP0_G0035070 | 595.1011 | 590.8975 | 662.9147 | 1151.145 | 991.0762 | 1061.251 | 751.7941 | 985.7483 | 882.7102 | 6.87E-08 | -0.79332801 | YLR014C | PPR1 | n |
| W303.asm02.HP0_G0035080 | 1475.046 | 1527.876 | 1590.995 | 1818.767 | 1781.681 | 1550.475 | 1217.19 | 1996.194 | 1700.832 | 0.445435395 | -0.165110922 | YLR015W | BRE2 | n |
| W303.asm02.HP0_G0035090 | 1608.467 | 1498.943 | 1492.906 | 450.8708 | 439.9962 | 261.3246 | 664.1465 | 605.6232 | 574.1205 | 1.09E-19 | 1.997525921 | YLR016C | PML1 | n |
| W303.asm02.HP0_G0035100 | 3391.653 | 3421.864 | 3626.09 | 3850.117 | 4296.688 | 3565.106 | 4083.637 | 4519.624 | 4694.631 | 0.171845343 | -0.165742503 | YLR017W | MEU1 | n |
| W303.asm02.HP0_G0035110 | 1149.964 | 1060.5 | 1076.832 | 3916.115 | 3634.524 | 3622.841 | 4181.16 | 4046.078 | 3877.705 | 8.50E-95 | -1.765103468 | YLR018C | POM34 | n |
| W303.asm02.HP0_G0035130 | 2908.795 | 2808.711 | 3012.759 | 2803.874 | 2611.338 | 3469.388 | 5826.713 | 3831.318 | 4351.355 | 0.913527841 | -0.025331774 | YLR020C | YEH2 | n |
| W303.asm02.HP0_G0035140 | 1690.002 | 1533.44 | 1689.085 | 1142.808 | 1417.187 | 1035.423 | 1324.59 | 908.4347 | 1022.652 | 0.024808931 | 0.450695885 | YLR021W | IRC25 | n |
| W303.asm02.HP0_G0035150 | 1268.561 | 1340.926 | 1338.764 | 1019.149 | 1011.904 | 409.4592 | 1090.04 | 607.7708 | 1102.79 | 0.074547535 | 0.694087627 | YLR022C | SDO1 | n |
| W303.asm02.HP0_G0035160 | 2687.485 | 2798.695 | 2773.463 | 2650.342 | 2281.558 | 1409.178 | 1715.918 | 1282.117 | 1648.204 | 0.144498028 | 0.38130358 | YLR023C | IZH3 | n |
| W303.asm02.HP0_G0035170 | 2465.116 | 2600.617 | 2447.934 | 2316.878 | 2150.514 | 2657.307 | 1645.553 | 1889.888 | 1596.773 | 0.591386521 | 0.076501656 | YLR024C | UBR2 | n |
| W303.asm02.HP0_G0035180 | 7819.967 | 8073.374 | 7407.398 | 3752.857 | 3751.683 | 4804.879 | 7273.515 | 5410.878 | 5968.461 | 3.02E-08 | 0.920585935 | YLR025W | SNF7 | n |
| W303.asm02.HP0_G0035190 | 1783.185 | 2070.923 | 1901.433 | 4402.416 | 4814.79 | 4916.549 | 4319.421 | 4313.454 | 4518.807 | 4.20E-40 | -1.296166144 | YLR026C | SED5 | n |
| W303.asm02.HP0_G0035200 | 8656.497 | 8121.224 | 9046.9 | 6784.598 | 7301.159 | 6365.23 | 7815.449 | 7075.268 | 7457.586 | 0.00079383 | 0.336679656 | YLR027C | AAT2 | n |
| W303.asm02.HP0_G0035220 | 21075.26 | 18796.33 | 21858.94 | 31433.13 | 35895.7 | 34089.95 | 37420.58 | 31320.6 | 31976.12 | 5.95E-09 | -0.716245182 | YLR029C | RPL15A | y |
| W303.asm02.HP0_G0035240 | 2164.389 | 2053.119 | 2120.249 | 1094.873 | 1004.962 | 798.4075 | 892.524 | 830.0473 | 1330.046 | 2.85E-07 | 1.128738677 | YLR032W | RAD5 | n |
| W303.asm02.HP0_G0035250 | 2270.279 | 2567.233 | 2585.906 | 3752.857 | 3435.789 | 3209.583 | 2512.153 | 2965.835 | 2516.561 | 0.000147087 | -0.486314665 | YLR033W | RSC58 | n |
| W303.asm02.HP0_G0035260 | 1505.754 | 1570.163 | 1530.632 | 2535.019 | 2718.951 | 3774.014 | 10230.08 | 6805.744 | 7755.411 | 8.94E-06 | -0.970729315 | YLR034C | SMF3 | n |
| W303.asm02.HP0_G0035270 | 1436.926 | 1332.023 | 1358.167 | 206.3307 | 203.943 | 171.6842 | 467.8653 | 366.1658 | 539.434 | 4.86E-58 | 2.826312161 | YLR035C | MLH2 | n |
| W303.asm02.HP0_G0035300 | 2821.965 | 2817.613 | 2707.71 | 7441.105 | 8650.654 | 7484.216 | 2773.861 | 6571.656 | 5677.812 | 3.61E-07 | -1.497922644 | YLR038C | COX12 | n |
| W303.asm02.HP0_G0035310 | 3064.453 | 2898.848 | 2824.125 | 1885.46 | 1843.298 | 2443.081 | 1234.473 | 2006.932 | 1577.635 | 0.019204298 | 0.509702098 | YLR039C | RIC1 | n |
| W303.asm02.HP0_G0035320 | 1047.251 | 1106.125 | 1068.209 | 45.15655 | 52.93839 | 25.8286 | 14.81368 | 61.2066 | 21.52952 | 6.57E-26 | 4.702133012 | YLR040C | AFB1 | n |
| W303.asm02.HP0_G0035350 | 72860.7 | 72010.49 | 82936.56 | 63828.43 | 73311 | 114855.2 | 184924.1 | 177260.7 | 183826.2 | 0.562322578 | -0.145575352 | YLR044C | PDC1 | n |

| | | | | | | | | | | | | | | |
|---|---|---|---|---|---|---|---|---|---|---|---|---|---|---|
| W303.asm02.HP0_G0035360 | 4157.236 | 4366.633 | 4392.484 | 2462.769 | 2143.571 | 2655.028 | 662.912 | 1596.74 | 1099.201 | 0.005369436 | 0.830837143 | YLR045C | STU2 | n |
| W303.asm02.HP0_G0035380 | 787.8206 | 807.8938 | 753.4592 | 830.1858 | 836.6002 | 847.026 | 1162.874 | 1398.088 | 897.0632 | 0.624353832 | -0.097739563 | YLR047C | FRE8 | n |
| W303.asm02.HP0_G0035410 | 2462.998 | 1845.025 | 2192.469 | 494.6379 | 503.3487 | 448.2021 | 435.769 | 590.59 | 714.0623 | 3.85E-25 | 2.168449097 | YLR050C | EMA19 | n |
| W303.asm02.HP0_G0035420 | 878.8859 | 1018.213 | 980.8982 | 2172.377 | 2115.8 | 862.979 | 4753.956 | 2286.12 | 3724.607 | 0.03414338 | -0.839826769 | YLR051C | FCF2 | n |
| W303.asm02.HP0_G0035430 | 1696.356 | 1783.821 | 1673.994 | 1137.945 | 1164.645 | 1273.958 | 1871.461 | 1772.844 | 1739.107 | 1.16E-08 | 0.5271332 | YLR052W | IES3 | n |
| W303.asm02.HP0_G0035450 | 1171.142 | 1332.023 | 928.0806 | 143.8062 | 122.3658 | 122.306 | 446.8793 | 510.055 | 531.0614 | 4.63E-63 | 3.140932389 | YLR054C | OSW2 | n |
| W303.asm02.HP0_G0035460 | 742.288 | 721.0953 | 718.9661 | 3524.99 | 3018.356 | 3272.635 | 2592.393 | 3004.492 | 2810.798 | 8.21E-87 | -2.169449571 | YLR055C | SPT8 | n |
| W303.asm02.HP0_G0035470 | 8764.505 | 10351.28 | 11047.5 | 7997.572 | 7649.164 | 11340.27 | 40538.86 | 24325.86 | 26751.62 | 0.567826193 | 0.160513821 | YLR056W | ERG3 | n |
| W303.asm02.HP0_G0035480 | 2270.279 | 2217.813 | 2341.221 | 2259.217 | 2022.073 | 2307.861 | 3056.555 | 2715.64 | 3010.544 | 0.651510892 | 0.051341348 | YLR057W | MNL2 | n |
| W303.asm02.HP0_G0035490 | 2105.091 | 1536.779 | 1987.666 | 15399.77 | 13546.15 | 20539.05 | 12300.29 | 31373.21 | 23286.57 | 6.51E-21 | -3.135878965 | YLR058C | SHM2 | n |
| W303.asm02.HP0_G0035500 | 3650.024 | 3414.075 | 3841.672 | 3332.553 | 3473.974 | 2996.117 | 2430.678 | 2356.991 | 2180.462 | 0.175416291 | 0.153991451 | YLR059C | REX2 | n |
| W303.asm02.HP0_G0035510 | 18695.92 | 18218.78 | 20029.73 | 16740.57 | 16735.48 | 18048.87 | 10520.18 | 13040.23 | 12257.47 | 0.240913962 | 0.144283952 | YLR060W | FRS1 | n |
| W303.asm02.HP0_G0035520 | 6846.839 | 6696.839 | 7095.882 | 9871.916 | 11666.41 | 9254.994 | 12008.95 | 8689.189 | 8636.925 | 0.001116146 | -0.577162544 | YLR061W | RPL22A | n |
| W303.asm02.HP0_G0035530 | 253.0768 | 272.6363 | 287.802 | 1140.724 | 1004.962 | 477.829 | 2560.297 | 1834.05 | 2462.738 | 1.56E-07 | -1.689324902 | YLR063W | BMT6 | n |
| W303.asm02.HP0_G0035540 | 3968.752 | 3662.229 | 3531.234 | 2665.626 | 2894.255 | 3470.908 | 2070.211 | 3440.455 | 2783.288 | 0.172576172 | 0.305724952 | YLR064W | PER33 | n |
| W303.asm02.HP0_G0035550 | 2097.678 | 1958.53 | 1940.238 | 2149.452 | 2280.69 | 2360.278 | 1608.518 | 2422.493 | 2258.207 | 0.338622182 | -0.17932999 | YLR065C | SND2 | n |
| W303.asm02.HP0_G0035560 | 2899.265 | 2954.488 | 2871.553 | 1474.188 | 1509.178 | 2074.644 | 1722.09 | 2406.386 | 1870.676 | 6.46E-05 | 0.786597116 | YLR066W | SPC3 | n |
| W303.asm02.HP0_G0035570 | 1693.179 | 1678.104 | 1570.515 | 4339.197 | 3820.243 | 3047.774 | 4291.028 | 3653.067 | 4585.787 | 2.30E-12 | -1.181375216 | YLR067C | PET309 | n |
| W303.asm02.HP0_G0035580 | 871.4736 | 834.601 | 933.4702 | 1460.988 | 1352.966 | 777.8966 | 1309.776 | 1136.08 | 1343.203 | 0.08459406 | -0.444419496 | YLR068W | FYV7 | n |
| W303.asm02.HP0_G0035590 | 23373.07 | 24330.29 | 23117.94 | 8281.711 | 7891.292 | 6920.545 | 2138.107 | 2281.825 | 2392.169 | 1.66E-55 | 1.61667158 | YLR069C | MEF1 | n |
| W303.asm02.HP0_G0035610 | 3144.929 | 3085.798 | 3141.03 | 5022.103 | 4616.054 | 5204.462 | 6588.383 | 5837.176 | 6495.934 | 1.52E-12 | -0.663474593 | YLR071C | RGR1 | n |
| W303.asm02.HP0_G0035630 | 309.1984 | 349.4196 | 363.2557 | 984.4128 | 899.0848 | 156.4909 | 815.9867 | 299.5902 | 662.6307 | 0.110904053 | -0.997340674 | YLR073C | RFU1 | n |
| W303.asm02.HP0_G0035640 | 1898.605 | 1991.914 | 2099.769 | 2387.045 | 2118.404 | 948.0614 | 3846.618 | 2287.194 | 3466.252 | 0.7275852 | 0.135430513 | YLR074C | BUD20 | n |
| W303.asm02.HP0_G0035650 | 26147.39 | 25488.72 | 27549.23 | 26447.84 | 31200.67 | 28546.68 | 26389.33 | 23935 | 22315.35 | 0.314730242 | -0.122343862 | YLR075W | RPL10 | n |
| W303.asm02.HP0_G0035660 | 4761.867 | 4415.596 | 4609.144 | 7597.416 | 7692.556 | 7729.587 | 4191.036 | 4474.524 | 4796.298 | 1.23E-18 | -0.739537222 | YLR077W | FMP25 | n |
| W303.asm02.HP0_G0035670 | 1589.407 | 1574.614 | 1372.18 | 2336.33 | 2342.307 | 2116.426 | 2444.257 | 2418.197 | 2577.562 | 3.06E-08 | -0.582974763 | YLR078C | BOS1 | n |
| W303.asm02.HP0_G0035680 | 5674.638 | 5412.666 | 5251.578 | 1513.786 | 1333.874 | 1267.88 | 1460.382 | 1376.612 | 1858.715 | 2.51E-40 | 1.988946056 | YLR079W | SIC1 | n |
| W303.asm02.HP0_G0035720 | 3391.653 | 3349.532 | 3745.738 | 6636.623 | 6283.18 | 6685.049 | 4858.886 | 7059.161 | 5694.557 | 1.31E-08 | -0.902634466 | YLR083C | EMP70 | n |
| W303.asm02.HP0_G0035730 | 703.1087 | 698.8393 | 720.044 | 8521.388 | 8026.675 | 8014.462 | 8148.757 | 7964.374 | 9333.046 | 3.73E-268 | -3.53298475 | YLR084C | RAX2 | n |
| W303.asm02.HP0_G0035740 | 2284.045 | 2208.911 | 2239.897 | 1795.147 | 1618.526 | 1163.806 | 3291.105 | 1551.641 | 2979.446 | 0.07270295 | 0.556642093 | YLR085C | ARP6 | n |
| W303.asm02.HP0_G0035750 | 2071.206 | 2005.268 | 2137.496 | 3656.986 | 3665.767 | 3437.482 | 6167.428 | 3877.492 | 5551.027 | 1.97E-05 | -0.79210459 | YLR086W | SMC4 | n |
| W303.asm02.HP0_G0035760 | 2481 | 2638.452 | 2357.389 | 3679.217 | 3493.934 | 4200.945 | 4352.752 | 5133.837 | 4871.651 | 1.13E-06 | -0.605370848 | YLR087C | CSF1 | n |
| W303.asm02.HP0_G0035770 | 1771.538 | 1654.736 | 1744.059 | 3336.722 | 3466.163 | 3805.16 | 2399.816 | 3801.252 | 3359.801 | 2.35E-08 | -1.036798041 | YLR088W | GAA1 | n |
| W303.asm02.HP0_G0035780 | 7847.498 | 7232.096 | 8174.871 | 4211.369 | 4375.662 | 5517.444 | 10144.9 | 5540.808 | 7708.763 | 0.002775765 | 0.721357289 | YLR089C | ALT1 | n |
| W303.asm02.HP0_G0035790 | 4061.935 | 4460.108 | 4040.007 | 1953.542 | 1673.2 | 1426.65 | 2096.135 | 1357.283 | 2066.834 | 3.90E-10 | 1.313631942 | YLR090W | XDJ1 | n |
| W303.asm02.HP0_G0035800 | 857.708 | 830.1498 | 779.329 | 1611.741 | 1504.839 | 742.1923 | 838.2072 | 425.2248 | 924.5732 | 0.091827075 | -0.645263138 | YLR091W | GEP5 | n |
| W303.asm02.HP0_G0035820 | 3957.104 | 4230.871 | 3781.309 | 838.5224 | 912.9703 | 933.6278 | 1067.819 | 1639.692 | 1150.633 | 3.90E-32 | 2.156409319 | YLR093C | NYV1 | n |
| W303.asm02.HP0_G0035830 | 892.6516 | 967.0244 | 861.2502 | 4231.516 | 3892.274 | 2136.177 | 6405.681 | 3035.632 | 4646.788 | 7.49E-09 | -1.914874882 | YLR094C | GIS3 | n |
| W303.asm02.HP0_G0035840 | 2234.276 | 2323.529 | 2484.583 | 8299.079 | 9174.831 | 10582.89 | 15124.76 | 11386.57 | 13752.58 | 3.36E-38 | -1.994203853 | YLR095C | IOC2 | n |
| W303.asm02.HP0_G0035860 | 1932.49 | 1898.439 | 1782.863 | 87.53423 | 85.91641 | 94.19841 | 70.36497 | 69.797 | 95.68675 | 1.34E-189 | 4.389879298 | YLR097C | HRT3 | n |
| W303.asm02.HP0_G0035870 | 1387.157 | 1495.605 | 1316.128 | 1376.233 | 1326.063 | 1600.613 | 1599.877 | 1359.431 | 1371.909 | 0.815829326 | -0.03553013 | YLR098C | CHA4 | n |
| W303.asm02.HP0_G0035890 | 354.7311 | 258.1699 | 309.3602 | 617.6026 | 696.8775 | 623.6847 | 545.6371 | 434.889 | 391.1196 | 2.47E-08 | -1.070580535 | YLR099W- | MIM2 | n |
| W303.asm02.HP0_G0035900 | 3830.037 | 3402.947 | 3891.256 | 2522.514 | 2491.576 | 3126.02 | 2797.316 | 2489.068 | 2784.484 | 0.001257119 | 0.450563107 | YLR100W | ERG27 | n |
| W303.asm02.HP0_G0035920 | 672.4007 | 693.2753 | 791.186 | 3008.121 | 3045.259 | 3744.387 | 1977.626 | 2087.467 | 1926.892 | 3.88E-58 | -2.183544308 | YLR103C | CDC45 | n |
| W303.asm02.HP0_G0035930 | 852.4135 | 885.7899 | 836.4583 | 796.8394 | 828.7896 | 695.8528 | 769.0767 | 647.5014 | 571.7283 | 0.360615683 | 0.149524102 | YLR104W | LCL2 | n |
| W303.asm02.HP0_G0035940 | 2526.532 | 2352.462 | 2406.973 | 1220.616 | 1311.31 | 1106.072 | 795.0007 | 1197.287 | 935.3379 | 8.00E-09 | 1.00215483 | YLR105C | SEN2 | n |
| W303.asm02.HP0_G0035950 | 1169.024 | 1225.194 | 1028.326 | 604.403 | 515.4985 | 447.4425 | 1036.957 | 590.59 | 909.0241 | 2.79E-06 | 1.12644962 | YLR107W | REX3 | n |
| W303.asm02.HP0_G0035970 | 17064.15 | 16775.48 | 15957.38 | 6021.799 | 8114.328 | 10715.83 | 4681.122 | 18622.91 | 11232.43 | 0.024765427 | 1.002707948 | YLR109W | AHP1 | n |
| W303.asm02.HP0_G0036000 | 12150.86 | 12063.88 | 11991.75 | 8684.646 | 8213.262 | 9208.654 | 7641.388 | 6734.873 | 6796.151 | 3.58E-07 | 0.471780282 | YLR113W | HOG1 | n |
| W303.asm02.HP0_G0036010 | 3114.221 | 2960.052 | 3218.64 | 2960.88 | 2915.951 | 2967.25 | 3287.402 | 3257.909 | 2999.779 | 0.416041481 | 0.071421747 | YLR114C | AVL9 | n |
| W303.asm02.HP0_G0036020 | 3283.645 | 3810.232 | 3397.573 | 3754.246 | 3718.705 | 3268.837 | 2676.338 | 2709.197 | 2679.229 | 0.776126775 | -0.03407877 | YLR115W | CFT2 | n |
| W303.asm02.HP0_G0036030 | 1951.55 | 2007.494 | 1932.693 | 1849.334 | 1695.764 | 1716.082 | 2778.799 | 2441.821 | 2358.678 | 0.134978123 | 0.163002964 | YLR116W | MSL5 | n |
| W303.asm02.HP0_G0036040 | 3274.115 | 3252.718 | 3130.251 | 2989.364 | 3132.044 | 3458.753 | 2889.902 | 2673.762 | 2478.287 | 0.927233999 | 0.011578747 | YLR117C | CLF1 | n |
| W303.asm02.HP0_G0036050 | 3382.122 | 3109.167 | 3116.238 | 1203.248 | 1462.315 | 1141.016 | 715.9944 | 980.3793 | 859.9846 | 2.29E-16 | 1.336095189 | YLR118C | TML25 | n |

| | | | | | | | | | | | | | |
|---|---|---|---|---|---|---|---|---|---|---|---|---|---|
| W303.asm02.HP0_G0036060 | 1954.727 | 1758.226 | 1541.412 | 698.8844 | 721.1771 | 739.9133 | 994.9853 | 1122.121 | 1003.515 | 1.33E-23 | 1.282673744 | YLR119W | SRN2 | n |
| W303.asm02.HP0_G0036080 | 3447.774 | 3195.966 | 3499.974 | 7650.214 | 8141.231 | 10114.93 | 9974.543 | 9046.764 | 9037.613 | 6.95E-23 | -1.352691377 | YLR121C | YPS3 | n |
| W303.asm02.HP0_G0036120 | 2165.448 | 2262.325 | 2044.796 | 696.8003 | 653.4854 | 816.6395 | 609.8297 | 781.7263 | 639.9051 | 2.98E-26 | 1.578329259 | YLR127C | APC2 | n |
| W303.asm02.HP0_G0036140 | 2215.216 | 2090.954 | 2398.35 | 5517.436 | 4957.984 | 1461.595 | 8582.057 | 3619.78 | 7407.35 | 0.094242251 | -0.832230631 | YLR129W | DIP2 | n |
| W303.asm02.HP0_G0036150 | 2579.477 | 2442.599 | 2597.763 | 4551.085 | 4972.738 | 5160.402 | 6485.922 | 8150.141 | 6304.56 | 2.66E-12 | -0.94636029 | YLR130C | ZRT2 | n |
| W303.asm02.HP0_G0036160 | 1980.141 | 1871.732 | 2137.496 | 6230.214 | 5893.519 | 6085.673 | 2423.271 | 1614.995 | 1713.989 | 6.37E-20 | -1.604203452 | YLR131C | ACE2 | n |
| W303.asm02.HP0_G0036170 | 1342.684 | 1471.123 | 1261.155 | 828.1016 | 925.1201 | 748.2696 | 124.6818 | 148.1844 | 114.8241 | 2.58E-06 | 0.704390839 | YLR132C | USB1 | n |
| W303.asm02.HP0_G0036180 | 3568.489 | 4023.89 | 3631.479 | 2385.655 | 2366.607 | 2050.335 | 839.4417 | 969.6413 | 515.5123 | 0.002273685 | 0.722406836 | YLR133W | CKI1 | n |
| W303.asm02.HP0_G0036200 | 1537.521 | 1676.992 | 1560.814 | 2141.115 | 1816.394 | 2139.216 | 3250.368 | 1494.73 | 2402.933 | 0.221013341 | -0.352522685 | YLR135W | SLX4 | n |
| W303.asm02.HP0_G0036210 | 220.2509 | 290.4412 | 205.8808 | 444.6183 | 280.3131 | 196.7531 | 541.9337 | 584.1472 | 620.7678 | 0.279219218 | -0.364357517 | YLR136C | TIS11 | n |
| W303.asm02.HP0_G0036220 | 937.1254 | 678.8088 | 798.7314 | 582.8668 | 554.5514 | 401.8626 | 87.64759 | 175.0294 | 117.2163 | 0.025556278 | 0.649660638 | YLR137W | RKM5 | n |
| W303.asm02.HP0_G0036240 | 1913.43 | 1969.658 | 1891.732 | 2239.765 | 2027.28 | 1514.012 | 1773.938 | 1675.128 | 1660.165 | 0.992921844 | -0.001637881 | YLR139C | SLS1 | n |
| W303.asm02.HP0_G0036250 | 1534.344 | 1277.496 | 1372.18 | 1269.941 | 1061.371 | 1065.809 | 1431.989 | 1598.888 | 1669.734 | 0.037244531 | 0.300225165 | YLR141W | RRN5 | n |
| W303.asm02.HP0_G0036260 | 929.7131 | 1254.127 | 920.5353 | 2290.479 | 2118.404 | 4314.895 | 7848.78 | 9466.62 | 3755.705 | 0.000171272 | -1.490715307 | YLR142W | PUT1 | n |
| W303.asm02.HP0_G0036270 | 1605.29 | 1622.464 | 1700.942 | 2507.231 | 2593.981 | 1911.316 | 1287.555 | 1546.272 | 1406.595 | 0.001364355 | -0.508638587 | YLR143W | DPH6 | n |
| W303.asm02.HP0_G0036280 | 1994.965 | 1850.589 | 1989.822 | 1583.258 | 1572.531 | 2473.468 | 2012.191 | 3256.835 | 2954.328 | 0.857783299 | 0.051850161 | YLR144C | ACF2 | n |
| W303.asm02.HP0_G0036290 | 959.3622 | 770.0586 | 893.5875 | 211.1937 | 218.6963 | 117.748 | 285.1633 | 227.6456 | 296.6289 | 5.05E-18 | 2.260592407 | YLR145W | RMP1 | n |
| W303.asm02.HP0_G0036300 | 386.498 | 408.3981 | 409.6059 | 1835.44 | 1813.791 | 1029.346 | 806.1109 | 777.4311 | 1119.535 | 1.25E-13 | -1.957624112 | YLR146C | SPE4 | n |
| W303.asm02.HP0_G0036330 | 284.8438 | 289.3284 | 256.5426 | 3004.647 | 3000.132 | 4491.137 | 3171.361 | 3964.469 | 3910 | 2.95E-70 | -3.659416272 | YLR148W | PEP3 | n |
| W303.asm02.HP0_G0036450 | 680.8719 | 751.1409 | 648.9019 | 300.8121 | 330.648 | 343.3684 | 235.7844 | 159.9962 | 175.8244 | 4.42E-10 | 1.094356715 | YLR151C | PCD1 | n |
| W303.asm02.HP0_G0036460 | 1327.859 | 1297.526 | 1190.013 | 1815.293 | 1637.619 | 2611.727 | 16061.73 | 9957.347 | 9840.186 | 0.012847447 | -0.668658698 | YLR152C | | n |
| W303.asm02.HP0_G0036470 | 13682.03 | 14496.46 | 15487.41 | 20207.21 | 20871.61 | 23617.97 | 18152.93 | 24366.67 | 26932.23 | 0.001257119 | -0.567187664 | YLR153C | ACS2 | n |
| W303.asm02.HP0_G0036520 | 8015.863 | 8511.818 | 8068.157 | 3286.702 | 3023.563 | 3132.097 | 8762.29 | 3066.773 | 7721.92 | 5.72E-05 | 1.381179641 | YLR163C | MAS1 | n |
| W303.asm02.HP0_G0036530 | 219.192 | 174.7098 | 183.2447 | 7671.75 | 8213.262 | 5229.531 | 9651.111 | 3503.809 | 9365.34 | 2.13E-44 | -5.192997975 | YLR164W | SHH4 | n |
| W303.asm02.HP0_G0036540 | 1571.406 | 1290.85 | 1228.818 | 1168.513 | 1011.904 | 1086.32 | 3467.635 | 1323.995 | 3305.977 | 0.379211348 | 0.324580391 | YLR165C | PUS5 | n |
| W303.asm02.HP0_G0036550 | 4431.491 | 4395.565 | 4142.409 | 4016.849 | 3945.212 | 3833.268 | 9893.067 | 4150.237 | 9250.516 | 0.685413869 | 0.136901562 | YLR166C | SEC10 | n |
| W303.asm02.HP0_G0036560 | 21338.93 | 21313.48 | 21699.41 | 20819.95 | 24514.82 | 22508.1 | 32474.05 | 16340.01 | 23999.43 | 0.778775524 | -0.076207979 | YLR167W | RPS31 | n |
| W303.asm02.HP0_G0036570 | 1485.635 | 1495.605 | 1580.216 | 1568.669 | 1479.671 | 1071.127 | 6589.617 | 2379.541 | 7016.231 | 0.726623769 | 0.147034312 | YLR168C | UPS2 | n |
| W303.asm02.HP0_G0036580 | 1386.098 | 1322.008 | 1362.478 | 1710.391 | 1828.544 | 1878.651 | 2115.887 | 2904.629 | 2491.444 | 0.004737393 | -0.412286815 | YLR170C | APS1 | n |
| W303.asm02.HP0_G0036590 | 7419.703 | 7700.586 | 8563.996 | 4249.579 | 4652.504 | 3277.953 | 2613.38 | 2474.035 | 2840.7 | 2.97E-09 | 0.959495638 | YLR172C | DPH5 | n |
| W303.asm02.HP0_G0036600 | 2192.979 | 2261.212 | 2084.678 | 1008.033 | 989.3405 | 968.5724 | 2068.977 | 1724.523 | 1821.636 | 2.20E-27 | 1.140439044 | YLR173W | TAG1 | n |
| W303.asm02.HP0_G0036610 | 688.2842 | 600.9127 | 628.4216 | 11470.46 | 12954.29 | 14155.59 | 10849.78 | 9407.561 | 7439.644 | 1.67E-137 | -4.330343063 | YLR174W | IDP2 | n |
| W303.asm02.HP0_G0036620 | 10925.72 | 10342.38 | 11363.33 | 10169.25 | 10475.73 | 5103.427 | 9169.666 | 4910.487 | 8468.277 | 0.331656428 | 0.341789576 | YLR175W | CBF5 | n |
| W303.asm02.HP0_G0036630 | 1405.159 | 1439.965 | 1313.972 | 5978.727 | 4949.306 | 4613.443 | 6164.959 | 4807.402 | 7715.94 | 2.51E-21 | -1.90184473 | YLR176C | RFX1 | n |
| W303.asm02.HP0_G0036640 | 5167.426 | 5511.705 | 4587.586 | 4683.081 | 4150.891 | 7344.438 | 6897.001 | 4382.178 | 4961.358 | 0.781164576 | -0.083714755 | YLR177W | | n |
| W303.asm02.HP0_G0036650 | 9045.113 | 7699.473 | 7438.658 | 528.679 | 597.9435 | 1241.292 | 3829.336 | 2193.773 | 2650.523 | 6.84E-22 | 3.352287736 | YLR178C | TFS1 | n |
| W303.asm02.HP0_G0036670 | 7218.512 | 4276.496 | 6590.343 | 20853.29 | 23487.29 | 18505.43 | 7572.258 | 7718.474 | 7126.27 | 2.01E-17 | -1.796953509 | YLR180W | SAM1 | y |
| W303.asm02.HP0_G0036680 | 3190.462 | 3522.016 | 3146.42 | 1330.381 | 1392.887 | 2116.426 | 1428.285 | 1703.047 | 1332.438 | 1.75E-06 | 1.026464052 | YLR181C | VTA1 | n |
| W303.asm02.HP0_G0036690 | 4517.262 | 4926.372 | 4484.106 | 4947.768 | 4796.566 | 5281.188 | 3063.962 | 3085.027 | 2746.21 | 0.296026973 | -0.109553552 | YLR182W | SWI6 | n |
| W303.asm02.HP0_G0036720 | 2084.972 | 2084.277 | 2339.065 | 2985.89 | 3140.722 | 1487.423 | 2447.96 | 1627.881 | 2139.795 | 0.490207473 | -0.226329759 | YLR186W | EMG1 | n |
| W303.asm02.HP0_G0036730 | 534.7438 | 568.6415 | 510.9294 | 5393.081 | 4888.557 | 4210.821 | 7042.669 | 4700.022 | 6194.521 | 2.25E-65 | -3.166409609 | YLR187W | SKG3 | n |
| W303.asm02.HP0_G0036740 | 2884.44 | 2464.855 | 2942.695 | 4650.43 | 4824.337 | 5323.73 | 3078.776 | 3405.02 | 3235.408 | 7.62E-13 | -0.835561133 | YLR188W | MDL1 | n |
| W303.asm02.HP0_G0036750 | 2955.386 | 3074.67 | 2700.165 | 1738.874 | 1705.311 | 5693.686 | 7774.712 | 10851.82 | 8353.453 | 0.898412541 | -0.065864733 | YLR189C | ATG26 | n |
| W303.asm02.HP0_G0036770 | 1175.378 | 1426.611 | 1210.493 | 1465.156 | 1464.918 | 1936.385 | 1970.219 | 1677.275 | 1594.38 | 0.054051329 | -0.352304405 | YLR191W | PEX13 | n |
| W303.asm02.HP0_G0036780 | 8067.749 | 7470.236 | 8096.183 | 5227.044 | 5818.884 | 4805.638 | 3340.484 | 2320.482 | 2884.955 | 0.000514361 | 0.576314968 | YLR192C | HCR1 | n |
| W303.asm02.HP0_G0036790 | 3970.87 | 3681.147 | 3574.35 | 3286.007 | 3526.044 | 3166.282 | 4708.28 | 3732.529 | 3463.86 | 0.282428538 | 0.170107993 | YLR193C | UPS1 | n |
| W303.asm02.HP0_G0036800 | 6267.621 | 6799.216 | 6128.997 | 3933.483 | 4040.675 | 3523.324 | 2759.047 | 4593.716 | 4683.866 | 0.000767187 | 0.739485951 | YLR194C | NCW2 | n |
| W303.asm02.HP0_G0036810 | 4779.869 | 4749.436 | 4770.83 | 5198.561 | 5169.738 | 4647.628 | 3078.776 | 3440.455 | 3754.509 | 0.561623312 | -0.0704501 | YLR195C | NMT1 | n |
| W303.asm02.HP0_G0036820 | 3447.774 | 3879.226 | 4063.721 | 9038.952 | 7916.459 | 3066.766 | 9032.64 | 4407.949 | 8275.707 | 0.054959086 | -0.813743377 | YLR196W | PWP1 | n |
| W303.asm02.HP0_G0036830 | 12028.03 | 12556.85 | 13070.74 | 16202.17 | 16074.18 | 10377.02 | 12942.22 | 9175.62 | 11862.76 | 0.451753393 | -0.179789782 | YLR197W | NOP56 | n |
| W303.asm02.HP0_G0036850 | 1696.356 | 1694.796 | 1754.838 | 2043.16 | 2395.245 | 1890.805 | 1360.389 | 2069.212 | 1757.048 | 0.125320995 | -0.298408045 | YLR200W | YKE2 | n |
| W303.asm02.HP0_G0036860 | 4040.757 | 3905.933 | 3676.751 | 2492.642 | 2533.232 | 3293.906 | 2950.391 | 3196.702 | 2833.524 | 0.001141976 | 0.482391598 | YLR201C | COQ9 | n |
| W303.asm02.HP0_G0036870 | 15268.26 | 14496.46 | 15364.53 | 4802.573 | 4416.451 | 3831.748 | 3824.398 | 2288.268 | 3602.606 | 1.37E-17 | 1.789911103 | YLR203C | MSS51 | n |
| W303.asm02.HP0_G0036890 | 3895.688 | 4359.956 | 4355.835 | 2503.062 | 2786.642 | 2085.279 | 4121.906 | 6417.028 | 5103.692 | 0.000127758 | 0.774110158 | YLR205C | HMX1 | n |

| | | | | | | | | | | | | | | |
|---|---|---|---|---|---|---|---|---|---|---|---|---|---|---|
| W303.asm02.HP0_G0036910 | 2987.153 | 2956.713 | 3027.85 | 2746.213 | 2679.898 | 4407.574 | 2697.324 | 3697.093 | 3095.466 | 0.608168678 | -0.132383607 | YLR207W | HRD3 | n |
| W303.asm02.HP0_G0036920 | 6805.542 | 6656.778 | 6963.3 | 6159.353 | 6981.793 | 6911.429 | 4600.881 | 5969.254 | 4877.632 | 0.860491682 | 0.026657479 | YLR208W | SEC13 | n |
| W303.asm02.HP0_G0036930 | 4037.581 | 3520.904 | 3870.775 | 4187.749 | 4496.292 | 5597.209 | 1434.458 | 4683.915 | 3451.899 | 0.430893313 | -0.321382391 | YLR209C | PNP1 | n |
| W303.asm02.HP0_G0036960 | 1045.133 | 1023.777 | 1197.558 | 1492.945 | 1439.751 | 1718.361 | 940.6685 | 929.9107 | 828.8864 | 0.000123595 | -0.509920905 | YLR212C | TUB4 | n |
| W303.asm02.HP0_G0036970 | 213.8975 | 185.8378 | 209.1146 | 193.8258 | 203.943 | 215.7448 | 351.8248 | 406.9702 | 336.0997 | 0.957562864 | -0.010011742 | YLR213C | CRR1 | n |
| W303.asm02.HP0_G0036980 | 1322.564 | 1487.815 | 1346.31 | 22725.55 | 23181.81 | 16611.59 | 1113.495 | 5257.324 | 12180.92 | 1.55E-10 | -3.910786469 | YLR214W | FRE1 | n |
| W303.asm02.HP0_G0036990 | 978.4224 | 1087.207 | 1089.767 | 4129.393 | 4240.279 | 3973.806 | 8275.908 | 6308.575 | 6418.188 | 2.92E-44 | -1.967908635 | YLR215C | CDC123 | n |
| W303.asm02.HP0_G0037000 | 11728.36 | 12817.25 | 11522.86 | 5841.868 | 6075.765 | 7632.35 | 2536.842 | 5144.575 | 4211.413 | 0.000842919 | 0.883567727 | YLR216C | CPR6 | n |
| W303.asm02.HP0_G0037010 | 1368.097 | 1112.801 | 1200.792 | 632.8864 | 684.7277 | 690.5351 | 774.0146 | 726.9626 | 755.9253 | 2.95E-13 | 0.875153868 | YLR218C | COA4 | n |
| W303.asm02.HP0_G0037020 | 2512.767 | 2893.284 | 2617.166 | 5217.318 | 4968.398 | 8903.269 | 6894.532 | 6389.11 | 5047.476 | 1.12E-06 | -1.250524174 | YLR219W | MSC3 | n |
| W303.asm02.HP0_G0037030 | 2749.96 | 2638.452 | 2527.699 | 6836.702 | 6791.736 | 7749.339 | 4662.605 | 5875.833 | 4210.217 | 6.59E-21 | -1.43324903 | YLR220W | CCC1 | n |
| W303.asm02.HP0_G0037040 | 2998.801 | 2823.177 | 3101.147 | 2837.221 | 3293.462 | 980.727 | 3276.292 | 1181.18 | 2753.386 | 0.528781555 | 0.327456434 | YLR221C | RSA3 | n |
| W303.asm02.HP0_G0037050 | 2061.676 | 2227.828 | 2549.257 | 7086.799 | 7001.753 | 1778.375 | 12347.2 | 4002.052 | 9927.5 | 0.025275097 | -1.214208826 | YLR222C | UTP13 | n |
| W303.asm02.HP0_G0037060 | 1040.897 | 1048.259 | 1147.974 | 6246.888 | 5558.531 | 3592.454 | 9211.638 | 4149.163 | 7415.723 | 1.04E-12 | -2.249947778 | YLR223C | IFH1 | n |
| W303.asm02.HP0_G0037090 | 891.5927 | 900.2563 | 854.7827 | 3324.911 | 3130.308 | 3623.6 | 3955.252 | 3496.293 | 3744.94 | 1.18E-77 | -1.929267305 | YLR226W | BUR2 | n |
| W303.asm02.HP0_G0037100 | 1000.659 | 924.7379 | 936.7039 | 280.6653 | 242.1281 | 221.8221 | 495.0237 | 405.8964 | 488.0024 | 2.51E-39 | 1.941238551 | YLR227C | ADY4 | n |
| W303.asm02.HP0_G0037120 | 5707.464 | 6508.775 | 6359.67 | 5147.152 | 5562.003 | 5992.994 | 6987.118 | 7032.316 | 6851.171 | 0.152872887 | 0.153422624 | YLR229C | CDC42 | n |
| W303.asm02.HP0_G0037150 | 340.9654 | 378.3525 | 376.1906 | 431.4187 | 362.7582 | 185.3582 | 649.3328 | 251.2692 | 711.6702 | 0.738066187 | 0.161300107 | YLR234W | TOP3 | n |
| W303.asm02.HP0_G0037180 | 1515.284 | 1561.26 | 1306.427 | 1637.446 | 1679.275 | 1834.59 | 5058.871 | 1990.825 | 5546.243 | 0.540815697 | -0.233034583 | YLR238W | FAR10 | n |
| W303.asm02.HP0_G0037190 | 4830.696 | 4808.415 | 4558.482 | 1830.577 | 1713.121 | 1363.598 | 1559.14 | 930.9845 | 2294.09 | 5.41E-07 | 1.532633902 | YLR239C | LIP2 | n |
| W303.asm02.HP0_G0037200 | 3591.784 | 3636.635 | 3589.441 | 1809.736 | 1645.429 | 2192.392 | 5341.565 | 2423.566 | 5606.047 | 0.002627708 | 0.937671024 | YLR240W | VPS34 | n |
| W303.asm02.HP0_G0037210 | 5812.295 | 5718.686 | 5710.768 | 2211.976 | 2117.536 | 2575.263 | 11599.11 | 3798.03 | 9994.481 | 0.000279537 | 1.320225949 | YLR241W | CSC1 | n |
| W303.asm02.HP0_G0037220 | 507.2125 | 469.6022 | 522.7864 | 1423.473 | 1424.13 | 1358.28 | 5229.228 | 1888.814 | 4697.023 | 1.36E-05 | -1.487797098 | YLR242C | ARV1 | n |
| W303.asm02.HP0_G0037230 | 2378.286 | 2492.675 | 2552.491 | 1642.309 | 1542.156 | 823.4764 | 6580.976 | 1590.298 | 5661.067 | 0.070578566 | 0.889232619 | YLR243W | GPN3 | n |
| W303.asm02.HP0_G0037240 | 5841.944 | 5508.367 | 6274.515 | 6694.285 | 6740.533 | 6407.771 | 23268.58 | 7343.718 | 19405.27 | 0.675544609 | -0.170987561 | YLR244C | MAP1 | n |
| W303.asm02.HP0_G0037250 | 488.1523 | 406.1725 | 448.4106 | 227.8669 | 284.6523 | 567.4695 | 2371.423 | 1131.785 | 2308.443 | 0.470663246 | 0.314225434 | YLR245C | CDD1 | n |
| W303.asm02.HP0_G0037260 | 795.2329 | 773.397 | 778.2511 | 2011.203 | 1973.474 | 1770.019 | 10625.11 | 3320.189 | 7950.373 | 0.000415485 | -1.293989418 | YLR246W | ERF2 | n |
| W303.asm02.HP0_G0037270 | 4496.084 | 4408.919 | 4287.927 | 2741.35 | 2652.127 | 3212.622 | 2796.082 | 3725.012 | 2778.504 | 0.000124202 | 0.61628437 | YLR247C | IRC20 | n |
| W303.asm02.HP0_G0037280 | 7837.968 | 8502.915 | 7601.422 | 5654.295 | 5319.007 | 6220.134 | 5073.684 | 4594.79 | 4569.042 | 1.02E-05 | 0.477610334 | YLR248W | RCK2 | n |
| W303.asm02.HP0_G0037290 | 86808.52 | 84399.31 | 96931.07 | 82010.55 | 91972.21 | 83740.11 | 100867.6 | 84464.03 | 90238.58 | 0.663558987 | 0.057167286 | YLR249W | YEF3 | n |
| W303.asm02.HP0_G0037300 | 4769.28 | 4909.68 | 4501.353 | 4250.968 | 4756.645 | 4739.548 | 4286.091 | 5512.889 | 4926.671 | 0.755021404 | 0.044828851 | YLR250W | SSP120 | n |
| W303.asm02.HP0_G0037310 | 1898.605 | 1971.884 | 1639.501 | 642.6124 | 719.4414 | 877.4126 | 797.4696 | 1586.002 | 1017.868 | 1.69E-06 | 1.298951565 | YLR251W | SYM1 | n |
| W303.asm02.HP0_G0037320 | 6922.021 | 6882.676 | 7123.908 | 2450.959 | 2463.805 | 2986.242 | 1320.886 | 1705.194 | 1429.321 | 2.39E-23 | 1.405337669 | YLR253W | CQD2 | n |
| W303.asm02.HP0_G0037340 | 3616.139 | 3386.255 | 3503.208 | 6318.443 | 6015.884 | 13066.99 | 34202.31 | 23453.94 | 26736.07 | 5.89E-05 | -1.273752959 | YLR256W | HAP1 | n |
| W303.asm02.HP0_G0037350 | 20781.95 | 22227.09 | 20062.06 | 2785.117 | 2554.061 | 4678.774 | 5795.851 | 6468.571 | 6814.092 | 3.73E-29 | 2.654336614 | YLR257W | | n |
| W303.asm02.HP0_G0037360 | 27980.34 | 29194.34 | 25426.82 | 4231.516 | 3698.745 | 10605.68 | 53537.86 | 26374.67 | 33277.46 | 2.85E-07 | 2.155828114 | YLR258W | GSY2 | n |
| W303.asm02.HP0_G0037370 | 41693.08 | 43236.78 | 42214.19 | 15136.48 | 15609.88 | 16218.08 | 8820.31 | 9773.727 | 8034.098 | 5.77E-48 | 1.436829997 | YLR259C | HSP60 | n |
| W303.asm02.HP0_G0037380 | 2943.738 | 3221.56 | 2867.241 | 4057.142 | 4380.869 | 5118.62 | 6343.957 | 6261.327 | 5805.793 | 3.29E-06 | -0.585803913 | YLR260W | LCB5 | n |
| W303.asm02.HP0_G0037400 | 1126.668 | 1049.372 | 1103.78 | 782.2504 | 721.1771 | 869.0563 | 1396.189 | 1991.899 | 1346.791 | 0.015730868 | 0.467030998 | YLR263W | RED1 | n |
| W303.asm02.HP0_G0037410 | 2845.261 | 2927.78 | 3108.693 | 11962.32 | 14552.85 | 12727.42 | 12792.84 | 9578.295 | 9618.91 | 1.40E-42 | -2.143471843 | YLR264W | RPS28B | y |
| W303.asm02.HP0_G0037420 | 2436.526 | 2195.557 | 2326.13 | 718.3365 | 708.1595 | 829.5538 | 582.6713 | 803.2023 | 644.6895 | 5.38E-26 | 1.624826952 | YLR265C | NEJ1 | n |
| W303.asm02.HP0_G0037450 | 3178.814 | 3153.679 | 3161.51 | 2536.409 | 2855.202 | 2699.088 | 2588.69 | 2984.09 | 2785.68 | 0.019790203 | 0.231049941 | YLR268W | SEC22 | n |
| W303.asm02.HP0_G0037460 | 7840.086 | 7650.509 | 7319.01 | 4535.802 | 4721.931 | 6092.51 | 7459.921 | 7263.183 | 7220.761 | 4.52E-05 | 0.571375041 | YLR270W | DCS1 | n |
| W303.asm02.HP0_G0037470 | 2069.088 | 2201.121 | 1814.123 | 830.1858 | 860.8998 | 1099.235 | 1444.334 | 1495.803 | 1563.282 | 2.06E-13 | 1.124598363 | YLR271W | CMG1 | n |
| W303.asm02.HP0_G0037480 | 1448.573 | 1424.386 | 1551.113 | 2877.514 | 2688.576 | 2389.905 | 3851.556 | 2646.917 | 3413.625 | 6.59E-07 | -0.84669133 | YLR272C | YCS4 | n |
| W303.asm02.HP0_G0037490 | 992.1881 | 945.8812 | 892.5096 | 503.6692 | 348.0048 | 831.0731 | 3816.991 | 4097.621 | 4847.73 | 0.015260132 | 0.749804239 | YLR273C | PIG1 | n |
| W303.asm02.HP0_G0037500 | 2558.299 | 2756.409 | 2658.126 | 2502.368 | 2617.413 | 2555.512 | 1070.288 | 1343.324 | 1065.711 | 0.692000143 | 0.054908241 | YLR274W | MCM5 | n |
| W303.asm02.HP0_G0037510 | 500.8591 | 596.4615 | 580.9936 | 688.4637 | 792.3402 | 418.5752 | 662.912 | 987.8959 | 818.1217 | 0.555339794 | -0.178260793 | YLR275W | SMD2 | n |
| W303.asm02.HP0_G0037520 | 5792.176 | 6488.745 | 6666.874 | 4140.508 | 3447.938 | 1009.594 | 7083.407 | 3065.699 | 6567.699 | 0.022327959 | 1.139942428 | YLR276C | DBP9 | n |
| W303.asm02.HP0_G0037530 | 1874.251 | 1812.753 | 1949.939 | 4234.99 | 3846.278 | 3890.243 | 4268.808 | 4468.082 | 3627.724 | 5.04E-21 | -1.086693059 | YLR277C | YSH1 | n |
| W303.asm02.HP0_G0037540 | 4559.618 | 4899.664 | 4779.454 | 2804.569 | 2506.329 | 3669.94 | 10749.79 | 5862.948 | 8089.118 | 0.009169923 | 0.664842965 | YLR278C | | n |
| W303.asm02.HP0_G0037550 | 823.8232 | 744.4641 | 745.9138 | 1199.08 | 1372.059 | 1655.309 | 2283.775 | 2004.784 | 1759.44 | 5.66E-07 | -0.868737676 | YLR281C | RSO55 | n |
| W303.asm02.HP0_G0037570 | 840.7656 | 983.7164 | 800.8872 | 302.8962 | 293.3308 | 335.0121 | 467.8653 | 483.21 | 398.2961 | 8.88E-24 | 1.494694903 | YLR284C | ECI1 | n |
| W303.asm02.HP0_G0037580 | 1610.585 | 1379.874 | 1442.244 | 2896.272 | 2945.457 | 1694.812 | 2477.587 | 1458.22 | 2265.384 | 0.007627425 | -0.765659381 | YLR285W | NNT1 | n |

| | | | | | | | | | | | | | | |
|---|---|---|---|---|---|---|---|---|---|---|---|---|---|---|
| W303.asm02.HP0_G0037600 | 31860.14 | 33142.56 | 34749.67 | 6100.302 | 5929.1 | 8041.81 | 4952.706 | 6020.796 | 6250.737 | 1.35E-45 | 2.313197793 | YLR286C | CTS1 | n |
| W303.asm02.HP0_G0037610 | 1891.193 | 2013.058 | 2007.069 | 2030.655 | 1891.897 | 937.4261 | 2271.43 | 1481.844 | 1934.068 | 0.391606552 | 0.282532794 | YLR287C | | n |
| W303.asm02.HP0_G0037620 | 1466.575 | 1431.063 | 1405.595 | 967.0449 | 977.1907 | 1108.351 | 749.3252 | 1283.191 | 820.5138 | 0.029117456 | 0.495378859 | YLR288C | MEC3 | n |
| W303.asm02.HP0_G0037630 | 2114.621 | 2262.325 | 2048.029 | 2338.415 | 2209.527 | 2245.569 | 1235.708 | 1282.117 | 1263.065 | 0.363218221 | -0.08080032 | YLR289W | GUF1 | n |
| W303.asm02.HP0_G0037640 | 1972.728 | 2116.548 | 1850.772 | 1709.002 | 1837.223 | 1991.841 | 1556.671 | 2062.77 | 1804.891 | 0.528115497 | 0.101130304 | YLR290C | COQ11 | n |
| W303.asm02.HP0_G0037650 | 2543.475 | 2707.446 | 2874.786 | 2837.221 | 2626.091 | 2273.676 | 2978.784 | 2712.419 | 2822.759 | 0.606471247 | 0.070617227 | YLR291C | GCD7 | n |
| W303.asm02.HP0_G0037660 | 2533.945 | 2175.527 | 2402.662 | 2297.426 | 2473.351 | 2303.303 | 1812.206 | 2483.699 | 2103.912 | 0.963746521 | 0.007892626 | YLR292C | SEC72 | n |
| W303.asm02.HP0_G0037670 | 13354.83 | 12590.23 | 13977.26 | 9161.916 | 11267.2 | 9080.271 | 8073.454 | 7455.393 | 7579.586 | 0.000466677 | 0.436125393 | YLR293C | GSP1 | y |
| W303.asm02.HP0_G0037680 | 4250.419 | 4130.719 | 4080.968 | 5887.025 | 6606.017 | 6456.39 | 3752.798 | 3952.658 | 4106.157 | 1.98E-12 | -0.604449229 | YLR295C | ATP14 | n |
| W303.asm02.HP0_G0037700 | 2499.001 | 2440.373 | 2389.727 | 862.8374 | 879.1245 | 783.9739 | 1398.658 | 1171.516 | 1326.458 | 5.80E-46 | 1.53703978 | YLR298C | YHC1 | n |
| W303.asm02.HP0_G0037710 | 7213.218 | 7121.929 | 7311.464 | 637.0547 | 605.7541 | 1077.204 | 1090.04 | 1181.18 | 802.5726 | 1.10E-36 | 3.221775189 | YLR299W | ECM38 | n |
| W303.asm02.HP0_G0037720 | 8166.227 | 7281.059 | 9272.183 | 5012.377 | 5062.993 | 4662.062 | 6332.847 | 5221.889 | 5663.459 | 3.78E-08 | 0.746196538 | YLR300W | EXG1 | n |
| W303.asm02.HP0_G0037730 | 2615.48 | 2270.115 | 2709.866 | 2804.569 | 3412.357 | 5869.928 | 14553.2 | 16880.13 | 13363.85 | 0.01946292 | -0.670206372 | YLR301W | HRI1 | n |
| W303.asm02.HP0_G0037740 | 21386.58 | 14287.26 | 18488.31 | 470.3228 | 443.4675 | 776.3772 | 2804.723 | 3545.687 | 3823.882 | 2.78E-79 | 5.001845811 | YLR303W | MET17 | n |
| W303.asm02.HP0_G0037750 | 24838.59 | 23802.82 | 24289.63 | 57392.58 | 59268.44 | 52143.38 | 14077.93 | 25906.5 | 16680.59 | 1.18E-07 | -1.210745525 | YLR304C | ACO1 | n |
| W303.asm02.HP0_G0037760 | 4509.85 | 4374.422 | 4333.199 | 2900.44 | 2738.911 | 2559.31 | 1383.844 | 1976.866 | 1783.362 | 7.58E-06 | 0.688949212 | YLR305C | STT4 | n |
| W303.asm02.HP0_G0037770 | 1237.853 | 1000.408 | 1117.793 | 1179.628 | 1086.539 | 1167.605 | 1208.549 | 1333.66 | 1247.516 | 0.808881517 | -0.033105846 | YLR306W | UBC12 | n |
| W303.asm02.HP0_G0037790 | 115.42 | 92.36251 | 85.1549 | 405.7142 | 446.0711 | 1327.894 | 12194.13 | 7402.777 | 8218.295 | 1.86E-10 | -2.895333245 | YLR307C-/ | DPA10 | n |
| W303.asm02.HP0_G0037800 | 26.47247 | 25.59443 | 15.09074 | 165.3424 | 159.683 | 109.3917 | 927.0893 | 597.0328 | 538.2379 | 4.38E-14 | -2.693653951 | YLR308W | CDA2 | n |
| W303.asm02.HP0_G0037810 | 5707.464 | 6310.697 | 5606.211 | 5956.496 | 5345.042 | 6452.591 | 4745.315 | 5598.793 | 5766.323 | 0.945940969 | -0.010680326 | YLR309C | IMH1 | n |
| W303.asm02.HP0_G0037820 | 7070.267 | 7178.682 | 7009.65 | 3703.532 | 3184.114 | 3508.891 | 7531.52 | 4108.359 | 5943.343 | 3.13E-06 | 1.031886738 | YLR310C | CDC25 | n |
| W303.asm02.HP0_G0037840 | 6613.881 | 6647.875 | 6264.814 | 5428.512 | 5231.355 | 4197.907 | 2871.384 | 2286.12 | 3107.427 | 0.020163278 | 0.394218186 | YLR312W- | MRPL15 | n |
| W303.asm02.HP0_G0037870 | 8676.616 | 8976.969 | 8666.398 | 9670.449 | 9664.294 | 10541.87 | 5925.471 | 7986.924 | 6954.034 | 0.198564796 | -0.182867693 | YLR314C | CDC3 | n |
| W303.asm02.HP0_G0037880 | 1544.933 | 1554.584 | 1571.593 | 1338.023 | 1345.156 | 1120.505 | 1193.735 | 891.2539 | 1040.593 | 0.055776201 | 0.296441156 | YLR316C | TAD3 | n |
| W303.asm02.HP0_G0037900 | 479.6811 | 499.6478 | 493.6828 | 1313.014 | 1352.966 | 1213.184 | 1413.472 | 1355.136 | 1220.006 | 2.21E-37 | -1.39689443 | YLR318W | EST2 | n |
| W303.asm02.HP0_G0037910 | 2390.993 | 2231.167 | 2233.43 | 3917.504 | 3795.943 | 4682.573 | 3503.435 | 3976.281 | 3735.371 | 2.63E-12 | -0.854551951 | YLR319C | BUD6 | n |
| W303.asm02.HP0_G0037920 | 1130.904 | 1129.493 | 1087.611 | 2649.647 | 2503.726 | 2601.851 | 2686.213 | 2470.814 | 2664.876 | 2.60E-51 | -1.212101502 | YLR320W | MMS22 | n |
| W303.asm02.HP0_G0037930 | 1572.465 | 1431.063 | 1475.659 | 3830.665 | 3770.776 | 3491.419 | 4249.056 | 3267.573 | 3886.078 | 5.58E-24 | -1.308258439 | YLR321C | SFH1 | n |
| W303.asm02.HP0_G0037940 | 943.4787 | 1044.92 | 1076.832 | 771.8296 | 764.5693 | 725.4797 | 846.8485 | 1119.973 | 1072.888 | 0.00301142 | 0.438398194 | YLR323C | CWC24 | n |
| W303.asm02.HP0_G0037950 | 5402.501 | 5032.088 | 4964.854 | 2934.481 | 2895.123 | 4456.952 | 5274.904 | 5219.741 | 4628.846 | 0.003490455 | 0.582083917 | YLR324W | PEX30 | n |
| W303.asm02.HP0_G0037960 | 7117.917 | 6840.39 | 6965.455 | 10748.65 | 13251.96 | 11316.72 | 9720.241 | 7565.994 | 8174.04 | 8.56E-08 | -0.755169647 | YLR325C | RPL38 | n |
| W303.asm02.HP0_G0037970 | 2118.856 | 2210.024 | 2105.159 | 780.8609 | 905.1597 | 775.6176 | 970.2958 | 897.6967 | 954.4753 | 3.76E-41 | 1.387150558 | YLR326W | | n |
| W303.asm02.HP0_G0037990 | 1740.829 | 1739.309 | 1755.916 | 4786.594 | 5172.341 | 4997.074 | 9588.152 | 7218.083 | 7628.626 | 9.13E-30 | -1.514119298 | YLR328W | NMA1 | n |
| W303.asm02.HP0_G0038010 | 15761.71 | 16923.48 | 16890.85 | 11209.24 | 11353.12 | 14087.98 | 5699.562 | 10196.8 | 7670.489 | 0.063142063 | 0.435812695 | YLR330W | CHS5 | n |
| W303.asm02.HP0_G0038020 | 2397.347 | 2885.494 | 2512.609 | 4696.976 | 4144.816 | 4300.461 | 7741.381 | 6264.549 | 7934.823 | 1.56E-07 | -0.75366209 | YLR332W | MID2 | n |
| W303.asm02.HP0_G0038030 | 3997.343 | 3632.184 | 3690.764 | 10438.8 | 13159.1 | 10575.29 | 11468.26 | 9047.838 | 8435.983 | 5.56E-22 | -1.593887755 | YLR333C | RPS25B | y |
| W303.asm02.HP0_G0038040 | 8272.117 | 8508.479 | 8983.303 | 12623.69 | 11389.57 | 12671.21 | 9628.89 | 11210.47 | 10079.4 | 7.26E-07 | -0.5098736 | YLR335W | NUP2 | n |
| W303.asm02.HP0_G0038050 | 1669.883 | 1786.046 | 1916.524 | 5829.363 | 4803.508 | 2109.589 | 5751.41 | 3407.167 | 5737.617 | 0.000854481 | -1.246005803 | YLR336C | SGD1 | n |
| W303.asm02.HP0_G0038060 | 5794.294 | 5724.25 | 6009.349 | 5395.166 | 4756.645 | 6864.329 | 4879.872 | 5698.656 | 5397.929 | 0.81695902 | 0.042682764 | YLR337C | VRP1 | n |
| W303.asm02.HP0_G0038070 | 29229.84 | 28568.95 | 31434.02 | 34441.94 | 40035.31 | 35291.74 | 34965.22 | 26755.87 | 26206.21 | 0.057869942 | -0.298811272 | YLR340W | RPP0 | n |
| W303.asm02.HP0_G0038090 | 36446.23 | 36159.37 | 37394.86 | 33739.58 | 34570.51 | 36810.31 | 43215.2 | 45935.01 | 47929.49 | 0.449796088 | 0.065471236 | YLR342W | FKS1 | n |
| W303.asm02.HP0_G0038110 | 1068.429 | 865.7595 | 1028.326 | 839.2171 | 795.8116 | 547.7182 | 522.1821 | 786.0215 | 734.3958 | 0.075368174 | 0.440754949 | YLR343W | GAS2 | n |
| W303.asm02.HP0_G0038130 | 7112.623 | 7661.637 | 6683.043 | 2905.303 | 2614.81 | 3176.158 | 2766.454 | 2694.164 | 1990.284 | 2.90E-15 | 1.3028963 | YLR345W | | n |
| W303.asm02.HP0_G0038150 | 3610.845 | 4120.703 | 3704.777 | 13105.82 | 12199.26 | 14338.67 | 11426.28 | 11798.91 | 12319.67 | 1.91E-65 | -1.793568129 | YLR347C | KAP95 | n |
| W303.asm02.HP0_G0038190 | 3751.678 | 3386.255 | 3558.181 | 2150.146 | 2370.946 | 2639.835 | 1385.079 | 1840.493 | 1229.575 | 0.001603578 | 0.578938024 | YLR351C | NIT3 | n |
| W303.asm02.HP0_G0038200 | 1243.147 | 1106.125 | 1220.194 | 1875.734 | 1634.147 | 2232.654 | 1444.334 | 2010.153 | 1539.361 | 0.000296395 | -0.686078861 | YLR352W | LUG1 | n |
| W303.asm02.HP0_G0038220 | 20624.17 | 19297.09 | 21307.05 | 10629.16 | 12890.06 | 13590.4 | 10462.16 | 15791.3 | 11454.9 | 0.000185896 | 0.722429286 | YLR354C | TAL1 | n |
| W303.asm02.HP0_G0038230 | 3029.509 | 2736.379 | 3437.455 | 9783.687 | 9751.079 | 13962.64 | 10899.16 | 9159.513 | 11448.92 | 5.70E-22 | -1.863807177 | YLR355C | ILV5 | n |
| W303.asm02.HP0_G0038240 | 2643.011 | 2576.135 | 2394.038 | 3633.365 | 4440.75 | 4918.069 | 2825.709 | 5886.571 | 3827.47 | 0.005453833 | -0.771040958 | YLR356W | ATG33 | n |
| W303.asm02.HP0_G0038260 | 9750.339 | 7287.736 | 9138.522 | 9686.427 | 8737.438 | 12373.42 | 8572.181 | 17986.15 | 15148.41 | 0.460314489 | -0.23452747 | YLR359W | ADE13 | n |
| W303.asm02.HP0_G0038270 | 1820.247 | 1889.537 | 2024.315 | 2002.172 | 2037.694 | 2541.078 | 3647.868 | 3554.278 | 3660.018 | 0.145200989 | -0.198777894 | YLR361C | DCR2 | n |
| W303.asm02.HP0_G0038290 | 3764.385 | 3970.475 | 3459.014 | 4297.514 | 3712.63 | 4140.172 | 5951.395 | 5231.553 | 5885.931 | 0.339490872 | -0.11848881 | YLR362W | STE11 | n |
| W303.asm02.HP0_G0038300 | 972.069 | 1024.89 | 889.2759 | 2712.172 | 2508.065 | 1419.054 | 5108.25 | 1703.047 | 4681.474 | 0.003772891 | -1.201849028 | YLR363C | NMD4 | n |
| W303.asm02.HP0_G0038310 | 441.5608 | 399.4957 | 443.0211 | 816.9862 | 804.49 | 211.1868 | 1825.786 | 791.3905 | 1578.831 | 0.332561466 | -0.513127765 | YLR363W-A | | n |

| | | | | | | | | | | | | | | |
|---|---|---|---|---|---|---|---|---|---|---|---|---|---|---|
| W303.asm02.HP0_G0038320 | 595.1011 | 357.2092 | 492.6049 | 795.45 | 963.3052 | 891.8463 | 569.0921 | 1195.139 | 986.7696 | 0.005267446 | -0.874984047 | YLR364W | GRX8 | n |
| W303.asm02.HP0_G0038340 | 1729.182 | 1936.274 | 1690.163 | 583.5616 | 466.8993 | 407.1802 | 596.2505 | 442.4056 | 686.5524 | 1.13E-18 | 1.876847322 | YLR368W | MDM30 | n |
| W303.asm02.HP0_G0038350 | 3271.997 | 3522.016 | 3027.85 | 4132.866 | 3958.23 | 5379.185 | 3483.683 | 4320.971 | 3890.862 | 0.007718575 | -0.455775342 | YLR369W | SSQ1 | n |
| W303.asm02.HP0_G0038360 | 6370.335 | 6479.842 | 6168.88 | 2091.096 | 2267.672 | 2835.068 | 2718.31 | 3646.625 | 3189.957 | 4.45E-17 | 1.402635398 | YLR370C | ARC18 | n |
| W303.asm02.HP0_G0038370 | 3217.993 | 3404.059 | 3203.549 | 8747.171 | 7900.838 | 11525.63 | 11678.12 | 12001.86 | 11560.15 | 5.80E-22 | -1.519780566 | YLR371W | ROM2 | n |
| W303.asm02.HP0_G0038380 | 5356.969 | 5312.514 | 6045.998 | 10716 | 10533 | 11949.52 | 7508.065 | 8141.551 | 7939.608 | 4.17E-22 | -0.989942798 | YLR372W | ELO3 | n |
| W303.asm02.HP0_G0038400 | 3804.623 | 3565.416 | 3744.66 | 8469.979 | 8890.179 | 6556.665 | 14144.59 | 14561.8 | 17522.64 | 3.35E-12 | -1.105513714 | YLR375W | STP3 | n |
| W303.asm02.HP0_G0038410 | 301.7861 | 199.1914 | 261.9322 | 944.8139 | 1001.49 | 1073.406 | 841.9106 | 984.6745 | 849.2199 | 8.02E-29 | -1.983864411 | YLR376C | PSY3 | n |
| W303.asm02.HP0_G0038420 | 422.5006 | 409.5109 | 465.6572 | 279.2759 | 157.9473 | 126.1043 | 5572.412 | 2370.95 | 1796.519 | 0.008717739 | 1.202967794 | YLR377C | FBP1 | n |
| W303.asm02.HP0_G0038430 | 7213.218 | 6817.021 | 7422.489 | 5201.34 | 5231.355 | 6969.163 | 4834.197 | 7104.26 | 6066.54 | 0.137303605 | 0.301911928 | YLR378C | SEC61 | n |
| W303.asm02.HP0_G0038440 | 3831.096 | 3626.62 | 3975.333 | 9249.451 | 9878.651 | 9486.692 | 4309.546 | 5204.708 | 4511.63 | 5.24E-36 | -1.323484393 | YLR380W | CSR1 | n |
| W303.asm02.HP0_G0038450 | 1313.034 | 1360.956 | 1381.881 | 1424.863 | 1378.134 | 1511.733 | 1249.287 | 1511.91 | 1373.105 | 0.456482294 | -0.089454037 | YLR381W | CTF3 | n |
| W303.asm02.HP0_G0038460 | 8568.608 | 8051.118 | 8256.792 | 2646.174 | 2423.884 | 2051.854 | 830.8004 | 1130.711 | 911.4163 | 3.55E-30 | 1.804393592 | YLR382C | NAM2 | n |
| W303.asm02.HP0_G0038470 | 1783.185 | 1971.884 | 1830.291 | 3245.019 | 2993.189 | 3555.99 | 2931.874 | 3257.909 | 2878.975 | 9.36E-13 | -0.810504613 | YLR383W | SMC6 | n |
| W303.asm02.HP0_G0038480 | 3812.035 | 3678.921 | 4215.707 | 10844.52 | 10823.73 | 6309.015 | 6816.76 | 4564.723 | 7047.329 | 1.88E-06 | -1.256912888 | YLR384C | IKI3 | n |
| W303.asm02.HP0_G0038490 | 70.94621 | 77.8961 | 95.934 | 394.5988 | 380.9829 | 315.2608 | 454.2861 | 328.5828 | 306.1976 | 1.02E-19 | -2.156047396 | YLR385C | SWC7 | n |
| W303.asm02.HP0_G0038510 | 5532.746 | 6025.819 | 5646.093 | 2070.949 | 2009.923 | 1932.587 | 3697.247 | 4198.558 | 3810.725 | 2.22E-62 | 1.516420537 | YLR387C | REH1 | n |
| W303.asm02.HP0_G0038520 | 2817.729 | 2508.254 | 2503.985 | 3040.078 | 3572.908 | 2926.228 | 3028.162 | 2605.039 | 2257.011 | 0.08128984 | -0.284681332 | YLR388W | RPS29A | n |
| W303.asm02.HP0_G0038530 | 6677.415 | 6506.55 | 6745.562 | 5336.115 | 4901.575 | 5471.105 | 12903.95 | 7438.212 | 12267.04 | 0.130578409 | 0.343313196 | YLR389C | STE23 | n |
| W303.asm02.HP0_G0038540 | 1412.571 | 1008.198 | 1130.728 | 888.5419 | 807.0935 | 764.2226 | 1544.326 | 1248.829 | 1578.831 | 0.003747511 | 0.529824757 | YLR390W | ECM19 | n |
| W303.asm02.HP0_G0038550 | 19685.99 | 19477.36 | 20333.7 | 14587.65 | 13471.52 | 15459.93 | 19852.8 | 16411.96 | 16802.59 | 9.58E-05 | 0.451140392 | YLR390W- | CCW14 | n |
| W303.asm02.HP0_G0038570 | 939.2432 | 783.4122 | 804.121 | 1473.493 | 1470.125 | 1120.505 | 235.7844 | 318.9186 | 278.6876 | 0.000215928 | -0.685476056 | YLR393W | ATP10 | n |
| W303.asm02.HP0_G0038580 | 1189.143 | 1107.237 | 1281.635 | 111.1546 | 102.4054 | 139.0186 | 80.24075 | 350.0588 | 117.2163 | 1.55E-12 | 3.342879041 | YLR394W | CST9 | n |
| W303.asm02.HP0_G0038590 | 2968.093 | 2902.186 | 2707.71 | 7609.921 | 8812.941 | 7634.629 | 4034.258 | 6309.648 | 6116.775 | 8.99E-15 | -1.487739782 | YLR395C | COX8 | n |
| W303.asm02.HP0_G0038600 | 938.1843 | 869.0979 | 939.9376 | 2229.344 | 2086.293 | 2677.818 | 639.4571 | 1409.899 | 1111.162 | 1.44E-06 | -1.34804403 | YLR396C | VPS33 | n |
| W303.asm02.HP0_G0038610 | 2576.301 | 2254.536 | 2648.425 | 4073.815 | 4204.697 | 1592.257 | 3704.654 | 1362.652 | 3588.253 | 0.409026703 | -0.400243823 | YLR397C | AFG2 | n |
| W303.asm02.HP0_G0038620 | 6275.034 | 6368.562 | 6074.024 | 6563.678 | 6092.254 | 6729.869 | 5490.936 | 7835.518 | 7735.077 | 0.780180244 | -0.050636496 | YLR398C | SKI2 | n |
| W303.asm02.HP0_G0038630 | 2544.534 | 2832.079 | 2352 | 7400.116 | 7373.19 | 7412.807 | 8564.774 | 7154.729 | 7137.035 | 8.20E-36 | -1.521443315 | YLR399C | BDF1 | n |
| W303.asm02.HP0_G0038640 | 3227.523 | 3291.666 | 3655.193 | 4197.475 | 4220.318 | 1710.765 | 3946.61 | 1322.922 | 2855.053 | 0.990172095 | 0.00652625 | YLR401C | DUS3 | n |
| W303.asm02.HP0_G0038650 | 1925.078 | 2282.356 | 2063.12 | 7429.989 | 6904.555 | 9002.026 | 18188.73 | 10675.72 | 14080.3 | 4.68E-18 | -1.895977348 | YLR403W | SFP1 | n |
| W303.asm02.HP0_G0038660 | 462.7387 | 444.0077 | 450.5664 | 540.4892 | 574.5117 | 948.0614 | 239.4878 | 857.9661 | 563.3557 | 0.183287457 | -0.604045905 | YLR404W | SEI1 | n |
| W303.asm02.HP0_G0038670 | 931.8309 | 999.2956 | 958.2621 | 1480.44 | 1444.09 | 854.6227 | 920.9169 | 446.7008 | 620.7678 | 0.243518176 | -0.387286666 | YLR405W | DUS4 | n |
| W303.asm02.HP0_G0038700 | 591.9244 | 597.5743 | 667.2264 | 3545.136 | 3860.164 | 4510.129 | 9051.157 | 6708.028 | 6395.463 | 7.69E-50 | -2.681964761 | YLR407W | GAG1 | n |
| W303.asm02.HP0_G0038710 | 258.3713 | 336.066 | 267.3217 | 427.9451 | 432.1856 | 421.6139 | 307.3838 | 443.4794 | 455.7081 | 0.006075515 | -0.573155534 | YLR408C | BLS1 | n |
| W303.asm02.HP0_G0038720 | 2826.201 | 3188.176 | 3493.507 | 6688.727 | 6371.7 | 2776.574 | 8548.726 | 4345.668 | 7572.41 | 0.054424298 | -0.736100921 | YLR409C | UTP21 | n |
| W303.asm02.HP0_G0038730 | 9088.528 | 8861.237 | 8983.303 | 6640.097 | 6657.22 | 8731.585 | 5879.795 | 6151.8 | 6829.641 | 0.055406217 | 0.289965904 | YLR410W | VIP1 | n |
| W303.asm02.HP0_G0038750 | 2002.377 | 2197.783 | 2110.548 | 407.1037 | 367.0974 | 132.9413 | 595.016 | 269.5238 | 608.8069 | 1.04E-10 | 2.798403654 | YLR412W | BER1 | n |
| W303.asm02.HP0_G0038780 | 4647.506 | 5313.627 | 4291.16 | 268.8551 | 317.6304 | 219.5431 | 449.3482 | 1068.431 | 639.9051 | 8.59E-41 | 4.144723746 | YLR414C | PUN1 | n |
| W303.asm02.HP0_G0038800 | 1552.346 | 1633.592 | 1571.593 | 1828.493 | 1805.98 | 2060.97 | 1661.601 | 1680.497 | 1751.067 | 0.007390767 | -0.259786608 | YLR417W | VPS36 | n |
| W303.asm02.HP0_G0038810 | 3118.457 | 3461.925 | 3206.783 | 3618.776 | 3704.82 | 2961.932 | 2813.364 | 3066.773 | 2660.092 | 0.625111207 | -0.071628788 | YLR418C | CDC73 | n |
| W303.asm02.HP0_G0038820 | 2476.764 | 2590.602 | 2643.036 | 3534.716 | 3313.423 | 3073.603 | 2376.361 | 2123.976 | 2745.014 | 0.006375828 | -0.363857636 | YLR419W | | n |
| W303.asm02.HP0_G0038830 | 3695.556 | 3271.636 | 3357.69 | 5073.512 | 4923.271 | 5817.512 | 3036.804 | 4486.336 | 3869.333 | 0.000431675 | -0.615103367 | YLR420W | URA4 | n |
| W303.asm02.HP0_G0038840 | 3993.107 | 4184.133 | 3840.594 | 3727.847 | 4130.063 | 5003.151 | 3175.065 | 5090.885 | 4387.237 | 0.674216182 | -0.097821522 | YLR421C | RPN13 | n |
| W303.asm02.HP0_G0038850 | 2610.185 | 2650.693 | 2549.257 | 10584.7 | 9972.378 | 11176.19 | 10160.95 | 13571.76 | 14473.82 | 1.05E-39 | -2.02259987 | YLR422W | DCK1 | n |
| W303.asm02.HP0_G0038860 | 1219.851 | 1271.932 | 1121.027 | 407.7984 | 425.2428 | 531.0056 | 582.6713 | 589.5162 | 355.237 | 7.37E-10 | 1.405201931 | YLR423C | ATG17 | n |
| W303.asm02.HP0_G0038870 | 891.5927 | 919.1739 | 854.7827 | 1594.374 | 1576.87 | 1632.519 | 1543.091 | 1462.516 | 1309.712 | 3.42E-17 | -0.849876146 | YLR424W | SPP382 | n |
| W303.asm02.HP0_G0038880 | 1835.071 | 2038.652 | 1727.89 | 2258.522 | 2331.893 | 2959.653 | 2271.43 | 2418.197 | 2498.62 | 0.004442677 | -0.430724002 | YLR425W | TUS1 | n |
| W303.asm02.HP0_G0038890 | 585.571 | 545.2727 | 654.2915 | 499.5009 | 518.9698 | 625.204 | 1751.717 | 1306.815 | 1605.145 | 0.536986899 | 0.119214838 | YLR426W | TDA5 | n |
| W303.asm02.HP0_G0038900 | 2909.854 | 2847.659 | 3033.239 | 3651.428 | 3524.308 | 4235.89 | 5356.379 | 4825.657 | 4697.023 | 0.001319739 | -0.376506106 | YLR427W | MAG2 | n |
| W303.asm02.HP0_G0038920 | 5131.423 | 5253.535 | 5434.823 | 3856.369 | 3664.031 | 3350.121 | 2583.752 | 3228.916 | 2985.426 | 8.08E-06 | 0.541264921 | YLR430W | SEN1 | n |
| W303.asm02.HP0_G0038930 | 2064.852 | 2075.375 | 1983.355 | 1565.195 | 1628.94 | 2308.621 | 860.4277 | 1580.633 | 932.9458 | 0.611326011 | 0.154206365 | YLR431C | ATG23 | n |
| W303.asm02.HP0_G0038950 | 2945.856 | 2743.055 | 2810.112 | 4759.5 | 4813.055 | 5206.741 | 4876.169 | 4661.365 | 4774.769 | 2.51E-24 | -0.798142651 | YLR433C | CNA1 | n |
| W303.asm02.HP0_G0038960 | 1239.97 | 1238.548 | 1343.076 | 1800.01 | 2077.615 | 769.5403 | 1267.804 | 599.1804 | 1151.829 | 0.517713161 | -0.282095886 | YLR435W | TSR2 | n |
| W303.asm02.HP0_G0038970 | 2383.581 | 2108.759 | 2304.572 | 2031.35 | 1816.394 | 2449.918 | 2807.192 | 2471.887 | 2424.463 | 0.507204961 | 0.109923205 | YLR436C | ECM30 | n |

| | | | | | | | | | | | | | | |
|---|---|---|---|---|---|---|---|---|---|---|---|---|---|---|
| W303.asm02.HP0_G0038980 | 813.2342 | 645.4248 | 689.8625 | 4765.753 | 4449.429 | 4731.191 | 10322.66 | 7422.105 | 9462.223 | 3.11E-61 | -2.698301736 | YLR437C | DIF1 | n |
| W303.asm02.HP0_G0038990 | 355.79 | 346.0812 | 356.7883 | 8822.895 | 9406.545 | 17013.45 | 10194.28 | 8069.606 | 5938.559 | 1.51E-64 | -5.057006652 | YLR438W | CAR2 | n |
| W303.asm02.HP0_G0039000 | 2371.933 | 2065.359 | 2063.12 | 1071.947 | 1248.825 | 818.1588 | 897.4619 | 743.0696 | 741.5723 | 3.50E-08 | 1.050647959 | YLR438C-/ | LSM3 | n |
| W303.asm02.HP0_G0039010 | 10619.7 | 9699.177 | 9969.591 | 5745.997 | 5647.051 | 4039.896 | 2507.215 | 3037.78 | 2750.994 | 1.15E-08 | 0.972774863 | YLR439W | MRPL4 | n |
| W303.asm02.HP0_G0039020 | 3891.453 | 4088.432 | 3888.022 | 2387.739 | 2240.769 | 2096.674 | 1961.578 | 2117.533 | 2249.835 | 9.22E-17 | 0.819256147 | YLR440C | SEC39 | n |
| W303.asm02.HP0_G0039050 | 1615.879 | 1586.855 | 1567.281 | 266.0763 | 203.943 | 248.4103 | 287.6322 | 211.5386 | 200.9422 | 4.02E-54 | 2.729070649 | YLR443W | ECM7 | n |
| W303.asm02.HP0_G0039060 | 19.06018 | 22.25603 | 23.71402 | 47.93541 | 45.99565 | 39.50256 | 103.6957 | 68.7232 | 63.39247 | 0.00446933 | -1.037889829 | YLR445W | GMC2 | n |
| W303.asm02.HP0_G0039080 | 12071.45 | 11797.92 | 12605.08 | 4143.982 | 4386.076 | 5306.257 | 4640.384 | 3796.957 | 4107.354 | 3.62E-24 | 1.398439391 | YLR447C | VMA6 | n |
| W303.asm02.HP0_G0039120 | 3144.929 | 3062.429 | 3296.249 | 2319.657 | 1697.5 | 1116.707 | 6179.772 | 2717.788 | 5914.637 | 0.015595589 | 0.888354132 | YLR451W | LEU3 | n |
| W303.asm02.HP0_G0039130 | 1247.383 | 1242.999 | 1309.661 | 432.1134 | 411.3574 | 454.2794 | 528.3545 | 1536.608 | 546.6105 | 9.59E-05 | 1.549949248 | YLR452C | SST2 | n |
| W303.asm02.HP0_G0039150 | 7312.754 | 7589.305 | 6838.262 | 5566.76 | 5627.091 | 6596.928 | 4468.793 | 8193.093 | 7238.702 | 0.241576904 | 0.28923769 | YLR454W | FMP27 | n |
| W303.asm02.HP0_G0039160 | 1063.134 | 1214.066 | 880.6526 | 3930.009 | 3513.894 | 3919.11 | 3707.123 | 3607.968 | 3050.015 | 7.45E-32 | -1.847539816 | YLR455W | PDP3 | n |
| W303.asm02.HP0_G0039180 | 1053.604 | 1080.53 | 1054.196 | 1121.967 | 1205.433 | 1356.761 | 949.3098 | 1009.372 | 743.9644 | 0.205146392 | -0.208465404 | YLR457C | NBP1 | n |
| W303.asm02.HP0_G0039190 | 945.5965 | 899.1435 | 963.6517 | 2368.982 | 2646.92 | 2813.038 | 2630.662 | 3377.101 | 2741.425 | 7.29E-26 | -1.478911335 | YLR459W | GAB1 | n |
| W303.asm02.HP0_G0039260 | 4834.931 | 4986.463 | 5012.282 | 5608.443 | 5456.994 | 8186.146 | 11473.19 | 11917.03 | 12310.1 | 0.041901227 | -0.376131895 | YML130C | ERO1 | n |
| W303.asm02.HP0_G0039270 | 4262.067 | 3993.844 | 3717.712 | 1836.829 | 1945.703 | 1368.916 | 1749.248 | 1364.8 | 1618.302 | 7.17E-12 | 1.216944097 | YML129C | COX14 | n |
| W303.asm02.HP0_G0039280 | 18019.28 | 17694.65 | 16207.46 | 114.6282 | 95.46268 | 385.9096 | 1306.073 | 1634.323 | 1879.048 | 1.24E-44 | 6.444411642 | YML128C | MSC1 | n |
| W303.asm02.HP0_G0039290 | 2574.183 | 2415.892 | 2676.451 | 4670.577 | 4407.772 | 4616.482 | 3789.832 | 3541.392 | 3631.312 | 2.57E-25 | -0.837029276 | YML127W | RSC9 | n |
| W303.asm02.HP0_G0039300 | 9931.411 | 8939.133 | 11158.53 | 1126.135 | 1109.971 | 1150.892 | 1057.943 | 919.1727 | 1022.652 | 5.74E-153 | 3.14823137 | YML126C | ERG13 | n |
| W303.asm02.HP0_G0039310 | 3217.993 | 2958.939 | 3346.911 | 2712.172 | 2725.893 | 3011.311 | 1615.925 | 3023.821 | 2304.854 | 0.489925416 | 0.172705544 | YML125C | PGA3 | n |
| W303.asm02.HP0_G0039330 | 1758.831 | 1924.034 | 1990.9 | 27247.46 | 28081.65 | 20921.16 | 3186.175 | 7976.186 | 4473.355 | 1.02E-32 | -3.748368469 | YML123C | PHO84 | n |
| W303.asm02.HP0_G0039350 | 2700.192 | 2447.05 | 2588.062 | 1367.896 | 1378.134 | 1347.645 | 1133.246 | 1245.608 | 1235.555 | 2.19E-26 | 0.918219488 | YML121W | GTR1 | n |
| W303.asm02.HP0_G0039360 | 10640.87 | 12021.59 | 10352.25 | 19078.99 | 20054.1 | 14811.94 | 2560.297 | 6164.685 | 4064.295 | 0.023979076 | -0.708379261 | YML120C | NDI1 | n |
| W303.asm02.HP0_G0039370 | 4372.193 | 4051.71 | 4356.913 | 853.8061 | 745.4767 | 829.5538 | 622.1744 | 1151.114 | 626.7482 | 8.83E-22 | 2.395483579 | YML119W | | n |
| W303.asm02.HP0_G0039390 | 15852.77 | 16817.77 | 15522.98 | 6748.473 | 5995.056 | 8354.791 | 7527.817 | 7931.086 | 6938.485 | 1.15E-15 | 1.191647894 | YML117W | NAB6 | n |
| W303.asm02.HP0_G0039410 | 1657.176 | 1666.976 | 1545.723 | 5534.109 | 5797.188 | 6155.562 | 3603.427 | 4128.761 | 3552.37 | 1.40E-70 | -1.844304027 | YML115C | VAN1 | n |
| W303.asm02.HP0_G0039420 | 2210.981 | 2299.048 | 2454.401 | 2015.372 | 1966.531 | 2020.708 | 2807.192 | 2394.574 | 2567.993 | 0.038496654 | 0.214314615 | YML114C | TAF8 | n |
| W303.asm02.HP0_G0039440 | 1563.993 | 1713.714 | 1661.06 | 1981.33 | 1970.87 | 1390.186 | 2557.828 | 2609.334 | 2268.972 | 0.550159307 | -0.113310883 | YML112W | CTK3 | n |
| W303.asm02.HP0_G0039450 | 2243.806 | 2286.807 | 2294.871 | 2698.972 | 2545.382 | 2751.505 | 3981.176 | 3418.979 | 3521.272 | 0.021865165 | -0.228501005 | YML111W | BUL2 | n |
| W303.asm02.HP0_G0039460 | 13257.41 | 12763.83 | 12877.79 | 4105.078 | 4524.931 | 3682.854 | 1960.343 | 3078.584 | 2779.7 | 3.97E-19 | 1.659624123 | YML110C | COQ5 | n |
| W303.asm02.HP0_G0039480 | 401.3226 | 357.2092 | 361.0999 | 1463.072 | 1387.68 | 755.1066 | 1902.323 | 1363.726 | 1782.166 | 8.17E-10 | -1.687226054 | YML108W | | n |
| W303.asm02.HP0_G0039490 | 996.4237 | 921.3995 | 951.7947 | 842.6907 | 951.1554 | 696.6125 | 1510.995 | 1488.287 | 1385.066 | 0.183963169 | 0.205121358 | YML107C | PML39 | n |
| W303.asm02.HP0_G0039500 | 6914.609 | 7050.71 | 7655.318 | 7223.658 | 8081.35 | 7744.021 | 10353.53 | 8226.381 | 8665.631 | 0.505213526 | -0.092241195 | YML106W | URA5 | n |
| W303.asm02.HP0_G0039510 | 5269.08 | 4722.729 | 5042.464 | 3559.725 | 3884.463 | 3246.047 | 3172.596 | 2945.433 | 2918.446 | 6.67E-06 | 0.49216267 | YML105C | SEC65 | n |
| W303.asm02.HP0_G0039520 | 1334.212 | 1337.587 | 1320.44 | 1208.111 | 945.9483 | 970.8514 | 1183.86 | 1200.508 | 1202.065 | 0.006834054 | 0.352519413 | YML104C | MDM1 | n |
| W303.asm02.HP0_G0039530 | 4543.734 | 4562.486 | 4439.912 | 9159.138 | 9101.064 | 7459.907 | 5418.102 | 6114.217 | 6108.403 | 1.02E-14 | -0.925005618 | YML103C | NUP188 | n |
| W303.asm02.HP0_G0039540 | 1567.17 | 1463.334 | 1780.708 | 4158.571 | 4882.482 | 4090.034 | 5776.1 | 3897.894 | 4499.669 | 2.99E-15 | -1.44840524 | YML102W | CAC2 | n |
| W303.asm02.HP0_G0039570 | 2278.75 | 2445.937 | 2273.312 | 1363.728 | 1374.663 | 525.6879 | 1709.745 | 631.3944 | 1178.143 | 0.012659705 | 1.100306749 | YML099C | ARG81 | n |
| W303.asm02.HP0_G0039580 | 1275.973 | 1229.646 | 1304.271 | 863.5322 | 934.6664 | 913.8765 | 1141.888 | 1045.881 | 1182.927 | 4.17E-07 | 0.490975129 | YML098W | TAF13 | n |
| W303.asm02.HP0_G0039590 | 2287.221 | 2314.627 | 2232.352 | 2825.411 | 2890.783 | 3149.57 | 2261.555 | 2846.644 | 2483.071 | 0.001994071 | -0.375474086 | YML097C | VPS9 | n |
| W303.asm02.HP0_G0039600 | 1189.143 | 1290.85 | 1122.104 | 652.3385 | 744.6089 | 734.5957 | 569.0921 | 755.9552 | 636.3169 | 3.71E-07 | 0.757336792 | YML095C | RAD10 | n |
| W303.asm02.HP0_G0039620 | 3279.409 | 3850.293 | 4125.162 | 6288.57 | 5233.958 | 1204.828 | 7622.871 | 2979.795 | 6589.229 | 0.769399269 | -0.177388207 | YML093W | UTP14 | n |
| W303.asm02.HP0_G0039630 | 9397.726 | 9257.395 | 9029.653 | 6161.437 | 6752.683 | 8480.136 | 4424.352 | 8469.06 | 6025.873 | 0.158938841 | 0.371874842 | YML092C | PRE8 | n |
| W303.asm02.HP0_G0039640 | 2701.251 | 3425.203 | 2755.138 | 6285.097 | 5551.589 | 3994.317 | 6066.201 | 4485.262 | 3815.509 | 0.000830169 | -0.833905582 | YML091C | RPM2 | n |
| W303.asm02.HP0_G0039660 | 2209.922 | 2152.158 | 2425.298 | 2952.544 | 2677.294 | 3533.96 | 4355.221 | 3830.244 | 3466.252 | 0.006587156 | -0.433183045 | YML088W | UFO1 | n |
| W303.asm02.HP0_G0039680 | 7917.386 | 6882.676 | 7387.996 | 9555.82 | 9932.458 | 10461.34 | 7326.598 | 8587.178 | 7770.96 | 7.25E-05 | -0.432709635 | YML086C | ALO1 | n |
| W303.asm02.HP0_G0039690 | 12529.95 | 12583.56 | 13404.89 | 9168.864 | 9428.241 | 10083.03 | 3950.314 | 6683.331 | 5901.48 | 0.040222787 | 0.425498685 | YML085C | TUB1 | n |
| W303.asm02.HP0_G0039720 | 1359.626 | 1185.133 | 1154.442 | 6264.255 | 7382.736 | 6729.109 | 2619.552 | 4361.775 | 3942.294 | 1.16E-32 | -2.461514563 | YML081C- | ATP18 | n |
| W303.asm02.HP0_G0039730 | 1617.997 | 1744.873 | 1731.124 | 5147.152 | 4466.785 | 4725.114 | 5041.588 | 4294.126 | 4633.631 | 6.78E-43 | -1.493293136 | YML081W | TDA9 | n |
| W303.asm02.HP0_G0039740 | 752.877 | 919.1739 | 1003.534 | 3893.884 | 3646.674 | 2228.096 | 3777.488 | 2541.684 | 3565.527 | 1.49E-12 | -1.868349126 | YML080W | DUS1 | n |
| W303.asm02.HP0_G0039750 | 3934.868 | 3877 | 3546.324 | 1010.117 | 998.8867 | 1050.616 | 838.2072 | 1293.929 | 1017.868 | 1.42E-28 | 1.892271611 | YML079W | CFF1 | n |
| W303.asm02.HP0_G0039760 | 6532.346 | 5685.302 | 6055.699 | 3430.508 | 4109.234 | 3932.024 | 2061.57 | 3041.001 | 2399.345 | 0.000137113 | 0.671758008 | YML078W | CPR3 | n |
| W303.asm02.HP0_G0039780 | 1953.668 | 1966.32 | 1965.03 | 2696.888 | 2387.435 | 2537.28 | 4678.653 | 3114.02 | 3423.193 | 0.038952684 | -0.373146709 | YML076C | WAR1 | n |
| W303.asm02.HP0_G0039790 | 5277.551 | 5462.742 | 6358.592 | 10789.64 | 10413.24 | 12172.87 | 26318.97 | 28354.76 | 31384.06 | 2.25E-13 | -0.964915473 | YML075C | HMG1 | n |

| | | | | | | | | | | | | | | |
|---|---|---|---|---|---|---|---|---|---|---|---|---|---|---|
| W303.asm02.HP0_G0039800 | 14137.36 | 12544.61 | 13768.15 | 14766.89 | 14493.84 | 7263.913 | 14739.61 | 10845.38 | 16354.06 | 0.651942442 | 0.14728107 | YML074C | FPR3 | n |
| W303.asm02.HP0_G0039810 | 7424.998 | 7255.465 | 8048.755 | 18394.69 | 21526.83 | 20108.32 | 17106.09 | 15086.89 | 13881.75 | 1.90E-30 | -1.401094032 | YML073C | RPL6A | n |
| W303.asm02.HP0_G0039820 | 19346.08 | 18494.76 | 18961.52 | 12228.39 | 12723.44 | 16882.79 | 14748.25 | 22294.23 | 17861.13 | 0.032841651 | 0.441252954 | YML072C | TCB3 | n |
| W303.asm02.HP0_G0039830 | 2926.796 | 2917.765 | 2997.668 | 2796.232 | 2567.946 | 3425.328 | 3304.684 | 3568.237 | 3517.684 | 0.957562864 | 0.008489876 | YML071C | COG8 | n |
| W303.asm02.HP0_G0039840 | 7739.491 | 7209.84 | 7138.999 | 4633.062 | 5233.958 | 12493.44 | 12475.59 | 32405.13 | 20201.86 | 0.973187472 | -0.017667787 | YML070W | DAK1 | n |
| W303.asm02.HP0_G0039850 | 8153.52 | 8082.276 | 7949.587 | 8114.285 | 8328.685 | 9492.769 | 3639.227 | 5279.874 | 3748.528 | 0.602155486 | -0.100805342 | YML069W | POB3 | n |
| W303.asm02.HP0_G0039860 | 1939.902 | 1978.561 | 1861.551 | 1555.469 | 1596.83 | 1817.118 | 2857.805 | 2744.633 | 2729.464 | 0.032958216 | 0.217971631 | YML068W | ITT1 | n |
| W303.asm02.HP0_G0039880 | 311.3162 | 292.6668 | 299.659 | 115.3229 | 83.31288 | 116.2287 | 188.8744 | 179.3246 | 116.0202 | 4.86E-10 | 1.518025473 | YML066C | SMA2 | n |
| W303.asm02.HP0_G0039890 | 2231.1 | 2373.605 | 2413.441 | 3688.248 | 3354.211 | 3439.002 | 4145.361 | 3763.669 | 3797.568 | 1.86E-10 | -0.578925753 | YML065W | ORC1 | n |
| W303.asm02.HP0_G0039900 | 2202.509 | 2189.993 | 2148.275 | 1098.346 | 1239.279 | 904.0009 | 878.9448 | 1038.365 | 784.6313 | 1.25E-09 | 1.013089618 | YML064C | TEM1 | n |
| W303.asm02.HP0_G0039910 | 16765.54 | 16536.23 | 17978.46 | 32966.36 | 37203.54 | 35907.07 | 28323.75 | 24994.84 | 23688.45 | 4.17E-22 | -1.048617201 | YML063W | RPS1B | y |
| W303.asm02.HP0_G0039920 | 2733.018 | 2601.73 | 2568.66 | 3802.876 | 3530.383 | 3627.399 | 2402.285 | 2996.976 | 3265.31 | 0.000761952 | -0.471836436 | YML062C | MFT1 | n |
| W303.asm02.HP0_G0039930 | 2613.362 | 2339.108 | 2553.569 | 2473.884 | 2424.752 | 2594.255 | 1301.135 | 1595.667 | 1261.869 | 0.986491869 | 0.002539326 | YML061C | PIF1 | n |
| W303.asm02.HP0_G0039940 | 333.5531 | 292.6668 | 357.8662 | 867.7005 | 862.6355 | 504.4173 | 403.6727 | 177.177 | 233.2364 | 0.000741288 | -1.18313382 | YML060W | OGG1 | n |
| W303.asm02.HP0_G0039950 | 1652.941 | 1360.956 | 1596.385 | 976.0762 | 905.1597 | 1343.087 | 983.8751 | 1322.922 | 1072.888 | 0.016001389 | 0.515747893 | YML059C | NTE1 | n |
| W303.asm02.HP0_G0039990 | 4630.564 | 3812.458 | 5147.021 | 4983.894 | 5102.046 | 4334.646 | 8057.406 | 8256.448 | 6938.485 | 0.626652844 | -0.085516465 | YML056C | IMD4 | n |
| W303.asm02.HP0_G0040000 | 1359.626 | 1111.689 | 1150.13 | 1399.158 | 1451.033 | 1517.81 | 1078.929 | 1300.372 | 1309.712 | 0.060254675 | -0.270192897 | YML055W | SPC2 | n |
| W303.asm02.HP0_G0040010 | 6680.592 | 8457.29 | 6406.02 | 3558.336 | 3044.392 | 4467.588 | 20246.59 | 10780.95 | 14831.45 | 0.000395288 | 0.960528441 | YML054C | CYB2 | n |
| W303.asm02.HP0_G0040040 | 2540.298 | 2188.88 | 2259.3 | 8079.549 | 8663.672 | 9280.823 | 8442.561 | 10930.21 | 10244.46 | 1.48E-42 | -1.896718967 | YML052W | SUR7 | n |
| W303.asm02.HP0_G0040050 | 2518.061 | 2262.325 | 2471.648 | 1394.99 | 1323.46 | 1672.022 | 10738.68 | 3268.647 | 6700.464 | 0.060776799 | 0.723990836 | YML051W | GAL80 | n |
| W303.asm02.HP0_G0040060 | 1847.778 | 1837.235 | 1627.644 | 902.4363 | 912.9703 | 1188.875 | 645.6294 | 1106.014 | 808.553 | 0.000382049 | 0.822368093 | YML050W | AIM32 | n |
| W303.asm02.HP0_G0040070 | 3111.044 | 3249.38 | 3158.277 | 2621.164 | 2406.527 | 2964.971 | 4986.037 | 4604.454 | 4217.393 | 0.047201697 | 0.25190203 | YML049C | RSE1 | n |
| W303.asm02.HP0_G0040080 | 13539.08 | 13896.66 | 13656.04 | 8489.431 | 8840.712 | 12652.97 | 22373.59 | 26188.91 | 22100.05 | 0.015603076 | 0.454692953 | YML048W | GSF2 | n |
| W303.asm02.HP0_G0040100 | 1118.197 | 1037.131 | 1140.429 | 1345.665 | 1198.491 | 957.9371 | 1592.47 | 1143.597 | 1425.733 | 0.679173943 | -0.087688629 | YML046W | PRP39 | n |
| W303.asm02.HP0_G0040110 | 178.9539 | 178.0482 | 198.3355 | 835.0488 | 573.6439 | 297.0289 | 1311.01 | 659.3132 | 1230.771 | 5.73E-05 | -1.619164871 | YML043C | RRN11 | n |
| W303.asm02.HP0_G0040120 | 2196.156 | 2285.694 | 2087.912 | 279.2759 | 247.3351 | 201.3111 | 962.889 | 565.8926 | 850.416 | 1.05E-44 | 3.173558624 | YML042W | CAT2 | n |
| W303.asm02.HP0_G0040130 | 1293.974 | 1366.52 | 1141.507 | 1165.039 | 1081.332 | 1096.956 | 1710.98 | 1270.305 | 1552.517 | 0.253884522 | 0.185244989 | YML041C | VPS71 | n |
| W303.asm02.HP0_G0040140 | 1239.97 | 1439.965 | 1244.986 | 1492.945 | 1352.966 | 1890.046 | 1435.692 | 1524.796 | 1437.693 | 0.108107915 | -0.271296104 | YML038C | YMD8 | n |
| W303.asm02.HP0_G0040150 | 376.9679 | 410.6237 | 370.8011 | 1442.231 | 1388.548 | 1472.99 | 1217.19 | 1285.339 | 1178.143 | 1.37E-81 | -1.894105439 | YML037C | LFT1 | n |
| W303.asm02.HP0_G0040160 | 1125.609 | 1020.439 | 1062.819 | 767.6613 | 861.7676 | 498.34 | 609.8297 | 861.1875 | 730.8075 | 0.015413604 | 0.593069827 | YML036W | CGI121 | n |
| W303.asm02.HP0_G0040170 | 6350.216 | 5953.487 | 6117.14 | 7767.621 | 7371.454 | 10269.91 | 9705.427 | 8289.735 | 8851.024 | 0.00413137 | -0.464020382 | YML035C | AMD1 | n |
| W303.asm02.HP0_G0040180 | 5866.299 | 5779.89 | 6000.726 | 5307.631 | 4755.777 | 4823.111 | 3366.408 | 4614.118 | 4360.923 | 0.106934986 | 0.245350982 | YML034W | SRC1 | n |
| W303.asm02.HP0_G0040190 | 2893.97 | 3024.594 | 2876.942 | 2635.753 | 2628.695 | 3738.31 | 1354.217 | 2002.637 | 1413.772 | 0.893767654 | -0.033634987 | YML032C | RAD52 | n |
| W303.asm02.HP0_G0040200 | 1787.421 | 1871.732 | 1890.654 | 4523.991 | 4426.865 | 5425.525 | 5968.677 | 5745.903 | 5528.302 | 9.84E-34 | -1.373273121 | YML031W | NDC1 | n |
| W303.asm02.HP0_G0040210 | 5220.371 | 5048.78 | 4644.715 | 2102.906 | 2329.289 | 1689.494 | 2023.301 | 2314.039 | 2686.405 | 5.46E-14 | 1.284810039 | YML030W | RCF1 | n |
| W303.asm02.HP0_G0040220 | 2575.242 | 2636.226 | 2655.971 | 6214.931 | 6810.828 | 10272.18 | 12064.51 | 12335.81 | 10913.07 | 1.13E-14 | -1.566240477 | YML029W | USA1 | n |
| W303.asm02.HP0_G0040230 | 19199.95 | 18902.04 | 19091.94 | 12243.68 | 14316.8 | 18424.15 | 5103.312 | 10682.16 | 8006.588 | 0.245570608 | 0.346435447 | YML028W | TSA1 | n |
| W303.asm02.HP0_G0040240 | 403.4404 | 381.6909 | 449.4885 | 967.7396 | 1405.037 | 1294.469 | 193.8123 | 992.1911 | 198.55 | 0.00699086 | -1.57056245 | YML027W | YOX1 | n |
| W303.asm02.HP0_G0040260 | 6620.235 | 7110.801 | 6529.98 | 3756.33 | 3891.406 | 2538.799 | 1885.04 | 2000.489 | 1607.537 | 2.93E-07 | 0.992086689 | YML025C | YML6 | n |
| W303.asm02.HP0_G0040280 | 1479.281 | 1416.596 | 1633.034 | 1268.552 | 1158.57 | 1096.196 | 1855.413 | 1549.493 | 1615.91 | 0.005703221 | 0.362032322 | YML023C | NSE5 | n |
| W303.asm02.HP0_G0040290 | 3840.626 | 3234.914 | 3786.698 | 2937.955 | 3397.603 | 1790.53 | 1413.472 | 1579.56 | 1563.282 | 0.096879144 | 0.418805589 | YML022W | APT1 | n |
| W303.asm02.HP0_G0040300 | 1267.502 | 1129.493 | 1188.935 | 1039.295 | 1081.332 | 1031.625 | 601.1884 | 801.0547 | 749.9449 | 0.205146392 | 0.186198419 | YML021C | UNG1 | n |
| W303.asm02.HP0_G0040310 | 2151.682 | 2346.898 | 2048.029 | 1537.407 | 1531.742 | 2525.885 | 1798.627 | 1693.382 | 1716.381 | 0.339770881 | 0.226519082 | YML020W | | n |
| W303.asm02.HP0_G0040320 | 2366.639 | 2284.581 | 2230.196 | 2276.585 | 2298.915 | 2329.891 | 1387.548 | 2763.961 | 2204.383 | 0.985973269 | -0.005010229 | YML019W | OST6 | n |
| W303.asm02.HP0_G0040330 | 516.7426 | 459.587 | 518.4748 | 3080.371 | 2586.171 | 3370.632 | 9011.654 | 7983.702 | 9605.753 | 9.49E-69 | -2.595898923 | YML018C | | n |
| W303.asm02.HP0_G0040340 | 1074.782 | 1150.637 | 1085.456 | 9175.116 | 8622.015 | 9574.053 | 8352.445 | 7778.607 | 8773.279 | 3.56E-233 | -3.047560657 | YML017W | PSP2 | n |
| W303.asm02.HP0_G0040350 | 5583.573 | 5811.049 | 5785.144 | 6132.259 | 5675.69 | 6521.721 | 7390.79 | 7403.851 | 6571.287 | 0.383656707 | -0.093578093 | YML016C | PPZ1 | n |
| W303.asm02.HP0_G0040360 | 2339.107 | 2201.121 | 2275.468 | 1197.691 | 1166.38 | 911.5975 | 1666.539 | 1887.74 | 1755.852 | 2.27E-14 | 1.057131815 | YML015C | TAF11 | n |
| W303.asm02.HP0_G0040370 | 1634.94 | 1594.644 | 1749.448 | 2152.925 | 2133.157 | 1148.613 | 2455.367 | 1561.305 | 2295.286 | 0.692000143 | -0.12629341 | YML014W | TRM9 | n |
| W303.asm02.HP0_G0040380 | 3930.632 | 3651.101 | 3879.399 | 2048.023 | 1870.201 | 2442.322 | 3134.327 | 3091.47 | 3012.936 | 1.28E-11 | 0.849331736 | YML013W | UBX2 | n |
| W303.asm02.HP0_G0040390 | 5637.577 | 5213.474 | 5693.521 | 5826.584 | 6363.889 | 6403.213 | 4076.23 | 5901.604 | 4923.083 | 0.32059328 | -0.168410748 | YML012W | ERV25 | n |
| W303.asm02.HP0_G0040400 | 759.2304 | 585.3335 | 602.5518 | 986.4969 | 1062.239 | 974.6497 | 982.6406 | 1146.818 | 862.3768 | 0.000199305 | -0.634392796 | YML011C | RAD33 | n |
| W303.asm02.HP0_G0040410 | 9902.821 | 9740.35 | 9902.76 | 11917.16 | 11594.38 | 11801.39 | 15260.56 | 11280.27 | 12415.36 | 0.062682932 | -0.257243168 | YML010W | SPT5 | n |
| W303.asm02.HP0_G0040420 | 2634.54 | 2460.404 | 2262.533 | 1380.401 | 1490.086 | 929.0698 | 648.0984 | 521.8668 | 575.3166 | 4.30E-06 | 0.953562492 | YML009C | MRPL39 | n |

| | | | | | | | | | | | | | | |
|---|---|---|---|---|---|---|---|---|---|---|---|---|---|---|
| W303.asm02.HP0_G0040430 | 10780.65 | 11733.38 | 12652.51 | 9677.396 | 10865.39 | 9780.682 | 7340.177 | 4986.727 | 5003.221 | 0.284188383 | 0.21378828 | YML008C | ERG6 | n |
| W303.asm02.HP0_G0040440 | 11233.86 | 12095.04 | 11724.43 | 14381.32 | 13409.03 | 20106.04 | 26826.33 | 27161.77 | 24970.65 | 0.011624015 | -0.450385744 | YML007W | YAP1 | n |
| W303.asm02.HP0_G0040450 | 973.1279 | 1356.505 | 1018.625 | 508.5322 | 540.6659 | 168.6455 | 932.0272 | 635.6896 | 1237.947 | 0.000888873 | 1.459229301 | YML006C | GIS4 | n |
| W303.asm02.HP0_G0040470 | 11904.14 | 11776.78 | 11249.07 | 9801.75 | 11011.19 | 12582.33 | 18256.62 | 21165.67 | 16394.73 | 0.696480847 | 0.064842325 | YML004C | GLO1 | n |
| W303.asm02.HP0_G0040480 | 2466.175 | 2560.556 | 2298.104 | 1850.029 | 1805.112 | 2223.538 | 2209.707 | 2033.777 | 2221.129 | 0.012590837 | 0.317160173 | YML002W | VRL1 | n |
| W303.asm02.HP0_G0040490 | 2809.258 | 2539.413 | 2605.309 | 2096.653 | 2471.615 | 3036.379 | 1692.463 | 1446.409 | 1342.007 | 0.741749615 | 0.064896395 | YML001W | YPT7 | n |
| W303.asm02.HP0_G0040500 | 1505.754 | 1547.907 | 1588.84 | 2715.645 | 2551.457 | 2279.754 | 1445.568 | 1771.77 | 1048.966 | 0.00065659 | -0.701004457 | YMR001C | CDC5 | n |
| W303.asm02.HP0_G0040520 | 6973.907 | 6886.015 | 6574.174 | 4887.328 | 5347.646 | 4039.137 | 1022.144 | 2154.043 | 1544.145 | 0.061147201 | 0.51760289 | YMR002W | MIX17 | n |
| W303.asm02.HP0_G0040530 | 438.3841 | 406.1725 | 410.6838 | 3000.479 | 3026.167 | 3623.6 | 7452.514 | 6598.501 | 7219.565 | 2.01E-121 | -2.942452679 | YMR003W | AIM34 | n |
| W303.asm02.HP0_G0040540 | 5414.149 | 5644.129 | 5341.045 | 2548.914 | 2660.805 | 3998.115 | 1876.399 | 3605.82 | 2404.129 | 0.003395516 | 0.832690635 | YMR004W | MVP1 | n |
| W303.asm02.HP0_G0040550 | 3994.166 | 4102.899 | 4071.267 | 4098.131 | 4019.847 | 3701.086 | 5711.907 | 4811.697 | 5048.672 | 0.717152345 | 0.041996918 | YMR005W | TAF4 | n |
| W303.asm02.HP0_G0040570 | 3406.477 | 3350.645 | 3394.339 | 1292.867 | 1418.923 | 1197.231 | 639.4571 | 1524.796 | 1205.653 | 2.13E-06 | 1.376875568 | YMR009W | ADI1 | n |
| W303.asm02.HP0_G0040580 | 937.1254 | 928.0763 | 937.7818 | 2366.898 | 2387.435 | 2074.644 | 2824.474 | 2374.172 | 2913.661 | 3.78E-25 | -1.284637727 | YMR010W | ANY1 | n |
| W303.asm02.HP0_G0040600 | 22749.38 | 24453.81 | 23487.66 | 21159.66 | 19648.82 | 20482.08 | 8519.099 | 9889.697 | 9166.79 | 0.029856642 | 0.205824701 | YMR012W | CLU1 | n |
| W303.asm02.HP0_G0040610 | 478.6222 | 388.3677 | 397.7488 | 1661.066 | 1648.901 | 2645.152 | 5505.75 | 5665.368 | 6190.932 | 3.22E-24 | -2.235173113 | YMR013C | SEC59 | n |
| W303.asm02.HP0_G0040620 | 1315.152 | 1399.904 | 1678.306 | 3742.436 | 3537.326 | 1503.376 | 2738.061 | 1588.15 | 2648.131 | 0.007139359 | -0.999403399 | YMR014W | BUD22 | n |
| W303.asm02.HP0_G0040630 | 2034.144 | 2688.528 | 3158.277 | 8803.443 | 8676.689 | 9758.652 | 15866.68 | 18518.75 | 19674.39 | 6.70E-22 | -1.789265735 | YMR015C | ERG5 | n |
| W303.asm02.HP0_G0040640 | 2547.71 | 3063.542 | 2873.708 | 6276.76 | 5892.651 | 5060.886 | 4262.636 | 3874.27 | 3619.351 | 1.34E-13 | -1.022049895 | YMR016C | SOK2 | n |
| W303.asm02.HP0_G0040680 | 3090.925 | 3121.408 | 2879.098 | 2942.818 | 3166.757 | 3232.373 | 1703.573 | 1343.324 | 1414.968 | 0.779030447 | -0.039145219 | YMR020W | FMS1 | n |
| W303.asm02.HP0_G0040690 | 1542.815 | 1862.829 | 1356.011 | 2212.671 | 1895.368 | 2241.011 | 3015.818 | 2082.098 | 2916.054 | 0.047357096 | -0.415268164 | YMR021C | MAC1 | n |
| W303.asm02.HP0_G0040700 | 3294.234 | 3102.49 | 2886.643 | 1790.283 | 1770.399 | 2389.145 | 1781.345 | 1936.061 | 1755.852 | 4.34E-05 | 0.641733777 | YMR022W | UBC7 | n |
| W303.asm02.HP0_G0040710 | 1585.171 | 1557.922 | 1469.192 | 1943.121 | 1905.782 | 1463.874 | 2834.35 | 2396.721 | 3230.624 | 0.270861097 | -0.203956683 | YMR023C | MSS1 | n |
| W303.asm02.HP0_G0040720 | 13570.85 | 14103.64 | 13407.05 | 4237.768 | 4082.331 | 3052.332 | 1902.323 | 1608.552 | 1740.303 | 2.23E-33 | 1.852951735 | YMR024W | MRPL3 | n |
| W303.asm02.HP0_G0040730 | 490.2701 | 514.1142 | 397.7488 | 372.3679 | 365.3617 | 397.3046 | 396.2659 | 580.9258 | 486.8063 | 0.131521808 | 0.304645752 | YMR025W | CSI1 | n |
| W303.asm02.HP0_G0040740 | 1363.862 | 1392.115 | 1280.557 | 685.6848 | 729.8556 | 765.7419 | 1756.655 | 1182.254 | 870.7494 | 0.000479557 | 0.887996379 | YMR026C | PEX12 | n |
| W303.asm02.HP0_G0040750 | 8260.469 | 8461.742 | 8026.119 | 4581.653 | 5069.068 | 6912.188 | 11220.13 | 11853.68 | 11958.45 | 0.000809523 | 0.579382383 | YMR027W | | n |
| W303.asm02.HP0_G0040760 | 1994.965 | 2128.789 | 2059.886 | 2543.356 | 2619.149 | 3040.178 | 3293.574 | 4032.119 | 3381.33 | 0.002383347 | -0.407673672 | YMR028W | TAP42 | n |
| W303.asm02.HP0_G0040770 | 5266.962 | 5816.613 | 4936.828 | 3089.403 | 3178.907 | 4352.119 | 2949.156 | 2931.474 | 2566.797 | 0.000721784 | 0.593035131 | YMR029C | FAR8 | n |
| W303.asm02.HP0_G0040790 | 11526.11 | 11548.65 | 10830.84 | 11286.36 | 11844.31 | 17527.74 | 15369.19 | 14871.06 | 13697.56 | 0.194733183 | -0.262034611 | YMR031C | EIS1 | n |
| W303.asm02.HP0_G0040810 | 2099.796 | 2008.606 | 2332.598 | 2924.755 | 2777.964 | 2718.08 | 814.7522 | 724.815 | 650.6699 | 0.001987913 | -0.386697956 | YMR032W | HOF1 | n |
| W303.asm02.HP0_G0040830 | 4501.378 | 4809.528 | 4657.65 | 4118.277 | 4133.534 | 4451.635 | 2844.226 | 3902.189 | 3546.39 | 0.369585315 | 0.136942567 | YMR033W | ARP9 | n |
| W303.asm02.HP0_G0040840 | 267.9014 | 367.2245 | 226.3611 | 2032.045 | 1943.967 | 1680.378 | 6236.558 | 4293.052 | 2657.699 | 2.59E-17 | -2.715239488 | YMR034C | RCH1 | n |
| W303.asm02.HP0_G0040850 | 946.6554 | 913.6099 | 780.4069 | 650.949 | 744.6089 | 644.9553 | 756.732 | 563.745 | 655.4542 | 0.026360881 | 0.372494309 | YMR035W | IMP2 | n |
| W303.asm02.HP0_G0040860 | 815.352 | 1000.408 | 835.3804 | 4789.373 | 4417.318 | 5969.445 | 3849.087 | 4139.499 | 3420.801 | 2.16E-51 | -2.51733472 | YMR036C | MIH1 | n |
| W303.asm02.HP0_G0040870 | 2006.613 | 1803.851 | 2157.976 | 2883.767 | 2511.536 | 4535.198 | 5435.385 | 4792.369 | 5260.379 | 0.00207894 | -0.734559658 | YMR037C | MSN2 | n |
| W303.asm02.HP0_G0040880 | 4333.014 | 4197.487 | 4162.889 | 1937.563 | 2242.505 | 2895.842 | 2070.211 | 3676.691 | 2376.62 | 0.000988945 | 0.843121807 | YMR038C | CCS1 | n |
| W303.asm02.HP0_G0040890 | 7557.36 | 8177.977 | 7350.269 | 5270.811 | 5482.161 | 6280.907 | 8416.638 | 7259.961 | 7597.528 | 0.000245968 | 0.438578365 | YMR039C | SUB1 | n |
| W303.asm02.HP0_G0040910 | 959.3622 | 1003.747 | 860.1723 | 1135.166 | 1222.79 | 983.7657 | 1009.799 | 790.3167 | 1021.456 | 0.154775204 | -0.24304363 | YMR041C | ARA2 | n |
| W303.asm02.HP0_G0040920 | 684.0486 | 714.4185 | 815.978 | 4933.179 | 5023.072 | 5908.671 | 5867.451 | 5340.007 | 5476.87 | 1.99E-112 | -2.840847592 | YMR042W | ARG80 | n |
| W303.asm02.HP0_G0040930 | 2723.487 | 2982.308 | 2563.27 | 3387.436 | 3261.352 | 3290.107 | 2182.548 | 2393.5 | 2462.738 | 0.010199441 | -0.265506237 | YMR044W | IOC4 | n |
| W303.asm02.HP0_G0040940 | 7165.568 | 6248.38 | 7020.429 | 18478.06 | 18056.33 | 20052.11 | 17030.79 | 18200.91 | 18162.54 | 2.45E-53 | -1.46943061 | YMR047C | NUP116 | n |
| W303.asm02.HP0_G0040950 | 1118.197 | 1015.988 | 1122.104 | 1171.986 | 1287.878 | 846.2664 | 596.2505 | 529.3834 | 516.7084 | 0.921112924 | -0.021595761 | YMR048W | CSM3 | n |
| W303.asm02.HP0_G0040960 | 3765.444 | 4146.298 | 4822.57 | 7761.369 | 6863.766 | 2229.616 | 9182.011 | 3494.145 | 7259.036 | 0.432721646 | -0.404434426 | YMR049C | ERB1 | n |
| W303.asm02.HP0_G0040970 | 697.8142 | 706.6289 | 681.2392 | 839.2171 | 988.4726 | 967.8127 | 936.9651 | 1017.962 | 904.2397 | 0.000355667 | -0.421861251 | YMR052W | FAR3 | n |
| W303.asm02.HP0_G0040990 | 5939.363 | 5698.656 | 5784.066 | 4395.469 | 4368.719 | 5618.48 | 5626.728 | 5754.494 | 5840.48 | 0.037469251 | 0.276556397 | YMR054W | STV1 | n |
| W303.asm02.HP0_G0041000 | 646.9871 | 738.9001 | 623.0321 | 232.7299 | 217.8285 | 315.2608 | 119.7439 | 131.0036 | 89.70632 | 1.06E-10 | 1.390669009 | YMR055C | BUB2 | n |
| W303.asm02.HP0_G0041020 | 886.2982 | 885.7899 | 1001.379 | 15541.49 | 16338.87 | 11739.1 | 18051.7 | 11689.39 | 11395.1 | 1.06E-69 | -3.975191194 | YMR058W | FET3 | n |
| W303.asm02.HP0_G0041030 | 630.0447 | 553.0623 | 526.0201 | 479.3541 | 536.3267 | 357.0424 | 614.7676 | 335.0256 | 580.1009 | 0.283785184 | 0.316590293 | YMR059W | SEN15 | n |
| W303.asm02.HP0_G0041040 | 1665.648 | 1660.3 | 1559.736 | 1650.646 | 1556.909 | 1381.83 | 1351.748 | 1017.962 | 1242.732 | 0.577305032 | 0.090183125 | YMR060C | SAM37 | n |
| W303.asm02.HP0_G0041050 | 6467.753 | 6475.391 | 6591.421 | 4491.34 | 4525.799 | 4558.747 | 4194.74 | 3792.661 | 3782.019 | 1.26E-12 | 0.524998315 | YMR061W | RNA14 | n |
| W303.asm02.HP0_G0041060 | 3022.097 | 2930.006 | 2870.475 | 5878.688 | 5820.62 | 7447.752 | 2728.186 | 4478.82 | 3014.132 | 2.51E-07 | -1.117857501 | YMR062C | ARG7 | n |
| W303.asm02.HP0_G0041070 | 273.1959 | 311.5844 | 296.4253 | 102.1233 | 106.7446 | 88.88076 | 454.2861 | 148.1844 | 172.2361 | 0.000212352 | 1.565586048 | YMR063W | RIM9 | n |
| W303.asm02.HP0_G0041080 | 3332.354 | 3367.337 | 3141.03 | 3517.348 | 3210.149 | 2345.844 | 1552.967 | 1489.361 | 2030.951 | 0.600279756 | 0.117103603 | YMR064W | AEP1 | n |
| W303.asm02.HP0_G0041090 | 459.562 | 481.843 | 472.1246 | 603.0136 | 630.0537 | 322.0978 | 374.0453 | 262.0072 | 297.825 | 0.665313434 | -0.137570185 | YMR065W | KAR5 | n |

| | | | | | | | | | | | | | | |
|---|---|---|---|---|---|---|---|---|---|---|---|---|---|---|
| W303.asm02.HP0_G0041100 | 1910.253 | 2089.841 | 1919.758 | 1467.24 | 1107.367 | 494.5417 | 2834.35 | 1176.885 | 2550.052 | 0.031343536 | 0.94764349 | YMR066W | SOV1 | n |
| W303.asm02.HP0_G0041110 | 4185.827 | 4149.636 | 4196.304 | 1406.105 | 1398.094 | 1674.301 | 814.7522 | 1522.648 | 979.5931 | 3.00E-10 | 1.484477145 | YMR067C | UBX4 | n |
| W303.asm02.HP0_G0041120 | 1279.15 | 1217.405 | 1164.143 | 2328.688 | 2632.166 | 2579.061 | 3024.459 | 2221.692 | 2517.758 | 1.00E-12 | -1.042271815 | YMR068W | AVO2 | n |
| W303.asm02.HP0_G0041130 | 183.1895 | 132.4234 | 147.6737 | 368.1996 | 395.7362 | 249.17 | 185.171 | 387.6418 | 299.0211 | 0.000647189 | -1.128191155 | YMR069W | NAT4 | n |
| W303.asm02.HP0_G0041140 | 781.4672 | 1112.801 | 851.549 | 2043.855 | 1805.112 | 1975.128 | 1602.346 | 2653.36 | 1460.419 | 4.11E-05 | -1.084972233 | YMR070W | MOT3 | n |
| W303.asm02.HP0_G0041150 | 6041.017 | 6327.389 | 5960.843 | 4405.195 | 5064.729 | 5827.387 | 5858.809 | 7014.061 | 6367.953 | 0.076452677 | 0.260918474 | YMR072W | ABF2 | n |
| W303.asm02.HP0_G0041160 | 872.5325 | 839.0522 | 838.6141 | 900.3521 | 1019.715 | 698.8914 | 793.7662 | 718.3722 | 619.5717 | 0.854871663 | -0.037955403 | YMR073C | IRC21 | n |
| W303.asm02.HP0_G0041170 | 2159.094 | 1952.966 | 1935.927 | 1916.027 | 2147.042 | 1722.16 | 1666.539 | 1258.494 | 1499.89 | 0.71145853 | 0.064315658 | YMR074C | SDD2 | n |
| W303.asm02.HP0_G0041180 | 2708.663 | 2914.427 | 2853.228 | 3053.972 | 2638.241 | 2342.046 | 2759.047 | 2175.519 | 2453.169 | 0.645639995 | 0.077123486 | YMR075W | RCO1 | n |
| W303.asm02.HP0_G0041190 | 2264.984 | 2459.291 | 2510.453 | 4673.355 | 4939.76 | 6060.604 | 1001.158 | 2141.157 | 1627.871 | 4.22E-05 | -1.11534196 | YMR076C | PDS5 | n |
| W303.asm02.HP0_G0041200 | 1327.859 | 1337.587 | 1324.752 | 855.1956 | 885.1994 | 1345.366 | 949.3098 | 783.8739 | 833.6708 | 0.085741466 | 0.370764933 | YMR077C | VPS20 | n |
| W303.asm02.HP0_G0041210 | 1251.618 | 1202.938 | 1308.583 | 3790.371 | 4309.706 | 4343.003 | 1278.914 | 2986.238 | 2078.795 | 1.04E-09 | -1.725303563 | YMR078C | CTF18 | n |
| W303.asm02.HP0_G0041220 | 3151.283 | 3580.995 | 3613.155 | 8331.731 | 8968.285 | 8625.992 | 5078.622 | 5507.52 | 4796.298 | 4.75E-38 | -1.325412371 | YMR079W | SEC14 | n |
| W303.asm02.HP0_G0041230 | 4587.149 | 4181.908 | 4804.245 | 8624.901 | 8635.901 | 7098.306 | 6015.587 | 5727.649 | 6120.363 | 2.37E-12 | -0.843641597 | YMR080C | NAM7 | n |
| W303.asm02.HP0_G0041250 | 14969.65 | 14488.67 | 15256.74 | 647.4754 | 825.3182 | 632.8006 | 304.9149 | 912.7299 | 309.7858 | 8.35E-29 | 4.408612277 | YMR083W | ADH3 | n |
| W303.asm02.HP0_G0041270 | 7007.792 | 7072.966 | 6951.442 | 9806.613 | 9391.792 | 12463.06 | 8196.901 | 9434.406 | 8077.157 | 5.39E-05 | -0.590153039 | YMR086W | SEG1 | n |
| W303.asm02.HP0_G0041280 | 1929.313 | 1767.129 | 1831.369 | 719.0312 | 814.9041 | 788.5319 | 1091.274 | 905.2133 | 1208.045 | 3.80E-19 | 1.251562011 | YMR087W | PDL32 | n |
| W303.asm02.HP0_G0041290 | 2066.97 | 2134.353 | 2023.237 | 1012.896 | 1256.636 | 1128.102 | 1083.867 | 1249.903 | 1176.947 | 5.19E-13 | 0.874199568 | YMR088C | VBA1 | n |
| W303.asm02.HP0_G0041300 | 16800.49 | 17576.7 | 17180.81 | 8770.097 | 8328.685 | 10140 | 5263.793 | 4839.616 | 4470.963 | 3.61E-15 | 0.920492547 | YMR089C | YTA12 | n |
| W303.asm02.HP0_G0041310 | 3198.933 | 2830.967 | 2749.749 | 1185.88 | 1355.57 | 1924.99 | 2987.425 | 3287.975 | 3014.132 | 1.16E-06 | 0.975098179 | YMR090W | | n |
| W303.asm02.HP0_G0041320 | 2701.251 | 2400.313 | 2440.389 | 2712.172 | 2484.633 | 2952.816 | 2851.633 | 2423.566 | 2524.934 | 0.405022517 | -0.111897931 | YMR091C | NPL6 | n |
| W303.asm02.HP0_G0041330 | 5846.18 | 5641.903 | 5646.093 | 5290.263 | 5614.941 | 6983.597 | 5721.783 | 8128.665 | 6683.719 | 0.753413188 | -0.062172277 | YMR092C | AIP1 | n |
| W303.asm02.HP0_G0041340 | 786.7617 | 758.9305 | 881.7305 | 5288.179 | 5001.376 | 2002.476 | 8536.381 | 3792.661 | 6425.365 | 4.53E-09 | -2.340210495 | YMR093W | UTP15 | n |
| W303.asm02.HP0_G0041350 | 199.073 | 208.0939 | 173.5435 | 557.1623 | 593.6043 | 470.2324 | 220.9707 | 200.8006 | 218.8834 | 6.57E-20 | -1.480773354 | YMR094W | CTF13 | n |
| W303.asm02.HP0_G0041380 | 2375.11 | 2465.968 | 2291.637 | 1162.955 | 1126.46 | 586.4611 | 1271.507 | 806.4237 | 1198.476 | 8.68E-06 | 1.310489186 | YMR097C | MTG1 | n |
| W303.asm02.HP0_G0041390 | 5028.71 | 4988.689 | 4988.568 | 2120.968 | 1787.756 | 1833.83 | 1111.026 | 955.6819 | 1148.241 | 6.07E-32 | 1.385409651 | YMR098C | ATP25 | n |
| W303.asm02.HP0_G0041400 | 8887.337 | 9122.746 | 9301.287 | 4508.013 | 5141.099 | 5837.263 | 3254.071 | 6288.172 | 4722.141 | 0.000875435 | 0.818520052 | YMR099C | | n |
| W303.asm02.HP0_G0041410 | 1448.573 | 1715.94 | 1449.789 | 1560.332 | 1555.174 | 2165.804 | 2065.273 | 1405.604 | 1642.224 | 0.400337563 | -0.194882324 | YMR100W | MUB1 | n |
| W303.asm02.HP0_G0041420 | 162.0115 | 198.0786 | 164.9203 | 92.39725 | 88.51994 | 101.7951 | 83.94417 | 109.5276 | 63.39247 | 0.000338734 | 0.891971345 | YMR101C | SRT1 | n |
| W303.asm02.HP0_G0041430 | 1694.238 | 1976.335 | 1667.527 | 5443.796 | 5343.306 | 7126.414 | 11215.19 | 7977.26 | 9109.378 | 2.29E-20 | -1.746720213 | YMR102C | LAF1 | n |
| W303.asm02.HP0_G0041450 | 29798.47 | 28853.83 | 26887.39 | 6684.559 | 6980.925 | 11505.88 | 50683.76 | 26196.42 | 39293.76 | 2.06E-09 | 1.76480611 | YMR105C | PGM2 | n |
| W303.asm02.HP0_G0041470 | 657.5761 | 708.8545 | 661.8368 | 1581.869 | 1656.711 | 1770.778 | 6169.897 | 2792.954 | 3649.253 | 3.63E-06 | -1.304392633 | YMR106C | YKU80 | n |
| W303.asm02.HP0_G0041480 | 1347.978 | 1555.696 | 1036.95 | 78.50292 | 77.23798 | 56.97485 | 1271.507 | 720.5198 | 435.3747 | 2.08E-28 | 4.211562985 | YMR107W | SPG4 | n |
| W303.asm02.HP0_G0041490 | 10491.57 | 11588.71 | 11451.72 | 8413.012 | 8324.345 | 8390.496 | 12534.84 | 5595.571 | 6420.581 | 0.180157932 | 0.416248776 | YMR108W | ILV2 | n |
| W303.asm02.HP0_G0041500 | 6307.86 | 5674.174 | 5987.791 | 11328.04 | 11058.92 | 15502.48 | 9716.538 | 11745.22 | 9988.5 | 3.24E-10 | -1.076220552 | YMR109W | MYO5 | n |
| W303.asm02.HP0_G0041510 | 11318.57 | 11538.64 | 11097.08 | 3411.056 | 3592.868 | 4241.208 | 2809.661 | 4051.447 | 3509.311 | 4.80E-22 | 1.59430117 | YMR110C | HFD1 | n |
| W303.asm02.HP0_G0041520 | 1217.734 | 1436.627 | 1185.701 | 1323.434 | 1349.495 | 1646.953 | 3060.259 | 1977.939 | 2221.129 | 0.453109767 | -0.169947243 | YMR111C | EUC1 | n |
| W303.asm02.HP0_G0041530 | 786.7617 | 718.8697 | 617.6425 | 935.7826 | 1113.442 | 967.0531 | 1394.955 | 1053.398 | 1015.476 | 0.006912996 | -0.506085633 | YMR112C | MED11 | n |
| W303.asm02.HP0_G0041540 | 1605.29 | 1565.712 | 1605.008 | 1686.771 | 1753.91 | 1781.414 | 1138.184 | 1573.117 | 1483.145 | 0.412094584 | -0.128750079 | YMR113W | FOL3 | n |
| W303.asm02.HP0_G0041550 | 4740.69 | 4582.516 | 4222.174 | 1292.867 | 1471.861 | 1830.032 | 2477.587 | 2907.85 | 2156.54 | 3.51E-18 | 1.559825062 | YMR114C | | n |
| W303.asm02.HP0_G0041560 | 3663.79 | 3909.271 | 3528 | 5029.05 | 4867.729 | 4783.608 | 2099.839 | 2859.529 | 2517.758 | 0.004022756 | -0.403232625 | YMR115W | MGR3 | n |
| W303.asm02.HP0_G0041570 | 17610.54 | 15342.19 | 18853.73 | 32067.4 | 35911.32 | 35299.34 | 23266.11 | 20666.35 | 19991.35 | 9.40E-16 | -0.995306404 | YMR116C | ASC1 | n |
| W303.asm02.HP0_G0041600 | 2048.969 | 2215.588 | 2063.12 | 3891.105 | 3926.12 | 5746.103 | 8041.358 | 6471.792 | 6147.873 | 3.92E-08 | -1.100002776 | YMR119W | ASI1 | n |
| W303.asm02.HP0_G0041610 | 3766.503 | 2000.817 | 3509.675 | 17176.86 | 16494.21 | 27467.95 | 14448.27 | 49258.42 | 25915.56 | 8.10E-10 | -2.720344998 | YMR120C | ADE17 | n |
| W303.asm02.HP0_G0041630 | 7130.624 | 7204.276 | 6987.014 | 3956.408 | 5194.037 | 3784.649 | 2065.273 | 2486.921 | 1521.419 | 0.000916124 | 0.721144423 | YMR122W | NCW1 | n |
| W303.asm02.HP0_G0041640 | 1440.102 | 1468.898 | 1536.022 | 845.4695 | 979.7942 | 817.3991 | 657.9742 | 746.291 | 613.5913 | 6.05E-09 | 0.750895321 | YMR123W | PKR1 | n |
| W303.asm02.HP0_G0041650 | 6736.714 | 7368.971 | 7363.204 | 6054.451 | 4907.649 | 5789.404 | 8470.954 | 5655.704 | 7294.918 | 0.054438942 | 0.357876043 | YMR124W | EPO1 | n |
| W303.asm02.HP0_G0041660 | 8158.815 | 8687.64 | 8521.958 | 8727.719 | 8674.086 | 9704.715 | 6598.259 | 7724.917 | 7316.448 | 0.391848094 | -0.095635423 | YMR125W | STO1 | n |
| W303.asm02.HP0_G0041670 | 411.9116 | 455.1358 | 411.7617 | 638.4441 | 620.5074 | 779.4159 | 1143.122 | 675.4202 | 968.8283 | 0.003163314 | -0.672827344 | YMR126C | DLT1 | n |
| W303.asm02.HP0_G0041680 | 785.7028 | 777.8482 | 851.549 | 1198.385 | 1145.552 | 717.8831 | 1343.107 | 1111.383 | 1218.81 | 0.136930734 | -0.342265577 | YMR127C | SAS2 | n |
| W303.asm02.HP0_G0041690 | 2585.831 | 2854.336 | 3193.848 | 5060.312 | 4175.19 | 919.9539 | 5778.569 | 1808.279 | 4973.319 | 0.717734912 | -0.234155553 | YMR128W | ECM16 | n |
| W303.asm02.HP0_G0041700 | 2420.642 | 2500.465 | 2511.531 | 7732.191 | 7087.67 | 6861.291 | 5273.669 | 6082.003 | 5708.91 | 1.32E-57 | -1.544586062 | YMR129W | POM152 | n |
| W303.asm02.HP0_G0041710 | 2781.727 | 2598.391 | 2777.774 | 980.9392 | 968.5122 | 1397.023 | 1472.726 | 1698.751 | 1849.146 | 2.55E-12 | 1.28545219 | YMR130W | DPI35 | n |
| W303.asm02.HP0_G0041720 | 3940.162 | 4149.636 | 4576.806 | 8947.249 | 8441.504 | 4834.506 | 11469.49 | 7351.234 | 10743.23 | 0.003703791 | -0.811039485 | YMR131C | RRB1 | n |

| | | | | | | | | | | | | | | |
|---|---|---|---|---|---|---|---|---|---|---|---|---|---|---|
| W303.asm02.HP0_G0041730 | 1254.795 | 1211.841 | 1240.675 | 345.2739 | 327.1766 | 252.2087 | 165.4194 | 302.8116 | 191.3735 | 2.44E-16 | 2.003342177 | YMR132C | JLP2 | n |
| W303.asm02.HP0_G0041740 | 2356.05 | 2518.27 | 2519.076 | 1410.968 | 1251.429 | 1490.462 | 2741.765 | 2321.555 | 2637.366 | 2.46E-12 | 0.831685857 | YMR133W | REC114 | n |
| W303.asm02.HP0_G0041750 | 5425.797 | 5529.51 | 5034.918 | 1251.878 | 1267.918 | 2377.75 | 1914.668 | 2508.397 | 1821.636 | 7.73E-10 | 1.707007463 | YMR135C | GID8 | n |
| W303.asm02.HP0_G0041770 | 1169.024 | 1115.027 | 1088.689 | 1567.974 | 1513.517 | 1842.187 | 2324.513 | 2143.305 | 1784.558 | 0.000265747 | -0.545895841 | YMR137C | PSO2 | n |
| W303.asm02.HP0_G0041780 | 964.6567 | 825.6986 | 861.2502 | 936.4774 | 911.2346 | 585.7014 | 1708.511 | 951.3867 | 1115.947 | 0.692482178 | 0.123964113 | YMR138W | CIN4 | n |
| W303.asm02.HP0_G0041800 | 3727.323 | 3981.603 | 3442.845 | 1679.129 | 1778.209 | 3942.659 | 5599.57 | 6224.818 | 4604.925 | 0.089131409 | 0.591647724 | YMR140W | SIP5 | n |
| W303.asm02.HP0_G0041820 | 9524.794 | 9389.818 | 9985.76 | 18118.2 | 22035.39 | 19058.47 | 16977.71 | 14232.14 | 13277.73 | 4.69E-14 | -1.034766292 | YMR143W | RPS16A | y |
| W303.asm02.HP0_G0041830 | 570.7464 | 598.6871 | 639.2007 | 1351.223 | 1259.239 | 1845.225 | 359.2317 | 472.472 | 276.2955 | 6.64E-08 | -1.300851088 | YMR144W | FDO1 | n |
| W303.asm02.HP0_G0041840 | 16203.27 | 18287.78 | 16942.59 | 27889.38 | 28547.68 | 23393.11 | 17015.98 | 30328.41 | 30766.88 | 0.010052644 | -0.634221707 | YMR145C | NDE1 | n |
| W303.asm02.HP0_G0041850 | 17011.21 | 19027.79 | 19117.81 | 13140.56 | 14289.03 | 11016.66 | 7875.938 | 8063.164 | 7735.077 | 5.25E-05 | 0.520727169 | YMR146C | TIF34 | n |
| W303.asm02.HP0_G0041880 | 4058.759 | 4128.493 | 4148.876 | 3099.823 | 3339.458 | 4146.249 | 1662.835 | 4206.074 | 3282.055 | 0.522706159 | 0.220800183 | YMR149W | SWP1 | n |
| W303.asm02.HP0_G0041890 | 1542.815 | 1397.679 | 1471.347 | 450.1761 | 492.9345 | 378.313 | 697.4773 | 693.6748 | 773.8666 | 1.00E-37 | 1.740279013 | YMR150C | IMP1 | n |
| W303.asm02.HP0_G0041920 | 1355.39 | 1301.978 | 1230.973 | 2644.09 | 2283.294 | 3012.07 | 2888.667 | 2840.201 | 2994.995 | 3.78E-15 | -1.030094794 | YMR154C | RIM13 | n |
| W303.asm02.HP0_G0041930 | 1205.027 | 1181.795 | 1040.183 | 605.7925 | 522.4412 | 755.1066 | 1888.744 | 2132.567 | 1459.223 | 3.56E-05 | 0.863237819 | YMR155W | | n |
| W303.asm02.HP0_G0041940 | 1134.081 | 1107.237 | 1004.612 | 915.6359 | 1002.358 | 912.3572 | 780.187 | 684.0106 | 741.5723 | 0.089552734 | 0.198188186 | YMR156C | TPP1 | n |
| W303.asm02.HP0_G0041950 | 3809.918 | 3851.406 | 3425.598 | 1636.751 | 1474.464 | 1395.504 | 1064.116 | 1123.195 | 1149.437 | 1.19E-33 | 1.298387251 | YMR157C | AIM36 | n |
| W303.asm02.HP0_G0041960 | 4936.586 | 4996.478 | 4736.337 | 2141.81 | 2289.369 | 1622.644 | 1341.872 | 1271.379 | 1221.202 | 2.63E-17 | 1.277046859 | YMR158W | MRPS8 | n |
| W303.asm02.HP0_G0041970 | 780.4083 | 855.7443 | 775.0174 | 270.2446 | 316.7625 | 202.8305 | 411.0795 | 413.413 | 380.3548 | 1.78E-18 | 1.611543823 | YMR159C | ATG16 | n |
| W303.asm02.HP0_G0041980 | 3945.457 | 4059.499 | 3432.066 | 1059.442 | 931.195 | 1443.363 | 1573.953 | 1259.567 | 1251.104 | 6.66E-18 | 1.735514864 | YMR160W | CVM1 | n |
| W303.asm02.HP0_G0041990 | 2758.431 | 3113.618 | 2825.202 | 2096.653 | 2043.769 | 1770.019 | 1498.65 | 907.3609 | 1094.417 | 0.007171025 | 0.557271794 | YMR161W | HLJ1 | n |
| W303.asm02.HP0_G0042000 | 1952.609 | 1931.823 | 1768.851 | 3113.023 | 3184.114 | 3334.928 | 3538 | 3440.455 | 3358.605 | 8.90E-21 | -0.768729494 | YMR162C | DNF3 | n |
| W303.asm02.HP0_G0042010 | 357.9078 | 317.1484 | 343.8533 | 889.9314 | 711.6309 | 834.8714 | 623.4089 | 659.3132 | 737.984 | 8.27E-18 | -1.258294865 | YMR163C | INP2 | n |
| W303.asm02.HP0_G0042020 | 1237.853 | 1360.956 | 1140.429 | 1298.424 | 1125.592 | 1438.805 | 902.3998 | 846.1543 | 834.8669 | 0.761511696 | -0.047415558 | YMR164C | MSS11 | n |
| W303.asm02.HP0_G0042030 | 3265.644 | 3453.023 | 3074.2 | 4699.755 | 4470.257 | 5254.6 | 3799.708 | 3981.65 | 3588.253 | 2.67E-07 | -0.558843875 | YMR165C | PAH1 | n |
| W303.asm02.HP0_G0042040 | 4030.168 | 3901.482 | 3999.047 | 4018.238 | 3994.679 | 3258.202 | 1645.553 | 2205.585 | 2100.324 | 0.637227231 | 0.082072655 | YMR166C | MME1 | n |
| W303.asm02.HP0_G0042050 | 2258.631 | 2472.645 | 2305.65 | 775.9979 | 750.6838 | 656.3502 | 527.12 | 546.5642 | 569.3361 | 2.26E-54 | 1.68846886 | YMR167W | MLH1 | n |
| W303.asm02.HP0_G0042060 | 1673.06 | 1702.586 | 1629.8 | 1021.927 | 1126.46 | 1031.625 | 511.0719 | 666.8298 | 509.5319 | 2.53E-06 | 0.654768706 | YMR168C | CEP3 | n |
| W303.asm02.HP0_G0042080 | 2069.088 | 2031.975 | 1869.096 | 915.6359 | 1004.962 | 1905.239 | 160.4815 | 390.8632 | 220.0795 | 0.106948169 | 0.641977282 | YMR170C | ALD2 | n |
| W303.asm02.HP0_G0042090 | 2928.914 | 2844.32 | 2916.825 | 2398.855 | 2297.179 | 3100.191 | 9105.474 | 6579.172 | 7798.47 | 0.419143701 | 0.156531186 | YMR171C | EAR1 | n |
| W303.asm02.HP0_G0042100 | 1602.114 | 1599.096 | 1655.67 | 2562.808 | 2642.58 | 3304.541 | 892.524 | 1346.545 | 813.3373 | 0.00026903 | -0.809120976 | YMR172W | HOT1 | n |
| W303.asm02.HP0_G0042110 | 22043.09 | 19135.73 | 20300.28 | 1731.927 | 2187.831 | 1787.491 | 2059.101 | 2679.131 | 2339.541 | 1.99E-112 | 3.429570075 | YMR173W | DDR48 | n |
| W303.asm02.HP0_G0042150 | 2338.048 | 2236.731 | 2258.222 | 1640.92 | 1410.244 | 1356.001 | 1204.846 | 1106.014 | 1074.084 | 3.75E-08 | 0.632258956 | YMR176W | ECM5 | n |
| W303.asm02.HP0_G0042160 | 1063.134 | 1007.085 | 1048.807 | 4594.853 | 4314.913 | 6677.452 | 1494.947 | 3503.809 | 2365.855 | 1.03E-13 | -2.321199656 | YMR177W | MMT1 | n |
| W303.asm02.HP0_G0042170 | 4334.072 | 4061.725 | 3921.437 | 843.3854 | 849.6178 | 851.584 | 1119.667 | 946.0177 | 863.5729 | 2.19E-77 | 2.275220067 | YMR178W | FPY1 | n |
| W303.asm02.HP0_G0042180 | 956.1855 | 991.506 | 874.1851 | 1482.524 | 1897.972 | 2079.202 | 253.067 | 275.9666 | 179.4126 | 3.91E-06 | -0.952036974 | YMR179W | SPT21 | n |
| W303.asm02.HP0_G0042230 | 1573.523 | 1421.047 | 1486.438 | 821.8492 | 925.988 | 1204.068 | 1539.388 | 1169.368 | 1095.613 | 0.003743062 | 0.602284826 | YMR184W | ADD37 | n |
| W303.asm02.HP0_G0042240 | 857.708 | 959.2348 | 1003.534 | 2350.225 | 1950.042 | 1156.969 | 4836.666 | 3413.61 | 4403.982 | 0.000626669 | -0.952313128 | YMR185W | RTP1 | n |
| W303.asm02.HP0_G0042250 | 40246.62 | 42093.94 | 41387.44 | 26458.96 | 26413.65 | 34801.76 | 5962.505 | 12554.87 | 10152.36 | 0.079493221 | 0.496943778 | YMR186W | HSC82 | y |
| W303.asm02.HP0_G0042260 | 1050.428 | 1087.207 | 1112.403 | 1469.325 | 1424.13 | 1822.435 | 904.8688 | 941.7225 | 767.8861 | 0.000372981 | -0.537215738 | YMR187C | | n |
| W303.asm02.HP0_G0042270 | 6940.022 | 7470.236 | 6887.846 | 4582.348 | 4786.151 | 3732.232 | 3138.031 | 2160.485 | 2820.367 | 9.37E-05 | 0.70110759 | YMR188C | MRPS17 | n |
| W303.asm02.HP0_G0042280 | 1173.26 | 972.5884 | 1100.546 | 14475.11 | 11378.28 | 21999.13 | 15642.01 | 29685.2 | 22080.91 | 8.97E-38 | -3.881660097 | YMR189W | GCV2 | n |
| W303.asm02.HP0_G0042290 | 1548.11 | 1545.681 | 1430.387 | 1956.321 | 2206.924 | 2920.151 | 2267.727 | 3198.85 | 2390.973 | 0.002285012 | -0.646760392 | YMR190C | SGS1 | n |
| W303.asm02.HP0_G0042320 | 6265.504 | 6696.839 | 5930.662 | 2739.266 | 2851.731 | 2105.79 | 1577.657 | 1752.441 | 1480.752 | 2.72E-17 | 1.295601139 | YMR193W | MRPL24 | n |
| W303.asm02.HP0_G0042360 | 10162.25 | 9681.372 | 9033.965 | 2145.283 | 1832.883 | 3655.506 | 9473.346 | 8259.669 | 8201.55 | 7.47E-14 | 1.919418622 | YMR196W | | n |
| W303.asm02.HP0_G0042370 | 3944.398 | 4003.859 | 3893.411 | 2848.336 | 3114.687 | 3547.634 | 3804.646 | 4354.259 | 3500.939 | 0.022317535 | 0.316309407 | YMR197C | VTI1 | n |
| W303.asm02.HP0_G0042400 | 5653.46 | 5796.582 | 5530.757 | 5186.751 | 5194.905 | 5938.298 | 7430.294 | 7499.419 | 7223.153 | 0.563386336 | 0.05722375 | YMR200W | ROT1 | n |
| W303.asm02.HP0_G0042420 | 6180.792 | 6130.423 | 6789.756 | 3585.43 | 4016.375 | 3619.042 | 2934.343 | 2972.278 | 3058.388 | 4.63E-17 | 0.767710715 | YMR202W | ERG2 | n |
| W303.asm02.HP0_G0042430 | 20026.95 | 19464.01 | 20064.22 | 10753.51 | 11271.54 | 10215.97 | 5778.569 | 5923.08 | 5926.598 | 1.43E-38 | 0.88540777 | YMR203W | TOM40 | n |
| W303.asm02.HP0_G0042440 | 549.5684 | 647.6504 | 557.2795 | 188.9628 | 180.5112 | 318.2995 | 206.157 | 612.066 | 148.3145 | 0.008233363 | 1.350886719 | YMR204C | INP1 | n |
| W303.asm02.HP0_G0042450 | 36704.61 | 36457.6 | 39362.04 | 44018.6 | 44585.41 | 83449.92 | 119222.9 | 118825.6 | 139562.7 | 0.023522809 | -0.612625873 | YMR205C | PFK2 | n |
| W303.asm02.HP0_G0042460 | 1133.022 | 1290.85 | 1093.001 | 233.4246 | 212.6214 | 307.6642 | 6882.187 | 2257.127 | 4601.336 | 1.55E-09 | 2.221946959 | YMR206W | | n |
| W303.asm02.HP0_G0042480 | 5868.417 | 6309.584 | 6121.452 | 3279.755 | 3679.652 | 4349.08 | 7310.55 | 6148.578 | 6521.052 | 1.06E-06 | 0.69443851 | YMR208W | ERG12 | n |
| W303.asm02.HP0_G0042490 | 1075.841 | 1001.521 | 998.1448 | 2284.921 | 1947.439 | 2387.626 | 1713.449 | 2264.644 | 2123.05 | 3.29E-13 | -1.106163933 | YMR209C | | n |
| W303.asm02.HP0_G0042500 | 4320.307 | 4215.292 | 3772.685 | 2073.728 | 2050.712 | 1289.151 | 453.0516 | 776.3573 | 566.944 | 5.57E-06 | 1.185001864 | YMR210W | MGL2 | n |

| | | | | | | | | | | | | | |
|---|---|---|---|---|---|---|---|---|---|---|---|---|---|
| W303.asm02.HP0_G0042510 | 1198.673 | 1239.661 | 1194.324 | 2084.843 | 2091.5 | 2189.353 | 927.0893 | 1678.349 | 1491.517 | 0.000272433 | -0.809307664 | YMR211W | DML1 | n |
| W303.asm02.HP0_G0042520 | 3084.572 | 2638.452 | 3098.992 | 4455.909 | 4408.64 | 5232.57 | 3615.772 | 4620.561 | 4222.178 | 5.46E-06 | -0.676193553 | YMR212C | EFR3 | n |
| W303.asm02.HP0_G0042530 | 2146.388 | 2422.569 | 2166.599 | 1389.432 | 1244.486 | 1195.712 | 1951.702 | 1705.194 | 1773.793 | 3.02E-12 | 0.81416869 | YMR213W | CEF1 | n |
| W303.asm02.HP0_G0042540 | 2154.859 | 2340.221 | 2194.625 | 1644.393 | 1718.328 | 2279.754 | 454.2861 | 937.4273 | 535.8458 | 0.42828276 | 0.245600115 | YMR214W | SCJ1 | n |
| W303.asm02.HP0_G0042550 | 3048.569 | 2561.669 | 2941.617 | 6972.866 | 6842.938 | 8291.739 | 17934.43 | 16396.92 | 17043.01 | 1.26E-27 | -1.370181192 | YMR215W | GAS3 | n |
| W303.asm02.HP0_G0042560 | 2600.655 | 3032.384 | 2591.296 | 5374.324 | 5014.394 | 5529.599 | 7246.357 | 5086.59 | 5290.281 | 2.87E-08 | -0.952788466 | YMR216C | SKY1 | n |
| W303.asm02.HP0_G0042570 | 8426.716 | 7957.643 | 9842.397 | 18289.79 | 18509.35 | 14627.34 | 16234.56 | 20304.48 | 14857.76 | 5.17E-08 | -0.971459962 | YMR217W | GUA1 | n |
| W303.asm02.HP0_G0042580 | 3302.705 | 3232.688 | 3063.421 | 1909.775 | 1906.65 | 1865.736 | 1361.624 | 1787.877 | 1789.342 | 1.97E-08 | 0.756423203 | YMR218C | TRS130 | n |
| W303.asm02.HP0_G0042600 | 3461.54 | 3420.751 | 3693.998 | 2475.274 | 2623.488 | 3134.376 | 2083.791 | 2446.116 | 2472.306 | 0.010199441 | 0.361349349 | YMR220W | ERG8 | n |
| W303.asm02.HP0_G0042610 | 2580.536 | 2783.116 | 2861.851 | 1270.636 | 1229.733 | 1406.899 | 1570.25 | 2483.699 | 1777.381 | 2.09E-08 | 1.073867001 | YMR221C | FMP42 | n |
| W303.asm02.HP0_G0042620 | 2880.204 | 2462.629 | 2730.346 | 1574.921 | 1824.205 | 1172.922 | 409.8451 | 487.5052 | 588.4735 | 0.000140146 | 0.820517356 | YMR222C | FSH2 | n |
| W303.asm02.HP0_G0042630 | 995.3648 | 1014.875 | 943.1714 | 1296.34 | 1294.821 | 976.9287 | 1099.916 | 1001.855 | 1027.436 | 0.079693253 | -0.27269646 | YMR223W | UBP8 | n |
| W303.asm02.HP0_G0042640 | 1273.855 | 1445.529 | 1263.311 | 1983.415 | 1821.601 | 2707.445 | 4394.724 | 2862.751 | 4076.255 | 0.002095554 | -0.709573398 | YMR224C | MRE11 | n |
| W303.asm02.HP0_G0042650 | 2040.498 | 1979.674 | 1926.225 | 964.266 | 967.6444 | 711.8057 | 509.8374 | 438.1104 | 471.2572 | 6.81E-14 | 1.169599064 | YMR225C | MRPL44 | n |
| W303.asm02.HP0_G0042660 | 9471.849 | 9622.393 | 9404.766 | 1251.184 | 1418.055 | 2648.191 | 843.1451 | 2205.585 | 1475.968 | 2.31E-10 | 2.422104223 | YMR226C | ORA1 | n |
| W303.asm02.HP0_G0042670 | 2194.038 | 2222.264 | 2189.235 | 5330.557 | 5212.262 | 5284.227 | 5002.085 | 4876.125 | 5220.908 | 9.88E-87 | -1.260738333 | YMR227C | TAF7 | n |
| W303.asm02.HP0_G0042680 | 2188.744 | 2233.392 | 2068.51 | 461.9862 | 369.7009 | 178.5212 | 170.3573 | 95.56819 | 161.4714 | 2.27E-14 | 2.683479704 | YMR228W | MTF1 | n |
| W303.asm02.HP0_G0042690 | 6667.885 | 6617.83 | 7421.411 | 11890.76 | 10961.72 | 3454.195 | 12513.85 | 6417.028 | 13289.69 | 0.477587954 | -0.345298297 | YMR229C | RRP5 | n |
| W303.asm02.HP0_G0042720 | 1991.788 | 1983.012 | 1845.382 | 1338.023 | 1232.336 | 1862.698 | 911.0411 | 1281.043 | 966.4361 | 0.077634573 | 0.39263477 | YMR231W | PEP5 | n |
| W303.asm02.HP0_G0042730 | 619.4557 | 695.5009 | 543.2667 | 93.09196 | 68.55956 | 85.8421 | 122.2128 | 120.2656 | 77.74548 | 3.09E-36 | 2.904631744 | YMR232W | FUS2 | n |
| W303.asm02.HP0_G0042740 | 1359.626 | 1299.752 | 1284.869 | 947.5928 | 868.7104 | 713.3251 | 812.2833 | 617.435 | 772.6705 | 0.000131219 | 0.640697338 | YMR233W | TRI1 | n |
| W303.asm02.HP0_G0042750 | 1665.648 | 1537.891 | 1639.501 | 1226.869 | 1173.323 | 1315.739 | 761.6699 | 802.1285 | 721.2388 | 0.000143166 | 0.381996645 | YMR234W | RNH1 | n |
| W303.asm02.HP0_G0042760 | 5814.413 | 5459.404 | 5977.012 | 7002.739 | 7744.627 | 7003.348 | 6632.824 | 5099.476 | 5784.264 | 0.01203432 | -0.334326183 | YMR235C | RNA1 | n |
| W303.asm02.HP0_G0042770 | 1938.844 | 1806.077 | 1865.862 | 1413.747 | 1595.095 | 1642.395 | 1233.239 | 1150.04 | 1010.691 | 0.030885265 | 0.270914169 | YMR236W | TAF9 | n |
| W303.asm02.HP0_G0042790 | 4541.617 | 4694.909 | 4375.237 | 3909.862 | 3642.335 | 3830.989 | 2971.377 | 4110.506 | 3427.978 | 0.081575336 | 0.257880932 | YMR238W | DFG5 | n |
| W303.asm02.HP0_G0042800 | 1563.993 | 1640.269 | 1886.343 | 4141.898 | 4002.49 | 3130.578 | 3867.604 | 3454.414 | 3618.155 | 4.66E-14 | -1.147216354 | YMR239C | RNT1 | n |
| W303.asm02.HP0_G0042810 | 1116.079 | 1073.853 | 1011.08 | 1485.998 | 1444.09 | 1457.796 | 2071.446 | 1913.511 | 2056.069 | 2.26E-07 | -0.455102212 | YMR240C | CUS1 | n |
| W303.asm02.HP0_G0042820 | 1409.394 | 1149.524 | 1416.374 | 3496.506 | 3663.163 | 4088.515 | 4168.816 | 3745.414 | 2943.564 | 4.66E-18 | -1.50044676 | YMR241W | YHM2 | n |
| W303.asm02.HP0_G0042840 | 6972.848 | 7127.493 | 7350.269 | 4155.792 | 4090.142 | 4459.231 | 4936.658 | 5528.996 | 5175.457 | 3.60E-19 | 0.755542014 | YMR243C | ZRC1 | n |
| W303.asm02.HP0_G0042850 | 96.35978 | 84.5729 | 104.5573 | 123.6595 | 96.33052 | 101.0354 | 153.0747 | 179.3246 | 132.7654 | 0.496232222 | -0.171512045 | YMR244W | | n |
| W303.asm02.HP0_G0042860 | 2649.365 | 2144.368 | 2163.366 | 1668.013 | 1762.588 | 1634.798 | 465.3964 | 751.66 | 637.5129 | 0.02404924 | 0.457912507 | YMR244C- | COA6 | n |
| W303.asm02.HP0_G0042880 | 3183.05 | 3178.161 | 3037.551 | 4795.626 | 4450.296 | 4432.643 | 5167.504 | 4608.749 | 4836.965 | 3.24E-10 | -0.5415066 | YMR247C | RKR1 | n |
| W303.asm02.HP0_G0042890 | 13069.99 | 11408.44 | 12405.67 | 1743.043 | 1674.936 | 5898.036 | 5646.48 | 14708.91 | 10603.29 | 0.000108435 | 1.985196547 | YMR250W | GAD1 | n |
| W303.asm02.HP0_G0042920 | 1245.265 | 1135.057 | 1223.428 | 1183.796 | 1337.345 | 1139.497 | 1340.638 | 1357.283 | 1288.183 | 0.858482049 | -0.021825784 | YMR252C | MLO1 | n |
| W303.asm02.HP0_G0042950 | 1450.691 | 1199.6 | 1252.532 | 5328.473 | 5920.422 | 5743.065 | 3087.417 | 5872.612 | 5470.89 | 4.09E-18 | -2.122200251 | YMR256C | COX7 | n |
| W303.asm02.HP0_G0042960 | 1057.84 | 980.378 | 1000.301 | 1700.665 | 1443.222 | 1301.305 | 1086.336 | 747.3648 | 1251.104 | 0.013363649 | -0.548977532 | YMR257C | PET111 | n |
| W303.asm02.HP0_G0042970 | 3455.186 | 2991.21 | 3196.004 | 3310.322 | 3243.127 | 3776.293 | 9849.861 | 7274.995 | 8605.827 | 0.555339794 | -0.099337085 | YMR258C | ROY1 | n |
| W303.asm02.HP0_G0042980 | 2272.397 | 2344.672 | 2497.518 | 3579.872 | 3320.365 | 3054.611 | 2420.802 | 2553.496 | 2661.288 | 3.44E-06 | -0.48473249 | YMR259C | TRM732 | n |
| W303.asm02.HP0_G0042990 | 8619.435 | 9389.818 | 8838.863 | 9357.132 | 10311.7 | 9359.068 | 7443.873 | 6561.991 | 6605.974 | 0.290922221 | -0.112569295 | YMR260C | TIF11 | n |
| W303.asm02.HP0_G0043000 | 10936.31 | 9551.174 | 10273.56 | 9203.599 | 10190.21 | 18768.27 | 53913.14 | 39085.24 | 42519.6 | 0.314120756 | -0.311034319 | YMR261C | TPS3 | n |
| W303.asm02.HP0_G0043010 | 2610.185 | 2183.316 | 2473.804 | 795.45 | 768.0406 | 828.7941 | 1038.192 | 1199.435 | 1422.144 | 2.57E-26 | 1.60295194 | YMR262W | | n |
| W303.asm02.HP0_G0043020 | 1866.838 | 1743.76 | 1665.371 | 790.587 | 844.4108 | 776.3772 | 1555.436 | 1036.217 | 1263.065 | 2.76E-11 | 1.12979694 | YMR263W | SAP30 | n |
| W303.asm02.HP0_G0043030 | 2648.306 | 2472.645 | 2543.868 | 3120.665 | 3564.229 | 3923.668 | 7140.192 | 5429.132 | 5920.617 | 0.002305857 | -0.468807893 | YMR264W | CUE1 | n |
| W303.asm02.HP0_G0043040 | 1373.392 | 1259.691 | 1303.193 | 980.9392 | 914.706 | 1049.857 | 1424.582 | 1072.726 | 1302.536 | 0.004064098 | 0.418118395 | YMR265C | | n |
| W303.asm02.HP0_G0043050 | 4136.058 | 4362.181 | 4124.084 | 19325.61 | 18395.66 | 21600.3 | 38340.27 | 28437.44 | 33204.5 | 4.35E-57 | -2.2326031 | YMR266W | RSN1 | n |
| W303.asm02.HP0_G0043060 | 6869.076 | 6957.234 | 6772.509 | 3248.493 | 3165.022 | 4181.954 | 2144.28 | 3982.724 | 2932.799 | 6.99E-05 | 0.959092602 | YMR267W | PPA2 | n |
| W303.asm02.HP0_G0043070 | 733.8168 | 755.5921 | 695.252 | 1201.859 | 1097.821 | 1169.124 | 2526.966 | 2056.327 | 2538.091 | 1.99E-07 | -0.667406383 | YMR268C | PRP24 | n |
| W303.asm02.HP0_G0043080 | 1417.865 | 1093.884 | 1252.532 | 1018.454 | 808.8292 | 183.0792 | 1507.292 | 725.8888 | 1307.32 | 0.105169074 | 0.904890652 | YMR269W | TMA23 | n |
| W303.asm02.HP0_G0043090 | 1768.361 | 1717.053 | 1737.591 | 783.6398 | 707.2917 | 637.3586 | 309.8527 | 344.6898 | 395.9039 | 6.94E-22 | 1.294822335 | YMR270C | RRN9 | n |
| W303.asm02.HP0_G0043110 | 6549.289 | 6316.261 | 6953.598 | 6039.862 | 6320.497 | 7592.848 | 32271.6 | 24158.35 | 22666.99 | 0.962479714 | -0.009718804 | YMR272C | SCS7 | n |
| W303.asm02.HP0_G0043120 | 2050.028 | 2008.606 | 1927.303 | 2976.859 | 2772.757 | 3216.42 | 2065.273 | 1429.228 | 1754.656 | 0.00030376 | -0.582960047 | YMR273C | ZDS1 | n |
| W303.asm02.HP0_G0043130 | 560.1574 | 559.7391 | 552.9679 | 797.5341 | 710.763 | 1057.453 | 630.8157 | 946.0177 | 905.4358 | 0.00665691 | -0.617297837 | YMR274C | RCE1 | n |
| W303.asm02.HP0_G0043150 | 12340.41 | 12525.69 | 13449.08 | 6028.747 | 6728.383 | 8942.772 | 4546.564 | 6946.412 | 5923.01 | 0.000143166 | 0.820235026 | YMR276W | DSK2 | n |
| W303.asm02.HP0_G0043160 | 3939.103 | 3185.95 | 4228.641 | 3579.178 | 3455.749 | 3044.736 | 2136.873 | 2103.574 | 2216.344 | 0.261559075 | 0.171760731 | YMR277W | FCP1 | n |

| | | | | | | | | | | | | | | |
|---|---|---|---|---|---|---|---|---|---|---|---|---|---|---|
| W303.asm02.HP0_G0043170 | 3923.22 | 3956.009 | 3662.739 | 4109.246 | 3914.838 | 6147.966 | 4957.644 | 7031.242 | 6077.304 | 0.208150032 | -0.296192522 | YMR278W | PRM15 | n |
| W303.asm02.HP0_G0043190 | 1945.197 | 2207.798 | 1676.15 | 7474.451 | 7111.102 | 11375.22 | 7366.101 | 7763.573 | 6952.838 | 6.10E-24 | -2.155031735 | YMR280C | CAT8 | n |
| W303.asm02.HP0_G0043200 | 1927.196 | 1786.046 | 1631.956 | 1320.655 | 1597.698 | 1653.03 | 2312.168 | 2146.526 | 1997.461 | 0.125652253 | 0.225976519 | YMR281W | GPI12 | n |
| W303.asm02.HP0_G0043210 | 1451.75 | 1384.325 | 1344.154 | 2589.207 | 2299.783 | 1961.454 | 1435.692 | 1321.848 | 1760.636 | 1.84E-05 | -0.712688044 | YMR282C | AEP2 | n |
| W303.asm02.HP0_G0043220 | 1686.826 | 1856.153 | 1696.631 | 2195.998 | 1877.143 | 2036.661 | 1475.195 | 1812.574 | 1931.676 | 0.13940968 | -0.221959656 | YMR283C | RIT1 | n |
| W303.asm02.HP0_G0043230 | 2670.543 | 3027.933 | 2405.895 | 1850.029 | 1712.253 | 2141.495 | 2822.005 | 2904.629 | 2980.642 | 0.00033233 | 0.506549898 | YMR284W | YKU70 | n |
| W303.asm02.HP0_G0043240 | 3022.097 | 2684.077 | 2852.15 | 1797.231 | 1844.165 | 2177.199 | 1923.309 | 2655.507 | 2140.991 | 0.000733507 | 0.556680487 | YMR285C | NGL2 | n |
| W303.asm02.HP0_G0043250 | 2646.188 | 2192.219 | 2145.041 | 758.63 | 726.3842 | 601.6544 | 295.0391 | 322.14 | 350.4527 | 2.50E-29 | 1.742748836 | YMR286W | MRPL33 | n |
| W303.asm02.HP0_G0043260 | 1175.378 | 1105.012 | 1165.221 | 3718.815 | 2997.528 | 2747.707 | 7164.882 | 3737.898 | 7324.82 | 6.96E-08 | -1.457735763 | YMR287C | DSS1 | y |
| W303.asm02.HP0_G0043280 | 1682.59 | 1828.333 | 1694.475 | 2430.117 | 2161.796 | 1886.247 | 3131.858 | 2193.773 | 2474.698 | 0.087167089 | -0.315693939 | YMR288W | HSH155 | n |
| W303.asm02.HP0_G0043290 | 3921.102 | 3637.748 | 3732.803 | 1253.268 | 1291.35 | 930.5892 | 1345.576 | 1006.151 | 1173.359 | 3.35E-22 | 1.700233558 | YMR289W | ABZ2 | n |
| W303.asm02.HP0_G0043300 | 4408.195 | 4295.413 | 5066.178 | 7709.96 | 7755.909 | 3167.801 | 9894.302 | 3816.285 | 7843.921 | 0.328166833 | -0.436395696 | YMR290C | HAS1 | n |
| W303.asm02.HP0_G0043310 | 2861.144 | 3792.427 | 2784.242 | 4411.447 | 3826.318 | 4510.888 | 23909.27 | 9956.273 | 21502.01 | 0.199483723 | -0.433861868 | YMR291W | TDA1 | n |
| W303.asm02.HP0_G0043330 | 1908.135 | 1931.823 | 1764.539 | 3716.731 | 3706.555 | 4681.813 | 3317.029 | 3673.47 | 3500.939 | 3.62E-17 | -1.111011278 | YMR293C | HER2 | n |
| W303.asm02.HP0_G0043340 | 1823.424 | 1810.528 | 1882.031 | 1240.763 | 1107.367 | 1248.889 | 791.2973 | 1122.121 | 923.3771 | 6.47E-05 | 0.61655678 | YMR294W | JNM1 | n |
| W303.asm02.HP0_G0043350 | 9101.234 | 9099.377 | 9361.65 | 2321.047 | 2567.078 | 2461.313 | 6261.248 | 4129.835 | 4998.436 | 4.81E-31 | 1.907077689 | YMR295C | GSR1 | n |
| W303.asm02.HP0_G0043360 | 7542.535 | 6735.787 | 7492.553 | 11683.04 | 12370.23 | 14881.07 | 15635.84 | 14648.78 | 15086.21 | 2.44E-11 | -0.838618788 | YMR296C | LCB1 | n |
| W303.asm02.HP0_G0043370 | 33443.2 | 31396.58 | 32425.69 | 5017.935 | 5898.726 | 11605.4 | 9536.305 | 16471.02 | 10466.93 | 6.56E-10 | 2.110588575 | YMR297W | PRC1 | n |
| W303.asm02.HP0_G0043380 | 2725.605 | 2542.751 | 2494.284 | 2282.837 | 2548.853 | 2864.695 | 4179.926 | 3519.916 | 3197.133 | 0.944851079 | 0.012465779 | YMR298W | LIP1 | n |
| W303.asm02.HP0_G0043390 | 1536.462 | 1537.891 | 1470.269 | 712.084 | 670.8423 | 772.5789 | 329.6043 | 424.151 | 336.0997 | 9.00E-16 | 1.075779042 | YMR299C | DYN3 | n |
| W303.asm02.HP0_G0043400 | 1607.408 | 1259.691 | 1672.917 | 6242.719 | 5903.065 | 7269.99 | 9673.331 | 12451.78 | 11270.7 | 6.63E-35 | -2.096377784 | YMR300C | ADE4 | n |
| W303.asm02.HP0_G0043410 | 3673.32 | 3231.575 | 3935.45 | 4594.158 | 4167.38 | 3162.484 | 7636.451 | 4679.62 | 7660.92 | 0.61131964 | -0.137461441 | YMR301C | ATM1 | n |
| W303.asm02.HP0_G0043420 | 6155.378 | 6247.267 | 5844.429 | 5772.396 | 5904.8 | 7924.062 | 4632.978 | 5424.837 | 4805.867 | 0.558555718 | -0.103296525 | YMR302C | YME2 | n |
| W303.asm02.HP0_G0043440 | 6641.413 | 7003.972 | 6230.321 | 4046.027 | 4094.481 | 6900.793 | 5911.892 | 8018.064 | 6400.247 | 0.119339818 | 0.402056229 | YMR304W | UBP15 | n |
| W303.asm02.HP0_G0043460 | 1332.095 | 1278.609 | 1334.453 | 1210.89 | 996.2832 | 966.2934 | 2299.823 | 3630.518 | 2288.109 | 0.174879438 | 0.313813777 | YMR306W | FKS3 | n |
| W303.asm02.HP0_G0043470 | 30031.43 | 33391.83 | 33459.41 | 26023.37 | 26437.95 | 27756.63 | 22153.85 | 29149.37 | 26075.83 | 0.044687772 | 0.272311099 | YMR307W | GAS1 | n |
| W303.asm02.HP0_G0043480 | 7530.888 | 7244.337 | 8377.518 | 16300.82 | 16347.55 | 17526.22 | 7236.481 | 9371.052 | 8392.924 | 9.53E-18 | -1.115766786 | YMR308C | PSE1 | n |
| W303.asm02.HP0_G0043490 | 9971.649 | 10364.63 | 10960.19 | 25199.44 | 24560.81 | 22081.17 | 16297.51 | 16634.23 | 16218.9 | 6.00E-43 | -1.198823467 | YMR309C | NIP1 | n |
| W303.asm02.HP0_G0043500 | 1451.75 | 1734.857 | 1883.109 | 2450.264 | 2153.985 | 696.6125 | 2554.125 | 806.4237 | 1570.459 | 0.913535645 | -0.064328821 | YMR310C | UPA2 | n |
| W303.asm02.HP0_G0043510 | 9135.119 | 9629.07 | 8497.166 | 3390.215 | 3317.762 | 4752.462 | 6248.903 | 5545.103 | 6188.54 | 2.42E-13 | 1.250146644 | YMR311C | GLC8 | n |
| W303.asm02.HP0_G0043520 | 1478.223 | 1532.327 | 1513.386 | 1425.558 | 1477.936 | 386.6693 | 1760.359 | 636.7634 | 1344.399 | 0.399873602 | 0.459465823 | YMR312W | ELP6 | n |
| W303.asm02.HP0_G0043530 | 2348.637 | 2359.139 | 2147.197 | 1500.587 | 1641.09 | 2569.186 | 3487.386 | 3085.027 | 3042.839 | 0.266973543 | 0.263430011 | YMR313C | TGL3 | n |
| W303.asm02.HP0_G0043540 | 10120.95 | 9769.283 | 9165.47 | 5239.549 | 6296.197 | 6749.62 | 5286.014 | 7898.872 | 6054.579 | 0.000381576 | 0.668176336 | YMR314W | PRE5 | n |
| W303.asm02.HP0_G0043550 | 7512.886 | 7039.581 | 7090.493 | 3368.679 | 3940.005 | 5569.861 | 19371.35 | 13561.02 | 11222.86 | 0.004781106 | 0.748943509 | YMR315W | | n |
| W303.asm02.HP0_G0043580 | 14631.86 | 13980.12 | 14963.55 | 5348.62 | 5837.109 | 3789.207 | 1313.479 | 6467.497 | 1917.323 | 0.002928985 | 1.5409776 | YMR318C | ADH6 | n |
| W303.asm02.HP0_G0043590 | 4812.695 | 4651.51 | 4782.687 | 3725.763 | 3396.736 | 5167.998 | 17494.95 | 6792.858 | 17874.28 | 0.587866157 | 0.213088148 | YMR319C | FET4 | n |
| W303.asm02.HP0_G0043660 | 717.9333 | 709.9673 | 642.4344 | 6324.696 | 6229.374 | 7842.018 | 4162.643 | 10331.03 | 8696.729 | 1.19E-26 | -3.300362708 | YNL334C | SNO2 | y |
| W303.asm02.HP0_G0043670 | 2130.504 | 2043.103 | 1982.277 | 22756.82 | 22286.2 | 27662.43 | 6751.333 | 25135.51 | 23320.06 | 2.83E-18 | -3.562022565 | YNL333W | SNZ2 | y |
| W303.asm02.HP0_G0043700 | 4002.637 | 4179.682 | 3940.839 | 4938.042 | 4899.839 | 4806.398 | 3512.076 | 3140.865 | 3349.036 | 0.000592073 | -0.27261908 | YNL330C | RPD3 | n |
| W303.asm02.HP0_G0043710 | 2421.701 | 2623.986 | 2502.907 | 1464.462 | 1390.284 | 1614.287 | 735.746 | 1083.464 | 803.7687 | 1.08E-05 | 0.756124639 | YNL329C | PEX6 | n |
| W303.asm02.HP0_G0043720 | 175.7772 | 189.1762 | 182.1668 | 761.4089 | 831.3931 | 520.3703 | 264.1772 | 513.2764 | 452.1199 | 2.83E-11 | -1.949393571 | YNL328C | MDJ2 | n |
| W303.asm02.HP0_G0043730 | 17038.74 | 19071.19 | 19215.9 | 16219.54 | 16771.06 | 15175.06 | 14391.49 | 21963.5 | 24297.26 | 0.36874924 | 0.199950725 | YNL327W | EGT2 | n |
| W303.asm02.HP0_G0043740 | 478.6222 | 452.9102 | 436.5536 | 477.9647 | 438.2605 | 450.4811 | 759.201 | 932.0583 | 574.1205 | 0.99544114 | 0.001208531 | YNL326C | PFA3 | n |
| W303.asm02.HP0_G0043750 | 1285.503 | 1267.481 | 1254.687 | 1194.912 | 1125.592 | 1400.822 | 1140.653 | 1154.335 | 1191.3 | 0.803969704 | 0.032779999 | YNL325C | FIG4 | n |
| W303.asm02.HP0_G0043770 | 2086.03 | 1985.238 | 1972.576 | 4405.89 | 4478.935 | 4347.561 | 4924.313 | 4902.97 | 4704.2 | 6.04E-65 | -1.130384382 | YNL323W | LEM3 | n |
| W303.asm02.HP0_G0043780 | 4730.101 | 4643.72 | 4694.299 | 7598.805 | 6827.317 | 7379.382 | 10833.74 | 8898.58 | 9575.851 | 4.23E-09 | -0.632336856 | YNL322C | KRE1 | n |
| W303.asm02.HP0_G0043790 | 4196.416 | 4007.198 | 4268.524 | 2958.796 | 2900.33 | 3838.585 | 3278.761 | 4545.395 | 3291.624 | 0.063515598 | 0.362964628 | YNL321W | VNX1 | n |
| W303.asm02.HP0_G0043800 | 876.7681 | 692.1625 | 776.0953 | 1734.011 | 1894.5 | 2193.152 | 4442.869 | 4435.868 | 3804.744 | 1.50E-17 | -1.311528547 | YNL320W | | n |
| W303.asm02.HP0_G0043810 | 56.12163 | 79.0089 | 46.35014 | 254.2661 | 312.4233 | 281.8356 | 897.4619 | 1034.069 | 507.1398 | 8.98E-12 | -2.225862784 | YNL318C | HXT14 | n |
| W303.asm02.HP0_G0043820 | 2683.249 | 2799.808 | 2714.178 | 1966.741 | 1682.747 | 1969.051 | 1429.52 | 1713.785 | 1717.577 | 1.24E-05 | 0.544570973 | YNL317W | PFS2 | n |
| W303.asm02.HP0_G0043830 | 1041.956 | 1002.634 | 1091.923 | 2280.058 | 2297.179 | 2562.349 | 1175.218 | 1555.936 | 1291.771 | 1.33E-17 | -1.186648651 | YNL316C | PHA2 | n |
| W303.asm02.HP0_G0043840 | 11561.06 | 11549.77 | 11644.66 | 4410.753 | 4802.64 | 4298.942 | 1883.806 | 2275.382 | 1954.402 | 2.71E-38 | 1.363080437 | YNL315C | ATP11 | n |
| W303.asm02.HP0_G0043850 | 1019.719 | 1000.408 | 939.9376 | 398.7671 | 394.8683 | 440.6055 | 769.0767 | 653.9442 | 826.4943 | 1.34E-21 | 1.261801119 | YNL314W | DAL82 | n |
| W303.asm02.HP0_G0043860 | 1458.104 | 1770.467 | 2034.016 | 4441.32 | 4193.415 | 1776.096 | 5905.719 | 1805.058 | 4987.672 | 0.040471005 | -0.984242335 | YNL313C | EMW1 | n |

| | | | | | | | | | | | | | | |
|---|---|---|---|---|---|---|---|---|---|---|---|---|---|---|
| W303.asm02.HP0_G0043870 | 2041.557 | 1980.786 | 2068.51 | 2293.258 | 2659.069 | 2902.679 | 1259.163 | 2258.201 | 1617.106 | 0.122438606 | -0.366908792 | YNL312W | RFA2 | n |
| W303.asm02.HP0_G0043880 | 649.1049 | 814.5706 | 635.967 | 1431.81 | 1342.552 | 1940.183 | 1777.641 | 2194.847 | 1715.185 | 1.62E-08 | -1.167247199 | YNL311C | SKP2 | n |
| W303.asm02.HP0_G0043890 | 1982.258 | 1744.873 | 1845.382 | 971.2132 | 932.9307 | 648.7536 | 1003.627 | 630.3206 | 769.0822 | 1.15E-06 | 1.126235723 | YNL310C | ZIM17 | n |
| W303.asm02.HP0_G0043900 | 566.5108 | 553.0623 | 617.6425 | 2209.197 | 2237.298 | 2376.231 | 1506.057 | 1316.479 | 952.0831 | 7.38E-27 | -1.973530923 | YNL309W | STB1 | n |
| W303.asm02.HP0_G0043910 | 4415.608 | 4898.552 | 5269.903 | 5152.015 | 4521.46 | 1499.578 | 6909.346 | 3058.182 | 5711.303 | 0.431447846 | 0.384367062 | YNL308C | KRI1 | n |
| W303.asm02.HP0_G0043920 | 10092.36 | 10699.59 | 10780.18 | 5725.85 | 5645.316 | 7670.334 | 5002.085 | 5697.582 | 5180.241 | 2.13E-06 | 0.729473981 | YNL307C | MCK1 | n |
| W303.asm02.HP0_G0043930 | 5834.532 | 5529.51 | 5335.655 | 2673.962 | 2823.092 | 1962.214 | 2325.747 | 1530.165 | 1980.716 | 2.09E-08 | 1.162783928 | YNL306W | MRPS18 | n |
| W303.asm02.HP0_G0043940 | 5017.062 | 5288.032 | 4704 | 956.6241 | 1009.301 | 1036.942 | 2601.035 | 2262.496 | 2768.935 | 1.92E-91 | 2.32159502 | YNL305C | BXI1 | n |
| W303.asm02.HP0_G0043950 | 774.055 | 810.1194 | 807.3547 | 3481.223 | 3485.256 | 3689.691 | 2802.254 | 3077.511 | 3290.428 | 1.98E-106 | -2.155790122 | YNL304W | YPT11 | n |
| W303.asm02.HP0_G0043970 | 4696.216 | 4129.606 | 4988.568 | 16110.47 | 19227.92 | 17114.48 | 17309.78 | 13979.8 | 12921.3 | 8.75E-35 | -1.924799812 | YOL120C | RPL18A | y |
| W303.asm02.HP0_G0043990 | 328.2586 | 334.9532 | 377.2685 | 4549.696 | 4009.432 | 2448.399 | 6983.414 | 3665.953 | 5924.206 | 1.41E-29 | -3.403176096 | YNL299W | TRF5 | n |
| W303.asm02.HP0_G0044000 | 1093.842 | 1081.643 | 1221.272 | 9539.842 | 9099.329 | 11607.68 | 7366.101 | 6000.394 | 5407.497 | 1.61E-80 | -3.154552527 | YNL298W | CLA4 | n |
| W303.asm02.HP0_G0044010 | 3394.829 | 3398.495 | 3063.421 | 5383.355 | 5471.747 | 6978.279 | 3326.905 | 5495.708 | 4798.69 | 7.93E-05 | -0.855405802 | YNL297C | MON2 | n |
| W303.asm02.HP0_G0044020 | 1983.317 | 2399.2 | 1862.629 | 2400.939 | 2210.395 | 2522.846 | 1975.157 | 1987.604 | 2095.54 | 0.170989874 | -0.19227439 | YNL295W | MRX6 | n |
| W303.asm02.HP0_G0044030 | 4432.55 | 4392.227 | 4477.639 | 349.4422 | 273.3704 | 328.9348 | 412.314 | 314.6234 | 349.2566 | 2.88E-152 | 3.802936622 | YNL294C | RIM21 | n |
| W303.asm02.HP0_G0044040 | 2664.189 | 2725.251 | 2496.44 | 3854.285 | 3769.908 | 4382.505 | 4863.824 | 4644.185 | 5313.007 | 3.59E-08 | -0.606575543 | YNL293W | MSB3 | n |
| W303.asm02.HP0_G0044050 | 2662.071 | 2660.708 | 2643.036 | 2960.88 | 3089.519 | 1130.381 | 3493.559 | 1189.77 | 2803.622 | 0.766435113 | 0.149706017 | YNL292W | PUS4 | n |
| W303.asm02.HP0_G0044060 | 744.4058 | 770.0586 | 785.7965 | 2557.25 | 2889.916 | 2813.038 | 3382.456 | 2670.54 | 2817.975 | 1.67E-46 | -1.844258388 | YNL291C | MID1 | n |
| W303.asm02.HP0_G0044070 | 2911.971 | 2636.226 | 3023.538 | 2789.285 | 2725.893 | 3215.66 | 2859.04 | 2116.46 | 2545.267 | 0.880926145 | -0.02654367 | YNL290W | RFC3 | n |
| W303.asm02.HP0_G0044080 | 195.8963 | 132.4234 | 238.2181 | 395.9882 | 615.3003 | 754.347 | 214.7983 | 115.9704 | 80.13765 | 9.25E-05 | -1.639516936 | YNL289W | PCL1 | n |
| W303.asm02.HP0_G0044090 | 3591.784 | 4151.862 | 3784.543 | 4865.792 | 4718.46 | 5320.691 | 6293.344 | 4920.151 | 5252.006 | 0.007756103 | -0.370697102 | YNL288W | CAF40 | n |
| W303.asm02.HP0_G0044100 | 13643.91 | 13536.12 | 14234.88 | 16139.65 | 16219.11 | 20156.18 | 13549.58 | 16396.92 | 13959.5 | 0.016391795 | -0.34258237 | YNL287W | SEC21 | n |
| W303.asm02.HP0_G0044120 | 3342.943 | 3311.697 | 3254.211 | 5931.486 | 6180.774 | 4329.329 | 2440.553 | 2455.78 | 2496.228 | 2.73E-06 | -0.730512845 | YNL284C | MRPL10 | n |
| W303.asm02.HP0_G0044130 | 1464.457 | 1462.221 | 1735.435 | 9327.259 | 8266.2 | 8266.67 | 14006.33 | 10562.97 | 11834.06 | 7.64E-62 | -2.471690234 | YNL283C | WSC2 | n |
| W303.asm02.HP0_G0044140 | 668.1651 | 676.5832 | 730.8231 | 1310.235 | 1342.552 | 864.4983 | 670.3189 | 737.7006 | 843.2394 | 0.000265509 | -0.760834596 | YNL282W | POP3 | n |
| W303.asm02.HP0_G0044150 | 3580.137 | 3309.471 | 3480.572 | 2433.591 | 2674.691 | 2314.698 | 1678.883 | 1680.497 | 1728.342 | 2.19E-07 | 0.48274711 | YNL281W | HCH1 | n |
| W303.asm02.HP0_G0044160 | 3255.055 | 2874.366 | 3250.977 | 1745.822 | 1709.65 | 1835.35 | 4370.035 | 2655.507 | 2817.975 | 9.67E-05 | 0.826151982 | YNL280C | ERG24 | n |
| W303.asm02.HP0_G0044170 | 390.7336 | 383.9165 | 392.3593 | 82.67122 | 85.04857 | 72.16814 | 67.89602 | 183.6198 | 39.47078 | 2.18E-06 | 2.282562739 | YNL279W | PRM1 | n |
| W303.asm02.HP0_G0044200 | 1623.292 | 1170.667 | 1381.881 | 631.497 | 617.036 | 769.5403 | 809.8143 | 1137.154 | 971.2205 | 1.57E-07 | 1.049070841 | YNL277W | MET2 | n |
| W303.asm02.HP0_G0044210 | 339.9065 | 349.4196 | 331.9963 | 1326.908 | 1341.685 | 1421.332 | 2335.623 | 1960.759 | 1961.578 | 8.87E-62 | -2.00174958 | YNL275W | BOR1 | n |
| W303.asm02.HP0_G0044220 | 7559.478 | 6998.408 | 6718.614 | 4512.876 | 5414.469 | 5028.22 | 2804.723 | 6650.043 | 4234.139 | 0.09605596 | 0.508606917 | YNL274C | GOR1 | n |
| W303.asm02.HP0_G0044230 | 807.9397 | 796.7658 | 846.1595 | 3155.401 | 3355.079 | 2566.907 | 1202.377 | 1379.833 | 1525.008 | 2.63E-35 | -1.888872103 | YNL273W | TOF1 | n |
| W303.asm02.HP0_G0044240 | 3066.571 | 3301.682 | 3204.627 | 4147.455 | 4010.3 | 4368.072 | 2804.723 | 3242.876 | 3030.878 | 5.25E-05 | -0.38800145 | YNL272C | SEC2 | n |
| W303.asm02.HP0_G0044250 | 4751.279 | 4986.463 | 4822.57 | 6251.751 | 6202.47 | 7167.436 | 6022.994 | 6871.246 | 5897.892 | 0.000124294 | -0.43044565 | YNL271C | BNI1 | n |
| W303.asm02.HP0_G0044280 | 4827.519 | 4874.07 | 4880.777 | 8770.791 | 8748.72 | 9454.786 | 20283.63 | 16458.13 | 16505.96 | 1.78E-14 | -0.887370574 | YNL268W | LYP1 | n |
| W303.asm02.HP0_G0044290 | 6294.094 | 6834.826 | 6409.254 | 6879.774 | 6785.661 | 8050.166 | 4176.222 | 5266.989 | 4643.199 | 0.287290844 | -0.15247047 | YNL267W | PIK1 | n |
| W303.asm02.HP0_G0044300 | 2731.959 | 2773.101 | 2598.841 | 1561.722 | 1649.769 | 2474.228 | 3329.374 | 3713.2 | 3506.919 | 0.011139818 | 0.511244477 | YNL265C | IST1 | n |
| W303.asm02.HP0_G0044320 | 3704.028 | 3900.369 | 3974.255 | 2712.867 | 2784.907 | 3088.796 | 3712.061 | 4033.193 | 3616.959 | 1.28E-05 | 0.431304623 | YNL263C | YIF1 | n |
| W303.asm02.HP0_G0044330 | 3100.455 | 2970.067 | 3052.642 | 4257.915 | 4624.733 | 3935.063 | 1473.961 | 2104.648 | 1697.244 | 0.002175676 | -0.490459204 | YNL262W | POL2 | n |
| W303.asm02.HP0_G0044340 | 1705.886 | 1606.885 | 1716.033 | 2398.16 | 2402.188 | 2415.733 | 2756.578 | 2347.327 | 2351.502 | 3.53E-07 | -0.520928042 | YNL261W | ORC5 | n |
| W303.asm02.HP0_G0044350 | 1108.667 | 1176.231 | 1109.17 | 578.6985 | 598.8113 | 493.782 | 703.6497 | 1263.863 | 1045.378 | 6.84E-06 | 1.02216734 | YNL260C | LTO1 | n |
| W303.asm02.HP0_G0044360 | 378.0268 | 304.9076 | 312.5939 | 1511.702 | 1803.377 | 1632.519 | 1388.782 | 2049.884 | 1965.167 | 1.80E-32 | -2.312729163 | YNL259C | ATX1 | n |
| W303.asm02.HP0_G0044370 | 3048.569 | 3016.805 | 2931.916 | 4071.731 | 4037.203 | 4431.124 | 3887.356 | 4664.587 | 4534.356 | 8.96E-06 | -0.479009342 | YNL258C | DSL1 | n |
| W303.asm02.HP0_G0044380 | 2997.742 | 3224.898 | 2904.968 | 5013.072 | 4824.337 | 6418.406 | 4652.729 | 5416.247 | 4163.57 | 5.69E-07 | -0.832700224 | YNL257C | SIP3 | n |
| W303.asm02.HP0_G0044390 | 3561.076 | 3795.765 | 4156.422 | 3230.43 | 2860.409 | 2256.964 | 2049.225 | 6181.866 | 4266.433 | 0.22672782 | 0.463812812 | YNL256W | FOL1 | n |
| W303.asm02.HP0_G0044400 | 1064.193 | 960.3476 | 963.6517 | 1249.794 | 1155.098 | 1220.781 | 1035.723 | 1056.619 | 932.9458 | 0.009676191 | -0.279198893 | YNL254C | RTC4 | n |
| W303.asm02.HP0_G0044410 | 1343.742 | 1483.364 | 1531.71 | 3091.487 | 3112.951 | 2825.952 | 2501.042 | 2878.858 | 3027.289 | 9.20E-19 | -1.050890823 | YNL253W | TEX1 | n |
| W303.asm02.HP0_G0044420 | 9195.476 | 8993.661 | 8812.993 | 3913.336 | 3614.564 | 3013.59 | 878.9448 | 1088.833 | 977.2009 | 1.99E-22 | 1.356952721 | YNL252C | MRPL17 | n |
| W303.asm02.HP0_G0044430 | 7655.838 | 7003.972 | 8026.119 | 4168.297 | 3925.252 | 2672.5 | 3471.338 | 2859.529 | 2181.658 | 5.00E-06 | 1.075312634 | YNL251C | NRD1 | n |
| W303.asm02.HP0_G0044440 | 2359.226 | 2550.541 | 2499.674 | 1929.921 | 1879.747 | 2394.463 | 1356.686 | 1896.331 | 1465.203 | 0.166639887 | 0.256054806 | YNL250W | RAD50 | n |
| W303.asm02.HP0_G0044450 | 860.8846 | 774.5098 | 810.5884 | 1519.344 | 1560.381 | 1287.632 | 1908.495 | 1693.382 | 1892.205 | 2.66E-11 | -0.836042246 | YNL249C | MPA43 | n |
| W303.asm02.HP0_G0044460 | 4647.506 | 4875.183 | 5460.693 | 6726.242 | 6838.599 | 3461.792 | 7883.345 | 4378.956 | 6816.485 | 0.596530913 | -0.184416897 | YNL248C | RPA49 | n |
| W303.asm02.HP0_G0044470 | 9418.904 | 9821.585 | 9896.293 | 7909.343 | 7762.851 | 6929.661 | 6072.373 | 6151.8 | 6730.366 | 0.000133842 | 0.36639295 | YNL247W | CRS1 | n |
| W303.asm02.HP0_G0044480 | 3192.58 | 3283.877 | 3274.691 | 3339.5 | 3435.789 | 2771.257 | 4010.803 | 3451.193 | 3978.176 | 0.831427366 | 0.030659383 | YNL246W | VPS75 | n |

| | | | | | | | | | | | | | |
|---|---|---|---|---|---|---|---|---|---|---|---|---|---|
| W303.asm02.HP0_G0044490 | 4389.135 | 4071.74 | 4195.226 | 7259.089 | 8369.473 | 7089.19 | 6811.823 | 6093.815 | 5842.872 | 1.55E-13 | -0.843861595 | YNL244C | SUI1 | n |
| W303.asm02.HP0_G0044500 | 8211.759 | 7489.153 | 8657.774 | 5720.293 | 5225.28 | 8014.462 | 6081.014 | 8647.311 | 7165.741 | 0.107259196 | 0.361449587 | YNL243W | SLA2 | n |
| W303.asm02.HP0_G0044510 | 2428.055 | 2460.404 | 1998.445 | 1433.894 | 1374.663 | 2381.549 | 7510.534 | 1849.083 | 5619.204 | 0.407776676 | 0.408068038 | YNL242W | ATG2 | n |
| W303.asm02.HP0_G0044520 | 22954.81 | 20975.19 | 23822.89 | 8678.394 | 8899.725 | 13342.75 | 17566.55 | 14518.85 | 20968.55 | 3.36E-07 | 1.131697382 | YNL241C | ZWF1 | n |
| W303.asm02.HP0_G0044530 | 2655.718 | 2257.874 | 2781.008 | 6028.747 | 5923.893 | 6502.729 | 18992.37 | 9460.177 | 19053.62 | 3.35E-06 | -1.262107799 | YNL240C | NAR1 | n |
| W303.asm02.HP0_G0044560 | 1453.868 | 1322.008 | 1327.985 | 814.902 | 964.173 | 1260.284 | 2466.477 | 3289.049 | 8718.259 | 0.354823206 | 0.433250469 | YNL237W | YTP1 | n |
| W303.asm02.HP0_G0044570 | 3637.317 | 3485.294 | 3538.779 | 4797.015 | 4256.768 | 5064.684 | 10680.66 | 4627.004 | 11519.49 | 0.227311825 | -0.405202842 | YNL236W | SIN4 | n |
| W303.asm02.HP0_G0044580 | 1177.495 | 1237.435 | 1142.585 | 5136.731 | 4683.746 | 5412.61 | 19056.56 | 7671.227 | 20424.34 | 1.97E-10 | -2.098286491 | YNL234W | | n |
| W303.asm02.HP0_G0044590 | 2846.32 | 2795.357 | 2982.577 | 4129.393 | 4097.952 | 4909.712 | 6999.462 | 3523.138 | 8843.847 | 0.057047535 | -0.607173597 | YNL233W | BNI4 | n |
| W303.asm02.HP0_G0044600 | 1776.832 | 1707.037 | 1924.07 | 2185.577 | 2225.148 | 1297.507 | 4611.991 | 1894.183 | 4603.729 | 0.849408677 | -0.077934389 | YNL232W | CSL4 | n |
| W303.asm02.HP0_G0044610 | 2451.351 | 2538.3 | 2766.995 | 7559.901 | 7751.569 | 8374.543 | 11862.05 | 8361.68 | 16101.69 | 4.57E-12 | -1.610530599 | YNL231C | PDR16 | n |
| W303.asm02.HP0_G0044620 | 1217.734 | 1119.478 | 1098.39 | 858.6691 | 858.2962 | 742.1923 | 2182.548 | 859.0399 | 2417.286 | 0.191387445 | 0.482423798 | YNL230C | ELA1 | n |
| W303.asm02.HP0_G0044630 | 6655.178 | 7663.863 | 7159.479 | 5705.704 | 5603.659 | 5279.669 | 15609.91 | 5671.811 | 13555.22 | 0.296800158 | 0.372661319 | YNL229C | URE2 | n |
| W303.asm02.HP0_G0044640 | 2380.404 | 2572.797 | 2655.971 | 3386.046 | 3174.568 | 2988.521 | 7473.5 | 2424.64 | 7506.625 | 0.425159521 | -0.327640243 | YNL227C | JJJ1 | n |
| W303.asm02.HP0_G0044650 | 1064.193 | 1147.298 | 1003.534 | 1092.094 | 932.0629 | 1198.751 | 1565.312 | 620.6564 | 1473.576 | 0.992697538 | -0.003642726 | YNL225C | CNM67 | n |
| W303.asm02.HP0_G0044660 | 2512.767 | 2607.294 | 2286.247 | 4363.512 | 3916.573 | 4491.137 | 10870.77 | 4839.616 | 12288.57 | 0.015474542 | -0.786091663 | YNL224C | SQS1 | n |
| W303.asm02.HP0_G0044670 | 1233.617 | 1306.429 | 1173.844 | 633.5811 | 684.7277 | 1263.322 | 3781.191 | 1675.128 | 2761.759 | 0.163280864 | 0.524607193 | YNL223W | ATG4 | n |
| W303.asm02.HP0_G0044680 | 989.0114 | 1047.146 | 1022.937 | 814.902 | 845.2786 | 604.693 | 2139.342 | 783.8739 | 1577.635 | 0.232338256 | 0.43372881 | YNL222W | SSU72 | n |
| W303.asm02.HP0_G0044690 | 2494.765 | 2362.477 | 2644.114 | 4488.561 | 3883.595 | 4420.488 | 14498.89 | 5920.933 | 18096.76 | 0.035536696 | -0.770103669 | YNL221C | POP1 | n |
| W303.asm02.HP0_G0044700 | 3359.886 | 3111.393 | 2885.565 | 11565.63 | 11401.71 | 15762.28 | 35201 | 27381.9 | 47160.41 | 1.35E-18 | -2.049342946 | YNL220W | ADE12 | n |
| W303.asm02.HP0_G0044710 | 4087.349 | 4252.014 | 4258.823 | 4052.974 | 4066.71 | 5027.46 | 22493.33 | 7955.784 | 19924.37 | 0.877497891 | -0.061538213 | YNL219C | ALG9 | n |
| W303.asm02.HP0_G0044720 | 626.868 | 646.5376 | 702.7974 | 1289.393 | 1249.693 | 1550.475 | 2810.895 | 1190.844 | 2487.855 | 0.000707522 | -1.049248695 | YNL218W | MGS1 | n |
| W303.asm02.HP0_G0044730 | 4848.697 | 5054.344 | 4753.584 | 2347.446 | 2246.844 | 2207.585 | 4765.066 | 2102.5 | 5470.89 | 0.000549025 | 1.107536536 | YNL217W | PPN2 | n |
| W303.asm02.HP0_G0044740 | 2222.628 | 2258.987 | 2340.143 | 4796.32 | 4617.79 | 5268.274 | 10337.48 | 4979.21 | 9724.166 | 3.74E-05 | -1.105881105 | YNL216W | RAP1 | n |
| W303.asm02.HP0_G0044750 | 4564.912 | 4454.544 | 4215.707 | 2752.465 | 2658.202 | 2588.937 | 8474.658 | 3294.418 | 7425.291 | 0.025088324 | 0.726369796 | YNL215W | IES2 | n |
| W303.asm02.HP0_G0044760 | 1054.663 | 1322.008 | 1067.131 | 672.4852 | 709.0273 | 620.646 | 1770.234 | 908.4347 | 1795.323 | 0.005083982 | 0.782478313 | YNL214W | PEX17 | n |
| W303.asm02.HP0_G0044770 | 1523.755 | 1478.913 | 1441.166 | 973.2973 | 791.4724 | 341.0894 | 2801.019 | 556.2284 | 2748.602 | 0.063622913 | 1.077360747 | YNL213C | RRG9 | n |
| W303.asm02.HP0_G0044780 | 7563.713 | 6977.265 | 7099.116 | 1012.896 | 967.6444 | 1377.272 | 3876.246 | 1418.49 | 3380.134 | 4.79E-15 | 2.688070432 | YNL212W | VID27 | n |
| W303.asm02.HP0_G0044790 | 463.7976 | 413.9621 | 390.2035 | 887.8472 | 731.5912 | 1027.067 | 3265.181 | 1195.139 | 2910.073 | 0.003228181 | -1.061663701 | YNL211C | MRX7 | n |
| W303.asm02.HP0_G0044800 | 133.4212 | 86.79851 | 114.2585 | 250.0978 | 222.1677 | 135.98 | 711.0565 | 294.2212 | 794.2 | 0.045790179 | -0.862429923 | YNL210W | MER1 | n |
| W303.asm02.HP0_G0044830 | 1278.091 | 1344.264 | 1463.802 | 2364.814 | 2271.144 | 766.5016 | 7493.252 | 899.8443 | 6162.226 | 0.572227073 | -0.402863078 | YNL207W | RIO2 | n |
| W303.asm02.HP0_G0044840 | 2171.801 | 2139.917 | 2313.195 | 2025.098 | 1838.09 | 2177.958 | 3579.972 | 1437.818 | 3660.018 | 0.717734912 | 0.133049621 | YNL206C | RTT106 | n |
| W303.asm02.HP0_G0044860 | 888.416 | 987.0548 | 785.7965 | 88.92367 | 72.03093 | 72.9278 | 380.2177 | 191.1364 | 325.3349 | 1.91E-37 | 3.506331789 | YNL202W | SPS19 | n |
| W303.asm02.HP0_G0044870 | 1128.786 | 1266.368 | 1112.403 | 2764.97 | 2109.725 | 1418.294 | 13397.74 | 3114.02 | 12409.37 | 0.092354376 | -0.843317108 | YNL201C | PSY2 | n |
| W303.asm02.HP0_G0044880 | 6537.641 | 6373.013 | 6148.399 | 709.3052 | 852.2214 | 1725.958 | 3389.863 | 4381.104 | 3626.528 | 1.48E-15 | 2.535414275 | YNL200C | NNR1 | n |
| W303.asm02.HP0_G0044890 | 3193.639 | 3465.263 | 3194.926 | 3156.095 | 3211.885 | 5266.755 | 5751.41 | 6612.46 | 7927.647 | 0.352518218 | -0.239712332 | YNL199C | GCR2 | n |
| W303.asm02.HP0_G0044950 | 11562.11 | 13050.93 | 11243.68 | 9834.402 | 8778.227 | 6741.264 | 13381.69 | 13887.45 | 14619.74 | 0.002913439 | 0.500008396 | YNL192W | CHS1 | n |
| W303.asm02.HP0_G0044960 | 2007.672 | 1396.566 | 1714.955 | 2079.285 | 2202.584 | 5626.076 | 15713.61 | 16342.16 | 19443.55 | 0.014844677 | -0.952658853 | YNL191W | DUG3 | n |
| W303.asm02.HP0_G0044980 | 8293.295 | 8468.418 | 8567.23 | 12204.77 | 11752.32 | 11479.29 | 9079.55 | 7778.607 | 8934.75 | 1.74E-07 | -0.484468072 | YNL189W | SRP1 | n |
| W303.asm02.HP0_G0044990 | 1579.877 | 1549.02 | 1592.073 | 223.6986 | 210.0179 | 197.5128 | 124.6818 | 107.38 | 138.7458 | 9.00E-122 | 2.902044524 | YNL188W | KAR1 | n |
| W303.asm02.HP0_G0045000 | 717.9333 | 699.9521 | 731.901 | 229.951 | 214.3571 | 126.864 | 167.8883 | 163.2176 | 105.2554 | 3.20E-12 | 1.912201933 | YNL187W | SWT21 | n |
| W303.asm02.HP0_G0045010 | 3588.608 | 3769.058 | 4134.863 | 4603.884 | 3924.384 | 1629.481 | 7005.635 | 2917.514 | 5180.241 | 0.702858886 | 0.178108289 | YNL186W | UBP10 | n |
| W303.asm02.HP0_G0045020 | 5943.598 | 5569.571 | 5350.746 | 1829.882 | 2067.201 | 1206.347 | 902.3998 | 760.2503 | 779.847 | 2.84E-16 | 1.724572618 | YNL185C | MRPL19 | n |
| W303.asm02.HP0_G0045030 | 3492.248 | 3960.46 | 3284.392 | 4061.311 | 4130.93 | 5076.079 | 4025.617 | 3620.853 | 3297.604 | 0.052832063 | -0.305430032 | YNL183C | NPR1 | n |
| W303.asm02.HP0_G0045040 | 1520.579 | 1471.123 | 1829.214 | 4089.794 | 3582.454 | 892.6059 | 5557.598 | 2173.371 | 4882.416 | 0.131905892 | -0.829117256 | YNL182C | IPI3 | n |
| W303.asm02.HP0_G0045050 | 5098.597 | 5355.913 | 5089.892 | 4621.252 | 4400.829 | 4630.156 | 5035.416 | 4987.801 | 5736.42 | 0.051125641 | 0.187149696 | YNL181W | PBR1 | n |
| W303.asm02.HP0_G0045080 | 17020.74 | 15707.19 | 17595.81 | 26008.78 | 31347.34 | 27380.59 | 24546.26 | 17915.28 | 17760.66 | 8.98E-06 | -0.751729112 | YNL178W | RPS3 | n |
| W303.asm02.HP0_G0045090 | 7942.799 | 8224.715 | 7413.866 | 7541.838 | 7938.155 | 5974.762 | 6601.962 | 4982.432 | 5503.184 | 0.453339085 | 0.136376314 | YNL177C | MRPL22 | n |
| W303.asm02.HP0_G0045100 | 4452.669 | 4792.836 | 4889.4 | 5684.862 | 4837.354 | 4265.517 | 9985.653 | 7410.293 | 8529.277 | 0.727464594 | -0.065210176 | YNL176C | TDA7 | n |
| W303.asm02.HP0_G0045110 | 3981.459 | 4160.764 | 4545.547 | 3718.121 | 3253.542 | 797.6478 | 4761.363 | 1802.91 | 4046.353 | 0.205163917 | 0.70758043 | YNL175C | NOP13 | n |
| W303.asm02.HP0_G0045130 | 1315.152 | 1352.054 | 1267.622 | 2483.61 | 2321.479 | 2306.342 | 1356.686 | 1901.7 | 1388.654 | 8.91E-08 | -0.853924227 | YNL172W | APC1 | n |
| W303.asm02.HP0_G0045140 | 3986.754 | 4071.74 | 4135.941 | 5205.508 | 4623.865 | 3595.493 | 2493.636 | 2099.279 | 2327.58 | 0.437012854 | -0.13872942 | YNL169C | PSD1 | n |
| W303.asm02.HP0_G0045150 | 3160.813 | 2974.518 | 2925.448 | 2819.158 | 3080.841 | 3040.178 | 965.358 | 2802.618 | 2078.795 | 0.960658745 | 0.019350579 | YNL168C | FMP41 | n |
| W303.asm02.HP0_G0045160 | 3335.531 | 3781.299 | 3198.159 | 3352.005 | 2893.387 | 2597.293 | 3097.293 | 1764.253 | 2516.561 | 0.365961435 | 0.22211824 | YNL167C | SKO1 | n |

| | | | | | | | | | | | | | | |
|---|---|---|---|---|---|---|---|---|---|---|---|---|---|---|
| W303.asm02.HP0_G0045170 | 2931.032 | 2674.062 | 2858.618 | 3181.8 | 2997.528 | 3186.033 | 3457.759 | 3007.714 | 3045.231 | 0.162980332 | -0.146109563 | YNL166C | BNI5 | n |
| W303.asm02.HP0_G0045180 | 1339.507 | 1424.386 | 1292.414 | 667.6222 | 643.0713 | 808.2832 | 1159.17 | 1025.479 | 842.0434 | 2.06E-08 | 0.936544585 | YNL165W | | n |
| W303.asm02.HP0_G0045190 | 648.046 | 652.1016 | 735.1347 | 1729.843 | 1647.165 | 1208.626 | 2330.685 | 1647.209 | 2065.638 | 6.57E-09 | -1.17188197 | YNL164C | IBD2 | n |
| W303.asm02.HP0_G0045200 | 4733.277 | 4655.961 | 4926.049 | 5213.845 | 4860.786 | 4585.336 | 7699.409 | 6188.309 | 7473.135 | 0.803750685 | -0.034366807 | YNL163C | RIA1 | n |
| W303.asm02.HP0_G0045210 | 99.53648 | 114.6185 | 113.1806 | 320.9589 | 283.7845 | 166.3666 | 918.448 | 500.3908 | 733.1997 | 0.000172399 | -1.236504876 | YNL162W-A | | n |
| W303.asm02.HP0_G0045230 | 4331.955 | 4292.075 | 4400.029 | 7280.625 | 6529.647 | 7353.553 | 6829.105 | 6549.106 | 5858.421 | 3.11E-12 | -0.700516387 | YNL161W | CBK1 | n |
| W303.asm02.HP0_G0045250 | 2432.29 | 2577.248 | 2222.651 | 411.272 | 471.2385 | 370.7163 | 603.6573 | 490.7266 | 417.4334 | 2.58E-47 | 2.529528991 | YNL159C | ASI2 | n |
| W303.asm02.HP0_G0045260 | 1265.384 | 1216.292 | 1261.155 | 249.4031 | 265.5598 | 232.4574 | 475.2721 | 466.0292 | 453.316 | 9.52E-112 | 2.32571434 | YNL158W | PGA1 | n |
| W303.asm02.HP0_G0045280 | 3375.769 | 3852.518 | 3094.68 | 3753.551 | 3933.93 | 5248.523 | 8256.156 | 7385.596 | 7322.428 | 0.072545271 | -0.325571104 | YNL156C | NSG2 | n |
| W303.asm02.HP0_G0045290 | 4387.017 | 4485.702 | 4213.551 | 1422.084 | 1446.693 | 1914.355 | 1809.738 | 1465.737 | 1480.752 | 5.16E-19 | 1.451973021 | YNL155W | CUZ1 | n |
| W303.asm02.HP0_G0045300 | 9120.295 | 9428.766 | 9988.993 | 11852.55 | 11121.4 | 12504.08 | 9405.45 | 9036.026 | 9628.479 | 0.000336778 | -0.31409698 | YNL154C | YCK2 | n |
| W303.asm02.HP0_G0045310 | 2098.737 | 1749.324 | 1927.303 | 2508.62 | 2773.625 | 2349.643 | 1312.245 | 1936.061 | 1920.911 | 0.038598527 | -0.401991175 | YNL153C | GIM3 | n |
| W303.asm02.HP0_G0045320 | 1128.786 | 1257.466 | 1308.583 | 1416.526 | 1398.094 | 1320.297 | 1443.099 | 1200.508 | 1291.771 | 0.200870434 | -0.162451748 | YNL152W | INN1 | n |
| W303.asm02.HP0_G0045330 | 1809.658 | 2126.563 | 2100.847 | 2234.902 | 2009.923 | 1059.732 | 2277.603 | 1186.549 | 1935.264 | 0.613802459 | 0.18659496 | YNL151C | RPC31 | n |
| W303.asm02.HP0_G0045340 | 3203.169 | 2872.14 | 3087.135 | 1879.207 | 2180.888 | 1684.176 | 1629.504 | 2067.065 | 1709.204 | 1.33E-05 | 0.673892274 | YNL149C | PGA2 | n |
| W303.asm02.HP0_G0045350 | 1736.594 | 1651.397 | 1601.774 | 1481.135 | 1494.425 | 1338.529 | 797.4696 | 1217.689 | 1127.908 | 0.271779723 | 0.209966939 | YNL148C | ALF1 | n |
| W303.asm02.HP0_G0045400 | 664.9884 | 758.9305 | 822.4454 | 696.8003 | 684.7277 | 752.068 | 841.9106 | 1068.431 | 641.1012 | 0.752849828 | 0.074187519 | YNL142W | MEP2 | n |
| W303.asm02.HP0_G0045410 | 1169.024 | 1091.658 | 1181.39 | 1312.319 | 1076.125 | 535.5636 | 2115.887 | 1225.206 | 1984.304 | 0.535081243 | 0.235271629 | YNL141W | AAH1 | n |
| W303.asm02.HP0_G0045420 | 4522.556 | 4572.501 | 4143.487 | 6162.132 | 5638.373 | 7194.024 | 4508.296 | 5672.885 | 5510.36 | 0.000556759 | -0.520896536 | YNL139C | THO2 | n |
| W303.asm02.HP0_G0045440 | 11591.76 | 10189.92 | 10698.26 | 8042.729 | 8287.896 | 12470.65 | 11791.69 | 15821.37 | 13713.11 | 0.456482294 | 0.173414516 | YNL138W | SRV2 | n |
| W303.asm02.HP0_G0045450 | 15956.54 | 17127.13 | 16310.94 | 8907.65 | 8517.006 | 7009.425 | 2533.139 | 2874.562 | 2920.838 | 2.87E-15 | 1.015447651 | YNL137C | NAM9 | n |
| W303.asm02.HP0_G0045460 | 3725.206 | 3678.921 | 3617.466 | 2899.05 | 2704.197 | 2277.475 | 2178.845 | 2455.78 | 2034.539 | 0.000368236 | 0.483881664 | YNL136W | EAF7 | n |
| W303.asm02.HP0_G0045470 | 8804.743 | 9048.188 | 8748.319 | 3759.109 | 4719.328 | 3678.296 | 3141.734 | 3378.175 | 3273.683 | 1.58E-19 | 1.129980409 | YNL135C | FPR1 | n |
| W303.asm02.HP0_G0045480 | 12251.46 | 12495.65 | 11734.13 | 322.3483 | 309.8198 | 753.5873 | 1757.89 | 9270.115 | 3642.077 | 5.10E-18 | 4.71832886 | YNL134C | | n |
| W303.asm02.HP0_G0045490 | 697.8142 | 661.004 | 566.9807 | 1788.199 | 1726.139 | 1827.753 | 1819.613 | 1503.32 | 1301.34 | 8.95E-20 | -1.471977646 | YNL133C | FYV6 | n |
| W303.asm02.HP0_G0045500 | 2030.968 | 2131.015 | 2603.153 | 7767.621 | 6882.859 | 2691.492 | 9399.278 | 3737.898 | 8458.708 | 0.00232367 | -1.358076958 | YNL132W | KRE33 | n |
| W303.asm02.HP0_G0045510 | 6997.203 | 7251.014 | 6822.093 | 4030.048 | 4448.561 | 3445.079 | 2304.761 | 2267.865 | 2150.56 | 1.10E-11 | 0.821517085 | YNL131W | TOM22 | n |
| W303.asm02.HP0_G0045530 | 551.6862 | 479.6174 | 493.6828 | 650.2543 | 671.7101 | 470.9921 | 441.9414 | 512.2026 | 560.9635 | 0.255227625 | -0.233156214 | YNL129W | NRK1 | n |
| W303.asm02.HP0_G0045540 | 289.0793 | 264.8467 | 264.088 | 440.45 | 469.5028 | 245.3717 | 234.5499 | 254.4906 | 211.7069 | 0.075242795 | -0.497647363 | YNL128W | TEP1 | n |
| W303.asm02.HP0_G0045550 | 3267.761 | 3520.904 | 2985.811 | 2576.702 | 2296.311 | 2482.584 | 1885.04 | 2006.932 | 2009.422 | 0.00012429 | 0.409859241 | YNL127W | FAR11 | n |
| W303.asm02.HP0_G0045560 | 874.6503 | 970.3628 | 977.6645 | 1224.09 | 1150.759 | 1451.719 | 1455.444 | 1345.471 | 1435.301 | 0.000970433 | -0.439353244 | YNL126W | SPC98 | n |
| W303.asm02.HP0_G0045570 | 1511.048 | 1634.705 | 1532.788 | 1526.986 | 1321.724 | 1515.531 | 5104.546 | 4062.185 | 4408.767 | 0.489925416 | 0.099895058 | YNL125C | ESBP6 | n |
| W303.asm02.HP0_G0045580 | 1121.374 | 1231.871 | 1395.894 | 1419.305 | 1198.491 | 598.6157 | 4221.898 | 1686.94 | 3705.469 | 0.620968255 | 0.221068038 | YNL124W | NAF1 | n |
| W303.asm02.HP0_G0045590 | 5093.303 | 5042.103 | 5377.694 | 7314.666 | 7098.952 | 4128.018 | 5939.05 | 4139.499 | 5597.675 | 0.342835538 | -0.257272251 | YNL123W | NMA111 | n |
| W303.asm02.HP0_G0045600 | 2881.263 | 2485.998 | 2392.961 | 1552.691 | 1759.985 | 1172.922 | 1232.004 | 929.9107 | 1040.593 | 8.05E-05 | 0.791022346 | YNL122C | MRP35 | n |
| W303.asm02.HP0_G0045610 | 23171.88 | 24822.15 | 24335.98 | 17024.71 | 15622.9 | 16068.43 | 7793.229 | 9380.716 | 9272.046 | 1.28E-07 | 0.570165712 | YNL121C | TOM70 | n |
| W303.asm02.HP0_G0045620 | 1735.535 | 2072.036 | 2062.042 | 1215.059 | 1208.037 | 281.0759 | 1557.905 | 434.889 | 1222.398 | 0.056537008 | 1.118110514 | YNL119W | NCS2 | n |
| W303.asm02.HP0_G0045630 | 8553.784 | 9196.191 | 8602.801 | 9284.881 | 8690.575 | 9667.492 | 6190.883 | 6613.534 | 6744.72 | 0.465361072 | -0.069048634 | YNL118C | DCP2 | n |
| W303.asm02.HP0_G0045640 | 294.3738 | 245.9291 | 273.7892 | 2907.387 | 3245.731 | 4001.154 | 2534.373 | 2020.891 | 1861.107 | 2.58E-81 | -3.64040844 | YNL117W | MLS1 | n |
| W303.asm02.HP0_G0045660 | 4934.468 | 4797.287 | 4546.625 | 3067.172 | 3046.127 | 5778.009 | 3665.151 | 5488.191 | 4798.69 | 0.395874407 | 0.263903215 | YNL115C | | n |
| W303.asm02.HP0_G0045670 | 1422.101 | 1265.255 | 1433.62 | 2475.968 | 3011.414 | 1239.773 | 2981.253 | 1502.246 | 2457.953 | 0.057513068 | -0.706919778 | YNL113W | RPC19 | n |
| W303.asm02.HP0_G0045680 | 1873.192 | 1537.891 | 2215.105 | 5815.469 | 5336.364 | 1744.95 | 9162.259 | 5944.556 | 8462.297 | 0.004573052 | -1.196763232 | YNL112W | DBP2 | n |
| W303.asm02.HP0_G0045690 | 2908.795 | 3336.179 | 3545.246 | 5629.98 | 6469.766 | 3875.049 | 3810.818 | 7028.021 | 6102.422 | 0.013540717 | -0.706357962 | YNL111C | CYB5 | n |
| W303.asm02.HP0_G0045700 | 2637.717 | 2877.704 | 3095.758 | 6158.659 | 6195.528 | 1809.521 | 5906.954 | 2629.736 | 5562.988 | 0.140610327 | -0.717906874 | YNL110C | NOP15 | n |
| W303.asm02.HP0_G0045720 | 975.2457 | 1080.53 | 976.5866 | 1950.068 | 2148.778 | 2499.297 | 5083.56 | 5378.664 | 5241.242 | 8.36E-19 | -1.121645333 | YNL107W | YAF9 | n |
| W303.asm02.HP0_G0045740 | 10026.71 | 10276.72 | 10574.3 | 6924.236 | 6862.899 | 6008.947 | 6356.302 | 6895.943 | 7040.152 | 4.13E-11 | 0.641361276 | YNL104C | LEU4 | n |
| W303.asm02.HP0_G0045750 | 1615.879 | 2003.042 | 1748.37 | 3502.759 | 3323.837 | 3508.131 | 4728.032 | 2864.898 | 3877.705 | 2.97E-06 | -0.945317178 | YNL103W | MET4 | n |
| W303.asm02.HP0_G0045760 | 1615.879 | 1722.617 | 1898.2 | 6158.659 | 6671.105 | 5822.07 | 1443.099 | 2208.806 | 1936.461 | 9.24E-25 | -1.832570175 | YNL102W | POL1 | n |
| W303.asm02.HP0_G0045770 | 3208.463 | 3700.065 | 3535.545 | 2101.516 | 2245.976 | 3252.884 | 2360.313 | 3639.108 | 2285.717 | 0.08061363 | 0.458524626 | YNL101W | AVT4 | n |
| W303.asm02.HP0_G0045780 | 3287.88 | 3378.465 | 2898.5 | 4958.884 | 5586.302 | 4206.263 | 1451.74 | 3193.481 | 2542.875 | 0.031992447 | -0.6250208 | YNL100W | MIC27 | n |
| W303.asm02.HP0_G0045790 | 1966.375 | 2169.963 | 1996.29 | 2286.311 | 2383.096 | 2241.011 | 2465.243 | 2188.404 | 1873.068 | 0.218653443 | -0.172248619 | YNL099C | OCA1 | n |
| W303.asm02.HP0_G0045810 | 5169.543 | 5222.377 | 4973.477 | 2175.156 | 2033.355 | 2288.869 | 3510.841 | 2392.426 | 2633.778 | 3.89E-14 | 1.241670322 | YNL097C | PHO23 | n |
| W303.asm02.HP0_G0045840 | 3518.72 | 3437.443 | 3717.712 | 1770.831 | 1933.553 | 2474.228 | 3088.652 | 2440.747 | 2819.171 | 2.96E-06 | 0.788767171 | YNL094W | APP1 | n |
| W303.asm02.HP0_G0045860 | 882.0626 | 880.2259 | 780.4069 | 1116.409 | 1074.389 | 1658.348 | 876.4759 | 1622.512 | 1200.869 | 0.033757217 | -0.598254398 | YNL092W | | n |

| | | | | | | | | | | | | | | |
|---|---|---|---|---|---|---|---|---|---|---|---|---|---|---|
| W303.asm02.HP0_G0045870 | 10396.27 | 10106.46 | 10516.09 | 6603.972 | 6034.109 | 5989.956 | 5846.465 | 5202.561 | 5664.655 | 3.61E-17 | 0.735570659 | YNL091W | NST1 | n |
| W303.asm02.HP0_G0045880 | 775.1139 | 656.5528 | 776.0953 | 1324.824 | 1444.09 | 1738.872 | 539.4647 | 1304.667 | 706.8858 | 0.001521578 | -1.029772511 | YNL090W | RHO2 | n |
| W303.asm02.HP0_G0045890 | 4261.008 | 4677.104 | 4692.143 | 7591.163 | 7960.719 | 8398.092 | 2486.229 | 3927.96 | 3046.427 | 8.02E-06 | -0.813213949 | YNL088W | TOP2 | n |
| W303.asm02.HP0_G0045910 | 532.626 | 502.9862 | 488.2933 | 260.5186 | 308.9519 | 316.7801 | 357.9972 | 444.5532 | 338.4919 | 2.09E-06 | 0.783169192 | YNL086W | SNN1 | n |
| W303.asm02.HP0_G0045920 | 6283.505 | 6986.167 | 6697.056 | 7830.84 | 8085.689 | 10782.68 | 7875.938 | 9456.956 | 8214.707 | 0.012952591 | -0.419216182 | YNL085W | MKT1 | n |
| W303.asm02.HP0_G0045930 | 6147.966 | 6601.138 | 6040.608 | 4756.027 | 4728.006 | 5541.753 | 3997.224 | 5174.642 | 4200.648 | 0.028139012 | 0.322470502 | YNL084C | END3 | n |
| W303.asm02.HP0_G0045940 | 2812.435 | 2875.479 | 2807.956 | 3326.996 | 2911.612 | 2354.201 | 2503.511 | 2040.22 | 2492.64 | 0.932995457 | -0.016454172 | YNL082W | PMS1 | n |
| W303.asm02.HP0_G0045950 | 4662.331 | 4480.138 | 4373.081 | 1222.006 | 1279.2 | 1039.981 | 551.8095 | 560.5236 | 589.6696 | 1.66E-67 | 1.932630722 | YNL081C | SWS2 | n |
| W303.asm02.HP0_G0045960 | 1736.594 | 1790.497 | 1809.811 | 3807.044 | 4168.248 | 4082.438 | 4451.51 | 2944.359 | 3272.487 | 8.73E-12 | -1.17584825 | YNL080C | EOS1 | n |
| W303.asm02.HP0_G0045980 | 1760.949 | 1927.372 | 1861.551 | 1715.949 | 1607.244 | 1947.78 | 2535.608 | 2780.068 | 3111.015 | 0.613618859 | 0.074151339 | YNL078W | NIS1 | n |
| W303.asm02.HP0_G0045990 | 4391.253 | 6004.676 | 3530.156 | 4917.201 | 4688.085 | 4949.975 | 9152.383 | 7962.226 | 9648.812 | 0.784992848 | -0.063806276 | YNL077W | APJ1 | n |
| W303.asm02.HP0_G0046000 | 3862.862 | 4375.535 | 4244.81 | 4314.882 | 4168.248 | 5424.005 | 6256.31 | 4978.136 | 5963.676 | 0.363280321 | -0.155897488 | YNL076W | MKS1 | n |
| W303.asm02.HP0_G0046010 | 1591.525 | 1521.199 | 1788.253 | 2892.103 | 2469.88 | 780.1756 | 4176.222 | 1474.327 | 3091.878 | 0.548014576 | -0.325678155 | YNL075W | IMP4 | n |
| W303.asm02.HP0_G0046030 | 5924.538 | 6002.451 | 5784.066 | 1572.143 | 1507.442 | 1477.548 | 1182.625 | 1014.741 | 1227.183 | 4.84E-86 | 1.958308463 | YNL073W | MSK1 | n |
| W303.asm02.HP0_G0046040 | 645.9282 | 607.5895 | 649.9798 | 1181.017 | 1298.292 | 1194.952 | 409.8451 | 511.1288 | 355.237 | 1.32E-09 | -0.948507043 | YNL072W | RNH201 | n |
| W303.asm02.HP0_G0046050 | 19438.2 | 17862.69 | 19367.89 | 13932.53 | 14932.97 | 17106.13 | 5263.793 | 11782.81 | 10233.7 | 0.322010709 | 0.301813977 | YNL071W | LAT1 | n |
| W303.asm02.HP0_G0046060 | 2625.01 | 2313.514 | 2184.924 | 1489.471 | 1859.787 | 1099.994 | 815.9867 | 661.4608 | 665.0229 | 0.002290609 | 0.679303489 | YNL070W | TOM7 | n |
| W303.asm02.HP0_G0046080 | 2658.895 | 2585.038 | 2803.644 | 5850.205 | 5478.69 | 4270.834 | 4149.064 | 4113.728 | 3809.529 | 3.91E-11 | -0.954896122 | YNL068C | FKH2 | n |
| W303.asm02.HP0_G0046090 | 4523.615 | 3973.814 | 4587.586 | 22008.61 | 24832.45 | 23134.83 | 17171.52 | 14992.39 | 14020.5 | 2.98E-87 | -2.418881563 | YNL067W | RPL9B | y |
| W303.asm02.HP0_G0046100 | 2634.54 | 2369.154 | 3197.081 | 15921.5 | 17135.55 | 17956.19 | 20081.17 | 22210.48 | 18888.56 | 4.63E-74 | -2.636983372 | YNL066W | SUN4 | n |
| W303.asm02.HP0_G0046110 | 151.4225 | 144.6642 | 163.8423 | 2356.477 | 2202.584 | 4266.276 | 22797.01 | 17077.71 | 22926.54 | 1.28E-51 | -4.262120607 | YNL065W | AQR1 | n |
| W303.asm02.HP0_G0046120 | 11318.57 | 12293.12 | 12129.72 | 13239.9 | 12868.37 | 11163.27 | 7912.973 | 8695.632 | 8383.355 | 0.607344681 | -0.060491096 | YNL064C | YDJ1 | n |
| W303.asm02.HP0_G0046130 | 1750.36 | 1757.113 | 1575.905 | 222.3092 | 207.4144 | 67.61015 | 245.6601 | 100.9372 | 246.3934 | 3.07E-13 | 3.353595588 | YNL063W | MTQ1 | n |
| W303.asm02.HP0_G0046140 | 1191.261 | 1401.017 | 1504.763 | 2596.154 | 2042.901 | 587.2207 | 4409.538 | 1574.191 | 3630.116 | 0.540868538 | -0.351226317 | YNL062C | GCD10 | n |
| W303.asm02.HP0_G0046150 | 9588.328 | 9407.623 | 10712.27 | 7559.901 | 6983.529 | 2286.59 | 6826.636 | 3241.802 | 5811.774 | 0.070317677 | 0.819829368 | YNL061W | NOP2 | n |
| W303.asm02.HP0_G0046160 | 4211.24 | 4441.19 | 4554.17 | 6474.06 | 6249.334 | 7028.417 | 4678.653 | 6296.763 | 5523.517 | 2.96E-05 | -0.580760306 | YNL059C | ARP5 | n |
| W303.asm02.HP0_G0046170 | 8690.382 | 8687.64 | 8885.213 | 2437.064 | 2436.902 | 2614.766 | 3001.004 | 2922.883 | 2596.699 | 6.41E-92 | 1.810218578 | YNL058C | | n |
| W303.asm02.HP0_G0046180 | 3046.452 | 2829.854 | 2892.033 | 941.3404 | 1015.376 | 1008.835 | 1414.706 | 1144.671 | 1392.242 | 1.27E-40 | 1.564336334 | YNL056W | OCA2 | n |
| W303.asm02.HP0_G0046190 | 17379.7 | 17426.47 | 17743.48 | 13540.02 | 14540.7 | 16072.22 | 7292.033 | 12231.65 | 11573.31 | 0.260318546 | 0.2511763 | YNL055C | POR1 | n |
| W303.asm02.HP0_G0046210 | 913.8296 | 951.4452 | 1017.547 | 3946.682 | 3476.577 | 3704.125 | 7678.423 | 5512.889 | 6790.171 | 6.40E-38 | -1.948669674 | YNL054W | VAC7 | n |
| W303.asm02.HP0_G0046220 | 1302.445 | 1304.203 | 1384.037 | 4844.95 | 4195.151 | 4792.724 | 8089.502 | 6178.645 | 6970.779 | 1.00E-39 | -1.793487383 | YNL053W | MSG5 | n |
| W303.asm02.HP0_G0046230 | 4990.59 | 5005.381 | 4978.867 | 11564.24 | 12900.48 | 11392.69 | 7057.483 | 9643.797 | 9642.832 | 5.84E-17 | -1.259703949 | YNL052W | COX5A | n |
| W303.asm02.HP0_G0046240 | 2226.864 | 2516.044 | 2164.444 | 1262.299 | 1182.869 | 1153.931 | 864.1312 | 1082.39 | 1082.456 | 2.90E-12 | 0.940274034 | YNL051W | COG5 | n |
| W303.asm02.HP0_G0046250 | 1717.534 | 1832.784 | 1934.849 | 2161.262 | 2107.989 | 1398.543 | 2998.535 | 2000.489 | 2814.386 | 0.85794513 | -0.047238351 | YNL050C | | n |
| W303.asm02.HP0_G0046270 | 2479.941 | 2595.053 | 2554.647 | 2791.369 | 2614.81 | 3123.741 | 1211.018 | 2367.729 | 2358.678 | 0.575049099 | -0.160949801 | YNL048W | ALG11 | n |
| W303.asm02.HP0_G0046280 | 837.5889 | 903.5947 | 882.8084 | 13945.73 | 13051.48 | 18594.31 | 18517.1 | 16578.4 | 17596.79 | 1.55E-147 | -4.119006162 | YNL047C | SLM2 | n |
| W303.asm02.HP0_G0046300 | 6597.998 | 5955.713 | 5978.09 | 7720.38 | 8506.592 | 8357.07 | 5985.96 | 5422.69 | 6187.344 | 6.01E-05 | -0.407622551 | YNL045W | LAP2 | n |
| W303.asm02.HP0_G0046330 | 1898.605 | 1929.598 | 1834.603 | 1033.738 | 968.5122 | 1086.32 | 514.7753 | 371.5348 | 421.0217 | 4.80E-10 | 0.874336278 | YNL042W | BOP3 | n |
| W303.asm02.HP0_G0046340 | 2909.854 | 3190.402 | 2867.241 | 2810.127 | 2625.224 | 2747.707 | 1871.461 | 2201.29 | 2029.755 | 0.23515979 | 0.131855598 | YNL041C | COG6 | n |
| W303.asm02.HP0_G0046350 | 3479.541 | 3450.797 | 3351.223 | 2976.164 | 2746.722 | 3368.353 | 1290.024 | 2459.002 | 1797.715 | 0.492482038 | 0.177472861 | YNL040W | | n |
| W303.asm02.HP0_G0046360 | 5037.181 | 5036.539 | 5178.28 | 3447.181 | 3542.533 | 3586.377 | 6636.527 | 5088.738 | 6586.836 | 5.79E-05 | 0.528216691 | YNL039W | BDP1 | n |
| W303.asm02.HP0_G0046370 | 1569.288 | 1609.111 | 1600.697 | 299.4227 | 289.8594 | 293.2305 | 209.8604 | 336.0994 | 202.1383 | 2.24E-33 | 2.436890311 | YNL038W | GPI15 | n |
| W303.asm02.HP0_G0046380 | 7704.547 | 7215.404 | 7004.26 | 13453.87 | 13274.52 | 11240.76 | 2339.327 | 3749.709 | 4412.355 | 0.000879808 | -0.792298474 | YNL037C | IDH1 | n |
| W303.asm02.HP0_G0046390 | 828.0588 | 851.2931 | 842.9257 | 1319.266 | 1234.94 | 1844.466 | 924.6203 | 1035.143 | 502.3554 | 0.006684968 | -0.802409231 | YNL036W | NCE103 | n |
| W303.asm02.HP0_G0046400 | 1862.603 | 1846.137 | 1940.238 | 955.2347 | 774.1155 | 159.5296 | 3773.784 | 1107.088 | 3137.329 | 0.00949334 | 1.580432167 | YNL035C | | n |
| W303.asm02.HP0_G0046430 | 2852.673 | 2974.518 | 2769.151 | 2145.978 | 2370.946 | 1928.029 | 3126.92 | 2464.371 | 2395.757 | 0.006952035 | 0.415702677 | YNL032W | SIW14 | n |
| W303.asm02.HP0_G0046460 | 1241.029 | 1136.17 | 1239.597 | 2091.79 | 2252.051 | 2623.122 | 846.8485 | 1432.449 | 1027.436 | 1.12E-05 | -0.9457632 | YNL029C | KTR5 | n |
| W303.asm02.HP0_G0046470 | 4236.654 | 5020.96 | 4367.692 | 4842.171 | 4772.266 | 4968.966 | 5920.533 | 3846.351 | 4993.652 | 0.622998311 | -0.098071085 | YNL027W | CRZ1 | n |
| W303.asm02.HP0_G0046480 | 5076.36 | 5268.002 | 4774.064 | 4346.839 | 4520.592 | 3574.222 | 2531.904 | 2803.692 | 2851.465 | 0.034972437 | 0.281185951 | YNL026W | SAM50 | n |
| W303.asm02.HP0_G0046490 | 1485.635 | 1695.909 | 1350.621 | 1439.452 | 1462.315 | 1614.287 | 1211.018 | 1025.479 | 917.3967 | 0.978596391 | 0.005044922 | YNL025C | SSN8 | n |
| W303.asm02.HP0_G0046500 | 1481.399 | 1315.331 | 1376.491 | 681.5165 | 820.979 | 545.4392 | 802.4075 | 587.3686 | 797.7882 | 5.36E-07 | 1.02757353 | YNL024C-, | KSH1 | n |
| W303.asm02.HP0_G0046510 | 879.9448 | 618.7176 | 784.7186 | 1622.857 | 1556.042 | 998.1993 | 1810.972 | 1369.095 | 2077.598 | 0.001049742 | -0.871186011 | YNL024C | EFM6 | n |
| W303.asm02.HP0_G0046520 | 1794.833 | 1916.244 | 2141.807 | 2925.45 | 2663.409 | 1533.763 | 1977.626 | 1358.357 | 1806.087 | 0.33109601 | -0.28327247 | YNL023C | FAP1 | n |
| W303.asm02.HP0_G0046530 | 2043.675 | 2201.121 | 2325.052 | 1902.828 | 1562.984 | 733.836 | 2699.793 | 1426.006 | 2569.189 | 0.095485523 | 0.645548439 | YNL022C | RCM1 | n |

| | | | | | | | | | | | | | | |
|---|---|---|---|---|---|---|---|---|---|---|---|---|---|---|
| W303.asm02.HP0_G0046540 | 2356.05 | 2657.37 | 2442.544 | 3839.001 | 3636.26 | 4248.804 | 5557.598 | 5415.173 | 6303.364 | 3.20E-08 | -0.653158791 | YNL021W | HDA1 | n |
| W303.asm02.HP0_G0046580 | 20813.71 | 21393.61 | 21436.4 | 21964.15 | 22648.09 | 20698.58 | 12171.9 | 13029.49 | 12857.91 | 0.647189673 | -0.037284371 | YNL016W | PUB1 | n |
| W303.asm02.HP0_G0046590 | 5325.202 | 5605.181 | 5035.996 | 653.0332 | 797.5473 | 1212.425 | 1339.403 | 1472.18 | 1297.751 | 1.06E-29 | 2.583998018 | YNL015W | PBI2 | n |
| W303.asm02.HP0_G0046620 | 1668.824 | 1576.84 | 1570.515 | 1271.331 | 1290.482 | 1829.272 | 1564.077 | 1335.807 | 1303.732 | 0.505213526 | 0.133267835 | YNL011C | | n |
| W303.asm02.HP0_G0046630 | 6257.032 | 5749.845 | 6200.139 | 2230.734 | 3014.885 | 5279.669 | 7358.694 | 9583.664 | 8630.944 | 0.01075205 | 0.790657183 | YNL010W | PYP1 | n |
| W303.asm02.HP0_G0046660 | 16746.48 | 17681.3 | 14570.11 | 4324.608 | 4288.878 | 5701.283 | 3571.331 | 3671.322 | 5144.359 | 1.38E-18 | 1.775195751 | YNL007C | SIS1 | n |
| W303.asm02.HP0_G0046670 | 3715.676 | 3919.286 | 3335.054 | 1377.622 | 1418.923 | 1726.718 | 2351.671 | 1184.401 | 1923.304 | 5.75E-07 | 1.27811828 | YNL006W | LST8 | n |
| W303.asm02.HP0_G0046680 | 12119.1 | 12087.25 | 11786.95 | 7102.083 | 7444.353 | 5365.511 | 2618.317 | 2493.363 | 2844.289 | 1.38E-08 | 0.854134174 | YNL005C | MRP7 | n |
| W303.asm02.HP0_G0046700 | 2638.776 | 2394.749 | 2396.194 | 4710.87 | 5397.113 | 4469.867 | 2112.183 | 2813.356 | 2528.522 | 5.33E-11 | -0.972274776 | YNL003C | PET8 | n |
| W303.asm02.HP0_G0046710 | 5060.477 | 5856.674 | 6015.816 | 5989.148 | 5353.72 | 1983.484 | 7232.778 | 2370.95 | 5829.715 | 0.503524202 | 0.345551774 | YNL002C | RLP7 | n |
| W303.asm02.HP0_G0046720 | 1430.572 | 1658.074 | 1510.152 | 3558.336 | 3447.07 | 4371.11 | 2694.855 | 3682.06 | 3535.625 | 3.92E-14 | -1.306828437 | YNL001W | DOM34 | n |
| W303.asm02.HP0_G0046730 | 13660.85 | 13736.42 | 12051.04 | 19477.76 | 21707.34 | 22438.97 | 23966.06 | 9513.867 | 6676.543 | 0.110904053 | -0.68960676 | YNR001C | CIT1 | n |
| W303.asm02.HP0_G0046750 | 3385.299 | 3288.328 | 3584.051 | 1098.346 | 1168.116 | 433.0088 | 1411.003 | 665.756 | 1227.183 | 1.27E-06 | 1.92604727 | YNR003C | RPC34 | n |
| W303.asm02.HP0_G0046770 | 6909.314 | 7266.593 | 7031.208 | 5534.109 | 5162.795 | 8873.642 | 6829.105 | 8906.097 | 7433.664 | 0.656266967 | 0.115836216 | YNR006W | VPS27 | n |
| W303.asm02.HP0_G0046780 | 3977.224 | 4324.346 | 3677.829 | 1458.904 | 1549.967 | 2551.713 | 3024.459 | 2826.241 | 2130.226 | 1.52E-05 | 1.107216447 | YNR007C | ATG3 | n |
| W303.asm02.HP0_G0046790 | 2321.106 | 1853.927 | 2064.198 | 6833.228 | 7062.502 | 9119.774 | 6090.89 | 7994.44 | 7084.407 | 4.65E-26 | -1.883107815 | YNR008W | LRO1 | n |
| W303.asm02.HP0_G0046810 | 1391.393 | 1334.249 | 1181.39 | 657.2015 | 798.4151 | 676.1015 | 527.12 | 603.4756 | 507.1398 | 9.35E-10 | 0.874945149 | YNR010W | CSE2 | n |
| W303.asm02.HP0_G0046820 | 2367.698 | 2471.532 | 2431.765 | 1481.83 | 1231.469 | 979.2077 | 2075.149 | 1157.556 | 1712.793 | 9.34E-05 | 0.977423263 | YNR011C | PRP2 | n |
| W303.asm02.HP0_G0046830 | 1119.256 | 1204.051 | 1364.634 | 2159.872 | 1759.985 | 1071.887 | 2094.901 | 893.4015 | 1615.91 | 0.249006261 | -0.436777272 | YNR012W | URK1 | n |
| W303.asm02.HP0_G0046840 | 3600.256 | 3296.118 | 3883.71 | 5222.876 | 5453.522 | 7675.651 | 6308.157 | 9251.86 | 7459.978 | 0.000419552 | -0.767565678 | YNR013C | PHO91 | n |
| W303.asm02.HP0_G0046860 | 1148.905 | 1170.667 | 1356.011 | 2803.874 | 2612.206 | 1501.857 | 5655.121 | 2987.311 | 3875.313 | 0.003634694 | -0.912364932 | YNR015W | SMM1 | n |
| W303.asm02.HP0_G0046870 | 37941.4 | 35764.32 | 38728.23 | 23045.12 | 24514.82 | 34611.08 | 12782.97 | 27612.77 | 28741.91 | 0.174545773 | 0.452376979 | YNR016C | ACC1 | n |
| W303.asm02.HP0_G0046880 | 11568.47 | 11345.01 | 11776.17 | 6033.61 | 6444.599 | 5471.105 | 3538 | 3649.846 | 3509.311 | 2.06E-26 | 0.950693709 | YNR017W | TIM23 | n |
| W303.asm02.HP0_G0046890 | 2191.92 | 2089.841 | 2230.196 | 8237.249 | 8882.368 | 13809.94 | 22919.23 | 23812.59 | 23097.58 | 6.96E-28 | -2.247814766 | YNR018W | RCF2 | n |
| W303.asm02.HP0_G0046900 | 5901.242 | 6012.466 | 5954.376 | 6032.915 | 5919.554 | 10136.96 | 15792.61 | 20369.98 | 17642.24 | 0.223755263 | -0.305988834 | YNR019W | ARE2 | n |
| W303.asm02.HP0_G0046920 | 807.9397 | 747.8025 | 690.9404 | 2143.894 | 2301.518 | 1732.035 | 1527.043 | 1519.427 | 1625.479 | 8.93E-24 | -1.458982477 | YNR020C | ATP23 | n |
| W303.asm02.HP0_G0046930 | 7605.011 | 7498.056 | 7926.951 | 7097.915 | 6897.612 | 8246.159 | 6574.804 | 8601.137 | 8007.785 | 0.751575039 | 0.050229393 | YNR021W | | n |
| W303.asm02.HP0_G0046940 | 4701.51 | 4460.108 | 4263.135 | 1112.935 | 1147.288 | 722.4411 | 462.9274 | 564.8188 | 582.4931 | 1.59E-26 | 2.17037306 | YNR022C | MRPL50 | n |
| W303.asm02.HP0_G0046950 | 2999.86 | 3007.902 | 3075.278 | 5031.829 | 4987.491 | 5188.509 | 6210.634 | 5787.782 | 5696.95 | 8.38E-26 | -0.743599993 | YNR023W | SNF12 | n |
| W303.asm02.HP0_G0046960 | 1364.92 | 1283.06 | 1318.284 | 959.403 | 879.1245 | 395.0256 | 1513.464 | 1043.734 | 1267.849 | 0.012793631 | 0.828456022 | YNR024W | MPP6 | n |
| W303.asm02.HP0_G0046980 | 1634.94 | 1427.724 | 1624.411 | 1583.258 | 1528.271 | 2140.735 | 2688.682 | 2421.419 | 2408.914 | 0.368572381 | -0.164297564 | YNR027W | BUD17 | n |
| W303.asm02.HP0_G0047000 | 2362.403 | 2342.447 | 2370.324 | 1781.252 | 1833.751 | 1450.2 | 2180.079 | 2015.522 | 2456.757 | 0.000566319 | 0.482276017 | YNR029C | ZNG1 | n |
| W303.asm02.HP0_G0047010 | 2625.01 | 2224.49 | 2347.688 | 3991.144 | 3841.939 | 4159.164 | 2809.661 | 4787 | 5125.221 | 0.001835172 | -0.736586242 | YNR030W | ALG12 | n |
| W303.asm02.HP0_G0047020 | 2449.233 | 2544.977 | 2237.741 | 3130.391 | 2915.951 | 3163.243 | 3361.47 | 2905.703 | 3017.721 | 0.001265849 | -0.348931815 | YNR031C | SSK2 | n |
| W303.asm02.HP0_G0047030 | 1190.202 | 1264.142 | 1188.935 | 1196.301 | 1136.874 | 1323.336 | 888.8206 | 1262.789 | 1057.339 | 0.977573304 | -0.00541265 | YNR032W | PPG1 | n |
| W303.asm02.HP0_G0047040 | 972.069 | 806.781 | 845.0816 | 329.9902 | 398.3397 | 231.6977 | 248.1291 | 231.9408 | 197.3539 | 1.83E-10 | 1.451672473 | YNR032C- | HUB1 | n |
| W303.asm02.HP0_G0047050 | 5711.7 | 5817.726 | 5568.484 | 1932.006 | 1601.169 | 933.6278 | 371.5764 | 410.1916 | 540.6301 | 1.79E-12 | 1.936434873 | YNR033W | ABZ1 | n |
| W303.asm02.HP0_G0047060 | 2729.841 | 2847.659 | 2308.884 | 6097.524 | 6527.911 | 8006.105 | 7125.379 | 7647.603 | 6716.013 | 8.12E-20 | -1.387431936 | YNR034W | SOL1 | n |
| W303.asm02.HP0_G0047080 | 8577.08 | 8462.854 | 8319.31 | 7275.067 | 8017.997 | 10196.22 | 9879.488 | 10833.57 | 10285.13 | 0.966356516 | -0.007365476 | YNR035C | ARC35 | n |
| W303.asm02.HP0_G0047090 | 6327.979 | 5392.635 | 5346.434 | 2478.052 | 2883.841 | 1866.496 | 1956.64 | 1368.021 | 1727.146 | 6.22E-09 | 1.239603703 | YNR036C | MRPS12 | n |
| W303.asm02.HP0_G0047100 | 2935.267 | 2572.797 | 2389.727 | 1640.225 | 1761.72 | 995.9203 | 1209.784 | 994.3387 | 1088.437 | 0.000384347 | 0.844784125 | YNR037C | RSM19 | n |
| W303.asm02.HP0_G0047110 | 642.7515 | 650.9888 | 728.6673 | 3206.81 | 2758.004 | 869.0563 | 3388.629 | 1388.423 | 3101.447 | 0.000267129 | -1.756631073 | YNR038W | DBP6 | n |
| W303.asm02.HP0_G0047120 | 1133.022 | 1015.988 | 1093.001 | 3950.851 | 3684.859 | 4409.093 | 5613.149 | 5448.461 | 4997.24 | 3.48E-62 | -1.893417404 | YNR039C | ZRG17 | n |
| W303.asm02.HP0_G0047130 | 2669.484 | 2681.851 | 2631.179 | 1491.556 | 1618.526 | 630.5216 | 1067.819 | 520.793 | 865.9651 | 0.004564583 | 1.093650183 | YNR040W | MRX15 | n |
| W303.asm02.HP0_G0047140 | 1480.34 | 1406.581 | 1477.815 | 2965.743 | 3313.423 | 2715.041 | 1088.805 | 1590.298 | 1203.261 | 1.59E-09 | -1.042981173 | YNR041C | COQ2 | n |
| W303.asm02.HP0_G0047150 | 8543.195 | 8679.851 | 9596.634 | 5707.093 | 6146.929 | 7004.867 | 6684.672 | 6383.741 | 5944.539 | 2.70E-05 | 0.508076521 | YNR043W | MVD1 | n |
| W303.asm02.HP0_G0047160 | 1099.137 | 1003.747 | 1333.375 | 1520.039 | 1464.05 | 1245.85 | 2257.851 | 1809.353 | 1929.284 | 0.091280122 | -0.299738987 | YNR044W | AGA1 | n |
| W303.asm02.HP0_G0047170 | 2991.389 | 3283.877 | 2701.243 | 1795.147 | 1685.35 | 1780.654 | 2393.643 | 1674.054 | 2309.639 | 6.76E-06 | 0.770668105 | YNR045W | PET494 | n |
| W303.asm02.HP0_G0047180 | 1316.211 | 1358.73 | 1481.049 | 2007.73 | 1799.905 | 1245.09 | 2729.42 | 1756.737 | 2474.698 | 0.272468429 | -0.281909506 | YNR046W | TRM112 | n |
| W303.asm02.HP0_G0047190 | 2421.701 | 2704.107 | 2246.365 | 3996.007 | 3814.168 | 5356.395 | 5621.79 | 5283.096 | 4949.397 | 9.02E-07 | -0.836802194 | YNR047W | FPK1 | n |
| W303.asm02.HP0_G0047210 | 2306.281 | 2322.416 | 2210.794 | 712.084 | 702.9524 | 700.4108 | 925.8548 | 682.9368 | 776.2587 | 9.13E-38 | 1.692865133 | YNR049C | MSO1 | n |
| W303.asm02.HP0_G0047220 | 1733.417 | 2018.622 | 2205.404 | 5661.242 | 4811.319 | 5589.612 | 3667.62 | 2475.109 | 1999.853 | 2.10E-09 | -1.430954225 | YNR050C | LYS9 | n |
| W303.asm02.HP0_G0047230 | 7148.625 | 7358.955 | 8113.43 | 9458.56 | 8999.527 | 9859.687 | 10060.96 | 8061.016 | 8679.984 | 0.009768777 | -0.324071511 | YNR051C | BRE5 | n |
| W303.asm02.HP0_G0047240 | 1810.717 | 1931.823 | 1697.708 | 7325.782 | 6877.652 | 7164.397 | 5068.747 | 5859.726 | 5650.302 | 2.17E-86 | -1.973834083 | YNR052C | POP2 | n |

| | | | | | | | | | | | | | | |
|---|---|---|---|---|---|---|---|---|---|---|---|---|---|---|
| W303.asm02.HP0_G0047250 | 1615.879 | 1818.317 | 2012.458 | 4118.972 | 3933.93 | 1094.677 | 11063.35 | 5561.21 | 10770.74 | 0.126199906 | -0.748012839 | YNR053C | NOG2 | n |
| W303.asm02.HP0_G0047260 | 1700.591 | 1670.315 | 1984.433 | 2620.469 | 2564.475 | 997.4396 | 5813.134 | 3177.374 | 4332.217 | 0.621711609 | -0.207159264 | YNR054C | ESF2 | n |
| W303.asm02.HP0_G0047270 | 4349.956 | 3889.241 | 4368.77 | 8330.341 | 8304.385 | 7017.782 | 13353.3 | 7419.958 | 10048.3 | 5.36E-05 | -0.907642308 | YNR055C | HOL1 | n |
| W303.asm02.HP0_G0047280 | 672.4007 | 602.0255 | 670.4601 | 419.6085 | 410.4895 | 785.4932 | 1154.232 | 930.9845 | 995.1422 | 0.371506598 | 0.267473547 | YNR056C | BIO5 | n |
| W303.asm02.HP0_G0047290 | 656.5172 | 598.6871 | 614.4088 | 31.26223 | 36.44939 | 41.78155 | 107.3992 | 102.011 | 93.29458 | 1.58E-106 | 4.097736324 | YNR057C | BIO4 | n |
| W303.asm02.HP0_G0047300 | 438.3841 | 399.4957 | 434.3978 | 880.9001 | 821.8469 | 1445.642 | 5448.964 | 2874.562 | 6297.384 | 9.67E-05 | -1.307240528 | YNR058W | BIO3 | n |
| W303.asm02.HP0_G0047330 | 281.6671 | 326.0508 | 343.8533 | 1861.145 | 1669.729 | 2196.95 | 2798.55 | 2533.094 | 2750.994 | 9.74E-65 | -2.590068809 | YNR062C | PUL3 | n |
| W303.asm02.HP0_G0047340 | 321.9052 | 353.8708 | 355.7103 | 2941.428 | 3069.559 | 2747.707 | 2102.308 | 2418.197 | 2326.384 | 1.57E-163 | -3.086178587 | YNR063W | PUL4 | n |
| W303.asm02.HP0_G0047380 | 8397.067 | 8688.753 | 9462.973 | 4863.708 | 5351.117 | 5262.197 | 2399.816 | 6740.242 | 6048.598 | 0.026183618 | 0.778528066 | YNR067C | DSE4 | n |
| W303.asm02.HP0_G0047650 | 1484.576 | 1537.891 | 1442.244 | 2160.567 | 2226.016 | 2316.977 | 2255.382 | 2344.105 | 2206.776 | 1.12E-13 | -0.586355055 | YOL149W | DCP1 | n |
| W303.asm02.HP0_G0047660 | 1623.292 | 1650.284 | 1559.736 | 1863.229 | 1669.729 | 1677.339 | 1567.781 | 1505.468 | 1655.381 | 0.288258083 | -0.108799342 | YOL148C | SPT20 | n |
| W303.asm02.HP0_G0047670 | 2622.892 | 2616.196 | 2796.099 | 690.5478 | 782.7939 | 1106.831 | 317.2596 | 496.0956 | 345.6684 | 6.85E-12 | 1.638910513 | YOL147C | PEX11 | n |
| W303.asm02.HP0_G0047680 | 525.2138 | 456.2486 | 530.3318 | 546.7416 | 622.2431 | 629.762 | 229.612 | 255.5644 | 180.6087 | 0.158145317 | -0.250100898 | YOL146W | PSF3 | n |
| W303.asm02.HP0_G0047690 | 8250.939 | 8726.588 | 8588.788 | 10262.35 | 10055.69 | 9689.522 | 6756.271 | 5720.132 | 6097.638 | 0.016984169 | -0.231104321 | YOL145C | CTR9 | n |
| W303.asm02.HP0_G0047700 | 1069.488 | 1275.27 | 1351.699 | 1960.489 | 1656.711 | 575.0661 | 2538.077 | 1323.995 | 2675.641 | 0.713927019 | -0.181613572 | YOL144W | NOP8 | n |
| W303.asm02.HP0_G0047710 | 4131.823 | 3964.911 | 4115.461 | 1756.937 | 1740.024 | 2458.275 | 1206.08 | 2370.95 | 1612.322 | 0.000116212 | 1.036065504 | YOL143C | RIB4 | n |
| W303.asm02.HP0_G0047720 | 925.4775 | 948.1068 | 1101.624 | 1076.115 | 1042.279 | 431.4895 | 795.0007 | 613.1398 | 692.5328 | 0.546931249 | 0.222615331 | YOL142W | RRP40 | n |
| W303.asm02.HP0_G0047740 | 10408.97 | 10010.76 | 10359.79 | 8790.243 | 9780.585 | 8600.923 | 5742.769 | 5896.235 | 5816.558 | 0.034381634 | 0.179998126 | YOL139C | CDC33 | n |
| W303.asm02.HP0_G0047750 | 1782.127 | 1676.992 | 1800.11 | 3870.958 | 3832.393 | 4691.689 | 6626.652 | 5308.867 | 5646.714 | 9.51E-19 | -1.23683579 | YOL138C | RTC1 | n |
| W303.asm02.HP0_G0047760 | 1088.548 | 1015.988 | 1105.936 | 1852.808 | 1247.09 | 1343.087 | 11073.22 | 9464.473 | 5700.538 | 0.100351603 | -0.468943647 | YOL137W | BSC6 | n |
| W303.asm02.HP0_G0047770 | 2962.799 | 3565.416 | 3185.224 | 4766.447 | 4268.049 | 4662.821 | 8006.792 | 6456.759 | 7751.823 | 0.000284225 | -0.495986219 | YOL136C | PFK27 | n |
| W303.asm02.HP0_G0047780 | 2002.377 | 1847.25 | 2090.068 | 1946.595 | 1965.663 | 1905.239 | 4149.064 | 3164.488 | 3492.566 | 0.843226213 | 0.030067584 | YOL135C | MED7 | n |
| W303.asm02.HP0_G0047790 | 1648.705 | 1729.293 | 1589.917 | 1524.207 | 1824.205 | 1768.499 | 2094.901 | 1762.106 | 1522.615 | 0.810577156 | -0.042410409 | YOL133W | HRT1 | n |
| W303.asm02.HP0_G0047800 | 378.0268 | 344.9684 | 343.8533 | 170.2055 | 158.8152 | 206.6288 | 243.1912 | 230.867 | 186.5892 | 3.81E-08 | 0.993116023 | YOL132W | GAS4 | n |
| W303.asm02.HP0_G0047820 | 3105.75 | 3227.124 | 3244.51 | 7359.823 | 7214.375 | 4747.904 | 8672.173 | 4008.495 | 5840.48 | 0.000787009 | -1.012545527 | YOL130W | ALR1 | n |
| W303.asm02.HP0_G0047830 | 4342.544 | 4252.014 | 4042.163 | 753.767 | 821.8469 | 1028.586 | 1493.712 | 1590.298 | 1404.203 | 7.73E-57 | 2.278820524 | YOL129W | VPS68 | n |
| W303.asm02.HP0_G0047850 | 9769.399 | 8874.591 | 9824.073 | 16423.09 | 19339 | 16927.61 | 15491.4 | 12991.91 | 12469.18 | 1.36E-11 | -0.888129148 | YOL127W | RPL25 | n |
| W303.asm02.HP0_G0047860 | 1507.872 | 1959.643 | 1408.829 | 10941.78 | 10716.12 | 11078.19 | 34841.77 | 14964.48 | 12063.71 | 1.10E-13 | -2.747036899 | YOL126C | MDH2 | n |
| W303.asm02.HP0_G0047870 | 1544.933 | 1275.27 | 1444.4 | 1951.458 | 1966.531 | 1112.909 | 4840.369 | 1182.254 | 3479.409 | 0.637410394 | -0.238373714 | YOL125W | TRM13 | n |
| W303.asm02.HP0_G0047880 | 201.1908 | 233.6883 | 301.8148 | 2585.733 | 2240.769 | 685.9771 | 3370.112 | 1480.77 | 2506.993 | 1.24E-09 | -2.903589022 | YOL124C | TRM11 | n |
| W303.asm02.HP0_G0047890 | 23571.09 | 24446.02 | 24764.99 | 6842.259 | 6732.722 | 5035.817 | 7119.206 | 6174.35 | 5879.951 | 2.34E-37 | 1.967454478 | YOL123W | HRP1 | n |
| W303.asm02.HP0_G0047900 | 2950.092 | 2874.366 | 2802.566 | 7219.49 | 7742.023 | 8665.494 | 8147.522 | 9350.65 | 6993.505 | 1.41E-24 | -1.453480092 | YOL122C | SMF1 | n |
| W303.asm02.HP0_G0047910 | 6014.545 | 5643.016 | 5844.429 | 21596.64 | 26076.93 | 23638.48 | 21458.85 | 18645.46 | 17278.63 | 1.18E-58 | -2.026583958 | YOL121C | RPS19A | y |
| W303.asm02.HP0_G0047940 | 1947.315 | 2031.975 | 1654.592 | 1060.832 | 921.6488 | 1569.467 | 429.5966 | 809.6451 | 583.6892 | 0.025782374 | 0.665386162 | YOL117W | RRI2 | n |
| W303.asm02.HP0_G0047950 | 2133.681 | 2321.304 | 2149.353 | 2302.984 | 2403.924 | 2196.19 | 1409.768 | 1103.866 | 1078.868 | 0.679491498 | -0.06379352 | YOL116W | MSN1 | n |
| W303.asm02.HP0_G0047970 | 294.3738 | 252.6059 | 282.4125 | 101.4286 | 112.8195 | 41.78155 | 66.66155 | 66.5756 | 55.01988 | 2.88E-06 | 1.697201813 | YOL114C | PTH4 | n |
| W303.asm02.HP0_G0048000 | 3596.02 | 3532.032 | 3720.946 | 1835.44 | 1936.157 | 1957.656 | 2219.583 | 2684.5 | 2337.149 | 9.55E-19 | 0.921289817 | YOL111C | MDY2 | n |
| W303.asm02.HP0_G0048010 | 1643.411 | 1313.106 | 1357.089 | 564.1095 | 525.0447 | 752.068 | 613.5331 | 649.649 | 556.1792 | 4.48E-11 | 1.227977554 | YOL110W | SHR5 | n |
| W303.asm02.HP0_G0048030 | 2914.089 | 3047.963 | 2740.048 | 289.0019 | 328.0445 | 301.5869 | 418.4864 | 422.0034 | 275.0994 | 1.94E-71 | 3.244522152 | YOL108C | INO4 | n |
| W303.asm02.HP0_G0048040 | 1129.845 | 1140.621 | 1084.378 | 1293.561 | 1374.663 | 1350.684 | 490.0858 | 1191.918 | 840.8473 | 0.411810945 | -0.260529155 | YOL107W | | n |
| W303.asm02.HP0_G0048080 | 1189.143 | 1293.075 | 1297.804 | 921.1936 | 856.5606 | 875.8933 | 782.6559 | 1013.667 | 685.3563 | 0.002753531 | 0.510202065 | YOL102C | TPT1 | n |
| W303.asm02.HP0_G0048100 | 3850.156 | 4089.545 | 3701.543 | 5545.224 | 5437.033 | 7401.412 | 7946.303 | 7569.216 | 7890.568 | 1.20E-05 | -0.65922717 | YOL100W | PKH2 | n |
| W303.asm02.HP0_G0048110 | 8936.046 | 8460.629 | 9251.703 | 7110.42 | 6831.656 | 6516.403 | 4199.677 | 4397.211 | 5254.398 | 0.001655468 | 0.381339911 | YOL098C | SDD3 | n |
| W303.asm02.HP0_G0048130 | 8530.488 | 8186.88 | 9047.978 | 9079.245 | 9529.779 | 9073.434 | 6053.856 | 5938.114 | 6524.64 | 0.248255282 | -0.103471394 | YOL097C | WRS1 | n |
| W303.asm02.HP0_G0048140 | 5090.126 | 4671.54 | 4548.781 | 2544.745 | 2661.673 | 2659.586 | 1678.883 | 2173.371 | 2074.01 | 1.45E-11 | 0.863442061 | YOL096C | COQ3 | n |
| W303.asm02.HP0_G0048150 | 622.6324 | 599.7999 | 548.6563 | 1015.675 | 954.6268 | 1194.952 | 632.0502 | 853.6709 | 650.6699 | 1.57E-06 | -0.837845165 | YOL095C | HMI1 | n |
| W303.asm02.HP0_G0048160 | 1015.484 | 911.3843 | 1020.781 | 1905.606 | 1943.967 | 2068.567 | 1060.412 | 1666.537 | 1343.203 | 3.61E-08 | -1.005504344 | YOL094C | RFC4 | n |
| W303.asm02.HP0_G0048170 | 3637.317 | 3576.544 | 3923.593 | 2439.148 | 2453.391 | 1774.577 | 1796.158 | 1386.276 | 1587.204 | 2.84E-05 | 0.740341058 | YOL093W | TRM10 | n |
| W303.asm02.HP0_G0048180 | 1861.544 | 1855.04 | 1947.784 | 2870.567 | 2562.739 | 2198.469 | 2724.482 | 2398.869 | 2711.523 | 0.001623461 | -0.430215497 | YOL092W | YPQ1 | n |
| W303.asm02.HP0_G0048190 | 231.8988 | 253.7187 | 205.8808 | 159.7847 | 174.4363 | 93.43875 | 335.7767 | 199.7268 | 143.5301 | 0.073349895 | 0.693771924 | YOL091W | SPO21 | n |
| W303.asm02.HP0_G0048200 | 1734.476 | 1663.638 | 1835.681 | 3374.236 | 3821.978 | 4333.127 | 1369.031 | 2019.818 | 1423.34 | 2.24E-09 | -1.13932635 | YOL090W | MSH2 | n |
| W303.asm02.HP0_G0048210 | 1318.329 | 1520.087 | 1288.103 | 4106.467 | 3764.701 | 4301.981 | 9830.109 | 5568.726 | 6238.776 | 1.01E-11 | -1.560776487 | YOL089C | HAL9 | n |
| W303.asm02.HP0_G0048220 | 2553.005 | 2519.382 | 2558.959 | 2559.334 | 2817.885 | 4230.572 | 2580.049 | 4318.823 | 2971.073 | 0.234070606 | -0.332265033 | YOL088C | MPD2 | n |
| W303.asm02.HP0_G0048230 | 6734.596 | 6403.059 | 6318.709 | 4191.917 | 4143.948 | 5277.39 | 4657.667 | 5209.003 | 4460.198 | 0.000203325 | 0.515200162 | YOL087C | DUF1 | n |

| | | | | | | | | | | | | | | |
|---|---|---|---|---|---|---|---|---|---|---|---|---|---|---|
| W303.asm02.HP0_G0048240 | 939.2432 | 682.1472 | 765.3162 | 796.8394 | 968.5122 | 705.7284 | 857.9588 | 961.0509 | 717.6506 | 0.831552997 | -0.04950364 | YOL086W- | MHF1 | n |
| W303.asm02.HP0_G0048250 | 73743.82 | 80496.71 | 79542.22 | 41758 | 49540.79 | 74053.63 | 130659.1 | 109085.2 | 118686.3 | 0.029574499 | 0.499625606 | YOL086C | ADH1 | n |
| W303.asm02.HP0_G0048270 | 407.676 | 417.3005 | 355.7103 | 3452.739 | 3973.851 | 6610.601 | 2968.908 | 6512.597 | 3651.645 | 4.19E-25 | -3.571574668 | YOL084W | PHM7 | n |
| W303.asm02.HP0_G0048280 | 2257.572 | 2353.575 | 1871.252 | 2102.906 | 2186.963 | 3919.11 | 3586.144 | 3851.72 | 3012.936 | 0.236894397 | -0.340708818 | YOL083W | ATG34 | n |
| W303.asm02.HP0_G0048310 | 498.7413 | 588.6719 | 659.681 | 1854.892 | 1372.059 | 353.244 | 3105.934 | 1106.014 | 2633.778 | 0.073004886 | -1.035130643 | YOL080C | REX4 | n |
| W303.asm02.HP0_G0048320 | 1037.721 | 1196.261 | 1028.326 | 4272.504 | 3958.23 | 5114.062 | 4226.836 | 3366.363 | 4064.295 | 3.49E-38 | -2.032480428 | YOL078W | AVO1 | n |
| W303.asm02.HP0_G0048330 | 1149.964 | 760.0433 | 798.7314 | 3821.633 | 4916.328 | 2959.653 | 774.0146 | 1853.379 | 1405.399 | 1.14E-09 | -2.110406069 | YOL077W- | ATP19 | n |
| W303.asm02.HP0_G0048340 | 2983.977 | 3446.346 | 3559.259 | 5011.682 | 4603.037 | 1580.102 | 5441.557 | 2215.249 | 4037.981 | 0.741811162 | -0.164334789 | YOL077C | BRX1 | n |
| W303.asm02.HP0_G0048350 | 983.7169 | 983.7164 | 1044.495 | 5027.661 | 5138.496 | 4926.425 | 11180.62 | 7237.412 | 10342.54 | 6.61E-40 | -2.325062489 | YOL076W | MDM20 | n |
| W303.asm02.HP0_G0048360 | 1551.287 | 1496.718 | 1468.114 | 1806.262 | 1617.658 | 1941.703 | 1161.639 | 1465.737 | 1218.81 | 0.080616088 | -0.248880889 | YOL075C | | n |
| W303.asm02.HP0_G0048370 | 2525.473 | 2389.185 | 2181.69 | 335.5479 | 305.4806 | 470.2324 | 993.7508 | 897.6967 | 849.2199 | 2.75E-46 | 2.674241977 | YOL073C | DSC2 | n |
| W303.asm02.HP0_G0048380 | 1742.947 | 1752.662 | 1854.005 | 1229.648 | 1136.874 | 1351.443 | 1182.625 | 1216.615 | 1137.476 | 5.17E-07 | 0.524488031 | YOL072W | THP1 | n |
| W303.asm02.HP0_G0048390 | 5106.01 | 4864.055 | 4423.743 | 2380.097 | 2401.32 | 2455.236 | 1513.464 | 1308.962 | 1566.87 | 2.25E-19 | 0.992071583 | YOL071W | SDH5 | n |
| W303.asm02.HP0_G0048400 | 3748.501 | 3601.025 | 3842.75 | 5365.987 | 5226.148 | 5526.56 | 2841.757 | 2232.43 | 2949.544 | 7.40E-05 | -0.526239161 | YOL070C | NBA1 | n |
| W303.asm02.HP0_G0048410 | 698.8731 | 713.3057 | 718.9661 | 978.855 | 814.9041 | 1119.746 | 801.173 | 796.7595 | 746.3566 | 0.002593537 | -0.451765474 | YOL069W | NUF2 | n |
| W303.asm02.HP0_G0048430 | 1459.162 | 1468.898 | 1229.895 | 477.9647 | 396.604 | 476.3097 | 356.7627 | 367.2396 | 370.7861 | 1.41E-32 | 1.620615204 | YOL067C | RTG1 | n |
| W303.asm02.HP0_G0048440 | 1772.596 | 1661.412 | 1699.864 | 3367.289 | 3496.537 | 2926.988 | 2344.264 | 2557.791 | 2355.09 | 5.08E-18 | -0.931216763 | YOL066C | RIB2 | n |
| W303.asm02.HP0_G0048450 | 1211.38 | 1308.654 | 1124.26 | 370.2837 | 339.3264 | 379.0726 | 774.0146 | 561.5974 | 944.9066 | 2.68E-17 | 1.742650272 | YOL065C | INP54 | n |
| W303.asm02.HP0_G0048460 | 4866.698 | 4194.148 | 4810.713 | 4462.162 | 4611.715 | 6795.2 | 6372.35 | 8824.488 | 7342.762 | 0.4086825 | -0.194084586 | YOL064C | MET22 | n |
| W303.asm02.HP0_G0048470 | 1734.476 | 1720.391 | 1763.461 | 1374.843 | 1404.169 | 1837.629 | 1423.347 | 1566.674 | 1497.498 | 0.258260188 | 0.17669601 | YOL063C | CRT10 | n |
| W303.asm02.HP0_G0048480 | 5536.981 | 5062.133 | 5281.76 | 4289.872 | 4136.137 | 5262.197 | 5950.16 | 5853.283 | 6204.089 | 0.101769218 | 0.2143153 | YOL062C | APM4 | n |
| W303.asm02.HP0_G0048490 | 6398.925 | 5767.65 | 6822.093 | 7870.439 | 7897.367 | 7863.288 | 11368.26 | 8375.639 | 9970.559 | 0.033408071 | -0.31552811 | YOL061W | PRS5 | n |
| W303.asm02.HP0_G0048500 | 5558.159 | 4998.704 | 5564.172 | 4753.248 | 4790.491 | 5276.63 | 7927.786 | 6803.596 | 7541.312 | 0.297429088 | 0.121387984 | YOL060C | MAM3 | n |
| W303.asm02.HP0_G0048520 | 959.3622 | 747.8025 | 959.34 | 6388.61 | 5613.205 | 6900.034 | 4383.614 | 3093.618 | 2362.267 | 3.25E-31 | -2.825437626 | YOL058W | ARG1 | n |
| W303.asm02.HP0_G0048530 | 5308.259 | 4973.109 | 5602.977 | 6985.371 | 7473.86 | 9091.666 | 5636.604 | 6953.928 | 5621.596 | 0.000302703 | -0.568156693 | YOL057W | | n |
| W303.asm02.HP0_G0048540 | 1185.967 | 1111.689 | 1179.234 | 3049.109 | 3385.454 | 5464.268 | 12591.63 | 10290.22 | 13231.08 | 1.15E-13 | -1.774933548 | YOL056W | GPM3 | n |
| W303.asm02.HP0_G0048550 | 922.3008 | 842.3906 | 740.5243 | 321.6536 | 405.2825 | 654.8309 | 2367.719 | 3634.813 | 2919.642 | 0.004068832 | 0.858558125 | YOL055C | THI20 | n |
| W303.asm02.HP0_G0048560 | 1389.275 | 1683.668 | 1605.008 | 1810.43 | 1937.024 | 1336.25 | 970.2958 | 633.542 | 813.3373 | 0.624353832 | -0.119952275 | YOL054W | PSH1 | n |
| W303.asm02.HP0_G0048570 | 3480.6 | 3763.494 | 3201.393 | 2330.078 | 2396.981 | 3628.158 | 1955.405 | 2591.079 | 1859.911 | 0.182718857 | 0.322097705 | YOL053W | AIM39 | n |
| W303.asm02.HP0_G0048590 | 3359.886 | 3148.115 | 3254.211 | 3121.36 | 3336.854 | 3248.326 | 2865.212 | 2310.817 | 1844.362 | 0.967025572 | 0.00830612 | YOL052C | SPE2 | n |
| W303.asm02.HP0_G0048600 | 9182.77 | 9395.382 | 9958.812 | 12509.75 | 11313.2 | 14245.23 | 15760.52 | 14380.33 | 13809.99 | 0.000869178 | -0.415794943 | YOL051W | GAL11 | n |
| W303.asm02.HP0_G0048610 | 6802.365 | 6447.571 | 6760.652 | 2658.679 | 2679.898 | 2626.161 | 1597.408 | 3362.068 | 2150.56 | 2.19E-07 | 1.329071558 | YOL049W | GSH2 | n |
| W303.asm02.HP0_G0048620 | 2228.982 | 2378.057 | 2046.951 | 83.36594 | 67.69172 | 78.24546 | 49.37892 | 92.34679 | 41.86295 | 2.78E-79 | 4.8566738 | YOL048C | RRT8 | n |
| W303.asm02.HP0_G0048650 | 1207.145 | 1347.602 | 1215.883 | 1006.644 | 934.6664 | 1247.369 | 802.4075 | 1341.176 | 826.4943 | 0.349086293 | 0.241700912 | YOL044W | PEX15 | n |
| W303.asm02.HP0_G0048660 | 1557.64 | 1651.397 | 1550.035 | 1381.096 | 1383.341 | 1325.615 | 2847.929 | 2263.57 | 2704.347 | 0.081575336 | 0.218532793 | YOL043C | NTG2 | n |
| W303.asm02.HP0_G0048670 | 2367.698 | 2497.126 | 2382.181 | 1118.493 | 1031.865 | 1188.115 | 1856.648 | 1958.611 | 1966.363 | 2.70E-34 | 1.117555957 | YOL042W | NGL1 | n |
| W303.asm02.HP0_G0048680 | 3646.847 | 3958.234 | 4191.993 | 4754.637 | 4058.032 | 1258.005 | 6885.891 | 2912.145 | 6297.384 | 0.668129074 | 0.228264493 | YOL041C | NOP12 | n |
| W303.asm02.HP0_G0048690 | 14302.54 | 14447.5 | 14663.89 | 20601.11 | 23285.95 | 23825.36 | 17782.58 | 15016.02 | 14693.9 | 4.23E-08 | -0.641241914 | YOL040C | RPS15 | n |
| W303.asm02.HP0_G0048700 | 8640.613 | 7633.817 | 8021.807 | 9729.5 | 11606.53 | 10641.38 | 10548.57 | 7445.729 | 8270.923 | 0.018947165 | -0.396271062 | YOL039W | RPP2A | n |
| W303.asm02.HP0_G0048720 | 2438.644 | 2480.434 | 2637.646 | 5441.712 | 4788.755 | 5467.306 | 12271.9 | 7943.972 | 10593.72 | 3.42E-09 | -1.054777728 | YOL036W | | n |
| W303.asm02.HP0_G0048730 | 1280.209 | 1486.703 | 1378.647 | 2799.011 | 2698.99 | 2704.406 | 1624.567 | 1572.043 | 1314.497 | 5.72E-16 | -0.984667502 | YOL034W | SMC5 | n |
| W303.asm02.HP0_G0048740 | 3502.837 | 3230.462 | 3236.964 | 1992.446 | 1828.544 | 1766.98 | 1170.28 | 1004.003 | 1041.789 | 4.40E-15 | 0.835130254 | YOL033W | MSE1 | n |
| W303.asm02.HP0_G0048750 | 3819.448 | 4438.965 | 3351.223 | 440.45 | 481.6526 | 514.2929 | 338.2456 | 221.2028 | 248.7855 | 3.77E-55 | 3.014991976 | YOL032W | OPI10 | n |
| W303.asm02.HP0_G0048760 | 2293.575 | 2661.821 | 2496.44 | 2041.076 | 2053.315 | 3195.149 | 1727.028 | 2656.581 | 2231.893 | 0.911893327 | 0.031716938 | YOL031C | SIL1 | n |
| W303.asm02.HP0_G0048770 | 17229.34 | 18272.2 | 18830.01 | 4877.602 | 5225.28 | 4775.252 | 3808.349 | 3182.743 | 2865.818 | 1.02E-46 | 1.868649782 | YOL030W | GAS5 | n |
| W303.asm02.HP0_G0048800 | 11605.53 | 12960.8 | 12189.01 | 4844.95 | 4728.874 | 6065.922 | 2055.398 | 3736.824 | 2350.306 | 3.75E-07 | 1.232721207 | YOL027C | MDM38 | n |
| W303.asm02.HP0_G0048810 | 816.4109 | 741.1257 | 701.7195 | 2363.424 | 2760.607 | 2817.596 | 6026.698 | 5142.428 | 5917.029 | 1.50E-43 | -1.813293375 | YOL026C | MIM1 | n |
| W303.asm02.HP0_G0048820 | 1368.097 | 1577.952 | 1306.427 | 2782.338 | 2703.329 | 5179.393 | 4689.763 | 4405.801 | 4313.08 | 6.72E-07 | -1.326582359 | YOL025W | LAG2 | n |
| W303.asm02.HP0_G0048840 | 2906.677 | 3143.664 | 2691.542 | 1898.659 | 1579.473 | 1317.258 | 493.7892 | 444.5532 | 455.7081 | 1.79E-07 | 0.866052628 | YOL023W | IFM1 | n |
| W303.asm02.HP0_G0048850 | 3666.966 | 3589.897 | 3912.814 | 3401.33 | 3246.599 | 2313.938 | 3467.635 | 2365.581 | 3352.624 | 0.158684994 | 0.317722446 | YOL022C | TSR4 | n |
| W303.asm02.HP0_G0048860 | 3282.586 | 3224.898 | 3665.972 | 7026.359 | 6844.674 | 4514.687 | 9249.907 | 5852.21 | 7388.213 | 0.000367128 | -0.85376472 | YOL021C | DIS3 | n |
| W303.asm02.HP0_G0048870 | 1452.809 | 1418.822 | 1557.58 | 5578.571 | 5570.681 | 5339.683 | 21008.26 | 15635.6 | 17929.3 | 1.01E-45 | -1.89634611 | YOL020W | TAT2 | n |
| W303.asm02.HP0_G0048890 | 2940.562 | 3554.288 | 3430.988 | 3703.532 | 3876.653 | 3584.857 | 1812.206 | 2179.814 | 1568.067 | 0.314986412 | -0.169731202 | YOL019W | TOS7 | n |
| W303.asm02.HP0_G0048900 | 4201.71 | 4469.01 | 4124.084 | 1536.712 | 1516.121 | 1391.706 | 2050.46 | 1807.205 | 1764.224 | 3.17E-50 | 1.52542512 | YOL018C | TLG2 | n |

| | | | | | | | | | | | | | | |
|---|---|---|---|---|---|---|---|---|---|---|---|---|---|---|
| W303.asm02.HP0_G0048920 | 12695.14 | 16794.4 | 11539.03 | 4509.402 | 4997.037 | 4921.867 | 4000.927 | 4881.494 | 4020.039 | 1.21E-17 | 1.507749936 | YOL016C | CMK2 | n |
| W303.asm02.HP0_G0048950 | 1684.708 | 1974.11 | 1582.372 | 1668.708 | 1794.698 | 2683.135 | 2576.345 | 3578.975 | 2639.758 | 0.375541958 | -0.229925784 | YOL013C | HRD1 | n |
| W303.asm02.HP0_G0048960 | 1637.057 | 1663.638 | 1693.397 | 2637.142 | 3232.713 | 2825.952 | 1596.174 | 2258.201 | 1650.596 | 3.69E-06 | -0.799948439 | YOL012C | HTZ1 | n |
| W303.asm02.HP0_G0048970 | 1298.21 | 1146.185 | 1246.064 | 14012.42 | 16517.65 | 15751.65 | 13113.81 | 22286.72 | 15952.18 | 4.59E-70 | -3.648511147 | YOL011W | PLB3 | n |
| W303.asm02.HP0_G0048980 | 979.4813 | 1084.981 | 1165.221 | 3354.784 | 3127.704 | 859.9403 | 4619.398 | 1717.006 | 3618.155 | 0.02129132 | -1.184861358 | YOL010W | RCL1 | n |
| W303.asm02.HP0_G0048990 | 1746.124 | 1680.33 | 1599.619 | 1174.07 | 1090.878 | 967.8127 | 1043.13 | 1102.793 | 1150.633 | 3.15E-08 | 0.636461786 | YOL009C | MDM12 | n |
| W303.asm02.HP0_G0049000 | 1848.837 | 1601.321 | 1611.476 | 1240.068 | 1333.006 | 947.3018 | 706.1186 | 680.7892 | 855.2003 | 0.004220165 | 0.524127493 | YOL008W | COQ10 | n |
| W303.asm02.HP0_G0049020 | 2951.151 | 3133.649 | 3241.276 | 3200.557 | 2981.907 | 1973.609 | 1947.999 | 2366.655 | 2986.623 | 0.452716446 | 0.193385531 | YOL006C | TOP1 | n |
| W303.asm02.HP0_G0049030 | 1963.198 | 1838.348 | 1871.252 | 1841.692 | 2074.144 | 1444.882 | 1369.031 | 1492.582 | 1285.791 | 0.651419997 | 0.081894572 | YOL005C | RPB11 | n |
| W303.asm02.HP0_G0049040 | 7427.116 | 7524.763 | 7519.501 | 11727.5 | 11549.25 | 11514.24 | 9363.478 | 10100.16 | 10448.99 | 4.92E-19 | -0.630645059 | YOL004W | SIN3 | n |
| W303.asm02.HP0_G0049050 | 1369.156 | 1253.014 | 1220.194 | 1216.448 | 1025.79 | 1469.951 | 2159.093 | 2495.511 | 2462.738 | 0.790902728 | 0.049398597 | YOL003C | PFA4 | n |
| W303.asm02.HP0_G0049060 | 3324.942 | 2940.021 | 3202.471 | 650.2543 | 706.4238 | 1103.793 | 2040.584 | 3114.02 | 2944.76 | 1.43E-14 | 1.944031503 | YOL002C | IZH2 | n |
| W303.asm02.HP0_G0049070 | 1000.659 | 985.942 | 1027.248 | 2086.927 | 1940.496 | 2316.217 | 2275.134 | 2222.766 | 2303.658 | 2.28E-26 | -1.073936463 | YOL001W | PHO80 | n |
| W303.asm02.HP0_G0049080 | 4734.336 | 4516.861 | 5226.786 | 4846.34 | 4267.182 | 2225.058 | 7443.873 | 4373.587 | 7026.995 | 0.304181713 | 0.352615857 | YOR001W | RRP6 | n |
| W303.asm02.HP0_G0049090 | 4376.428 | 4030.567 | 4487.34 | 2810.127 | 2686.84 | 3818.834 | 3614.537 | 4587.273 | 4398.002 | 0.01155016 | 0.468939699 | YOR002W | ALG6 | n |
| W303.asm02.HP0_G0049110 | 1199.732 | 1229.646 | 1315.05 | 2470.411 | 2322.347 | 828.7941 | 4141.657 | 1390.571 | 3328.703 | 0.241228728 | -0.586213225 | YOR004W | UTP23 | n |
| W303.asm02.HP0_G0049120 | 2717.134 | 2866.576 | 2414.519 | 1988.972 | 1734.817 | 2076.163 | 2713.372 | 2305.448 | 2650.523 | 0.000634864 | 0.463339088 | YOR005C | DNL4 | n |
| W303.asm02.HP0_G0049130 | 2190.861 | 2135.466 | 2043.718 | 2243.238 | 2270.276 | 1447.921 | 3047.914 | 2133.64 | 2785.68 | 0.703113218 | 0.095697705 | YOR006C | TSR3 | n |
| W303.asm02.HP0_G0049140 | 23519.2 | 25256.14 | 24399.57 | 6321.917 | 6880.255 | 7759.214 | 5198.366 | 6099.184 | 5650.302 | 1.55E-53 | 1.803643414 | YOR007C | SGT2 | n |
| W303.asm02.HP0_G0049220 | 10258.61 | 11091.29 | 10794.19 | 8762.455 | 9179.17 | 12253.39 | 17282.62 | 14824.88 | 15028.8 | 0.618879789 | 0.090235311 | YOR014W | RTS1 | n |
| W303.asm02.HP0_G0049230 | 2058.499 | 2186.655 | 2134.262 | 6459.471 | 7103.291 | 7358.871 | 3447.883 | 5285.243 | 4482.924 | 1.70E-23 | -1.713493282 | YOR016C | ERP4 | n |
| W303.asm02.HP0_G0049240 | 2600.655 | 2820.951 | 2493.206 | 1645.088 | 1362.513 | 1094.677 | 867.8346 | 704.4128 | 900.6515 | 5.20E-07 | 0.947911331 | YOR017W | PET127 | n |
| W303.asm02.HP0_G0049250 | 2038.38 | 1760.452 | 2095.457 | 4296.819 | 3219.696 | 5496.933 | 26732.51 | 10913.03 | 13341.12 | 0.001337708 | -1.142636582 | YOR018W | ROD1 | n |
| W303.asm02.HP0_G0049270 | 7062.854 | 6488.745 | 6214.152 | 1782.642 | 2203.452 | 1983.484 | 611.0642 | 697.97 | 773.8666 | 4.77E-36 | 1.727644095 | YOR020C | HSP10 | n |
| W303.asm02.HP0_G0049280 | 2008.731 | 1827.22 | 1559.736 | 1822.935 | 2200.849 | 1784.452 | 1375.203 | 1633.25 | 1601.557 | 0.546088097 | -0.105993703 | YOR020W | MCO10 | n |
| W303.asm02.HP0_G0049290 | 3384.24 | 3191.514 | 3380.326 | 3809.823 | 3861.899 | 2304.822 | 3220.74 | 1719.154 | 2933.995 | 0.992921844 | -0.002936747 | YOR021C | SFM1 | n |
| W303.asm02.HP0_G0049300 | 5207.664 | 5795.47 | 5208.462 | 3511.095 | 3381.982 | 6106.184 | 1759.124 | 3926.886 | 3268.898 | 0.384584719 | 0.318576698 | YOR022C | DDL1 | n |
| W303.asm02.HP0_G0049310 | 1048.31 | 973.7012 | 1188.935 | 2318.962 | 2252.919 | 2038.18 | 1267.804 | 918.0989 | 566.944 | 0.000203325 | -1.041645375 | YOR025W | HST3 | n |
| W303.asm02.HP0_G0049320 | 1223.028 | 1034.905 | 1205.104 | 1244.931 | 1293.085 | 1136.458 | 1282.618 | 1223.058 | 1180.535 | 0.509608404 | -0.085079017 | YOR026W | BUB3 | n |
| W303.asm02.HP0_G0049330 | 25755.59 | 28572.29 | 26028.3 | 11345.41 | 11425.15 | 15447.78 | 5016.899 | 8166.248 | 7690.822 | 1.90E-06 | 1.072138155 | YOR027W | STI1 | n |
| W303.asm02.HP0_G0049340 | 1754.595 | 2302.386 | 1740.825 | 4214.843 | 4045.882 | 6406.252 | 4894.686 | 5181.085 | 4292.747 | 3.12E-09 | -1.339079751 | YOR028C | CIN5 | n |
| W303.asm02.HP0_G0049360 | 1882.722 | 1883.973 | 1699.864 | 1064.305 | 1030.129 | 1035.423 | 1409.768 | 1319.7 | 1269.045 | 9.00E-18 | 0.804311192 | YOR030W | DFG16 | n |
| W303.asm02.HP0_G0049380 | 1375.509 | 2045.329 | 1317.206 | 13087.06 | 11294.97 | 16990.66 | 31384.01 | 22097.73 | 27208.53 | 7.55E-37 | -3.126418679 | YOR032C | HMS1 | n |
| W303.asm02.HP0_G0049400 | 2124.151 | 2240.069 | 2228.04 | 3124.138 | 3558.154 | 2892.043 | 2329.451 | 2598.596 | 2602.679 | 4.82E-06 | -0.538203025 | YOR033C | EXO1 | n |
| W303.asm02.HP0_G0049430 | 4717.394 | 5330.319 | 4498.119 | 3142.201 | 2686.84 | 3840.105 | 1141.888 | 2264.644 | 1632.655 | 0.031650838 | 0.589085944 | YOR035C | SHE4 | n |
| W303.asm02.HP0_G0049440 | 6748.361 | 6843.728 | 6036.297 | 1540.186 | 1609.848 | 2004.755 | 2175.142 | 2265.718 | 2002.245 | 1.19E-43 | 1.92895011 | YOR036W | PEP12 | n |
| W303.asm02.HP0_G0049450 | 3483.777 | 3468.602 | 3242.354 | 763.493 | 752.4195 | 679.8998 | 457.9895 | 417.7082 | 449.7277 | 1.34E-114 | 2.215003928 | YOR037W | CYC2 | n |
| W303.asm02.HP0_G0049460 | 1063.134 | 1018.213 | 1100.546 | 3932.788 | 3555.551 | 3503.573 | 3747.86 | 3478.038 | 3499.743 | 1.39E-81 | -1.788567224 | YOR038C | HIR2 | n |
| W303.asm02.HP0_G0049470 | 5138.835 | 5331.431 | 5353.98 | 4546.222 | 4864.257 | 4592.173 | 4966.285 | 4579.757 | 4752.043 | 0.021500721 | 0.176588271 | YOR039W | CKB2 | n |
| W303.asm02.HP0_G0049480 | 3008.331 | 2872.14 | 2800.411 | 1635.362 | 1827.676 | 2859.378 | 3384.925 | 2744.633 | 2628.993 | 0.061007691 | 0.457371899 | YOR040W | GLO4 | n |
| W303.asm02.HP0_G0049490 | 10414.27 | 10446.98 | 10388.9 | 3665.322 | 3731.723 | 5926.144 | 1973.922 | 3504.883 | 2443.6 | 5.75E-06 | 1.229910747 | YOR042W | CUE5 | n |
| W303.asm02.HP0_G0049500 | 4276.892 | 4014.987 | 4334.277 | 6054.451 | 5820.62 | 5624.557 | 5751.41 | 3777.628 | 4419.532 | 0.006916958 | -0.470908841 | YOR043W | WHI2 | n |
| W303.asm02.HP0_G0049520 | 1777.891 | 1495.605 | 1546.801 | 1351.918 | 1630.676 | 1122.025 | 662.912 | 746.291 | 619.5717 | 0.224727962 | 0.232285828 | YOR045W | TOM6 | n |
| W303.asm02.HP0_G0049530 | 5460.741 | 5517.269 | 5548.003 | 11121.02 | 11037.22 | 11044 | 7354.991 | 8768.65 | 8414.453 | 6.97E-28 | -1.00654961 | YOR046C | DBP5 | n |
| W303.asm02.HP0_G0049550 | 4161.472 | 4312.105 | 4588.663 | 6560.204 | 6041.052 | 4651.426 | 8098.143 | 5464.568 | 7965.922 | 0.062294714 | -0.401430735 | YOR048C | RAT1 | n |
| W303.asm02.HP0_G0049560 | 1804.363 | 1961.869 | 1774.24 | 1162.26 | 1119.517 | 1362.838 | 2749.172 | 2380.614 | 1696.048 | 0.003219786 | 0.60413583 | YOR049C | RSB1 | n |
| W303.asm02.HP0_G0049570 | 7526.652 | 7620.464 | 8181.338 | 4109.941 | 4183.869 | 4313.376 | 13011.35 | 10316 | 12021.84 | 1.19E-14 | 0.887857134 | YOR051C | ETT1 | n |
| W303.asm02.HP0_G0049580 | 8134.46 | 9284.102 | 7829.939 | 1588.121 | 1936.157 | 2445.36 | 7053.779 | 6001.468 | 6444.502 | 4.46E-30 | 2.080577727 | YOR052C | TMC1 | n |
| W303.asm02.HP0_G0049600 | 3249.76 | 3463.038 | 3556.026 | 3241.546 | 2902.933 | 751.3083 | 4877.403 | 1788.951 | 3776.038 | 0.297991781 | 0.574485951 | YOR056C | NOB1 | n |
| W303.asm02.HP0_G0049610 | 2366.639 | 2372.493 | 2372.48 | 1492.945 | 1375.53 | 1800.405 | 1719.621 | 2133.64 | 1819.244 | 4.92E-05 | 0.606894154 | YOR057W | SGT1 | n |
| W303.asm02.HP0_G0049620 | 2938.444 | 3284.99 | 2775.619 | 1332.466 | 1050.089 | 1412.976 | 559.2163 | 357.5754 | 330.1193 | 8.62E-08 | 1.245229281 | YOR058C | ASE1 | n |
| W303.asm02.HP0_G0049630 | 4571.266 | 4869.619 | 4234.031 | 2360.645 | 2416.074 | 3000.675 | 1767.765 | 3247.171 | 2314.423 | 0.000705093 | 0.81415938 | YOR059C | LPL1 | n |
| W303.asm02.HP0_G0049640 | 1280.209 | 1536.779 | 1263.311 | 1549.912 | 1784.284 | 1569.467 | 1804.8 | 1467.885 | 1462.811 | 0.094510981 | -0.265057505 | YOR060C | SLD7 | n |
| W303.asm02.HP0_G0049650 | 9147.826 | 9497.76 | 9400.454 | 4988.757 | 5382.359 | 5235.609 | 3255.306 | 4000.979 | 3210.29 | 3.46E-13 | 0.845687768 | YOR061W | CKA2 | n |

| | | | | | | | | | | | | | | |
|---|---|---|---|---|---|---|---|---|---|---|---|---|---|---|
| W303.asm02.HP0_G0049670 | 49870.95 | 47784.8 | 54214.57 | 51593.09 | 60735.96 | 51252.29 | 57062.28 | 45201.61 | 44440.51 | 0.510023869 | -0.107157524 | YOR063W | RPL3 | n |
| W303.asm02.HP0_G0049680 | 501.918 | 491.8582 | 534.6434 | 1057.358 | 1247.09 | 907.7992 | 781.4215 | 603.4756 | 641.1012 | 8.90E-10 | -1.071086738 | YOR064C | YNG1 | n |
| W303.asm02.HP0_G0049690 | 12428.29 | 13313.56 | 13179.61 | 20257.23 | 20789.17 | 19681.39 | 3610.834 | 11595.97 | 9658.381 | 0.092692413 | -0.64178996 | YOR065W | CYT1 | n |
| W303.asm02.HP0_G0049700 | 1902.841 | 2135.466 | 1819.512 | 3313.796 | 3352.476 | 3397.22 | 1872.696 | 2479.404 | 2179.266 | 1.40E-08 | -0.780758113 | YOR066W | MSA1 | n |
| W303.asm02.HP0_G0049710 | 2429.114 | 2192.219 | 2394.038 | 2589.207 | 2482.03 | 3303.781 | 3036.804 | 4339.226 | 3546.39 | 0.194880766 | -0.255601248 | YOR067C | ALG8 | n |
| W303.asm02.HP0_G0049730 | 2361.344 | 2666.272 | 2364.935 | 1345.665 | 1431.072 | 1606.691 | 1706.042 | 1633.25 | 1510.655 | 2.85E-10 | 0.754038516 | YOR070C | GYP1 | n |
| W303.asm02.HP0_G0049750 | 320.8463 | 327.1636 | 366.4894 | 602.3189 | 612.6968 | 691.2948 | 141.9644 | 273.819 | 131.5693 | 0.001670936 | -0.910017238 | YOR073W | SGO1 | n |
| W303.asm02.HP0_G0049760 | 1508.931 | 1402.13 | 1497.217 | 1863.923 | 2311.065 | 2069.326 | 608.5952 | 729.1102 | 592.0617 | 0.00034608 | -0.501937151 | YOR074C | CDC21 | n |
| W303.asm02.HP0_G0049770 | 1195.497 | 1280.834 | 1152.286 | 1925.058 | 1850.24 | 2050.335 | 1328.293 | 1626.807 | 1594.38 | 2.44E-08 | -0.683194646 | YOR075W | UFE1 | n |
| W303.asm02.HP0_G0049790 | 540.0383 | 523.0166 | 528.176 | 562.7201 | 471.2385 | 283.3549 | 1011.033 | 569.114 | 1005.907 | 0.428280515 | 0.272384304 | YOR077W | RTS2 | n |
| W303.asm02.HP0_G0049800 | 854.5313 | 842.3906 | 953.9505 | 3552.778 | 3145.061 | 1113.668 | 3708.357 | 1754.589 | 3256.938 | 0.000317213 | -1.55913004 | YOR078W | BUD21 | n |
| W303.asm02.HP0_G0049810 | 239.3111 | 193.6274 | 219.8937 | 897.5733 | 975.455 | 1281.554 | 2773.861 | 2460.076 | 2760.563 | 1.90E-35 | -2.272130106 | YOR079C | ATX2 | n |
| W303.asm02.HP0_G0049820 | 2108.267 | 2085.39 | 2066.354 | 1862.534 | 1834.619 | 2507.653 | 4105.857 | 4023.528 | 4348.963 | 0.944327747 | 0.012695669 | YOR080W | DIA2 | n |
| W303.asm02.HP0_G0049840 | 643.8104 | 644.312 | 611.175 | 1961.878 | 2094.104 | 2684.655 | 3741.688 | 2259.275 | 2640.954 | 3.30E-16 | -1.827419421 | YOR083W | WHI5 | n |
| W303.asm02.HP0_G0049850 | 591.9244 | 508.5502 | 643.5124 | 766.9666 | 797.5473 | 1029.346 | 1402.361 | 1136.08 | 1322.869 | 0.001921476 | -0.572564596 | YOR084W | LPX1 | n |
| W303.asm02.HP0_G0049860 | 3213.758 | 3254.944 | 3279.003 | 3101.213 | 3338.59 | 3722.357 | 1896.151 | 4982.432 | 3646.861 | 0.867606269 | -0.060064262 | YOR085W | OST3 | n |
| W303.asm02.HP0_G0049870 | 9329.957 | 8917.99 | 8883.057 | 16279.98 | 16547.15 | 21778.07 | 15607.44 | 20968.09 | 19152.9 | 4.32E-09 | -1.009096628 | YOR086C | TCB1 | n |
| W303.asm02.HP0_G0049880 | 1958.963 | 1929.598 | 1894.966 | 2276.585 | 2244.241 | 4048.253 | 4583.599 | 8064.237 | 5857.225 | 0.065290136 | -0.567217744 | YOR087W | YVC1 | n |
| W303.asm02.HP0_G0049890 | 6789.659 | 7158.651 | 6344.579 | 2418.307 | 2461.201 | 2992.319 | 3163.955 | 2598.596 | 2827.543 | 2.41E-23 | 1.366170972 | YOR089C | VPS21 | n |
| W303.asm02.HP0_G0049900 | 5270.139 | 5206.798 | 5396.018 | 4762.279 | 4925.874 | 7981.036 | 5936.581 | 8692.41 | 6542.581 | 0.571094167 | -0.154674271 | YOR090C | PTC5 | n |
| W303.asm02.HP0_G0049910 | 3284.704 | 3429.654 | 3916.048 | 5487.563 | 5383.227 | 3372.151 | 5095.905 | 4220.034 | 5401.517 | 0.067621841 | -0.422037933 | YOR091W | TMA46 | n |
| W303.asm02.HP0_G0049920 | 1339.507 | 1362.069 | 1292.414 | 2317.573 | 2233.827 | 4840.583 | 8325.287 | 6340.789 | 8522.101 | 8.49E-05 | -1.233635508 | YOR092W | ECM3 | n |
| W303.asm02.HP0_G0049930 | 2106.15 | 2171.075 | 2042.64 | 5031.134 | 4590.019 | 6716.195 | 4125.609 | 6767.087 | 5466.105 | 1.20E-09 | -1.370245841 | YOR093C | CMR2 | n |
| W303.asm02.HP0_G0049940 | 1190.202 | 1253.014 | 1107.014 | 1959.1 | 2008.188 | 1550.475 | 1765.297 | 1834.05 | 1431.713 | 0.000115876 | -0.636130489 | YOR094W | ARF3 | n |
| W303.asm02.HP0_G0049950 | 1001.718 | 932.5276 | 1213.727 | 4489.95 | 4447.693 | 2182.516 | 9833.813 | 4678.546 | 4498.473 | 1.97E-06 | -1.820657175 | YOR095C | RKI1 | n |
| W303.asm02.HP0_G0049960 | 8835.451 | 7580.403 | 8973.602 | 22790.86 | 25862.57 | 25381.15 | 21388.48 | 18910.69 | 19040.47 | 5.69E-40 | -1.54392239 | YOR096W | RPS7A | n |
| W303.asm02.HP0_G0049970 | 1657.176 | 1537.891 | 1471.347 | 296.6438 | 255.1457 | 429.9702 | 406.1416 | 444.5532 | 406.6687 | 1.28E-27 | 2.248046924 | YOR097C | | n |
| W303.asm02.HP0_G0049980 | 3099.397 | 3231.575 | 3261.756 | 8399.118 | 7935.552 | 8026.616 | 9716.538 | 7301.84 | 8214.707 | 6.22E-26 | -1.344614501 | YOR098C | NUP1 | n |
| W303.asm02.HP0_G0049990 | 9011.228 | 9084.91 | 9191.34 | 8308.11 | 8896.254 | 8756.654 | 6240.261 | 7795.787 | 5042.692 | 0.707685799 | 0.071908025 | YOR099W | KTR1 | n |
| W303.asm02.HP0_G0050000 | 369.5556 | 354.9836 | 271.6334 | 85.45009 | 91.99131 | 77.48579 | 959.1856 | 404.8226 | 305.0015 | 2.62E-06 | 1.966706592 | YOR100C | CRC1 | n |
| W303.asm02.HP0_G0050020 | 1667.765 | 1592.419 | 1576.983 | 805.8707 | 1002.358 | 760.4243 | 648.0984 | 1133.933 | 782.2391 | 8.52E-05 | 0.91350694 | YOR103C | OST2 | n |
| W303.asm02.HP0_G0050030 | 2202.509 | 2185.542 | 2043.718 | 2985.89 | 3255.277 | 2611.727 | 4155.236 | 4693.579 | 4967.338 | 0.000482587 | -0.460787697 | YOR104W | PIN2 | n |
| W303.asm02.HP0_G0050040 | 1471.869 | 1569.05 | 1474.581 | 607.8766 | 693.4062 | 815.8798 | 492.5548 | 579.852 | 377.9626 | 1.96E-08 | 1.09291903 | YOR106W | VAM3 | n |
| W303.asm02.HP0_G0050050 | 207.5441 | 202.5298 | 213.4262 | 3575.009 | 2876.898 | 3136.655 | 6214.338 | 3846.351 | 5411.085 | 1.24E-81 | -3.942892139 | YOR107W | RGS2 | n |
| W303.asm02.HP0_G0050070 | 6011.368 | 5552.879 | 5638.548 | 8069.823 | 7336.741 | 8636.627 | 6562.459 | 7454.319 | 7486.292 | 1.24E-05 | -0.483035493 | YOR109W | INP53 | n |
| W303.asm02.HP0_G0050080 | 2406.877 | 2386.959 | 2423.142 | 6232.298 | 6721.44 | 8665.494 | 22016.83 | 21132.38 | 20902.77 | 3.66E-28 | -1.582837556 | YOR110W | TFC7 | n |
| W303.asm02.HP0_G0050100 | 3694.498 | 3510.888 | 3628.246 | 4880.381 | 4845.165 | 5778.769 | 4956.409 | 6964.666 | 7327.213 | 0.004315527 | -0.517157604 | YOR112W | CEX1 | n |
| W303.asm02.HP0_G0050110 | 4044.993 | 4511.297 | 4119.773 | 9855.243 | 9726.779 | 12957.6 | 29589.09 | 19491.62 | 26988.45 | 1.19E-11 | -1.360112027 | YOR113W | AZF1 | n |
| W303.asm02.HP0_G0050120 | 645.9282 | 584.2207 | 558.3575 | 575.9197 | 644.807 | 645.7149 | 103.6957 | 178.2508 | 100.4711 | 0.812398642 | -0.061285292 | YOR114W | CIM1 | n |
| W303.asm02.HP0_G0050130 | 2908.795 | 2852.11 | 2783.164 | 1967.436 | 2136.628 | 2339.007 | 2307.23 | 3023.821 | 2814.386 | 0.004197871 | 0.407263234 | YOR115C | TRS33 | n |
| W303.asm02.HP0_G0050140 | 2268.161 | 2307.95 | 2444.7 | 7578.658 | 7387.075 | 5231.81 | 5219.352 | 4389.694 | 5216.124 | 2.01E-18 | -1.524462957 | YOR116C | RPO31 | n |
| W303.asm02.HP0_G0050150 | 9390.314 | 10433.63 | 9700.113 | 6967.308 | 7353.23 | 9249.676 | 4518.172 | 7717.4 | 5704.126 | 0.166882677 | 0.32492261 | YOR117W | RPT5 | n |
| W303.asm02.HP0_G0050160 | 3367.298 | 3549.836 | 3559.259 | 1521.428 | 1537.817 | 1566.428 | 2650.414 | 2070.286 | 2540.483 | 1.06E-22 | 1.179407336 | YOR118W | RTC5 | n |
| W303.asm02.HP0_G0050170 | 1740.829 | 1967.433 | 2000.601 | 1476.967 | 1232.336 | 312.9818 | 1912.199 | 626.0254 | 1473.576 | 0.107324796 | 0.917546679 | YOR119C | RIO1 | n |
| W303.asm02.HP0_G0050190 | 7536.182 | 7885.311 | 7584.176 | 3529.853 | 4214.243 | 4158.404 | 4241.65 | 5213.299 | 4093.001 | 4.68E-12 | 0.950848179 | YOR122C | PFY1 | n |
| W303.asm02.HP0_G0050200 | 7382.642 | 7880.859 | 7743.706 | 4668.492 | 4465.05 | 4049.772 | 3173.83 | 3086.101 | 3081.113 | 3.38E-20 | 0.80330453 | YOR123C | LEO1 | n |
| W303.asm02.HP0_G0050210 | 8257.292 | 8853.448 | 8042.288 | 5488.257 | 5145.438 | 7304.175 | 4235.477 | 6007.911 | 4454.218 | 0.01794752 | 0.487689839 | YOR124C | UBP2 | n |
| W303.asm02.HP0_G0050220 | 3165.048 | 3191.514 | 2834.904 | 2498.894 | 2533.232 | 2955.095 | 5671.169 | 4416.539 | 5432.615 | 0.189348658 | 0.202576495 | YOR125C | CAT5 | n |
| W303.asm02.HP0_G0050230 | 1433.749 | 1277.496 | 1339.842 | 378.6203 | 540.6659 | 875.1336 | 16.04815 | 83.75639 | 25.11777 | 0.027999922 | 1.174858884 | YOR126C | IAH1 | n |
| W303.asm02.HP0_G0050240 | 6026.193 | 6305.133 | 6446.981 | 7235.469 | 6356.946 | 6619.717 | 7910.504 | 6646.822 | 6650.229 | 0.394440445 | -0.106251094 | YOR127W | RGA1 | n |
| W303.asm02.HP0_G0050250 | 2593.243 | 1919.582 | 2280.858 | 7505.713 | 6346.532 | 9542.907 | 4692.232 | 14087.18 | 11358.02 | 1.92E-06 | -1.783936178 | YOR128C | ADE2 | n |
| W303.asm02.HP0_G0050260 | 511.4481 | 541.9343 | 566.9807 | 2715.645 | 2367.474 | 2348.883 | 2441.788 | 2451.485 | 2672.052 | 8.97E-86 | -2.19780549 | YOR129C | AFI1 | n |
| W303.asm02.HP0_G0050270 | 860.8846 | 786.7506 | 831.0687 | 1439.452 | 1225.394 | 472.5114 | 1901.089 | 856.8923 | 1417.36 | 0.460104115 | -0.339973632 | YOR130C | ORT1 | n |
| W303.asm02.HP0_G0050280 | 4044.993 | 3710.08 | 3528 | 585.6457 | 542.4016 | 539.3619 | 1412.237 | 1185.475 | 1392.242 | 3.70E-120 | 2.75800115 | YOR131C | | n |

| | | | | | | | | | | | | | | |
|---|---|---|---|---|---|---|---|---|---|---|---|---|---|---|
| W303.asm02.HP0_G0050290 | 6790.717 | 6916.061 | 6680.887 | 3446.487 | 3239.656 | 4665.86 | 6087.187 | 5532.217 | 6147.873 | 2.06E-07 | 0.844678012 | YOR132W | VPS17 | n |
| W303.asm02.HP0_G0050320 | 11281.51 | 10394.68 | 10815.75 | 19101.22 | 19522.12 | 16634.38 | 4592.24 | 7585.323 | 7028.191 | 0.000183041 | -0.766086535 | YOR136W | IDH2 | n |
| W303.asm02.HP0_G0050330 | 728.5223 | 913.6099 | 706.0311 | 5400.723 | 4822.601 | 6090.991 | 10458.46 | 9693.192 | 12088.82 | 2.39E-64 | -2.796811022 | YOR137C | SIA1 | n |
| W303.asm02.HP0_G0050340 | 3565.312 | 2911.088 | 3525.844 | 2865.009 | 2871.691 | 3975.325 | 2133.169 | 2755.371 | 2333.561 | 0.847117208 | 0.042484855 | YOR138C | RUP1 | n |
| W303.asm02.HP0_G0050350 | 1013.366 | 1196.261 | 1096.235 | 5463.942 | 4998.773 | 6871.926 | 5409.461 | 4736.531 | 5162.3 | 7.80E-53 | -2.390726324 | YOR140W | SFL1 | n |
| W303.asm02.HP0_G0050360 | 3442.48 | 4023.89 | 3538.779 | 4587.211 | 4234.204 | 5375.387 | 4118.202 | 2865.972 | 3875.313 | 0.053020065 | -0.367456172 | YOR141C | ARP8 | n |
| W303.asm02.HP0_G0050370 | 9008.051 | 9841.615 | 9320.689 | 10464.51 | 10809.85 | 13855.52 | 2362.781 | 7137.548 | 5567.773 | 0.411978927 | -0.318522044 | YOR142W | LSC1 | n |
| W303.asm02.HP0_G0050380 | 901.1228 | 795.653 | 889.2759 | 2388.434 | 2633.902 | 3448.877 | 5325.517 | 6236.63 | 6881.073 | 8.90E-21 | -1.711713213 | YOR143C | THI80 | n |
| W303.asm02.HP0_G0050390 | 587.6888 | 617.6048 | 670.4601 | 2141.115 | 1948.306 | 2105.79 | 969.0614 | 1044.807 | 1082.456 | 4.64E-60 | -1.724025134 | YOR144C | ELG1 | n |
| W303.asm02.HP0_G0050400 | 2861.144 | 2935.57 | 3179.835 | 3122.749 | 3037.449 | 706.4881 | 3640.461 | 1303.593 | 2887.348 | 0.50567447 | 0.386561214 | YOR145C | PNO1 | n |
| W303.asm02.HP0_G0050410 | 1419.983 | 1760.452 | 1325.829 | 1791.673 | 1635.015 | 1238.253 | 2097.37 | 1302.519 | 1725.95 | 0.854871663 | -0.050020597 | YOR147W | MDM32 | n |
| W303.asm02.HP0_G0050420 | 728.5223 | 604.2511 | 666.1485 | 398.7671 | 462.5601 | 381.3516 | 546.8716 | 508.9812 | 612.3952 | 5.66E-06 | 0.687127945 | YOR148C | SPP2 | n |
| W303.asm02.HP0_G0050430 | 1166.906 | 1314.218 | 1358.167 | 952.4558 | 807.9614 | 822.7168 | 633.2847 | 915.9513 | 850.416 | 0.001453677 | 0.571311474 | YOR149C | SMP3 | n |
| W303.asm02.HP0_G0050440 | 3404.359 | 3258.282 | 3315.652 | 2488.473 | 2396.113 | 1710.765 | 1070.288 | 922.3941 | 922.181 | 0.00062442 | 0.597364407 | YOR150W | MRPL23 | n |
| W303.asm02.HP0_G0050450 | 9531.147 | 9266.297 | 10014.86 | 12984.24 | 12283.44 | 14214.08 | 14919.84 | 12810.43 | 12710.79 | 6.35E-05 | -0.454522223 | YOR151C | RPB2 | n |
| W303.asm02.HP0_G0050470 | 13661.91 | 13873.29 | 13796.17 | 4813.688 | 4506.706 | 3963.93 | 2962.735 | 4111.58 | 2834.72 | 1.47E-21 | 1.637492204 | YOR153W | PDR5 | n |
| W303.asm02.HP0_G0050480 | 884.1804 | 1061.613 | 1061.741 | 971.2132 | 1016.244 | 925.2715 | 920.9169 | 1028.7 | 1063.319 | 0.746412691 | 0.04625291 | YOR154W | SLP1 | n |
| W303.asm02.HP0_G0050490 | 5204.487 | 4835.122 | 4913.114 | 1367.896 | 1416.319 | 3119.183 | 6485.922 | 6175.423 | 5824.931 | 2.13E-05 | 1.340755261 | YOR155C | ISN1 | n |
| W303.asm02.HP0_G0050510 | 5683.109 | 4834.009 | 5809.936 | 3767.446 | 4091.877 | 5081.397 | 3199.754 | 4726.867 | 4073.863 | 0.106439157 | 0.33537408 | YOR157C | PUP1 | n |
| W303.asm02.HP0_G0050520 | 6593.762 | 7261.029 | 6762.808 | 4797.71 | 4351.362 | 3917.59 | 2339.327 | 2194.847 | 2213.952 | 2.61E-09 | 0.657873444 | YOR158W | PET123 | n |
| W303.asm02.HP0_G0050530 | 1018.661 | 964.7988 | 969.0412 | 868.3952 | 990.2083 | 638.8779 | 1017.206 | 970.7151 | 1071.692 | 0.209107647 | 0.241957246 | YOR159C | SME1 | n |
| W303.asm02.HP0_G0050540 | 2578.418 | 2644.016 | 2655.971 | 6790.85 | 6570.436 | 6169.236 | 6005.712 | 6273.139 | 6254.325 | 1.43E-71 | -1.309819073 | YOR160W | MTR10 | n |
| W303.asm02.HP0_G0050550 | 1188.084 | 1277.496 | 1130.728 | 1806.262 | 1702.707 | 2918.631 | 3560.22 | 4536.805 | 3233.016 | 0.000841908 | -0.837852599 | YOR161C | PNS1 | n |
| W303.asm02.HP0_G0050560 | 2146.388 | 2550.541 | 2346.61 | 8113.59 | 8176.812 | 7349.755 | 6783.43 | 9610.509 | 5320.183 | 3.58E-15 | -1.746890878 | YOR162C | YRR1 | n |
| W303.asm02.HP0_G0050570 | 2801.846 | 2682.964 | 2714.178 | 1018.454 | 1155.966 | 1270.919 | 1815.91 | 1493.656 | 1745.087 | 1.48E-21 | 1.251097977 | YOR163W | DDP1 | n |
| W303.asm02.HP0_G0050580 | 3951.81 | 3753.479 | 3742.504 | 4200.254 | 4483.274 | 5275.871 | 4587.302 | 6190.457 | 5608.439 | 0.079493221 | -0.286142538 | YOR164C | GET4 | n |
| W303.asm02.HP0_G0050590 | 9027.111 | 9378.69 | 9702.269 | 6321.222 | 6238.92 | 7987.114 | 7326.598 | 8276.85 | 7436.056 | 0.000956633 | 0.452009323 | YOR165W | SEY1 | n |
| W303.asm02.HP0_G0050600 | 1893.311 | 1743.76 | 1800.11 | 1472.103 | 1471.861 | 1343.087 | 1472.726 | 1610.7 | 1507.066 | 0.000298517 | 0.343039923 | YOR166C | SWT1 | n |
| W303.asm02.HP0_G0050610 | 13328.36 | 13701.92 | 13887.79 | 13464.99 | 14202.24 | 13082.94 | 5542.784 | 6356.896 | 6304.56 | 0.956829779 | 0.005963288 | YOR168W | GLN4 | n |
| W303.asm02.HP0_G0050650 | 1811.776 | 1714.827 | 1803.344 | 1361.644 | 1469.257 | 1312.7 | 1380.141 | 1197.287 | 1303.732 | 0.000412798 | 0.36368616 | YOR174W | MED4 | n |
| W303.asm02.HP0_G0050660 | 8229.761 | 7966.545 | 8543.516 | 5347.23 | 5602.791 | 7250.999 | 6800.712 | 10237.61 | 6833.23 | 0.043130974 | 0.442816749 | YOR175C | ALE1 | n |
| W303.asm02.HP0_G0050670 | 8644.849 | 9195.078 | 8547.827 | 5406.281 | 5457.862 | 8003.067 | 1224.597 | 1649.357 | 1109.966 | 0.033731651 | 0.483972578 | YOR176W | HEM15 | n |
| W303.asm02.HP0_G0050720 | 2277.691 | 2035.314 | 2173.067 | 3920.283 | 3874.917 | 4782.848 | 2535.608 | 4389.694 | 3070.348 | 1.80E-05 | -0.955484258 | YOR181W | LAS17 | n |
| W303.asm02.HP0_G0050740 | 3217.993 | 2755.296 | 3134.563 | 5459.079 | 5342.439 | 7070.199 | 4041.665 | 10845.38 | 8178.825 | 0.003932473 | -0.972491644 | YOR184W | SER1 | n |
| W303.asm02.HP0_G0050770 | 22902.92 | 23983.1 | 23198.78 | 7625.899 | 8203.715 | 6964.605 | 1855.413 | 2599.67 | 2557.228 | 4.17E-25 | 1.620463173 | YOR187W | TUF1 | n |
| W303.asm02.HP0_G0050780 | 1448.573 | 1478.913 | 1679.384 | 6030.831 | 5686.972 | 6704.8 | 6998.228 | 5922.007 | 6239.972 | 8.21E-58 | -1.999662173 | YOR188W | MSB1 | n |
| W303.asm02.HP0_G0050810 | 3528.25 | 3684.485 | 3571.116 | 2523.904 | 2282.426 | 2546.396 | 5988.429 | 4082.587 | 5695.754 | 0.001167884 | 0.552416204 | YOR191W | ULS1 | n |
| W303.asm02.HP0_G0050840 | 4974.706 | 5170.075 | 5016.594 | 4833.14 | 4898.971 | 5539.474 | 3768.846 | 4395.063 | 4249.688 | 0.934110501 | -0.010472303 | YOR194C | TOA1 | n |
| W303.asm02.HP0_G0050850 | 1711.18 | 1947.402 | 1990.9 | 2307.152 | 2273.747 | 2967.25 | 2478.822 | 2825.168 | 2297.678 | 0.011705953 | -0.418100911 | YOR195W | SLK19 | n |
| W303.asm02.HP0_G0050860 | 2976.564 | 3011.241 | 3033.239 | 8702.709 | 8787.773 | 8527.995 | 5477.357 | 6413.807 | 6381.11 | 4.99E-62 | -1.528165014 | YOR196C | LIP5 | n |
| W303.asm02.HP0_G0050870 | 8069.867 | 8033.313 | 8236.311 | 3158.18 | 3355.947 | 4294.384 | 4881.107 | 4123.392 | 3869.333 | 2.25E-13 | 1.171136962 | YOR197W | MCA1 | n |
| W303.asm02.HP0_G0050880 | 16225.5 | 17171.64 | 17571.01 | 13727.59 | 14179.68 | 15720.5 | 7031.559 | 9606.214 | 7863.058 | 0.144038765 | 0.224349338 | YOR198C | BFR1 | n |
| W303.asm02.HP0_G0050890 | 1660.353 | 1727.068 | 1741.903 | 1647.172 | 1443.222 | 761.9436 | 880.1793 | 817.1617 | 790.6117 | 0.151371879 | 0.412992245 | YOR201C | MRM1 | n |
| W303.asm02.HP0_G0050900 | 867.238 | 667.6808 | 794.4198 | 1718.728 | 1779.077 | 1732.795 | 2015.895 | 1650.43 | 1308.516 | 1.04E-09 | -1.166846942 | YOR202W | HIS3 | n |
| W303.asm02.HP0_G0050910 | 9605.27 | 9683.598 | 9143.912 | 22746.4 | 23531.55 | 22598.5 | 23703.12 | 28209.8 | 25894.03 | 8.43E-43 | -1.276448258 | YOR204W | DED1 | n |
| W303.asm02.HP0_G0050920 | 3885.099 | 4066.176 | 4151.032 | 935.7826 | 767.1728 | 593.2981 | 627.1123 | 351.1326 | 657.8464 | 3.02E-19 | 2.397746485 | YOR205C | GEP3 | n |
| W303.asm02.HP0_G0050930 | 5570.866 | 6053.639 | 6542.915 | 8613.091 | 7706.442 | 2914.833 | 9484.457 | 4654.923 | 9514.851 | 0.862217886 | -0.082333 | YOR206W | NOC2 | n |
| W303.asm02.HP0_G0050940 | 5058.359 | 5150.045 | 5366.915 | 6162.132 | 5893.519 | 3429.886 | 7645.092 | 5052.229 | 7150.192 | 0.978596391 | 0.008345848 | YOR207C | RET1 | n |
| W303.asm02.HP0_G0050950 | 3225.405 | 3666.681 | 3244.51 | 2847.641 | 2336.232 | 2604.13 | 3882.418 | 4517.476 | 3566.723 | 0.009670669 | 0.37997925 | YOR208W | PTP2 | n |
| W303.asm02.HP0_G0050960 | 10803.94 | 10936.61 | 11226.43 | 7035.39 | 7215.243 | 8144.364 | 6763.678 | 10570.49 | 8060.412 | 0.002476922 | 0.557845882 | YOR209C | NPT1 | n |
| W303.asm02.HP0_G0050970 | 2001.319 | 1868.394 | 1977.965 | 1986.888 | 2377.021 | 1313.46 | 2056.632 | 1518.353 | 1716.381 | 0.876253583 | 0.042818362 | YOR210W | RPB10 | n |
| W303.asm02.HP0_G0050980 | 4583.973 | 4306.541 | 4466.86 | 3720.205 | 3123.365 | 3635.755 | 5242.807 | 3985.945 | 5401.517 | 0.028324805 | 0.349968121 | YOR211C | MGM1 | n |
| W303.asm02.HP0_G0050990 | 8055.042 | 8135.691 | 8263.259 | 1447.788 | 1474.464 | 1525.407 | 1678.883 | 1728.818 | 1438.889 | 2.33E-141 | 2.45899516 | YOR212W | STE4 | n |
| W303.asm02.HP0_G0051020 | 5546.511 | 5829.966 | 5306.552 | 2108.463 | 2063.73 | 2026.026 | 775.2491 | 1046.955 | 993.9461 | 5.91E-27 | 1.428416058 | YOR215C | AIM41 | n |

| | | | | | | | | | | | | | | |
|---|---|---|---|---|---|---|---|---|---|---|---|---|---|---|
| W303.asm02.HP0_G0051030 | 2410.053 | 2646.242 | 2600.997 | 3704.921 | 3312.555 | 3299.223 | 2491.167 | 2787.585 | 2829.936 | 5.20E-05 | -0.430302912 | YOR216C | RUD3 | n |
| W303.asm02.HP0_G0051040 | 3896.747 | 3737.9 | 4084.202 | 6936.046 | 6679.784 | 7874.683 | 5809.43 | 6906.681 | 7481.507 | 9.69E-11 | -0.874883046 | YOR217W | RFC1 | n |
| W303.asm02.HP0_G0051050 | 3351.414 | 3212.658 | 3103.303 | 1417.916 | 1224.526 | 1124.304 | 1506.057 | 1153.261 | 1460.419 | 3.74E-19 | 1.359550347 | YOR219C | STE13 | n |
| W303.asm02.HP0_G0051060 | 5178.015 | 6059.203 | 4880.777 | 3355.479 | 3842.807 | 3543.835 | 3767.612 | 3547.835 | 3485.39 | 1.91E-06 | 0.58550681 | YOR220W | RCN2 | n |
| W303.asm02.HP0_G0051070 | 3696.615 | 3831.375 | 3672.44 | 1428.336 | 1490.953 | 1293.709 | 1957.874 | 1583.855 | 1843.166 | 5.23E-33 | 1.410812232 | YOR221C | MCT1 | n |
| W303.asm02.HP0_G0051090 | 2378.286 | 2490.449 | 2184.924 | 523.1213 | 606.6219 | 894.8849 | 1150.529 | 1254.198 | 1327.654 | 2.99E-17 | 1.800787687 | YOR223W | DSC3 | n |
| W303.asm02.HP0_G0051100 | 2913.03 | 2519.382 | 2588.062 | 3445.792 | 3712.63 | 2164.284 | 2891.136 | 1905.995 | 2654.111 | 0.429285393 | -0.216994226 | YOR224C | RPB8 | n |
| W303.asm02.HP0_G0051130 | 1665.648 | 1835.009 | 1658.904 | 1194.912 | 1201.094 | 1004.277 | 1369.031 | 1617.143 | 1722.361 | 2.36E-05 | 0.601623236 | YOR228C | MCP1 | n |
| W303.asm02.HP0_G0051160 | 2511.708 | 2274.566 | 2548.18 | 3989.755 | 4040.675 | 4844.381 | 4397.193 | 4400.432 | 3875.313 | 1.64E-10 | -0.811755638 | YOR231W | MKK1 | n |
| W303.asm02.HP0_G0051170 | 11258.21 | 12116.18 | 11435.55 | 7471.672 | 7773.265 | 7055.005 | 3315.795 | 3177.374 | 3040.446 | 7.57E-15 | 0.642504034 | YOR232W | MGE1 | n |
| W303.asm02.HP0_G0051200 | 446.8553 | 359.4348 | 383.736 | 1175.46 | 1400.698 | 1340.808 | 817.2212 | 845.0805 | 779.847 | 1.23E-36 | -1.717804262 | YOR236W | DFR1 | n |
| W303.asm02.HP0_G0051240 | 5621.693 | 5488.336 | 5570.64 | 4078.678 | 4270.653 | 3303.781 | 1913.433 | 1887.74 | 2014.206 | 3.55E-05 | 0.517556161 | YOR239W | ABP140 | n |
| W303.asm02.HP0_G0051250 | 3008.331 | 2855.448 | 3142.108 | 3390.215 | 3537.326 | 3108.548 | 1738.138 | 2655.507 | 2327.58 | 0.412797256 | -0.156205817 | YOR241W | MET7 | n |
| W303.asm02.HP0_G0051270 | 5526.392 | 5290.258 | 6157.023 | 7052.758 | 6666.766 | 3524.084 | 6497.032 | 3873.196 | 6565.307 | 0.951250616 | -0.022756678 | YOR243C | PUS7 | n |
| W303.asm02.HP0_G0051280 | 1497.283 | 1808.302 | 1554.346 | 1561.722 | 1386.812 | 1538.321 | 2575.111 | 2173.371 | 2552.444 | 0.431875162 | 0.114809191 | YOR244W | ESA1 | n |
| W303.asm02.HP0_G0051290 | 2581.595 | 2736.379 | 2441.466 | 1819.462 | 1928.346 | 3461.032 | 6182.241 | 6068.043 | 5968.461 | 0.710819148 | 0.106151247 | YOR245C | DGA1 | n |
| W303.asm02.HP0_G0051300 | 2023.555 | 1719.278 | 2175.223 | 2179.325 | 2039.43 | 1741.911 | 3286.167 | 3442.603 | 4137.256 | 0.957562864 | -0.01032958 | YOR246C | ENV9 | n |
| W303.asm02.HP0_G0051330 | 2004.495 | 1988.576 | 1968.264 | 1727.759 | 1667.125 | 1520.089 | 1383.844 | 1790.024 | 1525.008 | 0.037663332 | 0.278411414 | YOR250C | CLP1 | n |
| W303.asm02.HP0_G0051340 | 3555.782 | 3083.573 | 3404.04 | 1710.391 | 1991.699 | 2833.549 | 1414.706 | 2475.109 | 1693.655 | 0.024965673 | 0.619880267 | YOR251C | TUM1 | n |
| W303.asm02.HP0_G0051350 | 1957.904 | 1738.196 | 1873.408 | 1170.597 | 1083.067 | 202.8305 | 1646.787 | 638.911 | 1338.418 | 0.043717669 | 1.180972633 | YOR252W | TMA16 | n |
| W303.asm02.HP0_G0051360 | 741.2291 | 685.4856 | 689.8625 | 1331.771 | 1506.575 | 1021.749 | 1211.018 | 1343.324 | 1495.105 | 1.63E-06 | -0.866562171 | YOR253W | NAT5 | n |
| W303.asm02.HP0_G0051370 | 8040.218 | 7972.109 | 8098.339 | 5896.751 | 5652.258 | 5978.561 | 3209.63 | 4179.229 | 3606.194 | 0.000140142 | 0.460012298 | YOR254C | SEC63 | n |
| W303.asm02.HP0_G0051380 | 114.3611 | 124.6338 | 91.62236 | 152.8376 | 165.7579 | 217.2641 | 181.4675 | 483.21 | 187.7852 | 0.112950442 | -0.696792892 | YOR255W | OSW1 | n |
| W303.asm02.HP0_G0051390 | 2827.26 | 2995.661 | 2657.048 | 2521.125 | 2492.444 | 2761.381 | 4165.112 | 3991.314 | 3823.882 | 0.225056764 | 0.125100104 | YOR256C | TRE2 | n |
| W303.asm02.HP0_G0051400 | 2361.344 | 2045.329 | 1898.2 | 1279.667 | 1415.451 | 1220.021 | 913.5101 | 1121.047 | 941.3184 | 3.01E-06 | 0.687717819 | YOR257W | CDC31 | n |
| W303.asm02.HP0_G0051410 | 658.635 | 677.696 | 645.6682 | 619.6868 | 537.1945 | 670.0242 | 192.5778 | 543.3428 | 498.7672 | 0.764980383 | 0.117413395 | YOR258W | HNT3 | n |
| W303.asm02.HP0_G0051420 | 12788.32 | 13218.97 | 12682.69 | 8042.729 | 8469.275 | 10581.37 | 5308.234 | 8532.414 | 6116.775 | 0.016120208 | 0.51401956 | YOR259C | RPT4 | n |
| W303.asm02.HP0_G0051430 | 5858.887 | 6312.922 | 6929.884 | 6819.334 | 6622.506 | 6248.241 | 6950.083 | 6464.276 | 7272.193 | 0.714526418 | -0.043795696 | YOR260W | GCD1 | n |
| W303.asm02.HP0_G0051440 | 7424.998 | 8087.84 | 7217.686 | 5409.755 | 5784.17 | 7583.732 | 4041.665 | 7009.766 | 5157.516 | 0.273294841 | 0.275610814 | YOR261C | RPN8 | n |
| W303.asm02.HP0_G0051450 | 712.6388 | 715.5313 | 751.3034 | 1984.109 | 2027.28 | 1516.291 | 2517.091 | 1570.969 | 2187.638 | 2.26E-10 | -1.342640052 | YOR262W | GPN2 | n |
| W303.asm02.HP0_G0051460 | 2393.111 | 2827.628 | 2534.167 | 4448.267 | 4892.028 | 5127.736 | 6377.288 | 6708.028 | 7573.606 | 4.83E-13 | -0.89970565 | YOR264W | DSE3 | n |
| W303.asm02.HP0_G0051480 | 1105.49 | 913.6099 | 963.6517 | 1065.695 | 1096.085 | 967.8127 | 756.732 | 1000.782 | 1058.535 | 0.717366993 | -0.069110632 | YOR265W | RBL2 | n |
| W303.asm02.HP0_G0051490 | 1322.564 | 1328.685 | 1188.935 | 1958.405 | 1894.5 | 1708.486 | 1539.388 | 1545.198 | 1669.734 | 6.70E-07 | -0.534347983 | YOR266W | PNT1 | n |
| W303.asm02.HP0_G0051500 | 7863.382 | 7611.561 | 7506.566 | 9141.77 | 9042.051 | 11135.92 | 23021.69 | 24777.93 | 32748.79 | 0.04929532 | -0.351406135 | YOR267C | HRK1 | n |
| W303.asm02.HP0_G0051520 | 286.9615 | 290.4412 | 302.8927 | 862.1427 | 1011.904 | 1163.806 | 1044.364 | 940.6487 | 1053.75 | 1.90E-31 | -1.786691232 | YOR269W | PAC1 | n |
| W303.asm02.HP0_G0051530 | 15369.91 | 14230.5 | 15420.58 | 9409.235 | 9891.669 | 12547.38 | 15282.78 | 17589.92 | 14541.99 | 0.001234489 | 0.499387258 | YOR270C | VPH1 | n |
| W303.asm02.HP0_G0051540 | 2435.467 | 2200.008 | 2770.229 | 3757.025 | 4169.983 | 3579.54 | 3920.687 | 3852.794 | 3697.097 | 4.40E-07 | -0.635554455 | YOR271C | FSF1 | n |
| W303.asm02.HP0_G0051550 | 1531.168 | 1811.641 | 2122.405 | 3458.297 | 3215.357 | 1043.779 | 3244.195 | 1521.574 | 2656.503 | 0.297991781 | -0.497841136 | YOR272W | YTM1 | n |
| W303.asm02.HP0_G0051560 | 2250.16 | 2344.672 | 2042.64 | 3106.076 | 2884.709 | 4994.795 | 6157.552 | 8764.355 | 7043.741 | 0.004122962 | -0.726955403 | YOR273C | TPO4 | n |
| W303.asm02.HP0_G0051570 | 2046.851 | 2237.844 | 2023.237 | 1579.09 | 1547.363 | 964.7741 | 1706.042 | 1046.955 | 1376.693 | 0.016666037 | 0.62465404 | YOR274W | MOD5 | n |
| W303.asm02.HP0_G0051580 | 4142.412 | 4407.806 | 4073.422 | 3108.855 | 2808.338 | 4198.666 | 3834.273 | 4127.687 | 3921.961 | 0.072710503 | 0.319403041 | YOR275C | RIM20 | n |
| W303.asm02.HP0_G0051590 | 4496.084 | 4543.568 | 4781.609 | 4018.238 | 4603.905 | 3975.325 | 2218.348 | 2216.323 | 2203.187 | 0.179662653 | 0.134010389 | YOR276W | CAF20 | n |
| W303.asm02.HP0_G0051600 | 1939.902 | 1882.86 | 1748.37 | 732.2308 | 736.7983 | 530.2459 | 127.1507 | 278.1142 | 244.0012 | 1.75E-07 | 1.478561727 | YOR278W | HEM4 | n |
| W303.asm02.HP0_G0051610 | 3711.44 | 3463.038 | 3270.379 | 1549.217 | 1574.266 | 1909.797 | 2260.32 | 2212.028 | 2211.56 | 2.11E-18 | 1.053195514 | YOR279C | RFM1 | n |
| W303.asm02.HP0_G0051620 | 3709.322 | 3873.662 | 3831.971 | 2778.17 | 2929.836 | 2213.663 | 1223.363 | 1595.667 | 1649.4 | 0.002237987 | 0.527121071 | YOR281C | PLP2 | n |
| W303.asm02.HP0_G0051630 | 3072.924 | 2568.346 | 3087.135 | 3426.34 | 3584.19 | 4604.327 | 8782.042 | 7835.518 | 7910.902 | 0.012262632 | -0.412136342 | YOR283W | | n |
| W303.asm02.HP0_G0051650 | 5584.632 | 5409.327 | 5098.515 | 187.5734 | 223.9034 | 303.8658 | 1304.838 | 689.3796 | 1239.143 | 2.32E-59 | 4.491878647 | YOR285W | RDL1 | n |
| W303.asm02.HP0_G0051660 | 5268.021 | 4901.89 | 4810.713 | 1859.755 | 2128.818 | 1616.566 | 587.6092 | 856.8923 | 1008.299 | 4.10E-11 | 1.418405815 | YOR286W | RDL2 | n |
| W303.asm02.HP0_G0051670 | 392.8514 | 392.8189 | 444.099 | 2613.522 | 2163.531 | 1440.324 | 1520.871 | 825.7521 | 1465.203 | 4.71E-15 | -2.337952509 | YOR287C | RRP36 | n |
| W303.asm02.HP0_G0051680 | 2323.224 | 2285.694 | 2134.262 | 2186.966 | 2435.166 | 2658.826 | 1446.802 | 1252.051 | 1327.654 | 0.397187861 | -0.110557654 | YOR288C | MPD1 | n |
| W303.asm02.HP0_G0051690 | 2926.796 | 3036.835 | 2770.229 | 491.859 | 591.0008 | 1152.411 | 1102.384 | 1296.077 | 1222.398 | 1.06E-10 | 1.966163996 | YOR289W | | n |
| W303.asm02.HP0_G0051700 | 7848.557 | 8190.218 | 8201.818 | 5705.009 | 5029.147 | 7358.111 | 11485.54 | 7971.891 | 8560.376 | 0.044705348 | 0.42200815 | YOR290C | SNF2 | n |
| W303.asm02.HP0_G0051710 | 2502.178 | 2088.728 | 2263.611 | 5049.197 | 5098.575 | 5397.417 | 5004.554 | 6787.489 | 6357.188 | 2.82E-15 | -1.181276535 | YOR291W | YPK9 | n |
| W303.asm02.HP0_G0051720 | 1903.9 | 1561.26 | 1692.319 | 715.5576 | 833.9967 | 735.3553 | 2956.563 | 1954.316 | 2373.031 | 3.20E-10 | 1.174964317 | YOR292C | | n |

| | | | | | | | | | | | | | |
|---|---|---|---|---|---|---|---|---|---|---|---|---|---|
| W303.asm02.HP0_G0051730 | 8525.193 | 8382.733 | 8879.824 | 16466.16 | 18697.67 | 19336.5 | 16199.99 | 11646.43 | 12491.9 | 3.55E-12 | -1.079561685 | YOR293W | RPS10A | y |
| W303.asm02.HP0_G0051750 | 1194.438 | 1428.837 | 1447.633 | 2753.16 | 2825.695 | 845.5067 | 3787.363 | 1602.109 | 2911.269 | 0.175126226 | -0.658186254 | YOR294W | RRS1 | n |
| W303.asm02.HP0_G0051770 | 1357.508 | 1334.249 | 1220.194 | 3418.698 | 3161.55 | 4037.617 | 3259.009 | 3282.606 | 3552.37 | 2.73E-29 | -1.440647666 | YOR296W | | n |
| W303.asm02.HP0_G0051780 | 1404.1 | 1453.319 | 1449.789 | 1384.569 | 1500.5 | 1079.483 | 1383.844 | 1116.752 | 1185.32 | 0.50749763 | 0.119810275 | YOR297C | TIM18 | n |
| W303.asm02.HP0_G0051790 | 195.8963 | 260.3955 | 178.9331 | 188.2681 | 201.3395 | 144.3363 | 214.7983 | 127.7822 | 160.2753 | 0.405022517 | 0.250710797 | YOR298W | MUM3 | n |
| W303.asm02.HP0_G0051800 | 9428.434 | 9637.973 | 8367.816 | 4648.346 | 5602.791 | 5785.606 | 6130.393 | 5851.136 | 5387.164 | 3.10E-09 | 0.774695634 | YOR298C- | MBF1 | n |
| W303.asm02.HP0_G0051810 | 2374.051 | 2506.029 | 2445.778 | 5063.091 | 5117.667 | 7205.419 | 4304.608 | 6606.017 | 5658.675 | 5.12E-09 | -1.246891965 | YOR299W | BUD7 | n |
| W303.asm02.HP0_G0051820 | 859.8258 | 973.7012 | 937.7818 | 2913.639 | 2977.568 | 2521.327 | 3080.01 | 2874.562 | 3148.094 | 3.86E-45 | -1.602066182 | YOR301W | RAX1 | n |
| W303.asm02.HP0_G0051830 | 2144.27 | 2019.734 | 2024.315 | 4084.931 | 4005.961 | 3215.66 | 10385.62 | 7749.614 | 8657.258 | 5.97E-08 | -0.869509756 | YOR303W | CPA1 | n |
| W303.asm02.HP0_G0051840 | 4519.38 | 4799.512 | 4507.82 | 3272.808 | 3107.744 | 3485.341 | 4184.864 | 4387.547 | 4516.414 | 1.59E-08 | 0.486736042 | YOR304W | ISW2 | n |
| W303.asm02.HP0_G0051860 | 1462.339 | 1411.032 | 1395.894 | 928.8355 | 847.8821 | 614.5687 | 614.7676 | 601.328 | 818.1217 | 5.24E-05 | 0.836128026 | YOR305W | RRG7 | n |
| W303.asm02.HP0_G0051870 | 2711.84 | 2665.159 | 2747.593 | 1418.61 | 1133.402 | 1194.193 | 8172.212 | 5127.395 | 5713.695 | 3.59E-08 | 1.11660951 | YOR306C | MCH5 | n |
| W303.asm02.HP0_G0051880 | 2543.475 | 2424.794 | 2368.169 | 1464.462 | 1234.072 | 1450.959 | 4715.687 | 2726.378 | 3770.058 | 0.000120475 | 0.821974347 | YOR307C | SLY41 | n |
| W303.asm02.HP0_G0051890 | 1614.821 | 1648.059 | 1585.606 | 1731.927 | 1566.456 | 1451.719 | 1688.759 | 1971.497 | 1917.323 | 0.825209285 | 0.029310342 | YOR308C | SNU66 | n |
| W303.asm02.HP0_G0051900 | 11841.66 | 12431.1 | 12960.79 | 19349.23 | 19234.86 | 12049.04 | 16977.71 | 12844.79 | 16398.32 | 0.050394567 | -0.44347409 | YOR310C | NOP58 | n |
| W303.asm02.HP0_G0051910 | 2365.58 | 2403.651 | 2526.621 | 2753.855 | 3102.537 | 2831.27 | 3661.447 | 3670.248 | 3421.997 | 0.006606759 | -0.251563162 | YOR311C | DGK1 | n |
| W303.asm02.HP0_G0051930 | 238.2522 | 258.1699 | 209.1146 | 263.9921 | 262.0884 | 198.2725 | 312.3217 | 656.0918 | 559.7675 | 0.914105155 | -0.037974056 | YOR313C | SPS4 | n |
| W303.asm02.HP0_G0051940 | 583.4532 | 446.2334 | 704.9532 | 614.1291 | 426.9785 | 464.1551 | 281.4599 | 396.2322 | 300.2172 | 0.459561017 | 0.20409 | YOR315W | SFG1 | n |
| W303.asm02.HP0_G0051970 | 19832.11 | 20158.4 | 19334.47 | 11804.62 | 12499.54 | 15691.63 | 5193.428 | 8677.377 | 6712.425 | 0.009213368 | 0.568792478 | YOR317W | FAA1 | n |
| W303.asm02.HP0_G0051990 | 1192.32 | 1484.477 | 1196.48 | 534.2367 | 570.1725 | 577.3451 | 511.0719 | 477.841 | 452.1199 | 8.99E-19 | 1.203712511 | YOR319W | HSH49 | n |
| W303.asm02.HP0_G0052000 | 2141.093 | 2248.972 | 2288.403 | 2802.485 | 2613.942 | 3394.941 | 3515.779 | 3835.613 | 3726.999 | 0.002486199 | -0.399998212 | YOR320C | GNT1 | n |
| W303.asm02.HP0_G0052020 | 2079.677 | 2168.85 | 2110.548 | 4578.179 | 4182.133 | 5140.65 | 5588.46 | 3282.606 | 4647.984 | 5.70E-08 | -1.128339546 | YOR322C | LDB19 | n |
| W303.asm02.HP0_G0052030 | 8371.653 | 7898.664 | 8615.736 | 4451.046 | 4780.077 | 4564.825 | 3349.125 | 3876.418 | 3075.133 | 2.06E-13 | 0.851176397 | YOR323C | PRO2 | n |
| W303.asm02.HP0_G0052040 | 744.4058 | 802.3298 | 716.8102 | 288.3072 | 326.3088 | 417.0559 | 317.2596 | 241.605 | 234.4325 | 1.46E-08 | 1.133801496 | YOR324C | FRT1 | n |
| W303.asm02.HP0_G0052050 | 12722.67 | 13215.63 | 13557.95 | 11899.1 | 12018.75 | 13174.1 | 5500.812 | 9091.864 | 7172.918 | 0.67402414 | 0.090607668 | YOR326W | MYO2 | n |
| W303.asm02.HP0_G0052060 | 2364.521 | 2122.112 | 2081.444 | 3901.526 | 4015.507 | 4716.758 | 2917.06 | 3460.857 | 2919.642 | 2.76E-12 | -0.943701555 | YOR327C | SNC2 | n |
| W303.asm02.HP0_G0052080 | 7830.556 | 7204.276 | 8095.105 | 2393.297 | 2206.056 | 3144.252 | 1538.153 | 2127.198 | 1509.458 | 3.13E-15 | 1.578616349 | YOR329C | SCD5 | n |
| W303.asm02.HP0_G0052090 | 1648.705 | 1747.098 | 1641.657 | 2696.888 | 2722.422 | 2547.915 | 2619.552 | 2879.931 | 3122.976 | 9.77E-11 | -0.661405291 | YOR330C | MIP1 | n |
| W303.asm02.HP0_G0052100 | 12609.37 | 11997.11 | 13137.57 | 7052.064 | 7883.481 | 8481.655 | 6956.256 | 9310.919 | 7147.8 | 8.44E-06 | 0.688712891 | YOR332W | VMA4 | n |
| W303.asm02.HP0_G0052120 | 597.2189 | 598.6871 | 576.6819 | 1387.348 | 1311.31 | 1410.697 | 848.083 | 860.1137 | 942.5144 | 9.79E-37 | -1.213412304 | YOR334W | MRS2 | n |
| W303.asm02.HP0_G0052130 | 12028.03 | 11150.27 | 12711.79 | 13870.01 | 14803.66 | 14198.13 | 10184.4 | 10412.64 | 9572.263 | 0.004715532 | -0.256387784 | YOR335C | ALA1 | n |
| W303.asm02.HP0_G0052140 | 1973.787 | 2001.93 | 1889.576 | 4011.986 | 3753.419 | 4810.196 | 2028.239 | 3688.503 | 3461.468 | 4.65E-06 | -1.100381312 | YOR336W | KRE5 | n |
| W303.asm02.HP0_G0052150 | 552.7451 | 577.5439 | 663.9926 | 1287.309 | 1228.865 | 1217.742 | 1497.416 | 1708.416 | 1851.539 | 1.38E-15 | -1.057420431 | YOR337W | TEA1 | n |
| W303.asm02.HP0_G0052160 | 638.5159 | 676.5832 | 574.5261 | 3443.708 | 2495.047 | 3987.48 | 15181.55 | 11428.45 | 19829.88 | 3.07E-21 | -2.393282801 | YOR338W | | n |
| W303.asm02.HP0_G0052180 | 1248.442 | 1287.511 | 1356.011 | 3627.808 | 3669.238 | 1522.368 | 3515.779 | 1944.652 | 2969.877 | 0.001050502 | -1.180163175 | YOR340C | RPA43 | n |
| W303.asm02.HP0_G0052190 | 8842.863 | 8775.552 | 9955.578 | 15178.85 | 14125.87 | 7099.826 | 14861.82 | 10317.07 | 15289.55 | 0.209580174 | -0.400808339 | YOR341W | RPA190 | n |
| W303.asm02.HP0_G0052200 | 2207.804 | 2480.434 | 2419.908 | 743.3463 | 718.5736 | 452.7601 | 1841.834 | 1795.393 | 2224.717 | 6.75E-19 | 1.89241783 | YOR342C | | n |
| W303.asm02.HP0_G0052220 | 3984.636 | 5855.561 | 3799.633 | 15728.37 | 17180.68 | 18416.55 | 16692.55 | 19112.56 | 25246.95 | 6.37E-17 | -1.911884275 | YOR344C | TYE7 | n |
| W303.asm02.HP0_G0052230 | 2089.207 | 2069.811 | 2168.755 | 1092.094 | 995.4154 | 1077.964 | 1917.137 | 1021.184 | 1325.261 | 1.44E-05 | 0.999172302 | YOR346W | REV1 | n |
| W303.asm02.HP0_G0052250 | 524.1549 | 513.0014 | 530.3318 | 2822.632 | 3431.449 | 3725.395 | 1219.659 | 1822.238 | 1033.417 | 6.44E-31 | -2.670439582 | YOR348C | PUT4 | n |
| W303.asm02.HP0_G0052260 | 1129.845 | 1062.725 | 1173.844 | 1470.019 | 1374.663 | 1327.134 | 2307.23 | 1877.002 | 2576.366 | 0.044853281 | -0.309523307 | YOR349W | CIN1 | n |
| W303.asm02.HP0_G0052270 | 1461.28 | 1561.26 | 1441.166 | 3327.69 | 3179.775 | 2933.065 | 2413.395 | 2628.662 | 2362.267 | 7.85E-28 | -1.080781288 | YOR350C | MNE1 | n |
| W303.asm02.HP0_G0052280 | 118.5967 | 110.1673 | 125.0376 | 242.4559 | 230.8461 | 208.9078 | 269.1151 | 693.6748 | 458.1003 | 0.006087415 | -0.947228977 | YOR351C | MEK1 | n |
| W303.asm02.HP0_G0052290 | 2062.735 | 2609.519 | 2082.522 | 1645.783 | 1578.606 | 2239.491 | 3973.769 | 3162.341 | 3234.212 | 0.145315819 | 0.305844827 | YOR352W | TFB6 | n |
| W303.asm02.HP0_G0052300 | 2163.33 | 2296.822 | 2184.924 | 6075.987 | 5775.492 | 7728.068 | 11878.1 | 10433.04 | 11043.45 | 5.25E-28 | -1.559058319 | YOR353C | SOG2 | n |
| W303.asm02.HP0_G0052310 | 14109.83 | 15246.49 | 14263.98 | 5272.201 | 4927.61 | 3837.826 | 2154.156 | 2288.268 | 2522.542 | 4.40E-28 | 1.635678337 | YOR354C | MSC6 | n |
| W303.asm02.HP0_G0052320 | 4518.321 | 4933.048 | 4983.179 | 4763.669 | 4493.689 | 2607.929 | 6511.846 | 4448.753 | 5486.439 | 0.305831868 | 0.282778762 | YOR355W | GDS1 | n |
| W303.asm02.HP0_G0052330 | 5096.479 | 5051.005 | 5089.892 | 7672.445 | 7059.031 | 7769.85 | 7806.808 | 9865 | 10541.09 | 4.99E-05 | -0.562433013 | YOR356W | CIR2 | n |
| W303.asm02.HP0_G0052340 | 1435.867 | 1191.81 | 1284.869 | 1254.657 | 1389.416 | 1623.403 | 1708.511 | 1977.939 | 1814.46 | 0.443079007 | -0.125036662 | YOR357C | SNX3 | n |
| W303.asm02.HP0_G0052350 | 3572.724 | 3238.252 | 3443.923 | 1604.1 | 1715.725 | 2108.829 | 3144.203 | 2899.26 | 3321.526 | 4.79E-11 | 0.917711785 | YOR358W | HAP5 | n |
| W303.asm02.HP0_G0052360 | 335.6709 | 360.5476 | 422.5408 | 3923.757 | 3041.788 | 2724.157 | 2657.821 | 1688.013 | 2284.521 | 7.13E-43 | -3.114664773 | YOR359W | VTS1 | n |
| W303.asm02.HP0_G0052370 | 919.1241 | 1022.664 | 928.0806 | 5777.954 | 4764.455 | 6032.497 | 13203.92 | 8548.521 | 10301.87 | 4.61E-40 | -2.530066562 | YOR360C | PDE2 | n |
| W303.asm02.HP0_G0052380 | 16519.88 | 15911.95 | 17932.11 | 15564.42 | 16283.33 | 13900.34 | 10168.35 | 10483.51 | 10309.05 | 0.191381025 | 0.138714178 | YOR361C | PRT1 | n |
| W303.asm02.HP0_G0052420 | 1054.663 | 997.07 | 955.0284 | 1465.851 | 1435.412 | 1672.781 | 1291.259 | 1885.593 | 1796.519 | 0.000697787 | -0.605291206 | YOR367W | SCP1 | n |

| | | | | | | | | | | | | | |
|---|---|---|---|---|---|---|---|---|---|---|---|---|---|
| W303.asm02.HP0_G0052430 | 1419.983 | 1471.123 | 1437.932 | 1381.096 | 1360.777 | 1959.175 | 2062.805 | 2627.588 | 2193.619 | 0.567699554 | -0.119033186 | YOR368W | RAD17 | n |
| W303.asm02.HP0_G0052440 | 10531.81 | 9664.68 | 10655.14 | 14281.28 | 17789.9 | 14766.36 | 11923.78 | 10402.97 | 9223.006 | 5.55E-05 | -0.602267695 | YOR369C | RPS12 | n |
| W303.asm02.HP0_G0052450 | 9360.665 | 9287.44 | 10282.18 | 7435.547 | 7158.833 | 7250.239 | 5498.343 | 5397.992 | 4528.375 | 0.000322359 | 0.40528202 | YOR370C | MRS6 | n |
| W303.asm02.HP0_G0052470 | 2697.015 | 3045.737 | 2896.345 | 3755.635 | 3580.718 | 4592.932 | 5345.268 | 4078.292 | 3390.899 | 0.024388826 | -0.465606779 | YOR372C | NDD1 | n |
| W303.asm02.HP0_G0052480 | 1469.751 | 1382.099 | 1534.944 | 2712.867 | 2849.127 | 3081.2 | 1062.881 | 1681.571 | 1155.417 | 5.08E-07 | -0.978354181 | YOR373W | NUD1 | n |
| W303.asm02.HP0_G0052490 | 9783.165 | 12828.37 | 9577.232 | 681.5165 | 678.6528 | 1106.831 | 180.2331 | 1432.449 | 306.1976 | 3.04E-09 | 3.705709427 | YOR374W | ALD4 | n |
| W303.asm02.HP0_G0052530 | 1855.191 | 1688.12 | 1894.966 | 1824.325 | 1653.24 | 1678.859 | 2041.818 | 2070.286 | 2047.696 | 0.45089137 | 0.076509995 | YOR377W | ATF1 | n |
| W303.asm02.HP0_G0052540 | 439.443 | 442.8949 | 443.0211 | 213.9726 | 211.7536 | 252.2087 | 662.912 | 387.6418 | 497.5711 | 1.58E-05 | 0.966955415 | YOR378W | AMF1 | n |
| W303.asm02.HP0_G0052560 | 612.0435 | 604.2511 | 554.0458 | 2173.072 | 2235.562 | 2629.199 | 3392.332 | 3554.278 | 2960.309 | 6.43E-50 | -1.991116792 | YOR381W | FRE3 | n |
| W303.asm02.HP0_G0052620 | 5521.098 | 5819.951 | 4770.83 | 1416.526 | 1289.614 | 1132.66 | 750.5596 | 1328.291 | 1475.968 | 5.04E-16 | 2.069341858 | YOR386W | PHR1 | n |
| W303.asm02.HP0_G0052650 | 2367.698 | 2102.082 | 2027.549 | 306.3698 | 262.9563 | 271.2003 | 477.7411 | 781.7263 | 620.7678 | 1.67E-49 | 2.949753361 | YOR389W | | n |
| W303.asm02.HP0_G0052730 | 169.4238 | 172.4842 | 170.3098 | 352.9158 | 342.7978 | 335.0121 | 1088.805 | 915.9513 | 734.3958 | 6.07E-08 | -1.009018966 | YPL279C | FEX2 | y |
| W303.asm02.HP0_G0052760 | 3440.362 | 1936.274 | 2582.673 | 4891.496 | 4115.309 | 1972.089 | 12342.26 | 7365.194 | 9443.086 | 0.243502176 | -0.463999573 | YPL274W | SAM3 | n |
| W303.asm02.HP0_G0052770 | 5791.117 | 5432.696 | 6592.498 | 8227.523 | 8452.786 | 8243.88 | 11663.3 | 6820.777 | 8516.12 | 0.024265006 | -0.484335738 | YPL273W | SAM4 | n |
| W303.asm02.HP0_G0052790 | 2897.147 | 2354.688 | 2186.002 | 5893.277 | 7037.335 | 6670.615 | 2715.841 | 4884.716 | 4032 | 3.21E-09 | -1.3979347 | YPL271W | ATP15 | n |
| W303.asm02.HP0_G0052800 | 3346.12 | 3118.069 | 3157.199 | 3493.033 | 3651.881 | 3344.044 | 3658.978 | 3594.008 | 3504.527 | 0.130737096 | -0.124324807 | YPL270W | MDL2 | n |
| W303.asm02.HP0_G0052810 | 433.0896 | 431.7669 | 463.5014 | 1715.254 | 1643.694 | 1796.607 | 486.3824 | 686.1582 | 636.3169 | 2.95E-37 | -1.956538514 | YPL269W | KAR9 | n |
| W303.asm02.HP0_G0052820 | 1704.827 | 1932.936 | 1667.527 | 1608.268 | 1676.672 | 1363.598 | 1414.706 | 1142.523 | 1298.948 | 0.218861822 | 0.190719649 | YPL268W | PLC1 | n |
| W303.asm02.HP0_G0052830 | 477.5633 | 466.2638 | 457.0339 | 779.4715 | 950.2876 | 748.2696 | 899.9309 | 339.3208 | 612.3952 | 0.014070972 | -0.822708225 | YPL267W | ACM1 | n |
| W303.asm02.HP0_G0052840 | 2599.596 | 2717.461 | 2847.839 | 5510.488 | 6031.505 | 1985.004 | 8626.498 | 3132.274 | 6523.444 | 0.126250012 | -0.728317898 | YPL266W | DIM1 | n |
| W303.asm02.HP0_G0052850 | 761.3482 | 914.7227 | 789.0302 | 16448.79 | 16589.68 | 18388.44 | 16133.33 | 12289.64 | 7071.251 | 2.60E-56 | -4.382861639 | YPL265W | DIP5 | n |
| W303.asm02.HP0_G0052870 | 2525.473 | 2516.044 | 2873.708 | 5195.087 | 5915.215 | 3928.226 | 4335.469 | 3318.042 | 4237.727 | 2.41E-06 | -0.925891301 | YPL263C | KEL3 | n |
| W303.asm02.HP0_G0052880 | 8906.397 | 8732.152 | 8986.537 | 18382.88 | 19763.38 | 20023.24 | 15693.86 | 21980.68 | 16987.99 | 2.22E-14 | -1.127465248 | YPL262W | FUM1 | n |
| W303.asm02.HP0_G0052890 | 12326.64 | 13273.49 | 12470.34 | 6783.903 | 7265.578 | 7759.974 | 14200.14 | 9650.24 | 11140.33 | 1.10E-06 | 0.803730103 | YPL260W | CUB1 | n |
| W303.asm02.HP0_G0052900 | 3929.573 | 4290.962 | 3881.554 | 5244.412 | 5292.104 | 6416.887 | 4616.929 | 5509.667 | 4616.885 | 0.000502644 | -0.486358181 | YPL259C | APM1 | n |
| W303.asm02.HP0_G0052930 | 2435.467 | 2625.098 | 2861.851 | 4701.144 | 5828.43 | 5756.738 | 2676.338 | 2178.74 | 1856.323 | 4.44E-09 | -1.039588411 | YPL256C | CLN2 | n |
| W303.asm02.HP0_G0052940 | 763.466 | 778.961 | 730.8231 | 1171.291 | 1211.508 | 1355.242 | 1566.546 | 833.2687 | 938.9262 | 0.004602053 | -0.717529841 | YPL255W | BBP1 | n |
| W303.asm02.HP0_G0052950 | 937.1254 | 915.8355 | 1055.274 | 2234.207 | 1942.232 | 2198.469 | 2731.889 | 2506.249 | 2608.66 | 3.69E-24 | -1.132534523 | YPL254W | HFI1 | n |
| W303.asm02.HP0_G0052960 | 735.9346 | 784.525 | 799.8093 | 1121.967 | 1183.737 | 1286.872 | 271.5841 | 382.2728 | 247.5895 | 0.000612601 | -0.630712708 | YPL253C | VIK1 | n |
| W303.asm02.HP0_G0052970 | 1598.937 | 1405.468 | 1545.723 | 3247.798 | 3546.005 | 1905.239 | 1391.251 | 1048.029 | 1084.848 | 0.000266241 | -0.934860005 | YPL252C | YAH1 | n |
| W303.asm02.HP0_G0052990 | 10566.75 | 10541.57 | 10475.13 | 15246.94 | 18275.9 | 16116.28 | 14513.7 | 11448.85 | 11329.31 | 3.58E-06 | -0.652272509 | YPL249C-A | RPL36B | y |
| W303.asm02.HP0_G0053000 | 5553.924 | 5575.135 | 5272.058 | 6483.786 | 6423.77 | 8162.596 | 11802.8 | 11310.33 | 12527.79 | 0.006246978 | -0.361435388 | YPL249C | GYP5 | n |
| W303.asm02.HP0_G0053010 | 1372.333 | 1377.648 | 1169.533 | 2000.782 | 1620.262 | 2342.046 | 7038.966 | 2364.507 | 2794.053 | 0.146010996 | -0.605435422 | YPL248C | GAL4 | n |
| W303.asm02.HP0_G0053020 | 8513.546 | 8292.596 | 7796.524 | 4924.148 | 4669.861 | 7352.034 | 5626.728 | 5495.708 | 6018.696 | 0.00653125 | 0.537832078 | YPL247C | | n |
| W303.asm02.HP0_G0053030 | 3161.872 | 2927.78 | 3040.785 | 2329.383 | 2620.017 | 2255.444 | 2793.613 | 3574.68 | 3561.939 | 0.01693917 | 0.341922496 | YPL246C | RBD2 | n |
| W303.asm02.HP0_G0053040 | 1634.94 | 1418.822 | 1759.149 | 540.4892 | 524.1769 | 692.8141 | 1704.807 | 1870.559 | 1808.479 | 2.24E-20 | 1.453186513 | YPL245W | | n |
| W303.asm02.HP0_G0053050 | 2219.452 | 2151.045 | 2245.287 | 1447.788 | 1481.407 | 1518.57 | 639.4571 | 1473.254 | 1209.241 | 0.048446196 | 0.57284471 | YPL244C | HUT1 | n |
| W303.asm02.HP0_G0053060 | 6480.46 | 7026.228 | 6838.262 | 9198.042 | 8804.262 | 8745.259 | 5656.356 | 5962.811 | 6194.521 | 4.45E-07 | -0.394817621 | YPL243W | SRP68 | n |
| W303.asm02.HP0_G0053070 | 4717.394 | 4641.495 | 4726.636 | 8039.255 | 7502.499 | 6274.07 | 1846.772 | 3255.761 | 3596.626 | 0.014621503 | -0.631165388 | YPL242C | IQG1 | n |
| W303.asm02.HP0_G0053100 | 2566.77 | 2978.969 | 2538.478 | 2267.553 | 2073.276 | 742.952 | 2265.258 | 855.8185 | 1861.107 | 0.16220184 | 0.669205787 | YPL239W | YAR1 | n |
| W303.asm02.HP0_G0053110 | 7487.473 | 7708.375 | 7452.671 | 15898.58 | 17037.48 | 14156.35 | 12560.76 | 11064.43 | 11381.94 | 5.39E-23 | -1.056052372 | YPL237W | SUI3 | n |
| W303.asm02.HP0_G0053140 | 3680.732 | 3193.74 | 3318.885 | 1765.968 | 2114.064 | 2078.442 | 3060.259 | 3478.038 | 2803.622 | 2.28E-08 | 0.774917073 | YPL234C | VMA11 | n |
| W303.asm02.HP0_G0053150 | 886.2982 | 940.3172 | 908.6782 | 432.8082 | 518.9698 | 461.8761 | 627.1123 | 374.7562 | 517.9045 | 3.65E-06 | 0.952815304 | YPL233W | NSL1 | n |
| W303.asm02.HP0_G0053160 | 10890.77 | 11220.38 | 10466.51 | 9332.817 | 9985.396 | 10568.45 | 11638.61 | 10799.21 | 11414.23 | 0.175109114 | 0.124413032 | YPL232W | SSO1 | n |
| W303.asm02.HP0_G0053170 | 72158.65 | 72935.23 | 76326.82 | 30429.96 | 33581.17 | 33808.11 | 14627.27 | 31028.52 | 28412.98 | 1.10E-05 | 1.178602729 | YPL231W | FAS2 | n |
| W303.asm02.HP0_G0053180 | 4624.211 | 4868.506 | 4418.354 | 443.2289 | 328.0445 | 458.8374 | 3141.734 | 1230.575 | 1874.264 | 1.07E-27 | 3.49893612 | YPL230W | USV1 | n |
| W303.asm02.HP0_G0053190 | 856.6491 | 969.25 | 820.2896 | 764.1878 | 755.8908 | 2218.221 | 5079.857 | 2330.146 | 3072.741 | 0.308074024 | -0.498529233 | YPL229W | | n |
| W303.asm02.HP0_G0053200 | 2099.796 | 2233.392 | 1824.902 | 5067.954 | 4705.442 | 5566.822 | 5055.167 | 4320.971 | 4685.062 | 7.27E-24 | -1.316879909 | YPL228W | CET1 | n |
| W303.asm02.HP0_G0053210 | 1130.904 | 1146.185 | 1255.765 | 3090.097 | 3421.903 | 3397.98 | 1991.205 | 2826.241 | 2424.463 | 5.87E-22 | -1.487970833 | YPL227C | ALG5 | n |
| W303.asm02.HP0_G0053220 | 4610.445 | 4264.255 | 5324.876 | 11177.98 | 10865.39 | 5049.491 | 10239.95 | 4309.159 | 8577.121 | 0.017886366 | -0.932057318 | YPL226W | NEW1 | n |
| W303.asm02.HP0_G0053230 | 6679.533 | 6391.931 | 6277.749 | 2011.203 | 2416.941 | 1700.889 | 1680.118 | 1357.283 | 1637.439 | 3.23E-24 | 1.65880982 | YPL225W | CHP1 | n |
| W303.asm02.HP0_G0053260 | 2210.981 | 1861.717 | 1949.939 | 2379.403 | 2252.919 | 2642.873 | 762.9044 | 2394.574 | 1623.086 | 0.488080016 | -0.272592469 | YPL222W | FMP40 | n |
| W303.asm02.HP0_G0053270 | 1922.96 | 1927.372 | 1808.733 | 4834.53 | 4426.865 | 4576.22 | 10527.59 | 8443.289 | 8969.436 | 1.86E-28 | -1.290053867 | YPL221W | FLC1 | n |
| W303.asm02.HP0_G0053300 | 3737.912 | 3565.416 | 3766.218 | 4440.626 | 5004.848 | 4868.691 | 3398.504 | 4049.3 | 3660.018 | 0.000625976 | -0.370694263 | YPL218W | SAR1 | n |

| | | | | | | | | | | | | | | |
|---|---|---|---|---|---|---|---|---|---|---|---|---|---|---|
| W303.asm02.HP0_G0053310 | 4139.235 | 4073.966 | 4605.91 | 8265.038 | 7344.551 | 2964.971 | 11353.45 | 4366.071 | 10413.11 | 0.250206351 | -0.535030752 | YPL217C | BMS1 | n |
| W303.asm02.HP0_G0053330 | 6268.68 | 6541.046 | 6049.232 | 3604.187 | 3419.3 | 2898.121 | 2199.831 | 2432.157 | 2645.739 | 5.99E-13 | 0.926566505 | YPL215W | CBP3 | n |
| W303.asm02.HP0_G0053340 | 1415.748 | 1394.34 | 1363.556 | 1328.992 | 1300.896 | 1540.6 | 1950.467 | 3894.672 | 2469.914 | 0.996795312 | 0.001061467 | YPL214C | THI6 | n |
| W303.asm02.HP0_G0053350 | 1185.967 | 1138.396 | 1124.26 | 934.3932 | 939.0056 | 824.2361 | 1083.867 | 1074.874 | 1072.888 | 0.000315183 | 0.354566776 | YPL213W | LEA1 | n |
| W303.asm02.HP0_G0053360 | 1915.548 | 1886.198 | 2173.067 | 9100.087 | 9228.637 | 6182.91 | 6056.325 | 5424.837 | 7276.977 | 4.06E-24 | -2.036477352 | YPL212C | PUS1 | n |
| W303.asm02.HP0_G0053370 | 1220.91 | 1170.667 | 1264.389 | 1594.374 | 1611.584 | 477.829 | 1501.119 | 775.2835 | 1252.3 | 0.983768722 | -0.010887571 | YPL211W | NIP7 | n |
| W303.asm02.HP0_G0053380 | 8487.073 | 9363.111 | 8908.927 | 10172.03 | 10052.22 | 10136.96 | 7741.381 | 9078.978 | 9272.046 | 0.094858252 | -0.182225925 | YPL210C | SRP72 | n |
| W303.asm02.HP0_G0053390 | 231.8988 | 243.7035 | 258.6984 | 587.7299 | 515.4985 | 756.626 | 634.5192 | 777.4311 | 545.4145 | 3.73E-10 | -1.341182867 | YPL209C | IPL1 | n |
| W303.asm02.HP0_G0053400 | 1799.069 | 1555.696 | 1903.589 | 2089.706 | 1875.408 | 1469.951 | 1166.577 | 1424.933 | 1499.89 | 0.821832107 | -0.047704535 | YPL208W | RKM1 | n |
| W303.asm02.HP0_G0053420 | 3782.386 | 3519.791 | 3523.688 | 851.0273 | 1023.186 | 2962.692 | 5527.97 | 5654.63 | 4866.867 | 0.007181046 | 1.162306709 | YPL206C | PGC1 | n |
| W303.asm02.HP0_G0053440 | 5438.504 | 5656.369 | 5817.481 | 9557.21 | 9037.712 | 10234.96 | 18430.68 | 13445.05 | 15950.98 | 6.47E-08 | -0.769514537 | YPL204W | HRR25 | n |
| W303.asm02.HP0_G0053460 | 321.9052 | 338.2916 | 282.4125 | 848.9431 | 641.3356 | 662.4275 | 12882.96 | 4105.137 | 7925.255 | 0.001599919 | -1.191625406 | YPL201C | YIG1 | n |
| W303.asm02.HP0_G0053480 | 3456.245 | 3361.773 | 3376.015 | 2388.434 | 2280.69 | 1928.788 | 1961.578 | 2245.316 | 1594.38 | 0.000120931 | 0.627629204 | YPL199C | | n |
| W303.asm02.HP0_G0053500 | 4390.194 | 4161.877 | 3871.853 | 502.9745 | 490.331 | 566.7098 | 2117.121 | 1496.877 | 2163.717 | 3.25E-67 | 2.993324269 | YPL196W | OXR1 | n |
| W303.asm02.HP0_G0053510 | 5883.241 | 5863.35 | 5770.053 | 5672.357 | 5185.359 | 6819.509 | 3691.075 | 5742.682 | 5150.339 | 0.955585057 | -0.01319846 | YPL195W | APL5 | n |
| W303.asm02.HP0_G0053520 | 1022.896 | 1137.283 | 973.3529 | 664.8433 | 619.6396 | 525.6879 | 324.6664 | 474.6196 | 387.5313 | 1.80E-05 | 0.791461803 | YPL194W | DDC1 | n |
| W303.asm02.HP0_G0053530 | 1547.051 | 1703.699 | 1668.605 | 572.4461 | 413.093 | 99.51606 | 1334.465 | 381.199 | 1209.241 | 0.000279616 | 2.180572433 | YPL193W | RSA1 | n |
| W303.asm02.HP0_G0053560 | 8081.515 | 8202.459 | 8472.374 | 6395.557 | 5346.778 | 5407.293 | 2959.032 | 4271.576 | 3078.721 | 0.00295428 | 0.529560927 | YPL190C | NAB3 | n |
| W303.asm02.HP0_G0053570 | 942.4198 | 918.0611 | 868.7956 | 1753.464 | 1847.637 | 1096.956 | 566.6231 | 1017.962 | 855.2003 | 0.005514596 | -0.783471104 | YPL189C-A | COA2 | n |
| W303.asm02.HP0_G0053590 | 4086.29 | 3417.413 | 3799.633 | 1945.9 | 1876.276 | 1880.93 | 1868.992 | 1779.286 | 1467.595 | 8.47E-14 | 0.986919382 | YPL188W | POS5 | n |
| W303.asm02.HP0_G0053600 | 40.23815 | 44.51205 | 44.19432 | 191.0469 | 222.1677 | 156.4909 | 125.9163 | 170.7342 | 80.13765 | 1.20E-10 | -2.14308872 | YPL187W | MF(ALPHA | n |
| W303.asm02.HP0_G0053610 | 4274.774 | 4717.165 | 3872.931 | 1160.176 | 1193.283 | 2492.46 | 2488.698 | 3881.787 | 3350.232 | 1.76E-05 | 1.408548426 | YPL186C | UIP4 | n |
| W303.asm02.HP0_G0053620 | 4102.174 | 3785.75 | 4244.81 | 9427.298 | 8456.257 | 7048.928 | 8789.448 | 6145.357 | 8054.432 | 1.15E-08 | -1.039126154 | YPL184C | MRN1 | n |
| W303.asm02.HP0_G0053630 | 1427.395 | 1230.758 | 1068.209 | 936.4774 | 1014.508 | 461.1164 | 346.8869 | 327.509 | 438.9629 | 0.050691549 | 0.627659934 | YPL183W- | RTC6 | n |
| W303.asm02.HP0_G0053640 | 2227.923 | 2235.618 | 2605.309 | 1859.06 | 1665.39 | 913.8765 | 7284.626 | 1439.966 | 6261.501 | 0.219369972 | 0.671455698 | YPL183C | RTT10 | n |
| W303.asm02.HP0_G0053650 | 6906.137 | 7366.745 | 7257.569 | 2539.882 | 2764.946 | 2367.115 | 2335.623 | 2843.422 | 2564.405 | 2.32E-38 | 1.488888266 | YPL181W | CTI6 | n |
| W303.asm02.HP0_G0053660 | 2622.892 | 2730.815 | 2570.816 | 3801.487 | 3437.524 | 4342.243 | 4382.379 | 4330.635 | 4576.219 | 2.43E-06 | -0.54757233 | YPL180W | TCO89 | n |
| W303.asm02.HP0_G0053670 | 6770.598 | 7549.245 | 7084.025 | 7590.469 | 7109.366 | 5773.451 | 10021.45 | 8287.588 | 7583.175 | 0.726988184 | 0.064106944 | YPL179W | PPQ1 | n |
| W303.asm02.HP0_G0053680 | 3464.717 | 3363.999 | 3231.575 | 2016.066 | 2220.809 | 1808.002 | 1340.638 | 1646.135 | 1568.067 | 1.93E-08 | 0.735118696 | YPL178W | CBC2 | n |
| W303.asm02.HP0_G0053690 | 2400.523 | 2487.111 | 2147.197 | 6026.663 | 6472.369 | 6359.153 | 3736.75 | 5357.188 | 4150.413 | 6.90E-19 | -1.42259626 | YPL177C | CUP9 | n |
| W303.asm02.HP0_G0053710 | 3090.925 | 2953.375 | 3037.551 | 975.3815 | 877.3888 | 826.5151 | 1290.024 | 1290.708 | 1225.986 | 5.67E-74 | 1.760545598 | YPL175W | SPT14 | n |
| W303.asm02.HP0_G0053720 | 1038.78 | 1052.71 | 995.989 | 1014.286 | 1094.349 | 1622.644 | 460.4585 | 722.6674 | 339.6879 | 0.409026703 | -0.273251179 | YPL174C | NIP100 | n |
| W303.asm02.HP0_G0053730 | 7842.204 | 8350.462 | 7855.809 | 4999.177 | 5010.055 | 3723.876 | 3078.776 | 2354.843 | 2825.151 | 1.82E-06 | 0.80830453 | YPL173W | MRPL40 | n |
| W303.asm02.HP0_G0053740 | 3011.508 | 3143.664 | 2930.838 | 5127.005 | 4926.742 | 4453.914 | 1403.596 | 1852.305 | 1745.087 | 9.17E-07 | -0.675128648 | YPL172C | COX10 | n |
| W303.asm02.HP0_G0053760 | 4942.939 | 4851.814 | 4172.59 | 2235.597 | 2654.73 | 2945.22 | 5442.792 | 4623.782 | 4622.866 | 6.00E-08 | 0.834077852 | YPL170W | DAP1 | n |
| W303.asm02.HP0_G0053770 | 5440.622 | 5665.272 | 5615.912 | 5522.993 | 5279.954 | 4728.912 | 3340.484 | 3014.156 | 3205.506 | 0.312253193 | 0.106454461 | YPL169C | MEX67 | n |
| W303.asm02.HP0_G0053780 | 1756.713 | 1926.259 | 1607.164 | 1835.44 | 1657.579 | 1581.622 | 1239.411 | 1653.652 | 1625.479 | 0.737988399 | 0.059818591 | YPL168W | MRX4 | n |
| W303.asm02.HP0_G0053790 | 1285.503 | 1249.676 | 1177.078 | 831.5752 | 750.6838 | 804.4848 | 1212.253 | 1186.549 | 1060.927 | 2.13E-09 | 0.636569793 | YPL167C | REV3 | n |
| W303.asm02.HP0_G0053800 | 1464.457 | 1645.833 | 1334.453 | 455.0391 | 485.9918 | 635.0796 | 1002.392 | 856.8923 | 769.0822 | 1.67E-15 | 1.495701887 | YPL166W | ATG29 | n |
| W303.asm02.HP0_G0053810 | 1875.31 | 1885.086 | 1737.591 | 1219.922 | 1097.821 | 994.401 | 746.8562 | 1025.479 | 734.3958 | 2.50E-05 | 0.73096957 | YPL164C | MLH3 | n |
| W303.asm02.HP0_G0053830 | 1023.955 | 1169.554 | 995.989 | 1349.833 | 1474.464 | 2285.831 | 850.552 | 2436.452 | 1774.989 | 0.081861119 | -0.680052587 | YPL162C | | n |
| W303.asm02.HP0_G0053840 | 1686.826 | 1849.476 | 1706.332 | 2537.103 | 2311.065 | 2553.233 | 1306.073 | 2038.072 | 1681.695 | 0.006543515 | -0.49761952 | YPL161C | BEM4 | n |
| W303.asm02.HP0_G0053850 | 14481.5 | 14307.29 | 15255.66 | 28541.72 | 28893.95 | 33829.38 | 48827.11 | 44323.24 | 47457.04 | 7.53E-24 | -1.051102489 | YPL160W | CDC60 | n |
| W303.asm02.HP0_G0053860 | 3241.289 | 3597.687 | 3478.416 | 3445.097 | 3162.418 | 3541.556 | 4162.643 | 3636.96 | 3576.292 | 0.851133105 | 0.023538031 | YPL159C | PET20 | n |
| W303.asm02.HP0_G0053870 | 2425.937 | 2893.284 | 2866.163 | 5022.103 | 4880.746 | 4674.976 | 3971.3 | 5614.9 | 5369.223 | 5.48E-07 | -0.832702173 | YPL158C | AIM44 | n |
| W303.asm02.HP0_G0053880 | 943.4787 | 1007.085 | 945.3272 | 1096.262 | 1051.825 | 550.7568 | 1586.298 | 1144.671 | 1581.223 | 0.74677942 | 0.101688779 | YPL157W | TGS1 | n |
| W303.asm02.HP0_G0053890 | 1424.219 | 1552.358 | 1287.025 | 1092.788 | 971.1158 | 843.9874 | 2360.313 | 1769.622 | 1465.203 | 0.011385527 | 0.551931999 | YPL156C | PRM4 | n |
| W303.asm02.HP0_G0053900 | 1711.18 | 1712.601 | 1885.265 | 5931.486 | 5531.628 | 5876.006 | 1982.564 | 3687.429 | 2856.249 | 1.22E-14 | -1.707517451 | YPL155C | KIP2 | n |
| W303.asm02.HP0_G0053910 | 18025.63 | 18195.42 | 17488.01 | 5909.95 | 6015.884 | 11123.77 | 14260.63 | 21444.86 | 16850.44 | 1.52E-05 | 1.220413736 | YPL154C | PEP4 | n |
| W303.asm02.HP0_G0053920 | 670.2829 | 627.62 | 704.9532 | 2234.902 | 2187.831 | 2949.778 | 2009.722 | 1822.238 | 1995.069 | 6.51E-34 | -1.880079887 | YPL153C | RAD53 | n |
| W303.asm02.HP0_G0053930 | 2378.286 | 2118.774 | 2174.145 | 1222.7 | 1386.812 | 1747.988 | 2507.215 | 2943.286 | 3266.506 | 0.000745933 | 0.614558364 | YPL152W | RRD2 | n |
| W303.asm02.HP0_G0053940 | 2083.913 | 2188.88 | 2001.679 | 1397.074 | 1235.808 | 1288.391 | 2315.872 | 1956.463 | 2290.501 | 3.83E-09 | 0.677706542 | YPL151C | PRP46 | n |
| W303.asm02.HP0_G0053950 | 2807.14 | 3012.353 | 2731.424 | 3813.992 | 3664.031 | 4324.771 | 2725.717 | 3088.249 | 2984.23 | 3.93E-05 | -0.46510024 | YPL150W | | n |
| W303.asm02.HP0_G0053960 | 1484.576 | 1702.586 | 1281.635 | 245.2348 | 263.8241 | 252.9683 | 448.1137 | 360.7968 | 403.0804 | 1.48E-58 | 2.552518897 | YPL149W | ATG5 | n |

| | | | | | | | | | | | | | | |
|---|---|---|---|---|---|---|---|---|---|---|---|---|---|---|
| W303.asm02.HP0_G0053970 | 349.4366 | 373.9013 | 376.1906 | 167.4266 | 189.1897 | 192.9548 | 88.88206 | 76.23979 | 47.84337 | 1.72E-06 | 1.001236841 | YPL148C | PPT2 | n |
| W303.asm02.HP0_G0053990 | 1575.641 | 1700.36 | 1366.79 | 171.5949 | 147.5332 | 164.0876 | 435.769 | 342.5422 | 167.4518 | 6.07E-25 | 3.263684587 | YPL147W | PXA1 | n |
| W303.asm02.HP0_G0054000 | 2420.642 | 2980.082 | 3027.85 | 3115.107 | 2485.501 | 743.7117 | 4304.608 | 1343.324 | 3087.094 | 0.476901998 | 0.409805588 | YPL146C | NOP53 | n |
| W303.asm02.HP0_G0054010 | 14923.06 | 16126.72 | 16705.45 | 10653.47 | 11282.82 | 13039.64 | 10917.68 | 13902.49 | 11743.16 | 0.001865545 | 0.449298922 | YPL145C | KES1 | n |
| W303.asm02.HP0_G0054020 | 640.6337 | 540.8215 | 573.4482 | 367.5048 | 407.886 | 297.0289 | 416.0174 | 322.14 | 453.316 | 0.000397536 | 0.711330145 | YPL144W | POC4 | n |
| W303.asm02.HP0_G0054040 | 2216.275 | 2394.749 | 2438.233 | 4731.017 | 4181.265 | 4453.154 | 4687.294 | 4107.285 | 3921.961 | 2.90E-16 | -0.923137116 | YPL141C | FRK1 | n |
| W303.asm02.HP0_G0054060 | 2422.76 | 2232.28 | 2286.247 | 2615.606 | 2673.823 | 3834.027 | 2968.908 | 4352.111 | 3701.881 | 0.075947311 | -0.394383187 | YPL139C | UME1 | n |
| W303.asm02.HP0_G0054070 | 1286.562 | 1409.919 | 1437.932 | 1188.659 | 1122.12 | 1026.307 | 1359.155 | 1345.471 | 1428.125 | 0.004260762 | 0.308793643 | YPL138C | SPP1 | n |
| W303.asm02.HP0_G0054080 | 1688.943 | 1904.003 | 1665.371 | 7951.721 | 8138.627 | 10294.97 | 16356.77 | 16584.84 | 15520.39 | 9.51E-65 | -2.327145357 | YPL137C | GIP3 | n |
| W303.asm02.HP0_G0054090 | 2388.875 | 2348.011 | 2127.795 | 3614.608 | 3532.119 | 2980.164 | 5231.697 | 3348.108 | 3662.41 | 0.005787684 | -0.560924966 | YPL135W | ISU1 | n |
| W303.asm02.HP0_G0054100 | 1863.662 | 1889.537 | 1803.344 | 4614.999 | 4675.935 | 4061.927 | 1667.773 | 1665.464 | 1227.183 | 7.25E-17 | -1.264877806 | YPL134C | ODC1 | n |
| W303.asm02.HP0_G0054110 | 1440.102 | 1353.166 | 1333.375 | 1181.712 | 1088.275 | 1201.03 | 912.2756 | 536.9 | 444.9434 | 0.379120995 | 0.249551405 | YPL133C | RDS2 | n |
| W303.asm02.HP0_G0054120 | 1840.366 | 1848.363 | 1678.306 | 2566.976 | 2615.677 | 2527.404 | 3500.966 | 2603.965 | 3115.8 | 0.000150595 | -0.522599074 | YPL132W | COX11 | n |
| W303.asm02.HP0_G0054130 | 34559.28 | 34033.92 | 36759.97 | 42734.77 | 48449.04 | 45691.55 | 34925.71 | 29128.97 | 28935.67 | 0.00146053 | -0.377615475 | YPL131W | RPL5 | n |
| W303.asm02.HP0_G0054150 | 6093.962 | 6173.822 | 6060.011 | 3198.473 | 3577.247 | 3170.08 | 3962.659 | 4314.528 | 3906.411 | 1.46E-22 | 0.882144417 | YPL129W | TAF14 | n |
| W303.asm02.HP0_G0054160 | 823.8232 | 774.5098 | 864.4839 | 1561.027 | 1079.596 | 594.0577 | 5578.584 | 2632.957 | 5195.79 | 0.344068766 | -0.393431417 | YPL128C | TBF1 | n |
| W303.asm02.HP0_G0054180 | 2812.435 | 3042.399 | 3419.131 | 6967.308 | 5835.373 | 1945.501 | 8235.17 | 4118.023 | 7292.526 | 0.149711672 | -0.669284351 | YPL126W | NAN1 | n |
| W303.asm02.HP0_G0054190 | 3617.198 | 3560.964 | 3748.971 | 7318.14 | 7072.917 | 8286.422 | 6710.596 | 7331.906 | 7055.701 | 2.81E-27 | -1.053380029 | YPL125W | KAP120 | n |
| W303.asm02.HP0_G0054210 | 5444.857 | 5434.922 | 4681.364 | 771.1349 | 801.0186 | 2898.121 | 855.4899 | 4034.266 | 1936.461 | 0.003615035 | 1.799489327 | YPL123C | RNY1 | n |
| W303.asm02.HP0_G0054220 | 1184.908 | 1157.313 | 1184.623 | 1613.826 | 1549.967 | 1656.069 | 1156.701 | 1322.922 | 1320.477 | 2.87E-06 | -0.450772922 | YPL122C | TFB2 | n |
| W303.asm02.HP0_G0054230 | 36.00256 | 32.27124 | 37.72685 | 38.20939 | 57.27761 | 44.82021 | 41.97209 | 105.2324 | 72.96114 | 0.371274209 | -0.400693791 | YPL121C | MEI5 | n |
| W303.asm02.HP0_G0054240 | 1574.582 | 1658.074 | 1606.086 | 764.1878 | 702.9524 | 935.9068 | 581.4368 | 815.0141 | 598.0422 | 4.85E-08 | 1.009486666 | YPL120W | VPS30 | n |
| W303.asm02.HP0_G0054270 | 12714.2 | 13026.45 | 12484.36 | 4736.575 | 4645.561 | 3890.243 | 1277.68 | 1485.065 | 1400.615 | 2.24E-39 | 1.526095807 | YPL118W | MRP51 | n |
| W303.asm02.HP0_G0054280 | 5632.282 | 5538.412 | 5792.689 | 6103.776 | 6107.008 | 5876.006 | 4492.248 | 3951.584 | 4338.198 | 0.296026973 | -0.09249009 | YPL117C | IDI1 | n |
| W303.asm02.HP0_G0054290 | 1443.279 | 1487.815 | 1393.738 | 5299.295 | 5528.157 | 7107.422 | 5060.105 | 5326.048 | 4313.08 | 2.31E-39 | -2.052075929 | YPL116W | HOS3 | n |
| W303.asm02.HP0_G0054300 | 2839.966 | 3014.579 | 2768.073 | 4544.138 | 4459.843 | 5864.611 | 4909.499 | 5070.483 | 5293.869 | 1.66E-08 | -0.786134558 | YPL115C | BEM3 | n |
| W303.asm02.HP0_G0054310 | 711.5799 | 673.2448 | 652.1356 | 3239.461 | 3519.101 | 2634.517 | 707.3531 | 1374.464 | 968.8283 | 1.08E-18 | -2.205119937 | YPL113C | | n |
| W303.asm02.HP0_G0054320 | 2100.855 | 1783.821 | 1965.03 | 1507.534 | 1543.024 | 2117.945 | 459.224 | 1124.269 | 782.2391 | 0.611419706 | 0.178621315 | YPL112C | PEX25 | n |
| W303.asm02.HP0_G0054330 | 1254.795 | 1632.48 | 1214.805 | 8379.666 | 8752.192 | 10810.03 | 5926.705 | 6991.511 | 5505.576 | 3.34E-52 | -2.768125689 | YPL111W | CAR1 | n |
| W303.asm02.HP0_G0054340 | 6532.346 | 6750.253 | 6404.942 | 3759.109 | 3338.59 | 5138.371 | 3579.972 | 7573.511 | 5352.477 | 0.020346271 | 0.686110757 | YPL110C | GDE1 | n |
| W303.asm02.HP0_G0054350 | 1313.034 | 1471.123 | 1280.557 | 1852.113 | 1732.214 | 2238.732 | 1935.654 | 2370.95 | 2162.52 | 0.000789697 | -0.518840817 | YPL109C | CQD1 | n |
| W303.asm02.HP0_G0054360 | 311.3162 | 284.8772 | 306.1265 | 1469.325 | 1485.746 | 607.7317 | 1352.983 | 558.376 | 934.1419 | 1.19E-06 | -1.981222206 | YPL108W | | n |
| W303.asm02.HP0_G0054370 | 745.4647 | 768.9457 | 736.2126 | 1051.8 | 1103.896 | 799.1672 | 727.1046 | 1060.914 | 814.5334 | 0.058668491 | -0.392631332 | YPL107W | DPC25 | n |
| W303.asm02.HP0_G0054380 | 26199.27 | 27292.57 | 27196.75 | 20030.75 | 21134.57 | 20940.91 | 11610.22 | 10436.26 | 11177.41 | 5.44E-07 | 0.377648425 | YPL106C | SSE1 | n |
| W303.asm02.HP0_G0054390 | 8437.305 | 8591.939 | 8673.943 | 10037.95 | 8632.43 | 7948.371 | 7766.07 | 9045.691 | 7947.98 | 0.721691129 | -0.050555519 | YPL105C | SYH1 | n |
| W303.asm02.HP0_G0054400 | 6276.093 | 6367.449 | 5991.025 | 2815.685 | 2601.792 | 2925.468 | 1751.717 | 1905.995 | 2051.285 | 2.04E-31 | 1.159151944 | YPL104W | MSD1 | n |
| W303.asm02.HP0_G0054410 | 4048.17 | 4119.591 | 4002.28 | 1151.145 | 1141.213 | 647.2343 | 670.3189 | 842.9329 | 723.631 | 2.15E-18 | 2.049726767 | YPL103C | FMP30 | n |
| W303.asm02.HP0_G0054420 | 2191.92 | 2514.931 | 2416.675 | 2747.602 | 2517.611 | 1122.025 | 2746.703 | 1444.261 | 2130.226 | 0.696480847 | 0.157393899 | YPL101W | ELP4 | n |
| W303.asm02.HP0_G0054430 | 3728.382 | 3899.256 | 3416.975 | 1137.25 | 1134.27 | 1593.776 | 2518.325 | 2236.725 | 2419.679 | 1.91E-19 | 1.514579895 | YPL100W | ATG21 | n |
| W303.asm02.HP0_G0054440 | 1712.239 | 1783.821 | 1575.905 | 1704.833 | 1710.518 | 1682.657 | 1569.015 | 1923.176 | 1853.931 | 0.957562864 | -0.007424752 | YPL099C | INA17 | n |
| W303.asm02.HP0_G0054450 | 3202.11 | 2808.711 | 2871.553 | 1103.209 | 1341.685 | 1033.904 | 701.1807 | 652.8704 | 495.1789 | 2.22E-14 | 1.352765698 | YPL098C | MGR2 | n |
| W303.asm02.HP0_G0054460 | 3902.042 | 3977.152 | 3548.48 | 4118.277 | 4248.957 | 4157.644 | 1387.548 | 1871.633 | 1703.224 | 0.370517771 | -0.132249618 | YPL097W | MSY1 | n |
| W303.asm02.HP0_G0054470 | 540.0383 | 431.7669 | 496.9166 | 194.5205 | 217.8285 | 219.5431 | 154.3091 | 280.2618 | 161.4714 | 2.93E-06 | 1.217373925 | YPL096C-/ | ERI1 | n |
| W303.asm02.HP0_G0054480 | 2160.153 | 2045.329 | 1846.46 | 1006.644 | 1057.9 | 1219.262 | 993.7508 | 1408.826 | 1038.201 | 8.30E-07 | 0.882072292 | YPL096W | PNG1 | n |
| W303.asm02.HP0_G0054500 | 4339.367 | 4065.063 | 4328.887 | 2855.978 | 2843.052 | 2947.499 | 3180.003 | 3806.621 | 3752.117 | 8.05E-08 | 0.558425008 | YPL094C | SEC62 | n |
| W303.asm02.HP0_G0054510 | 3887.217 | 4333.249 | 5108.216 | 8370.635 | 7604.904 | 2449.918 | 13917.45 | 5676.106 | 11045.84 | 0.349603115 | -0.467164972 | YPL093W | NOG1 | n |
| W303.asm02.HP0_G0054520 | 525.2138 | 564.1903 | 504.4619 | 3986.976 | 4042.41 | 3053.092 | 8699.332 | 7006.545 | 6544.973 | 4.96E-58 | -2.797742543 | YPL092W | SSU1 | n |
| W303.asm02.HP0_G0054530 | 8530.488 | 8565.232 | 8491.776 | 4178.718 | 4196.886 | 5228.012 | 3208.396 | 5678.254 | 4742.474 | 4.59E-05 | 0.911439397 | YPL091W | GLR1 | n |
| W303.asm02.HP0_G0054550 | 4684.568 | 5083.277 | 4889.4 | 10345.71 | 8561.266 | 10202.3 | 13308.85 | 13189.48 | 12630.65 | 2.30E-20 | -0.989999121 | YPL089C | RLM1 | n |
| W303.asm02.HP0_G0054570 | 7621.953 | 6960.573 | 6576.33 | 31.26223 | 29.50665 | 85.8421 | 1571.484 | 1493.656 | 1175.751 | 2.14E-86 | 7.171350002 | YPL087W | YDC1 | n |
| W303.asm02.HP0_G0054580 | 2733.018 | 2830.967 | 3104.381 | 4304.461 | 4097.085 | 2398.261 | 5814.368 | 3184.891 | 5402.713 | 0.320918225 | -0.317169031 | YPL086C | ELP3 | n |
| W303.asm02.HP0_G0054590 | 5688.404 | 5540.638 | 5842.273 | 14916.94 | 13701.5 | 17318.83 | 13261.94 | 16944.56 | 16721.26 | 1.62E-22 | -1.428106435 | YPL085W | SEC16 | n |
| W303.asm02.HP0_G0054600 | 6753.656 | 6663.455 | 6560.161 | 3557.641 | 3399.339 | 4167.52 | 3699.716 | 4322.045 | 3558.351 | 5.01E-11 | 0.844547919 | YPL084W | BRO1 | n |
| W303.asm02.HP0_G0054610 | 1495.165 | 1488.928 | 1414.218 | 1573.532 | 1431.072 | 957.1774 | 1291.259 | 1154.335 | 1291.771 | 0.494323684 | 0.150752062 | YPL083C | SEN54 | n |

| | | | | | | | | | | | | | | |
|---|---|---|---|---|---|---|---|---|---|---|---|---|---|---|
| W303.asm02.HP0_G0054620 | 2045.792 | 2015.283 | 2112.704 | 5818.942 | 6130.44 | 3682.854 | 5776.1 | 4442.31 | 5435.007 | 7.56E-10 | -1.340260541 | YPL082C | MOT1 | n |
| W303.asm02.HP0_G0054640 | 7792.436 | 7125.267 | 8191.039 | 15092.01 | 17902.72 | 15888.39 | 13140.97 | 11544.42 | 10732.46 | 4.70E-17 | -1.080842124 | YPL079W | RPL21B | y |
| W303.asm02.HP0_G0054650 | 6565.172 | 6137.1 | 6174.269 | 18788.6 | 20880.29 | 20271.65 | 8217.887 | 12242.39 | 10993.21 | 1.90E-24 | -1.666919081 | YPL078C | ATP4 | n |
| W303.asm02.HP0_G0054670 | 1712.239 | 1742.647 | 1610.398 | 1539.491 | 1616.791 | 2342.046 | 2350.437 | 2453.633 | 2335.953 | 0.560669202 | -0.118386997 | YPL076W | GPI2 | n |
| W303.asm02.HP0_G0054680 | 4839.167 | 4802.851 | 4988.568 | 8848.599 | 7910.385 | 21067.78 | 25046.22 | 23634.34 | 27497.98 | 0.000106987 | -1.370428067 | YPL075W | GCR1 | n |
| W303.asm02.HP0_G0054690 | 3602.373 | 3838.052 | 3534.467 | 325.1272 | 323.7053 | 491.503 | 527.12 | 691.5272 | 564.5518 | 4.29E-58 | 3.266394093 | YPL074W | YTA6 | n |
| W303.asm02.HP0_G0054700 | 1178.554 | 1032.68 | 1166.299 | 3856.369 | 3520.837 | 5029.739 | 4289.794 | 5688.992 | 3778.43 | 4.14E-19 | -1.877098939 | YPL072W | UBP16 | n |
| W303.asm02.HP0_G0054710 | 919.1241 | 928.0763 | 974.4308 | 526.5948 | 553.6835 | 675.3418 | 797.4696 | 558.376 | 549.0027 | 0.000517557 | 0.684573368 | YPL071C | | n |
| W303.asm02.HP0_G0054720 | 4491.848 | 4608.11 | 4470.093 | 1443.62 | 1376.398 | 2046.536 | 2018.364 | 1462.516 | 1581.223 | 2.25E-13 | 1.479353569 | YPL070W | MUK1 | n |
| W303.asm02.HP0_G0054730 | 522.0371 | 541.9343 | 499.0724 | 1157.397 | 1098.689 | 1258.005 | 978.9372 | 1326.143 | 1010.691 | 1.51E-13 | -1.169009952 | YPL069C | BTS1 | n |
| W303.asm02.HP0_G0054740 | 447.9142 | 436.2181 | 469.9688 | 808.6496 | 680.3885 | 527.2072 | 1706.042 | 1019.036 | 1603.949 | 0.026162222 | -0.574514636 | YPL068C | | n |
| W303.asm02.HP0_G0054750 | 1831.895 | 1632.48 | 1645.969 | 1367.896 | 1482.275 | 1811.8 | 2155.39 | 3892.525 | 2536.895 | 0.626652844 | 0.1325223 | YPL067C | HTC1 | n |
| W303.asm02.HP0_G0054760 | 2413.23 | 2687.415 | 2409.129 | 578.6985 | 499.0094 | 692.8141 | 582.6713 | 550.8594 | 458.1003 | 4.51E-36 | 2.083929745 | YPL066W | RGL1 | n |
| W303.asm02.HP0_G0054770 | 1427.395 | 1424.386 | 1388.348 | 573.8355 | 615.3003 | 814.3605 | 830.8004 | 1001.855 | 691.3367 | 5.14E-08 | 1.081606719 | YPL065W | VPS28 | n |
| W303.asm02.HP0_G0054790 | 10606.99 | 10122.04 | 10409.38 | 11927.58 | 11852.99 | 11712.51 | 6336.55 | 5494.634 | 5734.028 | 0.028960736 | -0.188833018 | YPL063W | TIM50 | n |
| W303.asm02.HP0_G0054800 | 2107.208 | 3424.09 | 2294.871 | 4599.716 | 3522.573 | 4924.906 | 23117.98 | 5670.737 | 8199.158 | 0.141089305 | -0.737395521 | YPL061W | ALD6 | n |
| W303.asm02.HP0_G0054810 | 1323.623 | 1452.206 | 1198.636 | 1004.56 | 946.8162 | 863.7387 | 1254.225 | 1171.516 | 1212.83 | 5.02E-05 | 0.497257383 | YPL060W | MFM1 | n |
| W303.asm02.HP0_G0054820 | 4286.422 | 4470.123 | 4329.965 | 2876.82 | 3181.511 | 2885.966 | 1614.691 | 2309.744 | 2095.54 | 0.000407071 | 0.549129182 | YPL059W | GRX5 | n |
| W303.asm02.HP0_G0054830 | 1670.942 | 1817.205 | 1693.397 | 2480.137 | 2481.162 | 2030.584 | 2049.225 | 1917.807 | 2363.463 | 0.002362231 | -0.43230549 | YPL058C | PDR12 | n |
| W303.asm02.HP0_G0054840 | 6828.838 | 9536.708 | 6427.578 | 3490.949 | 3762.965 | 4410.613 | 8510.457 | 4610.897 | 6839.21 | 0.00030725 | 0.966464956 | YPL057C | SUR1 | n |
| W303.asm02.HP0_G0054860 | 3013.626 | 2843.208 | 2783.164 | 543.9627 | 500.7451 | 526.4476 | 557.9818 | 493.948 | 611.1991 | 2.25E-105 | 2.458537157 | YPL055C | LGE1 | n |
| W303.asm02.HP0_G0054870 | 1460.221 | 1199.6 | 1213.727 | 1403.327 | 1439.751 | 2372.433 | 3167.658 | 3480.186 | 2587.13 | 0.099072729 | -0.429148429 | YPL054W | LEE1 | n |
| W303.asm02.HP0_G0054890 | 826.9999 | 847.9546 | 831.0687 | 509.9216 | 552.8157 | 417.0559 | 846.8485 | 612.066 | 450.9238 | 0.002337225 | 0.760252661 | YPL052W | OAZ1 | n |
| W303.asm02.HP0_G0054900 | 919.1241 | 835.7138 | 892.5096 | 1257.436 | 1289.614 | 1566.428 | 1698.635 | 2008.006 | 1952.01 | 6.21E-06 | -0.635732346 | YPL051W | ARL3 | n |
| W303.asm02.HP0_G0054910 | 3777.092 | 3564.303 | 3938.684 | 10858.41 | 11296.71 | 10241.04 | 4651.495 | 5912.342 | 4870.455 | 3.43E-34 | -1.522004291 | YPL050C | MNN9 | n |
| W303.asm02.HP0_G0054920 | 5786.881 | 6022.481 | 6038.453 | 1619.383 | 1821.601 | 2027.545 | 1591.236 | 1430.302 | 1216.418 | 2.74E-33 | 1.706651084 | YPL049C | DIG1 | n |
| W303.asm02.HP0_G0054930 | 14335.37 | 12651.44 | 13652.81 | 3235.988 | 4083.199 | 4943.897 | 6020.525 | 5967.106 | 5541.459 | 2.69E-24 | 1.728649763 | YPL048W | CAM1 | n |
| W303.asm02.HP0_G0054940 | 608.8668 | 600.9127 | 630.5774 | 345.9686 | 277.7096 | 294.7499 | 296.2735 | 277.0404 | 349.2566 | 4.50E-11 | 1.00098295 | YPL047W | SGF11 | n |
| W303.asm02.HP0_G0054950 | 718.9922 | 722.2081 | 632.7333 | 255.6555 | 325.4409 | 248.4103 | 388.859 | 356.5016 | 309.7858 | 5.59E-15 | 1.324053933 | YPL046C | ELC1 | n |
| W303.asm02.HP0_G0054960 | 2989.271 | 3041.286 | 2983.655 | 2550.998 | 2516.743 | 3391.143 | 1903.558 | 2802.618 | 2068.03 | 0.681028982 | 0.091696288 | YPL045W | VPS16 | n |
| W303.asm02.HP0_G0054970 | 2567.829 | 2556.105 | 2982.577 | 5909.95 | 5078.614 | 1380.311 | 7810.511 | 2276.456 | 5607.243 | 0.294216395 | -0.609560439 | YPL043W | NOP4 | n |
| W303.asm02.HP0_G0054980 | 1113.961 | 1165.103 | 1085.456 | 2371.761 | 2073.276 | 2640.594 | 2372.657 | 2888.522 | 2582.346 | 7.15E-15 | -1.074756518 | YPL042C | SSN3 | n |
| W303.asm02.HP0_G0054990 | 806.8808 | 674.3576 | 770.7058 | 525.2054 | 531.1196 | 542.4005 | 812.2833 | 610.9922 | 554.9831 | 0.006572726 | 0.494422834 | YPL041C | MRX11 | n |
| W303.asm02.HP0_G0055000 | 6844.721 | 6600.025 | 6509.499 | 3103.992 | 2900.33 | 3032.581 | 3998.458 | 3529.58 | 4609.709 | 7.14E-21 | 1.142708842 | YPL040C | ISM1 | n |
| W303.asm02.HP0_G0055010 | 681.9308 | 650.9888 | 649.9798 | 1839.608 | 1621.998 | 1916.634 | 11088.04 | 5642.819 | 6385.894 | 3.61E-08 | -1.439575185 | YPL039W | | n |
| W303.asm02.HP0_G0055040 | 7839.027 | 7589.305 | 7506.566 | 8608.922 | 9960.229 | 9646.221 | 5855.106 | 6890.574 | 6102.422 | 0.007139359 | -0.298854267 | YPL037C | EGD1 | n |
| W303.asm02.HP0_G0055060 | 803.7041 | 881.3387 | 850.4711 | 527.9843 | 556.2871 | 660.9082 | 1185.094 | 899.8443 | 798.9843 | 0.00513918 | 0.538891269 | YPL034W | | n |
| W303.asm02.HP0_G0055080 | 4202.769 | 4423.385 | 4504.586 | 20596.25 | 19938.68 | 19724.69 | 9238.797 | 14799.11 | 13274.14 | 5.67E-34 | -2.198250982 | YPL032C | SVL3 | n |
| W303.asm02.HP0_G0055090 | 4945.057 | 4844.024 | 4916.348 | 2577.397 | 2789.246 | 3217.18 | 2883.729 | 3959.1 | 3011.74 | 2.13E-06 | 0.776694893 | YPL031C | PHO85 | n |
| W303.asm02.HP0_G0055100 | 517.8015 | 520.791 | 696.33 | 3267.25 | 3167.625 | 2966.49 | 2050.46 | 2409.607 | 2217.54 | 4.59E-57 | -2.43794418 | YPL030W | TRM44 | n |
| W303.asm02.HP0_G0055110 | 3374.71 | 3392.931 | 3603.454 | 1193.522 | 1000.622 | 391.9869 | 1156.701 | 520.793 | 856.3964 | 1.48E-06 | 2.003664873 | YPL029W | SUV3 | n |
| W303.asm02.HP0_G0055120 | 13208.7 | 13447.09 | 14481.72 | 9394.646 | 10779.47 | 11103.26 | 9449.892 | 9148.775 | 7384.625 | 0.005117295 | 0.3953714 | YPL028W | ERG10 | n |
| W303.asm02.HP0_G0055140 | 673.4596 | 794.5402 | 628.4216 | 16282.76 | 16899.5 | 22401.75 | 28154.63 | 23611.79 | 35491.41 | 2.05E-112 | -4.728787746 | YPL026C | SKS1 | n |
| W303.asm02.HP0_G0055160 | 727.4634 | 608.7024 | 751.3034 | 1012.896 | 1090.01 | 1264.842 | 1271.507 | 766.6931 | 742.7684 | 0.006025464 | -0.68990675 | YPL024W | RMI1 | n |
| W303.asm02.HP0_G0055170 | 3799.329 | 3813.57 | 3787.776 | 5641.79 | 5604.527 | 4968.207 | 5013.195 | 3760.447 | 4535.552 | 0.000224044 | -0.508160361 | YPL023C | MET12 | n |
| W303.asm02.HP0_G0055180 | 3915.807 | 4068.402 | 3688.609 | 1777.084 | 1719.196 | 2171.881 | 999.9232 | 1422.785 | 973.6126 | 3.64E-08 | 1.042131522 | YPL022W | RAD1 | n |
| W303.asm02.HP0_G0055200 | 3586.49 | 4020.551 | 3605.609 | 2942.818 | 2934.176 | 2851.021 | 1889.978 | 2001.563 | 1911.343 | 2.71E-05 | 0.361319016 | YPL020C | ULP1 | n |
| W303.asm02.HP0_G0055240 | 2254.395 | 2208.911 | 2581.595 | 4341.281 | 4084.935 | 4270.075 | 3520.717 | 2955.097 | 2932.799 | 3.36E-12 | -0.849791073 | YPL016W | SWI1 | n |
| W303.asm02.HP0_G0055250 | 4981.06 | 4737.195 | 4766.519 | 1261.605 | 1314.781 | 1489.702 | 1227.066 | 1532.312 | 1254.692 | 1.10E-45 | 1.83287338 | YPL015C | HST2 | n |
| W303.asm02.HP0_G0055260 | 3455.186 | 4053.935 | 3207.861 | 4872.739 | 4951.042 | 6792.161 | 7632.747 | 10178.55 | 9705.028 | 0.001804298 | -0.632707606 | YPL014W | CIP1 | n |
| W303.asm02.HP0_G0055270 | 6276.093 | 5397.087 | 5371.226 | 1645.783 | 1673.2 | 1184.317 | 845.6141 | 832.1949 | 836.0629 | 1.05E-31 | 1.920364081 | YPL013C | MRPS16 | n |
| W303.asm02.HP0_G0055280 | 2501.119 | 2707.446 | 2931.916 | 8481.789 | 7269.917 | 2174.16 | 10621.41 | 4847.133 | 9614.126 | 0.018300973 | -1.13885934 | YPL012W | RRP12 | n |
| W303.asm02.HP0_G0055290 | 1156.317 | 1284.173 | 1252.532 | 1743.043 | 1635.883 | 1764.701 | 2172.673 | 1827.607 | 2076.402 | 1.88E-05 | -0.478313363 | YPL011C | TAF3 | n |
| W303.asm02.HP0_G0055300 | 5363.322 | 4960.869 | 5272.058 | 3672.964 | 3591.132 | 3612.965 | 3366.408 | 3226.769 | 3390.899 | 8.33E-14 | 0.519896628 | YPL010W | RET3 | n |

| | | | | | | | | | | | | | | |
|---|---|---|---|---|---|---|---|---|---|---|---|---|---|---|
| W303.asm02.HP0_G0055310 | 9879.525 | 9895.03 | 10440.64 | 5293.042 | 4993.566 | 5349.558 | 2262.789 | 3486.628 | 2994.995 | 1.97E-08 | 0.950361759 | YPL009C | RQC2 | n |
| W303.asm02.HP0_G0055320 | 2824.083 | 2702.995 | 2867.241 | 1404.021 | 1367.72 | 1214.704 | 791.2973 | 840.7853 | 1123.123 | 2.29E-11 | 1.074313027 | YPL008W | CHL1 | n |
| W303.asm02.HP0_G0055330 | 1332.095 | 1466.672 | 1271.934 | 1037.906 | 1045.75 | 1058.972 | 490.0858 | 472.472 | 340.884 | 0.023890527 | 0.37325178 | YPL007C | TFC8 | n |
| W303.asm02.HP0_G0055340 | 3138.576 | 3051.301 | 2810.112 | 1331.076 | 1332.138 | 1911.316 | 1606.049 | 1659.021 | 1368.32 | 1.18E-07 | 0.976228156 | YPL006W | NCR1 | n |
| W303.asm02.HP0_G0055350 | 1034.544 | 1170.667 | 888.198 | 1797.231 | 1624.601 | 1292.19 | 1434.458 | 1307.888 | 1417.36 | 0.000627149 | -0.607949029 | YPL005W | AEP3 | n |
| W303.asm02.HP0_G0055360 | 17204.99 | 18362.34 | 16571.79 | 8961.838 | 10060.9 | 13843.37 | 9321.506 | 17568.44 | 14966.6 | 0.015079929 | 0.665767416 | YPL004C | LSP1 | n |
| W303.asm02.HP0_G0055370 | 2158.036 | 1803.851 | 1839.993 | 1749.295 | 1760.852 | 2377.75 | 1883.806 | 2341.958 | 2011.814 | 0.919805869 | -0.021225943 | YPL003W | ULA1 | n |
| W303.asm02.HP0_G0055380 | 1721.769 | 1649.172 | 1680.462 | 684.2954 | 778.4547 | 720.9217 | 559.2163 | 935.2797 | 602.8265 | 9.98E-09 | 1.210186203 | YPL002C | SNF8 | n |
| W303.asm02.HP0_G0055390 | 730.6401 | 681.0344 | 717.8882 | 3010.9 | 3059.145 | 3338.726 | 1699.869 | 2392.426 | 2167.305 | 5.26E-45 | -2.143370279 | YPL001W | HAT1 | n |
| W303.asm02.HP0_G0055410 | 303.9039 | 348.3068 | 337.3859 | 398.0723 | 384.4542 | 227.1397 | 493.7892 | 522.9406 | 352.8449 | 0.928179465 | -0.028967709 | YPR002W | PDH1 | n |
| W303.asm02.HP0_G0055420 | 2129.445 | 2158.835 | 1870.174 | 1275.499 | 1283.539 | 1711.524 | 723.4012 | 1146.818 | 808.553 | 0.020025975 | 0.528094909 | YPR003C | | n |
| W303.asm02.HP0_G0055430 | 3755.914 | 3635.522 | 3441.767 | 4169.686 | 5001.376 | 4764.616 | 1569.015 | 4089.03 | 3100.251 | 0.280029689 | -0.363298293 | YPR004C | AIM45 | n |
| W303.asm02.HP0_G0055450 | 2300.987 | 2332.432 | 2067.432 | 544.6575 | 545.8729 | 271.9599 | 298.7425 | 539.0476 | 244.0012 | 1.23E-10 | 2.298202636 | YPR006C | ICL2 | n |
| W303.asm02.HP0_G0055460 | 583.4532 | 629.8456 | 590.6948 | 232.7299 | 219.5642 | 237.0154 | 381.4522 | 331.8042 | 251.1777 | 4.59E-15 | 1.387363021 | YPR007C | REC8 | n |
| W303.asm02.HP0_G0055470 | 4574.442 | 5133.353 | 5044.619 | 6877.69 | 6881.123 | 7034.494 | 9321.506 | 6627.493 | 8008.981 | 0.000961256 | -0.495188225 | YPR008W | HAA1 | n |
| W303.asm02.HP0_G0055490 | 6034.664 | 5849.997 | 7148.7 | 14991.97 | 14238.69 | 6135.051 | 14921.08 | 8591.473 | 14482.19 | 0.014754769 | -0.893816446 | YPR010C | RPA135 | n |
| W303.asm02.HP0_G0055500 | 1080.077 | 1088.32 | 920.5353 | 4054.363 | 4704.574 | 3727.674 | 2536.842 | 4222.181 | 4014.059 | 5.99E-20 | -2.015161364 | YPR010C-/ | MRA1 | n |
| W303.asm02.HP0_G0055510 | 3449.892 | 3069.106 | 3352.301 | 1774.305 | 1871.068 | 1076.445 | 162.9504 | 360.7968 | 333.7075 | 0.00115527 | 1.063957525 | YPR011C | MRX21 | n |
| W303.asm02.HP0_G0055520 | 2236.394 | 2694.092 | 2207.56 | 818.3756 | 833.9967 | 965.5337 | 1125.839 | 2358.065 | 1627.871 | 6.79E-08 | 1.447083832 | YPR013C | CMR3 | n |
| W303.asm02.HP0_G0055540 | 1921.901 | 1832.784 | 1877.719 | 5311.8 | 5840.58 | 2291.148 | 5768.693 | 3032.411 | 4793.906 | 0.000678347 | -1.255059457 | YPR016C | TIF6 | n |
| W303.asm02.HP0_G0055570 | 1785.303 | 2047.555 | 2150.431 | 2695.499 | 2618.281 | 2900.4 | 1365.327 | 1707.342 | 1253.496 | 0.004114965 | -0.457260047 | YPR018W | RLF2 | n |
| W303.asm02.HP0_G0055580 | 2681.132 | 2571.684 | 2892.033 | 2797.622 | 2635.638 | 2779.613 | 1191.267 | 1526.943 | 1344.399 | 0.934396795 | -0.012064556 | YPR019W | MCM4 | n |
| W303.asm02.HP0_G0055590 | 1739.771 | 1493.379 | 1356.011 | 8385.224 | 9676.444 | 7623.994 | 1886.275 | 4136.277 | 3576.292 | 2.71E-18 | -2.484625465 | YPR020W | ATP20 | n |
| W303.asm02.HP0_G0055600 | 3605.55 | 3500.873 | 3404.04 | 2665.626 | 2527.157 | 2135.417 | 3065.197 | 1627.881 | 2108.697 | 0.034198514 | 0.52028641 | YPR021C | AGC1 | n |
| W303.asm02.HP0_G0055610 | 2848.438 | 2740.83 | 2904.968 | 5409.755 | 5258.258 | 6524.759 | 10741.15 | 9736.144 | 11634.31 | 4.05E-15 | -1.017266863 | YPR022C | SDD4 | n |
| W303.asm02.HP0_G0055620 | 5412.031 | 5053.231 | 5248.344 | 1486.693 | 1412.848 | 1432.727 | 1430.754 | 1194.066 | 1441.282 | 8.41E-70 | 1.858664774 | YPR023C | EAF3 | n |
| W303.asm02.HP0_G0055630 | 8837.569 | 8951.374 | 8859.343 | 7846.124 | 8324.345 | 10084.55 | 6142.738 | 7132.179 | 5945.735 | 0.893767654 | 0.021454415 | YPR024W | YME1 | n |
| W303.asm02.HP0_G0055640 | 2888.676 | 3057.978 | 2926.526 | 2727.456 | 2492.444 | 2098.953 | 2101.073 | 2663.024 | 2054.873 | 0.096015606 | 0.277756538 | YPR025C | CCL1 | n |
| W303.asm02.HP0_G0055650 | 2000.26 | 2008.606 | 1778.552 | 793.3658 | 700.3489 | 1785.212 | 4597.178 | 1277.822 | 3079.917 | 0.103756854 | 0.819636109 | YPR026W | ATH1 | n |
| W303.asm02.HP0_G0055680 | 7708.783 | 7387.888 | 7835.329 | 8115.674 | 8049.239 | 11388.13 | 8287.018 | 9242.196 | 9054.358 | 0.128327648 | -0.264861457 | YPR029C | APL4 | n |
| W303.asm02.HP0_G0055700 | 639.5748 | 609.8152 | 672.6159 | 1047.632 | 932.0629 | 1040.741 | 1367.796 | 802.1285 | 1020.26 | 0.002242013 | -0.652276659 | YPR031W | NTO1 | n |
| W303.asm02.HP0_G0055720 | 7359.346 | 6596.687 | 7524.891 | 9994.186 | 10746.49 | 9793.596 | 7105.627 | 9281.927 | 7372.664 | 0.000355667 | -0.507331321 | YPR033C | HTS1 | n |
| W303.asm02.HP0_G0055730 | 4520.439 | 4382.212 | 4907.725 | 4856.761 | 5023.072 | 4893.759 | 2655.352 | 3544.614 | 3093.074 | 0.50567447 | -0.097231171 | YPR034W | ARP7 | n |
| W303.asm02.HP0_G0055740 | 5025.533 | 5787.68 | 5067.256 | 15418.53 | 16830.94 | 21240.98 | 8091.971 | 13930.41 | 7587.959 | 1.77E-11 | -1.752033898 | YPR035W | GLN1 | n |
| W303.asm02.HP0_G0055750 | 16040.2 | 14644.47 | 16652.63 | 8092.748 | 8431.09 | 10772.04 | 8190.729 | 9950.904 | 8392.924 | 5.46E-07 | 0.794296882 | YPR036W | VMA13 | n |
| W303.asm02.HP0_G0055760 | 2758.431 | 2771.988 | 2249.598 | 2077.201 | 1779.077 | 1814.079 | 9391.871 | 4049.3 | 3689.92 | 0.220122493 | 0.45629176 | YPR036W- | SPO24 | n |
| W303.asm02.HP0_G0055770 | 3348.238 | 3064.655 | 3113.004 | 1067.084 | 1136.874 | 1065.809 | 1834.427 | 1648.283 | 1919.715 | 7.54E-54 | 1.543084608 | YPR037C | ERV2 | n |
| W303.asm02.HP0_G0055780 | 1979.082 | 1991.914 | 1865.862 | 2114.716 | 2263.333 | 2490.181 | 1981.329 | 2696.312 | 2312.031 | 0.132279174 | -0.234700714 | YPR040W | TIP41 | n |
| W303.asm02.HP0_G0055790 | 9704.807 | 10260.03 | 10144.21 | 9673.228 | 9898.612 | 7796.438 | 6645.169 | 6056.232 | 6052.187 | 0.307248378 | 0.137712485 | YPR041W | TIF5 | n |
| W303.asm02.HP0_G0055820 | 2206.745 | 2332.432 | 2190.313 | 2980.332 | 2675.558 | 3471.667 | 5978.553 | 4919.077 | 5686.185 | 0.002340963 | -0.439895054 | YPR045C | THP3 | n |
| W303.asm02.HP0_G0055830 | 205.4263 | 179.161 | 171.3877 | 193.8258 | 177.0399 | 154.2119 | 317.2596 | 259.8596 | 344.4723 | 0.707981375 | 0.082154393 | YPR046W | MCM16 | n |
| W303.asm02.HP0_G0055840 | 8171.521 | 8796.695 | 7793.29 | 2232.123 | 2064.597 | 1687.975 | 1017.206 | 770.9883 | 920.9849 | 8.88E-38 | 2.04868789 | YPR047W | MSF1 | n |
| W303.asm02.HP0_G0055850 | 963.5978 | 955.8964 | 1126.416 | 1991.751 | 1811.187 | 1452.479 | 1344.341 | 1100.645 | 1310.908 | 4.65E-06 | -0.786980654 | YPR048W | TAH18 | n |
| W303.asm02.HP0_G0055860 | 1459.162 | 1632.48 | 1514.464 | 1260.215 | 1281.803 | 1657.588 | 1430.754 | 1144.671 | 1380.281 | 0.458672882 | 0.133190243 | YPR049C | ATG11 | n |
| W303.asm02.HP0_G0055870 | 1370.215 | 1212.953 | 1463.802 | 533.542 | 665.6352 | 567.4695 | 953.0132 | 671.125 | 831.2786 | 2.38E-11 | 1.196633425 | YPR051W | MAK3 | n |
| W303.asm02.HP0_G0055880 | 2476.764 | 2313.514 | 2326.13 | 8488.042 | 9447.334 | 8648.782 | 10981.87 | 7802.23 | 9417.968 | 4.77E-38 | -1.901287245 | YPR052C | NHP6A | n |
| W303.asm02.HP0_G0055890 | 316.6107 | 316.0356 | 355.7103 | 799.6183 | 870.446 | 952.6194 | 827.097 | 891.2539 | 784.6313 | 5.86E-27 | -1.407602451 | YPR054W | SMK1 | n |
| W303.asm02.HP0_G0055900 | 2475.705 | 2808.711 | 2426.376 | 4733.796 | 4378.265 | 5740.786 | 4994.678 | 5130.616 | 5320.183 | 7.11E-12 | -0.94591693 | YPR055W | SEC8 | n |
| W303.asm02.HP0_G0055910 | 1766.243 | 1749.324 | 1746.214 | 1469.325 | 1503.103 | 1718.361 | 2794.847 | 1966.128 | 2370.639 | 0.333566196 | 0.165719906 | YPR056W | TFB4 | n |
| W303.asm02.HP0_G0055920 | 1842.484 | 1767.129 | 1861.551 | 644.6966 | 512.8949 | 365.3987 | 1451.74 | 668.9774 | 1283.398 | 6.32E-09 | 1.844720033 | YPR057W | BRR1 | n |
| W303.asm02.HP0_G0055930 | 1998.142 | 1820.543 | 2064.198 | 1965.352 | 2173.078 | 1548.196 | 1130.777 | 1470.032 | 1644.616 | 0.825880393 | 0.049108448 | YPR058W | YMC1 | n |
| W303.asm02.HP0_G0055940 | 979.4813 | 1079.417 | 1043.417 | 2054.276 | 2110.593 | 1967.531 | 1014.737 | 1504.394 | 1130.3 | 5.66E-09 | -0.98312655 | YPR060C | ARO7 | n |
| W303.asm02.HP0_G0055950 | 178.9539 | 192.5146 | 183.2447 | 1303.287 | 1175.059 | 940.4648 | 1387.548 | 659.3132 | 1446.066 | 1.08E-17 | -2.62379946 | YPR061C | JID1 | n |
| W303.asm02.HP0_G0055960 | 3681.791 | 3377.352 | 3449.312 | 3117.191 | 3749.08 | 3243.768 | 2921.998 | 2511.618 | 2416.09 | 0.699281843 | 0.056007657 | YPR062W | FCY1 | n |

| | | | | | | | | | | | | | |
|---|---|---|---|---|---|---|---|---|---|---|---|---|---|
| W303.asm02.HP0_G0055970 | 4899.524 | 4832.896 | 5266.669 | 1761.8 | 2015.998 | 2053.373 | 2129.466 | 2073.508 | 1948.421 | 9.74E-41 | 1.36336727 | YPR063C | | n |
| W303.asm02.HP0_G0055980 | 984.7758 | 1187.359 | 1068.209 | 8224.05 | 7728.138 | 10064.8 | 5240.338 | 5894.088 | 5956.5 | 4.20E-93 | -3.00541206 | YPR065W | ROX1 | n |
| W303.asm02.HP0_G0055990 | 1083.253 | 964.7988 | 966.8854 | 279.2759 | 265.5598 | 291.7112 | 112.3371 | 192.2102 | 117.2163 | 2.31E-18 | 1.849373053 | YPR066W | UBA3 | n |
| W303.asm02.HP0_G0056000 | 2462.998 | 2599.504 | 2237.741 | 851.722 | 1033.6 | 627.483 | 509.8374 | 704.4128 | 537.0419 | 4.56E-12 | 1.539039393 | YPR067W | ISA2 | n |
| W303.asm02.HP0_G0056010 | 584.5121 | 616.492 | 616.5646 | 1003.17 | 921.6488 | 1235.215 | 1083.867 | 1738.482 | 1478.36 | 0.00019537 | -0.798121778 | YPR068C | HOS1 | n |
| W303.asm02.HP0_G0056020 | 5546.511 | 5148.932 | 5486.563 | 8353.267 | 9547.136 | 7978.757 | 6095.828 | 8011.621 | 7724.313 | 3.61E-06 | -0.677333933 | YPR069C | SPE3 | n |
| W303.asm02.HP0_G0056030 | 2027.791 | 2120.999 | 2107.314 | 2123.053 | 1926.61 | 2007.034 | 1545.56 | 1586.002 | 1651.792 | 0.618248593 | 0.046247149 | YPR070W | MED1 | n |
| W303.asm02.HP0_G0056050 | 6886.018 | 7690.57 | 7181.037 | 6137.817 | 5857.069 | 6068.961 | 5871.154 | 5315.31 | 5840.48 | 0.00274749 | 0.268312679 | YPR072W | NOT5 | n |
| W303.asm02.HP0_G0056060 | 2916.207 | 3027.933 | 2874.786 | 1374.843 | 1558.645 | 1642.395 | 745.6217 | 1747.072 | 1070.495 | 0.00127382 | 0.946598515 | YPR073C | LTP1 | n |
| W303.asm02.HP0_G0056070 | 14910.35 | 13596.21 | 15975.71 | 17055.98 | 17962.6 | 22340.22 | 13429.83 | 21570.49 | 15242.9 | 0.102037193 | -0.366780621 | YPR074C | TKL1 | n |
| W303.asm02.HP0_G0056090 | 4654.919 | 4638.156 | 4847.362 | 3044.246 | 3099.933 | 2832.789 | 2933.108 | 1332.586 | 1963.97 | 0.016684633 | 0.655536699 | YPR075C | OPY2 | n |
| W303.asm02.HP0_G0056110 | 2044.733 | 1921.808 | 1887.421 | 2197.387 | 2014.262 | 2468.15 | 2565.235 | 2435.378 | 2588.326 | 0.104096329 | -0.190594889 | YPR079W | MRL1 | n |
| W303.asm02.HP0_G0056140 | 655.4583 | 693.2753 | 681.2392 | 731.5361 | 697.7454 | 1070.367 | 636.9881 | 666.8298 | 619.5717 | 0.139922521 | -0.300528555 | YPR082C | DIB1 | n |
| W303.asm02.HP0_G0056150 | 367.4379 | 376.1269 | 336.308 | 3666.017 | 3744.74 | 4692.448 | 2346.733 | 4768.745 | 5183.829 | 4.20E-33 | -3.486484979 | YPR083W | MDM36 | n |
| W303.asm02.HP0_G0056160 | 1848.837 | 1833.897 | 1686.929 | 2256.438 | 2081.954 | 2447.639 | 3028.162 | 2168.002 | 2577.562 | 0.036510384 | -0.337850131 | YPR084W | | n |
| W303.asm02.HP0_G0056170 | 1740.829 | 1810.528 | 1765.617 | 602.3189 | 465.1636 | 489.224 | 393.7969 | 467.103 | 461.6885 | 1.10E-35 | 1.77070814 | YPR085C | ASA1 | n |
| W303.asm02.HP0_G0056180 | 5301.906 | 5426.019 | 5341.045 | 6394.167 | 6928.855 | 7006.387 | 11291.73 | 8411.075 | 9500.498 | 0.013379511 | -0.339259367 | YPR086W | SUA7 | n |
| W303.asm02.HP0_G0056190 | 7716.195 | 7616.013 | 7895.692 | 5898.835 | 6258.012 | 6663.018 | 5873.623 | 5985.361 | 5681.401 | 0.000162143 | 0.303674664 | YPR088C | SRP54 | n |
| W303.asm02.HP0_G0056200 | 5072.125 | 5463.855 | 5222.475 | 5611.917 | 5450.051 | 6252.799 | 4632.978 | 4358.554 | 4046.353 | 0.216454185 | -0.135952447 | YPR089W | | n |
| W303.asm02.HP0_G0056210 | 6182.91 | 5779.89 | 5770.053 | 2622.553 | 2308.461 | 4980.361 | 1987.502 | 2819.799 | 2474.698 | 0.006290854 | 0.839224212 | YPR091C | NVJ2 | n |
| W303.asm02.HP0_G0056220 | 1878.486 | 1908.454 | 1606.086 | 434.1976 | 446.9389 | 477.829 | 664.1465 | 497.1694 | 592.0617 | 1.81E-40 | 1.988648813 | YPR093C | ASR1 | n |
| W303.asm02.HP0_G0056230 | 396.0281 | 327.1636 | 321.2172 | 250.0978 | 239.5245 | 199.0321 | 272.8186 | 225.498 | 186.5892 | 0.004068832 | 0.600973331 | YPR094W | RDS3 | n |
| W303.asm02.HP0_G0056240 | 1338.448 | 1384.325 | 1329.063 | 1362.338 | 1254.9 | 1526.166 | 1382.61 | 1488.287 | 1435.301 | 0.791983059 | -0.032673142 | YPR095C | SYT1 | n |
| W303.asm02.HP0_G0056260 | 3072.924 | 3483.068 | 3068.81 | 1844.471 | 1829.412 | 1808.002 | 375.2798 | 1397.014 | 833.6708 | 0.045287547 | 0.812078811 | YPR097W | LEC1 | n |
| W303.asm02.HP0_G0056280 | 4085.231 | 3536.483 | 3546.324 | 1394.295 | 1477.936 | 880.4513 | 912.2756 | 644.28 | 645.8855 | 8.36E-11 | 1.573509478 | YPR100W | MRPL51 | n |
| W303.asm02.HP0_G0056290 | 1536.462 | 1459.995 | 1338.764 | 351.5264 | 337.5907 | 265.8826 | 232.0809 | 245.9002 | 183.0009 | 6.28E-37 | 2.182519965 | YPR101W | SNT309 | n |
| W303.asm02.HP0_G0056310 | 10554.04 | 10120.93 | 9810.06 | 5094.353 | 5466.54 | 6430.561 | 3696.012 | 5826.438 | 4530.767 | 1.29E-05 | 0.843302969 | YPR103W | PRE2 | n |
| W303.asm02.HP0_G0056320 | 1303.504 | 1349.828 | 1364.634 | 3531.937 | 2913.347 | 3170.84 | 6246.434 | 3839.909 | 5918.225 | 6.56E-10 | -1.259087643 | YPR104C | FHL1 | n |
| W303.asm02.HP0_G0056330 | 2162.271 | 1842.799 | 2021.082 | 3069.951 | 2928.101 | 3576.501 | 1946.764 | 2897.112 | 2108.697 | 0.000477511 | -0.667965529 | YPR105C | COG4 | n |
| W303.asm02.HP0_G0056350 | 1774.714 | 1835.009 | 1593.151 | 587.7299 | 624.8466 | 791.5705 | 1001.158 | 760.2503 | 806.1608 | 6.33E-15 | 1.376329965 | YPR107C | YTH1 | n |
| W303.asm02.HP0_G0056360 | 11166.09 | 11912.54 | 11390.28 | 9254.314 | 9751.079 | 12842.13 | 7895.69 | 16142.43 | 11405.86 | 0.7054447 | 0.114112603 | YPR108W | RPN7 | n |
| W303.asm02.HP0_G0056370 | 1965.316 | 2055.344 | 1805.499 | 786.4187 | 911.2346 | 925.2715 | 1220.894 | 1519.427 | 1284.595 | 5.00E-17 | 1.151798153 | YPR109W | GLD1 | n |
| W303.asm02.HP0_G0056380 | 5019.18 | 5243.52 | 5582.497 | 5636.232 | 5544.646 | 2924.709 | 7627.809 | 3546.761 | 6041.422 | 0.65432168 | 0.167787455 | YPR110C | RPC40 | n |
| W303.asm02.HP0_G0056400 | 2419.584 | 2710.784 | 2871.553 | 5629.98 | 5437.901 | 1469.191 | 7469.797 | 2492.29 | 6047.402 | 0.238037135 | -0.647776574 | YPR112C | MRD1 | n |
| W303.asm02.HP0_G0056410 | 7118.976 | 6625.619 | 7101.272 | 4095.352 | 4726.27 | 4764.616 | 6098.297 | 6240.925 | 6157.442 | 8.04E-11 | 0.617826796 | YPR113W | PIS1 | n |
| W303.asm02.HP0_G0056420 | 3666.966 | 3311.697 | 3812.568 | 2871.262 | 2970.625 | 3541.556 | 3599.724 | 3587.566 | 3524.861 | 0.114986154 | 0.201710951 | YPR114W | | n |
| W303.asm02.HP0_G0056440 | 493.4468 | 725.5465 | 429.0082 | 569.6672 | 564.0976 | 347.9264 | 392.5624 | 283.4832 | 374.3744 | 0.639255532 | 0.153365109 | YPR116W | RRG8 | n |
| W303.asm02.HP0_G0056450 | 3098.338 | 2942.247 | 2754.06 | 1245.626 | 1083.935 | 1079.483 | 4642.853 | 3386.765 | 5665.851 | 1.31E-11 | 1.367094248 | YPR117W | HOB2 | n |
| W303.asm02.HP0_G0056460 | 2674.778 | 2556.105 | 2888.799 | 4576.79 | 4503.235 | 4247.285 | 1134.481 | 2403.164 | 2437.62 | 0.012914206 | -0.71488513 | YPR118W | MRI1 | n |
| W303.asm02.HP0_G0056470 | 626.868 | 593.1231 | 623.0321 | 3682.69 | 3059.145 | 2961.173 | 2214.645 | 2248.537 | 2236.678 | 3.88E-89 | -2.396479488 | YPR119W | CLB2 | n |
| W303.asm02.HP0_G0056480 | 600.3956 | 563.0775 | 556.2016 | 586.3404 | 753.2873 | 932.8681 | 35.79972 | 61.2066 | 29.90211 | 0.146079151 | -0.401918384 | YPR120C | CLB5 | n |
| W303.asm02.HP0_G0056500 | 3740.03 | 3854.744 | 3591.597 | 320.2641 | 313.2911 | 249.9297 | 543.1682 | 608.8446 | 606.4148 | 3.59E-175 | 3.66249637 | YPR122W | AXL1 | n |
| W303.asm02.HP0_G0056510 | 11942.26 | 13895.55 | 12063.97 | 13231.56 | 14196.17 | 14078.1 | 4763.832 | 7375.932 | 16916.22 | 0.769781348 | -0.131047267 | YPR124W | CTR1 | n |
| W303.asm02.HP0_G0056520 | 5132.482 | 5971.292 | 5592.198 | 5871.046 | 5451.787 | 2625.401 | 1403.596 | 1474.327 | 1902.97 | 0.430465502 | 0.259415213 | YPR125W | YLH47 | n |
| W303.asm02.HP0_G0056530 | 4824.343 | 4100.673 | 4375.237 | 3955.019 | 4021.582 | 7532.075 | 1927.012 | 9725.406 | 4594.16 | 0.696480847 | -0.221622369 | YPR127W | | n |
| W303.asm02.HP0_G0056540 | 1037.721 | 1019.326 | 1043.417 | 653.0332 | 687.3313 | 1003.517 | 1275.211 | 1234.87 | 644.6895 | 0.19739852 | 0.403564911 | YPR128C | ANT1 | n |
| W303.asm02.HP0_G0056550 | 4891.053 | 5362.59 | 5578.185 | 3768.835 | 4073.653 | 4493.416 | 5520.564 | 5350.745 | 5092.927 | 0.00133282 | 0.360005818 | YPR129W | SCD6 | n |
| W303.asm02.HP0_G0056560 | 1795.892 | 1850.589 | 1705.254 | 931.6143 | 899.0848 | 888.8076 | 1035.723 | 1159.704 | 1184.123 | 3.01E-24 | 0.976392135 | YPR131C | NAT3 | n |
| W303.asm02.HP0_G0056580 | 7044.853 | 7611.561 | 7306.075 | 6909.647 | 7368.851 | 6222.413 | 5572.412 | 5036.122 | 4820.22 | 0.410319638 | 0.099403043 | YPR133C | SPN1 | n |
| W303.asm02.HP0_G0056590 | 3365.18 | 2940.021 | 2865.085 | 458.5127 | 610.0933 | 275.7583 | 258.0049 | 225.498 | 166.2557 | 7.20E-21 | 2.770708615 | YPR133W-TOM5 | | n |
| W303.asm02.HP0_G0056600 | 1554.463 | 1587.968 | 1453.023 | 1441.536 | 1562.984 | 1336.25 | 1227.066 | 1329.364 | 1166.182 | 0.502175181 | 0.082573448 | YPR134W | MSS18 | n |
| W303.asm02.HP0_G0056610 | 2587.948 | 2689.641 | 2516.92 | 3470.802 | 3628.45 | 4320.972 | 678.9602 | 1000.782 | 798.9843 | 0.001836143 | -0.551068085 | YPR135W | CTF4 | n |
| W303.asm02.HP0_G0056620 | 2600.655 | 2709.671 | 3011.681 | 3929.314 | 3259.616 | 1762.422 | 6148.91 | 3046.37 | 5707.714 | 0.78881036 | -0.105195205 | YPR137W | RRP9 | n |
| W303.asm02.HP0_G0056630 | 1381.863 | 1498.943 | 1359.245 | 2647.563 | 3293.462 | 4147.009 | 1101.15 | 2265.718 | 1257.085 | 3.38E-05 | -1.250477679 | YPR138C | MEP3 | n |

| | | | | | | | | | | | | | | |
|---|---|---|---|---|---|---|---|---|---|---|---|---|---|---|
| W303.asm02.HP0_G0056640 | 2331.695 | 2615.083 | 2240.975 | 1555.469 | 1532.61 | 1541.36 | 1582.595 | 1765.327 | 1798.911 | 1.91E-09 | 0.634554253 | YPR139C | LOA1 | n |
| W303.asm02.HP0_G0056650 | 4247.243 | 4204.164 | 3960.242 | 3345.753 | 3306.48 | 2643.633 | 1515.933 | 1415.268 | 1430.517 | 0.001096684 | 0.417062734 | YPR140W | TAZ1 | n |
| W303.asm02.HP0_G0056660 | 919.1241 | 1072.741 | 951.7947 | 763.493 | 647.4105 | 802.9655 | 746.8562 | 907.3609 | 831.2786 | 0.006379979 | 0.410200585 | YPR141C | KAR3 | n |
| W303.asm02.HP0_G0056670 | 1486.694 | 1560.148 | 1744.059 | 2325.215 | 2080.219 | 897.9236 | 1560.374 | 910.5823 | 1498.694 | 0.719054821 | -0.146614474 | YPR143W | RRP15 | n |
| W303.asm02.HP0_G0056680 | 2042.616 | 2439.261 | 2447.934 | 3242.935 | 2782.303 | 751.3083 | 4684.825 | 1421.711 | 3488.978 | 0.959210133 | 0.032259073 | YPR144C | NOC4 | n |
| W303.asm02.HP0_G0056720 | 12916.45 | 14332.88 | 13482.5 | 4900.528 | 4644.693 | 6144.927 | 3467.635 | 6526.556 | 5030.731 | 1.21E-08 | 1.37627672 | YPR148C | | n |
| W303.asm02.HP0_G0056730 | 6952.729 | 7632.705 | 6787.6 | 3280.45 | 3744.74 | 5288.025 | 3293.574 | 6016.501 | 4226.962 | 0.00358045 | 0.795589015 | YPR149W | NCE102 | n |
| W303.asm02.HP0_G0056740 | 1216.675 | 1333.136 | 1082.222 | 556.4676 | 566.7012 | 704.2091 | 4123.14 | 4518.55 | 5103.692 | 8.19E-10 | 0.990844481 | YPR151C | SUE1 | n |
| W303.asm02.HP0_G0056750 | 1101.255 | 1112.801 | 994.9111 | 3195.694 | 3099.066 | 3272.635 | 1044.364 | 2034.851 | 1747.479 | 4.36E-11 | -1.57603878 | YPR152C | URN1 | n |
| W303.asm02.HP0_G0056770 | 6126.788 | 7193.148 | 5365.837 | 3943.904 | 4317.517 | 4951.494 | 12369.42 | 8929.72 | 10624.82 | 0.00854298 | 0.499994643 | YPR154W | PIN3 | n |
| W303.asm02.HP0_G0056780 | 1319.388 | 1526.763 | 1216.961 | 6148.933 | 6585.189 | 6184.43 | 2698.558 | 5618.121 | 4289.158 | 6.76E-18 | -2.21917128 | YPR155C | NCA2 | n |
| W303.asm02.HP0_G0056790 | 2799.728 | 2491.562 | 2433.921 | 10884.81 | 11032.01 | 6833.183 | 12300.29 | 15330.64 | 19858.59 | 2.08E-14 | -1.895894391 | YPR156C | TPO3 | n |
| W303.asm02.HP0_G0056820 | 8290.118 | 8487.336 | 9213.976 | 13630.33 | 12442.26 | 11132.13 | 19656.52 | 16973.56 | 17214.05 | 3.19E-05 | -0.517472883 | YPR159W | KRE6 | n |
| W303.asm02.HP0_G0056830 | 51353.41 | 54737.59 | 47544.46 | 1906.301 | 1578.606 | 7851.893 | 30656.9 | 30462.63 | 28784.97 | 3.39E-13 | 3.760404519 | YPR160W | GPH1 | n |
| W303.asm02.HP0_G0056840 | 3772.856 | 4252.014 | 4028.15 | 6561.594 | 6346.532 | 6225.452 | 8527.74 | 7517.673 | 7525.763 | 6.79E-12 | -0.666798697 | YPR161C | SGV1 | n |
| W303.asm02.HP0_G0056850 | 2413.23 | 2378.057 | 2280.858 | 1770.137 | 1764.324 | 1416.775 | 1206.08 | 1509.763 | 1611.126 | 0.001412573 | 0.514407805 | YPR162C | ORC4 | n |
| W303.asm02.HP0_G0056860 | 18216.23 | 18621.62 | 19033.74 | 11344.02 | 11354.85 | 8704.997 | 11007.8 | 7961.153 | 11170.23 | 5.11E-06 | 0.83117999 | YPR163C | TIF3 | n |
| W303.asm02.HP0_G0056880 | 14329.02 | 15400.06 | 14440.76 | 8801.359 | 9866.502 | 10064.8 | 11017.67 | 10014.26 | 9151.241 | 3.51E-08 | 0.620412088 | YPR165W | RHO1 | n |
| W303.asm02.HP0_G0056890 | 1727.064 | 1682.556 | 1529.554 | 2093.874 | 2028.148 | 1704.687 | 1033.254 | 996.4863 | 1053.75 | 0.061480266 | -0.238414998 | YPR166C | MRP2 | n |
| W303.asm02.HP0_G0056900 | 1382.922 | 1125.042 | 1241.752 | 610.6555 | 628.318 | 627.483 | 776.4836 | 662.5346 | 928.1614 | 5.23E-10 | 1.006696494 | YPR167C | MET16 | n |
| W303.asm02.HP0_G0056910 | 507.2125 | 459.587 | 427.9303 | 757.2406 | 721.1771 | 752.068 | 495.0237 | 472.472 | 527.4732 | 7.63E-09 | -0.677479806 | YPR168W | NUT2 | n |
| W303.asm02.HP0_G0056920 | 3841.685 | 4160.764 | 4128.396 | 3310.322 | 3430.581 | 1985.004 | 2682.51 | 1689.087 | 2398.149 | 0.080005084 | 0.475336113 | YPR169W | JIP5 | n |
| W303.asm02.HP0_G0056940 | 1166.906 | 1087.207 | 1093.001 | 4418.395 | 4022.45 | 5584.295 | 4076.23 | 4986.727 | 4880.024 | 4.50E-37 | -2.067050506 | YPR171W | BSP1 | n |
| W303.asm02.HP0_G0056950 | 2402.641 | 2380.282 | 2091.146 | 3056.751 | 3097.33 | 4192.589 | 4760.128 | 5952.073 | 4767.592 | 0.001288774 | -0.589957283 | YPR172W | | n |
| W303.asm02.HP0_G0056960 | 6144.789 | 6239.477 | 6195.827 | 5160.351 | 4958.852 | 6409.29 | 7451.28 | 7473.648 | 7163.349 | 0.203089958 | 0.168743713 | YPR173C | VPS4 | n |
| W303.asm02.HP0_G0056980 | 756.0537 | 766.7201 | 689.8625 | 2211.281 | 2488.972 | 1921.951 | 461.6929 | 679.7154 | 466.4729 | 5.17E-16 | -1.581451043 | YPR175W | DPB2 | n |
| W303.asm02.HP0_G0056990 | 1969.552 | 1946.29 | 2072.821 | 1626.33 | 1681.879 | 2017.669 | 993.7508 | 1929.618 | 1525.008 | 0.529495974 | 0.169216747 | YPR176C | BET2 | n |
| W303.asm02.HP0_G0057000 | 2180.272 | 2433.697 | 2270.079 | 1153.229 | 1199.358 | 1133.42 | 1182.625 | 1360.505 | 1069.299 | 1.01E-15 | 0.981776061 | YPR178W | PRP4 | n |
| W303.asm02.HP0_G0057010 | 2613.362 | 2483.773 | 2427.454 | 3880.684 | 3488.727 | 4085.476 | 4029.32 | 3854.942 | 4126.491 | 5.17E-10 | -0.606450402 | YPR179C | HDA3 | n |
| W303.asm02.HP0_G0057020 | 2351.814 | 2424.794 | 2205.404 | 1467.24 | 1515.253 | 1608.97 | 943.1374 | 1743.851 | 1430.517 | 0.008148333 | 0.604696484 | YPR180W | AOS1 | n |
| W303.asm02.HP0_G0057030 | 11432.93 | 11417.34 | 13042.71 | 11936.61 | 12249.6 | 15178.1 | 7833.966 | 11316.78 | 8964.652 | 0.50655185 | -0.133186865 | YPR181C | SEC23 | n |
| W303.asm02.HP0_G0057050 | 11931.67 | 11783.45 | 12637.42 | 7727.328 | 8446.711 | 8736.903 | 8146.288 | 8776.167 | 8892.887 | 1.30E-09 | 0.545348727 | YPR183W | DPM1 | n |
| W303.asm02.HP0_G0057060 | 20947.13 | 21442.57 | 19000.32 | 3072.035 | 3079.973 | 7500.928 | 9667.159 | 10787.39 | 8979.005 | 1.92E-10 | 2.168774564 | YPR184W | GDB1 | n |
| W303.asm02.HP0_G0057070 | 4020.638 | 4038.356 | 4141.331 | 4662.24 | 4387.812 | 7472.821 | 5774.865 | 5795.298 | 4411.159 | 0.07472194 | -0.437572744 | YPR185W | ATG13 | n |
| W303.asm02.HP0_G0057080 | 273.1959 | 260.3955 | 339.5417 | 366.1154 | 284.6523 | 98.7564 | 744.3873 | 224.4242 | 592.0617 | 0.712635275 | 0.220093991 | YPR186C | PZF1 | n |
| W303.asm02.HP0_G0057090 | 4164.649 | 4105.124 | 4222.174 | 3878.6 | 3906.159 | 2802.403 | 3482.449 | 2745.706 | 3388.507 | 0.19839063 | 0.238719361 | YPR187W | RPO26 | n |
| W303.asm02.HP0_G0057100 | 1364.92 | 1073.853 | 1167.377 | 1326.908 | 1424.13 | 1600.613 | 1304.838 | 1583.855 | 1406.595 | 0.090442812 | -0.270867685 | YPR188C | MLC2 | n |
| W303.asm02.HP0_G0057110 | 2712.898 | 2806.485 | 2829.514 | 6516.437 | 6343.929 | 6678.212 | 5298.359 | 5748.051 | 6006.735 | 2.87E-51 | -1.226741951 | YPR189W | SKI3 | n |
| W303.asm02.HP0_G0057120 | 1069.488 | 962.5732 | 1321.518 | 6462.944 | 5406.659 | 4030.78 | 7550.037 | 4035.34 | 6299.776 | 1.17E-15 | -2.245276264 | YPR190C | RPC82 | n |
| W303.asm02.HP0_G0057130 | 12618.9 | 12739.35 | 12433.69 | 21880.08 | 23765 | 21741.6 | 6339.019 | 13574.98 | 11328.11 | 0.001654364 | -0.834381049 | YPR191W | QCR2 | n |
| W303.asm02.HP0_G0057140 | 477.5633 | 531.9191 | 483.9817 | 7466.809 | 6881.123 | 5174.835 | 10193.04 | 8837.373 | 8715.866 | 1.67E-105 | -3.708577829 | YPR192W | AQY1 | n |
| W303.asm02.HP0_G0057160 | 1634.94 | 1710.376 | 1469.192 | 864.9216 | 731.5912 | 995.9203 | 1149.294 | 969.6413 | 1084.848 | 1.52E-08 | 0.892504014 | YPR194C | OPT2 | n |
| W303.asm02.HP0_G0057180 | 694.6376 | 602.0255 | 685.5508 | 5479.921 | 6166.021 | 9887.795 | 10528.82 | 11070.88 | 13896.11 | 1.67E-44 | -3.44134461 | YPR196W | | n |
| W303.asm02.HP0_G0057220 | 337.7887 | 363.886 | 312.5939 | 1004.56 | 954.6268 | 1122.784 | 1867.758 | 2261.423 | 1914.931 | 1.02E-29 | -1.603888692 | YPR201W | ARR3 | n |

**NaCl_gw**

| | Normalized read counts (genome-wide) | | | | | | | | | | | | | |
|---|---|---|---|---|---|---|---|---|---|---|---|---|---|---|
| W303_gene | SC.RI.1_NaCl | SC.RI.2_NaCl | SC.RI.3_NaCl | Km.V.1_NaCl | Km.V.2_NaCl | Km.V.3_NaCl | KS.1.1_NaCl | KS.1.2_NaCl | KS.1.3_NaCl | padj.sc.v.km | log2FC.sc.v.km | locus_tag | gene | ambiguous |
| W303.asm02.HP0_G0000040 | 194.8290856 | 235.1251578 | 219.89293 | 317.1389421 | 225.5783322 | 369.3416899 | 387.1559335 | 581.5234251 | 677.6405358 | 0.059304298 | -0.49273412 | YAL067C | SEO1 | n |
| W303.asm02.HP0_G0000100 | 569.9477729 | 681.5773805 | 531.6522396 | 3593.592798 | 3870.658794 | 4176.819993 | 605.8185757 | 1928.412955 | 1468.903578 | 6.34E-19 | -2.706670517 | YAL063C | FLO9 | n |
| W303.asm02.HP0_G0000110 | 255.8949184 | 208.4712938 | 188.6192688 | 562.28929 | 675.4080652 | 557.6335318 | 441.1116504 | 485.7317128 | 525.1202339 | 2.75E-16 | -1.457161888 | YAL062W | GDH3 | n |
| W303.asm02.HP0_G0000120 | 344.1011214 | 217.0386072 | 258.9850064 | 603.1476813 | 295.9056946 | 526.8550576 | 246.1137964 | 194.4862037 | 258.9774253 | 0.004737546 | -0.800746038 | YAL061W | BDH2 | n |
| W303.asm02.HP0_G0000130 | 1550.878293 | 983.3371984 | 1584.206398 | 2752.104501 | 822.6974468 | 2483.098665 | 1083.847296 | 427.6761296 | 1416.698643 | 0.383071132 | -0.556947243 | YAL060W | BDH1 | n |
| W303.asm02.HP0_G0000140 | 1118.570969 | 1981.905176 | 1237.264219 | 4110.159603 | 3902.505147 | 4141.515273 | 2924.021219 | 2726.677225 | 2174.182021 | 1.90E-12 | -1.486411392 | YAL059W | ECM1 | n |
| W303.asm02.HP0_G0000150 | 337.3160288 | 369.3464017 | 335.2145555 | 922.2322611 | 778.9087117 | 878.997012 | 1882.770542 | 1790.047149 | 1951.031512 | 1.85E-23 | -1.309821072 | YAL058W | CNE1 | n |
| W303.asm02.HP0_G0000170 | 646.5223886 | 473.1060868 | 662.6106957 | 526.2949929 | 463.0990467 | 509.6553221 | 591.6197028 | 465.4122587 | 461.6554103 | 0.200036453 | 0.248408271 | YAL055W | PEX22 | n |
| W303.asm02.HP0_G0000180 | 424.5529328 | 415.0387401 | 378.2158396 | 100.2003406 | 86.25053878 | 87.80917627 | 158.0807846 | 106.4352359 | 211.8906207 | 3.63E-17 | 2.148959581 | YAL054C | ACS1 | n |
| W303.asm02.HP0_G0000190 | 1344.417621 | 1356.491295 | 1387.768714 | 1653.792029 | 1719.70305 | 1563.365437 | 1500.347566 | 2017.431516 | 1574.337076 | 0.065996683 | -0.271566332 | YAL053W | FLC2 | n |
| W303.asm02.HP0_G0000200 | 358.6406054 | 458.827231 | 321.5323287 | 655.6798987 | 867.8131132 | 654.4952005 | 616.2310825 | 795.3614899 | 764.6487617 | 2.91E-06 | -0.933128365 | YAL051W | OAF1 | n |
| W303.asm02.HP0_G0000210 | 1413.237845 | 1334.59705 | 1379.950298 | 2182.03266 | 1893.531059 | 2069.399762 | 1736.048856 | 1720.380449 | 1371.659091 | 2.66E-05 | -0.574559052 | YAL049C | AIM2 | n |
| W303.asm02.HP0_G0000220 | 203.552776 | 209.4232175 | 171.0278344 | 663.4624494 | 817.3897213 | 756.7883646 | 617.177674 | 762.4633261 | 668.4279001 | 7.94E-31 | -1.936491537 | YAL048C | GEM1 | n |
| W303.asm02.HP0_G0000230 | 450.724004 | 377.9137152 | 391.8980663 | 439.7141161 | 533.426409 | 493.3608358 | 427.8593691 | 527.3382141 | 462.6790365 | 0.119625812 | -0.263244231 | YAL047C | SPC72 | n |
| W303.asm02.HP0_G0000240 | 485.6187656 | 614.9427204 | 472.036823 | 363.8342465 | 410.021792 | 396.4991671 | 515.8923809 | 456.7039212 | 355.1982869 | 0.028947027 | 0.427150307 | YAL046C | BOL3 | n |
| W303.asm02.HP0_G0000250 | 850.0751646 | 831.9813276 | 911.822683 | 593.4194929 | 664.7926143 | 671.694936 | 849.0925975 | 669.574393 | 530.2383649 | 0.022350357 | 0.426832306 | YAL044W- | BOL1 | n |
| W303.asm02.HP0_G0000260 | 1064.290229 | 1153.731544 | 1226.513898 | 3578.027697 | 2338.053066 | 3406.45289 | 2289.804898 | 1981.630573 | 1736.070013 | 5.82E-15 | -1.437046317 | YAL044C | GCV3 | n |
| W303.asm02.HP0_G0000270 | 495.311755 | 394.0964183 | 327.3961402 | 1373.620203 | 1460.951434 | 1443.872538 | 872.7573856 | 1220.13484 | 1220.162415 | 2.55E-21 | -1.813835619 | YAL043C | PTA1 | n |
| W303.asm02.HP0_G0000280 | 2467.835085 | 1942.876304 | 2098.267203 | 2300.716559 | 2200.052204 | 2273.080841 | 2584.194862 | 2446.075239 | 2475.12812 | 0.670121867 | -0.057766711 | YAL042W | ERV46 | n |
| W303.asm02.HP0_G0000290 | 599.0267409 | 696.80816 | 574.6535237 | 1610.988 | 1572.413668 | 1598.670158 | 1162.414392 | 1310.120994 | 1539.533785 | 5.89E-20 | -1.354190483 | YAL041W | CDC24 | n |
| W303.asm02.HP0_G0000300 | 281.0966907 | 611.1350255 | 190.5738726 | 1241.316841 | 2205.35993 | 1410.378316 | 2069.249073 | 1178.528339 | 2057.488636 | 3.64E-05 | -2.16497417 | YAL040C | CLN3 | n |
| W303.asm02.HP0_G0000310 | 1423.900133 | 1165.154628 | 1238.241521 | 1568.183971 | 1775.434167 | 1615.869893 | 1517.386214 | 1597.496131 | 1447.407428 | 0.004412794 | -0.373203689 | YAL039C | CYC3 | n |
| W303.asm02.HP0_G0000320 | 19575.96126 | 17395.45398 | 15820.56334 | 41020.85206 | 33811.53813 | 38651.4268 | 17052.84631 | 17814.35571 | 17069.99029 | 1.62E-18 | -1.104139515 | YAL038W | CDC19 | n |
| W303.asm02.HP0_G0000340 | 1041.027055 | 1655.395342 | 1150.284349 | 3170.416602 | 3155.442788 | 3085.08941 | 2630.577847 | 2537.028986 | 2392.214399 | 9.21E-14 | -1.290587069 | YAL036C | RBG1 | n |
| W303.asm02.HP0_G0000350 | 2605.475533 | 2717.742209 | 2418.82223 | 8920.74877 | 8824.093582 | 8862.390059 | 5234.651131 | 6607.692962 | 6494.908156 | 3.56E-46 | -1.781017393 | YAL035W | FUN12 | n |
| W303.asm02.HP0_G0000360 | 399.3511606 | 414.0868164 | 417.307916 | 280.1718262 | 303.8672828 | 242.6067963 | 305.7490624 | 329.9492312 | 237.4812754 | 0.001858102 | 0.577424217 | YAL034W- | MTW1 | n |
| W303.asm02.HP0_G0000380 | 510.8205379 | 571.1542295 | 543.3798625 | 339.5137755 | 396.7524784 | 305.0689939 | 322.7877099 | 292.2131021 | 273.3081919 | 8.85E-05 | 0.646588809 | YAL033W | POP5 | n |
| W303.asm02.HP0_G0000390 | 214.2150643 | 195.1443617 | 186.664665 | 236.3949783 | 298.5595573 | 231.7438054 | 270.725176 | 295.1158813 | 341.8911465 | 0.072003542 | -0.358452624 | YAL032C | PRP45 | n |
| W303.asm02.HP0_G0000400 | 318.8993491 | 361.731012 | 277.5537427 | 665.4080871 | 688.6773788 | 685.2736746 | 659.7742926 | 717.9540456 | 743.1526117 | 2.73E-13 | -1.089234602 | YAL031C | GIP4 | n |
| W303.asm02.HP0_G0000440 | 702.7417267 | 690.144694 | 690.9524511 | 200.4006812 | 197.7127735 | 210.9230729 | 226.2353744 | 244.8010425 | 198.5834803 | 2.48E-31 | 1.773120265 | YAL027W | SAW1 | n |
| W303.asm02.HP0_G0000450 | 787.0707339 | 767.2505149 | 783.7961326 | 2305.580653 | 2340.706929 | 2252.260109 | 1594.060127 | 2447.042832 | 2294.969911 | 7.55E-21 | -1.560869806 | YAL026C | DRS2 | n |
| W303.asm02.HP0_G0000460 | 2629.708006 | 3984.752674 | 3102.933567 | 4926.35461 | 4565.97083 | 4938.134604 | 2977.976936 | 2508.968788 | 2163.945759 | 0.001635354 | -0.570536497 | YAL025C | MAK16 | n |
| W303.asm02.HP0_G0000470 | 169.6273133 | 324.6059871 | 178.8462497 | 1599.314174 | 1670.60659 | 1623.111887 | 1442.605483 | 1738.764717 | 1555.911804 | 9.95E-38 | -2.861249798 | YAL024C | LTE1 | n |
| W303.asm02.HP0_G0000480 | 2484.313166 | 2460.722805 | 2272.226943 | 7414.825205 | 7294.141718 | 7336.13984 | 7629.527688 | 9809.458376 | 9494.132884 | 3.18E-37 | -1.610934533 | YAL023C | PMT2 | n |
| W303.asm02.HP0_G0000490 | 789.9786307 | 629.2215761 | 708.5438855 | 243.2047102 | 272.02093 | 251.6592887 | 531.0378453 | 422.8381643 | 445.2773913 | 8.69E-19 | 1.474273469 | YAL022C | FUN26 | n |
| W303.asm02.HP0_G0000500 | 844.259371 | 986.1929696 | 641.1100536 | 3909.758921 | 3512.387325 | 3840.067276 | 1803.256854 | 2704.422584 | 2760.719826 | 1.45E-27 | -2.188001093 | YAL021C | CCR4 | n |
| W303.asm02.HP0_G0000510 | 368.3335947 | 378.8656389 | 316.6458192 | 133.2761812 | 147.2893816 | 104.1036626 | 267.8854014 | 218.6760301 | 235.434023 | 3.53E-13 | 1.475439944 | YAL020C | ATS1 | n |
| W303.asm02.HP0_G0000520 | 371.2414915 | 685.3850754 | 372.3520281 | 2066.267218 | 1974.473872 | 1946.285866 | 963.630172 | 1418.491416 | 1138.27232 | 5.31E-18 | -2.066633869 | YAL019W | FUN30 | n |
| W303.asm02.HP0_G0000540 | 401.2897584 | 466.4426207 | 355.7378956 | 1519.543029 | 1409.20111 | 1509.050483 | 638.0026875 | 1263.676528 | 915.1218112 | 2.55E-16 | -1.858918362 | YAL017W | PSK1 | n |
| W303.asm02.HP0_G0000550 | 3104.664484 | 2689.184497 | 3015.953697 | 3058.542436 | 3095.730876 | 2989.13299 | 2788.658631 | 3152.418168 | 3418.911465 | 0.689015526 | -0.053544325 | YAL016W | TPD3 | n |
| W303.asm02.HP0_G0000570 | 982.8691185 | 880.5294371 | 1033.00812 | 114.7926232 | 144.6355189 | 105.9141611 | 182.6921642 | 149.9769233 | 222.1268826 | 3.80E-51 | 2.996072126 | YAL014C | SYN8 | n |
| W303.asm02.HP0_G0000580 | 795.7944243 | 523.5580437 | 594.1995619 | 531.1590871 | 514.8493699 | 506.0343251 | 673.026574 | 568.9447154 | 553.7817672 | 0.115244134 | 0.301994838 | YAL013W | DEP1 | n |
| W303.asm02.HP0_G0000590 | 2424.216633 | 2030.453286 | 1888.147292 | 11094.99888 | 9746.310882 | 10286.34711 | 6203.960852 | 6019.396385 | 5389.391874 | 7.87E-67 | -2.295077198 | YAL012W | CYS3 | n |
| W303.asm02.HP0_G0000600 | 220.0308579 | 246.5482424 | 196.4376841 | 919.3138046 | 1000.50625 | 982.1954253 | 672.0799824 | 1006.296776 | 979.6102609 | 1.50E-29 | -2.129463918 | YAL011W | SWC3 | n |
| W303.asm02.HP0_G0000610 | 624.2285131 | 433.1252907 | 534.5841453 | 429.9859276 | 471.0606348 | 485.2135926 | 379.5832013 | 459.6067004 | 415.5922319 | 0.287086203 | 0.199605102 | YAL010C | MDM10 | n |
| W303.asm02.HP0_G0000620 | 723.0970043 | 456.9233836 | 713.430395 | 358.9701523 | 387.4639588 | 384.730927 | 923.873328 | 624.0975195 | 737.0108546 | 0.000762166 | 0.743384924 | YAL009W | SPO7 | n |
| W303.asm02.HP0_G0000630 | 1080.768311 | 742.5004983 | 1212.831672 | 449.4423045 | 282.6363809 | 458.0561154 | 748.7538959 | 505.0835739 | 567.0889076 | 7.12E-08 | 1.347946894 | YAL008W | FUN14 | n |
| W303.asm02.HP0_G0000660 | 14493.92695 | 13590.61489 | 13720.34153 | 8910.047763 | 9100.095306 | 8155.390402 | 9555.84144 | 7818.151872 | 7808.220554 | 3.23E-08 | 0.676108582 | YAL003W | EFB1 | n |
| W303.asm02.HP0_G0000670 | 430.3687264 | 431.2214433 | 403.6256892 | 469.8715001 | 567.9266245 | 444.4773768 | 512.1060148 | 655.0604972 | 733.9399761 | 0.235598752 | -0.225843215 | YAL002W | VPS8 | n |
| W303.asm02.HP0_G0000680 | 719.2198086 | 587.3369326 | 719.2942065 | 674.1634567 | 716.5429375 | 763.1251093 | 551.8628588 | 769.2364775 | 800.4756782 | 0.650967804 | -0.088617464 | YAL001C | TFC3 | n |
| W303.asm02.HP0_G0000700 | 1449.101905 | 1110.894976 | 1208.922464 | 1354.163826 | 1318.969778 | 1399.515325 | 1296.830389 | 1194.977421 | 1344.021184 | 0.446651358 | -0.112191999 | YAR002W | NUP60 | n |
| W303.asm02.HP0_G0000710 | 3180.269801 | 3248.915642 | 3403.942556 | 2927.211892 | 2872.806407 | 2898.608066 | 3541.198893 | 2836.982833 | 2527.333056 | 0.235584252 | 0.176819605 | YAR002C- | ERP1 | n |
| W303.asm02.HP0_G0000720 | 2109.194479 | 1855.299322 | 2157.882619 | 647.897348 | 697.9658984 | 740.4938783 | 726.9822909 | 781.8151872 | 796.3811735 | 9.95E-35 | 1.552867642 | YAR003W | SWD1 | n |
| W303.asm02.HP0_G0000730 | 1764.124059 | 1464.058675 | 1770.871063 | 2036.109834 | 1869.646294 | 2110.135978 | 1846.800064 | 2236.107546 | 2091.2683 | 0.057532638 | -0.267691404 | YAR007C | RFA1 | n |
| W303.asm02.HP0_G0000740 | 758.9610649 | 975.7218087 | 810.1832842 | 369.6711595 | 461.7721153 | 411.8884042 | 434.4855098 | 321.2408937 | 364.4109226 | 8.56E-09 | 1.035492059 | YAR008W | SEN34 | n |
| W303.asm02.HP0_G0000750 | 1060.413033 | 983.3371984 | 1059.395272 | 832.7329278 | 855.8707309 | 779.4195956 | 740.2345722 | 715.0512665 | 742.1289856 | 0.005054056 | 0.331412485 | YAR014C | BUD14 | n |
| W303.asm02.HP0_G0000760 | 1122.448165 | 1044.260316 | 1197.194841 | 3441.833059 | 1011.121701 | 2963.786012 | 983.508594 | 943.4032271 | 811.7355663 | 0.000466961 | -1.141264746 | YAR015W | ADE1 | n |
| W303.asm02.HP0_G0000770 | 602.9039366 | 507.3753405 | 572.6989198 | 350.2147827 | 457.7913212 | 372.9626868 | 580.2606046 | 490.5696781 | 509.7658411 | 0.002332873 | 0.514726154 | YAR018C | KIN3 | n |
| W303.asm02.HP0_G0000780 | 156.0571283 | 169.4424214 | 109.457814 | 197.4822247 | 298.5595573 | 238.0805501 | 252.7399371 | 319.3057076 | 352.1274083 | 0.003029134 | -0.751055992 | YAR019C | CDC15 | n |
| W303.asm02.HP0_G0000810 | 1296.921973 | 1128.981527 | 1249.969144 | 409.556732 | 341.021361 | 370.2469391 | 229.0751489 | 413.1622338 | 220.0796302 | 1.38E-13 | 1.712500112 | YAR027W | UIP3 | n |

| | | | | | | | | | | | | | | |
|---|---|---|---|---|---|---|---|---|---|---|---|---|---|---|
| W303.asm02.HP0_G0000840 | 103.7149859 | 178.0097349 | 114.3443236 | 57.39631161 | 76.96201921 | 84.18817931 | 244.2206133 | 93.85652618 | 228.2686397 | 0.027298299 | 0.858640421 | YAR035W | YAT1 | n |
| W303.asm02.HP0_G0000850 | 328.5923384 | 589.2407801 | 295.1451771 | 2224.836689 | 2115.128597 | 2197.945155 | 2408.128838 | 2899.876381 | 2864.106071 | 1.54E-26 | -2.430020974 | YAR042W | SWH1 | n |
| W303.asm02.HP0_G0000910 | 1431.654525 | 1540.212572 | 1544.137019 | 380.3721668 | 402.0602038 | 439.0458814 | 428.8059606 | 348.3334992 | 265.1191824 | 7.96E-26 | 1.885928793 | YBL107C | MIX23 | n |
| W303.asm02.HP0_G0000920 | 180.2896016 | 176.1058874 | 170.0505325 | 1075.937678 | 1054.910436 | 1073.625599 | 835.8403162 | 1006.296776 | 878.2712684 | 5.12E-76 | -2.605856934 | YBL106C | SRO77 | n |
| W303.asm02.HP0_G0000930 | 296.6054736 | 446.4522227 | 239.4389682 | 2244.293066 | 2314.168302 | 2335.543039 | 1531.585087 | 2156.764916 | 2131.189721 | 3.29E-37 | -2.810456231 | YBL105C | PKC1 | n |
| W303.asm02.HP0_G0000940 | 266.5572067 | 385.5291049 | 276.5764408 | 815.2221886 | 816.0627899 | 810.1980697 | 776.2050501 | 847.6115148 | 998.0355323 | 1.33E-14 | -1.394159367 | YBL104C | SEA4 | n |
| W303.asm02.HP0_G0000950 | 612.5969259 | 798.6639976 | 615.7002039 | 458.197674 | 553.3303795 | 414.6041519 | 605.8185757 | 715.0512665 | 701.1839381 | 0.005056604 | 0.51032282 | YBL103C | RTG3 | n |
| W303.asm02.HP0_G0000960 | 1138.926247 | 1435.500963 | 1232.37771 | 963.0906524 | 1003.160113 | 865.4182734 | 1342.266782 | 1334.310821 | 1128.036058 | 0.00423444 | 0.427929109 | YBL102W | SFT2 | n |
| W303.asm02.HP0_G0000970 | 146.3641389 | 178.9616586 | 108.4805121 | 319.0845798 | 297.2326259 | 317.7424832 | 582.1537876 | 745.0466511 | 844.4916043 | 7.12E-07 | -1.106711868 | YBL101C | ECM21 | n |
| W303.asm02.HP0_G0000980 | 3837.454477 | 3594.463951 | 3877.933983 | 28110.57322 | 27446.24837 | 26760.07278 | 17289.4942 | 22021.4503 | 21048.82528 | 3.33E-124 | -2.863617959 | YBL099W | ATP1 | n |
| W303.asm02.HP0_G0001000 | 81.42111041 | 179.9135823 | 68.41113377 | 922.2322611 | 966.0060343 | 909.7754861 | 973.0960872 | 985.0097284 | 856.7751185 | 5.79E-28 | -3.083789482 | YBL097W | BRN1 | n |
| W303.asm02.HP0_G0001010 | 513.7284347 | 391.2406472 | 406.557595 | 355.0788769 | 173.8280089 | 361.1944467 | 528.1980707 | 471.217817 | 474.9625508 | 0.036627904 | 0.55363677 | YBL095W | MRX3 | n |
| W303.asm02.HP0_G0001020 | 1529.553717 | 1468.818293 | 1699.528023 | 719.8859422 | 865.1592505 | 712.4311518 | 1014.746114 | 900.8291327 | 1157.721217 | 8.47E-12 | 1.033876769 | YBL093C | ROX3 | n |
| W303.asm02.HP0_G0001030 | 8919.488785 | 10834.79573 | 9223.775436 | 9063.75314 | 10787.952 | 8989.124952 | 10034.81675 | 7224.049737 | 6290.182919 | 0.972219063 | 0.006956603 | YBL092W | RPL32 | n |
| W303.asm02.HP0_G0001050 | 1171.882411 | 715.8466343 | 908.8907772 | 2982.662566 | 2943.133769 | 2999.090732 | 2840.721165 | 2933.742138 | 2871.271454 | 5.24E-23 | -1.674282273 | YBL091C | MAP2 | n |
| W303.asm02.HP0_G0001060 | 1363.803599 | 1694.424214 | 1462.043659 | 1205.322544 | 1202.199817 | 1106.214571 | 1617.724915 | 1121.440349 | 985.7520181 | 0.066074621 | 0.363644608 | YBL090W | MRP21 | n |
| W303.asm02.HP0_G0001080 | 379.9651819 | 603.5196358 | 412.4214064 | 722.8043987 | 846.5822114 | 748.6411214 | 977.8290449 | 805.0374205 | 818.9009496 | 0.000248596 | -0.730624513 | YBL088C | TEL1 | n |
| W303.asm02.HP0_G0001100 | 629.0750078 | 350.3079274 | 423.1717275 | 290.8728334 | 285.2902436 | 254.3750364 | 158.0807846 | 287.3751369 | 305.0406037 | 0.008389253 | 0.756379095 | YBL086C | | n |
| W303.asm02.HP0_G0001120 | 464.2941891 | 618.7504153 | 459.3318982 | 483.4909639 | 589.1575264 | 509.6553221 | 411.7673132 | 630.8706708 | 634.6482359 | 0.879436246 | -0.035476764 | YBL084C | CDC27 | n |
| W303.asm02.HP0_G0001130 | 432.3073243 | 387.4329523 | 463.2411058 | 501.001703 | 496.2723308 | 492.4555865 | 854.7721467 | 927.9217382 | 998.0355323 | 0.142416468 | -0.215711122 | YBL082C | ALG3 | n |
| W303.asm02.HP0_G0001140 | 280.1273918 | 628.2696524 | 244.3254777 | 728.6413118 | 793.5049567 | 790.2825865 | 1008.119974 | 496.3752364 | 506.6949626 | 0.004499296 | -1.00408882 | YBL081W | | n |
| W303.asm02.HP0_G0001150 | 376.0879862 | 362.6829357 | 389.9434625 | 553.5339204 | 644.8886438 | 562.159778 | 528.1980707 | 508.9539461 | 530.2383649 | 4.16E-06 | -0.639066088 | YBL080C | PET112 | n |
| W303.asm02.HP0_G0001160 | 965.4217377 | 1535.452954 | 1019.325893 | 4522.634791 | 4893.722877 | 4576.940157 | 2774.459758 | 3644.923032 | 3294.02907 | 5.13E-26 | -1.990823303 | YBL079W | NUP170 | n |
| W303.asm02.HP0_G0001170 | 576.7328654 | 505.4714931 | 541.4252587 | 62.26040581 | 79.61588195 | 62.46219755 | 108.8580253 | 163.523226 | 153.543928 | 1.18E-41 | 3.000895267 | YBL078C | ATG8 | n |
| W303.asm02.HP0_G0001180 | 5843.90327 | 5358.378596 | 5605.803761 | 8121.091683 | 8533.495613 | 8274.883302 | 6274.955216 | 5562.692464 | 5663.723693 | 2.44E-08 | -0.568626234 | YBL076C | ILS1 | n |
| W303.asm02.HP0_G0001190 | 457.5090966 | 1617.318393 | 490.6055593 | 400.8013624 | 765.639398 | 524.1393099 | 3997.456008 | 2404.468738 | 5281.911125 | 0.347228525 | 0.602171972 | YBL075C | SSA3 | n |
| W303.asm02.HP0_G0001200 | 103.7149859 | 87.57698185 | 98.70749301 | 129.3849058 | 136.6739307 | 114.0614042 | 262.2058523 | 185.7778663 | 226.2213873 | 0.108642419 | -0.387102653 | YBL074C | AAR2 | n |
| W303.asm02.HP0_G0001210 | 8462.948988 | 9502.102531 | 8695.055102 | 28858.67091 | 32115.71985 | 27244.38112 | 25513.48136 | 20748.09784 | 18296.29447 | 5.25E-32 | -1.726356975 | YER102W | RPS8B | y |
| W303.asm02.HP0_G0001220 | 1314.369354 | 1325.077812 | 1452.27064 | 778.2550727 | 617.0230851 | 736.8728813 | 413.6604962 | 334.7871965 | 272.2845657 | 9.82E-08 | 0.938860723 | YBL071W- | KTI11 | n |
| W303.asm02.HP0_G0001260 | 180.2896016 | 286.5290385 | 167.1186268 | 1083.720189 | 1450.335983 | 1160.529526 | 954.1642567 | 1433.005312 | 1333.784922 | 6.99E-27 | -2.541865544 | YBL066C | SEF1 | n |
| W303.asm02.HP0_G0001270 | 2708.22122 | 1873.385873 | 2373.866342 | 2317.254479 | 1541.894247 | 2320.153802 | 2569.995989 | 2203.209383 | 2485.364382 | 0.413195871 | 0.16981427 | YBL064C | PRX1 | n |
| W303.asm02.HP0_G0001290 | 938.2813676 | 1129.933451 | 921.5957021 | 1819.171232 | 1836.47301 | 1881.107921 | 1595.006719 | 2386.08447 | 2057.488636 | 1.53E-07 | -0.888955391 | YBL061C | SKT5 | n |
| W303.asm02.HP0_G0001300 | 443.9389115 | 563.5388398 | 479.8552383 | 564.2349277 | 609.0614969 | 538.6232978 | 779.0448247 | 773.1068497 | 879.2948946 | 0.202165198 | -0.201253517 | YBL060W | YEL1 | n |
| W303.asm02.HP0_G0001310 | 1010.009489 | 1105.183434 | 1027.144308 | 978.6557539 | 1073.487475 | 992.153167 | 1139.696196 | 865.9957828 | 743.1526117 | 0.803373302 | 0.046269614 | YBL059C-/ | CMC2 | n |
| W303.asm02.HP0_G0001330 | 3314.033053 | 2072.337929 | 2956.338281 | 1281.202413 | 1004.487044 | 1323.474389 | 1854.372797 | 1642.973005 | 1817.960108 | 2.12E-10 | 1.207787311 | YBL058W | SHP1 | n |
| W303.asm02.HP0_G0001340 | 1432.623824 | 1151.827696 | 1472.79398 | 933.9060872 | 979.2753479 | 945.0802065 | 1065.862057 | 793.4263038 | 812.7591925 | 0.002460651 | 0.505576069 | YBL057C | PTH2 | n |
| W303.asm02.HP0_G0001350 | 4605.139233 | 4030.445013 | 4772.165231 | 3425.295139 | 2610.073996 | 3293.296735 | 2123.2049 | 2741.19112 | 2977.728578 | 0.00223012 | 0.522832941 | YBL056W | PTC3 | n |
| W303.asm02.HP0_G0001360 | 1173.821008 | 1569.722207 | 1299.811542 | 1107.067841 | 1082.775994 | 1080.867592 | 1172.826899 | 1093.38015 | 1170.004732 | 0.033494723 | 0.305973858 | YBL055C | | n |
| W303.asm02.HP0_G0001380 | 600.9653387 | 774.8659047 | 683.1340358 | 588.5553987 | 701.9466925 | 636.3902157 | 587.8333367 | 578.620646 | 581.4196742 | 0.544651594 | 0.097345336 | YBL052C | SAS3 | n |
| W303.asm02.HP0_G0001390 | 596.1188441 | 663.4908299 | 563.9032026 | 2360.058508 | 2085.936107 | 2479.477668 | 1282.631516 | 1822.945313 | 1982.763924 | 4.95E-27 | -1.925490606 | YBL051C | PIN4 | n |
| W303.asm02.HP0_G0001410 | 234.5703419 | 272.2501827 | 228.6886472 | 15.56510145 | 14.59624502 | 24.44172948 | 35.97047793 | 43.5416874 | 46.06317841 | 1.20E-32 | 3.727451893 | YBL049W | MOH1 | n |
| W303.asm02.HP0_G0001420 | 1094.338496 | 1360.29899 | 997.8252511 | 3008.928675 | 2385.822596 | 2977.36475 | 2460.191372 | 2534.126207 | 2757.648948 | 7.04E-16 | -1.278424808 | YBL047C | EDE1 | n |
| W303.asm02.HP0_G0001430 | 824.8733923 | 633.029271 | 717.3396027 | 574.9359349 | 638.2539869 | 613.7589847 | 795.1368806 | 699.5697776 | 881.342147 | 0.125588162 | 0.252387325 | YBL046W | PSY4 | n |
| W303.asm02.HP0_G0001440 | 6439.052815 | 4885.272509 | 6956.435002 | 12735.17145 | 12329.84625 | 12413.68283 | 5948.38114 | 10574.82448 | 9241.297216 | 7.11E-07 | -1.035762031 | YBL045C | COR1 | n |
| W303.asm02.HP0_G0001460 | 250.0791248 | 398.8560369 | 267.7807236 | 43.77684784 | 69.00043102 | 22.631231 | 53.00912537 | 47.41205962 | 66.53570215 | 1.49E-14 | 2.782587226 | YBL043W | ECM13 | n |
| W303.asm02.HP0_G0001470 | 1159.281524 | 2048.539836 | 1235.309615 | 4191.876385 | 4958.742514 | 4076.337327 | 1896.969415 | 2030.977819 | 1868.117791 | 3.34E-14 | -1.573564449 | YBL042C | FUI1 | n |
| W303.asm02.HP0_G0001480 | 5394.148565 | 4542.579972 | 5312.613188 | 2038.055472 | 1736.953158 | 2056.726273 | 2487.642527 | 1939.056479 | 1871.18867 | 1.48E-19 | 1.386116154 | YBL041W | PRE7 | n |
| W303.asm02.HP0_G0001500 | 335.377431 | 272.2501827 | 340.101065 | 45.72248552 | 25.21169595 | 52.50445592 | 108.8580253 | 80.31022343 | 97.24448776 | 1.20E-25 | 2.914538733 | YBL039W- | MIN6 | n |
| W303.asm02.HP0_G0001510 | 2727.607199 | 3218.454083 | 2810.720296 | 12572.7107 | 13825.2979 | 12303.24242 | 7892.680132 | 9241.481253 | 7852.23648 | 3.37E-66 | -2.143778233 | YBL039C | URA7 | n |
| W303.asm02.HP0_G0001520 | 1692.395938 | 1331.741278 | 1556.841944 | 1531.216855 | 1571.086737 | 1547.070951 | 1824.081868 | 1534.602583 | 1373.706343 | 0.897825572 | -0.021330454 | YBL038W | MRPL16 | n |
| W303.asm02.HP0_G0001530 | 306.298463 | 454.0676124 | 283.4175542 | 1448.527254 | 1415.835767 | 1450.209282 | 987.2949601 | 1314.95896 | 1282.603612 | 2.84E-25 | -2.047138455 | YBL037W | APL3 | n |
| W303.asm02.HP0_G0001540 | 2405.799953 | 2209.414944 | 2458.891608 | 1133.33395 | 1129.218592 | 1093.541082 | 1416.100921 | 1127.245907 | 1142.366825 | 4.09E-16 | 1.075908021 | YBL036C | | n |
| W303.asm02.HP0_G0001550 | 370.2721926 | 335.077148 | 296.122479 | 1093.448377 | 1076.141338 | 1151.477033 | 901.1551314 | 1226.907992 | 1004.177289 | 1.63E-26 | -1.729765446 | YBL035C | POL12 | n |
| W303.asm02.HP0_G0001560 | 327.6230395 | 466.4426207 | 362.579009 | 677.0819132 | 704.6005552 | 705.1891579 | 409.8741301 | 635.7086361 | 634.6482359 | 5.87E-05 | -0.851146595 | YBL034C | STU1 | n |
| W303.asm02.HP0_G0001570 | 947.005058 | 858.6351916 | 726.1353199 | 793.8201741 | 902.3133287 | 752.2621184 | 1442.605483 | 1036.29216 | 1353.233819 | 0.79656316 | 0.049525503 | YBL033C | RIB1 | n |
| W303.asm02.HP0_G0001580 | 642.6451929 | 706.3273971 | 546.3117682 | 1612.933638 | 1414.508836 | 1581.470422 | 1464.377088 | 1274.320051 | 1945.913381 | 2.60E-12 | -1.282322154 | YBL032W | HEK2 | n |
| W303.asm02.HP0_G0001590 | 527.2986198 | 587.3369326 | 537.516051 | 708.2121161 | 663.4656829 | 759.5041123 | 841.5198653 | 799.2318621 | 824.0190805 | 0.004763091 | -0.368873347 | YBL031W | SHE1 | n |
| W303.asm02.HP0_G0001600 | 4798.02972 | 4887.176357 | 4622.638039 | 21532.37222 | 22458.31337 | 20359.96066 | 16846.48936 | 18275.89759 | 19046.61246 | 1.07E-103 | -2.169074744 | YBL030C | PET9 | n |
| W303.asm02.HP0_G0001610 | 962.5138409 | 675.8658382 | 926.4822116 | 48.64094204 | 45.11566644 | 54.3149544 | 81.40687111 | 58.05558321 | 54.25218791 | 9.08E-65 | 4.107650217 | YBL029C-A | | n |
| W303.asm02.HP0_G0001620 | 419.7064382 | 331.2694531 | 423.1717275 | 368.6983407 | 354.2906747 | 382.9204285 | 317.1081607 | 351.2362784 | 252.8356682 | 0.672433615 | 0.085318507 | YBL029W | | n |
| W303.asm02.HP0_G0001630 | 1478.180873 | 2022.837896 | 1708.32374 | 4224.952226 | 4072.352362 | 4328.901865 | 2625.844889 | 2356.089085 | 2180.323778 | 1.30E-18 | -1.277297359 | YBL028C | | n |
| W303.asm02.HP0_G0001640 | 532.1451145 | 389.3367998 | 558.0393912 | 13.61946377 | 10.61545093 | 14.48398784 | 28.39774574 | 24.18982634 | 30.70878561 | 1.70E-57 | 5.234987087 | NA | NA | NA |
| W303.asm02.HP0_G0001650 | 873.338339 | 930.981394 | 844.3888511 | 630.3866089 | 725.8314571 | 682.5579269 | 882.2233009 | 638.6114153 | 559.9235243 | 0.044638921 | 0.378330789 | YBL026W | LSM2 | n |

| | | | | | | | | | | | | | | |
|---|---|---|---|---|---|---|---|---|---|---|---|---|---|---|
| W303.asm02.HP0_G0001660 | 698.864531 | 432.173367 | 697.7935644 | 118.6838986 | 155.2509698 | 130.3558905 | 50.1693508 | 108.370422 | 102.3626187 | 9.99E-14 | 2.180518128 | YBL025W | RRN10 | n |
| W303.asm02.HP0_G0001670 | 1298.860571 | 1362.202837 | 1279.288201 | 5526.583835 | 6287.000811 | 5593.535053 | 3904.690039 | 4151.941792 | 4157.969572 | 1.88E-90 | -2.142898779 | YBL024W | NCL1 | n |
| W303.asm02.HP0_G0001680 | 641.6758939 | 691.0966177 | 634.2689402 | 1500.086653 | 1659.991139 | 1543.449954 | 1134.963238 | 1629.426702 | 1695.124966 | 1.19E-13 | -1.257329725 | YBL023C | MCM2 | n |
| W303.asm02.HP0_G0001690 | 1578.018664 | 1767.72234 | 1462.043659 | 2459.28603 | 2058.070548 | 2290.280577 | 2098.59341 | 2525.417869 | 2479.222625 | 0.000376422 | -0.50225551 | YBL022C | PIM1 | n |
| W303.asm02.HP0_G0001700 | 826.8119902 | 720.6062529 | 895.2085505 | 301.5738407 | 306.5211455 | 312.3109878 | 308.588837 | 371.5557325 | 417.6394843 | 4.43E-17 | 1.407881555 | YBL021C | HAP3 | n |
| W303.asm02.HP0_G0001710 | 1200.961379 | 1280.337398 | 1309.584561 | 719.8859422 | 705.9274866 | 719.6731457 | 886.9562585 | 979.2041701 | 1060.47673 | 3.71E-11 | 0.820926113 | YBL020W | RFT1 | n |
| W303.asm02.HP0_G0001720 | 529.2372177 | 497.8561034 | 566.8351084 | 542.8329132 | 613.042291 | 577.5490151 | 677.7595316 | 584.4262043 | 635.6718621 | 0.417861448 | -0.11982671 | YBL019W | APN2 | n |
| W303.asm02.HP0_G0001730 | 337.3160288 | 416.9425875 | 304.9181962 | 266.5523624 | 285.2902436 | 263.4275288 | 188.3717134 | 232.2223328 | 263.0719301 | 0.064279121 | 0.379046311 | YBL018C | POP8 | n |
| W303.asm02.HP0_G0001740 | 872.3690401 | 1086.14496 | 897.1631543 | 1302.604428 | 1353.469993 | 1310.800899 | 1810.829586 | 2774.089284 | 2106.622693 | 0.00870164 | -0.473952204 | YBL017C | PEP1 | n |
| W303.asm02.HP0_G0001750 | 955.7287484 | 891.9525217 | 808.2286804 | 36.96711595 | 38.48100961 | 29.87322492 | 132.5228134 | 69.66669985 | 121.8115163 | 4.88E-69 | 4.665543134 | YBL016W | FUS3 | n |
| W303.asm02.HP0_G0001760 | 402.2590574 | 400.7598843 | 295.1451771 | 627.4681523 | 664.7926143 | 628.2429725 | 845.3062314 | 1643.940598 | 1284.650865 | 0.000713005 | -0.805986186 | YBL015W | ACH1 | n |
| W303.asm02.HP0_G0001770 | 332.4695342 | 624.4619576 | 319.5777249 | 563.2621088 | 546.6957227 | 544.0547932 | 323.7343014 | 434.449281 | 370.5526797 | 0.159044549 | -0.373551261 | YBL014C | RRN6 | n |
| W303.asm02.HP0_G0001780 | 31.9868648 | 34.26925377 | 24.43254777 | 335.6225001 | 382.1562333 | 373.8679361 | 561.3287741 | 402.5187102 | 507.7185887 | 7.86E-50 | -3.587557187 | YBL013W | FMT1 | n |
| W303.asm02.HP0_G0001790 | 812.2725062 | 917.654462 | 749.5905657 | 1358.055102 | 1728.99157 | 1274.59093 | 1920.634203 | 2661.84849 | 2803.712126 | 1.71E-05 | -0.813716266 | YBL011W | SCT1 | n |
| W303.asm02.HP0_G0001800 | 270.4344024 | 237.0290052 | 257.0304026 | 739.342319 | 693.9851043 | 766.7461062 | 826.3744009 | 699.5697776 | 571.1834123 | 1.24E-18 | -1.525815175 | YBL010C | LAA2 | n |
| W303.asm02.HP0_G0001810 | 812.2725062 | 865.2986577 | 833.6385301 | 2024.436008 | 1853.723118 | 2074.831258 | 1364.984978 | 1551.051665 | 1675.676068 | 6.77E-23 | -1.245787607 | YBL009W | ALK2 | n |
| W303.asm02.HP0_G0001830 | 300.4826694 | 349.3560037 | 334.2372536 | 568.126203 | 570.5804873 | 621.9062278 | 673.9731655 | 820.5189093 | 755.436126 | 1.87E-08 | -0.839911268 | YBL008W | HIR1 | n |
| W303.asm02.HP0_G0001840 | 1251.364923 | 1178.48156 | 1022.257799 | 3454.479704 | 3359.790218 | 3593.839483 | 1961.337639 | 3610.089682 | 3607.258683 | 1.34E-13 | -1.592202104 | YBL007C | SLA1 | n |
| W303.asm02.HP0_G0001850 | 1367.680795 | 1452.63559 | 1376.041091 | 548.6698262 | 586.5036637 | 521.4235622 | 831.1073586 | 643.4493805 | 624.4119741 | 4.04E-19 | 1.342295379 | YBL006C | LDB7 | n |
| W303.asm02.HP0_G0001870 | 1026.48757 | 2017.126354 | 1182.535312 | 7085.039618 | 7607.29752 | 7376.876056 | 2909.822346 | 3881.015737 | 3096.469216 | 9.62E-26 | -2.384514771 | YBL004W | UTP20 | n |
| W303.asm02.HP0_G0001880 | 3587.375353 | 5440.244036 | 3665.859468 | 4565.43882 | 4209.026292 | 4422.142537 | 4630.725738 | 4987.94219 | 3598.046047 | 0.786792316 | -0.056120718 | YBL003C | HTA2 | y |
| W303.asm02.HP0_G0001920 | 949.9129548 | 855.7794205 | 889.344739 | 995.1936742 | 988.5638675 | 1057.331112 | 1056.396141 | 952.1115646 | 903.8619231 | 0.177828759 | -0.174837534 | YBR002C | RER2 | n |
| W303.asm02.HP0_G0001930 | 974.1454281 | 835.7890225 | 879.5717199 | 1592.504442 | 1452.989846 | 1591.428164 | 1215.423518 | 1241.421888 | 1317.406903 | 7.28E-11 | -0.786606125 | YBR003W | COQ1 | n |
| W303.asm02.HP0_G0001940 | 1159.281524 | 1125.173832 | 1189.376426 | 464.0345871 | 497.5992622 | 439.9511306 | 451.5241572 | 507.01876 | 573.2306647 | 8.08E-21 | 1.311350504 | YBR004C | GPI18 | n |
| W303.asm02.HP0_G0001950 | 863.6453497 | 616.8465678 | 879.5717199 | 518.5124422 | 510.8685758 | 525.9498084 | 969.3097211 | 846.6439218 | 843.4679781 | 0.000473517 | 0.601178939 | YBR005W | RCR1 | n |
| W303.asm02.HP0_G0001960 | 615.5048227 | 443.5964516 | 536.5387491 | 193.5909493 | 237.5207145 | 188.2918419 | 510.2128317 | 515.7270975 | 590.6323099 | 1.93E-13 | 1.370332463 | YBR006W | UGA2 | n |
| W303.asm02.HP0_G0001970 | 326.6537406 | 315.0867499 | 301.9862905 | 470.844319 | 508.2147131 | 498.7923312 | 692.904996 | 723.759604 | 1044.098711 | 0.000702261 | -0.646623778 | YBR007C | DSF2 | n |
| W303.asm02.HP0_G0001990 | 6490.425658 | 8225.572828 | 6890.955774 | 6751.362755 | 6805.830975 | 6787.558801 | 7584.091295 | 6222.590927 | 5631.991281 | 0.588059537 | 0.086857207 | YNL030W | HHF2 | y |
| W303.asm02.HP0_G0002000 | 3473.967377 | 4537.820353 | 3353.122857 | 4051.790472 | 4019.275107 | 3998.485893 | 4386.505125 | 2766.34854 | 3338.044996 | 0.669926302 | -0.086762025 | YNL031C | HHT2 | y |
| W303.asm02.HP0_G0002010 | 8127.571557 | 5994.263638 | 8091.082521 | 12817.86105 | 12256.86503 | 12206.38075 | 13647.01001 | 9955.564927 | 10565.8695 | 8.15E-06 | -0.747066736 | YBR011C | IPP1 | n |
| W303.asm02.HP0_G0002020 | 1965.738237 | 1564.010665 | 2095.335297 | 690.701377 | 686.0235161 | 689.7999208 | 1294.937206 | 978.236577 | 1031.815196 | 1.30E-18 | 1.44455874 | YBR014C | GRX7 | n |
| W303.asm02.HP0_G0002030 | 1185.452596 | 1171.818094 | 987.0749301 | 3286.182044 | 3316.001483 | 3267.044507 | 3154.989551 | 4326.108542 | 4413.876118 | 2.26E-23 | -1.561139826 | YBR015C | MNN2 | n |
| W303.asm02.HP0_G0002040 | 843.2900721 | 647.3081267 | 770.1139059 | 1549.700413 | 1549.855835 | 1541.639456 | 2063.569524 | 1446.551615 | 1434.100288 | 5.39E-09 | -1.037782462 | YBR016W | CPP1 | n |
| W303.asm02.HP0_G0002050 | 1710.812617 | 1717.270383 | 1754.25693 | 3942.834762 | 3509.733462 | 3869.940501 | 1796.630714 | 2829.242088 | 2428.041316 | 1.47E-11 | -1.127697507 | YBR017C | KAP104 | n |
| W303.asm02.HP0_G0002060 | 76.57461574 | 55.21157552 | 56.68351084 | 240.2862537 | 244.1553713 | 248.943541 | 212.0365015 | 295.1158813 | 282.5208276 | 3.36E-17 | -1.960581085 | YBR018C | GAL7 | n |
| W303.asm02.HP0_G0002070 | 108.5614805 | 133.2693202 | 92.84368154 | 266.5523624 | 287.9441064 | 294.206003 | 301.0161048 | 540.8845169 | 534.3328696 | 2.53E-07 | -1.341948745 | YBR019C | GAL10 | n |
| W303.asm02.HP0_G0002090 | 986.7463142 | 781.5293707 | 838.5250356 | 118.6838986 | 83.59667604 | 105.9141611 | 225.2887828 | 142.2361789 | 202.677985 | 1.22E-43 | 3.072555769 | YBR021W | FUR4 | n |
| W303.asm02.HP0_G0002110 | 1073.013919 | 1208.943119 | 1043.758441 | 3315.36661 | 3120.942572 | 3203.67706 | 1728.476124 | 2720.871666 | 2594.892384 | 2.89E-18 | -1.535395491 | YBR023C | CHS3 | n |
| W303.asm02.HP0_G0002130 | 13174.7111 | 9583.967971 | 11832.19424 | 10550.22033 | 11086.51156 | 10159.61222 | 8693.496561 | 8900.888499 | 7510.345334 | 0.429881061 | 0.121581636 | YBR025C | OLA1 | n |
| W303.asm02.HP0_G0002140 | 1960.891742 | 1383.145159 | 1775.757572 | 2290.015551 | 1317.642846 | 1727.21555 | 1481.415736 | 1504.607198 | 1488.352476 | 0.789012557 | -0.060230141 | YBR026C | ETR1 | n |
| W303.asm02.HP0_G0002150 | 1321.154446 | 1182.289255 | 1360.40426 | 581.7456668 | 643.5617124 | 592.0330029 | 701.4243197 | 751.8198025 | 624.4119741 | 8.68E-16 | 1.089547119 | YBR028C | YPK3 | n |
| W303.asm02.HP0_G0002160 | 2377.690284 | 2623.501761 | 2628.942141 | 3189.872979 | 3217.808562 | 3112.246887 | 2833.148433 | 3687.497127 | 3201.902713 | 0.015911515 | -0.319155506 | YBR029C | CDS1 | n |
| W303.asm02.HP0_G0002170 | 516.6363315 | 414.0868164 | 479.8552383 | 1391.130942 | 1252.623209 | 1398.610076 | 1031.784762 | 1123.375535 | 926.3816992 | 7.95E-26 | -1.519760537 | YBR030W | RKM3 | n |
| W303.asm02.HP0_G0002200 | 2827.444989 | 6161.802212 | 3196.754551 | 5592.735516 | 4027.236695 | 5342.781014 | 2162.015042 | 2777.959656 | 2264.261126 | 0.285275164 | -0.296271233 | YBR034C | HMT1 | n |
| W303.asm02.HP0_G0002230 | 860.7374529 | 575.9138481 | 816.0470957 | 1005.894681 | 1048.275779 | 1036.51038 | 1081.007521 | 1034.356974 | 1034.886075 | 0.006838871 | -0.45628776 | YBR037C | SCO1 | n |
| W303.asm02.HP0_G0002240 | 850.0751646 | 856.7313442 | 722.2261122 | 3971.046508 | 4901.684465 | 3898.003227 | 3245.862338 | 5083.733903 | 4666.711786 | 4.52E-38 | -2.394058798 | YBR038W | CHS2 | n |
| W303.asm02.HP0_G0002250 | 5044.23165 | 5318.3978 | 5400.57036 | 9709.70485 | 9116.018483 | 9366.613886 | 4962.979363 | 7464.980407 | 7616.802457 | 4.75E-07 | -0.838770533 | YBR039W | ATP3 | n |
| W303.asm02.HP0_G0002270 | 1116.632371 | 1093.760349 | 1191.33103 | 880.401051 | 761.658604 | 855.4605317 | 568.9015063 | 787.6207455 | 855.7514923 | 0.010783905 | 0.44484815 | YBR041W | FAT1 | n |
| W303.asm02.HP0_G0002280 | 1275.597396 | 1507.847166 | 1367.245373 | 355.0788769 | 436.5604193 | 401.9306625 | 786.6175569 | 667.6392069 | 765.6723879 | 1.22E-34 | 1.800590532 | YBR042C | CST26 | n |
| W303.asm02.HP0_G0002290 | 541.8381038 | 716.798558 | 503.3104842 | 231.5308841 | 337.0405669 | 257.0907841 | 294.3899641 | 399.6159311 | 432.9938771 | 1.51E-06 | 1.097624796 | YBR043C | QDR3 | n |
| W303.asm02.HP0_G0002300 | 947.005058 | 733.9331849 | 919.6410982 | 1138.198044 | 1260.584797 | 1231.138966 | 1008.119974 | 1078.866255 | 1119.847049 | 0.000821255 | -0.480841002 | YBR044C | TCM62 | n |
| W303.asm02.HP0_G0002460 | 1538.277407 | 973.8179613 | 1431.7473 | 439.7141161 | 327.7520473 | 481.5925956 | 1046.930226 | 991.7828798 | 774.8850235 | 4.80E-14 | 1.655841562 | YBR053C | | n |
| W303.asm02.HP0_G0002480 | 666.8776662 | 768.2024386 | 731.9991313 | 890.1292394 | 893.0248092 | 913.3964831 | 678.7061231 | 854.3846662 | 785.1212854 | 0.02662561 | -0.31545031 | YBR055C | PRP6 | n |
| W303.asm02.HP0_G0002490 | 1276.566695 | 659.683135 | 1161.03467 | 162.4607464 | 192.405048 | 167.4711094 | 247.0603879 | 270.926055 | 392.0488296 | 3.38E-22 | 2.569979243 | YBR056W | MRX18 | n |
| W303.asm02.HP0_G0002510 | 1096.277094 | 1152.77962 | 1058.41797 | 150.7869203 | 139.3277934 | 143.9346291 | 96.5523355 | 153.8472955 | 136.1422829 | 3.24E-57 | 2.928321751 | YBR057C | MUM2 | n |
| W303.asm02.HP0_G0002520 | 764.7768585 | 692.0485414 | 786.7280384 | 1204.349725 | 1198.219023 | 1258.296443 | 1258.020136 | 1390.431218 | 1572.289823 | 1.41E-07 | -0.706761627 | YBR058C | UBP14 | n |
| W303.asm02.HP0_G0002530 | 1183.513998 | 1288.904711 | 1287.106617 | 602.1748625 | 670.1003397 | 602.8959938 | 842.4664569 | 585.3937973 | 592.6795623 | 4.01E-09 | 1.004571154 | YBR058C-, | TSC3 | n |
| W303.asm02.HP0_G0002540 | 549.5924953 | 586.3850089 | 537.516051 | 930.9876307 | 934.1596815 | 842.7870424 | 827.3209925 | 1108.861639 | 1269.296472 | 8.15E-05 | -0.693850074 | YBR059C | AKL1 | n |
| W303.asm02.HP0_G0002550 | 296.6054736 | 350.3079274 | 251.1665911 | 682.9188263 | 630.2923987 | 675.315933 | 520.6253385 | 631.8382639 | 557.8762719 | 1.12E-11 | -1.147159968 | YBR060C | ORC2 | n |
| W303.asm02.HP0_G0002580 | 1084.645507 | 1249.875839 | 1107.283065 | 702.3752031 | 709.9082807 | 712.4311518 | 821.6414433 | 831.1624329 | 674.5696572 | 5.51E-07 | 0.695976311 | YBR061C | TRM7 | n |
| W303.asm02.HP0_G0002590 | 896.6015134 | 874.8178948 | 866.8667951 | 194.5637682 | 201.6935676 | 163.8501124 | 362.5445539 | 273.8288341 | 262.0483039 | 1.82E-36 | 2.239661174 | YBR062C | | n |
| W303.asm02.HP0_G0002610 | 247.171228 | 355.067546 | 276.5764408 | 438.7412972 | 293.2518318 | 442.6668783 | 125.8966728 | 291.2455091 | 232.3631444 | 0.176574709 | -0.420453871 | YBR065C | ECM2 | n |
| W303.asm02.HP0_G0002660 | 783.1935382 | 726.3177952 | 825.8201148 | 188.7268551 | 254.7708222 | 231.7438054 | 776.2050501 | 425.7409435 | 627.4828526 | 2.20E-15 | 1.792640381 | YBR070C | ALG14 | n |

| | | | | | | | | | | | | | | |
|---|---|---|---|---|---|---|---|---|---|---|---|---|---|---|
| W303.asm02.HP0_G0002670 | 1694.334536 | 1480.241378 | 1953.62652 | 252.9328986 | 232.212989 | 259.8065319 | 577.42083 | 349.3010923 | 369.5290535 | 7.77E-40 | 2.781687655 | YBR071W | | n |
| W303.asm02.HP0_G0002680 | 1079.799012 | 2219.886105 | 674.3383186 | 16.53792029 | 66.34656829 | 53.40970516 | 40.70343556 | 19.35186107 | 45.03955223 | 4.34E-13 | 4.87071622 | YBR072W | HSP26 | n |
| W303.asm02.HP0_G0002700 | 1109.847279 | 1464.058675 | 1119.010688 | 296.7097465 | 410.021792 | 310.5004893 | 308.588837 | 396.7131519 | 370.5526797 | 2.97E-22 | 1.864336672 | YBR073W | RDH54 | n |
| W303.asm02.HP0_G0002710 | 766.7154563 | 820.558243 | 850.2526626 | 2201.489037 | 2081.955313 | 2238.68137 | 1595.006719 | 2412.209849 | 2018.590841 | 2.10E-18 | -1.420139473 | YBR074W | PFF1 | n |
| W303.asm02.HP0_G0002730 | 1033.272663 | 793.904379 | 947.0055518 | 212.0745073 | 222.9244694 | 236.2700516 | 411.7673132 | 265.1204966 | 255.9065467 | 6.59E-22 | 2.046719018 | YBR077C | SLM4 | n |
| W303.asm02.HP0_G0002740 | 14908.7869 | 14652.96176 | 16125.48153 | 11951.07946 | 11219.2047 | 11184.35436 | 9652.393776 | 9128.272866 | 9366.179611 | 1.96E-05 | 0.411243349 | YBR078W | ECM33 | n |
| W303.asm02.HP0_G0002750 | 3121.142566 | 3059.482823 | 2790.196956 | 14320.86616 | 14613.49513 | 14174.3926 | 9694.043803 | 11091.51917 | 10326.34097 | 9.41E-106 | -2.264618064 | YBR079C | RPG1 | n |
| W303.asm02.HP0_G0002760 | 1347.325517 | 1056.635325 | 1225.536596 | 2130.473261 | 1791.357344 | 2014.179559 | 1868.571669 | 2119.028787 | 2185.441909 | 7.58E-07 | -0.710388399 | YBR080C | SEC18 | n |
| W303.asm02.HP0_G0002770 | 752.1759723 | 808.1832347 | 794.5464536 | 1385.294029 | 1552.509698 | 1416.71506 | 1125.497323 | 1646.843377 | 1709.455732 | 3.35E-07 | -0.886407168 | YBR081C | SPT7 | n |
| W303.asm02.HP0_G0002790 | 838.4435774 | 673.0100671 | 695.8389606 | 264.6067247 | 311.828871 | 259.8065319 | 239.4876557 | 325.111266 | 363.3872964 | 9.35E-13 | 1.403097204 | YBR083W | TEC1 | n |
| W303.asm02.HP0_G0002800 | 1867.839045 | 1928.597448 | 1796.280912 | 4742.491849 | 4282.007517 | 4700.054054 | 1732.26249 | 2388.987249 | 2650.168198 | 1.46E-15 | -1.295267066 | YBR084W | MIS1 | n |
| W303.asm02.HP0_G0002830 | 3733.739492 | 3021.405874 | 3583.766108 | 612.8758697 | 461.7721153 | 662.6424436 | 1291.150839 | 757.6253608 | 918.1926897 | 9.27E-34 | 2.571083879 | YBR085C-A | | n |
| W303.asm02.HP0_G0002840 | 2445.541209 | 3085.184763 | 2668.034217 | 3521.604204 | 3039.999759 | 3453.52585 | 2594.607369 | 3198.862635 | 3052.45329 | 0.04972198 | -0.289019425 | YBR086C | IST2 | n |
| W303.asm02.HP0_G0002850 | 1644.90029 | 1784.856967 | 1628.184984 | 1068.155087 | 1076.141338 | 1125.224805 | 1360.252021 | 1004.361589 | 1020.555308 | 3.03E-05 | 0.629308507 | YBR087W | RFC5 | n |
| W303.asm02.HP0_G0002860 | 1821.312696 | 2339.828493 | 2068.948146 | 2265.69508 | 2174.840508 | 2413.394474 | 2910.768938 | 2774.089284 | 2536.545691 | 0.328734884 | -0.137991423 | YBR088C | POL30 | n |
| W303.asm02.HP0_G0002880 | 739.5750862 | 630.1734999 | 724.180716 | 611.9030509 | 623.6577419 | 645.4427081 | 904.9414975 | 620.2271472 | 619.2938431 | 0.422599384 | 0.154518315 | YBR091C | TIM12 | n |
| W303.asm02.HP0_G0002920 | 853.9523603 | 814.8467007 | 757.408981 | 585.6369422 | 636.9270556 | 646.3479573 | 701.4243197 | 671.5095791 | 780.0031545 | 0.005326194 | 0.376856452 | YBR095C | RXT2 | n |
| W303.asm02.HP0_G0002930 | 1719.536308 | 1324.125889 | 1809.963139 | 598.2835871 | 506.8877817 | 597.4644984 | 624.7504062 | 586.3613904 | 629.530105 | 2.22E-21 | 1.509324425 | YBR096W | | n |
| W303.asm02.HP0_G0002940 | 330.5309363 | 561.6349923 | 357.6924994 | 483.4909639 | 477.6952917 | 486.1188418 | 466.6696216 | 566.0419363 | 464.7262889 | 0.354114835 | -0.211469348 | YBR097W | VPS15 | n |
| W303.asm02.HP0_G0002950 | 481.7415699 | 503.5676457 | 482.787144 | 266.5523624 | 221.5975381 | 227.2175592 | 236.6478811 | 270.926055 | 292.7570895 | 3.63E-10 | 1.035133359 | YBR098W | MMS4 | n |
| W303.asm02.HP0_G0002960 | 5976.697224 | 3981.896903 | 4696.912984 | 1231.588652 | 1316.315915 | 1224.802222 | 1718.063617 | 1339.148786 | 1342.997557 | 2.81E-29 | 1.958080867 | YBR101C | FES1 | n |
| W303.asm02.HP0_G0002970 | 351.8555128 | 435.0291381 | 347.9194803 | 931.9604495 | 847.9091427 | 930.5962186 | 830.160767 | 1003.393996 | 1149.532208 | 1.43E-13 | -1.256548734 | YBR102C | EXO84 | n |
| W303.asm02.HP0_G0002980 | 659.1232747 | 473.1060868 | 571.7216179 | 854.1349423 | 752.3700844 | 815.6295652 | 675.8663485 | 732.4679414 | 956.0668586 | 0.007195663 | -0.508121836 | YBR103W | SIF2 | n |
| W303.asm02.HP0_G0002990 | 1169.943813 | 1655.395342 | 1229.445804 | 598.2835871 | 943.4482011 | 602.8959938 | 1359.305429 | 1430.102533 | 1050.240468 | 4.34E-05 | 0.921330442 | YBR104W | YMC2 | n |
| W303.asm02.HP0_G0003000 | 536.9916091 | 893.8563691 | 425.1263313 | 104.091616 | 47.76952917 | 89.61967475 | 73.83413892 | 58.05558321 | 106.4571234 | 2.99E-17 | 2.932476246 | YBR105C | VID24 | n |
| W303.asm02.HP0_G0003010 | 6681.377548 | 7678.216692 | 6830.363056 | 4766.81232 | 4442.566213 | 4534.393443 | 7645.619744 | 5787.174053 | 6177.584038 | 5.99E-06 | 0.624531535 | YBR106W | SND3 | n |
| W303.asm02.HP0_G0003030 | 186.1053952 | 224.6539969 | 179.8235516 | 364.8070653 | 382.1562333 | 370.2469391 | 436.3786928 | 681.1855096 | 591.6559361 | 7.71E-06 | -0.918986655 | YBR108W | AIM3 | n |
| W303.asm02.HP0_G0003040 | 4726.301599 | 4273.18556 | 4796.597779 | 2791.017254 | 2748.074859 | 2744.715695 | 3349.040814 | 2202.24179 | 2268.35563 | 1.75E-05 | 0.735873652 | YBR109C | CMD1 | n |
| W303.asm02.HP0_G0003050 | 1180.606101 | 1009.991062 | 1266.583277 | 1124.57858 | 1179.641984 | 1110.740817 | 1734.155673 | 1818.107347 | 1964.338653 | 0.900095696 | 0.018130114 | YBR110W | ALG1 | n |
| W303.asm02.HP0_G0003060 | 2669.449263 | 2433.117018 | 2619.169121 | 1017.568508 | 928.851956 | 977.6691791 | 1278.84515 | 1045.000498 | 1072.760244 | 3.10E-28 | 1.400214261 | YBR111C | YSA1 | n |
| W303.asm02.HP0_G0003080 | 494.342456 | 763.4428201 | 377.2385376 | 3912.677378 | 3592.003207 | 3991.243899 | 2888.050741 | 2985.02457 | 2886.625847 | 9.90E-40 | -2.81358405 | YBR112C | CYC8 | n |
| W303.asm02.HP0_G0003090 | 185.1360963 | 456.9233836 | 183.7327593 | 784.0919857 | 598.446046 | 773.9881001 | 443.951425 | 611.5188098 | 510.7894673 | 1.34E-05 | -1.38513861 | YBR114W | RAD16 | n |
| W303.asm02.HP0_G0003100 | 1157.342927 | 1470.722141 | 1010.530176 | 791.8745364 | 954.063652 | 888.9547536 | 656.934518 | 886.3152369 | 885.4366517 | 0.015438152 | 0.466374487 | YBR115C | LYS2 | n |
| W303.asm02.HP0_G0003120 | 9693.958633 | 7303.158748 | 8499.59472 | 87158.73123 | 93462.41075 | 79904.53991 | 70165.15017 | 63827.27577 | 61329.53937 | 3.61E-134 | -3.353039037 | YPR080W | TEF1 | y |
| W303.asm02.HP0_G0003130 | 696.9259331 | 571.1542295 | 696.8162625 | 343.4050508 | 348.9829492 | 372.0574376 | 352.1320471 | 357.0418367 | 297.8752204 | 3.01E-08 | 0.883555263 | YBR119W | MUD1 | n |
| W303.asm02.HP0_G0003140 | 888.847122 | 1006.183368 | 963.6196842 | 1399.886312 | 1192.911298 | 1426.672802 | 1134.016646 | 980.1717631 | 763.6251355 | 0.004321975 | -0.492431445 | YBR120C | CBP6 | n |
| W303.asm02.HP0_G0003150 | 8645.177187 | 8656.794271 | 9438.781856 | 8255.340683 | 8443.26428 | 7889.247126 | 6395.17234 | 6168.405716 | 5911.44123 | 0.237630102 | 0.121203863 | YBR121C | GRS1 | n |
| W303.asm02.HP0_G0003160 | 1984.154917 | 2073.289853 | 2066.01624 | 1520.515848 | 1606.913884 | 1576.944176 | 1719.010209 | 1373.982136 | 1268.272846 | 0.007506158 | 0.380535755 | YBR122C | MRPL36 | n |
| W303.asm02.HP0_G0003170 | 451.693303 | 467.3945445 | 390.9207644 | 550.6154639 | 589.1575264 | 605.6117415 | 552.8094503 | 559.2687849 | 555.8290195 | 0.003326488 | -0.413703039 | YBR123C | TFC1 | n |
| W303.asm02.HP0_G0003180 | 786.101435 | 741.5485746 | 692.9070549 | 452.360761 | 496.2723308 | 467.1086078 | 701.4243197 | 639.5790083 | 665.3570215 | 7.24E-07 | 0.650930594 | YBR125C | PTC4 | n |
| W303.asm02.HP0_G0003190 | 1923.089084 | 856.7313442 | 1169.830387 | 1373.620203 | 1231.392307 | 1322.56914 | 1187.972363 | 1304.315436 | 1417.722269 | 0.975958725 | 0.007847007 | YBR126C | TPS1 | n |
| W303.asm02.HP0_G0003210 | 7634.1984 | 4656.810818 | 7137.235856 | 5054.766697 | 4295.276831 | 4866.619914 | 4190.560679 | 4266.117773 | 4004.425643 | 0.015057148 | 0.450397982 | YBR127C | VMA2 | n |
| W303.asm02.HP0_G0003220 | 299.5133704 | 181.8174297 | 300.0316867 | 29.18456523 | 27.86555868 | 30.77847416 | 71.94095587 | 74.50466511 | 99.29174014 | 4.06E-24 | 3.149133034 | YBR128C | ATG14 | n |
| W303.asm02.HP0_G0003230 | 913.0795953 | 975.7218087 | 859.0483798 | 268.4980001 | 290.5979691 | 245.322544 | 342.6661319 | 246.7362286 | 260.0010515 | 9.13E-25 | 1.775142709 | YBR129C | OPY1 | n |
| W303.asm02.HP0_G0003240 | 1097.246393 | 1092.808476 | 994.8933454 | 809.3852756 | 882.4093582 | 773.0828509 | 916.3005948 | 852.4494801 | 920.2399421 | 0.002953893 | 0.371446779 | YBR130C | SHE3 | n |
| W303.asm02.HP0_G0003250 | 245.2326302 | 235.1251578 | 221.8475338 | 311.3020291 | 311.828871 | 324.0792279 | 472.3491708 | 499.2780156 | 562.9944028 | 0.006891216 | -0.432293136 | YBR131W | CCZ1 | n |
| W303.asm02.HP0_G0003260 | 211.3071675 | 190.3847432 | 173.9597402 | 113.8198044 | 88.90440151 | 122.2086474 | 243.2740218 | 423.8057574 | 313.2296132 | 0.001652897 | 0.818561823 | YBR132C | AGP2 | n |
| W303.asm02.HP0_G0003270 | 796.7637233 | 706.3273971 | 702.680074 | 1556.510145 | 1766.145648 | 1606.817401 | 1190.812138 | 1548.148885 | 1498.588738 | 2.37E-15 | -1.15960354 | YBR133C | HSL7 | n |
| W303.asm02.HP0_G0003280 | 930.5269761 | 1117.558442 | 1033.985422 | 747.1248698 | 765.639398 | 682.5579269 | 761.0595857 | 716.0188595 | 639.7663669 | 0.00051973 | 0.490517743 | YBR135W | CKS1 | n |
| W303.asm02.HP0_G0003290 | 237.4782387 | 392.1925709 | 260.9396102 | 339.5137755 | 467.0798408 | 362.099696 | 315.2149777 | 402.5187102 | 357.2455393 | 0.106675233 | -0.38926504 | YBR136W | MEC1 | n |
| W303.asm02.HP0_G0003310 | 624.2285131 | 504.5195694 | 648.9284689 | 211.1016885 | 238.8476458 | 230.8385562 | 318.0547522 | 302.8566257 | 459.608158 | 2.24E-11 | 1.385774077 | YBR138C | | n |
| W303.asm02.HP0_G0003320 | 1658.470475 | 1026.173766 | 1444.452224 | 295.7369276 | 256.0977536 | 285.1535106 | 567.9549147 | 488.634492 | 642.8372454 | 4.11E-30 | 2.300762208 | YBR139W | ATG42 | n |
| W303.asm02.HP0_G0003330 | 855.8909582 | 1110.894976 | 937.2325326 | 1254.936305 | 1257.930935 | 1299.032659 | 1269.379234 | 1195.945014 | 1000.082785 | 0.010873041 | -0.392492768 | YBR140C | IRA1 | n |
| W303.asm02.HP0_G0003340 | 367.3642958 | 644.4523556 | 430.9901427 | 641.0876161 | 670.1003397 | 663.5476929 | 470.4559877 | 531.2085863 | 380.7889416 | 0.047540006 | -0.45245845 | YBR141C | BMT2 | n |
| W303.asm02.HP0_G0003350 | 1706.935422 | 2514.030533 | 1955.581124 | 4912.735146 | 4106.852577 | 4729.927279 | 1905.488739 | 2463.491914 | 1914.18097 | 8.24E-11 | -1.154651156 | YBR142W | MAK5 | n |
| W303.asm02.HP0_G0003360 | 8426.115628 | 9035.65991 | 8951.108203 | 8088.988662 | 8525.534025 | 8051.28674 | 6377.187101 | 6185.822391 | 5515.297895 | 0.371848037 | 0.098843969 | YBR143C | SUP45 | n |
| W303.asm02.HP0_G0003380 | 3420.655936 | 3957.146887 | 3570.083881 | 3297.85587 | 3377.040326 | 3250.750021 | 3030.986062 | 3145.645017 | 2649.144572 | 0.254903654 | 0.14156961 | YBR146W | MRPS9 | n |
| W303.asm02.HP0_G0003420 | 3678.489452 | 2694.896039 | 3536.855616 | 169.2704783 | 200.3666362 | 153.8923708 | 238.5410642 | 180.939901 | 223.1505088 | 5.25E-118 | 4.248647887 | YBR149W | ARA1 | n |
| W303.asm02.HP0_G0003440 | 2333.102533 | 2382.665061 | 2391.457776 | 527.2678117 | 595.7921832 | 525.9498084 | 935.2324263 | 780.8475941 | 705.2784428 | 2.08E-50 | 2.109592731 | YBR151W | APD1 | n |
| W303.asm02.HP0_G0003460 | 832.6277838 | 930.0294703 | 872.7306065 | 3324.121979 | 2332.745341 | 3013.57472 | 1962.28423 | 1369.144171 | 1501.659616 | 2.79E-21 | -1.718588249 | YBR153W | RIB7 | n |
| W303.asm02.HP0_G0003470 | 3132.774153 | 3778.185228 | 3427.397802 | 4582.949559 | 4851.261073 | 4637.591856 | 2796.231364 | 2940.515289 | 2400.403408 | 0.000581805 | -0.44466111 | YBR154C | RPB5 | n |
| W303.asm02.HP0_G0003480 | 1013.886684 | 1104.23151 | 1053.53146 | 2883.435044 | 3036.018965 | 2981.890996 | 1826.921642 | 1635.23226 | 1381.895352 | 4.05E-29 | -1.488655438 | YBR155W | CNS1 | n |
| W303.asm02.HP0_G0003490 | 220.0308579 | 227.5097681 | 249.2119873 | 361.8886088 | 408.6948607 | 335.847468 | 301.0161048 | 358.9770228 | 468.8207936 | 0.001310514 | -0.665217749 | YBR156C | SLI15 | n |

| | | | | | | | | | | | | | | |
|---|---|---|---|---|---|---|---|---|---|---|---|---|---|---|
| W303.asm02.HP0_G0003500 | 896.6015134 | 941.4525549 | 816.0470957 | 126.4664493 | 148.616313 | 127.6401428 | 186.4785303 | 141.2685858 | 163.7801899 | 4.54E-55 | 2.726175497 | YBR157C | ICS2 | n |
| W303.asm02.HP0_G0003510 | 3112.418875 | 3742.964051 | 3227.05091 | 1393.07658 | 1368.066238 | 1512.67148 | 1588.380578 | 1463.000697 | 1250.8712 | 4.52E-19 | 1.23782239 | YBR158W | AMN1 | n |
| W303.asm02.HP0_G0003520 | 3229.704046 | 2846.25191 | 3364.85048 | 2618.82832 | 2575.573781 | 2600.781066 | 3094.407694 | 3091.459806 | 2818.042893 | 0.016681711 | 0.276222442 | YBR159W | IFA38 | n |
| W303.asm02.HP0_G0003530 | 2036.497059 | 1436.452887 | 1849.055216 | 1446.581616 | 1267.219454 | 1472.840513 | 1238.141714 | 1118.53757 | 1224.25692 | 0.032384894 | 0.345393564 | YBR160W | CDC28 | n |
| W303.asm02.HP0_G0003550 | 10282.32309 | 8047.563093 | 10764.98055 | 5963.379494 | 5644.76603 | 5751.95367 | 5984.351618 | 5769.757378 | 6061.914279 | 2.03E-08 | 0.744917289 | YBR162C | TOS1 | n |
| W303.asm02.HP0_G0003560 | 1292.075478 | 1558.299123 | 1426.86079 | 818.1406451 | 733.7930453 | 699.7576625 | 1056.396141 | 835.0328051 | 968.3503729 | 4.26E-10 | 0.925577276 | YBR162W- | YSY6 | n |
| W303.asm02.HP0_G0003570 | 831.6584849 | 760.5870489 | 820.9336052 | 548.6698262 | 701.9466925 | 563.0650272 | 406.087764 | 656.9956833 | 471.8916722 | 0.042692904 | 0.413766559 | YBR163W | EXO5 | n |
| W303.asm02.HP0_G0003580 | 2424.216633 | 2409.318925 | 2416.867626 | 2016.653457 | 2012.954882 | 1997.885073 | 2177.160506 | 2015.49633 | 1800.558463 | 0.020177262 | 0.266530566 | YBR164C | ARL1 | n |
| W303.asm02.HP0_G0003590 | 1213.562265 | 816.7505482 | 1144.420538 | 319.0845798 | 313.1558023 | 353.9524528 | 234.7546981 | 243.8334495 | 289.6862109 | 4.95E-19 | 1.685381346 | YBR165W | UBS1 | n |
| W303.asm02.HP0_G0003600 | 2447.479807 | 2115.174496 | 2506.779402 | 997.1393119 | 948.7559265 | 957.7536958 | 865.1846534 | 975.3337979 | 981.6575133 | 7.55E-25 | 1.283409245 | YBR166C | TYR1 | n |
| W303.asm02.HP0_G0003610 | 841.3514742 | 731.0774137 | 727.1126218 | 319.0845798 | 351.6368119 | 314.1214863 | 401.3548064 | 252.5417869 | 268.190061 | 8.23E-10 | 1.224804771 | YBR167C | POP7 | n |
| W303.asm02.HP0_G0003620 | 825.8426913 | 608.2792544 | 762.2954906 | 87.55369568 | 126.0584797 | 103.1984134 | 166.6001083 | 145.138958 | 200.6307326 | 3.59E-38 | 2.801670944 | YBR168W | PEX32 | n |
| W303.asm02.HP0_G0003640 | 865.5839476 | 643.5004319 | 699.7481683 | 1015.62287 | 817.3897213 | 1018.405395 | 1090.473436 | 1157.241292 | 1495.517859 | 0.051537428 | -0.369616736 | YBR170C | NPL4 | n |
| W303.asm02.HP0_G0003650 | 1886.255724 | 1675.38574 | 1864.692046 | 1313.305435 | 1285.796493 | 1261.91744 | 1553.356692 | 1220.13484 | 1100.398151 | 0.00146183 | 0.490993593 | YBR171W | SEC66 | n |
| W303.asm02.HP0_G0003670 | 2867.186245 | 2473.097814 | 3068.728001 | 1534.135312 | 1470.239953 | 1557.933942 | 2196.092337 | 1839.394395 | 1501.659616 | 5.78E-08 | 0.881914438 | YBR173C | UMP1 | n |
| W303.asm02.HP0_G0003680 | 894.6629156 | 660.6350588 | 829.7293224 | 483.4909639 | 510.8685758 | 469.8243555 | 566.0617317 | 456.7039212 | 403.3087177 | 0.000118943 | 0.704623484 | YBR175W | SWD3 | n |
| W303.asm02.HP0_G0003690 | 780.2856414 | 559.7311449 | 803.3421708 | 633.3050654 | 740.4277021 | 659.0214467 | 589.7265198 | 627.0002986 | 492.3641959 | 0.698515502 | 0.077351528 | YBR176W | ECM31 | n |
| W303.asm02.HP0_G0003700 | 1917.27329 | 1181.337331 | 1554.88734 | 2977.798472 | 2444.207576 | 2903.134312 | 2379.731093 | 2419.950227 | 1959.220522 | 9.31E-06 | -0.839533947 | YBR177C | EHT1 | n |
| W303.asm02.HP0_G0003710 | 945.0664601 | 855.7794205 | 1002.711761 | 967.9547466 | 940.7943383 | 955.0379481 | 721.3027417 | 991.7828798 | 992.9174014 | 0.860084996 | -0.030872332 | YBR179C | FZO1 | n |
| W303.asm02.HP0_G0003720 | 94.02199654 | 93.28852415 | 91.86637963 | 2121.717892 | 1634.779443 | 2270.365094 | 4709.292835 | 5606.234152 | 9518.699913 | 7.54E-69 | -4.43258667 | YBR180W | DTR1 | n |
| W303.asm02.HP0_G0003730 | 7605.119432 | 10060.88175 | 7822.324496 | 32164.30933 | 35413.14429 | 30314.0813 | 26726.0651 | 22286.5708 | 20116.30183 | 1.40E-34 | -1.941305413 | YPL090C | RPS6A | y |
| W303.asm02.HP0_G0003780 | 1376.404485 | 1373.625922 | 1472.79398 | 1704.378609 | 1685.202835 | 1774.28851 | 1892.236458 | 1739.73231 | 1457.64369 | 0.031696726 | -0.290488438 | YBR185C | MBA1 | n |
| W303.asm02.HP0_G0003790 | 258.8028152 | 232.2693867 | 258.9850064 | 31.13020291 | 21.23090185 | 32.58897264 | 20.82501354 | 39.67131519 | 24.56702849 | 3.45E-24 | 3.123365713 | YBR186W | PCH2 | n |
| W303.asm02.HP0_G0003800 | 2000.632999 | 1600.183766 | 1911.602538 | 1205.322544 | 1277.834905 | 1200.360492 | 1941.459217 | 1802.625859 | 1645.990909 | 1.81E-05 | 0.582034752 | YBR187W | GDT1 | n |
| W303.asm02.HP0_G0003820 | 12056.14013 | 14044.6825 | 12334.52742 | 14454.14234 | 16289.40945 | 13509.03441 | 15412.4032 | 10379.37068 | 9048.855493 | 0.317571225 | -0.203270462 | YBR189W | RPS9B | y |
| W303.asm02.HP0_G0003840 | 1496.597553 | 1640.164562 | 1594.956719 | 1545.809138 | 1176.988121 | 1547.070951 | 970.2563127 | 1091.444964 | 795.3575473 | 0.404834645 | 0.147000582 | YBR192W | RIM2 | n |
| W303.asm02.HP0_G0003850 | 940.2199654 | 745.3562695 | 1035.940026 | 436.7956595 | 463.0990467 | 437.2353829 | 599.192435 | 492.5048642 | 543.5455053 | 7.93E-10 | 1.025959089 | YBR193C | MED8 | n |
| W303.asm02.HP0_G0003860 | 512.7591358 | 413.1348927 | 503.3104842 | 429.0131088 | 384.8100961 | 442.6668783 | 603.9253927 | 407.3566755 | 417.6394843 | 0.378450293 | 0.184345551 | YBR194W | AIM4 | n |
| W303.asm02.HP0_G0003870 | 714.3733139 | 546.4042129 | 629.3824307 | 559.3708335 | 656.8310261 | 614.6642339 | 620.0174486 | 608.6160306 | 571.1834123 | 0.778703425 | 0.047144033 | YBR195C | MSI1 | n |
| W303.asm02.HP0_G0003880 | 13096.19789 | 11149.88248 | 12832.95139 | 16728.59279 | 12592.57866 | 15887.12416 | 9364.629952 | 9832.680609 | 9041.69011 | 0.0461043 | -0.286136946 | YBR196C | PGI1 | n |
| W303.asm02.HP0_G0003900 | 323.7458438 | 308.4232839 | 297.0997809 | 168.2976595 | 163.212558 | 173.8078541 | 143.8819117 | 220.6112162 | 236.4576492 | 6.59E-05 | 0.87784587 | YBR197C | | n |
| W303.asm02.HP0_G0003910 | 761.8689617 | 794.8563027 | 637.200846 | 1840.573247 | 1914.761961 | 1860.287188 | 1529.691904 | 2281.58442 | 2320.560566 | 4.64E-14 | -1.355758089 | YBR198C | TAF5 | n |
| W303.asm02.HP0_G0003930 | 689.1715417 | 653.9715928 | 609.8363925 | 1365.837653 | 1482.182336 | 1465.598519 | 892.6358077 | 1198.847793 | 1082.996506 | 5.90E-15 | -1.142973757 | YBR200W | BEM1 | n |
| W303.asm02.HP0_G0003950 | 631.0136057 | 710.135092 | 604.9498829 | 231.5308841 | 218.9436754 | 197.3443343 | 444.8980165 | 387.0372214 | 416.6158581 | 4.19E-24 | 1.588316883 | YBR201W | DER1 | n |
| W303.asm02.HP0_G0003960 | 706.6189225 | 680.6254568 | 595.1768638 | 2273.477631 | 2303.552851 | 2287.564829 | 1644.229478 | 2210.950127 | 1932.606241 | 3.59E-35 | -1.791812473 | YBR202W | MCM7 | n |
| W303.asm02.HP0_G0003970 | 299.5133704 | 414.0868164 | 320.5550268 | 79.77114495 | 100.8467838 | 82.37768083 | 168.4932914 | 195.4537968 | 195.5126017 | 1.76E-19 | 1.98348702 | YBR203W | COS111 | n |
| W303.asm02.HP0_G0003980 | 426.4915307 | 355.067546 | 377.2385376 | 76.85268843 | 104.8275779 | 76.94618539 | 194.9978541 | 96.75930534 | 141.2604138 | 1.66E-15 | 2.171768136 | YBR204C | LDH1 | n |
| W303.asm02.HP0_G0003990 | 1725.352102 | 1533.549106 | 1837.327593 | 2549.758182 | 2506.57335 | 2657.811768 | 3865.879786 | 3079.848689 | 3054.500542 | 2.00E-05 | -0.598282229 | YBR205W | KTR3 | n |
| W303.asm02.HP0_G0004000 | 1066.228827 | 1090.904578 | 1127.806405 | 1266.610131 | 1151.776425 | 1283.643422 | 1520.225988 | 1440.746057 | 1569.218945 | 0.138113721 | -0.173624675 | YBR207W | FTH1 | n |
| W303.asm02.HP0_G0004010 | 301.4519683 | 384.5771812 | 256.0531007 | 879.4282321 | 962.0252402 | 770.3671032 | 1055.44955 | 991.7828798 | 896.6965398 | 5.06E-16 | -1.469651065 | YBR208C | DUR12 | n |
| W303.asm02.HP0_G0004030 | 1001.285798 | 734.8851086 | 835.5931339 | 122.5751739 | 156.5779012 | 139.4083829 | 263.1524438 | 190.6158315 | 227.2450135 | 4.80E-39 | 2.623787712 | YBR211C | AME1 | n |
| W303.asm02.HP0_G0004040 | 941.1892644 | 809.1351584 | 782.8188307 | 199.4278624 | 254.7708222 | 207.3020759 | 411.7673132 | 319.3057076 | 302.9933513 | 9.85E-26 | 1.942015748 | YBR212W | NGR1 | n |
| W303.asm02.HP0_G0004050 | 143.4562421 | 158.0193368 | 116.2989274 | 403.7198189 | 449.829733 | 389.2571732 | 731.7152485 | 689.8938471 | 729.8454713 | 1.78E-20 | -1.570010536 | YBR213W | MET8 | n |
| W303.asm02.HP0_G0004060 | 429.3994275 | 222.7501495 | 262.8942141 | 730.5869495 | 686.0235161 | 753.1673676 | 588.7799283 | 673.4447652 | 883.3893994 | 5.04E-07 | -1.246103708 | YBR214W | SDS24 | n |
| W303.asm02.HP0_G0004070 | 372.2107904 | 501.6637982 | 390.9207644 | 701.4023842 | 666.1195456 | 651.7794527 | 775.2584586 | 699.5697776 | 768.7432664 | 2.11E-05 | -0.674759323 | YBR215W | HPC2 | n |
| W303.asm02.HP0_G0004090 | 529.2372177 | 493.0964848 | 502.3331823 | 67.12450002 | 57.05804873 | 62.46219755 | 97.49892703 | 92.88893313 | 108.5043758 | 5.27E-54 | 3.023982308 | YBR217W | ATG12 | n |
| W303.asm02.HP0_G0004110 | 688.2022427 | 725.3658714 | 657.7241861 | 413.4480074 | 443.1950762 | 402.8359118 | 769.5789095 | 873.7365272 | 1038.98058 | 4.58E-06 | 0.71921827 | YBR220C | | n |
| W303.asm02.HP0_G0004120 | 4880.42013 | 3656.338992 | 4898.237178 | 13860.72284 | 12759.77201 | 13451.09846 | 4688.467821 | 9670.124976 | 9827.835021 | 9.87E-11 | -1.576608117 | YBR221C | PDB1 | n |
| W303.asm02.HP0_G0004140 | 126.9781603 | 139.9327862 | 103.5940026 | 208.1832319 | 144.6355189 | 183.7655957 | 206.3569524 | 185.7778663 | 321.4186227 | 0.056521422 | -0.537661084 | YBR223C | TDP1 | n |
| W303.asm02.HP0_G0004150 | 185.1360963 | 245.5963187 | 175.914344 | 258.7698117 | 171.1741462 | 252.5645379 | 133.469405 | 157.7176677 | 190.3944708 | 0.513414779 | -0.174366289 | YBR225W | | n |
| W303.asm02.HP0_G0004160 | 820.9961966 | 781.5293707 | 871.7533046 | 827.8688336 | 761.658604 | 848.2185378 | 695.7447705 | 803.1022343 | 902.8382969 | 0.897825572 | 0.020376994 | YBR227C | MCX1 | n |
| W303.asm02.HP0_G0004170 | 567.0398761 | 520.7022725 | 551.1982778 | 272.3892754 | 260.0785477 | 254.3750364 | 338.8797658 | 268.9908689 | 283.5444538 | 2.19E-11 | 1.058495183 | YBR228W | SLX1 | n |
| W303.asm02.HP0_G0004180 | 627.1364099 | 620.6542627 | 696.8162625 | 1944.664863 | 1914.761961 | 1949.001614 | 1521.17258 | 2274.811269 | 1753.471658 | 4.42E-22 | -1.57878068 | YBR229C | ROT2 | n |
| W303.asm02.HP0_G0004190 | 2028.742668 | 1546.876038 | 1905.738726 | 519.485261 | 331.7328414 | 478.8768479 | 543.3435351 | 372.5233256 | 466.7735413 | 1.68E-22 | 2.039750434 | YBR230C | OM14 | n |
| W303.asm02.HP0_G0004210 | 484.6494667 | 388.384876 | 368.4428204 | 279.1990073 | 345.0021551 | 315.9319847 | 336.9865827 | 313.5001493 | 318.3477441 | 0.024783285 | 0.403536172 | YBR231C | SWC5 | n |
| W303.asm02.HP0_G0004220 | 430.3687264 | 343.6444614 | 392.8753682 | 214.020145 | 220.2706067 | 200.060082 | 313.3217946 | 331.8844173 | 486.2224388 | 3.92E-05 | 0.880505478 | YBR233W | PBP2 | n |
| W303.asm02.HP0_G0004230 | 505.9740432 | 602.5677121 | 522.8565224 | 65.17886234 | 91.55826424 | 61.55694831 | 135.362588 | 81.27781649 | 91.10273064 | 1.08E-31 | 2.915067437 | YBR233W- | DAD3 | n |
| W303.asm02.HP0_G0004240 | 3575.743765 | 2663.482557 | 3438.148123 | 3901.976371 | 3500.444943 | 3967.707419 | 3395.423799 | 2926.001394 | 2629.695674 | 0.154480838 | -0.23281745 | YBR234C | ARC40 | n |
| W303.asm02.HP0_G0004250 | 411.9520467 | 493.0964848 | 405.5802931 | 425.1218334 | 448.5028016 | 458.0561154 | 600.1390266 | 776.0096289 | 800.4756782 | 0.902195849 | -0.022757527 | YBR235W | VHC1 | n |
| W303.asm02.HP0_G0004260 | 1495.628254 | 1313.654728 | 1479.635093 | 1183.920529 | 1234.04617 | 1212.128732 | 941.8585669 | 955.9819368 | 878.2712684 | 0.043171256 | 0.240833739 | YBR236C | ABD1 | n |
| W303.asm02.HP0_G0004270 | 468.1713848 | 651.1158216 | 463.2411058 | 668.3265437 | 658.1579574 | 614.6642339 | 528.1980707 | 697.6345915 | 715.5147047 | 0.137041069 | -0.294213176 | YBR237W | PRP5 | n |
| W303.asm02.HP0_G0004280 | 678.5092534 | 1359.347066 | 792.5918498 | 7825.354756 | 5887.59447 | 7562.45215 | 2357.959488 | 3866.501842 | 3672.770759 | 4.93E-29 | -2.910081298 | YBR238C | | n |
| W303.asm02.HP0_G0004290 | 166.7194165 | 283.6732673 | 161.2548153 | 248.0688044 | 298.5595573 | 249.8487902 | 408.9275386 | 398.648338 | 511.8130935 | 0.141251229 | -0.37839791 | YBR239C | ERT1 | n |

| | | | | | | | | | | | | | | |
|---|---|---|---|---|---|---|---|---|---|---|---|---|---|---|
| W303.asm02.HP0_G0004300 | 157.9957261 | 152.3077945 | 141.7087771 | 33.07584059 | 35.82714688 | 35.30472036 | 25.55797116 | 107.4028289 | 24.56702849 | 2.31E-05 | 2.117650008 | YBR240C | THI2 | n |
| W303.asm02.HP0_G0004310 | 270.4344024 | 212.2789886 | 224.7794395 | 130.3577247 | 98.19292107 | 119.4928997 | 128.7364473 | 191.5834246 | 239.5285278 | 0.00014291 | 1.018944131 | YBR241C | VVS1 | n |
| W303.asm02.HP0_G0004320 | 1643.930991 | 1681.097282 | 1613.525455 | 2006.925269 | 2357.957037 | 2067.589264 | 1931.993302 | 1792.949928 | 1489.376102 | 0.006678624 | -0.38062652 | YBR242W | | n |
| W303.asm02.HP0_G0004330 | 955.7287484 | 828.1736327 | 1081.873215 | 1076.910457 | 962.0252402 | 1026.552638 | 1765.393193 | 1618.783178 | 1720.71562 | 0.510738399 | -0.09805385 | YBR243C | ALG7 | n |
| W303.asm02.HP0_G0004350 | 548.6231963 | 618.7504153 | 450.536181 | 3084.808544 | 2905.979691 | 3112.246887 | 2158.228676 | 2825.371716 | 2551.900084 | 1.55E-56 | -2.492183871 | YBR245C | ISW1 | n |
| W303.asm02.HP0_G0004360 | 614.5355238 | 534.0292046 | 626.4505249 | 724.7500364 | 652.850232 | 780.3248448 | 576.4742385 | 534.1113655 | 524.0966077 | 0.053145023 | -0.283761651 | YBR246W | RRT2 | n |
| W303.asm02.HP0_G0004370 | 1748.615276 | 1917.174364 | 1769.893761 | 5284.351943 | 4447.873938 | 5126.426446 | 2311.576503 | 2647.334594 | 2411.663297 | 3.57E-33 | -1.451109191 | YBR247C | ENP1 | n |
| W303.asm02.HP0_G0004380 | 1280.443891 | 1107.087281 | 1234.332314 | 1951.474595 | 1723.683844 | 1894.686659 | 1973.643329 | 1676.838762 | 1896.779324 | 1.27E-06 | -0.621579355 | YBR248C | HIS7 | n |
| W303.asm02.HP0_G0004390 | 8570.541169 | 7607.774337 | 9096.726188 | 6721.205371 | 6362.635899 | 6759.496075 | 5246.956821 | 4409.321544 | 4601.19971 | 0.00484105 | 0.348939695 | YBR249C | ARO4 | n |
| W303.asm02.HP0_G0004400 | 260.7414131 | 160.875108 | 215.0064204 | 81.71678263 | 69.00043102 | 56.12545288 | 275.4581336 | 134.4954344 | 486.2224388 | 5.16E-05 | 1.622618814 | YBR250W | SPO23 | n |
| W303.asm02.HP0_G0004410 | 1563.47918 | 1135.644993 | 1455.202545 | 2941.804175 | 2773.286554 | 2799.935899 | 2728.076774 | 2818.598565 | 2738.20005 | 1.10E-13 | -1.035577647 | YBR251W | MRPS5 | n |
| W303.asm02.HP0_G0004420 | 894.6629156 | 730.12549 | 995.8706473 | 286.981558 | 372.8677138 | 347.6157081 | 491.2810012 | 358.0094298 | 378.7416892 | 1.52E-12 | 1.380995619 | YBR253W | SRB6 | n |
| W303.asm02.HP0_G0004430 | 933.4348729 | 883.3852083 | 993.9160435 | 357.0245146 | 370.213851 | 339.468465 | 520.6253385 | 384.1344422 | 356.2219131 | 8.16E-16 | 1.39882327 | YBR254C | TRS20 | n |
| W303.asm02.HP0_G0004450 | 302.4212672 | 371.2502492 | 288.3040637 | 244.177529 | 265.3862732 | 237.1753009 | 431.6457352 | 317.3705215 | 276.3790705 | 0.096281435 | 0.367157446 | YBR255C-[illegible] | RCF3 | n |
| W303.asm02.HP0_G0004460 | 3229.704046 | 2848.155758 | 3321.849195 | 764.6356089 | 609.0614969 | 688.8946716 | 874.6505687 | 618.2919611 | 585.514179 | 1.08E-34 | 2.186912537 | YBR256C | RIB5 | n |
| W303.asm02.HP0_G0004470 | 773.5005489 | 895.7602166 | 811.1605861 | 340.4865943 | 388.7908902 | 399.2149148 | 398.5150318 | 384.1344422 | 454.490027 | 2.99E-14 | 1.136820257 | YBR257W | POP4 | n |
| W303.asm02.HP0_G0004480 | 456.5397976 | 362.6829357 | 517.9700128 | 216.9386015 | 290.5979691 | 240.7962978 | 277.3513167 | 203.1945412 | 191.418097 | 0.000185178 | 0.840588521 | YBR258C | SHG1 | n |
| W303.asm02.HP0_G0004500 | 482.7108688 | 376.9617915 | 440.7631619 | 1868.784993 | 1957.223765 | 1867.529182 | 1466.270272 | 2146.121393 | 1978.669419 | 1.66E-35 | -2.130298851 | YBR260C | RGD1 | n |
| W303.asm02.HP0_G0004510 | 1132.141154 | 749.1639643 | 1131.715613 | 1140.143681 | 1061.545093 | 1058.236361 | 1322.38836 | 826.3244676 | 1012.366299 | 0.620697674 | -0.113802646 | YBR261C | TAE1 | n |
| W303.asm02.HP0_G0004520 | 1039.088457 | 1392.664396 | 1081.873215 | 545.7513697 | 532.0994777 | 508.7500728 | 792.2971061 | 619.2595542 | 523.0729815 | 1.47E-09 | 1.147258173 | YBR262C | MIC12 | n |
| W303.asm02.HP0_G0004530 | 3545.695498 | 2335.068875 | 3439.125425 | 5660.832835 | 5229.436512 | 5434.211187 | 3816.657027 | 4296.113157 | 3777.18063 | 1.49E-06 | -0.808756388 | YBR263W | SHM1 | n |
| W303.asm02.HP0_G0004540 | 617.4434206 | 481.6734002 | 666.5199033 | 140.0859131 | 195.0589108 | 154.79762 | 344.5593149 | 203.1945412 | 246.6939111 | 1.65E-14 | 1.854304567 | YBR264C | YPT10 | n |
| W303.asm02.HP0_G0004550 | 1582.865158 | 1057.587248 | 1464.975565 | 667.3537248 | 635.6001242 | 628.2429725 | 963.630172 | 860.1902245 | 865.9877542 | 1.68E-10 | 1.08785553 | YBR265W | TSC10 | n |
| W303.asm02.HP0_G0004560 | 940.2199654 | 1049.019935 | 896.1858524 | 5046.984146 | 4100.21792 | 4890.156394 | 2082.501354 | 2681.200351 | 2262.213873 | 5.27E-54 | -2.282670151 | YBR267W | REI1 | n |
| W303.asm02.HP0_G0004570 | 1291.106179 | 1320.318194 | 1449.338734 | 644.0060726 | 662.1387515 | 610.1379877 | 814.0687111 | 554.4308196 | 565.0416552 | 3.84E-10 | 1.08398913 | YBR268W | MRPL37 | n |
| W303.asm02.HP0_G0004580 | 1542.154603 | 1281.289321 | 1533.386698 | 403.7198189 | 427.2718998 | 402.8359118 | 637.056096 | 456.7039212 | 499.5295792 | 2.67E-28 | 1.820966806 | YBR269C | SDH8 | n |
| W303.asm02.HP0_G0004600 | 421.645036 | 511.1830354 | 378.2158396 | 563.2621088 | 591.8113891 | 587.5067567 | 468.5628047 | 567.9771224 | 411.4977272 | 0.025985247 | -0.410093794 | YBR271W | EFM2 | n |
| W303.asm02.HP0_G0004610 | 1098.215692 | 984.2891221 | 1225.536596 | 584.6641233 | 571.9074186 | 542.2442947 | 556.5958164 | 523.4678419 | 490.3169436 | 6.47E-12 | 0.961743453 | YBR272C | HSM3 | n |
| W303.asm02.HP0_G0004620 | 1119.540268 | 809.1351584 | 1013.462082 | 331.7312247 | 306.5211455 | 344.8999604 | 341.7195404 | 353.1714645 | 466.7735413 | 3.07E-17 | 1.579804447 | YBR273C | UBX7 | n |
| W303.asm02.HP0_G0004630 | 789.9786307 | 792.0005315 | 802.3648689 | 375.5080726 | 390.1178215 | 337.6579665 | 379.5832013 | 421.8705713 | 495.4350745 | 2.80E-13 | 1.113850173 | YBR274W | CHK1 | n |
| W303.asm02.HP0_G0004640 | 646.5223886 | 858.6351916 | 710.4984893 | 853.1621234 | 878.4285641 | 826.4925561 | 585.9401537 | 752.7873956 | 665.3570215 | 0.219124619 | -0.206811531 | YBR275C | RIF1 | n |
| W303.asm02.HP0_G0004650 | 548.6231963 | 772.0101335 | 591.2676562 | 753.9346016 | 690.0043102 | 769.4618539 | 562.2753656 | 752.7873956 | 795.3575473 | 0.288520447 | -0.211804173 | YBR276C | PPS1 | n |
| W303.asm02.HP0_G0004670 | 1517.92213 | 1257.491229 | 1485.498905 | 1492.304102 | 1609.567747 | 1482.798255 | 1384.8634 | 1444.616429 | 1331.737669 | 0.427062953 | -0.105119156 | YBR279W | PAF1 | n |
| W303.asm02.HP0_G0004680 | 349.916915 | 261.7790218 | 311.7593096 | 115.7654421 | 131.3662052 | 137.5978845 | 229.0751489 | 186.7454593 | 180.1582089 | 1.60E-09 | 1.263394438 | YBR280C | SAF1 | n |
| W303.asm02.HP0_G0004690 | 865.5839476 | 860.5390391 | 964.5969862 | 617.7399639 | 577.2151441 | 668.9791883 | 561.3287741 | 740.2086859 | 685.8295453 | 0.000653705 | 0.528185585 | YBR281C | DUG2 | n |
| W303.asm02.HP0_G0004700 | 1227.13245 | 1516.414479 | 1295.902334 | 1538.026587 | 1521.990277 | 1524.43972 | 1585.540804 | 1348.824716 | 1258.036584 | 0.217529148 | -0.182555955 | YBR282W | MRPL27 | n |
| W303.asm02.HP0_G0004710 | 4123.397663 | 3463.098478 | 4328.470164 | 5292.134494 | 4588.528663 | 5225.098613 | 5599.088868 | 6298.063185 | 6364.907631 | 0.011216023 | -0.342621776 | YBR283C | SSH1 | n |
| W303.asm02.HP0_G0004740 | 8477.488472 | 5883.840487 | 8831.87737 | 3814.422675 | 2932.518318 | 3910.676716 | 2726.183591 | 2787.635587 | 2539.61657 | 4.93E-10 | 1.121390262 | YBR286W | APE3 | n |
| W303.asm02.HP0_G0004750 | 1372.52729 | 1431.693269 | 1325.221391 | 1190.730261 | 1032.352603 | 1153.287532 | 1285.47129 | 1552.019258 | 1503.706869 | 0.029846814 | 0.289391106 | YBR287W | | n |
| W303.asm02.HP0_G0004760 | 1244.579831 | 1264.154695 | 1329.130599 | 1152.790326 | 1127.891661 | 1160.529526 | 1265.592868 | 1385.593253 | 1440.242045 | 0.179689421 | 0.157047482 | YBR288C | APM3 | n |
| W303.asm02.HP0_G0004770 | 387.7195734 | 436.9329856 | 291.2359695 | 1003.949044 | 951.4097893 | 946.890705 | 649.3617858 | 866.9633759 | 978.5866348 | 5.52E-12 | -1.378842013 | YBR289W | SNF5 | n |
| W303.asm02.HP0_G0004780 | 1935.68997 | 1747.731942 | 1878.374273 | 949.4711887 | 944.7751324 | 791.1878357 | 896.4221738 | 1197.8802 | 1090.161889 | 3.63E-12 | 1.051289776 | YBR290W | BSD2 | n |
| W303.asm02.HP0_G0004790 | 764.7768585 | 607.3273307 | 695.8389606 | 353.1332392 | 306.5211455 | 433.6143859 | 456.2571148 | 363.8149881 | 451.4191485 | 1.07E-06 | 0.91586284 | YBR291C | CTP1 | n |
| W303.asm02.HP0_G0004800 | 956.6980473 | 853.8755731 | 928.4368154 | 387.1818987 | 367.5599883 | 395.5939178 | 268.831993 | 592.1669487 | 292.7570895 | 2.43E-06 | 1.251157744 | YBR293W | VBA2 | n |
| W303.asm02.HP0_G0004810 | 210.3378686 | 250.3559373 | 248.2485654 | 250.0144421 | 281.3094495 | 240.7962978 | 385.2627505 | 313.5001493 | 369.5290535 | 0.53255927 | -0.120029803 | YBR294W | SUL1 | n |
| W303.asm02.HP0_G0004830 | 225.8466515 | 216.0866835 | 233.5751567 | 380.3721668 | 439.2142821 | 405.5516595 | 395.6752573 | 434.449281 | 407.4032224 | 2.13E-08 | -0.856970251 | YBR296C | PHO89 | n |
| W303.asm02.HP0_G0004980 | 648.4609865 | 417.8945112 | 480.8325402 | 447.4966668 | 410.021792 | 506.0343251 | 321.8411183 | 345.4307201 | 309.1351085 | 0.383893135 | 0.180511362 | YCL064C | CHA1 | n |
| W303.asm02.HP0_G0004990 | 546.6845985 | 665.3946773 | 586.3811466 | 523.3765364 | 586.5036637 | 548.5810394 | 445.8446081 | 508.9539461 | 486.2224388 | 0.446078917 | 0.118315041 | YCL063W | VAC17 | n |
| W303.asm02.HP0_G0005000 | 307.2677619 | 596.8561698 | 382.1250472 | 696.53829 | 778.9087117 | 777.6090971 | 716.5697841 | 808.9077927 | 682.7586667 | 0.000421945 | -0.808277055 | YCL061C | MRC1 | n |
| W303.asm02.HP0_G0005010 | 2265.251607 | 2286.520765 | 2383.639361 | 4460.374385 | 4584.547869 | 4501.80447 | 2613.539199 | 2921.163428 | 2141.425983 | 6.21E-13 | -0.965849791 | YCL059C | KRR1 | n |
| W303.asm02.HP0_G0005030 | 1848.453066 | 1870.530102 | 2030.833371 | 3774.537102 | 4151.968243 | 3615.565464 | 3013.000823 | 3068.237572 | 2688.042367 | 7.14E-18 | -1.004909318 | YCL057C-[illegible] | MIC10 | n |
| W303.asm02.HP0_G0005040 | 1485.935265 | 1092.808426 | 1187.421822 | 1481.603095 | 1192.911298 | 1439.346291 | 1404.741822 | 1985.500946 | 1460.714569 | 0.523446791 | -0.128141588 | YCL057W | PRD1 | n |
| W303.asm02.HP0_G0005060 | 1389.005372 | 1229.885441 | 1195.240237 | 59.34194929 | 51.75032327 | 49.7887082 | 37.86366098 | 39.67131519 | 47.0868046 | 2.11E-123 | 4.565826557 | YCL055W | KAR4 | n |
| W303.asm02.HP0_G0005070 | 1599.34324 | 2467.386271 | 1830.486479 | 7382.722183 | 6371.924418 | 7127.932515 | 2428.953852 | 3014.052361 | 2581.585244 | 1.53E-25 | -1.824240107 | YCL054W | SPB1 | n |
| W303.asm02.HP0_G0005080 | 1893.040817 | 1717.270383 | 1878.374273 | 458.197674 | 565.2727618 | 507.8448236 | 614.3378994 | 681.1855096 | 786.1449116 | 4.32E-36 | 1.843900493 | YCL052C | PBN1 | n |
| W303.asm02.HP0_G0005100 | 6985.737413 | 5348.859359 | 6021.157074 | 4853.393197 | 3684.888403 | 5000.596801 | 3081.155412 | 3110.811667 | 2590.797879 | 0.009074972 | 0.438706368 | YCL050C | APA1 | n |
| W303.asm02.HP0_G0005130 | 75.60531681 | 94.24044786 | 76.22954906 | 34.04865943 | 25.21169595 | 21.72598176 | 31.23752031 | 53.21761794 | 29.68515942 | 3.42E-05 | 1.602343103 | YCL048W | SPS22 | n |
| W303.asm02.HP0_G0005150 | 1468.487884 | 1229.885441 | 1387.768714 | 1409.6145 | 1337.546817 | 1350.631866 | 1152.001885 | 1531.699804 | 1526.226645 | 0.978603956 | -0.004295313 | YCL045C | EMC1 | n |
| W303.asm02.HP0_G0005160 | 386.7502744 | 267.4905641 | 285.372158 | 891.1020582 | 952.7367206 | 868.1340211 | 880.3301178 | 902.7643189 | 796.3811735 | 1.08E-19 | -1.528726221 | YCL044C | MGR1 | n |
| W303.asm02.HP0_G0005170 | 2626.80011 | 2624.453684 | 2132.47277 | 9166.871937 | 8208.397429 | 8624.309509 | 12438.21263 | 12272.95029 | 12579.34221 | 5.72E-52 | -1.81612714 | YCL043C | PDI1 | n |
| W303.asm02.HP0_G0005180 | 3225.82685 | 1267.962389 | 2270.272339 | 143.9771884 | 171.1741462 | 118.5876504 | 136.3091795 | 149.9769233 | 182.2054613 | 2.38E-42 | 3.967030857 | YCL040W | GLK1 | n |
| W303.asm02.HP0_G0005190 | 671.7241609 | 566.3946109 | 556.0847874 | 178.9986667 | 210.9820872 | 192.8180881 | 253.6865286 | 338.6575687 | 379.7653154 | 1.38E-16 | 1.624860224 | YCL039W | GID7 | n |
| W303.asm02.HP0_G0005200 | 429.3994275 | 323.6540634 | 401.6710854 | 310.3292102 | 298.5595573 | 342.1842127 | 623.8038147 | 673.4447652 | 666.3806477 | 0.102279277 | 0.277661185 | YCL038C | ATG22 | n |

| | | | | | | | | | | | | | | |
|---|---|---|---|---|---|---|---|---|---|---|---|---|---|---|
| W303.asm02.HP0_G0005210 | 43.618452 | 79.96159213 | 39.09207644 | 5693.908675 | 5306.398532 | 5586.29306 | 2960.938289 | 3864.566655 | 4180.489348 | 5.40E-189 | -6.669911358 | YCL037C | SRO9 | n |
| W303.asm02.HP0_G0005230 | 2447.479807 | 2330.309256 | 2435.436362 | 2261.803805 | 2361.937831 | 2169.882428 | 2261.407152 | 1619.750771 | 1549.770047 | 0.615982447 | 0.086674251 | YCL035C | GRX1 | n |
| W303.asm02.HP0_G0005240 | 1539.246706 | 777.7216758 | 1184.489916 | 833.7057466 | 419.3103116 | 774.8933494 | 754.4334451 | 678.2827305 | 639.7663669 | 0.005619738 | 0.786024635 | YCL034W | LSB5 | n |
| W303.asm02.HP0_G0005250 | 1140.864845 | 1120.414213 | 1248.991842 | 401.7741813 | 437.8873507 | 383.8256777 | 455.3105233 | 358.0094298 | 323.4658751 | 8.06E-21 | 1.522368073 | YCL033C | MXR2 | n |
| W303.asm02.HP0_G0005260 | 949.9129548 | 573.0580769 | 805.2967747 | 558.3980146 | 581.1959382 | 537.7180485 | 640.8424621 | 542.819703 | 551.7345148 | 0.01745024 | 0.473524753 | YCL032W | STE50 | n |
| W303.asm02.HP0_G0005270 | 2329.225337 | 3036.636653 | 2392.435078 | 2352.275957 | 2615.381722 | 2348.216528 | 1825.975051 | 1923.57499 | 1641.896404 | 0.58619567 | 0.085140732 | YCL031C | RRP7 | n |
| W303.asm02.HP0_G0005280 | 1841.667974 | 1856.251246 | 1571.501473 | 2574.078653 | 2145.648018 | 2564.571097 | 1909.275105 | 1844.23236 | 2137.331478 | 0.000737637 | -0.467797144 | YCL030C | HIS4 | n |
| W303.asm02.HP0_G0005290 | 440.0617158 | 476.9137816 | 374.3066319 | 717.9403045 | 725.8314571 | 665.3581913 | 586.8867452 | 595.0697279 | 523.0729815 | 2.51E-06 | -0.706952826 | YCL029C | BIK1 | n |
| W303.asm02.HP0_G0005310 | 213.2457654 | 177.0578111 | 135.8449656 | 448.4694856 | 463.0990467 | 494.266085 | 130.6296304 | 111.2732011 | 126.9296472 | 2.98E-12 | -1.418006429 | YCL027W | FUS1 | n |
| W303.asm02.HP0_G0005320 | 408.074851 | 286.5290385 | 364.5336128 | 644.9788915 | 489.637674 | 556.7282826 | 740.2345722 | 548.6252613 | 450.3955223 | 0.003029134 | -0.67654357 | YCL026C-I | HBN1 | n |
| W303.asm02.HP0_G0005340 | 1229.071048 | 2016.17443 | 1114.124179 | 2809.500812 | 4459.81632 | 2904.944811 | 2254.781011 | 3714.589732 | 2890.720352 | 4.73E-06 | -1.222319282 | YCL025C | AGP1 | n |
| W303.asm02.HP0_G0005370 | 965.4217377 | 1265.106618 | 932.3460231 | 165.3792029 | 200.3666362 | 198.2495835 | 413.6604962 | 315.4353354 | 388.9779511 | 7.15E-39 | 2.489286424 | YCL018W | LEU2 | n |
| W303.asm02.HP0_G0005380 | 1260.088613 | 1042.356469 | 1197.194841 | 3411.675675 | 3395.617365 | 3441.75761 | 2624.898298 | 4173.228839 | 4057.654205 | 1.20E-17 | -1.550264977 | YCL017C | NFS1 | n |
| W303.asm02.HP0_G0005390 | 663.0004705 | 558.7792212 | 625.473223 | 331.7312247 | 383.4831647 | 398.3096656 | 470.4559877 | 438.3196532 | 468.8207936 | 1.25E-06 | 0.73066024 | YCL016C | DCC1 | n |
| W303.asm02.HP0_G0005400 | 410.9827478 | 562.586916 | 360.6244052 | 1659.628942 | 1857.703912 | 1790.582997 | 1133.070055 | 1652.648935 | 1502.683242 | 1.10E-23 | -1.991771356 | YCL014W | BUD3 | n |
| W303.asm02.HP0_G0005430 | 365.4256979 | 297.952123 | 289.2813657 | 340.4865943 | 431.2526939 | 347.6157081 | 492.2275928 | 413.1622338 | 391.0252034 | 0.240219981 | -0.229454238 | YCL010C | SGF29 | n |
| W303.asm02.HP0_G0005440 | 1649.746785 | 1496.424081 | 1267.560579 | 2261.803805 | 1637.433305 | 2417.015471 | 2961.88488 | 1758.116578 | 2421.899558 | 0.019411666 | -0.51768315 | YCL009C | ILV6 | n |
| W303.asm02.HP0_G0005450 | 383.8423776 | 264.634793 | 293.1905733 | 54.47785509 | 45.11566644 | 47.07296048 | 14.19887287 | 69.66669985 | 74.72471165 | 6.99E-10 | 2.680957692 | YCL008C | STP22 | n |
| W303.asm02.HP0_G0005470 | 995.4700046 | 709.1831683 | 977.301911 | 88.52651452 | 100.8467838 | 91.43017323 | 156.1876016 | 126.75469 | 146.3785447 | 1.69E-57 | 3.259713571 | YCL005W | LDB16 | n |
| W303.asm02.HP0_G0005480 | 410.9827478 | 258.9232507 | 383.1023491 | 858.9990365 | 928.851956 | 938.7434618 | 534.8242114 | 862.1254106 | 827.0899591 | 1.26E-09 | -1.372699864 | YCL004W | PGS1 | n |
| W303.asm02.HP0_G0005500 | 2093.685696 | 1868.626254 | 2205.770413 | 1792.905124 | 1706.433736 | 1691.00558 | 2073.035439 | 1791.982335 | 1633.707394 | 0.072611681 | 0.248878289 | YCL001W | RER1 | n |
| W303.asm02.HP0_G0005530 | 3477.844573 | 2405.51123 | 3449.875746 | 2887.32632 | 3053.269073 | 2916.713051 | 3112.392933 | 3352.70993 | 3130.24888 | 0.658661427 | 0.075588338 | YCR002C | CDC10 | n |
| W303.asm02.HP0_G0005540 | 1676.887155 | 1536.404877 | 1612.548153 | 2156.73937 | 2027.551127 | 2129.146212 | 2273.712842 | 1926.477769 | 1774.967808 | 0.00321953 | -0.387936912 | YCR003W | MRPL32 | n |
| W303.asm02.HP0_G0005550 | 3028.089868 | 2811.030733 | 2985.657338 | 3266.725668 | 2319.476027 | 3024.437711 | 2752.688153 | 1734.894345 | 2440.32483 | 0.867336082 | 0.034957893 | YCR004C | YCP4 | n |
| W303.asm02.HP0_G0005580 | 499.1889507 | 197.0482092 | 433.9220485 | 115.7654421 | 165.8664207 | 133.9768875 | 244.2206133 | 0 | 192.4417232 | 0.348935932 | 1.443395026 | YCR007C | DFP3 | n |
| W303.asm02.HP0_G0005590 | 862.6760508 | 1275.577779 | 936.2552307 | 611.9030509 | 672.7542024 | 609.2327385 | 961.7369889 | 632.8058569 | 947.8778491 | 0.000936285 | 0.699720796 | YCR008W | SAT4 | n |
| W303.asm02.HP0_G0005600 | 3818.068499 | 3209.88677 | 3755.771244 | 1941.746406 | 1805.953589 | 1942.664869 | 3830.8559 | 2817.630972 | 2949.067045 | 1.15E-09 | 0.921949484 | YCR009C | RVS161 | n |
| W303.asm02.HP0_G0005610 | 63.97372961 | 62.82696524 | 60.59271848 | 220.8298769 | 207.0012931 | 210.9230729 | 242.3274303 | 271.893648 | 248.7411634 | 1.75E-18 | -1.769977952 | YCR010C | ADY2 | n |
| W303.asm02.HP0_G0005620 | 1391.913268 | 1175.625789 | 1374.086487 | 713.0762103 | 703.2736239 | 731.4413859 | 883.1698924 | 1228.843178 | 1278.509108 | 2.10E-07 | 0.875635564 | YCR011C | ADP1 | n |
| W303.asm02.HP0_G0005630 | 35291.20487 | 35699.04319 | 33476.49966 | 28906.33904 | 23817.09108 | 27591.09158 | 17410.65791 | 15314.09526 | 14041.08041 | 0.003561871 | 0.379246617 | YCR012W | PGK1 | n |
| W303.asm02.HP0_G0005640 | 469.1406838 | 518.7984251 | 499.4012765 | 279.1990073 | 289.2710377 | 315.0267355 | 513.0526063 | 478.9585614 | 514.883972 | 1.67E-07 | 0.750286585 | YCR014C | POL4 | n |
| W303.asm02.HP0_G0005650 | 564.1319793 | 545.4522892 | 597.1314676 | 92.41778988 | 78.28895058 | 77.85143463 | 54.90230842 | 150.9445163 | 128.9768996 | 5.23E-20 | 2.778327539 | YCR015C | CTO1 | n |
| W303.asm02.HP0_G0005660 | 1523.737923 | 2281.761147 | 1802.144724 | 2516.682341 | 2127.070979 | 2444.172948 | 1327.121317 | 1403.977521 | 1272.36735 | 0.046501449 | -0.33831925 | YCR016W | RBP95 | n |
| W303.asm02.HP0_G0005670 | 1365.742197 | 1239.404678 | 1268.53788 | 1556.510145 | 1747.568609 | 1659.321857 | 1509.813482 | 1878.098117 | 1695.124966 | 0.006818904 | -0.35714664 | YCR017C | CWH43 | n |
| W303.asm02.HP0_G0005690 | 676.5706555 | 577.8176955 | 617.6548077 | 398.8557247 | 479.022223 | 472.5401032 | 605.8185757 | 446.0603976 | 489.2933174 | 0.006752511 | 0.472140291 | YCR019W | MAK32 | n |
| W303.asm02.HP0_G0005700 | 306.298463 | 144.6924048 | 306.8728001 | 37.93993479 | 34.50021551 | 43.45196352 | 45.43639318 | 39.67131519 | 46.06317841 | 2.31E-15 | 2.702863597 | YCR020C | PET18 | n |
| W303.asm02.HP0_G0005710 | 456.5397976 | 524.509674 | 538.493353 | 90.4721522 | 72.98122512 | 91.43017323 | 115.484166 | 95.79171229 | 113.6225068 | 1.60E-38 | 2.566694965 | YCR020C-. | MAK31 | n |
| W303.asm02.HP0_G0005720 | 816.1497019 | 850.0678782 | 843.4115492 | 332.7040436 | 364.9061256 | 315.9319847 | 190.2648964 | 352.2038714 | 316.3004918 | 1.17E-09 | 1.309477361 | YCR020W- | HTL1 | n |
| W303.asm02.HP0_G0005760 | 609.6890291 | 425.509901 | 553.1528816 | 507.8114349 | 512.1955072 | 491.5503373 | 668.2936163 | 606.6808445 | 666.3806477 | 0.692513056 | 0.071684882 | YCR024C | SLM5 | n |
| W303.asm02.HP0_G0005780 | 531.1758155 | 616.8465678 | 449.5588791 | 202.3463189 | 232.212989 | 193.7233373 | 195.9444456 | 205.1297273 | 272.2845657 | 3.99E-11 | 1.349774631 | YCR026C | NPP1 | n |
| W303.asm02.HP0_G0005790 | 758.9610649 | 901.4717589 | 769.136604 | 1092.475558 | 854.5437996 | 1075.436097 | 722.2493332 | 604.7456584 | 704.2548166 | 0.051557363 | -0.316549331 | YCR027C | RHB1 | n |
| W303.asm02.HP0_G0005800 | 602.9039366 | 633.9811947 | 600.0633734 | 743.2335944 | 777.5817803 | 719.6731457 | 1255.180362 | 1344.954344 | 1561.029935 | 0.039960763 | -0.285587201 | YCR028C | FEN2 | n |
| W303.asm02.HP0_G0005810 | 2747.962476 | 2851.011529 | 2857.630788 | 2628.556508 | 2656.516594 | 2559.139601 | 2650.456269 | 2341.575189 | 2233.55234 | 0.351102051 | 0.108607085 | YCR028C-. | RIM1 | n |
| W303.asm02.HP0_G0005840 | 669.785563 | 890.0486743 | 686.0659415 | 344.3778697 | 368.8869197 | 368.4364406 | 504.5332826 | 668.6067999 | 480.0806817 | 2.18E-08 | 1.05444415 | YCR032W | BPH1 | n |
| W303.asm02.HP0_G0005850 | 423.5836339 | 669.2023722 | 377.2385376 | 1814.307138 | 1973.146941 | 1835.845459 | 1329.0145 | 1930.348142 | 2083.07929 | 4.86E-17 | -1.935220956 | YCR033W | SNT1 | n |
| W303.asm02.HP0_G0005860 | 4885.266624 | 4452.147219 | 4996.944671 | 5956.569762 | 6079.999518 | 6076.938148 | 4466.018813 | 5477.544275 | 4562.301915 | 0.006075611 | -0.337587935 | YCR034W | ELO2 | n |
| W303.asm02.HP0_G0005870 | 1807.742511 | 2115.174496 | 1872.510461 | 1866.839356 | 1750.222471 | 1843.992702 | 1447.338441 | 1375.917322 | 1128.036058 | 0.563411086 | 0.085479513 | YCR035C | RRP43 | n |
| W303.asm02.HP0_G0005880 | 1880.439931 | 1442.164429 | 1845.146008 | 917.3681669 | 828.0051722 | 901.628243 | 871.8107941 | 656.0280902 | 670.4751525 | 8.90E-09 | 0.964487421 | YCR036W | RBK1 | n |
| W303.asm02.HP0_G0005890 | 818.0882998 | 914.7986909 | 791.6145479 | 5321.319059 | 3484.521767 | 5235.961604 | 2128.884339 | 4788.618021 | 3826.314687 | 1.63E-21 | -2.475850322 | YCR037C | PHO87 | n |
| W303.asm02.HP0_G0005900 | 139.5790464 | 133.2693202 | 115.3216255 | 1434.90779 | 1625.490923 | 1477.36676 | 933.3392432 | 1416.55623 | 1108.58716 | 1.17E-79 | -3.546623958 | YCR038C | BUD5 | n |
| W303.asm02.HP0_G0005920 | 785.1321361 | 1020.462223 | 855.1391721 | 1395.995037 | 1567.105943 | 1505.429486 | 910.6210466 | 1443.648836 | 1329.690417 | 9.65E-05 | -0.74768297 | YCR042C | TAF2 | n |
| W303.asm02.HP0_G0005930 | 1428.746628 | 1877.193568 | 1508.954151 | 797.7114495 | 962.0252402 | 734.1571336 | 981.615411 | 789.5559316 | 667.4042739 | 6.09E-07 | 0.95060114 | YCR043C | | n |
| W303.asm02.HP0_G0005940 | 1180.606101 | 1048.068011 | 1170.807689 | 1546.781957 | 1965.185353 | 1554.312945 | 1571.341931 | 1965.181492 | 1787.251224 | 0.000178995 | -0.574514477 | YCR044C | PER1 | n |
| W303.asm02.HP0_G0005950 | 184.1667974 | 188.4808957 | 129.9811542 | 5.836913045 | 9.28851956 | 9.957741639 | 7.572732196 | 15.48148885 | 10.23626187 | 1.36E-22 | 4.330848028 | YCR045C | RRT12 | n |
| W303.asm02.HP0_G0005960 | 803.5488158 | 957.6352581 | 918.6637963 | 1648.927935 | 1744.914746 | 1613.154146 | 1905.488739 | 1606.204469 | 1363.470081 | 6.13E-09 | -0.901437976 | YCR046C | IMG1 | n |
| W303.asm02.HP0_G0005970 | 978.0226238 | 1138.500764 | 899.1177581 | 2907.755515 | 2704.286123 | 2890.460823 | 1662.214717 | 1569.435933 | 1292.839874 | 1.18E-24 | -1.495527251 | YCR047C | BUD23 | n |
| W303.asm02.HP0_G0005990 | 1778.663543 | 1895.280118 | 1918.443651 | 2011.789363 | 1851.069255 | 1941.75962 | 1638.549929 | 1284.963575 | 1054.334973 | 0.766749895 | -0.053940445 | YCR051W | | n |
| W303.asm02.HP0_G0006010 | 3031.967064 | 2714.886437 | 3049.181962 | 6287.328168 | 5541.265383 | 6190.094303 | 4496.309742 | 3773.612908 | 4018.75641 | 1.49E-17 | -1.034826032 | YCR053W | THR4 | n |
| W303.asm02.HP0_G0006020 | 536.0223102 | 821.5101667 | 535.5614472 | 698.4839277 | 627.638536 | 698.8524132 | 479.9219029 | 527.3382141 | 424.8048676 | 0.651448416 | -0.097686433 | YCR054C | CTR86 | n |
| W303.asm02.HP0_G0006030 | 971.2375313 | 1670.626121 | 1010.530176 | 6615.168118 | 5663.343069 | 6369.333652 | 2554.850525 | 3272.399707 | 2665.522591 | 1.34E-29 | -2.352078693 | YCR057C | PWP2 | n |
| W303.asm02.HP0_G0006040 | 1254.27282 | 1192.760416 | 1390.700619 | 542.8329132 | 566.5996932 | 517.8025652 | 603.9253927 | 525.403028 | 461.6554103 | 4.64E-17 | 1.238835939 | YCR059C | YIH1 | n |
| W303.asm02.HP0_G0006050 | 1012.917385 | 1002.375673 | 1087.737027 | 196.5094058 | 164.5394894 | 186.4813434 | 66.26140672 | 120.9491317 | 140.2367876 | 7.93E-28 | 2.500252472 | YCR060W | TAH1 | n |
| W303.asm02.HP0_G0006060 | 216.1536622 | 208.4712938 | 189.5965707 | 1419.342689 | 1571.086737 | 1163.245273 | 1362.145204 | 2742.158713 | 2066.701272 | 2.68E-31 | -2.756960142 | YCR061W | TVS1 | n |

| | | | | | | | | | | | | | | |
|---|---|---|---|---|---|---|---|---|---|---|---|---|---|---|
| W303.asm02.HP0_G0006070 | 70.75882214 | 104.7116087 | 84.04796435 | 141.0587319 | 74.30815648 | 124.9243951 | 26.50456269 | 51.28243183 | 55.2758141 | 0.295909729 | -0.396740869 | YCR063W | BUD31 | n |
| W303.asm02.HP0_G0006080 | 343.1318224 | 370.2983254 | 232.5978548 | 1031.187971 | 870.4669759 | 1039.226127 | 567.0083232 | 960.8199021 | 739.058107 | 1.49E-12 | -1.636680902 | YCR065W | HCM1 | n |
| W303.asm02.HP0_G0006090 | 214.2150643 | 229.4136155 | 198.3922879 | 768.5268843 | 724.5045257 | 752.2621184 | 723.1959248 | 706.342929 | 624.4119741 | 2.36E-36 | -1.806526471 | YCR066W | RAD18 | n |
| W303.asm02.HP0_G0006100 | 720.1891075 | 711.0870157 | 669.451809 | 1896.99674 | 1941.300588 | 1868.434431 | 1547.677143 | 2341.575189 | 2349.222099 | 5.76E-17 | -1.441649126 | YCR067C | SED4 | n |
| W303.asm02.HP0_G0006110 | 200.6448792 | 171.3462688 | 148.5498905 | 241.2590725 | 327.7520473 | 285.1535106 | 278.2979082 | 413.1622338 | 550.7108886 | 0.008295368 | -0.712567468 | YCR068W | ATG15 | n |
| W303.asm02.HP0_G0006120 | 675.6013566 | 603.5196358 | 649.9057708 | 1798.742037 | 1634.779443 | 1706.394817 | 2218.810534 | 2175.149184 | 2204.890807 | 2.17E-37 | -1.414510935 | YCR069W | CPR4 | n |
| W303.asm02.HP0_G0006130 | 1173.821008 | 1467.86637 | 1248.01454 | 1254.936305 | 1316.315915 | 1197.644744 | 1442.605483 | 1063.384766 | 971.4212514 | 0.809672587 | 0.045925037 | YCR071C | IMG2 | n |
| W303.asm02.HP0_G0006140 | 404.1976552 | 656.8273639 | 376.2612357 | 3711.303878 | 3180.654484 | 3728.721619 | 1931.04671 | 2220.626058 | 2070.795776 | 1.86E-46 | -2.88537145 | YCR072C | RSA4 | n |
| W303.asm02.HP0_G0006170 | 556.3775878 | 466.4426207 | 545.3344663 | 374.5352537 | 395.425547 | 401.0254133 | 605.8185757 | 471.217817 | 438.112008 | 0.017532453 | 0.421395915 | YCR075C | ERS1 | n |
| W303.asm02.HP0_G0006180 | 207.4299718 | 213.2309123 | 183.7327593 | 139.1130942 | 149.9432443 | 138.5031337 | 223.3955998 | 148.0417372 | 158.662059 | 0.029336636 | 0.501349352 | YCR075W- | EGO2 | n |
| W303.asm02.HP0_G0006190 | 658.1539758 | 494.0484085 | 680.2021301 | 84.63523915 | 71.65429375 | 85.99867779 | 131.5762219 | 115.1435734 | 138.1895352 | 2.34E-43 | 2.912009446 | YCR076C | FUB1 | n |
| W303.asm02.HP0_G0006200 | 1149.588535 | 1374.577846 | 966.55159 | 3914.623016 | 4173.199145 | 4148.757267 | 3031.932653 | 3451.404422 | 3658.439992 | 7.96E-33 | -1.809438272 | YCR077C | PAT1 | n |
| W303.asm02.HP0_G0006210 | 409.0441499 | 376.9617915 | 337.1691593 | 178.9986667 | 192.405048 | 191.9128389 | 280.1910913 | 319.3057076 | 306.0642299 | 3.43E-09 | 0.996454678 | YCR079W | PTC6 | n |
| W303.asm02.HP0_G0006220 | 446.8468083 | 534.9811283 | 497.4466727 | 474.7355943 | 517.5032327 | 505.1290759 | 366.33092 | 573.7826807 | 504.6477102 | 0.93449823 | -0.017013574 | YCR081W | SRB8 | n |
| W303.asm02.HP0_G0006230 | 1475.272977 | 1791.520433 | 1500.158433 | 600.2292248 | 514.8493699 | 586.6015075 | 822.5880348 | 695.6994054 | 749.2943689 | 1.18E-24 | 1.484253758 | YCR082W | AHC2 | n |
| W303.asm02.HP0_G0006240 | 783.1935382 | 960.4910292 | 825.8201148 | 843.433935 | 912.9287796 | 877.1865135 | 1056.396141 | 911.4726563 | 744.1762379 | 0.845479335 | -0.035033533 | YCR083W | TRX3 | n |
| W303.asm02.HP0_G0006250 | 1122.448165 | 1450.731743 | 914.7545887 | 4449.673378 | 4384.181233 | 4349.722598 | 3398.263573 | 4835.062488 | 5007.579307 | 1.14E-22 | -1.918205199 | YCR084C | TUP1 | n |
| W303.asm02.HP0_G0006260 | 515.6670326 | 453.1156887 | 604.9498829 | 696.53829 | 691.3312416 | 657.2109482 | 945.644933 | 768.2688844 | 720.6328356 | 0.028545249 | -0.377864566 | YCR086W | CSM1 | n |
| W303.asm02.HP0_G0006270 | 1013.886684 | 1537.356801 | 1256.810258 | 2398.971262 | 3305.386032 | 2386.236996 | 1719.9568 | 1538.472955 | 1225.280546 | 1.41E-07 | -1.08660491 | YCR087C-A | | n |
| W303.asm02.HP0_G0006280 | 763.8075595 | 681.5773805 | 528.7203338 | 2842.576653 | 2352.649312 | 2847.00886 | 2248.154871 | 2132.57509 | 1978.669419 | 8.00E-36 | -2.026788305 | YCR088W | ABP1 | n |
| W303.asm02.HP0_G0006300 | 1141.834144 | 1176.577713 | 1145.39784 | 917.3681669 | 958.0444461 | 903.4387414 | 1085.740479 | 815.680944 | 658.1916382 | 0.088249162 | 0.318194313 | YCR090C | | n |
| W303.asm02.HP0_G0006320 | 291.758979 | 450.2599176 | 297.0997809 | 589.5282175 | 667.446477 | 683.4631761 | 536.7173944 | 801.1670482 | 622.3647217 | 4.85E-05 | -0.90059503 | YCR092C | MSH3 | n |
| W303.asm02.HP0_G0006330 | 802.5795169 | 1555.443352 | 803.3421708 | 3981.747516 | 4224.949469 | 4085.38982 | 2796.231364 | 3468.821097 | 3315.52522 | 5.35E-17 | -1.958953628 | YCR093W | CDC39 | n |
| W303.asm02.HP0_G0006340 | 848.1365667 | 788.1928367 | 915.7318906 | 1002.003406 | 1052.256573 | 1023.83689 | 1481.415736 | 1443.648836 | 1365.517333 | 0.026412689 | -0.270045646 | YCR094W | CDC50 | n |
| W303.asm02.HP0_G0006350 | 863.6453497 | 937.6448601 | 786.7280384 | 447.4966668 | 415.3295175 | 443.5721276 | 463.829847 | 456.7039212 | 518.9784768 | 2.13E-12 | 0.985013509 | YCR095C | OCA4 | n |
| W303.asm02.HP0_G0006450 | 286.9124843 | 444.5483753 | 320.5550268 | 823.9775582 | 745.7354276 | 720.578395 | 584.9935622 | 1134.019059 | 954.0196063 | 7.48E-06 | -1.1222707 | YCR106W | RDS1 | n |
| W303.asm02.HP0_G0006500 | 92.08339868 | 121.8462356 | 111.4124179 | 34.04865943 | 23.88476458 | 26.25222796 | 58.68867452 | 32.89816382 | 52.20493554 | 3.49E-08 | 1.942769375 | YJR159W | SOR1 | y |
| W303.asm02.HP0_G0006530 | 121.1623667 | 99.95199016 | 129.0038523 | 19.45637682 | 34.50021551 | 28.06272644 | 20.82501354 | 29.0277916 | 33.77966417 | 2.88E-09 | 2.108511424 | YDL244W | THI13 | y |
| W303.asm02.HP0_G0006570 | 293.6975768 | 603.5196358 | 295.1451771 | 635.2507031 | 701.9466925 | 651.7794527 | 2415.701571 | 807.9401996 | 2473.080868 | 0.156430794 | -0.738032467 | YDL240W | LRG1 | n |
| W303.asm02.HP0_G0006590 | 82.39040934 | 85.67313442 | 66.45652995 | 111.8741667 | 108.808372 | 105.9141611 | 464.7764386 | 193.5186107 | 565.0416552 | 0.192457235 | -0.477503544 | YDL238C | GUD1 | n |
| W303.asm02.HP0_G0006600 | 1346.356219 | 1164.202704 | 1370.177279 | 68.09731886 | 61.03884283 | 53.40970516 | 270.725176 | 60.95836237 | 224.1741349 | 4.31E-31 | 4.410688131 | YDL237W | AIM6 | n |
| W303.asm02.HP0_G0006610 | 2433.909622 | 1801.03967 | 2256.590112 | 1840.573247 | 1580.375257 | 1786.962 | 3887.651391 | 1168.852409 | 3184.501068 | 0.345778509 | 0.317712513 | YDL236W | PHO13 | n |
| W303.asm02.HP0_G0006620 | 2062.66813 | 2254.155359 | 2220.429942 | 787.9832611 | 729.8122512 | 714.2416503 | 2622.058523 | 832.130026 | 2008.354579 | 2.75E-07 | 1.550284063 | YDL235C | YPD1 | n |
| W303.asm02.HP0_G0006630 | 567.0398761 | 471.2022393 | 512.1062014 | 173.1617537 | 147.2893816 | 151.1766231 | 443.951425 | 199.324169 | 607.0103289 | 8.78E-08 | 1.716040526 | YDL234C | GYP7 | n |
| W303.asm02.HP0_G0006640 | 273.3422992 | 251.307861 | 271.6899313 | 439.7141161 | 475.0414289 | 449.9088722 | 1301.563346 | 546.6900752 | 1785.20407 | 0.017273874 | -0.776868948 | YDL233W | MFG1 | n |
| W303.asm02.HP0_G0006650 | 727.943499 | 1203.231577 | 775.9777173 | 1816.252776 | 1669.279658 | 1870.24493 | 2740.382464 | 1308.185808 | 3181.430189 | 0.000701198 | -0.98437947 | YDL231C | BRE4 | n |
| W303.asm02.HP0_G0006660 | 553.469691 | 340.7886903 | 382.1520472 | 487.3822393 | 492.2915367 | 499.6975804 | 1310.08267 | 566.0419363 | 1432.053036 | 0.511304184 | -0.212992869 | YDL230W | PTP1 | n |
| W303.asm02.HP0_G0006670 | 8462.948988 | 8272.21709 | 8280.679092 | 49946.46493 | 53017.54272 | 46717.19752 | 77378.17758 | 38203.47653 | 76610.23109 | 9.49E-31 | -2.580971325 | YDL229W | SSB1 | y |
| W303.asm02.HP0_G0006690 | 1411.299247 | 1186.09695 | 1325.221391 | 2400.916899 | 2562.304467 | 2407.057729 | 6803.153287 | 2248.686256 | 5513.250643 | 0.002328518 | -0.909858133 | YDL226C | GCS1 | n |
| W303.asm02.HP0_G0006700 | 1733.106493 | 1320.318194 | 1752.302326 | 1990.387348 | 1949.262176 | 1930.896629 | 1949.978541 | 2009.690772 | 2116.858955 | 0.043908428 | -0.28883359 | YDL225W | SHS1 | n |
| W303.asm02.HP0_G0006710 | 348.947616 | 408.3752741 | 293.1905733 | 1912.561841 | 1825.857559 | 1996.074574 | 761.0595857 | 1702.963774 | 1926.464484 | 1.49E-18 | -2.448556053 | YDL224C | WHI4 | n |
| W303.asm02.HP0_G0006730 | 101.776388 | 82.81736327 | 103.5940026 | 13.61946377 | 39.80794097 | 12.67348936 | 35.97047793 | 33.86575687 | 24.56702849 | 1.79E-06 | 2.167215357 | YDL222C | FMP45 | n |
| W303.asm02.HP0_G0006740 | 135.7018507 | 166.5866503 | 100.6620968 | 478.6268697 | 496.2723308 | 464.3928601 | 365.3843285 | 472.1854101 | 459.608158 | 1.76E-18 | -1.835501228 | YDL220C | CDC13 | n |
| W303.asm02.HP0_G0006750 | 1024.548973 | 1058.539172 | 991.9614397 | 370.6439784 | 323.7712532 | 372.9626868 | 734.5550231 | 538.9493308 | 514.883972 | 6.18E-19 | 1.52438795 | YDL219W | DTD1 | n |
| W303.asm02.HP0_G0006770 | 925.6804814 | 1110.894976 | 864.9121912 | 689.7285582 | 761.658604 | 674.4106837 | 963.630172 | 756.6577678 | 721.6564618 | 0.008517291 | 0.449897934 | YDL217C | TIM22 | n |
| W303.asm02.HP0_G0006780 | 265.5879078 | 309.3752076 | 303.9408943 | 70.04295654 | 76.96201921 | 67.89369299 | 56.79549147 | 106.4352359 | 124.8823948 | 1.18E-12 | 2.035208024 | YDL216C | RRI1 | n |
| W303.asm02.HP0_G0006790 | 1228.101749 | 1234.645059 | 1196.217539 | 876.5097756 | 575.8882127 | 809.2928205 | 1358.358838 | 2021.301889 | 1972.527662 | 0.000651602 | 0.692038043 | YDL215C | GDH2 | n |
| W303.asm02.HP0_G0006810 | 1972.52333 | 2199.895707 | 2189.156281 | 2535.165899 | 2279.668086 | 2492.151158 | 1645.17607 | 1354.630275 | 1237.56406 | 0.170757401 | -0.200237836 | YDL213C | NOP6 | n |
| W303.asm02.HP0_G0006820 | 3368.313794 | 4008.550767 | 3491.899728 | 1157.654421 | 1338.873748 | 1235.665212 | 2755.527928 | 2365.765016 | 2206.938059 | 5.08E-29 | 1.542887895 | YDL212W | SHR3 | n |
| W303.asm02.HP0_G0006840 | 190.9518899 | 107.5673799 | 151.4817962 | 145.9228261 | 171.1741462 | 157.5133677 | 44.48980165 | 172.2315635 | 150.4730495 | 0.864576976 | -0.076184091 | YDL210W | UGA4 | n |
| W303.asm02.HP0_G0006850 | 465.263488 | 404.5675792 | 434.8993504 | 341.4594131 | 341.021361 | 382.0151792 | 274.5115421 | 307.694591 | 328.584006 | 0.070271161 | 0.292023814 | YDL209C | CWC2 | n |
| W303.asm02.HP0_G0006860 | 4315.318852 | 5030.916838 | 4799.529685 | 6478.97348 | 7352.526698 | 6406.448871 | 4947.833899 | 4508.983629 | 3883.637753 | 0.000148952 | -0.51648402 | YDL208W | NHP2 | n |
| W303.asm02.HP0_G0006870 | 713.404015 | 542.596518 | 719.2942065 | 890.1292394 | 889.0440151 | 881.7127597 | 752.540262 | 752.7873956 | 770.7905188 | 0.003938312 | -0.429978965 | YDL207W | GLE1 | n |
| W303.asm02.HP0_G0006880 | 574.7942675 | 408.3752741 | 557.0620893 | 248.0688044 | 273.3478613 | 233.5543039 | 351.1854556 | 309.6297771 | 343.9383988 | 3.16E-08 | 1.031143026 | YDL206W | YCX1 | n |
| W303.asm02.HP0_G0006890 | 873.338339 | 1623.981859 | 924.5276078 | 3578.027697 | 3891.889696 | 3624.617957 | 3473.044304 | 3574.288739 | 2877.413212 | 1.30E-14 | -1.696813253 | YDL205C | HEM3 | n |
| W303.asm02.HP0_G0006910 | 380.9344808 | 495.9522559 | 353.7832918 | 610.930232 | 582.5228696 | 592.0330029 | 562.2753656 | 610.5512167 | 679.6877882 | 0.001957963 | -0.536887984 | YDL203C | ACK1 | n |
| W303.asm02.HP0_G0006920 | 1185.452596 | 1037.59685 | 1271.469786 | 2173.27729 | 2140.340293 | 2187.082164 | 2482.909569 | 2127.737124 | 1999.141943 | 3.83E-11 | -0.895673462 | YDL202W | MRPL11 | n |
| W303.asm02.HP0_G0006930 | 1547.001098 | 1490.712539 | 1575.410681 | 1854.192711 | 1928.031274 | 1900.118155 | 1668.840858 | 1442.681243 | 1238.587686 | 0.030884578 | -0.300637398 | YDL201W | TRM8 | n |
| W303.asm02.HP0_G0006940 | 747.3294777 | 648.2600505 | 798.4556613 | 27.23892754 | 39.80794097 | 29.87322492 | 8.519323721 | 50.31483878 | 60.39394503 | 2.46E-26 | 4.509605617 | YDL200C | MGT1 | n |
| W303.asm02.HP0_G0006950 | 197.7369824 | 170.3943451 | 171.0278344 | 80.74396379 | 75.63508785 | 85.99867779 | 124.0034897 | 205.1297273 | 164.8038161 | 5.86E-06 | 1.150156657 | YDL199C | | n |
| W303.asm02.HP0_G0006960 | 2428.093828 | 2351.251578 | 2288.841076 | 4633.536139 | 4199.737773 | 4808.683962 | 4170.682257 | 3001.473652 | 2938.830783 | 1.53E-09 | -0.948829659 | YDL198C | GGC1 | n |
| W303.asm02.HP0_G0006980 | 1197.084183 | 1097.568044 | 907.9134753 | 5879.717074 | 5179.013121 | 5665.954993 | 3373.652194 | 5275.317327 | 5064.902373 | 7.10E-38 | -2.384743053 | YDL195W | SEC31 | n |
| W303.asm02.HP0_G0006990 | 107.5921816 | 228.4616918 | 100.6620968 | 179.9714856 | 197.7127735 | 178.3341003 | 102.2318847 | 215.7732509 | 209.8433683 | 0.365618836 | -0.347235491 | YDL194W | SNF3 | n |

| | | | | | | | | | | | | | | |
|---|---|---|---|---|---|---|---|---|---|---|---|---|---|---|
| W303.asm02.HP0_G0007000 | 990.62351 | 890.0486743 | 864.9121912 | 838.5698408 | 874.44777 | 847.3132886 | 1055.44955 | 1265.611714 | 1158.744844 | 0.46739865 | 0.101216037 | YDL193W | NUS1 | n |
| W303.asm02.HP0_G0007030 | 1640.053795 | 1411.702871 | 1669.231664 | 1424.206783 | 1387.970209 | 1436.630544 | 1252.340587 | 1880.033303 | 1836.385379 | 0.406641431 | 0.151864306 | YDL190C | UFD2 | n |
| W303.asm02.HP0_G0007040 | 549.5924953 | 599.711941 | 441.7404638 | 606.0661378 | 494.9453994 | 575.7385166 | 545.2367181 | 627.0002986 | 724.7273404 | 0.696895373 | -0.077140272 | YDL189W | RBS1 | n |
| W303.asm02.HP0_G0007060 | 121.1623667 | 82.81736327 | 105.5486064 | 47.6681232 | 47.76952917 | 46.16771124 | 51.11594233 | 62.89354847 | 61.41757122 | 0.00014783 | 1.128744467 | YDL186W | | n |
| W303.asm02.HP0_G0007070 | 3567.020075 | 3264.146421 | 3121.502304 | 4534.308617 | 3631.811148 | 4475.552242 | 4023.960571 | 4234.187202 | 3561.195504 | 0.01346962 | -0.345498428 | YDL185W | VMA1 | n |
| W303.asm02.HP0_G0007080 | 318.8993491 | 237.0290052 | 277.5537427 | 393.0188117 | 388.7908902 | 372.0574376 | 492.2275928 | 372.5233256 | 442.2065128 | 0.011520239 | -0.469107959 | YDL183C | MRX19 | n |
| W303.asm02.HP0_G0007090 | 4366.691695 | 3445.011927 | 4320.651749 | 597.3107683 | 516.1763013 | 716.0521488 | 150.5080524 | 263.1853105 | 167.8746947 | 8.64E-38 | 2.727689791 | YDL182W | LYS20 | n |
| W303.asm02.HP0_G0007100 | 2383.506077 | 2613.0306 | 2525.348138 | 4353.364313 | 3909.139804 | 4484.604735 | 4676.162131 | 4195.48348 | 3365.682903 | 1.95E-07 | -0.761243282 | YDL181W | INH1 | n |
| W303.asm02.HP0_G0007110 | 1146.680638 | 981.433351 | 957.7558728 | 216.9386015 | 207.0012931 | 207.3020759 | 98.44551855 | 195.4537968 | 218.0323778 | 2.52E-20 | 2.289173945 | YDL180W | AIT1 | n |
| W303.asm02.HP0_G0007130 | 2756.686167 | 2134.212971 | 2794.106164 | 1641.145384 | 1207.507543 | 1619.49089 | 1747.407954 | 1457.195138 | 1300.005257 | 2.16E-05 | 0.781329255 | YDL178W | DLD2 | n |
| W303.asm02.HP0_G0007140 | 613.5662249 | 561.6349923 | 666.5199033 | 168.2976595 | 222.9244694 | 157.5133677 | 309.5354285 | 223.5139953 | 203.7016112 | 4.83E-16 | 1.75334873 | YDL177C | | n |
| W303.asm02.HP0_G0007150 | 278.1887939 | 341.740614 | 318.600423 | 47.6681232 | 55.73111736 | 51.59920668 | 122.1103067 | 98.69449145 | 130.0005257 | 2.77E-29 | 2.603489986 | YDL176W | GID12 | n |
| W303.asm02.HP0_G0007160 | 992.5621078 | 831.9813276 | 819.9563033 | 1050.644348 | 875.7747014 | 986.7216715 | 1262.753094 | 1799.723079 | 1399.296998 | 0.445159696 | -0.14045184 | YDL174C | DLD1 | n |
| W303.asm02.HP0_G0007170 | 1995.786504 | 1501.1837 | 2012.264635 | 560.3436523 | 687.3504475 | 578.4542643 | 1167.14735 | 739.2410928 | 827.0899591 | 4.00E-15 | 1.59433255 | YDL173W | PAR32 | n |
| W303.asm02.HP0_G0007180 | 1417.115041 | 1679.193435 | 1224.559294 | 3302.719965 | 4486.354948 | 3196.435066 | 3766.487676 | 6223.55852 | 5130.414449 | 2.64E-10 | -1.345792397 | YDL171C | GLT1 | n |
| W303.asm02.HP0_G0007190 | 518.5749294 | 542.596518 | 468.1276154 | 454.3063987 | 542.7149286 | 440.8563798 | 501.693508 | 598.9401001 | 807.6410615 | 0.677499006 | 0.090867996 | YDL170W | UGA3 | n |
| W303.asm02.HP0_G0007200 | 305.329164 | 263.6828693 | 271.6899313 | 10.70100725 | 14.59624502 | 9.052492399 | 85.19323721 | 25.15741939 | 106.4571234 | 1.52E-22 | 4.635744493 | YDL169C | UGX2 | n |
| W303.asm02.HP0_G0007210 | 2262.343711 | 1680.145358 | 2006.400823 | 8299.117531 | 7223.814355 | 7662.934816 | 7440.209383 | 6679.294848 | 5856.165416 | 4.74E-39 | -1.962684683 | YDL168W | SFA1 | n |
| W303.asm02.HP0_G0007220 | 836.5049795 | 1370.770151 | 1014.439384 | 1316.223892 | 1239.353896 | 1379.599842 | 1048.823409 | 1052.741242 | 1124.965179 | 0.124769854 | -0.288868609 | YDL167C | NRP1 | n |
| W303.asm02.HP0_G0007230 | 1442.316813 | 1762.962722 | 1532.409396 | 1764.693377 | 1811.261314 | 1764.330769 | 2187.573013 | 1506.542384 | 1532.368402 | 0.338315567 | -0.172607384 | YDL166C | FAP7 | n |
| W303.asm02.HP0_G0007240 | 1076.891115 | 976.6737324 | 1190.353728 | 930.0148118 | 902.3133287 | 869.9445196 | 1275.058784 | 983.0745423 | 966.3031205 | 0.101729265 | 0.263591324 | YDL165W | CDC36 | n |
| W303.asm02.HP0_G0007250 | 488.5266624 | 459.7791547 | 431.9674447 | 772.4181596 | 671.4272711 | 795.7140819 | 375.7968352 | 673.4447652 | 986.7756442 | 0.013764216 | -0.698738486 | YDL164C | CDC9 | n |
| W303.asm02.HP0_G0007260 | 2825.506391 | 2369.338129 | 2805.833786 | 2313.363204 | 2152.282675 | 2229.628878 | 1436.925934 | 2595.084569 | 2839.539043 | 0.26963232 | 0.256886694 | YDL161W | ENT1 | n |
| W303.asm02.HP0_G0007270 | 419.7064382 | 433.1252907 | 416.3306141 | 37.93993479 | 53.07725463 | 40.7362158 | 105.0716592 | 53.21761794 | 82.91372115 | 3.00E-33 | 3.281108274 | YDL160C-, | MHF2 | n |
| W303.asm02.HP0_G0007280 | 631.9829046 | 737.7408797 | 682.1567339 | 4161.719001 | 4116.141097 | 4284.544653 | 1199.331462 | 3778.450874 | 3301.194453 | 7.96E-18 | -2.614091887 | YDL160C | DHH1 | n |
| W303.asm02.HP0_G0007290 | 426.4915307 | 408.3752741 | 401.6710854 | 976.7101162 | 977.9484166 | 971.3324344 | 896.4221738 | 1326.570076 | 1314.336024 | 7.19E-13 | -1.242586358 | YDL159W | STE7 | n |
| W303.asm02.HP0_G0007300 | 1042.965652 | 1349.827829 | 1135.624821 | 344.3778697 | 428.5988311 | 337.6579665 | 452.4707487 | 327.0464521 | 333.702137 | 9.93E-19 | 1.670756539 | YDL157C | DMO2 | n |
| W303.asm02.HP0_G0007310 | 89.17550188 | 117.086617 | 63.52462421 | 637.1963407 | 582.5228696 | 635.4849664 | 656.934518 | 711.1808943 | 748.2707427 | 2.50E-42 | -2.781180152 | YDL156W | CMR1 | n |
| W303.asm02.HP0_G0007320 | 723.0970043 | 840.548641 | 685.0886396 | 682.9188263 | 781.5625744 | 691.6104193 | 796.0834721 | 854.3846662 | 1075.831123 | 0.732311756 | 0.06196337 | YDL155W | CLB3 | n |
| W303.asm02.HP0_G0007330 | 512.7591358 | 585.4330852 | 492.5601631 | 81.71678263 | 112.7891661 | 81.47243159 | 117.377349 | 101.5972706 | 66.53570215 | 9.10E-26 | 2.536982374 | YDL154W | MSH5 | n |
| W303.asm02.HP0_G0007340 | 2036.497059 | 2742.492225 | 2345.524586 | 6773.737589 | 5821.247902 | 6680.739391 | 3307.390787 | 3811.349037 | 2942.925288 | 2.73E-20 | -1.436048919 | YDL153C | SAS10 | n |
| W303.asm02.HP0_G0007350 | 756.0531681 | 976.6737324 | 778.9096231 | 2688.871276 | 2023.570333 | 2661.432765 | 1254.23377 | 1634.264667 | 1309.217893 | 2.10E-18 | -1.554322538 | YDL150W | RPC53 | n |
| W303.asm02.HP0_G0007360 | 252.9870216 | 262.7309456 | 271.6899313 | 159.5422899 | 165.8664207 | 159.3238662 | 232.861515 | 358.9770228 | 314.2532394 | 0.000824561 | 0.700875937 | YDL149W | ATG9 | n |
| W303.asm02.HP0_G0007370 | 1868.808344 | 2551.155558 | 2120.745147 | 6057.742922 | 4934.857749 | 5919.42478 | 1941.459217 | 2966.640302 | 2472.057242 | 1.44E-13 | -1.370672012 | YDL148C | NOP14 | n |
| W303.asm02.HP0_G0007380 | 5558.929383 | 5312.686258 | 5410.343379 | 2723.892754 | 2202.706067 | 2568.192094 | 2615.432382 | 2530.255835 | 2532.451187 | 1.70E-22 | 1.118408718 | YDL147W | RPN5 | n |
| W303.asm02.HP0_G0007390 | 315.0221534 | 356.9713934 | 343.0329708 | 240.2862537 | 301.21342 | 240.7962978 | 342.6661319 | 273.8288341 | 460.6317841 | 0.096486286 | 0.378986057 | YDL146W | LDB17 | n |
| W303.asm02.HP0_G0007400 | 2399.01486 | 2316.030401 | 2133.450072 | 5572.50632 | 5404.591453 | 5725.701443 | 4655.337118 | 5627.521199 | 5571.597336 | 5.05E-27 | -1.286304858 | YDL145C | COP1 | n |
| W303.asm02.HP0_G0007410 | 1115.663072 | 890.0486743 | 946.0282498 | 1297.740334 | 1037.660328 | 1221.181225 | 1570.395339 | 1327.537669 | 1482.210719 | 0.084863361 | -0.27014258 | YDL144C | | n |
| W303.asm02.HP0_G0007420 | 3549.572694 | 3081.377068 | 3474.308294 | 5096.597907 | 4595.16332 | 5063.964248 | 4298.472113 | 4006.802834 | 3874.425118 | 1.68E-06 | -0.546474393 | YDL143W | CCT4 | n |
| W303.asm02.HP0_G0007430 | 715.3426129 | 540.6926706 | 637.200846 | 955.3081017 | 960.6983088 | 933.3119664 | 984.4551855 | 1095.315336 | 924.3344468 | 7.72E-05 | -0.589712806 | YDL142C | CRD1 | n |
| W303.asm02.HP0_G0007440 | 376.0879862 | 458.827231 | 386.0342548 | 1453.391348 | 1072.160544 | 1427.578051 | 1037.464311 | 1436.875684 | 1154.650339 | 5.49E-20 | -1.695991194 | YDL141W | BPL1 | n |
| W303.asm02.HP0_G0007450 | 953.7901505 | 1335.548973 | 727.1126218 | 6670.618792 | 6710.291917 | 7050.98633 | 2926.860994 | 6115.188098 | 6130.497234 | 2.29E-24 | -2.759835227 | YDL140C | RPO21 | n |
| W303.asm02.HP0_G0007460 | 497.2503528 | 213.2309123 | 447.6042752 | 114.7926232 | 102.1737152 | 117.6824012 | 88.03301178 | 129.6574692 | 160.7093114 | 5.08E-08 | 1.78902773 | YDL139C | SCM3 | n |
| W303.asm02.HP0_G0007480 | 5399.964358 | 5512.590238 | 5298.930961 | 8856.542727 | 8429.994967 | 8166.253393 | 5508.216081 | 6669.618917 | 5727.188516 | 9.36E-09 | -0.650800587 | YDL137W | ARF2 | y |
| W303.asm02.HP0_G0007500 | 2108.22518 | 2072.337929 | 2150.064204 | 1961.202783 | 1746.241677 | 1870.24493 | 2205.558252 | 1725.218414 | 1584.573337 | 0.238441679 | 0.182298359 | YDL135C | RDI1 | n |
| W303.asm02.HP0_G0007510 | 3408.05505 | 2709.174895 | 3397.101443 | 2489.443414 | 2454.823027 | 2373.563507 | 2957.151923 | 2901.811567 | 3103.634599 | 0.003222241 | 0.378716089 | YDL134C | PPH21 | n |
| W303.asm02.HP0_G0007520 | 1271.720201 | 837.6928699 | 1115.10148 | 187.7540363 | 226.9052635 | 198.2495835 | 335.0933997 | 289.310323 | 353.1510345 | 8.61E-35 | 2.398562717 | YDL133W | SRF1 | n |
| W303.asm02.HP0_G0007530 | 1433.593123 | 1407.895176 | 1471.816678 | 2055.566211 | 2087.263038 | 1954.433109 | 1836.387558 | 2051.297273 | 1925.440858 | 2.59E-06 | -0.499015765 | YDL132W | CDC53 | n |
| W303.asm02.HP0_G0007540 | 5401.902956 | 3403.127284 | 4681.276154 | 1793.877943 | 1304.373533 | 1937.233373 | 2043.691102 | 1348.824716 | 1806.70022 | 1.63E-10 | 1.420533273 | YDL131W | LYS21 | n |
| W303.asm02.HP0_G0007560 | 1780.602141 | 1899.087813 | 1680.959287 | 9626.04243 | 10822.45222 | 9130.343834 | 9129.875254 | 7297.586809 | 6606.483411 | 7.87E-62 | -2.46393243 | YDL130W | RPP1B | y |
| W303.asm02.HP0_G0007570 | 376.0879862 | 222.7501495 | 339.1237631 | 578.8272103 | 391.4447529 | 525.9498084 | 151.4546439 | 285.4399508 | 376.6944368 | 0.036096776 | -0.674988516 | YDL129W | | n |
| W303.asm02.HP0_G0007580 | 1465.579987 | 1103.279587 | 1307.629957 | 2672.333356 | 2400.418841 | 2757.389185 | 2955.25874 | 3085.654247 | 2812.924762 | 2.50E-13 | -1.01481109 | YDL128W | VCX1 | n |
| W303.asm02.HP0_G0007590 | 577.7021643 | 531.1734334 | 599.0860714 | 1021.459783 | 1009.794769 | 1101.688325 | 1123.60414 | 1157.241292 | 1554.888178 | 6.59E-08 | -0.875715605 | YDL127W | PCL2 | n |
| W303.asm02.HP0_G0007600 | 5237.122137 | 4393.127948 | 4458.451318 | 7818.545024 | 6184.827096 | 7453.822242 | 8298.827896 | 8576.744826 | 8293.419367 | 4.96E-06 | -0.60721198 | YDL126C | CDC48 | n |
| W303.asm02.HP0_G0007620 | 4402.555756 | 2798.655724 | 4084.144686 | 1630.444377 | 900.9863974 | 1326.190136 | 1236.248531 | 998.5560311 | 1122.917927 | 1.76E-11 | 1.547558165 | YDL124W | | n |
| W303.asm02.HP0_G0007630 | 413.8906446 | 192.2885906 | 301.0089886 | 183.8627609 | 195.0589108 | 194.6285866 | 289.6570065 | 199.324169 | 234.4103968 | 0.01954077 | 0.661559233 | YDL123W | SNA4 | n |
| W303.asm02.HP0_G0007640 | 1919.211888 | 2469.290119 | 1956.558426 | 3842.634421 | 3712.753961 | 3767.647337 | 3025.306512 | 3828.765712 | 4034.110803 | 7.42E-08 | -0.835550523 | YDL122W | UBP1 | n |
| W303.asm02.HP0_G0007650 | 1345.38692 | 1603.991461 | 1430.769998 | 720.8587611 | 828.0051722 | 761.3146108 | 1188.918955 | 815.680944 | 888.5075303 | 7.14E-08 | 0.923833678 | YDL121C | EXP1 | n |
| W303.asm02.HP0_G0007660 | 1866.869746 | 1841.020466 | 2020.08305 | 334.6496812 | 311.828871 | 295.1112522 | 714.676601 | 406.3890824 | 451.4191485 | 1.73E-36 | 2.604936993 | YDL120W | YFH1 | n |
| W303.asm02.HP0_G0007670 | 1004.193695 | 811.0390059 | 1027.144308 | 214.9929638 | 204.3474303 | 207.3020759 | 287.7638235 | 209.9676926 | 293.7807157 | 1.93E-32 | 2.18068929 | YDL119C | HEM25 | n |
| W303.asm02.HP0_G0007680 | 926.6497804 | 1006.183368 | 1008.575572 | 1440.744703 | 1516.682551 | 1416.71506 | 980.6688194 | 1320.764518 | 1317.406903 | 0.000141955 | -0.572135895 | YDL117W | CYK3 | n |
| W303.asm02.HP0_G0007690 | 1388.036073 | 1344.116287 | 1399.496337 | 1912.561841 | 1888.223333 | 1939.043872 | 1487.095285 | 1701.996181 | 1723.786499 | 2.33E-05 | -0.474475808 | YDL116W | NUP84 | n |
| W303.asm02.HP0_G0007700 | 880.1234316 | 871.0102 | 837.5477377 | 493.2191523 | 485.6568799 | 479.7820972 | 279.2444997 | 495.4076434 | 685.8295453 | 0.001660825 | 0.827591207 | YDL115C | IWR1 | n |

| | | | | | | | | | | | | | | |
|---|---|---|---|---|---|---|---|---|---|---|---|---|---|---|
| W303.asm02.HP0_G0007720 | 533.1144134 | 523.5580437 | 515.0381071 | 298.6553841 | 236.1937831 | 281.5325136 | 441.1116504 | 799.2318621 | 799.452052 | 5.98E-05 | 0.942803457 | YDL113C | ATG20 | n |
| W303.asm02.HP0_G0007730 | 878.1848337 | 1515.462556 | 1076.986706 | 2533.220262 | 2834.325397 | 2680.442999 | 1587.433987 | 2063.875983 | 1987.882065 | 1.34E-09 | -1.2132086 | YDL112W | TRM3 | n |
| W303.asm02.HP0_G0007740 | 1453.9484 | 1815.318526 | 1545.114321 | 1485.49437 | 1374.700895 | 1472.840513 | 1135.909829 | 1008.231962 | 830.1608376 | 0.357084667 | 0.151654568 | YDL111C | RRP42 | n |
| W303.asm02.HP0_G0007750 | 1666.224867 | 1129.933451 | 1427.838092 | 251.9600798 | 241.5015086 | 224.5018115 | 372.0104692 | 382.1992561 | 243.6230325 | 2.35E-33 | 2.557035121 | YDL110C | TMA17 | n |
| W303.asm02.HP0_G0007770 | 919.8646878 | 738.6928035 | 900.09506 | 424.1490146 | 471.0606348 | 444.4773768 | 460.0434809 | 425.7409435 | 367.4818011 | 3.70E-09 | 0.93455994 | YDL108W | KIN28 | n |
| W303.asm02.HP0_G0007780 | 936.3427697 | 907.1833012 | 906.9361734 | 468.8986813 | 538.7341345 | 476.1611002 | 569.8480978 | 562.171564 | 558.8998981 | 8.39E-13 | 0.893244569 | YDL107W | MSS2 | n |
| W303.asm02.HP0_G0007790 | 943.1278622 | 695.8562362 | 831.6839263 | 1035.079247 | 1058.89123 | 1070.909851 | 563.2219571 | 996.620845 | 1092.209142 | 0.135147304 | -0.357290224 | YDL106C | PHO2 | n |
| W303.asm02.HP0_G0007800 | 341.1932246 | 342.6925377 | 380.1704434 | 455.2792175 | 490.9646053 | 448.0983738 | 606.7651672 | 528.3058072 | 500.5532054 | 0.011380693 | -0.388627725 | YDL105W | NSE4 | n |
| W303.asm02.HP0_G0007810 | 513.7284347 | 558.7792212 | 561.9485988 | 496.1376088 | 506.8877817 | 474.3506017 | 558.4889995 | 484.7641198 | 509.7658411 | 0.305598638 | 0.146748664 | YDL104C | QRI7 | n |
| W303.asm02.HP0_G0007820 | 633.9215025 | 657.7792876 | 604.9498829 | 1147.926232 | 1126.56473 | 1129.751051 | 1368.771345 | 1460.097918 | 1369.611838 | 4.11E-14 | -0.843902229 | YDL103C | QRI1 | n |
| W303.asm02.HP0_G0007830 | 781.2549403 | 967.1544952 | 835.5931339 | 1375.565841 | 1531.278796 | 1539.828957 | 1171.880307 | 1519.121094 | 1397.249745 | 2.25E-07 | -0.782846808 | YDL102W | POL3 | n |
| W303.asm02.HP0_G0007840 | 1169.943813 | 931.9333178 | 1223.581993 | 475.7084132 | 484.3299485 | 452.62462 | 492.2275928 | 440.2548393 | 499.5295792 | 6.28E-16 | 1.235797451 | YDL101C | DUN1 | n |
| W303.asm02.HP0_G0007850 | 5983.482316 | 4858.618645 | 6002.588337 | 3167.498146 | 2720.2093 | 3218.161048 | 3317.803294 | 2975.348639 | 2750.483564 | 3.92E-10 | 0.886944782 | YDL100C | GET3 | n |
| W303.asm02.HP0_G0007860 | 1997.725102 | 1568.770284 | 2086.53958 | 972.8188408 | 944.7751324 | 1009.352903 | 1269.379234 | 1128.2135 | 993.9410276 | 1.28E-09 | 0.949183605 | YDL099W | BUG1 | n |
| W303.asm02.HP0_G0007870 | 532.1451145 | 517.8465014 | 608.8590905 | 99.22752177 | 118.0968916 | 120.3981489 | 159.9739676 | 114.1759803 | 128.9768996 | 8.06E-31 | 2.297767749 | YDL098C | SNU23 | n |
| W303.asm02.HP0_G0007880 | 5046.170247 | 3935.252641 | 4839.599063 | 2622.719595 | 2485.342448 | 2708.505726 | 2964.724655 | 3101.135736 | 3072.925813 | 1.08E-10 | 0.821919734 | YDL097C | RPN6 | n |
| W303.asm02.HP0_G0007890 | 1548.939696 | 1532.597182 | 1422.951582 | 6477.027842 | 5924.748548 | 6292.387467 | 6778.541907 | 9048.930236 | 8555.467671 | 1.12E-51 | -2.053243404 | YDL095W | PMT1 | n |
| W303.asm02.HP0_G0007910 | 1611.944126 | 1867.67433 | 1767.939157 | 1082.74737 | 1206.180611 | 1124.319556 | 1773.912517 | 1151.435734 | 1096.303646 | 0.00131919 | 0.620881868 | YDL092W | SRP14 | n |
| W303.asm02.HP0_G0007920 | 688.2022427 | 503.5676457 | 640.1327517 | 235.4221595 | 254.7708222 | 248.943541 | 566.0617317 | 481.8613406 | 618.2702169 | 3.53E-13 | 1.310202837 | YDL091C | UBX3 | n |
| W303.asm02.HP0_G0007930 | 1014.855983 | 835.7890225 | 999.7798549 | 327.8399494 | 330.4059101 | 385.6361762 | 425.0195945 | 328.0140451 | 405.35597 | 4.48E-18 | 1.447456993 | YDL090C | RAM1 | n |
| W303.asm02.HP0_G0007940 | 826.8119902 | 658.7312113 | 764.2500944 | 374.5352537 | 463.0990467 | 371.1521884 | 496.0139589 | 583.4586112 | 581.4196742 | 6.36E-08 | 0.899645155 | YDL089W | NUR1 | n |
| W303.asm02.HP0_G0007950 | 1494.658955 | 1643.020333 | 1521.659075 | 2243.320247 | 2555.66981 | 2316.532805 | 2039.904735 | 2237.07514 | 2084.102917 | 8.98E-08 | -0.610143031 | YDL088C | ASM4 | n |
| W303.asm02.HP0_G0007960 | 293.6975768 | 182.7693534 | 227.7113453 | 416.3664639 | 403.3871352 | 417.3198996 | 519.678747 | 420.9029782 | 459.608158 | 5.22E-05 | -0.813549214 | YDL087C | LUC7 | n |
| W303.asm02.HP0_G0007970 | 1393.851866 | 871.9621237 | 1325.221391 | 1295.794696 | 1007.140907 | 1302.653656 | 1152.001885 | 958.884716 | 1045.122337 | 0.974686121 | -0.006836123 | YDL086W | | n |
| W303.asm02.HP0_G0007980 | 439.0924168 | 598.7600172 | 516.9927109 | 667.3537248 | 794.8318881 | 665.3581913 | 1464.377088 | 952.1115646 | 819.9245758 | 0.053241487 | -0.451448985 | YDL085C-A | | n |
| W303.asm02.HP0_G0008000 | 9936.283367 | 7843.851418 | 10350.60454 | 9816.714923 | 9433.155079 | 9563.95822 | 9724.334732 | 8767.360657 | 8013.969418 | 0.820040973 | -0.034650157 | YDL084W | SUB2 | n |
| W303.asm02.HP0_G0008030 | 6790.908328 | 7657.27437 | 7171.441423 | 11280.80728 | 13191.02471 | 10796.00244 | 11085.53334 | 7948.776934 | 7438.691501 | 3.47E-05 | -0.705906443 | YDL081C | RPP1A | n |
| W303.asm02.HP0_G0008040 | 454.6011998 | 397.9041132 | 454.4453886 | 725.7228553 | 566.5996932 | 745.0201245 | 554.7026334 | 712.1484873 | 560.9471505 | 0.000301127 | -0.642754279 | YDL080C | THI3 | n |
| W303.asm02.HP0_G0008050 | 408.074851 | 252.2597847 | 388.9661606 | 216.9386015 | 230.8860576 | 203.681079 | 618.1242655 | 743.111465 | 881.342147 | 0.002469325 | 0.688974004 | YDL079C | MRK1 | n |
| W303.asm02.HP0_G0008070 | 3243.274231 | 2673.953718 | 3247.57425 | 1384.321211 | 1178.315053 | 1434.820045 | 2196.092337 | 1945.82963 | 1713.550237 | 2.77E-15 | 1.195968527 | YDL078C | MDH3 | n |
| W303.asm02.HP0_G0008080 | 462.3555912 | 526.4138148 | 481.8098421 | 345.3506885 | 345.0021551 | 339.468465 | 274.5115421 | 450.8983629 | 399.2142129 | 0.011093045 | 0.514170825 | YDL077C | VAM6 | n |
| W303.asm02.HP0_G0008090 | 1119.540268 | 770.1062861 | 1101.419254 | 515.5939856 | 459.1182526 | 495.1713342 | 300.0695133 | 552.4956335 | 755.436126 | 0.000323031 | 1.024509369 | YDL076C | RXT3 | n |
| W303.asm02.HP0_G0008110 | 2121.795365 | 2195.136089 | 2333.796963 | 1976.767885 | 1880.261745 | 1949.001614 | 1314.815628 | 1612.97762 | 1665.439806 | 0.143446369 | 0.195721499 | YDL074C | BRE1 | n |
| W303.asm02.HP0_G0008120 | 1246.518428 | 1611.606851 | 1412.201261 | 877.4825944 | 830.659035 | 908.8702369 | 658.8277011 | 865.9957828 | 777.9559021 | 7.32E-06 | 0.705843893 | YDL073W | AHK1 | n |
| W303.asm02.HP0_G0008130 | 2697.558932 | 1915.270516 | 2778.469333 | 506.8386161 | 435.233488 | 439.9511306 | 1330.907684 | 910.5050633 | 864.964128 | 1.09E-31 | 2.418462181 | YDL072C | YET3 | n |
| W303.asm02.HP0_G0008160 | 1752.492472 | 2483.568975 | 2057.220523 | 3899.057914 | 3516.368119 | 3496.977814 | 2472.497062 | 2518.644718 | 2359.458361 | 7.18E-08 | -0.79411451 | YDL067C | COX9 | n |
| W303.asm02.HP0_G0008170 | 1339.571126 | 1355.539371 | 1449.338734 | 1478.684638 | 1226.084582 | 1551.597197 | 1843.96029 | 1701.996181 | 1609.140366 | 0.786620608 | -0.03992721 | YDL066W | IDP1 | n |
| W303.asm02.HP0_G0008180 | 1775.755646 | 1672.529969 | 1928.21667 | 1247.153754 | 1101.353034 | 1203.07624 | 1327.121317 | 1160.144071 | 1007.248168 | 3.41E-05 | 0.59744606 | YDL065C | PEX19 | n |
| W303.asm02.HP0_G0008190 | 1248.457026 | 1378.38554 | 1296.879636 | 1061.345355 | 1153.103357 | 1008.447653 | 1189.865546 | 1090.477371 | 836.3025948 | 0.079889731 | 0.28466796 | YDL064W | UBC9 | n |
| W303.asm02.HP0_G0008200 | 1278.505293 | 2372.1939 | 1562.705756 | 4348.500219 | 4033.871352 | 4404.942801 | 1866.678486 | 2320.288142 | 1848.668894 | 8.91E-10 | -1.294429386 | YDL063C | SYO1 | n |
| W303.asm02.HP0_G0008220 | 1107.908681 | 1636.356867 | 1182.535312 | 9864.383046 | 9829.907558 | 9772.165545 | 4793.53948 | 6544.799413 | 5483.565484 | 9.81E-63 | -2.907558264 | YDL060W | TSR1 | n |
| W303.asm02.HP0_G0008230 | 297.5747726 | 321.7502159 | 338.1464612 | 138.1402754 | 119.4238229 | 137.5978845 | 212.983093 | 148.0417372 | 130.0005564 | 2.36E-08 | 1.273424981 | YDL059C | RAD59 | n |
| W303.asm02.HP0_G0008240 | 809.3646094 | 1200.375806 | 843.4115492 | 1954.393051 | 1754.203266 | 2035.905541 | 1396.222499 | 1712.639705 | 1460.714569 | 1.55E-08 | -1.009892856 | YDL058W | USO1 | n |
| W303.asm02.HP0_G0008250 | 261.710712 | 178.9616586 | 255.0757988 | 1011.731594 | 571.9074186 | 1044.657623 | 451.5241572 | 535.0789585 | 629.530105 | 5.51E-14 | -1.919404122 | YDL057W | | n |
| W303.asm02.HP0_G0008260 | 400.3204595 | 556.8753737 | 444.6723695 | 855.1077611 | 867.8131132 | 907.0597384 | 707.1038688 | 938.5652618 | 949.9251015 | 4.19E-07 | -0.90768098 | YDL056W | MBP1 | n |
| W303.asm02.HP0_G0008280 | 829.719887 | 646.356203 | 805.2967747 | 437.7684784 | 467.0798408 | 447.1931245 | 568.9015063 | 780.8475941 | 700.1603119 | 1.31E-05 | 0.755336963 | YDL054C | MCH1 | n |
| W303.asm02.HP0_G0008290 | 1600.312539 | 1519.27025 | 1792.371705 | 541.8600943 | 542.7149286 | 536.8127993 | 750.647079 | 541.8521099 | 534.3328696 | 3.02E-22 | 1.599122758 | YDL053C | PBP4 | n |
| W303.asm02.HP0_G0008300 | 704.6803246 | 461.6830022 | 661.6333937 | 2046.810841 | 1801.972795 | 1971.632845 | 2913.608713 | 2320.288142 | 2236.623219 | 7.00E-20 | -1.67130588 | YDL052C | SLC1 | n |
| W303.asm02.HP0_G0008310 | 3604.822733 | 4116.118147 | 4026.483873 | 3625.69582 | 4252.815027 | 3786.657571 | 4773.661058 | 3008.246803 | 3084.185701 | 0.95885471 | 0.010321759 | YDL051W | LHP1 | n |
| W303.asm02.HP0_G0008340 | 1135.049051 | 1017.606452 | 929.4141174 | 1704.378609 | 1719.70305 | 1566.081185 | 1151.055294 | 1535.570176 | 1411.580512 | 4.41E-06 | -0.694872292 | YDL047W | SIT4 | n |
| W303.asm02.HP0_G0008350 | 4501.424247 | 3558.29085 | 4686.162663 | 1040.91616 | 841.2744859 | 962.279942 | 1223.942841 | 804.0698274 | 1038.98058 | 3.68E-32 | 2.162919459 | YDL046W | NPC2 | n |
| W303.asm02.HP0_G0008360 | 1249.426325 | 1021.414147 | 1334.99441 | 898.8846089 | 956.7175147 | 904.3439907 | 1425.566836 | 1017.907892 | 974.49213 | 0.037922303 | 0.385980086 | YDL045W- | MRP10 | n |
| W303.asm02.HP0_G0008370 | 855.8909582 | 878.6255897 | 818.0016995 | 314.2204856 | 342.3482924 | 332.2264711 | 642.7356452 | 440.2548393 | 503.624084 | 8.18E-16 | 1.369090237 | YDL045C | FAD1 | n |
| W303.asm02.HP0_G0008380 | 978.0226238 | 845.3082596 | 881.5263237 | 895.9661524 | 906.2941228 | 927.8804709 | 1331.854275 | 1085.639406 | 1243.705817 | 0.925869961 | -0.013581768 | YDL044C | MTF2 | n |
| W303.asm02.HP0_G0008390 | 1028.426168 | 1109.943053 | 1038.871931 | 378.4265291 | 402.0602038 | 350.3314559 | 448.6843826 | 375.4261047 | 411.4977272 | 3.80E-27 | 1.492934025 | YDL043C | PRP11 | n |
| W303.asm02.HP0_G0008400 | 380.9344808 | 488.3368662 | 301.9862905 | 1688.813508 | 1977.127735 | 1798.73024 | 2245.315096 | 2606.695686 | 2430.088568 | 5.77E-36 | -2.221357095 | YDL042C | SIR2 | n |
| W303.asm02.HP0_G0008410 | 1549.908995 | 1230.837365 | 1610.593549 | 4756.111313 | 4891.069014 | 4624.918367 | 3721.997875 | 4322.23817 | 3918.441044 | 3.74E-36 | -1.70045941 | YDL040C | NAT1 | n |
| W303.asm02.HP0_G0008440 | 693.0487374 | 867.2025051 | 740.7948485 | 1350.272551 | 1458.297571 | 1327.095386 | 1016.639297 | 1413.653451 | 1715.597489 | 1.55E-05 | -0.845458169 | YDL035C | GPR1 | n |
| W303.asm02.HP0_G0008450 | 310.1756587 | 277.0098013 | 243.3481758 | 377.4537102 | 330.4059101 | 348.5209574 | 257.4728947 | 296.0834743 | 218.0323778 | 0.070128867 | -0.348140522 | YDL033C | SLM3 | n |
| W303.asm02.HP0_G0008460 | 996.4393036 | 1306.039338 | 1057.440668 | 5984.781509 | 5535.957658 | 6152.979084 | 2631.524438 | 3358.515488 | 2837.49179 | 7.84E-58 | -2.395146985 | YDL031W | DBP10 | n |
| W303.asm02.HP0_G0008470 | 595.1495451 | 674.9139145 | 585.4038447 | 409.556732 | 403.3871352 | 426.372392 | 372.0104691 | 349.3010923 | 345.9856512 | 3.35E-05 | 0.581490238 | YDL030W | PRP9 | n |
| W303.asm02.HP0_G0008480 | 3675.581556 | 2987.13662 | 3647.290732 | 2649.958522 | 2452.169164 | 2564.571097 | 2446.939091 | 2559.283626 | 2295.993537 | 0.000775396 | 0.427082513 | YDL029W | ARP2 | n |
| W303.asm02.HP0_G0008490 | 406.1362531 | 478.817629 | 415.3533122 | 537.968819 | 613.042291 | 574.8332674 | 544.2901266 | 606.6808445 | 707.3256952 | 0.012720226 | -0.407217833 | YDL028C | MPS1 | n |

| | | | | | | | | | | | | | | |
|---|---|---|---|---|---|---|---|---|---|---|---|---|---|---|
| W303.asm02.HP0_G0008510 | 411.9520467 | 369.3464017 | 351.828688 | 339.5137755 | 417.9833802 | 325.8897264 | 561.3287741 | 584.4262043 | 874.1767637 | 0.772044454 | 0.067423815 | YDL025C | RTK1 | n |
| W303.asm02.HP0_G0008520 | 170.5966123 | 86.62505814 | 115.3216255 | 660.5439929 | 481.6760858 | 730.5361366 | 118.3239406 | 371.5557325 | 65.51207597 | 0.000426237 | -2.329984955 | YDL024C | DIA3 | n |
| W303.asm02.HP0_G0008530 | 4427.757528 | 906.2313774 | 2764.787106 | 2160.630645 | 1932.012069 | 2144.535449 | 2442.206133 | 2542.834544 | 5091.516654 | 0.552143222 | 0.376776928 | YDL022W | GPD1 | n |
| W303.asm02.HP0_G0008550 | 2465.896487 | 1773.433883 | 2093.380693 | 1407.668863 | 1342.854542 | 1339.768875 | 1243.821263 | 1576.209084 | 2169.06389 | 0.002513287 | 0.630548169 | YDL020C | RPN4 | n |
| W303.asm02.HP0_G0008570 | 867.5225454 | 825.3178616 | 961.6650804 | 82.68960147 | 95.53905834 | 80.56718235 | 44.48980165 | 81.27781649 | 141.2604138 | 1.44E-27 | 3.361767949 | YDL018C | ERP3 | n |
| W303.asm02.HP0_G0008580 | 268.4958046 | 255.1155558 | 200.3468918 | 354.1060581 | 352.9637433 | 357.5734498 | 373.9036522 | 374.4585117 | 319.3713703 | 0.001857568 | -0.556333301 | YDL017W | CDC7 | n |
| W303.asm02.HP0_G0008590 | 5172.179109 | 4554.003056 | 5253.975074 | 2037.082653 | 1878.934814 | 2106.514981 | 2663.70855 | 2659.913304 | 2425.994063 | 3.84E-30 | 1.313933999 | YDL015C | TSC13 | n |
| W303.asm02.HP0_G0008600 | 3922.752784 | 4125.637384 | 3513.40037 | 14629.24973 | 16841.41289 | 14182.53984 | 10176.80548 | 10835.10701 | 10637.52334 | 2.57E-63 | -1.981188465 | YDL014W | NOP1 | n |
| W303.asm02.HP0_G0008610 | 509.851239 | 458.827231 | 416.3306141 | 475.7084132 | 404.7140666 | 426.372392 | 367.2775115 | 487.6668989 | 487.246065 | 0.66720531 | 0.082706554 | YDL013W | SLX5 | n |
| W303.asm02.HP0_G0008630 | 1002.255097 | 981.433351 | 1113.146877 | 187.7540363 | 200.3666362 | 209.1125744 | 350.2388641 | 190.6158315 | 265.1191824 | 7.25E-29 | 2.374312294 | YDL008W | APC11 | n |
| W303.asm02.HP0_G0008650 | 5492.047757 | 4254.147086 | 5123.016617 | 2659.686711 | 2139.013362 | 2529.266376 | 3104.820201 | 3009.214396 | 2932.689026 | 4.23E-13 | 1.020166284 | YDL007W | RPT2 | n |
| W303.asm02.HP0_G0008660 | 921.8032857 | 891.9525217 | 885.4355314 | 914.4497104 | 810.7550645 | 884.4285074 | 856.6653297 | 917.2782147 | 816.8536972 | 0.720770214 | 0.047333008 | YDL006W | PTC1 | n |
| W303.asm02.HP0_G0008670 | 1218.408759 | 1133.741146 | 1394.609827 | 332.7040436 | 387.4639588 | 362.099696 | 531.0378453 | 481.8613406 | 508.7422149 | 1.47E-35 | 1.793667892 | YDL005C | MED2 | n |
| W303.asm02.HP0_G0008680 | 3946.985257 | 4619.685793 | 4202.398217 | 4642.291508 | 4739.798839 | 4564.266668 | 3782.579732 | 3694.270278 | 3356.470267 | 0.277536285 | -0.127114975 | YDL004W | ATP16 | n |
| W303.asm02.HP0_G0008690 | 1063.32093 | 1231.789288 | 1000.757157 | 1601.259812 | 1678.568178 | 1614.964644 | 1518.332805 | 1621.685958 | 1570.242571 | 4.03E-06 | -0.570163966 | YDL003W | MCD1 | n |
| W303.asm02.HP0_G0008700 | 1616.790621 | 1387.904778 | 1646.75372 | 839.5426596 | 863.8323191 | 922.4489755 | 1079.114338 | 840.8383634 | 766.696014 | 3.33E-07 | 0.82461886 | YDL002C | NHP10 | n |
| W303.asm02.HP0_G0008710 | 1376.404485 | 1036.644927 | 1153.216255 | 603.1476813 | 682.042722 | 610.1379877 | 750.647079 | 796.329083 | 912.0509326 | 7.08E-09 | 0.913227371 | YDL001W | RMD1 | n |
| W303.asm02.HP0_G0008720 | 1029.395467 | 822.4620904 | 851.2299645 | 1847.382979 | 1525.971071 | 1803.256486 | 974.9892703 | 1302.38025 | 1155.673965 | 9.49E-09 | -0.938103296 | YDR001C | NTH1 | n |
| W303.asm02.HP0_G0008730 | 7202.860374 | 7510.678118 | 7493.951054 | 7323.380234 | 7859.414479 | 7181.34222 | 6111.194883 | 6103.576981 | 5020.886447 | 0.937840193 | -0.009993571 | YDR002W | YRB1 | n |
| W303.asm02.HP0_G0008750 | 1500.474749 | 1047.116087 | 1310.561863 | 350.2147827 | 317.1365964 | 365.7206929 | 345.5059065 | 387.0372214 | 394.096082 | 1.09E-28 | 1.898853515 | YDR004W | RAD57 | n |
| W303.asm02.HP0_G0008770 | 358.6406054 | 787.240913 | 331.3053478 | 757.825877 | 524.1378895 | 723.2941427 | 338.8797658 | 458.6391073 | 638.7427407 | 0.19922009 | -0.441554231 | YDR006C | SOK1 | n |
| W303.asm02.HP0_G0008780 | 426.4915307 | 418.8464349 | 381.1477453 | 804.5211814 | 650.1963692 | 859.9867779 | 1053.556367 | 163.523226 | 754.4124998 | 0.14522708 | -0.916447927 | YDR007W | TRP1 | n |
| W303.asm02.HP0_G0008790 | 105.6535837 | 112.3269985 | 131.935758 | 89.49933336 | 80.94281331 | 100.4826656 | 158.0807846 | 137.3982136 | 161.7329375 | 0.135076371 | 0.363120626 | YDR009W | GAL3 | n |
| W303.asm02.HP0_G0008800 | 2893.357316 | 2838.636521 | 2799.969975 | 6714.395639 | 5716.420324 | 5937.529765 | 3552.557992 | 8151.003882 | 5324.903425 | 4.21E-06 | -1.106313946 | YDR011W | SNQ2 | n |
| W303.asm02.HP0_G0008810 | 6778.307441 | 5123.253438 | 6202.935229 | 32327.7429 | 34453.77291 | 30353.91226 | 30307.96743 | 24282.71527 | 24311.12194 | 1.75E-60 | -2.4236379 | YDR012W | RPL4B | y |
| W303.asm02.HP0_G0008820 | 786.101435 | 652.0677453 | 801.387567 | 430.9587465 | 326.425116 | 429.0881397 | 494.1207758 | 401.5511172 | 399.2142129 | 3.81E-07 | 0.912914721 | YDR013W | PSF1 | n |
| W303.asm02.HP0_G0008830 | 357.6713064 | 560.6830686 | 443.6950676 | 251.9600798 | 285.2902436 | 257.9960334 | 347.3990895 | 410.2594547 | 370.5526797 | 0.000107542 | 0.778293129 | YDR014W | RAD61 | n |
| W303.asm02.HP0_G0008850 | 352.8248118 | 396.9521895 | 387.0115568 | 119.6567174 | 131.3662052 | 121.3033981 | 283.9774574 | 167.3935982 | 206.7724898 | 8.53E-13 | 1.612547546 | YDR016C | DAD1 | n |
| W303.asm02.HP0_G0008860 | 577.7021643 | 930.981394 | 570.744316 | 1113.877573 | 1098.699171 | 1101.688325 | 711.8368265 | 1042.097719 | 813.7828186 | 0.002392649 | -0.672370107 | YDR017C | KCS1 | n |
| W303.asm02.HP0_G0008880 | 308.2370608 | 326.5098345 | 352.8059899 | 3776.48274 | 1301.71967 | 3610.133969 | 1180.399631 | 1149.500547 | 1042.051458 | 2.00E-25 | -3.137731858 | YDR019C | GCV1 | n |
| W303.asm02.HP0_G0008890 | 820.9961966 | 1125.173832 | 904.0042677 | 1223.806102 | 1131.872455 | 1210.318234 | 843.4130484 | 613.4539959 | 524.0966077 | 0.113570989 | -0.323460774 | YDR020C | DAS2 | n |
| W303.asm02.HP0_G0008900 | 704.6803246 | 1031.885308 | 793.5691517 | 1288.984964 | 1216.796062 | 1347.916118 | 1032.731353 | 1027.583823 | 997.0119061 | 0.000172467 | -0.607357681 | YDR021W | FAL1 | n |
| W303.asm02.HP0_G0008910 | 484.6494667 | 375.057944 | 467.1503135 | 18.48355798 | 10.61545093 | 11.76824012 | 52.06253385 | 18.38426802 | 60.39394503 | 3.02E-35 | 5.006740215 | YDR022C | ATG31 | n |
| W303.asm02.HP0_G0008920 | 9479.743569 | 9128.948434 | 9691.903051 | 7159.946669 | 7202.583453 | 7053.702077 | 5799.766271 | 5683.641596 | 5306.478153 | 1.50E-05 | 0.402166294 | YDR023W | SES1 | n |
| W303.asm02.HP0_G0008950 | 856.8602571 | 953.8275632 | 912.7999849 | 999.0849495 | 1031.025671 | 961.3746928 | 879.3835263 | 1081.769034 | 1088.114637 | 0.348141577 | -0.134826339 | YDR027C | VPS54 | n |
| W303.asm02.HP0_G0008960 | 1210.654368 | 1470.722141 | 1351.608543 | 947.525551 | 1049.60271 | 912.4912338 | 1189.865546 | 1305.283029 | 1321.501407 | 0.000424641 | 0.472390751 | YDR028C | REG1 | n |
| W303.asm02.HP0_G0008990 | 3816.129901 | 3217.502159 | 3602.334844 | 1975.795066 | 1592.317639 | 1766.141267 | 1842.067107 | 1262.708953 | 1259.06021 | 2.22E-08 | 0.995132654 | YDR032C | PST2 | n |
| W303.asm02.HP0_G0009010 | 645.5530897 | 719.6543291 | 708.5438855 | 1047.725892 | 1126.56473 | 1120.698559 | 641.7890536 | 925.0189591 | 848.586109 | 4.03E-05 | -0.667822577 | YDR034C | LYS14 | n |
| W303.asm02.HP0_G0009030 | 4770.88935 | 4882.416738 | 4849.372082 | 2501.11724 | 1950.589108 | 2648.759276 | 2093.860452 | 1360.435833 | 1705.361228 | 8.49E-09 | 1.029689901 | YDR035W | ARO3 | n |
| W303.asm02.HP0_G0009040 | 625.1978121 | 602.5677121 | 574.6535237 | 2170.358834 | 2085.936107 | 2230.534127 | 1942.405808 | 1948.73241 | 2195.678171 | 5.32E-58 | -1.847916566 | YDR036C | EHD3 | n |
| W303.asm02.HP0_G0009050 | 6581.539758 | 5591.599907 | 6469.738651 | 13018.26173 | 13456.41098 | 12647.23713 | 9737.587013 | 9831.713016 | 8769.405544 | 1.98E-21 | -1.069314211 | YDR037W | KRS1 | n |
| W303.asm02.HP0_G0009090 | 14.539484 | 13.32693202 | 20.52334013 | 803.5483625 | 1041.641122 | 727.8203889 | 1116.031407 | 1552.986851 | 1456.620064 | 1.61E-89 | -5.732689653 | YDR040C | ENA1 | y |
| W303.asm02.HP0_G0009100 | 2172.19891 | 2034.260981 | 2131.495468 | 1348.326913 | 1426.451218 | 1470.124766 | 1805.150037 | 1346.88953 | 1265.201967 | 0.000216508 | 0.578279987 | YDR041W | RSM10 | n |
| W303.asm02.HP0_G0009120 | 707.5882214 | 570.2023058 | 607.8817886 | 107.9828913 | 145.9624502 | 140.3136322 | 127.7898558 | 139.3333997 | 181.1818351 | 1.89E-27 | 2.261421484 | YDR043C | NRG1 | n |
| W303.asm02.HP0_G0009140 | 943.1278622 | 1366.962456 | 1033.985422 | 1060.372537 | 1175.66119 | 1075.436097 | 1251.393995 | 866.9633759 | 713.4674523 | 0.951276273 | 0.0145197 | YDR045C | RPC11 | n |
| W303.asm02.HP0_G0009150 | 2505.637743 | 2634.924845 | 2327.933152 | 3749.243813 | 4117.468028 | 3735.963613 | 2217.863942 | 2432.528936 | 2763.790705 | 5.74E-07 | -0.635256923 | YDR046C | BAP3 | n |
| W303.asm02.HP0_G0009160 | 2746.023878 | 2200.847631 | 2611.350706 | 2541.975631 | 2494.630968 | 2691.30599 | 2406.235655 | 2360.92705 | 2270.402883 | 0.80867489 | -0.032333499 | YDR047W | HEM12 | n |
| W303.asm02.HP0_G0009170 | 582.548659 | 684.4331517 | 566.8351084 | 745.1792321 | 782.8895058 | 799.3350789 | 664.5072502 | 722.7920109 | 728.8218451 | 0.01322255 | -0.343655365 | YDR049W | VMS1 | n |
| W303.asm02.HP0_G0009180 | 20437.66801 | 19341.18606 | 19709.24764 | 20476.86378 | 17864.47698 | 19530.75235 | 12406.02852 | 11059.5886 | 9768.464702 | 0.770853498 | 0.03966864 | YDR050C | TPI1 | n |
| W303.asm02.HP0_G0009190 | 1665.255568 | 1584.952987 | 1584.206398 | 322.0030363 | 287.9441064 | 322.2687294 | 511.1594233 | 358.0094298 | 450.3955223 | 3.23E-50 | 2.372360419 | YDR051C | DET1 | n |
| W303.asm02.HP0_G0009200 | 807.4260115 | 898.6159877 | 798.4556613 | 1121.660123 | 1142.487906 | 1129.751051 | 527.2514792 | 1285.931168 | 1191.500882 | 0.111742836 | -0.438370892 | YDR052C | DBF4 | n |
| W303.asm02.HP0_G0009210 | 2922.436284 | 2926.213502 | 3091.205944 | 1629.471558 | 1419.816561 | 1651.174614 | 589.7265198 | 1267.5469 | 1111.658039 | 7.73E-05 | 0.92714513 | YDR054C | CDC34 | n |
| W303.asm02.HP0_G0009230 | 2000.632999 | 2112.318765 | 2118.790543 | 630.3866089 | 610.3884283 | 594.7487506 | 922.9267364 | 662.8012416 | 791.2630425 | 1.18E-33 | 1.763502353 | YDR056C | EMC10 | n |
| W303.asm02.HP0_G0009240 | 705.6496235 | 774.8659047 | 759.3635848 | 480.5725074 | 479.022223 | 510.5605713 | 707.1038688 | 633.77345 | 777.9559021 | 1.35E-05 | 0.606709462 | YDR057W | YOS9 | n |
| W303.asm02.HP0_G0009250 | 576.7328654 | 353.1636986 | 550.2209759 | 288.9271957 | 281.3094495 | 264.3327781 | 424.073003 | 413.1622338 | 498.5059531 | 9.98E-05 | 0.826819059 | YDR058C | TGL2 | n |
| W303.asm02.HP0_G0009270 | 2006.448792 | 3283.184896 | 2331.84236 | 7356.456074 | 6738.157475 | 6894.378211 | 3757.968352 | 3824.89534 | 3576.549897 | 5.64E-17 | -1.461478641 | YDR060W | MAK21 | n |
| W303.asm02.HP0_G0009280 | 1257.180717 | 1080.433417 | 1207.945162 | 130.3577247 | 168.5202835 | 144.8398784 | 294.3899641 | 250.6066008 | 361.340044 | 7.19E-59 | 3.004092436 | YDR061W | | n |
| W303.asm02.HP0_G0009290 | 2467.835085 | 2711.078743 | 2878.154128 | 2740.430675 | 2611.400928 | 2589.012826 | 2663.70855 | 2771.186505 | 2641.979189 | 0.860084996 | 0.020880364 | YDR062W | LCB2 | n |
| W303.asm02.HP0_G0009300 | 1308.55356 | 1546.876038 | 1343.790128 | 703.3480219 | 642.234781 | 601.0854953 | 1317.655402 | 908.5698772 | 1053.311346 | 5.74E-11 | 1.109206992 | YDR063W | AIM7 | n |
| W303.asm02.HP0_G0009320 | 569.9477729 | 650.1638979 | 551.1982778 | 468.8986813 | 477.6952917 | 459.8666139 | 441.1116504 | 392.8427797 | 386.9306987 | 0.022748614 | 0.333444596 | YDR065W | RRG1 | n |
| W303.asm02.HP0_G0009340 | 938.2813676 | 919.5583095 | 870.7760027 | 221.8026957 | 265.3862732 | 218.1650668 | 326.574076 | 253.50938 | 299.9224728 | 5.66E-34 | 1.956848022 | YDR067C | OCA6 | n |
| W303.asm02.HP0_G0009350 | 1751.523173 | 1531.645259 | 1852.964423 | 901.8030655 | 891.6978778 | 922.4489755 | 955.1108483 | 814.713351 | 810.7119401 | 6.17E-12 | 0.918892168 | YDR068W | DOS2 | n |
| W303.asm02.HP0_G0009360 | 707.5882214 | 602.5677121 | 712.4530931 | 410.5295508 | 352.9637433 | 401.9306625 | 407.0343556 | 422.8381643 | 432.9938771 | 7.33E-08 | 0.792547493 | YDR069C | DOA4 | n |

| | | | | | | | | | | | | | |
|---|---|---|---|---|---|---|---|---|---|---|---|---|---|
| W303.asm02.HP0_G0009370 | 326.6537406 | 161.8270317 | 292.2132714 | 17.51073914 | 25.21169595 | 9.957741639 | 29.34433726 | 20.31945412 | 22.51977611 | 1.70E-22 | 3.924540991 | YDR070C | FMP16 | n |
| W303.asm02.HP0_G0009380 | 3336.326929 | 3828.637185 | 3264.188383 | 2591.589392 | 2484.015517 | 2727.51596 | 2772.566575 | 2170.311219 | 1977.645793 | 0.007139992 | 0.418298493 | YDR071C | PAA1 | n |
| W303.asm02.HP0_G0009390 | 1159.281524 | 1084.241112 | 1075.032102 | 2113.935341 | 1792.684275 | 2123.714717 | 1306.296304 | 1799.723079 | 1942.842503 | 3.21E-07 | -0.862205158 | YDR072C | IPT1 | n |
| W303.asm02.HP0_G0009400 | 895.6322145 | 745.3562695 | 889.344739 | 218.8842392 | 200.3666362 | 192.8180881 | 260.3126993 | 177.0695288 | 218.0323778 | 1.64E-28 | 2.047291653 | YDR073W | SNF11 | n |
| W303.asm02.HP0_G0009410 | 1609.036229 | 874.8178948 | 1139.534028 | 1657.683305 | 1360.10465 | 1759.804522 | 903.0483144 | 1382.690473 | 1551.817299 | 0.12212036 | -0.399399872 | YDR074W | TPS2 | n |
| W303.asm02.HP0_G0009420 | 765.7461574 | 1338.404744 | 818.0016995 | 1352.218189 | 971.3137597 | 1345.200371 | 610.5515333 | 744.0790581 | 685.8295453 | 0.169095775 | -0.329005147 | YDR075W | PPH3 | n |
| W303.asm02.HP0_G0009430 | 260.7414131 | 227.5097681 | 219.89293 | 366.752703 | 360.9253315 | 367.5311914 | 128.7364473 | 436.3844671 | 510.7894673 | 0.099398831 | -0.629155798 | YDR076W | RAD55 | n |
| W303.asm02.HP0_G0009440 | 11347.58261 | 8416.909495 | 10692.66021 | 1686.86787 | 1032.352603 | 1171.392516 | 1098.99276 | 2983.089384 | 1645.990909 | 4.56E-23 | 2.968324513 | YDR077W | SED1 | n |
| W303.asm02.HP0_G0009450 | 344.1011214 | 268.4424879 | 294.1678752 | 311.3020291 | 382.1562333 | 318.6477325 | 327.5206675 | 217.708437 | 235.434023 | 0.49006203 | -0.156411971 | YDR078C | SHU2 | n |
| W303.asm02.HP0_G0009460 | 798.7023211 | 748.2120406 | 773.0458116 | 1078.856094 | 1176.988121 | 1105.309322 | 1249.500812 | 911.4726563 | 905.9091755 | 0.000879444 | -0.534409614 | YDR079W | PET100 | n |
| W303.asm02.HP0_G0009470 | 320.837947 | 321.7502159 | 316.6458192 | 30.15738407 | 47.76952917 | 52.50445592 | 45.43639318 | 49.34724572 | 70.6302069 | 7.39E-26 | 2.881678087 | YDR079C- | TFB5 | n |
| W303.asm02.HP0_G0009480 | 849.1058657 | 868.1544288 | 966.55159 | 444.5782103 | 483.0030171 | 462.5823616 | 402.3013979 | 648.2873458 | 622.3647217 | 8.90E-07 | 0.949596333 | YDR080W | VPS41 | n |
| W303.asm02.HP0_G0009490 | 487.5573635 | 676.8177619 | 477.9006345 | 821.0591017 | 724.5045257 | 814.7243159 | 602.0322096 | 702.4725568 | 598.8213194 | 0.003458778 | -0.523927583 | YDR081C | PDC2 | n |
| W303.asm02.HP0_G0009500 | 634.8908014 | 744.4043458 | 559.993995 | 103.1187971 | 107.4814406 | 119.4928997 | 138.2023626 | 116.1111664 | 143.3076662 | 3.45E-39 | 2.552706596 | YDR082W | STN1 | n |
| W303.asm02.HP0_G0009510 | 1481.08877 | 2435.972789 | 1711.255646 | 4476.912306 | 3902.505147 | 4434.816026 | 2053.157017 | 1977.760201 | 1911.110091 | 5.53E-11 | -1.187070876 | YDR083W | RRP8 | n |
| W303.asm02.HP0_G0009520 | 942.1585633 | 998.5679779 | 955.801269 | 648.8701668 | 640.9078497 | 670.7896868 | 982.5620025 | 771.1716636 | 726.7745928 | 0.000202693 | 0.562677131 | YDR084C | TVP23 | n |
| W303.asm02.HP0_G0009530 | 615.5048227 | 506.4234168 | 580.5173351 | 668.3265437 | 639.5809183 | 717.8626473 | 429.7525521 | 699.5697776 | 847.5624828 | 0.295494634 | -0.251365282 | YDR085C | AFR1 | n |
| W303.asm02.HP0_G0009540 | 2566.703576 | 3181.329058 | 2711.035501 | 1232.561471 | 1454.316777 | 1308.085152 | 1854.372797 | 1159.176478 | 1229.375051 | 2.05E-08 | 1.082804651 | YDR086C | SSS1 | n |
| W303.asm02.HP0_G0009550 | 2043.282152 | 2158.011064 | 2090.448788 | 2433.019921 | 2388.476458 | 2422.446966 | 2392.983374 | 1823.912906 | 1855.834277 | 0.149621306 | -0.203369564 | YDR087C | RRP1 | n |
| W303.asm02.HP0_G0009560 | 754.1145702 | 831.0294039 | 771.0912078 | 454.3063987 | 489.637674 | 423.6566443 | 625.6969977 | 447.9955837 | 593.7031884 | 2.44E-06 | 0.786531136 | YDR088C | SLU7 | n |
| W303.asm02.HP0_G0009570 | 742.482983 | 667.2985248 | 784.7734345 | 293.7912899 | 269.3670673 | 317.7424832 | 425.966186 | 618.2919611 | 607.0103289 | 1.33E-12 | 1.314562562 | YDR089W | VTC5 | n |
| W303.asm02.HP0_G0009580 | 944.0971612 | 1049.019935 | 969.4834957 | 24.32047102 | 29.19249005 | 38.02046808 | 85.19323721 | 60.95836237 | 101.3389925 | 3.73E-90 | 5.007949058 | YDR090C | ILT1 | n |
| W303.asm02.HP0_G0009590 | 4535.34971 | 4510.214565 | 4853.28129 | 8875.026285 | 8460.514388 | 8559.131563 | 4200.026594 | 5174.68765 | 4546.947523 | 5.24E-15 | -0.897739083 | YDR091C | RLI1 | n |
| W303.asm02.HP0_G0009600 | 1638.115197 | 1387.904778 | 1634.048795 | 537.968819 | 437.8873507 | 563.0650272 | 1030.83817 | 797.296676 | 884.4130255 | 6.17E-23 | 1.595547561 | YDR092W | UBC13 | n |
| W303.asm02.HP0_G0009630 | 683.3557481 | 792.9524553 | 740.7948485 | 1433.934971 | 1661.31807 | 1588.712416 | 1403.795231 | 1813.269382 | 1534.415654 | 1.16E-13 | -1.078540918 | YDR097C | MSH6 | n |
| W303.asm02.HP0_G0009650 | 5722.740903 | 6580.648647 | 5974.246582 | 8977.172263 | 7649.759324 | 8683.150709 | 8295.928121 | 8218.735396 | 8500.191857 | 3.63E-05 | -0.469845756 | YDR099W | BMH2 | n |
| W303.asm02.HP0_G0009660 | 822.9347945 | 694.9043125 | 804.3194727 | 841.4882973 | 565.2727618 | 846.4080393 | 1575.128297 | 987.9125076 | 1119.847049 | 0.861978865 | 0.041612462 | YDR100W | TVP15 | n |
| W303.asm02.HP0_G0009670 | 1580.92656 | 2789.136487 | 1750.347723 | 6481.891936 | 5920.767754 | 6489.731801 | 3417.195404 | 3346.904372 | 3284.816434 | 9.65E-17 | -1.626105474 | YDR101C | ARX1 | n |
| W303.asm02.HP0_G0009680 | 408.074851 | 584.4811615 | 375.2839338 | 63.23322465 | 67.67349965 | 59.74644984 | 68.15458977 | 87.08337481 | 114.6461329 | 2.75E-26 | 2.846496544 | YDR103W | STE5 | n |
| W303.asm02.HP0_G0009690 | 186.1053952 | 210.3751412 | 161.2548153 | 409.556732 | 387.4639588 | 439.0458814 | 301.0161048 | 399.6159311 | 443.230139 | 1.19E-08 | -1.148917271 | YDR104C | SPO71 | n |
| W303.asm02.HP0_G0009700 | 961.544542 | 800.567845 | 978.2792129 | 269.4708189 | 277.3286554 | 257.9960334 | 407.9809471 | 444.1252115 | 533.3092434 | 7.49E-27 | 1.768863758 | YDR105C | TMS1 | n |
| W303.asm02.HP0_G0009710 | 99.83779014 | 39.02887235 | 81.11605861 | 116.7382609 | 148.616313 | 116.777152 | 181.7455727 | 158.6852608 | 132.0477781 | 0.019606433 | -0.794157102 | YDR106W | ARP10 | n |
| W303.asm02.HP0_G0009730 | 708.5575203 | 711.0870157 | 714.4076969 | 756.8530582 | 736.446908 | 758.5988631 | 478.0287199 | 702.4725568 | 636.6954883 | 0.661696336 | -0.07780572 | YDR108W | TRS85 | n |
| W303.asm02.HP0_G0009740 | 268.4958046 | 316.9905974 | 278.5310446 | 261.6882682 | 177.808803 | 219.0703161 | 719.4095587 | 543.787296 | 864.964128 | 0.09318981 | 0.387742012 | YDR109C | | n |
| W303.asm02.HP0_G0009750 | 620.3513174 | 826.2697853 | 698.7708664 | 910.558435 | 928.851956 | 986.7216715 | 811.2289365 | 806.9726066 | 786.1449116 | 0.00643776 | -0.397748054 | YDR110W | FOB1 | n |
| W303.asm02.HP0_G0009770 | 297.5747726 | 217.9905309 | 273.6445351 | 442.6325726 | 552.0034482 | 457.1508662 | 664.5072502 | 583.4586112 | 674.5696572 | 1.02E-06 | -0.877139509 | YDR113C | PDS1 | n |
| W303.asm02.HP0_G0009790 | 1887.225023 | 1788.664662 | 1979.03637 | 1809.443044 | 1856.376981 | 1773.383261 | 1783.378432 | 1617.815585 | 1579.455207 | 0.6387039 | 0.056384461 | YDR116C | MRPL1 | n |
| W303.asm02.HP0_G0009800 | 955.7287484 | 851.9717256 | 1008.575572 | 1195.594355 | 1186.276641 | 1213.939231 | 1029.891579 | 1342.051565 | 1217.091536 | 0.015512984 | -0.35273715 | YDR117C | TMA64 | n |
| W303.asm02.HP0_G0009830 | 59.12723494 | 24.75001661 | 38.11477453 | 27.23892754 | 30.51942141 | 20.82073252 | 33.13070336 | 23.22223328 | 22.51977611 | 0.188971845 | 0.642268297 | YDR119W· | COX26 | n |
| W303.asm02.HP0_G0009840 | 1165.097318 | 1407.895176 | 1253.878352 | 2889.271957 | 3212.500837 | 3024.437711 | 1840.173924 | 2360.92705 | 2379.930885 | 5.86E-18 | -1.253580611 | YDR120C | TRM1 | n |
| W303.asm02.HP0_G0009850 | 925.6804814 | 962.3948767 | 981.2111186 | 520.4580798 | 541.3879972 | 528.6655561 | 564.1685486 | 507.9863531 | 483.1515603 | 1.90E-11 | 0.851782271 | YDR121W | DPB4 | n |
| W303.asm02.HP0_G0009860 | 429.3994275 | 453.1156887 | 388.9661606 | 1434.90779 | 1515.35562 | 1408.567817 | 1265.592868 | 1973.889829 | 1945.913381 | 5.49E-22 | -1.777095832 | YDR122W | KIN1 | n |
| W303.asm02.HP0_G0009870 | 321.8072459 | 308.4232839 | 272.6672332 | 253.9057175 | 290.5979691 | 247.1330425 | 573.6344639 | 367.6853603 | 445.2773913 | 0.37519577 | 0.192256149 | YDR123C | INO2 | n |
| W303.asm02.HP0_G0009890 | 230.6931462 | 214.1828361 | 217.9383262 | 658.5983552 | 601.0999087 | 642.7269603 | 708.9970519 | 629.9030778 | 733.9399761 | 3.29E-26 | -1.522324688 | YDR125C | ECM18 | n |
| W303.asm02.HP0_G0009900 | 663.0004705 | 384.5771812 | 587.3584485 | 406.6382755 | 402.0602038 | 355.7629513 | 784.7243739 | 594.1021348 | 560.9471505 | 0.034165398 | 0.490063278 | YDR126W | SWF1 | n |
| W303.asm02.HP0_G0009910 | 1863.961849 | 1980.001329 | 1547.068925 | 6235.76877 | 5926.07548 | 6070.601403 | 4288.059606 | 4687.020751 | 5011.673811 | 5.03E-42 | -1.757869568 | YDR127W | ARO1 | n |
| W303.asm02.HP0_G0009920 | 458.4783955 | 563.5388398 | 421.2171236 | 255.8513551 | 319.7904592 | 266.1432765 | 340.7729488 | 375.4261047 | 444.2537651 | 5.03E-05 | 0.781369773 | YDR128W | MTC5 | n |
| W303.asm02.HP0_G0009930 | 2320.501647 | 1812.462755 | 2071.880051 | 4795.996885 | 4027.236695 | 4648.454847 | 3772.167225 | 4305.789088 | 4267.497574 | 7.06E-16 | -1.118815297 | YDR129C | SAC6 | n |
| W303.asm02.HP0_G0009940 | 481.7415699 | 351.2598511 | 401.6710854 | 110.9013479 | 118.0968916 | 117.6824012 | 66.26140672 | 121.9167247 | 114.6461329 | 2.98E-13 | 1.832932312 | YDR130C | FIN1 | n |
| W303.asm02.HP0_G0009950 | 566.0705771 | 455.9714599 | 487.6736536 | 404.6926378 | 355.617606 | 365.7206929 | 216.7694591 | 252.5417869 | 301.9697252 | 0.021521194 | 0.422038376 | YDR131C | | n |
| W303.asm02.HP0_G0009960 | 699.8338299 | 689.1927702 | 647.951167 | 369.6711595 | 447.1758703 | 355.7629513 | 354.9718217 | 493.4724572 | 510.7894673 | 1.39E-05 | 0.799663921 | YDR132C | MRX16 | n |
| W303.asm02.HP0_G0009970 | 7168.934912 | 5499.263306 | 7369.833711 | 30615.58174 | 33880.53856 | 28443.83637 | 26144.85791 | 21622.80197 | 20956.69893 | 1.56E-46 | -2.213538652 | YLR110C | CCW12 | y |
| W303.asm02.HP0_G0009980 | 2065.576027 | 2198.943783 | 2136.381877 | 2024.436008 | 1998.358637 | 1935.422875 | 2740.382464 | 3255.950625 | 2718.751153 | 0.408237273 | 0.103505627 | YDR135C | YCF1 | n |
| W303.asm02.HP0_G0009990 | 923.7418836 | 1009.039139 | 1001.734459 | 331.7312247 | 313.1558023 | 314.1214863 | 405.1411725 | 386.0696283 | 443.230139 | 3.71E-33 | 1.612900939 | YDR137W | RGP1 | n |
| W303.asm02.HP0_G0010000 | 532.1451145 | 596.8561698 | 566.8351084 | 1081.774551 | 1198.219023 | 1132.466799 | 950.3778907 | 1061.44958 | 999.0591585 | 3.71E-16 | -1.007958365 | YDR138W | HPR1 | n |
| W303.asm02.HP0_G0010010 | 588.3644526 | 803.4236161 | 641.1100536 | 279.1990073 | 298.5595573 | 292.3955045 | 407.0343556 | 277.6992063 | 248.7411634 | 1.63E-08 | 1.224757497 | YDR139C | RUB1 | n |
| W303.asm02.HP0_G0010020 | 766.7154563 | 749.1639643 | 840.4796435 | 960.1721959 | 1078.7952 | 994.8689147 | 1565.662382 | 1071.12551 | 1143.390451 | 0.036903347 | -0.364034955 | YDR140W | MTQ2 | n |
| W303.asm02.HP0_G0010030 | 634.8908014 | 778.6735995 | 689.9751492 | 1846.41016 | 1843.107667 | 1882.01317 | 1588.380578 | 1902.287943 | 1910.086465 | 2.33E-25 | -1.405222792 | YDR141C | DOP1 | n |
| W303.asm02.HP0_G0010040 | 549.5924953 | 536.8849757 | 492.5601631 | 230.5580653 | 274.6747927 | 224.5018115 | 456.2571148 | 395.7455589 | 423.7812414 | 1.57E-12 | 1.118949102 | YDR142C | PEX7 | n |
| W303.asm02.HP0_G0010050 | 320.837947 | 400.7598843 | 329.350744 | 1334.70745 | 1506.0671 | 1383.220839 | 994.8676923 | 1312.05618 | 1427.958531 | 6.15E-31 | -2.006291228 | YDR143C | SAN1 | n |
| W303.asm02.HP0_G0010070 | 1845.545169 | 2153.251445 | 1960.467633 | 1346.381276 | 1398.58566 | 1396.799577 | 1154.84166 | 1492.028488 | 1602.998609 | 0.000555721 | 0.524996556 | YDR145W | TAF12 | n |
| W303.asm02.HP0_G0010100 | 2886.572224 | 2205.607249 | 2831.243636 | 2314.336022 | 2510.554144 | 2292.996325 | 1629.084014 | 1826.815685 | 1938.747998 | 0.309309852 | 0.154951516 | YDR148C | KGD2 | n |
| W303.asm02.HP0_G0010110 | 628.1057089 | 714.8947106 | 564.8805046 | 1939.800769 | 1699.79908 | 1972.538094 | 1760.660236 | 1669.098017 | 1700.243097 | 3.64E-31 | -1.557163465 | YDR150W | NUM1 | n |

| | | | | | | | | | | | | | | |
|---|---|---|---|---|---|---|---|---|---|---|---|---|---|---|
| W303.asm02.HP0_G0010130 | 2399.01486 | 2587.32866 | 2458.891608 | 2681.088725 | 2549.035154 | 2620.69655 | 2253.83442 | 2135.477869 | 2312.371556 | 0.47255753 | -0.07674875 | YDR152W | GIR2 | n |
| W303.asm02.HP0_G0010140 | 1175.759606 | 922.4140806 | 1076.986706 | 3110.101834 | 2851.575505 | 2931.197039 | 2460.191372 | 3109.844074 | 3303.241705 | 2.88E-22 | -1.486013005 | YDR153C | ENT5 | n |
| W303.asm02.HP0_G0010150 | 14646.10688 | 14540.63476 | 13805.36679 | 9192.165227 | 9281.884904 | 8679.529712 | 9609.797157 | 7296.619216 | 6885.93336 | 4.19E-06 | 0.662975072 | YDR155C | CPR1 | n |
| W303.asm02.HP0_G0010160 | 922.7725846 | 654.9235165 | 918.6637963 | 3038.11324 | 3016.114994 | 2995.469735 | 2326.721967 | 1792.949928 | 1791.345827 | 1.43E-27 | -1.858166338 | YDR156W | RPA14 | n |
| W303.asm02.HP0_G0010170 | 8461.979689 | 7986.639976 | 8250.382733 | 4903.979777 | 4482.374154 | 4749.842762 | 4271.96755 | 3106.941295 | 3384.108174 | 7.73E-09 | 0.804921314 | YDR158W | HOM2 | n |
| W303.asm02.HP0_G0010180 | 507.9126411 | 719.6543291 | 531.6522396 | 1209.213819 | 1269.873317 | 1257.391194 | 722.2493332 | 1248.195039 | 1255.989331 | 1.45E-06 | -1.086563626 | YDR159W | SAC3 | n |
| W303.asm02.HP0_G0010190 | 527.2986198 | 676.8177619 | 539.4706549 | 320.0573986 | 378.1754392 | 307.7847416 | 271.6717675 | 410.2594547 | 365.4345488 | 9.38E-05 | 0.796003791 | YDR160W | SSY1 | n |
| W303.asm02.HP0_G0010200 | 1728.259998 | 1606.847232 | 1738.6201 | 4609.215668 | 4389.488958 | 4550.687929 | 2940.113275 | 3033.404223 | 2409.616044 | 3.42E-30 | -1.417243419 | YDR161W | ACL4 | n |
| W303.asm02.HP0_G0010210 | 1278.505293 | 1206.087348 | 1331.085203 | 609.9574132 | 591.8113891 | 569.4017719 | 776.2050501 | 669.574393 | 690.9476762 | 1.12E-18 | 1.107426224 | YDR162C | NBP2 | n |
| W303.asm02.HP0_G0010220 | 1050.720044 | 901.4717589 | 1119.010688 | 190.6724928 | 232.212989 | 196.4390851 | 347.3990895 | 205.1297273 | 281.4972014 | 1.94E-28 | 2.313320431 | YDR163W | CWC15 | n |
| W303.asm02.HP0_G0010230 | 593.2109473 | 580.6734666 | 590.2903542 | 823.0047393 | 883.7362896 | 878.0917627 | 959.8438059 | 973.3986118 | 935.5943349 | 1.83E-06 | -0.55062431 | YDR164C | SEC1 | n |
| W303.asm02.HP0_G0010240 | 2398.045561 | 3220.357931 | 2755.014087 | 3710.331059 | 3836.158578 | 3798.425811 | 2521.719821 | 2743.126306 | 2370.718249 | 0.001688159 | -0.438073729 | YDR165W | TRM82 | n |
| W303.asm02.HP0_G0010250 | 917.92609 | 1133.741146 | 1021.280497 | 1135.279587 | 1299.065807 | 1180.445009 | 1111.29845 | 1359.46824 | 1176.146489 | 0.118182561 | -0.233420736 | YDR166C | SEC5 | n |
| W303.asm02.HP0_G0010260 | 1696.273133 | 1691.568443 | 1804.099328 | 799.6570872 | 899.659466 | 815.6295652 | 1308.189487 | 920.1809938 | 920.2399421 | 2.37E-10 | 1.04655928 | YDR167W | TAF10 | n |
| W303.asm02.HP0_G0010270 | 3250.059324 | 3689.656322 | 3297.416648 | 1481.603095 | 1445.028257 | 1458.356526 | 1453.01799 | 1428.167347 | 1485.281597 | 4.38E-31 | 1.223041297 | YDR168W | CDC37 | n |
| W303.asm02.HP0_G0010280 | 283.0352886 | 338.8848428 | 218.9156281 | 35.02147827 | 31.84635278 | 28.96797568 | 134.4159965 | 67.73151374 | 187.3235922 | 1.28E-17 | 3.134099753 | YDR169C | STB3 | n |
| W303.asm02.HP0_G0010300 | 1304.676364 | 1646.828028 | 1326.198693 | 2739.457856 | 2622.016379 | 2735.663203 | 2839.774574 | 2590.246604 | 2566.230851 | 1.74E-12 | -0.920651556 | YDR170C | SEC7 | n |
| W303.asm02.HP0_G0010310 | 3177.361904 | 1546.876038 | 1791.394403 | 235.4221595 | 293.2518318 | 236.2700516 | 1593.113536 | 675.3799513 | 1348.115688 | 7.61E-23 | 3.092009174 | YDR171W | HSP42 | n |
| W303.asm02.HP0_G0010320 | 3803.529015 | 3679.185162 | 3721.565677 | 7038.344313 | 7194.621865 | 7157.80574 | 5020.721446 | 5533.664673 | 5379.155613 | 5.79E-24 | -0.932910094 | YDR172W | SUP35 | n |
| W303.asm02.HP0_G0010330 | 761.8689617 | 398.8560369 | 678.2475262 | 325.8943117 | 335.7136355 | 286.9640091 | 360.6513709 | 327.0464521 | 326.5367536 | 4.81E-05 | 0.956030012 | YDR173C | ARG82 | n |
| W303.asm02.HP0_G0010340 | 1560.571283 | 1578.289521 | 1533.386698 | 1816.252776 | 1947.935245 | 1864.813434 | 5059.531699 | 1718.445263 | 3926.630053 | 0.374590702 | -0.268696503 | YDR174W | HMO1 | n |
| W303.asm02.HP0_G0010350 | 2131.488355 | 1475.481759 | 2103.153712 | 2995.309211 | 3189.943003 | 3108.62589 | 7531.082169 | 2894.070823 | 6497.979035 | 0.014383171 | -0.702750897 | YDR175C | RSM24 | n |
| W303.asm02.HP0_G0010360 | 858.798855 | 687.2889228 | 803.3421708 | 1026.323877 | 1114.622347 | 1041.941875 | 2445.045908 | 1051.773649 | 3285.84006 | 0.166220276 | -0.437911166 | YDR176W | NGG1 | n |
| W303.asm02.HP0_G0010370 | 2755.716868 | 2817.694199 | 2953.406375 | 1347.354095 | 1557.817423 | 1399.515325 | 1462.483905 | 1133.051466 | 1073.78387 | 2.49E-11 | 0.986791354 | YDR177W | UBC1 | n |
| W303.asm02.HP0_G0010390 | 297.5747726 | 250.3559373 | 310.7820077 | 147.8684638 | 159.2317639 | 152.9871215 | 584.0469706 | 169.3287844 | 384.8834463 | 0.011220023 | 0.900707749 | YDR179C | CSN9 | n |
| W303.asm02.HP0_G0010400 | 532.1451145 | 672.0581434 | 550.2209759 | 442.6325726 | 537.4072031 | 449.9088722 | 1417.994104 | 515.7270975 | 1282.603612 | 0.354114835 | 0.295856586 | YDR179W- | NVJ3 | n |
| W303.asm02.HP0_G0010410 | 352.8248118 | 570.2023058 | 342.0556688 | 840.5154785 | 780.2356431 | 905.2492399 | 1334.69405 | 710.2133012 | 1658.274423 | 0.000623457 | -0.997784071 | YDR180W | SCC2 | n |
| W303.asm02.HP0_G0010420 | 875.2769369 | 971.9141138 | 861.9802855 | 455.2792175 | 540.0610659 | 458.0561154 | 1602.579451 | 477.0233753 | 1336.8558 | 0.006547861 | 0.899035314 | YDR181C | SAS4 | n |
| W303.asm02.HP0_G0010430 | 1105.000784 | 1025.221842 | 1100.441952 | 1125.551399 | 1178.315053 | 1126.130054 | 3594.208019 | 1553.954444 | 4126.23716 | 0.775013135 | -0.08628884 | YDR182W | CDC1 | n |
| W303.asm02.HP0_G0010450 | 772.5312499 | 822.4620904 | 882.5036256 | 708.2121161 | 748.3892903 | 710.6206533 | 1500.347566 | 686.9910679 | 1902.921082 | 0.525103363 | 0.193195898 | YDR183W | PLP1 | n |
| W303.asm02.HP0_G0010460 | 1266.873706 | 1977.145558 | 1368.222675 | 2818.256182 | 2243.84094 | 2716.652969 | 2947.686007 | 1344.954344 | 1987.882055 | 0.004688915 | -0.754230779 | YDR184C | ATC1 | n |
| W303.asm02.HP0_G0010480 | 751.2066734 | 915.7506146 | 743.7267543 | 731.5597683 | 778.9087117 | 725.1046412 | 1379.183851 | 737.3059067 | 1537.486533 | 0.677954416 | 0.109130116 | YDR186C | SND1 | n |
| W303.asm02.HP0_G0010490 | 2793.519526 | 2572.09788 | 2429.572551 | 4802.806617 | 4705.298623 | 4638.497105 | 9584.239186 | 3869.404621 | 9689.645486 | 0.001659622 | -0.859797093 | YDR188W | CCT6 | n |
| W303.asm02.HP0_G0010500 | 949.9129548 | 921.4621569 | 959.7104766 | 2035.137015 | 2040.820441 | 2107.420231 | 4258.715269 | 2117.093601 | 5356.635836 | 1.54E-05 | -1.12707076 | YDR189W | SLY1 | n |
| W303.asm02.HP0_G0010510 | 3651.349082 | 3298.415675 | 3607.221353 | 3488.528363 | 3473.906316 | 3531.377285 | 7039.801168 | 2791.505959 | 6856.2482 | 0.975958725 | 0.008647511 | YDR190C | RVB1 | n |
| W303.asm02.HP0_G0010520 | 510.8205379 | 331.2694531 | 433.9220485 | 267.5251812 | 264.0593418 | 320.4582309 | 802.7096128 | 298.9862535 | 899.7674184 | 0.092169246 | 0.581854976 | YDR191W | HST4 | n |
| W303.asm02.HP0_G0010530 | 317.9300502 | 272.2501827 | 282.4402523 | 824.950937 | 752.3700844 | 840.0712947 | 1903.595556 | 867.9309689 | 2535.522065 | 7.86E-07 | -1.470311379 | YDR192C | NUP42 | n |
| W303.asm02.HP0_G0010540 | 4799.968318 | 6616.821748 | 5339.977642 | 3443.778697 | 2562.304467 | 3599.270978 | 4177.308398 | 2081.292658 | 5495.848998 | 0.005439761 | 0.80262525 | YDR194C | MSS116 | n |
| W303.asm02.HP0_G0010560 | 689.1715417 | 566.3946109 | 659.6787899 | 1141.1165 | 1068.179749 | 1118.888061 | 2293.591264 | 988.8801006 | 1893.708446 | 0.001917829 | -0.797379123 | YDR195W | REF2 | n |
| W303.asm02.HP0_G0010570 | 1475.272977 | 1324.125889 | 1492.340018 | 347.2963262 | 429.9257625 | 348.5209574 | 1312.922445 | 416.065013 | 1391.107988 | 6.20E-09 | 1.93161296 | YDR196C | CAB5 | n |
| W303.asm02.HP0_G0010580 | 881.0927305 | 758.6832015 | 849.2753607 | 769.4997031 | 808.1012018 | 759.5041123 | 1890.343275 | 686.9910679 | 1974.574915 | 0.783375583 | 0.09100458 | YDR197W | CBS2 | n |
| W303.asm02.HP0_G0010590 | 712.4347161 | 973.8179613 | 792.5918498 | 893.0476959 | 1074.814406 | 931.5014679 | 2630.577847 | 1038.227346 | 2322.607818 | 0.452596101 | -0.225647385 | YDR198C | RKM2 | n |
| W303.asm02.HP0_G0010610 | 571.8863707 | 571.1542295 | 579.5400332 | 241.2590725 | 276.0017241 | 238.0805501 | 1126.443914 | 321.2408937 | 870.0822589 | 0.00044951 | 1.190280145 | YDR201W | SPC19 | n |
| W303.asm02.HP0_G0010620 | 1115.663072 | 780.577447 | 990.9841377 | 160.5151087 | 225.5783322 | 199.1548328 | 875.5971602 | 257.3797522 | 765.6723879 | 4.05E-11 | 2.304069282 | YDR202C | RAV2 | n |
| W303.asm02.HP0_G0010630 | 684.325047 | 481.6734002 | 727.1126218 | 886.237964 | 820.043584 | 872.6602673 | 2779.192716 | 932.7597035 | 2518.12042 | 0.17659494 | -0.446218816 | YDR204W | COQ4 | n |
| W303.asm02.HP0_G0010640 | 536.9916091 | 479.7695528 | 527.7430319 | 1261.746037 | 1405.220316 | 1267.348936 | 1283.578107 | 1722.315635 | 1513.943131 | 1.46E-19 | -1.348375254 | YDR205W | MSC2 | n |
| W303.asm02.HP0_G0010650 | 1787.387233 | 1975.24171 | 1934.080482 | 3374.708559 | 3751.234971 | 3496.072565 | 2747.955196 | 4135.49271 | 3698.361414 | 2.46E-08 | -0.898681458 | YDR206W | EBS1 | n |
| W303.asm02.HP0_G0010660 | 229.7238472 | 246.5482424 | 185.6873631 | 491.2735146 | 618.3500164 | 551.2967871 | 381.4763844 | 734.4031276 | 715.5147047 | 1.76E-07 | -1.326171091 | YDR207C | UME6 | n |
| W303.asm02.HP0_G0010670 | 615.5048227 | 778.6735995 | 641.1100536 | 715.021848 | 866.4861818 | 681.6526777 | 695.7447705 | 1178.528339 | 1026.697066 | 0.502943036 | -0.15181512 | YDR208W | MSS4 | n |
| W303.asm02.HP0_G0010700 | 2097.562892 | 2396.943916 | 2253.658207 | 4265.810617 | 4447.873938 | 4373.259078 | 3681.294439 | 3684.594347 | 3215.209853 | 2.03E-17 | -0.955448401 | YDR211W | GCD6 | n |
| W303.asm02.HP0_G0010710 | 3225.82685 | 3425.021529 | 3251.483458 | 5072.277436 | 4714.587143 | 5140.910434 | 4069.396964 | 4226.446457 | 4208.127255 | 7.10E-10 | -0.592458387 | YDR212W | TCP1 | n |
| W303.asm02.HP0_G0010720 | 157.0264272 | 228.4616918 | 139.7541733 | 1510.78766 | 991.2177302 | 1427.578051 | 514.9457894 | 587.3289834 | 609.0575813 | 1.61E-38 | -2.903844487 | YDR213W | UPC2 | n |
| W303.asm02.HP0_G0010730 | 4951.178952 | 3432.636919 | 4026.483873 | 1072.046363 | 1134.526318 | 1084.489589 | 3258.168028 | 2287.389978 | 2542.687448 | 4.24E-27 | 1.915149238 | YDR214W | AHA1 | n |
| W303.asm02.HP0_G0010750 | 680.4478513 | 722.5101003 | 626.4505249 | 209.1560508 | 197.7127735 | 182.8603465 | 151.4546439 | 187.7130524 | 242.5994063 | 1.21E-19 | 1.783571242 | YDR216W | ADR1 | n |
| W303.asm02.HP0_G0010760 | 437.153819 | 653.019669 | 430.9901427 | 505.8657972 | 553.3303795 | 487.9293403 | 400.4082149 | 508.9539461 | 540.4746267 | 0.917804428 | -0.023287982 | YDR217C | RAD9 | n |
| W303.asm02.HP0_G0010770 | 78.51321361 | 59.01927038 | 55.70620893 | 115.7654421 | 156.5779012 | 98.67216715 | 149.5614609 | 126.75469 | 109.528002 | 0.001433026 | -0.933472491 | YDR218C | SPR28 | n |
| W303.asm02.HP0_G0010780 | 621.3206163 | 690.144694 | 553.1528816 | 681.9460074 | 822.6974468 | 659.9266959 | 829.2141755 | 709.2457082 | 775.9086497 | 0.194353998 | -0.212911353 | YDR219C | MFB1 | n |
| W303.asm02.HP0_G0010790 | 665.9083673 | 682.5293042 | 673.3610167 | 1271.474225 | 1187.603572 | 1232.044216 | 1300.616755 | 1731.991566 | 1778.038687 | 7.86E-09 | -0.86872396 | YDR221W | GTB1 | n |
| W303.asm02.HP0_G0010800 | 1469.457183 | 1240.356602 | 1304.698051 | 21.4020145 | 15.92317639 | 34.39947112 | 34.07729488 | 28.06019855 | 37.87416892 | 3.62E-116 | 5.765543824 | YDR222W | | n |
| W303.asm02.HP0_G0010820 | 4437.450517 | 6721.533357 | 4602.114699 | 5503.236183 | 5342.225679 | 5292.992306 | 6515.389463 | 3916.81668 | 4759.861769 | 0.882853312 | -0.034120479 | YDR224C | HTB1 | y |
| W303.asm02.HP0_G0010840 | 6140.508743 | 7449.755 | 6354.417025 | 6352.507031 | 6070.710998 | 6009.044455 | 7840.617598 | 5764.919412 | 5861.283547 | 0.485045444 | 0.113768719 | YDR226W | ADK1 | n |
| W303.asm02.HP0_G0010850 | 414.8599435 | 579.7215429 | 458.3545963 | 463.0617682 | 472.3875662 | 493.3608358 | 234.7546981 | 382.1992561 | 436.0647557 | 0.924855414 | 0.024116866 | YDR227W | SIR4 | n |
| W303.asm02.HP0_G0010860 | 994.5007057 | 853.8755731 | 1011.507478 | 915.4225292 | 1036.333397 | 989.4374192 | 715.6231926 | 878.5744925 | 847.5624828 | 0.799381187 | -0.039857123 | YDR228C | PCF11 | n |

| | | | | | | | | | | | | | | |
|---|---|---|---|---|---|---|---|---|---|---|---|---|---|---|
| W303.asm02.HP0_G0010870 | 745.3908798 | 472.154163 | 594.1995619 | 512.6755291 | 540.0610659 | 456.2456169 | 514.9457894 | 555.3984127 | 570.1597861 | 0.176557062 | 0.264934814 | YDR229W | IVY1 | n |
| W303.asm02.HP0_G0010880 | 890.7857198 | 728.2216426 | 915.7318906 | 679.0275509 | 736.446908 | 711.5259026 | 810.282345 | 719.8892318 | 665.3570215 | 0.101729265 | 0.253479231 | YDR231C | COX20 | n |
| W303.asm02.HP0_G0010890 | 614.5355238 | 1049.971859 | 558.0393912 | 2329.901124 | 1861.684706 | 2453.22544 | 764.8459518 | 1261.741342 | 1230.398677 | 1.39E-09 | -1.580184813 | YDR232W | HEM1 | n |
| W303.asm02.HP0_G0010900 | 5373.793287 | 4664.426207 | 5137.676146 | 8606.528285 | 7994.761479 | 8266.736059 | 6806.939653 | 7140.836734 | 6965.776202 | 3.33E-12 | -0.712629116 | YDR233C | RTN1 | n |
| W303.asm02.HP0_G0010910 | 1443.286112 | 1445.972124 | 1232.37771 | 636.2235219 | 720.5237316 | 808.3875713 | 422.17982 | 715.0512665 | 580.396048 | 3.58E-06 | 0.928571836 | YDR234W | LYS4 | n |
| W303.asm02.HP0_G0010920 | 1087.553403 | 1060.443019 | 988.052232 | 619.6856016 | 615.6961537 | 603.801243 | 338.8797658 | 515.7270975 | 580.396048 | 7.16E-05 | 0.76997275 | YDR235W | PRP42 | n |
| W303.asm02.HP0_G0010930 | 852.9830614 | 850.0678782 | 863.9348893 | 488.3550581 | 439.2142821 | 487.0240911 | 574.5810554 | 492.5048642 | 487.246065 | 2.42E-10 | 0.857680741 | YDR236C | FMN1 | n |
| W303.asm02.HP0_G0010940 | 1748.615276 | 1783.905043 | 1871.53316 | 2342.547769 | 2344.687723 | 2280.322835 | 2321.042418 | 2021.301889 | 2027.803476 | 0.00103326 | -0.366512111 | YDR237W | MRPL7 | n |
| W303.asm02.HP0_G0010950 | 2847.800266 | 2598.751744 | 2946.565262 | 5144.26603 | 4792.876093 | 5006.933546 | 3825.176351 | 4911.502339 | 4565.372794 | 1.17E-10 | -0.8324352 | YDR238C | SEC26 | n |
| W303.asm02.HP0_G0010960 | 374.1493883 | 476.9137816 | 396.7845759 | 346.3235073 | 406.0409979 | 392.8781701 | 375.7968352 | 497.3428295 | 493.3878221 | 0.524199554 | 0.125027162 | YDR239C | | n |
| W303.asm02.HP0_G0010970 | 658.1539758 | 706.3273971 | 642.0873555 | 613.8486886 | 642.234781 | 563.9702765 | 475.1889453 | 552.4956335 | 533.3092434 | 0.324104154 | 0.142361679 | YDR240C | SNU56 | n |
| W303.asm02.HP0_G0010980 | 304.3598651 | 409.3271978 | 345.9648765 | 222.7755146 | 268.0401359 | 174.7131033 | 193.104671 | 223.5139953 | 175.040078 | 0.002465671 | 0.676653794 | YDR242W | AMD2 | n |
| W303.asm02.HP0_G0010990 | 336.3467299 | 428.3656721 | 329.350744 | 664.4352683 | 589.1575264 | 621.9062278 | 503.5866911 | 550.5604474 | 540.4746267 | 9.46E-07 | -0.778198852 | YDR243C | PRP28 | n |
| W303.asm02.HP0_G0011000 | 775.4391467 | 673.0100671 | 760.3408868 | 406.6382755 | 423.2911057 | 391.9729209 | 701.4243197 | 648.2873458 | 881.342147 | 1.50E-07 | 0.855081909 | YDR244W | PEX5 | n |
| W303.asm02.HP0_G0011010 | 2735.36159 | 2441.684331 | 2740.354558 | 3519.658566 | 3850.754823 | 3459.862595 | 3470.204529 | 4387.066904 | 3779.227882 | 0.000613848 | -0.451694832 | YDR245W | MNN10 | n |
| W303.asm02.HP0_G0011020 | 610.6583281 | 406.4714266 | 572.6989198 | 313.2476667 | 282.6363809 | 337.6579665 | 113.5909829 | 184.8102732 | 205.7488636 | 0.002016648 | 0.766585006 | YDR246W | TRS23 | n |
| W303.asm02.HP0_G0011050 | 1123.417464 | 1031.885308 | 1073.077498 | 53.50503625 | 41.13487234 | 43.45196352 | 117.377349 | 71.60188595 | 68.58295453 | 6.43E-82 | 4.5407741 | YDR248C | | n |
| W303.asm02.HP0_G0011060 | 373.1800894 | 478.817629 | 315.6685173 | 250.0144421 | 286.617175 | 260.7117811 | 315.2149777 | 451.865956 | 363.3872964 | 0.011697065 | 0.552167469 | YDR249C | | n |
| W303.asm02.HP0_G0011100 | 443.9389115 | 423.6060535 | 464.2184077 | 320.0573986 | 294.5787632 | 310.5004893 | 386.209342 | 455.7363282 | 358.2691654 | 0.00110722 | 0.524147894 | YDR253C | MET32 | n |
| W303.asm02.HP0_G0011110 | 392.566068 | 335.077148 | 388.9661606 | 428.04029 | 338.3674983 | 397.4044163 | 362.5445539 | 312.5325563 | 319.3713703 | 0.732234991 | -0.062800283 | YDR254W | CHL4 | n |
| W303.asm02.HP0_G0011120 | 330.5309363 | 349.3560037 | 315.6685173 | 312.2748479 | 310.5019396 | 320.4582309 | 517.7855639 | 352.2038714 | 375.6708106 | 0.706308741 | 0.077612743 | YDR255C | RMD5 | n |
| W303.asm02.HP0_G0011130 | 330.5309363 | 279.8655724 | 311.7593096 | 346.3235073 | 538.7341345 | 366.6259422 | 629.4833638 | 739.2410928 | 978.5866348 | 0.062982799 | -0.437271926 | YDR256C | CTA1 | n |
| W303.asm02.HP0_G0011140 | 1247.487727 | 1096.616121 | 1232.37771 | 2246.238703 | 2127.070979 | 2296.617322 | 1402.848639 | 1579.111863 | 1493.470607 | 9.32E-15 | -0.899611955 | YDR257C | RKM4 | n |
| W303.asm02.HP0_G0011150 | 5121.775564 | 3722.021729 | 3629.699297 | 247.0959856 | 468.4067721 | 321.3634802 | 942.8051585 | 660.8660555 | 1017.48443 | 1.81E-48 | 3.592692218 | YDR258C | HSP78 | n |
| W303.asm02.HP0_G0011170 | 787.0707339 | 631.1254236 | 776.9550192 | 180.9443044 | 185.7703912 | 149.3661246 | 165.6535168 | 173.1991566 | 233.3867706 | 5.88E-26 | 2.09201715 | YDR260C | SWM1 | n |
| W303.asm02.HP0_G0011190 | 856.8602571 | 697.7600837 | 743.7267543 | 354.1060581 | 400.7332725 | 351.2367051 | 547.1299012 | 508.9539461 | 701.1839381 | 1.47E-09 | 1.05710628 | YDR262W | | n |
| W303.asm02.HP0_G0011210 | 2084.962006 | 1866.722407 | 2167.655639 | 1071.073544 | 1093.391445 | 1133.372048 | 1424.620244 | 1514.283129 | 1449.454681 | 4.43E-15 | 0.891611858 | YDR264C | AKR1 | n |
| W303.asm02.HP0_G0011220 | 529.2372177 | 673.0100671 | 499.4012765 | 236.3949783 | 244.1553713 | 229.9333069 | 275.4581336 | 324.1436729 | 392.0488296 | 1.09E-10 | 1.261135169 | YDR265W | PEX10 | n |
| W303.asm02.HP0_G0011230 | 440.0617158 | 581.6253904 | 423.1717275 | 1541.917863 | 1383.989414 | 1485.514003 | 1202.171236 | 1384.625659 | 1323.54866 | 5.52E-25 | -1.610483147 | YDR266C | HEL2 | n |
| W303.asm02.HP0_G0011240 | 1471.395781 | 1427.885574 | 1481.589697 | 495.16479 | 518.830164 | 499.6975804 | 525.3582961 | 569.9123085 | 650.0026287 | 1.79E-32 | 1.533932134 | YDR267C | CIA1 | n |
| W303.asm02.HP0_G0011250 | 679.4785523 | 756.7793541 | 809.2059823 | 466.9530436 | 477.6952917 | 435.4248844 | 455.3105233 | 488.634492 | 479.0570555 | 3.47E-07 | 0.703557901 | YDR268W | MSW1 | n |
| W303.asm02.HP0_G0011260 | 2303.054266 | 2124.693734 | 2220.429942 | 3062.433711 | 2518.515732 | 3007.237975 | 2129.83093 | 2821.501344 | 3802.771285 | 0.069267079 | -0.369688283 | YDR270W | CCC2 | n |
| W303.asm02.HP0_G0011290 | 3610.638527 | 3749.627517 | 3852.524133 | 2266.667899 | 2210.667655 | 2242.302367 | 1940.512625 | 1669.098017 | 1574.337076 | 3.07E-10 | 0.738583203 | YDR276C | PMP3 | n |
| W303.asm02.HP0_G0011300 | 286.9124843 | 220.8463021 | 223.8021376 | 450.4151233 | 492.2915367 | 467.1086078 | 463.829847 | 646.3521597 | 634.6482359 | 9.07E-07 | -0.945819175 | YDR277C | MTH1 | n |
| W303.asm02.HP0_G0011310 | 628.1057089 | 658.7312113 | 563.9032026 | 591.4738552 | 506.8877817 | 571.2122704 | 567.0083232 | 537.9817377 | 485.1988126 | 0.34038118 | 0.146808696 | YDR279W | RNH202 | n |
| W303.asm02.HP0_G0011320 | 1573.172169 | 2242.732274 | 1666.299758 | 1825.008145 | 1824.530628 | 1863.002936 | 1423.673653 | 1286.898761 | 1302.05251 | 0.961916637 | -0.007941675 | YDR280W | RRP45 | n |
| W303.asm02.HP0_G0011340 | 483.6801678 | 459.7791547 | 508.1969937 | 620.6584205 | 597.1191146 | 651.7794527 | 667.3470248 | 696.6669985 | 669.4515263 | 0.004583417 | -0.366350242 | YDR282C | MRX10 | n |
| W303.asm02.HP0_G0011350 | 473.9871784 | 630.1734999 | 514.0608052 | 641.0876161 | 704.6005552 | 706.0944071 | 697.6379536 | 846.6439218 | 777.9559021 | 0.037492129 | -0.341969023 | YDR283C | GCN2 | n |
| W303.asm02.HP0_G0011360 | 1655.562578 | 1540.212572 | 1629.162286 | 335.6225001 | 367.5599883 | 335.847468 | 639.8958706 | 581.5234251 | 613.152086 | 1.78E-72 | 2.217934583 | YDR284C | DPP1 | n |
| W303.asm02.HP0_G0011370 | 464.2941891 | 400.7598843 | 456.3999924 | 281.144645 | 310.5019396 | 257.0907841 | 277.3513167 | 350.2686853 | 256.9301729 | 0.000497122 | 0.641536836 | YDR285W | ZIP1 | n |
| W303.asm02.HP0_G0011380 | 271.4037014 | 295.0963519 | 271.6899313 | 185.8083986 | 209.6551558 | 175.6183525 | 229.0751489 | 161.5880399 | 141.2604138 | 0.013413178 | 0.556609116 | YDR286C | MGP12 | n |
| W303.asm02.HP0_G0011400 | 947.9743569 | 905.2794537 | 987.0749301 | 415.393645 | 419.3103116 | 406.4569087 | 636.1095045 | 587.3289834 | 777.9559021 | 1.23E-15 | 1.194752111 | YDR288W | NSE3 | n |
| W303.asm02.HP0_G0011410 | 826.8119902 | 960.4910292 | 885.4355314 | 852.1893046 | 863.8323191 | 816.5348144 | 838.6800908 | 716.9864526 | 706.302069 | 0.589792281 | 0.078260485 | YDR289C | RTT103 | n |
| W303.asm02.HP0_G0011420 | 451.693303 | 594.0003987 | 457.3772943 | 538.9416378 | 642.234781 | 528.6655561 | 609.6049418 | 723.759604 | 698.1130595 | 0.310907609 | -0.183713596 | YDR291W | HRQ1 | n |
| W303.asm02.HP0_G0011430 | 2218.725259 | 2259.866901 | 2388.52587 | 1739.400087 | 1876.280951 | 1711.826313 | 1320.495177 | 1679.741541 | 1641.896404 | 0.006210685 | 0.36673735 | YDR292C | SRP101 | n |
| W303.asm02.HP0_G0011440 | 845.2286699 | 1032.837232 | 751.5451696 | 4604.351574 | 4515.547438 | 4688.285814 | 2937.273501 | 4043.57137 | 3988.047624 | 1.87E-46 | -2.392529374 | YDR293C | SSD1 | y |
| W303.asm02.HP0_G0011460 | 972.2068302 | 951.9237158 | 899.1177581 | 485.4366016 | 404.7140666 | 468.9191063 | 668.2936163 | 856.3198523 | 646.9317502 | 4.56E-11 | 1.052468104 | YDR294C | DPL1 | n |
| W303.asm02.HP0_G0011470 | 550.5617942 | 573.0580769 | 575.6308256 | 607.0389567 | 611.7153596 | 541.3390455 | 526.3048877 | 606.6808445 | 590.6323099 | 0.739094733 | -0.049445856 | YDR295C | HDA2 | n |
| W303.asm02.HP0_G0011480 | 1348.294816 | 1160.39501 | 1342.812826 | 2679.143088 | 2734.805545 | 2846.10361 | 3001.641724 | 2593.149383 | 2419.852306 | 2.62E-17 | -1.100863833 | YDR296W | MHR1 | n |
| W303.asm02.HP0_G0011490 | 1373.496589 | 1528.789488 | 1480.612395 | 1685.895051 | 1472.893816 | 1607.72265 | 1747.407954 | 1925.510176 | 1560.006309 | 0.37817218 | -0.121711192 | YDR297W | SUR2 | n |
| W303.asm02.HP0_G0011500 | 5623.872412 | 5522.109475 | 5836.447012 | 9376.027988 | 9001.902385 | 8806.264606 | 7329.458175 | 7497.878571 | 6735.46031 | 2.77E-12 | -0.678737309 | YDR298C | ATP5 | n |
| W303.asm02.HP0_G0011510 | 1484.965966 | 2470.242042 | 1741.552005 | 3612.076356 | 2948.441495 | 3563.061008 | 1622.457873 | 1826.815685 | 1515.990383 | 2.08E-05 | -0.829716107 | YDR299W | BFR2 | n |
| W303.asm02.HP0_G0011520 | 2490.12896 | 2396.943916 | 2508.734006 | 1939.800769 | 1874.95402 | 1986.116832 | 1383.916809 | 1444.616429 | 1261.107462 | 0.001244613 | 0.350031784 | YDR300C | PRO1 | n |
| W303.asm02.HP0_G0011530 | 428.4301286 | 603.5196358 | 392.8753682 | 1153.763145 | 1245.988552 | 1128.845802 | 718.4629671 | 1053.708835 | 906.9328017 | 9.41E-11 | -1.307743965 | YDR301W | CFT1 | n |
| W303.asm02.HP0_G0011540 | 789.0093318 | 1018.558376 | 912.7999849 | 632.3322465 | 735.1199766 | 623.7167263 | 845.3062314 | 895.9911675 | 705.2784428 | 0.006353901 | 0.451885916 | YDR302W | GPI11 | n |
| W303.asm02.HP0_G0011560 | 3780.26584 | 2963.338527 | 4000.096722 | 1978.713522 | 1778.08803 | 2035.905541 | 2489.53571 | 1715.542484 | 1640.872778 | 2.69E-06 | 0.890841578 | YDR304C | CPR5 | n |
| W303.asm02.HP0_G0011580 | 1024.548973 | 731.0774137 | 964.5969862 | 544.7785509 | 577.2151441 | 515.9920668 | 452.4707487 | 558.3011918 | 456.5372794 | 3.03E-05 | 0.732681307 | YDR306C | PFU1 | n |
| W303.asm02.HP0_G0011590 | 1286.259685 | 1027.125689 | 1278.3109 | 402.7470001 | 380.829302 | 395.5939178 | 478.9753114 | 495.4076434 | 449.3718961 | 2.32E-28 | 1.605936071 | YDR307W | PMT7 | n |
| W303.asm02.HP0_G0011600 | 1041.027055 | 1282.241245 | 1142.465934 | 261.6882682 | 327.7520473 | 288.7745075 | 524.4117046 | 329.9492312 | 421.733989 | 3.88E-24 | 1.983036733 | YDR308C | SRB7 | n |
| W303.asm02.HP0_G0011610 | 937.3120686 | 824.3659379 | 1032.030818 | 1431.989334 | 1495.451649 | 1558.839191 | 1503.187341 | 1832.621243 | 1718.668368 | 1.10E-06 | -0.683516625 | YDR309C | GIC2 | n |
| W303.asm02.HP0_G0011620 | 540.8688049 | 591.1446275 | 502.3331823 | 3680.173675 | 3903.832078 | 3794.804814 | 2430.847035 | 3695.237871 | 3549.935616 | 2.80E-60 | -2.799410327 | YDR310C | SUM1 | n |
| W303.asm02.HP0_G0011630 | 813.2418051 | 847.2121071 | 794.5464536 | 676.1090944 | 639.5809183 | 666.2634406 | 488.4412267 | 577.6530529 | 615.1993384 | 0.028339298 | 0.308194403 | YDR311W | TFB1 | n |
| W303.asm02.HP0_G0011640 | 1326.000941 | 1632.549173 | 1455.202545 | 5895.282175 | 4961.396377 | 5766.437658 | 2898.463248 | 3158.223726 | 2546.781953 | 5.57E-42 | -1.913318856 | YDR312W | SSF2 | n |

| | | | | | | | | | | | | | | |
|---|---|---|---|---|---|---|---|---|---|---|---|---|---|---|
| W303.asm02.HP0_G0011650 | 704.6803246 | 711.0870157 | 645.9965632 | 326.8671305 | 298.5595573 | 300.5427477 | 302.9092879 | 318.3381146 | 315.2768656 | 8.56E-17 | 1.153899061 | YDR313C | PIB1 | n |
| W303.asm02.HP0_G0011670 | 343.1318224 | 421.7022061 | 365.5109147 | 135.2218189 | 147.2893816 | 112.2509058 | 185.5319388 | 165.4584121 | 180.1582789 | 3.11E-15 | 1.524377707 | YDR315C | IPK1 | n |
| W303.asm02.HP0_G0011680 | 1165.097318 | 1530.693335 | 1305.675353 | 1700.487334 | 1498.105512 | 1610.438398 | 1604.472634 | 1573.306305 | 1548.746421 | 0.059073532 | -0.265685601 | YDR316W | OMS1 | n |
| W303.asm02.HP0_G0011690 | 480.772271 | 426.4618247 | 453.4680867 | 116.7382609 | 138.000862 | 123.1138966 | 297.2297387 | 202.2269482 | 288.6625847 | 1.15E-19 | 1.852478969 | YDR317W | HIM1 | n |
| W303.asm02.HP0_G0011710 | 801.6102179 | 672.0581434 | 918.6637963 | 125.4936305 | 123.404617 | 112.2509058 | 201.6239947 | 172.2315635 | 188.3472184 | 1.62E-48 | 2.729531777 | YDR319C | YFT2 | n |
| W303.asm02.HP0_G0011720 | 250.0791248 | 290.3367333 | 257.0304026 | 739.342319 | 806.7742704 | 802.0508266 | 761.0595857 | 872.7689342 | 972.4448776 | 3.10E-24 | -1.557519298 | YDR320C | SWA2 | n |
| W303.asm02.HP0_G0011730 | 599.9960398 | 754.8755066 | 695.8389606 | 123.5479928 | 176.4818716 | 135.787386 | 276.4047252 | 222.5464023 | 239.5285278 | 1.50E-30 | 2.243157256 | YDR320C- | DAD4 | n |
| W303.asm02.HP0_G0011740 | 2744.085281 | 2596.847897 | 2797.038069 | 1544.836319 | 1718.376119 | 1705.489568 | 3132.271355 | 2548.640103 | 3148.674151 | 3.16E-08 | 0.71206552 | YDR321W | ASP1 | n |
| W303.asm02.HP0_G0011750 | 1567.356375 | 1396.472091 | 1579.319888 | 3700.602871 | 3545.560609 | 3649.964935 | 4109.153808 | 3452.372015 | 3714.739433 | 3.48E-27 | -1.262230095 | YDR322W | MRPL35 | n |
| W303.asm02.HP0_G0011760 | 2564.764978 | 2930.973121 | 2672.920727 | 2838.685378 | 3193.923797 | 2852.440355 | 3513.747739 | 2621.209582 | 2333.867706 | 0.495983798 | -0.121007879 | YDR322C- | TIM11 | n |
| W303.asm02.HP0_G0011770 | 525.3600219 | 466.4426207 | 572.6989198 | 333.6768624 | 372.8677138 | 314.1214863 | 389.0491166 | 418.0001991 | 436.0647557 | 9.34E-05 | 0.619134199 | YDR323C | PEP7 | n |
| W303.asm02.HP0_G0011780 | 1999.6637 | 2936.684663 | 2283.954566 | 6684.238255 | 5736.324294 | 6630.950682 | 3084.941779 | 3715.557325 | 3405.604324 | 1.11E-17 | -1.399881424 | YDR324C | UTP4 | n |
| W303.asm02.HP0_G0011800 | 767.6847553 | 1089.000731 | 782.8188307 | 1310.386979 | 1288.450356 | 1268.254185 | 1007.173382 | 1481.384965 | 1611.187618 | 0.008139697 | -0.550874571 | YDR326C | YSP2 | n |
| W303.asm02.HP0_G0011810 | 3955.708947 | 4091.36813 | 4326.51556 | 1481.603095 | 1433.085875 | 1453.830279 | 1956.604681 | 1284.963575 | 1395.202493 | 6.52E-20 | 1.501950125 | YDR328C | SKP1 | n |
| W303.asm02.HP0_G0011820 | 1512.106336 | 1338.404744 | 1568.569567 | 461.1161306 | 494.9453994 | 546.7705409 | 866.131245 | 812.7781649 | 962.2086158 | 1.93E-28 | 1.555314009 | YDR329C | PEX3 | n |
| W303.asm02.HP0_G0011830 | 633.9215025 | 508.3272642 | 570.744316 | 1209.213819 | 1042.968053 | 1175.918763 | 979.7222279 | 1295.607099 | 1181.26462 | 3.15E-10 | -1.001782479 | YDR330W | UBX5 | n |
| W303.asm02.HP0_G0011840 | 977.0533249 | 1502.135624 | 1060.372573 | 1106.095022 | 1101.353034 | 1069.099352 | 1040.304085 | 1136.921838 | 970.3976253 | 0.558208501 | 0.111589084 | YDR331W | GPI8 | n |
| W303.asm02.HP0_G0011850 | 429.3994275 | 510.2311117 | 472.036823 | 1050.644348 | 1058.89123 | 1042.847124 | 886.009667 | 1080.801441 | 1153.626713 | 5.09E-15 | -1.158848322 | YDR332W | IRC3 | n |
| W303.asm02.HP0_G0011860 | 1263.965809 | 1345.06821 | 1285.152013 | 1178.083616 | 1015.102495 | 1135.182547 | 702.3709112 | 943.4032271 | 901.8146707 | 0.14373942 | 0.225634496 | YDR333C | RQC1 | n |
| W303.asm02.HP0_G0011870 | 458.4783955 | 840.548641 | 468.1276154 | 916.3953481 | 874.44777 | 908.8702369 | 656.934518 | 924.051366 | 839.3734733 | 0.012100755 | -0.611336032 | YDR334W | SWR1 | n |
| W303.asm02.HP0_G0011880 | 1105.970083 | 1301.279719 | 1270.492484 | 1695.62324 | 1728.99157 | 1786.962 | 1531.585087 | 1923.57499 | 1744.259023 | 0.000205866 | -0.502956318 | YDR335W | MSN5 | n |
| W303.asm02.HP0_G0011890 | 603.8732355 | 573.0580769 | 554.1301835 | 320.0573986 | 359.5984001 | 352.1419543 | 427.8593691 | 348.3334992 | 322.4422489 | 4.50E-06 | 0.747550607 | YDR336W | MRX8 | n |
| W303.asm02.HP0_G0011900 | 2260.405113 | 2291.280384 | 2400.253493 | 2534.19308 | 2563.631399 | 2600.781066 | 2481.016386 | 2353.186306 | 2026.77985 | 0.228397141 | -0.14721782 | YDR337W | MRPS28 | n |
| W303.asm02.HP0_G0011910 | 388.6888723 | 458.827231 | 387.0115568 | 536.9960001 | 473.7144976 | 503.3185774 | 746.8607129 | 779.8800011 | 1098.350899 | 0.130022048 | -0.295152093 | YDR338C | | n |
| W303.asm02.HP0_G0011920 | 1745.707379 | 2058.059074 | 1884.238084 | 2467.06858 | 2575.573781 | 2495.772154 | 2633.417621 | 2053.232459 | 1876.306801 | 0.008709662 | -0.406202098 | YDR339C | FCF1 | n |
| W303.asm02.HP0_G0011940 | 5118.867667 | 4107.550834 | 5496.345947 | 7366.184263 | 7580.758893 | 7432.09626 | 5128.63288 | 5065.349635 | 4944.114483 | 3.63E-06 | -0.604098021 | YDR341C | | n |
| W303.asm02.HP0_G0011960 | 3932.445773 | 4915.734068 | 4232.694577 | 5181.233146 | 4830.030171 | 5059.438002 | 3010.161048 | 3773.612908 | 4212.221759 | 0.202439834 | -0.204354051 | YDR346C | SVF1 | n |
| W303.asm02.HP0_G0011970 | 1429.715927 | 1534.50103 | 1470.839376 | 2306.553472 | 2267.725704 | 2314.722306 | 2043.691102 | 1966.149085 | 1687.959582 | 7.14E-08 | -0.635427943 | YDR347W | MRP1 | n |
| W303.asm02.HP0_G0011980 | 1015.825282 | 1009.039139 | 946.0282498 | 880.401051 | 526.7917522 | 748.6411214 | 177.9592066 | 415.0974199 | 396.1433344 | 0.117047433 | 0.461495349 | YDR348C | PAL1 | n |
| W303.asm02.HP0_G0012000 | 1088.522702 | 1205.135424 | 1069.168291 | 1739.400087 | 2024.897264 | 1737.173291 | 920.0869619 | 1595.560945 | 1331.737669 | 0.000308357 | -0.709707283 | YDR349C | YPS7 | n |
| W303.asm02.HP0_G0012010 | 1037.149859 | 1231.789288 | 1115.10148 | 373.5624349 | 421.9641743 | 372.9626868 | 310.4820201 | 348.3334992 | 333.702137 | 1.72E-27 | 1.536788425 | YDR350C | ATP22 | n |
| W303.asm02.HP0_G0012030 | 1252.334222 | 1516.414479 | 1377.018393 | 603.1476813 | 579.8690068 | 623.7167263 | 1371.611119 | 1109.829232 | 1051.264094 | 2.13E-15 | 1.197542132 | YDR352W | YPQ2 | n |
| W303.asm02.HP0_G0012040 | 9145.335437 | 9067.073393 | 9854.135169 | 4795.024066 | 4175.853008 | 4796.915722 | 5522.414954 | 4890.215292 | 4928.76009 | 1.75E-19 | 1.027146052 | YDR353W | TRR1 | n |
| W303.asm02.HP0_G0012050 | 1547.970397 | 1272.722008 | 1601.797832 | 640.1147973 | 448.5028016 | 615.5694831 | 794.1902891 | 587.3289834 | 655.1207597 | 3.01E-13 | 1.373065438 | YDR354W | TRP4 | n |
| W303.asm02.HP0_G0012060 | 519.5442283 | 697.7600837 | 541.4252587 | 632.3322465 | 683.3696534 | 647.2532065 | 495.0673673 | 627.9678917 | 549.6872624 | 0.377768636 | -0.157719809 | YDR356W | SPC110 | n |
| W303.asm02.HP0_G0012070 | 721.1584065 | 794.8563027 | 791.6145479 | 47.6681232 | 31.84635278 | 53.40970516 | 76.67391349 | 46.44446656 | 42.99229985 | 6.59E-58 | 4.097526788 | YDR357C | CNL1 | n |
| W303.asm02.HP0_G0012090 | 352.8248118 | 384.5771812 | 302.9635924 | 1029.242334 | 977.9484166 | 1016.594986 | 752.542062 | 806.9726066 | 940.7124658 | 4.31E-24 | -1.539372219 | YDR359C | EAF1 | n |
| W303.asm02.HP0_G0012100 | 2493.036857 | 2638.73254 | 2534.143855 | 2505.981334 | 2486.669379 | 2465.89893 | 1618.671507 | 1291.736726 | 1067.642113 | 0.81591858 | 0.039576977 | YDR361C | BCP1 | n |
| W303.asm02.HP0_G0012110 | 569.9477729 | 570.2023058 | 605.9271848 | 535.0503625 | 597.1191146 | 652.684702 | 473.2957623 | 498.3104225 | 562.9944028 | 0.841290269 | -0.032026215 | YDR362C | TFC6 | n |
| W303.asm02.HP0_G0012120 | 389.6581712 | 504.5195694 | 434.8993504 | 547.6970074 | 610.3884283 | 555.8230333 | 475.1889453 | 359.9446159 | 337.7966417 | 0.052401722 | -0.365799542 | YDR363W | ESC2 | n |
| W303.asm02.HP0_G0012130 | 1797.080223 | 2261.770749 | 1924.307463 | 701.4023842 | 818.7166527 | 748.6411214 | 1092.366619 | 896.9587605 | 894.6492874 | 1.57E-20 | 1.400200203 | YDR363W- | SEM1 | n |
| W303.asm02.HP0_G0012140 | 739.5750862 | 888.1448268 | 729.0672256 | 437.7684784 | 433.9065566 | 450.8141215 | 361.5979624 | 570.8799015 | 384.8834463 | 2.73E-05 | 0.83335158 | YDR364C | CDC40 | n |
| W303.asm02.HP0_G0012150 | 1207.746471 | 1553.539504 | 1368.222675 | 6149.187893 | 5923.421617 | 6178.326062 | 2785.818857 | 3065.334793 | 2453.63197 | 6.22E-55 | -2.143932241 | YDR365C | ESF1 | n |
| W303.asm02.HP0_G0012170 | 3530.186716 | 2141.828361 | 3272.006798 | 56.42349277 | 47.76952917 | 47.07296048 | 42.5966186 | 49.34724572 | 46.06317841 | 1.52E-139 | 5.883323441 | YDR368W | YPR1 | n |
| W303.asm02.HP0_G0012180 | 485.6187656 | 594.9523224 | 587.3584485 | 765.6084277 | 903.6402601 | 805.6718235 | 638.0026875 | 875.6717134 | 774.8850235 | 0.000908667 | -0.568240109 | YDR369C | XRS2 | n |
| W303.asm02.HP0_G0012190 | 968.3296345 | 1027.125689 | 853.1845683 | 998.1121307 | 1011.121701 | 1000.30041 | 480.8684945 | 666.6716138 | 467.7971674 | 0.666482959 | -0.079134959 | YDR370C | DXO1 | n |
| W303.asm02.HP0_G0012200 | 1017.76388 | 922.4140806 | 1075.032102 | 41.83121016 | 33.17328414 | 34.39947112 | 81.40687111 | 66.76392069 | 95.19723539 | 1.61E-103 | 4.777824225 | YDR371W | CTS2 | n |
| W303.asm02.HP0_G0012210 | 3070.739021 | 3426.925377 | 3324.781101 | 2137.282993 | 1951.916039 | 2046.768531 | 2131.724113 | 1890.676826 | 1727.881004 | 4.23E-08 | 0.678445017 | YDR372C | VPS74 | n |
| W303.asm02.HP0_G0012220 | 1308.55356 | 1472.625988 | 1234.332314 | 564.2349277 | 644.8886438 | 586.6015075 | 768.6323179 | 587.3289834 | 640.789993 | 2.95E-14 | 1.162434418 | YDR373W | FRQ1 | n |
| W303.asm02.HP0_G0012230 | 136.6711496 | 49.50003322 | 87.95717199 | 5.836913045 | 6.634656829 | 3.62099696 | 5.679549147 | 13.54630275 | 5.118130935 | 6.07E-11 | 4.115537925 | YDR374C | PHO92 | n |
| W303.asm02.HP0_G0012240 | 297.5747726 | 272.2501827 | 267.7807236 | 165.3792029 | 196.3858421 | 163.8501124 | 183.6387558 | 130.6250622 | 142.28404 | 0.001037105 | 0.676512899 | YDR374W- | WIP1 | n |
| W303.asm02.HP0_G0012250 | 1076.891115 | 1073.769951 | 1056.463366 | 992.2752177 | 1074.814406 | 1048.27862 | 1162.414392 | 1216.264468 | 1093.232768 | 0.729428209 | 0.042497704 | YDR375C | BCS1 | n |
| W303.asm02.HP0_G0012260 | 989.654211 | 921.4621569 | 976.3246091 | 638.1691596 | 650.1963692 | 611.9484862 | 866.131245 | 855.3522592 | 813.7828186 | 2.31E-07 | 0.604408323 | YDR376W | ARH1 | n |
| W303.asm02.HP0_G0012280 | 1134.079752 | 1294.616253 | 1244.105333 | 886.237964 | 958.0444461 | 890.7652521 | 1248.554221 | 901.7967258 | 887.4839041 | 0.010821701 | 0.425882442 | YDR378C | LSM6 | n |
| W303.asm02.HP0_G0012300 | 323.7458438 | 303.6636653 | 340.101065 | 433.877203 | 475.0414289 | 437.2353829 | 373.9036522 | 303.8242188 | 242.5994063 | 0.013037674 | -0.475586059 | YDR379C- | SDH6 | n |
| W303.asm02.HP0_G0012310 | 2038.435657 | 1591.616453 | 2021.060352 | 2930.130349 | 5242.705826 | 3316.833215 | 8687.817012 | 7425.309092 | 7172.548692 | 1.94E-06 | -1.023198974 | YDR380W | ARO10 | n |
| W303.asm02.HP0_G0012320 | 4583.814656 | 5350.763206 | 4411.540826 | 2349.357501 | 1999.685568 | 2324.680048 | 1638.549929 | 2209.982534 | 2170.087516 | 1.93E-12 | 1.103624608 | YDR381W | YRA1 | n |
| W303.asm02.HP0_G0012330 | 749.2680755 | 766.2985912 | 758.3862829 | 437.7684784 | 436.5604193 | 391.9729209 | 508.3196487 | 308.662184 | 288.6625847 | 9.52E-05 | 0.845179767 | YDR381C- | COI1 | n |
| W303.asm02.HP0_G0012340 | 8977.646721 | 10276.96844 | 9262.867512 | 12053.22544 | 14069.45327 | 11609.8215 | 12384.25692 | 9405.004479 | 8517.593502 | 0.014813208 | -0.403856803 | YDR382W | RPP2B | n |
| W303.asm02.HP0_G0012350 | 733.7592926 | 445.500299 | 679.2248281 | 347.2963262 | 400.7332725 | 337.6579665 | 416.5002708 | 288.3427299 | 372.5999321 | 0.000646171 | 0.776967146 | YDR383C | NKP1 | n |
| W303.asm02.HP0_G0012360 | 630.0443067 | 582.5773141 | 642.0873555 | 36.96711595 | 41.13487234 | 42.54671428 | 17.03864744 | 41.6065013 | 49.13405697 | 9.55E-44 | 3.942862834 | YDR384C | ATO3 | n |
| W303.asm02.HP0_G0012370 | 8297.19887 | 7329.812612 | 7996.284236 | 59871.16274 | 63411.39611 | 56781.75857 | 39452.04156 | 41711.96894 | 38866.06269 | 1.65E-181 | -2.930215094 | YOR133W | EFT1 | y |
| W303.asm02.HP0_G0012380 | 576.7328654 | 496.9041796 | 610.8136944 | 200.4006812 | 212.3090185 | 214.5440699 | 239.4876557 | 237.0602981 | 221.1032564 | 9.19E-19 | 1.425408525 | YDR386W | MUS81 | n |

| | | | | | | | | | | | | | | |
|---|---|---|---|---|---|---|---|---|---|---|---|---|---|---|
| W303.asm02.HP0_G0012390 | 426.4915307 | 356.0194697 | 361.6017071 | 188.7268551 | 232.212989 | 210.0178237 | 281.1376828 | 316.4029285 | 410.474101 | 2.17E-05 | 0.861401083 | YDR387C | CIN10 | n |
| W303.asm02.HP0_G0012400 | 1430.685226 | 1114.702671 | 1110.214971 | 3479.772994 | 2904.65276 | 3452.620601 | 4022.067388 | 3336.260848 | 3346.234005 | 3.45E-20 | -1.428523038 | YDR388W | RVS167 | n |
| W303.asm02.HP0_G0012410 | 465.263488 | 579.7215429 | 453.4680867 | 1173.219522 | 1036.333397 | 1227.517969 | 514.9457894 | 985.9773214 | 1095.28002 | 1.80E-06 | -1.198007686 | YDR389W | SAC7 | n |
| W303.asm02.HP0_G0012420 | 1121.478866 | 938.5967838 | 1130.738311 | 1266.610131 | 1147.795631 | 1228.423219 | 1020.425663 | 1120.472756 | 1101.421777 | 0.155474084 | -0.191916189 | YDR390C | UBA2 | n |
| W303.asm02.HP0_G0012430 | 1232.948243 | 765.3466675 | 1191.33103 | 62.26040581 | 65.01963692 | 51.59920668 | 119.2705321 | 105.4676428 | 90.07910445 | 9.61E-66 | 4.162761372 | YDR391C | | n |
| W303.asm02.HP0_G0012440 | 1030.364766 | 686.3369991 | 922.573004 | 648.8701668 | 682.042722 | 706.0944071 | 866.131245 | 779.8800011 | 804.570183 | 0.032410403 | 0.373532348 | YDR392W | SPT3 | n |
| W303.asm02.HP0_G0012450 | 1793.203027 | 1320.318194 | 1777.712176 | 397.8829059 | 457.7913212 | 440.8563798 | 341.7195404 | 485.7317128 | 393.0724558 | 2.44E-25 | 1.91636211 | YDR393W | SHE9 | n |
| W303.asm02.HP0_G0012460 | 3769.603552 | 2669.194099 | 3487.013218 | 2675.251812 | 2197.398342 | 2474.046173 | 2870.065502 | 2592.18179 | 2712.609395 | 0.006384827 | 0.433626967 | YDR394W | RPT3 | n |
| W303.asm02.HP0_G0012470 | 2312.747255 | 2634.924845 | 2526.32544 | 4760.002588 | 5291.802287 | 4756.179507 | 4014.494656 | 4475.117872 | 3951.197082 | 1.26E-17 | -0.986107854 | YDR395W | SXM1 | n |
| W303.asm02.HP0_G0012480 | 1474.303678 | 1507.847166 | 1501.135735 | 2264.722261 | 2046.128166 | 2097.462489 | 2134.563888 | 2122.899159 | 1535.43928 | 0.000772192 | -0.515580319 | YDR397C | NCB2 | n |
| W303.asm02.HP0_G0012490 | 1506.290543 | 1811.510831 | 1489.408112 | 4417.570356 | 3745.927246 | 4261.913422 | 2334.2947 | 2555.413254 | 2291.899033 | 2.86E-25 | -1.370299133 | YDR398W | UTP5 | n |
| W303.asm02.HP0_G0012500 | 3001.918797 | 6288.408067 | 3495.808936 | 3810.5314 | 3323.963071 | 3706.090388 | 3665.202383 | 2950.19122 | 2341.03309 | 0.379446982 | 0.238073629 | YDR399W | HPT1 | n |
| W303.asm02.HP0_G0012510 | 1314.369354 | 1126.125756 | 1337.926316 | 2016.653457 | 1880.261745 | 1972.538094 | 481.815086 | 948.2411924 | 603.9394503 | 0.004347175 | -0.635575898 | YDR400W | URH1 | n |
| W303.asm02.HP0_G0012520 | 111.4693773 | 71.39427868 | 85.02526626 | 1707.297066 | 1231.392307 | 1900.118155 | 2056.943383 | 1489.125709 | 3349.304884 | 4.91E-47 | -4.17573488 | YDR402C | DIT2 | n |
| W303.asm02.HP0_G0012530 | 415.8292424 | 316.9905974 | 386.0342548 | 2354.221595 | 1269.873317 | 2683.158747 | 2345.653798 | 1240.454294 | 6235.930731 | 5.83E-05 | -2.495141037 | YDR403W | DIT1 | n |
| W303.asm02.HP0_G0012540 | 1747.645977 | 1437.404811 | 1766.961855 | 1137.225225 | 1150.449494 | 1094.446331 | 1243.821263 | 1034.356974 | 1025.673439 | 0.000119461 | 0.550306123 | YDR404C | RPB7 | n |
| W303.asm02.HP0_G0012550 | 585.4565558 | 391.2406472 | 446.6269733 | 3155.82432 | 3042.653622 | 3164.751343 | 3825.176351 | 3099.20055 | 2877.413212 | 3.47E-55 | -2.717934886 | YDR405W | MRP20 | n |
| W303.asm02.HP0_G0012570 | 734.7285915 | 798.6639976 | 702.680074 | 936.8245437 | 1054.910436 | 936.0277141 | 936.1790178 | 1129.181093 | 1043.075085 | 0.005134318 | -0.387533164 | YDR407C | TRS120 | n |
| W303.asm02.HP0_G0012580 | 1674.948557 | 1316.510499 | 1788.462497 | 627.4681523 | 526.7917522 | 619.1904801 | 795.1368806 | 588.2965765 | 542.5218791 | 5.39E-14 | 1.429064454 | YDR408C | ADE8 | n |
| W303.asm02.HP0_G0012590 | 596.1188441 | 492.1445611 | 539.4706549 | 566.1805654 | 582.5228696 | 499.6975804 | 412.7139047 | 488.634492 | 531.261991 | 0.92378617 | -0.01696074 | YDR409W | SIZ1 | n |
| W303.asm02.HP0_G0012600 | 1388.036073 | 944.3083261 | 1379.950298 | 520.4580798 | 587.830595 | 440.8563798 | 872.7573856 | 753.7549886 | 679.6877882 | 1.50E-10 | 1.262753355 | YDR410C | STE14 | n |
| W303.asm02.HP0_G0012610 | 737.6364883 | 431.2214433 | 770.1139059 | 123.5479928 | 123.404617 | 151.1766231 | 185.5319388 | 209.0000995 | 231.3395183 | 1.77E-20 | 2.281137148 | YDR411C | DFM1 | n |
| W303.asm02.HP0_G0012620 | 1849.422365 | 1945.732075 | 2048.424805 | 2413.563544 | 2429.611331 | 2462.277933 | 1401.902048 | 1497.834047 | 1275.438229 | 0.004950891 | -0.322168855 | YDR412W | RRP17 | n |
| W303.asm02.HP0_G0012630 | 613.5662249 | 471.2022393 | 626.4505249 | 660.5439929 | 679.3888593 | 563.0650272 | 744.0209383 | 924.051366 | 855.7514923 | 0.402850701 | -0.1521837 | YDR414C | ERD1 | n |
| W303.asm02.HP0_G0012640 | 987.7156132 | 713.9427868 | 921.5957021 | 1332.761812 | 899.659466 | 1259.201693 | 1318.601994 | 1190.139456 | 1216.06791 | 0.030923503 | -0.414091304 | YDR415C | | n |
| W303.asm02.HP0_G0012650 | 790.9479297 | 637.7888896 | 776.9550192 | 1094.421196 | 1106.660759 | 1094.446331 | 907.781272 | 1031.454195 | 932.5234563 | 2.31E-05 | -0.579318899 | YDR416W | SYF1 | n |
| W303.asm02.HP0_G0012660 | 894.6629156 | 626.365805 | 971.4380995 | 347.2963262 | 378.1754392 | 360.2891975 | 293.4433726 | 366.7177673 | 282.5208276 | 2.03E-09 | 1.199491978 | YDR419W | RAD30 | n |
| W303.asm02.HP0_G0012670 | 230.6931462 | 487.3849425 | 179.8235516 | 1281.202413 | 1188.930504 | 1209.412985 | 653.1481519 | 1027.583823 | 1033.862449 | 5.81E-11 | -2.034622376 | YDR420W | HKR1 | n |
| W303.asm02.HP0_G0012680 | 462.3555912 | 500.7118745 | 455.4226905 | 481.5453262 | 602.4268401 | 544.0547932 | 462.8832555 | 735.3707206 | 648.9790025 | 0.338069528 | -0.197413147 | YDR421W | ARO80 | n |
| W303.asm02.HP0_G0012720 | 506.9433422 | 323.6540634 | 429.0355389 | 511.7027103 | 514.8493699 | 516.897316 | 550.9162673 | 702.4725568 | 705.2784428 | 0.146137161 | -0.293340137 | YDR425W | SNX41 | n |
| W303.asm02.HP0_G0012730 | 4493.669855 | 4136.108545 | 4299.151106 | 1847.382979 | 1624.163992 | 1774.28851 | 2268.979884 | 2086.130623 | 1786.227696 | 3.66E-24 | 1.300738896 | YDR427W | RPN9 | n |
| W303.asm02.HP0_G0012750 | 4858.126254 | 5872.417403 | 5261.793489 | 6544.152342 | 6832.369602 | 6559.435992 | 5437.221717 | 5029.548692 | 4608.365094 | 0.009579352 | -0.317896596 | YDR429C | TIF35 | n |
| W303.asm02.HP0_G0012760 | 1274.628097 | 1377.433617 | 1169.830387 | 1728.69908 | 1642.741031 | 1748.941532 | 1245.714446 | 1826.815685 | 1549.770047 | 0.009594267 | -0.42215935 | YDR430C | CYM1 | n |
| W303.asm02.HP0_G0012770 | 2331.163935 | 2174.193767 | 1819.736158 | 5917.657009 | 5578.419462 | 5991.844719 | 3447.486332 | 4057.117673 | 4005.44927 | 3.22E-28 | -1.467285182 | YDR432W | NPL3 | n |
| W303.asm02.HP0_G0012780 | 1869.777643 | 1786.760815 | 1862.737442 | 2039.02829 | 1807.28052 | 1937.233373 | 1709.544293 | 2385.116877 | 2038.039738 | 0.669589365 | -0.067866979 | YDR434W | GPI17 | n |
| W303.asm02.HP0_G0012790 | 764.7768585 | 620.6542627 | 708.5438855 | 321.0302175 | 309.1750082 | 339.468465 | 328.467259 | 355.1066506 | 336.7730155 | 1.09E-13 | 1.108899764 | YDR435C | PPM1 | n |
| W303.asm02.HP0_G0012810 | 96.92989334 | 124.7020068 | 116.2989274 | 239.3134348 | 285.2902436 | 214.5440699 | 383.3695674 | 442.1900254 | 256.9301729 | 7.73E-06 | -1.125931759 | YDR437W | GPI19 | n |
| W303.asm02.HP0_G0012830 | 442.0003136 | 458.827231 | 462.2638039 | 411.5023697 | 407.3679293 | 414.6041519 | 522.5185216 | 406.3890824 | 408.4268486 | 0.386731962 | 0.143899564 | YDR439W | LRS4 | n |
| W303.asm02.HP0_G0012840 | 432.3073243 | 610.1831018 | 491.5828612 | 466.9530436 | 471.0606348 | 440.8563798 | 410.8207217 | 523.4678419 | 435.0411295 | 0.418238859 | 0.154574311 | YDR440W | DOT1 | n |
| W303.asm02.HP0_G0012860 | 245.2326302 | 310.3271313 | 201.3241937 | 680.9731886 | 648.8694379 | 688.8946716 | 543.3435351 | 850.514294 | 785.1212854 | 5.63E-11 | -1.415313684 | YDR443C | SSN2 | n |
| W303.asm02.HP0_G0012870 | 234.5703419 | 314.1348262 | 211.0972128 | 178.9986667 | 180.4626657 | 168.3763586 | 127.7898558 | 275.7640202 | 220.0796302 | 0.073442435 | 0.526292826 | YDR444W | | n |
| W303.asm02.HP0_G0012880 | 59.12723494 | 17.13462688 | 58.63811466 | 134.249 | 119.4238229 | 119.4928997 | 132.5228134 | 129.6574692 | 131.0241519 | 0.000100961 | -1.469775503 | YDR446W | ECM11 | n |
| W303.asm02.HP0_G0012900 | 725.0356022 | 620.6542627 | 702.680074 | 967.9547466 | 1009.794769 | 927.8804709 | 981.615411 | 967.5930534 | 993.9410276 | 4.85E-05 | -0.503636754 | YDR448W | ADA2 | n |
| W303.asm02.HP0_G0012910 | 1421.961535 | 2071.386006 | 1630.139588 | 3736.597168 | 3170.039033 | 3640.912443 | 2046.530876 | 2220.626058 | 1657.250797 | 3.26E-09 | -1.041896029 | YDR449C | UTP6 | n |
| W303.asm02.HP0_G0012920 | 8457.133194 | 9984.727855 | 8359.850547 | 16693.57131 | 19229.88935 | 16147.83594 | 15153.03713 | 11509.51397 | 9857.520181 | 3.14E-08 | -0.958098975 | YML026C | RPS18B | y |
| W303.asm02.HP0_G0012940 | 2705.313323 | 2215.126487 | 2849.812372 | 2038.055472 | 1897.511853 | 2084.789 | 2196.092337 | 2576.700301 | 2437.253951 | 0.008774807 | 0.367667293 | YDR452W | PPN1 | n |
| W303.asm02.HP0_G0012960 | 3484.629666 | 3580.185095 | 3707.88345 | 1998.169899 | 2059.39748 | 2038.621288 | 3491.976134 | 2258.362187 | 2293.946285 | 2.81E-06 | 0.821449046 | YDR454C | GUK1 | n |
| W303.asm02.HP0_G0012970 | 943.1278622 | 764.3947438 | 950.9147594 | 550.6154639 | 601.0999087 | 542.2442947 | 924.8199195 | 1206.588538 | 1261.107462 | 0.000128293 | 0.651174173 | YDR456W | NHX1 | n |
| W303.asm02.HP0_G0012980 | 1107.908681 | 1575.43375 | 1225.536596 | 2653.849798 | 3130.231092 | 2746.526194 | 2283.178757 | 2496.390078 | 2086.150169 | 3.37E-12 | -1.125451473 | YDR457W | TOM1 | n |
| W303.asm02.HP0_G0013000 | 783.1935382 | 553.0676789 | 768.159302 | 780.2007104 | 837.2936918 | 786.6615895 | 791.3505145 | 884.3800508 | 706.302069 | 0.291285805 | -0.191758406 | YDR459C | PFA5 | n |
| W303.asm02.HP0_G0013010 | 885.9392252 | 624.4619576 | 835.5931339 | 640.1147973 | 660.8118202 | 687.9894223 | 708.9970519 | 643.4493805 | 613.152086 | 0.167575636 | 0.237941227 | YDR460W | TFB3 | n |
| W303.asm02.HP0_G0013030 | 1372.52729 | 1007.135291 | 1350.631241 | 1568.183971 | 1421.143493 | 1578.754674 | 1736.995448 | 1465.903476 | 1323.54866 | 0.090026056 | -0.29280761 | YDR462W | MRPL28 | n |
| W303.asm02.HP0_G0013040 | 831.6584849 | 629.2215761 | 735.908339 | 2273.477631 | 2164.225058 | 2220.576386 | 1610.152183 | 2172.246405 | 2322.607818 | 5.34E-21 | -1.599929333 | YDR463W | STP1 | n |
| W303.asm02.HP0_G0013050 | 284.9738864 | 359.8271646 | 209.142609 | 763.6627901 | 822.6974468 | 741.3991275 | 531.0378453 | 689.8938471 | 672.5224048 | 3.50E-12 | -1.445840542 | YDR464W | SPP41 | n |
| W303.asm02.HP0_G0013060 | 1099.184991 | 1381.241312 | 1198.172143 | 2354.221595 | 2253.129459 | 2342.785033 | 1584.594212 | 1494.931268 | 1660.321675 | 3.39E-13 | -0.918000807 | YDR465C | RMT2 | n |
| W303.asm02.HP0_G0013070 | 370.2721926 | 473.1060868 | 331.3053478 | 556.452377 | 599.7729773 | 561.2545288 | 549.9696758 | 907.6022841 | 867.0113804 | 0.014435359 | -0.547378593 | YDR466W | PKH3 | n |
| W303.asm02.HP0_G0013080 | 1548.939696 | 1581.145292 | 1587.138303 | 293.7912899 | 228.2321949 | 267.0485258 | 93.71256093 | 323.1760798 | 412.5213534 | 2.16E-07 | 2.579200209 | YDR468C | TLG1 | n |
| W303.asm02.HP0_G0013090 | 904.3559049 | 1036.644927 | 988.052232 | 166.3520218 | 175.1549403 | 207.3020759 | 126.8432643 | 201.2593551 | 210.8669945 | 2.12E-31 | 2.414128519 | YDR469W | SDC1 | n |
| W303.asm02.HP0_G0013100 | 1010.978788 | 566.3946109 | 929.4141174 | 775.3366161 | 849.2360741 | 813.8190667 | 1187.972363 | 1003.393996 | 971.4212514 | 0.861978865 | 0.040091001 | YDR470C | UGO1 | n |
| W303.asm02.HP0_G0013120 | 1427.777329 | 1047.116086 | 1389.723317 | 248.0688044 | 248.1361654 | 244.4172948 | 190.2648964 | 303.8242188 | 406.3795962 | 4.27E-22 | 2.383588932 | YDR472W | TRS31 | n |
| W303.asm02.HP0_G0013130 | 675.6013566 | 592.0965512 | 651.8603746 | 522.4037175 | 483.0030171 | 576.6437658 | 655.041335 | 559.2687849 | 589.6086837 | 0.064913157 | 0.276712356 | YDR473C | PRP3 | n |
| W303.asm02.HP0_G0013140 | 457.5090966 | 408.3752741 | 403.6256892 | 241.2590725 | 360.9253315 | 246.2277933 | 409.8741301 | 467.3474448 | 557.8762719 | 0.005509075 | 0.587307607 | YDR475C | JIP4 | n |
| W303.asm02.HP0_G0013150 | 533.1144134 | 367.4425543 | 419.2625198 | 419.2849204 | 453.8105271 | 429.993389 | 678.7061231 | 534.1113655 | 694.0185548 | 0.925869961 | 0.0190347 | YDR476C | | n |

| | | | | | | | | | | | | | | |
|---|---|---|---|---|---|---|---|---|---|---|---|---|---|---|
| W303.asm02.HP0_G0013160 | 730.8513958 | 661.5869825 | 616.6775058 | 1098.312471 | 1102.679965 | 1139.708793 | 1169.987124 | 1272.384865 | 1474.021709 | 1.73E-07 | -0.733732045 | YDR477W | SNF1 | n |
| W303.asm02.HP0_G0013170 | 384.8116766 | 464.5387733 | 413.3987084 | 275.307732 | 276.0017241 | 258.9012826 | 208.2501354 | 214.8056579 | 300.946099 | 0.001066003 | 0.6409927 | YDR478W | SNM1 | n |
| W303.asm02.HP0_G0013180 | 555.4082889 | 508.3272642 | 493.5374651 | 368.6983407 | 403.3871352 | 391.9729209 | 506.4264656 | 499.2780156 | 520.002103 | 0.00232475 | 0.421068213 | YDR479C | PEX29 | n |
| W303.asm02.HP0_G0013200 | 1785.448635 | 1631.597249 | 1696.596117 | 2482.633682 | 1885.569471 | 2409.773447 | 1650.855619 | 1726.186007 | 1790.322201 | 0.003672453 | -0.407540184 | YDR481C | PHO8 | n |
| W303.asm02.HP0_G0013210 | 565.1012782 | 499.7599508 | 551.1982778 | 706.2664784 | 555.9842423 | 633.6744679 | 298.1763302 | 415.0974199 | 271.2609395 | 0.249996461 | -0.231744646 | YDR482C | CWC21 | n |
| W303.asm02.HP0_G0013220 | 4425.81893 | 3774.377533 | 4742.846174 | 2727.78403 | 2794.517456 | 2608.928309 | 2530.239145 | 3310.135836 | 2579.537991 | 7.74E-06 | 0.670765113 | YDR483W | KRE2 | n |
| W303.asm02.HP0_G0013230 | 875.2769369 | 803.4236161 | 961.6650804 | 746.1520509 | 725.8314571 | 695.2314163 | 568.9015063 | 636.6762292 | 612.1284598 | 0.035776645 | 0.285023727 | YDR484W | VPS52 | n |
| W303.asm02.HP0_G0013240 | 636.8293993 | 666.3466011 | 635.2462421 | 517.5396233 | 545.3687913 | 507.8448236 | 475.1889453 | 521.5326558 | 546.6163838 | 0.022675162 | 0.30472972 | YDR485C | VPS72 | n |
| W303.asm02.HP0_G0013250 | 1705.966123 | 1663.962655 | 1684.868495 | 429.9859276 | 379.5023706 | 403.741161 | 592.5662944 | 491.5372711 | 498.5059531 | 4.74E-54 | 2.056935672 | YDR486C | VPS60 | n |
| W303.asm02.HP0_G0013260 | 4254.253019 | 4209.406671 | 4626.547247 | 1762.74774 | 1787.37655 | 1759.804522 | 2177.160506 | 1879.06571 | 1764.731546 | 1.38E-27 | 1.301813838 | YDR487C | RIB3 | n |
| W303.asm02.HP0_G0013270 | 231.6624451 | 241.7886238 | 180.8008535 | 211.1016885 | 191.0781167 | 205.4915775 | 177.9592066 | 183.8426802 | 206.7724898 | 0.620697674 | 0.105019839 | YDR488C | PAC11 | n |
| W303.asm02.HP0_G0013280 | 547.6538974 | 497.8561034 | 516.9927109 | 303.5194783 | 285.2902436 | 318.6477325 | 316.1615692 | 260.2825314 | 231.3395183 | 3.89E-06 | 0.782105288 | YDR489W | SLD5 | n |
| W303.asm02.HP0_G0013310 | 1311.461457 | 1932.405143 | 1230.423106 | 652.7614422 | 566.5996932 | 619.1904801 | 559.435591 | 671.5095791 | 589.6086837 | 2.55E-11 | 1.282356179 | YDR492W | IZH1 | n |
| W303.asm02.HP0_G0013320 | 770.5926521 | 936.6929363 | 776.9550192 | 1026.323877 | 1020.41022 | 1037.415629 | 1123.60414 | 906.6346911 | 846.5388566 | 0.050697216 | -0.312062488 | YDR493W | MZM1 | n |
| W303.asm02.HP0_G0013330 | 2185.769095 | 1853.395475 | 2318.160133 | 2642.175972 | 2839.633123 | 2763.725929 | 2769.726801 | 2507.033601 | 2330.796828 | 0.005134318 | -0.375082686 | YDR494W | RSM28 | n |
| W303.asm02.HP0_G0013340 | 515.6670326 | 737.7408797 | 591.2676562 | 599.256406 | 559.9650364 | 648.1584558 | 549.0230842 | 760.52814 | 697.0894333 | 0.890306672 | 0.028619075 | YDR495C | VPS3 | n |
| W303.asm02.HP0_G0013350 | 2412.585045 | 3259.386803 | 2624.055631 | 9396.457184 | 8443.26428 | 9326.782919 | 3324.429434 | 4273.858517 | 3447.572998 | 4.15E-28 | -1.711382269 | YDR496C | PUF6 | n |
| W303.asm02.HP0_G0013360 | 2588.028152 | 2321.741943 | 2818.538711 | 1150.844689 | 1604.260021 | 1164.150523 | 1642.336295 | 1783.273997 | 2268.35563 | 9.33E-08 | 0.980937542 | YDR497C | ITR1 | n |
| W303.asm02.HP0_G0013370 | 631.9829046 | 387.4329523 | 576.6081275 | 394.9644494 | 406.0409979 | 419.1303981 | 556.5958164 | 424.7733505 | 477.0098031 | 0.069361113 | 0.387166305 | YDR498C | SEC20 | n |
| W303.asm02.HP0_G0013380 | 384.8116766 | 475.9618579 | 460.3092001 | 471.8171378 | 561.2919677 | 471.634854 | 377.6900183 | 526.3706211 | 453.4664008 | 0.331990101 | -0.18580436 | YDR499W | LCD1 | n |
| W303.asm02.HP0_G0013400 | 422.614335 | 285.5771147 | 385.0569529 | 1472.847725 | 1557.817423 | 1540.734206 | 1302.509938 | 1579.111863 | 1258.036584 | 1.82E-33 | -2.064158736 | YDR501W | PLM2 | n |
| W303.asm02.HP0_G0013420 | 792.8865275 | 667.2985248 | 759.3635848 | 311.3020291 | 207.0012931 | 285.1535106 | 458.1502979 | 335.7547895 | 381.8125677 | 1.73E-13 | 1.460716765 | YDR503C | LPP1 | n |
| W303.asm02.HP0_G0013450 | 176.4124059 | 154.211642 | 185.6873631 | 373.5624349 | 431.2526939 | 358.478699 | 853.8255551 | 875.6717134 | 948.9014753 | 4.17E-13 | -1.168945774 | YDR506C | GMC1 | n |
| W303.asm02.HP0_G0013460 | 850.0751646 | 1137.54884 | 921.5957021 | 1932.018218 | 2401.745772 | 2052.200027 | 1641.389704 | 2158.700102 | 2118.906207 | 4.15E-11 | -1.133654079 | YDR507C | GIN4 | n |
| W303.asm02.HP0_G0013480 | 1335.69393 | 1194.664263 | 1357.472354 | 1370.701747 | 1669.279658 | 1391.368082 | 2278.4458 | 1705.866553 | 1696.148592 | 0.270795941 | -0.187920333 | YDR510W | SMT3 | n |
| W303.asm02.HP0_G0013490 | 1056.535837 | 1134.693069 | 1045.713045 | 634.2778842 | 656.8310261 | 608.3274892 | 630.4299554 | 544.7548891 | 469.8444198 | 1.49E-07 | 0.769887622 | YDR511W | SDH7 | n |
| W303.asm02.HP0_G0013500 | 2357.335006 | 1950.491694 | 2289.818377 | 226.6667899 | 253.4438909 | 237.1753009 | 298.1763302 | 180.939901 | 191.418097 | 2.13E-59 | 3.202777264 | YDR512C | EMI1 | n |
| W303.asm02.HP0_G0013510 | 545.7152995 | 686.3369991 | 538.493353 | 747.1248698 | 701.9466925 | 668.0739391 | 656.934518 | 625.0651125 | 696.0658071 | 0.097615411 | -0.257661678 | YDR514C | | n |
| W303.asm02.HP0_G0013520 | 859.768154 | 1434.54904 | 965.5742881 | 23.34765218 | 25.21169595 | 27.1574772 | 16.09205592 | 20.31945412 | 10.23626187 | 5.42E-77 | 5.426692786 | YDR515W | SLF1 | n |
| W303.asm02.HP0_G0013540 | 975.114727 | 633.029271 | 825.8201148 | 666.380906 | 725.8314571 | 699.7576625 | 899.2619483 | 694.7318124 | 863.9405018 | 0.264728514 | 0.218750553 | YDR517W | GRH1 | n |
| W303.asm02.HP0_G0013560 | 1375.435187 | 1643.020333 | 1511.886056 | 1421.288326 | 1238.026964 | 1334.33738 | 1498.454383 | 1182.398711 | 1099.374525 | 0.268322661 | 0.181407177 | YDR519W | FPR2 | n |
| W303.asm02.HP0_G0013570 | 392.566068 | 452.163765 | 428.058237 | 473.7627755 | 417.9833802 | 439.0458814 | 305.7490624 | 350.2686853 | 313.2296132 | 0.688485488 | -0.065554746 | YDR520C | URC2 | n |
| W303.asm02.HP0_G0013590 | 115.3465731 | 86.62505814 | 119.2308331 | 49.61376088 | 69.00043102 | 65.17794527 | 67.20799824 | 83.2130026 | 70.6302069 | 0.005296363 | 0.810025968 | YDR523C | SPS1 | n |
| W303.asm02.HP0_G0013620 | 510.8205379 | 438.836833 | 539.4706549 | 272.3892754 | 237.5207145 | 252.5645379 | 303.8558794 | 297.0510674 | 211.8906207 | 5.89E-07 | 0.964024273 | YDR525W- | SNA2 | n |
| W303.asm02.HP0_G0013630 | 1414.207144 | 1638.260715 | 1421.97428 | 1270.501406 | 1223.430719 | 1263.727939 | 1009.066565 | 984.0421353 | 804.570183 | 0.075116997 | 0.251667562 | YDR527W | RBA50 | n |
| W303.asm02.HP0_G0013640 | 334.408132 | 185.6251246 | 218.9156281 | 574.9359349 | 699.2928298 | 597.4644984 | 739.2879807 | 612.4864028 | 571.1834123 | 3.81E-09 | -1.339962521 | YDR528W | HLR1 | n |
| W303.asm02.HP0_G0013650 | 5075.249215 | 5322.205495 | 5205.109978 | 7265.011103 | 7917.79946 | 6796.611293 | 6085.636911 | 5989.401001 | 5175.454001 | 2.56E-05 | -0.494131348 | YDR529C | QCR7 | n |
| W303.asm02.HP0_G0013670 | 2022.926874 | 1685.856901 | 1893.033802 | 724.7500364 | 802.7934763 | 669.8844375 | 784.7243739 | 779.8800011 | 844.4916043 | 4.32E-24 | 1.352045968 | YDR531W | CAB1 | n |
| W303.asm02.HP0_G0013680 | 698.864531 | 545.4522892 | 661.6333937 | 168.2976595 | 233.5399204 | 174.7131033 | 236.6478811 | 228.3519606 | 237.4812754 | 2.39E-20 | 1.732537778 | YDR532C | KRE28 | n |
| W303.asm02.HP0_G0013690 | 3478.813872 | 2473.097814 | 3490.922426 | 260.7154493 | 352.9637433 | 260.7117811 | 355.9184132 | 257.3797522 | 312.205987 | 2.56E-66 | 3.437336696 | YDR533C | HSP31 | n |
| W303.asm02.HP0_G0013710 | 585.4565558 | 121.8462356 | 392.8753682 | 525.3221741 | 831.9859663 | 452.62462 | 902.1017229 | 1132.083873 | 1770.873303 | 0.254116603 | -0.717773369 | YDR536W | STL1 | n |
| W303.asm02.HP0_G0013750 | 566.0705771 | 550.2119077 | 580.5173351 | 106.0372537 | 116.7699602 | 99.57741639 | 92.76596941 | 82.24540954 | 68.58295453 | 1.23E-38 | 2.401457007 | YDR541C | | n |
| W303.asm02.HP0_G0013840 | 354.7634096 | 505.4714931 | 307.850102 | 2576.997109 | 1033.679534 | 1830.413963 | 823.5346264 | 1864.551814 | 1956.149643 | 3.22E-06 | -2.219957536 | YEL065W | SIT1 | n |
| W303.asm02.HP0_G0013850 | 322.7765448 | 337.9329191 | 309.8047058 | 406.6382755 | 439.2142821 | 408.2674072 | 426.9127776 | 495.4076434 | 461.6554103 | 0.01360966 | -0.368225027 | YEL064C | AVT2 | n |
| W303.asm02.HP0_G0013870 | 473.9871784 | 660.6350588 | 506.2423899 | 382.3178044 | 394.0986156 | 346.7104589 | 367.2775115 | 342.5279409 | 463.7026627 | 0.005161443 | 0.548239583 | YEL062W | NPR2 | n |
| W303.asm02.HP0_G0013880 | 463.3248902 | 703.471626 | 495.4920689 | 787.9832611 | 708.5813493 | 751.3568691 | 573.6344639 | 739.2410928 | 720.6328356 | 0.027436733 | -0.435720086 | YEL061C | CIN8 | n |
| W303.asm02.HP0_G0013890 | 1260.088613 | 1195.616187 | 1009.552874 | 1057.45408 | 709.9082807 | 995.7741639 | 1003.387016 | 988.8801006 | 1099.374525 | 0.074473406 | 0.324609181 | YEL060C | PRB1 | n |
| W303.asm02.HP0_G0013900 | 301.4519683 | 432.173367 | 347.9194803 | 453.3335798 | 465.7529094 | 481.5925956 | 680.5993062 | 515.7270975 | 543.5455008 | 0.052752646 | -0.372958275 | YEL059C-/ | SOM1 | n |
| W303.asm02.HP0_G0013910 | 2829.383587 | 2114.222573 | 2798.015371 | 1097.339652 | 992.5446616 | 1079.057094 | 1578.914663 | 1367.208984 | 1418.745895 | 8.94E-18 | 1.287942339 | YEL058W | PCM1 | n |
| W303.asm02.HP0_G0013930 | 1255.242119 | 970.9621901 | 1219.672785 | 1433.934971 | 1325.604434 | 1397.704826 | 1340.373599 | 1319.796925 | 1330.714043 | 0.050903854 | -0.271300279 | YEL056W | HAT2 | n |
| W303.asm02.HP0_G0013940 | 1820.343397 | 2057.10715 | 1935.057784 | 5992.56406 | 5550.553903 | 6112.242868 | 2116.578649 | 2836.01524 | 2093.315552 | 1.03E-27 | -1.602980148 | YEL055C | POL5 | n |
| W303.asm02.HP0_G0013950 | 8642.26929 | 9731.516147 | 8800.603708 | 19366.87748 | 21885.07902 | 18641.7976 | 17345.3431 | 13707.89079 | 12148.39559 | 8.72E-14 | -1.140095306 | YEL054C | RPL12A | y |
| W303.asm02.HP0_G0013960 | 887.877823 | 1033.789155 | 959.7104766 | 1594.45008 | 1735.626226 | 1546.165702 | 1387.703175 | 1778.436082 | 1659.298049 | 7.93E-08 | -0.758367279 | YEL053C | MAK10 | n |
| W303.asm02.HP0_G0013970 | 1911.457497 | 1471.674065 | 1829.509177 | 1337.625906 | 1531.278796 | 1290.885416 | 1257.073545 | 1610.074841 | 1330.714043 | 0.047354729 | 0.326266345 | YEL052W | AFG1 | n |
| W303.asm02.HP0_G0013980 | 3170.576811 | 3861.002591 | 3248.551552 | 2066.267218 | 1997.031705 | 2091.125744 | 2628.684664 | 2199.33901 | 2027.803476 | 1.98E-07 | 0.740010928 | YEL051W | VMA8 | n |
| W303.asm02.HP0_G0013990 | 1467.518585 | 1563.058741 | 1502.113037 | 3767.727371 | 3743.273383 | 3842.783023 | 3532.67957 | 3116.617225 | 3052.45329 | 6.25E-35 | -1.324787375 | YEL050C | RML2 | n |
| W303.asm02.HP0_G0014010 | 482.7108688 | 426.4618247 | 550.2209759 | 232.503703 | 208.3282244 | 218.1650668 | 218.6626422 | 288.3427299 | 319.3713703 | 6.27E-09 | 1.145556527 | YEL048C | TCA17 | n |
| W303.asm02.HP0_G0014020 | 887.877823 | 630.1734999 | 723.2034141 | 629.41379 | 345.0021551 | 623.7167263 | 438.2718759 | 353.1714645 | 530.2383649 | 0.062910046 | 0.48513914 | YEL047C | FRD1 | n |
| W303.asm02.HP0_G0014030 | 2971.87053 | 3556.387002 | 2268.317735 | 7462.493328 | 6532.483114 | 7410.370278 | 1983.109244 | 4364.812264 | 5318.761668 | 8.82E-06 | -1.282979473 | YEL046C | GLY1 | n |
| W303.asm02.HP0_G0014050 | 1344.417621 | 1479.289454 | 1330.107901 | 709.184935 | 635.6001242 | 694.326167 | 787.5641484 | 588.2965765 | 605.9867027 | 2.07E-11 | 1.025502272 | YEL044W | IES6 | n |
| W303.asm02.HP0_G0014060 | 450.724004 | 587.3369326 | 433.9220485 | 1787.068211 | 1873.627088 | 1795.109243 | 1591.220353 | 2302.871467 | 2132.213347 | 3.38E-26 | -1.889702807 | YEL043W | GTA1 | n |
| W303.asm02.HP0_G0014070 | 847.1672678 | 1088.048807 | 811.1605861 | 2253.048435 | 2040.820441 | 2251.35486 | 2155.388901 | 1809.39901 | 2032.921607 | 8.95E-17 | -1.253130759 | YEL042W | GDA1 | n |
| W303.asm02.HP0_G0014090 | 3827.761488 | 5109.926506 | 4198.48901 | 4485.667675 | 5524.015276 | 4623.107868 | 6453.861014 | 8212.929838 | 6461.128492 | 0.371617996 | -0.155372195 | YEL040W | UTR2 | n |

| | | | | | | | | | | | | | | |
|---|---|---|---|---|---|---|---|---|---|---|---|---|---|---|
| W303.asm02.HP0_G0014110 | 829.719887 | 675.8658382 | 727.1126218 | 992.2752177 | 1003.160113 | 920.638477 | 1048.823409 | 895.9911675 | 911.0273064 | 0.00690727 | -0.384756861 | YEL038W | UTR4 | n |
| W303.asm02.HP0_G0014120 | 2121.795365 | 1871.482025 | 1808.008535 | 3404.865943 | 2710.92078 | 3372.053419 | 2286.018532 | 2744.0939 | 2809.853883 | 1.79E-06 | -0.710262788 | YEL037C | RAD23 | n |
| W303.asm02.HP0_G0014130 | 1133.110453 | 1060.443019 | 888.3674371 | 4222.033769 | 4425.316105 | 3925.160704 | 3876.292293 | 5105.988543 | 4973.799643 | 2.79E-40 | -2.028151766 | YEL036C | ANP1 | n |
| W303.asm02.HP0_G0014160 | 547.6538974 | 568.2984583 | 385.0569529 | 1793.877943 | 1702.452942 | 1707.300066 | 1324.281543 | 1859.713849 | 1652.132666 | 2.26E-22 | -1.793485177 | YEL032W | MCM3 | n |
| W303.asm02.HP0_G0014170 | 1893.040817 | 2009.510964 | 1905.738726 | 5349.530806 | 4344.373292 | 5117.373953 | 4873.99976 | 5881.998172 | 5948.291773 | 5.33E-24 | -1.350849682 | YEL031W | SPF1 | n |
| W303.asm02.HP0_G0014190 | 862.6760508 | 970.9621901 | 892.2766447 | 713.0762103 | 802.7934763 | 776.7038479 | 1203.117828 | 965.6578673 | 1009.29542 | 0.085628655 | 0.250573513 | YEL029C | BUD16 | n |
| W303.asm02.HP0_G0014210 | 6079.44291 | 7120.389394 | 6322.166062 | 4257.055247 | 3495.137217 | 3898.003227 | 4116.72654 | 3543.325762 | 3193.713703 | 1.90E-07 | 0.74444347 | YEL027W | VMA3 | n |
| W303.asm02.HP0_G0014220 | 4221.296855 | 4655.858894 | 4201.420915 | 7513.079908 | 9821.945969 | 7404.938783 | 7243.318346 | 5423.359064 | 4556.160158 | 9.83E-07 | -0.919450421 | YEL026W | SNU13 | n |
| W303.asm02.HP0_G0014240 | 5177.994902 | 5485.936374 | 5298.930961 | 10019.06124 | 10791.9328 | 9813.80701 | 5507.26949 | 8621.254106 | 7462.234903 | 8.17E-09 | -0.939919051 | YEL024W | RIP1 | n |
| W303.asm02.HP0_G0014250 | 244.2633312 | 267.4905641 | 248.2346854 | 186.7812174 | 252.1169595 | 190.1023404 | 307.6422455 | 254.4769731 | 229.2922659 | 0.189791477 | 0.279013566 | YEL023C | | n |
| W303.asm02.HP0_G0014260 | 1079.799012 | 1323.173965 | 1196.217539 | 2648.985704 | 2750.728721 | 2634.275288 | 2060.729749 | 2862.140252 | 2850.798931 | 2.73E-13 | -1.158290946 | YEL022W | GEA2 | n |
| W303.asm02.HP0_G0014270 | 1661.378372 | 1504.039471 | 1662.390551 | 4916.626422 | 3643.75353 | 4841.272935 | 982.5620025 | 704.4077429 | 743.1526117 | 1.27E-17 | -1.473362921 | YEL021W | URA3 | n |
| W303.asm02.HP0_G0014280 | 1928.904878 | 2684.424879 | 2030.833371 | 1814.307138 | 1932.012069 | 1802.351237 | 2289.804898 | 1908.093501 | 1640.872778 | 0.149744735 | 0.260209076 | YEL020W- | TIM9 | n |
| W303.asm02.HP0_G0014300 | 60.09653387 | 70.44235497 | 47.88779364 | 127.4392681 | 151.2701757 | 150.2713738 | 159.0273761 | 154.8148885 | 178.1109565 | 2.95E-07 | -1.263271215 | YEL019C | MMS21 | n |
| W303.asm02.HP0_G0014310 | 1019.702478 | 986.1929696 | 1031.053516 | 457.2248552 | 403.3871352 | 419.1303981 | 505.4798741 | 362.847395 | 368.5054273 | 5.17E-14 | 1.245745857 | YEL018W | EAF5 | n |
| W303.asm02.HP0_G0014320 | 866.5532465 | 528.3176623 | 873.7079084 | 1981.631979 | 1983.762392 | 1939.043872 | 1622.457873 | 2882.459706 | 2805.759379 | 1.24E-08 | -1.380088611 | YEL017W | GTT3 | n |
| W303.asm02.HP0_G0014340 | 2099.50149 | 1786.760815 | 2057.220523 | 1447.554435 | 1313.662052 | 1403.136322 | 1065.862057 | 862.1254106 | 1028.744318 | 0.00018197 | 0.512649008 | YEL015W | EDC3 | n |
| W303.asm02.HP0_G0014350 | 1672.04066 | 1739.164629 | 1810.940441 | 1914.507479 | 1991.72398 | 1884.728918 | 1238.141714 | 2163.538067 | 2268.35563 | 0.49869198 | -0.149053019 | YEL013W | VAC8 | n |
| W303.asm02.HP0_G0014360 | 964.4524388 | 846.2601833 | 976.3246091 | 876.5097756 | 853.2168682 | 842.7870424 | 1591.220353 | 950.1763785 | 1006.224542 | 0.594764018 | 0.115505171 | YEL012W | UBC8 | n |
| W303.asm02.HP0_G0014370 | 349.916915 | 213.2309123 | 303.9408943 | 288.9271957 | 242.8284399 | 295.1112522 | 209.1967269 | 307.694591 | 313.2296132 | 0.801661997 | 0.066330725 | YEL011W | GLC3 | n |
| W303.asm02.HP0_G0014380 | 8358.264703 | 10093.24716 | 9612.741597 | 8921.721589 | 10119.1786 | 8875.968797 | 5317.951185 | 4857.317128 | 4411.828866 | 0.956050043 | 0.007752538 | YEL009C | GCN4 | n |
| W303.asm02.HP0_G0014420 | 545.7152995 | 273.2021064 | 580.5173351 | 201.3735001 | 213.6359499 | 215.4493191 | 290.603598 | 228.3519606 | 283.5444538 | 2.30E-05 | 1.150355269 | YEL005C | VAB2 | n |
| W303.asm02.HP0_G0014430 | 609.6890291 | 536.8849757 | 652.8376765 | 657.6255364 | 609.0614969 | 616.4747324 | 940.9119754 | 782.7827802 | 789.2157902 | 0.680895339 | -0.066231221 | YEL004W | YEA4 | n |
| W303.asm02.HP0_G0014440 | 844.259371 | 1087.096883 | 910.845381 | 520.4580798 | 615.6961537 | 574.8332674 | 780.9380078 | 570.8799015 | 568.1125338 | 5.51E-05 | 0.733253284 | YEL003W | GIM4 | n |
| W303.asm02.HP0_G0014450 | 4054.577439 | 3597.319722 | 4175.033764 | 2079.886682 | 1897.511853 | 2027.758297 | 2380.677684 | 2273.843676 | 2196.701797 | 2.90E-18 | 0.977255841 | YEL002C | WBP1 | n |
| W303.asm02.HP0_G0014460 | 3030.028466 | 3343.15609 | 3306.212365 | 1460.20108 | 1253.950141 | 1409.473067 | 2457.351598 | 1630.394295 | 1891.661194 | 9.43E-14 | 1.230411719 | YEL001C | IRC22 | n |
| W303.asm02.HP0_G0014470 | 2352.488511 | 3467.858097 | 2828.31173 | 110.9013479 | 134.0200679 | 123.1138966 | 34.07729488 | 151.9121094 | 215.9851255 | 2.19E-14 | 4.555068498 | YER001W | MNN1 | n |
| W303.asm02.HP0_G0014480 | 3480.75247 | 3754.387135 | 3814.409359 | 4248.299878 | 4402.758272 | 4173.198996 | 2690.213113 | 2096.774147 | 1829.219996 | 0.176574709 | -0.214801458 | YER002W | NOP16 | n |
| W303.asm02.HP0_G0014490 | 2051.036543 | 1479.289454 | 1943.853501 | 4729.845204 | 4727.856456 | 4859.37792 | 4489.683601 | 4033.89544 | 3996.236634 | 4.39E-22 | -1.387123821 | YER003C | PMI40 | n |
| W303.asm02.HP0_G0014500 | 2336.01043 | 1591.616453 | 2138.336581 | 1210.186638 | 1086.756789 | 1215.749729 | 2005.827441 | 1087.574592 | 1320.477781 | 0.000605448 | 0.787809817 | YER004W | FMP52 | n |
| W303.asm02.HP0_G0014510 | 891.7550188 | 704.4235497 | 875.6625123 | 726.6956741 | 801.4665449 | 662.6424436 | 672.0799824 | 736.3383137 | 784.0976592 | 0.281026665 | 0.175620997 | YER005W | YND1 | n |
| W303.asm02.HP0_G0014520 | 3252.967221 | 3502.12735 | 3583.766108 | 4913.707965 | 4416.027585 | 4747.127014 | 2286.965123 | 2016.463923 | 2104.57544 | 5.79E-05 | -0.445515296 | YER006W | NUG1 | n |
| W303.asm02.HP0_G0014530 | 447.8161072 | 452.163765 | 417.307916 | 286.0087392 | 277.3286554 | 311.4057385 | 201.6239947 | 325.111266 | 366.4581749 | 0.008960935 | 0.58963096 | YER007W | PAC2 | n |
| W303.asm02.HP0_G0014540 | 1751.523173 | 2018.078277 | 1880.328877 | 2433.99274 | 2714.901574 | 2336.448288 | 1923.473978 | 1522.023873 | 1388.03711 | 0.009439398 | -0.40539202 | YER007C-/ | TMA20 | n |
| W303.asm02.HP0_G0014550 | 505.9740432 | 659.683135 | 485.7190498 | 912.5040727 | 995.1985243 | 926.9752217 | 996.7608754 | 1188.20427 | 1413.627764 | 1.77E-05 | -0.778884961 | YER008C | SEC3 | n |
| W303.asm02.HP0_G0014560 | 4197.064382 | 4378.849093 | 4272.763955 | 2080.859501 | 2457.476889 | 2070.305012 | 2661.815367 | 1808.431417 | 1680.794199 | 1.10E-07 | 0.959631737 | YER009W | NTF2 | n |
| W303.asm02.HP0_G0014570 | 1034.241962 | 879.5775134 | 1009.552874 | 47.6681232 | 37.15407824 | 43.45196352 | 129.6830389 | 76.43985122 | 119.7642639 | 1.66E-79 | 4.50080381 | YER010C | | n |
| W303.asm02.HP0_G0014590 | 3597.068342 | 4034.252708 | 4177.965669 | 1300.65879 | 1111.968485 | 1252.864948 | 1789.057981 | 1321.732111 | 1424.887652 | 2.63E-29 | 1.687026317 | YER012W | PRE1 | n |
| W303.asm02.HP0_G0014600 | 327.6230395 | 299.8559705 | 262.8942141 | 914.4497104 | 842.6014173 | 863.6077749 | 578.3674215 | 805.0374205 | 754.4124998 | 9.10E-20 | -1.557695525 | YER013W | PRP22 | n |
| W303.asm02.HP0_G0014610 | 387.7195734 | 401.7118081 | 360.6244052 | 397.8829059 | 414.0025861 | 376.5836838 | 578.3674215 | 596.0373209 | 713.4674523 | 0.785417766 | -0.045883218 | YER014W | HEM14 | n |
| W303.asm02.HP0_G0014620 | 264.6186088 | 270.3463353 | 323.4869325 | 429.0131088 | 542.7149286 | 537.7180485 | 566.0617317 | 508.9539461 | 504.6477102 | 1.84E-06 | -0.81324886 | YER015W | FAA2 | n |
| W303.asm02.HP0_G0014630 | 1069.136724 | 990.0006644 | 1048.64495 | 508.7842538 | 579.8690068 | 556.7282826 | 709.9436434 | 548.6252613 | 484.1751864 | 4.58E-08 | 0.918278598 | YER016W | BIM1 | n |
| W303.asm02.HP0_G0014640 | 1091.430599 | 1133.741146 | 1024.212403 | 2502.090059 | 2407.053498 | 2503.919398 | 2281.285574 | 2615.404023 | 2871.271454 | 2.65E-21 | -1.190035271 | YER017C | AFG3 | n |
| W303.asm02.HP0_G0014650 | 529.2372177 | 449.3079939 | 542.4025606 | 227.6396088 | 278.6555868 | 267.0485258 | 124.9500812 | 257.3797522 | 257.9537991 | 0.000206775 | 0.976742353 | YER018C | SPC25 | n |
| W303.asm02.HP0_G0014670 | 1590.61955 | 1671.578045 | 1808.008535 | 1535.108131 | 1756.857128 | 1524.43972 | 2088.180903 | 1441.71365 | 1445.360176 | 0.689595254 | 0.074619424 | YER019C-/ | SBH2 | n |
| W303.asm02.HP0_G0014680 | 1448.132607 | 980.4814273 | 1358.449656 | 1170.301066 | 1320.296709 | 1146.950787 | 876.5437517 | 1540.408141 | 1816.936482 | 0.8285647 | 0.058401821 | YER020W | GPA2 | n |
| W303.asm02.HP0_G0014690 | 5493.986355 | 4614.926174 | 5181.654732 | 3340.659899 | 2898.018103 | 3376.579665 | 3626.392131 | 3383.672908 | 3169.146675 | 1.21E-07 | 0.668719751 | YER021W | RPN3 | n |
| W303.asm02.HP0_G0014700 | 1128.263959 | 1052.82763 | 1182.535312 | 987.4111234 | 1085.429857 | 950.5117019 | 740.2345722 | 896.9587605 | 869.0586327 | 0.274179985 | 0.154987265 | YER022W | SRB4 | n |
| W303.asm02.HP0_G0014710 | 2827.444989 | 2016.17443 | 2582.031649 | 1524.407124 | 1215.469131 | 1505.429486 | 2256.674195 | 1545.246106 | 1861.976034 | 2.17E-05 | 0.805875869 | YER023W | PRO3 | n |
| W303.asm02.HP0_G0014720 | 410.0134488 | 444.5483753 | 378.2158396 | 814.2493698 | 786.8702999 | 782.1353433 | 751.5936705 | 730.5327553 | 613.152086 | 4.71E-11 | -0.95095674 | YER024W | YAT2 | n |
| W303.asm02.HP0_G0014730 | 3691.090338 | 3845.771812 | 3745.020923 | 11351.82305 | 11821.63154 | 11094.73468 | 8354.616796 | 8876.698672 | 8122.473794 | 7.87E-67 | -1.602752415 | YER025W | GCD11 | n |
| W303.asm02.HP0_G0014740 | 3147.313637 | 2118.030268 | 3032.56783 | 1939.800769 | 1594.971502 | 1852.139945 | 1784.325024 | 2371.570574 | 2369.694623 | 0.001081324 | 0.622800378 | YER026C | CHO1 | n |
| W303.asm02.HP0_G0014750 | 616.4741217 | 505.4714931 | 495.4920689 | 948.4983698 | 1029.69874 | 962.279942 | 908.7278636 | 1248.195039 | 1167.957479 | 2.36E-07 | -0.861861497 | YER027C | GAL83 | n |
| W303.asm02.HP0_G0014770 | 1014.855983 | 882.4332845 | 1137.579424 | 1080.801732 | 1056.237367 | 1100.783076 | 1053.556367 | 970.4958326 | 782.0504069 | 0.585335775 | -0.093704889 | YER029C | SMB1 | n |
| W303.asm02.HP0_G0014780 | 1796.110924 | 2118.982191 | 2179.383262 | 1909.643385 | 1865.6655 | 2034.095042 | 1314.815628 | 1197.8802 | 881.342147 | 0.706308741 | 0.068904411 | YER030W | CHZ1 | n |
| W303.asm02.HP0_G0014800 | 318.8993491 | 391.2406472 | 273.6445351 | 1526.352761 | 1698.472148 | 1509.955732 | 893.5823992 | 1536.537769 | 1468.903578 | 3.06E-25 | -2.26640029 | YER032W | FIR1 | n |
| W303.asm02.HP0_G0014810 | 122.1316656 | 138.9808625 | 123.1400408 | 572.0174784 | 634.2731928 | 599.2749968 | 487.4946351 | 563.1391571 | 552.758141 | 8.45E-45 | -2.230969825 | YER033C | ZRG8 | n |
| W303.asm02.HP0_G0014820 | 451.693303 | 256.0674795 | 453.4680867 | 363.8342465 | 317.1365964 | 391.0676716 | 359.7047793 | 395.7455589 | 307.0878561 | 0.662005052 | 0.113485357 | YER034W | | n |
| W303.asm02.HP0_G0014840 | 5331.144134 | 5311.734334 | 5372.228605 | 9628.960887 | 10220.02538 | 10005.71985 | 6865.628328 | 7600.443435 | 6536.87683 | 7.97E-19 | -0.898464304 | YER036C | ARB1 | n |
| W303.asm02.HP0_G0014860 | 394.5046659 | 485.4810951 | 433.9220485 | 247.0959856 | 264.0593418 | 248.0382917 | 239.4876557 | 268.9908689 | 246.6939111 | 1.18E-06 | 0.793062539 | YER038C | KRE29 | n |
| W303.asm02.HP0_G0014870 | 308.2370608 | 118.9904645 | 211.0972128 | 4402.978074 | 4494.316536 | 4325.280868 | 4892.93159 | 5687.511968 | 4903.169436 | 2.23E-69 | -4.372924138 | YER039C | HVG1 | n |
| W303.asm02.HP0_G0014890 | 357.6713064 | 412.1829689 | 280.4856485 | 1480.630276 | 1081.449063 | 1479.177258 | 619.0708571 | 1094.347743 | 898.7437922 | 1.31E-16 | -1.94451222 | YER040W | GLN3 | n |
| W303.asm02.HP0_G0014900 | 252.0177227 | 272.2501827 | 247.2573835 | 489.3278769 | 512.1955072 | 518.7078145 | 628.5367723 | 668.6067999 | 658.1916382 | 1.02E-12 | -0.978166609 | YER041W | YEN1 | n |

| | | | | | | | | | | | | | | |
|---|---|---|---|---|---|---|---|---|---|---|---|---|---|---|
| W303.asm02.HP0_G0014910 | 1396.759763 | 1421.222108 | 1309.584561 | 913.4768915 | 762.9855353 | 914.3017323 | 779.0448247 | 929.8569243 | 684.8059191 | 1.59E-05 | 0.670439759 | YER042W | MXR1 | n |
| W303.asm02.HP0_G0014920 | 9370.212789 | 8656.794271 | 8973.586147 | 19982.67181 | 20507.72426 | 18409.14854 | 8898.906922 | 12454.85778 | 10820.75242 | 1.12E-15 | -1.125231858 | YER043C | SAH1 | n |
| W303.asm02.HP0_G0014930 | 1768.001255 | 1839.116619 | 1502.113037 | 866.7815872 | 956.7175147 | 938.7434618 | 1027.051804 | 841.8059565 | 644.8844978 | 1.04E-06 | 0.887578885 | YER044C | ERG28 | n |
| W303.asm02.HP0_G0014960 | 398.3818616 | 372.2021729 | 384.079651 | 62.26040581 | 53.07725463 | 68.79894223 | 29.34433726 | 61.92595542 | 88.03185208 | 9.32E-17 | 2.643103991 | YER046W | SPO73 | n |
| W303.asm02.HP0_G0014970 | 186.1053952 | 275.1059539 | 171.0278344 | 729.6141306 | 654.1771633 | 716.957398 | 375.7968352 | 545.7224821 | 654.0971335 | 7.77E-13 | -1.732427566 | YER047C | SAP1 | n |
| W303.asm02.HP0_G0014980 | 2527.931618 | 1961.914778 | 2459.86891 | 1620.716189 | 1337.546817 | 1585.996668 | 1328.067909 | 1564.597967 | 1362.446455 | 6.44E-05 | 0.611990434 | YER048C | CAJ1 | n |
| W303.asm02.HP0_G0014990 | 1697.242432 | 1550.683733 | 1693.664212 | 707.2392973 | 676.7349965 | 638.2007141 | 1074.38138 | 849.5467009 | 885.4366517 | 9.21E-21 | 1.289302655 | YER048W- | ISD11 | n |
| W303.asm02.HP0_G0015000 | 3175.423306 | 3873.3776 | 3322.826497 | 4666.611979 | 2980.287848 | 4353.343595 | 1089.526845 | 1585.885015 | 1099.374525 | 0.334709785 | -0.210802515 | YER049W | TPA1 | n |
| W303.asm02.HP0_G0015010 | 1640.053795 | 1868.626254 | 1611.570851 | 1123.605761 | 1133.199386 | 1147.856036 | 1157.681435 | 978.236577 | 782.0504069 | 0.000296335 | 0.58870616 | YER050C | RSM18 | n |
| W303.asm02.HP0_G0015020 | 705.6496235 | 608.2792544 | 647.951167 | 476.681232 | 461.7721153 | 488.8345896 | 314.2683862 | 456.7039212 | 438.112008 | 0.009948302 | 0.4583443 | YER051W | JHD1 | n |
| W303.asm02.HP0_G0015030 | 4230.020545 | 4243.675925 | 4143.760103 | 3269.644124 | 2694.997604 | 3439.041862 | 1459.644131 | 1697.158216 | 1499.612364 | 0.001480281 | 0.423405774 | YER052C | HOM3 | n |
| W303.asm02.HP0_G0015050 | 235.5396408 | 139.9327862 | 182.7554574 | 102.1459783 | 132.6931366 | 91.43017323 | 99.39211008 | 160.6204469 | 166.8510685 | 0.010044618 | 0.780745138 | YER054C | GIP2 | n |
| W303.asm02.HP0_G0015060 | 6969.259331 | 7443.091534 | 7518.383601 | 1569.15679 | 1326.931366 | 1521.723972 | 1832.601192 | 1203.685758 | 1390.084362 | 4.46E-45 | 2.310918638 | YER055C | HIS1 | n |
| W303.asm02.HP0_G0015070 | 4401.586457 | 7190.831749 | 4632.411058 | 4897.170045 | 3503.098806 | 4744.411266 | 2482.909569 | 3611.057275 | 2750.483564 | 0.195172677 | 0.303500142 | YER056C | FCY2 | n |
| W303.asm02.HP0_G0015100 | 684.325047 | 621.6061864 | 683.1340358 | 402.7470001 | 407.3679293 | 458.0561154 | 587.8333367 | 422.8381643 | 415.5922319 | 0.00028548 | 0.648345552 | YER058W | PET117 | n |
| W303.asm02.HP0_G0015110 | 786.101435 | 518.7984251 | 693.8843568 | 636.2235219 | 559.9650364 | 629.1482217 | 368.2241031 | 448.9631768 | 477.0098031 | 0.524415396 | 0.129883748 | YER059W | PCL6 | n |
| W303.asm02.HP0_G0015130 | 378.026584 | 390.2887235 | 363.5563109 | 574.9359349 | 455.1374585 | 579.3595136 | 469.5093962 | 587.3289834 | 533.3092434 | 0.002293921 | -0.510610625 | YER061C | CEM1 | n |
| W303.asm02.HP0_G0015150 | 1012.917385 | 825.3178616 | 1088.714329 | 803.5483625 | 786.8702999 | 794.8088327 | 1228.675799 | 720.8568248 | 822.9954543 | 0.171750708 | 0.295137342 | YER063W | THO1 | n |
| W303.asm02.HP0_G0015170 | 341.1932246 | 238.9328527 | 301.0089886 | 19.45637682 | 19.90397049 | 4.5262462 | 10.41250677 | 5.805558321 | 11.25988806 | 3.87E-26 | 4.365104705 | YER065C | ICL1 | n |
| W303.asm02.HP0_G0015190 | 767.6847553 | 377.9137152 | 513.0835033 | 18.48355798 | 29.19249005 | 8.147243159 | 57.742083 | 29.0277916 | 52.20493554 | 9.83E-34 | 4.935983704 | YER067W | RGI1 | n |
| W303.asm02.HP0_G0015200 | 522.4521251 | 591.1446275 | 499.4012765 | 1041.888979 | 1141.160975 | 1062.762608 | 1063.968874 | 1045.968091 | 1116.77617 | 1.80E-15 | -1.007813459 | YER068W | MOT2 | n |
| W303.asm02.HP0_G0015210 | 1940.536465 | 1567.81836 | 1737.642798 | 1672.275587 | 1442.374395 | 1691.910829 | 1077.221155 | 916.3106216 | 1241.658565 | 0.454512399 | 0.125499126 | YER069W | ARG56 | n |
| W303.asm02.HP0_G0015220 | 1446.194009 | 1627.789554 | 1413.178563 | 8973.280988 | 8755.093151 | 9050.681901 | 5898.211789 | 8733.4949 | 8154.206205 | 4.16E-60 | -2.577218025 | YER070W | RNR1 | n |
| W303.asm02.HP0_G0015230 | 733.7592926 | 749.1639643 | 759.3635848 | 570.0718407 | 508.2147131 | 550.3915379 | 560.3821825 | 374.4585117 | 397.1669605 | 0.010877141 | 0.460329165 | YER071C | TDA2 | n |
| W303.asm02.HP0_G0015240 | 6855.851356 | 8413.1018 | 6908.547209 | 2863.005849 | 1844.434598 | 2789.072908 | 3156.882734 | 2006.787993 | 2297.017164 | 1.88E-13 | 1.564196929 | YER072W | VTC1 | n |
| W303.asm02.HP0_G0015250 | 1883.347828 | 2382.665061 | 1777.712176 | 10614.42637 | 8143.377792 | 10039.21407 | 10689.85809 | 11554.02865 | 14062.57656 | 4.52E-40 | -2.252502946 | YER073W | ALD5 | n |
| W303.asm02.HP0_G0015270 | 289.8203811 | 254.1636321 | 291.2359695 | 118.6838986 | 135.3469993 | 124.0191459 | 168.4932914 | 132.5602483 | 139.2131614 | 8.14E-09 | 1.146828229 | YER074W- | YOS1 | n |
| W303.asm02.HP0_G0015280 | 313.0835555 | 405.5195029 | 308.8274039 | 782.146348 | 579.8690068 | 765.840857 | 377.6900183 | 632.8058569 | 507.7185887 | 3.27E-06 | -1.051580923 | YER075C | PTP3 | n |
| W303.asm02.HP0_G0015290 | 362.5178011 | 217.0386072 | 312.7366115 | 223.7483334 | 270.6939986 | 261.6170303 | 342.6661319 | 311.5649632 | 288.6625847 | 0.299475547 | 0.240073578 | YER076C | | n |
| W303.asm02.HP0_G0015300 | 1199.022781 | 1377.433617 | 1337.926316 | 888.1836017 | 910.2749169 | 942.3644588 | 642.7356452 | 814.713351 | 733.9399761 | 0.000200579 | 0.514059744 | YER077C | MRX1 | n |
| W303.asm02.HP0_G0015310 | 726.9742001 | 686.3369991 | 682.1567339 | 694.5926524 | 818.7166527 | 715.1468995 | 835.8403162 | 865.9957828 | 747.2471165 | 0.550291403 | -0.086812077 | YER078C | ICP55 | n |
| W303.asm02.HP0_G0015330 | 477.8643742 | 202.7597515 | 400.6937835 | 12.64664493 | 11.94238229 | 11.76824012 | 37.86366098 | 38.70372214 | 50.15768316 | 1.19E-33 | 4.893614916 | YER079W | | n |
| W303.asm02.HP0_G0015340 | 1743.768781 | 1361.250914 | 1579.319888 | 3885.43845 | 3641.099668 | 3687.080154 | 2767.833618 | 3277.237672 | 3006.390111 | 2.02E-21 | -1.259479415 | YER080W | AIM9 | n |
| W303.asm02.HP0_G0015350 | 2974.778427 | 2870.050003 | 3342.372536 | 2731.675305 | 2087.263038 | 2398.910486 | 2357.959488 | 2406.403924 | 2639.931936 | 0.014301975 | 0.347339228 | YER081W | SER3 | n |
| W303.asm02.HP0_G0015360 | 1410.329948 | 1726.78962 | 1446.406828 | 3972.019327 | 3455.329276 | 3898.003227 | 2221.650308 | 2307.709432 | 1895.755698 | 7.48E-21 | -1.305270529 | YER082C | UTP7 | n |
| W303.asm02.HP0_G0015370 | 635.8601003 | 587.3369326 | 622.5413173 | 1425.179602 | 1435.739738 | 1441.15679 | 2019.079722 | 1862.616628 | 1880.401305 | 6.65E-29 | -1.220910605 | YER083C | GET2 | n |
| W303.asm02.HP0_G0015400 | 2256.527917 | 2288.424613 | 2359.206813 | 2696.653827 | 2014.281813 | 2746.526194 | 2394.876557 | 2040.65375 | 2269.379257 | 0.480696156 | -0.112163084 | YER086W | ILV1 | n |
| W303.asm02.HP0_G0015410 | 901.4480081 | 852.9236494 | 997.8252511 | 783.1191669 | 605.0807028 | 791.1878357 | 691.0118129 | 768.2688844 | 829.1372115 | 0.040118863 | 0.334165207 | YER087W | AIM10 | n |
| W303.asm02.HP0_G0015430 | 503.0661464 | 630.1734999 | 348.8967822 | 2502.090059 | 2404.399635 | 2511.161392 | 1349.839514 | 2219.658465 | 3009.46099 | 6.95E-18 | -2.32319033 | YER088C | DOT6 | n |
| W303.asm02.HP0_G0015450 | 4391.893467 | 3186.088677 | 4097.826913 | 2142.147088 | 1860.357775 | 2260.407352 | 1920.634203 | 1744.570275 | 2107.646319 | 1.66E-08 | 0.89801308 | YER090W | TRP2 | n |
| W303.asm02.HP0_G0015460 | 4534.380411 | 5020.445677 | 4571.81834 | 589.5282175 | 254.7708222 | 542.2442947 | 832.0539501 | 793.4263038 | 1778.038687 | 2.08E-25 | 3.346441095 | YER091C | MET6 | n |
| W303.asm02.HP0_G0015480 | 903.386606 | 722.5101003 | 969.4834957 | 121.6023551 | 123.404617 | 135.787386 | 106.9648423 | 105.4676428 | 123.8587686 | 4.83E-50 | 2.766802857 | YER092W | IES5 | n |
| W303.asm02.HP0_G0015490 | 271.4037014 | 407.4233504 | 296.122479 | 614.8215074 | 647.5425065 | 659.0214467 | 993.9211008 | 758.5929539 | 844.4916043 | 2.68E-07 | -0.978390652 | YER093C | TSC11 | n |
| W303.asm02.HP0_G0015500 | 365.4256979 | 339.8367665 | 369.4201224 | 453.3335798 | 486.9838112 | 527.7603069 | 639.8958706 | 464.4446656 | 412.5213534 | 0.022112481 | -0.450354264 | YER093C-/ | AIM11 | n |
| W303.asm02.HP0_G0015510 | 3871.37994 | 3247.963718 | 3534.901012 | 1650.873573 | 1555.163561 | 1618.585641 | 2422.327711 | 1762.954543 | 1786.227696 | 1.45E-13 | 1.142713154 | YER094C | PUP3 | n |
| W303.asm02.HP0_G0015520 | 1330.847436 | 698.7120074 | 1013.462082 | 1631.417196 | 1575.067531 | 1619.49089 | 1944.298991 | 1917.769432 | 1818.983734 | 0.001265511 | -0.665487504 | YER095W | RAD51 | n |
| W303.asm02.HP0_G0015540 | 184.1667974 | 242.7405475 | 172.9824382 | 762.6899712 | 850.5630055 | 681.6526777 | 711.8368265 | 963.7226812 | 1261.107462 | 4.03E-17 | -1.934523445 | YER098W | UBP9 | n |
| W303.asm02.HP0_G0015550 | 1533.430913 | 1306.039338 | 1441.520319 | 1614.879276 | 1721.029981 | 1610.438398 | 1670.734041 | 1671.033203 | 1528.273897 | 0.087836105 | -0.207955179 | YER099C | PRS2 | n |
| W303.asm02.HP0_G0015560 | 2119.856767 | 1958.107083 | 2316.205529 | 739.342319 | 931.5058188 | 777.6090971 | 1560.929424 | 1298.509878 | 1322.525034 | 4.84E-21 | 1.386852662 | YER100W | UBC6 | n |
| W303.asm02.HP0_G0015570 | 531.1758155 | 343.6444614 | 494.514767 | 1357.082283 | 1076.141338 | 1368.736851 | 1196.491687 | 1322.699704 | 1162.839348 | 3.11E-15 | -1.47441923 | YER101C | AST2 | n |
| W303.asm02.HP0_G0015630 | 1265.904407 | 1041.404545 | 1289.061221 | 1722.862167 | 1795.338138 | 1717.257808 | 1586.487395 | 1547.181292 | 1380.871726 | 4.63E-05 | -0.541610445 | YER107C | GLE2 | n |
| W303.asm02.HP0_G0015650 | 1789.325831 | 1988.568642 | 1959.490332 | 15351.08131 | 17486.30154 | 15232.62896 | 6951.768156 | 9221.161799 | 7880.898014 | 1.71E-111 | -3.066580122 | YER110C | KAP123 | n |
| W303.asm02.HP0_G0015660 | 523.4214241 | 694.9043125 | 587.3584485 | 958.2265582 | 1003.160113 | 1088.109586 | 720.3561502 | 1000.491267 | 1125.988806 | 9.32E-05 | -0.756113168 | YER111C | SWI4 | n |
| W303.asm02.HP0_G0015670 | 2319.532348 | 2815.790351 | 2565.417516 | 1997.19708 | 1994.377843 | 1970.727595 | 2016.239947 | 1662.324866 | 1531.344776 | 0.010853767 | 0.369181789 | YER112W | LSM4 | n |
| W303.asm02.HP0_G0015680 | 665.9083673 | 821.5101667 | 669.451809 | 1841.546066 | 1762.164854 | 1824.982468 | 1401.902048 | 2300.936281 | 2261.190247 | 1.25E-11 | -1.331689621 | YER113C | TMN3 | n |
| W303.asm02.HP0_G0015690 | 421.645036 | 511.1830354 | 342.0556688 | 1862.94808 | 1885.569471 | 1792.393495 | 1138.749604 | 1724.250821 | 1850.716146 | 1.14E-24 | -2.119484898 | YER114C | BOI2 | n |
| W303.asm02.HP0_G0015700 | 380.9344808 | 340.7886903 | 361.6017071 | 157.5966522 | 119.4238229 | 140.3136322 | 155.24101 | 149.9769233 | 137.1659091 | 2.30E-13 | 1.36987492 | YER115C | SPR6 | n |
| W303.asm02.HP0_G0015710 | 545.7152995 | 334.1252242 | 515.0381071 | 295.7369276 | 203.020499 | 231.7438054 | 147.6682778 | 262.2177175 | 294.8043418 | 0.001411555 | 0.931213274 | YER116C | SLX8 | n |
| W303.asm02.HP0_G0015730 | 2003.540895 | 1873.385873 | 2150.064204 | 1197.539993 | 1366.739307 | 1206.697237 | 1617.724915 | 1678.773948 | 1555.911804 | 1.48E-08 | 0.677729302 | YER118C | SHO1 | n |
| W303.asm02.HP0_G0015740 | 399.3511606 | 247.5001661 | 324.4642344 | 481.5453262 | 516.1763013 | 577.5490151 | 1226.782616 | 1083.70422 | 1323.54866 | 0.000366802 | -0.698281568 | YER119C | AVT6 | n |
| W303.asm02.HP0_G0015770 | 4055.546737 | 3229.877168 | 4111.50914 | 1852.247073 | 1986.416255 | 1887.444665 | 2282.232166 | 2161.602881 | 2178.276526 | 1.34E-14 | 0.99330803 | YER122C | GLO3 | n |
| W303.asm02.HP0_G0015780 | 1031.334065 | 1240.356602 | 1097.510046 | 1315.251073 | 1366.739307 | 1329.811133 | 776.2050501 | 1432.037719 | 1554.888178 | 0.292959984 | -0.251737122 | YER123W | YCK3 | n |
| W303.asm02.HP0_G0015790 | 4120.489766 | 3516.406206 | 3896.502719 | 792.8473553 | 975.2945538 | 931.5014679 | 1169.987124 | 1232.71355 | 998.0355323 | 1.72E-49 | 2.095950201 | YER124C | DSE1 | n |

| | | | | | | | | | | | | | | |
|---|---|---|---|---|---|---|---|---|---|---|---|---|---|---|
| W303.asm02.HP0_G0015800 | 1871.71624 | 1766.770417 | 1750.347723 | 3024.493776 | 2963.03774 | 2994.564486 | 2351.333347 | 2650.237373 | 2879.460464 | 2.36E-10 | -0.737144665 | YER125W | RSP5 | n |
| W303.asm02.HP0_G0015810 | 3205.471573 | 4360.762542 | 3495.808936 | 5966.297951 | 5813.286313 | 6114.958616 | 3623.552356 | 3524.941494 | 3426.076848 | 3.46E-07 | -0.693933025 | YER126C | NSA2 | n |
| W303.asm02.HP0_G0015820 | 1570.264272 | 2184.664928 | 1748.393119 | 2742.376312 | 2318.149096 | 2694.926987 | 1798.523897 | 1666.195238 | 1568.195318 | 0.00167305 | -0.495291487 | YER127W | LCP5 | n |
| W303.asm02.HP0_G0015830 | 381.9037798 | 397.9041152 | 399.7164816 | 279.1990073 | 281.3094495 | 283.3430121 | 352.1320471 | 210.9352856 | 223.1505088 | 0.027669775 | 0.483122115 | YER128W | VFA1 | n |
| W303.asm02.HP0_G0015840 | 509.851239 | 748.2120406 | 556.0847874 | 537.968819 | 575.8882127 | 515.9920668 | 467.6162131 | 604.7456584 | 676.6169096 | 0.466965399 | 0.155477167 | YER129W | SAK1 | n |
| W303.asm02.HP0_G0015850 | 806.4567126 | 1039.500698 | 825.8201148 | 529.2134494 | 627.638536 | 532.2865531 | 643.6822367 | 793.4263038 | 886.4602779 | 0.000225675 | 0.663287331 | YER130C | COM2 | n |
| W303.asm02.HP0_G0015860 | 8591.865746 | 11686.76746 | 9756.404977 | 15202.24003 | 18169.67119 | 14722.97364 | 15291.23949 | 11652.72314 | 10580.20027 | 0.000164906 | -0.679157884 | YER131W | RPS26B | y |
| W303.asm02.HP0_G0015880 | 645.5530897 | 653.9715928 | 647.951167 | 171.216116 | 147.2893816 | 208.2073252 | 209.1967269 | 196.4213898 | 222.1268826 | 1.19E-27 | 1.878484952 | YER139C | RTR1 | n |
| W303.asm02.HP0_G0015890 | 592.2416483 | 818.6543956 | 614.722902 | 452.360761 | 445.8489389 | 391.9729209 | 600.1390266 | 602.8104723 | 678.664162 | 0.000183959 | 0.651950208 | YER140W | EMP65 | n |
| W303.asm02.HP0_G0015900 | 2060.729532 | 1865.770483 | 1910.625236 | 1524.407124 | 1466.259159 | 1496.376994 | 1127.390506 | 1219.167247 | 1023.626187 | 0.001496726 | 0.379247724 | YER141W | COX15 | n |
| W303.asm02.HP0_G0015910 | 548.6231963 | 629.2215761 | 546.3117682 | 92.41778988 | 74.30815648 | 76.04093615 | 177.9592066 | 116.1111664 | 120.7878901 | 1.69E-36 | 2.824949509 | YER142C | MAG1 | n |
| W303.asm02.HP0_G0015920 | 1108.87798 | 840.548641 | 1046.690347 | 853.1621234 | 688.6773788 | 787.5668387 | 949.4312991 | 829.2272468 | 802.5229306 | 0.028995837 | 0.361667092 | YER143W | DDI1 | n |
| W303.asm02.HP0_G0015930 | 2069.453223 | 2276.049604 | 2123.677053 | 13696.31646 | 10310.25671 | 13053.69404 | 14755.46868 | 11911.07049 | 17336.1331 | 1.96E-51 | -2.51833135 | YER144C | UBP5 | n |
| W303.asm02.HP0_G0015940 | 931.496275 | 1352.6836 | 968.5061938 | 872.6185002 | 924.8711619 | 896.1967475 | 1086.68707 | 679.2503235 | 743.1526117 | 0.219161475 | 0.2723095 | YER146W | LSM5 | n |
| W303.asm02.HP0_G0015950 | 323.7458438 | 353.1636986 | 294.1678752 | 458.197674 | 420.6372429 | 476.1611002 | 412.7139047 | 419.9353852 | 434.0175033 | 0.001451552 | -0.482356486 | YER147C | SCC4 | n |
| W303.asm02.HP0_G0015960 | 3235.51984 | 3174.665592 | 3308.166969 | 1962.175602 | 1977.127735 | 1893.78141 | 2031.385412 | 1890.676826 | 1697.172218 | 3.48E-11 | 0.736663516 | YER148W | SPT15 | n |
| W303.asm02.HP0_G0015970 | 632.9522035 | 732.9812612 | 740.7948485 | 737.3966814 | 782.8895058 | 760.4093615 | 762.9527688 | 690.8614401 | 695.042181 | 0.41328041 | -0.113897774 | YER149C | PEA2 | n |
| W303.asm02.HP0_G0015990 | 755.0838691 | 1108.991129 | 672.3837148 | 3144.150494 | 3074.499974 | 3100.478647 | 1613.938549 | 2419.950227 | 2176.229274 | 1.30E-19 | -1.877264919 | YER151C | UBP3 | n |
| W303.asm02.HP0_G0016000 | 836.5049795 | 811.0390059 | 705.6119797 | 625.5225147 | 557.3111736 | 576.6437658 | 755.3800366 | 796.329083 | 852.6806138 | 0.002650334 | 0.418522742 | YER152C | | n |
| W303.asm02.HP0_G0016010 | 85.29830614 | 139.9327862 | 90.88907772 | 375.5080726 | 382.1562333 | 417.3198996 | 613.3913079 | 515.7270975 | 484.1751864 | 3.01E-18 | -1.893773459 | YER153C | PET122 | n |
| W303.asm02.HP0_G0016020 | 784.1628371 | 930.0294703 | 731.9991313 | 1955.36587 | 1841.780736 | 1894.686659 | 1889.396683 | 1859.713849 | 1612.211244 | 5.95E-19 | -1.218414785 | YER154W | OXA1 | n |
| W303.asm02.HP0_G0016030 | 1422.930834 | 2009.510964 | 1489.408112 | 2024.436008 | 2225.2639 | 2066.684015 | 1807.989812 | 2654.107746 | 2068.748524 | 0.05962115 | -0.359634275 | YER155C | BEM2 | n |
| W303.asm02.HP0_G0016040 | 1432.623824 | 1220.366204 | 1401.45094 | 3135.395124 | 3718.061687 | 3176.519583 | 2235.849181 | 2623.144768 | 2158.827628 | 4.62E-21 | -1.306326808 | YER156C | MYG1 | n |
| W303.asm02.HP0_G0016050 | 896.6015134 | 1049.019935 | 1002.711761 | 1317.19671 | 1299.065807 | 1356.968611 | 1245.714446 | 1305.283029 | 1182.288246 | 0.000354709 | -0.430676556 | YER157W | COG3 | n |
| W303.asm02.HP0_G0016070 | 2517.26933 | 2963.338527 | 2894.76826 | 1424.206783 | 1426.451218 | 1399.515325 | 1661.268126 | 1088.542185 | 1057.405851 | 9.54E-08 | 0.978685629 | YER159C | BUR6 | n |
| W303.asm02.HP0_G0016080 | 651.3688833 | 518.7984251 | 583.4492409 | 503.9201596 | 534.7533404 | 563.0650272 | 494.1207758 | 420.9029782 | 415.5922319 | 0.430120824 | 0.130423168 | YER161C | SPT2 | n |
| W303.asm02.HP0_G0016090 | 243.2940323 | 246.5482424 | 214.0291185 | 437.7684784 | 372.8677138 | 465.2981093 | 340.7729488 | 507.9863531 | 471.8916722 | 1.86E-05 | -0.860153708 | YER162C | RAD4 | n |
| W303.asm02.HP0_G0016100 | 1040.057756 | 996.6641304 | 1091.646235 | 552.5611016 | 506.8877817 | 544.0547932 | 443.951425 | 468.3150379 | 414.5686057 | 3.25E-14 | 0.962666049 | YER163C | GCG1 | n |
| W303.asm02.HP0_G0016110 | 393.535367 | 410.2791215 | 324.4642344 | 2225.809508 | 2202.706067 | 2188.892662 | 1649.909027 | 1956.473154 | 2063.630393 | 3.27E-70 | -2.551923187 | YER164W | CHD1 | n |
| W303.asm02.HP0_G0016120 | 4133.090652 | 3862.906439 | 3878.911285 | 15692.54072 | 16090.36974 | 15519.59297 | 11377.08353 | 12466.4689 | 12874.14655 | 4.41E-92 | -1.993966731 | YER165W | PAB1 | n |
| W303.asm02.HP0_G0016130 | 953.7901505 | 1072.818028 | 1008.575572 | 2754.050138 | 2945.787632 | 2812.609388 | 1864.785303 | 3321.746952 | 2997.177475 | 4.08E-14 | -1.487681027 | YER166W | DNF1 | n |
| W303.asm02.HP0_G0016140 | 386.7502744 | 610.1831018 | 389.9434625 | 1113.877573 | 1259.257866 | 1064.573106 | 745.9141213 | 1547.181292 | 1655.203544 | 3.36E-06 | -1.309165929 | YER167W | BCK2 | n |
| W303.asm02.HP0_G0016150 | 1181.5754 | 1082.337265 | 1271.469786 | 2013.735001 | 1926.704343 | 1943.570118 | 1176.613265 | 1423.329382 | 1281.579096 | 7.33E-09 | -0.735132461 | YER168C | CCA1 | n |
| W303.asm02.HP0_G0016160 | 307.2677619 | 495.0003322 | 259.9623083 | 1346.381276 | 979.2753479 | 1293.601164 | 628.5367723 | 663.7688347 | 575.2779171 | 4.68E-14 | -1.769046157 | YER169W | RPH1 | n |
| W303.asm02.HP0_G0016170 | 892.7243177 | 758.6832015 | 887.3901352 | 611.9030509 | 563.9458305 | 628.2429725 | 621.9106316 | 537.9817377 | 453.4664008 | 0.002620051 | 0.491709984 | YER170W | ADK2 | n |
| W303.asm02.HP0_G0016180 | 1011.948086 | 1143.260383 | 1055.486064 | 1324.979261 | 1443.701326 | 1354.252863 | 1162.414392 | 1242.389481 | 1116.77617 | 0.002623713 | -0.360037085 | YER171W | RAD3 | n |
| W303.asm02.HP0_G0016190 | 646.5223886 | 922.4140806 | 648.9284689 | 1897.969558 | 1963.858421 | 1908.265398 | 969.3097211 | 1488.158116 | 1269.296472 | 1.82E-12 | -1.379218599 | YER172C | BRR2 | n |
| W303.asm02.HP0_G0016200 | 298.5440715 | 315.0867499 | 283.4175542 | 383.2906233 | 403.3871352 | 372.9626868 | 291.5501896 | 416.065013 | 398.1905867 | 0.047619459 | -0.369346856 | YER173W | RAD24 | n |
| W303.asm02.HP0_G0016210 | 1215.500863 | 1557.347199 | 1392.655223 | 4094.594501 | 3820.235402 | 4009.348884 | 3527.00002 | 3703.946209 | 2980.799456 | 1.26E-27 | -1.517404314 | YER174C | GRX4 | n |
| W303.asm02.HP0_G0016220 | 991.5928089 | 757.7312778 | 1003.689063 | 8.755369568 | 11.94238229 | 8.147243159 | 8.519323721 | 6.773151374 | 19.44889755 | 1.61E-75 | 6.607384506 | YER175C | TMT1 | n |
| W303.asm02.HP0_G0016250 | 11298.14837 | 8607.294238 | 11130.49146 | 11108.61834 | 10144.39029 | 10951.7053 | 6853.322638 | 10264.22711 | 10120.59211 | 0.78600029 | -0.053381677 | YER178W | PDA1 | n |
| W303.asm02.HP0_G0016260 | 251.0484238 | 164.6828028 | 245.3027797 | 44.74966668 | 46.4425978 | 37.11521884 | 64.36822367 | 61.92595542 | 48.11043079 | 2.43E-16 | 2.371238842 | YER179W | DMC1 | n |
| W303.asm02.HP0_G0016270 | 462.3555912 | 395.0483421 | 383.1023491 | 272.3892754 | 310.5019396 | 266.1432765 | 444.8980165 | 320.2733007 | 371.5763059 | 0.003399303 | 0.549683912 | YER180C | ISC10 | n |
| W303.asm02.HP0_G0016280 | 201.6141782 | 171.3462688 | 181.7781554 | 62.26040581 | 99.51985243 | 77.85143463 | 53.9557169 | 82.24540954 | 92.12635683 | 1.82E-05 | 1.219749787 | YER180C-/ | SLO1 | n |
| W303.asm02.HP0_G0016290 | 1843.606571 | 1410.750947 | 1816.804263 | 517.5396233 | 464.425978 | 564.8775257 | 1184.185997 | 760.52814 | 915.1218112 | 8.94E-19 | 1.711514486 | YER182W | FMP10 | n |
| W303.asm02.HP0_G0016300 | 1106.939382 | 811.9909296 | 1051.576856 | 175.1073914 | 136.6739307 | 199.1548328 | 431.6457352 | 214.8056579 | 321.4186227 | 8.05E-23 | 2.534838689 | YER183C | FAU1 | n |
| W303.asm02.HP0_G0016320 | 65.91232747 | 90.432753 | 70.36573759 | 3930.188117 | 3300.078307 | 3625.523206 | 5005.575982 | 5490.122985 | 6025.063737 | 4.86E-242 | -5.580804372 | YER185W | PUG1 | n |
| W303.asm02.HP0_G0016410 | 961.544542 | 1082.337265 | 947.0055518 | 1147.926232 | 1074.814406 | 1094.446331 | 914.4074127 | 880.5096786 | 849.6097352 | 0.23135829 | -0.149695248 | YFL049W | SWP82 | n |
| W303.asm02.HP0_G0016420 | 2429.063127 | 2029.501362 | 2325.001246 | 2025.408827 | 1906.800373 | 2069.399762 | 1842.067107 | 1753.278613 | 1581.502459 | 0.176474248 | 0.176236437 | YFL048C | EMP47 | n |
| W303.asm02.HP0_G0016430 | 1501.444048 | 1243.212373 | 1480.612395 | 1201.431268 | 1109.314622 | 1113.456565 | 1083.847296 | 1258.838563 | 1122.917927 | 0.026737929 | 0.302929065 | YFL047W | RGD2 | n |
| W303.asm02.HP0_G0016440 | 1142.803443 | 1136.596917 | 1197.194841 | 139.1130942 | 177.808803 | 140.3136322 | 357.8115963 | 273.8288341 | 325.5131275 | 1.17E-67 | 2.935176615 | YFL046W | FMP32 | n |
| W303.asm02.HP0_G0016450 | 7785.409033 | 8061.841949 | 8340.294508 | 6901.176857 | 7064.582591 | 6661.729157 | 6975.432944 | 5675.900851 | 4938.996352 | 0.116428656 | 0.229755232 | YFL045C | SEC53 | n |
| W303.asm02.HP0_G0016460 | 1052.658642 | 891.000598 | 1044.735743 | 265.5795435 | 233.5399204 | 239.8910486 | 326.574076 | 390.9075936 | 317.324118 | 7.88E-37 | 2.01403431 | YFL044C | OTU1 | n |
| W303.asm02.HP0_G0016470 | 445.8775094 | 281.7694199 | 337.1691593 | 690.701377 | 648.8694379 | 716.0521488 | 593.5128859 | 884.3800508 | 797.4047997 | 7.66E-06 | -0.949615338 | YFL042C | LAM5 | n |
| W303.asm02.HP0_G0016500 | 845.2286699 | 841.5005648 | 872.7306065 | 1734.535993 | 1494.124718 | 1738.98379 | 1800.41708 | 1804.561045 | 1998.118317 | 2.57E-15 | -0.957846995 | YFL041W | FET5 | n |
| W303.asm02.HP0_G0016510 | 116.315872 | 124.7020068 | 106.5259083 | 3.891275363 | 11.94238229 | 9.052492399 | 16.09205592 | 22.25464023 | 26.61428086 | 2.99E-17 | 3.843032932 | YFL040W | | n |
| W303.asm02.HP0_G0016520 | 9634.831398 | 8037.091932 | 9406.530893 | 20388.33727 | 19561.62219 | 19514.45787 | 15409.56343 | 15421.49809 | 13960.21394 | 8.34E-24 | -1.134904413 | YFL039C | ACT1 | n |
| W303.asm02.HP0_G0016530 | 7160.211221 | 6540.667851 | 7202.715084 | 2921.374979 | 2729.497819 | 2866.019094 | 3637.751229 | 2900.843974 | 2352.292978 | 8.10E-16 | 1.295213362 | YFL038C | YPT1 | n |
| W303.asm02.HP0_G0016540 | 713.404015 | 976.6737324 | 627.4278269 | 3896.139458 | 2923.229799 | 3748.637103 | 1105.618901 | 2004.852807 | 1564.100814 | 9.30E-21 | -2.18919338 | YFL036W | RPO41 | n |
| W303.asm02.HP0_G0016570 | 367.3642958 | 379.8175626 | 366.4882166 | 511.7027103 | 630.2923987 | 526.8550576 | 623.8038147 | 764.3985122 | 778.9795283 | 0.000321322 | -0.580982838 | YFL034W | MIL1 | n |
| W303.asm02.HP0_G0016580 | 536.0223102 | 600.6638647 | 591.2676562 | 135.2218189 | 175.1549403 | 143.0293799 | 220.5558252 | 257.3797522 | 406.3795962 | 9.68E-16 | 1.934562931 | YFL033C | RIM15 | n |
| W303.asm02.HP0_G0016590 | 2521.146526 | 2716.790285 | 2738.399955 | 10543.4106 | 9601.675363 | 9888.037448 | 9162.059366 | 10020.39366 | 8759.169282 | 1.78E-72 | -1.912836793 | YFL031W | HAC1 | n |
| W303.asm02.HP0_G0016600 | 358.6406054 | 283.6732673 | 350.851386 | 49.61376088 | 39.80794097 | 48.88345896 | 57.742083 | 23.22223328 | 85.98459971 | 1.12E-14 | 2.839022191 | YFL030W | AGX1 | n |

| | | | | | | | | | | | | | | |
|---|---|---|---|---|---|---|---|---|---|---|---|---|---|---|
| W303.asm02.HP0_G0016610 | 759.9303638 | 478.817629 | 716.3623008 | 634.2778842 | 676.7349965 | 681.6526777 | 703.3175027 | 597.972507 | 531.261991 | 0.900095696 | -0.0274686 | YFL029C | CAK1 | n |
| W303.asm02.HP0_G0016620 | 1335.69393 | 1348.875905 | 1291.993126 | 533.1047248 | 425.9449684 | 481.5925956 | 611.4981249 | 431.5465018 | 524.0966077 | 4.09E-19 | 1.462782735 | YFL028C | CAF16 | n |
| W303.asm02.HP0_G0016630 | 688.2022427 | 806.2793873 | 582.471939 | 708.2121161 | 594.4652519 | 735.0623828 | 480.8684945 | 590.2317626 | 471.8916722 | 0.894667223 | 0.026047663 | YFL027C | GYP8 | n |
| W303.asm02.HP0_G0016640 | 7884.277524 | 8518.765333 | 7701.139059 | 135.2218189 | 188.4242539 | 143.9346291 | 327.5206675 | 209.9676926 | 165.8274423 | 1.83E-142 | 5.69409337 | YFL026W | STE2 | n |
| W303.asm02.HP0_G0016650 | 571.8863707 | 615.8946441 | 589.3130523 | 550.6154639 | 599.7729773 | 544.9600424 | 339.8263573 | 670.541986 | 642.8372454 | 0.788255854 | 0.068497415 | YFL025C | BST1 | n |
| W303.asm02.HP0_G0016670 | 410.9827478 | 392.1925709 | 382.1250472 | 1295.794696 | 1220.776857 | 1276.401428 | 1000.547241 | 1149.500547 | 1284.650865 | 9.76E-34 | -1.678447716 | YFL024C | EPL1 | y |
| W303.asm02.HP0_G0016680 | 502.0968475 | 484.5291713 | 486.6963517 | 1716.052435 | 1434.412806 | 1652.079863 | 1019.479072 | 1344.954344 | 1203.784396 | 1.20E-30 | -1.705527029 | YFL023W | BUD27 | n |
| W303.asm02.HP0_G0016690 | 2448.449106 | 1983.809024 | 1971.217954 | 7212.478886 | 7234.429806 | 7152.374245 | 5438.168309 | 5882.965765 | 5563.408326 | 5.79E-48 | -1.754055238 | YFL022C | FRS2 | n |
| W303.asm02.HP0_G0016700 | 611.627627 | 707.2793208 | 496.4693708 | 496.1376088 | 537.4072031 | 479.7820972 | 753.4868535 | 833.097619 | 1224.25692 | 0.240775271 | 0.263444378 | YFL021W | GAT1 | n |
| W303.asm02.HP0_G0016720 | 4941.485963 | 3285.088743 | 4857.190498 | 13636.97451 | 12643.00205 | 13061.84128 | 11131.91633 | 11506.61659 | 11701.07094 | 3.19E-23 | -1.588343295 | YFL018C | LPD1 | n |
| W303.asm02.HP0_G0016740 | 958.6366452 | 807.231311 | 942.1190422 | 986.4383046 | 846.5822114 | 857.2710302 | 1356.465655 | 932.7597035 | 950.9487277 | 0.962582754 | 0.009008747 | YFL017C | GNA1 | n |
| W303.asm02.HP0_G0016750 | 3357.651505 | 2178.001462 | 2235.08947 | 1646.982298 | 1394.604865 | 1560.64969 | 1739.835222 | 1613.945213 | 1670.557937 | 3.27E-05 | 0.755242887 | YFL016C | MDJ1 | n |
| W303.asm02.HP0_G0016770 | 2599.659739 | 2236.068808 | 2884.995241 | 183.8627609 | 268.0401359 | 124.9243951 | 790.403923 | 256.4121592 | 752.3652474 | 9.80E-15 | 3.7449971 | YFL014W | HSP12 | n |
| W303.asm02.HP0_G0016780 | 839.4128763 | 739.6447272 | 763.2727925 | 1275.3655 | 1314.988983 | 1251.959699 | 1371.611119 | 1637.167446 | 1730.951882 | 1.51E-07 | -0.713737261 | YFL013C | IES1 | n |
| W303.asm02.HP0_G0016810 | 2444.57191 | 1834.357 | 2506.779402 | 1044.807435 | 752.3700844 | 1017.500146 | 1836.387558 | 1927.445362 | 1270.320098 | 1.13E-09 | 1.268220853 | YFL010C | WWM1 | n |
| W303.asm02.HP0_G0016820 | 1003.224396 | 1046.164164 | 990.9841377 | 736.4238625 | 831.9859663 | 770.3671032 | 669.2402079 | 1044.032905 | 1317.406903 | 0.092052195 | 0.378965597 | YFL009W | CDC4 | n |
| W303.asm02.HP0_G0016830 | 600.9653387 | 838.6447936 | 642.0873555 | 1324.979261 | 1426.451218 | 1352.442364 | 1395.275907 | 1506.542384 | 1251.894827 | 8.56E-10 | -0.978738639 | YFL008W | SMC1 | n |
| W303.asm02.HP0_G0016840 | 932.465574 | 1191.808492 | 958.7331747 | 1884.350095 | 1703.779874 | 1908.265398 | 1959.444456 | 1763.922136 | 1710.479358 | 3.85E-09 | -0.834565584 | YFL007W | BLM10 | n |
| W303.asm02.HP0_G0016850 | 3378.976082 | 3879.089142 | 3521.218785 | 3398.056211 | 3694.176922 | 3216.350549 | 4989.483926 | 3746.520303 | 3801.747658 | 0.68811551 | 0.064715968 | YFL005W | SEC4 | n |
| W303.asm02.HP0_G0016860 | 2989.317911 | 2792.944182 | 2997.384961 | 3312.448153 | 2377.861007 | 3275.19175 | 2745.115421 | 3786.191618 | 3318.596098 | 0.869317284 | -0.030856679 | YFL004W | VTC2 | n |
| W303.asm02.HP0_G0016870 | 189.982591 | 164.6828028 | 194.4830803 | 407.6110943 | 467.0798408 | 394.6886686 | 503.5866911 | 511.8567253 | 505.6713364 | 3.61E-14 | -1.206306789 | YFL003C | MSH4 | n |
| W303.asm02.HP0_G0016880 | 1049.750745 | 1127.07768 | 1000.757157 | 1950.501776 | 1644.067962 | 1955.338358 | 972.1494957 | 1241.421888 | 979.6102609 | 7.20E-08 | -0.805225294 | YFL002C | SPB4 | n |
| W303.asm02.HP0_G0016890 | 563.1626803 | 468.3464682 | 602.9952791 | 1825.980964 | 1772.780305 | 1796.919741 | 1054.502958 | 1313.991367 | 1106.539908 | 9.53E-32 | -1.723228897 | YFL001W | DEG1 | n |
| W303.asm02.HP0_G0016900 | 1582.865158 | 2123.74181 | 1811.917743 | 3244.350834 | 2757.363378 | 3053.405686 | 1598.793085 | 1429.13494 | 1138.27232 | 2.69E-05 | -0.714668459 | YFR001W | LOC1 | n |
| W303.asm02.HP0_G0016910 | 2716.94491 | 2475.953585 | 2931.905733 | 3388.328023 | 3592.003207 | 3424.557875 | 2210.29121 | 3016.955141 | 2793.475864 | 0.015977327 | -0.35675373 | YFR002W | NIC96 | n |
| W303.asm02.HP0_G0016920 | 910.1716985 | 829.1255565 | 936.2552307 | 168.2976595 | 200.3666362 | 187.3865927 | 246.1137964 | 232.2223328 | 188.3472184 | 3.33E-42 | 2.269687714 | YFR003C | YPI1 | n |
| W303.asm02.HP0_G0016930 | 3864.594848 | 3778.185228 | 3776.294584 | 2283.205819 | 2091.243832 | 2343.690282 | 3112.392933 | 2673.459607 | 2728.987414 | 1.20E-11 | 0.764673463 | YFR004W | RPN11 | n |
| W303.asm02.HP0_G0016940 | 1507.259841 | 1327.933584 | 1404.382846 | 429.0131088 | 465.7529094 | 486.1188418 | 457.2037064 | 467.3474448 | 363.3872964 | 6.73E-28 | 1.618315462 | YFR005C | SAD1 | n |
| W303.asm02.HP0_G0016950 | 3378.976082 | 2404.559306 | 3294.484742 | 2631.474964 | 2652.5358 | 2505.729896 | 1382.970217 | 2522.51509 | 1417.722269 | 0.36652338 | 0.220834494 | YFR006W | | n |
| W303.asm02.HP0_G0016960 | 794.8251254 | 463.5868496 | 660.6560918 | 169.2704783 | 165.8664207 | 184.6708449 | 298.1763302 | 243.8334495 | 217.0087516 | 2.36E-16 | 1.882864034 | YFR007W | YFH7 | n |
| W303.asm02.HP0_G0016970 | 878.1848337 | 491.1926374 | 787.7053403 | 76.85268843 | 90.23133287 | 72.41993919 | 35.97047793 | 109.338015 | 75.74833784 | 5.52E-21 | 3.174549486 | YFR008W | FAR7 | n |
| W303.asm02.HP0_G0016980 | 2130.519056 | 1782.001196 | 2157.882619 | 3434.050508 | 3435.425306 | 3410.073887 | 2354.173122 | 3132.098714 | 2874.342333 | 1.35E-07 | -0.759943669 | YFR009W | GCN20 | n |
| W303.asm02.HP0_G0016990 | 4451.990001 | 3013.790484 | 4168.19265 | 2008.870906 | 1588.336845 | 1901.928653 | 2704.411986 | 2210.950127 | 2258.119368 | 1.32E-09 | 1.08045026 | YFR010W | UBP6 | n |
| W303.asm02.HP0_G0017000 | 1463.641389 | 1860.058941 | 1662.390551 | 1979.686341 | 2316.822165 | 2021.421553 | 2356.066305 | 2201.274197 | 1882.448558 | 0.024859515 | -0.340928993 | YFR011C | MIC19 | n |
| W303.asm02.HP0_G0017020 | 739.5750862 | 776.7697521 | 746.65866 | 1650.873573 | 1394.604865 | 1493.661246 | 1044.090452 | 1458.162732 | 1384.966231 | 2.00E-10 | -1.004659225 | YFR013W | IOC3 | n |
| W303.asm02.HP0_G0017050 | 509.851239 | 402.6637318 | 499.4012765 | 1226.724558 | 1318.969778 | 1226.61272 | 1846.800064 | 1571.371119 | 1560.006309 | 1.79E-22 | -1.417481708 | YFR016C | AIP5 | n |
| W303.asm02.HP0_G0017070 | 921.8032857 | 750.115888 | 829.7293224 | 516.5668045 | 512.1955072 | 507.8448236 | 606.7651672 | 537.0141447 | 526.1438601 | 7.98E-07 | 0.703096072 | YFR018C | | n |
| W303.asm02.HP0_G0017080 | 478.8336731 | 576.8657718 | 428.058237 | 771.4453408 | 813.4089272 | 773.0828509 | 706.1572773 | 903.7319119 | 784.0976592 | 5.40E-05 | -0.667617199 | YFR019W | FAB1 | n |
| W303.asm02.HP0_G0017100 | 279.1580928 | 249.4040135 | 199.3695898 | 699.4567466 | 639.5809183 | 700.6629117 | 709.9436434 | 775.0420358 | 784.0976592 | 3.31E-20 | -1.487238339 | YFR021W | ATG18 | n |
| W303.asm02.HP0_G0017120 | 135.7018507 | 138.9808625 | 105.5486064 | 50.58657972 | 43.78873507 | 59.74644984 | 118.3239406 | 86.11578176 | 128.9768996 | 7.52E-06 | 1.29402633 | YFR023W | PES4 | n |
| W303.asm02.HP0_G0017140 | 1474.303678 | 1115.654595 | 1541.205114 | 285.0359204 | 212.3090185 | 270.6695227 | 274.5115421 | 185.7778663 | 295.827968 | 3.27E-29 | 2.423842872 | YFR025C | HIS2 | n |
| W303.asm02.HP0_G0017160 | 495.311755 | 302.7117416 | 411.4441045 | 66.15168118 | 62.36577419 | 46.16771124 | 120.2171236 | 79.34263038 | 118.7406377 | 1.57E-23 | 2.796608354 | YFR027W | ECO1 | n |
| W303.asm02.HP0_G0017170 | 1294.983375 | 1103.279587 | 1231.400408 | 1667.411493 | 1939.973657 | 1705.489568 | 1476.682778 | 1713.607298 | 1513.943131 | 3.81E-05 | -0.548892009 | YFR028C | CDC14 | n |
| W303.asm02.HP0_G0017180 | 169.6273133 | 183.7212771 | 115.3216255 | 381.3449856 | 439.2142821 | 368.4364406 | 375.7968352 | 518.6298766 | 415.5922319 | 1.40E-09 | -1.34103178 | YFR029W | PTR3 | n |
| W303.asm02.HP0_G0017190 | 755.0836891 | 735.8370323 | 782.8188307 | 1058.426809 | 927.5250247 | 1052.804866 | 720.3561502 | 804.0698274 | 813.7828186 | 0.001083181 | -0.419848334 | YFR030W | MET10 | n |
| W303.asm02.HP0_G0017200 | 541.8381038 | 755.8274303 | 592.2449581 | 1297.740334 | 1397.258728 | 1277.306678 | 1045.983635 | 1174.657967 | 1128.036058 | 3.89E-12 | -1.070986034 | YFR031C | SMC2 | n |
| W303.asm02.HP0_G0017220 | 60.09653387 | 17.13462688 | 43.00128408 | 59.34194929 | 67.67349965 | 57.93595136 | 53.9557169 | 89.01856092 | 53.22856172 | 0.182538469 | -0.619991085 | YFR032C | RRT5 | n |
| W303.asm02.HP0_G0017250 | 4768.950752 | 5797.215429 | 5603.849918 | 6520.80469 | 6881.466063 | 6267.945737 | 4465.072221 | 5458.192414 | 4995.295792 | 0.033907867 | -0.282554436 | YFR033C | QCR6 | n |
| W303.asm02.HP0_G0017260 | 624.2285131 | 591.1446275 | 546.3117682 | 1970.930972 | 1756.857128 | 1983.401085 | 1924.420569 | 2519.612311 | 2690.089619 | 2.46E-27 | -1.697260735 | YFR034C | PHO4 | n |
| W303.asm02.HP0_G0017280 | 275.2808971 | 179.9135823 | 242.3708739 | 106.0372537 | 114.1160975 | 84.18817931 | 130.6296304 | 116.1111664 | 102.3626187 | 1.71E-06 | 1.202062044 | YFR036W | CDC26 | n |
| W303.asm02.HP0_G0017290 | 2069.453223 | 1418.366337 | 1785.530591 | 2069.185674 | 2166.87892 | 2127.335714 | 1834.494375 | 1927.445362 | 1930.558989 | 0.081741273 | -0.271031329 | YFR037C | RSC8 | n |
| W303.asm02.HP0_G0017300 | 500.1582496 | 562.586916 | 444.6723695 | 323.948674 | 334.3867042 | 323.1739787 | 327.5206675 | 426.7085366 | 441.1828866 | 0.000529276 | 0.619705951 | YFR038W | IRC5 | n |
| W303.asm02.HP0_G0017320 | 805.4874137 | 1253.683534 | 830.7006243 | 1242.28966 | 1492.797786 | 1295.411662 | 1130.23028 | 1558.792409 | 1199.689891 | 0.022789046 | -0.479328015 | YFR040W | SAP155 | n |
| W303.asm02.HP0_G0017330 | 1237.794738 | 1288.904711 | 1299.811542 | 854.1349423 | 801.4665449 | 850.0290363 | 1452.071399 | 1234.648736 | 1192.524508 | 1.74E-06 | 0.610083503 | YFR041C | ERJ5 | n |
| W303.asm02.HP0_G0017340 | 820.9961966 | 697.7600837 | 764.2500944 | 258.7698117 | 278.6555868 | 328.6054741 | 559.435591 | 437.3520602 | 402.2850915 | 1.58E-14 | 1.396905606 | YFR042W | KEG1 | n |
| W303.asm02.HP0_G0017350 | 772.5312499 | 707.2793208 | 803.3421708 | 163.4335653 | 217.616744 | 186.4813434 | 310.4820201 | 244.8010425 | 300.946099 | 1.74E-30 | 2.013840328 | YFR043C | IRC6 | n |
| W303.asm02.HP0_G0017360 | 3267.506705 | 1924.789753 | 2875.222222 | 3889.329726 | 4499.624261 | 3865.414254 | 4406.383547 | 4717.016135 | 3417.887838 | 0.003485492 | -0.60296917 | YFR044C | DUG1 | n |
| W303.asm02.HP0_G0017370 | 749.2680755 | 560.6830686 | 701.7027721 | 1289.957783 | 1353.469993 | 1362.400106 | 1287.364473 | 1330.440448 | 1202.76077 | 3.15E-12 | -0.993790065 | YFR045W | MRX20 | n |
| W303.asm02.HP0_G0017380 | 654.2767801 | 499.7599508 | 681.179432 | 111.8741667 | 118.0968916 | 117.6824012 | 215.8228676 | 176.1019357 | 171.9691994 | 1.88E-33 | 2.400796154 | YFR046C | CNN1 | n |
| W303.asm02.HP0_G0017400 | 482.7108688 | 435.9810618 | 322.5096306 | 526.2949929 | 520.1570954 | 506.9395744 | 435.4321013 | 634.7410431 | 645.908124 | 0.136765693 | -0.323486492 | YFR048W | RMD8 | n |
| W303.asm02.HP0_G0017410 | 1384.158877 | 1567.81836 | 1295.902334 | 2039.02829 | 2279.668086 | 2016.895307 | 2847.347306 | 2442.204867 | 2328.749575 | 3.22E-05 | -0.576124501 | YFR049W | KGD4 | n |
| W303.asm02.HP0_G0017420 | 2997.072302 | 3169.905974 | 3077.523718 | 2346.439044 | 1963.858421 | 2249.544361 | 2580.408496 | 2329.964073 | 2291.899033 | 3.43E-05 | 0.494091317 | YFR050C | PRE4 | n |
| W303.asm02.HP0_G0017430 | 1283.351788 | 980.4814273 | 940.1644384 | 3908.786102 | 3814.927677 | 4011.159382 | 3471.151121 | 3847.14998 | 3520.250457 | 2.05E-39 | -1.872963931 | YFR051C | RET2 | n |

| | | | | | | | | | | | | | | |
|---|---|---|---|---|---|---|---|---|---|---|---|---|---|---|
| W303.asm02.HP0_G0017440 | 2640.370295 | 2229.405342 | 2627.964839 | 1840.573247 | 1592.317639 | 1767.046516 | 2400.556106 | 1970.019457 | 1898.826577 | 0.000282128 | 0.527351307 | YFR052W | RPN12 | n |
| W303.asm02.HP0_G0017570 | 806.4567126 | 871.9621237 | 778.9096231 | 160.5151087 | 173.8280089 | 176.5236018 | 109.8046168 | 149.9769233 | 163.7801899 | 4.64E-36 | 2.266611577 | YGL256W | ADH4 | n |
| W303.asm02.HP0_G0017600 | 12884.89072 | 11684.86361 | 13072.39036 | 6617.113755 | 6561.675604 | 6621.89819 | 6434.929184 | 8488.693858 | 8331.293536 | 4.67E-12 | 0.926780274 | YGL253W | HXK2 | n |
| W303.asm02.HP0_G0017610 | 1801.926717 | 1707.751146 | 1909.647934 | 1247.153754 | 1403.893385 | 1281.832924 | 1410.421372 | 1492.996081 | 1792.369453 | 0.000783225 | 0.463247795 | YGL252C | RTG2 | n |
| W303.asm02.HP0_G0017620 | 288.8510822 | 369.3464017 | 316.6458192 | 155.6510145 | 183.1165285 | 180.1445987 | 262.2058523 | 247.7038217 | 317.324118 | 3.17E-06 | 0.911555543 | YGL251C | HFM1 | n |
| W303.asm02.HP0_G0017630 | 490.4652603 | 336.0290717 | 451.5134829 | 107.9828913 | 107.4814406 | 107.7246596 | 185.5319388 | 154.8148885 | 227.2450135 | 2.86E-18 | 1.983213503 | YGL250W | RMR1 | n |
| W303.asm02.HP0_G0017640 | 297.5747726 | 209.4232175 | 309.8047058 | 19.45637682 | 10.61545093 | 19.91548328 | 13.25228134 | 23.22223328 | 23.5434023 | 2.74E-29 | 3.998725868 | YGL249W | ZIP2 | n |
| W303.asm02.HP0_G0017650 | 1357.018507 | 1179.433484 | 1354.540449 | 142.0315508 | 167.1933521 | 170.1868571 | 255.5797116 | 221.5788092 | 273.3081919 | 4.67E-78 | 3.022288782 | YGL248W | PDE1 | n |
| W303.asm02.HP0_G0017660 | 436.18452 | 418.8464349 | 393.8526701 | 112.8469855 | 130.0392738 | 98.67216715 | 232.861515 | 111.2732011 | 160.7093114 | 1.43E-12 | 1.875210427 | YGL247W | BRR6 | n |
| W303.asm02.HP0_G0017670 | 1404.514155 | 1613.510698 | 1394.609827 | 1339.571544 | 1374.700895 | 1351.537115 | 1304.403121 | 1168.852409 | 1131.106937 | 0.36362425 | 0.118340673 | YGL246C | RAI1 | n |
| W303.asm02.HP0_G0017680 | 4008.05109 | 3095.655924 | 3995.210212 | 12669.99258 | 13086.19713 | 12649.95288 | 8570.439663 | 10181.9817 | 8764.287413 | 8.91E-39 | -1.790927151 | YGL245W | GUS1 | n |
| W303.asm02.HP0_G0017690 | 3384.791876 | 2536.876703 | 3575.947692 | 2550.731001 | 2940.479907 | 2713.031972 | 2283.178757 | 2806.987448 | 2464.891858 | 0.208304381 | 0.211424385 | YGL244W | RTF1 | n |
| W303.asm02.HP0_G0017700 | 726.9742001 | 756.7793541 | 701.7027721 | 680.0003697 | 682.042722 | 660.8319451 | 555.6492249 | 628.9354847 | 551.7345148 | 0.404076612 | 0.111931609 | YGL243W | TAD1 | n |
| W303.asm02.HP0_G0017710 | 1200.961379 | 1390.760549 | 1374.086487 | 768.5268843 | 755.0239471 | 792.0930849 | 1132.123463 | 988.8801006 | 800.4756782 | 8.23E-07 | 0.775873861 | YGL242C | ANK1 | n |
| W303.asm02.HP0_G0017720 | 1220.347357 | 1357.443219 | 1332.062505 | 1398.913493 | 1297.738876 | 1370.547349 | 529.1446622 | 1268.514493 | 1075.831123 | 0.837268246 | -0.057042852 | YGL241W | KAP114 | n |
| W303.asm02.HP0_G0017730 | 59.12723494 | 101.8558376 | 86.97987008 | 420.2577392 | 425.9449684 | 390.1624224 | 439.2184674 | 405.4214894 | 311.1823608 | 8.43E-24 | -2.316845204 | YGL240W | DOC1 | n |
| W303.asm02.HP0_G0017740 | 2095.624294 | 2189.424546 | 2405.140003 | 4301.804914 | 4526.162889 | 4366.922333 | 2794.33818 | 4037.765812 | 3395.368062 | 8.12E-11 | -0.979797805 | YGL238W | CSE1 | n |
| W303.asm02.HP0_G0017750 | 534.0837123 | 356.9713934 | 549.243674 | 280.1718262 | 327.7520473 | 278.8167659 | 548.0764927 | 390.9075936 | 466.7735413 | 0.001366642 | 0.701674169 | YGL237C | HAP2 | n |
| W303.asm02.HP0_G0017760 | 523.4214241 | 578.7696192 | 468.1276154 | 1645.03666 | 1325.604434 | 1662.942854 | 744.0209383 | 1024.681044 | 787.1685378 | 4.85E-19 | -1.561785213 | YGL236C | MTO1 | n |
| W303.asm02.HP0_G0017770 | 1534.400212 | 1282.241245 | 1330.107901 | 6620.032212 | 3327.943865 | 6046.159673 | 2704.411986 | 4763.460602 | 3264.34391 | 5.68E-14 | -1.947773738 | YGL234W | ADE57 | n |
| W303.asm02.HP0_G0017780 | 897.5708124 | 1085.193036 | 969.4834957 | 1129.442674 | 1219.449925 | 1175.918763 | 855.7187382 | 1036.29216 | 897.720166 | 0.071559349 | -0.255243071 | YGL233W | SEC15 | n |
| W303.asm02.HP0_G0017790 | 1504.351945 | 1518.318327 | 1527.522887 | 1193.648718 | 1220.776857 | 1222.086474 | 1062.075691 | 992.7504728 | 968.3503729 | 0.002432827 | 0.323468563 | YGL232W | TAN1 | n |
| W303.asm02.HP0_G0017800 | 2714.037014 | 2431.21317 | 2772.605521 | 410.5295508 | 412.6756548 | 417.3198996 | 602.0322096 | 379.2964769 | 412.5213534 | 4.35E-50 | 2.674078307 | YGL231C | EMC4 | n |
| W303.asm02.HP0_G0017830 | 2737.300188 | 1683.00113 | 2460.846212 | 1596.395718 | 1531.278796 | 1609.533149 | 1394.329316 | 1767.792509 | 1517.014009 | 0.00454169 | 0.53841503 | YGL228W | SHE10 | n |
| W303.asm02.HP0_G0017840 | 673.6627587 | 491.1926374 | 602.0179772 | 136.1946377 | 172.5010776 | 131.2611398 | 308.588837 | 313.5001493 | 424.8048676 | 4.96E-21 | 2.012141426 | YGL227W | VID30 | n |
| W303.asm02.HP0_G0017850 | 1185.452596 | 1204.1835 | 1217.718181 | 201.3735001 | 213.6359499 | 204.5863282 | 284.9240489 | 216.740844 | 158.662059 | 2.94E-37 | 2.542513903 | YGL226C-/ | OST5 | n |
| W303.asm02.HP0_G0017860 | 506.9433422 | 539.7407469 | 575.6308256 | 641.0876161 | 693.9851043 | 706.0944071 | 820.6948518 | 553.4632266 | 552.758141 | 0.082922302 | -0.331227772 | YGL226W | MTC3 | n |
| W303.asm02.HP0_G0017890 | 1688.518742 | 1435.500963 | 1762.075346 | 178.9986667 | 216.2898126 | 162.9448632 | 418.3934539 | 421.8705713 | 333.702137 | 1.43E-72 | 3.136697744 | YGL224C | SDT1 | n |
| W303.asm02.HP0_G0017900 | 1900.795208 | 1227.981593 | 1894.988405 | 511.7027103 | 502.9069876 | 514.1815683 | 651.2549689 | 572.8150876 | 579.3724218 | 1.44E-20 | 1.716105655 | YGL223C | COG1 | n |
| W303.asm02.HP0_G0017920 | 2852.646761 | 1674.433816 | 2659.2385 | 1200.45845 | 1166.372671 | 1182.255507 | 1266.53946 | 1051.773649 | 999.0591585 | 3.68E-07 | 1.017687987 | YGL221C | NIF3 | n |
| W303.asm02.HP0_G0017930 | 2021.957575 | 1996.184032 | 2231.180263 | 542.8329132 | 567.9266245 | 519.6130637 | 694.798179 | 567.0095293 | 443.230139 | 6.78E-30 | 1.939269131 | YGL220W | BOL2 | n |
| W303.asm02.HP0_G0017940 | 1472.36508 | 975.7218087 | 1423.928884 | 1067.182268 | 1265.892523 | 1067.288854 | 820.6948518 | 1151.435734 | 1000.082785 | 0.36652338 | 0.188042788 | YGL219C | MDM34 | n |
| W303.asm02.HP0_G0017950 | 240.3861355 | 247.5001661 | 181.7781554 | 343.4050508 | 410.021792 | 310.5004893 | 425.966186 | 492.5048642 | 498.5059531 | 0.000705815 | -0.664212025 | YGL216W | KIP3 | n |
| W303.asm02.HP0_G0017960 | 1301.768468 | 1765.818493 | 1378.972996 | 8283.55243 | 8392.840888 | 8242.29433 | 3935.927559 | 6250.651125 | 5391.439127 | 7.91E-42 | -2.486402964 | YGL215W | CLG1 | n |
| W303.asm02.HP0_G0017970 | 1865.900447 | 1554.491428 | 1844.168706 | 768.5268843 | 778.9087117 | 796.6193311 | 612.4447164 | 830.1948398 | 855.7514923 | 4.27E-13 | 1.16718772 | YGL213C | SKI8 | n |
| W303.asm02.HP0_G0017980 | 167.6887155 | 158.9712605 | 139.7541733 | 528.2406306 | 545.3687913 | 562.159778 | 769.5789095 | 749.8846164 | 702.2075643 | 6.78E-34 | -1.810213963 | YGL212W | VAM7 | n |
| W303.asm02.HP0_G0017990 | 1167.035916 | 1581.145292 | 1223.581993 | 1300.65879 | 1166.372671 | 1325.284887 | 826.3744009 | 818.5837232 | 699.1366857 | 0.703009523 | 0.06611846 | YGL211W | NCS6 | n |
| W303.asm02.HP0_G0018010 | 168.6580144 | 363.6348594 | 119.2308331 | 838.5698408 | 1179.641984 | 869.9445196 | 935.2324263 | 1195.945014 | 799.452052 | 8.60E-11 | -2.147235761 | YGL209W | MIG2 | n |
| W303.asm02.HP0_G0018030 | 1700.150329 | 1737.260781 | 1667.27706 | 3759.94482 | 3870.658794 | 3788.468069 | 3222.19755 | 3488.172958 | 3053.476916 | 4.75E-30 | -1.161419309 | YGL207W | SPT16 | n |
| W303.asm02.HP0_G0018040 | 1458.794895 | 1980.001329 | 1670.208966 | 8982.036357 | 7073.871111 | 8794.496366 | 4663.856441 | 6767.345816 | 6274.828526 | 2.42E-36 | -2.282260764 | YGL206C | CHC1 | n |
| W303.asm02.HP0_G0018050 | 50.40354454 | 69.49043125 | 54.72890702 | 12.64664493 | 7.961588195 | 10.86299088 | 13.25228134 | 14.5138958 | 21.49614993 | 3.18E-07 | 2.45404956 | YGL205W | POX1 | n |
| W303.asm02.HP0_G0018070 | 844.259371 | 764.3947438 | 861.9802855 | 566.1805654 | 727.1583884 | 564.8755257 | 1105.618901 | 706.342929 | 808.6646877 | 0.039965262 | 0.412887159 | YGL203C | KEX1 | n |
| W303.asm02.HP0_G0018080 | 3070.739021 | 2257.01113 | 2799.969975 | 3979.801878 | 4523.509026 | 4161.430756 | 5577.317263 | 4571.877177 | 5306.478153 | 2.25E-05 | -0.639737381 | YGL202W | ARO8 | n |
| W303.asm02.HP0_G0018090 | 446.8468083 | 584.4811615 | 460.3092001 | 1032.16079 | 1025.717946 | 1018.405395 | 1001.493833 | 1000.491217 | 1147.484956 | 2.60E-12 | -1.044051382 | YGL201C | MCM6 | n |
| W303.asm02.HP0_G0018100 | 4923.069283 | 3899.07954 | 5054.605484 | 1859.056805 | 1902.819579 | 1829.508714 | 3703.066044 | 2515.741939 | 2258.119368 | 9.31E-12 | 1.311465061 | YGL200C | EMP24 | n |
| W303.asm02.HP0_G0018110 | 1843.606571 | 1350.779753 | 1855.896329 | 455.2792175 | 453.8105271 | 458.0561154 | 925.766511 | 714.0836734 | 923.3108207 | 2.30E-27 | 1.885050081 | YGL198W | YIP4 | n |
| W303.asm02.HP0_G0018120 | 397.4125627 | 461.6830022 | 339.1237631 | 379.3993479 | 407.3679293 | 385.6361762 | 473.2957623 | 451.865956 | 549.6872624 | 0.862156239 | 0.032598774 | YGL197W | MDS3 | n |
| W303.asm02.HP0_G0018130 | 1432.623824 | 1284.145093 | 1424.906186 | 691.6741958 | 522.8109581 | 729.6308874 | 828.267584 | 748.9170234 | 869.0586327 | 6.34E-12 | 1.08790265 | YGL196W | DSD1 | n |
| W303.asm02.HP0_G0018140 | 1662.347671 | 1938.116685 | 1724.937873 | 6836.970813 | 7233.102875 | 7188.584214 | 4531.333628 | 5220.164523 | 4619.624982 | 3.43E-67 | -1.99698925 | YGL195W | GCN1 | n |
| W303.asm02.HP0_G0018150 | 213.2457654 | 262.7309456 | 239.4389682 | 48.64094204 | 62.36577419 | 48.88345896 | 55.84889995 | 30.96297771 | 26.61428086 | 5.48E-13 | 2.169957316 | YGL194C-A | | n |
| W303.asm02.HP0_G0018160 | 368.3335947 | 450.2599176 | 393.8526701 | 654.7070799 | 739.1007707 | 701.5681609 | 800.8164298 | 812.7781649 | 868.0350066 | 2.28E-08 | -0.788064118 | YGL194C | HOS2 | n |
| W303.asm02.HP0_G0018180 | 384.8116766 | 340.7886903 | 308.8274039 | 55.45067393 | 71.65429375 | 46.16771124 | 45.43639318 | 106.4352359 | 54.25218791 | 9.77E-16 | 2.587884423 | YGL192W | IME4 | n |
| W303.asm02.HP0_G0018190 | 3337.296228 | 3840.06027 | 3321.849195 | 8290.362162 | 8622.400015 | 7880.194634 | 7727.026615 | 6287.419661 | 5584.904476 | 1.47E-17 | -1.239556818 | YGL191W | COX13 | n |
| W303.asm02.HP0_G0018200 | 1629.391507 | 1566.866436 | 1543.159717 | 1360.973558 | 1476.87461 | 1491.850747 | 1682.093139 | 1780.371218 | 1764.731546 | 0.239357828 | 0.130693683 | YGL190C | CDC55 | n |
| W303.asm02.HP0_G0018220 | 3946.985257 | 4471.185693 | 3823.205076 | 7396.341647 | 7463.988932 | 7101.680287 | 5962.580013 | 6009.720455 | 5248.131461 | 2.12E-13 | -0.843151781 | YGL187C | COX4 | n |
| W303.asm02.HP0_G0018230 | 1260.088613 | 1148.020001 | 1204.035954 | 920.2866234 | 878.4285641 | 968.6166867 | 1057.342733 | 1495.898861 | 1142.366825 | 0.017289503 | 0.383733981 | YGL186C | TPN1 | n |
| W303.asm02.HP0_G0018240 | 789.9786307 | 600.6638647 | 700.7254702 | 421.2305581 | 439.2142821 | 433.6143859 | 482.7616775 | 397.680745 | 458.5845318 | 1.75E-05 | 0.692809186 | YGL185C | | n |
| W303.asm02.HP0_G0018250 | 221.9694558 | 222.7501495 | 207.1880051 | 23.34765218 | 41.13487234 | 28.96797568 | 120.2171236 | 41.6065013 | 234.4103968 | 5.54E-09 | 2.812660466 | YGL184C | STR3 | n |
| W303.asm02.HP0_G0018260 | 44.58775094 | 39.98079606 | 29.31905733 | 6.809731886 | 6.634656829 | 9.052492399 | 13.25228134 | 13.54630275 | 8.189009496 | 4.27E-05 | 2.327846141 | YGL183C | MND1 | n |
| W303.asm02.HP0_G0018270 | 1115.663072 | 773.9139809 | 988.052232 | 688.7557393 | 733.7930453 | 678.9369299 | 1003.387016 | 909.5374702 | 1236.540434 | 0.013495677 | 0.453960852 | YGL181W | GTS1 | n |
| W303.asm02.HP0_G0018280 | 184.1667974 | 180.865506 | 148.5498905 | 239.3134348 | 266.7132045 | 228.1228085 | 408.9275386 | 478.9585614 | 589.6086837 | 0.013804736 | -0.512762771 | YGL180W | ATG1 | n |
| W303.asm02.HP0_G0018310 | 411.9520467 | 714.8947106 | 442.7177657 | 2135.337356 | 1851.069255 | 2170.787677 | 513.0526063 | 1470.741441 | 1639.849152 | 2.16E-05 | -1.971921762 | YGL178W | MPT5 | n |
| W303.asm02.HP0_G0018330 | 71.72812107 | 66.63466011 | 59.61541657 | 139.1130942 | 140.6547248 | 159.3238662 | 195.9444456 | 185.7778663 | 166.8510685 | 3.84E-07 | -1.151048332 | YGL175C | SAE2 | n |

| | | | | | | | | | | | | | | |
|---|---|---|---|---|---|---|---|---|---|---|---|---|---|---|
| W303.asm02.HP0_G0018340 | 318.8993491 | 283.6732673 | 307.850102 | 209.1560508 | 208.3282244 | 248.0382917 | 168.4932914 | 166.4260052 | 96.22086158 | 0.060960913 | 0.450335192 | YGL174W | BUD13 | n |
| W303.asm02.HP0_G0018350 | 1326.000941 | 1427.885574 | 1187.421822 | 6003.265067 | 5825.228696 | 6149.358087 | 4635.458696 | 5349.821992 | 4956.397997 | 1.32E-77 | -2.18948951 | YGL173C | XRN1 | n |
| W303.asm02.HP0_G0018360 | 834.5663817 | 974.769885 | 745.6813581 | 2940.831356 | 2836.97926 | 2984.606744 | 2174.320732 | 2717.001294 | 2562.136346 | 5.11E-35 | -1.777997841 | YGL172W | NUP49 | n |
| W303.asm02.HP0_G0018370 | 1056.535837 | 1218.462356 | 1237.264219 | 4404.923711 | 3763.177353 | 4273.681662 | 2041.797918 | 2411.241889 | 2099.457309 | 8.85E-44 | -1.825052391 | YGL171W | ROK1 | n |
| W303.asm02.HP0_G0018390 | 1112.755176 | 1128.029603 | 1226.513898 | 1632.390015 | 1057.564299 | 1509.955732 | 840.5732738 | 820.5189093 | 778.9795283 | 0.118895653 | -0.278107744 | YGL169W | SUA5 | n |
| W303.asm02.HP0_G0018400 | 995.4700046 | 1113.750747 | 1076.986706 | 3519.658566 | 3373.059532 | 3468.915087 | 2893.730291 | 3843.279608 | 3183.477441 | 5.14E-36 | -1.701408481 | YGL167C | PMR1 | n |
| W303.asm02.HP0_G0018430 | 267.5265056 | 412.1829689 | 231.6205529 | 477.6540509 | 447.1758703 | 477.9715987 | 417.4468623 | 505.0835739 | 514.883972 | 0.005970061 | -0.622408172 | YGL163C | RAD54 | n |
| W303.asm02.HP0_G0018440 | 157.9957261 | 141.8366337 | 115.3216255 | 253.9057175 | 326.425116 | 297.8269999 | 407.0343556 | 419.9353852 | 595.7504408 | 2.00E-06 | -1.078756162 | YGL162W | SUT1 | n |
| W303.asm02.HP0_G0018450 | 1705.966123 | 1347.923982 | 1737.642798 | 402.7470001 | 521.4840267 | 437.2353829 | 730.768657 | 796.329083 | 764.6487617 | 3.13E-30 | 1.818374029 | YGL161C | YIP5 | n |
| W303.asm02.HP0_G0018460 | 1015.825282 | 875.7698185 | 1041.803837 | 608.9845944 | 591.8113891 | 592.0330029 | 620.9640401 | 760.52814 | 701.1839381 | 5.57E-07 | 0.710095019 | YGL160W | AIM14 | n |
| W303.asm02.HP0_G0018470 | 829.719887 | 935.7410126 | 902.0496638 | 1059.399718 | 857.1976623 | 1114.361814 | 900.2085398 | 662.8012416 | 778.9795283 | 0.287456955 | -0.18577013 | YGL159W | | n |
| W303.asm02.HP0_G0018510 | 798.7023211 | 803.4236161 | 800.4102651 | 335.6225001 | 374.1946451 | 360.2891975 | 723.1959248 | 508.9539461 | 700.1603119 | 3.99E-12 | 1.167878699 | YGL155W | CDC43 | n |
| W303.asm02.HP0_G0018520 | 544.7460006 | 410.2791215 | 521.8792205 | 131.3305435 | 99.51985243 | 118.5876504 | 260.3126693 | 154.8148885 | 206.7724898 | 2.98E-18 | 2.074486691 | YGL154C | LYS5 | n |
| W303.asm02.HP0_G0018530 | 586.4258547 | 424.5579772 | 599.0860714 | 418.3121016 | 387.4639588 | 380.2046808 | 744.9675298 | 644.4169736 | 815.830071 | 0.017556078 | 0.440635202 | YGL153W | PEX14 | n |
| W303.asm02.HP0_G0018540 | 978.9919228 | 1130.885374 | 1129.761009 | 1336.653087 | 1271.200248 | 1402.231073 | 1045.037043 | 1307.218215 | 1294.887127 | 0.032828801 | -0.308328068 | YGL151W | NUT1 | n |
| W303.asm02.HP0_G0018550 | 827.7812891 | 1204.1835 | 884.4582294 | 1952.447414 | 1914.761961 | 1875.676425 | 1483.308919 | 2205.144569 | 2100.480936 | 4.47E-07 | -0.977468178 | YGL150C | INO80 | n |
| W303.asm02.HP0_G0018560 | 1963.799639 | 1975.24171 | 1767.939157 | 4084.866313 | 3958.236264 | 3935.118446 | 3615.979624 | 2983.089384 | 3034.028018 | 1.26E-18 | -1.06960857 | YGL148W | ARO2 | n |
| W303.asm02.HP0_G0018580 | 181.2589006 | 84.72121071 | 154.4137019 | 100.2003406 | 134.0200679 | 107.7246596 | 272.6183591 | 135.4630275 | 200.6307326 | 0.404116399 | 0.30104623 | YGL146C | RRT6 | n |
| W303.asm02.HP0_G0018590 | 1267.843005 | 1453.587514 | 1333.039807 | 887.2107828 | 806.7742704 | 839.1660454 | 537.6639859 | 852.4494801 | 858.8223709 | 0.000346828 | 0.678247746 | YGL145W | TIP20 | n |
| W303.asm02.HP0_G0018600 | 265.5879078 | 331.2694531 | 283.4175542 | 566.1805654 | 623.6577419 | 599.2749968 | 527.2514792 | 550.5604474 | 486.2224388 | 1.70E-11 | -1.022142987 | YGL144C | ROG1 | n |
| W303.asm02.HP0_G0018610 | 809.3646094 | 885.2890557 | 839.5023415 | 777.2822538 | 728.4853198 | 743.209626 | 815.9618942 | 737.3059067 | 613.152086 | 0.260677485 | 0.171866919 | YGL143C | MRF1 | n |
| W303.asm02.HP0_G0018620 | 1637.145899 | 1635.404944 | 1687.8004 | 737.3966814 | 693.9851043 | 706.9996564 | 855.7187382 | 960.8199021 | 860.8696232 | 1.89E-26 | 1.213305897 | YGL142C | GPI10 | n |
| W303.asm02.HP0_G0018630 | 349.916915 | 476.9137816 | 359.6471032 | 210.1288696 | 171.1741462 | 220.8808145 | 105.0716592 | 209.0000995 | 187.3235922 | 0.000287385 | 0.975750331 | YGL141W | HUL5 | n |
| W303.asm02.HP0_G0018640 | 752.1759723 | 891.9525217 | 697.7935644 | 831.7601089 | 729.8122512 | 830.113553 | 662.6140672 | 806.0050135 | 616.2229646 | 0.861978865 | -0.031277106 | YGL140C | | n |
| W303.asm02.HP0_G0018660 | 11.6315872 | 7.615389726 | 16.61413249 | 57.39631161 | 70.32736239 | 61.55694831 | 87.08642026 | 85.1481887 | 81.89009496 | 7.98E-10 | -2.39852131 | YGL138C | | n |
| W303.asm02.HP0_G0018670 | 1971.554031 | 1758.203103 | 1837.327593 | 2782.261885 | 2843.613917 | 2688.590243 | 3052.757667 | 2680.232758 | 2575.443486 | 8.63E-07 | -0.578517062 | YGL137W | SEC27 | n |
| W303.asm02.HP0_G0018680 | 747.3294777 | 687.2889228 | 715.3849988 | 449.4423045 | 459.1182526 | 451.7193707 | 478.0287199 | 445.0928046 | 423.7812414 | 4.06E-07 | 0.660793372 | YGL136C | MRM2 | n |
| W303.asm02.HP0_G0018690 | 8138.233845 | 8940.467539 | 8089.127917 | 21251.22758 | 22507.40983 | 19328.88177 | 21490.46738 | 15979.79928 | 15851.87513 | 3.38E-19 | -1.325722638 | YPL220W | RPL1A | y |
| W303.asm02.HP0_G0018700 | 520.5135273 | 326.5098345 | 478.8779364 | 255.8513551 | 282.6363809 | 254.3750364 | 309.5354285 | 381.2316631 | 356.2219131 | 0.000417211 | 0.743215292 | YGL134W | PCL10 | n |
| W303.asm02.HP0_G0018710 | 512.7591358 | 712.0389394 | 595.1768638 | 1306.495703 | 1340.200679 | 1282.738173 | 984.4551855 | 1255.935783 | 1117.799796 | 5.58E-12 | -1.110065428 | YGL133W | ITC1 | n |
| W303.asm02.HP0_G0018720 | 273.3422992 | 546.4042129 | 317.6231211 | 808.4124567 | 806.7742704 | 773.0828509 | 726.9822909 | 912.4402494 | 777.9559021 | 7.93E-06 | -1.069872325 | YGL131C | SNT2 | n |
| W303.asm02.HP0_G0018730 | 844.259371 | 694.9043125 | 841.4569454 | 1221.860464 | 1293.758082 | 1228.423219 | 1142.53597 | 1313.991367 | 1178.193741 | 1.27E-06 | -0.652947192 | YGL130W | CEG1 | n |
| W303.asm02.HP0_G0018740 | 1790.29513 | 1735.356934 | 1762.075346 | 3335.795805 | 3146.154268 | 3354.853683 | 2552.01075 | 3172.737622 | 2833.397286 | 4.45E-14 | -0.895789964 | YGL129C | RSM23 | n |
| W303.asm02.HP0_G0018750 | 373.1800894 | 198.9520566 | 336.1918574 | 177.053029 | 205.6743617 | 175.6183525 | 305.7490624 | 190.6158315 | 263.0719301 | 0.011195213 | 0.703838222 | YGL128C | CWC23 | n |
| W303.asm02.HP0_G0018760 | 553.469691 | 331.2694531 | 503.3104842 | 350.2147827 | 322.4443219 | 324.9844771 | 475.1889453 | 328.0140451 | 324.4895013 | 0.047997363 | 0.475962995 | YGL127C | SOH1 | n |
| W303.asm02.HP0_G0018770 | 1304.676364 | 1085.193036 | 1288.083919 | 205.2647754 | 197.7127735 | 180.1445987 | 411.7673132 | 360.9122089 | 341.8911465 | 1.78E-63 | 2.658353048 | YGL126W | SCS3 | n |
| W303.asm02.HP0_G0018780 | 602.9039366 | 430.2695195 | 465.1957096 | 426.0946523 | 445.8489389 | 472.5401032 | 265.9922184 | 309.6297771 | 266.1428086 | 0.408720514 | 0.155986047 | YGL125W | MET13 | n |
| W303.asm02.HP0_G0018790 | 1416.145742 | 958.5871818 | 1263.651371 | 108.9557102 | 122.0776857 | 107.7246596 | 88.97960331 | 209.0000995 | 279.449949 | 1.45E-26 | 3.42670103 | YGL124C | MON1 | n |
| W303.asm02.HP0_G0018800 | 21393.39676 | 23235.50598 | 21279.77181 | 21204.53227 | 23391.14612 | 20302.92995 | 22625.43062 | 15619.85466 | 15358.48731 | 0.901187812 | 0.022321626 | YGL123W | RPS2 | n |
| W303.asm02.HP0_G0018810 | 786.101435 | 931.9333178 | 599.0860714 | 1472.847725 | 1512.701757 | 1547.9762 | 1967.96378 | 2015.49633 | 2764.814331 | 6.24E-07 | -0.968187655 | YGL122C | NAB2 | n |
| W303.asm02.HP0_G0018830 | 1838.760077 | 1782.95312 | 1983.922879 | 7728.072872 | 6605.464339 | 7599.567369 | 3086.834962 | 3762.001792 | 3175.288432 | 9.92E-53 | -1.968399097 | YGL120C | PRP43 | n |
| W303.asm02.HP0_G0018850 | 2193.523486 | 2161.818759 | 1880.328877 | 886.237964 | 628.9654674 | 900.7229937 | 463.829847 | 338.6575687 | 415.5922319 | 6.35E-14 | 1.365818398 | YGL117W | ARO5 | n |
| W303.asm02.HP0_G0018860 | 829.719887 | 731.0774137 | 720.2715084 | 841.4882973 | 854.5437996 | 819.2505621 | 943.75175 | 802.1346413 | 898.7437922 | 0.310967173 | -0.140625245 | YGL116W | CDC20 | n |
| W303.asm02.HP0_G0018870 | 2367.027995 | 2672.04987 | 2373.866342 | 745.1792321 | 777.5817803 | 697.0419147 | 1154.84166 | 1045.968091 | 1221.186041 | 2.72E-44 | 1.740883733 | YGL115W | SNF4 | n |
| W303.asm02.HP0_G0018880 | 754.1145702 | 736.788956 | 618.6321097 | 352.1604204 | 395.425547 | 320.4852309 | 611.4981249 | 600.8752862 | 734.9636022 | 2.47E-09 | 0.985153692 | YGL114W | | n |
| W303.asm02.HP0_G0018890 | 456.5397976 | 581.6253904 | 473.9914268 | 408.5839132 | 441.8681448 | 413.6989026 | 303.8558794 | 430.5789088 | 373.6235582 | 0.171050068 | 0.259473125 | YGL113W | SLD3 | n |
| W303.asm02.HP0_G0018900 | 986.7463142 | 871.0102 | 889.344739 | 1551.646051 | 1588.336845 | 1584.18617 | 1345.106556 | 1392.366404 | 1411.580512 | 1.52E-12 | -0.782091657 | YGL112C | TAF6 | n |
| W303.asm02.HP0_G0018910 | 975.114727 | 1094.712273 | 1068.190989 | 3037.140421 | 2702.959192 | 2986.417243 | 1842.067107 | 1812.301789 | 1782.133192 | 1.73E-40 | -1.476049765 | YGL111W | NSA1 | n |
| W303.asm02.HP0_G0018920 | 571.8863707 | 571.1542295 | 566.8351084 | 592.4466741 | 537.4072031 | 600.1802461 | 509.2662402 | 468.3150379 | 555.8290195 | 0.900548629 | -0.018620089 | YGL110C | CUE3 | n |
| W303.asm02.HP0_G0018930 | 632.9522035 | 692.0485414 | 748.6132638 | 301.5738407 | 281.3094495 | 271.574772 | 312.3752031 | 202.2269482 | 196.5362279 | 3.42E-10 | 1.278990516 | YGL108C | | n |
| W303.asm02.HP0_G0018950 | 3333.419032 | 2851.963453 | 3446.94384 | 2624.665233 | 2815.748358 | 2672.295756 | 3692.653537 | 2673.459607 | 2626.624796 | 0.138614527 | 0.247831866 | YGL106W | MLC1 | n |
| W303.asm02.HP0_G0018960 | 5349.560814 | 4394.079872 | 5470.936098 | 6768.873495 | 7126.948366 | 6879.894223 | 6447.234874 | 4953.108841 | 4911.358445 | 0.003043626 | -0.449413389 | YGL105W | ARC1 | n |
| W303.asm02.HP0_G0018990 | 1789.325831 | 1283.193169 | 1776.734874 | 1341.517182 | 1329.585229 | 1367.831602 | 2181.893464 | 1453.324766 | 1792.369453 | 0.174188172 | 0.263651946 | YGL100W | SEH1 | n |
| W303.asm02.HP0_G0019000 | 522.4521251 | 683.4812279 | 446.6269733 | 5670.561023 | 5169.724601 | 5568.188075 | 2596.500552 | 3465.918317 | 3111.823608 | 1.92E-79 | -3.311531085 | YGL099W | LSG1 | n |
| W303.asm02.HP0_G0019010 | 945.0664601 | 784.3851418 | 864.9121912 | 397.8829059 | 414.0025861 | 364.8154437 | 550.9162673 | 520.5650627 | 424.8048676 | 7.65E-13 | 1.142190824 | YGL098W | USE1 | n |
| W303.asm02.HP0_G0019020 | 1088.522702 | 1251.779686 | 1072.100196 | 2746.267588 | 2757.363378 | 2757.389185 | 1907.381922 | 2131.607497 | 2161.898507 | 2.08E-27 | -1.275430541 | YGL097W | SRM1 | n |
| W303.asm02.HP0_G0019040 | 1286.259685 | 1386.952854 | 1385.81411 | 628.4409712 | 632.9462615 | 646.3479573 | 628.5367723 | 697.6345915 | 667.4042739 | 9.74E-21 | 1.089072522 | YGL095C | VPS45 | n |
| W303.asm02.HP0_G0019050 | 423.5836339 | 666.3466011 | 488.6509555 | 658.5983552 | 700.6197611 | 701.5681609 | 567.9549147 | 796.329083 | 876.2240161 | 0.083202808 | -0.384294088 | YGL094C | PAN2 | n |
| W303.asm02.HP0_G0019060 | 317.9300502 | 426.4618247 | 334.2372536 | 533.1047248 | 618.3500164 | 549.4862886 | 508.3196487 | 585.3937973 | 605.9867027 | 0.000145803 | -0.65543426 | YGL093W | SPC105 | n |
| W303.asm02.HP0_G0019070 | 951.8515526 | 1275.577779 | 891.2993428 | 3418.485407 | 3568.118443 | 3599.270978 | 2039.904735 | 2746.029086 | 2294.969911 | 4.36E-25 | -1.76297814 | YGL092W | NUP145 | n |
| W303.asm02.HP0_G0019080 | 650.3995843 | 559.7311449 | 550.2209759 | 277.2533696 | 303.8672828 | 277.9115167 | 407.9809471 | 393.8103727 | 466.7735413 | 3.21E-11 | 1.037233591 | YGL091C | NBP35 | n |
| W303.asm02.HP0_G0019110 | 1041.027055 | 993.8083593 | 1112.169575 | 1018.541326 | 825.3513095 | 971.3324344 | 1067.75524 | 839.8707704 | 740.0817332 | 0.373127098 | 0.159541468 | YGL087C | MMS2 | n |
| W303.asm02.HP0_G0019120 | 825.8426913 | 928.1256229 | 899.1177581 | 572.0174784 | 634.2731928 | 626.432474 | 572.6878724 | 596.0373209 | 504.6477102 | 9.07E-05 | 0.534413596 | YGL086W | MAD1 | n |

| | | | | | | | | | | | | | | |
|---|---|---|---|---|---|---|---|---|---|---|---|---|---|---|
| W303.asm02.HP0_G0019130 | 688.2022427 | 584.4811615 | 617.6548077 | 283.0902827 | 313.1558023 | 298.7322492 | 393.7820742 | 334.7871965 | 298.8988466 | 5.76E-11 | 1.080061357 | YGL085W | LCL3 | n |
| W303.asm02.HP0_G0019140 | 1346.356219 | 1196.568111 | 1292.970428 | 1002.003406 | 1151.776425 | 1058.236361 | 1463.430497 | 1894.547199 | 1648.038161 | 0.078298308 | 0.256936502 | YGL084C | GUP1 | n |
| W303.asm02.HP0_G0019150 | 403.2283563 | 335.077148 | 335.2145555 | 619.6856016 | 717.8698689 | 695.2314163 | 503.5866911 | 732.4679414 | 651.0262549 | 4.46E-07 | -0.92053536 | YGL083W | SCY1 | n |
| W303.asm02.HP0_G0019160 | 2042.312853 | 1641.116486 | 1948.740011 | 974.7644785 | 735.1199766 | 888.9547536 | 966.4699466 | 1048.87087 | 1112.681665 | 5.80E-13 | 1.113986207 | YGL082W | | n |
| W303.asm02.HP0_G0019170 | 94.02199654 | 46.64426207 | 85.02526626 | 26.2661087 | 26.53862732 | 27.1574772 | 70.04777282 | 52.25002489 | 74.72471165 | 0.000108961 | 1.495958619 | YGL081W | | n |
| W303.asm02.HP0_G0019180 | 1168.974514 | 1170.86617 | 1092.623536 | 2356.167233 | 2477.38086 | 2411.583975 | 3309.28397 | 2744.0939 | 2368.670997 | 7.66E-14 | -1.077615992 | YGL080W | MPC1 | n |
| W303.asm02.HP0_G0019190 | 694.9873353 | 868.1544288 | 845.366153 | 95.3362464 | 108.808372 | 109.535158 | 121.1637151 | 121.9167247 | 110.5516282 | 2.85E-59 | 2.942489695 | YGL079W | KXD1 | n |
| W303.asm02.HP0_G0019200 | 2368.966593 | 3345.059937 | 2687.580255 | 7275.712111 | 6600.156613 | 6942.356421 | 2791.498406 | 3121.45519 | 2764.814331 | 6.39E-19 | -1.309147878 | YGL078C | DBP3 | n |
| W303.asm02.HP0_G0019210 | 3655.226278 | 2984.280849 | 4002.051326 | 883.3195075 | 1349.489199 | 952.3222004 | 1196.491687 | 1729.088786 | 1592.762347 | 2.60E-17 | 1.741828914 | YGL077C | HNM1 | n |
| W303.asm02.HP0_G0019230 | 298.5440715 | 269.3944116 | 246.2800816 | 121.6023551 | 114.1160975 | 94.14592095 | 100.3387016 | 139.3333997 | 106.4571234 | 6.82E-09 | 1.306254254 | YGL075C | MPS2 | n |
| W303.asm02.HP0_G0019240 | 612.5969259 | 623.5100338 | 526.76573 | 1570.129609 | 1476.87461 | 1546.165702 | 975.9358618 | 1941.959258 | 2149.614993 | 2.16E-08 | -1.381624967 | YGL073W | HSF1 | n |
| W303.asm02.HP0_G0019250 | 223.9080536 | 284.625191 | 215.9837223 | 1531.216855 | 1706.433736 | 1536.20796 | 1476.682778 | 2010.658365 | 2254.024864 | 3.08E-47 | -2.719405166 | YGL071W | AFT1 | n |
| W303.asm02.HP0_G0019260 | 1278.505293 | 1667.77035 | 1406.33745 | 821.0591017 | 915.5826424 | 817.4400637 | 1256.126953 | 945.3384132 | 869.0586327 | 1.59E-05 | 0.769874339 | YGL070C | RPB9 | n |
| W303.asm02.HP0_G0019270 | 3118.234669 | 3405.031131 | 3134.207229 | 2463.177305 | 2619.362516 | 2488.530161 | 3500.495458 | 2478.973403 | 2422.923185 | 0.031696726 | 0.351314432 | YGL068W | MNP1 | n |
| W303.asm02.HP0_G0019280 | 1007.101592 | 930.981394 | 1124.8745 | 597.3107683 | 485.6568799 | 525.9498084 | 778.0982332 | 608.6160306 | 762.6015093 | 9.98E-09 | 0.927245253 | YGL067W | NPY1 | n |
| W303.asm02.HP0_G0019290 | 523.4214241 | 561.6349923 | 522.8565224 | 820.0862828 | 838.6206232 | 886.2390059 | 992.0279177 | 818.5837232 | 834.2553424 | 1.32E-06 | -0.662694765 | YGL066W | SGF73 | n |
| W303.asm02.HP0_G0019300 | 951.8515526 | 693.9523888 | 851.2299645 | 1788.041029 | 1531.278796 | 1721.784054 | 2024.759271 | 2037.750971 | 2407.568792 | 1.01E-10 | -1.01419077 | YGL065C | ALG2 | n |
| W303.asm02.HP0_G0019310 | 589.3337515 | 582.5773141 | 559.993995 | 751.0161451 | 825.3513095 | 836.4502977 | 525.3582961 | 640.5466014 | 548.6636362 | 0.000621112 | -0.478039794 | YGL064C | MRH4 | n |
| W303.asm02.HP0_G0019330 | 1595.466044 | 1797.231975 | 1370.177279 | 4007.040805 | 4182.487665 | 3914.297713 | 4168.789074 | 4260.312214 | 4447.655782 | 3.55E-26 | -1.345308002 | YGL062W | PYC1 | n |
| W303.asm02.HP0_G0019340 | 757.9917659 | 826.2697853 | 765.2273963 | 248.0688044 | 248.1361654 | 277.0062674 | 369.1706946 | 264.1529036 | 310.1587347 | 1.39E-21 | 1.601852316 | YGL061C | DUO1 | n |
| W303.asm02.HP0_G0019350 | 414.8599435 | 396.0002658 | 422.1944255 | 715.9946669 | 782.8895058 | 749.5463707 | 760.1129942 | 812.7781649 | 814.8064448 | 4.81E-12 | -0.865813945 | YGL060W | YBP2 | n |
| W303.asm02.HP0_G0019360 | 860.7374529 | 733.9331849 | 865.8894931 | 343.4050508 | 290.5979691 | 313.216237 | 501.693508 | 463.4770726 | 560.9471505 | 7.41E-19 | 1.374842763 | YGL059W | PKP2 | n |
| W303.asm02.HP0_G0019370 | 1414.207144 | 1514.510632 | 1340.858222 | 1744.264182 | 1402.566454 | 1649.364115 | 1839.227332 | 1516.218315 | 1534.415654 | 0.260677485 | -0.168730896 | YGL058W | RAD6 | n |
| W303.asm02.HP0_G0019400 | 8901.072106 | 10837.6515 | 9026.36045 | 8322.465183 | 6987.620572 | 8165.348144 | 6296.726821 | 8270.985421 | 9268.935123 | 0.090648529 | 0.293044394 | YGL055W | OLE1 | n |
| W303.asm02.HP0_G0019410 | 2838.107277 | 2947.155824 | 2961.22479 | 2544.894088 | 2418.99588 | 2331.016793 | 2534.972103 | 2558.316033 | 2300.088042 | 0.013998797 | 0.26184222 | YGL054C | ERV14 | n |
| W303.asm02.HP0_G0019420 | 721.1584065 | 345.5483088 | 601.0406753 | 258.7698117 | 345.0021551 | 277.9115167 | 955.1108483 | 778.912408 | 883.3893994 | 0.000300107 | 0.921787284 | YGL053W | PRM8 | n |
| W303.asm02.HP0_G0019430 | 369.3028936 | 267.4905641 | 358.6698013 | 390.1003552 | 550.6765168 | 353.9524528 | 550.9162673 | 459.6067004 | 425.8284938 | 0.101252487 | -0.375750905 | YGL051W | MST27 | y |
| W303.asm02.HP0_G0019460 | 3812.252705 | 2423.59778 | 3518.28688 | 2749.186044 | 2409.70736 | 2732.042206 | 4453.713123 | 3686.529534 | 3846.787211 | 0.098678078 | 0.305516843 | YGL048C | RPT6 | n |
| W303.asm02.HP0_G0019470 | 684.325047 | 668.2504485 | 772.0685097 | 189.699674 | 254.7708222 | 205.4915775 | 496.0139589 | 352.2038714 | 371.5763059 | 9.30E-19 | 1.713884345 | YGL047W | ALG13 | n |
| W303.asm02.HP0_G0019480 | 270.4344024 | 240.8367001 | 227.7113453 | 328.8127682 | 321.1173905 | 337.6579665 | 316.1615692 | 385.1020353 | 440.1592604 | 0.024603397 | -0.419019812 | YGL045W | RIM8 | n |
| W303.asm02.HP0_G0019490 | 982.8691185 | 829.1255565 | 922.573004 | 511.7027103 | 526.7917522 | 511.4658206 | 742.1277552 | 564.1067502 | 591.6559361 | 1.38E-07 | 0.819436745 | YGL044C | RNA15 | n |
| W303.asm02.HP0_G0019500 | 2292.391978 | 1789.616586 | 2235.08947 | 1193.648718 | 1248.642415 | 1166.86627 | 1272.219009 | 1195.945014 | 1141.343198 | 3.07E-09 | 0.808015815 | YGL043W | DST1 | n |
| W303.asm02.HP0_G0019520 | 2462.98859 | 1702.991528 | 2142.245789 | 3433.077689 | 3314.674552 | 3453.52585 | 2141.190029 | 2357.056678 | 1900.873829 | 1.31E-05 | -0.693571436 | YGL040C | HEM2 | n |
| W303.asm02.HP0_G0019540 | 880.1234316 | 789.1447604 | 819.9563033 | 649.8429857 | 488.3107426 | 641.8217111 | 478.9753114 | 761.4957331 | 765.6723879 | 0.022845395 | 0.481985822 | YGL038C | OCH1 | n |
| W303.asm02.HP0_G0019550 | 6704.640723 | 3150.867499 | 5636.100121 | 222.7755146 | 163.212558 | 217.2598176 | 407.0343556 | 386.0696283 | 350.0801559 | 9.14E-80 | 4.678864734 | YGL037C | PNC1 | n |
| W303.asm02.HP0_G0019560 | 1113.724475 | 847.2121071 | 1076.009404 | 31.13020291 | 41.13487234 | 21.72598176 | 83.30005416 | 84.18059565 | 111.5752544 | 1.33E-86 | 5.04533724 | YGL036W | VIR1 | n |
| W303.asm02.HP0_G0019570 | 599.0267409 | 717.7504817 | 431.9674447 | 927.0963553 | 1435.739738 | 964.0904405 | 854.7721467 | 1219.167247 | 1224.25692 | 0.000148907 | -0.926522031 | YGL035C | MIG1 | n |
| W303.asm02.HP0_G0019600 | 7919.172286 | 10375.9685 | 8171.221278 | 15899.75113 | 17937.4582 | 15283.32292 | 15466.35892 | 11881.0751 | 10531.06621 | 3.35E-07 | -0.892102729 | YGL031C | RPL24A | y |
| W303.asm02.HP0_G0019610 | 7133.070851 | 8200.822812 | 7343.446559 | 10941.2935 | 12759.77201 | 10836.73865 | 13363.97914 | 9653.675894 | 8917.831341 | 0.000374504 | -0.60682895 | YGL030W | RPL30 | n |
| W303.asm02.HP0_G0019620 | 997.4086025 | 1694.424214 | 1270.492484 | 1864.893718 | 2233.225489 | 2011.463811 | 1758.767053 | 1380.755287 | 1181.26462 | 0.003946714 | -0.624400686 | YGL029W | CGR1 | n |
| W303.asm02.HP0_G0019630 | 1436.501019 | 1684.904977 | 1289.061221 | 2330.873943 | 2162.898126 | 2497.582653 | 1739.835222 | 2624.112361 | 2342.056716 | 0.000212457 | -0.664875626 | YGL028C | SCW11 | n |
| W303.asm02.HP0_G0019640 | 1098.215692 | 935.7410126 | 1075.032102 | 997.1393119 | 934.1596815 | 945.9854557 | 1003.387016 | 1253.033004 | 1163.862975 | 0.452302703 | 0.111361631 | YGL027C | CWH41 | n |
| W303.asm02.HP0_G0019650 | 2267.190205 | 1718.222307 | 2113.904033 | 3006.983037 | 2622.016379 | 2932.102288 | 3023.413329 | 2975.348639 | 3582.691654 | 0.001136189 | -0.489535502 | YGL026C | TRP5 | n |
| W303.asm02.HP0_G0019660 | 449.7547051 | 405.5195029 | 372.3520281 | 333.6768624 | 356.9445374 | 339.468465 | 524.4117046 | 290.277916 | 380.7889416 | 0.273914413 | 0.253726253 | YGL025C | PGD1 | n |
| W303.asm02.HP0_G0019670 | 554.4389899 | 635.8850422 | 456.3999924 | 810.3580944 | 776.254849 | 776.7038479 | 345.5059065 | 720.8568248 | 856.7751185 | 0.067892742 | -0.521194787 | YGL023C | PIB2 | n |
| W303.asm02.HP0_G0019680 | 1547.970397 | 1589.712605 | 1360.40426 | 3356.225001 | 2988.249436 | 3372.053419 | 4766.088326 | 4196.451073 | 3889.779511 | 2.23E-17 | -1.111460928 | YGL022W | STT3 | n |
| W303.asm02.HP0_G0019700 | 1604.189735 | 1728.693468 | 1702.459929 | 1794.850761 | 1792.684275 | 1805.972234 | 3673.721707 | 2651.204966 | 2587.727001 | 0.554402091 | -0.099147637 | YGL020C | GET1 | n |
| W303.asm02.HP0_G0019710 | 1736.983689 | 1328.885507 | 1773.802968 | 1975.795066 | 2223.936969 | 2069.399762 | 2787.71204 | 1887.774047 | 2024.732598 | 0.046827287 | -0.373171632 | YGL019W | CKB1 | n |
| W303.asm02.HP0_G0019720 | 476.8950752 | 386.4810286 | 474.9687287 | 439.7141161 | 329.0789787 | 408.2674072 | 712.783418 | 548.6252613 | 517.9548506 | 0.374721516 | 0.182388895 | YGL018C | JAC1 | n |
| W303.asm02.HP0_G0019730 | 348.947616 | 325.5579108 | 317.6231211 | 342.432232 | 366.233057 | 376.5836838 | 339.8263573 | 356.0742437 | 410.474101 | 0.436311509 | -0.129313211 | YGL017W | ATE1 | n |
| W303.asm02.HP0_G0019740 | 1205.807873 | 1402.183633 | 1373.109185 | 1702.432971 | 2050.10896 | 1743.510036 | 1296.830389 | 1612.010027 | 1215.044284 | 0.003142472 | -0.464489197 | YGL016W | KAP122 | n |
| W303.asm02.HP0_G0019760 | 710.4961182 | 1174.673865 | 644.0419593 | 2790.044436 | 2815.748358 | 2900.418565 | 1193.651912 | 1580.079456 | 1610.163992 | 2.04E-15 | -1.749702746 | YGL014W | PUF4 | n |
| W303.asm02.HP0_G0019770 | 667.8469651 | 840.548641 | 709.5211874 | 1034.106428 | 1050.929642 | 1076.341346 | 717.5163756 | 1373.982136 | 1479.13984 | 0.037440684 | -0.51133135 | YGL013C | PDR1 | n |
| W303.asm02.HP0_G0019780 | 5109.174678 | 4009.502691 | 4974.466727 | 2853.27766 | 2587.516163 | 2780.925665 | 4658.176892 | 4146.136234 | 4039.228934 | 8.44E-09 | 0.777168753 | YGL012W | ERG4 | n |
| W303.asm02.HP0_G0019790 | 3875.257136 | 3687.752475 | 3783.135697 | 2001.088356 | 1697.145217 | 1939.949121 | 2944.846233 | 2330.931666 | 2120.953459 | 1.34E-11 | 1.008296796 | YGL011C | SCL1 | n |
| W303.asm02.HP0_G0019800 | 655.246079 | 366.4906306 | 555.1074854 | 46.69530436 | 65.01963692 | 49.7887082 | 202.5705863 | 147.0741441 | 154.5675542 | 6.50E-33 | 3.29821936 | YGL010W | MPO1 | n |
| W303.asm02.HP0_G0019810 | 9277.160092 | 8117.053525 | 9470.055518 | 378.4265291 | 502.9069876 | 407.362158 | 465.7230301 | 695.6994054 | 690.9476762 | 2.05E-129 | 4.384599896 | YGL009C | LEU1 | n |
| W303.asm02.HP0_G0019820 | 43415.86853 | 33491.53209 | 42726.66225 | 47647.69401 | 51183.72357 | 46250.08892 | 17573.47165 | 48069.0553 | 43522.53822 | 0.352461679 | -0.278231247 | YGL008C | PMA1 | n |
| W303.asm02.HP0_G0019830 | 532.1451145 | 638.7408133 | 534.5841453 | 803.5483625 | 1020.41022 | 845.5027901 | 875.5971602 | 1669.098017 | 1381.895352 | 0.005154056 | -0.645469265 | YGL006W | PMC1 | n |
| W303.asm02.HP0_G0019840 | 1031.334065 | 1007.135291 | 993.9160435 | 499.0560653 | 513.5224386 | 478.8768479 | 602.0322096 | 470.250224 | 470.868046 | 2.06E-12 | 1.024620576 | YGL005C | COG7 | n |
| W303.asm02.HP0_G0019850 | 837.4742785 | 481.6734002 | 774.0231135 | 257.7969928 | 286.617175 | 294.206003 | 336.0399912 | 295.1158813 | 403.3087177 | 8.44E-09 | 1.319716556 | YGL004C | RPN14 | n |
| W303.asm02.HP0_G0019860 | 371.2414915 | 333.1733005 | 264.8488179 | 151.7597392 | 200.3666362 | 182.8603465 | 237.5944727 | 216.740844 | 313.2296132 | 0.000161273 | 0.86048968 | YGL003C | CDH1 | n |
| W303.asm02.HP0_G0019880 | 3381.883979 | 2668.242175 | 2979.793527 | 2550.731001 | 2480.034723 | 2662.338015 | 2280.338983 | 2092.903775 | 1901.897455 | 0.09465697 | 0.23090697 | YGL001C | ERG26 | n |

| | | | | | | | | | | | | | | |
|---|---|---|---|---|---|---|---|---|---|---|---|---|---|---|
| W303.asm02.HP0_G0019890 | 1355.079909 | 1114.702671 | 1514.817962 | 2136.310174 | 2286.302743 | 2101.083486 | 2059.783157 | 1851.973104 | 1481.187093 | 2.32E-05 | -0.711090955 | YGR001C | EFM5 | n |
| W303.asm02.HP0_G0019900 | 890.7857198 | 479.7695528 | 847.3207568 | 876.5097756 | 873.1208387 | 832.8293007 | 719.4095587 | 813.7457579 | 798.4284258 | 0.337044045 | -0.219587846 | YGR002C | SWC4 | n |
| W303.asm02.HP0_G0019910 | 982.8691185 | 781.5293707 | 956.7785709 | 572.0174784 | 621.0038792 | 572.1175196 | 531.0378453 | 598.9401001 | 537.4037482 | 2.20E-05 | 0.625489926 | YGR003W | CUL3 | n |
| W303.asm02.HP0_G0019930 | 2120.826066 | 1509.751013 | 2010.310031 | 1485.49437 | 1559.144355 | 1475.556261 | 1704.811356 | 1670.06561 | 1548.746421 | 0.038900497 | 0.319748541 | YGR005C | TFG2 | n |
| W303.asm02.HP0_G0019940 | 180.2896016 | 119.9423882 | 172.9824382 | 190.6724928 | 214.9628813 | 210.9230729 | 171.3330659 | 294.1482882 | 296.8515942 | 0.168944242 | -0.381361475 | YGR006W | PRP18 | n |
| W303.asm02.HP0_G0019950 | 1569.294973 | 1237.500831 | 1508.954151 | 667.3537248 | 655.5040947 | 717.8626473 | 985.4017771 | 783.7503733 | 713.4674523 | 8.28E-11 | 1.079883728 | YGR007W | ECT1 | n |
| W303.asm02.HP0_G0019960 | 1245.549129 | 1073.769951 | 1124.8745 | 38.91275363 | 102.1737152 | 45.262462 | 200.6774032 | 68.69910679 | 251.812042 | 6.16E-14 | 4.216500099 | YGR008C | STF2 | n |
| W303.asm02.HP0_G0019970 | 781.2549403 | 743.452422 | 743.7267543 | 1350.272551 | 1102.679965 | 1303.558905 | 1205.957602 | 1227.875585 | 1233.469555 | 7.68E-09 | -0.729297093 | YGR009C | SEC9 | n |
| W303.asm02.HP0_G0019990 | 1104.031485 | 762.4908964 | 936.2552307 | 1064.263812 | 1098.699171 | 1132.466799 | 1038.410902 | 1059.514394 | 991.8937752 | 0.147507754 | -0.233762985 | YGR012W | MCY1 | n |
| W303.asm02.HP0_G0020000 | 541.8381038 | 489.2887899 | 466.1730115 | 431.9315653 | 489.637674 | 420.9408966 | 520.6253385 | 467.3474448 | 514.883972 | 0.300855384 | 0.160065298 | YGR013W | SNU71 | n |
| W303.asm02.HP0_G0020010 | 980.9305206 | 1346.972058 | 951.8920613 | 2793.935711 | 3004.172612 | 2771.873173 | 2849.240489 | 3610.089682 | 3707.574049 | 4.91E-16 | -1.385406908 | YGR014W | MSB2 | n |
| W303.asm02.HP0_G0020020 | 358.6406054 | 246.5482424 | 355.7378956 | 229.5852464 | 343.6752237 | 278.8167659 | 606.7651672 | 510.8891322 | 551.7345148 | 0.440994716 | 0.177113606 | YGR015C | EAT1 | n |
| W303.asm02.HP0_G0020030 | 255.8949184 | 266.5386404 | 261.9169121 | 337.5681378 | 399.4063411 | 338.5632157 | 274.5115421 | 463.4770726 | 318.3477441 | 0.042414225 | -0.453646829 | YGR016W | | n |
| W303.asm02.HP0_G0020050 | 1353.141311 | 807.231311 | 1068.190989 | 103.1187971 | 43.78873507 | 86.90392703 | 70.04777282 | 86.11578176 | 82.91372115 | 1.10E-40 | 3.771991316 | YGR019W | UGA1 | n |
| W303.asm02.HP0_G0020060 | 3813.222004 | 4078.041198 | 4005.960533 | 807.4396379 | 874.44777 | 833.73455 | 1386.756583 | 950.1763785 | 904.8855493 | 1.19E-39 | 2.242066074 | YGR020C | VMA7 | n |
| W303.asm02.HP0_G0020070 | 1252.334222 | 1010.942986 | 1062.327177 | 410.5295508 | 419.3103116 | 433.6143859 | 771.4720925 | 621.1947403 | 582.4433004 | 4.95E-18 | 1.395996939 | YGR021W | DPC29 | n |
| W303.asm02.HP0_G0020090 | 1307.584261 | 1216.558503 | 1246.059937 | 968.9275655 | 863.8323191 | 998.4899116 | 482.7616775 | 446.0603976 | 437.0883818 | 0.001023877 | 0.411593325 | YGR024C | THG1 | n |
| W303.asm02.HP0_G0020100 | 3431.318224 | 2848.155758 | 3490.922426 | 1966.066877 | 1689.183629 | 1913.696893 | 1535.371453 | 1165.949629 | 1296.934379 | 8.78E-08 | 0.810434692 | YGR026W | | n |
| W303.asm02.HP0_G0020120 | 1376.404485 | 1518.318327 | 1363.336166 | 1593.477261 | 1342.854542 | 1571.512681 | 1241.92808 | 1288.833947 | 1171.028358 | 0.535924379 | -0.083384025 | YGR028W | MSP1 | n |
| W303.asm02.HP0_G0020140 | 757.022467 | 777.7216758 | 711.4757912 | 334.6496812 | 396.7524784 | 341.2789635 | 320.8945268 | 339.6251618 | 288.6625847 | 4.37E-13 | 1.069790469 | YGR030C | POP6 | n |
| W303.asm02.HP0_G0020170 | 1833.913582 | 2129.453352 | 2063.084334 | 1024.378239 | 1167.699602 | 1015.689647 | 1103.725718 | 1057.579207 | 1001.106411 | 3.93E-13 | 0.911194606 | YGR033C | TIM21 | n |
| W303.asm02.HP0_G0020210 | 705.6496235 | 856.7313442 | 733.9537352 | 404.6926378 | 498.9261935 | 444.4773768 | 746.8607129 | 619.2595542 | 621.3410955 | 1.77E-06 | 0.770782634 | YGR036C | CAX4 | n |
| W303.asm02.HP0_G0020220 | 4161.200321 | 4012.358462 | 4210.216633 | 2934.021624 | 2920.575936 | 2883.218829 | 3582.84892 | 2597.019755 | 2158.827628 | 0.004846258 | 0.503115922 | YGR037C | ACB1 | n |
| W303.asm02.HP0_G0020230 | 1335.69393 | 1168.010399 | 1347.699335 | 1058.426899 | 983.256142 | 1020.215893 | 1079.114338 | 1411.718265 | 1521.108514 | 0.040965961 | 0.33056324 | YGR038W | ORM1 | n |
| W303.asm02.HP0_G0020240 | 1073.013919 | 1270.818161 | 1094.57814 | 517.5396233 | 634.2731928 | 509.6553221 | 867.0778365 | 675.3799513 | 985.7520181 | 4.77E-09 | 1.051482179 | YGR040W | KSS1 | n |
| W303.asm02.HP0_G0020250 | 374.1493883 | 398.8560369 | 305.8954981 | 986.4383046 | 1190.257435 | 1053.710115 | 1047.876818 | 1326.570076 | 1452.525559 | 2.98E-19 | -1.581139757 | YGR041W | BUD9 | n |
| W303.asm02.HP0_G0020260 | 797.7330222 | 783.4332181 | 763.2727925 | 109.928529 | 124.7315484 | 110.4404073 | 33.13070336 | 113.2083873 | 121.8115163 | 1.82E-16 | 2.765789354 | YGR042W | MTE1 | n |
| W303.asm02.HP0_G0020280 | 1678.825753 | 1659.203037 | 1636.980701 | 239.3134348 | 238.8476458 | 244.4172948 | 330.3604421 | 336.7223826 | 384.8834463 | 1.76E-100 | 2.782921998 | YGR044C | RME1 | n |
| W303.asm02.HP0_G0020290 | 1590.61955 | 1542.11642 | 1473.771282 | 741.2879567 | 866.4861818 | 731.4413859 | 884.1164839 | 930.8245174 | 961.1849896 | 2.90E-15 | 0.980063094 | YGR046W | TAM41 | n |
| W303.asm02.HP0_G0020300 | 468.1713848 | 522.60612 | 494.514767 | 442.6325726 | 534.7533404 | 462.5823616 | 129.6830389 | 460.5742934 | 497.4823269 | 0.907716975 | 0.045203863 | YGR047C | TFC4 | n |
| W303.asm02.HP0_G0020310 | 1153.465731 | 1000.471825 | 1056.463366 | 537.968819 | 476.3683603 | 553.1072856 | 806.4959789 | 598.9401001 | 627.4828526 | 1.51E-10 | 1.032583757 | YGR048W | UFD1 | n |
| W303.asm02.HP0_G0020340 | 565.1012782 | 191.3366669 | 222.8248357 | 366.752703 | 362.2522629 | 340.3737142 | 297.2297387 | 367.6853603 | 350.0801559 | 0.732234991 | -0.126940969 | YGR052W | FMP48 | n |
| W303.asm02.HP0_G0020350 | 167.6887155 | 154.211642 | 148.5498905 | 214.020145 | 199.0397049 | 228.1228085 | 111.6977999 | 215.7732509 | 252.8356682 | 0.128278904 | -0.447859784 | YGR053C | MCO32 | n |
| W303.asm02.HP0_G0020360 | 1214.531564 | 1397.424015 | 984.1430244 | 6476.055023 | 6210.038792 | 6181.04181 | 3924.568461 | 4982.136632 | 4641.121132 | 1.01E-53 | -2.391285137 | YGR054W | | n |
| W303.asm02.HP0_G0020370 | 5246.815127 | 3817.2141 | 4831.780648 | 1747.182638 | 1308.354327 | 1904.644401 | 2289.804898 | 2605.728093 | 3010.484616 | 1.64E-15 | 1.485168005 | YGR055W | MUP1 | n |
| W303.asm02.HP0_G0020380 | 321.8072459 | 435.9810618 | 337.1691593 | 1559.428602 | 1673.260452 | 1652.985112 | 1280.738333 | 1461.065511 | 1710.479358 | 2.78E-38 | -2.157287218 | YGR056W | RSC1 | n |
| W303.asm02.HP0_G0020390 | 883.0313284 | 968.106419 | 942.1190422 | 613.8486886 | 640.9078497 | 679.8421792 | 722.2493332 | 575.7178668 | 471.8916722 | 0.002428295 | 0.529717324 | YGR057C | LST7 | n |
| W303.asm02.HP0_G0020400 | 503.0661464 | 381.72141 | 464.2184077 | 295.7369276 | 233.5399204 | 251.6592887 | 372.9570607 | 280.6019855 | 290.7098371 | 7.50E-05 | 0.786286743 | YGR058W | PEF1 | n |
| W303.asm02.HP0_G0020410 | 91.11409974 | 38.07694863 | 74.27494524 | 15.56510145 | 15.92317639 | 7.241993919 | 17.03864744 | 17.41667496 | 17.40164518 | 2.66E-06 | 2.412339166 | YGR059W | SPR3 | n |
| W303.asm02.HP0_G0020420 | 7692.356336 | 7947.611103 | 7754.890664 | 7974.196038 | 7142.871542 | 7611.335609 | 6352.575721 | 6160.664971 | 5524.510531 | 0.719401442 | 0.041528803 | YGR060W | ERG25 | n |
| W303.asm02.HP0_G0020430 | 1541.185304 | 1620.174164 | 1275.378994 | 4536.254255 | 2368.572488 | 4045.558853 | 2203.665069 | 2638.626257 | 2221.268826 | 4.26E-09 | -1.303897222 | YGR061C | ADE6 | n |
| W303.asm02.HP0_G0020440 | 797.7330222 | 746.3081932 | 781.8415288 | 407.6110943 | 425.9449684 | 388.3519239 | 576.4742385 | 524.435435 | 553.7817672 | 4.91E-13 | 0.93044878 | YGR062C | COX18 | n |
| W303.asm02.HP0_G0020450 | 896.6015134 | 1196.568111 | 934.3006269 | 639.1419784 | 693.9851043 | 591.1277537 | 825.4278094 | 491.5372711 | 537.4037482 | 0.002753232 | 0.65466525 | YGR063C | SPT4 | n |
| W303.asm02.HP0_G0020460 | 1394.821165 | 1381.241312 | 1538.273208 | 878.4554133 | 701.9466925 | 852.744784 | 790.403923 | 1334.310821 | 956.0668586 | 3.46E-05 | 0.82519701 | YGR065C | VHT1 | n |
| W303.asm02.HP0_G0020480 | 102.7456869 | 93.28852415 | 64.50192613 | 306.4379349 | 400.7332725 | 267.953775 | 587.8333367 | 720.8568248 | 525.1202339 | 5.76E-15 | -1.899951049 | YGR067C | | n |
| W303.asm02.HP0_G0020490 | 144.4255411 | 166.5866503 | 99.68479492 | 75.87986959 | 59.71191146 | 63.36744679 | 28.39774574 | 78.37503733 | 138.1895352 | 0.025501327 | 1.044204886 | YGR068C | ART5 | n |
| W303.asm02.HP0_G0020520 | 744.4215809 | 787.240913 | 773.0458116 | 1418.36987 | 1556.490492 | 1478.272009 | 1299.670163 | 1672.968389 | 1350.162941 | 6.95E-12 | -0.949785549 | YGR072W | UPF3 | n |
| W303.asm02.HP0_G0020530 | 955.7287484 | 864.3467339 | 998.802553 | 405.6654566 | 415.3295175 | 410.0779057 | 582.1537876 | 424.7733505 | 337.7966417 | 1.77E-09 | 1.195347542 | YGR075C | PRP38 | n |
| W303.asm02.HP0_G0020540 | 1003.224396 | 1157.539238 | 1101.419254 | 1516.624573 | 1592.317639 | 1566.986434 | 1570.395339 | 1304.315436 | 1105.516282 | 0.0008343 | -0.519248902 | YGR076C | MRPL25 | n |
| W303.asm02.HP0_G0020550 | 836.5049795 | 751.0678118 | 729.0672256 | 286.0087392 | 301.21342 | 282.4377629 | 331.3070336 | 303.8242188 | 363.3872964 | 8.41E-22 | 1.415051425 | YGR077C | PEX8 | n |
| W303.asm02.HP0_G0020560 | 659.1232747 | 506.4234168 | 553.1528816 | 1395.022218 | 1378.681689 | 1427.578051 | 1457.750948 | 1166.917222 | 983.7047657 | 1.81E-13 | -1.289691659 | YGR078C | PAC10 | n |
| W303.asm02.HP0_G0020570 | 2008.38739 | 2153.251445 | 1816.804253 | 555.4795581 | 522.8109581 | 536.8127993 | 819.7482603 | 776.9772219 | 832.20809 | 3.33E-53 | 1.88742805 | YGR079W | | n |
| W303.asm02.HP0_G0020580 | 1307.584261 | 1806.751213 | 1373.109185 | 1706.324247 | 1594.971502 | 1634.880127 | 1332.800867 | 1069.190324 | 883.3893994 | 0.487187265 | -0.137569816 | YGR081C | SLX9 | n |
| W303.asm02.HP0_G0020590 | 1745.707379 | 1725.837697 | 1738.6201 | 4863.121385 | 4983.95421 | 4771.568744 | 3328.2158 | 3366.256233 | 3024.815383 | 6.18E-52 | -1.488271693 | YGR082W | TOM20 | n |
| W303.asm02.HP0_G0020600 | 1237.794738 | 1607.799156 | 1290.038523 | 4091.676045 | 4195.756979 | 3996.675394 | 3007.321274 | 3802.6407 | 3126.154375 | 2.26E-26 | -1.57045545 | YGR083C | GCD2 | n |
| W303.asm02.HP0_G0020610 | 1610.974827 | 1378.38554 | 1607.661644 | 3028.385052 | 2941.806838 | 2956.544018 | 2593.660777 | 2583.473453 | 2057.488636 | 1.97E-12 | -0.957530482 | YGR084C | MRP13 | n |
| W303.asm02.HP0_G0020620 | 6768.614452 | 7264.129875 | 6810.817018 | 16129.33638 | 18389.9418 | 15592.01291 | 14782.91984 | 11775.60746 | 10519.80632 | 1.22E-17 | -1.265453119 | YGR085C | RPL11B | y |
| W303.asm02.HP0_G0020630 | 5212.889664 | 3719.165958 | 4506.339112 | 3536.196486 | 2827.69074 | 3368.432422 | 2104.272959 | 3081.783875 | 3078.043944 | 0.016764062 | 0.465208126 | YGR086C | PIL1 | n |
| W303.asm02.HP0_G0020650 | 565.1012782 | 177.0578111 | 367.4655185 | 377.4537102 | 347.6560178 | 335.847468 | 335.0933997 | 715.0512665 | 365.4345488 | 0.873723702 | 0.064544916 | YGR088W | CTT1 | n |
| W303.asm02.HP0_G0020660 | 841.3514742 | 1144.212306 | 975.3473072 | 1614.879276 | 1551.182767 | 1538.923708 | 2662.761959 | 2638.626257 | 2823.161074 | 1.57E-06 | -0.668104693 | YGR089W | NNF2 | n |
| W303.asm02.HP0_G0020670 | 1495.628254 | 2255.107283 | 1602.775134 | 7612.30743 | 7501.143011 | 7420.32802 | 4863.587253 | 5855.873159 | 6100.812074 | 4.69E-34 | -2.073459802 | YGR090W | UTP22 | n |
| W303.asm02.HP0_G0020680 | 600.9653387 | 588.2888564 | 573.6762218 | 518.5124422 | 541.3879972 | 522.3288114 | 701.4243197 | 566.0419363 | 662.286143 | 0.285022525 | 0.156694091 | YGR091W | PRP31 | n |
| W303.asm02.HP0_G0020690 | 1018.733179 | 842.4524885 | 948.9601556 | 629.41379 | 636.9270556 | 630.053471 | 957.0040313 | 599.9076931 | 1061.500356 | 0.006838871 | 0.56742757 | YGR092W | DBF2 | n |

| | | | | | | | | | | | | | | |
|---|---|---|---|---|---|---|---|---|---|---|---|---|---|---|
| W303.asm02.HP0_G0020710 | 6105.613982 | 4763.426274 | 5756.308256 | 6512.049321 | 6285.67388 | 6512.363032 | 4891.984999 | 4842.803232 | 4764.9799 | 0.084860773 | -0.216060488 | YGR094W | VAS1 | n |
| W303.asm02.HP0_G0020720 | 1455.886998 | 1731.549239 | 1642.844512 | 1583.749073 | 1767.472579 | 1643.02737 | 1483.308919 | 1222.070026 | 1198.666265 | 0.754847578 | -0.047642564 | YGR095C | RRP46 | n |
| W303.asm02.HP0_G0020730 | 172.5352102 | 175.1539637 | 170.0505325 | 301.5738407 | 313.1558023 | 270.6695227 | 368.2241031 | 329.9492312 | 364.4109226 | 3.69E-06 | -0.771609475 | YGR096W | TPC1 | n |
| W303.asm02.HP0_G0020740 | 540.8688049 | 692.0485414 | 538.493353 | 1394.049399 | 1074.814406 | 1369.6421 | 596.3526605 | 937.5976588 | 1073.78387 | 7.47E-07 | -1.116229045 | YGR097W | ASK10 | n |
| W303.asm02.HP0_G0020750 | 510.8205379 | 875.7698185 | 548.2663721 | 672.217819 | 797.4857508 | 682.5579269 | 633.2697299 | 757.6253608 | 710.3965738 | 0.506924136 | -0.15254777 | YGR098C | ESP1 | n |
| W303.asm02.HP0_G0020760 | 645.5530897 | 678.7216094 | 720.2715084 | 611.9030509 | 733.7930453 | 675.315933 | 599.192435 | 702.4725568 | 631.5773574 | 0.905374137 | 0.018279685 | YGR099W | TEL2 | n |
| W303.asm02.HP0_G0020770 | 963.4831398 | 909.0871486 | 1046.690347 | 1465.065174 | 1410.528042 | 1471.935264 | 1169.987124 | 1391.398811 | 1294.887127 | 4.63E-06 | -0.574997599 | YGR100W | MDR1 | n |
| W303.asm02.HP0_G0020780 | 970.2682324 | 942.4044786 | 1009.552874 | 1090.529921 | 1061.545093 | 1160.529526 | 1003.387016 | 1058.5468 | 955.0432324 | 0.129220154 | -0.181751739 | YGR101W | PCP1 | n |
| W303.asm02.HP0_G0020790 | 797.7330222 | 718.7024054 | 778.9096231 | 284.0631015 | 303.8672828 | 299.6374984 | 347.3990895 | 288.3427299 | 266.1428086 | 1.52E-18 | 1.371487933 | YGR102C | GTF1 | n |
| W303.asm02.HP0_G0020800 | 4557.643585 | 6088.504086 | 5293.06715 | 10749.64819 | 11251.05105 | 10508.13318 | 5115.380599 | 5159.206161 | 4578.679934 | 1.35E-14 | -1.028176078 | YGR103W | NOP7 | n |
| W303.asm02.HP0_G0020810 | 781.2549403 | 531.1734334 | 768.159302 | 434.8500219 | 494.9453994 | 483.4030941 | 573.6344639 | 517.6622836 | 526.1438601 | 0.00234906 | 0.558578477 | YGR104C | SRB5 | n |
| W303.asm02.HP0_G0020820 | 1421.961535 | 1311.75088 | 1350.631241 | 382.3178044 | 355.617606 | 388.3519239 | 451.5241572 | 318.3381146 | 307.0878561 | 2.24E-27 | 1.857290004 | YGR105W | VMA21 | n |
| W303.asm02.HP0_G0020830 | 5282.679187 | 4738.676257 | 5459.208475 | 2209.271588 | 2007.647156 | 2206.092398 | 3087.781553 | 2433.496529 | 2416.781427 | 3.51E-20 | 1.26872021 | YGR106C | VOA1 | n |
| W303.asm02.HP0_G0020860 | 315.9914523 | 194.192438 | 311.7593096 | 860.9446741 | 1190.257435 | 891.6705013 | 1377.290668 | 1357.533054 | 1390.084362 | 6.25E-19 | -1.839065198 | YGR110W | CLD1 | n |
| W303.asm02.HP0_G0020870 | 1564.448479 | 1072.818028 | 1527.522887 | 449.4423045 | 644.8886438 | 423.6566443 | 1484.25551 | 1368.176578 | 1174.099236 | 4.22E-12 | 1.459294856 | YGR111W | | n |
| W303.asm02.HP0_G0020880 | 827.7812891 | 785.3370655 | 765.2273963 | 918.3409857 | 833.3128977 | 817.4400637 | 744.0209383 | 744.0790581 | 624.4119741 | 0.436720616 | -0.111576125 | YGR112W | SHY1 | n |
| W303.asm02.HP0_G0020890 | 504.0354454 | 364.5867831 | 422.1944255 | 351.1876015 | 330.4059101 | 350.3314559 | 373.9036522 | 355.1066506 | 468.8207936 | 0.094873216 | 0.321966811 | YGR113W | DAM1 | n |
| W303.asm02.HP0_G0020900 | 2169.291013 | 2251.299588 | 2356.274907 | 4704.551914 | 4832.684034 | 4651.170595 | 3190.013438 | 4510.918815 | 4221.434395 | 2.43E-13 | -1.065966529 | YGR116W | SPT6 | n |
| W303.asm02.HP0_G0020910 | 1260.088613 | 1311.75088 | 1325.221391 | 777.2822538 | 740.4277021 | 781.2300941 | 738.3413892 | 797.296676 | 683.7822929 | 1.89E-10 | 0.760610828 | YGR117C | | n |
| W303.asm02.HP0_G0020920 | 6061.99553 | 8019.005382 | 6010.406753 | 47.6681232 | 61.03884283 | 68.79894223 | 48.27616775 | 42.57409435 | 72.67745928 | 3.28E-210 | 6.822278023 | YPR132W | RPS23B | y |
| W303.asm02.HP0_G0020930 | 2272.0367 | 2146.587979 | 2214.56613 | 2978.771291 | 2887.402652 | 3002.711729 | 2073.035439 | 2695.714247 | 2415.757801 | 0.001085552 | -0.41919716 | YGR119C | NUP57 | n |
| W303.asm02.HP0_G0020940 | 1280.443891 | 1189.904645 | 1242.150729 | 241.2590725 | 268.0401359 | 229.9333069 | 331.3070336 | 273.8288341 | 233.3867706 | 1.09E-45 | 2.331567426 | YGR120C | COG2 | n |
| W303.asm02.HP0_G0020950 | 973.1761292 | 674.9139145 | 808.2286804 | 267.5251812 | 283.9633123 | 292.3955045 | 466.6696216 | 356.0742437 | 371.5763059 | 3.01E-16 | 1.541341552 | YGR122W | | n |
| W303.asm02.HP0_G0020980 | 1829.067087 | 2367.434281 | 1906.716028 | 2115.880979 | 2007.647156 | 2169.882428 | 1680.199956 | 1973.889829 | 1673.628816 | 0.777463247 | -0.04449915 | YGR123C | PPT1 | n |
| W303.asm02.HP0_G0020990 | 7292.035876 | 5398.359392 | 7104.007591 | 11819.74892 | 10940.54911 | 11449.59239 | 9958.142838 | 9176.652519 | 9630.275167 | 1.07E-08 | -0.789389009 | YGR124W | ASN2 | n |
| W303.asm02.HP0_G0021000 | 966.3910366 | 1172.770018 | 1087.737027 | 1042.861797 | 1332.239091 | 1195.834246 | 836.7869077 | 1675.871169 | 1113.705291 | 0.558208501 | -0.145676751 | YGR125W | VSB1 | n |
| W303.asm02.HP0_G0021010 | 108.5614805 | 151.3558708 | 98.70749301 | 116.7382609 | 80.94281331 | 117.6824012 | 97.49892703 | 118.0463525 | 72.67745928 | 0.577751035 | 0.179460949 | YGR126W | | n |
| W303.asm02.HP0_G0021020 | 351.8555128 | 290.3367333 | 324.4642344 | 241.2590725 | 203.020499 | 247.1330425 | 310.4820201 | 198.356576 | 231.3395183 | 0.02828844 | 0.480383394 | YGR127W | | n |
| W303.asm02.HP0_G0021030 | 2196.431383 | 2695.847963 | 2477.460344 | 4983.750922 | 4534.124477 | 4996.975804 | 2221.650308 | 2776.992063 | 2350.245725 | 8.04E-13 | -0.978009006 | YGR128C | UTP8 | n |
| W303.asm02.HP0_G0021040 | 818.0882998 | 828.1736327 | 809.2059823 | 234.4493406 | 236.1937831 | 186.4813434 | 211.08991 | 186.7454593 | 191.418097 | 1.05E-33 | 1.906055353 | YGR129W | SYF2 | n |
| W303.asm02.HP0_G0021050 | 1273.658799 | 1168.962323 | 1105.328461 | 1125.551399 | 894.3517405 | 1064.573106 | 980.6688194 | 1257.870969 | 1246.776696 | 0.221343599 | 0.200647236 | YGR130C | | n |
| W303.asm02.HP0_G0021070 | 4173.801207 | 4327.445212 | 4075.348969 | 2188.842392 | 1866.992432 | 2156.30369 | 3466.418163 | 2663.783676 | 2773.003341 | 2.90E-13 | 1.016946051 | YGR132C | PHB1 | n |
| W303.asm02.HP0_G0021080 | 315.9914523 | 308.4232839 | 330.3280459 | 372.589616 | 228.2321949 | 362.099696 | 597.299252 | 548.6252613 | 529.2147387 | 0.934523012 | -0.018115915 | YGR133W | PEX4 | n |
| W303.asm02.HP0_G0021090 | 451.693303 | 559.7311449 | 488.6509555 | 296.7097465 | 370.213851 | 307.7847416 | 260.3126693 | 370.5881395 | 370.5526797 | 0.001767115 | 0.624902079 | YGR134W | CAF130 | n |
| W303.asm02.HP0_G0021100 | 5240.030034 | 4808.166688 | 5254.952375 | 2793.935711 | 2250.475596 | 2752.862939 | 3636.804637 | 2841.820798 | 2671.664348 | 3.40E-10 | 0.972208977 | YGR135W | PRE9 | n |
| W303.asm02.HP0_G0021130 | 353.7941107 | 476.9137816 | 359.6471032 | 524.3493552 | 549.3495854 | 540.4337962 | 370.1172861 | 475.0881892 | 474.9625508 | 0.017539627 | -0.4388578 | YGR140W | CBF2 | n |
| W303.asm02.HP0_G0021140 | 482.7108688 | 370.2983254 | 491.5828612 | 419.2849204 | 345.0021551 | 410.9831549 | 398.5150318 | 405.4214894 | 382.8361939 | 0.29089119 | 0.191317892 | YGR141W | VPS62 | n |
| W303.asm02.HP0_G0021150 | 4490.761959 | 2975.713536 | 2477.460344 | 190.6724928 | 406.0409979 | 328.6054741 | 2174.320732 | 772.1392566 | 1828.19637 | 2.02E-12 | 3.426804754 | YGR142W | BTN2 | n |
| W303.asm02.HP0_G0021170 | 132.7939539 | 75.20197355 | 128.0265503 | 273.3620943 | 276.0017241 | 287.8692583 | 290.603598 | 353.1714645 | 302.9933513 | 1.68E-08 | -1.318106316 | YGR144W | THI4 | n |
| W303.asm02.HP0_G0021180 | 1484.965966 | 1983.809024 | 1750.347723 | 5791.190559 | 5029.069876 | 5812.60537 | 2426.114077 | 3199.830228 | 2372.765501 | 4.83E-24 | -1.672288954 | YGR145W | ENP2 | n |
| W303.asm02.HP0_G0021190 | 943.1278622 | 951.9237158 | 806.2740766 | 170.2432971 | 126.0584797 | 193.7233373 | 458.1502979 | 238.0278911 | 431.9702509 | 5.98E-22 | 2.457153419 | YGR146C | ECL1 | n |
| W303.asm02.HP0_G0021210 | 1809.681109 | 1689.664596 | 1791.394403 | 805.4940002 | 768.2932608 | 870.8497688 | 1112.245041 | 856.3198523 | 928.4289516 | 1.26E-15 | 1.112776941 | YGR147C | NAT2 | n |
| W303.asm02.HP0_G0021230 | 1104.031485 | 707.2793208 | 1022.257799 | 306.4379349 | 281.3094495 | 282.4377629 | 230.968332 | 408.3242685 | 390.0015772 | 2.02E-12 | 1.702815086 | YGR149W | GPC1 | n |
| W303.asm02.HP0_G0021240 | 1806.773212 | 2439.780484 | 2020.08305 | 1357.082283 | 1138.507112 | 1261.91744 | 576.4742385 | 1013.069927 | 762.6015093 | 0.000441335 | 0.737521554 | YGR150C | CCM1 | n |
| W303.asm02.HP0_G0021250 | 1681.733649 | 1922.885906 | 1733.73359 | 942.6614568 | 1025.717946 | 946.890705 | 2025.705863 | 1489.125709 | 2024.732598 | 1.59E-08 | 0.873422116 | YGR152C | RSR1 | n |
| W303.asm02.HP0_G0021280 | 10665.19616 | 9022.332978 | 10797.23151 | 9175.627307 | 9881.657881 | 8926.662755 | 6096.049418 | 6695.74393 | 6025.063737 | 0.32443209 | 0.123603627 | YGR155W | CYS4 | n |
| W303.asm02.HP0_G0021290 | 366.3949968 | 306.5194365 | 356.7151975 | 1067.182268 | 1025.717946 | 1014.784398 | 1035.571128 | 950.1763785 | 1002.130037 | 2.33E-34 | -1.594038296 | YGR156W | PTI1 | n |
| W303.asm02.HP0_G0021300 | 4249.406524 | 4587.320386 | 4739.914268 | 10479.20455 | 7936.376499 | 8225.094594 | 5691.854837 | 7796.864825 | 8590.270961 | 4.19E-08 | -0.972586519 | YGR157W | CHO2 | n |
| W303.asm02.HP0_G0021310 | 1587.711653 | 2141.828361 | 1733.73359 | 1274.392681 | 1314.988983 | 1335.242629 | 977.8290449 | 907.6022841 | 801.4993044 | 0.001858102 | 0.477279047 | YGR158C | MTR3 | n |
| W303.asm02.HP0_G0021320 | 4490.761959 | 8947.131005 | 5166.017902 | 16320.98169 | 16590.62287 | 16359.66426 | 8480.513468 | 8168.420557 | 7067.115195 | 3.01E-10 | -1.405116585 | YGR159C | NSR1 | n |
| W303.asm02.HP0_G0021330 | 848.1365667 | 532.1253571 | 553.1528816 | 112.8469855 | 157.9048325 | 127.6401428 | 731.7152485 | 254.4769731 | 1095.28002 | 1.87E-05 | 2.280162468 | YGR161C | RTS3 | n |
| W303.asm02.HP0_G0021360 | 2296.269173 | 2678.713336 | 1922.352859 | 8938.25951 | 8676.804201 | 8738.370913 | 5066.157839 | 4852.479163 | 5283.958377 | 1.48E-45 | -1.933830522 | YGR162W | TIF4631 | n |
| W303.asm02.HP0_G0021370 | 1431.654525 | 1140.404612 | 1506.999547 | 599.256406 | 686.0235161 | 562.159778 | 941.8585669 | 832.130026 | 913.0745588 | 2.66E-13 | 1.144680291 | YGR163W | GTR2 | n |
| W303.asm02.HP0_G0021380 | 2606.444832 | 2063.770616 | 2765.764408 | 2024.436008 | 2008.974088 | 1974.348592 | 2017.186539 | 2173.213998 | 2215.127069 | 0.029226556 | 0.307676893 | YGR165W | MRPS35 | n |
| W303.asm02.HP0_G0021390 | 915.9874921 | 1069.962257 | 929.4141174 | 393.0188117 | 432.5796252 | 401.9306625 | 368.2241031 | 349.3010923 | 423.7812414 | 1.62E-17 | 1.249560198 | YGR166W | TRS65 | n |
| W303.asm02.HP0_G0021400 | 2831.322185 | 2683.472955 | 3035.499736 | 2459.28603 | 2064.705205 | 2331.922042 | 2646.669903 | 2146.121393 | 1987.882055 | 0.034009624 | 0.318104201 | YGR167W | CLC1 | n |
| W303.asm02.HP0_G0021410 | 1109.847279 | 689.1927702 | 1027.144308 | 224.7211522 | 232.212989 | 253.4697872 | 296.2831472 | 269.9584619 | 317.324118 | 9.24E-23 | 1.991265304 | YGR168C | PEX35 | n |
| W303.asm02.HP0_G0021420 | 926.6497804 | 601.6157884 | 805.2967747 | 1811.388682 | 1364.085444 | 1664.753352 | 1055.44955 | 867.9309689 | 768.7432664 | 1.53E-07 | -1.053252709 | YGR169C | PUS6 | n |
| W303.asm02.HP0_G0021430 | 2242.957732 | 2501.655525 | 2532.189251 | 1321.087986 | 1410.528042 | 1455.640778 | 1107.512084 | 1011.134741 | 898.7437922 | 4.77E-10 | 0.797325721 | YGR169C- | LSO2 | n |
| W303.asm02.HP0_G0021440 | 698.864531 | 764.3947438 | 669.451809 | 585.6369422 | 627.638536 | 591.1277537 | 450.5775657 | 733.4355345 | 742.1289856 | 0.247356107 | 0.241690035 | YGR170W | PSD2 | n |
| W303.asm02.HP0_G0021450 | 776.4084457 | 665.3946773 | 731.9991313 | 704.3208408 | 652.850232 | 763.1251093 | 567.9549147 | 675.3799513 | 756.4597522 | 0.840158856 | 0.034613601 | YGR171C | MSM1 | n |
| W303.asm02.HP0_G0021460 | 1416.145742 | 1072.818028 | 1449.338734 | 1054.535623 | 1224.757651 | 1152.382282 | 1727.529532 | 1556.857223 | 1592.762347 | 0.211447991 | 0.199204132 | YGR172C | YIP1 | n |
| W303.asm02.HP0_G0021470 | 2321.470946 | 1676.337664 | 2331.84236 | 1440.744703 | 1452.989846 | 1489.135 | 1375.397485 | 1278.190424 | 1207.878901 | 0.000702659 | 0.530124375 | YGR173W | RBG2 | n |

| | | | | | | | | | | | | | | |
|---|---|---|---|---|---|---|---|---|---|---|---|---|---|---|
| W303.asm02.HP0_G0021480 | 631.0136057 | 592.0965512 | 633.2916383 | 622.6040581 | 757.6778099 | 676.2211822 | 840.5732738 | 729.5651623 | 746.2234903 | 0.323504604 | -0.145774993 | YGR174C | CBP4 | n |
| W303.asm02.HP0_G0021490 | 5271.0476 | 6078.984849 | 5109.334391 | 5317.427784 | 4438.585418 | 5350.928257 | 2126.044564 | 2372.538167 | 2040.086991 | 0.386052098 | 0.123360427 | YGR175C | ERG1 | n |
| W303.asm02.HP0_G0021510 | 1797.080223 | 1425.029803 | 1537.295906 | 2583.806841 | 2349.995449 | 2563.665847 | 2024.759271 | 2691.843875 | 2503.789653 | 1.75E-05 | -0.655929174 | YGR178C | PBP1 | n |
| W303.asm02.HP0_G0021520 | 518.5749294 | 400.7598843 | 476.9233326 | 242.2318914 | 221.5975381 | 235.3648024 | 248.953571 | 240.9306703 | 251.812042 | 3.00E-09 | 0.99590215 | YGR179C | OKP1 | n |
| W303.asm02.HP0_G0021550 | 806.4567126 | 805.3274636 | 820.9336052 | 913.4768915 | 962.0252402 | 945.0802065 | 1042.197269 | 719.8892318 | 611.1048336 | 0.291470014 | -0.213271544 | YGR181W | TIM13 | n |
| W303.asm02.HP0_G0021560 | 3930.507175 | 3911.454548 | 3970.777664 | 3865.009255 | 4069.698499 | 3629.144203 | 2690.213113 | 3120.487597 | 2661.428086 | 0.801310231 | 0.031053487 | YGR183C | QCR9 | n |
| W303.asm02.HP0_G0021570 | 996.4393036 | 1226.077746 | 1053.53146 | 1848.355798 | 1999.685568 | 1961.675103 | 1861.945529 | 2318.352956 | 2022.685345 | 3.11E-09 | -0.826233379 | YGR184C | UBR1 | n |
| W303.asm02.HP0_G0021580 | 3519.524427 | 3141.348262 | 3478.217501 | 6324.295284 | 6450.213369 | 6299.629461 | 5847.095847 | 4951.173654 | 4735.294741 | 4.33E-14 | -0.911666721 | YGR185C | TYS1 | n |
| W303.asm02.HP0_G0021590 | 1388.036073 | 1365.058608 | 1378.972996 | 1961.202783 | 2027.551127 | 2093.841492 | 1766.339785 | 2049.362087 | 1975.598541 | 5.06E-07 | -0.557893457 | YGR186W | TFG1 | n |
| W303.asm02.HP0_G0021600 | 1877.532034 | 2621.597913 | 1965.354143 | 3222.94882 | 3313.34762 | 3278.812747 | 2210.29121 | 1937.121293 | 1527.250271 | 0.000783225 | -0.602393151 | YGR187C | HGH1 | n |
| W303.asm02.HP0_G0021610 | 171.5659112 | 307.4713602 | 160.2775134 | 609.9574132 | 679.3888593 | 615.5694831 | 573.6344639 | 717.9540456 | 603.9394503 | 6.30E-11 | -1.57407693 | YGR188C | BUB1 | n |
| W303.asm02.HP0_G0021620 | 2173.168209 | 2338.87657 | 2361.161417 | 3298.828689 | 3266.905023 | 3321.359461 | 4039.106035 | 4794.42358 | 4307.418995 | 2.69E-06 | -0.524622304 | YGR189C | CRH1 | n |
| W303.asm02.HP0_G0021640 | 49909.20208 | 41240.19114 | 46497.09302 | 57106.41159 | 56175.63937 | 53190.63484 | 39811.74634 | 32420.17285 | 32508.32045 | 0.040418988 | -0.274313918 | YGR192C | TDH3 | y |
| W303.asm02.HP0_G0021650 | 1685.610845 | 1386.00093 | 1677.050079 | 3267.698486 | 2594.15082 | 3390.158404 | 2731.86314 | 2176.116777 | 2891.743978 | 7.79E-09 | -0.962802749 | YGR193C | PDX1 | n |
| W303.asm02.HP0_G0021660 | 360.5792032 | 244.644395 | 304.9181962 | 71.01577538 | 92.8851956 | 50.69395744 | 266.9388099 | 174.1667496 | 367.4818011 | 4.49E-11 | 2.095445778 | YGR194C | XKS1 | n |
| W303.asm02.HP0_G0021670 | 1180.606101 | 1230.837365 | 1219.672785 | 1392.103761 | 1488.816992 | 1434.820045 | 1229.62239 | 1101.120895 | 939.6888396 | 0.070355362 | -0.248847301 | YGR195W | SKI6 | n |
| W303.asm02.HP0_G0021680 | 560.2547835 | 570.2023058 | 524.8111262 | 945.5799133 | 1062.872024 | 1068.194103 | 894.5289907 | 1134.986652 | 1129.059684 | 1.09E-09 | -0.893916674 | YGR196C | FYV8 | n |
| W303.asm02.HP0_G0021690 | 473.9871784 | 554.0196026 | 470.0822192 | 496.1376088 | 735.1199766 | 472.5401032 | 861.3982873 | 1707.801739 | 1509.848626 | 0.50840739 | -0.183457648 | YGR197C | SNG1 | n |
| W303.asm02.HP0_G0021700 | 1079.799012 | 1654.443418 | 1224.559294 | 1033.133609 | 1113.295416 | 1041.941875 | 954.1642567 | 1185.30149 | 1082.996506 | 0.090267151 | 0.312698092 | YGR198W | YPP1 | n |
| W303.asm02.HP0_G0021710 | 919.8646878 | 785.3370655 | 958.7331747 | 729.6141306 | 704.6005552 | 723.2941427 | 679.6527146 | 836.0003982 | 813.7828186 | 0.042118867 | 0.303754602 | YGR199W | PMT6 | n |
| W303.asm02.HP0_G0021720 | 1238.764037 | 1403.135557 | 1209.899766 | 2572.133015 | 2639.266487 | 2527.455878 | 1715.223842 | 2171.278812 | 1989.929307 | 2.07E-14 | -1.006373551 | YGR200C | ELP2 | n |
| W303.asm02.HP0_G0021740 | 1010.978788 | 776.7697521 | 888.3674371 | 2079.886682 | 2059.39748 | 2119.188471 | 1783.378432 | 1850.037918 | 1685.91233 | 5.68E-21 | -1.225879751 | YGR202C | PCT1 | n |
| W303.asm02.HP0_G0021750 | 588.3644526 | 542.596518 | 571.7216179 | 214.9929638 | 213.6359499 | 211.8283221 | 264.0990354 | 238.0278911 | 207.796116 | 8.14E-19 | 1.410684903 | YGR203W | YCH1 | n |
| W303.asm02.HP0_G0021770 | 564.1319793 | 413.1348927 | 501.3558803 | 239.3134348 | 184.4434598 | 226.31231 | 469.5093962 | 363.8149881 | 458.5845318 | 2.33E-09 | 1.181239511 | YGR205W | TDA10 | n |
| W303.asm02.HP0_G0021780 | 389.6581712 | 341.740614 | 369.4201224 | 107.9828913 | 102.1737152 | 109.535158 | 233.8081066 | 119.9815386 | 154.5675542 | 2.01E-12 | 1.782274243 | YGR206W | MVB12 | n |
| W303.asm02.HP0_G0021790 | 2139.242746 | 1775.33773 | 2074.811957 | 955.3081017 | 886.3901523 | 934.2172156 | 1876.144402 | 1173.690374 | 1195.595386 | 1.01E-08 | 1.109117218 | YGR207C | CIR1 | n |
| W303.asm02.HP0_G0021800 | 1274.628097 | 1432.645192 | 1331.085203 | 1834.736334 | 1507.394032 | 1832.224462 | 2368.371994 | 1761.98695 | 1854.810651 | 0.025744667 | -0.358386394 | YGR208W | SER2 | n |
| W303.asm02.HP0_G0021810 | 2442.633312 | 2665.386404 | 2387.548569 | 4295.968001 | 4641.605917 | 4220.271957 | 4690.361004 | 3806.511072 | 3259.225779 | 8.02E-08 | -0.811632202 | YGR209C | TRX2 | n |
| W303.asm02.HP0_G0021820 | 1522.768624 | 1073.769951 | 1202.081351 | 2163.543189 | 2291.610469 | 2249.544361 | 2611.646016 | 2206.112162 | 2560.089094 | 1.35E-07 | -0.819637572 | YGR210C | | n |
| W303.asm02.HP0_G0021830 | 3560.234982 | 2657.771015 | 2585.940856 | 5692.935857 | 5778.786098 | 5640.608014 | 4100.634484 | 4881.506955 | 4641.121132 | 1.98E-10 | -0.958791762 | YGR211W | ZPR1 | n |
| W303.asm02.HP0_G0021840 | 317.9300502 | 278.9136487 | 297.0997809 | 239.3134348 | 201.6935676 | 226.31231 | 291.5501896 | 286.4075438 | 323.4658751 | 0.014932473 | 0.41819472 | YGR212W | SLI1 | n |
| W303.asm02.HP0_G0021860 | 9302.361864 | 9525.900624 | 9771.064506 | 375.5080726 | 380.829302 | 350.3314559 | 206.3569524 | 271.893648 | 338.8202679 | 1.61E-180 | 4.692625419 | YGR214W | RPS0A | n |
| W303.asm02.HP0_G0021870 | 1100.154289 | 1276.529703 | 1110.214971 | 920.2866234 | 1032.352603 | 945.9854557 | 1016.639297 | 801.1670482 | 734.9636022 | 0.10325014 | 0.267352335 | YGR215W | RSM27 | n |
| W303.asm02.HP0_G0021880 | 856.8602571 | 828.1736327 | 953.8466651 | 592.4466741 | 562.6188991 | 626.432474 | 484.6548606 | 691.8290332 | 515.9075982 | 0.000975684 | 0.565896224 | YGR216C | GPI1 | n |
| W303.asm02.HP0_G0021890 | 300.4826694 | 514.9907302 | 332.2826497 | 1329.843355 | 1451.662914 | 1278.211927 | 1290.204248 | 1704.89896 | 1593.785973 | 2.40E-18 | -1.821921304 | YGR217W | CCH1 | n |
| W303.asm02.HP0_G0021900 | 1799.988119 | 1933.357067 | 1906.716028 | 5826.212038 | 6237.90435 | 5947.487506 | 3394.477207 | 5288.86363 | 4444.584904 | 3.24E-25 | -1.675074314 | YGR218W | CRM1 | n |
| W303.asm02.HP0_G0021910 | 1941.505764 | 2265.578444 | 2117.813241 | 2504.035696 | 2692.343741 | 2797.220151 | 2479.123203 | 2136.445462 | 1914.18097 | 0.015949073 | -0.337776938 | YGR220C | MRPL9 | n |
| W303.asm02.HP0_G0021930 | 1066.228827 | 738.6928035 | 992.9387416 | 535.0503625 | 457.7913212 | 500.6028297 | 586.8867452 | 516.6946905 | 588.5850575 | 2.20E-07 | 0.904295244 | YGR222W | PET54 | n |
| W303.asm02.HP0_G0021940 | 974.1454281 | 748.2120406 | 855.1391721 | 185.8083986 | 181.7895971 | 197.3443343 | 372.0104691 | 252.5417869 | 437.0883818 | 4.46E-24 | 2.188632922 | YGR223C | HSV2 | n |
| W303.asm02.HP0_G0021960 | 22.29387547 | 47.59618579 | 37.13747262 | 3.891275363 | 5.307725463 | 7.241993919 | 9.465915246 | 20.31945412 | 28.66153324 | 6.07E-05 | 2.69520481 | YGR225W | AMA1 | n |
| W303.asm02.HP0_G0021970 | 471.0792816 | 450.2599176 | 443.6950676 | 738.3695002 | 774.9279176 | 782.1353433 | 1064.915465 | 1099.185709 | 1014.413551 | 5.31E-10 | -0.749640487 | YGR227W | DIE2 | n |
| W303.asm02.HP0_G0021980 | 3261.690911 | 3900.983387 | 3516.332276 | 4167.555914 | 3631.811148 | 4078.147826 | 3120.912256 | 3718.460104 | 3564.266383 | 0.25868297 | -0.153763375 | YGR229C | SMI1 | n |
| W303.asm02.HP0_G0021990 | 482.7108688 | 505.4714931 | 472.036823 | 858.9990365 | 845.25528 | 839.1660454 | 841.5198653 | 801.1670482 | 730.8690975 | 1.68E-10 | -0.800522763 | YGR230W | BNS1 | n |
| W303.asm02.HP0_G0022000 | 4356.998706 | 3721.069805 | 4202.398217 | 2194.679305 | 1881.588677 | 2109.230729 | 2415.701571 | 1951.635189 | 1873.235922 | 5.05E-12 | 0.988861744 | YGR231C | PHB2 | n |
| W303.asm02.HP0_G0022010 | 974.1454281 | 1009.991062 | 1024.212403 | 465.9802248 | 555.9842423 | 510.5605713 | 771.4720925 | 660.8660555 | 571.1834123 | 2.87E-10 | 0.974761519 | YGR232W | NAS6 | n |
| W303.asm02.HP0_G0022020 | 1014.855983 | 1422.174031 | 1065.259083 | 1448.527254 | 1228.738445 | 1358.779109 | 773.3652756 | 1195.945014 | 877.2476422 | 0.329711169 | -0.204985299 | YGR233C | PHO81 | n |
| W303.asm02.HP0_G0022030 | 26114.85186 | 24748.11276 | 25208.52549 | 13775.11479 | 7989.453753 | 11989.12093 | 3426.661319 | 3321.746952 | 3455.762007 | 8.70E-11 | 1.172107641 | YGR234W | YHB1 | n |
| W303.asm02.HP0_G0022040 | 2103.378686 | 2019.982125 | 2166.678337 | 1137.225225 | 1460.951434 | 1151.477033 | 1516.439622 | 1364.306205 | 1101.421777 | 4.39E-06 | 0.747665449 | YGR235C | MIC26 | n |
| W303.asm02.HP0_G0022060 | 489.4959614 | 430.2695195 | 525.7884281 | 1165.436971 | 989.8907989 | 1112.551316 | 947.5381161 | 1159.176478 | 1294.887127 | 6.87E-13 | -1.177684558 | YGR237C | | n |
| W303.asm02.HP0_G0022080 | 399.3511606 | 414.0868164 | 359.6471032 | 643.0332538 | 471.0606348 | 687.9894223 | 348.345681 | 327.0464521 | 247.7175372 | 0.002317512 | -0.621774316 | YGR239C | PEX21 | n |
| W303.asm02.HP0_G0022090 | 6295.596573 | 5930.484749 | 5959.587053 | 10732.13745 | 6957.101151 | 10039.21407 | 3715.371734 | 3690.399906 | 4319.702509 | 0.000368249 | -0.608812836 | YGR240C | PFK1 | n |
| W303.asm02.HP0_G0022130 | 4190.279289 | 3155.627118 | 4168.19265 | 9082.236698 | 7734.682931 | 9196.427028 | 11907.17479 | 8218.735396 | 10745.00408 | 2.31E-11 | -1.175982543 | YGR244C | LSC2 | n |
| W303.asm02.HP0_G0022140 | 1432.623824 | 2123.74181 | 1572.478775 | 5912.792915 | 4848.607211 | 5713.027953 | 3799.61838 | 3321.746952 | 3736.235582 | 1.10E-23 | -1.68365223 | YGR245C | SDA1 | n |
| W303.asm02.HP0_G0022150 | 763.8075595 | 704.4235497 | 751.5451696 | 631.3594277 | 574.5612814 | 653.5899512 | 818.8016687 | 726.6623831 | 703.2311905 | 0.065601308 | 0.253636021 | YGR246C | BRF1 | n |
| W303.asm02.HP0_G0022160 | 392.566068 | 376.0098677 | 356.7151975 | 486.4094204 | 433.9065566 | 515.9920668 | 958.8972144 | 667.6392069 | 740.0817332 | 0.053241487 | -0.3538055 | YGR247W | CPD1 | n |
| W303.asm02.HP0_G0022180 | 36.83335947 | 55.21157552 | 29.31905733 | 789.9288988 | 593.1383205 | 721.4836442 | 278.2979082 | 464.4446656 | 456.5372794 | 4.03E-49 | -4.115444604 | YGR249W | MGA1 | n |
| W303.asm02.HP0_G0022190 | 1411.299247 | 1528.789488 | 1352.585845 | 362.8614276 | 408.6948607 | 392.8781701 | 325.6274844 | 568.9447154 | 610.0812074 | 3.83E-18 | 1.882914747 | YGR250C | RIE1 | n |
| W303.asm02.HP0_G0022200 | 623.2592142 | 660.6350588 | 610.8136944 | 1314.278254 | 1147.795631 | 1383.220839 | 655.9879265 | 609.5836237 | 427.8757462 | 1.01E-08 | -1.021817474 | YGR251W | NOP19 | n |
| W303.asm02.HP0_G0022210 | 1357.018507 | 1025.221842 | 1235.309615 | 1180.029254 | 1387.970209 | 1231.138966 | 1201.224645 | 1419.459009 | 1336.8558 | 0.671551953 | -0.069870888 | YGR252W | GCN5 | n |
| W303.asm02.HP0_G0022220 | 4347.305716 | 4157.050867 | 4274.718559 | 2620.773957 | 2226.590832 | 2562.760598 | 3173.921382 | 2750.867051 | 2579.537991 | 5.72E-10 | 0.785560181 | YGR253C | PUP2 | n |
| W303.asm02.HP0_G0022240 | 1148.619236 | 1027.125689 | 1081.873215 | 1194.621537 | 1261.911729 | 1184.066006 | 1292.097431 | 1250.130225 | 1367.564586 | 0.178915494 | -0.159711996 | YGR255C | COQ6 | n |
| W303.asm02.HP0_G0022260 | 1224.224553 | 1244.164297 | 1194.262935 | 348.269145 | 435.233488 | 420.9408966 | 531.9844368 | 463.4770726 | 515.9075982 | 3.71E-30 | 1.606610232 | YGR257C | MTM1 | n |
| W303.asm02.HP0_G0022270 | 247.171228 | 410.2791215 | 216.9610242 | 333.6768624 | 315.8096651 | 293.3007537 | 229.0751489 | 244.8010425 | 241.5757801 | 0.683648695 | -0.108010104 | YGR258C | RAD2 | n |

| | | | | | | | | | | | | | | |
|---|---|---|---|---|---|---|---|---|---|---|---|---|---|---|
| W303.asm02.HP0_G0022290 | 1809.681109 | 1577.337597 | 1699.528023 | 2829.930008 | 2681.72829 | 2845.198361 | 2962.831472 | 7361.447951 | 2702.373134 | 0.019783434 | -0.716319621 | YGR260W | TNA1 | n |
| W303.asm02.HP0_G0022300 | 1326.000941 | 1243.212373 | 1444.452224 | 1251.045029 | 1316.315915 | 1286.35917 | 1265.592868 | 1456.227545 | 1494.494233 | 0.667837349 | 0.058899956 | YGR261C | APL6 | n |
| W303.asm02.HP0_G0022310 | 1521.799325 | 1399.327862 | 1556.841944 | 890.1292394 | 1020.41022 | 887.1442551 | 1135.909829 | 960.8199021 | 912.0509326 | 1.08E-06 | 0.67996093 | YGR262C | BUD32 | n |
| W303.asm02.HP0_G0022320 | 428.4301286 | 440.7406804 | 485.7190498 | 229.5852464 | 229.5591263 | 214.5440699 | 429.7525521 | 329.9492263 | 392.0488296 | 2.16E-09 | 1.009070563 | YGR263C | SAY1 | n |
| W303.asm02.HP0_G0022330 | 5115.959771 | 4721.54163 | 5287.203338 | 6078.172118 | 5835.844147 | 6324.976439 | 4814.364494 | 5072.122786 | 5040.335345 | 0.008070434 | -0.270333141 | YGR264C | MES1 | n |
| W303.asm02.HP0_G0022340 | 1035.211261 | 757.7312778 | 977.301911 | 1854.192711 | 1790.030412 | 1949.001614 | 1158.628026 | 2049.362087 | 1659.298049 | 1.25E-06 | -1.013878092 | YGR266W | | n |
| W303.asm02.HP0_G0022350 | 3032.936363 | 2843.396139 | 3174.276607 | 1674.221225 | 1642.741031 | 1735.362793 | 2324.828784 | 1938.088886 | 1871.18867 | 2.75E-11 | 0.84077268 | YGR267C | FOL2 | n |
| W303.asm02.HP0_G0022360 | 214.2150643 | 155.1635657 | 164.186721 | 96.30906524 | 80.94281331 | 95.95641943 | 192.1580795 | 119.9815386 | 172.9928256 | 0.000289473 | 0.961217228 | YGR268C | HUA1 | n |
| W303.asm02.HP0_G0022370 | 668.8162641 | 732.0293374 | 583.4492409 | 2908.728334 | 2923.229799 | 3023.532461 | 2827.468884 | 1672.968389 | 3813.007546 | 5.81E-19 | -2.157916774 | YGR270W | YTA7 | n |
| W303.asm02.HP0_G0022380 | 529.2372177 | 719.6543291 | 525.7884281 | 1381.402754 | 1556.490492 | 1364.210605 | 1371.611119 | 1452.357173 | 1347.092062 | 3.96E-16 | -1.276593838 | YGR271W | SLH1 | n |
| W303.asm02.HP0_G0022390 | 817.1190009 | 1617.318393 | 1020.303195 | 2874.679675 | 3012.1342 | 2953.82827 | 1938.619442 | 1603.30169 | 1505.754121 | 1.49E-09 | -1.355441845 | YGR271C- | EFG1 | n |
| W303.asm02.HP0_G0022410 | 590.3030505 | 880.5294371 | 575.6308256 | 1359.027921 | 1297.738876 | 1331.621632 | 1282.631516 | 1369.144171 | 1517.014009 | 1.07E-07 | -0.96261678 | YGR274C | TAF1 | n |
| W303.asm02.HP0_G0022420 | 969.2989334 | 1098.519968 | 1079.918612 | 722.8043987 | 716.5429375 | 722.3888935 | 948.4847076 | 674.4123582 | 695.042181 | 0.001028573 | 0.542063036 | YGR275W | RTT102 | n |
| W303.asm02.HP0_G0022430 | 938.2813676 | 1041.404545 | 1022.257799 | 3898.085095 | 4060.409979 | 3963.181172 | 2912.662121 | 3173.705215 | 3296.076322 | 7.22E-76 | -1.989466969 | YGR276C | RNH70 | n |
| W303.asm02.HP0_G0022440 | 1204.838574 | 1007.135291 | 1136.602122 | 494.1919711 | 489.637674 | 476.1611002 | 602.9788011 | 567.9771224 | 571.1834123 | 3.56E-20 | 1.197804857 | YGR277C | CAB4 | n |
| W303.asm02.HP0_G0022450 | 513.7284347 | 470.2503156 | 511.1288994 | 569.0990219 | 599.7729773 | 559.4440303 | 533.8776198 | 581.5234251 | 593.7031884 | 0.127656459 | -0.208071869 | YGR278W | CWC22 | n |
| W303.asm02.HP0_G0022460 | 3703.691225 | 3245.107947 | 3588.652617 | 14391.88193 | 13489.58426 | 13482.78218 | 12534.76497 | 12073.62612 | 12885.40644 | 5.27E-85 | -1.97291666 | YGR279C | SCW4 | n |
| W303.asm02.HP0_G0022470 | 904.3559049 | 1346.020134 | 1119.010688 | 2310.444747 | 2319.476027 | 2374.468756 | 1825.028459 | 1518.153501 | 1344.021184 | 1.69E-09 | -1.055737413 | YGR280C | PXR1 | n |
| W303.asm02.HP0_G0022480 | 597.088143 | 858.6351916 | 626.4505249 | 871.6456814 | 887.7170837 | 813.8190667 | 955.1108483 | 1515.250722 | 1161.815722 | 0.153214759 | -0.305039408 | YGR281W | YOR1 | n |
| W303.asm02.HP0_G0022490 | 16770.81015 | 12810.98937 | 16223.21172 | 9510.276988 | 8780.304847 | 8896.78953 | 8862.936444 | 9311.147953 | 9244.368095 | 6.62E-09 | 0.752515158 | YGR282C | BGL2 | n |
| W303.asm02.HP0_G0022510 | 4113.704673 | 3267.954116 | 3743.066319 | 2002.061174 | 1768.799511 | 1918.223139 | 3698.333086 | 3039.209781 | 2956.232428 | 1.92E-11 | 0.967053671 | YGR284C | ERV29 | n |
| W303.asm02.HP0_G0022520 | 2826.47569 | 2111.366802 | 2513.620515 | 6564.581538 | 7134.909954 | 6351.228667 | 6100.782376 | 5561.724871 | 5464.116586 | 1.04E-25 | -1.427945577 | YGR285C | ZUO1 | n |
| W303.asm02.HP0_G0022530 | 2100.470789 | 2068.530234 | 2238.021376 | 836.6242031 | 467.0798408 | 816.5348144 | 886.9562585 | 1000.491217 | 860.8696232 | 8.05E-15 | 1.592451791 | YGR286C | BIO2 | n |
| W303.asm02.HP0_G0022660 | 244.2633312 | 342.6925377 | 280.4856485 | 122.5751739 | 55.73111736 | 113.156155 | 1671.680632 | 355.1066506 | 2981.823083 | 0.035372134 | 1.571306012 | YHL047C | ARN2 | n |
| W303.asm02.HP0_G0022710 | 227.7852494 | 388.384876 | 222.8248357 | 4627.699226 | 2144.321087 | 3451.715352 | 7993.018833 | 8393.869739 | 9932.244892 | 4.90E-40 | -3.607216301 | YHL040C | ARN1 | n |
| W303.asm02.HP0_G0022720 | 1609.036229 | 1659.203037 | 1772.825667 | 2374.65079 | 2433.592125 | 2359.079519 | 1836.387558 | 1994.209283 | 1665.439806 | 1.38E-05 | -0.50756391 | YHL039W | EFM1 | n |
| W303.asm02.HP0_G0022750 | 234.5703419 | 205.6155226 | 183.7327593 | 102.1459783 | 115.4430288 | 101.3879149 | 146.7216863 | 163.523226 | 174.0164518 | 4.44E-06 | 0.972375612 | YHL036W | MUP3 | n |
| W303.asm02.HP0_G0022770 | 4325.011841 | 3158.482889 | 4332.379371 | 3052.705523 | 2712.247712 | 3001.80648 | 3737.143339 | 2769.251319 | 2056.46501 | 0.049501086 | 0.430451053 | YHL034C | SBP1 | n |
| W303.asm02.HP0_G0022790 | 329.5616374 | 370.2983254 | 296.122479 | 297.6825653 | 423.2911057 | 265.2380273 | 408.9275386 | 548.6252613 | 543.5455053 | 0.93610248 | 0.019284026 | YHL032C | GUT1 | n |
| W303.asm02.HP0_G0022800 | 1131.171855 | 949.0679446 | 1170.807689 | 594.3923118 | 602.4268401 | 604.7064923 | 749.7004874 | 596.0373209 | 512.8367197 | 7.48E-07 | 0.851647311 | YHL031C | GOS1 | n |
| W303.asm02.HP0_G0022810 | 638.7679971 | 821.5101667 | 598.1087695 | 816.1950075 | 770.9471235 | 917.01748 | 944.6983415 | 1138.857024 | 1020.555308 | 0.100874848 | -0.283658553 | YHL030W | ECM29 | n |
| W303.asm02.HP0_G0022820 | 687.2329438 | 902.4236826 | 638.1781479 | 761.7171524 | 834.6398291 | 764.0303585 | 789.4573315 | 882.4448647 | 925.358073 | 0.648511374 | -0.082463956 | YHL029C | OCA5 | n |
| W303.asm02.HP0_G0022830 | 206.4606728 | 188.4808957 | 135.8449656 | 191.6453116 | 209.6551558 | 187.3865927 | 249.9001625 | 485.7317128 | 495.4350745 | 0.629333216 | -0.148016045 | YHL028W | WSC4 | n |
| W303.asm02.HP0_G0022840 | 663.0004705 | 791.0486078 | 609.8363925 | 1498.141015 | 1975.800804 | 1540.734206 | 2239.635547 | 2213.852906 | 2525.285803 | 3.35E-15 | -1.27954548 | YHL027W | RIM101 | n |
| W303.asm02.HP0_G0022860 | 946.035759 | 997.6160542 | 798.4556613 | 661.5168118 | 700.6197611 | 685.2736746 | 1122.657548 | 847.6115148 | 972.4448776 | 0.008112491 | 0.421938338 | YHL025W | SNF6 | n |
| W303.asm02.HP0_G0022870 | 403.2283563 | 392.1925709 | 365.5109147 | 211.1016885 | 207.0012931 | 203.681079 | 70.99436434 | 253.50938 | 271.2609395 | 0.015949073 | 0.900825821 | YHL024W | RIM4 | n |
| W303.asm02.HP0_G0022880 | 398.3818616 | 413.1348927 | 351.828688 | 321.0302175 | 541.3879972 | 395.5939178 | 596.3526605 | 817.6161301 | 1057.405851 | 0.683648695 | -0.109914703 | YHL023C | NPR3 | n |
| W303.asm02.HP0_G0022900 | 1910.488198 | 863.3948102 | 1460.089055 | 6.809731886 | 7.961588195 | 5.43149544 | 11.35909829 | 17.41667496 | 17.40164518 | 1.98E-70 | 7.736139978 | YHL021C | AIM17 | n |
| W303.asm02.HP0_G0022910 | 626.167111 | 580.6734666 | 577.5854294 | 449.4423045 | 449.829733 | 448.0983738 | 570.7946893 | 579.588239 | 598.8213194 | 0.001589015 | 0.405365637 | YHL020C | OPI1 | n |
| W303.asm02.HP0_G0022920 | 277.219495 | 232.2693867 | 235.5297605 | 317.1389421 | 380.829302 | 321.3634802 | 414.6070878 | 402.5187102 | 390.0015772 | 0.007920563 | -0.448871365 | YHL019C | APM2 | n |
| W303.asm02.HP0_G0022930 | 67.85092534 | 66.63466011 | 58.63811466 | 141.0587319 | 167.1933521 | 156.6081185 | 298.1763302 | 184.8102732 | 179.1345827 | 3.97E-06 | -1.265342773 | YHL018W | MCO14 | n |
| W303.asm02.HP0_G0022950 | 273.3422992 | 152.3077945 | 252.143893 | 256.824174 | 197.7127735 | 238.0805501 | 188.3717134 | 293.1806952 | 256.9301729 | 0.907716975 | -0.033702322 | YHL016C | DUR3 | n |
| W303.asm02.HP0_G0022960 | 13892.96161 | 16600.59768 | 14675.1655 | 15918.23469 | 17458.43598 | 14862.38202 | 12997.64822 | 11328.57947 | 9840.118535 | 0.535172222 | -0.094796731 | YHL015W | RPS20 | n |
| W303.asm02.HP0_G0022970 | 1031.334065 | 734.8851086 | 1001.734459 | 437.7684784 | 464.425978 | 468.013857 | 524.4117046 | 546.6900752 | 463.7026627 | 2.73E-09 | 1.014522221 | YHL014C | YLF2 | n |
| W303.asm02.HP0_G0022980 | 1334.724631 | 1475.481759 | 1239.218823 | 874.5641379 | 943.4482011 | 868.1340211 | 1070.595014 | 995.653252 | 1037.956954 | 1.81E-06 | 0.593294443 | YHL013C | OTU2 | n |
| W303.asm02.HP0_G0023000 | 3363.467299 | 3110.886703 | 3542.719427 | 6003.265067 | 6670.483976 | 5951.108503 | 5097.39536 | 5321.761794 | 4721.987601 | 2.41E-15 | -0.894538896 | YHL011C | PRS3 | n |
| W303.asm02.HP0_G0023010 | 303.3905662 | 232.2693867 | 270.7126293 | 178.9986667 | 179.1357344 | 176.5236018 | 257.4728947 | 237.0602981 | 321.4186227 | 0.003573014 | 0.592894826 | YHL010C | ETP1 | n |
| W303.asm02.HP0_G0023020 | 442.9696126 | 370.2983254 | 470.0822192 | 981.5742104 | 930.1788874 | 972.2376837 | 1401.902048 | 810.8429788 | 1004.177289 | 1.40E-08 | -1.168483086 | YHL009C | YAP3 | n |
| W303.asm02.HP0_G0023030 | 254.9256195 | 139.9327862 | 191.5511746 | 586.609761 | 712.5621434 | 644.5374588 | 506.4264656 | 676.3475443 | 598.8213194 | 1.18E-14 | -1.728359928 | YHL008C | | n |
| W303.asm02.HP0_G0023040 | 710.4961182 | 961.442953 | 599.0860714 | 2370.759515 | 2489.323242 | 2445.983446 | 1738.888631 | 2208.047348 | 2409.616044 | 1.10E-18 | -1.685534477 | YHL007C | STE20 | n |
| W303.asm02.HP0_G0023050 | 312.1142566 | 246.5482424 | 282.4402523 | 56.42349277 | 92.8851956 | 58.8412006 | 178.9057981 | 77.40744427 | 105.4334973 | 8.00E-10 | 2.025630101 | YHL006C | SHU1 | n |
| W303.asm02.HP0_G0023060 | 1410.329948 | 1091.856502 | 1212.831672 | 3400.974668 | 3472.579384 | 3407.358139 | 2995.015584 | 3048.885711 | 2921.429138 | 1.03E-32 | -1.46852607 | YHL004W | MRP4 | n |
| W303.asm02.HP0_G0023070 | 3052.322341 | 2519.742076 | 3000.316867 | 1056.481261 | 1064.198955 | 1131.56155 | 1233.408756 | 1294.639505 | 1197.642639 | 7.81E-30 | 1.397838742 | YHL003C | LAG1 | n |
| W303.asm02.HP0_G0023080 | 1572.20287 | 613.038873 | 1278.3109 | 927.0963553 | 690.0043102 | 878.0917627 | 780.9380078 | 931.7921105 | 1123.941553 | 0.113817088 | 0.472393304 | YHL002W | HSE1 | n |
| W303.asm02.HP0_G0023090 | 3693.028936 | 3855.291049 | 3493.854332 | 2838.685378 | 3195.250729 | 2630.654291 | 2378.784501 | 2198.371417 | 2038.039738 | 0.005814504 | 0.350507975 | YHL001W | RPL14B | y |
| W303.asm02.HP0_G0023100 | 2091.747098 | 1351.731676 | 1982.945577 | 1750.101095 | 1719.70305 | 1840.371705 | 1535.371453 | 2179.987149 | 1808.747472 | 0.882928234 | 0.031052144 | YHR001W | OSH7 | n |
| W303.asm02.HP0_G0023120 | 985.7770153 | 502.6157219 | 806.2740766 | 273.3620943 | 325.0981846 | 261.6170303 | 273.5649506 | 376.3936978 | 482.1279341 | 1.87E-07 | 1.417378865 | YHR002W | LEU5 | n |
| W303.asm02.HP0_G0023140 | 851.0444635 | 583.5292378 | 796.5010575 | 350.2147827 | 335.7136355 | 353.9524528 | 407.0343556 | 491.5372711 | 581.4196742 | 2.66E-08 | 1.100592826 | YHR004C | NEM1 | n |
| W303.asm02.HP0_G0023150 | 3174.454007 | 2803.415343 | 2859.585392 | 15.56510145 | 25.21169595 | 12.67348936 | 20.82501354 | 30.96297771 | 34.80329036 | 4.82E-177 | 7.424148044 | YHR005C | GPA1 | n |
| W303.asm02.HP0_G0023160 | 3179.300502 | 4001.887301 | 3586.698013 | 2017.626276 | 2300.898988 | 2042.242285 | 2501.841399 | 1857.778663 | 1759.613415 | 5.80E-06 | 0.759859947 | YHR005C- | TIM10 | n |
| W303.asm02.HP0_G0023180 | 6263.609708 | 6107.542561 | 6062.203754 | 5981.863052 | 5112.666552 | 5795.405634 | 3492.922726 | 4151.941792 | 3910.252034 | 0.309842828 | 0.125846242 | YHR007C | ERG11 | n |
| W303.asm02.HP0_G0023210 | 3802.559716 | 2707.271048 | 3890.638908 | 3117.884385 | 3144.827337 | 3099.573398 | 3906.583222 | 2936.644917 | 2944.97254 | 0.416502984 | 0.151708019 | YHR008C | SOD2 | n |
| W303.asm02.HP0_G0023220 | 1051.689343 | 1184.193102 | 935.2779288 | 3292.991776 | 2856.883231 | 3229.929288 | 1820.295502 | 2793.441145 | 2900.956614 | 1.09E-16 | -1.564697964 | YHR009C | TDA3 | n |

| | | | | | | | | | | | | | | |
|---|---|---|---|---|---|---|---|---|---|---|---|---|---|---|
| W303.asm02.HP0_G0023240 | 494.342456 | 318.8944448 | 421.2171236 | 738.3695002 | 639.5809183 | 691.6104193 | 818.8016687 | 819.5513163 | 777.9559021 | 2.88E-05 | -0.746512102 | YHR011W | DIA4 | n |
| W303.asm02.HP0_G0023260 | 2188.676992 | 1784.856967 | 2195.020092 | 1007.840319 | 891.6978778 | 997.5846624 | 1457.750948 | 1009.199555 | 1088.114637 | 2.39E-10 | 1.089577737 | YHR013C | ARD1 | n |
| W303.asm02.HP0_G0023300 | 873.338339 | 781.5293707 | 775.9777173 | 893.0476959 | 911.6018483 | 925.1647232 | 955.1108483 | 942.435634 | 1134.177815 | 0.231262694 | -0.167380503 | YHR017W | YSC83 | n |
| W303.asm02.HP0_G0023310 | 5539.543405 | 3869.569905 | 4765.324118 | 1617.797732 | 1406.547248 | 1744.415285 | 2569.049398 | 1564.597967 | 2536.545691 | 7.18E-14 | 1.571129822 | YHR018C | ARG4 | n |
| W303.asm02.HP0_G0023320 | 3358.620804 | 2666.338328 | 2693.444067 | 8181.406451 | 8664.861818 | 8073.012722 | 7271.716092 | 7172.767305 | 6563.491111 | 9.68E-33 | -1.515050084 | YHR019C | DED81 | n |
| W303.asm02.HP0_G0023330 | 3174.454007 | 2891.944249 | 2917.246204 | 6609.331205 | 6831.042671 | 6575.730479 | 5231.811356 | 5820.072216 | 5677.030833 | 2.25E-30 | -1.155700826 | YHR020W | | n |
| W303.asm02.HP0_G0023350 | 273.3422992 | 121.8462356 | 223.8021376 | 2093.506145 | 1942.627519 | 2034.095042 | 666.4004333 | 804.0698274 | 581.4196742 | 1.89E-41 | -3.294261851 | YHR022C | | n |
| W303.asm02.HP0_G0023360 | 727.943499 | 1148.020001 | 890.3220409 | 1920.344392 | 2215.975381 | 2008.748063 | 1700.078378 | 1874.227744 | 1652.132666 | 3.83E-11 | -1.151023792 | YHR023W | MYO1 | n |
| W303.asm02.HP0_G0023370 | 1261.057912 | 1254.635457 | 1071.122894 | 1422.261145 | 1320.296709 | 1406.757319 | 1201.224645 | 1261.741342 | 1355.281072 | 0.103598316 | -0.210658401 | YHR024C | MAS2 | n |
| W303.asm02.HP0_G0023380 | 1060.413033 | 1121.366137 | 873.7079084 | 2649.958522 | 2393.784184 | 2635.180537 | 3101.033834 | 1831.65365 | 2362.52924 | 7.09E-12 | -1.329631173 | YHR025W | THR1 | n |
| W303.asm02.HP0_G0023390 | 3283.984786 | 2857.674995 | 3232.914722 | 1600.286993 | 1536.586522 | 1570.607431 | 2668.441508 | 2225.464023 | 2060.559514 | 1.77E-13 | 0.993599878 | YHR026W | VMA16 | n |
| W303.asm02.HP0_G0023400 | 3415.809441 | 2943.348129 | 3082.410227 | 4893.278769 | 4284.66138 | 4690.096312 | 4299.418705 | 5485.28502 | 5672.936328 | 0.000117808 | -0.554874522 | YHR027C | RPN1 | n |
| W303.asm02.HP0_G0023410 | 2085.931305 | 1669.674198 | 2052.334013 | 690.701377 | 451.1566644 | 654.4952005 | 697.6379536 | 881.4772717 | 902.8382969 | 1.75E-18 | 1.690116829 | YHR028C | DAP2 | n |
| W303.asm02.HP0_G0023430 | 1355.079909 | 1048.068011 | 1334.99441 | 1057.45408 | 700.6197611 | 942.3644588 | 1055.44955 | 1292.704319 | 1361.422829 | 0.017024084 | 0.46737937 | YHR030C | SLT2 | n |
| W303.asm02.HP0_G0023440 | 423.5836339 | 432.173367 | 364.5336128 | 958.2265582 | 956.7175147 | 986.7216715 | 621.9106316 | 711.1808943 | 804.570183 | 6.69E-17 | -1.249689466 | YHR031C | RRM3 | n |
| W303.asm02.HP0_G0023450 | 703.7110257 | 850.0678782 | 642.0873555 | 4558.629088 | 2781.248143 | 4280.923656 | 1095.206394 | 1941.959258 | 1332.761295 | 1.43E-23 | -2.404119823 | YHR032W | ERC1 | n |
| W303.asm02.HP0_G0023470 | 321.8072459 | 243.6924712 | 269.7353274 | 520.4580798 | 575.8882127 | 541.3390455 | 570.7946893 | 517.6622836 | 501.5768316 | 1.75E-09 | -0.970568495 | YHR034C | PIH1 | n |
| W303.asm02.HP0_G0023480 | 252.9870216 | 309.3752076 | 248.2346854 | 401.7741813 | 536.0802718 | 365.7206929 | 542.3969436 | 706.342929 | 576.3015433 | 0.000976712 | -0.681564489 | YHR035W | NEL1 | n |
| W303.asm02.HP0_G0023490 | 561.2240825 | 549.259984 | 542.4025606 | 635.2507031 | 635.6001242 | 612.8537354 | 508.3196487 | 823.4216885 | 830.1608376 | 0.363438018 | -0.188409475 | YHR036W | BRL1 | n |
| W303.asm02.HP0_G0023500 | 2063.637429 | 1506.895242 | 1677.050079 | 600.2292248 | 688.6773788 | 548.5810394 | 1393.382724 | 2027.107447 | 1708.432106 | 4.43E-16 | 1.515507982 | YHR037W | PUT2 | n |
| W303.asm02.HP0_G0023510 | 694.9873353 | 561.6349923 | 781.8415288 | 873.5913191 | 808.1012018 | 915.2069816 | 752.540262 | 748.9170234 | 576.3015433 | 0.052438903 | -0.35016475 | YHR038W | RRF1 | n |
| W303.asm02.HP0_G0023520 | 2130.519056 | 2073.289853 | 1875.442367 | 2351.303138 | 2392.457252 | 2315.627556 | 2081.554762 | 2689.908689 | 2453.63197 | 0.113817088 | -0.215509797 | YHR039C | MSC7 | n |
| W303.asm02.HP0_G0023530 | 2152.812931 | 2529.261313 | 2286.886472 | 1142.089319 | 1157.084151 | 1156.908529 | 1642.336295 | 1054.676428 | 987.7992704 | 1.66E-07 | 1.01184013 | YHR039C- | VMA10 | n |
| W303.asm02.HP0_G0023540 | 333.4388331 | 329.3656057 | 262.8942141 | 961.1450147 | 818.7166527 | 988.53217 | 612.4447164 | 685.0558818 | 536.380122 | 7.78E-21 | -1.581484441 | YHR040W | BCD1 | n |
| W303.asm02.HP0_G0023560 | 827.7812891 | 974.769885 | 627.4278269 | 4357.255588 | 3631.811148 | 4095.347561 | 3500.495458 | 4510.918815 | 4198.914619 | 2.71E-38 | -2.314188339 | YHR042W | NCP1 | n |
| W303.asm02.HP0_G0023570 | 1514.044934 | 1138.500764 | 1520.681774 | 2.918456523 | 7.961588195 | 5.43149544 | 76.67391349 | 14.5138958 | 89.05547827 | 3.82E-27 | 8.028018046 | YHR043C | DOG2 | n |
| W303.asm02.HP0_G0023590 | 964.4524388 | 991.9045119 | 940.1644384 | 555.4795581 | 595.7921832 | 574.8332674 | 482.7616775 | 620.2271472 | 560.9471505 | 8.68E-08 | 0.74757767 | YHR045W | DDE1 | n |
| W303.asm02.HP0_G0023640 | 3643.594691 | 3477.377334 | 3661.95026 | 573.9631161 | 598.446046 | 607.42224 | 411.7673132 | 457.6715143 | 280.4735752 | 9.73E-53 | 2.598984803 | YHR049W | FSH1 | n |
| W303.asm02.HP0_G0023650 | 1119.540268 | 1128.981527 | 1138.556726 | 760.7443335 | 743.0815648 | 740.4938783 | 657.8811096 | 966.6254604 | 1181.26462 | 0.003018307 | 0.593722651 | YHR050W | SMF2 | n |
| W303.asm02.HP0_G0023670 | 3035.844259 | 3473.569639 | 3071.659906 | 8510.21922 | 8541.457201 | 8193.410871 | 7373.001385 | 5819.104623 | 5362.777594 | 1.33E-22 | -1.397695301 | YHR051W | COX6 | n |
| W303.asm02.HP0_G0023680 | 3296.585673 | 4769.137816 | 3527.082597 | 8484.92593 | 7961.588195 | 8226.905092 | 3665.202383 | 3900.367598 | 3157.886787 | 1.50E-11 | -1.089731695 | YHR052W | CIC1 | n |
| W303.asm02.HP0_G0023860 | 668.8162641 | 396.9521895 | 612.7682982 | 488.3550581 | 498.9261935 | 522.3288114 | 771.4720925 | 621.1947403 | 670.4751525 | 0.48606381 | 0.152670447 | YHR058C | MED6 | n |
| W303.asm02.HP0_G0023870 | 1223.255254 | 999.5199016 | 1262.674069 | 1053.562805 | 1038.987259 | 1077.246596 | 1222.99625 | 965.6578673 | 812.7591925 | 0.465405692 | 0.136738442 | YHR059W | FYV4 | n |
| W303.asm02.HP0_G0023880 | 814.2111041 | 845.3082596 | 861.9802855 | 283.0902827 | 307.8480769 | 343.0894619 | 409.8741301 | 289.310323 | 270.2373134 | 2.13E-14 | 1.43207475 | YHR060W | VMA22 | n |
| W303.asm02.HP0_G0023900 | 1096.277094 | 848.1640308 | 1055.486064 | 835.6513843 | 931.5058188 | 806.5770728 | 979.7222279 | 931.7921105 | 851.6569876 | 0.153248036 | 0.222200786 | YHR062C | RPP1 | n |
| W303.asm02.HP0_G0023910 | 2591.905348 | 2385.520832 | 2454.005099 | 1231.588652 | 858.5245937 | 1289.980167 | 851.9323721 | 622.1623334 | 689.92405 | 6.69E-10 | 1.134878538 | YHR063C | PAN5 | n |
| W303.asm02.HP0_G0023920 | 7858.106453 | 6894.783474 | 7864.348478 | 10121.20722 | 10579.62378 | 9966.794132 | 8706.748843 | 8790.58289 | 8416.254509 | 1.85E-05 | -0.439217334 | YHR064C | SSZ1 | n |
| W303.asm02.HP0_G0023930 | 2519.207928 | 2148.491827 | 2426.640645 | 4162.69182 | 3569.445374 | 4226.608701 | 1663.161309 | 2126.769531 | 1683.865078 | 2.91E-07 | -0.753686266 | YHR065C | RRP3 | n |
| W303.asm02.HP0_G0023950 | 595.1495451 | 463.5868496 | 627.4278269 | 182.8899421 | 161.8856266 | 182.8603465 | 163.7603337 | 169.3287844 | 225.1977611 | 1.71E-16 | 1.673332705 | YHR067W | HTD2 | n |
| W303.asm02.HP0_G0023960 | 3774.450047 | 4058.0508 | 3940.481305 | 3791.075023 | 4075.006224 | 3708.806136 | 3366.079461 | 2964.705116 | 2817.019267 | 0.842930897 | 0.024797653 | YHR068W | DYS1 | n |
| W303.asm02.HP0_G0023980 | 560.2547835 | 743.452422 | 426.1036332 | 2097.397421 | 2153.609607 | 2226.007881 | 2059.783157 | 1955.505561 | 2162.922133 | 8.48E-25 | -1.904500936 | YHR070W | TRM5 | n |
| W303.asm02.HP0_G0023990 | 1795.141625 | 2132.309123 | 2029.856069 | 1219.914826 | 1106.660759 | 1194.928997 | 1083.847296 | 423.8057574 | 688.9004238 | 0.004597548 | 0.758290862 | YHR071W | PCL5 | n |
| W303.asm02.HP0_G0024000 | 1829.067087 | 1552.58758 | 1438.588413 | 1226.724558 | 1138.507112 | 1261.012191 | 1001.493833 | 1131.116279 | 1112.681665 | 0.003290451 | 0.409928037 | YHR072W | ERG7 | n |
| W303.asm02.HP0_G0024010 | 2069.453223 | 2648.251777 | 2189.156281 | 2272.504812 | 2725.517025 | 2320.153802 | 2042.74451 | 1375.917322 | 1196.619013 | 0.708999439 | -0.083122588 | YHR072W- | NOP10 | n |
| W303.asm02.HP0_G0024020 | 824.8733923 | 959.5391055 | 846.3434549 | 1882.404457 | 1787.37655 | 1902.833902 | 1868.571669 | 2549.607696 | 2468.986363 | 1.04E-12 | -1.083058634 | YHR073W | OSH3 | n |
| W303.asm02.HP0_G0024030 | 1115.663072 | 940.5006312 | 987.0749301 | 2249.15716 | 1896.184922 | 2223.292133 | 1772.965925 | 1780.371218 | 1871.18867 | 9.23E-17 | -1.066244093 | YHR074W | QNS1 | n |
| W303.asm02.HP0_G0024040 | 303.3905662 | 231.3174629 | 287.3267618 | 271.4164566 | 269.3670673 | 292.3955045 | 323.7343014 | 353.1714645 | 491.3405467 | 0.929381711 | -0.020399426 | YHR075C | PPE1 | n |
| W303.asm02.HP0_G0024050 | 1028.426168 | 791.0486078 | 987.0749301 | 798.6842683 | 858.5245937 | 816.5348144 | 967.4165381 | 1132.083873 | 998.0355323 | 0.24016668 | 0.182571317 | YHR076W | PTC7 | n |
| W303.asm02.HP0_G0024060 | 1258.150016 | 1707.751146 | 1427.838092 | 1763.720558 | 2048.782029 | 1872.960677 | 1427.460019 | 1693.287844 | 1420.793148 | 0.018751855 | -0.371312401 | YHR077C | NMD2 | n |
| W303.asm02.HP0_G0024070 | 682.3864491 | 646.356203 | 739.8175466 | 468.8986813 | 533.426409 | 492.4555865 | 697.6379536 | 800.1994552 | 700.1603119 | 0.0008801 | 0.470371637 | YHR078W | | n |
| W303.asm02.HP0_G0024080 | 309.2063598 | 360.7790883 | 298.0770829 | 870.6728625 | 922.2172992 | 878.997012 | 829.2141755 | 1077.898662 | 1051.264094 | 1.90E-20 | -1.464032593 | YHR079C | IRE1 | n |
| W303.asm02.HP0_G0024090 | 9.692989334 | 7.615389726 | 4.886509555 | 9.728188408 | 9.28851956 | 15.38923708 | 36.91706946 | 39.67131519 | 22.51977611 | 0.335513829 | -0.645939556 | YHR079C- | SAE3 | n |
| W303.asm02.HP0_G0024110 | 1326.000941 | 1687.760748 | 1366.268072 | 1083.720189 | 943.4482011 | 1072.720349 | 789.4573315 | 648.2873458 | 596.774067 | 0.002541209 | 0.497944069 | YHR081W | LRP1 | n |
| W303.asm02.HP0_G0024120 | 1164.128019 | 1163.250781 | 1104.351159 | 1449.500073 | 1658.664207 | 1471.935264 | 1380.130443 | 1967.116678 | 1672.60519 | 0.008159152 | -0.415793695 | YHR082C | KSP1 | n |
| W303.asm02.HP0_G0024130 | 599.9960398 | 469.2983919 | 543.3798625 | 374.5352537 | 360.9253315 | 383.8256777 | 456.2571148 | 417.032606 | 583.4669266 | 0.004038947 | 0.526029505 | YHR083W | SAM35 | n |
| W303.asm02.HP0_G0024140 | 446.8468083 | 617.7984916 | 483.7644459 | 575.9087538 | 850.5630055 | 582.9805105 | 1048.823409 | 1114.667198 | 1431.029409 | 0.091194559 | -0.373517327 | YHR084W | STE12 | n |
| W303.asm02.HP0_G0024150 | 1186.421895 | 1823.885839 | 1173.739595 | 2017.626276 | 1409.20111 | 1952.622611 | 645.5754197 | 822.4540954 | 596.774067 | 0.113674901 | -0.36314033 | YHR085W | IPI1 | n |
| W303.asm02.HP0_G0024160 | 693.0487374 | 635.8850422 | 634.2689402 | 352.1604204 | 293.2518318 | 321.3634802 | 69.10118129 | 323.1760798 | 424.8048676 | 0.092113888 | 1.021757854 | YHR086W | NAM8 | n |
| W303.asm02.HP0_G0024180 | 2048.128646 | 446.4522227 | 1437.611111 | 30.15738407 | 31.84635278 | 19.01023404 | 166.6001083 | 48.37965267 | 131.0241519 | 3.45E-16 | 5.605276509 | YHR087W | RTC3 | n |
| W303.asm02.HP0_G0024190 | 1808.71181 | 2194.184165 | 1895.965707 | 3107.183378 | 2622.016379 | 3156.6041 | 1459.644131 | 1529.764617 | 1240.634939 | 5.12E-05 | -0.591533293 | YHR088W | RPF1 | n |
| W303.asm02.HP0_G0024200 | 4246.498627 | 5206.070802 | 4660.752814 | 7813.68093 | 8629.034672 | 7612.240859 | 5641.685486 | 4864.09028 | 4320.726135 | 6.31E-08 | -0.76913047 | YHR089C | GAR1 | n |
| W303.asm02.HP0_G0024210 | 923.7418836 | 718.7024054 | 807.2513785 | 461.1161306 | 518.830164 | 471.634854 | 868.9710195 | 564.1067502 | 644.8844978 | 8.94E-05 | 0.755973959 | YHR090C | YNG2 | n |
| W303.asm02.HP0_G0024230 | 1586.742354 | 1561.154894 | 1527.522887 | 275.307732 | 453.8105271 | 315.9319847 | 336.9865827 | 323.1760798 | 267.1664348 | 2.67E-29 | 2.167233887 | YHR092C | HXT4 | n |

| | | | | | | | | | | | | | | |
|---|---|---|---|---|---|---|---|---|---|---|---|---|---|---|
| W303.asm02.HP0_G0024260 | 146.3641389 | 144.6924048 | 133.8903618 | 3210.302175 | 4109.50644 | 3383.821659 | 1536.318044 | 3529.779459 | 2399.379782 | 2.34E-74 | -4.654435767 | YHR096C | HXT5 | n |
| W303.asm02.HP0_G0024280 | 1036.18056 | 1374.577846 | 1068.190989 | 1617.797732 | 1674.587384 | 1640.311623 | 1443.552075 | 1528.797024 | 1494.494233 | 0.000382701 | -0.503462787 | YHR098C | SFB3 | n |
| W303.asm02.HP0_G0024290 | 596.1188441 | 947.1640972 | 615.7002039 | 2824.093095 | 3192.596866 | 3155.69885 | 2352.279939 | 2210.950127 | 2319.53694 | 7.68E-31 | -2.086660032 | YHR099W | TRA1 | n |
| W303.asm02.HP0_G0024300 | 488.5266624 | 576.8657718 | 534.5841453 | 590.5010364 | 658.1579574 | 679.8421792 | 873.7039772 | 651.190125 | 694.0185548 | 0.113552999 | -0.269185088 | YHR100C | GEP4 | n |
| W303.asm02.HP0_G0024320 | 277.219495 | 480.7214765 | 236.5070625 | 841.4882973 | 943.4482011 | 854.5552825 | 974.0426788 | 984.0421353 | 1160.792096 | 3.56E-09 | -1.407532463 | YHR102W | KIC1 | n |
| W303.asm02.HP0_G0024330 | 888.847122 | 1176.577713 | 864.9121912 | 2223.86387 | 2381.841802 | 2266.744097 | 2411.915205 | 2160.635288 | 2624.577543 | 1.34E-15 | -1.229434366 | YHR103W | SBE22 | n |
| W303.asm02.HP0_G0024340 | 2928.252078 | 1880.049339 | 2520.461628 | 513.648348 | 358.2714688 | 482.4978449 | 1472.896412 | 903.7319119 | 1159.76847 | 1.13E-26 | 2.433667268 | YHR104W | GRE3 | n |
| W303.asm02.HP0_G0024370 | 1296.921973 | 1408.847099 | 1170.807689 | 4652.992516 | 4434.604624 | 4472.836494 | 4293.739155 | 4417.062289 | 4350.411295 | 1.18E-61 | -1.806526052 | YHR107C | CDC12 | n |
| W303.asm02.HP0_G0024380 | 2029.711967 | 1807.703136 | 1968.286049 | 2750.158863 | 2103.186215 | 2750.147191 | 2112.792283 | 2519.612311 | 2403.474287 | 0.009560698 | -0.390045906 | YHR108W | GGA2 | n |
| W303.asm02.HP0_G0024390 | 191.9211888 | 242.7405475 | 175.914344 | 461.1161306 | 455.1374585 | 429.0881397 | 330.3604421 | 409.2918616 | 328.584006 | 1.01E-09 | -1.138926684 | YHR109W | CTM1 | n |
| W303.asm02.HP0_G0024400 | 1204.838574 | 1214.654661 | 1142.465934 | 146.895645 | 169.8472148 | 143.0293799 | 219.6092337 | 205.1297273 | 200.6307326 | 2.15E-89 | 2.960725352 | YHR110W | ERP5 | n |
| W303.asm02.HP0_G0024410 | 1285.290386 | 944.3083261 | 1269.515182 | 1207.268181 | 1068.179749 | 1225.707471 | 505.4798741 | 899.8615397 | 808.6646877 | 0.995773659 | -0.00131093 | YHR111W | UBA4 | n |
| W303.asm02.HP0_G0024420 | 670.7548619 | 487.3849425 | 615.7002039 | 864.8359495 | 887.7170837 | 984.0059238 | 1035.571128 | 896.9587605 | 941.736092 | 9.03E-05 | -0.626100436 | YHR112C | | n |
| W303.asm02.HP0_G0024430 | 1215.500863 | 936.6929363 | 1168.853086 | 2682.061544 | 2513.208007 | 2731.136957 | 2051.263834 | 2346.413155 | 2017.567215 | 2.65E-19 | -1.255392634 | YHR113W | APE4 | n |
| W303.asm02.HP0_G0024440 | 654.2767801 | 574.9619243 | 685.0886396 | 674.1634567 | 672.7542024 | 611.0432369 | 774.3118671 | 772.1392566 | 763.6251355 | 0.825690881 | -0.031738435 | YHR114W | BZZ1 | n |
| W303.asm02.HP0_G0024450 | 546.6845985 | 535.933052 | 501.3558803 | 1388.212486 | 1520.663345 | 1394.989079 | 1116.031407 | 1547.181292 | 1746.306275 | 1.77E-16 | -1.441684287 | YHR115C | DMA1 | n |
| W303.asm02.HP0_G0024460 | 623.2592142 | 537.8368994 | 637.200846 | 441.6597537 | 360.9253315 | 378.3941823 | 444.8980165 | 328.0140451 | 333.702137 | 0.000951675 | 0.60517612 | YHR116W | COX23 | n |
| W303.asm02.HP0_G0024480 | 542.8074027 | 765.3466675 | 600.0633734 | 214.020145 | 203.020499 | 223.5965623 | 28.39774574 | 177.0695288 | 183.2290875 | 0.013032271 | 1.574485075 | YHR118C | ORC6 | n |
| W303.asm02.HP0_G0024490 | 404.1976552 | 564.4907635 | 409.4895007 | 936.8245437 | 956.7175147 | 937.8382126 | 673.9731655 | 925.0189591 | 857.7987447 | 1.67E-08 | -1.038412171 | YHR119W | SET1 | n |
| W303.asm02.HP0_G0024500 | 408.074851 | 644.4523556 | 468.1276154 | 1066.20945 | 1129.218592 | 1059.141611 | 760.1129942 | 1017.907892 | 767.7196402 | 3.14E-08 | -1.097321942 | YHR120W | MSH1 | n |
| W303.asm02.HP0_G0024510 | 1995.786504 | 2161.818759 | 2073.834655 | 1392.103761 | 1297.738876 | 1320.758641 | 1313.869036 | 1109.829232 | 891.5784089 | 4.60E-05 | 0.635553133 | YHR121W | LSM12 | n |
| W303.asm02.HP0_G0024520 | 1647.808187 | 1374.577846 | 1574.433379 | 465.9802248 | 440.5412134 | 461.6771124 | 674.919757 | 572.8150876 | 618.2702169 | 1.72E-37 | 1.747351358 | YHR122W | CIA2 | n |
| W303.asm02.HP0_G0024540 | 322.7765448 | 253.2117084 | 308.8274039 | 67.12450002 | 62.36577419 | 85.99867779 | 21.77160506 | 79.34263038 | 49.13405697 | 2.01E-08 | 2.033582591 | YHR124W | NDT80 | n |
| W303.asm02.HP0_G0024560 | 960.575243 | 810.0870821 | 913.7772868 | 828.8416524 | 859.851525 | 800.2403281 | 1040.304085 | 924.051366 | 888.5075303 | 0.440994716 | 0.109722719 | YHR127W | | n |
| W303.asm02.HP0_G0024570 | 3555.388488 | 3504.983122 | 3671.72328 | 3275.481037 | 4309.873076 | 3474.346583 | 3133.217946 | 2880.52452 | 2064.654019 | 0.822868078 | -0.04299538 | YHR128W | FUR1 | n |
| W303.asm02.HP0_G0024580 | 260.7414131 | 231.3174629 | 219.89293 | 286.981558 | 322.4443219 | 296.9217507 | 361.5979624 | 330.9168243 | 322.4422489 | 0.040613376 | -0.346155781 | YHR129C | ARP1 | n |
| W303.asm02.HP0_G0024590 | 388.6888723 | 412.1829689 | 358.6698013 | 480.5725074 | 465.7529094 | 471.634854 | 645.5754197 | 493.4724572 | 507.7185887 | 0.083033329 | -0.290422565 | YHR131C | | n |
| W303.asm02.HP0_G0024600 | 2266.220906 | 1686.808824 | 2136.381977 | 898.8846089 | 1040.314191 | 945.0802065 | 1350.786106 | 1420.426602 | 1353.233819 | 1.46E-13 | 1.079030483 | YHR132C | ECM14 | n |
| W303.asm02.HP0_G0024610 | 1188.360492 | 888.1448268 | 1102.396556 | 412.4751885 | 394.0986156 | 430.8968382 | 592.5662944 | 448.9631768 | 508.7422149 | 1.32E-15 | 1.360023047 | YHR132W- | IGO2 | n |
| W303.asm02.HP0_G0024630 | 510.8205379 | 491.1926374 | 555.1074854 | 278.2261885 | 281.3094495 | 312.3109878 | 184.5853473 | 240.9306703 | 257.9537991 | 2.32E-06 | 0.835457167 | YHR134W | WSS1 | n |
| W303.asm02.HP0_G0024650 | 682.3864491 | 1023.317994 | 660.6560918 | 12.64664493 | 2.653862732 | 15.38923708 | 17.03864744 | 12.57870969 | 17.40164518 | 5.50E-63 | 6.186773474 | YHR136C | SPL2 | n |
| W303.asm02.HP0_G0024660 | 1250.395624 | 681.5773805 | 994.8933454 | 573.9631161 | 683.3696534 | 591.1277537 | 992.0279177 | 1051.773649 | 959.1377372 | 0.001771637 | 0.663994346 | YHR137W | ARO9 | n |
| W303.asm02.HP0_G0024670 | 757.022467 | 748.2120406 | 725.158018 | 121.6023551 | 141.9816561 | 107.7246596 | 289.6570065 | 166.4260052 | 231.3395183 | 3.96E-32 | 2.592829915 | YHR138C | | n |
| W303.asm02.HP0_G0024690 | 221.0001568 | 140.8847099 | 185.6873631 | 595.3651306 | 514.8493699 | 575.7385166 | 462.8832555 | 529.2734002 | 528.1911125 | 1.30E-17 | -1.623980265 | YHR140W | | n |
| W303.asm02.HP0_G0024710 | 2238.111237 | 1528.789488 | 2157.882619 | 1440.744703 | 1276.507974 | 1381.41034 | 1556.196466 | 1547.181292 | 1410.556886 | 0.001431024 | 0.531098856 | YHR142W | CHS7 | n |
| W303.asm02.HP0_G0024720 | 5594.793444 | 4704.407003 | 5960.564355 | 2227.755146 | 2262.417979 | 2197.039905 | 2313.469686 | 2348.348341 | 1869.141417 | 9.18E-20 | 1.281905987 | YHR143W | DSE2 | n |
| W303.asm02.HP0_G0024730 | 1780.602141 | 2239.876503 | 1879.351575 | 1560.401421 | 1674.587384 | 1520.818753 | 1638.549929 | 1132.083873 | 1018.508056 | 0.111924612 | 0.311289211 | YHR143W- | RPC10 | n |
| W303.asm02.HP0_G0024740 | 1039.088457 | 969.0583427 | 979.2565148 | 954.3352829 | 1041.641122 | 976.7639299 | 1418.940695 | 1184.333897 | 1289.768996 | 0.953089235 | 0.007930706 | YHR144C | DCD1 | n |
| W303.asm02.HP0_G0024750 | 1748.615276 | 1170.86617 | 1859.805537 | 5426.383494 | 5163.089944 | 5322.865531 | 5048.1726 | 5548.178568 | 5141.674337 | 1.46E-24 | -1.735410962 | YHR146W | CRP1 | n |
| W303.asm02.HP0_G0024760 | 1376.404485 | 1664.914579 | 1411.223959 | 2166.467559 | 1925.377412 | 2194.324158 | 2481.962977 | 1964.213898 | 1994.023812 | 0.00085845 | -0.497986564 | YHR147C | MRPL6 | n |
| W303.asm02.HP0_G0024770 | 1226.163151 | 1903.847432 | 1350.631241 | 1750.101095 | 1587.009913 | 1720.878805 | 1116.031407 | 1073.060696 | 867.0113804 | 0.374590702 | -0.175048232 | YHR148W | IMP3 | n |
| W303.asm02.HP0_G0024780 | 757.022467 | 862.4428865 | 782.8188307 | 1582.776254 | 1508.720963 | 1651.174614 | 1295.883797 | 1604.269283 | 2137.331478 | 1.00E-07 | -0.981507008 | YHR149C | SKG6 | n |
| W303.asm02.HP0_G0024790 | 357.6713064 | 330.3175294 | 337.1691593 | 225.6939711 | 276.0017241 | 248.943541 | 417.4468623 | 348.3334992 | 364.4109226 | 0.007517546 | 0.453091139 | YHR150W | PEX28 | n |
| W303.asm02.HP0_G0024800 | 379.9651819 | 565.4426872 | 399.7164816 | 318.111761 | 309.1750082 | 302.3532461 | 133.469405 | 340.5927548 | 341.8911465 | 0.098240651 | 0.533103198 | YHR151C | MTC6 | n |
| W303.asm02.HP0_G0024830 | 929.5576772 | 1121.366187 | 985.1203263 | 675.1362765 | 617.0230851 | 717.8626473 | 536.7173494 | 594.1021348 | 535.3564958 | 2.57E-05 | 0.593373685 | YHR154W | RTT107 | n |
| W303.asm02.HP0_G0024850 | 629.0750078 | 604.4715595 | 600.0633734 | 458.197674 | 577.2151441 | 533.1918023 | 525.3582961 | 440.2548393 | 435.0411295 | 0.156866705 | 0.227133947 | YHR156C | LIN1 | n |
| W303.asm02.HP0_G0024860 | 407.105552 | 607.3273307 | 351.828688 | 1506.896384 | 1354.796924 | 1530.776465 | 1223.942841 | 1519.121094 | 1501.659616 | 9.84E-17 | -1.684870327 | YHR158C | KEL1 | n |
| W303.asm02.HP0_G0024870 | 282.0659896 | 276.0578776 | 266.8034217 | 336.5953189 | 391.4447529 | 361.1944467 | 542.3969436 | 434.449281 | 518.9784768 | 0.016131338 | -0.398913983 | YHR159W | TDA11 | n |
| W303.asm02.HP0_G0024890 | 891.7550188 | 675.8658382 | 801.387567 | 2079.886682 | 1645.394894 | 2062.157769 | 1712.384068 | 1823.912906 | 2451.584718 | 3.17E-12 | -1.289313009 | YHR161C | YAP1801 | n |
| W303.asm02.HP0_G0024900 | 5168.301913 | 5599.215296 | 5238.338243 | 2494.307508 | 2968.345465 | 2570.907841 | 5631.27298 | 3445.598863 | 3916.393791 | 5.68E-08 | 0.994810971 | YHR162W | MPC2 | n |
| W303.asm02.HP0_G0024910 | 2940.852964 | 2999.511628 | 3132.252625 | 1112.904754 | 889.0440151 | 1071.8151 | 1350.786106 | 826.3244676 | 988.8228966 | 3.19E-17 | 1.560543189 | YHR163W | SOL3 | n |
| W303.asm02.HP0_G0024920 | 244.2633312 | 342.6925377 | 228.6886472 | 614.8215074 | 624.9846733 | 582.9805105 | 598.2458435 | 654.0929041 | 672.5224048 | 1.40E-10 | -1.159225958 | YHR164C | DNA2 | n |
| W303.asm02.HP0_G0024930 | 543.7767017 | 855.7794205 | 559.993995 | 1207.268181 | 1393.277934 | 1259.201693 | 974.0426788 | 1231.745957 | 1197.642639 | 7.99E-07 | -0.97737357 | YHR165C | PRP8 | n |
| W303.asm02.HP0_G0024940 | 382.8730787 | 419.7983587 | 411.4441045 | 321.0302175 | 356.9445374 | 319.5529817 | 351.1854556 | 350.2686853 | 340.8675203 | 0.055101305 | 0.286240911 | YHR166C | CDC23 | n |
| W303.asm02.HP0_G0024950 | 487.5573635 | 581.6253904 | 516.9927109 | 525.3221741 | 488.3107426 | 573.9280181 | 628.5367723 | 502.1807947 | 536.380122 | 0.986511404 | -0.002876446 | YHR167W | THP2 | n |
| W303.asm02.HP0_G0024960 | 357.6713064 | 407.4233504 | 309.8047058 | 352.1604204 | 343.6752237 | 292.3955045 | 216.7694591 | 300.9214396 | 366.4581749 | 0.610869733 | 0.122397715 | YHR168W | MTG2 | n |
| W303.asm02.HP0_G0024970 | 488.5266624 | 636.8369659 | 483.7644459 | 2376.596428 | 1881.588677 | 2292.091075 | 1114.138224 | 1235.616329 | 974.49213 | 6.20E-33 | -2.025692668 | YHR169W | DBP8 | n |
| W303.asm02.HP0_G0024980 | 2968.962633 | 3898.127616 | 3195.777249 | 6257.170784 | 5725.708843 | 6404.638372 | 3360.399912 | 3544.293355 | 2884.578595 | 1.97E-09 | -0.869796861 | YHR170W | NMD3 | n |
| W303.asm02.HP0_G0024990 | 415.8292424 | 418.8464349 | 387.9888587 | 74.90705074 | 90.23133287 | 83.28293007 | 182.6921642 | 174.1667496 | 193.4653493 | 1.79E-35 | 2.305033724 | YHR171W | ATG7 | n |
| W303.asm02.HP0_G0025000 | 312.1142566 | 410.2791215 | 386.0342548 | 337.5681378 | 352.9637433 | 343.9947112 | 98.44551855 | 345.4307201 | 448.3482699 | 0.812501068 | 0.099679504 | YHR172W | SPC97 | n |
| W303.asm02.HP0_G0025020 | 34518.67362 | 28626.24998 | 29874.16482 | 48698.33835 | 41637.77933 | 45044.29693 | 25450.05973 | 22900.99239 | 22921.03758 | 1.23E-05 | -0.541449601 | YHR174W | ENO2 | n |
| W303.asm02.HP0_G0025030 | 423.5836339 | 321.7502159 | 358.6698013 | 296.7097465 | 249.4630968 | 283.3430121 | 677.7595316 | 396.7131519 | 628.5064788 | 0.080308609 | 0.41061312 | YHR175W | CTR2 | n |
| W303.asm02.HP0_G0025050 | 212.2764664 | 186.5770483 | 193.5057784 | 203.3191377 | 136.6739307 | 173.8078541 | 497.9071419 | 259.3149383 | 510.7894673 | 0.49869198 | 0.201522909 | YHR176W | FMO1 | n |

| | | | | | | | | | | | | | | |
|---|---|---|---|---|---|---|---|---|---|---|---|---|---|---|
| W303.asm02.HP0_G0025060 | 70.75882214 | 103.759685 | 61.57002039 | 65.17886234 | 78.28895058 | 68.79894223 | 83.30005416 | 80.31022343 | 85.98459971 | 0.620916912 | 0.158443703 | YHR177W | ROF1 | n |
| W303.asm02.HP0_G0025070 | 240.3861355 | 229.4136155 | 237.4843644 | 795.7658118 | 882.4093582 | 841.8817931 | 1074.38138 | 971.4634256 | 1175.122863 | 9.98E-39 | -1.832362953 | YHR178W | STB5 | n |
| W303.asm02.HP0_G0025080 | 5310.788856 | 5057.570702 | 5311.635886 | 3061.460892 | 1131.872455 | 1651.174614 | 665.4538418 | 537.9817377 | 853.7042399 | 1.15E-06 | 1.423191312 | YHR179W | OYE2 | n |
| W303.asm02.HP0_G0025100 | 1429.715927 | 1336.500897 | 1487.453509 | 993.2480365 | 992.5446616 | 970.4271852 | 2262.353744 | 1498.80164 | 1659.298049 | 0.002025051 | 0.525019645 | YHR181W | SVP26 | n |
| W303.asm02.HP0_G0025110 | 442.0003136 | 321.7502159 | 414.3760103 | 797.7114495 | 812.0819959 | 813.8190667 | 703.3175027 | 715.0512665 | 1018.508056 | 7.59E-08 | -1.040800047 | YHR182W | RGD3 | n |
| W303.asm02.HP0_G0025120 | 5909.815597 | 5441.195959 | 5857.947654 | 13167.10301 | 11209.91618 | 12284.23219 | 10743.8138 | 9106.985819 | 11232.25015 | 2.67E-18 | -1.091211754 | YHR183W | GND1 | n |
| W303.asm02.HP0_G0025130 | 248.140527 | 271.298259 | 235.5297605 | 36.96711595 | 55.73111736 | 53.40970516 | 55.84889995 | 59.02317626 | 64.48844978 | 2.10E-21 | 2.377164707 | YHR184W | SSP1 | n |
| W303.asm02.HP0_G0025140 | 22.29387547 | 12.37500831 | 24.43254777 | 23.34765218 | 26.53862732 | 15.38923708 | 29.34433726 | 18.38426802 | 20.47252374 | 0.815470348 | -0.133110875 | YHR185C | PFS1 | n |
| W303.asm02.HP0_G0025150 | 429.3994275 | 599.711941 | 483.7644459 | 883.3195075 | 1004.487044 | 848.2185378 | 675.8663485 | 830.1948398 | 964.2558681 | 5.96E-06 | -0.853753235 | YHR186C | KOG1 | n |
| W303.asm02.HP0_G0025160 | 1041.027055 | 1100.423815 | 1076.009404 | 1282.175232 | 1228.738445 | 1276.401428 | 1225.836024 | 996.620845 | 990.870149 | 0.080300109 | -0.235566313 | YHR187W | IKI1 | n |
| W303.asm02.HP0_G0025170 | 774.4698478 | 780.577447 | 682.1567339 | 1612.933638 | 1640.087168 | 1610.438398 | 1558.089649 | 1861.649035 | 1799.534837 | 7.95E-19 | -1.120076451 | YHR188C | GPI16 | n |
| W303.asm02.HP0_G0025180 | 79.48251254 | 67.58658382 | 63.52462421 | 184.8355798 | 228.2321949 | 181.049848 | 212.0365015 | 163.523226 | 150.4730495 | 9.58E-10 | -1.492377758 | YHR189W | PTH1 | n |
| W303.asm02.HP0_G0025190 | 4907.5605 | 3763.906372 | 4414.472732 | 3990.502885 | 3833.504716 | 4029.264367 | 2791.498406 | 3059.529235 | 2892.767604 | 0.289082142 | 0.142579875 | YHR190W | ERG9 | n |
| W303.asm02.HP0_G0025200 | 644.5837907 | 595.9042461 | 729.0672256 | 231.5308841 | 249.4630968 | 224.5018115 | 336.9865827 | 282.5371716 | 244.6466587 | 5.47E-17 | 1.483189295 | YHR191C | CTF8 | n |
| W303.asm02.HP0_G0025210 | 1191.268389 | 917.654462 | 1228.468502 | 912.5040727 | 919.5634365 | 974.0481822 | 1355.519063 | 1024.681044 | 984.7283919 | 0.171030441 | 0.249854485 | YHR192W | LNP1 | n |
| W303.asm02.HP0_G0025220 | 8394.128763 | 7036.620107 | 8660.849535 | 6296.083538 | 6601.483545 | 6193.7153 | 7249.944487 | 5095.345019 | 4579.703561 | 0.067758561 | 0.335660528 | YHR193C | EGD2 | n |
| W303.asm02.HP0_G0025230 | 357.6713064 | 321.7502159 | 277.5537427 | 266.5523624 | 265.3862732 | 235.3648024 | 247.0603879 | 265.1204966 | 274.3318181 | 0.075050171 | 0.320397602 | YHR194W | MDM31 | n |
| W303.asm02.HP0_G0025250 | 2736.330889 | 4040.916174 | 2920.17811 | 4372.82069 | 3842.793235 | 4042.843106 | 2324.828784 | 2359.959457 | 1884.49581 | 0.058377514 | -0.338182391 | YHR196W | UTP9 | n |
| W303.asm02.HP0_G0025260 | 1418.08434 | 2249.39574 | 1423.928884 | 5390.389197 | 4333.757841 | 5282.129315 | 2215.970759 | 2816.663379 | 2483.31713 | 4.48E-15 | -1.559537669 | YHR197W | RIX1 | n |
| W303.asm02.HP0_G0025270 | 1599.34324 | 1156.587315 | 1550.978133 | 407.6110943 | 386.1370274 | 396.4991671 | 498.8537334 | 478.9585614 | 474.9625508 | 2.13E-31 | 1.854713497 | YHR198C | AIM18 | n |
| W303.asm02.HP0_G0025300 | 1842.637272 | 1403.135557 | 1615.480059 | 1686.86787 | 1553.836629 | 1715.44731 | 2906.982572 | 2126.769531 | 2145.520488 | 0.877985635 | -0.028280534 | YHR200W | RPN10 | n |
| W303.asm02.HP0_G0025310 | 1939.567166 | 1765.818493 | 1915.511746 | 733.505406 | 816.0627899 | 763.1251093 | 680.5993062 | 627.9678917 | 647.9553764 | 2.97E-28 | 1.282520393 | YHR201C | PPX1 | n |
| W303.asm02.HP0_G0025320 | 292.7282779 | 218.9424546 | 272.6672332 | 137.1674566 | 165.8664207 | 143.0293799 | 284.9240489 | 336.7223826 | 349.0565298 | 5.38E-05 | 0.81862353 | YHR202W | SMN1 | n |
| W303.asm02.HP0_G0025340 | 485.6187656 | 436.9329856 | 382.1250472 | 702.3752031 | 646.2155751 | 667.1686898 | 870.8642026 | 996.620845 | 882.3657732 | 2.14E-05 | -0.62818577 | YHR204W | MNL1 | n |
| W303.asm02.HP0_G0025350 | 799.6716201 | 1010.942986 | 724.180716 | 5067.413342 | 4158.6029 | 5136.384187 | 1364.984978 | 2986.959756 | 2716.7039 | 6.01E-23 | -2.502414224 | YHR205W | SCH9 | n |
| W303.asm02.HP0_G0025360 | 633.9215025 | 910.0390723 | 678.2475262 | 1477.711819 | 1535.25959 | 1520.818723 | 708.9970519 | 1334.310821 | 1278.509108 | 1.28E-05 | -1.028600193 | YHR206W | SKN7 | n |
| W303.asm02.HP0_G0025370 | 671.7241609 | 500.7118745 | 677.2702243 | 463.0617682 | 443.1950762 | 424.5618935 | 699.5311366 | 563.1391571 | 609.0575813 | 0.006769952 | 0.474962462 | YHR207C | SET5 | n |
| W303.asm02.HP0_G0025380 | 6406.096651 | 5668.705728 | 6069.044867 | 7951.821205 | 7003.543749 | 7901.920615 | 10385.05562 | 7305.327553 | 8126.568298 | 0.031984932 | -0.333303318 | YHR208W | BAT1 | n |
| W303.asm02.HP0_G0025580 | 10.66228827 | 15.23077945 | 5.863811466 | 31.13020291 | 57.05804873 | 39.83096656 | 52.06253385 | 77.40744427 | 156.6148066 | 0.000295753 | -2.001128838 | YOL156W | HXT11 | y |
| W303.asm02.HP0_G0025620 | 280.1273918 | 374.1060203 | 289.2813657 | 180.9443044 | 180.4626657 | 169.2816079 | 325.6274844 | 233.1899259 | 323.4658751 | 9.68E-05 | 0.831226083 | YCL064C | CHA1 | y |
| W303.asm02.HP0_G0025630 | 261.710712 | 237.0290052 | 224.7794395 | 1972.876609 | 1399.912591 | 2332.827291 | 2937.273501 | 3098.232957 | 5542.935802 | 5.19E-31 | -2.979885235 | YIL166C | SOA1 | n |
| W303.asm02.HP0_G0025670 | 938.2813676 | 924.317928 | 922.573004 | 157.5966522 | 123.404617 | 120.3981489 | 111.6977999 | 120.9491317 | 166.8510685 | 1.56E-46 | 2.792369651 | YIL162W | SUC2 | n |
| W303.asm02.HP0_G0025680 | 331.5002352 | 415.9906638 | 413.3987084 | 181.9171232 | 207.0012931 | 219.0703161 | 210.1433185 | 179.9723079 | 188.3472184 | 4.92E-07 | 0.933307792 | YIL161W | SMU2 | n |
| W303.asm02.HP0_G0025690 | 74.63601787 | 51.40388065 | 60.59271848 | 97.28188408 | 124.7315484 | 108.6299088 | 142.9353202 | 106.4352359 | 91.10273056 | 0.005439761 | -0.821749777 | YIL160C | POT1 | n |
| W303.asm02.HP0_G0025700 | 189.013292 | 339.8367665 | 226.7340434 | 214.020145 | 236.1937831 | 200.060082 | 213.9296845 | 246.7362286 | 288.6625847 | 0.406463376 | 0.218750725 | YIL159W | BNR1 | n |
| W303.asm02.HP0_G0025720 | 1941.505764 | 1826.741611 | 2141.268487 | 932.9332684 | 784.2164372 | 898.007246 | 1148.215519 | 780.8475941 | 810.7119401 | 2.19E-11 | 1.175169812 | YIL157C | COA1 | n |
| W303.asm02.HP0_G0025740 | 245.2326302 | 345.5483088 | 244.3254777 | 357.9973334 | 351.6368119 | 330.4159726 | 330.3604421 | 405.4214894 | 409.4504748 | 0.119625812 | -0.315848218 | YIL156W | UBP7 | n |
| W303.asm02.HP0_G0025750 | 541.8381038 | 403.6156555 | 405.5802931 | 958.2265582 | 882.4093582 | 1025.647389 | 933.3392432 | 1027.583823 | 966.3031205 | 8.86E-12 | -1.086119682 | YIL155C | GUT2 | n |
| W303.asm02.HP0_G0025760 | 1350.233414 | 836.7409462 | 1188.399124 | 687.7829205 | 762.9855353 | 690.7051701 | 759.1664027 | 732.4679414 | 825.0427067 | 0.000438267 | 0.657099102 | YIL154C | IMP21 | n |
| W303.asm02.HP0_G0025770 | 530.2065166 | 385.5291049 | 328.3734421 | 637.1963407 | 721.850663 | 594.7487506 | 903.9949059 | 814.713351 | 767.7196402 | 0.001043904 | -0.649897461 | YIL153W | RRD1 | n |
| W303.asm02.HP0_G0025800 | 63.97372961 | 160.875108 | 50.81969937 | 278.2261885 | 321.1173905 | 283.3430121 | 166.6001083 | 400.5835241 | 341.8911465 | 3.96E-05 | -1.677701803 | YIL150C | MCM10 | n |
| W303.asm02.HP0_G0025820 | 590.3030505 | 802.4716924 | 611.7909963 | 1261.746037 | 1050.929642 | 1204.886738 | 672.0799824 | 1015.972706 | 880.3185208 | 3.22E-05 | -0.811785838 | YIL147C | SLN1 | n |
| W303.asm02.HP0_G0025830 | 162.8422208 | 182.7693534 | 157.3456077 | 520.4580798 | 534.7533404 | 529.5708054 | 457.2037064 | 597.972507 | 442.2065128 | 1.28E-20 | -1.655343281 | YIL146C | ATG32 | n |
| W303.asm02.HP0_G0025840 | 1141.834144 | 945.2602498 | 1043.758441 | 1006.8675 | 709.9082807 | 967.7114375 | 1146.322336 | 947.2735993 | 1024.649813 | 0.219480839 | 0.219803499 | YIL145C | PAN6 | n |
| W303.asm02.HP0_G0025850 | 708.5575203 | 737.7408797 | 693.8843568 | 730.5869495 | 793.5049567 | 743.209626 | 692.904996 | 750.8522095 | 677.6405358 | 0.532012975 | -0.082047527 | YIL144W | NDC80 | n |
| W303.asm02.HP0_G0025860 | 748.2987766 | 920.5102332 | 652.8376765 | 1899.915196 | 2047.455097 | 1930.896629 | 1184.185997 | 2109.352856 | 1896.779324 | 4.67E-10 | -1.340023319 | YIL143C | SSL2 | n |
| W303.asm02.HP0_G0025870 | 4156.353827 | 3267.002193 | 3843.728416 | 5224.037175 | 4834.010965 | 5127.331695 | 4209.49251 | 4128.719559 | 4058.677831 | 0.000463733 | -0.430756024 | YIL142W | CCT2 | n |
| W303.asm02.HP0_G0025880 | 892.7243177 | 874.8178948 | 965.5742881 | 1787.068211 | 1754.203266 | 1919.128389 | 1809.882995 | 2265.135338 | 2053.394131 | 1.46E-14 | -0.998890799 | YIL140W | AXL2 | n |
| W303.asm02.HP0_G0025890 | 596.1188441 | 517.8465014 | 544.3571644 | 173.1617537 | 193.7319794 | 197.3443343 | 278.2979082 | 186.7454593 | 199.6071065 | 2.49E-15 | 1.555922354 | YIL139C | REV7 | n |
| W303.asm02.HP0_G0025900 | 3340.204125 | 3384.08881 | 3703.974243 | 8463.523915 | 8428.668035 | 8458.648898 | 7880.374442 | 7824.925023 | 7299.478339 | 7.76E-40 | -1.28156299 | YIL138C | TPM2 | n |
| W303.asm02.HP0_G0025910 | 691.1101395 | 572.1061532 | 558.0393912 | 1923.262848 | 1840.453804 | 1969.822346 | 1113.191633 | 1530.732211 | 1281.579986 | 8.01E-26 | -1.654734848 | YIL137C | TMA108 | n |
| W303.asm02.HP0_G0025920 | 1224.224553 | 582.5773141 | 991.9614397 | 85.60805799 | 144.6355189 | 51.59920668 | 169.4398829 | 92.88893313 | 222.1268826 | 5.85E-18 | 3.322287291 | YIL136W | OM45 | n |
| W303.asm02.HP0_G0025930 | 790.9479297 | 481.6734002 | 622.5413173 | 536.0231813 | 486.9838112 | 544.9600424 | 374.8502437 | 616.356775 | 821.9718281 | 0.339370662 | 0.272875029 | YIL135C | VHS2 | n |
| W303.asm02.HP0_G0025950 | 518.5749294 | 628.2696524 | 547.2890702 | 346.3235073 | 388.7908902 | 363.9101944 | 440.1650589 | 380.26407 | 424.8048676 | 4.96E-05 | 0.626139214 | YIL134W | FLX1 | n |
| W303.asm02.HP0_G0025960 | 5869.105042 | 6583.504418 | 5920.494977 | 21989.59708 | 24321.325 | 20877.76322 | 21358.89116 | 16361.99853 | 14979.74562 | 1.88E-34 | -1.870568974 | YIL133C | RPL16A | n |
| W303.asm02.HP0_G0025970 | 114.3772741 | 105.6635325 | 109.457814 | 215.9657827 | 168.5202835 | 194.6285866 | 185.5319388 | 208.0325065 | 140.2367876 | 0.000432368 | -0.816838033 | YIL132C | CSM2 | n |
| W303.asm02.HP0_G0025990 | 409.0441499 | 804.3755398 | 437.8312561 | 2363.949783 | 1702.452942 | 2200.660902 | 757.2732196 | 1096.28293 | 854.7278661 | 9.45E-14 | -1.924398114 | YIL130W | ASG1 | n |
| W303.asm02.HP0_G0026000 | 757.9917659 | 1004.27952 | 757.408981 | 1074.964819 | 1202.199817 | 1007.542404 | 981.615411 | 1122.407942 | 1025.673439 | 0.019343941 | -0.381381031 | YIL129C | TAO3 | n |
| W303.asm02.HP0_G0026010 | 1197.084183 | 1633.501096 | 1310.561863 | 1045.780254 | 1137.18018 | 1072.720349 | 823.5346264 | 983.0745423 | 884.4130255 | 0.027968206 | 0.347619721 | YIL128W | MET18 | n |
| W303.asm02.HP0_G0026020 | 953.7901505 | 1845.780085 | 1267.560579 | 1659.628942 | 1593.64457 | 1648.458866 | 1090.473436 | 1015.972706 | 772.8377712 | 0.259419866 | -0.269289335 | YIL127C | RRT14 | n |
| W303.asm02.HP0_G0026030 | 1226.163151 | 1366.962456 | 1269.515182 | 2977.798472 | 2956.403083 | 2908.565808 | 1716.170434 | 2276.746455 | 2232.528714 | 9.43E-18 | -1.194853888 | YIL126W | STH1 | n |
| W303.asm02.HP0_G0026040 | 2504.668444 | 1833.405077 | 2256.590112 | 12617.46037 | 14305.64705 | 12876.26519 | 7971.247228 | 14386.17352 | 12164.77361 | 2.24E-36 | -2.593360479 | YIL125W | KGD1 | n |
| W303.asm02.HP0_G0026050 | 2453.295601 | 2220.838029 | 2276.136151 | 1109.013479 | 1048.275779 | 1102.593574 | 1382.023626 | 1277.222831 | 1076.854749 | 9.11E-17 | 1.091756241 | YIL124W | AYR1 | n |

| | | | | | | | | | | | | | |
|---|---|---|---|---|---|---|---|---|---|---|---|---|---|
| W303.asm02.HP0_G0026080 | 514.6977336 | 355.067546 | 566.8351084 | 627.4681523 | 423.2911057 | 608.3274892 | 219.6092337 | 298.0186605 | 305.0406037 | 0.404076612 | -0.20989331 | YIL121W | QDR2 | n |
| W303.asm02.HP0_G0026100 | 609.6890291 | 819.6063193 | 648.9284689 | 106.0372537 | 92.8851956 | 87.80917627 | 29.34433726 | 175.1343427 | 99.29174014 | 8.62E-12 | 2.857431193 | YIL119C | RPI1 | n |
| W303.asm02.HP0_G0026110 | 1932.782073 | 2624.453684 | 2015.19654 | 1796.796399 | 1848.415393 | 1698.247574 | 2446.939091 | 1923.57499 | 2110.717198 | 0.067892742 | 0.298960252 | YIL118W | RHO3 | n |
| W303.asm02.HP0_G0026130 | 2197.400682 | 1593.5203 | 2149.086902 | 877.4825944 | 898.3325346 | 912.4912338 | 1139.696196 | 864.0605967 | 1089.138263 | 1.47E-11 | 1.14370642 | YIL116W | HIS5 | n |
| W303.asm02.HP0_G0026140 | 920.8339868 | 1163.250781 | 909.8680791 | 2256.939711 | 2404.399635 | 2299.333069 | 1475.736187 | 1947.764817 | 1850.716146 | 6.86E-15 | -1.216907773 | YIL115C | NUP159 | n |
| W303.asm02.HP0_G0026170 | 595.1495451 | 799.6159213 | 576.6081275 | 742.2607756 | 822.6974468 | 723.2941427 | 633.2697299 | 777.944815 | 931.4998301 | 0.291285805 | -0.214140226 | YIL112W | HOS4 | n |
| W303.asm02.HP0_G0026190 | 1176.728905 | 1735.356934 | 1329.130599 | 1825.008145 | 1654.683413 | 1855.760942 | 1264.646277 | 1233.681143 | 1138.27232 | 0.047248921 | -0.331459523 | YIL110W | HPM1 | n |
| W303.asm02.HP0_G0026200 | 1569.294973 | 1729.645392 | 1284.174711 | 3919.48711 | 3570.772305 | 3838.256777 | 3651.950102 | 3774.580501 | 3936.866315 | 4.42E-22 | -1.305677259 | YIL109C | SEC24 | n |
| W303.asm02.HP0_G0026210 | 460.4169934 | 562.586916 | 428.058237 | 549.6426451 | 465.7529094 | 529.5708054 | 434.4855098 | 652.157718 | 805.5938092 | 0.714574478 | -0.091286725 | YIL108W | | n |
| W303.asm02.HP0_G0026220 | 663.9697694 | 494.0484085 | 755.4543772 | 497.1104277 | 453.8105271 | 499.6975804 | 501.693508 | 624.0975195 | 759.5306307 | 0.06300049 | 0.398602036 | YIL107C | PFK26 | n |
| W303.asm02.HP0_G0026240 | 722.1277054 | 939.5487075 | 808.2286804 | 1393.07658 | 1129.218592 | 1375.073595 | 681.5458977 | 793.4263038 | 744.1762379 | 2.89E-05 | -0.65906241 | YIL104C | SHQ1 | n |
| W303.asm02.HP0_G0026250 | 1361.865001 | 1738.212705 | 1365.29077 | 1590.558805 | 1415.835767 | 1623.111887 | 857.6119212 | 1020.810671 | 826.0663329 | 0.757290583 | -0.052623117 | YIL103W | DPH1 | n |
| W303.asm02.HP0_G0026260 | 213.2457654 | 235.1251578 | 223.8021376 | 27.23892754 | 23.88476458 | 19.01023404 | 12.30568982 | 25.15741939 | 34.80329036 | 7.78E-21 | 3.264817687 | YIL102C-A | | n |
| W303.asm02.HP0_G0026280 | 158.9650251 | 165.6347265 | 170.0505325 | 140.0859131 | 149.9432443 | 156.6081185 | 76.67391349 | 193.5186107 | 137.1659091 | 0.656983113 | 0.147241443 | YIL101C | XBP1 | n |
| W303.asm02.HP0_G0026290 | 97.89919228 | 93.28852415 | 93.82098346 | 475.7084132 | 513.5224386 | 486.1188418 | 691.0118129 | 715.0512665 | 554.8053933 | 5.49E-41 | -2.371137511 | YIL099W | SGA1 | n |
| W303.asm02.HP0_G0026300 | 924.7111825 | 1101.375739 | 1018.348591 | 882.3466886 | 859.851525 | 869.9445196 | 1171.880307 | 729.5651623 | 770.7905188 | 0.273698966 | 0.220874719 | YIL098C | FMC1 | n |
| W303.asm02.HP0_G0026310 | 784.1628371 | 570.2023058 | 715.3849988 | 83.66242031 | 87.57747014 | 66.98844375 | 187.4251219 | 186.7454593 | 198.5834803 | 3.78E-53 | 3.12689193 | YIL097W | FYV10 | n |
| W303.asm02.HP0_G0026320 | 642.6451929 | 1100.423815 | 701.7027721 | 1346.381276 | 1628.144786 | 1475.556261 | 1168.093941 | 1095.315336 | 1010.319047 | 2.71E-05 | -0.863590046 | YIL096C | BMT5 | n |
| W303.asm02.HP0_G0026330 | 318.8993491 | 349.3560037 | 328.3734421 | 991.3023988 | 931.5058188 | 1019.310644 | 895.4755822 | 1006.296776 | 1104.492656 | 2.64E-29 | -1.562415725 | YIL095W | PRK1 | n |
| W303.asm02.HP0_G0026340 | 5700.447027 | 5276.513157 | 6049.498829 | 722.8043987 | 528.1186836 | 804.7665743 | 687.2254468 | 406.3890824 | 568.1125338 | 3.11E-49 | 3.047966171 | YIL094C | LYS12 | n |
| W303.asm02.HP0_G0026350 | 1637.145899 | 1406.943252 | 1724.937873 | 2494.307508 | 2558.323673 | 2539.224118 | 2664.655142 | 2191.598266 | 2031.897981 | 3.50E-06 | -0.67076521 | YIL093C | RSM25 | n |
| W303.asm02.HP0_G0026370 | 1268.812304 | 1806.751213 | 1404.382846 | 2724.865573 | 2306.206714 | 2619.7913 | 1252.340587 | 1558.792409 | 1549.770047 | 5.50E-06 | -0.772417033 | YIL091C | UTP25 | n |
| W303.asm02.HP0_G0026380 | 803.5488158 | 1063.298791 | 812.137888 | 987.4111234 | 911.6018483 | 867.2287718 | 1219.209884 | 1102.088488 | 1286.698117 | 0.787624368 | -0.046107461 | YIL090W | ICE2 | n |
| W303.asm02.HP0_G0026400 | 1650.716084 | 1480.241378 | 1728.847081 | 477.6540509 | 575.8882127 | 504.2238266 | 1064.915465 | 1165.949629 | 1239.611312 | 1.04E-32 | 1.644000888 | YIL088C | AVT7 | n |
| W303.asm02.HP0_G0026410 | 548.6231963 | 384.5771812 | 517.9700128 | 196.5094058 | 161.8856266 | 233.5543039 | 297.2297387 | 175.1343427 | 200.6307326 | 3.30E-07 | 1.289551467 | YIL087C | AIM19 | n |
| W303.asm02.HP0_G0026440 | 188.0439931 | 237.0290052 | 158.3229096 | 550.6154639 | 567.9266245 | 525.0445592 | 785.6709654 | 509.9215392 | 816.8536972 | 2.22E-11 | -1.493516144 | YIL084C | SDS3 | n |
| W303.asm02.HP0_G0026450 | 1099.184991 | 945.2602498 | 1094.57814 | 674.1634567 | 704.6005552 | 709.7154041 | 1056.396141 | 809.8753857 | 1165.910227 | 0.000393237 | 0.587917619 | YIL083C | CAB2 | n |
| W303.asm02.HP0_G0026460 | 7609.965926 | 6201.783008 | 7848.711647 | 8874.053466 | 9183.691982 | 8778.20188 | 7346.496822 | 7410.795196 | 6404.829052 | 0.01954077 | -0.309071099 | YIL078W | THS1 | n |
| W303.asm02.HP0_G0026470 | 679.4785523 | 334.1252242 | 538.493353 | 227.6396088 | 201.6935676 | 240.7962978 | 369.1706946 | 345.4307201 | 351.1037821 | 8.79E-07 | 1.209589666 | YIL077C | RCI37 | n |
| W303.asm02.HP0_G0026480 | 3871.37994 | 3717.26211 | 3964.913853 | 1719.943711 | 1685.202835 | 1738.98379 | 2227.329857 | 1592.658166 | 1741.188144 | 4.08E-16 | 1.16719439 | YIL076W | SEC28 | n |
| W303.asm02.HP0_G0026490 | 3554.419189 | 3244.156023 | 3346.281743 | 4939.001255 | 4600.471045 | 5015.080789 | 4358.107379 | 5661.386956 | 5200.02103 | 6.31E-05 | -0.520907342 | YIL075C | RPN2 | n |
| W303.asm02.HP0_G0026510 | 491.4345592 | 481.6734002 | 517.9700128 | 42.804029 | 26.53862732 | 46.16771124 | 53.00912537 | 56.1203971 | 44.01592604 | 5.43E-53 | 3.663402433 | YIL073C | SPO22 | n |
| W303.asm02.HP0_G0026520 | 183.1974984 | 151.3558708 | 157.3456077 | 7.782550727 | 10.61545093 | 12.67348936 | 27.45115421 | 45.47687351 | 26.61428086 | 3.56E-23 | 3.981928065 | YIL072W | HOP1 | n |
| W303.asm02.HP0_G0026530 | 632.9522035 | 656.8273639 | 639.1554498 | 259.7426305 | 238.8476458 | 250.7540395 | 372.0104691 | 367.6853603 | 283.5444538 | 3.54E-17 | 1.362621334 | YIL071C | PCI8 | n |
| W303.asm02.HP0_G0026540 | 2224.541052 | 2019.030201 | 2198.9293 | 3461.289436 | 2895.36424 | 3458.052097 | 2542.544835 | 2695.714247 | 2688.042367 | 4.12E-07 | -0.608016861 | YIL070C | MAM33 | n |
| W303.asm02.HP0_G0026560 | 895.6322145 | 943.3564024 | 901.0723619 | 1193.648718 | 1203.526749 | 1119.79331 | 1276.005375 | 1276.255237 | 1442.289297 | 0.002539704 | -0.359546284 | YIL068C | SEC6 | n |
| W303.asm02.HP0_G0026570 | 264.6186088 | 351.2598511 | 252.143893 | 427.0674711 | 485.6568799 | 403.741161 | 526.3048877 | 685.0558818 | 746.2234903 | 0.003533504 | -0.598888615 | YIL067C | | n |
| W303.asm02.HP0_G0026590 | 817.1190009 | 795.8082264 | 806.2740766 | 287.9543769 | 318.4635278 | 283.3430121 | 602.0322096 | 458.6391073 | 452.4427746 | 4.91E-19 | 1.445414042 | YIL065C | FIS1 | n |
| W303.asm02.HP0_G0026600 | 933.4348729 | 1238.452754 | 1114.124179 | 955.3081017 | 1037.660328 | 1021.121143 | 1240.034897 | 1128.2135 | 1076.854749 | 0.418238859 | 0.125019469 | YIL064W | EFM4 | n |
| W303.asm02.HP0_G0026610 | 1818.404799 | 1660.15496 | 1861.76014 | 1253.963486 | 1272.52718 | 1239.286209 | 1396.222499 | 1283.028389 | 1405.438755 | 6.53E-06 | 0.504179918 | YIL063C | YRB2 | n |
| W303.asm02.HP0_G0026620 | 2807.089711 | 2445.492026 | 2831.243636 | 1070.100725 | 997.8523871 | 1074.530848 | 1617.724915 | 1116.602384 | 1100.398151 | 1.42E-15 | 1.362703207 | YIL062C | ARC15 | n |
| W303.asm02.HP0_G0026630 | 233.601043 | 161.8270317 | 182.7554574 | 229.5852464 | 222.9244694 | 254.3750364 | 389.0491166 | 315.4353354 | 344.962025 | 0.165189124 | -0.29192985 | YIL061C | SNP1 | n |
| W303.asm02.HP0_G0026660 | 426.4915307 | 410.2791215 | 349.8740841 | 1360.973558 | 1015.102495 | 1318.948143 | 967.4165381 | 1164.982036 | 886.4602779 | 1.46E-20 | -1.639856872 | YIL056W | VHR1 | n |
| W303.asm02.HP0_G0026670 | 179.3203027 | 125.6539305 | 172.0051363 | 84.63523915 | 78.28895058 | 84.18817931 | 131.5762219 | 134.4954344 | 227.2450135 | 0.001242464 | 0.946748337 | YIL055C | | n |
| W303.asm02.HP0_G0026690 | 6719.180207 | 7820.053325 | 6622.197749 | 4431.18982 | 2779.921211 | 4332.522862 | 2133.617296 | 1774.56566 | 3641.038347 | 0.000758146 | 0.874049354 | YIL053W | GPP1 | n |
| W303.asm02.HP0_G0026700 | 5594.793444 | 6271.27344 | 5519.801193 | 18436.86267 | 21002.66966 | 17699.43314 | 14659.86294 | 12783.83942 | 11121.69852 | 6.15E-35 | -1.716471647 | YIL052C | RPL34B | y |
| W303.asm02.HP0_G0026710 | 2588.997451 | 2021.885972 | 2631.874046 | 3536.196486 | 3212.500837 | 3681.648659 | 2792.444997 | 2000.982434 | 1922.369979 | 0.003571985 | -0.52640483 | YIL051C | MMF1 | n |
| W303.asm02.HP0_G0026730 | 832.6277838 | 471.2022393 | 710.4984893 | 712.1033915 | 631.6193301 | 668.0739391 | 811.2289365 | 730.5327553 | 655.1207597 | 0.995773659 | 0.001196223 | YIL049W | DFG10 | n |
| W303.asm02.HP0_G0026740 | 1268.812304 | 1109.943053 | 1380.9276 | 1421.288326 | 1429.105081 | 1489.135 | 1274.112192 | 1894.547199 | 1864.023286 | 0.26059571 | -0.207076566 | YIL048W | NEO1 | n |
| W303.asm02.HP0_G0026750 | 725.0356022 | 1420.270184 | 728.0899237 | 951.4168263 | 963.3521715 | 957.7536958 | 859.5051043 | 1066.287545 | 957.0904848 | 0.998255 | 0.000591116 | YIL047C | SYG1 | n |
| W303.asm02.HP0_G0026760 | 480.772271 | 520.7022725 | 358.6698013 | 414.4208262 | 394.0986156 | 386.5414254 | 308.588837 | 368.6529534 | 387.9543249 | 0.344721032 | 0.186783139 | YIL046W | MET30 | n |
| W303.asm02.HP0_G0026780 | 4507.24004 | 3651.579374 | 4219.989652 | 1155.708783 | 1031.025671 | 1125.224805 | 989.1881432 | 876.6393064 | 835.2789686 | 3.99E-46 | 1.901265518 | YIL044C | AGE2 | y |
| W303.asm02.HP0_G0026790 | 346.0397192 | 217.9905309 | 215.0064204 | 324.9214928 | 331.7328414 | 351.2367051 | 499.800325 | 503.1483878 | 520.002103 | 0.083378702 | -0.372029486 | YIL042C | PKP1 | n |
| W303.asm02.HP0_G0026800 | 2059.760234 | 1485.95292 | 2077.743863 | 3922.405566 | 3308.039895 | 3829.204285 | 2938.220092 | 3133.066307 | 2849.775305 | 5.31E-10 | -0.976143054 | YIL041W | GVP36 | n |
| W303.asm02.HP0_G0026810 | 982.8691185 | 1113.750747 | 1043.758441 | 84.63523915 | 91.55826424 | 90.52492399 | 307.6422455 | 197.3889829 | 244.6466587 | 2.64E-71 | 3.558970954 | YIL040W | APQ12 | n |
| W303.asm02.HP0_G0026820 | 1980.277721 | 1666.818426 | 1993.695898 | 1307.468522 | 1370.720101 | 1268.254185 | 1608.259 | 1819.07494 | 1692.054087 | 4.72E-05 | 0.515854917 | YIL039W | TED1 | n |
| W303.asm02.HP0_G0026830 | 1302.737767 | 1381.241312 | 1360.40426 | 2511.818247 | 2641.920349 | 2484.909164 | 1957.551273 | 2367.700202 | 2256.072116 | 6.25E-15 | -0.917086828 | YIL038C | NOT3 | n |
| W303.asm02.HP0_G0026840 | 192.8904878 | 131.3654728 | 162.2321172 | 186.7812174 | 265.3862732 | 161.1343647 | 241.3808388 | 250.6066008 | 190.3944708 | 0.220953749 | -0.328144746 | YIL037C | PRM2 | n |
| W303.asm02.HP0_G0026850 | 445.8775094 | 447.4041464 | 379.1931415 | 869.7000437 | 806.7742704 | 862.7025256 | 502.6400995 | 842.7735495 | 1069.689365 | 3.67E-05 | -0.996931249 | YIL036W | CST6 | n |
| W303.asm02.HP0_G0026860 | 2549.256195 | 2538.78055 | 2650.442783 | 2684.980001 | 2810.440633 | 2616.170303 | 2422.327711 | 2669.589234 | 2584.656122 | 0.532424688 | -0.067539322 | YIL035C | CKA1 | n |
| W303.asm02.HP0_G0026870 | 1784.479336 | 1371.722074 | 1908.670632 | 1491.331283 | 1399.912591 | 1519.008225 | 2117.52524 | 1635.23226 | 1597.880478 | 0.266447978 | 0.19929264 | YIL034C | CAP2 | n |
| W303.asm02.HP0_G0026880 | 1771.87845 | 1190.856568 | 1394.609827 | 3052.705523 | 2093.897695 | 2811.704139 | 2221.650308 | 2310.612212 | 2373.789128 | 3.37E-06 | -0.869649152 | YIL033C | BCY1 | n |
| W303.asm02.HP0_G0026900 | 268.4958046 | 451.2118413 | 286.3494599 | 781.1735292 | 822.6974468 | 725.1046412 | 506.4264656 | 724.727197 | 621.3410955 | 6.25E-08 | -1.210217811 | YIL031W | ULP2 | n |

| | | | | | | | | | | | | | | |
|---|---|---|---|---|---|---|---|---|---|---|---|---|---|---|
| W303.asm02.HP0_G0026910 | 1261.057912 | 1467.86637 | 1149.307047 | 1347.354095 | 1409.20111 | 1393.17858 | 1206.904194 | 1643.940598 | 1367.564586 | 0.564455729 | -0.097423525 | YIL030C | SSM4 | n |
| W303.asm02.HP0_G0026940 | 789.9786307 | 732.0293374 | 796.5010575 | 652.7614422 | 627.638536 | 649.063705 | 1116.977999 | 801.1670482 | 725.7509066 | 0.153214759 | 0.26465609 | YIL027C | EMC5 | n |
| W303.asm02.HP0_G0026950 | 826.8119902 | 869.1063525 | 887.3901352 | 969.9003843 | 985.9100048 | 985.8164223 | 856.6653297 | 1159.176478 | 900.7910445 | 0.228943872 | -0.187347199 | YIL026C | IRR1 | n |
| W303.asm02.HP0_G0026960 | 151.2106336 | 172.2981926 | 160.2775141 | 40.85839132 | 47.76952917 | 50.69395744 | 56.79549147 | 77.40744427 | 67.55932834 | 1.25E-11 | 1.796534116 | YIL024C | | n |
| W303.asm02.HP0_G0026970 | 606.7811323 | 581.6253904 | 563.9032026 | 505.8657972 | 459.1182526 | 434.5196352 | 919.1403703 | 627.0002986 | 771.814145 | 0.068665752 | 0.324313566 | YIL023C | YKE4 | n |
| W303.asm02.HP0_G0026980 | 1826.159191 | 1662.058808 | 1425.883488 | 3679.200856 | 3779.10053 | 3598.365729 | 3178.654339 | 3575.256332 | 3536.628476 | 4.46E-20 | -1.169760265 | YIL022W | TIM44 | n |
| W303.asm02.HP0_G0026990 | 2398.045561 | 2255.107283 | 2437.390966 | 2972.934378 | 3231.077876 | 3123.109878 | 3836.535449 | 3238.53395 | 3347.257631 | 0.000693548 | -0.395328925 | YIL021W | RPB3 | n |
| W303.asm02.HP0_G0027000 | 978.0226238 | 1015.702605 | 1023.235101 | 1040.91616 | 968.659897 | 1031.984134 | 1475.736187 | 945.3384132 | 1145.437703 | 0.948142427 | -0.012112003 | YIL020C | HIS6 | n |
| W303.asm02.HP0_G0027010 | 881.0927305 | 1087.096883 | 897.1631543 | 2860.087392 | 2413.688154 | 2787.26241 | 2288.858306 | 1561.695188 | 1642.92003 | 4.23E-17 | -1.492505916 | YIL019W | FAF1 | n |
| W303.asm02.HP0_G0027030 | 439.0924168 | 347.4521563 | 413.3987084 | 161.4879276 | 169.8472148 | 148.4608753 | 215.8228676 | 295.1158813 | 247.7175372 | 1.44E-11 | 1.3251089 | YIL017C | VID28 | n |
| W303.asm02.HP0_G0027040 | 1036.18056 | 1048.068011 | 1144.420538 | 748.0976886 | 772.2740549 | 718.7678965 | 1207.850785 | 861.1578176 | 836.3025948 | 0.001962803 | 0.528452552 | YIL016W | SNL1 | n |
| W303.asm02.HP0_G0027050 | 3117.26537 | 2862.434613 | 2611.350706 | 103.1187971 | 100.8467838 | 115.8719027 | 185.5319388 | 131.5926553 | 152.5203019 | 1.02E-166 | 4.743195381 | YIL015W | BAR1 | n |
| W303.asm02.HP0_G0027070 | 911.1409974 | 1101.375739 | 1025.189705 | 403.7198189 | 406.0409979 | 374.7731853 | 375.7968352 | 510.8891322 | 457.5609056 | 4.20E-17 | 1.359558777 | YIL014W | MNT3 | n |
| W303.asm02.HP0_G0027110 | 631.9829046 | 535.933052 | 605.9271848 | 1272.447044 | 939.467407 | 1186.781754 | 1425.566836 | 1125.310721 | 1233.469555 | 1.78E-08 | -0.939607677 | YIL010W | DOT5 | n |
| W303.asm02.HP0_G0027120 | 639.7372961 | 508.3272642 | 605.9271848 | 86.58087683 | 132.6931366 | 114.9666535 | 133.469405 | 112.2407942 | 186.299966 | 2.89E-23 | 2.39777882 | YIL009C-A | EST3 | n |
| W303.asm02.HP0_G0027140 | 1156.373628 | 1436.452887 | 1291.993126 | 485.4366016 | 610.3884283 | 521.4235622 | 456.2571148 | 400.5835241 | 383.8598201 | 5.40E-16 | 1.266806888 | YIL008W | URM1 | n |
| W303.asm02.HP0_G0027150 | 1199.99208 | 916.7025383 | 1110.214971 | 93.39060872 | 67.67349965 | 87.80917627 | 110.7512084 | 72.56947901 | 93.14998301 | 3.28E-66 | 3.686702437 | YIL007C | NAS2 | n |
| W303.asm02.HP0_G0027160 | 261.710712 | 153.2597182 | 161.2548153 | 227.6396088 | 318.4635278 | 279.7220151 | 545.2367181 | 451.865956 | 632.6009836 | 0.04535221 | -0.517139161 | YIL006W | YIA6 | n |
| W303.asm02.HP0_G0027170 | 813.2418051 | 752.9716592 | 906.9361734 | 983.5198481 | 1145.141769 | 964.9956898 | 1015.692706 | 1240.454294 | 1302.05251 | 0.042670332 | -0.321813368 | YIL005W | EPS1 | n |
| W303.asm02.HP0_G0027180 | 851.0444635 | 814.8467007 | 901.0723619 | 743.2335944 | 821.3705154 | 738.6833798 | 757.2732196 | 628.9354847 | 613.152086 | 0.28619635 | 0.157551752 | YIL004C | BET1 | n |
| W303.asm02.HP0_G0027190 | 710.4961182 | 496.9041796 | 711.4757912 | 527.2678117 | 443.1950762 | 511.4658206 | 382.4229759 | 471.217817 | 509.7658411 | 0.064289121 | 0.371234194 | YIL003W | CFD1 | n |
| W303.asm02.HP0_G0027210 | 528.2679187 | 541.6445943 | 507.2196918 | 574.9359349 | 552.0034482 | 641.8217111 | 833.0005416 | 732.4679414 | 821.9718281 | 0.242433523 | -0.167179235 | YIL002C | INP51 | n |
| W303.asm02.HP0_G0027220 | 1126.325361 | 705.3754734 | 1109.237669 | 416.3664639 | 465.7529094 | 444.4773768 | 562.2753656 | 552.4956335 | 551.7345148 | 2.66E-09 | 1.149183161 | YIL001W | | n |
| W303.asm02.HP0_G0027230 | 1397.729062 | 872.9140474 | 1369.199977 | 2361.031327 | 2339.379998 | 2423.352215 | 2156.335493 | 2389.954842 | 2147.56774 | 7.42E-08 | -0.968928489 | YIR001C | SGN1 | n |
| W303.asm02.HP0_G0027240 | 1013.886684 | 833.885175 | 1131.715613 | 452.360761 | 488.3107426 | 439.9511306 | 410.8207217 | 470.250224 | 566.0652814 | 3.94E-10 | 1.110791749 | YIR002C | MPH1 | n |
| W303.asm02.HP0_G0027250 | 699.8338299 | 498.8080271 | 466.1730115 | 377.4537102 | 276.0017241 | 364.8154437 | 374.8502437 | 267.0556827 | 394.096082 | 0.002524612 | 0.706555361 | YIR003W | AIM21 | n |
| W303.asm02.HP0_G0027260 | 1959.922443 | 2285.568842 | 1966.331445 | 1810.415863 | 1865.6655 | 1749.846781 | 1803.256854 | 1789.079556 | 1840.479884 | 0.095786051 | 0.195600565 | YIR004W | DJP1 | n |
| W303.asm02.HP0_G0027270 | 489.4959614 | 425.509901 | 466.1730115 | 171.216116 | 189.7511853 | 163.8501124 | 229.0751489 | 162.555633 | 169.921947 | 4.54E-13 | 1.399265618 | YIR005W | IST3 | n |
| W303.asm02.HP0_G0027280 | 605.8118334 | 843.4044122 | 457.3772943 | 1977.740703 | 1459.624502 | 1796.014492 | 1039.357494 | 1302.38025 | 1696.148592 | 2.57E-09 | -1.45700511 | YIR006C | PAN1 | n |
| W303.asm02.HP0_G0027290 | 169.6273133 | 118.9904645 | 140.7314752 | 146.895645 | 165.8664207 | 190.1023404 | 150.5080524 | 207.0649134 | 207.796116 | 0.368407833 | -0.229195515 | YIR007W | EGH1 | n |
| W303.asm02.HP0_G0027300 | 894.6629156 | 709.1831683 | 886.4128333 | 597.3107683 | 585.1767323 | 613.7589847 | 690.0652214 | 691.8290332 | 771.814145 | 0.001410911 | 0.470665442 | YIR008C | PRI1 | n |
| W303.asm02.HP0_G0027310 | 532.1451145 | 433.1252907 | 505.265088 | 90.4721522 | 116.7699602 | 103.1984134 | 15.14546439 | 97.7268984 | 113.6225068 | 0.000471613 | 2.244695238 | YIR009W | MSL1 | n |
| W303.asm02.HP0_G0027320 | 278.1887939 | 290.3367333 | 230.643251 | 270.4436378 | 327.7520473 | 286.9640091 | 279.2444997 | 291.2455091 | 436.0647557 | 0.540741858 | -0.145330006 | YIR010W | DSN1 | n |
| W303.asm02.HP0_G0027330 | 2394.168366 | 2278.905376 | 2294.704887 | 1577.91216 | 1342.854542 | 1504.524237 | 1025.158621 | 1351.727496 | 1383.942605 | 1.12E-05 | 0.654220598 | YIR011C | STS1 | n |
| W303.asm02.HP0_G0027340 | 1471.395781 | 2071.386006 | 1503.090339 | 5668.615386 | 5468.284158 | 5851.531087 | 3574.329597 | 3848.117573 | 3376.942791 | 2.00E-30 | -1.751368264 | YIR012W | SQT1 | n |
| W303.asm02.HP0_G0027350 | 28.10966907 | 8.567313442 | 20.52334013 | 64.2060435 | 61.03884283 | 58.8412006 | 178.9057981 | 118.0463525 | 130.0005257 | 2.34E-05 | -1.688258572 | YIR013C | GAT4 | n |
| W303.asm02.HP0_G0027360 | 47.49564774 | 37.12502492 | 19.54603822 | 83.66242031 | 107.4814406 | 81.47243159 | 169.4398529 | 171.2639705 | 266.1428086 | 0.000100961 | -1.382453532 | YIR014W | VLD1 | n |
| W303.asm02.HP0_G0027370 | 138.6097475 | 98.04814273 | 63.52462421 | 339.5137755 | 392.7716843 | 336.7527173 | 785.6709654 | 485.7317128 | 607.0103289 | 3.00E-11 | -1.8310882 | YIR015W | RPR2 | n |
| W303.asm02.HP0_G0027390 | 377.0572851 | 277.0098013 | 380.1704434 | 31.13020291 | 14.59624502 | 19.01023404 | 52.06253385 | 31.93057076 | 67.55932834 | 9.88E-30 | 3.978173496 | YIR017C | MET28 | n |
| W303.asm02.HP0_G0027400 | 107.5921816 | 150.4039471 | 86.97987008 | 708.2121161 | 339.6944296 | 668.0739391 | 294.3899641 | 237.0602981 | 445.2773913 | 1.42E-12 | -2.315905915 | YIR018W | YAP5 | n |
| W303.asm02.HP0_G0027410 | 62.03513174 | 55.21157552 | 41.04668026 | 324.9214928 | 184.4434598 | 266.1432765 | 62.47504062 | 105.4676428 | 182.2054613 | 1.76E-09 | -2.294373855 | YIR019C | FLO11 | n |
| W303.asm02.HP0_G0027460 | 1439.408916 | 1145.16423 | 1475.725886 | 901.8030655 | 895.6786719 | 902.5334922 | 1658.428351 | 1299.477471 | 1245.75307 | 0.000325949 | 0.588496045 | YIR022W | SEC11 | n |
| W303.asm02.HP0_G0027470 | 315.0221534 | 364.5867831 | 244.3254777 | 2092.533327 | 2204.032999 | 2171.692927 | 1034.624536 | 1846.167546 | 1831.267249 | 6.46E-35 | -2.80726634 | YIR023W | DAL81 | n |
| W303.asm02.HP0_G0027480 | 631.0136057 | 779.6255232 | 670.4291109 | 1034.106428 | 958.0444461 | 1019.310644 | 1216.370109 | 847.6115148 | 898.7437922 | 0.002382964 | -0.533458813 | YIR024C | INA22 | n |
| W303.asm02.HP0_G0027490 | 232.631744 | 237.0290052 | 224.7794395 | 143.0043696 | 177.808803 | 162.9448632 | 389.0491166 | 225.4491814 | 299.9224728 | 0.028048497 | 0.524289809 | YIR025W | MND2 | n |
| W303.asm02.HP0_G0027500 | 924.7111825 | 923.3660043 | 915.7318906 | 2104.207153 | 1999.685568 | 2122.809468 | 1391.489541 | 1283.028389 | 1223.233293 | 7.84E-27 | -1.172217572 | YIR026C | YVH1 | n |
| W303.asm02.HP0_G0027510 | 68.82022427 | 91.38467672 | 60.59271848 | 99.22752177 | 120.7507543 | 87.80917627 | 162.8137422 | 158.6852608 | 111.5752544 | 0.112344513 | -0.472537273 | YIR027C | DAL1 | n |
| W303.asm02.HP0_G0027530 | 187.0746942 | 190.3847432 | 151.4817962 | 391.073174 | 440.5412134 | 403.741161 | 513.9991978 | 531.2085863 | 631.5773574 | 2.62E-12 | -1.221918048 | YIR029W | DAL2 | n |
| W303.asm02.HP0_G0027540 | 520.5135273 | 610.1831018 | 665.5426014 | 199.4278624 | 149.9432443 | 199.1548328 | 401.3548064 | 270.926055 | 320.3949965 | 3.82E-16 | 1.705846171 | YIR030C | DCG1 | n |
| W303.asm02.HP0_G0027560 | 411.9520467 | 356.0194697 | 401.6710854 | 104.091616 | 104.8275779 | 96.86166867 | 145.7750948 | 140.3009927 | 94.1736092 | 6.36E-19 | 1.937111394 | YIR032C | DAL3 | n |
| W303.asm02.HP0_G0027570 | 911.1409974 | 1177.529636 | 950.9147594 | 1700.487334 | 1693.164423 | 1698.247574 | 1129.283689 | 1687.482285 | 2087.173795 | 0.000479999 | -0.744286333 | YIR033W | MGA2 | n |
| W303.asm02.HP0_G0027580 | 1197.084183 | 879.5775134 | 998.802553 | 273.3620943 | 278.6555868 | 318.6477325 | 431.6457352 | 438.3196532 | 466.7735413 | 2.02E-28 | 1.818868741 | YIR034C | LYS1 | n |
| W303.asm02.HP0_G0027590 | 1161.220122 | 1338.404744 | 1210.877068 | 433.877203 | 317.1365964 | 420.0356473 | 218.6626422 | 190.6158315 | 175.040078 | 8.26E-23 | 1.659700507 | YIR035C | NRE1 | n |
| W303.asm02.HP0_G0027610 | 2293.361276 | 2332.213104 | 2303.500604 | 2204.407493 | 2318.149096 | 2138.198705 | 2779.192716 | 3233.695985 | 2018.590841 | 0.759105722 | 0.057176953 | YIR037W | HYR1 | n |
| W303.asm02.HP0_G0027620 | 1153.465731 | 979.5295036 | 1078.94131 | 1647.955116 | 1139.834043 | 1682.858337 | 1308.189487 | 813.7457579 | 994.9646537 | 0.0259514 | -0.478044477 | YIR038C | GTT1 | n |
| W303.asm02.HP0_G0027720 | 498.2196518 | 483.5772476 | 371.3747262 | 20.42919566 | 23.88476458 | 23.53648024 | 92.76596941 | 41.6065013 | 80.86646877 | 9.84E-40 | 4.321242127 | YJL218W | | n |
| W303.asm02.HP0_G0027770 | 575.7635665 | 724.4139477 | 545.3344663 | 835.6513843 | 744.4084962 | 881.7127597 | 289.6570065 | 730.5327553 | 278.4263229 | 0.207406603 | -0.415954446 | YJL212C | OPT1 | n |
| W303.asm02.HP0_G0027780 | 709.5268193 | 524.5099674 | 625.473223 | 453.3335798 | 391.4447529 | 473.4453525 | 373.9036522 | 258.3473453 | 280.4735752 | 0.013623278 | 0.494485273 | YJL210W | PEX2 | n |
| W303.asm02.HP0_G0027800 | 1318.246549 | 2251.299588 | 1498.20383 | 2465.122943 | 2936.499112 | 2510.256142 | 2583.248271 | 2295.130723 | 1935.67712 | 0.002194762 | -0.642340099 | YJL208C | NUC1 | n |
| W303.asm02.HP0_G0027810 | 897.5708124 | 1189.904645 | 906.9361734 | 421.2305581 | 460.4451839 | 474.3506017 | 154.2944185 | 504.1159808 | 485.1988126 | 0.000702659 | 1.142957958 | YJL207C | LAA1 | n |
| W303.asm02.HP0_G0027820 | 296.6054736 | 332.2213768 | 285.372158 | 607.0389567 | 621.0038792 | 539.528547 | 467.6162131 | 513.7919114 | 535.3564958 | 2.51E-10 | -0.949631803 | YJL206C | | n |
| W303.asm02.HP0_G0027840 | 634.8908014 | 757.7312778 | 679.2248281 | 674.1634567 | 822.6974468 | 711.5259026 | 584.9935622 | 804.0698274 | 712.4438261 | 0.618965543 | -0.090547062 | YJL204C | RCY1 | n |
| W303.asm02.HP0_G0027850 | 850.0751646 | 878.6255897 | 892.2766447 | 359.9429711 | 360.9253315 | 338.5632157 | 361.5979624 | 322.2084868 | 335.7493893 | 1.97E-23 | 1.307913976 | YJL203W | PRP21 | n |

| | | | | | | | | | | | | | | |
|---|---|---|---|---|---|---|---|---|---|---|---|---|---|---|
| W303.asm02.HP0_G0027860 | 605.8118334 | 367.4425543 | 675.3156205 | 686.8101016 | 828.0051722 | 699.7576625 | 614.3378994 | 699.5697776 | 627.4828526 | 0.058923641 | -0.425090851 | YJL201W | ECM25 | n |
| W303.asm02.HP0_G0027870 | 1529.553717 | 1655.395342 | 1430.769998 | 848.2980292 | 871.7939073 | 1003.016158 | 747.8073044 | 664.7364277 | 971.4212514 | 7.39E-06 | 0.76077062 | YJL200C | ACO2 | n |
| W303.asm02.HP0_G0027900 | 429.3994275 | 630.1734999 | 420.2398217 | 2242.347428 | 2600.785477 | 2352.742775 | 1602.579451 | 2277.714048 | 2131.189721 | 2.32E-31 | -2.281303529 | YJL197W | UBP12 | n |
| W303.asm02.HP0_G0027920 | 241.3554344 | 228.4616918 | 186.664665 | 648.8701668 | 798.8126822 | 752.2621184 | 680.5993062 | 769.2364775 | 626.4592264 | 1.62E-24 | -1.743415999 | YJL194W | CDC6 | n |
| W303.asm02.HP0_G0027930 | 663.9697694 | 566.3946109 | 659.6787899 | 286.981558 | 276.0017241 | 262.5222796 | 279.2444997 | 425.7409435 | 329.6076322 | 2.60E-10 | 1.195288901 | YJL193W | | n |
| W303.asm02.HP0_G0027940 | 1801.926717 | 1280.337398 | 1859.805537 | 903.7487031 | 837.2936918 | 936.9329633 | 1311.975853 | 1021.778264 | 965.2794943 | 2.05E-06 | 0.883344556 | YJL192C | SOP4 | n |
| W303.asm02.HP0_G0027960 | 7648.737884 | 9647.74686 | 7726.548908 | 8181.406451 | 9401.308726 | 7991.54029 | 8358.403162 | 5956.502837 | 5283.958377 | 0.877985635 | -0.031330736 | YJL190C | RPS22A | y |
| W303.asm02.HP0_G0027980 | 710.4961182 | 572.1061532 | 724.180716 | 777.2822538 | 752.3700844 | 786.6615895 | 779.9914162 | 849.5467009 | 771.814145 | 0.159639338 | -0.207571982 | YJL187C | SWE1 | n |
| W303.asm02.HP0_G0027990 | 1571.233571 | 1442.164429 | 1624.275776 | 528.2406306 | 460.4451839 | 458.9613646 | 957.0040313 | 1587.820201 | 1129.059684 | 2.28E-18 | 1.679195345 | YJL186W | MNN5 | n |
| W303.asm02.HP0_G0028010 | 925.6804814 | 1168.010399 | 1109.237669 | 381.3449856 | 483.0030171 | 415.5094011 | 586.8867452 | 418.9677921 | 385.9070725 | 1.65E-11 | 1.325398296 | YJL184W | GON7 | n |
| W303.asm02.HP0_G0028020 | 1834.882881 | 1725.837697 | 1872.510461 | 3933.106573 | 3837.48551 | 3784.847072 | 4324.030084 | 4208.062189 | 4167.182207 | 1.16E-30 | -1.088737548 | YJL183W | MNN11 | n |
| W303.asm02.HP0_G0028040 | 1315.338653 | 1204.1835 | 1432.724602 | 1693.677602 | 1654.683413 | 1652.079863 | 1494.668017 | 1603.30169 | 1535.43928 | 0.003810754 | -0.339530494 | YJL180C | ATP12 | n |
| W303.asm02.HP0_G0028050 | 959.6059441 | 1094.712273 | 1081.873215 | 967.9547466 | 1073.487475 | 1001.205659 | 1115.084816 | 910.5050633 | 861.8932494 | 0.780797929 | 0.0443977 | YJL179W | PFD1 | n |
| W303.asm02.HP0_G0028060 | 2092.716397 | 1732.501163 | 2206.747715 | 1152.790326 | 1243.33469 | 1196.739495 | 2106.166142 | 1793.917521 | 2081.032038 | 7.83E-08 | 0.747835189 | YJL178C | ATG27 | n |
| W303.asm02.HP0_G0028070 | 6659.083673 | 7122.293242 | 6690.608883 | 13161.2661 | 13354.23727 | 11719.35666 | 10911.3605 | 9392.42577 | 9013.028576 | 4.66E-14 | -0.901145349 | YJL177W | RPL17B | y |
| W303.asm02.HP0_G0028080 | 912.1102964 | 746.3081932 | 775.0004154 | 2517.65516 | 2448.18837 | 2367.226762 | 1825.028459 | 2451.880797 | 2640.955562 | 2.10E-22 | -1.591435651 | YJL176C | SWI3 | n |
| W303.asm02.HP0_G0028090 | 3683.335947 | 2800.559572 | 3944.390513 | 1528.298399 | 1491.470855 | 1533.492212 | 1867.625078 | 1900.352757 | 1580.478833 | 1.53E-14 | 1.195365759 | YJL174W | KRE9 | n |
| W303.asm02.HP0_G0028100 | 1041.996353 | 1049.019935 | 1125.851801 | 474.7355943 | 468.4067721 | 461.6771124 | 747.8073044 | 457.6715143 | 379.7653154 | 1.32E-07 | 1.195292627 | YJL173C | RFA3 | n |
| W303.asm02.HP0_G0028110 | 536.9916091 | 633.9811947 | 534.5841453 | 257.7969928 | 295.9056946 | 284.2482613 | 371.0638776 | 503.1483878 | 549.6872624 | 6.36E-08 | 1.026686295 | YJL172W | CPS1 | n |
| W303.asm02.HP0_G0028130 | 260.7414131 | 111.3750747 | 188.6192688 | 4.864094204 | 3.980794097 | 6.336744679 | 7.572732196 | 12.57870969 | 17.40164518 | 2.54E-21 | 5.18050635 | YJL170C | ASG7 | n |
| W303.asm02.HP0_G0028140 | 911.1409974 | 831.0294039 | 944.073646 | 1426.152421 | 1423.797355 | 1454.735529 | 1125.497323 | 1314.95896 | 1333.784922 | 6.25E-08 | -0.6805144 | YJL168C | SET2 | n |
| W303.asm02.HP0_G0028150 | 9061.00643 | 7713.437869 | 8941.335184 | 7264.038285 | 6911.985484 | 7299.024622 | 7841.564189 | 5617.845268 | 5328.997929 | 0.1235616 | 0.259936513 | YJL167W | ERG20 | n |
| W303.asm02.HP0_G0028160 | 2430.032426 | 2519.742076 | 2432.504456 | 8257.286321 | 8818.785857 | 7891.962874 | 7464.820763 | 7729.133311 | 6976.012464 | 5.85E-68 | -1.757728355 | YJL166W | QCR8 | n |
| W303.asm02.HP0_G0028180 | 474.9564774 | 183.7212771 | 415.3533122 | 684.8644639 | 682.042722 | 738.6833798 | 797.9766552 | 1045.000498 | 940.7124658 | 0.000634604 | -0.971599528 | YJL164C | TPK1 | n |
| W303.asm02.HP0_G0028190 | 437.153819 | 235.1251578 | 340.101065 | 575.9087538 | 583.8498009 | 560.3492795 | 666.4004333 | 835.0328051 | 780.0031545 | 0.000483927 | -0.764769434 | YJL163C | | n |
| W303.asm02.HP0_G0028200 | 777.3777446 | 642.5485082 | 687.0432434 | 626.4953335 | 736.446908 | 576.6437658 | 649.3617858 | 664.7364277 | 617.2465907 | 0.449287013 | 0.121995651 | YJL162C | JJJ2 | n |
| W303.asm02.HP0_G0028210 | 395.4739648 | 213.2309123 | 416.3306141 | 71.98859422 | 65.01963692 | 39.83096656 | 90.87278636 | 74.50466511 | 89.05547827 | 1.90E-15 | 2.541074688 | YJL161W | FMP33 | n |
| W303.asm02.HP0_G0028220 | 167.6887155 | 73.29812612 | 125.0946446 | 5048.929784 | 3443.386894 | 4470.120747 | 3585.688695 | 4587.358666 | 3907.181156 | 1.89E-96 | -5.146840136 | YJL160C | PIR5 | n |
| W303.asm02.HP0_G0028250 | 1266.873706 | 1570.674131 | 1205.990558 | 141.0587319 | 95.53905834 | 130.3558905 | 80.46027959 | 128.6898761 | 144.3312924 | 1.50E-49 | 3.454617185 | YJL157C | FAR1 | n |
| W303.asm02.HP0_G0028260 | 1065.259528 | 1137.54884 | 1119.98799 | 623.576877 | 483.0030171 | 581.170012 | 612.4447164 | 1028.551416 | 852.6806138 | 9.13E-07 | 0.975695367 | YJL156C | SSY5 | n |
| W303.asm02.HP0_G0028270 | 948.9436558 | 888.1448268 | 968.5061938 | 135.2218189 | 260.0785477 | 158.418617 | 525.3582961 | 469.2826309 | 480.0806817 | 7.46E-27 | 2.35215006 | YJL155C | FBP26 | n |
| W303.asm02.HP0_G0028280 | 760.8996627 | 1063.298791 | 754.4770753 | 834.6785654 | 739.1007707 | 832.8293007 | 738.3413892 | 856.3198523 | 913.0745588 | 0.604885204 | 0.098923025 | YJL154C | VPS35 | n |
| W303.asm02.HP0_G0028290 | 336.3467299 | 238.9328527 | 367.4655185 | 1679.085319 | 1333.566023 | 1667.4691 | 3083.048595 | 2920.195835 | 2421.899558 | 1.13E-34 | -2.312660201 | YJL153C | INO1 | n |
| W303.asm02.HP0_G0028310 | 445.8775094 | 336.9809954 | 447.6042752 | 260.7154493 | 350.3098806 | 307.7847416 | 281.1376828 | 450.8983629 | 521.0257292 | 0.097810546 | 0.423099362 | YJL149W | DAS1 | n |
| W303.asm02.HP0_G0028320 | 3074.616217 | 4291.272111 | 3444.011934 | 3742.434081 | 4339.065566 | 3682.553908 | 3270.473717 | 2393.825214 | 2315.442435 | 0.531447129 | -0.121783232 | YJL148W | RPA34 | n |
| W303.asm02.HP0_G0028330 | 711.4654171 | 592.0965512 | 677.2702243 | 419.2849204 | 415.3295175 | 411.8884042 | 574.5810554 | 448.9631768 | 747.2471165 | 0.000823651 | 0.668184408 | YJL147C | SMT1 | n |
| W303.asm02.HP0_G0028340 | 864.6146486 | 603.5196358 | 883.4809275 | 250.9872609 | 201.6935676 | 274.2905197 | 363.4911454 | 333.8196034 | 400.2378391 | 7.76E-17 | 1.68915095 | YJL146W | IDS2 | n |
| W303.asm02.HP0_G0028350 | 2195.462084 | 1899.087813 | 2182.315167 | 582.7184857 | 516.1763013 | 573.0227689 | 751.5936705 | 543.787296 | 694.0185548 | 2.18E-34 | 1.90716936 | YJL145W | SFH5 | n |
| W303.asm02.HP0_G0028370 | 2870.094142 | 3033.780882 | 3290.575534 | 2660.65953 | 2832.998466 | 2654.190771 | 3289.405548 | 2693.779061 | 2386.072642 | 0.248888757 | 0.174542603 | YJL143W | TIM17 | n |
| W303.asm02.HP0_G0028380 | 487.5573635 | 350.3079274 | 349.8740841 | 691.6741958 | 825.3513095 | 735.9676321 | 860.4516958 | 1132.083873 | 1358.35195 | 1.15E-05 | -0.92288344 | YJL141C | YAK1 | n |
| W303.asm02.HP0_G0028390 | 2893.357316 | 2779.61725 | 3042.340849 | 1041.888979 | 1105.333828 | 993.9636654 | 1363.091795 | 1193.042235 | 1370.635464 | 2.23E-36 | 1.473285447 | YJL140W | RPB4 | n |
| W303.asm02.HP0_G0028400 | 726.0049011 | 610.1831018 | 709.5211874 | 721.8315799 | 768.2932608 | 714.2416503 | 1329.961092 | 882.4448647 | 965.2794943 | 0.581182998 | -0.107305524 | YJL139C | YUR1 | n |
| W303.asm02.HP0_G0028410 | 8243.887429 | 7812.437936 | 8631.530478 | 22343.70314 | 23365.93442 | 21782.10721 | 22417.18048 | 17814.35571 | 18098.73461 | 1.65E-31 | -1.450884799 | YKR059W | TIF1 | y |
| W303.asm02.HP0_G0028420 | 439.0924168 | 202.7597515 | 453.4680867 | 59.34194929 | 62.36577419 | 48.88345896 | 102.2318847 | 97.7268984 | 145.3549186 | 4.93E-16 | 2.686898309 | YJL137C | GLG2 | n |
| W303.asm02.HP0_G0028450 | 758.9610649 | 596.8561698 | 741.7721504 | 838.5698408 | 1004.487044 | 832.8293007 | 1274.112192 | 1292.704319 | 1340.950305 | 0.021656806 | -0.349685554 | YJL134W | LCB3 | n |
| W303.asm02.HP0_G0028460 | 883.0313284 | 554.9715263 | 776.9550192 | 28.21174638 | 54.404186 | 38.02046808 | 23.66478811 | 53.21761794 | 11.25988806 | 5.62E-27 | 4.211116056 | YJL133C-A | DPI8 | n |
| W303.asm02.HP0_G0028490 | 700.8031289 | 741.5485746 | 736.8856409 | 704.3208408 | 656.8310261 | 697.0419147 | 502.6400995 | 591.1993556 | 574.2542909 | 0.55963147 | 0.081508082 | YJL131C | AIM23 | n |
| W303.asm02.HP0_G0028500 | 2402.892056 | 3216.550236 | 2525.348138 | 9504.440075 | 9936.062067 | 9433.602329 | 2874.79846 | 5717.507353 | 4624.743113 | 1.42E-16 | -1.825787733 | YJL130C | URA2 | n |
| W303.asm02.HP0_G0028520 | 519.5442283 | 711.0870157 | 469.1049173 | 2737.512218 | 2656.516594 | 2718.463467 | 1757.820461 | 2657.010525 | 2351.269351 | 4.78E-31 | -2.254697592 | YJL128C | PBS2 | n |
| W303.asm02.HP0_G0028530 | 607.7504313 | 727.2697189 | 602.0179772 | 334.6496812 | 333.0597728 | 326.7949756 | 306.695654 | 282.5371716 | 332.6785108 | 4.66E-10 | 0.962153391 | YJL127C-B | MCO6 | n |
| W303.asm02.HP0_G0028540 | 142.4869432 | 157.0674131 | 113.3670217 | 511.7027103 | 590.4844578 | 561.2545288 | 544.2901266 | 660.8660555 | 479.0570555 | 4.62E-25 | -2.00896616 | YJL127C | SPT10 | n |
| W303.asm02.HP0_G0028550 | 608.7197302 | 403.6156555 | 533.6068434 | 393.0188117 | 265.3862732 | 437.2353829 | 339.8263573 | 226.4167745 | 319.3713703 | 0.053169431 | 0.493274069 | YJL126W | NIT2 | n |
| W303.asm02.HP0_G0028560 | 1815.496902 | 1737.260781 | 1892.0565 | 2859.114573 | 2346.014655 | 2828.903875 | 1394.329316 | 1487.190523 | 1185.359125 | 6.26E-05 | -0.561904352 | YJL125C | GCD14 | n |
| W303.asm02.HP0_G0028570 | 2477.528074 | 2415.982391 | 2628.942141 | 1606.123906 | 1678.568178 | 1567.891684 | 1706.704519 | 1556.857223 | 1482.210719 | 1.50E-08 | 0.632949474 | YJL124C | LSM1 | n |
| W303.asm02.HP0_G0028580 | 2915.651192 | 2435.972789 | 3049.181962 | 1953.420232 | 1776.761099 | 1901.023404 | 1872.358036 | 1877.130524 | 1706.384854 | 8.40E-06 | 0.576602173 | YJL123C | MTC1 | n |
| W303.asm02.HP0_G0028590 | 1676.887155 | 3129.925178 | 2119.767845 | 4472.048211 | 4891.069014 | 4492.751978 | 3351.880588 | 2640.561443 | 2336.938585 | 6.09E-06 | -1.000167934 | YJL122W | ALB1 | n |
| W303.asm02.HP0_G0028600 | 1586.742354 | 1305.087414 | 1769.893761 | 1789.013848 | 1442.374395 | 1785.151501 | 2466.817513 | 1680.709134 | 1547.722795 | 0.629367761 | -0.106407992 | YJL121C | RPE1 | n |
| W303.asm02.HP0_G0028620 | 1726.3214 | 1485.000997 | 1772.825667 | 3010.874312 | 2460.130752 | 3100.478647 | 3294.138505 | 3252.080253 | 3138.437889 | 7.90E-09 | -0.782986215 | YJL117W | PHO86 | n |
| W303.asm02.HP0_G0028640 | 710.4961182 | 732.9812612 | 728.0899237 | 1340.544363 | 1346.835336 | 1430.293799 | 1823.135276 | 1515.250722 | 1290.792622 | 8.97E-10 | -0.923301175 | YJL115W | ASF1 | n |
| W303.asm02.HP0_G0028650 | 1116.632371 | 1000.471825 | 1150.284349 | 885.2651452 | 935.4866129 | 856.365781 | 756.3266281 | 920.1809938 | 771.814145 | 0.040319636 | 0.288206113 | YJL112W | MDV1 | n |
| W303.asm02.HP0_G0028660 | 4943.42456 | 4623.493488 | 5124.971221 | 5143.293211 | 4729.183388 | 4982.491817 | 3830.8559 | 3966.163926 | 3597.022421 | 0.886814968 | -0.01615783 | YJL111W | CCT7 | n |
| W303.asm02.HP0_G0028670 | 1757.338966 | 2811.030733 | 1911.602538 | 7371.048357 | 6608.118202 | 7561.546901 | 2779.192716 | 3122.422783 | 2793.475864 | 3.46E-22 | -1.733041237 | YJL109C | UTP10 | n |
| W303.asm02.HP0_G0028680 | 449.7547051 | 161.8270317 | 207.1880051 | 672.217819 | 792.1780254 | 678.9369299 | 850.9857806 | 1389.463625 | 1316.383276 | 1.78E-05 | -1.388050031 | YJL107C | | n |

| | | | | | | | | | | | | | | |
|---|---|---|---|---|---|---|---|---|---|---|---|---|---|---|
| W303.asm02.HP0_G0028690 | 120.1930677 | 99.00006644 | 117.2762293 | 47.6681232 | 39.80794097 | 57.93595136 | 59.63526605 | 120.9491317 | 94.1736092 | 0.000324166 | 1.201204478 | YJL106W | IME2 | n |
| W303.asm02.HP0_G0028700 | 254.9256195 | 202.7597515 | 297.0997809 | 871.6456814 | 1107.98769 | 849.123787 | 785.6709654 | 1040.162532 | 843.4679781 | 4.06E-22 | -1.904995402 | YJL105W | SET4 | n |
| W303.asm02.HP0_G0028710 | 2494.006156 | 2272.24191 | 2799.969975 | 1358.055102 | 1374.700895 | 1282.738173 | 1716.170434 | 1322.699704 | 1170.004732 | 3.26E-08 | 0.91420215 | YJL104W | PAM16 | n |
| W303.asm02.HP0_G0028720 | 346.0397192 | 450.2599176 | 396.7845759 | 57.39631161 | 67.67349965 | 61.55694831 | 77.62050501 | 54.18521099 | 69.60658071 | 6.63E-31 | 2.681752208 | YJL103C | GSM1 | n |
| W303.asm02.HP0_G0028730 | 314.0528544 | 376.9617915 | 269.7353274 | 1230.615834 | 1163.718808 | 1189.497501 | 776.2050501 | 1255.935783 | 1215.044284 | 3.39E-20 | -1.899181749 | YJL102W | MEF2 | n |
| W303.asm02.HP0_G0028740 | 1136.01835 | 1245.11622 | 956.7785709 | 5356.340538 | 4864.530387 | 5158.110169 | 3386.904475 | 4125.81678 | 4239.859666 | 1.66E-52 | -2.203970312 | YJL101C | GSH1 | n |
| W303.asm02.HP0_G0028750 | 319.868648 | 362.6829357 | 314.6912153 | 433.877203 | 268.0401359 | 372.0574376 | 392.8354827 | 371.5557325 | 624.4119741 | 0.685871998 | -0.109727109 | YJL100W | LSB6 | n |
| W303.asm02.HP0_G0028760 | 542.8074027 | 505.4714931 | 513.0835033 | 728.6413118 | 737.7738394 | 764.9356077 | 1209.743968 | 1116.602384 | 1304.099762 | 5.40E-05 | -0.515428955 | YJL099W | CHS6 | n |
| W303.asm02.HP0_G0028780 | 1589.650251 | 1180.385408 | 1875.442367 | 757.825877 | 683.3696534 | 732.3466351 | 1419.887287 | 1139.824617 | 1008.271794 | 2.90E-08 | 1.095205526 | YJL097W | PHS1 | n |
| W303.asm02.HP0_G0028790 | 1041.027055 | 1651.587647 | 1051.576856 | 1622.661827 | 1710.41453 | 1653.890361 | 2063.569524 | 1504.607198 | 1412.604138 | 0.05794996 | -0.413323162 | YJL096W | MRPL49 | n |
| W303.asm02.HP0_G0028800 | 206.4606728 | 276.0578776 | 223.8021376 | 564.2349277 | 583.8498009 | 540.4337962 | 489.3878182 | 614.4215889 | 660.2388906 | 4.51E-12 | -1.25644916 | YJL095W | BCK1 | n |
| W303.asm02.HP0_G0028810 | 413.8906446 | 501.6637982 | 472.036823 | 734.4782248 | 642.234781 | 763.1251093 | 1027.998396 | 1159.176478 | 1096.303646 | 1.63E-05 | -0.626716754 | YJL094C | KHA1 | n |
| W303.asm02.HP0_G0028820 | 757.022467 | 750.115888 | 746.65866 | 379.3993479 | 555.9842423 | 416.4146504 | 424.073003 | 518.6298766 | 463.7026627 | 2.84E-05 | 0.741727832 | YJL093C | TOK1 | n |
| W303.asm02.HP0_G0028830 | 342.1625235 | 451.2118413 | 330.3280459 | 546.7241885 | 619.6769478 | 534.0970516 | 438.2718759 | 662.8012416 | 595.7504408 | 0.003946714 | -0.596392018 | YJL092W | SRS2 | n |
| W303.asm02.HP0_G0028840 | 940.2199654 | 946.2121735 | 1025.189705 | 628.4409712 | 624.9846733 | 605.6117415 | 954.1642567 | 901.7967258 | 887.4839041 | 4.08E-08 | 0.647575658 | YJL091C | GWT1 | n |
| W303.asm02.HP0_G0028850 | 524.390723 | 703.471626 | 593.22226 | 626.4953335 | 623.6577419 | 638.2007141 | 372.9570607 | 557.3335988 | 530.2383649 | 0.800599336 | -0.052365142 | YJL090C | DPB11 | n |
| W303.asm02.HP0_G0028860 | 253.9563206 | 554.0196026 | 282.4402523 | 130.3577247 | 124.7315484 | 132.166389 | 118.3239406 | 143.2037719 | 145.3549186 | 5.95E-07 | 1.492989735 | YJL089W | SIP4 | n |
| W303.asm02.HP0_G0028870 | 1042.965652 | 817.7024719 | 835.5931339 | 1302.604428 | 1117.27621 | 1315.327146 | 1010.013157 | 740.2086859 | 868.0350066 | 0.0062034 | -0.47104282 | YJL088W | ARG3 | n |
| W303.asm02.HP0_G0028880 | 611.627627 | 1142.308459 | 693.8843568 | 1789.013848 | 1794.011207 | 1711.826313 | 1173.77349 | 1426.232161 | 1205.831648 | 3.40E-07 | -1.112879903 | YJL087C | TRL1 | n |
| W303.asm02.HP0_G0028890 | 970.2682324 | 1084.241112 | 1123.897198 | 592.4466741 | 627.638536 | 611.0432369 | 595.4060689 | 658.9308694 | 705.2784428 | 2.48E-09 | 0.796172117 | YJL085W | EXO70 | n |
| W303.asm02.HP0_G0028910 | 59.12723494 | 115.1827696 | 37.13747262 | 167.3248406 | 165.8664207 | 168.3763586 | 156.1876016 | 216.740844 | 370.5526797 | 0.002183995 | -1.24534631 | YJL083W | TAX4 | n |
| W303.asm02.HP0_G0028930 | 1145.711339 | 1240.356602 | 1178.626105 | 1087.611464 | 1069.506681 | 1089.014836 | 1038.410902 | 1102.088488 | 1218.115163 | 0.277805398 | 0.134885726 | YJL081C | ARP4 | n |
| W303.asm02.HP0_G0028940 | 1990.940009 | 2071.386006 | 1740.574703 | 5238.629458 | 4803.491544 | 5407.05371 | 4054.2515 | 3873.274993 | 3321.666977 | 1.83E-27 | -1.412858302 | YJL080C | SCP160 | n |
| W303.asm02.HP0_G0028990 | 662.0311715 | 887.1929031 | 702.680074 | 883.3195075 | 958.0444461 | 975.8586806 | 1027.998396 | 1098.218116 | 817.8773234 | 0.066882277 | -0.322898755 | YJL074C | SMC3 | n |
| W303.asm02.HP0_G0029000 | 447.8161072 | 563.5388398 | 455.4226905 | 659.5711741 | 610.3884283 | 646.3479573 | 666.4004333 | 809.8753857 | 631.5773574 | 0.020998207 | -0.386181899 | YJL073W | JEM1 | n |
| W303.asm02.HP0_G0029010 | 878.1848337 | 999.5199016 | 961.6650804 | 577.8543915 | 448.5028016 | 538.6232978 | 246.1137964 | 405.4214894 | 309.1351085 | 1.58E-05 | 0.857714596 | YJL072C | PSF2 | n |
| W303.asm02.HP0_G0029020 | 910.1716985 | 870.0582762 | 971.4380995 | 413.4480074 | 229.5591263 | 372.0574376 | 270.725176 | 234.1575189 | 289.6862109 | 1.91E-11 | 1.433536092 | YJL071W | ARG2 | n |
| W303.asm02.HP0_G0029040 | 890.7857198 | 1135.644993 | 812.137888 | 4167.555914 | 3463.290865 | 4028.359118 | 1745.514771 | 2398.663179 | 2092.291926 | 1.19E-31 | -2.038368386 | YJL069C | UTP18 | n |
| W303.asm02.HP0_G0029050 | 1577.049365 | 1046.164164 | 1527.522887 | 1526.352761 | 1295.085013 | 1515.387228 | 1502.240749 | 1360.435833 | 1008.271794 | 0.77428441 | -0.063805157 | YJL068C | | n |
| W303.asm02.HP0_G0029060 | 805.4874137 | 484.5291713 | 682.1567339 | 821.0591017 | 875.7747014 | 796.6193311 | 1626.244239 | 1125.310721 | 1086.067384 | 0.136435388 | -0.338082941 | YJL066C | MPM1 | n |
| W303.asm02.HP0_G0029070 | 664.9390683 | 465.490697 | 665.5426014 | 365.7798842 | 387.4639588 | 427.2776412 | 296.2831472 | 435.416874 | 431.9702509 | 0.005148767 | 0.604779858 | YJL065C | DLS1 | n |
| W303.asm02.HP0_G0029080 | 1404.514155 | 1363.154761 | 1466.930168 | 1512.733297 | 1450.335983 | 1487.324501 | 1713.330659 | 1440.746057 | 1273.390977 | 0.632150239 | -0.071927946 | YJL063C | MRPL8 | n |
| W303.asm02.HP0_G0029090 | 1634.238002 | 1706.799222 | 1760.120742 | 722.8043987 | 740.4277021 | 735.9676321 | 1034.624536 | 746.9818372 | 663.3097692 | 1.88E-12 | 1.213944737 | YJL062W-/ | COA3 | n |
| W303.asm02.HP0_G0029100 | 694.9873353 | 641.5965844 | 660.6560918 | 718.9131234 | 731.1391825 | 747.7358722 | 520.6253385 | 952.1115646 | 906.9328017 | 0.547892466 | -0.138101643 | YJL062W | LAS21 | n |
| W303.asm02.HP0_G0029110 | 1761.216162 | 1944.780151 | 1969.263351 | 1842.518885 | 1823.203697 | 1814.119477 | 1559.982832 | 1839.394395 | 1597.880478 | 0.696070643 | 0.05055133 | YJL061W | NUP82 | n |
| W303.asm02.HP0_G0029120 | 1149.588535 | 717.7504817 | 1195.240237 | 1034.106428 | 650.1963692 | 1200.360492 | 1095.206394 | 2583.473453 | 1245.75307 | 0.809672587 | 0.085404212 | YJL060W | BNA3 | n |
| W303.asm02.HP0_G0029130 | 595.1495451 | 398.8560369 | 639.1554498 | 186.7812174 | 242.8284399 | 156.6081185 | 327.5206675 | 308.662184 | 311.1823608 | 1.24E-10 | 1.484299043 | YJL059W | YHC3 | n |
| W303.asm02.HP0_G0029140 | 287.8817832 | 292.2405807 | 287.3267618 | 163.4335653 | 157.9048325 | 161.1343647 | 229.0751489 | 219.6436231 | 258.9774253 | 8.81E-07 | 0.845496135 | YJL058C | BIT61 | n |
| W303.asm02.HP0_G0029150 | 737.6364883 | 472.154163 | 604.9498829 | 277.2533696 | 148.616313 | 305.0689939 | 296.2831472 | 473.1530031 | 465.7499151 | 6.78E-06 | 1.306750117 | YJL057C | IKS1 | n |
| W303.asm02.HP0_G0029160 | 270.4344024 | 247.5001661 | 190.5738726 | 1638.226928 | 1491.470855 | 1559.74444 | 913.4608212 | 1349.79231 | 1272.36735 | 1.83E-46 | -2.726588391 | YJL056C | ZAP1 | n |
| W303.asm02.HP0_G0029180 | 1167.035916 | 1012.846834 | 1033.985422 | 1981.631979 | 1874.95402 | 1945.380617 | 1665.054492 | 1813.269382 | 1791.345827 | 9.97E-14 | -0.852573564 | YJL054W | TIM54 | n |
| W303.asm02.HP0_G0029190 | 1612.913425 | 933.8371652 | 1442.497621 | 350.2147827 | 326.425116 | 321.3634802 | 439.2184674 | 417.032606 | 448.3482627 | 5.40E-23 | 1.998720125 | YJL053W | PEP8 | n |
| W303.asm02.HP0_G0029210 | 584.4872569 | 605.4234832 | 553.1528816 | 634.2778842 | 769.6201921 | 617.3799816 | 640.8424621 | 664.7364277 | 885.4366517 | 0.24716399 | -0.211692902 | YJL051W | IRC8 | n |
| W303.asm02.HP0_G0029220 | 1096.277094 | 1582.097216 | 1153.216255 | 7187.185596 | 6845.638916 | 7420.32802 | 2332.401516 | 3223.052461 | 2672.687974 | 5.24E-46 | -2.485144851 | YJL050W | MTR4 | n |
| W303.asm02.HP0_G0029230 | 1111.785877 | 1116.606519 | 1153.216255 | 302.5466595 | 319.7904592 | 283.3430121 | 385.2627505 | 350.2686853 | 383.8598201 | 5.61E-48 | 1.903562983 | YJL049W | CHM7 | n |
| W303.asm02.HP0_G0029250 | 605.8118334 | 671.1062196 | 616.6775058 | 402.7470001 | 467.0798408 | 445.382626 | 376.7434268 | 426.7085366 | 324.4895013 | 0.000974844 | 0.527240153 | YJL047C | RTT101 | n |
| W303.asm02.HP0_G0029260 | 914.0488942 | 1019.5103 | 1033.00812 | 694.5926524 | 654.1771633 | 621.9062278 | 647.4686028 | 648.2873458 | 639.7663669 | 2.31E-06 | 0.59026571 | YJL046W | AIM22 | n |
| W303.asm02.HP0_G0029280 | 1085.614805 | 886.2409794 | 1038.871931 | 253.9057175 | 169.8472148 | 238.9857993 | 384.316159 | 264.1529036 | 354.1746607 | 5.38E-26 | 2.177708293 | YJL044C | GYP6 | n |
| W303.asm02.HP0_G0029300 | 1241.671934 | 1560.20297 | 1239.218823 | 3949.644494 | 4049.794528 | 3859.982759 | 2477.23002 | 3280.140451 | 3098.516468 | 1.01E-24 | -1.55306186 | YJL042W | MHP1 | n |
| W303.asm02.HP0_G0029310 | 1263.965809 | 1759.155027 | 1380.9276 | 3792.047842 | 4271.392066 | 3961.370674 | 2008.667215 | 3130.163528 | 2801.664874 | 1.56E-14 | -1.448942133 | YJL039C | NUP192 | n |
| W303.asm02.HP0_G0029340 | 1147.649937 | 1209.895043 | 1204.035954 | 269.4708189 | 268.0401359 | 271.574772 | 406.087764 | 380.26407 | 379.7653154 | 9.62E-64 | 2.137805673 | YJL036W | SNX4 | n |
| W303.asm02.HP0_G0029350 | 467.2020859 | 407.4233504 | 434.8993504 | 302.5466595 | 352.9637433 | 296.0165015 | 245.1672049 | 266.0880897 | 219.056004 | 0.005671255 | 0.464095095 | YJL035C | TAD2 | n |
| W303.asm02.HP0_G0029360 | 4885.266624 | 4919.541763 | 4042.120704 | 7584.095683 | 6634.656829 | 7403.128284 | 6881.720383 | 11321.80632 | 11915.00882 | 0.001449236 | -0.642990881 | YJL034W | KAR2 | n |
| W303.asm02.HP0_G0029370 | 631.9829046 | 1159.443086 | 660.6560918 | 5314.509327 | 4544.739928 | 5221.477616 | 2330.508333 | 2852.464322 | 2242.764976 | 2.64E-32 | -2.620580767 | YJL033W | HCA4 | n |
| W303.asm02.HP0_G0029390 | 875.2769369 | 859.5871154 | 918.6637963 | 337.5681378 | 289.2710377 | 315.9319847 | 123.0568982 | 275.7640202 | 197.5598541 | 8.19E-10 | 1.49189103 | YJL030W | MAD2 | n |
| W303.asm02.HP0_G0029400 | 492.4038582 | 495.0003322 | 552.1755797 | 1085.665826 | 1121.257004 | 1071.8151 | 856.6653297 | 1268.514493 | 1090.161889 | 5.08E-11 | -1.090415346 | YJL029C | VPS53 | n |
| W303.asm02.HP0_G0029430 | 2834.230081 | 2488.328593 | 2873.267618 | 6248.415415 | 5573.111736 | 6139.400345 | 5072.78398 | 4773.136533 | 4173.323964 | 2.04E-19 | -1.132124478 | YJL026W | RNR2 | n |
| W303.asm02.HP0_G0029440 | 220.0308579 | 211.3270649 | 179.8235516 | 662.4896306 | 687.3504475 | 676.2211822 | 619.0708571 | 675.3799513 | 684.8059191 | 1.91E-32 | -1.728450466 | YJL025W | RRN7 | n |
| W303.asm02.HP0_G0029460 | 505.0047443 | 471.2022393 | 498.4239746 | 561.3164712 | 492.2915367 | 486.1188418 | 650.3083774 | 806.9726066 | 614.1757122 | 0.718369634 | -0.06292875 | YJL023C | PET130 | n |
| W303.asm02.HP0_G0029470 | 681.4171502 | 595.9042461 | 511.1288994 | 1298.713153 | 1069.506681 | 1408.567817 | 886.009667 | 1306.250622 | 1396.226119 | 1.45E-07 | -1.079237385 | YJL020C | BBC1 | n |
| W303.asm02.HP0_G0029480 | 226.8159504 | 297.0001993 | 204.2560994 | 175.1073914 | 236.1937831 | 179.2393495 | 137.2557711 | 262.2177175 | 328.584006 | 0.349772517 | 0.305020954 | YJL019W | MPS3 | n |
| W303.asm02.HP0_G0029490 | 632.9522035 | 688.2408465 | 642.0873555 | 613.8486886 | 563.9458305 | 631.8639695 | 1175.666673 | 1249.162632 | 1731.975508 | 0.53341775 | 0.116602599 | YJL016W | TPH3 | n |
| W303.asm02.HP0_G0029500 | 4145.691538 | 3639.204365 | 4215.103142 | 4862.148566 | 4495.643467 | 4704.5803 | 3502.388641 | 3404.959955 | 3459.856512 | 0.036096457 | -0.229029532 | YJL014W | CCT3 | n |

| | | | | | | | | | | | | | | |
|---|---|---|---|---|---|---|---|---|---|---|---|---|---|---|
| W303.asm02.HP0_G0029520 | 6030.977964 | 5315.542029 | 6014.31596 | 4287.212632 | 2672.439771 | 4213.935212 | 2033.278595 | 3892.626854 | 2740.247303 | 0.009156013 | 0.635384709 | YJL012C | VTC4 | n |
| W303.asm02.HP0_G0029530 | 694.0180363 | 771.0582098 | 748.6132638 | 798.6842683 | 861.1784564 | 774.8933494 | 685.3322638 | 607.6484376 | 556.8526457 | 0.349338422 | -0.136177529 | YJL011C | RPC17 | n |
| W303.asm02.HP0_G0029540 | 1345.38692 | 1795.328128 | 1642.844512 | 3957.427045 | 3317.328414 | 3928.781701 | 1673.573815 | 1866.487 | 1838.432632 | 4.68E-17 | -1.22813482 | YJL010C | NOP9 | n |
| W303.asm02.HP0_G0029550 | 4763.134959 | 4511.166489 | 4842.530969 | 4699.67892 | 4186.468459 | 4487.320482 | 4117.673132 | 3957.455589 | 3931.748184 | 0.467488007 | 0.07768528 | YJL008C | CCT8 | n |
| W303.asm02.HP0_G0029560 | 829.719887 | 770.1062861 | 793.5691517 | 805.4940002 | 749.7162217 | 854.5552825 | 571.7412808 | 646.3521597 | 523.0729815 | 0.940431674 | -0.011129404 | YJL006C | CTK2 | n |
| W303.asm02.HP0_G0029570 | 597.088143 | 867.2025051 | 670.4291109 | 1093.448377 | 1131.872455 | 1118.888061 | 1158.628026 | 1332.375635 | 1237.56406 | 6.71E-05 | -0.647333219 | YJL005W | CYR1 | n |
| W303.asm02.HP0_G0029580 | 1223.255254 | 965.2506478 | 1304.698051 | 339.5137755 | 334.3867042 | 335.847468 | 585.9401537 | 476.0557823 | 431.9702509 | 2.67E-24 | 1.790343871 | YJL004C | SYS1 | n |
| W303.asm02.HP0_G0029590 | 598.0574419 | 543.5484417 | 585.4038447 | 404.6926378 | 388.7908902 | 401.9306625 | 455.3105233 | 368.6529534 | 372.5999321 | 0.000466108 | 0.530023465 | YJL003W | COX16 | n |
| W303.asm02.HP0_G0029600 | 3321.787445 | 2939.540434 | 3510.468464 | 2433.019921 | 2282.321949 | 2362.700516 | 2401.502698 | 2609.598465 | 2521.191299 | 5.37E-05 | 0.46497507 | YJL002C | OST1 | n |
| W303.asm02.HP0_G0029610 | 3206.440872 | 2519.742076 | 3198.709155 | 136.1946377 | 114.1160975 | 133.0716383 | 109.8046168 | 131.5926553 | 147.4021709 | 2.81E-147 | 4.535294574 | YJL001W | PRE3 | n |
| W303.asm02.HP0_G0029620 | 1391.913268 | 951.9237158 | 1212.831672 | 1325.95208 | 1287.123425 | 1338.863626 | 1199.331462 | 1652.648935 | 1530.32115 | 0.421041186 | -0.152254753 | YJR001W | AVT1 | n |
| W303.asm02.HP0_G0029630 | 1371.557991 | 1596.376071 | 1363.336166 | 5871.934523 | 5035.704533 | 5751.048421 | 2265.193518 | 3044.047746 | 2144.496862 | 2.74E-33 | -1.943541036 | YJR002W | MPP10 | n |
| W303.asm02.HP0_G0029640 | 1313.400055 | 1440.260582 | 1490.385414 | 453.3335798 | 628.9654674 | 477.0663494 | 757.2732196 | 688.926254 | 814.8064448 | 2.83E-19 | 1.448375212 | YJR003C | MRX12 | n |
| W303.asm02.HP0_G0029650 | 288.8510822 | 249.4040135 | 280.4856485 | 184.8355798 | 192.405048 | 209.1125744 | 159.9739676 | 224.4815884 | 241.5757801 | 0.024133861 | 0.480710134 | YJR004C | SAG1 | n |
| W303.asm02.HP0_G0029660 | 1001.285798 | 1148.971925 | 1013.462082 | 840.5154785 | 895.6786719 | 858.1762794 | 800.8164298 | 860.1902245 | 951.9723539 | 0.032469755 | 0.286915521 | YJR005W | APL1 | n |
| W303.asm02.HP0_G0029680 | 806.4567126 | 615.8946441 | 818.0016995 | 787.0104422 | 760.3316726 | 802.9560758 | 724.1425163 | 754.7225817 | 734.9636022 | 0.66720531 | -0.069764624 | YJR006W | POL31 | n |
| W303.asm02.HP0_G0029690 | 4240.682834 | 5189.888099 | 4685.185361 | 7855.51214 | 8392.840888 | 7615.861855 | 7061.572773 | 5890.706509 | 5977.976932 | 5.36E-09 | -0.757409201 | YJR007W | SUI2 | n |
| W303.asm02.HP0_G0029700 | 786.101435 | 524.5099674 | 751.5451696 | 58.36913045 | 69.00043102 | 64.27269603 | 169.4398829 | 106.4352359 | 153.543928 | 8.17E-43 | 3.431462914 | YJR008W | MHO1 | n |
| W303.asm02.HP0_G0029720 | 835.5356806 | 633.029271 | 805.2967747 | 288.9271957 | 199.0397049 | 298.7322492 | 298.1763302 | 228.3519606 | 258.9774253 | 2.21E-13 | 1.525753238 | YJR010W | MET3 | n |
| W303.asm02.HP0_G0029730 | 805.4874137 | 702.5197023 | 778.9096231 | 471.8171378 | 468.4067721 | 456.2456169 | 826.3744009 | 539.9169238 | 528.1911125 | 0.000277921 | 0.711721114 | YJR010C-A | SPC1 | n |
| W303.asm02.HP0_G0029750 | 1236.825439 | 886.2409794 | 1253.878352 | 489.3278769 | 570.5804873 | 461.6771124 | 757.2732196 | 857.2874453 | 758.5070046 | 6.48E-11 | 1.152192168 | YJR013W | GPI14 | n |
| W303.asm02.HP0_G0029760 | 2521.146526 | 2152.299521 | 2458.891608 | 2442.748109 | 2879.441064 | 2437.836203 | 2233.955998 | 1748.440648 | 1515.990383 | 0.497873337 | -0.121281388 | YJR014W | TMA22 | n |
| W303.asm02.HP0_G0029800 | 5466.845985 | 4076.137351 | 4244.422199 | 6547.070799 | 5669.977726 | 7146.0375 | 3658.576242 | 3809.413851 | 4238.83604 | 0.00188335 | -0.490191526 | YJR016C | ILV3 | n |
| W303.asm02.HP0_G0029810 | 1505.321244 | 1218.462356 | 1587.138303 | 727.6684929 | 639.5809183 | 749.5463707 | 919.1403703 | 564.1067502 | 713.4674523 | 1.87E-07 | 1.025131387 | YJR017C | ESS1 | n |
| W303.asm02.HP0_G0029820 | 645.5530897 | 302.7117416 | 529.6976358 | 277.2533696 | 249.4630968 | 334.0369695 | 448.6843826 | 344.463127 | 428.8993723 | 0.004013925 | 0.777865059 | YJR019C | TES1 | n |
| W303.asm02.HP0_G0029830 | 208.3992707 | 90.432753 | 149.5271924 | 17.51073914 | 14.59624502 | 12.67348936 | 39.75684403 | 20.31945412 | 26.61428086 | 6.23E-14 | 3.324248711 | YJR021C | REC107 | y |
| W303.asm02.HP0_G0029840 | 479.802972 | 391.2406472 | 556.0847874 | 277.2533696 | 355.617606 | 250.7540395 | 377.6900183 | 315.4353354 | 343.9383988 | 0.000705815 | 0.696080028 | YJR022W | LSM8 | n |
| W303.asm02.HP0_G0029850 | 1610.005528 | 1677.289587 | 1695.618816 | 471.8171378 | 563.9458305 | 527.7603069 | 703.3175027 | 577.6530529 | 616.2229646 | 5.21E-35 | 1.674059592 | YJR024C | MDE1 | n |
| W303.asm02.HP0_G0029870 | 402.2590574 | 434.0772144 | 388.9661606 | 2054.593392 | 2092.570764 | 2164.450933 | 1278.84515 | 1757.148985 | 1642.92003 | 5.40E-56 | -2.364826716 | YJR030C | RBH2 | n |
| W303.asm02.HP0_G0029890 | 920.8339868 | 1378.38554 | 1109.237669 | 791.8745364 | 851.8899368 | 818.3453129 | 625.6969977 | 656.0280902 | 598.8213194 | 0.0054995 | 0.46972286 | YJR032W | CPR7 | n |
| W303.asm02.HP0_G0029900 | 257.8335163 | 343.6444614 | 264.8488179 | 455.2792175 | 441.8681448 | 442.6668783 | 381.4763844 | 523.4678419 | 498.5059531 | 0.001033407 | -0.628924183 | YJR033C | RAV1 | n |
| W303.asm02.HP0_G0029910 | 531.1758155 | 632.0773473 | 547.2890702 | 550.6154639 | 563.9458305 | 624.6219755 | 701.4243197 | 583.4586112 | 507.7185887 | 0.894667223 | -0.024492494 | YJR034W | PET191 | n |
| W303.asm02.HP0_G0029920 | 283.0352886 | 356.9713934 | 257.0304026 | 339.5137755 | 420.6372429 | 362.099696 | 120.2171236 | 471.217817 | 376.6944368 | 0.411671606 | -0.322447527 | YJR035W | RAD26 | n |
| W303.asm02.HP0_G0029930 | 163.8115197 | 150.4039471 | 158.3229096 | 58.36913045 | 69.00043102 | 45.262462 | 51.11594233 | 70.6342929 | 70.6302069 | 9.02E-08 | 1.464772332 | YJR036C | HUL4 | n |
| W303.asm02.HP0_G0029950 | 496.2810539 | 582.5773141 | 528.7203338 | 1033.133609 | 1050.929642 | 1070.909851 | 958.8972144 | 1137.889431 | 1068.665739 | 1.97E-13 | -0.972680485 | YJR040W | GEF1 | n |
| W303.asm02.HP0_G0029960 | 874.307638 | 1247.020068 | 1027.144308 | 4256.082429 | 4108.179508 | 4288.16565 | 1483.308919 | 2368.667795 | 1908.039213 | 3.11E-26 | -2.006675534 | YJR041C | URB2 | n |
| W303.asm02.HP0_G0029970 | 1690.45734 | 1701.08768 | 1802.144724 | 2397.998443 | 2464.111546 | 2433.309957 | 2125.097973 | 2204.176976 | 1915.204596 | 6.26E-06 | -0.490137967 | YJR042W | NUP85 | n |
| W303.asm02.HP0_G0029980 | 635.8601003 | 717.7504817 | 693.8843568 | 556.452377 | 452.4835957 | 564.8755257 | 530.0912537 | 507.9863531 | 479.0570555 | 0.012449241 | 0.376467662 | YJR043C | POL32 | n |
| W303.asm02.HP0_G0029990 | 1568.325674 | 1602.087614 | 1719.074061 | 322.0030363 | 278.6555868 | 356.6682005 | 756.3266281 | 552.4956335 | 506.6949626 | 2.15E-38 | 2.349376301 | YJR044C | VPS55 | n |
| W303.asm02.HP0_G0030000 | 7331.777132 | 7370.745331 | 5158.199486 | 13090.25032 | 12633.71353 | 12858.1602 | 11622.25074 | 12056.20945 | 11560.83416 | 1.50E-10 | -0.95801191 | YJR045C | SSC1 | n |
| W303.asm02.HP0_G0030010 | 950.8822537 | 724.4139477 | 566.8351084 | 842.4611162 | 826.6782409 | 851.8395348 | 839.6266823 | 824.3892815 | 822.9954543 | 0.388776187 | -0.169235981 | YJR046W | TAH11 | n |
| W303.asm02.HP0_G0030020 | 5567.653074 | 5501.167154 | 6094.454717 | 8721.320908 | 10425.69974 | 8264.020311 | 8593.15786 | 8184.869639 | 7539.006867 | 1.70E-07 | -0.675180634 | YJR048W | CYC1 | n |
| W303.asm02.HP0_G0030030 | 906.2945028 | 925.2698518 | 1022.257799 | 312.2748479 | 372.8677138 | 324.9844771 | 488.4412267 | 547.6576682 | 564.018029 | 2.65E-25 | 1.501921374 | YJR049C | UTR1 | n |
| W303.asm02.HP0_G0030040 | 916.956791 | 855.7794205 | 892.2766447 | 571.0446596 | 488.3107426 | 541.3390455 | 567.0083232 | 553.4632266 | 492.3641959 | 5.49E-08 | 0.733305021 | YJR050W | ISY1 | n |
| W303.asm02.HP0_G0030060 | 645.5530897 | 693.9523888 | 657.7241861 | 680.0003697 | 732.4661139 | 656.3056989 | 478.9753114 | 667.6392069 | 553.7817672 | 0.774255337 | -0.049720164 | YJR052W | RAD7 | n |
| W303.asm02.HP0_G0030070 | 127.9474592 | 176.1058874 | 155.3910038 | 390.1003552 | 382.1562333 | 351.2367051 | 333.2002166 | 470.250224 | 436.0647557 | 2.33E-10 | -1.289222548 | YJR053W | BFA1 | n |
| W303.asm02.HP0_G0030080 | 561.2240825 | 1196.568111 | 668.4745071 | 463.0617682 | 303.8672828 | 363.9101944 | 297.2297387 | 434.449281 | 370.5526797 | 0.000130262 | 1.099972939 | YJR054W | KCH1 | n |
| W303.asm02.HP0_G0030090 | 555.4082889 | 538.7888231 | 507.2196918 | 568.126203 | 440.5412134 | 526.8550576 | 272.6183591 | 242.8658564 | 191.418097 | 0.765847798 | 0.058494358 | YJR055W | HIT1 | n |
| W303.asm02.HP0_G0030110 | 1164.128019 | 935.7410126 | 1149.307047 | 499.0560653 | 455.1374585 | 479.7820972 | 743.0743468 | 513.7919114 | 463.7026627 | 2.13E-09 | 1.179381146 | YJR057W | CDC8 | n |
| W303.asm02.HP0_G0030120 | 987.7156132 | 751.0678118 | 941.1417403 | 285.0359204 | 335.7136355 | 278.8167659 | 534.8242114 | 407.3566755 | 377.718063 | 1.63E-16 | 1.577619771 | YJR058C | APS2 | n |
| W303.asm02.HP0_G0030140 | 1053.627941 | 804.3755398 | 968.5061938 | 1465.065174 | 1502.086306 | 1590.522915 | 2058.836566 | 1339.148786 | 1653.156292 | 0.000152095 | -0.689440628 | YJR060W | CBF1 | n |
| W303.asm02.HP0_G0030150 | 433.2766232 | 426.4618247 | 451.5134829 | 220.8298769 | 244.1553713 | 232.6490547 | 306.695654 | 423.8057574 | 479.0570555 | 3.16E-06 | 0.911675935 | YJR061W | MNN14 | n |
| W303.asm02.HP0_G0030160 | 813.2418051 | 726.3177952 | 826.7974167 | 308.3835725 | 305.1942141 | 285.1535106 | 501.693508 | 448.9631768 | 587.5614313 | 5.43E-19 | 1.397503481 | YJR062C | NTA1 | n |
| W303.asm02.HP0_G0030180 | 373.1800894 | 707.2793208 | 460.3092001 | 1498.141015 | 1775.434167 | 1700.058073 | 1351.732697 | 854.3846662 | 944.8069706 | 4.65E-12 | -1.690317984 | YJR063W | RPA12 | n |
| W303.asm02.HP0_G0030190 | 2461.049992 | 2543.540169 | 2466.710023 | 3492.419639 | 3435.425306 | 3567.587255 | 2858.706404 | 3148.547796 | 3178.359311 | 8.55E-07 | -0.490474257 | YJR064W | CCT5 | n |
| W303.asm02.HP0_G0030200 | 1938.597867 | 1319.36627 | 1678.027381 | 2866.897124 | 2448.18837 | 2748.336692 | 3388.797658 | 3104.038515 | 3277.651051 | 7.90E-06 | -0.708459806 | YJR065C | ARP3 | n |
| W303.asm02.HP0_G0030220 | 430.3687264 | 629.2215761 | 447.6042752 | 334.6496812 | 402.0602038 | 319.5529817 | 429.7525521 | 364.7825811 | 352.1274083 | 0.012068015 | 0.515766947 | YJR067C | YAE1 | n |
| W303.asm02.HP0_G0030230 | 1652.654681 | 1692.520367 | 1745.461213 | 874.5641379 | 959.3713775 | 947.7959542 | 1285.47129 | 990.8152867 | 1058.429477 | 1.93E-10 | 0.872271606 | YJR068W | RFC2 | n |
| W303.asm02.HP0_G0030240 | 1746.676678 | 1784.856567 | 1869.578556 | 1026.323877 | 1183.622778 | 1095.35158 | 1103.725718 | 824.3892815 | 890.5547827 | 1.10E-06 | 0.709253915 | YJR069C | HAM1 | n |
| W303.asm02.HP0_G0030250 | 2570.580771 | 2257.963054 | 2643.601669 | 4597.541842 | 4872.491975 | 4719.064288 | 3704.012636 | 2965.672709 | 2793.475864 | 4.65E-11 | -0.925136914 | YJR070C | LIA1 | n |
| W303.asm02.HP0_G0030260 | 2444.57191 | 1774.385806 | 2352.3657 | 2934.021624 | 2904.65276 | 2870.54534 | 2083.447946 | 2040.65375 | 2023.708972 | 0.004765147 | -0.406409707 | YJR072C | NPA3 | n |
| W303.asm02.HP0_G0030270 | 1185.452596 | 876.7217422 | 1040.826535 | 1546.781957 | 1471.566885 | 1448.398784 | 2584.194862 | 2040.65375 | 1736.070013 | 0.003636942 | -0.525651387 | YJR073C | OPI3 | n |
| W303.asm02.HP0_G0030280 | 836.5049795 | 667.2985248 | 885.4355314 | 782.146348 | 805.447339 | 762.21986 | 1350.786106 | 768.2688844 | 808.6646877 | 0.92110327 | 0.024150928 | YJR074W | MOG1 | n |

| | | | | | | | | | | | | | | |
|---|---|---|---|---|---|---|---|---|---|---|---|---|---|---|
| W303.asm02.HP0_G0030290 | 2055.883038 | 1646.828028 | 2247.794395 | 3247.269291 | 3250.981846 | 3120.39413 | 3354.720363 | 3417.538665 | 3386.155427 | 3.36E-07 | -0.692817415 | YJR075W | HOC1 | n |
| W303.asm02.HP0_G0030300 | 3031.967064 | 2251.299588 | 3050.159264 | 3124.694117 | 3070.51918 | 3360.285179 | 3549.718217 | 3470.756283 | 3380.013669 | 0.184625785 | -0.197648263 | YJR076C | CDC11 | n |
| W303.asm02.HP0_G0030310 | 3217.10316 | 3105.175161 | 3041.363547 | 18087.62071 | 17787.51496 | 16921.82404 | 10243.06689 | 14453.90503 | 13472.96787 | 2.01E-69 | -2.495299265 | YJR077C | MIR1 | n |
| W303.asm02.HP0_G0030330 | 1424.869432 | 1233.693136 | 1397.541733 | 1023.405421 | 1028.371808 | 1099.877827 | 1039.357494 | 1094.347743 | 1222.209667 | 0.005159535 | 0.363501107 | YJR080C | AIM24 | n |
| W303.asm02.HP0_G0030340 | 1171.882411 | 1192.760416 | 1251.923748 | 273.3620943 | 319.7904592 | 264.3327781 | 283.0308658 | 221.5788092 | 202.677985 | 3.59E-36 | 2.080202824 | YJR082C | EAF6 | n |
| W303.asm02.HP0_G0030350 | 420.6757371 | 231.3174629 | 400.6937835 | 441.6597537 | 513.5224386 | 493.3608358 | 559.435591 | 553.4632266 | 409.4504748 | 0.060566571 | -0.460237035 | YJR083C | ACF4 | n |
| W303.asm02.HP0_G0030360 | 1889.163621 | 1456.443285 | 1893.033802 | 1199.485631 | 1430.432012 | 1134.277298 | 1790.951164 | 1324.63489 | 1110.634413 | 0.019518398 | 0.477590216 | YJR085C | TMH11 | n |
| W303.asm02.HP0_G0030370 | 847.1672678 | 589.2407801 | 853.1845683 | 7.782550727 | 6.634656829 | 7.241993919 | 4.732957623 | 5.805558321 | 6.141757122 | 3.40E-66 | 6.714573633 | YJR086W | STE18 | n |
| W303.asm02.HP0_G0030380 | 3044.56795 | 2303.655392 | 3049.181962 | 625.5225147 | 557.3111736 | 587.5067567 | 893.5823992 | 656.0280902 | 663.3097692 | 8.25E-38 | 2.245179818 | YJR088C | EMC2 | n |
| W303.asm02.HP0_G0030390 | 609.6890291 | 583.5292378 | 653.8149785 | 784.0919857 | 862.5053878 | 789.3773372 | 540.5037605 | 716.9864526 | 701.1839381 | 0.011307104 | -0.398491674 | YJR089W | BIR1 | n |
| W303.asm02.HP0_G0030400 | 603.8732355 | 904.32753 | 591.2676562 | 819.113464 | 882.4093582 | 819.2505621 | 597.299252 | 1072.093103 | 891.5784089 | 0.28658253 | -0.263435108 | YJR090C | GRR1 | n |
| W303.asm02.HP0_G0030410 | 390.6274702 | 544.5003654 | 377.2385376 | 1239.371203 | 1277.834905 | 1228.423219 | 774.3118671 | 1499.769233 | 1477.092588 | 8.28E-10 | -1.51281095 | YJR091C | JSN1 | n |
| W303.asm02.HP0_G0030420 | 516.6363315 | 657.7792876 | 474.9687287 | 2668.44208 | 3016.114994 | 2687.684993 | 1382.970217 | 2191.598266 | 2191.583666 | 2.37E-31 | -2.34335631 | YJR092W | BUD4 | n |
| W303.asm02.HP0_G0030430 | 396.4432638 | 361.731012 | 366.4882166 | 799.6570872 | 932.8327501 | 877.1865135 | 896.4221738 | 936.6300757 | 925.358073 | 9.57E-21 | -1.213192626 | YJR093C | FIP1 | n |
| W303.asm02.HP0_G0030440 | 114.3772741 | 67.58658382 | 97.7301911 | 114.7926232 | 79.61588195 | 114.0614042 | 67.20799824 | 128.6898761 | 128.9768996 | 0.691429421 | -0.146675327 | YJR094C | IME1 | n |
| W303.asm02.HP0_G0030460 | 83.35970827 | 47.59618579 | 67.43383186 | 2817.283363 | 2872.806407 | 2979.175249 | 2198.932112 | 2980.186605 | 2538.592944 | 9.16E-171 | -5.450606606 | YJR095W | SFC1 | n |
| W303.asm02.HP0_G0030470 | 1083.676208 | 840.548641 | 1066.236385 | 21.4020145 | 31.84635278 | 17.19973556 | 172.2796575 | 120.9491317 | 168.8983209 | 3.51E-89 | 5.445963012 | YJR096W | | n |
| W303.asm02.HP0_G0030480 | 504.0354454 | 534.0292046 | 537.516051 | 351.1876015 | 368.8869197 | 314.1214863 | 279.2444997 | 261.2501244 | 176.0637042 | 0.001995226 | 0.608983679 | YJR097W | JJJ3 | n |
| W303.asm02.HP0_G0030490 | 347.9783171 | 241.7886238 | 347.9194803 | 406.6382755 | 453.8105271 | 383.8256777 | 399.4616234 | 616.356775 | 403.3087177 | 0.08500654 | -0.407095366 | YJR098C | | n |
| W303.asm02.HP0_G0030500 | 1062.351631 | 710.135092 | 1002.711761 | 208.1832319 | 152.5971071 | 229.0280577 | 224.3421913 | 247.7038217 | 292.7570895 | 5.47E-24 | 2.228204041 | YJR099W | YUH1 | n |
| W303.asm02.HP0_G0030510 | 1474.303678 | 1043.308393 | 1437.611111 | 493.2191523 | 404.7140666 | 456.2456169 | 488.4412267 | 430.5789088 | 447.3246437 | 6.58E-19 | 1.544472389 | YJR100C | AIM25 | n |
| W303.asm02.HP0_G0030520 | 1874.624137 | 2259.866901 | 2000.537012 | 1969.958153 | 1863.011638 | 1980.685337 | 1896.022824 | 1610.074841 | 1479.13984 | 0.606507831 | 0.077378196 | YJR101W | RSM26 | n |
| W303.asm02.HP0_G0030530 | 997.4086025 | 672.0581434 | 940.1644384 | 110.9013479 | 96.8659897 | 91.43017323 | 133.469405 | 133.5278414 | 112.5988806 | 2.30E-49 | 3.124088788 | YJR102C | VPS25 | n |
| W303.asm02.HP0_G0030550 | 8348.571714 | 7397.399195 | 7883.894516 | 1573.048066 | 2031.531921 | 1453.830279 | 2324.828784 | 1905.190722 | 1284.650866 | 2.28E-26 | 2.224609252 | YJR104C | SOD1 | n |
| W303.asm02.HP0_G0030560 | 4938.578066 | 3401.223437 | 4831.780648 | 5842.749958 | 6094.595763 | 5676.817984 | 4883.465675 | 4187.742735 | 3770.015247 | 0.014989731 | -0.419267877 | YJR105W | ADO1 | n |
| W303.asm02.HP0_G0030570 | 245.2326302 | 270.3463353 | 228.6886472 | 453.3335798 | 489.637674 | 485.2135926 | 556.5958164 | 626.0327056 | 607.0103289 | 2.74E-10 | -0.939430318 | YJR106W | ECM27 | n |
| W303.asm02.HP0_G0030580 | 257.8335163 | 257.971327 | 279.5083465 | 7.782550727 | 13.26931366 | 15.38923708 | 19.87842202 | 20.31945412 | 24.56702849 | 2.85E-38 | 4.450427507 | YJR107W | LIH1 | n |
| W303.asm02.HP0_G0030610 | 3528.248118 | 4028.541165 | 3462.580671 | 2999.200486 | 2346.014655 | 3179.235331 | 1944.298991 | 1690.385064 | 2519.144046 | 0.046369601 | 0.369851876 | YJR109C | CPA2 | n |
| W303.asm02.HP0_G0030620 | 430.3687264 | 455.0195362 | 433.9220485 | 419.2849204 | 497.5992622 | 429.993389 | 466.6696216 | 643.4493805 | 662.286143 | 0.886814968 | -0.027804545 | YJR110W | YMR1 | n |
| W303.asm02.HP0_G0030630 | 703.7110257 | 846.2601833 | 763.2727925 | 305.465116 | 351.6368119 | 286.0587598 | 450.5775657 | 391.8751866 | 448.3482699 | 1.39E-16 | 1.298387349 | YJR111C | PXP2 | n |
| W303.asm02.HP0_G0030640 | 580.6100611 | 544.5003654 | 557.0620893 | 201.3735001 | 268.0401359 | 212.7335714 | 280.1910913 | 261.2501244 | 266.1428086 | 1.73E-15 | 1.309052863 | YJR112W | NNF1 | n |
| W303.asm02.HP0_G0030650 | 705.6496235 | 588.2888564 | 674.3383186 | 151.7597392 | 188.4242539 | 145.7451276 | 233.8081066 | 174.1667496 | 168.8983209 | 1.14E-24 | 2.02438235 | YJR112W-A | | n |
| W303.asm02.HP0_G0030660 | 2009.356689 | 1513.558708 | 1925.284765 | 2281.260182 | 2117.78246 | 2135.482957 | 2350.386755 | 2132.57509 | 1982.763924 | 0.078723961 | -0.262494227 | YJR113C | RSM7 | n |
| W303.asm02.HP0_G0030680 | 347.9783171 | 214.1828361 | 266.8034217 | 405.6654566 | 414.0025861 | 407.362158 | 977.8290449 | 912.4402494 | 904.8855493 | 0.002871393 | -0.565821246 | YJR116W | TDA4 | n |
| W303.asm02.HP0_G0030690 | 2485.282465 | 1776.289654 | 2159.837223 | 2057.511848 | 1902.819579 | 2033.189793 | 2355.119713 | 2569.92715 | 2433.159446 | 0.523446791 | 0.099139709 | YJR117W | STE24 | n |
| W303.asm02.HP0_G0030700 | 1145.711339 | 1188.000797 | 1161.03467 | 960.1721959 | 1060.218161 | 1007.542404 | 1304.403121 | 1104.991267 | 882.3657732 | 0.21374988 | 0.207346397 | YJR118C | ILM1 | n |
| W303.asm02.HP0_G0030710 | 424.5529328 | 516.8945777 | 423.1717275 | 492.2463335 | 611.7153596 | 529.5708054 | 472.3491708 | 657.9632763 | 607.0103289 | 0.176900153 | -0.257628738 | YJR119C | JHD2 | n |
| W303.asm02.HP0_G0030730 | 6025.16217 | 5907.63858 | 5559.870572 | 29044.47931 | 28989.46955 | 27524.10314 | 17553.59323 | 23459.29358 | 22438.90964 | 1.08E-67 | -2.290137423 | YJR121W | ATP2 | n |
| W303.asm02.HP0_G0030740 | 1222.285955 | 1055.683401 | 1231.400408 | 464.0345871 | 525.4648208 | 448.0983738 | 511.1594233 | 752.7873956 | 577.3251695 | 2.38E-13 | 1.289242735 | YJR122W | IBA57 | n |
| W303.asm02.HP0_G0030750 | 21623.12061 | 24727.17044 | 22002.97522 | 32938.67313 | 34455.09984 | 30139.36819 | 22075.46094 | 22301.0847 | 20601.50064 | 3.68E-06 | -0.512831589 | YJR123W | RPS5 | n |
| W303.asm02.HP0_G0030760 | 1197.084183 | 1000.471825 | 1209.899766 | 1315.251073 | 1012.448632 | 1301.748407 | 491.2810012 | 832.130026 | 622.3647217 | 0.681566654 | -0.09195033 | YJR124C | | n |
| W303.asm02.HP0_G0030770 | 1463.641389 | 1066.154562 | 1405.360148 | 1142.089319 | 975.2945538 | 1184.971255 | 1499.400975 | 1413.653451 | 1363.470081 | 0.126061158 | 0.251710966 | YJR125C | ENT3 | n |
| W303.asm02.HP0_G0030780 | 820.9961966 | 807.231311 | 715.3849988 | 333.6768624 | 305.1942141 | 285.1535106 | 393.7820742 | 583.4586112 | 596.774067 | 1.89E-12 | 1.342882321 | YJR126C | VPS70 | n |
| W303.asm02.HP0_G0030800 | 443.9389115 | 473.1060868 | 496.4693708 | 668.3265437 | 716.5429375 | 717.8626473 | 678.7061231 | 578.620646 | 514.883972 | 0.000184219 | -0.572732161 | YJR129C | EFM3 | n |
| W303.asm02.HP0_G0030810 | 915.0181932 | 758.6832015 | 872.7306065 | 741.2879567 | 824.0243781 | 752.2621184 | 1165.254167 | 1128.2135 | 1262.131089 | 0.331735668 | 0.137022023 | YJR130C | STR2 | n |
| W303.asm02.HP0_G0030820 | 307.2677619 | 351.2598511 | 264.8488179 | 844.4067538 | 857.1976623 | 817.4400637 | 1086.68707 | 1230.778364 | 1182.288246 | 5.44E-23 | -1.447162926 | YJR131W | MNS1 | n |
| W303.asm02.HP0_G0030830 | 783.1935382 | 1069.010333 | 895.2085505 | 5124.809654 | 4852.588005 | 5058.532753 | 2083.447946 | 2924.066207 | 2535.522065 | 1.07E-50 | -2.452246131 | YJR132W | NMD5 | n |
| W303.asm02.HP0_G0030840 | 901.4480081 | 978.5775798 | 1029.098912 | 4702.606277 | 6208.71186 | 4862.998917 | 3866.826378 | 3423.344223 | 2666.546217 | 2.01E-45 | -2.438619293 | YJR133W | XPT1 | n |
| W303.asm02.HP0_G0030850 | 859.768154 | 1043.308393 | 905.9588715 | 1213.105095 | 1244.661621 | 1255.580696 | 939.9653839 | 1092.412557 | 850.6333614 | 0.00607493 | -0.402538885 | YJR134C | SGM1 | n |
| W303.asm02.HP0_G0030860 | 331.5002352 | 351.2598511 | 336.1918574 | 223.7483334 | 276.0017241 | 235.3648024 | 163.7603337 | 287.3751369 | 238.5049016 | 0.03859339 | 0.473696827 | YJR135C | MCM22 | n |
| W303.asm02.HP0_G0030870 | 736.6671894 | 781.5293707 | 707.5665836 | 893.0476959 | 963.3521715 | 936.9329633 | 901.1551314 | 742.143872 | 653.0735073 | 0.036185573 | -0.327253865 | YJR135W- | TIM8 | n |
| W303.asm02.HP0_G0030880 | 688.2022427 | 562.586916 | 612.7682982 | 362.8614276 | 386.1370274 | 360.2891975 | 373.9036522 | 411.2270477 | 354.1746607 | 7.52E-07 | 0.749704552 | YJR136C | TTI2 | n |
| W303.asm02.HP0_G0030890 | 933.4348729 | 1232.741212 | 962.6423823 | 1507.869203 | 1532.605727 | 1656.606109 | 1193.651912 | 1422.361789 | 1460.714569 | 0.000174831 | -0.586255325 | YJR137C | MET5 | n |
| W303.asm02.HP0_G0030900 | 253.9563206 | 331.2694531 | 304.9181962 | 432.9043842 | 558.638105 | 420.9408966 | 439.2184674 | 615.389182 | 610.0812074 | 0.001458194 | -0.663425176 | YJR138W | IML1 | n |
| W303.asm02.HP0_G0030910 | 5277.832693 | 4541.628048 | 5264.725395 | 2991.417936 | 2515.861869 | 3015.385218 | 2959.991697 | 2186.760301 | 2243.788602 | 1.80E-07 | 0.82320894 | YJR139C | HOM6 | n |
| W303.asm02.HP0_G0030920 | 552.5003921 | 837.6928699 | 597.1314676 | 770.4725219 | 895.6786719 | 825.5873068 | 772.418684 | 1028.551416 | 905.9091755 | 0.102550708 | -0.325536652 | YJR140C | HIR3 | n |
| W303.asm02.HP0_G0030930 | 601.9346377 | 760.5870489 | 631.3370345 | 588.5553987 | 643.5617124 | 607.42224 | 573.6344639 | 598.9401001 | 551.7345148 | 0.447950689 | 0.117327904 | YJR141W | IPA1 | n |
| W303.asm02.HP0_G0030940 | 1220.347357 | 1089.952655 | 1271.469786 | 662.4896306 | 690.0043102 | 649.9689543 | 947.5381161 | 800.1994552 | 848.586109 | 2.21E-10 | 0.839610387 | YJR142W | | n |
| W303.asm02.HP0_G0030950 | 2290.45338 | 1885.760881 | 2144.200393 | 3722.004885 | 3576.080031 | 3574.829248 | 3898.063898 | 5229.840454 | 5060.807868 | 1.05E-07 | -0.782711365 | YJR143C | PMT4 | n |
| W303.asm02.HP0_G0030960 | 2361.212202 | 2379.809289 | 2448.141287 | 1122.632942 | 1162.391876 | 1150.571784 | 1273.165601 | 1172.722781 | 1115.752544 | 2.75E-23 | 1.065504377 | YJR144W | MGM101 | n |
| W303.asm02.HP0_G0030990 | 413.8906446 | 392.1925709 | 418.2852179 | 31.13020291 | 34.50021551 | 35.30472036 | 123.0568982 | 144.171365 | 135.1186567 | 4.80E-57 | 3.601706324 | YJR149W | | n |
| W303.asm02.HP0_G0031020 | 220.0308579 | 301.7598179 | 259.9623083 | 56.42349277 | 53.07725463 | 38.02046808 | 82.35346264 | 122.8843178 | 95.19723539 | 1.50E-18 | 2.412838457 | YJR152W | DAL5 | n |
| W303.asm02.HP0_G0031030 | 126.9781603 | 207.51937 | 150.5044943 | 107.0100725 | 92.8851956 | 101.3879149 | 111.6977999 | 111.2732011 | 124.8823948 | 0.008139697 | 0.684564934 | YJR153W | PGU1 | n |

| | | | | | | | | | | | | | | |
|---|---|---|---|---|---|---|---|---|---|---|---|---|---|---|
| W303.asm02.HP0_G0031140 | 337.3160288 | 436.9329856 | 384.079651 | 488.3550581 | 383.4831647 | 496.0765835 | 291.5501896 | 448.9631768 | 365.4345488 | 0.269038597 | -0.242019968 | YKL222C | | n |
| W303.asm02.HP0_G0031150 | 73.66671894 | 82.81736327 | 102.6167007 | 99.22752177 | 94.21212697 | 124.0191459 | 94.65915246 | 141.2685858 | 98.26811395 | 0.318084136 | -0.297435693 | YKL221W | MCH2 | n |
| W303.asm02.HP0_G0031190 | 211.3071675 | 283.6732673 | 168.0959287 | 178.0258479 | 234.8668517 | 161.1343647 | 312.3752031 | 216.740844 | 361.340044 | 0.469527589 | 0.212672947 | YKL217W | JEN1 | n |
| W303.asm02.HP0_G0031200 | 2910.804697 | 2977.617383 | 3012.04449 | 2188.842492 | 960.6983088 | 2002.411319 | 446.7911996 | 281.5695785 | 378.7416892 | 0.003629711 | 0.787809485 | YKL216W | URA1 | n |
| W303.asm02.HP0_G0031210 | 483.6801678 | 520.7022725 | 456.3999924 | 320.0573986 | 228.2321949 | 274.2905197 | 176.0660236 | 275.7640202 | 189.3708446 | 0.000160257 | 0.82550392 | YKL215C | OXP1 | n |
| W303.asm02.HP0_G0031230 | 1515.983532 | 1401.23171 | 1499.181131 | 701.4023842 | 624.9846733 | 674.4106837 | 995.8142838 | 1015.005113 | 1292.839874 | 4.02E-15 | 1.141189955 | YKL213C | DOA1 | n |
| W303.asm02.HP0_G0031240 | 1884.317127 | 1915.270516 | 1970.240653 | 4222.033769 | 3391.636571 | 3976.759911 | 3022.466738 | 3692.335092 | 3785.369639 | 3.58E-13 | -1.006765273 | YKL212W | SAC1 | n |
| W303.asm02.HP0_G0031250 | 1258.150016 | 1018.558376 | 1024.212403 | 2126.581986 | 1855.050049 | 2206.092398 | 1941.459217 | 1816.172161 | 2150.638619 | 3.28E-10 | -0.907222489 | YKL211C | TRP3 | n |
| W303.asm02.HP0_G0031260 | 4020.651976 | 3151.819423 | 3661.95026 | 4295.968001 | 3651.715119 | 4068.190084 | 3574.329597 | 5169.849684 | 4746.554629 | 0.400313207 | -0.149535886 | YKL210W | UBA1 | n |
| W303.asm02.HP0_G0031270 | 2015.172483 | 2539.732474 | 2110.972128 | 300.6010218 | 358.2714688 | 304.1637446 | 265.0456269 | 336.7223826 | 299.9224728 | 8.47E-68 | 2.794526314 | YKL209C | STE6 | n |
| W303.asm02.HP0_G0031280 | 878.1848337 | 475.0099342 | 894.2312486 | 369.6711595 | 352.9637433 | 370.2469391 | 357.8115963 | 363.8149881 | 359.2927916 | 7.11E-06 | 1.039569619 | YKL208W | CBT1 | n |
| W303.asm02.HP0_G0031290 | 1636.1766 | 1247.020068 | 1624.275776 | 674.1634567 | 628.9654674 | 665.3581913 | 1191.758729 | 1064.352359 | 1162.839348 | 5.33E-16 | 1.194427175 | YKL207W | EMC3 | n |
| W303.asm02.HP0_G0031300 | 1226.163151 | 996.6641304 | 1151.261651 | 491.2735146 | 436.5604193 | 476.1611002 | 755.3800366 | 515.7270975 | 630.5537312 | 3.80E-13 | 1.263748238 | YKL206C | ADD66 | n |
| W303.asm02.HP0_G0031310 | 1357.018507 | 1845.780085 | 1547.068925 | 3593.592798 | 3739.292589 | 3554.008516 | 1660.321534 | 3249.177473 | 2896.862109 | 1.08E-07 | -1.196559673 | YKL205W | LOS1 | n |
| W303.asm02.HP0_G0031320 | 483.6801678 | 512.1349591 | 322.5096306 | 1622.661827 | 1571.086737 | 1786.962 | 1379.183851 | 1451.38958 | 1637.801899 | 5.03E-26 | -1.917729434 | YKL204W | EAP1 | n |
| W303.asm02.HP0_G0031330 | 405.1669542 | 546.4042129 | 383.1023491 | 1641.145384 | 1711.741462 | 1666.563851 | 1869.518261 | 1703.931367 | 1671.581563 | 1.36E-34 | -1.910584067 | YKL203C | TOR2 | n |
| W303.asm02.HP0_G0031350 | 346.0397192 | 326.5098345 | 258.9850064 | 1047.725892 | 1316.315915 | 1076.341346 | 1250.447404 | 1450.421987 | 1611.187618 | 3.35E-27 | -1.883562018 | YKL198C | PTK1 | n |
| W303.asm02.HP0_G0031360 | 270.4344024 | 312.2309788 | 240.4162701 | 123.5479928 | 115.4430288 | 140.3136322 | 27.45115421 | 157.7176677 | 218.0323778 | 0.024158386 | 1.116832452 | YKL197C | PEX1 | n |
| W303.asm02.HP0_G0031370 | 4188.340691 | 4491.176091 | 4275.695861 | 1801.660493 | 1916.088892 | 1923.654635 | 2892.783699 | 1901.32035 | 1742.21177 | 8.26E-11 | 1.199465659 | YKL196C | YKT6 | n |
| W303.asm02.HP0_G0031380 | 3312.094456 | 2916.694265 | 3082.410227 | 2478.742406 | 2460.130752 | 2308.385562 | 2950.525782 | 3380.770627 | 3233.635125 | 0.001594746 | 0.361727627 | YKL195W | MIA40 | n |
| W303.asm02.HP0_G0031390 | 757.022467 | 600.6638647 | 684.1113377 | 1006.8675 | 959.3713775 | 958.6589451 | 877.4903433 | 947.2735993 | 855.7514923 | 0.000147142 | -0.518836634 | YKL194C | MST1 | n |
| W303.asm02.HP0_G0031400 | 1577.049365 | 987.1448933 | 1350.631241 | 584.6641233 | 621.0038792 | 554.9177841 | 669.2402079 | 647.3197527 | 636.6954883 | 1.65E-10 | 1.153675784 | YKL193C | SDS22 | n |
| W303.asm02.HP0_G0031410 | 2426.15523 | 1979.049405 | 2258.544716 | 2170.358834 | 2534.438909 | 2187.082164 | 2731.86314 | 2145.153799 | 1926.464484 | 0.787870339 | -0.048135792 | YKL192C | ACP1 | n |
| W303.asm02.HP0_G0031420 | 645.5530897 | 810.0870821 | 538.493353 | 1363.892015 | 1346.835336 | 1379.599842 | 1333.747458 | 1457.195138 | 1376.777221 | 1.17E-10 | -1.036344931 | YKL191W | DPH2 | n |
| W303.asm02.HP0_G0031430 | 2345.703419 | 2107.559107 | 2279.068056 | 519.485261 | 545.3687913 | 512.3710698 | 676.8129401 | 520.5650627 | 543.5455053 | 4.23E-50 | 2.09461483 | YKL190W | CNB1 | n |
| W303.asm02.HP0_G0031440 | 675.6013566 | 419.7983587 | 599.0860714 | 367.7255218 | 471.0606348 | 372.9626868 | 616.2310825 | 592.1669487 | 650.0026287 | 0.018631176 | 0.486248215 | YKL189W | HYM1 | n |
| W303.asm02.HP0_G0031450 | 505.0047443 | 456.9233836 | 462.2638039 | 23.34765218 | 22.55783322 | 20.82073252 | 42.5966186 | 36.76853603 | 55.2758141 | 3.28E-61 | 4.418546079 | YKL188C | PXA2 | n |
| W303.asm02.HP0_G0031460 | 106.6228827 | 133.2693202 | 110.4351159 | 9936.37164 | 10112.54394 | 10144.22298 | 8982.206976 | 8855.411625 | 9284.289516 | 0 | -6.428603837 | YKL187C | FAT3 | n |
| W303.asm02.HP0_G0031470 | 1015.825282 | 1173.721942 | 975.3473072 | 888.1836017 | 1074.814406 | 941.4592095 | 1151.055294 | 776.9772219 | 783.074033 | 0.533046339 | 0.124778819 | YKL186C | MTR2 | n |
| W303.asm02.HP0_G0031480 | 619.3820185 | 1157.599238 | 604.9498829 | 507.8114349 | 550.6765168 | 532.2865531 | 341.7195404 | 478.9585614 | 436.0647557 | 0.024569592 | 0.582773588 | YKL185W | ASH1 | n |
| W303.asm02.HP0_G0031490 | 903.386606 | 981.433351 | 1064.281781 | 1242.28966 | 1421.143493 | 1270.064684 | 1623.404465 | 1688.449878 | 1336.8558 | 0.00457699 | -0.414741297 | YKL184W | SPE1 | n |
| W303.asm02.HP0_G0031510 | 742.482983 | 670.1542959 | 731.9991313 | 481.5453262 | 544.04186 | 505.1290759 | 705.2106858 | 511.8567253 | 544.5691315 | 0.003421036 | 0.487667031 | YKL183W | LOT5 | n |
| W303.asm02.HP0_G0031520 | 6650.359982 | 7591.591633 | 6829.385754 | 12505.5862 | 9167.768806 | 12553.09121 | 5614.234332 | 11718.51947 | 10230.12011 | 0.00381855 | -0.699908888 | YKL182W | FAS1 | n |
| W303.asm02.HP0_G0031530 | 6101.736786 | 5589.696059 | 6616.333937 | 6653.108052 | 6742.138269 | 6445.374588 | 5173.122682 | 4987.94219 | 4660.570029 | 0.316744258 | -0.11594619 | YKL181W | PRS1 | n |
| W303.asm02.HP0_G0031550 | 1089.492001 | 1180.385408 | 1040.826535 | 751.988964 | 755.0239471 | 755.8831153 | 907.781272 | 973.3986118 | 1072.760244 | 1.59E-05 | 0.549046023 | YKL179C | COY1 | n |
| W303.asm02.HP0_G0031560 | 214.2150643 | 219.8943783 | 171.0278344 | 42.804029 | 51.75032327 | 40.7362158 | 76.67391349 | 105.4676428 | 84.96097352 | 3.54E-16 | 2.171676527 | YKL178C | STE3 | n |
| W303.asm02.HP0_G0031570 | 671.7241609 | 963.3468004 | 697.7935644 | 405.6654566 | 459.1182526 | 356.6682005 | 464.7764586 | 653.1253111 | 667.4042739 | 1.03E-05 | 0.935619099 | YKL176C | LST4 | n |
| W303.asm02.HP0_G0031580 | 853.9523603 | 760.5870489 | 807.2513785 | 400.8013624 | 414.0025861 | 390.1624224 | 385.2627505 | 534.1113655 | 590.6323099 | 1.87E-08 | 1.007720217 | YKL175W | ZRT3 | n |
| W303.asm02.HP0_G0031590 | 1114.693773 | 1150.875772 | 1259.742163 | 445.5510291 | 493.6184681 | 462.5823616 | 718.4629671 | 819.5513163 | 961.1849896 | 1.07E-18 | 1.331755657 | YKL174C | TPO5 | n |
| W303.asm02.HP0_G0031600 | 417.7678403 | 556.8753737 | 373.32933 | 899.8574278 | 869.1400446 | 816.5348144 | 727.9288824 | 852.4494801 | 883.3893994 | 1.39E-07 | -0.939232843 | YKL173W | SNU114 | n |
| W303.asm02.HP0_G0031610 | 3334.388331 | 5667.753804 | 4087.076592 | 5278.51503 | 4822.068583 | 5042.238266 | 2980.816711 | 2566.056778 | 2788.357733 | 0.279256548 | -0.210291939 | YKL172W | EBP2 | n |
| W303.asm02.HP0_G0031620 | 296.6054736 | 301.7598179 | 242.3708739 | 159.5422899 | 220.2706067 | 174.7131033 | 230.968332 | 271.893648 | 457.5609056 | 0.031161071 | 0.603981709 | YKL171W | NNK1 | n |
| W303.asm02.HP0_G0031630 | 1611.944126 | 1801.991594 | 1689.755004 | 1264.664493 | 1316.315915 | 1324.379638 | 1499.400975 | 1266.579307 | 1179.217367 | 0.003946714 | 0.38619446 | YKL170W | MRPL38 | n |
| W303.asm02.HP0_G0031640 | 230.6931462 | 216.0866835 | 198.3922879 | 746.1520509 | 800.1396136 | 710.6206533 | 744.9675298 | 892.1207953 | 967.3267467 | 1.16E-29 | -1.805488329 | YKL168C | KKQ8 | n |
| W303.asm02.HP0_G0031650 | 1144.74204 | 1017.606452 | 1149.307047 | 935.8517249 | 1046.948848 | 970.4271852 | 1061.129099 | 920.1809938 | 818.9009496 | 0.269414567 | 0.166009289 | YKL167C | MRP49 | n |
| W303.asm02.HP0_G0031670 | 1180.606101 | 1503.087547 | 1367.245373 | 2004.979631 | 1605.586953 | 2011.463811 | 2069.249073 | 2920.195835 | 3043.240654 | 0.01143156 | -0.473422345 | YKL165C | MCD4 | n |
| W303.asm02.HP0_G0031680 | 5741.157583 | 4854.810951 | 4920.715122 | 9652.308539 | 6693.041809 | 8337.3455 | 5653.044585 | 7674.9481 | 9565.786717 | 0.001158112 | -0.669819041 | YKL164C | PIR1 | n |
| W303.asm02.HP0_G0031710 | 238.4475376 | 229.4136155 | 215.0064204 | 285.0359204 | 264.0593418 | 262.5222796 | 445.8446081 | 391.8751866 | 465.7499151 | 0.141443463 | -0.24972202 | YKL162C | | n |
| W303.asm02.HP0_G0031730 | 2227.448949 | 2392.184298 | 2364.093323 | 2557.540733 | 2543.727428 | 2547.371361 | 2963.778063 | 2267.070524 | 1808.747472 | 0.478534236 | -0.131209191 | YKL160W | ELF1 | n |
| W303.asm02.HP0_G0031740 | 464.2941891 | 269.3944116 | 403.6256892 | 201.3735001 | 229.5591263 | 210.9230729 | 316.1615692 | 262.2177175 | 252.8356682 | 0.000341439 | 0.826701423 | YKL159C | RCN1 | n |
| W303.asm02.HP0_G0031760 | 1455.886998 | 1163.250781 | 1353.563147 | 3667.52703 | 2407.053498 | 3549.48227 | 2741.329055 | 3813.284224 | 2994.106597 | 8.06E-11 | -1.277097707 | YKL157W | APE2 | n |
| W303.asm02.HP0_G0031770 | 4737.933187 | 6624.437138 | 4889.441461 | 7544.210111 | 9459.693707 | 7540.726169 | 7271.716092 | 6261.294649 | 5397.580884 | 0.001099975 | -0.594627928 | YKL156W | RPS27A | y |
| W303.asm02.HP0_G0031780 | 1169.943813 | 1025.221842 | 1051.576856 | 1311.359797 | 1210.161406 | 1349.726617 | 829.2141755 | 1120.472756 | 1002.130037 | 0.097383791 | -0.254441275 | YKL155C | RSM22 | n |
| W303.asm02.HP0_G0031790 | 1088.522702 | 1164.202704 | 1139.534028 | 1250.07221 | 1253.950141 | 1327.095386 | 2170.534366 | 1628.459109 | 1579.455207 | 0.27023663 | -0.175722024 | YKL154W | SRP102 | n |
| W303.asm02.HP0_G0031800 | 19611.82532 | 15360.24108 | 19129.70761 | 17897.92103 | 14892.15072 | 16436.61045 | 12177.89996 | 10558.3754 | 10729.64969 | 0.358555632 | 0.136158292 | YKL152C | GPM1 | n |
| W303.asm02.HP0_G0031810 | 994.5007057 | 531.1734334 | 837.5477377 | 168.2976595 | 149.9432443 | 159.3238662 | 277.3513167 | 227.3843676 | 245.6702849 | 1.32E-22 | 2.305417216 | YKL151C | NNR2 | n |
| W303.asm02.HP0_G0031820 | 6384.772074 | 4430.252973 | 6028.975489 | 5667.642567 | 4522.182095 | 5418.82195 | 5572.584305 | 4924.081049 | 5115.060056 | 0.534378224 | 0.109626307 | YKL150W | MCR1 | n |
| W303.asm02.HP0_G0031830 | 775.4391467 | 549.259984 | 678.2475262 | 422.2033769 | 455.1374585 | 411.8884042 | 439.2184674 | 475.0881892 | 416.6158581 | 0.000182658 | 0.636800146 | YKL149C | DBR1 | n |
| W303.asm02.HP0_G0031840 | 1834.882881 | 1468.818293 | 1347.699335 | 8936.313872 | 10870.22175 | 8942.051992 | 5895.372015 | 10528.38001 | 10569.96401 | 1.04E-32 | -2.627675 | YKL148C | SDH1 | n |
| W303.asm02.HP0_G0031850 | 1159.281524 | 859.5871154 | 997.8252511 | 652.7614422 | 609.0614969 | 598.3697476 | 1015.692706 | 852.4494801 | 1158.744844 | 4.71E-05 | 0.697311299 | YKL146W | AVT3 | n |
| W303.asm02.HP0_G0031860 | 5403.841554 | 3946.675726 | 5078.06073 | 1788.041029 | 1499.432443 | 1721.784054 | 2931.593952 | 2095.806554 | 2602.057767 | 1.06E-18 | 1.525720829 | YKL145W | RPT1 | n |
| W303.asm02.HP0_G0031870 | 1227.13245 | 1308.895109 | 1336.949014 | 1781.231298 | 1661.31807 | 1742.604787 | 1761.606827 | 1275.287644 | 1339.926679 | 0.005785848 | -0.421201453 | YKL144C | RPC25 | n |
| W303.asm02.HP0_G0031880 | 1279.474592 | 1518.318327 | 1385.81411 | 2918.456523 | 2365.918625 | 2859.682349 | 1716.170434 | 1176.593153 | 1381.895352 | 1.38E-08 | -0.961416099 | YKL143W | LTV1 | n |

| | | | | | | | | | | | | | |
|---|---|---|---|---|---|---|---|---|---|---|---|---|---|
| W303.asm02.HP0_G0031890 | 3330.511135 | 2349.347731 | 3158.639776 | 455.2792175 | 380.829302 | 416.4146504 | 623.8038147 | 494.4400503 | 562.9944028 | 1.39E-59 | 2.817353309 | YKL142W | MRP8 | n |
| W303.asm02.HP0_G0031900 | 3430.348925 | 3445.011927 | 3770.430773 | 3104.264921 | 3901.178215 | 3013.57472 | 3753.235395 | 3733.941593 | 3088.280206 | 0.576696367 | 0.088177854 | YKL141W | SDH3 | n |
| W303.asm02.HP0_G0031920 | 657.1846769 | 738.6928035 | 597.1314676 | 586.609761 | 659.4848888 | 577.5490151 | 695.7447705 | 637.6438222 | 721.6564618 | 0.399489103 | 0.130041616 | YKL139W | CTK1 | n |
| W303.asm02.HP0_G0031930 | 1073.983218 | 1190.856568 | 1121.942594 | 147.8684638 | 241.5015086 | 160.2291155 | 253.6865286 | 161.5880399 | 127.9532734 | 1.67E-24 | 2.629949838 | YKL138C-/ | HSK3 | n |
| W303.asm02.HP0_G0031940 | 1112.755176 | 1085.193036 | 1108.260367 | 1478.684638 | 1568.432874 | 1558.839191 | 1660.321534 | 1332.375635 | 1203.784396 | 0.001122535 | -0.478189725 | YKL138C | MRPL31 | n |
| W303.asm02.HP0_G0031950 | 951.8515526 | 1145.16423 | 945.0509479 | 373.5624349 | 391.4447529 | 368.4364406 | 527.2514792 | 403.4863033 | 412.5213534 | 2.80E-18 | 1.425343481 | YKL137W | CMC1 | n |
| W303.asm02.HP0_G0031960 | 1930.843475 | 1591.616453 | 1909.647934 | 1659.628942 | 1588.336845 | 1554.312945 | 1717.117026 | 1924.542583 | 2117.882581 | 0.212586094 | 0.17772268 | YKL135C | APL2 | n |
| W303.asm02.HP0_G0031970 | 623.2592142 | 616.8465678 | 615.7002039 | 1039.943341 | 915.5826424 | 1041.036626 | 850.9857806 | 1136.921838 | 1032.838823 | 3.61E-06 | -0.692293342 | YKL134C | OCT1 | n |
| W303.asm02.HP0_G0032000 | 1226.163151 | 1057.587248 | 1208.922464 | 958.2265582 | 962.0252402 | 885.3337566 | 1367.824753 | 929.8569243 | 962.2086158 | 0.078119148 | 0.316328456 | YKL130C | SHE2 | n |
| W303.asm02.HP0_G0032020 | 1461.702792 | 1441.212506 | 1481.589697 | 674.1634567 | 623.6577419 | 693.4209178 | 1027.998396 | 525.403028 | 805.5938092 | 1.09E-07 | 1.138191457 | YKL128C | PMU1 | n |
| W303.asm02.HP0_G0032040 | 1855.238159 | 1683.953053 | 1840.259498 | 1590.558805 | 1413.181905 | 1560.64969 | 1864.785303 | 1526.861838 | 2044.181495 | 0.109414731 | 0.236411707 | YKL126W | YPK1 | n |
| W303.asm02.HP0_G0032050 | 795.7944243 | 950.0198684 | 836.5704358 | 3035.194783 | 2373.880213 | 3118.583632 | 1078.167746 | 1637.167446 | 1226.304172 | 1.57E-20 | -1.723863544 | YKL125W | RRN3 | n |
| W303.asm02.HP0_G0032060 | 454.6011998 | 243.6924712 | 384.079651 | 137.1674566 | 228.2321949 | 219.9755653 | 437.3252843 | 366.7177673 | 550.7108886 | 0.001921659 | 0.888822621 | YKL124W | SSH4 | n |
| W303.asm02.HP0_G0032070 | 1886.255724 | 1772.481959 | 2118.790543 | 1220.887645 | 1324.277503 | 1345.200371 | 1621.511282 | 1194.009828 | 1190.477255 | 0.000385192 | 0.570618653 | YKL122C | SRP21 | n |
| W303.asm02.HP0_G0032090 | 4825.170091 | 5601.119144 | 4680.298852 | 820.0862828 | 862.5053878 | 811.103319 | 1123.60414 | 1374.949729 | 1285.674491 | 3.13E-87 | 2.599516096 | YKL120W | OAC1 | n |
| W303.asm02.HP0_G0032100 | 890.7857198 | 811.0390059 | 896.1858524 | 302.5466595 | 363.5791942 | 313.216237 | 566.0617317 | 576.6854598 | 520.002103 | 2.13E-23 | 1.411570604 | YKL119C | VPH2 | n |
| W303.asm02.HP0_G0032110 | 6003.837594 | 5912.398199 | 6233.231588 | 1509.814841 | 1313.662052 | 1517.197726 | 2558.636891 | 1778.436032 | 1712.526611 | 4.82E-35 | 2.06333417 | YKL117W | SBA1 | n |
| W303.asm02.HP0_G0032120 | 529.2372177 | 569.250382 | 485.7190498 | 843.433935 | 887.7170837 | 894.386249 | 784.7243739 | 875.6717134 | 714.4910785 | 2.03E-07 | -0.728574471 | YKL116C | PRR1 | n |
| W303.asm02.HP0_G0032140 | 2274.944597 | 2109.462954 | 2228.248357 | 686.8101016 | 647.5425065 | 619.1904801 | 704.2640943 | 678.2827305 | 674.5696572 | 1.70E-55 | 1.75923482 | YKL114C | APN1 | n |
| W303.asm02.HP0_G0032150 | 1855.238159 | 1408.847099 | 1788.462497 | 1109.013479 | 1060.218161 | 1241.096708 | 1415.154329 | 1300.445064 | 1050.240468 | 0.000831545 | 0.566400076 | YKL113C | RAD27 | n |
| W303.asm02.HP0_G0032160 | 962.5138409 | 885.2890557 | 971.4380995 | 1087.611464 | 1224.757651 | 1054.615365 | 1010.013157 | 904.699505 | 983.7047657 | 0.049129877 | -0.254786362 | YKL112W | ABF1 | n |
| W303.asm02.HP0_G0032180 | 881.0927305 | 1204.1835 | 872.7306065 | 176.0802102 | 192.405048 | 143.0293799 | 166.6001083 | 272.8612411 | 309.1351085 | 1.09E-24 | 2.535326071 | YKL109W | HAP4 | n |
| W303.asm02.HP0_G0032190 | 435.2152211 | 431.2214433 | 405.5802931 | 209.1560508 | 213.6359499 | 190.1023404 | 274.5115421 | 220.6112162 | 218.0323778 | 8.21E-10 | 1.055668051 | YKL108W | SLD2 | n |
| W303.asm02.HP0_G0032210 | 1048.781446 | 1577.337597 | 1225.536596 | 394.9644494 | 384.8100961 | 390.1624224 | 475.1889453 | 404.4538963 | 470.868046 | 3.47E-22 | 1.718876058 | YKL106W | AAT1 | n |
| W303.asm02.HP0_G0032220 | 3126.958359 | 2605.41521 | 2945.58796 | 6715.368458 | 6196.769478 | 6194.620549 | 5171.229499 | 6711.225419 | 6717.035039 | 1.09E-15 | -1.138718691 | YKL104C | GFA1 | n |
| W303.asm02.HP0_G0032230 | 1088.522702 | 835.7890225 | 958.7331747 | 1481.603095 | 817.3897213 | 1417.62031 | 1026.105213 | 791.4911177 | 1238.587686 | 0.143452499 | -0.367675845 | YKL103C | APE1 | n |
| W303.asm02.HP0_G0032240 | 689.1715417 | 1017.606452 | 688.9978473 | 1515.651754 | 1590.990708 | 1520.818723 | 1311.975853 | 1751.343427 | 1798.511211 | 8.06E-07 | -0.949439806 | YKL101W | HSL1 | n |
| W303.asm02.HP0_G0032260 | 1328.908838 | 1923.83783 | 1520.681774 | 2465.122943 | 2385.822596 | 2422.446966 | 2022.866088 | 1593.625759 | 1662.368928 | 0.000275863 | -0.607597598 | YKL099C | UTP11 | n |
| W303.asm02.HP0_G0032270 | 554.4389899 | 396.9521895 | 432.9447466 | 133.2761812 | 119.4238229 | 125.8296443 | 280.1910913 | 170.2963774 | 204.7252374 | 1.86E-15 | 1.868993308 | YKL098W | MTC2 | n |
| W303.asm02.HP0_G0032300 | 3857.809755 | 2327.453485 | 4116.395649 | 5325.210335 | 4398.777478 | 4981.586567 | 5335.936424 | 6741.220803 | 4466.081054 | 0.023974284 | -0.513623814 | YKL096W | CWP1 | n |
| W303.asm02.HP0_G0032310 | 213.2457654 | 249.4040135 | 204.2560994 | 189.699674 | 226.9052635 | 182.8603465 | 346.452498 | 332.8520104 | 318.3477441 | 0.411329662 | 0.159470644 | YKL095W | YJU2 | n |
| W303.asm02.HP0_G0032320 | 2561.857081 | 2093.280251 | 2510.688609 | 560.3436523 | 555.9842423 | 557.6335318 | 946.5915246 | 747.9494303 | 887.4839041 | 6.01E-48 | 2.097761968 | YKL094W | YJU3 | n |
| W303.asm02.HP0_G0032330 | 554.4389899 | 513.0868828 | 504.2877861 | 244.177529 | 246.809234 | 262.5222796 | 265.9922184 | 315.4353354 | 278.4263229 | 5.12E-12 | 1.059728808 | YKL093W | MBR1 | n |
| W303.asm02.HP0_G0032340 | 619.3820185 | 760.5870489 | 651.8603746 | 858.9990365 | 927.5250247 | 871.755018 | 1000.547241 | 866.9633759 | 742.1289856 | 0.016551541 | -0.387057288 | YKL092C | BUD2 | n |
| W303.asm02.HP0_G0032360 | 739.5750862 | 453.1156887 | 684.1113377 | 237.3677972 | 260.0785477 | 278.8167659 | 373.9036522 | 312.5325563 | 302.9933513 | 2.69E-09 | 1.273313031 | YKL090W | CUE2 | n |
| W303.asm02.HP0_G0032370 | 526.3293209 | 565.4426872 | 489.6282574 | 201.3735001 | 226.9052635 | 214.5440699 | 249.9001625 | 176.1019357 | 281.4972014 | 9.47E-11 | 1.300321555 | YKL089W | MIF2 | n |
| W303.asm02.HP0_G0032380 | 665.9083673 | 676.8177619 | 606.9014867 | 913.4768915 | 1089.410651 | 965.900939 | 1053.556367 | 1008.231962 | 1036.933327 | 2.79E-06 | -0.604987892 | YKL088W | CAB3 | n |
| W303.asm02.HP0_G0032390 | 1410.329948 | 1610.654927 | 1458.134451 | 1138.198044 | 1165.045739 | 1111.646067 | 1337.533824 | 1061.44958 | 1047.169589 | 0.006112997 | 0.391716767 | YKL087C | CYT2 | n |
| W303.asm02.HP0_G0032400 | 98.86849121 | 66.63466011 | 99.68479492 | 226.6667899 | 229.5591263 | 226.31231 | 196.8910371 | 261.2501244 | 290.7098371 | 1.55E-08 | -1.364364726 | YKL086W | SRX1 | n |
| W303.asm02.HP0_G0032410 | 2660.725572 | 1751.539637 | 2599.623083 | 4065.409936 | 4370.911919 | 3974.044163 | 5832.896974 | 4474.150279 | 4023.874541 | 2.27E-05 | -0.823609501 | YKL085W | MDH1 | n |
| W303.asm02.HP0_G0032420 | 297.5747726 | 325.5579108 | 395.807274 | 35.99429711 | 25.21169595 | 28.06272644 | 52.06253385 | 28.06019855 | 38.89779511 | 3.79E-32 | 3.503256993 | YKL084W | HOT13 | n |
| W303.asm02.HP0_G0032430 | 1794.172326 | 2901.463486 | 2187.201677 | 3454.479704 | 3355.809424 | 3479.778078 | 2086.28772 | 2428.658564 | 2362.52924 | 0.001061225 | -0.580185081 | YKL082C | RRP14 | n |
| W303.asm02.HP0_G0032450 | 5864.258547 | 5638.244169 | 6154.070134 | 1288.984964 | 1068.179749 | 1304.464155 | 1459.644131 | 1121.440349 | 1117.799796 | 6.29E-53 | 2.26842993 | YKL080W | VMA5 | n |
| W303.asm02.HP0_G0032470 | 222.9387547 | 457.8753073 | 200.3468918 | 3109.129015 | 2763.998035 | 3133.067619 | 1896.022824 | 1647.81097 | 1754.495284 | 2.65E-39 | -3.353126782 | YKL078W | DHR2 | n |
| W303.asm02.HP0_G0032480 | 3885.919424 | 3765.81022 | 4085.121988 | 2216.081319 | 2266.398773 | 2193.418908 | 2261.407152 | 2041.621343 | 1913.157343 | 1.48E-13 | 0.814185973 | YKL077W | PSG1 | n |
| W303.asm02.HP0_G0032490 | 1085.614805 | 1090.904578 | 1045.713045 | 859.9718553 | 866.4861818 | 868.1340211 | 730.768657 | 755.6901747 | 809.6883139 | 0.006457722 | 0.312602166 | YKL075C | AAN1 | n |
| W303.asm02.HP0_G0032500 | 841.3514742 | 992.8564356 | 836.5704358 | 1230.615834 | 1008.467838 | 1330.716383 | 863.2914704 | 834.0652121 | 945.8305968 | 0.007099464 | -0.420047816 | YKL074C | MUD2 | n |
| W303.asm02.HP0_G0032510 | 1039.088457 | 1011.89491 | 1114.124179 | 1415.451413 | 1361.431581 | 1448.398784 | 1876.144402 | 2088.065809 | 2362.52924 | 0.001496726 | -0.41724411 | YKL073W | LHS1 | n |
| W303.asm02.HP0_G0032520 | 330.5309363 | 518.7984251 | 382.1250472 | 342.432232 | 330.4059101 | 293.3007537 | 353.0786387 | 459.6067004 | 508.7422149 | 0.122205275 | 0.350925991 | YKL072W | STB6 | n |
| W303.asm02.HP0_G0032540 | 390.6274702 | 375.057944 | 396.7845759 | 202.3463189 | 134.0200679 | 127.6401428 | 666.4004333 | 384.1344422 | 689.92405 | 4.58E-07 | 1.322812032 | YKL070W | | n |
| W303.asm02.HP0_G0032550 | 832.6277838 | 738.6928035 | 878.594418 | 605.093319 | 662.1387515 | 615.5694831 | 801.7630213 | 665.7040208 | 467.7971674 | 0.061970893 | 0.380382033 | YKL069W | | n |
| W303.asm02.HP0_G0032560 | 982.8691185 | 1751.539637 | 1028.12161 | 171.216116 | 127.3854111 | 175.6183525 | 224.3421913 | 170.2963774 | 197.5598541 | 9.68E-33 | 2.983302253 | YKL068W-A | | n |
| W303.asm02.HP0_G0032580 | 3110.480277 | 2678.713336 | 2945.58796 | 7742.665154 | 8790.920298 | 7549.778661 | 8263.744009 | 6141.31311 | 5477.423727 | 1.64E-18 | -1.463088333 | YKL067W | YNK1 | n |
| W303.asm02.HP0_G0032610 | 3365.405897 | 2894.80002 | 3455.739557 | 509.7570726 | 330.4059101 | 448.0983738 | 1004.333608 | 797.296676 | 890.5547827 | 1.65E-60 | 2.910905307 | YKL065C | YET1 | n |
| W303.asm02.HP0_G0032620 | 1249.426325 | 1228.933517 | 1294.925032 | 815.2221886 | 989.8907989 | 898.9124952 | 1089.526845 | 1146.597768 | 1462.761821 | 0.002042588 | 0.481815684 | YKL064W | MNR2 | n |
| W303.asm02.HP0_G0032630 | 1067.198126 | 1475.481759 | 1232.37771 | 1267.58295 | 1264.565592 | 1253.770197 | 1573.235114 | 1257.870969 | 1239.611312 | 0.981150021 | -0.00405282 | YKL063C | | n |
| W303.asm02.HP0_G0032650 | 670.7548619 | 595.9042461 | 660.6560918 | 44.74966668 | 43.78873507 | 38.92571732 | 44.48980165 | 66.76392069 | 55.2758141 | 7.80E-69 | 3.922334838 | YKL061W | BLI1 | n |
| W303.asm02.HP0_G0032660 | 49121.16205 | 42158.79753 | 49069.35165 | 47463.83124 | 42208.35981 | 44239.53036 | 26377.71942 | 21555.07045 | 22323.23989 | 0.625189805 | 0.067720063 | YKL060C | FBA1 | n |
| W303.asm02.HP0_G0032670 | 519.5442283 | 323.6540634 | 406.557595 | 653.734261 | 668.7734083 | 601.9907446 | 907.781272 | 742.143872 | 865.9877542 | 0.001271594 | -0.622422349 | YKL059C | MPE1 | n |
| W303.asm02.HP0_G0032680 | 1493.689656 | 1373.625922 | 1429.792696 | 883.3195075 | 1056.237367 | 976.7639299 | 1395.275907 | 876.6393064 | 856.7751185 | 0.005212708 | 0.559762061 | YKL058W | TOA2 | n |
| W303.asm02.HP0_G0032690 | 1964.768938 | 2358.866968 | 2077.743863 | 2484.579374 | 2576.900712 | 2484.909164 | 1592.166944 | 2238.042733 | 1897.802951 | 0.130268636 | -0.237259232 | YKL057C | NUP120 | n |
| W303.asm02.HP0_G0032700 | 12515.58783 | 12205.56588 | 12402.93855 | 15082.58331 | 17546.01345 | 14583.56526 | 14227.27061 | 11219.24145 | 9992.638837 | 0.025679303 | -0.346715218 | YKL056C | TMA19 | n |
| W303.asm02.HP0_G0032710 | 90.14480081 | 91.38467672 | 81.11605861 | 273.3620943 | 200.3666362 | 290.585006 | 272.6183591 | 295.1158813 | 282.5208276 | 4.35E-13 | -1.546428112 | YKL055C | OAR1 | n |

| | | | | | | | | | | | | | | |
|---|---|---|---|---|---|---|---|---|---|---|---|---|---|---|
| W303.asm02.HP0_G0032720 | 3078.493413 | 3073.761678 | 3594.516429 | 14782.95511 | 13252.06355 | 14012.35298 | 8450.22254 | 12385.19108 | 12411.46752 | 1.81E-37 | -2.109067837 | YKL054C | DEF1 | n |
| W303.asm02.HP0_G0032730 | 1093.369197 | 1199.423882 | 1079.918612 | 537.968819 | 575.8882127 | 580.2647628 | 997.7074669 | 591.1993556 | 532.2856172 | 8.69E-06 | 0.993515547 | YKL053C-/ | MDM35 | n |
| W303.asm02.HP0_G0032740 | 1323.093044 | 932.8852415 | 1112.169575 | 129.3849058 | 169.8472148 | 152.0818723 | 200.6774032 | 161.5880399 | 249.7647896 | 1.05E-41 | 2.90365578 | YKL052C | ASK1 | n |
| W303.asm02.HP0_G0032750 | 840.3821753 | 1009.039139 | 729.0672256 | 949.4711887 | 1252.623209 | 880.8075104 | 1164.307575 | 1467.838662 | 1347.092062 | 0.193358607 | -0.256000658 | YKL051W | SFK1 | n |
| W303.asm02.HP0_G0032770 | 830.6891859 | 822.4620904 | 823.865511 | 327.8399494 | 342.3482924 | 354.8577021 | 400.4082149 | 369.6205464 | 347.0092774 | 1.51E-21 | 1.272693211 | YKL049C | CSE4 | n |
| W303.asm02.HP0_G0032780 | 309.2063598 | 398.8560369 | 258.9850064 | 272.3892754 | 290.5979691 | 261.6170303 | 282.0842743 | 330.9168243 | 344.962025 | 0.277094169 | 0.231609283 | YKL048C | ELM1 | n |
| W303.asm02.HP0_G0032790 | 1232.948243 | 930.0294703 | 1304.698051 | 286.981558 | 290.5979691 | 249.8487902 | 319.0013438 | 290.277916 | 343.9383988 | 1.18E-31 | 2.069051724 | YKL047W | ANR2 | n |
| W303.asm02.HP0_G0032800 | 1502.413347 | 1628.741478 | 1680.959287 | 1899.915196 | 1790.030412 | 1853.045194 | 2112.792283 | 2537.028986 | 2499.695149 | 0.108119283 | -0.204276947 | YKL046C | DCW1 | n |
| W303.asm02.HP0_G0032810 | 1299.82987 | 1176.577713 | 1218.695483 | 1294.821877 | 1199.545955 | 1358.779109 | 1231.515573 | 1320.764518 | 1137.248694 | 0.64634861 | -0.061422595 | YKL045W | PRI2 | n |
| W303.asm02.HP0_G0032840 | 386.7502744 | 279.8655724 | 431.9674447 | 227.6396088 | 233.5399204 | 235.3648024 | 294.3899641 | 242.8658564 | 226.2213873 | 0.002095533 | 0.657181179 | YKL042W | SPC42 | n |
| W303.asm02.HP0_G0032850 | 1117.60167 | 996.6641304 | 1150.284349 | 405.6654566 | 453.8105271 | 379.2994315 | 726.0356993 | 563.1391571 | 612.1284598 | 1.97E-19 | 1.400563357 | YKL041W | VPS24 | n |
| W303.asm02.HP0_G0032860 | 2311.777956 | 2345.540036 | 2360.184115 | 1040.91616 | 1203.526749 | 1070.004602 | 1506.027116 | 1541.375734 | 1340.950305 | 5.11E-20 | 1.083641164 | YKL040C | NFU1 | n |
| W303.asm02.HP0_G0032870 | 2486.251764 | 2186.568775 | 2565.417516 | 1842.518885 | 1746.241677 | 1792.393495 | 2346.600389 | 2773.121691 | 2307.253425 | 0.000948203 | 0.427465689 | YKL039W | PTM1 | n |
| W303.asm02.HP0_G0032880 | 211.3071675 | 328.4136819 | 195.4603822 | 64.2060435 | 92.8851956 | 48.88345896 | 305.7490624 | 245.7686356 | 322.4422489 | 3.56E-10 | 1.851124486 | YKL038W | RGT1 | n |
| W303.asm02.HP0_G0032900 | 3924.691381 | 2431.21317 | 3558.356258 | 5122.864016 | 3832.177784 | 5017.796537 | 2818.002969 | 4468.344721 | 4272.615704 | 0.031696726 | -0.495271153 | YKL035W | UGP1 | n |
| W303.asm02.HP0_G0032910 | 286.9124843 | 300.8078942 | 299.0543848 | 284.0631015 | 278.6555868 | 262.5222796 | 455.3105233 | 604.7456584 | 618.2702169 | 0.583062944 | 0.104392001 | YKL034W | TUL1 | n |
| W303.asm02.HP0_G0032920 | 2577.365864 | 2733.924912 | 2726.672332 | 815.2221886 | 630.2923987 | 767.6513555 | 868.9710195 | 722.7920109 | 602.9158241 | 1.57E-29 | 1.858925891 | YKL033W-A | | n |
| W303.asm02.HP0_G0032930 | 651.3688833 | 778.6735995 | 703.6573759 | 137.1674566 | 144.6355189 | 133.9768875 | 33.13070336 | 122.8843178 | 154.5675542 | 2.54E-10 | 2.360092673 | YKL033W | TTI1 | n |
| W303.asm02.HP0_G0032940 | 239.4168366 | 353.1636986 | 208.165307 | 1857.111167 | 1559.144355 | 1905.54965 | 656.934518 | 1485.255337 | 1656.227171 | 8.68E-20 | -2.732503579 | YKL032C | IXR1 | y |
| W303.asm02.HP0_G0032960 | 656.2153779 | 993.8083593 | 614.722902 | 5893.336538 | 4368.258056 | 5740.18543 | 1121.710957 | 1875.195338 | 1446.383802 | 4.49E-34 | -2.820862039 | YKL029C | MAE1 | n |
| W303.asm02.HP0_G0032970 | 2659.756273 | 2379.809289 | 2567.37212 | 1535.108131 | 1438.3936 | 1457.451276 | 1568.502156 | 1462.033104 | 1360.399202 | 9.13E-12 | 0.779382315 | YKL028W | TFA1 | n |
| W303.asm02.HP0_G0032980 | 1834.882881 | 1524.981793 | 1771.848365 | 4154.909269 | 3931.697637 | 4079.958324 | 2095.753635 | 2536.061393 | 1886.543063 | 5.14E-18 | -1.245544403 | YKL027W | TCD2 | n |
| W303.asm02.HP0_G0033000 | 519.5442283 | 520.7022725 | 420.2398217 | 888.1836017 | 875.7747014 | 876.2812642 | 675.8663485 | 784.7179663 | 812.7591925 | 7.51E-09 | -0.854124904 | YKL025C | PAN3 | n |
| W303.asm02.HP0_G0033010 | 1114.693773 | 1106.135358 | 1215.763577 | 2674.278993 | 2726.843957 | 2590.823325 | 3091.567919 | 2759.575388 | 2524.262177 | 7.55E-24 | -1.217427447 | YKL024C | URA6 | n |
| W303.asm02.HP0_G0033040 | 474.9564774 | 521.6541963 | 446.6269733 | 531.1590871 | 626.3116046 | 499.6975804 | 558.4889995 | 586.3613904 | 701.1839381 | 0.253966057 | -0.196791167 | YKL022C | CDC16 | n |
| W303.asm02.HP0_G0033050 | 1777.694244 | 2022.837896 | 1905.738726 | 4400.059617 | 3877.293451 | 4418.52154 | 2495.215259 | 2130.639904 | 1976.622167 | 3.94E-18 | -1.154021802 | YKL021C | MAK11 | n |
| W303.asm02.HP0_G0033070 | 1611.944126 | 1228.933517 | 1679.004683 | 459.1704929 | 414.0025861 | 449.9088722 | 676.8129401 | 511.8567253 | 576.3015433 | 1.16E-24 | 1.771176307 | YKL019W | RAM2 | n |
| W303.asm02.HP0_G0033080 | 789.0093318 | 772.9620572 | 804.3194727 | 164.4063841 | 200.3666362 | 186.4813434 | 354.0252302 | 236.092705 | 240.5521539 | 4.51E-27 | 2.104545173 | YKL018C-/ | MCO12 | n |
| W303.asm02.HP0_G0033090 | 897.5708124 | 607.3273307 | 895.2085505 | 616.7671451 | 651.5233006 | 588.412006 | 701.4243197 | 566.0419363 | 679.6877882 | 0.052045648 | 0.370933218 | YKL018W | SWD2 | n |
| W303.asm02.HP0_G0033100 | 561.2240825 | 467.3945445 | 508.1969937 | 439.7141161 | 475.0414289 | 486.1188418 | 560.3821825 | 527.3382141 | 515.9075982 | 0.37519577 | 0.133742179 | YKL017C | HCS1 | n |
| W303.asm02.HP0_G0033110 | 4774.766546 | 5175.609243 | 5074.151522 | 7447.901045 | 7507.777668 | 7145.132251 | 7124.047814 | 5621.71564 | 5055.689737 | 0.000115205 | -0.556735548 | YKL016C | ATP7 | n |
| W303.asm02.HP0_G0033120 | 1306.614962 | 1630.645325 | 1367.245373 | 752.9617828 | 780.2356431 | 730.5361366 | 600.1390266 | 748.9170234 | 727.7982189 | 4.71E-10 | 0.927778324 | YKL015W | PUT3 | n |
| W303.asm02.HP0_G0033130 | 765.7461574 | 1352.6836 | 813.1151899 | 4425.352907 | 4438.585418 | 4662.938835 | 1828.814825 | 2157.732509 | 1948.98426 | 1.80E-27 | -2.205989472 | YKL014C | URB1 | n |
| W303.asm02.HP0_G0033140 | 3080.43201 | 3086.136687 | 3032.56783 | 1528.298399 | 1350.81613 | 1485.514003 | 2150.655944 | 1551.051665 | 1546.699169 | 3.39E-12 | 1.075115362 | YKL013C | ARC19 | n |
| W303.asm02.HP0_G0033150 | 1011.948086 | 859.5871154 | 955.801269 | 622.6040581 | 651.5233006 | 562.159778 | 567.9549147 | 646.3521597 | 777.9559021 | 0.000123675 | 0.623817445 | YKL012W | PRP40 | n |
| W303.asm02.HP0_G0033160 | 768.6540542 | 661.5869825 | 679.2248281 | 83.66242031 | 71.65429375 | 90.52492399 | 101.2852931 | 128.6898761 | 134.0950305 | 3.24E-56 | 3.092239778 | YKL011C | CCE1 | n |
| W303.asm02.HP0_G0033170 | 1031.334065 | 1348.875905 | 1039.849303 | 1324.006442 | 1381.335552 | 1329.811133 | 1132.123463 | 1567.500747 | 1787.251322 | 0.22822612 | -0.23838505 | YKL010C | UFD4 | n |
| W303.asm02.HP0_G0033180 | 3696.906132 | 5425.96518 | 3971.754966 | 7223.179893 | 7514.412324 | 7088.101549 | 4667.642808 | 3959.390775 | 4131.355291 | 5.00E-06 | -0.736972013 | YKL009W | MRT4 | n |
| W303.asm02.HP0_G0033200 | 3150.221534 | 2424.549704 | 3199.686457 | 1051.617167 | 835.9667604 | 969.521936 | 1294.937206 | 985.9773214 | 955.0432324 | 5.79E-20 | 1.617720823 | YKL007W | CAP1 | n |
| W303.asm02.HP0_G0033240 | 1007.101592 | 1007.135291 | 1005.643666 | 1656.710486 | 1318.969778 | 1672.900595 | 739.2879807 | 1169.820002 | 882.3657732 | 0.000847499 | -0.623069544 | YKL004W | AUR1 | n |
| W303.asm02.HP0_G0033250 | 1351.202713 | 1319.36627 | 1330.107901 | 1398.913493 | 1439.720532 | 1409.473067 | 1760.660236 | 1371.079357 | 1174.099236 | 0.616139554 | -0.086458566 | YKL003C | MRP17 | n |
| W303.asm02.HP0_G0033270 | 1422.930834 | 1222.270051 | 1500.158433 | 515.5939856 | 460.4451839 | 483.4030941 | 749.7004874 | 516.6946905 | 518.9784768 | 8.04E-17 | 1.505267746 | YKL001C | MET14 | n |
| W303.asm02.HP0_G0033280 | 3074.616217 | 2095.184098 | 3034.522434 | 4591.704929 | 3662.33057 | 4366.922333 | 3681.294439 | 4468.344721 | 4180.489348 | 0.000483927 | -0.62166487 | YKR001C | VPS1 | n |
| W303.asm02.HP0_G0033290 | 752.1759723 | 595.9042461 | 645.0192613 | 1826.953783 | 1778.08803 | 1965.2961 | 1175.666673 | 1855.843476 | 1534.415654 | 3.38E-16 | -1.483017141 | YKR002W | PAP1 | n |
| W303.asm02.HP0_G0033310 | 676.5706555 | 423.6060535 | 663.5879267 | 230.5580653 | 254.7708222 | 186.4813434 | 267.8854014 | 259.3149883 | 237.4812754 | 3.43E-10 | 1.395977466 | YKR004C | ECM9 | n |
| W303.asm02.HP0_G0033320 | 361.5485022 | 236.0770815 | 328.3734421 | 13.61946377 | 17.25010776 | 15.38923708 | 20.82501354 | 23.22223328 | 29.68515942 | 2.35E-37 | 4.333661626 | YKR005C | | n |
| W303.asm02.HP0_G0033330 | 2507.576341 | 2587.32866 | 2637.737858 | 2512.791066 | 2656.516594 | 2600.781066 | 2299.270813 | 2180.954742 | 1899.850203 | 0.956050043 | -0.006792462 | YKR006C | MRPL13 | n |
| W303.asm02.HP0_G0033340 | 1051.689343 | 1081.385341 | 1016.393987 | 393.9916305 | 443.1950762 | 395.5939178 | 648.4151943 | 565.0743432 | 643.8608716 | 1.02E-24 | 1.355894134 | YKR007W | MEH1 | n |
| W303.asm02.HP0_G0033350 | 1758.308265 | 1550.683733 | 1732.756288 | 1149.87187 | 1175.66119 | 1103.498823 | 907.781272 | 1069.190324 | 1001.106411 | 6.87E-06 | 0.556568942 | YKR008W | RSC4 | n |
| W303.asm02.HP0_G0033360 | 561.2240825 | 415.0387401 | 522.8565224 | 194.5637682 | 298.5595573 | 222.691313 | 304.8024709 | 290.277916 | 366.4581749 | 6.24E-07 | 1.072002035 | YKR009C | FOX2 | n |
| W303.asm02.HP0_G0033370 | 284.0045875 | 255.1155558 | 247.2573835 | 2591.589392 | 2687.036016 | 2558.234352 | 1798.523897 | 2099.676926 | 1833.314501 | 2.89E-147 | -3.316772933 | YKR010C | TOF2 | n |
| W303.asm02.HP0_G0033380 | 1816.466201 | 1273.673932 | 1600.82053 | 164.4063841 | 184.4434598 | 181.9550972 | 459.0968894 | 376.3936978 | 479.0570555 | 5.63E-66 | 3.144869049 | YKR011C | FPT1 | n |
| W303.asm02.HP0_G0033390 | 1792.233728 | 1797.231975 | 1722.005967 | 1051.617167 | 886.3901523 | 972.2376837 | 937.1256093 | 832.130026 | 984.7283919 | 8.91E-12 | 0.866671641 | YKR013W | PRY2 | n |
| W303.asm02.HP0_G0033400 | 3053.29164 | 2791.040335 | 3123.456908 | 1059.399718 | 950.0828579 | 1037.415629 | 1354.572472 | 1090.477371 | 1217.091536 | 1.03E-33 | 1.556436625 | YKR014C | YPT52 | n |
| W303.asm02.HP0_G0033420 | 1837.790778 | 2008.55904 | 1885.215386 | 3835.824689 | 3749.90804 | 3850.930267 | 3512.801148 | 3535.585017 | 3139.461515 | 2.16E-21 | -0.99674035 | YKR016W | MIC60 | n |
| W303.asm02.HP0_G0033430 | 470.1099827 | 497.8561034 | 420.2398217 | 284.0631015 | 302.5403514 | 246.2277933 | 290.603598 | 299.9538466 | 368.5054273 | 2.89E-05 | 0.740281661 | YKR017C | HEL1 | n |
| W303.asm02.HP0_G0033440 | 1777.694244 | 1564.010665 | 1803.122026 | 1131.388312 | 1175.66119 | 1194.023747 | 1125.497323 | 1196.912607 | 1188.430003 | 1.63E-06 | 0.555308081 | YKR018C | | n |
| W303.asm02.HP0_G0033460 | 540.8688049 | 465.490697 | 517.9700128 | 147.8684638 | 167.1933521 | 162.0396139 | 354.9718217 | 220.6112162 | 307.0878561 | 6.71E-16 | 1.677273289 | YKR020W | VPS51 | n |
| W303.asm02.HP0_G0033470 | 536.9916091 | 663.4908299 | 467.1503135 | 714.0490292 | 662.1387515 | 734.1571336 | 496.9605504 | 837.9355843 | 910.0036802 | 0.157111612 | -0.340009602 | YKR021W | ALY1 | n |
| W303.asm02.HP0_G0033480 | 880.1234316 | 908.1352249 | 875.6625123 | 116.7382609 | 57.05804873 | 105.0089118 | 27.45115421 | 65.79632763 | 52.20493554 | 1.74E-29 | 3.244694374 | YKR022C | NTR2 | n |
| W303.asm02.HP0_G0033490 | 750.2373745 | 860.5390391 | 794.5464536 | 541.8600943 | 647.5425065 | 518.7078145 | 737.3947976 | 791.4911177 | 977.5630086 | 0.003031858 | 0.496278444 | YKR023W | RQT4 | n |
| W303.asm02.HP0_G0033500 | 477.8643742 | 892.9044454 | 485.7190498 | 3991.475704 | 3375.713395 | 3997.580644 | 1792.844348 | 2267.070524 | 2151.662245 | 3.29E-30 | -2.613872828 | YKR024C | DBP7 | n |
| W303.asm02.HP0_G0033510 | 1029.395467 | 1035.693003 | 978.2792129 | 2534.19308 | 2357.957037 | 2534.697872 | 1959.444456 | 1694.255437 | 1653.156292 | 1.25E-27 | -1.287425942 | YKR025W | RPC37 | n |

| | | | | | | | | | | | | | | |
|---|---|---|---|---|---|---|---|---|---|---|---|---|---|---|
| W303.asm02.HP0_G0033520 | 1996.755803 | 1924.789753 | 2109.017524 | 1907.697747 | 1837.799942 | 1945.380617 | 1990.681976 | 1511.380349 | 1451.501933 | 0.600089999 | 0.083406887 | YKR026C | GCN3 | n |
| W303.asm02.HP0_G0033540 | 960.575243 | 1174.673865 | 1000.757157 | 4330.989479 | 4741.12577 | 4357.869841 | 3419.088587 | 4811.840255 | 4652.38102 | 1.30E-39 | -2.098254043 | YKR028W | SAP190 | n |
| W303.asm02.HP0_G0033560 | 1381.25098 | 1405.991328 | 1303.720749 | 454.3063987 | 623.6577419 | 496.0765835 | 820.6948518 | 699.5697776 | 651.0262549 | 1.43E-17 | 1.381371566 | YKR030W | GMH1 | n |
| W303.asm02.HP0_G0033570 | 482.7108688 | 705.3754734 | 459.3318982 | 291.8456523 | 301.21342 | 297.8269999 | 188.3717134 | 464.4446656 | 458.5845318 | 0.004380615 | 0.887099875 | YKR031C | SPO14 | n |
| W303.asm02.HP0_G0033590 | 146.3641389 | 158.9712605 | 179.8235516 | 1279.256776 | 1231.392307 | 1289.980167 | 986.3483686 | 1134.986652 | 993.9410276 | 1.37E-94 | -2.970262703 | YKR034W | DAL80 | n |
| W303.asm02.HP0_G0033600 | 1880.439931 | 2315.078477 | 1919.420953 | 900.8302466 | 879.7554955 | 902.5334922 | 1239.088306 | 954.0467507 | 882.3657732 | 2.63E-13 | 1.188280965 | YKR035W- | DID2 | n |
| W303.asm02.HP0_G0033620 | 694.9873353 | 648.2600505 | 704.6346778 | 379.3993479 | 456.4643898 | 391.0676716 | 505.4798741 | 447.9955837 | 415.5922319 | 9.63E-07 | 0.742152868 | YKR037C | SPC34 | n |
| W303.asm02.HP0_G0033630 | 1542.154603 | 1010.942986 | 1375.063789 | 1548.727595 | 1663.971933 | 1585.091419 | 1499.400975 | 1454.292359 | 1331.737669 | 0.093438313 | -0.288364664 | YKR038C | KAE1 | n |
| W303.asm02.HP0_G0033640 | 2591.905348 | 3321.261844 | 2470.619231 | 378.4265291 | 346.3290865 | 383.8256777 | 198.7842202 | 437.3520602 | 589.6086837 | 2.99E-23 | 2.918376515 | YKR039W | GAP1 | n |
| W303.asm02.HP0_G0033660 | 5560.867981 | 5174.657319 | 4626.547247 | 2334.765218 | 1784.722687 | 2347.311279 | 606.7651672 | 1709.736925 | 1731.975508 | 4.05E-05 | 1.248013829 | YKR042W | UTH1 | n |
| W303.asm02.HP0_G0033670 | 2777.041444 | 2559.722872 | 2914.314299 | 1741.345725 | 1573.7406 | 1759.804522 | 1440.7123 | 1251.097818 | 1180.240994 | 4.89E-08 | 0.700503951 | YKR043C | SHB17 | n |
| W303.asm02.HP0_G0033700 | 1789.325831 | 1474.529836 | 1641.86721 | 1315.251073 | 640.9078497 | 1371.452598 | 539.557169 | 484.7641198 | 855.7514923 | 0.053968213 | 0.558665999 | YKR046C | PLN1 | n |
| W303.asm02.HP0_G0033710 | 1350.233414 | 1054.731477 | 924.5276078 | 3947.698856 | 3768.485079 | 3758.594844 | 3922.675278 | 3499.784074 | 3101.587347 | 2.97E-28 | -1.785109124 | YKR048C | NAP1 | n |
| W303.asm02.HP0_G0033720 | 462.3555912 | 409.3271978 | 479.8552383 | 64.2060435 | 55.73111736 | 47.97820972 | 119.2705321 | 108.370422 | 87.00822589 | 4.19E-41 | 3.010534272 | YKR049C | FMP46 | n |
| W303.asm02.HP0_G0033730 | 143.4562421 | 160.875108 | 103.5940026 | 1196.567174 | 1169.026533 | 1189.497501 | 745.9141213 | 1154.338513 | 921.2635683 | 3.62E-51 | -3.123044401 | YKR050W | TRK2 | n |
| W303.asm02.HP0_G0033740 | 696.9259331 | 693.0004651 | 569.7670141 | 196.5094058 | 224.2514008 | 175.6183525 | 437.3252843 | 372.5233256 | 473.9389246 | 7.83E-22 | 1.721607122 | YKR051W | HFL1 | n |
| W303.asm02.HP0_G0033750 | 783.1935382 | 647.3081267 | 700.7254702 | 3136.367943 | 2409.70736 | 3117.678382 | 2062.622932 | 2152.894544 | 1938.747998 | 1.33E-42 | -2.024084686 | YKR052C | MRS4 | n |
| W303.asm02.HP0_G0033770 | 455.5704987 | 660.6350588 | 400.6937835 | 769.4997031 | 843.9283486 | 830.113553 | 880.3301178 | 818.5837232 | 734.9636022 | 0.000518082 | -0.687341238 | YKR054C | DYN1 | n |
| W303.asm02.HP0_G0033780 | 656.2153779 | 661.5869825 | 647.951167 | 404.6926378 | 469.7337035 | 459.8666139 | 565.1151402 | 432.5140949 | 396.1433344 | 0.001216289 | 0.559921473 | YKR055W | RHO4 | n |
| W303.asm02.HP0_G0033790 | 874.307638 | 930.981394 | 880.5490218 | 2553.649457 | 2814.421427 | 2683.158747 | 1878.984176 | 2152.894544 | 1975.598541 | 1.07E-44 | -1.583431483 | YKR056W | TRM2 | n |
| W303.asm02.HP0_G0033830 | 861.7067518 | 910.0390723 | 930.3914193 | 1514.678935 | 1609.567747 | 1545.260453 | 1285.47129 | 1164.982036 | 1130.08331 | 6.95E-12 | -0.788744411 | YKR060W | UTP30 | n |
| W303.asm02.HP0_G0033850 | 1091.430599 | 1135.644993 | 1124.8745 | 867.754406 | 1013.775563 | 865.4182734 | 1332.800867 | 1029.519009 | 1215.044284 | 0.050631612 | 0.288692548 | YKR062W | TFA2 | n |
| W303.asm02.HP0_G0033860 | 484.6494667 | 701.5677785 | 535.5614472 | 947.525551 | 858.5245937 | 932.4067171 | 1000.547241 | 905.667098 | 918.1926897 | 5.38E-05 | -0.66990701 | YKR063C | LAS1 | n |
| W303.asm02.HP0_G0033870 | 660.0925737 | 681.5773805 | 677.2702243 | 520.4580798 | 427.2718998 | 521.4235622 | 472.3491708 | 524.435435 | 521.0257292 | 0.001589905 | 0.455288085 | YKR064W | OAF3 | n |
| W303.asm02.HP0_G0033880 | 1532.461614 | 1182.289255 | 1508.954151 | 1986.496073 | 1853.723118 | 1970.727595 | 2694.94607 | 1714.574891 | 1954.102391 | 0.015530868 | -0.46048222 | YKR065C | PAM17 | n |
| W303.asm02.HP0_G0033890 | 1673.009959 | 985.2410458 | 1484.521603 | 8941.177966 | 5756.228265 | 8083.875713 | 4449.926757 | 4939.562538 | 4472.222811 | 4.75E-31 | -2.459423842 | YKR066C | CCP1 | n |
| W303.asm02.HP0_G0033900 | 513.7284347 | 435.0291381 | 532.6295415 | 882.3466886 | 1098.699171 | 873.5655165 | 1286.417882 | 1746.505461 | 1619.376628 | 6.05E-08 | -0.944985646 | YKR067W | GPT2 | n |
| W303.asm02.HP0_G0033910 | 2715.006313 | 2082.80909 | 2639.692462 | 617.7399639 | 674.0811338 | 645.4427081 | 941.8585669 | 616.356775 | 648.9790025 | 4.87E-25 | 1.941227321 | YKR068C | BET3 | n |
| W303.asm02.HP0_G0033920 | 576.7328654 | 420.7502824 | 449.5588791 | 233.4765218 | 258.7516163 | 286.0587598 | 240.4342472 | 323.1760798 | 364.4109226 | 3.11E-05 | 0.894216365 | YKR069W | MET1 | n |
| W303.asm02.HP0_G0033930 | 951.8515526 | 740.5966509 | 903.0269658 | 466.9530436 | 508.2147131 | 505.1290759 | 804.6027959 | 652.157718 | 725.7509666 | 2.43E-07 | 0.810505923 | YKR070W | | n |
| W303.asm02.HP0_G0033940 | 7208.676168 | 7801.014851 | 7714.821285 | 2300.716559 | 1881.588677 | 2102.893984 | 2106.166142 | 1804.561045 | 1725.833751 | 1.66E-46 | 1.853527554 | YKR071C | DRE2 | n |
| W303.asm02.HP0_G0033950 | 499.1889507 | 739.6447272 | 465.1957096 | 1458.255442 | 1441.047463 | 1432.104298 | 1492.774834 | 1473.64422 | 1537.486533 | 3.73E-14 | -1.34563358 | YKR072C | SIS2 | n |
| W303.asm02.HP0_G0033970 | 1385.128176 | 1306.039338 | 1397.541733 | 828.8416524 | 954.063652 | 876.2812642 | 903.9949059 | 842.7735495 | 629.530105 | 0.000115717 | 0.621480006 | YKR074W | AIM29 | n |
| W303.asm02.HP0_G0033980 | 868.4918444 | 1011.89491 | 747.6359619 | 346.3235073 | 587.830595 | 382.0151792 | 736.4482061 | 771.1716636 | 926.3816692 | 5.96E-06 | 1.001836189 | YKR075C | | n |
| W303.asm02.HP0_G0033990 | 433.2766232 | 298.9040468 | 388.9661606 | 749.0705074 | 293.2518318 | 583.8857598 | 812.1755281 | 453.8011421 | 588.5850575 | 0.099686519 | -0.538764522 | YKR076W | ECM4 | n |
| W303.asm02.HP0_G0034010 | 517.6056305 | 613.038873 | 554.1301835 | 947.525551 | 903.6402601 | 916.1122308 | 992.0279177 | 1170.787595 | 1188.430003 | 2.20E-07 | -0.716073245 | YKR078W | VPS501 | n |
| W303.asm02.HP0_G0034020 | 1164.128019 | 1564.962589 | 1327.175995 | 2384.378979 | 2312.841371 | 2506.635145 | 1407.581597 | 1820.422534 | 1636.778273 | 8.46E-08 | -0.828751148 | YKR079C | TRZ1 | n |
| W303.asm02.HP0_G0034040 | 2676.234355 | 3528.781214 | 3102.933567 | 4874.795211 | 4966.704102 | 4893.777391 | 3074.529272 | 2597.019755 | 2239.694097 | 2.18E-05 | -0.662691555 | YKR081C | RPF2 | n |
| W303.asm02.HP0_G0034050 | 1558.632685 | 1861.962788 | 1685.845796 | 3223.921639 | 3200.558454 | 3299.63348 | 1985.949019 | 3007.27921 | 2756.625322 | 2.32E-08 | -0.929267631 | YKR082W | NUP133 | n |
| W303.asm02.HP0_G0034060 | 409.0441499 | 554.0196026 | 431.9674447 | 217.9114203 | 228.2321949 | 210.9230729 | 256.5263032 | 248.6714147 | 228.2686397 | 1.95E-09 | 1.087937012 | YKR083C | DAD2 | n |
| W303.asm02.HP0_G0034080 | 10.66228827 | 14.27885574 | 23.45524586 | 397.8829059 | 362.2522629 | 360.2891975 | 141.0421372 | 429.6113157 | 276.3790705 | 2.84E-29 | -4.534301679 | YKR084C | HBS1 | y |
| W303.asm02.HP0_G0034090 | 1222.285955 | 1561.154894 | 1276.356296 | 1579.857798 | 1629.471717 | 1542.544705 | 1862.89212 | 1415.588637 | 1491.423354 | 0.162197948 | -0.226805902 | YKR085C | MRPL20 | n |
| W303.asm02.HP0_G0034100 | 391.5967691 | 501.6637982 | 417.307916 | 634.2778842 | 658.1579574 | 614.6642339 | 453.4173403 | 723.759604 | 648.9790025 | 0.007189566 | -0.540651505 | YKR086W | PRP16 | n |
| W303.asm02.HP0_G0034110 | 859.768154 | 682.5293042 | 740.7948485 | 660.5439929 | 654.1771633 | 675.315933 | 781.8845993 | 667.6392069 | 637.7191145 | 0.206834097 | 0.197847471 | YKR087C | OMA1 | n |
| W303.asm02.HP0_G0034120 | 1161.220122 | 1011.89491 | 1234.332314 | 214.020145 | 248.1361654 | 189.1970911 | 368.2241031 | 308.662184 | 371.5763059 | 7.85E-46 | 2.393012946 | YKR088C | TVP38 | n |
| W303.asm02.HP0_G0034130 | 654.2767801 | 490.2407136 | 536.5387491 | 448.4694856 | 388.7908902 | 426.372392 | 375.7968352 | 524.435435 | 577.3251695 | 0.046300551 | 0.410605384 | YKR089C | TGL4 | n |
| W303.asm02.HP0_G0034140 | 706.6189225 | 579.7215429 | 710.4984893 | 443.6053914 | 432.5796252 | 476.1611002 | 459.0968894 | 465.4122587 | 544.5691315 | 0.000335114 | 0.561001053 | YKR090W | PXL1 | n |
| W303.asm02.HP0_G0034160 | 1886.255724 | 2307.463087 | 2057.220523 | 1957.311508 | 1825.857559 | 2021.421553 | 1334.69405 | 1014.03752 | 1005.200916 | 0.51791511 | 0.106556954 | YKR092C | SRP40 | y |
| W303.asm02.HP0_G0034170 | 760.8996627 | 1839.116619 | 770.1139059 | 452.360761 | 1097.372239 | 459.8666139 | 1588.380578 | 1322.699704 | 1190.477255 | 0.163206035 | 0.746902079 | YKR093W | PTR2 | n |
| W303.asm02.HP0_G0034180 | 1071.075321 | 1283.193169 | 1128.783707 | 2377.569247 | 2693.670673 | 2431.499458 | 2077.768396 | 2183.857522 | 1874.259548 | 1.10E-17 | -1.106503152 | YKR095W | MLP1 | n |
| W303.asm02.HP0_G0034190 | 332.4695342 | 455.9714599 | 289.2813657 | 357.0245146 | 378.1754392 | 370.2469391 | 518.7321555 | 252.5417869 | 338.8202679 | 0.901187812 | -0.036239541 | YKR095W- | PCC1 | n |
| W303.asm02.HP0_G0034200 | 508.88194 | 622.5581101 | 519.9246166 | 1166.40979 | 1317.642846 | 1166.86627 | 1103.725718 | 1451.38958 | 1485.281597 | 3.21E-12 | -1.143782244 | YKR096W | ESL2 | n |
| W303.asm02.HP0_G0034210 | 217.1229611 | 194.192438 | 199.3695898 | 109.928529 | 108.808372 | 85.99867779 | 281.1376828 | 100.6296977 | 278.4263229 | 0.004332903 | 1.004816977 | YKR097W | PCK1 | n |
| W303.asm02.HP0_G0034230 | 359.6099043 | 564.4907635 | 355.7378956 | 1795.82358 | 1277.834905 | 1640.311623 | 582.1537876 | 963.7226812 | 948.9014753 | 2.95E-14 | -1.88130575 | YKR099W | BAS1 | n |
| W303.asm02.HP0_G0034240 | 1440.378215 | 1417.414413 | 1485.498905 | 2261.803805 | 2251.802528 | 2267.649346 | 1843.96029 | 2431.561343 | 2286.780902 | 1.37E-06 | -0.642806767 | YKR100C | SKG1 | n |
| W303.asm02.HP0_G0034420 | 240.3861355 | 194.192438 | 238.4616663 | 135.2218189 | 127.3854111 | 133.9768875 | 150.5080524 | 145.138958 | 124.8823948 | 0.000204261 | 0.761237028 | YLL057C | JLP1 | n |
| W303.asm02.HP0_G0034440 | 315.0221534 | 442.6445278 | 258.9850064 | 216.9386015 | 180.4626657 | 217.2598176 | 207.3035439 | 166.4260052 | 194.4889755 | 0.002428295 | 0.723519004 | YLL055W | YCT1 | n |
| W303.asm02.HP0_G0034500 | 1405.483453 | 1912.414745 | 1603.752436 | 3442.805878 | 3268.231954 | 3389.253154 | 2831.25525 | 4307.724274 | 3138.437889 | 1.15E-08 | -1.037211547 | YLL048C | YBT1 | n |
| W303.asm02.HP0_G0034520 | 14525.91382 | 14001.84594 | 14853.03444 | 14000.80876 | 16045.25407 | 13406.74124 | 13719.89756 | 10947.34781 | 10506.49918 | 0.987679331 | -0.002286238 | YLL045C | RPL8B | y |
| W303.asm02.HP0_G0034530 | 843.2900721 | 957.6352581 | 824.8428129 | 912.5040727 | 740.4277021 | 1003.016158 | 833.0005416 | 1184.333897 | 905.9091755 | 0.927849195 | -0.018012497 | YLL043W | FPS1 | n |
| W303.asm02.HP0_G0034540 | 121.1623667 | 144.6924048 | 151.4817962 | 130.3577247 | 143.3085875 | 153.8923708 | 166.6001083 | 225.4491814 | 113.6225068 | 0.906838556 | -0.03514697 | YLL042C | ATG10 | n |
| W303.asm02.HP0_G0034550 | 2741.177384 | 2386.472755 | 2977.838923 | 5039.201596 | 7253.006845 | 5041.333017 | 4271.96755 | 6867.975493 | 5663.723693 | 1.16E-07 | -1.096330424 | YLL041C | SDH2 | n |
| W303.asm02.HP0_G0034560 | 634.8908014 | 654.9235165 | 613.7456001 | 649.8429857 | 553.3303795 | 662.6424436 | 828.267584 | 874.7041203 | 750.3179951 | 0.86211993 | 0.026481148 | YLL040C | VPS13 | n |

| | | | | | | | | | | | | | | |
|---|---|---|---|---|---|---|---|---|---|---|---|---|---|---|
| W303.asm02.HP0_G0034570 | 196.7676835 | 203.7116752 | 155.3910038 | 369.6711595 | 481.6760858 | 363.0049452 | 332.2536251 | 408.3242685 | 364.4109226 | 1.67E-08 | -1.12372027 | YLL038C | ENT4 | n |
| W303.asm02.HP0_G0034580 | 1038.119158 | 958.5871818 | 1029.098912 | 861.917493 | 920.8903678 | 831.0188023 | 674.919757 | 761.4957331 | 722.680088 | 0.092304996 | 0.212333255 | YLL036C | PRP19 | n |
| W303.asm02.HP0_G0034590 | 248.140527 | 388.384876 | 284.3948561 | 1103.176566 | 1171.680396 | 1102.593574 | 713.7300095 | 1054.676428 | 851.6569876 | 2.65E-20 | -1.874268502 | YLL035W | GRC3 | n |
| W303.asm02.HP0_G0034600 | 789.0093318 | 1157.539238 | 834.615832 | 3458.370979 | 2862.190956 | 3307.780723 | 1381.077034 | 1718.445263 | 1402.367876 | 7.10E-24 | -1.791757987 | YLL034C | RIX7 | n |
| W303.asm02.HP0_G0034610 | 312.1142566 | 316.0386736 | 322.5096306 | 189.699674 | 210.9820872 | 178.3341003 | 158.0807846 | 135.4630275 | 112.5988806 | 0.000188836 | 0.718866973 | YLL033W | IRC19 | n |
| W303.asm02.HP0_G0034620 | 540.8688049 | 686.3369991 | 604.9498829 | 770.4725219 | 873.1208387 | 813.8190667 | 590.6731113 | 773.1068497 | 750.3179951 | 0.011175549 | -0.422744375 | YLL032C | | n |
| W303.asm02.HP0_G0034630 | 695.9566342 | 768.2024386 | 710.4984893 | 1045.780254 | 1040.314191 | 1097.162079 | 804.6027959 | 1090.477371 | 882.3657732 | 0.000260143 | -0.549938986 | YLL031C | GPI13 | n |
| W303.asm02.HP0_G0034640 | 997.4086025 | 812.9428533 | 780.8642269 | 2807.555175 | 2782.575074 | 2830.714373 | 3116.179299 | 3332.390476 | 3993.165755 | 4.07E-31 | -1.700387514 | YLL029W | FRA1 | n |
| W303.asm02.HP0_G0034650 | 980.9305206 | 1043.308393 | 880.5490218 | 3853.335429 | 5644.76603 | 3640.007194 | 3686.973988 | 4838.93286 | 4503.955223 | 1.37E-30 | -2.176706201 | YLL028W | TPO1 | n |
| W303.asm02.HP0_G0034660 | 2052.975141 | 1625.885707 | 1978.059068 | 612.8758697 | 1121.257004 | 697.947164 | 1432.192977 | 1365.273798 | 1479.13984 | 1.31E-08 | 1.220480368 | YLL027W | ISA1 | n |
| W303.asm02.HP0_G0034670 | 6355.693106 | 5409.782477 | 4268.854747 | 1046.753073 | 869.1400446 | 1109.835568 | 1099.939352 | 560.2363779 | 1047.169589 | 2.75E-23 | 2.405171352 | YLL026W | HSP104 | n |
| W303.asm02.HP0_G0034690 | 10306.55556 | 10709.1418 | 7701.139059 | 26173.69091 | 27302.93978 | 24932.37457 | 20616.7634 | 23500.90008 | 20905.51762 | 3.28E-23 | -1.449086643 | YLL024C | SSA2 | y |
| W303.asm02.HP0_G0034700 | 770.5926521 | 643.5004319 | 724.180716 | 1084.693008 | 772.2740549 | 1020.215893 | 1368.771345 | 2176.116777 | 1381.895352 | 0.051498331 | -0.429408877 | YLL023C | POM33 | n |
| W303.asm02.HP0_G0034710 | 439.0924168 | 324.6059871 | 349.8740841 | 1828.899421 | 1695.818285 | 1826.792966 | 1600.686268 | 1484.287744 | 1399.296998 | 1.08E-53 | -2.265251484 | YLL022C | HIF1 | n |
| W303.asm02.HP0_G0034720 | 568.009175 | 787.240913 | 559.993995 | 3601.375349 | 3070.51918 | 3616.470714 | 1514.546439 | 2585.408639 | 2320.560566 | 4.76E-30 | -2.425479327 | YLL021W | SPA2 | n |
| W303.asm02.HP0_G0034730 | 568.009175 | 599.711941 | 433.9220485 | 77.82550727 | 124.7315484 | 72.41993919 | 135.362588 | 216.740844 | 230.3158921 | 3.07E-20 | 2.554857076 | YLL019C | KNS1 | n |
| W303.asm02.HP0_G0034740 | 182.2281995 | 272.2501827 | 195.4603822 | 436.7956595 | 459.1182526 | 462.5823616 | 502.6400995 | 428.6437227 | 359.2927916 | 2.94E-07 | -1.063126709 | YLL018C-A | COX19 | n |
| W303.asm02.HP0_G0034750 | 2543.440401 | 1824.837763 | 1932.125878 | 9036.514213 | 9162.461081 | 9115.859846 | 6778.541907 | 7060.526511 | 6178.607665 | 2.65E-48 | -2.116185646 | YLL018C | DPS1 | n |
| W303.asm02.HP0_G0034780 | 836.5049795 | 1069.962257 | 966.55159 | 563.2621088 | 720.5237316 | 631.8639695 | 719.4095587 | 1135.954245 | 737.0108546 | 0.005594146 | 0.585915415 | YLL015W | BPT1 | n |
| W303.asm02.HP0_G0034790 | 1220.347357 | 1216.558509 | 1243.128031 | 384.2634421 | 379.5023706 | 387.4466747 | 476.1355368 | 329.9492312 | 332.6785108 | 2.01E-23 | 1.67628712 | YLL014W | EMC6 | n |
| W303.asm02.HP0_G0034810 | 116.315872 | 285.5771147 | 119.2308331 | 1414.478595 | 1383.989414 | 1456.546027 | 746.8607129 | 1124.343128 | 1176.146489 | 1.50E-23 | -3.028950548 | YLL013C | PUF3 | n |
| W303.asm02.HP0_G0034830 | 1896.918013 | 2603.511363 | 1950.694614 | 3901.003552 | 3354.482493 | 3869.035251 | 1939.566034 | 1910.028687 | 1721.739246 | 4.28E-07 | -0.78634969 | YLL011W | SOF1 | n |
| W303.asm02.HP0_G0034840 | 155.0878293 | 275.1059539 | 135.8449656 | 1545.809138 | 1726.337707 | 1594.143912 | 1532.531678 | 2052.264866 | 2451.584718 | 8.05E-35 | -3.103124585 | YLL010C | PSR1 | n |
| W303.asm02.HP0_G0034850 | 1032.303364 | 1082.337265 | 1092.623536 | 834.6785654 | 926.1980933 | 912.4912338 | 1048.823409 | 809.8753857 | 712.4438261 | 0.120245738 | 0.263040643 | YLL009C | COX17 | n |
| W303.asm02.HP0_G0034860 | 1229.071048 | 1325.077812 | 1047.667649 | 5339.802617 | 5352.84113 | 5460.463415 | 2087.234312 | 2742.158713 | 2287.804528 | 3.87E-51 | -2.164957786 | YLL008W | DRS1 | n |
| W303.asm02.HP0_G0034870 | 405.1669542 | 480.7214765 | 419.2625198 | 786.0376234 | 956.7175147 | 875.376015 | 929.5528771 | 918.2458077 | 1008.271794 | 2.75E-12 | -1.002818349 | YLL007C | LMO1 | n |
| W303.asm02.HP0_G0034880 | 389.6581712 | 316.9905974 | 411.4441045 | 253.9057175 | 282.6363809 | 276.1010182 | 90.87278636 | 335.7547895 | 344.962025 | 0.237748437 | 0.460558482 | YLL006W | MMM1 | n |
| W303.asm02.HP0_G0034890 | 321.8072459 | 325.5579108 | 323.4869325 | 400.8013624 | 544.04186 | 401.0254133 | 509.2662402 | 597.004914 | 448.3482699 | 0.01419567 | -0.467507104 | YLL005C | SPO75 | n |
| W303.asm02.HP0_G0034900 | 602.9039366 | 977.6256561 | 646.9738651 | 647.897348 | 692.6581729 | 661.7371944 | 530.0912537 | 559.2687849 | 539.4510005 | 0.454581969 | 0.154326197 | YLL004W | ORC3 | n |
| W303.asm02.HP0_G0034910 | 403.2283563 | 481.6734002 | 385.0569529 | 421.2305581 | 483.0030171 | 462.5823616 | 474.2423538 | 496.3752364 | 462.6790365 | 0.526652476 | -0.104539649 | YLL003W | SFI1 | n |
| W303.asm02.HP0_G0034920 | 565.1012782 | 347.4521563 | 442.7177657 | 388.1547175 | 388.7908902 | 388.3519239 | 443.0048335 | 399.6159311 | 437.0883818 | 0.285859475 | 0.217738545 | YLL002W | RTT109 | n |
| W303.asm02.HP0_G0034930 | 820.9961966 | 801.5197687 | 775.9777173 | 2074.049769 | 2131.051773 | 2180.745419 | 1522.119171 | 1787.14437 | 1690.006835 | 5.74E-35 | -1.41276432 | YLL001W | DNM1 | n |
| W303.asm02.HP0_G0034940 | 773.5005489 | 519.7503488 | 778.9096231 | 342.432232 | 407.3679293 | 330.4159726 | 450.5775657 | 636.6762292 | 745.1998641 | 5.23E-05 | 0.941553132 | YLR001C | | n |
| W303.asm02.HP0_G0034950 | 1471.395781 | 1759.155027 | 1498.20383 | 4682.177081 | 4120.121891 | 4589.613646 | 1966.070596 | 2683.135537 | 2227.410583 | 1.72E-23 | -1.501984879 | YLR002C | NOC3 | n |
| W303.asm02.HP0_G0034960 | 1581.895859 | 2112.318725 | 1780.644082 | 2663.577986 | 2469.419272 | 2554.613355 | 1208.797377 | 1254.96819 | 1066.618487 | 0.00070218 | -0.489827748 | YLR003C | CMS1 | n |
| W303.asm02.HP0_G0034970 | 234.5703419 | 224.6539969 | 200.3468918 | 28.21174638 | 25.21169595 | 29.87322492 | 10.41250677 | 43.5416874 | 18.42527137 | 1.29E-13 | 2.980335731 | YLR004C | THI73 | n |
| W303.asm02.HP0_G0034980 | 771.561951 | 571.1542295 | 693.8843568 | 651.7886234 | 611.7153596 | 661.7371944 | 83.30005416 | 521.5326558 | 729.8454713 | 0.913094188 | 0.081075781 | YLR005W | SSL1 | n |
| W303.asm02.HP0_G0034990 | 377.0572851 | 327.4617582 | 311.7593096 | 268.4980001 | 311.828871 | 256.1855349 | 247.0603879 | 346.3983131 | 416.6158581 | 0.216961429 | 0.283251673 | YLR006C | SSK1 | n |
| W303.asm02.HP0_G0035000 | 1114.693773 | 1223.221975 | 1132.692915 | 470.844319 | 498.9261935 | 515.0868175 | 781.8845993 | 667.6392069 | 696.0658071 | 4.57E-21 | 1.224900443 | YLR007W | NSE1 | n |
| W303.asm02.HP0_G0035010 | 1774.786347 | 2276.049604 | 1805.07663 | 1413.505776 | 1464.932228 | 1381.41034 | 1897.916007 | 1257.870969 | 1266.225593 | 0.016442558 | 0.459309225 | YLR008C | PAM18 | n |
| W303.asm02.HP0_G0035020 | 2238.111237 | 3083.280915 | 2596.691178 | 7534.481922 | 7203.910385 | 7341.571336 | 4718.75875 | 4015.511172 | 3623.636702 | 4.95E-22 | -1.479510163 | YLR009W | RLP24 | n |
| W303.asm02.HP0_G0035070 | 283.0352886 | 441.6926041 | 240.4162701 | 268.4980001 | 347.6560178 | 280.6272644 | 258.4194862 | 334.7871965 | 415.5922319 | 0.710230148 | 0.108260532 | YLR014C | PPR1 | n |
| W303.asm02.HP0_G0035080 | 393.535367 | 415.9906638 | 407.5348969 | 557.4251958 | 534.7533404 | 547.6757902 | 524.4117046 | 540.8845169 | 482.1279341 | 0.001344699 | -0.430692289 | YLR015W | BRE2 | n |
| W303.asm02.HP0_G0035090 | 977.0533249 | 1070.91418 | 980.2338167 | 432.9043842 | 362.2522629 | 391.0676716 | 288.710415 | 304.7918118 | 185.2763398 | 3.22E-12 | 1.350820348 | YLR016C | PML1 | n |
| W303.asm02.HP0_G0035100 | 2134.396251 | 2482.617051 | 2438.368268 | 2633.420602 | 2592.823889 | 2749.241942 | 1840.173924 | 1613.945213 | 1506.777747 | 0.178724844 | -0.177017568 | YLR017W | MEU1 | n |
| W303.asm02.HP0_G0035110 | 1440.378215 | 1431.693269 | 1394.609827 | 1453.391348 | 1452.989846 | 1418.525559 | 1298.723572 | 1480.417372 | 1427.958531 | 0.870791904 | -0.019385721 | YLR018C | POM34 | n |
| W303.asm02.HP0_G0035130 | 1047.812147 | 1167.058476 | 1147.352444 | 959.1993771 | 977.9484166 | 936.0277141 | 1025.158621 | 1033.389381 | 967.3267467 | 0.052752646 | 0.227288221 | YLR020C | YEH2 | n |
| W303.asm02.HP0_G0035140 | 984.8077164 | 880.5294371 | 1041.803837 | 580.772848 | 537.4072031 | 565.780775 | 737.3947976 | 549.5928543 | 532.2856172 | 2.39E-06 | 0.787005749 | YLR021W | IRC25 | n |
| W303.asm02.HP0_G0035150 | 1359.926404 | 1761.058874 | 1550.000831 | 970.8732032 | 916.9095737 | 1015.689647 | 760.1129942 | 694.7318124 | 628.5064788 | 3.16E-06 | 0.685274564 | YLR022C | SDO1 | n |
| W303.asm02.HP0_G0035160 | 1448.132607 | 1853.395475 | 1431.7473 | 572.0174784 | 650.1963692 | 568.4965227 | 452.4707487 | 719.8892318 | 762.6015093 | 1.61E-11 | 1.403260014 | YLR023C | IZH3 | n |
| W303.asm02.HP0_G0035170 | 272.3730003 | 478.817629 | 259.9623083 | 502.9473407 | 449.829733 | 484.3083434 | 401.3548064 | 695.6994054 | 645.908124 | 0.069881302 | -0.507330602 | YLR024C | UBR2 | n |
| W303.asm02.HP0_G0035180 | 2929.221377 | 2304.607316 | 2972.952413 | 981.5742104 | 952.7367206 | 1013.879149 | 1987.842202 | 1265.611714 | 1573.313449 | 5.01E-16 | 1.47669033 | YLR025W | SNF7 | n |
| W303.asm02.HP0_G0035190 | 771.561951 | 626.365805 | 679.2248281 | 1350.272551 | 1393.277934 | 1398.610076 | 2144.976395 | 1583.949828 | 1887.566689 | 6.90E-11 | -0.995788947 | YLR026C | SED5 | n |
| W303.asm02.HP0_G0035200 | 3942.138762 | 3212.742541 | 3603.312146 | 2324.064211 | 2217.302312 | 2432.404708 | 2562.423257 | 2399.630773 | 2602.057767 | 2.10E-07 | 0.624999848 | YLR027C | AAT2 | n |
| W303.asm02.HP0_G0035220 | 19559.48318 | 15634.39511 | 19289.00782 | 20760.92688 | 23619.37831 | 19601.36179 | 24004.61447 | 19822.11129 | 19651.57554 | 0.125678828 | -0.231793596 | YLR029C | RPL15A | y |
| W303.asm02.HP0_G0035240 | 433.2766232 | 527.3657386 | 437.8312561 | 609.9574132 | 573.23435 | 589.3172552 | 334.1468082 | 445.0928046 | 452.4427746 | 0.052125676 | -0.34221403 | YLR032W | RAD5 | n |
| W303.asm02.HP0_G0035250 | 1562.509881 | 1690.616519 | 1614.502757 | 1339.571544 | 1381.335552 | 1362.400106 | 1148.215519 | 1266.579307 | 1218.115163 | 0.020593907 | 0.2537366 | YLR033W | RSC58 | n |
| W303.asm02.HP0_G0035260 | 1034.241962 | 1102.327663 | 1187.421822 | 969.9003843 | 690.0043102 | 1017.500146 | 1036.517719 | 1292.704319 | 1378.824474 | 0.102190031 | 0.310007318 | YLR034C | SMF3 | n |
| W303.asm02.HP0_G0035270 | 217.1229611 | 230.3655392 | 210.1199109 | 30.15738407 | 31.84635278 | 40.7362158 | 56.79549147 | 34.83334992 | 38.89779511 | 2.12E-20 | 2.670989935 | YLR035C | MLH2 | n |
| W303.asm02.HP0_G0035300 | 2547.317597 | 2752.011462 | 2585.940856 | 5928.358016 | 6117.153596 | 5734.753935 | 4661.963258 | 3930.362983 | 3470.092774 | 1.01E-18 | -1.172940282 | YLR038C | COX12 | n |
| W303.asm02.HP0_G0035310 | 619.3820185 | 699.6639311 | 565.8578065 | 508.7842538 | 578.5420755 | 514.1815683 | 623.8038147 | 770.2040705 | 682.7586667 | 0.141565205 | 0.236893074 | YLR039C | RIC1 | n |
| W303.asm02.HP0_G0035320 | 582.548659 | 433.1252907 | 631.3370345 | 9.728188408 | 13.26931366 | 10.86299088 | 10.41250677 | 20.31945412 | 17.40164518 | 1.64E-57 | 5.62295907 | YLR040C | AFB1 | n |
| W303.asm02.HP0_G0035350 | 20774.98404 | 18355.94501 | 16481.21943 | 48456.10646 | 33824.80744 | 44966.44549 | 10854.56501 | 13845.289 | 11690.83468 | 2.01E-12 | -1.194205682 | YLR044C | PDC1 | n |

| | | | | | | | | | | | | | | |
|---|---|---|---|---|---|---|---|---|---|---|---|---|---|---|
| W303.asm02.HP0_G0035360 | 574.7942675 | 533.0772808 | 478.8779364 | 378.4265291 | 347.6560178 | 332.2264711 | 67.20799824 | 306.7269979 | 418.6631105 | 0.3479966 | 0.584311413 | YLR045C | STU2 | n |
| W303.asm02.HP0_G0035380 | 163.8115197 | 182.7693534 | 184.7100612 | 271.4164566 | 240.1745772 | 242.6067963 | 308.588837 | 353.1714645 | 349.0565298 | 0.004048826 | -0.506632435 | YLR047C | FRE8 | n |
| W303.asm02.HP0_G0035410 | 1300.799169 | 1428.837497 | 1276.356296 | 435.8228407 | 431.2526939 | 440.8563798 | 779.9914162 | 545.7224821 | 653.0735073 | 2.42E-24 | 1.614634305 | YLR050C | EMA19 | n |
| W303.asm02.HP0_G0035420 | 1279.474592 | 1948.587846 | 1249.969144 | 1821.11687 | 1527.298002 | 1673.805845 | 1632.87038 | 1478.482184 | 1491.423354 | 0.400376636 | -0.165745013 | YLR051C | FCF2 | n |
| W303.asm02.HP0_G0035430 | 849.1058657 | 873.8659711 | 979.2565148 | 433.877203 | 437.8873507 | 422.751395 | 594.4594774 | 430.5789088 | 459.608158 | 7.68E-11 | 1.062082831 | YLR052W | IES3 | n |
| W303.asm02.HP0_G0035450 | 171.5659112 | 169.4424214 | 150.5044943 | 34.04865943 | 31.84635278 | 20.82073252 | 36.91706946 | 25.15741939 | 26.61428086 | 3.07E-15 | 2.513587618 | YLR054C | OSW2 | n |
| W303.asm02.HP0_G0035460 | 505.9740432 | 497.8561034 | 381.1477453 | 1026.323877 | 1062.872024 | 1091.730583 | 690.0652214 | 1060.481987 | 1013.389925 | 6.47E-10 | -1.199496388 | YLR055C | SPT8 | n |
| W303.asm02.HP0_G0035470 | 7974.422325 | 7817.197554 | 7958.169461 | 3838.743146 | 3743.273383 | 3758.594844 | 3454.112473 | 4274.82611 | 3655.369114 | 6.09E-21 | 1.066371705 | YLR056W | ERG3 | n |
| W303.asm02.HP0_G0035480 | 803.5488158 | 755.8274303 | 862.9575874 | 720.8587611 | 747.0623589 | 654.4952005 | 611.4981249 | 641.5141944 | 571.1834123 | 0.164166471 | 0.191978659 | YLR057W | MNL2 | n |
| W303.asm02.HP0_G0035490 | 634.8908014 | 550.2119077 | 565.8578065 | 3326.067617 | 1723.683844 | 2829.809124 | 1339.427007 | 1965.181492 | 1535.43928 | 1.09E-20 | -2.170610142 | YLR058C | SHM2 | n |
| W303.asm02.HP0_G0035500 | 889.8164209 | 762.4908964 | 882.5036256 | 1376.53866 | 1313.662052 | 1390.462833 | 1249.500812 | 1254.96819 | 1158.744844 | 1.50E-08 | -0.68751858 | YLR059C | REX2 | n |
| W303.asm02.HP0_G0035510 | 3427.441029 | 2947.155824 | 3214.345985 | 9144.497104 | 9470.309157 | 9088.702369 | 6657.378192 | 7518.198025 | 6374.120266 | 1.08E-40 | -1.53058984 | YLR060W | FRS1 | n |
| W303.asm02.HP0_G0035520 | 5342.775721 | 6297.927304 | 5454.321965 | 7005.268473 | 8360.994536 | 6953.219412 | 7022.762521 | 5037.289436 | 4134.426169 | 0.051262548 | -0.384596285 | YLR061W | RPL22A | n |
| W303.asm02.HP0_G0035530 | 365.4256979 | 472.154163 | 417.307916 | 1297.740334 | 992.5446616 | 1328.905884 | 599.192435 | 616.356775 | 459.608158 | 2.26E-16 | -1.529184733 | YLR063W | BMT6 | n |
| W303.asm02.HP0_G0035540 | 2367.997294 | 2102.799488 | 2355.297605 | 1234.507109 | 1308.354327 | 1207.602486 | 2364.585628 | 2165.473254 | 1769.849677 | 1.00E-09 | 0.864479381 | YLR064W | PER33 | n |
| W303.asm02.HP0_G0035550 | 1325.031642 | 1392.664396 | 1498.20383 | 795.7658118 | 817.3897213 | 767.6513555 | 1676.41359 | 1283.028389 | 1272.36735 | 3.14E-08 | 0.824869821 | YLR065C | SND2 | n |
| W303.asm02.HP0_G0035560 | 1452.009802 | 1486.904844 | 1515.795264 | 739.342319 | 679.3888593 | 745.0201245 | 1243.821263 | 878.5744925 | 900.7910445 | 1.04E-10 | 1.040973251 | YLR066W | SPC3 | n |
| W303.asm02.HP0_G0035570 | 695.9566342 | 1043.308393 | 737.8629428 | 1921.317211 | 1882.915608 | 1919.128389 | 966.4699466 | 1489.125709 | 1255.989331 | 1.62E-09 | -1.208126903 | YLR067C | PET309 | n |
| W303.asm02.HP0_G0035580 | 575.7635665 | 1180.385408 | 754.4770753 | 1190.730261 | 1234.04617 | 1058.236361 | 992.9745093 | 795.3614899 | 703.2311905 | 0.060366181 | -0.471891328 | YLR068W | FYV7 | n |
| W303.asm02.HP0_G0035590 | 2138.273447 | 1963.818626 | 2162.769129 | 3927.26966 | 3972.832509 | 4017.496127 | 3383.118109 | 3596.54338 | 3375.919165 | 5.54E-21 | -0.927796746 | YLR069C | MEF1 | n |
| W303.asm02.HP0_G0035610 | 604.8425345 | 803.4236161 | 677.2702243 | 1318.169529 | 1427.77815 | 1369.6421 | 1122.657548 | 1371.079357 | 1249.847574 | 6.55E-11 | -0.980221206 | YLR071C | RGR1 | n |
| W303.asm02.HP0_G0035630 | 503.0661464 | 1120.414213 | 635.2462421 | 1838.627609 | 1878.934814 | 1817.740474 | 1472.896412 | 1271.417272 | 1050.240468 | 4.49E-07 | -1.292971066 | YLR073C | RFU1 | n |
| W303.asm02.HP0_G0035640 | 1561.540582 | 1898.135889 | 1598.865926 | 3325.094798 | 2949.768426 | 3216.350549 | 2127.937747 | 1742.635089 | 1431.029409 | 5.21E-08 | -0.908032539 | YLR074C | BUD20 | n |
| W303.asm02.HP0_G0035650 | 22163.98941 | 20558.69649 | 21648.21463 | 24522.81734 | 27486.05631 | 23030.44591 | 23996.09515 | 20148.19015 | 18774.3279 | 0.105178014 | -0.22117466 | YLR075W | RPL10 | n |
| W303.asm02.HP0_G0035660 | 637.7986982 | 481.6734002 | 550.2209759 | 3083.835725 | 2847.594711 | 3200.961312 | 2677.907423 | 2971.478267 | 2815.99564 | 1.67E-72 | -2.451887295 | YLR077W | FMP25 | n |
| W303.asm02.HP0_G0035670 | 1062.351631 | 866.2505814 | 1014.439384 | 933.9060872 | 980.6022793 | 945.9854557 | 1009.066565 | 845.6763287 | 751.3416212 | 0.805687201 | 0.041297022 | YLR078C | BOS1 | n |
| W303.asm02.HP0_G0035680 | 942.1585633 | 744.4043458 | 717.3396027 | 396.9100871 | 557.3111736 | 445.382626 | 709.9436434 | 631.8382639 | 715.5147047 | 1.60E-05 | 0.783546638 | YLR079W | SIC1 | n |
| W303.asm02.HP0_G0035720 | 2730.515095 | 2841.492292 | 3116.615794 | 3332.877349 | 3412.867473 | 3239.88703 | 2650.456269 | 4284.502041 | 3784.346013 | 0.273336657 | -0.200658862 | YLR083C | EMP70 | n |
| W303.asm02.HP0_G0035730 | 915.0181932 | 1683.00113 | 1059.395272 | 3089.672638 | 3473.906316 | 3052.500437 | 2782.032491 | 3885.853703 | 3453.714755 | 2.68E-10 | -1.394387234 | YLR084C | RAX2 | n |
| W303.asm02.HP0_G0035740 | 796.7637233 | 642.5485082 | 818.0016995 | 644.0060726 | 635.6001242 | 642.7269603 | 498.8537334 | 664.7364277 | 774.8850235 | 0.237146512 | 0.231567555 | YLR085C | ARP6 | n |
| W303.asm02.HP0_G0035750 | 313.0835555 | 529.269586 | 312.7366115 | 966.0091089 | 1060.218161 | 983.1006746 | 1007.173382 | 940.5004479 | 863.9405018 | 4.06E-11 | -1.380763695 | YLR086W | SMC4 | n |
| W303.asm02.HP0_G0035760 | 345.0704203 | 524.5099674 | 378.2158396 | 618.7127828 | 577.2151441 | 615.5694831 | 519.678747 | 601.8428792 | 678.664162 | 0.006105919 | -0.53809249 | YLR087C | CSF1 | n |
| W303.asm02.HP0_G0035770 | 839.4128763 | 848.1640308 | 946.0282498 | 1273.419863 | 1183.622778 | 1256.485945 | 1222.049658 | 1631.361888 | 1605.045861 | 0.001080663 | -0.496209673 | YLR088W | GAA1 | n |
| W303.asm02.HP0_G0035780 | 822.9347945 | 748.2120406 | 675.3156205 | 857.0533988 | 1068.179749 | 913.3964831 | 1719.010209 | 903.7319119 | 1379.8481 | 0.143685076 | -0.336804613 | YLR089C | ALT1 | n |
| W303.asm02.HP0_G0035790 | 462.3555912 | 599.711941 | 387.9888587 | 835.6513843 | 688.6773788 | 762.21986 | 488.4412267 | 655.0604972 | 616.2229646 | 0.001103248 | -0.657647127 | YLR090W | XDJ1 | n |
| W303.asm02.HP0_G0035800 | 150.2413347 | 190.3847432 | 160.2775134 | 849.270848 | 812.0819959 | 797.5245804 | 686.2788553 | 729.5651623 | 700.1603119 | 5.23E-55 | -2.294998222 | YLR091W | GEP5 | n |
| W303.asm02.HP0_G0035820 | 1919.211888 | 1776.289654 | 1848.077914 | 169.2704783 | 188.4242539 | 181.9550972 | 301.0161048 | 243.8334495 | 243.6230325 | 6.61E-115 | 3.36294616 | YLR093C | NYV1 | n |
| W303.asm02.HP0_G0035830 | 190.9518899 | 221.7982258 | 135.8449656 | 793.8201741 | 851.8899368 | 784.851091 | 753.4868535 | 770.2040705 | 786.1449116 | 9.26E-33 | -2.146281452 | YLR094C | GIS3 | n |
| W303.asm02.HP0_G0035840 | 404.1976552 | 324.6059871 | 287.3267618 | 1770.53029 | 1827.184491 | 1721.784054 | 2027.599046 | 1791.982335 | 2082.055664 | 6.54E-56 | -2.387935071 | YLR095C | IOC2 | n |
| W303.asm02.HP0_G0035860 | 797.7330222 | 764.3947438 | 823.865511 | 31.13020291 | 58.38498009 | 42.54671428 | 19.87842202 | 33.86575687 | 46.06317841 | 1.94E-54 | 4.195002416 | YLR097C | HRT3 | n |
| W303.asm02.HP0_G0035870 | 308.2370608 | 348.40408 | 344.0102727 | 372.589616 | 468.4067721 | 414.6041519 | 333.2002166 | 432.5140949 | 399.2142129 | 0.06942481 | -0.325048735 | YLR098C | CHA4 | n |
| W303.asm02.HP0_G0035890 | 410.9827478 | 413.1348927 | 471.0595211 | 175.1073914 | 220.2706067 | 196.4390851 | 401.3548064 | 268.0232758 | 248.7411634 | 3.04E-07 | 1.132703964 | YLR099W- | MIM2 | n |
| W303.asm02.HP0_G0035900 | 4110.796777 | 3345.059937 | 3917.026059 | 1190.730261 | 1255.277072 | 1248.338702 | 1182.292814 | 1049.838463 | 933.5470825 | 4.16E-32 | 1.622353149 | YLR100W | ERG27 | n |
| W303.asm02.HP0_G0035920 | 700.8031289 | 566.3946109 | 701.7027721 | 866.7815872 | 810.7550645 | 877.1865135 | 758.2198112 | 897.9263536 | 897.720166 | 0.012635465 | -0.376628337 | YLR103C | CDC45 | n |
| W303.asm02.HP0_G0035930 | 964.4524388 | 854.8274968 | 1056.463366 | 373.5624349 | 468.4067721 | 360.2891975 | 648.4151943 | 452.833549 | 466.7735413 | 6.20E-11 | 1.261125804 | YLR104W | LCL2 | n |
| W303.asm02.HP0_G0035940 | 689.1715417 | 544.5003654 | 667.4972052 | 586.609761 | 570.5804873 | 542.2442947 | 570.7946893 | 511.8567253 | 479.0570555 | 0.31677068 | 0.161980835 | YLR105C | SEN2 | n |
| W303.asm02.HP0_G0035950 | 921.8032857 | 1442.164429 | 851.2299645 | 472.7899566 | 378.1754392 | 505.1290759 | 349.2922726 | 334.7871965 | 265.1191824 | 5.95E-08 | 1.243662645 | YLR107W | REX3 | n |
| W303.asm02.HP0_G0035970 | 16122.34916 | 13971.38438 | 15754.10681 | 3818.31395 | 2716.228506 | 3720.574376 | 2427.060669 | 2179.019556 | 2021.661719 | 1.94E-43 | 2.159866437 | YLR109W | AHP1 | n |
| W303.asm02.HP0_G0036000 | 3760.879862 | 3390.752276 | 3914.094154 | 2381.460522 | 2530.458115 | 2386.236996 | 2375.944727 | 2550.575289 | 2474.104494 | 2.87E-08 | 0.600838204 | YLR113W | HOG1 | n |
| W303.asm02.HP0_G0036010 | 791.9172286 | 1060.443019 | 838.5250396 | 773.3909785 | 788.1972313 | 755.8831153 | 691.9584044 | 799.2318621 | 821.9718281 | 0.184052235 | 0.215965792 | YLR114C | AVL9 | n |
| W303.asm02.HP0_G0036020 | 309.2063598 | 415.0387401 | 300.0316867 | 1061.345355 | 1222.103788 | 1087.204337 | 1090.473436 | 1253.033004 | 1259.06021 | 1.91E-25 | -1.717301454 | YLR115W | CFT2 | n |
| W303.asm02.HP0_G0036030 | 190.9518899 | 357.9233171 | 139.7541733 | 555.4795581 | 457.7913212 | 513.276319 | 440.1650589 | 452.833549 | 503.624084 | 7.26E-05 | -1.148615455 | YLR116W | MSL5 | n |
| W303.asm02.HP0_G0036040 | 436.18452 | 548.3080603 | 426.1036332 | 589.5282215 | 606.4076342 | 615.5694831 | 572.6878724 | 623.1299264 | 816.8536972 | 0.054503277 | -0.360716907 | YLR117C | CLF1 | n |
| W303.asm02.HP0_G0036050 | 1129.233257 | 936.6929363 | 1090.668933 | 451.3879421 | 428.5988311 | 440.8563798 | 589.7265198 | 398.648338 | 386.9306987 | 2.51E-11 | 1.256463109 | YLR118C | TML25 | n |
| W303.asm02.HP0_G0036060 | 402.2590574 | 444.5483753 | 384.079651 | 142.0315508 | 156.5779012 | 148.4608753 | 202.5705863 | 199.324169 | 175.040078 | 3.27E-17 | 1.463661018 | YLR119W | SRN2 | n |
| W303.asm02.HP0_G0036080 | 383.8423776 | 425.509901 | 378.2158396 | 1562.347058 | 1261.911729 | 1412.188814 | 2064.516115 | 1775.533253 | 2245.835854 | 1.36E-34 | -1.835580885 | YLR121C | YPS3 | n |
| W303.asm02.HP0_G0036120 | 433.2766232 | 541.6445943 | 381.1477453 | 190.6724928 | 222.9244694 | 200.9653313 | 300.0695133 | 253.50938 | 376.6944368 | 1.00E-07 | 1.144222685 | YLR127C | APC2 | n |
| W303.asm02.HP0_G0036140 | 1262.027211 | 2168.482225 | 1302.743447 | 7284.46748 | 6528.50232 | 7409.465029 | 2877.638235 | 4114.205663 | 3507.966943 | 9.07E-24 | -2.164676502 | YLR129W | DIP2 | n |
| W303.asm02.HP0_G0036150 | 1768.970554 | 1731.549239 | 1824.622668 | 1382.375573 | 1402.566454 | 1339.768875 | 1324.281543 | 1624.588737 | 1503.706869 | 0.002493375 | 0.368815651 | YLR130C | ZRT2 | n |
| W303.asm02.HP0_G0036160 | 216.1536622 | 230.3655392 | 164.186721 | 1600.286993 | 1791.357344 | 1571.512681 | 1410.421372 | 1722.315635 | 2058.512262 | 4.16E-59 | -3.021941536 | YLR131C | ACE2 | n |
| W303.asm02.HP0_G0036170 | 312.1142566 | 310.3271313 | 333.2599516 | 392.0459929 | 544.04186 | 390.1624224 | 751.5936705 | 533.1437724 | 601.8921979 | 0.021238028 | -0.469179087 | YLR132C | USB1 | n |
| W303.asm02.HP0_G0036180 | 286.9124843 | 258.9232507 | 232.5978548 | 420.2577392 | 626.3116046 | 401.0254133 | 873.7039772 | 906.6346911 | 786.1449116 | 1.26E-05 | -0.890751591 | YLR133W | CKI1 | n |
| W303.asm02.HP0_G0036200 | 443.9389115 | 415.0387401 | 470.0822192 | 620.6584205 | 632.9462615 | 649.063705 | 572.6878724 | 697.6345915 | 809.6883139 | 0.001817852 | -0.517806801 | YLR135W | SLX4 | n |

| | | | | | | | | | | | | | | |
|---|---|---|---|---|---|---|---|---|---|---|---|---|---|---|
| W303.asm02.HP0_G0036210 | 157.9957261 | 201.8078277 | 157.3456077 | 1897.969558 | 973.9676225 | 1827.698215 | 1098.046168 | 1270.449679 | 3124.107123 | 1.16E-10 | -3.184103199 | YLR136C | TIS11 | n |
| W303.asm02.HP0_G0036220 | 292.7282779 | 183.7212771 | 241.393572 | 218.8842392 | 283.9633123 | 252.5645379 | 299.1229218 | 334.7871965 | 302.9933513 | 0.762974275 | -0.071205921 | YLR137W | RKM5 | n |
| W303.asm02.HP0_G0036240 | 179.3203027 | 180.865506 | 129.9811542 | 1325.95208 | 1212.815268 | 1323.474389 | 719.4095587 | 965.6578673 | 923.3108207 | 9.34E-61 | -2.97809747 | YLR139C | SLS1 | n |
| W303.asm02.HP0_G0036250 | 510.8205379 | 494.0484865 | 545.3344663 | 326.8671305 | 301.21342 | 340.3737142 | 143.8819117 | 233.1899259 | 261.0246777 | 0.002024526 | 0.677441178 | YLR141W | RRN5 | n |
| W303.asm02.HP0_G0036260 | 236.5089398 | 219.8943783 | 251.1665911 | 301.5738407 | 532.0994777 | 331.3212218 | 972.1494957 | 1169.820002 | 986.7756442 | 0.001614659 | -0.714314175 | YLR142W | PUT1 | n |
| W303.asm02.HP0_G0036270 | 528.2679187 | 668.2504485 | 526.76573 | 930.0148118 | 973.9676225 | 875.376015 | 931.4460602 | 1173.690374 | 1155.673965 | 1.99E-05 | -0.688780034 | YLR143W | DPH6 | n |
| W303.asm02.HP0_G0036280 | 404.1976552 | 475.9618579 | 376.2612357 | 237.3677972 | 216.2898126 | 261.6170303 | 308.588837 | 351.2362784 | 322.4422489 | 3.33E-06 | 0.809358434 | YLR144C | ACF2 | n |
| W303.asm02.HP0_G0036290 | 847.1672678 | 833.885175 | 853.1845683 | 174.1345725 | 176.4818716 | 181.049848 | 206.3569524 | 182.8750871 | 159.6856852 | 2.19E-47 | 2.252531711 | YLR145W | RMP1 | n |
| W303.asm02.HP0_G0036300 | 318.8993491 | 325.5579108 | 298.0770829 | 1475.766182 | 1531.278796 | 1595.049161 | 1125.497323 | 1064.352359 | 906.9328017 | 7.04E-63 | -2.287650225 | YLR146C | SPE4 | n |
| W303.asm02.HP0_G0036330 | 41.67985414 | 66.63466011 | 56.68351084 | 534.0775436 | 579.8690068 | 585.6962582 | 507.3730572 | 757.6253608 | 763.6251355 | 6.98E-47 | -3.363893815 | YLR148W | PEP3 | n |
| W303.asm02.HP0_G0036450 | 163.8115197 | 136.1250914 | 200.3468918 | 60.31476813 | 67.67349965 | 66.08319451 | 136.3091795 | 112.2407942 | 145.3549186 | 1.03E-07 | 1.367862489 | YLR151C | PCD1 | n |
| W303.asm02.HP0_G0036460 | 403.2283563 | 280.8174962 | 365.5109147 | 435.8228407 | 396.7524784 | 468.013857 | 481.815086 | 406.3890824 | 482.1279341 | 0.102224498 | -0.311282599 | YLR152C | | n |
| W303.asm02.HP0_G0036470 | 7144.702438 | 5290.792012 | 6042.657716 | 9948.045466 | 8222.993674 | 10056.41381 | 5755.276469 | 7087.619116 | 7974.047997 | 0.000358107 | -0.611414774 | YLR153C | ACS2 | n |
| W303.asm02.HP0_G0036520 | 2374.782387 | 1574.481826 | 2319.137435 | 1413.505776 | 1360.10465 | 1401.325823 | 4069.396964 | 1607.172062 | 4433.325016 | 0.057517167 | 0.58627728 | YLR163C | MAS1 | n |
| W303.asm02.HP0_G0036530 | 391.5967691 | 107.5673799 | 295.1451771 | 3242.405197 | 4703.971692 | 3334.9382 | 10382.21584 | 4337.719659 | 8280.112226 | 1.03E-12 | -3.828127024 | YLR164W | SHH4 | n |
| W303.asm02.HP0_G0036540 | 560.2547835 | 375.057944 | 465.1957096 | 492.2463335 | 483.0030171 | 523.2340607 | 730.768657 | 504.1159808 | 1168.981106 | 0.744245018 | -0.097897164 | YLR165C | PUS5 | n |
| W303.asm02.HP0_G0036550 | 1845.545169 | 1675.38574 | 1834.395687 | 1140.143681 | 1285.796493 | 1191.308 | 2809.483645 | 1394.30159 | 3959.386091 | 0.049395331 | 0.566254547 | YLR166C | SEC10 | n |
| W303.asm02.HP0_G0036560 | 17233.16574 | 20728.13891 | 18330.27464 | 21868.96754 | 25190.46505 | 20932.98342 | 34304.47685 | 15229.91466 | 24669.39111 | 0.276233337 | -0.272428651 | YLR167W | RPS31 | n |
| W303.asm02.HP0_G0036570 | 2827.444989 | 2126.597581 | 2733.513445 | 1070.100725 | 780.2356431 | 993.9636654 | 1677.360182 | 966.6254604 | 2104.57544 | 1.51E-08 | 1.433686615 | YLR168C | UPS2 | n |
| W303.asm02.HP0_G0036580 | 1167.035916 | 955.7314107 | 1172.762293 | 656.6527176 | 659.4848888 | 649.9689543 | 831.1073586 | 663.7688347 | 666.3806477 | 1.17E-06 | 0.745214014 | YLR170C | APS1 | n |
| W303.asm02.HP0_G0036590 | 2428.093828 | 2261.770749 | 2533.166553 | 2493.334689 | 2844.940848 | 2466.804179 | 2711.038126 | 2135.477869 | 1796.463958 | 0.5279912 | -0.111462222 | YLR172C | DPH5 | n |
| W303.asm02.HP0_G0036600 | 430.3687264 | 308.4232839 | 364.5336128 | 211.1016885 | 260.0785477 | 227.2175592 | 263.1524438 | 370.5881395 | 251.812042 | 0.002990131 | 0.662192534 | YLR173W | TAG1 | n |
| W303.asm02.HP0_G0036610 | 275.2808971 | 152.3077945 | 234.5524586 | 5348.557987 | 4782.260642 | 5399.811716 | 6673.470248 | 4457.701197 | 4922.618333 | 6.95E-99 | -4.552291564 | YLR174W | IDP2 | n |
| W303.asm02.HP0_G0036620 | 4101.103787 | 4446.435676 | 4436.950676 | 8321.492365 | 8920.959572 | 8082.065214 | 7652.245884 | 5469.803531 | 5421.124286 | 4.33E-10 | -0.96366347 | YLR175W | CBF5 | n |
| W303.asm02.HP0_G0036630 | 108.5614805 | 159.9231843 | 130.9584561 | 1965.094058 | 1321.62364 | 1996.074574 | 308.588837 | 531.2085863 | 503.624084 | 3.49E-50 | -3.725614267 | YLR176C | RFX1 | n |
| W303.asm02.HP0_G0036640 | 494.342456 | 271.298259 | 410.4668026 | 248.0688044 | 348.9829492 | 287.8692583 | 543.3435351 | 858.2550384 | 937.6415873 | 0.140362575 | 0.412388332 | YLR177W | | n |
| W303.asm02.HP0_G0036650 | 1015.825282 | 421.7022061 | 822.8882091 | 138.1402754 | 152.5971071 | 125.8296443 | 363.4911454 | 234.1575189 | 306.0642299 | 2.45E-16 | 2.442068794 | YLR178C | TFS1 | n |
| W303.asm02.HP0_G0036670 | 1137.956948 | 1162.298857 | 946.0282498 | 16031.08168 | 15473.34666 | 15659.9066 | 13908.26927 | 12600.96433 | 12452.41256 | 3.86E-235 | -3.860753206 | YLR180W | SAM1 | y |
| W303.asm02.HP0_G0036680 | 1621.637116 | 1805.799289 | 1820.71346 | 272.3892754 | 257.424685 | 268.8590243 | 535.7708029 | 345.4307201 | 397.1669605 | 1.11E-52 | 2.715221765 | YLR181C | VTA1 | n |
| W303.asm02.HP0_G0036690 | 761.8689617 | 879.5775134 | 762.2954906 | 1331.788993 | 1455.643708 | 1254.675447 | 1161.467801 | 1683.611913 | 1704.337601 | 2.08E-05 | -0.749201669 | YLR182W | SWI6 | n |
| W303.asm02.HP0_G0036720 | 1575.110767 | 1907.655126 | 1737.642798 | 2420.373276 | 2506.57335 | 2363.605765 | 2073.98203 | 1694.255437 | 1509.848626 | 0.001496726 | -0.481668104 | YLR186W | EMG1 | n |
| W303.asm02.HP0_G0036730 | 95.96059441 | 85.67313442 | 58.63811466 | 2473.878312 | 2201.379136 | 2427.878461 | 1423.673653 | 1987.436132 | 1820.00736 | 5.81E-134 | -4.885363615 | YLR187W | SKG3 | n |
| W303.asm02.HP0_G0036740 | 695.9566342 | 733.9331849 | 609.8363925 | 1526.352761 | 1725.010776 | 1614.059395 | 1432.192977 | 1769.727695 | 1597.880478 | 2.49E-19 | -1.253565174 | YLR188W | MDL1 | n |
| W303.asm02.HP0_G0036750 | 686.2636449 | 739.6447272 | 730.0445275 | 370.6439784 | 342.3482924 | 352.1419543 | 327.5206675 | 405.4214894 | 427.8757462 | 3.84E-11 | 1.016379608 | YLR189C | ATG26 | n |
| W303.asm02.HP0_G0036770 | 889.8164209 | 792.9524553 | 911.822683 | 370.6439784 | 440.5412134 | 371.1521884 | 526.3048877 | 614.4215889 | 831.1844638 | 3.69E-09 | 1.135919025 | YLR191W | PEX13 | n |
| W303.asm02.HP0_G0036780 | 2720.822106 | 2021.885972 | 2679.76184 | 1896.023921 | 2014.281813 | 1858.47669 | 2004.880849 | 1245.29226 | 1312.288772 | 0.078330329 | 0.36377781 | YLR192C | HCR1 | n |
| W303.asm02.HP0_G0036790 | 539.8995059 | 457.8753075 | 559.0166931 | 774.3637973 | 869.1400446 | 784.851091 | 855.7187382 | 795.3614899 | 847.5624828 | 4.15E-06 | -0.640306349 | YLR193C | UPS1 | n |
| W303.asm02.HP0_G0036800 | 2367.997294 | 1637.308791 | 2562.485611 | 860.9446741 | 816.0627899 | 939.648711 | 955.1108483 | 983.0745423 | 1106.539908 | 1.73E-13 | 1.326953721 | YLR194C | NCW2 | n |
| W303.asm02.HP0_G0036810 | 2841.984473 | 2333.165027 | 2771.62822 | 3386.382385 | 2912.614348 | 3311.40172 | 2851.133672 | 2728.612411 | 2593.868758 | 0.036714766 | -0.274721563 | YLR195C | NMT1 | n |
| W303.asm02.HP0_G0036820 | 2057.821636 | 2680.617184 | 2209.679621 | 7032.5074 | 6272.404566 | 6828.295017 | 2966.617838 | 3513.330377 | 3359.541146 | 1.79E-28 | -1.534965924 | YLR196W | PWP1 | n |
| W303.asm02.HP0_G0036830 | 8252.611119 | 9333.612033 | 9101.612697 | 11614.48414 | 12222.36481 | 11400.70893 | 7138.246687 | 7269.52661 | 7015.933886 | 5.49E-05 | -0.400826212 | YLR197W | NOP56 | n |
| W303.asm02.HP0_G0036850 | 1343.448322 | 1552.58758 | 1578.342586 | 1021.459783 | 1050.929642 | 1024.74214 | 1341.32019 | 897.9263536 | 851.6569876 | 0.00474749 | 0.53089231 | YLR200W | YKE2 | n |
| W303.asm02.HP0_G0036860 | 993.5314068 | 629.2215761 | 951.8920613 | 536.9960001 | 590.4844578 | 559.4440303 | 873.7039772 | 623.1299264 | 708.3493214 | 0.003458778 | 0.610391452 | YLR201C | COQ9 | n |
| W303.asm02.HP0_G0036870 | 1084.645507 | 920.5102332 | 768.159302 | 2239.428972 | 2132.378705 | 2213.334392 | 2502.787991 | 2439.302088 | 2746.38906 | 1.74E-17 | -1.247718084 | YLR203C | MSS51 | n |
| W303.asm02.HP0_G0036890 | 1369.619393 | 979.5295036 | 1098.487348 | 1584.721892 | 1690.51056 | 1628.543383 | 2253.83442 | 1330.440448 | 2193.630919 | 0.018098191 | -0.508196099 | YLR205C | HMX1 | n |
| W303.asm02.HP0_G0036910 | 444.9082104 | 396.9521895 | 418.2852179 | 474.7355943 | 414.0025861 | 448.0983738 | 552.8094503 | 802.1346413 | 802.5229306 | 0.666612081 | -0.086443612 | YLR207W | HRD3 | n |
| W303.asm02.HP0_G0036920 | 3374.129587 | 3059.482823 | 3527.082597 | 3398.056211 | 3297.424444 | 3339.464446 | 3703.066044 | 3209.506158 | 2861.035193 | 0.939355072 | -0.010797249 | YLR208W | SEC13 | n |
| W303.asm02.HP0_G0036930 | 632.9522035 | 488.3368662 | 582.471939 | 2080.859501 | 1569.759806 | 1920.033638 | 1670.734041 | 1582.982235 | 1455.596438 | 5.89E-27 | -1.710105485 | YLR209C | PNP1 | n |
| W303.asm02.HP0_G0036960 | 839.4128763 | 662.5389062 | 816.0470957 | 641.0876161 | 744.4084962 | 612.8537354 | 698.5845451 | 743.111465 | 752.3652474 | 0.172494493 | 0.216049626 | YLR212C | TUB4 | n |
| W303.asm02.HP0_G0036970 | 386.7502744 | 385.5291049 | 308.8274039 | 45.72248552 | 30.51942141 | 45.262462 | 70.04777282 | 76.43985122 | 53.22856172 | 2.16E-33 | 3.135724493 | YLR213C | CRR1 | n |
| W303.asm02.HP0_G0036980 | 8872.962437 | 11576.34431 | 10155.14416 | 23764.01864 | 23380.53066 | 23005.09893 | 22097.23255 | 27772.82341 | 32212.49248 | 1.04E-13 | -1.196686726 | YLR214W | FRE1 | n |
| W303.asm02.HP0_G0036990 | 724.0663033 | 701.5677785 | 763.2727925 | 1601.259812 | 1621.510129 | 1638.501124 | 1090.473436 | 971.4634256 | 974.49213 | 1.00E-23 | -1.151212856 | YLR215C | CDC123 | n |
| W303.asm02.HP0_G0037000 | 10594.43734 | 9735.323841 | 8636.416987 | 1685.895051 | 1786.049618 | 1760.709772 | 3779.739598 | 3181.44596 | 3245.918659 | 9.97E-85 | 2.468944672 | YLR216C | CPR6 | n |
| W303.asm02.HP0_G0037010 | 436.18452 | 347.4521563 | 445.6496714 | 114.7926232 | 123.404617 | 109.535158 | 208.2501354 | 156.7500747 | 135.1186567 | 2.19E-16 | 1.824511821 | YLR218C | COA4 | n |
| W303.asm02.HP0_G0037020 | 492.4038582 | 483.5772476 | 380.1704434 | 644.0060726 | 719.1968002 | 619.1904801 | 868.9710195 | 1019.843078 | 1326.619538 | 0.005407947 | -0.546610671 | YLR219W | MSC3 | n |
| W303.asm02.HP0_G0037030 | 1735.045091 | 1368.866303 | 1731.778986 | 1270.501406 | 1762.164854 | 1161.434775 | 2287.911715 | 2085.16303 | 2146.544114 | 0.275396043 | 0.206902402 | YLR220W | CCC1 | n |
| W303.asm02.HP0_G0037040 | 2082.054109 | 2481.665127 | 2442.277476 | 2896.081689 | 3148.808131 | 2894.08182 | 2469.657288 | 1824.880499 | 2042.134243 | 0.019756857 | -0.351270708 | YLR221C | RSA3 | n |
| W303.asm02.HP0_G0037050 | 1352.172012 | 1851.491627 | 1416.110469 | 6104.438226 | 5058.262366 | 5714.838452 | 3420.035178 | 3556.872064 | 3886.708632 | 1.72E-34 | -1.869326795 | YLR222C | UTP13 | n |
| W303.asm02.HP0_G0037060 | 403.2283563 | 1004.27952 | 399.7164816 | 3559.544139 | 2851.575505 | 3378.390163 | 2187.573013 | 2687.973502 | 3016.626373 | 4.38E-16 | -2.437391575 | YLR223C | IFH1 | n |
| W303.asm02.HP0_G0037090 | 626.167111 | 878.6255897 | 576.6081275 | 543.805732 | 586.5036637 | 578.4542643 | 433.5389182 | 582.4910182 | 523.0729815 | 0.163112275 | 0.285054874 | YLR226W | BUR2 | n |
| W303.asm02.HP0_G0037100 | 528.2679187 | 637.7888896 | 514.0608052 | 44.74966668 | 58.38498009 | 63.36744679 | 65.31481519 | 41.6065013 | 59.37031884 | 4.46E-44 | 3.336240002 | YLR227C | ADY4 | n |
| W303.asm02.HP0_G0037120 | 2880.75643 | 2838.636521 | 3111.729285 | 2105.179972 | 2020.91647 | 1952.622611 | 2219.757125 | 2014.528737 | 2150.638619 | 4.13E-07 | 0.538854083 | YLR229C | CDC42 | n |
| W303.asm02.HP0_G0037150 | 80.45181147 | 99.00006644 | 52.77430319 | 349.2419639 | 379.5023706 | 355.7629513 | 323.7343014 | 399.6159311 | 472.9152984 | 1.20E-19 | -2.221783182 | YLR234W | TOP3 | n |

| | | | | | | | | | | | | | | |
|---|---|---|---|---|---|---|---|---|---|---|---|---|---|---|
| W303.asm02.HP0_G0037180 | 347.0090182 | 446.4522227 | 349.8740841 | 322.9758552 | 432.5796252 | 341.2789635 | 1505.080524 | 463.4770726 | 1444.33655 | 0.871076009 | 0.061075425 | YLR238W | FAR10 | n |
| W303.asm02.HP0_G0037190 | 705.6496235 | 672.0581434 | 663.5879976 | 773.3909785 | 727.1583884 | 687.0841731 | 2042.74451 | 788.5883385 | 2274.497387 | 0.759848814 | -0.099853941 | YLR239C | LIP2 | n |
| W303.asm02.HP0_G0037200 | 397.4125627 | 454.0676124 | 410.4668026 | 339.5137755 | 417.9833802 | 363.9101944 | 969.3097211 | 447.0279907 | 1381.895352 | 0.613397115 | 0.171195198 | YLR240W | VPS34 | n |
| W303.asm02.HP0_G0037210 | 939.2506665 | 789.1447604 | 887.3901352 | 966.9819278 | 1072.160544 | 939.648711 | 2917.395079 | 1469.773948 | 3806.865789 | 0.518077943 | -0.187194366 | YLR241W | CSC1 | n |
| W303.asm02.HP0_G0037220 | 321.8072459 | 361.731012 | 346.9421784 | 783.1191669 | 958.0444461 | 848.2185378 | 2367.425403 | 856.3198523 | 1993.000186 | 5.72E-06 | -1.328884059 | YLR242C | ARV1 | n |
| W303.asm02.HP0_G0037230 | 1298.860571 | 1624.933783 | 1495.271924 | 1962.175602 | 1831.165285 | 1887.444665 | 3111.446341 | 1170.787595 | 2679.853358 | 0.210763479 | -0.362393233 | YLR243W | GPN3 | n |
| W303.asm02.HP0_G0037240 | 1127.29466 | 1059.491096 | 1094.57814 | 4257.055247 | 4287.315243 | 4259.197674 | 7486.592368 | 3034.371816 | 6549.160344 | 1.76E-14 | -1.964178856 | YLR244C | MAP1 | n |
| W303.asm02.HP0_G0037250 | 109.5307795 | 107.5673799 | 97.7301911 | 143.0043696 | 74.30815648 | 125.8296443 | 424.073003 | 89.98615397 | 285.5917062 | 0.791896027 | -0.128200171 | YLR245C | CDD1 | n |
| W303.asm02.HP0_G0037260 | 774.4698478 | 619.702339 | 765.2273963 | 774.3637973 | 817.3897213 | 817.4400637 | 2366.478811 | 999.5236242 | 2374.812754 | 0.59337678 | -0.157898613 | YLR246W | ERF2 | n |
| W303.asm02.HP0_G0037270 | 697.8952321 | 975.7218087 | 678.2475262 | 802.5755437 | 643.5617124 | 735.9676321 | 571.7412808 | 644.4169736 | 721.6564618 | 0.598230707 | 0.107102813 | YLR247C | IRC20 | n |
| W303.asm02.HP0_G0037280 | 851.0444635 | 831.9813276 | 760.3408868 | 1225.751739 | 1146.4687 | 1108.02507 | 1270.325826 | 1246.259853 | 1544.651916 | 0.000245309 | -0.510310985 | YLR248W | RCK2 | n |
| W303.asm02.HP0_G0037290 | 21790.80932 | 20423.52332 | 22232.64117 | 48783.94641 | 51957.32456 | 45783.88556 | 29559.21354 | 34336.00709 | 35186.12655 | 4.46E-25 | -1.184937627 | YLR249W | YEF3 | n |
| W303.asm02.HP0_G0037300 | 2393.199067 | 2281.761147 | 2354.320304 | 2709.300472 | 2460.130752 | 2691.30599 | 3785.419507 | 3138.871865 | 3054.500542 | 0.216666234 | -0.161695432 | YLR250W | SSP120 | n |
| W303.asm02.HP0_G0037310 | 1006.132293 | 464.5387733 | 775.9777173 | 445.5510291 | 352.9637433 | 440.8563798 | 477.0821284 | 406.3890824 | 343.9383988 | 0.001261209 | 0.856719662 | YLR251W | SYM1 | n |
| W303.asm02.HP0_G0037320 | 1189.329791 | 1108.039205 | 1187.421822 | 679.0275509 | 739.1007707 | 728.7256381 | 772.418684 | 803.1022343 | 848.586109 | 3.32E-09 | 0.699485016 | YLR253W | CQD2 | n |
| W303.asm02.HP0_G0037340 | 523.4214241 | 598.7600172 | 510.1515975 | 1406.696044 | 1268.546386 | 1424.862304 | 693.8515875 | 1051.773649 | 993.9410276 | 9.06E-14 | -1.32909786 | YLR256W | HAP1 | n |
| W303.asm02.HP0_G0037350 | 5013.214084 | 3772.473686 | 4711.572513 | 415.393645 | 435.233488 | 393.7834194 | 140.0955456 | 530.2409933 | 537.4037482 | 1.29E-12 | 3.439277948 | YLR257W | | n |
| W303.asm02.HP0_G0037360 | 913.0795953 | 660.6350588 | 750.5678676 | 552.5611016 | 313.1558023 | 522.3288114 | 485.6014521 | 630.8706708 | 648.9790025 | 0.001809334 | 0.740376322 | YLR258W | GSY2 | n |
| W303.asm02.HP0_G0037370 | 9350.826811 | 7028.052794 | 6804.953206 | 5236.68382 | 5570.457873 | 5340.065266 | 7464.820763 | 7135.998769 | 7399.793706 | 0.000282243 | 0.52191419 | YLR259C | HSP60 | n |
| W303.asm02.HP0_G0037380 | 994.5007057 | 730.12549 | 954.823967 | 1303.577247 | 1027.044877 | 1264.633188 | 1253.287179 | 1373.014543 | 1260.083836 | 0.010953288 | -0.425441797 | YLR260W | LCB5 | n |
| W303.asm02.HP0_G0037400 | 192.8904878 | 178.0097349 | 188.6192688 | 215.9657827 | 225.5783322 | 218.1650668 | 133.469405 | 344.463127 | 228.2686397 | 0.454702471 | -0.237333871 | YLR263W | RED1 | n |
| W303.asm02.HP0_G0037410 | 4725.3323 | 5170.849624 | 4958.829896 | 13757.60405 | 15661.77091 | 12703.36258 | 11201.9641 | 9543.370286 | 8814.445096 | 1.03E-28 | -1.503516112 | YLR264W | RPS28B | y |
| W303.asm02.HP0_G0037420 | 881.0927305 | 779.6255232 | 908.8907772 | 165.3792029 | 179.1357344 | 168.3763586 | 199.7308117 | 175.1343427 | 159.6856852 | 2.04E-46 | 2.327055537 | YLR265C | NEJ1 | n |
| W303.asm02.HP0_G0037450 | 1385.128176 | 1363.154761 | 1459.111753 | 884.2923263 | 890.3709464 | 835.5450485 | 1775.8057 | 1242.389481 | 1348.115688 | 1.31E-05 | 0.689094695 | YLR268W | SEC22 | n |
| W303.asm02.HP0_G0037460 | 1466.549286 | 909.0871486 | 1305.675353 | 851.2164857 | 1196.892092 | 850.9342855 | 1126.443914 | 1230.778364 | 1700.243097 | 0.150484385 | 0.34583314 | YLR270W | DCS1 | n |
| W303.asm02.HP0_G0037470 | 331.5002352 | 336.9809954 | 313.7139134 | 171.216116 | 167.1933521 | 152.9871215 | 259.3660777 | 195.4537968 | 270.2373134 | 1.88E-07 | 1.000456096 | YLR271W | CMG1 | n |
| W303.asm02.HP0_G0037480 | 760.8996627 | 1010.942986 | 856.116474 | 917.3681669 | 944.7751324 | 1003.016158 | 615.284491 | 748.9170234 | 810.7119401 | 0.474779924 | -0.124784834 | YLR272C | YCS4 | n |
| W303.asm02.HP0_G0037490 | 245.2326302 | 206.5674463 | 212.0745147 | 14.59228261 | 19.90397049 | 18.1049848 | 5.679549147 | 41.6065013 | 29.68515942 | 2.44E-14 | 3.664049756 | YLR273C | PIG1 | n |
| W303.asm02.HP0_G0037500 | 1588.680952 | 1550.683733 | 1672.16357 | 786.0376234 | 890.3709464 | 871.755018 | 795.1368806 | 1020.810671 | 883.3893994 | 2.43E-11 | 0.917652639 | YLR274W | MCM5 | n |
| W303.asm02.HP0_G0037510 | 466.232787 | 733.9331849 | 498.4239746 | 361.8886088 | 396.7524784 | 355.7629513 | 482.7616775 | 331.8844173 | 358.2691654 | 0.007130281 | 0.60922003 | YLR275W | SMD2 | n |
| W303.asm02.HP0_G0037520 | 3476.875274 | 5255.570835 | 4048.961817 | 5173.450596 | 4494.316536 | 5197.035886 | 2370.265177 | 2757.640202 | 2351.269351 | 0.216355828 | -0.218007459 | YLR276C | DBP9 | n |
| W303.asm02.HP0_G0037530 | 477.8643742 | 574.9619243 | 475.9460307 | 1701.460153 | 1957.223765 | 1704.584319 | 1574.181705 | 1698.125809 | 1402.367876 | 3.37E-36 | -1.809858155 | YLR277C | YSH1 | n |
| W303.asm02.HP0_G0037540 | 819.0575987 | 1310.798957 | 974.3700053 | 399.8285436 | 464.425978 | 364.8154437 | 408.9275386 | 489.602085 | 501.5768316 | 4.09E-11 | 1.33893584 | YLR278C | | n |
| W303.asm02.HP0_G0037550 | 208.3992707 | 160.875108 | 215.0064204 | 236.3949783 | 242.8284399 | 221.7860638 | 564.1685486 | 380.26407 | 410.474101 | 0.248563215 | -0.261962502 | YLR281C | RSO55 | n |
| W303.asm02.HP0_G0037570 | 425.5222318 | 301.7598179 | 425.1263313 | 57.39631161 | 62.36577419 | 80.56718235 | 136.3091795 | 108.370422 | 120.7878901 | 5.86E-25 | 2.518510985 | YLR284C | ECI1 | n |
| W303.asm02.HP0_G0037580 | 1392.882567 | 1580.193368 | 1432.724602 | 1506.896384 | 1711.741462 | 1568.796933 | 1151.055294 | 980.1717631 | 983.7047657 | 0.383677617 | -0.119104459 | YLR285W | NNT1 | n |
| W303.asm02.HP0_G0037600 | 10160.19142 | 6207.494551 | 9972.3887 | 2303.635015 | 2449.515301 | 2508.445644 | 1688.71298 | 2711.195736 | 2116.858955 | 5.58E-18 | 1.85890134 | YLR286C | CTS1 | n |
| W303.asm02.HP0_G0037610 | 1682.702948 | 1875.28972 | 1791.394403 | 1782.204116 | 1794.011207 | 1925.465133 | 1506.973707 | 1315.926553 | 1171.028358 | 0.778932338 | -0.040720068 | YLR287C | | n |
| W303.asm02.HP0_G0037620 | 679.4785523 | 512.1349591 | 670.4291109 | 192.6181305 | 230.8860576 | 219.0703161 | 253.6865286 | 306.7269979 | 322.4422489 | 3.12E-16 | 1.536982436 | YLR288C | MEC3 | n |
| W303.asm02.HP0_G0037630 | 505.9740432 | 410.2791215 | 408.5121988 | 784.0919857 | 773.6009862 | 735.9676321 | 655.041335 | 823.4216885 | 761.5778831 | 4.22E-07 | -0.791697197 | YLR289W | GUF1 | n |
| W303.asm02.HP0_G0037640 | 833.5970827 | 623.5100338 | 728.0899237 | 381.3449856 | 421.9641743 | 433.6143859 | 675.8663485 | 460.5742934 | 527.1674863 | 1.72E-05 | 0.821149309 | YLR290C | COQ11 | n |
| W303.asm02.HP0_G0037650 | 1983.185618 | 1968.578244 | 1978.059068 | 919.3138046 | 962.0252402 | 994.8689147 | 882.2233009 | 878.5744925 | 836.3025948 | 2.25E-23 | 1.04372762 | YLR291C | GCD7 | n |
| W303.asm02.HP0_G0037660 | 1713.720514 | 2072.337929 | 1904.761425 | 969.9003843 | 1025.717946 | 997.5846624 | 1255.180362 | 1064.352359 | 935.5943349 | 3.61E-10 | 0.927282282 | YLR292C | SEC72 | n |
| W303.asm02.HP0_G0037670 | 7785.409033 | 7380.264569 | 7846.757043 | 7585.068502 | 8987.30614 | 7434.812008 | 8173.817815 | 5975.854698 | 6134.591739 | 0.715709499 | -0.060875134 | YLR293C | GSP1 | y |
| W303.asm02.HP0_G0037680 | 4356.998706 | 4204.647053 | 4515.134829 | 3409.730037 | 3523.002776 | 3366.621923 | 3516.587514 | 2783.765215 | 2384.025389 | 0.026208387 | 0.344540862 | YLR295C | ATP14 | n |
| W303.asm02.HP0_G0037700 | 861.7067518 | 886.2409794 | 955.801269 | 335.6225001 | 315.8096651 | 329.5107233 | 310.4820201 | 266.0880897 | 264.0955562 | 4.59E-25 | 1.461490718 | YLR298C | YHC1 | n |
| W303.asm02.HP0_G0037710 | 519.5442283 | 504.5195694 | 471.0595211 | 106.0372537 | 134.0200679 | 100.4826656 | 206.3569524 | 210.9352856 | 245.6702849 | 7.38E-31 | 2.145012093 | YLR299W | ECM38 | n |
| W303.asm02.HP0_G0037720 | 7459.724592 | 9110.861884 | 8508.390437 | 2296.825283 | 1970.493078 | 2466.804179 | 1212.583743 | 2473.167845 | 1608.11674 | 4.80E-17 | 1.896619462 | YLR300W | EXG1 | n |
| W303.asm02.HP0_G0037730 | 2705.313323 | 2615.886371 | 2633.82865 | 2472.905493 | 1540.567316 | 2364.511015 | 1819.34891 | 1325.602483 | 1193.548134 | 0.141818903 | 0.317973739 | YLR301W | HRI1 | n |
| W303.asm02.HP0_G0037740 | 6678.469651 | 5039.484151 | 6947.639285 | 136.1946377 | 123.404617 | 144.8398784 | 173.226249 | 104.5000498 | 148.4257971 | 1.94E-167 | 5.524548898 | YLR303W | MET17 | n |
| W303.asm02.HP0_G0037750 | 7177.658602 | 6083.744468 | 6752.178903 | 24247.50961 | 30369.47817 | 23302.02068 | 21016.22503 | 28763.6387 | 26895.77806 | 4.14E-34 | -1.960933659 | YLR304C | ACO1 | n |
| W303.asm02.HP0_G0037760 | 966.3910366 | 1346.020134 | 1142.465934 | 475.7084132 | 542.7149286 | 559.4440303 | 238.5410642 | 360.9122089 | 350.0801559 | 1.27E-08 | 1.130929331 | YLR305C | STT4 | n |
| W303.asm02.HP0_G0037770 | 443.9389115 | 419.7983587 | 380.1704434 | 258.7698117 | 294.5787632 | 252.5645379 | 187.4251219 | 364.7825811 | 340.8675203 | 0.0116717 | 0.627830816 | YLR306W | UBC12 | n |
| W303.asm02.HP0_G0037790 | 70.75882214 | 58.06734666 | 66.45652995 | 128.412087 | 151.2701757 | 110.4404073 | 212.0365015 | 160.6204469 | 178.1109565 | 8.83E-05 | -0.992498694 | YLR307C-/ | DPA10 | n |
| W303.asm02.HP0_G0037800 | 5.815793601 | 7.615389726 | 5.863811466 | 18.48355798 | 21.23090185 | 29.87322492 | 17.98523897 | 15.48148885 | 17.40164518 | 0.00232475 | -1.854844311 | YLR308W | CDA2 | n |
| W303.asm02.HP0_G0037810 | 1301.768468 | 1841.97239 | 1567.592265 | 1917.425935 | 1997.031705 | 1842.182203 | 1305.349712 | 1707.801739 | 1466.856326 | 0.088673613 | -0.288810235 | YLR309C | IMH1 | n |
| W303.asm02.HP0_G0037820 | 899.5094102 | 1046.164164 | 843.4115492 | 630.3866089 | 705.9274866 | 650.8742035 | 815.9618942 | 876.6393064 | 1028.744318 | 0.001530309 | 0.490235041 | YLR310C | CDC25 | n |
| W303.asm02.HP0_G0037840 | 1998.694401 | 2738.68453 | 2125.631656 | 3263.807211 | 2957.730014 | 3224.497793 | 2428.953852 | 2721.839259 | 2445.442961 | 0.002070666 | -0.461046822 | YLR312W- | MRPL15 | n |
| W303.asm02.HP0_G0037870 | 2115.010273 | 1908.60705 | 2179.383262 | 4001.203892 | 4017.948176 | 4067.284835 | 4271.020959 | 3907.14075 | 3760.802611 | 1.30E-18 | -0.962423013 | YLR314C | CDC3 | n |
| W303.asm02.HP0_G0037880 | 412.9213456 | 432.173367 | 339.1237631 | 770.4725219 | 812.0819959 | 731.4413859 | 776.2050501 | 728.5975692 | 634.6482359 | 6.07E-10 | -0.965408698 | YLR316C | TAD3 | n |
| W303.asm02.HP0_G0037900 | 362.5178011 | 455.9714599 | 359.6471032 | 458.197674 | 536.0802718 | 468.013857 | 487.4946351 | 509.9215392 | 483.1515603 | 0.060613158 | -0.309371216 | YLR318W | EST2 | n |
| W303.asm02.HP0_G0037910 | 312.1142566 | 422.6541298 | 303.9408943 | 1199.485631 | 1151.776425 | 1174.108264 | 1057.342733 | 1232.71355 | 1147.484956 | 2.38E-28 | -1.762793696 | YLR319C | BUD6 | n |
| W303.asm02.HP0_G0037920 | 238.4475376 | 468.3464682 | 241.393572 | 630.3866089 | 701.9466925 | 633.6744679 | 466.6696216 | 626.0327056 | 598.8213194 | 5.19E-05 | -1.051243427 | YLR320W | MMS22 | n |

| | | | | | | | | | | | | | | |
|---|---|---|---|---|---|---|---|---|---|---|---|---|---|---|
| W303.asm02.HP0_G0037930 | 769.6233531 | 617.7984916 | 785.7507364 | 1261.746037 | 1523.317208 | 1284.548671 | 1359.305429 | 1275.287644 | 1467.879952 | 2.21E-09 | -0.904147155 | YLR321C | SFH1 | n |
| W303.asm02.HP0_G0037940 | 549.5924953 | 591.1446275 | 572.6989198 | 165.3792029 | 180.4626657 | 178.3341003 | 206.3569524 | 222.5464023 | 195.5126017 | 4.39E-28 | 1.710180228 | YLR323C | CWC24 | n |
| W303.asm02.HP0_G0037950 | 385.7809755 | 336.9809954 | 344.0102727 | 492.2463335 | 492.2915367 | 522.3288114 | 768.6323179 | 796.329083 | 939.6888396 | 0.000998427 | -0.498968351 | YLR324W | PEX30 | n |
| W303.asm02.HP0_G0037960 | 6219.021957 | 6991.879693 | 6232.254286 | 11373.22507 | 13477.64188 | 10846.69639 | 10851.72524 | 8228.411326 | 7273.887685 | 2.01E-07 | -0.876446582 | YLR325C | RPL38 | n |
| W303.asm02.HP0_G0037970 | 624.2285131 | 536.8849757 | 663.5879976 | 244.177529 | 265.3862732 | 239.8910486 | 613.3913079 | 345.4307201 | 454.490027 | 4.53E-09 | 1.28504732 | YLR326W | | n |
| W303.asm02.HP0_G0037990 | 1068.167425 | 834.8370988 | 988.052232 | 1961.202783 | 2091.243832 | 1977.06434 | 1593.113536 | 1794.885114 | 1493.470607 | 2.43E-14 | -1.060256891 | YLR328W | NMA1 | n |
| W303.asm02.HP0_G0038010 | 1942.475063 | 1457.395209 | 1816.804253 | 2024.436008 | 1685.202835 | 1766.141267 | 2110.8991 | 2245.783477 | 2677.806105 | 0.688092069 | -0.070351654 | YLR330W | CHS5 | n |
| W303.asm02.HP0_G0038020 | 871.3997412 | 977.6256561 | 928.4368154 | 929.041993 | 756.3508785 | 910.6807354 | 494.1207758 | 793.4263038 | 933.5470825 | 0.684086745 | 0.096638631 | YLR332W | MID2 | n |
| W303.asm02.HP0_G0038030 | 4152.476631 | 5553.522958 | 4237.581086 | 11204.92741 | 13302.48694 | 10896.4851 | 12194.93861 | 8870.893114 | 8194.127627 | 4.82E-13 | -1.344203471 | YLR333C | RPS25B | y |
| W303.asm02.HP0_G0038040 | 826.8119902 | 910.990996 | 694.8616587 | 3722.004885 | 3615.887972 | 3615.565464 | 2464.92433 | 3333.358069 | 2903.003866 | 1.78E-45 | -2.170684309 | YLR335W | NUP2 | n |
| W303.asm02.HP0_G0038050 | 744.4215809 | 1246.068144 | 866.8667951 | 4723.035472 | 4171.872214 | 4622.202619 | 1474.789595 | 2629.917919 | 2092.291926 | 2.07E-22 | -2.242032802 | YLR336C | SGD1 | n |
| W303.asm02.HP0_G0038060 | 365.4256979 | 374.1060203 | 269.7353274 | 523.3765364 | 493.6184681 | 487.9293403 | 400.4082149 | 482.8289337 | 640.789993 | 0.007524375 | -0.57633828 | YLR337C | VRP1 | n |
| W303.asm02.HP0_G0038070 | 9036.773956 | 8038.99578 | 9517.943311 | 26818.6698 | 29865.24425 | 25627.60598 | 27629.11342 | 20926.13497 | 20373.232 | 2.42E-27 | -1.629976723 | YLR340W | RPP0 | n |
| W303.asm02.HP0_G0038090 | 5324.359041 | 6808.158415 | 5628.281705 | 19746.27683 | 19071.98452 | 20034.97618 | 8415.198653 | 16389.09114 | 15013.52528 | 2.11E-15 | -1.728425892 | YLR342W | FKS1 | n |
| W303.asm02.HP0_G0038110 | 214.2150643 | 114.2308459 | 201.3241937 | 316.1661233 | 459.1182526 | 335.847468 | 299.1229218 | 360.9122089 | 289.6862109 | 5.04E-05 | -1.066422945 | YLR343W | GAS2 | n |
| W303.asm02.HP0_G0038130 | 1548.939696 | 1127.07768 | 1513.84066 | 356.0516957 | 541.3879972 | 360.2891975 | 722.2493332 | 892.1207953 | 1002.130037 | 6.30E-16 | 1.739922653 | YLR345W | | n |
| W303.asm02.HP0_G0038150 | 1453.9484 | 1559.251046 | 1575.410681 | 4333.907936 | 4086.948607 | 4173.198996 | 2207.451435 | 3336.260848 | 3907.181156 | 1.32E-14 | -1.456654035 | YLR347C | KAP95 | n |
| W303.asm02.HP0_G0038190 | 2762.50196 | 2323.64579 | 2725.69503 | 1106.095022 | 1013.775563 | 1184.066006 | 1632.87038 | 1311.088587 | 1096.303646 | 1.70E-13 | 1.240730734 | YLR351C | NIT3 | n |
| W303.asm02.HP0_G0038200 | 473.9871784 | 269.3944116 | 430.0128408 | 196.5094058 | 229.5591263 | 219.9755653 | 227.1819659 | 243.8334495 | 327.5603798 | 0.000587088 | 0.861900852 | YLR352W | LUG1 | n |
| W303.asm02.HP0_G0038220 | 6139.539444 | 4140.868164 | 6226.390475 | 4298.886458 | 4088.275538 | 4148.757267 | 6559.879265 | 4741.205962 | 4125.213534 | 0.058189647 | 0.396947418 | YLR354C | TAL1 | n |
| W303.asm02.HP0_G0038230 | 5297.218671 | 5959.042461 | 4667.593927 | 8023.809799 | 5531.976864 | 8019.603016 | 4378.932393 | 3842.312015 | 4838.680986 | 0.015840643 | -0.438458524 | YLR355C | ILV5 | n |
| W303.asm02.HP0_G0038240 | 522.4521251 | 432.173367 | 607.8817886 | 2191.760848 | 1713.068393 | 2272.175592 | 1969.856963 | 1419.459009 | 1337.879426 | 4.56E-23 | -1.983748463 | YLR356W | ATG33 | n |
| W303.asm02.HP0_G0038260 | 2427.124529 | 2257.01113 | 2411.003814 | 4181.175378 | 2226.590832 | 3727.81637 | 1970.803554 | 2309.644619 | 1834.338127 | 0.017317164 | -0.515120502 | YLR359W | ADE13 | n |
| W303.asm02.HP0_G0038270 | 586.4258547 | 514.9907302 | 607.8817886 | 487.3822393 | 440.5412134 | 493.3608358 | 540.5037605 | 574.7502737 | 582.4433004 | 0.068045775 | 0.264084952 | YLR361C | DCR2 | n |
| W303.asm02.HP0_G0038290 | 681.4171502 | 579.7215429 | 566.8351084 | 855.1077611 | 814.7358586 | 841.8817931 | 741.1811637 | 990.8152867 | 1036.933327 | 0.006733509 | -0.458743073 | YLR362W | STE11 | n |
| W303.asm02.HP0_G0038300 | 1136.01835 | 742.5004983 | 1032.030818 | 1737.45445 | 1581.702188 | 1707.300066 | 1165.254167 | 1139.824617 | 1240.634939 | 2.79E-06 | -0.788681433 | YLR363C | NMD4 | n |
| W303.asm02.HP0_G0038310 | 739.5750862 | 1027.125689 | 832.6612282 | 1069.127906 | 1085.429857 | 1117.982811 | 823.5346264 | 634.7410431 | 597.7976932 | 0.067892742 | -0.332233192 | YLR363W-A | | n |
| W303.asm02.HP0_G0038320 | 469.1406838 | 321.7502159 | 500.3785784 | 554.5067393 | 480.3491544 | 626.432474 | 539.557169 | 455.7363282 | 403.3087177 | 0.094159711 | -0.365296919 | YLR364W | GRX8 | n |
| W303.asm02.HP0_G0038340 | 617.4434206 | 563.5388398 | 600.0633734 | 166.3520218 | 192.405048 | 192.8180881 | 184.5853473 | 155.7824816 | 232.3631444 | 2.59E-19 | 1.692278795 | YLR368W | MDM30 | n |
| W303.asm02.HP0_G0038350 | 916.956791 | 820.558243 | 937.2325326 | 804.5211814 | 847.9091427 | 878.997012 | 867.0778365 | 983.0745423 | 942.7597182 | 0.561307017 | 0.07930204 | YLR369W | SSQ1 | n |
| W303.asm02.HP0_G0038360 | 2090.777799 | 2018.078277 | 2171.564846 | 1052.589986 | 887.7170837 | 951.4169512 | 1147.268928 | 946.3060063 | 939.6888396 | 1.16E-16 | 1.117885794 | YLR370C | ARC18 | n |
| W303.asm02.HP0_G0038370 | 420.6757371 | 587.3369326 | 428.058237 | 1587.640348 | 1504.740169 | 1595.049161 | 878.4369348 | 1582.014642 | 1454.572812 | 2.29E-14 | -1.706664384 | YLR371W | ROM2 | n |
| W303.asm02.HP0_G0038380 | 6039.701654 | 7200.350986 | 6800.066697 | 6408.930523 | 6249.846733 | 6215.441281 | 3738.08993 | 5447.548891 | 4729.152984 | 0.602584146 | 0.086472557 | YLR372W | ELO3 | n |
| W303.asm02.HP0_G0038400 | 1176.728905 | 1011.89491 | 1142.465934 | 787.0104422 | 980.6022793 | 724.1993919 | 740.2345722 | 900.8291327 | 949.9251015 | 0.013733451 | 0.420906396 | YLR375W | STP3 | n |
| W303.asm02.HP0_G0038410 | 543.7767017 | 297.0001993 | 541.4252587 | 196.5094058 | 214.9628813 | 201.8705805 | 234.7546981 | 241.8982634 | 197.5598541 | 2.10E-06 | 1.173025853 | YLR376C | PSY3 | n |
| W303.asm02.HP0_G0038420 | 164.7808187 | 72.3462024 | 155.3910038 | 15.56510145 | 29.19249005 | 23.53648024 | 51.11594233 | 41.6065013 | 37.87416892 | 1.14E-09 | 2.536770481 | YLR377C | FBP1 | n |
| W303.asm02.HP0_G0038430 | 1583.834457 | 1571.626055 | 1272.447088 | 3253.106204 | 2908.633554 | 3153.888352 | 3893.33094 | 3935.200948 | 4005.44927 | 1.10E-17 | -1.073298567 | YLR378C | SEC61 | n |
| W303.asm02.HP0_G0038440 | 440.0617158 | 473.1060868 | 367.4655185 | 4587.813653 | 4592.509457 | 4532.582944 | 3574.329597 | 4121.946408 | 3613.40044 | 8.17E-150 | -3.420328558 | YLR380W | CSR1 | n |
| W303.asm02.HP0_G0038450 | 447.8161072 | 395.0483421 | 401.6710854 | 574.9359349 | 652.850232 | 624.6219755 | 503.5866911 | 678.2827305 | 544.5691315 | 0.000548358 | -0.573007338 | YLR381W | CTF3 | n |
| W303.asm02.HP0_G0038460 | 482.7108688 | 581.6253904 | 394.829972 | 1159.600058 | 1203.526749 | 1191.308 | 782.8311908 | 1084.671813 | 955.0432324 | 1.49E-12 | -1.284172905 | YLR382C | NAM2 | n |
| W303.asm02.HP0_G0038470 | 349.916915 | 508.3272642 | 398.7391797 | 899.8574278 | 966.0060343 | 940.5539603 | 660.7208841 | 895.0235744 | 694.0185548 | 4.07E-10 | -1.158322763 | YLR383W | SMC6 | n |
| W303.asm02.HP0_G0038480 | 1275.597396 | 1609.703003 | 1313.493768 | 5385.525103 | 5956.594901 | 5589.914057 | 3499.548866 | 4333.849286 | 3866.236108 | 4.05E-47 | -2.011502938 | YLR384C | IKI3 | n |
| W303.asm02.HP0_G0038490 | 12.60088613 | 33.31733005 | 16.61413249 | 175.1073914 | 189.7511853 | 184.6708449 | 230.0217405 | 195.4537968 | 157.6384328 | 1.98E-21 | -3.132057708 | YLR385C | SWC7 | n |
| W303.asm02.HP0_G0038510 | 2603.536935 | 2259.866901 | 2628.942141 | 633.3050654 | 599.7729773 | 543.149544 | 553.7560419 | 542.819703 | 543.5455053 | 3.48E-60 | 2.077258482 | YLR387C | REH1 | n |
| W303.asm02.HP0_G0038520 | 1937.628568 | 2253.203435 | 2017.151144 | 1932.018218 | 2602.112408 | 2058.536772 | 2106.166142 | 1760.051764 | 1397.249745 | 0.66526898 | -0.086053311 | YLR388W | RPS29A | n |
| W303.asm02.HP0_G0038530 | 1442.316813 | 1131.837298 | 1539.25051 | 2281.260182 | 1869.646294 | 2133.672459 | 1270.325826 | 2108.385263 | 1751.424406 | 0.002728905 | -0.611857931 | YLR389C | STE23 | n |
| W303.asm02.HP0_G0038540 | 501.1275486 | 473.1060868 | 499.4012765 | 1134.306768 | 735.1199766 | 1049.183869 | 520.6253385 | 521.5326558 | 351.1037821 | 3.27E-06 | -0.987416412 | YLR390W | ECM19 | n |
| W303.asm02.HP0_G0038550 | 2891.418718 | 2670.146023 | 2685.625651 | 6745.525842 | 6440.924849 | 6619.182442 | 4838.029282 | 5022.77554 | 5007.579307 | 4.57E-42 | -1.264052237 | YLR390W- | CCW14 | n |
| W303.asm02.HP0_G0038570 | 127.9474592 | 122.7981593 | 109.457814 | 810.3580944 | 835.9667604 | 753.1673676 | 885.0630755 | 897.9263536 | 698.1130595 | 2.01E-58 | -2.734951301 | YLR393W | ATP10 | n |
| W303.asm02.HP0_G0038580 | 177.3817048 | 205.6155226 | 163.2094191 | 15.56510145 | 33.17328414 | 25.34697872 | 31.23752031 | 44.50928046 | 37.87416892 | 2.20E-18 | 2.905412012 | YLR394W | CST9 | n |
| W303.asm02.HP0_G0038590 | 3036.813558 | 3530.685062 | 3100.001662 | 5979.917415 | 6111.845871 | 5546.462093 | 4554.051825 | 4568.974398 | 4172.300338 | 1.81E-14 | -0.867292415 | YLR395C | COX8 | n |
| W303.asm02.HP0_G0038600 | 631.0136057 | 409.3271978 | 597.1314676 | 314.2204856 | 333.0597728 | 339.468465 | 378.6366098 | 374.4585117 | 411.4977272 | 0.000155952 | 0.730653162 | YLR396C | VPS33 | n |
| W303.asm02.HP0_G0038610 | 639.7372961 | 796.7601501 | 647.951167 | 3021.57532 | 3243.020258 | 3028.058708 | 1878.037585 | 2454.783577 | 2168.040264 | 4.70E-48 | -2.156100927 | YLR397C | AFG2 | n |
| W303.asm02.HP0_G0038620 | 684.325047 | 851.9717256 | 651.8603746 | 1745.237 | 1954.569902 | 1831.319212 | 1266.53946 | 1595.560945 | 1718.668368 | 4.14E-16 | -1.337419478 | YLR398C | SKI2 | n |
| W303.asm02.HP0_G0038630 | 973.1761292 | 863.3948102 | 831.6839263 | 3113.99311 | 3432.771443 | 3063.363428 | 3172.97479 | 3182.413553 | 3266.391163 | 1.69E-58 | -1.848179915 | YLR399C | BDF1 | n |
| W303.asm02.HP0_G0038640 | 602.9039366 | 805.3274636 | 554.1301835 | 1973.849428 | 1898.838784 | 1942.664869 | 1807.989812 | 1225.940399 | 1565.12444 | 5.40E-17 | -1.567196634 | YLR401C | DUS3 | n |
| W303.asm02.HP0_G0038650 | 616.4741217 | 856.7313442 | 591.2676562 | 1804.57895 | 1565.779012 | 1682.858337 | 588.7799283 | 1289.80154 | 1541.581038 | 8.05E-06 | -1.291487398 | YLR403W | SFP1 | n |
| W303.asm02.HP0_G0038660 | 605.8118334 | 512.1349591 | 675.3156205 | 160.5151087 | 136.6739307 | 153.8923708 | 262.2058523 | 201.2593551 | 232.3631444 | 1.60E-25 | 1.987283175 | YLR404W | SEI1 | n |
| W303.asm02.HP0_G0038670 | 888.847122 | 663.4908299 | 829.7293224 | 724.7500364 | 834.6398291 | 731.4413859 | 998.6540584 | 889.2180161 | 920.2399421 | 0.733760175 | 0.057573241 | YLR405W | DUS4 | n |
| W303.asm02.HP0_G0038700 | 1085.614805 | 1229.885441 | 1081.873215 | 1003.949044 | 964.6791029 | 923.3542247 | 1431.246385 | 1140.79221 | 1241.658565 | 0.101158357 | 0.232543125 | YLR407W | GAG1 | n |
| W303.asm02.HP0_G0038710 | 319.868648 | 241.7886238 | 321.5323287 | 97.28188408 | 114.1160975 | 112.2509058 | 90.87278636 | 126.75469 | 111.5752544 | 2.07E-10 | 1.450011729 | YLR408C | BLS1 | n |
| W303.asm02.HP0_G0038720 | 1216.470161 | 1653.491494 | 1189.376426 | 6665.754697 | 6237.90435 | 6773.074813 | 2665.601733 | 3836.506457 | 3097.492842 | 3.40E-38 | -2.277162858 | YLR409C | UTP21 | n |
| W303.asm02.HP0_G0038730 | 1015.825282 | 1188.952721 | 949.9374575 | 2004.979631 | 1792.684275 | 2094.746741 | 1845.853473 | 2001.950028 | 1719.691994 | 9.01E-11 | -0.901900466 | YLR410W | VIP1 | n |

| | | | | | | | | | | | | | | |
|---|---|---|---|---|---|---|---|---|---|---|---|---|---|---|
| W303.asm02.HP0_G0038750 | 930.5269761 | 900.5198351 | 1068.190989 | 627.4681523 | 617.0230851 | 665.3581913 | 473.2957623 | 548.6252613 | 403.3087177 | 0.00014042 | 0.601546334 | YLR412W | BER1 | n |
| W303.asm02.HP0_G0038780 | 1459.764194 | 1479.289454 | 1505.044943 | 48.64094204 | 35.82714688 | 48.88345896 | 78.56709654 | 114.1759803 | 121.8115163 | 1.38E-123 | 5.041408547 | YLR414C | PUN1 | n |
| W303.asm02.HP0_G0038800 | 315.9914523 | 296.0482756 | 319.5777249 | 475.7084132 | 583.8498009 | 444.4773768 | 601.0856181 | 550.5604474 | 597.7976932 | 1.86E-05 | -0.687441864 | YLR417W | VPS36 | n |
| W303.asm02.HP0_G0038810 | 1675.917856 | 1523.077945 | 1732.756288 | 1237.425566 | 1353.469993 | 1312.611398 | 1504.133933 | 1186.269084 | 1246.776696 | 0.015312954 | 0.337676468 | YLR418C | CDC73 | n |
| W303.asm02.HP0_G0038820 | 443.9389115 | 731.0774137 | 462.2638039 | 1934.936674 | 2120.436322 | 2008.748063 | 863.2914704 | 1707.801739 | 1484.257971 | 1.38E-13 | -1.888764681 | YLR419W | | n |
| W303.asm02.HP0_G0038830 | 1935.68997 | 1820.078145 | 2105.108316 | 1891.159827 | 1705.106805 | 1939.949121 | 2240.582139 | 2171.278812 | 1924.417232 | 0.544931451 | 0.0814476 | YLR420W | URA4 | n |
| W303.asm02.HP0_G0038840 | 2289.484081 | 2323.64579 | 2236.066772 | 1962.175602 | 1626.817854 | 1800.540738 | 1980.269469 | 2061.940797 | 2024.732598 | 0.002701388 | 0.344786587 | YLR421C | RPN13 | n |
| W303.asm02.HP0_G0038850 | 520.5135273 | 542.596518 | 452.4907848 | 2055.566211 | 2265.071841 | 2234.155124 | 2043.691102 | 2640.561443 | 2404.497913 | 5.18E-49 | -2.112344888 | YLR422W | DCK1 | n |
| W303.asm02.HP0_G0038860 | 401.2897584 | 477.8657053 | 381.1477453 | 64.2060435 | 91.55826424 | 70.60944071 | 158.0807846 | 141.2685858 | 185.2763398 | 4.00E-29 | 2.489270365 | YLR423C | ATG17 | n |
| W303.asm02.HP0_G0038870 | 286.9124843 | 358.8752409 | 276.5764408 | 374.5352537 | 443.1950762 | 385.6361762 | 376.7434268 | 446.0603976 | 432.9938771 | 0.034103544 | -0.381150404 | YLR424W | SPP382 | n |
| W303.asm02.HP0_G0038880 | 201.6141782 | 273.2021064 | 198.3922879 | 366.752703 | 428.5988311 | 364.8154437 | 403.2479895 | 404.4538963 | 428.8993723 | 2.28E-05 | -0.78247476 | YLR425W | TUS1 | n |
| W303.asm02.HP0_G0038890 | 212.2764664 | 223.7020732 | 246.2800816 | 184.8355798 | 205.6743617 | 197.3443343 | 277.3513167 | 196.4213898 | 206.7724898 | 0.313313164 | 0.216190348 | YLR426W | TDA5 | n |
| W303.asm02.HP0_G0038900 | 750.2373745 | 791.0486078 | 754.4770753 | 880.401051 | 839.9475545 | 900.7229937 | 764.8459518 | 823.4216885 | 761.5778831 | 0.112714298 | -0.19204851 | YLR427W | MAG2 | n |
| W303.asm02.HP0_G0038920 | 638.7679971 | 761.5389726 | 527.7430319 | 1696.596058 | 1813.915177 | 1764.330769 | 1140.642787 | 1595.560945 | 1272.36735 | 3.22E-16 | -1.451690857 | YLR430W | SEN1 | n |
| W303.asm02.HP0_G0038930 | 424.5529328 | 493.0964848 | 445.6496714 | 88.52651452 | 95.53905834 | 111.3456565 | 142.9353202 | 148.0417372 | 223.1505088 | 4.37E-22 | 2.204017789 | YLR431C | ATG23 | n |
| W303.asm02.HP0_G0038950 | 654.2767801 | 754.8755066 | 689.9751492 | 1294.821877 | 1251.296278 | 1245.622954 | 1272.219009 | 1451.38958 | 1459.690943 | 7.98E-12 | -0.853068673 | YLR433C | CNA1 | n |
| W303.asm02.HP0_G0038960 | 1769.939852 | 2751.059539 | 2025.946861 | 1995.251443 | 2433.592125 | 2015.084808 | 2209.344618 | 1714.574891 | 1574.337076 | 0.913863938 | 0.023346426 | YLR435W | TSR2 | n |
| W303.asm02.HP0_G0038970 | 576.7328654 | 693.9523888 | 592.2449581 | 416.3664639 | 477.6952917 | 420.0356473 | 399.4616234 | 618.2919611 | 677.6405358 | 0.018128161 | 0.504807684 | YLR436C | ECM30 | n |
| W303.asm02.HP0_G0038980 | 682.3864491 | 1036.644927 | 750.5678676 | 2251.102798 | 2543.727428 | 2109.230729 | 2745.115421 | 2407.371517 | 3121.036244 | 5.37E-16 | -1.48264741 | YLR437C | DIF1 | n |
| W303.asm02.HP0_G0038990 | 114.3772741 | 102.8077613 | 86.97987008 | 2243.320247 | 1452.989846 | 1887.444665 | 4501.042699 | 5016.002389 | 3927.653679 | 4.65E-98 | -4.1989634 | YLR438W | CAR2 | n |
| W303.asm02.HP0_G0039000 | 1769.939852 | 1889.568576 | 1953.62652 | 634.2778842 | 725.8314571 | 666.2634406 | 739.2879807 | 512.8243183 | 478.0334293 | 1.11E-16 | 1.470767027 | YLR438C-/ | LSM3 | n |
| W303.asm02.HP0_G0039010 | 435.2152211 | 409.3271978 | 335.2145555 | 3087.727001 | 3120.942572 | 2920.334048 | 3132.271355 | 2889.232858 | 2889.696726 | 1.34E-113 | -2.951620802 | YLR439W | MRPL4 | n |
| W303.asm02.HP0_G0039020 | 547.6538974 | 557.8272975 | 585.4038447 | 1129.442674 | 1131.872455 | 1050.994368 | 717.5163756 | 1010.167148 | 923.3108207 | 5.24E-10 | -0.969761328 | YLR440C | SEC39 | n |
| W303.asm02.HP0_G0039050 | 994.5007057 | 1085.193036 | 1007.59827 | 255.8513551 | 260.0785477 | 212.7335714 | 126.8432643 | 278.6667994 | 349.0565298 | 3.84E-13 | 2.084151347 | YLR443W | ECM7 | n |
| W303.asm02.HP0_G0039060 | 11.6315872 | 1.903847432 | 10.75032102 | 28.21174638 | 41.13487234 | 17.19973556 | 50.1693508 | 36.76853603 | 27.63790705 | 0.004770652 | -1.823822853 | YLR445W | GMC2 | n |
| W303.asm02.HP0_G0039080 | 6134.69295 | 4855.762874 | 5923.426883 | 1954.393051 | 2048.782029 | 2054.915775 | 2086.28772 | 1865.519407 | 1699.21947 | 1.75E-27 | 1.481325123 | YLR447C | VMA6 | n |
| W303.asm02.HP0_G0039120 | 337.3160288 | 483.5772476 | 346.9421784 | 1125.551399 | 859.851525 | 1021.121143 | 398.5150318 | 697.6345915 | 587.5614313 | 4.48E-09 | -1.364767987 | YLR451W | LEU3 | n |
| W303.asm02.HP0_G0039130 | 1885.286426 | 2340.780417 | 1707.346439 | 89.49933336 | 100.8467838 | 97.76691791 | 84.24664569 | 164.4908191 | 68.58295453 | 6.76E-58 | 4.365571075 | YLR452C | SST2 | n |
| W303.asm02.HP0_G0039150 | 827.7812891 | 1060.443019 | 945.0509479 | 630.3866089 | 717.8698689 | 685.2736746 | 630.4299554 | 902.7643189 | 683.7822929 | 0.007905028 | 0.479182086 | YLR454W | FMP27 | n |
| W303.asm02.HP0_G0039160 | 923.7418836 | 1014.750681 | 873.7079084 | 909.5856162 | 831.9859663 | 840.9765439 | 649.3617858 | 822.4540954 | 770.7905188 | 0.418453026 | 0.122576598 | YLR455W | PDP3 | n |
| W303.asm02.HP0_G0039180 | 371.2414915 | 306.5194365 | 383.1023491 | 246.1231667 | 277.3286554 | 277.0062674 | 365.3843285 | 307.694591 | 381.8125677 | 0.024560396 | 0.407032053 | YLR457C | NBP1 | n |
| W303.asm02.HP0_G0039190 | 1296.921973 | 1211.79889 | 1296.879636 | 1346.381276 | 1361.431581 | 1311.706149 | 1823.135276 | 1792.949928 | 1711.502985 | 0.482534219 | -0.078610953 | YLR459W | GAB1 | n |
| W303.asm02.HP0_G0039260 | 3890.765919 | 4030.445013 | 3545.651333 | 2223.86387 | 1829.838353 | 1987.927331 | 2961.88488 | 2712.163329 | 3572.455393 | 2.16E-10 | 0.923967395 | YML130C | ERO1 | n |
| W303.asm02.HP0_G0039270 | 956.6980473 | 848.1640308 | 884.4582294 | 1410.587319 | 1370.720101 | 1333.43213 | 1252.340587 | 1101.120895 | 979.6102609 | 6.43E-06 | -0.613546793 | YML129C | COX14 | n |
| W303.asm02.HP0_G0039280 | 1109.847279 | 495.0003322 | 904.0042677 | 58.36913045 | 151.2701757 | 41.64146504 | 147.6682778 | 76.43985122 | 187.3235922 | 8.85E-08 | 3.326416976 | YML128C | MSC1 | n |
| W303.asm02.HP0_G0039290 | 1013.886684 | 883.3852083 | 986.0976282 | 1309.41416 | 1295.085013 | 1266.443687 | 908.7278636 | 1029.519009 | 1159.76847 | 0.002119216 | -0.424916573 | YML127W | RSC9 | n |
| W303.asm02.HP0_G0039300 | 6403.188754 | 4380.75294 | 5549.120251 | 983.5198481 | 895.6786719 | 948.7012034 | 238.5410642 | 645.3845666 | 484.1751864 | 5.06E-20 | 2.529814887 | YML126C | ERG13 | n |
| W303.asm02.HP0_G0039310 | 1751.523173 | 1989.520566 | 1812.895045 | 1635.308471 | 1598.952296 | 1610.438398 | 1896.969415 | 1774.56566 | 1686.935956 | 0.08980047 | 0.197037279 | YML125C | PGA3 | n |
| W303.asm02.HP0_G0039330 | 11847.74086 | 16080.84733 | 11675.82593 | 4513.879421 | 1575.067531 | 4252.860929 | 2123.20479 | 3664.274893 | 3394.344436 | 1.38E-05 | 1.936960083 | YML123C | PHO84 | n |
| W303.asm02.HP0_G0039350 | 1344.417621 | 1271.770084 | 1363.336166 | 410.5295508 | 477.6952917 | 428.1828905 | 560.3821825 | 588.2965765 | 543.5455053 | 6.18E-37 | 1.598910922 | YML121W | GTR1 | n |
| W303.asm02.HP0_G0039360 | 2244.89633 | 1645.876105 | 1936.035086 | 7289.331574 | 8587.899799 | 7461.969485 | 7327.564992 | 9290.828499 | 7921.843061 | 5.53E-38 | -2.001869383 | YML120C | NDI1 | n |
| W303.asm02.HP0_G0039370 | 1455.886998 | 965.2506478 | 1358.449656 | 444.5782103 | 566.5996932 | 399.2149148 | 774.3118671 | 567.9771224 | 605.9867027 | 1.97E-11 | 1.424718515 | YML119W | | n |
| W303.asm02.HP0_G0039390 | 736.6671894 | 787.240913 | 603.972581 | 641.0876161 | 663.4656829 | 626.432474 | 739.2879807 | 1197.8802 | 1205.831648 | 0.524415396 | 0.140475319 | YML117W | NAB6 | n |
| W303.asm02.HP0_G0039410 | 1237.794738 | 1045.21224 | 1232.37771 | 1866.839356 | 1930.685137 | 1924.559884 | 2201.771886 | 2583.473453 | 2535.522065 | 3.42E-08 | -0.702857261 | YML115C | VAN1 | n |
| W303.asm02.HP0_G0039420 | 348.947616 | 355.067546 | 260.9396102 | 614.8215074 | 642.234781 | 639.1059634 | 573.6344639 | 521.5326558 | 601.8921979 | 3.13E-09 | -0.974085951 | YML114C | TAF8 | n |
| W303.asm02.HP0_G0039440 | 315.9914523 | 311.2790551 | 299.0543848 | 602.1748625 | 607.7345655 | 559.4440303 | 741.1811637 | 758.5929539 | 756.4597522 | 6.10E-13 | -0.932529694 | YML112W | CTK3 | n |
| W303.asm02.HP0_G0039450 | 587.3951537 | 901.4717589 | 643.0646574 | 587.5825799 | 670.1003397 | 602.8959938 | 549.0230842 | 755.6901747 | 767.7196402 | 0.365011865 | 0.19733601 | YML111W | BUL2 | n |
| W303.asm02.HP0_G0039460 | 4102.073086 | 3271.761811 | 4190.670594 | 2502.090059 | 2651.208869 | 2603.496814 | 1981.216061 | 2012.593551 | 1665.439806 | 5.03E-05 | 0.576254841 | YML110C | COQ5 | n |
| W303.asm02.HP0_G0039480 | 752.1759723 | 1005.231444 | 838.5250396 | 1496.195377 | 1417.162699 | 1439.346291 | 1087.633662 | 783.7503733 | 728.8218451 | 5.41E-05 | -0.745696623 | YML108W | | n |
| W303.asm02.HP0_G0039490 | 667.8469651 | 584.4811615 | 645.9965632 | 310.3292102 | 327.7520473 | 321.3634802 | 505.4798741 | 328.9816382 | 535.3564958 | 8.32E-07 | 0.984854408 | YML107C | PML39 | n |
| W303.asm02.HP0_G0039500 | 4645.849788 | 5889.55203 | 5410.343379 | 3598.456892 | 3799.0045 | 3595.649981 | 3435.180643 | 2456.718763 | 2640.955562 | 0.000774445 | 0.536690036 | YML106W | URA5 | n |
| W303.asm02.HP0_G0039510 | 2198.369981 | 2469.290119 | 2318.160133 | 1202.404087 | 1402.566454 | 1227.517969 | 1501.294158 | 1526.861838 | 1389.060736 | 6.35E-13 | 0.867526054 | YML105C | SEC65 | n |
| W303.asm02.HP0_G0039520 | 519.5442283 | 630.1734999 | 627.4278269 | 411.5023697 | 449.829733 | 426.372392 | 417.4468623 | 563.1391571 | 559.9235243 | 0.008174722 | 0.465701359 | YML104C | MDM1 | n |
| W303.asm02.HP0_G0039530 | 1164.128019 | 1605.895309 | 1406.33745 | 2922.347798 | 3319.982277 | 3083.278911 | 1845.853473 | 2593.149383 | 2337.962211 | 1.11E-11 | -1.158716487 | YML103C | NUP188 | n |
| W303.asm02.HP0_G0039540 | 597.088143 | 451.2118413 | 459.3318982 | 1051.617167 | 1107.98769 | 1180.445009 | 1702.918153 | 1503.639605 | 1716.621116 | 5.17E-14 | -1.147809228 | YML102W | CAC2 | n |
| W303.asm02.HP0_G0039570 | 390.6274702 | 518.7984251 | 393.8526701 | 565.2077465 | 598.446046 | 580.2647628 | 409.8741301 | 509.9215392 | 540.4746267 | 0.021062864 | -0.419579111 | YML099C | ARG81 | n |
| W303.asm02.HP0_G0039580 | 1143.772741 | 1253.683534 | 1307.629957 | 276.2805508 | 305.1942141 | 251.6592887 | 386.209342 | 290.277916 | 306.0642299 | 1.40E-39 | 2.156359661 | YML098W | TAF13 | n |
| W303.asm02.HP0_G0039590 | 1300.799169 | 1792.472357 | 1465.952866 | 922.2322611 | 869.1400446 | 948.7012034 | 797.9766552 | 937.5976688 | 815.830071 | 2.63E-06 | 0.734029609 | YML097C | VPS9 | n |
| W303.asm02.HP0_G0039600 | 351.8555128 | 415.0387401 | 322.5096306 | 176.0802102 | 205.6743617 | 183.7655957 | 139.1489541 | 222.5464023 | 235.434023 | 5.06E-05 | 0.948217278 | YML095C | RAD10 | n |
| W303.asm02.HP0_G0039620 | 1126.325361 | 1487.856768 | 1155.170859 | 5551.877125 | 4564.643898 | 5188.888643 | 2205.558252 | 3086.62184 | 2810.877509 | 5.42E-32 | -2.021760035 | YML093W | UTP14 | n |
| W303.asm02.HP0_G0039630 | 3344.08132 | 3095.655924 | 3515.354974 | 2118.799435 | 1893.531059 | 2160.829936 | 2564.31644 | 2126.769531 | 2267.332004 | 5.01E-08 | 0.688738947 | YML092C | PRE8 | n |
| W303.asm02.HP0_G0039640 | 1151.527133 | 2194.184165 | 1376.041091 | 700.4295654 | 1125.237798 | 775.7985986 | 819.7482603 | 1880.033303 | 1740.164518 | 0.007332446 | 0.860939058 | YML091C | RPM2 | n |
| W303.asm02.HP0_G0039660 | 1293.044777 | 1363.154761 | 1288.083919 | 743.2335944 | 790.851094 | 700.6629117 | 661.6674757 | 772.1392566 | 644.8844978 | 2.62E-10 | 0.821056359 | YML088W | UFO1 | n |

| | | | | | | | | | | | | | | |
|---|---|---|---|---|---|---|---|---|---|---|---|---|---|---|
| W303.asm02.HP0_G0039680 | 1293.044777 | 1126.125756 | 1209.899766 | 3522.577023 | 3505.752668 | 3595.649981 | 3135.111129 | 3467.853503 | 3407.651576 | 1.16E-46 | -1.549787407 | YML086C | ALO1 | n |
| W303.asm02.HP0_G0039690 | 1899.82591 | 1396.472091 | 1531.432095 | 4849.501922 | 4778.279848 | 4557.929923 | 4594.75526 | 4616.386458 | 4828.444724 | 1.16E-30 | -1.555015751 | YML085C | TUB1 | n |
| W303.asm02.HP0_G0039720 | 1352.172012 | 1739.164629 | 1485.498905 | 4909.81669 | 5397.956796 | 4674.707075 | 4622.206414 | 3895.529633 | 3655.369114 | 2.35E-31 | -1.710586173 | YML081C- | ATP18 | n |
| W303.asm02.HP0_G0039730 | 496.2810539 | 788.1928367 | 592.2449581 | 1136.252406 | 1145.141769 | 1145.140289 | 735.5016146 | 1172.722781 | 1073.78387 | 5.92E-05 | -0.868408797 | YML081W | TDA9 | n |
| W303.asm02.HP0_G0039740 | 1349.264115 | 1452.63559 | 1424.906186 | 3450.588428 | 3001.518749 | 3403.737142 | 1955.65809 | 2201.274197 | 1708.432106 | 5.93E-20 | -1.221792914 | YML080W | DUS1 | n |
| W303.asm02.HP0_G0039750 | 1459.764194 | 1474.529836 | 1608.638945 | 409.556732 | 419.3103116 | 453.5298692 | 473.2957623 | 465.4122587 | 358.2691654 | 4.49E-34 | 1.824026221 | YML079W | CFF1 | n |
| W303.asm02.HP0_G0039760 | 3291.739178 | 2991.896239 | 3218.255193 | 2449.557841 | 2230.571626 | 2498.487902 | 2672.227874 | 1870.357372 | 1735.046387 | 0.021082832 | 0.404233693 | YML078W | CPR3 | n |
| W303.asm02.HP0_G0039780 | 353.7941107 | 361.731012 | 352.8059899 | 1328.870537 | 1151.776425 | 1232.949465 | 1090.473436 | 1032.421788 | 1040.004206 | 2.96E-48 | -1.798376221 | YML076C | WAR1 | n |
| W303.asm02.HP0_G0039790 | 1814.527603 | 1707.751146 | 1522.636377 | 7520.862458 | 6052.133959 | 7539.820919 | 2249.101462 | 3976.80745 | 3309.383462 | 4.99E-25 | -2.065340765 | YML075C | HMG1 | n |
| W303.asm02.HP0_G0039800 | 3169.607512 | 2342.684265 | 3207.504872 | 12424.84224 | 13774.87451 | 12118.57157 | 6117.821023 | 6970.540357 | 6082.386803 | 1.38E-46 | -2.135634674 | YML074C | FPR3 | n |
| W303.asm02.HP0_G0039810 | 6309.166758 | 7159.418266 | 6222.481267 | 15723.67092 | 17327.06977 | 14863.28727 | 15593.20218 | 11883.01029 | 10913.90241 | 1.95E-16 | -1.282835933 | YML073C | RPL6A | n |
| W303.asm02.HP0_G0039820 | 1869.777643 | 1980.001329 | 1846.12331 | 4187.98511 | 3723.369412 | 4115.263045 | 2758.367703 | 3700.075836 | 2926.547269 | 1.97E-14 | -1.078453879 | YML072C | TCB3 | n |
| W303.asm02.HP0_G0039830 | 936.3427697 | 1073.769951 | 1086.759725 | 1058.426899 | 996.5254557 | 1001.205659 | 780.9380078 | 950.1763785 | 917.1690635 | 0.899316223 | 0.018863432 | YML071C | COG8 | n |
| W303.asm02.HP0_G0039840 | 1246.518428 | 730.12549 | 1024.212403 | 554.5067393 | 484.3299485 | 500.6028297 | 841.5198653 | 1158.208885 | 997.0119061 | 5.33E-06 | 0.96234727 | YML070W | DAK1 | n |
| W303.asm02.HP0_G0039850 | 2141.181344 | 1869.578178 | 2115.858637 | 3333.850168 | 3094.403945 | 3355.758932 | 3015.840597 | 3224.987647 | 3029.933513 | 8.81E-10 | -0.675739473 | YML069W | POB3 | n |
| W303.asm02.HP0_G0039860 | 631.0136057 | 656.8273639 | 630.3597326 | 422.2033769 | 441.8681448 | 401.9306625 | 451.5241572 | 434.449281 | 438.112008 | 3.41E-06 | 0.601378535 | YML068W | ITT1 | n |
| W303.asm02.HP0_G0039880 | 168.6580144 | 128.5097016 | 132.9130599 | 189.699674 | 214.9628813 | 213.6388206 | 321.8411183 | 276.7316133 | 260.0010515 | 0.011556938 | -0.522809339 | YML066C | SMA2 | n |
| W303.asm02.HP0_G0039890 | 472.0485806 | 682.5293042 | 513.0835033 | 859.9718553 | 1001.833181 | 848.2185378 | 760.1129942 | 914.3754355 | 770.7905188 | 0.000103214 | -0.699256974 | YML065W | ORC1 | n |
| W303.asm02.HP0_G0039900 | 927.6190793 | 755.8274303 | 980.2338167 | 317.1389421 | 360.9253315 | 326.7949756 | 536.7173944 | 423.8057574 | 366.4581749 | 4.84E-14 | 1.407979655 | YML064C | TEM1 | n |
| W303.asm02.HP0_G0039910 | 11920.43828 | 14071.33637 | 12298.36725 | 29867.48405 | 33072.43736 | 28031.94796 | 26546.21271 | 21634.41308 | 20014.96283 | 4.41E-18 | -1.248400931 | YML063W | RPS1B | y |
| W303.asm02.HP0_G0039920 | 1249.426325 | 1367.91438 | 1258.764861 | 1215.050732 | 1163.718808 | 1199.455243 | 1110.351858 | 1134.986652 | 1180.240994 | 0.317571225 | 0.11498748 | YML062C | MFT1 | n |
| W303.asm02.HP0_G0039930 | 678.5092534 | 721.5581766 | 650.8830727 | 866.7815872 | 875.7747014 | 919.7332278 | 833.9471331 | 1039.194939 | 828.1135853 | 0.009192441 | -0.376586527 | YML061C | PIF1 | n |
| W303.asm02.HP0_G0039940 | 493.3731571 | 703.471626 | 526.76573 | 553.5339204 | 506.8877817 | 592.0330029 | 600.1390266 | 597.972507 | 518.9784768 | 0.75911897 | 0.059673605 | YML060W | OGG1 | n |
| W303.asm02.HP0_G0039950 | 586.4258547 | 645.4042793 | 538.493353 | 242.2318914 | 281.3094495 | 226.31231 | 212.983093 | 341.5603479 | 285.5917062 | 1.36E-09 | 1.242594909 | YML059C | NTE1 | n |
| W303.asm02.HP0_G0039990 | 3377.037484 | 4195.127816 | 3451.83035 | 3540.087762 | 3184.635278 | 3321.359461 | 1539.157819 | 2375.440946 | 2194.654545 | 0.477506605 | 0.133920082 | YML056C | IMD4 | n |
| W303.asm02.HP0_G0040000 | 1050.720044 | 1150.875772 | 1118.033386 | 830.7872901 | 792.1780254 | 800.2403281 | 968.3631296 | 668.6067999 | 636.6954883 | 0.01137742 | 0.453948645 | YML055W | SPC2 | n |
| W303.asm02.HP0_G0040010 | 503.0661464 | 284.625191 | 417.307916 | 205.2647754 | 220.2706067 | 237.1753009 | 498.8537334 | 496.3752364 | 640.789993 | 0.000199596 | 0.862042124 | YML054C | CYB2 | n |
| W303.asm02.HP0_G0040040 | 3301.432167 | 2391.232374 | 3483.104011 | 2990.445117 | 3396.944296 | 3179.235331 | 5039.653277 | 5098.247799 | 5119.154561 | 0.723212084 | -0.060015312 | YML052W | SUR7 | n |
| W303.asm02.HP0_G0040050 | 868.4918444 | 713.9427868 | 818.0016995 | 133.2761812 | 203.020499 | 149.3661246 | 256.5263032 | 277.6992063 | 331.6548846 | 4.96E-31 | 2.314283326 | YML051W | GAL80 | n |
| W303.asm02.HP0_G0040060 | 612.5969259 | 571.1542295 | 584.4265428 | 189.699674 | 179.1357344 | 180.1445987 | 249.9001625 | 218.6760301 | 287.6389585 | 2.36E-24 | 1.686669135 | YML050W | AIM32 | n |
| W303.asm02.HP0_G0040070 | 413.8906446 | 656.8273639 | 406.557595 | 605.093319 | 782.8895058 | 680.7474284 | 599.192435 | 732.4679414 | 673.546031 | 0.026941111 | -0.484502205 | YML049C | RSE1 | n |
| W303.asm02.HP0_G0040080 | 4114.673972 | 3315.550302 | 4232.694577 | 3267.698486 | 2257.110253 | 3160.225097 | 4286.166423 | 3604.284124 | 3610.329561 | 0.017843836 | 0.42464968 | YML048W | GSF2 | n |
| W303.asm02.HP0_G0040100 | 932.465574 | 1194.664263 | 992.9387416 | 480.5725074 | 492.2915367 | 465.2981093 | 389.0491166 | 460.5742934 | 449.3718961 | 2.12E-13 | 1.118124208 | YML046W | PRP39 | n |
| W303.asm02.HP0_G0040110 | 356.7020075 | 1008.087215 | 426.1036332 | 956.2809205 | 761.658604 | 989.4374192 | 444.8980165 | 562.171564 | 329.6076322 | 0.086698155 | -0.596491119 | YML043C | RRN11 | n |
| W303.asm02.HP0_G0040120 | 174.473808 | 162.7789554 | 161.2548153 | 92.41778988 | 122.0776857 | 95.95641943 | 210.1433185 | 150.9445163 | 192.4417232 | 0.00313183 | 0.691242846 | YML042W | CAT2 | n |
| W303.asm02.HP0_G0040130 | 950.8822537 | 752.9716592 | 873.7079084 | 248.0688044 | 230.8860576 | 213.6388206 | 132.5228134 | 307.694591 | 291.7334633 | 2.87E-12 | 1.896104948 | YML041C | VPS71 | n |
| W303.asm02.HP0_G0040140 | 1029.395467 | 1153.731544 | 1064.281781 | 581.7456668 | 621.0038792 | 600.1802461 | 1048.823049 | 986.9449145 | 1001.106411 | 8.20E-13 | 0.849916653 | YML038C | YMD8 | n |
| W303.asm02.HP0_G0040150 | 106.6228827 | 101.8558376 | 91.86637963 | 201.3735001 | 213.6359499 | 206.3968267 | 254.6331201 | 282.5371716 | 321.4186227 | 1.28E-07 | -1.047722667 | YML037C | LFT1 | n |
| W303.asm02.HP0_G0040160 | 606.7811323 | 903.3756063 | 669.451809 | 393.0188117 | 408.6948607 | 361.1944467 | 356.8650048 | 350.2686853 | 288.6625847 | 2.26E-06 | 0.907599688 | YML036W | CGI121 | n |
| W303.asm02.HP0_G0040170 | 829.719887 | 994.760283 | 802.3648689 | 2513.763885 | 2403.072703 | 2533.792623 | 1367.824753 | 2023.237075 | 1581.502459 | 3.00E-19 | -1.504076232 | YML035C | AMD1 | n |
| W303.asm02.HP0_G0040180 | 565.1012782 | 794.8563027 | 540.4479568 | 1594.45008 | 1682.548972 | 1529.871215 | 973.0960872 | 1935.186107 | 1976.622167 | 1.45E-07 | -1.338553546 | YML034W | SRC1 | n |
| W303.asm02.HP0_G0040190 | 749.2680755 | 676.8177619 | 598.1087695 | 681.9460074 | 704.6005552 | 683.4631761 | 709.9436434 | 859.2226314 | 799.452052 | 0.841290269 | -0.031860789 | YML032C | RAD52 | n |
| W303.asm02.HP0_G0040200 | 707.5882214 | 821.5101667 | 610.8136944 | 1819.171232 | 2024.897264 | 1826.792966 | 1934.833076 | 2319.320549 | 2179.300152 | 2.65E-21 | -1.405386999 | YML031W | NDC1 | n |
| W303.asm02.HP0_G0040210 | 1370.588692 | 1335.548973 | 1472.79398 | 638.1691596 | 686.0235161 | 686.1789239 | 1018.53248 | 905.667098 | 919.2163159 | 2.38E-18 | 1.056072419 | YML030W | RCF1 | n |
| W303.asm02.HP0_G0040220 | 440.0617158 | 337.9329191 | 363.5563109 | 618.7127828 | 692.6581729 | 653.5899512 | 1006.226791 | 1038.227346 | 1563.077188 | 0.000130941 | -0.783039658 | YML029W | USA1 | n |
| W303.asm02.HP0_G0040230 | 12777.29854 | 11575.39238 | 12701.01564 | 9044.296763 | 7637.816941 | 8773.675633 | 9411.959529 | 8757.684727 | 8626.097878 | 2.12E-06 | 0.541379636 | YML028W | TSA1 | n |
| W303.asm02.HP0_G0040240 | 360.5792032 | 285.5771147 | 252.143893 | 436.7956595 | 512.1955072 | 480.6873464 | 625.6969977 | 691.8290332 | 671.4987787 | 0.000138147 | -0.669124185 | YML027W | YOX1 | n |
| W303.asm02.HP0_G0040260 | 943.1278622 | 697.7600837 | 798.4556613 | 2718.055841 | 2663.151251 | 2830.714373 | 2048.424059 | 2181.922335 | 1960.244148 | 4.05E-37 | -1.751489693 | YML025C | YML6 | n |
| W303.asm02.HP0_G0040280 | 1130.202556 | 1139.452688 | 1067.213687 | 525.3221741 | 501.5800563 | 493.3608358 | 615.284491 | 560.2363779 | 561.9707767 | 1.12E-20 | 1.134064948 | YML023C | NSE5 | n |
| W303.asm02.HP0_G0040290 | 3538.910406 | 4124.685461 | 3641.42692 | 3189.872979 | 3590.676276 | 3207.298057 | 2392.036783 | 2241.913105 | 1943.866129 | 0.188971845 | 0.179109333 | YML022W | APT1 | n |
| W303.asm02.HP0_G0040300 | 897.5708124 | 805.3274636 | 845.366153 | 575.9087538 | 520.1570954 | 579.3595136 | 729.8220654 | 658.9308694 | 589.6086837 | 2.07E-05 | 0.603333173 | YML021C | UNG1 | n |
| W303.asm02.HP0_G0040310 | 470.1099827 | 446.4522227 | 396.7845759 | 259.7426305 | 277.3286554 | 247.1330425 | 448.6843826 | 591.1993556 | 610.0812074 | 3.28E-05 | 0.746007673 | YML020W | | n |
| W303.asm02.HP0_G0040320 | 1525.676521 | 1575.43375 | 1527.522887 | 1112.904754 | 1173.007327 | 1084.488589 | 2195.145745 | 2189.66308 | 2064.654019 | 1.22E-05 | 0.458679726 | YML019W | OST6 | n |
| W303.asm02.HP0_G0040330 | 1248.457026 | 1870.530102 | 1387.768714 | 951.4168263 | 712.5621434 | 980.3849268 | 607.7117588 | 638.6114153 | 620.3174693 | 5.75E-05 | 0.767596827 | YML018C | | n |
| W303.asm02.HP0_G0040340 | 258.8028152 | 351.2598511 | 240.4162701 | 3334.822986 | 3166.058239 | 3533.187783 | 1556.196466 | 2532.191021 | 2447.490213 | 7.44E-65 | -3.560316933 | YML017W | PSP2 | n |
| W303.asm02.HP0_G0040350 | 1722.444205 | 1616.366469 | 1732.756288 | 1354.163826 | 1395.931797 | 1295.411662 | 816.9084857 | 1521.05628 | 1393.15524 | 0.128954863 | 0.326283396 | YML016C | PPZ1 | n |
| W303.asm02.HP0_G0040360 | 881.0927305 | 761.5389726 | 821.9109071 | 353.1332392 | 395.425547 | 401.0254133 | 373.9036522 | 442.1900254 | 411.4977272 | 4.11E-14 | 1.10077907 | YML015C | TAF11 | n |
| W303.asm02.HP0_G0040370 | 530.2065166 | 411.2310452 | 511.1288994 | 1271.474225 | 1218.122994 | 1310.800899 | 1276.005375 | 1145.630175 | 1040.004206 | 1.34E-20 | -1.388139726 | YML014W | TRM9 | n |
| W303.asm02.HP0_G0040380 | 410.0134488 | 353.1636986 | 314.6912153 | 389.1275363 | 398.0794097 | 325.8897264 | 523.4651131 | 570.8799015 | 494.4114483 | 0.812737864 | -0.044190506 | YML013W | UBX2 | n |
| W303.asm02.HP0_G0040390 | 4330.827635 | 3986.656522 | 4431.086864 | 3292.991776 | 3266.905023 | 3363.906176 | 4658.176892 | 3869.404621 | 3733.164704 | 0.004066567 | 0.361279447 | YML012W | ERV25 | n |
| W303.asm02.HP0_G0040400 | 608.7197302 | 502.6157219 | 597.1314676 | 500.0288842 | 521.4840267 | 516.897316 | 639.8958706 | 470.250224 | 502.6004578 | 0.395783242 | 0.151410944 | YML011C | RAD33 | n |
| W303.asm02.HP0_G0040410 | 1420.022937 | 1500.231776 | 1206.96786 | 3524.52266 | 3664.984432 | 3589.313236 | 2688.31993 | 3611.057275 | 3736.235582 | 8.24E-19 | -1.384847074 | YML010W | SPT5 | n |
| W303.asm02.HP0_G0040420 | 842.3207731 | 1339.356668 | 972.4154014 | 956.2809205 | 966.0060343 | 978.5744284 | 1241.92808 | 864.0605967 | 810.7119401 | 0.60133775 | 0.120803955 | YML009C | MRPL39 | n |

| | | | | | | | | | | | | | |
|---|---|---|---|---|---|---|---|---|---|---|---|---|---|
| W303.asm02.HP0_G0040430 | 5282.679187 | 4082.800817 | 4247.354105 | 4052.763291 | 4258.122753 | 4090.821315 | 5015.041897 | 4321.270577 | 4059.701458 | 0.372276502 | 0.134513754 | YML008C | ERG6 | n |
| W303.asm02.HP0_G0040440 | 2678.172953 | 2695.847963 | 2694.421369 | 1486.467189 | 1340.200679 | 1336.147878 | 760.1129942 | 1959.375933 | 2358.434735 | 0.00162725 | 0.954677111 | YML007W | YAP1 | n |
| W303.asm02.HP0_G0040450 | 337.3160288 | 476.9137816 | 304.9181962 | 241.2590725 | 302.5403514 | 165.6606109 | 194.0512625 | 207.0649134 | 227.2450135 | 0.011650678 | 0.662965643 | YML006C | GIS4 | n |
| W303.asm02.HP0_G0040470 | 3125.98906 | 1998.087879 | 2810.720296 | 3301.747146 | 2965.691602 | 3130.351872 | 3833.695674 | 3030.501843 | 3105.681857 | 0.193331679 | -0.244296336 | YML004C | GLO1 | n |
| W303.asm02.HP0_G0040480 | 821.9654955 | 644.4523556 | 801.387567 | 421.2305581 | 455.1374585 | 429.0881397 | 428.8059606 | 553.4632266 | 549.6872624 | 2.51E-06 | 0.797569176 | YML002W | VRL1 | n |
| W303.asm02.HP0_G0040490 | 1983.185618 | 2057.10715 | 1866.64665 | 670.2721813 | 713.8890748 | 691.6104193 | 1076.274563 | 944.3708201 | 924.3344468 | 1.01E-35 | 1.509505419 | YML001W | YPT7 | n |
| W303.asm02.HP0_G0040500 | 934.4041718 | 890.0486743 | 992.9387416 | 1645.03666 | 2006.320225 | 1777.909507 | 951.3244822 | 1778.436032 | 1853.787025 | 3.40E-05 | -0.946117392 | YMR001C | CDC5 | n |
| W303.asm02.HP0_G0040520 | 5146.977336 | 4491.176091 | 5633.168215 | 2554.622276 | 2754.709515 | 2537.41362 | 3306.444195 | 3261.756183 | 2795.523117 | 3.27E-13 | 0.960950065 | YMR002W | MIX17 | n |
| W303.asm02.HP0_G0040530 | 693.0487374 | 737.7408797 | 736.8856409 | 1058.426899 | 1158.411082 | 995.7741639 | 1082.900704 | 981.1393562 | 1098.350899 | 7.42E-06 | -0.566182973 | YMR003W | AIM34 | n |
| W303.asm02.HP0_G0040540 | 950.8822537 | 760.5870489 | 902.0496638 | 811.3309133 | 609.0614969 | 754.0726169 | 804.6027959 | 920.1809938 | 882.3657732 | 0.115104333 | 0.263106002 | YMR004W | MVP1 | n |
| W303.asm02.HP0_G0040550 | 1526.64582 | 1303.183567 | 1629.162286 | 1322.060805 | 1240.680827 | 1263.727939 | 1208.797377 | 1377.852508 | 1268.272846 | 0.106036489 | 0.220344551 | YMR005W | TAF4 | n |
| W303.asm02.HP0_G0040570 | 1618.729219 | 1510.702937 | 1590.070209 | 411.5023697 | 416.6564489 | 395.5939178 | 598.2458435 | 473.1530031 | 454.490027 | 6.92E-41 | 1.948015142 | YMR009W | ADI1 | n |
| W303.asm02.HP0_G0040580 | 1756.369667 | 2217.030334 | 1788.462497 | 1062.318174 | 1242.007758 | 1135.182547 | 1701.02497 | 1701.028588 | 1695.124966 | 5.72E-08 | 0.745350152 | YMR010W | ANY1 | n |
| W303.asm02.HP0_G0040600 | 2956.361747 | 3162.290584 | 2989.566546 | 7180.375864 | 7590.047412 | 7433.001509 | 4546.479092 | 7146.642293 | 5517.345148 | 3.04E-15 | -1.285502941 | YMR012W | CLU1 | n |
| W303.asm02.HP0_G0040610 | 687.2329438 | 683.4812279 | 710.4984893 | 596.3379494 | 740.4277021 | 582.0752613 | 1104.672309 | 1142.727396 | 1373.706343 | 0.469764715 | 0.11993261 | YMR013C | SEC59 | n |
| W303.asm02.HP0_G0040620 | 1559.601984 | 2161.818759 | 1680.959287 | 2812.419269 | 2659.170457 | 2897.702817 | 1820.295502 | 1767.792509 | 1617.329375 | 2.40E-05 | -0.631658422 | YMR014W | BUD22 | n |
| W303.asm02.HP0_G0040630 | 5351.499411 | 4583.512692 | 4845.462875 | 4030.388458 | 3825.543128 | 3831.014783 | 3034.772428 | 3920.687053 | 3775.133378 | 0.013482018 | 0.33873442 | YMR015C | ERG5 | n |
| W303.asm02.HP0_G0040640 | 328.5923384 | 564.4907635 | 286.3494599 | 2074.049769 | 2434.919056 | 2097.462489 | 1688.71928 | 1939.056479 | 1913.157343 | 2.13E-28 | -2.48519088 | YMR016C | SOK2 | n |
| W303.asm02.HP0_G0040680 | 655.246079 | 715.8466343 | 667.4972052 | 616.7671451 | 705.9274866 | 654.4952005 | 1140.642787 | 698.6021846 | 858.8223709 | 0.826834438 | 0.044906537 | YMR020W | FMS1 | n |
| W303.asm02.HP0_G0040690 | 937.3120686 | 867.2025051 | 852.2072664 | 401.7741813 | 460.4451839 | 463.4876108 | 471.4025792 | 447.9955837 | 501.5768316 | 1.66E-13 | 1.003771712 | YMR021C | MAC1 | n |
| W303.asm02.HP0_G0040700 | 1422.930834 | 1463.106751 | 1432.724602 | 406.6382755 | 309.1750082 | 358.478699 | 430.6991437 | 282.5371716 | 402.2850915 | 7.15E-28 | 2.004644766 | YMR022W | UBC7 | n |
| W303.asm02.HP0_G0040710 | 640.706595 | 582.5773141 | 600.0633734 | 730.5869495 | 768.2932608 | 748.6411214 | 852.8789636 | 715.0512665 | 720.6328356 | 0.028900854 | -0.30132138 | YMR023C | MSS1 | n |
| W303.asm02.HP0_G0040720 | 2531.808814 | 2669.194099 | 2644.578971 | 2833.821283 | 2794.517456 | 2880.503081 | 2744.16883 | 2621.209582 | 2610.246777 | 0.234804985 | -0.117247573 | YMR024W | MRPL3 | n |
| W303.asm02.HP0_G0040730 | 296.6054736 | 248.4520898 | 301.9862905 | 82.68960147 | 98.19292107 | 91.43017323 | 71.94095587 | 108.370422 | 107.4807496 | 3.39E-12 | 1.640438954 | YMR025W | CSI1 | n |
| W303.asm02.HP0_G0040740 | 520.5135273 | 463.5868496 | 458.3545963 | 199.4278624 | 240.1745772 | 231.7438054 | 424.073003 | 294.1482882 | 315.2768656 | 1.11E-08 | 1.105193657 | YMR026C | PEX12 | n |
| W303.asm02.HP0_G0040750 | 1854.26886 | 1326.029736 | 1741.552005 | 2156.73937 | 1693.164423 | 2179.84017 | 2094.807044 | 1750.375834 | 2441.348456 | 0.128332209 | -0.293559359 | YMR027W | | n |
| W303.asm02.HP0_G0040760 | 1064.290229 | 1050.923782 | 1015.416686 | 791.8745364 | 749.7162217 | 787.5668387 | 770.525501 | 762.4633261 | 803.5465568 | 0.000164589 | 0.425763255 | YMR028W | TAP42 | n |
| W303.asm02.HP0_G0040770 | 1187.391193 | 1223.221975 | 1165.92118 | 725.7228553 | 699.2928298 | 616.4747324 | 601.0856181 | 789.5559316 | 886.4602779 | 1.32E-06 | 0.80953793 | YMR029C | FAR8 | n |
| W303.asm02.HP0_G0040790 | 1222.285955 | 1053.779553 | 1186.44452 | 1647.955116 | 1556.490492 | 1632.16438 | 1476.682778 | 1559.760002 | 1724.810125 | 0.00010333 | -0.482619141 | YMR031C | EIS1 | n |
| W303.asm02.HP0_G0040810 | 517.6056305 | 506.4234168 | 503.3104842 | 1646.982298 | 1833.819147 | 1544.355203 | 1317.655402 | 1386.560846 | 1651.10904 | 1.26E-34 | -1.717297495 | YMR032W | HOF1 | n |
| W303.asm02.HP0_G0040830 | 2023.896173 | 2029.501362 | 2146.154997 | 1572.075247 | 1583.029119 | 1606.817401 | 1538.211227 | 1485.255337 | 1414.65139 | 0.000214975 | 0.380524873 | YMR033W | ARP9 | n |
| W303.asm02.HP0_G0040840 | 192.8904878 | 201.8078277 | 166.1413249 | 597.3107683 | 384.8100961 | 549.4862886 | 314.2683862 | 767.3012914 | 477.0098031 | 1.31E-06 | -1.450737136 | YMR034C | RCH1 | n |
| W303.asm02.HP0_G0040850 | 348.947616 | 364.5867831 | 319.5777249 | 221.8026957 | 258.7516163 | 230.8385562 | 331.3070336 | 209.0000995 | 204.7252374 | 0.017273874 | 0.540302288 | YMR035W | IMP2 | n |
| W303.asm02.HP0_G0040860 | 360.5792032 | 502.6157219 | 322.5096306 | 749.0705074 | 821.3705154 | 743.209626 | 679.6527146 | 704.4077429 | 763.6251355 | 1.57E-07 | -0.96340907 | YMR036C | MIH1 | n |
| W303.asm02.HP0_G0040870 | 736.6671894 | 755.8274303 | 679.2248281 | 359.9429711 | 386.1370274 | 337.6579665 | 503.5866911 | 573.7826807 | 747.2471165 | 1.76E-08 | 1.004538154 | YMR037C | MSN2 | n |
| W303.asm02.HP0_G0040880 | 4393.832065 | 4555.906904 | 4496.566092 | 592.4466741 | 436.5604193 | 547.6757902 | 996.7608754 | 731.5003484 | 654.0971335 | 3.37E-66 | 3.089848703 | YMR038C | CCS1 | n |
| W303.asm02.HP0_G0040890 | 2801.271918 | 2619.694066 | 3124.434209 | 2863.005849 | 2599.458546 | 2841.577364 | 3226.930507 | 2195.468638 | 2268.35563 | 0.823041539 | 0.041109921 | YMR039C | SUB1 | n |
| W303.asm02.HP0_G0040910 | 964.4524388 | 616.8465678 | 944.073646 | 886.237964 | 745.7354276 | 895.2914983 | 1181.346223 | 846.6439218 | 1022.602561 | 0.992725792 | -0.00206429 | YMR041C | ARA2 | n |
| W303.asm02.HP0_G0040920 | 520.5135273 | 418.8464349 | 538.493353 | 1752.046732 | 1601.606158 | 1845.8032 | 755.3800366 | 1807.463824 | 1777.015061 | 3.63E-11 | -1.81508848 | YMR042W | ARG80 | n |
| W303.asm02.HP0_G0040930 | 686.2636449 | 675.8658382 | 618.6321097 | 825.9231959 | 914.255711 | 860.8920272 | 854.7721467 | 895.9911675 | 935.5943349 | 0.001764947 | -0.391949093 | YMR044W | IOC4 | n |
| W303.asm02.HP0_G0040940 | 991.5928089 | 1316.510499 | 893.2539466 | 6202.692929 | 5912.806166 | 6257.987996 | 3345.254448 | 5212.423779 | 4736.318367 | 5.55E-39 | -2.520819956 | YMR047C | NUP116 | n |
| W303.asm02.HP0_G0040950 | 659.1232747 | 496.9041796 | 676.2929224 | 431.9315653 | 395.425547 | 435.4248844 | 615.284491 | 469.2826309 | 420.7103628 | 0.007163505 | 0.53594209 | YMR048W | CSM3 | n |
| W303.asm02.HP0_G0040960 | 1239.733336 | 1648.731876 | 1214.786275 | 8589.017546 | 7713.452029 | 8460.459396 | 2610.699425 | 3759.099013 | 3175.288432 | 4.32E-51 | -2.593338404 | YMR049C | ERB1 | n |
| W303.asm02.HP0_G0040970 | 445.8775094 | 338.8848428 | 424.1490294 | 156.6238334 | 163.212558 | 159.3238662 | 243.2740218 | 209.0000995 | 273.3081919 | 5.38E-12 | 1.335788579 | YMR052W | FAR3 | n |
| W303.asm02.HP0_G0040990 | 1080.768311 | 1223.221975 | 1162.011972 | 732.5325871 | 760.3316726 | 747.7358722 | 733.6084315 | 1066.287545 | 1044.098711 | 0.0001812 | 0.629615878 | YMR054W | STV1 | n |
| W303.asm02.HP0_G0041000 | 284.0045875 | 250.3559373 | 242.3708739 | 161.4879276 | 183.1165285 | 171.0921063 | 308.588837 | 220.6112162 | 427.8757462 | 0.020128278 | 0.592351084 | YMR055C | BUB2 | n |
| W303.asm02.HP0_G0041020 | 1417.115041 | 1484.049073 | 1230.423106 | 11180.60694 | 9902.888783 | 10701.85651 | 6502.137182 | 8114.235346 | 8550.34954 | 3.26E-95 | -2.943618418 | YMR058W | FET3 | n |
| W303.asm02.HP0_G0041030 | 459.4476944 | 542.596518 | 459.3318982 | 184.8355798 | 209.6551558 | 180.1445987 | 272.6183591 | 198.356576 | 223.1505088 | 6.30E-13 | 1.350241114 | YMR059W | SEN15 | n |
| W303.asm02.HP0_G0041040 | 1350.233414 | 1034.741079 | 1369.199977 | 747.1248698 | 701.9466925 | 729.6308874 | 837.7334992 | 722.7920109 | 888.5075303 | 6.90E-07 | 0.784483967 | YMR060C | SAM37 | n |
| W303.asm02.HP0_G0041050 | 2093.685696 | 1920.030135 | 2150.064204 | 1663.520218 | 1695.818285 | 1824.982468 | 1302.509938 | 1491.060895 | 1556.93543 | 0.048420127 | 0.249349463 | YMR061W | RNA14 | n |
| W303.asm02.HP0_G0041060 | 3945.046659 | 2925.261579 | 3940.481305 | 1910.616203 | 1133.199386 | 1864.813434 | 1575.128297 | 981.1393562 | 1517.014009 | 2.25E-06 | 1.13819363 | YMR062C | ARG7 | n |
| W303.asm02.HP0_G0041070 | 118.2544699 | 121.8462356 | 139.7541733 | 50.58657972 | 61.03884283 | 57.93595136 | 120.2171236 | 68.69910679 | 92.12635683 | 6.71E-05 | 1.166909758 | YMR063W | RIM9 | n |
| W303.asm02.HP0_G0041080 | 1256.211418 | 1193.71234 | 1196.217576 | 740.3151379 | 679.3888593 | 649.063705 | 579.314013 | 722.7920109 | 923.3108207 | 4.77E-06 | 0.817559524 | YMR064W | AEP1 | n |
| W303.asm02.HP0_G0041090 | 493.3731571 | 474.0580105 | 448.5815771 | 230.5580653 | 254.7708222 | 218.1650668 | 155.24101 | 233.1899259 | 220.0796302 | 1.68E-07 | 1.0118991 | YMR065W | KAR5 | n |
| W303.asm02.HP0_G0041100 | 479.802972 | 589.2407801 | 552.1755797 | 919.3138046 | 825.3513095 | 932.4067171 | 498.8537334 | 711.1808943 | 526.1438601 | 4.46E-05 | -0.724271587 | YMR066W | SOV1 | n |
| W303.asm02.HP0_G0041110 | 573.8249686 | 444.5483753 | 567.8124103 | 279.1990073 | 258.7516163 | 254.3750364 | 417.4468623 | 337.6899756 | 376.6944368 | 1.02E-08 | 1.000881653 | YMR067C | UBX4 | n |
| W303.asm02.HP0_G0041120 | 349.916915 | 197.0482092 | 266.8034217 | 484.4637827 | 502.9069876 | 486.1188418 | 666.4004333 | 589.2641695 | 705.2784428 | 4.46E-05 | -0.856510093 | YMR068W | AVO2 | n |
| W303.asm02.HP0_G0041130 | 65.91232747 | 64.73081267 | 56.68351084 | 223.7483334 | 245.4823027 | 200.9653313 | 96.5523355 | 231.2547398 | 203.7016112 | 8.25E-09 | -1.837498211 | YMR069W | NAT4 | n |
| W303.asm02.HP0_G0041140 | 607.7504313 | 648.2600505 | 526.76573 | 478.6268697 | 639.5809183 | 452.62462 | 142.9353202 | 762.4633261 | 639.7663669 | 0.771670214 | 0.182962489 | YMR070W | MOT3 | n |
| W303.asm02.HP0_G0041150 | 3539.879705 | 2953.81929 | 3569.106579 | 2711.246109 | 2526.47732 | 2621.601799 | 2673.174465 | 2466.394693 | 1846.621641 | 0.030317435 | 0.356404289 | YMR072W | ABF2 | n |
| W303.asm02.HP0_G0041160 | 949.9129548 | 732.9812612 | 968.5061938 | 312.2748479 | 347.6560178 | 324.9844771 | 526.3048877 | 454.7687351 | 420.7103628 | 4.61E-17 | 1.430034466 | YMR073C | IRC21 | n |
| W303.asm02.HP0_G0041170 | 1403.544856 | 1407.895176 | 1463.020961 | 1001.030587 | 1139.834043 | 1058.236361 | 1495.614609 | 947.2735993 | 846.5388566 | 0.045353962 | 0.418461775 | YMR074C | SDD2 | n |
| W303.asm02.HP0_G0041180 | 753.1452713 | 890.0486743 | 833.6385301 | 820.0862828 | 797.4857508 | 783.9458418 | 585.9401537 | 764.3985122 | 824.0190805 | 0.799981989 | 0.044595284 | YMR075W | RCO1 | n |

| | | | | | | | | | | | | | | |
|---|---|---|---|---|---|---|---|---|---|---|---|---|---|---|
| W303.asm02.HP0_G0041190 | 630.0443067 | 737.7408797 | 681.179432 | 1338.598725 | 1378.681689 | 1453.830279 | 1074.38138 | 1525.894245 | 1195.595386 | 1.64E-10 | -1.025601775 | YMR076C | PDS5 | n |
| W303.asm02.HP0_G0041200 | 702.7417267 | 816.7505482 | 718.3169046 | 318.111761 | 262.7324104 | 317.7424832 | 431.6457352 | 341.5603479 | 453.4664008 | 2.13E-14 | 1.312961313 | YMR077C | VPS20 | n |
| W303.asm02.HP0_G0041210 | 501.1275486 | 547.3561366 | 480.8325402 | 746.1520509 | 744.4084962 | 728.7256381 | 1170.933716 | 1014.03752 | 1185.359125 | 5.94E-05 | -0.536900366 | YMR078C | CTF18 | n |
| W303.asm02.HP0_G0041220 | 1785.448635 | 1825.789687 | 1800.19012 | 4941.919711 | 4936.184681 | 4939.945102 | 4017.33443 | 4243.863132 | 3669.69988 | 1.69E-46 | -1.453269103 | YMR079W | SEC14 | n |
| W303.asm02.HP0_G0041230 | 911.1409974 | 1279.385474 | 861.0029836 | 3561.489776 | 3684.888403 | 3482.493826 | 1953.764907 | 3334.325662 | 2914.263754 | 1.14E-17 | -1.813753538 | YMR080C | NAM7 | n |
| W303.asm02.HP0_G0041250 | 3912.090495 | 3383.136886 | 4253.217917 | 153.7053769 | 132.6931366 | 148.4608753 | 742.1277552 | 562.171564 | 572.2070385 | 2.18E-168 | 4.72685849 | YMR083W | ADH3 | n |
| W303.asm02.HP0_G0041270 | 454.6011998 | 569.250382 | 404.6029912 | 1010.758776 | 1028.371808 | 941.4592095 | 975.9358618 | 1352.695089 | 1515.990383 | 7.54E-08 | -1.060677066 | YMR086W | SEG1 | n |
| W303.asm02.HP0_G0041280 | 409.0441499 | 331.2694531 | 342.0556688 | 110.9013479 | 127.3854111 | 124.0191459 | 224.3421913 | 121.9167247 | 180.1582089 | 1.26E-10 | 1.580385765 | YMR087W | PDL32 | n |
| W303.asm02.HP0_G0041290 | 728.9127979 | 889.0967506 | 796.5010575 | 316.1661233 | 313.1558023 | 346.7104589 | 377.6900183 | 398.648338 | 447.3246437 | 1.05E-17 | 1.305298421 | YMR088C | VBA1 | n |
| W303.asm02.HP0_G0041300 | 1088.522702 | 1464.058675 | 961.6650804 | 2094.478964 | 1908.127304 | 2031.379294 | 1797.577305 | 2327.061294 | 2189.536414 | 1.42E-05 | -0.779994279 | YMR089C | YTA12 | n |
| W303.asm02.HP0_G0041310 | 1267.843005 | 851.0198019 | 1122.919896 | 250.0144421 | 257.424685 | 257.0907841 | 572.6878724 | 411.2270477 | 559.9235243 | 8.30E-26 | 2.084261001 | YMR090W | | n |
| W303.asm02.HP0_G0041320 | 1155.404329 | 1179.433484 | 1181.55801 | 712.1033915 | 762.9855353 | 754.9778661 | 753.4868535 | 718.9216387 | 764.6487617 | 4.53E-09 | 0.657592818 | YMR091C | NPL6 | n |
| W303.asm02.HP0_G0041330 | 1718.567009 | 1233.693136 | 1547.068925 | 1623.634645 | 1342.854542 | 1634.880127 | 1807.04322 | 1500.736826 | 1561.029935 | 0.854055053 | -0.033219075 | YMR092C | AIP1 | n |
| W303.asm02.HP0_G0041340 | 931.496275 | 1116.606519 | 998.802553 | 4728.872385 | 3832.177784 | 4707.296048 | 2494.268667 | 2764.413354 | 2649.144572 | 1.19E-55 | -2.122954309 | YMR093W | UTP15 | n |
| W303.asm02.HP0_G0041350 | 71.72812107 | 133.2693202 | 106.5259083 | 193.5909493 | 208.3282244 | 220.8808145 | 255.5797116 | 273.8288341 | 239.5285278 | 4.20E-05 | -0.999109052 | YMR094W | CTF13 | n |
| W303.asm02.HP0_G0041380 | 979.9612217 | 841.5005648 | 912.7999849 | 690.701377 | 647.5425065 | 736.8728813 | 524.4117046 | 511.8567253 | 522.0493554 | 0.002570025 | 0.396322188 | YMR097C | MTG1 | n |
| W303.asm02.HP0_G0041390 | 1161.220122 | 1026.173766 | 1172.762293 | 1035.079247 | 1082.775994 | 1022.026392 | 893.5823992 | 853.4170731 | 824.0190805 | 0.438963138 | 0.098374074 | YMR098C | ATP25 | n |
| W303.asm02.HP0_G0041400 | 2801.273918 | 2128.501429 | 2892.813657 | 2777.397791 | 2478.707791 | 2809.893641 | 2737.542689 | 2506.066008 | 2212.05619 | 0.793619285 | -0.044578764 | YMR099C | | n |
| W303.asm02.HP0_G0041410 | 485.6187656 | 768.2024386 | 505.265088 | 170.2432971 | 200.3666362 | 197.3443343 | 70.04777282 | 290.277916 | 434.0175033 | 0.00848795 | 1.631176597 | YMR100W | MUB1 | n |
| W303.asm02.HP0_G0041420 | 41.67985414 | 59.9711941 | 59.61541657 | 36.96711595 | 53.07725463 | 38.02046808 | 92.76596941 | 71.60188595 | 85.98459971 | 0.326225521 | 0.345541048 | YMR101C | SRT1 | n |
| W303.asm02.HP0_G0041430 | 382.8730787 | 462.6349259 | 356.7151975 | 780.2007104 | 724.5045257 | 781.2300941 | 747.8073044 | 894.0559814 | 1079.925627 | 2.36E-07 | -0.927445527 | YMR102C | LAF1 | n |
| W303.asm02.HP0_G0041450 | 520.5135273 | 433.1252907 | 467.1503135 | 1219.914826 | 874.44777 | 1200.360492 | 1460.590722 | 1028.551416 | 1534.415654 | 1.29E-09 | -1.214776798 | YMR105C | PGM2 | n |
| W303.asm02.HP0_G0041470 | 148.3027368 | 176.1058874 | 133.8903618 | 286.0087392 | 272.02093 | 330.4159726 | 381.4763844 | 279.6343924 | 388.9779511 | 6.59E-06 | -0.95664462 | YMR106C | YKU80 | n |
| W303.asm02.HP0_G0041480 | 24.23247334 | 67.58658382 | 39.09207644 | 11.67382609 | 18.57703912 | 9.052492399 | 34.07729488 | 22.25464023 | 24.56702849 | 0.001561392 | 1.762284493 | YMR107W | SPG4 | n |
| W303.asm02.HP0_G0041490 | 6523.381822 | 6775.793009 | 5945.904826 | 909.5856162 | 999.1793184 | 907.0597384 | 1429.353202 | 867.9309689 | 1474.021709 | 5.19E-48 | 2.773358743 | YMR108W | ILV2 | n |
| W303.asm02.HP0_G0041500 | 706.6189225 | 642.5485082 | 553.1528816 | 1894.078283 | 1606.913884 | 1751.657279 | 1205.957602 | 1819.07494 | 2307.253425 | 2.27E-11 | -1.465500714 | YMR109W | MYO5 | n |
| W303.asm02.HP0_G0041510 | 2372.843789 | 1784.856967 | 2312.296321 | 803.5483625 | 740.4277021 | 850.0290363 | 762.9527688 | 907.6022841 | 986.7756442 | 1.22E-18 | 1.433343467 | YMR110C | HFD1 | n |
| W303.asm02.HP0_G0041520 | 427.4608296 | 293.1925045 | 376.2612357 | 191.6453116 | 207.0012931 | 205.4915775 | 299.1229218 | 258.3473453 | 435.0411295 | 0.000342367 | 0.86086242 | YMR111C | EUC1 | n |
| W303.asm02.HP0_G0041530 | 536.9916091 | 523.5580437 | 537.516051 | 278.2261885 | 314.4827337 | 254.3750364 | 425.0195945 | 317.3705215 | 349.0565298 | 8.68E-08 | 0.919453133 | YMR112C | MED11 | n |
| W303.asm02.HP0_G0041540 | 707.5882214 | 593.0484749 | 697.7935644 | 426.0946523 | 441.8681448 | 448.0983738 | 610.5515333 | 543.787296 | 670.4751525 | 6.72E-05 | 0.602599607 | YMR113W | FOL3 | n |
| W303.asm02.HP0_G0041550 | 598.0574419 | 465.490697 | 611.7909963 | 269.4708189 | 261.4054791 | 243.5120455 | 541.450352 | 424.7733505 | 464.7262889 | 4.49E-10 | 1.113844931 | YMR114C | | n |
| W303.asm02.HP0_G0041560 | 962.5138409 | 956.6833344 | 1061.349875 | 1346.381276 | 1438.3936 | 1381.41034 | 1057.342733 | 1524.926652 | 1526.226645 | 0.003934976 | -0.482939058 | YMR115W | MGR3 | n |
| W303.asm02.HP0_G0041570 | 16934.62167 | 16674.84773 | 16996.25753 | 25292.31704 | 26060.93202 | 23111.0131 | 21352.26502 | 18142.36975 | 18535.82299 | 5.16E-07 | -0.557196942 | YMR116C | ASC1 | n |
| W303.asm02.HP0_G0041600 | 874.307638 | 799.6159213 | 955.801269 | 387.1818987 | 403.3871352 | 351.2367051 | 637.056096 | 711.1808943 | 765.6723879 | 3.71E-16 | 1.205565933 | YMR119W | ASI1 | n |
| W303.asm02.HP0_G0041610 | 548.6231963 | 447.4041464 | 488.6509555 | 6580.146639 | 3093.077014 | 5639.702765 | 3981.363952 | 4341.590031 | 4130.331664 | 1.01E-47 | -3.36694081 | YMR120C | ADE17 | n |
| W303.asm02.HP0_G0041630 | 3158.945224 | 3230.829091 | 3015.953697 | 2259.858167 | 2331.41841 | 2215.14489 | 3531.732978 | 3825.862933 | 3055.524168 | 0.000123173 | 0.466894022 | YMR122W | NCW1 | n |
| W303.asm02.HP0_G0041640 | 1475.272977 | 1425.029803 | 1521.659075 | 440.6869349 | 395.425547 | 376.5836838 | 857.6119252 | 520.5650627 | 510.7894673 | 1.36E-19 | 1.866346233 | YMR123W | PKR1 | n |
| W303.asm02.HP0_G0041650 | 366.3949968 | 576.8657718 | 352.8059899 | 1083.720189 | 897.0056033 | 1126.130054 | 658.8277011 | 813.7457579 | 980.6338871 | 2.70E-08 | -1.261798531 | YMR124W | EPO1 | n |
| W303.asm02.HP0_G0041660 | 2811.936206 | 3678.233238 | 3087.296737 | 3392.219298 | 3239.039464 | 3248.939522 | 2310.629911 | 2717.968887 | 2596.939636 | 0.764579359 | -0.044917495 | YMR125W | STO1 | n |
| W303.asm02.HP0_G0041670 | 352.8248118 | 495.0003322 | 344.0102727 | 350.2147827 | 303.8672828 | 326.7949756 | 347.3990895 | 274.7964272 | 282.5208276 | 0.181214251 | 0.279998894 | YMR126C | DLT1 | n |
| W303.asm02.HP0_G0041680 | 668.8162641 | 738.6928035 | 712.4530931 | 566.1805654 | 525.4648208 | 640.0112126 | 400.4082149 | 538.9493308 | 377.718063 | 0.108992061 | 0.290393722 | YMR127C | SAS2 | n |
| W303.asm02.HP0_G0041690 | 528.2679187 | 787.240913 | 497.4466727 | 4168.528733 | 3469.925521 | 3955.033929 | 1580.807846 | 2314.482584 | 2198.74905 | 2.54E-38 | -2.676885473 | YMR128W | ECM16 | n |
| W303.asm02.HP0_G0041700 | 735.6978905 | 1031.885308 | 813.1151899 | 2856.196117 | 2949.768426 | 2946.586276 | 1863.838712 | 2764.413354 | 2324.655071 | 6.26E-23 | -1.761820927 | YMR129W | POM152 | n |
| W303.asm02.HP0_G0041710 | 1573.172169 | 1578.289521 | 1750.347723 | 168.2976595 | 189.7511853 | 199.1548328 | 235.7012896 | 163.523226 | 182.2054613 | 3.15E-75 | 3.137264006 | YMR130W | DPI35 | n |
| W303.asm02.HP0_G0041720 | 2788.673031 | 3601.127417 | 3024.749415 | 7473.194335 | 7132.256091 | 7249.235913 | 3299.818055 | 4394.807669 | 3882.614127 | 2.03E-16 | -1.214980777 | YMR131C | RRB1 | n |
| W303.asm02.HP0_G0041730 | 872.3690401 | 1049.019935 | 945.0509479 | 192.6181305 | 164.5394894 | 207.3020759 | 136.3091795 | 113.2083873 | 85.98459971 | 7.96E-34 | 2.339845402 | YMR132C | JLP2 | n |
| W303.asm02.HP0_G0041740 | 1421.961535 | 1283.193169 | 1397.541733 | 686.8101016 | 631.6193301 | 678.0316807 | 671.1333909 | 636.6762292 | 495.4350745 | 4.00E-12 | 1.038234773 | YMR133W | REC114 | n |
| W303.asm02.HP0_G0041750 | 1057.505136 | 746.3081932 | 949.9374575 | 107.0100725 | 94.21212697 | 95.95641943 | 134.4159965 | 162.555633 | 220.0796302 | 1.94E-45 | 3.210235331 | YMR135C | GID8 | n |
| W303.asm02.HP0_G0041770 | 289.8203811 | 401.7118081 | 299.0543848 | 375.5080726 | 416.6564489 | 375.6784346 | 137.2557711 | 518.6298766 | 523.0729815 | 0.557978418 | -0.237077059 | YMR137C | PSO2 | n |
| W303.asm02.HP0_G0041780 | 429.3994275 | 554.0196026 | 497.4466727 | 363.8342465 | 347.6560178 | 373.8679361 | 469.5093962 | 400.5835241 | 418.6631105 | 0.006243222 | 0.44723563 | YMR138W | CIN4 | n |
| W303.asm02.HP0_G0041800 | 338.2853278 | 243.6924712 | 236.5070625 | 119.6567174 | 112.7891661 | 128.5453921 | 239.4876557 | 316.4029285 | 268.190061 | 1.49E-07 | 1.178128388 | YMR140W | SIP5 | n |
| W303.asm02.HP0_G0041820 | 8276.843593 | 9241.275433 | 8135.061107 | 16650.76728 | 19344.00545 | 16309.87556 | 16020.11496 | 12464.53371 | 11127.84028 | 6.77E-11 | -1.027725779 | YMR143W | RPS16A | y |
| W303.asm02.HP0_G0041830 | 1005.162994 | 682.5293042 | 953.8466651 | 492.2463385 | 419.3103116 | 465.2981093 | 508.3196487 | 630.8706708 | 644.8844978 | 7.34E-07 | 0.938605231 | YMR144W | FDO1 | n |
| W303.asm02.HP0_G0041840 | 7109.807677 | 6503.542826 | 7199.783178 | 14023.18359 | 15381.78839 | 13445.66696 | 7163.804658 | 8237.119664 | 7357.825032 | 4.53E-20 | -1.041715693 | YMR145C | NDE1 | n |
| W303.asm02.HP0_G0041850 | 7054.557637 | 5936.196292 | 7034.619155 | 7937.228922 | 8203.089703 | 7920.930849 | 6942.302241 | 5867.484276 | 5494.825372 | 0.050770298 | -0.264851874 | YMR146C | TIF34 | n |
| W303.asm02.HP0_G0041880 | 2592.874647 | 2167.530301 | 2647.510877 | 1658.656124 | 1492.797786 | 1585.091419 | 2982.709894 | 2421.885413 | 2243.788602 | 2.06E-05 | 0.644822108 | YMR149W | SWP1 | n |
| W303.asm02.HP0_G0041890 | 878.1848337 | 1014.750681 | 923.5503059 | 141.0587319 | 148.616313 | 122.2086474 | 235.7012896 | 139.3333997 | 167.8746947 | 8.01E-40 | 2.777303895 | YMR150C | IMP1 | n |
| W303.asm02.HP0_G0041920 | 307.2677619 | 257.0194033 | 333.2599516 | 649.8429857 | 680.7157906 | 695.2314163 | 454.3639318 | 769.2364775 | 699.1366817 | 3.00E-08 | -1.174576355 | YMR154C | RIM13 | n |
| W303.asm02.HP0_G0041930 | 779.3163425 | 532.1253571 | 673.3610167 | 87.55369568 | 87.57747014 | 79.66193311 | 195.9444456 | 205.1297273 | 251.812042 | 1.61E-43 | 2.963757378 | YMR155W | | n |
| W303.asm02.HP0_G0041940 | 410.9827478 | 238.9328527 | 456.3999924 | 419.2849204 | 420.6372429 | 463.4876108 | 563.2219571 | 382.1992561 | 434.0175033 | 0.36737042 | -0.237184645 | YMR156C | TPP1 | n |
| W303.asm02.HP0_G0041950 | 1329.878137 | 1150.875772 | 1268.53788 | 1012.704413 | 894.3517405 | 926.9752217 | 957.9506228 | 886.3152369 | 797.4047997 | 0.003031858 | 0.403010801 | YMR157C | AIM36 | n |
| W303.asm02.HP0_G0041960 | 1247.487727 | 1499.279852 | 1340.858222 | 1328.870537 | 1377.354758 | 1260.106942 | 1440.7123 | 1240.454294 | 1096.303646 | 0.783099184 | 0.043948964 | YMR158W | MRPS8 | n |
| W303.asm02.HP0_G0041970 | 177.3817048 | 135.1731676 | 164.186721 | 44.74966668 | 54.404186 | 34.39947112 | 114.5375745 | 49.34724572 | 47.0868046 | 1.54E-06 | 1.844863941 | YMR159C | ATG16 | n |

| | | | | | | | | | | | | | | |
|---|---|---|---|---|---|---|---|---|---|---|---|---|---|---|
| W303.asm02.HP0_G0041980 | 373.1800894 | 303.6636653 | 276.5764408 | 153.7053769 | 164.5394894 | 136.6926352 | 226.2353744 | 244.8010425 | 266.1428086 | 6.53E-08 | 1.071422142 | YMR160W | CVM1 | n |
| W303.asm02.HP0_G0041990 | 1562.509881 | 1446.924048 | 1454.225244 | 788.9560799 | 846.5822114 | 771.2723524 | 981.615411 | 961.7874951 | 939.6888396 | 1.88E-15 | 0.892534885 | YMR161W | HLJ1 | n |
| W303.asm02.HP0_G0042000 | 457.5090966 | 579.7215429 | 470.0822192 | 715.9946669 | 893.0248092 | 685.2736746 | 704.2640943 | 767.3012914 | 910.0036802 | 0.000837365 | -0.604017689 | YMR162C | DNF3 | n |
| W303.asm02.HP0_G0042010 | 200.6448792 | 216.0866835 | 201.3241937 | 336.5953189 | 403.3871352 | 334.0369695 | 373.9036522 | 378.3288839 | 468.8207936 | 9.15E-06 | -0.7939799 | YMR163C | INP2 | n |
| W303.asm02.HP0_G0042020 | 188.0439931 | 217.9905309 | 172.9824382 | 306.4379349 | 322.4443219 | 353.0472036 | 212.983093 | 238.0278911 | 320.3949965 | 0.000327845 | -0.762297282 | YMR164C | MSS11 | n |
| W303.asm02.HP0_G0042030 | 398.3818616 | 382.6733337 | 362.579009 | 914.4497104 | 1037.660328 | 902.5334922 | 851.9323721 | 1118.53757 | 956.0668586 | 5.40E-18 | -1.318497473 | YMR165C | PAH1 | n |
| W303.asm02.HP0_G0042040 | 640.706595 | 777.7216758 | 688.0205453 | 1447.554435 | 1524.644139 | 1477.36676 | 1572.288522 | 1522.991466 | 1642.92003 | 1.12E-18 | -1.078397137 | YMR166C | MME1 | n |
| W303.asm02.HP0_G0042050 | 448.7854062 | 623.5100338 | 472.036823 | 221.8026957 | 232.212989 | 243.5120455 | 139.1489541 | 229.3195537 | 211.8906207 | 4.07E-07 | 1.146568791 | YMR167W | MLH1 | n |
| W303.asm02.HP0_G0042060 | 469.1406838 | 513.0868828 | 510.1515975 | 475.7084132 | 510.8685758 | 484.3083434 | 404.194581 | 480.8937476 | 447.3246437 | 0.886814968 | 0.022044293 | YMR168C | CEP3 | n |
| W303.asm02.HP0_G0042080 | 515.6670326 | 330.3175294 | 499.4012765 | 305.465116 | 252.1169595 | 305.0689939 | 653.1481519 | 435.416874 | 558.8998981 | 0.005769698 | 0.638838434 | YMR170C | ALD2 | n |
| W303.asm02.HP0_G0042090 | 640.706595 | 743.452422 | 550.2209759 | 250.0144421 | 360.9253315 | 250.7540395 | 676.8129401 | 656.0280902 | 752.3652474 | 2.03E-09 | 1.172741219 | YMR171C | EAR1 | n |
| W303.asm02.HP0_G0042100 | 367.3642958 | 392.1925709 | 308.8274039 | 513.648348 | 565.2727618 | 493.3608358 | 584.0469706 | 763.4309192 | 759.5306307 | 0.001647288 | -0.555840892 | YMR172W | HOT1 | n |
| W303.asm02.HP0_G0042110 | 5525.003921 | 5022.349525 | 6067.090263 | 1107.067841 | 1321.62364 | 1130.656301 | 692.904996 | 667.6392069 | 513.8603459 | 1.25E-47 | 2.22378064 | YMR173W | DDR48 | n |
| W303.asm02.HP0_G0042150 | 365.4256979 | 584.4811615 | 397.7618778 | 636.2235219 | 723.1775943 | 659.0214467 | 452.4707487 | 685.0558818 | 554.8053933 | 0.009495187 | -0.5819671 | YMR176W | ECM5 | n |
| W303.asm02.HP0_G0042160 | 556.3775878 | 710.135092 | 586.3811466 | 1189.757442 | 634.2731928 | 1089.920085 | 1220.156475 | 1005.329183 | 1128.036058 | 0.004403875 | -0.654974948 | YMR177W | MMT1 | n |
| W303.asm02.HP0_G0042170 | 1249.426325 | 1362.202837 | 1246.059937 | 284.0631015 | 351.6368119 | 308.6899908 | 535.7708029 | 477.0233753 | 573.2306647 | 2.99E-44 | 2.034240868 | YMR178W | FPY1 | n |
| W303.asm02.HP0_G0042180 | 326.6537406 | 432.173367 | 317.6231211 | 1037.997703 | 988.5638675 | 1061.857358 | 960.7903974 | 1084.671813 | 1082.996506 | 2.75E-22 | -1.520714135 | YMR179W | SPT21 | n |
| W303.asm02.HP0_G0042230 | 854.9216593 | 930.0294703 | 860.0256817 | 212.0745073 | 272.02093 | 218.1650668 | 613.3913079 | 479.9261545 | 440.1592604 | 1.21E-26 | 1.91836521 | YMR184W | ADD37 | n |
| W303.asm02.HP0_G0042240 | 787.0707339 | 1183.241179 | 831.6839263 | 1793.877943 | 1394.604865 | 1835.845459 | 914.4074127 | 1174.657967 | 890.5547827 | 3.33E-05 | -0.843051329 | YMR185W | RTP1 | n |
| W303.asm02.HP0_G0042250 | 29541.32359 | 19522.05156 | 25299.41457 | 9934.426003 | 10013.02409 | 9537.705992 | 12798.864 | 10939.60706 | 10839.17769 | 7.61E-17 | 1.334611718 | YMR186W | HSC82 | y |
| W303.asm02.HP0_G0042260 | 550.5617942 | 537.8368994 | 541.4252587 | 261.6882682 | 287.9441064 | 307.7847416 | 364.437737 | 371.5557325 | 396.1433344 | 1.46E-10 | 0.926314711 | YMR187C | | n |
| W303.asm02.HP0_G0042270 | 2106.286582 | 1986.664795 | 2317.182831 | 2654.822617 | 2563.631399 | 2677.727252 | 2487.642527 | 2196.436231 | 2185.441909 | 0.011583901 | -0.301046804 | YMR188C | MRPS17 | n |
| W303.asm02.HP0_G0042280 | 241.3554344 | 239.8847764 | 233.5751567 | 3524.52266 | 916.9095737 | 2689.495492 | 949.4312991 | 1213.361689 | 1493.470607 | 4.33E-12 | -3.318733526 | YMR189W | GCV2 | n |
| W303.asm02.HP0_G0042290 | 275.2808971 | 408.3752741 | 251.1665911 | 485.4366016 | 447.1758703 | 465.2981093 | 423.1264115 | 473.1530031 | 429.9229985 | 0.004688804 | -0.580652647 | YMR190C | SGS1 | n |
| W303.asm02.HP0_G0042320 | 1760.246863 | 1404.087481 | 1712.232948 | 2380.487704 | 2421.649743 | 2405.24723 | 2429.900444 | 2209.014941 | 1895.755698 | 9.42E-05 | -0.563629033 | YMR193W | MRPL24 | n |
| W303.asm02.HP0_G0042360 | 525.3600219 | 495.0003322 | 473.0141249 | 213.0473261 | 218.9436754 | 214.5440699 | 261.2592608 | 236.092705 | 344.962025 | 6.58E-11 | 1.208302852 | YMR196W | | n |
| W303.asm02.HP0_G0042370 | 1462.672091 | 1280.337398 | 1433.701903 | 842.4611162 | 814.7358586 | 802.9560758 | 1317.655402 | 1040.162532 | 1132.130563 | 2.59E-08 | 0.763538335 | YMR197C | VTI1 | n |
| W303.asm02.HP0_G0042400 | 2918.559089 | 2482.617051 | 2922.132714 | 1773.448747 | 1721.029981 | 1675.616343 | 2091.967269 | 2058.070425 | 1937.724372 | 3.52E-09 | 0.686965989 | YMR200W | ROT1 | n |
| W303.asm02.HP0_G0042420 | 5508.525839 | 5370.753605 | 5053.628182 | 2683.034363 | 2608.747065 | 2576.339337 | 3069.796314 | 2763.445761 | 2322.607818 | 3.51E-15 | 1.017912744 | YMR202W | ERG2 | n |
| W303.asm02.HP0_G0042430 | 5335.02133 | 5235.580437 | 5517.846589 | 5196.798248 | 5192.282434 | 5086.595479 | 3890.491166 | 4829.25693 | 4521.356868 | 0.654593748 | 0.056049855 | YMR203W | TOM40 | n |
| W303.asm02.HP0_G0042440 | 187.0746942 | 111.3750747 | 152.4590981 | 77.82550727 | 116.7699602 | 73.32518843 | 125.8966728 | 162.555633 | 142.28404 | 0.00979103 | 0.76142589 | YMR204C | INP1 | n |
| W303.asm02.HP0_G0042450 | 4202.880175 | 4752.955113 | 3926.799078 | 15052.42592 | 10586.25844 | 14152.66662 | 4986.644151 | 6944.415344 | 6902.311379 | 2.45E-19 | -1.627135829 | YMR205C | PFK2 | n |
| W303.asm02.HP0_G0042460 | 101.776388 | 77.10582098 | 81.11605861 | 38.91275363 | 53.07725463 | 41.64146504 | 49.22275928 | 40.63890824 | 63.46482359 | 0.003299955 | 0.969694072 | YMR206W | | n |
| W303.asm02.HP0_G0042480 | 1847.483767 | 1511.654861 | 1860.782839 | 2362.004146 | 2073.993725 | 2331.922042 | 1830.708008 | 1854.875883 | 1601.974983 | 0.00715018 | -0.37525234 | YMR208W | ERG12 | n |
| W303.asm02.HP0_G0042490 | 1010.009489 | 968.106419 | 1020.303195 | 604.1205002 | 650.1963692 | 624.6219755 | 724.1425163 | 859.2226314 | 732.9163849 | 1.94E-07 | 0.67511879 | YMR209C | | n |
| W303.asm02.HP0_G0042500 | 964.4524388 | 936.6929363 | 848.2980587 | 905.6943408 | 992.5446616 | 924.259474 | 744.0209383 | 863.0930037 | 724.7273404 | 0.799706101 | -0.03693662 | YMR210W | MGL2 | n |
| W303.asm02.HP0_G0042510 | 920.8339868 | 1089.000731 | 875.6625123 | 792.8473553 | 904.9671915 | 824.6820576 | 677.7595316 | 738.2734998 | 631.5773574 | 0.196125485 | 0.19517581 | YMR211W | DML1 | n |
| W303.asm02.HP0_G0042520 | 1743.768781 | 2059.792921 | 2042.560994 | 1182.94771 | 1264.565592 | 1323.474389 | 1159.574618 | 1226.907992 | 1189.453629 | 2.54E-07 | 0.632538388 | YMR212C | EFR3 | n |
| W303.asm02.HP0_G0042530 | 175.443107 | 198.9520566 | 152.4590981 | 442.6325726 | 400.7332725 | 408.2674072 | 320.8945268 | 375.4261047 | 332.6785108 | 6.65E-13 | -1.248648152 | YMR213W | CEF1 | n |
| W303.asm02.HP0_G0042540 | 697.8952321 | 594.9523224 | 637.200846 | 595.3651306 | 615.6961537 | 573.9280181 | 1128.337097 | 889.2180161 | 1061.500356 | 0.471309221 | 0.113440717 | YMR214W | SCJ1 | n |
| W303.asm02.HP0_G0042550 | 2592.874647 | 2028.549438 | 2571.281328 | 3939.916305 | 3947.620813 | 4028.359118 | 2279.392391 | 3472.691469 | 2861.035193 | 2.69E-05 | -0.728331273 | YMR215W | GAS3 | n |
| W303.asm02.HP0_G0042560 | 701.7724278 | 1195.616187 | 695.8389606 | 1010.758776 | 1159.738014 | 1091.730583 | 817.8550772 | 1247.227446 | 1326.619538 | 0.193723284 | -0.330745305 | YMR216C | SKY1 | n |
| W303.asm02.HP0_G0042570 | 3373.160288 | 3442.156156 | 3139.093738 | 13030.90837 | 13175.10153 | 12360.27312 | 6912.011312 | 8521.592022 | 6899.2405 | 1.23E-57 | -1.953876257 | YMR217W | GUA1 | n |
| W303.asm02.HP0_G0042580 | 674.6320577 | 907.1833012 | 743.7267543 | 649.8429857 | 666.1195456 | 678.0316807 | 469.5093962 | 639.5790083 | 611.1048336 | 0.224126269 | 0.221983065 | YMR218C | TRS130 | n |
| W303.asm02.HP0_G0042600 | 2602.567636 | 1933.357067 | 2436.413664 | 1191.70308 | 1232.719239 | 1254.675447 | 1165.254167 | 1209.491317 | 1040.004206 | 2.61E-10 | 0.922249037 | YMR220W | ERG8 | n |
| W303.asm02.HP0_G0042610 | 2075.269016 | 1955.251312 | 2212.611526 | 874.5641379 | 781.5625744 | 842.7870424 | 942.8051585 | 916.3106216 | 884.4130255 | 5.72E-30 | 1.319622311 | YMR221C | FMP42 | n |
| W303.asm02.HP0_G0042620 | 705.6496235 | 519.7503488 | 618.6321097 | 1209.213819 | 1182.295847 | 1177.729261 | 1043.14386 | 908.5698772 | 851.6569876 | 6.76E-10 | -0.952918248 | YMR222C | FSH2 | n |
| W303.asm02.HP0_G0042630 | 864.6146486 | 1050.923782 | 843.4115492 | 510.7298914 | 492.2915367 | 488.8345896 | 367.2775115 | 460.5742934 | 453.4664008 | 2.04E-08 | 0.88698403 | YMR223W | UBP8 | n |
| W303.asm02.HP0_G0042640 | 354.7634096 | 326.5098345 | 298.0770829 | 506.8386161 | 522.8109581 | 452.62462 | 321.8411183 | 401.5511172 | 409.4504748 | 0.00043461 | -0.596523748 | YMR224C | MRE11 | n |
| W303.asm02.HP0_G0042650 | 765.7461574 | 1020.462223 | 819.9563033 | 703.3480219 | 759.0047412 | 721.4836442 | 781.8845993 | 613.4539959 | 554.8053933 | 0.167101514 | 0.255623313 | YMR225C | MRPL44 | n |
| W303.asm02.HP0_G0042660 | 4804.814813 | 4115.166223 | 5015.513407 | 814.2493698 | 356.9445374 | 828.3030545 | 764.8459518 | 528.3058072 | 575.2779171 | 2.32E-25 | 2.798513594 | YMR226C | ORA1 | n |
| W303.asm02.HP0_G0042670 | 215.1843632 | 327.4617582 | 212.0745147 | 945.5799133 | 967.3329656 | 955.0379481 | 1080.06093 | 1005.329183 | 976.5393824 | 2.14E-26 | -1.92539388 | YMR227C | TAF7 | n |
| W303.asm02.HP0_G0042680 | 710.4961182 | 828.1736327 | 763.2727925 | 800.629906 | 695.3120357 | 817.4400637 | 502.6400995 | 542.819703 | 417.6394843 | 0.95885471 | -0.008550642 | YMR228W | MTF1 | n |
| W303.asm02.HP0_G0042690 | 1566.387076 | 2158.011064 | 1566.614963 | 14924.01384 | 14596.24502 | 15503.29848 | 4370.413069 | 6065.840852 | 5336.163313 | 4.33E-75 | -3.089018771 | YMR229C | RRP5 | n |
| W303.asm02.HP0_G0042720 | 529.2372177 | 532.1253571 | 591.2676562 | 307.4107537 | 374.1946451 | 333.1317203 | 356.8650048 | 397.680745 | 495.4350745 | 6.83E-05 | 0.706059231 | YMR231W | PEP5 | n |
| W303.asm02.HP0_G0042730 | 130.855356 | 98.04814273 | 151.4817962 | 29.18456523 | 35.82714688 | 24.44172948 | 36.91706946 | 43.5416874 | 36.85054273 | 2.37E-10 | 2.102504357 | YMR232W | FUS2 | n |
| W303.asm02.HP0_G0042740 | 599.9960398 | 653.019669 | 567.8124103 | 438.7412972 | 421.9641743 | 420.0356473 | 425.0195945 | 308.662184 | 291.7334633 | 0.005032573 | 0.507536401 | YMR233W | TRI1 | n |
| W303.asm02.HP0_G0042750 | 887.877823 | 898.6159877 | 887.3901352 | 218.8842392 | 192.405048 | 215.4493191 | 211.08991 | 177.0695288 | 197.5598541 | 2.09E-45 | 2.089168997 | YMR234W | RNH1 | n |
| W303.asm02.HP0_G0042760 | 4011.928285 | 3646.819755 | 4165.260745 | 4141.289805 | 4254.141959 | 4046.464102 | 3400.156756 | 3420.441444 | 3115.918113 | 0.520893202 | -0.073354945 | YMR235C | RNA1 | n |
| W303.asm02.HP0_G0042770 | 1874.624137 | 1401.23171 | 1880.328877 | 731.5597683 | 781.5625744 | 726.0098904 | 992.9745093 | 782.7827802 | 734.9636022 | 3.09E-12 | 1.203873257 | YMR236W | TAF9 | n |
| W303.asm02.HP0_G0042790 | 1396.759763 | 1432.645192 | 1463.998263 | 1824.035327 | 1616.202404 | 1792.393495 | 1607.312409 | 1796.8203 | 1731.975508 | 0.012964687 | -0.286291244 | YMR238W | DFG5 | n |
| W303.asm02.HP0_G0042800 | 1188.360492 | 1870.530102 | 1296.879636 | 1716.052435 | 1792.684275 | 1695.531826 | 1752.140912 | 1766.824916 | 1589.691468 | 0.163112275 | -0.256504662 | YMR239C | RNT1 | n |
| W303.asm02.HP0_G0042810 | 531.1758155 | 353.1636986 | 469.1049173 | 361.8886088 | 351.6368119 | 399.2149148 | 378.6366098 | 356.0742437 | 383.8598201 | 0.142796613 | 0.281058523 | YMR240C | CUS1 | n |

| | | | | | | | | | | | | | | |
|---|---|---|---|---|---|---|---|---|---|---|---|---|---|---|
| W303.asm02.HP0_G0042820 | 3062.015331 | 3238.444481 | 3335.531422 | 1517.597392 | 989.8907989 | 1494.566495 | 1759.713644 | 1215.296875 | 1560.006309 | 7.03E-11 | 1.266333222 | YMR241W | YHM2 | n |
| W303.asm02.HP0_G0042840 | 4230.989844 | 4427.397202 | 4390.040184 | 2376.596428 | 2016.935676 | 2353.648024 | 1946.192174 | 2226.431616 | 1970.48041 | 1.24E-15 | 0.950672303 | YMR243C | ZRC1 | n |
| W303.asm02.HP0_G0042850 | 34.8947616 | 48.54810951 | 18.56873631 | 60.31476813 | 63.69270556 | 59.74644984 | 25.55797116 | 51.28243183 | 47.0868046 | 0.05768518 | -0.846894908 | YMR244W | | n |
| W303.asm02.HP0_G0042860 | 880.1234316 | 836.7409462 | 814.0924919 | 890.1292394 | 749.7162217 | 831.9240515 | 855.7187382 | 693.7642193 | 647.9553764 | 0.842651113 | 0.032971611 | YMR244C- | COA6 | n |
| W303.asm02.HP0_G0042880 | 1051.689343 | 1199.423882 | 1116.078782 | 2497.225964 | 2546.381291 | 2548.27661 | 1424.620244 | 1923.57499 | 1518.037635 | 3.07E-16 | -1.172852032 | YMR247C | RKR1 | n |
| W303.asm02.HP0_G0042890 | 490.4652603 | 257.971327 | 393.8526701 | 311.3020291 | 118.0968916 | 232.6490547 | 171.3330659 | 193.5186107 | 296.8515942 | 0.030066325 | 0.782641512 | YMR250W | GAD1 | n |
| W303.asm02.HP0_G0042920 | 254.9256195 | 245.5963187 | 291.2359695 | 445.5510291 | 468.4067721 | 442.6668783 | 496.9605504 | 492.5048642 | 553.7817672 | 2.91E-07 | -0.776135662 | YMR252C | MLO1 | n |
| W303.asm02.HP0_G0042950 | 1425.838731 | 1503.087547 | 1399.496337 | 3966.182414 | 4019.275107 | 3804.762555 | 2834.095025 | 3028.566257 | 2901.98024 | 2.94E-49 | -1.445440544 | YMR256C | COX7 | n |
| W303.asm02.HP0_G0042960 | 417.7678403 | 726.3177952 | 550.2209759 | 879.4282321 | 916.9095737 | 880.8075104 | 628.5367723 | 793.4263038 | 586.5378051 | 0.001972092 | -0.65966735 | YMR257C | PET111 | n |
| W303.asm02.HP0_G0042970 | 1038.119158 | 902.4236826 | 1137.579424 | 825.9231959 | 828.0051722 | 799.3350789 | 696.6913621 | 627.9678917 | 651.0262549 | 0.018264664 | 0.327508398 | YMR258C | ROY1 | n |
| W303.asm02.HP0_G0042980 | 985.7770153 | 1478.337531 | 981.2111186 | 1255.909124 | 1449.009051 | 1319.853392 | 858.5585128 | 1095.315336 | 1030.79157 | 0.262348126 | -0.223738247 | YMR259C | TRM732 | n |
| W303.asm02.HP0_G0042990 | 5481.385469 | 6527.340919 | 6013.338658 | 5953.651306 | 6391.828389 | 6015.381199 | 5782.727623 | 4988.909783 | 4630.88487 | 0.849088072 | -0.026742352 | YMR260C | TIF11 | n |
| W303.asm02.HP0_G0043000 | 1014.855983 | 930.0294703 | 908.8907772 | 911.5312539 | 822.6974468 | 821.9663099 | 1070.595014 | 1126.278314 | 1194.57176 | 0.221919834 | 0.158368931 | YMR261C | TPS3 | n |
| W303.asm02.HP0_G0043010 | 426.4915307 | 229.4136155 | 339.1237631 | 265.5795435 | 254.7708222 | 256.1855349 | 253.6865286 | 235.125112 | 270.2373134 | 0.135963444 | 0.357225013 | YMR262W | | n |
| W303.asm02.HP0_G0043020 | 1073.013919 | 983.3371984 | 1048.64495 | 176.0802102 | 205.6743617 | 171.0921063 | 416.5002708 | 213.8380648 | 397.1669605 | 4.64E-26 | 2.492492997 | YMR263W | SAP30 | n |
| W303.asm02.HP0_G0043030 | 1487.873863 | 1447.875972 | 1544.137019 | 1350.272551 | 1354.796924 | 1421.241307 | 2539.70506 | 1558.792409 | 1668.510685 | 0.553073097 | 0.118487923 | YMR264W | CUE1 | n |
| W303.asm02.HP0_G0043040 | 583.5179579 | 594.0003987 | 525.7884281 | 113.8198044 | 134.0200679 | 131.2611398 | 54.90230842 | 149.0093302 | 228.2686397 | 2.80E-09 | 2.168438666 | YMR265C | | n |
| W303.asm02.HP0_G0043050 | 1105.970083 | 1762.010798 | 1222.604691 | 14761.55309 | 12750.48349 | 14245.00204 | 5010.308939 | 6783.794898 | 7175.619571 | 4.61E-65 | -3.351604251 | YMR266W | RSN1 | n |
| W303.asm02.HP0_G0043060 | 1861.053952 | 1887.664728 | 2138.336581 | 634.2778842 | 516.1763013 | 659.0214467 | 888.8494416 | 829.2272468 | 886.4602779 | 2.61E-34 | 1.69877649 | YMR267W | PPA2 | n |
| W303.asm02.HP0_G0043070 | 433.2766232 | 588.2888564 | 431.9674447 | 293.7912899 | 276.0017241 | 270.6695227 | 248.953571 | 284.4723577 | 320.3949965 | 3.78E-05 | 0.790314638 | YMR268C | PRP24 | n |
| W303.asm02.HP0_G0043080 | 875.2769369 | 990.9525881 | 1033.985422 | 1463.119537 | 1272.52718 | 1513.576729 | 791.3505145 | 631.8382639 | 573.2306647 | 0.000724345 | -0.551813983 | YMR269W | TMA23 | n |
| W303.asm02.HP0_G0043090 | 1264.935108 | 1274.625855 | 1219.672785 | 215.9657827 | 290.5979691 | 255.2802857 | 310.4820201 | 314.4677424 | 359.2927916 | 3.65E-51 | 2.307992444 | YMR270C | RRN9 | n |
| W303.asm02.HP0_G0043110 | 1611.944126 | 1148.971925 | 1492.340018 | 2632.447783 | 2253.129459 | 2727.51596 | 1865.731895 | 2243.848291 | 2009.378205 | 2.45E-07 | -0.840452768 | YMR272C | SCS7 | n |
| W303.asm02.HP0_G0043120 | 130.855356 | 209.4232175 | 96.75288919 | 802.5755437 | 853.2168682 | 775.7985986 | 620.9640401 | 846.6439218 | 904.8855493 | 2.32E-22 | -2.47507978 | YMR273C | ZDS1 | n |
| W303.asm02.HP0_G0043130 | 632.9522035 | 522.60612 | 698.7708664 | 323.948674 | 307.8480769 | 290.585006 | 510.2128317 | 396.7131519 | 455.5136532 | 1.00E-08 | 1.007620587 | YMR274C | RCE1 | n |
| W303.asm02.HP0_G0043150 | 1958.953144 | 1337.452821 | 1583.229096 | 2254.994073 | 1844.434598 | 2188.892662 | 3254.381661 | 2637.658664 | 2799.617621 | 0.037464431 | -0.366458063 | YMR276W | DSK2 | n |
| W303.asm02.HP0_G0043160 | 541.8381038 | 489.2887899 | 432.9447466 | 841.4882973 | 874.44777 | 810.1980697 | 636.1095045 | 741.1762789 | 786.1449116 | 1.80E-07 | -0.786282243 | YMR277W | FCP1 | n |
| W303.asm02.HP0_G0043170 | 488.5266624 | 407.4233504 | 448.5815771 | 730.5869495 | 508.2147131 | 764.0303585 | 235.7012896 | 548.6252613 | 766.696014 | 0.092052195 | -0.575863722 | YMR278W | PRM15 | n |
| W303.asm02.HP0_G0043190 | 184.1667974 | 201.8078277 | 154.4137019 | 560.3436523 | 457.7913212 | 522.3288114 | 687.2254468 | 1398.171962 | 1264.178341 | 3.07E-09 | -1.511962948 | YMR280C | CAT8 | n |
| W303.asm02.HP0_G0043200 | 511.7898368 | 455.0195362 | 553.1528816 | 452.360761 | 431.2526939 | 530.4760546 | 708.9970519 | 751.8198025 | 517.9548506 | 0.613854021 | 0.102457625 | YMR281W | GPI12 | n |
| W303.asm02.HP0_G0043210 | 962.5138409 | 1029.029537 | 960.6877785 | 948.4983698 | 1048.275779 | 965.900939 | 881.2767094 | 887.28283 | 844.4916043 | 0.975375674 | -0.00382422 | YMR282C | AEP2 | n |
| W303.asm02.HP0_G0043220 | 540.8688049 | 632.0773473 | 582.471939 | 708.2121161 | 707.254418 | 668.9791883 | 693.8515875 | 617.3243681 | 600.8685718 | 0.078298308 | -0.247275882 | YMR283C | RIT1 | n |
| W303.asm02.HP0_G0043230 | 713.404015 | 716.798558 | 715.3849988 | 424.1490146 | 428.5988311 | 439.0458814 | 409.8741301 | 380.26407 | 386.9306987 | 6.75E-09 | 0.731710063 | YMR284W | YKU70 | n |
| W303.asm02.HP0_G0043240 | 724.0663033 | 654.9235165 | 621.5640154 | 511.7027103 | 508.2147131 | 499.6975804 | 447.7377911 | 560.2363779 | 437.0883818 | 0.01143368 | 0.396823575 | YMR285C | NGL2 | n |
| W303.asm02.HP0_G0043250 | 897.5708124 | 1186.09695 | 887.3901352 | 584.6641233 | 656.8310261 | 582.0752613 | 677.7595316 | 485.7317128 | 464.7262889 | 0.000314239 | 0.705233617 | YMR286W | MRPL33 | n |
| W303.asm02.HP0_G0043260 | 345.0704203 | 496.9041796 | 375.2839338 | 2120.745073 | 1869.646294 | 1997.885073 | 733.6084315 | 1261.741342 | 1024.649813 | 2.20E-27 | -2.298516941 | YMR287C | DSS1 | y |
| W303.asm02.HP0_G0043280 | 379.9651819 | 654.9235165 | 437.8312561 | 746.1520509 | 749.7162217 | 758.5988631 | 454.3639318 | 691.8290332 | 697.0894533 | 0.009826799 | -0.614155343 | YMR288W | HSH155 | n |
| W303.asm02.HP0_G0043290 | 1484.965966 | 1536.404877 | 1610.593549 | 298.6553841 | 342.3482924 | 313.216237 | 441.1116504 | 379.2964769 | 453.4664008 | 2.24E-62 | 2.282051684 | YMR289W | ABZ2 | n |
| W303.asm02.HP0_G0043300 | 1638.115197 | 2049.49176 | 1685.845796 | 8112.336314 | 8350.379085 | 8347.303241 | 5138.098795 | 5161.141347 | 4944.114483 | 1.16E-77 | -2.206905023 | YMR290C | HAS1 | n |
| W303.asm02.HP0_G0043310 | 237.4782387 | 224.6539969 | 198.3922879 | 372.589616 | 548.0226541 | 424.5618935 | 759.1664027 | 624.0975195 | 1004.177289 | 5.67E-06 | -1.023435016 | YMR291W | TDA1 | n |
| W303.asm02.HP0_G0043330 | 536.0223102 | 395.0483421 | 453.4680867 | 1004.921863 | 1008.467838 | 1011.163401 | 994.8676923 | 1085.639406 | 1016.460804 | 8.67E-15 | -1.127482758 | YMR293C | HER2 | n |
| W303.asm02.HP0_G0043340 | 858.798855 | 835.7890225 | 888.3674371 | 224.7211522 | 246.809234 | 240.7962978 | 288.710415 | 279.6343924 | 295.827968 | 3.43E-42 | 1.860063153 | YMR294W | JNM1 | n |
| W303.asm02.HP0_G0043350 | 3813.222004 | 3846.723736 | 4004.983231 | 1169.328247 | 1230.065376 | 1222.991723 | 1520.225988 | 1403.009927 | 1598.904104 | 1.05E-57 | 1.687434757 | YMR295C | GSR1 | n |
| W303.asm02.HP0_G0043360 | 1216.470161 | 1074.721875 | 999.7798549 | 4398.113979 | 4242.199576 | 4417.616291 | 5112.540824 | 5501.734102 | 5444.667689 | 1.56E-68 | -1.988449505 | YMR296C | LCB1 | n |
| W303.asm02.HP0_G0043370 | 6781.215338 | 4362.666389 | 6752.178903 | 3115.938747 | 2456.149958 | 3079.657914 | 3037.612202 | 3390.446059 | 2519.144046 | 1.35E-07 | 1.048212379 | YMR297W | PRC1 | n |
| W303.asm02.HP0_G0043380 | 1483.996667 | 1302.231643 | 1456.179847 | 1020.486964 | 971.3137597 | 1028.363137 | 1795.684122 | 1267.5469 | 1246.776696 | 0.003951162 | 0.489888291 | YMR298W | LIP1 | n |
| W303.asm02.HP0_G0043390 | 806.4567126 | 831.9813276 | 797.4783594 | 146.895645 | 138.000862 | 152.0818723 | 18.93183049 | 142.2361789 | 176.0637042 | 0.000332527 | 2.478613328 | YMR299C | DYN3 | n |
| W303.asm02.HP0_G0043400 | 1087.553403 | 1169.914247 | 1194.262935 | 3737.569986 | 2166.87892 | 3479.778078 | 1816.509136 | 1766.824916 | 1794.416706 | 3.99E-14 | -1.44363231 | YMR300C | ADE4 | n |
| W303.asm02.HP0_G0043410 | 597.088143 | 663.4908299 | 537.516051 | 2043.892385 | 1890.877196 | 1975.253842 | 1415.154329 | 1546.213699 | 1333.784922 | 1.93E-39 | -1.716827659 | YMR301C | ATM1 | n |
| W303.asm02.HP0_G0043420 | 988.6849121 | 1015.702605 | 922.573004 | 1289.957783 | 1264.565592 | 1300.843158 | 1346.053148 | 1534.602583 | 1608.11674 | 0.001325757 | -0.397646193 | YMR302C | YME2 | n |
| W303.asm02.HP0_G0043440 | 608.7197302 | 650.1638979 | 614.722902 | 308.3835725 | 364.9061256 | 358.478699 | 396.6218488 | 529.2734002 | 493.3878221 | 1.72E-07 | 0.862041212 | YMR304W | UBP15 | n |
| W303.asm02.HP0_G0043460 | 215.1843632 | 264.634793 | 212.0745147 | 156.6238334 | 169.8472148 | 159.3238662 | 147.6682778 | 218.6760301 | 156.6148066 | 0.024633516 | 0.511920134 | YMR306W | FKS3 | n |
| W303.asm02.HP0_G0043470 | 15369.20389 | 15976.13572 | 17206.37744 | 12962.81105 | 11983.51716 | 12069.68812 | 10386.00221 | 13744.65932 | 13108.55695 | 0.004051598 | 0.391343582 | YMR307W | GAS1 | n |
| W303.asm02.HP0_G0043480 | 2043.282152 | 2217.030334 | 2065.038938 | 6176.42682 | 6321.501027 | 6099.569379 | 4195.293637 | 5513.345218 | 5092.54028 | 9.83E-34 | -1.555815308 | YMR308C | PSE1 | n |
| W303.asm02.HP0_G0043490 | 5143.100141 | 6986.16815 | 5630.236309 | 12725.44326 | 12372.30805 | 12453.51379 | 6316.605243 | 9142.786762 | 8615.861616 | 2.18E-10 | -1.080262875 | YMR309C | NIP1 | n |
| W303.asm02.HP0_G0043500 | 476.8950752 | 550.2119077 | 492.5601631 | 1794.850761 | 1685.202835 | 1683.763586 | 1585.540804 | 1469.773848 | 1545.675542 | 1.50E-50 | -1.764704531 | YMR310C | UPA2 | n |
| W303.asm02.HP0_G0043510 | 2164.444518 | 1652.539571 | 2159.837223 | 903.7487031 | 719.1968002 | 846.4080393 | 1031.784762 | 963.7226812 | 935.5943349 | 4.01E-16 | 1.27364523 | YMR311C | GLC8 | n |
| W303.asm02.HP0_G0043520 | 1005.162994 | 1104.23151 | 1112.169575 | 1300.65879 | 1231.392307 | 1303.558905 | 1191.758729 | 1120.472756 | 1033.862449 | 0.039410874 | -0.252234248 | YMR312W | ELP6 | n |
| W303.asm02.HP0_G0043530 | 619.3820185 | 631.1254236 | 601.0406753 | 147.8684638 | 135.3469993 | 137.5978845 | 226.2353744 | 262.2177175 | 316.3004918 | 5.90E-33 | 2.136087515 | YMR313C | TGL3 | n |
| W303.asm02.HP0_G0043540 | 3100.787288 | 2413.12662 | 2836.130146 | 2000.115537 | 1853.723118 | 1981.590586 | 3554.451175 | 2661.84849 | 2920.405511 | 0.000835466 | 0.516645591 | YMR314W | PRE5 | n |
| W303.asm02.HP0_G0043550 | 1965.738237 | 1441.212506 | 1894.988405 | 1056.481261 | 683.3696534 | 1012.06865 | 910.6210466 | 784.7179663 | 831.1844638 | 1.51E-06 | 0.944140783 | YMR315W | | n |
| W303.asm02.HP0_G0043580 | 6089.1359 | 5589.696059 | 6370.053856 | 4251.218334 | 3769.81201 | 3079.657914 | 3268.580534 | 3323.682139 | 3341.115874 | 6.16E-07 | 0.701422948 | YMR318C | ADH6 | n |
| W303.asm02.HP0_G0043590 | 1690.45734 | 1701.08768 | 1814.849649 | 1725.780624 | 1340.200679 | 1633.069629 | 3307.390787 | 1623.621144 | 3843.716332 | 0.596200698 | 0.147548388 | YMR319C | FET4 | n |

| | | | | | | | | | | | | | | |
|---|---|---|---|---|---|---|---|---|---|---|---|---|---|---|
| W303.asm02.HP0_G0043660 | 521.4828262 | 533.0772808 | 571.7216179 | 4511.933784 | 2249.148665 | 3945.981437 | 2350.386755 | 2415.112261 | 2666.546217 | 6.26E-35 | -2.719490048 | YNL334C | SNO2 | y |
| W303.asm02.HP0_G0043670 | 796.7637233 | 769.1543624 | 825.8201148 | 30604.88073 | 18667.27045 | 27322.23256 | 13783.31919 | 17613.09635 | 18939.13171 | 3.93E-151 | -5.001201735 | YNL333W | SNZ2 | y |
| W303.asm02.HP0_G0043700 | 1254.27282 | 1024.269918 | 1178.626105 | 1884.350095 | 1959.877627 | 1901.023404 | 1805.150037 | 1961.311119 | 2156.780365 | 2.01E-08 | -0.732663882 | YNL330C | RPD3 | n |
| W303.asm02.HP0_G0043710 | 369.3028936 | 484.5291713 | 345.9648765 | 279.1990073 | 311.828871 | 251.6592887 | 302.9092879 | 387.0372214 | 462.6790365 | 0.021102035 | 0.512257883 | YNL329C | PEX6 | n |
| W303.asm02.HP0_G0043720 | 41.67985414 | 74.25004983 | 42.02398217 | 248.0688044 | 240.1745772 | 238.9857993 | 361.5979624 | 221.5788092 | 235.434023 | 3.75E-14 | -2.201773762 | YNL328C | MDJ2 | n |
| W303.asm02.HP0_G0043730 | 3227.765448 | 3622.069739 | 2660.215802 | 3887.384088 | 3917.101392 | 4103.494805 | 4094.954935 | 5849.100008 | 5099.705664 | 0.058444702 | -0.324417797 | YNL327W | EGT2 | n |
| W303.asm02.HP0_G0043740 | 755.0838691 | 707.2793208 | 711.4757912 | 256.824174 | 287.9441064 | 229.0280577 | 479.9219029 | 354.1390576 | 362.3636702 | 2.32E-17 | 1.494117006 | YNL326C | PFA3 | n |
| W303.asm02.HP0_G0043750 | 594.1802462 | 597.8080935 | 649.9057708 | 267.5251812 | 306.5211455 | 310.5004893 | 327.5206675 | 379.2964769 | 390.0015772 | 4.64E-12 | 1.059060736 | YNL325C | FIG4 | n |
| W303.asm02.HP0_G0043770 | 1088.522702 | 1089.952655 | 1188.399124 | 941.6886379 | 975.2945538 | 879.9022612 | 1290.204248 | 1205.620945 | 1379.8481 | 0.031122466 | 0.268547105 | YNL323W | LEM3 | n |
| W303.asm02.HP0_G0043780 | 894.6629156 | 671.1062196 | 721.2488103 | 2623.692414 | 2320.802959 | 2490.340659 | 2525.506188 | 2816.663379 | 2752.530817 | 2.60E-33 | -1.701230088 | YNL322C | KRE1 | n |
| W303.asm02.HP0_G0043790 | 495.311755 | 434.0772144 | 372.3520281 | 824.950377 | 907.6210542 | 833.73455 | 918.1937788 | 1325.602483 | 1090.161895 | 5.06E-08 | -0.978592557 | YNL321W | VNX1 | n |
| W303.asm02.HP0_G0043800 | 573.8249686 | 521.6541963 | 595.1768638 | 543.805732 | 542.7149286 | 567.5912734 | 309.5354285 | 704.4077434 | 435.0411295 | 0.909704543 | 0.031345717 | YNL320W | | n |
| W303.asm02.HP0_G0043810 | 16.47808187 | 21.89424546 | 13.68222675 | 143.9771884 | 163.212558 | 137.5978845 | 148.6148694 | 182.8750871 | 142.28404 | 1.26E-23 | -3.090030299 | YNL318C | HXT14 | n |
| W303.asm02.HP0_G0043820 | 311.1449576 | 277.961725 | 236.5070625 | 720.8587611 | 729.8122512 | 672.6001853 | 656.934518 | 847.6115148 | 667.4042739 | 1.70E-15 | -1.36213872 | YNL317W | PFS2 | n |
| W303.asm02.HP0_G0043830 | 721.1584065 | 578.7696192 | 758.3862829 | 723.7772176 | 571.9074186 | 742.3043767 | 824.4812179 | 810.8429788 | 893.6256612 | 0.947683276 | 0.011896568 | YNL316C | PHA2 | n |
| W303.asm02.HP0_G0043840 | 1754.43107 | 1818.174297 | 1864.692046 | 2309.471928 | 2152.282675 | 2378.089753 | 2437.473176 | 2328.028887 | 2157.804002 | 0.00276361 | -0.331663051 | YNL315C | ATP11 | n |
| W303.asm02.HP0_G0043850 | 223.9080536 | 183.7212771 | 249.2119873 | 108.9557102 | 149.9432443 | 114.0614042 | 240.4342472 | 170.2963774 | 192.4417232 | 0.000658775 | 0.823499295 | YNL314W | DAL82 | n |
| W303.asm02.HP0_G0043860 | 852.9830614 | 1168.962323 | 867.844097 | 3621.804544 | 3630.484217 | 3499.693562 | 2092.913861 | 2498.325264 | 2369.694623 | 1.14E-35 | -1.895377698 | YNL313C | EMW1 | n |
| W303.asm02.HP0_G0043870 | 2247.804227 | 1721.078078 | 2103.153712 | 1255.909124 | 1080.122132 | 1199.455243 | 1721.849983 | 1435.908091 | 1247.800322 | 2.96E-06 | 0.779585601 | YNL312W | RFA2 | n |
| W303.asm02.HP0_G0043880 | 853.9523603 | 885.2890557 | 835.5931339 | 181.9171232 | 226.9052635 | 184.6708449 | 152.4012355 | 299.9538466 | 351.1037821 | 7.61E-16 | 2.119898662 | YNL311C | SKP2 | n |
| W303.asm02.HP0_G0043890 | 431.3380254 | 377.9137152 | 399.7164816 | 718.9131234 | 667.446477 | 731.4413859 | 713.7300095 | 559.2687849 | 552.758141 | 2.95E-07 | -0.809740517 | YNL310C | ZIM17 | n |
| W303.asm02.HP0_G0043900 | 268.4958046 | 200.855904 | 167.1186268 | 753.9346016 | 679.3888593 | 799.3350789 | 752.540262 | 691.8290332 | 678.664162 | 4.35E-22 | -1.811533767 | YNL309W | STB1 | n |
| W303.asm02.HP0_G0043910 | 1912.426796 | 1672.529969 | 1889.124594 | 5616.083168 | 5314.36012 | 5696.733467 | 2925.914402 | 2992.765314 | 2284.733649 | 2.24E-31 | -1.602997525 | YNL308C | KRI1 | n |
| W303.asm02.HP0_G0043920 | 3376.068185 | 2756.771081 | 3394.169537 | 1278.283957 | 1175.66119 | 1283.643422 | 979.7222279 | 1467.838662 | 1641.896404 | 1.62E-12 | 1.349566936 | YNL307C | MCK1 | n |
| W303.asm02.HP0_G0043930 | 2227.448949 | 2036.164828 | 2350.411096 | 1591.531624 | 1844.434598 | 1751.657279 | 2008.667215 | 1710.704518 | 1582.526085 | 0.012733086 | 0.35089866 | YNL306W | MRPS18 | n |
| W303.asm02.HP0_G0043940 | 894.6629156 | 404.5675792 | 848.2980587 | 375.5080726 | 339.6944296 | 344.8999604 | 681.5458977 | 437.3520602 | 496.4587007 | 0.000335896 | 1.018065636 | YNL305C | BXI1 | n |
| W303.asm02.HP0_G0043950 | 476.8950752 | 257.971327 | 382.1250472 | 901.8030655 | 922.2172992 | 922.4489755 | 943.75175 | 902.7643189 | 1045.122337 | 2.55E-10 | -1.298162482 | YNL304W | YPT11 | n |
| W303.asm02.HP0_G0043970 | 3104.664484 | 3284.136819 | 3054.068472 | 13487.16041 | 15022.18999 | 12775.78252 | 13770.06691 | 9663.351825 | 8841.059377 | 1.35E-35 | -2.128253343 | YOL120C | RPL18A | y |
| W303.asm02.HP0_G0043990 | 298.5440715 | 440.7406804 | 318.600423 | 1696.596058 | 1671.933521 | 1664.753352 | 1378.23726 | 1433.005312 | 1624.494759 | 7.95E-42 | -2.249984714 | YNL299W | TRF5 | n |
| W303.asm02.HP0_G0044000 | 378.995883 | 433.1252907 | 285.372158 | 2334.765218 | 2422.976674 | 2192.513659 | 1809.882995 | 2482.843775 | 2830.326407 | 1.03E-40 | -2.662473991 | YNL298W | CLA4 | n |
| W303.asm02.HP0_G0044010 | 567.0398761 | 784.3851418 | 602.0179772 | 1382.375573 | 1442.374395 | 1349.726617 | 995.8142838 | 1242.389481 | 1199.689891 | 2.16E-11 | -1.095128079 | YNL297C | MON2 | n |
| W303.asm02.HP0_G0044020 | 321.8072459 | 455.0195362 | 285.372158 | 499.0560653 | 501.5800563 | 520.518313 | 462.8832555 | 485.7317128 | 597.7976932 | 0.012610831 | -0.517994182 | YNL295W | MRX6 | n |
| W303.asm02.HP0_G0044030 | 1286.259685 | 1111.8469 | 1283.197409 | 127.4392681 | 131.3662052 | 114.9666535 | 142.9353202 | 138.3658066 | 224.1741349 | 3.26E-57 | 3.302691252 | YNL294C | RIM21 | n |
| W303.asm02.HP0_G0044040 | 390.6274702 | 362.6829357 | 340.101065 | 906.6671597 | 919.5634365 | 865.4182734 | 918.1937788 | 979.2041701 | 1161.815722 | 4.39E-19 | -1.299188245 | YNL293W | MSB3 | n |
| W303.asm02.HP0_G0044050 | 1513.075635 | 1442.164429 | 1518.72717 | 2276.396088 | 1908.127304 | 2189.797911 | 1412.314555 | 1198.847793 | 1469.927204 | 0.000126779 | -0.51145302 | YNL292W | PUS4 | n |
| W303.asm02.HP0_G0044060 | 434.2459222 | 455.0195362 | 439.7858599 | 674.1634567 | 700.6197611 | 674.4106837 | 904.9414975 | 971.4634256 | 1072.760244 | 2.05E-06 | -0.624010845 | YNL291C | MID1 | n |
| W303.asm02.HP0_G0044070 | 1564.448479 | 1227.02967 | 1531.432095 | 537.968819 | 504.233919 | 566.6860242 | 715.6231926 | 511.8567253 | 705.2784428 | 3.90E-16 | 1.424874992 | YNL290W | RFC3 | n |
| W303.asm02.HP0_G0044080 | 503.0661464 | 804.3755398 | 574.6535237 | 576.8815726 | 489.637674 | 628.2429725 | 1072.488197 | 755.6901747 | 835.2789686 | 0.518433225 | 0.149961147 | YNL289W | PCL1 | n |
| W303.asm02.HP0_G0044090 | 959.6059441 | 793.904379 | 1027.144308 | 1359.027921 | 1474.220747 | 1355.158112 | 1694.398829 | 1482.352558 | 1525.203019 | 2.84E-05 | -0.590546933 | YNL288W | CAF40 | n |
| W303.asm02.HP0_G0044100 | 3682.366648 | 3969.521895 | 4150.601216 | 5036.283139 | 4649.567506 | 4819.546953 | 4979.071419 | 5440.775739 | 6380.262023 | 0.023228026 | -0.297619052 | YNL287W | SEC21 | n |
| W303.asm02.HP0_G0044120 | 2308.870059 | 2070.434082 | 2321.092039 | 2752.104501 | 2940.479907 | 2877.787334 | 3036.665611 | 2800.214297 | 2547.805579 | 0.003031954 | -0.354973568 | YNL284C | MRPL10 | n |
| W303.asm02.HP0_G0044130 | 561.2240825 | 638.7408133 | 445.6496714 | 2755.022957 | 2647.228075 | 2598.970568 | 1427.460019 | 2053.232459 | 1996.071065 | 3.75E-36 | -2.281440091 | YNL283C | WSC2 | n |
| W303.asm02.HP0_G0044140 | 342.1625235 | 326.5098345 | 300.0316867 | 577.8543915 | 490.9646053 | 599.2749968 | 335.0933997 | 486.6993059 | 450.3955223 | 2.60E-05 | -0.785749664 | YNL282W | POP3 | n |
| W303.asm02.HP0_G0044150 | 4005.143193 | 4044.723868 | 3649.245336 | 1467.010812 | 1588.336845 | 1461.072273 | 1882.770542 | 1265.611714 | 1094.256394 | 4.96E-13 | 1.373424735 | YNL281W | HCH1 | n |
| W303.asm02.HP0_G0044160 | 1793.203027 | 1379.337464 | 1712.232948 | 1815.279957 | 1606.913864 | 1777.004258 | 1716.170434 | 1739.73231 | 1502.683242 | 0.558208501 | -0.090593821 | YNL280C | ERG24 | n |
| W303.asm02.HP0_G0044170 | 330.5309363 | 189.4328194 | 271.6899313 | 31.13020291 | 29.19249005 | 35.30472036 | 36.91706946 | 67.73151374 | 53.22856172 | 2.21E-20 | 3.043222354 | YNL279W | PRM1 | n |
| W303.asm02.HP0_G0044200 | 466.232787 | 394.0964183 | 405.5802931 | 144.9500073 | 159.2317639 | 143.9346291 | 248.953571 | 150.9445163 | 235.434023 | 1.34E-11 | 1.500436634 | YNL277W | MET2 | n |
| W303.asm02.HP0_G0044210 | 419.7064382 | 554.0196026 | 427.0809351 | 351.1876015 | 444.5220075 | 385.6361762 | 442.058242 | 447.0279907 | 641.8136192 | 0.258870084 | 0.247836708 | YNL275W | BOR1 | n |
| W303.asm02.HP0_G0044220 | 1275.597396 | 804.3755398 | 1170.807689 | 1293.849058 | 1401.239522 | 1363.305355 | 1646.122661 | 1577.176677 | 1510.872252 | 0.07630127 | -0.320025864 | YNL274C | GOR1 | n |
| W303.asm02.HP0_G0044230 | 315.0221534 | 455.0195362 | 401.6710854 | 1359.027921 | 1471.566885 | 1386.841836 | 1296.830389 | 1238.519108 | 1044.098711 | 7.07E-29 | -1.847259786 | YNL273W | TOF1 | n |
| W303.asm02.HP0_G0044240 | 899.5094102 | 1080.433417 | 994.8933454 | 1352.218189 | 1398.58566 | 1323.474389 | 1116.977999 | 1306.250622 | 1353.233819 | 0.000846871 | -0.453323034 | YNL272C | SEC2 | n |
| W303.asm02.HP0_G0044250 | 402.2590574 | 624.4619576 | 364.5336128 | 800.629906 | 972.6406911 | 900.7229937 | 1024.21203 | 1064.352359 | 1207.878901 | 6.90E-06 | -0.941769231 | YNL271C | BNI1 | n |
| W303.asm02.HP0_G0044280 | 2323.409543 | 2475.953585 | 2298.614095 | 4629.644864 | 4349.681017 | 4613.150127 | 2962.831472 | 3901.335191 | 4085.292112 | 4.99E-11 | -0.937430178 | YNL268W | LYP1 | n |
| W303.asm02.HP0_G0044290 | 719.2198086 | 764.3947438 | 700.7254702 | 2090.587689 | 2190.763685 | 1993.358826 | 1762.553419 | 2101.612112 | 2055.441383 | 6.41E-35 | -1.5218355 | YNL267W | PIK1 | n |
| W303.asm02.HP0_G0044300 | 1348.294816 | 1314.606652 | 1436.633809 | 323.948674 | 306.5211455 | 333.1317203 | 526.3048877 | 426.7085366 | 489.2933174 | 1.86E-51 | 2.087066615 | YNL265C | IST1 | n |
| W303.asm02.HP0_G0044320 | 1703.058226 | 1725.837697 | 1817.781554 | 1478.684638 | 1584.356051 | 1465.598519 | 1575.128297 | 1668.130424 | 1464.809074 | 0.06365118 | 0.213023082 | YNL263C | YIF1 | n |
| W303.asm02.HP0_G0044330 | 571.8863707 | 703.471626 | 615.7002039 | 1231.588652 | 1414.508836 | 1332.526881 | 986.3483686 | 1353.662682 | 1097.327272 | 3.65E-11 | -1.072513124 | YNL262W | POL2 | n |
| W303.asm02.HP0_G0044340 | 1003.224396 | 1110.894976 | 1102.396556 | 1461.173899 | 1674.587384 | 1547.9762 | 1144.429153 | 1333.343228 | 1187.406377 | 1.98E-05 | -0.541403824 | YNL261W | ORC5 | n |
| W303.asm02.HP0_G0044350 | 926.6497804 | 709.1831683 | 990.0068358 | 239.3134348 | 294.5787632 | 263.4275288 | 331.3070336 | 320.2733007 | 229.2922659 | 2.51E-17 | 1.721758207 | YNL260C | LTO1 | n |
| W303.asm02.HP0_G0044360 | 329.5616374 | 389.3367998 | 321.5323287 | 1184.893348 | 1139.834043 | 1142.424541 | 1541.051002 | 1128.2135 | 1270.320098 | 1.34E-27 | -1.736468283 | YNL259C | ATX1 | n |
| W303.asm02.HP0_G0044370 | 1227.13245 | 1498.327929 | 1306.652655 | 1463.119537 | 1450.335983 | 1480.987757 | 1262.753094 | 1292.704319 | 1278.509108 | 0.326115118 | -0.124204058 | YNL258C | DSL1 | n |
| W303.asm02.HP0_G0044380 | 485.6187656 | 704.4235497 | 517.9700128 | 1284.12087 | 1545.875041 | 1334.33738 | 1006.226791 | 1492.996081 | 1308.194267 | 9.39E-11 | -1.285064083 | YNL257C | SIP3 | n |
| W303.asm02.HP0_G0044390 | 1404.514155 | 1454.539438 | 1537.295906 | 1845.437341 | 1352.143062 | 1689.195082 | 1011.90634 | 1216.264468 | 965.2794943 | 0.358358687 | -0.153656619 | YNL256W | FOL1 | n |

| | | | | | | | | | | | | | | |
|---|---|---|---|---|---|---|---|---|---|---|---|---|---|---|
| W303.asm02.HP0_G0044400 | 620.3513174 | 392.1925709 | 555.1074854 | 210.1288696 | 257.424685 | 231.7438054 | 201.6239947 | 269.9584619 | 265.1191824 | 1.35E-07 | 1.166656852 | YNL254C | RTC4 | n |
| W303.asm02.HP0_G0044410 | 1295.952674 | 1020.462223 | 1275.378994 | 1203.376906 | 1281.815699 | 1207.602486 | 1108.458675 | 1173.690374 | 1095.28002 | 0.788255854 | -0.039616864 | YNL253W | TEX1 | n |
| W303.asm02.HP0_G0044420 | 1885.286426 | 1745.828095 | 2108.040222 | 2027.354464 | 2071.339862 | 2055.821024 | 1843.96029 | 1732.959159 | 1436.14754 | 0.495069487 | -0.100793399 | YNL252C | MRPL17 | n |
| W303.asm02.HP0_G0044430 | 916.956791 | 574.9619243 | 747.6359619 | 2018.599095 | 2550.362085 | 2140.914452 | 1565.662382 | 2070.649134 | 1948.98424 | 3.58E-16 | -1.582732943 | YNL251C | NRD1 | n |
| W303.asm02.HP0_G0044440 | 426.4915307 | 752.0197355 | 455.4226905 | 528.2406306 | 579.8690068 | 627.3377233 | 442.058242 | 559.2687849 | 494.4114483 | 0.726479432 | -0.086875669 | YNL250W | RAD50 | n |
| W303.asm02.HP0_G0044450 | 422.614335 | 640.6446607 | 467.1503135 | 612.8758697 | 617.0230851 | 624.6219755 | 465.7230301 | 504.1159808 | 469.8444198 | 0.13477842 | -0.276957854 | YNL249C | MPA43 | n |
| W303.asm02.HP0_G0044460 | 2875.909935 | 3906.69493 | 3309.144271 | 7102.550357 | 7648.432392 | 6870.841731 | 4215.172059 | 4541.881793 | 3677.88889 | 9.03E-14 | -1.099173621 | YNL248C | RPA49 | n |
| W303.asm02.HP0_G0044470 | 2089.8085 | 1990.47249 | 2020.08305 | 3908.786102 | 4004.678862 | 3771.268334 | 2610.699425 | 3134.0339 | 2549.852832 | 4.65E-15 | -0.937512812 | YNL247W | CRS1 | n |
| W303.asm02.HP0_G0044480 | 1461.702792 | 1247.020068 | 1543.159717 | 1464.092355 | 1841.780736 | 1544.355203 | 1802.310263 | 1442.681243 | 1263.154715 | 0.29918692 | -0.189244224 | YNL246W | VPS75 | n |
| W303.asm02.HP0_G0044490 | 4025.498471 | 4910.97445 | 4051.893723 | 6390.446965 | 7283.526267 | 6110.432369 | 5427.755802 | 4508.983629 | 3610.329561 | 0.000437173 | -0.606975125 | YNL244C | SUI1 | n |
| W303.asm02.HP0_G0044500 | 1274.628097 | 1192.760416 | 1078.94131 | 2402.862537 | 2084.609176 | 2236.870872 | 1257.073545 | 2351.25112 | 2329.773202 | 2.94E-05 | -0.923215643 | YNL243W | SLA2 | n |
| W303.asm02.HP0_G0044510 | 318.8993491 | 413.1348927 | 289.2813657 | 310.3292102 | 311.828871 | 303.2584954 | 1114.138224 | 388.9724075 | 1237.56406 | 0.697672759 | 0.142443256 | YNL242W | ATG2 | n |
| W303.asm02.HP0_G0044520 | 3204.502274 | 2358.866968 | 2337.706171 | 4732.763661 | 3289.462856 | 4089.916066 | 8881.868275 | 4087.113058 | 9508.463651 | 0.02849412 | -0.616488465 | YNL241C | ZWF1 | n |
| W303.asm02.HP0_G0044530 | 543.7767017 | 625.4138813 | 463.2411058 | 1998.169899 | 2103.186215 | 2003.316568 | 5543.239968 | 2243.848291 | 6172.465907 | 5.05E-11 | -1.902780255 | YNL240C | NAR1 | n |
| W303.asm02.HP0_G0044560 | 238.4475376 | 173.2501163 | 172.9824382 | 240.2862537 | 120.7507543 | 228.1228085 | 1315.762219 | 388.9724075 | 1194.57176 | 0.972312312 | -0.014500645 | YNL237W | YTP1 | n |
| W303.asm02.HP0_G0044570 | 618.4127195 | 782.4812944 | 705.6119797 | 1265.637312 | 1281.815699 | 1256.485945 | 1869.518261 | 1309.153401 | 3378.990043 | 0.002283167 | -0.852583543 | YNL236W | SIN4 | n |
| W303.asm02.HP0_G0044580 | 277.219495 | 584.4811615 | 275.5991389 | 1136.252406 | 1219.449925 | 1174.108264 | 3285.619182 | 1352.695089 | 3912.299287 | 0.000465143 | -1.633840653 | YNL234W | | n |
| W303.asm02.HP0_G0044590 | 267.5265056 | 415.0387401 | 266.8034217 | 937.7973626 | 950.0828579 | 880.8075104 | 2478.176611 | 1060.481987 | 3192.690077 | 1.38E-06 | -1.543916293 | YNL233W | BNI4 | n |
| W303.asm02.HP0_G0044600 | 1743.768781 | 1795.328128 | 1735.688194 | 1290.930602 | 1447.68212 | 1348.821367 | 3491.976134 | 1381.72288 | 3485.447167 | 0.200036453 | 0.368073704 | YNL232W | CSL4 | n |
| W303.asm02.HP0_G0044610 | 2584.150957 | 3100.415542 | 2561.508309 | 2768.642421 | 2456.149958 | 2617.075553 | 6353.522313 | 3164.996878 | 6901.287753 | 0.785411228 | 0.072425005 | YNL231C | PDR16 | n |
| W303.asm02.HP0_G0044620 | 192.8904878 | 126.6058542 | 137.7995695 | 305.465116 | 250.7900281 | 268.8590243 | 852.8789636 | 242.8658564 | 726.7745928 | 0.023495826 | -0.852099904 | YNL230C | ELA1 | n |
| W303.asm02.HP0_G0044630 | 2480.435971 | 2515.934381 | 2653.374688 | 2885.380682 | 2813.094495 | 2826.188127 | 6379.080284 | 2638.626257 | 6970.894333 | 0.596496919 | -0.156243363 | YNL229C | URE2 | n |
| W303.asm02.HP0_G0044640 | 1136.987649 | 1022.366071 | 1338.903618 | 1281.202413 | 1320.296709 | 1234.759963 | 3256.274844 | 1160.144071 | 2976.704952 | 0.678620717 | -0.132999175 | YNL227C | JJJ1 | n |
| W303.asm02.HP0_G0044650 | 926.6497804 | 1041.404545 | 867.844097 | 392.0459929 | 533.426409 | 416.4146504 | 1280.738333 | 496.3752364 | 1448.431055 | 0.000584595 | 1.080725346 | YNL225C | CNM67 | n |
| W303.asm02.HP0_G0044660 | 506.9433422 | 534.9811283 | 501.3558803 | 1004.921863 | 1082.775994 | 1037.415629 | 2182.840056 | 1045.968091 | 2843.633547 | 0.000281864 | -1.017767491 | YNL224C | SQS1 | n |
| W303.asm02.HP0_G0044670 | 275.2808971 | 240.8367001 | 257.0304026 | 115.7654421 | 96.8659897 | 132.166389 | 456.2571148 | 232.2223328 | 557.8762719 | 0.000120383 | 1.160759914 | YNL223W | ATG4 | n |
| W303.asm02.HP0_G0044680 | 645.5530897 | 617.7984916 | 726.1353199 | 465.0074059 | 624.9846733 | 476.1611002 | 1897.916007 | 556.3660057 | 1159.76847 | 0.318330281 | 0.346084928 | YNL222W | SSU72 | n |
| W303.asm02.HP0_G0044690 | 492.4038582 | 910.990996 | 536.5387491 | 1986.496073 | 1748.89554 | 1960.769854 | 3360.399912 | 1555.88963 | 3719.857563 | 2.35E-07 | -1.554005381 | YNL221C | POP1 | n |
| W303.asm02.HP0_G0044700 | 1396.759763 | 1465.962522 | 1322.289486 | 7892.479256 | 4770.31826 | 7221.173187 | 10325.42035 | 6501.257726 | 9767.441076 | 1.66E-25 | -2.248525983 | YNL220W | ADE12 | n |
| W303.asm02.HP0_G0044710 | 2164.444518 | 1722.981926 | 2211.634225 | 1793.877943 | 1908.127304 | 1807.782732 | 5414.50352 | 2242.880698 | 5447.738567 | 0.61881171 | 0.146651273 | YNL219C | ALG9 | n |
| W303.asm02.HP0_G0044720 | 473.9871784 | 565.4426872 | 494.514767 | 524.3493552 | 597.1191146 | 583.8857598 | 1463.430497 | 604.7456584 | 1422.8404 | 0.613028733 | -0.152534775 | YNL218W | MGS1 | n |
| W303.asm02.HP0_G0044730 | 1578.987963 | 1850.539704 | 1592.024813 | 1417.397051 | 1337.546817 | 1429.38855 | 3817.603619 | 1433.005312 | 3337.02137 | 0.369697531 | 0.263042863 | YNL217W | PPN2 | n |
| W303.asm02.HP0_G0044740 | 762.8382606 | 685.3850754 | 675.3156205 | 1738.427269 | 1866.992432 | 1825.887717 | 3784.472915 | 1576.209084 | 4141.591553 | 1.11E-06 | -1.354777863 | YNL216W | RAP1 | n |
| W303.asm02.HP0_G0044750 | 963.4831398 | 725.3658714 | 875.6625123 | 636.2235219 | 668.7734083 | 650.8742035 | 1789.057981 | 705.375336 | 1593.785973 | 0.17965132 | 0.390969009 | YNL215W | IES2 | n |
| W303.asm02.HP0_G0044760 | 742.482983 | 459.7791547 | 625.473223 | 214.9929638 | 240.1745772 | 230.8385562 | 1071.541606 | 328.9816382 | 805.5938092 | 3.38E-05 | 1.414162868 | YNL214W | PEX17 | n |
| W303.asm02.HP0_G0044770 | 1129.233257 | 1138.500764 | 1120.965292 | 839.5426596 | 603.7537714 | 785.7563403 | 1280.738333 | 387.0372214 | 965.2794943 | 0.067004516 | 0.603660967 | YNL213C | RRG9 | n |
| W303.asm02.HP0_G0044780 | 1855.238159 | 1596.376071 | 2006.400823 | 277.2533696 | 282.6363809 | 279.7220151 | 1167.14735 | 341.5603479 | 1152.603087 | 1.05E-15 | 2.70064716 | YNL212W | VID27 | n |
| W303.asm02.HP0_G0044790 | 343.1318224 | 351.2598511 | 356.7151975 | 273.3620943 | 242.8284399 | 281.5325136 | 549.0230842 | 284.4723577 | 794.3339211 | 0.203286626 | 0.396943403 | YNL211C | MRX7 | n |
| W303.asm02.HP0_G0044800 | 126.0088613 | 80.91351584 | 123.1400408 | 137.1674566 | 130.0392738 | 128.5453921 | 185.5319388 | 76.43985122 | 169.921947 | 0.468707766 | -0.262312824 | YNL210W | MER1 | n |
| W303.asm02.HP0_G0044830 | 1496.597553 | 1690.616519 | 1540.227812 | 2690.816914 | 2579.554575 | 2681.348249 | 5609.501374 | 1944.862037 | 4692.302441 | 0.009944177 | -0.750232106 | YNL207W | RIO2 | n |
| W303.asm02.HP0_G0044840 | 1587.711653 | 1643.972257 | 1679.981985 | 716.9674857 | 715.2160061 | 706.9996564 | 1732.26249 | 641.5141944 | 1514.966757 | 1.95E-05 | 1.199172421 | YNL206C | RTT106 | n |
| W303.asm02.HP0_G0044860 | 156.0571283 | 158.0193368 | 156.3683058 | 30.15738407 | 23.88476458 | 26.25222796 | 103.1784762 | 22.25464023 | 90.07910445 | 3.80E-08 | 2.546801888 | YNL202W | SPS19 | n |
| W303.asm02.HP0_G0044870 | 560.2547835 | 1088.048807 | 576.6081275 | 1870.730631 | 1564.45208 | 1862.097687 | 2155.388901 | 929.8569243 | 2160.874881 | 8.06E-05 | -1.251586052 | YNL201C | PSY2 | n |
| W303.asm02.HP0_G0044880 | 715.3426129 | 554.0196026 | 700.7254702 | 59.34194929 | 76.96201921 | 61.55694831 | 168.4932914 | 123.8519108 | 141.2604138 | 2.83E-52 | 3.326492263 | YNL200C | NNR1 | n |
| W303.asm02.HP0_G0044890 | 597.088143 | 753.9235859 | 468.1276154 | 338.5490566 | 261.4054791 | 320.4582309 | 265.9922184 | 269.9584619 | 331.6548846 | 5.46E-06 | 0.980455707 | YNL199C | GCR2 | n |
| W303.asm02.HP0_G0044950 | 1634.238002 | 1912.414745 | 1500.158433 | 2429.128646 | 2024.897264 | 2269.459844 | 1507.920299 | 2621.209582 | 2069.77215 | 0.044138625 | -0.414075805 | YNL192W | CHS1 | n |
| W303.asm02.HP0_G0044960 | 802.5795169 | 982.3852747 | 825.8201148 | 514.6211668 | 408.6948607 | 570.3070212 | 692.904996 | 575.7178668 | 597.7976932 | 4.27E-06 | 0.802460003 | YNL191W | DUG3 | n |
| W303.asm02.HP0_G0044980 | 3220.011057 | 2766.290318 | 3291.552836 | 6298.029176 | 6156.961537 | 6300.53471 | 3955.805981 | 5088.571868 | 4754.743639 | 2.90E-14 | -1.015518746 | YNL189W | SRP1 | n |
| W303.asm02.HP0_G0044990 | 738.6057873 | 1010.942986 | 864.9121912 | 122.5751739 | 128.7123425 | 133.0716383 | 113.5909829 | 128.6898761 | 131.0241519 | 1.06E-50 | 2.765894714 | YNL188W | KAR1 | n |
| W303.asm02.HP0_G0045000 | 122.1316656 | 191.3366669 | 117.2762293 | 118.6838986 | 132.6931366 | 98.67216715 | 177.9592066 | 113.2083873 | 99.29174014 | 0.354199837 | 0.303248678 | YNL187W | SWT21 | n |
| W303.asm02.HP0_G0045010 | 927.6190793 | 1575.43375 | 898.1404562 | 3463.235073 | 2763.998035 | 3506.030306 | 1308.189487 | 1746.505461 | 1719.691994 | 2.13E-11 | -1.517025699 | YNL186W | UBP10 | n |
| W303.asm02.HP0_G0045020 | 1304.676364 | 1550.683733 | 1433.701903 | 1231.588652 | 1240.680827 | 1209.412985 | 1394.329316 | 1134.019059 | 934.5707087 | 0.198180304 | 0.22042557 | YNL185C | MRPL19 | n |
| W303.asm02.HP0_G0045030 | 1094.338496 | 954.7794869 | 1023.235101 | 563.2621088 | 607.7345655 | 568.4965227 | 885.0630755 | 856.3198523 | 972.4448776 | 1.57E-10 | 0.821891073 | YNL183C | NPR1 | n |
| W303.asm02.HP0_G0045040 | 1588.680952 | 2640.636388 | 1847.100612 | 4008.986443 | 3501.771874 | 4154.188762 | 2162.961634 | 2494.454892 | 2135.284226 | 7.21E-07 | -0.941029418 | YNL182C | IPI3 | n |
| W303.asm02.HP0_G0045050 | 1097.246393 | 1165.154628 | 1249.969144 | 1882.404457 | 1616.202404 | 1980.685337 | 1560.929424 | 1617.815585 | 1421.816774 | 1.28E-06 | -0.642586593 | YNL181W | PBR1 | n |
| W303.asm02.HP0_G0045080 | 18618.29391 | 17761.94461 | 17824.03225 | 22079.09641 | 25128.09927 | 20952.89891 | 21236.78085 | 16029.14652 | 14938.80057 | 0.036089496 | -0.330465169 | YNL178W | RPS3 | n |
| W303.asm02.HP0_G0045090 | 2352.488511 | 2138.020666 | 2377.775549 | 3899.057914 | 3771.138942 | 3870.84575 | 3851.680913 | 3514.29797 | 3216.233479 | 6.92E-11 | -0.748887299 | YNL177C | MRPL22 | n |
| W303.asm02.HP0_G0045100 | 694.0180363 | 625.4138813 | 598.1087695 | 1213.105095 | 873.1208387 | 1156.003279 | 738.3413892 | 946.3060063 | 802.5229306 | 1.31E-05 | -0.759299183 | YNL176C | TDA7 | n |
| W303.asm02.HP0_G0045110 | 1921.150486 | 2281.761147 | 2108.040222 | 4712.334465 | 4320.488527 | 4400.416555 | 2233.955998 | 2052.264866 | 2002.212822 | 6.68E-21 | -1.089972283 | YNL175C | NOP13 | n |
| W303.asm02.HP0_G0045130 | 321.8072459 | 506.4234168 | 303.9408943 | 597.3107683 | 712.5621434 | 604.7064923 | 482.7616775 | 609.5836237 | 582.4433004 | 0.000584595 | -0.756598577 | YNL172W | APC1 | n |
| W303.asm02.HP0_G0045140 | 894.6629156 | 688.2408465 | 733.9537352 | 2178.141385 | 2158.917332 | 2149.061696 | 1753.087503 | 2187.727894 | 2096.386431 | 3.73E-24 | -1.485248601 | YNL169C | PSD1 | n |
| W303.asm02.HP0_G0045150 | 1452.979101 | 1125.173832 | 1366.268072 | 1607.096725 | 1350.81613 | 1512.67148 | 1771.072742 | 1346.88953 | 1384.966231 | 0.294062627 | -0.181296031 | YNL168C | FMP41 | n |
| W303.asm02.HP0_G0045160 | 480.772271 | 572.1061532 | 494.514767 | 1010.758776 | 1167.699602 | 1041.036626 | 1206.904194 | 1271.417272 | 1587.644216 | 2.48E-11 | -1.055999026 | YNL167C | SKO1 | n |

| | | | | | | | | | | | | | | |
|---|---|---|---|---|---|---|---|---|---|---|---|---|---|---|
| W303.asm02.HP0_G0045170 | 1359.926404 | 1041.404545 | 1275.378994 | 793.8201741 | 697.9658984 | 693.4209178 | 824.4812179 | 827.2920607 | 971.4212514 | 1.36E-06 | 0.750131066 | YNL166C | BNI5 | n |
| W303.asm02.HP0_G0045180 | 527.2986198 | 554.0196026 | 561.9485988 | 194.5637682 | 237.5207145 | 174.7131033 | 299.1229218 | 273.8288341 | 379.7653154 | 3.90E-14 | 1.443443546 | YNL165W | | n |
| W303.asm02.HP0_G0045190 | 706.6189225 | 1162.298857 | 727.1126218 | 545.7513697 | 545.3687913 | 511.4658206 | 564.1685486 | 517.6622836 | 482.1279341 | 0.000840531 | 0.696357185 | YNL164C | IBD2 | n |
| W303.asm02.HP0_G0045200 | 678.5092534 | 860.5390391 | 662.6106957 | 3758.972001 | 3383.674983 | 3855.456513 | 2112.792283 | 2256.427001 | 1918.275474 | 2.35E-58 | -2.32071382 | YNL163C | RIA1 | n |
| W303.asm02.HP0_G0045210 | 362.5178011 | 562.586916 | 365.5109147 | 440.6869349 | 390.1178215 | 401.0254133 | 236.6478811 | 243.8334495 | 179.1345827 | 0.782131451 | 0.066924139 | YNL162W-A | | n |
| W303.asm02.HP0_G0045230 | 708.5575203 | 666.3466011 | 571.7216179 | 2001.088356 | 2160.244263 | 2052.200027 | 1651.80221 | 2418.015041 | 2273.473761 | 3.17E-23 | -1.6741966 | YNL161W | CBK1 | n |
| W303.asm02.HP0_G0045250 | 879.1541326 | 566.3946109 | 789.6599441 | 235.4221595 | 276.0017241 | 233.5543039 | 192.1580795 | 309.6297771 | 278.4263229 | 5.90E-12 | 1.587232112 | YNL159C | ASI2 | n |
| W303.asm02.HP0_G0045260 | 566.0705771 | 342.6925377 | 580.5173351 | 207.2104131 | 244.1553713 | 201.8705805 | 372.9570607 | 305.7594049 | 270.2373134 | 6.29E-07 | 1.19134114 | YNL158W | PGA1 | n |
| W303.asm02.HP0_G0045280 | 2863.309049 | 2529.261313 | 2632.851348 | 1702.432971 | 1519.336414 | 1704.584319 | 1900.755781 | 1913.89906 | 1786.227696 | 6.56E-10 | 0.703111352 | YNL156C | NSG2 | n |
| W303.asm02.HP0_G0045290 | 1562.509881 | 1919.078211 | 1467.90747 | 305.465116 | 313.1558023 | 305.0689939 | 430.6991437 | 323.1760798 | 411.4977272 | 6.94E-47 | 2.422338841 | YNL155W | CUZ1 | n |
| W303.asm02.HP0_G0045300 | 1774.786347 | 1723.933849 | 1711.255646 | 4139.344168 | 4607.105702 | 4349.722598 | 2075.875213 | 4013.575986 | 3778.204254 | 5.36E-10 | -1.329726953 | YNL154C | YCK2 | n |
| W303.asm02.HP0_G0045310 | 2094.654995 | 2100.895641 | 2328.910454 | 1120.687305 | 1204.85368 | 1067.288854 | 1498.454383 | 1050.806056 | 1093.232768 | 5.39E-09 | 0.943980613 | YNL153C | GIM3 | n |
| W303.asm02.HP0_G0045320 | 376.0879862 | 237.9809289 | 286.3494599 | 418.3121016 | 457.7913212 | 422.751395 | 512.1060148 | 385.1020353 | 434.0175033 | 0.011546265 | -0.527904977 | YNL152W | INN1 | n |
| W303.asm02.HP0_G0045330 | 1232.948243 | 1808.65506 | 1511.886056 | 1727.726261 | 1752.876334 | 1775.193759 | 1401.902048 | 1212.394096 | 1005.200916 | 0.265989401 | -0.206896318 | YNL151C | RPC31 | n |
| W303.asm02.HP0_G0045340 | 2283.668287 | 2428.357399 | 2347.47919 | 1093.448377 | 1240.680827 | 1076.341346 | 1597.846493 | 1134.019059 | 967.3267467 | 1.20E-08 | 1.050243652 | YNL149C | PGA2 | n |
| W303.asm02.HP0_G0045350 | 660.0925737 | 540.6926706 | 670.4291109 | 620.6584205 | 582.5228696 | 564.8755257 | 731.7152485 | 562.171564 | 517.9548506 | 0.67046536 | 0.081659048 | YNL148C | ALF1 | n |
| W303.asm02.HP0_G0045400 | 773.5005489 | 813.894777 | 685.0886396 | 206.2375943 | 258.7516163 | 243.5120455 | 202.5705863 | 268.0232758 | 226.2213873 | 9.47E-22 | 1.684197983 | YNL142W | MEP2 | n |
| W303.asm02.HP0_G0045410 | 1063.32093 | 1626.83763 | 1152.238953 | 4769.730777 | 3813.600745 | 4623.107868 | 1032.731353 | 1146.597768 | 786.1449116 | 6.53E-19 | -1.781289165 | YNL141W | AAH1 | n |
| W303.asm02.HP0_G0045420 | 604.8425345 | 858.6351916 | 626.4505249 | 1753.99237 | 1886.896402 | 1796.014492 | 1412.314555 | 1637.167446 | 1461.738195 | 9.22E-18 | -1.378920178 | YNL139C | THO2 | n |
| W303.asm02.HP0_G0045440 | 1229.071048 | 1051.875706 | 948.9601556 | 3449.61561 | 2564.95833 | 3239.88703 | 2514.147089 | 2756.672609 | 2752.530817 | 1.99E-22 | -1.5192299 | YNL138W | SRV2 | n |
| W303.asm02.HP0_G0045450 | 3721.138605 | 3614.454349 | 4023.551968 | 4702.606277 | 4463.797114 | 4713.632792 | 3838.428632 | 4403.515986 | 4053.5597 | 0.008449495 | -0.289355346 | YNL137C | NAM9 | n |
| W303.asm02.HP0_G0045460 | 1030.364766 | 870.0582762 | 1058.41797 | 519.485261 | 550.6765168 | 529.5708054 | 557.542408 | 649.2549389 | 701.1839381 | 3.10E-09 | 0.887784941 | YNL136W | EAF7 | n |
| W303.asm02.HP0_G0045470 | 4230.020545 | 3998.079606 | 4215.103147 | 3540.087762 | 4012.64045 | 3457.146847 | 3552.557992 | 2620.241989 | 2366.623744 | 0.304773953 | 0.176817855 | YNL135C | FPR1 | n |
| W303.asm02.HP0_G0045480 | 7385.088574 | 5484.98445 | 5986.951507 | 167.3248406 | 120.7507543 | 98.67216715 | 192.1580795 | 186.7454593 | 191.418097 | 2.34E-180 | 5.607004306 | YNL134C | | n |
| W303.asm02.HP0_G0045490 | 572.8556697 | 385.5291049 | 584.4265428 | 975.7372974 | 1037.660328 | 1003.921407 | 721.3027417 | 676.3475443 | 648.9790025 | 4.14E-08 | -0.967682175 | YNL133C | FYV6 | n |
| W303.asm02.HP0_G0045500 | 568.9784739 | 973.8179613 | 625.473223 | 8190.161821 | 7989.453753 | 8296.609284 | 2906.982572 | 3956.487995 | 3363.63565 | 3.47E-68 | -3.496618473 | YNL132W | KRE33 | n |
| W303.asm02.HP0_G0045510 | 3008.703889 | 3911.454548 | 3145.934851 | 2608.127312 | 2681.72829 | 2750.147191 | 3022.466738 | 2396.727993 | 2097.410057 | 0.058444702 | 0.324247042 | YNL131W | TOM22 | n |
| W303.asm02.HP0_G0045530 | 746.3601787 | 752.0197355 | 847.3207568 | 627.4681523 | 571.9074186 | 648.1584558 | 233.8081066 | 385.1020353 | 334.7257631 | 0.079287265 | 0.343491465 | YNL129W | NRK1 | n |
| W303.asm02.HP0_G0045540 | 164.7808187 | 97.09621901 | 126.0719465 | 475.7084132 | 456.4643898 | 482.4978449 | 374.8502437 | 339.6251618 | 265.1191824 | 7.62E-17 | -1.86734358 | YNL128W | TEP1 | n |
| W303.asm02.HP0_G0045550 | 347.9783171 | 492.1445611 | 384.079651 | 659.5711741 | 701.9466925 | 715.1468995 | 534.8242114 | 717.9540456 | 750.3179951 | 7.03E-05 | -0.762162115 | YNL127W | FAR11 | n |
| W303.asm02.HP0_G0045560 | 429.3994275 | 586.3850089 | 486.6963517 | 456.2520364 | 468.4067721 | 523.2340607 | 400.4082149 | 424.7733505 | 388.9779511 | 0.772044454 | 0.052711899 | YNL126W | SPC98 | n |
| W303.asm02.HP0_G0045570 | 433.2766232 | 516.8945777 | 362.579009 | 1048.69871 | 1035.006465 | 1027.457887 | 491.2810012 | 866.9633759 | 735.9872284 | 1.02E-08 | -1.244711887 | YNL125C | ESBP6 | n |
| W303.asm02.HP0_G0045580 | 320.837947 | 695.8562362 | 271.6899313 | 1365.837653 | 1200.872886 | 1423.051805 | 620.0174486 | 746.9818372 | 709.3729476 | 1.84E-08 | -1.630724212 | YNL124W | NAF1 | n |
| W303.asm02.HP0_G0045590 | 1683.672247 | 1995.232108 | 1783.575988 | 4451.619016 | 4140.025861 | 4409.469048 | 2352.279939 | 2960.834743 | 2735.129122 | 1.14E-21 | -1.251126772 | YNL123W | NMA111 | n |
| W303.asm02.HP0_G0045600 | 757.022467 | 978.5775798 | 777.9323212 | 944.6070945 | 1013.775563 | 948.7012034 | 1060.182507 | 846.6439218 | 666.3806477 | 0.295658717 | -0.209423245 | YNL122C | MRP35 | n |
| W303.asm02.HP0_G0045610 | 4024.529172 | 4034.252708 | 4041.143402 | 6605.439929 | 6383.866801 | 6517.794527 | 3081.155412 | 5553.984127 | 4655.451898 | 0.00031566 | -0.689020922 | YNL121C | TOM70 | n |
| W303.asm02.HP0_G0045620 | 1553.78619 | 2276.049604 | 1654.572135 | 1932.018218 | 1893.531059 | 1824.982468 | 1189.865546 | 1283.028389 | 1006.224542 | 0.819133716 | -0.04293619 | YNL119W | NCS2 | n |
| W303.asm02.HP0_G0045630 | 1581.895859 | 1617.318393 | 1639.912607 | 2553.649457 | 2478.707791 | 2443.267699 | 1797.577305 | 2221.593651 | 2401.427035 | 3.16E-06 | -0.627457766 | YNL118C | DCP2 | n |
| W303.asm02.HP0_G0045640 | 105.6535837 | 77.10582098 | 85.02526626 | 219.857058 | 172.5010776 | 224.5018115 | 594.4594774 | 358.9770228 | 540.4746267 | 2.10E-06 | -1.207604419 | YNL117W | MLS1 | n |
| W303.asm02.HP0_G0045660 | 720.1891075 | 498.8080271 | 709.5211874 | 739.342319 | 505.5608504 | 754.9778661 | 875.5971602 | 914.3754355 | 933.5470825 | 0.803089073 | -0.054941475 | YNL115C | | n |
| W303.asm02.HP0_G0045670 | 2178.984002 | 3019.502027 | 2317.182831 | 2750.158863 | 3297.424444 | 2939.344282 | 2723.343816 | 1865.519407 | 1595.833225 | 0.240820993 | -0.25770342 | YNL113W | RPC19 | n |
| W303.asm02.HP0_G0045680 | 1707.904721 | 4122.781613 | 1851.987121 | 9599.776321 | 6767.349965 | 9402.823855 | 2275.606025 | 2985.992163 | 3112.847235 | 2.49E-09 | -1.746045459 | YNL112W | DBP2 | n |
| W303.asm02.HP0_G0045690 | 2872.03274 | 3682.040933 | 2902.586676 | 5207.499255 | 5444.399394 | 4968.913078 | 3952.966207 | 3602.348938 | 3266.391163 | 3.24E-07 | -0.723897719 | YNL111C | CYB5 | n |
| W303.asm02.HP0_G0045700 | 2567.672875 | 3618.262044 | 2901.609374 | 6078.172118 | 5806.651657 | 6212.725534 | 4076.969696 | 3273.3673 | 3377.966417 | 2.10E-10 | -0.993869367 | YNL110C | NOP15 | n |
| W303.asm02.HP0_G0045720 | 983.8384174 | 905.2794537 | 972.4154014 | 561.3164712 | 591.8113891 | 517.8025652 | 373.9036522 | 529.2734002 | 536.380122 | 4.88E-06 | 0.777253075 | YNL107W | YAF9 | n |
| W303.asm02.HP0_G0045740 | 4354.090809 | 4333.156754 | 3967.845759 | 1360.973558 | 1035.006465 | 1351.537115 | 1138.749604 | 1485.255337 | 1698.195844 | 9.23E-24 | 1.754540645 | YNL104C | LEU4 | n |
| W303.asm02.HP0_G0045750 | 578.6714633 | 703.471626 | 487.6736536 | 715.9946669 | 679.3888593 | 733.2518843 | 515.8923809 | 513.7919114 | 724.7273404 | 0.189329931 | -0.266629227 | YNL103W | MET4 | n |
| W303.asm02.HP0_G0045760 | 416.7985414 | 691.0966177 | 437.8312561 | 1614.879276 | 1547.201972 | 1625.827635 | 1361.198612 | 1899.385164 | 1659.298049 | 4.27E-15 | -1.631016238 | YNL102W | POL1 | n |
| W303.asm02.HP0_G0045770 | 1027.456869 | 1049.019935 | 1090.668933 | 249.0416233 | 179.1357344 | 203.681079 | 173.226249 | 420.9029782 | 306.0642299 | 8.67E-18 | 2.323358069 | YNL101W | AVT4 | n |
| W303.asm02.HP0_G0045780 | 2296.269173 | 2221.789953 | 2203.815809 | 1881.431638 | 2079.30145 | 1813.214228 | 2742.275647 | 2359.959457 | 2277.568266 | 0.083419022 | 0.22007331 | YNL100W | MIC27 | n |
| W303.asm02.HP0_G0045790 | 742.482983 | 569.250382 | 643.0646574 | 620.6584205 | 690.0043102 | 661.7371944 | 1023.265438 | 785.6855594 | 803.5465568 | 0.946066083 | -0.012375607 | YNL099C | OCA1 | n |
| W303.asm02.HP0_G0045810 | 771.561951 | 455.9714599 | 651.8603746 | 588.5553987 | 639.5809183 | 583.8857598 | 853.8255551 | 833.097619 | 919.2163159 | 0.800599336 | 0.053223792 | YNL097C | PHO23 | n |
| W303.asm02.HP0_G0045840 | 575.7635665 | 689.1927702 | 558.0393912 | 280.1718262 | 261.4054791 | 268.8590243 | 286.8172839 | 298.0186605 | 295.827968 | 5.00E-14 | 1.168623965 | YNL094W | APP1 | n |
| W303.asm02.HP0_G0045860 | 128.9167581 | 83.76928699 | 129.9811542 | 220.8298769 | 201.6935676 | 191.0075896 | 298.1763302 | 319.3057076 | 296.8515942 | 0.000136969 | -0.841037492 | YNL092W | | n |
| W303.asm02.HP0_G0045870 | 652.3381822 | 798.6639976 | 598.1087695 | 1591.531624 | 1792.684275 | 1604.101653 | 1029.891579 | 1514.283129 | 1620.400254 | 3.22E-11 | -1.283107963 | YNL091W | NST1 | n |
| W303.asm02.HP0_G0045880 | 1072.04462 | 1005.231444 | 1031.053516 | 585.6369422 | 573.23435 | 601.0854953 | 772.418684 | 679.2503235 | 666.3806477 | 5.64E-11 | 0.819876567 | YNL090W | RHO2 | n |
| W303.asm02.HP0_G0045890 | 751.2066734 | 1124.221908 | 849.2753607 | 2409.672269 | 2400.418841 | 2617.980802 | 2226.383266 | 2398.663179 | 1947.960634 | 8.71E-18 | -1.446911982 | YNL088W | TOP2 | n |
| W303.asm02.HP0_G0045910 | 314.0528544 | 373.1540966 | 373.32933 | 78.79832611 | 71.65429375 | 74.23043767 | 119.2705321 | 105.4676428 | 82.91372115 | 3.72E-24 | 2.236622729 | YNL086W | SNN1 | n |
| W303.asm02.HP0_G0045920 | 1798.049522 | 1900.039737 | 2074.811957 | 2490.416233 | 2461.457684 | 2477.66717 | 2271.819659 | 2597.987348 | 2242.764976 | 0.002227087 | -0.364043381 | YNL085W | MKT1 | n |
| W303.asm02.HP0_G0045930 | 2701.436127 | 2713.934514 | 2647.510877 | 1535.108131 | 1429.105081 | 1530.776465 | 1493.721426 | 1522.023873 | 1487.32885 | 6.88E-18 | 0.842157406 | YNL084C | END3 | n |
| W303.asm02.HP0_G0045940 | 1276.566695 | 1353.635524 | 1197.194841 | 1598.341355 | 1332.239091 | 1600.480656 | 806.4959789 | 1175.62556 | 899.7674184 | 0.164070534 | -0.24419897 | YNL082W | PMS1 | n |
| W303.asm02.HP0_G0045950 | 1958.953144 | 2360.770815 | 2011.287333 | 1070.100725 | 1082.775994 | 1076.341346 | 1185.132589 | 885.3476439 | 781.0267807 | 1.50E-08 | 0.971324529 | YNL081C | SWS2 | n |
| W303.asm02.HP0_G0045960 | 648.4609865 | 472.154163 | 581.494637 | 1300.65879 | 1460.951434 | 1390.462833 | 1733.209081 | 1324.63489 | 1322.525034 | 5.41E-14 | -1.286296969 | YNL080C | EOS1 | n |

| | | | | | | | | | | | | | |
|---|---|---|---|---|---|---|---|---|---|---|---|---|---|
| W303.asm02.HP0_G0045980 | 803.5488158 | 595.9042461 | 518.9473147 | 264.6067247 | 253.4438909 | 267.953775 | 281.1376828 | 364.7825811 | 292.7570895 | 3.30E-10 | 1.286518614 | YNL078W | NIS1 | n |
| W303.asm02.HP0_G0045990 | 1297.891272 | 568.2984583 | 702.680074 | 1242.28966 | 1507.394032 | 1292.695915 | 1332.800867 | 1794.885114 | 1502.683242 | 0.015656439 | -0.653790184 | YNL077W | APJ1 | n |
| W303.asm02.HP0_G0046000 | 757.022467 | 749.1639643 | 762.2954906 | 761.7171524 | 751.043153 | 809.2928205 | 563.2219571 | 654.0929041 | 623.3883479 | 0.800175764 | -0.034466645 | YNL076W | MKS1 | n |
| W303.asm02.HP0_G0046010 | 1695.303835 | 2159.914911 | 1877.396971 | 2460.258848 | 2109.820872 | 2330.111544 | 1366.878161 | 1489.125709 | 1250.8712 | 0.065744573 | -0.267856404 | YNL075W | IMP4 | n |
| W303.asm02.HP0_G0046030 | 1121.478866 | 852.9236494 | 1007.59827 | 442.6325726 | 428.5988311 | 454.4351184 | 397.5684403 | 477.9909684 | 434.0175033 | 6.81E-14 | 1.168717662 | YNL073W | MSK1 | n |
| W303.asm02.HP0_G0046040 | 316.9607512 | 174.20204 | 247.2573835 | 362.8614276 | 352.9637433 | 369.3416899 | 711.8368265 | 565.0743432 | 493.3878221 | 0.020024243 | -0.556020506 | YNL072W | RNH201 | n |
| W303.asm02.HP0_G0046050 | 2559.918483 | 1954.299389 | 2030.833371 | 8117.200408 | 7551.566403 | 8008.740026 | 8737.986363 | 6881.521796 | 7318.927237 | 3.21E-36 | -1.855124324 | YNL071W | LAT1 | n |
| W303.asm02.HP0_G0046060 | 1070.106023 | 1478.337531 | 1098.487348 | 1184.893348 | 1515.35562 | 1311.706149 | 1354.572472 | 1056.611614 | 963.2322419 | 0.502943036 | -0.136856544 | YNL070W | TOM7 | n |
| W303.asm02.HP0_G0046080 | 415.8292424 | 618.7504153 | 342.0556688 | 2360.058508 | 2064.705205 | 2277.607088 | 1424.620244 | 1799.723079 | 1628.589263 | 1.57E-28 | -2.283456136 | YNL068C | FKH2 | n |
| W303.asm02.HP0_G0046090 | 6963.443538 | 7789.591766 | 7055.142495 | 18975.80431 | 21490.9804 | 18083.25882 | 16551.15281 | 12775.13108 | 11423.66825 | 8.85E-20 | -1.424735989 | YNL067W | RPL9B | y |
| W303.asm02.HP0_G0046100 | 1570.264272 | 2122.789886 | 1694.641514 | 11488.99051 | 11131.62723 | 11108.31342 | 6279.688174 | 9682.703686 | 8533.971521 | 2.12E-50 | -2.646207855 | YNL066W | SUN4 | n |
| W303.asm02.HP0_G0046110 | 401.2897584 | 1919.078211 | 419.2625198 | 1235.479928 | 813.4089272 | 1222.086474 | 453.4173403 | 635.7086361 | 524.0966077 | 0.719968667 | -0.255842759 | YNL065W | AQR1 | n |
| W303.asm02.HP0_G0046120 | 7026.447968 | 6136.100272 | 5540.324533 | 9560.863568 | 10003.73557 | 9519.601007 | 3385.011292 | 7328.549787 | 6102.859327 | 0.007163505 | -0.636957417 | YNL064C | YDJ1 | n |
| W303.asm02.HP0_G0046130 | 753.1452713 | 381.72141 | 751.5451696 | 148.8412826 | 200.3666362 | 178.3341003 | 198.7842202 | 154.8148885 | 167.8746947 | 1.03E-11 | 1.840620471 | YNL063W | MTQ1 | n |
| W303.asm02.HP0_G0046140 | 1210.654368 | 1261.298923 | 1190.353728 | 3667.52703 | 3045.307484 | 3513.2723 | 1519.279397 | 1745.537868 | 1420.793148 | 1.87E-29 | -1.4818801 | YNL062C | GCD10 | n |
| W303.asm02.HP0_G0046150 | 1891.102219 | 2298.895774 | 1982.945577 | 9473.309872 | 9003.229317 | 9669.872381 | 3761.754719 | 4860.219907 | 4140.567926 | 6.02E-58 | -2.188944296 | YNL061W | NOP2 | n |
| W303.asm02.HP0_G0046160 | 927.6190793 | 925.2698518 | 1002.711761 | 1322.060805 | 1350.81613 | 1278.211927 | 1321.441768 | 1492.996081 | 1546.699169 | 0.000107169 | -0.46809138 | YNL059C | ARP5 | n |
| W303.asm02.HP0_G0046170 | 1034.241962 | 914.7986909 | 1066.236385 | 753.9346016 | 778.9087117 | 716.0521488 | 763.8993603 | 720.8568248 | 789.2157902 | 0.000958863 | 0.423932665 | YNL058C | | n |
| W303.asm02.HP0_G0046180 | 1388.036073 | 1058.539172 | 1372.131883 | 220.8298769 | 217.616744 | 201.8705805 | 342.6661319 | 200.2917621 | 233.3867706 | 3.09E-33 | 2.576875662 | YNL056W | OCA2 | n |
| W303.asm02.HP0_G0046190 | 10306.55556 | 8069.457339 | 10816.77755 | 5203.60798 | 5327.629434 | 5083.879731 | 5965.419788 | 5038.257029 | 5024.980952 | 3.24E-10 | 0.902719486 | YNL055C | POR1 | n |
| W303.asm02.HP0_G0046210 | 153.1492315 | 253.2117084 | 151.4817962 | 601.2020436 | 480.3491544 | 561.2545288 | 332.2536251 | 451.865956 | 574.2542909 | 2.19E-09 | -1.558689175 | YNL054W | VAC7 | n |
| W303.asm02.HP0_G0046220 | 525.3600219 | 468.3464682 | 455.4226905 | 974.7644785 | 796.1588195 | 938.7434618 | 1302.509938 | 1183.366304 | 1198.666265 | 1.25E-10 | -0.90470638 | YNL053W | MSG5 | n |
| W303.asm02.HP0_G0046230 | 3900.458908 | 4277.945179 | 4007.915137 | 8541.349423 | 8420.706447 | 8101.980697 | 4833.296324 | 6624.142044 | 5888.921454 | 1.97E-14 | -1.040324845 | YNL052W | COX5A | n |
| W303.asm02.HP0_G0046240 | 1199.99208 | 1003.327596 | 1138.556726 | 561.3164712 | 621.0038792 | 609.2327385 | 527.2514792 | 477.9909684 | 515.9075982 | 2.17E-11 | 0.900152609 | YNL051W | COG5 | n |
| W303.asm02.HP0_G0046250 | 1494.658955 | 1704.895375 | 1522.636377 | 907.6399785 | 1011.121701 | 1017.500146 | 853.8255551 | 872.7689342 | 754.4124998 | 1.09E-07 | 0.68589321 | YNL050C | | n |
| W303.asm02.HP0_G0046270 | 1253.303521 | 991.9045119 | 1229.445804 | 978.6557539 | 1017.756358 | 941.4592095 | 1133.070055 | 1447.519208 | 1293.8635 | 0.122528037 | 0.242525836 | YNL048W | ALG11 | n |
| W303.asm02.HP0_G0046280 | 263.6493099 | 358.8752409 | 213.0518166 | 2745.294769 | 2550.362085 | 2756.483936 | 1527.798721 | 3066.302386 | 2750.483564 | 2.63E-38 | -3.268351257 | YNL047C | SLM2 | n |
| W303.asm02.HP0_G0046300 | 739.5750862 | 572.1061532 | 635.2462421 | 3122.748479 | 3055.922935 | 3146.646358 | 3610.300075 | 3516.233156 | 3499.777933 | 3.85E-75 | -2.260182002 | YNL045W | LAP2 | n |
| W303.asm02.HP0_G0046330 | 182.2281995 | 203.7116752 | 109.457814 | 322.9758552 | 314.4827337 | 319.5529817 | 381.4763844 | 366.7177673 | 521.0257292 | 0.000146278 | -0.949683589 | YNL042W | BOP3 | n |
| W303.asm02.HP0_G0046340 | 1246.518428 | 1249.875839 | 1356.495052 | 800.629906 | 755.0239471 | 794.8088327 | 652.2015604 | 812.7781649 | 726.7745928 | 6.59E-08 | 0.712281786 | YNL041C | COG6 | n |
| W303.asm02.HP0_G0046350 | 1586.742354 | 1436.452887 | 1564.66036 | 566.1805654 | 636.9270556 | 586.6015075 | 802.7096128 | 746.9818372 | 819.9245758 | 5.32E-29 | 1.359869229 | YNL040W | | n |
| W303.asm02.HP0_G0046360 | 578.6714633 | 540.6926706 | 516.9927109 | 830.7872901 | 893.0248092 | 821.0610606 | 740.2345722 | 886.3152369 | 862.9168756 | 3.16E-06 | -0.636149457 | YNL039W | BDP1 | n |
| W303.asm02.HP0_G0046370 | 773.5005489 | 901.4717589 | 854.1618702 | 139.1130942 | 128.7123425 | 141.2188814 | 105.0716592 | 162.555633 | 135.1186567 | 2.22E-43 | 2.625695643 | YNL038W | GPI15 | n |
| W303.asm02.HP0_G0046380 | 5783.806736 | 4254.147086 | 5863.811466 | 8776.771582 | 9604.329225 | 8524.732092 | 8662.259041 | 7910.073212 | 7952.551847 | 2.84E-07 | -0.75865567 | YNL037C | IDH1 | n |
| W303.asm02.HP0_G0046390 | 10540.1566 | 6886.21616 | 11127.55956 | 573.9631161 | 1316.315915 | 618.2852309 | 3865.879786 | 2524.450276 | 3167.099423 | 1.42E-32 | 3.510812741 | YNL036W | NCE103 | n |
| W303.asm02.HP0_G0046400 | 1311.461457 | 918.6063857 | 1278.3109 | 2018.599095 | 1567.105943 | 2014.179559 | 925.766511 | 864.0605967 | 889.5311565 | 0.000120445 | -0.675448463 | YNL035C | | n |
| W303.asm02.HP0_G0046430 | 1539.246706 | 1557.347199 | 1561.728454 | 415.393645 | 486.9838112 | 451.7193707 | 893.5823992 | 683.1206957 | 722.680088 | 4.79E-34 | 1.784307322 | YNL032W | SIW14 | n |
| W303.asm02.HP0_G0046460 | 336.3467299 | 296.0482756 | 327.3961402 | 491.2735146 | 505.5608504 | 509.6553221 | 873.7039772 | 655.0604972 | 899.7674184 | 0.000135194 | -0.650443541 | YNL029C | KTR5 | n |
| W303.asm02.HP0_G0046470 | 544.7460006 | 658.7312113 | 553.1528816 | 480.5725074 | 512.1955072 | 497.887082 | 594.4594774 | 423.8057574 | 646.9317502 | 0.224528247 | 0.237379603 | YNL027W | CRZ1 | n |
| W303.asm02.HP0_G0046480 | 1186.421895 | 1069.010333 | 1259.742163 | 1088.584283 | 1131.872455 | 1076.341346 | 1356.465655 | 1474.611813 | 1503.706869 | 0.472768498 | 0.092955422 | YNL026W | SAM50 | n |
| W303.asm02.HP0_G0046490 | 328.5923384 | 278.9136487 | 347.9194803 | 169.2704783 | 187.0973226 | 155.7028693 | 180.7989812 | 189.6482385 | 281.4972014 | 7.23E-05 | 0.902698859 | YNL025C | SSN8 | n |
| W303.asm02.HP0_G0046500 | 1223.255254 | 1311.75088 | 1262.674069 | 338.5409566 | 408.6948607 | 366.6259422 | 489.3878182 | 362.847395 | 365.4345488 | 1.62E-27 | 1.771930027 | YNL024C- | KSH1 | n |
| W303.asm02.HP0_G0046510 | 274.3115982 | 493.0964848 | 234.5524586 | 416.3664639 | 415.3295175 | 420.0356473 | 300.0695133 | 284.4723577 | 314.2532394 | 0.232669814 | -0.320813987 | YNL024C | EFM6 | n |
| W303.asm02.HP0_G0046520 | 255.8949184 | 489.2887899 | 244.3254777 | 781.1735292 | 842.6014173 | 816.5348144 | 787.5641484 | 830.1948398 | 796.3811735 | 7.69E-08 | -1.301721074 | YNL023C | FAP1 | n |
| W303.asm02.HP0_G0046530 | 1141.834144 | 1438.356735 | 1126.829103 | 2775.452153 | 2349.995449 | 2671.390507 | 1243.821263 | 1419.459009 | 1116.77617 | 4.96E-12 | -1.072947372 | YNL022C | RCM1 | n |
| W303.asm02.HP0_G0046540 | 474.9564774 | 699.6639311 | 490.6055593 | 2440.802472 | 1986.416255 | 2463.183182 | 1179.45304 | 1627.491516 | 1462.761821 | 3.84E-26 | -2.049124801 | YNL021W | HDA1 | n |
| W303.asm02.HP0_G0046580 | 4359.906603 | 4069.473885 | 4631.433756 | 8408.073241 | 8444.591212 | 8343.682244 | 7230.066065 | 7286.943285 | 7641.369486 | 1.97E-22 | -0.947976277 | YNL016W | PUB1 | n |
| W303.asm02.HP0_G0046590 | 1559.601984 | 2001.895574 | 1690.732306 | 147.8684638 | 127.3854111 | 158.418617 | 360.6513709 | 276.7316133 | 396.1433344 | 4.39E-81 | 3.592780115 | YNL015W | PBI2 | n |
| W303.asm02.HP0_G0046620 | 1002.255097 | 568.2984583 | 905.9588715 | 386.2090798 | 335.7136355 | 416.4146504 | 459.0968894 | 474.1205962 | 502.6004578 | 2.20E-07 | 1.119452948 | YNL011C | | n |
| W303.asm02.HP0_G0046630 | 3077.524114 | 2965.242375 | 3170.367399 | 1853.219892 | 1474.220747 | 1903.739152 | 2287.911715 | 1724.250821 | 1496.541485 | 4.61E-06 | 0.81578038 | YNL010W | PYP1 | n |
| W303.asm02.HP0_G0046660 | 5400.933657 | 3563.050468 | 4246.376803 | 1385.294029 | 1540.567316 | 1458.356526 | 2673.174465 | 2221.593651 | 2875.365959 | 5.24E-20 | 1.59156023 | YNL007C | SIS1 | n |
| W303.asm02.HP0_G0046670 | 3311.125157 | 2305.55924 | 2938.746846 | 584.6641233 | 562.6188991 | 540.4337962 | 893.5823992 | 683.1206957 | 784.0976592 | 2.23E-43 | 2.341848853 | YNL006W | LST8 | n |
| W303.asm02.HP0_G0046680 | 3289.80058 | 2539.732474 | 3333.576818 | 3388.328023 | 3098.384739 | 3167.46709 | 3208.945268 | 3070.172759 | 3217.257106 | 0.60645308 | -0.075504776 | YNL005C | MRP7 | n |
| W303.asm02.HP0_G0046700 | 1011.948086 | 741.5485746 | 1000.757157 | 1354.163826 | 1495.451649 | 1445.683036 | 2054.103608 | 1730.05638 | 1829.219996 | 4.70E-05 | -0.640920186 | YNL003C | PET8 | n |
| W303.asm02.HP0_G0046710 | 4751.503372 | 5275.561233 | 5143.539958 | 5875.825799 | 5314.36012 | 5606.208543 | 2779.192716 | 2746.029086 | 2470.009989 | 0.195718405 | -0.147055414 | YNL002C | RLP7 | n |
| W303.asm02.HP0_G0046720 | 1873.654838 | 1520.222174 | 1774.78027 | 1146.953413 | 1311.008189 | 1191.308 | 1259.913319 | 1360.435833 | 1075.831123 | 0.000671037 | 0.502902948 | YNL001W | DOM34 | n |
| W303.asm02.HP0_G0046730 | 6758.921463 | 4342.675991 | 6417.941649 | 3995.366979 | 8358.340673 | 4097.15806 | 8101.876859 | 9310.18036 | 8290.348488 | 0.756844209 | 0.091153765 | YNR001C | CIT1 | n |
| W303.asm02.HP0_G0046750 | 1357.987806 | 1029.98146 | 1397.541733 | 1332.761812 | 1308.354327 | 1253.770197 | 1106.565492 | 851.481887 | 793.3102949 | 0.829486618 | -0.041077528 | YNR003C | RPC34 | n |
| W303.asm02.HP0_G0046770 | 505.0047443 | 398.8560369 | 430.9901427 | 930.9876307 | 877.1016328 | 1026.552638 | 689.1186299 | 1137.889431 | 1084.020132 | 1.34E-07 | -1.087098266 | YNR006W | VPS27 | n |
| W303.asm02.HP0_G0046780 | 971.2375313 | 700.6158548 | 942.1190422 | 129.3849058 | 141.9816561 | 127.6401428 | 364.437737 | 289.310323 | 400.2378391 | 1.78E-41 | 2.714617584 | YNR007C | ATG3 | n |
| W303.asm02.HP0_G0046790 | 442.9696126 | 541.6445943 | 478.8779364 | 1686.86787 | 1410.528042 | 1665.658601 | 1971.750146 | 2616.371616 | 2268.35563 | 4.07E-27 | -1.703129139 | YNR008W | LRO1 | n |
| W303.asm02.HP0_G0046810 | 500.1582496 | 446.4522227 | 511.1288994 | 257.7969928 | 359.5984001 | 283.3430121 | 537.6639859 | 372.5233256 | 452.4427746 | 0.000492275 | 0.698676585 | YNR010W | CSE2 | n |
| W303.asm02.HP0_G0046820 | 469.1406838 | 437.8849093 | 417.307916 | 721.8315799 | 717.8698689 | 739.588629 | 398.5150318 | 613.4539959 | 577.3251695 | 8.58E-05 | -0.718736885 | YNR011C | PRP2 | n |

| | | | | | | | | | | | | | | |
|---|---|---|---|---|---|---|---|---|---|---|---|---|---|---|
| W303.asm02.HP0_G0046830 | 1409.360649 | 1650.635723 | 1447.38413 | 2045.838022 | 2265.071841 | 2140.914452 | 1010.959748 | 1188.20427 | 994.9646537 | 9.00E-05 | -0.517000782 | YNR012W | URK1 | n |
| W303.asm02.HP0_G0046840 | 675.6013566 | 757.7312778 | 606.9044867 | 923.20508 | 733.7930453 | 902.5334922 | 931.4460602 | 1401.074741 | 1203.784396 | 0.089669383 | -0.328305558 | YNR013C | PHO91 | n |
| W303.asm02.HP0_G0046860 | 476.8950752 | 546.4042129 | 481.8098421 | 2389.243073 | 2336.726135 | 2456.846437 | 1464.377088 | 1705.866553 | 1272.36735 | 2.70E-55 | -2.254713238 | YNR015W | SMM1 | n |
| W303.asm02.HP0_G0046870 | 5662.644369 | 5678.224965 | 5252.997772 | 8082.17893 | 7176.044826 | 8381.702712 | 7817.899401 | 8096.818571 | 8696.728085 | 4.85E-06 | -0.510824199 | YNR016C | ACC1 | n |
| W303.asm02.HP0_G0046880 | 2025.834771 | 1638.260715 | 1988.809389 | 3759.94482 | 3585.36855 | 3639.101944 | 2926.860994 | 3415.603479 | 3028.909887 | 9.62E-14 | -0.958533384 | YNR017W | TIM23 | n |
| W303.asm02.HP0_G0046890 | 873.338339 | 1013.798757 | 818.0016995 | 3959.372682 | 3142.173474 | 4011.159382 | 2844.507531 | 3370.126605 | 2986.941214 | 7.57E-42 | -2.038836983 | YNR018W | RCF2 | n |
| W303.asm02.HP0_G0046900 | 1409.360649 | 1200.375806 | 1046.690347 | 1574.020884 | 1161.064945 | 1695.531826 | 1098.046168 | 1387.528439 | 1406.462381 | 0.158503613 | -0.278186005 | YNR019W | ARE2 | n |
| W303.asm02.HP0_G0046920 | 622.2899153 | 788.1928367 | 673.3610167 | 1228.670196 | 1127.891661 | 1193.118498 | 1088.580253 | 1062.417173 | 968.3503729 | 2.49E-08 | -0.768844639 | YNR020C | ATP23 | n |
| W303.asm02.HP0_G0046930 | 3223.888253 | 2566.386338 | 3275.916006 | 2870.788399 | 2939.152975 | 2928.481291 | 4418.689237 | 3963.261147 | 3666.629002 | 0.723367507 | 0.053119094 | YNR021W | | n |
| W303.asm02.HP0_G0046940 | 1280.443891 | 1658.251113 | 1438.588413 | 844.4067538 | 845.25528 | 847.3132886 | 1102.779126 | 750.8522095 | 747.2471165 | 2.04E-05 | 0.786954936 | YNR022C | MRPL50 | n |
| W303.asm02.HP0_G0046950 | 574.7942675 | 474.0580105 | 477.9006345 | 1252.990667 | 1243.33469 | 1223.896972 | 1255.180362 | 1288.833947 | 1402.367876 | 9.12E-23 | -1.284913915 | YNR023W | SNF12 | n |
| W303.asm02.HP0_G0046960 | 698.864531 | 987.1448933 | 820.9336052 | 670.2721813 | 587.830595 | 761.3146108 | 496.9605504 | 487.6668989 | 347.0092774 | 0.135076371 | 0.310687117 | YNR024W | MPP6 | n |
| W303.asm02.HP0_G0046980 | 445.8775094 | 493.0964848 | 413.3987084 | 303.5194783 | 329.0789787 | 288.7745075 | 620.0174486 | 473.1530031 | 503.624084 | 0.001312047 | 0.555948869 | YNR027W | BUD17 | n |
| W303.asm02.HP0_G0047000 | 864.6146486 | 787.240913 | 845.366153 | 664.4352683 | 660.8118202 | 631.8639695 | 710.8902349 | 755.6901747 | 767.7196402 | 0.00406012 | 0.352016235 | YNR029C | ZNG1 | n |
| W303.asm02.HP0_G0047010 | 936.3427697 | 993.8083593 | 1073.077498 | 1753.99237 | 1840.453804 | 1784.246252 | 1706.704519 | 2078.389879 | 2047.252374 | 6.77E-11 | -0.840540195 | YNR030W | ALG12 | n |
| W303.asm02.HP0_G0047020 | 334.408132 | 607.3273307 | 358.6698013 | 499.0560653 | 504.233919 | 515.0868175 | 387.1559335 | 566.0419363 | 533.3092434 | 0.388975326 | -0.223402106 | YNR031C | SSK2 | n |
| W303.asm02.HP0_G0047030 | 799.6716201 | 775.8178284 | 797.4783594 | 306.4379349 | 368.8869197 | 295.1112522 | 390.9422996 | 372.5233256 | 379.7653154 | 1.61E-19 | 1.295263183 | YNR032W | PPG1 | n |
| W303.asm02.HP0_G0047040 | 510.8205379 | 455.9714599 | 490.6055593 | 108.9557102 | 159.2317639 | 115.8719027 | 198.7842202 | 112.2407942 | 112.5988806 | 9.97E-14 | 1.930864059 | YNR032C- | HUB1 | n |
| W303.asm02.HP0_G0047050 | 733.7592926 | 672.0581434 | 583.4492409 | 1164.464152 | 1088.08372 | 1096.25683 | 1065.862057 | 1271.417272 | 1097.327272 | 1.43E-07 | -0.751587347 | YNR033W | ABZ1 | n |
| W303.asm02.HP0_G0047060 | 570.9170718 | 211.3270649 | 368.4428204 | 1563.319877 | 1565.779012 | 1566.081185 | 1801.363671 | 1892.612013 | 1808.747472 | 2.23E-14 | -2.028872446 | YNR034W | SOL1 | n |
| W303.asm02.HP0_G0047080 | 4209.665268 | 3019.502027 | 4132.03248 | 2439.829653 | 2141.667224 | 2301.143568 | 2834.095025 | 2218.609872 | 2115.835328 | 3.30E-05 | 0.722815074 | YNR035C | ARC35 | n |
| W303.asm02.HP0_G0047090 | 1861.053952 | 1557.347199 | 1984.900181 | 1491.331283 | 1487.490061 | 1481.893006 | 2073.98203 | 1579.111863 | 1556.93543 | 0.089762803 | 0.276523673 | YNR036C | MRPS12 | n |
| W303.asm02.HP0_G0047100 | 977.0533249 | 941.4525549 | 940.1644384 | 931.9604495 | 1029.69874 | 945.0802065 | 1182.292814 | 890.1856092 | 763.6251355 | 0.900825573 | -0.02348508 | YNR037C | RSM19 | n |
| W303.asm02.HP0_G0047110 | 799.6716201 | 792.0005315 | 760.3408868 | 2227.755146 | 2186.782891 | 2243.207617 | 1331.854275 | 1676.838762 | 1510.872252 | 2.10E-33 | -1.501236247 | YNR038W | DBP6 | n |
| W303.asm02.HP0_G0047120 | 509.851239 | 473.1060868 | 453.4680867 | 934.878906 | 1009.794769 | 911.5859846 | 859.5051043 | 1308.185808 | 1108.58716 | 1.27E-08 | -0.991033893 | YNR039C | ZRG17 | n |
| W303.asm02.HP0_G0047130 | 712.4347161 | 671.1062196 | 736.8856409 | 708.2121161 | 753.6970158 | 719.6731457 | 1052.609775 | 782.7827802 | 887.4839041 | 0.802962964 | -0.040494211 | YNR040W | MRX15 | n |
| W303.asm02.HP0_G0047140 | 761.8689617 | 535.933052 | 625.473223 | 946.5527321 | 956.7175147 | 979.4796776 | 1260.859911 | 1192.074642 | 1340.950305 | 0.000185285 | -0.584087504 | YNR041C | COQ2 | n |
| W303.asm02.HP0_G0047150 | 2810.966907 | 2351.251578 | 2356.274907 | 5516.855646 | 4857.89573 | 5207.898877 | 2816.109786 | 3265.626555 | 2800.641248 | 6.05E-16 | -1.051617951 | YNR043W | MVD1 | n |
| W303.asm02.HP0_G0047160 | 196.7676835 | 310.3271313 | 137.7995695 | 626.4953335 | 621.0038792 | 697.947164 | 440.1650589 | 689.8938471 | 657.168012 | 7.70E-09 | -1.59281859 | YNR044W | AGA1 | n |
| W303.asm02.HP0_G0047170 | 445.8775094 | 336.0290717 | 323.4869325 | 587.5825799 | 611.7153596 | 576.6437658 | 528.1980707 | 522.5002489 | 550.7108886 | 3.40E-05 | -0.683458406 | YNR045W | PET494 | n |
| W303.asm02.HP0_G0047180 | 1312.430756 | 988.096817 | 1098.487348 | 1990.387348 | 1851.069255 | 1922.749386 | 1259.913319 | 1030.486602 | 707.3256952 | 0.000229436 | -0.762146785 | YNR046W | TRM112 | n |
| W303.asm02.HP0_G0047190 | 247.171228 | 234.1732341 | 208.165307 | 836.6242031 | 930.1788874 | 783.0405925 | 951.3244822 | 952.1115646 | 1024.649813 | 4.24E-39 | -1.884959334 | YNR047W | FPK1 | n |
| W303.asm02.HP0_G0047210 | 643.6144918 | 466.4426207 | 635.2462421 | 264.6067247 | 294.5787632 | 250.7540395 | 400.4082149 | 338.6575687 | 457.5609056 | 1.32E-08 | 1.109839185 | YNR049C | MSO1 | n |
| W303.asm02.HP0_G0047220 | 3953.770349 | 3160.386736 | 3908.230342 | 755.8802393 | 1130.545524 | 963.1851913 | 1166.200758 | 1045.000498 | 1088.114637 | 9.34E-31 | 1.953210027 | YNR050C | LYS9 | n |
| W303.asm02.HP0_G0047230 | 1096.277094 | 1181.337331 | 1085.782423 | 4119.887791 | 4189.122322 | 4178.630491 | 2946.739416 | 3630.409136 | 3709.621302 | 2.96E-51 | -1.892443857 | YNR051C | BRE5 | n |
| W303.asm02.HP0_G0047240 | 1092.399898 | 1029.029537 | 999.7798549 | 1902.833653 | 1898.838784 | 1977.06434 | 1313.869036 | 1780.371218 | 1730.951882 | 9.25E-10 | -0.888763687 | YNR052C | POP2 | n |
| W303.asm02.HP0_G0047250 | 1732.137194 | 3980.944979 | 1901.829519 | 5368.014364 | 4520.855163 | 5074.827239 | 3448.432924 | 3106.941295 | 2575.443486 | 0.000346827 | -0.974588229 | YNR053C | NOG2 | n |
| W303.asm02.HP0_G0047260 | 1071.075321 | 971.9141138 | 1110.214971 | 2279.314544 | 2146.97495 | 2238.68137 | 1760.660236 | 1715.542484 | 1553.864552 | 1.14E-20 | -1.080156434 | YNR054C | ESF2 | n |
| W303.asm02.HP0_G0047270 | 1003.224396 | 922.4140806 | 1002.711761 | 2664.550805 | 2758.690309 | 2808.988391 | 2959.991697 | 2967.607895 | 3445.525745 | 7.33E-38 | -1.491249647 | YNR055C | HOL1 | n |
| W303.asm02.HP0_G0047280 | 119.2237688 | 125.6539305 | 129.0038523 | 62.26040581 | 53.07725463 | 65.17794527 | 112.6443914 | 80.31022343 | 131.0241519 | 0.000169311 | 1.04407025 | YNR056C | BIO5 | n |
| W303.asm02.HP0_G0047290 | 213.2457654 | 176.1058874 | 218.9156281 | 6.809731886 | 9.28851956 | 4.5262462 | 6.626140672 | 2.90277916 | 6.141757122 | 1.80E-28 | 4.928880549 | YNR057C | BIO4 | n |
| W303.asm02.HP0_G0047300 | 191.9211888 | 158.9712605 | 193.5057784 | 242.2318914 | 168.5202835 | 233.5543039 | 450.5775657 | 222.5464023 | 636.6954883 | 0.484898336 | -0.245634045 | YNR058W | BIO3 | n |
| W303.asm02.HP0_G0047330 | 139.5790464 | 169.4424214 | 164.186721 | 642.060435 | 614.3692224 | 653.5899512 | 562.2753656 | 513.7919114 | 574.2542909 | 7.76E-40 | -2.013634571 | YNR062C | PUL3 | n |
| W303.asm02.HP0_G0047340 | 50.40354454 | 88.52890557 | 51.79700128 | 1183.920529 | 1107.98769 | 1116.172313 | 812.1755281 | 871.8013411 | 886.4602779 | 9.12E-98 | -4.157708039 | YNR063W | PUL4 | n |
| W303.asm02.HP0_G0047380 | 1643.930991 | 2118.982191 | 1408.292054 | 1825.008145 | 1608.240815 | 2051.294778 | 1209.743968 | 2567.024371 | 2169.06389 | 0.759848814 | -0.085138211 | YNR067C | DSE4 | n |
| W303.asm02.HP0_G0047650 | 1365.742197 | 1668.722274 | 1360.40426 | 892.074877 | 849.2360741 | 900.7229937 | 985.4017771 | 794.3938969 | 752.3652474 | 2.30E-06 | 0.733772962 | YOL149W | DCP1 | n |
| W303.asm02.HP0_G0047660 | 392.566068 | 379.8175626 | 405.5802931 | 559.3708335 | 642.234781 | 592.9382522 | 710.8902349 | 702.4725568 | 719.6092094 | 4.25E-06 | -0.60557747 | YOL148C | SPT20 | n |
| W303.asm02.HP0_G0047670 | 1195.145585 | 1109.943053 | 1193.285633 | 228.6124276 | 212.3090185 | 238.0805501 | 437.3252843 | 389.9400005 | 371.5763059 | 7.33E-63 | 2.361878514 | YOL147C | PEX11 | n |
| W303.asm02.HP0_G0047680 | 410.9827478 | 459.7791547 | 463.2411058 | 340.4865943 | 338.3674983 | 336.7527173 | 496.9605504 | 360.9122089 | 295.827968 | 0.060960913 | 0.393439366 | YOL146W | PSF3 | n |
| W303.asm02.HP0_G0047690 | 1947.321557 | 2243.684198 | 2018.128446 | 3527.441117 | 3816.254608 | 3719.669127 | 2587.034637 | 3143.709831 | 2734.105545 | 1.94E-11 | -0.833145574 | YOL145C | CTR9 | n |
| W303.asm02.HP0_G0047700 | 436.18452 | 554.0196026 | 438.808558 | 2693.73537 | 2233.225489 | 2641.517282 | 1066.808648 | 1262.708935 | 1066.618487 | 1.94E-54 | -2.405305257 | YOL144W | NOP8 | n |
| W303.asm02.HP0_G0047710 | 1739.891586 | 1446.924048 | 1848.077914 | 1237.425566 | 952.7367206 | 1090.825334 | 1187.972363 | 782.7827802 | 812.7591925 | 0.001861067 | 0.617046553 | YOL143C | RIB4 | n |
| W303.asm02.HP0_G0047720 | 1389.974671 | 1456.443285 | 1461.066357 | 811.3309133 | 891.6978778 | 803.8613251 | 759.1664027 | 706.342929 | 595.7504408 | 7.08E-09 | 0.782117267 | YOL142W | RRP40 | n |
| W303.asm02.HP0_G0047740 | 6783.153936 | 8344.563293 | 7162.645706 | 6850.590277 | 7491.854491 | 6750.443582 | 7444.942341 | 5736.859214 | 5122.22544 | 0.650467565 | 0.079771073 | YOL139C | CDC33 | n |
| W303.asm02.HP0_G0047750 | 231.6624451 | 319.8463685 | 195.4603822 | 1002.003406 | 1038.987259 | 980.3849268 | 1122.657548 | 1255.935783 | 1085.043758 | 4.24E-28 | -2.015287125 | YOL138C | RTC1 | n |
| W303.asm02.HP0_G0047760 | 906.2945028 | 828.1736327 | 988.052232 | 263.6339059 | 269.3670673 | 229.0280577 | 142.9353202 | 272.8612411 | 292.7570895 | 3.40E-14 | 1.838245545 | YOL137W | BSC6 | n |
| W303.asm02.HP0_G0047770 | 1015.825282 | 1551.635657 | 924.5276078 | 425.1218334 | 376.8485079 | 378.3941823 | 281.1376828 | 443.1576185 | 440.1592604 | 5.31E-11 | 1.564526674 | YOL136C | PFK27 | n |
| W303.asm02.HP0_G0047780 | 385.7809755 | 203.7116752 | 266.8034217 | 443.6053914 | 437.8873507 | 435.4248844 | 1010.013157 | 583.4586112 | 675.5932834 | 0.020436899 | -0.621147963 | YOL135C | MED7 | n |
| W303.asm02.HP0_G0047790 | 1900.795208 | 1618.270317 | 1851.987121 | 742.2607756 | 805.447339 | 759.5041123 | 1010.959748 | 932.7597035 | 811.7355663 | 5.32E-19 | 1.219818851 | YOL133W | HRT1 | n |
| W303.asm02.HP0_G0047800 | 113.4079752 | 83.76928699 | 87.95717199 | 37.93993479 | 43.78873507 | 43.45196352 | 159.9739676 | 71.60188595 | 70.6302069 | 0.002196935 | 1.189032328 | YOL132W | GAS4 | n |
| W303.asm02.HP0_G0047820 | 888.847122 | 1356.491295 | 868.8213989 | 2864.951486 | 2448.18837 | 2852.440355 | 1808.936403 | 1882.936082 | 1969.456784 | 2.73E-14 | -1.390880324 | YOL130W | ALR1 | n |
| W303.asm02.HP0_G0047830 | 2114.040974 | 1989.520566 | 1927.239368 | 415.393645 | 391.4447529 | 426.372392 | 721.3027417 | 552.4956335 | 520.002103 | 1.75E-49 | 2.28868101 | YOL129W | VPS68 | n |
| W303.asm02.HP0_G0047850 | 9162.782818 | 10796.71878 | 8957.949316 | 13898.66278 | 16039.94635 | 13582.3596 | 14277.43996 | 11135.06086 | 10332.48273 | 0.000194747 | -0.589680535 | YOL127W | RPL25 | n |

| | | | | | | | | | | | | | | |
|---|---|---|---|---|---|---|---|---|---|---|---|---|---|---|
| W303.asm02.HP0_G0047860 | 676.5706555 | 405.5195029 | 489.6282574 | 177.053029 | 684.6965847 | 206.3968267 | 657.8811096 | 957.9171229 | 1257.012958 | 0.370077504 | 0.55922683 | YOL126C | MDH2 | n |
| W303.asm02.HP0_G0047870 | 712.4347161 | 691.0966177 | 694.8616587 | 1317.19671 | 1174.334259 | 1334.33738 | 986.3483686 | 1007.264369 | 920.2399421 | 3.42E-13 | -0.867727422 | YOL125W | TRM13 | n |
| W303.asm02.HP0_G0047880 | 1392.882567 | 1955.251312 | 1475.725886 | 3461.289436 | 2622.016379 | 3334.9382 | 1427.460019 | 1781.338811 | 1581.502459 | 6.16E-08 | -0.96565082 | YOL124C | TRM11 | n |
| W303.asm02.HP0_G0047890 | 3615.485022 | 2734.876835 | 4023.551968 | 6362.235219 | 6482.059722 | 6181.04181 | 3345.254448 | 4415.127103 | 3550.959243 | 2.56E-07 | -0.874959462 | YOL123W | HRP1 | n |
| W303.asm02.HP0_G0047900 | 1409.360649 | 1349.827829 | 1462.043659 | 2638.284696 | 1865.6655 | 2326.490547 | 1329.961092 | 2537.028986 | 1648.038161 | 0.002409511 | -0.694719246 | YOL122C | SMF1 | n |
| W303.asm02.HP0_G0047910 | 5254.569518 | 5999.023257 | 5376.137812 | 19867.87919 | 22859.04664 | 19186.75764 | 18237.9789 | 14632.90975 | 13311.23494 | 1.46E-37 | -1.896418543 | YOL121C | RPS19A | y |
| W303.asm02.HP0_G0047940 | 417.7678403 | 453.1156887 | 455.4226905 | 121.6023551 | 107.4814406 | 116.777152 | 63.42163215 | 127.7222831 | 197.5598541 | 1.65E-09 | 1.937790419 | YOL117W | RRI2 | n |
| W303.asm02.HP0_G0047950 | 820.9961966 | 697.7600837 | 818.9790014 | 380.3721668 | 333.0597728 | 392.8781701 | 348.345681 | 396.7131519 | 564.018029 | 8.62E-08 | 1.077562075 | YOL116W | MSN1 | n |
| W303.asm02.HP0_G0047970 | 118.2544699 | 77.10582098 | 97.7301911 | 71.98859422 | 78.28895058 | 66.08319451 | 125.8966728 | 77.40744427 | 73.70108546 | 0.178650865 | 0.440976743 | YOL114C | PTH4 | n |
| W303.asm02.HP0_G0048000 | 459.4476944 | 306.5194365 | 399.7164816 | 821.0591017 | 752.3700844 | 904.3439907 | 1115.084816 | 654.0929041 | 691.9713024 | 3.61E-06 | -1.088587743 | YOL111C | MDY2 | n |
| W303.asm02.HP0_G0048010 | 394.5046659 | 355.067546 | 335.2145555 | 66.15168118 | 53.07725463 | 54.3149544 | 17.98523897 | 73.53707206 | 90.07910445 | 9.02E-11 | 2.642399043 | YOL110W | SHR5 | n |
| W303.asm02.HP0_G0048030 | 968.3296345 | 600.6638647 | 990.0068358 | 119.6567174 | 122.0776857 | 100.4826656 | 142.9353202 | 89.01856092 | 108.5043758 | 5.83E-31 | 2.905361622 | YOL108C | INO4 | n |
| W303.asm02.HP0_G0048040 | 560.2547835 | 445.500299 | 642.0873555 | 651.7886234 | 687.3504475 | 622.8114771 | 1171.880307 | 946.3060063 | 954.0196063 | 0.16674525 | -0.251158021 | YOL107W | | n |
| W303.asm02.HP0_G0048080 | 756.0531681 | 881.4813608 | 866.8667951 | 342.432232 | 411.3487234 | 434.5196352 | 509.2662402 | 363.8149881 | 415.5922319 | 1.52E-09 | 1.075941081 | YOL102C | TPT1 | n |
| W303.asm02.HP0_G0048100 | 377.0572851 | 477.8657053 | 337.1691593 | 874.5641379 | 939.467407 | 831.9240515 | 834.8937247 | 1121.440349 | 1016.460804 | 1.58E-10 | -1.149367723 | YOL100W | PKH2 | n |
| W303.asm02.HP0_G0048110 | 1381.25098 | 1471.674065 | 1468.884772 | 3048.814247 | 3111.654053 | 3095.047151 | 1971.750146 | 2354.153899 | 2229.457835 | 1.78E-22 | -1.098619221 | YOL098C | SDD3 | n |
| W303.asm02.HP0_G0048130 | 3788.989531 | 3354.579174 | 4061.666742 | 4411.733443 | 4767.664397 | 4410.374297 | 4117.673132 | 3577.191519 | 3512.061448 | 0.030286955 | -0.278177681 | YOL097C | WRS1 | n |
| W303.asm02.HP0_G0048140 | 879.1541326 | 530.2215097 | 775.0004154 | 1053.562805 | 996.5254557 | 1101.688325 | 966.4699466 | 892.1207953 | 822.9954543 | 0.006013762 | -0.529482756 | YOL096C | COQ3 | n |
| W303.asm02.HP0_G0048150 | 229.7238472 | 266.5386404 | 221.8475338 | 200.4006812 | 232.212989 | 218.1650668 | 216.7694591 | 313.5001493 | 312.205987 | 0.520317838 | 0.143961815 | YOL095C | HMI1 | n |
| W303.asm02.HP0_G0048160 | 725.0356022 | 653.019669 | 729.0672256 | 862.8903118 | 818.7166527 | 905.2492399 | 876.5437517 | 809.8753857 | 686.8531715 | 0.043210774 | -0.296840555 | YOL094C | RFC4 | n |
| W303.asm02.HP0_G0048170 | 1236.825439 | 1132.789222 | 1324.244089 | 1431.989334 | 1484.836198 | 1457.451276 | 1132.123463 | 969.5282395 | 859.8459971 | 0.094078181 | -0.243815924 | YOL093W | TRM10 | n |
| W303.asm02.HP0_G0048180 | 1991.909308 | 1958.107083 | 2232.157565 | 1087.611464 | 1038.987259 | 1104.404073 | 938.0722008 | 1147.565361 | 1036.933327 | 2.50E-13 | 0.935556462 | YOL092W | YPQ1 | n |
| W303.asm02.HP0_G0048190 | 201.6141782 | 209.4232175 | 173.9597402 | 111.8741667 | 127.3854111 | 114.9666535 | 173.226249 | 120.9491317 | 89.05547827 | 0.007665343 | 0.726180946 | YOL091W | SPO21 | n |
| W303.asm02.HP0_G0048200 | 671.7241609 | 793.904379 | 703.6573759 | 1163.491334 | 1113.295416 | 1193.118498 | 1025.158621 | 1164.014443 | 964.2558681 | 5.85E-07 | -0.678088915 | YOL090W | MSH2 | n |
| W303.asm02.HP0_G0048210 | 501.1275486 | 740.5966509 | 575.6308256 | 1084.693008 | 979.2753479 | 1110.740817 | 757.2732196 | 975.3337979 | 839.3734733 | 7.90E-06 | -0.805249142 | YOL089C | HAL9 | n |
| W303.asm02.HP0_G0048220 | 1349.264115 | 1143.260383 | 1453.247942 | 1360.973558 | 997.8523871 | 1290.885416 | 1857.212571 | 1518.153501 | 1421.816774 | 0.555784029 | 0.111217831 | YOL088C | MPD2 | n |
| W303.asm02.HP0_G0048230 | 546.6845985 | 536.8849757 | 509.1742956 | 465.0074059 | 536.0802718 | 489.7398388 | 488.4412267 | 619.2595542 | 653.0735073 | 0.574537815 | 0.09697543 | YOL087C | DUF1 | n |
| W303.asm02.HP0_G0048240 | 367.3642958 | 471.2022393 | 373.32933 | 354.1060581 | 399.4063411 | 368.4364406 | 550.9162673 | 415.0974199 | 367.4818011 | 0.601648522 | 0.112568205 | YOL086W- | MHF1 | n |
| W303.asm02.HP0_G0048250 | 27069.61131 | 22900.42883 | 26061.71006 | 32911.4342 | 17718.51453 | 30294.16581 | 6036.414152 | 3208.538565 | 5938.055511 | 0.75314567 | -0.090017516 | YOL086C | ADH1 | n |
| W303.asm02.HP0_G0048270 | 56.21933814 | 51.40388065 | 38.11477453 | 285.0359204 | 363.5791942 | 247.1330425 | 423.1264115 | 744.0790581 | 568.1125338 | 3.41E-21 | -2.616130975 | YOL084W | PHM7 | n |
| W303.asm02.HP0_G0048280 | 514.6977336 | 351.2598511 | 498.4239746 | 202.3463189 | 249.4630968 | 208.2073252 | 359.7047793 | 461.5418865 | 534.3328696 | 2.39E-06 | 1.050641674 | YOL083W | ATG34 | n |
| W303.asm02.HP0_G0048310 | 535.0530113 | 1058.539172 | 636.2235441 | 2757.941414 | 2465.438478 | 2829.809124 | 1568.502156 | 1449.454394 | 1437.171167 | 5.35E-17 | -1.852648809 | YOL080C | REX4 | n |
| W303.asm02.HP0_G0048320 | 223.9080536 | 376.9617915 | 244.3254777 | 875.5369568 | 712.5621434 | 873.5655165 | 674.919757 | 925.0189591 | 1029.767944 | 3.95E-11 | -1.54278396 | YOL078W | AVO1 | n |
| W303.asm02.HP0_G0048330 | 1102.092887 | 1623.029935 | 1125.851801 | 3160.688414 | 3613.234109 | 3201.866562 | 3515.640922 | 2879.556927 | 2612.294029 | 8.13E-14 | -1.372911917 | YOL077W- | ATP19 | n |
| W303.asm02.HP0_G0048340 | 3439.072616 | 5024.253372 | 4045.05261 | 4892.305951 | 4522.182095 | 4735.358774 | 3314.963519 | 2738.288341 | 2774.026967 | 0.290797185 | -0.177952452 | YOL077C | BRX1 | n |
| W303.asm02.HP0_G0048350 | 948.9436558 | 1308.895109 | 1143.443236 | 2527.383348 | 2044.801235 | 2497.582653 | 1961.337639 | 1913.89906 | 1918.275474 | 6.13E-12 | -1.056159228 | YOL076W | MDM20 | n |
| W303.asm02.HP0_G0048360 | 368.3335947 | 518.7984251 | 402.6483873 | 374.5352537 | 456.4643898 | 403.741161 | 326.574076 | 496.3752364 | 534.3328696 | 0.7924182 | 0.064384119 | YOL075C | | n |
| W303.asm02.HP0_G0048370 | 600.9653387 | 385.5291049 | 505.265088 | 192.6181305 | 160.5586953 | 196.4390851 | 466.6696216 | 417.032606 | 461.6554103 | 1.39E-12 | 1.436271959 | YOL073C | DSC2 | n |
| W303.asm02.HP0_G0048380 | 396.4432638 | 282.7213436 | 289.2813657 | 520.4580798 | 521.4840267 | 537.7180485 | 474.2423538 | 545.7224821 | 529.2147387 | 6.01E-05 | -0.706177513 | YOL072W | THP1 | n |
| W303.asm02.HP0_G0048390 | 1333.755332 | 1296.520101 | 1387.768714 | 939.7430002 | 959.3713775 | 976.7639299 | 1278.84515 | 733.4355345 | 808.6646877 | 0.018446571 | 0.48246154 | YOL071W | SDH5 | n |
| W303.asm02.HP0_G0048400 | 1391.913268 | 1287.952787 | 1336.949014 | 1379.457116 | 1421.143493 | 1357.87386 | 1403.795231 | 1105.95886 | 1456.620064 | 0.742905162 | -0.049750324 | YOL070C | NBA1 | n |
| W303.asm02.HP0_G0048410 | 485.6187656 | 538.7888231 | 501.3558803 | 458.197674 | 480.3491544 | 470.7296048 | 437.3252843 | 446.0603976 | 343.9383988 | 0.48890373 | 0.115110307 | YOL069W | NUF2 | n |
| W303.asm02.HP0_G0048430 | 214.2150643 | 246.5482424 | 167.1186268 | 59.34194929 | 74.30815648 | 51.59920668 | 19.87842202 | 76.43985122 | 72.67745928 | 1.86E-05 | 1.766381834 | YOL067C | RTG1 | n |
| W303.asm02.HP0_G0048440 | 569.9477729 | 363.6348594 | 404.6029912 | 1373.620203 | 1390.624071 | 1335.242629 | 1022.318847 | 1061.44958 | 1008.271794 | 9.36E-21 | -1.615152802 | YOL066C | RIB2 | n |
| W303.asm02.HP0_G0048450 | 305.329164 | 286.5290385 | 316.6458192 | 195.536587 | 172.5010776 | 186.4813434 | 193.104671 | 186.7454593 | 225.1977611 | 6.48E-05 | 0.709481415 | YOL065C | INP54 | n |
| W303.asm02.HP0_G0048460 | 2926.31348 | 3450.72347 | 3024.749415 | 1670.32995 | 1313.662052 | 1719.068307 | 1425.566836 | 1472.676627 | 1233.469555 | 3.19E-11 | 0.998121404 | YOL064C | MET22 | n |
| W303.asm02.HP0_G0048470 | 410.9827478 | 529.269586 | 421.2171236 | 494.1919711 | 492.2915367 | 554.9177841 | 436.3786928 | 569.9123085 | 528.1911125 | 0.329048093 | -0.179803286 | YOL063C | CRT10 | n |
| W303.asm02.HP0_G0048480 | 2144.089241 | 1798.183899 | 2109.994826 | 753.9346016 | 717.8698689 | 687.0841731 | 715.6231926 | 778.912408 | 907.9564278 | 1.92E-25 | 1.487251409 | YOL062C | APM4 | n |
| W303.asm02.HP0_G0048490 | 2667.510665 | 2707.271048 | 2867.403807 | 2441.77529 | 2505.246419 | 2438.741452 | 2420.434528 | 2328.028887 | 2202.843554 | 0.127135038 | 0.158456573 | YOL061W | PRS5 | n |
| W303.asm02.HP0_G0048500 | 715.3426129 | 658.7312113 | 633.2916383 | 1241.316841 | 1145.141769 | 1194.023747 | 1324.281543 | 1829.718464 | 1725.833751 | 6.80E-08 | -0.835199307 | YOL060C | MAM3 | n |
| W303.asm02.HP0_G0048520 | 2150.874333 | 1699.183833 | 1734.710892 | 1992.332986 | 1662.645001 | 1995.169325 | 1541.997593 | 738.2734998 | 1706.384854 | 0.951400755 | -0.01704691 | YOL058W | ARG1 | n |
| W303.asm02.HP0_G0048530 | 970.2682324 | 827.221709 | 888.3674371 | 1615.852095 | 1476.87461 | 1602.291155 | 1695.34542 | 1822.945313 | 1620.400254 | 4.81E-11 | -0.806456965 | YOL057W | | n |
| W303.asm02.HP0_G0048540 | 568.009175 | 640.6446607 | 662.6106957 | 674.1634567 | 541.3879972 | 746.8306229 | 492.2275928 | 497.3428295 | 525.1202339 | 0.683648695 | -0.071469324 | YOL056W | GPM3 | n |
| W303.asm02.HP0_G0048550 | 748.2987766 | 482.6253239 | 631.3370345 | 61.28758697 | 39.80794097 | 64.27269603 | 28.39774574 | 27.0926055 | 38.89779511 | 1.60E-39 | 3.4775354 | YOL055C | THI20 | n |
| W303.asm02.HP0_G0048560 | 1132.141154 | 1030.933384 | 1045.713045 | 697.5111089 | 701.9466925 | 714.2416503 | 764.8459518 | 646.3521597 | 775.9086497 | 4.12E-06 | 0.602112053 | YOL054W | PSH1 | n |
| W303.asm02.HP0_G0048570 | 786.101435 | 675.8658382 | 784.7734345 | 257.7969928 | 221.5975381 | 252.5645379 | 385.2627505 | 369.6205464 | 394.096082 | 3.48E-27 | 1.614035868 | YOL053W | AIM39 | n |
| W303.asm02.HP0_G0048590 | 1093.369197 | 885.2890557 | 1021.280497 | 1409.6145 | 2024.897264 | 1540.734206 | 2382.570867 | 2075.4871 | 1594.809599 | 0.000216835 | -0.728952585 | YOL052C | SPE2 | n |
| W303.asm02.HP0_G0048600 | 583.5179579 | 831.9813276 | 479.8552383 | 2165.49474 | 2312.841371 | 2207.902896 | 1956.604681 | 2471.232658 | 2440.32483 | 2.98E-20 | -1.818408394 | YOL051W | GAL11 | n |
| W303.asm02.HP0_G0048610 | 2705.313323 | 2202.751478 | 2960.247488 | 771.4453408 | 751.043153 | 732.3466351 | 835.8403162 | 872.7689342 | 923.3108207 | 6.51E-37 | 1.803046819 | YOL049W | GSH2 | n |
| W303.asm02.HP0_G0048620 | 779.3163425 | 692.0485414 | 735.908339 | 25.29328986 | 25.21169595 | 31.6837234 | 38.81025251 | 37.73612908 | 27.63790705 | 1.16E-86 | 4.736121304 | YOL048C | RRT8 | n |
| W303.asm02.HP0_G0048650 | 581.5793601 | 608.2792544 | 558.0393912 | 108.9557102 | 138.000862 | 111.3456565 | 150.5080524 | 127.7222831 | 159.6856852 | 2.23E-36 | 2.295057951 | YOL044W | PEX15 | n |
| W303.asm02.HP0_G0048660 | 573.8249686 | 408.3752741 | 498.4239746 | 474.7355943 | 521.4840267 | 476.1611002 | 620.0174486 | 539.9169238 | 567.0889076 | 0.959361164 | 0.009105535 | YOL043C | NTG2 | n |
| W303.asm02.HP0_G0048670 | 1254.27282 | 1365.058608 | 1255.832956 | 324.9214928 | 338.3674983 | 361.1944467 | 121.1637151 | 358.9770228 | 334.7257631 | 1.98E-10 | 1.919210564 | YOL042W | NGL1 | n |

| | | | | | | | | | | | | | | |
|---|---|---|---|---|---|---|---|---|---|---|---|---|---|---|
| W303.asm02.HP0_G0048680 | 2363.1508 | 2722.501827 | 2519.484327 | 5632.621088 | 4782.260642 | 5295.708054 | 2667.494916 | 2929.871766 | 2704.420386 | 8.16E-19 | -1.04696485 | YOL041C | NOP12 | n |
| W303.asm02.HP0_G0048690 | 7607.05803 | 8343.611369 | 7709.934776 | 18139.18011 | 19357.27476 | 17008.72797 | 13428.34737 | 13335.36746 | 11883.2764 | 3.39E-28 | -1.203810303 | YOL040C | RPS15 | n |
| W303.asm02.HP0_G0048700 | 7312.391154 | 7974.264967 | 7163.623008 | 7741.692335 | 8935.555817 | 7385.928549 | 8315.806543 | 5992.30378 | 5860.25992 | 0.566424608 | -0.099936248 | YOL039W | RPP2A | n |
| W303.asm02.HP0_G0048720 | 753.1452713 | 914.7986909 | 624.4959211 | 1319.142348 | 1158.411082 | 1290.885416 | 1021.372255 | 1126.278314 | 1360.399202 | 6.71E-05 | -0.717458938 | YOL036W | | n |
| W303.asm02.HP0_G0048730 | 484.6494667 | 598.7600172 | 471.0595211 | 784.0919857 | 1001.833181 | 847.3132886 | 826.3744009 | 985.0097284 | 938.6652135 | 4.66E-06 | -0.758699125 | YOL034W | SMC5 | n |
| W303.asm02.HP0_G0048740 | 906.2945028 | 861.4909628 | 813.1151899 | 717.9403045 | 680.7157906 | 752.2621184 | 641.7890536 | 660.8660555 | 756.4597522 | 0.054503277 | 0.26179422 | YOL033W | MSE1 | n |
| W303.asm02.HP0_G0048750 | 1822.281995 | 746.3081932 | 1437.611111 | 114.7926232 | 136.6739307 | 143.0293799 | 536.7173944 | 236.092705 | 341.8911465 | 7.54E-24 | 3.344459147 | YOL032W | OPI10 | n |
| W303.asm02.HP0_G0048760 | 1303.707065 | 1423.125955 | 1320.334882 | 537.968819 | 467.0798408 | 515.9920668 | 913.4608212 | 648.2873458 | 782.0504069 | 4.26E-19 | 1.410375064 | YOL031C | SIL1 | n |
| W303.asm02.HP0_G0048770 | 2673.326458 | 1986.664795 | 2474.528439 | 3193.764254 | 3288.135924 | 3184.666826 | 3745.662663 | 3721.362883 | 3412.769707 | 0.001479989 | -0.438143407 | YOL030W | GAS5 | n |
| W303.asm02.HP0_G0048800 | 1016.794581 | 947.1640972 | 988.052232 | 1144.034957 | 996.5254557 | 1134.277298 | 1187.025772 | 1465.903476 | 1630.636516 | 0.346947477 | -0.150674222 | YOL027C | MDM38 | n |
| W303.asm02.HP0_G0048810 | 649.4302854 | 482.6253239 | 678.2475262 | 484.4637827 | 524.1378895 | 453.5298692 | 855.7187382 | 645.3845666 | 771.814145 | 0.103958452 | 0.309308669 | YOL026C | MIM1 | n |
| W303.asm02.HP0_G0048820 | 225.8466515 | 232.2693867 | 206.2107032 | 221.8026957 | 203.020499 | 218.1650668 | 258.4194862 | 238.9954842 | 292.7570895 | 0.814500028 | 0.045269096 | YOL025W | LAG2 | n |
| W303.asm02.HP0_G0048840 | 292.7282779 | 447.4041464 | 263.871516 | 784.0919857 | 768.2932608 | 771.2723524 | 548.0764927 | 766.3336983 | 747.2471165 | 3.94E-08 | -1.210362561 | YOL023W | IFM1 | n |
| W303.asm02.HP0_G0048850 | 1570.264272 | 1484.049073 | 1782.598686 | 2232.61924 | 2474.726997 | 2402.531483 | 1774.859109 | 1890.676826 | 1620.400254 | 1.40E-05 | -0.555529064 | YOL022C | TSR4 | n |
| W303.asm02.HP0_G0048860 | 709.5268193 | 991.9045119 | 602.0179772 | 4518.743516 | 4161.256763 | 4438.437023 | 2130.777522 | 2787.635587 | 2273.473761 | 2.97E-40 | -2.509642204 | YOL021C | DIS3 | n |
| W303.asm02.HP0_G0048870 | 927.6190793 | 1406.943252 | 851.2299645 | 2245.265885 | 2184.129028 | 2268.554595 | 1612.991958 | 2136.445462 | 2263.237499 | 1.89E-07 | -1.072020514 | YOL020W | TAT2 | n |
| W303.asm02.HP0_G0048890 | 1100.154289 | 1269.866237 | 1320.334882 | 766.5812466 | 940.7943383 | 798.4298296 | 1033.677945 | 1335.278414 | 1568.195318 | 0.002221849 | 0.559692973 | YOL019W | TOS7 | n |
| W303.asm02.HP0_G0048900 | 1497.566852 | 1416.462489 | 1455.202545 | 246.1231667 | 242.8284399 | 248.0382917 | 407.9809471 | 344.463127 | 379.7653154 | 7.15E-83 | 2.567020309 | YOL018C | TLG2 | n |
| W303.asm02.HP0_G0048920 | 1046.842848 | 710.135092 | 848.2980587 | 1098.312471 | 781.5625744 | 1036.51038 | 841.5198653 | 956.9495298 | 927.4053254 | 0.412624966 | -0.164292291 | YOL016C | CMK2 | n |
| W303.asm02.HP0_G0048950 | 885.9392252 | 745.3562695 | 927.4595135 | 290.8728334 | 315.8096651 | 264.3327781 | 493.1741843 | 526.3706211 | 591.6559361 | 2.58E-22 | 1.557734824 | YOL013C | HRD1 | n |
| W303.asm02.HP0_G0048960 | 1383.189578 | 1299.375872 | 1332.062505 | 1559.428602 | 1748.89554 | 1582.375671 | 1772.965925 | 1501.704419 | 1359.375576 | 0.04329161 | -0.284153763 | YOL012C | HTZ1 | n |
| W303.asm02.HP0_G0048970 | 489.4959614 | 466.4426207 | 388.9661606 | 2261.803805 | 2360.6109 | 2204.281899 | 4176.361806 | 4007.770427 | 5170.33587 | 8.47E-56 | -2.3433487 | YOL011W | PLB3 | n |
| W303.asm02.HP0_G0048980 | 1458.794895 | 1848.635856 | 1418.065073 | 3672.391124 | 3074.499974 | 3578.450245 | 2098.59341 | 2163.538067 | 1929.535362 | 1.18E-14 | -1.127958758 | YOL010W | RCL1 | n |
| W303.asm02.HP0_G0048990 | 704.6803246 | 477.8657053 | 647.951167 | 309.3563914 | 337.0405669 | 325.8897264 | 385.2627505 | 313.5001493 | 375.6708106 | 1.42E-06 | 0.913428786 | YOL009C | MDM12 | n |
| W303.asm02.HP0_G0049000 | 808.3953105 | 771.0582098 | 711.4757912 | 784.0919857 | 828.0051722 | 768.5566047 | 695.7447705 | 543.787296 | 613.152086 | 0.72607439 | -0.05464247 | YOL008W | COQ10 | n |
| W303.asm02.HP0_G0049020 | 989.654211 | 968.106419 | 872.7306065 | 2075.995406 | 2142.994156 | 2082.978501 | 1503.187341 | 1671.033203 | 1362.446455 | 3.17E-20 | -1.154537475 | YOL006C | TOP1 | n |
| W303.asm02.HP0_G0049030 | 882.0620294 | 851.0198019 | 969.4834957 | 703.3480219 | 871.7939073 | 749.5463707 | 898.3153568 | 658.9308694 | 521.0257292 | 0.305788223 | 0.218276104 | YOL005C | RPB11 | n |
| W303.asm02.HP0_G0049040 | 847.1672678 | 1038.548774 | 757.408981 | 3580.946153 | 3952.928539 | 3740.489859 | 2981.763302 | 3555.904471 | 3403.557072 | 7.27E-46 | -2.092420178 | YOL004W | SIN3 | n |
| W303.asm02.HP0_G0049050 | 939.2506665 | 681.5773805 | 848.2980587 | 449.4423045 | 417.9833802 | 412.7936534 | 481.815086 | 491.5372711 | 537.4037482 | 4.97E-09 | 0.947209404 | YOL003C | PFA4 | n |
| W303.asm02.HP0_G0049060 | 1905.641703 | 1802.943518 | 1808.985837 | 120.6295363 | 79.61588195 | 124.0191459 | 173.226249 | 247.7038217 | 161.7329375 | 1.18E-84 | 4.076399855 | YOL002C | IZH2 | n |
| W303.asm02.HP0_G0049070 | 1023.579674 | 566.3946109 | 1013.462082 | 580.772848 | 605.0807028 | 583.8857598 | 369.1706946 | 476.0557823 | 510.7894673 | 0.01868631 | 0.556981604 | YOL001W | PHO80 | n |
| W303.asm02.HP0_G0049080 | 980.9305206 | 920.5102332 | 918.6637963 | 3785.23811 | 3396.944296 | 3646.343938 | 1908.328513 | 2239.977919 | 2075.913907 | 5.18E-63 | -1.941340696 | YOR001W | RRP6 | n |
| W303.asm02.HP0_G0049090 | 2027.773369 | 1776.289654 | 2119.767845 | 958.2265582 | 954.063652 | 949.6064527 | 1182.292814 | 1234.648736 | 1308.194267 | 3.23E-18 | 1.049517937 | YOR002W | ALG6 | n |
| W303.asm02.HP0_G0049110 | 491.4345592 | 642.5485082 | 528.7203338 | 2585.752479 | 2304.879782 | 2466.804179 | 1978.376286 | 1492.028488 | 1463.785447 | 5.37E-38 | -2.145762521 | YOR004W | UTP23 | n |
| W303.asm02.HP0_G0049120 | 770.5926521 | 441.6926041 | 690.9524511 | 429.0131088 | 536.0802718 | 450.8141215 | 404.194581 | 463.4770726 | 437.0883818 | 0.053669325 | 0.42818132 | YOR005C | DNL4 | n |
| W303.asm02.HP0_G0049130 | 1543.123902 | 1506.895242 | 1587.138303 | 1241.316841 | 1224.757651 | 1182.255507 | 1478.575961 | 1129.181093 | 1145.437703 | 0.015518384 | 0.346117285 | YOR006C | TSR3 | n |
| W303.asm02.HP0_G0049140 | 10216.41076 | 8381.688379 | 9251.139889 | 3377.627015 | 2936.499112 | 3219.971546 | 3554.451175 | 2573.797522 | 2764.814331 | 6.19E-24 | 1.546210059 | YOR007C | SGT2 | n |
| W303.asm02.HP0_G0049220 | 959.6059441 | 1205.135424 | 1034.962724 | 1730.644718 | 1784.722687 | 1668.374349 | 2486.695935 | 2423.820599 | 2692.136872 | 6.38E-08 | -0.695611691 | YOR014W | RTS1 | n |
| W303.asm02.HP0_G0049230 | 1237.794738 | 1332.693202 | 1309.584561 | 3180.144791 | 3038.672828 | 3076.942167 | 5802.606046 | 3904.237971 | 3791.511397 | 2.30E-13 | -1.260529952 | YOR016C | ERP4 | n |
| W303.asm02.HP0_G0049240 | 528.2679187 | 700.6158548 | 526.76573 | 763.6627901 | 732.4661139 | 743.209626 | 616.2310825 | 702.4725568 | 643.8608716 | 0.029984429 | -0.351102616 | YOR017W | PET127 | n |
| W303.asm02.HP0_G0049250 | 441.0310147 | 469.2983919 | 376.2612357 | 369.6711595 | 315.8096651 | 323.1739787 | 394.7286657 | 506.0511669 | 419.6867367 | 0.051769407 | 0.35004875 | YOR018W | ROD1 | n |
| W303.asm02.HP0_G0049270 | 5634.5347 | 5910.494351 | 5238.338243 | 1483.548732 | 1600.279227 | 1472.840513 | 3022.466738 | 1900.352757 | 1831.267249 | 3.00E-23 | 1.881248862 | YOR020C | HSP10 | n |
| W303.asm02.HP0_G0049280 | 909.2023996 | 750.115888 | 804.3194727 | 961.1450147 | 1038.987259 | 966.8061882 | 1341.32019 | 904.699505 | 853.7042399 | 0.174999382 | -0.267835033 | YOR020W | MCO10 | n |
| W303.asm02.HP0_G0049290 | 2608.38343 | 3002.3674 | 2891.836355 | 1667.411493 | 1859.030843 | 1589.617665 | 1620.56469 | 1432.037719 | 1392.131614 | 9.30E-09 | 0.733783757 | YOR021C | SFM1 | n |
| W303.asm02.HP0_G0049300 | 1150.557834 | 1002.375673 | 953.8466651 | 345.3506885 | 303.8672828 | 343.9947112 | 549.9696758 | 567.0095293 | 778.9795283 | 3.75E-20 | 1.643310952 | YOR022C | DDL1 | n |
| W303.asm02.HP0_G0049310 | 757.9917659 | 424.5579772 | 624.4959211 | 1162.518515 | 1391.951003 | 1119.79331 | 1186.07918 | 1401.074741 | 1692.054087 | 4.19E-06 | -1.023312822 | YOR025W | HST3 | n |
| W303.asm02.HP0_G0049320 | 1106.939382 | 668.2504485 | 943.0963441 | 418.3121016 | 416.6564489 | 409.1726564 | 513.0526063 | 481.8613406 | 545.5927577 | 5.51E-09 | 1.127555342 | YOR026W | BUB3 | n |
| W303.asm02.HP0_G0049330 | 8735.321988 | 5785.792345 | 6464.852141 | 3360.116276 | 2859.537093 | 3317.738464 | 3661.416017 | 3501.71926 | 4327.891519 | 7.38E-11 | 1.137475278 | YOR027W | STI1 | n |
| W303.asm02.HP0_G0049340 | 664.9390683 | 323.6540634 | 526.76573 | 558.3980146 | 662.1387515 | 528.6655561 | 796.0834721 | 929.8569243 | 1355.281072 | 0.47391007 | -0.206267687 | YOR028C | CIN5 | n |
| W303.asm02.HP0_G0049360 | 300.4826694 | 406.4714266 | 313.7139134 | 223.7483334 | 264.0593418 | 232.6490547 | 319.9479353 | 335.7547895 | 326.5367536 | 0.006601267 | 0.505855213 | YOR030W | DFG16 | n |
| W303.asm02.HP0_G0049380 | 122.1316656 | 204.6635989 | 135.8449656 | 1968.012515 | 1475.547679 | 1896.497158 | 313.3217946 | 779.8800011 | 1043.075085 | 7.19E-25 | -3.52886899 | YOR032C | HMS1 | n |
| W303.asm02.HP0_G0049400 | 495.311755 | 594.0003987 | 481.8098421 | 1378.484297 | 1086.756789 | 1348.821367 | 1189.865546 | 1057.579207 | 1031.815196 | 2.31E-16 | -1.280647083 | YOR033C | EXO1 | n |
| W303.asm02.HP0_G0049430 | 348.947616 | 407.4233504 | 321.5323287 | 436.7956595 | 370.213851 | 392.8781701 | 395.6752573 | 507.9863531 | 637.7191145 | 0.481045251 | -0.155541465 | YOR035C | SHE4 | n |
| W303.asm02.HP0_G0049440 | 1452.009802 | 1162.298857 | 1253.878352 | 417.3392827 | 465.7529094 | 447.1931245 | 543.3435351 | 430.5789088 | 422.7576152 | 5.73E-22 | 1.540871961 | YOR036W | PEP12 | n |
| W303.asm02.HP0_G0049450 | 864.6146486 | 632.0773473 | 798.4556613 | 351.1876015 | 338.3674983 | 371.1521884 | 416.5002708 | 320.2733007 | 406.3795962 | 5.94E-10 | 1.112380382 | YOR037W | CYC2 | n |
| W303.asm02.HP0_G0049460 | 437.153819 | 526.4138148 | 431.9674447 | 1151.817508 | 1212.815268 | 1118.888061 | 1003.387016 | 1328.505262 | 1224.25692 | 2.74E-17 | -1.319098578 | YOR038C | HIR2 | n |
| W303.asm02.HP0_G0049470 | 2629.708006 | 2904.319257 | 2778.469333 | 1807.497406 | 1730.318501 | 1830.413963 | 2125.097973 | 1940.991665 | 1792.369453 | 4.51E-08 | 0.630468714 | YOR039W | CKB2 | n |
| W303.asm02.HP0_G0049480 | 759.9303638 | 649.2119742 | 829.7293224 | 427.0674711 | 337.0405669 | 381.10993 | 598.2458435 | 440.2548393 | 554.8053933 | 1.13E-07 | 0.964878633 | YOR040W | GLO4 | n |
| W303.asm02.HP0_G0049490 | 1166.066617 | 778.6735995 | 1047.667649 | 781.1735292 | 639.5809183 | 759.5041123 | 1407.581597 | 973.3986118 | 1102.445403 | 0.026234993 | 0.455658899 | YOR042W | CUE5 | n |
| W303.asm02.HP0_G0049500 | 1030.364766 | 1089.952655 | 992.9387416 | 1647.955116 | 1905.473441 | 1656.606109 | 1771.072742 | 1892.612013 | 2746.38906 | 4.24E-05 | -0.742375541 | YOR043W | WHI2 | n |
| W303.asm02.HP0_G0049520 | 1078.829713 | 1242.260449 | 1140.51133 | 1029.242334 | 1139.834043 | 1042.847124 | 1352.679289 | 863.0930037 | 859.8459971 | 0.599665015 | 0.108464239 | YOR045W | TOM6 | n |
| W303.asm02.HP0_G0049530 | 2871.063441 | 2429.309323 | 3049.181962 | 4655.910972 | 4437.258487 | 4732.643026 | 3453.165882 | 3752.325861 | 3677.88889 | 2.34E-09 | -0.727796486 | YOR046C | DBP5 | n |
| W303.asm02.HP0_G0049550 | 753.1452713 | 1029.98146 | 691.929753 | 3327.040436 | 3244.347189 | 3281.528495 | 1362.145204 | 2355.121492 | 2187.489162 | 2.83E-20 | -1.993008971 | YOR048C | RAT1 | n |

| | | | | | | | | | | | | | | |
|---|---|---|---|---|---|---|---|---|---|---|---|---|---|---|
| W303.asm02.HP0_G0049560 | 267.5265056 | 251.307861 | 211.0972128 | 438.7412972 | 403.3871352 | 446.2878753 | 457.2037064 | 587.3289834 | 514.883972 | 2.31E-06 | -0.82095582 | YOR049C | RSB1 | n |
| W303.asm02.HP0_G0049570 | 3299.493569 | 3964.762276 | 3234.869325 | 3237.541102 | 2741.440202 | 3124.015127 | 1684.932914 | 1741.667496 | 1366.54096 | 0.185178447 | 0.205507773 | YOR051C | ETT1 | n |
| W303.asm02.HP0_G0049580 | 2689.80454 | 2968.098146 | 2213.588828 | 251.9600798 | 192.405048 | 260.7117811 | 765.7925434 | 627.9678917 | 721.6564618 | 9.34E-88 | 3.474795924 | YOR052C | TMC1 | n |
| W303.asm02.HP0_G0049600 | 1767.031956 | 1955.251312 | 1895.965707 | 3147.06895 | 2838.306191 | 3401.926264 | 1910.221697 | 1484.287744 | 1462.761821 | 4.55E-07 | -0.740983479 | YOR056C | NOB1 | n |
| W303.asm02.HP0_G0049610 | 619.3820185 | 474.0580105 | 553.1528816 | 298.6553841 | 327.7520473 | 298.7322492 | 316.1615692 | 332.8520104 | 324.4895013 | 2.36E-07 | 0.833609004 | YOR057W | SGT1 | n |
| W303.asm02.HP0_G0049620 | 895.6322145 | 835.7890225 | 937.2325326 | 505.8657972 | 630.2923987 | 481.5925956 | 311.4286116 | 624.0975195 | 639.7663669 | 0.004346503 | 0.723479722 | YOR058C | ASE1 | n |
| W303.asm02.HP0_G0049630 | 785.1321361 | 825.3178616 | 760.3408868 | 275.307732 | 262.7324104 | 282.4377629 | 389.9957081 | 469.2826309 | 485.1988126 | 8.04E-25 | 1.529225401 | YOR059C | LPL1 | n |
| W303.asm02.HP0_G0049640 | 157.9957261 | 292.2405807 | 166.1413249 | 593.4194929 | 598.446046 | 579.3595136 | 711.8368265 | 588.2965765 | 529.2147387 | 1.90E-10 | -1.522221838 | YOR060C | SLD7 | n |
| W303.asm02.HP0_G0049650 | 2476.558775 | 2259.866901 | 2485.27876 | 1971.90379 | 2015.608745 | 1958.054106 | 2058.836566 | 2067.746355 | 1934.653493 | 0.007407749 | 0.280712251 | YOR061W | CKA2 | n |
| W303.asm02.HP0_G0049670 | 13920.10198 | 11153.69018 | 12119.521 | 34931.97894 | 36733.441 | 32798.08521 | 35005.90117 | 27810.55954 | 26929.55773 | 4.36E-25 | -1.489865494 | YOR063W | RPL3 | n |
| W303.asm02.HP0_G0049680 | 867.5225454 | 532.1253571 | 815.0697938 | 284.0631015 | 363.5791942 | 311.4057385 | 535.7708029 | 337.6899756 | 331.6548846 | 1.36E-06 | 1.209073218 | YOR064C | YNG1 | n |
| W303.asm02.HP0_G0049690 | 2713.067715 | 2332.213104 | 2744.263766 | 11619.34823 | 11350.5709 | 11185.25961 | 5590.569544 | 9199.874752 | 8546.255035 | 2.27E-32 | -2.132511025 | YOR065W | CYT1 | n |
| W303.asm02.HP0_G0049700 | 227.7852494 | 311.2790551 | 183.7327593 | 702.3752031 | 690.0043102 | 717.8626473 | 599.192435 | 779.8800011 | 819.9245758 | 2.16E-13 | -1.545498025 | YOR066W | MSA1 | n |
| W303.asm02.HP0_G0049710 | 1333.755332 | 1429.789421 | 1465.952866 | 824.950377 | 834.6398291 | 791.1878357 | 939.0187924 | 1280.12561 | 1101.421777 | 6.30E-08 | 0.787631647 | YOR067C | ALG8 | n |
| W303.asm02.HP0_G0049730 | 221.9694558 | 277.961725 | 210.1199109 | 216.9386015 | 248.1361654 | 223.5965623 | 412.7139047 | 259.3149383 | 385.9070725 | 0.852202478 | 0.046107568 | YOR070C | GYP1 | n |
| W303.asm02.HP0_G0049750 | 364.456399 | 422.6541298 | 384.079651 | 288.9271957 | 302.5403514 | 251.6592887 | 225.2887828 | 269.9584619 | 289.6862109 | 0.007562684 | 0.476898485 | YOR073W | SGO1 | n |
| W303.asm02.HP0_G0049760 | 710.4961182 | 507.3753405 | 675.3156205 | 1080.801732 | 989.8907989 | 1057.331112 | 1598.793085 | 1271.417272 | 1074.807496 | 0.000141298 | -0.724912642 | YOR074C | CDC21 | n |
| W303.asm02.HP0_G0049770 | 1046.842848 | 1025.221842 | 998.802553 | 688.7557393 | 644.8886438 | 701.5681609 | 866.131245 | 628.9354847 | 695.042181 | 0.000105214 | 0.592612722 | YOR075W | UFE1 | n |
| W303.asm02.HP0_G0049790 | 187.0746942 | 118.9904645 | 159.3002115 | 263.6339059 | 244.1553713 | 264.3327781 | 239.4876557 | 202.2269482 | 209.8433683 | 0.00071056 | -0.732083403 | YOR077W | RTS2 | n |
| W303.asm02.HP0_G0049800 | 1471.395781 | 1738.212705 | 1595.934021 | 2410.645088 | 2067.359068 | 2373.563507 | 1398.115682 | 1265.611714 | 1341.973931 | 5.44E-05 | -0.512391082 | YOR078W | BUD21 | n |
| W303.asm02.HP0_G0049810 | 814.2111041 | 882.4332845 | 800.4102651 | 335.6225001 | 364.9061256 | 322.2687294 | 738.3413892 | 562.171564 | 614.1757122 | 1.01E-16 | 1.289994494 | YOR079C | ATX2 | n |
| W303.asm02.HP0_G0049820 | 627.1364099 | 820.558243 | 693.8843568 | 470.844319 | 460.4451839 | 469.8243555 | 364.437737 | 504.1159808 | 469.8444198 | 0.000562189 | 0.611938936 | YOR080W | DIA2 | n |
| W303.asm02.HP0_G0049840 | 100.8070891 | 124.7020068 | 78.18415288 | 326.8671305 | 390.1178215 | 393.7834194 | 454.3639318 | 449.9307698 | 590.6323099 | 2.26E-17 | -1.869509668 | YOR083W | WHI5 | n |
| W303.asm02.HP0_G0049850 | 136.6711496 | 200.855904 | 118.2535312 | 195.536587 | 169.8472148 | 201.8705805 | 216.7694591 | 128.6898761 | 206.7724898 | 0.28086068 | -0.317133387 | YOR084W | LPX1 | n |
| W303.asm02.HP0_G0049860 | 1216.470161 | 1524.981793 | 1394.609827 | 1198.512812 | 1131.872455 | 1127.940553 | 2396.76974 | 2236.107546 | 2504.81328 | 0.052053695 | 0.258062918 | YOR085W | OST3 | n |
| W303.asm02.HP0_G0049870 | 1208.71577 | 1424.077879 | 1213.808973 | 6129.731516 | 5683.24704 | 6105.906123 | 2817.056377 | 4611.548493 | 3754.660854 | 6.46E-36 | -2.219853488 | YOR086C | TCB1 | n |
| W303.asm02.HP0_G0049880 | 1046.842848 | 810.0870821 | 1065.259083 | 1447.554435 | 1218.122994 | 1449.304033 | 743.0743468 | 1113.699604 | 771.814145 | 0.012509614 | -0.494493934 | YOR087W | YVC1 | n |
| W303.asm02.HP0_G0049890 | 2966.054736 | 2299.847697 | 2841.016655 | 787.0104422 | 854.5437996 | 749.5463707 | 1242.874672 | 958.884716 | 963.2322419 | 4.72E-28 | 1.762408496 | YOR089C | VPS21 | n |
| W303.asm02.HP0_G0049900 | 783.1935382 | 665.3946773 | 751.5451696 | 1358.055102 | 1069.506681 | 1462.882772 | 1358.358838 | 1280.12561 | 1224.25692 | 1.07E-07 | -0.824019141 | YOR090C | PTC5 | n |
| W303.asm02.HP0_G0049910 | 919.8646878 | 770.1062861 | 765.2273963 | 3414.594131 | 3550.868335 | 3520.514294 | 3043.291751 | 2609.598465 | 2489.458887 | 3.25E-56 | -2.094513821 | YOR091W | TMA46 | n |
| W303.asm02.HP0_G0049920 | 310.1756587 | 289.3848096 | 324.4642344 | 488.3550581 | 313.1558023 | 496.0765835 | 244.2206133 | 388.9724075 | 349.0565298 | 0.041949287 | -0.493008616 | YOR092W | ECM3 | n |
| W303.asm02.HP0_G0049930 | 775.4391467 | 874.8178948 | 818.0016995 | 772.4181596 | 814.7358586 | 753.1673676 | 695.7447705 | 1036.29216 | 858.8223709 | 0.670643327 | 0.077395329 | YOR093C | CMR2 | n |
| W303.asm02.HP0_G0049940 | 988.6849121 | 879.5775134 | 1002.711761 | 528.2406306 | 650.1963692 | 480.6873464 | 951.3244822 | 1006.296776 | 1223.233293 | 2.58E-06 | 0.794149322 | YOR094W | ARF3 | n |
| W303.asm02.HP0_G0049950 | 2080.115511 | 3137.540567 | 2187.201677 | 2842.576653 | 2762.671104 | 2792.693905 | 3091.567919 | 2114.190822 | 2456.702849 | 0.373005477 | -0.181550652 | YOR095C | RKI1 | n |
| W303.asm02.HP0_G0049960 | 6645.513488 | 8265.553624 | 6325.097968 | 20007.9651 | 22492.81358 | 19060.928 | 18114.92201 | 14669.67828 | 13654.14971 | 1.88E-22 | -1.535442216 | YOR096W | RPS7A | n |
| W303.asm02.HP0_G0049970 | 769.6233531 | 574.0100006 | 804.3194727 | 122.5751739 | 104.8275779 | 80.56718235 | 147.6682778 | 133.5278414 | 123.8587686 | 4.85E-37 | 2.804760309 | YOR097C | | n |
| W303.asm02.HP0_G0049980 | 770.5926521 | 888.1448268 | 664.5652995 | 1934.936674 | 1801.972795 | 1882.918419 | 959.8438059 | 1528.797024 | 1589.691468 | 2.23E-10 | -1.274395967 | YOR098C | NUP1 | n |
| W303.asm02.HP0_G0049990 | 3220.011057 | 3371.713801 | 3425.443198 | 2957.369276 | 2831.671535 | 2965.59651 | 3936.874151 | 4764.428195 | 3683.007021 | 0.143021551 | 0.194185619 | YOR099W | KTR1 | n |
| W303.asm02.HP0_G0050000 | 88.20620294 | 50.45195694 | 79.16145479 | 8.755369568 | 19.90397049 | 11.76824012 | 258.4194862 | 136.4306205 | 138.1895352 | 2.68E-08 | 2.465775394 | YOR100C | CRC1 | n |
| W303.asm02.HP0_G0050020 | 1498.536151 | 1574.481826 | 1558.796548 | 539.9144567 | 554.6573109 | 580.2647628 | 735.5016146 | 593.1345418 | 526.1438601 | 1.75E-22 | 1.467300681 | YOR103C | OST2 | n |
| W303.asm02.HP0_G0050030 | 635.8601003 | 792.0005315 | 669.451809 | 1002.976225 | 992.5446616 | 1013.879149 | 1217.316701 | 1193.042235 | 1426.934905 | 0.000289513 | -0.520958846 | YOR104W | PIN2 | n |
| W303.asm02.HP0_G0050040 | 852.0137625 | 1071.866104 | 886.4128333 | 170.2432971 | 193.7319794 | 187.3865927 | 311.4286116 | 216.740844 | 201.6543588 | 3.61E-31 | 2.351160953 | YOR106W | VAM3 | n |
| W303.asm02.HP0_G0050050 | 323.7458438 | 500.7118745 | 272.6672332 | 661.5168118 | 561.2919677 | 669.8844375 | 395.6752573 | 440.2548393 | 541.4982529 | 0.000961454 | -0.787294511 | YOR107W | RGS2 | n |
| W303.asm02.HP0_G0050070 | 756.0531681 | 794.8563027 | 668.4745071 | 1470.902087 | 1529.951865 | 1480.987757 | 1245.714446 | 1346.88953 | 1376.777221 | 7.90E-17 | -1.01351355 | YOR109W | INP53 | n |
| W303.asm02.HP0_G0050080 | 622.2899153 | 533.0772808 | 586.3811466 | 2080.859501 | 1779.414961 | 2147.251197 | 2082.501354 | 1586.852608 | 1900.873829 | 4.88E-32 | -1.786992483 | YOR110W | TFC7 | n |
| W303.asm02.HP0_G0050100 | 373.1800894 | 325.5579108 | 292.2132714 | 1180.029254 | 1118.603141 | 1218.465477 | 880.3301178 | 1552.019258 | 1155.673965 | 1.20E-18 | -1.827735034 | YOR112W | CEX1 | n |
| W303.asm02.HP0_G0050110 | 358.6406054 | 367.4425543 | 346.9421784 | 3681.146494 | 3116.961778 | 3636.386197 | 1334.69405 | 2452.84839 | 2651.191824 | 4.28E-48 | -3.281738116 | YOR113W | AZF1 | n |
| W303.asm02.HP0_G0050120 | 326.6537406 | 280.8174962 | 365.5109147 | 222.7755146 | 208.3282244 | 251.6592887 | 347.3990895 | 281.5695785 | 223.1505088 | 0.020677461 | 0.508497329 | YOR114W | CIM1 | n |
| W303.asm02.HP0_G0050130 | 1465.579987 | 1605.895309 | 1550.978133 | 629.41379 | 626.3116046 | 639.1059634 | 843.4130484 | 710.2133012 | 707.3256952 | 4.96E-24 | 1.286368541 | YOR115C | TRS33 | n |
| W303.asm02.HP0_G0050140 | 703.7110257 | 1176.577713 | 715.3849988 | 4260.946523 | 4891.069014 | 4576.940157 | 2834.095025 | 3482.367399 | 3169.146675 | 1.59E-33 | -2.402796195 | YOR116C | RPO31 | n |
| W303.asm02.HP0_G0050150 | 3941.169463 | 3503.079274 | 3729.384092 | 3187.927341 | 2610.073996 | 3022.627212 | 3162.562284 | 3193.057076 | 3002.295606 | 0.00526377 | 0.34045057 | YOR117W | RPT5 | n |
| W303.asm02.HP0_G0050160 | 1372.52729 | 1494.520234 | 1473.771282 | 862.8903118 | 853.2168682 | 920.638477 | 771.4720925 | 619.2595542 | 611.1048336 | 1.88E-07 | 0.718665503 | YOR118W | RTC5 | n |
| W303.asm02.HP0_G0050170 | 616.4741217 | 1030.933384 | 648.9284689 | 2147.984001 | 1707.760668 | 2180.745419 | 630.4299554 | 1238.519108 | 1086.067384 | 1.25E-07 | -1.394579581 | YOR119C | RIO1 | n |
| W303.asm02.HP0_G0050190 | 5803.192714 | 6342.667718 | 6033.861998 | 3355.252182 | 3400.92509 | 3254.371018 | 3444.646558 | 2718.93648 | 2543.711075 | 3.76E-10 | 0.860907472 | YOR122C | PFY1 | n |
| W303.asm02.HP0_G0050200 | 2046.190048 | 1713.462688 | 2122.699751 | 1238.398384 | 1135.853249 | 1165.961021 | 1105.618901 | 1097.250523 | 1195.595386 | 1.68E-08 | 0.732070295 | YOR123C | LEO1 | n |
| W303.asm02.HP0_G0050210 | 1254.27282 | 1373.625922 | 1305.675353 | 1046.753073 | 1012.448632 | 1012.973899 | 766.7391349 | 1211.426503 | 1199.689891 | 0.055878973 | 0.356527274 | YOR124C | UBP2 | n |
| W303.asm02.HP0_G0050220 | 880.1234316 | 1050.923782 | 853.1845683 | 570.0718407 | 667.446477 | 565.780775 | 773.3652756 | 687.958661 | 656.1443859 | 4.21E-05 | 0.628821081 | YOR125C | CAT5 | n |
| W303.asm02.HP0_G0050230 | 299.5133704 | 310.3271313 | 304.9181962 | 465.0074059 | 516.1763013 | 459.8666139 | 684.3856723 | 596.0373209 | 511.8130935 | 6.41E-05 | -0.653993724 | YOR126C | IAH1 | n |
| W303.asm02.HP0_G0050240 | 572.8556697 | 684.4331517 | 575.6308256 | 1360.000739 | 1474.220747 | 1266.443687 | 1277.898558 | 1617.815585 | 1577.407954 | 1.67E-14 | -1.160847375 | YOR127W | RGA1 | n |
| W303.asm02.HP0_G0050250 | 367.3642958 | 339.8367665 | 258.0077045 | 2813.392088 | 1507.394032 | 2475.856671 | 1813.669361 | 1833.588836 | 1811.818351 | 1.98E-36 | -2.816587902 | YOR128C | ADE2 | n |
| W303.asm02.HP0_G0050260 | 339.2546267 | 423.6060535 | 392.8753682 | 584.6641233 | 611.7153596 | 580.2647628 | 400.4082149 | 496.3752364 | 514.883972 | 0.000167491 | -0.619655313 | YOR129C | AFI1 | n |
| W303.asm02.HP0_G0050270 | 1453.9484 | 1073.769951 | 1421.97428 | 1923.262848 | 1356.123856 | 1897.402407 | 848.146006 | 1003.393996 | 674.5696572 | 0.064249253 | -0.391158477 | YOR130C | ORT1 | n |
| W303.asm02.HP0_G0050280 | 1096.277094 | 950.0198684 | 1252.90105 | 203.3191377 | 191.0781167 | 184.6708449 | 328.467259 | 257.3797522 | 239.5285278 | 6.12E-43 | 2.51020203 | YOR131C | | n |

| | | | | | | | | | | | | | | |
|---|---|---|---|---|---|---|---|---|---|---|---|---|---|---|
| W303.asm02.HP0_G0050290 | 727.943499 | 549.259984 | 623.5186192 | 458.197674 | 412.6756548 | 470.7296048 | 532.9310283 | 526.3706211 | 574.2542909 | 0.001686356 | 0.500615017 | YOR132W | VPS17 | n |
| W303.asm02.HP0_G0050320 | 5634.5347 | 5108.022659 | 5770.967784 | 10244.75521 | 11394.35964 | 9924.247417 | 10528.93753 | 10297.12527 | 9784.842721 | 1.19E-17 | -0.934436069 | YOR136W | IDH2 | n |
| W303.asm02.HP0_G0050330 | 331.5002352 | 462.6349259 | 308.8274039 | 823.0047393 | 548.0226541 | 753.1673676 | 636.1095045 | 1076.931068 | 977.5630086 | 0.000151926 | -0.946721305 | YOR137C | SIA1 | n |
| W303.asm02.HP0_G0050340 | 821.9654955 | 844.3563359 | 740.7948485 | 421.2305581 | 400.7332725 | 433.6143859 | 765.7925434 | 581.5234251 | 744.1762379 | 2.09E-09 | 0.937882617 | YOR138C | RUP1 | n |
| W303.asm02.HP0_G0050350 | 191.9211888 | 321.7502159 | 148.5498905 | 1356.109464 | 1231.392307 | 1340.674124 | 825.4278094 | 1278.190424 | 1275.438229 | 1.45E-22 | -2.568191774 | YOR140W | SFL1 | n |
| W303.asm02.HP0_G0050360 | 619.3820185 | 568.2984583 | 553.1528816 | 974.7644785 | 1125.237798 | 1038.320878 | 961.7369889 | 1070.157917 | 1197.642639 | 1.76E-09 | -0.849286994 | YOR141C | ARP8 | n |
| W303.asm02.HP0_G0050370 | 4320.165346 | 3668.714001 | 4415.450034 | 3224.894457 | 2679.074427 | 3095.952401 | 3463.578388 | 2323.190921 | 3554.030121 | 0.011757888 | 0.462560216 | YOR142W | LSC1 | n |
| W303.asm02.HP0_G0050380 | 1127.29466 | 1556.395275 | 1140.51133 | 528.2406306 | 363.5791942 | 562.159778 | 727.9288824 | 566.0419363 | 679.6877882 | 2.84E-11 | 1.392060794 | YOR143C | THI80 | n |
| W303.asm02.HP0_G0050390 | 387.7195734 | 519.7503488 | 437.8312561 | 377.4537102 | 431.2526939 | 420.0356473 | 410.8207217 | 450.8983629 | 537.4037482 | 0.492565946 | 0.131766014 | YOR144C | ELG1 | n |
| W303.asm02.HP0_G0050400 | 2665.572067 | 3074.713602 | 2849.812372 | 4523.60761 | 4458.489389 | 4485.509984 | 2466.817513 | 2374.473353 | 2191.583666 | 2.39E-09 | -0.648751053 | YOR145C | PNO1 | n |
| W303.asm02.HP0_G0050410 | 1059.443734 | 1394.568244 | 981.2111186 | 487.3822393 | 475.0414289 | 497.887082 | 365.3843285 | 482.8289337 | 457.5609056 | 8.29E-12 | 1.233798013 | YOR147W | MDM32 | n |
| W303.asm02.HP0_G0050420 | 564.1319793 | 510.2311117 | 613.7456001 | 121.6023551 | 136.6739307 | 114.9666535 | 188.3717134 | 139.3333997 | 145.3549186 | 2.28E-29 | 2.182211712 | YOR148C | SPP2 | n |
| W303.asm02.HP0_G0050430 | 565.1012782 | 576.8657718 | 615.7002039 | 532.1319059 | 626.3116046 | 544.9600424 | 741.1811637 | 950.1763785 | 884.4130255 | 0.777560747 | 0.047246851 | YOR149C | SMP3 | n |
| W303.asm02.HP0_G0050440 | 1679.795052 | 2050.443684 | 1672.16357 | 1576.939341 | 1663.971933 | 1601.385905 | 1592.166944 | 1367.208984 | 1312.288772 | 0.278000607 | 0.158225606 | YOR150W | MRPL23 | n |
| W303.asm02.HP0_G0050450 | 2386.413974 | 2828.16536 | 2495.051779 | 4627.699226 | 4913.626847 | 4891.061643 | 3366.079461 | 3883.918516 | 3494.659802 | 2.11E-14 | -0.904471669 | YOR151C | RPB2 | n |
| W303.asm02.HP0_G0050470 | 1716.628411 | 1617.318393 | 1564.66036 | 873.5913191 | 1231.392307 | 809.2928205 | 1052.609775 | 1661.357273 | 1549.770047 | 0.000395429 | 0.750952597 | YOR153W | PDR5 | n |
| W303.asm02.HP0_G0050480 | 844.259371 | 1181.337331 | 908.8907772 | 373.5624349 | 518.830164 | 428.1828905 | 494.1207758 | 527.3382141 | 495.4350745 | 3.49E-10 | 1.154953917 | YOR154W | SLP1 | n |
| W303.asm02.HP0_G0050490 | 802.5795169 | 636.8369659 | 732.9764332 | 142.0315508 | 98.19292107 | 155.7028693 | 149.5614609 | 101.5972706 | 161.7329375 | 9.54E-26 | 2.44713648 | YOR155C | ISN1 | n |
| W303.asm02.HP0_G0050510 | 2567.672875 | 2002.847498 | 2468.664627 | 1832.790696 | 1556.490492 | 1754.373027 | 2144.029803 | 1807.463824 | 1723.786499 | 0.003361263 | 0.451977592 | YOR157C | PUP1 | n |
| W303.asm02.HP0_G0050520 | 2307.90076 | 2514.030533 | 2399.276191 | 2350.330319 | 2322.12989 | 2321.059051 | 2342.814023 | 1943.894444 | 1844.574389 | 0.738596027 | 0.046216094 | YOR158W | PET123 | n |
| W303.asm02.HP0_G0050530 | 988.6849121 | 1141.356535 | 1038.871931 | 378.4265291 | 465.7529094 | 417.3198996 | 519.678747 | 473.1530031 | 377.718063 | 1.27E-15 | 1.33105574 | YOR159C | SME1 | n |
| W303.asm02.HP0_G0050540 | 426.4915307 | 460.7310784 | 369.4201224 | 2732.648124 | 2888.729583 | 2656.00127 | 1754.980687 | 2678.297572 | 2095.362805 | 3.47E-55 | -2.719295674 | YOR160W | MTR10 | n |
| W303.asm02.HP0_G0050550 | 1037.149859 | 423.6060535 | 659.6787899 | 322.9758552 | 399.4063411 | 355.7629513 | 445.8446081 | 567.0095293 | 479.0570555 | 0.000501726 | 0.976715601 | YOR161C | PNS1 | n |
| W303.asm02.HP0_G0050560 | 606.7811323 | 573.0580769 | 610.8136944 | 1649.900754 | 1547.201972 | 1400.420574 | 2349.440164 | 4664.766111 | 4007.496522 | 7.98E-10 | -1.360304538 | YOR162C | YRR1 | n |
| W303.asm02.HP0_G0050570 | 1293.044777 | 1025.221842 | 1334.99441 | 365.7798842 | 362.2522629 | 360.2891975 | 637.056096 | 400.5835241 | 571.1834123 | 3.13E-19 | 1.747024241 | YOR163W | DDP1 | n |
| W303.asm02.HP0_G0050580 | 2054.913739 | 1797.231975 | 2045.4929 | 1612.933638 | 1445.028257 | 1609.533149 | 1877.090993 | 1536.537769 | 1562.053561 | 0.014056409 | 0.336777585 | YOR164C | GET4 | n |
| W303.asm02.HP0_G0050590 | 2677.203654 | 2461.674729 | 2857.630788 | 1717.025254 | 1585.682982 | 1629.448632 | 1463.430497 | 1986.468539 | 1855.834277 | 1.80E-06 | 0.696891377 | YOR165W | SEY1 | n |
| W303.asm02.HP0_G0050600 | 916.956791 | 1009.039139 | 824.8428129 | 662.4896306 | 680.7157906 | 646.3479573 | 583.1003791 | 597.004914 | 591.6559361 | 0.000402054 | 0.468201092 | YOR166C | SWT1 | n |
| W303.asm02.HP0_G0050610 | 3629.055207 | 3487.848495 | 3797.795226 | 6398.229516 | 7016.813062 | 6333.123683 | 5329.310283 | 6482.873458 | 5536.794045 | 9.30E-13 | -0.855268942 | YOR168W | GLN4 | n |
| W303.asm02.HP0_G0050650 | 1156.373628 | 838.6447936 | 1100.441952 | 444.5782103 | 586.5036637 | 457.1508662 | 717.5163756 | 604.7456584 | 603.9394503 | 8.48E-09 | 1.059290451 | YOR174W | MED4 | n |
| W303.asm02.HP0_G0050660 | 2128.580458 | 1701.08768 | 2309.364416 | 1308.441341 | 1527.298002 | 1322.56914 | 2172.427549 | 3012.117175 | 2436.230325 | 0.001304246 | 0.562624288 | YOR175C | ALE1 | n |
| W303.asm02.HP0_G0050670 | 1841.667974 | 1236.548907 | 1939.944293 | 1398.913493 | 1857.703912 | 1527.155468 | 3013.947414 | 3791.029583 | 3195.760956 | 0.74602291 | 0.069642452 | YOR176W | HEM15 | n |
| W303.asm02.HP0_G0050720 | 252.9870216 | 237.9809289 | 162.2321172 | 567.1533842 | 594.4652519 | 589.3172552 | 442.058242 | 597.972507 | 700.1603119 | 1.51E-10 | -1.422153346 | YOR181W | LAS17 | n |
| W303.asm02.HP0_G0050740 | 3504.984943 | 2808.174962 | 3418.602085 | 2781.289066 | 1763.491785 | 2564.571097 | 1990.681976 | 1801.658265 | 1895.755698 | 0.014231316 | 0.452102127 | YOR184W | SER1 | n |
| W303.asm02.HP0_G0050770 | 5461.030191 | 4862.42634 | 5539.347232 | 4981.805284 | 4873.818906 | 5015.080789 | 5389.892141 | 4713.145763 | 4371.907445 | 0.471700657 | 0.093094179 | YOR187W | TUF1 | n |
| W303.asm02.HP0_G0050780 | 249.1098259 | 362.6829357 | 249.2119873 | 1067.182268 | 1143.814837 | 1131.56155 | 754.4334451 | 1037.259753 | 1067.642113 | 5.06E-23 | -1.956391129 | YOR188W | MSB1 | n |
| W303.asm02.HP0_G0050810 | 466.232787 | 649.2119742 | 438.808558 | 332.7040436 | 363.5791942 | 354.8577021 | 478.0287199 | 397.680745 | 427.8757462 | 0.003441271 | 0.565139028 | YOR191W | ULS1 | n |
| W303.asm02.HP0_G0050840 | 1610.974827 | 1632.549173 | 1678.027381 | 1840.573247 | 1847.088461 | 1870.24493 | 1882.770542 | 1700.060995 | 1561.029935 | 0.140813319 | -0.175485899 | YOR194C | TOA1 | n |
| W303.asm02.HP0_G0050850 | 754.1145702 | 1388.856701 | 959.7104766 | 563.2621088 | 675.4080652 | 563.0650272 | 570.7946893 | 625.0651125 | 507.7185887 | 0.000438267 | 0.785352865 | YOR195W | SLK19 | n |
| W303.asm02.HP0_G0050860 | 1486.904564 | 1364.106685 | 1489.408112 | 3046.868609 | 3005.499543 | 3052.500437 | 2923.074628 | 3060.496828 | 3096.469216 | 4.19E-27 | -1.068951867 | YOR196C | LIP5 | n |
| W303.asm02.HP0_G0050870 | 1417.115041 | 1184.193102 | 1222.604691 | 866.7815872 | 808.1012018 | 858.1762794 | 822.5880348 | 853.4170731 | 879.2948946 | 3.01E-06 | 0.593324776 | YOR197W | MCA1 | n |
| W303.asm02.HP0_G0050880 | 5239.060735 | 5476.417137 | 5322.386207 | 7665.812466 | 7373.7576 | 7357.865822 | 6175.563106 | 6636.720753 | 6564.514737 | 1.29E-07 | -0.48188241 | YOR198C | BFR1 | n |
| W303.asm02.HP0_G0050890 | 1034.241962 | 1080.433417 | 984.1430244 | 985.4654858 | 975.2945538 | 1044.657623 | 625.6969977 | 805.0374205 | 757.4833784 | 0.772715577 | 0.043749179 | YOR201C | MRM1 | n |
| W303.asm02.HP0_G0050900 | 703.7110257 | 552.1157552 | 700.7254702 | 538.9416378 | 497.5992622 | 515.0868175 | 691.9584044 | 530.2409933 | 613.152086 | 0.050959677 | 0.333780793 | YOR202W | HIS3 | n |
| W303.asm02.HP0_G0050910 | 1454.917596 | 2732.972988 | 1386.791412 | 10103.69648 | 8230.955262 | 10244.70565 | 2658.975592 | 6545.767006 | 6290.182919 | 8.65E-14 | -2.358012017 | YOR204W | DED1 | n |
| W303.asm02.HP0_G0050920 | 481.7415699 | 619.702339 | 501.3558803 | 732.5325871 | 695.3120357 | 796.6193311 | 465.7230301 | 603.7780653 | 506.6949626 | 0.005509075 | -0.473621941 | YOR205C | GEP3 | n |
| W303.asm02.HP0_G0050930 | 3105.633783 | 4648.243504 | 3462.580671 | 8367.21485 | 7763.875421 | 8331.914004 | 3667.095566 | 4102.594547 | 3424.029595 | 6.03E-12 | -1.125009764 | YOR206W | NOC2 | n |
| W303.asm02.HP0_G0050940 | 1186.421895 | 1346.972058 | 1223.581993 | 5591.762697 | 5590.361844 | 5655.092002 | 2437.473176 | 3488.172958 | 2572.372608 | 6.11E-45 | -2.163985622 | YOR207C | RET1 | n |
| W303.asm02.HP0_G0050950 | 230.6931462 | 428.3656721 | 230.643251 | 492.2463335 | 435.233488 | 514.1815683 | 394.7286657 | 605.7132514 | 423.7812414 | 0.010727694 | -0.696805872 | YOR208W | PTP2 | n |
| W303.asm02.HP0_G0050960 | 3857.809755 | 3500.223503 | 3961.004645 | 2991.417936 | 2397.764978 | 2900.418565 | 2072.088847 | 3190.154297 | 2081.032038 | 0.018663742 | 0.449005975 | YOR209C | NPT1 | n |
| W303.asm02.HP0_G0050970 | 1420.022937 | 1879.097415 | 1518.72717 | 1725.780624 | 2070.012931 | 1836.750708 | 1896.022824 | 1431.070126 | 1164.886601 | 0.273698966 | -0.224991388 | YOR210W | RPB10 | n |
| W303.asm02.HP0_G0050980 | 753.1452713 | 940.5006312 | 754.4770753 | 1586.667529 | 1411.854973 | 1613.154146 | 936.1790178 | 1408.815486 | 1106.539908 | 3.84E-07 | -0.913987097 | YOR211C | MGM1 | n |
| W303.asm02.HP0_G0050990 | 2313.716554 | 2202.751478 | 2297.636793 | 570.0718407 | 570.5804873 | 588.412006 | 501.693508 | 663.7688347 | 528.1911125 | 3.62E-46 | 1.978232524 | YOR212W | STE4 | n |
| W303.asm02.HP0_G0051020 | 1685.610845 | 1691.568443 | 1747.415817 | 1319.142348 | 1230.065376 | 1355.158112 | 1232.462165 | 1010.167148 | 929.4525778 | 0.005299371 | 0.391727346 | YOR215C | AIM41 | n |
| W303.asm02.HP0_G0051030 | 1196.114884 | 1541.164496 | 1209.899766 | 1358.055102 | 1389.29714 | 1410.378316 | 1010.013157 | 1134.019059 | 1006.224542 | 0.626878846 | -0.074887438 | YOR216C | RUD3 | n |
| W303.asm02.HP0_G0051040 | 938.2813676 | 933.8371652 | 909.8680791 | 1654.764848 | 1564.45208 | 1611.343647 | 1216.370109 | 1630.394295 | 1528.273897 | 1.34E-08 | -0.796289835 | YOR217W | RFC1 | n |
| W303.asm02.HP0_G0051050 | 434.2459222 | 475.0099342 | 432.9447466 | 687.7829205 | 773.6009862 | 776.7038479 | 639.8958706 | 942.435634 | 880.3185208 | 2.91E-05 | -0.737285098 | YOR219C | STE13 | n |
| W303.asm02.HP0_G0051060 | 1652.654681 | 913.8467672 | 1375.063789 | 195.536587 | 173.8280089 | 197.3443343 | 326.574076 | 383.1668492 | 470.868046 | 2.97E-33 | 2.796091139 | YOR220W | RCN2 | n |
| W303.asm02.HP0_G0051070 | 622.2899153 | 547.3561366 | 519.9246166 | 237.3677972 | 273.3478613 | 237.1753009 | 367.2775115 | 325.111266 | 268.190061 | 4.25E-11 | 1.17880069 | YOR221C | MCT1 | n |
| W303.asm02.HP0_G0051090 | 816.1497019 | 319.8463685 | 675.3156205 | 129.3849058 | 149.9432443 | 107.7246596 | 272.6183591 | 218.6760301 | 289.6862109 | 5.45E-13 | 2.229972314 | YOR223W | DSC3 | n |
| W303.asm02.HP0_G0051100 | 1960.891742 | 2404.559306 | 2251.703603 | 2903.86424 | 3166.058239 | 2963.786012 | 2577.568721 | 1990.338911 | 1769.849677 | 0.006629161 | -0.448921028 | YOR224C | RPB8 | n |
| W303.asm02.HP0_G0051130 | 436.18452 | 299.8559705 | 415.3533122 | 324.9214928 | 376.8485079 | 308.6899908 | 372.9570607 | 331.8844173 | 383.8598201 | 0.349938929 | 0.190748128 | YOR228C | MCP1 | n |
| W303.asm02.HP0_G0051160 | 345.0704203 | 376.9617915 | 264.8488179 | 683.8916451 | 650.1963692 | 669.8844375 | 886.009667 | 882.4448647 | 1065.594861 | 2.17E-09 | -1.021978608 | YOR231W | MKK1 | n |

| | | | | | | | | | | | | | | |
|---|---|---|---|---|---|---|---|---|---|---|---|---|---|---|
| W303.asm02.HP0_G0051170 | 3167.668914 | 4845.291713 | 3447.921142 | 3308.556878 | 3369.078738 | 3258.897264 | 4076.023105 | 3190.154297 | 3028.909887 | 0.287078383 | 0.205977937 | YOR232W | MGE1 | n |
| W303.asm02.HP0_G0051200 | 436.18452 | 399.8079606 | 440.7631619 | 512.6755291 | 496.2723308 | 549.4862886 | 851.9323721 | 523.4678419 | 533.3092434 | 0.181977827 | -0.288282531 | YOR236W | DFR1 | n |
| W303.asm02.HP0_G0051240 | 835.5356806 | 729.1735663 | 858.0710779 | 1988.441711 | 1989.070117 | 1910.075896 | 1794.737531 | 1822.945313 | 1648.038161 | 3.33E-27 | -1.280971586 | YOR239W | ABP140 | n |
| W303.asm02.HP0_G0051250 | 1879.470632 | 1978.097481 | 1957.535728 | 1753.019551 | 1519.336414 | 1786.962 | 1400.008865 | 1450.421987 | 1501.659616 | 0.093152802 | 0.19966374 | YOR241W | MET7 | n |
| W303.asm02.HP0_G0051270 | 1636.1766 | 2133.261047 | 1892.0565 | 5747.413712 | 5297.110012 | 5817.131616 | 2426.114077 | 3079.848689 | 2684.971488 | 1.02E-27 | -1.574587467 | YOR243C | PUS7 | n |
| W303.asm02.HP0_G0051280 | 971.2375313 | 1038.548774 | 922.573004 | 463.0617682 | 508.2147131 | 470.7296048 | 365.3843285 | 409.2918616 | 386.9306987 | 6.05E-15 | 1.025738366 | YOR244W | ESA1 | n |
| W303.asm02.HP0_G0051290 | 1100.154289 | 928.1256229 | 1048.64495 | 562.28929 | 447.1758703 | 537.7180485 | 512.1060148 | 566.0419363 | 629.530105 | 2.40E-10 | 0.989161064 | YOR245C | DGA1 | n |
| W303.asm02.HP0_G0051300 | 1730.198596 | 1432.645192 | 1831.463781 | 1441.717522 | 1161.064945 | 1378.694592 | 680.5993062 | 872.7689342 | 734.9636022 | 0.049952037 | 0.325995034 | YOR246C | ENV9 | n |
| W303.asm02.HP0_G0051330 | 608.7197302 | 467.3945445 | 563.9032026 | 594.3923118 | 524.1378895 | 630.053471 | 556.5958164 | 632.8058569 | 537.4037482 | 0.591408631 | -0.09444981 | YOR250C | CLP1 | n |
| W303.asm02.HP0_G0051340 | 1290.13688 | 894.8082928 | 1303.720749 | 665.4080871 | 479.022223 | 667.1686898 | 631.3765469 | 662.8012416 | 477.0098031 | 1.09E-05 | 0.943349782 | YOR251C | TUM1 | n |
| W303.asm02.HP0_G0051350 | 1156.373628 | 1365.058608 | 1327.175995 | 1870.730631 | 1787.37655 | 1793.298744 | 1083.847296 | 891.1532022 | 714.4910785 | 0.002854344 | -0.502343383 | YOR252W | TMA16 | n |
| W303.asm02.HP0_G0051360 | 1133.110453 | 1330.789355 | 1268.53788 | 940.7158191 | 1024.391014 | 922.4489755 | 1044.090452 | 786.6531524 | 783.074033 | 0.020547712 | 0.371038393 | YOR253W | NAT5 | n |
| W303.asm02.HP0_G0051370 | 3499.16915 | 4172.281646 | 4094.895007 | 1959.257145 | 1832.492216 | 1881.107921 | 1867.625078 | 2303.83906 | 2236.623219 | 1.85E-15 | 1.052309427 | YOR254C | SEC63 | n |
| W303.asm02.HP0_G0051380 | 28.10966907 | 7.615389726 | 16.61413249 | 40.85839132 | 34.50021551 | 46.16771124 | 32.18411183 | 68.69910679 | 22.51977611 | 0.03804857 | -1.219818238 | YOR255W | OSW1 | n |
| W303.asm02.HP0_G0051390 | 742.482983 | 793.904379 | 755.4543772 | 694.5926524 | 789.5241626 | 715.1468995 | 804.6027959 | 946.3060063 | 900.7910445 | 0.669926302 | 0.061092803 | YOR256C | TRE2 | n |
| W303.asm02.HP0_G0051400 | 961.544542 | 999.5199016 | 1124.8745 | 420.2577392 | 524.1378895 | 447.1931245 | 634.2163215 | 550.5604474 | 512.8367197 | 1.44E-13 | 1.15177092 | YOR257W | CDC31 | n |
| W303.asm02.HP0_G0051410 | 317.9300502 | 330.3175294 | 323.4869325 | 246.1231667 | 200.3666362 | 212.7335714 | 217.7160506 | 297.0510674 | 260.0010515 | 0.003176057 | 0.557343919 | YOR258W | HNT3 | n |
| W303.asm02.HP0_G0051420 | 4966.687735 | 3732.49289 | 4770.210628 | 2846.467928 | 2405.726566 | 2749.241942 | 3076.422455 | 3244.339508 | 3184.501068 | 1.45E-07 | 0.750857601 | YOR259C | RPT4 | n |
| W303.asm02.HP0_G0051430 | 1577.049365 | 1810.558907 | 1683.891193 | 3787.183747 | 3860.043343 | 3850.930267 | 2669.388099 | 2941.482882 | 2513.002289 | 3.67E-25 | -1.180848348 | YOR260W | GCD1 | n |
| W303.asm02.HP0_G0051440 | 4270.731101 | 4647.291581 | 4111.50914 | 1685.895051 | 1406.547248 | 1691.00558 | 2042.74451 | 1861.649035 | 1959.220522 | 7.34E-32 | 1.444410217 | YOR261C | RPN8 | n |
| W303.asm02.HP0_G0051450 | 1569.294973 | 1007.135291 | 1428.815394 | 704.3208408 | 719.1968002 | 735.9676321 | 638.0026875 | 836.0003982 | 971.4212514 | 1.55E-05 | 0.891100366 | YOR262W | GPN2 | n |
| W303.asm02.HP0_G0051460 | 1020.671777 | 933.8371652 | 925.5049097 | 1222.833283 | 1401.239522 | 1341.579374 | 1498.454383 | 1676.838762 | 1345.04481 | 0.000746146 | -0.460904018 | YOR264W | DSE3 | n |
| W303.asm02.HP0_G0051480 | 807.4260115 | 972.8660375 | 866.8667951 | 518.5124422 | 476.3683603 | 556.7282826 | 748.7538959 | 446.0603976 | 436.0647557 | 0.000490943 | 0.76979362 | YOR265W | RBL2 | n |
| W303.asm02.HP0_G0051490 | 779.3163425 | 762.4908964 | 770.1139059 | 605.093319 | 642.234781 | 645.4427081 | 651.2549689 | 575.7178668 | 560.9471505 | 0.026603037 | 0.288890769 | YOR266W | PNT1 | n |
| W303.asm02.HP0_G0051500 | 557.3468867 | 752.9716592 | 471.0595211 | 870.6728625 | 1035.006465 | 803.8613251 | 653.1481519 | 929.8569243 | 890.5547827 | 0.005620852 | -0.603894484 | YOR267C | HRK1 | n |
| W303.asm02.HP0_G0051520 | 274.3115982 | 311.2790551 | 251.1665911 | 284.0631015 | 322.4443219 | 272.4800212 | 309.5354285 | 284.4723577 | 282.5208276 | 0.71802952 | -0.067563523 | YOR269W | PAC1 | n |
| W303.asm02.HP0_G0051530 | 6814.171502 | 5460.234434 | 6966.208022 | 2606.181675 | 2744.094064 | 2517.498136 | 4161.216342 | 5152.43301 | 4858.129883 | 5.61E-20 | 1.290360043 | YOR270C | VPH1 | n |
| W303.asm02.HP0_G0051540 | 1270.750902 | 1378.38554 | 1284.174711 | 1348.326913 | 1322.950572 | 1479.177258 | 1720.903392 | 1348.824716 | 1569.218945 | 0.591331612 | -0.078055092 | YOR271C | FSF1 | n |
| W303.asm02.HP0_G0051550 | 2014.203184 | 1993.328261 | 2230.202961 | 5443.894233 | 4696.010103 | 5343.686263 | 2519.826638 | 3062.432014 | 2531.42756 | 1.15E-23 | -1.311940969 | YOR272W | YTM1 | n |
| W303.asm02.HP0_G0051560 | 707.5882214 | 221.7982258 | 494.514767 | 717.9403045 | 797.4857508 | 756.7883646 | 578.3674215 | 865.9957828 | 655.1207597 | 0.040623483 | -0.674217247 | YOR273C | TPO4 | n |
| W303.asm02.HP0_G0051570 | 775.4391467 | 806.2793873 | 819.9563033 | 930.0148118 | 1060.218161 | 980.3849268 | 1037.464311 | 944.3708201 | 767.7196402 | 0.045503117 | -0.305909365 | YOR274W | MOD5 | n |
| W303.asm02.HP0_G0051580 | 1136.987649 | 848.1640308 | 1095.555442 | 801.6027248 | 755.0239471 | 765.840857 | 543.3435351 | 826.3244676 | 659.2152644 | 0.033431173 | 0.407329461 | YOR275C | RIM20 | n |
| W303.asm02.HP0_G0051590 | 2901.111708 | 3528.781214 | 3018.885603 | 1489.385645 | 1670.60659 | 1479.177258 | 2230.169632 | 1404.945114 | 1237.56406 | 1.03E-06 | 1.026594417 | YOR276W | CAF20 | n |
| W303.asm02.HP0_G0051600 | 724.0663033 | 441.6926041 | 718.3169046 | 369.6711595 | 402.0602038 | 420.9408966 | 458.1502979 | 418.0001991 | 392.0488296 | 0.001804757 | 0.659490507 | YOR278W | HEM4 | n |
| W303.asm02.HP0_G0051610 | 1848.453066 | 1212.750814 | 1790.417101 | 804.5211814 | 769.6201921 | 846.4080393 | 792.2971061 | 605.7132514 | 635.6718621 | 1.32E-07 | 1.00260463 | YOR279C | RFM1 | n |
| W303.asm02.HP0_G0051620 | 2485.282465 | 2641.588311 | 2605.486895 | 1152.790326 | 1227.411513 | 1159.624276 | 1203.117828 | 1181.431118 | 1087.091011 | 3.65E-26 | 1.127994979 | YOR281C | PLP2 | n |
| W303.asm02.HP0_G0051630 | 1893.040817 | 1971.434015 | 1911.602538 | 1005.894681 | 924.8711619 | 1031.984134 | 1164.307575 | 995.653252 | 925.358073 | 1.26E-13 | 0.962135204 | YOR283W | | n |
| W303.asm02.HP0_G0051650 | 2714.037014 | 2399.799688 | 2460.846212 | 117.7110797 | 99.51985243 | 103.1984134 | 117.377349 | 47.41205962 | 85.98459971 | 2.53E-81 | 4.56101125 | YOR285W | RDL1 | n |
| W303.asm02.HP0_G0051660 | 2364.120099 | 1965.722473 | 2401.230795 | 1307.468522 | 1325.604434 | 1336.147878 | 1463.430497 | 977.268984 | 818.9009496 | 0.000203081 | 0.761960345 | YOR286W | RDL2 | n |
| W303.asm02.HP0_G0051670 | 730.8513958 | 991.9045119 | 729.0672256 | 1672.275587 | 1612.221609 | 1699.152823 | 992.0279177 | 897.9263536 | 824.0190805 | 1.19E-10 | -1.023396965 | YOR287C | RRP36 | n |
| W303.asm02.HP0_G0051680 | 997.4086025 | 752.9716592 | 988.052232 | 436.7956595 | 468.4067721 | 353.0472036 | 1103.725718 | 839.8707704 | 947.8778491 | 1.41E-09 | 1.124272422 | YOR288C | MPD1 | n |
| W303.asm02.HP0_G0051690 | 397.4125627 | 364.5867831 | 390.9207644 | 83.66242031 | 78.28895058 | 86.90392703 | 216.7694591 | 114.1759803 | 189.3708446 | 1.65E-18 | 2.209232123 | YOR289W | | n |
| W303.asm02.HP0_G0051700 | 586.4258547 | 805.3274636 | 559.0166931 | 2110.044066 | 1836.47301 | 2063.063018 | 1146.322336 | 1623.621144 | 1969.456784 | 3.16E-14 | -1.62334079 | YOR290C | SNF2 | n |
| W303.asm02.HP0_G0051710 | 328.5923384 | 507.3753405 | 369.4201224 | 1503.977928 | 1401.239522 | 1510.860981 | 1207.850785 | 1743.602682 | 1465.8327 | 7.38E-22 | -1.873278621 | YOR291W | YPK9 | n |
| W303.asm02.HP0_G0051720 | 799.6716201 | 446.4522227 | 796.5010575 | 292.8184711 | 228.2321949 | 313.216237 | 302.9092879 | 358.0094298 | 359.2927916 | 1.19E-07 | 1.288771234 | YOR292C | | n |
| W303.asm02.HP0_G0051730 | 5990.267409 | 8242.707455 | 6168.729662 | 18755.94725 | 20794.34143 | 17476.74183 | 16667.58356 | 13876.25198 | 12760.52405 | 2.87E-19 | -1.482887856 | YOR293W | RPS10A | y |
| W303.asm02.HP0_G0051750 | 1570.264272 | 2583.520965 | 1760.120742 | 3517.712928 | 3136.865749 | 3361.190428 | 2599.340326 | 1932.283328 | 1785.20407 | 0.000293806 | -0.760144574 | YOR294W | RRS1 | n |
| W303.asm02.HP0_G0051770 | 425.5222318 | 450.2599176 | 467.1503135 | 509.7570726 | 545.3687913 | 538.6232978 | 515.8923809 | 586.3613904 | 696.0658071 | 0.131508037 | -0.246612363 | YOR296W | | n |
| W303.asm02.HP0_G0051780 | 872.3690401 | 1029.029537 | 880.5490218 | 616.7671451 | 782.8895058 | 704.2839087 | 755.3800366 | 656.9956833 | 603.9394503 | 0.012276223 | 0.40465483 | YOR297C | TIM18 | n |
| W303.asm02.HP0_G0051790 | 290.78968 | 140.8847099 | 235.5297605 | 51.55939856 | 87.57747014 | 78.75668387 | 140.0955456 | 82.24540954 | 74.72471165 | 6.75E-06 | 1.618974583 | YOR298W | MUM3 | n |
| W303.asm02.HP0_G0051800 | 10372.46789 | 6678.69679 | 9254.071795 | 2804.636718 | 2882.094926 | 2789.072908 | 3706.85241 | 3009.214396 | 2568.278103 | 2.13E-18 | 1.633973197 | YOR298C- | MBF1 | n |
| W303.asm02.HP0_G0051810 | 1047.812147 | 832.9332513 | 1118.033386 | 1381.402754 | 1249.969347 | 1436.630544 | 1454.911173 | 1462.033104 | 1774.967808 | 0.006280205 | -0.440702348 | YOR299W | BUD7 | n |
| W303.asm02.HP0_G0051820 | 367.3642958 | 365.5387069 | 338.1464612 | 996.166493 | 999.1793184 | 948.7012034 | 1160.521209 | 1210.45891 | 1170.004732 | 1.44E-34 | -1.458204292 | YOR301W | RAX1 | n |
| W303.asm02.HP0_G0051830 | 2847.800266 | 3142.300186 | 3079.478322 | 3391.246479 | 2675.093633 | 3571.208252 | 2321.042418 | 1585.885015 | 1740.164518 | 0.643871712 | -0.088099795 | YOR303W | CPA1 | n |
| W303.asm02.HP0_G0051840 | 619.3820185 | 851.9717256 | 606.9044867 | 985.4654858 | 1001.833181 | 1007.542404 | 917.2471873 | 1077.898662 | 951.9723539 | 0.001661107 | -0.526905739 | YOR304W | ISW2 | n |
| W303.asm02.HP0_G0051860 | 1036.18056 | 1192.760416 | 1175.694199 | 402.7470001 | 395.425547 | 374.7731853 | 450.5775657 | 387.0372214 | 351.1037821 | 3.64E-25 | 1.537897823 | YOR305W | RRG7 | n |
| W303.asm02.HP0_G0051870 | 555.4082889 | 1302.231643 | 657.7241861 | 1661.57458 | 1300.392738 | 1599.575407 | 663.5606587 | 1011.134741 | 765.6723879 | 0.003154661 | -0.858946913 | YOR306C | MCH5 | n |
| W303.asm02.HP0_G0051880 | 1091.430599 | 1285.097016 | 1098.487348 | 843.433935 | 934.1596815 | 829.2083038 | 935.2324263 | 1107.894046 | 1106.539908 | 0.003452505 | 0.416000058 | YOR307C | SLY41 | n |
| W303.asm02.HP0_G0051890 | 171.5659112 | 253.2117084 | 173.9597402 | 553.5339204 | 558.638105 | 525.0445592 | 442.058242 | 511.8567253 | 379.7653154 | 6.58E-13 | -1.450426863 | YOR308C | SNU66 | n |
| W303.asm02.HP0_G0051900 | 2878.817832 | 3036.636653 | 2780.423937 | 16511.65419 | 17450.47439 | 16235.64512 | 10396.41471 | 9925.569542 | 9128.698335 | 1.08E-134 | -2.52912656 | YOR310C | NOP58 | n |
| W303.asm02.HP0_G0051910 | 1420.022937 | 1378.38554 | 1516.772566 | 1695.62324 | 1683.875903 | 1642.122121 | 1179.45304 | 1313.023774 | 1229.375051 | 0.05192883 | -0.218660292 | YOR311C | DGK1 | n |
| W303.asm02.HP0_G0051930 | 313.0835555 | 466.4426207 | 375.2839338 | 82.68960147 | 88.90440151 | 79.66193311 | 73.83413892 | 140.3009927 | 96.22086158 | 5.10E-16 | 2.20303572 | YOR313C | SPS4 | n |
| W303.asm02.HP0_G0051940 | 3686.243844 | 4378.849093 | 4187.738689 | 430.9587465 | 449.829733 | 383.8256777 | 317.1081607 | 544.7548891 | 585.514179 | 2.09E-55 | 3.277413916 | YOR315W | SFG1 | n |

| | | | | | | | | | | | | | | |
|---|---|---|---|---|---|---|---|---|---|---|---|---|---|---|
| W303.asm02.HP0_G0051970 | 3618.392918 | 2508.318991 | 3512.423068 | 4761.948226 | 4442.566213 | 4534.393443 | 3178.654339 | 4852.479163 | 4731.200236 | 0.009851797 | -0.511357784 | YOR317W | FAA1 | n |
| W303.asm02.HP0_G0051990 | 722.1277054 | 759.6351252 | 745.6813581 | 191.6453116 | 250.7900281 | 250.7540395 | 312.3752031 | 250.6066008 | 330.6312584 | 1.40E-21 | 1.68615447 | YOR319W | HSH49 | n |
| W303.asm02.HP0_G0052000 | 1830.036386 | 1966.674397 | 2003.468918 | 1060.372537 | 1031.025671 | 1116.172313 | 910.6210466 | 1194.977421 | 979.6102609 | 6.72E-10 | 0.854097752 | YOR320C | GNT1 | n |
| W303.asm02.HP0_G0052020 | 454.6011998 | 603.5196358 | 394.829972 | 1721.889348 | 1449.009051 | 1690.100331 | 1413.261146 | 1592.658166 | 1884.49581 | 1.29E-20 | -1.742541499 | YOR322C | LDB19 | n |
| W303.asm02.HP0_G0052030 | 1994.817205 | 1424.077879 | 1880.328877 | 2368.813877 | 2343.360792 | 2380.805501 | 2596.500552 | 2058.070425 | 1852.763398 | 0.017375585 | -0.420697434 | YOR323C | PRO2 | n |
| W303.asm02.HP0_G0052040 | 203.552776 | 348.40408 | 209.142609 | 175.1073914 | 191.0781167 | 179.2393495 | 258.4194862 | 222.5464023 | 335.7493893 | 0.073442698 | 0.482063575 | YOR324C | FRT1 | n |
| W303.asm02.HP0_G0052050 | 3736.647388 | 3099.463619 | 4024.529269 | 4855.338835 | 5021.108288 | 4882.9144 | 3480.617036 | 4797.326359 | 4512.144232 | 0.005514042 | -0.442502534 | YOR326W | MYO2 | n |
| W303.asm02.HP0_G0052060 | 1476.242276 | 2211.318792 | 1516.772566 | 2333.792399 | 2190.763685 | 2286.65958 | 1640.443112 | 1876.162931 | 1502.683242 | 0.032795253 | -0.388250945 | YOR327C | SNC2 | n |
| W303.asm02.HP0_G0052080 | 404.1976552 | 453.1156887 | 349.8740841 | 393.9916305 | 355.617606 | 435.4248844 | 358.7581878 | 436.3844671 | 456.5372794 | 0.900095696 | 0.02460549 | YOR329C | SCD5 | n |
| W303.asm02.HP0_G0052090 | 357.6713064 | 554.0196026 | 376.2612357 | 1106.095022 | 1244.661621 | 1112.551316 | 801.7630213 | 1192.074642 | 1012.366299 | 7.86E-12 | -1.426425108 | YOR330C | MIP1 | n |
| W303.asm02.HP0_G0052100 | 2932.129274 | 2053.299455 | 2706.148992 | 3488.528363 | 2997.537955 | 3308.685972 | 3538.359119 | 2988.894942 | 2709.538517 | 0.043912738 | -0.348991839 | YOR332W | VMA4 | n |
| W303.asm02.HP0_G0052120 | 329.5616374 | 435.9810618 | 354.7605937 | 466.9530436 | 406.0409979 | 445.382626 | 452.4707487 | 449.9307698 | 420.7103628 | 0.166499473 | -0.236265071 | YOR334W | MRS2 | n |
| W303.asm02.HP0_G0052130 | 6547.614295 | 6487.360123 | 7102.052987 | 8058.831277 | 8579.938211 | 8141.811664 | 5319.844368 | 5750.405517 | 4821.279341 | 0.00816446 | -0.299267546 | YOR335C | ALA1 | n |
| W303.asm02.HP0_G0052140 | 653.3074811 | 673.0100671 | 697.7935644 | 1673.248406 | 1577.721394 | 1634.880127 | 1343.213373 | 1596.528538 | 1338.903053 | 1.89E-24 | -1.271658518 | YOR336W | KRE5 | n |
| W303.asm02.HP0_G0052150 | 260.7414131 | 530.2215097 | 234.5524586 | 249.0416233 | 221.5975381 | 275.1957689 | 21.77160506 | 169.3287844 | 213.9378731 | 0.577301157 | 0.459245559 | YOR337W | TEA1 | n |
| W303.asm02.HP0_G0052160 | 282.0659896 | 1306.991262 | 224.7794395 | 568.126203 | 337.0405669 | 519.6130637 | 180.7989812 | 261.2501244 | 446.3010175 | 0.669964337 | 0.348073661 | YOR338W | | n |
| W303.asm02.HP0_G0052180 | 1838.760077 | 2348.395807 | 1978.059068 | 3859.172342 | 4003.351931 | 3796.615312 | 2888.997333 | 2344.477968 | 2171.111143 | 1.09E-09 | -0.919099754 | YOR340C | RPA43 | n |
| W303.asm02.HP0_G0052190 | 2966.054736 | 3826.733337 | 3187.958834 | 15675.02998 | 15158.86392 | 16056.40577 | 6531.481519 | 8358.068796 | 7421.289856 | 1.19E-56 | -2.232067162 | YOR341W | RPA190 | n |
| W303.asm02.HP0_G0052200 | 1712.751215 | 2231.30919 | 1890.101896 | 480.5725074 | 432.5796252 | 468.9191063 | 460.0434809 | 471.217817 | 441.1828866 | 2.93E-47 | 2.076231772 | YOR342C | | n |
| W303.asm02.HP0_G0052220 | 948.9436558 | 2674.905641 | 1044.735743 | 1336.653087 | 821.3705154 | 1342.484623 | 450.5775657 | 582.4910182 | 1065.514981 | 0.471092925 | 0.41520872 | YOR344C | TYE7 | n |
| W303.asm02.HP0_G0052230 | 843.2900721 | 1018.558376 | 1035.940026 | 128.412087 | 100.8467838 | 106.8194103 | 31.23752031 | 71.60188595 | 51.18130935 | 1.08E-38 | 3.105094788 | YOR346W | REV1 | n |
| W303.asm02.HP0_G0052250 | 642.6451929 | 344.5963851 | 490.6055593 | 130.3577247 | 122.0776857 | 150.2713738 | 428.8059606 | 235.125112 | 425.8284938 | 8.54E-11 | 1.873251619 | YOR348C | PUT4 | n |
| W303.asm02.HP0_G0052260 | 249.1098259 | 223.7020732 | 227.7113453 | 530.1862683 | 479.022223 | 582.0752613 | 388.1025251 | 520.5650627 | 466.7735413 | 1.03E-11 | -1.185577532 | YOR349W | CIN1 | n |
| W303.asm02.HP0_G0052270 | 478.8336731 | 574.0100006 | 477.9006345 | 783.1191669 | 874.44777 | 755.8831153 | 600.1390266 | 907.6022841 | 750.3179951 | 0.000379319 | -0.655839279 | YOR350C | MNE1 | n |
| W303.asm02.HP0_G0052280 | 33.92546267 | 48.54810951 | 34.20556688 | 62.26040581 | 55.73111736 | 57.03070212 | 44.48980165 | 57.08799015 | 40.94504748 | 0.097736474 | -0.585668216 | YOR351C | MEK1 | n |
| W303.asm02.HP0_G0052290 | 604.8425345 | 547.3561366 | 604.9498829 | 182.8899421 | 216.2898126 | 199.1548328 | 368.2241031 | 330.9168243 | 319.3713703 | 3.26E-23 | 1.557790065 | YOR352W | TFB6 | n |
| W303.asm02.HP0_G0052300 | 868.4918444 | 878.6255897 | 943.0963441 | 1393.07658 | 1354.796924 | 1309.89565 | 926.7131025 | 1254.000597 | 1240.634939 | 8.28E-05 | -0.592928468 | YOR353C | SOG2 | n |
| W303.asm02.HP0_G0052310 | 2417.43154 | 2662.530633 | 2593.759272 | 3047.841428 | 2963.03774 | 3105.004893 | 2067.35589 | 2666.686455 | 2211.032564 | 0.062135 | -0.248587336 | YOR354C | MSC6 | n |
| W303.asm02.HP0_G0052320 | 1901.764507 | 2441.684331 | 2291.772981 | 4083.893494 | 4039.179077 | 3945.981437 | 2546.331201 | 3577.191519 | 2958.27968 | 4.52E-08 | -0.863057492 | YOR355W | GDS1 | n |
| W303.asm02.HP0_G0052330 | 1063.32093 | 1022.366071 | 1000.757157 | 2457.340392 | 2376.534076 | 2509.350893 | 1550.516917 | 1934.218514 | 1844.574389 | 7.07E-24 | -1.250687299 | YOR356W | CIR2 | n |
| W303.asm02.HP0_G0052340 | 957.6673462 | 773.9139809 | 893.2539466 | 621.6312393 | 607.7345655 | 620.0957293 | 728.8754739 | 615.389182 | 502.6004578 | 0.003431463 | 0.504852807 | YOR357C | SNX3 | n |
| W303.asm02.HP0_G0052350 | 499.1889507 | 364.5867831 | 424.1490294 | 383.2906233 | 436.5604193 | 404.6464102 | 543.3435351 | 442.1900254 | 374.6471844 | 0.72845947 | 0.073868818 | YOR358W | HAP5 | n |
| W303.asm02.HP0_G0052360 | 272.3730003 | 676.8177619 | 265.8261198 | 3305.638421 | 3114.307915 | 3200.961312 | 1971.750146 | 2532.191021 | 2446.466587 | 2.08E-24 | -2.984940176 | YOR359W | VTS1 | n |
| W303.asm02.HP0_G0052370 | 1836.821479 | 3580.185095 | 1699.528023 | 2120.745073 | 1476.87461 | 1983.401085 | 686.2788553 | 1157.241292 | 860.8696232 | 0.240820993 | 0.350329021 | YOR360C | PDE2 | n |
| W303.asm02.HP0_G0052380 | 6113.368373 | 5612.542228 | 6209.776342 | 8076.342017 | 8491.03381 | 8208.800108 | 6839.123765 | 6616.401299 | 5974.906053 | 1.11E-05 | -0.46605594 | YOR361C | PRT1 | n |
| W303.asm02.HP0_G0052420 | 885.9392252 | 630.1734999 | 859.0483798 | 598.2835871 | 638.2539869 | 660.8319451 | 796.0834721 | 663.7688347 | 556.8526457 | 0.097196342 | 0.323923148 | YOR367W | SCP1 | n |
| W303.asm02.HP0_G0052430 | 901.4480081 | 741.5485746 | 894.2312486 | 633.3050654 | 571.9074186 | 640.0112126 | 655.9879265 | 653.1253111 | 529.2147387 | 0.003596868 | 0.458098428 | YOR368W | RAD17 | n |
| W303.asm02.HP0_G0052440 | 9975.055324 | 11797.19061 | 9668.447805 | 15948.39208 | 18585.00071 | 15446.26778 | 16222.68555 | 12301.97808 | 11460.51879 | 5.19E-05 | -0.668629408 | YOR369C | RPS12 | n |
| W303.asm02.HP0_G0052450 | 1506.290543 | 1640.164562 | 1431.7473 | 1625.580283 | 1797.992001 | 1668.374349 | 2228.276449 | 2656.042932 | 2436.230325 | 0.241543615 | -0.152797466 | YOR370C | MRS6 | n |
| W303.asm02.HP0_G0052470 | 303.3905662 | 410.2791215 | 273.6445351 | 936.8245437 | 1049.60271 | 995.7741639 | 900.2085398 | 1147.565361 | 1122.917927 | 3.91E-18 | -1.594038318 | YOR372C | NDD1 | n |
| W303.asm02.HP0_G0052480 | 207.4299718 | 258.9232507 | 170.0505325 | 701.4023842 | 749.7162217 | 731.4413859 | 771.4720925 | 909.5374702 | 846.5388566 | 3.57E-24 | -1.777127918 | YOR373W | NUD1 | n |
| W303.asm02.HP0_G0052490 | 2036.497059 | 1317.462423 | 1593.979417 | 284.0631015 | 258.7516163 | 161.1343647 | 1735.102265 | 3553.001692 | 2900.956614 | 7.54E-24 | 2.815357903 | YOR374W | ALD4 | n |
| W303.asm02.HP0_G0052530 | 411.9520467 | 534.9811283 | 519.9246166 | 650.8158045 | 488.3107426 | 604.7064923 | 618.1242655 | 646.3521597 | 575.2779171 | 0.161159365 | -0.251848281 | YOR377W | ATF1 | n |
| W303.asm02.HP0_G0052540 | 80.45181147 | 118.9904645 | 86.00256817 | 21.4020145 | 30.51942141 | 33.49422188 | 80.46027959 | 69.66669985 | 104.4098711 | 5.52E-07 | 1.742926032 | YOR378W | AMF1 | n |
| W303.asm02.HP0_G0052560 | 157.9957261 | 195.1443617 | 138.7768714 | 1229.643015 | 675.4080652 | 1145.140289 | 965.523355 | 1251.097818 | 1484.257971 | 1.79E-25 | -2.633555978 | YOR381W | FRE3 | n |
| W303.asm02.HP0_G0052620 | 428.4301286 | 434.0772144 | 444.6723695 | 255.8513551 | 216.2898126 | 268.8590243 | 231.9149235 | 328.0140451 | 319.3713703 | 1.77E-05 | 0.815173256 | YOR386W | PHR1 | n |
| W303.asm02.HP0_G0052650 | 285.9431854 | 372.2021729 | 288.3040637 | 92.41778988 | 65.01963692 | 79.66193311 | 226.2353744 | 169.3287844 | 245.6702849 | 1.95E-16 | 1.98921506 | YOR389W | | n |
| W303.asm02.HP0_G0052730 | 325.6844416 | 494.0484085 | 400.6937835 | 288.9271957 | 270.6939986 | 261.6170303 | 238.5410642 | 250.6066008 | 229.2922659 | 0.003541445 | 0.571645671 | YPL279C | FEX2 | y |
| W303.asm02.HP0_G0052760 | 1863.961849 | 1520.222174 | 1569.546869 | 3660.717298 | 3053.269073 | 3642.722941 | 1408.528189 | 2214.820499 | 2022.685345 | 7.97E-09 | -1.064257281 | YPL274W | SAM3 | n |
| W303.asm02.HP0_G0052770 | 5656.828575 | 6572.081334 | 5890.198618 | 2940.831356 | 2911.287416 | 3262.518261 | 3771.220634 | 2704.422584 | 2997.177475 | 4.00E-11 | 0.991063872 | YPL273W | SAM4 | n |
| W303.asm02.HP0_G0052790 | 2360.242903 | 2600.655592 | 2428.595249 | 5034.337501 | 5075.512474 | 4757.084756 | 4459.392672 | 4063.890824 | 3498.754307 | 5.96E-16 | -1.008438043 | YPL271W | ATP15 | n |
| W303.asm02.HP0_G0052800 | 805.4874137 | 901.4717589 | 729.0672256 | 1012.704413 | 1019.083289 | 1076.341346 | 899.2619483 | 1169.820002 | 1317.406903 | 0.04329161 | -0.351633519 | YPL270W | MDL2 | n |
| W303.asm02.HP0_G0052810 | 103.7149859 | 142.7885574 | 101.6393987 | 481.5453262 | 504.233919 | 514.1815683 | 375.7968352 | 495.4076434 | 451.4191485 | 6.46E-27 | -2.1064396 | YPL269W | KAR9 | n |
| W303.asm02.HP0_G0052820 | 603.8732355 | 952.8756395 | 692.9070549 | 511.7027103 | 561.2919677 | 489.7398388 | 629.4833638 | 576.6854598 | 703.2311905 | 0.007251308 | 0.526879365 | YPL268W | PLC1 | n |
| W303.asm02.HP0_G0052830 | 563.1626803 | 693.0004651 | 563.9032026 | 613.8486886 | 648.8694379 | 673.5054345 | 1116.977999 | 795.3614899 | 829.1372115 | 0.643158302 | -0.089180921 | YPL267W | ACM1 | n |
| W303.asm02.HP0_G0052840 | 1679.795052 | 2151.347598 | 1829.509177 | 5907.92882 | 5813.286313 | 5891.362053 | 5415.450112 | 4105.497326 | 4217.33989 | 8.74E-28 | -1.637542215 | YPL266W | DIM1 | n |
| W303.asm02.HP0_G0052850 | 2199.33928 | 2277.001528 | 2034.742579 | 569.0990219 | 1403.893385 | 683.4631761 | 1304.403121 | 906.6346911 | 1545.675542 | 1.38E-05 | 1.295391074 | YPL265W | DIP5 | n |
| W303.asm02.HP0_G0052870 | 1141.834144 | 1017.606452 | 1117.056084 | 1883.377276 | 2144.321087 | 2025.947799 | 2420.434528 | 2066.778762 | 2062.606767 | 3.05E-12 | -0.885229936 | YPL263C | KEL3 | n |
| W303.asm02.HP0_G0052880 | 3235.51984 | 2416.934314 | 3176.231211 | 8387.644046 | 8906.363327 | 8322.861512 | 9999.792865 | 9801.717631 | 8583.105578 | 2.36E-27 | -1.536798276 | YPL262W | FUM1 | n |
| W303.asm02.HP0_G0052890 | 3085.278505 | 3019.502027 | 3047.227358 | 962.1178336 | 1247.315484 | 994.8689147 | 1532.531678 | 1653.616528 | 2173.158395 | 1.02E-19 | 1.515534385 | YPL260W | CUB1 | n |
| W303.asm02.HP0_G0052900 | 797.7330222 | 627.3177287 | 688.0205453 | 1430.043696 | 1552.509698 | 1452.019781 | 1428.406611 | 1656.519307 | 1655.203544 | 1.66E-14 | -1.069070756 | YPL259C | APM1 | n |
| W303.asm02.HP0_G0052930 | 1071.075321 | 1538.308725 | 1302.743447 | 1239.371203 | 1417.162699 | 1364.210605 | 1795.684122 | 1965.181492 | 2177.2529 | 0.824991661 | -0.03906972 | YPL256C | CLN2 | n |
| W303.asm02.HP0_G0052940 | 405.1669542 | 458.827231 | 400.6937835 | 244.177529 | 268.0401359 | 230.8385562 | 270.725176 | 228.3519606 | 309.1351085 | 1.83E-05 | 0.77014782 | YPL255W | BBP1 | n |

| | | | | | | | | | | | | | | |
|---|---|---|---|---|---|---|---|---|---|---|---|---|---|---|
| W303.asm02.HP0_G0052950 | 1012.917385 | 1203.231577 | 1123.897198 | 634.2778842 | 686.0235161 | 616.4747324 | 645.5754197 | 567.9771224 | 648.9790025 | 7.07E-09 | 0.787673279 | YPL254W | HFI1 | n |
| W303.asm02.HP0_G0052960 | 520.5135273 | 401.7118081 | 475.9460307 | 596.3379494 | 656.8310261 | 553.1072856 | 544.2901266 | 578.620646 | 531.261991 | 0.020730918 | -0.367825548 | YPL253C | VIK1 | n |
| W303.asm02.HP0_G0052970 | 583.5179579 | 507.3753405 | 562.9259007 | 1752.046732 | 2083.282244 | 1821.361471 | 2057.889974 | 1890.676826 | 1572.289823 | 1.67E-32 | -1.773573082 | YPL252C | YAH1 | n |
| W303.asm02.HP0_G0052990 | 7982.176717 | 9427.852481 | 8275.792582 | 13974.54265 | 15571.53958 | 13213.92316 | 12795.07764 | 10249.71321 | 9423.502677 | 8.95E-07 | -0.7351991 | YPL249C-/ | RPL36B | y |
| W303.asm02.HP0_G0053000 | 493.3731571 | 392.1925709 | 475.9460307 | 1896.99674 | 1779.414961 | 1832.224462 | 1579.861254 | 2029.042633 | 1795.440332 | 2.66E-42 | -2.01685972 | YPL249C | GYP5 | n |
| W303.asm02.HP0_G0053010 | 315.9914523 | 335.077148 | 304.9181962 | 93.39060872 | 135.3469993 | 99.57741639 | 133.469405 | 233.1899259 | 257.9537991 | 1.15E-08 | 1.549152628 | YPL248C | GAL4 | n |
| W303.asm02.HP0_G0053020 | 574.7942675 | 276.0578776 | 367.4655185 | 534.0775436 | 537.4072031 | 513.276319 | 404.194581 | 590.2317626 | 717.5619571 | 0.185766284 | -0.379342585 | YPL247C | | n |
| W303.asm02.HP0_G0053030 | 1006.132293 | 912.8948434 | 903.0269658 | 1340.544363 | 1165.045739 | 1261.91744 | 1609.205592 | 1492.028488 | 1463.785447 | 0.000713972 | -0.417839293 | YPL246C | RBD2 | n |
| W303.asm02.HP0_G0053040 | 870.4304422 | 629.2215761 | 846.3434549 | 417.3392827 | 262.7324104 | 365.7206929 | 107.9114338 | 135.4630275 | 109.528002 | 1.13E-07 | 1.16188865 | YPL245W | | n |
| W303.asm02.HP0_G0053050 | 1367.680795 | 1154.683467 | 1347.699335 | 610.930232 | 586.5036637 | 535.90755 | 1029.891579 | 874.7041203 | 827.0899591 | 2.98E-15 | 1.159297285 | YPL244C | HUT1 | n |
| W303.asm02.HP0_G0053060 | 4546.981297 | 4241.772078 | 4623.615341 | 2674.278993 | 2564.95833 | 2434.215206 | 2469.657288 | 2424.788192 | 2502.766027 | 6.45E-16 | 0.805713237 | YPL243W | SRP68 | n |
| W303.asm02.HP0_G0053070 | 1363.803599 | 1428.837497 | 1408.292054 | 2397.998443 | 2603.43934 | 2307.480313 | 1593.113536 | 2037.750971 | 1965.362279 | 1.99E-09 | -0.798485023 | YPL242C | IQG1 | n |
| W303.asm02.HP0_G0053100 | 2688.835241 | 2540.684397 | 2712.990105 | 2498.198783 | 2433.592125 | 2437.836203 | 1991.628568 | 1676.838762 | 1448.431055 | 0.463608772 | 0.107949393 | YPL239W | YAR1 | n |
| W303.asm02.HP0_G0053110 | 5567.653074 | 6339.811947 | 6126.70568 | 8874.053466 | 9552.578902 | 8577.236548 | 8772.063658 | 7691.397182 | 7435.620622 | 8.39E-07 | -0.582287046 | YPL237W | SUI3 | n |
| W303.asm02.HP0_G0053140 | 1869.777643 | 1653.491494 | 1999.55971 | 1352.218189 | 1194.238229 | 1338.863626 | 2109.952508 | 1627.491516 | 1620.400254 | 0.001069016 | 0.506655191 | YPL234C | VMA11 | n |
| W303.asm02.HP0_G0053150 | 380.9344808 | 369.3464017 | 307.850102 | 150.7869203 | 132.6931366 | 143.0293799 | 230.0217405 | 135.4630275 | 224.1741349 | 3.58E-08 | 1.309018009 | YPL233W | NSL1 | n |
| W303.asm02.HP0_G0053160 | 2809.028309 | 2317.934248 | 2710.058199 | 5002.23448 | 4347.027154 | 4919.12437 | 4268.181184 | 4711.210577 | 4728.129358 | 5.98E-12 | -0.86478744 | YPL232W | SSO1 | n |
| W303.asm02.HP0_G0053170 | 5246.815127 | 4468.329922 | 4548.363094 | 11730.24958 | 12301.98069 | 12704.26783 | 10754.22631 | 9576.26845 | 8770.42917 | 3.75E-28 | -1.364894999 | YPL231W | FAS2 | n |
| W303.asm02.HP0_G0053180 | 276.250196 | 159.9231843 | 258.0077045 | 39.88557247 | 29.19249005 | 40.7362158 | 61.5284491 | 87.08337481 | 103.3862449 | 4.46E-16 | 2.648533821 | YPL230W | USV1 | n |
| W303.asm02.HP0_G0053190 | 785.1321361 | 613.038873 | 723.2034141 | 128.412087 | 96.8659897 | 103.1984134 | 187.4251219 | 190.6158315 | 257.9537991 | 1.06E-37 | 2.686688777 | YPL229W | | n |
| W303.asm02.HP0_G0053200 | 951.8515526 | 782.4812944 | 889.344739 | 1353.191008 | 1316.315915 | 1291.790665 | 1111.29845 | 1447.519208 | 1548.746421 | 0.000203603 | -0.594436256 | YPL228W | CET1 | n |
| W303.asm02.HP0_G0053210 | 1877.532034 | 2244.636122 | 2052.334013 | 1739.400087 | 1735.626226 | 1763.425519 | 2667.494916 | 2413.177075 | 2354.34023 | 0.052813901 | 0.23714267 | YPL227C | ALG5 | n |
| W303.asm02.HP0_G0053220 | 2022.926874 | 2675.857565 | 1876.419669 | 11889.79187 | 10308.92978 | 11827.98657 | 4138.498145 | 6489.646609 | 5862.307173 | 8.55E-35 | -2.371575529 | YPL226W | NEW1 | n |
| W303.asm02.HP0_G0053230 | 3740.524584 | 3344.108014 | 3830.046189 | 1470.902087 | 1609.567747 | 1541.639456 | 1977.429695 | 1391.398811 | 1165.910227 | 3.07E-11 | 1.239838776 | YPL225W | CHP1 | n |
| W303.asm02.HP0_G0053260 | 272.3730003 | 167.538574 | 247.2573835 | 407.6110943 | 410.021792 | 393.7834194 | 468.5628047 | 429.6113157 | 577.3251695 | 0.000134905 | -0.818028754 | YPL222W | FMP40 | n |
| W303.asm02.HP0_G0053270 | 818.0882998 | 1060.443019 | 727.1126218 | 3447.669972 | 2937.826044 | 3503.314559 | 2337.134474 | 3164.029285 | 2493.553391 | 1.54E-26 | -1.924312359 | YPL221W | FLC1 | n |
| W303.asm02.HP0_G0053300 | 3334.388331 | 3826.733337 | 3411.760971 | 3967.155233 | 3842.793235 | 3928.781701 | 3804.351337 | 3009.214396 | 2723.869284 | 0.329529202 | -0.150946227 | YPL218W | SAR1 | n |
| W303.asm02.HP0_G0053310 | 972.2068302 | 1495.472158 | 1005.643666 | 8543.29506 | 8246.878438 | 8603.488776 | 3509.014782 | 4339.654845 | 4019.780036 | 3.14E-60 | -2.869991296 | YPL217C | BMS1 | n |
| W303.asm02.HP0_G0053330 | 1742.799482 | 1482.145225 | 1766.961855 | 2334.765218 | 2066.032137 | 2243.207617 | 1747.407954 | 1678.773948 | 1624.494759 | 0.000882356 | -0.413044241 | YPL215W | CBP3 | n |
| W303.asm02.HP0_G0053340 | 819.0575987 | 749.1639643 | 765.2273963 | 310.3292102 | 341.021361 | 305.9742431 | 322.7877099 | 375.4261047 | 364.4109226 | 1.24E-19 | 1.288004515 | YPL214C | THI6 | n |
| W303.asm02.HP0_G0053350 | 999.3472004 | 1115.654595 | 1034.962724 | 334.6496812 | 371.5407824 | 378.3941823 | 403.2479895 | 386.0696283 | 383.8598201 | 2.69E-31 | 1.538860447 | YPL213W | LEA1 | n |
| W303.asm02.HP0_G0053360 | 2079.146212 | 1911.462821 | 2135.404676 | 4826.154269 | 5063.570092 | 4798.726221 | 3116.179299 | 3688.46472 | 3148.674151 | 2.30E-27 | -1.26155666 | YPL212C | PUS1 | n |
| W303.asm02.HP0_G0053370 | 1641.023094 | 2530.213237 | 1833.418385 | 2556.567914 | 2879.441064 | 2684.969246 | 1717.117026 | 1521.05628 | 1283.627238 | 0.020914366 | -0.43534251 | YPL211W | NIP7 | n |
| W303.asm02.HP0_G0053380 | 4047.792346 | 4394.079872 | 4363.653033 | 3838.743146 | 3925.06298 | 3915.202963 | 2931.593952 | 3668.145266 | 3194.73733 | 0.28995973 | 0.13288413 | YPL210C | SRP72 | n |
| W303.asm02.HP0_G0053390 | 135.7018507 | 136.1250914 | 143.6633809 | 168.2976595 | 176.4818716 | 169.2816079 | 251.7933455 | 272.8612411 | 222.1268826 | 0.121204289 | -0.305942494 | YPL209C | IPL1 | n |
| W303.asm02.HP0_G0053400 | 378.026584 | 371.2502492 | 314.6912153 | 756.8530582 | 801.4665449 | 837.3555469 | 359.7047793 | 853.4170731 | 733.9399761 | 1.26E-05 | -1.170955422 | YPL208W | RKM1 | n |
| W303.asm02.HP0_G0053420 | 2451.357003 | 1657.299189 | 2259.522018 | 86.58087683 | 30.51942141 | 87.80917627 | 163.7603337 | 103.5324567 | 218.0323778 | 1.94E-50 | 4.940529261 | YPL206C | PGC1 | n |
| W303.asm02.HP0_G0053440 | 1326.000941 | 1322.222041 | 1175.694199 | 3918.514291 | 3690.196128 | 3918.82396 | 2470.603879 | 2930.839359 | 2747.412686 | 1.55E-41 | -1.592114523 | YPL204W | HRR25 | n |
| W303.asm02.HP0_G0053460 | 40.7105552 | 59.01927038 | 35.1828688 | 5.836913045 | 14.59624502 | 10.86299088 | 32.18411183 | 14.5138958 | 28.66153324 | 0.000114637 | 2.133051121 | YPL201C | YIG1 | n |
| W303.asm02.HP0_G0053480 | 1600.312539 | 1529.741411 | 1686.823098 | 1097.339652 | 1085.429857 | 1086.299088 | 1091.420028 | 1034.356974 | 985.7520181 | 4.78E-07 | 0.559115476 | YPL199C | | n |
| W303.asm02.HP0_G0053500 | 888.847122 | 597.8080935 | 807.2513785 | 64.2060435 | 88.90440151 | 59.74644984 | 148.6148694 | 92.88893313 | 125.906021 | 4.76E-42 | 3.442614947 | YPL196W | OXR1 | n |
| W303.asm02.HP0_G0053510 | 1325.031642 | 1573.529902 | 1446.406828 | 1613.906457 | 1575.067531 | 1657.511358 | 1198.38487 | 1487.190523 | 1324.572286 | 0.256070192 | -0.157812821 | YPL195W | APL5 | n |
| W303.asm02.HP0_G0053520 | 214.2150643 | 297.0001993 | 251.1665911 | 215.9657827 | 276.0017241 | 230.8385562 | 441.1116504 | 300.9214396 | 361.340044 | 0.736723896 | 0.080317644 | YPL194W | DDC1 | n |
| W303.asm02.HP0_G0053530 | 597.088143 | 693.0004651 | 727.1126218 | 1277.311138 | 971.3137597 | 1126.130054 | 357.8115963 | 645.3845666 | 509.7658411 | 0.000654711 | -0.743079796 | YPL193W | RSA1 | n |
| W303.asm02.HP0_G0053560 | 911.1409974 | 885.2890557 | 719.2942065 | 1561.37424 | 1808.607452 | 1587.807167 | 1090.473436 | 1581.047049 | 1468.903578 | 2.50E-08 | -0.978165574 | YPL190C | NAB3 | n |
| W303.asm02.HP0_G0053570 | 845.2286699 | 765.3466675 | 800.4102651 | 1165.436971 | 1308.354327 | 1172.297766 | 828.267584 | 1046.935684 | 893.6256612 | 2.06E-05 | -0.59588073 | YPL189C-/ | COA2 | n |
| W303.asm02.HP0_G0053590 | 997.4086025 | 712.0389394 | 1143.443236 | 351.1876015 | 395.425547 | 359.3839483 | 528.1980707 | 518.6298766 | 577.3251695 | 6.42E-13 | 1.368309332 | YPL188W | POS5 | n |
| W303.asm02.HP0_G0053600 | 21.32457654 | 20.94232175 | 20.52334013 | 44.74966668 | 87.57747014 | 67.89369299 | 85.19323721 | 60.95836237 | 56.29944028 | 2.84E-05 | -1.66457689 | YPL187W | MF(ALPHA | n |
| W303.asm02.HP0_G0053610 | 268.4958046 | 196.0962855 | 256.0531007 | 386.2090798 | 220.2706067 | 383.8256777 | 308.588837 | 232.2223328 | 273.3081919 | 0.075253206 | -0.463019567 | YPL186C | UIP4 | n |
| W303.asm02.HP0_G0053620 | 769.6233531 | 864.3467339 | 835.5931339 | 1974.822247 | 1694.491354 | 1912.791644 | 482.7616775 | 1030.486602 | 1075.831123 | 3.04E-06 | -1.176743847 | YPL184C | MRN1 | n |
| W303.asm02.HP0_G0053630 | 679.4785523 | 716.798558 | 752.5224715 | 716.9674857 | 866.4861818 | 763.1251093 | 797.0300637 | 623.1299264 | 595.7504408 | 0.464994433 | -0.125680054 | YPL183W- | RTC6 | n |
| W303.asm02.HP0_G0053640 | 1011.948086 | 1591.616453 | 1041.803837 | 1379.457116 | 1437.066669 | 1385.936586 | 932.3926517 | 1138.857024 | 1009.29542 | 0.3030567 | -0.20492649 | YPL183C | RTT10 | n |
| W303.asm02.HP0_G0053650 | 1192.237688 | 1228.933517 | 1185.467218 | 715.9946669 | 762.9855353 | 679.8421792 | 982.5620025 | 864.0605967 | 802.5229306 | 1.36E-08 | 0.741803347 | YPL181W | CTI6 | n |
| W303.asm02.HP0_G0053660 | 303.3905662 | 307.4713602 | 279.5083465 | 727.6684929 | 709.9082807 | 697.947164 | 571.7412808 | 746.9818372 | 785.1212854 | 1.02E-14 | -1.262005773 | YPL180W | TCO89 | n |
| W303.asm02.HP0_G0053670 | 3020.335477 | 2994.75201 | 3188.936136 | 3086.754182 | 3051.942141 | 2959.259765 | 1462.483905 | 3233.695985 | 3402.533446 | 0.951139584 | 0.016734135 | YPL179W | PPQ1 | n |
| W303.asm02.HP0_G0053680 | 2365.089398 | 2212.270716 | 2240.953282 | 1052.589986 | 1093.391445 | 1070.004602 | 1381.077034 | 959.852309 | 874.1767637 | 5.48E-10 | 1.084300608 | YPL178W | CBC2 | n |
| W303.asm02.HP0_G0053690 | 924.7111825 | 814.8467007 | 922.573004 | 643.0332538 | 814.7358586 | 675.315933 | 761.0595857 | 1314.95896 | 1453.549186 | 0.175667887 | 0.320651932 | YPL177C | CUP9 | n |
| W303.asm02.HP0_G0053710 | 753.1452713 | 906.2313774 | 779.886925 | 535.0503625 | 553.3303795 | 582.0752613 | 566.0617317 | 593.1345418 | 443.230139 | 0.000818085 | 0.546029879 | YPL175W | SPT14 | n |
| W303.asm02.HP0_G0053720 | 352.8248118 | 588.2888564 | 391.8980663 | 345.3506885 | 406.0409979 | 353.9524528 | 375.7968352 | 485.7317128 | 397.1669605 | 0.241543615 | 0.271879465 | YPL174C | NIP100 | n |
| W303.asm02.HP0_G0053730 | 1471.395781 | 1623.981859 | 1493.31732 | 2180.087022 | 2177.494371 | 2280.322835 | 2361.745854 | 2116.126008 | 2101.504562 | 1.79E-06 | -0.532795422 | YPL173W | MRPL40 | n |
| W303.asm02.HP0_G0053740 | 800.640919 | 929.0775466 | 781.8415288 | 1735.508812 | 1702.452942 | 1876.581674 | 1259.913319 | 1691.352657 | 1612.211244 | 1.32E-12 | -1.081581443 | YPL172C | COX10 | n |
| W303.asm02.HP0_G0053760 | 1511.137037 | 1709.654994 | 1539.25051 | 1000.057768 | 1042.968053 | 935.1224648 | 1057.342733 | 850.514294 | 666.3806477 | 0.000126206 | 0.677076227 | YPL170W | DAP1 | n |
| W303.asm02.HP0_G0053770 | 1277.535994 | 1159.443086 | 1308.607259 | 2576.997109 | 2588.843095 | 2606.212562 | 1333.747458 | 2055.167645 | 1711.502985 | 1.39E-10 | -1.053143958 | YPL169C | MEX67 | n |

| | | | | | | | | | | | | | | |
|---|---|---|---|---|---|---|---|---|---|---|---|---|---|---|
| W303.asm02.HP0_G0053780 | 553.469691 | 633.029271 | 510.1515975 | 447.4966668 | 472.3875662 | 441.7616291 | 420.2866369 | 515.7270975 | 447.3246437 | 0.047151602 | 0.318479718 | YPL168W | MRX4 | n |
| W303.asm02.HP0_G0053790 | 228.7545483 | 450.2599176 | 237.4843644 | 177.053029 | 175.1549403 | 177.428851 | 200.6774032 | 168.3611913 | 211.8906207 | 0.004786434 | 0.791283122 | YPL167C | REV3 | n |
| W303.asm02.HP0_G0053800 | 169.6273133 | 138.9808625 | 129.9811542 | 173.1617537 | 200.3666362 | 202.7758297 | 109.8046168 | 123.8519108 | 123.8587686 | 0.069953286 | -0.393046283 | YPL166W | ATG29 | n |
| W303.asm02.HP0_G0053810 | 223.9080536 | 249.4040135 | 182.7554574 | 560.3436523 | 729.8122512 | 521.4235622 | 381.4763844 | 607.6484376 | 445.2773913 | 1.54E-10 | -1.463210695 | YPL164C | MLH3 | n |
| W303.asm02.HP0_G0053830 | 556.3775878 | 574.9619243 | 497.4466727 | 402.7470001 | 363.5791942 | 390.1624224 | 429.7525521 | 537.0141447 | 399.2142129 | 0.003573014 | 0.492818745 | YPL162C | | n |
| W303.asm02.HP0_G0053840 | 1038.119158 | 926.2217755 | 1120.965292 | 746.1520509 | 732.4661139 | 757.6936138 | 534.8242114 | 714.0836734 | 576.3015433 | 0.003043626 | 0.463940988 | YPL161C | BEM4 | n |
| W303.asm02.HP0_G0053850 | 4327.919738 | 4400.743338 | 4456.496714 | 13271.19463 | 12692.09851 | 13337.9423 | 7556.640141 | 9071.184876 | 8417.278136 | 6.28E-50 | -1.575718203 | YPL160W | CDC60 | n |
| W303.asm02.HP0_G0053860 | 1130.202556 | 1146.116154 | 1076.986706 | 1082.74737 | 1066.852818 | 983.1006746 | 1242.874672 | 1083.70422 | 1078.902001 | 0.448789716 | 0.098724728 | YPL159C | PET20 | n |
| W303.asm02.HP0_G0053870 | 735.6978905 | 815.7986244 | 711.4757912 | 1665.465856 | 1759.510991 | 1810.49848 | 1482.362327 | 2098.709333 | 1486.305223 | 2.52E-13 | -1.210034328 | YPL158C | AIM44 | n |
| W303.asm02.HP0_G0053880 | 510.8205379 | 745.3562695 | 540.4479568 | 810.3580944 | 725.8314571 | 710.6206533 | 687.2254468 | 718.9216387 | 551.7345148 | 0.097383791 | -0.322617387 | YPL157W | TGS1 | n |
| W303.asm02.HP0_G0053890 | 241.3554344 | 217.0386072 | 252.143893 | 220.8298769 | 214.9628813 | 200.060082 | 378.6366098 | 364.7825811 | 319.3713703 | 0.378579587 | 0.160976957 | YPL156C | PRM4 | n |
| W303.asm02.HP0_G0053900 | 333.4388331 | 329.3656057 | 287.3267618 | 1395.022218 | 1620.183198 | 1368.736851 | 1394.329316 | 1591.690573 | 1635.754647 | 8.58E-54 | -2.204824217 | YPL155C | KIP2 | n |
| W303.asm02.HP0_G0053910 | 2924.374882 | 2439.780484 | 2914.314299 | 5621.920081 | 4374.892713 | 5184.362397 | 2215.024167 | 4007.770427 | 2995.130223 | 2.55E-05 | -0.875022682 | YPL154C | PEP4 | n |
| W303.asm02.HP0_G0053920 | 145.39484 | 215.1347598 | 116.2989274 | 713.0762103 | 806.7742704 | 727.8203889 | 719.4095587 | 909.5374702 | 805.5938092 | 5.97E-25 | -2.235561551 | YPL153C | RAD53 | n |
| W303.asm02.HP0_G0053930 | 591.2723494 | 374.1060203 | 548.2663721 | 259.7426305 | 289.2710377 | 254.3750364 | 319.9479353 | 280.6019855 | 305.0406037 | 7.41E-06 | 0.915587835 | YPL152W | RRD2 | n |
| W303.asm02.HP0_G0053940 | 481.7415699 | 681.5773805 | 367.4655185 | 628.4409712 | 471.0606348 | 622.8114771 | 350.2388641 | 455.7363282 | 359.2927916 | 0.50484048 | -0.171342731 | YPL151C | PRP46 | n |
| W303.asm02.HP0_G0053950 | 503.0661464 | 598.7600172 | 478.8779364 | 735.4510437 | 716.5429375 | 754.0726169 | 681.5458977 | 791.4911177 | 879.2948946 | 0.002456234 | -0.481162474 | YPL150W | | n |
| W303.asm02.HP0_G0053960 | 640.706595 | 732.0293374 | 673.3610167 | 60.31476813 | 72.98122512 | 74.23043767 | 145.7750948 | 96.75930534 | 113.6225068 | 1.07E-54 | 3.303871161 | YPL149W | ATG5 | n |
| W303.asm02.HP0_G0053970 | 272.3730003 | 255.1155558 | 273.6445351 | 51.55939856 | 71.65429375 | 47.07296048 | 39.75684403 | 46.44446656 | 81.89009496 | 9.92E-14 | 2.246342147 | YPL148C | PPT2 | n |
| W303.asm02.HP0_G0053990 | 471.0792816 | 467.3945445 | 501.3558803 | 13.61946377 | 19.90397049 | 23.53648024 | 51.11594233 | 62.89354847 | 41.96867367 | 4.04E-60 | 4.656096239 | YPL147W | PXA1 | n |
| W303.asm02.HP0_G0054000 | 2213.878764 | 3595.415875 | 2616.237216 | 3977.85634 | 3229.750944 | 3904.339972 | 1890.343275 | 1842.297174 | 1496.541485 | 0.044015484 | -0.399483638 | YPL146C | NOP53 | n |
| W303.asm02.HP0_G0054010 | 4737.933187 | 4310.310585 | 5379.069718 | 3986.61161 | 3680.907609 | 3918.82396 | 3724.837649 | 4283.534448 | 3642.061973 | 0.016873571 | 0.316164855 | YPL145C | KES1 | n |
| W303.asm02.HP0_G0054020 | 563.1626803 | 556.8753737 | 632.3143364 | 133.2761812 | 122.0776857 | 147.5556261 | 255.5797116 | 171.2639705 | 159.6856852 | 1.05E-22 | 2.116900522 | YPL144W | POC4 | n |
| W303.asm02.HP0_G0054040 | 474.9564774 | 648.2600505 | 392.8753682 | 1455.336986 | 1541.894247 | 1349.726617 | 1200.278053 | 1849.070325 | 1568.195318 | 1.04E-12 | -1.519156014 | YPL141C | FRK1 | n |
| W303.asm02.HP0_G0054060 | 1052.658642 | 881.4813608 | 993.9160435 | 854.1349423 | 806.7742704 | 851.8395348 | 957.9506228 | 875.6717134 | 835.2789686 | 0.104277824 | 0.219926019 | YPL139C | UME1 | n |
| W303.asm02.HP0_G0054070 | 568.9784739 | 655.8754402 | 593.22226 | 297.6825653 | 339.6944296 | 293.3007537 | 375.7968352 | 356.0742437 | 310.1587347 | 6.91E-10 | 0.969302992 | YPL138C | SPP1 | n |
| W303.asm02.HP0_G0054080 | 622.2899153 | 1034.741079 | 662.6106957 | 781.1735292 | 968.659897 | 759.5041123 | 1387.703175 | 1602.334096 | 1431.029409 | 0.619048018 | -0.112061556 | YPL137C | GIP3 | n |
| W303.asm02.HP0_G0054090 | 3451.673502 | 2947.155824 | 3785.090301 | 2258.885348 | 3012.1342 | 2217.860638 | 3250.595295 | 3039.209781 | 3235.682377 | 0.006024367 | 0.444333304 | YPL135W | ISU1 | n |
| W303.asm02.HP0_G0054100 | 447.8161072 | 430.2695195 | 489.6282574 | 2026.381645 | 1772.780305 | 2083.88375 | 2404.342472 | 1920.672211 | 2215.127069 | 2.60E-50 | -2.105615424 | YPL134C | ODC1 | n |
| W303.asm02.HP0_G0054110 | 319.868648 | 290.3367333 | 230.643251 | 83.66242031 | 156.5779012 | 88.71442551 | 253.6865286 | 305.7594049 | 378.7416892 | 1.12E-06 | 1.366575394 | YPL133C | RDS2 | n |
| W303.asm02.HP0_G0054120 | 979.9612217 | 1142.308459 | 948.9601556 | 832.7329278 | 806.7742704 | 820.1558114 | 1208.797377 | 951.1439715 | 907.9564278 | 0.044791866 | 0.32023373 | YPL132W | COX11 | n |
| W303.asm02.HP0_G0054130 | 15007.65539 | 12047.54655 | 14510.00147 | 30001.73305 | 32073.25804 | 28270.93376 | 27406.66441 | 22632.00152 | 22315.05088 | 4.78E-16 | -1.120057172 | YPL131W | RPL5 | n |
| W303.asm02.HP0_G0054150 | 1953.137351 | 1689.664596 | 2002.491616 | 1453.391348 | 1514.028688 | 1418.525559 | 1534.424861 | 1326.570076 | 1298.981631 | 0.005464763 | 0.364522799 | YPL129W | TAF14 | n |
| W303.asm02.HP0_G0054160 | 313.0835555 | 452.163765 | 280.4856485 | 3536.196486 | 2484.015517 | 3328.601455 | 1053.556367 | 1214.329282 | 780.0031545 | 8.03E-44 | -3.160356534 | YPL128C | TBF1 | n |
| W303.asm02.HP0_G0054180 | 1697.242432 | 2439.780484 | 1826.577272 | 7736.828241 | 6832.369602 | 7629.440594 | 2761.207477 | 4385.131718 | 3268.438415 | 2.67E-22 | -1.89624576 | YPL126W | NAN1 | n |
| W303.asm02.HP0_G0054190 | 1092.399898 | 1128.981527 | 1171.784991 | 2452.476298 | 2515.861869 | 2482.193416 | 1512.653256 | 2201.274197 | 1869.141417 | 5.24E-14 | -1.134661816 | YPL125W | KAP120 | n |
| W303.asm02.HP0_G0054210 | 967.3603356 | 723.462024 | 896.1858524 | 209.1560508 | 151.2701757 | 198.2495835 | 178.9057981 | 181.907494 | 222.1268826 | 1.82E-29 | 2.205026673 | YPL123C | RNY1 | n |
| W303.asm02.HP0_G0054220 | 641.6758939 | 732.9812612 | 632.3143364 | 748.0976886 | 715.2160061 | 750.4516199 | 690.0652214 | 635.7086361 | 527.1674863 | 0.372414407 | -0.141884587 | YPL122C | TFB2 | n |
| W303.asm02.HP0_G0054230 | 20.3552776 | 12.37500831 | 18.56873631 | 12.64664493 | 5.307725463 | 18.1049848 | 12.30568982 | 5.805558321 | 16.37801899 | 0.509564177 | 0.481716041 | YPL121C | MEI5 | n |
| W303.asm02.HP0_G0054240 | 610.6583281 | 565.4426872 | 628.4051288 | 226.6667899 | 238.8476458 | 260.7117811 | 229.0751489 | 247.7038217 | 227.2450135 | 3.91E-18 | 1.311880065 | YPL120W | VPS30 | n |
| W303.asm02.HP0_G0054270 | 3203.532975 | 3588.752409 | 3508.51386 | 3014.765588 | 2787.882799 | 2969.217507 | 2794.33818 | 2804.084669 | 2562.136346 | 0.035872482 | 0.231452982 | YPL118W | MRP51 | n |
| W303.asm02.HP0_G0054280 | 5627.749607 | 6513.062063 | 5606.781063 | 3172.36224 | 3339.886248 | 3071.510671 | 2326.721967 | 2699.584619 | 2353.316604 | 1.82E-13 | 0.88925323 | YPL117C | IDI1 | n |
| W303.asm02.HP0_G0054290 | 789.0093318 | 676.8177619 | 697.7935644 | 1118.741667 | 1247.315484 | 1151.477033 | 1080.06093 | 1349.79231 | 1364.493707 | 1.74E-06 | -0.700482159 | YPL116W | HOS3 | n |
| W303.asm02.HP0_G0054300 | 373.1800894 | 618.7504153 | 384.079651 | 951.4168263 | 987.2369361 | 952.3222004 | 906.8346805 | 1206.588538 | 1131.106937 | 4.12E-07 | -1.070660731 | YPL115C | BEM3 | n |
| W303.asm02.HP0_G0054310 | 291.758979 | 157.0674131 | 232.5978548 | 838.5698408 | 761.658604 | 866.3235226 | 1519.279397 | 1492.028488 | 1352.210193 | 6.83E-20 | -1.856892309 | YPL113C | | n |
| W303.asm02.HP0_G0054320 | 2376.720985 | 1919.078211 | 2387.548569 | 501.001703 | 473.7144976 | 513.276319 | 594.4594774 | 604.7456584 | 527.1674863 | 5.80E-53 | 2.165984806 | YPL112C | PEX25 | n |
| W303.asm02.HP0_G0054330 | 694.9873353 | 658.7312113 | 676.2929224 | 1719.943711 | 2284.975812 | 1662.942854 | 4168.789074 | 5451.419263 | 3631.825711 | 1.44E-15 | -1.480438002 | YPL111W | CAR1 | n |
| W303.asm02.HP0_G0054340 | 549.5924953 | 619.702339 | 468.1276154 | 825.9231959 | 678.0619279 | 841.8817931 | 460.0434809 | 882.4448647 | 729.8454713 | 0.029799276 | -0.5194708 | YPL110C | GDE1 | n |
| W303.asm02.HP0_G0054350 | 484.6494667 | 431.2214433 | 512.1062014 | 344.3778697 | 355.617606 | 353.9524528 | 363.4911454 | 411.2270477 | 446.3010175 | 0.005520957 | 0.438323982 | YPL109C | CQD1 | n |
| W303.asm02.HP0_G0054360 | 538.930207 | 527.3657386 | 528.7203338 | 1266.610131 | 1340.200679 | 1248.338702 | 1198.38487 | 1188.20427 | 863.9405018 | 3.71E-16 | -1.272832624 | YPL108W | | n |
| W303.asm02.HP0_G0054370 | 937.3120686 | 887.1929031 | 985.1203263 | 337.5681378 | 382.1562333 | 305.0689939 | 513.9991978 | 388.9724075 | 393.0724558 | 4.15E-18 | 1.458323466 | YPL107W | DPC25 | n |
| W303.asm02.HP0_G0054380 | 8391.220867 | 8390.255631 | 6591.90139 | 9087.100792 | 9819.292107 | 9216.342512 | 9680.791522 | 7084.716337 | 6760.027339 | 0.124962505 | -0.266814703 | YPL106C | SSE1 | n |
| W303.asm02.HP0_G0054390 | 805.4874137 | 1486.904844 | 726.1353199 | 2997.254849 | 2961.710808 | 2941.154781 | 2029.492299 | 2469.297472 | 2297.017164 | 3.76E-11 | -1.559849428 | YPL105C | SYH1 | n |
| W303.asm02.HP0_G0054400 | 562.1933814 | 534.0292046 | 461.286502 | 1178.083616 | 1090.737583 | 1213.939231 | 869.9176111 | 1028.551416 | 962.2086158 | 8.56E-17 | -1.161669737 | YPL104W | MSD1 | n |
| W303.asm02.HP0_G0054410 | 611.627627 | 526.4138148 | 606.9044867 | 1394.049399 | 1553.836629 | 1452.019781 | 1022.318847 | 1067.255138 | 905.9091755 | 1.11E-23 | -1.333823477 | YPL103C | FMP30 | n |
| W303.asm02.HP0_G0054420 | 577.7021643 | 577.8176955 | 542.4025606 | 1520.515848 | 1559.144355 | 1524.43972 | 1305.349712 | 1585.885015 | 1187.406377 | 3.00E-24 | -1.438966879 | YPL101W | ELP4 | n |
| W303.asm02.HP0_G0054430 | 543.7767017 | 328.4136819 | 409.4895007 | 357.0245146 | 398.0794097 | 366.6259422 | 530.0912537 | 441.2224324 | 608.0339551 | 0.39628726 | 0.193141391 | YPL100W | ATG21 | n |
| W303.asm02.HP0_G0054440 | 733.7592926 | 581.6253904 | 696.8162625 | 711.1305727 | 707.254418 | 705.1891579 | 807.4425704 | 633.77345 | 585.514179 | 0.670643327 | -0.077818364 | YPL099C | INA17 | n |
| W303.asm02.HP0_G0054450 | 1436.501019 | 1181.337331 | 1479.635093 | 821.0591017 | 983.256142 | 895.2914983 | 1303.456529 | 903.7319119 | 854.7278661 | 0.001727982 | 0.602647067 | YPL098C | MGR2 | n |
| W303.asm02.HP0_G0054460 | 1227.13245 | 1295.568177 | 1317.402976 | 1607.096725 | 1610.894678 | 1626.732884 | 1739.835222 | 1805.528638 | 1623.471133 | 0.00193385 | -0.335322945 | YPL097W | MSY1 | n |
| W303.asm02.HP0_G0054470 | 221.9694558 | 286.5290385 | 247.6672332 | 107.0100725 | 114.1160975 | 114.9666535 | 194.0512625 | 127.7222831 | 124.8823948 | 5.06E-07 | 1.217285697 | YPL096C-/ | ERI1 | n |
| W303.asm02.HP0_G0054480 | 979.9612217 | 824.3659379 | 881.5263237 | 330.7584059 | 359.5984001 | 313.216237 | 470.4559877 | 389.9400005 | 420.7103628 | 3.82E-21 | 1.422774559 | YPL096W | PNG1 | n |
| W303.asm02.HP0_G0054500 | 3176.392605 | 3562.098545 | 3474.308294 | 1433.934971 | 1308.354327 | 1308.085152 | 1835.440966 | 1716.510077 | 1682.841451 | 2.99E-33 | 1.333914645 | YPL094C | SEC62 | n |

| | | | | | | | | | | | | | | |
|---|---|---|---|---|---|---|---|---|---|---|---|---|---|---|
| W303.asm02.HP0_G0054510 | 2281.729689 | 3873.3776 | 2618.19182 | 14388.96347 | 12865.92652 | 14105.59366 | 5742.024188 | 6783.794898 | 5686.243469 | 2.87E-32 | -2.237054323 | YPL093W | NOG1 | n |
| W303.asm02.HP0_G0054520 | 661.0618726 | 509.279188 | 623.5186192 | 2009.843725 | 2369.899419 | 2065.778766 | 1834.494375 | 2902.77916 | 2257.095742 | 1.09E-22 | -1.844960982 | YPL092W | SSU1 | n |
| W303.asm02.HP0_G0054530 | 3379.945381 | 2394.088145 | 3141.048342 | 2586.725298 | 2166.87892 | 2444.172948 | 2037.064961 | 2187.727894 | 2019.614467 | 0.052752646 | 0.308229477 | YPL091W | GLR1 | n |
| W303.asm02.HP0_G0054550 | 385.7809755 | 590.1927038 | 356.7151975 | 1313.305435 | 1016.429426 | 1170.487267 | 1016.639297 | 1559.760002 | 1659.298049 | 5.61E-09 | -1.393395199 | YPL089C | RLM1 | n |
| W303.asm02.HP0_G0054570 | 1233.917542 | 940.5006312 | 1068.190989 | 6.809731886 | 14.59624502 | 11.76824012 | 12.30568982 | 23.22223328 | 16.37801899 | 4.17E-90 | 6.644827353 | YPL087W | YDC1 | n |
| W303.asm02.HP0_G0054580 | 1326.000941 | 1224.173899 | 1307.629957 | 3820.259588 | 3765.831216 | 3908.866218 | 2257.620786 | 3054.69127 | 2547.805579 | 4.78E-31 | -1.575250029 | YPL086C | ELP3 | n |
| W303.asm02.HP0_G0054590 | 317.9300502 | 527.3657386 | 301.0089886 | 2782.261885 | 2729.497819 | 2793.599154 | 1667.894266 | 2310.612212 | 2359.458361 | 5.60E-39 | -2.856778978 | YPL085W | SEC16 | n |
| W303.asm02.HP0_G0054600 | 662.0311715 | 585.4330852 | 556.0847874 | 564.2349277 | 668.7734083 | 556.7282826 | 676.8129401 | 744.0790581 | 704.2548166 | 0.931823509 | 0.013652694 | YPL084W | BRO1 | n |
| W303.asm02.HP0_G0054610 | 445.8775094 | 542.596518 | 481.8098421 | 906.6671597 | 1074.814406 | 972.2376837 | 790.403923 | 954.0467507 | 712.4438261 | 7.31E-10 | -1.005432482 | YPL083C | SEN54 | n |
| W303.asm02.HP0_G0054620 | 411.9520467 | 787.240913 | 361.6017071 | 4543.063987 | 4503.605055 | 4557.929923 | 2589.874411 | 3028.566257 | 2441.348456 | 1.59E-37 | -3.123501259 | YPL082C | MOT1 | n |
| W303.asm02.HP0_G0054640 | 6413.851042 | 8564.457671 | 6608.515522 | 13781.92452 | 15677.69409 | 13255.56462 | 13895.01699 | 10419.042 | 9303.738413 | 5.77E-08 | -0.984525399 | YPL079W | RPL21B | y |
| W303.asm02.HP0_G0054650 | 6189.942989 | 6448.331251 | 6315.324949 | 11241.89452 | 11570.84151 | 11056.71422 | 10729.61493 | 10438.39386 | 9783.819095 | 5.24E-20 | -0.837432922 | YPL078C | ATP4 | n |
| W303.asm02.HP0_G0054670 | 389.6581712 | 291.288657 | 393.8526701 | 536.0231813 | 561.2919677 | 557.6335318 | 574.5810554 | 657.9632763 | 600.8685718 | 0.000169731 | -0.622749667 | YPL076W | GPI2 | n |
| W303.asm02.HP0_G0054680 | 981.8998196 | 1050.923782 | 916.7091925 | 507.8114349 | 630.2923987 | 537.7180485 | 502.6400995 | 626.0327056 | 703.2311905 | 1.36E-06 | 0.81770073 | YPL075W | GCR1 | n |
| W303.asm02.HP0_G0054690 | 672.6934598 | 566.3946109 | 537.516051 | 26.2661087 | 33.17328414 | 31.6837234 | 50.1693508 | 56.1203971 | 52.20493554 | 6.94E-71 | 4.292415911 | YPL074W | YTA6 | n |
| W303.asm02.HP0_G0054700 | 415.8292424 | 356.9713934 | 363.5563109 | 642.060435 | 752.3700844 | 624.6219755 | 570.7946893 | 865.0281898 | 738.0344808 | 1.09E-05 | -0.827895578 | YPL072W | UBP16 | n |
| W303.asm02.HP0_G0054710 | 684.325047 | 655.8754402 | 738.8402447 | 162.4607464 | 229.5591263 | 194.6285866 | 317.1081607 | 217.708437 | 246.6939111 | 3.84E-20 | 1.830331955 | YPL071C | | n |
| W303.asm02.HP0_G0054720 | 1016.794581 | 1215.606585 | 1096.532744 | 188.7268551 | 242.8284399 | 226.31231 | 462.8832555 | 373.4909186 | 509.7658411 | 5.13E-40 | 2.342408487 | YPL070W | MUK1 | n |
| W303.asm02.HP0_G0054730 | 725.0356022 | 752.0197355 | 677.2702243 | 344.3778697 | 334.3867042 | 357.5734498 | 354.9718217 | 379.2964769 | 324.4895013 | 5.88E-14 | 1.054519608 | YPL069C | BTS1 | n |
| W303.asm02.HP0_G0054740 | 161.8729219 | 158.0193368 | 164.186721 | 498.0832465 | 346.3290865 | 507.8448236 | 304.8024709 | 343.495534 | 239.5285278 | 7.62E-12 | -1.485208562 | YPL068C | | n |
| W303.asm02.HP0_G0054750 | 349.916915 | 266.5386404 | 342.0556688 | 250.0144421 | 238.8476458 | 226.31231 | 401.3548064 | 571.8474946 | 392.0488296 | 0.050386378 | 0.422598368 | YPL067C | HTC1 | n |
| W303.asm02.HP0_G0054760 | 537.9609081 | 502.6157219 | 440.7631619 | 128.412087 | 132.6931366 | 144.8398784 | 113.5909829 | 178.0371218 | 191.418097 | 6.86E-17 | 1.866484816 | YPL066W | RGL1 | n |
| W303.asm02.HP0_G0054770 | 761.8689617 | 655.8754402 | 759.3635848 | 75.87986959 | 94.21212697 | 66.98844375 | 162.8137422 | 149.0093302 | 174.0164518 | 1.43E-62 | 3.213288733 | YPL065W | VPS28 | n |
| W303.asm02.HP0_G0054790 | 1270.750902 | 1220.366204 | 1157.125463 | 3845.552878 | 3572.099237 | 4004.822637 | 4203.812961 | 4113.23807 | 4430.254137 | 3.12E-54 | -1.646996834 | YPL063W | TIM50 | n |
| W303.asm02.HP0_G0054800 | 12139.49984 | 8426.428732 | 11259.49532 | 762.6899712 | 743.0815648 | 649.9689543 | 3495.7625 | 4961.817178 | 5034.193588 | 1.08E-95 | 3.884404432 | YPL061W | ALD6 | n |
| W303.asm02.HP0_G0054810 | 443.9389115 | 431.2214433 | 414.3760103 | 173.1617537 | 191.0781167 | 175.6183525 | 126.8432643 | 165.4584121 | 171.9691994 | 3.43E-12 | 1.258716515 | YPL060W | MFM1 | n |
| W303.asm02.HP0_G0054820 | 1561.540582 | 1401.23171 | 1523.613679 | 1391.130942 | 1506.0671 | 1373.263097 | 1591.220353 | 978.236577 | 1118.823422 | 0.719968667 | 0.071481309 | YPL059W | GRX5 | n |
| W303.asm02.HP0_G0054830 | 633.9215025 | 859.5871154 | 626.4505249 | 458.197674 | 782.8895058 | 565.780775 | 80.46027959 | 661.8336485 | 397.1669605 | 0.759364845 | 0.230829213 | YPL058C | PDR12 | n |
| W303.asm02.HP0_G0054840 | 820.9961966 | 852.9236494 | 857.0937759 | 1756.910827 | 1499.432443 | 1721.784054 | 977.8290449 | 1186.269084 | 1582.526085 | 7.73E-08 | -0.976358924 | YPL057C | SUR1 | n |
| W303.asm02.HP0_G0054860 | 428.4301286 | 382.6733337 | 315.6685173 | 271.4164566 | 287.9441064 | 278.8167659 | 293.4433726 | 236.092705 | 211.8906207 | 0.032104975 | 0.427642523 | YPL055C | LGE1 | n |
| W303.asm02.HP0_G0054870 | 182.2281995 | 118.0385408 | 161.2548153 | 399.8285436 | 277.3286554 | 362.099696 | 383.3695674 | 398.648338 | 403.3087177 | 6.53E-08 | -1.174754883 | YPL054W | LEE1 | n |
| W303.asm02.HP0_G0054890 | 489.4959614 | 681.5773805 | 461.286502 | 530.1862683 | 522.8109581 | 492.4555865 | 334.1468082 | 276.7316133 | 241.5757801 | 0.715709499 | 0.07939265 | YPL052W | OAZ1 | n |
| W303.asm02.HP0_G0054900 | 744.4215809 | 854.8274968 | 722.2261122 | 449.4423045 | 394.0986156 | 407.362158 | 421.2332284 | 332.8520104 | 343.9383988 | 2.39E-08 | 0.891055081 | YPL051W | ARL3 | n |
| W303.asm02.HP0_G0054910 | 3485.598965 | 4358.858695 | 3765.544263 | 4766.81232 | 4954.76172 | 4881.103902 | 4381.772167 | 4944.400503 | 4464.033801 | 0.008267688 | -0.330777613 | YPL050C | MNN9 | n |
| W303.asm02.HP0_G0054920 | 992.5621078 | 1140.404612 | 814.0924919 | 439.7141161 | 480.3491544 | 410.0779057 | 777.1516417 | 599.9076931 | 792.2866687 | 2.46E-10 | 1.149376949 | YPL049C | DIG1 | n |
| W303.asm02.HP0_G0054930 | 2105.317283 | 1554.491428 | 1655.549437 | 1706.324247 | 1305.700464 | 1636.690626 | 2224.490083 | 1651.681342 | 1657.250797 | 0.326036652 | 0.192543112 | YPL048W | CAM1 | n |
| W303.asm02.HP0_G0054940 | 740.5443851 | 668.2504485 | 736.8856409 | 81.71678263 | 95.53905834 | 93.24067171 | 39.75864803 | 59.02317626 | 75.74833784 | 3.60E-43 | 2.990182708 | YPL047W | SGF11 | n |
| W303.asm02.HP0_G0054950 | 406.1362531 | 491.1926374 | 443.6950676 | 76.85268843 | 100.8467838 | 107.7246596 | 119.2705321 | 109.338015 | 84.96097352 | 2.68E-24 | 2.233642047 | YPL046C | ELC1 | n |
| W303.asm02.HP0_G0054960 | 869.4611433 | 965.2506478 | 882.5036256 | 511.7027103 | 544.04186 | 527.7603069 | 528.1980707 | 638.6114153 | 661.2625168 | 4.13E-08 | 0.779738128 | YPL045W | VPS16 | n |
| W303.asm02.HP0_G0054970 | 1007.101592 | 2081.857166 | 1214.786275 | 7566.584944 | 6320.174095 | 7271.867144 | 3096.300877 | 4051.312115 | 3619.542197 | 1.88E-22 | -2.297571882 | YPL043W | NOP4 | n |
| W303.asm02.HP0_G0054980 | 429.3994275 | 480.7214765 | 367.4655185 | 460.1433117 | 485.6568799 | 438.1406321 | 368.2241031 | 433.4816879 | 497.4823269 | 0.547892466 | -0.114123263 | YPL042C | SSN3 | n |
| W303.asm02.HP0_G0054990 | 227.7852494 | 230.3655392 | 186.664665 | 286.981558 | 278.6555868 | 262.5222796 | 281.1376828 | 220.6112162 | 211.8906207 | 0.070339926 | -0.360364178 | YPL041C | MRX11 | n |
| W303.asm02.HP0_G0055000 | 422.614335 | 504.5195694 | 437.8312561 | 1021.459783 | 996.5254557 | 1036.51038 | 564.1685486 | 838.9031773 | 865.9877542 | 2.89E-10 | -1.162152642 | YPL040C | ISM1 | n |
| W303.asm02.HP0_G0055010 | 591.2723494 | 528.3176623 | 592.2449581 | 364.8070653 | 307.8480769 | 380.2046808 | 324.6808929 | 337.6899756 | 286.6153324 | 1.20E-05 | 0.697478315 | YPL039W | | n |
| W303.asm02.HP0_G0055040 | 6869.421541 | 7357.418399 | 7208.578895 | 6946.899342 | 7498.489148 | 6989.429381 | 6181.242655 | 5040.195812 | 4525.451373 | 0.999148516 | 0.000142375 | YPL037C | EGD1 | n |
| W303.asm02.HP0_G0055060 | 576.7328654 | 502.6157219 | 516.9927109 | 255.8513551 | 221.5975381 | 288.7745075 | 153.347827 | 276.7316133 | 196.5362279 | 3.57E-06 | 1.056084431 | YPL034W | | n |
| W303.asm02.HP0_G0055080 | 777.3777446 | 997.6160542 | 689.9751492 | 4411.733443 | 3617.214903 | 4249.239932 | 2156.335493 | 4141.298269 | 3395.368062 | 3.52E-24 | -2.316519316 | YPL032C | SVL3 | n |
| W303.asm02.HP0_G0055090 | 1047.812147 | 757.7312778 | 961.6650804 | 641.0876161 | 644.8886438 | 670.7896868 | 1186.07918 | 933.7272966 | 946.854223 | 0.003614604 | 0.499649126 | YPL031C | PHO85 | n |
| W303.asm02.HP0_G0055100 | 296.6054736 | 415.9906638 | 261.9169121 | 929.041993 | 943.4482011 | 938.7434618 | 778.0982332 | 972.4310187 | 934.5707087 | 5.74E-16 | -1.528029791 | YPL030W | TRM44 | n |
| W303.asm02.HP0_G0055110 | 532.1451145 | 568.2984583 | 586.3811466 | 1297.740334 | 1273.854111 | 1261.012191 | 644.6288282 | 903.7319119 | 890.5547827 | 8.44E-14 | -1.184026035 | YPL029W | SUV3 | n |
| W303.asm02.HP0_G0055120 | 9513.669032 | 7559.226227 | 8843.604993 | 5187.070059 | 4914.953779 | 5313.813038 | 4260.608452 | 4183.872363 | 3561.195504 | 1.50E-08 | 0.749309791 | YPL028W | ERG10 | n |
| W303.asm02.HP0_G0055140 | 987.7156132 | 2100.895641 | 1116.078782 | 1149.87187 | 874.44777 | 1147.856036 | 344.5593149 | 512.8243183 | 536.380122 | 0.161372949 | 0.406025116 | YPL026C | SKS1 | n |
| W303.asm02.HP0_G0055160 | 1204.838574 | 1089.000731 | 1247.037238 | 271.4164566 | 293.2518318 | 272.4800212 | 283.0308658 | 306.7269979 | 246.6939111 | 1.30E-45 | 2.082476556 | YPL024W | RMI1 | n |
| W303.asm02.HP0_G0055170 | 793.8558265 | 862.4428865 | 707.5665836 | 1972.876609 | 2221.283106 | 2170.787677 | 1856.26598 | 2168.376033 | 2102.528188 | 3.25E-27 | -1.428595027 | YPL023C | MET12 | n |
| W303.asm02.HP0_G0055180 | 572.8556697 | 792.0005315 | 556.0847874 | 419.2849204 | 489.637674 | 409.1726564 | 297.2297387 | 544.7548891 | 374.6471844 | 0.026469965 | 0.544687916 | YPL022W | RAD1 | n |
| W303.asm02.HP0_G0055200 | 609.6890291 | 955.7314107 | 625.473223 | 1034.106428 | 955.3905834 | 1027.457887 | 897.3687653 | 1119.505163 | 1014.413551 | 0.019577289 | -0.46176341 | YPL020C | ULP1 | n |
| W303.asm02.HP0_G0055240 | 286.9124843 | 356.9713934 | 223.8021376 | 1098.312471 | 1257.930935 | 1173.203015 | 805.5493874 | 1113.699604 | 1115.752544 | 7.99E-24 | -2.023491397 | YPL016W | SWI1 | n |
| W303.asm02.HP0_G0055250 | 566.0705771 | 558.7792212 | 459.3318982 | 158.5694711 | 132.6931366 | 144.8398784 | 25.55797116 | 149.0093302 | 151.4966757 | 1.94E-05 | 1.860251325 | YPL015C | HST2 | n |
| W303.asm02.HP0_G0055260 | 210.3378686 | 203.7116752 | 170.0505325 | 446.5238479 | 354.2906747 | 439.0458814 | 375.7968352 | 596.0373209 | 759.5306307 | 2.04E-05 | -1.08725799 | YPL014W | CIP1 | n |
| W303.asm02.HP0_G0055270 | 2176.076106 | 1916.22244 | 2283.954566 | 1260.773218 | 1275.181043 | 1334.33738 | 1683.986322 | 1279.158017 | 1179.217367 | 7.48E-06 | 0.720102449 | YPL013C | MRPS16 | n |
| W303.asm02.HP0_G0055280 | 1879.470632 | 3119.454017 | 2089.471486 | 9396.457184 | 7916.472528 | 9499.685524 | 3182.440706 | 4198.386259 | 3459.856512 | 4.77E-22 | -1.91942049 | YPL012W | RRP12 | n |
| W303.asm02.HP0_G0055290 | 1105.000784 | 1141.356535 | 1081.873215 | 413.4480074 | 448.5028016 | 441.7616291 | 588.7799283 | 519.5974697 | 547.64001 | 2.89E-27 | 1.353087396 | YPL011C | TAF3 | n |
| W303.asm02.HP0_G0055300 | 2563.795679 | 2785.328792 | 2623.078329 | 1584.721892 | 1653.356482 | 1607.72265 | 2228.276449 | 1636.199853 | 1608.11674 | 2.05E-06 | 0.718416959 | YPL010W | RET3 | n |

| | | | | | | | | | | | | | | |
|---|---|---|---|---|---|---|---|---|---|---|---|---|---|---|
| W303.asm02.HP0_G0055310 | 1096.277094 | 875.7698185 | 1060.372573 | 1417.397051 | 1401.239522 | 1475.556261 | 1115.084816 | 1303.347843 | 1344.021184 | 0.000415312 | -0.502247818 | YPL009C | RQC2 | n |
| W303.asm02.HP0_G0055320 | 504.0354454 | 621.6061864 | 521.8792205 | 813.2765509 | 691.3312416 | 764.0303585 | 543.3435351 | 813.7457579 | 636.6954883 | 0.013936498 | -0.46227428 | YPL008W | CHL1 | n |
| W303.asm02.HP0_G0055330 | 713.404015 | 661.5869825 | 681.179432 | 315.1933044 | 390.1178215 | 353.9524528 | 319.9479353 | 340.5927548 | 354.1746607 | 5.52E-11 | 0.960011757 | YPL007C | TFC8 | n |
| W303.asm02.HP0_G0055340 | 657.1846769 | 658.7312113 | 681.179432 | 197.4822247 | 278.6555868 | 249.8487902 | 357.8115963 | 505.0835739 | 526.1438601 | 1.97E-13 | 1.463334889 | YPL006W | NCR1 | n |
| W303.asm02.HP0_G0055350 | 211.3071675 | 431.2214433 | 189.5965707 | 353.1332392 | 379.5023706 | 365.7206929 | 280.1910913 | 362.847395 | 349.0565298 | 0.182404079 | -0.399804599 | YPL005W | AEP3 | n |
| W303.asm02.HP0_G0055360 | 3631.963104 | 2166.578377 | 2764.787106 | 5101.462001 | 4222.295606 | 5160.825917 | 3831.802491 | 3628.47395 | 3214.186227 | 7.12E-05 | -0.758499186 | YPL004C | LSP1 | n |
| W303.asm02.HP0_G0055370 | 890.7857198 | 396.9521895 | 864.9121912 | 220.8298769 | 209.6551558 | 222.691313 | 419.3400454 | 384.1344422 | 378.7416892 | 2.02E-10 | 1.719826023 | YPL003W | ULA1 | n |
| W303.asm02.HP0_G0055380 | 1508.22914 | 814.8467007 | 1437.611111 | 193.5909493 | 207.0012931 | 190.1023404 | 277.3513167 | 217.708437 | 278.4263229 | 1.34E-29 | 2.671630665 | YPL002C | SNF8 | n |
| W303.asm02.HP0_G0055390 | 2122.764664 | 1113.750747 | 1984.900181 | 1472.847725 | 1406.547248 | 1501.808489 | 1176.613265 | 1060.481987 | 1022.602561 | 0.266447978 | 0.252898243 | YPL001W | HAT1 | n |
| W303.asm02.HP0_G0055410 | 676.5706555 | 520.7022725 | 630.3597326 | 157.5966522 | 167.1933521 | 146.6503769 | 117.377349 | 125.7870969 | 157.6384328 | 2.70E-24 | 1.957602679 | YPR002W | PDH1 | n |
| W303.asm02.HP0_G0055420 | 639.7372961 | 546.4042129 | 659.6787899 | 366.752703 | 432.5796252 | 348.5209574 | 612.4447164 | 713.1160804 | 775.9086497 | 4.18E-05 | 0.688499982 | YPR003C | | n |
| W303.asm02.HP0_G0055430 | 2614.199223 | 2374.097747 | 2660.215802 | 2241.374609 | 1946.608314 | 2258.596854 | 2466.817513 | 1965.181492 | 1970.48041 | 0.088517614 | 0.246016429 | YPR004C | AIM45 | n |
| W303.asm02.HP0_G0055450 | 631.0136057 | 735.8370323 | 656.7468842 | 43.77684784 | 62.36577419 | 61.55694831 | 31.23752031 | 59.99076931 | 52.20493554 | 2.70E-48 | 3.598250481 | YPR006C | ICL2 | n |
| W303.asm02.HP0_G0055460 | 64.94302854 | 81.86543956 | 62.5473223 | 174.1345725 | 183.1165285 | 158.418617 | 175.119432 | 146.1065511 | 148.4257971 | 2.05E-08 | -1.2974539 | YPR007C | REC8 | n |
| W303.asm02.HP0_G0055470 | 535.0530113 | 823.4140142 | 505.265088 | 1226.724558 | 1292.43115 | 1207.602486 | 993.9211008 | 1206.588538 | 1226.304172 | 4.56E-07 | -0.999287486 | YPR008W | HAA1 | n |
| W303.asm02.HP0_G0055490 | 1995.786504 | 2684.424879 | 1948.740011 | 15040.7521 | 14011.06829 | 14566.36552 | 6022.215279 | 7693.332368 | 6905.382257 | 9.51E-68 | -2.718043548 | YPR010C | RPA135 | n |
| W303.asm02.HP0_G0055500 | 1624.545012 | 1791.520433 | 1456.179847 | 2896.081689 | 3095.730876 | 2936.628534 | 3108.606567 | 2297.065909 | 2042.134243 | 3.74E-07 | -0.873661984 | YPR010C-/ | MRA1 | n |
| W303.asm02.HP0_G0055510 | 933.4348729 | 780.577447 | 857.0937759 | 643.0332538 | 801.4665449 | 678.0316807 | 1060.182507 | 801.1670482 | 843.4679781 | 0.105861796 | 0.278304693 | YPR011C | MRX21 | n |
| W303.asm02.HP0_G0055520 | 170.5966123 | 181.8174297 | 136.8222675 | 111.8741667 | 120.7507543 | 122.2086474 | 70.04777282 | 132.5602483 | 70.6302069 | 0.125603792 | 0.46389031 | YPR013C | CMR3 | n |
| W303.asm02.HP0_G0055540 | 3441.011214 | 3300.319523 | 3495.808936 | 5627.756994 | 5829.20949 | 5591.724555 | 4344.855098 | 3639.117474 | 3627.731207 | 1.11E-10 | -0.73576366 | YPR016C | TIF6 | n |
| W303.asm02.HP0_G0055570 | 490.4652603 | 595.9042461 | 553.1528816 | 829.8144712 | 768.2932608 | 790.2825865 | 919.1403703 | 893.0883883 | 882.3657732 | 3.71E-05 | -0.543262976 | YPR018W | RLF2 | n |
| W303.asm02.HP0_G0055580 | 481.7415699 | 553.0676789 | 352.8059899 | 924.1778988 | 997.8523871 | 941.4592095 | 915.3540042 | 906.6346911 | 744.1762379 | 1.50E-08 | -1.044563505 | YPR019W | MCM4 | n |
| W303.asm02.HP0_G0055590 | 1535.369511 | 2079.001395 | 1687.8004 | 4645.209965 | 4873.818906 | 4585.992649 | 5124.846514 | 4548.654944 | 4062.772336 | 2.85E-21 | -1.411396337 | YPR020W | ATP20 | n |
| W303.asm02.HP0_G0055600 | 888.847122 | 1009.991062 | 1025.189705 | 654.7070799 | 931.5058188 | 688.8946716 | 973.0960872 | 1021.778264 | 1044.098711 | 0.030063855 | 0.364952965 | YPR021C | AGC1 | n |
| W303.asm02.HP0_G0055610 | 437.153819 | 655.8754402 | 392.8753682 | 1170.301066 | 1086.756789 | 1122.509058 | 1231.515573 | 1404.945114 | 1482.210719 | 1.26E-09 | -1.185434068 | YPR022C | SDD4 | n |
| W303.asm02.HP0_G0055620 | 896.6015134 | 772.9620572 | 759.3635848 | 293.7912899 | 301.21342 | 301.4479969 | 357.8115963 | 210.9352856 | 369.5290535 | 1.91E-11 | 1.438121135 | YPR023C | EAF3 | n |
| W303.asm02.HP0_G0055630 | 1337.632528 | 1226.077746 | 1021.280497 | 3067.297805 | 2742.767133 | 2975.554252 | 2436.526584 | 2950.19122 | 3238.753256 | 7.72E-17 | -1.293369753 | YPR024W | YME1 | n |
| W303.asm02.HP0_G0055640 | 1090.4613 | 990.9525881 | 1035.940026 | 752.9617828 | 883.7362896 | 789.3773372 | 873.7039772 | 1006.296776 | 921.2635683 | 0.006112997 | 0.363370115 | YPR025C | CCL1 | n |
| W303.asm02.HP0_G0055650 | 229.7238472 | 211.3270649 | 176.8916459 | 91.44497104 | 116.7699602 | 71.51468995 | 229.0751489 | 284.4723577 | 440.1592604 | 9.38E-05 | 1.152127411 | YPR026W | ATH1 | n |
| W303.asm02.HP0_G0055680 | 2369.935892 | 2200.847631 | 2551.73529 | 1784.149754 | 1799.318932 | 1754.373027 | 1717.117026 | 1818.107347 | 1958.196896 | 0.000330372 | 0.416233806 | YPR029C | APL4 | n |
| W303.asm02.HP0_G0055700 | 352.8248118 | 476.9137816 | 357.6924994 | 257.7969928 | 322.4443219 | 257.9960334 | 359.7047793 | 323.1760798 | 343.9383988 | 0.008808881 | 0.506605495 | YPR031W | NTO1 | n |
| W303.asm02.HP0_G0055720 | 5069.433422 | 4996.647584 | 5091.742956 | 4839.773733 | 4960.069445 | 4735.358774 | 4557.838191 | 4364.812264 | 3793.558649 | 0.607231254 | 0.060628867 | YPR033C | HTS1 | n |
| W303.asm02.HP0_G0055730 | 1261.057912 | 1013.798757 | 1070.145593 | 1482.575913 | 1480.855404 | 1447.493535 | 1471.949821 | 1507.509977 | 1574.337076 | 0.001730091 | -0.399014921 | YPR034W | ARP7 | n |
| W303.asm02.HP0_G0055740 | 3307.247961 | 5860.042394 | 3364.85048 | 2760.85987 | 4280.680586 | 2710.316224 | 5093.608994 | 4675.409634 | 3641.038347 | 0.171030441 | 0.362317484 | YPR035W | GLN1 | n |
| W303.asm02.HP0_G0055750 | 3718.230709 | 2717.742209 | 3476.262897 | 3425.295139 | 3176.67369 | 3378.390163 | 3420.98177 | 3124.35797 | 3012.531868 | 0.948272369 | -0.010090673 | YPR036W | VMA13 | n |
| W303.asm02.HP0_G0055760 | 4230.020545 | 5111.830354 | 4228.785369 | 108.9557102 | 414.0025861 | 105.0089118 | 510.2128317 | 298.9862535 | 482.1279341 | 2.11E-14 | 4.43736188 | YPR036W- | SPO24 | n |
| W303.asm02.HP0_G0055770 | 880.1234316 | 861.4909628 | 931.3687212 | 433.877203 | 453.8105271 | 408.2674072 | 738.3413892 | 679.2503235 | 615.1993384 | 1.28E-14 | 1.046155315 | YPR037C | ERV2 | n |
| W303.asm02.HP0_G0055780 | 287.8817832 | 218.9424546 | 243.3481758 | 473.7627755 | 431.2526939 | 448.0983738 | 511.1594233 | 513.7919114 | 667.4042739 | 3.30E-06 | -0.851861686 | YPR040W | TIP41 | n |
| W303.asm02.HP0_G0055790 | 4781.551639 | 4459.762609 | 4953.943387 | 5225.009994 | 5677.939314 | 5361.791248 | 4375.146026 | 3778.450874 | 3582.691654 | 0.112250107 | -0.196194774 | YPR041W | TIF5 | n |
| W303.asm02.HP0_G0055820 | 769.6233531 | 687.2889228 | 749.5905657 | 609.9574132 | 628.9654674 | 641.8217111 | 620.9640401 | 509.9215392 | 578.3487956 | 0.103223466 | 0.230360194 | YPR045C | THP3 | n |
| W303.asm02.HP0_G0055830 | 610.6583281 | 559.7311449 | 629.3824307 | 43.77684784 | 46.4425978 | 52.50445592 | 60.58185757 | 51.28243183 | 75.74833784 | 1.13E-61 | 3.652926883 | YPR046W | MCM16 | n |
| W303.asm02.HP0_G0055840 | 1124.386763 | 908.1352249 | 1092.623536 | 1191.70308 | 1133.199386 | 1193.118498 | 1355.519063 | 1288.833947 | 1344.021184 | 0.198879643 | -0.171421983 | YPR047W | MSF1 | n |
| W303.asm02.HP0_G0055850 | 973.1761292 | 961.442953 | 1048.64495 | 824.950377 | 842.6014173 | 878.0917627 | 516.8389724 | 815.680944 | 849.6097352 | 0.249869943 | 0.228730777 | YPR048W | TAH18 | n |
| W303.asm02.HP0_G0055860 | 256.8642174 | 355.067546 | 299.0543848 | 376.4808914 | 433.9065566 | 388.3519239 | 383.3695674 | 507.01876 | 492.3641959 | 0.043908428 | -0.394267653 | YPR049C | ATG11 | n |
| W303.asm02.HP0_G0055870 | 939.2506665 | 1058.539172 | 1037.894629 | 357.0245146 | 350.3098806 | 383.8256777 | 531.9844368 | 352.2038714 | 373.6235582 | 4.85E-16 | 1.475252222 | YPR051W | MAK3 | n |
| W303.asm02.HP0_G0055880 | 1747.645977 | 2493.088212 | 1908.670632 | 5212.363349 | 5299.763875 | 5123.710698 | 4154.590201 | 4054.214894 | 4463.010175 | 6.79E-20 | -1.346227691 | YPR052C | NHP6A | n |
| W303.asm02.HP0_G0055890 | 164.7808187 | 103.759685 | 104.5713045 | 330.7584059 | 358.2714688 | 302.3532461 | 367.2775115 | 381.2316631 | 358.2691654 | 6.35E-11 | -1.408292407 | YPR054W | SMK1 | n |
| W303.asm02.HP0_G0055900 | 661.0618726 | 707.2793208 | 632.3143364 | 975.7372974 | 950.0828579 | 992.153167 | 1086.68707 | 1162.079257 | 1296.934379 | 2.34E-05 | -0.54482355 | YPR055W | SEC8 | n |
| W303.asm02.HP0_G0055910 | 1129.233257 | 1232.741212 | 1135.624821 | 536.9960001 | 570.5804873 | 522.3288114 | 620.0174486 | 513.7919114 | 550.7108886 | 9.53E-17 | 1.103009407 | YPR056W | TFB4 | n |
| W303.asm02.HP0_G0055920 | 1256.211418 | 920.5102332 | 1150.284349 | 483.4909639 | 382.1562333 | 470.7296048 | 256.5263032 | 326.078859 | 344.962025 | 1.82E-12 | 1.313511139 | YPR057W | BRR1 | n |
| W303.asm02.HP0_G0055930 | 1052.658642 | 957.6352581 | 1123.897198 | 1686.86787 | 1604.260021 | 1685.574085 | 1358.358838 | 1261.741342 | 1148.508582 | 1.14E-07 | -0.667525087 | YPR058W | YMC1 | n |
| W303.asm02.HP0_G0055940 | 2105.317283 | 2016.17443 | 2114.881335 | 1086.638645 | 1035.006465 | 1083.58334 | 1265.592868 | 1101.120895 | 1151.57946 | 6.12E-18 | 0.959659171 | YPR060C | ARO7 | n |
| W303.asm02.HP0_G0055950 | 472.0485806 | 237.0792052 | 417.307916 | 338.5409566 | 352.9637433 | 353.0472036 | 378.6366098 | 208.0325065 | 466.7735413 | 0.746673926 | 0.108754291 | YPR061C | JID1 | n |
| W303.asm02.HP0_G0055960 | 2355.396408 | 2512.126686 | 2538.053063 | 2372.705153 | 2322.12989 | 2397.099987 | 3213.678226 | 2056.135239 | 2247.883107 | 0.738830333 | 0.062360757 | YPR062W | FCY1 | n |
| W303.asm02.HP0_G0055970 | 1056.535837 | 697.7600837 | 1054.508762 | 743.2335944 | 740.4277021 | 734.1571336 | 1370.664528 | 1004.361589 | 1025.673439 | 0.095786051 | 0.340735136 | YPR063C | | n |
| W303.asm02.HP0_G0055980 | 591.2723494 | 576.8657718 | 450.536181 | 1646.009479 | 1805.953589 | 1785.151501 | 1249.500812 | 2669.589234 | 2452.608344 | 7.54E-12 | -1.693836131 | YPR065W | ROX1 | n |
| W303.asm02.HP0_G0055990 | 457.5090966 | 311.2790551 | 436.8539542 | 33.07584059 | 37.15407824 | 50.69395744 | 100.3387016 | 67.73151374 | 112.5988806 | 4.52E-30 | 3.309985286 | YPR066W | UBA3 | n |
| W303.asm02.HP0_G0056000 | 1183.513998 | 941.4525549 | 1293.94773 | 283.0902827 | 351.6368119 | 296.0165015 | 625.6969977 | 409.2918616 | 434.0175033 | 1.62E-19 | 1.879479825 | YPR067W | ISA2 | n |
| W303.asm02.HP0_G0056010 | 752.1759723 | 421.7022061 | 637.200846 | 188.7268551 | 176.4818716 | 182.8603465 | 214.8762761 | 269.9584619 | 241.5757801 | 5.43E-14 | 1.723479922 | YPR068C | HOS1 | n |
| W303.asm02.HP0_G0056020 | 4466.529485 | 3857.194896 | 4454.54211 | 4331.962298 | 4877.799701 | 4527.151449 | 5483.604702 | 6053.262142 | 4617.577729 | 0.481127731 | -0.104192826 | YPR069C | SPE3 | n |
| W303.asm02.HP0_G0056030 | 1029.395467 | 1284.145093 | 1027.144308 | 553.5339204 | 598.446046 | 578.4542643 | 817.8550772 | 811.8105718 | 788.192164 | 1.22E-11 | 0.949879202 | YPR070W | MED1 | n |
| W303.asm02.HP0_G0056050 | 1695.303835 | 1561.154894 | 1797.258214 | 1703.40579 | 1750.222471 | 1643.02737 | 1161.467801 | 1454.292359 | 1477.092588 | 0.935862265 | -0.011962262 | YPR072W | NOT5 | n |
| W303.asm02.HP0_G0056060 | 1818.404799 | 1946.683999 | 1900.852217 | 752.9617828 | 780.2356431 | 834.6397992 | 846.2528229 | 757.6253608 | 610.0812074 | 1.11E-17 | 1.258439546 | YPR073C | LTP1 | n |

| | | | | | | | | | | | | | | |
|---|---|---|---|---|---|---|---|---|---|---|---|---|---|---|
| W303.asm02.HP0_G0056070 | 5366.038895 | 6353.138879 | 5354.63717 | 7643.437632 | 6598.829682 | 7351.529077 | 5570.691122 | 5421.423878 | 5133.485328 | 0.00607493 | -0.339013952 | YPR074C | TKL1 | n |
| W303.asm02.HP0_G0056090 | 1326.97024 | 827.221709 | 1385.81411 | 1438.799066 | 1544.54811 | 1452.019781 | 2606.913059 | 1629.426702 | 2787.334107 | 0.18348139 | -0.325225943 | YPR075C | OPY2 | n |
| W303.asm02.HP0_G0056110 | 663.9697694 | 395.0483421 | 563.9032026 | 659.5711741 | 540.0610659 | 660.8319451 | 1026.105213 | 838.9031773 | 890.5547827 | 0.359265241 | -0.198564847 | YPR079W | MRL1 | n |
| W303.asm02.HP0_G0056140 | 597.088143 | 670.1542959 | 678.2475262 | 166.3520218 | 172.5010776 | 163.8501124 | 79.51368806 | 193.5186107 | 226.2213873 | 7.82E-11 | 1.952881975 | YPR082C | DIB1 | n |
| W303.asm02.HP0_G0056150 | 517.6056305 | 560.6830686 | 493.5374651 | 640.1147973 | 597.1191146 | 609.2327385 | 520.6253385 | 551.5280405 | 540.4746267 | 0.082641193 | -0.232842958 | YPR083W | MDM36 | n |
| W303.asm02.HP0_G0056160 | 258.8028152 | 355.067546 | 304.9181962 | 454.3063987 | 463.0990467 | 446.2878753 | 584.0469706 | 560.2363779 | 676.6169096 | 0.000926694 | -0.569071132 | YPR084W | | n |
| W303.asm02.HP0_G0056170 | 847.1672678 | 763.4428201 | 805.2967747 | 249.0416233 | 236.1937831 | 218.1650668 | 239.4876557 | 250.6066008 | 196.5362279 | 2.57E-29 | 1.780912166 | YPR085C | ASA1 | n |
| W303.asm02.HP0_G0056180 | 2641.339594 | 2666.338328 | 2866.426505 | 2192.733667 | 2387.149527 | 2196.134656 | 2011.50699 | 2573.797522 | 2585.679748 | 0.050861858 | 0.270982999 | YPR086W | SUA7 | n |
| W303.asm02.HP0_G0056190 | 2405.799953 | 2056.155226 | 2342.592681 | 2037.082653 | 2173.513577 | 2155.39844 | 2338.081066 | 2025.172261 | 1953.078765 | 0.475082743 | 0.096230856 | YPR088C | SRP54 | n |
| W303.asm02.HP0_G0056200 | 655.246079 | 773.9139809 | 681.179432 | 1279.256776 | 1252.623209 | 1260.106942 | 996.7608754 | 1341.083972 | 1291.816248 | 3.25E-08 | -0.845466237 | YPR089W | | n |
| W303.asm02.HP0_G0056210 | 601.9346377 | 493.0964848 | 560.9712969 | 280.1718262 | 273.3478613 | 287.8692583 | 470.4559877 | 656.9956833 | 621.3410955 | 4.58E-08 | 0.975982778 | YPR091C | NVJ2 | n |
| W303.asm02.HP0_G0056220 | 378.026584 | 342.6925377 | 386.0342548 | 125.4936305 | 144.6355189 | 124.0191459 | 212.0365015 | 160.6204469 | 204.7252374 | 9.48E-15 | 1.494355552 | YPR093C | ASR1 | n |
| W303.asm02.HP0_G0056230 | 351.8555128 | 362.6829357 | 336.1918574 | 150.7869203 | 164.5394894 | 162.0396139 | 158.0807846 | 122.8843178 | 159.6856852 | 6.88E-10 | 1.139502036 | YPR094W | RDS3 | n |
| W303.asm02.HP0_G0056240 | 232.631744 | 327.4617582 | 231.6205529 | 351.1876015 | 384.8100961 | 373.8679361 | 527.2514792 | 530.2409933 | 551.7345148 | 0.008139697 | -0.486140354 | YPR095C | SYT1 | n |
| W303.asm02.HP0_G0056260 | 722.1277054 | 1318.414346 | 845.366153 | 175.1073914 | 183.1165285 | 210.9230729 | 227.1819659 | 312.5325563 | 272.2845657 | 3.74E-22 | 2.34108107 | YPR097W | LEC1 | n |
| W303.asm02.HP0_G0056280 | 1814.527603 | 1739.164629 | 1854.919027 | 703.3480219 | 849.2360741 | 766.7461062 | 1109.405267 | 805.0374205 | 727.7982189 | 2.65E-12 | 1.222553982 | YPR100W | MRPL51 | n |
| W303.asm02.HP0_G0056290 | 1732.137194 | 968.106419 | 1623.298474 | 117.7110797 | 126.0584797 | 124.9243951 | 282.0842743 | 177.0695288 | 250.7884158 | 7.80E-46 | 3.552291397 | YPR101W | SNT309 | n |
| W303.asm02.HP0_G0056310 | 6340.184324 | 5110.87843 | 6243.004607 | 2429.128646 | 2089.916901 | 2307.480313 | 3161.615692 | 2802.149483 | 2722.845657 | 2.59E-24 | 1.373536675 | YPR103W | PRE2 | n |
| W303.asm02.HP0_G0056320 | 377.0572851 | 508.3272642 | 358.6698013 | 1109.013479 | 883.7362896 | 1023.83689 | 533.8776198 | 791.4911177 | 751.3416212 | 4.91E-10 | -1.278409009 | YPR104C | FHL1 | n |
| W303.asm02.HP0_G0056330 | 1265.904407 | 1206.087348 | 1350.631241 | 861.917493 | 931.5058188 | 821.0610606 | 842.4664569 | 948.2411924 | 856.7751185 | 1.20E-05 | 0.549398148 | YPR105C | COG4 | n |
| W303.asm02.HP0_G0056350 | 894.6629156 | 837.6928699 | 833.6385301 | 166.3520218 | 209.6551558 | 168.3763586 | 179.8523897 | 231.2547398 | 248.7411634 | 4.03E-36 | 2.243088964 | YPR107C | YTH1 | n |
| W303.asm02.HP0_G0056360 | 5855.534857 | 5465.945976 | 5620.46329 | 2455.394754 | 2007.647156 | 2346.40603 | 2656.135818 | 2847.626356 | 2820.090145 | 8.60E-30 | 1.314032478 | YPR108W | RPN7 | n |
| W303.asm02.HP0_G0056370 | 1092.399898 | 1037.59685 | 1020.303195 | 214.020145 | 208.3282244 | 237.1753009 | 431.6457352 | 361.879802 | 363.3872964 | 1.02E-52 | 2.25302559 | YPR109W | GLD1 | n |
| W303.asm02.HP0_G0056380 | 3932.445773 | 3893.367998 | 4247.354105 | 3814.422675 | 3722.042481 | 3846.40402 | 3408.67608 | 1983.56576 | 1997.094691 | 0.697672759 | 0.084900761 | YPR110C | RPC40 | n |
| W303.asm02.HP0_G0056400 | 651.3688833 | 1218.462356 | 665.5426014 | 5245.43919 | 4328.450115 | 4920.029619 | 2844.507531 | 2977.283825 | 2592.845132 | 1.59E-29 | -2.515135804 | YPR112C | MRD1 | n |
| W303.asm02.HP0_G0056410 | 3057.168836 | 3038.540501 | 3049.181962 | 1984.550435 | 2022.243401 | 2129.146212 | 3422.874953 | 2480.908589 | 2707.491265 | 5.31E-05 | 0.575566278 | YPR113W | PIS1 | n |
| W303.asm02.HP0_G0056420 | 1027.456869 | 1062.346867 | 1050.599554 | 908.6127973 | 958.0444461 | 937.8382126 | 1836.387558 | 1361.403426 | 1586.62059 | 0.271226645 | 0.163530837 | YPR114W | | n |
| W303.asm02.HP0_G0056440 | 46.5263488 | 90.432753 | 32.25096306 | 245.1503479 | 236.1937831 | 238.9857993 | 309.5354285 | 256.4121592 | 256.9301729 | 1.92E-11 | -2.088399826 | YPR116W | RRG8 | n |
| W303.asm02.HP0_G0056450 | 365.4256979 | 661.5869825 | 368.4428204 | 194.5637682 | 179.1357344 | 196.4390851 | 192.1580795 | 247.7038217 | 242.5994063 | 5.62E-07 | 1.290372154 | YPR117W | HOB2 | n |
| W303.asm02.HP0_G0056460 | 2734.392291 | 3108.982856 | 3017.908301 | 2067.240037 | 2202.706067 | 2044.958033 | 2361.745854 | 2099.676926 | 2138.355105 | 1.83E-05 | 0.489234872 | YPR118W | MRI1 | n |
| W303.asm02.HP0_G0056470 | 1341.509724 | 1535.452954 | 1489.408112 | 1578.884979 | 1502.086306 | 1444.777787 | 635.162913 | 1338.181193 | 1099.374525 | 0.835640352 | -0.051702563 | YPR119W | CLB2 | n |
| W303.asm02.HP0_G0056480 | 774.4698478 | 902.4236826 | 803.3421708 | 751.0161451 | 776.254849 | 793.9035834 | 604.8719842 | 720.8568248 | 650.0026287 | 0.508562017 | 0.095674436 | YPR120C | CLB5 | n |
| W303.asm02.HP0_G0056500 | 762.8382606 | 1046.164164 | 773.0458116 | 79.77114495 | 104.8275779 | 88.71442551 | 92.76596941 | 142.2361789 | 144.3312924 | 4.29E-44 | 3.246458387 | YPR122W | AXL1 | n |
| W303.asm02.HP0_G0056510 | 8920.458084 | 9726.756528 | 9594.17286 | 12132.02376 | 12482.44336 | 11940.23747 | 18094.09699 | 15695.32692 | 20056.93151 | 0.002459962 | -0.372204745 | YPR124W | CTR1 | n |
| W303.asm02.HP0_G0056520 | 1662.347671 | 2255.107283 | 1795.30361 | 2742.376312 | 3111.654053 | 2631.55954 | 2552.957342 | 2557.34844 | 2817.019267 | 0.000147994 | -0.570319317 | YPR125W | YLH47 | n |
| W303.asm02.HP0_G0056530 | 793.8558265 | 490.2407136 | 655.7695823 | 3312.448153 | 2123.090185 | 2685.874495 | 1978.376286 | 2594.116976 | 2413.710549 | 7.20E-22 | -2.066296175 | YPR127W | | n |
| W303.asm02.HP0_G0056540 | 1474.303678 | 1067.106485 | 1362.358864 | 91.44497104 | 102.1737152 | 76.04093615 | 241.3808388 | 217.708437 | 261.0246777 | 1.25E-85 | 3.864942131 | YPR128C | ANT1 | n |
| W303.asm02.HP0_G0056550 | 929.5576772 | 725.3658714 | 732.9764332 | 1575.966522 | 1539.240384 | 1652.079863 | 1694.398829 | 1457.195138 | 1504.730495 | 3.40E-12 | -0.997762216 | YPR129W | SCD6 | n |
| W303.asm02.HP0_G0056560 | 1766.062657 | 974.769885 | 1770.871063 | 565.2077465 | 508.2147131 | 538.6232978 | 552.8094503 | 507.01876 | 396.1433344 | 1.68E-10 | 1.484260426 | YPR131C | NAT3 | n |
| W303.asm02.HP0_G0056580 | 2627.769409 | 2303.655392 | 2713.967407 | 2832.848465 | 2920.575936 | 2732.042206 | 2993.122401 | 2971.478267 | 3139.461515 | 0.19552683 | -0.150176267 | YPR133C | SPN1 | n |
| W303.asm02.HP0_G0056590 | 1164.128019 | 1388.856701 | 1212.831672 | 313.2476667 | 374.1946451 | 362.099696 | 510.2128317 | 280.6019855 | 292.7570895 | 6.03E-16 | 1.844036742 | YPR133W- | TOM5 | n |
| W303.asm02.HP0_G0056600 | 1105.970083 | 861.4909628 | 1131.715613 | 416.3664639 | 436.5604193 | 416.4146504 | 710.8902349 | 552.4956335 | 607.0103289 | 8.34E-15 | 1.288406165 | YPR134W | MSS18 | n |
| W303.asm02.HP0_G0056610 | 626.167111 | 613.9907967 | 589.3130523 | 1319.142348 | 1215.469131 | 1399.515325 | 946.5915246 | 1173.690374 | 1056.382225 | 2.63E-16 | -1.10549738 | YPR135W | CTF4 | n |
| W303.asm02.HP0_G0056620 | 1389.974671 | 2192.280317 | 1505.044943 | 5035.31032 | 4118.794959 | 4983.397066 | 1896.022824 | 2275.778862 | 1841.50351 | 7.68E-15 | -1.474686829 | YPR137W | RRP9 | n |
| W303.asm02.HP0_G0056630 | 692.0794385 | 1406.943252 | 739.8175466 | 202.3463189 | 322.4443219 | 199.1548583 | 243.2740218 | 198.356576 | 246.6939111 | 1.15E-11 | 1.975752088 | YPR138C | MEP3 | n |
| W303.asm02.HP0_G0056640 | 1526.64582 | 1429.789421 | 1513.84066 | 958.2265582 | 952.7367206 | 930.5962186 | 994.8676923 | 1254.96819 | 1060.47673 | 5.35E-07 | 0.65379669 | YPR139C | LOA1 | n |
| W303.asm02.HP0_G0056650 | 661.0618726 | 650.1638979 | 673.3610167 | 2006.925269 | 2117.78246 | 2078.452255 | 2090.074086 | 2079.357472 | 1741.188144 | 1.62E-41 | -1.644003017 | YPR140W | TAZ1 | n |
| W303.asm02.HP0_G0056660 | 507.9126411 | 670.1542959 | 571.7216179 | 379.3993479 | 351.6368119 | 429.0881397 | 280.1910913 | 389.9400005 | 325.5131275 | 0.001929346 | 0.591039249 | YPR141C | KAR3 | n |
| W303.asm02.HP0_G0056670 | 2147.966436 | 2739.636454 | 2491.142571 | 3484.637088 | 2986.922504 | 3390.158404 | 1724.689758 | 1695.22303 | 1393.15524 | 0.005152991 | -0.418776741 | YPR143W | RRP15 | n |
| W303.asm02.HP0_G0056680 | 1026.48757 | 1081.385341 | 1136.602122 | 3035.194783 | 2835.652329 | 3065.173926 | 1811.776178 | 1990.338911 | 1851.739772 | 2.12E-41 | -1.46205062 | YPR144C | NOC4 | n |
| W303.asm02.HP0_G0056720 | 2475.589476 | 1746.780018 | 2405.140003 | 1851.274254 | 1511.374826 | 1716.352559 | 1297.77698 | 1472.676627 | 1067.642113 | 0.040518804 | 0.383400671 | YPR148C | | n |
| W303.asm02.HP0_G0056730 | 11844.83297 | 8179.88049 | 11763.7831 | 937.7973626 | 989.8907989 | 882.6180089 | 673.026574 | 1751.343427 | 1275.438229 | 1.37E-37 | 3.499934532 | YPR149W | NCE102 | n |
| W303.asm02.HP0_G0056740 | 400.3204595 | 233.2213104 | 321.5323287 | 44.74966668 | 33.17328414 | 57.93595136 | 170.3864744 | 113.2083873 | 281.4972014 | 9.07E-15 | 2.801819337 | YPR151C | SUE1 | n |
| W303.asm02.HP0_G0056750 | 187.0746942 | 240.8367001 | 202.3014956 | 897.9117901 | 761.658604 | 859.9867779 | 801.7630213 | 904.699505 | 892.602035 | 1.95E-37 | -2.000177414 | YPR152C | URN1 | n |
| W303.asm02.HP0_G0056770 | 308.2370608 | 344.5963851 | 210.1199109 | 1492.304102 | 1494.124718 | 1533.492212 | 1789.057981 | 1479.449779 | 1709.455732 | 1.36E-39 | -2.388619716 | YPR154W | PIN3 | n |
| W303.asm02.HP0_G0056780 | 832.6277838 | 712.9908631 | 813.1151899 | 1141.1165 | 1024.391014 | 1086.299088 | 1164.307575 | 1362.371019 | 1094.256394 | 0.001168939 | -0.463946187 | YPR155C | NCA2 | n |
| W303.asm02.HP0_G0056790 | 675.6013566 | 645.4042793 | 579.5400332 | 2404.808175 | 2720.2093 | 2481.288167 | 472.3491708 | 1550.084072 | 849.6097352 | 1.02E-10 | -2.000691157 | YPR156C | TPO3 | n |
| W303.asm02.HP0_G0056820 | 2139.242746 | 1990.47249 | 2195.020092 | 4012.877718 | 3308.039895 | 3814.720297 | 1942.405808 | 3406.895141 | 3066.784056 | 3.83E-05 | -0.816314514 | YPR159W | KRE6 | n |
| W303.asm02.HP0_G0056830 | 629.0750078 | 564.4907635 | 614.722902 | 342.432232 | 249.4630968 | 294.206003 | 354.0252302 | 225.4491814 | 485.1988126 | 6.50E-05 | 1.027141495 | YPR160W | GPH1 | n |
| W303.asm02.HP0_G0056840 | 611.627627 | 654.9235165 | 531.6522396 | 1261.746037 | 1271.200248 | 1301.748407 | 965.523355 | 1282.060796 | 1535.43928 | 1.99E-09 | -1.092521941 | YPR161C | SGV1 | n |
| W303.asm02.HP0_G0056850 | 433.2766232 | 441.6926041 | 456.3999924 | 730.5869495 | 805.447339 | 745.9253737 | 748.7538959 | 806.0050135 | 726.7745928 | 1.08E-09 | -0.776110914 | YPR162C | ORC4 | n |
| W303.asm02.HP0_G0056860 | 4580.906759 | 4237.964383 | 4609.933114 | 4748.328762 | 4887.08822 | 5002.4073 | 3677.508073 | 3197.895042 | 3734.18833 | 0.282128592 | -0.124398912 | YPR163C | TIF3 | n |
| W303.asm02.HP0_G0056880 | 6011.591985 | 5932.388597 | 6184.366493 | 6656.999328 | 6468.790408 | 6387.438637 | 7242.371754 | 5433.034995 | 4962.539754 | 0.510907329 | -0.106210928 | YPR165W | RHO1 | n |

| | | | | | | | | | | | | | | |
|---|---|---|---|---|---|---|---|---|---|---|---|---|---|---|
| W303.asm02.HP0_G0056890 | 783.1935382 | 917.654462 | 764.2500944 | 1260.773218 | 1383.989414 | 1261.012191 | 1421.78047 | 1102.088488 | 1046.145963 | 3.64E-05 | -0.663359977 | YPR166C | MRP2 | n |
| W303.asm02.HP0_G0056900 | 1175.759606 | 1121.366137 | 1194.262935 | 294.7641088 | 230.8860576 | 279.7220151 | 458.1502979 | 309.6297771 | 384.8834463 | 2.28E-32 | 2.112214276 | YPR167C | MET16 | n |
| W303.asm02.HP0_G0056910 | 162.8422208 | 145.6443285 | 178.8462497 | 259.7426305 | 218.9436754 | 266.1432765 | 93.71256093 | 250.6066008 | 292.7570895 | 0.083682045 | -0.613216265 | YPR168W | NUT2 | n |
| W303.asm02.HP0_G0056920 | 1877.532034 | 1780.097349 | 1883.260782 | 3813.449856 | 3511.060394 | 3634.575698 | 2627.738072 | 2481.876182 | 2288.828154 | 4.74E-20 | -0.984185206 | YPR169W | JIP5 | n |
| W303.asm02.HP0_G0056940 | 344.1011214 | 237.9809289 | 248.2346854 | 966.9819278 | 1045.621916 | 1036.51038 | 911.5676381 | 966.6254604 | 975.5157562 | 4.71E-30 | -1.876562413 | YPR171W | BSP1 | n |
| W303.asm02.HP0_G0056950 | 931.496275 | 672.0581434 | 1004.666365 | 1307.468522 | 1066.852818 | 1165.055772 | 1271.272417 | 1260.773749 | 1062.523982 | 0.016201469 | -0.441163171 | YPR172W | | n |
| W303.asm02.HP0_G0056960 | 1660.409073 | 1683.953053 | 1639.912607 | 1422.261145 | 1169.026533 | 1336.147878 | 1240.034897 | 1262.708935 | 1221.186041 | 0.004319441 | 0.342336985 | YPR173C | VPS4 | n |
| W303.asm02.HP0_G0056980 | 432.3073243 | 477.8657053 | 349.8740841 | 1021.459783 | 962.0252402 | 1012.06865 | 857.6119212 | 1155.306106 | 959.1377372 | 2.38E-13 | -1.249478511 | YPR175W | DPB2 | n |
| W303.asm02.HP0_G0056990 | 1452.009802 | 984.2891221 | 1419.042375 | 553.5339204 | 566.5996932 | 608.3274892 | 729.8220654 | 646.3521597 | 618.2702169 | 6.71E-11 | 1.156921485 | YPR176C | BET2 | n |
| W303.asm02.HP0_G0057000 | 713.404015 | 481.6734002 | 625.473223 | 354.1060581 | 396.7524784 | 363.0049452 | 402.3013979 | 445.0928046 | 435.0411295 | 8.21E-05 | 0.710112063 | YPR178W | PRP4 | n |
| W303.asm02.HP0_G0057010 | 807.4260115 | 716.798558 | 776.9550192 | 960.1721959 | 987.2369361 | 941.4592095 | 927.6596941 | 922.1161799 | 835.2789686 | 0.008064055 | -0.327694832 | YPR179C | HDA3 | n |
| W303.asm02.HP0_G0057020 | 1372.52729 | 1176.577713 | 1291.015824 | 632.3322465 | 674.0811338 | 611.0432369 | 739.2879807 | 609.5836237 | 634.6482359 | 3.67E-13 | 1.003131904 | YPR180W | AOS1 | n |
| W303.asm02.HP0_G0057030 | 2458.142095 | 2679.66526 | 2126.608958 | 4821.290175 | 4600.471045 | 4760.705753 | 3931.194601 | 4612.516086 | 4441.514025 | 4.44E-14 | -0.965230564 | YPR181C | SEC23 | n |
| W303.asm02.HP0_G0057050 | 2117.91817 | 1386.00093 | 2219.45264 | 3722.977704 | 3479.214041 | 3861.793258 | 4312.670986 | 3739.747152 | 3387.179053 | 3.32E-07 | -0.951125508 | YPR183W | DPM1 | n |
| W303.asm02.HP0_G0057060 | 394.5046659 | 342.6925377 | 308.8274039 | 410.5295508 | 362.2522629 | 396.4991671 | 499.800325 | 555.3984127 | 635.6718621 | 0.368981816 | -0.16224519 | YPR184W | GDB1 | n |
| W303.asm02.HP0_G0057070 | 454.6011998 | 493.0964848 | 457.3772943 | 545.7513697 | 591.8113891 | 517.8025652 | 499.800325 | 775.0420358 | 883.3893994 | 0.283754668 | -0.235600324 | YPR185W | ATG13 | n |
| W303.asm02.HP0_G0057080 | 772.5312499 | 826.2697853 | 817.0243976 | 355.0788769 | 282.6363809 | 359.3839483 | 194.9978541 | 251.5741939 | 174.0164518 | 1.65E-12 | 1.273182501 | YPR186C | PZF1 | n |
| W303.asm02.HP0_G0057090 | 3650.379783 | 4756.762808 | 3997.164816 | 3238.513921 | 3326.616934 | 3132.16237 | 2755.527928 | 2350.283527 | 1988.905681 | 0.027012589 | 0.355315738 | YPR187W | RPO26 | n |
| W303.asm02.HP0_G0057100 | 1081.73761 | 1111.8469 | 1129.761009 | 547.6970074 | 460.4451839 | 556.7282826 | 694.798179 | 485.7317128 | 569.13616 | 6.12E-11 | 1.084448304 | YPR188C | MLC2 | n |
| W303.asm02.HP0_G0057110 | 606.7811323 | 898.6159877 | 621.5640154 | 1570.129609 | 1728.99157 | 1578.754674 | 1363.091795 | 1580.079456 | 1390.084362 | 5.57E-12 | -1.196927953 | YPR189W | SKI3 | n |
| W303.asm02.HP0_G0057120 | 397.4125627 | 652.0677453 | 376.2612357 | 3913.650197 | 3369.078738 | 3838.256777 | 1929.153527 | 2594.116976 | 2046.228748 | 5.18E-44 | -2.963393227 | YPR190C | RPC82 | n |
| W303.asm02.HP0_G0057130 | 3727.923698 | 2695.847963 | 3494.831634 | 12839.26306 | 12795.59916 | 12334.92614 | 8277.942882 | 10157.79187 | 9405.077406 | 1.93E-39 | -1.936663598 | YPR191W | QCR2 | n |
| W303.asm02.HP0_G0057140 | 58.15793601 | 100.9039139 | 65.47922804 | 9785.58472 | 11310.76296 | 10166.85421 | 2161.068451 | 2437.366902 | 1563.077188 | 8.64E-230 | -7.120178013 | YPR192W | AQY1 | n |
| W303.asm02.HP0_G0057160 | 620.3513174 | 671.1062196 | 604.9498829 | 210.1288696 | 258.7516163 | 200.9653313 | 215.8228676 | 244.8010425 | 254.8829206 | 1.84E-19 | 1.507751661 | YPR194C | OPT2 | n |
| W303.asm02.HP0_G0057180 | 306.298463 | 296.0482756 | 300.0316867 | 460.1433117 | 453.8105271 | 432.7091367 | 225.2887828 | 368.6529534 | 279.449949 | 0.00389615 | -0.577341018 | YPR196W | | n |
| W303.asm02.HP0_G0057220 | 303.3905662 | 185.6251246 | 283.4175542 | 378.4265291 | 399.4063411 | 364.8154437 | 592.5662944 | 536.0465516 | 637.7191145 | 0.006333752 | -0.564365881 | YPR201W | ARR3 | n |

**TM_gw**

| | Normalized read counts (genome-wide) | | | | | | | | | | | | | |
|---|---|---|---|---|---|---|---|---|---|---|---|---|---|---|
| W303_gene | SC.RI.1_TM | SC.RI.2_TM | SC.RI.3_TM | Km.V.1_TM | Km.V.2_TM | Km.V.3_TM | KS.1.1_TM | KS.1.2_TM | KS.1.3_TM | padj.sc.v.km | log2FC.sc.v.km | locus_tag | gene | ambiguous |
| W303.asm02.HP0_G0000040 | 284.45615 | 216.952082 | 279.1523362 | 781.707498 | 682.23665 | 334.696399 | 777.54407 | 688.84366 | 838.45097 | 0.000208605 | -1.204692402 | YAL067C | SEO1 | n |
| W303.asm02.HP0_G0000100 | 796.47722 | 854.313703 | 802.6837071 | 634.199793 | 663.36054 | 772.740977 | 649.85355 | 736.07866 | 663.88389 | 0.011329803 | 0.246100188 | YAL063C | FLO9 | n |
| W303.asm02.HP0_G0000110 | 239.15388 | 294.805701 | 305.2323122 | 1366.73806 | 1246.7223 | 1316.97818 | 858.49075 | 612.74284 | 875.48035 | 1.10E-62 | -2.226625881 | YAL062W | GDH3 | n |
| W303.asm02.HP0_G0000120 | 272.8672 | 247.055481 | 258.8679104 | 290.848525 | 311.9053 | 274.014899 | 578.02763 | 405.4337 | 561.61227 | 0.231686743 | -0.171426244 | YAL061W | BDH2 | n |
| W303.asm02.HP0_G0000130 | 1161.0025 | 1226.97302 | 1516.502311 | 477.524943 | 433.25174 | 420.977906 | 1727.2423 | 940.76363 | 1532.311 | 1.28E-10 | 1.551730343 | YAL060W | BDH1 | n |
| W303.asm02.HP0_G0000140 | 1864.7681 | 1619.35526 | 2108.614359 | 2402.6255 | 2561.7582 | 2473.71927 | 1773.9862 | 2255.471 | 1773.8836 | 0.000226416 | -0.411056346 | YAL059W | ECM1 | n |
| W303.asm02.HP0_G0000150 | 396.13153 | 378.887608 | 342.9033887 | 983.384699 | 971.67039 | 999.348453 | 2047.6087 | 1701.7719 | 1998.7049 | 4.83E-48 | -1.403143287 | YAL058W | CNE1 | n |
| W303.asm02.HP0_G0000170 | 568.9123 | 581.307015 | 640.4083006 | 637.533301 | 694.82073 | 659.911313 | 678.3559 | 612.74284 | 648.89581 | 0.090394433 | -0.152772205 | YAL055W | PEX22 | n |
| W303.asm02.HP0_G0000180 | 515.1817 | 481.654384 | 563.1342975 | 343.351268 | 364.03932 | 288.237125 | 409.29373 | 300.46705 | 343.84425 | 5.86E-07 | 0.647304076 | YAL054C | ACS1 | n |
| W303.asm02.HP0_G0000190 | 1747.825 | 1646.34451 | 1821.734623 | 3370.17604 | 3242.1971 | 3539.43812 | 3289.171 | 3442.9062 | 3199.5148 | 4.61E-46 | -0.960354951 | YAL053W | FLC2 | n |
| W303.asm02.HP0_G0000200 | 597.35792 | 602.06798 | 636.5446005 | 1048.38809 | 1119.0838 | 1060.9781 | 977.06052 | 856.79031 | 972.46206 | 7.71E-23 | -0.813704802 | YAL051W | OAF1 | n |
| W303.asm02.HP0_G0000210 | 1552.9199 | 1610.01282 | 1632.413315 | 2074.27501 | 2110.5292 | 2107.73398 | 1864.0536 | 2057.3464 | 1812.6763 | 9.97E-09 | -0.391761335 | YAL049C | AIM2 | n |
| W303.asm02.HP0_G0000220 | 246.52866 | 252.245723 | 262.7316105 | 870.878823 | 838.63873 | 849.541 | 827.70821 | 913.20989 | 855.20235 | 1.45E-82 | -1.748115894 | YAL048C | GEM1 | n |
| W303.asm02.HP0_G0000230 | 376.11424 | 386.153946 | 370.9152148 | 575.030036 | 560.89021 | 602.074258 | 633.89224 | 673.09867 | 628.61781 | 4.59E-12 | -0.616905775 | YAL047C | SPC72 | n |
| W303.asm02.HP0_G0000240 | 725.88996 | 758.813265 | 761.1489304 | 546.695223 | 547.40728 | 511.052008 | 583.7281 | 579.94076 | 567.78383 | 2.69E-09 | 0.484115446 | YAL046C | BOL3 | n |
| W303.asm02.HP0_G0000250 | 994.54299 | 1047.39068 | 1058.653842 | 633.366417 | 740.66272 | 729.126149 | 614.51064 | 665.22617 | 597.76 | 1.58E-10 | 0.561225178 | YAL044W- | BOL1 | n |
| W303.asm02.HP0_G0000260 | 1540.2774 | 1429.39243 | 1721.278419 | 2469.29565 | 2438.614 | 2622.57858 | 2141.0964 | 2507.3909 | 2255.2656 | 3.92E-15 | -0.682319875 | YAL044C | GCV3 | n |
| W303.asm02.HP0_G0000270 | 575.23355 | 727.671817 | 742.7963547 | 1336.73649 | 1345.5972 | 1429.80784 | 1354.4316 | 1308.1469 | 1258.9989 | 7.63E-26 | -1.006172699 | YAL043C | PTA1 | n |
| W303.asm02.HP0_G0000280 | 2717.083 | 2635.60449 | 3026.243146 | 3441.84645 | 3470.5082 | 3481.60106 | 3917.3628 | 3820.7862 | 3842.239 | 2.12E-06 | -0.310510525 | YAL042W | ERV46 | n |
| W303.asm02.HP0_G0000290 | 834.40471 | 943.585851 | 891.5488106 | 2097.60957 | 2212.9995 | 2054.63766 | 2074.971 | 1909.081 | 2020.7462 | 3.42E-65 | -1.253537293 | YAL041W | CDC24 | n |
| W303.asm02.HP0_G0000300 | 816.49451 | 932.167321 | 1023.880541 | 2198.44817 | 2267.8301 | 2109.63027 | 2305.27 | 2357.8135 | 2671.4053 | 1.87E-41 | -1.245278431 | YAL040C | CLN3 | n |
| W303.asm02.HP0_G0000310 | 1359.0683 | 1370.22368 | 1826.564248 | 1329.2361 | 1311.4404 | 1377.65968 | 1242.7024 | 1289.7778 | 1206.0998 | 0.095200631 | 0.181849888 | YAL039C | CYC3 | n |
| W303.asm02.HP0_G0000320 | 24216.7 | 27022.4718 | 28566.26709 | 20178.554 | 18938.134 | 20412.6877 | 18142.315 | 15846.028 | 16966.509 | 4.88E-08 | 0.422915919 | YAL038W | CDC19 | n |
| W303.asm02.HP0_G0000340 | 1665.6488 | 1485.44703 | 1718.380644 | 2668.47272 | 2485.3549 | 2643.43784 | 2106.8936 | 2393.2397 | 2176.7986 | 7.81E-18 | -0.678990787 | YAL036C | RBG1 | n |
| W303.asm02.HP0_G0000350 | 2890.9173 | 3165.00909 | 3405.851686 | 7092.87049 | 6723.4917 | 7275.14296 | 5426.8472 | 5640.6455 | 5526.1942 | 4.36E-61 | -1.156156616 | YAL035W | FUN12 | n |
| W303.asm02.HP0_G0000360 | 466.7188 | 485.806577 | 433.7003423 | 346.684775 | 357.74728 | 359.348258 | 286.16358 | 349.01412 | 342.08094 | 0.000331792 | 0.381697716 | YAL034W- | MTW1 | n |
| W303.asm02.HP0_G0000380 | 684.80184 | 651.894296 | 838.4229335 | 450.856883 | 483.58804 | 412.44457 | 388.77204 | 422.49078 | 402.03327 | 8.79E-10 | 0.691905719 | YAL033W | POP5 | n |
| W303.asm02.HP0_G0000390 | 233.88617 | 324.9091 | 268.5271607 | 421.688693 | 395.49951 | 337.540844 | 368.25035 | 387.06453 | 360.59563 | 0.000299691 | -0.482296397 | YAL032C | PRP45 | n |
| W303.asm02.HP0_G0000400 | 419.30944 | 493.072915 | 447.2232929 | 951.716379 | 926.72726 | 852.385446 | 957.67892 | 877.78364 | 957.47398 | 3.15E-28 | -1.00673237 | YAL031C | GIP4 | n |
| W303.asm02.HP0_G0000440 | 853.36845 | 740.128396 | 822.0022079 | 387.520242 | 384.71316 | 353.659367 | 307.82537 | 349.01412 | 333.26442 | 7.48E-30 | 1.100432885 | YAL027W | SAW1 | n |
| W303.asm02.HP0_G0000450 | 1016.6674 | 1142.89111 | 1185.190022 | 2692.64065 | 2554.5673 | 2689.89712 | 2909.5198 | 2891.8313 | 2882.1201 | 5.26E-70 | -1.246065869 | YAL026C | DRS2 | n |
| W303.asm02.HP0_G0000460 | 3383.9747 | 3145.28617 | 3209.768903 | 2280.1191 | 2334.346 | 2440.53408 | 1340.7505 | 1682.0906 | 1424.7495 | 2.63E-08 | 0.465247104 | YAL025C | MAK16 | n |
| W303.asm02.HP0_G0000470 | 329.75843 | 364.354933 | 346.7670889 | 1462.5764 | 1413.9107 | 1536.00047 | 1639.4551 | 1535.1373 | 1606.3698 | 4.60E-144 | -2.083390713 | YAL024C | LTE1 | n |
| W303.asm02.HP0_G0000480 | 4748.3106 | 5648.02048 | 5365.71359 | 10929.7376 | 10933.764 | 11669.811 | 13869.243 | 13572.188 | 12880.053 | 3.33E-50 | -1.08902684 | YAL023C | PMT2 | n |
| W303.asm02.HP0_G0000490 | 832.29763 | 860.541992 | 881.8895603 | 436.689477 | 397.29724 | 408.651977 | 661.25449 | 665.22617 | 660.35728 | 5.65E-37 | 1.050399086 | YAL022C | FUN26 | n |
| W303.asm02.HP0_G0000500 | 1287.4275 | 1437.69681 | 1647.868116 | 3726.86134 | 3549.6081 | 3723.37892 | 3706.4454 | 3440.2821 | 3599.7848 | 9.40E-55 | -1.330113622 | YAL021C | CCR4 | n |
| W303.asm02.HP0_G0000510 | 464.61171 | 422.485634 | 510.0084204 | 275.847742 | 252.58037 | 253.155633 | 279.32302 | 276.84955 | 284.77357 | 2.83E-14 | 0.837809047 | YAL020C | ATS1 | n |
| W303.asm02.HP0_G0000520 | 820.70867 | 906.216115 | 1020.016841 | 1295.06765 | 1401.3267 | 1477.21527 | 1184.5576 | 1100.8378 | 1135.5677 | 5.79E-11 | -0.60217409 | YAL019W | FUN30 | n |
| W303.asm02.HP0_G0000540 | 624.74999 | 767.11765 | 719.6141538 | 1330.90285 | 1380.6528 | 1373.86709 | 1195.9586 | 1381.6236 | 1294.265 | 2.89E-26 | -0.951726681 | YAL017W | PSK1 | n |
| W303.asm02.HP0_G0000550 | 3219.6222 | 2966.74187 | 2868.797365 | 4116.88171 | 4169.8232 | 3937.66046 | 4171.6038 | 3963.8033 | 4253.0888 | 3.52E-11 | -0.433279697 | YAL016W | TPD3 | n |
| W303.asm02.HP0_G0000570 | 1030.3634 | 959.156575 | 1002.63019 | 455.023768 | 435.04946 | 404.859383 | 425.25504 | 404.12162 | 409.08649 | 1.39E-45 | 1.206997839 | YAL014C | SYN8 | n |
| W303.asm02.HP0_G0000580 | 548.89502 | 624.905041 | 558.3046723 | 664.201361 | 675.94462 | 641.896492 | 694.31722 | 684.90741 | 728.24448 | 0.025907346 | -0.194965296 | YAL013W | DEP1 | n |
| W303.asm02.HP0_G0000590 | 2805.5805 | 3726.59319 | 3489.887165 | 6542.00838 | 6576.0782 | 6727.11317 | 5316.2581 | 6141.8613 | 5199.1013 | 8.77E-23 | -0.98548092 | YAL012W | CYS3 | n |
| W303.asm02.HP0_G0000600 | 253.90345 | 286.501315 | 292.6752867 | 1365.90468 | 1316.8336 | 1332.14856 | 1206.2194 | 1158.5694 | 1246.6558 | 9.04E-154 | -2.26764067 | YAL011W | SWC3 | n |
| W303.asm02.HP0_G0000610 | 474.09358 | 502.415349 | 453.9847682 | 508.359887 | 568.08111 | 657.066867 | 568.90688 | 642.92075 | 585.41687 | 0.012743966 | -0.276073655 | YAL010C | MDM10 | n |
| W303.asm02.HP0_G0000620 | 509.91399 | 504.491445 | 563.1342975 | 616.698879 | 611.22652 | 603.970555 | 890.41338 | 805.61907 | 877.24365 | 0.014930036 | -0.214922263 | YAL009W | SPO7 | n |
| W303.asm02.HP0_G0000630 | 1041.9523 | 936.319514 | 1157.178196 | 875.879084 | 805.38082 | 806.874321 | 1648.5759 | 1556.1306 | 1684.8368 | 0.00023522 | 0.33373729 | YAL008W | FUN14 | n |
| W303.asm02.HP0_G0000660 | 15836.833 | 16104.2804 | 14275.40615 | 7432.88825 | 7001.2402 | 8150.28397 | 7635.2092 | 7135.1083 | 7105.2328 | 5.04E-43 | 1.033113857 | YAL003W | EFB1 | n |
| W303.asm02.HP0_G0000670 | 496.21795 | 547.051423 | 588.2483485 | 1075.88953 | 1106.4998 | 1037.27439 | 1279.1854 | 1083.7807 | 1218.4429 | 2.17E-26 | -0.979858611 | YAL002W | VPS8 | n |
| W303.asm02.HP0_G0000680 | 645.82082 | 629.057234 | 700.295653 | 899.213636 | 924.92954 | 934.87436 | 1046.6062 | 868.59906 | 1009.4914 | 9.40E-08 | -0.481355196 | YAL001C | TFC3 | n |
| W303.asm02.HP0_G0000700 | 1135.7175 | 1185.45109 | 907.9695363 | 1123.39201 | 1194.5883 | 1177.60036 | 1085.3694 | 1107.3982 | 1145.2658 | 0.250579704 | -0.115008398 | YAR002W | NUP60 | n |
| W303.asm02.HP0_G0000710 | 4796.7735 | 4951.49011 | 5398.555041 | 3150.99792 | 3215.2312 | 3331.79361 | 3380.3786 | 3322.1946 | 3179.2368 | 3.79E-23 | 0.643594402 | YAR002C- | ERP1 | n |
| W303.asm02.HP0_G0000720 | 2549.5699 | 2396.85339 | 2494.984375 | 1130.8924 | 1066.9498 | 1101.74848 | 1016.9638 | 989.31071 | 1004.2015 | 2.40E-75 | 1.173035673 | YAR003W | SWD1 | n |
| W303.asm02.HP0_G0000730 | 1780.4848 | 1690.98059 | 1689.402893 | 2593.4688 | 2547.3764 | 2508.80077 | 2763.5877 | 2552.0018 | 2905.0431 | 5.07E-17 | -0.568047708 | YAR007C | RFA1 | n |
| W303.asm02.HP0_G0000740 | 1007.1855 | 971.613154 | 1103.086394 | 589.197443 | 551.90159 | 535.703867 | 430.95551 | 418.55453 | 490.19846 | 2.35E-21 | 0.877800829 | YAR008W | SEN34 | n |
| W303.asm02.HP0_G0000750 | 1036.6846 | 1011.05899 | 1059.619767 | 1379.23871 | 1364.4733 | 1285.68928 | 1294.0066 | 1173.0024 | 1283.6852 | 2.55E-07 | -0.375064455 | YAR014C | BUD14 | n |
| W303.asm02.HP0_G0000760 | 1063.0232 | 969.537058 | 935.9813624 | 1860.09716 | 1707.8388 | 1536.00047 | 1518.6051 | 1560.0669 | 1646.0441 | 8.30E-19 | -0.782866316 | YAR015W | ADE1 | n |

| | | | | | | | | | | | | | | |
|---|---|---|---|---|---|---|---|---|---|---|---|---|---|---|
| W303.asm02.HP0_G0000770 | 507.80691 | 534.594844 | 507.1106453 | 448.356753 | 444.93695 | 452.266805 | 462.87814 | 481.53452 | 459.34065 | 0.022139933 | 0.203506668 | YAR018C | KIN3 | n |
| W303.asm02.HP0_G0000780 | 219.13659 | 267.816446 | 269.4930858 | 452.523637 | 482.68917 | 453.214953 | 501.64134 | 484.15869 | 506.94985 | 2.89E-16 | -0.87460949 | YAR019C | CDC15 | n |
| W303.asm02.HP0_G0000810 | 1550.8128 | 1402.40317 | 1620.822215 | 282.514757 | 353.25297 | 273.06675 | 281.60321 | 367.38329 | 305.93321 | 1.62E-86 | 2.331491571 | YAR027W | UIP3 | n |
| W303.asm02.HP0_G0000840 | 207.54764 | 207.609648 | 227.9583091 | 126.673283 | 124.94189 | 109.985219 | 210.91738 | 162.69831 | 208.06985 | 2.42E-08 | 0.829689219 | YAR035W | YAT1 | n |
| W303.asm02.HP0_G0000850 | 624.74999 | 820.058111 | 803.6496321 | 3565.18622 | 3519.0468 | 3644.6826 | 4555.8154 | 4037.2799 | 4468.2119 | 1.22E-124 | -2.253754075 | YAR042W | SWH1 | n |
| W303.asm02.HP0_G0000910 | 2310.4161 | 2336.64659 | 2585.781328 | 495.859234 | 526.73344 | 440.889024 | 383.07157 | 455.29286 | 398.50666 | 1.05E-135 | 2.304548305 | YBL107C | MIX23 | n |
| W303.asm02.HP0_G0000920 | 245.47512 | 264.702302 | 255.0042102 | 1435.90834 | 1457.0561 | 1471.52638 | 1303.1274 | 1305.5228 | 1316.3063 | 6.71E-208 | -2.511626609 | YBL106C | SRO77 | n |
| W303.asm02.HP0_G0000930 | 508.86045 | 526.290458 | 533.1906213 | 2804.31315 | 2703.7785 | 2756.26751 | 2849.0948 | 2538.8809 | 2759.5705 | 4.24E-234 | -2.397362924 | YBL105C | PKC1 | n |
| W303.asm02.HP0_G0000940 | 431.95193 | 494.110963 | 473.3032689 | 1275.89998 | 1133.4656 | 1164.32628 | 1300.8472 | 1256.9757 | 1244.0108 | 3.99E-57 | -1.352785276 | YBL104C | SEA4 | n |
| W303.asm02.HP0_G0000950 | 820.70867 | 759.851313 | 778.5355811 | 930.881957 | 805.38082 | 862.815078 | 947.41807 | 919.7703 | 943.36755 | 0.092241115 | -0.140563652 | YBL103C | RTG3 | n |
| W303.asm02.HP0_G0000960 | 1916.3916 | 1798.9376 | 2036.169981 | 1098.39071 | 1171.2179 | 1171.91147 | 1442.2188 | 1553.5065 | 1394.7733 | 2.27E-22 | 0.741557529 | YBL102W | SFT2 | n |
| W303.asm02.HP0_G0000970 | 268.65303 | 334.251534 | 292.6752867 | 1145.05981 | 1145.1509 | 1145.36331 | 1861.7734 | 1466.909 | 1779.1736 | 4.94E-71 | -1.939641077 | YBL101C | ECM21 | n |
| W303.asm02.HP0_G0000980 | 4105.6504 | 4495.78693 | 4792.920042 | 21325.2806 | 20788.892 | 23430.6442 | 19250.486 | 18916.303 | 17848.161 | 1.04E-207 | -2.290653224 | YBL099W | ATP1 | n |
| W303.asm02.HP0_G0001000 | 154.87057 | 143.250657 | 128.4680301 | 1124.22539 | 1078.635 | 1155.79295 | 1070.5482 | 1092.9653 | 1112.6447 | 3.78E-191 | -2.979562098 | YBL097W | BRN1 | n |
| W303.asm02.HP0_G0001010 | 464.61171 | 514.871928 | 534.1565464 | 445.023245 | 423.36425 | 431.407539 | 587.14838 | 649.48117 | 634.78938 | 0.021611757 | 0.220216062 | YBL095W | MRX3 | n |
| W303.asm02.HP0_G0001020 | 1914.2845 | 1943.22631 | 1738.66507 | 2126.77776 | 1984.6885 | 1894.40058 | 2188.9804 | 2129.511 | 2231.461 | 0.168436852 | -0.102734825 | YBL093C | ROX3 | n |
| W303.asm02.HP0_G0001030 | 12193.687 | 12211.5995 | 12560.8892 | 7672.90079 | 8180.5478 | 8204.32843 | 6678.6704 | 7755.7236 | 6339.9589 | 7.00E-16 | 0.619774097 | YBL092W | RPL32 | n |
| W303.asm02.HP0_G0001050 | 800.69139 | 1083.72236 | 956.2657882 | 2604.3027 | 2419.7379 | 2575.17116 | 2013.4059 | 2385.3672 | 2245.5674 | 7.60E-43 | -1.419443525 | YBL091C | MAP2 | n |
| W303.asm02.HP0_G0001060 | 2064.9409 | 2127.99889 | 2765.443386 | 995.885352 | 1051.6692 | 1074.25218 | 962.23929 | 1041.794 | 974.22536 | 1.52E-27 | 1.157032008 | YBL090W | MRP21 | n |
| W303.asm02.HP0_G0001080 | 701.65851 | 693.416225 | 717.6823037 | 1224.23061 | 1278.1825 | 1302.75595 | 1160.6156 | 1238.6065 | 1147.9108 | 2.90E-31 | -0.848191188 | YBL088C | TEL1 | n |
| W303.asm02.HP0_G0001100 | 306.58052 | 357.088595 | 245.3449598 | 549.195353 | 564.48566 | 482.607555 | 535.84416 | 440.85994 | 494.60672 | 1.24E-09 | -0.814511139 | YBL086C | | n |
| W303.asm02.HP0_G0001120 | 601.57208 | 586.497256 | 607.5668493 | 730.87151 | 676.84348 | 657.066867 | 810.6068 | 716.39741 | 735.29769 | 0.022009053 | -0.202204406 | YBL084C | CDC27 | n |
| W303.asm02.HP0_G0001130 | 612.1075 | 669.541116 | 673.2497519 | 711.703842 | 715.49457 | 626.726117 | 943.99779 | 987.99863 | 970.69876 | 0.42439168 | -0.071750549 | YBL082C | ALG3 | n |
| W303.asm02.HP0_G0001140 | 807.01264 | 828.362497 | 870.2984598 | 599.197965 | 679.54007 | 641.896492 | 606.52998 | 680.97116 | 663.00224 | 9.59E-06 | 0.384627328 | YBL081W | | n |
| W303.asm02.HP0_G0001150 | 362.41821 | 401.724669 | 363.1878145 | 829.20998 | 906.95229 | 750.933563 | 710.27853 | 721.64574 | 748.52247 | 1.20E-31 | -1.142322151 | YBL080C | PET112 | n |
| W303.asm02.HP0_G0001160 | 1734.129 | 1743.92105 | 1949.236728 | 3912.70438 | 4097.0154 | 4151.94201 | 3598.1365 | 3528.1916 | 3654.4472 | 3.25E-70 | -1.163382334 | YBL079W | NUP170 | n |
| W303.asm02.HP0_G0001170 | 789.10243 | 771.269843 | 931.1517372 | 268.34735 | 262.46786 | 227.555625 | 371.67063 | 328.02079 | 356.18737 | 9.04E-53 | 1.715084675 | YBL078C | ATG8 | n |
| W303.asm02.HP0_G0001180 | 5670.1593 | 5693.6946 | 6274.649051 | 6326.16377 | 6512.259 | 6941.39471 | 4820.3172 | 4526.6869 | 4708.0212 | 0.013591999 | -0.1650319 | YBL076C | ILS1 | n |
| W303.asm02.HP0_G0001190 | 374.00716 | 2137.34133 | 428.8707171 | 2540.96606 | 2648.049 | 3172.50467 | 8602.0089 | 4635.5898 | 6906.8611 | 0.044386071 | -1.507871946 | YBL075C | SSA3 | n |
| W303.asm02.HP0_G0001200 | 90.604552 | 119.375548 | 113.9791546 | 335.017499 | 308.30985 | 402.014938 | 351.14894 | 293.90663 | 341.19929 | 1.39E-28 | -1.687568969 | YBL074C | AAR2 | n |
| W303.asm02.HP0_G0001210 | 11267.624 | 10703.3154 | 12677.76613 | 21861.9753 | 21823.483 | 24208.1259 | 19470.525 | 21226.882 | 17380.004 | 1.35E-27 | -0.97039765 | YER102W | RPS8B | y |
| W303.asm02.HP0_G0001220 | 1163.1096 | 1251.88618 | 1249.907 | 411.688171 | 430.55515 | 428.563094 | 274.76264 | 250.60789 | 263.61392 | 4.47E-75 | 1.528748211 | YBL071W- | KTI11 | n |
| W303.asm02.HP0_G0001260 | 311.84822 | 411.067104 | 362.2218895 | 1400.07313 | 1373.4619 | 1445.92637 | 1445.6391 | 1295.0261 | 1467.9504 | 1.17E-88 | -1.95871457 | YBL066C | SEF1 | n |
| W303.asm02.HP0_G0001270 | 2460.0189 | 2596.15865 | 2495.9503 | 2195.11466 | 2192.3257 | 2225.30438 | 2812.6118 | 2798.6734 | 2768.387 | 0.00061525 | 0.191684699 | YBL064C | PRX1 | n |
| W303.asm02.HP0_G0001290 | 1191.5552 | 1268.49495 | 1253.7707 | 2921.81928 | 2770.2943 | 2998.99351 | 2655.2788 | 2423.4176 | 2679.3402 | 5.57E-70 | -1.226416346 | YBL061C | SKT5 | n |
| W303.asm02.HP0_G0001300 | 616.32166 | 556.393857 | 591.1461236 | 921.714811 | 893.46935 | 874.19286 | 801.48605 | 889.59239 | 803.18489 | 2.33E-13 | -0.608953812 | YBL060W | YEL1 | n |
| W303.asm02.HP0_G0001310 | 1300.07 | 1379.56611 | 1718.380644 | 1031.72056 | 1075.0396 | 1078.04477 | 995.30202 | 1056.227 | 987.45014 | 3.96E-06 | 0.46647042 | YBL059C-/ | CMC2 | n |
| W303.asm02.HP0_G0001330 | 2387.3246 | 2448.7558 | 2410.948896 | 2011.77175 | 1945.1385 | 1978.78579 | 2239.1445 | 2015.3597 | 2223.5261 | 6.55E-06 | 0.287881531 | YBL058W | SHP1 | n |
| W303.asm02.HP0_G0001340 | 1271.6243 | 1225.93497 | 1222.861099 | 918.381304 | 832.3467 | 891.259532 | 787.80492 | 906.64947 | 947.77581 | 6.90E-09 | 0.493496517 | YBL057C | PTH2 | n |
| W303.asm02.HP0_G0001350 | 4934.7874 | 4621.39077 | 4551.438782 | 3167.66546 | 2953.6622 | 3081.48242 | 3431.6828 | 3187.05 | 3336.1708 | 3.71E-23 | 0.616102122 | YBL056W | PTC3 | n |
| W303.asm02.HP0_G0001360 | 1685.6661 | 1489.59923 | 1618.890365 | 1155.89371 | 1102.0055 | 1121.6596 | 1061.4275 | 1077.2203 | 1050.9291 | 2.08E-12 | 0.504227306 | YBL055C | | n |
| W303.asm02.HP0_G0001380 | 745.90724 | 728.709866 | 760.1830054 | 682.535652 | 707.40481 | 803.081727 | 738.78088 | 719.02158 | 826.10784 | 0.77273065 | 0.028402612 | YBL052C | SAS3 | n |
| W303.asm02.HP0_G0001390 | 727.99704 | 783.726422 | 873.1962349 | 2600.13582 | 2345.1323 | 2399.7637 | 2470.5836 | 1964.1885 | 2517.1162 | 2.11E-54 | -1.62234368 | YBL051C | PIN4 | n |
| W303.asm02.HP0_G0001410 | 217.02951 | 282.349122 | 217.3331337 | 33.3350746 | 26.965876 | 34.1333438 | 58.144792 | 44.610828 | 49.372507 | 2.31E-42 | 2.923495201 | YBL049W | MOH1 | n |
| W303.asm02.HP0_G0001420 | 1847.9114 | 2057.41161 | 2062.249957 | 2738.47637 | 2798.159 | 2987.61573 | 3137.5386 | 2717.3242 | 3220.6744 | 5.01E-10 | -0.514067297 | YBL047C | EDE1 | n |
| W303.asm02.HP0_G0001430 | 651.08852 | 676.807453 | 559.2705974 | 883.379476 | 889.8739 | 829.629883 | 847.08981 | 850.22989 | 864.90053 | 1.52E-07 | -0.465378324 | YBL046W | PSY4 | n |
| W303.asm02.HP0_G0001440 | 5593.2508 | 5239.02947 | 5944.302688 | 9578.0003 | 9459.6292 | 10447.6476 | 8938.3366 | 8570.5273 | 8022.1508 | 1.98E-29 | -0.813377532 | YBL045C | COR1 | n |
| W303.asm02.HP0_G0001460 | 414.04173 | 358.126643 | 359.3241144 | 66.6701491 | 73.706727 | 61.6296485 | 94.627798 | 82.66124 | 104.91658 | 4.45E-57 | 2.484175672 | YBL043W | ECM13 | n |
| W303.asm02.HP0_G0001470 | 1753.0927 | 1667.10548 | 1841.053124 | 1015.05302 | 958.18745 | 1121.6596 | 719.39928 | 976.18988 | 844.62253 | 2.46E-13 | 0.765959597 | YBL042C | FUI1 | n |
| W303.asm02.HP0_G0001480 | 5326.7048 | 5251.48605 | 5253.666285 | 2877.65031 | 2930.2918 | 2928.83052 | 3077.1136 | 3088.6438 | 3010.8413 | 5.09E-67 | 0.857730134 | YBL041W | PRE7 | n |
| W303.asm02.HP0_G0001500 | 418.2559 | 420.409538 | 521.5995209 | 85.837817 | 76.403314 | 84.385211 | 82.086764 | 86.597489 | 74.940413 | 3.76E-59 | 2.464060368 | YBL039W- | MIN6 | n |
| W303.asm02.HP0_G0001510 | 2778.1884 | 2467.44067 | 2877.49069 | 6896.19355 | 6714.503 | 7642.07641 | 4676.6654 | 5481.8835 | 4583.7083 | 1.31E-53 | -1.387352945 | YBL039C | URA7 | n |
| W303.asm02.HP0_G0001520 | 1355.9077 | 1444.96315 | 1398.659456 | 1491.74459 | 1547.8413 | 1546.4301 | 1504.924 | 1615.1744 | 1470.5954 | 0.066490341 | -0.126754797 | YBL038W | MRPL16 | n |
| W303.asm02.HP0_G0001530 | 544.68085 | 615.562607 | 641.3742256 | 1804.26091 | 1762.6694 | 1727.52645 | 1735.223 | 1679.4665 | 1798.5699 | 3.15E-89 | -1.554330649 | YBL037W | APL3 | n |
| W303.asm02.HP0_G0001540 | 2539.0345 | 2386.47291 | 2927.718792 | 1161.72735 | 1160.4315 | 1169.06702 | 965.65958 | 1161.1936 | 1035.941 | 5.30E-37 | 1.169858637 | YBL036C | | n |
| W303.asm02.HP0_G0001550 | 389.81028 | 446.360744 | 435.6321924 | 1200.06268 | 1208.9701 | 1115.02256 | 1054.5869 | 1174.3144 | 1171.7154 | 1.51E-62 | -1.470122273 | YBL035C | POL12 | n |
| W303.asm02.HP0_G0001560 | 518.34232 | 554.317761 | 595.9757488 | 823.376342 | 797.29106 | 803.081727 | 593.98895 | 628.48784 | 610.10312 | 1.59E-10 | -0.537760953 | YBL034C | STU1 | n |
| W303.asm02.HP0_G0001570 | 1217.8937 | 1468.83826 | 1398.659456 | 1158.39384 | 1174.8133 | 1112.17812 | 1263.2241 | 1200.5561 | 1383.3119 | 0.005974373 | 0.245879396 | YBL033C | RIB1 | n |
| W303.asm02.HP0_G0001580 | 870.22511 | 908.292211 | 1002.63019 | 1810.92793 | 1853.4545 | 1962.66727 | 1710.1409 | 1401.3048 | 1820.6112 | 2.57E-24 | -1.016044245 | YBL032W | HEK2 | n |
| W303.asm02.HP0_G0001590 | 574.18001 | 507.60559 | 446.2573678 | 690.86942 | 602.23789 | 698.785399 | 733.08041 | 637.67242 | 776.73533 | 0.000950776 | -0.383536189 | YBL031W | SHE1 | n |

| | | | | | | | | | | | | | | |
|---|---|---|---|---|---|---|---|---|---|---|---|---|---|---|
| W303.asm02.HP0_G0001600 | 5739.693 | 6282.26796 | 6865.795175 | 14856.6094 | 14255.061 | 17407.0572 | 14307.039 | 12913.523 | 13208.909 | 1.55E-46 | -1.30021118 | YBL030C | PET9 | n |
| W303.asm02.HP0_G0001610 | 1110.4325 | 1109.67357 | 1177.462622 | 51.6693656 | 33.257913 | 24.6518594 | 70.685825 | 60.355826 | 59.070678 | 1.55E-191 | 4.936789035 | YBL029C-A | | n |
| W303.asm02.HP0_G0001620 | 330.81197 | 320.756907 | 348.6989389 | 135.007052 | 118.64985 | 142.222266 | 114.0094 | 161.38623 | 148.11752 | 4.76E-21 | 1.338144503 | YBL029W | | n |
| W303.asm02.HP0_G0001630 | 2199.7942 | 2173.67302 | 2858.172189 | 2020.10552 | 1852.5557 | 1953.18578 | 1295.1467 | 1456.4123 | 1330.4127 | 0.003031012 | 0.31216468 | YBL028C | | n |
| W303.asm02.HP0_G0001640 | 508.86045 | 477.502191 | 415.3477666 | 28.3348134 | 34.156776 | 29.3926016 | 55.864604 | 38.050412 | 40.555988 | 5.25E-91 | 3.931761111 | NA | NA | NA |
| W303.asm02.HP0_G0001650 | 1274.785 | 1209.3262 | 1644.970341 | 677.53539 | 778.41494 | 711.111328 | 614.51064 | 703.27658 | 681.51693 | 1.66E-15 | 0.930871697 | YBL026W | LSM2 | n |
| W303.asm02.HP0_G0001660 | 524.66357 | 469.197805 | 524.497296 | 116.672761 | 98.874877 | 133.68893 | 143.65184 | 91.845822 | 116.37805 | 3.10E-45 | 2.121909434 | YBL025W | RRN10 | n |
| W303.asm02.HP0_G0001670 | 1558.1876 | 1510.36019 | 1549.343762 | 3443.5132 | 3385.1163 | 3775.52708 | 2564.0713 | 2848.5326 | 2667.8787 | 1.17E-65 | -1.199091177 | YBL024W | NCL1 | n |
| W303.asm02.HP0_G0001680 | 760.65682 | 806.563484 | 905.0717612 | 1823.42858 | 1798.6239 | 1655.46717 | 1543.6872 | 1560.0669 | 1617.8313 | 3.33E-41 | -1.093386177 | YBL023C | MCM2 | n |
| W303.asm02.HP0_G0001690 | 2130.2605 | 2633.52839 | 2419.642222 | 3146.83104 | 3083.0985 | 3074.84538 | 3379.2385 | 2991.5496 | 3206.568 | 7.86E-06 | -0.373242632 | YBL022C | PIM1 | n |
| W303.asm02.HP0_G0001700 | 785.94181 | 783.726422 | 899.276211 | 653.367461 | 642.6867 | 582.163141 | 632.75214 | 644.23284 | 564.25722 | 3.83E-05 | 0.394473078 | YBL021C | HAP3 | n |
| W303.asm02.HP0_G0001710 | 1756.2533 | 1534.2353 | 1588.946689 | 1245.06503 | 1242.228 | 1209.83741 | 1759.165 | 1566.6273 | 1726.2744 | 2.76E-07 | 0.399961057 | YBL020W | RFT1 | n |
| W303.asm02.HP0_G0001720 | 594.19729 | 555.355809 | 527.3950711 | 748.372424 | 725.38206 | 667.4965 | 579.16773 | 646.857 | 622.44625 | 0.000105727 | -0.354110691 | YBL019W | APN2 | n |
| W303.asm02.HP0_G0001730 | 502.5392 | 461.931467 | 661.6586515 | 255.846697 | 252.58037 | 223.763031 | 205.21691 | 179.75539 | 198.37168 | 3.64E-16 | 1.151348917 | YBL018C | POP8 | n |
| W303.asm02.HP0_G0001740 | 1369.6037 | 1456.38168 | 1442.126083 | 2432.62707 | 2433.2208 | 2294.51922 | 3166.0409 | 3110.9492 | 3130.7459 | 1.70E-34 | -0.74650524 | YBL017C | PEP1 | n |
| W303.asm02.HP0_G0001750 | 1044.0594 | 1123.1682 | 1225.758874 | 108.338992 | 146.51459 | 127.051891 | 199.51644 | 99.718321 | 186.91021 | 2.61E-71 | 3.152900623 | YBL016W | FUS3 | n |
| W303.asm02.HP0_G0001760 | 403.50632 | 485.806577 | 477.1669691 | 1155.89371 | 1416.6073 | 1202.25222 | 1069.4081 | 942.07572 | 1065.9172 | 1.03E-42 | -1.465135298 | YBL015W | ACH1 | n |
| W303.asm02.HP0_G0001770 | 548.89502 | 516.948024 | 583.4187233 | 500.026118 | 547.40728 | 493.985336 | 354.56922 | 440.85994 | 370.2938 | 0.361393811 | 0.098062584 | YBL014C | RRN6 | n |
| W303.asm02.HP0_G0001780 | 47.409358 | 64.358991 | 51.19402705 | 485.858712 | 512.35164 | 503.46682 | 409.29373 | 414.61828 | 440.82596 | 1.78E-104 | -3.20453317 | YBL013W | FMT1 | n |
| W303.asm02.HP0_G0001790 | 1096.7365 | 1257.07642 | 1323.317303 | 1925.10056 | 2015.2498 | 2040.41544 | 2768.1481 | 2374.8705 | 2625.5594 | 6.30E-16 | -0.701062688 | YBL011W | SCT1 | n |
| W303.asm02.HP0_G0001800 | 271.81366 | 285.463266 | 243.4131097 | 1179.22826 | 1163.1281 | 1170.96332 | 989.60155 | 1167.754 | 1107.3548 | 1.08E-109 | -2.135056452 | YBL010C | LAA2 | n |
| W303.asm02.HP0_G0001810 | 853.36845 | 784.764471 | 730.2393292 | 1765.0922 | 1756.3774 | 1581.51159 | 1805.9088 | 1596.8052 | 1693.6533 | 3.53E-38 | -1.108516461 | YBL009W | ALK2 | n |
| W303.asm02.HP0_G0001830 | 475.14713 | 537.708989 | 501.315095 | 1020.05328 | 930.32271 | 951.941032 | 1042.0459 | 1011.6161 | 1095.0117 | 9.98E-29 | -0.939092089 | YBL008W | HIR1 | n |
| W303.asm02.HP0_G0001840 | 1589.7938 | 1765.72006 | 1736.733219 | 3806.86551 | 3809.3794 | 3816.29746 | 3822.735 | 3403.5437 | 3799.0381 | 7.70E-63 | -1.166560182 | YBL007C | SLA1 | n |
| W303.asm02.HP0_G0001850 | 1870.0358 | 1723.16008 | 1876.79235 | 665.034737 | 702.91049 | 688.355766 | 698.87759 | 740.01491 | 695.62336 | 2.21E-81 | 1.411977026 | YBL006C | LDB7 | n |
| W303.asm02.HP0_G0001870 | 1629.8284 | 1563.30065 | 2008.158155 | 3135.16376 | 3076.8064 | 3496.77144 | 1940.4399 | 2516.5755 | 1970.492 | 1.15E-14 | -0.899969111 | YBL004W | UTP20 | n |
| W303.asm02.HP0_G0001880 | 7692.9586 | 7232.0821 | 8985.034709 | 3915.20451 | 4012.5223 | 3686.40113 | 4396.2023 | 6330.8013 | 4444.4073 | 1.74E-16 | 1.041756245 | YBL003C | HTA2 | y |
| W303.asm02.HP0_G0001920 | 1215.7867 | 1313.13103 | 1221.895174 | 1174.228 | 1191.8917 | 1210.78556 | 1050.0265 | 1191.3715 | 1243.1292 | 0.40888363 | 0.068546177 | YBR002C | RER2 | n |
| W303.asm02.HP0_G0001930 | 1074.6121 | 1265.38081 | 1233.486274 | 1540.08044 | 1453.4607 | 1479.11156 | 1401.1755 | 1469.5331 | 1472.3587 | 3.76E-05 | -0.32362755 | YBR003W | COQ1 | n |
| W303.asm02.HP0_G0001940 | 1302.177 | 1201.02182 | 1498.149735 | 570.029775 | 550.10386 | 520.533492 | 709.13844 | 633.73617 | 685.92519 | 1.08E-39 | 1.286104103 | YBR004C | GPI18 | n |
| W303.asm02.HP0_G0001950 | 723.78287 | 711.063045 | 687.7386275 | 801.708543 | 723.58433 | 745.244672 | 954.25864 | 843.66948 | 1012.1364 | 0.292108992 | -0.098019431 | YBR005W | RCR1 | n |
| W303.asm02.HP0_G0001960 | 602.62562 | 610.372366 | 680.9771522 | 685.869159 | 613.02424 | 603.022406 | 696.5974 | 687.53158 | 685.92519 | 0.948712426 | -0.006249997 | YBR006W | UGA2 | n |
| W303.asm02.HP0_G0001970 | 434.05902 | 416.257345 | 430.8025672 | 858.37817 | 882.683 | 860.918782 | 845.94971 | 768.88074 | 971.58041 | 1.20E-25 | -1.022116585 | YBR007C | DSF2 | n |
| W303.asm02.HP0_G0001990 | 10165.62 | 9823.05051 | 10963.24919 | 6516.1737 | 6686.6383 | 6703.40946 | 7923.653 | 8153.2848 | 7460.5385 | 2.12E-24 | 0.636933858 | YNL030W | HHF2 | y |
| W303.asm02.HP0_G0002000 | 5695.4443 | 5303.38847 | 5754.015455 | 3942.70594 | 3826.4578 | 3972.74195 | 4407.6032 | 4160.6157 | 4384.455 | 1.47E-18 | 0.512820895 | YNL031C | HHT2 | y |
| W303.asm02.HP0_G0002010 | 6907.0168 | 6425.51861 | 5744.356205 | 11535.6025 | 11221.4 | 12119.2333 | 10479.744 | 10247.37 | 10003.223 | 9.87E-33 | -0.87055367 | YBR011C | IPP1 | n |
| W303.asm02.HP0_G0002020 | 2204.0084 | 2236.99396 | 2770.273011 | 869.212069 | 823.35807 | 852.385446 | 1138.9539 | 1048.3545 | 1105.5915 | 1.44E-56 | 1.502900807 | YBR014C | GRX7 | n |
| W303.asm02.HP0_G0002030 | 1571.8836 | 1727.31227 | 1768.608746 | 2885.1507 | 2781.0806 | 2858.66754 | 3397.48 | 3131.9425 | 3248.0056 | 8.34E-29 | -0.749982396 | YBR015C | MNN2 | n |
| W303.asm02.HP0_G0002040 | 902.88489 | 942.547803 | 1027.744241 | 2147.61218 | 2022.4407 | 2048.00063 | 2055.5894 | 2021.9202 | 2100.0949 | 4.67E-56 | -1.113223922 | YBR016W | CPP1 | n |
| W303.asm02.HP0_G0002050 | 1774.1635 | 1714.85569 | 1854.576074 | 2523.46514 | 2558.1627 | 2694.63786 | 2049.8889 | 2196.4272 | 2118.6095 | 4.40E-17 | -0.540753275 | YBR017C | KAP104 | n |
| W303.asm02.HP0_G0002060 | 61.105395 | 61.2448462 | 63.75105255 | 286.681641 | 266.96217 | 260.74082 | 249.68058 | 229.61455 | 233.63776 | 1.26E-43 | -2.129846324 | YBR018C | GAL7 | n |
| W303.asm02.HP0_G0002070 | 145.3887 | 161.935526 | 167.1050317 | 604.198226 | 573.47429 | 575.526102 | 695.45731 | 600.93409 | 659.47563 | 2.05E-65 | -1.884651534 | YBR019C | GAL10 | n |
| W303.asm02.HP0_G0002090 | 689.01601 | 637.36162 | 644.2720008 | 245.012798 | 184.26682 | 160.237086 | 289.58386 | 223.05414 | 264.49557 | 5.36E-34 | 1.737078245 | YBR021W | FUR4 | n |
| W303.asm02.HP0_G0002110 | 1433.8697 | 1361.91929 | 1536.786736 | 3231.83548 | 3163.9961 | 3336.53435 | 3232.1664 | 3162.1204 | 3185.4084 | 9.23E-74 | -1.166972704 | YBR023C | CHS3 | n |
| W303.asm02.HP0_G0002130 | 10347.883 | 10078.4104 | 8009.45042 | 8277.09901 | 8273.1307 | 8735.29156 | 6987.6358 | 7953.8482 | 6907.7427 | 0.085278722 | 0.169322362 | YBR025C | OLA1 | n |
| W303.asm02.HP0_G0002140 | 1589.7938 | 1599.63234 | 1590.878539 | 1818.42832 | 1869.634 | 1958.87467 | 2336.0525 | 2412.9209 | 2161.8105 | 0.000452428 | -0.240031981 | YBR026C | ETR1 | n |
| W303.asm02.HP0_G0002150 | 1320.0872 | 1236.31546 | 1072.176793 | 852.544532 | 884.48072 | 833.422477 | 857.35065 | 843.66948 | 853.43905 | 1.44E-08 | 0.496557953 | YBR028C | YPK3 | n |
| W303.asm02.HP0_G0002160 | 2820.3301 | 2545.29429 | 2801.182612 | 2049.27371 | 2106.9338 | 2227.20068 | 2576.6123 | 2628.1026 | 2522.4061 | 1.20E-07 | 0.355866195 | YBR029C | CDS1 | n |
| W303.asm02.HP0_G0002170 | 462.50463 | 435.980261 | 424.041092 | 1049.22147 | 950.99655 | 1016.41513 | 721.67947 | 793.81032 | 704.43988 | 1.07E-41 | -1.19049772 | YBR030W | RKM3 | n |
| W303.asm02.HP0_G0002200 | 6228.5362 | 5611.68879 | 5827.425758 | 2739.30975 | 2584.2298 | 2443.37852 | 1750.0442 | 2300.0818 | 1914.948 | 4.92E-36 | 1.185484932 | YBR034C | HMT1 | n |
| W303.asm02.HP0_G0002230 | 694.28372 | 666.426971 | 783.3652063 | 1148.39332 | 1186.4985 | 1139.67442 | 1152.635 | 1272.7207 | 1240.4842 | 5.80E-17 | -0.695537767 | YBR037C | SCO1 | n |
| W303.asm02.HP0_G0002240 | 1015.6138 | 1112.78771 | 1283.714376 | 3556.85246 | 3362.6447 | 3812.50487 | 4743.9309 | 4656.5832 | 4671.8735 | 6.07E-82 | -1.652256853 | YBR038W | CHS2 | n |
| W303.asm02.HP0_G0002250 | 5828.1905 | 6030.02223 | 5671.911827 | 7218.71039 | 7108.2048 | 7633.54307 | 6085.8215 | 6022.4617 | 5661.9686 | 3.53E-08 | -0.325059337 | YBR039W | ATP3 | n |
| W303.asm02.HP0_G0002270 | 1274.785 | 1220.74473 | 1180.360397 | 1170.06112 | 1131.6679 | 1225.00778 | 1262.084 | 1149.3849 | 1260.7622 | 0.437210651 | 0.059680946 | YBR041W | FAT1 | n |
| W303.asm02.HP0_G0002280 | 2032.2812 | 1825.92686 | 2015.885556 | 1038.38757 | 1004.9283 | 832.474328 | 1503.7839 | 1363.2544 | 1552.589 | 1.87E-27 | 1.029562047 | YBR042C | CST26 | n |
| W303.asm02.HP0_G0002290 | 786.99535 | 724.557673 | 750.523755 | 255.846697 | 257.07468 | 264.533414 | 380.79138 | 276.84955 | 348.25251 | 4.45E-43 | 1.541080694 | YBR043C | QDR3 | n |
| W303.asm02.HP0_G0002300 | 819.65513 | 881.302957 | 793.9903817 | 1120.89188 | 1048.9726 | 1128.29664 | 1101.3308 | 948.63613 | 1013.018 | 1.43E-06 | -0.403001661 | YBR044C | TCM62 | n |
| W303.asm02.HP0_G0002460 | 1207.3583 | 1187.52719 | 1203.542598 | 833.376864 | 814.36945 | 806.874321 | 1094.4902 | 1031.2974 | 1123.2245 | 4.89E-15 | 0.551575334 | YBR053C | | n |
| W303.asm02.HP0_G0002480 | 816.49451 | 779.574229 | 816.2066576 | 1432.57483 | 1460.6516 | 1401.36339 | 1388.6344 | 1358.0061 | 1436.211 | 4.63E-34 | -0.832103838 | YBR055C | PRP6 | n |
| W303.asm02.HP0_G0002490 | 909.20614 | 851.199558 | 866.4347596 | 305.849309 | 257.07468 | 245.570445 | 264.5018 | 304.4033 | 272.43044 | 1.06E-58 | 1.697257947 | YBR056W | MRX18 | n |

| | | | | | | | | | | | | | | |
|---|---|---|---|---|---|---|---|---|---|---|---|---|---|---|
| W303.asm02.HP0_G0002510 | 1257.9283 | 1248.77203 | 1160.075971 | 210.844347 | 204.04179 | 215.229695 | 229.15888 | 146.95331 | 221.29463 | 1.09E-97 | 2.540813085 | YBR057C | MUM2 | n |
| W303.asm02.HP0_G0002520 | 782.78119 | 759.851313 | 761.1489304 | 1796.76052 | 1837.275 | 1775.88202 | 1852.6527 | 1708.3323 | 1784.4635 | 2.65E-74 | -1.231810197 | YBR058C | UBP14 | n |
| W303.asm02.HP0_G0002530 | 1696.2015 | 1695.13278 | 2114.40991 | 872.545576 | 861.1103 | 790.755797 | 873.31197 | 918.45822 | 909.86477 | 3.13E-31 | 1.125043485 | YBR058C-[illegible] | TSC3 | n |
| W303.asm02.HP0_G0002540 | 754.33557 | 913.482452 | 801.7177821 | 1565.08175 | 1528.9652 | 1405.15598 | 1574.4697 | 1403.929 | 1528.7844 | 3.07E-22 | -0.865848065 | YBR059C | AKL1 | n |
| W303.asm02.HP0_G0002550 | 397.18507 | 483.730481 | 430.8025672 | 668.368245 | 650.77647 | 588.80018 | 584.8682 | 621.92742 | 587.18017 | 5.45E-08 | -0.541326149 | YBR060C | ORC2 | n |
| W303.asm02.HP0_G0002580 | 1267.4102 | 1199.98377 | 1168.769297 | 675.03526 | 671.4503 | 717.748367 | 645.29318 | 742.63907 | 655.94902 | 2.42E-23 | 0.816991188 | YBR061C | TRM7 | n |
| W303.asm02.HP0_G0002590 | 1119.9144 | 1081.64627 | 1253.7707 | 420.855316 | 426.9597 | 368.829742 | 388.77204 | 423.80286 | 441.70761 | 2.21E-53 | 1.505287667 | YBR062C | | n |
| W303.asm02.HP0_G0002610 | 368.73945 | 301.03399 | 294.6071368 | 251.679813 | 263.36672 | 260.74082 | 159.61315 | 234.86289 | 150.76248 | 0.046306086 | 0.313433889 | YBR065C | ECM2 | n |
| W303.asm02.HP0_G0002660 | 1025.0957 | 1012.09704 | 1155.246346 | 538.361454 | 527.6323 | 515.79275 | 769.56342 | 650.79325 | 737.061 | 1.77E-28 | 1.013298438 | YBR070C | ALG14 | n |
| W303.asm02.HP0_G0002670 | 1679.3448 | 1641.15427 | 1672.016242 | 466.691044 | 525.83458 | 470.281625 | 608.81017 | 573.38034 | 644.48755 | 9.63E-107 | 1.771259164 | YBR071W | | n |
| W303.asm02.HP0_G0002680 | 1079.8798 | 2981.27455 | 627.8512751 | 33.3350746 | 68.313552 | 41.7185313 | 18.241503 | 15.744998 | 15.869734 | 4.16E-16 | 5.031941945 | YBR072W | HSP26 | n |
| W303.asm02.HP0_G0002700 | 1598.2222 | 1562.2626 | 1622.754065 | 491.69235 | 542.0141 | 548.977946 | 533.56397 | 467.10161 | 514.00306 | 4.73E-87 | 1.596852473 | YBR073W | RDH54 | n |
| W303.asm02.HP0_G0002710 | 912.36677 | 842.895172 | 1024.846466 | 1622.58475 | 1580.2003 | 1688.65237 | 2088.6521 | 1965.5006 | 2089.515 | 1.82E-23 | -0.81415565 | YBR074W | PFF1 | n |
| W303.asm02.HP0_G0002730 | 936.59822 | 903.10197 | 904.1058361 | 395.85401 | 412.5779 | 377.363078 | 435.51589 | 415.93036 | 414.3764 | 6.80E-47 | 1.209688977 | YBR077C | SLM4 | n |
| W303.asm02.HP0_G0002740 | 16387.835 | 15240.6243 | 12870.95114 | 9858.8483 | 9600.7506 | 10681.8403 | 9889.1749 | 9769.7713 | 9498.0361 | 7.78E-11 | 0.562008354 | YBR078W | ECM33 | n |
| W303.asm02.HP0_G0002750 | 3666.3237 | 3940.43112 | 4236.547219 | 10112.1949 | 10004.34 | 10732.0922 | 7826.745 | 8283.181 | 7762.9451 | 7.57E-92 | -1.380856958 | YBR079C | RPG1 | n |
| W303.asm02.HP0_G0002760 | 1278.9991 | 1419.01195 | 1249.907 | 2640.1379 | 2688.4978 | 2646.28229 | 2805.7712 | 2691.0826 | 2722.5411 | 1.21E-52 | -1.014713657 | YBR080C | SEC18 | n |
| W303.asm02.HP0_G0002770 | 880.76053 | 894.797584 | 858.7073593 | 1689.2549 | 1733.0069 | 1810.96352 | 1823.0102 | 1640.104 | 1820.6112 | 3.78E-42 | -0.990204454 | YBR081C | SPT7 | n |
| W303.asm02.HP0_G0002790 | 706.92621 | 901.025874 | 832.6273833 | 1009.21938 | 1057.0623 | 1050.54847 | 703.43797 | 617.99117 | 791.72342 | 0.001206603 | -0.352453534 | YBR083W | TEC1 | n |
| W303.asm02.HP0_G0002800 | 1919.5522 | 2107.23793 | 2198.445388 | 2169.27998 | 1974.801 | 2234.78587 | 1487.8226 | 1441.9794 | 1490.8734 | 0.664730872 | -0.034798578 | YBR084W | MIS1 | n |
| W303.asm02.HP0_G0002830 | 3805.3912 | 3854.27312 | 4371.776725 | 729.204756 | 707.40481 | 732.918742 | 1031.785 | 990.62279 | 1053.574 | 3.18E-228 | 2.471606 | YBR085C-A | | n |
| W303.asm02.HP0_G0002840 | 3636.8246 | 3805.48485 | 3969.951909 | 2037.60643 | 2183.3371 | 2408.29703 | 2762.4476 | 2453.5955 | 2709.3163 | 7.02E-23 | 0.78423364 | YBR086C | IST2 | n |
| W303.asm02.HP0_G0002850 | 2113.4038 | 2005.5092 | 2267.026066 | 1261.73257 | 1260.2053 | 1227.85223 | 1223.3208 | 1313.3953 | 1252.8274 | 1.22E-27 | 0.768251708 | YBR087W | RFC5 | n |
| W303.asm02.HP0_G0002860 | 2614.8895 | 2727.99078 | 2469.870324 | 2464.29539 | 2488.0515 | 2384.59332 | 2235.7242 | 2348.6289 | 2357.5372 | 0.162845052 | 0.090281711 | YBR088C | POL30 | n |
| W303.asm02.HP0_G0002880 | 1010.3461 | 1027.66776 | 1231.554424 | 699.203189 | 718.19116 | 636.207602 | 584.8682 | 740.01491 | 652.42242 | 6.39E-10 | 0.671114241 | YBR091C | TIM12 | n |
| W303.asm02.HP0_G0002920 | 1308.4983 | 1349.46271 | 1160.075971 | 914.21442 | 938.41247 | 782.222461 | 980.4808 | 969.62946 | 964.52719 | 4.55E-09 | 0.533916947 | YBR095C | RXT2 | n |
| W303.asm02.HP0_G0002930 | 1775.2171 | 1780.25273 | 1936.679702 | 651.700708 | 804.48196 | 716.800219 | 889.27328 | 719.02158 | 841.09592 | 2.12E-41 | 1.33843151 | YBR096W | | n |
| W303.asm02.HP0_G0002940 | 712.19392 | 716.253287 | 822.9681329 | 799.208412 | 830.54897 | 902.637313 | 924.61619 | 830.54865 | 962.76389 | 0.080397622 | -0.168251177 | YBR097W | VPS15 | n |
| W303.asm02.HP0_G0002950 | 516.23524 | 561.584099 | 423.0751669 | 388.353619 | 366.73591 | 337.540844 | 407.01354 | 326.70871 | 346.4892 | 0.000353986 | 0.45609285 | YBR098W | MMS4 | n |
| W303.asm02.HP0_G0002960 | 3703.1977 | 4320.35678 | 2786.693736 | 1277.56673 | 1222.453 | 1238.28186 | 1306.5477 | 1287.1536 | 1258.9989 | 9.18E-36 | 1.531661135 | YBR101C | FES1 | n |
| W303.asm02.HP0_G0002970 | 510.96753 | 568.850436 | 503.2469451 | 1262.56595 | 1179.3076 | 1175.70406 | 1156.0553 | 1044.4182 | 1125.8695 | 2.99E-46 | -1.193173945 | YBR102C | EXO84 | n |
| W303.asm02.HP0_G0002980 | 459.34401 | 512.795831 | 410.5181414 | 1060.05537 | 973.46811 | 1007.88179 | 1024.9445 | 894.84072 | 1020.9529 | 4.93E-31 | -1.138645017 | YBR103W | SIF2 | n |
| W303.asm02.HP0_G0002990 | 1806.8233 | 1666.06743 | 1926.054527 | 447.523376 | 391.0052 | 494.933485 | 557.50594 | 703.27658 | 662.12059 | 7.55E-77 | 2.018273338 | YBR104W | YMC2 | n |
| W303.asm02.HP0_G0003000 | 843.88658 | 913.482452 | 989.1072395 | 94.1715856 | 97.077152 | 66.3703906 | 112.8693 | 108.9029 | 123.43127 | 2.70E-134 | 3.409399182 | YBR105C | VID24 | n |
| W303.asm02.HP0_G0003010 | 9442.8907 | 9444.1629 | 9085.490913 | 6469.50459 | 6192.2639 | 6712.89094 | 7945.3147 | 8230.6977 | 7775.2882 | 9.34E-22 | 0.52985368 | YBR106W | SND3 | n |
| W303.asm02.HP0_G0003030 | 303.41989 | 349.822257 | 356.4263393 | 635.866547 | 623.81059 | 655.170571 | 636.17242 | 614.05492 | 664.76554 | 2.01E-22 | -0.921980602 | YBR108W | AIM3 | n |
| W303.asm02.HP0_G0003040 | 5306.6875 | 5383.31818 | 6113.339569 | 3158.49831 | 3147.8166 | 3322.31213 | 3101.0555 | 3134.5667 | 3042.5807 | 2.06E-31 | 0.803659511 | YBR109C | CMD1 | n |
| W303.asm02.HP0_G0003050 | 1289.5346 | 1199.98377 | 1191.951498 | 1633.41865 | 1614.3571 | 1628.91902 | 1998.5847 | 1839.5406 | 1944.9241 | 1.79E-09 | -0.405978997 | YBR110W | ALG1 | n |
| W303.asm02.HP0_G0003060 | 3188.016 | 3211.72126 | 3794.153552 | 1180.06164 | 1185.5997 | 1152.00035 | 1170.8765 | 1154.6332 | 1188.4668 | 3.44E-82 | 1.53521382 | YBR111C | YSA1 | n |
| W303.asm02.HP0_G0003080 | 858.63616 | 1071.26579 | 1039.335342 | 4061.04546 | 4009.8257 | 4138.66793 | 3527.4507 | 3263.1508 | 3615.6545 | 3.34E-130 | -2.039313407 | YBR112C | CYC8 | n |
| W303.asm02.HP0_G0003090 | 501.48566 | 531.4807 | 605.6349992 | 647.533823 | 629.20377 | 666.548352 | 506.20171 | 561.5716 | 476.97368 | 0.015325074 | -0.244759997 | YBR114W | RAD16 | n |
| W303.asm02.HP0_G0003100 | 1498.1357 | 1667.10548 | 1741.562845 | 1217.5636 | 1130.7691 | 1204.14852 | 1286.026 | 1121.8311 | 1342.7559 | 1.44E-07 | 0.466204344 | YBR115C | LYS2 | n |
| W303.asm02.HP0_G0003120 | 8020.6099 | 8838.98078 | 8822.759303 | 67131.8399 | 65198.993 | 75457.4453 | 66765.042 | 67610.334 | 58041.79 | 4.34E-298 | -3.01621863 | YPR080W | TEF1 | y |
| W303.asm02.HP0_G0003130 | 795.42368 | 767.11765 | 735.0689544 | 430.022462 | 394.60065 | 416.237164 | 393.33241 | 367.38329 | 416.1397 | 8.13E-23 | 0.887941073 | YBR119W | MUD1 | n |
| W303.asm02.HP0_G0003140 | 1301.1235 | 1304.82664 | 1685.539192 | 893.379998 | 948.29996 | 920.652133 | 772.9837 | 827.92448 | 781.14359 | 7.13E-10 | 0.636377707 | YBR120C | CBP6 | n |
| W303.asm02.HP0_G0003150 | 9195.3085 | 8089.50995 | 8313.716807 | 6231.99219 | 6108.6697 | 6528.00199 | 5093.9398 | 5096.131 | 4990.1498 | 7.22E-12 | 0.440074106 | YBR121C | GRS1 | n |
| W303.asm02.HP0_G0003160 | 2509.5354 | 2457.06019 | 2736.465634 | 1374.23845 | 1483.1232 | 1414.63747 | 1287.1661 | 1441.9794 | 1362.1522 | 5.55E-31 | 0.850947783 | YBR122C | MRPL36 | n |
| W303.asm02.HP0_G0003170 | 500.43212 | 524.214362 | 583.4187233 | 791.708021 | 761.33656 | 750.933563 | 819.72755 | 830.54865 | 863.13722 | 1.71E-09 | -0.518005354 | YBR123C | TFC1 | n |
| W303.asm02.HP0_G0003180 | 757.49619 | 761.927409 | 793.0244567 | 768.373468 | 806.27968 | 724.385407 | 861.91103 | 880.40781 | 887.82348 | 0.920018626 | 0.008306535 | YBR125C | PTC4 | n |
| W303.asm02.HP0_G0003190 | 1132.5569 | 1572.64309 | 1412.182406 | 1360.07104 | 1203.5769 | 1225.95593 | 1587.0108 | 1592.869 | 1732.446 | 0.289627877 | 0.119585387 | YBR126C | TPS1 | n |
| W303.asm02.HP0_G0003210 | 4639.7959 | 4815.50579 | 3643.469246 | 5536.95588 | 5317.6707 | 5450.90537 | 5117.8817 | 4955.7381 | 5123.2793 | 0.000728509 | -0.316179256 | YBR127C | VMA2 | n |
| W303.asm02.HP0_G0003220 | 260.2247 | 252.245723 | 257.9019853 | 165.008619 | 128.53734 | 122.311148 | 178.99475 | 154.82581 | 202.77994 | 4.62E-09 | 0.885800998 | YBR128C | ATG14 | n |
| W303.asm02.HP0_G0003230 | 1386.4604 | 1266.41885 | 1478.831234 | 383.353357 | 371.23022 | 360.296406 | 329.48715 | 338.51746 | 325.32956 | 9.40E-98 | 1.889520723 | YBR129C | OPY1 | n |
| W303.asm02.HP0_G0003240 | 1232.6433 | 1183.375 | 1008.42574 | 749.205801 | 714.59571 | 786.015055 | 832.26858 | 839.73323 | 862.25557 | 2.19E-11 | 0.605596559 | YBR130C | SHE3 | n |
| W303.asm02.HP0_G0003250 | 310.79468 | 285.463266 | 284.9478864 | 696.703058 | 656.16964 | 643.792789 | 839.10915 | 671.78658 | 808.4748 | 1.77E-28 | -1.181074607 | YBR131W | CCZ1 | n |
| W303.asm02.HP0_G0003260 | 251.79637 | 289.615459 | 310.0619374 | 253.346567 | 233.70426 | 198.163024 | 460.59796 | 385.75245 | 425.83787 | 0.019574114 | 0.312276695 | YBR132C | AGP2 | n |
| W303.asm02.HP0_G0003270 | 799.63785 | 924.900983 | 838.4229335 | 1240.06477 | 1213.4644 | 1379.55598 | 1254.1033 | 1200.5561 | 1209.6264 | 3.87E-12 | -0.580264721 | YBR133C | HSL7 | n |
| W303.asm02.HP0_G0003280 | 1716.2188 | 1658.80109 | 1773.438371 | 740.872032 | 766.72973 | 789.807649 | 849.36999 | 719.02158 | 873.71705 | 4.59E-44 | 1.164716923 | YBR135W | CKS1 | n |
| W303.asm02.HP0_G0003290 | 397.18507 | 379.925656 | 393.1314907 | 541.694962 | 620.21514 | 589.748328 | 492.52059 | 564.19576 | 516.64802 | 2.45E-09 | -0.581152356 | YBR136W | MEC1 | n |
| W303.asm02.HP0_G0003310 | 571.01938 | 563.660195 | 427.9047921 | 541.694962 | 491.6778 | 466.489031 | 601.96961 | 532.70577 | 543.97923 | 0.653954053 | 0.056971644 | YBR138C | | n |

| | | | | | | | | | | | | | | |
|---|---|---|---|---|---|---|---|---|---|---|---|---|---|---|
| W303.asm02.HP0_G0003320 | 1359.0683 | 1366.07149 | 1210.304073 | 549.195353 | 523.13799 | 502.518672 | 933.73695 | 787.2499 | 911.62808 | 9.65E-47 | 1.32017564 | YBR139W | ATG42 | n |
| W303.asm02.HP0_G0003330 | 1218.9473 | 1256.03837 | 1449.853483 | 1756.75843 | 1782.4444 | 1797.68944 | 2074.971 | 2010.1114 | 2134.4793 | 1.11E-08 | -0.442526114 | YBR140C | IRA1 | n |
| W303.asm02.HP0_G0003340 | 615.26812 | 599.991884 | 554.4409722 | 505.859756 | 559.99135 | 516.740899 | 340.88809 | 527.45743 | 394.0984 | 0.233317615 | 0.161009094 | YBR141C | BMT2 | n |
| W303.asm02.HP0_G0003350 | 2169.2415 | 1870.56293 | 1879.690125 | 2109.27684 | 1996.3737 | 2254.69699 | 1162.8958 | 1518.0802 | 1302.1999 | 0.319546106 | -0.103689168 | YBR142W | MAK5 | n |
| W303.asm02.HP0_G0003360 | 11114.861 | 10047.2689 | 11410.47248 | 5556.95693 | 5732.0463 | 5855.76475 | 4534.1536 | 4960.9865 | 4380.9284 | 5.03E-39 | 0.926002866 | YBR143C | SUP45 | n |
| W303.asm02.HP0_G0003380 | 4664.0273 | 4359.80261 | 4953.263598 | 2779.31184 | 2590.5218 | 2797.98604 | 2588.0133 | 2894.4555 | 2587.6484 | 1.55E-25 | 0.775162778 | YBR146W | MRPS9 | n |
| W303.asm02.HP0_G0003420 | 3357.6361 | 3580.22838 | 2979.878744 | 154.17472 | 168.98615 | 159.288938 | 245.1202 | 213.86956 | 283.89192 | 0 | 4.361988011 | YBR149W | ARA1 | n |
| W303.asm02.HP0_G0003440 | 2798.2057 | 2592.00646 | 3009.82242 | 600.864719 | 548.30614 | 531.911274 | 562.06632 | 620.61534 | 625.09121 | 1.18E-158 | 2.320155586 | YBR151W | APD1 | n |
| W303.asm02.HP0_G0003460 | 1045.113 | 1006.90679 | 974.618364 | 1610.91748 | 1514.5834 | 1588.14863 | 1858.3531 | 1735.886 | 1750.079 | 1.85E-20 | -0.639641077 | YBR153W | RIB7 | n |
| W303.asm02.HP0_G0003470 | 4034.0096 | 3769.15316 | 4188.250967 | 2727.64248 | 2684.9024 | 2753.42306 | 2158.1978 | 2532.3205 | 2161.8105 | 4.27E-13 | 0.554421636 | YBR154C | RPB5 | n |
| W303.asm02.HP0_G0003480 | 1100.9507 | 1052.58092 | 888.6510355 | 1510.07888 | 1516.3811 | 1628.91902 | 970.21995 | 1254.3515 | 1042.1126 | 9.36E-09 | -0.614216133 | YBR155W | CNS1 | n |
| W303.asm02.HP0_G0003490 | 215.97597 | 235.636951 | 228.9242342 | 633.366417 | 583.36178 | 563.200172 | 591.70876 | 544.51452 | 542.21593 | 1.67E-41 | -1.38767952 | YBR156C | SLI15 | n |
| W303.asm02.HP0_G0003500 | 1986.9789 | 2076.09648 | 1221.895174 | 213.344477 | 229.20994 | 220.918586 | 174.43437 | 173.19498 | 178.09369 | 3.52E-76 | 2.993641659 | YBR157C | ICS2 | n |
| W303.asm02.HP0_G0003510 | 4536.5488 | 4544.5752 | 4309.957522 | 1385.07235 | 1510.089 | 1475.31897 | 1492.383 | 1456.4123 | 1542.0092 | 1.30E-145 | 1.615721694 | YBR158W | AMN1 | n |
| W303.asm02.HP0_G0003520 | 3707.4118 | 3485.76599 | 3798.017252 | 2450.96136 | 2517.7139 | 2401.65999 | 2866.1962 | 2843.2842 | 2784.2567 | 1.67E-22 | 0.576606511 | YBR159W | IFA38 | n |
| W303.asm02.HP0_G0003530 | 1498.1357 | 1556.03431 | 1231.554424 | 1567.58188 | 1532.5606 | 1507.55602 | 1486.6825 | 1382.9357 | 1449.4357 | 0.252639991 | -0.105269508 | YBR160W | CDC28 | n |
| W303.asm02.HP0_G0003550 | 8980.386 | 8065.63484 | 8227.749479 | 5377.7809 | 5231.3799 | 5552.35725 | 6503.0959 | 6363.6034 | 6390.2131 | 4.88E-27 | 0.645073239 | YBR162C | TOS1 | n |
| W303.asm02.HP0_G0003560 | 2005.9426 | 2016.92773 | 2353.959319 | 1646.75268 | 1551.4367 | 1645.03754 | 1784.247 | 1603.3656 | 1609.8964 | 1.88E-06 | 0.397252157 | YBR162W- | YSY6 | n |
| W303.asm02.HP0_G0003570 | 852.31491 | 786.840567 | 734.1030294 | 380.853227 | 397.29724 | 499.674227 | 432.09561 | 431.67536 | 393.21675 | 4.21E-15 | 0.894556856 | YBR163W | EXO5 | n |
| W303.asm02.HP0_G0003580 | 3085.8225 | 3074.69889 | 3691.765498 | 2669.3061 | 2755.9125 | 2879.52681 | 2572.052 | 2448.3472 | 2540.0392 | 0.002511205 | 0.247000184 | YBR164C | ARL1 | n |
| W303.asm02.HP0_G0003590 | 887.08177 | 872.998571 | 967.8568887 | 436.689477 | 496.17211 | 425.718649 | 457.17767 | 450.04453 | 446.99752 | 3.44E-28 | 1.006111118 | YBR165W | UBS1 | n |
| W303.asm02.HP0_G0003600 | 2504.2677 | 2376.09242 | 2435.097022 | 1068.38914 | 1051.6692 | 1024.00031 | 1013.5435 | 1012.9282 | 1016.5447 | 8.18E-85 | 1.2179118 | YBR166C | TYR1 | n |
| W303.asm02.HP0_G0003610 | 883.92115 | 904.140018 | 1115.64342 | 352.518413 | 400.89269 | 351.76307 | 323.78668 | 325.39663 | 320.03964 | 1.83E-36 | 1.394485225 | YBR167C | POP7 | n |
| W303.asm02.HP0_G0003620 | 820.70867 | 771.269843 | 814.2748076 | 207.510839 | 219.32246 | 205.748211 | 294.14424 | 297.84288 | 284.77357 | 5.57E-88 | 1.927529178 | YBR168W | PEX32 | n |
| W303.asm02.HP0_G0003640 | 821.76221 | 938.39561 | 852.9118091 | 2407.62576 | 2451.1981 | 2360.88961 | 2525.3081 | 2490.3339 | 2558.5538 | 1.06E-102 | -1.466358947 | YBR170C | NPL4 | n |
| W303.asm02.HP0_G0003650 | 2606.4612 | 2543.21819 | 2381.971145 | 1531.74668 | 1603.5707 | 1546.4301 | 1561.9287 | 1619.1106 | 1542.0092 | 1.67E-26 | 0.68571015 | YBR171W | SEC66 | n |
| W303.asm02.HP0_G0003670 | 3200.6585 | 3217.94955 | 3130.56305 | 2252.61766 | 2452.9958 | 2289.77848 | 2378.236 | 2613.6697 | 2500.3648 | 1.90E-12 | 0.448999361 | YBR173C | UMP1 | n |
| W303.asm02.HP0_G0003680 | 770.13869 | 649.818199 | 743.7622797 | 631.699663 | 604.93448 | 546.1335 | 567.76679 | 591.74951 | 602.16826 | 0.003994655 | 0.278513289 | YBR175W | SWD3 | n |
| W303.asm02.HP0_G0003690 | 751.17495 | 788.916664 | 683.8749273 | 452.523637 | 422.46539 | 454.163102 | 445.77673 | 494.65535 | 457.57734 | 1.71E-15 | 0.741940776 | YBR176W | ECM31 | n |
| W303.asm02.HP0_G0003700 | 1611.9182 | 1638.04013 | 1719.346569 | 2280.1191 | 2269.6279 | 2399.7637 | 2121.7148 | 2204.2997 | 2103.6215 | 8.62E-15 | -0.483324242 | YBR177C | EHT1 | n |
| W303.asm02.HP0_G0003710 | 946.08009 | 965.384865 | 924.390262 | 904.213897 | 914.14319 | 918.755836 | 825.42802 | 871.22322 | 864.90053 | 0.490243427 | 0.05103981 | YBR179C | FZO1 | n |
| W303.asm02.HP0_G0003720 | 106.40767 | 100.690679 | 115.9110046 | 3084.32777 | 3451.6321 | 3419.97142 | 7452.7942 | 6464.6338 | 6277.3616 | 0 | -4.943922158 | YBR180W | DTR1 | n |
| W303.asm02.HP0_G0003730 | 11458.315 | 11168.361 | 12730.89201 | 24104.5924 | 24627.035 | 27108.512 | 21504.452 | 23415.434 | 19255.278 | 4.07E-38 | -1.100869195 | YPL090C | RPS6A | y |
| W303.asm02.HP0_G0003780 | 1685.6661 | 1678.52401 | 1783.097621 | 1428.40794 | 1539.7515 | 1498.07453 | 1320.2288 | 1537.7615 | 1361.2706 | 0.007941941 | 0.205201128 | YBR185C | MBA1 | n |
| W303.asm02.HP0_G0003790 | 233.88617 | 229.408661 | 225.060534 | 59.1697573 | 77.302177 | 50.2518672 | 54.72451 | 45.922911 | 49.372507 | 8.47E-26 | 1.880424298 | YBR186W | PCH2 | n |
| W303.asm02.HP0_G0003800 | 1899.535 | 1666.06743 | 1889.349376 | 1638.41891 | 1640.4241 | 1554.96344 | 1927.8989 | 2088.8364 | 1956.3856 | 0.020992861 | 0.174336315 | YBR187W | GDT1 | n |
| W303.asm02.HP0_G0003820 | 15917.955 | 15915.3556 | 16866.98302 | 12029.795 | 12192.171 | 13366.0485 | 9689.6585 | 10176.517 | 9209.7359 | 1.15E-08 | 0.373763613 | YBR189W | RPS9B | y |
| W303.asm02.HP0_G0003840 | 1640.3638 | 1547.72993 | 1793.722797 | 755.039439 | 748.75248 | 795.496539 | 527.8635 | 674.41075 | 572.19209 | 7.47E-32 | 1.116033152 | YBR192W | RIM2 | n |
| W303.asm02.HP0_G0003850 | 800.69139 | 740.128396 | 615.2942496 | 488.358842 | 563.5868 | 560.355727 | 547.2451 | 635.04825 | 574.83705 | 0.000236652 | 0.419190684 | YBR193C | MED8 | n |
| W303.asm02.HP0_G0003860 | 543.62731 | 618.676752 | 735.0689544 | 395.020633 | 459.31875 | 417.185313 | 373.95082 | 384.44037 | 431.12779 | 2.14E-06 | 0.578880166 | YBR194W | AIM4 | n |
| W303.asm02.HP0_G0003870 | 489.8967 | 504.491445 | 452.0529181 | 557.529122 | 558.19363 | 628.622414 | 609.95026 | 671.78658 | 612.74808 | 0.004966157 | -0.269988808 | YBR195C | MSI1 | n |
| W303.asm02.HP0_G0003880 | 12939.594 | 12302.9478 | 11726.32997 | 16212.5135 | 15579.984 | 16772.7459 | 16737.719 | 18438.705 | 15591.132 | 9.61E-08 | -0.393643485 | YBR196C | PGI1 | n |
| W303.asm02.HP0_G0003900 | 372.95362 | 423.523683 | 425.972942 | 202.510578 | 211.23269 | 194.37043 | 210.91738 | 198.12456 | 210.71481 | 1.40E-18 | 1.007572949 | YBR197C | | n |
| W303.asm02.HP0_G0003910 | 842.83304 | 1033.89605 | 1018.084991 | 2277.61897 | 2230.9768 | 2368.4748 | 2410.1586 | 2399.8001 | 2319.6262 | 9.75E-53 | -1.247642447 | YBR198C | TAF5 | n |
| W303.asm02.HP0_G0003930 | 775.4064 | 788.916664 | 864.5029096 | 1575.91565 | 1440.8766 | 1327.40781 | 1379.5137 | 1462.9727 | 1399.1816 | 6.08E-23 | -0.838907414 | YBR200W | BEM1 | n |
| W303.asm02.HP0_G0003950 | 2040.7095 | 2325.22806 | 2602.202054 | 1086.72343 | 1079.5339 | 1027.79291 | 1251.8232 | 1343.5732 | 1355.9806 | 4.43E-35 | 1.125544383 | YBR201W | DER1 | n |
| W303.asm02.HP0_G0003960 | 766.97807 | 951.890237 | 902.1739861 | 2030.93942 | 1973.9021 | 1852.68205 | 2012.2658 | 1916.9535 | 1943.1608 | 1.73E-42 | -1.159990163 | YBR202W | MCM7 | n |
| W303.asm02.HP0_G0003970 | 352.93634 | 333.213486 | 371.8811399 | 248.346305 | 221.12018 | 227.555625 | 327.20696 | 221.74206 | 282.12861 | 1.02E-05 | 0.601677413 | YBR203W | COS111 | n |
| W303.asm02.HP0_G0003980 | 409.82757 | 478.540239 | 510.0084204 | 132.506921 | 152.80663 | 147.911156 | 166.45372 | 149.57748 | 162.22395 | 1.44E-38 | 1.693043116 | YBR204C | LDH1 | n |
| W303.asm02.HP0_G0003990 | 1834.2154 | 1849.80197 | 1669.118467 | 2560.9671 | 2542.8821 | 2499.31928 | 3014.4084 | 2853.7809 | 2933.2559 | 2.47E-15 | -0.506768894 | YBR205W | KTR3 | n |
| W303.asm02.HP0_G0004000 | 1391.7281 | 1434.58267 | 1497.18381 | 1353.40403 | 1397.7312 | 1508.50416 | 1731.8027 | 1676.8423 | 1728.9194 | 0.763594376 | 0.022397439 | YBR207W | FTH1 | n |
| W303.asm02.HP0_G0004010 | 467.77234 | 513.83388 | 579.5550232 | 1507.57875 | 1584.6946 | 1192.77073 | 1159.4755 | 1166.4419 | 1274.8687 | 2.34E-36 | -1.456247022 | YBR208C | DUR12 | n |
| W303.asm02.HP0_G0004030 | 756.44265 | 768.155699 | 985.2435394 | 323.350223 | 338.87117 | 310.044539 | 321.50649 | 303.09121 | 296.23504 | 1.88E-31 | 1.368815095 | YBR211C | AME1 | n |
| W303.asm02.HP0_G0004040 | 1040.8988 | 1163.65208 | 1241.213675 | 862.545054 | 854.81826 | 848.592852 | 558.64604 | 841.04531 | 766.15551 | 0.000691122 | 0.425561022 | YBR212W | NGR1 | n |
| W303.asm02.HP0_G0004050 | 215.97597 | 269.892543 | 255.0042102 | 781.707498 | 755.94338 | 747.140969 | 725.09975 | 808.24323 | 870.19044 | 1.16E-51 | -1.624154611 | YBR213W | MET8 | n |
| W303.asm02.HP0_G0004060 | 309.74114 | 425.599779 | 410.5181414 | 755.872816 | 798.18992 | 822.992844 | 767.28323 | 726.89408 | 848.14914 | 9.83E-20 | -1.051377873 | YBR214W | SDS24 | n |
| W303.asm02.HP0_G0004070 | 552.05564 | 593.763594 | 631.7149753 | 1013.38627 | 989.64764 | 979.437336 | 922.33601 | 981.43821 | 1045.6392 | 9.97E-19 | -0.745967761 | YBR215W | HPC2 | n |
| W303.asm02.HP0_G0004090 | 631.07124 | 578.19287 | 717.6823037 | 113.339253 | 126.73962 | 93.8666953 | 158.47306 | 146.95331 | 139.301 | 2.05E-74 | 2.527036055 | YBR217W | ATG12 | n |
| W303.asm02.HP0_G0004110 | 952.40134 | 1019.36337 | 1086.665668 | 819.209457 | 870.09892 | 898.844719 | 1176.577 | 1174.3144 | 1183.1769 | 0.00218763 | 0.242196538 | YBR220C | | n |
| W303.asm02.HP0_G0004120 | 4170.97 | 4195.79099 | 3881.086805 | 9273.81774 | 9069.5229 | 10486.5217 | 8423.0141 | 6915.9904 | 7151.9603 | 4.91E-45 | -1.235055481 | YBR221C | PDB1 | n |

| | | | | | | | | | | | | | | |
|---|---|---|---|---|---|---|---|---|---|---|---|---|---|---|
| W303.asm02.HP0_G0004140 | 198.06576 | 198.267214 | 213.4694335 | 250.013059 | 242.69288 | 255.05193 | 254.24095 | 175.81914 | 250.38914 | 0.042487342 | -0.29345791 | YBR223C | TDP1 | n |
| W303.asm02.HP0_G0004150 | 284.45615 | 311.414472 | 339.0396886 | 375.019589 | 313.70302 | 290.133422 | 425.25504 | 387.06453 | 462.86725 | 0.616210338 | -0.067260694 | YBR225W | | n |
| W303.asm02.HP0_G0004160 | 776.45994 | 799.297146 | 746.6600549 | 1069.22252 | 969.87266 | 978.489188 | 959.95911 | 926.33072 | 906.33817 | 1.25E-06 | -0.378778958 | YBR227C | MCX1 | n |
| W303.asm02.HP0_G0004170 | 556.26981 | 512.795831 | 638.4764505 | 394.187257 | 339.77003 | 354.607516 | 331.76734 | 322.77246 | 297.11669 | 8.03E-09 | 0.649418041 | YBR228W | SLX1 | n |
| W303.asm02.HP0_G0004180 | 741.69308 | 817.982014 | 825.865908 | 2290.953 | 2242.662 | 2302.10441 | 2440.9411 | 2342.0685 | 2371.6436 | 7.61E-113 | -1.518133265 | YBR229C | ROT2 | n |
| W303.asm02.HP0_G0004190 | 2003.8356 | 2272.2876 | 2375.20967 | 425.855577 | 400.89269 | 382.10382 | 758.16248 | 574.69243 | 750.28578 | 1.42E-109 | 2.459408436 | YBR230C | OM14 | n |
| W303.asm02.HP0_G0004210 | 386.64966 | 456.741226 | 432.7344173 | 459.190652 | 475.49827 | 438.044578 | 475.41918 | 388.37662 | 427.60118 | 0.340302002 | -0.105228535 | YBR231C | SWC5 | n |
| W303.asm02.HP0_G0004220 | 448.80859 | 448.43684 | 423.0751669 | 662.534607 | 657.0685 | 529.066828 | 709.13844 | 624.55159 | 751.16743 | 1.43E-05 | -0.487074671 | YBR233W | PBP2 | n |
| W303.asm02.HP0_G0004230 | 711.14038 | 693.416225 | 804.6155572 | 105.838862 | 110.56009 | 102.400031 | 133.39099 | 120.71165 | 118.14136 | 2.01E-110 | 2.79295757 | YBR233W- | DAD3 | n |
| W303.asm02.HP0_G0004240 | 3027.8777 | 3088.19352 | 2827.262588 | 3232.66886 | 3178.3779 | 3265.42322 | 3048.6112 | 3159.4963 | 2920.9128 | 0.069805375 | -0.113879169 | YBR234C | ARC40 | n |
| W303.asm02.HP0_G0004250 | 622.64291 | 652.932344 | 692.5682527 | 765.873338 | 754.14566 | 714.903922 | 1254.1033 | 1016.8645 | 1142.6209 | 0.056434303 | -0.183245991 | YBR235W | VHC1 | n |
| W303.asm02.HP0_G0004260 | 1331.6762 | 1294.44616 | 1306.896577 | 1072.55602 | 1067.8487 | 1098.90404 | 965.65958 | 970.94155 | 949.53911 | 1.93E-05 | 0.280085256 | YBR236C | ABD1 | n |
| W303.asm02.HP0_G0004270 | 580.50126 | 650.856247 | 579.5550232 | 808.375558 | 804.48196 | 767.052086 | 761.58276 | 778.06532 | 757.33899 | 1.69E-06 | -0.394945254 | YBR237W | PRP5 | n |
| W303.asm02.HP0_G0004280 | 1611.9182 | 1350.50076 | 1533.888961 | 4326.0593 | 4039.4882 | 3901.63082 | 3567.354 | 3478.3325 | 3429.6259 | 6.93E-80 | -1.448132264 | YBR238C | | n |
| W303.asm02.HP0_G0004290 | 301.31281 | 326.985196 | 359.3241144 | 617.532256 | 570.7777 | 520.533492 | 588.28848 | 591.74951 | 607.45817 | 2.42E-14 | -0.790803969 | YBR239C | ERT1 | n |
| W303.asm02.HP0_G0004300 | 242.3145 | 225.256468 | 260.7997604 | 48.3358581 | 61.122652 | 52.1481641 | 30.782537 | 98.406238 | 34.384425 | 5.82E-07 | 2.172606764 | YBR240C | THI2 | n |
| W303.asm02.HP0_G0004310 | 286.56323 | 323.871051 | 305.2323122 | 257.513451 | 292.13032 | 251.259336 | 388.77204 | 346.38996 | 381.75528 | 0.098111115 | 0.193264219 | YBR241C | VVS1 | n |
| W303.asm02.HP0_G0004320 | 2033.3347 | 1906.89462 | 1970.487079 | 1540.08044 | 1655.7048 | 1652.62273 | 1243.8425 | 1436.7311 | 1192.875 | 0.000411619 | 0.286070183 | YBR242W | | n |
| W303.asm02.HP0_G0004330 | 1256.8748 | 1182.33695 | 1181.326322 | 2649.30505 | 2709.1716 | 2816.94901 | 4171.6038 | 3694.8262 | 3876.6235 | 2.07E-63 | -1.175130425 | YBR243C | ALG7 | n |
| W303.asm02.HP0_G0004350 | 896.56365 | 1123.1682 | 1187.121872 | 3806.03214 | 3480.3957 | 3645.63074 | 3404.3205 | 3247.4058 | 3258.5855 | 1.58E-80 | -1.76867725 | YBR245C | ISW1 | n |
| W303.asm02.HP0_G0004360 | 680.58768 | 674.731357 | 712.8526785 | 654.200838 | 597.74358 | 611.555742 | 449.19702 | 494.65535 | 478.73699 | 0.082758885 | 0.150049775 | YBR246W | RRT2 | n |
| W303.asm02.HP0_G0004370 | 1474.9578 | 1429.39243 | 1232.520349 | 2101.77645 | 1983.7896 | 2176.00066 | 1206.2194 | 1578.4361 | 1351.5724 | 5.54E-09 | -0.598336457 | YBR247C | ENP1 | n |
| W303.asm02.HP0_G0004380 | 1076.7192 | 1112.78771 | 1137.859695 | 2375.95744 | 2332.5482 | 2365.63035 | 2358.8544 | 1913.0173 | 2202.3665 | 1.12E-40 | -1.087986325 | YBR248C | HIS7 | n |
| W303.asm02.HP0_G0004390 | 8490.4893 | 7548.68681 | 7060.912032 | 5179.43721 | 5044.4165 | 5166.46084 | 4138.541 | 4125.1895 | 4080.2851 | 2.65E-16 | 0.585736602 | YBR249C | ARO4 | n |
| W303.asm02.HP0_G0004400 | 120.10371 | 127.679934 | 108.1836043 | 131.673544 | 141.12142 | 91.9703985 | 857.35065 | 543.20243 | 824.34454 | 0.854354162 | -0.038727835 | YBR250W | SPO23 | n |
| W303.asm02.HP0_G0004410 | 1235.8039 | 1181.2989 | 1221.895174 | 2276.78559 | 2239.0665 | 2230.99327 | 2128.5554 | 2263.3435 | 2119.4912 | 1.45E-46 | -0.890741074 | YBR251W | MRPS5 | n |
| W303.asm02.HP0_G0004420 | 959.77612 | 958.118527 | 1195.815198 | 456.690521 | 495.27325 | 462.696438 | 522.16303 | 524.83327 | 570.42879 | 1.49E-27 | 1.139095645 | YBR253W | SRB6 | n |
| W303.asm02.HP0_G0004430 | 1009.2926 | 985.107781 | 1182.292247 | 547.5286 | 591.45154 | 594.489071 | 510.76209 | 558.94743 | 543.09758 | 1.59E-20 | 0.875175657 | YBR254C | TRS20 | n |
| W303.asm02.HP0_G0004450 | 443.54089 | 448.43684 | 528.3609961 | 489.192219 | 479.09372 | 438.992727 | 407.01354 | 478.91036 | 463.74891 | 0.904669364 | 0.013719859 | YBR255C-, | RCF3 | n |
| W303.asm02.HP0_G0004460 | 3484.0611 | 3499.26062 | 4224.956119 | 813.375819 | 828.75125 | 899.792868 | 793.50539 | 814.80365 | 761.74725 | 9.07E-125 | 2.141344372 | YBR256C | RIB5 | n |
| W303.asm02.HP0_G0004470 | 946.08009 | 984.069733 | 1106.950094 | 500.859495 | 505.16074 | 467.43718 | 473.13899 | 522.2091 | 515.76637 | 5.93E-30 | 1.043592546 | YBR257W | POP4 | n |
| W303.asm02.HP0_G0004480 | 501.48566 | 506.567542 | 629.7831252 | 335.017499 | 408.98245 | 299.614906 | 262.22161 | 268.97705 | 287.41852 | 1.08E-06 | 0.650638701 | YBR258C | SHG1 | n |
| W303.asm02.HP0_G0004500 | 453.02276 | 541.861182 | 529.3269212 | 3423.51216 | 3458.823 | 3468.32699 | 2926.6212 | 3156.8721 | 2999.3798 | 2.11E-288 | -2.762670166 | YBR260C | RGD1 | n |
| W303.asm02.HP0_G0004510 | 755.38911 | 843.93322 | 760.1830054 | 800.875166 | 756.84224 | 796.444688 | 621.3512 | 658.66575 | 620.68295 | 0.974021079 | 0.002951164 | YBR261C | TAE1 | n |
| W303.asm02.HP0_G0004520 | 1588.7403 | 1515.55043 | 1943.441178 | 533.361193 | 553.69931 | 508.207563 | 572.32716 | 568.13201 | 572.19209 | 4.81E-61 | 1.662077966 | YBR262C | MIC12 | n |
| W303.asm02.HP0_G0004530 | 2206.1155 | 2310.69539 | 2248.67349 | 4037.71091 | 3913.6474 | 4315.97169 | 3505.7889 | 3832.5949 | 3405.8213 | 1.04E-35 | -0.858419796 | YBR263W | SHM1 | n |
| W303.asm02.HP0_G0004540 | 638.44603 | 644.627958 | 592.1120487 | 342.517891 | 313.70302 | 292.977867 | 348.86875 | 363.44704 | 365.88554 | 1.96E-23 | 0.9798834 | YBR264C | YPT10 | n |
| W303.asm02.HP0_G0004550 | 1197.8765 | 1186.48914 | 1070.244943 | 965.883785 | 915.94091 | 891.259532 | 883.57281 | 1039.1699 | 923.9712 | 0.000262631 | 0.31610018 | YBR265W | TSC10 | n |
| W303.asm02.HP0_G0004560 | 903.93844 | 955.004382 | 878.0258601 | 2170.11335 | 2026.935 | 2064.11915 | 1239.2821 | 1545.634 | 1349.8091 | 9.27E-43 | -1.194234602 | YBR267W | REI1 | n |
| W303.asm02.HP0_G0004570 | 1726.7542 | 1624.5455 | 2045.829232 | 640.033431 | 603.13675 | 564.148321 | 608.81017 | 557.63535 | 650.65911 | 3.08E-53 | 1.578017211 | YBR268W | MRPL37 | n |
| W303.asm02.HP0_G0004580 | 1622.4536 | 1495.82752 | 1755.085795 | 663.367984 | 699.31504 | 667.4965 | 599.68942 | 678.347 | 648.01416 | 1.03E-49 | 1.263754124 | YBR269C | SDH8 | n |
| W303.asm02.HP0_G0004600 | 484.629 | 465.045612 | 468.4736437 | 586.697312 | 578.86746 | 586.903883 | 514.18237 | 548.45076 | 499.89663 | 0.000442083 | -0.305580845 | YBR271W | EFM2 | n |
| W303.asm02.HP0_G0004610 | 1172.5915 | 1131.47258 | 1190.985573 | 790.041267 | 863.80688 | 869.452117 | 747.90163 | 833.17281 | 782.9069 | 1.45E-09 | 0.470897174 | YBR272C | HSM3 | n |
| W303.asm02.HP0_G0004620 | 870.22511 | 885.45515 | 771.7741059 | 765.873338 | 742.46044 | 627.674266 | 828.8483 | 898.77697 | 835.80601 | 0.014626676 | 0.241118941 | YBR273C | UBX7 | n |
| W303.asm02.HP0_G0004630 | 978.73987 | 843.93322 | 1002.63019 | 420.021939 | 433.25174 | 409.600125 | 432.09561 | 393.62495 | 433.77274 | 6.03E-34 | 1.161856452 | YBR274W | CHK1 | n |
| W303.asm02.HP0_G0004640 | 823.8693 | 869.884426 | 907.0036113 | 1030.0538 | 1045.3771 | 1048.65217 | 923.4761 | 848.91781 | 923.9712 | 0.000521128 | -0.26382504 | YBR275C | RIF1 | n |
| W303.asm02.HP0_G0004650 | 1022.9886 | 1003.79265 | 1184.224097 | 851.711155 | 909.64887 | 851.437297 | 867.6115 | 884.34406 | 910.74643 | 0.000500476 | 0.298216557 | YBR276C | PPS1 | n |
| W303.asm02.HP0_G0004670 | 1493.9216 | 1558.11041 | 1171.667072 | 1573.41552 | 1705.1422 | 1530.31158 | 1482.1221 | 1556.1306 | 1542.8908 | 0.068078006 | -0.187847479 | YBR279W | PAF1 | n |
| W303.asm02.HP0_G0004680 | 313.95531 | 339.441775 | 358.3581893 | 217.511361 | 197.74975 | 196.266727 | 290.72396 | 254.54413 | 304.16991 | 1.79E-09 | 0.725851195 | YBR280C | SAF1 | n |
| W303.asm02.HP0_G0004690 | 955.56196 | 904.140018 | 1025.812391 | 879.212591 | 866.50347 | 894.103977 | 1101.3308 | 1026.049 | 1065.9172 | 0.100739697 | 0.129085295 | YBR281C | DUG2 | n |
| W303.asm02.HP0_G0004700 | 1904.8027 | 1863.29659 | 2137.59211 | 1324.23584 | 1389.6415 | 1373.86709 | 1238.142 | 1402.6169 | 1356.8623 | 1.66E-11 | 0.531360223 | YBR282W | MRPL27 | n |
| W303.asm02.HP0_G0004710 | 4262.6281 | 4098.21446 | 4360.185624 | 4181.05173 | 3936.119 | 4333.03836 | 5342.4803 | 4854.7077 | 4930.1975 | 0.652164497 | 0.031183221 | YBR283C | SSH1 | n |
| W303.asm02.HP0_G0004740 | 6569.8835 | 6568.76927 | 6304.592727 | 2558.46697 | 2589.6229 | 2549.57115 | 3175.1617 | 3093.8921 | 3180.1184 | 4.36E-149 | 1.336675442 | YBR286W | APE3 | n |
| W303.asm02.HP0_G0004750 | 1728.8613 | 1583.02357 | 1642.072566 | 1605.91722 | 1565.8185 | 1542.63751 | 2424.9798 | 2306.6422 | 2391.9216 | 0.283892146 | 0.071156074 | YBR287W | | n |
| W303.asm02.HP0_G0004760 | 1415.9595 | 1374.37587 | 1515.536386 | 1343.4035 | 1335.7097 | 1334.04485 | 1397.7552 | 1411.8015 | 1478.5303 | 0.142991136 | 0.101957525 | YBR288C | APM3 | n |
| W303.asm02.HP0_G0004770 | 489.8967 | 579.230919 | 520.6335958 | 1093.39045 | 1075.0396 | 1100.80034 | 1032.9251 | 956.50863 | 1048.2841 | 5.44E-35 | -1.040135152 | YBR289W | SNF5 | n |
| W303.asm02.HP0_G0004780 | 2129.207 | 2049.10723 | 2421.574072 | 1575.08227 | 1493.9095 | 1614.69679 | 1617.7933 | 1650.6006 | 1659.2689 | 1.56E-10 | 0.495284003 | YBR290W | BSD2 | n |
| W303.asm02.HP0_G0004790 | 557.32335 | 532.518748 | 608.5327743 | 408.354663 | 398.1961 | 405.807531 | 527.8635 | 497.27952 | 543.09758 | 1.94E-07 | 0.487059613 | YBR291C | CTP1 | n |
| W303.asm02.HP0_G0004800 | 913.42031 | 862.618089 | 892.5147357 | 400.854272 | 449.43126 | 431.407539 | 460.59796 | 520.89702 | 441.70761 | 9.54E-31 | 1.058791968 | YBR293W | VBA2 | n |
| W303.asm02.HP0_G0004810 | 241.26096 | 266.778398 | 286.8797365 | 449.19013 | 477.296 | 433.303836 | 493.66068 | 414.61828 | 476.97368 | 4.41E-12 | -0.773332636 | YBR294W | SUL1 | n |

| | | | | | | | | | | | | | | |
|---|---|---|---|---|---|---|---|---|---|---|---|---|---|---|
| W303.asm02.HP0_G0004830 | 208.60118 | 217.990131 | 255.9701352 | 1093.39045 | 1197.2849 | 992.711414 | 909.79497 | 661.28992 | 858.72896 | 2.86E-66 | -2.264796166 | YBR296C | PHO89 | n |
| W303.asm02.HP0_G0004980 | 507.80691 | 425.599779 | 399.892966 | 228.345261 | 293.02918 | 180.148203 | 367.11025 | 283.40996 | 405.55988 | 9.34E-08 | 0.9250282 | YCL064C | CHA1 | n |
| W303.asm02.HP0_G0004990 | 651.08852 | 622.828945 | 632.6809003 | 643.366939 | 721.78661 | 505.363117 | 613.37055 | 667.85033 | 657.71233 | 0.831681877 | 0.026596898 | YCL063W | VAC17 | n |
| W303.asm02.HP0_G0005000 | 599.465 | 570.926533 | 616.2601746 | 970.05067 | 1050.7703 | 1006.93364 | 1003.2827 | 981.43821 | 1039.4676 | 1.58E-22 | -0.760268442 | YCL061C | MRC1 | n |
| W303.asm02.HP0_G0005010 | 2256.6855 | 2043.91699 | 2114.40991 | 2595.96893 | 2746.9239 | 2820.7416 | 1667.9575 | 2271.216 | 1802.0965 | 0.000752069 | -0.347693015 | YCL059C | KRR1 | n |
| W303.asm02.HP0_G0005030 | 2098.6543 | 2103.08574 | 2303.731217 | 3288.5051 | 3484.89 | 3571.67517 | 3130.698 | 3324.8188 | 2945.599 | 1.61E-20 | -0.668735775 | YCL057C-/ | MIC10 | n |
| W303.asm02.HP0_G0005040 | 1539.2238 | 1727.31227 | 1845.882749 | 2030.10604 | 2194.1234 | 2041.36359 | 2801.2108 | 3198.8588 | 2755.1622 | 0.000920954 | -0.293028 | YCL057W | PRD1 | n |
| W303.asm02.HP0_G0005060 | 1603.4899 | 1568.49089 | 1470.137909 | 85.837817 | 73.706727 | 46.4592735 | 55.864604 | 52.483327 | 52.899115 | 6.21E-219 | 4.482999966 | YCL055W | KAR4 | n |
| W303.asm02.HP0_G0005070 | 2111.2968 | 1908.97072 | 2094.125484 | 2808.48003 | 2930.2918 | 2897.54163 | 1758.0249 | 2191.1789 | 1838.2442 | 8.17E-09 | -0.498080407 | YCL054W | SPB1 | n |
| W303.asm02.HP0_G0005080 | 1992.2466 | 1823.85076 | 1679.743642 | 1015.05302 | 1081.3316 | 911.170649 | 1081.9492 | 1123.1432 | 1106.4731 | 4.21E-23 | 0.868934581 | YCL052C | PBN1 | n |
| W303.asm02.HP0_G0005100 | 6377.0855 | 6541.78002 | 5612.990399 | 3726.02796 | 3874.0975 | 3698.72706 | 3522.8903 | 3726.3162 | 3521.3177 | 4.04E-24 | 0.713665932 | YCL050C | APA1 | n |
| W303.asm02.HP0_G0005130 | 106.40767 | 119.375548 | 115.9110046 | 22.5011753 | 22.471563 | 17.0666719 | 20.521691 | 48.547077 | 19.396342 | 4.21E-16 | 2.457605324 | YCL048W | SPS22 | n |
| W303.asm02.HP0_G0005150 | 1543.438 | 1590.28991 | 1620.822215 | 1552.5811 | 1675.4797 | 1437.39303 | 2065.8502 | 1882.8393 | 1953.7406 | 0.736574743 | 0.027168186 | YCL045C | EMC1 | n |
| W303.asm02.HP0_G0005160 | 670.05227 | 835.628834 | 697.3978779 | 831.71011 | 1054.3657 | 884.622492 | 879.01244 | 949.94821 | 919.56294 | 0.003050263 | -0.330620363 | YCL044C | MGR1 | n |
| W303.asm02.HP0_G0005170 | 5802.9055 | 7566.33363 | 6689.996817 | 23477.893 | 24934.446 | 26089.2524 | 26592.691 | 27456.652 | 24908.43 | 6.76E-97 | -1.892958899 | YCL043C | PDI1 | n |
| W303.asm02.HP0_G0005180 | 2342.0223 | 2924.1819 | 2807.944087 | 177.509272 | 181.57023 | 183.940797 | 178.99475 | 187.62789 | 208.9515 | 6.30E-269 | 3.894867769 | YCL040W | GLK1 | n |
| W303.asm02.HP0_G0005190 | 633.17832 | 726.633769 | 722.5119289 | 586.697312 | 614.82197 | 528.11868 | 956.53883 | 789.87407 | 891.35008 | 0.00982771 | 0.267542025 | YCL039W | GID7 | n |
| W303.asm02.HP0_G0005200 | 410.88111 | 416.257345 | 440.4618176 | 567.529644 | 660.66395 | 512.948305 | 883.57281 | 829.23656 | 893.99504 | 1.41E-05 | -0.457918829 | YCL038C | ATG22 | n |
| W303.asm02.HP0_G0005210 | 77.962056 | 101.728728 | 114.9450796 | 3351.84175 | 3483.0923 | 3382.99363 | 2232.304 | 2432.6022 | 2400.7382 | 0 | -5.110401517 | YCL037C | SRO9 | n |
| W303.asm02.HP0_G0005230 | 3029.9848 | 3160.8569 | 3717.845474 | 2267.61845 | 2485.3549 | 2287.88218 | 2313.2506 | 2569.0588 | 2204.1298 | 7.56E-08 | 0.493234271 | YCL035C | GRX1 | n |
| W303.asm02.HP0_G0005240 | 860.74324 | 984.069733 | 876.09401 | 863.378431 | 914.14319 | 751.881711 | 1335.05 | 1125.7674 | 1251.9457 | 0.311673805 | 0.104607129 | YCL034W | LSB5 | n |
| W303.asm02.HP0_G0005250 | 1496.0286 | 1452.22949 | 1734.801369 | 530.027685 | 623.81059 | 550.874242 | 523.30312 | 513.02452 | 551.03245 | 1.16E-51 | 1.458714528 | YCL033C | MXR2 | n |
| W303.asm02.HP0_G0005260 | 624.74999 | 585.459208 | 465.5758686 | 1188.39541 | 1294.362 | 1012.62253 | 918.91572 | 1020.8007 | 1042.9942 | 1.04E-18 | -1.062054514 | YCL032W | STE50 | n |
| W303.asm02.HP0_G0005270 | 3285.9953 | 3005.14966 | 3372.04431 | 1955.9355 | 2296.5937 | 1997.74876 | 1663.3971 | 2003.551 | 1765.9488 | 4.25E-12 | 0.628738138 | YCL031C | RRP7 | n |
| W303.asm02.HP0_G0005280 | 2082.8511 | 2233.87982 | 2009.12408 | 2245.11727 | 2473.6697 | 2325.80812 | 2215.2025 | 1777.8727 | 2187.3784 | 0.093552491 | -0.155295708 | YCL030C | HIS4 | n |
| W303.asm02.HP0_G0005290 | 592.09021 | 603.106028 | 568.9298477 | 790.041267 | 852.12167 | 822.044696 | 776.40398 | 737.39074 | 771.44542 | 1.42E-09 | -0.482228897 | YCL029C | BIK1 | n |
| W303.asm02.HP0_G0005310 | 205.44055 | 224.21842 | 127.5021051 | 142.507444 | 145.61573 | 114.725961 | 125.41033 | 116.7754 | 163.1056 | 0.028361584 | 0.464478449 | YCL027W | FUS1 | n |
| W303.asm02.HP0_G0005320 | 398.23861 | 416.257345 | 291.7093617 | 1047.55472 | 1163.1281 | 1059.08181 | 1052.3067 | 1145.4486 | 1006.8465 | 1.69E-40 | -1.565498911 | YCL026C-I | HBN1 | n |
| W303.asm02.HP0_G0005340 | 2495.8393 | 2658.44155 | 2391.630396 | 2540.96606 | 2707.3739 | 2449.06741 | 2489.9652 | 2961.3717 | 2646.719 | 0.736322915 | -0.028885047 | YCL025C | AGP1 | n |
| W303.asm02.HP0_G0005370 | 1414.906 | 1487.52313 | 1299.169177 | 456.690521 | 426.9597 | 394.42975 | 625.91158 | 476.28619 | 618.03799 | 3.25E-55 | 1.715700986 | YCL018W | LEU2 | n |
| W303.asm02.HP0_G0005380 | 1255.8212 | 1379.56611 | 1447.921633 | 3275.17108 | 3111.8621 | 3262.57877 | 3102.1956 | 3183.1138 | 3194.2249 | 7.21E-75 | -1.240170273 | YCL017C | NFS1 | n |
| W303.asm02.HP0_G0005390 | 574.18001 | 616.600655 | 517.7358207 | 636.699924 | 674.14689 | 597.333516 | 645.29318 | 658.66575 | 737.061 | 0.112148065 | -0.160597386 | YCL016C | DCC1 | n |
| W303.asm02.HP0_G0005400 | 497.27149 | 639.437717 | 577.6231731 | 1724.25673 | 1697.9513 | 1863.11168 | 1781.9668 | 1730.6377 | 1756.2506 | 8.59E-74 | -1.623747318 | YCL014W | BUD3 | n |
| W303.asm02.HP0_G0005430 | 430.89839 | 426.637827 | 448.1892179 | 444.189868 | 423.36425 | 426.666797 | 446.91683 | 480.22244 | 442.58926 | 0.897206229 | 0.012709208 | YCL010C | SGF29 | n |
| W303.asm02.HP0_G0005440 | 2011.2103 | 2401.00558 | 2499.814 | 2292.61975 | 2349.6266 | 2335.2896 | 2650.7184 | 1969.4368 | 2589.4117 | 0.910807574 | -0.013390081 | YCL009C | ILV6 | n |
| W303.asm02.HP0_G0005450 | 301.31281 | 363.316885 | 404.7225912 | 140.007313 | 123.14417 | 129.896336 | 114.0094 | 111.52707 | 126.95788 | 6.56E-23 | 1.444133938 | YCL008C | STP22 | n |
| W303.asm02.HP0_G0005470 | 671.10581 | 690.302081 | 610.4646244 | 168.342127 | 176.17705 | 146.014859 | 245.1202 | 192.87623 | 265.37723 | 8.28E-55 | 2.0052087 | YCL005W | LDB16 | n |
| W303.asm02.HP0_G0005480 | 328.70489 | 339.441775 | 315.8574876 | 1150.06007 | 1106.4998 | 1083.73366 | 1071.6883 | 847.60573 | 963.64554 | 2.01E-70 | -1.763983564 | YCL004W | PGS1 | n |
| W303.asm02.HP0_G0005500 | 2582.2297 | 2494.42992 | 2873.62699 | 2565.96736 | 2657.9365 | 2744.88973 | 2927.7613 | 2893.1434 | 2808.0613 | 0.971185302 | -0.002808984 | YCL001W | RER1 | n |
| W303.asm02.HP0_G0005530 | 2791.8844 | 2821.41512 | 2570.326528 | 3976.04102 | 4053.87 | 4029.63086 | 3812.4742 | 3890.3266 | 3840.4757 | 3.83E-22 | -0.559647265 | YCR002C | CDC10 | n |
| W303.asm02.HP0_G0005540 | 1924.82 | 1980.59604 | 2328.845268 | 1720.08985 | 1700.6479 | 1601.42271 | 1548.2476 | 1701.7719 | 1500.5716 | 0.000415255 | 0.312125515 | YCR003W | MRPL32 | n |
| W303.asm02.HP0_G0005550 | 3576.7727 | 3364.31435 | 3586.479668 | 2734.30949 | 2639.9592 | 2811.26012 | 3509.2092 | 2740.9417 | 3435.7975 | 6.42E-05 | 0.363096782 | YCR004C | YCP4 | n |
| W303.asm02.HP0_G0005580 | 228.61846 | 237.713047 | 249.20866 | 154.17472 | 121.34644 | 143.170414 | 204.07682 | 69.540408 | 221.29463 | 0.059319742 | 0.773046761 | YCR007C | DFP3 | n |
| W303.asm02.HP0_G0005590 | 1343.2652 | 1394.09879 | 1636.277015 | 869.212069 | 773.02177 | 747.140969 | 1175.4369 | 847.60573 | 1094.13 | 5.56E-13 | 0.872011045 | YCR008W | SAT4 | n |
| W303.asm02.HP0_G0005600 | 4386.946 | 4088.87202 | 4457.744053 | 2522.63177 | 2517.7139 | 2614.04524 | 2529.8685 | 2751.4384 | 2693.4466 | 1.41E-33 | 0.756978335 | YCR009C | RVS161 | n |
| W303.asm02.HP0_G0005610 | 60.051854 | 58.1307015 | 64.71697759 | 311.682947 | 328.08482 | 365.037149 | 344.30837 | 396.24912 | 304.16991 | 1.32E-51 | -2.455241704 | YCR010C | ADY2 | n |
| W303.asm02.HP0_G0005620 | 1529.742 | 1583.02357 | 1597.640014 | 1030.0538 | 1016.6135 | 954.785477 | 1762.5852 | 1619.1106 | 1780.9369 | 5.59E-20 | 0.649794223 | YCR011C | ADP1 | n |
| W303.asm02.HP0_G0005630 | 45854.332 | 40960.3456 | 40961.98311 | 19771.0327 | 18340.391 | 20379.5025 | 21320.897 | 21052.374 | 19624.69 | 5.41E-59 | 1.127335983 | YCR012W | PGK1 | n |
| W303.asm02.HP0_G0005640 | 492.00379 | 526.290458 | 518.7017457 | 601.698096 | 570.7777 | 580.266844 | 617.93092 | 675.72283 | 660.35728 | 0.029749278 | -0.189623989 | YCR014C | POL4 | n |
| W303.asm02.HP0_G0005650 | 644.76728 | 661.23673 | 761.1489304 | 190.843302 | 187.86227 | 151.70375 | 171.01409 | 149.57748 | 167.51386 | 7.03E-57 | 1.960517082 | YCR015C | CTO1 | n |
| W303.asm02.HP0_G0005660 | 2257.739 | 2116.58036 | 2565.496903 | 1176.72813 | 1104.702 | 1202.25222 | 564.34651 | 713.77324 | 638.31598 | 7.86E-25 | 0.994601786 | YCR016W | RBP95 | n |
| W303.asm02.HP0_G0005670 | 1373.8179 | 1518.66458 | 1643.038491 | 1641.75242 | 1599.9753 | 1682.01533 | 1942.7201 | 1752.9431 | 1884.0901 | 0.151616345 | -0.117877539 | YCR017C | CWH43 | n |
| W303.asm02.HP0_G0005690 | 643.71373 | 586.497256 | 543.8157967 | 474.191436 | 472.80169 | 486.400149 | 495.94087 | 497.27952 | 504.30489 | 0.000961673 | 0.306828785 | YCR019W | MAK32 | n |
| W303.asm02.HP0_G0005700 | 161.19182 | 156.745284 | 181.5939073 | 65.8367722 | 49.437439 | 49.3037188 | 45.603758 | 61.667909 | 47.609203 | 4.09E-16 | 1.597775518 | YCR020C | PET18 | n |
| W303.asm02.HP0_G0005710 | 616.32166 | 588.573353 | 780.4674312 | 93.3382088 | 123.14417 | 104.296328 | 109.44902 | 119.39957 | 97.863362 | 7.40E-67 | 2.632191445 | YCR020C-. | MAK31 | n |
| W303.asm02.HP0_G0005720 | 1106.2184 | 999.640457 | 1319.453603 | 280.848003 | 250.78264 | 297.718609 | 319.22631 | 234.86289 | 242.45428 | 1.11E-52 | 2.046943917 | YCR020W- | HTL1 | n |
| W303.asm02.HP0_G0005760 | 551.0021 | 606.220173 | 592.1120487 | 736.705148 | 771.22404 | 664.652055 | 710.27853 | 691.46783 | 715.90135 | 0.000435093 | -0.3129736 | YCR024C | SLM5 | n |
| W303.asm02.HP0_G0005780 | 713.24746 | 744.280589 | 797.8540819 | 226.678507 | 226.51336 | 217.125992 | 263.3617 | 203.37289 | 256.56071 | 6.91E-57 | 1.750409169 | YCR026C | NPP1 | n |
| W303.asm02.HP0_G0005790 | 1009.2926 | 948.776093 | 1052.858292 | 689.202666 | 687.62983 | 687.407617 | 595.12904 | 611.43076 | 588.06183 | 1.75E-12 | 0.544981807 | YCR027C | RHB1 | n |
| W303.asm02.HP0_G0005800 | 642.66019 | 626.981138 | 669.3860518 | 1400.90651 | 1473.2357 | 1384.29672 | 1895.9762 | 1892.0239 | 1864.6938 | 3.52E-59 | -1.134522908 | YCR028C | FEN2 | n |

| | | | | | | | | | | | | | | |
|---|---|---|---|---|---|---|---|---|---|---|---|---|---|---|
| W303.asm02.HP0_G0005810 | 3954.994 | 3926.9365 | 4976.445799 | 2745.97677 | 2806.2488 | 2856.77124 | 2430.6803 | 2891.8313 | 2466.8621 | 1.00E-09 | 0.612962353 | YCR028C- | RIM1 | n |
| W303.asm02.HP0_G0005840 | 1081.9869 | 925.939031 | 1165.871522 | 855.878039 | 880.88527 | 862.815078 | 1121.8524 | 1075.9082 | 1137.331 | 0.001513352 | 0.288553933 | YCR032W | BPH1 | n |
| W303.asm02.HP0_G0005850 | 690.06955 | 723.519624 | 766.9444807 | 2099.27632 | 2142.8883 | 1948.44504 | 2113.7342 | 1919.5777 | 2127.4261 | 1.68E-82 | -1.504978913 | YCR033W | SNT1 | n |
| W303.asm02.HP0_G0005860 | 5119.1572 | 4518.624 | 4690.531988 | 4721.91331 | 4678.5794 | 4764.4459 | 5009.5728 | 5119.7485 | 4886.9965 | 0.803832416 | 0.016472117 | YCR034W | ELO2 | n |
| W303.asm02.HP0_G0005870 | 2839.2938 | 2861.899 | 2597.372429 | 1294.23427 | 1392.338 | 1337.83745 | 865.33131 | 1005.0557 | 903.69321 | 6.07E-43 | 1.044057862 | YCR035C | RRP43 | n |
| W303.asm02.HP0_G0005880 | 1744.6644 | 1642.19232 | 1667.186617 | 834.210241 | 842.23418 | 819.20025 | 778.68417 | 782.00157 | 764.39221 | 2.71E-51 | 1.017749018 | YCR036W | RBK1 | n |
| W303.asm02.HP0_G0005890 | 921.84864 | 875.074668 | 941.7769127 | 4033.54402 | 3900.1645 | 3687.34927 | 3671.1025 | 3811.6016 | 3838.7124 | 1.88E-218 | -2.085094615 | YCR037C | PHO87 | n |
| W303.asm02.HP0_G0005900 | 166.45953 | 150.516995 | 175.798357 | 1445.90886 | 1567.6162 | 1613.74864 | 1545.9674 | 1566.6273 | 1633.701 | 6.08E-238 | -3.229169291 | YCR038C | BUD5 | n |
| W303.asm02.HP0_G0005920 | 1031.4169 | 1093.0648 | 1183.258172 | 1705.92244 | 1897.4988 | 1763.55609 | 1665.6773 | 1478.7177 | 1576.3936 | 8.37E-18 | -0.697556773 | YCR042C | TAF2 | n |
| W303.asm02.HP0_G0005930 | 2023.8528 | 1928.69363 | 2426.403697 | 950.049625 | 1022.0067 | 980.385485 | 872.17187 | 1171.6903 | 898.4033 | 1.51E-21 | 1.111777189 | YCR043C | | n |
| W303.asm02.HP0_G0005940 | 1174.6985 | 1099.29309 | 1068.313093 | 1575.08227 | 1643.1207 | 1574.87456 | 1815.0296 | 1936.6348 | 1774.7653 | 1.90E-13 | -0.520532056 | YCR044C | PER1 | n |
| W303.asm02.HP0_G0005950 | 218.08305 | 221.104275 | 252.1064351 | 7.50039177 | 8.0897627 | 9.48148438 | 11.40094 | 5.2483327 | 10.579823 | 1.68E-50 | 4.792093655 | YCR045C | RRT12 | n |
| W303.asm02.HP0_G0005960 | 1221.0544 | 1088.91261 | 1269.225501 | 1466.74328 | 1557.7288 | 1521.77824 | 1364.6925 | 1558.7548 | 1486.4651 | 2.86E-05 | -0.34452796 | YCR046C | IMG1 | n |
| W303.asm02.HP0_G0005970 | 1052.4878 | 1072.30383 | 1294.339552 | 1373.40507 | 1311.4404 | 1429.80784 | 921.19591 | 1087.7169 | 877.24365 | 0.010839866 | -0.266438175 | YCR047C | BUD23 | n |
| W303.asm02.HP0_G0005990 | 1852.1256 | 1670.21962 | 2053.556632 | 1185.89528 | 1128.9713 | 1220.26704 | 839.10915 | 1056.227 | 967.17215 | 3.28E-11 | 0.657806763 | YCR051W | | n |
| W303.asm02.HP0_G0006010 | 3371.3322 | 3478.49966 | 3786.426151 | 5782.80206 | 6034.963 | 6217.00931 | 5158.9251 | 4814.0331 | 5137.3857 | 6.42E-31 | -0.76143496 | YCR053W | THR4 | n |
| W303.asm02.HP0_G0006020 | 810.17326 | 718.329383 | 779.5015062 | 674.201883 | 684.03438 | 676.029836 | 519.88284 | 502.52785 | 486.67186 | 0.029517736 | 0.182192011 | YCR054C | CTR86 | n |
| W303.asm02.HP0_G0006030 | 1605.5969 | 1522.81677 | 1877.758275 | 2606.80283 | 2508.7253 | 2746.78602 | 1526.5858 | 1767.376 | 1460.8972 | 1.32E-11 | -0.650750213 | YCR057C | PWP2 | n |
| W303.asm02.HP0_G0006040 | 1281.1062 | 1019.36337 | 1145.587096 | 535.027947 | 537.51979 | 485.452 | 492.52059 | 539.26618 | 514.88472 | 1.34E-30 | 1.144483929 | YCR059C | YIH1 | n |
| W303.asm02.HP0_G0006050 | 1103.0577 | 1068.15164 | 1275.021051 | 222.511623 | 194.1543 | 210.488953 | 156.19287 | 140.3929 | 130.48448 | 9.05E-107 | 2.457757728 | YCR060W | TAH1 | n |
| W303.asm02.HP0_G0006060 | 328.70489 | 399.648573 | 373.8129899 | 1220.89711 | 1182.9031 | 1639.34865 | 2295.0091 | 2157.0647 | 1902.6048 | 6.33E-47 | -1.874304579 | YCR061W | TVS1 | n |
| W303.asm02.HP0_G0006070 | 119.05017 | 124.565789 | 132.3317303 | 66.6701491 | 56.628339 | 54.9926094 | 43.32357 | 57.731659 | 43.200944 | 5.33E-08 | 1.07398875 | YCR063W | BUD31 | n |
| W303.asm02.HP0_G0006080 | 426.68423 | 498.263156 | 447.2232929 | 1077.55629 | 1066.051 | 1019.25957 | 957.67892 | 1103.4619 | 1121.4612 | 2.30E-38 | -1.205122042 | YCR065W | HCM1 | n |
| W303.asm02.HP0_G0006090 | 265.49241 | 281.311073 | 302.3345371 | 653.367461 | 658.86623 | 642.844641 | 628.19177 | 680.97116 | 644.48755 | 7.76E-38 | -1.201900161 | YCR066W | RAD18 | n |
| W303.asm02.HP0_G0006100 | 912.36677 | 997.56436 | 1026.778316 | 3677.6921 | 3784.2112 | 4021.09752 | 3851.2374 | 3738.1249 | 3801.683 | 1.57E-181 | -1.9664095 | YCR067C | SED4 | n |
| W303.asm02.HP0_G0006110 | 231.77909 | 299.995942 | 235.6857094 | 769.206845 | 781.11153 | 718.696516 | 1058.0072 | 817.42781 | 977.75197 | 2.76E-37 | -1.564815439 | YCR068W | ATG15 | n |
| W303.asm02.HP0_G0006120 | 887.08177 | 1019.36337 | 908.9354613 | 2386.79134 | 2324.4585 | 2299.25996 | 2952.8433 | 2788.1767 | 2809.8246 | 1.16E-76 | -1.316517588 | YCR069W | CPR4 | n |
| W303.asm02.HP0_G0006130 | 1682.5055 | 1729.38837 | 1878.7242 | 1041.72108 | 1122.6793 | 1131.14109 | 923.4761 | 1128.3915 | 1029.7694 | 2.56E-15 | 0.683603431 | YCR071C | IMG2 | n |
| W303.asm02.HP0_G0006140 | 616.32166 | 700.682563 | 703.1934281 | 1756.75843 | 1688.9627 | 1806.22277 | 1031.785 | 1344.8852 | 1069.4438 | 1.44E-40 | -1.37790521 | YCR072C | RSA4 | n |
| W303.asm02.HP0_G0006170 | 636.33895 | 628.019186 | 735.0689544 | 655.034215 | 711.89912 | 701.629844 | 702.29787 | 686.2195 | 689.4518 | 0.61245603 | -0.047478241 | YCR075C | ERS1 | n |
| W303.asm02.HP0_G0006180 | 285.50969 | 294.805701 | 317.7893377 | 252.51319 | 296.62463 | 244.622297 | 217.75794 | 261.10455 | 258.32401 | 0.161731352 | 0.178841088 | YCR075W- | EGO2 | n |
| W303.asm02.HP0_G0006190 | 627.91061 | 619.7148 | 655.8631012 | 272.514234 | 257.97354 | 207.644508 | 293.00415 | 326.70871 | 313.86808 | 3.35E-34 | 1.364048256 | YCR076C | FUB1 | n |
| W303.asm02.HP0_G0006200 | 1782.5919 | 1801.0137 | 2233.218689 | 3845.20085 | 3883.985 | 4321.66058 | 3617.5181 | 3369.4296 | 3771.7069 | 1.18E-28 | -1.050237966 | YCR077C | PAT1 | n |
| W303.asm02.HP0_G0006210 | 481.46837 | 490.996818 | 559.2705974 | 324.1836 | 302.91667 | 301.511203 | 381.93147 | 406.74578 | 369.41215 | 3.21E-12 | 0.722197341 | YCR079W | PTC6 | n |
| W303.asm02.HP0_G0006220 | 603.67916 | 548.089471 | 558.3046723 | 886.712983 | 903.35684 | 864.711375 | 900.67422 | 831.86073 | 859.61061 | 2.49E-15 | -0.635106892 | YCR081W | SRB8 | n |
| W303.asm02.HP0_G0006230 | 2172.4022 | 2260.86907 | 2465.040699 | 1034.22069 | 1087.6237 | 1040.11884 | 1184.5576 | 1137.5761 | 1151.4374 | 1.99E-55 | 1.125860471 | YCR082W | AHC2 | n |
| W303.asm02.HP0_G0006240 | 1186.2875 | 1190.64133 | 1502.013435 | 1045.88796 | 1037.2874 | 1100.80034 | 873.31197 | 947.32405 | 887.82348 | 0.003983874 | 0.285732059 | YCR083W | TRX3 | n |
| W303.asm02.HP0_G0006250 | 1599.2757 | 1901.70438 | 2025.544806 | 5080.26536 | 5074.0789 | 5113.36453 | 5057.4568 | 4543.744 | 4864.9553 | 4.99E-67 | -1.465577587 | YCR084C | TUP1 | n |
| W303.asm02.HP0_G0006260 | 540.46669 | 486.844625 | 395.0633408 | 864.211808 | 817.06603 | 832.474328 | 688.61675 | 767.56865 | 720.30961 | 2.32E-14 | -0.823374207 | YCR086W | CSM1 | n |
| W303.asm02.HP0_G0006270 | 1304.2841 | 1322.47346 | 1578.321513 | 1132.55916 | 1091.2191 | 1212.68185 | 531.28378 | 917.14614 | 605.69486 | 0.06092989 | 0.291514863 | YCR087C-A | | n |
| W303.asm02.HP0_G0006280 | 986.11466 | 1186.48914 | 1152.348571 | 3130.99688 | 3051.6383 | 2980.03054 | 2935.7419 | 2789.4888 | 2905.9247 | 1.96E-82 | -1.462103507 | YCR088W | ABP1 | n |
| W303.asm02.HP0_G0006300 | 1528.6884 | 1471.95241 | 1960.827828 | 1043.38783 | 1109.1964 | 1054.34106 | 840.24924 | 967.0053 | 844.62253 | 8.12E-09 | 0.630136794 | YCR090C | | n |
| W303.asm02.HP0_G0006320 | 482.52192 | 482.692432 | 565.0661476 | 953.383132 | 1072.343 | 1046.75588 | 1043.186 | 974.87779 | 1020.0713 | 1.94E-28 | -1.003830058 | YCR092C | MSH3 | n |
| W303.asm02.HP0_G0006330 | 1536.0632 | 1690.98059 | 1867.1331 | 5570.29096 | 5459.691 | 5417.72017 | 5125.8624 | 4723.4994 | 5057.1554 | 5.65E-111 | -1.690428702 | YCR093W | CDC39 | n |
| W303.asm02.HP0_G0006340 | 973.47216 | 897.911729 | 845.1844088 | 1527.57979 | 1475.9323 | 1401.36339 | 1614.373 | 1680.7785 | 1693.6533 | 5.45E-20 | -0.698458566 | YCR094W | CDC50 | n |
| W303.asm02.HP0_G0006350 | 1191.5552 | 1185.45109 | 1423.773507 | 610.031864 | 556.3959 | 573.629805 | 647.57354 | 591.74951 | 617.15634 | 8.87E-33 | 1.127297679 | YCR095C | OCA4 | n |
| W303.asm02.HP0_G0006450 | 466.7188 | 408.991007 | 449.155143 | 1530.9133 | 1561.3242 | 1660.20791 | 1451.3396 | 1591.5569 | 1549.0624 | 3.77E-111 | -1.842400307 | YCR106W | RDS1 | n |
| W303.asm02.HP0_G0006500 | 144.33516 | 174.392105 | 193.1850077 | 50.8359887 | 66.515827 | 41.7185313 | 103.74855 | 72.164574 | 82.87528 | 2.02E-15 | 1.685705729 | YJR159W | SOR1 | y |
| W303.asm02.HP0_G0006530 | 122.21079 | 103.804824 | 119.7747048 | 32.5016977 | 32.359051 | 25.6000078 | 34.202819 | 34.114162 | 27.331209 | 2.40E-16 | 1.930826988 | YDL244W | THI13 | y |
| W303.asm02.HP0_G0006570 | 639.49957 | 610.372366 | 550.577272 | 997.552106 | 1040.8828 | 1105.54108 | 3223.0456 | 1780.4969 | 3157.1955 | 0.000665256 | -0.804305777 | YDL240W | LRG1 | n |
| W303.asm02.HP0_G0006590 | 97.979341 | 139.098464 | 120.7406298 | 271.680858 | 366.73591 | 313.837133 | 1210.7798 | 679.65908 | 1336.5843 | 2.39E-06 | -1.41166834 | YDL238C | GUD1 | n |
| W303.asm02.HP0_G0006600 | 1440.191 | 1380.60416 | 1280.816601 | 90.8380782 | 91.683977 | 74.9037266 | 458.31777 | 240.11122 | 488.43516 | 1.85E-47 | 3.992967608 | YDL237W | AIM6 | n |
| W303.asm02.HP0_G0006610 | 2205.0619 | 2174.71107 | 2099.921034 | 2188.44764 | 2209.4041 | 1902.93391 | 4856.8002 | 3194.9225 | 4941.659 | 0.777263875 | 0.040217927 | YDL236W | PHO13 | n |
| W303.asm02.HP0_G0006620 | 3233.3182 | 3275.0422 | 3897.507531 | 1084.2233 | 1190.094 | 1077.09663 | 3194.5433 | 2032.4168 | 3115.7579 | 1.50E-31 | 1.634662398 | YDL235C | YPD1 | n |
| W303.asm02.HP0_G0006630 | 539.41315 | 605.182125 | 660.6927264 | 615.032126 | 555.49704 | 495.881633 | 1788.8074 | 1001.1195 | 1809.1497 | 0.658949581 | 0.115398567 | YDL234C | GYP7 | n |
| W303.asm02.HP0_G0006640 | 294.99156 | 289.615459 | 336.1419134 | 804.208674 | 765.83087 | 712.059477 | 2501.3661 | 1399.9927 | 2464.2171 | 3.17E-08 | -1.30930505 | YDL233W | MFG1 | n |
| W303.asm02.HP0_G0006650 | 1327.462 | 1165.72818 | 1312.692127 | 957.550017 | 999.53513 | 1163.37813 | 3514.9097 | 1986.4939 | 3269.1653 | 0.214298565 | 0.286633381 | YDL231C | BRE4 | n |
| W303.asm02.HP0_G0006660 | 324.49072 | 385.115898 | 361.2559644 | 623.365894 | 622.01287 | 567.940914 | 1720.4018 | 1106.0861 | 1749.1974 | 5.20E-07 | -0.759936919 | YDL230W | PTP1 | n |
| W303.asm02.HP0_G0006670 | 9010.9387 | 9091.2265 | 9831.185043 | 35803.5368 | 34429.131 | 38678.7674 | 63196.548 | 47454.112 | 56816.294 | 1.23E-93 | -1.963050728 | YDL229W | SSB1 | y |
| W303.asm02.HP0_G0006690 | 1371.7108 | 1462.60997 | 1362.920229 | 2502.63072 | 2591.4207 | 2551.46745 | 6736.8152 | 4277.3911 | 6481.0232 | 5.46E-11 | -0.865196901 | YDL226C | GCS1 | n |

| | | | | | | | | | | | | | | |
|---|---|---|---|---|---|---|---|---|---|---|---|---|---|---|
| W303.asm02.HP0_G0006700 | 1446.5122 | 1501.01776 | 1303.032877 | 2162.61296 | 2183.3371 | 2116.26731 | 2022.5267 | 1974.6852 | 2073.6453 | 5.14E-18 | -0.605029659 | YDL225W | SHS1 | n |
| W303.asm02.HP0_G0006710 | 522.55648 | 660.198682 | 628.8172001 | 2460.1285 | 2310.0767 | 2206.34141 | 2413.5789 | 2225.2931 | 2454.5189 | 1.10E-100 | -1.94490621 | YDL224C | WHI4 | n |
| W303.asm02.HP0_G0006730 | 97.979341 | 94.46239 | 134.2635804 | 31.6683208 | 36.853363 | 32.2370469 | 45.603758 | 52.483327 | 47.609203 | 1.02E-12 | 1.700246705 | YDL222C | FMP45 | n |
| W303.asm02.HP0_G0006740 | 213.86888 | 238.751096 | 226.026459 | 638.366678 | 684.03438 | 653.274274 | 582.58801 | 591.74951 | 582.77191 | 2.24E-57 | -1.541178167 | YDL220C | CDC13 | n |
| W303.asm02.HP0_G0006750 | 1251.6071 | 1171.95646 | 1509.740835 | 682.535652 | 684.93324 | 742.400227 | 749.04173 | 816.11573 | 759.98395 | 4.96E-19 | 0.899630919 | YDL219W | DTD1 | n |
| W303.asm02.HP0_G0006770 | 1300.07 | 1160.53793 | 1385.136505 | 735.871771 | 788.30243 | 704.474289 | 625.91158 | 757.07199 | 688.57014 | 3.18E-16 | 0.787243789 | YDL217C | TIM22 | n |
| W303.asm02.HP0_G0006780 | 288.67032 | 289.615459 | 244.3790348 | 207.510839 | 200.44634 | 164.02968 | 222.31832 | 187.62789 | 199.25333 | 0.000172998 | 0.52083721 | YDL216C | RRI1 | n |
| W303.asm02.HP0_G0006790 | 1371.7108 | 1385.7944 | 1342.635804 | 1593.41656 | 1544.2458 | 1468.68193 | 2621.076 | 2537.5688 | 2555.9089 | 0.008934696 | -0.168629762 | YDL215C | GDH2 | n |
| W303.asm02.HP0_G0006810 | 1808.9304 | 1910.00876 | 1798.552422 | 978.384438 | 981.55787 | 1148.20776 | 563.20641 | 665.22617 | 593.35174 | 4.18E-21 | 0.828630379 | YDL213C | NOP6 | n |
| W303.asm02.HP0_G0006820 | 5110.7288 | 4978.47937 | 6464.936283 | 1666.75373 | 1698.8502 | 1617.54123 | 2424.9798 | 2693.7067 | 2398.9749 | 2.13E-69 | 1.732177735 | YDL212W | SHR3 | n |
| W303.asm02.HP0_G0006840 | 149.60286 | 131.832127 | 134.2635804 | 88.3379476 | 88.088527 | 91.02225 | 83.226858 | 97.094155 | 82.87528 | 0.000156789 | 0.636032256 | YDL210W | UGA4 | n |
| W303.asm02.HP0_G0006850 | 365.57883 | 434.942213 | 437.5640425 | 554.195614 | 486.28462 | 481.659406 | 519.88284 | 506.4641 | 506.94985 | 0.004658688 | -0.298212994 | YDL209C | CWC2 | n |
| W303.asm02.HP0_G0006860 | 5650.142 | 5285.74165 | 6253.3987 | 4180.21835 | 4389.1457 | 4568.17917 | 3283.4706 | 4035.9678 | 3356.4488 | 2.53E-05 | 0.388005235 | YDL208W | NHP2 | n |
| W303.asm02.HP0_G0006870 | 639.49957 | 602.06798 | 566.0320726 | 1148.39332 | 1151.4429 | 1080.88922 | 1054.5869 | 1053.6028 | 1038.586 | 2.01E-31 | -0.904526939 | YDL207W | GLE1 | n |
| W303.asm02.HP0_G0006880 | 637.39249 | 607.258221 | 618.1920247 | 447.523376 | 469.20624 | 434.251985 | 584.8682 | 509.08827 | 598.64165 | 6.74E-07 | 0.463179791 | YDL206W | YCX1 | n |
| W303.asm02.HP0_G0006890 | 1562.4017 | 1455.34363 | 1581.219288 | 2597.63568 | 2734.3398 | 2918.40089 | 2541.2694 | 2709.4517 | 2556.7905 | 6.26E-33 | -0.84263563 | YDL205C | HEM3 | n |
| W303.asm02.HP0_G0006910 | 584.71542 | 676.807453 | 667.4542017 | 695.036304 | 721.78661 | 705.422438 | 706.85825 | 705.90075 | 713.2564 | 0.112449272 | -0.137013789 | YDL203C | ACK1 | n |
| W303.asm02.HP0_G0006920 | 1205.2512 | 1209.3262 | 1266.327726 | 1563.415 | 1676.3786 | 1698.13385 | 1557.3683 | 1737.1981 | 1559.6422 | 1.01E-08 | -0.423203576 | YDL202W | MRPL11 | n |
| W303.asm02.HP0_G0006930 | 1421.2272 | 1446.0012 | 1190.985573 | 1373.40507 | 1361.7767 | 1355.85227 | 1008.9831 | 1171.6903 | 1041.2309 | 0.89859595 | -0.012244712 | YDL201W | TRM8 | n |
| W303.asm02.HP0_G0006940 | 940.81238 | 949.814141 | 1120.473045 | 55.8362499 | 53.931751 | 50.2518672 | 44.463664 | 49.85916 | 59.070678 | 1.22E-170 | 4.232301926 | YDL200C | MGT1 | n |
| W303.asm02.HP0_G0006950 | 192.79806 | 236.674999 | 213.4694335 | 180.842779 | 203.14293 | 182.0445 | 320.3664 | 292.59455 | 299.76165 | 0.163683898 | 0.18428559 | YDL199C | | n |
| W303.asm02.HP0_G0006960 | 2724.4578 | 2719.68639 | 2710.385658 | 2664.30583 | 2651.6444 | 2958.22313 | 2334.9124 | 2376.1826 | 2320.5078 | 0.759338641 | -0.020668383 | YDL198C | GGC1 | n |
| W303.asm02.HP0_G0006980 | 1545.5451 | 2249.45054 | 2041.965532 | 8749.62369 | 8576.9462 | 9086.10648 | 7847.2667 | 6246.828 | 7427.0357 | 1.02E-74 | -2.177768132 | YDL195W | SEC31 | n |
| W303.asm02.HP0_G0006990 | 128.53204 | 193.076973 | 143.9228308 | 167.50875 | 147.41345 | 162.133383 | 232.57917 | 183.69164 | 201.01664 | 0.84086896 | -0.036091631 | YDL194W | SNF3 | n |
| W303.asm02.HP0_G0007000 | 1370.6572 | 1632.84988 | 1167.803372 | 1214.23009 | 1170.319 | 1123.5559 | 1235.8618 | 1300.2744 | 1206.0998 | 0.022142443 | 0.249098754 | YDL193W | NUS1 | n |
| W303.asm02.HP0_G0007030 | 1609.8111 | 1611.05087 | 1467.240134 | 2323.4547 | 2284.0097 | 2241.42291 | 2527.5883 | 2378.8068 | 2492.43 | 4.93E-17 | -0.547496428 | YDL190C | UFD2 | n |
| W303.asm02.HP0_G0007040 | 650.03498 | 722.481576 | 776.6037311 | 1158.39384 | 1082.2305 | 979.437336 | 897.25394 | 907.96155 | 953.94737 | 4.10E-10 | -0.583285458 | YDL189W | RBS1 | n |
| W303.asm02.HP0_G0007060 | 72.69435 | 66.4350875 | 56.98957728 | 39.1687126 | 53.931751 | 44.5629766 | 33.062725 | 68.228325 | 37.02938 | 0.070877019 | 0.510597341 | YDL186W | | n |
| W303.asm02.HP0_G0007070 | 3786.4274 | 4349.42213 | 4592.007634 | 4974.4265 | 4957.2268 | 5044.14969 | 5049.4761 | 4951.8019 | 4797.0681 | 0.001766687 | -0.234404841 | YDL185W | VMA1 | n |
| W303.asm02.HP0_G0007080 | 194.90514 | 241.86524 | 175.798357 | 497.525988 | 511.45278 | 438.992727 | 575.74745 | 531.39368 | 601.2866 | 3.03E-23 | -1.243524763 | YDL183C | MRX19 | n |
| W303.asm02.HP0_G0007090 | 3790.6416 | 3393.3797 | 3169.200052 | 220.011492 | 226.51336 | 207.644508 | 163.03344 | 196.81248 | 202.77994 | 0 | 3.983415145 | YDL182W | LYS20 | n |
| W303.asm02.HP0_G0007100 | 3133.2318 | 3180.57981 | 3661.821821 | 4314.39202 | 4839.4758 | 4637.39401 | 4982.2106 | 5199.7856 | 4728.2992 | 4.21E-09 | -0.466902979 | YDL181W | INH1 | n |
| W303.asm02.HP0_G0007110 | 1256.8748 | 1147.04331 | 1021.948691 | 217.511361 | 225.61449 | 261.688969 | 266.78198 | 217.80581 | 257.44236 | 1.35E-85 | 2.282609515 | YDL180W | AIT1 | n |
| W303.asm02.HP0_G0007130 | 2761.3317 | 2659.47959 | 2644.702756 | 1653.4197 | 1732.1081 | 1504.71157 | 1545.9674 | 1545.634 | 1585.2101 | 9.21E-26 | 0.721370631 | YDL178W | DLD2 | n |
| W303.asm02.HP0_G0007140 | 671.10581 | 686.149888 | 731.2052542 | 247.512929 | 226.51336 | 211.437102 | 240.55982 | 237.48705 | 245.09923 | 3.13E-55 | 1.605540273 | YDL177C | | n |
| W303.asm02.HP0_G0007150 | 453.02276 | 412.105152 | 412.4499915 | 246.679552 | 246.28833 | 198.163024 | 272.48245 | 268.97705 | 305.93321 | 1.66E-13 | 0.883792659 | YDL176W | GID12 | n |
| W303.asm02.HP0_G0007160 | 952.40134 | 1177.14671 | 1278.884751 | 1791.76026 | 1674.5809 | 1725.63016 | 1743.2037 | 2132.1352 | 1816.2029 | 2.89E-08 | -0.606686263 | YDL174C | DLD1 | n |
| W303.asm02.HP0_G0007170 | 1939.5695 | 1825.92686 | 1921.224902 | 720.03761 | 737.96613 | 757.570602 | 758.16248 | 761.00824 | 831.39775 | 1.27E-78 | 1.360428733 | YDL173W | PAR32 | n |
| W303.asm02.HP0_G0007180 | 1862.661 | 1863.29659 | 2349.129694 | 6511.17344 | 6591.3589 | 7085.51328 | 6158.7875 | 5525.1822 | 5996.1147 | 3.88E-78 | -1.731983229 | YDL171C | GLT1 | n |
| W303.asm02.HP0_G0007190 | 556.26981 | 564.698243 | 550.577272 | 823.376342 | 782.01039 | 766.103938 | 868.75159 | 829.23656 | 883.41522 | 2.15E-10 | -0.505393743 | YDL170W | UGA3 | n |
| W303.asm02.HP0_G0007200 | 246.52866 | 294.805701 | 262.7316105 | 14.1674067 | 15.280663 | 12.3259297 | 34.202819 | 26.241663 | 36.147728 | 1.38E-60 | 4.263729947 | YDL169C | UGX2 | n |
| W303.asm02.HP0_G0007210 | 1904.8027 | 1869.52488 | 2011.05593 | 7327.88276 | 7303.258 | 8059.26172 | 6202.1111 | 6132.6767 | 5942.3339 | 4.33E-208 | -1.971212885 | YDL168W | SFA1 | n |
| W303.asm02.HP0_G0007220 | 1257.9283 | 1196.86962 | 1368.71578 | 1085.89005 | 1028.2987 | 1015.46698 | 824.28793 | 918.45822 | 802.30324 | 0.000556554 | 0.288951882 | YDL167C | NRP1 | n |
| W303.asm02.HP0_G0007230 | 1907.9633 | 1914.16096 | 2113.443985 | 1353.40403 | 1222.453 | 1432.65229 | 1005.5629 | 1237.2944 | 1136.4493 | 9.32E-10 | 0.566702537 | YDL166C | FAP7 | n |
| W303.asm02.HP0_G0007240 | 1277.9456 | 1244.61984 | 1506.84306 | 1088.39018 | 1067.8487 | 1105.54108 | 1145.7944 | 1120.519 | 1128.5144 | 0.000298812 | 0.305671088 | YDL165W | CDC36 | n |
| W303.asm02.HP0_G0007250 | 602.62562 | 672.65526 | 647.1697759 | 697.536435 | 693.023 | 795.499539 | 611.09036 | 467.10161 | 613.62973 | 0.105633921 | -0.184688118 | YDL164C | CDC9 | n |
| W303.asm02.HP0_G0007260 | 2285.1311 | 2284.74418 | 1932.816002 | 2728.47585 | 2722.6546 | 2545.77856 | 2739.6458 | 2242.3501 | 2618.5062 | 0.001363734 | -0.298835063 | YDL161W | ENT1 | n |
| W303.asm02.HP0_G0007270 | 580.50126 | 556.393857 | 734.1030294 | 148.341082 | 174.37933 | 120.414852 | 194.95607 | 157.44998 | 193.08177 | 1.62E-41 | 2.077149743 | YDL160C-. | MHF2 | n |
| W303.asm02.HP0_G0007280 | 969.258 | 859.503944 | 826.8318331 | 3558.51921 | 3341.9709 | 3689.24557 | 2999.5872 | 2626.7905 | 2772.7953 | 6.72E-129 | -1.996146217 | YDL160C | DHH1 | n |
| W303.asm02.HP0_G0007290 | 428.79131 | 449.474889 | 411.4840665 | 2033.43955 | 1858.8477 | 1813.80796 | 1610.9528 | 1844.7889 | 1696.2983 | 1.83E-135 | -2.146351967 | YDL159W | STE7 | n |
| W303.asm02.HP0_G0007300 | 1625.6142 | 1517.62653 | 2020.715181 | 382.519981 | 364.03932 | 395.377899 | 311.24565 | 286.03413 | 312.10478 | 1.02E-83 | 2.177654315 | YDL157C | DMO2 | n |
| W303.asm02.HP0_G0007310 | 133.79975 | 152.593091 | 192.2190827 | 803.375297 | 764.03314 | 814.459508 | 787.80492 | 764.94449 | 831.39775 | 3.11E-90 | -2.310754046 | YDL156W | CMR1 | n |
| W303.asm02.HP0_G0007320 | 827.02992 | 822.134207 | 752.4556051 | 945.049364 | 906.95229 | 936.770657 | 1099.0506 | 1024.737 | 1143.5025 | 0.006927011 | -0.216352757 | YDL155W | CLB3 | n |
| W303.asm02.HP0_G0007330 | 639.49957 | 665.388923 | 762.1148555 | 80.8375558 | 59.324926 | 78.6963203 | 68.405637 | 69.540408 | 52.017463 | 1.04E-100 | 3.238940411 | YDL154W | MSH5 | n |
| W303.asm02.HP0_G0007340 | 1957.4797 | 1906.89462 | 1788.893172 | 2895.15123 | 2648.049 | 3066.31205 | 1557.3683 | 2056.0343 | 1704.2331 | 1.21E-09 | -0.606931602 | YDL153C | SAS10 | n |
| W303.asm02.HP0_G0007350 | 959.77612 | 951.890237 | 1011.323515 | 1442.57535 | 1361.7767 | 1499.97083 | 906.37469 | 1119.2069 | 947.77581 | 3.26E-10 | -0.557968289 | YDL150W | RPC53 | n |
| W303.asm02.HP0_G0007360 | 357.1505 | 357.088595 | 368.9833648 | 471.691305 | 426.9597 | 430.459391 | 613.37055 | 540.57827 | 618.91964 | 0.003329596 | -0.29574819 | YDL149W | ATG9 | n |
| W303.asm02.HP0_G0007370 | 2217.7044 | 1966.06337 | 1836.223498 | 2422.62654 | 2318.1664 | 2534.40077 | 1210.7798 | 1540.3856 | 1312.7797 | 0.005686535 | -0.273430115 | YDL148C | NOP14 | n |
| W303.asm02.HP0_G0007380 | 6810.091 | 6640.3946 | 6797.214497 | 3477.68165 | 3537.9229 | 3293.86767 | 3501.2285 | 3635.7825 | 3581.2701 | 1.25E-73 | 0.973613422 | YDL147W | RPN5 | n |
| W303.asm02.HP0_G0007390 | 579.44771 | 617.638704 | 673.2497519 | 608.365111 | 637.29353 | 477.866813 | 623.63139 | 551.07493 | 610.98478 | 0.327185898 | 0.11741961 | YDL146W | LDB17 | n |

| | | | | | | | | | | | | | | |
|---|---|---|---|---|---|---|---|---|---|---|---|---|---|---|
| W303.asm02.HP0_G0007400 | 2991.0038 | 3499.26062 | 3516.933066 | 7850.41006 | 7612.4667 | 8231.82474 | 7935.0539 | 7167.9104 | 7416.4559 | 8.60E-59 | -1.243304824 | YDL145C | COP1 | n |
| W303.asm02.HP0_G0007410 | 1048.2736 | 1113.82576 | 1236.384049 | 1530.07992 | 1575.706 | 1532.20788 | 2008.8455 | 1809.3627 | 1966.0838 | 1.79E-08 | -0.447874785 | YDL144C | | n |
| W303.asm02.HP0_G0007420 | 3016.2887 | 2962.58968 | 2782.830036 | 5182.77072 | 5002.1699 | 4994.84597 | 5062.0171 | 4964.9227 | 4743.2873 | 1.38E-41 | -0.793211645 | YDL143W | CCT4 | n |
| W303.asm02.HP0_G0007430 | 633.17832 | 619.7148 | 569.8957728 | 997.552106 | 988.74877 | 1023.05216 | 1014.6836 | 998.49529 | 984.80519 | 2.55E-21 | -0.724040292 | YDL142C | CRD1 | n |
| W303.asm02.HP0_G0007440 | 465.66525 | 579.230919 | 489.7239946 | 1573.41552 | 1606.2673 | 1611.85234 | 1803.6286 | 1564.0031 | 1703.3515 | 3.40E-72 | -1.642769296 | YDL141W | BPL1 | n |
| W303.asm02.HP0_G0007450 | 1545.5451 | 1831.1171 | 1867.1331 | 7152.87362 | 6858.321 | 7106.37254 | 6606.8444 | 6629.9563 | 6780.7849 | 7.50E-164 | -2.009320782 | YDL140C | RPO21 | n |
| W303.asm02.HP0_G0007460 | 202.27993 | 240.827192 | 183.5257573 | 210.844347 | 185.16568 | 195.318578 | 205.21691 | 177.13123 | 182.50195 | 0.582625133 | 0.081406075 | YDL139C | SCM3 | n |
| W303.asm02.HP0_G0007480 | 8079.6082 | 8406.11466 | 9858.230944 | 8298.76681 | 8322.5681 | 9148.68428 | 7477.8762 | 6832.0171 | 6600.9279 | 0.72781794 | 0.03194096 | YDL137W | ARF2 | y |
| W303.asm02.HP0_G0007500 | 2663.3524 | 2528.68552 | 2964.423944 | 3300.17238 | 3403.9924 | 3389.63067 | 3387.2191 | 3461.2754 | 3150.1423 | 3.22E-05 | -0.307077207 | YDL135C | RDI1 | n |
| W303.asm02.HP0_G0007510 | 3239.6395 | 3085.07937 | 3013.68612 | 3308.50615 | 3204.4449 | 3116.56392 | 3746.3487 | 3610.8529 | 3768.1803 | 0.463646575 | -0.04468837 | YDL134C | PPH21 | n |
| W303.asm02.HP0_G0007520 | 913.42031 | 824.210304 | 695.4660278 | 277.514496 | 327.18596 | 283.496383 | 465.15833 | 397.5612 | 448.76082 | 2.51E-33 | 1.453380521 | YDL133W | SRF1 | n |
| W303.asm02.HP0_G0007530 | 1569.7765 | 1756.37762 | 1733.835444 | 2225.11623 | 2129.4053 | 2400.71184 | 2421.5596 | 2272.528 | 2392.8033 | 2.19E-08 | -0.416423426 | YDL132W | CDC53 | n |
| W303.asm02.HP0_G0007540 | 3580.9869 | 3654.96786 | 2927.718792 | 1921.76705 | 1999.0703 | 2119.11176 | 2127.4153 | 1697.8356 | 2199.7215 | 4.79E-12 | 0.750788217 | YDL131W | LYS21 | n |
| W303.asm02.HP0_G0007560 | 2401.0206 | 2220.38519 | 2615.725005 | 7391.21941 | 7230.4501 | 8005.21726 | 6246.5748 | 5921.4313 | 5719.276 | 2.43E-104 | -1.644188639 | YDL130W | RPP1B | y |
| W303.asm02.HP0_G0007570 | 211.7618 | 211.761841 | 222.1627589 | 563.36276 | 497.07098 | 488.296445 | 506.20171 | 372.63162 | 405.55988 | 5.41E-23 | -1.262538084 | YDL129W | | n |
| W303.asm02.HP0_G0007580 | 1688.8267 | 1672.29572 | 1987.873729 | 2982.6558 | 2853.8885 | 2834.01568 | 3571.9143 | 3576.7387 | 3488.6966 | 3.79E-20 | -0.696398774 | YDL128W | VCX1 | n |
| W303.asm02.HP0_G0007590 | 662.67748 | 652.932344 | 550.577272 | 1724.25673 | 1676.3786 | 1517.98565 | 1683.9188 | 1746.3827 | 1958.1489 | 5.77E-48 | -1.399518446 | YDL127W | PCL2 | n |
| W303.asm02.HP0_G0007600 | 5480.5218 | 6248.01237 | 6408.912631 | 11808.1168 | 11536.002 | 12046.2259 | 13024.433 | 12139.393 | 12207.352 | 1.31E-43 | -0.964240917 | YDL126C | CDC48 | n |
| W303.asm02.HP0_G0007620 | 5971.4721 | 7355.60984 | 5492.24977 | 1380.90546 | 1393.2369 | 1357.74856 | 2256.2459 | 1909.081 | 2115.9646 | 1.21E-99 | 2.18721675 | YDL124W | | n |
| W303.asm02.HP0_G0007630 | 292.88448 | 324.9091 | 320.6871128 | 235.012276 | 269.65876 | 221.866734 | 308.96546 | 284.72205 | 320.9213 | 0.002110384 | 0.369330903 | YDL123W | SNA4 | n |
| W303.asm02.HP0_G0007640 | 2335.7011 | 2334.5705 | 2116.34176 | 2953.48761 | 2876.3601 | 3149.74911 | 3057.732 | 2695.0188 | 3061.9771 | 1.37E-07 | -0.40408401 | YDL122W | UBP1 | n |
| W303.asm02.HP0_G0007650 | 1936.4089 | 1913.12291 | 2496.916225 | 724.204495 | 695.71959 | 671.289094 | 788.94501 | 838.42115 | 761.74725 | 7.33E-52 | 1.601679347 | YDL121C | EXP1 | n |
| W303.asm02.HP0_G0007660 | 2426.3056 | 2502.73431 | 2703.624183 | 470.857928 | 509.65505 | 437.09643 | 491.38049 | 388.37662 | 421.42961 | 1.98E-145 | 2.428418061 | YDL120W | YFH1 | n |
| W303.asm02.HP0_G0007670 | 943.973 | 940.471707 | 963.0272635 | 364.18569 | 375.72453 | 355.555664 | 336.32772 | 342.45371 | 354.42407 | 1.31E-60 | 1.378021784 | YDL119C | HEM25 | n |
| W303.asm02.HP0_G0007680 | 1090.4152 | 1019.36337 | 1162.973746 | 1498.4116 | 1351.8892 | 1408.94858 | 1317.9486 | 1411.8015 | 1208.7448 | 9.19E-06 | -0.379968298 | YDL117W | CYK3 | n |
| W303.asm02.HP0_G0007690 | 1392.7816 | 1301.71249 | 1178.428547 | 1936.76783 | 1919.0715 | 1810.96352 | 1737.5032 | 1756.8794 | 1711.2864 | 3.00E-13 | -0.55002998 | YDL116W | NUP84 | n |
| W303.asm02.HP0_G0007700 | 946.08009 | 926.97708 | 963.9931885 | 618.365633 | 575.27201 | 596.385367 | 573.46726 | 497.27952 | 549.26914 | 6.91E-16 | 0.66417438 | YDL115C | IWR1 | n |
| W303.asm02.HP0_G0007720 | 642.66019 | 571.964581 | 583.4187233 | 1028.38705 | 1074.1407 | 1043.91143 | 1406.8759 | 1275.3448 | 1363.0339 | 6.28E-24 | -0.807664716 | YDL113C | ATG20 | n |
| W303.asm02.HP0_G0007730 | 1388.5674 | 1331.81589 | 1605.367414 | 1905.93289 | 1980.1941 | 2124.80065 | 1460.4604 | 1512.8319 | 1482.9385 | 1.28E-08 | -0.473738881 | YDL112W | TRM3 | n |
| W303.asm02.HP0_G0007740 | 1972.2293 | 1906.89462 | 1614.06074 | 1040.8877 | 1030.0965 | 1032.53365 | 799.20586 | 980.12613 | 898.4033 | 6.85E-18 | 0.823296059 | YDL111C | RRP42 | n |
| W303.asm02.HP0_G0007750 | 1663.5417 | 1819.69857 | 1579.287438 | 225.84513 | 248.08606 | 256.948227 | 321.50649 | 352.95037 | 322.6846 | 4.48E-183 | 2.793716037 | YDL110C | TMA17 | n |
| W303.asm02.HP0_G0007770 | 839.67242 | 780.612278 | 808.4792573 | 502.526249 | 488.98121 | 470.281625 | 436.65598 | 516.96077 | 461.10395 | 2.61E-16 | 0.731785272 | YDL108W | KIN28 | n |
| W303.asm02.HP0_G0007780 | 1147.3065 | 1108.63552 | 1239.281825 | 825.043095 | 810.774 | 750.933563 | 703.43797 | 751.82366 | 790.84177 | 3.46E-11 | 0.550131745 | YDL107W | MSS2 | n |
| W303.asm02.HP0_G0007790 | 764.87098 | 746.356686 | 623.9875749 | 1507.57875 | 1441.7755 | 1305.6004 | 1358.992 | 1250.4153 | 1342.7559 | 3.57E-26 | -0.996247573 | YDL106C | PHO2 | n |
| W303.asm02.HP0_G0007800 | 358.20404 | 338.403727 | 283.0160363 | 615.865502 | 597.74358 | 585.955735 | 559.78613 | 524.83327 | 550.15079 | 6.51E-18 | -0.879726427 | YDL105W | NSE4 | n |
| W303.asm02.HP0_G0007810 | 702.71205 | 679.921598 | 782.3992813 | 456.690521 | 459.31875 | 444.681617 | 413.8541 | 468.41369 | 448.76082 | 6.00E-13 | 0.670560449 | YDL104C | QRI7 | n |
| W303.asm02.HP0_G0007820 | 864.95741 | 919.710742 | 1056.721992 | 2189.28102 | 2249.8529 | 2260.38588 | 2207.2219 | 2297.4576 | 2286.1234 | 4.51E-56 | -1.236309983 | YDL103C | QRI1 | n |
| W303.asm02.HP0_G0007830 | 991.38236 | 864.694185 | 915.6969366 | 1890.09873 | 1846.2636 | 1849.8376 | 1610.9528 | 1663.7215 | 1650.4524 | 2.80E-47 | -1.011406455 | YDL102W | POL3 | n |
| W303.asm02.HP0_G0007840 | 1058.809 | 966.422913 | 869.3325348 | 648.3672 | 661.56282 | 615.348336 | 591.70876 | 631.112 | 633.02607 | 1.45E-10 | 0.587193879 | YDL101C | DUN1 | n |
| W303.asm02.HP0_G0007850 | 6041.0058 | 6432.78495 | 5177.358207 | 4437.7318 | 4485.324 | 4645.92735 | 4690.3465 | 4702.5061 | 4369.4669 | 3.52E-06 | 0.379348691 | YDL100C | GET3 | n |
| W303.asm02.HP0_G0007860 | 1625.6142 | 1830.07905 | 1474.001609 | 1275.89998 | 1386.046 | 1428.8597 | 1103.6109 | 1230.734 | 1207.8631 | 0.003774755 | 0.269142793 | YDL099W | BUG1 | n |
| W303.asm02.HP0_G0007870 | 677.42706 | 691.340129 | 737.9667295 | 150.841212 | 173.48047 | 171.614867 | 178.99475 | 169.25873 | 177.21203 | 3.96E-81 | 2.089362863 | YDL098C | SNU23 | n |
| W303.asm02.HP0_G0007880 | 4758.8461 | 4792.66873 | 4170.864317 | 3993.54193 | 4015.2189 | 3918.69749 | 4157.9226 | 4230.1561 | 3958.6171 | 0.003615246 | 0.201986198 | YDL097C | RPN6 | n |
| W303.asm02.HP0_G0007890 | 3061.591 | 3566.73376 | 3547.842667 | 9933.01884 | 9808.3878 | 10564.2699 | 12580.937 | 12483.159 | 11715.391 | 5.30E-104 | -1.574173563 | YDL095W | PMT1 | n |
| W303.asm02.HP0_G0007910 | 2342.0223 | 2363.63585 | 2840.785539 | 1262.56595 | 1297.0586 | 1315.08188 | 1254.1033 | 1361.9423 | 1327.7678 | 2.85E-29 | 0.962268155 | YDL092W | SRP14 | n |
| W303.asm02.HP0_G0007920 | 597.35792 | 683.035743 | 547.6794969 | 684.202405 | 684.03438 | 617.244633 | 941.7176 | 837.10906 | 948.65746 | 0.248817238 | -0.120480136 | YDL091C | UBX3 | n |
| W303.asm02.HP0_G0007930 | 919.74155 | 865.732233 | 744.7282048 | 449.19013 | 427.85856 | 439.940875 | 457.17767 | 495.96744 | 491.96177 | 1.47E-21 | 0.940922311 | YDL090C | RAM1 | n |
| W303.asm02.HP0_G0007940 | 1014.5603 | 1060.8853 | 1020.982766 | 720.870987 | 707.40481 | 657.066867 | 859.63084 | 809.55532 | 836.68766 | 7.39E-14 | 0.569424161 | YDL089W | NUR1 | n |
| W303.asm02.HP0_G0007950 | 1712.0046 | 1710.7035 | 1612.128889 | 2492.6302 | 2446.7038 | 2450.01556 | 2307.5502 | 2452.2834 | 2316.0996 | 9.72E-20 | -0.553915047 | YDL088C | ASM4 | n |
| W303.asm02.HP0_G0007960 | 214.92243 | 177.506249 | 211.5375835 | 650.033954 | 616.61969 | 516.740899 | 517.60265 | 574.69243 | 501.65994 | 1.16E-36 | -1.562697517 | YDL087C | LUC7 | n |
| W303.asm02.HP0_G0007970 | 997.70361 | 1243.58179 | 1025.812391 | 1376.73858 | 1344.6983 | 1310.34114 | 1337.3302 | 1343.5732 | 1332.176 | 0.000601649 | -0.303829494 | YDL086W | | n |
| W303.asm02.HP0_G0007980 | 723.78287 | 739.090348 | 844.2184838 | 1045.05459 | 1052.568 | 1073.30403 | 822.00774 | 905.33739 | 844.62253 | 4.57E-08 | -0.457712777 | YDL085C-A | | n |
| W303.asm02.HP0_G0008000 | 9556.6731 | 8914.7583 | 8629.574295 | 9137.14394 | 9407.4952 | 9789.63262 | 8559.8254 | 8584.9602 | 8343.9537 | 0.290839381 | -0.064209002 | YDL084W | SUB2 | n |
| W303.asm02.HP0_G0008030 | 9262.7351 | 9164.92792 | 10428.12672 | 8791.29254 | 9184.5773 | 9654.04739 | 7408.3305 | 7372.5953 | 6771.0867 | 0.401939202 | 0.06279587 | YDL081C | RPP1A | n |
| W303.asm02.HP0_G0008040 | 488.84316 | 471.273902 | 565.0661476 | 785.874383 | 772.12291 | 835.318774 | 767.28323 | 825.30031 | 758.22064 | 2.75E-12 | -0.648412293 | YDL080C | THI3 | n |
| W303.asm02.HP0_G0008050 | 259.17116 | 233.560854 | 281.0841862 | 570.029775 | 573.47429 | 590.696477 | 825.42802 | 679.65908 | 850.7941 | 8.85E-25 | -1.162822747 | YDL079C | MRK1 | n |
| W303.asm02.HP0_G0008070 | 3213.301 | 3129.71545 | 3137.324525 | 2544.29957 | 2393.6709 | 2726.87491 | 2900.399 | 2864.2776 | 2974.6936 | 2.14E-06 | 0.306869382 | YDL078C | MDH3 | n |
| W303.asm02.HP0_G0008080 | 485.68254 | 570.926533 | 584.3846484 | 659.201099 | 675.04575 | 605.866852 | 640.7328 | 671.78658 | 678.87197 | 0.012765617 | -0.241213312 | YDL077C | VAM6 | n |
| W303.asm02.HP0_G0008090 | 871.27865 | 848.085413 | 792.0585317 | 561.696006 | 569.87884 | 539.496461 | 529.00359 | 406.74578 | 528.1095 | 1.11E-08 | 0.587134572 | YDL076C | RXT3 | n |
| W303.asm02.HP0_G0008110 | 2226.1328 | 2303.42905 | 2129.86471 | 1760.09194 | 1854.3534 | 2071.70434 | 1856.073 | 1695.2115 | 1801.2149 | 0.003777719 | 0.228342732 | YDL074C | BRE1 | n |

| | | | | | | | | | | | | | | |
|---|---|---|---|---|---|---|---|---|---|---|---|---|---|---|
| W303.asm02.HP0_G0008120 | 1860.5539 | 1559.14846 | 1803.382047 | 729.204756 | 719.09002 | 835.318774 | 743.34126 | 762.32032 | 716.783 | 1.51E-38 | 1.194411999 | YDL073W | AHK1 | n |
| W303.asm02.HP0_G0008130 | 2723.4043 | 2577.47378 | 2959.594318 | 1640.08567 | 1644.9184 | 1657.36347 | 2445.5015 | 2621.5422 | 2519.7612 | 1.12E-26 | 0.741364772 | YDL072C | YET3 | n |
| W303.asm02.HP0_G0008160 | 2769.7601 | 2751.86589 | 3194.314103 | 2666.80596 | 2671.4194 | 3044.50463 | 2655.2788 | 2233.1656 | 2380.4602 | 0.554095405 | 0.056675648 | YDL067C | COX9 | n |
| W303.asm02.HP0_G0008170 | 1667.7559 | 1812.43223 | 1832.359798 | 1715.08959 | 1671.8843 | 1707.61534 | 1722.682 | 1613.8623 | 1716.5763 | 0.379291638 | 0.060696685 | YDL066W | IDP1 | n |
| W303.asm02.HP0_G0008180 | 1843.6973 | 1745.99714 | 1758.949495 | 1169.22774 | 1201.7792 | 1255.34853 | 1034.0652 | 1119.2069 | 1119.6979 | 3.70E-16 | 0.560997853 | YDL065C | PEX19 | n |
| W303.asm02.HP0_G0008190 | 1512.8853 | 1380.60416 | 1473.035684 | 1150.06007 | 1250.3178 | 1195.61518 | 1065.9878 | 1243.8548 | 1089.7218 | 0.000590296 | 0.280336056 | YDL064W | UBC9 | n |
| W303.asm02.HP0_G0008200 | 2189.2588 | 1789.59517 | 2007.19223 | 1876.7647 | 1903.7908 | 2063.171 | 1207.3595 | 1369.8148 | 1208.7448 | 0.712230811 | 0.034903865 | YDL063C | SYO1 | n |
| W303.asm02.HP0_G0008220 | 1383.2997 | 1412.78366 | 1556.105237 | 5040.26327 | 4692.0624 | 5120.00156 | 3099.9155 | 3694.8262 | 3106.9413 | 1.41E-95 | -1.77050302 | YDL060W | TSR1 | n |
| W303.asm02.HP0_G0008230 | 427.73777 | 387.191994 | 393.1314907 | 146.674328 | 189.65999 | 199.111172 | 174.43437 | 186.31581 | 182.50195 | 1.98E-19 | 1.177141214 | YDL059C | RAD59 | n |
| W303.asm02.HP0_G0008240 | 1238.9646 | 1156.38574 | 1159.110046 | 3386.0102 | 3640.3932 | 3555.55664 | 2995.0268 | 2760.623 | 2877.7118 | 2.73E-125 | -1.574027796 | YDL058W | USO1 | n |
| W303.asm02.HP0_G0008250 | 245.47512 | 216.952082 | 235.6857094 | 800.041789 | 773.92063 | 786.015055 | 816.30727 | 688.84366 | 752.93073 | 1.12E-68 | -1.757343227 | YDL057W | | n |
| W303.asm02.HP0_G0008260 | 566.80522 | 606.220173 | 715.7504536 | 1275.0666 | 1235.0371 | 1177.60036 | 1176.577 | 1148.0728 | 1208.7448 | 3.72E-26 | -0.964213247 | YDL056W | MBP1 | n |
| W303.asm02.HP0_G0008280 | 702.71205 | 652.932344 | 604.6690742 | 849.211024 | 924.03067 | 844.800258 | 1105.8911 | 1127.0794 | 1144.3842 | 6.06E-07 | -0.418310714 | YDL054C | MCH1 | n |
| W303.asm02.HP0_G0008290 | 1752.0392 | 1714.85569 | 1850.712374 | 718.370857 | 708.30367 | 658.015016 | 686.33656 | 743.95116 | 715.90135 | 9.64E-73 | 1.350430173 | YDL053C | PBP4 | n |
| W303.asm02.HP0_G0008300 | 772.24577 | 861.58004 | 842.2866337 | 2085.94229 | 2002.6657 | 2161.77844 | 2524.168 | 2462.7801 | 2473.0336 | 2.25E-82 | -1.335359446 | YDL052C | SLC1 | n |
| W303.asm02.HP0_G0008310 | 4306.8768 | 4132.47005 | 3973.815609 | 2837.64822 | 3050.7394 | 3063.4676 | 2574.3321 | 2785.5526 | 2672.2869 | 1.64E-13 | 0.47172104 | YDL051W | LHP1 | n |
| W303.asm02.HP0_G0008340 | 1321.1408 | 1348.42467 | 1558.037087 | 1640.08567 | 1592.7844 | 1627.02272 | 1688.4791 | 1578.4361 | 1570.2221 | 0.012021017 | -0.200467148 | YDL047W | SIT4 | n |
| W303.asm02.HP0_G0008350 | 4120.4 | 3827.28387 | 3657.958121 | 1033.38731 | 1003.1306 | 1099.85219 | 1186.8378 | 885.65614 | 1184.0585 | 4.18E-75 | 1.887767993 | YDL046W | NPC2 | n |
| W303.asm02.HP0_G0008360 | 1407.5312 | 1410.70756 | 1411.216481 | 957.550017 | 926.72726 | 938.666953 | 1014.6836 | 1066.7236 | 1009.4914 | 1.47E-18 | 0.583077353 | YDL045W- | MRP10 | n |
| W303.asm02.HP0_G0008370 | 981.90049 | 943.585851 | 866.4347596 | 546.695223 | 496.17211 | 540.44461 | 522.16303 | 476.28619 | 536.04436 | 9.10E-20 | 0.817567518 | YDL045C | FAD1 | n |
| W303.asm02.HP0_G0008380 | 1027.2028 | 1071.26579 | 1016.153141 | 843.377386 | 807.17855 | 870.400266 | 725.09975 | 712.46116 | 785.55185 | 7.79E-05 | 0.305133891 | YDL044C | MTF2 | n |
| W303.asm02.HP0_G0008390 | 1258.9819 | 1149.1194 | 1239.281825 | 518.360409 | 569.87884 | 473.12607 | 427.53523 | 541.89035 | 472.56542 | 1.16E-33 | 1.223605954 | YDL043C | PRP11 | n |
| W303.asm02.HP0_G0008400 | 632.12478 | 681.997695 | 663.5905015 | 2314.28755 | 2159.9666 | 2256.59328 | 2268.787 | 2205.6118 | 2236.7509 | 1.97E-148 | -1.766888148 | YDL042C | SIR2 | n |
| W303.asm02.HP0_G0008410 | 1445.4587 | 1509.32214 | 1424.739432 | 4081.87988 | 3963.0849 | 4103.58644 | 3883.16 | 3710.5712 | 3895.1381 | 4.88E-142 | -1.472132033 | YDL040C | NAT1 | n |
| W303.asm02.HP0_G0008440 | 1027.2028 | 973.68925 | 1039.335342 | 2785.97886 | 2695.6887 | 2419.67481 | 2885.5778 | 2777.6801 | 2794.8366 | 1.43E-78 | -1.378135937 | YDL035C | GPR1 | n |
| W303.asm02.HP0_G0008450 | 297.09865 | 274.044736 | 288.8115865 | 270.847481 | 290.33259 | 277.807492 | 216.61785 | 295.21871 | 245.09923 | 0.786371834 | 0.035897339 | YDL033C | SLM3 | n |
| W303.asm02.HP0_G0008460 | 1076.7192 | 1111.74967 | 1176.496697 | 2508.46436 | 2485.3549 | 2823.58605 | 1392.0547 | 1750.3189 | 1407.9981 | 5.68E-36 | -1.215653602 | YDL031W | DBP10 | n |
| W303.asm02.HP0_G0008470 | 719.56871 | 625.94309 | 671.3179018 | 517.527032 | 565.38453 | 492.089039 | 515.32247 | 505.15202 | 510.47646 | 0.000130598 | 0.356283244 | YDL030W | PRP9 | n |
| W303.asm02.HP0_G0008480 | 3879.1391 | 3630.0547 | 3703.356598 | 2700.97442 | 2682.2058 | 2692.74156 | 2630.1967 | 2761.9351 | 2612.3346 | 8.06E-17 | 0.473328486 | YDL029W | ARP2 | n |
| W303.asm02.HP0_G0008490 | 499.37858 | 510.719735 | 503.2469451 | 665.868114 | 680.43893 | 713.955774 | 809.46671 | 732.14241 | 729.12613 | 2.16E-07 | -0.444514789 | YDL028C | MPS1 | n |
| W303.asm02.HP0_G0008510 | 527.82419 | 640.475765 | 640.4083006 | 914.21442 | 802.68423 | 823.940992 | 877.87234 | 753.13574 | 1025.3612 | 5.08E-05 | -0.490235519 | YDL025C | RTK1 | n |
| W303.asm02.HP0_G0008520 | 89.55101 | 94.46239 | 96.59250386 | 292.515279 | 287.63601 | 240.829703 | 103.74855 | 175.81914 | 89.046843 | 6.40E-14 | -1.549039103 | YDL024C | DIA3 | n |
| W303.asm02.HP0_G0008530 | 1703.5763 | 1978.51995 | 1965.657454 | 5021.0956 | 4778.3532 | 4377.60134 | 4366.5598 | 4594.9153 | 4920.4993 | 1.99E-57 | -1.327740385 | YDL022W | GPD1 | n |
| W303.asm02.HP0_G0008550 | 1699.3621 | 2081.28672 | 1371.613555 | 3294.33874 | 3297.0277 | 3298.60842 | 3794.2327 | 3318.2583 | 3689.7133 | 1.69E-15 | -0.941161501 | YDL020C | RPN4 | n |
| W303.asm02.HP0_G0008570 | 1114.6467 | 1134.58673 | 1344.567654 | 182.509533 | 174.37933 | 149.807453 | 165.31362 | 174.50706 | 177.21203 | 5.90E-129 | 2.824276709 | YDL018C | ERP3 | n |
| W303.asm02.HP0_G0008580 | 251.79637 | 257.435964 | 241.4812597 | 532.527816 | 475.49827 | 457.007547 | 540.40453 | 485.47077 | 513.12141 | 1.03E-19 | -0.966462881 | YDL017W | CDC7 | n |
| W303.asm02.HP0_G0008590 | 5512.1281 | 5247.33386 | 5990.66709 | 2145.94542 | 2125.8099 | 2249.00809 | 2819.4523 | 2946.9388 | 2744.5824 | 1.83E-88 | 1.36136963 | YDL015C | TSC13 | n |
| W303.asm02.HP0_G0008600 | 4085.6332 | 4242.50316 | 4759.112665 | 9421.32545 | 8537.3962 | 9738.4326 | 6563.5209 | 7305.6791 | 6411.3727 | 3.10E-39 | -1.081391384 | YDL014W | NOP1 | n |
| W303.asm02.HP0_G0008610 | 406.66694 | 484.768529 | 429.8366422 | 602.531473 | 535.72206 | 441.837172 | 598.54932 | 528.76952 | 548.38749 | 0.036897531 | -0.259682607 | YDL013W | SLX5 | n |
| W303.asm02.HP0_G0008630 | 1340.1045 | 1421.08804 | 1717.414719 | 270.847481 | 282.24283 | 282.548234 | 318.08621 | 321.46038 | 342.08094 | 7.98E-110 | 2.423109539 | YDL008W | APC11 | n |
| W303.asm02.HP0_G0008650 | 5077.0155 | 4796.82092 | 4694.395688 | 3371.00941 | 3417.4753 | 3429.4529 | 3527.4507 | 3437.6579 | 3403.1764 | 2.48E-20 | 0.511643306 | YDL007W | RPT2 | n |
| W303.asm02.HP0_G0008660 | 1105.1648 | 1099.29309 | 1165.871522 | 1328.40272 | 1399.5289 | 1408.94858 | 1176.577 | 1255.6636 | 1240.4842 | 2.22E-05 | -0.294911879 | YDL006W | PTC1 | n |
| W303.asm02.HP0_G0008670 | 1225.2685 | 1017.28728 | 1068.313093 | 762.53983 | 721.78661 | 736.711336 | 744.48135 | 734.76657 | 759.1023 | 3.25E-11 | 0.575370487 | YDL005C | MED2 | n |
| W303.asm02.HP0_G0008680 | 5624.857 | 5337.64406 | 6363.514154 | 4444.39882 | 4464.6502 | 4671.52735 | 4053.034 | 4180.297 | 3725.861 | 7.18E-06 | 0.351614663 | YDL004W | ATP16 | n |
| W303.asm02.HP0_G0008690 | 1337.9975 | 1285.10372 | 1253.7707 | 2103.4432 | 2150.0792 | 1988.26727 | 1984.9036 | 2066.531 | 2069.237 | 5.91E-26 | -0.687617056 | YDL003W | MCD1 | n |
| W303.asm02.HP0_G0008700 | 1618.2394 | 1625.58355 | 1799.518347 | 1324.23584 | 1385.1471 | 1399.46709 | 1166.3161 | 1216.3011 | 1304.8448 | 9.50E-05 | 0.296335895 | YDL002C | NHP10 | n |
| W303.asm02.HP0_G0008710 | 1207.3583 | 1048.42872 | 1200.644823 | 1099.22408 | 1039.0851 | 1059.08181 | 1154.9152 | 1062.7874 | 1131.1594 | 0.171314676 | 0.112362411 | YDL001W | RMD1 | n |
| W303.asm02.HP0_G0008720 | 1075.6657 | 1244.61984 | 1186.155947 | 1840.09612 | 1706.9399 | 1854.57834 | 1925.6187 | 1911.7052 | 1817.9662 | 2.12E-16 | -0.623159477 | YDR001C | NTH1 | n |
| W303.asm02.HP0_G0008730 | 8962.4758 | 8446.59854 | 10910.12331 | 5976.97887 | 5927.0995 | 6295.70563 | 4877.3219 | 5467.4506 | 4876.4167 | 9.19E-12 | 0.637994967 | YDR002W | YRB1 | n |
| W303.asm02.HP0_G0008750 | 1177.8592 | 1141.85307 | 1147.518946 | 578.363544 | 538.41865 | 505.363117 | 493.66068 | 410.68203 | 478.73699 | 9.68E-35 | 1.094664424 | YDR004W | RAD57 | n |
| W303.asm02.HP0_G0008770 | 916.58093 | 1050.50482 | 1141.723396 | 921.714811 | 871.89665 | 839.111367 | 881.29262 | 747.88741 | 934.55103 | 0.023995557 | 0.239970376 | YDR006C | SOK1 | n |
| W303.asm02.HP0_G0008780 | 418.2559 | 428.713924 | 450.121068 | 1390.07261 | 1354.5858 | 1296.11891 | 1324.7892 | 742.63907 | 1283.6852 | 2.04E-24 | -1.639232192 | YDR007W | TRP1 | n |
| W303.asm02.HP0_G0008790 | 144.33516 | 131.832127 | 124.60433 | 185.009664 | 159.09867 | 182.0445 | 299.84471 | 238.79914 | 287.41852 | 0.008436843 | -0.394388046 | YDR009W | GAL3 | n |
| W303.asm02.HP0_G0008800 | 3389.2424 | 3700.64198 | 3515.001216 | 6039.48213 | 5906.4256 | 6353.54268 | 5911.3871 | 8893.2997 | 5623.1759 | 4.90E-09 | -0.787051877 | YDR011W | SNQ2 | n |
| W303.asm02.HP0_G0008810 | 5413.0952 | 5584.69954 | 5559.864522 | 24542.1153 | 24293.557 | 27142.6453 | 20381.46 | 21590.329 | 19532.116 | 1.80E-251 | -2.198029651 | YDR012W | RPL4B | y |
| W303.asm02.HP0_G0008820 | 845.99366 | 788.916664 | 1038.369417 | 498.359365 | 532.12661 | 516.740899 | 464.01824 | 467.10161 | 482.2636 | 1.65E-13 | 0.790066387 | YDR013W | PSF1 | n |
| W303.asm02.HP0_G0008830 | 755.38911 | 690.302081 | 753.4215301 | 308.34944 | 331.68027 | 347.970477 | 434.3758 | 383.12829 | 435.53604 | 1.65E-32 | 1.155888992 | YDR014W | RAD61 | n |
| W303.asm02.HP0_G0008850 | 559.43043 | 533.556796 | 658.7608763 | 153.341343 | 176.17705 | 140.325969 | 194.95607 | 209.93331 | 209.83316 | 1.16E-49 | 1.898580059 | YDR016C | DAD1 | n |
| W303.asm02.HP0_G0008860 | 1100.9507 | 1187.52719 | 1133.995995 | 838.377125 | 874.59323 | 924.444727 | 820.86764 | 856.79031 | 861.37392 | 5.76E-07 | 0.376682039 | YDR017C | KCS1 | n |
| W303.asm02.HP0_G0008880 | 383.48903 | 397.572476 | 407.6203663 | 1316.73544 | 1177.5099 | 1211.7337 | 1455.9 | 1246.479 | 1571.9854 | 6.01E-60 | -1.640433339 | YDR019C | GCV1 | n |

| | | | | | | | | | | | | | | |
|---|---|---|---|---|---|---|---|---|---|---|---|---|---|---|
| W303.asm02.HP0_G0008890 | 1065.1303 | 948.776093 | 1187.121872 | 611.698618 | 631.00149 | 634.311305 | 352.28903 | 414.61828 | 349.13416 | 2.96E-14 | 0.770843284 | YDR020C | DAS2 | n |
| W303.asm02.HP0_G0008900 | 949.24071 | 861.58004 | 957.2317133 | 796.708282 | 671.4503 | 814.459508 | 503.92153 | 574.69243 | 509.59481 | 0.005047632 | 0.27838639 | YDR021W | FAL1 | n |
| W303.asm02.HP0_G0008910 | 520.4494 | 510.719735 | 503.2469451 | 114.17263 | 110.56009 | 93.8666953 | 163.03344 | 140.3929 | 178.09369 | 8.08E-68 | 2.264587862 | YDR022C | ATG31 | n |
| W303.asm02.HP0_G0008920 | 10521.717 | 9791.90906 | 10121.92848 | 5361.94674 | 5104.6403 | 5579.85356 | 4213.7872 | 4188.1695 | 4054.7171 | 4.60E-57 | 0.923562029 | YDR023W | SES1 | n |
| W303.asm02.HP0_G0008950 | 1354.8541 | 1389.9466 | 1368.71578 | 1779.2596 | 1721.3217 | 1714.25238 | 1747.764 | 1623.0469 | 1754.4873 | 1.10E-07 | -0.342463573 | YDR027C | VPS54 | n |
| W303.asm02.HP0_G0008960 | 1542.3845 | 1678.52401 | 1683.607342 | 1925.10056 | 1801.3205 | 1669.6894 | 2147.937 | 2046.8497 | 2101.8582 | 0.068732957 | -0.138150314 | YDR028C | REG1 | n |
| W303.asm02.HP0_G0008990 | 3953.9405 | 3782.64779 | 4269.388671 | 1449.24237 | 1378.8551 | 1369.12634 | 1682.7787 | 1508.8956 | 1578.1569 | 2.59E-98 | 1.516115766 | YDR032C | PST2 | n |
| W303.asm02.HP0_G0009010 | 831.24409 | 800.335194 | 780.4674312 | 871.7122 | 899.76139 | 927.289172 | 761.58276 | 736.07866 | 754.69404 | 0.032352817 | -0.161928454 | YDR034C | LYS14 | n |
| W303.asm02.HP0_G0009030 | 5555.3233 | 4961.87059 | 5158.039706 | 2343.45574 | 2354.1209 | 2432.94889 | 2261.9464 | 1760.8156 | 2010.1664 | 5.26E-39 | 1.136450567 | YDR035W | ARO3 | n |
| W303.asm02.HP0_G0009040 | 714.301 | 855.351751 | 812.3429575 | 1891.76548 | 1814.8034 | 1863.11168 | 1739.7834 | 1734.5739 | 1728.0377 | 1.18E-58 | -1.225066584 | YDR036C | EHD3 | n |
| W303.asm02.HP0_G0009050 | 6162.1631 | 6202.33824 | 6553.801387 | 10034.6908 | 10038.497 | 10412.5661 | 8005.7397 | 8498.3627 | 7842.2938 | 3.27E-34 | -0.688203271 | YDR037W | KRS1 | n |
| W303.asm02.HP0_G0009090 | 12.642496 | 11.4185307 | 5.795550232 | 1645.91931 | 1583.7958 | 1819.49685 | 2756.7472 | 2860.3413 | 3126.3377 | 7.53E-153 | -7.417500218 | YDR040C | ENA1 | y |
| W303.asm02.HP0_G0009100 | 2536.9274 | 2452.90799 | 2882.320315 | 1273.39985 | 1276.3848 | 1296.11891 | 1292.8665 | 1327.8282 | 1248.4191 | 5.35E-42 | 1.033874987 | YDR041W | RSM10 | n |
| W303.asm02.HP0_G0009120 | 614.21458 | 706.910852 | 640.4083006 | 291.681902 | 225.61449 | 231.348219 | 229.15888 | 190.25206 | 226.58454 | 3.04E-29 | 1.387187245 | YDR043C | NRG1 | n |
| W303.asm02.HP0_G0009140 | 1559.2411 | 1550.84407 | 2116.34176 | 861.711677 | 906.05342 | 939.615102 | 695.45731 | 838.42115 | 721.19126 | 3.64E-15 | 0.949649918 | YDR045C | RPC11 | n |
| W303.asm02.HP0_G0009150 | 3297.5843 | 3161.89494 | 3420.340562 | 3252.6699 | 3157.704 | 2812.20827 | 2173.0191 | 1930.0743 | 2102.7398 | 0.207339751 | 0.099107904 | YDR046C | BAP3 | n |
| W303.asm02.HP0_G0009160 | 2938.3267 | 2752.90394 | 3026.243146 | 2752.64378 | 2653.4422 | 2739.20084 | 2570.9119 | 2571.683 | 2451.874 | 0.118873136 | 0.098087575 | YDR047W | HEM12 | n |
| W303.asm02.HP0_G0009170 | 640.55311 | 609.334318 | 611.4305494 | 1029.22043 | 1006.726 | 933.926211 | 881.29262 | 915.83405 | 852.5574 | 1.92E-17 | -0.674964691 | YDR049W | VMS1 | n |
| W303.asm02.HP0_G0009180 | 25648.463 | 24892.3968 | 26755.15764 | 13338.1967 | 12798.005 | 13952.0043 | 12668.724 | 13107.711 | 11798.266 | 1.39E-51 | 0.947264692 | YDR050C | TPI1 | n |
| W303.asm02.HP0_G0009190 | 1911.1239 | 1763.64396 | 2009.12408 | 453.357014 | 447.63354 | 414.340867 | 503.92153 | 482.84661 | 514.88472 | 2.37E-139 | 2.11077177 | YDR051C | DET1 | n |
| W303.asm02.HP0_G0009200 | 907.09906 | 981.993636 | 905.0717612 | 1323.40246 | 1266.4973 | 1333.0967 | 1209.6397 | 1153.3211 | 1166.4255 | 5.77E-12 | -0.489750273 | YDR052C | DBF4 | n |
| W303.asm02.HP0_G0009210 | 3174.3199 | 2921.06775 | 2812.773712 | 1516.74589 | 1427.3937 | 1438.34118 | 1287.1661 | 1184.8111 | 1263.4072 | 1.22E-46 | 1.022864929 | YDR054C | CDC34 | n |
| W303.asm02.HP0_G0009230 | 2563.266 | 2574.35964 | 3050.391272 | 717.53748 | 697.51732 | 668.444649 | 850.51009 | 863.35073 | 906.33817 | 3.26E-115 | 1.974365617 | YDR056C | EMC10 | n |
| W303.asm02.HP0_G0009240 | 809.11972 | 814.86787 | 766.9444807 | 803.375297 | 838.63873 | 801.18543 | 989.60155 | 925.01863 | 960.11893 | 0.691268777 | -0.031620821 | YDR057W | YOS9 | n |
| W303.asm02.HP0_G0009250 | 525.71711 | 449.474889 | 566.9979977 | 540.861585 | 592.3504 | 467.43718 | 591.70876 | 587.81326 | 619.80129 | 0.654512868 | -0.053654489 | YDR058C | TGL2 | n |
| W303.asm02.HP0_G0009270 | 2921.47 | 2688.54495 | 2768.341161 | 2906.8185 | 3014.7849 | 3006.5787 | 1388.6344 | 1815.9231 | 1591.3817 | 0.313722831 | -0.091680843 | YDR060W | MAK21 | n |
| W303.asm02.HP0_G0009280 | 1252.6606 | 1169.88037 | 1117.57527 | 404.187779 | 402.69041 | 400.118641 | 584.8682 | 429.0512 | 560.73062 | 7.17E-47 | 1.55204222 | YDR061W | | n |
| W303.asm02.HP0_G0009290 | 2949.9156 | 2718.64834 | 2700.726408 | 1571.74877 | 1674.5809 | 1670.63755 | 2141.0964 | 2037.6652 | 2096.5682 | 8.75E-32 | 0.767492849 | YDR062W | LCB2 | n |
| W303.asm02.HP0_G0009300 | 2100.7614 | 2089.59111 | 2294.071967 | 1047.55472 | 1056.1635 | 1027.79291 | 1113.8718 | 1107.3982 | 1165.5438 | 1.12E-52 | 1.050334556 | YDR063W | AIM7 | n |
| W303.asm02.HP0_G0009320 | 788.04889 | 694.454274 | 845.1844088 | 435.022723 | 427.85856 | 398.222344 | 370.53053 | 427.73911 | 389.69015 | 4.85E-18 | 0.884049813 | YDR065W | RRG1 | n |
| W303.asm02.HP0_G0009340 | 1173.645 | 1201.02182 | 1272.123276 | 269.180727 | 350.55638 | 330.903805 | 347.72866 | 321.46038 | 331.50112 | 3.20E-80 | 1.94208495 | YDR067C | OCA6 | n |
| W303.asm02.HP0_G0009350 | 1704.6298 | 1719.00789 | 1875.826425 | 1025.05354 | 1041.7817 | 1017.36327 | 961.0992 | 1012.9282 | 963.64554 | 7.59E-29 | 0.781320564 | YDR068W | DOS2 | n |
| W303.asm02.HP0_G0009360 | 994.54299 | 925.939031 | 814.2748076 | 933.382088 | 889.8739 | 930.133618 | 868.75159 | 805.61907 | 908.10147 | 0.911996505 | -0.010667366 | YDR069C | DOA4 | n |
| W303.asm02.HP0_G0009370 | 210.70826 | 265.74035 | 236.6516345 | 7.50039177 | 11.685213 | 6.63703906 | 21.661785 | 22.305414 | 22.92295 | 7.26E-52 | 4.786713929 | YDR070C | FMP16 | n |
| W303.asm02.HP0_G0009380 | 4606.0826 | 4660.8366 | 5017.980576 | 2657.63882 | 2692.0933 | 2793.2453 | 2578.8925 | 2625.4784 | 2552.3823 | 1.04E-42 | 0.811239586 | YDR071C | PAA1 | n |
| W303.asm02.HP0_G0009390 | 1329.5691 | 1191.67938 | 1277.918826 | 2050.94046 | 1942.4419 | 1804.32648 | 1860.6333 | 1699.1477 | 1934.3443 | 5.34E-14 | -0.610213103 | YDR072C | IPT1 | n |
| W303.asm02.HP0_G0009400 | 816.49451 | 748.432782 | 823.9340579 | 259.180205 | 221.12018 | 241.777852 | 231.43907 | 205.99706 | 219.53133 | 8.78E-63 | 1.725175767 | YDR073W | SNF11 | n |
| W303.asm02.HP0_G0009410 | 1285.3204 | 1476.1046 | 1403.489081 | 2115.11048 | 2048.5077 | 2059.37841 | 2489.9652 | 2181.9943 | 2446.5841 | 7.81E-14 | -0.579263744 | YDR074W | TPS2 | n |
| W303.asm02.HP0_G0009420 | 1468.6366 | 1390.98464 | 1456.614958 | 1067.55576 | 923.13181 | 890.311383 | 633.89224 | 751.82366 | 689.4518 | 1.53E-10 | 0.582316199 | YDR075W | PPH3 | n |
| W303.asm02.HP0_G0009430 | 296.04511 | 315.566665 | 382.5063153 | 469.191174 | 461.11647 | 438.044578 | 499.36115 | 392.31287 | 454.93239 | 0.000183437 | -0.459669926 | YDR076W | RAD55 | n |
| W303.asm02.HP0_G0009440 | 10603.893 | 11500.5365 | 14050.34561 | 3049.32594 | 1928.959 | 2419.67481 | 2995.0268 | 4814.0331 | 2945.599 | 1.14E-16 | 2.288938061 | YDR077W | SED1 | n |
| W303.asm02.HP0_G0009450 | 347.66863 | 363.316885 | 398.927041 | 421.688693 | 471.90282 | 441.837172 | 323.78668 | 325.39663 | 327.09286 | 0.009729537 | -0.265227185 | YDR078C | SHU2 | n |
| W303.asm02.HP0_G0009460 | 981.90049 | 1034.9341 | 1143.655246 | 1134.22591 | 1198.1837 | 1099.85219 | 1036.3454 | 1075.9082 | 1067.6805 | 0.146192289 | -0.118410519 | YDR079W | PET100 | n |
| W303.asm02.HP0_G0009470 | 458.29047 | 408.991007 | 632.6809003 | 55.8362499 | 50.336301 | 68.2666875 | 46.743852 | 64.292075 | 59.070678 | 4.88E-56 | 3.107728037 | YDR079C- | TFB5 | n |
| W303.asm02.HP0_G0009480 | 880.76053 | 862.618089 | 1021.948691 | 934.215464 | 900.66025 | 855.229891 | 1093.3501 | 1053.6028 | 1044.7575 | 0.658140929 | 0.040115271 | YDR080W | VPS41 | n |
| W303.asm02.HP0_G0009490 | 637.39249 | 722.481576 | 727.3415541 | 799.208412 | 762.23542 | 833.422477 | 715.979 | 737.39074 | 678.87197 | 0.024135392 | -0.197536826 | YDR081C | PDC2 | n |
| W303.asm02.HP0_G0009500 | 792.26306 | 690.302081 | 732.1711793 | 137.507183 | 132.13279 | 104.296328 | 147.07212 | 132.5204 | 119.90466 | 1.37E-87 | 2.562310329 | YDR082W | STN1 | n |
| W303.asm02.HP0_G0009510 | 2034.3882 | 1854.99221 | 2221.627589 | 1746.75791 | 1661.9968 | 1917.15614 | 970.21995 | 1275.3448 | 993.62171 | 0.073417468 | 0.198729029 | YDR083W | RRP8 | n |
| W303.asm02.HP0_G0009520 | 1349.5864 | 1335.96809 | 1722.244344 | 943.38261 | 950.09769 | 877.985453 | 1210.7798 | 1186.1232 | 1277.5136 | 4.44E-11 | 0.669769335 | YDR084C | TVP23 | n |
| W303.asm02.HP0_G0009530 | 492.00379 | 482.692432 | 456.8825433 | 1001.71899 | 949.19882 | 914.963243 | 893.83366 | 792.49823 | 929.26112 | 5.13E-29 | -1.00246379 | YDR085C | AFR1 | n |
| W303.asm02.HP0_G0009540 | 4294.2343 | 4338.0036 | 5292.303287 | 1660.92009 | 1660.1991 | 1650.72643 | 1581.3103 | 2032.4168 | 1627.5294 | 2.75E-45 | 1.485915799 | YDR086C | SSS1 | n |
| W303.asm02.HP0_G0009550 | 2322.005 | 2226.61348 | 2164.638012 | 1122.55864 | 1131.6679 | 1298.96336 | 700.01769 | 883.03197 | 782.02525 | 1.04E-23 | 0.918334664 | YDR087C | RRP1 | n |
| W303.asm02.HP0_G0009560 | 817.54805 | 778.536181 | 701.261578 | 432.522592 | 407.18472 | 374.518633 | 425.25504 | 426.42703 | 506.94985 | 5.58E-18 | 0.918182532 | YDR088C | SLU7 | n |
| W303.asm02.HP0_G0009570 | 788.04889 | 761.927409 | 745.6941298 | 1332.56961 | 1275.4859 | 1128.29664 | 1462.7405 | 1161.1936 | 1521.7312 | 4.87E-11 | -0.703347304 | YDR089W | VTC5 | n |
| W303.asm02.HP0_G0009580 | 1223.1614 | 1081.64627 | 1066.381243 | 78.3374252 | 78.201039 | 58.7852031 | 111.72921 | 82.66124 | 89.046843 | 1.10E-175 | 3.963449296 | YDR090C | ILT1 | n |
| W303.asm02.HP0_G0009590 | 4821.005 | 4254.95974 | 4024.043711 | 5568.6242 | 5478.5671 | 5791.29066 | 4493.1103 | 4799.6002 | 4281.3017 | 3.88E-06 | -0.362337179 | YDR091C | RLI1 | n |
| W303.asm02.HP0_G0009600 | 1749.9321 | 1830.07905 | 1729.005819 | 1298.40115 | 1423.7982 | 1280.94854 | 1558.5084 | 1570.5636 | 1688.3634 | 1.73E-08 | 0.407181519 | YDR092W | UBC13 | n |
| W303.asm02.HP0_G0009630 | 789.10243 | 799.297146 | 900.242136 | 2013.4385 | 2109.6303 | 1985.42283 | 2029.3672 | 1918.2656 | 2006.6398 | 5.80E-69 | -1.294547023 | YDR097C | MSH6 | n |
| W303.asm02.HP0_G0009650 | 8118.5893 | 8279.47277 | 7511.999025 | 9072.97392 | 8619.1927 | 9574.40293 | 10150.256 | 9237.0655 | 9572.0948 | 0.005581836 | -0.189552898 | YDR099W | BMH2 | n |
| W303.asm02.HP0_G0009660 | 1010.3461 | 1026.62971 | 1173.598922 | 895.880129 | 902.45797 | 877.037305 | 1445.6391 | 1318.6436 | 1433.566 | 0.001503558 | 0.2637102 | YDR100W | TVP15 | n |

| | | | | | | | | | | | | | |
|---|---|---|---|---|---|---|---|---|---|---|---|---|---|
| W303.asm02.HP0_G0009670 | 2331.4869 | 2084.40087 | 2264.128291 | 2584.30165 | 2507.8264 | 2784.71196 | 1459.3203 | 1759.5035 | 1537.6009 | 0.005168956 | -0.237594822 | YDR101C | ARX1 | n |
| W303.asm02.HP0_G0009680 | 760.65682 | 800.335194 | 883.8214103 | 111.6725 | 85.39194 | 81.5407657 | 131.1108 | 107.59082 | 142.82761 | 1.00E-108 | 3.128393252 | YDR103W | STE5 | n |
| W303.asm02.HP0_G0009690 | 245.47512 | 243.941337 | 264.6634606 | 822.542965 | 818.86376 | 809.718766 | 814.02708 | 683.59533 | 788.19681 | 2.86E-66 | -1.699790667 | YDR104C | SPO71 | n |
| W303.asm02.HP0_G0009700 | 888.13532 | 940.471707 | 961.0954134 | 510.026641 | 513.2505 | 487.348297 | 768.42332 | 746.57532 | 816.40967 | 1.82E-27 | 0.884879552 | YDR105C | TMS1 | n |
| W303.asm02.HP0_G0009710 | 56.89123 | 53.9785086 | 92.72880371 | 144.174197 | 180.67137 | 139.37782 | 159.61315 | 182.37956 | 171.04047 | 6.51E-09 | -1.183026902 | YDR106W | ARP10 | n |
| W303.asm02.HP0_G0009730 | 705.87267 | 713.139142 | 689.6704776 | 755.872816 | 851.22281 | 857.126188 | 865.33131 | 747.88741 | 829.63445 | 0.012033167 | -0.224143133 | YDR108W | TRS85 | n |
| W303.asm02.HP0_G0009740 | 342.40092 | 353.97445 | 355.4604142 | 991.718468 | 935.71589 | 949.096586 | 1678.2183 | 1309.459 | 1664.5588 | 1.26E-42 | -1.451409546 | YDR109C | | n |
| W303.asm02.HP0_G0009750 | 797.53076 | 681.997695 | 759.2170804 | 958.383393 | 910.54774 | 946.252141 | 832.26858 | 817.42781 | 846.38584 | 5.35E-05 | -0.330625373 | YDR110W | FOB1 | n |
| W303.asm02.HP0_G0009770 | 272.8672 | 294.805701 | 235.6857094 | 894.213375 | 918.6375 | 897.896571 | 893.83366 | 961.75696 | 927.49781 | 4.66E-77 | -1.756633906 | YDR113C | PDS1 | n |
| W303.asm02.HP0_G0009790 | 1785.7525 | 1902.74243 | 1721.278419 | 1512.57901 | 1503.797 | 1519.88195 | 1429.6778 | 1609.926 | 1515.5596 | 0.00038509 | 0.253848428 | YDR116C | MRPL1 | n |
| W303.asm02.HP0_G0009800 | 1214.7331 | 1323.51151 | 998.7664899 | 1285.06712 | 1310.5416 | 1358.69671 | 1170.8765 | 1347.5094 | 1200.8099 | 0.122073213 | -0.16136779 | YDR117C | TMA64 | n |
| W303.asm02.HP0_G0009830 | 49.516441 | 40.4838814 | 38.63700154 | 12.500653 | 16.179525 | 20.8592656 | 15.961315 | 20.993331 | 20.277994 | 5.62E-05 | 1.383777482 | YDR119W | COX26 | n |
| W303.asm02.HP0_G0009840 | 1218.9473 | 1195.83157 | 1373.545405 | 2115.94386 | 2015.2498 | 2222.45994 | 1558.5084 | 1550.8823 | 1593.145 | 3.69E-23 | -0.745253387 | YDR120C | TRM1 | n |
| W303.asm02.HP0_G0009850 | 830.19054 | 924.900983 | 855.8095842 | 614.198749 | 580.66519 | 598.281664 | 493.66068 | 582.56493 | 573.95539 | 1.87E-09 | 0.541760252 | YDR121W | DPB4 | n |
| W303.asm02.HP0_G0009860 | 597.35792 | 776.460085 | 831.6614583 | 1959.26901 | 2109.6303 | 2027.14136 | 2163.8983 | 1890.7118 | 2071.882 | 9.58E-43 | -1.465769086 | YDR122W | KIN1 | n |
| W303.asm02.HP0_G0009870 | 418.2559 | 432.866117 | 396.0292658 | 713.370595 | 691.22528 | 613.452039 | 799.20586 | 649.48117 | 710.61144 | 2.65E-11 | -0.695654967 | YDR123C | INO2 | n |
| W303.asm02.HP0_G0009890 | 229.672 | 216.952082 | 284.9478864 | 952.549755 | 949.19882 | 841.007664 | 946.27798 | 934.20322 | 941.60424 | 6.40E-72 | -1.904401434 | YDR125C | ECM18 | n |
| W303.asm02.HP0_G0009900 | 389.81028 | 386.153946 | 360.2900394 | 512.526771 | 533.02548 | 539.496461 | 685.19646 | 628.48784 | 619.80129 | 2.32E-07 | -0.480523705 | YDR126W | SWF1 | n |
| W303.asm02.HP0_G0009910 | 2441.0552 | 2640.79473 | 3065.846073 | 5149.43564 | 5278.1207 | 5407.29054 | 4873.9016 | 4138.3103 | 4785.6066 | 1.03E-25 | -0.958275436 | YDR127W | ARO1 | n |
| W303.asm02.HP0_G0009920 | 725.88996 | 714.17719 | 788.1948315 | 554.195614 | 555.49704 | 540.44461 | 559.78613 | 502.52785 | 562.49392 | 5.93E-07 | 0.433717332 | YDR128W | MTC5 | n |
| W303.asm02.HP0_G0009930 | 2078.637 | 2121.77061 | 2485.325124 | 4914.42337 | 4765.7691 | 4898.13483 | 4945.7276 | 4922.9361 | 4858.7837 | 2.02E-53 | -1.124162827 | YDR129C | SAC6 | n |
| W303.asm02.HP0_G0009940 | 404.55986 | 448.43684 | 374.778915 | 134.173675 | 127.63848 | 111.881516 | 114.0094 | 131.20832 | 126.07622 | 6.46E-37 | 1.712922725 | YDR130C | FIN1 | n |
| W303.asm02.HP0_G0009950 | 529.93127 | 509.681687 | 515.8039706 | 601.698096 | 611.22652 | 493.985336 | 477.69937 | 486.78286 | 465.51221 | 0.188779468 | -0.135425184 | YDR131C | | n |
| W303.asm02.HP0_G0009960 | 954.50842 | 839.781027 | 789.1607566 | 525.860801 | 531.22775 | 479.76311 | 653.27383 | 659.97783 | 603.04991 | 9.78E-15 | 0.747976062 | YDR132C | MRX16 | n |
| W303.asm02.HP0_G0009970 | 6478.2255 | 5542.13956 | 5179.290057 | 23578.7316 | 22348.419 | 26284.571 | 25274.743 | 24715.711 | 21745.944 | 9.20E-104 | -2.069949391 | YLR110C | CCW12 | y |
| W303.asm02.HP0_G0009980 | 2728.672 | 2846.32828 | 3141.188226 | 4316.05878 | 4319.0344 | 4421.21617 | 5413.1661 | 4964.9227 | 5003.3746 | 4.52E-17 | -0.582609433 | YDR135C | YCF1 | n |
| W303.asm02.HP0_G0009990 | 1074.6121 | 923.862935 | 963.0272635 | 686.702536 | 703.80936 | 601.12611 | 679.49599 | 659.97783 | 685.92519 | 4.67E-10 | 0.571289383 | YDR137W | RGP1 | n |
| W303.asm02.HP0_G0010000 | 622.64291 | 633.209427 | 645.2379258 | 1312.56856 | 1361.7767 | 1262.93372 | 1211.9199 | 1142.8244 | 1165.5438 | 9.89E-47 | -1.050290144 | YDR138W | HPR1 | n |
| W303.asm02.HP0_G0010010 | 1044.0594 | 1014.17313 | 1227.690724 | 337.51763 | 330.78141 | 326.163063 | 310.10555 | 350.32621 | 341.19929 | 1.66E-64 | 1.724602386 | YDR139C | RUB1 | n |
| W303.asm02.HP0_G0010020 | 741.69308 | 798.259098 | 658.7608763 | 940.049102 | 891.67162 | 870.400266 | 912.07516 | 969.62946 | 849.91244 | 0.001350726 | -0.298646619 | YDR140W | MTQ2 | n |
| W303.asm02.HP0_G0010030 | 969.258 | 915.558549 | 1122.404895 | 2215.94908 | 2225.5836 | 2301.15626 | 2312.1105 | 2078.3397 | 2240.2775 | 5.52E-44 | -1.163962757 | YDR141C | DOP1 | n |
| W303.asm02.HP0_G0010040 | 576.28709 | 590.649449 | 613.3623995 | 543.361715 | 547.40728 | 472.177922 | 652.13374 | 674.41075 | 640.07929 | 0.042319935 | 0.187076741 | YDR142C | PEX7 | n |
| W303.asm02.HP0_G0010050 | 462.50463 | 524.214362 | 569.8957728 | 1910.09977 | 1940.6442 | 1919.05244 | 1987.1838 | 1797.5539 | 1892.025 | 1.51E-113 | -1.888656838 | YDR143C | SAN1 | n |
| W303.asm02.HP0_G0010070 | 2118.6716 | 1999.28091 | 1696.164368 | 1567.58188 | 1590.9867 | 1623.23013 | 1579.0301 | 1582.3723 | 1682.1918 | 0.001116434 | 0.281617714 | YDR145W | TAF12 | n |
| W303.asm02.HP0_G0010100 | 2385.2175 | 2251.52664 | 2022.647031 | 1825.09533 | 2078.1702 | 2132.38584 | 1640.5952 | 1525.9527 | 1661.9139 | 0.107372421 | 0.141981288 | YDR148C | KGD2 | n |
| W303.asm02.HP0_G0010110 | 696.3908 | 941.509755 | 899.276211 | 1893.43223 | 1830.0841 | 1777.77832 | 2053.3092 | 1817.2352 | 2041.0242 | 3.95E-27 | -1.116156795 | YDR150W | NUM1 | n |
| W303.asm02.HP0_G0010130 | 2433.6804 | 2493.39188 | 2335.606743 | 2214.28233 | 2131.203 | 2301.15626 | 1816.1697 | 1932.6985 | 1954.6223 | 0.051159888 | 0.1278411 | YDR152W | GIR2 | n |
| W303.asm02.HP0_G0010140 | 1021.9351 | 1041.16239 | 1056.721992 | 4896.08908 | 4442.1786 | 4811.85332 | 4613.9602 | 4461.0828 | 4565.1936 | 1.34E-255 | -2.181140161 | YDR153C | ENT5 | n |
| W303.asm02.HP0_G0010150 | 17700.547 | 17549.2436 | 15879.80763 | 8810.4602 | 8554.4746 | 9347.79545 | 8587.1876 | 9125.5384 | 7837.0038 | 2.83E-36 | 0.936634381 | YDR155C | CPR1 | n |
| W303.asm02.HP0_G0010160 | 637.39249 | 703.796708 | 606.6009243 | 1462.5764 | 1371.6642 | 1538.84491 | 937.15723 | 1091.6532 | 999.79327 | 8.92E-39 | -1.167194751 | YDR156W | RPA14 | n |
| W303.asm02.HP0_G0010170 | 9041.4914 | 8659.39843 | 7625.012255 | 3805.19876 | 3808.4805 | 3991.70492 | 3026.9494 | 2702.8913 | 3009.078 | 6.24E-48 | 1.125785876 | YDR158W | HOM2 | n |
| W303.asm02.HP0_G0010180 | 723.78287 | 677.845502 | 736.0348794 | 1556.74798 | 1567.6162 | 1524.62269 | 1501.5037 | 1368.5027 | 1516.4413 | 1.43E-51 | -1.120642427 | YDR159W | SAC3 | n |
| W303.asm02.HP0_G0010190 | 935.54467 | 883.379053 | 1022.914616 | 489.192219 | 504.26188 | 460.800141 | 529.00359 | 472.34994 | 535.16271 | 1.44E-25 | 0.966618411 | YDR160W | SSY1 | n |
| W303.asm02.HP0_G0010200 | 1386.4604 | 1316.24517 | 1227.690724 | 2512.63124 | 2541.0844 | 2676.62304 | 1694.1796 | 2105.8935 | 1717.4579 | 1.19E-26 | -0.976067604 | YDR161W | ACL4 | n |
| W303.asm02.HP0_G0010210 | 1374.8714 | 1313.13103 | 1368.71578 | 990.051714 | 957.28859 | 929.185469 | 936.01713 | 998.49529 | 1048.2841 | 1.35E-11 | 0.495571953 | YDR162C | NBP2 | n |
| W303.asm02.HP0_G0010220 | 1032.4705 | 1026.62971 | 1048.994592 | 528.360932 | 556.3959 | 495.881633 | 505.06162 | 448.73244 | 503.42324 | 2.51E-31 | 0.975102165 | YDR163W | CWC15 | n |
| W303.asm02.HP0_G0010230 | 653.19561 | 603.106028 | 686.7727025 | 1043.38783 | 1133.4656 | 1190.87444 | 1238.142 | 1115.2707 | 1257.2356 | 4.57E-19 | -0.792274078 | YDR164C | SEC1 | n |
| W303.asm02.HP0_G0010240 | 3389.2424 | 3027.98672 | 3067.777923 | 2325.12145 | 2306.4812 | 2415.88222 | 1631.4744 | 2020.6081 | 1688.3634 | 7.63E-07 | 0.428525634 | YDR165W | TRM82 | n |
| W303.asm02.HP0_G0010250 | 1295.8558 | 1134.58673 | 1153.314496 | 2212.61557 | 2256.1449 | 2377.00813 | 2315.5308 | 2309.2664 | 2321.3895 | 9.27E-40 | -0.933888509 | YDR166C | SEC5 | n |
| W303.asm02.HP0_G0010260 | 2022.7993 | 2009.6614 | 2476.631799 | 1190.89554 | 1220.6553 | 1226.90408 | 1238.142 | 1313.3953 | 1221.0879 | 4.49E-21 | 0.839706668 | YDR167W | TAF10 | n |
| W303.asm02.HP0_G0010270 | 5449.9691 | 5361.51917 | 6255.33055 | 1744.25778 | 1748.2876 | 1688.65237 | 1672.5178 | 1671.594 | 1651.334 | 2.97E-125 | 1.719898384 | YDR168W | CDC37 | n |
| W303.asm02.HP0_G0010280 | 509.91399 | 612.448462 | 582.4527983 | 319.183339 | 318.19733 | 280.651938 | 607.67008 | 394.93703 | 549.26914 | 4.57E-10 | 0.892483453 | YDR169C | STB3 | n |
| W303.asm02.HP0_G0010300 | 1723.5936 | 1984.74824 | 2024.578881 | 3631.85637 | 3626.0114 | 3582.1048 | 3270.9295 | 3131.9425 | 3404.9397 | 2.17E-36 | -0.918673421 | YDR170C | SEC7 | n |
| W303.asm02.HP0_G0010310 | 1085.1475 | 2403.08168 | 703.1934281 | 945.882741 | 1125.3759 | 1008.82994 | 3460.1851 | 1999.6147 | 3010.8413 | 0.370475456 | 0.44441252 | YDR171W | HSP42 | n |
| W303.asm02.HP0_G0010320 | 3544.1129 | 3592.68496 | 3554.604142 | 6708.68375 | 6709.1099 | 6713.83909 | 5845.2617 | 6052.6397 | 5922.9375 | 4.68E-82 | -0.913019313 | YDR172W | SUP35 | n |
| W303.asm02.HP0_G0010330 | 417.20235 | 429.751972 | 442.3936677 | 385.020111 | 445.83581 | 403.911235 | 407.01354 | 482.84661 | 401.15162 | 0.573058313 | 0.063257019 | YDR173C | ARG82 | n |
| W303.asm02.HP0_G0010340 | 1871.0893 | 2006.54725 | 2069.011433 | 2189.28102 | 2308.279 | 2255.64513 | 4750.7715 | 3562.3058 | 4932.8424 | 0.095069388 | -0.183326543 | YDR174W | HMO1 | n |
| W303.asm02.HP0_G0010350 | 1499.1893 | 1540.46359 | 1431.500907 | 2413.4594 | 2567.1514 | 2578.96375 | 6389.0865 | 4357.4282 | 6283.5332 | 3.37E-10 | -0.757674862 | YDR175C | RSM24 | n |
| W303.asm02.HP0_G0010360 | 726.9435 | 790.99276 | 686.7727025 | 1377.57196 | 1349.1926 | 1271.46706 | 4108.8986 | 2525.7601 | 4016.8061 | 4.14E-09 | -0.85907867 | YDR176W | NGG1 | n |
| W303.asm02.HP0_G0010370 | 3051.0556 | 3172.27543 | 3660.855896 | 1773.42597 | 1971.2055 | 1769.24498 | 1549.3877 | 1579.7481 | 1665.4405 | 1.31E-23 | 0.842352014 | YDR177W | UBC1 | n |

| | | | | | | | | | | | | | | |
|---|---|---|---|---|---|---|---|---|---|---|---|---|---|---|
| W303.asm02.HP0_G0010390 | 274.97428 | 273.006688 | 394.0974158 | 197.510317 | 229.20994 | 189.629688 | 558.64604 | 317.52413 | 550.15079 | 0.002476165 | 0.613121811 | YDR179C | CSN9 | n |
| W303.asm02.HP0_G0010400 | 769.08515 | 740.128396 | 855.8095842 | 542.528338 | 590.55268 | 573.629805 | 1531.1462 | 1027.3611 | 1603.7248 | 0.000860432 | 0.471137555 | YDR179W- | NVJ3 | n |
| W303.asm02.HP0_G0010410 | 564.69814 | 485.806577 | 525.463221 | 1421.74093 | 1453.4607 | 1424.11895 | 3403.1804 | 1924.826 | 3184.5267 | 2.46E-11 | -1.447898893 | YDR180W | SCC2 | n |
| W303.asm02.HP0_G0010420 | 1047.2201 | 985.107781 | 978.4820641 | 694.202928 | 723.58433 | 698.785399 | 2161.6181 | 1262.224 | 2146.8224 | 0.001137948 | 0.508374315 | YDR181C | SAS4 | n |
| W303.asm02.HP0_G0010430 | 1122.0215 | 1088.91261 | 1113.71157 | 1375.07183 | 1371.6642 | 1311.28929 | 4300.4344 | 2705.5155 | 4194.0181 | 0.037825302 | -0.287642838 | YDR182W | CDC1 | n |
| W303.asm02.HP0_G0010450 | 1099.8971 | 1067.11359 | 1313.658053 | 630.032909 | 702.91049 | 700.681696 | 1664.5372 | 1145.4486 | 1758.8956 | 3.31E-08 | 0.775943514 | YDR183W | PLP1 | n |
| W303.asm02.HP0_G0010460 | 1776.2706 | 1699.28497 | 1519.400086 | 1199.22931 | 1155.0383 | 1197.51148 | 1898.2564 | 1419.674 | 2011.048 | 2.10E-05 | 0.491687856 | YDR184C | ATC1 | n |
| W303.asm02.HP0_G0010480 | 977.68633 | 1175.07061 | 1179.394472 | 628.366155 | 692.12414 | 775.585422 | 2000.8649 | 1116.5828 | 1821.4929 | 0.004454379 | 0.669051343 | YDR186C | SND1 | n |
| W303.asm02.HP0_G0010490 | 3027.8777 | 3115.18277 | 3605.798169 | 4560.2382 | 4730.7135 | 4790.04591 | 11391.819 | 7265.0045 | 10849.608 | 7.74E-05 | -0.530315182 | YDR188W | CCT6 | n |
| W303.asm02.HP0_G0010500 | 1302.177 | 1472.99045 | 1478.831234 | 2627.63725 | 2668.7228 | 2713.60083 | 7061.7419 | 4363.9886 | 6920.9675 | 8.79E-11 | -0.912846644 | YDR189W | SLY1 | n |
| W303.asm02.HP0_G0010510 | 3293.3701 | 3129.71545 | 3299.599932 | 3307.67277 | 3386.0151 | 3310.93434 | 7948.735 | 5052.8323 | 7476.4082 | 0.761085064 | -0.041223859 | YDR190C | RVB1 | n |
| W303.asm02.HP0_G0010520 | 349.77571 | 397.572476 | 391.1996406 | 370.852704 | 331.68027 | 265.481563 | 1005.5629 | 620.61534 | 1058.8639 | 0.199149726 | 0.233042304 | YDR191W | HST4 | n |
| W303.asm02.HP0_G0010530 | 301.31281 | 323.871051 | 343.8693137 | 1040.05433 | 970.77152 | 994.607711 | 2797.7906 | 1693.8994 | 2786.02 | 6.76E-26 | -1.632544032 | YDR192C | NUP42 | n |
| W303.asm02.HP0_G0010540 | 7467.5007 | 7063.91828 | 7367.11027 | 2455.96162 | 2426.9288 | 1975.94134 | 5101.9204 | 3679.0812 | 5324.2959 | 2.39E-42 | 1.674577216 | YDR194C | MSS116 | n |
| W303.asm02.HP0_G0010560 | 548.89502 | 604.144077 | 550.577272 | 677.53539 | 664.2594 | 828.681735 | 2072.6908 | 1204.4923 | 1894.67 | 0.033116456 | -0.349071126 | YDR195W | REF2 | n |
| W303.asm02.HP0_G0010570 | 1689.8802 | 1678.52401 | 1930.884152 | 698.369812 | 768.52746 | 601.12611 | 1950.7008 | 1238.6065 | 2109.793 | 3.83E-10 | 1.357458555 | YDR196C | CAB5 | n |
| W303.asm02.HP0_G0010580 | 817.54805 | 832.51469 | 738.9326545 | 765.873338 | 809.87513 | 798.340985 | 2074.971 | 1247.7911 | 2077.1719 | 0.956785856 | 0.008918348 | YDR197W | CBS2 | n |
| W303.asm02.HP0_G0010590 | 1036.6846 | 872.998571 | 926.322112 | 1253.3988 | 1247.6212 | 1283.79298 | 3232.1664 | 2113.766 | 3127.2193 | 0.00201744 | -0.416439443 | YDR198C | RKM2 | n |
| W303.asm02.HP0_G0010610 | 733.26474 | 752.584975 | 925.356187 | 396.687387 | 438.64491 | 408.651977 | 1257.5236 | 835.79698 | 1251.0641 | 6.86E-11 | 0.955449416 | YDR201W | SPC19 | n |
| W303.asm02.HP0_G0010620 | 984.00757 | 1002.7546 | 959.1635634 | 275.014365 | 324.48937 | 310.044539 | 954.25864 | 648.16909 | 1047.4025 | 1.02E-31 | 1.696166741 | YDR202C | RAV2 | n |
| W303.asm02.HP0_G0010630 | 672.15935 | 718.329383 | 747.6259799 | 1009.21938 | 903.35684 | 906.429907 | 2515.0473 | 1647.9765 | 2435.1226 | 0.003462826 | -0.398887279 | YDR204W | COQ4 | n |
| W303.asm02.HP0_G0010640 | 576.28709 | 507.60559 | 538.9861716 | 2028.43929 | 2053.9009 | 2033.7784 | 2668.9599 | 2411.6089 | 2517.9979 | 6.84E-143 | -1.914280649 | YDR205W | MSC2 | n |
| W303.asm02.HP0_G0010650 | 1993.3001 | 1729.38837 | 2057.420332 | 3426.01229 | 3471.4071 | 3582.1048 | 3616.378 | 3412.7283 | 3641.2224 | 9.47E-31 | -0.858044477 | YDR206W | EBS1 | n |
| W303.asm02.HP0_G0010660 | 289.72386 | 372.659319 | 334.2100634 | 945.882741 | 924.92954 | 940.56325 | 940.57751 | 763.6324 | 908.98312 | 1.67E-43 | -1.495861439 | YDR207C | UME6 | n |
| W303.asm02.HP0_G0010670 | 1000.8642 | 1137.70087 | 1143.655246 | 1398.40638 | 1332.1143 | 1249.65964 | 1371.533 | 1389.4961 | 1395.655 | 0.00045738 | -0.278259789 | YDR208W | MSS4 | n |
| W303.asm02.HP0_G0010700 | 2185.0447 | 2106.19988 | 2165.603937 | 3505.18309 | 3442.6435 | 3693.98631 | 3010.9881 | 3192.2984 | 3005.5514 | 2.60E-33 | -0.720667662 | YDR211W | GCD6 | n |
| W303.asm02.HP0_G0010710 | 3832.7832 | 3609.29374 | 3478.296064 | 4889.42206 | 4857.4531 | 4888.65335 | 4847.6795 | 4975.4194 | 4737.9974 | 1.95E-13 | -0.422678615 | YDR212W | TCP1 | n |
| W303.asm02.HP0_G0010720 | 206.49409 | 222.142324 | 199.946483 | 625.866025 | 625.60832 | 629.570563 | 644.15308 | 644.23284 | 557.20401 | 1.14E-51 | -1.582077064 | YDR213W | UPC2 | n |
| W303.asm02.HP0_G0010730 | 3595.7365 | 3807.56095 | 3039.766097 | 2548.46645 | 2442.2095 | 2747.73417 | 3316.5333 | 2969.2442 | 3046.1074 | 2.18E-06 | 0.432332809 | YDR214W | AHA1 | n |
| W303.asm02.HP0_G0010750 | 681.64122 | 785.802519 | 779.5015062 | 221.678246 | 236.40084 | 236.088961 | 180.13484 | 169.25873 | 174.56708 | 2.42E-58 | 1.695890186 | YDR216W | ADR1 | n |
| W303.asm02.HP0_G0010760 | 665.8381 | 671.617212 | 707.0571283 | 665.034737 | 723.58433 | 640.000196 | 714.83891 | 673.09867 | 680.63528 | 0.898029373 | 0.011470762 | YDR217C | RAD9 | n |
| W303.asm02.HP0_G0010770 | 84.283304 | 74.7394734 | 71.47845286 | 120.839645 | 133.03165 | 120.414852 | 111.72921 | 131.20832 | 138.41935 | 7.78E-05 | -0.70093061 | YDR218C | SPR28 | n |
| W303.asm02.HP0_G0010780 | 776.45994 | 781.650326 | 792.0585317 | 808.375558 | 716.39343 | 768.000235 | 835.68887 | 848.91781 | 913.39138 | 0.681924476 | 0.035303653 | YDR219C | MFB1 | n |
| W303.asm02.HP0_G0010790 | 730.10412 | 738.0523 | 719.6141538 | 2656.80544 | 2495.2424 | 2557.15634 | 2843.3943 | 2558.5622 | 2772.7953 | 8.27E-147 | -1.817433236 | YDR221W | GTB1 | n |
| W303.asm02.HP0_G0010800 | 1435.9768 | 1244.61984 | 1191.951498 | 35.8352051 | 27.864738 | 19.9111172 | 29.642443 | 19.681248 | 25.567905 | 6.82E-195 | 5.519148641 | YDR222W | | n |
| W303.asm02.HP0_G0010820 | 7951.0762 | 7952.48758 | 7674.274432 | 5345.2792 | 5327.5582 | 5196.80159 | 5928.4885 | 5861.0755 | 6005.8128 | 2.09E-32 | 0.571006749 | YDR224C | HTB1 | y |
| W303.asm02.HP0_G0010840 | 9186.8801 | 8499.539 | 9478.622404 | 5051.09717 | 4862.8462 | 5116.20897 | 4723.4092 | 4596.2273 | 4439.999 | 4.15E-44 | 0.853972921 | YDR226W | ADK1 | n |
| W303.asm02.HP0_G0010850 | 667.94518 | 700.682563 | 734.1030294 | 550.862107 | 643.58557 | 558.45943 | 570.04698 | 488.09494 | 561.61227 | 0.008062403 | 0.263137894 | YDR227W | SIR4 | n |
| W303.asm02.HP0_G0010860 | 788.04889 | 749.47083 | 724.443779 | 1005.88587 | 994.14195 | 1047.70402 | 1027.2247 | 987.99863 | 936.31433 | 2.43E-08 | -0.430411385 | YDR228C | PCF11 | n |
| W303.asm02.HP0_G0010870 | 564.69814 | 622.828945 | 567.9639227 | 835.876994 | 862.90802 | 783.17061 | 799.20586 | 795.1224 | 813.76472 | 1.42E-09 | -0.500286494 | YDR229W | IVY1 | n |
| W303.asm02.HP0_G0010880 | 1044.0594 | 1126.28234 | 1084.733818 | 582.530428 | 522.23913 | 634.311305 | 510.76209 | 505.15202 | 519.29298 | 1.19E-24 | 0.905017018 | YDR231C | COX20 | n |
| W303.asm02.HP0_G0010890 | 1165.2167 | 1465.72412 | 1454.683108 | 1127.5589 | 1037.2874 | 1141.57072 | 990.74164 | 839.73323 | 904.57486 | 0.002968751 | 0.305648858 | YDR232W | HEM1 | n |
| W303.asm02.HP0_G0010900 | 5651.1955 | 5558.74833 | 4321.548623 | 6984.5315 | 6963.488 | 7443.91339 | 7056.0415 | 6706.0571 | 6693.5013 | 9.36E-07 | -0.462030497 | YDR233C | RTN1 | n |
| W303.asm02.HP0_G0010910 | 1480.2255 | 1571.60504 | 1822.700548 | 706.703581 | 663.36054 | 707.318735 | 662.39459 | 633.73617 | 674.46371 | 9.28E-42 | 1.231004015 | YDR234W | LYS4 | n |
| W303.asm02.HP0_G0010920 | 1435.9768 | 1283.02763 | 1522.297861 | 725.037872 | 753.24679 | 694.044656 | 674.93562 | 658.66575 | 675.34536 | 2.07E-30 | 0.96537988 | YDR235W | PRP42 | n |
| W303.asm02.HP0_G0010930 | 1157.8419 | 1181.2989 | 1389.000206 | 629.199532 | 673.24803 | 722.48911 | 590.56867 | 649.48117 | 639.19764 | 3.09E-20 | 0.882012746 | YDR236C | FMN1 | n |
| W303.asm02.HP0_G0010940 | 2257.739 | 2169.52082 | 2368.448195 | 1885.93184 | 1866.0386 | 1947.49689 | 1800.2083 | 1913.0173 | 1892.025 | 7.48E-05 | 0.254196303 | YDR237W | MRPL7 | n |
| W303.asm02.HP0_G0010950 | 3188.016 | 3299.95536 | 3104.483074 | 6988.69838 | 6978.7686 | 7278.93556 | 6994.4764 | 6719.1779 | 6570.9517 | 4.45E-95 | -1.147309911 | YDR238C | SEC26 | n |
| W303.asm02.HP0_G0010960 | 466.7188 | 459.855371 | 455.9166182 | 551.695484 | 529.43003 | 490.192742 | 558.64604 | 586.50118 | 613.62973 | 0.045634415 | -0.185952644 | YDR239C | | n |
| W303.asm02.HP0_G0010970 | 686.90893 | 625.94309 | 572.7935479 | 745.038916 | 828.75125 | 731.022446 | 653.27383 | 733.45449 | 657.71233 | 0.003116792 | -0.290499989 | YDR240C | SNU56 | n |
| W303.asm02.HP0_G0010980 | 481.46837 | 447.398792 | 499.383245 | 118.339515 | 124.04303 | 140.325969 | 134.53109 | 146.95331 | 129.60283 | 8.65E-52 | 1.902986692 | YDR242W | AMD2 | n |
| W303.asm02.HP0_G0010990 | 450.91568 | 392.382235 | 379.6085402 | 625.032648 | 642.6867 | 630.518711 | 565.4866 | 636.36034 | 593.35174 | 3.49E-11 | -0.635380388 | YDR243C | PRP28 | n |
| W303.asm02.HP0_G0011000 | 705.87267 | 671.617212 | 661.6586515 | 916.71455 | 885.37958 | 886.518789 | 1140.094 | 997.18321 | 1109.9998 | 8.86E-07 | -0.3995119 | YDR244W | PEX5 | n |
| W303.asm02.HP0_G0011010 | 2569.5872 | 2404.11973 | 2135.66026 | 3222.66833 | 3172.9847 | 3350.75658 | 3665.4021 | 3680.3933 | 3419.0461 | 3.19E-09 | -0.455479772 | YDR245W | MNN10 | n |
| W303.asm02.HP0_G0011020 | 424.57714 | 476.464143 | 370.9152148 | 265.84722 | 263.36672 | 239.881555 | 282.7433 | 236.17497 | 283.01026 | 7.83E-09 | 0.723928285 | YDR246W | TRS23 | n |
| W303.asm02.HP0_G0011050 | 1383.2997 | 1342.19638 | 1502.97936 | 135.840429 | 166.28957 | 104.296328 | 120.84996 | 123.33582 | 162.22395 | 1.30E-126 | 3.376904067 | YDR248C | | n |
| W303.asm02.HP0_G0011060 | 565.75168 | 521.100217 | 554.4409722 | 934.215464 | 911.4466 | 867.555821 | 727.37994 | 821.36406 | 737.061 | 1.51E-17 | -0.725572338 | YDR249C | | n |
| W303.asm02.HP0_G0011100 | 479.36129 | 462.969516 | 505.1787952 | 378.353096 | 359.54501 | 327.111211 | 362.54988 | 485.47077 | 412.61309 | 0.000163212 | 0.441946475 | YDR253C | MET32 | n |
| W303.asm02.HP0_G0011110 | 403.50632 | 371.62127 | 394.0974158 | 376.686342 | 374.82567 | 367.881594 | 364.83006 | 313.58788 | 337.67268 | 0.552434478 | 0.062641125 | YDR254W | CHL4 | n |
| W303.asm02.HP0_G0011120 | 406.66694 | 384.077849 | 327.4485881 | 497.525988 | 541.11524 | 497.77793 | 583.7281 | 530.0816 | 581.89026 | 1.25E-05 | -0.459940944 | YDR255C | RMD5 | n |

| | | | | | | | | | | | | | | |
|---|---|---|---|---|---|---|---|---|---|---|---|---|---|---|
| W303.asm02.HP0_G0011130 | 308.6876 | 268.854495 | 280.1182612 | 1040.8877 | 1063.3544 | 1039.17069 | 1101.3308 | 816.11573 | 1066.7988 | 1.44E-58 | -1.874264097 | YDR256C | CTA1 | n |
| W303.asm02.HP0_G0011140 | 1172.5915 | 1048.42872 | 993.9368647 | 1990.93733 | 1878.6227 | 2230.99327 | 1594.9914 | 1810.6748 | 1544.6542 | 4.55E-22 | -0.924353779 | YDR257C | RKM4 | n |
| W303.asm02.HP0_G0011150 | 3493.543 | 4808.23945 | 2702.658258 | 1082.55655 | 1062.4555 | 1072.35588 | 2191.2606 | 1448.5398 | 1744.7891 | 4.07E-13 | 1.774069498 | YDR258C | HSP78 | n |
| W303.asm02.HP0_G0011170 | 738.53245 | 684.073791 | 680.9771522 | 196.67694 | 188.76113 | 201.007469 | 230.29898 | 195.50039 | 210.71481 | 2.48E-68 | 1.842709192 | YDR260C | SWM1 | n |
| W303.asm02.HP0_G0011190 | 824.92284 | 852.237606 | 763.0807805 | 748.372424 | 752.34793 | 651.377977 | 910.93507 | 835.79698 | 940.72259 | 0.052085551 | 0.179960075 | YDR262W | | n |
| W303.asm02.HP0_G0011210 | 2240.8823 | 2065.716 | 1996.567055 | 2010.93837 | 1798.6239 | 1588.14863 | 2036.2078 | 1906.4568 | 1972.2553 | 0.01457819 | 0.222912431 | YDR264C | AKR1 | n |
| W303.asm02.HP0_G0011220 | 739.58599 | 746.356686 | 688.7045525 | 485.858712 | 434.1506 | 425.718649 | 527.8635 | 498.5916 | 524.58289 | 2.43E-14 | 0.690509031 | YDR265W | PEX10 | n |
| W303.asm02.HP0_G0011230 | 659.51685 | 778.536181 | 681.9430773 | 1563.415 | 1539.7515 | 1481.95601 | 1405.7358 | 1381.6236 | 1486.4651 | 1.28E-43 | -1.113332434 | YDR266C | HEL2 | n |
| W303.asm02.HP0_G0011240 | 1579.2584 | 1423.16414 | 1453.717183 | 906.714028 | 906.95229 | 857.126188 | 800.34595 | 869.91114 | 801.42159 | 3.05E-22 | 0.737935308 | YDR267C | CIA1 | n |
| W303.asm02.HP0_G0011250 | 839.67242 | 768.155699 | 817.1725827 | 591.697573 | 601.33903 | 602.074258 | 551.80547 | 519.58494 | 561.61227 | 9.55E-08 | 0.434115226 | YDR268W | MSW1 | n |
| W303.asm02.HP0_G0011260 | 2655.9776 | 2664.66984 | 2824.364813 | 1858.43041 | 1802.2194 | 2036.62284 | 2108.0337 | 1893.336 | 2085.9884 | 2.28E-12 | 0.516123161 | YDR270W | CCC2 | n |
| W303.asm02.HP0_G0011290 | 4651.3848 | 4420.00941 | 4753.317115 | 1982.60356 | 2021.5418 | 1985.42283 | 2007.7054 | 1777.8727 | 1825.9011 | 2.60E-76 | 1.206816104 | YDR276C | PMP3 | n |
| W303.asm02.HP0_G0011300 | 161.19182 | 205.533552 | 204.7761082 | 674.201883 | 700.21391 | 663.703906 | 688.61675 | 661.28992 | 626.85451 | 5.54E-65 | -1.832560226 | YDR277C | MTH1 | n |
| W303.asm02.HP0_G0011310 | 723.78287 | 637.36162 | 710.9208284 | 572.529905 | 667.85485 | 627.674266 | 519.88284 | 560.25951 | 581.00861 | 0.12233287 | 0.150377839 | YDR279W | RNH202 | n |
| W303.asm02.HP0_G0011320 | 2556.9447 | 2334.5705 | 2752.88636 | 1343.4035 | 1310.5416 | 1346.37078 | 1064.8478 | 1255.6636 | 1087.9585 | 3.18E-28 | 0.934507024 | YDR280W | RRP45 | n |
| W303.asm02.HP0_G0011340 | 553.10918 | 532.518748 | 456.8825433 | 1141.7263 | 1139.7577 | 1101.74848 | 1214.2001 | 1309.459 | 1247.5375 | 5.26E-40 | -1.134795969 | YDR282C | MRX10 | n |
| W303.asm02.HP0_G0011350 | 758.54974 | 735.976203 | 886.7191855 | 1023.38679 | 1111.8929 | 1144.41516 | 1274.625 | 1128.3915 | 1292.5017 | 1.40E-06 | -0.460537973 | YDR283C | GCN2 | n |
| W303.asm02.HP0_G0011360 | 2115.5109 | 2089.59111 | 2205.206863 | 730.038133 | 675.94462 | 649.48168 | 981.62089 | 915.83405 | 1000.6749 | 4.15E-107 | 1.639984568 | YDR284C | DPP1 | n |
| W303.asm02.HP0_G0011370 | 458.29047 | 401.724669 | 419.2114668 | 270.847481 | 321.79278 | 290.133422 | 245.1202 | 253.23205 | 205.4249 | 7.01E-06 | 0.53561628 | YDR285W | ZIP1 | n |
| W303.asm02.HP0_G0011380 | 333.97259 | 385.115898 | 503.2469451 | 164.175242 | 178.87364 | 135.585227 | 142.51174 | 132.5204 | 158.69734 | 3.54E-16 | 1.35263066 | YDR286C | MGP12 | n |
| W303.asm02.HP0_G0011400 | 1049.3271 | 1012.09704 | 1254.736625 | 540.028208 | 576.17088 | 564.148321 | 631.61205 | 579.94076 | 628.61781 | 8.23E-24 | 0.981802437 | YDR288W | NSE3 | n |
| W303.asm02.HP0_G0011410 | 956.6155 | 911.406356 | 855.8095842 | 880.045968 | 895.26707 | 925.392875 | 853.93037 | 881.71989 | 886.94182 | 0.882128698 | 0.012039138 | YDR289C | RTT103 | n |
| W303.asm02.HP0_G0011420 | 578.39417 | 576.116774 | 566.9979977 | 808.375558 | 808.07741 | 881.778047 | 850.51009 | 855.47823 | 870.19044 | 1.25E-11 | -0.536725979 | YDR291W | HRQ1 | n |
| W303.asm02.HP0_G0011430 | 2293.5594 | 2172.63497 | 2361.686719 | 1625.91826 | 1590.0878 | 1670.63755 | 1608.6726 | 1590.2448 | 1631.056 | 1.37E-14 | 0.482939203 | YDR292C | SRP101 | n |
| W303.asm02.HP0_G0011440 | 1165.2167 | 1437.69681 | 1383.204655 | 5574.45784 | 5422.8376 | 5763.79435 | 5356.1614 | 5316.561 | 5414.2244 | 3.63E-159 | -2.071635052 | YDR293C | SSD1 | y |
| W303.asm02.HP0_G0011460 | 1190.5017 | 1258.11447 | 1337.806178 | 699.203189 | 747.85362 | 675.081688 | 1099.0506 | 1016.8645 | 1024.4795 | 3.52E-25 | 0.83556808 | YDR294C | DPL1 | n |
| W303.asm02.HP0_G0011470 | 600.51854 | 624.905041 | 642.3401507 | 792.541398 | 754.14566 | 679.82243 | 672.65543 | 704.58866 | 689.4518 | 0.003796238 | -0.254163265 | YDR295C | HDA2 | n |
| W303.asm02.HP0_G0011480 | 1276.8921 | 1265.38081 | 1387.068355 | 2111.77697 | 2263.3358 | 2345.71924 | 2068.1304 | 2473.2768 | 2189.1417 | 2.89E-20 | -0.773748724 | YDR296W | MHR1 | n |
| W303.asm02.HP0_G0011490 | 1607.704 | 1420.04999 | 1452.751258 | 1241.73153 | 1273.6882 | 1269.57076 | 1501.5037 | 1763.4398 | 1527.0211 | 0.003855598 | 0.243268428 | YDR297W | SUR2 | n |
| W303.asm02.HP0_G0011500 | 6756.3604 | 6517.90491 | 7483.987199 | 8020.41894 | 7955.8322 | 8884.15086 | 7333.0843 | 7275.5012 | 6597.4013 | 0.000831774 | -0.25998266 | YDR298C | ATP5 | n |
| W303.asm02.HP0_G0011510 | 2186.0982 | 2071.94429 | 1778.267996 | 1434.24158 | 1410.3153 | 1435.49673 | 777.54407 | 1102.1499 | 831.39775 | 2.51E-05 | 0.495794084 | YDR299W | BFR2 | n |
| W303.asm02.HP0_G0011520 | 2512.696 | 2242.1842 | 2504.643625 | 1697.58867 | 1708.7377 | 1653.57088 | 1347.5911 | 1460.3486 | 1352.454 | 1.85E-13 | 0.52076796 | YDR300C | PRO1 | n |
| W303.asm02.HP0_G0011530 | 692.17663 | 727.671817 | 685.8067774 | 1532.58005 | 1618.8514 | 1541.68936 | 1390.9146 | 1369.8148 | 1370.0871 | 8.45E-63 | -1.156472722 | YDR301W | CFT1 | n |
| W303.asm02.HP0_G0011540 | 1201.0371 | 1237.3535 | 1559.968937 | 675.03526 | 707.40481 | 695.940953 | 877.87234 | 856.79031 | 792.60507 | 1.02E-19 | 0.944792932 | YDR302W | GPI11 | n |
| W303.asm02.HP0_G0011560 | 3927.602 | 3924.8604 | 3758.414325 | 2174.28024 | 2354.1209 | 2264.17847 | 2403.318 | 2307.9543 | 2238.5142 | 7.19E-37 | 0.773539399 | YDR304C | CPR5 | n |
| W303.asm02.HP0_G0011580 | 895.5101 | 909.33026 | 930.1858122 | 690.036043 | 675.94462 | 707.318735 | 624.77149 | 762.32032 | 603.04991 | 2.16E-05 | 0.399915252 | YDR306C | PFU1 | n |
| W303.asm02.HP0_G0011590 | 1195.7694 | 1112.78771 | 1246.0433 | 538.361454 | 581.56405 | 573.629805 | 768.42332 | 747.88741 | 789.07846 | 1.98E-40 | 1.07064383 | YDR307W | PMT7 | n |
| W303.asm02.HP0_G0011600 | 1690.9338 | 1568.49089 | 1995.60113 | 735.871771 | 768.52746 | 689.303914 | 818.58746 | 755.75991 | 827.87115 | 9.10E-38 | 1.260481022 | YDR308C | SRB7 | n |
| W303.asm02.HP0_G0011610 | 1046.1665 | 1010.02094 | 943.7087627 | 2170.11335 | 2046.71 | 1914.3117 | 2078.3913 | 2206.9239 | 2212.9463 | 1.79E-42 | -1.032220192 | YDR309C | GIC2 | n |
| W303.asm02.HP0_G0011620 | 830.19054 | 1000.6785 | 909.9013864 | 3266.00393 | 3168.4904 | 3335.5862 | 2842.2542 | 3025.6638 | 2983.5101 | 3.41E-131 | -1.83361802 | YDR310C | SUM1 | n |
| W303.asm02.HP0_G0011630 | 977.68633 | 881.302957 | 860.6392094 | 851.711155 | 865.60461 | 744.296524 | 801.48605 | 747.88741 | 840.21427 | 0.128351247 | 0.142673449 | YDR311W | TFB1 | n |
| W303.asm02.HP0_G0011640 | 1394.8887 | 1409.66951 | 1250.872925 | 2810.14679 | 2795.4624 | 2815.05271 | 1724.9621 | 2107.2056 | 1750.9607 | 5.17E-34 | -1.05445398 | YDR312W | SSF2 | n |
| W303.asm02.HP0_G0011650 | 675.31997 | 854.313703 | 631.7149753 | 338.351007 | 333.478 | 344.177883 | 328.34706 | 347.70204 | 322.6846 | 4.86E-20 | 1.088567828 | YDR313C | PIB1 | n |
| W303.asm02.HP0_G0011670 | 566.80522 | 516.948024 | 681.9430773 | 183.34291 | 176.17705 | 141.274117 | 155.05278 | 200.74872 | 186.02855 | 6.78E-36 | 1.816302038 | YDR315C | IPK1 | n |
| W303.asm02.HP0_G0011680 | 1505.5105 | 1593.40405 | 1437.296457 | 1498.4116 | 1427.3937 | 1511.34861 | 1290.5864 | 1233.3582 | 1299.5549 | 0.666292378 | 0.031544056 | YDR316W | OMS1 | n |
| W303.asm02.HP0_G0011690 | 559.43043 | 509.681687 | 553.4750471 | 339.184354 | 337.07345 | 299.614906 | 363.68997 | 372.63162 | 386.16354 | 1.40E-13 | 0.732354982 | YDR317W | HIM1 | n |
| W303.asm02.HP0_G0011710 | 1045.113 | 931.129273 | 1102.120469 | 138.340559 | 153.70549 | 164.02968 | 230.29898 | 278.16163 | 250.38914 | 5.22E-123 | 2.757700903 | YDR319C | YFT2 | n |
| W303.asm02.HP0_G0011720 | 541.52023 | 650.856247 | 627.8512751 | 1060.88875 | 1004.9283 | 1023.05216 | 1119.5723 | 1026.049 | 1117.053 | 2.43E-18 | -0.76260667 | YDR320C | SWA2 | n |
| W303.asm02.HP0_G0011730 | 1095.683 | 1111.74967 | 1306.896577 | 370.019328 | 403.58927 | 319.526024 | 440.07627 | 524.83327 | 448.76082 | 7.31E-50 | 1.684310073 | YDR320C- | DAD4 | n |
| W303.asm02.HP0_G0011740 | 3260.7103 | 2773.6649 | 3239.71258 | 2146.7788 | 1988.2839 | 2400.71184 | 1999.7248 | 1785.7452 | 2086.8701 | 1.24E-07 | 0.505115876 | YDR321W | ASP1 | n |
| W303.asm02.HP0_G0011750 | 1459.1547 | 1487.52313 | 1281.782526 | 3020.15775 | 2687.5989 | 3053.03797 | 2900.399 | 3139.815 | 3017.8945 | 1.30E-38 | -1.05142949 | YDR322W | MRPL35 | n |
| W303.asm02.HP0_G0011760 | 3480.9005 | 3330.05876 | 4216.262794 | 2539.2993 | 2636.3638 | 2671.8823 | 2409.0185 | 2601.8609 | 2341.6675 | 5.76E-08 | 0.491112602 | YDR322C- | TIM11 | n |
| W303.asm02.HP0_G0011770 | 518.34232 | 503.453397 | 601.7712991 | 513.360148 | 536.62093 | 486.400149 | 490.2404 | 460.54119 | 493.72507 | 0.432026501 | 0.080400258 | YDR323C | PEP7 | n |
| W303.asm02.HP0_G0011780 | 2582.2297 | 2234.91786 | 2429.301472 | 2830.98121 | 2767.5977 | 3036.91945 | 1551.6679 | 1952.3798 | 1640.7542 | 0.006746466 | -0.252920034 | YDR324C | UTP4 | n |
| W303.asm02.HP0_G0011800 | 1203.1442 | 1288.21787 | 1417.977957 | 2055.10735 | 1999.9691 | 2084.97841 | 2295.0091 | 2025.8564 | 2347.839 | 4.43E-15 | -0.650739632 | YDR326C | YSP2 | n |
| W303.asm02.HP0_G0011810 | 5100.1934 | 4721.0434 | 5592.705974 | 1753.42492 | 1665.5923 | 1664.94866 | 1523.1655 | 1495.7748 | 1715.6946 | 8.53E-86 | 1.600160884 | YDR328C | SKP1 | n |
| W303.asm02.HP0_G0011820 | 1525.5278 | 1464.68607 | 1355.192829 | 851.711155 | 764.93201 | 820.148399 | 1191.3982 | 1040.482 | 1144.3842 | 2.13E-23 | 0.833809957 | YDR329C | PEX3 | n |
| W303.asm02.HP0_G0011830 | 545.73439 | 660.198682 | 580.5209482 | 1620.918 | 1500.2016 | 1426.01525 | 1381.7939 | 1529.889 | 1401.8265 | 1.16E-49 | -1.348372773 | YDR330W | UBX5 | n |
| W303.asm02.HP0_G0011840 | 1876.3571 | 1796.86151 | 1904.804176 | 1708.42257 | 1718.6251 | 1700.03015 | 1830.9909 | 1751.631 | 1864.6938 | 0.051850301 | 0.121750247 | YDR331W | GPI8 | n |
| W303.asm02.HP0_G0011850 | 679.53414 | 616.600655 | 637.5105255 | 859.211547 | 793.69561 | 887.466938 | 761.58276 | 692.77991 | 723.83622 | 3.97E-06 | -0.393784058 | YDR332W | IRC3 | n |

| | | | | | | | | | | | | | | |
|---|---|---|---|---|---|---|---|---|---|---|---|---|---|---|
| W303.asm02.HP0_G0011860 | 1468.6366 | 1380.60416 | 1353.260979 | 1100.89084 | 1054.3657 | 1103.64478 | 951.97845 | 1091.6532 | 1053.574 | 1.58E-06 | 0.366577441 | YDR333C | RQC1 | n |
| W303.asm02.HP0_G0011870 | 904.99198 | 941.509755 | 917.6287867 | 1010.05276 | 1048.0737 | 970.904 | 1077.3888 | 1060.1632 | 1055.3373 | 0.066554614 | -0.132296312 | YDR334W | SWR1 | n |
| W303.asm02.HP0_G0011880 | 1468.6366 | 1562.2626 | 1670.084392 | 2543.46619 | 2571.6457 | 2757.21566 | 2736.2255 | 2432.6022 | 2614.9796 | 6.83E-23 | -0.743194354 | YDR335W | MSN5 | n |
| W303.asm02.HP0_G0011890 | 679.53414 | 641.513813 | 765.0126306 | 361.685559 | 338.87117 | 354.607516 | 290.72396 | 337.20537 | 311.22313 | 1.01E-21 | 0.98381785 | YDR336W | MRX8 | n |
| W303.asm02.HP0_G0011900 | 2634.9068 | 2642.87082 | 2806.012237 | 1948.43511 | 1994.5759 | 2058.43026 | 1780.8268 | 2098.021 | 1801.2149 | 1.71E-08 | 0.430052629 | YDR337W | MRPS28 | n |
| W303.asm02.HP0_G0011910 | 440.38026 | 428.713924 | 417.2796167 | 980.051192 | 916.83977 | 987.970672 | 1190.2581 | 944.69988 | 1191.9934 | 4.57E-30 | -1.165512689 | YDR338C | | n |
| W303.asm02.HP0_G0011920 | 2202.9549 | 2167.44473 | 2137.59211 | 1424.24106 | 1405.821 | 1497.12638 | 1148.0746 | 1519.3923 | 1270.4604 | 1.78E-10 | 0.588881524 | YDR339C | FCF1 | n |
| W303.asm02.HP0_G0011940 | 4124.6142 | 3543.8967 | 3566.195243 | 4871.92115 | 4892.5087 | 5241.36456 | 3915.0826 | 3908.6958 | 3781.4051 | 1.05E-08 | -0.417621709 | YDR341C | | n |
| W303.asm02.HP0_G0011960 | 5597.4649 | 5166.3661 | 5183.153757 | 3711.86055 | 3751.8522 | 3929.12713 | 4031.3722 | 3360.245 | 3647.394 | 1.37E-10 | 0.48519677 | YDR346C | SVF1 | n |
| W303.asm02.HP0_G0011970 | 1730.9684 | 1814.50833 | 1859.405699 | 1977.6033 | 2150.978 | 2032.83025 | 1732.9428 | 1923.5139 | 1767.7121 | 0.007880166 | -0.188641656 | YDR347W | MRP1 | n |
| W303.asm02.HP0_G0011980 | 1396.9958 | 1367.10953 | 1401.557231 | 434.189346 | 402.69041 | 470.281625 | 328.34706 | 360.82287 | 360.59563 | 1.62E-86 | 1.672927087 | YDR348C | PAL1 | n |
| W303.asm02.HP0_G0012000 | 1214.7331 | 1365.03344 | 1274.055126 | 1722.58998 | 1758.1751 | 1718.04497 | 1649.7159 | 1796.2419 | 1592.2634 | 6.04E-09 | -0.431871301 | YDR349C | YPS7 | n |
| W303.asm02.HP0_G0012010 | 1543.438 | 1327.6637 | 1482.694934 | 611.698618 | 616.61969 | 554.666836 | 518.74275 | 502.52785 | 442.58926 | 1.33E-43 | 1.287373355 | YDR350C | ATP22 | n |
| W303.asm02.HP0_G0012030 | 1694.0944 | 1536.3114 | 1686.505117 | 793.374774 | 749.65134 | 820.148399 | 1167.4562 | 1108.7103 | 1307.4898 | 1.30E-35 | 1.057294884 | YDR352W | YPQ2 | n |
| W303.asm02.HP0_G0012040 | 11065.344 | 10161.4542 | 11123.59274 | 5624.46045 | 5457.8932 | 5802.66844 | 6403.9077 | 6586.6575 | 6414.0177 | 5.00E-59 | 0.938129931 | YDR353W | TRR1 | n |
| W303.asm02.HP0_G0012050 | 1422.2808 | 1492.71337 | 1533.888961 | 790.874644 | 803.5831 | 834.370625 | 852.79028 | 863.35073 | 944.2492 | 1.72E-31 | 0.87388013 | YDR354W | TRP4 | n |
| W303.asm02.HP0_G0012060 | 757.49619 | 742.204493 | 651.9994011 | 865.878562 | 912.34546 | 806.874321 | 784.38464 | 761.00824 | 780.26194 | 0.002609855 | -0.266007265 | YDR356W | SPC110 | n |
| W303.asm02.HP0_G0012070 | 1059.8625 | 1042.20043 | 1321.385453 | 80.8375558 | 77.302177 | 87.2296563 | 70.685825 | 83.973323 | 93.455103 | 1.22E-152 | 3.804120433 | YDR357C | CNL1 | n |
| W303.asm02.HP0_G0012090 | 439.32672 | 507.60559 | 512.9061955 | 957.550017 | 1084.0282 | 1087.52626 | 1039.7657 | 738.70282 | 989.21345 | 6.72E-19 | -1.099136386 | YDR359C | EAF1 | n |
| W303.asm02.HP0_G0012100 | 2752.9034 | 2610.69133 | 3236.814804 | 1388.40586 | 1327.6199 | 1335.94115 | 787.80492 | 1033.9215 | 891.35008 | 8.77E-26 | 1.085898068 | YDR361C | BCP1 | n |
| W303.asm02.HP0_G0012110 | 693.23018 | 785.802519 | 708.9889783 | 752.539308 | 811.67286 | 718.696516 | 746.76154 | 734.76657 | 724.71787 | 0.491043919 | -0.061649139 | YDR362C | TFC6 | n |
| W303.asm02.HP0_G0012120 | 554.16272 | 479.578288 | 474.269194 | 506.693133 | 549.205 | 498.726078 | 407.01354 | 435.61161 | 426.71953 | 0.664730872 | -0.044757443 | YDR363W | ESC2 | n |
| W303.asm02.HP0_G0012130 | 2855.0969 | 2772.62685 | 3258.065155 | 1127.5589 | 1124.477 | 1096.05959 | 1063.7077 | 1159.8815 | 1113.5264 | 4.54E-72 | 1.40831818 | YDR363W- | SEM1 | n |
| W303.asm02.HP0_G0012140 | 1002.9713 | 915.558549 | 898.3102859 | 492.525727 | 558.19363 | 519.585344 | 494.80077 | 581.25284 | 520.17463 | 1.59E-19 | 0.843174127 | YDR364C | CDC40 | n |
| W303.asm02.HP0_G0012150 | 1265.3031 | 1187.52719 | 1192.917423 | 2863.4829 | 2885.3487 | 3179.14171 | 1622.3537 | 2098.021 | 1810.913 | 1.00E-43 | -1.292078836 | YDR365C | ESF1 | n |
| W303.asm02.HP0_G0012170 | 3085.8225 | 2954.2853 | 2943.173593 | 44.1689738 | 44.044264 | 42.6666797 | 58.144792 | 76.100824 | 69.650501 | 0 | 6.099996311 | YDR368W | YPR1 | n |
| W303.asm02.HP0_G0012180 | 656.35623 | 670.579164 | 822.9681329 | 807.542181 | 819.76262 | 870.400266 | 788.94501 | 776.75324 | 789.07846 | 0.027993979 | -0.214741802 | YDR369C | XRS2 | n |
| W303.asm02.HP0_G0012190 | 1043.0059 | 1012.09704 | 1042.233117 | 615.865502 | 742.46044 | 714.903922 | 506.20171 | 516.96077 | 432.00944 | 2.16E-09 | 0.580102955 | YDR370C | DXO1 | n |
| W303.asm02.HP0_G0012200 | 1162.0561 | 1114.86381 | 1204.508523 | 263.347089 | 231.90653 | 221.866734 | 242.84001 | 207.30914 | 272.43044 | 1.08E-99 | 2.277424483 | YDR371W | CTS2 | n |
| W303.asm02.HP0_G0012210 | 4234.1825 | 3885.41457 | 4704.054938 | 2363.45679 | 2408.0527 | 2292.62292 | 2260.8063 | 2271.216 | 2334.6143 | 6.75E-30 | 0.86035546 | YDR372C | VPS74 | n |
| W303.asm02.HP0_G0012220 | 1925.8735 | 1870.56293 | 2379.07337 | 980.051192 | 1022.9056 | 914.963243 | 1081.9492 | 1032.6095 | 1109.9998 | 1.88E-28 | 1.081813212 | YDR373W | FRQ1 | n |
| W303.asm02.HP0_G0012230 | 34.766863 | 78.8916664 | 43.46662674 | 10.0005224 | 12.584075 | 11.3777813 | 10.260846 | 19.681248 | 14.106431 | 1.10E-07 | 2.211467607 | YDR374C | PHO92 | n |
| W303.asm02.HP0_G0012240 | 414.04173 | 429.751972 | 491.6558447 | 163.341865 | 207.63724 | 167.822273 | 134.53109 | 160.07445 | 151.64413 | 6.75E-23 | 1.310894642 | YDR374W- | WIP1 | n |
| W303.asm02.HP0_G0012250 | 1119.9144 | 1096.17894 | 1076.040493 | 1081.72317 | 1093.0168 | 1147.25961 | 1069.4081 | 1119.2069 | 1056.219 | 0.862375039 | -0.012818626 | YDR375C | BCS1 | n |
| W303.asm02.HP0_G0012260 | 1051.4342 | 1186.48914 | 1063.483468 | 938.382349 | 954.592 | 921.600282 | 877.87234 | 918.45822 | 916.03634 | 0.002887052 | 0.229867736 | YDR376W | ARH1 | n |
| W303.asm02.HP0_G0012280 | 1627.7213 | 1597.55624 | 1881.621975 | 1060.05537 | 1117.2861 | 1005.98549 | 991.88174 | 1171.6903 | 1119.6979 | 1.01E-13 | 0.682130737 | YDR378C | LSM6 | n |
| W303.asm02.HP0_G0012300 | 423.5236 | 391.344187 | 542.8498717 | 382.519981 | 435.04946 | 366.933445 | 328.34706 | 379.19204 | 337.67268 | 0.141879689 | 0.198107913 | YDR379C- | SDH6 | n |
| W303.asm02.HP0_G0012310 | 1940.6231 | 1975.4058 | 1711.619168 | 7533.72685 | 7921.6754 | 8536.18039 | 9307.727 | 8371.0906 | 8853.5485 | 2.87E-158 | -2.092206708 | YDR380W | ARO10 | n |
| W303.asm02.HP0_G0012320 | 6964.9615 | 6586.41609 | 7423.133922 | 1832.59572 | 1694.3559 | 1816.65241 | 2004.2852 | 1829.0439 | 1862.0488 | 8.36E-174 | 1.97277708 | YDR381W | YRA1 | n |
| W303.asm02.HP0_G0012330 | 852.31491 | 858.465896 | 1108.881944 | 427.522331 | 450.33012 | 412.44457 | 416.13429 | 440.85994 | 405.55988 | 2.05E-23 | 1.128494456 | YDR381C- | COI1 | n |
| W303.asm02.HP0_G0012340 | 12749.957 | 12350.698 | 14179.77957 | 9578.83367 | 9748.1641 | 10417.3069 | 8144.8312 | 8800.1418 | 7487.8697 | 3.77E-07 | 0.40130349 | YDR382W | RPP2B | n |
| W303.asm02.HP0_G0012350 | 637.39249 | 613.486511 | 699.329728 | 376.686342 | 352.35411 | 323.318617 | 311.24565 | 295.21871 | 279.48366 | 3.38E-18 | 0.889110668 | YDR383C | NKP1 | n |
| W303.asm02.HP0_G0012360 | 409.82757 | 414.181248 | 526.429146 | 31.6683208 | 41.347676 | 34.1333438 | 38.763194 | 36.738329 | 39.674336 | 6.09E-81 | 3.658731693 | YDR384C | ATO3 | n |
| W303.asm02.HP0_G0012370 | 7736.1538 | 7498.8605 | 8457.639638 | 43533.1072 | 42128.788 | 47485.1701 | 35276.787 | 37474.407 | 32828.309 | 5.77E-240 | -2.490415539 | YOR133W | EFT1 | y |
| W303.asm02.HP0_G0012380 | 484.629 | 486.844625 | 424.041092 | 272.514234 | 268.75989 | 264.533414 | 243.98011 | 299.15496 | 255.67905 | 1.94E-12 | 0.791090493 | YDR386W | MUS81 | n |
| W303.asm02.HP0_G0012390 | 401.39924 | 374.735415 | 370.9152148 | 404.187779 | 351.45525 | 342.281586 | 478.83946 | 453.98078 | 484.0269 | 0.576783852 | 0.060901894 | YDR387C | CIN10 | n |
| W303.asm02.HP0_G0012400 | 1352.747 | 1624.5455 | 1804.347972 | 3662.69132 | 3522.6422 | 3697.77891 | 3685.9237 | 3704.0108 | 3890.7299 | 2.83E-38 | -1.185924954 | YDR388W | RVS167 | n |
| W303.asm02.HP0_G0012410 | 560.48397 | 644.627958 | 677.1134521 | 1385.90572 | 1286.2723 | 1292.32632 | 1213.06 | 1005.0557 | 1156.7273 | 6.82E-29 | -1.074154215 | YDR389W | SAC7 | n |
| W303.asm02.HP0_G0012420 | 948.18717 | 852.237606 | 847.1162589 | 1393.40612 | 1333.912 | 1381.45227 | 1348.7311 | 1319.9557 | 1322.4779 | 1.72E-18 | -0.634687719 | YDR390C | UBA2 | n |
| W303.asm02.HP0_G0012430 | 1015.6138 | 1070.22774 | 1022.914616 | 91.671455 | 97.077152 | 98.6074375 | 117.42968 | 111.52707 | 106.67988 | 1.74E-192 | 3.436943458 | YDR391C | | n |
| W303.asm02.HP0_G0012440 | 653.19561 | 720.40548 | 608.5327743 | 955.883263 | 972.56925 | 996.504008 | 956.53883 | 936.82738 | 940.72259 | 6.53E-12 | -0.561903284 | YDR392W | SPT3 | n |
| W303.asm02.HP0_G0012450 | 1288.481 | 1408.63146 | 1293.373627 | 637.533301 | 680.43893 | 616.296485 | 584.8682 | 537.9541 | 546.62419 | 2.65E-38 | 1.044438789 | YDR393W | SHE9 | n |
| W303.asm02.HP0_G0012460 | 3054.2162 | 3258.43343 | 2507.5414 | 3892.70333 | 3739.2681 | 3889.30489 | 3932.184 | 3802.417 | 3724.9793 | 1.54E-05 | -0.385752195 | YDR394W | RPT3 | n |
| W303.asm02.HP0_G0012470 | 2468.4473 | 2314.84758 | 2527.825826 | 3661.85794 | 3783.3124 | 4056.17902 | 3626.6389 | 3679.0812 | 3605.0747 | 2.77E-24 | -0.653218342 | YDR395W | SXM1 | n |
| W303.asm02.HP0_G0012480 | 1797.3415 | 1741.84495 | 2069.011433 | 1839.26274 | 1813.9046 | 1909.57095 | 1673.6579 | 1878.9031 | 1738.6176 | 0.887635494 | 0.012304409 | YDR397C | NCB2 | n |
| W303.asm02.HP0_G0012490 | 1817.3587 | 1928.69363 | 2082.534383 | 2210.11544 | 2164.461 | 2347.61553 | 1417.1368 | 1873.6548 | 1549.9441 | 0.04087872 | -0.205462717 | YDR398W | UTP5 | n |
| W303.asm02.HP0_G0012500 | 6903.8561 | 6015.48956 | 7869.39129 | 2568.46749 | 2796.3613 | 2618.78599 | 2394.1973 | 2806.5459 | 2532.1043 | 1.51E-45 | 1.380830466 | YDR399W | HPT1 | n |
| W303.asm02.HP0_G0012510 | 2182.9376 | 2163.29254 | 2293.106042 | 799.208412 | 727.17978 | 849.541 | 521.02294 | 682.28325 | 495.48837 | 5.21E-44 | 1.482839434 | YDR400W | URH1 | n |
| W303.asm02.HP0_G0012520 | 87.443928 | 87.1960523 | 87.89917851 | 1548.41421 | 1802.2194 | 1778.72647 | 3547.9724 | 2524.448 | 2969.4036 | 1.77E-192 | -4.287840668 | YDR402C | DIT2 | n |
| W303.asm02.HP0_G0012530 | 359.25758 | 406.914911 | 396.9951909 | 1817.59494 | 1927.1612 | 2364.6822 | 5691.349 | 3156.8721 | 5068.6168 | 7.31E-24 | -2.39280288 | YDR403W | DIT1 | n |

| | | | | | | | | | | | | | | |
|---|---|---|---|---|---|---|---|---|---|---|---|---|---|---|
| W303.asm02.HP0_G0012540 | 1506.5641 | 1595.48015 | 1463.376434 | 1287.56725 | 1282.6768 | 1293.27447 | 1283.7458 | 1325.204 | 1295.1467 | 0.000271471 | 0.240583278 | YDR404C | RPB7 | n |
| W303.asm02.HP0_G0012550 | 481.46837 | 634.247476 | 653.9312511 | 2532.63229 | 2489.8492 | 2616.88969 | 2446.6416 | 2634.663 | 2388.395 | 7.16E-100 | -2.10881376 | YDR405W | MRP20 | n |
| W303.asm02.HP0_G0012570 | 1092.5223 | 1230.08717 | 1229.622574 | 1576.74903 | 1600.8742 | 1664.00051 | 1640.5952 | 1640.104 | 1665.4405 | 3.27E-10 | -0.446061085 | YDR407C | TRS120 | n |
| W303.asm02.HP0_G0012580 | 1678.2913 | 1559.14846 | 1751.222095 | 822.542965 | 874.59323 | 672.237242 | 829.9884 | 909.27364 | 916.91799 | 4.88E-27 | 1.073472122 | YDR408C | ADE8 | n |
| W303.asm02.HP0_G0012590 | 565.75168 | 628.019186 | 701.261578 | 840.043879 | 748.75248 | 628.622414 | 711.41863 | 627.17575 | 700.03162 | 0.060037928 | -0.22702959 | YDR409W | SIZ1 | n |
| W303.asm02.HP0_G0012600 | 1371.7108 | 1291.33201 | 1580.253363 | 658.367722 | 727.17978 | 713.955774 | 847.08981 | 879.09572 | 902.81156 | 8.40E-29 | 1.016223828 | YDR410C | STE14 | n |
| W303.asm02.HP0_G0012610 | 1144.1459 | 1245.65789 | 1252.804775 | 294.182033 | 245.38947 | 274.963047 | 404.73335 | 293.90663 | 402.03327 | 2.44E-74 | 2.160286721 | YDR411C | DFM1 | n |
| W303.asm02.HP0_G0012620 | 1693.0409 | 1859.1444 | 1674.914017 | 1450.07574 | 1406.7198 | 1481.00786 | 948.55817 | 1128.3915 | 950.42076 | 0.001371523 | 0.268979297 | YDR412W | RRP17 | n |
| W303.asm02.HP0_G0012630 | 569.96584 | 559.508002 | 651.9994011 | 815.87595 | 879.08755 | 863.763227 | 1130.9732 | 1157.2574 | 1109.9998 | 8.55E-10 | -0.52077665 | YDR414C | ERD1 | n |
| W303.asm02.HP0_G0012640 | 755.38911 | 730.785962 | 769.8422558 | 1120.05851 | 1159.5327 | 1092.267 | 1715.8414 | 1532.5131 | 1733.3277 | 1.12E-13 | -0.579655505 | YDR415C | | n |
| W303.asm02.HP0_G0012650 | 666.89164 | 629.057234 | 668.4201267 | 1027.55367 | 1057.9612 | 1029.6892 | 987.32136 | 963.06905 | 983.04188 | 7.57E-20 | -0.665000139 | YDR416W | SYF1 | n |
| W303.asm02.HP0_G0012660 | 665.8381 | 603.106028 | 597.9075989 | 360.852182 | 347.8598 | 384.000117 | 293.00415 | 343.76579 | 297.99835 | 1.76E-14 | 0.772810262 | YDR419W | RAD30 | n |
| W303.asm02.HP0_G0012670 | 547.84148 | 561.584099 | 655.8631012 | 1781.75973 | 1765.366 | 1848.88945 | 1462.7405 | 1455.1002 | 1603.7248 | 2.58E-80 | -1.610537236 | YDR420W | HKR1 | n |
| W303.asm02.HP0_G0012680 | 537.30606 | 515.909976 | 532.2246963 | 934.215464 | 893.46935 | 933.926211 | 970.21995 | 955.19655 | 943.36755 | 7.41E-26 | -0.800614565 | YDR421W | ARO80 | n |
| W303.asm02.HP0_G0012720 | 396.13153 | 404.838814 | 435.6321924 | 1594.24994 | 1512.7856 | 1540.74121 | 1585.8707 | 1540.3856 | 1482.0569 | 9.56E-134 | -1.909304807 | YDR425W | SNX41 | n |
| W303.asm02.HP0_G0012730 | 5415.2023 | 5422.76401 | 6302.660877 | 2724.30897 | 2855.6862 | 2737.30454 | 2891.2783 | 2865.5896 | 2724.3044 | 9.64E-46 | 1.043440314 | YDR427W | RPN9 | n |
| W303.asm02.HP0_G0012750 | 6404.4776 | 5953.20666 | 6311.354202 | 5261.10814 | 4973.4063 | 5163.61639 | 4570.6367 | 4807.4727 | 4495.5431 | 1.98E-06 | 0.277858226 | YDR429C | TIF35 | n |
| W303.asm02.HP0_G0012760 | 1494.9751 | 1617.27916 | 1857.473849 | 1880.93158 | 1901.0942 | 1834.66723 | 1752.3244 | 1851.3494 | 1712.168 | 0.039430502 | -0.176030304 | YDR430C | CYM1 | n |
| W303.asm02.HP0_G0012770 | 2662.2989 | 2875.39363 | 3200.109653 | 4493.56805 | 4184.205 | 4320.71243 | 3625.4988 | 3609.5408 | 3621.8261 | 7.62E-15 | -0.57271926 | YDR432W | NPL3 | n |
| W303.asm02.HP0_G0012780 | 2344.1294 | 2441.48946 | 2380.039295 | 3303.50589 | 3149.6143 | 3189.57134 | 3332.4946 | 3197.5467 | 3269.1653 | 1.49E-14 | -0.428490217 | YDR434W | GPI17 | n |
| W303.asm02.HP0_G0012790 | 620.53583 | 657.084537 | 560.2365224 | 644.200316 | 698.41618 | 629.570563 | 580.30782 | 670.4745 | 685.04354 | 0.315777553 | -0.102565674 | YDR435C | PPM1 | n |
| W303.asm02.HP0_G0012810 | 127.4785 | 102.766776 | 133.2976553 | 376.686342 | 437.74605 | 394.42975 | 422.97486 | 536.64202 | 409.08649 | 2.66E-32 | -1.731711814 | YDR437W | GPI19 | n |
| W303.asm02.HP0_G0012830 | 447.75505 | 413.1432 | 424.041092 | 618.365633 | 634.59694 | 592.592774 | 565.4866 | 616.67909 | 633.02607 | 8.25E-09 | -0.522921299 | YDR439W | LRS4 | n |
| W303.asm02.HP0_G0012840 | 613.16104 | 611.410414 | 583.4187233 | 835.876994 | 898.86252 | 792.652094 | 761.58276 | 792.49823 | 779.38029 | 4.35E-09 | -0.4839697 | YDR440W | DOT1 | n |
| W303.asm02.HP0_G0012860 | 360.31112 | 399.648573 | 391.1996406 | 1104.22434 | 1167.6224 | 1171.91147 | 1209.6397 | 1073.284 | 1234.3127 | 4.62E-71 | -1.580127522 | YDR443C | SSN2 | n |
| W303.asm02.HP0_G0012870 | 289.72386 | 339.441775 | 310.0619374 | 332.517369 | 357.74728 | 362.192703 | 427.53523 | 410.68203 | 428.48283 | 0.128935525 | -0.163201937 | YDR444W | | n |
| W303.asm02.HP0_G0012880 | 16.856661 | 15.5707236 | 17.3866507 | 100.838601 | 136.6271 | 127.051891 | 115.14949 | 97.094155 | 101.38997 | 3.59E-26 | -2.868032229 | YDR446W | ECM11 | n |
| W303.asm02.HP0_G0012900 | 582.60834 | 598.953835 | 525.463221 | 1266.73283 | 1301.5529 | 1231.64482 | 1205.0793 | 1133.6399 | 1229.0228 | 1.34E-47 | -1.155611563 | YDR448W | ADA2 | n |
| W303.asm02.HP0_G0012910 | 1988.0324 | 1694.09473 | 1877.758275 | 1762.59207 | 1905.5885 | 1770.19313 | 962.23929 | 1462.9727 | 1113.5264 | 0.809040485 | 0.031892278 | YDR449C | UTP6 | n |
| W303.asm02.HP0_G0012920 | 12217.918 | 11801.5705 | 13836.87618 | 12885.6731 | 13263.615 | 14504.7748 | 11530.91 | 12483.159 | 10362.937 | 0.253590832 | -0.102766854 | YML026C | RPS18B | y |
| W303.asm02.HP0_G0012940 | 2419.9844 | 2317.96172 | 2321.117868 | 2070.10813 | 1919.9703 | 2035.6747 | 2502.5062 | 2268.5918 | 2462.4538 | 0.000624068 | 0.228135126 | YDR452W | PPN1 | n |
| W303.asm02.HP0_G0012960 | 4066.6694 | 4148.04077 | 4260.695345 | 2136.77828 | 2106.9338 | 2224.35624 | 2043.0484 | 2025.8564 | 2003.9948 | 4.72E-65 | 0.948003445 | YDR454C | GUK1 | n |
| W303.asm02.HP0_G0012970 | 945.02655 | 969.537058 | 1025.812391 | 1007.55263 | 1037.2874 | 961.422516 | 1644.0155 | 1549.5702 | 1527.0211 | 0.684679697 | -0.031774814 | YDR456W | NHX1 | n |
| W303.asm02.HP0_G0012980 | 1761.5211 | 1840.45953 | 1959.861903 | 2865.14966 | 2898.8316 | 2953.48238 | 2593.7137 | 2659.5926 | 2676.6952 | 1.10E-25 | -0.647785326 | YDR457W | TOM1 | n |
| W303.asm02.HP0_G0013000 | 674.26643 | 607.258221 | 589.2142736 | 1571.74877 | 1581.998 | 1344.47448 | 1250.6831 | 1310.7711 | 1237.8393 | 3.27E-45 | -1.266901628 | YDR459C | PFA5 | n |
| W303.asm02.HP0_G0013010 | 755.38911 | 827.324448 | 635.5786754 | 1157.56046 | 1139.7577 | 1055.28921 | 930.31666 | 998.49529 | 1019.1896 | 1.66E-09 | -0.597119081 | YDR460W | TFB3 | n |
| W303.asm02.HP0_G0013030 | 1107.2719 | 1176.10866 | 1130.132295 | 1276.73336 | 1325.8222 | 1243.97075 | 1344.1708 | 1414.4257 | 1334.821 | 0.013131486 | -0.172445039 | YDR462W | MRPL28 | n |
| W303.asm02.HP0_G0013040 | 761.71036 | 958.118527 | 928.2539621 | 2179.2805 | 2144.686 | 2194.01549 | 2317.811 | 2210.8601 | 2163.5738 | 1.15E-52 | -1.298886483 | YDR463W | STP1 | n |
| W303.asm02.HP0_G0013050 | 375.0607 | 459.855371 | 464.6099436 | 1050.88823 | 1036.3885 | 935.822508 | 831.12849 | 867.28697 | 880.77026 | 4.65E-36 | -1.217452755 | YDR464W | SPP41 | n |
| W303.asm02.HP0_G0013060 | 1202.0906 | 1146.00526 | 1217.065549 | 1515.91252 | 1510.9879 | 1759.7635 | 1016.9638 | 1144.1365 | 1095.8933 | 4.63E-07 | -0.424407406 | YDR465C | RMT2 | n |
| W303.asm02.HP0_G0013070 | 572.07293 | 648.780151 | 708.9889783 | 965.883785 | 981.55787 | 1066.66699 | 918.91572 | 783.31365 | 934.55103 | 2.87E-10 | -0.641973557 | YDR466W | PKH3 | n |
| W303.asm02.HP0_G0013080 | 2033.3347 | 2065.716 | 2341.402294 | 428.355708 | 413.47676 | 438.044578 | 485.68002 | 392.31287 | 493.72507 | 8.01E-126 | 2.3315835 | YDR468C | TLG1 | n |
| W303.asm02.HP0_G0013090 | 1281.1062 | 1319.35931 | 1586.048913 | 227.511884 | 239.09743 | 258.844524 | 223.45841 | 224.36622 | 211.59646 | 4.66E-117 | 2.530776602 | YDR469W | SDC1 | n |
| W303.asm02.HP0_G0013100 | 696.3908 | 614.524559 | 668.4201267 | 1397.573 | 1371.6642 | 1342.57819 | 1428.5377 | 1503.6473 | 1508.5064 | 1.86E-45 | -1.054970823 | YDR470C | UGO1 | n |
| W303.asm02.HP0_G0013120 | 1090.4152 | 1174.03256 | 1051.892367 | 435.022723 | 478.19486 | 404.859383 | 446.91683 | 442.17203 | 474.32873 | 3.85E-48 | 1.330360471 | YDR472W | TRS31 | n |
| W303.asm02.HP0_G0013130 | 537.30606 | 536.670941 | 426.9388671 | 451.69026 | 451.22899 | 458.903844 | 373.95082 | 372.63162 | 341.19929 | 0.212099022 | 0.139051176 | YDR473C | PRP3 | n |
| W303.asm02.HP0_G0013140 | 552.05564 | 573.002629 | 581.4868732 | 459.190652 | 413.47676 | 473.12607 | 588.28848 | 568.13201 | 622.44625 | 0.000217832 | 0.343105498 | YDR475C | JIP4 | n |
| W303.asm02.HP0_G0013150 | 461.45109 | 601.029932 | 516.7698957 | 1505.91199 | 1330.3165 | 1176.65221 | 1381.7939 | 1422.2982 | 1416.8146 | 1.97E-34 | -1.345982752 | YDR476C | | n |
| W303.asm02.HP0_G0013160 | 955.56196 | 975.765347 | 984.2776144 | 1879.26483 | 1788.7364 | 1699.082 | 1895.9762 | 1873.6548 | 1974.9003 | 2.03E-37 | -0.880684188 | YDR477W | SNF1 | n |
| W303.asm02.HP0_G0013170 | 440.38026 | 468.159757 | 534.1565464 | 332.517369 | 337.97231 | 304.355649 | 241.69992 | 259.79247 | 260.96897 | 2.70E-07 | 0.565890057 | YDR478W | SNM1 | n |
| W303.asm02.HP0_G0013180 | 506.75337 | 565.736292 | 576.6572481 | 686.702536 | 583.36178 | 607.763149 | 760.44267 | 666.53825 | 741.46926 | 0.064100119 | -0.187656629 | YDR479C | PEX29 | n |
| W303.asm02.HP0_G0013200 | 2273.5421 | 2252.56468 | 2422.539997 | 1377.57196 | 1240.4303 | 1185.18555 | 1777.4065 | 1760.8156 | 1733.3277 | 2.77E-32 | 0.868873259 | YDR481C | PHO8 | n |
| W303.asm02.HP0_G0013210 | 508.86045 | 590.649449 | 498.4173199 | 370.019328 | 398.1961 | 417.185313 | 311.24565 | 338.51746 | 283.01026 | 8.81E-05 | 0.431372623 | YDR482C | CWC21 | n |
| W303.asm02.HP0_G0013220 | 4333.2154 | 4420.00941 | 3841.483879 | 2503.4641 | 2468.2765 | 2585.60079 | 2735.0854 | 3037.4725 | 2615.8612 | 5.17E-21 | 0.736763622 | YDR483W | KRE2 | n |
| W303.asm02.HP0_G0013230 | 961.88321 | 992.374119 | 972.6865139 | 841.710633 | 864.70575 | 859.022485 | 768.42332 | 802.9949 | 801.42159 | 0.00800963 | 0.190389805 | YDR484W | VPS52 | n |
| W303.asm02.HP0_G0013240 | 645.82082 | 663.312826 | 648.1357009 | 662.534607 | 736.16841 | 640.000196 | 676.07571 | 709.83699 | 686.80684 | 0.505401652 | -0.059019846 | YDR485C | VPS72 | n |
| W303.asm02.HP0_G0013250 | 1798.395 | 1783.36688 | 1962.759678 | 608.365111 | 620.21514 | 601.12611 | 622.4913 | 606.18242 | 653.30407 | 2.13E-98 | 1.599709138 | YDR486C | VPS60 | n |
| W303.asm02.HP0_G0013260 | 5095.9793 | 4842.49505 | 5480.658669 | 3090.99479 | 3280.8482 | 3143.11207 | 2818.3122 | 3172.6171 | 2873.3036 | 2.19E-22 | 0.696632261 | YDR487C | RIB3 | n |
| W303.asm02.HP0_G0013270 | 281.29553 | 264.702302 | 344.8352388 | 335.017499 | 337.07345 | 300.563055 | 294.14424 | 305.71538 | 318.27634 | 0.32160459 | -0.125746584 | YDR488C | PAC11 | n |
| W303.asm02.HP0_G0013280 | 483.57546 | 475.426095 | 395.0633408 | 399.187518 | 367.63477 | 335.644547 | 299.84471 | 282.09788 | 319.15799 | 0.012245845 | 0.294044129 | YDR489W | SLD5 | n |

| | | | | | | | | | | | | | | |
|---|---|---|---|---|---|---|---|---|---|---|---|---|---|---|
| W303.asm02.HP0_G0013310 | 1863.7146 | 1548.76798 | 1622.754065 | 614.198749 | 598.64244 | 665.600203 | 661.25449 | 814.80365 | 730.00778 | 2.95E-46 | 1.422792245 | YDR492W | IZH1 | n |
| W303.asm02.HP0_G0013320 | 1125.1821 | 1206.21206 | 1341.669879 | 944.215987 | 989.64764 | 1005.98549 | 964.51948 | 1068.0357 | 1012.1364 | 0.000157713 | 0.322240564 | YDR493W | MZM1 | n |
| W303.asm02.HP0_G0013330 | 1997.5143 | 1801.0137 | 1866.167175 | 2510.13111 | 2513.2196 | 2509.74891 | 2371.3954 | 2418.1693 | 2251.739 | 1.72E-10 | -0.41139209 | YDR494W | RSM28 | n |
| W303.asm02.HP0_G0013340 | 816.49451 | 770.231795 | 793.0244567 | 981.717946 | 1117.2861 | 1037.27439 | 1023.8044 | 921.08238 | 953.06572 | 1.27E-06 | -0.397927299 | YDR495C | VPS3 | n |
| W303.asm02.HP0_G0013350 | 2941.4873 | 2565.0172 | 2320.151943 | 3638.52339 | 3602.641 | 3958.51973 | 2174.1592 | 2649.0959 | 2178.5619 | 3.87E-07 | -0.517157016 | YDR496C | PUF6 | n |
| W303.asm02.HP0_G0013360 | 2818.223 | 2672.97422 | 2691.067158 | 2442.62759 | 2557.2639 | 2306.84515 | 3912.8024 | 3393.0471 | 3576.8618 | 0.027916413 | 0.163057625 | YDR497C | ITR1 | n |
| W303.asm02.HP0_G0013370 | 428.79131 | 442.208551 | 378.6426151 | 528.360932 | 512.35164 | 526.222383 | 506.20171 | 486.78286 | 545.74253 | 0.000818145 | -0.327658819 | YDR498C | SEC20 | n |
| W303.asm02.HP0_G0013380 | 538.3596 | 489.95877 | 447.2232929 | 515.860279 | 613.02424 | 497.77793 | 543.82481 | 549.76285 | 573.95539 | 0.201790504 | -0.14168671 | YDR499W | LCD1 | n |
| W303.asm02.HP0_G0013400 | 323.43718 | 360.20274 | 339.0396886 | 1585.0828 | 1776.1523 | 1683.91163 | 1633.7546 | 1611.2381 | 1755.369 | 1.08E-157 | -2.302188758 | YDR501W | PLM2 | n |
| W303.asm02.HP0_G0013420 | 1409.6383 | 1614.16502 | 1562.866712 | 222.511623 | 273.25421 | 243.674149 | 304.40508 | 297.84288 | 354.42407 | 3.59E-135 | 2.634294699 | YDR503C | LPP1 | n |
| W303.asm02.HP0_G0013450 | 199.11931 | 180.620394 | 131.3658053 | 627.532779 | 702.01163 | 709.215031 | 801.48605 | 761.00824 | 843.74088 | 3.19E-56 | -1.999421099 | YDR506C | GMC1 | n |
| W303.asm02.HP0_G0013460 | 1269.5173 | 1434.58267 | 1331.044703 | 3088.49466 | 3295.23 | 2942.1046 | 2568.6317 | 2592.6763 | 2799.2448 | 4.25E-58 | -1.208692469 | YDR507C | GIN4 | n |
| W303.asm02.HP0_G0013480 | 1680.3984 | 1738.7308 | 1546.445987 | 1736.75738 | 1885.8136 | 1638.4005 | 1910.7975 | 2069.1552 | 1976.6636 | 0.298960606 | -0.083867268 | YDR510W | SMT3 | n |
| W303.asm02.HP0_G0013490 | 1457.0476 | 1430.43048 | 1811.109447 | 659.201099 | 664.2594 | 598.281664 | 543.82481 | 602.24617 | 587.18017 | 8.71E-36 | 1.289854773 | YDR511W | SDH7 | n |
| W303.asm02.HP0_G0013500 | 2727.6184 | 2644.94692 | 3005.95872 | 299.182294 | 391.0052 | 310.044539 | 270.20227 | 276.84955 | 297.99835 | 1.40E-192 | 3.067082891 | YDR512C | EMI1 | n |
| W303.asm02.HP0_G0013510 | 646.87436 | 577.154822 | 642.3401507 | 588.364066 | 628.3049 | 526.222383 | 441.21636 | 498.5916 | 486.67186 | 0.337414019 | 0.098293108 | YDR514C | | n |
| W303.asm02.HP0_G0013520 | 1669.863 | 1523.85482 | 1719.346569 | 15.8341604 | 25.168151 | 12.3259297 | 6.8405637 | 24.92958 | 15.869734 | 2.53E-208 | 6.522543668 | YDR515W | SLF1 | n |
| W303.asm02.HP0_G0013540 | 935.54467 | 1045.31458 | 1067.347168 | 1078.38966 | 1162.2292 | 1017.36327 | 1094.4902 | 1221.5494 | 1167.3071 | 0.278743694 | -0.095807848 | YDR517W | GRH1 | n |
| W303.asm02.HP0_G0013560 | 3068.9658 | 3152.55251 | 3478.296064 | 1797.5939 | 1923.5658 | 1728.4746 | 1743.2037 | 1809.3627 | 1727.1561 | 1.82E-30 | 0.831937043 | YDR519W | FPR2 | n |
| W303.asm02.HP0_G0013570 | 509.91399 | 507.60559 | 480.0647442 | 466.691044 | 479.99259 | 407.703828 | 422.97486 | 405.4337 | 408.20484 | 0.148221669 | 0.143422431 | YDR520C | URC2 | n |
| W303.asm02.HP0_G0013590 | 111.67538 | 86.158004 | 119.7747048 | 40.0020895 | 57.527201 | 50.2518672 | 64.985355 | 82.66124 | 58.189026 | 1.33E-06 | 1.108249771 | YDR523C | SPS1 | n |
| W303.asm02.HP0_G0013620 | 528.87773 | 492.034866 | 592.1120487 | 228.345261 | 279.54624 | 222.814883 | 297.56452 | 309.65163 | 318.27634 | 8.78E-22 | 1.143158276 | YDR525W- | SNA2 | n |
| W303.asm02.HP0_G0013630 | 1516.0459 | 1512.43629 | 1317.521753 | 878.379215 | 919.53636 | 882.726196 | 694.31722 | 801.68282 | 697.38666 | 1.07E-15 | 0.6966975 | YDR527W | RBA50 | n |
| W303.asm02.HP0_G0013640 | 509.91399 | 638.399668 | 524.497296 | 750.872554 | 830.54897 | 861.86693 | 990.74164 | 969.62946 | 952.18407 | 4.75E-08 | -0.546190888 | YDR528W | HLR1 | n |
| W303.asm02.HP0_G0013650 | 6176.9126 | 6025.87004 | 6835.851498 | 5646.12825 | 6186.8707 | 6325.09823 | 5414.3062 | 5618.3401 | 5023.6526 | 0.380581307 | 0.068561103 | YDR529C | QCR7 | n |
| W303.asm02.HP0_G0013670 | 1910.0704 | 1698.24692 | 1871.962725 | 1010.88614 | 1061.5566 | 910.2225 | 900.67422 | 970.94155 | 900.1666 | 3.84E-26 | 0.877208097 | YDR531W | CAB1 | n |
| W303.asm02.HP0_G0013680 | 582.60834 | 575.078726 | 571.8276229 | 300.015671 | 305.61326 | 303.4075 | 302.1249 | 305.71538 | 297.99835 | 1.36E-23 | 0.928015986 | YDR532C | KRE28 | n |
| W303.asm02.HP0_G0013690 | 3796.9628 | 3905.13748 | 4016.316311 | 215.844608 | 217.52473 | 201.955617 | 271.34236 | 258.48038 | 248.62584 | 0 | 4.204111063 | YDR533C | HSP31 | n |
| W303.asm02.HP0_G0013710 | 163.2989 | 156.745284 | 130.3998802 | 1134.22591 | 916.83977 | 998.400305 | 588.28848 | 631.112 | 725.59952 | 1.33E-101 | -2.76195934 | YDR536W | STL1 | n |
| W303.asm02.HP0_G0013750 | 593.14375 | 604.144077 | 440.4618176 | 161.675112 | 169.88502 | 127.051891 | 118.56977 | 76.100824 | 108.44319 | 8.57E-28 | 1.833901372 | YDR541C | | n |
| W303.asm02.HP0_G0013840 | 448.80859 | 649.818199 | 473.3032689 | 1424.24106 | 1298.8563 | 1558.75603 | 1520.8853 | 1813.2989 | 1826.7828 | 1.08E-28 | -1.445908187 | YEL065W | SIT1 | n |
| W303.asm02.HP0_G0013850 | 505.69982 | 434.942213 | 455.9166182 | 590.03082 | 710.10139 | 653.274274 | 858.49075 | 726.89408 | 782.9069 | 4.89E-06 | -0.483835107 | YEL064C | AVT2 | n |
| W303.asm02.HP0_G0013870 | 816.49451 | 664.350875 | 680.9771522 | 620.865764 | 667.85485 | 559.407578 | 570.04698 | 531.39368 | 612.74808 | 0.040323978 | 0.225219312 | YEL062W | NPR2 | n |
| W303.asm02.HP0_G0013880 | 758.54974 | 693.416225 | 710.9208284 | 848.377647 | 811.67286 | 804.978024 | 864.19122 | 795.1224 | 882.53356 | 0.020965382 | -0.189313637 | YEL061C | CIN8 | n |
| W303.asm02.HP0_G0013890 | 1428.602 | 1912.08486 | 1698.096218 | 1248.39854 | 1123.5782 | 1171.91147 | 1498.0835 | 1238.6065 | 1479.4119 | 4.54E-06 | 0.507652459 | YEL060C | PRB1 | n |
| W303.asm02.HP0_G0013900 | 553.10918 | 548.089471 | 608.5327743 | 494.19248 | 472.80169 | 490.192742 | 501.64134 | 358.19871 | 446.11587 | 0.045538446 | 0.231007076 | YEL059C-A | SOM1 | n |
| W303.asm02.HP0_G0013910 | 2650.7099 | 2541.1421 | 3089.994199 | 1598.41682 | 1604.4696 | 1594.78567 | 1581.3103 | 1531.2011 | 1576.3936 | 5.38E-23 | 0.787975673 | YEL058W | PCM1 | n |
| W303.asm02.HP0_G0013930 | 1145.1994 | 993.412167 | 1036.437566 | 1467.57666 | 1428.2925 | 1498.07453 | 1195.9586 | 1211.0528 | 1220.2062 | 2.02E-10 | -0.46896284 | YEL056W | HAT2 | n |
| W303.asm02.HP0_G0013940 | 1893.2137 | 1806.20394 | 1922.190827 | 3108.4957 | 3003.0997 | 3333.68991 | 1732.9428 | 2218.7326 | 1775.647 | 1.27E-15 | -0.748448063 | YEL055C | POL5 | n |
| W303.asm02.HP0_G0013950 | 11091.683 | 10762.4842 | 11616.21451 | 14774.9384 | 15215.046 | 16505.368 | 13470.21 | 14768.808 | 12298.163 | 1.09E-09 | -0.47412469 | YEL054C | RPL12A | y |
| W303.asm02.HP0_G0013960 | 1139.9317 | 1109.67357 | 909.9013864 | 1786.76 | 1751.8831 | 1756.91906 | 1736.3631 | 1726.7015 | 1739.4992 | 1.57E-18 | -0.746096834 | YEL053C | MAK10 | n |
| W303.asm02.HP0_G0013970 | 1539.2238 | 1591.32795 | 1408.318706 | 1391.73936 | 1374.3608 | 1397.5708 | 1177.7171 | 1287.1536 | 1156.7273 | 0.103406508 | 0.124064533 | YEL052W | AFG1 | n |
| W303.asm02.HP0_G0013980 | 4799.9342 | 4842.49505 | 5600.433374 | 2450.96136 | 2407.1538 | 2455.70445 | 2398.7577 | 2454.9076 | 2363.7088 | 1.76E-49 | 1.059655804 | YEL051W | VMA8 | n |
| W303.asm02.HP0_G0013990 | 1786.806 | 1722.12203 | 2110.546209 | 3105.99557 | 3107.3677 | 3218.0158 | 3058.8721 | 3177.8654 | 3037.2908 | 6.73E-21 | -0.746415345 | YEL050C | RML2 | n |
| W303.asm02.HP0_G0014010 | 494.11087 | 434.942213 | 522.5654459 | 333.350746 | 343.36548 | 342.281586 | 347.72866 | 335.89329 | 372.93876 | 1.06E-06 | 0.511533925 | YEL048C | TCA17 | n |
| W303.asm02.HP0_G0014020 | 847.04721 | 903.10197 | 1017.119066 | 1757.59181 | 1558.6276 | 1576.77085 | 2531.0086 | 2014.0477 | 2392.8033 | 9.49E-16 | -0.821942352 | YEL047C | FRD1 | n |
| W303.asm02.HP0_G0014030 | 4430.1412 | 4810.31555 | 5042.128702 | 4665.24368 | 4371.1684 | 5120.00156 | 3856.9378 | 3398.2954 | 3657.0921 | 0.88147023 | 0.013024415 | YEL046C | GLY1 | n |
| W303.asm02.HP0_G0014050 | 1670.9165 | 1567.45284 | 1816.904998 | 1043.38783 | 1032.793 | 1130.19294 | 1130.9732 | 1152.009 | 1168.1888 | 2.93E-17 | 0.657691002 | YEL044W | IES6 | n |
| W303.asm02.HP0_G0014060 | 650.03498 | 772.307892 | 719.6141538 | 1705.08906 | 1615.256 | 1654.51902 | 1992.8842 | 1844.7889 | 1902.6048 | 4.61E-54 | -1.21560223 | YEL043W | GTA1 | n |
| W303.asm02.HP0_G0014070 | 1358.0147 | 1457.41973 | 1485.592709 | 1720.92322 | 1690.7604 | 1883.0228 | 1834.4112 | 1697.8356 | 1857.6406 | 7.12E-05 | -0.299236565 | YEL042W | GDA1 | n |
| W303.asm02.HP0_G0014090 | 5537.4131 | 5165.32805 | 5242.075185 | 6812.02249 | 6541.9214 | 6678.7576 | 7821.0445 | 8151.9727 | 7900.4828 | 8.70E-10 | -0.329388867 | YEL040W | UTR2 | n |
| W303.asm02.HP0_G0014110 | 858.63616 | 799.297146 | 835.5251584 | 1058.38862 | 1078.635 | 1128.29664 | 1062.5676 | 1058.8511 | 1119.6979 | 1.26E-07 | -0.388649222 | YEL038W | UTR4 | n |
| W303.asm02.HP0_G0014120 | 2242.9894 | 2427.99484 | 2526.859901 | 3465.181 | 3520.8445 | 3492.0307 | 3275.4899 | 3099.1404 | 3174.8285 | 8.59E-18 | -0.541372086 | YEL037C | RAD23 | n |
| W303.asm02.HP0_G0014130 | 1226.3221 | 1389.9466 | 1250.872925 | 3225.16846 | 3031.8633 | 3092.8602 | 3606.1172 | 3669.8966 | 3532.7792 | 8.83E-80 | -1.27397095 | YEL036C | ANP1 | n |
| W303.asm02.HP0_G0014160 | 674.26643 | 754.661072 | 909.9013864 | 1527.57979 | 1475.0334 | 1480.05971 | 1388.6344 | 1468.2211 | 1405.3531 | 8.40E-22 | -0.937440282 | YEL032W | MCM3 | n |
| W303.asm02.HP0_G0014170 | 3447.1871 | 4039.04571 | 3689.833648 | 8452.10815 | 8303.692 | 8668.92117 | 10363.454 | 9508.6667 | 9607.3609 | 5.56E-64 | -1.185781209 | YEL031W | SPF1 | n |
| W303.asm02.HP0_G0014190 | 1166.2702 | 1042.20043 | 1214.167774 | 995.885352 | 959.08631 | 1035.37809 | 1092.21 | 1065.4115 | 1013.018 | 0.017680902 | 0.195388297 | YEL029C | BUD16 | n |
| W303.asm02.HP0_G0014210 | 8399.8848 | 8094.70019 | 9498.90683 | 3480.18178 | 3725.7852 | 3944.2975 | 4176.1641 | 3854.9004 | 3777.8784 | 6.00E-53 | 1.221367869 | YEL027W | VMA3 | n |
| W303.asm02.HP0_G0014220 | 5374.1142 | 5089.55053 | 5922.086412 | 5043.59678 | 5135.2016 | 5501.15724 | 3895.701 | 5052.8323 | 3909.2446 | 0.550060117 | 0.063658973 | YEL026W | SNU13 | n |
| W303.asm02.HP0_G0014240 | 6269.6243 | 5558.74833 | 5985.837464 | 8089.58922 | 8136.5036 | 8679.3508 | 7483.5767 | 7098.3699 | 6693.5013 | 3.05E-12 | -0.483371598 | YEL024W | RIP1 | n |

| | | | | | | | | | | | | | | |
|---|---|---|---|---|---|---|---|---|---|---|---|---|---|---|
| W303.asm02.HP0_G0014250 | 271.81366 | 270.930591 | 289.7775116 | 268.34735 | 279.54624 | 323.318617 | 342.02819 | 359.51079 | 387.04519 | 0.607318466 | -0.063033204 | YEL023C | | n |
| W303.asm02.HP0_G0014260 | 1431.7626 | 1419.01195 | 1721.278419 | 5687.7971 | 5826.4269 | 5928.77218 | 5543.1368 | 5391.3497 | 5520.0226 | 1.23E-139 | -1.930954839 | YEL022W | GEA2 | n |
| W303.asm02.HP0_G0014270 | 1430.7091 | 1462.60997 | 1392.863906 | 4189.38549 | 2437.7152 | 4427.8532 | 844.80962 | 943.3878 | 908.10147 | 1.06E-09 | -1.366941039 | YEL021W | URA3 | n |
| W303.asm02.HP0_G0014280 | 3602.0577 | 3502.37477 | 4447.118878 | 1506.74537 | 1717.7263 | 1584.35604 | 1582.4504 | 1632.2315 | 1500.5716 | 2.35E-39 | 1.264716049 | YEL020W- | TIM9 | n |
| W303.asm02.HP0_G0014300 | 132.7462 | 97.5765347 | 122.6724799 | 243.346044 | 241.79402 | 216.177844 | 250.82067 | 220.42997 | 262.73227 | 2.50E-11 | -0.99114803 | YEL019C | MMS21 | n |
| W303.asm02.HP0_G0014310 | 1217.8937 | 1232.16326 | 1168.769297 | 580.030297 | 533.92434 | 565.096469 | 560.92622 | 558.94743 | 582.77191 | 5.77E-47 | 1.107256857 | YEL018W | EAF5 | n |
| W303.asm02.HP0_G0014320 | 588.92959 | 644.627958 | 533.1906213 | 1718.42309 | 1653.0082 | 1711.40793 | 2566.3515 | 2393.2397 | 2563.8438 | 7.21E-81 | -1.5256609 | YEL017W | GTT3 | n |
| W303.asm02.HP0_G0014340 | 1892.1602 | 1723.16008 | 1666.220692 | 1181.72839 | 1078.635 | 1132.08923 | 995.30202 | 907.96155 | 1114.408 | 8.53E-13 | 0.638163811 | YEL015W | EDC3 | n |
| W303.asm02.HP0_G0014350 | 1980.6576 | 1841.49758 | 1995.60113 | 2130.94464 | 2230.0779 | 2478.46002 | 2468.3034 | 2050.786 | 2432.4776 | 0.009902915 | -0.233020262 | YEL013W | VAC8 | n |
| W303.asm02.HP0_G0014360 | 1109.379 | 1170.91842 | 1318.487678 | 972.5508 | 938.41247 | 950.044735 | 1238.142 | 1187.4353 | 1358.6256 | 0.000151428 | 0.331479263 | YEL012W | UBC8 | n |
| W303.asm02.HP0_G0014370 | 242.3145 | 312.452521 | 249.20866 | 227.511884 | 213.92928 | 239.881555 | 902.95441 | 543.20243 | 758.22064 | 0.169118702 | 0.238764761 | YEL011W | GLC3 | n |
| W303.asm02.HP0_G0014380 | 11711.165 | 11244.1386 | 11577.57751 | 6585.34398 | 6523.0453 | 7445.80968 | 5744.9334 | 5365.1081 | 5341.0473 | 8.77E-29 | 0.748687649 | YEL009C | GCN4 | n |
| W303.asm02.HP0_G0014420 | 306.58052 | 317.642762 | 266.5953107 | 280.014626 | 311.9053 | 280.651938 | 272.48245 | 305.71538 | 341.19929 | 0.828042791 | 0.028810891 | YEL005C | VAB2 | n |
| W303.asm02.HP0_G0014430 | 612.1075 | 607.258221 | 715.7504536 | 494.19248 | 559.09249 | 582.163141 | 631.61205 | 612.74284 | 583.65357 | 0.016736586 | 0.244811409 | YEL004W | YEA4 | n |
| W303.asm02.HP0_G0014440 | 1254.7677 | 1203.09791 | 1577.355588 | 709.203711 | 855.71712 | 687.407617 | 632.75214 | 729.51824 | 708.84814 | 1.93E-12 | 0.841691088 | YEL003W | GIM4 | n |
| W303.asm02.HP0_G0014450 | 4852.6112 | 4905.81599 | 4975.479874 | 2970.98852 | 2958.1566 | 2983.82313 | 3355.2965 | 3395.6712 | 3129.8643 | 3.58E-39 | 0.725229418 | YEL002C | WBP1 | n |
| W303.asm02.HP0_G0014460 | 3966.583 | 3927.97455 | 4334.105648 | 3694.35964 | 3790.5033 | 3803.02338 | 4163.6231 | 4211.787 | 4110.2612 | 0.054653214 | 0.115818536 | YEL001C | IRC22 | n |
| W303.asm02.HP0_G0014470 | 4130.9354 | 3598.91325 | 3536.251566 | 505.02638 | 431.45401 | 434.251985 | 293.00415 | 246.67164 | 291.82678 | 8.21E-216 | 3.037534081 | YER001W | MNN1 | n |
| W303.asm02.HP0_G0014480 | 3152.1956 | 3136.98179 | 3082.266798 | 2035.93968 | 1970.3066 | 2177.89696 | 1207.3595 | 1502.3352 | 1206.0998 | 4.99E-12 | 0.599821146 | YER002W | NOP16 | n |
| W303.asm02.HP0_G0014490 | 2326.2192 | 2547.37038 | 2137.59211 | 6391.16717 | 6359.4523 | 6535.58718 | 6271.6568 | 5707.5618 | 5788.0448 | 1.43E-87 | -1.460129235 | YER003C | PMI40 | n |
| W303.asm02.HP0_G0014500 | 1927.9806 | 1962.94922 | 1948.270803 | 1200.06268 | 1217.9587 | 1247.76334 | 1366.9726 | 1281.9053 | 1422.9862 | 1.54E-24 | 0.671942265 | YER004W | FMP52 | n |
| W303.asm02.HP0_G0014510 | 794.37014 | 848.085413 | 776.6037311 | 1062.5555 | 1105.6009 | 994.607711 | 1047.7463 | 1049.6665 | 1069.4438 | 4.73E-07 | -0.3874557 | YER005W | YND1 | n |
| W303.asm02.HP0_G0014520 | 3105.8398 | 2667.78398 | 2828.228513 | 2143.44529 | 1985.5873 | 2054.63766 | 1237.0019 | 1418.3619 | 1320.7146 | 1.89E-09 | 0.475951025 | YER006W | NUG1 | n |
| W303.asm02.HP0_G0014530 | 500.43212 | 413.1432 | 500.34917 | 514.193525 | 525.83458 | 464.592735 | 453.75739 | 427.73911 | 453.16908 | 0.405267632 | -0.089908253 | YER007W | PAC2 | n |
| W303.asm02.HP0_G0014540 | 2387.3246 | 2323.15196 | 2654.362006 | 1920.10029 | 1958.6214 | 1971.2006 | 1549.3877 | 1768.6881 | 1592.2634 | 1.46E-05 | 0.3326306 | YER007C-/ | TMA20 | n |
| W303.asm02.HP0_G0014550 | 857.58262 | 922.824887 | 830.6955332 | 1465.9099 | 1503.797 | 1597.63012 | 1487.8226 | 1322.5798 | 1495.2816 | 1.10E-23 | -0.806685889 | YER008C | SEC3 | n |
| W303.asm02.HP0_G0014560 | 5738.6395 | 5676.04778 | 6851.306299 | 1891.76548 | 1893.9033 | 1961.71912 | 1682.7787 | 1743.7585 | 1657.5056 | 6.13E-96 | 1.668464331 | YER009W | NTF2 | n |
| W303.asm02.HP0_G0014570 | 1068.2909 | 1021.43947 | 882.8554853 | 135.007052 | 149.21118 | 140.325969 | 240.55982 | 174.50706 | 238.92767 | 2.32E-102 | 2.807885493 | YER010C | | n |
| W303.asm02.HP0_G0014590 | 5360.4181 | 5499.57958 | 5924.984187 | 2040.10656 | 2238.1677 | 2048.00063 | 2305.27 | 2319.763 | 2270.2537 | 1.51E-101 | 1.407937842 | YER012W | PRE1 | n |
| W303.asm02.HP0_G0014600 | 390.86382 | 450.512937 | 412.4499915 | 1153.39358 | 1152.3418 | 1147.25961 | 974.78033 | 942.07572 | 946.0125 | 6.26E-74 | -1.461539148 | YER013W | PRP22 | n |
| W303.asm02.HP0_G0014610 | 454.0763 | 474.388046 | 493.5876947 | 750.872554 | 750.55021 | 654.222422 | 802.62614 | 788.56198 | 865.78218 | 9.32E-11 | -0.600452318 | YER014W | HEM14 | n |
| W303.asm02.HP0_G0014620 | 308.6876 | 356.050547 | 362.2218895 | 536.6947 | 524.03685 | 522.429789 | 508.4819 | 510.40035 | 518.41132 | 7.99E-11 | -0.623548787 | YER015W | FAA2 | n |
| W303.asm02.HP0_G0014630 | 930.27697 | 888.569295 | 871.2643848 | 731.704886 | 735.26954 | 697.83725 | 723.95966 | 698.02825 | 698.26831 | 4.06E-05 | 0.312658951 | YER016W | BIM1 | n |
| W303.asm02.HP0_G0014640 | 1514.9924 | 1685.79034 | 1948.270803 | 3336.84096 | 3293.4323 | 3317.57138 | 3684.7837 | 3364.1812 | 3343.2241 | 4.49E-27 | -0.949514707 | YER017C | AFG3 | n |
| W303.asm02.HP0_G0014650 | 626.85707 | 573.002629 | 661.6586515 | 273.347611 | 283.14169 | 269.274156 | 282.7433 | 244.04747 | 230.11115 | 2.61E-27 | 1.173047916 | YER018C | SPC25 | n |
| W303.asm02.HP0_G0014670 | 2222.9721 | 2168.48278 | 2511.4051 | 2215.1157 | 2338.8403 | 2224.35624 | 2216.3426 | 2288.273 | 2145.9408 | 0.730861792 | 0.026693097 | YER019C-/ | SBH2 | n |
| W303.asm02.HP0_G0014680 | 1211.5725 | 1258.11447 | 1044.164967 | 1668.42048 | 1526.2686 | 1495.23009 | 2131.9757 | 1667.6577 | 1823.2562 | 6.20E-05 | -0.417229407 | YER020W | GPA2 | n |
| W303.asm02.HP0_G0014690 | 5266.653 | 4948.37597 | 4776.499316 | 4377.72867 | 4438.5831 | 4400.3569 | 4421.2843 | 4487.3244 | 4497.3064 | 0.001361776 | 0.181632315 | YER021W | RPN3 | n |
| W303.asm02.HP0_G0014700 | 1067.2373 | 1135.62478 | 817.1725827 | 1230.06425 | 1200.8803 | 1120.71145 | 1059.1473 | 1094.2774 | 1091.4851 | 0.033140988 | -0.235009361 | YER022W | SRB4 | n |
| W303.asm02.HP0_G0014710 | 2312.5232 | 2311.73343 | 2183.956512 | 1778.42623 | 1737.5013 | 1868.80057 | 1783.1069 | 1611.2381 | 1840.0075 | 2.27E-06 | 0.338417275 | YER023W | PRO3 | n |
| W303.asm02.HP0_G0014720 | 441.4338 | 408.991007 | 423.0751669 | 490.025596 | 538.41865 | 563.200172 | 556.36585 | 625.86367 | 567.78383 | 0.001062485 | -0.32069286 | YER024W | YAT2 | n |
| W303.asm02.HP0_G0014730 | 3767.4637 | 3712.06051 | 3669.549222 | 8349.6028 | 8078.0775 | 8543.76557 | 7066.3023 | 7680.9349 | 6936.8372 | 1.28E-85 | -1.163385532 | YER025W | GCD11 | n |
| W303.asm02.HP0_G0014740 | 2864.5788 | 2580.58793 | 2870.729215 | 1882.59834 | 1782.4444 | 1887.76354 | 2031.6474 | 2042.9135 | 2108.0297 | 3.38E-18 | 0.582756799 | YER026C | CHO1 | n |
| W303.asm02.HP0_G0014750 | 860.74324 | 1053.61897 | 981.3798392 | 1182.56177 | 1105.6009 | 1171.91147 | 1222.1807 | 1205.8044 | 1243.1292 | 0.002573834 | -0.256667956 | YER027C | GAL83 | n |
| W303.asm02.HP0_G0014770 | 888.13532 | 901.025874 | 931.1517372 | 916.71455 | 896.16594 | 870.400266 | 843.66952 | 915.83405 | 921.32625 | 0.805752935 | 0.019665595 | YER029C | SMB1 | n |
| W303.asm02.HP0_G0014780 | 2303.0413 | 2417.61435 | 2820.501113 | 1561.74824 | 1672.7832 | 1619.43753 | 1272.3448 | 1221.5494 | 1258.9989 | 2.21E-14 | 0.636161864 | YER030W | CHZ1 | n |
| W303.asm02.HP0_G0014800 | 478.30775 | 462.969516 | 479.0988192 | 1344.23688 | 1341.1029 | 1353.00782 | 1475.2816 | 1389.4961 | 1486.4651 | 1.20E-92 | -1.507295746 | YER032W | FIR1 | n |
| W303.asm02.HP0_G0014810 | 191.74452 | 232.522806 | 196.0827828 | 649.200577 | 679.54007 | 718.696516 | 612.23045 | 625.86367 | 607.45817 | 4.88E-62 | -1.722627062 | YER033C | ZRG8 | n |
| W303.asm02.HP0_G0014820 | 269.70657 | 239.789144 | 237.6175595 | 440.022984 | 424.26311 | 417.185313 | 323.78668 | 394.93703 | 354.42407 | 3.21E-12 | -0.779760312 | YER034W | | n |
| W303.asm02.HP0_G0014840 | 5376.2213 | 5024.15349 | 4693.429763 | 7870.4111 | 7606.1747 | 8151.23212 | 6427.8497 | 7104.9304 | 6160.9836 | 5.76E-18 | -0.646643354 | YER036C | ARB1 | n |
| W303.asm02.HP0_G0014860 | 509.91399 | 459.855371 | 521.5995209 | 335.850876 | 352.35411 | 329.955656 | 340.88809 | 342.45371 | 350.89746 | 2.21E-08 | 0.551018835 | YER038C | KRE29 | n |
| W303.asm02.HP0_G0014870 | 132.7462 | 124.565789 | 139.0932056 | 4693.5785 | 4717.2305 | 4942.69781 | 5853.2423 | 5934.5522 | 5648.7438 | 0 | -5.176769874 | YER039C | HVG1 | n |
| W303.asm02.HP0_G0014890 | 591.03667 | 614.524559 | 666.4882766 | 1315.06869 | 1203.5769 | 1321.71892 | 1213.06 | 1216.3011 | 1177.887 | 4.88E-39 | -1.035744452 | YER040W | GLN3 | n |
| W303.asm02.HP0_G0014900 | 280.24199 | 255.359867 | 272.3908609 | 670.868375 | 786.50471 | 673.185391 | 738.78088 | 666.53825 | 694.74171 | 1.71E-41 | -1.398653075 | YER041W | YEN1 | n |
| W303.asm02.HP0_G0014910 | 1799.4485 | 1769.87225 | 2222.593514 | 775.87386 | 805.38082 | 768.948383 | 582.58801 | 784.62574 | 603.93156 | 1.38E-28 | 1.301553748 | YER042W | MXR1 | n |
| W303.asm02.HP0_G0014920 | 9895.9134 | 8609.57211 | 9257.42557 | 12668.9951 | 12569.694 | 13275.0263 | 9264.4034 | 10445.494 | 8924.9623 | 7.67E-10 | -0.472203407 | YER043C | SAH1 | n |
| W303.asm02.HP0_G0014930 | 2476.8756 | 2353.25536 | 2290.208267 | 799.208412 | 828.75125 | 794.548391 | 770.70351 | 894.84072 | 718.54631 | 5.55E-76 | 1.555295995 | YER044C | ERG28 | n |
| W303.asm02.HP0_G0014960 | 423.5236 | 495.149011 | 405.6885162 | 193.343432 | 159.09867 | 175.407461 | 169.874 | 148.2654 | 148.11752 | 3.17E-24 | 1.325187128 | YER046W | SPO73 | n |
| W303.asm02.HP0_G0014970 | 317.11593 | 358.126643 | 368.9833648 | 1140.05955 | 965.37835 | 988.918821 | 928.03648 | 683.59533 | 830.5161 | 1.22E-38 | -1.566957791 | YER047C | SAP1 | n |
| W303.asm02.HP0_G0014980 | 2020.6922 | 2080.24868 | 1771.506521 | 2172.61348 | 2100.6417 | 2076.44508 | 2139.9563 | 2146.5681 | 1969.6104 | 0.15077017 | -0.113302976 | YER048C | CAJ1 | n |

| | | | | | | | | | | | | | |
|---|---|---|---|---|---|---|---|---|---|---|---|---|---|
| W303.asm02.HP0_G0014990 | 2204.0084 | 2108.27598 | 2545.212477 | 1160.06059 | 1261.1041 | 1146.31146 | 1040.9058 | 1280.5932 | 1140.8576 | 1.25E-22 | 0.943056118 | YER048W- | ISD11 | n |
| W303.asm02.HP0_G0015000 | 3836.9974 | 3837.66435 | 3456.079788 | 1290.06739 | 1207.1724 | 1360.59301 | 876.73225 | 1140.2003 | 906.33817 | 1.67E-55 | 1.528720072 | YER049W | TPA1 | n |
| W303.asm02.HP0_G0015010 | 2391.5388 | 2530.76161 | 2976.980969 | 946.716117 | 1012.1192 | 943.407696 | 863.05112 | 1064.0995 | 927.49781 | 2.12E-47 | 1.444856852 | YER050C | RSM18 | n |
| W303.asm02.HP0_G0015020 | 701.65851 | 619.7148 | 697.3978779 | 532.527816 | 559.09249 | 580.266844 | 527.8635 | 480.22244 | 515.76637 | 0.002818637 | 0.272985311 | YER051W | JHD1 | n |
| W303.asm02.HP0_G0015030 | 4854.7183 | 4752.18485 | 3715.913624 | 1877.59807 | 1877.7238 | 1895.34873 | 1539.1268 | 1529.889 | 1548.1808 | 9.80E-40 | 1.237191887 | YER052C | HOM3 | n |
| W303.asm02.HP0_G0015050 | 158.03119 | 164.011622 | 145.8546808 | 114.17263 | 116.85213 | 117.570406 | 171.01409 | 143.01707 | 166.63221 | 0.005253761 | 0.424081027 | YER054C | GIP2 | n |
| W303.asm02.HP0_G0015060 | 8949.8333 | 8169.43966 | 8407.411536 | 1553.41447 | 1626.9412 | 1668.74125 | 1504.924 | 1216.3011 | 1410.6431 | 3.38E-199 | 2.396456977 | YER055C | HIS1 | n |
| W303.asm02.HP0_G0015070 | 7391.6458 | 6608.21511 | 7515.862726 | 4759.41527 | 4098.8131 | 2361.83776 | 3274.3498 | 4046.4645 | 3813.1445 | 0.000127166 | 0.939204543 | YER056C | FCY2 | n |
| W303.asm02.HP0_G0015100 | 866.01095 | 869.884426 | 929.2198872 | 470.857928 | 437.74605 | 445.629766 | 451.4772 | 435.61161 | 428.48283 | 3.64E-31 | 0.976473894 | YER058W | PET117 | n |
| W303.asm02.HP0_G0015110 | 584.71542 | 602.06798 | 494.5536198 | 704.20345 | 655.27078 | 582.163141 | 718.25919 | 677.03492 | 686.80684 | 0.046921307 | -0.209759186 | YER059W | PCL6 | n |
| W303.asm02.HP0_G0015130 | 348.72217 | 324.9091 | 365.1196646 | 554.195614 | 579.76633 | 551.822391 | 614.51064 | 606.18242 | 625.09121 | 2.29E-14 | -0.697742787 | YER061C | CEM1 | n |
| W303.asm02.HP0_G0015150 | 1055.6484 | 1130.43453 | 1048.994592 | 934.215464 | 898.86252 | 897.896571 | 831.12849 | 716.39741 | 853.43905 | 0.003782568 | 0.243974762 | YER063W | THO1 | n |
| W303.asm02.HP0_G0015170 | 273.92074 | 328.023244 | 304.2663872 | 9.1671455 | 27.864738 | 6.63703906 | 6.8405637 | 10.496665 | 2.6449557 | 6.06E-14 | 4.374957984 | YER065C | ICL1 | n |
| W303.asm02.HP0_G0015190 | 302.36635 | 445.322696 | 338.0737635 | 35.8352051 | 30.561326 | 20.8592656 | 54.72451 | 40.674578 | 82.87528 | 4.68E-47 | 3.629119201 | YER067W | RGI1 | n |
| W303.asm02.HP0_G0015200 | 741.69308 | 880.264909 | 795.9222318 | 1601.75033 | 1607.1662 | 1449.71896 | 1641.7353 | 1653.2248 | 1732.446 | 2.05E-30 | -0.946495195 | YER068W | MOT2 | n |
| W303.asm02.HP0_G0015210 | 1909.0168 | 1925.57949 | 2079.636608 | 1503.41186 | 1477.73 | 1592.88938 | 1433.0981 | 1087.7169 | 1428.2761 | 0.000223604 | 0.371019731 | YER069W | ARG56 | n |
| W303.asm02.HP0_G0015220 | 2086.0118 | 2361.55975 | 2450.551823 | 8696.28757 | 8588.6314 | 9436.9214 | 9146.9738 | 9113.7297 | 9669.9582 | 4.53E-155 | -1.953358069 | YER070W | RNR1 | n |
| W303.asm02.HP0_G0015230 | 961.88321 | 886.493198 | 1241.213675 | 683.369028 | 683.13552 | 693.096508 | 603.1097 | 645.54492 | 627.73616 | 3.17E-07 | 0.585926843 | YER071C | TDA2 | n |
| W303.asm02.HP0_G0015240 | 10005.482 | 10008.8611 | 10920.74849 | 3232.66886 | 3321.297 | 3327.05287 | 3224.1857 | 2549.3776 | 3204.8047 | 2.35E-79 | 1.646599259 | YER072W | VTC1 | n |
| W303.asm02.HP0_G0015250 | 2364.1467 | 2429.03289 | 2547.144327 | 5863.63961 | 3945.1076 | 6978.3725 | 10540.169 | 10281.484 | 11191.689 | 1.62E-16 | -1.193340119 | YER073W | ALD5 | n |
| W303.asm02.HP0_G0015270 | 359.25758 | 322.833003 | 426.9388671 | 189.176548 | 216.62587 | 175.407461 | 189.2556 | 170.57081 | 162.22395 | 2.98E-11 | 0.932837541 | YER074W- | YOS1 | n |
| W303.asm02.HP0_G0015280 | 515.1817 | 549.12752 | 626.8853501 | 1033.38731 | 1039.0851 | 1002.1929 | 727.37994 | 755.75991 | 778.49864 | 1.06E-22 | -0.86117837 | YER075C | PTP3 | n |
| W303.asm02.HP0_G0015290 | 231.77909 | 243.941337 | 234.7197844 | 347.518152 | 323.59051 | 326.163063 | 332.90743 | 397.5612 | 357.95068 | 1.56E-05 | -0.490038345 | YER076C | | n |
| W303.asm02.HP0_G0015300 | 1492.868 | 1427.31633 | 1361.954304 | 857.544793 | 888.97503 | 833.422477 | 664.67477 | 775.44115 | 662.12059 | 7.55E-19 | 0.730552234 | YER077C | MRX1 | n |
| W303.asm02.HP0_G0015310 | 720.62225 | 782.688374 | 725.409704 | 1138.3928 | 1215.2621 | 1105.54108 | 970.21995 | 1081.1565 | 984.80519 | 4.62E-15 | -0.634498375 | YER078C | ICP55 | n |
| W303.asm02.HP0_G0015330 | 285.50969 | 286.501315 | 289.7775116 | 76.6706715 | 62.920377 | 53.0963125 | 102.60846 | 129.89623 | 154.28908 | 5.08E-34 | 2.155361041 | YER079W | | n |
| W303.asm02.HP0_G0015340 | 1589.7938 | 1682.6762 | 1495.25196 | 3333.50746 | 3467.8116 | 3571.67517 | 3209.3645 | 3141.1271 | 3064.622 | 2.85E-65 | -1.121666695 | YER080W | AIM9 | n |
| W303.asm02.HP0_G0015350 | 2766.5995 | 2572.28354 | 2277.651241 | 4146.0499 | 4141.0596 | 3757.51226 | 3648.3006 | 3370.7417 | 3696.7665 | 1.38E-15 | -0.661654563 | YER081W | SER3 | n |
| W303.asm02.HP0_G0015360 | 1448.6193 | 1464.68607 | 1303.998802 | 1876.7647 | 1832.7807 | 2027.14136 | 1146.9345 | 1363.2544 | 1191.9934 | 2.20E-07 | -0.444058469 | YER082C | UTP7 | n |
| W303.asm02.HP0_G0015370 | 694.28372 | 817.982014 | 817.1725827 | 1560.91487 | 1594.5821 | 1495.23009 | 1881.155 | 1876.2789 | 1927.2911 | 6.84E-37 | -0.996935578 | YER083C | GET2 | n |
| W303.asm02.HP0_G0015400 | 2481.0898 | 2651.17521 | 2794.421137 | 2277.61897 | 2274.1222 | 2311.58589 | 2242.5648 | 2071.7793 | 2205.0114 | 0.00197363 | 0.208161198 | YER086W | ILV1 | n |
| W303.asm02.HP0_G0015410 | 921.84864 | 930.091224 | 850.979959 | 935.048841 | 908.75001 | 915.911391 | 958.81901 | 820.05198 | 916.91799 | 0.725261498 | -0.03063076 | YER087W | AIM10 | n |
| W303.asm02.HP0_G0015430 | 639.49957 | 793.068856 | 787.2289065 | 2402.6255 | 2257.0438 | 2322.96367 | 2013.4059 | 1784.4331 | 1943.1608 | 5.02E-76 | -1.652727799 | YER088C | DOT6 | n |
| W303.asm02.HP0_G0015450 | 3432.4376 | 3385.07532 | 2890.047716 | 2581.80152 | 2474.5685 | 2511.64521 | 2416.9992 | 2133.4472 | 2411.318 | 1.50E-05 | 0.358854572 | YER090W | TRP2 | n |
| W303.asm02.HP0_G0015460 | 5957.7761 | 5700.96094 | 5915.324937 | 1934.2677 | 1880.4204 | 1505.65972 | 3744.0685 | 2582.1797 | 3670.3169 | 1.13E-41 | 1.723548704 | YER091C | MET6 | n |
| W303.asm02.HP0_G0015480 | 845.99366 | 853.275654 | 886.7191855 | 150.841212 | 142.91914 | 136.533375 | 141.37165 | 95.782071 | 145.47257 | 2.04E-99 | 2.586015381 | YER092W | IES5 | n |
| W303.asm02.HP0_G0015490 | 420.36298 | 472.31195 | 473.3032689 | 1161.72735 | 1146.9486 | 1138.72627 | 1306.5477 | 1187.4353 | 1354.2173 | 1.60E-55 | -1.334958068 | YER093C | TSC11 | n |
| W303.asm02.HP0_G0015500 | 397.18507 | 421.447586 | 371.8811399 | 499.192741 | 509.65505 | 540.44461 | 486.82012 | 502.52785 | 480.50029 | 5.73E-05 | -0.380148562 | YER093C-/ | AIM11 | n |
| W303.asm02.HP0_G0015510 | 4062.4553 | 3978.83891 | 3685.969947 | 2262.61819 | 2299.2903 | 2211.08216 | 2419.2794 | 2482.4614 | 2453.6373 | 5.41E-39 | 0.791680685 | YER094C | PUP3 | n |
| W303.asm02.HP0_G0015520 | 686.90893 | 708.986949 | 658.7608763 | 1736.75738 | 1648.5139 | 1645.03754 | 1640.5952 | 1448.5398 | 1648.6891 | 6.41E-61 | -1.292398986 | YER095W | RAD51 | n |
| W303.asm02.HP0_G0015540 | 257.06408 | 284.425218 | 299.436762 | 1291.73414 | 1294.362 | 1167.17073 | 1475.2816 | 1339.6369 | 1489.1101 | 2.03E-116 | -2.157146268 | YER098W | UBP9 | n |
| W303.asm02.HP0_G0015550 | 1232.6433 | 1051.54287 | 1173.598922 | 1704.25569 | 1632.3343 | 1755.02276 | 1617.7933 | 1667.6577 | 1588.7367 | 2.12E-13 | -0.558148075 | YER099C | PRS2 | n |
| W303.asm02.HP0_G0015560 | 2039.656 | 2129.03694 | 1876.79235 | 1543.41395 | 1590.0878 | 1484.80045 | 1680.4985 | 1772.6244 | 1719.2212 | 4.27E-08 | 0.387974387 | YER100W | UBC6 | n |
| W303.asm02.HP0_G0015570 | 379.27487 | 386.153946 | 400.858891 | 2015.93863 | 2152.7757 | 2268.91921 | 2349.7336 | 2238.4139 | 2243.8041 | 2.43E-211 | -2.463575503 | YER101C | AST2 | n |
| W303.asm02.HP0_G0015630 | 1125.1821 | 1018.32532 | 1060.585692 | 1765.0922 | 1785.141 | 1829.92649 | 1612.0928 | 1675.5302 | 1645.1625 | 2.67E-29 | -0.747722389 | YER107C | GLE2 | n |
| W303.asm02.HP0_G0015650 | 1757.3069 | 1856.03026 | 2066.113658 | 7453.72267 | 7180.1138 | 8549.45446 | 4629.9215 | 6077.5692 | 4780.3167 | 7.45E-77 | -2.028980856 | YER110C | KAP123 | n |
| W303.asm02.HP0_G0015660 | 739.58599 | 769.193747 | 728.3074791 | 1102.55759 | 1152.3418 | 1204.14852 | 1050.0265 | 910.58572 | 1147.0291 | 1.06E-11 | -0.628545161 | YER111C | SWI4 | n |
| W303.asm02.HP0_G0015670 | 3328.137 | 3270.89001 | 4039.498512 | 1959.26901 | 2002.6657 | 1955.08208 | 1748.9041 | 1788.3694 | 1905.2498 | 1.11E-22 | 0.846643402 | YER112W | LSM4 | n |
| W303.asm02.HP0_G0015680 | 979.79341 | 966.422913 | 1082.801968 | 2814.31367 | 2745.1261 | 2832.11938 | 3448.7842 | 3261.8388 | 3316.7745 | 1.38E-107 | -1.469300753 | YER113C | TMN3 | n |
| W303.asm02.HP0_G0015690 | 471.9865 | 603.106028 | 606.6009243 | 2235.95013 | 2296.5937 | 2319.17108 | 2442.0812 | 2101.9572 | 2395.4482 | 1.63E-95 | -2.025519436 | YER114C | BOI2 | n |
| W303.asm02.HP0_G0015700 | 377.16779 | 356.050547 | 396.0292658 | 213.344477 | 270.55762 | 204.800063 | 204.07682 | 228.30247 | 214.24141 | 2.52E-08 | 0.71376532 | YER115C | SPR6 | n |
| W303.asm02.HP0_G0015710 | 330.81197 | 332.175437 | 278.1864111 | 554.195614 | 505.16074 | 442.78532 | 421.83476 | 443.48411 | 391.45345 | 1.06E-08 | -0.67714532 | YER116C | SLX8 | n |
| W303.asm02.HP0_G0015730 | 2334.6475 | 2173.67302 | 2214.866114 | 1582.58266 | 1556.8299 | 1491.43749 | 1731.8027 | 1851.3494 | 1754.4873 | 1.04E-16 | 0.537416398 | YER118C | SHO1 | n |
| W303.asm02.HP0_G0015740 | 329.75843 | 424.561731 | 346.7670889 | 1223.39724 | 1159.5327 | 1144.41516 | 1865.1937 | 1659.7852 | 1745.6708 | 1.39E-66 | -1.680355453 | YER119C | AVT6 | n |
| W303.asm02.HP0_G0015770 | 3755.8747 | 3718.2888 | 3345.964334 | 3028.49152 | 3063.3235 | 3181.98616 | 2952.8433 | 2904.9521 | 2924.4394 | 0.000563658 | 0.222362594 | YER122C | GLO3 | n |
| W303.asm02.HP0_G0015780 | 1188.3946 | 1198.94572 | 1055.756067 | 1707.58919 | 1599.0764 | 1686.75607 | 1935.8795 | 1508.8956 | 1817.0846 | 2.78E-08 | -0.536765087 | YER123W | YCK3 | n |
| W303.asm02.HP0_G0015790 | 3831.7297 | 3450.47235 | 3644.435171 | 960.050147 | 1053.4669 | 1138.72627 | 1301.9873 | 1251.7273 | 1303.9632 | 1.31E-118 | 1.794355615 | YER124C | DSE1 | n |
| W303.asm02.HP0_G0015800 | 1975.3899 | 2185.09155 | 2387.766695 | 3060.15984 | 3005.7963 | 3069.15649 | 3477.2866 | 2878.7105 | 3359.9754 | 7.27E-08 | -0.47999032 | YER125W | RSP5 | n |
| W303.asm02.HP0_G0015810 | 4701.9548 | 4231.08463 | 4875.989595 | 3706.86029 | 3338.3754 | 4001.18641 | 2509.3468 | 3138.5029 | 2622.9144 | 0.001200988 | 0.322165902 | YER126C | NSA2 | n |
| W303.asm02.HP0_G0015820 | 2128.1534 | 2097.8955 | 1883.553825 | 1269.23296 | 1166.7236 | 1380.50413 | 720.53938 | 926.33072 | 792.60507 | 9.07E-12 | 0.678863736 | YER127W | LCP5 | n |

| | | | | | | | | | | | | | |
|---|---|---|---|---|---|---|---|---|---|---|---|---|---|
| W303.asm02.HP0_G0015830 | 428.79131 | 426.637827 | 341.9374637 | 250.846436 | 265.16444 | 283.496383 | 184.69522 | 208.62122 | 178.97534 | 3.64E-06 | 0.582477383 | YER128W | VFA1 | n |
| W303.asm02.HP0_G0015840 | 805.95909 | 916.596597 | 917.6287867 | 1527.57979 | 1528.0663 | 1363.43745 | 1451.3396 | 1339.6369 | 1456.489 | 7.81E-19 | -0.74307381 | YER129W | SAK1 | n |
| W303.asm02.HP0_G0015850 | 1234.7504 | 1232.16326 | 1109.847869 | 1829.26222 | 1873.2295 | 1698.13385 | 1581.3103 | 1573.1877 | 1579.0386 | 2.36E-16 | -0.595337637 | YER130C | COM2 | n |
| W303.asm02.HP0_G0015860 | 12973.308 | 12630.971 | 13894.83168 | 12453.1505 | 13223.167 | 13769.9598 | 11942.484 | 13603.678 | 10658.29 | 0.983463244 | 0.001998451 | YER131W | RPS26B | y |
| W303.asm02.HP0_G0015880 | 869.17157 | 808.63958 | 878.0258601 | 318.349962 | 337.07345 | 318.577875 | 320.3664 | 312.27579 | 326.21121 | 4.87E-54 | 1.39211035 | YER139C | RTR1 | n |
| W303.asm02.HP0_G0015890 | 938.7053 | 824.210304 | 816.2066576 | 600.864719 | 618.41742 | 595.437219 | 798.06577 | 743.95116 | 743.23256 | 4.19E-09 | 0.506526507 | YER140W | EMP65 | n |
| W303.asm02.HP0_G0015900 | 2213.4903 | 2226.61348 | 2089.295859 | 1420.07418 | 1345.5972 | 1403.25969 | 1441.0788 | 1447.2277 | 1336.5843 | 3.02E-22 | 0.646940313 | YER141W | COX15 | n |
| W303.asm02.HP0_G0015910 | 830.19054 | 887.531246 | 996.8346399 | 245.846175 | 257.07468 | 210.488953 | 320.3664 | 254.54413 | 266.25888 | 1.40E-58 | 1.927058945 | YER142C | MAG1 | n |
| W303.asm02.HP0_G0015920 | 1001.9178 | 929.053176 | 943.7087627 | 1224.23061 | 1224.2508 | 1132.08923 | 1244.9826 | 1132.3278 | 1128.5144 | 4.32E-05 | -0.317415653 | YER143W | DDI1 | n |
| W303.asm02.HP0_G0015930 | 2856.1505 | 3084.04133 | 2982.776519 | 7152.04025 | 6368.441 | 7862.04685 | 12806.675 | 10996.569 | 12763.675 | 2.15E-45 | -1.260754758 | YER144C | UBP5 | n |
| W303.asm02.HP0_G0015940 | 1708.844 | 1757.41567 | 2137.59211 | 1038.38757 | 995.04081 | 1011.67438 | 873.31197 | 904.0253 | 929.26112 | 7.32E-22 | 0.880321274 | YER146W | LSM5 | n |
| W303.asm02.HP0_G0015950 | 359.25758 | 387.191994 | 441.4277426 | 572.529905 | 548.30614 | 625.777969 | 564.34651 | 519.58494 | 579.24531 | 1.32E-07 | -0.55419543 | YER147C | SCC4 | n |
| W303.asm02.HP0_G0015960 | 3774.8385 | 3524.17378 | 3698.526973 | 1926.76731 | 2024.2384 | 2011.97099 | 1713.5612 | 1810.6748 | 1754.4873 | 4.56E-49 | 0.883326137 | YER148W | SPT15 | n |
| W303.asm02.HP0_G0015970 | 824.92284 | 796.183001 | 691.6023277 | 959.21677 | 1000.434 | 978.489188 | 827.70821 | 910.58572 | 831.39775 | 7.66E-05 | -0.346157987 | YER149C | PEA2 | n |
| W303.asm02.HP0_G0015990 | 1240.0181 | 1328.70175 | 1592.810389 | 2776.81171 | 2764.0023 | 2969.60091 | 2410.1586 | 2449.6593 | 2496.8382 | 2.55E-31 | -1.031257016 | YER151C | UBP3 | n |
| W303.asm02.HP0_G0016000 | 1075.6657 | 1142.89111 | 1329.112853 | 1165.06086 | 1164.027 | 1129.24479 | 1215.3402 | 1238.6065 | 1231.6677 | 0.676648578 | 0.037476139 | YER152C | | n |
| W303.asm02.HP0_G0016010 | 220.19013 | 204.495504 | 237.6175595 | 374.186212 | 422.46539 | 418.133461 | 388.77204 | 438.23578 | 424.07457 | 1.33E-14 | -0.872942426 | YER153C | PET122 | n |
| W303.asm02.HP0_G0016020 | 1212.626 | 1299.6364 | 1480.763084 | 1603.41709 | 1531.6617 | 1628.91902 | 1537.9867 | 1594.1811 | 1490.8734 | 0.002394214 | -0.253953531 | YER154W | OXA1 | n |
| W303.asm02.HP0_G0016030 | 2438.9481 | 2531.79966 | 2760.61376 | 3280.17134 | 3207.1415 | 3281.54174 | 3306.2725 | 3732.8766 | 3345.869 | 3.55E-06 | -0.337174214 | YER155C | BEM2 | n |
| W303.asm02.HP0_G0016040 | 1547.6522 | 1658.80109 | 1590.878539 | 2498.46384 | 2539.2866 | 2404.50444 | 1656.5565 | 2288.273 | 1720.9845 | 3.27E-09 | -0.633551426 | YER156C | MYG1 | n |
| W303.asm02.HP0_G0016050 | 1220.0008 | 1179.2228 | 1043.199042 | 1426.74119 | 1487.6175 | 1508.50416 | 1296.2868 | 1338.3248 | 1350.6907 | 2.47E-06 | -0.362032317 | YER157W | COG3 | n |
| W303.asm02.HP0_G0016070 | 3178.5341 | 3208.60711 | 3569.093018 | 1304.23479 | 1377.0574 | 1368.1782 | 1284.8859 | 1376.3752 | 1339.2293 | 1.66E-75 | 1.298406281 | YER159C | BUR6 | n |
| W303.asm02.HP0_G0016080 | 659.51685 | 733.900107 | 662.6245765 | 762.53983 | 853.9194 | 742.400227 | 584.8682 | 711.14908 | 697.38666 | 0.051471421 | -0.198461681 | YER161C | SPT2 | n |
| W303.asm02.HP0_G0016090 | 271.81366 | 242.903289 | 250.174585 | 515.026902 | 481.79031 | 493.985336 | 509.622 | 476.28619 | 504.30489 | 5.62E-22 | -0.963748397 | YER162C | RAD4 | n |
| W303.asm02.HP0_G0016100 | 1087.2546 | 1211.4023 | 1012.28944 | 588.364066 | 588.75495 | 587.852031 | 492.52059 | 551.07493 | 491.96177 | 1.98E-23 | 0.907140816 | YER163C | GCG1 | n |
| W303.asm02.HP0_G0016110 | 534.14544 | 691.340129 | 634.6127504 | 2300.12014 | 2212.1007 | 2308.74145 | 2043.0484 | 1872.3427 | 2086.8701 | 8.47E-95 | -1.874139529 | YER164W | CHD1 | n |
| W303.asm02.HP0_G0016120 | 4460.6939 | 5091.62662 | 5137.75528 | 9712.17397 | 9548.6166 | 10243.7957 | 8050.2034 | 8165.0936 | 7618.3542 | 4.58E-47 | -1.005918742 | YER165W | PAB1 | n |
| W303.asm02.HP0_G0016130 | 1311.6589 | 1287.17982 | 1294.339552 | 3325.17369 | 3199.0517 | 3229.39358 | 3373.538 | 3269.7113 | 3395.2415 | 3.23E-119 | -1.325170827 | YER166W | DNF1 | n |
| W303.asm02.HP0_G0016140 | 742.74662 | 729.747914 | 869.3325348 | 1561.74824 | 1547.8413 | 1558.75603 | 1677.0782 | 1460.3486 | 1572.867 | 6.84E-31 | -0.994300183 | YER167W | BCK2 | n |
| W303.asm02.HP0_G0016150 | 1243.1787 | 1179.2228 | 1230.588499 | 1270.89972 | 1352.7881 | 1370.07449 | 1072.8284 | 1203.1803 | 1085.3135 | 0.090138569 | -0.128204372 | YER168C | CCA1 | n |
| W303.asm02.HP0_G0016160 | 476.20067 | 534.594844 | 566.0320726 | 473.358059 | 467.40851 | 598.281664 | 552.94557 | 486.78286 | 604.81321 | 0.780110266 | 0.036743954 | YER169W | RPH1 | n |
| W303.asm02.HP0_G0016170 | 959.77612 | 909.33026 | 973.6524389 | 608.365111 | 612.12538 | 568.889063 | 522.16303 | 612.74284 | 551.9141 | 6.36E-15 | 0.667483119 | YER170W | ADK2 | n |
| W303.asm02.HP0_G0016180 | 1264.2496 | 1182.33695 | 1306.896577 | 1468.41003 | 1447.1687 | 1383.34857 | 1301.9873 | 1420.9861 | 1425.6311 | 0.008335234 | -0.195744408 | YER171W | RAD3 | n |
| W303.asm02.HP0_G0016190 | 939.75884 | 994.450215 | 1080.870118 | 2065.94125 | 2023.3395 | 2056.53396 | 1742.0636 | 1731.9498 | 1689.2451 | 3.10E-49 | -1.026605378 | YER172C | BRR2 | n |
| W303.asm02.HP0_G0016200 | 440.38026 | 490.996818 | 497.4513949 | 387.520242 | 437.74605 | 356.503813 | 381.93147 | 393.62495 | 398.50666 | 0.010500134 | 0.273910962 | YER173W | RAD24 | n |
| W303.asm02.HP0_G0016210 | 1847.9114 | 1548.76798 | 2107.648434 | 2992.65632 | 3067.8178 | 3133.63059 | 2767.008 | 2848.5326 | 2674.9319 | 1.51E-14 | -0.73965954 | YER174C | GRX4 | n |
| W303.asm02.HP0_G0016220 | 901.83135 | 990.298022 | 849.0481089 | 11.6672761 | 12.584075 | 5.68889063 | 22.801879 | 10.496665 | 18.51469 | 9.27E-126 | 6.500806367 | YER175C | TMT1 | n |
| W303.asm02.HP0_G0016250 | 11135.932 | 11629.2544 | 12326.16942 | 8897.1314 | 8840.3129 | 9997.27713 | 9133.2926 | 7549.7266 | 8128.8306 | 5.16E-05 | 0.339530027 | YER178W | PDA1 | n |
| W303.asm02.HP0_G0016260 | 187.53035 | 222.142324 | 213.4694335 | 49.169235 | 47.639714 | 50.2518672 | 58.144792 | 66.916242 | 65.242242 | 1.30E-30 | 2.083956431 | YER179W | DMC1 | n |
| W303.asm02.HP0_G0016270 | 433.00547 | 404.838814 | 337.1078385 | 502.526249 | 520.4414 | 457.007547 | 521.02294 | 497.27952 | 510.47646 | 0.002310163 | -0.335211305 | YER180C | ISC10 | n |
| W303.asm02.HP0_G0016280 | 185.42327 | 145.326754 | 128.4680301 | 143.340821 | 149.21118 | 135.585227 | 134.53109 | 110.21499 | 131.36613 | 0.575080637 | 0.098263873 | YER180C-/ | SLO1 | n |
| W303.asm02.HP0_G0016290 | 1568.723 | 1526.96896 | 1598.605939 | 810.875689 | 808.97627 | 730.074297 | 859.63084 | 942.07572 | 834.92436 | 1.65E-37 | 0.99765139 | YER182W | FMP10 | n |
| W303.asm02.HP0_G0016300 | 1016.6674 | 1030.7819 | 1155.246346 | 238.345783 | 229.20994 | 241.777852 | 303.26499 | 247.98372 | 275.0754 | 2.15E-99 | 2.175365176 | YER183C | FAU1 | n |
| W303.asm02.HP0_G0016320 | 58.998313 | 59.1687498 | 84.03547836 | 9005.47039 | 8476.2736 | 8177.78028 | 11065.752 | 11462.359 | 10772.023 | 0 | -6.97978123 | YER185W | PUG1 | n |
| W303.asm02.HP0_G0016410 | 1011.3996 | 977.841443 | 1017.119066 | 1182.56177 | 1205.3746 | 1147.25961 | 1189.118 | 1259.5998 | 1258.1173 | 0.000749269 | -0.233833947 | YFL049W | SWP82 | n |
| W303.asm02.HP0_G0016420 | 2353.6113 | 2227.65153 | 2267.991991 | 2125.94438 | 2177.9439 | 2044.28049 | 2198.1011 | 2294.8335 | 2170.627 | 0.079060067 | 0.109376417 | YFL048C | EMP47 | n |
| W303.asm02.HP0_G0016430 | 1314.8195 | 1280.95153 | 1304.964727 | 2282.61923 | 2295.6949 | 2158.93399 | 1974.6427 | 2020.6081 | 2002.2315 | 3.13E-38 | -0.788660497 | YFL047W | RGD2 | n |
| W303.asm02.HP0_G0016440 | 1402.2635 | 1528.00701 | 1686.505117 | 495.859234 | 515.94709 | 414.340867 | 524.44322 | 549.76285 | 573.07374 | 1.32E-61 | 1.694001694 | YFL046W | FMP32 | n |
| W303.asm02.HP0_G0016450 | 11098.004 | 10809.1963 | 11505.13314 | 11746.4469 | 12033.073 | 13147.0262 | 10858.255 | 11399.379 | 10188.369 | 0.035178703 | -0.144166839 | YFL045C | SEC53 | n |
| W303.asm02.HP0_G0016460 | 961.88321 | 929.053176 | 777.5696561 | 515.026902 | 538.41865 | 584.059438 | 616.79083 | 576.00451 | 596.87834 | 1.77E-12 | 0.704571566 | YFL044C | OTU1 | n |
| W303.asm02.HP0_G0016470 | 317.11593 | 398.610525 | 335.1759884 | 1558.41474 | 1598.1776 | 1531.25973 | 1209.6397 | 1474.7815 | 1249.3008 | 7.85E-97 | -2.157603705 | YFL042C | LAM5 | n |
| W303.asm02.HP0_G0016500 | 984.00757 | 952.928286 | 902.1739861 | 1433.40821 | 1338.4063 | 1428.8597 | 1788.8074 | 1718.829 | 1798.5699 | 1.01E-15 | -0.565788582 | YFL041W | FET5 | n |
| W303.asm02.HP0_G0016510 | 144.33516 | 125.603837 | 140.0591306 | 25.0013059 | 14.3818 | 18.9629688 | 21.661785 | 15.744998 | 22.041298 | 1.71E-26 | 2.805607204 | YFL040W | | n |
| W303.asm02.HP0_G0016520 | 9442.8907 | 8949.01389 | 8219.056154 | 18693.4764 | 18693.644 | 20542.5841 | 17396.694 | 16776.295 | 15480.926 | 1.30E-51 | -1.122335463 | YFL039C | ACT1 | n |
| W303.asm02.HP0_G0016530 | 8037.4666 | 8033.45534 | 8992.76211 | 3494.34919 | 3626.0114 | 3876.03081 | 3476.1465 | 3713.1954 | 3479.8801 | 1.88E-64 | 1.188887374 | YFL038C | YPT1 | n |
| W303.asm02.HP0_G0016540 | 1055.6484 | 1241.5057 | 1339.738029 | 1694.25516 | 1748.2876 | 1780.62277 | 1287.1661 | 1314.7073 | 1191.1117 | 7.58E-09 | -0.521427748 | YFL036W | RPO41 | n |
| W303.asm02.HP0_G0016570 | 384.54257 | 452.589033 | 408.5862913 | 1010.88614 | 1010.3215 | 989.866969 | 1071.6883 | 1036.5457 | 1074.7337 | 3.25E-53 | -1.27344526 | YFL034W | MIL1 | n |
| W303.asm02.HP0_G0016580 | 798.58431 | 737.014251 | 890.5828856 | 1013.38627 | 981.55787 | 824.889141 | 1076.2487 | 754.44782 | 963.64554 | 0.097232575 | -0.217040028 | YFL033C | RIM15 | n |
| W303.asm02.HP0_G0016590 | 4586.0653 | 4710.66292 | 5368.611365 | 6420.33536 | 6251.5888 | 6734.69835 | 8746.8008 | 7236.1387 | 8091.8013 | 2.51E-06 | -0.403949311 | YFL031W | HAC1 | n |
| W303.asm02.HP0_G0016600 | 324.49072 | 341.517871 | 384.4381654 | 41.6688432 | 35.954501 | 33.1851953 | 54.72451 | 40.674578 | 54.662419 | 1.46E-67 | 3.240709611 | YFL030W | AGX1 | n |

| | | | | | | | | | | | | | | |
|---|---|---|---|---|---|---|---|---|---|---|---|---|---|---|
| W303.asm02.HP0_G0016610 | 554.16272 | 586.497256 | 565.0661476 | 697.536435 | 786.50471 | 681.718727 | 648.71346 | 637.67242 | 643.6059 | 8.84E-05 | -0.344556224 | YFL029C | CAK1 | n |
| W303.asm02.HP0_G0016620 | 1530.7955 | 1512.43629 | 1656.561441 | 612.531995 | 618.41742 | 607.763149 | 660.1144 | 610.11867 | 591.58843 | 2.01E-67 | 1.354193275 | YFL028C | CAF16 | n |
| W303.asm02.HP0_G0016630 | 804.90555 | 745.318637 | 785.2970564 | 703.370073 | 728.07864 | 708.266883 | 786.66483 | 700.65241 | 729.12613 | 0.125188669 | 0.126427795 | YFL027C | GYP8 | n |
| W303.asm02.HP0_G0016640 | 10492.218 | 10266.2971 | 10637.73245 | 188.343171 | 202.24407 | 253.155633 | 238.27964 | 234.86289 | 272.43044 | 0 | 5.612431754 | YFL026W | STE2 | n |
| W303.asm02.HP0_G0016650 | 694.28372 | 680.959646 | 680.0112272 | 1006.71925 | 959.08631 | 972.800297 | 1130.9732 | 1081.1565 | 1173.4787 | 4.06E-12 | -0.516235041 | YFL025C | BST1 | n |
| W303.asm02.HP0_G0016670 | 382.43549 | 378.887608 | 353.5285641 | 1640.08567 | 1651.2105 | 1512.29676 | 1445.6391 | 1436.7311 | 1456.489 | 4.24E-155 | -2.108567462 | YFL024C | EPL1 | y |
| W303.asm02.HP0_G0016680 | 487.78962 | 549.12752 | 634.6127504 | 1077.55629 | 967.17607 | 1109.33367 | 736.50069 | 784.62574 | 748.52247 | 3.32E-19 | -0.914657786 | YFL023W | BUD27 | n |
| W303.asm02.HP0_G0016690 | 2581.1762 | 3052.89988 | 3169.200052 | 5471.95249 | 5360.8161 | 5812.14992 | 4666.4045 | 4568.6736 | 4762.6836 | 9.12E-31 | -0.918633143 | YFL022C | FRS2 | n |
| W303.asm02.HP0_G0016700 | 822.81576 | 1070.22774 | 978.4820641 | 1245.06503 | 1266.4973 | 1276.2078 | 1751.1843 | 1319.9557 | 1780.0552 | 0.000958801 | -0.399291993 | YFL021W | GAT1 | n |
| W303.asm02.HP0_G0016720 | 3827.5155 | 3762.92488 | 3981.543009 | 10254.7023 | 10565.23 | 11300.9812 | 9777.4457 | 9935.0938 | 9697.2894 | 4.22E-144 | -1.472644151 | YFL018C | LPD1 | n |
| W303.asm02.HP0_G0016740 | 948.18717 | 976.803395 | 911.8332365 | 1327.56934 | 1354.5858 | 1383.34857 | 1479.8419 | 1531.2011 | 1505.8615 | 1.01E-14 | -0.519272488 | YFL017C | GNA1 | n |
| W303.asm02.HP0_G0016750 | 2313.5767 | 2835.9478 | 2357.823019 | 2011.77175 | 1970.3066 | 1949.39319 | 1876.5946 | 1725.3894 | 1877.9186 | 7.87E-05 | 0.339713713 | YFL016C | MDJ1 | n |
| W303.asm02.HP0_G0016770 | 2713.9224 | 3188.8842 | 4075.237738 | 320.850093 | 184.26682 | 225.659328 | 1387.4943 | 922.39447 | 1349.8091 | 6.17E-41 | 3.770473173 | YFL014W | HSP12 | n |
| W303.asm02.HP0_G0016780 | 821.76221 | 945.661948 | 809.4451824 | 1975.93654 | 1834.5784 | 1845.09686 | 1950.7008 | 2021.9202 | 2027.7994 | 2.15E-48 | -1.134763689 | YFL013C | IES1 | n |
| W303.asm02.HP0_G0016810 | 2059.6732 | 2173.67302 | 2205.206863 | 2490.96345 | 2369.4016 | 2482.25261 | 2988.1862 | 3008.6067 | 2875.9485 | 0.001915026 | -0.189372558 | YFL010C | WWM1 | n |
| W303.asm02.HP0_G0016820 | 1151.5206 | 1127.32039 | 1173.598922 | 941.715856 | 929.42385 | 1034.42995 | 1157.1954 | 914.52197 | 1101.1832 | 0.008253367 | 0.249309824 | YFL009W | CDC4 | n |
| W303.asm02.HP0_G0016830 | 943.973 | 931.129273 | 764.0467055 | 1535.08018 | 1646.7161 | 1674.43014 | 1455.9 | 1577.124 | 1574.6303 | 1.71E-22 | -0.880447307 | YFL008W | SMC1 | n |
| W303.asm02.HP0_G0016840 | 1224.215 | 1381.64221 | 1514.570461 | 2698.47429 | 2708.2728 | 2573.27486 | 2658.6991 | 2667.4651 | 2648.4823 | 7.63E-34 | -0.953013232 | YFL007W | BLM10 | n |
| W303.asm02.HP0_G0016850 | 5002.2141 | 4629.69516 | 5701.855503 | 3980.2079 | 3591.8546 | 4128.2383 | 4126 | 4130.4378 | 4119.0777 | 4.05E-06 | 0.390366967 | YFL005W | SEC4 | n |
| W303.asm02.HP0_G0016860 | 2851.9363 | 2756.01808 | 2692.033083 | 5106.10005 | 5155.8754 | 4802.37184 | 5299.1567 | 4605.4119 | 5271.3968 | 1.30E-33 | -0.860160247 | YFL004W | VTC2 | n |
| W303.asm02.HP0_G0016870 | 153.81703 | 177.506249 | 192.2190827 | 939.215726 | 965.37835 | 925.392875 | 1060.2874 | 967.0053 | 1057.1006 | 3.51E-127 | -2.431883623 | YFL003C | MSH4 | n |
| W303.asm02.HP0_G0016880 | 1056.7019 | 996.526312 | 1009.391665 | 1383.40559 | 1412.113 | 1324.56337 | 868.75159 | 1134.9519 | 876.362 | 1.95E-05 | -0.428109377 | YFL002C | SPB4 | n |
| W303.asm02.HP0_G0016890 | 481.46837 | 507.60559 | 496.4854699 | 1218.39697 | 1185.5997 | 1145.36331 | 859.63084 | 949.94821 | 812.88306 | 6.50E-50 | -1.256771889 | YFL001W | DEG1 | n |
| W303.asm02.HP0_G0016900 | 2240.8823 | 2187.16764 | 2858.172189 | 1303.40142 | 1412.113 | 1451.61526 | 806.04642 | 1056.227 | 924.85286 | 4.32E-12 | 0.806634142 | YFR001W | LOC1 | n |
| W303.asm02.HP0_G0016910 | 2858.2575 | 2652.21326 | 2656.293856 | 3091.82817 | 3045.3462 | 3197.15653 | 2998.4471 | 2910.2005 | 2886.5284 | 0.001308495 | -0.192846587 | YFR002W | NIC96 | n |
| W303.asm02.HP0_G0016920 | 1187.341 | 1209.3262 | 1208.372223 | 315.016455 | 310.10757 | 308.148242 | 352.28903 | 343.76579 | 352.66076 | 2.46E-123 | 1.949363506 | YFR003C | YPI1 | n |
| W303.asm02.HP0_G0016930 | 4432.2482 | 4334.88946 | 4441.323328 | 3542.68505 | 3424.6662 | 3461.68995 | 3731.5275 | 3554.4333 | 3575.9802 | 8.31E-11 | 0.340782005 | YFR004W | RPN11 | n |
| W303.asm02.HP0_G0016940 | 1447.5657 | 1449.11535 | 1445.023858 | 508.359887 | 482.68917 | 545.185352 | 445.77673 | 586.50118 | 492.84342 | 6.85E-55 | 1.499336522 | YFR005C | SAD1 | n |
| W303.asm02.HP0_G0016950 | 2688.6374 | 2649.09911 | 2189.752063 | 1714.25621 | 1533.4595 | 1686.75607 | 1563.0688 | 2629.4147 | 1570.2221 | 0.010560744 | 0.609211181 | YFR006W | | n |
| W303.asm02.HP0_G0016960 | 514.12815 | 532.518748 | 425.972942 | 434.189346 | 411.67904 | 396.326047 | 448.05692 | 392.31287 | 435.53604 | 0.02858534 | 0.24356477 | YFR007W | YFH7 | n |
| W303.asm02.HP0_G0016970 | 604.73271 | 588.573353 | 631.7149753 | 115.839384 | 114.15554 | 119.466703 | 149.35231 | 104.96665 | 132.24779 | 3.73E-79 | 2.385433206 | YFR008W | FAR7 | n |
| W303.asm02.HP0_G0016980 | 1785.7525 | 1713.81765 | 1533.888961 | 4190.21887 | 4159.9358 | 4375.70504 | 3598.1365 | 3753.8699 | 3692.3582 | 1.11E-84 | -1.338638228 | YFR009W | GCN20 | n |
| W303.asm02.HP0_G0016990 | 3792.7487 | 3648.73957 | 3481.193839 | 3002.65684 | 2952.7634 | 3040.71204 | 3551.3927 | 3188.3621 | 3271.8102 | 1.64E-05 | 0.279810992 | YFR010W | UBP6 | n |
| W303.asm02.HP0_G0017000 | 2273.5421 | 2235.95591 | 2834.024063 | 1952.60199 | 2124.0121 | 2028.08951 | 1799.0683 | 1956.316 | 1827.6644 | 0.004966687 | 0.267040998 | YFR011C | MIC19 | n |
| W303.asm02.HP0_G0017020 | 810.17326 | 808.63958 | 855.8095842 | 1464.24315 | 1421.1016 | 1451.61526 | 1323.6491 | 1162.5057 | 1418.5779 | 5.01E-22 | -0.809191437 | YFR013W | IOC3 | n |
| W303.asm02.HP0_G0017050 | 444.59443 | 610.372366 | 547.6794969 | 1863.43067 | 2028.7327 | 1902.93391 | 2074.971 | 1986.4939 | 2153.8756 | 7.79E-78 | -1.853802362 | YFR016C | AIP5 | n |
| W303.asm02.HP0_G0017070 | 932.38405 | 898.949777 | 966.8909637 | 770.040222 | 728.97751 | 659.911313 | 760.44267 | 732.14241 | 797.89498 | 1.23E-05 | 0.373348266 | YFR018C | | n |
| W303.asm02.HP0_G0017080 | 658.46331 | 699.644515 | 851.9458841 | 1319.23558 | 1369.8665 | 1360.59301 | 1205.0793 | 1241.2307 | 1282.8035 | 1.27E-20 | -0.872328589 | YFR019W | FAB1 | n |
| W303.asm02.HP0_G0017100 | 321.3301 | 423.523683 | 348.6989389 | 1125.05877 | 1148.7463 | 1152.9485 | 1190.2581 | 1110.0224 | 1221.9696 | 6.44E-64 | -1.64794531 | YFR021W | ATG18 | n |
| W303.asm02.HP0_G0017120 | 146.44224 | 165.04967 | 162.2754065 | 140.84069 | 142.91914 | 127.051891 | 186.97541 | 153.51373 | 192.20012 | 0.181776371 | 0.205112572 | YFR023W | PES4 | n |
| W303.asm02.HP0_G0017140 | 1408.5847 | 1230.08717 | 1394.795756 | 330.017238 | 329.88255 | 346.07418 | 337.46781 | 272.9133 | 326.21121 | 2.09E-92 | 2.004023252 | YFR025C | HIS2 | n |
| W303.asm02.HP0_G0017160 | 290.7774 | 315.566665 | 276.254561 | 152.507966 | 148.31232 | 127.051891 | 184.69522 | 177.13123 | 189.55516 | 2.13E-15 | 1.041480041 | YFR027W | ECO1 | n |
| W303.asm02.HP0_G0017170 | 1212.626 | 1203.09791 | 1061.551617 | 1135.89267 | 1137.0611 | 1169.06702 | 1142.3741 | 1148.0728 | 1109.9998 | 0.860309202 | 0.014168666 | YFR028C | CDC14 | n |
| W303.asm02.HP0_G0017180 | 181.2091 | 268.854495 | 257.9019853 | 516.693656 | 481.79031 | 521.481641 | 494.80077 | 570.75618 | 537.80767 | 3.93E-17 | -1.100708685 | YFR029W | PTR3 | n |
| W303.asm02.HP0_G0017190 | 851.26137 | 920.74879 | 938.8791375 | 824.209718 | 891.67162 | 887.466938 | 930.31666 | 847.60573 | 873.71705 | 0.476765947 | 0.059382113 | YFR030W | MET10 | n |
| W303.asm02.HP0_G0017200 | 716.40808 | 797.221049 | 797.8540819 | 1300.90128 | 1310.5416 | 1286.63743 | 1256.3835 | 1275.3448 | 1216.6796 | 5.15E-25 | -0.753446085 | YFR031C | SMC2 | n |
| W303.asm02.HP0_G0017220 | 6.3212478 | 20.7609648 | 9.659250386 | 66.6701491 | 60.223789 | 43.6148281 | 34.202819 | 74.788741 | 53.780767 | 2.99E-08 | -2.219026757 | YFR032C | RRT5 | n |
| W303.asm02.HP0_G0017250 | 5608.0003 | 5755.9775 | 5137.75528 | 5296.10997 | 5179.2459 | 5830.16474 | 4476.0089 | 4014.9745 | 4119.0777 | 0.828604637 | 0.017263235 | YFR033C | QCR6 | n |
| W303.asm02.HP0_G0017260 | 664.78456 | 741.166444 | 751.48968 | 3925.20503 | 4015.2189 | 3857.06785 | 3545.6922 | 3015.1671 | 3546.004 | 3.58E-190 | -2.450577656 | YFR034C | PHO4 | n |
| W303.asm02.HP0_G0017280 | 223.35076 | 213.837938 | 176.7642821 | 159.174981 | 170.78388 | 145.066711 | 166.45372 | 164.0104 | 179.85699 | 0.011540984 | 0.367346468 | YFR036W | CDC26 | n |
| W303.asm02.HP0_G0017290 | 1556.0805 | 1781.29078 | 1807.245747 | 2222.6161 | 2233.6734 | 2213.9266 | 2097.7729 | 2101.9572 | 2028.6811 | 1.53E-07 | -0.374273312 | YFR037C | RSC8 | n |
| W303.asm02.HP0_G0017300 | 607.89333 | 662.274778 | 668.4201267 | 469.191174 | 458.41989 | 474.074219 | 457.17767 | 410.68203 | 484.90855 | 5.41E-07 | 0.468339934 | YFR038W | IRC5 | n |
| W303.asm02.HP0_G0017320 | 1342.2116 | 1354.65296 | 1402.523156 | 2085.10891 | 2173.4496 | 2110.57842 | 2060.1498 | 2006.1752 | 2007.5214 | 4.34E-26 | -0.63532193 | YFR040W | SAP155 | n |
| W303.asm02.HP0_G0017330 | 2723.4043 | 2826.60536 | 3112.210474 | 2114.2771 | 2409.8504 | 2302.10441 | 2418.1393 | 2562.4984 | 2398.0932 | 7.82E-06 | 0.344176631 | YFR041C | ERJ5 | n |
| W303.asm02.HP0_G0017340 | 1383.2997 | 1532.1592 | 1723.210269 | 962.550278 | 1081.3316 | 982.281782 | 1289.4463 | 1209.7407 | 1210.5081 | 1.90E-11 | 0.616886327 | YFR042W | KEG1 | n |
| W303.asm02.HP0_G0017350 | 807.01264 | 794.106905 | 947.5724629 | 385.020111 | 432.35287 | 356.503813 | 448.05692 | 425.11495 | 462.86725 | 4.05E-26 | 1.118705905 | YFR043C | IRC6 | n |
| W303.asm02.HP0_G0017360 | 2402.0742 | 2738.37126 | 2624.41833 | 3514.35023 | 3224.2199 | 3413.33438 | 3810.194 | 4455.8344 | 3787.5766 | 3.54E-06 | -0.386700291 | YFR044C | DUG1 | n |
| W303.asm02.HP0_G0017370 | 635.2854 | 662.274778 | 623.9875749 | 1329.2361 | 1396.8324 | 1335.94115 | 1278.0453 | 1474.7815 | 1375.377 | 1.63E-42 | -1.080143757 | YFR045W | MRX20 | n |
| W303.asm02.HP0_G0017380 | 558.37689 | 643.58991 | 496.4854699 | 221.678246 | 226.51336 | 198.163024 | 223.45841 | 250.60789 | 221.29463 | 5.13E-29 | 1.392006586 | YFR046C | CNN1 | n |
| W303.asm02.HP0_G0017400 | 480.41483 | 453.627082 | 525.463221 | 907.547405 | 863.80688 | 808.770617 | 1018.1039 | 1077.2203 | 963.64554 | 1.97E-19 | -0.821716979 | YFR048W | RMD8 | n |

| | | | | | | | | | | | | | | |
|---|---|---|---|---|---|---|---|---|---|---|---|---|---|---|
| W303.asm02.HP0_G0017410 | 1902.6956 | 1790.63322 | 2363.61857 | 2393.45835 | 2355.9187 | 2415.88222 | 2429.5402 | 2334.196 | 2443.0574 | 0.01127588 | -0.241915536 | YFR049W | KGD4 | n |
| W303.asm02.HP0_G0017420 | 3751.6606 | 3612.40788 | 3376.873935 | 3068.49361 | 3094.7837 | 3101.39354 | 3421.4219 | 3416.6646 | 3314.1295 | 0.000391545 | 0.213087697 | YFR050C | PRE4 | n |
| W303.asm02.HP0_G0017430 | 1378.032 | 1894.43804 | 1812.075372 | 6108.65241 | 6040.3562 | 6353.54268 | 5899.9862 | 6224.5226 | 5851.5237 | 1.11E-78 | -1.863200155 | YFR051C | RET2 | n |
| W303.asm02.HP0_G0017440 | 2876.1677 | 2876.43168 | 2854.308489 | 2437.62733 | 2692.9921 | 2512.59336 | 2597.134 | 2855.093 | 2690.8016 | 0.009175026 | 0.171431126 | YFR052W | RPN12 | n |
| W303.asm02.HP0_G0017570 | 1037.7382 | 995.488264 | 944.6746878 | 181.676156 | 175.27819 | 208.592656 | 180.13484 | 157.44998 | 178.97534 | 1.84E-113 | 2.398196153 | YGL256W | ADH4 | n |
| W303.asm02.HP0_G0017600 | 11949.265 | 11826.4836 | 10942.96476 | 9049.63936 | 8820.5379 | 9052.92128 | 9621.2529 | 8180.8386 | 9137.4404 | 2.22E-07 | 0.366814761 | YGL253W | HXK2 | n |
| W303.asm02.HP0_G0017610 | 1781.5383 | 1721.08398 | 1702.925843 | 2075.94177 | 1928.959 | 2047.05248 | 2046.4686 | 1869.7185 | 1956.3856 | 0.001064226 | -0.217602247 | YGL252C | RTG2 | n |
| W303.asm02.HP0_G0017620 | 387.7032 | 373.697367 | 402.7907411 | 304.182555 | 319.99506 | 291.08157 | 399.03288 | 331.95704 | 352.66076 | 0.00139628 | 0.347124869 | YGL251C | HFM1 | n |
| W303.asm02.HP0_G0017630 | 464.61171 | 490.996818 | 449.155143 | 363.352313 | 344.26435 | 301.511203 | 450.33711 | 415.93036 | 435.53604 | 4.89E-06 | 0.4749205 | YGL250W | RMR1 | n |
| W303.asm02.HP0_G0017640 | 298.15219 | 248.09353 | 311.9937875 | 50.8359887 | 51.235164 | 35.0814922 | 47.883946 | 36.738329 | 44.964248 | 5.89E-43 | 2.640449954 | YGL249W | ZIP2 | n |
| W303.asm02.HP0_G0017650 | 1312.7125 | 1435.62072 | 1209.338148 | 405.021156 | 337.97231 | 343.229734 | 369.39044 | 408.05787 | 414.3764 | 5.30E-73 | 1.86336655 | YGL248W | PDE1 | n |
| W303.asm02.HP0_G0017660 | 499.37858 | 494.110963 | 583.4187233 | 168.342127 | 147.41345 | 169.71857 | 140.23156 | 149.57748 | 138.41935 | 3.29E-43 | 1.700429976 | YGL247W | BRR6 | n |
| W303.asm02.HP0_G0017670 | 1768.8958 | 1556.03431 | 1762.813195 | 1208.39645 | 1155.9372 | 1096.05959 | 942.8577 | 1082.4686 | 1024.4795 | 2.51E-11 | 0.555752257 | YGL246C | RAI1 | n |
| W303.asm02.HP0_G0017680 | 3270.1922 | 3382.99922 | 3502.44419 | 10256.3691 | 10021.418 | 10731.144 | 8213.2368 | 8506.2352 | 7727.679 | 2.98E-152 | -1.610205652 | YGL245W | GUS1 | n |
| W303.asm02.HP0_G0017690 | 2608.5683 | 2566.05525 | 2343.334144 | 3236.00236 | 3212.5347 | 3185.77875 | 2698.6024 | 2810.4821 | 2708.4347 | 8.23E-09 | -0.358260621 | YGL244W | RTF1 | n |
| W303.asm02.HP0_G0017700 | 880.76053 | 803.449339 | 1023.880541 | 595.864458 | 638.19239 | 612.503891 | 515.32247 | 519.58494 | 470.80212 | 4.84E-08 | 0.553397057 | YGL243W | TAD1 | n |
| W303.asm02.HP0_G0017710 | 1707.7904 | 1736.65471 | 2131.79656 | 1011.71951 | 1065.1521 | 1017.36327 | 990.74164 | 1112.6465 | 991.8584 | 8.16E-19 | 0.850235172 | YGL242C | ANK1 | n |
| W303.asm02.HP0_G0017720 | 1579.2584 | 1470.91436 | 1523.263786 | 1598.41682 | 1629.6378 | 1636.5042 | 1319.0887 | 1386.8719 | 1299.5549 | 0.179735048 | -0.088950882 | YGL241W | KAP114 | n |
| W303.asm02.HP0_G0017730 | 103.24705 | 101.728728 | 83.06955332 | 452.523637 | 532.12661 | 456.059399 | 417.27439 | 459.22911 | 437.29935 | 6.57E-67 | -2.325463797 | YGL240W | DOC1 | n |
| W303.asm02.HP0_G0017740 | 2522.1779 | 2304.4671 | 2522.030276 | 4008.54272 | 4153.6437 | 4518.87545 | 3837.5562 | 4084.5149 | 3902.1914 | 1.63E-29 | -0.786795646 | YGL238W | CSE1 | n |
| W303.asm02.HP0_G0017750 | 383.48903 | 400.686621 | 368.0174397 | 436.689477 | 444.93695 | 427.614945 | 456.03758 | 473.66202 | 475.21038 | 0.051115392 | -0.18513118 | YGL237C | HAP2 | n |
| W303.asm02.HP0_G0017760 | 754.33557 | 790.99276 | 970.7546638 | 964.217032 | 889.8739 | 952.88918 | 581.44792 | 711.14908 | 613.62973 | 0.160374218 | -0.157037394 | YGL236C | MTO1 | n |
| W303.asm02.HP0_G0017770 | 1775.2171 | 2016.92773 | 2172.365412 | 5131.93473 | 4777.4543 | 4588.09029 | 4480.5692 | 5172.2319 | 4658.6487 | 4.37E-48 | -1.281091812 | YGL234W | ADE57 | n |
| W303.asm02.HP0_G0017780 | 1157.8419 | 1202.05986 | 1294.339552 | 1353.40403 | 1503.797 | 1460.14859 | 1307.6878 | 1376.3752 | 1337.466 | 0.001543015 | -0.239656067 | YGL233W | SEC15 | n |
| W303.asm02.HP0_G0017790 | 2002.782 | 1815.54637 | 2369.41412 | 1251.73205 | 1355.4847 | 1282.84484 | 1038.6256 | 1081.1565 | 1034.1777 | 2.02E-12 | 0.670135247 | YGL232W | TAN1 | n |
| W303.asm02.HP0_G0017800 | 3221.7293 | 2915.87751 | 3668.583297 | 527.527555 | 560.89021 | 511.052008 | 564.34651 | 587.81326 | 577.482 | 4.82E-172 | 2.616168056 | YGL231C | EMC4 | n |
| W303.asm02.HP0_G0017830 | 1998.5678 | 2020.04188 | 1489.45641 | 1667.5871 | 1709.6365 | 1808.11907 | 2055.5894 | 2066.531 | 2018.1012 | 0.423605484 | 0.086762338 | YGL228W | SHE10 | n |
| W303.asm02.HP0_G0017840 | 572.07293 | 590.649449 | 625.919425 | 602.531473 | 603.13675 | 524.326086 | 857.35065 | 736.07866 | 882.53356 | 0.652164497 | 0.047694743 | YGL227W | VID30 | n |
| W303.asm02.HP0_G0017850 | 1623.5071 | 1564.3387 | 2009.12408 | 385.853488 | 465.61079 | 403.911235 | 380.79138 | 331.95704 | 365.88554 | 5.92E-72 | 2.05035619 | YGL226C-/ | OST5 | n |
| W303.asm02.HP0_G0017860 | 684.80184 | 694.454274 | 765.0126306 | 886.712983 | 915.94091 | 866.607672 | 735.3606 | 831.86073 | 790.84177 | 0.000175622 | -0.3153179 | YGL226W | MTC3 | n |
| W303.asm02.HP0_G0017890 | 1765.7352 | 1813.47028 | 1759.91542 | 622.532517 | 680.43893 | 640.948344 | 886.99309 | 876.47156 | 896.63999 | 2.62E-96 | 1.457974623 | YGL224C | SDT1 | n |
| W303.asm02.HP0_G0017900 | 1437.0303 | 1373.33782 | 1392.863906 | 780.040745 | 787.40357 | 741.452078 | 723.95966 | 703.27658 | 714.13805 | 8.40E-35 | 0.863761164 | YGL223C | COG1 | n |
| W303.asm02.HP0_G0017920 | 1803.6627 | 1901.70438 | 1571.560038 | 1335.90311 | 1331.2154 | 1267.67446 | 1209.6397 | 1178.2507 | 1214.9163 | 1.99E-07 | 0.422702352 | YGL221C | NIF3 | n |
| W303.asm02.HP0_G0017930 | 2406.2883 | 2352.21732 | 2896.809191 | 541.694962 | 651.67533 | 620.089078 | 511.90218 | 659.97783 | 551.9141 | 2.74E-75 | 2.078620075 | YGL220W | BOL2 | n |
| W303.asm02.HP0_G0017940 | 966.09737 | 966.422913 | 866.4347596 | 944.215987 | 942.90679 | 942.459547 | 1043.186 | 1023.4249 | 974.22536 | 0.84086896 | -0.016420484 | YGL219C | MDM34 | n |
| W303.asm02.HP0_G0017950 | 296.04511 | 349.822257 | 352.5626391 | 512.526771 | 520.4414 | 525.274235 | 589.42857 | 579.94076 | 627.73616 | 9.32E-11 | -0.640825137 | YGL216W | KIP3 | n |
| W303.asm02.HP0_G0017960 | 2064.9409 | 2090.62916 | 2464.074774 | 5346.11258 | 5351.8275 | 5896.53513 | 5119.0218 | 4576.5461 | 4648.9505 | 2.06E-55 | -1.32538844 | YGL215W | CLG1 | n |
| W303.asm02.HP0_G0017970 | 1632.989 | 1579.90942 | 1611.162964 | 1301.73466 | 1159.5327 | 1139.67442 | 1076.2487 | 1163.8178 | 1087.9585 | 2.48E-08 | 0.421072541 | YGL213C | SKI8 | n |
| W303.asm02.HP0_G0017980 | 162.24536 | 174.392105 | 154.5480062 | 697.536435 | 735.26954 | 651.377977 | 835.68887 | 835.79698 | 812.00141 | 3.56E-89 | -2.086724593 | YGL212W | VAM7 | n |
| W303.asm02.HP0_G0017990 | 1702.5227 | 1545.65383 | 1709.687318 | 800.875166 | 805.38082 | 792.652094 | 601.96961 | 753.13574 | 628.61781 | 5.86E-32 | 1.047397001 | YGL211W | NCS6 | n |
| W303.asm02.HP0_G0018010 | 361.36467 | 414.181248 | 450.121068 | 853.377909 | 924.92954 | 982.281782 | 492.52059 | 555.01118 | 530.75445 | 3.75E-30 | -1.1694643 | YGL209W | MIG2 | n |
| W303.asm02.HP0_G0018030 | 1836.3225 | 2114.50427 | 2076.738833 | 3806.86551 | 3915.4451 | 3810.60857 | 3494.388 | 3448.1546 | 3526.6076 | 2.42E-45 | -0.935825034 | YGL207W | SPT16 | n |
| W303.asm02.HP0_G0018040 | 2253.5248 | 2333.53245 | 2507.5414 | 9155.47823 | 9255.5874 | 9676.80296 | 9609.8519 | 8832.9439 | 9243.2386 | 1.29E-214 | -1.984755535 | YGL206C | CHC1 | n |
| W303.asm02.HP0_G0018050 | 76.908515 | 63.3209427 | 84.03547836 | 19.1676679 | 21.572701 | 17.0666719 | 22.801879 | 27.553747 | 29.094513 | 1.22E-11 | 1.955518888 | YGL205W | POX1 | n |
| W303.asm02.HP0_G0018070 | 830.19054 | 813.829821 | 813.3088825 | 1974.26979 | 1971.2055 | 1633.65976 | 1811.6093 | 1805.4264 | 1972.2553 | 8.63E-41 | -1.183520773 | YGL203C | KEX1 | n |
| W303.asm02.HP0_G0018080 | 2865.6323 | 3274.00415 | 3544.944892 | 5497.78717 | 5685.3055 | 5886.1055 | 4416.724 | 4059.5853 | 4375.6384 | 6.51E-24 | -0.81728365 | YGL202W | ARO8 | n |
| W303.asm02.HP0_G0018090 | 591.03667 | 747.394734 | 792.0585317 | 916.71455 | 972.56925 | 1089.42256 | 844.80962 | 851.54198 | 973.34371 | 3.32E-05 | -0.482214908 | YGL201C | MCM6 | n |
| W303.asm02.HP0_G0018100 | 5577.4476 | 5775.70042 | 5509.63642 | 2625.13712 | 2681.3069 | 2663.34896 | 3033.79 | 3450.7787 | 3046.989 | 1.66E-61 | 1.081243329 | YGL200C | EMP24 | n |
| W303.asm02.HP0_G0018110 | 1707.7904 | 1536.3114 | 1656.561441 | 758.372946 | 797.29106 | 717.748367 | 1010.1232 | 926.33072 | 986.56849 | 1.18E-45 | 1.107782153 | YGL198W | YIP4 | n |
| W303.asm02.HP0_G0018120 | 528.87773 | 603.106028 | 623.0216499 | 826.709849 | 840.43646 | 777.481719 | 824.28793 | 758.38407 | 862.25557 | 2.11E-07 | -0.47766226 | YGL197W | MDS3 | n |
| W303.asm02.HP0_G0018130 | 1625.6142 | 1577.83333 | 1655.595516 | 1103.39097 | 1015.7147 | 1127.34849 | 762.72285 | 673.09867 | 804.06654 | 6.49E-13 | 0.581978464 | YGL196W | DSD1 | n |
| W303.asm02.HP0_G0018140 | 2029.1205 | 2431.10898 | 2855.274414 | 5673.62969 | 5695.1929 | 5916.44625 | 4833.9984 | 5254.8931 | 4841.1507 | 4.86E-34 | -1.240094764 | YGL195W | GCN1 | n |
| W303.asm02.HP0_G0018150 | 455.12984 | 507.60559 | 619.1579498 | 40.8354663 | 55.729476 | 52.1481641 | 42.183476 | 38.050412 | 43.200944 | 1.98E-80 | 3.416457922 | YGL194C-A | | n |
| W303.asm02.HP0_G0018160 | 460.39755 | 422.485634 | 409.5522164 | 1139.22617 | 1121.7804 | 1121.6596 | 1197.0986 | 1220.2373 | 1193.7567 | 3.00E-71 | -1.389201643 | YGL194C | HOS2 | n |
| W303.asm02.HP0_G0018180 | 346.61509 | 347.746161 | 372.8470649 | 64.1700185 | 47.639714 | 60.6815 | 57.004698 | 78.72499 | 63.478938 | 3.70E-57 | 2.627938905 | YGL192W | IME4 | n |
| W303.asm02.HP0_G0018190 | 4646.1171 | 4348.38408 | 5113.607154 | 6579.51034 | 6595.8532 | 6903.46878 | 6069.8602 | 6200.9051 | 5752.7787 | 6.30E-13 | -0.508909847 | YGL191W | COX13 | n |
| W303.asm02.HP0_G0018200 | 1817.3587 | 1735.61666 | 1859.405699 | 2145.11205 | 2015.2498 | 1810.01537 | 2268.787 | 2195.1151 | 2213.828 | 0.064395817 | -0.142005545 | YGL190C | CDC55 | n |
| W303.asm02.HP0_G0018220 | 5231.8861 | 5096.81687 | 5959.757488 | 5521.12172 | 5376.0967 | 5931.61663 | 5131.5629 | 5100.0673 | 4837.624 | 0.550060117 | -0.04682964 | YGL187C | COX4 | n |
| W303.asm02.HP0_G0018230 | 1266.3566 | 1220.74473 | 1324.283228 | 827.543226 | 811.67286 | 749.037266 | 1112.7317 | 1235.9823 | 1113.5264 | 2.42E-17 | 0.673917941 | YGL186C | TPN1 | n |
| W303.asm02.HP0_G0018240 | 658.46331 | 679.921598 | 665.5223516 | 514.193525 | 526.73344 | 570.78536 | 584.8682 | 439.54786 | 521.05628 | 0.002697453 | 0.314903366 | YGL185C | | n |

| | | | | | | | | | | | | | | |
|---|---|---|---|---|---|---|---|---|---|---|---|---|---|---|
| W303.asm02.HP0_G0018250 | 318.16947 | 332.175437 | 270.4590108 | 78.3374252 | 99.77374 | 71.1111328 | 344.30837 | 203.37289 | 347.37085 | 5.29E-21 | 1.884316172 | YGL184C | STR3 | n |
| W303.asm02.HP0_G0018260 | 56.89123 | 52.9404603 | 39.60292658 | 11.6672761 | 16.179525 | 20.8592656 | 11.40094 | 13.120832 | 15.869734 | 3.45E-06 | 1.624366046 | YGL183C | MND1 | n |
| W303.asm02.HP0_G0018270 | 823.8693 | 947.738044 | 832.6273833 | 1274.23322 | 1208.9701 | 1218.37074 | 1501.5037 | 1361.9423 | 1441.5009 | 2.84E-10 | -0.507782727 | YGL181W | GTS1 | n |
| W303.asm02.HP0_G0018280 | 234.93971 | 278.196929 | 250.174585 | 746.70567 | 764.03314 | 756.622453 | 1050.0265 | 834.4849 | 1019.1896 | 4.41E-46 | -1.5706078 | YGL180W | ATG1 | n |
| W303.asm02.HP0_G0018310 | 719.56871 | 775.422036 | 813.3088825 | 2240.11701 | 2128.5065 | 2251.85254 | 1784.247 | 1443.2915 | 1783.5818 | 2.20E-60 | -1.519616252 | YGL178W | MPT5 | n |
| W303.asm02.HP0_G0018330 | 85.336845 | 91.3482452 | 70.51252782 | 175.009141 | 213.03042 | 187.733391 | 213.19757 | 190.25206 | 213.35976 | 4.40E-14 | -1.221921652 | YGL175C | SAE2 | n |
| W303.asm02.HP0_G0018340 | 327.65134 | 366.431029 | 212.5035085 | 250.846436 | 226.51336 | 219.970438 | 204.07682 | 229.61455 | 190.43681 | 0.033140988 | 0.376118164 | YGL174W | BUD13 | n |
| W303.asm02.HP0_G0018350 | 1933.2483 | 2271.24955 | 2369.41412 | 5937.81016 | 5986.4244 | 6195.20189 | 5807.6386 | 5505.501 | 5446.8455 | 1.83E-79 | -1.462287173 | YGL173C | XRN1 | n |
| W303.asm02.HP0_G0018360 | 992.4359 | 1139.77697 | 1198.712973 | 2781.81197 | 2717.2614 | 2882.37125 | 2730.525 | 2656.9684 | 2602.6364 | 8.08E-68 | -1.330474861 | YGL172W | NUP49 | n |
| W303.asm02.HP0_G0018370 | 1007.1855 | 904.140018 | 1001.664265 | 1985.93707 | 1872.3306 | 1980.68209 | 1078.5289 | 1378.9994 | 1154.0824 | 8.72E-27 | -1.003097313 | YGL171W | ROK1 | n |
| W303.asm02.HP0_G0018390 | 1143.0923 | 989.259974 | 959.1635634 | 898.380259 | 835.04328 | 873.244711 | 631.61205 | 686.2195 | 694.74171 | 0.006416406 | 0.245345709 | YGL169W | SUA5 | n |
| W303.asm02.HP0_G0018400 | 1278.9991 | 1386.83245 | 1518.434161 | 2937.65345 | 3075.9076 | 3089.06761 | 3536.5714 | 3713.1954 | 3565.4003 | 6.86E-54 | -1.120502999 | YGL167C | PMR1 | n |
| W303.asm02.HP0_G0018430 | 349.77571 | 433.904165 | 412.4499915 | 592.53095 | 650.77647 | 666.548352 | 617.93092 | 576.00451 | 660.35728 | 1.65E-10 | -0.673743048 | YGL163C | RAD54 | n |
| W303.asm02.HP0_G0018440 | 181.2091 | 147.40285 | 153.5820811 | 606.698357 | 508.75619 | 342.281586 | 619.07102 | 646.857 | 615.39303 | 2.28E-20 | -1.59832101 | YGL162W | SUT1 | n |
| W303.asm02.HP0_G0018450 | 1684.6125 | 1594.4421 | 1588.946689 | 750.872554 | 807.17855 | 695.940953 | 1034.0652 | 938.13947 | 1041.2309 | 3.34E-43 | 1.110241919 | YGL161C | YIP5 | n |
| W303.asm02.HP0_G0018460 | 1122.0215 | 1075.41798 | 949.504313 | 749.205801 | 794.59447 | 801.18543 | 863.05112 | 850.22989 | 864.01887 | 4.23E-07 | 0.424105904 | YGL160W | AIM14 | n |
| W303.asm02.HP0_G0018470 | 1207.3583 | 1218.66864 | 1445.989783 | 698.369812 | 691.22528 | 762.311344 | 778.68417 | 547.13868 | 658.59398 | 9.03E-13 | 0.848159264 | YGL159W | | n |
| W303.asm02.HP0_G0018510 | 810.17326 | 798.259098 | 731.2052542 | 777.540614 | 713.69684 | 763.259492 | 745.62144 | 759.69616 | 849.03079 | 0.564951113 | 0.052710889 | YGL155W | CDC43 | n |
| W303.asm02.HP0_G0018520 | 489.8967 | 474.388046 | 558.3046723 | 211.677723 | 229.20994 | 194.37043 | 247.40039 | 225.67831 | 224.82124 | 2.63E-27 | 1.26115127 | YGL154C | LYS5 | n |
| W303.asm02.HP0_G0018530 | 482.52192 | 503.453397 | 445.2914428 | 693.369551 | 730.77523 | 696.889102 | 817.44736 | 814.80365 | 838.45097 | 1.72E-11 | -0.56831863 | YGL153W | PEX14 | n |
| W303.asm02.HP0_G0018540 | 1267.4102 | 1195.83157 | 1198.712973 | 1733.42388 | 1787.8376 | 1773.03758 | 1705.5806 | 1567.9394 | 1699.8249 | 2.04E-15 | -0.531922143 | YGL151W | NUT1 | n |
| W303.asm02.HP0_G0018550 | 1423.3343 | 1605.86063 | 1727.073969 | 2329.28833 | 2408.9516 | 2354.25257 | 2415.8591 | 2414.233 | 2413.9629 | 2.19E-14 | -0.575833362 | YGL150C | INO80 | n |
| W303.asm02.HP0_G0018560 | 2433.6804 | 2749.78979 | 2825.330738 | 3233.50223 | 3155.0075 | 3428.50475 | 2769.2882 | 2692.3947 | 2677.5769 | 5.03E-05 | -0.293298343 | YGL148W | ARO2 | n |
| W303.asm02.HP0_G0018580 | 85.336845 | 93.4243417 | 83.06955332 | 148.341082 | 137.52597 | 142.222266 | 221.17823 | 191.56414 | 253.0341 | 2.02E-05 | -0.710644712 | YGL146C | RRT6 | n |
| W303.asm02.HP0_G0018590 | 1528.6884 | 1469.87631 | 1560.934862 | 954.216509 | 1004.0294 | 964.266961 | 946.27798 | 994.55904 | 906.33817 | 8.76E-20 | 0.641958624 | YGL145W | TIP20 | n |
| W303.asm02.HP0_G0018600 | 379.27487 | 355.012499 | 332.2782133 | 624.199271 | 583.36178 | 641.896492 | 542.68472 | 614.05492 | 561.61227 | 3.20E-16 | -0.795036167 | YGL144C | ROG1 | n |
| W303.asm02.HP0_G0018610 | 906.04552 | 826.2864 | 919.5606368 | 705.870204 | 725.38206 | 770.84468 | 690.89693 | 641.60867 | 581.89026 | 0.003088614 | 0.268957786 | YGL143C | MRF1 | n |
| W303.asm02.HP0_G0018620 | 2118.6716 | 1999.28091 | 2092.193634 | 816.709327 | 828.75125 | 838.163219 | 1044.3261 | 1037.8578 | 1076.497 | 3.61E-90 | 1.322474096 | YGL142C | GPI10 | n |
| W303.asm02.HP0_G0018630 | 638.44603 | 651.894296 | 697.3978779 | 512.526771 | 496.17211 | 494.933485 | 548.38519 | 490.71911 | 481.38194 | 7.58E-06 | 0.402899797 | YGL141W | HUL5 | n |
| W303.asm02.HP0_G0018640 | 1093.5759 | 956.04243 | 1153.314496 | 720.03761 | 812.57172 | 757.570602 | 833.40868 | 759.69616 | 818.17297 | 1.60E-07 | 0.484699032 | YGL140C | | n |
| W303.asm02.HP0_G0018660 | 7.3747891 | 4.15219297 | 12.5570255 | 56.6696267 | 55.729476 | 46.4592735 | 35.342912 | 74.788741 | 68.768849 | 8.49E-11 | -2.708703412 | YGL138C | | n |
| W303.asm02.HP0_G0018670 | 2398.9135 | 2795.46391 | 2691.067158 | 4356.06087 | 4355.8878 | 4358.63837 | 4375.6806 | 4159.3036 | 4280.42 | 2.04E-27 | -0.728888257 | YGL137W | SEC27 | n |
| W303.asm02.HP0_G0018680 | 658.46331 | 716.253287 | 707.0571283 | 415.021678 | 453.92557 | 441.837172 | 337.46781 | 447.42036 | 381.75528 | 2.10E-10 | 0.668111803 | YGL136C | MRM2 | n |
| W303.asm02.HP0_G0018690 | 10279.402 | 10156.264 | 10286.13574 | 15757.4897 | 15952.113 | 17533.1609 | 13088.279 | 14023.545 | 12232.039 | 1.74E-23 | -0.680591525 | YPL220W | RPL1A | y |
| W303.asm02.HP0_G0018700 | 285.50969 | 314.528617 | 315.8574876 | 569.196398 | 522.23913 | 432.355688 | 476.55927 | 452.66869 | 435.53604 | 4.62E-10 | -0.735486261 | YGL134W | PCL10 | n |
| W303.asm02.HP0_G0018710 | 642.66019 | 768.155699 | 774.671881 | 1775.09272 | 1801.3205 | 1877.33391 | 1678.2183 | 1545.634 | 1590.5 | 2.02E-56 | -1.318321425 | YGL133W | ITC1 | n |
| W303.asm02.HP0_G0018720 | 526.77065 | 509.681687 | 524.497296 | 1259.23244 | 1238.6326 | 1263.88187 | 973.64023 | 1111.3344 | 1106.4731 | 1.01E-55 | -1.268898496 | YGL131C | SNT2 | n |
| W303.asm02.HP0_G0018730 | 764.87098 | 718.329383 | 732.1711793 | 1693.42179 | 1601.773 | 1559.70418 | 1418.2769 | 1512.8319 | 1489.9917 | 1.58E-55 | -1.132513773 | YGL130W | CEG1 | n |
| W303.asm02.HP0_G0018740 | 1924.82 | 1951.53069 | 1792.756872 | 2703.47455 | 2746.025 | 2746.78602 | 2345.1733 | 2596.6126 | 2376.9336 | 1.09E-15 | -0.532109686 | YGL129C | RSM23 | n |
| W303.asm02.HP0_G0018750 | 288.67032 | 296.881797 | 322.6189629 | 480.85845 | 525.83458 | 447.526063 | 403.59326 | 429.0512 | 487.55351 | 1.80E-09 | -0.678696128 | YGL128C | CWC23 | n |
| W303.asm02.HP0_G0018760 | 830.19054 | 896.873681 | 806.5474072 | 447.523376 | 482.68917 | 483.555703 | 471.9989 | 527.45743 | 534.28106 | 1.04E-20 | 0.842105943 | YGL127C | SOH1 | n |
| W303.asm02.HP0_G0018770 | 2705.4941 | 2798.57806 | 3069.709773 | 438.35623 | 479.09372 | 431.407539 | 671.51534 | 612.74284 | 671.81876 | 3.00E-222 | 2.668390177 | YGL126W | SCS3 | n |
| W303.asm02.HP0_G0018780 | 543.62731 | 577.154822 | 652.9653261 | 430.022462 | 404.48814 | 398.222344 | 290.72396 | 321.46038 | 325.32956 | 3.46E-07 | 0.525233152 | YGL125W | MET13 | n |
| W303.asm02.HP0_G0018790 | 978.73987 | 935.281466 | 963.9931885 | 320.850093 | 321.79278 | 303.4075 | 396.7527 | 325.39663 | 390.5718 | 1.95E-64 | 1.604462081 | YGL124C | MON1 | n |
| W303.asm02.HP0_G0018800 | 26069.879 | 24698.2818 | 26846.92052 | 16750.875 | 16588.508 | 18017.6648 | 13863.542 | 14691.395 | 12824.509 | 2.79E-18 | 0.595823374 | YGL123W | RPS2 | n |
| W303.asm02.HP0_G0018810 | 942.91946 | 1157.42379 | 1233.486274 | 2216.78246 | 2168.0564 | 2057.48211 | 2097.7729 | 2267.2797 | 2130.071 | 3.10E-24 | -0.949866737 | YGL122C | NAB2 | n |
| W303.asm02.HP0_G0018830 | 1949.0514 | 2056.37357 | 2103.784734 | 3010.15723 | 2908.7191 | 3152.59356 | 1766.0055 | 2313.2026 | 1835.5993 | 3.00E-09 | -0.570181616 | YGL120C | PRP43 | n |
| W303.asm02.HP0_G0018850 | 3501.9713 | 3699.60393 | 3275.451806 | 362.518936 | 393.70178 | 344.177883 | 510.76209 | 421.1787 | 581.89026 | 5.77E-207 | 3.250722994 | YGL117W | ARO5 | n |
| W303.asm02.HP0_G0018860 | 627.91061 | 639.437717 | 636.5446005 | 573.363282 | 509.65505 | 605.866852 | 747.90163 | 721.64574 | 721.19126 | 0.054641356 | 0.173312955 | YGL116W | CDC20 | n |
| W303.asm02.HP0_G0018870 | 3221.7293 | 3000.99747 | 3504.37604 | 1032.55393 | 1095.7134 | 998.400305 | 1386.3542 | 1268.7844 | 1288.0934 | 4.61E-95 | 1.637454801 | YGL115W | SNF4 | n |
| W303.asm02.HP0_G0018880 | 995.59653 | 1020.40142 | 1182.292247 | 551.695484 | 513.2505 | 482.607555 | 795.78558 | 670.4745 | 761.74725 | 3.20E-25 | 1.046951912 | YGL114W | | n |
| W303.asm02.HP0_G0018890 | 508.86045 | 551.203616 | 472.3373439 | 585.030558 | 506.95846 | 442.78532 | 458.31777 | 469.72577 | 433.77274 | 0.971972377 | -0.004444174 | YGL113W | SLD3 | n |
| W303.asm02.HP0_G0018900 | 894.45656 | 913.482452 | 878.0258601 | 1649.25281 | 1618.8514 | 1568.23752 | 1535.7066 | 1511.5198 | 1498.8083 | 1.10E-38 | -0.849012498 | YGL112C | TAF6 | n |
| W303.asm02.HP0_G0018910 | 995.59653 | 1049.46677 | 1078.938268 | 1559.24811 | 1473.2357 | 1536.94862 | 892.69356 | 1125.7674 | 961.00058 | 5.85E-10 | -0.548419775 | YGL111W | NSA1 | n |
| W303.asm02.HP0_G0018920 | 598.41146 | 591.687498 | 668.4201267 | 790.874644 | 794.59447 | 714.903922 | 664.67477 | 610.11867 | 638.31598 | 0.00061725 | -0.307509456 | YGL110C | CUE3 | n |
| W303.asm02.HP0_G0018930 | 654.24915 | 731.82401 | 666.4882766 | 299.182294 | 343.36548 | 283.496383 | 263.3617 | 283.40996 | 285.65522 | 5.21E-28 | 1.14780786 | YGL108C | | n |
| W303.asm02.HP0_G0018950 | 3295.4772 | 3436.97773 | 3429.033887 | 2583.46828 | 2448.5015 | 2506.90447 | 2451.202 | 2662.2168 | 2510.9446 | 6.32E-13 | 0.430617682 | YGL106W | MLC1 | n |
| W303.asm02.HP0_G0018960 | 4472.2828 | 4579.86884 | 3780.630601 | 5343.61245 | 5103.7414 | 5749.57213 | 4216.0674 | 4551.6165 | 4246.0356 | 6.55E-05 | -0.336021548 | YGL105W | ARC1 | n |
| W303.asm02.HP0_G0018990 | 1669.863 | 1598.59429 | 1535.820811 | 1511.74563 | 1502.8981 | 1589.09678 | 1780.8268 | 1675.5302 | 1720.9845 | 0.372780485 | 0.061441786 | YGL100W | SEH1 | n |
| W303.asm02.HP0_G0019000 | 738.53245 | 776.460085 | 920.5265618 | 2564.30061 | 2402.6595 | 2518.28225 | 1478.7019 | 1926.1381 | 1530.5477 | 1.34E-49 | -1.619118904 | YGL099W | LSG1 | n |

| | | | | | | | | | | | | | | |
|---|---|---|---|---|---|---|---|---|---|---|---|---|---|---|
| W303.asm02.HP0_G0019010 | 854.42199 | 853.275654 | 909.9013864 | 645.86707 | 657.96737 | 560.355727 | 631.61205 | 666.53825 | 642.72424 | 2.22E-08 | 0.489019469 | YGL098W | USE1 | n |
| W303.asm02.HP0_G0019020 | 1233.6969 | 1277.83739 | 1467.240134 | 2048.44033 | 1943.3408 | 2070.75619 | 2004.2852 | 1979.9335 | 1913.1846 | 5.99E-15 | -0.606861851 | YGL097W | SRM1 | n |
| W303.asm02.HP0_G0019040 | 1721.4865 | 1593.40405 | 1663.322917 | 1056.72186 | 1131.6679 | 969.955852 | 1002.1426 | 1056.227 | 975.10702 | 6.70E-17 | 0.65601572 | YGL095C | VPS45 | n |
| W303.asm02.HP0_G0019050 | 753.28203 | 770.231795 | 786.2629814 | 986.718207 | 971.67039 | 1054.34106 | 1143.5142 | 1027.3611 | 1137.331 | 1.21E-06 | -0.382646161 | YGL094C | PAN2 | n |
| W303.asm02.HP0_G0019060 | 476.20067 | 507.60559 | 510.9743454 | 772.540353 | 747.85362 | 809.718766 | 769.56342 | 732.14241 | 835.80601 | 3.10E-13 | -0.6397495 | YGL093W | SPC105 | n |
| W303.asm02.HP0_G0019070 | 1431.7626 | 1431.46852 | 1736.733219 | 2939.3202 | 3125.345 | 3169.66023 | 2849.0948 | 2801.2976 | 2779.8485 | 1.98E-34 | -1.004563289 | YGL092W | NUP145 | n |
| W303.asm02.HP0_G0019080 | 593.14375 | 691.340129 | 639.4423756 | 515.860279 | 536.62093 | 512.000156 | 636.17242 | 632.42409 | 579.24531 | 0.001214703 | 0.298464617 | YGL091C | NBP35 | n |
| W303.asm02.HP0_G0019110 | 1328.5156 | 1272.64714 | 1287.578076 | 964.217032 | 1020.209 | 960.474368 | 1129.8331 | 1024.737 | 1094.13 | 2.24E-08 | 0.401011583 | YGL087C | MMS2 | n |
| W303.asm02.HP0_G0019120 | 1052.4878 | 945.661948 | 1032.573866 | 549.195353 | 622.01287 | 565.096469 | 557.50594 | 560.25951 | 501.65994 | 2.42E-19 | 0.804043739 | YGL086W | MAD1 | n |
| W303.asm02.HP0_G0019130 | 643.71373 | 612.448462 | 660.6927264 | 470.857928 | 465.61079 | 450.370508 | 503.92153 | 528.76952 | 516.64802 | 4.88E-08 | 0.466812868 | YGL085W | LCL3 | n |
| W303.asm02.HP0_G0019140 | 1685.6661 | 1596.5182 | 1624.685915 | 1759.25856 | 1791.433 | 1751.23016 | 2453.4822 | 2528.3843 | 2451.874 | 0.065220641 | -0.111837601 | YGL084C | GUP1 | n |
| W303.asm02.HP0_G0019150 | 457.23692 | 498.263156 | 517.7358207 | 764.206584 | 806.27968 | 775.585422 | 733.08041 | 719.02158 | 716.783 | 1.07E-15 | -0.670196961 | YGL083W | SCY1 | n |
| W303.asm02.HP0_G0019160 | 2107.0826 | 1963.98727 | 1827.530173 | 1032.55393 | 1036.3885 | 1052.44477 | 1128.693 | 1095.5894 | 1230.7861 | 7.51E-32 | 0.917929543 | YGL082W | | n |
| W303.asm02.HP0_G0019170 | 62.158937 | 70.5872804 | 49.26217697 | 125.00653 | 110.56009 | 77.7481719 | 167.59381 | 127.27207 | 205.4249 | 0.000982949 | -0.789447929 | YGL081W | | n |
| W303.asm02.HP0_G0019180 | 1868.9823 | 1841.49758 | 2211.968338 | 1990.10395 | 2066.4849 | 2105.83768 | 1989.4639 | 1981.2456 | 1931.6993 | 0.496755698 | -0.056618197 | YGL080W | MPC1 | n |
| W303.asm02.HP0_G0019190 | 976.63278 | 951.890237 | 1068.313093 | 151.674589 | 177.97478 | 179.200055 | 233.71926 | 188.93998 | 230.11115 | 1.30E-112 | 2.560960975 | YGL079W | KXD1 | n |
| W303.asm02.HP0_G0019200 | 2984.6825 | 2603.42499 | 2318.220093 | 2815.98042 | 2667.824 | 2978.13424 | 1458.1802 | 1861.846 | 1557.8789 | 0.376102677 | -0.098161694 | YGL078C | DBP3 | n |
| W303.asm02.HP0_G0019210 | 3216.4616 | 2669.86008 | 2871.69514 | 1135.89267 | 1211.6667 | 1239.23001 | 1618.9334 | 1389.4961 | 1587.8551 | 1.58E-47 | 1.288178632 | YGL077C | HNM1 | n |
| W303.asm02.HP0_G0019230 | 321.3301 | 271.968639 | 255.0042102 | 142.507444 | 135.72824 | 163.081531 | 150.4924 | 162.69831 | 148.11752 | 2.66E-11 | 0.943101917 | YGL075C | MPS2 | n |
| W303.asm02.HP0_G0019240 | 822.81576 | 1075.41798 | 907.0036113 | 2615.1366 | 2605.8025 | 2799.88234 | 2737.3656 | 2353.8772 | 2714.6062 | 3.71E-53 | -1.51554729 | YGL073W | HSF1 | n |
| W303.asm02.HP0_G0019250 | 338.18676 | 494.110963 | 372.8470649 | 1847.59651 | 1822.8932 | 1739.85238 | 1858.3531 | 1632.2315 | 1881.4452 | 1.96E-78 | -2.166937072 | YGL071W | AFT1 | n |
| W303.asm02.HP0_G0019260 | 1910.0704 | 1922.46534 | 2298.901592 | 1055.88849 | 1058.8601 | 964.266961 | 1031.785 | 1141.5124 | 1123.2245 | 4.73E-28 | 0.993796813 | YGL070C | RPB9 | n |
| W303.asm02.HP0_G0019270 | 4229.9683 | 3944.58332 | 4825.761493 | 2059.27423 | 2112.3269 | 2152.29695 | 2014.546 | 2147.8801 | 2035.7343 | 5.92E-39 | 1.040002518 | YGL068W | MNP1 | n |
| W303.asm02.HP0_G0019280 | 1071.4515 | 1045.31458 | 885.7532604 | 880.045968 | 873.69437 | 771.792828 | 889.27328 | 806.93115 | 837.56932 | 0.010307251 | 0.24825395 | YGL067W | NPY1 | n |
| W303.asm02.HP0_G0019290 | 559.43043 | 599.991884 | 548.6454219 | 1066.72239 | 1091.2191 | 1100.80034 | 1213.06 | 1071.9719 | 1106.4731 | 1.19E-30 | -0.932147067 | YGL066W | SGF73 | n |
| W303.asm02.HP0_G0019300 | 904.99198 | 827.324448 | 757.2852303 | 3143.49753 | 3058.8292 | 3208.53431 | 4170.4637 | 3801.1049 | 3834.3042 | 4.99E-136 | -1.919320613 | YGL065C | ALG2 | n |
| W303.asm02.HP0_G0019310 | 592.09021 | 558.469954 | 574.725398 | 878.379215 | 927.62612 | 747.140969 | 701.15778 | 673.09867 | 662.12059 | 3.45E-09 | -0.566250604 | YGL064C | MRH4 | n |
| W303.asm02.HP0_G0019330 | 2295.6665 | 3059.12817 | 2802.148537 | 4340.22671 | 4497.9081 | 4570.07547 | 4137.4009 | 3721.0679 | 4143.764 | 8.22E-14 | -0.716841896 | YGL062W | PYC1 | n |
| W303.asm02.HP0_G0019340 | 1068.2909 | 1081.64627 | 1189.053723 | 464.190913 | 414.37562 | 435.200133 | 451.4772 | 471.03786 | 480.50029 | 2.00E-53 | 1.345408767 | YGL061C | DUO1 | n |
| W303.asm02.HP0_G0019350 | 389.81028 | 423.523683 | 367.0515147 | 1090.05694 | 1148.7463 | 1071.40773 | 1222.1807 | 1162.5057 | 1274.8687 | 1.83E-67 | -1.488678199 | YGL060W | YBP2 | n |
| W303.asm02.HP0_G0019360 | 895.5101 | 859.503944 | 855.8095842 | 625.032648 | 583.36178 | 572.681656 | 795.78558 | 715.08533 | 782.02525 | 2.00E-11 | 0.550695672 | YGL059W | PKP2 | n |
| W303.asm02.HP0_G0019370 | 1597.1686 | 1694.09473 | 1423.773507 | 1772.59259 | 1767.1637 | 1801.48203 | 1797.9282 | 1730.6377 | 1750.079 | 0.017284404 | -0.180385002 | YGL058W | RAD6 | n |
| W303.asm02.HP0_G0019400 | 10125.585 | 8353.1742 | 9456.406128 | 8289.59967 | 7902.7993 | 8223.2914 | 10769.327 | 10311.662 | 10444.93 | 0.009002094 | 0.194259739 | YGL055W | OLE1 | n |
| W303.asm02.HP0_G0019410 | 3708.4654 | 3693.37564 | 4705.986788 | 2105.10996 | 2272.3245 | 2195.91178 | 2637.0373 | 2533.6326 | 2450.9923 | 8.16E-21 | 0.881587235 | YGL054C | ERV14 | n |
| W303.asm02.HP0_G0019420 | 469.87942 | 475.426095 | 401.8248161 | 526.694178 | 516.84595 | 500.622375 | 874.45206 | 934.20322 | 956.59232 | 0.039319952 | -0.198737041 | YGL053W | PRM8 | n |
| W303.asm02.HP0_G0019430 | 287.61677 | 288.577411 | 280.1182612 | 257.513451 | 238.19857 | 296.770461 | 283.88339 | 402.80953 | 325.32956 | 0.434454333 | 0.112611713 | YGL051W | MST27 | y |
| W303.asm02.HP0_G0019460 | 2776.0813 | 3098.574 | 2385.834845 | 4020.20999 | 3790.5033 | 3717.69002 | 4350.5985 | 4416.4719 | 4204.598 | 5.91E-08 | -0.481304668 | YGL048C | RPT6 | n |
| W303.asm02.HP0_G0019470 | 946.08009 | 885.45515 | 1020.982766 | 362.518936 | 371.23022 | 348.918625 | 477.69937 | 459.22911 | 453.16908 | 1.43E-53 | 1.397793355 | YGL047W | ALG13 | n |
| W303.asm02.HP0_G0019480 | 238.10033 | 305.186183 | 230.8560842 | 669.201622 | 572.57543 | 467.43718 | 540.40453 | 514.3366 | 522.81958 | 5.70E-16 | -1.144774341 | YGL045W | RIM8 | n |
| W303.asm02.HP0_G0019490 | 896.56365 | 832.51469 | 822.9681329 | 627.532779 | 595.04699 | 592.592774 | 633.89224 | 661.28992 | 709.72979 | 5.11E-09 | 0.490630951 | YGL044C | RNA15 | n |
| W303.asm02.HP0_G0019500 | 2181.884 | 2360.5217 | 1863.2694 | 1229.23087 | 1250.3178 | 1225.95593 | 1065.9878 | 1140.2003 | 1214.0347 | 1.11E-17 | 0.789312113 | YGL043W | DST1 | n |
| W303.asm02.HP0_G0019520 | 1783.6454 | 1814.50833 | 1789.859097 | 2226.78298 | 2221.9882 | 2305.897 | 1779.6867 | 1928.7623 | 1773.8836 | 1.66E-07 | -0.325990793 | YGL040C | HEM2 | n |
| W303.asm02.HP0_G0019540 | 1124.1286 | 1040.12434 | 794.9563068 | 1872.59781 | 1756.3774 | 1732.2672 | 1622.3537 | 1533.8252 | 1578.1569 | 1.71E-15 | -0.858445834 | YGL038C | OCH1 | n |
| W303.asm02.HP0_G0019550 | 4372.1964 | 4757.37509 | 3945.803783 | 627.532779 | 686.73097 | 651.377977 | 655.55402 | 704.58866 | 696.50501 | 1.34E-226 | 2.733937895 | YGL037C | PNC1 | n |
| W303.asm02.HP0_G0019560 | 966.09737 | 1007.94484 | 1066.381243 | 65.8367722 | 70.111277 | 54.9926094 | 93.487704 | 93.157905 | 99.626666 | 2.38E-193 | 3.990024959 | YGL036W | VIR1 | n |
| W303.asm02.HP0_G0019570 | 805.95909 | 884.417102 | 918.5947117 | 1459.24289 | 1466.9436 | 1197.51148 | 734.2205 | 867.28697 | 823.46289 | 8.77E-11 | -0.660592182 | YGL035C | MIG1 | n |
| W303.asm02.HP0_G0019600 | 11929.248 | 11670.7764 | 13185.8427 | 12308.9763 | 11937.793 | 13429.5745 | 10505.966 | 11011.002 | 9484.8113 | 0.679787094 | -0.034413592 | YGL031C | RPL24A | y |
| W303.asm02.HP0_G0019610 | 9972.8219 | 9578.07112 | 11279.10668 | 9474.66157 | 9860.5219 | 10081.6623 | 8324.966 | 9143.9076 | 7573.3899 | 0.437897702 | 0.067797905 | YGL030W | RPL30 | n |
| W303.asm02.HP0_G0019620 | 1538.1703 | 1594.4421 | 2032.306281 | 1062.5555 | 1031.8942 | 1059.08181 | 645.29318 | 944.69988 | 752.04908 | 5.08E-08 | 0.712110251 | YGL029W | CGR1 | n |
| W303.asm02.HP0_G0019630 | 1916.3916 | 2177.82521 | 2245.775715 | 2293.45313 | 2382.8845 | 2337.1859 | 3122.7173 | 2901.0159 | 3067.267 | 0.052088691 | -0.145237634 | YGL028C | SCW11 | n |
| W303.asm02.HP0_G0019640 | 1126.2356 | 1171.95646 | 1158.144121 | 1385.90572 | 1335.7097 | 1323.61522 | 1405.7358 | 1420.9861 | 1418.5779 | 0.000468079 | -0.227225836 | YGL027C | CWH41 | n |
| W303.asm02.HP0_G0019650 | 1917.4452 | 2202.73837 | 2228.389064 | 4010.20947 | 3875.8952 | 4094.10495 | 4128.2802 | 3334.0033 | 3999.1731 | 4.39E-24 | -0.915926059 | YGL026C | TRP5 | n |
| W303.asm02.HP0_G0019660 | 466.7188 | 541.861182 | 540.9180216 | 501.692872 | 471.90282 | 455.11125 | 494.80077 | 408.05787 | 526.34619 | 0.311252244 | 0.116950046 | YGL025C | PGD1 | n |
| W303.asm02.HP0_G0019670 | 694.28372 | 857.427847 | 798.8200069 | 1064.22226 | 1084.0282 | 1013.57068 | 1265.5043 | 1110.0224 | 1178.7686 | 3.63E-06 | -0.427654838 | YGL023C | PIB2 | n |
| W303.asm02.HP0_G0019680 | 2257.739 | 2397.89144 | 2567.428753 | 5380.28103 | 5225.0878 | 5680.35729 | 6165.6281 | 5826.9614 | 5788.0448 | 4.64E-67 | -1.172506764 | YGL022W | STT3 | n |
| W303.asm02.HP0_G0019700 | 2016.478 | 1994.09067 | 2356.857094 | 1900.09925 | 1735.7035 | 1829.92649 | 2680.3609 | 2737.0055 | 2469.507 | 0.008936936 | 0.220561751 | YGL020C | GET1 | n |
| W303.asm02.HP0_G0019710 | 1518.153 | 1443.9251 | 1303.998802 | 1730.09037 | 1662.8957 | 1757.8672 | 1735.223 | 1787.0573 | 1806.5048 | 0.000222969 | -0.272466972 | YGL019W | CKB1 | n |
| W303.asm02.HP0_G0019720 | 416.14881 | 459.855371 | 390.2337156 | 944.215987 | 852.12167 | 838.163219 | 1148.0746 | 1103.4619 | 1184.9402 | 7.17E-31 | -1.058637196 | YGL018C | JAC1 | n |
| W303.asm02.HP0_G0019730 | 445.64797 | 439.094406 | 499.383245 | 476.691566 | 515.04823 | 507.259414 | 494.80077 | 461.85328 | 488.43516 | 0.243810747 | -0.113379368 | YGL017W | ATE1 | n |
| W303.asm02.HP0_G0019740 | 1514.9924 | 1353.61491 | 1624.685915 | 1610.91748 | 1732.1081 | 1847.94131 | 1556.2282 | 1468.2211 | 1453.844 | 0.016864252 | -0.207434086 | YGL016W | KAP122 | n |

| | | | | | | | | | | | | | | |
|---|---|---|---|---|---|---|---|---|---|---|---|---|---|---|
| W303.asm02.HP0_G0019760 | 1218.9473 | 1345.31052 | 1499.11566 | 2086.77567 | 2081.7656 | 2145.65991 | 1903.9569 | 1608.614 | 1893.7883 | 4.04E-12 | -0.63532033 | YGL014W | PUF4 | n |
| W303.asm02.HP0_G0019770 | 873.38574 | 966.422913 | 888.6510355 | 1555.08123 | 1607.1662 | 1650.72643 | 1506.0641 | 1369.8148 | 1637.2276 | 9.12E-23 | -0.818726977 | YGL013C | PDR1 | n |
| W303.asm02.HP0_G0019780 | 4728.2934 | 4285.06314 | 4440.357403 | 2166.77985 | 2159.9666 | 2147.55621 | 3086.2343 | 3054.5296 | 2903.2797 | 1.81E-66 | 1.055102629 | YGL012W | ERG4 | n |
| W303.asm02.HP0_G0019790 | 4297.395 | 4272.60656 | 3763.24395 | 2780.14522 | 2611.1956 | 2763.8527 | 2935.7419 | 3075.5229 | 2935.9009 | 1.41E-17 | 0.596493487 | YGL011C | SCL1 | n |
| W303.asm02.HP0_G0019800 | 463.55817 | 442.208551 | 488.7580695 | 166.675373 | 166.28957 | 171.614867 | 287.30368 | 234.86289 | 252.15245 | 2.89E-36 | 1.467504055 | YGL010W | MPO1 | n |
| W303.asm02.HP0_G0019810 | 7848.8827 | 8472.54975 | 8248.99983 | 549.195353 | 567.18225 | 508.207563 | 742.20116 | 650.79325 | 670.93711 | 0 | 3.918074354 | YGL009C | LEU1 | n |
| W303.asm02.HP0_G0019820 | 37080.44 | 34907.4863 | 39507.3 | 33088.395 | 31447.604 | 36434.5 | 32303.422 | 31301.056 | 28462.369 | 0.072747199 | 0.143079025 | YGL008C | PMA1 | n |
| W303.asm02.HP0_G0019830 | 892.34948 | 883.379053 | 993.9368647 | 2337.6221 | 2224.6847 | 2255.64513 | 2429.5402 | 2474.5889 | 2614.9796 | 1.18E-73 | -1.298979691 | YGL006W | PMC1 | n |
| W303.asm02.HP0_G0019840 | 1267.4102 | 1269.533 | 1430.534982 | 727.538002 | 846.7285 | 776.533571 | 743.34126 | 722.95783 | 694.74171 | 2.96E-18 | 0.756014673 | YGL005C | COG7 | n |
| W303.asm02.HP0_G0019850 | 492.00379 | 508.643638 | 381.5403903 | 481.691827 | 449.43126 | 491.140891 | 436.65598 | 464.47744 | 477.85534 | 0.727701374 | -0.042644975 | YGL004C | RPN14 | n |
| W303.asm02.HP0_G0019860 | 379.27487 | 450.512937 | 441.4277426 | 373.352835 | 388.30861 | 274.963047 | 437.79608 | 324.08454 | 341.19929 | 0.053721652 | 0.293308634 | YGL003C | CDH1 | n |
| W303.asm02.HP0_G0019880 | 3368.1715 | 3103.76424 | 3141.188226 | 2049.27371 | 2172.5507 | 2133.33399 | 1808.189 | 1881.5273 | 1772.1203 | 2.60E-21 | 0.597155081 | YGL001C | ERG26 | n |
| W303.asm02.HP0_G0019890 | 1236.8575 | 1135.62478 | 1162.973746 | 1586.74955 | 1578.4026 | 1712.35608 | 1172.0166 | 1238.6065 | 1160.2539 | 1.11E-10 | -0.464076829 | YGR001C | EFM5 | n |
| W303.asm02.HP0_G0019900 | 608.94687 | 602.06798 | 602.7372241 | 1095.0572 | 1047.1748 | 1020.20772 | 972.50014 | 972.25363 | 1014.7814 | 7.46E-27 | -0.802652845 | YGR002C | SWC4 | n |
| W303.asm02.HP0_G0019910 | 808.06618 | 794.106905 | 675.181602 | 888.379737 | 891.67162 | 791.703946 | 818.58746 | 797.74657 | 841.97758 | 0.060329963 | -0.17690475 | YGR003W | CUL3 | n |
| W303.asm02.HP0_G0019930 | 1556.0805 | 1575.75723 | 1264.395876 | 1727.59024 | 1705.1422 | 1817.60056 | 1613.2329 | 1674.2181 | 1624.0028 | 0.00381617 | -0.256647691 | YGR005C | TFG2 | n |
| W303.asm02.HP0_G0019940 | 131.69266 | 152.593091 | 128.4680301 | 275.847742 | 290.33259 | 270.222305 | 189.2556 | 228.30247 | 215.12307 | 3.32E-14 | -1.020033476 | YGR006W | PRP18 | n |
| W303.asm02.HP0_G0019950 | 1314.8195 | 1328.70175 | 1223.827024 | 945.049364 | 967.17607 | 933.926211 | 820.86764 | 907.96155 | 804.9482 | 6.98E-09 | 0.441901011 | YGR007W | ECT1 | n |
| W303.asm02.HP0_G0019960 | 1819.4658 | 2162.25449 | 2091.227709 | 80.8375558 | 83.594215 | 57.8370547 | 264.5018 | 164.0104 | 302.40661 | 3.42E-78 | 4.769963825 | YGR008C | STF2 | n |
| W303.asm02.HP0_G0019970 | 873.38574 | 959.156575 | 970.7546638 | 1615.91774 | 1514.5834 | 1514.19306 | 1654.2763 | 1666.3456 | 1668.9671 | 1.10E-24 | -0.728316053 | YGR009C | SEC9 | n |
| W303.asm02.HP0_G0019990 | 908.1526 | 908.292211 | 927.2880371 | 1210.89658 | 1205.3746 | 1238.28186 | 1152.635 | 1208.4286 | 1105.5915 | 3.44E-09 | -0.413246846 | YGR012W | MCY1 | n |
| W303.asm02.HP0_G0020000 | 493.05733 | 552.241664 | 522.5654459 | 685.869159 | 658.86623 | 595.437219 | 614.51064 | 629.79992 | 581.89026 | 0.000966092 | -0.308274997 | YGR013W | SNU71 | n |
| W303.asm02.HP0_G0020010 | 1575.0442 | 1938.03607 | 1841.053124 | 4079.37975 | 3984.6576 | 4128.2383 | 4509.0716 | 4542.4319 | 4653.3588 | 1.85E-54 | -1.186971604 | YGR014W | MSB2 | n |
| W303.asm02.HP0_G0020020 | 310.79468 | 339.441775 | 311.9937875 | 855.878039 | 876.39096 | 767.052086 | 1013.5435 | 948.63613 | 1057.1006 | 1.04E-47 | -1.37799888 | YGR015C | EAT1 | n |
| W303.asm02.HP0_G0020030 | 291.83094 | 269.892543 | 350.630789 | 313.349701 | 309.20871 | 373.570485 | 326.06687 | 330.64496 | 293.59009 | 0.354184817 | -0.123962345 | YGR016W | | n |
| W303.asm02.HP0_G0020050 | 1083.0405 | 1170.91842 | 1260.532175 | 136.673806 | 137.52597 | 99.555586 | 95.767892 | 124.6479 | 94.336755 | 9.56E-132 | 3.229471909 | YGR019W | UGA1 | n |
| W303.asm02.HP0_G0020060 | 5211.8688 | 5096.81687 | 5852.539809 | 1190.06216 | 1183.8019 | 1145.36331 | 1303.1274 | 1277.969 | 1318.9513 | 5.13E-210 | 2.1991653 | YGR020C | VMA7 | n |
| W303.asm02.HP0_G0020070 | 929.22343 | 1003.79265 | 870.2984598 | 447.523376 | 443.13922 | 520.533492 | 509.622 | 561.5716 | 528.99115 | 1.12E-24 | 0.991033332 | YGR021W | DPC29 | n |
| W303.asm02.HP0_G0020090 | 1490.7609 | 1382.68026 | 1808.211672 | 447.523376 | 515.94709 | 522.429789 | 295.28433 | 384.44037 | 303.28826 | 1.68E-41 | 1.656829529 | YGR024C | THG1 | n |
| W303.asm02.HP0_G0020100 | 3692.6623 | 3707.90832 | 3623.18482 | 1212.56334 | 1280.8791 | 1304.65225 | 1697.5999 | 1490.5265 | 1635.4643 | 2.04E-114 | 1.53768952 | YGR026W | | n |
| W303.asm02.HP0_G0020120 | 2738.1538 | 2934.56238 | 3226.189629 | 1802.59416 | 1711.4342 | 1702.87459 | 1760.3051 | 1844.7889 | 1695.4166 | 4.24E-24 | 0.770511528 | YGR028W | MSP1 | n |
| W303.asm02.HP0_G0020140 | 875.49282 | 830.438593 | 1041.267192 | 284.181511 | 292.13032 | 292.029719 | 300.9848 | 244.04747 | 244.21758 | 1.91E-46 | 1.662531498 | YGR030C | POP6 | n |
| W303.asm02.HP0_G0020170 | 2185.0447 | 2312.77148 | 2193.615763 | 1135.05929 | 1166.7236 | 1129.24479 | 962.23929 | 1096.9015 | 1053.574 | 2.96E-43 | 0.963603527 | YGR033C | TIM21 | n |
| W303.asm02.HP0_G0020210 | 1058.809 | 888.569295 | 992.0050147 | 499.192741 | 547.40728 | 482.607555 | 663.53468 | 699.34033 | 694.74171 | 2.39E-24 | 0.942524898 | YGR036C | CAX4 | n |
| W303.asm02.HP0_G0020220 | 6056.8089 | 6022.7559 | 6074.702568 | 4283.55708 | 4355.8878 | 4388.03097 | 4277.6325 | 4580.4823 | 4197.5448 | 1.33E-18 | 0.478835132 | YGR037C | ACB1 | n |
| W303.asm02.HP0_G0020230 | 1450.7264 | 1465.72412 | 1543.548212 | 1732.5905 | 1698.8502 | 1641.24495 | 1981.4833 | 1931.3864 | 1965.2021 | 0.003427472 | -0.185672192 | YGR038W | ORM1 | n |
| W303.asm02.HP0_G0020240 | 1336.9439 | 1127.32039 | 1087.631593 | 1352.57065 | 1320.429 | 1178.54851 | 989.60155 | 1124.4553 | 1132.9227 | 0.246531385 | -0.117726381 | YGR040W | KSS1 | n |
| W303.asm02.HP0_G0020250 | 489.8967 | 515.909976 | 533.1906213 | 1039.22095 | 1015.7147 | 1004.0892 | 1323.6491 | 1300.2744 | 1434.4477 | 2.24E-35 | -0.990663703 | YGR041W | BUD9 | n |
| W303.asm02.HP0_G0020260 | 850.20783 | 879.226861 | 939.8450626 | 97.5050931 | 123.14417 | 125.155594 | 96.907986 | 86.597489 | 94.336755 | 6.93E-129 | 2.953118589 | YGR042W | MTE1 | n |
| W303.asm02.HP0_G0020280 | 1854.2327 | 1877.82927 | 2006.226305 | 339.184384 | 360.44387 | 350.814922 | 470.8588 | 394.93703 | 457.57734 | 5.75E-175 | 2.450322576 | YGR044C | RME1 | n |
| W303.asm02.HP0_G0020290 | 1857.3933 | 1675.40986 | 1924.122677 | 1003.38574 | 1095.7134 | 888.415086 | 1078.5289 | 1047.0424 | 1023.5979 | 3.81E-22 | 0.868853722 | YGR046W | TAM41 | n |
| W303.asm02.HP0_G0020300 | 588.92959 | 573.002629 | 517.7358207 | 610.031864 | 591.45154 | 634.311305 | 574.60735 | 534.01785 | 536.04436 | 0.158580414 | -0.128874994 | YGR047C | TFC4 | n |
| W303.asm02.HP0_G0020310 | 1337.9975 | 1253.96228 | 1231.554424 | 1107.55785 | 1229.6439 | 1024.00031 | 1162.8958 | 1043.1061 | 1054.4557 | 0.036090654 | 0.185418391 | YGR048W | UFD1 | n |
| W303.asm02.HP0_G0020340 | 229.672 | 225.256468 | 231.8220093 | 304.182555 | 297.5235 | 344.177883 | 295.28433 | 305.71538 | 326.21121 | 7.17E-05 | -0.460486783 | YGR052W | FMP48 | n |
| W303.asm02.HP0_G0020350 | 174.88786 | 176.468201 | 206.7079583 | 222.511623 | 233.70426 | 234.192664 | 231.43907 | 219.11789 | 228.34785 | 0.017675109 | -0.30447611 | YGR053C | MCO32 | n |
| W303.asm02.HP0_G0020360 | 1702.5227 | 2058.44966 | 2134.694335 | 5230.2732 | 5247.5594 | 5232.83123 | 4281.0528 | 4556.8648 | 4292.7632 | 5.21E-67 | -1.413604448 | YGR054W | | n |
| W303.asm02.HP0_G0020370 | 5739.693 | 5128.99636 | 5495.147545 | 1772.59259 | 1678.1763 | 1432.65229 | 3313.113 | 3416.6646 | 3553.0572 | 3.29E-96 | 1.743874047 | YGR055W | MUP1 | n |
| W303.asm02.HP0_G0020380 | 544.68085 | 614.524559 | 596.9416739 | 1895.93237 | 1925.3635 | 1848.88945 | 1833.2711 | 1725.3894 | 1853.2323 | 8.64E-116 | -1.690628229 | YGR056W | RSC1 | n |
| W303.asm02.HP0_G0020390 | 1194.7158 | 1150.15745 | 1336.840253 | 597.531211 | 651.67533 | 692.14836 | 582.58801 | 593.06159 | 577.482 | 2.67E-25 | 0.924887966 | YGR057C | LST7 | n |
| W303.asm02.HP0_G0020400 | 495.16441 | 426.637827 | 461.7121685 | 594.197704 | 617.51855 | 579.318696 | 445.77673 | 510.40035 | 496.37003 | 0.000108403 | -0.372763166 | YGR058W | PEF1 | n |
| W303.asm02.HP0_G0020410 | 58.998313 | 42.5599779 | 52.15995209 | 10.8338992 | 15.280663 | 20.8592656 | 9.1207516 | 14.432915 | 15.869734 | 1.27E-06 | 1.720839478 | YGR059W | SPR3 | n |
| W303.asm02.HP0_G0020420 | 8824.4619 | 8371.85907 | 7017.445406 | 3721.86107 | 3568.4842 | 3807.76413 | 4395.0622 | 4535.8715 | 4313.0412 | 1.48E-41 | 1.125407892 | YGR060W | ERG25 | n |
| W303.asm02.HP0_G0020430 | 1740.4502 | 1894.43804 | 2134.694335 | 2644.30479 | 2622.8808 | 2716.44527 | 2774.9887 | 2650.408 | 2809.8246 | 2.75E-09 | -0.467987493 | YGR061C | ADE6 | n |
| W303.asm02.HP0_G0020440 | 1065.1303 | 1072.30383 | 1048.028667 | 640.033431 | 621.114 | 581.214992 | 630.47196 | 628.48784 | 644.48755 | 3.77E-25 | 0.788859823 | YGR062C | COX18 | n |
| W303.asm02.HP0_G0020450 | 1522.3672 | 1481.29484 | 1789.859097 | 727.538002 | 728.97751 | 703.526141 | 739.92097 | 847.60573 | 782.9069 | 3.56E-36 | 1.150335391 | YGR063C | SPT4 | n |
| W303.asm02.HP0_G0020460 | 1538.1703 | 1304.82664 | 1347.465429 | 675.03526 | 666.05713 | 708.266883 | 1048.8864 | 1187.4353 | 1041.2309 | 1.29E-30 | 1.031977052 | YGR065C | VHT1 | n |
| W303.asm02.HP0_G0020480 | 104.30059 | 95.5004382 | 78.23992813 | 540.861585 | 608.52993 | 527.170531 | 637.31252 | 720.33366 | 654.18572 | 7.19E-86 | -2.596260404 | YGR067C | | n |
| W303.asm02.HP0_G0020490 | 179.10202 | 215.914034 | 212.5035085 | 319.183339 | 263.36672 | 247.466742 | 290.72396 | 178.44331 | 267.14053 | 0.009349391 | -0.450963997 | YGR068C | ART5 | n |
| W303.asm02.HP0_G0020520 | 700.60496 | 676.807453 | 754.3874552 | 1425.07444 | 1540.6504 | 1498.07453 | 1197.0986 | 1229.4219 | 1175.242 | 4.63E-46 | -1.065190611 | YGR072W | UPF3 | n |
| W303.asm02.HP0_G0020530 | 1036.6846 | 1005.86875 | 1136.89377 | 619.19901 | 613.9231 | 562.252024 | 563.20641 | 669.16242 | 622.44625 | 1.91E-19 | 0.824261322 | YGR075C | PRP38 | n |

| | | | | | | | | | | | | | | |
|---|---|---|---|---|---|---|---|---|---|---|---|---|---|---|
| W303.asm02.HP0_G0020540 | 1395.9422 | 1327.6637 | 1572.525963 | 1381.73884 | 1525.3697 | 1470.57823 | 1279.1854 | 1460.3486 | 1345.4008 | 0.773503103 | -0.026407144 | YGR076C | MRPL25 | n |
| W303.asm02.HP0_G0020550 | 811.2268 | 891.683439 | 804.6155572 | 475.024812 | 471.90282 | 464.592735 | 495.94087 | 474.97411 | 558.96731 | 1.78E-19 | 0.828580699 | YGR077C | PEX8 | n |
| W303.asm02.HP0_G0020560 | 389.81028 | 537.708989 | 444.3255178 | 1300.90128 | 1295.2609 | 1354.90412 | 1127.5529 | 1195.3078 | 1205.2182 | 2.72E-51 | -1.525975885 | YGR078C | PAC10 | n |
| W303.asm02.HP0_G0020570 | 2392.5923 | 2487.16359 | 2214.866114 | 1030.0538 | 1035.4896 | 962.370664 | 1094.4902 | 1116.5828 | 1098.5383 | 2.85E-66 | 1.227668637 | YGR079W | | n |
| W303.asm02.HP0_G0020580 | 1954.3191 | 1914.16096 | 2186.854287 | 706.703581 | 700.21391 | 804.978024 | 478.83946 | 646.857 | 501.65994 | 1.31E-40 | 1.453652141 | YGR081C | SLX9 | n |
| W303.asm02.HP0_G0020590 | 1926.927 | 1810.35613 | 1632.413315 | 3504.34971 | 3408.4867 | 3657.95667 | 2969.9447 | 3005.9825 | 2875.0669 | 1.20E-41 | -0.977609901 | YGR082W | TOM20 | n |
| W303.asm02.HP0_G0020600 | 1617.1859 | 1654.6489 | 1764.745046 | 3499.34945 | 3514.5525 | 3634.25296 | 2938.0221 | 3142.4392 | 2937.6642 | 6.33E-66 | -1.079571016 | YGR083C | GCD2 | n |
| W303.asm02.HP0_G0020610 | 1537.1168 | 1605.86063 | 1348.431354 | 2056.7741 | 2103.3383 | 2144.71177 | 1986.0437 | 2325.0114 | 2026.0361 | 1.53E-08 | -0.489634403 | YGR084C | MRP13 | n |
| W303.asm02.HP0_G0020620 | 8212.3544 | 8234.8367 | 9106.741264 | 12248.1398 | 12666.771 | 13532.9227 | 10309.87 | 11522.714 | 9459.2434 | 7.85E-13 | -0.589246901 | YGR085C | RPL11B | y |
| W303.asm02.HP0_G0020630 | 4990.6251 | 5373.97575 | 6236.012049 | 2643.47141 | 2594.1172 | 2647.23044 | 3209.3645 | 2555.938 | 2993.2082 | 2.40E-28 | 1.074226202 | YGR086C | PIL1 | n |
| W303.asm02.HP0_G0020650 | 254.95699 | 247.055481 | 255.0042102 | 522.527294 | 525.83458 | 478.814961 | 517.60265 | 907.96155 | 525.46454 | 1.27E-08 | -1.01262881 | YGR088W | CTT1 | n |
| W303.asm02.HP0_G0020660 | 1336.9439 | 1197.90767 | 1352.295054 | 1693.42179 | 1641.323 | 1627.02272 | 2416.9992 | 2134.7593 | 2432.4776 | 8.67E-06 | -0.352083632 | YGR089W | NNF2 | n |
| W303.asm02.HP0_G0020670 | 2236.6682 | 2103.08574 | 2361.686719 | 5070.26484 | 4941.0473 | 5558.04614 | 3978.9279 | 3794.5445 | 3974.4868 | 5.81E-67 | -1.215786699 | YGR090W | UTP22 | n |
| W303.asm02.HP0_G0020680 | 546.78793 | 634.247476 | 530.2928462 | 562.529383 | 642.6867 | 700.681696 | 476.55927 | 465.78953 | 531.6361 | 0.173610266 | -0.154653369 | YGR091W | PRP31 | n |
| W303.asm02.HP0_G0020690 | 790.15597 | 889.607343 | 892.5147357 | 749.205801 | 793.69561 | 885.570641 | 864.19122 | 837.10906 | 1025.3612 | 0.432772683 | 0.084064686 | YGR092W | DBF2 | n |
| W303.asm02.HP0_G0020710 | 5196.0657 | 5416.53572 | 5736.628804 | 4931.92428 | 5015.6529 | 5020.44598 | 3921.9232 | 3694.8262 | 3817.5528 | 0.031588757 | 0.127547568 | YGR094W | VAS1 | n |
| W303.asm02.HP0_G0020720 | 1708.844 | 1692.01863 | 1613.094814 | 1250.89867 | 1227.8462 | 1348.26708 | 1133.2534 | 1313.3953 | 1104.7098 | 2.05E-06 | 0.389876185 | YGR095C | RRP46 | n |
| W303.asm02.HP0_G0020730 | 185.42327 | 166.087719 | 219.2649838 | 400.854272 | 394.60065 | 369.777891 | 355.70931 | 439.54786 | 378.22867 | 1.50E-15 | -1.027967861 | YGR096W | TPC1 | n |
| W303.asm02.HP0_G0020740 | 732.2112 | 920.74879 | 936.9472875 | 1220.06373 | 1275.4859 | 1339.73374 | 1296.2868 | 1049.6665 | 1359.5072 | 8.59E-07 | -0.565709537 | YGR097W | ASK10 | n |
| W303.asm02.HP0_G0020750 | 893.40302 | 825.248352 | 807.5133323 | 967.550539 | 949.19882 | 974.696594 | 1024.9445 | 1035.2336 | 1039.4676 | 0.008610697 | -0.195371252 | YGR098C | ESP1 | n |
| W303.asm02.HP0_G0020760 | 651.08852 | 595.839691 | 592.1120487 | 661.70123 | 725.38206 | 768.000235 | 640.7328 | 669.16242 | 679.75362 | 0.011795281 | -0.228190027 | YGR099W | TEL2 | n |
| W303.asm02.HP0_G0020770 | 985.06112 | 1005.86875 | 818.1385077 | 1865.9308 | 1924.4647 | 2084.03027 | 1948.4206 | 1797.5539 | 1825.9011 | 7.15E-33 | -1.064926872 | YGR100W | MDR1 | n |
| W303.asm02.HP0_G0020780 | 1197.8765 | 1126.28234 | 1265.361801 | 1248.39854 | 1170.319 | 1170.96332 | 1149.2147 | 1184.8111 | 1124.1062 | 0.999458016 | -4.96E-05 | YGR101W | PCP1 | n |
| W303.asm02.HP0_G0020790 | 757.49619 | 775.422036 | 759.2170804 | 336.684253 | 381.11771 | 353.659367 | 350.00884 | 363.44704 | 353.54242 | 1.24E-34 | 1.097798102 | YGR102C | GTF1 | n |
| W303.asm02.HP0_G0020800 | 5138.1209 | 4324.50897 | 5187.017457 | 5704.46463 | 5412.9501 | 6174.34263 | 3445.3639 | 4104.1962 | 3338.8158 | 0.015723291 | -0.239112559 | YGR103W | NOP7 | n |
| W303.asm02.HP0_G0020810 | 640.55311 | 619.7148 | 591.1461236 | 748.372424 | 747.85362 | 697.83725 | 725.09975 | 770.19282 | 754.69404 | 0.002747155 | -0.246117578 | YGR104C | SRB5 | n |
| W303.asm02.HP0_G0020820 | 1689.8802 | 1631.81184 | 2078.670683 | 452.523637 | 474.59941 | 451.318656 | 445.77673 | 476.28619 | 474.32873 | 6.76E-83 | 1.970517123 | YGR105W | VMA21 | n |
| W303.asm02.HP0_G0020830 | 5834.5117 | 5773.62432 | 7237.676314 | 2177.61375 | 2039.5191 | 2105.83768 | 2496.8058 | 2448.3472 | 2402.5015 | 1.91E-71 | 1.575651185 | YGR106C | VOA1 | n |
| W303.asm02.HP0_G0020860 | 254.95699 | 271.968639 | 235.6857094 | 1211.72996 | 1140.6565 | 1250.60779 | 1348.7311 | 1204.4923 | 1314.543 | 9.62E-127 | -2.241347886 | YGR110W | CLD1 | n |
| W303.asm02.HP0_G0020870 | 1327.462 | 1257.07642 | 1187.121872 | 1339.23662 | 1395.0346 | 1188.02999 | 1974.6427 | 2366.998 | 2025.1544 | 0.561483702 | -0.057163358 | YGR111W | | n |
| W303.asm02.HP0_G0020880 | 837.56533 | 889.607343 | 756.3193052 | 967.550539 | 964.47949 | 1023.05216 | 1026.0846 | 1086.4049 | 1044.7575 | 0.002342882 | -0.251271563 | YGR112W | SHY1 | n |
| W303.asm02.HP0_G0020890 | 358.20404 | 394.458332 | 362.2218895 | 396.687387 | 358.64615 | 351.76307 | 446.91683 | 400.18537 | 455.81404 | 0.938753429 | 0.008658495 | YGR113W | DAM1 | n |
| W303.asm02.HP0_G0020900 | 2590.6581 | 2405.15778 | 2639.873131 | 5366.947 | 5365.3104 | 5512.53502 | 5001.5922 | 5182.7285 | 4994.5581 | 1.68E-79 | -1.088905176 | YGR116W | SPT6 | n |
| W303.asm02.HP0_G0020910 | 1626.6678 | 1710.7035 | 1894.179001 | 767.540092 | 773.02177 | 775.585422 | 650.99365 | 695.40408 | 659.47563 | 1.66E-49 | 1.176067901 | YGR117C | | n |
| W303.asm02.HP0_G0020920 | 9311.198 | 8792.26861 | 10457.10447 | 139.173936 | 142.02028 | 95.7629922 | 174.43437 | 99.718321 | 95.218407 | 4.29E-110 | 6.242282035 | YPR132W | RPS23B | y |
| W303.asm02.HP0_G0020930 | 2143.9565 | 2092.70525 | 2353.959319 | 2443.46096 | 2391.8732 | 2541.03781 | 2138.8163 | 2172.8097 | 2052.4857 | 0.018655883 | -0.162003463 | YGR119C | NUP57 | n |
| W303.asm02.HP0_G0020940 | 1335.8904 | 1330.77785 | 1474.001609 | 397.520764 | 392.80292 | 347.970477 | 357.9895 | 372.63162 | 380.87363 | 4.70E-96 | 1.861982058 | YGR120C | COG2 | n |
| W303.asm02.HP0_G0020950 | 979.79341 | 874.036619 | 791.0926066 | 665.034737 | 691.22528 | 568.889063 | 546.105 | 638.9845 | 666.52885 | 5.73E-05 | 0.457038401 | YGR122W | | n |
| W303.asm02.HP0_G0020980 | 2427.3592 | 2012.77554 | 2377.14152 | 3077.66076 | 3101.9746 | 3069.15649 | 2690.6217 | 2851.1567 | 2750.754 | 1.10E-08 | -0.439959542 | YGR123C | PPT1 | n |
| W303.asm02.HP0_G0020990 | 5865.0644 | 5908.57059 | 6094.021069 | 10868.0677 | 10777.362 | 11580.685 | 8274.8019 | 7991.8986 | 8128.8306 | 1.05E-63 | -0.894815091 | YGR124W | ASN2 | n |
| W303.asm02.HP0_G0021000 | 1270.5708 | 1180.26085 | 1324.283228 | 1274.23322 | 1207.1724 | 1115.02256 | 1187.9779 | 1184.8111 | 1235.1943 | 0.393007467 | 0.069571604 | YGR125W | VSB1 | n |
| W303.asm02.HP0_G0021010 | 144.33516 | 157.783333 | 148.7524559 | 178.342649 | 177.97478 | 148.859305 | 134.53109 | 178.44331 | 166.63221 | 0.293139488 | -0.165830131 | YGR126W | | n |
| W303.asm02.HP0_G0021020 | 342.40092 | 350.860306 | 337.1078385 | 300.849048 | 345.16321 | 367.881594 | 375.09091 | 366.0712 | 352.66076 | 0.828900743 | 0.025068725 | YGR127W | | n |
| W303.asm02.HP0_G0021030 | 2482.1433 | 2133.18914 | 2071.909208 | 2245.95065 | 2306.4812 | 2421.57111 | 1492.383 | 1780.4969 | 1556.1156 | 0.519731286 | -0.060676871 | YGR128C | UTP8 | n |
| W303.asm02.HP0_G0021040 | 813.33388 | 817.982014 | 903.1399111 | 300.849048 | 270.55762 | 274.014899 | 224.59851 | 261.10455 | 215.12307 | 4.24E-53 | 1.58324866 | YGR129W | SYF2 | n |
| W303.asm02.HP0_G0021050 | 1386.4604 | 1614.16502 | 1500.081585 | 1249.23192 | 1298.8563 | 1201.30407 | 1693.0395 | 1338.3248 | 1759.7772 | 0.011845222 | 0.263449042 | YGR130C | | n |
| W303.asm02.HP0_G0021070 | 5624.857 | 5274.32311 | 6157.772121 | 2671.80623 | 2680.408 | 2610.25265 | 3145.5192 | 3089.9559 | 3106.0597 | 7.94E-60 | 1.099169655 | YGR132C | PHB1 | n |
| W303.asm02.HP0_G0021080 | 268.65303 | 315.566665 | 279.1523362 | 1195.8958 | 1081.3316 | 1116.91886 | 1034.0652 | 1012.9282 | 1034.1777 | 2.87E-106 | -1.975694863 | YGR133W | PEX4 | n |
| W303.asm02.HP0_G0021090 | 591.03667 | 620.752848 | 599.839449 | 715.870726 | 683.13552 | 610.607594 | 573.46726 | 530.0816 | 604.81321 | 0.112251808 | -0.15072691 | YGR134W | CAF130 | n |
| W303.asm02.HP0_G0021100 | 5915.6344 | 5972.92958 | 6419.537807 | 3755.19615 | 3593.6524 | 3766.99374 | 4057.5944 | 4363.9886 | 4017.6878 | 3.00E-31 | 0.719985699 | YGR135W | PRE9 | n |
| W303.asm02.HP0_G0021130 | 497.27149 | 479.578288 | 433.7003423 | 568.363021 | 570.7777 | 578.370547 | 566.62669 | 499.90369 | 492.84342 | 0.003135512 | -0.284922545 | YGR140W | CBF2 | n |
| W303.asm02.HP0_G0021140 | 463.55817 | 451.550985 | 534.1565464 | 474.191436 | 477.296 | 399.170492 | 529.00359 | 560.25951 | 558.08566 | 0.357012693 | 0.101469776 | YGR141W | VPS62 | n |
| W303.asm02.HP0_G0021150 | 2090.2259 | 3645.62542 | 1068.313093 | 304.182555 | 453.02671 | 398.222344 | 1734.0829 | 926.33072 | 1279.2769 | 4.33E-08 | 2.558060129 | YGR142W | BTN2 | n |
| W303.asm02.HP0_G0021170 | 89.55101 | 91.3482452 | 96.59250386 | 323.350223 | 388.30861 | 393.481602 | 410.43382 | 381.8162 | 329.73782 | 1.02E-41 | -1.991382718 | YGR144W | THI4 | n |
| W303.asm02.HP0_G0021180 | 1535.0097 | 1419.01195 | 1338.772104 | 2437.62733 | 2337.9414 | 2669.986 | 1337.3302 | 1788.3694 | 1400.0632 | 1.21E-13 | -0.794577488 | YGR145W | ENP2 | n |
| W303.asm02.HP0_G0021190 | 1114.6467 | 1353.61491 | 924.390262 | 952.549755 | 884.48072 | 922.54843 | 1354.4316 | 967.0053 | 1159.3723 | 0.032625135 | 0.297460348 | YGR146C | ECL1 | n |
| W303.asm02.HP0_G0021210 | 1851.0721 | 1762.60591 | 1921.224902 | 903.38052 | 889.8739 | 861.86693 | 850.51009 | 915.83405 | 886.06017 | 3.29E-51 | 1.059630164 | YGR147C | NAT2 | n |
| W303.asm02.HP0_G0021230 | 901.83135 | 871.960523 | 810.4111074 | 495.859234 | 489.88007 | 407.703828 | 535.84416 | 527.45743 | 566.02053 | 5.15E-20 | 0.889090313 | YGR149W | GPC1 | n |
| W303.asm02.HP0_G0021240 | 2719.1901 | 2502.73431 | 2498.848075 | 885.879606 | 856.61598 | 850.489149 | 660.1144 | 787.2499 | 670.93711 | 5.61E-86 | 1.573723996 | YGR150C | CCM1 | n |
| W303.asm02.HP0_G0021250 | 2227.1863 | 2092.70525 | 2199.411313 | 1859.26378 | 1697.9513 | 1845.09686 | 1972.3625 | 1772.6244 | 2078.9352 | 0.000463548 | 0.271107002 | YGR152C | RSR1 | n |

| | | | | | | | | | | | | | | |
|---|---|---|---|---|---|---|---|---|---|---|---|---|---|---|
| W303.asm02.HP0_G0021280 | 9748.4176 | 9367.34733 | 8642.131321 | 5834.47142 | 5827.3257 | 6205.63153 | 4375.6806 | 5185.3527 | 4420.6027 | 1.35E-15 | 0.635562065 | YGR155W | CYS4 | n |
| W303.asm02.HP0_G0021290 | 333.97259 | 427.675875 | 452.0529181 | 1102.55759 | 1188.2963 | 1117.86701 | 945.13789 | 821.36406 | 1030.6511 | 1.14E-37 | -1.488484917 | YGR156W | PTI1 | n |
| W303.asm02.HP0_G0021300 | 5982.0075 | 5796.46138 | 6201.238748 | 6827.02327 | 7142.3616 | 7327.29113 | 7974.9572 | 6901.5575 | 7462.3018 | 0.000363363 | -0.244110605 | YGR157W | CHO2 | n |
| W303.asm02.HP0_G0021310 | 2327.2727 | 2248.41249 | 2603.167979 | 840.043879 | 872.79551 | 924.444727 | 628.19177 | 809.55532 | 671.81876 | 4.38E-50 | 1.445338849 | YGR158C | MTR3 | n |
| W303.asm02.HP0_G0021320 | 8109.1074 | 6818.9389 | 7785.355811 | 6598.67801 | 6182.3764 | 7088.35772 | 3549.1125 | 5163.0473 | 3769.0619 | 0.12294778 | 0.193025851 | YGR159C | NSR1 | n |
| W303.asm02.HP0_G0021330 | 928.16989 | 1034.9341 | 784.3311314 | 3731.8616 | 3293.4323 | 2523.97114 | 4179.5844 | 3610.8529 | 4405.6146 | 3.75E-40 | -1.797974171 | YGR161C | RTS3 | n |
| W303.asm02.HP0_G0021360 | 3268.0851 | 3507.56501 | 3799.949102 | 5729.46594 | 5734.7429 | 6295.70563 | 4160.2028 | 4568.6736 | 4458.5137 | 2.47E-23 | -0.747550208 | YGR162W | TIF4631 | n |
| W303.asm02.HP0_G0021370 | 1296.9093 | 1468.83826 | 1205.474448 | 1184.22852 | 1160.4315 | 1095.11145 | 1207.3595 | 1211.0528 | 1272.2237 | 0.016627564 | 0.20647122 | YGR163W | GTR2 | n |
| W303.asm02.HP0_G0021380 | 2354.6648 | 2248.41249 | 2228.389064 | 1739.25751 | 1685.3672 | 1748.38572 | 1857.213 | 1747.6948 | 1869.1021 | 1.04E-10 | 0.401038024 | YGR165W | MRPS35 | n |
| W303.asm02.HP0_G0021390 | 1122.0215 | 996.526312 | 1104.052319 | 497.525988 | 515.94709 | 533.807571 | 503.92153 | 489.40702 | 494.60672 | 9.20E-37 | 1.059325722 | YGR166W | TRS65 | n |
| W303.asm02.HP0_G0021400 | 3191.1766 | 3094.42181 | 2629.247955 | 2470.9624 | 2410.7493 | 2548.623 | 2358.8544 | 2408.9847 | 2355.7739 | 0.000915377 | 0.26249916 | YGR167W | CLC1 | n |
| W303.asm02.HP0_G0021410 | 929.22343 | 991.336071 | 1103.086394 | 329.183861 | 357.74728 | 318.577875 | 353.42912 | 371.31954 | 351.77911 | 4.16E-59 | 1.588779699 | YGR168C | PEX35 | n |
| W303.asm02.HP0_G0021420 | 577.34063 | 589.611401 | 599.839449 | 760.873077 | 621.114 | 718.696516 | 386.49185 | 501.21577 | 431.12779 | 0.023848736 | -0.249944893 | YGR169C | PUS6 | n |
| W303.asm02.HP0_G0021430 | 2566.4266 | 2464.32653 | 3060.050522 | 1149.2267 | 1048.0737 | 1237.33371 | 856.21056 | 985.37446 | 953.06572 | 2.11E-35 | 1.23661432 | YGR169C- | LSO2 | n |
| W303.asm02.HP0_G0021440 | 850.20783 | 964.346816 | 924.390262 | 1415.90729 | 1297.9575 | 1264.83002 | 1219.9005 | 1152.009 | 1224.6145 | 1.65E-11 | -0.539025002 | YGR170W | PSD2 | n |
| W303.asm02.HP0_G0021450 | 827.02992 | 808.63958 | 820.0703578 | 910.047535 | 935.71589 | 915.911391 | 808.32661 | 863.35073 | 844.62253 | 0.020799564 | -0.169313214 | YGR171C | MSM1 | n |
| W303.asm02.HP0_G0021460 | 1576.0978 | 1442.88706 | 1500.081585 | 1323.40246 | 1300.6541 | 1233.54112 | 1654.2763 | 1789.6814 | 1713.9313 | 0.001274523 | 0.227806583 | YGR172C | YIP1 | n |
| W303.asm02.HP0_G0021470 | 1630.8819 | 1464.68607 | 1800.484272 | 1451.7425 | 1424.6971 | 1380.50413 | 1263.2241 | 1276.6569 | 1281.0402 | 0.01475009 | 0.202064884 | YGR173W | RBG2 | n |
| W303.asm02.HP0_G0021480 | 655.30269 | 625.94309 | 752.4556051 | 811.709065 | 770.32518 | 772.740977 | 727.37994 | 868.59906 | 799.65828 | 0.032176203 | -0.210881981 | YGR174C | CBP4 | n |
| W303.asm02.HP0_G0021490 | 6952.319 | 6045.59296 | 6320.047528 | 2160.11283 | 2110.5292 | 2187.37845 | 1680.4985 | 1750.3189 | 1590.5 | 1.25E-111 | 1.580756676 | YGR175C | ERG1 | n |
| W303.asm02.HP0_G0021510 | 1856.3398 | 2004.47115 | 2257.366815 | 2927.65292 | 2851.1919 | 2972.44535 | 2936.882 | 2603.173 | 2843.3274 | 3.09E-10 | -0.515923334 | YGR178C | PBP1 | n |
| W303.asm02.HP0_G0021520 | 396.13153 | 394.458332 | 281.0841862 | 330.850615 | 339.77003 | 323.318617 | 307.82537 | 333.26912 | 320.03964 | 0.434738365 | 0.106620277 | YGR179C | OKP1 | n |
| W303.asm02.HP0_G0021550 | 970.31154 | 1030.7819 | 1073.142718 | 857.544793 | 948.29996 | 930.133618 | 750.18182 | 864.66281 | 862.25557 | 0.053219041 | 0.169053738 | YGR181W | TIM13 | n |
| W303.asm02.HP0_G0021560 | 4507.0497 | 4415.85722 | 5451.680918 | 3151.8313 | 3287.1402 | 3572.62331 | 3003.0075 | 2887.8951 | 2643.1924 | 1.84E-08 | 0.522198675 | YGR183C | QCR9 | n |
| W303.asm02.HP0_G0021570 | 1443.3516 | 1431.46852 | 1552.241537 | 2561.80048 | 2628.274 | 2689.89712 | 2940.3023 | 2883.9588 | 3068.1487 | 5.25E-39 | -0.831212234 | YGR184C | UBR1 | n |
| W303.asm02.HP0_G0021580 | 3484.0611 | 3281.27049 | 2892.945491 | 5250.27424 | 5083.0676 | 5531.49799 | 4200.1061 | 4283.9515 | 4190.4915 | 1.18E-21 | -0.7162489 | YGR185C | TYS1 | n |
| W303.asm02.HP0_G0021590 | 1347.4793 | 1397.21293 | 1513.604536 | 2091.77593 | 2058.3952 | 2045.15618 | 1987.1838 | 1907.7689 | 1981.0718 | 3.78E-16 | -0.540425198 | YGR186W | TFG1 | n |
| W303.asm02.HP0_G0021600 | 2543.2487 | 2176.78716 | 2539.416927 | 1802.59416 | 1826.4886 | 2010.07469 | 1094.4902 | 1355.3819 | 1170.8337 | 0.0001306 | 0.364715127 | YGR187C | HGH1 | n |
| W303.asm02.HP0_G0021610 | 305.52698 | 388.230042 | 321.6530379 | 721.704364 | 660.66395 | 712.059477 | 790.08511 | 800.37073 | 759.1023 | 4.46E-25 | -1.04500803 | YGR188C | BUB1 | n |
| W303.asm02.HP0_G0021620 | 2993.1108 | 3094.42181 | 3557.501917 | 6988.69838 | 6470.0124 | 6704.3576 | 6385.6662 | 6772.9733 | 6698.7912 | 2.49E-46 | -1.063582319 | YGR189C | CRH1 | n |
| W303.asm02.HP0_G0021640 | 49545.94 | 49200.3725 | 49042.91199 | 43619.7784 | 41058.242 | 46036.3993 | 42036.404 | 43515.238 | 37767.323 | 0.010914757 | 0.177130625 | YGR192C | TDH3 | y |
| W303.asm02.HP0_G0021650 | 1531.849 | 1621.43135 | 1389.000206 | 2127.61113 | 2139.2928 | 2666.19341 | 1990.604 | 1561.379 | 2148.5857 | 6.43E-07 | -0.610010334 | YGR193C | PDX1 | n |
| W303.asm02.HP0_G0021660 | 304.47344 | 380.963705 | 371.8811399 | 285.848264 | 245.38947 | 241.777852 | 336.32772 | 230.92664 | 367.64885 | 0.005265848 | 0.451265117 | YGR194C | XKS1 | n |
| W303.asm02.HP0_G0021670 | 1173.645 | 1199.98377 | 1049.960517 | 1105.05772 | 1226.9473 | 1206.99296 | 841.38934 | 1100.8378 | 911.62808 | 0.657589612 | -0.0478881 | YGR195W | SKI6 | n |
| W303.asm02.HP0_G0021680 | 553.10918 | 587.535305 | 475.235119 | 2078.4419 | 2109.6303 | 2101.09694 | 1881.155 | 1910.3931 | 1882.3268 | 3.75E-135 | -1.962050153 | YGR196C | FYV8 | n |
| W303.asm02.HP0_G0021690 | 793.3166 | 897.911729 | 897.3443609 | 1087.55681 | 1110.9941 | 996.504008 | 1818.4499 | 2000.9268 | 1907.8947 | 0.000266022 | -0.303662028 | YGR197C | SNG1 | n |
| W303.asm02.HP0_G0021700 | 1759.414 | 1688.90449 | 1668.152542 | 1409.24028 | 1490.3141 | 1485.7486 | 1439.9387 | 1424.9223 | 1444.1458 | 0.000468143 | 0.222635613 | YGR198W | YPP1 | n |
| W303.asm02.HP0_G0021710 | 936.59822 | 905.178067 | 953.3680131 | 1111.72474 | 1120.8816 | 1124.50405 | 1322.509 | 1274.0328 | 1328.6494 | 0.000110196 | -0.26395804 | YGR199W | PMT6 | n |
| W303.asm02.HP0_G0021720 | 1284.2668 | 1214.51644 | 1491.38826 | 2200.9483 | 2089.8554 | 2425.3637 | 1797.9282 | 1927.4502 | 1789.7534 | 1.24E-16 | -0.750426567 | YGR200C | ELP2 | n |
| W303.asm02.HP0_G0021740 | 961.88321 | 1020.40142 | 1069.279018 | 2270.11858 | 2143.7871 | 2202.54882 | 1795.648 | 1802.8023 | 1773.002 | 5.97E-63 | -1.116039251 | YGR202C | PCT1 | n |
| W303.asm02.HP0_G0021750 | 672.15935 | 613.486511 | 719.6141538 | 288.348395 | 322.69165 | 290.133422 | 269.06217 | 322.77246 | 290.94513 | 6.70E-27 | 1.154625118 | YGR203W | YCH1 | n |
| W303.asm02.HP0_G0021770 | 396.13153 | 344.632016 | 344.8352388 | 587.530689 | 565.38453 | 555.614985 | 793.50539 | 649.48117 | 778.49864 | 6.38E-10 | -0.655405277 | YGR205W | TDA10 | n |
| W303.asm02.HP0_G0021780 | 471.9865 | 388.230042 | 590.1801986 | 215.844608 | 210.33383 | 217.125992 | 200.65654 | 238.79914 | 241.57262 | 3.45E-15 | 1.174005109 | YGR206W | MVB12 | n |
| W303.asm02.HP0_G0021790 | 2287.2382 | 2175.74911 | 2332.708968 | 1226.73074 | 1164.9258 | 1225.95593 | 1065.9878 | 1116.5828 | 1230.7861 | 4.52E-35 | 0.909665153 | YGR207C | CIR1 | n |
| W303.asm02.HP0_G0021800 | 1657.2205 | 1485.44703 | 1563.832638 | 2010.93837 | 1901.9931 | 2106.78583 | 1898.2564 | 2078.3397 | 2009.2847 | 1.44E-06 | -0.354978182 | YGR208W | SER2 | n |
| W303.asm02.HP0_G0021810 | 4097.2221 | 4002.71402 | 4905.933271 | 4331.89294 | 4458.3581 | 4527.40489 | 4350.5985 | 4867.8286 | 4345.6623 | 0.709585397 | -0.033902762 | YGR209C | TRX2 | n |
| W303.asm02.HP0_G0021820 | 1099.8971 | 1232.16326 | 1169.735222 | 1645.08593 | 1532.5606 | 1830.87463 | 1512.9047 | 1444.6036 | 1480.2936 | 5.26E-10 | -0.515854122 | YGR210C | | n |
| W303.asm02.HP0_G0021830 | 2339.9152 | 2932.48628 | 1862.303474 | 3711.02717 | 3566.6865 | 3949.03824 | 3209.3645 | 3352.3725 | 3189.8166 | 1.37E-07 | -0.654261699 | YGR211W | ZPR1 | n |
| W303.asm02.HP0_G0021840 | 252.84991 | 306.224231 | 248.2427349 | 509.193264 | 498.8687 | 454.163102 | 570.04698 | 489.40702 | 550.15079 | 5.09E-14 | -0.858548893 | YGR212W | SLI1 | n |
| W303.asm02.HP0_G0021860 | 9640.9564 | 10126.1606 | 8349.456034 | 586.697312 | 487.18349 | 573.629805 | 531.28378 | 469.72577 | 487.55351 | 0 | 4.092705033 | YGR214W | RPS0A | n |
| W303.asm02.HP0_G0021870 | 1659.3275 | 1676.44791 | 1971.453004 | 837.543748 | 905.15456 | 864.711375 | 768.42332 | 928.95488 | 849.03079 | 1.64E-27 | 1.025935342 | YGR215W | RSM27 | n |
| W303.asm02.HP0_G0021880 | 946.08009 | 909.33026 | 999.732415 | 612.531995 | 679.54007 | 713.955774 | 713.69881 | 772.81699 | 700.91327 | 6.29E-09 | 0.510726657 | YGR216C | GPI1 | n |
| W303.asm02.HP0_G0021890 | 580.50126 | 557.431906 | 655.8631012 | 1801.76078 | 1841.7693 | 1884.91909 | 2004.2852 | 1899.8964 | 2114.2013 | 1.18E-89 | -1.622545878 | YGR217W | CCH1 | n |
| W303.asm02.HP0_G0021900 | 2213.4903 | 2231.80372 | 2609.929454 | 4750.24812 | 4972.5075 | 5460.38685 | 4718.8489 | 4613.2844 | 4537.8624 | 1.72E-42 | -1.105161905 | YGR218W | CRM1 | n |
| W303.asm02.HP0_G0021910 | 2698.1193 | 2605.50109 | 2997.265395 | 2036.77306 | 2099.7429 | 2168.41548 | 2062.43 | 2143.9439 | 1970.492 | 5.33E-08 | 0.397302273 | YGR220C | MRPL9 | n |
| W303.asm02.HP0_G0021930 | 948.18717 | 1033.89605 | 1077.972343 | 579.19692 | 570.7777 | 571.733508 | 567.76679 | 565.50785 | 553.6774 | 1.86E-24 | 0.830139125 | YGR222W | PET54 | n |
| W303.asm02.HP0_G0021940 | 738.53245 | 758.813265 | 704.1593532 | 690.036043 | 589.65381 | 585.955735 | 747.90163 | 640.29659 | 762.6289 | 0.018838427 | 0.237422002 | YGR223C | HSV2 | n |
| W303.asm02.HP0_G0021960 | 33.713322 | 39.4458332 | 33.80737635 | 15.0007835 | 17.97725 | 9.48148438 | 17.101409 | 27.553747 | 25.567905 | 0.000316136 | 1.32650094 | YGR225W | AMA1 | n |
| W303.asm02.HP0_G0021970 | 626.85707 | 643.58991 | 716.7163787 | 870.045446 | 944.70451 | 812.563211 | 1164.0359 | 1239.9186 | 1161.1356 | 6.46E-06 | -0.402350072 | YGR227W | DIE2 | n |
| W303.asm02.HP0_G0021980 | 4089.8473 | 3759.81073 | 3604.832244 | 3534.35128 | 3329.3868 | 3354.54917 | 3437.3833 | 3152.9359 | 3328.236 | 0.018056216 | 0.164427378 | YGR229C | SMI1 | n |

| | | | | | | | | | | | | | | |
|---|---|---|---|---|---|---|---|---|---|---|---|---|---|---|
| W303.asm02.HP0_G0021990 | 610.00041 | 650.856247 | 772.7400309 | 561.696006 | 583.36178 | 596.385367 | 701.15778 | 745.26324 | 740.58761 | 0.024971693 | 0.225368481 | YGR230W | BNS1 | n |
| W303.asm02.HP0_G0022000 | 4531.2811 | 4550.80349 | 4670.247562 | 2590.96867 | 2526.7026 | 2381.74888 | 2741.926 | 2767.1834 | 2819.5228 | 2.09E-53 | 0.87451443 | YGR231C | PHB2 | n |
| W303.asm02.HP0_G0022010 | 1412.7989 | 1498.94166 | 1709.687318 | 862.545054 | 832.3467 | 875.141008 | 815.16718 | 858.10239 | 857.84731 | 2.49E-23 | 0.847197263 | YGR232W | NAS6 | n |
| W303.asm02.HP0_G0022020 | 1427.5485 | 1499.97971 | 1310.760277 | 2327.62158 | 2342.4357 | 2036.62284 | 1500.3636 | 1665.0335 | 1594.9083 | 2.05E-15 | -0.662797567 | YGR233C | PHO81 | n |
| W303.asm02.HP0_G0022030 | 28229.639 | 27059.8416 | 26383.2765 | 2506.79761 | 2442.2095 | 2081.18582 | 2304.1299 | 2268.5918 | 2671.4053 | 0 | 3.537827736 | YGR234W | YHB1 | n |
| W303.asm02.HP0_G0022040 | 2574.8549 | 2477.82115 | 2778.000411 | 1392.57274 | 1418.4051 | 1348.26708 | 1246.1227 | 1470.8452 | 1318.0696 | 2.11E-31 | 0.912900482 | YGR235C | MIC26 | n |
| W303.asm02.HP0_G0022060 | 595.25083 | 561.584099 | 566.9979977 | 1935.93445 | 1771.658 | 1695.28941 | 1699.8801 | 1487.9023 | 1771.2387 | 1.99E-75 | -1.648717259 | YGR237C | | n |
| W303.asm02.HP0_G0022080 | 462.50463 | 459.855371 | 471.3714188 | 535.027947 | 405.387 | 377.363078 | 344.30837 | 360.82287 | 395.86171 | 0.558758475 | 0.078935868 | YGR239C | PEX21 | n |
| W303.asm02.HP0_G0022090 | 7240.9894 | 7736.57354 | 8363.944909 | 5028.596 | 4801.7236 | 4726.51996 | 4933.1865 | 4077.9545 | 4821.7543 | 4.48E-16 | 0.681209155 | YGR240C | PFK1 | n |
| W303.asm02.HP0_G0022130 | 4039.2773 | 3973.64867 | 3816.369828 | 6375.33301 | 6279.4536 | 7216.35776 | 7303.4419 | 5762.6693 | 7320.3558 | 4.55E-15 | -0.74828317 | YGR244C | LSC2 | n |
| W303.asm02.HP0_G0022140 | 1636.1496 | 1487.52313 | 1699.062143 | 2152.61244 | 2097.0463 | 2277.45255 | 1224.4609 | 1480.0298 | 1401.8265 | 3.49E-07 | -0.436242311 | YGR245C | SDA1 | n |
| W303.asm02.HP0_G0022150 | 751.17495 | 700.682563 | 610.4646244 | 720.03761 | 698.41618 | 690.252063 | 786.66483 | 709.83699 | 790.84177 | 0.738425245 | -0.03341281 | YGR246C | BRF1 | n |
| W303.asm02.HP0_G0022160 | 454.0763 | 384.077849 | 447.2232929 | 763.373207 | 696.61846 | 728.178 | 981.62089 | 909.27364 | 971.58041 | 9.40E-17 | -0.767527547 | YGR247W | CPD1 | n |
| W303.asm02.HP0_G0022180 | 72.69435 | 68.5111839 | 68.58067774 | 650.867331 | 605.83334 | 657.066867 | 484.53993 | 534.01785 | 555.4407 | 2.72E-123 | -3.190117656 | YGR249W | MGA1 | n |
| W303.asm02.HP0_G0022190 | 1518.153 | 1673.33377 | 1390.932056 | 645.033693 | 614.82197 | 650.429828 | 795.78558 | 602.24617 | 828.7528 | 1.12E-28 | 1.262107796 | YGR250C | RIE1 | n |
| W303.asm02.HP0_G0022200 | 600.51854 | 634.247476 | 584.3846484 | 553.362238 | 594.14813 | 588.80018 | 251.96076 | 358.19871 | 291.82678 | 0.556347304 | 0.067381531 | YGR251W | NOP19 | n |
| W303.asm02.HP0_G0022210 | 1198.93 | 1121.0921 | 1024.846466 | 1502.57849 | 1511.8868 | 1504.71157 | 1408.016 | 1436.7311 | 1399.1816 | 3.30E-09 | -0.434867329 | YGR252W | GCN5 | n |
| W303.asm02.HP0_G0022220 | 5145.4957 | 5290.93189 | 5603.331149 | 3611.85533 | 3414.7787 | 3503.40848 | 3673.3827 | 3616.1012 | 3427.8626 | 1.17E-23 | 0.607178862 | YGR253C | PUP2 | n |
| W303.asm02.HP0_G0022240 | 1682.5055 | 1652.5728 | 1747.358395 | 2072.60826 | 2172.5507 | 2044.20803 | 2128.5554 | 2125.5747 | 2093.0416 | 4.81E-07 | -0.307185225 | YGR255C | COQ6 | n |
| W303.asm02.HP0_G0022260 | 1463.3689 | 1295.48421 | 1319.453603 | 814.209196 | 679.54007 | 667.4965 | 798.06577 | 678.347 | 789.96011 | 1.52E-19 | 0.914853693 | YGR257C | MTM1 | n |
| W303.asm02.HP0_G0022270 | 567.85876 | 645.666006 | 665.5223516 | 310.016193 | 293.02918 | 320.474172 | 242.84001 | 200.74872 | 241.57262 | 3.94E-21 | 1.025641279 | YGR258C | RAD2 | n |
| W303.asm02.HP0_G0022290 | 2263.0067 | 2070.90624 | 2472.768099 | 1550.08097 | 1484.9209 | 1615.64494 | 1911.9376 | 4594.9153 | 1818.8479 | 0.150288945 | 0.549541439 | YGR260W | TNA1 | n |
| W303.asm02.HP0_G0022300 | 1331.6762 | 1324.54956 | 1154.280421 | 1951.76862 | 1864.2409 | 1847.94131 | 1908.5173 | 1797.5539 | 1915.8296 | 3.03E-14 | -0.572710411 | YGR261C | APL6 | n |
| W303.asm02.HP0_G0022310 | 1552.9199 | 1504.1319 | 1637.24294 | 940.049102 | 948.29996 | 952.88918 | 973.64023 | 968.31738 | 1011.2547 | 4.54E-26 | 0.724838612 | YGR262C | BUD32 | n |
| W303.asm02.HP0_G0022320 | 549.94856 | 529.404603 | 544.7817218 | 370.019328 | 414.37562 | 360.296406 | 420.69467 | 366.0712 | 428.48283 | 5.62E-07 | 0.504626106 | YGR263C | SAY1 | n |
| W303.asm02.HP0_G0022330 | 4505.9961 | 4082.64373 | 3940.974158 | 5341.11232 | 5169.3584 | 5426.25351 | 4575.197 | 4724.8115 | 4479.6734 | 1.23E-07 | -0.347191387 | YGR264C | MES1 | n |
| W303.asm02.HP0_G0022340 | 744.8537 | 732.862058 | 775.637806 | 3288.5051 | 2941.977 | 3173.45282 | 2957.4037 | 2729.133 | 3065.5037 | 3.15E-161 | -2.060959747 | YGR266W | | n |
| W303.asm02.HP0_G0022350 | 3235.4253 | 3176.42762 | 3098.687524 | 2340.95561 | 2140.1917 | 2197.80808 | 2330.352 | 2319.763 | 2399.8565 | 3.71E-17 | 0.50945744 | YGR267C | FOL2 | n |
| W303.asm02.HP0_G0022360 | 192.79806 | 221.104275 | 176.7642821 | 465.857667 | 478.19486 | 442.78532 | 310.10555 | 299.15496 | 362.35894 | 4.30E-24 | -1.23312672 | YGR268C | HUA1 | n |
| W303.asm02.HP0_G0022370 | 825.97638 | 1015.21118 | 972.6865139 | 4237.72135 | 4255.2152 | 4657.30513 | 4265.0915 | 2506.0789 | 4202.8347 | 8.61E-28 | -2.224361704 | YGR270W | YTA7 | n |
| W303.asm02.HP0_G0022380 | 952.40134 | 969.537058 | 1093.427144 | 1888.43197 | 1905.5885 | 1898.19317 | 1747.764 | 1603.3656 | 1827.6644 | 2.61E-31 | -0.915900053 | YGR271W | SLH1 | n |
| W303.asm02.HP0_G0022390 | 1667.7559 | 1543.57774 | 2128.898785 | 1314.23531 | 1167.6224 | 1417.48191 | 701.15778 | 1026.049 | 751.16743 | 0.001743108 | 0.45404921 | YGR271C- | EFG1 | n |
| W303.asm02.HP0_G0022410 | 923.95572 | 967.460961 | 860.6392094 | 1654.25307 | 1681.7718 | 1838.45982 | 1843.5319 | 1595.4931 | 1922.8828 | 2.00E-24 | -0.911002322 | YGR274C | TAF1 | n |
| W303.asm02.HP0_G0022420 | 1443.3516 | 1443.9251 | 1674.914017 | 863.378431 | 799.98765 | 806.874321 | 757.02238 | 800.37073 | 807.59315 | 7.48E-26 | 0.885140638 | YGR275W | RTT102 | n |
| W303.asm02.HP0_G0022430 | 1097.79 | 1025.59166 | 973.6524389 | 2887.65083 | 2952.7634 | 3104.23799 | 2418.1393 | 2495.5822 | 2563.8438 | 2.71E-108 | -1.530556124 | YGR276C | RNH70 | n |
| W303.asm02.HP0_G0022440 | 1213.6796 | 1176.10866 | 1194.849273 | 692.536174 | 682.23665 | 710.16318 | 631.61205 | 763.6324 | 700.91327 | 1.12E-21 | 0.781983124 | YGR277C | CAB4 | n |
| W303.asm02.HP0_G0022450 | 534.14544 | 520.062169 | 445.2914428 | 714.203972 | 766.72973 | 707.318735 | 730.80022 | 662.602 | 640.07929 | 2.44E-08 | -0.546520308 | YGR278W | CWC22 | n |
| W303.asm02.HP0_G0022460 | 4278.4312 | 4633.84735 | 5017.980576 | 10863.0674 | 10175.124 | 11394.8479 | 11848.996 | 11014.938 | 10798.473 | 2.89E-58 | -1.219040429 | YGR279C | SCW4 | n |
| W303.asm02.HP0_G0022470 | 938.7053 | 991.336071 | 952.4020881 | 999.21886 | 968.9738 | 1121.6596 | 609.95026 | 802.9949 | 664.76554 | 0.343498837 | -0.099855446 | YGR280C | PXR1 | n |
| W303.asm02.HP0_G0022480 | 1186.2875 | 1243.58179 | 1268.259576 | 1286.73388 | 1214.3633 | 1278.10409 | 1544.8273 | 1731.9498 | 1640.7542 | 0.684679697 | -0.031082979 | YGR281W | YOR1 | n |
| W303.asm02.HP0_G0022490 | 15926.384 | 15149.276 | 15361.10589 | 9522.99742 | 9061.4331 | 10469.4551 | 10832.033 | 10887.666 | 10234.215 | 1.26E-24 | 0.676607622 | YGR282C | BGL2 | n |
| W303.asm02.HP0_G0022510 | 5448.9156 | 5736.25458 | 5565.660073 | 3178.49936 | 3142.4234 | 3209.48246 | 4455.4872 | 4476.8278 | 4289.2366 | 1.69E-54 | 0.813664862 | YGR284C | ERV29 | n |
| W303.asm02.HP0_G0022520 | 2721.2972 | 2974.00821 | 3311.191032 | 5142.76863 | 5116.3255 | 5493.57205 | 3705.3053 | 4371.8611 | 3842.239 | 3.02E-20 | -0.806236986 | YGR285C | ZUO1 | n |
| W303.asm02.HP0_G0022530 | 2065.9945 | 1886.13365 | 2101.852884 | 1170.89449 | 1095.7134 | 1094.1633 | 1486.6825 | 1649.2885 | 1466.1871 | 3.74E-28 | 0.848840952 | YGR286C | BIO2 | n |
| W303.asm02.HP0_G0022660 | 378.22133 | 466.08366 | 325.516738 | 137.507183 | 103.36919 | 194.37043 | 3512.6295 | 1993.0543 | 4190.4915 | 4.16E-05 | 1.426985395 | YHL047C | ARN2 | n |
| W303.asm02.HP0_G0022710 | 414.04173 | 599.991884 | 424.041092 | 3867.70203 | 3556.799 | 4412.68283 | 6669.5496 | 6860.8829 | 7252.4686 | 2.62E-136 | -3.041567993 | YHL040C | ARN1 | n |
| W303.asm02.HP0_G0022720 | 1706.7369 | 1406.55537 | 1535.820811 | 1972.60304 | 2045.8111 | 2113.42287 | 1746.6239 | 1666.3456 | 1590.5 | 1.15E-06 | -0.399303265 | YHL039W | EFM1 | n |
| W303.asm02.HP0_G0022750 | 228.61846 | 279.234977 | 170.0028068 | 225.011753 | 215.72701 | 182.992649 | 270.20227 | 250.60789 | 293.59009 | 0.503337706 | 0.116763384 | YHL036W | MUP3 | n |
| W303.asm02.HP0_G0022770 | 3628.3962 | 3873.99604 | 3434.829437 | 2400.95874 | 2482.6583 | 2609.3045 | 3083.9541 | 2969.2442 | 3003.7881 | 2.36E-16 | 0.545758683 | YHL034C | SBP1 | n |
| W303.asm02.HP0_G0022790 | 411.93465 | 528.366555 | 496.4854699 | 276.681119 | 278.64738 | 239.881555 | 239.41973 | 271.60122 | 275.0754 | 6.35E-12 | 0.852722939 | YHL032C | GUT1 | n |
| W303.asm02.HP0_G0022800 | 991.38236 | 1057.77116 | 880.9236352 | 908.380782 | 933.91816 | 850.489149 | 754.7422 | 846.29364 | 836.68766 | 0.19412365 | 0.120904613 | YHL031C | GOS1 | n |
| W303.asm02.HP0_G0022810 | 1137.8246 | 1307.94078 | 1416.046107 | 2527.63203 | 2498.8378 | 2488.88965 | 2736.2255 | 2426.0418 | 2586.7667 | 7.56E-30 | -0.959962966 | YHL030W | ECM29 | n |
| W303.asm02.HP0_G0022820 | 868.11803 | 894.797584 | 940.8109876 | 1238.39802 | 1270.0927 | 1114.07441 | 1372.6731 | 1226.7978 | 1317.188 | 2.79E-07 | -0.422135384 | YHL029C | OCA5 | n |
| W303.asm02.HP0_G0022830 | 284.45615 | 330.099341 | 294.6071368 | 599.197965 | 570.7777 | 585.955735 | 681.77618 | 560.25951 | 691.2151 | 2.13E-18 | -0.9499835 | YHL028W | WSC4 | n |
| W303.asm02.HP0_G0022840 | 805.95909 | 841.857124 | 909.9013864 | 2269.2852 | 2244.4597 | 2306.84515 | 2431.8204 | 2462.7801 | 2487.14 | 9.07E-100 | -1.414070284 | YHL027W | RIM101 | n |
| W303.asm02.HP0_G0022860 | 1050.3807 | 1139.77697 | 1209.338148 | 1180.89502 | 1055.2646 | 1021.15587 | 1008.9831 | 1158.5694 | 1053.574 | 0.514861014 | 0.061402393 | YHL025W | SNF6 | n |
| W303.asm02.HP0_G0022870 | 433.00547 | 407.952959 | 360.2900394 | 355.018544 | 330.78141 | 347.022328 | 388.77204 | 346.38996 | 363.24059 | 0.047068616 | 0.216374926 | YHL024W | RIM4 | n |
| W303.asm02.HP0_G0022880 | 498.32503 | 602.06798 | 594.0438988 | 1039.22095 | 1024.7033 | 1080.88922 | 1112.7317 | 936.82738 | 1087.0768 | 4.68E-20 | -0.891352817 | YHL023C | NPR3 | n |
| W303.asm02.HP0_G0022900 | 920.7951 | 1016.24923 | 751.48968 | 5.83363805 | 8.0897627 | 8.53333594 | 12.541033 | 13.120832 | 7.0532153 | 8.80E-104 | 6.913707708 | YHL021C | AIM17 | n |
| W303.asm02.HP0_G0022910 | 583.66188 | 697.568418 | 596.9416739 | 585.863935 | 551.00273 | 556.563133 | 603.1097 | 561.5716 | 569.54714 | 0.123806976 | 0.148660425 | YHL020C | OPI1 | n |

| | | | | | | | | | | | | | | |
|---|---|---|---|---|---|---|---|---|---|---|---|---|---|---|
| W303.asm02.HP0_G0022920 | 308.6876 | 333.213486 | 358.3581893 | 551.695484 | 527.6323 | 544.237203 | 593.98895 | 564.19576 | 585.41687 | 1.62E-13 | -0.697597677 | YHL019C | APM2 | n |
| W303.asm02.HP0_G0022930 | 102.19351 | 66.4350875 | 56.98957728 | 198.343694 | 228.31108 | 221.866734 | 225.7386 | 233.5508 | 223.05793 | 1.09E-17 | -1.527897404 | YHL018W | MCO14 | n |
| W303.asm02.HP0_G0022950 | 147.49578 | 138.060416 | 167.1050317 | 140.84069 | 152.80663 | 189.629688 | 189.2556 | 190.25206 | 201.01664 | 0.579727225 | -0.089789448 | YHL016C | DUR3 | n |
| W303.asm02.HP0_G0022960 | 19179.719 | 18655.803 | 21411.66033 | 11921.456 | 12514.863 | 13615.4116 | 10420.459 | 10928.341 | 9429.2672 | 3.98E-15 | 0.638879808 | YHL015W | RPS20 | n |
| W303.asm02.HP0_G0022970 | 820.70867 | 699.644515 | 770.8081808 | 544.195092 | 583.36178 | 561.303875 | 522.16303 | 665.22617 | 601.2866 | 2.02E-05 | 0.440221663 | YHL014C | YLF2 | n |
| W303.asm02.HP0_G0022980 | 1607.704 | 1574.71918 | 1706.789543 | 790.041267 | 781.11153 | 877.037305 | 606.52998 | 733.45449 | 694.74171 | 1.25E-30 | 0.998703392 | YHL013C | OTU2 | n |
| W303.asm02.HP0_G0023000 | 3117.4287 | 2877.46973 | 2895.843266 | 4119.38184 | 4159.0369 | 4435.43839 | 3411.1611 | 3890.3266 | 3470.1819 | 6.65E-13 | -0.516002428 | YHL011C | PRS3 | n |
| W303.asm02.HP0_G0023010 | 257.06408 | 239.789144 | 258.8679104 | 434.189346 | 396.39837 | 375.466781 | 401.31307 | 408.05787 | 421.42961 | 3.09E-10 | -0.675423414 | YHL010C | ETP1 | n |
| W303.asm02.HP0_G0023020 | 368.73945 | 379.925656 | 370.9152148 | 1079.22304 | 1066.9498 | 1003.14105 | 1055.727 | 910.58572 | 1182.2952 | 9.77E-48 | -1.492399753 | YHL009C | YAP3 | n |
| W303.asm02.HP0_G0023030 | 159.08474 | 159.859429 | 146.8206059 | 544.195092 | 543.81183 | 597.333516 | 518.74275 | 476.28619 | 540.45262 | 3.16E-61 | -1.855867448 | YHL008C | | n |
| W303.asm02.HP0_G0023040 | 1151.5206 | 1297.5603 | 1307.862502 | 2634.30427 | 2544.6798 | 2596.03042 | 2576.6123 | 2432.6022 | 2689.92 | 1.24E-47 | -1.048906838 | YHL007C | STE20 | n |
| W303.asm02.HP0_G0023050 | 337.13322 | 282.349122 | 341.9374637 | 120.006268 | 89.886252 | 84.385211 | 94.627798 | 106.27874 | 100.50832 | 2.18E-26 | 1.703384672 | YHL006C | SHU1 | n |
| W303.asm02.HP0_G0023060 | 1360.1218 | 1551.88212 | 1568.662263 | 2790.97912 | 2696.5876 | 2773.33418 | 2615.3755 | 2706.8276 | 2566.4887 | 3.26E-36 | -0.882175164 | YHL004W | MRP4 | n |
| W303.asm02.HP0_G0023070 | 3081.6083 | 2775.741 | 2940.275818 | 1331.73623 | 1191.8917 | 1281.89669 | 1702.1603 | 1637.4798 | 1610.778 | 1.03E-64 | 1.2087111 | YHL003C | LAG1 | n |
| W303.asm02.HP0_G0023080 | 711.14038 | 768.155699 | 601.7712991 | 1277.56673 | 1071.4441 | 1122.60775 | 1094.4902 | 944.69988 | 1086.1952 | 8.53E-12 | -0.739527083 | YHL002W | HSE1 | n |
| W303.asm02.HP0_G0023090 | 4102.4898 | 4167.76369 | 4288.707171 | 2205.11518 | 2033.227 | 2332.44516 | 2097.7729 | 2234.4776 | 1845.2975 | 2.34E-30 | 0.934755132 | YHL001W | RPL14B | y |
| W303.asm02.HP0_G0023100 | 1503.4034 | 1307.94078 | 1276.952901 | 2166.77985 | 2183.3371 | 2121.00806 | 2216.3426 | 2218.7326 | 2213.828 | 1.12E-19 | -0.663136354 | YHR001W | OSH7 | n |
| W303.asm02.HP0_G0023120 | 567.85876 | 563.660195 | 540.9180216 | 539.194831 | 485.38576 | 456.059399 | 619.07102 | 552.38701 | 589.82513 | 0.073339149 | 0.173903731 | YHR002W | LEU5 | n |
| W303.asm02.HP0_G0023140 | 692.17663 | 669.541116 | 662.6245765 | 515.026902 | 521.34026 | 414.340867 | 673.79553 | 682.28325 | 677.99032 | 1.22E-06 | 0.478913505 | YHR004C | NEM1 | n |
| W303.asm02.HP0_G0023150 | 2895.1315 | 2884.73606 | 2539.416927 | 36.668582 | 51.235164 | 40.7703828 | 52.444322 | 40.674578 | 47.609203 | 0 | 6.017745321 | YHR005C | GPA1 | n |
| W303.asm02.HP0_G0023160 | 5167.6201 | 4971.21303 | 6012.883365 | 1873.43119 | 2108.7315 | 2018.60802 | 1808.189 | 1987.806 | 1767.7121 | 1.88E-60 | 1.428822313 | YHR005C- | TIM10 | n |
| W303.asm02.HP0_G0023180 | 6553.0269 | 5977.08177 | 5195.710783 | 3517.68374 | 3556.799 | 3602.01592 | 3672.2426 | 3810.2895 | 3529.2526 | 9.26E-19 | 0.731274834 | YHR007C | ERG11 | n |
| W303.asm02.HP0_G0023210 | 3240.693 | 3010.3399 | 2875.55884 | 3146.83104 | 3148.7154 | 3306.1936 | 3387.2191 | 3635.7825 | 3280.6268 | 0.30546632 | -0.073328165 | YHR008C | SOD2 | n |
| W303.asm02.HP0_G0023220 | 1205.2512 | 1253.96228 | 1216.099624 | 2584.30165 | 2320.863 | 2764.80084 | 2785.2495 | 2340.7564 | 2769.2687 | 3.47E-32 | -1.061258927 | YHR009C | TDA3 | n |
| W303.asm02.HP0_G0023240 | 335.02613 | 399.648573 | 354.4944892 | 813.375819 | 863.80688 | 802.133578 | 758.16248 | 902.71322 | 797.89498 | 2.10E-32 | -1.186840951 | YHR011W | DIA4 | n |
| W303.asm02.HP0_G0023260 | 1927.9806 | 1868.48683 | 1804.347972 | 1073.3894 | 1143.3531 | 1091.31885 | 929.17657 | 1064.0995 | 1050.9291 | 2.03E-24 | 0.759587271 | YHR013C | ARD1 | n |
| W303.asm02.HP0_G0023300 | 1125.1821 | 1163.65208 | 1115.64342 | 1099.22408 | 1046.276 | 1095.11145 | 1195.9586 | 1106.0861 | 1142.6209 | 0.324384342 | 0.070966423 | YHR017W | YSC83 | n |
| W303.asm02.HP0_G0023310 | 4302.6627 | 4359.80261 | 3812.506127 | 2221.78272 | 2149.1803 | 2307.7933 | 2492.2454 | 1798.866 | 2571.7786 | 3.56E-15 | 0.901347511 | YHR018C | ARG4 | n |
| W303.asm02.HP0_G0023320 | 3100.572 | 3807.56095 | 3847.279429 | 6715.35077 | 6484.3942 | 7098.78735 | 5497.533 | 5620.9643 | 5296.083 | 2.79E-27 | -0.916079603 | YHR019C | DED81 | n |
| W303.asm02.HP0_G0023330 | 2918.3094 | 2871.24144 | 3053.289047 | 5058.59756 | 4914.9803 | 5193.00899 | 4167.0434 | 4119.9412 | 3948.0373 | 1.36E-42 | -0.778070931 | YHR020W | | n |
| W303.asm02.HP0_G0023350 | 180.15556 | 124.565789 | 180.6279822 | 606.698357 | 604.93448 | 551.822391 | 359.12959 | 404.12162 | 335.02773 | 3.55E-43 | -1.860390256 | YHR022C | | n |
| W303.asm02.HP0_G0023360 | 1165.2167 | 1197.90767 | 1312.692127 | 1784.25987 | 1705.1422 | 1855.52649 | 1930.1791 | 1817.2352 | 1868.2204 | 1.13E-13 | -0.539361155 | YHR023W | MYO1 | n |
| W303.asm02.HP0_G0023370 | 1491.8145 | 1649.45866 | 1761.84727 | 1543.41395 | 1395.9335 | 1464.88934 | 1313.3882 | 1251.7273 | 1203.4549 | 0.061861114 | 0.155002006 | YHR024C | MAS2 | n |
| W303.asm02.HP0_G0023380 | 1458.1012 | 1752.22543 | 1763.779121 | 2197.61479 | 2103.3383 | 2178.84511 | 2004.2852 | 1655.849 | 2011.048 | 8.13E-05 | -0.381252738 | YHR025W | THR1 | n |
| W303.asm02.HP0_G0023390 | 3704.2512 | 3376.77093 | 3981.543009 | 1995.10421 | 1939.7453 | 1905.77836 | 2550.3902 | 2441.7868 | 2482.7318 | 1.18E-37 | 0.921523078 | YHR026W | VMA16 | n |
| W303.asm02.HP0_G0023400 | 3402.9384 | 3585.41863 | 3859.836454 | 6642.01361 | 6602.1452 | 6660.74278 | 6642.1874 | 5935.8643 | 6283.5332 | 6.43E-39 | -0.875430882 | YHR027C | RPN1 | n |
| W303.asm02.HP0_G0023410 | 1980.6576 | 1914.16096 | 1989.80558 | 574.196659 | 577.9686 | 573.629805 | 1008.9831 | 814.80365 | 966.2905 | 1.78E-96 | 1.769734466 | YHR028C | DAP2 | n |
| W303.asm02.HP0_G0023430 | 1176.8056 | 1313.13103 | 1238.3159 | 2558.46697 | 2526.7026 | 2570.43041 | 2146.7969 | 2078.3397 | 2382.2235 | 1.27E-44 | -1.03795434 | YHR030C | SLT2 | n |
| W303.asm02.HP0_G0023440 | 385.59612 | 401.724669 | 364.1537396 | 1190.89554 | 1115.4884 | 1039.17069 | 882.43272 | 883.03197 | 1003.3199 | 1.20E-59 | -1.540038304 | YHR031C | RRM3 | n |
| W303.asm02.HP0_G0023450 | 1040.8988 | 1003.79265 | 1185.190022 | 1611.75085 | 1640.4241 | 1704.77089 | 1995.1644 | 2134.7593 | 1988.1251 | 4.73E-15 | -0.616986532 | YHR032W | ERC1 | n |
| W303.asm02.HP0_G0023470 | 247.58221 | 268.854495 | 240.5153346 | 771.706976 | 792.79674 | 749.985414 | 685.19646 | 772.81699 | 701.79492 | 3.81E-63 | -1.613334207 | YHR034C | PIH1 | n |
| W303.asm02.HP0_G0023480 | 290.7774 | 279.234977 | 323.5848879 | 542.528338 | 579.76633 | 611.555742 | 554.08566 | 621.92742 | 502.54159 | 7.53E-18 | -0.954427399 | YHR035W | NEL1 | n |
| W303.asm02.HP0_G0023490 | 539.41315 | 538.747037 | 425.972942 | 826.709849 | 735.26954 | 836.266922 | 819.72755 | 778.06532 | 817.29132 | 9.09E-11 | -0.674486803 | YHR036W | BRL1 | n |
| W303.asm02.HP0_G0023500 | 1842.6437 | 2117.61841 | 2128.898785 | 1545.08071 | 1457.0561 | 1336.8893 | 2698.6024 | 3303.8254 | 2856.5522 | 3.72E-07 | 0.488637878 | YHR037W | PUT2 | n |
| W303.asm02.HP0_G0023510 | 736.42537 | 770.231795 | 704.1593532 | 950.883002 | 1026.501 | 972.800297 | 860.77093 | 957.82071 | 845.50418 | 4.84E-07 | -0.416465339 | YHR038W | RRF1 | n |
| W303.asm02.HP0_G0023520 | 2560.1054 | 2628.33815 | 2646.634606 | 2385.12458 | 2310.9755 | 2468.97853 | 2695.1821 | 2628.1026 | 2585.0034 | 0.027801389 | 0.129213103 | YHR039C | MSC7 | n |
| W303.asm02.HP0_G0023530 | 2989.9502 | 3067.43255 | 3424.204262 | 1426.74119 | 1468.7414 | 1474.37082 | 1478.7019 | 1616.4865 | 1547.2991 | 2.63E-53 | 1.117997253 | YHR039C- | VMA10 | n |
| W303.asm02.HP0_G0023540 | 278.1349 | 308.300328 | 340.0056136 | 675.03526 | 667.85485 | 660.859461 | 411.57392 | 577.31659 | 449.64248 | 6.15E-19 | -1.111873143 | YHR040W | BCD1 | n |
| W303.asm02.HP0_G0023560 | 1146.2529 | 1308.97883 | 1427.637207 | 3451.01359 | 3407.5878 | 3620.97888 | 3663.1219 | 3791.9204 | 3781.4051 | 7.82E-73 | -1.431579153 | YHR042W | NCP1 | n |
| W303.asm02.HP0_G0023570 | 2053.352 | 2166.40668 | 2075.772908 | 130.840168 | 107.8635 | 109.985219 | 399.03288 | 224.36622 | 372.05711 | 2.05E-151 | 4.172215594 | YHR043C | DOG2 | n |
| W303.asm02.HP0_G0023590 | 1686.7196 | 1520.74067 | 1483.660859 | 809.208935 | 793.69561 | 841.007664 | 743.34126 | 799.05865 | 797.89498 | 4.75E-33 | 0.940634391 | YHR045W | DDE1 | n |
| W303.asm02.HP0_G0023640 | 3776.9456 | 3498.22257 | 3339.202859 | 465.02429 | 533.92434 | 531.911274 | 368.25035 | 494.65535 | 388.80849 | 3.77E-164 | 2.794414341 | YHR049W | FSH1 | n |
| W303.asm02.HP0_G0023650 | 1348.5329 | 1247.73399 | 1277.918826 | 1538.41369 | 1622.4469 | 1627.97087 | 2241.4247 | 1876.2789 | 2033.971 | 0.000146815 | -0.305665538 | YHR050W | SMF2 | n |
| W303.asm02.HP0_G0023670 | 4174.1306 | 4280.91095 | 5067.242753 | 6521.17396 | 6976.9709 | 7202.13553 | 5846.4018 | 6067.0726 | 5550.8804 | 1.21E-13 | -0.614006744 | YHR051W | COX6 | n |
| W303.asm02.HP0_G0023680 | 5078.0691 | 4580.90689 | 5692.196253 | 3435.17943 | 3323.0947 | 3629.51222 | 1999.7248 | 2872.1501 | 2154.7573 | 2.95E-06 | 0.563576356 | YHR052W | CIC1 | n |
| W303.asm02.HP0_G0023860 | 431.95193 | 428.713924 | 343.8693137 | 775.87386 | 760.43769 | 683.615024 | 689.75684 | 810.8674 | 825.22619 | 4.79E-15 | -0.884015624 | YHR058C | MED6 | n |
| W303.asm02.HP0_G0023870 | 1151.5206 | 1239.4296 | 1423.773507 | 875.045707 | 1075.9384 | 1051.49662 | 817.44736 | 883.03197 | 902.81156 | 0.000982949 | 0.346597776 | YHR059W | FYV4 | n |
| W303.asm02.HP0_G0023880 | 980.84695 | 974.727299 | 1201.610748 | 594.197704 | 684.93324 | 578.370547 | 637.31252 | 617.99117 | 603.93156 | 2.37E-13 | 0.76591444 | YHR060W | VMA22 | n |
| W303.asm02.HP0_G0023900 | 833.35117 | 835.628834 | 657.7949513 | 827.543226 | 827.85238 | 795.496539 | 814.02708 | 995.87113 | 896.63999 | 0.49911809 | -0.076041885 | YHR062C | RPP1 | n |
| W303.asm02.HP0_G0023910 | 2869.8465 | 2771.5888 | 3203.007428 | 873.378953 | 879.08755 | 923.496578 | 898.39403 | 758.38407 | 886.06017 | 1.41E-90 | 1.725237815 | YHR063C | PAN5 | n |

| | | | | | | | | | | | | | |
|---|---|---|---|---|---|---|---|---|---|---|---|---|---|
| W303.asm02.HP0_G0023920 | 7026.0669 | 6573.95951 | 6429.197057 | 8657.95224 | 8735.146 | 9200.83244 | 6682.0906 | 7296.4945 | 6666.1701 | 1.35E-10 | -0.409012538 | YHR064C | SSZ1 | n |
| W303.asm02.HP0_G0023930 | 2165.0274 | 2031.46041 | 1898.042701 | 1735.92401 | 1630.5366 | 1884.91909 | 1007.8431 | 1277.969 | 1001.5566 | 0.036899177 | 0.214793579 | YHR065C | RRP3 | n |
| W303.asm02.HP0_G0023950 | 630.0177 | 696.53037 | 622.0557249 | 272.514234 | 266.96217 | 266.429711 | 259.94142 | 217.80581 | 237.16436 | 4.01E-35 | 1.273175716 | YHR067W | HTD2 | n |
| W303.asm02.HP0_G0023960 | 4755.6854 | 4313.09044 | 4612.292059 | 3068.49361 | 2844.8999 | 3036.91945 | 2474.0039 | 2712.0759 | 2486.2584 | 1.50E-19 | 0.612127048 | YHR068W | DYS1 | n |
| W303.asm02.HP0_G0023980 | 689.01601 | 862.618089 | 879.9577102 | 1446.74224 | 1350.0915 | 1485.7486 | 1081.9492 | 1087.7169 | 1071.2071 | 5.21E-18 | -0.815752902 | YHR070W | TRM5 | n |
| W303.asm02.HP0_G0023990 | 2280.9169 | 2235.95591 | 2288.276416 | 860.8783 | 835.04328 | 918.755836 | 709.13844 | 450.04453 | 707.96649 | 4.16E-27 | 1.380164559 | YHR071W | PCL5 | n |
| W303.asm02.HP0_G0024000 | 2062.8339 | 2176.78716 | 2310.492692 | 1455.90938 | 1444.4721 | 1354.90412 | 1523.1655 | 1358.0061 | 1477.6486 | 6.62E-17 | 0.622221698 | YHR072W | ERG7 | n |
| W303.asm02.HP0_G0024010 | 3199.6049 | 2925.21994 | 3563.297467 | 1615.08436 | 1784.2421 | 1568.23752 | 1229.0213 | 1628.2952 | 1316.3063 | 4.40E-19 | 0.96378392 | YHR072W- | NOP10 | n |
| W303.asm02.HP0_G0024020 | 1267.4102 | 1381.64221 | 1546.445987 | 3678.52548 | 3599.0455 | 3804.91968 | 3805.6336 | 3456.0271 | 3888.0849 | 1.60E-65 | -1.400694462 | YHR073W | OSH3 | n |
| W303.asm02.HP0_G0024030 | 1145.1994 | 1316.24517 | 1473.035684 | 2024.2724 | 1963.1157 | 1968.35616 | 1867.4739 | 1770.0002 | 1738.6176 | 1.39E-11 | -0.597483154 | YHR074W | QNS1 | n |
| W303.asm02.HP0_G0024040 | 280.24199 | 290.653508 | 291.7093617 | 610.031864 | 591.45154 | 549.926094 | 718.25919 | 644.23284 | 661.23893 | 5.40E-26 | -1.02227293 | YHR075C | PPE1 | n |
| W303.asm02.HP0_G0024050 | 946.08009 | 915.558549 | 944.6746878 | 1010.05276 | 1039.9839 | 1076.14848 | 892.69356 | 905.33739 | 900.1666 | 0.02916036 | -0.155147181 | YHR076W | PTC7 | n |
| W303.asm02.HP0_G0024060 | 1650.8992 | 1680.6001 | 1570.594113 | 2044.27345 | 2085.3611 | 2373.21554 | 1858.3531 | 1894.6481 | 1894.67 | 4.92E-08 | -0.407402811 | YHR077C | NMD2 | n |
| W303.asm02.HP0_G0024070 | 836.51179 | 735.976203 | 737.0008045 | 581.697051 | 608.52993 | 666.548352 | 855.07046 | 868.59906 | 879.88861 | 0.000422609 | 0.31534209 | YHR078W | | n |
| W303.asm02.HP0_G0024080 | 404.55986 | 431.828068 | 390.2337156 | 1297.56778 | 1170.319 | 1260.08927 | 1258.6637 | 1229.4219 | 1287.2118 | 1.68E-84 | -1.604575464 | YHR079C | IRE1 | n |
| W303.asm02.HP0_G0024090 | 10.535413 | 14.5326754 | 5.795550232 | 14.1674067 | 27.864738 | 20.8592656 | 13.681127 | 30.177913 | 25.567905 | 0.034462698 | -1.031281981 | YHR079C- | SAE3 | n |
| W303.asm02.HP0_G0024110 | 1699.3621 | 1702.39912 | 2218.729814 | 682.535652 | 734.37068 | 756.622453 | 468.57861 | 581.25284 | 521.05628 | 1.19E-32 | 1.371480814 | YHR081W | LRP1 | n |
| W303.asm02.HP0_G0024120 | 1286.3739 | 1436.65877 | 1298.203252 | 2107.61009 | 2338.8403 | 2397.8674 | 2348.5935 | 2197.7393 | 2162.6921 | 1.15E-22 | -0.767018372 | YHR082C | KSP1 | n |
| W303.asm02.HP0_G0024130 | 540.46669 | 623.866993 | 554.4409722 | 510.860018 | 503.36301 | 480.711258 | 499.36115 | 545.8266 | 566.90218 | 0.038554423 | 0.200594824 | YHR083W | SAM35 | n |
| W303.asm02.HP0_G0024140 | 755.38911 | 843.93322 | 886.7191855 | 2430.96031 | 2400.8618 | 2655.76377 | 2106.8936 | 2111.1418 | 2184.7334 | 1.10E-92 | -1.589622992 | YHR084W | STE12 | n |
| W303.asm02.HP0_G0024150 | 1762.5746 | 1562.2626 | 1736.733219 | 681.702275 | 668.75372 | 777.481719 | 373.95082 | 469.72577 | 414.3764 | 1.77E-38 | 1.250904481 | YHR085W | IPI1 | n |
| W303.asm02.HP0_G0024160 | 684.80184 | 671.617212 | 672.2838269 | 1178.39489 | 1084.9271 | 1054.34106 | 1115.0119 | 1035.2336 | 1128.5144 | 2.90E-19 | -0.71040516 | YHR086W | NAM8 | n |
| W303.asm02.HP0_G0024180 | 615.26812 | 793.068856 | 733.1371043 | 72.5037872 | 71.010139 | 70.1629844 | 624.77149 | 289.97038 | 428.48283 | 2.43E-30 | 3.324954303 | YHR087W | RTC3 | n |
| W303.asm02.HP0_G0024190 | 2015.4245 | 1844.61173 | 2010.090005 | 1394.23949 | 1402.2255 | 1380.50413 | 852.79028 | 1123.1432 | 916.91799 | 1.60E-07 | 0.490967873 | YHR088W | RPF1 | n |
| W303.asm02.HP0_G0024200 | 5010.6424 | 4458.4172 | 5286.507736 | 4665.24368 | 4298.3606 | 4724.62367 | 3444.2238 | 3696.1383 | 3464.0104 | 0.167836084 | 0.108451137 | YHR089C | GAR1 | n |
| W303.asm02.HP0_G0024210 | 719.56871 | 706.910852 | 555.4068972 | 712.537219 | 678.6412 | 611.555742 | 652.13374 | 720.33366 | 691.2151 | 0.886240294 | -0.01690903 | YHR090C | YNG2 | n |
| W303.asm02.HP0_G0024230 | 872.3322 | 933.205369 | 983.3116893 | 342.517891 | 337.07345 | 351.76307 | 209.77729 | 268.97705 | 240.69097 | 1.19E-47 | 1.435804245 | YHR092C | HXT4 | n |
| W303.asm02.HP0_G0024260 | 148.54932 | 174.392105 | 167.1050317 | 3150.99792 | 2962.6509 | 2750.57862 | 1788.8074 | 2387.9914 | 1981.9535 | 3.22E-257 | -4.1765082 | YHR096C | HXT5 | n |
| W303.asm02.HP0_G0024280 | 2107.0826 | 2395.81534 | 2474.699949 | 3267.67068 | 3315.9038 | 3422.81586 | 3026.9494 | 3021.7275 | 3035.5275 | 1.31E-13 | -0.519654467 | YHR098C | SFB3 | n |
| W303.asm02.HP0_G0024290 | 989.27528 | 1124.20625 | 1218.997399 | 3207.66755 | 3154.1086 | 3098.54909 | 3187.7027 | 3164.7446 | 3314.1295 | 7.11E-85 | -1.504539273 | YHR099W | TRA1 | n |
| W303.asm02.HP0_G0024300 | 594.19729 | 584.42116 | 593.0779737 | 536.6947 | 485.38576 | 533.807571 | 446.91683 | 544.51452 | 515.76637 | 0.055397468 | 0.187291581 | YHR100C | GEP4 | n |
| W303.asm02.HP0_G0024320 | 709.03329 | 811.753725 | 873.1962349 | 1382.57222 | 1457.0561 | 1397.5708 | 1400.0354 | 1209.7407 | 1560.5239 | 2.54E-15 | -0.823010819 | YHR102W | KIC1 | n |
| W303.asm02.HP0_G0024330 | 1122.0215 | 1127.32039 | 1133.03007 | 2565.13399 | 2556.365 | 2899.43792 | 2882.1575 | 2355.1893 | 2884.7651 | 1.57E-42 | -1.2454873 | YHR103W | SBE22 | n |
| W303.asm02.HP0_G0024340 | 2402.0742 | 2463.28848 | 2150.149136 | 1033.38731 | 959.08631 | 1003.14105 | 1553.9481 | 1178.2507 | 1636.3459 | 7.61E-29 | 1.227430029 | YHR104W | GRE3 | n |
| W303.asm02.HP0_G0024370 | 1526.5813 | 1530.08311 | 1580.253363 | 4956.92559 | 4889.8121 | 5105.77934 | 4849.9597 | 4808.7848 | 4787.3699 | 9.47E-215 | -1.688825506 | YHR107C | CDC12 | n |
| W303.asm02.HP0_G0024380 | 1899.535 | 1990.97653 | 1795.654647 | 2676.80649 | 2737.0364 | 2582.75634 | 2829.7132 | 2520.5118 | 2689.0383 | 2.03E-12 | -0.492301799 | YHR108W | GGA2 | n |
| W303.asm02.HP0_G0024390 | 259.17116 | 284.425218 | 275.288636 | 466.691044 | 487.18349 | 504.414969 | 383.07157 | 421.1787 | 392.3351 | 1.85E-16 | -0.831536232 | YHR109W | CTM1 | n |
| W303.asm02.HP0_G0024400 | 1490.7609 | 1570.56699 | 1626.617765 | 259.180205 | 235.50198 | 251.259336 | 308.96546 | 276.84955 | 282.12861 | 5.76E-187 | 2.651458807 | YHR110W | ERP5 | n |
| W303.asm02.HP0_G0024410 | 930.27697 | 878.188812 | 793.0244567 | 894.213375 | 869.20006 | 916.859539 | 733.08041 | 780.68949 | 749.40413 | 0.614745267 | -0.043733392 | YHR111W | UBA4 | n |
| W303.asm02.HP0_G0024420 | 678.4806 | 741.166444 | 632.6809003 | 1419.2408 | 1469.6402 | 1450.66711 | 1304.2675 | 1221.5494 | 1221.9696 | 2.67E-43 | -1.08095096 | YHR112C | | n |
| W303.asm02.HP0_G0024430 | 1201.0371 | 1130.43453 | 1217.065549 | 2500.13059 | 2581.5332 | 2486.0452 | 2440.9411 | 2601.8609 | 2421.0162 | 7.72E-64 | -1.092405825 | YHR113W | APE4 | n |
| W303.asm02.HP0_G0024440 | 603.67916 | 712.101094 | 648.1357009 | 902.547144 | 822.45921 | 854.281742 | 848.2299 | 860.72656 | 883.41522 | 5.01E-06 | -0.393585982 | YHR114W | BZZ1 | n |
| W303.asm02.HP0_G0024450 | 680.58768 | 676.807453 | 795.9222318 | 2019.27214 | 1872.3306 | 1882.07465 | 2082.9516 | 1780.4969 | 2060.4205 | 3.79E-55 | -1.422212765 | YHR115C | DMA1 | n |
| W303.asm02.HP0_G0024460 | 520.4494 | 561.584099 | 529.3269212 | 324.1836 | 312.80416 | 334.696399 | 303.26499 | 305.71538 | 300.6433 | 1.90E-14 | 0.729966365 | YHR116W | COX23 | n |
| W303.asm02.HP0_G0024480 | 693.23018 | 649.818199 | 589.2142736 | 207.510839 | 186.9634 | 192.474133 | 186.97541 | 158.76206 | 146.35422 | 3.52E-47 | 1.717316904 | YHR118C | ORC6 | n |
| W303.asm02.HP0_G0024490 | 596.30438 | 756.737168 | 732.1711769 | 1065.05563 | 1108.2975 | 1170.01517 | 1030.6449 | 1083.7807 | 1078.2603 | 6.16E-13 | -0.680088173 | YHR119W | SET1 | n |
| W303.asm02.HP0_G0024500 | 716.40808 | 658.122585 | 691.6023277 | 978.384438 | 976.1647 | 995.55586 | 806.04642 | 890.90447 | 871.07209 | 3.82E-11 | -0.513764643 | YHR120W | MSH1 | n |
| W303.asm02.HP0_G0024510 | 2531.6597 | 2444.60361 | 3019.481671 | 1230.89763 | 1387.8437 | 1252.50409 | 1135.5336 | 1242.5428 | 1169.0704 | 8.41E-30 | 1.046848916 | YHR121W | LSM12 | n |
| W303.asm02.HP0_G0024520 | 1524.4743 | 1454.30559 | 1592.810389 | 640.866808 | 658.86623 | 607.763149 | 623.63139 | 720.33366 | 692.09675 | 4.53E-54 | 1.260857558 | YHR122W | CIA2 | n |
| W303.asm02.HP0_G0024540 | 257.06408 | 234.598903 | 285.9138114 | 49.169235 | 62.920377 | 45.511125 | 29.642443 | 40.674578 | 25.567905 | 1.59E-33 | 2.30286845 | YHR124W | NDT80 | n |
| W303.asm02.HP0_G0024560 | 839.67242 | 828.362497 | 708.9889783 | 1070.88927 | 1028.2987 | 981.333633 | 1010.1232 | 978.81404 | 986.56849 | 1.11E-05 | -0.375478148 | YHR127W | | n |
| W303.asm02.HP0_G0024570 | 3739.0181 | 3364.31435 | 3170.165977 | 2570.96763 | 2646.2513 | 2733.51195 | 1549.3877 | 2336.8201 | 1687.4818 | 0.003261008 | 0.369738974 | YHR128W | FUR1 | n |
| W303.asm02.HP0_G0024580 | 240.20742 | 276.120832 | 225.060534 | 525.027424 | 504.26188 | 471.229774 | 498.22106 | 502.52785 | 520.17463 | 1.21E-21 | -1.019302659 | YHR129C | ARP1 | n |
| W303.asm02.HP0_G0024590 | 540.46669 | 483.730481 | 533.1906213 | 605.031603 | 624.70945 | 556.563133 | 665.81487 | 653.41742 | 744.99587 | 0.041071006 | -0.198214921 | YHR131C | | n |
| W303.asm02.HP0_G0024600 | 1822.6264 | 1828.00295 | 1695.198443 | 1210.89658 | 1177.5099 | 1113.12627 | 1477.5618 | 1356.694 | 1431.8027 | 1.07E-17 | 0.609595804 | YHR132C | ECM14 | n |
| W303.asm02.HP0_G0024610 | 1173.645 | 1162.61403 | 1279.850676 | 683.369028 | 694.82073 | 644.740938 | 859.63084 | 785.93782 | 916.03634 | 4.13E-23 | 0.837989765 | YHR132W- | IGO2 | n |
| W303.asm02.HP0_G0024630 | 681.64122 | 607.258221 | 672.2838269 | 357.518675 | 330.78141 | 342.281586 | 305.54518 | 247.98372 | 292.70843 | 1.80E-19 | 0.927908096 | YHR134W | WSS1 | n |
| W303.asm02.HP0_G0024650 | 1149.4136 | 1137.70087 | 1243.145525 | 206.677462 | 186.06454 | 94.8148438 | 196.09616 | 116.7754 | 211.59646 | 2.37E-17 | 2.854784014 | YHR136C | SPL2 | n |
| W303.asm02.HP0_G0024660 | 893.40302 | 985.107781 | 984.2776144 | 1341.73675 | 1206.2735 | 931.081766 | 1081.9492 | 1451.164 | 1228.1411 | 0.03621121 | -0.281837506 | YHR137W | ARO9 | n |
| W303.asm02.HP0_G0024670 | 978.73987 | 1012.09704 | 1158.144121 | 251.679813 | 260.67013 | 245.570445 | 307.82537 | 337.20537 | 369.41215 | 1.45E-83 | 2.055123476 | YHR138C | | n |

| | | | | | | | | | | | | | | |
|---|---|---|---|---|---|---|---|---|---|---|---|---|---|---|
| W303.asm02.HP0_G0024690 | 194.90514 | 225.256468 | 239.5494096 | 329.183861 | 338.87117 | 402.963086 | 501.64134 | 472.34994 | 544.86088 | 3.04E-08 | -0.696055907 | YHR140W | | n |
| W303.asm02.HP0_G0024710 | 2076.5299 | 1878.86732 | 1924.122677 | 1180.06164 | 1065.1521 | 1125.4522 | 1298.567 | 1522.0165 | 1368.3238 | 7.16E-23 | 0.802257775 | YHR142W | CHS7 | n |
| W303.asm02.HP0_G0024720 | 4719.865 | 4355.65042 | 3908.132706 | 2205.94856 | 2347.8289 | 2299.25996 | 2336.0525 | 2298.7697 | 2329.3244 | 3.13E-34 | 0.921776784 | YHR143W | DSE2 | n |
| W303.asm02.HP0_G0024730 | 2846.6686 | 2725.91468 | 3289.940682 | 1127.5589 | 1156.8361 | 1199.40777 | 912.07516 | 1090.3411 | 989.21345 | 2.12E-50 | 1.347553532 | YHR143W-A | RPC10 | n |
| W303.asm02.HP0_G0024740 | 1084.094 | 1107.59747 | 1134.96192 | 1030.88718 | 956.38972 | 1014.51883 | 883.57281 | 980.12613 | 863.13722 | 0.058416905 | 0.14830428 | YHR144C | DCD1 | n |
| W303.asm02.HP0_G0024750 | 1255.8212 | 1147.04331 | 1188.087798 | 5610.29305 | 5724.8554 | 6120.29817 | 5172.6063 | 5139.4298 | 5169.1252 | 4.55E-285 | -2.281239367 | YHR146W | CRP1 | n |
| W303.asm02.HP0_G0024760 | 1942.7302 | 1846.68782 | 2297.935667 | 1645.91931 | 1641.323 | 1680.11903 | 1537.9867 | 1640.104 | 1539.3642 | 0.000678848 | 0.293867533 | YHR147C | MRPL6 | n |
| W303.asm02.HP0_G0024770 | 1746.7715 | 1618.31721 | 1885.485675 | 989.218337 | 941.10906 | 980.385485 | 611.09036 | 837.10906 | 676.22702 | 3.84E-16 | 0.851258323 | YHR148W | IMP3 | n |
| W303.asm02.HP0_G0024780 | 833.35117 | 851.199558 | 774.671881 | 1678.421 | 1657.5025 | 1621.33383 | 1673.6579 | 1434.1069 | 1661.9139 | 7.61E-36 | -1.011970735 | YHR149C | SKG6 | n |
| W303.asm02.HP0_G0024790 | 406.66694 | 378.887608 | 384.4381654 | 404.187779 | 425.16197 | 377.363078 | 446.91683 | 396.24912 | 454.05073 | 0.672705521 | -0.04541883 | YHR150W | PEX28 | n |
| W303.asm02.HP0_G0024800 | 617.3752 | 652.932344 | 628.8172001 | 260.013582 | 263.36672 | 254.103781 | 270.20227 | 223.05414 | 240.69097 | 2.77E-37 | 1.288174036 | YHR151C | MTC6 | n |
| W303.asm02.HP0_G0024830 | 1182.0733 | 1103.44528 | 1189.053723 | 710.870465 | 696.61846 | 731.022446 | 639.59271 | 608.80659 | 676.22702 | 1.56E-19 | 0.700609746 | YHR154W | RTT107 | n |
| W303.asm02.HP0_G0024850 | 662.67748 | 700.682563 | 712.8526785 | 622.532517 | 709.20253 | 607.763149 | 556.36585 | 528.76952 | 561.61227 | 0.287489995 | 0.098556534 | YHR156C | LIN1 | n |
| W303.asm02.HP0_G0024860 | 825.97638 | 924.900983 | 973.6524389 | 2486.79656 | 2360.413 | 2455.70445 | 2361.1346 | 2196.4272 | 2347.839 | 3.91E-79 | -1.421727431 | YHR158C | KEL1 | n |
| W303.asm02.HP0_G0024870 | 303.41989 | 328.023244 | 268.5271607 | 635.03317 | 593.24926 | 629.570563 | 608.81017 | 541.89035 | 599.5233 | 1.17E-24 | -1.047296796 | YHR159W | TDA11 | n |
| W303.asm02.HP0_G0024890 | 855.47554 | 893.759536 | 865.4688346 | 2083.44216 | 2098.844 | 2381.74888 | 2053.3092 | 1646.6644 | 2056.8939 | 1.20E-42 | -1.327691705 | YHR161C | YAP1801 | n |
| W303.asm02.HP0_G0024900 | 6760.5745 | 6597.83462 | 7940.869743 | 2481.7963 | 2461.0856 | 2689.89712 | 2705.4429 | 2494.2701 | 2832.7476 | 8.78E-68 | 1.480807707 | YHR162W | MPC2 | n |
| W303.asm02.HP0_G0024910 | 3460.8832 | 3491.99428 | 3370.11246 | 1056.72186 | 1011.2203 | 1101.74848 | 1077.3888 | 883.03197 | 1167.3071 | 4.19E-78 | 1.703560901 | YHR163W | SOL3 | n |
| W303.asm02.HP0_G0024920 | 352.93634 | 375.773463 | 379.6085402 | 830.876733 | 879.98641 | 869.452117 | 786.66483 | 788.56198 | 868.42713 | 6.81E-44 | -1.218283178 | YHR164C | DNA2 | n |
| W303.asm02.HP0_G0024930 | 980.84695 | 1171.95646 | 1174.564847 | 1665.08697 | 1702.4456 | 1810.96352 | 1584.7306 | 1624.359 | 1646.9258 | 1.16E-14 | -0.637341641 | YHR165C | PRP8 | n |
| W303.asm02.HP0_G0024940 | 425.63069 | 435.980261 | 413.4159165 | 425.855577 | 427.85856 | 407.703828 | 418.41448 | 402.80953 | 422.31127 | 0.882128698 | 0.014658389 | YHR166C | CDC23 | n |
| W303.asm02.HP0_G0024950 | 716.40808 | 653.970392 | 797.8540819 | 706.703581 | 663.36054 | 618.192781 | 663.53468 | 686.2195 | 633.90772 | 0.208113845 | 0.124939469 | YHR167W | THP2 | n |
| W303.asm02.HP0_G0024960 | 434.05902 | 414.181248 | 493.5876947 | 354.185167 | 329.88255 | 351.76307 | 321.50649 | 289.97038 | 301.52495 | 0.00047596 | 0.37437069 | YHR168W | MTG2 | n |
| W303.asm02.HP0_G0024970 | 583.66188 | 605.182125 | 592.1120487 | 1082.55655 | 1093.0168 | 1102.69663 | 632.75214 | 862.03864 | 603.04991 | 3.72E-14 | -0.880225676 | YHR169W | DBP8 | n |
| W303.asm02.HP0_G0024980 | 3481.954 | 3066.39451 | 3482.159764 | 2903.48499 | 2835.9113 | 3125.09725 | 1856.073 | 2264.6556 | 1928.1727 | 0.046789319 | 0.178464382 | YHR170W | NMD3 | n |
| W303.asm02.HP0_G0024990 | 464.61171 | 467.121709 | 448.1892179 | 283.348134 | 288.53487 | 300.563055 | 438.93617 | 360.82287 | 373.82041 | 4.15E-10 | 0.661828926 | YHR171W | ATG7 | n |
| W303.asm02.HP0_G0025000 | 369.793 | 370.583222 | 385.4040904 | 360.018805 | 362.2416 | 384.000117 | 334.04753 | 267.66497 | 280.36531 | 0.82063758 | 0.026151762 | YHR172W | SPC97 | n |
| W303.asm02.HP0_G0025020 | 38371.028 | 39989.7705 | 42947.92499 | 31160.7943 | 29429.658 | 33237.3435 | 31870.186 | 30625.333 | 28703.06 | 1.88E-07 | 0.370628168 | YHR174W | ENO2 | n |
| W303.asm02.HP0_G0025030 | 500.43212 | 520.062169 | 539.9520966 | 823.376342 | 793.69561 | 800.237282 | 1072.8284 | 904.0253 | 1090.6034 | 9.46E-12 | -0.631153609 | YHR175W | CTR2 | n |
| W303.asm02.HP0_G0025050 | 224.4043 | 268.854495 | 255.0042102 | 725.037872 | 624.70945 | 663.703906 | 1034.0652 | 728.20616 | 1009.4914 | 1.54E-25 | -1.427988704 | YHR176W | FMO1 | n |
| W303.asm02.HP0_G0025060 | 116.94308 | 162.973574 | 139.0932056 | 105.838862 | 86.290802 | 82.4889141 | 72.966013 | 77.412907 | 82.87528 | 0.001209044 | 0.606759598 | YHR177W | ROF1 | n |
| W303.asm02.HP0_G0025070 | 280.24199 | 298.957894 | 296.5389869 | 1135.89267 | 1085.8259 | 1275.25965 | 1182.2774 | 956.50863 | 1161.1356 | 2.40E-76 | -1.996861907 | YHR178W | STB5 | n |
| W303.asm02.HP0_G0025080 | 6461.3688 | 6580.1878 | 6245.6713 | 1376.73858 | 1360.8779 | 1199.40777 | 1574.4697 | 1600.7415 | 1373.6137 | 2.37E-185 | 2.291851932 | YHR179W | OYE2 | n |
| W303.asm02.HP0_G0025100 | 1683.559 | 1631.81184 | 1883.553825 | 1081.72317 | 1066.9498 | 1087.52626 | 1491.2429 | 1529.889 | 1616.9496 | 2.83E-19 | 0.684403161 | YHR181W | SVP26 | n |
| W303.asm02.HP0_G0025110 | 401.39924 | 365.392981 | 301.368612 | 1110.89136 | 1008.5238 | 1130.19294 | 1056.8671 | 831.86073 | 1020.9529 | 1.73E-41 | -1.606714798 | YHR182W | RGD3 | n |
| W303.asm02.HP0_G0025120 | 6584.6331 | 7150.07629 | 7569.954528 | 11328.9251 | 10852.866 | 12681.4854 | 10600.594 | 8762.0914 | 10240.387 | 3.35E-15 | -0.710425298 | YHR183W | GND1 | n |
| W303.asm02.HP0_G0025130 | 401.39924 | 340.479823 | 483.9284443 | 65.0033954 | 68.313552 | 73.9555782 | 41.043382 | 57.731659 | 57.307374 | 1.54E-47 | 2.56692801 | YHR184W | SSP1 | n |
| W303.asm02.HP0_G0025140 | 18.963743 | 17.6468201 | 25.114051 | 34.1684514 | 35.055638 | 24.6518594 | 31.922631 | 27.553747 | 37.02938 | 0.078087796 | -0.604035755 | YHR185C | PFS1 | n |
| W303.asm02.HP0_G0025150 | 666.89164 | 750.508879 | 765.0126306 | 1500.07835 | 1484.9209 | 1400.41524 | 1583.5905 | 1430.1707 | 1556.9973 | 9.94E-36 | -1.006420642 | YHR186C | KOG1 | n |
| W303.asm02.HP0_G0025160 | 1065.1303 | 962.27072 | 1020.016841 | 1110.89136 | 1146.0497 | 1128.29664 | 873.31197 | 1001.1195 | 943.36755 | 0.055252874 | -0.151612585 | YHR187W | IKI1 | n |
| W303.asm02.HP0_G0025170 | 1172.5915 | 1370.22368 | 1316.555828 | 2195.94804 | 2175.2473 | 2144.71177 | 2437.5209 | 2481.1493 | 2365.4721 | 2.48E-26 | -0.755385521 | YHR188C | GPI16 | n |
| W303.asm02.HP0_G0025180 | 114.836 | 135.98432 | 117.8428547 | 208.344216 | 227.41222 | 219.022289 | 198.37635 | 230.92664 | 208.9515 | 3.99E-09 | -0.828601036 | YHR189W | PTH1 | n |
| W303.asm02.HP0_G0025190 | 4266.8423 | 4359.80261 | 3587.445593 | 2925.98617 | 3043.5485 | 3026.48981 | 2707.7231 | 2786.8647 | 2686.3934 | 2.08E-08 | 0.441007082 | YHR190W | ERG9 | n |
| W303.asm02.HP0_G0025200 | 670.05227 | 596.877739 | 691.6023277 | 269.180727 | 281.34397 | 267.377859 | 247.40039 | 313.58788 | 291.82678 | 2.00E-30 | 1.26012984 | YHR191C | CTF8 | n |
| W303.asm02.HP0_G0025210 | 1088.3082 | 1065.0375 | 981.3798392 | 977.551061 | 987.84991 | 1026.84476 | 1093.3501 | 1102.1499 | 1099.4199 | 0.373886217 | 0.066851357 | YHR192W | LNP1 | n |
| W303.asm02.HP0_G0025220 | 7794.0985 | 8110.27091 | 7745.752885 | 5471.11911 | 5482.1625 | 5762.84621 | 5027.8143 | 5351.9872 | 4892.2865 | 4.36E-18 | 0.500683071 | YHR193C | EGD2 | n |
| W303.asm02.HP0_G0025230 | 313.95531 | 359.164692 | 455.9166182 | 310.016193 | 305.61326 | 300.563055 | 314.66593 | 310.96371 | 336.79103 | 0.024022861 | 0.303012679 | YHR194W | MDM31 | n |
| W303.asm02.HP0_G0025250 | 4317.4122 | 3812.75119 | 4617.121685 | 1922.60042 | 1954.1271 | 1946.54874 | 1181.1373 | 1511.5198 | 1288.0934 | 1.18E-32 | 1.130424163 | YHR196W | UTP9 | n |
| W303.asm02.HP0_G0025260 | 1825.7871 | 1621.43135 | 1730.937669 | 2330.95509 | 2352.3232 | 2420.62296 | 1406.8759 | 1637.4798 | 1403.5898 | 7.57E-09 | -0.45611478 | YHR197W | RIX1 | n |
| W303.asm02.HP0_G0025270 | 1531.849 | 1468.83826 | 1591.844464 | 668.368245 | 636.39467 | 599.229813 | 728.52003 | 701.9645 | 746.75917 | 4.11E-61 | 1.269467444 | YHR198C | AIM18 | n |
| W303.asm02.HP0_G0025300 | 1640.3638 | 1814.50833 | 1585.082988 | 2759.3108 | 2753.2159 | 2633.00821 | 2965.3844 | 3021.7275 | 2981.7468 | 6.25E-25 | -0.693066474 | YHR200W | RPN10 | n |
| W303.asm02.HP0_G0025310 | 2076.5299 | 1906.89462 | 2048.727007 | 680.868898 | 727.17978 | 679.82243 | 501.64134 | 514.3366 | 544.86088 | 4.93E-92 | 1.530747619 | YHR201C | PPX1 | n |
| W303.asm02.HP0_G0025320 | 235.99325 | 255.359867 | 234.7197844 | 391.687126 | 391.0052 | 356.503813 | 563.20641 | 503.83994 | 610.98478 | 9.14E-09 | -0.650986728 | YHR202W | SMN1 | n |
| W303.asm02.HP0_G0025340 | 553.10918 | 604.144077 | 589.2142736 | 576.69679 | 552.80045 | 686.459469 | 864.19122 | 754.44782 | 766.15551 | 0.613966065 | -0.055085206 | YHR204W | MNL1 | n |
| W303.asm02.HP0_G0025350 | 1197.8765 | 1401.36513 | 1352.295054 | 2986.82268 | 2809.8442 | 2750.57862 | 2338.3327 | 2516.5755 | 2297.5849 | 5.42E-47 | -1.112979557 | YHR205W | SCH9 | n |
| W303.asm02.HP0_G0025360 | 971.36508 | 1063.99945 | 1062.517542 | 1439.24184 | 1348.2938 | 1347.31893 | 1390.9146 | 1289.7778 | 1305.7265 | 1.83E-08 | -0.416586554 | YHR206W | SKN7 | n |
| W303.asm02.HP0_G0025370 | 554.16272 | 470.235853 | 463.6440185 | 664.201361 | 686.73097 | 676.029836 | 698.87759 | 631.112 | 655.94902 | 2.29E-06 | -0.44672368 | YHR207C | SET5 | n |
| W303.asm02.HP0_G0025380 | 6464.5294 | 6846.9662 | 7484.953124 | 6782.8543 | 6532.034 | 7459.08376 | 6479.1539 | 6274.3817 | 6400.7929 | 0.982605216 | 0.001776568 | YHR208W | BAT1 | n |
| W303.asm02.HP0_G0025580 | 26.338532 | 35.2936402 | 28.97775116 | 46.6691044 | 40.448814 | 26.5481563 | 34.202819 | 26.241663 | 32.621121 | 0.300597864 | -0.333238811 | YOL156W | HXT11 | y |
| W303.asm02.HP0_G0025620 | 429.84485 | 373.697367 | 450.121068 | 563.36276 | 540.21638 | 506.311266 | 680.63609 | 636.36034 | 688.57014 | 0.000297074 | -0.36082516 | YCL064C | CHA1 | y |

| | | | | | | | | | | | | | | |
|---|---|---|---|---|---|---|---|---|---|---|---|---|---|---|
| W303.asm02.HP0_G0025630 | 289.72386 | 266.778398 | 311.9937875 | 2587.63516 | 2748.7216 | 3105.18613 | 5206.8091 | 3982.1724 | 4230.1659 | 7.23E-174 | -3.279869088 | YIL166C | SOA1 | n |
| W303.asm02.HP0_G0025670 | 786.99535 | 937.357562 | 829.7296082 | 96.6717162 | 134.82938 | 77.7481719 | 82.086764 | 107.59082 | 115.4964 | 1.69E-76 | 3.044391069 | YIL162W | SUC2 | n |
| W303.asm02.HP0_G0025680 | 520.4494 | 494.110963 | 541.8839467 | 230.845391 | 193.25544 | 222.814883 | 164.17353 | 179.75539 | 189.55516 | 9.38E-29 | 1.26628853 | YIL161W | SMU2 | n |
| W303.asm02.HP0_G0025690 | 58.998313 | 80.9677628 | 87.89917851 | 57.5030036 | 80.897627 | 92.9185469 | 59.284885 | 65.604158 | 67.887197 | 0.953193612 | -0.014743821 | YIL160C | POT1 | n |
| W303.asm02.HP0_G0025700 | 344.50801 | 343.593968 | 468.4736437 | 353.35179 | 384.71316 | 356.503813 | 346.58856 | 274.22538 | 357.06902 | 0.582631169 | 0.081119598 | YIL159W | BNR1 | n |
| W303.asm02.HP0_G0025720 | 2354.6648 | 2175.74911 | 2428.335547 | 925.048319 | 877.28982 | 892.20768 | 904.0945 | 1001.1195 | 990.97675 | 6.07E-75 | 1.368672564 | YIL157C | COA1 | n |
| W303.asm02.HP0_G0025740 | 435.11256 | 502.415349 | 503.2469451 | 875.879084 | 829.65011 | 780.326164 | 863.05112 | 779.3774 | 872.83539 | 5.65E-17 | -0.786839918 | YIL156W | UBP7 | n |
| W303.asm02.HP0_G0025750 | 478.30775 | 585.459208 | 614.3283246 | 1881.76496 | 1914.5772 | 1869.74872 | 1698.74 | 1728.0135 | 1669.8487 | 2.77E-91 | -1.754205293 | YIL155C | GUT2 | n |
| W303.asm02.HP0_G0025760 | 1153.6277 | 1131.47258 | 1213.201849 | 1180.06164 | 1186.4985 | 1157.68924 | 1168.5963 | 1092.9653 | 1128.5144 | 0.887061872 | -0.010407919 | YIL154C | IMP21 | n |
| W303.asm02.HP0_G0025770 | 538.3596 | 567.812388 | 654.8971762 | 698.369812 | 744.25817 | 802.133578 | 758.16248 | 732.14241 | 748.52247 | 0.000312368 | -0.348114347 | YIL153W | RRD1 | n |
| W303.asm02.HP0_G0025800 | 184.36973 | 154.669188 | 173.866507 | 430.022462 | 432.35287 | 403.911235 | 384.21166 | 366.0712 | 369.41215 | 8.90E-31 | -1.304353138 | YIL150C | MCM10 | n |
| W303.asm02.HP0_G0025820 | 859.6897 | 935.281466 | 947.5724629 | 1082.55655 | 1111.8929 | 1116.91886 | 1032.9251 | 1111.3344 | 1081.7869 | 0.000262723 | -0.271222735 | YIL147C | SLN1 | n |
| W303.asm02.HP0_G0025830 | 197.01222 | 230.44671 | 216.3672087 | 340.01776 | 408.98245 | 452.266805 | 542.68472 | 467.10161 | 544.86088 | 7.58E-12 | -0.897698626 | YIL146C | ATG32 | n |
| W303.asm02.HP0_G0025840 | 1046.1665 | 1084.76041 | 1176.496697 | 755.039439 | 716.39343 | 855.229891 | 710.27853 | 600.93409 | 693.86005 | 1.45E-07 | 0.508456774 | YIL145C | PAN6 | n |
| W303.asm02.HP0_G0025850 | 673.21289 | 605.182125 | 577.6231731 | 840.877256 | 836.84101 | 846.696555 | 849.36999 | 851.54198 | 873.71705 | 6.36E-08 | -0.444648584 | YIL144W | NDC80 | n |
| W303.asm02.HP0_G0025860 | 1037.7382 | 1121.0921 | 1272.123276 | 2342.62236 | 2182.4382 | 2180.74141 | 2027.087 | 2088.8364 | 2089.515 | 7.20E-33 | -0.966191016 | YIL143C | SSL2 | n |
| W303.asm02.HP0_G0025870 | 3485.1146 | 3553.23913 | 2582.883553 | 5311.11075 | 5199.0208 | 5203.43863 | 4851.0998 | 4984.604 | 4640.134 | 6.59E-12 | -0.708037223 | YIL142W | CCT2 | n |
| W303.asm02.HP0_G0025880 | 1145.1994 | 1161.57598 | 1215.133699 | 2775.97833 | 2783.7772 | 2834.01568 | 3115.8768 | 2902.328 | 2974.6936 | 1.60E-91 | -1.252479657 | YIL140W | AXL2 | n |
| W303.asm02.HP0_G0025890 | 808.06618 | 822.134207 | 1005.527965 | 216.677985 | 248.08606 | 225.659328 | 230.29898 | 241.4233 | 250.38914 | 3.87E-62 | 1.933990564 | YIL139C | REV7 | n |
| W303.asm02.HP0_G0025900 | 4158.3275 | 3923.82235 | 4989.002824 | 9492.16248 | 9494.6848 | 10973.87 | 10000.904 | 10079.423 | 9051.0385 | 1.88E-35 | -1.196440322 | YIL138C | TPM2 | n |
| W303.asm02.HP0_G0025910 | 660.57039 | 853.275654 | 893.4806607 | 2002.6046 | 1905.5885 | 1879.2302 | 1469.5811 | 1692.5873 | 1561.4055 | 6.51E-34 | -1.264809876 | YIL137C | TMA108 | n |
| W303.asm02.HP0_G0025920 | 598.41146 | 727.671817 | 758.2511553 | 83.3376864 | 73.706727 | 51.2000156 | 204.07682 | 137.76873 | 236.28271 | 1.13E-66 | 3.317927297 | YIL136W | OM45 | n |
| W303.asm02.HP0_G0025930 | 685.85539 | 711.063045 | 680.9771522 | 961.716901 | 963.58062 | 1049.60032 | 814.02708 | 699.34033 | 987.45014 | 4.12E-06 | -0.517523032 | YIL135C | VHS2 | n |
| W303.asm02.HP0_G0025950 | 716.40808 | 681.997695 | 763.0807805 | 508.359887 | 549.205 | 469.333477 | 519.88284 | 532.70577 | 489.31681 | 8.49E-08 | 0.501358026 | YIL134W | FLX1 | n |
| W303.asm02.HP0_G0025960 | 6929.1411 | 6771.18868 | 6078.566268 | 16618.368 | 17001.985 | 18572.3316 | 14685.55 | 15990.358 | 13498.091 | 1.37E-66 | -1.399947069 | YIL133C | RPL16A | n |
| W303.asm02.HP0_G0025970 | 77.962056 | 119.375548 | 76.30807805 | 173.342388 | 179.7725 | 169.71857 | 133.39099 | 205.99706 | 172.80377 | 2.41E-06 | -0.935824871 | YIL132C | CSM2 | n |
| W303.asm02.HP0_G0025990 | 884.97469 | 1013.13508 | 1041.267192 | 1339.23662 | 1482.2243 | 1473.42267 | 1014.6836 | 1062.7874 | 1051.8107 | 5.27E-11 | -0.546116037 | YIL130W | ASG1 | n |
| W303.asm02.HP0_G0026000 | 1128.3427 | 1146.00526 | 1253.7707 | 1686.75477 | 1675.4797 | 1634.60791 | 1843.5319 | 1802.8023 | 1889.38 | 1.82E-13 | -0.501635095 | YIL129C | TAO3 | n |
| W303.asm02.HP0_G0026010 | 1764.6817 | 1699.28497 | 1555.139312 | 1474.24367 | 1486.7186 | 1387.14116 | 1405.7358 | 1485.2781 | 1413.288 | 0.005082731 | 0.206298659 | YIL128W | MET18 | n |
| W303.asm02.HP0_G0026020 | 1839.4831 | 1898.59023 | 2239.01424 | 833.376864 | 880.88527 | 882.726196 | 536.98425 | 736.07866 | 565.13888 | 2.09E-26 | 1.203084298 | YIL127C | RRT14 | n |
| W303.asm02.HP0_G0026030 | 1523.4207 | 1655.68695 | 1526.161561 | 3438.51294 | 3456.1264 | 3434.19364 | 3342.7555 | 3047.9692 | 3257.7038 | 9.01E-70 | -1.134478531 | YIL126W | STH1 | n |
| W303.asm02.HP0_G0026040 | 2043.8701 | 2284.74418 | 2486.291049 | 9677.17214 | 10045.688 | 10754.8477 | 9634.934 | 9830.1271 | 9230.8955 | 5.54E-169 | -2.160528375 | YIL125W | KGD1 | n |
| W303.asm02.HP0_G0026050 | 2878.2748 | 2904.45898 | 3327.611758 | 1339.23662 | 1343.7995 | 1236.38556 | 1589.291 | 1583.6844 | 1586.0918 | 1.29E-57 | 1.21680895 | YIL124W | AYR1 | n |
| W303.asm02.HP0_G0026080 | 503.59274 | 474.388046 | 598.8735239 | 265.84722 | 229.20994 | 275.911195 | 205.21691 | 212.55747 | 197.49003 | 3.99E-17 | 1.03350518 | YIL121W | QDR2 | n |
| W303.asm02.HP0_G0026100 | 858.63616 | 777.498133 | 768.8763307 | 198.343694 | 199.54748 | 182.0445 | 226.8787 | 333.26912 | 229.2295 | 3.10E-52 | 2.051088267 | YIL119C | RPI1 | n |
| W303.asm02.HP0_G0026110 | 2938.3267 | 2871.24144 | 3228.121479 | 2925.98617 | 2795.4624 | 2871.94162 | 3057.732 | 3190.9863 | 3200.3964 | 0.276075959 | 0.072950841 | YIL118W | RHO3 | n |
| W303.asm02.HP0_G0026130 | 1932.1947 | 1612.08892 | 1673.948092 | 1429.24132 | 1410.3153 | 1345.42263 | 1415.9967 | 1190.0594 | 1419.4596 | 0.000685217 | 0.317948358 | YIL116W | HIS5 | n |
| W303.asm02.HP0_G0026140 | 1214.7331 | 1353.61491 | 1477.865309 | 1985.93707 | 2043.1145 | 2085.92656 | 1870.8942 | 1790.9935 | 1903.4865 | 5.32E-14 | -0.594957543 | YIL115C | NUP159 | n |
| W303.asm02.HP0_G0026170 | 682.69476 | 934.243417 | 773.7059559 | 835.043618 | 839.5376 | 896.948422 | 785.52473 | 663.91408 | 821.69958 | 0.386783877 | -0.104987492 | YIL112W | HOS4 | n |
| W303.asm02.HP0_G0026190 | 1882.6783 | 1673.33377 | 1994.635205 | 1035.88744 | 1026.501 | 1061.92625 | 718.25919 | 974.87779 | 751.16743 | 2.23E-14 | 0.829358666 | YIL110W | HPM1 | n |
| W303.asm02.HP0_G0026200 | 2859.3111 | 3454.62455 | 3515.001216 | 6488.67226 | 6502.3715 | 6510.93532 | 6123.4446 | 5479.2593 | 6198.8946 | 1.42E-31 | -0.988304517 | YIL109C | SEC24 | n |
| W303.asm02.HP0_G0026210 | 946.08009 | 999.640457 | 1166.837447 | 998.385483 | 948.29996 | 943.407696 | 1060.2874 | 823.98823 | 1045.6392 | 0.340935785 | 0.107353625 | YIL108W | | n |
| W303.asm02.HP0_G0026220 | 474.09358 | 437.01831 | 497.4513949 | 686.702536 | 599.5413 | 607.763149 | 849.36999 | 749.19949 | 874.5987 | 2.25E-05 | -0.427508678 | YIL107C | PFK26 | n |
| W303.asm02.HP0_G0026240 | 862.85032 | 807.601532 | 881.8895603 | 755.872816 | 804.48196 | 779.378016 | 454.89749 | 578.62868 | 477.85534 | 0.19412365 | 0.125893841 | YIL104C | SHQ1 | n |
| W303.asm02.HP0_G0026250 | 1729.9148 | 1517.62653 | 1971.453004 | 957.550017 | 948.29996 | 937.718805 | 632.75214 | 766.25657 | 587.18017 | 2.84E-15 | 0.876316066 | YIL103W | DPH1 | n |
| W303.asm02.HP0_G0026260 | 357.1505 | 377.84956 | 383.4722403 | 32.5016977 | 34.156776 | 37.9259375 | 44.463664 | 30.177913 | 40.555988 | 5.59E-78 | 3.422498986 | YIL102C-A | | n |
| W303.asm02.HP0_G0026280 | 174.88786 | 244.979385 | 245.3449598 | 77.5040483 | 100.6726 | 119.466703 | 109.44902 | 132.5204 | 93.455103 | 4.77E-09 | 1.165120943 | YIL101C | XBP1 | n |
| W303.asm02.HP0_G0026290 | 142.22808 | 135.98432 | 185.4576074 | 409.18804 | 449.43126 | 484.503852 | 637.31252 | 564.19576 | 621.5646 | 4.84E-32 | -1.529896218 | YIL099W | SGA1 | n |
| W303.asm02.HP0_G0026300 | 1283.2133 | 1277.83739 | 1578.321513 | 778.373991 | 790.99902 | 818.252102 | 661.25449 | 692.77991 | 637.43433 | 6.32E-17 | 0.794741527 | YIL098C | FMC1 | n |
| W303.asm02.HP0_G0026310 | 743.80016 | 745.318637 | 674.215677 | 288.348395 | 312.80416 | 265.481563 | 353.42912 | 391.00078 | 385.28189 | 4.50E-39 | 1.318440304 | YIL097W | FYV10 | n |
| W303.asm02.HP0_G0026320 | 1019.828 | 917.634645 | 1183.258172 | 998.385483 | 997.7374 | 1080.88922 | 761.58276 | 940.76363 | 813.76472 | 0.853205786 | 0.021189034 | YIL096C | BMT5 | n |
| W303.asm02.HP0_G0026330 | 422.47006 | 497.225108 | 447.2232929 | 1410.90703 | 1363.5744 | 1356.80041 | 1331.6297 | 1203.1803 | 1368.3238 | 6.83E-76 | -1.595883923 | YIL095W | PRK1 | n |
| W303.asm02.HP0_G0026340 | 6079.9868 | 5721.72191 | 6061.179617 | 755.039439 | 741.56158 | 810.666914 | 722.81957 | 646.857 | 733.53439 | 0 | 2.953313137 | YIL094C | LYS12 | n |
| W303.asm02.HP0_G0026350 | 1584.5261 | 1630.77379 | 1367.749855 | 1971.76966 | 2010.7555 | 2054.63766 | 1925.6187 | 1922.2018 | 1922.0012 | 1.55E-07 | -0.398039061 | YIL093C | RSM25 | n |
| W303.asm02.HP0_G0026370 | 1487.6003 | 1331.81589 | 1382.23873 | 1250.0653 | 1200.8803 | 1298.96336 | 657.83421 | 831.86073 | 675.34536 | 0.082720996 | 0.164115877 | YIL091C | UTP25 | n |
| W303.asm02.HP0_G0026380 | 1225.2685 | 1228.01107 | 1362.920229 | 1191.72892 | 1019.3101 | 998.400305 | 1362.4123 | 1283.2173 | 1333.0577 | 0.004850951 | 0.249323568 | YIL090W | ICE2 | n |
| W303.asm02.HP0_G0026400 | 1903.7491 | 1849.80197 | 1776.336146 | 735.871771 | 687.62983 | 701.629844 | 1352.1514 | 1318.6436 | 1293.3834 | 1.25E-87 | 1.378886739 | YIL088C | AVT7 | n |
| W303.asm02.HP0_G0026410 | 458.29047 | 422.485634 | 448.1892179 | 317.516585 | 317.29847 | 296.770461 | 355.70931 | 312.27579 | 398.50666 | 3.98E-06 | 0.511953179 | YIL087C | AIM19 | n |
| W303.asm02.HP0_G0026440 | 213.86888 | 260.550109 | 246.3108848 | 711.703842 | 704.70822 | 781.274313 | 698.87759 | 628.48784 | 743.23256 | 7.87E-50 | -1.607249101 | YIL084C | SDS3 | n |
| W303.asm02.HP0_G0026450 | 1190.5017 | 1181.2989 | 1013.255366 | 1218.39697 | 1253.9132 | 1206.04481 | 1210.7798 | 1263.5361 | 1231.6677 | 0.137010123 | -0.120694281 | YIL083C | CAB2 | n |

| | | | | | | | | | | | | | | |
|---|---|---|---|---|---|---|---|---|---|---|---|---|---|---|
| W303.asm02.HP0_G0026460 | 6534.0631 | 6048.7071 | 6195.443198 | 6992.86526 | 6932.9266 | 7369.95781 | 5537.4363 | 5917.4951 | 5505.9162 | 0.002666353 | -0.18147118 | YIL078W | THS1 | n |
| W303.asm02.HP0_G0026470 | 414.04173 | 442.208551 | 379.6085402 | 344.184645 | 355.0507 | 330.903805 | 453.75739 | 381.8162 | 452.28743 | 0.017545938 | 0.261533624 | YIL077C | RCI37 | n |
| W303.asm02.HP0_G0026480 | 5287.7238 | 5216.19241 | 6183.852097 | 2172.61348 | 2141.0905 | 2266.07477 | 2243.7049 | 2398.488 | 2287.8867 | 9.10E-68 | 1.342977906 | YIL076W | SEC28 | n |
| W303.asm02.HP0_G0026490 | 3638.9316 | 3945.62137 | 3852.109054 | 7235.37793 | 7012.0265 | 7477.09858 | 8065.0246 | 8066.6873 | 7639.5138 | 2.36E-56 | -0.925537305 | YIL075C | RPN2 | n |
| W303.asm02.HP0_G0026510 | 544.68085 | 501.377301 | 547.6794969 | 38.3353357 | 33.257913 | 29.3926016 | 27.362255 | 41.986661 | 37.911032 | 2.04E-109 | 3.974094419 | YIL073C | SPO22 | n |
| W303.asm02.HP0_G0026520 | 267.59949 | 305.186183 | 242.4471847 | 30.0015671 | 29.662463 | 26.5481563 | 25.082067 | 47.234994 | 21.159646 | 3.76E-45 | 3.238534547 | YIL072W | HOP1 | n |
| W303.asm02.HP0_G0026530 | 830.19054 | 756.737168 | 809.4451824 | 416.688432 | 460.21761 | 447.526063 | 483.39984 | 476.28619 | 486.67186 | 5.16E-23 | 0.856368441 | YIL071C | PCI8 | n |
| W303.asm02.HP0_G0026540 | 2429.4662 | 2273.32565 | 2063.215882 | 2179.2805 | 2295.6949 | 2205.39327 | 1832.131 | 2065.2189 | 1739.4992 | 0.84093667 | 0.018106643 | YIL070C | MAM33 | n |
| W303.asm02.HP0_G0026560 | 1117.8073 | 1096.17894 | 940.8109876 | 1716.75634 | 1698.8502 | 1734.16349 | 1681.6386 | 1575.8119 | 1729.8011 | 1.07E-18 | -0.707755275 | YIL068C | SEC6 | n |
| W303.asm02.HP0_G0026570 | 322.38364 | 353.97445 | 356.4263393 | 562.529383 | 538.41865 | 498.726078 | 695.45731 | 638.9845 | 702.67657 | 9.90E-11 | -0.631573491 | YIL067C | | n |
| W303.asm02.HP0_G0026590 | 952.40134 | 919.710742 | 1087.631593 | 928.381826 | 944.70451 | 929.185469 | 972.50014 | 1016.8645 | 1000.6749 | 0.344869965 | 0.079713398 | YIL065C | FIS1 | n |
| W303.asm02.HP0_G0026600 | 1230.5362 | 1073.34188 | 1457.580883 | 1269.23296 | 1259.3064 | 1306.54855 | 1134.3935 | 1121.8311 | 1115.2897 | 0.797820842 | -0.027226767 | YIL064W | EFM4 | n |
| W303.asm02.HP0_G0026610 | 1921.6593 | 1898.59023 | 1986.907804 | 1772.59259 | 1821.9943 | 1767.34869 | 1648.5759 | 1749.0069 | 1687.4818 | 0.061671194 | 0.115272682 | YIL063C | YRB2 | n |
| W303.asm02.HP0_G0026620 | 3100.572 | 3081.96523 | 3256.133305 | 1408.4069 | 1561.3242 | 1496.17823 | 1592.7112 | 1691.2752 | 1568.4588 | 1.13E-58 | 1.080139225 | YIL062C | ARC15 | n |
| W303.asm02.HP0_G0026630 | 207.54764 | 219.028179 | 213.4694335 | 365.852443 | 347.8598 | 374.518633 | 361.40978 | 355.57454 | 387.04519 | 2.30E-12 | -0.765450707 | YIL061C | SNP1 | n |
| W303.asm02.HP0_G0026660 | 510.96753 | 526.290458 | 537.0543215 | 742.538786 | 767.62859 | 1006.93364 | 925.75629 | 805.61907 | 847.26749 | 9.62E-09 | -0.675780817 | YIL056W | VHR1 | n |
| W303.asm02.HP0_G0026670 | 188.58389 | 172.316008 | 212.5035085 | 150.841212 | 138.42483 | 139.37782 | 253.10086 | 153.51373 | 213.35976 | 0.01654595 | 0.420023188 | YIL055C | | n |
| W303.asm02.HP0_G0026690 | 10144.549 | 9809.55588 | 8807.304502 | 6011.14732 | 5535.1954 | 5609.24616 | 4237.7292 | 3903.4474 | 4760.0387 | 1.04E-18 | 0.745316642 | YIL053W | GPP1 | n |
| W303.asm02.HP0_G0026700 | 7242.0429 | 7145.92409 | 8000.757095 | 14243.244 | 14799.771 | 15633.0714 | 12805.535 | 13552.507 | 11391.824 | 9.20E-37 | -0.996608908 | YIL052C | RPL34B | y |
| W303.asm02.HP0_G0026710 | 2631.7462 | 2559.82696 | 2317.254168 | 2084.27554 | 2207.6064 | 2243.3192 | 2065.8502 | 1944.5073 | 1935.2259 | 0.00544919 | 0.200290401 | YIL051C | MMF1 | n |
| W303.asm02.HP0_G0026730 | 632.12478 | 620.752848 | 575.691323 | 522.527294 | 558.19363 | 522.429789 | 558.64604 | 657.35367 | 557.20401 | 0.051115392 | 0.189292516 | YIL049W | DFG10 | n |
| W303.asm02.HP0_G0026740 | 1348.5329 | 1300.67445 | 1380.30688 | 3138.49727 | 3196.3551 | 3147.85281 | 3497.8082 | 3623.9737 | 3572.4535 | 5.14E-103 | -1.234402169 | YIL048W | NEO1 | n |
| W303.asm02.HP0_G0026750 | 1359.0683 | 1420.04999 | 1460.478658 | 934.215464 | 968.07494 | 917.807688 | 1120.7124 | 1096.9015 | 1168.1888 | 3.82E-17 | 0.588384251 | YIL047C | SYG1 | n |
| W303.asm02.HP0_G0026760 | 541.52023 | 626.981138 | 624.9535 | 442.523115 | 399.99382 | 389.689008 | 410.43382 | 393.62495 | 424.07457 | 7.25E-08 | 0.540833854 | YIL046W | MET30 | n |
| W303.asm02.HP0_G0026780 | 4500.7284 | 4468.79768 | 4528.256581 | 1604.25046 | 1570.3128 | 1652.62273 | 1713.5612 | 1712.2685 | 1690.1267 | 2.75E-162 | 1.483689801 | YIL044C | AGE2 | y |
| W303.asm02.HP0_G0026790 | 338.18676 | 394.458332 | 355.4604142 | 680.035521 | 661.56282 | 665.600203 | 775.26389 | 758.38407 | 855.20235 | 8.04E-21 | -0.88361934 | YIL042C | PKP1 | n |
| W303.asm02.HP0_G0026800 | 1637.2032 | 1805.16589 | 1592.810389 | 4418.56413 | 4360.3821 | 4513.18656 | 3855.7977 | 3975.612 | 3850.1739 | 1.08E-107 | -1.400722489 | YIL041W | GVP36 | n |
| W303.asm02.HP0_G0026810 | 1435.9768 | 1427.31633 | 1986.907804 | 306.682686 | 319.99506 | 310.992688 | 690.89693 | 596.99784 | 631.26277 | 6.07E-77 | 2.371530244 | YIL040W | APQ12 | n |
| W303.asm02.HP0_G0026820 | 1845.8044 | 1693.05668 | 1506.84306 | 1651.75294 | 1598.1776 | 1687.70422 | 1862.9135 | 1865.7823 | 1858.5222 | 0.7165802 | 0.030727991 | YIL039W | TED1 | n |
| W303.asm02.HP0_G0026830 | 1464.4224 | 1296.52225 | 1387.068355 | 2325.12145 | 2339.7391 | 2189.27474 | 1964.3819 | 2216.1085 | 2160.0472 | 6.72E-22 | -0.724785883 | YIL038C | NOT3 | n |
| W303.asm02.HP0_G0026840 | 183.31619 | 182.69649 | 192.2190827 | 179.176026 | 245.38947 | 244.622297 | 217.75794 | 236.17497 | 224.82124 | 0.080107273 | -0.258370056 | YIL037C | PRM2 | n |
| W303.asm02.HP0_G0026850 | 589.98313 | 708.986949 | 755.3533802 | 1216.73022 | 1132.5668 | 1054.34106 | 1221.0406 | 1018.1765 | 1124.1062 | 8.70E-12 | -0.728063726 | YIL036W | CST6 | n |
| W303.asm02.HP0_G0026860 | 2725.5113 | 2483.01139 | 2795.387062 | 2826.81432 | 2852.0908 | 2787.55641 | 2512.7671 | 2612.3576 | 2716.3695 | 0.235127397 | -0.080913493 | YIL035C | CKA1 | n |
| W303.asm02.HP0_G0026870 | 1653.0063 | 1666.06743 | 1516.502311 | 1940.93472 | 1921.7681 | 1986.37098 | 1862.9135 | 1843.4769 | 1900.8415 | 2.16E-05 | -0.274880045 | YIL034C | CAP2 | n |
| W303.asm02.HP0_G0026880 | 1702.5227 | 1982.67214 | 2117.307685 | 3004.32359 | 3050.7394 | 3008.47499 | 3882.0199 | 3358.9329 | 3695.0032 | 1.39E-13 | -0.642997166 | YIL033C | BCY1 | n |
| W303.asm02.HP0_G0026900 | 545.73439 | 581.307015 | 592.1120487 | 811.709065 | 868.3012 | 866.607672 | 733.08041 | 707.21283 | 737.94265 | 2.13E-12 | -0.565819896 | YIL031W | ULP2 | n |
| W303.asm02.HP0_G0026910 | 1880.5712 | 1985.78629 | 2176.229112 | 1434.24158 | 1453.4607 | 1466.78563 | 1786.5272 | 1838.2285 | 1812.6763 | 2.27E-11 | 0.473254339 | YIL030C | SSM4 | n |
| W303.asm02.HP0_G0026940 | 994.54299 | 979.91754 | 1309.794352 | 729.204756 | 793.69561 | 736.711336 | 943.99779 | 978.81404 | 970.69876 | 8.67E-07 | 0.540392855 | YIL027C | EMC5 | n |
| W303.asm02.HP0_G0026950 | 966.09737 | 941.509755 | 819.1044328 | 1075.05615 | 1161.3304 | 1132.08923 | 1073.9685 | 1075.9082 | 1129.3961 | 0.000247988 | -0.305502578 | YIL026C | IRR1 | n |
| W303.asm02.HP0_G0026960 | 173.83431 | 209.685745 | 198.980558 | 119.172892 | 108.76237 | 108.088922 | 163.03344 | 157.44998 | 148.11752 | 8.79E-08 | 0.792764552 | YIL024C | | n |
| W303.asm02.HP0_G0026970 | 619.48228 | 746.356686 | 731.2052542 | 614.198749 | 525.83458 | 612.503891 | 788.94501 | 624.55159 | 762.6289 | 0.025752244 | 0.259200406 | YIL023C | YKE4 | n |
| W303.asm02.HP0_G0026980 | 1873.1964 | 2283.70613 | 2285.378641 | 3194.33352 | 3145.12 | 3225.60099 | 2919.7806 | 2815.7305 | 2833.6292 | 1.19E-12 | -0.569866298 | YIL022W | TIM44 | n |
| W303.asm02.HP0_G0026990 | 2403.1277 | 2322.11392 | 2255.434965 | 2900.15149 | 2836.8101 | 2955.37868 | 2946.0028 | 3190.9863 | 3131.6276 | 2.09E-07 | -0.316529262 | YIL021W | RPB3 | n |
| W303.asm02.HP0_G0027000 | 1057.7555 | 920.74879 | 1001.664265 | 1140.05955 | 1084.9271 | 1049.60032 | 925.75629 | 952.57238 | 953.94737 | 0.08653656 | -0.136467587 | YIL020C | HIS6 | n |
| W303.asm02.HP0_G0027010 | 809.11972 | 721.443528 | 694.5001028 | 1235.89789 | 1133.4656 | 1361.54116 | 774.12379 | 872.53531 | 837.56932 | 5.79E-15 | -0.745936008 | YIL019W | FAF1 | n |
| W303.asm02.HP0_G0027030 | 466.7188 | 428.713924 | 436.5981175 | 446.689999 | 452.12785 | 434.251985 | 636.17242 | 543.20243 | 580.12696 | 0.98713375 | -0.001696093 | YIL017C | VID28 | n |
| W303.asm02.HP0_G0027040 | 1263.196 | 1210.36425 | 1405.420931 | 820.876211 | 808.07741 | 858.074336 | 997.58221 | 1020.8007 | 1017.4263 | 4.63E-16 | 0.64213414 | YIL016W | SNL1 | n |
| W303.asm02.HP0_G0027050 | 3672.645 | 3875.03409 | 2903.570666 | 180.009403 | 151.90777 | 141.274117 | 223.45841 | 196.81248 | 196.60838 | 5.78E-266 | 4.461859001 | YIL015W | BAR1 | n |
| W303.asm02.HP0_G0027070 | 1183.1269 | 1087.87456 | 1158.144121 | 658.367722 | 653.47305 | 635.259453 | 584.8682 | 603.55826 | 588.94348 | 3.18E-27 | 0.816226138 | YIL014W | MNT3 | n |
| W303.asm02.HP0_G0027110 | 586.8225 | 579.230919 | 535.1224714 | 1426.74119 | 1377.0574 | 1408.94858 | 1493.5231 | 1459.0365 | 1469.7137 | 1.22E-73 | -1.309248164 | YIL010W | DOT5 | n |
| W303.asm02.HP0_G0027120 | 667.94518 | 637.36162 | 662.6245765 | 125.00653 | 141.12142 | 134.637078 | 119.70986 | 94.469988 | 119.90466 | 5.31E-84 | 2.297252777 | YIL009C-A | EST3 | n |
| W303.asm02.HP0_G0027140 | 1664.5953 | 1503.09385 | 2049.692932 | 425.022201 | 425.16197 | 409.600125 | 296.42443 | 397.5612 | 315.63138 | 5.71E-60 | 2.050317339 | YIL008W | URM1 | n |
| W303.asm02.HP0_G0027150 | 1120.9679 | 1044.27653 | 1191.951498 | 106.672239 | 88.98739 | 96.7111407 | 139.09146 | 128.58415 | 144.59091 | 8.53E-186 | 3.519412278 | YIL007C | NAS2 | n |
| W303.asm02.HP0_G0027160 | 162.24536 | 212.799889 | 171.9346569 | 698.369812 | 623.81059 | 579.318696 | 779.82426 | 770.19282 | 780.26194 | 1.75E-53 | -1.799225803 | YIL006W | YIA6 | n |
| W303.asm02.HP0_G0027170 | 879.70699 | 867.80833 | 843.2525587 | 1461.74302 | 1442.6743 | 1265.77816 | 1504.924 | 1451.164 | 1571.9854 | 7.53E-18 | -0.687600974 | YIL005W | EPS1 | n |
| W303.asm02.HP0_G0027180 | 1007.1855 | 916.596597 | 1093.427144 | 848.377647 | 832.3467 | 826.785438 | 743.34126 | 763.6324 | 731.77109 | 0.001428878 | 0.267404065 | YIL004C | BET1 | n |
| W303.asm02.HP0_G0027190 | 621.58937 | 522.138265 | 579.5550232 | 492.525727 | 471.90282 | 494.933485 | 474.27908 | 410.68203 | 488.43516 | 0.019472779 | 0.239752826 | YIL003W | CFD1 | n |
| W303.asm02.HP0_G0027210 | 642.66019 | 716.253287 | 726.375629 | 1163.3941 | 1170.319 | 1211.7337 | 1355.5717 | 1258.2878 | 1296.0283 | 3.46E-23 | -0.764810515 | YIL002C | INP51 | n |
| W303.asm02.HP0_G0027220 | 1145.1994 | 1211.4023 | 1033.539791 | 673.368506 | 711.89912 | 750.933563 | 774.12379 | 772.81699 | 770.56377 | 4.16E-15 | 0.666546197 | YIL001W | | n |
| W303.asm02.HP0_G0027230 | 934.49113 | 874.036619 | 800.751857 | 2280.95248 | 2307.3801 | 2442.43038 | 2215.2025 | 2137.3835 | 2164.4554 | 6.64E-85 | -1.430707162 | YIR001C | SGN1 | n |

| | | | | | | | | | | | | | | |
|---|---|---|---|---|---|---|---|---|---|---|---|---|---|---|
| W303.asm02.HP0_G0027240 | 1007.1855 | 1096.17894 | 959.1635634 | 525.860801 | 540.21638 | 557.511281 | 513.04228 | 438.23578 | 518.41132 | 2.17E-23 | 0.915660349 | YIR002C | MPH1 | n |
| W303.asm02.HP0_G0027250 | 652.14206 | 739.090348 | 766.9444807 | 585.863935 | 578.86746 | 561.303875 | 489.10031 | 354.26246 | 516.64802 | 0.009517458 | 0.32255785 | YIR003W | AIM21 | n |
| W303.asm02.HP0_G0027260 | 2527.4456 | 2459.13628 | 2611.861304 | 2166.77985 | 2066.4849 | 2054.63766 | 1804.7687 | 1977.3093 | 1954.6223 | 2.46E-05 | 0.27302487 | YIR004W | DJP1 | n |
| W303.asm02.HP0_G0027270 | 458.29047 | 481.654384 | 678.0793771 | 222.511623 | 197.74975 | 201.007469 | 192.67588 | 204.68497 | 204.54324 | 2.56E-20 | 1.381351536 | YIR005W | IST3 | n |
| W303.asm02.HP0_G0027280 | 1135.7175 | 1432.50657 | 1313.658053 | 1684.25464 | 1703.3445 | 1902.93391 | 1574.4697 | 1314.7073 | 1668.0854 | 3.65E-05 | -0.446289676 | YIR006C | PAN1 | n |
| W303.asm02.HP0_G0027290 | 205.45055 | 275.082784 | 230.8560842 | 255.01332 | 240.89516 | 240.829703 | 352.28903 | 300.46705 | 357.95068 | 0.716593616 | -0.051083449 | YIR007W | EGH1 | n |
| W303.asm02.HP0_G0027300 | 737.47891 | 685.111839 | 678.0793771 | 697.536435 | 672.34917 | 642.844641 | 644.15308 | 635.04825 | 686.80684 | 0.486340654 | 0.060572794 | YIR008C | PRI1 | n |
| W303.asm02.HP0_G0027310 | 468.82588 | 443.246599 | 344.8352388 | 93.3382088 | 88.088527 | 117.570406 | 78.666483 | 47.234994 | 50.254159 | 2.94E-30 | 2.073180978 | YIR009W | MSL1 | n |
| W303.asm02.HP0_G0027320 | 271.81366 | 325.947148 | 340.9715386 | 330.850615 | 349.65752 | 365.037149 | 352.28903 | 299.15496 | 331.50112 | 0.205800631 | -0.153352232 | YIR010W | DSN1 | n |
| W303.asm02.HP0_G0027330 | 2579.0691 | 2306.54319 | 2531.689526 | 1509.2455 | 1451.663 | 1442.13377 | 1573.3297 | 1327.8282 | 1418.5779 | 1.47E-21 | 0.752244251 | YIR011C | STS1 | n |
| W303.asm02.HP0_G0027340 | 1906.9098 | 1724.19813 | 1993.66928 | 2959.32124 | 2714.5648 | 3079.58613 | 2154.7776 | 2643.8476 | 2097.4499 | 8.22E-11 | -0.637854746 | YIR012W | SQT1 | n |
| W303.asm02.HP0_G0027350 | 16.856661 | 22.8370613 | 16.42072566 | 119.172892 | 122.2453 | 134.637078 | 181.27494 | 181.06748 | 196.60838 | 2.09E-29 | -2.746043325 | YIR013C | GAT4 | n |
| W303.asm02.HP0_G0027360 | 35.820404 | 46.7121709 | 57.95550232 | 265.013843 | 212.13156 | 240.829703 | 345.44847 | 325.39663 | 380.87363 | 1.09E-38 | -2.347770368 | YIR014W | VLD1 | n |
| W303.asm02.HP0_G0027370 | 168.56661 | 228.370613 | 235.6857094 | 525.027424 | 568.08111 | 541.392758 | 574.60735 | 633.73617 | 633.90772 | 3.03E-30 | -1.36695999 | YIR015W | RPR2 | n |
| W303.asm02.HP0_G0027390 | 482.52192 | 511.757783 | 492.6217697 | 42.5022201 | 48.538576 | 37.9259375 | 71.825919 | 45.922911 | 79.348672 | 7.57E-91 | 3.525518161 | YIR017C | MET28 | n |
| W303.asm02.HP0_G0027400 | 173.83431 | 208.647697 | 174.832432 | 386.686865 | 311.9053 | 368.829742 | 310.10555 | 270.28913 | 309.45982 | 1.19E-12 | -0.938690918 | YIR018W | YAP5 | n |
| W303.asm02.HP0_G0027410 | 62.158937 | 79.9297146 | 78.23992813 | 176.675895 | 175.27819 | 202.903766 | 280.46311 | 169.25873 | 249.50749 | 1.40E-11 | -1.330122499 | YIR019C | FLO11 | n |
| W303.asm02.HP0_G0027460 | 1673.0236 | 1731.46447 | 1783.097621 | 1188.39541 | 1162.2292 | 1185.18555 | 1560.7886 | 1510.2077 | 1527.0211 | 2.09E-18 | 0.553262975 | YIR022W | SEC11 | n |
| W303.asm02.HP0_G0027470 | 444.59443 | 523.176314 | 534.1565464 | 2249.28416 | 2225.5836 | 2285.03774 | 1862.9135 | 1818.5473 | 1937.8709 | 2.69E-166 | -2.168940066 | YIR023W | DAL81 | n |
| W303.asm02.HP0_G0027480 | 980.84695 | 1013.13508 | 1169.735222 | 955.883263 | 921.33409 | 964.266961 | 848.2299 | 872.53531 | 861.37392 | 0.067189436 | 0.155834971 | YIR024C | INA22 | n |
| W303.asm02.HP0_G0027490 | 214.92243 | 252.245723 | 214.4353586 | 403.354402 | 391.0052 | 351.76307 | 428.67533 | 456.60494 | 392.3351 | 2.02E-10 | -0.751481058 | YIR025W | MND2 | n |
| W303.asm02.HP0_G0027500 | 799.63785 | 764.003506 | 680.9771522 | 1209.22983 | 1142.4543 | 1129.24479 | 712.55872 | 904.0253 | 744.11421 | 3.13E-10 | -0.633904977 | YIR026C | YVH1 | n |
| W303.asm02.HP0_G0027510 | 114.836 | 108.995065 | 85.96732844 | 163.341865 | 206.73838 | 151.70375 | 182.41503 | 164.0104 | 180.73864 | 1.19E-05 | -0.755259562 | YIR027C | DAL1 | n |
| W303.asm02.HP0_G0027530 | 170.67369 | 177.506249 | 172.9005819 | 434.189346 | 406.28586 | 376.41493 | 498.22106 | 572.06826 | 465.51221 | 7.05E-24 | -1.224757312 | YIR029W | DAL2 | n |
| W303.asm02.HP0_G0027540 | 776.45994 | 701.720611 | 856.7755093 | 352.518413 | 403.58927 | 394.42975 | 428.67533 | 368.69537 | 406.44153 | 7.33E-22 | 1.022685751 | YIR030C | DCG1 | n |
| W303.asm02.HP0_G0027560 | 446.70151 | 437.01831 | 510.0084204 | 142.507444 | 161.79525 | 163.081531 | 134.53109 | 179.75539 | 147.23587 | 6.50E-33 | 1.578757288 | YIR032C | DAL3 | n |
| W303.asm02.HP0_G0027570 | 1425.4414 | 1505.16995 | 1651.731816 | 2021.77227 | 2024.2384 | 2116.26731 | 2090.9323 | 1751.631 | 2196.1949 | 3.83E-06 | -0.426919425 | YIR033W | MGA2 | n |
| W303.asm02.HP0_G0027580 | 1066.1838 | 1202.05986 | 1183.258172 | 687.535913 | 657.96737 | 644.740938 | 1030.6449 | 791.18615 | 981.27858 | 5.97E-15 | 0.794104116 | YIR034C | LYS1 | n |
| W303.asm02.HP0_G0027590 | 1657.2205 | 1704.47521 | 1794.688722 | 748.372424 | 854.81826 | 676.977985 | 286.16358 | 338.51746 | 310.34147 | 9.80E-35 | 1.176987229 | YIR035C | NRE1 | n |
| W303.asm02.HP0_G0027610 | 3164.8381 | 3234.55832 | 3481.193839 | 2103.4432 | 2062.8895 | 2168.41548 | 2551.5303 | 3415.3525 | 2454.5189 | 4.30E-09 | 0.641435303 | YIR037W | HYR1 | n |
| W303.asm02.HP0_G0027620 | 1207.3583 | 1209.3262 | 1022.914616 | 1496.74485 | 1544.2458 | 1431.70414 | 1769.4258 | 1486.5902 | 1827.6644 | 9.64E-05 | -0.37955153 | YIR038C | GTT1 | n |
| W303.asm02.HP0_G0027720 | 560.48397 | 585.459208 | 490.6899196 | 58.3363805 | 49.437439 | 56.8889063 | 134.53109 | 77.412907 | 119.90466 | 5.94E-79 | 3.311933466 | YJL218W | | n |
| W303.asm02.HP0_G0027770 | 1028.2563 | 1115.90186 | 1172.632997 | 759.206323 | 734.37068 | 660.859461 | 522.16303 | 656.04158 | 477.85534 | 3.12E-08 | 0.622182179 | YJL212C | OPT1 | n |
| W303.asm02.HP0_G0027780 | 694.28372 | 722.481576 | 716.7163787 | 458.357275 | 462.01534 | 447.526063 | 356.84941 | 284.72205 | 379.99197 | 2.61E-10 | 0.64116361 | YJL210W | PEX2 | n |
| W303.asm02.HP0_G0027800 | 2360.9861 | 2101.00964 | 2323.049718 | 1011.71951 | 1066.9498 | 1228.80038 | 645.29318 | 991.93488 | 632.14442 | 2.38E-13 | 1.036986489 | YJL208C | NUC1 | n |
| W303.asm02.HP0_G0027810 | 1372.7643 | 1312.09298 | 1496.217885 | 1668.42048 | 1671.8843 | 1741.74868 | 1506.0641 | 1361.9423 | 1490.8734 | 0.000205775 | -0.280892799 | YJL207C | LAA1 | n |
| W303.asm02.HP0_G0027820 | 389.81028 | 384.077849 | 316.8234127 | 443.356492 | 465.61079 | 534.755719 | 587.14838 | 486.78286 | 571.31044 | 0.000851458 | -0.404729137 | YJL206C | | n |
| W303.asm02.HP0_G0027840 | 809.11972 | 715.215238 | 886.7191855 | 888.379737 | 988.74877 | 954.785477 | 938.29732 | 934.20322 | 917.79964 | 0.012171604 | -0.230972901 | YJL204C | RCY1 | n |
| W303.asm02.HP0_G0027850 | 824.92284 | 839.781027 | 789.1607566 | 496.692611 | 513.2505 | 469.333477 | 433.2357 | 434.29953 | 454.93239 | 3.48E-18 | 0.729305901 | YJL203W | PRP21 | n |
| W303.asm02.HP0_G0027860 | 381.38195 | 387.191994 | 402.7907411 | 645.033693 | 649.8776 | 661.80761 | 662.39459 | 708.52491 | 691.2151 | 5.72E-18 | -0.73944713 | YJL201W | ECM25 | n |
| W303.asm02.HP0_G0027870 | 1883.7318 | 1917.2751 | 2334.640818 | 789.20789 | 751.44907 | 856.178039 | 824.28793 | 680.97116 | 879.88861 | 2.44E-34 | 1.356690031 | YJL200C | ACO2 | n |
| W303.asm02.HP0_G0027900 | 701.65851 | 713.139142 | 820.0703578 | 1816.76156 | 1841.7693 | 2071.70434 | 2128.5554 | 1982.5577 | 2142.4141 | 1.74E-55 | -1.357105255 | YJL197W | UBP12 | n |
| W303.asm02.HP0_G0027920 | 245.47512 | 264.702302 | 291.7093617 | 530.861062 | 539.31751 | 520.533492 | 535.84416 | 600.93409 | 644.48755 | 5.38E-20 | -0.986792469 | YJL194W | CDC6 | n |
| W303.asm02.HP0_G0027930 | 654.24915 | 581.307015 | 644.2720008 | 345.851399 | 361.34273 | 326.163063 | 360.26969 | 425.11495 | 369.41215 | 2.63E-17 | 0.862911228 | YJL193W | | n |
| W303.asm02.HP0_G0027940 | 1442.298 | 1491.67532 | 1370.64763 | 990.885091 | 1031.8942 | 999.348453 | 1331.6297 | 1398.6807 | 1390.3651 | 1.22E-13 | 0.510075953 | YJL192C | SOP4 | n |
| W303.asm02.HP0_G0027960 | 11405.638 | 11110.2303 | 12895.09927 | 6386.16691 | 6920.3426 | 7057.06882 | 5670.8273 | 6086.7538 | 5161.1903 | 3.16E-22 | 0.798352187 | YJL190C | RPS22A | y |
| W303.asm02.HP0_G0027980 | 668.99873 | 732.862058 | 684.8408524 | 1467.57666 | 1564.9197 | 1413.68932 | 1555.0881 | 1435.419 | 1521.7312 | 2.10E-45 | -1.091560884 | YJL187C | SWE1 | n |
| W303.asm02.HP0_G0027990 | 1785.7525 | 1574.71918 | 1794.688722 | 1990.93733 | 2014.3509 | 1958.87467 | 2263.0865 | 2672.7134 | 2302.8748 | 0.013060666 | -0.210196511 | YJL186W | MNN5 | n |
| W303.asm02.HP0_G0028010 | 1221.0544 | 1248.77203 | 1478.831234 | 503.359626 | 515.94709 | 537.600164 | 500.50124 | 539.26618 | 515.76637 | 2.56E-46 | 1.343824538 | YJL184W | GON7 | n |
| W303.asm02.HP0_G0028020 | 2149.2243 | 1853.95416 | 1879.690125 | 3836.0337 | 3715.8977 | 4017.30493 | 4136.2609 | 4193.4178 | 4105.853 | 4.00E-43 | -0.975910592 | YJL183W | MNN11 | n |
| W303.asm02.HP0_G0028040 | 1410.6918 | 1382.68026 | 1304.964727 | 1670.08724 | 1536.1561 | 1533.15602 | 1300.8472 | 1499.7111 | 1258.1173 | 0.012646956 | -0.210262609 | YJL180C | ATP12 | n |
| W303.asm02.HP0_G0028050 | 1274.785 | 1289.25592 | 1585.082988 | 1003.38574 | 1032.793 | 975.644743 | 880.15253 | 1086.4049 | 969.8171 | 8.39E-06 | 0.462634148 | YJL179W | PFD1 | n |
| W303.asm02.HP0_G0028060 | 2015.4245 | 2005.5092 | 2113.443985 | 1509.2455 | 1433.6857 | 1406.10413 | 2274.4874 | 2166.2493 | 2213.828 | 4.93E-15 | 0.495966904 | YJL178C | ATG27 | n |
| W303.asm02.HP0_G0028070 | 7819.3835 | 8200.58111 | 7633.70558 | 9152.14472 | 9009.2991 | 10249.4846 | 8396.792 | 7997.1469 | 7408.521 | 0.00033338 | -0.264332166 | YJL177W | RPL17B | y |
| W303.asm02.HP0_G0028080 | 814.38742 | 917.634645 | 960.1294884 | 2201.78167 | 2248.0552 | 2403.55629 | 1972.3625 | 1654.5369 | 1936.9893 | 2.72E-50 | -1.347304181 | YJL176C | SWI3 | n |
| W303.asm02.HP0_G0028090 | 3294.4236 | 3010.3399 | 2848.512939 | 1773.42597 | 1762.6694 | 1866.90427 | 1883.4352 | 2003.551 | 1950.214 | 2.46E-26 | 0.760461178 | YJL174W | KRE9 | n |
| W303.asm02.HP0_G0028100 | 1492.868 | 1372.29978 | 1488.490485 | 603.364849 | 586.05836 | 546.1335 | 549.52528 | 690.15575 | 610.98478 | 3.94E-47 | 1.326194834 | YJL173C | RFA3 | n |
| W303.asm02.HP0_G0028110 | 863.90387 | 958.118527 | 1070.244943 | 865.878562 | 886.27845 | 697.83725 | 879.01244 | 716.39741 | 797.89498 | 0.04372587 | 0.239241824 | YJL172W | CPS1 | n |
| W303.asm02.HP0_G0028130 | 146.44224 | 147.40285 | 98.52435394 | 12.500653 | 7.1909002 | 6.63703906 | 14.821221 | 17.057081 | 13.224779 | 3.88E-28 | 3.8795837 | YJL170C | ASG7 | n |
| W303.asm02.HP0_G0028140 | 811.2268 | 775.422036 | 707.0571283 | 1434.24158 | 1482.2243 | 1604.26716 | 1377.2335 | 1397.3686 | 1450.3174 | 1.56E-34 | -0.979212655 | YJL168C | SET2 | n |

| | | | | | | | | | | | | | | |
|---|---|---|---|---|---|---|---|---|---|---|---|---|---|---|
| W303.asm02.HP0_G0028150 | 9111.0252 | 9546.92968 | 7981.438594 | 6203.65737 | 6187.7696 | 6778.31318 | 5397.2048 | 5628.8368 | 5210.5628 | 1.07E-09 | 0.47471857 | YJL167W | ERG20 | n |
| W303.asm02.HP0_G0028160 | 2876.1677 | 2821.41512 | 3088.062348 | 6747.85247 | 6861.9165 | 7468.56524 | 6694.6317 | 6822.8325 | 6083.3982 | 1.47E-69 | -1.262246404 | YJL166W | QCR8 | n |
| W303.asm02.HP0_G0028180 | 298.15219 | 336.32763 | 261.7656855 | 2196.78141 | 2191.4268 | 2086.87471 | 2548.11 | 2759.3109 | 2495.9566 | 6.73E-217 | -2.855044708 | YJL164C | TPK1 | n |
| W303.asm02.HP0_G0028190 | 280.24199 | 330.099341 | 287.8456615 | 944.215987 | 972.56925 | 1056.23736 | 1054.5869 | 1154.6332 | 1067.6805 | 1.35E-73 | -1.726638339 | YJL163C | | n |
| W303.asm02.HP0_G0028200 | 672.15935 | 598.953835 | 561.2024474 | 914.21442 | 947.4011 | 950.992883 | 915.49544 | 835.79698 | 901.92991 | 1.33E-12 | -0.619024572 | YJL162C | JJJ2 | n |
| W303.asm02.HP0_G0028210 | 299.20573 | 285.463266 | 349.664864 | 114.17263 | 110.56009 | 114.725961 | 163.03344 | 131.20832 | 167.51386 | 9.66E-24 | 1.461873757 | YJL161W | FMP33 | n |
| W303.asm02.HP0_G0028220 | 66.373102 | 76.8155699 | 49.26217697 | 7063.7023 | 7068.6549 | 7297.89853 | 7229.3357 | 6021.1497 | 7375.0182 | 0 | -6.805227812 | YJL160C | PIR5 | n |
| W303.asm02.HP0_G0028250 | 1563.4553 | 1618.31721 | 1663.322917 | 170.842257 | 148.31232 | 159.288938 | 147.07212 | 154.82581 | 149.88083 | 5.92E-253 | 3.338715008 | YJL157C | FAR1 | n |
| W303.asm02.HP0_G0028260 | 1261.0889 | 1165.72818 | 1270.191426 | 723.371118 | 700.21391 | 702.577992 | 1063.7077 | 1162.5057 | 1060.6273 | 2.01E-25 | 0.798013074 | YJL156C | SSY5 | n |
| W303.asm02.HP0_G0028270 | 1115.7002 | 1025.59166 | 1098.256769 | 489.192219 | 409.88131 | 414.340867 | 596.26914 | 555.01118 | 608.33982 | 2.75E-44 | 1.300828081 | YJL155C | FBP26 | n |
| W303.asm02.HP0_G0028280 | 1409.6383 | 1654.6489 | 1699.062143 | 1205.89632 | 1180.2065 | 1235.43741 | 1422.8373 | 1401.3048 | 1445.0275 | 1.18E-06 | 0.39592373 | YJL154C | VPS35 | n |
| W303.asm02.HP0_G0028290 | 263.38532 | 349.822257 | 266.5953107 | 4711.91279 | 5803.0564 | 4937.00892 | 3823.8751 | 3214.6038 | 3481.6434 | 9.64E-273 | -4.135152106 | YJL153C | INO1 | n |
| W303.asm02.HP0_G0028310 | 353.98988 | 422.485634 | 363.1878145 | 605.86498 | 559.09249 | 679.82243 | 832.26858 | 686.2195 | 728.24448 | 1.39E-09 | -0.694563424 | YJL149W | DAS1 | n |
| W303.asm02.HP0_G0028320 | 4167.8094 | 3984.02915 | 4856.671094 | 2108.44347 | 1843.567 | 2048.94877 | 1412.5764 | 1590.2448 | 1522.6129 | 1.87E-35 | 1.116276061 | YJL148W | RPA34 | n |
| W303.asm02.HP0_G0028330 | 772.24577 | 708.986949 | 744.7282048 | 654.200838 | 654.37192 | 648.533531 | 623.63139 | 729.51824 | 722.07292 | 0.036186076 | 0.185598316 | YJL147C | SMT1 | n |
| W303.asm02.HP0_G0028340 | 691.12309 | 651.894296 | 650.067551 | 451.69026 | 403.58927 | 393.481602 | 640.7328 | 497.27952 | 616.27469 | 5.35E-10 | 0.673188801 | YJL146W | IDS2 | n |
| W303.asm02.HP0_G0028350 | 1980.6576 | 1858.10635 | 1841.053124 | 633.366417 | 657.0685 | 682.666875 | 588.28848 | 645.54492 | 619.80129 | 9.61E-93 | 1.52590356 | YJL145W | SFH5 | n |
| W303.asm02.HP0_G0028370 | 3290.2095 | 2964.66578 | 3639.605546 | 2291.78638 | 2222.887 | 2304.94885 | 2236.8643 | 2244.9743 | 2181.2068 | 6.06E-12 | 0.537196976 | YJL143W | TIM17 | n |
| W303.asm02.HP0_G0028380 | 364.52529 | 530.442651 | 484.8943694 | 1405.90677 | 1321.3279 | 1304.65225 | 1724.9621 | 1439.3552 | 1626.6478 | 4.28E-39 | -1.54641053 | YJL141C | YAK1 | n |
| W303.asm02.HP0_G0028390 | 3442.973 | 3622.78836 | 3807.676502 | 1352.57065 | 1241.3291 | 1282.84484 | 1355.5717 | 1247.7911 | 1341.8742 | 2.11E-100 | 1.487722307 | YJL140W | RPB4 | n |
| W303.asm02.HP0_G0028400 | 678.4806 | 723.519624 | 723.4778539 | 1035.88744 | 957.28859 | 936.770657 | 1026.0846 | 993.24696 | 1011.2547 | 2.45E-09 | -0.463500284 | YJL139C | YUR1 | n |
| W303.asm02.HP0_G0028410 | 8672.752 | 8424.79953 | 8039.394096 | 17754.2607 | 17426.248 | 18907.028 | 14911.289 | 15750.246 | 14355.938 | 8.54E-72 | -1.105525428 | YKR059W | TIF1 | y |
| W303.asm02.HP0_G0028420 | 230.72554 | 260.550109 | 220.2309088 | 183.34291 | 157.30094 | 117.570406 | 225.7386 | 182.37956 | 268.02218 | 0.000426537 | 0.631067922 | YJL137C | GLG2 | n |
| W303.asm02.HP0_G0028450 | 757.49619 | 653.970392 | 731.2052542 | 1501.74511 | 1375.2597 | 1412.74117 | 1762.5852 | 1746.3827 | 1836.4809 | 2.75E-37 | -1.00169777 | YJL134W | LCB3 | n |
| W303.asm02.HP0_G0028460 | 721.67579 | 789.954712 | 652.9653261 | 100.005224 | 119.54872 | 71.1111328 | 30.782537 | 48.547077 | 46.727551 | 9.17E-70 | 2.893159629 | YJL133C-A | DPI8 | n |
| W303.asm02.HP0_G0028490 | 907.09906 | 908.292211 | 887.6851105 | 701.703319 | 714.59571 | 664.652055 | 617.93092 | 627.17575 | 640.07929 | 6.95E-07 | 0.376729719 | YJL131C | AIM23 | n |
| W303.asm02.HP0_G0028500 | 3310.2268 | 3619.67422 | 3973.815609 | 8157.09274 | 7886.6198 | 8724.86192 | 5165.7657 | 5636.7093 | 5472.4134 | 2.57E-52 | -1.1833962 | YJL130C | URA2 | n |
| W303.asm02.HP0_G0028520 | 962.93675 | 1201.02182 | 1127.23452 | 2447.62785 | 2367.6039 | 2432.94889 | 2340.6129 | 2322.3872 | 2396.3299 | 1.25E-45 | -1.138683319 | YJL128C | PBS2 | n |
| W303.asm02.HP0_G0028530 | 917.63447 | 889.607343 | 1011.323515 | 261.680335 | 266.96217 | 286.340828 | 272.48245 | 185.00373 | 246.86254 | 6.80E-52 | 1.79127253 | YJL127C-B | MCO6 | n |
| W303.asm02.HP0_G0028540 | 132.7462 | 166.087719 | 177.7302071 | 495.025857 | 540.21638 | 589.748328 | 524.44322 | 564.19576 | 562.49392 | 8.22E-47 | -1.766047579 | YJL127C | SPT10 | n |
| W303.asm02.HP0_G0028550 | 469.87942 | 444.284647 | 466.5417937 | 440.856361 | 415.27449 | 440.889024 | 565.4866 | 402.80953 | 502.54159 | 0.457476936 | 0.090237896 | YJL126W | NIT2 | n |
| W303.asm02.HP0_G0028560 | 1552.9199 | 1496.86556 | 1438.262383 | 1251.73205 | 1235.0371 | 1234.48927 | 745.62144 | 900.08905 | 788.19681 | 0.000675668 | 0.269973604 | YJL125C | GCD14 | n |
| W303.asm02.HP0_G0028570 | 2829.8119 | 2879.54582 | 2582.883553 | 1788.42675 | 1847.1625 | 1762.60795 | 1754.6046 | 1894.6481 | 1819.7295 | 6.12E-20 | 0.618956561 | YJL124C | LSM1 | n |
| W303.asm02.HP0_G0028580 | 2501.107 | 2539.066 | 2354.925244 | 2253.45104 | 2185.1348 | 2321.06738 | 2344.0332 | 2370.9343 | 2217.3546 | 0.044591845 | 0.129492764 | YJL123C | MTC1 | n |
| W303.asm02.HP0_G0028590 | 2957.2904 | 2896.15459 | 3209.768903 | 2321.78794 | 2261.5381 | 2420.62296 | 1365.8326 | 1940.571 | 1444.1458 | 0.000904428 | 0.371994243 | YJL122W | ALB1 | n |
| W303.asm02.HP0_G0028600 | 1404.3706 | 1419.01195 | 1278.884751 | 2517.63151 | 2414.3447 | 2452.86001 | 2541.2694 | 2566.4347 | 2551.5006 | 3.76E-40 | -0.848846006 | YJL121C | RPE1 | n |
| W303.asm02.HP0_G0028620 | 1745.7179 | 1848.76392 | 1678.777717 | 5327.77829 | 5195.4254 | 5211.02381 | 5292.3161 | 6001.4684 | 5227.3142 | 3.81E-107 | -1.577397889 | YJL117W | PHO86 | n |
| W303.asm02.HP0_G0028640 | 781.72764 | 784.764471 | 921.4924868 | 1740.09089 | 1705.1422 | 1751.23016 | 1821.8701 | 1855.2856 | 1892.025 | 7.90E-43 | -1.061409467 | YJL115W | ASF1 | n |
| W303.asm02.HP0_G0028650 | 1086.2011 | 1087.87456 | 1035.471641 | 992.551845 | 993.24309 | 1085.62996 | 1060.2874 | 963.06905 | 1005.0832 | 0.417832829 | 0.063760111 | YJL112W | MDV1 | n |
| W303.asm02.HP0_G0028660 | 5007.4818 | 4823.81018 | 4729.168989 | 5001.92794 | 5004.8665 | 5147.49787 | 4463.4678 | 4533.2473 | 4496.4248 | 0.273636178 | -0.057713142 | YJL111W | CCT7 | n |
| W303.asm02.HP0_G0028670 | 2697.0657 | 2588.89231 | 2974.083194 | 3255.17003 | 3182.8722 | 3508.14922 | 2163.8983 | 2332.8839 | 2145.0591 | 0.000388554 | -0.267577235 | YJL109C | UTP10 | n |
| W303.asm02.HP0_G0028680 | 198.06576 | 169.201863 | 190.2872326 | 2299.28677 | 1911.8806 | 1938.01541 | 2475.144 | 2365.686 | 2695.2099 | 5.45E-220 | -3.463447068 | YJL107C | | n |
| W303.asm02.HP0_G0028690 | 109.5683 | 122.489692 | 108.1836043 | 141.674067 | 133.93052 | 133.68893 | 174.43437 | 164.0104 | 171.04047 | 0.081975562 | -0.268118962 | YJL106W | IME2 | n |
| W303.asm02.HP0_G0028700 | 190.69098 | 208.647697 | 211.5375835 | 855.044662 | 964.47949 | 1002.1929 | 1054.5869 | 1047.0424 | 1082.6685 | 2.33E-106 | -2.205744371 | YJL105W | SET4 | n |
| W303.asm02.HP0_G0028710 | 2440.0017 | 2499.62017 | 2518.166576 | 1037.5542 | 988.74877 | 1059.08181 | 1005.5629 | 1136.264 | 980.39693 | 3.33E-68 | 1.273463901 | YJL104W | PAM16 | n |
| W303.asm02.HP0_G0028720 | 396.13153 | 346.708113 | 358.3581893 | 92.5048319 | 76.403314 | 92.9185469 | 68.405637 | 62.979992 | 68.768849 | 2.27E-45 | 2.071771588 | YJL103C | GSM1 | n |
| W303.asm02.HP0_G0028730 | 377.16779 | 449.474889 | 458.8143933 | 1085.05668 | 1056.1635 | 1207.94111 | 917.77563 | 956.50863 | 954.82902 | 5.33E-48 | -1.379976463 | YJL102W | MEF2 | n |
| W303.asm02.HP0_G0028740 | 1636.1496 | 1816.58442 | 1982.078179 | 3486.8488 | 3384.2174 | 3721.48262 | 3239.0069 | 3142.4392 | 3381.1351 | 9.92E-34 | -0.962163411 | YJL101C | GSH1 | n |
| W303.asm02.HP0_G0028750 | 455.12984 | 522.138265 | 516.7698957 | 483.358581 | 444.03809 | 432.355688 | 573.46726 | 518.27285 | 619.80129 | 0.204058591 | 0.135452869 | YJL100W | LSB6 | n |
| W303.asm02.HP0_G0028760 | 622.64291 | 592.725546 | 707.0571283 | 1297.56778 | 1178.4088 | 1158.63739 | 1376.0934 | 1193.9957 | 1389.4834 | 8.40E-22 | -0.918458795 | YJL099W | CHS6 | n |
| W303.asm02.HP0_G0028780 | 1404.3706 | 1337.00613 | 1176.496697 | 836.710371 | 866.50347 | 828.681735 | 1138.9539 | 1149.3849 | 1102.9465 | 1.80E-14 | 0.629150925 | YJL097W | PHS1 | n |
| W303.asm02.HP0_G0028790 | 1968.0151 | 1947.3785 | 2415.778522 | 1499.24498 | 1554.1333 | 1430.75599 | 1374.9533 | 1561.379 | 1503.2165 | 1.35E-07 | 0.497949128 | YJL096W | MRPL49 | n |
| W303.asm02.HP0_G0028800 | 329.75843 | 394.458332 | 398.927041 | 1002.55237 | 1028.2987 | 987.970672 | 978.20061 | 957.82071 | 1061.5089 | 6.63E-54 | -1.425286445 | YJL095W | BCK1 | n |
| W303.asm02.HP0_G0028810 | 594.19729 | 594.801642 | 613.3623995 | 2650.13843 | 2808.0465 | 2993.30462 | 3388.3592 | 3259.2146 | 3113.9946 | 4.43E-192 | -2.228679156 | YJL094C | KHA1 | n |
| W303.asm02.HP0_G0028820 | 902.88489 | 881.302957 | 854.8436592 | 394.187257 | 414.37562 | 409.600125 | 497.08096 | 440.85994 | 423.19292 | 1.26E-35 | 1.115494796 | YJL093C | TOK1 | n |
| W303.asm02.HP0_G0028830 | 515.1817 | 563.660195 | 544.7817218 | 594.197704 | 586.05836 | 632.415008 | 603.1097 | 566.81993 | 621.5646 | 0.07592622 | -0.158190841 | YJL092W | SRS2 | n |
| W303.asm02.HP0_G0028840 | 1806.8233 | 1677.48596 | 1777.302071 | 790.041267 | 827.85238 | 819.20025 | 1199.3788 | 1103.4619 | 1186.7035 | 5.92E-55 | 1.110727769 | YJL091C | GWT1 | n |
| W303.asm02.HP0_G0028850 | 848.10075 | 728.709866 | 721.5460038 | 743.372163 | 677.74234 | 681.718727 | 612.23045 | 553.6991 | 618.03799 | 0.188415302 | 0.127090714 | YJL090C | DPB11 | n |
| W303.asm02.HP0_G0028860 | 228.61846 | 674.731357 | 250.174585 | 140.84069 | 145.61573 | 174.459313 | 194.95607 | 148.2654 | 176.33038 | 0.003163397 | 1.323571953 | YJL089W | SIP4 | n |
| W303.asm02.HP0_G0028870 | 1152.5742 | 996.526312 | 992.9709397 | 1275.0666 | 1279.0814 | 1371.97079 | 1195.9586 | 873.84739 | 1091.4851 | 0.003556482 | -0.321492318 | YJL088W | ARG3 | n |

| | | | | | | | | | | | | | | |
|---|---|---|---|---|---|---|---|---|---|---|---|---|---|---|
| W303.asm02.HP0_G0028880 | 1152.5742 | 949.814141 | 1004.56204 | 1540.08044 | 1581.998 | 1657.36347 | 1143.5142 | 1263.5361 | 1171.7154 | 4.04E-13 | -0.621426632 | YJL087C | TRL1 | n |
| W303.asm02.HP0_G0028890 | 1504.457 | 1428.35438 | 1288.544002 | 973.384177 | 1013.0181 | 976.592891 | 1078.5289 | 1073.284 | 1035.0593 | 5.07E-11 | 0.510106004 | YJL085W | EXO70 | n |
| W303.asm02.HP0_G0028910 | 139.06745 | 176.468201 | 147.7865309 | 582.530428 | 512.35164 | 526.222383 | 574.60735 | 506.4641 | 555.4407 | 1.52E-51 | -1.808041087 | YJL083W | TAX4 | n |
| W303.asm02.HP0_G0028930 | 1270.5708 | 1369.18563 | 1125.30267 | 1426.74119 | 1297.9575 | 1298.96336 | 1474.1415 | 1302.8986 | 1348.9274 | 0.297691891 | -0.09671267 | YJL081C | ARP4 | n |
| W303.asm02.HP0_G0028940 | 2571.6943 | 2990.61698 | 2996.29947 | 8005.41815 | 7954.0345 | 8102.87655 | 7355.8862 | 7313.5516 | 7297.4329 | 5.78E-110 | -1.491053338 | YJL080C | SCP160 | n |
| W303.asm02.HP0_G0028990 | 1216.8402 | 1348.42467 | 1217.065549 | 1165.06086 | 1178.4088 | 1229.74852 | 1281.4656 | 1237.2944 | 1306.6081 | 0.272182667 | 0.082180923 | YJL074C | SMC3 | n |
| W303.asm02.HP0_G0029000 | 1546.5986 | 1717.96984 | 1546.445987 | 2048.44033 | 2172.5507 | 2409.24518 | 2198.1011 | 2262.0314 | 2072.7636 | 1.58E-08 | -0.462407525 | YJL073W | JEM1 | n |
| W303.asm02.HP0_G0029010 | 1205.2512 | 1195.83157 | 1319.453603 | 290.015149 | 337.07345 | 413.392719 | 323.78668 | 341.14162 | 331.50112 | 2.51E-58 | 1.841249746 | YJL072C | PSF2 | n |
| W303.asm02.HP0_G0029020 | 860.74324 | 759.851313 | 764.0467055 | 336.684253 | 358.64615 | 388.74086 | 476.55927 | 370.00745 | 479.61864 | 1.06E-23 | 1.138149486 | YJL071W | ARG2 | n |
| W303.asm02.HP0_G0029040 | 1181.0198 | 1175.07061 | 1254.736625 | 1917.60016 | 1882.2181 | 2019.55617 | 1167.4562 | 1514.144 | 1352.454 | 4.80E-14 | -0.688192798 | YJL069C | UTP18 | n |
| W303.asm02.HP0_G0029050 | 1354.8541 | 1344.27247 | 1334.908403 | 1456.74276 | 1540.6504 | 1534.10417 | 1514.0448 | 1468.2211 | 1464.4238 | 0.009051435 | -0.16748128 | YJL068C | | n |
| W303.asm02.HP0_G0029060 | 571.01938 | 646.704054 | 627.8512751 | 1047.55472 | 1019.3101 | 1068.56329 | 1193.6784 | 1284.5294 | 1222.8512 | 2.68E-22 | -0.764060605 | YJL066C | MPM1 | n |
| W303.asm02.HP0_G0029070 | 454.0763 | 564.698243 | 359.3241144 | 357.518675 | 361.34273 | 367.881594 | 410.43382 | 309.65163 | 321.80295 | 0.028361584 | 0.341805432 | YJL065C | DLS1 | n |
| W303.asm02.HP0_G0029080 | 1487.6003 | 1508.28409 | 1426.671282 | 1297.56778 | 1303.3507 | 1309.39299 | 1227.8812 | 1363.2544 | 1271.3421 | 0.009983534 | 0.17740751 | YJL063C | MRPL8 | n |
| W303.asm02.HP0_G0029090 | 2171.3486 | 2197.54813 | 2346.231919 | 869.212069 | 835.04328 | 818.252102 | 810.6068 | 928.95488 | 869.30879 | 5.73E-80 | 1.412313314 | YJL062W-/ | COA3 | n |
| W303.asm02.HP0_G0029100 | 922.90218 | 988.221926 | 1033.539791 | 873.378953 | 808.07741 | 867.555821 | 940.57751 | 844.98156 | 934.55103 | 0.012116504 | 0.208558415 | YJL062W | LAS21 | n |
| W303.asm02.HP0_G0029110 | 2086.0118 | 1971.25361 | 1741.562845 | 1693.42179 | 1804.9159 | 1788.20795 | 1598.4117 | 1712.2685 | 1652.2157 | 0.098551464 | 0.133167373 | YJL061W | NUP82 | n |
| W303.asm02.HP0_G0029120 | 1107.2719 | 1114.86381 | 974.618364 | 1224.23061 | 1199.9815 | 1141.57072 | 1274.625 | 2283.0247 | 1318.0696 | 0.54213599 | -0.157718727 | YJL060W | BNA3 | n |
| W303.asm02.HP0_G0029130 | 509.91399 | 508.643638 | 506.1447202 | 274.180988 | 270.55762 | 298.666758 | 407.01354 | 391.00078 | 384.40023 | 1.15E-18 | 0.855274048 | YJL059W | YHC3 | n |
| W303.asm02.HP0_G0029140 | 350.82925 | 312.452521 | 358.3581893 | 246.679552 | 233.70426 | 172.563016 | 230.29898 | 237.48705 | 271.54879 | 6.85E-06 | 0.643386772 | YJL058C | BIT61 | n |
| W303.asm02.HP0_G0029150 | 679.53414 | 737.014251 | 618.1920247 | 1451.7425 | 1479.5277 | 1597.63012 | 1658.8367 | 1514.144 | 1495.2816 | 1.08E-40 | -1.154759324 | YJL057C | IKS1 | n |
| W303.asm02.HP0_G0029160 | 451.96922 | 553.279713 | 497.4513949 | 1389.23923 | 1546.9424 | 1596.68197 | 1240.4222 | 1283.2173 | 1216.6796 | 2.86E-68 | -1.59222724 | YJL056C | ZAP1 | n |
| W303.asm02.HP0_G0029180 | 1068.2909 | 1102.40723 | 943.7087627 | 1710.08932 | 1695.2547 | 1742.69683 | 1392.0547 | 1565.3152 | 1439.7376 | 7.64E-20 | -0.725694561 | YJL054W | TIM54 | n |
| W303.asm02.HP0_G0029190 | 1378.032 | 1372.29978 | 1198.712973 | 523.360671 | 550.10386 | 558.45943 | 615.65073 | 606.18242 | 621.5646 | 1.73E-51 | 1.274974948 | YJL053W | PEP8 | n |
| W303.asm02.HP0_G0029210 | 633.17832 | 631.133331 | 634.6127504 | 412.521548 | 383.8143 | 431.407539 | 665.81487 | 627.17575 | 710.61144 | 4.63E-12 | 0.6295957 | YJL051W | IRC8 | n |
| W303.asm02.HP0_G0029220 | 1218.9473 | 1231.12521 | 1406.386856 | 3147.66441 | 3092.0871 | 3426.60845 | 1697.5999 | 2016.6718 | 1665.4405 | 3.47E-47 | -1.325120213 | YJL050W | MTR4 | n |
| W303.asm02.HP0_G0029230 | 1332.7297 | 1305.86469 | 1439.228308 | 820.876211 | 902.45797 | 715.852071 | 717.1191 | 834.4849 | 814.64637 | 4.47E-14 | 0.741094853 | YJL049W | CHM7 | n |
| W303.asm02.HP0_G0029250 | 774.35286 | 819.020063 | 602.7372241 | 491.69235 | 517.74481 | 569.837211 | 461.73805 | 464.47744 | 500.77829 | 6.18E-05 | 0.475525825 | YJL047C | RTT101 | n |
| W303.asm02.HP0_G0029260 | 1260.0354 | 1127.32039 | 1271.157351 | 723.371118 | 728.97751 | 650.429828 | 617.93092 | 708.52491 | 668.29215 | 1.45E-19 | 0.798478405 | YJL046W | AIM22 | n |
| W303.asm02.HP0_G0029280 | 882.86761 | 891.683439 | 855.8095842 | 348.351529 | 310.10757 | 379.259375 | 486.82012 | 347.70204 | 512.23976 | 1.55E-25 | 1.342226229 | YJL044C | GYP6 | n |
| W303.asm02.HP0_G0029300 | 1902.6956 | 2208.96666 | 2097.023259 | 3483.51529 | 3572.0797 | 3532.80108 | 3207.0843 | 3238.2213 | 3232.1359 | 1.58E-31 | -0.769923013 | YJL042W | MHP1 | n |
| W303.asm02.HP0_G0029310 | 1764.6817 | 1754.30153 | 1963.725604 | 3493.51581 | 3572.9785 | 3641.83815 | 3465.8856 | 3256.5904 | 3164.2487 | 4.03E-45 | -0.965224121 | YJL039C | NUP192 | n |
| W303.asm02.HP0_G0029340 | 1634.0426 | 1538.38749 | 1553.207462 | 658.367722 | 756.84224 | 609.659446 | 726.23985 | 751.82366 | 799.65828 | 1.17E-42 | 1.222225337 | YJL036W | SNX4 | n |
| W303.asm02.HP0_G0029350 | 871.27865 | 998.602408 | 950.470238 | 410.854794 | 411.67904 | 416.237164 | 303.26499 | 364.75912 | 338.55433 | 1.96E-36 | 1.187259026 | YJL035C | TAD2 | n |
| W303.asm02.HP0_G0029360 | 13167.159 | 15438.8915 | 13334.59516 | 28370.6486 | 26986.55 | 31265.1947 | 33787.824 | 30314.37 | 29131.542 | 1.81E-33 | -1.04639014 | YJL034W | KAR2 | n |
| W303.asm02.HP0_G0029370 | 879.70699 | 819.020063 | 934.0495123 | 2190.1144 | 2226.4825 | 2385.54147 | 1237.0019 | 1574.4998 | 1205.2182 | 1.56E-40 | -1.368938325 | YJL033W | HCA4 | n |
| W303.asm02.HP0_G0029390 | 1276.8921 | 1213.47839 | 1451.785333 | 346.684775 | 356.84842 | 342.281586 | 290.72396 | 228.30247 | 239.80932 | 7.26E-75 | 1.914702179 | YJL030W | MAD2 | n |
| W303.asm02.HP0_G0029400 | 556.26981 | 599.991884 | 571.8276229 | 1549.24759 | 1541.5492 | 1488.59305 | 1556.2282 | 1451.164 | 1515.5596 | 9.99E-84 | -1.406199271 | YJL029C | VPS53 | n |
| W303.asm02.HP0_G0029430 | 2838.2403 | 2574.35964 | 2400.323721 | 5857.80598 | 5677.2157 | 5780.86103 | 5337.9199 | 5955.5455 | 5589.6731 | 7.55E-55 | -1.14850933 | YJL026W | RNR2 | n |
| W303.asm02.HP0_G0029440 | 221.24367 | 203.457455 | 230.8560842 | 560.862629 | 596.84472 | 595.437219 | 549.52528 | 569.4441 | 563.37557 | 5.00E-46 | -1.41759984 | YJL025W | RRN7 | n |
| W303.asm02.HP0_G0029460 | 501.48566 | 464.007564 | 430.8025672 | 734.205017 | 749.65134 | 770.84468 | 747.90163 | 931.57905 | 804.9482 | 7.52E-12 | -0.691952808 | YJL023C | PET130 | n |
| W303.asm02.HP0_G0029470 | 814.38742 | 1044.27653 | 968.8228137 | 1735.92401 | 1778.8489 | 1732.2672 | 1572.1896 | 1238.6065 | 1606.3698 | 6.81E-16 | -0.891746911 | YJL020C | BBC1 | n |
| W303.asm02.HP0_G0029480 | 336.07967 | 342.55592 | 379.6085402 | 423.355447 | 414.37562 | 493.985336 | 485.68002 | 482.84661 | 499.89663 | 0.002012817 | -0.329335487 | YJL019W | MPS3 | n |
| W303.asm02.HP0_G0029490 | 797.53076 | 780.612278 | 851.9458841 | 1891.76548 | 1921.7681 | 1889.65984 | 3508.0691 | 2852.4688 | 3327.3543 | 4.43E-46 | -1.230342676 | YJL016W | TPH3 | n |
| W303.asm02.HP0_G0029500 | 3985.5467 | 3647.70152 | 3799.949102 | 5076.93185 | 4705.5453 | 4842.19407 | 4682.3659 | 4697.2577 | 4625.1459 | 1.94E-09 | -0.355345969 | YJL014W | CCT3 | n |
| W303.asm02.HP0_G0029520 | 5947.2406 | 5585.73759 | 5370.543215 | 5384.44792 | 5515.4204 | 5217.66085 | 4652.7234 | 4437.4653 | 4458.5137 | 0.268410452 | 0.068495902 | YJL012C | VTC4 | n |
| W303.asm02.HP0_G0029530 | 852.31491 | 901.025874 | 1133.03007 | 567.529644 | 562.68794 | 658.015016 | 448.05692 | 384.44037 | 415.25805 | 5.96E-09 | 0.692127738 | YJL011C | RPC17 | n |
| W303.asm02.HP0_G0029540 | 1587.6867 | 1503.09385 | 1384.17058 | 1730.92375 | 1599.0764 | 1755.97091 | 853.93037 | 1052.2907 | 820.81793 | 0.062003371 | -0.184931799 | YJL010C | NOP9 | n |
| W303.asm02.HP0_G0029550 | 4863.1466 | 4308.93825 | 4309.957522 | 4643.57589 | 4338.8094 | 4606.10511 | 4076.976 | 4004.4778 | 4123.486 | 0.871480805 | -0.011518379 | YJL008C | CCT8 | n |
| W303.asm02.HP0_G0029560 | 884.97469 | 877.150764 | 1029.676091 | 650.033954 | 691.22528 | 672.237242 | 554.08566 | 583.87701 | 522.81958 | 2.23E-07 | 0.472520466 | YJL006C | CTK2 | n |
| W303.asm02.HP0_G0029570 | 996.65007 | 1205.17401 | 1072.176793 | 1548.41421 | 1534.3583 | 1573.92641 | 1815.0296 | 1592.869 | 1795.9249 | 3.34E-09 | -0.508195744 | YJL005W | CYR1 | n |
| W303.asm02.HP0_G0029580 | 1208.4119 | 1157.42379 | 1570.594113 | 418.355186 | 434.1506 | 381.155672 | 624.77149 | 637.67242 | 680.63528 | 1.16E-45 | 1.674037913 | YJL004C | SYS1 | n |
| W303.asm02.HP0_G0029590 | 795.42368 | 752.584975 | 932.1176623 | 495.025857 | 462.01534 | 451.318656 | 441.21636 | 568.13201 | 483.14525 | 6.01E-13 | 0.816453763 | YJL003W | COX16 | n |
| W303.asm02.HP0_G0029600 | 3702.1441 | 3401.68409 | 3300.565857 | 3531.01777 | 3483.0923 | 3556.50479 | 3807.9138 | 3892.9508 | 3605.9563 | 0.736038112 | -0.023065331 | YJL002C | OST1 | n |
| W303.asm02.HP0_G0029610 | 3106.8933 | 3007.22576 | 2724.874534 | 240.845914 | 205.83952 | 190.577836 | 223.45841 | 245.35955 | 189.55516 | 5.99E-268 | 3.79072403 | YJL001W | PRE3 | n |
| W303.asm02.HP0_G0029620 | 1288.481 | 1268.49495 | 1348.431354 | 1313.40194 | 1281.778 | 1279.05224 | 1633.7546 | 1624.359 | 1714.813 | 0.867631969 | 0.011694754 | YJR001W | AVT1 | n |
| W303.asm02.HP0_G0029630 | 1310.6054 | 1243.58179 | 1418.943882 | 2520.96501 | 2433.2208 | 2595.08227 | 1555.0881 | 1802.8023 | 1534.956 | 2.34E-29 | -0.92561493 | YJR002W | MPP10 | n |
| W303.asm02.HP0_G0029640 | 1500.2428 | 1441.84901 | 1400.591306 | 777.540614 | 826.95352 | 846.696555 | 888.13319 | 780.68949 | 911.62808 | 1.66E-24 | 0.82549307 | YJR003C | MRX12 | n |
| W303.asm02.HP0_G0029650 | 326.5978 | 316.604714 | 350.630789 | 230.845391 | 236.40084 | 241.777852 | 240.55982 | 265.0408 | 225.70289 | 2.10E-05 | 0.488471564 | YJR004C | SAG1 | n |
| W303.asm02.HP0_G0029660 | 1234.7504 | 1106.55943 | 1002.63019 | 1145.05981 | 1039.0851 | 1037.27439 | 970.21995 | 991.93488 | 1040.3493 | 0.577893477 | 0.052708599 | YJR005W | APL1 | n |

| | | | | | | | | | | | | | | |
|---|---|---|---|---|---|---|---|---|---|---|---|---|---|---|
| W303.asm02.HP0_G0029680 | 685.85539 | 743.242541 | 765.9785556 | 920.048058 | 911.4466 | 833.422477 | 822.00774 | 827.92448 | 912.50973 | 0.001226522 | -0.279868482 | YJR006W | POL31 | n |
| W303.asm02.HP0_G0029690 | 5638.553 | 5227.61094 | 4825.761493 | 6285.32831 | 5895.6393 | 6589.63164 | 5461.05 | 5690.5047 | 5344.5739 | 0.000476761 | -0.258568513 | YJR007W | SUI2 | n |
| W303.asm02.HP0_G0029700 | 584.71542 | 588.573353 | 568.9298477 | 111.6725 | 115.0544 | 106.192625 | 144.79193 | 160.07415 | 165.75056 | 3.22E-88 | 2.38669785 | YJR008W | MHO1 | n |
| W303.asm02.HP0_G0029720 | 874.43928 | 839.781027 | 796.8881569 | 299.182294 | 328.98368 | 299.614906 | 302.1249 | 216.49372 | 303.28826 | 2.49E-35 | 1.436292337 | YJR010W | MET3 | n |
| W303.asm02.HP0_G0029730 | 862.85032 | 754.661072 | 992.9709397 | 652.534084 | 622.91173 | 562.252024 | 734.2205 | 806.93115 | 767.91882 | 2.90E-06 | 0.506344708 | YJR010C-A | SPC1 | n |
| W303.asm02.HP0_G0029750 | 1011.3996 | 947.738044 | 884.7873354 | 718.370857 | 775.71836 | 689.303914 | 941.7176 | 961.75696 | 1034.1777 | 1.13E-05 | 0.380523436 | YJR013W | GPI14 | n |
| W303.asm02.HP0_G0029760 | 2242.9894 | 2495.46797 | 1895.144926 | 2152.61244 | 2261.5381 | 2306.84515 | 1445.6391 | 1701.7719 | 1463.5422 | 0.86031159 | -0.019077887 | YJR014W | TMA22 | n |
| W303.asm02.HP0_G0029800 | 4782.024 | 5214.11632 | 5735.662879 | 4712.74617 | 4665.0965 | 4826.07555 | 4937.7469 | 4029.4074 | 4876.4167 | 0.110360673 | 0.147546512 | YJR016C | ILV3 | n |
| W303.asm02.HP0_G0029810 | 1263.196 | 1292.37006 | 1263.429951 | 825.876472 | 824.25693 | 785.066907 | 814.02708 | 690.15575 | 780.26194 | 1.27E-16 | 0.64875183 | YJR017C | ESS1 | n |
| W303.asm02.HP0_G0029820 | 438.27318 | 483.730481 | 445.2914428 | 443.356492 | 457.52102 | 496.829781 | 649.85355 | 443.48411 | 609.22147 | 0.821358813 | -0.031262083 | YJR019C | TES1 | n |
| W303.asm02.HP0_G0029830 | 125.37141 | 122.489692 | 109.1495294 | 31.6683208 | 27.864738 | 30.34075 | 30.782537 | 14.432915 | 34.384425 | 2.05E-15 | 1.988222787 | YJR021C | REC107 | y |
| W303.asm02.HP0_G0029840 | 664.78456 | 739.090348 | 874.16216 | 340.01776 | 355.94956 | 351.76307 | 320.3664 | 364.75912 | 352.66076 | 4.96E-23 | 1.121778098 | YJR022W | LSM8 | n |
| W303.asm02.HP0_G0029850 | 2248.2571 | 2063.6399 | 2710.385658 | 561.696006 | 553.69931 | 577.422399 | 554.08566 | 534.01785 | 559.84896 | 1.10E-91 | 2.0530373 | YJR024C | MDE1 | n |
| W303.asm02.HP0_G0029870 | 415.09527 | 504.491445 | 490.6899196 | 3115.16272 | 2993.2122 | 2985.71943 | 2938.0221 | 2679.2738 | 2783.3751 | 1.56E-240 | -2.68815836 | YJR030C | RBH2 | n |
| W303.asm02.HP0_G0029890 | 1636.1496 | 1589.25186 | 1764.745046 | 670.868375 | 693.023 | 727.229852 | 494.80077 | 572.06826 | 499.89663 | 6.65E-52 | 1.255710659 | YJR032W | CPR7 | n |
| W303.asm02.HP0_G0029900 | 377.16779 | 441.170503 | 409.5522164 | 685.035782 | 736.16841 | 695.940953 | 834.54877 | 745.26324 | 856.96566 | 1.89E-16 | -0.785583028 | YJR033C | RAV1 | n |
| W303.asm02.HP0_G0029910 | 830.19054 | 851.199558 | 1031.607941 | 733.37164 | 804.48196 | 711.111328 | 751.32191 | 704.58866 | 726.48118 | 0.006799551 | 0.271413622 | YJR034W | PET191 | n |
| W303.asm02.HP0_G0029920 | 309.74114 | 370.583222 | 352.5626391 | 672.535129 | 644.48443 | 621.985375 | 598.54932 | 599.62201 | 646.25085 | 6.89E-21 | -0.908592358 | YJR035W | RAD26 | n |
| W303.asm02.HP0_G0029930 | 214.92243 | 193.076973 | 212.5035085 | 140.84069 | 139.32369 | 131.792633 | 220.03813 | 204.68497 | 257.44236 | 3.16E-05 | 0.590329193 | YJR036C | HUL4 | n |
| W303.asm02.HP0_G0029950 | 678.4806 | 732.862058 | 735.0689544 | 992.551845 | 1014.8158 | 993.659563 | 1164.0359 | 1092.9653 | 1090.6034 | 1.25E-10 | -0.483062384 | YJR040W | GEF1 | n |
| W303.asm02.HP0_G0029960 | 1300.07 | 1161.57598 | 1301.101027 | 2188.44764 | 2295.6949 | 2472.77113 | 1376.0934 | 1666.3456 | 1350.6907 | 4.24E-21 | -0.886295474 | YJR041C | URB2 | n |
| W303.asm02.HP0_G0029970 | 1675.1307 | 1548.76798 | 1442.126083 | 2336.78873 | 2352.3232 | 2412.08963 | 2316.6709 | 2313.2026 | 2285.2418 | 9.52E-19 | -0.606333251 | YJR042W | NUP85 | n |
| W303.asm02.HP0_G0029980 | 855.47554 | 741.166444 | 572.7935479 | 561.696006 | 587.85609 | 563.200172 | 506.20171 | 465.78953 | 515.76637 | 0.007862228 | 0.339967029 | YJR043C | POL32 | n |
| W303.asm02.HP0_G0029990 | 1998.5678 | 1960.87313 | 1850.712374 | 551.695484 | 551.00273 | 517.689047 | 844.80962 | 754.44782 | 879.88861 | 8.72E-114 | 1.841435259 | YJR044C | VPS55 | n |
| W303.asm02.HP0_G0030000 | 9480.8182 | 11872.1577 | 11112.96757 | 12255.6402 | 12383.629 | 13859.0857 | 13288.935 | 11483.352 | 11790.331 | 0.00919766 | -0.245799344 | YJR045C | SSC1 | n |
| W303.asm02.HP0_G0030010 | 723.78287 | 1094.10285 | 811.3770324 | 688.36929 | 711.00026 | 749.985414 | 559.78613 | 658.66575 | 552.79575 | 0.033325077 | 0.290894536 | YJR046W | TAH11 | n |
| W303.asm02.HP0_G0030020 | 6576.2048 | 6158.74022 | 6803.010047 | 7513.7258 | 7793.1381 | 8373.09885 | 7471.0357 | 7537.9178 | 6801.9445 | 9.07E-05 | -0.277214797 | YJR048W | CYC1 | n |
| W303.asm02.HP0_G0030030 | 1030.3634 | 1000.6785 | 1031.607941 | 596.697835 | 587.85609 | 511.052008 | 617.93092 | 594.37368 | 610.98478 | 2.90E-24 | 0.851696046 | YJR049C | UTR1 | n |
| W303.asm02.HP0_G0030040 | 1022.9886 | 1057.77116 | 1020.016841 | 602.531473 | 586.95723 | 576.47425 | 509.622 | 568.13201 | 575.7187 | 2.95E-24 | 0.811648559 | YJR050W | ISY1 | n |
| W303.asm02.HP0_G0030060 | 750.12141 | 742.204493 | 663.5905015 | 642.533562 | 649.8776 | 747.140969 | 717.1191 | 725.58199 | 678.87197 | 0.40265779 | 0.080361844 | YJR052W | RAD7 | n |
| W303.asm02.HP0_G0030070 | 208.60118 | 180.620394 | 170.0028068 | 685.035782 | 693.023 | 621.037227 | 680.63609 | 682.28325 | 692.9784 | 9.02E-68 | -1.840605478 | YJR053W | BFA1 | n |
| W303.asm02.HP0_G0030080 | 2216.6509 | 2052.22137 | 2239.01424 | 938.382349 | 894.36821 | 856.178039 | 799.20586 | 932.89113 | 819.05463 | 7.80E-57 | 1.274674233 | YJR054W | KCH1 | n |
| W303.asm02.HP0_G0030090 | 699.55142 | 776.460085 | 749.55783 | 299.182294 | 304.7144 | 287.288977 | 174.43437 | 220.42997 | 150.76248 | 1.81E-30 | 1.320172323 | YJR055W | HIT1 | n |
| W303.asm02.HP0_G0030110 | 1187.341 | 1222.82083 | 1460.478658 | 538.361454 | 515.94709 | 531.911274 | 468.57861 | 464.47744 | 501.65994 | 8.60E-41 | 1.287515802 | YJR057W | CDC8 | n |
| W303.asm02.HP0_G0030120 | 1007.1855 | 959.156575 | 1183.258172 | 425.855577 | 393.70178 | 384.000117 | 458.31777 | 486.78286 | 509.59481 | 7.36E-42 | 1.387609006 | YJR058C | APS2 | n |
| W303.asm02.HP0_G0030140 | 823.8693 | 853.275654 | 867.4006847 | 1329.2361 | 1249.4189 | 1334.04485 | 1360.1321 | 1144.1365 | 1386.8385 | 7.81E-13 | -0.620554249 | YJR060W | CBF1 | n |
| W303.asm02.HP0_G0030150 | 482.52192 | 492.034866 | 475.235119 | 205.010709 | 213.92928 | 201.955617 | 224.59851 | 203.37289 | 253.0341 | 5.72E-28 | 1.22314808 | YJR061W | MNN14 | n |
| W303.asm02.HP0_G0030160 | 952.40134 | 825.248352 | 764.0467055 | 638.366678 | 597.74358 | 575.526102 | 800.34595 | 737.39074 | 721.19126 | 9.60E-07 | 0.487057477 | YJR062C | NTA1 | n |
| W303.asm02.HP0_G0030180 | 759.60328 | 732.862058 | 860.6392094 | 839.210502 | 788.30243 | 860.918782 | 465.15833 | 532.70577 | 548.38749 | 0.41696973 | -0.079854409 | YJR063W | RPA12 | n |
| W303.asm02.HP0_G0030190 | 2867.7394 | 2686.46885 | 3022.379446 | 3876.86917 | 3567.5854 | 3733.80855 | 3516.0497 | 3419.2887 | 3474.5902 | 4.69E-09 | -0.382107206 | YJR064W | CCT5 | n |
| W303.asm02.HP0_G0030200 | 1820.5194 | 1956.72094 | 1962.759678 | 3009.32386 | 2844.001 | 2762.90455 | 3034.9301 | 2851.1567 | 2982.6284 | 2.70E-19 | -0.586084199 | YJR065C | ARP3 | n |
| W303.asm02.HP0_G0030220 | 724.83641 | 695.492322 | 830.6955332 | 376.686342 | 371.23022 | 377.363078 | 307.82537 | 413.3062 | 335.90938 | 1.88E-18 | 1.000908839 | YJR067C | YAE1 | n |
| W303.asm02.HP0_G0030230 | 2090.2259 | 1960.87313 | 2182.024662 | 1078.38966 | 1054.3657 | 1019.25957 | 989.60155 | 1045.7303 | 1027.1245 | 1.39E-44 | 0.98353007 | YJR068W | RFC2 | n |
| W303.asm02.HP0_G0030240 | 2057.5662 | 1936.99802 | 2194.581688 | 734.205017 | 798.18992 | 773.689125 | 493.66068 | 615.36701 | 543.09758 | 1.23E-58 | 1.424853971 | YJR069C | HAM1 | n |
| W303.asm02.HP0_G0030250 | 2134.4747 | 2101.00964 | 1933.781927 | 2480.12955 | 2346.93 | 2725.92676 | 1815.0296 | 2289.5851 | 1921.1195 | 0.0023518 | -0.291973251 | YJR070C | LIA1 | n |
| W303.asm02.HP0_G0030260 | 1643.5244 | 1558.11041 | 1389.966131 | 2197.61479 | 2278.6165 | 2239.52661 | 1491.2429 | 1874.9668 | 1580.8019 | 4.72E-09 | -0.548859759 | YJR072C | NPA3 | n |
| W303.asm02.HP0_G0030270 | 1274.785 | 1410.70756 | 1390.932056 | 1545.91408 | 1525.3697 | 1689.60052 | 1872.0343 | 2086.2122 | 1841.7708 | 0.005807924 | -0.223386638 | YJR073C | OPI3 | n |
| W303.asm02.HP0_G0030280 | 828.08346 | 822.134207 | 709.9549034 | 816.709327 | 817.9649 | 823.940992 | 791.2252 | 841.04531 | 776.73533 | 0.499420168 | -0.059829717 | YJR074W | MOG1 | n |
| W303.asm02.HP0_G0030290 | 1947.9979 | 1901.70438 | 1729.005819 | 3169.33221 | 2992.3133 | 3116.56392 | 3325.6541 | 3639.7187 | 3425.2177 | 2.93E-26 | -0.734401372 | YJR075W | HOC1 | n |
| W303.asm02.HP0_G0030300 | 2745.5286 | 2626.26205 | 2399.357796 | 3604.35494 | 3526.2377 | 3691.14187 | 3253.8281 | 3406.1679 | 3340.5791 | 3.78E-13 | -0.478050518 | YJR076C | CDC11 | n |
| W303.asm02.HP0_G0030310 | 3921.2807 | 4154.26906 | 4792.920042 | 12654.8277 | 12379.135 | 14421.3377 | 12173.923 | 11829.742 | 11114.986 | 5.57E-76 | -1.61613144 | YJR077C | MIR1 | n |
| W303.asm02.HP0_G0030330 | 1329.5691 | 1326.62565 | 1420.875732 | 1230.89763 | 1216.161 | 1191.82259 | 1081.9492 | 1162.5057 | 1120.5796 | 0.017850399 | 0.164207701 | YJR080C | AIM24 | n |
| W303.asm02.HP0_G0030340 | 1448.6193 | 1422.12609 | 1752.18802 | 270.014104 | 283.14169 | 288.237125 | 264.5018 | 287.34621 | 267.14053 | 1.25E-120 | 2.45918164 | YJR082C | EAF6 | n |
| W303.asm02.HP0_G0030350 | 305.52698 | 320.756907 | 270.4590108 | 480.025074 | 507.85733 | 457.007547 | 502.78143 | 523.52118 | 524.58289 | 1.48E-11 | -0.689978974 | YJR083C | ACF4 | n |
| W303.asm02.HP0_G0030360 | 2131.314 | 2015.88968 | 2416.744447 | 1415.07391 | 1404.0233 | 1293.27447 | 1372.6731 | 1545.634 | 1333.0577 | 3.85E-14 | 0.674609224 | YJR085C | TMH11 | n |
| W303.asm02.HP0_G0030370 | 754.33557 | 797.221049 | 928.2539621 | 8.33376864 | 5.3931751 | 12.3259297 | 12.541033 | 11.808749 | 9.698171 | 3.22E-110 | 6.584148982 | YJR086W | STE18 | n |
| W303.asm02.HP0_G0030380 | 2977.3077 | 2790.27367 | 3302.497707 | 974.217554 | 916.83977 | 877.037305 | 906.37469 | 897.46489 | 928.37946 | 2.13E-99 | 1.711835823 | YJR088C | EMC2 | n |
| W303.asm02.HP0_G0030390 | 642.66019 | 616.600655 | 569.8957728 | 900.88039 | 855.71712 | 840.059516 | 828.8483 | 797.74657 | 832.27941 | 5.03E-10 | -0.506888107 | YJR089W | BIR1 | n |
| W303.asm02.HP0_G0030400 | 939.75884 | 1076.45603 | 1020.016841 | 1305.06817 | 1297.0586 | 1353.00782 | 1433.0981 | 1339.6369 | 1407.9981 | 3.49E-07 | -0.381062286 | YJR090C | GRR1 | n |
| W303.asm02.HP0_G0030410 | 719.56871 | 797.221049 | 776.6037311 | 2241.78376 | 2191.4268 | 2187.37845 | 2159.3379 | 1947.1314 | 2270.2537 | 5.25E-85 | -1.529290677 | YJR091C | JSN1 | n |

| | | | | | | | | | | | | | |
|---|---|---|---|---|---|---|---|---|---|---|---|---|---|
| W303.asm02.HP0_G0030420 | 684.80184 | 749.47083 | 793.9903817 | 2330.12171 | 2337.0426 | 2351.40813 | 2622.2161 | 2349.941 | 2537.3942 | 4.96E-106 | -1.654349579 | YJR092W | BUD4 | n |
| W303.asm02.HP0_G0030430 | 487.78962 | 537.708989 | 563.1342975 | 1160.06059 | 1132.5668 | 1106.48923 | 1080.8091 | 1062.7874 | 1065.9172 | 6.09E-44 | -1.09661313 | YJR093C | FIP1 | n |
| W303.asm02.HP0_G0030440 | 97.979341 | 106.918969 | 87.89917851 | 121.673022 | 97.077152 | 125.155594 | 158.47306 | 116.7754 | 148.99917 | 0.214576769 | -0.233499854 | YJR094C | IME1 | n |
| W303.asm02.HP0_G0030460 | 34.766863 | 43.5980261 | 50.22810201 | 2799.31289 | 2892.5396 | 3170.60838 | 2472.8638 | 2317.1389 | 2218.2362 | 0 | -6.10033604 | YJR095W | SFC1 | n |
| W303.asm02.HP0_G0030470 | 1165.2167 | 1128.35844 | 1232.520349 | 131.673544 | 140.22255 | 116.622258 | 434.3758 | 272.9133 | 422.31127 | 1.23E-100 | 3.180975135 | YJR096W | | n |
| W303.asm02.HP0_G0030480 | 575.23355 | 580.268967 | 667.4542017 | 238.345783 | 320.89392 | 274.014899 | 231.43907 | 232.23872 | 224.82124 | 1.06E-20 | 1.13146924 | YJR097W | JJJ3 | n |
| W303.asm02.HP0_G0030490 | 311.84822 | 267.816446 | 283.9819614 | 523.360671 | 551.90159 | 549.926094 | 519.88284 | 530.0816 | 442.58926 | 2.25E-17 | -0.912140004 | YJR098C | | n |
| W303.asm02.HP0_G0030500 | 871.27865 | 839.781027 | 876.09401 | 370.019328 | 327.18596 | 354.607516 | 493.66068 | 356.88662 | 424.95622 | 4.95E-32 | 1.298095539 | YJR099W | YUH1 | n |
| W303.asm02.HP0_G0030510 | 1261.0889 | 1197.90767 | 1337.806178 | 590.864196 | 563.5868 | 566.992766 | 544.96491 | 520.89702 | 501.65994 | 1.67E-45 | 1.141112803 | YJR100C | AIM25 | n |
| W303.asm02.HP0_G0030520 | 2663.3524 | 2448.7558 | 3083.232723 | 1684.25464 | 1696.1536 | 1631.76346 | 1498.0835 | 1714.8927 | 1653.979 | 4.93E-15 | 0.709584152 | YJR101W | RSM26 | n |
| W303.asm02.HP0_G0030530 | 982.95403 | 968.499009 | 1140.757471 | 163.341865 | 150.11004 | 200.05932 | 266.78198 | 166.63456 | 230.9928 | 1.66E-68 | 2.592237942 | YJR102C | VPS25 | n |
| W303.asm02.HP0_G0030550 | 9374.4105 | 9668.38132 | 9616.749685 | 1553.41447 | 1444.4721 | 1580.56345 | 1549.3877 | 1743.7585 | 1779.1736 | 0 | 2.64614559 | YJR104C | SOD1 | n |
| W303.asm02.HP0_G0030560 | 3669.4843 | 3572.96205 | 3567.161168 | 5262.7749 | 5136.1005 | 5531.49799 | 4354.0188 | 4491.2607 | 4117.3144 | 4.92E-21 | -0.559420946 | YJR105W | ADO1 | n |
| W303.asm02.HP0_G0030570 | 514.12815 | 554.317761 | 538.0202465 | 850.877778 | 891.67162 | 827.733586 | 1030.6449 | 948.63613 | 1009.4914 | 6.33E-17 | -0.678026135 | YJR106W | ECM27 | n |
| W303.asm02.HP0_G0030580 | 374.00716 | 440.132454 | 433.7003423 | 24.1679291 | 21.572701 | 18.0148203 | 25.082067 | 36.738329 | 26.449557 | 7.71E-87 | 4.284149489 | YJR107W | LIH1 | n |
| W303.asm02.HP0_G0030610 | 4831.5404 | 5159.09976 | 5558.898597 | 1876.7647 | 1929.8578 | 2111.52657 | 2285.8884 | 1430.1707 | 2190.0234 | 4.80E-25 | 1.39382421 | YJR109C | CPA2 | n |
| W303.asm02.HP0_G0030620 | 513.07461 | 508.643638 | 532.2246963 | 808.375558 | 743.35931 | 744.296524 | 904.0945 | 844.98156 | 902.81156 | 1.12E-11 | -0.563543812 | YJR110W | YMR1 | n |
| W303.asm02.HP0_G0030630 | 749.06786 | 701.720611 | 704.1593532 | 450.856883 | 465.61079 | 441.837172 | 535.84416 | 516.96077 | 530.75445 | 2.27E-15 | 0.665370924 | YJR111C | PXP2 | n |
| W303.asm02.HP0_G0030640 | 490.95025 | 478.540239 | 411.4840665 | 340.851137 | 372.12908 | 310.044539 | 302.1249 | 376.56787 | 353.54242 | 0.000344321 | 0.431322177 | YJR112W | NNF1 | n |
| W303.asm02.HP0_G0030650 | 697.44434 | 713.139142 | 876.09401 | 125.839906 | 104.26805 | 112.829664 | 148.21221 | 169.25873 | 149.88083 | 1.93E-90 | 2.736104248 | YJR112W-A | | n |
| W303.asm02.HP0_G0030660 | 1839.4831 | 1710.7035 | 1859.405699 | 1839.26274 | 1792.3319 | 1848.88945 | 1610.9528 | 1608.614 | 1578.1569 | 0.781362188 | -0.018585442 | YJR113C | RSM7 | n |
| W303.asm02.HP0_G0030680 | 287.61677 | 326.985196 | 294.6071368 | 1453.40925 | 1469.6402 | 1474.37082 | 2289.3087 | 2233.1656 | 2189.1417 | 3.02E-180 | -2.274231782 | YJR116W | TDA4 | n |
| W303.asm02.HP0_G0030690 | 2120.7786 | 2208.96666 | 1581.219288 | 2693.47402 | 2797.2602 | 2691.79342 | 3233.3064 | 3349.7483 | 3244.479 | 8.19E-06 | -0.469630368 | YJR117W | STE24 | n |
| W303.asm02.HP0_G0030700 | 1811.0375 | 1724.19813 | 2214.866114 | 798.375036 | 921.33409 | 787.911352 | 1097.9105 | 1073.284 | 1087.0768 | 1.23E-31 | 1.197704137 | YJR118C | ILM1 | n |
| W303.asm02.HP0_G0030710 | 611.05395 | 565.736292 | 562.1683725 | 596.697835 | 586.05836 | 531.911274 | 608.81017 | 598.30993 | 531.6361 | 0.850023122 | 0.0190398 | YJR119C | JHD2 | n |
| W303.asm02.HP0_G0030730 | 7244.15 | 8197.46696 | 8896.169606 | 22472.8405 | 21918.763 | 24689.7853 | 20631.14 | 19258.757 | 18199.94 | 1.29E-69 | -1.504980356 | YJR121W | ATP2 | n |
| W303.asm02.HP0_G0030740 | 1268.4637 | 1341.15833 | 1106.950094 | 495.859234 | 584.26064 | 554.666836 | 571.18707 | 628.48784 | 559.84896 | 1.86E-32 | 1.185206931 | YJR122W | IBA57 | n |
| W303.asm02.HP0_G0030750 | 27456.34 | 27640.1105 | 27584.88725 | 22767.8559 | 22682.796 | 25604.7486 | 19698.543 | 19824.265 | 17203.674 | 0.003325013 | 0.218652676 | YJR123W | RPS5 | n |
| W303.asm02.HP0_G0030760 | 1084.094 | 936.319514 | 1019.050916 | 545.028469 | 604.03562 | 640.000196 | 481.11965 | 558.94743 | 461.10395 | 3.37E-14 | 0.765572248 | YJR124C | | n |
| W303.asm02.HP0_G0030770 | 1202.0906 | 1233.20131 | 1188.087798 | 1545.91408 | 1514.5834 | 1404.20784 | 1633.7546 | 1588.9327 | 1717.4579 | 1.98E-05 | -0.301861509 | YJR125C | ENT3 | n |
| W303.asm02.HP0_G0030780 | 869.17157 | 948.776093 | 939.8450626 | 648.3672 | 618.41742 | 635.259453 | 817.44736 | 835.79698 | 866.66383 | 7.30E-12 | 0.536031993 | YJR126C | VPS70 | n |
| W303.asm02.HP0_G0030800 | 487.78962 | 428.713924 | 498.4173199 | 445.023245 | 434.1506 | 452.266805 | 311.24565 | 363.44704 | 346.4892 | 0.399419452 | 0.088351316 | YJR129C | EFM3 | n |
| W303.asm02.HP0_G0030810 | 713.24746 | 724.557673 | 701.261578 | 1600.08358 | 1526.2686 | 1559.70418 | 1727.2423 | 1814.611 | 1777.4103 | 5.17E-63 | -1.131799145 | YJR130C | STR2 | n |
| W303.asm02.HP0_G0030820 | 396.13153 | 435.980261 | 429.8366422 | 1101.72421 | 1067.8487 | 1040.11884 | 1303.1274 | 1272.7207 | 1272.2237 | 1.10E-65 | -1.346650169 | YJR131W | MNS1 | n |
| W303.asm02.HP0_G0030830 | 1252.6606 | 1056.73311 | 1210.304073 | 2525.1319 | 2517.7139 | 2852.97865 | 1722.682 | 2045.5377 | 1761.5405 | 3.13E-35 | -1.165354123 | YJR132W | NMD5 | n |
| W303.asm02.HP0_G0030840 | 1094.6294 | 1069.18969 | 1142.689321 | 2496.79708 | 2677.7115 | 2868.14902 | 1441.0788 | 2099.3331 | 1437.0926 | 2.66E-24 | -1.28212928 | YJR133W | XPT1 | n |
| W303.asm02.HP0_G0030850 | 1097.79 | 1078.53212 | 991.0390896 | 1285.06712 | 1349.1926 | 1257.24483 | 1016.9638 | 1027.3611 | 1025.3612 | 4.25E-05 | -0.297742798 | YJR134C | SGM1 | n |
| W303.asm02.HP0_G0030860 | 389.81028 | 321.794955 | 370.9152148 | 225.011753 | 269.65876 | 274.014899 | 235.99945 | 209.93331 | 184.26525 | 0.000211303 | 0.495706717 | YJR135C | MCM22 | n |
| W303.asm02.HP0_G0030870 | 1020.8815 | 1035.97214 | 1325.249153 | 820.876211 | 959.08631 | 869.452117 | 731.94032 | 877.78364 | 793.48672 | 0.002256754 | 0.35313253 | YJR135W- | TIM8 | n |
| W303.asm02.HP0_G0030880 | 543.62731 | 592.725546 | 524.497296 | 436.689477 | 467.40851 | 452.266805 | 424.11495 | 387.06453 | 409.08649 | 0.001963843 | 0.292061102 | YJR136C | TTI2 | n |
| W303.asm02.HP0_G0030890 | 1500.2428 | 1565.37675 | 1733.835444 | 1692.58841 | 1720.4229 | 1743.64498 | 1755.7447 | 1545.634 | 1725.3928 | 0.201788826 | -0.102974023 | YJR137C | MET5 | n |
| W303.asm02.HP0_G0030900 | 396.13153 | 441.170503 | 432.7344173 | 910.880912 | 911.4466 | 898.844719 | 979.3407 | 831.86073 | 999.79327 | 1.11E-31 | -1.099003643 | YJR138W | IML1 | n |
| W303.asm02.HP0_G0030910 | 6241.1787 | 5809.95601 | 6861.931474 | 2748.4769 | 2826.9226 | 2968.65276 | 2375.9558 | 2259.4072 | 2416.6079 | 1.92E-54 | 1.146709466 | YJR139C | HOM6 | n |
| W303.asm02.HP0_G0030920 | 950.29425 | 904.140018 | 877.0599351 | 1411.74041 | 1370.7653 | 1314.13373 | 1441.0788 | 1359.3182 | 1389.4834 | 1.84E-16 | -0.585643139 | YJR140C | HIR3 | n |
| W303.asm02.HP0_G0030930 | 1041.9523 | 1076.45603 | 1175.530772 | 609.198487 | 617.51855 | 600.177961 | 465.15833 | 549.76285 | 422.31127 | 4.26E-18 | 0.850734561 | YJR141W | IPA1 | n |
| W303.asm02.HP0_G0030940 | 1315.8731 | 1380.60416 | 1319.453603 | 885.879606 | 893.46935 | 885.570641 | 885.553 | 883.03197 | 916.91799 | 3.02E-18 | 0.591558266 | YJR142W | | n |
| W303.asm02.HP0_G0030950 | 2349.3971 | 2350.14122 | 2340.436369 | 7763.73886 | 7672.6905 | 8035.55801 | 8998.7616 | 8903.7964 | 8476.2015 | 1.02E-230 | -1.737257754 | YJR143C | PMT4 | n |
| W303.asm02.HP0_G0030960 | 2845.6151 | 2740.44736 | 2818.569263 | 1210.06321 | 1237.7337 | 1250.60779 | 1132.1133 | 1237.2944 | 1156.7273 | 3.23E-79 | 1.184545417 | YJR144W | MGM101 | n |
| W303.asm02.HP0_G0030990 | 467.77234 | 456.741226 | 451.086993 | 163.341865 | 159.99753 | 145.066711 | 303.26499 | 221.74206 | 272.43044 | 1.76E-34 | 1.552550257 | YJR149W | | n |
| W303.asm02.HP0_G0031020 | 304.47344 | 282.349122 | 259.8338354 | 99.1718468 | 113.25668 | 109.985219 | 128.83062 | 170.57081 | 138.41935 | 6.43E-21 | 1.393541922 | YJR152W | DAL5 | n |
| W303.asm02.HP0_G0031030 | 217.02951 | 195.153069 | 173.866507 | 80.8375558 | 110.56009 | 110.933367 | 123.13015 | 107.59082 | 111.08814 | 1.63E-08 | 0.957530129 | YJR153W | PGU1 | n |
| W303.asm02.HP0_G0031140 | 475.14713 | 430.79002 | 453.0188431 | 489.192219 | 532.12661 | 494.933485 | 542.68472 | 522.2091 | 514.00306 | 0.09016111 | -0.157946443 | YKL222C | | n |
| W303.asm02.HP0_G0031150 | 102.19351 | 101.728728 | 119.7747048 | 129.173414 | 135.72824 | 129.896336 | 135.67118 | 112.83915 | 116.37805 | 0.086229654 | -0.284362411 | YKL221W | MCH2 | n |
| W303.asm02.HP0_G0031190 | 278.1349 | 227.332565 | 255.9701352 | 277.514496 | 259.77127 | 293.926016 | 304.40508 | 226.99039 | 253.91575 | 0.361917681 | -0.126208858 | YKL217W | JEN1 | n |
| W303.asm02.HP0_G0031200 | 2638.0674 | 2454.98409 | 2448.619973 | 850.877778 | 678.6412 | 585.955735 | 761.58276 | 596.99784 | 818.17297 | 9.43E-45 | 1.832671299 | YKL216W | URA1 | n |
| W303.asm02.HP0_G0031210 | 636.33895 | 650.856247 | 725.409704 | 600.864719 | 574.37315 | 550.874242 | 497.08096 | 455.29286 | 443.47091 | 0.017899675 | 0.221644506 | YKL215C | OXP1 | n |
| W303.asm02.HP0_G0031230 | 1492.868 | 1504.1319 | 1486.558634 | 1841.76287 | 1708.7377 | 1834.66723 | 2151.3573 | 1793.6177 | 2030.4444 | 0.000839092 | -0.264413975 | YKL213C | DOA1 | n |
| W303.asm02.HP0_G0031240 | 2032.2812 | 2007.5853 | 2012.021855 | 4421.89764 | 4467.3467 | 4348.20874 | 4800.9356 | 4546.3682 | 4689.5065 | 4.07E-99 | -1.129250756 | YKL212W | SAC1 | n |
| W303.asm02.HP0_G0031250 | 1286.3739 | 1313.13103 | 1412.182406 | 3202.66729 | 2975.235 | 3138.37133 | 3218.4852 | 2906.2642 | 3144.8524 | 3.56E-67 | -1.215169925 | YKL211C | TRP3 | n |
| W303.asm02.HP0_G0031260 | 3653.6812 | 3654.96786 | 4112.908814 | 6692.84959 | 6736.0757 | 6975.52806 | 7079.9834 | 6014.5892 | 6602.6912 | 4.99E-28 | -0.83689071 | YKL210W | UBA1 | n |

| | | | | | | | | | | | | | | |
|---|---|---|---|---|---|---|---|---|---|---|---|---|---|---|
| W303.asm02.HP0_G0031270 | 2859.3111 | 2807.92049 | 3153.745251 | 432.522592 | 426.06084 | 434.251985 | 417.27439 | 387.06453 | 398.50666 | 1.64E-263 | 2.770675688 | YKL209C | STE6 | n |
| W303.asm02.HP0_G0031280 | 540.46669 | 564.698243 | 519.6676708 | 541.694962 | 581.56405 | 497.77793 | 474.27908 | 532.70577 | 514.88472 | 0.982008514 | 0.002414605 | YKL208W | CBT1 | n |
| W303.asm02.HP0_G0031290 | 1473.9043 | 1466.76217 | 1535.820811 | 884.212853 | 850.32395 | 847.644703 | 1355.5717 | 1247.7911 | 1329.5311 | 2.80E-30 | 0.793610027 | YKL207W | EMC3 | n |
| W303.asm02.HP0_G0031300 | 1681.4519 | 1737.69276 | 1504.91121 | 755.039439 | 801.78537 | 766.103938 | 856.21056 | 818.7399 | 875.48035 | 4.14E-42 | 1.083608604 | YKL206C | ADD66 | n |
| W303.asm02.HP0_G0031310 | 2156.599 | 1896.51414 | 2225.491289 | 2615.1366 | 2601.3081 | 3162.07504 | 2370.2553 | 2300.0818 | 2264.9638 | 6.33E-06 | -0.415779347 | YKL205W | LOS1 | n |
| W303.asm02.HP0_G0031320 | 663.73102 | 752.584975 | 760.1830054 | 1755.92505 | 1742.8944 | 1783.46721 | 1597.2716 | 1372.439 | 1661.9139 | 4.16E-47 | -1.278586031 | YKL204W | EAP1 | n |
| W303.asm02.HP0_G0031330 | 596.30438 | 696.53037 | 751.48968 | 2194.28128 | 2224.6847 | 2214.87475 | 2301.8497 | 2208.236 | 2380.4602 | 1.98E-92 | -1.696983417 | YKL203C | TOR2 | n |
| W303.asm02.HP0_G0031350 | 355.04342 | 420.409538 | 406.6544413 | 1480.91069 | 1438.18 | 1529.36343 | 1452.4797 | 1393.4323 | 1633.701 | 1.12E-94 | -1.911180283 | YKL198C | PTK1 | n |
| W303.asm02.HP0_G0031360 | 377.16779 | 442.208551 | 377.6766901 | 320.850093 | 291.23146 | 270.222305 | 308.96546 | 211.24539 | 284.77357 | 0.001648039 | 0.438669487 | YKL197C | PEX1 | n |
| W303.asm02.HP0_G0031370 | 5767.0851 | 5881.58134 | 7064.775732 | 1916.76679 | 1928.959 | 2008.17839 | 2056.7295 | 1978.6214 | 2063.0655 | 4.93E-92 | 1.676908526 | YKL196C | YKT6 | n |
| W303.asm02.HP0_G0031380 | 3304.9591 | 3559.46742 | 3152.779326 | 2533.46567 | 2486.2537 | 2757.21566 | 2759.0274 | 2590.0522 | 2600.8731 | 3.55E-07 | 0.36528675 | YKL195W | MIA40 | n |
| W303.asm02.HP0_G0031390 | 622.64291 | 657.084537 | 567.9639227 | 952.549755 | 1056.1635 | 1013.57068 | 1047.7463 | 1073.284 | 980.39693 | 1.67E-16 | -0.710341895 | YKL194C | MST1 | n |
| W303.asm02.HP0_G0031400 | 1358.0147 | 1320.39736 | 1323.317303 | 1105.8911 | 1143.3531 | 977.541039 | 1035.2053 | 1081.1565 | 1112.6447 | 8.55E-05 | 0.309808682 | YKL193C | SDS22 | n |
| W303.asm02.HP0_G0031410 | 2404.1812 | 2422.8046 | 2103.784734 | 2199.28154 | 2371.1993 | 2276.5044 | 1991.7441 | 2387.9914 | 2102.7398 | 0.850023122 | 0.017356412 | YKL192C | ACP1 | n |
| W303.asm02.HP0_G0031420 | 726.9435 | 821.096159 | 993.9368647 | 1022.55341 | 1014.8158 | 1005.03734 | 855.07046 | 978.81404 | 902.81156 | 0.017490347 | -0.258309751 | YKL191W | DPH2 | n |
| W303.asm02.HP0_G0031430 | 2476.8756 | 2413.46216 | 2664.021257 | 1123.39201 | 1166.7236 | 979.437336 | 1015.8237 | 997.18321 | 987.45014 | 2.17E-51 | 1.20776617 | YKL190W | CNB1 | n |
| W303.asm02.HP0_G0031440 | 569.96584 | 538.747037 | 494.5536198 | 715.870726 | 768.52746 | 697.83725 | 794.64548 | 779.3774 | 760.8656 | 3.69E-07 | -0.445926842 | YKL189W | HYM1 | n |
| W303.asm02.HP0_G0031450 | 529.93127 | 449.474889 | 525.463221 | 66.6701491 | 55.729476 | 70.1629844 | 46.743852 | 49.85916 | 44.082596 | 5.47E-82 | 2.967040284 | YKL188C | PXA2 | n |
| W303.asm02.HP0_G0031460 | 212.81534 | 197.229166 | 205.7420332 | 3365.17578 | 3060.6269 | 3382.99363 | 6508.7964 | 6917.3025 | 6514.526 | 0 | -3.993696808 | YKL187C | FAT3 | n |
| W303.asm02.HP0_G0031470 | 1327.462 | 1355.691 | 1313.658053 | 681.702275 | 737.96613 | 780.326164 | 689.75684 | 793.81032 | 753.81239 | 5.96E-26 | 0.86230013 | YKL186C | MTR2 | n |
| W303.asm02.HP0_G0031480 | 1356.9612 | 1650.4967 | 1867.1331 | 749.205801 | 723.58433 | 567.940914 | 785.52473 | 831.86073 | 854.3207 | 9.13E-24 | 1.255648637 | YKL185W | ASH1 | n |
| W303.asm02.HP0_G0031490 | 1165.2167 | 1123.1682 | 1304.964727 | 1336.73649 | 1337.5074 | 1458.2523 | 1251.8232 | 1409.1773 | 1231.6677 | 0.021850398 | -0.200839982 | YKL184W | SPE1 | n |
| W303.asm02.HP0_G0031510 | 778.56702 | 703.796708 | 827.7977581 | 768.373468 | 747.85362 | 723.437258 | 832.26858 | 851.54198 | 830.5161 | 0.610291054 | 0.044915906 | YKL183W | LOT5 | n |
| W303.asm02.HP0_G0031520 | 8269.2457 | 8002.31389 | 9759.70659 | 14509.9246 | 14201.129 | 15129.6046 | 13085.998 | 13780.81 | 13185.986 | 3.88E-22 | -0.751881605 | YKL182W | FAS1 | n |
| W303.asm02.HP0_G0031530 | 5747.0678 | 5346.98649 | 5237.245559 | 4545.23742 | 4550.0421 | 4900.97928 | 3806.7737 | 4316.7536 | 3857.2271 | 0.0020088 | 0.222629459 | YKL181W | PRS1 | n |
| W303.asm02.HP0_G0031550 | 1418.0666 | 1394.09879 | 1166.837447 | 1271.73309 | 1217.9587 | 1203.20037 | 1172.0166 | 1193.9957 | 1299.5549 | 0.240916912 | 0.106782259 | YKL179C | COY1 | n |
| W303.asm02.HP0_G0031560 | 267.59949 | 363.316885 | 255.9701352 | 94.1715856 | 110.56009 | 97.6592891 | 107.16883 | 103.65457 | 96.100058 | 3.54E-20 | 1.552169322 | YKL178C | STE3 | n |
| W303.asm02.HP0_G0031570 | 895.5101 | 1038.04824 | 1069.279018 | 860.8783 | 764.93201 | 773.689125 | 769.56342 | 674.41075 | 832.27941 | 0.001716648 | 0.323485848 | YKL176C | LST4 | n |
| W303.asm02.HP0_G0031580 | 960.82967 | 923.862935 | 942.7428377 | 780.040745 | 771.22404 | 669.392797 | 920.05582 | 867.28697 | 886.94182 | 3.02E-05 | 0.347388291 | YKL175W | ZRT3 | n |
| W303.asm02.HP0_G0031590 | 1597.1686 | 1469.87631 | 1574.457813 | 951.716379 | 956.38972 | 969.007703 | 1614.373 | 1347.5094 | 1527.9028 | 1.76E-17 | 0.690129511 | YKL174C | TPO5 | n |
| W303.asm02.HP0_G0031600 | 634.23186 | 661.23673 | 737.9667295 | 1072.55602 | 1024.7033 | 1090.3707 | 1002.1426 | 892.21656 | 1049.1658 | 6.41E-13 | -0.647638658 | YKL173W | SNU114 | n |
| W303.asm02.HP0_G0031610 | 5144.4422 | 5035.57202 | 4695.361613 | 2356.78977 | 2214.7973 | 2353.30442 | 1240.4222 | 1609.926 | 1392.1284 | 1.52E-36 | 1.1029664 | YKL172W | EBP2 | n |
| W303.asm02.HP0_G0031620 | 331.86551 | 325.947148 | 380.5744652 | 549.195353 | 533.92434 | 489.244594 | 622.4913 | 528.76952 | 585.41687 | 1.78E-08 | -0.59811361 | YKL171W | NNK1 | n |
| W303.asm02.HP0_G0031630 | 2103.922 | 2071.94429 | 2218.729814 | 1298.40115 | 1288.9689 | 1247.76334 | 1143.5142 | 1188.7474 | 1213.153 | 4.38E-30 | 0.737557345 | YKL170W | MRPL38 | n |
| W303.asm02.HP0_G0031640 | 291.83094 | 304.148135 | 370.9152148 | 1095.89058 | 1000.434 | 1056.23736 | 1065.9878 | 1082.4686 | 1132.0411 | 1.16E-68 | -1.702954409 | YKL168C | KKQ8 | n |
| W303.asm02.HP0_G0031650 | 1387.5139 | 1400.32708 | 1407.352781 | 845.877517 | 936.61475 | 805.926172 | 837.96905 | 851.54198 | 845.50418 | 4.05E-20 | 0.696468206 | YKL167C | MRP49 | n |
| W303.asm02.HP0_G0031670 | 5246.6357 | 5109.27344 | 4838.318518 | 7220.37715 | 7213.3717 | 7347.20224 | 8703.4772 | 8624.3227 | 7941.9204 | 5.04E-18 | -0.519657444 | YKL165C | MCD4 | n |
| W303.asm02.HP0_G0031680 | 6241.1787 | 7115.8207 | 8295.364232 | 17950.1043 | 16433.005 | 18412.0945 | 14997.936 | 13484.279 | 15875.024 | 1.17E-37 | -1.28575912 | YKL164C | PIR1 | n |
| W303.asm02.HP0_G0031710 | 215.97597 | 201.381359 | 158.4117063 | 372.519458 | 426.9597 | 376.41493 | 430.95551 | 384.44037 | 431.12779 | 1.18E-15 | -1.032808504 | YKL162C | | n |
| W303.asm02.HP0_G0031730 | 2741.3145 | 2744.59955 | 2931.582492 | 2325.12145 | 2446.7038 | 2527.76374 | 2054.4493 | 2116.3902 | 2143.2958 | 0.001165987 | 0.206139653 | YKL160W | ELF1 | n |
| W303.asm02.HP0_G0031740 | 317.11593 | 324.9091 | 311.9937875 | 200.010447 | 200.44634 | 209.540805 | 228.01879 | 229.61455 | 238.92767 | 1.06E-08 | 0.645606346 | YKL159C | RCN1 | n |
| W303.asm02.HP0_G0031760 | 1410.6918 | 1594.4421 | 1674.914017 | 3867.70203 | 3908.2542 | 3912.06045 | 4965.1092 | 4908.5031 | 4611.0395 | 1.66E-74 | -1.319877186 | YKL157W | APE2 | n |
| W303.asm02.HP0_G0031770 | 7786.7237 | 7579.82826 | 8532.015866 | 6294.49545 | 6731.5814 | 6815.29097 | 5784.8367 | 7056.3833 | 5190.2848 | 0.00843085 | 0.268506596 | YKL156W | RPS27A | y |
| W303.asm02.HP0_G0031780 | 1059.8625 | 1129.39649 | 1072.176793 | 1066.72239 | 1027.3999 | 1080.88922 | 807.18652 | 868.59906 | 801.42159 | 0.611522614 | 0.038735699 | YKL155C | RSM22 | n |
| W303.asm02.HP0_G0031790 | 1546.5986 | 1469.87631 | 1666.220692 | 1291.73414 | 1359.979 | 1312.23744 | 1559.6485 | 1756.8794 | 1683.9552 | 0.001755387 | 0.240881242 | YKL154W | SRP102 | n |
| W303.asm02.HP0_G0031800 | 21049.755 | 21025.6671 | 21082.27989 | 10727.227 | 10091.53 | 11081.0108 | 11353.056 | 11252.425 | 10891.046 | 1.09E-76 | 0.985447159 | YKL152C | GPM1 | n |
| W303.asm02.HP0_G0031810 | 684.80184 | 689.264032 | 666.4882766 | 286.681641 | 266.06331 | 230.40007 | 422.97486 | 328.02079 | 342.08094 | 3.53E-32 | 1.379372558 | YKL151C | NNR2 | n |
| W303.asm02.HP0_G0031820 | 6088.4152 | 5821.37454 | 6160.669896 | 6055.31629 | 6218.3309 | 6412.32789 | 6158.7875 | 5858.4514 | 5901.7779 | 0.394481743 | -0.048145155 | YKL150W | MCR1 | n |
| W303.asm02.HP0_G0031830 | 655.30269 | 597.915787 | 632.6809003 | 544.195092 | 533.02548 | 543.289055 | 568.90688 | 524.83327 | 521.93793 | 0.011854845 | 0.218741425 | YKL149C | DBR1 | n |
| W303.asm02.HP0_G0031840 | 1817.3587 | 2210.00471 | 2283.446791 | 9280.48476 | 9456.0337 | 10036.1512 | 8672.6947 | 8043.0698 | 8009.8076 | 7.39E-146 | -2.188404542 | YKL148C | SDH1 | n |
| W303.asm02.HP0_G0031850 | 987.1682 | 1060.8853 | 1146.553021 | 770.873599 | 815.26831 | 892.20768 | 1047.7463 | 909.27364 | 1029.7694 | 8.55E-05 | 0.36754697 | YKL146W | AVT3 | n |
| W303.asm02.HP0_G0031860 | 4320.5729 | 4307.9002 | 3836.654253 | 2715.9752 | 2693.891 | 2674.72674 | 3014.4084 | 2541.5051 | 2988.8 | 7.03E-15 | 0.624458103 | YKL145W | RPT1 | n |
| W303.asm02.HP0_G0031870 | 1512.8853 | 1488.56118 | 1807.245747 | 957.550017 | 864.70575 | 992.711414 | 692.03703 | 800.37073 | 741.46926 | 2.90E-15 | 0.773093343 | YKL144C | RPC25 | n |
| W303.asm02.HP0_G0031880 | 1128.3427 | 1016.24923 | 1185.190022 | 1088.39018 | 1117.2861 | 1414.63747 | 612.23045 | 585.18909 | 662.12059 | 0.289481795 | -0.119740567 | YKL143W | LTV1 | n |
| W303.asm02.HP0_G0031890 | 3333.4047 | 3330.05876 | 3486.989389 | 1103.39097 | 1145.1509 | 1164.32628 | 1019.244 | 1157.2574 | 1122.3429 | 1.64E-118 | 1.572976374 | YKL142W | MRP8 | n |
| W303.asm02.HP0_G0031900 | 3606.2719 | 3621.75031 | 3406.817611 | 3388.51033 | 3560.3945 | 3742.34188 | 3309.6927 | 3959.867 | 3243.5974 | 0.932922501 | -0.007560121 | YKL141W | SDH3 | n |
| W303.asm02.HP0_G0031920 | 813.33388 | 734.938155 | 824.899983 | 838.377125 | 817.06603 | 746.192821 | 785.52473 | 785.93782 | 816.40967 | 0.843965449 | -0.01766954 | YKL139W | CTK1 | n |
| W303.asm02.HP0_G0031930 | 1410.6918 | 1471.95241 | 1759.91542 | 164.175242 | 205.83952 | 194.37043 | 125.41033 | 288.6583 | 164.86891 | 1.02E-23 | 3.040605692 | YKL138C-A | HSK3 | n |
| W303.asm02.HP0_G0031940 | 1336.9439 | 1264.34276 | 1297.237327 | 1350.9039 | 1361.7767 | 1241.12631 | 1294.0066 | 1405.2411 | 1410.6431 | 0.788805068 | -0.020814103 | YKL138C | MRPL31 | n |
| W303.asm02.HP0_G0031950 | 1466.5295 | 1413.8217 | 1862.303474 | 328.350484 | 411.67904 | 301.511203 | 299.84471 | 414.61828 | 391.45345 | 1.63E-51 | 2.187033841 | YKL137W | CMC1 | n |

| | | | | | | | | | | | | | | |
|---|---|---|---|---|---|---|---|---|---|---|---|---|---|---|
| W303.asm02.HP0_G0031960 | 1862.661 | 1809.31808 | 1568.662263 | 2288.45287 | 2186.9325 | 2230.99327 | 2212.9224 | 2277.7764 | 2196.1949 | 2.18E-06 | -0.356454356 | YKL135C | APL2 | n |
| W303.asm02.HP0_G0031970 | 643.71373 | 662.274778 | 629.7831252 | 1260.8992 | 1240.4303 | 1211.7337 | 1219.9005 | 1141.5124 | 1224.6145 | 3.88E-38 | -0.940269162 | YKL134C | OCT1 | n |
| W303.asm02.HP0_G0032000 | 1271.6243 | 1301.71249 | 1324.283228 | 1105.05772 | 1069.6464 | 1099.85219 | 1056.8671 | 1029.9853 | 1061.5089 | 0.00014369 | 0.251439917 | YKL130C | SHE2 | n |
| W303.asm02.HP0_G0032020 | 1801.5556 | 1939.07412 | 1700.028068 | 760.873077 | 789.20129 | 788.8595 | 745.62144 | 585.18909 | 789.96011 | 4.57E-33 | 1.217938788 | YKL128C | PMU1 | n |
| W303.asm02.HP0_G0032040 | 1882.6783 | 1761.56787 | 1637.24294 | 1585.0828 | 1592.7844 | 1774.93388 | 1724.9621 | 1523.3286 | 1893.7883 | 0.337414019 | 0.092728066 | YKL126W | YPK1 | n |
| W303.asm02.HP0_G0032050 | 823.8693 | 808.63958 | 823.9340579 | 1305.90155 | 1227.8462 | 1434.54859 | 787.80492 | 1024.737 | 828.7528 | 1.34E-11 | -0.69166575 | YKL125W | RRN3 | n |
| W303.asm02.HP0_G0032060 | 356.09696 | 392.382235 | 325.516738 | 400.854272 | 410.78017 | 393.481602 | 596.26914 | 506.4641 | 589.82513 | 0.129882141 | -0.167341771 | YKL124W | SSH4 | n |
| W303.asm02.HP0_G0032070 | 2312.5232 | 2554.63672 | 2720.044909 | 1313.40194 | 1235.0371 | 1201.30407 | 1157.1954 | 1287.1536 | 1253.709 | 4.31E-36 | 1.016643524 | YKL122C | SRP21 | n |
| W303.asm02.HP0_G0032090 | 5847.1542 | 5934.5218 | 5595.603749 | 1264.2327 | 1195.4872 | 1232.59297 | 1075.1086 | 1015.5524 | 1072.0887 | 1.10E-294 | 2.234283278 | YKL120W | OAC1 | n |
| W303.asm02.HP0_G0032100 | 1140.9852 | 1139.77697 | 1484.626784 | 618.365633 | 596.84472 | 636.207602 | 589.42857 | 594.37368 | 652.42242 | 7.99E-21 | 1.024975606 | YKL119C | VPH2 | n |
| W303.asm02.HP0_G0032110 | 7313.6837 | 7403.36006 | 7150.743061 | 2235.11675 | 2308.279 | 2280.29699 | 3664.262 | 2983.6771 | 3178.3551 | 3.21E-108 | 1.680222134 | YKL117W | SBA1 | n |
| W303.asm02.HP0_G0032120 | 632.12478 | 664.350875 | 711.8867535 | 927.54845 | 960.88404 | 962.370664 | 904.0945 | 938.13947 | 957.47398 | 1.26E-10 | -0.504221098 | YKL116C | PRR1 | n |
| W303.asm02.HP0_G0032140 | 2464.2331 | 2398.92949 | 2404.187421 | 701.703319 | 676.84348 | 747.140969 | 652.13374 | 709.83699 | 700.91327 | 3.21E-138 | 1.774003597 | YKL114C | APN1 | n |
| W303.asm02.HP0_G0032150 | 1419.1201 | 1372.29978 | 1372.57948 | 1175.89475 | 1163.1281 | 1119.76331 | 1008.9831 | 1107.3982 | 1042.1126 | 0.000107943 | 0.267242906 | YKL113C | RAD27 | n |
| W303.asm02.HP0_G0032160 | 1086.2011 | 1195.83157 | 1245.077375 | 1094.22382 | 1051.6692 | 1256.29668 | 1283.7458 | 1235.9823 | 1244.0108 | 0.563483936 | 0.053034038 | YKL112W | ABF1 | n |
| W303.asm02.HP0_G0032180 | 1438.0839 | 1520.74067 | 1672.982167 | 409.18804 | 397.29724 | 420.029758 | 452.6173 | 381.8162 | 476.97368 | 4.54E-86 | 1.917575659 | YKL109W | HAP4 | n |
| W303.asm02.HP0_G0032190 | 422.47006 | 445.322696 | 429.8366422 | 245.846175 | 248.08606 | 249.363039 | 230.29898 | 278.16163 | 274.19374 | 1.43E-13 | 0.804036015 | YKL108W | SLD2 | n |
| W303.asm02.HP0_G0032210 | 2108.1361 | 1969.17751 | 1884.51975 | 524.194047 | 508.75619 | 489.244594 | 535.84416 | 520.89702 | 540.45262 | 1.52E-139 | 1.968554812 | YKL106W | AAT1 | n |
| W303.asm02.HP0_G0032220 | 4080.3655 | 4557.03178 | 4814.170392 | 9686.33929 | 9302.3282 | 10414.4624 | 9021.5634 | 8990.3939 | 8976.9798 | 2.88E-54 | -1.127956218 | YKL104C | GFA1 | n |
| W303.asm02.HP0_G0032230 | 1305.3377 | 1533.19725 | 1463.376434 | 1395.07287 | 1325.8222 | 1373.86709 | 1966.6621 | 1355.3819 | 1960.7939 | 0.583271609 | 0.071250009 | YKL103C | APE1 | n |
| W303.asm02.HP0_G0032240 | 1449.6728 | 1538.38749 | 1563.832638 | 2681.80675 | 2661.5319 | 2556.20819 | 2451.202 | 2289.5851 | 2567.3704 | 1.83E-31 | -0.795191344 | YKL101W | HSL1 | n |
| W303.asm02.HP0_G0032260 | 1735.1825 | 1659.83914 | 1966.623379 | 1575.91565 | 1433.6857 | 1518.9338 | 978.20061 | 1140.2003 | 1034.1777 | 0.005823109 | 0.243864381 | YKL099C | UTP11 | n |
| W303.asm02.HP0_G0032270 | 470.93296 | 485.806577 | 502.2810201 | 139.173936 | 136.6271 | 168.770422 | 217.75794 | 208.62122 | 182.50195 | 1.53E-43 | 1.717530337 | YKL098W | MTC2 | n |
| W303.asm02.HP0_G0032300 | 2873.0071 | 2569.1694 | 2825.330738 | 8794.62604 | 8025.0446 | 8781.75083 | 8784.4239 | 9663.4925 | 8931.1339 | 2.23E-117 | -1.630653945 | YKL096W | CWP1 | n |
| W303.asm02.HP0_G0032310 | 200.17285 | 186.848683 | 155.5139312 | 315.016455 | 298.42236 | 320.474172 | 331.76734 | 498.5916 | 418.78466 | 2.88E-07 | -0.78527606 | YKL095W | YJU2 | n |
| W303.asm02.HP0_G0032320 | 2238.7753 | 2357.40756 | 2198.445388 | 642.533562 | 632.79922 | 651.377977 | 847.08981 | 796.43448 | 850.7941 | 6.37E-148 | 1.818241658 | YKL094W | YJU3 | n |
| W303.asm02.HP0_G0032330 | 572.07293 | 473.349998 | 506.1447202 | 225.011753 | 254.37809 | 259.792672 | 239.41973 | 244.04747 | 232.7561 | 4.85E-21 | 1.071017345 | YKL093W | MBR1 | n |
| W303.asm02.HP0_G0032340 | 849.15429 | 785.802519 | 850.014034 | 1202.56281 | 1194.5883 | 1207.94111 | 1194.8185 | 1258.2878 | 1223.7329 | 2.99E-14 | -0.5365468 | YKL092C | BUD2 | n |
| W303.asm02.HP0_G0032360 | 518.34232 | 539.785086 | 508.0765703 | 282.514757 | 273.25421 | 280.651938 | 247.40039 | 247.98372 | 265.37723 | 4.74E-20 | 0.904567282 | YKL090W | CUE2 | n |
| W303.asm02.HP0_G0032370 | 527.82419 | 524.214362 | 384.4381654 | 265.013843 | 253.47923 | 281.600086 | 277.04283 | 217.80581 | 275.0754 | 2.51E-09 | 0.843494043 | YKL089W | MIF2 | n |
| W303.asm02.HP0_G0032380 | 701.65851 | 722.481576 | 701.261578 | 1337.56987 | 1184.7008 | 1120.71145 | 1175.4369 | 1159.8815 | 1320.7146 | 9.27E-18 | -0.778208039 | YKL088W | CAB3 | n |
| W303.asm02.HP0_G0032390 | 1828.9477 | 1830.07905 | 2036.169981 | 922.548188 | 955.49086 | 954.785477 | 833.40868 | 801.68282 | 804.06654 | 1.36E-43 | 1.008183491 | YKL087C | CYT2 | n |
| W303.asm02.HP0_G0032400 | 87.443928 | 108.995065 | 86.93325348 | 274.180988 | 235.50198 | 236.088961 | 329.48715 | 244.04747 | 350.01581 | 1.24E-16 | -1.398129181 | YKL086W | SRX1 | n |
| W303.asm02.HP0_G0032410 | 1930.0877 | 1965.02532 | 2028.442581 | 4727.74695 | 4925.7666 | 4879.17186 | 3992.609 | 3948.0583 | 4171.0952 | 1.05E-116 | -1.294441245 | YKL085W | MDH1 | n |
| W303.asm02.HP0_G0032420 | 381.38195 | 357.088595 | 375.74484 | 48.3358581 | 62.021514 | 49.3037188 | 49.02404 | 55.107493 | 40.555988 | 2.36E-64 | 2.803933214 | YKL084W | HOT13 | n |
| W303.asm02.HP0_G0032430 | 2560.1054 | 2350.14122 | 2232.252764 | 2026.77253 | 1876.8249 | 1978.78579 | 1119.5723 | 1528.5769 | 1290.7384 | 0.00657847 | 0.279811613 | YKL082C | RRP14 | n |
| W303.asm02.HP0_G0032450 | 7340.0222 | 6588.49219 | 8119.565875 | 1454.24263 | 1450.7641 | 1452.56341 | 1455.9 | 1381.6236 | 1482.9385 | 1.33E-185 | 2.339165028 | YKL080W | VMA5 | n |
| W303.asm02.HP0_G0032470 | 298.15219 | 252.245723 | 340.0056136 | 1063.38888 | 1058.8601 | 1354.90412 | 615.65073 | 610.11867 | 670.05545 | 2.11E-54 | -1.96346651 | YKL078W | DHR2 | n |
| W303.asm02.HP0_G0032480 | 4896.86 | 4485.40645 | 5121.334555 | 1872.59781 | 1860.6454 | 1913.36355 | 1949.5607 | 2010.1114 | 1924.6461 | 1.34E-93 | 1.36118012 | YKL077W | PSG1 | n |
| W303.asm02.HP0_G0032490 | 1197.8765 | 1144.96721 | 1241.213675 | 867.545315 | 874.59323 | 835.318774 | 777.54407 | 802.9949 | 807.59315 | 4.28E-11 | 0.475662984 | YKL075C | AAN1 | n |
| W303.asm02.HP0_G0032500 | 1500.2428 | 1383.71831 | 1350.363204 | 1124.22539 | 1072.343 | 1235.43741 | 942.8577 | 901.40114 | 916.91799 | 0.000150093 | 0.30323638 | YKL074C | MUD2 | n |
| W303.asm02.HP0_G0032510 | 2674.9414 | 2950.1331 | 2696.862708 | 6767.85351 | 6812.4791 | 7264.71333 | 7564.5234 | 7005.212 | 6984.4464 | 3.36E-89 | -1.324700536 | YKL073W | LHS1 | n |
| W303.asm02.HP0_G0032520 | 481.46837 | 445.322696 | 414.3818416 | 579.19692 | 545.60955 | 538.548313 | 584.8682 | 514.3366 | 558.96731 | 0.00132278 | -0.312227658 | YKL072W | STB6 | n |
| W303.asm02.HP0_G0032540 | 398.23861 | 401.724669 | 364.1537396 | 467.524421 | 457.52102 | 532.859422 | 718.25919 | 589.12534 | 817.29132 | 0.010129813 | -0.324516682 | YKL070W | | n |
| W303.asm02.HP0_G0032550 | 863.90387 | 832.51469 | 864.5029096 | 673.368506 | 727.17978 | 722.48911 | 635.03233 | 862.03864 | 700.91327 | 0.010700991 | 0.27095704 | YKL069W | | n |
| W303.asm02.HP0_G0032560 | 2036.4953 | 1999.28091 | 2295.037892 | 128.340037 | 127.63848 | 124.207445 | 200.65654 | 259.79247 | 245.98088 | 3.81E-277 | 4.057329779 | YKL068W-A | | n |
| W303.asm02.HP0_G0032580 | 3498.8107 | 3427.63529 | 3782.562451 | 6096.98514 | 6702.8178 | 6995.43917 | 5370.9826 | 5845.3305 | 5079.1967 | 7.64E-30 | -0.886051176 | YKL067W | YNK1 | n |
| W303.asm02.HP0_G0032610 | 4097.2221 | 4183.33441 | 4819.965943 | 1515.07914 | 1518.1788 | 1534.10417 | 1951.8408 | 1936.6348 | 1845.2975 | 2.84E-91 | 1.520486177 | YKL065C | YET1 | n |
| W303.asm02.HP0_G0032620 | 1485.4932 | 1452.22949 | 1621.78814 | 1899.26587 | 1980.1941 | 1844.14871 | 2236.8643 | 1928.7623 | 2157.4022 | 3.34E-05 | -0.327754915 | YKL064W | MNR2 | n |
| W303.asm02.HP0_G0032630 | 1768.8958 | 1639.07817 | 1662.356991 | 2070.10813 | 1967.6101 | 1979.73394 | 2145.6568 | 2197.7393 | 2199.7215 | 8.27E-05 | -0.247511699 | YKL063C | | n |
| W303.asm02.HP0_G0032650 | 822.81576 | 870.922475 | 1077.006418 | 65.0033954 | 70.111277 | 60.6815 | 72.966013 | 93.157905 | 81.111976 | 2.66E-134 | 3.822546374 | YKL061W | BLI1 | n |
| W303.asm02.HP0_G0032660 | 50754.352 | 48889.9961 | 46832.8755 | 25935.5214 | 24470.633 | 26025.7265 | 24697.855 | 24764.258 | 23072.831 | 5.00E-59 | 0.938414273 | YKL060C | FBA1 | n |
| W303.asm02.HP0_G0032670 | 386.64966 | 482.692432 | 448.1892179 | 831.71011 | 717.29229 | 788.8595 | 757.02238 | 799.05865 | 787.31516 | 1.31E-16 | -0.827244768 | YKL059C | MPE1 | n |
| W303.asm02.HP0_G0032680 | 1789.9667 | 1741.84495 | 2098.955109 | 1028.38705 | 1026.501 | 974.696594 | 1085.3694 | 1161.1936 | 1092.3667 | 8.01E-26 | 0.894428738 | YKL058W | TOA2 | n |
| W303.asm02.HP0_G0032690 | 2877.2213 | 2622.10986 | 2764.477461 | 2388.45809 | 2465.5799 | 2581.8082 | 2288.1686 | 2285.6489 | 2167.9821 | 0.020476398 | 0.152575426 | YKL057C | NUP120 | n |
| W303.asm02.HP0_G0032700 | 16256.142 | 15668.3002 | 18749.57092 | 11533.9358 | 12047.454 | 12651.1446 | 9746.6632 | 11008.378 | 9140.967 | 4.42E-08 | 0.484042982 | YKL056C | TMA19 | n |
| W303.asm02.HP0_G0032710 | 108.51475 | 86.158004 | 100.456204 | 310.84957 | 281.34397 | 266.429711 | 302.1249 | 308.33954 | 245.98088 | 1.75E-24 | -1.541894382 | YKL055C | OAR1 | n |
| W303.asm02.HP0_G0032720 | 2963.6117 | 2948.05701 | 3167.268202 | 12177.3027 | 11819.143 | 12534.5223 | 10373.715 | 9925.9092 | 10179.553 | 4.22E-284 | -2.00824841 | YKL054C | DEF1 | n |
| W303.asm02.HP0_G0032730 | 1640.3638 | 1602.74648 | 2190.717988 | 734.205017 | 675.04575 | 674.133539 | 655.55402 | 749.19949 | 673.58206 | 4.41E-32 | 1.383152456 | YKL053C-/ | MDM35 | n |
| W303.asm02.HP0_G0032740 | 1032.4705 | 932.167321 | 1075.074568 | 230.845391 | 248.98492 | 196.266727 | 207.4971 | 181.06748 | 221.29463 | 9.64E-81 | 2.167327465 | YKL052C | ASK1 | n |

| | | | | | | | | | | | | | | |
|---|---|---|---|---|---|---|---|---|---|---|---|---|---|---|
| W303.asm02.HP0_G0032750 | 1181.0198 | 1183.375 | 1170.701147 | 2565.96736 | 2405.3561 | 2326.75627 | 2467.1633 | 2373.5585 | 2354.8923 | 1.22E-57 | -1.046237061 | YKL051W | SFK1 | n |
| W303.asm02.HP0_G0032770 | 907.09906 | 912.444404 | 967.8568887 | 637.533301 | 700.21391 | 641.896492 | 635.03233 | 800.37073 | 609.22147 | 2.33E-06 | 0.49399782 | YKL049C | CSE4 | n |
| W303.asm02.HP0_G0032780 | 443.54089 | 476.464143 | 406.6544413 | 479.191697 | 425.16197 | 451.318656 | 372.81072 | 427.73911 | 407.32318 | 0.766880238 | -0.032634018 | YKL048C | ELM1 | n |
| W303.asm02.HP0_G0032790 | 1017.7209 | 950.852189 | 958.1976383 | 446.689999 | 464.71192 | 461.748289 | 482.25974 | 495.96744 | 489.31681 | 7.60E-42 | 1.092011073 | YKL047W | ANR2 | n |
| W303.asm02.HP0_G0032800 | 1949.0514 | 1736.65471 | 1980.146329 | 1450.90912 | 1528.0663 | 1402.31154 | 1735.223 | 1697.8356 | 1706.8781 | 4.23E-07 | 0.371013743 | YKL046C | DCW1 | n |
| W303.asm02.HP0_G0032810 | 1181.0198 | 1108.63552 | 1054.790142 | 1380.07209 | 1318.6313 | 1341.63004 | 1321.3689 | 1323.8919 | 1326.8861 | 9.96E-05 | -0.273488876 | YKL045W | PRI2 | n |
| W303.asm02.HP0_G0032840 | 361.36467 | 330.099341 | 291.7093617 | 567.529644 | 557.29476 | 552.770539 | 557.50594 | 472.34994 | 532.51776 | 3.03E-13 | -0.772685984 | YKL042W | SPC42 | n |
| W303.asm02.HP0_G0032850 | 1173.645 | 1057.77116 | 1187.121872 | 814.209196 | 892.57049 | 853.333594 | 869.89168 | 902.71322 | 904.57486 | 1.19E-07 | 0.417811789 | YKL041W | VPS24 | n |
| W303.asm02.HP0_G0032860 | 2884.5961 | 2657.4035 | 3295.736232 | 1308.40168 | 1341.1029 | 1248.71149 | 1268.9246 | 1524.6406 | 1310.1347 | 3.46E-35 | 1.180961719 | YKL040C | NFU1 | n |
| W303.asm02.HP0_G0032870 | 2449.4835 | 2247.37444 | 2096.057334 | 2241.78376 | 2148.2814 | 2237.63031 | 2755.6071 | 2866.9017 | 2590.2933 | 0.65666207 | 0.035149399 | YKL039W | PTM1 | n |
| W303.asm02.HP0_G0032880 | 454.0763 | 475.426095 | 427.9047921 | 228.345261 | 204.04179 | 207.644508 | 311.24565 | 225.67831 | 312.10478 | 8.30E-17 | 1.083209464 | YKL038W | RGT1 | n |
| W303.asm02.HP0_G0032900 | 2719.1901 | 2830.75755 | 2323.049718 | 5025.26249 | 4896.1042 | 5010.01635 | 5267.2341 | 4836.3386 | 5045.6939 | 1.73E-32 | -0.923738938 | YKL035W | UGP1 | n |
| W303.asm02.HP0_G0032910 | 470.93296 | 529.404603 | 477.1669691 | 524.194047 | 539.31751 | 460.800141 | 975.92042 | 814.80365 | 896.63999 | 0.674678429 | -0.046039736 | YKL034W | TUL1 | n |
| W303.asm02.HP0_G0032920 | 3430.3305 | 3143.21008 | 3935.178607 | 866.711938 | 937.51361 | 915.911391 | 843.66952 | 768.88074 | 877.24365 | 1.60E-98 | 1.950335976 | YKL033W-A | | n |
| W303.asm02.HP0_G0032930 | 713.24746 | 687.187936 | 803.6496321 | 318.349962 | 321.79278 | 322.370469 | 275.90274 | 237.48705 | 240.69097 | 1.50E-31 | 1.196144924 | YKL033W | TTI1 | n |
| W303.asm02.HP0_G0032940 | 394.02445 | 527.328507 | 466.5417937 | 1689.2549 | 1610.7616 | 1453.51156 | 1457.0401 | 1169.0661 | 1354.2173 | 2.78E-54 | -1.776104616 | YKL032C | IXR1 | y |
| W303.asm02.HP0_G0032960 | 973.47216 | 1040.12434 | 1194.849273 | 1910.09977 | 1841.7693 | 1925.68948 | 1021.5242 | 1195.3078 | 1105.5915 | 3.49E-20 | -0.822631397 | YKL029C | MAE1 | n |
| W303.asm02.HP0_G0032970 | 2685.4768 | 2625.224 | 2558.735427 | 1758.42518 | 1645.8173 | 1663.05236 | 1682.7787 | 1741.1344 | 1756.2506 | 2.64E-25 | 0.634527348 | YKL028W | TFA1 | n |
| W303.asm02.HP0_G0032980 | 1801.5556 | 1548.76798 | 1758.949495 | 1663.42022 | 1683.5695 | 2018.60802 | 1271.2048 | 1448.5398 | 1272.2237 | 0.492198081 | -0.070120367 | YKL027W | TCD2 | n |
| W303.asm02.HP0_G0033000 | 641.60665 | 726.633769 | 754.3874552 | 1388.40586 | 1413.9107 | 1279.05224 | 1419.417 | 1255.6636 | 1349.8091 | 2.58E-27 | -0.942770188 | YKL025C | PAN3 | n |
| W303.asm02.HP0_G0033010 | 1499.1893 | 1515.55043 | 1488.490485 | 2392.62498 | 2399.0641 | 2577.06745 | 2147.937 | 2161.001 | 2107.1481 | 5.29E-31 | -0.710207247 | YKL024C | URA6 | n |
| W303.asm02.HP0_G0033040 | 600.51854 | 702.758659 | 660.6927264 | 830.876733 | 844.03191 | 793.600242 | 824.28793 | 831.86073 | 815.52802 | 8.12E-05 | -0.329888029 | YKL022C | CDC16 | n |
| W303.asm02.HP0_G0033050 | 1850.0185 | 1668.14352 | 1997.53298 | 2234.28337 | 2255.2461 | 2364.6822 | 1657.6966 | 1871.0306 | 1682.1918 | 0.000151428 | -0.312970816 | YKL021C | MAK11 | n |
| W303.asm02.HP0_G0033070 | 1576.0978 | 1360.88124 | 1407.352781 | 656.700969 | 631.90035 | 646.637235 | 775.26389 | 770.19282 | 795.25002 | 1.57E-47 | 1.166195322 | YKL019W | RAM2 | n |
| W303.asm02.HP0_G0033080 | 923.95572 | 808.63958 | 1015.187216 | 330.017238 | 374.82567 | 291.08157 | 352.28903 | 384.44037 | 380.87363 | 1.93E-36 | 1.463841974 | YKL018C-/ | MCO12 | n |
| W303.asm02.HP0_G0033090 | 675.31997 | 692.378177 | 627.8512751 | 732.538263 | 695.71959 | 633.363156 | 661.25449 | 674.41075 | 623.3279 | 0.604647694 | -0.048579523 | YKL018W | SWD2 | n |
| W303.asm02.HP0_G0033100 | 516.23524 | 468.159757 | 441.4277426 | 655.034215 | 637.29353 | 581.214992 | 575.74745 | 555.01118 | 577.482 | 2.98E-05 | -0.39583007 | YKL017C | HCS1 | n |
| W303.asm02.HP0_G0033110 | 6278.0526 | 6399.56741 | 6680.337567 | 6433.66939 | 6603.943 | 7074.13549 | 5708.4504 | 6004.0926 | 5422.1593 | 0.410969843 | -0.054904153 | YKL016C | ATP7 | n |
| W303.asm02.HP0_G0033120 | 1815.2517 | 1721.08398 | 1606.333339 | 890.046491 | 869.20006 | 804.029875 | 761.58276 | 733.45449 | 767.03716 | 3.06E-38 | 1.003446373 | YKL015W | PUT3 | n |
| W303.asm02.HP0_G0033130 | 1208.4119 | 1102.40723 | 1423.773507 | 2114.2771 | 2301.0881 | 2401.65999 | 1377.2335 | 1720.141 | 1447.6724 | 1.17E-15 | -0.867505592 | YKL014C | URB1 | n |
| W303.asm02.HP0_G0033140 | 4077.2048 | 4041.1218 | 4648.997211 | 1735.92401 | 1715.0297 | 1745.54127 | 1746.6239 | 1798.866 | 1796.8066 | 4.14E-76 | 1.297108319 | YKL013C | ARC19 | n |
| W303.asm02.HP0_G0033150 | 984.00757 | 1062.9614 | 790.1266816 | 689.202666 | 667.85485 | 669.392797 | 646.43327 | 582.56493 | 658.59398 | 8.98E-06 | 0.484530577 | YKL012W | PRP40 | n |
| W303.asm02.HP0_G0033160 | 814.38742 | 810.715677 | 977.5161391 | 186.676418 | 177.97478 | 159.288938 | 181.27494 | 175.81914 | 189.55516 | 4.72E-85 | 2.311156343 | YKL011C | CCE1 | n |
| W303.asm02.HP0_G0033170 | 1479.172 | 1561.22456 | 1528.093411 | 1784.25987 | 1719.524 | 1920.94874 | 1745.4838 | 1586.3086 | 1865.5754 | 0.001919819 | -0.247543935 | YKL010C | UFD4 | n |
| W303.asm02.HP0_G0033180 | 5829.244 | 5398.8889 | 6547.039912 | 3315.17316 | 3121.7495 | 3434.19364 | 1786.5272 | 2372.2464 | 1939.6342 | 2.36E-16 | 0.848694797 | YKL009W | MRT4 | n |
| W303.asm02.HP0_G0033200 | 2943.5944 | 2983.35065 | 2866.865515 | 1311.73518 | 1391.4392 | 1315.08188 | 1362.4123 | 1396.0565 | 1487.3468 | 3.85E-69 | 1.129877829 | YKL007W | CAP1 | n |
| W303.asm02.HP0_G0033240 | 1092.5223 | 1116.93991 | 1166.837447 | 876.712461 | 844.03191 | 855.229891 | 905.2346 | 1089.029 | 893.11339 | 1.02E-05 | 0.390410979 | YKL004W | AUR1 | n |
| W303.asm02.HP0_G0033250 | 1605.5969 | 1577.83333 | 1823.666473 | 1212.56334 | 1319.5302 | 1303.7041 | 1160.6156 | 1285.8415 | 1195.52 | 2.48E-06 | 0.385249332 | YKL003C | MRP17 | n |
| W303.asm02.HP0_G0033270 | 1779.4313 | 1736.65471 | 2134.694335 | 665.034737 | 724.48319 | 717.748367 | 782.10445 | 743.95116 | 764.39221 | 1.81E-53 | 1.42403302 | YKL001C | MET14 | n |
| W303.asm02.HP0_G0033280 | 2549.5699 | 2449.79385 | 2430.267397 | 5035.26301 | 4962.62 | 5122.84601 | 5279.7751 | 5163.0473 | 5003.3746 | 8.30E-80 | -1.025282231 | YKR001C | VPS1 | n |
| W303.asm02.HAP0_G0033290 | 612.1075 | 703.796708 | 724.443779 | 1947.60173 | 1951.4305 | 1934.22281 | 1780.8268 | 1713.5806 | 1831.191 | 3.14E-87 | -1.51456526 | YKR002W | PAP1 | n |
| W303.asm02.HP0_G0033310 | 510.96753 | 533.556796 | 669.3860518 | 274.180988 | 333.478 | 305.303797 | 316.94612 | 346.38996 | 320.03964 | 3.52E-13 | 0.910709213 | YKR004C | ECM9 | n |
| W303.asm02.HP0_G0033320 | 271.81366 | 268.854495 | 247.2768099 | 26.6680596 | 36.853363 | 27.4963047 | 35.342912 | 20.993331 | 36.147728 | 3.24E-50 | 3.114785843 | YKR005C | | n |
| W303.asm02.HP0_G0033330 | 2904.6134 | 2825.56731 | 2948.969143 | 1991.7707 | 2057.4963 | 2044.20803 | 1759.165 | 2084.9002 | 1840.8892 | 1.40E-12 | 0.510451857 | YKR006C | MRPL13 | n |
| W303.asm02.HP0_G0033340 | 1265.3031 | 1232.16326 | 1349.397279 | 752.539308 | 759.53883 | 679.82243 | 795.78558 | 758.38407 | 801.42159 | 9.91E-25 | 0.81109619 | YKR007W | MEH1 | n |
| W303.asm02.HP0_G0033350 | 1777.3242 | 1717.96984 | 1498.149735 | 1211.72996 | 1235.936 | 1170.96332 | 1192.5383 | 1163.8178 | 1190.2301 | 4.25E-09 | 0.463882038 | YKR008W | RSC4 | n |
| W303.asm02.HP0_G0033360 | 506.75337 | 534.594844 | 482.9625193 | 204.177332 | 189.65999 | 212.38525 | 293.00415 | 226.99039 | 283.01026 | 2.66E-29 | 1.330225756 | YKR009C | FOX2 | n |
| W303.asm02.HP0_G0033370 | 312.90177 | 284.425218 | 293.6412117 | 2525.1319 | 2461.0856 | 2552.41559 | 2184.42 | 2220.0447 | 2277.3069 | 0 | -3.081287362 | YKR010C | TOF2 | n |
| W303.asm02.FHP0_G0033380 | 2205.0619 | 2431.10898 | 2328.845268 | 908.380782 | 919.53636 | 821.096547 | 1121.8524 | 1146.7607 | 1181.4136 | 3.62E-79 | 1.394038302 | YKR011C | FPT1 | n |
| W303.asm02.HP0_G0033390 | 2131.314 | 2441.48946 | 2404.187421 | 1590.08306 | 1532.5606 | 1586.25234 | 1324.7892 | 1060.1632 | 1327.7678 | 1.01E-09 | 0.567327788 | YKR013W | PRY2 | n |
| W303.asm02.HP0_G0033400 | 3514.6138 | 3338.36314 | 3776.766901 | 1530.07992 | 1675.4797 | 1552.11899 | 1527.7259 | 1562.6911 | 1569.3404 | 1.62E-61 | 1.160082215 | YKR014C | YPT52 | n |
| W303.asm02.HP0_G0033420 | 2150.2778 | 2164.33058 | 2030.374431 | 3688.526 | 3620.6182 | 3722.43077 | 3460.1851 | 3257.9025 | 3389.9516 | 2.63E-43 | -0.798215302 | YKR016W | MIC60 | n |
| W303.asm02.HP0_G0033430 | 547.84148 | 487.882673 | 538.0202465 | 670.034999 | 729.87637 | 619.14093 | 731.94032 | 644.23284 | 700.91327 | 0.000231449 | -0.359726307 | YKR017C | HEL1 | n |
| W303.asm02.HP0_G0033440 | 1564.5088 | 1509.32214 | 1429.569057 | 1795.92714 | 1801.3205 | 1706.66719 | 1710.1409 | 1623.0469 | 1768.5937 | 0.000636233 | -0.236658397 | YKR018C | | n |
| W303.asm02.HP0_G0033460 | 567.85876 | 516.948024 | 443.3595927 | 410.021417 | 401.79155 | 365.037149 | 466.29843 | 432.98745 | 456.69569 | 0.000712826 | 0.374701639 | YKR020W | VPS51 | n |
| W303.asm02.HP0_G0033470 | 741.69308 | 836.666883 | 764.0467055 | 1026.7203 | 1000.434 | 922.54843 | 1237.0019 | 1002.4315 | 1133.8044 | 0.000499569 | -0.333123837 | YKR021W | ALY1 | n |
| W303.asm02.HP0_G0033480 | 916.58093 | 965.384865 | 843.2525587 | 115.839384 | 131.23393 | 116.622258 | 107.16883 | 110.21499 | 95.218407 | 7.51E-132 | 2.905525834 | YKR022C | NTR2 | n |
| W303.asm02.HP0_G0033490 | 872.3322 | 826.2864 | 847.1162589 | 784.207629 | 770.32518 | 769.896532 | 700.01769 | 784.62574 | 749.40413 | 0.096591289 | 0.130966058 | YKR023W | RQT4 | n |
| W303.asm02.HP0_G0033500 | 703.76559 | 713.139142 | 738.9326545 | 2148.44556 | 1974.801 | 2196.85993 | 1297.4269 | 1634.8556 | 1238.7209 | 6.95E-52 | -1.551474836 | YKR024C | DBP7 | n |
| W303.asm02.HP0_G0033510 | 912.36677 | 988.221926 | 824.899983 | 1404.24002 | 1303.3507 | 1504.71157 | 910.93507 | 1070.6599 | 853.43905 | 1.10E-09 | -0.628390251 | YKR025W | RPC37 | n |

| | | | | | | | | | | | | | | |
|---|---|---|---|---|---|---|---|---|---|---|---|---|---|---|
| W303.asm02.HP0_G0033520 | 2185.0447 | 1881.98146 | 2224.525364 | 1469.24341 | 1486.7186 | 1601.42271 | 1258.6637 | 1438.0432 | 1296.0283 | 6.91E-08 | 0.465664441 | YKR026C | GCN3 | n |
| W303.asm02.HP0_G0033540 | 1370.6572 | 1493.75142 | 1530.025261 | 4316.05878 | 4161.7335 | 4443.97173 | 4169.3236 | 4009.7262 | 4222.231 | 9.30E-128 | -1.555567192 | YKR028W | SAP190 | n |
| W303.asm02.HP0_G0033560 | 1724.6471 | 1705.51326 | 2064.181808 | 720.03761 | 782.01039 | 689.303914 | 982.76099 | 1012.9282 | 1003.3199 | 1.59E-47 | 1.326408454 | YKR030W | GMH1 | n |
| W303.asm02.HP0_G0033570 | 835.45825 | 914.520501 | 1015.187216 | 518.360409 | 518.64368 | 471.229774 | 536.98425 | 413.3062 | 474.32873 | 1.27E-15 | 0.87451494 | YKR031C | SPO14 | n |
| W303.asm02.HP0_G0033590 | 175.9414 | 210.723793 | 131.3658053 | 1160.06059 | 1148.7463 | 1288.53373 | 1083.0893 | 1123.1432 | 1102.9465 | 1.97E-116 | -2.799299423 | YKR034W | DAL80 | n |
| W303.asm02.HP0_G0033600 | 2685.4768 | 2670.89813 | 2850.444789 | 1392.57274 | 1473.2357 | 1388.08931 | 1322.509 | 1464.2848 | 1424.7495 | 8.48E-45 | 0.948241756 | YKR035W- | DID2 | n |
| W303.asm02.HP0_G0033620 | 758.54974 | 776.460085 | 804.6155572 | 472.524682 | 471.00396 | 479.76311 | 481.11965 | 579.94076 | 529.8728 | 1.12E-15 | 0.717496735 | YKR037C | SPC34 | n |
| W303.asm02.HP0_G0033630 | 1107.2719 | 1096.17894 | 1142.689321 | 1711.75608 | 1867.8363 | 1736.05979 | 1658.8367 | 1642.7281 | 1620.4762 | 1.58E-23 | -0.667390255 | YKR038C | KAE1 | n |
| W303.asm02.HP0_G0033640 | 4015.0459 | 4329.69922 | 4296.434572 | 684.202405 | 678.6412 | 640.948344 | 702.29787 | 561.5716 | 643.6059 | 2.17E-232 | 2.656899995 | YKR039W | GAP1 | n |
| W303.asm02.HP0_G0033660 | 6773.217 | 7590.20874 | 8013.31412 | 1971.76966 | 1815.7023 | 1849.8376 | 1898.2564 | 1720.141 | 2022.5095 | 8.89E-126 | 1.988812445 | YKR042W | UTH1 | n |
| W303.asm02.HP0_G0033670 | 2645.4422 | 2549.44648 | 2742.261185 | 1910.93315 | 1912.7794 | 1912.4154 | 1706.7206 | 1637.4798 | 1632.8193 | 4.04E-15 | 0.468742978 | YKR043C | SHB17 | n |
| W303.asm02.HP0_G0033700 | 2082.8511 | 2178.86326 | 2187.820212 | 443.356492 | 408.08359 | 342.281586 | 1175.4369 | 883.03197 | 1189.3484 | 3.75E-93 | 2.432186915 | YKR046C | PLN1 | n |
| W303.asm02.HP0_G0033710 | 1303.2306 | 1548.76798 | 1682.641417 | 3716.02744 | 3590.9558 | 3758.46041 | 3379.2385 | 3604.2925 | 3368.792 | 1.37E-49 | -1.286402564 | YKR048C | NAP1 | n |
| W303.asm02.HP0_G0033720 | 476.20067 | 500.339252 | 461.7121685 | 163.341865 | 160.89639 | 161.185234 | 136.81127 | 152.20165 | 146.35422 | 7.30E-43 | 1.56643435 | YKR049C | FMP46 | n |
| W303.asm02.HP0_G0033730 | 283.40261 | 349.822257 | 340.0056136 | 1040.8877 | 1030.0965 | 1060.02995 | 1048.8864 | 1146.7607 | 1010.3731 | 4.55E-70 | -1.684635858 | YKR050W | TRK2 | n |
| W303.asm02.HP0_G0033740 | 880.76053 | 779.574229 | 825.865908 | 395.85401 | 399.09496 | 368.829742 | 593.98895 | 540.57827 | 605.69486 | 8.84E-33 | 1.094286131 | YKR051W | HFL1 | n |
| W303.asm02.HP0_G0033750 | 867.06449 | 862.618089 | 812.3429575 | 2037.60643 | 1848.0613 | 2002.4895 | 1787.6673 | 1822.4835 | 1763.3038 | 4.03E-66 | -1.212443128 | YKR052C | MRS4 | n |
| W303.asm02.HP0_G0033770 | 660.57039 | 687.187936 | 769.8422558 | 1287.56725 | 1208.0712 | 1158.63739 | 1189.118 | 1267.4723 | 1246.6558 | 4.08E-21 | -0.786709507 | YKR054C | DYN1 | n |
| W303.asm02.HP0_G0033780 | 769.08515 | 696.53037 | 677.1134521 | 560.862629 | 537.51979 | 510.10386 | 552.94557 | 627.17575 | 614.51138 | 8.79E-06 | 0.412387033 | YKR055W | RHO4 | n |
| W303.asm02.HP0_G0033790 | 851.26137 | 889.607343 | 849.0481089 | 1780.92636 | 1699.749 | 1754.07461 | 1385.2142 | 1594.1811 | 1484.7018 | 1.22E-42 | -1.015429938 | YKR056W | TRM2 | n |
| W303.asm02.HP0_G0033830 | 763.81744 | 731.82401 | 705.1252782 | 874.21233 | 861.1103 | 928.237321 | 557.50594 | 732.14241 | 559.84896 | 0.009611025 | -0.275402074 | YKR060W | UTP30 | n |
| W303.asm02.HP0_G0033850 | 1136.7711 | 1153.2716 | 1027.744241 | 1344.23688 | 1306.0472 | 1315.08188 | 1403.4557 | 1512.8319 | 1525.2578 | 0.000557704 | -0.257971921 | YKR062W | TFA2 | n |
| W303.asm02.HP0_G0033860 | 553.10918 | 628.019186 | 595.9757488 | 695.869681 | 714.59571 | 782.222461 | 508.4819 | 656.04158 | 561.61227 | 0.004268224 | -0.302442637 | YKR063C | LAS1 | n |
| W303.asm02.HP0_G0033870 | 667.94518 | 724.557673 | 646.2038508 | 833.376864 | 824.25693 | 814.459508 | 880.15253 | 900.08905 | 897.52165 | 0.000383034 | -0.278792458 | YKR064W | OAF3 | n |
| W303.asm02.HP0_G0033880 | 1532.9026 | 1521.77872 | 1547.411912 | 1558.41474 | 1508.2913 | 1574.87456 | 1716.9815 | 1864.4702 | 1926.4094 | 0.865955336 | -0.012226373 | YKR065C | PAM17 | n |
| W303.asm02.HP0_G0033890 | 1306.3912 | 1578.87138 | 1526.161561 | 4812.75139 | 4232.7436 | 4203.14202 | 5727.832 | 6162.8546 | 5850.6421 | 5.54E-77 | -1.586441305 | YKR066C | CCP1 | n |
| W303.asm02.HP0_G0033900 | 595.25083 | 645.666006 | 712.8526785 | 1365.0713 | 1281.778 | 1420.32636 | 1856.073 | 1578.4361 | 1682.1918 | 2.96E-30 | -1.056720985 | YKR067W | GPT2 | n |
| W303.asm02.HP0_G0033910 | 2795.0451 | 2858.78486 | 3157.608951 | 820.042834 | 840.43646 | 767.052086 | 753.6021 | 872.53531 | 823.46289 | 1.03E-115 | 1.859722003 | YKR068C | BET3 | n |
| W303.asm02.HP0_G0033920 | 429.84485 | 472.31195 | 448.1892179 | 353.35179 | 367.63477 | 322.370469 | 395.6126 | 338.51746 | 373.82041 | 0.000398451 | 0.37133451 | YKR069W | MET1 | n |
| W303.asm02.HP0_G0033930 | 856.52908 | 802.411291 | 801.7177821 | 896.713506 | 871.89665 | 864.711375 | 962.23929 | 974.87779 | 1027.1245 | 0.199196894 | -0.098484979 | YKR070W | | n |
| W303.asm02.HP0_G0033940 | 8892.9421 | 8489.15852 | 8599.630619 | 1468.41003 | 1359.0801 | 1426.01525 | 1362.4123 | 1298.9623 | 1389.4834 | 0 | 2.610500224 | YKR071C | DRE2 | n |
| W303.asm02.HP0_G0033950 | 735.37183 | 951.890237 | 883.8214103 | 1572.58214 | 1578.4026 | 1572.97826 | 1715.8414 | 1347.5094 | 1676.9019 | 9.28E-16 | -0.877333955 | YKR072C | SIS2 | n |
| W303.asm02.HP0_G0033970 | 1740.4502 | 1685.79034 | 2173.331337 | 855.044662 | 955.49086 | 912.118797 | 803.76624 | 860.72656 | 867.54548 | 1.17E-24 | 1.041058586 | YKR074W | AIM29 | n |
| W303.asm02.HP0_G0033980 | 523.61003 | 569.888485 | 495.5195448 | 752.539308 | 713.69684 | 517.689047 | 566.62669 | 675.72283 | 611.86643 | 0.016911016 | -0.321723114 | YKR075C | | n |
| W303.asm02.HP0_G0033990 | 382.43549 | 510.719735 | 432.7344173 | 445.856622 | 448.5324 | 379.259375 | 928.03648 | 759.69616 | 999.79327 | 0.683070319 | 0.057257739 | YKR076W | ECM4 | n |
| W303.asm02.HP0_G0034010 | 591.03667 | 585.459208 | 562.1683725 | 1421.74093 | 1377.0574 | 1342.57819 | 1600.6919 | 1415.7377 | 1549.9441 | 2.66E-58 | -1.252863644 | YKR078W | VPS501 | n |
| W303.asm02.HP0_G0034020 | 1457.0476 | 1230.08717 | 1465.308284 | 1557.58136 | 1565.8185 | 1501.86713 | 1107.0312 | 1288.4657 | 1128.5144 | 0.086628113 | -0.155475276 | YKR079C | TRZ1 | n |
| W303.asm02.HP0_G0034040 | 3149.0349 | 2942.86676 | 2952.832843 | 2195.94804 | 2162.6632 | 2464.23779 | 1333.9099 | 1826.4198 | 1378.9036 | 0.000181271 | 0.406825042 | YKR081C | RPF2 | n |
| W303.asm02.HP0_G0034050 | 1812.091 | 1579.90942 | 1530.991186 | 2913.48552 | 2960.8531 | 3017.95648 | 2664.3996 | 2642.5355 | 2600.8731 | 4.29E-32 | -0.853417419 | YKR082W | NUP133 | n |
| W303.asm02.HP0_G0034060 | 681.64122 | 672.65526 | 801.7177821 | 275.847742 | 266.96217 | 280.651938 | 266.78198 | 301.77913 | 276.8387 | 1.02E-38 | 1.389593089 | YKR083C | DAD2 | n |
| W303.asm02.HP0_G0034080 | 15.803119 | 12.4565789 | 11.59110046 | 440.022984 | 400.89269 | 425.718649 | 370.53053 | 372.63162 | 355.30572 | 3.56E-87 | -4.995982599 | YKR084C | HBS1 | y |
| W303.asm02.HP0_G0034090 | 2045.9772 | 2115.54232 | 2590.610954 | 1453.40925 | 1376.1585 | 1481.95601 | 1335.05 | 1430.1707 | 1359.5072 | 2.84E-12 | 0.64762818 | YKR085C | MRPL20 | n |
| W303.asm02.HP0_G0034100 | 532.03836 | 533.556796 | 552.5091221 | 743.372163 | 731.67409 | 768.000235 | 611.09036 | 625.86367 | 572.19209 | 1.12E-08 | -0.470561468 | YKR086W | PRP16 | n |
| W303.asm02.HP0_G0034110 | 727.99704 | 707.948901 | 722.5119289 | 629.199532 | 625.60832 | 676.977985 | 643.01299 | 699.34033 | 629.49947 | 0.05620228 | 0.160706582 | YKR087C | OMA1 | n |
| W303.asm02.HP0_G0034120 | 1332.7297 | 1241.5057 | 1285.646226 | 454.190391 | 462.9142 | 416.237164 | 598.54932 | 532.70577 | 566.02053 | 7.13E-75 | 1.532585638 | YKR088C | TVP38 | n |
| W303.asm02.HP0_G0034130 | 585.76896 | 702.758659 | 658.7608763 | 949.216248 | 950.99655 | 888.415086 | 1028.3647 | 985.37446 | 1074.7337 | 3.32E-09 | -0.518115541 | YKR089C | TGL4 | n |
| W303.asm02.HP0_G0034140 | 580.50126 | 630.095283 | 649.101626 | 423.355447 | 447.63354 | 523.377938 | 460.59796 | 383.12829 | 504.30489 | 0.000441646 | 0.417070717 | YKR090W | PXL1 | n |
| W303.asm02.HP0_G0034160 | 2098.6543 | 1927.65558 | 2289.242342 | 597.531211 | 671.4503 | 881.778047 | 393.33241 | 457.91703 | 449.64248 | 1.29E-34 | 1.555461201 | YKR092C | SRP40 | y |
| W303.asm02.HP0_G0034170 | 2312.5232 | 2707.22981 | 2573.224303 | 1105.05772 | 963.58062 | 887.466938 | 2402.178 | 2129.511 | 2749.8723 | 9.27E-36 | 1.360304404 | YKR093W | PTR2 | n |
| W303.asm02.HP0_G0034180 | 1331.6762 | 1295.48421 | 1176.496697 | 2913.48552 | 3022.8747 | 3011.31944 | 2729.3849 | 2725.1967 | 2759.5705 | 2.98E-80 | -1.234707741 | YKR095W | MLP1 | n |
| W303.asm02.HP0_G0034190 | 596.30438 | 571.964581 | 666.4882766 | 490.025596 | 532.12661 | 439.940875 | 449.19702 | 481.53452 | 478.73699 | 0.001456237 | 0.327627428 | YKR095W- | PCC1 | n |
| W303.asm02.HP0_G0034200 | 737.47891 | 839.781027 | 721.5460038 | 2232.61662 | 2149.1803 | 2047.05248 | 2445.5015 | 2141.3197 | 2410.4363 | 1.15E-68 | -1.484327623 | YKR096W | ESL2 | n |
| W303.asm02.HP0_G0034210 | 257.06408 | 229.408661 | 225.060534 | 251.679813 | 262.46786 | 242.726 | 470.8588 | 312.27579 | 474.32873 | 0.577893477 | -0.089911928 | YKR097W | PCK1 | n |
| W303.asm02.HP0_G0034230 | 437.21964 | 432.866117 | 405.6885162 | 690.036043 | 690.32642 | 763.259492 | 438.93617 | 453.98078 | 456.69569 | 9.77E-17 | -0.748628394 | YKR099W | BAS1 | n |
| W303.asm02.HP0_G0034240 | 1591.9009 | 1481.29484 | 1637.24294 | 1844.263 | 1742.8944 | 1938.96356 | 1998.5847 | 1914.3293 | 2045.4324 | 0.001503354 | -0.229984322 | YKR100C | SKG1 | n |
| W303.asm02.HP0_G0034420 | 233.88617 | 229.408661 | 224.094609 | 119.172892 | 129.4362 | 124.207445 | 135.67118 | 118.08749 | 148.11752 | 3.68E-10 | 0.883033623 | YLL057C | JLP1 | n |
| W303.asm02.HP0_G0034440 | 507.80691 | 452.589033 | 476.201044 | 237.512406 | 208.53611 | 274.014899 | 282.7433 | 249.2958 | 281.24696 | 9.75E-17 | 0.9976904 | YLL055W | YCT1 | n |
| W303.asm02.HP0_G0034500 | 2158.7061 | 2246.33639 | 2644.702756 | 2955.15436 | 2901.5282 | 2926.93423 | 3814.7544 | 4320.6899 | 3915.4161 | 0.000211195 | -0.316853715 | YLL048C | YBT1 | n |
| W303.asm02.HP0_G0034520 | 14495.675 | 14662.4314 | 12721.23276 | 12017.2944 | 11864.086 | 12935.5891 | 9766.0448 | 10206.695 | 9045.7486 | 0.015562603 | 0.185847192 | YLL045C | RPL8B | y |
| W303.asm02.HP0_G0034530 | 1015.6138 | 1067.11359 | 1247.97515 | 782.540875 | 799.98765 | 717.748367 | 940.57751 | 1010.304 | 890.46843 | 3.36E-08 | 0.534291434 | YLL043W | FPS1 | n |

| | | | | | | | | | | | | | | |
|---|---|---|---|---|---|---|---|---|---|---|---|---|---|---|
| W303.asm02.HP0_G0034540 | 176.99494 | 145.326754 | 220.2309088 | 216.677985 | 237.29971 | 209.540805 | 171.01409 | 212.55747 | 186.91021 | 0.073705366 | -0.28819608 | YLL042C | ATG10 | n |
| W303.asm02.HP0_G0034550 | 2197.6872 | 2173.67302 | 1923.156752 | 5786.13557 | 6285.7456 | 6363.02417 | 5259.2534 | 5215.5306 | 5046.5755 | 4.50E-107 | -1.550560799 | YLL041C | SDH2 | n |
| W303.asm02.HP0_G0034560 | 845.99366 | 949.814141 | 951.436163 | 1238.39802 | 1279.0814 | 1240.17816 | 1607.5325 | 1452.4761 | 1631.9377 | 1.33E-08 | -0.451371008 | YLL040C | VPS13 | n |
| W303.asm02.HP0_G0034570 | 218.08305 | 229.408661 | 295.5730618 | 662.534607 | 696.61846 | 583.111289 | 441.21636 | 544.51452 | 506.94985 | 2.55E-26 | -1.38432773 | YLL038C | ENT4 | n |
| W303.asm02.HP0_G0034580 | 933.43759 | 920.74879 | 812.3429575 | 1044.22121 | 1060.6578 | 953.837328 | 888.13319 | 967.0053 | 995.38501 | 0.024608484 | -0.199080711 | YLL036C | PRP19 | n |
| W303.asm02.HP0_G0034590 | 522.55648 | 539.785086 | 538.9861716 | 750.039177 | 802.68423 | 865.659524 | 522.16303 | 586.50118 | 529.8728 | 5.21E-11 | -0.593708887 | YLL035W | GRC3 | n |
| W303.asm02.HP0_G0034600 | 1099.8971 | 952.928286 | 1077.972343 | 1703.42231 | 1605.3685 | 1832.77093 | 1053.4468 | 1239.9186 | 1117.053 | 1.71E-15 | -0.715360356 | YLL034C | RIX7 | n |
| W303.asm02.HP0_G0034610 | 347.66863 | 386.153946 | 429.8366422 | 160.008358 | 189.65999 | 181.096352 | 102.60846 | 133.83248 | 121.66796 | 7.90E-17 | 1.134939141 | YLL033W | IRC19 | n |
| W303.asm02.HP0_G0034620 | 660.57039 | 641.513813 | 580.5209482 | 825.043095 | 815.26831 | 877.985453 | 985.04117 | 969.62946 | 1017.4263 | 3.49E-07 | -0.420239784 | YLL032C | | n |
| W303.asm02.HP0_G0034630 | 1039.8453 | 1017.28728 | 1047.062742 | 1067.55576 | 1053.4669 | 1108.38552 | 1161.7557 | 1154.6332 | 1104.7098 | 0.426256266 | -0.056668148 | YLL031C | GPI13 | n |
| W303.asm02.HP0_G0034640 | 939.75884 | 1250.84813 | 1133.995995 | 3851.86786 | 3797.6942 | 3782.16412 | 4673.2451 | 3868.0212 | 4290.9999 | 6.17E-71 | -1.781575449 | YLL029W | FRA1 | n |
| W303.asm02.HP0_G0034650 | 1163.1096 | 1250.84813 | 1381.272805 | 2626.80387 | 2571.6457 | 2920.29719 | 3633.4794 | 3618.7254 | 3559.2288 | 3.18E-43 | -1.0961926 | YLL028W | TPO1 | n |
| W303.asm02.HP0_G0034660 | 1651.9528 | 1642.19232 | 1565.764488 | 1340.07 | 1335.7097 | 1324.56337 | 1363.5524 | 1535.1373 | 1454.7257 | 4.80E-05 | 0.280488966 | YLL027W | ISA1 | n |
| W303.asm02.HP0_G0034670 | 4941.1087 | 7341.07716 | 4790.988192 | 803.375297 | 862.90802 | 865.659524 | 1724.9621 | 993.24696 | 1399.1816 | 8.34E-28 | 2.753452311 | YLL026W | HSP104 | n |
| W303.asm02.HP0_G0034690 | 13452.669 | 15053.7756 | 13717.10147 | 28916.5104 | 28183.834 | 31884.3357 | 29730.23 | 28382.983 | 26175.364 | 2.07E-46 | -1.075497265 | YLL024C | SSA2 | y |
| W303.asm02.HP0_G0034700 | 900.77781 | 919.710742 | 958.1976383 | 1243.39828 | 1172.1167 | 1113.12627 | 1764.8654 | 2196.4272 | 1758.0139 | 0.000444707 | -0.344889124 | YLL023C | POM33 | n |
| W303.asm02.HP0_G0034710 | 346.61509 | 395.49638 | 419.2114668 | 1536.74694 | 1499.3027 | 1580.56345 | 1306.5477 | 1266.1603 | 1267.8154 | 1.31E-127 | -1.989346083 | YLL022C | HIF1 | n |
| W303.asm02.HP0_G0034720 | 869.17157 | 1079.57017 | 1072.176793 | 4083.54663 | 4203.98 | 4121.60126 | 3615.2379 | 3441.5942 | 3665.9087 | 6.28E-137 | -2.037697647 | YLL021W | SPA2 | n |
| W303.asm02.HP0_G0034730 | 714.301 | 861.58004 | 852.9118091 | 351.685037 | 362.2416 | 326.163063 | 625.91158 | 617.99117 | 647.1325 | 1.47E-34 | 1.223374026 | YLL019C | KNS1 | n |
| W303.asm02.HP0_G0034740 | 244.42158 | 211.761841 | 218.2990587 | 423.355447 | 476.39714 | 438.992927 | 435.51589 | 482.84661 | 469.03882 | 1.65E-19 | -0.989391981 | YLL018C-/ | COX19 | n |
| W303.asm02.HP0_G0034750 | 1993.3001 | 2502.73431 | 2423.505922 | 8152.92586 | 8207.5137 | 8704.95081 | 6625.086 | 6959.2891 | 6184.7882 | 4.28E-101 | -1.856692291 | YLL018C | DPS1 | n |
| W303.asm02.HP0_G0034780 | 1226.3221 | 1176.10866 | 1396.727606 | 612.531995 | 618.41742 | 683.615024 | 922.33601 | 921.08238 | 949.53911 | 3.44E-29 | 0.990032997 | YLL015W | BPT1 | n |
| W303.asm02.HP0_G0034790 | 1505.5105 | 1548.76798 | 1929.918227 | 448.356753 | 493.47552 | 422.874203 | 461.73805 | 463.16536 | 465.51221 | 1.24E-67 | 1.868980792 | YLL014W | EMC6 | n |
| W303.asm02.HP0_G0034810 | 300.25927 | 326.985196 | 373.8129899 | 1489.24446 | 1561.3242 | 1520.83009 | 1507.2042 | 1322.5798 | 1401.8265 | 6.57E-124 | -2.188996651 | YLL013C | PUF3 | n |
| W303.asm02.HP0_G0034830 | 2556.9447 | 2158.10229 | 2232.252764 | 1357.57091 | 1440.8766 | 1582.45974 | 819.72755 | 1029.9853 | 797.89498 | 3.96E-10 | 0.665469051 | YLL011W | SOF1 | n |
| W303.asm02.HP0_G0034840 | 267.59949 | 356.050547 | 341.9374637 | 2281.78585 | 2251.6506 | 2024.29691 | 2420.4195 | 2429.978 | 2423.6611 | 5.79E-181 | -2.762664539 | YLL010C | PSR1 | n |
| W303.asm02.HP0_G0034850 | 1466.5295 | 1420.04999 | 1756.05172 | 825.043095 | 848.52622 | 822.044696 | 782.10445 | 932.89113 | 804.9482 | 1.36E-19 | 0.896033127 | YLL009C | COX17 | n |
| W303.asm02.HP0_G0034860 | 1155.7348 | 1211.4023 | 1337.806178 | 2353.45626 | 2241.7631 | 2529.66003 | 1401.1755 | 1745.0706 | 1398.2999 | 1.35E-21 | -0.942916275 | YLL008W | DRS1 | n |
| W303.asm02.HP0_G0034870 | 557.32335 | 537.708989 | 513.8721205 | 1224.23061 | 1245.8235 | 1163.37813 | 1238.142 | 1246.479 | 1349.8091 | 4.44E-50 | -1.176177101 | YLL007C | LMO1 | n |
| W303.asm02.HP0_G0034880 | 375.0607 | 340.479823 | 287.8456615 | 391.687126 | 416.17335 | 376.41493 | 396.7527 | 375.25579 | 401.15162 | 0.03726099 | -0.241182678 | YLL006W | MMM1 | n |
| W303.asm02.HP0_G0034890 | 377.16779 | 348.784209 | 409.5522164 | 448.356753 | 524.93571 | 572.681656 | 578.02763 | 561.5716 | 567.78383 | 7.12E-05 | -0.442738491 | YLL005C | SPO75 | n |
| W303.asm02.HP0_G0034900 | 1155.7348 | 1022.47752 | 1191.951498 | 813.375819 | 801.78537 | 865.659524 | 644.15308 | 638.9845 | 665.64719 | 1.15E-07 | 0.442749457 | YLL004W | ORC3 | n |
| W303.asm02.HP0_G0034910 | 465.66525 | 532.518748 | 479.0988192 | 680.868898 | 618.41742 | 697.83725 | 632.75214 | 629.79992 | 644.48755 | 2.11E-06 | -0.43504862 | YLL003W | SFI1 | n |
| W303.asm02.HP0_G0034920 | 437.21964 | 446.360744 | 429.8366422 | 520.86054 | 534.8232 | 484.503852 | 517.60265 | 457.91703 | 535.16271 | 0.01804105 | -0.230547609 | YLL002W | RTT109 | n |
| W303.asm02.HP0_G0034930 | 960.82967 | 999.640457 | 1091.495294 | 2024.2724 | 2008.9577 | 2056.53396 | 1921.0583 | 1825.1077 | 1750.079 | 4.83E-43 | -0.995865604 | YLL001W | DNM1 | n |
| W303.asm02.HP0_G0034940 | 676.37351 | 640.475765 | 599.839449 | 710.870465 | 745.15703 | 628.622414 | 1056.8671 | 918.45822 | 950.42076 | 0.217677883 | -0.122448345 | YLR001C | | n |
| W303.asm02.HP0_G0034950 | 1665.6488 | 1339.08223 | 1464.342359 | 2187.61427 | 2143.7871 | 2279.34884 | 1294.0066 | 1577.124 | 1388.6018 | 3.06E-09 | -0.564913703 | YLR002C | NOC3 | n |
| W303.asm02.HP0_G0034960 | 2127.0999 | 2068.83015 | 2480.495499 | 1413.40716 | 1407.6187 | 1482.90416 | 702.29787 | 911.8978 | 744.99587 | 3.40E-10 | 0.633816086 | YLR003C | CMS1 | n |
| W303.asm02.HP0_G0034970 | 220.19013 | 198.267214 | 162.2754065 | 20.0010447 | 13.482938 | 28.4444531 | 10.260846 | 43.298745 | 16.751386 | 2.82E-10 | 3.231335664 | YLR004C | THI73 | n |
| W303.asm02.HP0_G0034980 | 635.2854 | 688.225984 | 600.805374 | 628.366155 | 579.76633 | 713.007625 | 476.55927 | 358.19871 | 457.57734 | 0.983923089 | 0.002572903 | YLR005W | SSL1 | n |
| W303.asm02.HP0_G0034990 | 433.00547 | 484.768529 | 453.9847682 | 475.858189 | 462.9142 | 439.940875 | 580.30782 | 506.4641 | 498.13333 | 0.940653187 | -0.007932649 | YLR006C | SSK1 | n |
| W303.asm02.HP0_G0035000 | 1328.5156 | 1248.77203 | 1321.385453 | 664.201361 | 613.9231 | 641.896492 | 601.96961 | 674.41075 | 649.77746 | 6.06E-39 | 1.021557736 | YLR007W | NSE1 | n |
| W303.asm02.HP0_G0035010 | 2581.1762 | 2338.72269 | 2882.320315 | 1160.89397 | 1217.9587 | 1109.33367 | 1060.2874 | 1224.1736 | 1124.1062 | 1.49E-37 | 1.161520427 | YLR008C | PAM18 | n |
| W303.asm02.HP0_G0035020 | 2907.774 | 2764.32247 | 2876.524765 | 3321.84018 | 3205.3438 | 3685.45298 | 1952.9809 | 2735.6934 | 2163.5738 | 0.018697628 | -0.256485569 | YLR009W | RLP24 | n |
| W303.asm02.HP0_G0035070 | 461.45109 | 375.773463 | 457.8484683 | 515.026902 | 506.0596 | 488.296445 | 567.76679 | 506.4641 | 566.90218 | 0.036746872 | -0.220843507 | YLR014C | PPR1 | n |
| W303.asm02.HP0_G0035080 | 402.45278 | 410.029055 | 411.4840665 | 576.69679 | 538.41865 | 531.911274 | 444.63664 | 481.53452 | 465.51221 | 2.55E-06 | -0.428935428 | YLR015W | BRE2 | n |
| W303.asm02.HP0_G0035090 | 1246.3394 | 1289.25592 | 1438.262383 | 183.34291 | 201.34521 | 203.851914 | 180.13484 | 230.92664 | 160.46065 | 1.76E-118 | 2.756921028 | YLR016C | PML1 | n |
| W303.asm02.HP0_G0035100 | 2450.5371 | 2207.92861 | 2363.61857 | 1844.263 | 1706.0411 | 1875.43761 | 1434.2382 | 1428.8586 | 1412.4064 | 6.82E-08 | 0.372117722 | YLR017W | MEU1 | n |
| W303.asm02.HP0_G0035110 | 1528.6884 | 1521.77872 | 1417.977957 | 1751.75817 | 1659.3002 | 1627.02272 | 1734.0829 | 1859.2219 | 1824.1378 | 0.012079559 | -0.173863376 | YLR018C | POM34 | n |
| W303.asm02.HP0_G0035130 | 1247.3929 | 1180.26085 | 1353.260979 | 735.038394 | 764.03314 | 803.081727 | 930.31666 | 938.13947 | 971.58041 | 2.77E-19 | 0.716912728 | YLR020C | YEH2 | n |
| W303.asm02.HP0_G0035140 | 1120.9679 | 1122.13015 | 1288.544002 | 694.202928 | 710.10139 | 680.770578 | 598.54932 | 665.22617 | 660.35728 | 1.06E-18 | 0.760738918 | YLR021W | IRC25 | n |
| W303.asm02.HP0_G0035150 | 1862.661 | 1838.38344 | 2049.692932 | 743.372163 | 747.85362 | 726.281703 | 477.69937 | 638.9845 | 516.64802 | 1.07E-45 | 1.37490995 | YLR022C | SDO1 | n |
| W303.asm02.HP0_G0035160 | 1680.3984 | 1422.12609 | 1494.286035 | 1031.72056 | 902.45797 | 843.85211 | 850.51009 | 880.40781 | 834.92436 | 1.31E-14 | 0.725440667 | YLR023C | IZH3 | n |
| W303.asm02.HP0_G0035170 | 551.0021 | 537.708989 | 644.2720008 | 1310.90181 | 1411.2142 | 1327.40781 | 1496.9434 | 1284.5294 | 1430.0394 | 6.22E-40 | -1.223283095 | YLR024C | UBR2 | n |
| W303.asm02.HP0_G0035180 | 2505.3212 | 2609.65328 | 2384.86892 | 1355.90416 | 1406.7198 | 1307.4967 | 1384.0741 | 1478.7177 | 1608.1331 | 1.96E-31 | 0.881477388 | YLR025W | SNF7 | n |
| W303.asm02.HP0_G0035190 | 654.24915 | 683.035743 | 614.3283246 | 2295.95326 | 2226.4825 | 2318.22293 | 2464.8831 | 2199.0514 | 2514.4713 | 6.61E-119 | -1.810069004 | YLR026C | SED5 | n |
| W303.asm02.HP0_G0035200 | 3830.6762 | 4080.56764 | 4164.102841 | 2150.11231 | 1933.4533 | 2102.04509 | 2003.1451 | 1835.6044 | 1972.2553 | 3.46E-44 | 0.965068649 | YLR027C | AAT2 | n |
| W303.asm02.HP0_G0035220 | 17219.079 | 16971.0507 | 17622.3364 | 15674.9854 | 15987.169 | 17796.7462 | 14304.759 | 15704.323 | 13711.451 | 0.365456074 | 0.067138436 | YLR029C | RPL15A | y |
| W303.asm02.HP0_G0035240 | 545.73439 | 603.106028 | 664.5564266 | 600.031342 | 579.76633 | 600.177961 | 384.21166 | 421.1787 | 409.96814 | 0.784238913 | 0.027875857 | YLR032W | RAD5 | n |
| W303.asm02.HP0_G0035250 | 1984.8718 | 1824.88881 | 2050.658857 | 1521.74615 | 1522.6731 | 1503.76342 | 1343.0307 | 1478.7177 | 1455.6073 | 3.07E-07 | 0.365890367 | YLR033W | RSC58 | n |

| | | | | | | | | | | | | | |
|---|---|---|---|---|---|---|---|---|---|---|---|---|---|
| W303.asm02.HP0_G0035260 | 1322.1943 | 1301.71249 | 1273.089201 | 1007.55263 | 959.98517 | 996.504008 | 1766.0055 | 1657.161 | 1713.9313 | 4.14E-09 | 0.394454744 | YLR034C | SMF3 | n |
| W303.asm02.HP0_G0035270 | 262.33178 | 240.827192 | 240.5153346 | 58.3363805 | 57.527201 | 48.3555703 | 57.004698 | 53.79541 | 58.189026 | 2.32E-37 | 2.175367405 | YLR035C | MLH2 | n |
| W303.asm02.HP0_G0035300 | 3444.0265 | 3258.43343 | 4204.671693 | 4482.73415 | 4708.2419 | 4934.16447 | 4188.7052 | 4462.3949 | 4043.2557 | 7.25E-05 | -0.372641883 | YLR038C | COX12 | n |
| W303.asm02.HP0_G0035310 | 784.88827 | 806.563484 | 828.7636831 | 955.049886 | 956.38972 | 1020.20772 | 949.69826 | 1020.8007 | 999.79327 | 0.000254278 | -0.275753369 | YLR039C | RIC1 | n |
| W303.asm02.HP0_G0035320 | 510.96753 | 434.942213 | 429.8366422 | 18.334291 | 14.3818 | 9.48148438 | 10.260846 | 5.2483327 | 11.461475 | 2.03E-88 | 5.01036803 | YLR040C | AFB1 | n |
| W303.asm02.HP0_G0035350 | 23205.301 | 27387.8648 | 28409.78724 | 14562.4273 | 13587.206 | 13309.1596 | 17178.936 | 16954.739 | 16665.866 | 1.94E-32 | 0.930217624 | YLR044C | PDC1 | n |
| W303.asm02.HP0_G0035360 | 543.62731 | 605.182125 | 553.4750471 | 439.189607 | 417.07221 | 534.755719 | 373.95082 | 272.9133 | 327.09286 | 0.022083315 | 0.292258276 | YLR045C | STU2 | n |
| W303.asm02.HP0_G0035380 | 267.59949 | 261.588157 | 226.026459 | 346.684775 | 324.48937 | 364.089 | 458.31777 | 463.16536 | 523.70124 | 8.39E-05 | -0.456256457 | YLR047C | FRE8 | n |
| W303.asm02.HP0_G0035410 | 1931.1412 | 2000.31896 | 1922.190827 | 494.19248 | 562.68794 | 502.518672 | 660.1144 | 746.57532 | 748.52247 | 5.49E-119 | 1.908580286 | YLR050C | EMA19 | n |
| W303.asm02.HP0_G0035420 | 2268.2744 | 2270.2115 | 2966.355794 | 1037.5542 | 915.04205 | 1035.37809 | 658.9743 | 888.28031 | 797.89498 | 5.96E-27 | 1.328888369 | YLR051C | FCF2 | n |
| W303.asm02.HP0_G0035430 | 980.84695 | 969.537058 | 841.3207086 | 553.362238 | 549.205 | 520.533492 | 533.56397 | 544.51452 | 557.20401 | 1.33E-18 | 0.781122862 | YLR052W | IES3 | n |
| W303.asm02.HP0_G0035450 | 221.24367 | 170.239912 | 193.1850077 | 30.834944 | 46.740851 | 23.7037109 | 37.6231 | 35.426246 | 42.319292 | 7.86E-28 | 2.526603876 | YLR054C | OSW2 | n |
| W303.asm02.HP0_G0035460 | 602.62562 | 555.355809 | 674.215677 | 1210.89658 | 1100.2077 | 1137.77813 | 1044.3261 | 1061.4753 | 1074.7337 | 6.52E-26 | -0.911839064 | YLR055C | SPT8 | n |
| W303.asm02.HP0_G0035470 | 8035.3595 | 7459.41466 | 7084.094233 | 2366.79029 | 2325.3573 | 2439.58593 | 2748.7665 | 3213.2917 | 2808.943 | 1.36E-104 | 1.662639317 | YLR056W | ERG3 | n |
| W303.asm02.HP0_G0035480 | 716.40808 | 758.813265 | 753.4215301 | 640.033431 | 681.33779 | 694.044656 | 637.31252 | 774.12907 | 725.59952 | 0.114608942 | 0.145855996 | YLR057W | MNL2 | n |
| W303.asm02.HP0_G0035490 | 894.45656 | 1034.9341 | 1161.041896 | 6311.99637 | 5652.0475 | 3957.57158 | 4582.0376 | 5113.1881 | 4871.1268 | 1.53E-63 | -2.364909499 | YLR058C | SHM2 | n |
| W303.asm02.HP0_G0035500 | 813.33388 | 726.633769 | 605.6349992 | 1452.57587 | 1369.8665 | 1487.6449 | 1213.06 | 1386.8719 | 1278.3953 | 1.06E-22 | -1.007416193 | YLR059C | REX2 | n |
| W303.asm02.HP0_G0035510 | 3265.978 | 3403.76018 | 3822.165378 | 7137.87284 | 7451.5703 | 7774.81719 | 6293.3186 | 6725.7383 | 6054.3037 | 1.03E-47 | -1.091579309 | YLR060W | FRS1 | n |
| W303.asm02.HP0_G0035520 | 7421.1449 | 7517.54536 | 7494.612375 | 6400.33431 | 6643.4929 | 6991.64658 | 5275.2147 | 5842.7064 | 5157.6637 | 0.013257609 | 0.163213362 | YLR061W | RPL22A | n |
| W303.asm02.HP0_G0035530 | 384.54257 | 366.431029 | 395.0633408 | 630.032909 | 619.31628 | 576.47425 | 385.35176 | 430.36328 | 361.47728 | 9.00E-12 | -0.672108956 | YLR063W | BMT6 | n |
| W303.asm02.HP0_G0035540 | 2649.6564 | 2620.03376 | 2458.279223 | 1656.75321 | 1715.9286 | 1767.34869 | 2443.2213 | 2386.6793 | 2346.9574 | 4.19E-21 | 0.588422988 | YLR064W | PER33 | n |
| W303.asm02.HP0_G0035550 | 1766.7888 | 1754.30153 | 2116.34176 | 1767.59233 | 1859.7466 | 1693.39311 | 1886.8555 | 2084.9002 | 1978.4269 | 0.352686111 | 0.083791548 | YLR065C | SND2 | n |
| W303.asm02.HP0_G0035560 | 2062.8339 | 1961.91118 | 2219.695739 | 1575.91565 | 1674.5809 | 1611.85234 | 1857.213 | 1815.9231 | 1895.5516 | 1.69E-07 | 0.361400946 | YLR066W | SPC3 | n |
| W303.asm02.HP0_G0035570 | 1094.6294 | 870.922475 | 998.7664899 | 1470.91016 | 1545.1447 | 1523.67454 | 1274.625 | 1291.0898 | 1256.354 | 5.38E-13 | -0.614843587 | YLR067C | PET309 | n |
| W303.asm02.HP0_G0035580 | 1043.0059 | 990.298022 | 1448.887558 | 918.381304 | 1037.2874 | 882.726196 | 692.03703 | 804.30698 | 707.96649 | 0.027931343 | 0.295520663 | YLR068W | FYV7 | n |
| W303.asm02.HP0_G0035590 | 2218.758 | 2274.3637 | 2610.895379 | 3246.83626 | 3207.1415 | 3361.18621 | 2980.2056 | 2862.9655 | 2958.8238 | 1.54E-10 | -0.465855183 | YLR069C | MEF1 | n |
| W303.asm02.HP0_G0035610 | 992.4359 | 1045.31458 | 963.9931885 | 1659.25334 | 1732.1081 | 1557.80788 | 1641.7353 | 1630.9194 | 1653.0973 | 2.10E-24 | -0.721979459 | YLR071C | RGR1 | n |
| W303.asm02.HP0_G0035630 | 1030.3634 | 1074.37993 | 911.8332365 | 953.383132 | 1073.2419 | 1099.85219 | 544.96491 | 846.29364 | 689.4518 | 0.71007267 | -0.051585792 | YLR073C | RFU1 | n |
| W303.asm02.HP0_G0035640 | 1815.2517 | 1739.76885 | 1967.589304 | 1685.9214 | 1772.5569 | 1687.70422 | 1191.3982 | 1436.7311 | 1224.6145 | 0.24005517 | 0.102135602 | YLR074C | BUD20 | n |
| W303.asm02.HP0_G0035650 | 23709.947 | 23069.5841 | 25148.82431 | 18911.8212 | 18802.406 | 20676.273 | 17612.171 | 18831.018 | 15882.959 | 7.60E-05 | 0.300875677 | YLR075W | RPL10 | n |
| W303.asm02.HP0_G0035660 | 533.0919 | 634.247476 | 629.7831252 | 3047.65919 | 3143.3222 | 3077.68983 | 2722.5444 | 2953.4992 | 2680.2218 | 9.21E-195 | -2.365812154 | YLR077W | FMP25 | n |
| W303.asm02.HP0_G0035670 | 1432.8162 | 1342.19638 | 1539.684512 | 1162.56073 | 1153.2406 | 1113.12627 | 1010.1232 | 1054.9149 | 1077.3786 | 1.08E-05 | 0.331689279 | YLR078C | BOS1 | n |
| W303.asm02.HP0_G0035680 | 1097.79 | 1105.52138 | 1288.544002 | 736.705148 | 889.87439 | 753.778008 | 940.57751 | 1005.0557 | 1044.7575 | 2.70E-08 | 0.553533463 | YLR079W | SIC1 | n |
| W303.asm02.HP0_G0035720 | 2965.7188 | 2638.71863 | 2746.124885 | 3027.65815 | 3068.7167 | 3231.28988 | 3684.7837 | 3408.7921 | 3404.9397 | 0.021721089 | -0.159572059 | YLR083C | EMP70 | n |
| W303.asm02.HP0_G0035730 | 1547.6522 | 1501.01776 | 1475.933459 | 2546.7997 | 2473.6697 | 2643.43784 | 2944.8627 | 2966.62 | 2849.499 | 3.36E-36 | -0.76036774 | YLR084C | RAX2 | n |
| W303.asm02.HP0_G0035740 | 772.24577 | 744.280589 | 619.1579498 | 615.032126 | 591.45154 | 628.622414 | 532.42388 | 499.90369 | 529.8728 | 0.02858534 | 0.217871391 | YLR085C | ARP6 | n |
| W303.asm02.HP0_G0035750 | 560.48397 | 626.981138 | 557.3387473 | 1176.72813 | 1241.3291 | 1188.02999 | 997.58221 | 1024.737 | 1034.1777 | 3.81E-41 | -1.047710017 | YLR086W | SMC4 | n |
| W303.asm02.HP0_G0035760 | 604.73271 | 623.866993 | 680.0112272 | 1093.39045 | 1023.8044 | 940.56325 | 1076.2487 | 1044.4182 | 1071.2071 | 1.86E-15 | -0.680004468 | YLR087C | CSF1 | n |
| W303.asm02.HP0_G0035770 | 1168.3773 | 1096.17894 | 1045.130892 | 1687.58815 | 1619.7503 | 1649.77828 | 2057.8696 | 1897.2723 | 1857.6406 | 1.75E-15 | -0.583443679 | YLR088W | GAA1 | n |
| W303.asm02.HP0_G0035780 | 854.42199 | 931.129273 | 1037.403491 | 1379.23871 | 1371.6642 | 1278.10409 | 1454.7599 | 1360.6302 | 1453.844 | 1.44E-09 | -0.512664501 | YLR089C | ALT1 | n |
| W303.asm02.HP0_G0035790 | 611.05395 | 718.329383 | 766.9444807 | 889.213114 | 808.07741 | 713.007625 | 715.979 | 690.15575 | 664.76554 | 0.066461524 | -0.201542751 | YLR090W | XDJ1 | n |
| W303.asm02.HP0_G0035800 | 208.60118 | 200.343311 | 240.5153346 | 753.372685 | 762.23542 | 722.48911 | 560.92622 | 615.36701 | 566.02053 | 4.67E-69 | -1.783092427 | YLR091W | GEP5 | n |
| W303.asm02.HP0_G0035820 | 2179.7769 | 2095.8194 | 2544.246552 | 358.352051 | 348.75866 | 351.76307 | 383.07157 | 362.13495 | 432.89109 | 6.62E-159 | 2.687331456 | YLR093C | NYV1 | n |
| W303.asm02.HP0_G0035830 | 227.56492 | 270.930591 | 273.3567859 | 1865.09742 | 1697.9513 | 1675.37829 | 1165.176 | 1361.9423 | 1196.4016 | 3.83E-164 | -2.761749299 | YLR094C | GIS3 | n |
| W303.asm02.HP0_G0035840 | 565.75168 | 748.432782 | 738.9326545 | 2008.43824 | 1944.2396 | 2108.68213 | 1870.8942 | 1692.5873 | 1966.0838 | 2.20E-51 | -1.561012366 | YLR095C | IOC2 | n |
| W303.asm02.HP0_G0035860 | 966.09737 | 930.091224 | 928.2539621 | 72.5037872 | 56.628339 | 53.0963125 | 55.864604 | 44.610828 | 66.123893 | 4.53E-169 | 3.946981298 | YLR097C | HRT3 | n |
| W303.asm02.HP0_G0035870 | 368.73945 | 368.507126 | 373.8129899 | 475.858189 | 469.20624 | 495.881633 | 425.25504 | 414.61828 | 414.3764 | 5.04E-05 | -0.374530508 | YLR098C | CHA4 | n |
| W303.asm02.HP0_G0035890 | 504.64628 | 485.806577 | 581.4868732 | 322.516846 | 356.84842 | 367.881594 | 350.00884 | 322.77246 | 348.25251 | 4.59E-08 | 0.588202697 | YLR099W- | MIM2 | n |
| W303.asm02.HP0_G0035900 | 4546.0307 | 4390.94406 | 4287.741246 | 1276.73336 | 1333.912 | 1287.58558 | 1105.8911 | 1158.5694 | 1139.0943 | 3.87E-190 | 1.762325745 | YLR100W | ERG27 | n |
| W303.asm02.HP0_G0035920 | 608.94687 | 527.328507 | 514.8380456 | 1502.57849 | 1565.8185 | 1555.91159 | 1585.8707 | 1490.5265 | 1475.8853 | 1.32E-74 | -1.486670879 | YLR103C | CDC45 | n |
| W303.asm02.HP0_G0035930 | 1735.1825 | 1877.82927 | 1982.078179 | 1292.56752 | 1389.6415 | 1329.30411 | 1361.2722 | 1502.3352 | 1411.5247 | 2.35E-10 | 0.480546315 | YLR104W | LCL2 | n |
| W303.asm02.HP0_G0035940 | 685.85539 | 711.063045 | 671.3179018 | 763.373207 | 728.07864 | 626.726117 | 623.63139 | 711.14908 | 698.26831 | 0.72741578 | -0.035757167 | YLR105C | SEN2 | n |
| W303.asm02.HP0_G0035950 | 1603.4899 | 1570.56699 | 1722.244344 | 490.025596 | 469.20624 | 439.940875 | 361.40978 | 394.93703 | 356.18737 | 3.78E-103 | 1.806290937 | YLR107W | REX3 | n |
| W303.asm02.HP0_G0035970 | 16958.854 | 17409.1071 | 14996.95215 | 4906.92297 | 4921.2723 | 5035.61635 | 5288.8958 | 6253.3884 | 5166.4802 | 4.84E-94 | 1.731668959 | YLR109W | AHP1 | n |
| W303.asm02.HP0_G0036000 | 4467.0151 | 4187.48661 | 4190.182818 | 2702.64117 | 2682.2058 | 2871.94162 | 2572.052 | 2621.5422 | 2671.4053 | 8.58E-27 | 0.637644949 | YLR113W | HOG1 | n |
| W303.asm02.HP0_G0036010 | 1172.5915 | 1176.10866 | 1355.192829 | 1076.72291 | 1135.2634 | 1042.96328 | 1061.4275 | 1111.3344 | 1118.8163 | 0.023396731 | 0.186932551 | YLR114C | AVL9 | n |
| W303.asm02.HP0_G0036020 | 427.73777 | 461.931467 | 443.3595927 | 1308.40168 | 1188.2963 | 1267.67446 | 1307.6878 | 1314.7073 | 1363.0339 | 2.22E-79 | -1.497860011 | YLR115W | CFT2 | n |
| W303.asm02.HP0_G0036030 | 409.82757 | 519.024121 | 510.0084204 | 680.868898 | 640.88898 | 617.244633 | 603.1097 | 644.23284 | 646.25085 | 3.34E-05 | -0.430115869 | YLR116W | MSL5 | n |
| W303.asm02.HP0_G0036040 | 561.53751 | 550.165568 | 518.7017457 | 829.20998 | 817.9649 | 745.244672 | 819.72755 | 742.63907 | 780.26194 | 1.38E-10 | -0.554561759 | YLR117C | CLF1 | n |
| W303.asm02.HP0_G0036050 | 1195.7694 | 1199.98377 | 1204.508523 | 461.690783 | 420.66766 | 446.577914 | 476.55927 | 406.74578 | 461.9856 | 1.06E-64 | 1.437393244 | YLR118C | TML25 | n |

| | | | | | | | | | | | | | | |
|---|---|---|---|---|---|---|---|---|---|---|---|---|---|---|
| W303.asm02.HP0_G0036060 | 605.78625 | 611.410414 | 546.7135719 | 262.513712 | 286.73714 | 279.703789 | 282.7433 | 291.28246 | 284.77357 | 1.68E-27 | 1.089568759 | YLR119W | SRN2 | n |
| W303.asm02.HP0_G0036080 | 640.55311 | 735.976203 | 776.6037311 | 3710.1938 | 3634.1012 | 3755.61596 | 3004.1476 | 2508.703 | 3152.7872 | 1.49E-132 | -2.365323255 | YLR121C | YPS3 | n |
| W303.asm02.HP0_G0036120 | 641.60665 | 618.676752 | 574.725398 | 501.692872 | 444.93695 | 406.75568 | 519.88284 | 410.68203 | 442.58926 | 0.000120257 | 0.437336151 | YLR127C | APC2 | n |
| W303.asm02.HP0_G0036140 | 1973.2829 | 1815.54637 | 2224.525364 | 3246.00288 | 3170.2881 | 3582.1048 | 1866.3338 | 2401.1122 | 1925.5278 | 4.75E-12 | -0.733183227 | YLR129W | DIP2 | n |
| W303.asm02.HP0_G0036150 | 1976.4435 | 1821.77466 | 1866.167175 | 1291.73414 | 1172.1167 | 1116.91886 | 1283.7458 | 1527.2648 | 1439.7376 | 1.38E-14 | 0.660921826 | YLR130C | ZRT2 | n |
| W303.asm02.HP0_G0036160 | 291.83094 | 361.240788 | 369.9492898 | 1351.73727 | 1286.2723 | 1435.49673 | 1875.4545 | 1646.6644 | 1773.002 | 5.22E-90 | -1.991797541 | YLR131C | ACE2 | n |
| W303.asm02.HP0_G0036170 | 344.50801 | 318.68081 | 297.5049119 | 532.527816 | 535.72206 | 578.370547 | 556.36585 | 503.83994 | 588.94348 | 4.19E-14 | -0.777913101 | YLR132C | USB1 | n |
| W303.asm02.HP0_G0036180 | 288.67032 | 389.268091 | 377.6766901 | 934.215464 | 1037.2874 | 1033.4818 | 1064.8478 | 1001.1195 | 1059.7456 | 1.02E-45 | -1.507640413 | YLR133W | CKI1 | n |
| W303.asm02.HP0_G0036200 | 427.73777 | 404.838814 | 374.778915 | 801.708543 | 768.52746 | 786.963203 | 787.80492 | 628.48784 | 765.27386 | 3.45E-21 | -0.966219991 | YLR135W | SLX4 | n |
| W303.asm02.HP0_G0036210 | 243.36804 | 344.632016 | 161.3094814 | 2336.78873 | 2038.6202 | 2125.7488 | 2725.9646 | 2453.5955 | 2921.7944 | 2.01E-32 | -3.118045386 | YLR136C | TIS11 | n |
| W303.asm02.HP0_G0036220 | 254.95699 | 286.501315 | 307.1641623 | 350.85166 | 391.0052 | 323.318617 | 253.10086 | 363.44704 | 317.39469 | 0.020632612 | -0.327338065 | YLR137W | RKM5 | n |
| W303.asm02.HP0_G0036240 | 210.70826 | 232.522806 | 281.0841862 | 1017.55315 | 1012.1192 | 1002.1929 | 898.39403 | 998.49529 | 878.1253 | 5.33E-86 | -2.062670323 | YLR139C | SLS1 | n |
| W303.asm02.HP0_G0036250 | 529.93127 | 543.937279 | 647.1697759 | 407.521286 | 365.83705 | 408.651977 | 302.1249 | 295.21871 | 292.70843 | 4.29E-07 | 0.543084496 | YLR141W | RRN5 | n |
| W303.asm02.HP0_G0036260 | 237.04679 | 256.397916 | 223.1286839 | 815.042573 | 770.32518 | 697.83725 | 1477.5618 | 1956.316 | 1615.1863 | 3.32E-43 | -1.673278156 | YLR142W | PUT1 | n |
| W303.asm02.HP0_G0036270 | 675.31997 | 659.160633 | 722.5119289 | 1490.91121 | 1516.3811 | 1467.73378 | 1380.6538 | 1428.8586 | 1485.5835 | 4.04E-54 | -1.120734523 | YLR143W | DPH6 | n |
| W303.asm02.HP0_G0036280 | 571.01938 | 537.708989 | 673.2497519 | 1049.22147 | 1065.1521 | 1042.96328 | 972.50014 | 837.10906 | 1009.4914 | 3.47E-16 | -0.824156421 | YLR144C | ACF2 | n |
| W303.asm02.HP0_G0036290 | 922.90218 | 920.74879 | 1053.824217 | 307.516063 | 352.35411 | 301.511203 | 221.17823 | 255.85622 | 278.602 | 7.33E-48 | 1.592062728 | YLR145W | RMP1 | n |
| W303.asm02.HP0_G0036300 | 392.9709 | 313.490569 | 435.6321924 | 939.215726 | 1008.5238 | 922.54843 | 668.09506 | 796.43448 | 692.09675 | 2.96E-30 | -1.328338155 | YLR146C | SPE4 | n |
| W303.asm02.HP0_G0036330 | 67.426643 | 63.3209427 | 82.10362828 | 1005.88587 | 1018.4112 | 934.87436 | 1186.8378 | 1116.5828 | 1138.2126 | 3.14E-198 | -3.793063416 | YLR148W | PEP3 | n |
| W303.asm02.HP0_G0036450 | 134.85329 | 145.326754 | 133.2976553 | 197.510317 | 177.97478 | 180.148203 | 269.06217 | 217.80581 | 250.38914 | 0.002730539 | -0.427964671 | YLR151C | PCD1 | n |
| W303.asm02.HP0_G0036460 | 339.2403 | 303.110087 | 322.6189629 | 355.018544 | 340.6689 | 380.207524 | 547.2451 | 404.12162 | 539.57097 | 0.228130581 | -0.156620788 | YLR152C | | n |
| W303.asm02.HP0_G0036470 | 6016.7744 | 6269.81138 | 6102.714394 | 8492.11024 | 8392.6794 | 9461.57326 | 6610.2647 | 5540.9272 | 6720.8325 | 3.71E-10 | -0.5186694 | YLR153C | ACS2 | n |
| W303.asm02.HP0_G0036520 | 1762.5746 | 1572.64309 | 1713.551019 | 1755.92505 | 1742.8944 | 1649.77828 | 5003.8724 | 3354.9967 | 5013.0728 | 0.83519534 | -0.028317623 | YLR163C | MAS1 | n |
| W303.asm02.HP0_G0036530 | 178.04848 | 150.516995 | 144.8887558 | 4155.21704 | 4546.4466 | 4726.51996 | 9801.3877 | 7089.1854 | 9443.3736 | 5.73E-297 | -4.827377738 | YLR164W | SHH4 | n |
| W303.asm02.HP0_G0036540 | 485.68254 | 421.447586 | 486.8262195 | 480.85845 | 490.77894 | 490.192742 | 1166.3161 | 682.28325 | 1074.7337 | 0.681061019 | -0.06836153 | YLR165C | PUS5 | n |
| W303.asm02.HP0_G0036550 | 2486.3575 | 2396.85339 | 2807.944087 | 1551.74772 | 1544.2458 | 1495.23009 | 4641.3225 | 2893.1434 | 4655.1221 | 1.76E-07 | 0.744374161 | YLR166C | SEC10 | n |
| W303.asm02.HP0_G0036560 | 24168.237 | 23464.0424 | 27774.20856 | 17006.7217 | 17734.558 | 19212.3318 | 26241.542 | 21299.046 | 22385.142 | 2.28E-07 | 0.483025114 | YLR167W | RPS31 | n |
| W303.asm02.HP0_G0036570 | 1975.3899 | 2022.11797 | 2107.648434 | 889.213114 | 850.32395 | 787.911352 | 2106.8936 | 1355.3819 | 2169.7454 | 3.50E-20 | 1.272118518 | YLR168C | UPS2 | n |
| W303.asm02.HP0_G0036580 | 1274.785 | 1308.97883 | 1489.45641 | 1244.23166 | 1405.821 | 1240.17816 | 1182.2774 | 1249.1032 | 1272.2237 | 0.454820469 | 0.066890689 | YLR170C | APS1 | n |
| W303.asm02.HP0_G0036590 | 2174.5092 | 2019.00383 | 1925.088602 | 1976.76992 | 2033.227 | 2206.34141 | 1623.4938 | 2075.7156 | 1808.2681 | 0.820240137 | -0.02282154 | YLR172C | DPH5 | n |
| W303.asm02.HP0_G0036600 | 330.81197 | 290.653508 | 367.0515147 | 361.685559 | 368.53363 | 372.622336 | 315.80602 | 402.80953 | 349.13416 | 0.206745783 | -0.156525789 | YLR173W | TAG1 | n |
| W303.asm02.HP0_G0036610 | 185.42327 | 159.859429 | 188.3553825 | 5174.43695 | 5439.916 | 5572.26837 | 4740.5106 | 4101.572 | 4549.3239 | 0 | -4.921644666 | YLR174W | IDP2 | n |
| W303.asm02.HP0_G0036620 | 4209.951 | 4143.88858 | 4327.344173 | 4881.08829 | 4756.7805 | 4953.12744 | 3156.9202 | 3938.8737 | 3573.3352 | 0.010700991 | -0.202326935 | YLR175W | CBF5 | n |
| W303.asm02.HP0_G0036630 | 183.31619 | 202.419407 | 248.2427349 | 685.035782 | 631.00149 | 711.111328 | 442.35645 | 430.36328 | 499.89663 | 9.98E-44 | -1.674039255 | YLR176C | RFX1 | n |
| W303.asm02.HP0_G0036640 | 424.57714 | 400.686621 | 452.0529181 | 903.38052 | 968.07494 | 969.007703 | 1588.1509 | 1247.7911 | 1510.2697 | 6.58E-29 | -1.152081627 | YLR177W | | n |
| W303.asm02.HP0_G0036650 | 648.98144 | 663.312826 | 618.1920247 | 214.177854 | 212.13156 | 182.992649 | 365.97016 | 245.35955 | 367.64885 | 8.66E-33 | 1.662244528 | YLR178C | TFS1 | n |
| W303.asm02.HP0_G0036670 | 1366.4431 | 1540.46359 | 1498.149735 | 13435.7018 | 13412.827 | 13738.6709 | 12111.218 | 13005.368 | 12039.839 | 0 | -3.203518949 | YLR180W | SAM1 | y |
| W303.asm02.HP0_G0036680 | 2100.7614 | 2003.43311 | 2117.307685 | 500.026118 | 496.17211 | 450.370508 | 580.30782 | 539.26618 | 571.31044 | 8.01E-165 | 2.103566551 | YLR181C | VTA1 | n |
| W303.asm02.HP0_G0036690 | 919.74155 | 902.063922 | 937.9132125 | 1882.59834 | 2076.3724 | 1807.17092 | 2033.9276 | 1931.3864 | 1892.9067 | 6.97E-45 | -1.063019753 | YLR182W | SWI6 | n |
| W303.asm02.HP0_G0036720 | 1966.9616 | 1807.24199 | 1628.549615 | 1633.41865 | 1743.7933 | 1616.59309 | 1173.1567 | 1419.674 | 1236.9576 | 0.233206001 | 0.113176364 | YLR186W | EMG1 | n |
| W303.asm02.HP0_G0036730 | 123.26433 | 147.40285 | 99.49027898 | 2168.4466 | 2291.2006 | 2252.80069 | 2226.6035 | 2394.5518 | 2392.8033 | 0 | -4.185122661 | YLR187W | SKG3 | n |
| W303.asm02.HP0_G0036740 | 993.48945 | 1142.89111 | 1101.154544 | 1670.92061 | 1804.9159 | 1726.57831 | 1738.6433 | 1750.3189 | 1718.3396 | 8.42E-21 | -0.683814561 | YLR188W | MDL1 | n |
| W303.asm02.HP0_G0036750 | 798.58431 | 796.183001 | 828.7636831 | 333.350746 | 355.0507 | 351.76307 | 712.55872 | 494.65535 | 683.28023 | 1.09E-23 | 1.220829643 | YLR189C | ATG26 | n |
| W303.asm02.HP0_G0036770 | 740.63953 | 732.862058 | 726.375629 | 855.044662 | 738.86499 | 696.889102 | 1052.3067 | 923.70655 | 1021.8346 | 0.547945141 | -0.059745369 | YLR191W | PEX13 | n |
| W303.asm02.HP0_G0036780 | 2317.7909 | 2475.74506 | 2518.166576 | 1600.91696 | 1651.2105 | 1972.14875 | 1380.6538 | 1339.6369 | 1445.0275 | 2.47E-08 | 0.485701046 | YLR192C | HCR1 | n |
| W303.asm02.HP0_G0036790 | 515.1817 | 477.502191 | 532.2246963 | 1075.05615 | 1063.3544 | 992.711414 | 1080.8091 | 1104.774 | 1144.3842 | 9.93E-38 | -1.03786521 | YLR193C | UPS1 | n |
| W303.asm02.HP0_G0036800 | 2032.2812 | 1984.74824 | 2261.230515 | 896.713506 | 861.1103 | 788.8595 | 972.50014 | 1031.2974 | 1092.3667 | 6.14E-54 | 1.301284921 | YLR194C | NCW2 | n |
| W303.asm02.HP0_G0036810 | 3026.8242 | 2714.49615 | 2929.650642 | 2855.14914 | 2843.1022 | 2879.52681 | 2759.0274 | 2689.7705 | 2581.4768 | 0.817399712 | 0.015631361 | YLR195C | NMT1 | n |
| W303.asm02.HP0_G0036820 | 2285.1311 | 2061.56381 | 2057.420332 | 2865.98303 | 2673.2171 | 2842.54902 | 1671.3777 | 2073.0914 | 1788.8717 | 7.32E-06 | -0.388437052 | YLR196W | PWP1 | n |
| W303.asm02.HP0_G0036830 | 10312.062 | 9341.39612 | 10865.69076 | 6385.33353 | 6107.7708 | 6959.40953 | 4547.8348 | 5068.5773 | 4703.613 | 1.18E-16 | 0.649887738 | YLR197W | NOP56 | n |
| W303.asm02.HP0_G0036850 | 1953.2656 | 1949.4546 | 2178.160962 | 1055.05511 | 1046.276 | 1064.7707 | 1078.5289 | 1095.5894 | 1040.3493 | 1.14E-39 | 0.942036218 | YLR200W | YKE2 | n |
| W303.asm02.HP0_G0036860 | 821.76221 | 725.595721 | 754.3874552 | 738.371901 | 773.02177 | 710.16318 | 765.00304 | 743.95116 | 785.55185 | 0.559275286 | 0.050615325 | YLR201C | COQ9 | n |
| W303.asm02.HP0_G0036870 | 1285.3204 | 1558.11041 | 1633.37924 | 2007.60487 | 1934.3521 | 1981.63024 | 1883.4352 | 1792.3056 | 1940.5159 | 5.67E-06 | -0.403528533 | YLR203C | MSS51 | n |
| W303.asm02.HP0_G0036890 | 1302.177 | 1372.29978 | 1063.483468 | 2425.96005 | 2386.48 | 2359.94146 | 2493.3855 | 2399.8001 | 2470.3887 | 7.56E-27 | -0.941031785 | YLR205C | HMX1 | n |
| W303.asm02.HP0_G0036910 | 632.12478 | 762.965457 | 682.9090023 | 1500.91173 | 1518.1788 | 1518.9338 | 1903.9569 | 1780.4969 | 1850.5874 | 5.10E-46 | -1.126768901 | YLR207W | HRD3 | n |
| W303.asm02.HP0_G0036920 | 4111.9717 | 3848.04483 | 3631.878145 | 3734.36173 | 3605.3376 | 3809.66042 | 3408.8809 | 3688.2658 | 3636.8141 | 0.418974888 | 0.05598264 | YLR208W | SEC13 | n |
| W303.asm02.HP0_G0036930 | 486.73608 | 559.508002 | 488.7580695 | 1941.76809 | 2055.6986 | 1966.45986 | 2033.9276 | 1855.2856 | 1996.9416 | 9.26E-133 | -1.958404383 | YLR209C | PNP1 | n |
| W303.asm02.HP0_G0036960 | 711.14038 | 650.856247 | 641.3742256 | 760.0397 | 764.93201 | 749.037266 | 787.80492 | 808.24323 | 826.98949 | 0.022655059 | -0.183525095 | YLR212C | TUB4 | n |
| W303.asm02.HP0_G0036970 | 482.52192 | 468.159757 | 483.9284443 | 41.6688432 | 51.235164 | 48.3555703 | 61.565073 | 70.852491 | 41.43764 | 1.60E-85 | 3.347489676 | YLR213C | CRR1 | n |
| W303.asm02.HP0_G0036980 | 13522.203 | 12089.1098 | 13433.11951 | 17966.7718 | 18609.151 | 20679.1174 | 26149.195 | 25220.863 | 22837.429 | 3.24E-12 | -0.552226133 | YLR214W | FRE1 | n |

| | | | | | | | | | | | | | | |
|---|---|---|---|---|---|---|---|---|---|---|---|---|---|---|
| W303.asm02.HP0_G0036990 | 678.4806 | 695.492322 | 627.8512751 | 1395.90625 | 1339.3052 | 1341.63004 | 1150.3548 | 1149.3849 | 1168.1888 | 2.57E-45 | -1.027180458 | YLR215C | CDC123 | n |
| W303.asm02.HP0_G0037000 | 11098.004 | 11833.75 | 9697.887388 | 3881.86943 | 3919.0406 | 4049.54198 | 5960.4112 | 5081.6981 | 5278.45 | 2.24E-66 | 1.461225601 | YLR216C | CPR6 | n |
| W303.asm02.HP0_G0037010 | 439.32672 | 497.225108 | 490.6899196 | 489.192219 | 485.38576 | 499.674227 | 380.79138 | 430.36328 | 406.44153 | 0.645713887 | -0.046038986 | YLR218C | COA4 | n |
| W303.asm02.HP0_G0037020 | 646.87436 | 754.661072 | 701.261578 | 904.213897 | 878.18868 | 954.785477 | 1227.8812 | 913.20989 | 1290.7384 | 0.001668257 | -0.380150982 | YLR219W | MSC3 | n |
| W303.asm02.HP0_G0037030 | 1428.602 | 1286.14177 | 1472.069759 | 1457.57613 | 1442.6743 | 1463.94119 | 1542.5471 | 1532.5131 | 1601.0799 | 0.421716702 | -0.059503483 | YLR220W | CCC1 | n |
| W303.asm02.HP0_G0037040 | 2730.779 | 2605.50109 | 3073.573473 | 1352.57065 | 1283.5757 | 1438.34118 | 843.66952 | 994.55904 | 858.72896 | 1.16E-31 | 1.045883844 | YLR221C | RSA3 | n |
| W303.asm02.HP0_G0037050 | 1727.8077 | 1578.87138 | 1730.937669 | 2591.80205 | 2595.0161 | 2942.1046 | 1682.7787 | 1724.0773 | 1825.9011 | 3.11E-19 | -0.689894768 | YLR222C | UTP13 | n |
| W303.asm02.HP0_G0037060 | 878.65344 | 997.56436 | 978.4820641 | 2185.11414 | 2159.0678 | 2465.18594 | 1850.3725 | 1709.6444 | 2021.6278 | 1.78E-44 | -1.253648323 | YLR223C | IFH1 | n |
| W303.asm02.HP0_G0037090 | 1021.9351 | 868.846378 | 841.3207086 | 781.707498 | 776.61722 | 712.059477 | 725.09975 | 722.95783 | 740.58761 | 0.004215442 | 0.265876978 | YLR226W | BUR2 | n |
| W303.asm02.HP0_G0037100 | 750.12141 | 780.612278 | 967.8568887 | 95.8383393 | 96.17829 | 83.4370625 | 80.946671 | 76.100824 | 86.401887 | 6.26E-109 | 3.17994934 | YLR227C | ADY4 | n |
| W303.asm02.HP0_G0037120 | 3251.2285 | 3222.10174 | 3237.780729 | 2883.48395 | 2757.7102 | 2830.22309 | 3142.0989 | 3054.5296 | 3071.6753 | 0.00021829 | 0.196918028 | YLR229C | CDC42 | n |
| W303.asm02.HP0_G0037150 | 91.658093 | 112.10921 | 118.8087798 | 304.182555 | 322.69165 | 369.777891 | 331.76734 | 282.09788 | 280.36531 | 1.33E-28 | -1.623705623 | YLR234W | TOP3 | n |
| W303.asm02.HP0_G0037180 | 450.91568 | 461.931467 | 447.2232929 | 598.364588 | 552.80045 | 558.45943 | 1827.5706 | 1002.4315 | 1732.446 | 0.053560359 | -0.330229297 | YLR238W | FAR10 | n |
| W303.asm02.HP0_G0037190 | 811.2268 | 746.356686 | 807.5133323 | 785.041006 | 758.63997 | 769.896532 | 2364.5549 | 1434.1069 | 2443.9391 | 0.845811435 | 0.031774153 | YLR239C | LIP2 | n |
| W303.asm02.HP0_G0037200 | 522.55648 | 581.307015 | 679.0453022 | 689.202666 | 683.13552 | 575.526102 | 2116.0144 | 1279.2811 | 2093.9233 | 0.456618487 | -0.127525175 | YLR240W | VPS34 | n |
| W303.asm02.HP0_G0037210 | 1047.2201 | 1107.59747 | 1100.188619 | 1218.39697 | 1200.8803 | 1283.79298 | 5091.6596 | 2876.0863 | 4728.2992 | 0.395531474 | -0.18600467 | YLR241W | CSC1 | n |
| W303.asm02.HP0_G0037220 | 523.61003 | 469.197805 | 549.611347 | 657.534346 | 742.46044 | 753.778008 | 1995.1644 | 1167.754 | 1765.0671 | 0.002186752 | -0.481096909 | YLR242C | ARV1 | n |
| W303.asm02.HP0_G0037230 | 1864.7681 | 1859.1444 | 1786.961321 | 1096.72395 | 1103.8032 | 1060.9781 | 2176.4394 | 1540.3856 | 2177.6802 | 3.08E-11 | 0.756595253 | YLR243W | GPN3 | n |
| W303.asm02.HP0_G0037240 | 1126.2356 | 1231.12521 | 1305.930652 | 2875.98356 | 2811.642 | 2971.4972 | 6073.2805 | 4110.7566 | 6045.4872 | 4.98E-23 | -1.240840178 | YLR244C | MAP1 | n |
| W303.asm02.HP0_G0037250 | 138.01391 | 149.478947 | 152.6161561 | 135.840429 | 159.09867 | 144.118563 | 771.84361 | 339.82954 | 704.43988 | 0.990861908 | 0.003820594 | YLR245C | CDD1 | n |
| W303.asm02.HP0_G0037260 | 737.47891 | 690.302081 | 697.3978779 | 693.369551 | 731.67409 | 723.437258 | 2230.0238 | 1398.6807 | 2121.2545 | 0.916170964 | -0.015691179 | YLR246W | ERF2 | n |
| W303.asm02.HP0_G0037270 | 1022.9886 | 1086.83651 | 1139.791546 | 960.050147 | 937.51361 | 1024.00031 | 1138.9539 | 925.01863 | 1187.5851 | 0.124789799 | 0.154082885 | YLR247C | IRC20 | n |
| W303.asm02.HP0_G0037280 | 1116.7538 | 1274.72324 | 1154.280421 | 1585.91617 | 1528.9652 | 1638.4005 | 1678.2183 | 1529.889 | 1697.1799 | 5.79E-08 | -0.422790466 | YLR248W | RCK2 | n |
| W303.asm02.HP0_G0037290 | 20355.471 | 20688.3015 | 22664.46511 | 37769.4728 | 37142.797 | 40696.4272 | 31228.313 | 31619.892 | 29959.414 | 2.04E-41 | -0.859637539 | YLR249W | YEF3 | n |
| W303.asm02.HP0_G0037300 | 3137.446 | 3018.64429 | 3486.023464 | 3001.82346 | 2823.3272 | 3143.11207 | 3418.0017 | 3781.4237 | 3564.5187 | 0.182326157 | 0.104842706 | YLR250W | SSP120 | n |
| W303.asm02.HP0_G0037310 | 736.42537 | 774.383988 | 815.2407326 | 333.350746 | 315.50075 | 336.592695 | 503.92153 | 430.36328 | 394.0984 | 4.83E-32 | 1.239461072 | YLR251W | SYM1 | n |
| W303.asm02.HP0_G0037320 | 1169.4308 | 1221.78278 | 1212.235923 | 1005.0525 | 959.98517 | 926.341024 | 1023.8044 | 1010.304 | 1024.4795 | 5.90E-06 | 0.317067014 | YLR253W | CQD2 | n |
| W303.asm02.HP0_G0037340 | 646.87436 | 678.88355 | 684.8408524 | 1124.22539 | 1210.7678 | 1216.47445 | 1365.8326 | 1170.3782 | 1348.9274 | 3.51E-22 | -0.820107007 | YLR256W | HAP1 | n |
| W303.asm02.HP0_G0037350 | 4293.1808 | 4427.27575 | 3993.13411 | 625.032648 | 529.43003 | 532.859422 | 587.14838 | 536.64202 | 565.13888 | 7.50E-265 | 2.911956755 | YLR257W | | n |
| W303.asm02.HP0_G0037360 | 824.92284 | 950.852189 | 950.470238 | 304.182555 | 319.99506 | 311.940836 | 808.32661 | 583.87701 | 765.27386 | 1.67E-37 | 1.542646717 | YLR258W | GSY2 | n |
| W303.asm02.HP0_G0037370 | 7896.292 | 9560.4243 | 8857.532604 | 6325.3304 | 6181.4776 | 6599.11313 | 8074.1454 | 7199.4003 | 7174.0016 | 1.31E-08 | 0.461887334 | YLR259C | HSP60 | n |
| W303.asm02.HP0_G0037380 | 803.85201 | 867.80833 | 699.329728 | 1646.75268 | 1496.6061 | 1536.00047 | 1624.6339 | 1642.7281 | 1526.1395 | 7.33E-29 | -0.981982627 | YLR260W | LCB5 | n |
| W303.asm02.HP0_G0037400 | 163.2989 | 153.63114 | 149.718381 | 196.67694 | 249.88378 | 217.125992 | 206.35701 | 257.1683 | 196.60838 | 0.000784474 | -0.507829825 | YLR263W | RED1 | n |
| W303.asm02.HP0_G0037410 | 6621.5071 | 6115.14219 | 7090.855709 | 10936.4046 | 10566.129 | 12033.9 | 10155.957 | 9656.9321 | 8547.6153 | 5.70E-19 | -0.758108667 | YLR264W | RPS28B | y |
| W303.asm02.HP0_G0037420 | 903.93844 | 821.096159 | 944.6746878 | 189.176548 | 222.91791 | 201.007469 | 192.67588 | 211.24539 | 208.9515 | 1.32E-86 | 2.123832156 | YLR265C | NEJ1 | n |
| W303.asm02.HP0_G0037450 | 1452.8335 | 1523.85482 | 1344.567654 | 1545.08071 | 1395.0346 | 1412.74117 | 1540.2669 | 1630.9194 | 1575.512 | 0.887810605 | -0.011377371 | YLR268W | SEC22 | n |
| W303.asm02.HP0_G0037460 | 1254.7677 | 1123.1682 | 1090.529369 | 1718.42309 | 1608.0651 | 1622.28198 | 1841.2517 | 1821.1714 | 1870.8654 | 3.04E-12 | -0.513623661 | YLR270W | DCS1 | n |
| W303.asm02.HP0_G0037470 | 391.91736 | 401.724669 | 407.6203663 | 250.846436 | 238.19857 | 234.192664 | 217.75794 | 209.93331 | 237.16436 | 1.23E-11 | 0.731339146 | YLR271W | CMG1 | n |
| W303.asm02.HP0_G0037480 | 909.20614 | 994.450215 | 766.9444807 | 1033.38731 | 1018.4112 | 1105.54108 | 1014.6836 | 1047.0424 | 1025.3612 | 0.012193979 | -0.242133722 | YLR272C | YCS4 | n |
| W303.asm02.HP0_G0037490 | 238.10033 | 243.941337 | 198.980558 | 25.8346828 | 30.561326 | 33.1851953 | 46.743852 | 35.426246 | 43.200944 | 2.54E-43 | 2.930580267 | YLR273C | PIG1 | n |
| W303.asm02.HP0_G0037500 | 1875.3035 | 1536.3114 | 1809.177597 | 658.367722 | 616.61969 | 602.074258 | 668.09506 | 684.90741 | 724.71787 | 8.60E-58 | 1.475244032 | YLR274W | MCM5 | n |
| W303.asm02.HP0_G0037510 | 876.54636 | 901.025874 | 1010.35759 | 395.020633 | 469.20624 | 428.563094 | 387.63194 | 471.03786 | 450.52413 | 3.14E-26 | 1.109929175 | YLR275W | SMD2 | n |
| W303.asm02.HP0_G0037520 | 5000.107 | 4383.67772 | 5108.777529 | 2333.45522 | 2260.6392 | 2451.91186 | 1185.6977 | 1686.0269 | 1393.01 | 8.40E-22 | 1.040541584 | YLR276C | DBP9 | n |
| W303.asm02.HP0_G0037530 | 574.18001 | 605.182125 | 614.3283246 | 1681.75451 | 1705.1422 | 1745.54127 | 1681.6386 | 1670.2819 | 1582.5652 | 9.02E-102 | -1.516009738 | YLR277C | YSH1 | n |
| W303.asm02.HP0_G0037540 | 1451.7799 | 1486.48508 | 1573.491888 | 626.699402 | 646.28215 | 660.859461 | 654.41393 | 570.75618 | 677.10867 | 9.24E-50 | 1.22293902 | YLR278C | | n |
| W303.asm02.HP0_G0037550 | 186.47681 | 179.582346 | 201.8783331 | 430.022462 | 391.0052 | 406.75568 | 440.07627 | 494.65535 | 478.73699 | 2.45E-23 | -1.111609684 | YLR281C | RSO55 | n |
| W303.asm02.HP0_G0037570 | 385.59612 | 363.316885 | 374.778915 | 110.005746 | 125.84075 | 115.674109 | 142.51174 | 137.76873 | 164.86891 | 1.46E-37 | 1.677501144 | YLR284C | ECI1 | n |
| W303.asm02.HP0_G0037580 | 1934.3018 | 1686.82839 | 1996.567055 | 1055.88849 | 1092.118 | 1061.92625 | 737.64079 | 825.30031 | 756.45734 | 2.01E-22 | 0.807714047 | YLR285W | NNT1 | n |
| W303.asm02.HP0_G0037600 | 6784.806 | 6426.55666 | 6267.887576 | 1872.59781 | 1831.8818 | 1863.11168 | 2260.8063 | 2121.6385 | 2299.3482 | 7.04E-195 | 1.806679177 | YLR286C | CTS1 | n |
| W303.asm02.HP0_G0037610 | 2107.0826 | 1898.59023 | 2129.86471 | 1241.73153 | 1372.5631 | 1346.37078 | 965.65958 | 1098.2136 | 932.78772 | 5.96E-14 | 0.631918555 | YLR287C | | n |
| W303.asm02.HP0_G0037620 | 521.50294 | 517.986072 | 503.2469451 | 443.356492 | 408.08359 | 370.726039 | 401.31307 | 492.03119 | 443.47091 | 0.001713602 | 0.334322361 | YLR288C | MEC3 | n |
| W303.asm02.HP0_G0037630 | 479.36129 | 464.007564 | 470.4054938 | 831.71011 | 874.59323 | 745.244672 | 812.88699 | 872.53531 | 812.88306 | 7.75E-19 | -0.794828428 | YLR289W | GUF1 | n |
| W303.asm02.HP0_G0037640 | 940.81238 | 1019.36337 | 1171.667072 | 718.370857 | 605.83334 | 624.829821 | 649.85355 | 646.857 | 661.23893 | 1.45E-11 | 0.684050081 | YLR290C | COQ11 | n |
| W303.asm02.HP0_G0037650 | 2121.8322 | 2032.49846 | 1846.848674 | 1165.89423 | 1117.2861 | 1171.91147 | 985.04117 | 1074.5961 | 1012.1364 | 6.69E-26 | 0.7961277 | YLR291C | GCD7 | n |
| W303.asm02.HP0_G0037660 | 3257.5497 | 3401.68409 | 3571.024868 | 1434.24158 | 1552.3356 | 1464.88934 | 1522.0254 | 1592.869 | 1409.7614 | 3.54E-63 | 1.200791153 | YLR292C | SEC72 | n |
| W303.asm02.HP0_G0037670 | 8043.7878 | 8134.14602 | 6965.285453 | 6111.98592 | 6452.9341 | 6495.76495 | 5339.06 | 5981.7872 | 5293.4381 | 0.000395233 | 0.279941865 | YLR293C | GSP1 | y |
| W303.asm02.HP0_G0037680 | 5065.4266 | 5042.83836 | 5346.395089 | 3272.67094 | 3372.5322 | 3429.4529 | 2987.0462 | 2936.4421 | 2897.9898 | 1.49E-29 | 0.617625547 | YLR295C | ATP14 | n |
| W303.asm02.HP0_G0037700 | 928.16989 | 957.080479 | 875.128085 | 351.685037 | 325.38823 | 363.140852 | 319.22631 | 309.65163 | 320.03964 | 4.71E-55 | 1.407882808 | YLR298C | YHC1 | n |
| W303.asm02.HP0_G0037710 | 816.49451 | 863.656137 | 929.2198872 | 219.178115 | 204.94066 | 162.133383 | 299.84471 | 279.47372 | 267.14053 | 2.02E-75 | 2.151016884 | YLR299W | ECM38 | n |
| W303.asm02.HP0_G0037720 | 11930.302 | 10405.3956 | 12126.22293 | 1522.57953 | 1456.1573 | 1807.17092 | 1811.6093 | 1853.9735 | 1992.5333 | 1.33E-211 | 2.848621843 | YLR300W | EXG1 | n |

| | | | | | | | | | | | | | |
|---|---|---|---|---|---|---|---|---|---|---|---|---|---|
| W303.asm02.HP0_G0037730 | 3678.9662 | 3555.31523 | 4378.5382 | 2364.29016 | 2548.2753 | 2520.17855 | 3010.9881 | 2939.0663 | 2776.3219 | 3.92E-14 | 0.644146979 | YLR301W | HRI1 | n |
| W303.asm02.HP0_G0037740 | 7512.803 | 7571.52387 | 6830.055948 | 239.17916 | 198.64862 | 222.814883 | 223.45841 | 165.32248 | 196.60838 | 0 | 5.050494662 | YLR303W | MET17 | n |
| W303.asm02.HP0_G0037750 | 6003.0783 | 6171.1968 | 6686.133117 | 22939.5316 | 24031.09 | 25211.267 | 21063.236 | 21058.935 | 20068.161 | 2.02E-206 | -1.936098362 | YLR304C | ACO1 | n |
| W303.asm02.HP0_G0037760 | 1460.2082 | 1422.12609 | 1642.072566 | 748.372424 | 737.96613 | 694.992805 | 681.77618 | 587.81326 | 612.74808 | 2.65E-33 | 1.052474565 | YLR305C | STT4 | n |
| W303.asm02.HP0_G0037770 | 523.61003 | 464.007564 | 630.7490502 | 453.357014 | 384.71316 | 402.963086 | 444.63664 | 436.9237 | 421.42961 | 0.001619655 | 0.383266299 | YLR306W | UBC12 | n |
| W303.asm02.HP0_G0037790 | 89.55101 | 82.0058111 | 86.93325348 | 187.509794 | 209.43497 | 192.474133 | 206.35701 | 191.56414 | 255.67905 | 2.81E-13 | -1.188833349 | YLR307C-/ | DPA10 | n |
| W303.asm02.HP0_G0037800 | 6.3212478 | 5.19024121 | 5.795550232 | 25.8346828 | 17.97725 | 19.9111172 | 31.922631 | 34.114162 | 22.041298 | 0.000137403 | -1.884688019 | YLR308W | CDA2 | n |
| W303.asm02.HP0_G0037810 | 2104.9755 | 1949.4546 | 1989.80558 | 1755.09168 | 1697.9513 | 1736.05979 | 1523.1655 | 1525.9527 | 1512.033 | 0.000391399 | 0.21985001 | YLR309C | IMH1 | n |
| W303.asm02.HP0_G0037820 | 1204.1977 | 1220.74473 | 1306.896577 | 1199.22931 | 1199.0826 | 1275.25965 | 1458.1802 | 1413.1136 | 1519.0862 | 0.756268441 | 0.023525363 | YLR310C | CDC25 | n |
| W303.asm02.HP0_G0037840 | 3221.7293 | 3357.04801 | 3893.643831 | 2199.28154 | 2148.2814 | 2150.40066 | 2054.4493 | 2087.5243 | 1916.7113 | 7.22E-18 | 0.688763533 | YLR312W- | MRPL15 | n |
| W303.asm02.HP0_G0037870 | 2259.8461 | 2188.20569 | 2212.934263 | 4357.72762 | 4258.8106 | 4374.75689 | 4135.1208 | 4379.7336 | 4116.4328 | 6.31E-68 | -0.963781276 | YLR314C | CDC3 | n |
| W303.asm02.HP0_G0037880 | 468.82588 | 522.138265 | 550.577272 | 911.714289 | 904.2557 | 954.785477 | 744.48135 | 858.10239 | 804.06654 | 9.25E-21 | -0.844686843 | YLR316C | TAD3 | n |
| W303.asm02.HP0_G0037900 | 510.96753 | 510.719735 | 529.3269212 | 580.863674 | 614.82197 | 602.074258 | 530.14369 | 620.61534 | 556.32236 | 0.019852439 | -0.212353777 | YLR318W | EST2 | n |
| W303.asm02.HP0_G0037910 | 494.11087 | 511.757783 | 572.7935479 | 1332.56961 | 1395.0346 | 1426.01525 | 1319.0887 | 1262.224 | 1277.5136 | 2.19E-68 | -1.394010349 | YLR319C | BUD6 | n |
| W303.asm02.HP0_G0037920 | 440.38026 | 394.458332 | 459.7803184 | 865.045185 | 818.86376 | 963.318813 | 823.14783 | 808.24323 | 838.45097 | 1.02E-27 | -1.03068891 | YLR320W | MMS22 | n |
| W303.asm02.HP0_G0037930 | 665.8381 | 664.350875 | 604.6690742 | 1505.07862 | 1414.8096 | 1422.22266 | 1499.2235 | 1587.6206 | 1647.8074 | 1.36E-51 | -1.167273902 | YLR321C | SFH1 | n |
| W303.asm02.HP0_G0037940 | 613.16104 | 665.388923 | 498.4173199 | 240.012537 | 248.08606 | 247.466742 | 205.21691 | 241.4233 | 208.06985 | 4.56E-24 | 1.271835304 | YLR323C | CWC24 | n |
| W303.asm02.HP0_G0037950 | 449.86214 | 521.100217 | 485.8602944 | 1046.72134 | 1006.726 | 920.652133 | 1361.2722 | 1131.0157 | 1303.9632 | 1.50E-25 | -1.030026581 | YLR324W | PEX30 | n |
| W303.asm02.HP0_G0037960 | 8857.1217 | 8567.01214 | 10086.18925 | 8732.12278 | 9536.9314 | 9844.62523 | 8092.3869 | 9002.2026 | 7521.3725 | 0.737721258 | -0.031158285 | YLR325C | RPL38 | n |
| W303.asm02.HP0_G0037970 | 632.12478 | 633.209427 | 544.7817218 | 595.864458 | 578.86746 | 570.78536 | 581.44797 | 537.9541 | 604.81321 | 0.602650824 | 0.051081398 | YLR326W | | n |
| W303.asm02.HP0_G0037990 | 1030.3634 | 1018.32532 | 963.0272635 | 1695.08854 | 1640.4241 | 1689.60052 | 1555.0881 | 1625.671 | 1579.0386 | 1.71E-29 | -0.739099219 | YLR328W | NMA1 | n |
| W303.asm02.HP0_G0038010 | 1610.8646 | 1795.82346 | 1750.25617 | 2467.62889 | 2546.4775 | 2349.51183 | 2267.6469 | 2161.001 | 2406.9097 | 1.35E-12 | -0.513864403 | YLR330W | CHS5 | n |
| W303.asm02.HP0_G0038020 | 993.48945 | 864.694185 | 871.2643848 | 1246.73179 | 1186.4985 | 1281.89669 | 1216.4802 | 1065.4115 | 1268.6971 | 8.15E-07 | -0.44511442 | YLR332W | MID2 | n |
| W303.asm02.HP0_G0038030 | 6367.6036 | 6095.41927 | 7550.636027 | 9196.31369 | 9725.6925 | 10066.492 | 8386.5311 | 9662.1805 | 7956.0269 | 2.00E-08 | -0.534339368 | YLR333C | RPS25B | y |
| W303.asm02.HP0_G0038040 | 1150.4671 | 1342.19638 | 1448.887558 | 3288.5051 | 3258.3766 | 3415.23067 | 2845.6745 | 2953.4992 | 2705.7897 | 3.56E-57 | -1.336975153 | YLR335W | NUP2 | n |
| W303.asm02.HP0_G0038050 | 1043.0059 | 903.10197 | 996.8346399 | 2310.95404 | 2333.4471 | 2480.35631 | 1496.9434 | 1726.7015 | 1584.3285 | 4.74E-56 | -1.275376179 | YLR336C | SGD1 | n |
| W303.asm02.HP0_G0038060 | 408.77402 | 571.964581 | 557.3387473 | 775.87386 | 689.42755 | 789.807649 | 710.27853 | 557.63535 | 773.20873 | 9.37E-05 | -0.551432011 | YLR337C | VRP1 | n |
| W303.asm02.HP0_G0038070 | 8666.4307 | 9527.20676 | 8664.347596 | 21646.964 | 21522.364 | 23468.5701 | 19218.564 | 20267.749 | 17789.091 | 2.67E-76 | -1.31099176 | YLR340W | RPP0 | n |
| W303.asm02.HP0_G0038090 | 7938.4337 | 7879.8242 | 8618.94912 | 19476.0173 | 18904.877 | 20966.4064 | 19755.548 | 18659.135 | 19237.645 | 6.94E-91 | -1.279975218 | YLR342W | FKS1 | n |
| W303.asm02.HP0_G0038110 | 134.85329 | 135.98432 | 146.8206059 | 276.681119 | 308.30985 | 312.888984 | 285.02349 | 304.4033 | 259.20566 | 2.29E-17 | -1.101913921 | YLR343W | GAS2 | n |
| W303.asm02.HP0_G0038130 | 1415.9595 | 1422.12609 | 1341.669879 | 1066.72239 | 1093.9157 | 977.541039 | 1185.6977 | 1066.7236 | 1202.5732 | 1.52E-07 | 0.412783755 | YLR345W | | n |
| W303.asm02.HP0_G0038150 | 1654.0598 | 1764.68201 | 1651.731816 | 3806.86551 | 3874.9963 | 4072.29754 | 3925.3435 | 3268.3992 | 4034.4392 | 7.38E-47 | -1.212954638 | YLR347C | KAP95 | n |
| W303.asm02.HP0_G0038190 | 2799.2592 | 2751.86589 | 2707.487883 | 1270.89972 | 1331.2154 | 1408.00043 | 1333.9099 | 1260.9119 | 1288.0934 | 3.14E-58 | 1.042838124 | YLR351C | NIT3 | n |
| W303.asm02.HP0_G0038200 | 379.27487 | 398.610525 | 397.9611159 | 420.021939 | 444.03809 | 421.926055 | 424.11495 | 426.42703 | 485.7902 | 0.204557486 | -0.128799014 | YLR352W | LUG1 | n |
| W303.asm02.HP0_G0038220 | 5628.0176 | 5912.72278 | 5244.007035 | 4773.58268 | 4641.7261 | 4903.82372 | 4665.2644 | 5077.7619 | 4727.4176 | 0.000571369 | 0.229120935 | YLR354C | TAL1 | n |
| W303.asm02.HP0_G0038230 | 7627.639 | 8972.889 | 8853.668904 | 7456.2228 | 7154.0468 | 7607.94307 | 7368.4272 | 5845.3305 | 7270.1017 | 0.042319935 | 0.196211894 | YLR355C | ILV5 | n |
| W303.asm02.HP0_G0038240 | 588.92959 | 582.345063 | 552.5091221 | 2044.27345 | 2041.3168 | 2033.7784 | 2241.4247 | 2234.4776 | 2307.2831 | 4.79E-165 | -1.828524884 | YLR356W | ATG33 | n |
| W303.asm02.HP0_G0038260 | 2604.3541 | 2219.34714 | 2269.923841 | 3811.86578 | 3646.6853 | 2983.82313 | 2875.3169 | 3264.4629 | 3218.9111 | 3.86E-08 | -0.558234751 | YLR359W | ADE13 | n |
| W303.asm02.HP0_G0038270 | 559.43043 | 560.54605 | 606.6009243 | 750.872554 | 695.71959 | 694.044656 | 779.82426 | 747.88741 | 776.73533 | 0.00020916 | -0.310116415 | YLR361C | DCR2 | n |
| W303.asm02.HP0_G0038290 | 640.55311 | 634.247476 | 654.8971762 | 1734.25725 | 1615.256 | 1549.27455 | 1511.7646 | 1381.6236 | 1525.2578 | 6.84E-66 | -1.344341774 | YLR362W | STE11 | n |
| W303.asm02.HP0_G0038300 | 973.47216 | 995.488264 | 1162.973746 | 969.217293 | 1000.434 | 1076.14848 | 815.16718 | 801.68282 | 804.9482 | 0.653462657 | 0.041543319 | YLR363C | NMD4 | n |
| W303.asm02.HP0_G0038310 | 1035.6311 | 966.422913 | 1115.64342 | 547.5286 | 568.08111 | 555.614985 | 299.84471 | 499.90369 | 346.4892 | 9.17E-11 | 0.899824989 | YLR363W-A | | n |
| W303.asm02.HP0_G0038320 | 473.04004 | 451.550985 | 544.7817218 | 799.208412 | 858.41371 | 796.444688 | 721.67947 | 764.94449 | 690.33345 | 4.42E-15 | -0.738686956 | YLR364W | GRX8 | n |
| W303.asm02.HP0_G0038340 | 527.82419 | 508.643638 | 539.9520966 | 283.348134 | 303.81553 | 256.948227 | 266.78198 | 220.42997 | 231.87445 | 1.32E-16 | 0.900491526 | YLR368W | MDM30 | n |
| W303.asm02.HP0_G0038350 | 873.38574 | 910.368308 | 787.2289065 | 1410.07365 | 1385.1471 | 1286.63743 | 1411.4363 | 1440.6673 | 1482.0569 | 2.32E-17 | -0.668088243 | YLR369W | SSQ1 | n |
| W303.asm02.HP0_G0038360 | 2519.0172 | 2604.46304 | 2324.981568 | 1085.89005 | 1124.477 | 1044.85958 | 1022.6643 | 1132.3278 | 1083.5502 | 8.93E-59 | 1.193729174 | YLR370C | ARC18 | n |
| W303.asm02.HP0_G0038370 | 710.08684 | 688.225984 | 751.48968 | 1765.92557 | 1720.4229 | 1786.31166 | 1848.0923 | 1720.141 | 1871.747 | 6.92E-73 | -1.293683521 | YLR371W | ROM2 | n |
| W303.asm02.HP0_G0038380 | 8279.7811 | 7623.42629 | 8709.746073 | 4596.90678 | 4554.5364 | 4941.74966 | 4880.7422 | 4686.7611 | 4466.4486 | 1.18E-30 | 0.804608668 | YLR372W | ELO3 | n |
| W303.asm02.HP0_G0038400 | 1293.7487 | 1383.71831 | 1435.364607 | 1171.72787 | 1084.0282 | 1072.35588 | 1128.693 | 1183.499 | 1247.5375 | 8.51E-05 | 0.305239878 | YLR375W | STP3 | n |
| W303.asm02.HP0_G0038410 | 353.98988 | 386.153946 | 339.0396886 | 190.843302 | 186.06454 | 269.274156 | 273.62255 | 219.11789 | 231.87445 | 8.68E-07 | 0.742240648 | YLR376C | PSY3 | n |
| W303.asm02.HP0_G0038420 | 73.747891 | 89.2721488 | 76.30807805 | 45.8357275 | 53.931751 | 35.0814922 | 53.584416 | 43.298745 | 50.254159 | 0.00047848 | 0.824311358 | YLR377C | FBP1 | n |
| W303.asm02.HP0_G0038430 | 3041.5737 | 3678.84297 | 3675.344772 | 7494.55814 | 7496.5134 | 8062.10617 | 9036.3847 | 9080.9276 | 8517.6391 | 1.58E-46 | -1.14872713 | YLR378C | SEC61 | n |
| W303.asm02.HP0_G0038440 | 729.05058 | 835.628834 | 803.6496321 | 4109.38132 | 4044.8814 | 4252.44574 | 3530.871 | 3879.8299 | 3531.0159 | 3.99E-232 | -2.388772302 | YLR380W | CSR1 | n |
| W303.asm02.HP0_G0038450 | 399.29215 | 426.637827 | 443.3595927 | 610.865241 | 715.49457 | 734.815039 | 662.39459 | 627.17575 | 618.91964 | 2.28E-12 | -0.697616471 | YLR381W | CTF3 | n |
| W303.asm02.HP0_G0038460 | 743.80016 | 882.341005 | 851.9458841 | 1041.72108 | 1045.3771 | 1090.3707 | 1100.1907 | 1022.1128 | 982.16023 | 3.45E-05 | -0.358012466 | YLR382C | NAM2 | n |
| W303.asm02.HP0_G0038470 | 524.66357 | 536.670941 | 541.8839467 | 1010.88614 | 953.69314 | 952.88918 | 923.4761 | 884.34406 | 889.58678 | 3.32E-29 | -0.86397671 | YLR383W | SMC6 | n |
| W303.asm02.HP0_G0038480 | 1478.1184 | 1435.62072 | 1634.345165 | 3377.67643 | 3451.6321 | 3758.46041 | 2703.1628 | 2901.0159 | 2681.9851 | 5.60E-59 | -1.218400869 | YLR384C | IKI3 | n |
| W303.asm02.HP0_G0038490 | 25.284991 | 24.9131578 | 23.18220093 | 165.008619 | 180.67137 | 198.163024 | 190.39569 | 198.12456 | 185.1469 | 1.48E-40 | -2.889529506 | YLR385C | SWC7 | n |
| W303.asm02.HP0_G0038510 | 2684.4232 | 2477.82115 | 2406.119271 | 918.381304 | 922.23295 | 793.600242 | 848.2299 | 987.99863 | 845.50418 | 9.45E-65 | 1.521752921 | YLR387C | REH1 | n |
| W303.asm02.HP0_G0038520 | 2626.4785 | 2638.71863 | 2756.75006 | 1890.09873 | 2045.8111 | 1877.33391 | 1690.7593 | 2129.511 | 1518.2046 | 1.15E-05 | 0.46466974 | YLR388W | RPS29A | n |

| | | | | | | | | | | | | | | |
|---|---|---|---|---|---|---|---|---|---|---|---|---|---|---|
| W303.asm02.HP0_G0038530 | 1123.075 | 1190.64133 | 1112.745644 | 2490.96345 | 2495.2424 | 2488.88965 | 2352.0138 | 2562.4984 | 2344.3124 | 2.76E-65 | -1.125577322 | YLR389C | STE23 | n |
| W303.asm02.HP0_G0038540 | 561.53751 | 544.975327 | 492.6217697 | 558.362499 | 647.18102 | 554.666836 | 433.2357 | 502.52785 | 469.03882 | 0.185114379 | -0.139148704 | YLR390W | ECM19 | n |
| W303.asm02.HP0_G0038550 | 3251.2285 | 3580.22838 | 3832.790553 | 6818.6895 | 6530.2362 | 7126.28366 | 6620.5256 | 6059.2001 | 6433.414 | 5.66E-37 | -0.940817434 | YLR390W- | CCW14 | n |
| W303.asm02.HP0_G0038570 | 138.01391 | 146.364802 | 157.4457813 | 918.381304 | 972.56925 | 925.392875 | 793.50539 | 1002.4315 | 849.03079 | 2.07E-115 | -2.670681294 | YLR393W | ATP10 | n |
| W303.asm02.HP0_G0038580 | 228.61846 | 226.294517 | 257.9019853 | 29.1681902 | 30.561326 | 26.5481563 | 28.502349 | 53.79541 | 29.976165 | 1.48E-42 | 3.045517318 | YLR394W | CST9 | n |
| W303.asm02.HP0_G0038590 | 4336.376 | 4212.39976 | 4932.979172 | 4299.39124 | 4521.2785 | 4847.88296 | 4171.6038 | 4293.1361 | 3896.0198 | 0.818818244 | -0.019507225 | YLR395C | COX8 | n |
| W303.asm02.HP0_G0038600 | 504.64628 | 501.377301 | 537.0543215 | 760.0397 | 763.13428 | 676.977985 | 822.00774 | 754.44782 | 762.6289 | 8.51E-09 | -0.512018799 | YLR396C | VPS33 | n |
| W303.asm02.HP0_G0038610 | 769.08515 | 764.003506 | 1007.459815 | 1870.93106 | 1885.8136 | 2035.6747 | 1289.4463 | 1632.2315 | 1357.7439 | 2.21E-24 | -1.188040301 | YLR397C | AFG2 | n |
| W303.asm02.HP0_G0038620 | 1088.3082 | 1159.49989 | 1214.167774 | 2127.61113 | 2200.4155 | 2225.30438 | 1906.2371 | 1922.2018 | 2091.2783 | 3.71E-38 | -0.919955155 | YLR398C | SKI2 | n |
| W303.asm02.HP0_G0038630 | 1090.4152 | 1190.64133 | 1288.544002 | 4666.07706 | 4777.4543 | 5494.5202 | 5508.934 | 5079.0739 | 5457.4253 | 1.16E-127 | -2.064450365 | YLR399C | BDF1 | n |
| W303.asm02.HP0_G0038640 | 862.85032 | 896.873681 | 942.7428377 | 1446.74224 | 1471.4379 | 1617.54123 | 1010.1232 | 1036.5457 | 1036.8226 | 5.90E-23 | -0.74603087 | YLR401C | DUS3 | n |
| W303.asm02.HP0_G0038650 | 967.15091 | 980.955588 | 1130.132295 | 1625.08488 | 1558.6276 | 1575.8227 | 1625.774 | 1376.3752 | 1597.5533 | 1.31E-12 | -0.62815386 | YLR403W | SFP1 | n |
| W303.asm02.HP0_G0038660 | 659.51685 | 659.160633 | 638.4764505 | 322.516846 | 357.74728 | 344.177883 | 457.17767 | 438.23578 | 510.47646 | 2.18E-22 | 0.934526473 | YLR404W | SEI1 | n |
| W303.asm02.HP0_G0038670 | 706.92621 | 619.7148 | 676.147527 | 823.376342 | 808.07741 | 855.229891 | 906.37469 | 910.58572 | 928.37946 | 0.000109433 | -0.311939674 | YLR405W | DUS4 | n |
| W303.asm02.HP0_G0038700 | 1098.8436 | 1074.37993 | 992.0050147 | 970.05067 | 917.73864 | 1017.36327 | 1122.9925 | 1023.4249 | 1194.6383 | 0.156265886 | 0.12348202 | YLR407W | GAG1 | n |
| W303.asm02.HP0_G0038710 | 396.13153 | 377.84956 | 476.201044 | 125.839906 | 145.61573 | 128.000039 | 119.70986 | 145.64123 | 130.48448 | 8.25E-32 | 1.647524063 | YLR408C | BLS1 | n |
| W303.asm02.HP0_G0038720 | 1425.4414 | 1459.49583 | 1603.435564 | 3102.66206 | 2968.044 | 3399.11215 | 1990.604 | 2448.3472 | 1915.8296 | 1.25E-27 | -1.076766539 | YLR409C | UTP21 | n |
| W303.asm02.HP0_G0038730 | 1294.8023 | 1485.44703 | 1429.569057 | 2601.80257 | 2529.3991 | 2429.1563 | 2266.5068 | 2199.0514 | 2245.5674 | 5.86E-34 | -0.844684817 | YLR410W | VIP1 | n |
| W303.asm02.HP0_G0038750 | 949.24071 | 1042.20043 | 860.6392094 | 510.860018 | 524.03685 | 530.014977 | 287.30368 | 406.74578 | 330.61947 | 3.90E-14 | 0.865752963 | YLR412W | BER1 | n |
| W303.asm02.HP0_G0038780 | 1728.8613 | 1680.6001 | 1754.11987 | 481.691827 | 434.1506 | 465.540883 | 296.42443 | 362.13495 | 308.57817 | 4.01E-110 | 1.901885904 | YLR414C | PUN1 | n |
| W303.asm02.HP0_G0038800 | 316.06239 | 336.32763 | 321.6530379 | 770.873599 | 773.02177 | 765.155789 | 766.14314 | 817.42781 | 818.17297 | 2.37E-47 | -1.245366116 | YLR417W | VPS36 | n |
| W303.asm02.HP0_G0038810 | 1861.6075 | 1768.8342 | 1757.98357 | 1606.75059 | 1649.4127 | 1571.08196 | 1551.6679 | 1609.926 | 1719.2212 | 0.020584201 | 0.158389795 | YLR418C | CDC73 | n |
| W303.asm02.HP0_G0038820 | 842.83304 | 878.188812 | 925.356187 | 1700.0888 | 1690.7604 | 1717.09682 | 1560.7886 | 1428.8586 | 1421.2229 | 9.27E-40 | -0.948099517 | YLR419W | | n |
| W303.asm02.HP0_G0038830 | 1892.1602 | 1723.16008 | 1872.92865 | 2205.11518 | 2247.1563 | 2249.95624 | 2120.5747 | 2252.8468 | 2167.1004 | 7.14E-06 | -0.288095403 | YLR420W | URA4 | n |
| W303.asm02.HP0_G0038840 | 3042.6273 | 3025.91062 | 3622.218895 | 2525.1319 | 2513.2196 | 2603.61561 | 2756.7472 | 2531.0084 | 2498.6015 | 3.21E-05 | 0.343035592 | YLR421C | RPN13 | n |
| W303.asm02.HP0_G0038850 | 563.6446 | 652.932344 | 719.6141538 | 2565.96736 | 2572.5445 | 2632.06006 | 2654.1387 | 2624.1663 | 2734.8842 | 5.27E-131 | -2.003328627 | YLR422W | DCK1 | n |
| W303.asm02.HP0_G0038860 | 1201.0371 | 1437.69681 | 1496.217885 | 274.180988 | 304.7144 | 218.074141 | 342.02819 | 335.89329 | 343.84425 | 1.05E-84 | 2.373937814 | YLR423C | ATG17 | n |
| W303.asm02.HP0_G0038870 | 372.95362 | 406.914911 | 406.6544413 | 512.526771 | 546.50841 | 540.44461 | 515.32247 | 490.71911 | 515.76637 | 2.97E-06 | -0.429768968 | YLR424W | SPP382 | n |
| W303.asm02.HP0_G0038880 | 345.56155 | 373.697367 | 368.0174397 | 682.535652 | 728.97751 | 744.296524 | 829.9884 | 778.06532 | 824.34454 | 5.54E-29 | -0.98641028 | YLR425W | TUS1 | n |
| W303.asm02.HP0_G0038890 | 285.50969 | 287.539363 | 243.4131097 | 350.85166 | 318.19733 | 301.511203 | 279.32302 | 328.02079 | 325.32956 | 0.04127313 | -0.252140118 | YLR426W | TDA5 | n |
| W303.asm02.HP0_G0038900 | 975.57924 | 989.259974 | 986.2094644 | 1188.39541 | 1217.9587 | 1136.82998 | 995.30202 | 1001.1195 | 1022.7162 | 0.00012983 | -0.26406053 | YLR427W | MAG2 | n |
| W303.asm02.HP0_G0038920 | 831.24409 | 946.699996 | 1064.449393 | 1565.91513 | 1718.6251 | 1901.98577 | 1612.0928 | 1632.2315 | 1639.8726 | 2.26E-18 | -0.866437392 | YLR430W | SEN1 | n |
| W303.asm02.HP0_G0038930 | 657.40977 | 628.019186 | 611.4305494 | 327.517108 | 317.29847 | 238.933406 | 359.12959 | 335.89329 | 366.7672 | 8.46E-21 | 1.099076922 | YLR431C | ATG23 | n |
| W303.asm02.HP0_G0038950 | 744.8537 | 657.084537 | 630.7490502 | 2120.94412 | 2019.7441 | 2121.9562 | 2089.7922 | 2014.0477 | 2022.5095 | 7.67E-102 | -1.624403708 | YLR433C | CNA1 | n |
| W303.asm02.HP0_G0038960 | 2797.1522 | 2629.3762 | 3324.713983 | 1445.07548 | 1430.0903 | 1517.98565 | 1101.3308 | 1344.8852 | 1087.0768 | 6.28E-22 | 0.994583281 | YLR435W | TSR2 | n |
| W303.asm02.HP0_G0038970 | 876.54636 | 898.949777 | 895.4125108 | 969.217293 | 908.75001 | 822.992844 | 1010.1232 | 858.10239 | 976.87032 | 0.860309202 | -0.01695833 | YLR436C | ECM30 | n |
| W303.asm02.HP0_G0038980 | 1058.809 | 1102.40723 | 1135.927845 | 2789.31236 | 2792.7659 | 2706.01564 | 3159.2003 | 3407.48 | 3437.5608 | 2.85E-92 | -1.329534207 | YLR437C | DIF1 | n |
| W303.asm02.HP0_G0038990 | 88.497469 | 125.603837 | 90.79695363 | 2395.95848 | 2740.6318 | 2121.00806 | 3915.0826 | 4488.6365 | 4238.9824 | 2.43E-254 | -4.57480069 | YLR438W | CAR2 | n |
| W303.asm02.HP0_G0039000 | 2212.4367 | 2235.95591 | 2819.535188 | 638.366678 | 713.69684 | 659.911313 | 641.87289 | 686.2195 | 648.01416 | 1.32E-73 | 1.853525742 | YLR438C-/ | LSM3 | n |
| W303.asm02.HP0_G0039010 | 532.03836 | 663.312826 | 648.1357009 | 2600.13582 | 2675.9137 | 2608.35635 | 2406.7383 | 2645.1597 | 2470.3887 | 8.78E-135 | -2.095736677 | YLR439W | MRPL4 | n |
| W303.asm02.HP0_G0039020 | 627.91061 | 569.888485 | 581.4868732 | 1260.8992 | 1319.5302 | 1211.7337 | 1125.2727 | 1216.3011 | 1197.2833 | 9.70E-44 | -1.092286104 | YLR440C | SEC39 | n |
| W303.asm02.HP0_G0039050 | 1126.2356 | 1135.62478 | 1115.64342 | 450.856883 | 471.90282 | 445.629766 | 547.2451 | 411.99412 | 412.61309 | 1.48E-35 | 1.303395435 | YLR443W | ECM7 | n |
| W303.asm02.HP0_G0039060 | 10.535413 | 7.26633769 | 18.35257573 | 26.6680596 | 21.572701 | 26.5481563 | 27.362255 | 22.305414 | 17.633038 | 0.014815057 | -1.038237661 | YLR445W | GMC2 | n |
| W303.asm02.HP0_G0039080 | 5858.7432 | 5841.09745 | 6052.486292 | 2419.29304 | 2492.5458 | 2382.69702 | 2365.6949 | 2525.7601 | 2335.4959 | 3.85E-112 | 1.283133079 | YLR447C | VMA6 | n |
| W303.asm02.HP0_G0039120 | 428.79131 | 477.502191 | 473.3032689 | 748.372424 | 678.6412 | 775.585422 | 627.05167 | 736.07866 | 633.90772 | 1.14E-11 | -0.674368553 | YLR451W | LEU3 | n |
| W303.asm02.HP0_G0039130 | 2469.5008 | 2412.42411 | 1943.441178 | 113.339253 | 97.077152 | 85.33333594 | 98.04808 | 132.5204 | 86.401887 | 2.62E-212 | 4.52410088 | YLR452C | SST2 | n |
| W303.asm02.HP0_G0039150 | 1293.7487 | 1447.03925 | 1591.844464 | 1372.57169 | 1386.9449 | 1237.33371 | 1181.1373 | 1270.0965 | 1309.2531 | 0.2112084 | 0.116553548 | YLR454W | FMP27 | n |
| W303.asm02.HP0_G0039160 | 1116.7538 | 1026.62971 | 1133.995995 | 1027.55367 | 1043.5794 | 961.422516 | 954.25864 | 965.69321 | 968.0538 | 0.143142898 | 0.1118555 | YLR455W | PDP3 | n |
| W303.asm02.HP0_G0039180 | 374.00716 | 370.583222 | 323.5848879 | 362.518936 | 351.45525 | 374.518633 | 457.17767 | 346.38996 | 409.08649 | 0.824595462 | -0.027960891 | YLR457C | NBP1 | n |
| W303.asm02.HP0_G0039190 | 1513.9388 | 1523.85482 | 1409.284631 | 1894.26561 | 1906.4874 | 2057.48211 | 2624.4963 | 2618.918 | 2351.3657 | 9.74E-08 | -0.397624499 | YLR459W | GAB1 | n |
| W303.asm02.HP0_G0039260 | 12136.796 | 13392.8984 | 11179.6164 | 8437.10737 | 8167.9637 | 8814.93603 | 12297.053 | 10063.678 | 10370.871 | 4.50E-09 | 0.53016209 | YML130C | ERO1 | n |
| W303.asm02.HP0_G0039270 | 966.09737 | 1035.97214 | 883.8214103 | 1098.39071 | 1155.9372 | 1088.47441 | 1013.5435 | 1074.5961 | 1023.5979 | 0.008640202 | -0.212703516 | YML129C | COX14 | n |
| W303.asm02.HP0_G0039280 | 585.76896 | 722.481576 | 673.2497519 | 44.1689738 | 35.954501 | 29.3926016 | 196.09616 | 127.27207 | 215.12307 | 1.18E-94 | 4.169931865 | YML128C | MSC1 | n |
| W303.asm02.HP0_G0039290 | 1136.7711 | 1185.45109 | 1144.621171 | 1358.40429 | 1332.1143 | 1249.65964 | 1270.0647 | 1270.0965 | 1333.0577 | 0.008231864 | -0.185209033 | YML127W | RSC9 | n |
| W303.asm02.HP0_G0039300 | 5204.494 | 5181.93682 | 4592.007634 | 644.200316 | 565.38453 | 639.052047 | 499.36115 | 481.53452 | 410.84979 | 3.45E-243 | 3.018089066 | YML126C | ERG13 | n |
| W303.asm02.HP0_G0039310 | 2320.9515 | 2330.4183 | 2117.307685 | 1869.26431 | 2032.3282 | 1881.1265 | 2117.1545 | 2233.1656 | 2054.249 | 0.001537262 | 0.226888517 | YML125C | PGA3 | n |
| W303.asm02.HP0_G0039330 | 17220.133 | 15790.7898 | 14370.0668 | 12083.1311 | 11820.042 | 9435.0251 | 11632.379 | 9151.7801 | 11631.634 | 1.06E-05 | 0.507044698 | YML123C | PHO84 | n |
| W303.asm02.HP0_G0039350 | 1518.153 | 1357.7671 | 1446.955708 | 772.540353 | 841.33532 | 727.229852 | 703.43797 | 779.3774 | 713.2564 | 1.39E-25 | 0.884458013 | YML121W | GTR1 | n |
| W303.asm02.HP0_G0039360 | 1735.1825 | 2059.48771 | 1946.338953 | 7137.87284 | 7212.4729 | 7752.06163 | 6505.3761 | 6587.9696 | 6234.1607 | 7.25E-156 | -1.944576941 | YML120C | NDI1 | n |
| W303.asm02.HP0_G0039370 | 927.11634 | 761.927409 | 760.1830054 | 537.528077 | 539.31751 | 556.563133 | 791.2252 | 766.25657 | 768.80047 | 7.96E-10 | 0.584021978 | YML119W | | n |

| | | | | | | | | | | | | | | |
|---|---|---|---|---|---|---|---|---|---|---|---|---|---|---|
| W303.asm02.HP0_G0039390 | 958.72258 | 1193.75548 | 1157.178196 | 1897.59912 | 1919.9703 | 1846.04501 | 1760.3051 | 1637.4798 | 1944.9241 | 1.92E-16 | -0.774728538 | YML117W | NAB6 | n |
| W303.asm02.HP0_G0039410 | 1334.8368 | 1312.09298 | 1411.216481 | 2858.48264 | 2761.3057 | 2998.99351 | 3188.8428 | 3076.835 | 3267.402 | 1.31E-62 | -1.086146401 | YML115C | VAN1 | n |
| W303.asm02.HP0_G0039420 | 435.11256 | 460.893419 | 505.1787952 | 688.36929 | 677.74234 | 680.770578 | 663.53468 | 661.28992 | 715.90135 | 9.78E-10 | -0.545684339 | YML114C | TAF8 | n |
| W303.asm02.HP0_G0039440 | 350.82925 | 375.773463 | 375.74484 | 739.205278 | 672.34917 | 698.785399 | 647.57336 | 750.51157 | 722.07292 | 3.69E-23 | -0.936917386 | YML112W | CTK3 | n |
| W303.asm02.HP0_G0039450 | 971.36508 | 987.183878 | 1109.847869 | 986.718207 | 981.55787 | 929.185469 | 965.65958 | 944.69988 | 976.87032 | 0.306174528 | 0.083062512 | YML111W | BUL2 | n |
| W303.asm02.HP0_G0039460 | 4115.1323 | 3992.33354 | 4247.172395 | 2265.95169 | 2413.4459 | 2443.37852 | 1983.7635 | 2154.4406 | 1996.0599 | 4.65E-36 | 0.794944247 | YML110C | COQ5 | n |
| W303.asm02.HP0_G0039480 | 1246.3394 | 1245.65789 | 1582.185213 | 760.0397 | 800.88651 | 865.659524 | 571.18707 | 699.34033 | 604.81321 | 2.69E-11 | 0.748605681 | YML108W | | n |
| W303.asm02.HP0_G0039490 | 737.47891 | 638.399668 | 707.0571283 | 530.027685 | 524.93571 | 426.666797 | 451.4772 | 524.83327 | 496.37003 | 7.37E-06 | 0.490156179 | YML107C | PML39 | n |
| W303.asm02.HP0_G0039500 | 6621.5071 | 6263.58309 | 6692.894593 | 3057.65971 | 2983.3247 | 3135.52688 | 2851.375 | 2760.623 | 2823.9311 | 5.75E-87 | 1.0933587 | YML106W | URA5 | n |
| W303.asm02.HP0_G0039510 | 2949.9156 | 2800.65416 | 2867.83144 | 1551.74772 | 1511.8868 | 1484.80045 | 1592.7112 | 1712.2685 | 1646.9258 | 1.90E-51 | 0.921721296 | YML105C | SEC65 | n |
| W303.asm02.HP0_G0039520 | 667.94518 | 615.562607 | 583.4187233 | 542.528338 | 501.56529 | 595.437219 | 630.47196 | 604.87034 | 575.7187 | 0.057766674 | 0.187362151 | YML104C | MDM1 | n |
| W303.asm02.HP0_G0039530 | 1793.1273 | 1759.49177 | 2002.362605 | 2719.30871 | 2743.3284 | 2744.88973 | 2617.6557 | 2586.1159 | 2677.5769 | 2.11E-17 | -0.562617083 | YML103C | NUP188 | n |
| W303.asm02.HP0_G0039540 | 601.57208 | 714.17719 | 756.3193052 | 1325.90259 | 1282.6768 | 1385.24487 | 1345.3109 | 1243.8548 | 1436.211 | 2.73E-23 | -0.945656893 | YML102W | CAC2 | n |
| W303.asm02.HP0_G0039570 | 525.71711 | 510.719735 | 579.5550232 | 560.029253 | 542.0141 | 516.740899 | 364.83006 | 391.00078 | 370.2938 | 0.982008514 | -0.002323467 | YML099C | ARG81 | n |
| W303.asm02.HP0_G0039580 | 1210.519 | 1315.20712 | 1457.580883 | 391.687126 | 352.35411 | 407.703828 | 424.11495 | 432.98745 | 412.61309 | 9.86E-76 | 1.790931658 | YML098W | TAF13 | n |
| W303.asm02.HP0_G0039590 | 2352.5577 | 2233.87982 | 2251.571265 | 1327.56934 | 1439.0789 | 1347.31893 | 1258.6637 | 1251.7273 | 1221.9696 | 5.57E-30 | 0.733089573 | YML097C | VPS9 | n |
| W303.asm02.HP0_G0039600 | 478.30775 | 441.170503 | 396.9951909 | 282.514757 | 280.44511 | 252.207484 | 250.82067 | 266.35288 | 247.74419 | 9.26E-10 | 0.6892782 | YML095C | RAD10 | n |
| W303.asm02.HP0_G0039620 | 1234.7504 | 1220.74473 | 1195.815198 | 2445.12772 | 2321.7619 | 2582.75634 | 1370.3929 | 1695.2115 | 1497.9266 | 4.99E-33 | -1.009258188 | YML093W | UTP14 | n |
| W303.asm02.HP0_G0039630 | 3988.7074 | 3624.86446 | 3482.159764 | 3432.6793 | 3422.8685 | 3385.83807 | 3779.4114 | 3718.4437 | 3574.2169 | 0.085324339 | 0.115326202 | YML092C | PRE8 | n |
| W303.asm02.HP0_G0039640 | 2488.4646 | 2381.28267 | 2730.670084 | 1450.90912 | 1673.682 | 1812.85981 | 2104.6134 | 1926.1381 | 1945.8058 | 9.53E-12 | 0.623130835 | YML091C | RPM2 | n |
| W303.asm02.HP0_G0039660 | 1449.6728 | 1568.49089 | 1389.000206 | 974.217554 | 917.73864 | 989.866969 | 1005.5629 | 1054.9149 | 986.56849 | 9.06E-16 | 0.61262437 | YML088W | UFO1 | n |
| W303.asm02.HP0_G0039680 | 1317.9802 | 1471.95241 | 1490.422335 | 3481.01516 | 3429.1605 | 3620.97888 | 3756.6096 | 3584.6112 | 3753.1922 | 5.83E-84 | -1.298329392 | YML086C | ALO1 | n |
| W303.asm02.HP0_G0039690 | 1689.8802 | 1857.0683 | 1965.657454 | 3872.70229 | 3971.1746 | 4136.77163 | 4136.2609 | 4186.8574 | 4105.853 | 6.65E-61 | -1.11927856 | YML085C | TUB1 | n |
| W303.asm02.HP0_G0039720 | 2054.4055 | 2066.75405 | 2410.948896 | 4193.55238 | 4527.5705 | 4682.90513 | 3952.7057 | 3917.8803 | 3534.5425 | 9.32E-36 | -1.036522422 | YML081C- | ATP18 | n |
| W303.asm02.HP0_G0039730 | 845.99366 | 925.939031 | 1020.982766 | 1415.07391 | 1369.8665 | 1330.25226 | 1214.2001 | 1241.2307 | 1293.3834 | 9.97E-12 | -0.558588705 | YML081W | TDA9 | n |
| W303.asm02.HP0_G0039740 | 1251.6071 | 1182.33695 | 1128.200445 | 1694.25516 | 1734.8047 | 1846.99316 | 1281.4656 | 1629.6073 | 1367.4421 | 1.28E-09 | -0.566774724 | YML080W | DUS1 | n |
| W303.asm02.HP0_G0039750 | 1909.0168 | 1921.42729 | 2294.071967 | 404.187779 | 399.99382 | 430.459391 | 413.8541 | 432.98745 | 355.30572 | 7.21E-113 | 2.311337673 | YML079W | CFF1 | n |
| W303.asm02.HP0_G0039760 | 3874.9249 | 3679.88102 | 4411.379651 | 2219.28259 | 2274.1222 | 2255.64513 | 1927.8989 | 2045.5377 | 1943.1608 | 1.13E-27 | 0.82650338 | YML078W | CPR3 | n |
| W303.asm02.HP0_G0039780 | 476.20067 | 484.768529 | 499.383245 | 1102.55759 | 1156.8361 | 1189.92629 | 1110.4515 | 991.93488 | 1078.2603 | 1.77E-52 | -1.239128018 | YML076C | WAR1 | n |
| W303.asm02.HP0_G0039790 | 1835.2689 | 2013.81359 | 1824.632398 | 2264.28494 | 2230.0779 | 2430.10445 | 2203.8016 | 2373.5585 | 2183.8518 | 6.63E-05 | -0.287327818 | YML075C | HMG1 | n |
| W303.asm02.HP0_G0039800 | 2466.3402 | 2634.56644 | 1835.257573 | 4673.57745 | 4515.8853 | 5286.87569 | 2776.1288 | 3533.44 | 2883.0018 | 8.11E-17 | -1.061712636 | YML074C | FPR3 | n |
| W303.asm02.HP0_G0039810 | 8054.3232 | 8472.54975 | 8329.171608 | 11842.2852 | 12318.911 | 13395.4411 | 10590.333 | 11253.737 | 9561.515 | 1.73E-15 | -0.595390853 | YML073C | RPL6A | n |
| W303.asm02.HP0_G0039820 | 2298.8271 | 2797.54001 | 2437.028872 | 2714.30845 | 2660.6331 | 2622.57858 | 3844.3968 | 3599.0441 | 3779.6417 | 0.282966894 | -0.086373816 | YML072C | TCB3 | n |
| W303.asm02.HP0_G0039830 | 1296.9093 | 1154.30964 | 1021.948691 | 940.882479 | 897.0648 | 879.88175 | 875.59215 | 863.35073 | 892.23174 | 0.000136908 | 0.352717816 | YML071C | COG8 | n |
| W303.asm02.HP0_G0039840 | 946.08009 | 1056.73311 | 1136.89377 | 3147.66441 | 2774.7886 | 2794.19345 | 2839.974 | 3298.5771 | 3096.3615 | 9.68E-60 | -1.472787268 | YML070W | DAK1 | n |
| W303.asm02.HP0_G0039850 | 2288.2917 | 2119.69451 | 2302.765292 | 3448.51346 | 3401.2958 | 3413.33438 | 3405.4606 | 3299.8892 | 3315.8928 | 3.93E-26 | -0.612811084 | YML069W | POB3 | n |
| W303.asm02.HP0_G0039860 | 817.54805 | 724.557673 | 733.1371043 | 500.859495 | 525.83458 | 540.44461 | 494.80077 | 459.22911 | 481.38194 | 1.29E-09 | 0.538171046 | YML068W | ITT1 | n |
| W303.asm02.HP0_G0039880 | 115.88954 | 132.870175 | 95.62657882 | 255.01332 | 300.22008 | 274.963047 | 315.80602 | 308.33954 | 334.14607 | 3.85E-19 | -1.271718811 | YML066C | SMA2 | n |
| W303.asm02.HP0_G0039890 | 625.80353 | 662.274778 | 662.6245765 | 867.545315 | 843.13305 | 933.926211 | 855.07046 | 879.09572 | 886.06017 | 3.36E-08 | -0.438230879 | YML065W | ORC1 | n |
| W303.asm02.HP0_G0039900 | 744.8537 | 677.845502 | 706.0912032 | 475.024812 | 431.45401 | 458.903844 | 418.41448 | 540.57827 | 530.75445 | 1.89E-09 | 0.64030972 | YML064C | TEM1 | n |
| W303.asm02.HP0_G0039910 | 16857.714 | 16464.4832 | 17946.88722 | 21204.4409 | 21978.986 | 24444.2149 | 19125.076 | 20645.629 | 17243.348 | 1.03E-06 | -0.399464128 | YML063W | RPS1B | y |
| W303.asm02.HP0_G0039920 | 1403.317 | 1417.9739 | 1290.475852 | 1480.91069 | 1491.2129 | 1395.6745 | 1237.0019 | 1323.8919 | 1367.4421 | 0.246652387 | -0.087840419 | YML062C | MFT1 | n |
| W303.asm02.HP0_G0039930 | 681.64122 | 715.215238 | 638.4764505 | 820.876211 | 867.40233 | 970.904 | 857.35065 | 972.25363 | 872.83539 | 4.39E-05 | -0.385326609 | YML061C | PIF1 | n |
| W303.asm02.HP0_G0039940 | 709.03329 | 622.828945 | 615.2942496 | 495.025857 | 494.37439 | 499.674227 | 493.66068 | 497.27952 | 465.51221 | 2.43E-05 | 0.386361559 | YML060W | OGG1 | n |
| W303.asm02.HP0_G0039950 | 848.10075 | 928.015128 | 983.3116893 | 350.018283 | 357.74728 | 434.251985 | 527.8635 | 438.23578 | 461.10395 | 1.39E-30 | 1.274832263 | YML059C | NTE1 | n |
| W303.asm02.HP0_G0039990 | 4653.4919 | 4348.38408 | 4656.724611 | 2544.29957 | 2385.5811 | 2584.65264 | 2298.4294 | 2415.5451 | 2231.461 | 1.31E-41 | 0.862136843 | YML056C | IMD4 | n |
| W303.asm02.HP0_G0040000 | 1662.4882 | 1583.02357 | 2093.159559 | 902.547144 | 1029.1976 | 908.326203 | 980.4808 | 775.44115 | 894.87669 | 8.49E-15 | 0.910998646 | YML055W | SPC2 | n |
| W303.asm02.HP0_G0040010 | 378.22133 | 430.79002 | 401.8248161 | 635.03317 | 624.70945 | 649.48168 | 866.4714 | 653.41742 | 866.66383 | 1.34E-08 | -0.656808257 | YML054C | CYB2 | n |
| W303.asm02.HP0_G0040040 | 2726.5649 | 2579.54988 | 2562.599127 | 2380.12432 | 2147.3826 | 2376.05999 | 4255.9707 | 3959.867 | 4069.7052 | 0.005628705 | 0.188611385 | YML052W | SUR7 | n |
| W303.asm02.HP0_G0040050 | 838.61887 | 794.106905 | 766.9444807 | 343.351268 | 321.79278 | 317.629727 | 329.48715 | 271.60122 | 338.55433 | 2.04E-37 | 1.286691312 | YML051W | GAL80 | n |
| W303.asm02.HP0_G0040060 | 672.15935 | 640.475765 | 696.4319528 | 275.014365 | 267.86103 | 291.08157 | 376.231 | 307.02746 | 373.82041 | 3.48E-34 | 1.269382672 | YML050W | AIM32 | n |
| W303.asm02.HP0_G0040070 | 770.13869 | 756.737168 | 832.6273833 | 859.211547 | 835.94215 | 965.21511 | 877.87234 | 852.85406 | 834.04271 | 0.046748511 | -0.171908883 | YML049C | RSE1 | n |
| W303.asm02.HP0_G0040080 | 3972.9042 | 3629.01665 | 3806.710577 | 4239.38811 | 4170.7221 | 3968.00121 | 5785.9768 | 5640.6455 | 5736.0273 | 0.04874054 | -0.117875479 | YML048W | GSF2 | n |
| W303.asm02.HP0_G0040100 | 1553.9734 | 1370.22368 | 1473.035684 | 553.362238 | 574.37315 | 573.629805 | 514.18237 | 548.45076 | 527.22784 | 1.22E-65 | 1.370226577 | YML046W | PRP39 | n |
| W303.asm02.HP0_G0040110 | 956.6155 | 865.732233 | 925.356187 | 420.021939 | 429.65629 | 421.926055 | 249.68058 | 351.63829 | 267.14053 | 1.88E-24 | 1.111629592 | YML043C | RRN11 | n |
| W303.asm02.HP0_G0040120 | 143.28162 | 181.658442 | 160.3435564 | 103.338731 | 105.16692 | 104.296328 | 200.65654 | 190.25206 | 201.89829 | 3.21E-05 | 0.633957159 | YML042W | CAT2 | n |
| W303.asm02.HP0_G0040130 | 875.49282 | 773.34594 | 823.9340579 | 565.029514 | 492.57666 | 529.066828 | 554.08566 | 447.42036 | 509.59481 | 1.64E-10 | 0.639475579 | YML041C | VPS71 | n |
| W303.asm02.HP0_G0040140 | 1192.6088 | 1160.53793 | 1105.984169 | 933.382088 | 919.53636 | 904.53361 | 1244.9826 | 1255.6636 | 1281.0402 | 2.52E-06 | 0.326358153 | YML038C | YMD8 | n |
| W303.asm02.HP0_G0040150 | 97.979341 | 115.223355 | 114.9450796 | 402.521025 | 405.387 | 371.674188 | 411.57392 | 334.58121 | 372.93876 | 9.22E-44 | -1.844981513 | YML037C | LFT1 | n |
| W303.asm02.HP0_G0040160 | 1118.8609 | 1080.60822 | 1334.908403 | 430.022462 | 495.27325 | 456.059399 | 351.14894 | 476.28619 | 378.22867 | 1.75E-29 | 1.35618662 | YML036W | CGI121 | n |

| | | | | | | | | | | | | | | |
|---|---|---|---|---|---|---|---|---|---|---|---|---|---|---|
| W303.asm02.HP0_G0040170 | 1069.3444 | 1115.90186 | 1311.726202 | 2500.13059 | 2510.523 | 2505.95632 | 2434.1006 | 2377.4947 | 2396.3299 | 7.96E-45 | -1.103048258 | YML035C | AMD1 | n |
| W303.asm02.HP0_G0040180 | 1011.3996 | 1191.67938 | 1208.372223 | 1495.91147 | 1442.6743 | 1553.06714 | 1675.9381 | 1491.8386 | 1638.1093 | 4.61E-06 | -0.396251156 | YML034W | SRC1 | n |
| W303.asm02.HP0_G0040190 | 763.81744 | 820.058111 | 780.4674312 | 1125.05877 | 1126.2747 | 1152.9485 | 1216.4802 | 1100.8378 | 1202.5732 | 1.91E-12 | -0.52579068 | YML032C | RAD52 | n |
| W303.asm02.HP0_G0040200 | 1007.1855 | 1031.81995 | 1013.255366 | 3416.84514 | 3362.6447 | 3330.84546 | 3997.1694 | 4084.5149 | 3940.1024 | 2.14E-198 | -1.728012114 | YML031W | NDC1 | n |
| W303.asm02.HP0_G0040210 | 1880.5712 | 1833.19319 | 2386.80077 | 1075.88953 | 981.55787 | 975.644743 | 1047.7463 | 1111.3344 | 1085.3135 | 4.77E-23 | 1.00823173 | YML030W | RCF1 | n |
| W303.asm02.HP0_G0040220 | 675.31997 | 792.030808 | 774.671881 | 2327.62158 | 2145.5848 | 2401.65999 | 2821.7325 | 2281.7126 | 2771.9136 | 1.91E-60 | -1.61612806 | YML029W | USA1 | n |
| W303.asm02.HP0_G0040230 | 17755.332 | 18667.2215 | 18050.2412 | 8993.80312 | 8818.7402 | 9277.63246 | 10194.72 | 9494.2338 | 9565.9232 | 3.24E-78 | 1.007806187 | YML028W | TSA1 | n |
| W303.asm02.HP0_G0040240 | 403.50632 | 454.66513 | 458.8143933 | 1050.05485 | 1022.0067 | 1070.45959 | 1356.7118 | 1316.0194 | 1348.0458 | 1.81E-55 | -1.253634396 | YML027W | YOX1 | n |
| W303.asm02.HP0_G0040260 | 812.28034 | 948.776093 | 1004.56204 | 2168.4466 | 2104.2372 | 2126.69695 | 1962.1017 | 2225.2931 | 1975.7819 | 2.10E-43 | -1.209634669 | YML025C | YML6 | n |
| W303.asm02.HP0_G0040280 | 1218.9473 | 1046.35263 | 1058.653842 | 615.032126 | 650.77647 | 631.46686 | 693.17712 | 596.99784 | 608.33982 | 2.07E-18 | 0.808825879 | YML023C | NSE5 | n |
| W303.asm02.HP0_G0040290 | 4833.6475 | 4614.12443 | 6128.79437 | 2221.78272 | 2241.7631 | 2296.41552 | 1715.8414 | 1936.6348 | 1751.8423 | 4.53E-33 | 1.204529355 | YML022W | APT1 | n |
| W303.asm02.HP0_G0040300 | 1060.9161 | 969.537058 | 1031.607941 | 640.866808 | 660.66395 | 581.214992 | 660.1144 | 740.01491 | 709.72979 | 2.63E-16 | 0.700972839 | YML021C | UNG1 | n |
| W303.asm02.HP0_G0040310 | 517.28878 | 516.948024 | 551.5431971 | 775.87386 | 771.22404 | 811.615063 | 1117.2921 | 1040.482 | 1095.0117 | 9.11E-13 | -0.571798033 | YML020W | | n |
| W303.asm02.HP0_G0040320 | 2010.1568 | 1852.91611 | 2008.158155 | 1565.91513 | 1478.6288 | 1619.43753 | 2211.7823 | 2444.4109 | 2147.7041 | 1.25E-05 | 0.3323097 | YML019W | OST6 | n |
| W303.asm02.HP0_G0040330 | 1813.1446 | 1742.883 | 1657.527366 | 710.037088 | 684.93324 | 559.407578 | 672.65543 | 728.20616 | 789.07846 | 3.18E-46 | 1.414105404 | YML018C | | n |
| W303.asm02.HP0_G0040340 | 392.9709 | 494.110963 | 492.6217697 | 2952.65423 | 2839.5067 | 2874.78606 | 2451.202 | 2265.9676 | 2416.6079 | 2.35E-208 | -2.649991404 | YML017W | PSP2 | n |
| W303.asm02.HP0_G0040350 | 1803.6627 | 1551.88212 | 1712.585093 | 2073.44164 | 1917.2738 | 2009.12654 | 2150.2172 | 2178.0581 | 2061.3022 | 0.001064226 | -0.243614917 | YML016C | PPZ1 | n |
| W303.asm02.HP0_G0040360 | 788.04889 | 796.183001 | 674.215677 | 525.860801 | 577.06974 | 487.348297 | 483.39984 | 599.62201 | 521.93793 | 4.26E-06 | 0.505052784 | YML015C | TAF11 | n |
| W303.asm02.HP0_G0040370 | 438.27318 | 451.550985 | 508.0765703 | 1400.07313 | 1327.6199 | 1421.27451 | 1069.4081 | 1390.8082 | 1146.1475 | 3.36E-50 | -1.56857208 | YML014W | TRM9 | n |
| W303.asm02.HP0_G0040380 | 510.96753 | 633.209427 | 537.0543215 | 830.876733 | 870.09892 | 875.141008 | 897.25394 | 931.57905 | 999.79327 | 8.57E-11 | -0.615604877 | YML013W | UBX2 | n |
| W303.asm02.HP0_G0040390 | 5630.1247 | 5672.93364 | 6011.91744 | 4020.20999 | 4218.3618 | 4234.43092 | 4405.323 | 4669.704 | 4357.1238 | 6.54E-16 | 0.473474625 | YML012W | ERV25 | n |
| W303.asm02.HP0_G0040400 | 705.87267 | 702.758659 | 729.2734042 | 434.189346 | 455.7233 | 448.474211 | 473.13899 | 457.91703 | 501.65994 | 2.12E-15 | 0.676366216 | YML011C | RAD33 | n |
| W303.asm02.HP0_G0040410 | 1665.6488 | 1723.16008 | 2173.331337 | 3605.18831 | 3462.4184 | 3806.81598 | 3649.4407 | 3196.2346 | 3661.5004 | 1.05E-21 | -0.966680933 | YML010W | SPT5 | n |
| W303.asm02.HP0_G0040420 | 1594.008 | 1558.11041 | 1853.610149 | 890.879867 | 859.31257 | 821.096547 | 816.30727 | 957.82071 | 815.52802 | 1.74E-24 | 0.961100449 | YML009C | MRPL39 | n |
| W303.asm02.HP0_G0040430 | 4769.3815 | 4769.83167 | 4941.672498 | 3493.51581 | 3486.6877 | 3241.71951 | 2878.7372 | 3217.2279 | 3077.8468 | 2.89E-15 | 0.502352141 | YML008C | ERG6 | n |
| W303.asm02.HP0_G0040440 | 2785.5632 | 2876.43168 | 2747.09081 | 1906.76626 | 1958.6214 | 2054.63766 | 2163.8983 | 1847.4131 | 2182.0885 | 2.98E-11 | 0.506538492 | YML007W | YAP1 | n |
| W303.asm02.HP0_G0040450 | 467.77234 | 514.871928 | 485.8602944 | 305.849309 | 322.69165 | 360.296406 | 218.89804 | 237.48705 | 256.56071 | 1.35E-07 | 0.572145407 | YML006C | GIS4 | n |
| W303.asm02.HP0_G0040470 | 2549.5699 | 2552.56063 | 2240.94609 | 3202.66729 | 3011.1895 | 3104.23799 | 3518.3299 | 3277.5838 | 3411.1112 | 1.16E-06 | -0.34411578 | YML004C | GLO1 | n |
| W303.asm02.HP0_G0040480 | 774.35286 | 812.791773 | 755.3533802 | 682.535652 | 726.28092 | 753.778008 | 800.34595 | 703.27658 | 752.04908 | 0.18077193 | 0.115781242 | YML002W | VRL1 | n |
| W303.asm02.HP0_G0040490 | 2758.1711 | 2718.64834 | 3239.71258 | 1448.40899 | 1501.1004 | 1408.00043 | 1642.8754 | 1541.6977 | 1707.7598 | 1.76E-33 | 1.000480524 | YML001W | YPT7 | n |
| W303.asm02.HP0_G0040500 | 900.77781 | 850.16151 | 846.1503338 | 1265.89946 | 1260.2053 | 1428.8597 | 1805.9088 | 1605.9898 | 1701.5882 | 1.36E-13 | -0.606474468 | YMR001C | CDC5 | n |
| W303.asm02.HP0_G0040520 | 5238.2073 | 4851.83748 | 5109.743454 | 3075.994 | 3146.0188 | 3126.0454 | 3040.6306 | 3250.03 | 2905.0431 | 4.55E-29 | 0.701370604 | YMR002W | MIX17 | n |
| W303.asm02.HP0_G0040530 | 818.60159 | 798.259098 | 950.470238 | 985.88483 | 945.60337 | 1039.17069 | 1185.6977 | 1207.1165 | 1265.1705 | 0.017341434 | -0.209362494 | YMR003W | AIM34 | n |
| W303.asm02.HP0_G0040540 | 805.95909 | 876.112716 | 896.3784358 | 1304.23479 | 1344.6983 | 1243.0226 | 1463.8806 | 1405.2411 | 1443.2642 | 7.16E-16 | -0.593678984 | YMR004W | MVP1 | n |
| W303.asm02.HP0_G0040550 | 1284.2668 | 1368.14758 | 1104.052319 | 1471.74354 | 1371.6642 | 1334.993 | 1459.3203 | 1550.8823 | 1401.8265 | 0.09282777 | -0.154521001 | YMR005W | TAF4 | n |
| W303.asm02.HP0_G0040570 | 2005.9426 | 1852.91611 | 1859.405699 | 543.361715 | 568.08111 | 567.940914 | 497.08096 | 485.47077 | 511.35811 | 1.10E-122 | 1.76796205 | YMR009W | ADI1 | n |
| W303.asm02.HP0_G0040580 | 2672.8343 | 2426.95679 | 2423.505922 | 1564.24837 | 1383.3494 | 1402.31154 | 1903.9569 | 1987.806 | 1966.0838 | 1.43E-26 | 0.789638654 | YMR010W | ANY1 | n |
| W303.asm02.HP0_G0040600 | 3378.7069 | 3658.082 | 3515.001216 | 7612.06427 | 7776.9585 | 7905.66167 | 6622.8058 | 6782.1579 | 6523.3425 | 6.85E-99 | -1.14241023 | YMR012W | CLU1 | n |
| W303.asm02.HP0_G0040610 | 1165.2167 | 1235.27741 | 1230.588499 | 1319.23558 | 1474.1345 | 1215.5263 | 2302.9898 | 2049.4739 | 2242.0408 | 0.104335536 | -0.142921828 | YMR013C | SEC59 | n |
| W303.asm02.HP0_G0040620 | 1924.82 | 1738.7308 | 1608.265189 | 1887.5986 | 1884.9147 | 1965.51171 | 1202.7991 | 1350.1336 | 1275.7503 | 0.137752002 | -0.122579811 | YMR014W | BUD22 | n |
| W303.asm02.HP0_G0040630 | 5513.1816 | 4918.27257 | 4417.175202 | 2500.13059 | 2542.8821 | 2702.22305 | 3023.5292 | 3023.0396 | 2949.1256 | 1.94E-30 | 0.938919754 | YMR015C | ERG5 | n |
| W303.asm02.HP0_G0040640 | 616.32166 | 672.65526 | 702.2275031 | 2759.3108 | 2669.6217 | 2913.66015 | 2659.8392 | 2563.8105 | 2776.3219 | 4.18E-166 | -2.065882509 | YMR016C | SOK2 | n |
| W303.asm02.HP0_G0040680 | 857.58262 | 795.144953 | 808.4792573 | 837.543748 | 862.90802 | 799.289133 | 750.18182 | 715.08533 | 813.76472 | 0.793587011 | -0.02291191 | YMR020W | FMS1 | n |
| W303.asm02.HP0_G0040690 | 828.08346 | 883.379053 | 765.0126306 | 610.031864 | 615.72083 | 560.355727 | 624.77149 | 535.32993 | 545.74253 | 7.31E-07 | 0.470400794 | YMR021C | MAC1 | n |
| W303.asm02.HP0_G0040700 | 2067.048 | 2164.33058 | 2345.265994 | 905.047274 | 880.88527 | 914.963243 | 918.91572 | 717.70949 | 929.26112 | 2.01E-40 | 1.284153279 | YMR022W | UBC7 | n |
| W303.asm02.HP0_G0040710 | 627.91061 | 679.921598 | 685.8067774 | 696.703058 | 739.76386 | 727.229852 | 633.89224 | 616.67909 | 641.84259 | 0.157007281 | -0.11721148 | YMR023C | MSS1 | n |
| W303.asm02.HP0_G0040720 | 3017.3423 | 2806.88244 | 2966.355794 | 2195.11466 | 2251.6506 | 2171.25992 | 2174.1592 | 2297.4576 | 2124.7811 | 3.56E-11 | 0.409549135 | YMR024W | MRPL3 | n |
| W303.asm02.HP0_G0040730 | 286.56323 | 315.566665 | 360.2900394 | 190.843302 | 197.74975 | 190.577836 | 160.75325 | 164.0104 | 148.99917 | 1.02E-08 | 0.734174561 | YMR025W | CSI1 | n |
| W303.asm02.HP0_G0040740 | 499.37858 | 557.431906 | 476.201044 | 445.023245 | 453.92557 | 375.466781 | 438.93617 | 413.3062 | 437.29935 | 0.014151558 | 0.26474932 | YMR026C | PEX12 | n |
| W303.asm02.HP0_G0040750 | 1655.1134 | 1623.50745 | 1588.946689 | 5062.76445 | 4932.0587 | 5152.23861 | 5050.6162 | 4695.9457 | 4886.9965 | 2.16E-176 | -1.637934176 | YMR027W | | n |
| W303.asm02.HP0_G0040760 | 1303.2306 | 1372.29978 | 1583.151138 | 1014.21964 | 1011.2203 | 910.2225 | 869.89168 | 953.88446 | 881.65191 | 4.60E-09 | 0.536805208 | YMR028W | TAP42 | n |
| W303.asm02.HP0_G0040770 | 1585.5797 | 1520.74067 | 1553.207462 | 1070.88927 | 1051.6692 | 1017.36327 | 1027.2247 | 1099.5257 | 1111.7631 | 6.02E-17 | 0.5690293 | YMR029C | FAR8 | n |
| W303.asm02.HP0_G0040790 | 1181.0198 | 1220.74473 | 1361.954304 | 1522.57953 | 1520.8754 | 1429.80784 | 2225.4634 | 1957.6281 | 2260.5555 | 0.003342488 | -0.248857333 | YMR031C | EIS1 | n |
| W303.asm02.HP0_G0040810 | 498.32503 | 567.812388 | 599.839449 | 891.713244 | 888.97503 | 961.422516 | 1252.9633 | 1169.0661 | 1339.2293 | 1.22E-14 | -0.717555211 | YMR032W | HOF1 | n |
| W303.asm02.HP0_G0040830 | 2373.6285 | 2265.02126 | 1975.316704 | 1769.25908 | 1718.6251 | 1779.67462 | 1704.4405 | 1718.829 | 1655.7423 | 2.37E-05 | 0.32793605 | YMR033W | ARP9 | n |
| W303.asm02.HP0_G0040840 | 172.78077 | 181.658442 | 176.7642821 | 523.360671 | 463.81306 | 424.7705 | 436.65598 | 631.112 | 495.48837 | 1.05E-22 | -1.411280658 | YMR034C | RCH1 | n |
| W303.asm02.HP0_G0040850 | 470.93296 | 414.181248 | 453.9847682 | 382.519981 | 328.98368 | 336.592695 | 342.02819 | 372.63162 | 313.86808 | 0.001564428 | 0.352266189 | YMR035W | IMP2 | n |
| W303.asm02.HP0_G0040860 | 498.32503 | 495.149011 | 406.6544413 | 865.045185 | 823.35807 | 793.600242 | 958.81901 | 876.47156 | 978.63362 | 2.16E-17 | -0.827891249 | YMR036C | MIH1 | n |
| W303.asm02.HP0_G0040870 | 1029.3098 | 1068.15164 | 1010.35759 | 1033.38731 | 1023.8044 | 877.037305 | 1203.9392 | 1145.4486 | 1096.775 | 0.350954096 | 0.081911333 | YMR037C | MSN2 | n |
| W303.asm02.HP0_G0040880 | 5250.8498 | 5026.22959 | 5123.266405 | 810.042312 | 879.98641 | 859.022485 | 1075.1086 | 1120.519 | 1086.1952 | 0 | 2.595544293 | YMR038C | CCS1 | n |

| | | | | | | | | | | | | | | |
|---|---|---|---|---|---|---|---|---|---|---|---|---|---|---|
| W303.asm02.HP0_G0040890 | 3230.1576 | 3120.37301 | 3370.11246 | 2246.78403 | 2113.2258 | 2275.55625 | 2288.1686 | 2420.7934 | 2479.2052 | 1.65E-17 | 0.55100878 | YMR039C | SUB1 | n |
| W303.asm02.HP0_G0040910 | 691.12309 | 702.758659 | 657.7949513 | 1268.39959 | 1187.3974 | 1176.65221 | 1255.2434 | 1268.7844 | 1181.4136 | 1.09E-27 | -0.82509377 | YMR041C | ARA2 | n |
| W303.asm02.HP0_G0040920 | 423.5236 | 412.105152 | 450.121068 | 1864.26404 | 1767.1637 | 1869.74872 | 1810.4692 | 1575.8119 | 1640.7542 | 1.51E-140 | -2.096403195 | YMR042W | ARG80 | n |
| W303.asm02.HP0_G0040930 | 616.32166 | 712.101094 | 566.0320726 | 1065.05563 | 1077.7362 | 1062.8744 | 978.20061 | 1003.7436 | 1028.8878 | 1.44E-17 | -0.759757646 | YMR044W | IOC4 | n |
| W303.asm02.HP0_G0040940 | 1760.4675 | 1863.29659 | 2076.738833 | 4781.08307 | 4846.6667 | 5398.75721 | 4578.6173 | 4655.2711 | 4535.2174 | 1.11E-73 | -1.39773452 | YMR047C | NUP116 | n |
| W303.asm02.HP0_G0040950 | 546.78793 | 502.415349 | 502.2810201 | 456.690521 | 478.19486 | 457.955695 | 411.57392 | 471.03786 | 441.70761 | 0.102993045 | 0.155242929 | YMR048W | CSM3 | n |
| W303.asm02.HP0_G0040960 | 1458.1012 | 1497.90361 | 1629.51554 | 3448.51346 | 3373.431 | 3750.87522 | 1992.8842 | 2590.0522 | 2153.8756 | 8.89E-36 | -1.204851991 | YMR049C | ERB1 | n |
| W303.asm02.HP0_G0040970 | 429.84485 | 406.914911 | 437.5640425 | 227.511884 | 253.47923 | 218.074141 | 222.31832 | 220.42997 | 237.16436 | 4.04E-15 | 0.866192574 | YMR052W | FAR3 | n |
| W303.asm02.HP0_G0040990 | 1745.7179 | 1940.11216 | 2008.158155 | 1518.41265 | 1436.3823 | 1444.97822 | 1736.3631 | 1566.6273 | 1699.8249 | 1.28E-06 | 0.372106548 | YMR054W | STV1 | n |
| W303.asm02.HP0_G0041000 | 318.16947 | 320.756907 | 290.7434366 | 173.342388 | 180.67137 | 222.814883 | 207.4971 | 204.68497 | 242.45428 | 2.74E-07 | 0.690616155 | YMR055C | BUB2 | n |
| W303.asm02.HP0_G0041020 | 1650.8992 | 2375.05438 | 1724.176194 | 2720.97546 | 2408.0527 | 3080.53427 | 3981.2081 | 3349.7483 | 4474.3835 | 0.000279392 | -0.513718732 | YMR058W | FET3 | n |
| W303.asm02.HP0_G0041030 | 628.96416 | 613.486511 | 665.5223516 | 302.515802 | 282.24283 | 257.896375 | 230.29898 | 257.1683 | 240.69097 | 1.15E-30 | 1.177538534 | YMR059W | SEN15 | n |
| W303.asm02.HP0_G0041040 | 1203.1442 | 1238.39155 | 1342.635804 | 712.537219 | 599.5413 | 721.540961 | 604.24979 | 627.17575 | 674.46371 | 8.28E-22 | 0.896245167 | YMR060C | SAM37 | n |
| W303.asm02.HP0_G0041050 | 2131.314 | 2103.08574 | 2108.614359 | 1779.2596 | 1769.8603 | 1809.06722 | 1735.223 | 1697.8356 | 1728.9194 | 1.92E-05 | 0.243495422 | YMR061W | RNA14 | n |
| W303.asm02.HP0_G0041060 | 3323.9228 | 3074.69889 | 3033.004621 | 1461.74302 | 1469.6402 | 1635.55606 | 1649.7159 | 1196.6199 | 1615.1863 | 4.27E-22 | 1.046449466 | YMR062C | ARG7 | n |
| W303.asm02.HP0_G0041070 | 116.94308 | 113.147258 | 100.456204 | 116.672761 | 112.35782 | 87.2296563 | 93.487704 | 108.9029 | 92.573451 | 0.756268441 | 0.059181345 | YMR063W | RIM9 | n |
| W303.asm02.HP0_G0041080 | 1262.1425 | 1315.20712 | 1077.972343 | 864.211808 | 853.02053 | 811.615063 | 751.32191 | 616.67909 | 722.07292 | 8.10E-08 | 0.530713248 | YMR064W | AEP1 | n |
| W303.asm02.HP0_G0041090 | 603.67916 | 585.459208 | 568.9298477 | 211.677723 | 202.24407 | 224.71118 | 182.41503 | 166.63456 | 214.24141 | 1.05E-38 | 1.46133313 | YMR065W | KAR5 | n |
| W303.asm02.HP0_G0041100 | 701.65851 | 605.182125 | 653.9312511 | 772.540353 | 755.94338 | 753.778008 | 483.39984 | 553.6991 | 506.0682 | 0.014414092 | -0.2193577 | YMR066W | SOV1 | n |
| W303.asm02.HP0_G0041110 | 579.44771 | 610.372366 | 485.8602944 | 473.358059 | 541.11524 | 470.281625 | 552.94557 | 535.32993 | 562.49392 | 0.110841869 | 0.173499883 | YMR067C | UBX4 | n |
| W303.asm02.HP0_G0041120 | 246.52866 | 265.74035 | 251.14051 | 726.704625 | 735.26954 | 747.140969 | 754.7422 | 788.56198 | 876.362 | 8.02E-56 | -1.532859748 | YMR068W | AVO2 | n |
| W303.asm02.HP0_G0041130 | 76.908515 | 87.1960523 | 85.96732844 | 118.339515 | 118.64985 | 152.651898 | 164.17353 | 108.9029 | 119.02301 | 0.001363734 | -0.636976517 | YMR069W | NAT4 | n |
| W303.asm02.HP0_G0041140 | 780.6741 | 867.80833 | 867.4006847 | 709.203711 | 628.3049 | 665.600203 | 685.19646 | 675.72283 | 689.4518 | 0.000112963 | 0.328627964 | YMR070W | MOT3 | n |
| W303.asm02.HP0_G0041150 | 3871.7643 | 4054.61643 | 3980.577084 | 2610.13634 | 2934.7861 | 2795.14159 | 2683.7812 | 2911.5126 | 2666.997 | 1.98E-14 | 0.51392804 | YMR072W | ABF2 | n |
| W303.asm02.HP0_G0041160 | 979.79341 | 914.520501 | 1032.573866 | 442.523115 | 449.43126 | 439.940875 | 461.73805 | 543.20243 | 515.76637 | 3.20E-36 | 1.136235218 | YMR073C | IRC21 | n |
| W303.asm02.HP0_G0041170 | 1813.1446 | 1755.33958 | 2242.87794 | 1159.22722 | 1165.8247 | 1190.87444 | 1061.4275 | 1131.0157 | 1041.2309 | 2.44E-14 | 0.725557037 | YMR074C | SDD2 | n |
| W303.asm02.HP0_G0041180 | 1059.8625 | 829.400545 | 860.6392094 | 951.716379 | 1063.3544 | 941.511399 | 913.21525 | 885.65614 | 963.64554 | 0.318478306 | -0.10502817 | YMR075W | RCO1 | n |
| W303.asm02.HP0_G0041190 | 807.01264 | 799.297146 | 807.5133323 | 1710.9227 | 1663.7945 | 1659.25977 | 1745.4838 | 1686.0269 | 1718.3396 | 9.16E-61 | -1.060543201 | YMR076C | PDS5 | n |
| W303.asm02.HP0_G0041200 | 1028.2563 | 1019.36337 | 1229.622574 | 485.025335 | 480.89145 | 512.000156 | 617.93092 | 565.50785 | 572.19209 | 7.74E-33 | 1.15004036 | YMR077C | VPS20 | n |
| W303.asm02.HP0_G0041210 | 722.72933 | 793.068856 | 702.2275031 | 795.041528 | 799.08878 | 895.052125 | 908.65488 | 918.45822 | 1004.2015 | 0.066354013 | -0.166064534 | YMR078C | CTF18 | n |
| W303.asm02.HP0_G0041220 | 2150.2778 | 2085.43892 | 2057.420332 | 4558.57145 | 4352.2923 | 4677.21624 | 3884.3001 | 3886.3903 | 3717.9261 | 2.78E-81 | -1.110594953 | YMR079W | SEC14 | n |
| W303.asm02.HP0_G0041230 | 1227.3756 | 1310.01688 | 1354.226904 | 2970.15514 | 2846.6976 | 3119.40836 | 2444.3614 | 2592.6763 | 2257.9105 | 2.92E-56 | -1.198852374 | YMR080C | NAM7 | n |
| W303.asm02.HP0_G0041250 | 4026.6348 | 3839.74045 | 4048.191837 | 480.025074 | 391.0052 | 444.681617 | 453.75739 | 478.91036 | 548.38749 | 1.99E-267 | 3.177933587 | YMR083W | ADH3 | n |
| W303.asm02.HP0_G0041270 | 834.40471 | 901.025874 | 821.0362828 | 1350.9039 | 1355.4847 | 1282.84484 | 1594.9914 | 1410.4894 | 1686.6001 | 3.26E-14 | -0.642416784 | YMR086W | SEG1 | n |
| W303.asm02.HP0_G0041280 | 394.02445 | 398.610525 | 412.4499915 | 220.844869 | 200.44634 | 222.814883 | 269.06217 | 182.37956 | 226.58454 | 4.48E-12 | 0.903988088 | YMR087W | PDL32 | n |
| W303.asm02.HP0_G0041290 | 1001.9178 | 931.129273 | 1114.677495 | 608.365111 | 682.23665 | 609.659446 | 538.12435 | 448.73244 | 513.12141 | 8.89E-12 | 0.682120649 | YMR088C | VBA1 | n |
| W303.asm02.HP0_G0041300 | 1632.989 | 1824.88881 | 2099.921034 | 2595.96893 | 2488.0515 | 2523.97114 | 2630.1967 | 2646.4718 | 2599.9915 | 1.42E-07 | -0.452504418 | YMR089C | YTA12 | n |
| W303.asm02.HP0_G0041310 | 1431.7626 | 1748.07324 | 1623.71999 | 641.700185 | 628.3049 | 544.237203 | 760.44267 | 747.88741 | 807.59315 | 9.92E-48 | 1.403861676 | YMR090W | | n |
| W303.asm02.HP0_G0041320 | 1108.3254 | 1107.59747 | 952.4020881 | 837.543748 | 831.44783 | 849.541 | 872.17187 | 817.42781 | 831.39775 | 7.31E-05 | 0.330471596 | YMR091C | NPL6 | n |
| W303.asm02.HP0_G0041330 | 1487.6003 | 1518.66458 | 1582.185213 | 2670.13947 | 2542.8821 | 2851.08235 | 3012.1282 | 2641.2234 | 2973.8119 | 9.04E-27 | -0.813135784 | YMR092C | AIP1 | n |
| W303.asm02.HP0_G0041340 | 1122.0215 | 1120.05405 | 1237.349974 | 2380.12432 | 2253.4483 | 2396.91925 | 1467.3009 | 1668.9698 | 1471.477 | 2.57E-39 | -1.014278377 | YMR093W | UTP15 | n |
| W303.asm02.HP0_G0041350 | 143.28162 | 154.669188 | 182.5598323 | 235.845652 | 246.28833 | 272.118602 | 257.66123 | 242.73539 | 246.86254 | 1.50E-06 | -0.646573696 | YMR094W | CTF13 | n |
| W303.asm02.HP0_G0041380 | 1100.9507 | 1154.30964 | 1056.721992 | 680.868898 | 656.16964 | 593.540922 | 500.50124 | 545.8266 | 558.08566 | 1.68E-19 | 0.777354626 | YMR097C | MTG1 | n |
| W303.asm02.HP0_G0041390 | 1099.8971 | 1107.59747 | 1181.326322 | 992.551845 | 1086.7248 | 995.55586 | 1005.5629 | 963.06905 | 979.51527 | 0.062437408 | 0.140769092 | YMR098C | ATP25 | n |
| W303.asm02.HP0_G0041400 | 2640.1745 | 2466.40262 | 2351.061544 | 3942.70594 | 4260.6084 | 4315.97169 | 3966.3869 | 4003.1657 | 3790.2216 | 2.27E-28 | -0.747449586 | YMR099C | | n |
| W303.asm02.HP0_G0041410 | 785.94181 | 786.840567 | 810.4111074 | 611.699658 | 566.28339 | 550.874242 | 717.1191 | 558.94743 | 695.62336 | 4.39E-06 | 0.462501804 | YMR100W | MUB1 | n |
| W303.asm02.HP0_G0041420 | 62.158937 | 80.9677628 | 85.0014034 | 60.8365111 | 56.628339 | 61.6296485 | 79.806577 | 64.292075 | 59.070678 | 0.116170485 | 0.351364336 | YMR101C | SRT1 | n |
| W303.asm02.HP0_G0041430 | 481.46837 | 547.051423 | 543.8157967 | 990.885091 | 976.1647 | 1067.61514 | 1119.5723 | 930.26697 | 1080.9052 | 2.94E-23 | -0.947779211 | YMR102C | LAF1 | n |
| W303.asm02.HP0_G0041450 | 465.66525 | 601.029932 | 470.4054938 | 1575.91565 | 1518.1788 | 1655.46717 | 1572.1896 | 1234.6703 | 1682.1918 | 7.77E-40 | -1.627876698 | YMR105C | PGM2 | n |
| W303.asm02.HP0_G0041470 | 205.44055 | 224.21842 | 210.5716584 | 398.354141 | 391.90406 | 388.74086 | 387.63194 | 303.09121 | 391.45345 | 3.83E-13 | -0.881169231 | YMR106C | YKU80 | n |
| W303.asm02.HP0_G0041480 | 37.927487 | 73.7014251 | 40.56885162 | 25.0013059 | 12.584075 | 21.8074141 | 27.362255 | 28.86583 | 28.212861 | 0.000158991 | 1.353208972 | YMR107W | SPG4 | n |
| W303.asm02.HP0_G0041490 | 6866.9822 | 7299.55523 | 7622.11448 | 1395.90625 | 1512.7856 | 1598.57827 | 1591.5712 | 1200.5561 | 1675.1386 | 2.67E-100 | 2.273586164 | YMR108W | ILV2 | n |
| W303.asm02.HP0_G0041500 | 911.31322 | 1057.77116 | 1104.052319 | 2290.953 | 2309.1778 | 2710.75638 | 2281.328 | 1776.5606 | 2268.4904 | 1.39E-28 | -1.249750674 | YMR109W | MYO5 | n |
| W303.asm02.HP0_G0041510 | 2440.0017 | 2175.74911 | 2196.513538 | 1105.8911 | 1093.9157 | 1133.03738 | 1143.5142 | 1096.9015 | 1118.8163 | 2.08E-48 | 1.031306656 | YMR110C | HFD1 | n |
| W303.asm02.HP0_G0041520 | 323.43718 | 342.55592 | 287.8456615 | 507.52651 | 491.6778 | 475.022367 | 583.7281 | 514.3366 | 546.62419 | 8.61E-10 | -0.63003404 | YMR111C | EUC1 | n |
| W303.asm02.HP0_G0041530 | 855.47554 | 947.738044 | 926.322112 | 377.519719 | 364.03932 | 325.214914 | 453.75739 | 384.44037 | 383.51858 | 3.29E-41 | 1.354174502 | YMR112C | MED11 | n |
| W303.asm02.HP0_G0041540 | 643.71373 | 598.953835 | 580.5209482 | 765.873338 | 663.36054 | 681.718727 | 701.15778 | 764.94449 | 672.70041 | 0.025424765 | -0.212785292 | YMR113W | FOL3 | n |
| W303.asm02.HP0_G0041550 | 471.9865 | 514.871928 | 490.6899196 | 325.016977 | 312.80416 | 358.40011 | 413.8541 | 469.72577 | 382.63693 | 1.40E-07 | 0.569604892 | YMR114C | | n |
| W303.asm02.HP0_G0041560 | 1132.5569 | 1101.36918 | 972.6865139 | 1599.2502 | 1538.8526 | 1502.81527 | 1618.9334 | 1637.4798 | 1584.3285 | 1.52E-12 | -0.534450162 | YMR115W | MGR3 | n |
| W303.asm02.HP0_G0041570 | 19149.167 | 18130.5506 | 19764.75814 | 17327.5718 | 17259.958 | 19341.28 | 14923.83 | 15413.041 | 13813.722 | 0.262694803 | 0.081097825 | YMR116C | ASC1 | n |

| | | | | | | | | | | | | | | |
|---|---|---|---|---|---|---|---|---|---|---|---|---|---|---|
| W303.asm02.HP0_G0041600 | 1099.8971 | 993.412167 | 1076.040493 | 744.205539 | 736.16841 | 659.911313 | 767.28323 | 759.69616 | 810.23811 | 9.94E-12 | 0.565578041 | YMR119W | ASI1 | n |
| W303.asm02.HP0_G0041610 | 676.37351 | 623.866993 | 642.3401507 | 5386.94805 | 5211.6049 | 4912.35706 | 5186.2874 | 5169.6077 | 5237.894 | 0 | -2.997668827 | YMR120C | ADE17 | n |
| W303.asm02.HP0_G0041630 | 4024.5278 | 4121.05152 | 4133.19324 | 5806.13661 | 5231.3799 | 5208.17937 | 5383.5236 | 6473.8184 | 5746.6072 | 2.99E-07 | -0.404001464 | YMR122W | NCW1 | n |
| W303.asm02.HP0_G0041640 | 1734.129 | 1803.0898 | 1932.816002 | 620.032387 | 663.36054 | 629.570563 | 733.08041 | 881.71989 | 822.58123 | 2.29E-70 | 1.516185854 | YMR123W | PKR1 | n |
| W303.asm02.HP0_G0041650 | 720.62225 | 761.927409 | 843.2525587 | 1016.71977 | 1084.0282 | 1177.60036 | 837.96905 | 856.79031 | 971.58041 | 2.60E-07 | -0.493955502 | YMR124W | EPO1 | n |
| W303.asm02.HP0_G0041660 | 4095.115 | 3733.85952 | 3882.05273 | 3143.49753 | 3146.9177 | 3207.58617 | 3130.698 | 3114.8854 | 3042.5807 | 1.65E-07 | 0.302182439 | YMR125W | STO1 | n |
| W303.asm02.HP0_G0041670 | 587.87605 | 515.909976 | 564.1002226 | 373.352835 | 318.19733 | 329.955656 | 383.07157 | 363.44704 | 338.55433 | 2.27E-11 | 0.705987587 | YMR126C | DLT1 | n |
| W303.asm02.HP0_G0041680 | 873.38574 | 817.982014 | 996.8346399 | 489.192219 | 488.08235 | 479.76311 | 371.67063 | 425.11495 | 345.60755 | 1.12E-17 | 0.884106517 | YMR127C | SAS2 | n |
| W303.asm02.HP0_G0041690 | 681.64122 | 674.731357 | 822.9681329 | 1782.59311 | 1732.1081 | 1793.89684 | 962.23929 | 1053.6028 | 969.8171 | 1.26E-48 | -1.283240474 | YMR128W | ECM16 | n |
| W303.asm02.HP0_G0041700 | 1149.4136 | 1185.45109 | 1139.791546 | 2592.63542 | 2601.3081 | 2797.03789 | 2710.0033 | 2700.2672 | 2644.0741 | 4.77E-81 | -1.201344821 | YMR129W | POM152 | n |
| W303.asm02.HP0_G0041710 | 2024.9064 | 2012.77554 | 2162.706161 | 309.182817 | 326.2871 | 283.496383 | 308.96546 | 322.77246 | 363.24059 | 1.61E-204 | 2.753523605 | YMR130W | DPI35 | n |
| W303.asm02.HP0_G0041720 | 2936.2196 | 2780.93124 | 2636.975355 | 3215.16794 | 3066.9189 | 3472.11958 | 2085.2318 | 2607.1093 | 2162.6921 | 0.01586834 | -0.223568614 | YMR131C | RRB1 | n |
| W303.asm02.HP0_G0041730 | 1206.3048 | 1109.67357 | 1332.010628 | 272.514234 | 211.23269 | 213.333399 | 166.45372 | 205.99706 | 173.68543 | 4.94E-85 | 2.385371189 | YMR132C | JLP2 | n |
| W303.asm02.HP0_G0041740 | 1584.5261 | 1576.79528 | 1533.888961 | 615.032126 | 624.70945 | 693.096508 | 714.83891 | 708.52491 | 675.34536 | 8.36E-64 | 1.281450339 | YMR133W | REC114 | n |
| W303.asm02.HP0_G0041750 | 853.36845 | 947.738044 | 850.014034 | 291.681902 | 308.30985 | 267.377859 | 450.33711 | 381.8162 | 437.29935 | 1.28E-56 | 1.610850832 | YMR135C | GID8 | n |
| W303.asm02.HP0_G0041770 | 264.43887 | 458.817323 | 291.7093617 | 615.032126 | 611.22652 | 685.511321 | 646.43327 | 631.112 | 615.39303 | 6.75E-09 | -0.9137228 | YMR137C | PSO2 | n |
| W303.asm02.HP0_G0041780 | 597.35792 | 538.747037 | 596.9416739 | 527.527555 | 463.81306 | 450.370508 | 394.47251 | 408.05787 | 439.9443 | 0.008944126 | 0.264407014 | YMR138W | CIN4 | n |
| W303.asm02.HP0_G0041800 | 282.34907 | 350.860306 | 278.1864111 | 318.349962 | 327.18596 | 358.40011 | 519.88284 | 476.28619 | 536.92601 | 0.252639991 | -0.13929633 | YMR140W | SIP5 | n |
| W303.asm02.HP0_G0041820 | 11062.184 | 10895.3543 | 13537.43942 | 13334.8632 | 13873.044 | 14673.5452 | 12045.093 | 13236.295 | 10953.643 | 0.014097802 | -0.238595995 | YMR143W | RPS16A | y |
| W303.asm02.HP0_G0041830 | 719.56871 | 711.063045 | 676.147527 | 545.861846 | 610.32765 | 575.526102 | 633.89224 | 528.76952 | 625.09121 | 0.002924537 | 0.282956288 | YMR144W | FDO1 | n |
| W303.asm02.HP0_G0041840 | 7352.6647 | 7432.42541 | 8607.358019 | 9097.97522 | 9138.7353 | 9788.68447 | 6931.7712 | 7662.5657 | 7115.8126 | 0.000740665 | -0.260516304 | YMR145C | NDE1 | n |
| W303.asm02.HP0_G0041850 | 6147.4135 | 6006.14713 | 5401.452816 | 6646.18049 | 6620.1225 | 6784.95022 | 5903.4065 | 6317.6805 | 5847.1155 | 0.003940689 | -0.191952296 | YMR146C | TIF34 | n |
| W303.asm02.HP0_G0041880 | 2963.6117 | 3070.5467 | 2689.135308 | 2094.27606 | 2148.2814 | 2174.10437 | 2538.9892 | 2767.1834 | 2559.4355 | 3.96E-10 | 0.442916019 | YMR149W | SWP1 | n |
| W303.asm02.HP0_G0041890 | 1224.215 | 1166.76622 | 1523.263786 | 237.512406 | 238.19857 | 210.488953 | 212.05747 | 233.5508 | 208.9515 | 1.77E-96 | 2.511450852 | YMR150C | IMP1 | n |
| W303.asm02.HP0_G0041920 | 293.93802 | 321.794955 | 354.4944892 | 808.375558 | 890.77276 | 750.933563 | 771.84361 | 701.9645 | 715.0197 | 3.56E-38 | -1.335278561 | YMR154C | RIM13 | n |
| W303.asm02.HP0_G0041930 | 571.01938 | 510.719735 | 543.8157967 | 209.177593 | 207.63724 | 194.37043 | 322.64659 | 303.09121 | 281.24696 | 9.07E-39 | 1.410197274 | YMR155W | | n |
| W303.asm02.HP0_G0041940 | 288.67032 | 273.006688 | 254.0382852 | 440.022984 | 462.9142 | 462.696438 | 440.07627 | 413.3062 | 460.2223 | 4.42E-13 | -0.744165353 | YMR156C | TPP1 | n |
| W303.asm02.HP0_G0041950 | 1359.0683 | 1358.80515 | 1612.128889 | 941.715856 | 999.53513 | 1027.79291 | 905.2346 | 1024.737 | 901.04825 | 3.19E-09 | 0.545309962 | YMR157C | AIM36 | n |
| W303.asm02.HP0_G0041960 | 1996.4608 | 1971.25361 | 2514.302876 | 1133.39253 | 1265.5984 | 1209.83741 | 1045.4662 | 1144.1365 | 1065.9172 | 2.34E-17 | 0.845559784 | YMR158W | MRPS8 | n |
| W303.asm02.HP0_G0041970 | 162.24536 | 152.593091 | 109.1495294 | 102.505354 | 116.85213 | 79.6444688 | 103.74855 | 148.2654 | 151.64413 | 0.018955517 | 0.499996933 | YMR159C | ATG16 | n |
| W303.asm02.HP0_G0041980 | 369.793 | 410.029055 | 420.1773918 | 362.518936 | 423.36425 | 356.503813 | 474.27908 | 463.16536 | 484.0269 | 0.515526261 | 0.071731745 | YMR160W | CVM1 | n |
| W303.asm02.HP0_G0041990 | 1902.6956 | 1875.75317 | 1876.79235 | 1218.39697 | 1151.4429 | 1149.15591 | 1192.5383 | 1083.7807 | 1126.7511 | 1.13E-24 | 0.683939271 | YMR161W | HLJ1 | n |
| W303.asm02.HP0_G0042000 | 702.71205 | 778.536181 | 807.5133323 | 1242.5649 | 1208.0712 | 1170.01517 | 1227.8812 | 1207.1165 | 1327.7678 | 2.40E-16 | -0.661298398 | YMR162C | DNF3 | n |
| W303.asm02.HP0_G0042010 | 304.47344 | 299.995942 | 272.3908609 | 355.851921 | 402.69041 | 407.703828 | 580.30782 | 426.42703 | 490.19846 | 0.001232944 | -0.411302656 | YMR163C | INP2 | n |
| W303.asm02.HP0_G0042020 | 330.81197 | 357.088595 | 357.3922643 | 438.35623 | 441.3415 | 410.548274 | 445.77673 | 486.78286 | 484.90855 | 0.002122614 | -0.30384346 | YMR164C | MSS11 | n |
| W303.asm02.HP0_G0042030 | 488.84316 | 540.823134 | 558.3046723 | 1365.90468 | 1359.0801 | 1315.08188 | 1243.8425 | 1214.989 | 1219.3246 | 1.07E-69 | -1.346425295 | YMR165C | PAH1 | n |
| W303.asm02.HP0_G0042040 | 843.88658 | 802.411291 | 774.671881 | 1404.24002 | 1284.4745 | 1342.57819 | 1439.9387 | 1301.5865 | 1363.0339 | 7.24E-22 | -0.736420872 | YMR166C | MME1 | n |
| W303.asm02.HP0_G0042050 | 606.83979 | 553.279713 | 516.7698957 | 320.850093 | 310.10757 | 314.785281 | 274.76264 | 270.28913 | 258.32401 | 4.63E-16 | 0.825158477 | YMR167W | MLH1 | n |
| W303.asm02.HP0_G0042060 | 555.21626 | 505.529494 | 520.6335958 | 524.194047 | 547.40728 | 525.274235 | 597.40923 | 501.21577 | 535.16271 | 0.887287522 | -0.014257326 | YMR168C | CEP3 | n |
| W303.asm02.HP0_G0042080 | 403.50632 | 427.675875 | 377.6766901 | 619.19901 | 645.38329 | 691.200211 | 948.55817 | 793.81032 | 1003.3199 | 9.94E-11 | -0.694117459 | YMR170C | ALD2 | n |
| W303.asm02.HP0_G0042090 | 749.06786 | 840.819076 | 895.4125108 | 929.215203 | 895.26707 | 825.837289 | 988.46146 | 1031.2974 | 1026.2428 | 0.306544915 | -0.092653018 | YMR171C | EAR1 | n |
| W303.asm02.HP0_G0042100 | 389.81028 | 415.219297 | 421.1433168 | 1779.2596 | 1730.3104 | 1655.46717 | 1382.934 | 1381.6236 | 1347.1641 | 2.45E-163 | -2.074307027 | YMR172W | HOT1 | n |
| W303.asm02.HP0_G0042110 | 8672.752 | 9183.61279 | 8999.523585 | 815.87595 | 865.60461 | 740.50393 | 1021.5242 | 1330.4523 | 1090.6034 | 1.12E-297 | 3.470572401 | YMR173W | DDR48 | n |
| W303.asm02.HP0_G0042150 | 614.21458 | 588.573353 | 680.9771522 | 629.199532 | 609.42879 | 690.252063 | 605.38989 | 616.67909 | 579.24531 | 0.737721258 | -0.032691067 | YMR176W | ECM5 | n |
| W303.asm02.HP0_G0042160 | 679.53414 | 613.486511 | 623.9875749 | 1714.25621 | 1637.7275 | 1697.1857 | 2076.1111 | 1683.4027 | 2026.9177 | 1.63E-52 | -1.397542145 | YMR177W | MMT1 | n |
| W303.asm02.HP0_G0042170 | 1727.8077 | 1547.72993 | 1910.599726 | 690.036043 | 701.11277 | 623.881672 | 779.82426 | 770.19282 | 774.09038 | 2.86E-50 | 1.363641931 | YMR178W | FPY1 | n |
| W303.asm02.HP0_G0042180 | 427.73777 | 445.322696 | 420.1773918 | 1351.73727 | 1393.2369 | 1404.20784 | 1340.7505 | 1283.2173 | 1464.4238 | 1.37E-93 | -1.681983031 | YMR179W | SPT21 | n |
| W303.asm02.HP0_G0042230 | 2457.9119 | 2727.99078 | 2745.15896 | 641.700185 | 666.05713 | 683.615024 | 801.48605 | 923.70655 | 842.85923 | 5.27E-136 | 1.994442711 | YMR184W | ADD37 | n |
| W303.asm02.HP0_G0042240 | 1193.6623 | 1114.86381 | 1128.200445 | 944.215987 | 1021.1078 | 1105.54108 | 794.64548 | 902.71322 | 762.6289 | 0.073485172 | 0.163015647 | YMR185W | RTP1 | n |
| W303.asm02.HP0_G0042250 | 21294.177 | 20503.5289 | 15154.39793 | 14242.4106 | 14437.53 | 15544.8936 | 16253.179 | 15061.403 | 14568.416 | 0.000906877 | 0.364861822 | YMR186W | HSC82 | y |
| W303.asm02.HP0_G0042260 | 722.72933 | 718.329383 | 666.4882766 | 671.701752 | 598.64244 | 605.866852 | 601.96961 | 591.74951 | 599.5233 | 0.056095638 | 0.1662251 | YMR187C | | n |
| W303.asm02.HP0_G0042270 | 2475.8221 | 2371.94023 | 2696.862708 | 2285.11936 | 2235.4711 | 2345.71924 | 2069.2705 | 2134.7593 | 2079.8169 | 0.044609781 | 0.136336131 | YMR188C | MRPS17 | n |
| W303.asm02.HP0_G0042280 | 359.25758 | 391.344187 | 406.6544413 | 1242.5649 | 1028.2987 | 1036.32624 | 1320.2288 | 1060.1632 | 1482.9385 | 2.97E-33 | -1.514863327 | YMR189W | GCV2 | n |
| W303.asm02.HP0_G0042290 | 446.70151 | 464.007564 | 464.6099436 | 538.361454 | 564.48566 | 598.281664 | 529.00359 | 534.01785 | 566.02053 | 0.000719333 | -0.305524566 | YMR190C | SGS1 | n |
| W303.asm02.HP0_G0042320 | 1679.3448 | 1633.88793 | 1637.24294 | 1903.43276 | 1837.275 | 1861.21538 | 1650.856 | 1994.3664 | 1736.8543 | 0.023074536 | -0.178500918 | YMR193W | MRPL24 | n |
| W303.asm02.HP0_G0042360 | 532.03836 | 591.687498 | 711.8867535 | 307.516063 | 312.80416 | 289.185274 | 334.04753 | 287.34621 | 372.05711 | 3.56E-15 | 1.013680186 | YMR196W | | n |
| W303.asm02.HP0_G0042370 | 1487.6003 | 1459.49583 | 1338.772104 | 1995.10421 | 1989.1828 | 1956.97838 | 2039.6281 | 1969.4368 | 2025.1544 | 4.43E-13 | -0.471833971 | YMR197C | VTI1 | n |
| W303.asm02.HP0_G0042400 | 3000.4856 | 2971.93212 | 2662.089406 | 2496.79708 | 2398.1652 | 2431.05259 | 2564.0713 | 2760.623 | 2669.642 | 0.000552659 | 0.236669333 | YMR200W | ROT1 | n |
| W303.asm02.HP0_G0042420 | 6382.3532 | 6632.09021 | 5750.151755 | 2120.11074 | 2221.0893 | 2122.90435 | 2134.2559 | 2301.3939 | 2011.9297 | 1.71E-92 | 1.537388264 | YMR202W | ERG2 | n |
| W303.asm02.HP0_G0042430 | 6228.5362 | 5507.88397 | 5764.64063 | 4035.21077 | 4065.5552 | 4244.86056 | 3691.6242 | 3814.2258 | 3654.4472 | 1.69E-15 | 0.503502429 | YMR203W | TOM40 | n |

| | | | | | | | | | | | | | | |
|---|---|---|---|---|---|---|---|---|---|---|---|---|---|---|
| W303.asm02.HP0_G0042440 | 153.81703 | 128.717982 | 133.2976553 | 255.846697 | 298.42236 | 189.629688 | 280.46311 | 314.89996 | 297.11669 | 3.79E-07 | -0.841348729 | YMR204C | INP1 | n |
| W303.asm02.HP0_G0042450 | 5793.4236 | 6289.53429 | 6989.433579 | 7865.41084 | 7292.4716 | 7372.80225 | 7570.2238 | 7038.0141 | 7182.8181 | 0.001815293 | -0.240311329 | YMR205C | PFK2 | n |
| W303.asm02.HP0_G0042460 | 60.051854 | 85.1199558 | 85.0014034 | 48.3358581 | 42.246539 | 47.4074219 | 55.864604 | 44.610828 | 52.899115 | 0.001750891 | 0.739633965 | YMR206W | | n |
| W303.asm02.HP0_G0042480 | 1966.9616 | 1736.65471 | 1892.247151 | 1311.73518 | 1320.429 | 1408.94858 | 1363.5524 | 1298.9623 | 1286.3301 | 1.28E-10 | 0.469994411 | YMR208W | ERG12 | n |
| W303.asm02.HP0_G0042490 | 1210.519 | 1172.99451 | 1217.065549 | 714.203972 | 798.18992 | 755.674305 | 853.93037 | 817.42781 | 884.29687 | 1.77E-18 | 0.667402579 | YMR209C | | n |
| W303.asm02.HP0_G0042500 | 763.81744 | 1132.51063 | 897.3443609 | 1015.05302 | 993.24309 | 987.970672 | 991.88174 | 984.06238 | 953.94737 | 0.404230015 | -0.101077 | YMR210W | MGL2 | n |
| W303.asm02.HP0_G0042510 | 1125.1821 | 1054.65701 | 1093.427144 | 690.86942 | 676.84348 | 784.118758 | 672.65543 | 736.07866 | 712.37475 | 5.59E-13 | 0.606045829 | YMR211W | DML1 | n |
| W303.asm02.HP0_G0042520 | 2353.6113 | 2132.15109 | 2407.085196 | 1547.58084 | 1596.3798 | 1543.58566 | 1707.8607 | 1682.0906 | 1760.6589 | 3.88E-16 | 0.55647007 | YMR212C | EFR3 | n |
| W303.asm02.HP0_G0042530 | 172.78077 | 250.169626 | 231.8220093 | 498.359365 | 482.68917 | 408.651977 | 369.39044 | 423.80286 | 438.181 | 9.40E-15 | -1.085488352 | YMR213W | CEF1 | n |
| W303.asm02.HP0_G0042540 | 1231.5898 | 1505.16995 | 1398.659456 | 2582.6349 | 2568.9491 | 2697.48231 | 2834.2736 | 2662.2168 | 2744.5824 | 3.27E-32 | -0.924175044 | YMR214W | SCJ1 | n |
| W303.asm02.HP0_G0042550 | 2173.4557 | 2040.80284 | 1770.540596 | 2230.94986 | 2333.4471 | 2379.85258 | 2637.0373 | 2670.0892 | 2523.2878 | 0.008641955 | -0.214836335 | YMR215W | GAS3 | n |
| W303.asm02.HP0_G0042560 | 1288.481 | 1219.70668 | 1491.38826 | 1536.74694 | 1481.3254 | 1426.01525 | 1567.6292 | 1361.9423 | 1488.2284 | 0.098111544 | -0.151719437 | YMR216C | SKY1 | n |
| W303.asm02.HP0_G0042570 | 3471.4186 | 3721.40295 | 4324.446398 | 8282.93265 | 7945.0458 | 8909.75087 | 5630.924 | 7139.0445 | 5516.496 | 1.41E-25 | -1.12586837 | YMR217W | GUA1 | n |
| W303.asm02.HP0_G0042580 | 1181.0198 | 1128.35844 | 1194.849273 | 896.713506 | 925.8284 | 872.296563 | 832.26858 | 830.54865 | 797.01333 | 1.13E-07 | 0.378884166 | YMR218C | TRS130 | n |
| W303.asm02.HP0_G0042600 | 2078.637 | 2026.27017 | 1879.690125 | 1441.74197 | 1567.6162 | 1589.09678 | 1452.4797 | 1472.1573 | 1371.8504 | 1.79E-07 | 0.380266704 | YMR220W | ERG8 | n |
| W303.asm02.HP0_G0042610 | 2138.6888 | 2001.35701 | 1919.293052 | 794.208151 | 806.27968 | 767.052086 | 1056.8671 | 1003.7436 | 1069.4438 | 1.20E-77 | 1.355193567 | YMR221C | FMP42 | n |
| W303.asm02.HP0_G0042620 | 567.85876 | 580.268967 | 537.0543215 | 962.550278 | 927.62612 | 954.785477 | 755.88229 | 911.8978 | 765.27386 | 1.13E-16 | -0.756023059 | YMR222C | FSH2 | n |
| W303.asm02.HP0_G0042630 | 1225.2685 | 1130.43453 | 1314.623978 | 632.53304 | 572.57543 | 553.718688 | 530.14369 | 562.88368 | 580.12696 | 7.52E-32 | 1.060622829 | YMR223W | UBP8 | n |
| W303.asm02.HP0_G0042640 | 365.57883 | 345.670064 | 309.0960124 | 599.197965 | 573.47429 | 530.963125 | 498.22106 | 463.16536 | 475.21038 | 1.99E-15 | -0.741848172 | YMR224C | MRE11 | n |
| W303.asm02.HP0_G0042650 | 1196.8229 | 1164.69013 | 1400.591306 | 611.698618 | 649.8776 | 662.755758 | 635.03233 | 666.53825 | 600.40495 | 1.48E-25 | 0.968340167 | YMR225C | MRPL44 | n |
| W303.asm02.HP0_G0042660 | 5122.3178 | 5020.0013 | 5103.947904 | 1031.72056 | 1080.4328 | 1097.95589 | 1461.6004 | 1158.5694 | 1408.8798 | 1.94E-162 | 2.248079776 | YMR226C | ORA1 | n |
| W303.asm02.HP0_G0042670 | 306.58052 | 410.029055 | 364.1537396 | 1323.40246 | 1257.5087 | 1210.78556 | 1129.8331 | 1058.8511 | 1191.1117 | 4.10E-71 | -1.810619877 | YMR227C | TAF7 | n |
| W303.asm02.HP0_G0042680 | 974.5257 | 934.243417 | 1114.677495 | 617.532256 | 696.61846 | 585.955735 | 482.25974 | 516.96077 | 468.15717 | 1.60E-11 | 0.670480801 | YMR228W | MTF1 | n |
| W303.asm02.HP0_G0042690 | 1714.1117 | 1910.00876 | 1977.248554 | 5650.29514 | 5530.7011 | 6337.42416 | 3390.6394 | 4054.337 | 3387.3066 | 1.23E-71 | -1.644702397 | YMR229C | RRP5 | n |
| W303.asm02.HP0_G0042720 | 556.26981 | 560.54605 | 572.7935479 | 713.370595 | 795.49333 | 699.733547 | 840.24924 | 675.72283 | 783.78855 | 9.54E-05 | -0.386305276 | YMR231W | PEP5 | n |
| W303.asm02.HP0_G0042730 | 114.836 | 115.223355 | 126.5361801 | 34.1684514 | 32.359051 | 37.9259375 | 36.483006 | 48.547077 | 26.449557 | 4.20E-14 | 1.773726367 | YMR232W | FUS2 | n |
| W303.asm02.HP0_G0042740 | 853.36845 | 803.449339 | 873.1962349 | 430.855839 | 442.24036 | 447.526063 | 361.40978 | 354.26246 | 371.17546 | 1.51E-28 | 0.938660915 | YMR233W | TRI1 | n |
| W303.asm02.HP0_G0042750 | 1319.0337 | 1369.18563 | 1373.545405 | 310.016193 | 346.96093 | 348.918625 | 243.98011 | 217.80581 | 259.20566 | 7.28E-111 | 2.015225009 | YMR234W | RNH1 | n |
| W303.asm02.HP0_G0042760 | 3764.3031 | 3787.83803 | 3590.343369 | 3396.8441 | 3315.005 | 3792.59375 | 2826.2929 | 2853.7809 | 2784.2567 | 0.215839687 | 0.085227479 | YMR235C | RNA1 | n |
| W303.asm02.HP0_G0042770 | 1608.7576 | 1666.06743 | 1694.232518 | 985.051453 | 1050.7703 | 1073.30403 | 1071.6883 | 1068.0357 | 1095.0117 | 1.90E-23 | 0.677345359 | YMR236W | TAF9 | n |
| W303.asm02.HP0_G0042790 | 1632.989 | 1482.33289 | 1683.607342 | 1739.25751 | 1724.9172 | 1789.1561 | 2161.6181 | 1964.1885 | 2089.515 | 0.077691461 | -0.130126291 | YMR238W | DFG5 | n |
| W303.asm02.HP0_G0042800 | 1732.0219 | 1444.96315 | 1841.053124 | 1655.91983 | 1673.682 | 1604.26716 | 1412.5764 | 1617.7985 | 1527.9028 | 0.804890632 | 0.024638865 | YMR239C | RNT1 | n |
| W303.asm02.HP0_G0042810 | 337.13322 | 370.583222 | 293.6412117 | 539.194831 | 494.37439 | 457.007547 | 445.77473 | 492.03119 | 441.70761 | 4.25E-07 | -0.576325876 | YMR240C | CUS1 | n |
| W303.asm02.HP0_G0042820 | 3594.6829 | 3308.25975 | 3533.353791 | 1628.41839 | 1547.8413 | 1653.57088 | 1982.6234 | 1624.359 | 1938.7526 | 1.13E-41 | 1.111618183 | YMR241W | YHM2 | n |
| W303.asm02.HP0_G0042840 | 4540.763 | 4111.70908 | 4140.920641 | 2063.44111 | 1901.9931 | 2031.8821 | 2413.5789 | 2281.7126 | 2253.5023 | 2.64E-58 | 1.09280588 | YMR243C | ZRC1 | n |
| W303.asm02.HP0_G0042850 | 31.606239 | 65.3970392 | 21.25035085 | 26.6680596 | 32.359051 | 25.6000078 | 35.342912 | 31.489996 | 32.621121 | 0.210044655 | 0.478597682 | YMR244W | | n |
| W303.asm02.HP0_G0042860 | 1054.5948 | 1025.59166 | 1099.222694 | 785.874383 | 837.73987 | 814.459508 | 822.00774 | 793.81032 | 802.30324 | 1.73E-07 | 0.383711422 | YMR244C- | COA6 | n |
| W303.asm02.HP0_G0042880 | 1463.3689 | 1410.70756 | 1481.729009 | 2271.78533 | 2338.8403 | 2459.49705 | 2013.4059 | 2054.7222 | 1996.0599 | 7.24E-29 | -0.698290253 | YMR247C | RKR1 | n |
| W303.asm02.HP0_G0042890 | 377.16779 | 596.877739 | 529.3269212 | 758.372946 | 613.02424 | 626.726117 | 1140.094 | 841.04531 | 1054.4557 | 0.009562867 | -0.410526305 | YMR250W | GAD1 | n |
| W303.asm02.HP0_G0042920 | 403.50632 | 368.507126 | 365.1196646 | 510.860018 | 503.36301 | 498.726078 | 586.00829 | 541.89035 | 587.18017 | 9.57E-06 | -0.412912143 | YMR252C | MLO1 | n |
| W303.asm02.HP0_G0042950 | 1924.82 | 1925.57949 | 2366.516345 | 2912.65214 | 2832.3158 | 3179.14171 | 2861.6358 | 2674.0255 | 2561.1988 | 1.78E-08 | -0.520947217 | YMR256C | COX7 | n |
| W303.asm02.HP0_G0042960 | 862.85032 | 759.851313 | 878.0258601 | 675.868637 | 642.6867 | 762.311344 | 670.37524 | 686.2195 | 651.54076 | 0.004951659 | 0.266203492 | YMR257C | PET111 | n |
| W303.asm02.HP0_G0042970 | 1146.2529 | 1048.42872 | 1069.279018 | 958.383393 | 996.83854 | 933.926211 | 803.76624 | 856.79031 | 841.09592 | 0.019951314 | 0.175598361 | YMR258C | ROY1 | n |
| W303.asm02.HP0_G0042980 | 1326.4085 | 1089.95065 | 1420.875732 | 1272.56647 | 1297.0586 | 1352.05967 | 1199.3788 | 1096.9015 | 1090.6034 | 0.765936859 | -0.030797379 | YMR259C | TRM732 | n |
| W303.asm02.HP0_G0042990 | 7702.4404 | 7173.9514 | 9133.787165 | 4847.75322 | 4798.1281 | 5047.94228 | 4051.8939 | 4358.7403 | 3954.2088 | 5.83E-16 | 0.708595807 | YMR260C | TIF11 | n |
| W303.asm02.HP0_G0043000 | 1300.07 | 1465.72412 | 1389.966131 | 1205.06295 | 1235.936 | 1171.91147 | 1653.1362 | 1420.9861 | 1794.1616 | 0.030219694 | 0.201929588 | YMR261C | TPS3 | n |
| W303.asm02.HP0_G0043010 | 304.47344 | 309.338376 | 264.6634606 | 276.681119 | 265.16444 | 299.614906 | 256.52114 | 225.67831 | 251.2708 | 0.619858508 | 0.061592559 | YMR262W | | n |
| W303.asm02.HP0_G0043020 | 1093.5759 | 1051.54287 | 1246.0433 | 471.691305 | 451.22899 | 388.74086 | 473.13899 | 469.72577 | 518.41132 | 8.65E-41 | 1.369420099 | YMR263W | SAP30 | n |
| W303.asm02.HP0_G0043030 | 1884.7854 | 1728.35032 | 1870.9968 | 2199.28154 | 2087.1588 | 2189.27474 | 2518.4675 | 2470.6526 | 2509.1813 | 0.000154252 | -0.239708294 | YMR264W | CUE1 | n |
| W303.asm02.HP0_G0043040 | 647.9279 | 590.649449 | 566.0320726 | 187.509794 | 195.05317 | 219.970438 | 172.15419 | 160.07415 | 181.62029 | 6.12E-44 | 1.584056897 | YMR265C | | n |
| W303.asm02.HP0_G0043050 | 1927.9806 | 1861.2205 | 2054.522557 | 4034.3774 | 3697.9204 | 4276.14945 | 4872.7615 | 4353.492 | 4742.4056 | 3.51E-41 | -1.03877504 | YMR266W | RSN1 | n |
| W303.asm02.HP0_G0043060 | 2268.2744 | 2169.52082 | 2310.492692 | 755.039439 | 738.86499 | 771.792828 | 811.74689 | 823.98823 | 819.93628 | 1.06E-122 | 1.574898155 | YMR267W | PPA2 | n |
| W303.asm02.HP0_G0043070 | 607.89333 | 565.736292 | 555.4068972 | 498.359365 | 435.04946 | 376.41493 | 367.11025 | 380.50412 | 383.51858 | 0.00032503 | 0.398206601 | YMR268C | PRP24 | n |
| W303.asm02.HP0_G0043080 | 877.5999 | 832.51469 | 1021.948691 | 680.868898 | 639.99012 | 642.844641 | 393.33241 | 494.65535 | 429.36448 | 5.85E-06 | 0.476736373 | YMR269W | TMA23 | n |
| W303.asm02.HP0_G0043090 | 1485.4932 | 1351.53881 | 1361.954304 | 490.025596 | 507.85733 | 442.78532 | 418.41448 | 443.48411 | 444.35256 | 1.57E-72 | 1.54220342 | YMR270C | RRN9 | n |
| W303.asm02.HP0_G0043110 | 1372.7643 | 1382.68026 | 1336.840253 | 2000.93785 | 1884.9147 | 1802.43018 | 1805.9088 | 2132.1352 | 2026.9177 | 1.72E-09 | -0.475569358 | YMR272C | SCS7 | n |
| W303.asm02.HP0_G0043120 | 257.06408 | 289.615459 | 292.6752867 | 1187.56203 | 1237.7337 | 1147.25961 | 1182.2774 | 1054.9149 | 1143.5025 | 3.07E-116 | -2.0886751 | YMR273C | ZDS1 | n |
| W303.asm02.HP0_G0043130 | 701.65851 | 675.769405 | 705.1252782 | 266.680596 | 303.81553 | 253.155633 | 500.50124 | 431.67536 | 468.15717 | 1.12E-39 | 1.337965487 | YMR274C | RCE1 | n |
| W303.asm02.HP0_G0043150 | 1757.3069 | 2040.80284 | 2086.398083 | 3325.17369 | 3019.2792 | 3287.23063 | 3673.3827 | 3168.6809 | 3449.0223 | 1.18E-16 | -0.710614536 | YMR276W | DSK2 | n |
| W303.asm02.HP0_G0043160 | 659.51685 | 910.368308 | 888.6510355 | 1155.06033 | 1156.8361 | 1062.8744 | 1015.8237 | 968.31738 | 1089.7218 | 5.81E-05 | -0.456627021 | YMR277W | FCP1 | n |

| | | | | | | | | | | | | | | |
|---|---|---|---|---|---|---|---|---|---|---|---|---|---|---|
| W303.asm02.HP0_G0043170 | 482.52192 | 432.866117 | 494.5536198 | 745.872293 | 787.40357 | 845.748407 | 737.64079 | 536.64202 | 706.20318 | 1.94E-10 | -0.753935314 | YMR278W | PRM15 | n |
| W303.asm02.HP0_G0043190 | 206.49409 | 235.636951 | 211.5375835 | 1463.40977 | 1448.9664 | 1526.51898 | 1978.063 | 1613.8623 | 1970.492 | 3.51E-151 | -2.763741618 | YMR280C | CAT8 | n |
| W303.asm02.HP0_G0043200 | 512.02107 | 503.453397 | 477.1669691 | 668.368245 | 640.88898 | 659.911313 | 698.87759 | 754.44782 | 701.79492 | 2.11E-06 | -0.400492748 | YMR281W | GPI12 | n |
| W303.asm02.HP0_G0043210 | 1290.5881 | 1166.76622 | 1296.271402 | 979.217815 | 1004.0294 | 970.904 | 930.31666 | 901.40114 | 954.82902 | 3.07E-06 | 0.345718489 | YMR282C | AEP2 | n |
| W303.asm02.HP0_G0043220 | 774.35286 | 568.850436 | 697.3978779 | 700.869943 | 717.29229 | 696.889102 | 669.23515 | 652.10534 | 707.96649 | 0.646284928 | -0.051738423 | YMR283C | RIT1 | n |
| W303.asm02.HP0_G0043230 | 884.97469 | 816.946966 | 1013.255366 | 535.861323 | 553.69931 | 544.237203 | 505.06162 | 514.3366 | 504.30489 | 7.92E-15 | 0.73374693 | YMR284W | YKU70 | n |
| W303.asm02.HP0_G0043240 | 714.301 | 699.644515 | 678.0793771 | 650.867331 | 640.88898 | 658.963164 | 536.98425 | 543.20243 | 541.33427 | 0.21094573 | 0.100617498 | YMR285C | NGL2 | n |
| W303.asm02.HP0_G0043250 | 1458.1012 | 1365.03344 | 1731.903594 | 480.025074 | 551.00273 | 526.222383 | 478.83946 | 547.13868 | 497.25168 | 4.61E-49 | 1.549551454 | YMR286W | MRPL33 | n |
| W303.asm02.HP0_G0043260 | 553.10918 | 568.850436 | 567.9639227 | 1105.8911 | 1150.544 | 1225.95593 | 820.86764 | 946.01197 | 737.061 | 1.07E-26 | -1.042626015 | YMR287C | DSS1 | y |
| W303.asm02.HP0_G0043280 | 639.49957 | 615.562607 | 592.1120487 | 991.718468 | 967.17607 | 977.541039 | 826.56811 | 905.33739 | 874.5987 | 5.24E-18 | -0.669439617 | YMR288W | HSH155 | n |
| W303.asm02.HP0_G0043290 | 1718.3259 | 1573.68113 | 1702.925843 | 368.352574 | 380.21885 | 466.489031 | 363.68997 | 351.63829 | 399.38832 | 1.34E-92 | 2.041496672 | YMR289W | ABZ2 | n |
| W303.asm02.HP0_G0043300 | 1872.1429 | 1812.43223 | 1965.657454 | 3800.1985 | 3395.0037 | 4004.03085 | 2330.352 | 2933.818 | 2361.0638 | 4.62E-24 | -0.986850625 | YMR290C | HAS1 | n |
| W303.asm02.HP0_G0043310 | 270.76011 | 296.881797 | 304.2663872 | 495.025857 | 486.28462 | 539.496461 | 778.68417 | 568.13201 | 725.59952 | 1.00E-10 | -0.801510189 | YMR291W | TDA1 | n |
| W303.asm02.HP0_G0043330 | 530.98482 | 439.094406 | 477.1669691 | 1275.0666 | 1164.027 | 1226.90408 | 1178.8571 | 1208.4286 | 1195.52 | 1.04E-55 | -1.341525225 | YMR293C | HER2 | n |
| W303.asm02.HP0_G0043340 | 831.24409 | 757.775216 | 791.0926066 | 431.689215 | 427.85856 | 429.511242 | 373.95082 | 423.80286 | 424.07457 | 3.34E-23 | 0.884540851 | YMR294W | JNM1 | n |
| W303.asm02.HP0_G0043350 | 4508.1032 | 4259.11193 | 4672.179412 | 1443.40873 | 1386.9449 | 1497.12638 | 1726.1022 | 1663.7215 | 1765.9488 | 2.79E-139 | 1.635169817 | YMR295C | GSR1 | n |
| W303.asm02.HP0_G0043360 | 1513.9388 | 1781.29078 | 1735.767294 | 3392.67721 | 3406.689 | 3492.97884 | 4072.4156 | 3890.3266 | 4015.9245 | 1.75E-48 | -1.03227633 | YMR296C | LCB1 | n |
| W303.asm02.HP0_G0043370 | 5717.5686 | 5488.16105 | 5472.931269 | 737.538525 | 713.69684 | 753.778008 | 1470.7212 | 1279.2811 | 1327.7678 | 0 | 2.919260753 | YMR297W | PRC1 | n |
| W303.asm02.HP0_G0043380 | 1611.9182 | 1567.45284 | 1729.971744 | 1322.56908 | 1344.6983 | 1381.45227 | 1397.7552 | 1501.0231 | 1425.6311 | 6.12E-05 | 0.278709202 | YMR298W | LIP1 | n |
| W303.asm02.HP0_G0043390 | 1046.1665 | 1051.54287 | 1218.997399 | 228.345261 | 234.60312 | 239.881555 | 137.95157 | 166.63456 | 190.43681 | 6.16E-86 | 2.239509421 | YMR299C | DYN3 | n |
| W303.asm02.HP0_G0043400 | 1390.6745 | 1205.17401 | 1470.137909 | 2576.80126 | 2454.7935 | 2238.57846 | 2204.9417 | 2275.1522 | 2186.4967 | 1.39E-22 | -0.83832001 | YMR300C | ADE4 | n |
| W303.asm02.HP0_G0043410 | 725.88996 | 834.590786 | 912.7991615 | 928.381826 | 972.56925 | 1059.08181 | 690.89693 | 754.44782 | 653.30407 | 0.011722029 | -0.257905114 | YMR301C | ATM1 | n |
| W303.asm02.HP0_G0043420 | 1312.7125 | 1463.64802 | 1488.490485 | 1867.59755 | 1893.9033 | 1765.45239 | 1939.2998 | 1857.9098 | 1884.9718 | 1.79E-07 | -0.373791651 | YMR302C | YME2 | n |
| W303.asm02.HP0_G0043440 | 799.63785 | 864.694185 | 953.3680131 | 872.545576 | 807.17855 | 836.266922 | 1020.3841 | 943.3878 | 1067.6805 | 0.537407175 | 0.057661347 | YMR304W | UBP15 | n |
| W303.asm02.HP0_G0043460 | 317.11593 | 317.642762 | 313.9256376 | 466.691044 | 439.54377 | 448.474211 | 404.73335 | 440.85994 | 405.55988 | 1.29E-07 | -0.514555767 | YMR306W | FKS3 | n |
| W303.asm02.HP0_G0043470 | 17496.16 | 15628.8543 | 15228.77416 | 11903.9551 | 11298.702 | 12463.4112 | 12507.971 | 12531.706 | 12235.565 | 1.17E-10 | 0.439056607 | YMR307W | GAS1 | n |
| W303.asm02.HP0_G0043480 | 2413.6631 | 2443.56556 | 2727.772309 | 5475.286 | 5589.1272 | 5687.94248 | 4844.2592 | 4866.5165 | 4699.2047 | 6.86E-73 | -1.142663306 | YMR308C | PSE1 | n |
| W303.asm02.HP0_G0043490 | 7704.5475 | 7232.0821 | 6997.16098 | 9938.0191 | 9718.5016 | 10010.5512 | 8448.0962 | 8327.7919 | 8099.7361 | 4.67E-15 | -0.435818279 | YMR309C | NIP1 | n |
| W303.asm02.HP0_G0043500 | 582.60834 | 557.431906 | 579.5550232 | 1429.24132 | 1233.2394 | 1366.2819 | 931.45676 | 1158.5694 | 1010.3731 | 5.21E-37 | -1.228402394 | YMR310C | UPA2 | n |
| W303.asm02.HP0_G0043510 | 2079.6905 | 2105.16183 | 1885.485675 | 1135.05929 | 1215.2621 | 1141.57072 | 1346.451 | 1186.1232 | 1393.8917 | 1.58E-22 | 0.797550108 | YMR311C | GLC8 | n |
| W303.asm02.HP0_G0043520 | 1124.1286 | 1111.74967 | 1094.393069 | 878.379215 | 839.5376 | 913.066946 | 608.81017 | 787.2499 | 674.46371 | 0.000235313 | 0.340121628 | YMR312W | ELP6 | n |
| W303.asm02.HP0_G0043530 | 751.17495 | 643.58991 | 634.6127504 | 415.855055 | 372.12908 | 385.896414 | 521.02294 | 495.96744 | 515.76637 | 1.76E-15 | 0.788264681 | YMR313C | TGL3 | n |
| W303.asm02.HP0_G0043540 | 3118.4822 | 3080.92718 | 2948.969143 | 2889.31759 | 2858.3828 | 2995.20092 | 3308.5526 | 3240.8454 | 3216.2662 | 0.255706423 | 0.065353993 | YMR314W | PRE5 | n |
| W303.asm02.HP0_G0043550 | 2081.7976 | 2140.45547 | 1654.629591 | 1215.89684 | 1165.8247 | 1192.77073 | 1378.3736 | 1136.264 | 1226.3778 | 6.74E-12 | 0.716850096 | YMR315W | | n |
| W303.asm02.HP0_G0043580 | 6686.8266 | 6059.08759 | 6389.59413 | 3887.70307 | 3726.684 | 4142.46052 | 3072.5532 | 4067.4578 | 3205.6864 | 1.14E-12 | 0.702803188 | YMR318C | ADH6 | n |
| W303.asm02.HP0_G0043590 | 1801.5556 | 1676.44791 | 1829.462023 | 2239.28363 | 2309.1778 | 2203.49697 | 4862.5007 | 2889.2071 | 4189.6099 | 0.013117372 | -0.34725023 | YMR319C | FET4 | n |
| W303.asm02.HP0_G0043660 | 663.73102 | 699.644515 | 787.2289065 | 3981.04128 | 4241.7322 | 3931.97157 | 3688.2039 | 3652.8395 | 3683.5417 | 2.36E-252 | -2.497476342 | YNL334C | SNO2 | y |
| W303.asm02.HP0_G0043670 | 820.70867 | 860.541992 | 798.8200069 | 20990.2631 | 21449.556 | 24034.6148 | 20747.43 | 19807.208 | 17477.868 | 0 | -4.744546364 | YNL333W | SNZ2 | y |
| W303.asm02.HP0_G0043700 | 1031.4169 | 1107.59747 | 1025.812391 | 2695.97415 | 2534.7923 | 2551.46745 | 2573.192 | 2578.2434 | 2676.6952 | 3.02E-88 | -1.298517335 | YNL330C | RPD3 | n |
| W303.asm02.HP0_G0043710 | 642.66019 | 691.340129 | 722.5119289 | 646.700446 | 604.93448 | 650.429828 | 717.1191 | 661.28992 | 720.30961 | 0.199345238 | 0.113274803 | YNL329C | PEX6 | n |
| W303.asm02.HP0_G0043720 | 86.390387 | 79.9297146 | 98.52435394 | 315.849831 | 344.26435 | 271.170453 | 307.82537 | 313.58788 | 302.40661 | 6.11E-35 | -1.812653117 | YNL328C | MDJ2 | n |
| W303.asm02.HP0_G0043730 | 4067.723 | 4913.08233 | 5421.737242 | 3711.02717 | 3783.3124 | 3825.77895 | 4316.3957 | 4804.8486 | 4804.1213 | 0.000313292 | 0.347651143 | YNL327W | EGT2 | n |
| W303.asm02.HP0_G0043740 | 798.58431 | 820.058111 | 815.2407326 | 338.351007 | 399.99382 | 384.000117 | 470.8588 | 480.22244 | 488.43516 | 4.00E-35 | 1.118360758 | YNL326C | PFA3 | n |
| W303.asm02.HP0_G0043750 | 696.3908 | 688.225984 | 684.8408524 | 645.033693 | 635.4958 | 635.259453 | 768.42332 | 737.39074 | 756.45734 | 0.157459983 | 0.111032018 | YNL325C | FIG4 | n |
| W303.asm02.HP0_G0043770 | 1190.5017 | 1137.70087 | 1223.827024 | 1525.91304 | 1502.8981 | 1389.98561 | 1460.4604 | 1466.909 | 1653.0973 | 6.66E-05 | -0.315192388 | YNL323W | LEM3 | n |
| W303.asm02.HP0_G0043780 | 1059.8625 | 1150.15745 | 1194.849273 | 3115.99609 | 3003.9986 | 3241.71951 | 3396.3399 | 3532.1279 | 3387.3066 | 8.25E-103 | -1.458495325 | YNL322C | KRE1 | n |
| W303.asm02.HP0_G0043790 | 536.25252 | 646.704054 | 651.033476 | 1085.05668 | 1045.3771 | 1084.68181 | 1551.6679 | 1364.5665 | 1487.3468 | 7.38E-19 | -0.809080412 | YNL321W | VNX1 | n |
| W303.asm02.HP0_G0043800 | 657.40977 | 602.06798 | 583.4187233 | 606.698357 | 551.00273 | 574.577953 | 562.06632 | 544.51452 | 531.6361 | 0.336984387 | 0.088092098 | YNL320W | | n |
| W303.asm02.HP0_G0043810 | 16.856661 | 11.4185307 | 21.25035085 | 185.009664 | 196.85089 | 205.748211 | 186.97541 | 226.99039 | 180.73864 | 1.26E-45 | -3.55941315 | YNL318C | HXT14 | n |
| W303.asm02.HP0_G0043820 | 338.18676 | 422.485634 | 396.0292658 | 756.706192 | 814.36945 | 764.207641 | 707.99834 | 754.44782 | 731.77109 | 4.44E-26 | -1.012817516 | YNL317W | PFS2 | n |
| W303.asm02.HP0_G0043830 | 650.03498 | 578.19287 | 498.4173199 | 930.881957 | 994.14195 | 1002.1929 | 1030.6449 | 1031.2974 | 1017.4263 | 1.59E-15 | -0.762625039 | YNL316C | PHA2 | n |
| W303.asm02.HP0_G0043840 | 2104.9755 | 1890.28585 | 2092.193634 | 2018.43876 | 2058.3952 | 2014.81543 | 1860.6333 | 1985.1818 | 1855.8773 | 0.990525378 | -0.000869966 | YNL315C | ATP11 | n |
| W303.asm02.HP0_G0043850 | 200.17285 | 212.799889 | 177.7302071 | 211.677723 | 229.20994 | 226.607477 | 243.98011 | 223.05414 | 243.33593 | 0.174267651 | -0.17700287 | YNL314W | DAL82 | n |
| W303.asm02.HP0_G0043860 | 1080.9334 | 1271.6091 | 1273.089201 | 2125.94438 | 2165.3598 | 2385.54147 | 1420.5571 | 1460.3486 | 1342.7559 | 2.52E-25 | -0.88023403 | YNL313C | EMW1 | n |
| W303.asm02.HP0_G0043870 | 2067.048 | 1914.16096 | 1980.146329 | 1288.40063 | 1419.3039 | 1325.51152 | 1394.3349 | 1422.2982 | 1452.9624 | 1.43E-16 | 0.563864805 | YNL312W | RFA2 | n |
| W303.asm02.HP0_G0043880 | 836.51179 | 743.242541 | 620.1238748 | 401.687648 | 375.72453 | 363.140852 | 500.50124 | 431.67536 | 492.84342 | 1.22E-15 | 0.946032779 | YNL311C | SKP2 | n |
| W303.asm02.HP0_G0043890 | 446.70151 | 485.806577 | 419.2114668 | 567.529644 | 550.10386 | 559.407578 | 581.44792 | 545.8266 | 546.62419 | 0.000717778 | -0.312122474 | YNL310C | ZIM17 | n |
| W303.asm02.HP0_G0043900 | 259.17116 | 284.425218 | 279.1523362 | 748.372424 | 705.60708 | 690.252063 | 853.93037 | 785.93782 | 869.30879 | 1.62E-48 | -1.382071912 | YNL309W | STB1 | n |
| W303.asm02.HP0_G0043910 | 1248.4464 | 1394.09879 | 1205.474448 | 2205.11518 | 2247.1563 | 2487.9415 | 1213.06 | 1588.9327 | 1245.7742 | 8.80E-16 | -0.850862991 | YNL308C | KRI1 | n |
| W303.asm02.HP0_G0043920 | 3323.9228 | 3023.83453 | 3106.414924 | 1767.59233 | 1731.2092 | 1683.91163 | 1866.3338 | 1636.1677 | 1898.1966 | 1.70E-31 | 0.866976701 | YNL307C | MCK1 | n |

| | | | | | | | | | | | | | | |
|---|---|---|---|---|---|---|---|---|---|---|---|---|---|---|
| W303.asm02.HP0_G0043930 | 2491.6252 | 2348.06512 | 2906.468441 | 1651.75294 | 1791.433 | 1572.03011 | 1542.5471 | 1663.7215 | 1544.6542 | 3.92E-12 | 0.627378901 | YNL306W | MRPS18 | n |
| W303.asm02.HP0_G0043940 | 509.91399 | 576.116774 | 571.8276229 | 415.855055 | 391.90406 | 363.140852 | 551.80547 | 511.71244 | 576.60035 | 5.53E-07 | 0.500863537 | YNL305C | BXI1 | n |
| W303.asm02.HP0_G0043950 | 264.43887 | 310.376424 | 309.0960124 | 1542.58058 | 1363.5744 | 1413.68932 | 1509.4844 | 1305.5228 | 1470.5954 | 1.90E-119 | -2.288339215 | YNL304W | YPT11 | n |
| W303.asm02.HP0_G0043970 | 3492.4894 | 3497.18453 | 3318.918433 | 11068.0781 | 11168.367 | 12346.789 | 9197.1379 | 9646.4355 | 8485.018 | 7.75E-135 | -1.746255273 | YOL120C | RPL18A | y |
| W303.asm02.HP0_G0043990 | 385.59612 | 393.420284 | 344.8352388 | 1355.90416 | 1269.1939 | 1311.28929 | 923.4761 | 982.75029 | 1029.7694 | 4.35E-97 | -1.80987552 | YNL299W | TRF5 | n |
| W303.asm02.HP0_G0044000 | 498.32503 | 652.932344 | 603.7031491 | 3487.68218 | 3359.9481 | 3200.94913 | 3357.5767 | 2966.62 | 3191.5799 | 2.45E-159 | -2.517068546 | YNL298W | CLA4 | n |
| W303.asm02.HP0_G0044010 | 1005.0784 | 1056.73311 | 1130.132295 | 1908.43302 | 1926.2624 | 1821.39315 | 2014.546 | 1814.611 | 1943.1608 | 4.06E-29 | -0.824951017 | YNL297C | MON2 | n |
| W303.asm02.HP0_G0044020 | 448.80859 | 494.110963 | 546.7135719 | 618.365633 | 574.37315 | 588.80018 | 533.56397 | 535.32993 | 543.97923 | 0.007431979 | -0.257298007 | YNL295W | MRX6 | n |
| W303.asm02.HP0_G0044030 | 1185.234 | 1208.28815 | 1108.881944 | 384.186734 | 348.75866 | 272.118602 | 345.44847 | 310.96371 | 310.34147 | 1.39E-55 | 1.798227855 | YNL294C | RIM21 | n |
| W303.asm02.HP0_G0044040 | 407.72048 | 477.502191 | 472.3373439 | 1532.58005 | 1472.3368 | 1366.2819 | 1417.1368 | 1228.1098 | 1329.5311 | 1.64E-74 | -1.686694571 | YNL293W | MSB3 | n |
| W303.asm02.HP0_G0044050 | 1479.172 | 1475.06655 | 1395.761681 | 775.040483 | 764.03314 | 913.066946 | 474.27908 | 505.15202 | 516.64802 | 2.11E-21 | 0.827745163 | YNL292W | PUS4 | n |
| W303.asm02.HP0_G0044060 | 525.71711 | 506.567542 | 499.383245 | 980.884569 | 927.62612 | 909.274352 | 1132.1133 | 1016.8645 | 1136.4493 | 3.87E-26 | -0.8803368 | YNL291C | MID1 | n |
| W303.asm02.HP0_G0044070 | 1244.2323 | 1241.5057 | 1143.655246 | 563.36276 | 617.51855 | 648.533531 | 533.56397 | 461.85328 | 578.36365 | 6.00E-25 | 0.989005234 | YNL290W | RFC3 | n |
| W303.asm02.HP0_G0044080 | 1089.3617 | 1115.90186 | 1057.687917 | 890.046491 | 896.16594 | 940.56325 | 1272.3448 | 1313.3953 | 1456.489 | 0.001024459 | 0.259147362 | YNL289W | PCL1 | n |
| W303.asm02.HP0_G0044090 | 773.29931 | 902.063922 | 797.8540819 | 1680.92113 | 1655.7048 | 1752.17831 | 1762.5852 | 1779.1848 | 1908.7764 | 3.15E-40 | -1.040954143 | YNL288W | CAF40 | n |
| W303.asm02.HP0_G0044100 | 4792.5594 | 4725.1956 | 4422.970752 | 7297.04782 | 7268.2024 | 7438.22449 | 7276.0796 | 6472.5063 | 7210.1493 | 2.01E-24 | -0.65853918 | YNL287W | SEC21 | n |
| W303.asm02.HP0_G0044120 | 2373.6285 | 2418.6524 | 2522.996201 | 2283.45261 | 2337.0426 | 2368.4748 | 2233.4441 | 2655.6563 | 2372.5253 | 0.386907414 | 0.066097227 | YNL284C | MRPL10 | n |
| W303.asm02.HP0_G0044130 | 795.42368 | 925.939031 | 913.7650865 | 2506.79761 | 2377.4914 | 2538.19337 | 2640.4576 | 2387.9914 | 2580.5951 | 3.49E-81 | -1.493506557 | YNL283C | WSC2 | n |
| W303.asm02.HP0_G0044140 | 304.47344 | 271.968639 | 261.7656855 | 597.531211 | 559.09249 | 496.829781 | 464.01824 | 459.22911 | 454.05073 | 6.85E-20 | -0.982479433 | YNL282W | POP3 | n |
| W303.asm02.HP0_G0044150 | 5179.209 | 5230.72509 | 5819.698358 | 1375.9052 | 1417.5062 | 1469.63008 | 1535.7066 | 1472.1573 | 1436.211 | 1.01E-168 | 1.929188999 | YNL281W | HCH1 | n |
| W303.asm02.HP0_G0044160 | 1730.9684 | 1597.55624 | 1680.709567 | 1373.40507 | 1420.2028 | 1460.14859 | 1873.1744 | 1846.101 | 1779.1736 | 0.000383863 | 0.236233429 | YNL280C | ERG24 | n |
| W303.asm02.HP0_G0044170 | 204.38701 | 210.723793 | 172.9005819 | 84.1710633 | 83.594215 | 103.34818 | 58.144792 | 68.228325 | 52.017463 | 9.32E-11 | 1.118356984 | YNL279W | PRM1 | n |
| W303.asm02.HP0_G0044200 | 417.20235 | 489.95877 | 461.7121685 | 288.348395 | 322.69165 | 244.622297 | 385.35176 | 316.21204 | 388.80849 | 1.29E-07 | 0.677010311 | YNL277W | MET2 | n |
| W303.asm02.HP0_G0044210 | 685.85539 | 670.579164 | 576.6572481 | 526.694178 | 524.93571 | 524.326086 | 735.3606 | 610.11867 | 767.03716 | 0.006262529 | 0.293836549 | YNL275W | BOR1 | n |
| W303.asm02.HP0_G0044220 | 1099.8971 | 1130.43453 | 957.2317133 | 2590.96867 | 2625.5774 | 2344.77109 | 2247.1252 | 2410.2968 | 2242.0408 | 5.84E-52 | -1.247091511 | YNL274C | GOR1 | n |
| W303.asm02.HP0_G0044230 | 455.12984 | 447.398792 | 480.0647442 | 1609.25072 | 1875.9261 | 1776.83017 | 1511.7646 | 1573.1877 | 1563.1688 | 6.30E-118 | -1.927252188 | YNL273W | TOF1 | n |
| W303.asm02.HP0_G0044240 | 1083.0405 | 1024.55361 | 932.1176623 | 2097.60957 | 2171.6519 | 1955.08208 | 2002.005 | 2229.2293 | 2058.6572 | 1.81E-37 | -1.034836904 | YNL272C | SEC2 | n |
| W303.asm02.HP0_G0044250 | 751.17495 | 858.465896 | 905.0717612 | 1453.40925 | 1539.7515 | 1477.21527 | 1543.6872 | 1326.5161 | 1552.589 | 9.28E-20 | -0.829249079 | YNL271C | BNI1 | n |
| W303.asm02.HP0_G0044280 | 2587.4974 | 2788.19758 | 2695.896783 | 2865.98303 | 3057.9303 | 2774.28233 | 3041.7707 | 3080.7713 | 2946.4807 | 0.09672858 | -0.107897035 | YNL268W | LYP1 | n |
| W303.asm02.HP0_G0044290 | 828.08346 | 971.613154 | 899.276211 | 2800.14626 | 2993.2122 | 2821.68975 | 2763.5877 | 2793.4251 | 2786.9017 | 1.27E-120 | -1.674252623 | YNL267W | PIK1 | n |
| W303.asm02.HP0_G0044300 | 1712.0046 | 1641.15427 | 1891.281226 | 905.047274 | 926.72726 | 765.155789 | 990.74164 | 1049.6665 | 1012.1364 | 2.98E-29 | 1.013489271 | YNL265C | IST1 | n |
| W303.asm02.HP0_G0044320 | 2031.2276 | 1799.97565 | 1662.356991 | 1519.24602 | 1663.7945 | 1497.12638 | 1861.7734 | 2035.041 | 1909.658 | 0.008602033 | 0.230751876 | YNL263C | YIF1 | n |
| W303.asm02.HP0_G0044330 | 709.03329 | 740.128396 | 915.6969366 | 1455.076 | 1575.706 | 1528.41528 | 1428.5377 | 1439.3552 | 1452.0807 | 8.21E-24 | -0.945637086 | YNL262W | POL2 | n |
| W303.asm02.HP0_G0044340 | 1387.5139 | 1333.89199 | 1429.569057 | 1214.23009 | 1380.6528 | 1267.67446 | 1187.9779 | 1132.3278 | 1115.2897 | 0.1741208 | 0.10438753 | YNL261W | ORC5 | n |
| W303.asm02.HP0_G0044350 | 868.11803 | 774.383988 | 823.9340579 | 400.854272 | 444.93695 | 358.40011 | 319.22631 | 457.91703 | 344.7259 | 4.92E-16 | 1.033812873 | YNL260C | LTO1 | n |
| W303.asm02.HP0_G0044360 | 518.34232 | 533.556796 | 652.9653261 | 1479.24393 | 1713.232 | 1528.41528 | 1674.798 | 1949.7556 | 1590.5 | 7.76E-41 | -1.468182522 | YNL259C | ATX1 | n |
| W303.asm02.HP0_G0044370 | 1814.1981 | 1578.87138 | 1732.869519 | 1630.91852 | 1898.3976 | 1704.77089 | 1614.373 | 1775.2485 | 1670.7304 | 0.736021761 | -0.029937356 | YNL258C | DSL1 | n |
| W303.asm02.HP0_G0044380 | 903.93844 | 1023.51557 | 1034.505716 | 1739.25751 | 1970.3066 | 2037.57099 | 1714.7013 | 1605.9898 | 1660.1506 | 2.15E-29 | -0.955382984 | YNL257C | SIP3 | n |
| W303.asm02.HP0_G0044390 | 1471.7972 | 1508.28409 | 1627.58369 | 1326.73597 | 1504.6959 | 1364.3856 | 1032.9251 | 1358.0061 | 1108.2365 | 0.192138611 | 0.135403785 | YNL256W | FOL1 | n |
| W303.asm02.HP0_G0044400 | 391.91736 | 344.632016 | 348.6989389 | 360.852182 | 332.57913 | 299.614906 | 344.30837 | 262.41663 | 308.57817 | 0.323643928 | 0.126179381 | YNL254C | RTC4 | n |
| W303.asm02.HP0_G0044410 | 1172.5915 | 1075.41798 | 1266.327726 | 1397.573 | 1460.6516 | 1415.58562 | 1150.3548 | 1275.3448 | 1187.5851 | 0.00046474 | -0.281675537 | YNL253W | TEX1 | n |
| W303.asm02.HP0_G0044420 | 1898.4814 | 2042.87894 | 1908.667876 | 1707.58919 | 1902.892 | 1773.98573 | 1553.9481 | 1662.4094 | 1601.0799 | 0.093362648 | 0.11976577 | YNL252C | MRPL17 | n |
| W303.asm02.HP0_G0044430 | 563.6446 | 671.617212 | 648.1357009 | 1398.40638 | 1541.5492 | 1511.34861 | 1552.808 | 1695.2115 | 1557.8789 | 5.06E-47 | -1.240051066 | YNL251C | NRD1 | n |
| W303.asm02.HP0_G0044440 | 691.12309 | 739.090348 | 821.0362828 | 931.715334 | 1112.7918 | 969.007703 | 954.25864 | 907.96155 | 980.39693 | 1.29E-05 | -0.419716118 | YNL250W | RAD50 | n |
| W303.asm02.HP0_G0044450 | 864.95741 | 851.199558 | 854.8436592 | 602.531473 | 718.19116 | 580.266844 | 552.94557 | 548.45076 | 532.51776 | 3.34E-06 | 0.435448511 | YNL249C | MPA43 | n |
| W303.asm02.HP0_G0044460 | 3777.9991 | 3317.60218 | 3590.343369 | 3366.00915 | 3688.9318 | 3706.31224 | 2387.3567 | 2919.3851 | 2310.8097 | 0.920018626 | -0.010015791 | YNL248C | RPA49 | n |
| W303.asm02.HP0_G0044470 | 2052.2985 | 2154.98815 | 2474.699949 | 3280.17134 | 3572.0797 | 3439.88253 | 2992.7466 | 3295.9529 | 3058.4505 | 1.92E-14 | -0.622682026 | YNL247W | CRS1 | n |
| W303.asm02.HP0_G0044480 | 1396.9958 | 1298.59835 | 1397.693531 | 1491.74459 | 1719.524 | 1525.57084 | 1370.3929 | 1507.5836 | 1482.0569 | 0.009327075 | -0.210419322 | YNL246W | VPS75 | n |
| W303.asm02.HP0_G0044490 | 5995.7035 | 5787.11895 | 6992.331355 | 5329.44504 | 6096.9845 | 5604.50542 | 4425.8447 | 5118.4364 | 4270.7219 | 0.144243645 | 0.140825105 | YNL244C | SUI1 | n |
| W303.asm02.HP0_G0044500 | 1536.0632 | 1919.3512 | 1733.835444 | 3585.18727 | 3817.4691 | 3811.55672 | 3676.803 | 3310.3858 | 3356.4488 | 4.63E-39 | -1.111536418 | YNL243W | SLA2 | n |
| W303.asm02.HP0_G0044510 | 498.32503 | 462.969516 | 558.3046723 | 612.531995 | 719.09002 | 781.274313 | 2452.3421 | 1277.969 | 2321.3895 | 0.077022122 | -0.47518831 | YNL242W | ATG2 | n |
| W303.asm02.HP0_G0044520 | 3722.1614 | 4486.4445 | 4563.995807 | 4975.25988 | 5175.6504 | 5209.12752 | 13905.726 | 8857.8735 | 12916.201 | 0.051134729 | -0.266045475 | YNL241C | ZWF1 | n |
| W303.asm02.HP0_G0044530 | 697.44434 | 696.53037 | 765.9785556 | 3653.52417 | 3735.6726 | 3903.52712 | 9262.1233 | 6110.3713 | 8929.3706 | 2.69E-78 | -2.385996839 | YNL240C | NAR1 | n |
| W303.asm02.HP0_G0044560 | 234.93971 | 263.664253 | 262.7316105 | 338.351007 | 312.80416 | 329.007508 | 2297.2893 | 1317.3315 | 2300.2298 | 0.039763646 | -0.364312999 | YNL237W | YTP1 | n |
| W303.asm02.HP0_G0044570 | 903.93844 | 877.150764 | 870.2984598 | 1700.0888 | 1800.4216 | 1846.99316 | 5020.9738 | 2923.3213 | 4723.0093 | 1.07E-11 | -1.012082568 | YNL236W | SIN4 | n |
| W303.asm02.HP0_G0044580 | 636.33895 | 706.910852 | 542.8498717 | 1624.25151 | 1753.6808 | 1626.07457 | 4567.2164 | 2876.0863 | 4486.7266 | 2.40E-21 | -1.408145724 | YNL234W | | n |
| W303.asm02.HP0_G0044590 | 493.05733 | 625.94309 | 529.3269212 | 1714.25621 | 1861.5443 | 1675.37829 | 4727.9696 | 2851.1567 | 4827.9259 | 4.20E-14 | -1.671670698 | YNL233W | BNI4 | n |
| W303.asm02.HP0_G0044600 | 1715.1652 | 1608.97477 | 1628.549615 | 1288.40063 | 1313.2381 | 1239.23001 | 3150.0796 | 2234.4776 | 3274.4552 | 0.00228106 | 0.366658645 | YNL232W | CSL4 | n |
| W303.asm02.HP0_G0044610 | 3811.7124 | 3690.2615 | 3931.314907 | 3281.00471 | 3334.78 | 3436.08994 | 7665.9917 | 5281.1348 | 7603.3661 | 0.117418121 | 0.185845695 | YNL231C | PDR16 | n |
| W303.asm02.HP0_G0044620 | 172.78077 | 183.734539 | 180.6279822 | 558.362499 | 534.8232 | 504.414969 | 1320.2288 | 926.33072 | 1398.2999 | 3.47E-27 | -1.572742879 | YNL230C | ELA1 | n |

| | | | | | | | | | | | | | | |
|---|---|---|---|---|---|---|---|---|---|---|---|---|---|---|
| W303.asm02.HP0_G0044630 | 2807.6876 | 2732.14297 | 2738.397484 | 2779.31184 | 2629.1729 | 2815.05271 | 7087.9641 | 4625.0932 | 7144.0254 | 0.945139018 | 0.009556985 | YNL229C | URE2 | n |
| W303.asm02.HP0_G0044640 | 823.8693 | 892.721488 | 813.3088825 | 1633.41865 | 1709.6365 | 1663.05236 | 3918.5029 | 2529.6964 | 3910.1262 | 3.19E-13 | -0.98467662 | YNL227C | JJJ1 | n |
| W303.asm02.HP0_G0044650 | 1249.5 | 1130.43453 | 1113.71157 | 676.702013 | 823.35807 | 694.044656 | 2469.4435 | 1394.7444 | 2262.3188 | 0.002891726 | 0.671095277 | YNL225C | CNM67 | n |
| W303.asm02.HP0_G0044660 | 566.80522 | 579.230919 | 566.9979977 | 1340.90337 | 1366.271 | 1300.85966 | 3915.0826 | 2322.3872 | 3854.5822 | 4.19E-16 | -1.226396372 | YNL224C | SQS1 | n |
| W303.asm02.HP0_G0044670 | 269.70657 | 271.968639 | 299.436762 | 244.179421 | 248.08606 | 202.903766 | 1227.8812 | 602.24617 | 1107.3548 | 0.334841562 | 0.274737351 | YNL223W | ATG4 | n |
| W303.asm02.HP0_G0044680 | 811.2268 | 821.096159 | 1040.301267 | 508.359887 | 579.76633 | 537.600164 | 1265.5043 | 889.59239 | 1341.8742 | 7.16E-07 | 0.717896802 | YNL222W | SSU72 | n |
| W303.asm02.HP0_G0044690 | 906.04552 | 945.661948 | 1060.585692 | 1551.74772 | 1530.7629 | 1536.94862 | 3994.8892 | 2443.0989 | 3931.2859 | 5.59E-06 | -0.665329995 | YNL221C | POP1 | n |
| W303.asm02.HP0_G0044700 | 1740.4502 | 1591.32795 | 1872.92865 | 6898.69368 | 6958.0948 | 6829.5132 | 16391.131 | 11625.057 | 15348.678 | 1.38E-72 | -1.990613813 | YNL220W | ADE12 | n |
| W303.asm02.HP0_G0044710 | 2057.5662 | 1755.33958 | 1774.404296 | 1999.2711 | 2039.5191 | 2039.46729 | 6959.1335 | 4415.1599 | 6857.4886 | 0.39495389 | -0.121579223 | YNL219C | ALG9 | n |
| W303.asm02.HP0_G0044720 | 791.20952 | 702.758659 | 652.9653261 | 540.028208 | 564.48566 | 618.192781 | 1690.7593 | 1014.2403 | 1637.2276 | 0.046929267 | 0.317659767 | YNL218W | MGS1 | n |
| W303.asm02.HP0_G0044730 | 2291.4523 | 2324.19001 | 2220.661664 | 1216.73022 | 1260.2053 | 1090.3707 | 3124.9975 | 2023.2322 | 3160.7221 | 4.56E-12 | 0.938176963 | YNL217W | PPN2 | n |
| W303.asm02.HP0_G0044740 | 870.22511 | 891.683439 | 945.6406128 | 2025.93916 | 2104.2372 | 1974.04505 | 5120.1619 | 3251.3421 | 5149.7288 | 1.98E-17 | -1.172686388 | YNL216W | RAP1 | n |
| W303.asm02.HP0_G0044750 | 636.33895 | 718.329383 | 602.7372241 | 998.385483 | 953.69314 | 903.585461 | 2284.7483 | 1506.2715 | 2397.2115 | 0.000169391 | -0.545338596 | YNL215W | IES2 | n |
| W303.asm02.HP0_G0044760 | 677.42706 | 679.921598 | 853.8777341 | 381.686604 | 423.36425 | 389.689008 | 1154.9152 | 768.88074 | 1178.7686 | 3.34E-09 | 0.888813019 | YNL214W | PEX17 | n |
| W303.asm02.HP0_G0044770 | 1244.2323 | 1310.01688 | 1206.440373 | 582.530428 | 663.36054 | 566.992766 | 1116.152 | 712.46116 | 1189.3484 | 1.08E-12 | 1.052611758 | YNL213C | RRG9 | n |
| W303.asm02.HP0_G0044780 | 2151.3313 | 2204.81446 | 2139.523961 | 500.026118 | 550.10386 | 451.318656 | 1429.6778 | 977.50196 | 1581.6835 | 6.80E-52 | 2.112795747 | YNL212W | VID27 | n |
| W303.asm02.HP0_G0044790 | 522.55648 | 495.149011 | 622.0557249 | 305.015932 | 300.22008 | 305.303797 | 857.35065 | 583.87701 | 863.13722 | 5.04E-09 | 0.849344694 | YNL211C | MRX7 | n |
| W303.asm02.HP0_G0044800 | 92.711634 | 77.8536181 | 85.96732844 | 68.3369028 | 83.594215 | 85.3333594 | 132.2509 | 95.782071 | 136.65605 | 0.583709193 | 0.115276624 | YNL210W | MER1 | n |
| W303.asm02.HP0_G0044830 | 1533.9561 | 1372.29978 | 1467.240134 | 1439.24184 | 1430.9891 | 1514.19306 | 2762.4476 | 1981.2456 | 2914.7412 | 0.979082453 | -0.003521221 | YNL207W | RIO2 | n |
| W303.asm02.HP0_G0044840 | 1964.8545 | 1863.29659 | 1850.712374 | 864.211808 | 876.39096 | 822.044926 | 2177.5794 | 1405.2411 | 2373.4069 | 1.04E-15 | 1.147827831 | YNL206C | RTT106 | n |
| W303.asm02.HP0_G0044860 | 181.2091 | 204.495504 | 243.4131097 | 28.3348134 | 19.774975 | 24.6518594 | 102.60846 | 53.79541 | 85.520235 | 8.15E-35 | 3.109717122 | YNL202W | SPS19 | n |
| W303.asm02.HP0_G0044870 | 1169.4308 | 997.56436 | 1225.758874 | 1109.22461 | 1166.7236 | 1100.80034 | 2145.6568 | 1554.8186 | 2300.2298 | 0.959599367 | 0.007020201 | YNL201C | PSY2 | n |
| W303.asm02.HP0_G0044880 | 738.53245 | 746.356686 | 771.7741059 | 182.509533 | 193.25544 | 183.940797 | 275.90274 | 244.04747 | 264.49557 | 1.13E-87 | 2.011926518 | YNL200C | NNR1 | n |
| W303.asm02.HP0_G0044890 | 800.69139 | 837.704931 | 927.2880371 | 420.855316 | 372.12908 | 385.896414 | 422.97486 | 432.98745 | 434.65439 | 1.29E-32 | 1.121615965 | YNL199C | GCR2 | n |
| W303.asm02.HP0_G0044950 | 2290.3988 | 2183.01545 | 2896.809191 | 4771.91592 | 4552.7387 | 4766.3422 | 4178.4443 | 4438.7774 | 4139.3557 | 1.67E-22 | -0.934584122 | YNL192W | CHS1 | n |
| W303.asm02.HP0_G0044960 | 954.50842 | 937.357562 | 816.2066576 | 813.375819 | 760.43769 | 814.459508 | 948.55817 | 838.42115 | 945.13085 | 0.048386441 | 0.180531337 | YNL191W | DUG3 | n |
| W303.asm02.HP0_G0044980 | 3146.9279 | 2883.69801 | 2760.61376 | 4349.39385 | 4171.621 | 4477.15692 | 3970.9472 | 3576.7387 | 3704.7013 | 2.37E-15 | -0.564365802 | YNL189W | SRP1 | n |
| W303.asm02.HP0_G0044990 | 1134.664 | 1037.01019 | 951.436163 | 122.506399 | 125.84075 | 152.651898 | 145.93203 | 149.57748 | 159.579 | 1.05E-132 | 2.964215833 | YNL188W | KAR1 | n |
| W303.asm02.HP0_G0045000 | 188.58389 | 228.370613 | 182.5598323 | 125.839906 | 149.21118 | 110.933367 | 152.77259 | 167.94665 | 186.02855 | 8.80E-05 | 0.633482321 | YNL187W | SWT21 | n |
| W303.asm02.HP0_G0045010 | 1435.9768 | 1608.97477 | 1625.65184 | 1969.26953 | 1852.5557 | 1698.13385 | 1284.8859 | 1373.7511 | 1289.8567 | 0.003273205 | -0.241243087 | YNL186W | UBP10 | n |
| W303.asm02.HP0_G0045020 | 1754.1463 | 1680.6001 | 1811.109447 | 1125.89214 | 1174.8133 | 1126.40034 | 1117.2921 | 1295.0261 | 1146.1475 | 4.99E-16 | 0.614392161 | YNL185C | MRPL19 | n |
| W303.asm02.HP0_G0045030 | 1118.8609 | 1222.82083 | 1063.483468 | 1176.72813 | 1185.5997 | 1229.74852 | 1686.199 | 1550.8823 | 1731.5644 | 0.343402881 | -0.07727856 | YNL183C | NPR1 | n |
| W303.asm02.HP0_G0045040 | 2247.2036 | 2004.47115 | 2097.023259 | 1930.93419 | 1823.7921 | 1844.14871 | 1202.7991 | 1535.1373 | 1177.0053 | 0.069119591 | 0.18116023 | YNL182C | IPI3 | n |
| W303.asm02.HP0_G0045050 | 1322.1943 | 1223.85888 | 1188.087798 | 2033.43955 | 1981.093 | 1880.17835 | 1861.7734 | 1764.7519 | 1797.6882 | 3.88E-21 | -0.659426228 | YNL181W | PBR1 | n |
| W303.asm02.HP0_G0045080 | 20735.8 | 20031.2169 | 23122.31357 | 17289.2364 | 17345.35 | 19148.8059 | 15241.916 | 16507.318 | 14204.294 | 0.002111636 | 0.248481106 | YNL178W | RPS3 | n |
| W303.asm02.HP0_G0045090 | 2535.8739 | 2446.67971 | 2501.74585 | 3185.99975 | 3187.3665 | 3244.56395 | 2928.9014 | 3169.9929 | 2999.3798 | 2.79E-10 | -0.361796505 | YNL177C | MRPL22 | n |
| W303.asm02.HP0_G0045100 | 762.7639 | 851.199558 | 834.5592334 | 1635.08541 | 1512.7856 | 1687.70422 | 1506.0641 | 1620.4227 | 1414.1697 | 3.14E-32 | -0.981390728 | YNL176C | TDA7 | n |
| W303.asm02.HP0_G0045110 | 1865.8216 | 1678.52401 | 1609.231114 | 1733.42388 | 1620.6491 | 1754.07461 | 900.67422 | 1061.4753 | 938.07763 | 0.889890293 | 0.012465979 | YNL175C | NOP13 | n |
| W303.asm02.HP0_G0045130 | 616.32166 | 651.894296 | 680.0112272 | 1070.88927 | 1044.4783 | 1012.62253 | 1093.3501 | 964.38113 | 1056.219 | 4.45E-17 | -0.682795393 | YNL172W | APC1 | n |
| W303.asm02.HP0_G0045140 | 824.92284 | 788.916664 | 854.8436592 | 2042.60669 | 2035.9236 | 1985.42283 | 1843.5319 | 2036.3531 | 1920.2379 | 7.82E-77 | -1.296216552 | YNL169C | PSD1 | n |
| W303.asm02.HP0_G0045150 | 1190.5017 | 1234.23936 | 1012.28944 | 1509.2455 | 1585.5935 | 1506.60787 | 1498.0835 | 1447.2277 | 1442.3825 | 3.63E-07 | -0.421677288 | YNL168C | FMP41 | n |
| W303.asm02.HP0_G0045160 | 680.58768 | 674.731357 | 781.4333562 | 1660.92009 | 1508.2913 | 1462.04489 | 1647.4358 | 1502.3352 | 1562.2872 | 5.10E-38 | -1.115557714 | YNL167C | SKO1 | n |
| W303.asm02.HP0_G0045170 | 998.75715 | 1028.70581 | 851.9458841 | 928.381826 | 916.83977 | 840.059516 | 757.02238 | 802.9949 | 792.60507 | 0.285940397 | 0.099424703 | YNL166C | BNI5 | n |
| W303.asm02.HP0_G0045180 | 539.41315 | 567.812388 | 467.5077187 | 514.193525 | 519.54254 | 445.629766 | 600.82951 | 493.34327 | 520.17463 | 0.451296667 | 0.088460231 | YNL165W | | n |
| W303.asm02.HP0_G0045190 | 1198.93 | 1117.97796 | 1214.167774 | 639.200055 | 604.93448 | 659.911313 | 550.66538 | 568.13201 | 599.5233 | 6.34E-29 | 0.891460684 | YNL164C | IBD2 | n |
| W303.asm02.HP0_G0045200 | 729.05058 | 808.63958 | 917.6287867 | 2140.11179 | 2219.2916 | 2381.74888 | 1851.5126 | 1813.2989 | 1792.3983 | 3.67E-63 | -1.455757976 | YNL163C | RIA1 | n |
| W303.asm02.HP0_G0045210 | 544.68085 | 548.089471 | 711.8867535 | 185.009664 | 184.26682 | 195.318578 | 114.0094 | 115.46332 | 96.98171 | 2.42E-35 | 1.677984302 | YNL162W-A | | n |
| W303.asm02.HP0_G0045230 | 917.63447 | 1086.83651 | 978.4820641 | 2439.29408 | 2465.5799 | 2450.01556 | 2422.6996 | 2496.8943 | 2464.2171 | 1.57E-74 | -1.301942062 | YNL161W | CBK1 | n |
| W303.asm02.HP0_G0045250 | 671.10581 | 700.682563 | 591.1461236 | 292.515279 | 280.44511 | 309.096391 | 346.58856 | 304.4033 | 320.9213 | 5.61E-28 | 1.153837711 | YNL159C | ASI2 | n |
| W303.asm02.HP0_G0045260 | 331.86551 | 311.414472 | 288.8115865 | 290.015149 | 295.72577 | 284.444531 | 329.48715 | 355.57454 | 319.15799 | 0.386800008 | 0.097836643 | YNL158W | PGA1 | n |
| W303.asm02.HP0_G0045280 | 3191.1766 | 2990.61698 | 3236.814804 | 1844.263 | 1881.3193 | 1752.17831 | 2268.787 | 2179.3701 | 2268.4904 | 1.45E-35 | 0.781799463 | YNL156C | NSG2 | n |
| W303.asm02.HP0_G0045290 | 2501.107 | 2548.40843 | 2596.406504 | 603.364849 | 557.29476 | 589.748328 | 713.69881 | 646.857 | 700.03162 | 9.93E-193 | 2.126756737 | YNL155W | CUZ1 | n |
| W303.asm02.HP0_G0045300 | 1989.086 | 1959.83508 | 2156.910611 | 3794.36486 | 3734.7738 | 4249.6013 | 3688.2039 | 3511.1346 | 3529.2526 | 2.95E-39 | -0.94742394 | YNL154C | YCK2 | n |
| W303.asm02.HP0_G0045310 | 2371.5215 | 2315.88563 | 2750.95451 | 1025.88692 | 1102.9043 | 1078.04477 | 1058.0072 | 1083.7807 | 1072.0887 | 6.98E-51 | 1.214493516 | YNL153C | GIM3 | n |
| W303.asm02.HP0_G0045320 | 291.83094 | 359.164692 | 347.7330139 | 535.027947 | 462.01534 | 477.866813 | 550.66538 | 540.57827 | 531.6361 | 1.73E-07 | -0.562557224 | YNL152W | INN1 | n |
| W303.asm02.HP0_G0045330 | 1695.148 | 1703.43716 | 1793.722797 | 1213.39671 | 1202.6781 | 1280.94854 | 777.54407 | 985.37446 | 852.5574 | 1.08E-08 | 0.490391923 | YNL151C | RPC31 | n |
| W303.asm02.HP0_G0045340 | 3068.9658 | 3067.43255 | 3628.98037 | 1270.06634 | 1404.9221 | 1319.82263 | 1126.4128 | 1364.5665 | 1147.0291 | 8.85E-42 | 1.289907515 | YNL149C | PGA2 | n |
| W303.asm02.HP0_G0045350 | 740.63953 | 714.17719 | 727.3415541 | 730.87151 | 761.33656 | 772.740977 | 715.979 | 784.62574 | 764.39221 | 0.514604747 | -0.053269347 | YNL148C | ALF1 | n |
| W303.asm02.HP0_G0045400 | 946.08009 | 995.488264 | 935.9813624 | 204.177332 | 231.00767 | 256.000078 | 245.1202 | 240.11122 | 229.2295 | 2.02E-90 | 2.060635568 | YNL142W | MEP2 | n |
| W303.asm02.HP0_G0045410 | 1229.4827 | 1068.15164 | 1163.939672 | 730.87151 | 728.97751 | 917.807688 | 323.78668 | 503.83994 | 336.79103 | 0.00018592 | 0.542406451 | YNL141W | AAH1 | n |

| | | | | | | | | | | | | | | |
|---|---|---|---|---|---|---|---|---|---|---|---|---|---|---|
| W303.asm02.HP0_G0045420 | 996.65007 | 992.374119 | 982.3457643 | 1999.2711 | 2002.6657 | 2087.82286 | 1788.8074 | 1668.9698 | 1842.6525 | 1.91E-53 | -1.035158984 | YNL139C | THO2 | n |
| W303.asm02.HP0_G0045440 | 1432.8162 | 1700.32302 | 1782.131696 | 4491.9013 | 4440.3809 | 4507.49767 | 4545.5546 | 4252.4616 | 4415.3128 | 1.80E-74 | -1.450647072 | YNL138W | SRV2 | n |
| W303.asm02.HP0_G0045450 | 4445.9443 | 4366.0309 | 4265.524971 | 3518.51712 | 3534.3274 | 3618.13444 | 3235.5866 | 3478.3325 | 3249.7689 | 1.62E-07 | 0.293395668 | YNL137C | NAM9 | n |
| W303.asm02.HP0_G0045460 | 852.31491 | 837.704931 | 641.3742256 | 657.534346 | 620.21514 | 626.726117 | 589.42857 | 615.36701 | 615.39303 | 0.008149042 | 0.290461763 | YNL136W | EAF7 | n |
| W303.asm02.HP0_G0045470 | 5382.5425 | 4931.7672 | 5675.775527 | 2770.97807 | 2852.0908 | 2917.45274 | 2510.4869 | 2646.4718 | 2368.9987 | 1.53E-36 | 0.905030829 | YNL135C | FPR1 | n |
| W303.asm02.HP0_G0045480 | 8169.1592 | 9069.42749 | 9325.040323 | 250.013059 | 219.32246 | 244.622297 | 679.49599 | 450.04453 | 481.38194 | 0 | 5.217193261 | YNL134C | | n |
| W303.asm02.HP0_G0045490 | 426.68423 | 482.692432 | 515.8039706 | 822.542965 | 853.9194 | 905.481758 | 815.16718 | 776.75324 | 730.88944 | 1.08E-19 | -0.855625116 | YNL133C | FYV6 | n |
| W303.asm02.HP0_G0045500 | 851.26137 | 999.640457 | 1149.450796 | 3367.67591 | 3166.6927 | 3552.7122 | 1876.5946 | 2422.1055 | 2003.9948 | 2.57E-52 | -1.748617186 | YNL132W | KRE33 | n |
| W303.asm02.HP0_G0045510 | 4519.6922 | 4672.25513 | 5295.201062 | 2430.12694 | 2513.2196 | 2475.61557 | 2085.2318 | 2342.0685 | 2219.1179 | 3.88E-37 | 0.96573791 | YNL131W | TOM22 | n |
| W303.asm02.HP0_G0045530 | 925.00926 | 892.721488 | 1079.904193 | 349.184906 | 317.29847 | 360.296406 | 304.40508 | 295.21871 | 280.36531 | 4.89E-47 | 1.497610647 | YNL129W | NRK1 | n |
| W303.asm02.HP0_G0045540 | 121.15725 | 95.5004382 | 111.0813794 | 235.012276 | 282.24283 | 329.955656 | 243.98011 | 250.60789 | 235.40106 | 3.49E-18 | -1.367977651 | YNL128W | TEP1 | n |
| W303.asm02.HP0_G0045550 | 503.59274 | 534.594844 | 506.1447202 | 991.718468 | 985.15332 | 859.970633 | 950.83836 | 919.7703 | 908.10147 | 3.65E-24 | -0.878254576 | YNL127W | FAR11 | n |
| W303.asm02.HP0_G0045560 | 653.19561 | 567.812388 | 642.3401507 | 489.192219 | 561.78908 | 549.926094 | 482.25974 | 446.10828 | 507.8315 | 0.028909265 | 0.220107004 | YNL126W | SPC98 | n |
| W303.asm02.HP0_G0045570 | 483.57546 | 587.535305 | 557.3387473 | 536.6947 | 527.6323 | 586.903883 | 521.02294 | 583.87701 | 493.72507 | 0.864711927 | -0.019075019 | YNL125C | ESBP6 | n |
| W303.asm02.HP0_G0045580 | 645.82082 | 822.134207 | 855.8095842 | 704.20345 | 675.94462 | 739.555782 | 424.11495 | 450.04453 | 408.20484 | 0.233162064 | 0.133556726 | YNL124W | NAF1 | n |
| W303.asm02.HP0_G0045590 | 2045.9772 | 1995.12872 | 2036.169981 | 3005.15697 | 3101.0757 | 3311.88249 | 2429.5402 | 2603.173 | 2353.129 | 2.55E-21 | -0.63171952 | YNL123W | NMA111 | n |
| W303.asm02.HP0_G0045600 | 1280.0527 | 1154.30964 | 1469.171984 | 971.717423 | 1102.9043 | 934.87436 | 926.89638 | 1117.8949 | 981.27858 | 0.000568393 | 0.375619099 | YNL122C | MRP35 | n |
| W303.asm02.HP0_G0045610 | 4290.0202 | 4339.04165 | 3834.722403 | 5318.61115 | 5146.8868 | 5493.57205 | 4473.7287 | 4480.764 | 4194.0181 | 1.40E-07 | -0.356804119 | YNL121C | TOM70 | n |
| W303.asm02.HP0_G0045620 | 2338.8617 | 2156.0262 | 2696.862708 | 1094.22382 | 1066.9498 | 1175.70406 | 669.23515 | 809.55532 | 692.09675 | 5.47E-29 | 1.108355015 | YNL119W | NCS2 | n |
| W303.asm02.HP0_G0045630 | 1984.8718 | 1932.84583 | 2035.204056 | 2637.63777 | 2588.7241 | 2663.34896 | 2801.2108 | 2635.9751 | 2739.2925 | 3.24E-12 | -0.406159288 | YNL118C | DCP2 | n |
| W303.asm02.HP0_G0045640 | 92.711634 | 74.7394734 | 89.83102859 | 455.857145 | 495.27325 | 453.214953 | 434.3758 | 350.32621 | 520.17463 | 7.29E-53 | -2.447914309 | YNL117W | MLS1 | n |
| W303.asm02.HP0_G0045660 | 614.21458 | 628.019186 | 747.6259799 | 547.5286 | 555.49704 | 594.489071 | 1132.1133 | 850.22989 | 1097.6566 | 0.057997746 | 0.230381201 | YNL115C | | n |
| W303.asm02.HP0_G0045670 | 3284.9418 | 3152.55251 | 3652.162571 | 1527.57979 | 1528.0663 | 1546.4301 | 1061.4275 | 1316.0194 | 1002.4382 | 1.69E-31 | 1.132694896 | YNL113W | RPC19 | n |
| W303.asm02.HP0_G0045680 | 3599.9506 | 3061.20426 | 4138.98879 | 3887.70307 | 3519.9456 | 3389.63067 | 2073.8309 | 3030.9121 | 2240.2775 | 0.997759008 | 0.000410782 | YNL112W | DBP2 | n |
| W303.asm02.HP0_G0045690 | 6245.3928 | 6214.79482 | 7093.753484 | 4286.05721 | 4385.5502 | 4468.62359 | 3690.4841 | 4535.8715 | 3584.7967 | 5.52E-10 | 0.573599067 | YNL111C | CYB5 | n |
| W303.asm02.HP0_G0045700 | 3946.5657 | 3704.79417 | 4414.277426 | 2823.48081 | 2430.5243 | 2943.05275 | 1644.0155 | 1872.3427 | 1729.8011 | 2.05E-09 | 0.557916899 | YNL110C | NOP15 | n |
| W303.asm02.HP0_G0045720 | 1230.5362 | 1283.02763 | 1252.804775 | 566.696267 | 573.47429 | 569.837211 | 558.64604 | 606.18242 | 527.22784 | 7.65E-48 | 1.139252238 | YNL107W | YAF9 | n |
| W303.asm02.HP0_G0045740 | 5426.7912 | 6121.37048 | 6481.357009 | 1973.43641 | 1839.9716 | 1833.71908 | 1809.3291 | 1482.654 | 1706.8781 | 6.62E-79 | 1.674680233 | YNL104C | LEU4 | n |
| W303.asm02.HP0_G0045750 | 916.58093 | 1069.18969 | 1098.256769 | 686.702536 | 639.09125 | 712.059477 | 658.9743 | 545.8266 | 660.35728 | 4.72E-09 | 0.598350855 | YNL103W | MET4 | n |
| W303.asm02.HP0_G0045760 | 683.7483 | 769.193747 | 739.8985796 | 1860.09716 | 1805.8148 | 1804.32648 | 1803.6286 | 1768.6881 | 1836.4809 | 8.30E-80 | -1.31864877 | YNL102W | POL1 | n |
| W303.asm02.HP0_G0045770 | 1209.4654 | 1178.18475 | 1185.190022 | 394.187257 | 355.0507 | 323.318617 | 636.17242 | 656.04158 | 633.02607 | 2.38E-87 | 1.733513961 | YNL101W | AVT4 | n |
| W303.asm02.HP0_G0045780 | 2780.2955 | 2711.38201 | 3217.496304 | 2105.94334 | 2174.3484 | 2194.96363 | 2248.2653 | 2405.0484 | 2325.7977 | 2.52E-08 | 0.428070264 | YNL100W | MIC27 | n |
| W303.asm02.HP0_G0045790 | 589.98313 | 551.203616 | 514.8380456 | 954.216509 | 988.74877 | 912.118797 | 1026.0846 | 986.68654 | 1072.9704 | 6.65E-21 | -0.787009318 | YNL099C | OCA1 | n |
| W303.asm02.HP0_G0045810 | 536.25252 | 541.861182 | 560.2365224 | 746.70567 | 711.00026 | 769.896532 | 845.94971 | 829.23656 | 879.88861 | 4.68E-08 | -0.442649091 | YNL097C | PHO23 | n |
| W303.asm02.HP0_G0045840 | 772.24577 | 841.857124 | 835.5251584 | 272.514234 | 289.43373 | 316.681578 | 298.70462 | 288.6583 | 305.05156 | 4.01E-53 | 1.481364769 | YNL094W | APP1 | n |
| W303.asm02.HP0_G0045860 | 123.26433 | 137.022368 | 115.9110046 | 311.682947 | 356.84842 | 336.592695 | 378.51119 | 354.26246 | 335.02773 | 4.99E-28 | -1.418439972 | YNL092W | | n |
| W303.asm02.HP0_G0045870 | 864.95741 | 1050.50482 | 930.1858122 | 1671.75399 | 1673.682 | 1725.63016 | 1498.0835 | 1457.7244 | 1558.7606 | 3.13E-25 | -0.833422222 | YNL091W | NST1 | n |
| W303.asm02.HP0_G0045880 | 1175.7521 | 1020.40142 | 1193.883348 | 1124.22539 | 1063.3544 | 1026.84476 | 999.8624 | 980.12613 | 1009.4914 | 0.366831941 | 0.076546172 | YNL090W | RHO2 | n |
| W303.asm02.HP0_G0045890 | 1035.6311 | 1068.15164 | 1058.653842 | 2647.6383 | 2618.3865 | 2626.37117 | 2223.1832 | 2058.6585 | 2301.1115 | 4.74E-89 | -1.319382411 | YNL088W | TOP2 | n |
| W303.asm02.HP0_G0045910 | 412.98819 | 395.49638 | 423.0751669 | 110.005746 | 125.84075 | 126.103742 | 161.89334 | 132.5204 | 134.89274 | 4.98E-42 | 1.769115334 | YNL086W | SNN1 | n |
| W303.asm02.HP0_G0045920 | 2365.2002 | 2191.31984 | 2367.48227 | 3090.99479 | 3176.5802 | 3172.50467 | 3278.9102 | 3156.8721 | 3153.6689 | 3.84E-14 | -0.446970201 | YNL085W | MKT1 | n |
| W303.asm02.HP0_G0045930 | 3568.3444 | 3513.7933 | 4200.807993 | 2061.77436 | 2115.9224 | 2095.40805 | 2035.0677 | 2197.7393 | 2069.237 | 7.93E-27 | 0.847216298 | YNL084C | END3 | n |
| W303.asm02.HP0_G0045940 | 1197.8765 | 1321.43541 | 1205.474448 | 955.883263 | 973.46811 | 994.607711 | 825.42802 | 959.1328 | 839.33262 | 2.07E-05 | 0.349340076 | YNL082W | PMS1 | n |
| W303.asm02.HP0_G0045950 | 3090.0366 | 3064.31841 | 3619.32112 | 879.212591 | 944.70451 | 888.415086 | 810.6068 | 877.78364 | 853.43905 | 1.75E-108 | 1.849745371 | YNL081C | SWS2 | n |
| W303.asm02.HP0_G0045960 | 532.03836 | 522.138265 | 466.5417937 | 1732.5905 | 1604.4696 | 1679.17088 | 1644.0155 | 1590.2448 | 1528.7844 | 1.15E-104 | -1.723213134 | YNL080C | EOS1 | n |
| W303.asm02.HP0_G0045980 | 889.18886 | 1004.8307 | 1058.653842 | 553.362238 | 587.85609 | 511.052008 | 535.84416 | 598.30993 | 548.38749 | 2.97E-18 | 0.837630113 | YNL078W | NIS1 | n |
| W303.asm02.HP0_G0045990 | 425.63069 | 746.356686 | 406.6544413 | 1025.88692 | 1031.8942 | 1415.58562 | 1468.441 | 1222.8615 | 1337.466 | 3.89E-05 | -1.137733418 | YNL077W | APJ1 | n |
| W303.asm02.HP0_G0046000 | 848.10075 | 774.383988 | 795.9222318 | 939.215726 | 903.35684 | 828.681735 | 900.67422 | 833.17281 | 857.84731 | 0.085597258 | -0.144481052 | YNL076W | MKS1 | n |
| W303.asm02.HP0_G0046010 | 1906.9098 | 1754.30153 | 1984.010029 | 1334.23636 | 1387.8437 | 1303.7041 | 836.82896 | 1078.5324 | 870.19044 | 3.09E-07 | 0.487854642 | YNL075W | IMP4 | n |
| W303.asm02.HP0_G0046030 | 955.56196 | 940.471707 | 830.6955332 | 502.526249 | 515.04823 | 498.726078 | 555.22575 | 547.13868 | 535.16271 | 1.30E-22 | 0.845822255 | YNL073W | MSK1 | n |
| W303.asm02.HP0_G0046040 | 208.60118 | 276.120832 | 242.4471847 | 492.525727 | 460.21761 | 456.059399 | 484.53993 | 543.20243 | 517.52967 | 6.33E-17 | -0.954288027 | YNL072W | RNH201 | n |
| W303.asm02.HP0_G0046050 | 2475.8221 | 2855.67071 | 2942.207668 | 6740.35208 | 7010.2288 | 7235.32073 | 6700.3322 | 5678.696 | 6484.5498 | 7.55E-57 | -1.342544259 | YNL071W | LAT1 | n |
| W303.asm02.HP0_G0046060 | 1883.7318 | 1852.91611 | 2260.26459 | 1295.90102 | 1453.4607 | 1205.09666 | 1268.9246 | 1411.8015 | 1209.6264 | 4.35E-09 | 0.600930224 | YNL070W | TOM7 | n |
| W303.asm02.HP0_G0046080 | 651.08852 | 693.416225 | 773.7059559 | 1340.90337 | 1347.3949 | 1437.39303 | 1340.7505 | 1277.969 | 1423.8678 | 9.79E-30 | -0.960493446 | YNL068C | FKH2 | n |
| W303.asm02.HP0_G0046090 | 9870.6284 | 9505.40775 | 11303.2548 | 14365.7504 | 14747.637 | 15915.6197 | 12299.334 | 13922.515 | 11313.357 | 1.21E-09 | -0.553492096 | YNL067W | RPL9B | y |
| W303.asm02.HP0_G0046100 | 2280.9169 | 2438.37532 | 2716.181209 | 8562.94728 | 8152.6831 | 9242.55097 | 9124.1719 | 8989.0818 | 8306.0427 | 2.85E-112 | -1.803306929 | YNL066W | SUN4 | n |
| W303.asm02.HP0_G0046110 | 1800.5021 | 1571.60504 | 2143.387661 | 526.694178 | 488.08235 | 534.755719 | 536.98425 | 469.72577 | 503.42324 | 4.20E-60 | 1.832083733 | YNL065W | AQR1 | n |
| W303.asm02.HP0_G0046120 | 7275.7562 | 7707.50819 | 7319.779943 | 7005.36592 | 6745.9632 | 7435.38005 | 6020.8362 | 5792.8472 | 5125.9242 | 0.328322362 | 0.074124099 | YNL064C | YDJ1 | n |
| W303.asm02.HP0_G0046130 | 444.59443 | 498.263156 | 382.5063153 | 236.679029 | 222.91791 | 231.348219 | 132.2509 | 162.69831 | 145.47257 | 2.32E-13 | 0.938475461 | YNL063W | MTQ1 | n |
| W303.asm02.HP0_G0046140 | 1418.0666 | 1368.14758 | 1340.703954 | 1350.07052 | 1304.2495 | 1383.34857 | 752.46201 | 848.91781 | 734.41604 | 0.6908002 | 0.031444019 | YNL062C | GCD10 | n |

| | | | | | | | | | | | | | | |
|---|---|---|---|---|---|---|---|---|---|---|---|---|---|---|
| W303.asm02.HP0_G0046150 | 1863.7146 | 2045.99308 | 1930.884152 | 4566.07184 | 4332.5174 | 4745.48293 | 2551.5303 | 3353.6846 | 2641.4291 | 1.02E-35 | -1.224055718 | YNL061W | NOP2 | n |
| W303.asm02.HP0_G0046160 | 956.6155 | 997.56436 | 1120.473045 | 1655.91983 | 1731.2092 | 1584.35604 | 1780.8268 | 1587.6206 | 1690.1267 | 4.26E-17 | -0.692659532 | YNL059C | ARP5 | n |
| W303.asm02.HP0_G0046170 | 1109.379 | 1207.2501 | 1258.600325 | 1400.90651 | 1301.5529 | 1246.8152 | 1427.3976 | 1436.7311 | 1347.1641 | 0.073339149 | -0.143669299 | YNL058C | | n |
| W303.asm02.HP0_G0046180 | 1156.7883 | 1172.99451 | 1130.132295 | 370.852704 | 372.12908 | 384.000117 | 379.65129 | 360.82287 | 405.55988 | 3.62E-84 | 1.618555632 | YNL056W | OCA2 | n |
| W303.asm02.HP0_G0046190 | 9138.4172 | 9066.31334 | 8297.296082 | 5486.11989 | 5456.9944 | 5673.72025 | 5573.9193 | 5529.1185 | 5698.998 | 1.96E-31 | 0.673408936 | YNL055C | POR1 | n |
| W303.asm02.HP0_G0046210 | 355.04342 | 419.37149 | 395.0633408 | 631.699663 | 665.15827 | 669.392797 | 588.28848 | 522.2091 | 610.98478 | 3.89E-14 | -0.748798983 | YNL054W | VAC7 | n |
| W303.asm02.HP0_G0046220 | 448.80859 | 418.333441 | 495.5195448 | 2014.27188 | 1965.8123 | 2054.63766 | 1811.6093 | 1844.7889 | 1840.0075 | 2.77E-173 | -2.145365744 | YNL053W | MSG5 | n |
| W303.asm02.HP0_G0046230 | 4961.126 | 4616.20053 | 5134.857505 | 6018.64771 | 5714.9679 | 6373.4538 | 5576.1995 | 5304.7523 | 5106.5279 | 1.57E-05 | -0.299377847 | YNL052W | COX5A | n |
| W303.asm02.HP0_G0046240 | 1195.7694 | 1229.04912 | 1262.464025 | 880.045968 | 939.31134 | 877.037305 | 868.75159 | 886.96822 | 901.92991 | 2.06E-10 | 0.451827515 | YNL051W | COG5 | n |
| W303.asm02.HP0_G0046250 | 1854.2327 | 1825.92686 | 1874.8605 | 735.871771 | 764.03314 | 876.089157 | 607.67008 | 733.45449 | 652.42242 | 3.20E-43 | 1.226317499 | YNL050C | | n |
| W303.asm02.HP0_G0046270 | 1144.1459 | 1103.44528 | 1078.938268 | 1264.2327 | 1288.07 | 1227.85223 | 1547.1075 | 1525.9527 | 1584.3285 | 0.005868993 | -0.184944968 | YNL048W | ALG11 | n |
| W303.asm02.HP0_G0046280 | 438.27318 | 496.187059 | 531.2587712 | 2420.95979 | 2390.9743 | 2370.37109 | 2830.8533 | 2748.8142 | 2865.3687 | 5.12E-200 | -2.291343756 | YNL047C | SLM2 | n |
| W303.asm02.HP0_G0046300 | 702.71205 | 804.487387 | 900.242136 | 2689.30714 | 2566.2525 | 2595.08227 | 2935.7419 | 3083.3954 | 2942.9541 | 4.22E-91 | -1.704231555 | YNL045W | LAP2 | n |
| W303.asm02.HP0_G0046330 | 209.65472 | 237.713047 | 219.2649838 | 691.702797 | 700.21391 | 702.577992 | 737.64079 | 801.68282 | 774.97203 | 7.72E-67 | -1.65154035 | YNL042W | BOP3 | n |
| W303.asm02.HP0_G0046340 | 1620.3465 | 1484.40899 | 1530.991186 | 1025.05354 | 999.53513 | 1022.10402 | 993.02183 | 867.28697 | 986.56849 | 2.00E-15 | 0.605413015 | YNL041C | COG6 | n |
| W303.asm02.HP0_G0046350 | 1722.54 | 1641.15427 | 1742.52877 | 997.552106 | 952.79427 | 980.385485 | 965.65958 | 922.39447 | 1019.1896 | 3.06E-30 | 0.800959832 | YNL040W | | n |
| W303.asm02.HP0_G0046360 | 639.49957 | 667.465019 | 694.5001028 | 1020.88666 | 992.34423 | 997.452157 | 912.07516 | 889.59239 | 941.60424 | 4.26E-15 | -0.588615817 | YNL039W | BDP1 | n |
| W303.asm02.HP0_G0046370 | 1188.3946 | 1077.49407 | 1440.194233 | 103.338731 | 109.66123 | 138.429672 | 137.95137 | 143.01707 | 139.301 | 4.39E-130 | 3.402974928 | YNL038W | GPI15 | n |
| W303.asm02.HP0_G0046380 | 4789.3987 | 4233.16073 | 4011.486685 | 7042.0345 | 7301.4603 | 7884.80241 | 6489.4148 | 6553.8554 | 6403.4378 | 4.00E-24 | -0.770199299 | YNL037C | IDH1 | n |
| W303.asm02.HP0_G0046390 | 9665.1879 | 9992.25237 | 9182.083417 | 2238.45026 | 2231.8756 | 2249.95624 | 4348.3183 | 4563.4253 | 4378.2834 | 2.64E-302 | 2.101399216 | YNL036W | NCE103 | n |
| W303.asm02.HP0_G0046400 | 1007.1855 | 904.140018 | 911.8332365 | 995.885352 | 939.31134 | 1143.46702 | 508.4819 | 615.36701 | 556.32236 | 0.22369404 | -0.124721504 | YNL035C | | n |
| W303.asm02.HP0_G0046430 | 1851.0721 | 1623.50745 | 1924.122677 | 678.368767 | 658.86623 | 676.977985 | 798.06577 | 753.13574 | 782.9069 | 6.06E-66 | 1.42262796 | YNL032W | SIW14 | n |
| W303.asm02.HP0_G0046460 | 269.70657 | 293.767652 | 307.1641623 | 745.038916 | 766.72973 | 644.740938 | 822.00774 | 717.70949 | 828.7528 | 3.43E-34 | -1.308308569 | YNL029C | KTR5 | n |
| W303.asm02.HP0_G0046470 | 674.26643 | 707.948901 | 574.725398 | 957.550017 | 893.46935 | 790.755797 | 802.62614 | 606.18242 | 767.03716 | 0.000369994 | -0.434129069 | YNL027W | CRZ1 | n |
| W303.asm02.HP0_G0046480 | 1520.2601 | 1408.63146 | 1404.455006 | 1278.40011 | 1177.5099 | 1261.98557 | 1093.3501 | 1166.4419 | 1176.1237 | 0.002655114 | 0.220617978 | YNL026W | SAM50 | n |
| W303.asm02.HP0_G0046490 | 364.52529 | 323.871051 | 262.7316105 | 382.519981 | 361.34273 | 326.163063 | 352.28903 | 279.47372 | 416.1397 | 0.277931249 | -0.171992144 | YNL025C | SSN8 | n |
| W303.asm02.HP0_G0046500 | 1872.1429 | 1705.51326 | 2229.354989 | 460.024029 | 540.21638 | 434.251985 | 411.57392 | 505.15202 | 485.7902 | 1.31E-63 | 2.017372587 | YNL024C- | KSH1 | n |
| W303.asm02.HP0_G0046510 | 456.18338 | 417.295393 | 452.0529181 | 552.528861 | 522.23913 | 437.09643 | 307.82537 | 411.99412 | 360.59563 | 0.128174847 | -0.190824048 | YNL024C | EFM6 | n |
| W303.asm02.HP0_G0046520 | 473.04004 | 469.197805 | 561.2024474 | 987.551584 | 1047.1748 | 953.837328 | 806.04642 | 855.47823 | 783.78855 | 1.09E-26 | -0.989933103 | YNL023C | FAP1 | n |
| W303.asm02.HP0_G0046530 | 1247.3929 | 1255.00032 | 1132.064145 | 1031.72056 | 977.06356 | 1134.93368 | 588.28848 | 753.13574 | 569.54714 | 0.0539106 | 0.209301311 | YNL022C | RCM1 | n |
| W303.asm02.HP0_G0046540 | 743.80016 | 672.65526 | 735.0689544 | 1503.41186 | 1470.5391 | 1501.86713 | 1124.1326 | 1331.7644 | 1156.7273 | 2.20E-36 | -1.056650241 | YNL021W | HDA1 | n |
| W303.asm02.HP0_G0046580 | 4602.9219 | 4274.68266 | 4258.763495 | 6176.98932 | 6078.1084 | 6776.41689 | 5291.176 | 4921.624 | 5393.9464 | 7.13E-15 | -0.534796879 | YNL016W | PUB1 | n |
| W303.asm02.HP0_G0046590 | 2972.04 | 3255.31929 | 3677.276622 | 476.691566 | 502.46415 | 411.496422 | 628.19177 | 591.74951 | 663.00224 | 2.31E-172 | 2.831625529 | YNL015W | PBI2 | n |
| W303.asm02.HP0_G0046620 | 746.96078 | 806.563484 | 777.5696561 | 598.364588 | 540.21638 | 548.977946 | 558.64604 | 539.26618 | 590.70678 | 6.38E-08 | 0.465280784 | YNL011C | | n |
| W303.asm02.HP0_G0046630 | 4122.5071 | 4023.47498 | 4449.050728 | 3878.53592 | 3854.3225 | 4145.30497 | 3852.3775 | 3520.3191 | 3798.1564 | 0.219379262 | 0.084847394 | YNL010W | PYP1 | n |
| W303.asm02.HP0_G0046660 | 4263.6816 | 4969.13693 | 4061.714787 | 2455.12824 | 2415.2436 | 2816.00086 | 3060.0122 | 2423.4176 | 2699.6182 | 2.53E-14 | 0.790566661 | YNL007C | SIS1 | n |
| W303.asm02.HP0_G0046670 | 2010.1568 | 2670.89813 | 1735.767294 | 765.039961 | 779.31381 | 780.326164 | 835.68887 | 805.61907 | 840.21427 | 4.47E-31 | 1.46463019 | YNL006W | LST8 | n |
| W303.asm02.HP0_G0046680 | 2918.3094 | 2980.2365 | 2933.514342 | 2559.30035 | 2439.5129 | 2525.86744 | 2198.1011 | 2247.5985 | 2300.2298 | 3.77E-05 | 0.231022628 | YNL005C | MRP7 | n |
| W303.asm02.HP0_G0046700 | 912.36677 | 866.770282 | 966.8909637 | 1709.25595 | 1628.7389 | 1587.20048 | 1674.798 | 1654.5369 | 1666.3221 | 5.44E-33 | -0.842654349 | YNL003C | PET8 | n |
| W303.asm02.HP0_G0046710 | 5060.1589 | 4822.77213 | 5369.57729 | 2297.62001 | 2068.2827 | 2366.5785 | 1150.3548 | 1503.6473 | 1275.7503 | 8.57E-36 | 1.179935923 | YNL002C | RLP7 | n |
| W303.asm02.HP0_G0046720 | 1801.5556 | 1573.68113 | 1916.395277 | 1630.08515 | 1723.1195 | 1651.67458 | 1467.3009 | 1654.5369 | 1594.0267 | 0.359350324 | 0.080751228 | YNL001W | DOM34 | n |
| W303.asm02.HP0_G0046730 | 4271.0564 | 4221.7422 | 4208.535393 | 8929.6331 | 9857.8253 | 9680.59555 | 7606.7068 | 8760.7793 | 7612.1826 | 5.90E-59 | -1.164312651 | YNR001C | CIT1 | n |
| W303.asm02.HP0_G0046750 | 911.31322 | 851.199558 | 774.671881 | 763.373207 | 811.67286 | 815.407657 | 597.40923 | 743.95116 | 634.78938 | 0.400262171 | 0.085688006 | YNR003C | RPC34 | n |
| W303.asm02.HP0_G0046770 | 475.14713 | 567.812388 | 520.6335958 | 1719.25647 | 1688.9627 | 1630.81531 | 1629.1943 | 1575.8119 | 1648.6891 | 7.80E-102 | -1.6882848 | YNR006W | VPS27 | n |
| W303.asm02.HP0_G0046780 | 919.74155 | 866.770282 | 900.242136 | 496.692611 | 440.44264 | 478.814961 | 655.55402 | 586.50118 | 698.26831 | 8.40E-24 | 0.923642605 | YNR007C | ATG3 | n |
| W303.asm02.HP0_G0046790 | 621.58937 | 636.323572 | 680.9771522 | 2120.11074 | 2107.8326 | 2178.84511 | 2788.6698 | 2549.3776 | 2732.2393 | 1.26E-128 | -1.723469391 | YNR008W | LRO1 | n |
| W303.asm02.HP0_G0046810 | 488.84316 | 482.692432 | 471.3714188 | 383.353357 | 381.11771 | 353.659367 | 403.59326 | 453.98078 | 432.89109 | 0.000131242 | 0.366755335 | YNR010W | CSE2 | n |
| W303.asm02.HP0_G0046820 | 439.32672 | 465.045612 | 473.3032689 | 694.202928 | 689.42755 | 679.82243 | 526.72341 | 523.52118 | 570.42879 | 1.35E-11 | -0.582463918 | YNR011C | PRP2 | n |
| W303.asm02.HP0_G0046830 | 1448.6193 | 1207.2501 | 1394.795756 | 902.547144 | 1008.5238 | 1023.05216 | 721.67947 | 806.93115 | 680.63528 | 1.62E-06 | 0.465839551 | YNR012W | URK1 | n |
| W303.asm02.HP0_G0046840 | 947.13363 | 1104.48333 | 1049.960517 | 1477.57718 | 1493.0107 | 1411.79302 | 2025.947 | 1901.2085 | 2085.1068 | 1.63E-10 | -0.498653811 | YNR013C | PHO91 | n |
| W303.asm02.HP0_G0046860 | 558.37689 | 609.334318 | 616.2601746 | 1769.25908 | 1744.6922 | 1810.01537 | 1100.1907 | 1531.2011 | 1217.5613 | 3.32E-47 | -1.577050881 | YNR015W | SMM1 | n |
| W303.asm02.HP0_G0046870 | 5898.7777 | 6097.49537 | 7067.673508 | 11969.7919 | 11841.615 | 12002.6111 | 14006.054 | 13852.974 | 14408.837 | 3.34E-37 | -0.90948598 | YNR016C | ACC1 | n |
| W303.asm02.HP0_G0046880 | 1984.8718 | 2252.56468 | 2478.563649 | 2861.81615 | 2674.116 | 2842.54902 | 2819.4523 | 2662.2168 | 2622.0328 | 0.00015737 | -0.318705794 | YNR017W | TIM23 | n |
| W303.asm02.HP0_G0046890 | 1068.2909 | 1195.83157 | 1243.145525 | 1858.43041 | 1853.4545 | 1953.18578 | 2489.9652 | 2847.2205 | 2600.8731 | 9.37E-18 | -0.691079291 | YNR018W | RCF2 | n |
| W303.asm02.HP0_G0046900 | 1569.7765 | 1704.47521 | 1595.708164 | 1500.07835 | 1465.1459 | 1400.41524 | 2411.2987 | 2113.766 | 2301.9931 | 0.036746872 | 0.157329972 | YNR019W | ARE2 | n |
| W303.asm02.HP0_G0046920 | 778.56702 | 739.090348 | 895.4125108 | 1195.8958 | 1320.429 | 1210.78556 | 1053.4468 | 1152.009 | 1033.296 | 6.73E-12 | -0.626255183 | YNR020C | ATP23 | n |
| W303.asm02.HP0_G0046930 | 2860.3646 | 2696.84933 | 2262.19644 | 2921.81928 | 2966.2463 | 3066.31205 | 3651.7209 | 3504.5741 | 3575.0985 | 0.021356051 | -0.195851598 | YNR021W | | n |
| W303.asm02.HP0_G0046940 | 2060.7268 | 2239.07006 | 2316.288243 | 726.704625 | 785.60584 | 725.333555 | 631.61205 | 725.58199 | 661.23893 | 4.40E-83 | 1.564280285 | YNR022C | MRPL50 | n |
| W303.asm02.HP0_G0046950 | 536.25252 | 558.469954 | 504.2128702 | 1518.41265 | 1449.8652 | 1415.58562 | 1461.6004 | 1443.2915 | 1478.5303 | 1.13E-84 | -1.456286981 | YNR023W | SNF12 | n |
| W303.asm02.HP0_G0046960 | 911.31322 | 911.406356 | 1095.358994 | 490.858973 | 513.2505 | 454.163102 | 290.72396 | 478.91036 | 370.2938 | 1.86E-12 | 1.000858108 | YNR024W | MPP6 | n |

| | | | | | | | | | | | | | | |
|---|---|---|---|---|---|---|---|---|---|---|---|---|---|---|
| W303.asm02.HP0_G0046980 | 702.71205 | 708.986949 | 841.3207086 | 780.874121 | 715.49457 | 759.466899 | 973.64023 | 788.56198 | 907.21982 | 0.991388614 | -0.001159555 | YNR027W | BUD17 | n |
| W303.asm02.HP0_G0047000 | 716.40808 | 788.916664 | 701.261578 | 922.548188 | 904.2557 | 850.489149 | 792.3653 | 922.39447 | 829.63445 | 0.001627895 | -0.279768643 | YNR029C | ZNG1 | n |
| W303.asm02.HP0_G0047010 | 1517.0995 | 1567.45284 | 1522.297861 | 1891.76548 | 1903.7908 | 1967.40801 | 2084.0917 | 2143.9439 | 1990.77 | 2.44E-07 | -0.322872612 | YNR030W | ALG12 | n |
| W303.asm02.HP0_G0047020 | 689.01601 | 724.557673 | 895.4125108 | 712.537219 | 699.31504 | 717.748367 | 811.74689 | 696.71616 | 777.61699 | 0.283010866 | 0.117752474 | YNR031C | SSK2 | n |
| W303.asm02.HP0_G0047030 | 877.5999 | 863.656137 | 847.1162589 | 595.864456 | 553.69931 | 579.318696 | 612.23045 | 589.12534 | 598.64165 | 1.06E-13 | 0.581618026 | YNR032W | PPG1 | n |
| W303.asm02.HP0_G0047040 | 646.87436 | 613.486511 | 759.2170804 | 181.676156 | 192.35658 | 161.185234 | 166.45372 | 181.06748 | 163.1056 | 1.67E-54 | 1.915375718 | YNR032C- | HUB1 | n |
| W303.asm02.HP0_G0047050 | 815.44097 | 956.04243 | 985.2435394 | 1493.41134 | 1497.505 | 1517.0375 | 1235.8618 | 1393.4323 | 1193.7567 | 2.53E-15 | -0.708849719 | YNR033W | ABZ1 | n |
| W303.asm02.HP0_G0047060 | 273.92074 | 270.930591 | 255.0042102 | 1924.26718 | 1830.983 | 1875.43761 | 1959.8215 | 2095.3968 | 2027.7994 | 7.96E-264 | -2.81666561 | YNR034W | SOL1 | n |
| W303.asm02.HP0_G0047080 | 3796.9628 | 3594.76106 | 3525.626391 | 2571.801 | 2544.6798 | 2458.5489 | 2708.8632 | 2807.858 | 2713.7246 | 1.34E-19 | 0.526938817 | YNR035C | ARC35 | n |
| W303.asm02.HP0_G0047090 | 2179.7769 | 2153.9501 | 2423.505922 | 1275.0666 | 1274.5871 | 1223.11148 | 1316.8085 | 1430.1707 | 1296.0283 | 1.05E-29 | 0.840935189 | YNR036C | MRPS12 | n |
| W303.asm02.HP0_G0047100 | 1208.4119 | 1225.93497 | 1265.361801 | 886.712983 | 963.58062 | 923.496578 | 817.44736 | 967.0053 | 856.08401 | 2.98E-07 | 0.416036623 | YNR037C | RSM19 | n |
| W303.asm02.HP0_G0047110 | 774.35286 | 766.079602 | 772.7400309 | 1476.7438 | 1463.3482 | 1454.4597 | 963.37939 | 1119.2069 | 984.80519 | 1.23E-33 | -0.925906259 | YNR038W | DBP6 | n |
| W303.asm02.HP0_G0047120 | 644.76728 | 717.291335 | 627.8512751 | 1243.39828 | 1260.2053 | 1188.02999 | 1434.2382 | 1449.8519 | 1429.1577 | 1.04E-31 | -0.892363443 | YNR039C | ZRG17 | n |
| W303.asm02.HP0_G0047130 | 828.08346 | 868.846378 | 800.751857 | 790.041267 | 749.65134 | 827.733586 | 779.82426 | 796.43448 | 777.61699 | 0.339779443 | 0.077239702 | YNR040W | MRX15 | n |
| W303.asm02.HP0_G0047140 | 729.05058 | 690.302081 | 855.8095842 | 1315.06869 | 1178.4088 | 1284.74113 | 1205.0793 | 1162.5057 | 1210.5081 | 9.23E-16 | -0.730849331 | YNR041C | COQ2 | n |
| W303.asm02.HP0_G0047150 | 2474.7685 | 2557.75087 | 2171.399487 | 3211.00106 | 3124.4461 | 3387.73437 | 2572.052 | 2566.4347 | 2414.8446 | 1.20E-08 | -0.432900358 | YNR043W | MVD1 | n |
| W303.asm02.HP0_G0047160 | 300.25927 | 389.268091 | 290.7434366 | 536.6947 | 578.86746 | 658.015016 | 526.72341 | 502.52785 | 513.12141 | 3.40E-12 | -0.855322159 | YNR044W | AGA1 | n |
| W303.asm02.HP0_G0047170 | 389.81028 | 397.572476 | 358.3581893 | 580.030297 | 576.17088 | 579.318696 | 502.78143 | 596.99784 | 532.51776 | 6.19E-10 | -0.599947658 | YNR045W | PET494 | n |
| W303.asm02.HP0_G0047180 | 1022.9886 | 1035.97214 | 782.3992813 | 1035.05406 | 1030.0965 | 971.852149 | 710.27853 | 808.24323 | 725.59952 | 0.38581224 | -0.097111838 | YNR046W | TRM112 | n |
| W303.asm02.HP0_G0047190 | 306.58052 | 350.860306 | 375.74484 | 1364.23793 | 1304.2495 | 1394.72635 | 1487.8226 | 1375.0632 | 1527.0211 | 6.60E-106 | -1.973781779 | YNR047W | FPK1 | n |
| W303.asm02.HP0_G0047210 | 444.59443 | 503.453397 | 419.2114668 | 642.533562 | 615.72083 | 553.718688 | 616.79083 | 619.30326 | 641.84259 | 4.53E-05 | -0.408059176 | YNR049C | MSO1 | n |
| W303.asm02.HP0_G0047220 | 3972.9042 | 3724.51709 | 3779.664676 | 1575.91565 | 1430.9891 | 1470.57823 | 1626.9141 | 1573.1877 | 1685.7185 | 5.90E-95 | 1.357494042 | YNR050C | LYS9 | n |
| W303.asm02.HP0_G0047230 | 1248.4464 | 1320.39736 | 1347.465429 | 3486.01542 | 3314.1061 | 3709.15669 | 3098.7754 | 2915.4488 | 2957.0605 | 1.57E-95 | -1.42364902 | YNR051C | BRE5 | n |
| W303.asm02.HP0_G0047240 | 1139.9317 | 1154.30964 | 1171.667072 | 2080.10865 | 1898.3976 | 1919.05244 | 1867.4739 | 2024.5443 | 1904.3681 | 1.57E-28 | -0.767126087 | YNR052C | POP2 | n |
| W303.asm02.HP0_G0047250 | 3844.3722 | 3212.75931 | 4010.52076 | 2545.96632 | 2549.1741 | 2590.34153 | 1699.8801 | 2082.276 | 1721.8662 | 8.53E-08 | 0.526277051 | YNR053C | NOG2 | n |
| W303.asm02.HP0_G0047260 | 858.63616 | 963.308768 | 781.4333562 | 1292.56752 | 1243.1269 | 1397.5708 | 892.69356 | 1163.8178 | 933.66938 | 8.82E-08 | -0.595598651 | YNR054C | ESF2 | n |
| W303.asm02.HP0_G0047270 | 942.91946 | 922.824887 | 992.9709397 | 1813.42806 | 1748.2876 | 2084.97841 | 2111.454 | 1876.2789 | 2253.5023 | 5.13E-26 | -0.981451259 | YNR055C | HOL1 | n |
| W303.asm02.HP0_G0047280 | 126.42496 | 133.908223 | 122.6724799 | 109.172369 | 83.594215 | 116.622258 | 137.95137 | 80.037073 | 141.06431 | 0.149503902 | 0.307974099 | YNR056C | BIO5 | n |
| W303.asm02.HP0_G0047290 | 178.04848 | 165.04967 | 153.5820811 | 5.83363805 | 8.9886252 | 7.5851875 | 9.1207516 | 3.9362495 | 11.461475 | 5.89E-38 | 4.476797225 | YNR057C | BIO4 | n |
| W303.asm02.HP0_G0047300 | 190.69098 | 184.772587 | 184.4916824 | 316.683208 | 351.45525 | 314.785281 | 974.78033 | 547.13868 | 928.37946 | 6.03E-06 | -0.8118743 | YNR058W | BIO3 | n |
| W303.asm02.HP0_G0047330 | 227.56492 | 238.751096 | 222.1627589 | 592.53095 | 710.10139 | 675.081688 | 648.71346 | 498.5916 | 586.29852 | 1.42E-36 | -1.522085205 | YNR062C | PUL3 | n |
| W303.asm02.HP0_G0047340 | 97.979341 | 94.46239 | 110.1154544 | 1052.55498 | 1094.8146 | 1141.57072 | 1040.9058 | 810.8674 | 941.60424 | 1.95E-160 | -3.439943263 | YNR063W | PUL4 | n |
| W303.asm02.HP0_G0047380 | 2411.556 | 2708.26786 | 2814.705563 | 1176.72813 | 1054.3657 | 1243.97075 | 1344.1708 | 1590.2448 | 1595.79 | 6.61E-36 | 1.191440359 | YNR067C | DSE4 | n |
| W303.asm02.HP0_G0047650 | 1814.1981 | 1715.89374 | 2268.957916 | 1430.0747 | 1460.6516 | 1321.71892 | 1257.5236 | 1518.0802 | 1375.377 | 2.41E-05 | 0.461397736 | YOL149W | DCP1 | n |
| W303.asm02.HP0_G0047660 | 535.19898 | 524.214362 | 519.6676708 | 799.208412 | 815.26831 | 787.911352 | 871.03178 | 860.72656 | 893.11339 | 4.76E-15 | -0.605719227 | YOL148C | SPT20 | n |
| W303.asm02.HP0_G0047670 | 1182.0733 | 1220.74473 | 1045.130892 | 458.357275 | 471.00396 | 468.385328 | 489.10031 | 514.3366 | 528.1095 | 4.51E-49 | 1.302281122 | YOL147C | PEX11 | n |
| W303.asm02.HP0_G0047680 | 598.41146 | 631.133331 | 691.6023277 | 391.687126 | 390.10633 | 364.089 | 386.49185 | 389.6887 | 420.54796 | 1.25E-14 | 0.745617041 | YOL146W | PSF3 | n |
| W303.asm02.HP0_G0047690 | 2296.72 | 2281.63003 | 2138.558036 | 3637.69001 | 3744.6613 | 4007.82345 | 3310.8328 | 3532.1279 | 3512.5012 | 2.69E-31 | -0.761932356 | YOL145C | CTR9 | n |
| W303.asm02.HP0_G0047700 | 453.02276 | 522.138265 | 555.4068972 | 1260.8992 | 1297.0586 | 1325.51152 | 689.75684 | 876.47156 | 759.98395 | 3.70E-39 | -1.342175466 | YOL144W | NOP8 | n |
| W303.asm02.HP0_G0047710 | 2054.4055 | 2020.04188 | 1860.371624 | 2260.95143 | 2435.9174 | 2475.61557 | 2240.2846 | 2180.6822 | 2107.1481 | 8.58E-05 | -0.273291764 | YOL143C | RIB4 | n |
| W303.asm02.HP0_G0047720 | 1511.8318 | 1466.76217 | 1484.626784 | 722.537741 | 762.23542 | 673.185391 | 578.02763 | 673.09867 | 574.83705 | 3.63E-36 | 1.047954336 | YOL142W | RRP40 | n |
| W303.asm02.HP0_G0047740 | 10112.943 | 9690.18033 | 11730.19367 | 5010.26171 | 5114.5278 | 5405.39424 | 4263.9514 | 4854.7077 | 4160.5154 | 3.27E-32 | 1.021954217 | YOL139C | CDC33 | n |
| W303.asm02.HP0_G0047750 | 312.90177 | 370.583222 | 404.7225912 | 1417.57405 | 1522.6731 | 1517.98565 | 1508.3443 | 1575.8119 | 1508.5064 | 1.38E-109 | -2.032267829 | YOL138C | RTC1 | n |
| W303.asm02.HP0_G0047760 | 984.00757 | 909.33026 | 953.3680131 | 195.843563 | 227.41222 | 208.592656 | 235.99945 | 270.28913 | 186.91021 | 1.82E-72 | 2.1724387 | YOL137W | BSC6 | n |
| W303.asm02.HP0_G0047770 | 1542.3845 | 1414.85975 | 1509.740835 | 680.868898 | 648.07988 | 533.807571 | 560.92622 | 561.5716 | 654.18572 | 4.46E-35 | 1.260566359 | YOL136C | PFK27 | n |
| W303.asm02.HP0_G0047780 | 308.6876 | 358.126643 | 340.9715386 | 691.702797 | 675.94462 | 704.474289 | 757.02228 | 775.44115 | 779.38029 | 2.60E-31 | -1.039283608 | YOL135C | MED7 | n |
| W303.asm02.HP0_G0047790 | 1886.8925 | 1970.21556 | 2080.602533 | 1064.22226 | 1215.2621 | 1013.57068 | 1189.118 | 1232.0461 | 1164.6622 | 4.44E-25 | 0.850559432 | YOL133W | HRT1 | n |
| W303.asm02.HP0_G0047800 | 86.390387 | 97.5765347 | 84.03547836 | 94.1715856 | 89.886252 | 89.1259532 | 115.14949 | 91.845822 | 102.27162 | 0.88022123 | -0.029279555 | YOL132W | GAS4 | n |
| W303.asm02.HP0_G0047820 | 1336.9439 | 1283.02763 | 1359.056529 | 1975.93654 | 2175.2473 | 2250.90439 | 1492.383 | 1480.0298 | 1659.2689 | 1.47E-18 | -0.685578111 | YOL130W | ALR1 | n |
| W303.asm02.HP0_G0047830 | 2792.938 | 2716.57225 | 3173.063752 | 648.3672 | 710.10139 | 651.377977 | 996.44211 | 946.01197 | 876.362 | 8.75E-131 | 2.111390081 | YOL129W | VPS68 | n |
| W303.asm02.HP0_G0047850 | 13193.498 | 12610.21 | 14833.71082 | 11008.9084 | 11262.747 | 11788.3295 | 10039.667 | 11611.936 | 9259.99 | 0.005168956 | 0.254804824 | YOL127W | RPL25 | n |
| W303.asm02.HP0_G0047860 | 369.793 | 401.724669 | 351.5967141 | 1685.08802 | 1811.208 | 1331.20041 | 1124.1326 | 1127.0794 | 1147.9108 | 2.86E-77 | -2.104735445 | YOL126C | MDH2 | n |
| W303.asm02.HP0_G0047870 | 595.25083 | 622.828945 | 647.1697759 | 756.706192 | 692.12414 | 827.733586 | 475.41918 | 551.07493 | 524.58289 | 0.003161373 | -0.286607934 | YOL125W | TRM13 | n |
| W303.asm02.HP0_G0047880 | 1933.2483 | 1701.36107 | 1903.838251 | 1300.06791 | 1279.9802 | 1397.5708 | 929.17657 | 1243.8548 | 1013.018 | 3.27E-06 | 0.47782301 | YOL124C | TRM11 | n |
| W303.asm02.HP0_G0047890 | 2966.7723 | 2821.41512 | 2787.659661 | 2903.48499 | 2902.4271 | 3485.39366 | 3034.9301 | 2978.4288 | 2972.0486 | 0.158557297 | -0.115332738 | YOL123W | HRP1 | n |
| W303.asm02.HP0_G0047900 | 1845.8044 | 1686.82839 | 1772.472446 | 3475.18152 | 3403.0935 | 3156.38615 | 2346.3134 | 2851.1567 | 2372.5253 | 1.28E-26 | -0.919714068 | YOL122C | SMF1 | n |
| W303.asm02.HP0_G0047910 | 7308.416 | 7061.84219 | 8532.015866 | 15293.2988 | 15965.596 | 16861.8718 | 13832.76 | 15246.406 | 12791.006 | 7.64E-34 | -1.071034948 | YOL121C | RPS19A | y |
| W303.asm02.HP0_G0047940 | 686.90893 | 713.139142 | 739.8985796 | 203.343955 | 193.25544 | 199.111172 | 216.61785 | 158.76206 | 201.89829 | 1.34E-60 | 1.844845662 | YOL117W | RRI2 | n |
| W303.asm02.HP0_G0047950 | 731.15766 | 801.373242 | 661.6586515 | 649.200577 | 574.37315 | 586.903883 | 703.43797 | 581.25284 | 646.25085 | 0.008618139 | 0.276051202 | YOL116W | MSN1 | n |
| W303.asm02.HP0_G0047970 | 85.336845 | 104.842872 | 94.66065378 | 101.671977 | 79.998765 | 92.9185469 | 60.424979 | 59.043743 | 80.230324 | 0.805788176 | 0.051815737 | YOL114C | PTH4 | n |

| | | | | | | | | | | | | | | |
|---|---|---|---|---|---|---|---|---|---|---|---|---|---|---|
| W303.asm02.HP0_G0048000 | 395.07799 | 414.181248 | 425.972942 | 1050.05485 | 1077.7362 | 1069.51144 | 1007.8431 | 1083.7807 | 1007.7281 | 1.03E-65 | -1.371253276 | YOL111C | MDY2 | n |
| W303.asm02.HP0_G0048010 | 420.36298 | 439.094406 | 468.4736437 | 144.174197 | 127.63848 | 129.896336 | 147.07212 | 91.845822 | 111.08814 | 2.38E-33 | 1.723987485 | YOL110W | SHR5 | n |
| W303.asm02.HP0_G0048030 | 741.69308 | 808.63958 | 920.5265618 | 256.680074 | 263.36672 | 275.911195 | 183.55513 | 208.62122 | 219.53133 | 8.09E-48 | 1.63572516 | YOL108C | INO4 | n |
| W303.asm02.HP0_G0048040 | 683.7483 | 750.508879 | 698.3638029 | 804.208674 | 803.5831 | 870.400266 | 1083.0893 | 1108.7103 | 1117.053 | 0.006472646 | -0.216154916 | YOL107W | | n |
| W303.asm02.HP0_G0048080 | 1081.9869 | 1002.7546 | 1314.623978 | 543.361715 | 536.62093 | 494.933485 | 453.75739 | 486.78286 | 469.03882 | 2.71E-25 | 1.110118231 | YOL102C | TPT1 | n |
| W303.asm02.HP0_G0048100 | 587.87605 | 595.839691 | 684.8408524 | 1327.56934 | 1271.8905 | 1280.00039 | 1449.0594 | 1289.7778 | 1446.7908 | 8.79E-35 | -1.053007164 | YOL100W | PKH2 | n |
| W303.asm02.HP0_G0048110 | 1654.0598 | 1695.13278 | 1903.838251 | 3280.17134 | 3211.6358 | 3346.01584 | 2520.7477 | 2534.9447 | 2429.8327 | 1.21E-39 | -0.9045814 | YOL098C | SDD3 | n |
| W303.asm02.HP0_G0048130 | 3405.0455 | 3237.67247 | 3056.186822 | 3618.52234 | 3752.751 | 3664.59371 | 3289.171 | 3385.1746 | 3306.1947 | 0.002131543 | -0.18653508 | YOL097C | WRS1 | n |
| W303.asm02.HP0_G0048140 | 530.98482 | 611.410414 | 567.9639227 | 971.717423 | 1119.0838 | 1042.01513 | 1026.0846 | 993.24696 | 1009.4914 | 7.09E-24 | -0.872621429 | YOL096C | COQ3 | n |
| W303.asm02.HP0_G0048150 | 336.07967 | 324.9091 | 353.5285641 | 390.853749 | 462.9142 | 389.689008 | 430.95551 | 411.99412 | 444.35256 | 0.006927011 | -0.29285691 | YOL095C | HMI1 | n |
| W303.asm02.HP0_G0048160 | 768.03161 | 787.878615 | 682.9090023 | 999.21886 | 961.7829 | 987.022524 | 793.50539 | 917.14614 | 833.16106 | 4.64E-06 | -0.397873603 | YOL094C | RFC4 | n |
| W303.asm02.HP0_G0048170 | 1131.5034 | 1211.4023 | 924.390262 | 1162.56073 | 1233.2394 | 1215.5263 | 848.2299 | 917.14614 | 914.27303 | 0.149035055 | -0.14501066 | YOL093W | TRM10 | n |
| W303.asm02.HP0_G0048180 | 2204.0084 | 2037.6887 | 2054.522557 | 928.381826 | 957.28859 | 921.600282 | 1008.9831 | 1078.5324 | 1127.6328 | 4.14E-59 | 1.165137067 | YOL092W | YPQ1 | n |
| W303.asm02.HP0_G0048190 | 279.18844 | 225.256468 | 291.7093617 | 85.837817 | 87.189665 | 96.7111407 | 93.487704 | 95.782071 | 105.79823 | 9.09E-23 | 1.563963881 | YOL091W | SPO21 | n |
| W303.asm02.HP0_G0048200 | 987.1682 | 1043.23848 | 1043.199042 | 1690.08828 | 1806.7137 | 1791.0524 | 1728.3824 | 1621.7348 | 1639.8726 | 1.75E-30 | -0.782187011 | YOL090W | MSH2 | n |
| W303.asm02.HP0_G0048210 | 796.47722 | 773.34594 | 722.5119289 | 1242.5649 | 1205.3746 | 1279.05224 | 1152.635 | 1285.8415 | 1160.2539 | 3.98E-19 | -0.701679465 | YOL089C | HAL9 | n |
| W303.asm02.HP0_G0048220 | 1565.5624 | 1723.16008 | 1621.78814 | 1316.73544 | 1303.3507 | 1334.04485 | 1962.1017 | 1978.6214 | 1887.6167 | 2.40E-06 | 0.312589544 | YOL088C | MPD2 | n |
| W303.asm02.HP0_G0048230 | 617.3752 | 683.035743 | 700.295653 | 906.714028 | 829.65011 | 787.911352 | 890.41338 | 854.16614 | 886.94182 | 0.000130864 | -0.335661708 | YOL087C | DUF1 | n |
| W303.asm02.HP0_G0048240 | 626.85707 | 661.23673 | 785.2970564 | 424.188824 | 437.74605 | 432.355688 | 429.81542 | 409.36995 | 419.66631 | 2.68E-11 | 0.681122816 | YOL086W- | MHF1 | n |
| W303.asm02.HP0_G0048250 | 30207.136 | 29020.7147 | 33507.93959 | 8568.78091 | 6600.3475 | 7357.63188 | 10575.511 | 8073.2477 | 10146.932 | 7.43E-74 | 2.041424902 | YOL086C | ADH1 | n |
| W303.asm02.HP0_G0048270 | 124.31787 | 144.288706 | 166.1391066 | 497.525988 | 440.44264 | 373.570485 | 1213.06 | 1079.8444 | 1264.2888 | 2.99E-31 | -1.591967027 | YOL084W | PHM7 | n |
| W303.asm02.HP0_G0048280 | 459.34401 | 460.893419 | 468.4736437 | 621.69914 | 621.114 | 557.511281 | 936.01713 | 827.92448 | 921.32625 | 4.46E-05 | -0.375189448 | YOL083W | ATG34 | n |
| W303.asm02.HP0_G0048310 | 845.99366 | 774.383988 | 795.9222318 | 1085.05668 | 1016.6135 | 1150.10406 | 599.68942 | 772.81699 | 650.65911 | 1.44E-05 | -0.428369589 | YOL080C | REX4 | n |
| W303.asm02.HP0_G0048320 | 390.86382 | 396.534428 | 422.1092419 | 1150.06007 | 1089.4214 | 1073.30403 | 1197.0986 | 1099.5257 | 1224.6145 | 9.73E-67 | -1.453265467 | YOL078W | AVO1 | n |
| W303.asm02.HP0_G0048330 | 2021.7458 | 1922.46534 | 2399.357796 | 2705.1413 | 3054.3349 | 2965.80831 | 2595.9939 | 2981.053 | 2511.8263 | 3.07E-06 | -0.459425046 | YOL077W- | ATP19 | n |
| W303.asm02.HP0_G0048340 | 5294.045 | 4854.95163 | 5941.404913 | 2394.29173 | 2217.4938 | 2492.68224 | 1499.2235 | 1952.3798 | 1601.9615 | 3.61E-30 | 1.179546648 | YOL077C | BRX1 | n |
| W303.asm02.HP0_G0048350 | 1417.013 | 1220.74473 | 1356.158754 | 1720.92322 | 1774.3546 | 1938.96356 | 1511.7646 | 1494.4627 | 1525.2578 | 1.60E-08 | -0.443818205 | YOL076W | MDM20 | n |
| W303.asm02.HP0_G0048360 | 532.03836 | 531.4807 | 620.1238748 | 624.199271 | 632.79922 | 650.429828 | 799.20586 | 724.26991 | 779.38029 | 0.055444889 | -0.178552104 | YOL075C | | n |
| W303.asm02.HP0_G0048370 | 454.0763 | 529.404603 | 511.9402705 | 299.182294 | 374.82567 | 342.281586 | 631.61205 | 648.16909 | 670.05545 | 2.31E-07 | 0.559037078 | YOL073C | DSC2 | n |
| W303.asm02.HP0_G0048380 | 293.93802 | 341.517871 | 285.9138114 | 716.704103 | 794.59447 | 727.229852 | 760.44267 | 770.19282 | 736.17935 | 6.65E-40 | -1.281431282 | YOL072W | THP1 | n |
| W303.asm02.HP0_G0048390 | 1755.1998 | 1760.52982 | 2149.183211 | 919.214681 | 861.1103 | 943.407696 | 804.90633 | 797.74657 | 843.74088 | 2.55E-30 | 1.057043823 | YOL071W | SDH5 | n |
| W303.asm02.HP0_G0048400 | 1198.93 | 1225.93497 | 1196.781123 | 1089.22356 | 973.46811 | 1078.04477 | 1168.5963 | 930.26697 | 1296.91 | 0.061309833 | 0.20544973 | YOL070C | NBA1 | n |
| W303.asm02.HP0_G0048410 | 530.98482 | 457.779274 | 508.0765703 | 368.352574 | 410.78017 | 472.177922 | 458.31777 | 425.11495 | 446.11587 | 0.020811995 | 0.260391809 | YOL069W | NUF2 | n |
| W303.asm02.HP0_G0048430 | 326.5978 | 285.463266 | 353.5285641 | 85.0044401 | 84.493077 | 79.6444688 | 95.767892 | 51.171244 | 73.177109 | 3.40E-29 | 1.954213852 | YOL067C | RTG1 | n |
| W303.asm02.HP0_G0048440 | 419.30944 | 441.170503 | 400.858891 | 1305.06817 | 1295.2609 | 1236.38556 | 1150.3548 | 1187.4353 | 1231.6677 | 3.85E-91 | -1.605945767 | YOL066C | RIB2 | n |
| W303.asm02.HP0_G0048450 | 342.40092 | 344.632016 | 386.3700154 | 328.350484 | 363.14046 | 310.992688 | 368.25035 | 318.83621 | 343.84425 | 0.389899111 | 0.099234835 | YOL065C | INP54 | n |
| W303.asm02.HP0_G0048460 | 4773.5956 | 4514.4718 | 5183.153757 | 1833.4291 | 1910.0829 | 1941.808 | 1885.7154 | 1979.9335 | 1803.8598 | 1.67E-79 | 1.348257245 | YOL064C | MET22 | n |
| W303.asm02.HP0_G0048470 | 585.76896 | 573.002629 | 639.4423756 | 946.716117 | 1008.5238 | 919.703985 | 941.7176 | 946.01197 | 918.68129 | 6.54E-17 | -0.676270865 | YOL063C | CRT10 | n |
| W303.asm02.HP0_G0048480 | 1899.535 | 1816.58442 | 1708.721393 | 1176.72813 | 1172.1167 | 1011.67438 | 1101.3308 | 1045.7303 | 1060.6273 | 1.49E-17 | 0.689886459 | YOL062C | APM4 | n |
| W303.asm02.HP0_G0048490 | 2817.1694 | 2520.38113 | 2906.468441 | 2275.95222 | 2120.4167 | 2189.27474 | 1975.7828 | 2075.7156 | 2007.5214 | 6.28E-06 | 0.324053536 | YOL061W | PRS5 | n |
| W303.asm02.HP0_G0048500 | 852.31491 | 1010.02094 | 903.1399111 | 1000.05224 | 975.26584 | 1173.80777 | 1184.5576 | 1117.8949 | 1231.6677 | 0.05756465 | -0.18681549 | YOL060C | MAM3 | n |
| W303.asm02.HP0_G0048520 | 2210.3296 | 2439.41337 | 2396.460021 | 1760.09194 | 1832.7807 | 1767.34869 | 1979.2031 | 1171.6903 | 1883.2085 | 0.006313559 | 0.394595903 | YOL058W | ARG1 | n |
| W303.asm02.HP0_G0048530 | 1203.1442 | 1444.96315 | 1536.786736 | 1661.75347 | 1564.0208 | 1716.14867 | 1971.2224 | 2212.1722 | 1956.3856 | 0.013829375 | -0.239317393 | YOL057W | | n |
| W303.asm02.HP0_G0048540 | 689.01601 | 618.676752 | 655.8631012 | 1000.88561 | 931.22157 | 882.726196 | 1138.9539 | 1112.6465 | 1076.497 | 3.41E-10 | -0.520499728 | YOL056W | GPM3 | n |
| W303.asm02.HP0_G0048550 | 749.06786 | 720.40548 | 604.6690742 | 40.0020875 | 55.729476 | 48.3555703 | 26.222161 | 32.802079 | 29.094513 | 9.13E-117 | 3.850981555 | YOL055C | THI20 | n |
| W303.asm02.HP0_G0048560 | 1007.1855 | 1043.23848 | 1035.471641 | 782.540875 | 654.37192 | 707.318735 | 689.75644 | 772.81699 | 758.22064 | 2.52E-09 | 0.524478282 | YOL054W | PSH1 | n |
| W303.asm02.HP0_G0048570 | 687.96247 | 650.856247 | 706.0912032 | 594.197704 | 590.55268 | 532.859422 | 678.3559 | 670.4745 | 780.26194 | 0.008283773 | 0.251127231 | YOL053W | AIM39 | n |
| W303.asm02.HP0_G0048590 | 937.65176 | 984.069733 | 839.3888586 | 1043.38783 | 1147.8474 | 1298.96336 | 1193.6784 | 1413.1136 | 1231.6677 | 0.00127763 | -0.337813392 | YOL052C | SPE2 | n |
| W303.asm02.HP0_G0048600 | 966.09737 | 1235.27741 | 1149.450796 | 2535.13242 | 2533.8935 | 2527.76374 | 2675.8005 | 2630.7268 | 2871.5403 | 2.74E-42 | -1.180558337 | YOL051W | GAL11 | n |
| W303.asm02.HP0_G0048610 | 2537.981 | 2446.67971 | 2435.097022 | 961.716901 | 971.67039 | 898.844719 | 998.7223 | 1011.6161 | 981.27858 | 2.91E-100 | 1.388778388 | YOL049W | GSH2 | n |
| W303.asm02.HP0_G0048620 | 1067.2373 | 1147.04331 | 1197.747048 | 55.002873 | 53.931751 | 37.9259375 | 46.743852 | 49.85916 | 57.307374 | 1.73E-202 | 4.530840006 | YOL048C | RRT8 | n |
| W303.asm02.HP0_G0048650 | 705.87267 | 610.372366 | 714.7845286 | 224.178376 | 241.79402 | 209.540805 | 251.96076 | 220.42997 | 277.72035 | 6.82E-42 | 1.587846569 | YOL044W | PEX15 | n |
| W303.asm02.HP0_G0048660 | 442.48735 | 432.866117 | 403.7566661 | 590.03082 | 627.40604 | 571.733508 | 576.88754 | 595.68576 | 608.33982 | 7.10E-08 | -0.485167347 | YOL043C | NTG2 | n |
| W303.asm02.HP0_G0048670 | 1852.1256 | 1685.79034 | 1906.736026 | 348.351529 | 345.16321 | 363.140852 | 376.231 | 383.12829 | 366.7672 | 6.90E-170 | 2.366038695 | YOL042W | NGL1 | n |
| W303.asm02.HP0_G0048680 | 2337.8081 | 2255.67883 | 2108.614359 | 2530.96554 | 2376.5925 | 2709.80824 | 1395.475 | 1675.5302 | 1419.4596 | 0.035748361 | -0.18474448 | YOL041C | NOP12 | n |
| W303.asm02.HP0_G0048690 | 8887.6744 | 9187.76499 | 7963.086018 | 13487.3712 | 12983.17 | 14991.175 | 11923.103 | 11937.333 | 10578.06 | 3.16E-16 | -0.67116297 | YOL040C | RPS15 | n |
| W303.asm02.HP0_G0048700 | 9534.5488 | 9305.06444 | 10864.72483 | 6546.17527 | 6617.4259 | 6893.03914 | 5915.9475 | 6115.6197 | 5571.1584 | 6.15E-15 | 0.566748721 | YOL039W | RPP2A | n |
| W303.asm02.HP0_G0048720 | 829.137 | 1015.21118 | 819.1044328 | 1431.74145 | 1389.6415 | 1411.79302 | 1411.4363 | 1322.5798 | 1491.755 | 3.90E-13 | -0.668898049 | YOL036W | | n |
| W303.asm02.HP0_G0048730 | 720.62225 | 722.481576 | 711.8867535 | 1119.22513 | 1170.319 | 1213.63 | 1320.2288 | 1228.1098 | 1308.3714 | 1.88E-21 | -0.70054709 | YOL034W | SMC5 | n |
| W303.asm02.HP0_G0048740 | 1052.4878 | 976.803395 | 1111.779719 | 818.37608 | 783.80812 | 785.066907 | 733.08041 | 750.51157 | 770.56377 | 5.02E-07 | 0.396033261 | YOL033W | MSE1 | n |

| | | | | | | | | | | | | | | |
|---|---|---|---|---|---|---|---|---|---|---|---|---|---|---|
| W303.asm02.HP0_G0048750 | 820.70867 | 935.281466 | 738.9326545 | 323.350223 | 396.39837 | 320.474172 | 486.82012 | 419.86661 | 435.53604 | 3.73E-26 | 1.26172285 | YOL032W | OPI10 | n |
| W303.asm02.HP0_G0048760 | 5262.4388 | 5994.72859 | 4717.577889 | 1360.07104 | 1416.6073 | 1404.20784 | 1430.8179 | 1410.4894 | 1455.6073 | 2.55E-112 | 1.933891036 | YOL031C | SIL1 | n |
| W303.asm02.HP0_G0048770 | 2849.8292 | 3044.59549 | 2976.980969 | 2467.62889 | 2488.9503 | 2569.48227 | 3317.6734 | 3522.9433 | 3177.4735 | 0.000198261 | 0.237480992 | YOL030W | GAS5 | n |
| W303.asm02.HP0_G0048800 | 1142.0388 | 1504.1319 | 1385.136505 | 1910.93315 | 1843.567 | 1863.11168 | 2112.5941 | 2117.7022 | 2252.6206 | 2.78E-07 | -0.478517027 | YOL027C | MDM38 | n |
| W303.asm02.HP0_G0048810 | 628.96416 | 661.23673 | 715.7504536 | 885.879606 | 872.79551 | 840.059516 | 866.4714 | 1018.1765 | 938.07763 | 2.57E-05 | -0.373080606 | YOL026C | MIM1 | n |
| W303.asm02.HP0_G0048820 | 301.31281 | 335.289582 | 307.1641623 | 409.18804 | 364.93818 | 371.674188 | 416.13429 | 346.38996 | 409.08649 | 0.012967252 | -0.280948143 | YOL025W | LAG2 | n |
| W303.asm02.HP0_G0048840 | 508.86045 | 574.040678 | 580.5209482 | 1020.88666 | 946.50224 | 884.622492 | 932.59685 | 931.57905 | 890.46843 | 2.01E-18 | -0.777967888 | YOL023W | IFM1 | n |
| W303.asm02.HP0_G0048850 | 1437.0303 | 1424.20219 | 1429.569057 | 1631.7519 | 1658.4014 | 1737.00794 | 1217.6203 | 1445.9157 | 1204.3365 | 0.005021894 | -0.228278471 | YOL022C | TSR4 | n |
| W303.asm02.HP0_G0048860 | 1102.0042 | 1209.3262 | 1286.612151 | 2585.96841 | 2733.4409 | 2779.97122 | 1955.2611 | 2243.6622 | 2050.7223 | 4.95E-48 | -1.169925751 | YOL021C | DIS3 | n |
| W303.asm02.HP0_G0048870 | 1749.9321 | 1713.81765 | 1961.793753 | 2060.94098 | 1962.2169 | 1862.16353 | 2247.1252 | 2176.746 | 2312.573 | 0.126841464 | -0.11727903 | YOL020W | TAT2 | n |
| W303.asm02.HP0_G0048890 | 2017.5316 | 2110.35207 | 1938.611553 | 2515.96475 | 2295.6949 | 1831.82278 | 2495.6657 | 2474.5889 | 2626.441 | 0.209754634 | -0.131688537 | YOL019W | TOS7 | n |
| W303.asm02.HP0_G0048900 | 1490.7609 | 1468.83826 | 1423.773507 | 478.35832 | 462.01534 | 471.229774 | 475.41918 | 532.70577 | 522.81958 | 4.44E-97 | 1.63430139 | YOL018C | TLG2 | n |
| W303.asm02.HP0_G0048920 | 1030.3634 | 917.634645 | 827.7977581 | 2098.44294 | 1951.4305 | 1802.43018 | 1670.2376 | 1557.4427 | 1770.357 | 1.26E-29 | -1.077175468 | YOL016C | CMK2 | n |
| W303.asm02.HP0_G0048950 | 1647.7386 | 1655.68695 | 1377.409105 | 1355.90416 | 1362.6756 | 1159.58554 | 1809.3291 | 1634.8556 | 1811.7947 | 0.005524696 | 0.270481262 | YOL013C | HRD1 | n |
| W303.asm02.HP0_G0048960 | 1644.578 | 1612.08892 | 1781.165771 | 1631.7519 | 1554.1333 | 1571.08196 | 1577.89 | 1590.2448 | 1643.3992 | 0.230111841 | 0.08295527 | YOL012C | HTZ1 | n |
| W303.asm02.HP0_G0048970 | 543.62731 | 637.36162 | 675.181602 | 4217.72031 | 4000.8371 | 4033.42345 | 6724.2741 | 5992.2838 | 6483.6682 | 1.65E-232 | -2.721534414 | YOL011W | PLB3 | n |
| W303.asm02.HP0_G0048980 | 1892.1602 | 1639.07817 | 1766.676896 | 1880.0982 | 1740.1978 | 1784.41536 | 1190.2581 | 1504.9594 | 1242.2475 | 0.77273065 | -0.028982486 | YOL010W | RCL1 | n |
| W303.asm02.HP0_G0048990 | 540.46669 | 520.062169 | 557.3387473 | 455.023768 | 497.07098 | 425.718649 | 422.97486 | 472.34994 | 453.16908 | 0.017609778 | 0.231577704 | YOL009C | MDM12 | n |
| W303.asm02.HP0_G0049000 | 995.59653 | 918.672694 | 1051.892367 | 763.373207 | 756.84224 | 693.096508 | 611.09036 | 652.10534 | 660.35728 | 7.56E-07 | 0.422172389 | YOL008W | COQ10 | n |
| W303.asm02.HP0_G0049020 | 1021.9351 | 1216.59254 | 1158.144121 | 1402.57326 | 1454.3596 | 1799.58574 | 1493.5231 | 1483.9661 | 1382.4302 | 1.44E-05 | -0.454372884 | YOL006C | TOP1 | n |
| W303.asm02.HP0_G0049030 | 954.50842 | 952.928286 | 824.899983 | 974.217554 | 930.32271 | 851.437297 | 893.83366 | 997.18321 | 929.26112 | 0.88663544 | -0.013808715 | YOL005C | RPB11 | n |
| W303.asm02.HP0_G0049040 | 1185.234 | 1413.8217 | 1463.376434 | 4380.2288 | 4508.6944 | 4548.26806 | 4535.2937 | 4450.5861 | 4761.802 | 8.03E-105 | -1.725181178 | YOL004W | SIN3 | n |
| W303.asm02.HP0_G0049050 | 868.11803 | 838.742979 | 891.5488106 | 486.692089 | 444.03809 | 426.666797 | 648.71346 | 573.38034 | 633.90772 | 1.06E-25 | 0.935638826 | YOL003C | PFA4 | n |
| W303.asm02.HP0_G0049060 | 1927.9806 | 1740.8069 | 2246.74164 | 230.845391 | 226.51336 | 170.666719 | 350.00884 | 443.48411 | 384.40023 | 5.37E-133 | 3.233319104 | YOL002C | IZH2 | n |
| W303.asm02.HP0_G0049070 | 607.89333 | 561.584099 | 582.4527983 | 1085.89005 | 1073.2419 | 904.53361 | 786.66483 | 755.75991 | 840.21427 | 1.33E-17 | -0.807194103 | YOL001W | PHO80 | n |
| W303.asm02.HP0_G0049080 | 934.49113 | 952.928286 | 1022.914616 | 2222.6161 | 2141.9894 | 2155.1414 | 1482.1221 | 1797.5539 | 1583.4468 | 1.77E-45 | -1.163288211 | YOR001W | RRP6 | n |
| W303.asm02.HP0_G0049090 | 2874.0607 | 2643.90887 | 2662.089406 | 1151.72683 | 1217.0599 | 1139.67442 | 1550.5278 | 1508.8956 | 1535.8376 | 1.06E-76 | 1.221089122 | YOR002W | ALG6 | n |
| W303.asm02.HP0_G0049110 | 553.10918 | 529.404603 | 621.0897998 | 1018.38653 | 882.683 | 1063.82255 | 562.06632 | 659.97783 | 596.87834 | 1.12E-14 | -0.798508081 | YOR004W | UTP23 | n |
| W303.asm02.HP0_G0049120 | 518.34232 | 561.584099 | 510.9743454 | 877.545838 | 870.09892 | 923.496578 | 806.04642 | 688.84366 | 763.51056 | 1.93E-17 | -0.747668406 | YOR005C | DNL4 | n |
| W303.asm02.HP0_G0049130 | 1406.4776 | 1290.29396 | 1497.18381 | 1156.72709 | 1138.8588 | 1215.5263 | 1022.6643 | 1162.5057 | 1023.5979 | 0.002326938 | 0.256958703 | YOR006C | TSR3 | n |
| W303.asm02.HP0_G0049140 | 9241.6643 | 9607.13647 | 7156.538611 | 7121.2053 | 6910.4551 | 7337.72076 | 6077.8409 | 6106.4351 | 5909.7128 | 0.003597184 | 0.283162477 | YOR007C | SGT2 | n |
| W303.asm02.HP0_G0049220 | 1429.6555 | 1563.30065 | 1794.688722 | 2446.79447 | 2457.4901 | 2488.88965 | 2867.3363 | 2819.6667 | 2967.6403 | 2.03E-14 | -0.626179875 | YOR014W | RTS1 | n |
| W303.asm02.HP0_G0049230 | 1630.8819 | 1497.90361 | 1817.870923 | 3787.69785 | 3859.7157 | 4017.30493 | 4616.2404 | 4794.3519 | 4578.4184 | 6.84E-61 | -1.236992372 | YOR016C | ERP4 | n |
| W303.asm02.HP0_G0049240 | 733.26474 | 740.128396 | 684.8408524 | 707.536957 | 692.12414 | 671.289094 | 532.42388 | 603.55826 | 566.02053 | 0.501202245 | 0.058618145 | YOR017W | PET127 | n |
| W303.asm02.HP0_G0049250 | 528.87773 | 568.850436 | 542.8498717 | 431.689215 | 462.01534 | 413.392719 | 570.04698 | 539.26618 | 633.02607 | 0.000824866 | 0.327529151 | YOR018W | ROD1 | n |
| W303.asm02.HP0_G0049270 | 7530.7132 | 7142.80995 | 8546.504742 | 1811.7613 | 1991.8794 | 2022.40062 | 2279.0478 | 2163.6251 | 2100.9765 | 1.07E-133 | 1.995219676 | YOR020C | HSP10 | n |
| W303.asm02.HP0_G0049280 | 998.75715 | 1080.60822 | 1316.555828 | 990.051714 | 1072.343 | 931.081766 | 995.30202 | 926.33072 | 947.77581 | 0.086664387 | 0.182559792 | YOR020W | MCO10 | n |
| W303.asm02.HP0_G0049290 | 3199.6049 | 2930.41019 | 3848.245354 | 1119.22513 | 1179.3076 | 1234.48927 | 777.54407 | 868.59906 | 807.59315 | 4.97E-51 | 1.49845271 | YOR021C | SFM1 | n |
| W303.asm02.HP0_G0049300 | 1157.8419 | 1194.79353 | 1159.110046 | 1406.74015 | 1339.3052 | 1409.89673 | 1803.6286 | 1431.4827 | 1742.1442 | 0.007135786 | -0.243035845 | YOR022C | DDL1 | n |
| W303.asm02.HP0_G0049310 | 497.27149 | 555.355809 | 566.9979977 | 961.716901 | 908.75001 | 999.348453 | 1450.1995 | 1525.9527 | 1453.844 | 1.39E-23 | -0.824266665 | YOR025W | HST3 | n |
| W303.asm02.HP0_G0049320 | 672.15935 | 686.149888 | 661.6586515 | 545.028469 | 509.65505 | 493.985336 | 573.46726 | 518.27285 | 547.50584 | 1.02E-05 | 0.382120966 | YOR026W | BUB3 | n |
| W303.asm02.HP0_G0049330 | 5875.5998 | 6699.56335 | 5130.02788 | 4057.71195 | 4149.1494 | 4058.07531 | 4310.6952 | 3730.2525 | 4018.5694 | 1.51E-08 | 0.529510317 | YOR027W | STI1 | n |
| W303.asm02.HP0_G0049340 | 392.9709 | 316.604714 | 384.4381654 | 1280.06686 | 1218.8576 | 1164.32628 | 1552.808 | 1314.7073 | 1543.7725 | 9.11E-65 | -1.74339754 | YOR028C | CIN5 | n |
| W303.asm02.HP0_G0049360 | 446.70151 | 472.31195 | 486.8262195 | 443.356492 | 406.28586 | 384.948266 | 453.75739 | 480.22244 | 484.90855 | 0.059124359 | 0.186490968 | YOR030W | DFG16 | n |
| W303.asm02.HP0_G0049380 | 237.04679 | 222.142324 | 240.5153346 | 885.046229 | 786.50471 | 903.585461 | 978.20061 | 869.91114 | 1034.1777 | 1.08E-70 | -1.879349003 | YOR032C | HMS1 | n |
| W303.asm02.HP0_G0049400 | 605.78625 | 633.209427 | 556.3728222 | 1091.72369 | 1003.1306 | 1046.75588 | 947.41807 | 799.05865 | 974.22536 | 1.90E-17 | -0.808067658 | YOR033C | EXO1 | n |
| W303.asm02.HP0_G0049430 | 501.48566 | 609.334318 | 552.5091221 | 812.542442 | 740.66272 | 662.755758 | 824.28793 | 663.91408 | 784.6702 | 0.000301192 | -0.414604888 | YOR035C | SHE4 | n |
| W303.asm02.HP0_G0049440 | 1360.1218 | 1518.66458 | 1396.727606 | 635.866547 | 641.78784 | 586.903883 | 658.9743 | 633.73617 | 651.54076 | 3.29E-51 | 1.196529255 | YOR036W | PEP12 | n |
| W303.asm02.HP0_G0049450 | 926.0628 | 926.97708 | 761.1489304 | 488.358842 | 485.38576 | 447.526063 | 515.32247 | 474.97411 | 471.68377 | 3.77E-18 | 0.877732564 | YOR037W | CYC2 | n |
| W303.asm02.HP0_G0049460 | 492.00379 | 493.072915 | 590.1801986 | 1454.24263 | 1476.8311 | 1457.30415 | 1483.2622 | 1503.6473 | 1444.1458 | 1.13E-71 | -1.476423413 | YOR038C | HIR2 | n |
| W303.asm02.HP0_G0049470 | 3408.2061 | 3254.28124 | 3481.193839 | 1617.58449 | 1573.0094 | 1623.23013 | 1576.7499 | 1780.4969 | 1693.6533 | 8.12E-60 | 1.075425553 | YOR039W | CKB2 | n |
| W303.asm02.HP0_G0049480 | 785.94181 | 670.579164 | 832.6273833 | 590.03082 | 633.69808 | 583.111289 | 776.40398 | 715.08533 | 815.52802 | 0.000661795 | 0.3418811 | YOR040W | GLO4 | n |
| W303.asm02.HP0_G0049490 | 891.29594 | 1117.97796 | 999.732415 | 971.717423 | 981.55787 | 986.074375 | 1240.4222 | 1053.6028 | 1215.798 | 0.739612089 | 0.033961937 | YOR042W | CUE5 | n |
| W303.asm02.HP0_G0049500 | 1313.766 | 1349.46271 | 1373.545405 | 2493.46358 | 2389.1766 | 2413.98592 | 2765.8679 | 2309.2664 | 2681.1035 | 3.84E-28 | -0.85397527 | YOR043W | WHI2 | n |
| W303.asm02.HP0_G0049520 | 1561.3482 | 1534.2353 | 1944.407103 | 985.051453 | 1121.7804 | 987.022524 | 961.0992 | 1036.5457 | 975.10702 | 4.02E-12 | 0.704569681 | YOR045W | TOM6 | n |
| W303.asm02.HP0_G0049530 | 2790.8309 | 2543.21819 | 2320.151943 | 3499.34945 | 3519.0468 | 3609.6011 | 3070.273 | 3252.6542 | 3095.4799 | 1.59E-10 | -0.473867548 | YOR046C | DBP5 | n |
| W303.asm02.HP0_G0049550 | 1190.5017 | 1261.22861 | 1235.418124 | 2393.45835 | 2482.6583 | 2517.3341 | 2126.2752 | 2183.3064 | 2201.4848 | 4.07E-60 | -1.003391792 | YOR048C | RAT1 | n |
| W303.asm02.HP0_G0049560 | 223.35076 | 214.875986 | 273.3567859 | 556.695745 | 556.3959 | 587.852031 | 583.7281 | 602.24617 | 578.36365 | 3.51E-32 | -1.254246596 | YOR049C | RSB1 | n |
| W303.asm02.HP0_G0049570 | 4200.4692 | 4041.1218 | 3921.655657 | 1243.39828 | 1237.7337 | 1379.55598 | 897.25394 | 1142.8244 | 881.65191 | 2.57E-68 | 1.655867235 | YOR051C | ETT1 | n |
| W303.asm02.HP0_G0049580 | 3719.0008 | 3782.64779 | 3890.746056 | 1121.72526 | 1030.0965 | 1062.8744 | 1689.6192 | 1507.5836 | 1618.7129 | 3.10E-159 | 1.824965191 | YOR052C | TMC1 | n |

| | | | | | | | | | | | | | | |
|---|---|---|---|---|---|---|---|---|---|---|---|---|---|---|
| W303.asm02.HP0_G0049600 | 1550.8128 | 1470.91436 | 1276.952901 | 1368.40481 | 1280.8791 | 1358.69671 | 661.25449 | 846.29364 | 787.31516 | 0.331135719 | 0.100544755 | YOR056C | NOB1 | n |
| W303.asm02.HP0_G0049610 | 535.19898 | 560.54605 | 540.9180216 | 357.518675 | 384.71316 | 359.348258 | 400.17298 | 354.26246 | 350.89746 | 3.08E-09 | 0.571277946 | YOR057W | SGT1 | n |
| W303.asm02.HP0_G0049620 | 989.27528 | 915.558549 | 842.2866337 | 458.357275 | 472.80169 | 471.229774 | 635.03233 | 532.70577 | 622.44625 | 1.03E-23 | 0.96968507 | YOR058C | ASE1 | n |
| W303.asm02.HP0_G0049630 | 929.22343 | 957.080479 | 877.0599351 | 785.874383 | 753.24679 | 730.074297 | 934.87704 | 894.84072 | 874.5987 | 0.000290245 | 0.283159178 | YOR059C | LPL1 | n |
| W303.asm02.HP0_G0049640 | 325.54426 | 325.947148 | 300.402687 | 513.360148 | 523.13799 | 569.837211 | 567.76679 | 552.38701 | 521.05628 | 1.10E-14 | -0.754884598 | YOR060C | SLD7 | n |
| W303.asm02.HP0_G0049650 | 2612.7824 | 2456.02214 | 2591.576879 | 1997.60434 | 1900.1954 | 1990.16357 | 1988.3239 | 2059.9706 | 1902.6048 | 1.87E-09 | 0.379653969 | YOR061W | CKA2 | n |
| W303.asm02.HP0_G0049670 | 12150.492 | 13400.1647 | 13991.42418 | 26363.0437 | 25631.964 | 28654.9421 | 21714.229 | 23141.211 | 20127.232 | 4.35E-40 | -1.02821786 | YOR063W | RPL3 | n |
| W303.asm02.HP0_G0049680 | 576.28709 | 608.296269 | 600.805374 | 422.52207 | 412.5779 | 440.889024 | 381.93147 | 432.98745 | 383.51858 | 1.46E-07 | 0.485191288 | YOR064C | YNG1 | n |
| W303.asm02.HP0_G0049690 | 2826.6513 | 3075.73694 | 3355.623584 | 8149.59235 | 7756.2847 | 8946.72866 | 7549.7021 | 7051.135 | 6727.8857 | 3.07E-66 | -1.424324624 | YOR065W | CYT1 | n |
| W303.asm02.HP0_G0049700 | 345.56155 | 399.648573 | 435.6321924 | 770.873599 | 764.03314 | 783.17061 | 938.29732 | 880.40781 | 933.66938 | 4.11E-25 | -0.971328309 | YOR066W | MSA1 | n |
| W303.asm02.HP0_G0049710 | 1923.7664 | 1710.7035 | 1759.91542 | 937.548972 | 976.1647 | 942.459547 | 1324.7892 | 1187.4353 | 1244.8925 | 9.84E-34 | 0.917257179 | YOR067C | ALG8 | n |
| W303.asm02.HP0_G0049730 | 425.63069 | 425.599779 | 453.9847682 | 466.691044 | 426.06084 | 413.392719 | 514.18237 | 501.21577 | 571.31044 | 0.987711079 | -0.001654314 | YOR070C | GYP1 | n |
| W303.asm02.HP0_G0049750 | 365.57883 | 372.659319 | 297.5049119 | 229.178638 | 237.29971 | 219.970438 | 234.85935 | 213.86956 | 230.9928 | 1.77E-06 | 0.591580107 | YOR073W | SGO1 | n |
| W303.asm02.HP0_G0049760 | 746.96078 | 728.709866 | 751.48968 | 1110.89136 | 1083.1293 | 1138.72627 | 1042.0459 | 1028.6732 | 1148.7924 | 3.00E-14 | -0.58125195 | YOR074C | CDC21 | n |
| W303.asm02.HP0_G0049770 | 1335.8904 | 1325.5876 | 1638.208866 | 715.037349 | 713.69684 | 680.770578 | 827.70821 | 848.91781 | 892.23174 | 2.17E-27 | 1.027680632 | YOR075W | UFE1 | n |
| W303.asm02.HP0_G0049790 | 105.35413 | 147.40285 | 86.93325348 | 361.685559 | 324.48937 | 308.148242 | 255.38105 | 267.66497 | 270.66714 | 2.76E-22 | -1.553599072 | YOR077W | RTS2 | n |
| W303.asm02.HP0_G0049800 | 1786.806 | 1861.2205 | 1597.640014 | 1285.06712 | 1278.1825 | 1308.44484 | 761.58276 | 877.78364 | 849.03079 | 8.33E-08 | 0.437844509 | YOR078W | BUD21 | n |
| W303.asm02.HP0_G0049810 | 1138.8781 | 1039.08629 | 1277.918826 | 577.530167 | 548.30614 | 474.074219 | 808.32661 | 720.33366 | 804.9482 | 2.57E-26 | 1.110318086 | YOR079C | ATX2 | n |
| W303.asm02.HP0_G0049820 | 1033.524 | 930.091224 | 1044.164967 | 525.860801 | 541.11524 | 516.740899 | 535.84416 | 474.97411 | 533.39941 | 2.19E-26 | 0.925504565 | YOR080W | DIA2 | n |
| W303.asm02.HP0_G0049840 | 143.28182 | 161.935526 | 162.2754065 | 577.530167 | 585.1595 | 552.770539 | 729.66013 | 648.16909 | 786.43351 | 1.31E-59 | -1.874747097 | YOR083W | WHI5 | n |
| W303.asm02.HP0_G0049850 | 252.84991 | 263.664253 | 283.9819614 | 212.5111 | 186.06454 | 200.05932 | 231.43907 | 183.69164 | 263.61392 | 0.003309223 | 0.419231602 | YOR084W | LPX1 | n |
| W303.asm02.HP0_G0049860 | 1850.0185 | 1743.92105 | 1532.923036 | 2510.96449 | 2449.4004 | 2525.86744 | 3526.3106 | 3734.1887 | 3630.6426 | 4.54E-13 | -0.546747928 | YOR085W | OST3 | n |
| W303.asm02.HP0_G0049870 | 1820.5194 | 1835.26929 | 1941.509328 | 3356.84201 | 3478.598 | 3326.10472 | 4718.8489 | 4362.6765 | 4669.2285 | 4.28E-44 | -0.860033421 | YOR086C | TCB1 | n |
| W303.asm02.HP0_G0049880 | 1071.4515 | 933.205369 | 840.3547836 | 767.540092 | 718.19116 | 786.963203 | 1286.026 | 1277.969 | 1208.7448 | 0.000781191 | 0.323313062 | YOR087W | YVC1 | n |
| W303.asm02.HP0_G0049890 | 2646.4957 | 2609.65328 | 2657.259781 | 1285.06712 | 1328.5188 | 1226.90408 | 1351.0113 | 1213.6769 | 1311.898 | 1.46E-55 | 1.042778944 | YOR089C | VPS21 | n |
| W303.asm02.HP0_G0049900 | 807.01264 | 897.911729 | 962.0613385 | 1702.58893 | 1575.706 | 1605.21531 | 1861.7734 | 1792.3056 | 1789.7534 | 1.73E-28 | -0.872125986 | YOR090C | PTC5 | n |
| W303.asm02.HP0_G0049910 | 797.53076 | 928.015128 | 949.504313 | 2468.46227 | 2586.9263 | 2684.20823 | 2103.4733 | 2382.743 | 2152.1123 | 2.13E-73 | -1.531879001 | YOR091W | TMA46 | n |
| W303.asm02.HP0_G0049920 | 367.68591 | 383.039801 | 412.4499915 | 404.187779 | 358.64615 | 320.474172 | 445.77673 | 452.66869 | 444.35256 | 0.371964424 | 0.101290184 | YOR092W | ECM3 | n |
| W303.asm02.HP0_G0049930 | 1001.9178 | 932.167321 | 1022.914616 | 1242.5649 | 1216.161 | 1134.93368 | 1218.7604 | 1211.0528 | 1230.7861 | 0.000136441 | -0.281563406 | YOR093C | CMR2 | n |
| W303.asm02.HP0_G0049940 | 1006.1319 | 939.433659 | 1075.074568 | 1025.88692 | 911.4466 | 922.54843 | 1181.1373 | 1119.2069 | 1107.3548 | 0.362029879 | 0.078689595 | YOR094W | ARF3 | n |
| W303.asm02.HP0_G0049950 | 3135.3389 | 3027.98672 | 3528.524166 | 1970.93628 | 1973.9021 | 2136.17843 | 1601.832 | 1733.2619 | 1734.2093 | 3.41E-18 | 0.67295995 | YOR095C | RKI1 | n |
| W303.asm02.HP0_G0049960 | 10230.94 | 9929.96948 | 12098.21111 | 14777.4386 | 15695.937 | 16919.7089 | 13953.61 | 14713.701 | 12416.304 | 2.24E-09 | -0.554858341 | YOR096W | RPS7A | n |
| W303.asm02.HP0_G0049970 | 759.60328 | 742.204493 | 795.9222318 | 227.511884 | 239.99629 | 204.800063 | 235.99945 | 221.74206 | 243.33593 | 5.78E-68 | 1.772056623 | YOR097C | | n |
| W303.asm02.HP0_G0049980 | 948.18717 | 1111.74967 | 1140.757471 | 1478.41056 | 1502.8981 | 1605.21531 | 1335.05 | 1184.8111 | 1407.9981 | 2.55E-08 | -0.518285706 | YOR098C | NUP1 | n |
| W303.asm02.HP0_G0049990 | 5398.3456 | 5631.41171 | 5066.276828 | 2886.81746 | 2676.8126 | 2764.80084 | 3242.4272 | 3892.9508 | 3310.6029 | 8.66E-32 | 0.950404687 | YOR099W | KTR1 | n |
| W303.asm02.HP0_G0050000 | 55.837689 | 70.5872804 | 35.73922643 | 36.668582 | 26.965876 | 36.0296406 | 132.2509 | 86.597489 | 102.27162 | 0.016188194 | 0.697420261 | YOR100C | CRC1 | n |
| W303.asm02.HP0_G0050020 | 2489.5181 | 2576.43574 | 3140.222301 | 970.05067 | 1049.8714 | 929.185469 | 986.18127 | 1146.7607 | 939.84094 | 1.14E-44 | 1.476602056 | YOR103C | OST2 | n |
| W303.asm02.HP0_G0050030 | 911.31322 | 826.2864 | 865.4688346 | 2448.46123 | 2347.8289 | 2105.83768 | 2064.7101 | 2344.6926 | 2146.8224 | 5.20E-63 | -1.407304055 | YOR104W | PIN2 | n |
| W303.asm02.HP0_G0050040 | 1055.6484 | 1087.87456 | 1028.710166 | 283.348134 | 324.48937 | 298.666758 | 365.97016 | 356.88662 | 372.93876 | 3.13E-91 | 1.807791797 | YOR106W | VAM3 | n |
| W303.asm02.HP0_G0050050 | 610.00041 | 516.948024 | 504.2128702 | 642.533562 | 626.50718 | 641.896492 | 571.18707 | 418.55453 | 616.27469 | 0.081866277 | -0.228922097 | YOR107W | RGS2 | n |
| W303.asm02.HP0_G0050070 | 981.90049 | 977.841443 | 1221.895174 | 1695.08854 | 1652.1093 | 1627.02272 | 1467.3009 | 1321.2678 | 1542.0092 | 3.21E-11 | -0.64400328 | YOR109W | INP53 | n |
| W303.asm02.HP0_G0050080 | 577.34063 | 642.551861 | 549.611347 | 1889.26535 | 1854.3534 | 1846.04501 | 1833.2711 | 1947.1314 | 1892.025 | 6.54E-108 | -1.660335981 | YOR110W | TFC7 | n |
| W303.asm02.HP0_G0050100 | 400.34569 | 440.132454 | 477.1669691 | 1555.08123 | 1582.8969 | 1514.19306 | 1563.0688 | 1609.926 | 1424.7495 | 1.12E-98 | -1.818680208 | YOR112W | CEX1 | n |
| W303.asm02.HP0_G0050110 | 400.34569 | 452.589033 | 514.8380456 | 2242.61714 | 2363.1096 | 2530.60818 | 2209.5021 | 1865.7823 | 2093.0416 | 3.50E-124 | -2.381673185 | YOR113W | AZF1 | n |
| W303.asm02.HP0_G0050120 | 349.77571 | 368.507126 | 316.8234127 | 304.182555 | 341.56776 | 292.977867 | 329.48715 | 355.57454 | 335.02773 | 0.221521805 | 0.139804521 | YOR114W | CIM1 | n |
| W303.asm02.HP0_G0050130 | 2106.0291 | 1986.82433 | 2336.572668 | 826.709849 | 793.69561 | 786.963203 | 823.14783 | 886.96822 | 882.53356 | 8.75E-69 | 1.417230122 | YOR115C | TRS33 | n |
| W303.asm02.HP0_G0050140 | 1223.1614 | 1278.87543 | 1484.626784 | 3470.18126 | 3546.0127 | 3755.61596 | 2936.882 | 2960.0596 | 2935.9009 | 1.78E-75 | -1.433093483 | YOR116C | RPO31 | n |
| W303.asm02.HP0_G0050150 | 4282.6454 | 3975.72476 | 4065.578488 | 3939.37244 | 3999.0394 | 4004.03085 | 3874.0392 | 4047.7766 | 3914.5345 | 0.427725929 | 0.045345497 | YOR117W | RPT5 | n |
| W303.asm02.HP0_G0050160 | 1537.1168 | 1438.73486 | 1579.287438 | 784.207629 | 774.81949 | 793.600242 | 766.14314 | 770.19282 | 795.25002 | 2.46E-40 | 0.953565621 | YOR118W | RTC5 | n |
| W303.asm02.HP0_G0050170 | 824.92284 | 789.954712 | 828.7636831 | 1051.7216 | 1048.0737 | 1205.09666 | 682.91628 | 745.26324 | 655.94902 | 6.02E-07 | -0.434806998 | YOR119C | RIO1 | n |
| W303.asm02.HP0_G0050190 | 8307.1731 | 8136.22212 | 9682.432587 | 3486.01542 | 3839.9407 | 3846.63821 | 3356.4366 | 3377.3021 | 3216.2662 | 1.45E-54 | 1.225791741 | YOR122C | PFY1 | n |
| W303.asm02.HP0_G0050200 | 1663.5417 | 1824.88881 | 1390.932056 | 1534.24681 | 1501.9993 | 1404.20784 | 1183.4175 | 1276.6569 | 1359.5072 | 0.188770061 | 0.135308564 | YOR123C | LEO1 | n |
| W303.asm02.HP0_G0050210 | 1518.153 | 1462.60997 | 1601.503714 | 1678.421 | 1794.1296 | 1737.95609 | 1827.5706 | 1708.3323 | 1849.7057 | 0.008198997 | -0.18484054 | YOR124C | UBP2 | n |
| W303.asm02.HP0_G0050220 | 1316.9266 | 1342.19638 | 1559.968937 | 726.704625 | 744.25817 | 780.326164 | 816.30727 | 801.68282 | 758.22064 | 1.46E-25 | 0.907239863 | YOR125C | CAT5 | n |
| W303.asm02.HP0_G0050230 | 438.27318 | 387.191994 | 471.3714188 | 1100.89084 | 1154.1395 | 1205.09666 | 1046.6062 | 1274.0328 | 1028.8878 | 9.77E-41 | -1.414717122 | YOR126C | IAH1 | n |
| W303.asm02.HP0_G0050240 | 793.3166 | 897.911729 | 914.7310116 | 2363.45679 | 2347.8289 | 2403.55629 | 2180.9997 | 2138.6956 | 2180.3252 | 6.13E-96 | -1.448214794 | YOR127W | RGA1 | n |
| W303.asm02.HP0_G0050250 | 437.21964 | 393.420284 | 364.1537396 | 2854.31576 | 2875.4612 | 2624.47488 | 2643.8779 | 2597.9247 | 2868.8953 | 1.08E-233 | -2.80739501 | YOR128C | ADE2 | n |
| W303.asm02.HP0_G0050260 | 401.39924 | 399.648573 | 449.155143 | 538.361454 | 570.7777 | 647.585383 | 572.32716 | 489.40702 | 519.29298 | 4.99E-06 | -0.48873522 | YOR129C | AFI1 | n |
| W303.asm02.HP0_G0050270 | 1525.5278 | 1337.00613 | 1602.469639 | 672.535129 | 631.00149 | 722.48911 | 416.13429 | 558.94743 | 435.53604 | 2.15E-24 | 1.140489126 | YOR130C | ORT1 | n |
| W303.asm02.HP0_G0050280 | 1075.6657 | 1075.41798 | 1018.084991 | 267.513973 | 254.37809 | 269.274156 | 316.94612 | 284.72205 | 321.80295 | 1.08E-105 | 2.00178353 | YOR131C | | n |

| | | | | | | | | | | | | | | |
|---|---|---|---|---|---|---|---|---|---|---|---|---|---|---|
| W303.asm02.HP0_G0050290 | 578.39417 | 604.144077 | 445.2914428 | 703.370073 | 698.41618 | 610.607594 | 677.21581 | 694.092 | 722.95457 | 0.008275455 | -0.307858647 | YOR132W | VPS17 | n |
| W303.asm02.HP0_G0050320 | 5575.3406 | 5390.58452 | 4752.35119 | 8974.63545 | 9320.3055 | 9970.72897 | 8956.5781 | 8655.8127 | 8716.0108 | 1.78E-31 | -0.846711969 | YOR136W | IDH2 | n |
| W303.asm02.HP0_G0050330 | 690.06955 | 608.296269 | 597.9075989 | 1636.75216 | 1402.2255 | 1587.20048 | 1396.6151 | 1377.6873 | 1438.8559 | 4.85E-49 | -1.28750189 | YOR137C | SIA1 | n |
| W303.asm02.HP0_G0050340 | 950.29425 | 963.308768 | 1028.710166 | 933.382088 | 852.12167 | 851.437297 | 967.93976 | 829.23656 | 974.22536 | 0.076023109 | 0.15787815 | YOR138C | RUP1 | n |
| W303.asm02.HP0_G0050350 | 381.38195 | 485.806577 | 485.8602944 | 1495.91147 | 1473.2357 | 1558.75603 | 1575.6098 | 1380.3115 | 1546.4175 | 1.11E-70 | -1.741404033 | YOR140W | SFL1 | n |
| W303.asm02.HP0_G0050360 | 621.58937 | 722.481576 | 752.4556051 | 1706.75582 | 1860.6454 | 1622.28198 | 1723.8221 | 1605.9898 | 1763.3038 | 9.27E-48 | -1.306956967 | YOR141C | ARP8 | n |
| W303.asm02.HP0_G0050370 | 4318.4658 | 4195.79099 | 3993.13411 | 1950.10186 | 2158.1689 | 2398.81555 | 2307.5502 | 1788.3694 | 2466.8621 | 1.99E-17 | 0.942947599 | YOR142W | LSC1 | n |
| W303.asm02.HP0_G0050380 | 1925.8735 | 1670.21962 | 2069.977358 | 871.7122 | 753.24679 | 811.615063 | 910.93507 | 900.08905 | 936.31433 | 3.04E-39 | 1.217332474 | YOR143C | THI80 | n |
| W303.asm02.HP0_G0050390 | 494.11087 | 476.464143 | 347.7330139 | 581.697051 | 619.31628 | 544.237203 | 549.52528 | 561.5716 | 595.11504 | 0.001137948 | -0.406665691 | YOR144C | ELG1 | n |
| W303.asm02.HP0_G0050400 | 2583.2833 | 2365.71194 | 2501.74585 | 2092.60931 | 2145.5848 | 2204.44512 | 1231.3015 | 1594.1811 | 1365.6788 | 0.019111303 | 0.209850483 | YOR145C | PNO1 | n |
| W303.asm02.HP0_G0050410 | 1594.008 | 1432.50657 | 1528.093411 | 573.363282 | 531.22775 | 525.274235 | 494.80077 | 509.08827 | 521.05628 | 9.14E-76 | 1.481632203 | YOR147W | MDM32 | n |
| W303.asm02.HP0_G0050420 | 715.35454 | 667.465019 | 821.0362828 | 198.343694 | 227.41222 | 194.37043 | 208.63719 | 188.93998 | 242.45428 | 1.50E-48 | 1.829970897 | YOR148C | SPP2 | n |
| W303.asm02.HP0_G0050430 | 653.19561 | 578.19287 | 592.1120487 | 608.365111 | 649.8776 | 605.866852 | 886.99309 | 850.22989 | 902.81156 | 0.722774113 | -0.032201773 | YOR149C | SMP3 | n |
| W303.asm02.HP0_G0050440 | 2672.8343 | 2630.41424 | 3255.16738 | 1610.91748 | 1586.4924 | 1629.86716 | 1583.5905 | 1756.8794 | 1536.7193 | 1.68E-19 | 0.826485009 | YOR150W | MRPL23 | n |
| W303.asm02.HP0_G0050450 | 2947.8086 | 3175.38957 | 3242.610355 | 4797.75061 | 4800.8247 | 4863.05334 | 4849.9597 | 4820.5936 | 4700.0863 | 3.86E-28 | -0.626483717 | YOR151C | RPB2 | n |
| W303.asm02.HP0_G0050470 | 2024.9064 | 2158.10229 | 2332.708968 | 1631.7519 | 1534.3583 | 1567.28937 | 1948.4206 | 2100.6452 | 1976.6636 | 3.22E-10 | 0.461221328 | YOR153W | PDR5 | n |
| W303.asm02.HP0_G0050480 | 1145.1994 | 1050.50482 | 1014.221291 | 496.692611 | 584.26064 | 527.170531 | 625.91158 | 653.41742 | 640.96094 | 4.49E-29 | 0.997359909 | YOR154W | SLP1 | n |
| W303.asm02.HP0_G0050490 | 773.29931 | 760.889361 | 749.55783 | 295.848787 | 293.02918 | 252.207484 | 445.77673 | 316.21204 | 417.02135 | 7.37E-33 | 1.439686492 | YOR155C | ISN1 | n |
| W303.asm02.HP0_G0050510 | 2481.0898 | 2498.58212 | 2255.434965 | 2410.95927 | 2416.1425 | 2377.00813 | 2583.4529 | 2616.2938 | 2508.2997 | 0.932039158 | 0.005776997 | YOR157C | PUP1 | n |
| W303.asm02.HP0_G0050520 | 2834.0261 | 2796.50196 | 3064.880148 | 2020.10552 | 2164.461 | 2083.08212 | 1844.672 | 1920.8898 | 1925.5278 | 2.74E-13 | 0.472742769 | YOR158W | PET123 | n |
| W303.asm02.HP0_G0050530 | 1245.2858 | 1242.54375 | 1506.84306 | 515.026902 | 525.83458 | 430.459391 | 420.69467 | 490.71911 | 464.63056 | 3.62E-39 | 1.440441046 | YOR159C | SME1 | n |
| W303.asm02.HP0_G0050540 | 556.26981 | 549.12752 | 612.3964745 | 2625.13712 | 2733.4409 | 2887.11199 | 2494.5256 | 2659.5926 | 2532.9859 | 1.16E-198 | -2.26182804 | YOR160W | MTR10 | n |
| W303.asm02.HP0_G0050550 | 671.10581 | 671.617212 | 703.1934281 | 398.354141 | 382.01657 | 419.08161 | 921.19591 | 771.5049 | 908.10147 | 9.25E-16 | 0.771109036 | YOR161C | PNS1 | n |
| W303.asm02.HP0_G0050560 | 678.4806 | 618.676752 | 733.1371043 | 3105.99557 | 3066.0201 | 3653.21593 | 3779.4114 | 3730.2525 | 3465.7737 | 1.19E-135 | -2.273910152 | YOR162C | YRR1 | n |
| W303.asm02.HP0_G0050570 | 1220.0008 | 1308.97883 | 971.7205889 | 505.02638 | 488.08235 | 473.12607 | 511.90218 | 505.15202 | 544.86088 | 4.70E-30 | 1.25447034 | YOR163W | DDP1 | n |
| W303.asm02.HP0_G0050580 | 2330.4334 | 2163.29254 | 2596.406504 | 2004.27136 | 2035.0248 | 2070.75619 | 2070.4106 | 2103.2693 | 2030.4444 | 0.004602882 | 0.215115955 | YOR164C | GET4 | n |
| W303.asm02.HP0_G0050590 | 2851.9363 | 2542.18014 | 2788.625586 | 2416.79291 | 2400.8618 | 2328.65256 | 2430.6803 | 2340.7564 | 2358.4189 | 0.002736454 | 0.195300163 | YOR165W | SEY1 | n |
| W303.asm02.HP0_G0050600 | 1115.7002 | 1069.18969 | 935.0154374 | 542.528338 | 591.45154 | 619.14093 | 580.30782 | 547.13868 | 556.32236 | 5.22E-19 | 0.831891561 | YOR166C | SWT1 | n |
| W303.asm02.HP0_G0050610 | 3765.3566 | 3313.44999 | 3871.427555 | 5109.43355 | 5258.3458 | 5633.89802 | 4063.2948 | 4444.0257 | 4073.2318 | 1.19E-12 | -0.547019215 | YOR168W | GLN4 | n |
| W303.asm02.HP0_G0050650 | 1029.3098 | 1063.99945 | 919.5606368 | 782.540875 | 734.37068 | 727.229852 | 723.95966 | 728.20616 | 783.78855 | 6.88E-07 | 0.423807014 | YOR174W | MED4 | n |
| W303.asm02.HP0_G0050660 | 1739.3967 | 1757.41567 | 1521.331936 | 2103.4432 | 2026.0361 | 2139.97102 | 2834.2736 | 3373.3658 | 2888.2917 | 0.000243321 | -0.321532225 | YOR175C | ALE1 | n |
| W303.asm02.HP0_G0050670 | 1963.801 | 1826.9649 | 1788.893172 | 2765.97781 | 2672.3183 | 3002.7861 | 3577.6148 | 4009.7262 | 3597.1398 | 5.23E-15 | -0.597284058 | YOR176W | HEM15 | n |
| W303.asm02.HP0_G0050720 | 268.65303 | 323.871051 | 341.9374637 | 725.871248 | 675.94462 | 751.881711 | 741.06107 | 699.34033 | 796.13168 | 6.88E-30 | -1.202871686 | YOR181W | LAS17 | n |
| W303.asm02.HP0_G0050740 | 3202.7656 | 2951.17115 | 2802.148537 | 2453.46149 | 2419.7379 | 2263.23032 | 2143.3766 | 2052.0981 | 2212.0646 | 5.47E-06 | 0.327169775 | YOR184W | SER1 | n |
| W303.asm02.HP0_G0050770 | 5503.6998 | 5612.72684 | 5492.24977 | 4073.54611 | 3839.9407 | 4401.30505 | 4038.2128 | 3761.7424 | 3791.1032 | 5.88E-11 | 0.431685088 | YOR187W | TUF1 | n |
| W303.asm02.HP0_G0050780 | 470.93296 | 533.556796 | 526.429146 | 1143.39306 | 1204.4758 | 1265.77816 | 1104.751 | 1082.4686 | 1228.1411 | 2.10E-44 | -1.238058269 | YOR188W | MSB1 | n |
| W303.asm02.HP0_G0050810 | 681.64122 | 748.432782 | 734.1030294 | 664.201361 | 597.74358 | 674.133539 | 644.15308 | 611.43076 | 644.48755 | 0.066607783 | 0.160981775 | YOR191W | ULS1 | n |
| W303.asm02.HP0_G0050840 | 1704.6298 | 1824.88881 | 1567.696338 | 2017.60539 | 2102.4394 | 2144.71177 | 1899.3965 | 2083.5881 | 2081.5802 | 0.000103039 | -0.29772826 | YOR194C | TOA1 | n |
| W303.asm02.HP0_G0050850 | 1431.7626 | 1430.43048 | 1425.705357 | 760.0397 | 773.92063 | 742.400227 | 711.41863 | 742.63907 | 772.32708 | 9.63E-39 | 0.913324328 | YOR195W | SLK19 | n |
| W303.asm02.HP0_G0050860 | 1511.8318 | 1268.49495 | 1492.354185 | 2789.31236 | 2764.9011 | 2710.75638 | 2536.709 | 2603.173 | 2586.7667 | 1.73E-37 | -0.951736336 | YOR196C | LIP5 | n |
| W303.asm02.HP0_G0050870 | 1427.5485 | 1595.48015 | 1852.644224 | 1510.07888 | 1440.8766 | 1368.1782 | 1351.0113 | 1326.5161 | 1440.6192 | 0.072088024 | 0.175117316 | YOR197W | MCA1 | n |
| W303.asm02.HP0_G0050880 | 8448.3477 | 8780.85007 | 7725.468459 | 9159.64511 | 8977.8389 | 9682.49185 | 7457.3545 | 7124.6116 | 7158.1319 | 0.017187141 | -0.15686383 | YOR198C | BFR1 | n |
| W303.asm02.HP0_G0050890 | 1152.5742 | 1257.07642 | 1150.416721 | 987.551584 | 968.9738 | 876.089157 | 749.04173 | 759.69616 | 760.8656 | 3.18E-05 | 0.328844759 | YOR201C | MRM1 | n |
| W303.asm02.HP0_G0050900 | 604.73271 | 607.258221 | 517.7358207 | 722.537741 | 738.86499 | 776.533571 | 769.56342 | 663.91408 | 789.07846 | 0.00016868 | -0.372183816 | YOR202W | HIS3 | n |
| W303.asm02.HP0_G0050910 | 3254.3891 | 3098.574 | 3706.254373 | 8060.42103 | 7566.6247 | 7602.25417 | 7361.5866 | 6877.94 | 7041.7538 | 3.11E-59 | -1.207198376 | YOR204W | DED1 | n |
| W303.asm02.HP0_G0050920 | 667.94518 | 610.372366 | 639.4423756 | 735.871771 | 728.07864 | 673.185391 | 558.64604 | 625.86367 | 618.03799 | 0.076347152 | -0.156983717 | YOR205C | GEP3 | n |
| W303.asm02.HP0_G0050930 | 4709.3296 | 4081.60569 | 4310.923447 | 3906.03736 | 3896.569 | 4184.17906 | 2407.8784 | 2904.9521 | 2534.7492 | 0.133698211 | 0.12837155 | YOR206W | NOC2 | n |
| W303.asm02.HP0_G0050940 | 1266.3566 | 1197.90767 | 1297.237327 | 2766.81119 | 2799.9568 | 3073.89724 | 1854.9329 | 2159.6889 | 1916.7113 | 3.23E-51 | -1.199413182 | YOR207C | RET1 | n |
| W303.asm02.HP0_G0050950 | 504.64628 | 610.372366 | 572.7935479 | 1685.9214 | 1665.5923 | 1845.09686 | 1369.2528 | 1431.4827 | 1420.3412 | 2.04E-81 | -1.621682276 | YOR208W | PTP2 | n |
| W303.asm02.HP0_G0050960 | 3615.7537 | 3523.13573 | 2991.469845 | 3129.33012 | 3160.4006 | 3328.94917 | 2886.7179 | 3395.6712 | 2626.441 | 0.473895063 | 0.074659935 | YOR209C | NPT1 | n |
| W303.asm02.HP0_G0050970 | 2343.0759 | 2172.63497 | 2821.467038 | 1344.23688 | 1411.2142 | 1303.7041 | 1084.2293 | 1518.0802 | 1201.6916 | 2.23E-12 | 0.854223793 | YOR210W | RPB10 | n |
| W303.asm02.HP0_G0050980 | 932.38405 | 981.993636 | 976.550214 | 1267.56621 | 1312.3393 | 1375.76338 | 1254.1033 | 1306.8348 | 1262.5255 | 1.14E-10 | -0.45166522 | YOR211C | MGM1 | n |
| W303.asm02.HP0_G0050990 | 2220.8651 | 2207.92861 | 1810.143522 | 663.367984 | 626.50718 | 673.185391 | 648.71346 | 586.50118 | 651.54076 | 7.43E-69 | 1.667845957 | YOR212W | STE4 | n |
| W303.asm02.HP0_G0051020 | 2196.6336 | 2256.71688 | 2713.283433 | 1246.73179 | 1450.7641 | 1285.68928 | 1062.5676 | 1174.3144 | 1091.4851 | 1.57E-18 | 0.847850726 | YOR215C | AIM41 | n |
| W303.asm02.HP0_G0051030 | 1517.0995 | 1519.70263 | 1283.714376 | 2045.10682 | 2108.7315 | 1974.9932 | 1664.5372 | 1718.829 | 1707.7598 | 1.18E-10 | -0.505154749 | YOR216C | RUD3 | n |
| W303.asm02.HP0_G0051040 | 1037.7382 | 1011.05899 | 1137.859695 | 2045.9402 | 2105.136 | 2083.08212 | 1933.5993 | 1935.3227 | 1977.5452 | 5.66E-47 | -0.967340945 | YOR217W | RFC1 | n |
| W303.asm02.HP0_G0051050 | 585.76896 | 521.100217 | 542.8498717 | 748.372424 | 772.12291 | 842.903961 | 881.29262 | 938.13947 | 871.95374 | 3.46E-09 | -0.518122041 | YOR219C | STE13 | n |
| W303.asm02.HP0_G0051060 | 1265.3031 | 1270.57105 | 1430.534982 | 1350.07052 | 1283.5757 | 1084.68181 | 1193.6784 | 1193.9957 | 1268.6971 | 0.337684294 | 0.092810176 | YOR220W | RCN2 | n |
| W303.asm02.HP0_G0051070 | 673.21289 | 765.041554 | 721.5460038 | 353.35179 | 331.68027 | 391.585305 | 345.44847 | 379.19204 | 378.22867 | 8.97E-24 | 1.005464008 | YOR221C | MCT1 | n |
| W303.asm02.HP0_G0051090 | 409.82757 | 379.925656 | 423.0751669 | 375.019589 | 418.86994 | 378.311227 | 482.25974 | 489.40702 | 487.55351 | 0.628308038 | 0.04992232 | YOR223W | DSC3 | n |

| | | | | | | | | | | | | | | |
|---|---|---|---|---|---|---|---|---|---|---|---|---|---|---|
| W303.asm02.HP0_G0051100 | 2510.5889 | 2470.55481 | 2309.526767 | 2336.78873 | 2416.1425 | 2448.11927 | 1900.5366 | 2221.3568 | 1936.1076 | 0.824510745 | 0.017778553 | YOR224C | RPB8 | n |
| W303.asm02.HP0_G0051130 | 415.09527 | 385.115898 | 370.9152148 | 405.854533 | 415.27449 | 409.600125 | 392.19232 | 467.10161 | 496.37003 | 0.526347313 | -0.072185525 | YOR228C | MCP1 | n |
| W303.asm02.HP0_G0051160 | 439.32672 | 483.730481 | 539.9520966 | 1465.9099 | 1353.687 | 1355.85227 | 1585.8707 | 1367.1907 | 1601.9615 | 3.95E-55 | -1.512025582 | YOR231W | MKK1 | n |
| W303.asm02.HP0_G0051170 | 6204.3047 | 6237.63188 | 7224.153364 | 3695.19301 | 3716.7965 | 3845.69006 | 3408.8809 | 3627.91 | 3393.4782 | 1.53E-28 | 0.805038382 | YOR232W | MGE1 | n |
| W303.asm02.HP0_G0051200 | 648.98144 | 734.938155 | 630.7490502 | 752.539308 | 749.65134 | 768.000235 | 782.10445 | 842.35739 | 870.19044 | 0.05209048 | -0.172643303 | YOR236W | DFR1 | n |
| W303.asm02.HP0_G0051240 | 982.95403 | 1172.99451 | 993.9368647 | 1431.74145 | 1515.4822 | 1579.6153 | 1337.3302 | 1461.6607 | 1404.4715 | 1.42E-09 | -0.523091288 | YOR239W | ABP140 | n |
| W303.asm02.HP0_G0051250 | 2155.5455 | 2027.30822 | 2117.307685 | 2448.46123 | 2236.37 | 2454.75631 | 2141.0964 | 2212.1722 | 2198.8399 | 0.005860494 | -0.180462917 | YOR241W | MET7 | n |
| W303.asm02.HP0_G0051270 | 2159.7597 | 2104.12379 | 2049.692932 | 2807.64665 | 2684.0035 | 2867.20088 | 1602.9721 | 2073.0914 | 1722.7478 | 5.99E-06 | -0.404910493 | YOR243C | PUS7 | n |
| W303.asm02.HP0_G0051280 | 1083.0405 | 1044.27653 | 913.7650865 | 545.028469 | 567.18225 | 543.289055 | 573.46726 | 569.4441 | 542.21593 | 5.72E-23 | 0.876591386 | YOR244W | ESA1 | n |
| W303.asm02.HP0_G0051290 | 1147.3065 | 1137.70087 | 1193.883348 | 905.047274 | 844.03191 | 813.51136 | 983.90108 | 880.40781 | 942.48589 | 1.61E-08 | 0.44044312 | YOR245C | DGA1 | n |
| W303.asm02.HP0_G0051300 | 1505.5105 | 1436.65877 | 1644.970341 | 968.383916 | 979.76015 | 969.955852 | 753.6021 | 768.88074 | 796.13168 | 4.53E-18 | 0.653047846 | YOR246C | ENV9 | n |
| W303.asm02.HP0_G0051330 | 584.71542 | 696.53037 | 555.4068972 | 704.20345 | 738.86499 | 731.022446 | 681.77618 | 678.347 | 709.72979 | 0.011888722 | -0.243803812 | YOR250C | CLP1 | n |
| W303.asm02.HP0_G0051340 | 950.29425 | 905.178067 | 921.4924868 | 1110.05798 | 1155.9372 | 1027.79291 | 1138.9539 | 1127.0794 | 1154.964 | 0.000964982 | -0.246784315 | YOR251C | TUM1 | n |
| W303.asm02.HP0_G0051350 | 1284.2668 | 1192.71743 | 1539.684512 | 853.377909 | 963.58062 | 824.889141 | 445.77673 | 650.79325 | 541.33427 | 3.38E-06 | 0.604755205 | YOR252W | TMA16 | n |
| W303.asm02.HP0_G0051360 | 1569.7765 | 1549.80602 | 1861.337549 | 991.718468 | 1108.2975 | 959.526219 | 841.38934 | 968.31738 | 886.94182 | 3.19E-13 | 0.703452796 | YOR253W | NAT5 | n |
| W303.asm02.HP0_G0051370 | 6514.0459 | 6045.59296 | 6469.765909 | 3993.54193 | 3962.186 | 3763.20115 | 4159.0627 | 4084.5149 | 4104.9713 | 4.43E-35 | 0.699250318 | YOR254C | SEC63 | n |
| W303.asm02.HP0_G0051380 | 9.4818717 | 12.4565789 | 14.48887558 | 48.3358581 | 52.134026 | 45.511125 | 35.342912 | 43.298745 | 32.621121 | 7.34E-09 | -1.997269635 | YOR255W | OSW1 | n |
| W303.asm02.HP0_G0051390 | 896.56365 | 826.2864 | 904.1058361 | 1212.56334 | 1132.5668 | 1181.39295 | 1263.2241 | 1295.0261 | 1258.1173 | 5.00E-09 | -0.424788864 | YOR256C | TRE2 | n |
| W303.asm02.HP0_G0051400 | 1087.2546 | 1100.33114 | 1166.837447 | 613.365372 | 599.5413 | 594.489071 | 609.95026 | 633.73617 | 633.90772 | 4.29E-32 | 0.892221843 | YOR257W | CDC31 | n |
| W303.asm02.HP0_G0051410 | 372.95362 | 342.55592 | 475.235119 | 279.181249 | 291.23146 | 293.926016 | 246.26029 | 284.72205 | 253.91575 | 0.000471034 | 0.464146324 | YOR258W | HNT3 | n |
| W303.asm02.HP0_G0051420 | 4174.1306 | 4111.70908 | 3528.524166 | 3849.36773 | 3803.9862 | 3840.94932 | 3883.16 | 3726.3162 | 3831.6592 | 0.603922434 | 0.039335889 | YOR259C | RPT4 | n |
| W303.asm02.HP0_G0051430 | 1956.4262 | 2045.99308 | 2494.01845 | 3317.6733 | 3167.5915 | 3384.88992 | 2698.6024 | 3092.58 | 2757.8072 | 1.94E-10 | -0.602972039 | YOR260W | GCD1 | n |
| W303.asm02.HP0_G0051440 | 6352.854 | 6298.87673 | 7729.332159 | 2900.15149 | 2771.1932 | 2645.33414 | 3070.273 | 2969.2442 | 3041.6991 | 5.98E-53 | 1.293127755 | YOR261C | RPN8 | n |
| W303.asm02.HP0_G0051450 | 969.258 | 947.738044 | 954.3339382 | 1108.39123 | 1020.209 | 1078.04477 | 1024.9445 | 948.63613 | 1045.6392 | 0.034676519 | -0.159638429 | YOR262W | GPN2 | n |
| W303.asm02.HP0_G0051460 | 876.54636 | 792.030808 | 769.8422558 | 1110.05798 | 1105.6009 | 1120.71145 | 1270.0647 | 1394.7444 | 1307.4898 | 6.73E-09 | -0.452942007 | YOR264W | DSE3 | n |
| W303.asm02.HP0_G0051480 | 1148.36 | 1257.07642 | 1508.77491 | 711.703842 | 852.12167 | 697.83725 | 655.55402 | 742.63907 | 715.0197 | 4.85E-12 | 0.791877392 | YOR265W | RBL2 | n |
| W303.asm02.HP0_G0051490 | 842.83304 | 780.612278 | 750.523755 | 680.868898 | 690.32642 | 692.14836 | 663.53468 | 671.78658 | 621.5646 | 0.014891928 | 0.201848344 | YOR266W | PNT1 | n |
| W303.asm02.HP0_G0051500 | 798.58431 | 1072.30383 | 913.7650865 | 1038.38757 | 1014.8158 | 1012.62253 | 1036.3454 | 1032.6095 | 1072.0887 | 0.170181541 | -0.138807597 | YOR267C | HRK1 | n |
| W303.asm02.HP0_G0051520 | 379.27487 | 325.947148 | 286.8797365 | 203.343955 | 175.27819 | 213.333399 | 312.38574 | 259.79247 | 277.72035 | 8.21E-08 | 0.743966764 | YOR269W | PAC1 | n |
| W303.asm02.HP0_G0051530 | 5911.4202 | 5691.61851 | 5588.842273 | 2825.14757 | 2675.9137 | 2815.05271 | 5092.7997 | 4806.1606 | 4837.624 | 1.18E-75 | 1.047623522 | YOR270C | VPH1 | n |
| W303.asm02.HP0_G0051540 | 1422.2808 | 1669.18157 | 1672.016242 | 1590.91643 | 1599.0764 | 1543.58566 | 1453.6198 | 1292.4019 | 1512.9147 | 0.919676677 | 0.009301739 | YOR271C | FSF1 | n |
| W303.asm02.HP0_G0051550 | 1833.1619 | 1567.45284 | 1677.811792 | 2261.78481 | 2240.8643 | 2365.63035 | 1430.8179 | 1746.3827 | 1428.2761 | 2.15E-06 | -0.435569907 | YOR272W | YTM1 | n |
| W303.asm02.HP0_G0051560 | 619.48228 | 678.88355 | 643.3060757 | 654.200838 | 648.97874 | 624.829821 | 681.77618 | 617.99117 | 654.18572 | 0.912014777 | 0.009824852 | YOR273C | TPO4 | n |
| W303.asm02.HP0_G0051570 | 904.99198 | 823.172255 | 843.2525587 | 829.20998 | 892.57049 | 914.015094 | 701.15778 | 823.98823 | 755.57569 | 0.700328607 | -0.03528758 | YOR274W | MOD5 | n |
| W303.asm02.HP0_G0051580 | 920.7951 | 978.879492 | 802.6837071 | 761.706454 | 840.43646 | 845.748407 | 811.74689 | 797.74657 | 743.23256 | 0.136319144 | 0.142559972 | YOR275C | RIM20 | n |
| W303.asm02.HP0_G0051590 | 4416.4451 | 4343.19384 | 5371.50914 | 1320.90233 | 1243.1269 | 1243.0226 | 1004.4228 | 1099.5257 | 1117.053 | 1.59E-96 | 1.892104834 | YOR276W | CAF20 | n |
| W303.asm02.HP0_G0051600 | 568.9123 | 508.643638 | 502.2810201 | 502.526249 | 515.04823 | 525.274235 | 433.2357 | 498.5916 | 494.60672 | 0.738425245 | 0.033915965 | YOR278W | HEM4 | n |
| W303.asm02.HP0_G0051610 | 1458.1012 | 1298.59835 | 1373.545405 | 608.365111 | 628.3049 | 627.674266 | 656.69412 | 635.04825 | 600.40495 | 1.74E-47 | 1.147771152 | YOR279C | RFM1 | n |
| W303.asm02.HP0_G0051620 | 2974.1471 | 2825.56731 | 3213.632603 | 2379.29095 | 2364.9073 | 2285.03774 | 1943.8602 | 2268.5918 | 2075.4086 | 2.78E-06 | 0.358787662 | YOR281C | PLP2 | n |
| W303.asm02.HP0_G0051630 | 2181.884 | 2057.41161 | 2540.382852 | 1266.73283 | 1271.8905 | 1243.0226 | 1035.2053 | 1140.2003 | 1102.0649 | 1.84E-22 | 0.842524042 | YOR283W | | n |
| W303.asm02.HP0_G0051650 | 2862.4717 | 3086.11742 | 3540.115267 | 132.506921 | 117.75099 | 105.244477 | 150.4924 | 104.96665 | 140.18265 | 3.57E-303 | 4.734928752 | YOR285W | RDL1 | n |
| W303.asm02.HP0_G0051660 | 2160.8132 | 2335.60854 | 2279.583091 | 1260.8992 | 1258.4075 | 1234.48927 | 996.44211 | 1174.3144 | 1066.7988 | 8.47E-31 | 0.852049191 | YOR286W | RDL2 | n |
| W303.asm02.HP0_G0051670 | 980.84695 | 987.183878 | 1028.710166 | 1047.55472 | 1005.8272 | 1042.01513 | 677.21581 | 886.96822 | 829.63445 | 0.639376501 | -0.046584021 | YOR287C | RRP36 | n |
| W303.asm02.HP0_G0051680 | 2313.5767 | 2734.21907 | 2211.968338 | 1002.55237 | 1014.8158 | 1018.31142 | 1157.1954 | 1190.0594 | 1162.8989 | 5.86E-49 | 1.257762152 | YOR288C | MPD1 | n |
| W303.asm02.HP0_G0051690 | 614.21458 | 683.035743 | 716.7163787 | 343.351268 | 332.57913 | 310.992688 | 556.36585 | 434.29953 | 605.69486 | 5.23E-17 | 1.028751229 | YOR289W | | n |
| W303.asm02.HP0_G0051700 | 841.7795 | 987.183878 | 974.618364 | 1940.10134 | 2147.3826 | 2230.99327 | 2178.7195 | 1898.5843 | 2159.1655 | 7.05E-38 | -1.171589789 | YOR290C | SNF2 | n |
| W303.asm02.HP0_G0051710 | 561.53751 | 589.611401 | 684.8408524 | 1512.57901 | 1569.414 | 1651.67458 | 1946.1404 | 1734.5739 | 1927.2911 | 3.95E-52 | -1.364867396 | YOR291W | YPK9 | n |
| W303.asm02.HP0_G0051720 | 518.34232 | 546.013375 | 457.8484683 | 269.180727 | 297.5235 | 282.548234 | 367.11025 | 402.80953 | 361.47728 | 6.24E-15 | 0.841592074 | YOR292C | | n |
| W303.asm02.HP0_G0051730 | 9747.3641 | 9484.64678 | 10879.21371 | 13759.052 | 13786.753 | 15763.9159 | 12509.111 | 12203.686 | 11092.944 | 1.35E-10 | -0.52426906 | YOR293W | RPS10A | y |
| W303.asm02.HP0_G0051750 | 2353.6113 | 2288.89637 | 2400.323721 | 1634.25203 | 1525.3697 | 1695.28941 | 1087.6496 | 1426.2344 | 1102.0649 | 4.94E-08 | 0.536831771 | YOR294W | RRS1 | n |
| W303.asm02.HP0_G0051770 | 562.59105 | 591.687498 | 631.7149753 | 1083.38992 | 1075.9384 | 1214.57815 | 1045.4662 | 812.17948 | 1029.7694 | 6.22E-18 | -0.916828016 | YOR296W | | n |
| W303.asm02.HP0_G0051780 | 1169.4308 | 1132.51063 | 1224.792949 | 821.709588 | 786.50471 | 745.244672 | 710.27853 | 778.06532 | 768.80047 | 1.01E-13 | 0.583089713 | YOR297C | TIM18 | n |
| W303.asm02.HP0_G0051790 | 146.44224 | 132.870175 | 137.1613555 | 93.3382088 | 83.594215 | 80.5926172 | 67.265543 | 72.164574 | 72.295457 | 5.18E-05 | 0.690942363 | YOR298W | MUM3 | n |
| W303.asm02.HP0_G0051800 | 7866.7929 | 7612.00775 | 7253.131115 | 3369.34266 | 3357.2515 | 3471.17143 | 3260.6687 | 3662.0241 | 3117.5212 | 1.85E-60 | 1.156463472 | YOR298C- | MBF1 | n |
| W303.asm02.HP0_G0051810 | 974.5257 | 1016.24923 | 1015.187216 | 2932.65318 | 2827.8215 | 2783.76381 | 2759.0274 | 2587.428 | 2842.4458 | 3.53E-115 | -1.507121474 | YOR299W | BUD7 | n |
| W303.asm02.HP0_G0051820 | 418.2559 | 500.339252 | 425.972942 | 1269.23296 | 1231.4417 | 1262.93372 | 1484.4023 | 1309.459 | 1442.3825 | 3.13E-62 | -1.485441996 | YOR301W | RAX1 | n |
| W303.asm02.HP0_G0051830 | 3349.2078 | 3497.18453 | 3364.31691 | 1756.75843 | 1760.8717 | 1953.18578 | 1813.8895 | 1561.379 | 1760.6589 | 4.33E-33 | 0.900612773 | YOR303W | CPA1 | n |
| W303.asm02.HP0_G0051840 | 926.0628 | 980.955588 | 1073.142718 | 1447.57561 | 1481.3254 | 1486.69675 | 1426.2575 | 1478.7177 | 1504.9798 | 2.20E-14 | -0.566264137 | YOR304W | ISW2 | n |
| W303.asm02.HP0_G0051860 | 1280.0527 | 1243.58179 | 1423.773507 | 553.362238 | 470.1051 | 430.459391 | 410.43382 | 502.52785 | 491.08012 | 1.28E-38 | 1.439645953 | YOR305W | RRG7 | n |
| W303.asm02.HP0_G0051870 | 1118.8609 | 1140.81502 | 1216.099624 | 423.355447 | 387.40975 | 412.44457 | 558.64604 | 688.84366 | 551.03245 | 4.31E-52 | 1.506625663 | YOR306C | MCH5 | n |

| | | | | | | | | | | | | | | |
|---|---|---|---|---|---|---|---|---|---|---|---|---|---|---|
| W303.asm02.HP0_G0051880 | 1310.6054 | 1317.28322 | 1270.191426 | 669.201622 | 703.80936 | 746.192821 | 1351.0113 | 1253.0394 | 1328.6494 | 2.61E-32 | 0.880083878 | YOR307C | SLY41 | n |
| W303.asm02.HP0_G0051890 | 277.08136 | 265.74035 | 273.3567859 | 660.034476 | 703.80936 | 657.066867 | 573.46726 | 661.28992 | 578.36365 | 3.22E-40 | -1.308015049 | YOR308C | SNU66 | n |
| W303.asm02.HP0_G0051900 | 3065.8052 | 3009.30185 | 3397.158361 | 9170.47901 | 9109.9717 | 9994.43268 | 6603.4242 | 7906.6132 | 6630.0224 | 4.35E-76 | -1.577521928 | YOR310C | NOP58 | n |
| W303.asm02.HP0_G0051910 | 1791.0202 | 1962.94922 | 1667.186617 | 1704.25569 | 1693.457 | 1691.49681 | 1807.0489 | 1874.9668 | 1866.4571 | 0.224528908 | 0.090735193 | YOR311C | DGK1 | n |
| W303.asm02.HP0_G0051930 | 599.465 | 580.268967 | 613.3623995 | 120.006268 | 121.34644 | 106.192625 | 119.70986 | 191.56414 | 119.90466 | 2.58E-54 | 2.365930403 | YOR313C | SPS4 | n |
| W303.asm02.HP0_G0051940 | 4811.5231 | 4504.09132 | 4892.410321 | 495.025857 | 488.98121 | 576.47425 | 653.27383 | 603.55826 | 649.77746 | 0 | 3.18814487 | YOR315W | SFG1 | n |
| W303.asm02.HP0_G0051970 | 3297.5843 | 3429.71139 | 3764.209876 | 4582.73937 | 4630.0409 | 5082.07563 | 5244.4322 | 4238.0286 | 4708.0212 | 4.82E-07 | -0.446008756 | YOR317W | FAA1 | n |
| W303.asm02.HP0_G0051990 | 889.18886 | 822.134207 | 962.0613385 | 324.1836 | 358.64615 | 291.08157 | 399.03288 | 370.00745 | 391.45345 | 6.48E-45 | 1.456405526 | YOR319W | HSH49 | n |
| W303.asm02.HP0_G0052000 | 2474.7685 | 2376.09242 | 2652.430156 | 1311.73518 | 1361.7767 | 1276.2078 | 1330.4896 | 1367.1907 | 1346.2825 | 1.72E-42 | 0.925918533 | YOR320C | GNT1 | n |
| W303.asm02.HP0_G0052020 | 704.81913 | 827.324448 | 774.671881 | 1341.73675 | 1408.5176 | 1479.11156 | 1681.6386 | 1453.7882 | 1679.5469 | 1.47E-22 | -0.873991034 | YOR322C | LDB19 | n |
| W303.asm02.HP0_G0052030 | 1570.8301 | 1742.883 | 1769.574671 | 2296.78664 | 2357.7164 | 2506.90447 | 2048.7488 | 2094.0847 | 2018.1012 | 2.15E-12 | -0.4938432 | YOR323C | PRO2 | n |
| W303.asm02.HP0_G0052040 | 478.30775 | 529.404603 | 583.4187233 | 975.884308 | 887.17731 | 719.644664 | 819.72755 | 728.20616 | 793.48672 | 4.58E-09 | -0.699053373 | YOR324C | FRT1 | n |
| W303.asm02.HP0_G0052050 | 3380.814 | 3572.96205 | 3640.571471 | 5651.12851 | 5827.3257 | 5889.8981 | 5854.3824 | 5212.9064 | 5441.5556 | 2.83E-29 | -0.712959339 | YOR326W | MYO2 | n |
| W303.asm02.HP0_G0052060 | 2598.0328 | 2443.56556 | 2995.333545 | 2823.48081 | 2832.3158 | 2993.30462 | 2663.2595 | 2353.8772 | 2388.395 | 0.232613132 | -0.105492443 | YOR327C | SNC2 | n |
| W303.asm02.HP0_G0052080 | 589.98313 | 655.00844 | 661.6586515 | 766.706715 | 702.91049 | 719.644664 | 685.19646 | 704.58866 | 731.77109 | 0.02115926 | -0.19936169 | YOR329C | SCD5 | n |
| W303.asm02.HP0_G0052090 | 606.83979 | 692.378177 | 699.329728 | 1326.73597 | 1298.8563 | 1331.20041 | 1324.7892 | 1271.4086 | 1216.6796 | 1.54E-35 | -0.984636024 | YOR330C | MIP1 | n |
| W303.asm02.HP0_G0052100 | 2291.4523 | 2655.3274 | 2121.171385 | 3546.01856 | 3332.9822 | 3496.77144 | 3282.3305 | 3351.0604 | 3280.6268 | 1.21E-11 | -0.554177435 | YOR332W | VMA4 | n |
| W303.asm02.HP0_G0052120 | 494.11087 | 465.045612 | 450.121068 | 629.199532 | 575.27201 | 570.78536 | 568.90688 | 585.18909 | 559.84896 | 0.000210844 | -0.334630882 | YOR334W | MRS2 | n |
| W303.asm02.HP0_G0052130 | 6501.4034 | 5708.22728 | 6357.718604 | 6747.85247 | 6578.7748 | 7150.93552 | 5453.0694 | 5446.4572 | 5269.6335 | 0.037350509 | -0.14117701 | YOR335C | ALA1 | n |
| W303.asm02.HP0_G0052140 | 928.16989 | 975.765347 | 960.1294884 | 1484.24419 | 1509.1902 | 1636.5042 | 1555.0881 | 1481.3419 | 1464.4238 | 4.73E-22 | -0.692317983 | YOR336W | KRE5 | n |
| W303.asm02.HP0_G0052150 | 529.93127 | 568.850436 | 636.5446005 | 361.685559 | 367.63477 | 415.289016 | 304.40508 | 240.11122 | 322.6846 | 7.11E-07 | 0.601982902 | YOR337W | TEA1 | n |
| W303.asm02.HP0_G0052160 | 1195.7694 | 1191.67938 | 1241.213675 | 389.186995 | 382.91543 | 402.014938 | 380.79138 | 343.76579 | 408.20484 | 4.81E-80 | 1.628388025 | YOR338W | | n |
| W303.asm02.HP0_G0052180 | 2031.2276 | 2093.7433 | 1842.984974 | 2024.2724 | 2049.4066 | 2158.93399 | 1287.1661 | 1661.0973 | 1385.9568 | 0.523379468 | -0.062683028 | YOR340C | RPA43 | n |
| W303.asm02.HP0_G0052190 | 3029.9848 | 3000.99747 | 3444.488688 | 7109.53803 | 6995.847 | 7902.81723 | 4780.4139 | 5384.7893 | 4779.435 | 7.96E-50 | -1.215465619 | YOR341W | RPA190 | n |
| W303.asm02.HP0_G0052200 | 2076.5299 | 2064.67795 | 1828.496098 | 428.355708 | 381.11771 | 441.837172 | 384.21166 | 469.72577 | 391.45345 | 8.37E-116 | 2.254120498 | YOR342C | | n |
| W303.asm02.HP0_G0052220 | 4279.4848 | 3823.13167 | 3521.762691 | 1536.74694 | 1256.6098 | 1156.74109 | 1864.0536 | 1626.9831 | 1750.9607 | 5.61E-47 | 1.556341466 | YOR344C | TYE7 | n |
| W303.asm02.HP0_G0052230 | 1092.5223 | 980.955588 | 1106.950094 | 130.006791 | 121.34644 | 119.466703 | 103.74855 | 94.469988 | 107.56153 | 1.65E-161 | 3.099042937 | YOR346W | REV1 | n |
| W303.asm02.HP0_G0052250 | 537.30606 | 529.404603 | 675.181602 | 572.529905 | 534.8232 | 368.829742 | 722.81957 | 536.64202 | 843.74088 | 0.186858223 | 0.238245002 | YOR348C | PUT4 | n |
| W303.asm02.HP0_G0052260 | 276.02782 | 297.919845 | 311.0278624 | 780.874121 | 764.03314 | 808.770617 | 768.42332 | 770.19282 | 789.07846 | 1.24E-56 | -1.409768464 | YOR349W | CIN1 | n |
| W303.asm02.HP0_G0052270 | 630.0177 | 596.877739 | 599.839449 | 1229.23087 | 1233.2394 | 1186.1337 | 1172.0166 | 1145.4486 | 1157.609 | 3.98E-44 | -0.998624385 | YOR350C | MNE1 | n |
| W303.asm02.HP0_G0052280 | 49.516441 | 62.2828945 | 35.73922643 | 44.1689738 | 37.752226 | 41.7185313 | 33.062725 | 51.171244 | 44.082596 | 0.38581224 | 0.250426272 | YOR351C | MEK1 | n |
| W303.asm02.HP0_G0052290 | 628.96416 | 661.23673 | 614.3283246 | 453.357014 | 430.55515 | 408.651977 | 499.36115 | 426.42703 | 492.84342 | 5.31E-09 | 0.557926674 | YOR352W | TFB6 | n |
| W303.asm02.HP0_G0052300 | 893.40302 | 859.503944 | 793.9903817 | 1665.08697 | 1563.1219 | 1595.73382 | 1564.2089 | 1591.5569 | 1628.4111 | 1.15E-37 | -0.922497298 | YOR353C | SOG2 | n |
| W303.asm02.HP0_G0052310 | 3133.2318 | 2923.14385 | 2913.229916 | 2241.78376 | 2343.3346 | 2408.29703 | 2072.6908 | 2149.1922 | 2123.8995 | 6.72E-09 | 0.359206342 | YOR354C | MSC6 | n |
| W303.asm02.HP0_G0052320 | 2693.9051 | 2488.20163 | 2389.698546 | 1935.10108 | 2035.0248 | 2491.73409 | 2279.0478 | 2171.4976 | 2309.928 | 0.01761588 | 0.229044821 | YOR355W | GDS1 | n |
| W303.asm02.HP0_G0052330 | 1190.5017 | 1220.74473 | 1379.340955 | 2386.79134 | 2292.0994 | 2387.43777 | 1937.0196 | 1964.1885 | 1929.936 | 1.66E-36 | -0.897798855 | YOR356W | CIR2 | n |
| W303.asm02.HP0_G0052340 | 1021.9351 | 1042.20043 | 1240.24775 | 683.369028 | 749.65134 | 696.889102 | 694.31722 | 767.56865 | 733.53439 | 2.15E-11 | 0.634480314 | YOR357C | SNX3 | n |
| W303.asm02.HP0_G0052350 | 415.09527 | 398.610525 | 419.2114668 | 618.365633 | 661.56282 | 538.548313 | 538.12435 | 623.23951 | 631.26277 | 1.43E-07 | -0.561191084 | YOR358W | HAP5 | n |
| W303.asm02.HP0_G0052360 | 681.64122 | 765.041554 | 809.4451824 | 2504.29748 | 2577.9377 | 2591.28968 | 2347.4534 | 2496.8943 | 2197.0766 | 1.58E-106 | -1.76507751 | YOR359W | VTS1 | n |
| W303.asm02.HP0_G0052370 | 4119.3465 | 3583.34253 | 4243.308695 | 1294.23427 | 1260.2053 | 1342.57819 | 1156.0553 | 1268.7844 | 1139.9759 | 1.93E-88 | 1.616340889 | YOR360C | PDE2 | n |
| W303.asm02.HP0_G0052380 | 5855.5825 | 5391.62257 | 5594.637824 | 6786.1878 | 6343.2728 | 7013.45399 | 5921.648 | 6231.083 | 5757.187 | 6.62E-05 | -0.258212208 | YOR361C | PRT1 | n |
| W303.asm02.HP0_G0052420 | 861.79678 | 857.427847 | 855.8095842 | 752.539308 | 824.25693 | 815.407657 | 736.50069 | 879.09572 | 870.19044 | 0.238119191 | 0.106760475 | YOR367W | SCP1 | n |
| W303.asm02.HP0_G0052430 | 940.81238 | 926.97708 | 1048.994592 | 753.372685 | 815.26831 | 735.763188 | 679.49599 | 730.83033 | 725.59952 | 6.40E-05 | 0.340507296 | YOR368W | RAD17 | n |
| W303.asm02.HP0_G0052440 | 14346.072 | 13708.4651 | 16741.41277 | 12508.1534 | 12979.575 | 14167.234 | 10889.037 | 11803.5 | 9969.7198 | 0.057844547 | 0.175962332 | YOR369C | RPS12 | n |
| W303.asm02.HP0_G0052450 | 1812.091 | 1916.23705 | 2057.420332 | 2825.14757 | 2852.9896 | 2914.6083 | 3326.7941 | 3253.9663 | 3356.4488 | 5.95E-19 | -0.570032465 | YOR370C | MRS6 | n |
| W303.asm02.HP0_G0052470 | 509.91399 | 538.747037 | 604.6690742 | 1373.40507 | 1375.2597 | 1373.86769 | 1486.6825 | 1417.0494 | 1467.0688 | 8.86E-63 | -1.316721903 | YOR372C | NDD1 | n |
| W303.asm02.HP0_G0052480 | 298.15219 | 335.289582 | 338.0737635 | 1033.38731 | 1073.2419 | 1128.29664 | 1099.0506 | 1052.2907 | 1125.8695 | 2.62E-87 | -1.733912857 | YOR373W | NUD1 | n |
| W303.asm02.HP0_G0052490 | 1246.3394 | 1512.43629 | 1553.207462 | 403.354402 | 479.09372 | 392.533453 | 1516.325 | 1678.1544 | 1534.956 | 1.13E-60 | 1.758187886 | YOR374W | ALD4 | n |
| W303.asm02.HP0_G0052530 | 583.66188 | 470.235853 | 498.4173199 | 482.525204 | 435.04946 | 476.918664 | 502.78143 | 558.94743 | 546.62419 | 0.152842121 | 0.153959835 | YOR377W | ATF1 | n |
| W303.asm02.HP0_G0052540 | 140.12099 | 117.299451 | 146.8206059 | 56.6696267 | 57.527201 | 66.3703906 | 111.72921 | 72.164574 | 90.810147 | 6.86E-09 | 1.165738825 | YOR378W | AMF1 | n |
| W303.asm02.HP0_G0052560 | 221.24367 | 221.104275 | 232.7879343 | 1093.39045 | 1081.3316 | 1184.2374 | 1564.2089 | 1401.3048 | 1631.056 | 1.05E-123 | -2.313703552 | YOR381W | FRE3 | n |
| W303.asm02.HP0_G0052620 | 605.78625 | 520.062169 | 475.235119 | 362.518936 | 326.2871 | 296.770461 | 337.46781 | 299.15496 | 358.83233 | 1.66E-08 | 0.697615041 | YOR386W | PHR1 | n |
| W303.asm02.HP0_G0052650 | 377.16779 | 371.62127 | 419.2114668 | 314.183078 | 287.63601 | 352.711219 | 700.01769 | 480.22244 | 709.72979 | 0.04935417 | 0.292288423 | YOR389W | | n |
| W303.asm02.HP0_G0052730 | 579.44771 | 555.355809 | 611.4305494 | 147.507705 | 162.69412 | 132.740781 | 194.95607 | 152.20165 | 190.43681 | 5.25E-57 | 1.978225636 | YPL279C | FEX2 | y |
| W303.asm02.HP0_G0052760 | 2161.8667 | 1884.05756 | 2036.169981 | 1037.5542 | 1018.4112 | 1007.88179 | 943.99779 | 951.2603 | 987.45014 | 1.27E-41 | 0.988977673 | YPL274W | SAM3 | n |
| W303.asm02.HP0_G0052770 | 9101.5433 | 8450.75073 | 10662.8465 | 2914.31889 | 2705.5762 | 2935.46756 | 2616.5156 | 2395.8639 | 2644.9557 | 9.00E-85 | 1.721663973 | YPL273W | SAM4 | n |
| W303.asm02.HP0_G0052790 | 3239.6395 | 3238.71051 | 3675.344772 | 4064.37897 | 4258.8106 | 4687.64588 | 3845.5369 | 3875.8937 | 3446.3773 | 1.67E-05 | -0.35731876 | YPL271W | ATP15 | n |
| W303.asm02.HP0_G0052800 | 969.258 | 1082.68432 | 1017.119066 | 1376.73858 | 1308.7438 | 1448.77081 | 1417.1368 | 1141.5124 | 1380.6669 | 4.47E-06 | -0.429649302 | YPL270W | MDL2 | n |
| W303.asm02.HP0_G0052810 | 182.26264 | 198.267214 | 224.094609 | 580.863674 | 550.10386 | 551.822391 | 565.4866 | 532.70577 | 560.73062 | 1.43E-44 | -1.474790698 | YPL269W | KAR9 | n |
| W303.asm02.HP0_G0052820 | 1173.645 | 1075.41798 | 1010.35759 | 773.37373 | 729.87637 | 811.615063 | 728.52003 | 725.58199 | 767.03716 | 5.81E-09 | 0.49340662 | YPL268W | PLC1 | n |

| | | | | | | | | | | | | | | |
|---|---|---|---|---|---|---|---|---|---|---|---|---|---|---|
| W303.asm02.HP0_G0052830 | 763.81744 | 803.449339 | 808.4792573 | 871.7122 | 743.35931 | 846.696555 | 871.03178 | 831.86073 | 936.31433 | 0.580822992 | -0.051370266 | YPL267W | ACM1 | n |
| W303.asm02.HP0_G0052840 | 1884.7854 | 1693.05668 | 1834.291648 | 2751.8104 | 2548.2753 | 2874.78606 | 1824.1503 | 2365.686 | 1903.4865 | 1.82E-09 | -0.594925023 | YPL266W | DIM1 | n |
| W303.asm02.HP0_G0052850 | 2588.551 | 2729.02883 | 1868.099025 | 1361.7378 | 1361.7767 | 1332.14856 | 900.67422 | 810.8674 | 1167.3071 | 5.69E-09 | 0.824910343 | YPL265W | DIP5 | n |
| W303.asm02.HP0_G0052870 | 818.60159 | 924.900983 | 787.2289065 | 1970.10291 | 1940.6442 | 2082.13397 | 1592.7112 | 1680.7785 | 1700.7065 | 3.85E-57 | -1.244046818 | YPL263C | KEL3 | n |
| W303.asm02.HP0_G0052880 | 2560.1054 | 2682.31666 | 2610.895379 | 8797.95955 | 9003.9059 | 9472.95104 | 7742.378 | 8238.5702 | 7570.745 | 3.11E-197 | -1.796094 | YPL262W | FUM1 | n |
| W303.asm02.HP0_G0052890 | 3816.9801 | 3551.16303 | 3350.793959 | 1733.42388 | 1647.615 | 1591.94123 | 1914.2177 | 1713.5806 | 1973.137 | 1.40E-45 | 1.107480547 | YPL260W | CUB1 | n |
| W303.asm02.HP0_G0052900 | 670.05227 | 647.742103 | 650.067551 | 2193.44791 | 2145.5848 | 2069.80804 | 2146.7969 | 1962.8764 | 2041.9058 | 2.82E-130 | -1.703862072 | YPL259C | APM1 | n |
| W303.asm02.HP0_G0052930 | 1653.0063 | 1792.70931 | 1928.952302 | 2222.6161 | 2292.0994 | 2100.14879 | 2586.8732 | 2504.7668 | 2715.4879 | 8.62E-05 | -0.2992541 | YPL256C | CLN2 | n |
| W303.asm02.HP0_G0052940 | 486.73608 | 464.007564 | 430.8025672 | 498.359365 | 445.83581 | 426.666797 | 444.63664 | 497.27952 | 506.0682 | 0.934191337 | 0.009121682 | YPL255W | BBP1 | n |
| W303.asm02.HP0_G0052950 | 1223.1614 | 1107.59747 | 1269.225501 | 923.381565 | 955.49086 | 920.652133 | 936.01713 | 774.12907 | 895.75834 | 4.45E-05 | 0.363098127 | YPL254W | HFI1 | n |
| W303.asm02.HP0_G0052960 | 434.05902 | 394.458332 | 328.4145131 | 787.541136 | 773.02177 | 746.192821 | 849.36999 | 771.5049 | 810.23811 | 2.25E-22 | -0.997787059 | YPL253C | VIK1 | n |
| W303.asm02.HP0_G0052970 | 653.19561 | 667.465019 | 681.9430773 | 1383.40559 | 1433.6857 | 1474.37082 | 1272.3448 | 1512.8319 | 1399.1816 | 5.77E-41 | -1.099045139 | YPL252C | YAH1 | n |
| W303.asm02.HP0_G0052990 | 10656.57 | 10621.3096 | 10460.00224 | 11290.5898 | 11128.817 | 12176.1222 | 9877.774 | 10064.99 | 8936.4238 | 0.058801095 | -0.124335388 | YPL249C-/ | RPL36B | y |
| W303.asm02.HP0_G0053000 | 408.77402 | 492.034866 | 461.7121685 | 1944.26822 | 1927.1612 | 1899.14132 | 1862.9135 | 1802.8023 | 1880.5635 | 5.88E-158 | -2.081977966 | YPL249C | GYP5 | n |
| W303.asm02.HP0_G0053010 | 318.16947 | 335.289582 | 324.550813 | 250.013059 | 245.38947 | 257.896375 | 323.78668 | 213.86956 | 307.69652 | 0.007178493 | 0.376854716 | YPL248C | GAL4 | n |
| W303.asm02.HP0_G0053020 | 389.81028 | 510.719735 | 470.4054938 | 804.208674 | 750.55021 | 797.392836 | 730.80022 | 549.76285 | 793.48672 | 4.96E-09 | -0.778509317 | YPL247C | | n |
| W303.asm02.HP0_G0053030 | 975.57924 | 1104.48333 | 1084.733818 | 1081.72317 | 1079.5339 | 1014.51883 | 1380.6538 | 1514.144 | 1363.9155 | 0.953645378 | -0.005031198 | YPL246C | RBD2 | n |
| W303.asm02.HP0_G0053040 | 656.35623 | 567.812388 | 589.2142736 | 139.173936 | 109.66123 | 134.637078 | 100.32827 | 107.59082 | 96.98171 | 9.37E-68 | 2.240320862 | YPL245W | | n |
| W303.asm02.HP0_G0053050 | 1853.1791 | 1796.86151 | 1838.155348 | 584.197182 | 614.82197 | 573.629805 | 753.6021 | 751.82366 | 755.57569 | 9.14E-121 | 1.630305501 | YPL244C | HUT1 | n |
| W303.asm02.HP0_G0053060 | 4888.4316 | 4507.20546 | 4489.61958 | 2735.97624 | 2740.6318 | 2632.06006 | 2365.6949 | 2528.3843 | 2611.453 | 4.17E-33 | 0.775786139 | YPL243W | SRP68 | n |
| W303.asm02.HP0_G0053070 | 1568.723 | 1538.38749 | 1602.469639 | 1817.59494 | 1749.1865 | 1867.85242 | 2116.0144 | 2015.3597 | 2167.1004 | 0.001421562 | -0.206305655 | YPL242C | IQG1 | n |
| W303.asm02.HP0_G0053100 | 2525.3385 | 2834.90975 | 2346.231919 | 1138.3928 | 1108.2975 | 1307.4967 | 731.94032 | 999.80737 | 777.61699 | 4.43E-22 | 1.116753923 | YPL239W | YAR1 | n |
| W303.asm02.HP0_G0053110 | 6997.6213 | 6877.0696 | 6958.523978 | 6741.18545 | 6462.8215 | 7094.04661 | 5497.533 | 5779.7264 | 5588.7915 | 0.5190166 | 0.037630199 | YPL237W | SUI3 | n |
| W303.asm02.HP0_G0053140 | 1975.3899 | 1819.69857 | 2059.352182 | 1365.0713 | 1380.6528 | 1363.43745 | 1860.6333 | 1937.9468 | 1850.5874 | 8.27E-14 | 0.510977921 | YPL234C | VMA11 | n |
| W303.asm02.HP0_G0053150 | 344.50801 | 305.186183 | 295.5730618 | 201.677201 | 231.90653 | 228.503774 | 215.47776 | 220.42997 | 262.73227 | 6.48E-05 | 0.514249528 | YPL233W | NSL1 | n |
| W303.asm02.HP0_G0053160 | 2255.6319 | 2286.82028 | 2008.158155 | 5243.60723 | 4772.96 | 5241.36456 | 4886.4427 | 4874.389 | 4804.1213 | 2.62E-68 | -1.220258352 | YPL232W | SSO1 | n |
| W303.asm02.HP0_G0053170 | 5306.6875 | 6162.89241 | 6596.302089 | 14592.4289 | 14504.945 | 16544.2421 | 12851.139 | 13940.884 | 13812.841 | 5.58E-53 | -1.336913588 | YPL231W | FAS2 | n |
| W303.asm02.HP0_G0053180 | 176.99494 | 165.04967 | 211.5375835 | 110.839123 | 107.8635 | 92.9185469 | 125.41033 | 120.71165 | 130.48448 | 2.39E-07 | 0.828530308 | YPL230W | USV1 | n |
| W303.asm02.HP0_G0053190 | 658.46331 | 668.503067 | 602.7372241 | 253.346567 | 266.96217 | 265.481563 | 469.71871 | 367.38329 | 492.84342 | 8.81E-29 | 1.296235051 | YPL229W | | n |
| W303.asm02.HP0_G0053200 | 902.88489 | 813.829821 | 812.3429575 | 1802.59416 | 1790.5341 | 1673.48199 | 1575.6098 | 1611.2381 | 1607.2514 | 1.64E-48 | -1.059127131 | YPL228W | CET1 | n |
| W303.asm02.HP0_G0053210 | 2820.3301 | 2610.69133 | 3143.120076 | 4076.87962 | 4001.736 | 4080.83088 | 4595.7187 | 5039.7115 | 4430.3009 | 2.68E-10 | -0.503740029 | YPL227C | ALG5 | n |
| W303.asm02.HP0_G0053220 | 2910.9346 | 3003.07356 | 3363.350984 | 5929.47639 | 5743.7315 | 6510.93532 | 3477.2866 | 3965.1153 | 3709.9912 | 3.86E-33 | -0.97056894 | YPL226W | NEW1 | n |
| W303.asm02.HP0_G0053230 | 4238.3966 | 4232.12268 | 5127.130105 | 1250.0653 | 1409.4164 | 1364.3856 | 958.81901 | 1176.9386 | 1050.0474 | 1.86E-71 | 1.757138351 | YPL225W | CHP1 | n |
| W303.asm02.HP0_G0053260 | 232.83263 | 271.968639 | 268.5271607 | 717.53748 | 675.94462 | 659.911313 | 799.20586 | 705.90075 | 782.02525 | 2.27E-44 | -1.408460474 | YPL222W | FMP40 | n |
| W303.asm02.HP0_G0053270 | 1395.9422 | 1415.8978 | 1541.616362 | 2225.9496 | 2219.2916 | 2397.8674 | 2515.0473 | 2818.3546 | 2488.0217 | 1.88E-17 | -0.651882243 | YPL221W | FLC1 | n |
| W303.asm02.HP0_G0053300 | 5534.2524 | 5397.85086 | 6428.231132 | 3769.36356 | 3915.4451 | 4032.47531 | 3554.8129 | 3474.3962 | 3171.3019 | 1.76E-12 | 0.567419893 | YPL218W | SAR1 | n |
| W303.asm02.HP0_G0053310 | 1270.5708 | 1272.64714 | 1527.127486 | 3305.17264 | 3163.9961 | 3710.10484 | 2012.2658 | 2256.7831 | 1994.2966 | 2.49E-44 | -1.321725924 | YPL217C | BMS1 | n |
| W303.asm02.HP0_G0053330 | 1809.984 | 1835.26929 | 1714.516944 | 1726.75686 | 1757.2762 | 1650.72643 | 1531.1462 | 1654.5369 | 1555.234 | 0.371942125 | 0.061393595 | YPL215W | CBP3 | n |
| W303.asm02.HP0_G0053340 | 908.1526 | 870.922475 | 850.014034 | 345.018022 | 374.82567 | 315.73343 | 342.02819 | 293.90663 | 297.99835 | 1.21E-42 | 1.343131958 | YPL214C | THI6 | n |
| W303.asm02.HP0_G0053350 | 1294.8023 | 1245.65789 | 1235.418124 | 551.695484 | 565.38453 | 473.12607 | 498.22106 | 526.14535 | 507.8315 | 1.42E-47 | 1.246199308 | YPL213W | LEA1 | n |
| W303.asm02.HP0_G0053360 | 1833.1619 | 1728.35032 | 1817.870923 | 3017.65762 | 2969.8418 | 3053.03797 | 2043.0484 | 2553.3138 | 2129.1894 | 5.43E-19 | -0.748915433 | YPL212C | PUS1 | n |
| W303.asm02.HP0_G0053370 | 2738.1538 | 2442.52751 | 3304.429557 | 1554.24785 | 1601.773 | 1619.43753 | 987.32136 | 1406.5532 | 1049.1658 | 1.27E-10 | 0.829541007 | YPL211W | NIP7 | n |
| W303.asm02.HP0_G0053380 | 4922.145 | 4341.11775 | 4526.324731 | 4223.55395 | 4327.1242 | 4290.37168 | 3790.8124 | 4017.5987 | 3990.3566 | 0.114058244 | 0.10277476 | YPL210C | SRP72 | n |
| W303.asm02.HP0_G0053390 | 136.96037 | 135.98432 | 102.3880541 | 340.851137 | 340.6689 | 319.526024 | 326.06687 | 411.99412 | 346.4892 | 1.65E-23 | -1.419039207 | YPL209C | IPL1 | n |
| W303.asm02.HP0_G0053400 | 457.23692 | 453.627082 | 537.0543215 | 935.882218 | 913.24432 | 932.978063 | 847.08981 | 858.10239 | 752.04908 | 7.94E-24 | -0.940898754 | YPL208W | RKM1 | n |
| W303.asm02.HP0_G0053420 | 2040.7095 | 1962.94922 | 1955.998203 | 378.353096 | 350.55638 | 291.08157 | 489.10031 | 384.44037 | 461.9856 | 2.15E-128 | 2.544350827 | YPL206C | PGC1 | n |
| W303.asm02.HP0_G0053440 | 1501.2964 | 1689.94254 | 1585.082988 | 2360.95666 | 2239.9654 | 2369.42295 | 2320.0912 | 2348.6289 | 2297.5849 | 1.64E-16 | -0.545284859 | YPL204W | HRR25 | n |
| W303.asm02.HP0_G0053460 | 47.409358 | 58.1307015 | 43.46662674 | 13.3340298 | 10.78635 | 13.2740781 | 25.082067 | 14.432915 | 22.041298 | 1.92E-08 | 1.99261331 | YPL201C | YIG1 | n |
| W303.asm02.HP0_G0053480 | 1657.2205 | 1573.68113 | 1474.001609 | 890.879867 | 924.03067 | 888.415086 | 916.63554 | 994.55904 | 958.35563 | 7.38E-27 | 0.799012258 | YPL199C | | n |
| W303.asm02.HP0_G0053500 | 716.40808 | 683.035743 | 740.8645046 | 193.343432 | 195.05317 | 189.629688 | 223.45841 | 204.68497 | 235.40106 | 5.88E-74 | 1.888649276 | YPL196W | OXR1 | n |
| W303.asm02.HP0_G0053510 | 1580.3119 | 1590.28991 | 1617.92444 | 1872.59781 | 1997.2725 | 1882.07465 | 1914.2177 | 1844.7889 | 1822.3745 | 2.48E-05 | -0.264284992 | YPL195W | APL5 | n |
| W303.asm02.HP0_G0053520 | 343.45446 | 353.97445 | 314.8915626 | 410.021417 | 434.1506 | 397.274195 | 378.51119 | 362.13495 | 403.79658 | 0.004395673 | -0.295511255 | YPL194W | DDC1 | n |
| W303.asm02.HP0_G0053530 | 581.5548 | 515.909976 | 570.8616978 | 401.687648 | 369.4325 | 427.614945 | 204.07682 | 266.35288 | 171.92212 | 0.000298016 | 0.477278115 | YPL193W | RSA1 | n |
| W303.asm02.HP0_G0053560 | 1027.2028 | 1300.67445 | 1302.066952 | 1371.73832 | 1292.5643 | 1442.13377 | 1146.9345 | 1281.9053 | 1322.4779 | 0.087022577 | -0.177395855 | YPL190C | NAB3 | n |
| W303.asm02.HP0_G0053570 | 867.06449 | 836.666883 | 1087.631593 | 1185.89528 | 1296.1598 | 1191.82259 | 1072.8284 | 1137.5761 | 1039.4676 | 0.000109433 | -0.395369316 | YPL189C-/ | COA2 | n |
| W303.asm02.HP0_G0053590 | 855.47554 | 811.753725 | 831.6614583 | 610.865241 | 597.74358 | 543.289055 | 753.6021 | 753.13574 | 859.61061 | 9.47E-09 | 0.511388216 | YPL188W | POS5 | n |
| W303.asm02.HP0_G0053600 | 18.963743 | 33.2175437 | 19.31850077 | 49.169235 | 62.021514 | 38.874086 | 55.864604 | 56.419576 | 50.254159 | 0.000479368 | -1.074116097 | YPL187W | MF(ALPHA | n |
| W303.asm02.HP0_G0053610 | 224.4043 | 198.267214 | 227.9583091 | 190.009925 | 191.45772 | 181.096352 | 427.53523 | 317.52413 | 466.39386 | 0.173516758 | 0.209857031 | YPL186C | UIP4 | n |
| W303.asm02.HP0_G0053620 | 1194.7158 | 1213.47839 | 1258.600325 | 1575.91565 | 1595.481 | 1680.11903 | 1286.026 | 1165.1299 | 1312.7797 | 2.44E-08 | -0.403355508 | YPL184C | MRN1 | n |
| W303.asm02.HP0_G0053630 | 922.90218 | 908.292211 | 900.242136 | 689.202666 | 805.38082 | 669.392797 | 603.1097 | 771.5049 | 698.26831 | 0.001262514 | 0.33585288 | YPL183W- | RTC6 | n |

| | | | | | | | | | | | | | | |
|---|---|---|---|---|---|---|---|---|---|---|---|---|---|---|
| W303.asm02.HP0_G0053640 | 1513.9388 | 1316.24517 | 1565.764488 | 997.552106 | 1004.9283 | 1123.5559 | 902.95441 | 951.2603 | 911.62808 | 1.66E-08 | 0.492681799 | YPL183C | RTT10 | n |
| W303.asm02.HP0_G0053650 | 1332.7297 | 1230.08717 | 1324.283228 | 1021.72004 | 1002.2317 | 1005.03734 | 1046.6062 | 1039.1699 | 1045.6392 | 1.65E-07 | 0.3598466 | YPL181W | CTI6 | n |
| W303.asm02.HP0_G0053660 | 340.29384 | 438.056358 | 376.7107651 | 1196.72918 | 1253.9132 | 1103.64478 | 1128.693 | 1132.3278 | 1216.6796 | 1.70E-60 | -1.621831597 | YPL180W | TCO89 | n |
| W303.asm02.HP0_G0053670 | 3418.7415 | 3274.00415 | 3171.131902 | 2853.48238 | 2802.6533 | 2862.46013 | 3297.1517 | 3071.5867 | 3148.379 | 0.000380137 | 0.211353986 | YPL179W | PPQ1 | n |
| W303.asm02.HP0_G0053680 | 2773.9742 | 2727.99078 | 3152.779326 | 1240.06477 | 1287.1711 | 1243.0226 | 1205.0793 | 1304.2107 | 1193.7567 | 5.38E-55 | 1.199161553 | YPL178W | CBC2 | n |
| W303.asm02.HP0_G0053690 | 1047.2201 | 1120.05405 | 1164.905597 | 1530.9133 | 1452.5618 | 1394.72635 | 1875.4545 | 1860.5339 | 1949.3324 | 5.70E-08 | -0.393877098 | YPL177C | CUP9 | n |
| W303.asm02.HP0_G0053710 | 930.27697 | 844.971269 | 816.2066576 | 464.190913 | 479.09372 | 533.807571 | 471.9989 | 494.65535 | 491.96177 | 1.12E-18 | 0.811715834 | YPL175W | SPT14 | n |
| W303.asm02.HP0_G0053720 | 722.72933 | 612.448462 | 716.7163787 | 590.864196 | 577.06974 | 592.592774 | 612.23045 | 561.5716 | 543.09758 | 0.020150469 | 0.221248578 | YPL174C | NIP100 | n |
| W303.asm02.HP0_G0053730 | 1843.6973 | 1894.43804 | 1842.019049 | 1873.43119 | 1718.6251 | 1764.50424 | 1674.798 | 1621.7348 | 1710.4047 | 0.376680943 | 0.058586833 | YPL173W | MRPL40 | n |
| W303.asm02.HP0_G0053740 | 1107.2719 | 1060.8853 | 1029.676091 | 1877.59807 | 1751.8831 | 1740.80053 | 1812.7494 | 1922.2018 | 1817.9662 | 2.42E-27 | -0.748674891 | YPL172C | COX10 | n |
| W303.asm02.HP0_G0053760 | 1982.7647 | 2292.01052 | 2108.614359 | 1239.2314 | 1288.07 | 1326.45966 | 1239.2821 | 1329.1403 | 1266.9338 | 2.39E-22 | 0.728438695 | YPL170W | DAP1 | n |
| W303.asm02.HP0_G0053770 | 1271.6243 | 1346.34857 | 1313.658053 | 1860.09716 | 1867.8363 | 2038.51914 | 1483.2622 | 1532.5131 | 1407.1165 | 5.78E-15 | -0.552116222 | YPL169C | MEX67 | n |
| W303.asm02.HP0_G0053780 | 727.99704 | 740.128396 | 697.3978779 | 622.532517 | 579.76633 | 540.44461 | 542.68472 | 557.63535 | 566.90218 | 0.00033256 | 0.311751188 | YPL168W | MRX4 | n |
| W303.asm02.HP0_G0053790 | 493.05733 | 460.893419 | 511.9402705 | 316.683208 | 378.42112 | 412.44457 | 448.05692 | 363.44704 | 397.62501 | 0.000738443 | 0.406669233 | YPL167C | REV3 | n |
| W303.asm02.HP0_G0053800 | 199.11931 | 189.962828 | 166.1391066 | 130.840168 | 125.84075 | 140.325969 | 108.30893 | 89.221656 | 88.165191 | 0.001710917 | 0.483258736 | YPL166W | ATG29 | n |
| W303.asm02.HP0_G0053810 | 243.36804 | 213.837938 | 198.980558 | 492.525727 | 474.59941 | 507.259414 | 351.14894 | 359.51079 | 350.89746 | 2.46E-27 | -1.169623611 | YPL164C | MLH3 | n |
| W303.asm02.HP0_G0053830 | 625.80353 | 635.285524 | 624.9535 | 600.031342 | 592.3504 | 547.081649 | 768.42332 | 730.83033 | 789.07846 | 0.174479858 | 0.115797447 | YPL162C | | n |
| W303.asm02.HP0_G0053840 | 1220.0008 | 1131.47258 | 1018.084991 | 960.883524 | 1128.0725 | 972.800297 | 959.95911 | 981.43821 | 877.24365 | 0.162041709 | 0.137782434 | YPL161C | BEM4 | n |
| W303.asm02.HP0_G0053850 | 4329.0012 | 4463.60744 | 4508.93808 | 10168.8645 | 9913.5548 | 10597.4551 | 8466.3377 | 7914.4857 | 8080.3398 | 1.26E-105 | -1.205573624 | YPL160W | CDC60 | n |
| W303.asm02.HP0_G0053860 | 1227.3756 | 1288.21787 | 1021.948691 | 1405.07339 | 1446.2698 | 1396.62265 | 1346.451 | 1312.0832 | 1378.0219 | 0.002862642 | -0.264791092 | YPL159C | PET20 | n |
| W303.asm02.HP0_G0053870 | 797.53076 | 899.987825 | 895.4125108 | 1225.06399 | 1316.8336 | 1478.16341 | 1583.5905 | 1586.3086 | 1634.5826 | 5.87E-13 | -0.63151717 | YPL158C | AIM44 | n |
| W303.asm02.HP0_G0053880 | 704.81913 | 664.350875 | 731.2052542 | 576.69679 | 622.01287 | 613.452039 | 492.52059 | 581.25284 | 527.22784 | 0.019532303 | 0.213831398 | YPL157W | TGS1 | n |
| W303.asm02.HP0_G0053890 | 250.74283 | 301.03399 | 255.9701352 | 766.706715 | 673.24803 | 564.148321 | 492.52059 | 549.76285 | 514.00306 | 3.17E-25 | -1.312576025 | YPL156C | PRM4 | n |
| W303.asm02.HP0_G0053900 | 313.95531 | 380.963705 | 387.3359405 | 1442.57535 | 1384.2483 | 1428.8597 | 1760.3051 | 1733.2619 | 1768.5937 | 5.01E-120 | -1.973839565 | YPL155C | KIP2 | n |
| W303.asm02.HP0_G0053910 | 3047.895 | 3316.56413 | 3283.179206 | 1750.09141 | 1629.6378 | 1696.23756 | 3405.4606 | 3154.2479 | 2943.8357 | 3.18E-36 | 0.926431904 | YPL154C | PEP4 | n |
| W303.asm02.HP0_G0053920 | 249.68929 | 275.082784 | 263.6975355 | 884.212853 | 872.79551 | 870.400266 | 983.90108 | 847.60573 | 1033.296 | 4.18E-67 | -1.73639029 | YPL153C | RAD53 | n |
| W303.asm02.HP0_G0053930 | 482.52192 | 457.779274 | 455.9166182 | 554.195614 | 578.86746 | 452.266805 | 590.56867 | 587.81326 | 630.38112 | 0.087174274 | -0.184733996 | YPL152W | RRD2 | n |
| W303.asm02.HP0_G0053940 | 529.93127 | 712.101094 | 599.839449 | 531.694439 | 486.28462 | 498.726078 | 408.15363 | 451.35661 | 438.181 | 0.014166291 | 0.279837613 | YPL151C | PRP46 | n |
| W303.asm02.HP0_G0053950 | 639.49957 | 658.122585 | 655.8631012 | 950.883002 | 921.33409 | 980.385485 | 949.69826 | 902.71322 | 965.40884 | 5.00E-13 | -0.545887571 | YPL150W | | n |
| W303.asm02.HP0_G0053960 | 843.88658 | 767.11765 | 845.1844088 | 202.510578 | 178.87364 | 178.251906 | 212.05747 | 216.49372 | 203.66159 | 1.17E-90 | 2.131948576 | YPL149W | ATG5 | n |
| W303.asm02.HP0_G0053970 | 278.1349 | 296.881797 | 304.2663872 | 74.1705409 | 82.695352 | 74.9037266 | 69.545731 | 86.597489 | 77.585368 | 7.60E-37 | 1.924494851 | YPL148C | PPT2 | n |
| W303.asm02.HP0_G0053990 | 749.06786 | 656.046489 | 715.7504536 | 85.0044401 | 112.35782 | 138.429672 | 135.67118 | 171.8829 | 149.88083 | 1.09E-71 | 2.665249062 | YPL147W | PXA1 | n |
| W303.asm02.HP0_G0054000 | 3737.9645 | 3334.21095 | 4060.748862 | 1773.42597 | 1678.1763 | 1671.5857 | 972.50014 | 1146.7607 | 1035.0593 | 1.13E-37 | 1.119701434 | YPL146C | NOP53 | n |
| W303.asm02.HP0_G0054010 | 4814.6837 | 4476.06402 | 4205.637618 | 4468.56674 | 4522.1774 | 4595.67548 | 4032.5123 | 4247.2132 | 3947.1556 | 0.889240146 | -0.009738502 | YPL145C | KES1 | n |
| W303.asm02.HP0_G0054020 | 644.76728 | 607.258221 | 773.7059559 | 190.009925 | 205.83952 | 210.488953 | 182.41503 | 242.73539 | 200.13498 | 1.23E-40 | 1.742054052 | YPL144W | POC4 | n |
| W303.asm02.HP0_G0054040 | 709.03329 | 835.628834 | 834.5592334 | 1777.59285 | 1795.9273 | 1883.97095 | 1832.131 | 1894.6481 | 1944.0425 | 5.01E-52 | -1.196852652 | YPL141C | FRK1 | n |
| W303.asm02.HP0_G0054060 | 978.73987 | 879.226861 | 874.16216 | 1130.8924 | 1112.7918 | 1133.03738 | 1101.3308 | 1148.0728 | 1122.3429 | 3.74E-05 | -0.306132736 | YPL139C | UME1 | n |
| W303.asm02.HP0_G0054070 | 748.01432 | 756.737168 | 733.1371043 | 489.192219 | 524.93571 | 457.955695 | 430.95551 | 460.54119 | 435.53604 | 4.81E-12 | 0.603590015 | YPL138C | SPP1 | n |
| W303.asm02.HP0_G0054080 | 1503.4034 | 1551.88212 | 1842.984974 | 1834.26248 | 1795.9273 | 1700.9783 | 1943.8602 | 2225.2931 | 2213.828 | 0.195229053 | -0.121854833 | YPL137C | GIP3 | n |
| W303.asm02.HP0_G0054090 | 3603.1112 | 3151.51446 | 3233.917029 | 2830.98121 | 2845.7987 | 3034.075 | 3310.8328 | 3381.2383 | 3373.2002 | 0.004272092 | 0.19750671 | YPL135W | ISU1 | n |
| W303.asm02.HP0_G0054100 | 388.75674 | 430.79002 | 432.7344173 | 2405.95901 | 2342.4357 | 2537.24522 | 2301.8497 | 2183.3064 | 2274.6619 | 3.45E-242 | -2.539497766 | YPL134C | ODC1 | n |
| W303.asm02.HP0_G0054110 | 195.95868 | 261.588157 | 189.3213076 | 351.685037 | 331.68027 | 345.126031 | 461.73805 | 408.05787 | 439.9443 | 2.26E-07 | -0.670478462 | YPL133C | RDS2 | n |
| W303.asm02.HP0_G0054120 | 1276.8921 | 1308.97883 | 1282.748451 | 985.051453 | 962.68176 | 919.703985 | 966.79967 | 1092.9653 | 1102.9465 | 1.56E-08 | 0.43156839 | YPL132W | COX11 | n |
| W303.asm02.HP0_G0054130 | 12531.874 | 12095.3381 | 11286.83408 | 22151.157 | 21001.024 | 24147.4444 | 18169.677 | 19599.898 | 17171.053 | 5.04E-33 | -0.906073586 | YPL131W | RPL5 | n |
| W303.asm02.HP0_G0054150 | 1894.2673 | 2105.16183 | 1576.389663 | 1670.92061 | 1715.0297 | 1717.09682 | 1632.6145 | 1734.5739 | 1611.6597 | 0.196859854 | 0.127448248 | YPL129W | TAF14 | n |
| W303.asm02.HP0_G0054160 | 507.80691 | 550.165568 | 613.3623995 | 833.376864 | 865.60461 | 1005.03734 | 482.25974 | 617.99117 | 464.63056 | 1.50E-08 | -0.692858926 | YPL128C | TBF1 | n |
| W303.asm02.HP0_G0054180 | 2228.2398 | 2011.73749 | 2253.503115 | 3462.68087 | 3519.9456 | 3858.01599 | 2145.6568 | 2716.0122 | 2178.5619 | 1.49E-14 | -0.739176147 | YPL126W | NAN1 | n |
| W303.asm02.HP0_G0054190 | 1245.2858 | 1152.23355 | 1423.773507 | 2205.11518 | 2182.4382 | 2288.83033 | 2054.4493 | 1934.0106 | 2007.5214 | 5.65E-23 | -0.804226567 | YPL125W | KAP120 | n |
| W303.asm02.HP0_G0054210 | 1055.6484 | 1034.9341 | 1153.314496 | 393.35388 | 424.26311 | 323.318617 | 692.03703 | 619.30326 | 665.64719 | 2.08E-47 | 1.506409495 | YPL123C | RNY1 | n |
| W303.asm02.HP0_G0054220 | 815.44097 | 715.215238 | 777.5696561 | 852.544532 | 864.70575 | 892.20768 | 784.38464 | 825.30031 | 780.26194 | 0.029348039 | -0.176657705 | YPL122C | TFB2 | n |
| W303.asm02.HP0_G0054230 | 9.4818717 | 19.7229166 | 15.45480062 | 17.5009141 | 30.561326 | 7.5851875 | 18.241503 | 11.808749 | 7.9348672 | 0.564276671 | -0.319493622 | YPL121C | MEI5 | n |
| W303.asm02.HP0_G0054240 | 628.96416 | 636.323572 | 592.1120487 | 332.517369 | 355.0507 | 329.955656 | 353.42912 | 295.21871 | 335.02773 | 1.14E-18 | 0.867805144 | YPL120W | VPS30 | n |
| W303.asm02.HP0_G0054270 | 4057.1875 | 3883.33847 | 4193.080593 | 2478.46279 | 2407.1538 | 2336.23775 | 2326.9318 | 2503.4547 | 2312.573 | 6.32E-33 | 0.74844267 | YPL118W | MRP51 | n |
| W303.asm02.HP0_G0054280 | 7287.3452 | 6654.92728 | 7810.469862 | 2745.14339 | 2672.3183 | 2959.17127 | 2231.1639 | 2368.3101 | 2183.8518 | 5.75E-74 | 1.377013279 | YPL117C | IDI1 | n |
| W303.asm02.HP0_G0054290 | 648.98144 | 719.367431 | 658.7608763 | 1589.24968 | 1481.3254 | 1637.45235 | 1742.0636 | 1434.1069 | 1656.6239 | 4.59E-41 | -1.215762493 | YPL116W | HOS3 | n |
| W303.asm02.HP0_G0054300 | 757.49619 | 786.840567 | 763.0807805 | 1975.10317 | 2080.8667 | 2116.26731 | 2325.7917 | 1968.1248 | 2277.3069 | 6.93E-69 | -1.419332351 | YPL115C | BEM3 | n |
| W303.asm02.HP0_G0054310 | 169.62015 | 175.430153 | 143.9228308 | 2030.93942 | 2097.9451 | 1904.83021 | 2082.9516 | 2136.0714 | 2197.9582 | 0 | -3.628290901 | YPL113C | | n |
| W303.asm02.HP0_G0054320 | 2303.0413 | 2211.04275 | 2324.981568 | 700.869943 | 684.93324 | 714.903922 | 661.25449 | 661.28992 | 707.96649 | 6.78E-134 | 1.70318468 | YPL112C | PEX25 | n |
| W303.asm02.HP0_G0054330 | 525.71711 | 609.334318 | 659.7268014 | 3672.69184 | 4015.2189 | 3367.82325 | 5038.0752 | 6637.8288 | 5488.2832 | 9.10E-116 | -2.622237395 | YPL111W | CAR1 | n |
| W303.asm02.HP0_G0054340 | 799.63785 | 890.645391 | 973.6524389 | 1055.05511 | 1099.3089 | 1093.21515 | 1092.21 | 1074.5961 | 1066.7988 | 0.000672652 | -0.28468041 | YPL110C | GDE1 | n |

| | | | | | | | | | | | | | | |
|---|---|---|---|---|---|---|---|---|---|---|---|---|---|---|
| W303.asm02.HP0_G0054350 | 505.69982 | 495.149011 | 495.5195448 | 810.875689 | 816.16717 | 731.022446 | 820.86764 | 838.42115 | 881.65191 | 9.63E-15 | -0.656983911 | YPL109C | CQD1 | n |
| W303.asm02.HP0_G0054360 | 564.69814 | 633.209427 | 657.7949513 | 660.867853 | 774.81949 | 743.348375 | 432.09561 | 629.79992 | 495.48837 | 0.07269133 | -0.230894731 | YPL108W | | n |
| W303.asm02.HP0_G0054370 | 911.31322 | 960.194623 | 857.7414343 | 530.861062 | 594.14813 | 433.303836 | 562.06632 | 528.76952 | 552.79575 | 1.82E-13 | 0.807246995 | YPL107W | DPC25 | n |
| W303.asm02.HP0_G0054380 | 9237.4501 | 10550.7223 | 9262.255195 | 9410.49155 | 9282.5533 | 9830.403 | 8230.3382 | 7810.8311 | 7994.8195 | 0.710442527 | 0.026410482 | YPL106C | SSE1 | n |
| W303.asm02.HP0_G0054390 | 1660.3811 | 1903.78047 | 1920.258977 | 3057.65971 | 3088.4916 | 3036.91945 | 2804.6311 | 2925.9455 | 2788.665 | 1.24E-26 | -0.743278915 | YPL105C | SYH1 | n |
| W303.asm02.HP0_G0054400 | 610.00041 | 690.302081 | 668.4201267 | 1173.39462 | 1203.5769 | 1099.85219 | 1083.0893 | 1108.7103 | 1117.9346 | 2.14E-25 | -0.820481163 | YPL104W | MSD1 | n |
| W303.asm02.HP0_G0054410 | 687.96247 | 639.437717 | 640.4083006 | 652.534084 | 718.19116 | 861.86693 | 527.8635 | 558.94743 | 478.73699 | 0.109662734 | -0.181124786 | YPL103C | FMP30 | n |
| W303.asm02.HP0_G0054420 | 671.10581 | 742.204493 | 812.3429575 | 1101.72421 | 1086.7248 | 1152.00035 | 843.66952 | 1123.1432 | 911.62808 | 9.58E-08 | -0.585066215 | YPL101W | ELP4 | n |
| W303.asm02.HP0_G0054430 | 418.2559 | 411.067104 | 409.5522164 | 1789.26013 | 1810.3091 | 1675.37829 | 1931.3192 | 1834.2923 | 1773.8836 | 6.34E-159 | -2.090584072 | YPL100W | ATG21 | n |
| W303.asm02.HP0_G0054440 | 831.24409 | 817.982014 | 931.1517372 | 624.199271 | 687.62983 | 675.081688 | 632.75214 | 641.60867 | 633.90772 | 1.04E-05 | 0.378437089 | YPL099C | INA17 | n |
| W303.asm02.HP0_G0054450 | 1449.6728 | 1440.81096 | 1433.432757 | 721.704364 | 794.59447 | 835.318774 | 717.1191 | 741.32699 | 750.28578 | 1.02E-30 | 0.879836725 | YPL098C | MGR2 | n |
| W303.asm02.HP0_G0054460 | 1470.7437 | 1291.33201 | 1461.444583 | 1582.58266 | 1608.0651 | 1504.71157 | 1549.3877 | 1519.3923 | 1542.0092 | 0.039181175 | -0.152823246 | YPL097W | MSY1 | n |
| W303.asm02.HP0_G0054470 | 289.72386 | 303.110087 | 326.4826631 | 212.5111 | 222.01904 | 190.577836 | 204.07682 | 242.73539 | 239.80932 | 8.17E-06 | 0.556197676 | YPL096C-/ | ERI1 | n |
| W303.asm02.HP0_G0054480 | 980.84695 | 901.025874 | 1029.676091 | 875.045707 | 994.14195 | 926.341024 | 974.78033 | 980.12613 | 1046.5208 | 0.499242532 | 0.059474226 | YPL096W | PNG1 | n |
| W303.asm02.HP0_G0054500 | 5851.3684 | 6135.90316 | 6134.58992 | 2501.79735 | 2532.9946 | 2439.58593 | 2800.0707 | 2839.348 | 2881.2384 | 7.34E-127 | 1.277654823 | YPL094C | SEC62 | n |
| W303.asm02.HP0_G0054510 | 3764.3031 | 3626.94056 | 3673.412922 | 5803.63648 | 5694.2941 | 6386.72788 | 3415.7215 | 4635.5898 | 3516.0278 | 3.17E-11 | -0.69271842 | YPL093W | NOG1 | n |
| W303.asm02.HP0_G0054520 | 622.64291 | 549.12752 | 537.0543215 | 2152.61244 | 2372.9971 | 2280.29699 | 2489.9652 | 2378.8068 | 2357.5372 | 1.65E-141 | -1.994498149 | YPL092W | SSU1 | n |
| W303.asm02.HP0_G0054530 | 3025.7706 | 2753.94198 | 3162.438576 | 2683.4735 | 2448.5015 | 2692.74156 | 2719.1241 | 2698.9551 | 2663.4704 | 0.007800858 | 0.192754422 | YPL091W | GLR1 | n |
| W303.asm02.HP0_G0054550 | 687.96247 | 746.356686 | 775.637806 | 2655.97207 | 2603.1059 | 2747.73417 | 1984.9036 | 1884.1514 | 2063.0655 | 3.35E-144 | -1.856370946 | YPL089C | RLM1 | n |
| W303.asm02.HP0_G0054570 | 1439.1374 | 1421.08804 | 1488.490485 | 4.16688432 | 20.673838 | 10.4296328 | 34.202819 | 31.489996 | 27.331209 | 1.98E-171 | 6.966033153 | YPL087W | YDC1 | n |
| W303.asm02.HP0_G0054580 | 970.31154 | 916.596597 | 992.9709397 | 2689.30714 | 2662.4308 | 2834.96383 | 2120.5747 | 2250.2226 | 2059.5389 | 5.13E-106 | -1.506794874 | YPL086C | ELP3 | n |
| W303.asm02.HP0_G0054590 | 741.69308 | 991.336071 | 857.7414343 | 3281.83809 | 3448.9355 | 3419.02327 | 2899.2589 | 2510.0151 | 2922.6761 | 1.23E-87 | -1.969818792 | YPL085W | SEC16 | n |
| W303.asm02.HP0_G0054600 | 671.10581 | 772.307892 | 866.4347596 | 1039.22095 | 984.25446 | 927.289172 | 980.4808 | 1037.8578 | 1001.5566 | 0.000314902 | -0.352779332 | YPL084W | BRO1 | n |
| W303.asm02.HP0_G0054610 | 537.30606 | 512.795831 | 454.9506932 | 806.708804 | 996.83854 | 940.56325 | 738.78088 | 763.6324 | 763.51056 | 1.19E-17 | -0.866776056 | YPL083C | SEN54 | n |
| W303.asm02.HP0_G0054620 | 823.8693 | 892.721488 | 925.356187 | 3179.33274 | 3259.2755 | 3636.14926 | 3138.6786 | 3166.0567 | 3110.4679 | 1.20E-143 | -1.93021004 | YPL082C | MOT1 | n |
| W303.asm02.HP0_G0054640 | 9935.948 | 9611.28867 | 10871.48631 | 10602.2205 | 11311.286 | 11717.2184 | 9811.6485 | 10599.008 | 8674.5732 | 0.089563933 | -0.144716608 | YPL079W | RPL21B | y |
| W303.asm02.HP0_G0054650 | 8056.4303 | 7725.15501 | 9363.677324 | 9406.32466 | 9192.667 | 10120.5364 | 9017.0031 | 9302.6697 | 8473.5565 | 0.020155046 | -0.191591306 | YPL078C | ATP4 | n |
| W303.asm02.HP0_G0054670 | 355.04342 | 331.137389 | 366.0855896 | 435.8561 | 453.02671 | 496.829781 | 490.2404 | 505.15202 | 477.85534 | 7.32E-05 | -0.395370028 | YPL076W | GPI2 | n |
| W303.asm02.HP0_G0054680 | 1199.9835 | 1202.05986 | 1329.112853 | 748.372424 | 766.72973 | 808.770617 | 791.2252 | 787.2499 | 877.24365 | 8.63E-17 | 0.684106593 | YPL075W | GCR1 | n |
| W303.asm02.HP0_G0054690 | 638.44603 | 613.486511 | 536.0883964 | 56.6696267 | 59.324926 | 48.3555703 | 59.284885 | 38.050412 | 56.425722 | 3.88E-101 | 3.440242571 | YPL074W | YTA6 | n |
| W303.asm02.HP0_G0054700 | 337.13322 | 360.20274 | 302.3345371 | 810.042312 | 839.5376 | 804.978024 | 958.81901 | 926.33072 | 848.14914 | 5.98E-43 | -1.297325803 | YPL072W | UBP16 | n |
| W303.asm02.HP0_G0054710 | 839.67242 | 823.172255 | 977.5161391 | 277.514496 | 295.72577 | 321.42232 | 283.88339 | 253.23205 | 312.98643 | 1.64E-44 | 1.563303262 | YPL071C | | n |
| W303.asm02.HP0_G0054720 | 1500.2428 | 1383.71831 | 1421.841657 | 722.537741 | 673.24803 | 589.748328 | 826.56811 | 801.68282 | 844.62253 | 4.38E-38 | 1.115195743 | YPL070W | MUK1 | n |
| W303.asm02.HP0_G0054730 | 902.88489 | 888.569295 | 891.5488106 | 859.211547 | 888.97503 | 857.126188 | 743.34126 | 802.9949 | 793.48672 | 0.581231754 | 0.042351274 | YPL069C | BTS1 | n |
| W303.asm02.HP0_G0054740 | 134.85329 | 165.04967 | 119.7747048 | 440.856361 | 392.80292 | 386.844563 | 287.30368 | 368.69537 | 349.13416 | 3.47E-27 | -1.542927122 | YPL068C | | n |
| W303.asm02.HP0_G0054750 | 302.36635 | 384.077849 | 348.6989389 | 599.197965 | 586.95723 | 547.081649 | 601.96961 | 795.1224 | 611.86643 | 4.25E-09 | -0.743726033 | YPL067C | HTC1 | n |
| W303.asm02.HP0_G0054760 | 660.57039 | 645.666006 | 665.5223516 | 297.51554 | 293.92804 | 263.585266 | 319.22631 | 259.79247 | 313.86808 | 3.16E-31 | 1.204214901 | YPL066W | RGL1 | n |
| W303.asm02.HP0_G0054770 | 892.34948 | 747.394734 | 903.1399111 | 313.349701 | 329.88255 | 259.792672 | 359.12959 | 354.26246 | 377.34702 | 4.51E-39 | 1.492423081 | YPL065W | VPS28 | n |
| W303.asm02.HP0_G0054790 | 1373.8179 | 1731.46447 | 1785.029471 | 3461.01412 | 3449.8344 | 3584.0011 | 3463.6054 | 3459.9633 | 3562.7554 | 6.34E-35 | -1.101166344 | YPL063W | TIM50 | n |
| W303.asm02.HP0_G0054800 | 10823.03 | 11328.2205 | 10796.14416 | 1536.74694 | 1182.0042 | 1801.48203 | 5057.4568 | 6780.8458 | 5118.871 | 5.61E-100 | 2.865892022 | YPL061W | ALD6 | n |
| W303.asm02.HP0_G0054810 | 463.55817 | 434.942213 | 514.8380456 | 288.348395 | 289.43373 | 276.859344 | 307.82537 | 310.96371 | 299.76165 | 4.82E-12 | 0.726252419 | YPL060W | MFM1 | n |
| W303.asm02.HP0_G0054820 | 1696.2015 | 1622.4694 | 1907.701951 | 1342.57013 | 1457.955 | 1351.11152 | 1175.4369 | 1254.3515 | 1214.0347 | 3.47E-05 | 0.332642821 | YPL059W | GRX5 | n |
| W303.asm02.HP0_G0054830 | 1013.5067 | 1055.69506 | 1116.609345 | 724.204495 | 628.3049 | 695.940953 | 523.30312 | 442.17203 | 432.89109 | 2.13E-11 | 0.637120074 | YPL058C | PDR12 | n |
| W303.asm02.HP0_G0054840 | 791.20952 | 710.024997 | 680.9771522 | 3478.51503 | 3101.0757 | 2525.86744 | 3088.5145 | 2563.8105 | 3017.0128 | 2.16E-69 | -2.061630419 | YPL057C | SUR1 | n |
| W303.asm02.HP0_G0054860 | 598.41146 | 737.014251 | 653.9312511 | 346.684775 | 328.98368 | 338.488992 | 306.68527 | 314.89996 | 328.85616 | 7.26E-21 | 0.971784574 | YPL055C | LGE1 | n |
| W303.asm02.HP0_G0054870 | 112.72892 | 132.870175 | 117.8428547 | 452.523637 | 467.40851 | 431.407539 | 503.92153 | 447.42036 | 506.0682 | 1.36E-54 | -1.895214024 | YPL054W | LEE1 | n |
| W303.asm02.HP0_G0054890 | 808.06618 | 800.335194 | 1004.56204 | 229.178638 | 259.77127 | 285.39268 | 215.47776 | 217.80581 | 223.93959 | 1.26E-47 | 1.757393953 | YPL052W | OAZ1 | n |
| W303.asm02.HP0_G0054900 | 1075.6657 | 1028.70581 | 1223.827024 | 452.523637 | 443.13922 | 411.496422 | 395.6126 | 349.01412 | 426.71953 | 4.07E-40 | 1.348080968 | YPL051W | ARL3 | n |
| W303.asm02.HP0_G0054910 | 5146.5492 | 4783.3263 | 5356.054339 | 4556.90469 | 4317.2367 | 4599.46807 | 4832.8583 | 5350.6752 | 4839.3873 | 0.009313823 | 0.182174191 | YPL050C | MNN9 | n |
| W303.asm02.HP0_G0054920 | 1217.8937 | 1441.84901 | 1498.149735 | 1100.89084 | 1031.8942 | 995.55586 | 1121.8524 | 905.33739 | 1123.2245 | 0.000101248 | 0.410455688 | YPL049C | DIG1 | n |
| W303.asm02.HP0_G0054930 | 1843.6973 | 2228.68957 | 2242.87794 | 1601.75033 | 1579.3015 | 1727.52645 | 2022.5267 | 1827.7319 | 1975.7819 | 4.30E-05 | 0.364033383 | YPL048W | CAM1 | n |
| W303.asm02.HP0_G0054940 | 779.62056 | 756.737168 | 727.3415541 | 121.673022 | 84.493077 | 93.8666953 | 93.487704 | 80.037073 | 89.928495 | 2.68E-105 | 2.91017033 | YPL047W | SGF11 | n |
| W303.asm02.HP0_G0054950 | 563.6446 | 582.345063 | 725.409704 | 125.00653 | 121.34644 | 97.6592891 | 111.72921 | 87.909572 | 106.67988 | 4.11E-61 | 2.441454455 | YPL046C | ELC1 | n |
| W303.asm02.HP0_G0054960 | 1002.9713 | 1076.45603 | 944.6746878 | 884.212853 | 915.04205 | 870.400266 | 908.65488 | 855.47823 | 916.91799 | 0.024461042 | 0.179280887 | YPL045W | VPS16 | n |
| W303.asm02.HP0_G0054970 | 1589.7938 | 1538.38749 | 1570.594113 | 2959.32124 | 2804.4511 | 3171.55652 | 1420.5571 | 1940.571 | 1685.7185 | 1.79E-20 | -0.927152818 | YPL043W | NOP4 | n |
| W303.asm02.HP0_G0054980 | 549.94856 | 525.25241 | 586.3164984 | 668.368245 | 715.49457 | 625.777969 | 637.31252 | 611.43076 | 663.00224 | 0.00269892 | -0.274200662 | YPL042C | SSN3 | n |
| W303.asm02.HP0_G0054990 | 328.70489 | 320.756907 | 385.4040904 | 226.678507 | 244.49061 | 250.311188 | 238.27964 | 223.05414 | 230.11115 | 1.09E-05 | 0.522906875 | YPL041C | MRX11 | n |
| W303.asm02.HP0_G0055000 | 606.83979 | 657.084537 | 660.6927264 | 1046.72134 | 1028.2987 | 961.422516 | 970.21995 | 913.20989 | 999.79327 | 3.59E-16 | -0.65788228 | YPL040C | ISM1 | n |
| W303.asm02.HP0_G0055010 | 589.98313 | 575.078726 | 603.7031491 | 380.01985 | 316.39961 | 340.385289 | 441.21636 | 503.83994 | 473.44708 | 2.08E-14 | 0.76945408 | YPL039W | | n |
| W303.asm02.HP0_G0055040 | 8799.1769 | 8483.96828 | 9513.395705 | 5726.96581 | 6028.6709 | 6223.64635 | 5310.5576 | 5554.0481 | 4936.3691 | 2.05E-16 | 0.575898853 | YPL037C | EGD1 | n |

| | | | | | | | | | | | | | | |
|---|---|---|---|---|---|---|---|---|---|---|---|---|---|---|
| W303.asm02.HP0_G0055060 | 623.69645 | 590.649449 | 677.1134521 | 255.846697 | 267.86103 | 266.429711 | 250.82067 | 270.28913 | 209.83316 | 1.78E-29 | 1.260315711 | YPL034W | | n |
| W303.asm02.HP0_G0055080 | 1292.6952 | 1432.50657 | 1394.795756 | 5351.11284 | 5202.6163 | 5119.05342 | 4657.2838 | 4631.6536 | 4594.2881 | 1.71E-228 | -1.927399847 | YPL032C | SVL3 | n |
| W303.asm02.HP0_G0055090 | 914.47385 | 886.493198 | 861.6051344 | 1787.59337 | 1752.7819 | 1803.37833 | 1924.4786 | 1931.3864 | 1865.5754 | 5.17E-55 | -1.005424809 | YPL031C | PHO85 | n |
| W303.asm02.HP0_G0055100 | 445.64797 | 457.779274 | 479.0988192 | 1205.89632 | 1229.6439 | 1191.82259 | 1195.9586 | 1127.0794 | 1228.1411 | 1.52E-71 | -1.390969233 | YPL030W | TRM44 | n |
| W303.asm02.HP0_G0055110 | 642.66019 | 666.426971 | 694.5001028 | 914.21442 | 901.55911 | 977.541039 | 701.15778 | 778.06532 | 678.87197 | 1.56E-08 | -0.478526001 | YPL029W | SUV3 | n |
| W303.asm02.HP0_G0055120 | 10011.803 | 9566.65259 | 9719.137739 | 4466.06661 | 4342.4048 | 4546.37176 | 3701.8851 | 4236.7166 | 3742.6124 | 3.10E-68 | 1.133436324 | YPL028W | ERG10 | n |
| W303.asm02.HP0_G0055140 | 1824.7335 | 1676.44791 | 1589.912614 | 1224.23061 | 1067.8487 | 1034.42995 | 1029.5048 | 1213.6769 | 1150.5557 | 5.79E-11 | 0.612997564 | YPL026C | SKS1 | n |
| W303.asm02.HP0_G0055160 | 1029.3098 | 1020.40142 | 1172.632997 | 309.182817 | 275.05193 | 310.992688 | 266.78198 | 296.5308 | 292.70843 | 4.81E-76 | 1.848240445 | YPL024W | RMI1 | n |
| W303.asm02.HP0_G0055170 | 1051.4342 | 1142.89111 | 1157.178196 | 1744.25778 | 1651.2105 | 1735.11164 | 1800.2083 | 1726.7015 | 1679.5469 | 1.71E-18 | -0.61395326 | YPL023C | MET12 | n |
| W303.asm02.HP0_G0055180 | 1109.379 | 1175.07061 | 1177.462622 | 555.028991 | 549.205 | 610.607594 | 624.77149 | 600.93409 | 601.2866 | 2.56E-37 | 1.014613746 | YPL022W | RAD1 | n |
| W303.asm02.HP0_G0055200 | 1028.2563 | 962.27072 | 1117.57527 | 1046.72134 | 1040.8828 | 1108.38552 | 1144.6543 | 1120.519 | 1154.0824 | 0.63201206 | -0.039268232 | YPL020C | ULP1 | n |
| W303.asm02.HP0_G0055240 | 467.77234 | 472.31195 | 538.9861716 | 1267.56621 | 1260.2053 | 1334.993 | 1150.3548 | 1169.0661 | 1235.1943 | 5.39E-63 | -1.383355559 | YPL016W | SWI1 | n |
| W303.asm02.HP0_G0055250 | 693.23018 | 729.747914 | 769.8422558 | 308.34944 | 285.83828 | 331.851953 | 316.94612 | 236.17497 | 276.8387 | 5.11E-29 | 1.244626133 | YPL015C | HST2 | n |
| W303.asm02.HP0_G0055260 | 246.52866 | 301.03399 | 307.1641623 | 912.547666 | 822.45921 | 750.933563 | 1229.0213 | 969.62946 | 1138.2126 | 7.75E-37 | -1.539850364 | YPL014W | CIP1 | n |
| W303.asm02.HP0_G0055270 | 2503.2141 | 2512.07674 | 2877.49069 | 1187.56203 | 1297.0586 | 1146.31146 | 1214.2001 | 1245.1669 | 1120.5796 | 7.66E-41 | 1.120376944 | YPL013C | MRPS16 | n |
| W303.asm02.HP0_G0055280 | 2570.6408 | 2352.21732 | 2677.544207 | 3924.37165 | 3797.6942 | 4050.49013 | 2348.5935 | 2929.8817 | 2385.7501 | 3.24E-12 | -0.631120266 | YPL012W | RRP12 | n |
| W303.asm02.HP0_G0055290 | 1238.9646 | 1222.82083 | 1109.847869 | 652.534084 | 691.22528 | 699.733547 | 647.57336 | 644.23284 | 689.4518 | 1.39E-23 | 0.805636781 | YPL011C | TAF3 | n |
| W303.asm02.HP0_G0055300 | 4431.1947 | 4554.95568 | 5464.237943 | 2295.95326 | 2497.939 | 2315.37849 | 2336.0525 | 2604.4851 | 2371.6436 | 9.00E-31 | 1.023566237 | YPL010W | RET3 | n |
| W303.asm02.HP0_G0055310 | 916.58093 | 970.575106 | 987.1753895 | 2094.27606 | 2154.5735 | 2136.17843 | 1844.672 | 1725.3894 | 1904.3681 | 3.03E-63 | -1.150981321 | YPL009C | RQC2 | n |
| W303.asm02.HP0_G0055320 | 760.65682 | 671.617212 | 765.0126306 | 634.199793 | 698.41618 | 764.207641 | 723.95966 | 770.19282 | 645.3692 | 0.521777566 | 0.068747494 | YPL008W | CHL1 | n |
| W303.asm02.HP0_G0055330 | 771.19223 | 690.302081 | 708.0230533 | 468.357798 | 481.79031 | 406.75568 | 525.58331 | 430.36328 | 507.8315 | 1.19E-10 | 0.675871889 | YPL007C | TFC8 | n |
| W303.asm02.HP0_G0055340 | 823.8693 | 799.297146 | 804.6155572 | 617.532256 | 570.7777 | 531.911274 | 998.7223 | 804.30698 | 992.74005 | 9.15E-07 | 0.49599604 | YPL006W | NCR1 | n |
| W303.asm02.HP0_G0055350 | 251.79637 | 495.149011 | 279.1523362 | 548.361976 | 563.5868 | 493.037188 | 481.11965 | 452.66869 | 492.84342 | 0.000611599 | -0.646112477 | YPL005W | AEP3 | n |
| W303.asm02.HP0_G0055360 | 3264.9245 | 3723.47904 | 3958.360808 | 3841.03397 | 3672.7523 | 3895.94193 | 4574.0569 | 4112.0687 | 4357.1238 | 0.473761377 | -0.059517289 | YPL004C | LSP1 | n |
| W303.asm02.HP0_G0055370 | 487.78962 | 522.138265 | 507.1106453 | 396.687387 | 346.06207 | 378.311227 | 515.32247 | 488.09494 | 525.46454 | 5.73E-06 | 0.435823591 | YPL003W | ULA1 | n |
| W303.asm02.HP0_G0055380 | 988.22174 | 1010.02094 | 1007.459815 | 350.018283 | 388.30861 | 347.970477 | 329.48715 | 352.95037 | 357.06902 | 9.96E-64 | 1.46842563 | YPL002C | SNF8 | n |
| W303.asm02.HP0_G0055390 | 1501.2964 | 1546.69188 | 1694.232518 | 1521.74615 | 1591.8855 | 1496.17823 | 1247.2628 | 1183.499 | 1225.4962 | 0.582693716 | 0.041297785 | YPL001W | HAT1 | n |
| W303.asm02.HP0_G0055410 | 543.62731 | 475.426095 | 402.7907411 | 120.839645 | 151.90777 | 118.518555 | 165.31362 | 124.6479 | 151.64413 | 3.53E-33 | 1.860734302 | YPR002W | PDH1 | n |
| W303.asm02.HP0_G0055420 | 706.92621 | 663.312826 | 714.7845286 | 807.542181 | 761.33656 | 729.126149 | 1022.6643 | 934.20322 | 998.91162 | 0.092846787 | -0.140871469 | YPR003C | | n |
| W303.asm02.HP0_G0055430 | 2756.064 | 2487.16359 | 3010.788345 | 2093.44268 | 1990.0816 | 2197.80808 | 2094.3526 | 2014.0477 | 2013.693 | 8.91E-07 | 0.394414212 | YPR004C | AIM45 | n |
| W303.asm02.HP0_G0055450 | 896.56365 | 925.939031 | 1103.086394 | 106.672239 | 126.73962 | 86.2815078 | 69.545731 | 81.349156 | 61.715634 | 2.08E-107 | 3.192220866 | YPR006C | ICL2 | n |
| W303.asm02.HP0_G0055460 | 84.283304 | 73.7014251 | 77.27400309 | 155.841474 | 161.79525 | 142.222266 | 155.05278 | 129.89623 | 136.65605 | 7.05E-09 | -0.968418343 | YPR007C | REC8 | n |
| W303.asm02.HP0_G0055470 | 886.02823 | 869.884426 | 920.5265618 | 1443.40873 | 1424.6971 | 1321.71892 | 1309.968 | 1393.4323 | 1376.2586 | 3.24E-19 | -0.646727521 | YPR008W | HAA1 | n |
| W303.asm02.HP0_G0055490 | 2364.1467 | 2463.28848 | 2815.671488 | 6382.00002 | 6271.3638 | 7017.24659 | 4385.9414 | 5212.9064 | 4463.8036 | 3.25E-51 | -1.363467939 | YPR010C | RPA135 | n |
| W303.asm02.HP0_G0055500 | 2120.7786 | 2103.08574 | 2519.132501 | 2410.95927 | 2706.4751 | 2555.26004 | 2248.2653 | 2405.0484 | 2214.7096 | 0.032158807 | -0.185809207 | YPR010C-/ | MRA1 | n |
| W303.asm02.HP0_G0055510 | 886.02823 | 798.259098 | 936.9472875 | 538.361454 | 540.21638 | 590.696477 | 614.51064 | 602.24617 | 556.32236 | 1.62E-12 | 0.652218097 | YPR011C | MRX21 | n |
| W303.asm02.HP0_G0055520 | 170.67369 | 158.821381 | 155.5139312 | 88.3379476 | 80.897627 | 95.7629922 | 69.545731 | 103.65457 | 89.046843 | 5.28E-07 | 0.872416062 | YPR013C | CMR3 | n |
| W303.asm02.HP0_G0055540 | 3377.6534 | 2979.19845 | 3069.709773 | 3336.84096 | 3296.1289 | 3620.97888 | 2580.0326 | 2966.62 | 2664.3521 | 0.130568528 | -0.121328693 | YPR016C | TIF6 | n |
| W303.asm02.HP0_G0055570 | 598.41146 | 606.220173 | 411.4840665 | 829.20998 | 854.81826 | 840.059516 | 795.78558 | 881.71989 | 912.50973 | 3.98E-07 | -0.644618906 | YPR018W | RLF2 | n |
| W303.asm02.HP0_G0055580 | 745.90724 | 880.264909 | 850.014034 | 1055.88849 | 1065.1521 | 1114.07441 | 1063.7077 | 1082.4686 | 1037.7043 | 2.19E-06 | -0.384955799 | YPR019W | MCM4 | n |
| W303.asm02.HP0_G0055590 | 2460.0189 | 2387.51096 | 2957.662468 | 4041.04441 | 4221.0584 | 4338.72725 | 3866.0586 | 4396.7907 | 3905.718 | 1.31E-14 | -0.690584251 | YPR020W | ATP20 | n |
| W303.asm02.HP0_G0055600 | 1179.9663 | 1044.27653 | 1105.018244 | 936.715595 | 1013.9169 | 1071.40773 | 955.39873 | 936.82738 | 955.71067 | 0.087728601 | 0.14036893 | YPR021C | AGC1 | n |
| W303.asm02.HP0_G0055610 | 799.63785 | 884.417102 | 887.6851105 | 1555.08123 | 1597.2787 | 1698.13385 | 1727.2423 | 1493.1506 | 1581.6835 | 6.68E-29 | -0.914655941 | YPR022C | SDD4 | n |
| W303.asm02.HP0_G0055620 | 866.01095 | 937.357562 | 984.2776144 | 355.851921 | 377.52226 | 399.170492 | 364.83006 | 304.4033 | 372.05711 | 2.40E-38 | 1.300980239 | YPR023C | EAF3 | n |
| W303.asm02.HP0_G0055630 | 1673.0236 | 1857.0683 | 1932.816002 | 4102.7143 | 3932.5235 | 4077.03828 | 4546.6947 | 4116.0049 | 4130.5392 | 4.46E-56 | -1.148365977 | YPR024W | YME1 | n |
| W303.asm02.HP0_G0055640 | 1222.1079 | 1319.35931 | 1143.655246 | 1039.22095 | 1014.8158 | 1012.62253 | 888.13319 | 959.1328 | 914.27303 | 0.000693755 | 0.264410441 | YPR025C | CCL1 | n |
| W303.asm02.HP0_G0055650 | 272.8672 | 291.691556 | 324.550813 | 205.010709 | 216.62587 | 239.881555 | 379.65129 | 301.77913 | 368.5305 | 0.001044102 | 0.429322482 | YPR026W | ATH1 | n |
| W303.asm02.HP0_G0055680 | 2541.1416 | 2451.86995 | 2250.60534 | 2541.79943 | 2588.7241 | 2742.04528 | 2510.4869 | 2252.8468 | 2568.252 | 0.121497999 | -0.120207515 | YPR029C | APL4 | n |
| W303.asm02.HP0_G0055700 | 447.75505 | 395.49638 | 427.9047921 | 479.191697 | 526.73344 | 527.170531 | 510.76209 | 511.71244 | 540.45262 | 0.005047426 | -0.269443228 | YPR031W | NTO1 | n |
| W303.asm02.HP0_G0055720 | 5359.3646 | 4921.38671 | 4945.536198 | 4131.04911 | 3960.3883 | 4136.77163 | 3267.5093 | 3858.8366 | 3367.9103 | 2.51E-05 | 0.316280859 | YPR033C | HTS1 | n |
| W303.asm02.HP0_G0055730 | 1140.9852 | 1245.65789 | 1172.632997 | 1392.57274 | 1288.9689 | 1377.65968 | 1428.5377 | 1258.2878 | 1382.4302 | 0.0148978 | -0.189727838 | YPR034W | ARP7 | n |
| W303.asm02.HP0_G0055740 | 5784.9953 | 5313.76895 | 5357.986189 | 5326.11154 | 5356.3218 | 4448.71247 | 3187.7027 | 4562.1132 | 3699.4114 | 0.311123547 | 0.121038894 | YPR035W | GLN1 | n |
| W303.asm02.HP0_G0055750 | 2961.5046 | 3004.11161 | 2847.547014 | 3632.68975 | 3994.5451 | 3680.71224 | 3775.9912 | 3701.3866 | 3750.5472 | 3.28E-09 | -0.359688156 | YPR036W | VMA13 | n |
| W303.asm02.HP0_G0055760 | 6621.5071 | 7121.01094 | 6378.00303 | 655.867592 | 751.44907 | 568.889063 | 551.80547 | 578.62868 | 561.61227 | 4.32E-288 | 3.347127961 | YPR036W- | SPO24 | n |
| W303.asm02.HP0_G0055770 | 1027.2028 | 963.308768 | 950.470238 | 655.034215 | 612.12538 | 631.46686 | 747.90163 | 758.38407 | 796.13168 | 7.88E-16 | 0.630581413 | YPR037C | ERV2 | n |
| W303.asm02.HP0_G0055780 | 308.6876 | 413.1432 | 397.9611159 | 893.379998 | 913.24432 | 841.007664 | 865.33131 | 864.66281 | 884.29687 | 2.14E-33 | -1.240581531 | YPR040W | TIP41 | n |
| W303.asm02.HP0_G0055790 | 4935.841 | 4309.9763 | 5335.769913 | 4221.88719 | 4165.3289 | 4239.17167 | 3646.0205 | 3921.8166 | 3747.9023 | 0.00943282 | 0.207863088 | YPR041W | TIF5 | n |
| W303.asm02.HP0_G0055820 | 734.31829 | 719.367431 | 787.2289065 | 810.875689 | 810.774 | 774.637274 | 800.34595 | 726.89408 | 834.92436 | 0.262637666 | -0.096489896 | YPR045C | THP3 | n |
| W303.asm02.HP0_G0055830 | 736.42537 | 744.280589 | 914.7310116 | 113.339253 | 88.088527 | 87.2296563 | 78.666483 | 101.0304 | 84.638584 | 1.19E-95 | 3.049254566 | YPR046W | MCM16 | n |
| W303.asm02.HP0_G0055840 | 1053.5413 | 1098.25504 | 1030.642016 | 1255.06556 | 1211.6667 | 1218.37074 | 1254.1033 | 1182.1869 | 1206.9815 | 0.002113747 | -0.212061066 | YPR047W | MSF1 | n |

| | | | | | | | | | | | | | | |
|---|---|---|---|---|---|---|---|---|---|---|---|---|---|---|
| W303.asm02.HP0_G0055850 | 1044.0594 | 958.118527 | 831.6614583 | 880.879345 | 885.37958 | 898.844719 | 726.23985 | 758.38407 | 754.69404 | 0.347241854 | 0.087765654 | YPR048W | TAH18 | n |
| W303.asm02.HP0_G0055860 | 317.11593 | 371.62127 | 400.858891 | 665.034737 | 675.94462 | 692.14836 | 749.04173 | 640.29659 | 666.52885 | 7.21E-18 | -0.898186678 | YPR049C | ATG11 | n |
| W303.asm02.HP0_G0055870 | 1286.3739 | 1259.15252 | 1484.626784 | 413.354924 | 377.52226 | 362.192703 | 381.93147 | 377.87995 | 394.98006 | 9.88E-79 | 1.804537371 | YPR051W | MAK3 | n |
| W303.asm02.HP0_G0055880 | 3113.2145 | 2995.80722 | 3139.256376 | 4565.23846 | 4548.2444 | 4523.6162 | 4963.9691 | 4702.5061 | 4735.3524 | 9.46E-26 | -0.56022035 | YPR052C | NHP6A | n |
| W303.asm02.HP0_G0055890 | 135.90683 | 151.555043 | 181.5939073 | 354.185167 | 371.23022 | 413.392719 | 454.89749 | 436.9237 | 491.96177 | 4.50E-23 | -1.275492316 | YPR054W | SMK1 | n |
| W303.asm02.HP0_G0055900 | 883.92115 | 812.791773 | 819.1044328 | 1493.41134 | 1442.6743 | 1375.76338 | 1557.3683 | 1577.124 | 1568.4588 | 6.17E-28 | -0.778128505 | YPR055W | SEC8 | n |
| W303.asm02.HP0_G0055910 | 1458.1012 | 1378.52806 | 1477.865309 | 680.868898 | 655.27078 | 609.659446 | 653.27383 | 695.40408 | 682.39858 | 3.71E-50 | 1.14797253 | YPR056W | TFB4 | n |
| W303.asm02.HP0_G0055920 | 1014.5603 | 990.298022 | 902.1739861 | 343.351268 | 346.06207 | 322.370469 | 240.55982 | 233.5508 | 268.90383 | 3.19E-57 | 1.521493089 | YPR057W | BRR1 | n |
| W303.asm02.HP0_G0055930 | 1128.3427 | 934.243417 | 1128.200445 | 1137.55942 | 1113.6907 | 1225.00778 | 985.04117 | 987.99863 | 971.58041 | 0.173208043 | -0.123031513 | YPR058W | YMC1 | n |
| W303.asm02.HP0_G0055940 | 2467.3937 | 2355.33146 | 2522.996201 | 1055.05511 | 1132.5668 | 1148.20776 | 1177.7171 | 986.68654 | 1087.0768 | 3.07E-48 | 1.139432865 | YPR060C | ARO7 | n |
| W303.asm02.HP0_G0055950 | 332.91905 | 328.023244 | 259.8338354 | 397.520764 | 450.33012 | 461.748289 | 486.82012 | 400.18537 | 520.17463 | 0.000116453 | -0.508896599 | YPR061C | JID1 | n |
| W303.asm02.HP0_G0055960 | 2866.6859 | 2706.19177 | 3134.42675 | 2343.45574 | 2426.0299 | 2487.9415 | 2346.3134 | 2612.3576 | 2277.3069 | 0.000860996 | 0.263156142 | YPR062W | FCY1 | n |
| W303.asm02.HP0_G0055970 | 870.22511 | 949.814141 | 879.9577102 | 1196.72918 | 1168.5213 | 1118.81516 | 1413.7165 | 1300.2744 | 1343.6375 | 8.61E-07 | -0.368379574 | YPR063C | | n |
| W303.asm02.HP0_G0055980 | 844.94012 | 916.596597 | 960.1294884 | 2791.81249 | 2685.8012 | 2816.94901 | 2842.2542 | 2826.2271 | 2613.2163 | 8.97E-111 | -1.606973896 | YPR065W | ROX1 | n |
| W303.asm02.HP0_G0055990 | 366.63237 | 304.148135 | 378.6426151 | 141.674067 | 149.21118 | 127.051891 | 148.21221 | 128.58415 | 138.41935 | 8.35E-22 | 1.327644295 | YPR066W | UBA3 | n |
| W303.asm02.HP0_G0056000 | 1184.1804 | 1283.02763 | 1180.360397 | 532.527816 | 536.62093 | 545.185352 | 564.34651 | 530.0816 | 529.8728 | 2.17E-50 | 1.17607161 | YPR067W | ISA2 | n |
| W303.asm02.HP0_G0056010 | 554.16272 | 516.948024 | 538.9861716 | 332.517369 | 287.63601 | 329.007508 | 332.90743 | 360.82287 | 314.74973 | 1.40E-13 | 0.762114079 | YPR068C | HOS1 | n |
| W303.asm02.HP0_G0056020 | 4613.4574 | 4347.34604 | 5123.266405 | 3282.67147 | 3390.5094 | 4080.83088 | 3519.47 | 4484.7003 | 3511.6196 | 0.000589774 | 0.389454029 | YPR069C | SPE3 | n |
| W303.asm02.HP0_G0056030 | 1524.4743 | 1253.96228 | 1321.385453 | 808.375558 | 811.67286 | 770.84468 | 848.2299 | 875.15947 | 837.56932 | 2.93E-19 | 0.77744609 | YPR070W | MED1 | n |
| W303.asm02.HP0_G0056050 | 1623.5071 | 1640.11622 | 1344.567654 | 2072.60826 | 1988.2839 | 1876.38576 | 1848.0923 | 1840.8527 | 1907.8947 | 1.96E-05 | -0.366465965 | YPR072W | NOT5 | n |
| W303.asm02.HP0_G0056060 | 2808.7411 | 2704.11567 | 3496.64864 | 1659.25334 | 1824.6909 | 1753.12646 | 1499.2235 | 1661.0973 | 1482.9385 | 2.97E-15 | 0.783123583 | YPR073C | LTP1 | n |
| W303.asm02.HP0_G0056070 | 7799.3662 | 8090.54799 | 8626.67652 | 7602.06375 | 7773.3631 | 7765.33571 | 6938.6118 | 6670.6308 | 6730.5307 | 0.147985149 | 0.083481806 | YPR074C | TKL1 | n |
| W303.asm02.HP0_G0056090 | 956.6155 | 1013.13508 | 896.3784358 | 2280.1191 | 2241.7631 | 2278.4007 | 2843.3943 | 2353.8772 | 2879.4751 | 2.46E-45 | -1.246882816 | YPR075C | OPY2 | n |
| W303.asm02.HP0_G0056110 | 677.42706 | 704.834756 | 625.919425 | 1135.05929 | 1020.209 | 1011.67438 | 1325.9293 | 1350.1336 | 1362.1522 | 7.06E-16 | -0.658632486 | YPR079W | MRL1 | n |
| W303.asm02.HP0_G0056140 | 781.72764 | 823.172255 | 1026.778316 | 189.176548 | 160.89639 | 187.733391 | 223.45841 | 182.37956 | 193.08177 | 1.49E-66 | 2.291280697 | YPR082C | DIB1 | n |
| W303.asm02.HP0_G0056150 | 563.6446 | 478.540239 | 447.2232929 | 527.527555 | 526.73344 | 464.592735 | 555.22575 | 524.83327 | 577.482 | 0.793040192 | -0.030131244 | YPR083W | MDM36 | n |
| W303.asm02.HP0_G0056160 | 339.2403 | 349.822257 | 341.9374637 | 860.044924 | 790.10016 | 748.089117 | 734.2205 | 768.88074 | 784.6702 | 1.83E-40 | -1.218937605 | YPR084W | | n |
| W303.asm02.HP0_G0056170 | 783.83473 | 759.851313 | 792.0585317 | 275.847742 | 288.53487 | 259.792672 | 245.1202 | 305.71538 | 294.47174 | 1.50E-49 | 1.502363888 | YPR085C | ASA1 | n |
| W303.asm02.HP0_G0056180 | 3179.5876 | 2926.25799 | 3348.862109 | 2305.95378 | 2289.4028 | 2413.98592 | 2607.3949 | 2439.1626 | 2468.6254 | 4.46E-10 | 0.432104206 | YPR086W | SUA7 | n |
| W303.asm02.HP0_G0056190 | 2348.3436 | 2154.98815 | 2563.565053 | 2204.2818 | 2070.0804 | 2233.83772 | 2188.9804 | 2277.7764 | 2213.828 | 0.123502606 | 0.11916485 | YPR088C | SRP54 | n |
| W303.asm02.HP0_G0056200 | 875.49282 | 908.292211 | 949.504313 | 1791.76026 | 1811.208 | 1760.71165 | 1682.7787 | 1687.339 | 1707.7598 | 1.42E-50 | -0.972088756 | YPR089W | | n |
| W303.asm02.HP0_G0056210 | 673.21289 | 703.796708 | 701.261578 | 862.545054 | 759.53883 | 785.066907 | 1176.577 | 1054.9149 | 1076.497 | 0.013542717 | -0.212504168 | YPR091C | NVJ2 | n |
| W303.asm02.HP0_G0056220 | 421.41652 | 385.115898 | 378.6426151 | 290.015149 | 299.32122 | 243.674149 | 291.86405 | 261.10455 | 294.47174 | 1.37E-05 | 0.506587491 | YPR093C | ASR1 | n |
| W303.asm02.HP0_G0056230 | 487.78962 | 465.045612 | 634.6127504 | 197.510317 | 209.43497 | 219.022289 | 145.93203 | 124.6479 | 171.04047 | 2.99E-20 | 1.34442072 | YPR094W | RDS3 | n |
| W303.asm02.HP0_G0056240 | 400.34569 | 437.01831 | 471.3714188 | 735.038394 | 763.13428 | 765.155789 | 942.8577 | 859.41448 | 846.38584 | 4.53E-18 | -0.788763418 | YPR095C | SYT1 | n |
| W303.asm02.HP0_G0056260 | 1333.7833 | 1239.4296 | 1183.258172 | 328.350484 | 307.41098 | 311.940836 | 318.08621 | 358.19871 | 359.71398 | 9.36E-103 | 1.985795551 | YPR097W | LEC1 | n |
| W303.asm02.HP0_G0056280 | 2220.8651 | 2246.33639 | 2666.919032 | 736.705148 | 782.01039 | 781.274313 | 721.67947 | 858.10239 | 797.89498 | 4.81E-68 | 1.633777847 | YPR100W | MRPL51 | n |
| W303.asm02.HP0_G0056290 | 1229.4827 | 1271.6091 | 1431.500907 | 300.849048 | 324.48937 | 297.718609 | 394.47251 | 354.26246 | 358.83233 | 1.86E-104 | 2.09147838 | YPR101W | SNT309 | n |
| W303.asm02.HP0_G0056310 | 6531.9561 | 6284.34405 | 6584.710988 | 3686.85925 | 3554.1024 | 3516.68256 | 3934.4642 | 4159.3036 | 3882.795 | 9.92E-53 | 0.850642703 | YPR103W | PRE2 | n |
| W303.asm02.HP0_G0056320 | 477.25421 | 478.540239 | 482.9625193 | 915.881173 | 945.60337 | 945.303993 | 725.09975 | 780.68949 | 823.46289 | 1.62E-31 | -0.963758179 | YPR104C | FHL1 | n |
| W303.asm02.HP0_G0056330 | 1311.6589 | 1329.7398 | 1285.646226 | 970.884046 | 982.45674 | 1024.00031 | 907.51479 | 894.84072 | 874.5987 | 3.66E-09 | 0.399705615 | YPR105C | COG4 | n |
| W303.asm02.HP0_G0056350 | 1151.5206 | 1116.93991 | 1261.4981 | 256.680074 | 266.06331 | 256.948227 | 288.44377 | 249.2958 | 280.36531 | 1.95E-113 | 2.179169957 | YPR107C | YTH1 | n |
| W303.asm02.HP0_G0056360 | 6785.8595 | 6672.5741 | 7094.719409 | 3177.66598 | 3161.2995 | 3217.06765 | 3510.3493 | 3610.8529 | 3526.6076 | 1.65E-99 | 1.105026289 | YPR108W | RPN7 | n |
| W303.asm02.HP0_G0056370 | 1452.8335 | 1503.09385 | 1873.894575 | 430.855839 | 360.44387 | 366.933445 | 566.62669 | 509.08827 | 574.83705 | 2.31E-71 | 2.059286491 | YPR109W | GLD1 | n |
| W303.asm02.HP0_G0056380 | 3865.443 | 3559.46742 | 3419.374637 | 1937.60121 | 1854.3534 | 2037.57099 | 1130.9732 | 1241.2307 | 1230.7861 | 2.48E-34 | 0.895469381 | YPR110C | RPC40 | n |
| W303.asm02.HP0_G0056400 | 963.99029 | 998.602408 | 1113.71157 | 2166.77985 | 2017.0475 | 2324.85997 | 1100.1907 | 1511.5198 | 1204.3365 | 3.27E-22 | -1.080705912 | YPR112C | MRD1 | n |
| W303.asm02.HP0_G0056410 | 3652.6277 | 3425.5592 | 3990.236335 | 1973.43641 | 1999.9691 | 2058.43026 | 2562.9312 | 2708.1397 | 2639.6658 | 1.00E-35 | 0.87617029 | YPR113W | PIS1 | n |
| W303.asm02.HP0_G0056420 | 1374.8714 | 1371.26173 | 1438.262383 | 1406.74015 | 1368.0688 | 1402.31154 | 1763.7253 | 1545.634 | 1815.3213 | 0.974473732 | 0.002688705 | YPR114W | | n |
| W303.asm02.HP0_G0056440 | 158.03119 | 140.136513 | 153.5820811 | 326.683731 | 329.88255 | 347.970477 | 239.41973 | 320.14829 | 269.78549 | 6.02E-18 | -1.152300096 | YPR116W | RRG8 | n |
| W303.asm02.HP0_G0056450 | 872.3322 | 908.292211 | 1089.563444 | 300.015671 | 316.39961 | 268.326008 | 299.84471 | 286.03413 | 304.16991 | 3.34E-52 | 1.697633966 | YPR117W | HOB2 | n |
| W303.asm02.HP0_G0056460 | 3525.1492 | 3150.47641 | 3684.038097 | 2025.10578 | 2093.4508 | 2262.28217 | 1990.604 | 1793.6177 | 1890.2617 | 1.52E-18 | 0.699613374 | YPR118W | MRI1 | n |
| W303.asm02.HP0_G0056470 | 1608.7576 | 1437.69681 | 1592.810389 | 878.379215 | 849.42508 | 840.059516 | 896.11385 | 871.22322 | 826.98949 | 3.48E-29 | 0.853106013 | YPR119W | CLB2 | n |
| W303.asm02.HP0_G0056480 | 883.92115 | 788.916664 | 781.4333562 | 942.549233 | 960.88404 | 953.837328 | 949.69826 | 867.28697 | 950.42076 | 0.007105467 | -0.219813832 | YPR120C | CLB5 | n |
| W303.asm02.HP0_G0056500 | 1207.3583 | 1202.05986 | 1127.23452 | 100.005224 | 116.85213 | 105.244477 | 110.58911 | 115.46332 | 120.78631 | 1.85E-204 | 3.457827841 | YPR122W | AXL1 | n |
| W303.asm02.HP0_G0056510 | 11057.969 | 11102.964 | 9773.229541 | 10049.6916 | 10056.474 | 10970.0774 | 18097.851 | 17722.307 | 16635.008 | 0.601224904 | 0.039269711 | YPR124W | CTR1 | n |
| W303.asm02.HP0_G0056520 | 2403.1277 | 2516.22894 | 2118.27361 | 2754.31054 | 2587.8252 | 2627.31932 | 2127.4153 | 2042.9135 | 2158.2839 | 0.018557071 | -0.179901855 | YPR125W | YLH47 | n |
| W303.asm02.HP0_G0056530 | 592.09021 | 637.36162 | 651.9994011 | 3207.66755 | 3219.7256 | 3435.14179 | 4631.0616 | 4462.3949 | 4206.3613 | 4.17E-234 | -2.389298869 | YPR127W | | n |
| W303.asm02.HP0_G0056540 | 1275.8385 | 1235.27741 | 1340.703954 | 302.515802 | 251.68151 | 243.674149 | 372.81072 | 326.70871 | 364.12224 | 6.75E-112 | 2.268724619 | YPR128C | ANT1 | n |
| W303.asm02.HP0_G0056550 | 1020.8815 | 1186.48914 | 1234.452199 | 1700.92218 | 1648.5139 | 1665.89681 | 1679.3584 | 1985.1818 | 1797.6882 | 1.17E-09 | -0.542719602 | YPR129W | SCD6 | n |
| W303.asm02.HP0_G0056560 | 987.1682 | 977.841443 | 912.7991615 | 552.528861 | 556.3959 | 544.237203 | 506.20171 | 586.50118 | 574.83705 | 5.00E-21 | 0.799205206 | YPR131C | NAT3 | n |
