## Supplementary material for "Intergeneric chromosomal transfer in yeast results in improved phenotypes and widespread transcriptional responses": TableS7

Table S7: Strains used in this study.

| Name | Species | Genotype | Sources |
|---|---|---|---|
| FIM-1ΔU | *Kluyveromyces marxianus* | *ura3* Δ | *Biotechnol Biofuels* 2018, 11:235. doi:10.1186/s13068-018-1232-7 |
| W303-1A | *Saccharomyces cerevisiae* | *MAT* **a** *leu2-3,112 trp1-1 can1-100 ura3-1 ade2-1 his3-11,15* | |
| W303-1B | *Saccharomyces cerevisiae* | *MATα leu2-3,112 trp1-1 can1-100 ura3-1 ade2-1 his3-11,15* | |
| LHP560[1] | *Kluyveromyces marxianus* | *MATα HMR* **a**Δ *his3* Δ | *Front Microbiol* 2022, 13:865829. doi: 10.3389/fmicb.2022.865829 |
| LHP508[1] | *Kluyveromyces marxianus* | *MAT* **a** *HMLα* Δ *trp1* Δ | *Front Microbiol* 2022, 13:865829. doi: 10.3389/fmicb.2022.865829 |
| Sc-Ring1[2] | *Saccharomyces cerevisiae* | chrI (TELΔ::*KmURA3* ) | This study |
| Sc-Ring1L[2] | *Saccharomyces cerevisiae* | chrI (TELΔ::*KmURA3 KmARS1 -hyg* $^R$ ) | This study |
| Sc-R1[2] | *Saccharomyces cerevisiae* | chrI (TELΔ::*KmURA3 KmARS1 -hyg* $^R$ *KmARS1 /KmCEN5 -kan* $^R$ )[3] | This study |
| Sc-Ring3[2] | *Saccharomyces cerevisiae* | chrIII (TELΔ::*KmURA3* ) | This study |
| Sc-Ring3L[2] | *Saccharomyces cerevisiae* | chrIII (TELΔ::*KmURA3 KmARS1 /KmCEN5 -hyg* $^R$ ) | This study |
| Sc-R3[2] | *Saccharomyces cerevisiae* | chrIII (TELΔ::*KmURA3 KmARS1/KmCEN5 -hyg* $^R$ , *KmARS1 -kan* $^R$ )[4] | This study |
| KS-R1 | Hybrid | FIM-1ΔU, R1 | This study |
| KS-R3 | Hybrid | FIM-1ΔU, R3 | This study |

1. Isogenic relative to FIM-1ΔU
2. Isogenic relative to W303-1A
3. Engineered Sc chrI was designated as R1
4. Engineered Sc chrIII was designated as R3
