## Supplementary material for "Intergeneric chromosomal transfer in yeast results in improved phenotypes and widespread transcriptional responses": TableS8

Table S8: Plasmids used in this study, and sequences of LHZ626, LHZ1495 and pRS425-Cas9-2xSapI.

**Plasmids**

| Name | Essential features | application | Backbone | source |
|---|---|---|---|---|
| LHZ626 | *KmARS1, KmCEN5, ScARSH4, ScCEN6, HPHMX4* (*hyg* $^{R}$), *KANMX6* (*kan* $^{R}$), *KmURA3* | a vector control of circular chromosomes | | *Biotechnol J* . 2021 Dec;16(12):e2100122. doi:10.1002/biot.202100122 |
| LHZ882 | *KmARS1, KmCEN5, HPHMX4* | | | *Microorganisms* . 2022 Jun; 10(6): 1240. doi:10.3390/microorganisms10061240 |
| LHZ1493 | *KmARS1* , *KmCEN5* , *ScARSH4* , *ScCEN6* , | selelect diploid or hybrid | LHZ882 | This study |
| LHZ1494 | *KmARS1* , *KmCEN5* , *ScARSH4* , *ScCEN6* , *KANMX6* | selelect diploid or hybrid | LHZ1493 | This study |
| LHZ1495 | *KmARS1* , *KmCEN5* , *HPHMX4* | evaluate functionality of ARS and CEN in *K.marxianus* and *S. cerevisiae* | LHZ882; a BamHI site was introduced between KmARS1 and KmCEN5 of LHZ882 | This study |
| LHZ1496 | *KmARS1* , *ScCEN6* , *HPHMX4* | evaluate functionality of ARS and CEN in *K.marxianus* and *S. cerevisiae* | LHZ1495; ScCEN6 was inserted between BamHI and SalI | This study |
| LHZ1497 | *ScARSH4* , *KmCEN5* , *HPHMX4* | evaluate functionality of ARS and CEN in *K.marxianus* and *S. cerevisiae* | LHZ1495; ScARSH4 was inserted between BamHI and HindIII | This study |
| LHZ1498 | *ScARSH4* , *ScCEN6* , *HPHMX4* | evaluate functionality of ARS and CEN in *K.marxianus* and *S. cerevisiae* | LHZ1495; ScARSH4 and ScCEN6 were inserted between HindIII and SalI | This study |
| LHZ1783 | *KmARS18* , *KmCEN3* , *HPHMX4* | evaluate functionality of ARS and CEN in *K.marxianus* and *S. cerevisiae* | LHZ1495; KmARS18 and KmCEN3 were inserted between HindIII and SalI | This study |
| LHZ1784 | *ScARS1, ScCEN4, HPHMX4* | evaluate functionality of ARS and CEN in *K.marxianus* and *S. cerevisiae* | LHZ1783; ScARS1 and ScCEN4 were inserted between HindIII and SalI | This study |
| LHZ1785 | *ScARS1* , *KmCEN3* , *HPHMX4* | evaluate functionality of ARS and CEN in *K.marxianus* and *S. cerevisiae* | LHZ1783; ScARS1 was inserted between BamHI and HindIII | This study |
| LHZ1786 | *KmARS18, ScCEN4, HPHMX4* | evaluate functionality of ARS and CEN in *K.marxianus* and *S. cerevisiae* | LHZ1783; ScCEN4 was inserted between BamHI and SalI | This study |
| LHZ1787 | *KmARS18* , *KmCEN3* , *ScARS1* , *ScCEN4* , *HPHMX4* | evaluate functionality of ARS and CEN in *K.marxianus* and *S. cerevisiae* | LHZ1783; ScARS1 and ScCEN4 were inserted into SalI | This study |
| pRS425-Cas9-2xSapI | *TEF1* promoter-*SpCas9* -*CYC1* terminator, *SNR52* promoter-(2xSapI site)-gRNA-*SUP4* terminator, *LEU2* , 2μm | CRISPR vector | pRS425 | Bruce Futcher, Stony Brook University |
| LHZ1499A | gRNA-A1F/A1R | Intemediate CRISPR plasmid to delete telomeres of *S. cerevisiae* chr1 | pRS425-Cas9-2xSapI | This study |
| LHZ1499B | gRNA-A2F/A2R | Intemediate CRISPR plasmid to delete telomeres of *S. cerevisiae* chr1 | pRS425-Cas9-2xSapI | This study |
| LHZ1500A | gRNA-C1F/C1R | Intemediate CRISPR plasmid to delete telomeres of *S. cerevisiae* chr3 | pRS425-Cas9-2xSapI | This study |
| LHZ1500B | gRNA-C2F/C2R | Intemediate CRISPR plasmid to delete telomeres of *S. cerevisiae* chr3 | pRS425-Cas9-2xSapI | This study |
| LHZ1499 | gRNA-A1F/A1R, gRNA-A2F/A2R | CRISPR plasmid to delete telomeres and circularize S. cerevisiae chr1, expressing two gRNAs | LHZ1499B, with gRNA-A1F/A1R inserted into NotI | This study |
| LHZ1500 | gRNA-C1F/C1R, gRNA-C2F/C2R | CRISPR plasmid to delete telomeres and circularize S. cerevisiae chr3, expressing two gRNAs | LHZ1500B, with gRNA-C1F/C1R inserted into NotI | This study |
| LHZ1501 | gRNA-ALF/ALR | CRISPR plasmid to insert *KmARS1* into the left arm of *S. cerevisiae* chr1 | pRS425-Cas9-2xSapI | This study |

| | | | | |
|---|---|---|---|---|
| LHZ1502 | gRNA-ARF/ARR | CRISPR plasmid to insert *KmARS1* &*KmCEN5* into the right arm of *S. cerevisiae* chr1 | pRS425-Cas9-2xSapI | This study |
| LHZ1503 | gRNA-CLF/CLR | CRISPR plasmid to insert *KmARS1* &*KmCEN5* into the left arm of *S. cerevisiae* chr3 | pRS425-Cas9-2xSapI | This study |
| LHZ1504 | gRNA-CRF/CRR | CRISPR plasmid to insert *KmARS1* into the right arm of *S. cerevisiae* chr3 | pRS425-Cas9-2xSapI | This study |
| LHZ1505 | $P_{ScFLO9}$-*ScFLO9*-$T_{ScFLO9}$ | centromeric plasmid to express *ScFLO9* driven by *ScFLO9* promoter in *K.marxianus* | LHZ626 | This study |
| LHZ1506 | $P_{ScSPS22}$-*ScSPS22*-$T_{ScSPS22}$ | centromeric plasmid to express *ScSPS22* driven by *ScSPS22* promoter in *K.marxianus* | LHZ626 | This study |
| LHZ1507 | $P_{KmSPS22}$-*ScSPS22*-$T_{ScSPS22}$ | centromeric plasmid to express *ScSPS22* driven by KmSPS22 promoter in *K.marxianus* | LHZ626 | This study |
| LHZ1508 | $P_{KmSPS22}$-*KmSPS22*-$T_{KmSPS22}$ | centromeric plasmid to express *KmSPS22* driven by *KmSPS22* promoter in *K.marxianus* | LHZ626 | This study |
| pRS425 | *LEU2* , 2 micron | | | Rolf Sternglanz, Stony Brook University |
| pRS315 | *LEU2* , *ScARSH4* , *ScCEN6* | | | Rolf Sternglanz, Stony Brook University |
| LHZ1509 | $P_{ScFLO9}$-*ScFLO9*-$T_{ScFLO9}$ | centromeric plasmid to express *ScFLO9* driven by *ScFLO9* promoter in *S.cerevisiae* | pRS315 | This study |
| LHZ1510 | $P_{ScFLO9}$-*ScFLO9*-$T_{ScFLO9}$ | 2 micron plasmid to express *ScFLO9* driven by *ScFLO9* promoter in *S.cerevisiae* | pRS425 | This study |
| LHZ1511 | $P_{ScSPS22}$-*ScSPS22*-$T_{ScSPS22}$ | centromeric plasmid to express *ScSPS22* driven by *ScSPS22* promoter in *S.cerevisiae* | pRS315 | This study |
| LHZ1512 | $P_{ScSPS22}$-*ScSPS22*-$T_{ScSPS22}$ | 2 micron plasmid to express *ScSPS22* driven by *ScSPS22* promoter in *S.cerevisiae* | pRS425 | This study |

**LHZ626 in genbank format.**

```
LOCUS       Exported                9916 bp ds-DNA     circular SYN 15-APR-2024
DEFINITION  synthetic circular DNA.
ACCESSION   .
VERSION     .
KEYWORDS    LHZ626
SOURCE      synthetic DNA construct
  ORGANISM  synthetic DNA construct
REFERENCE   1  (bases 1 to 9916)
  AUTHORS   Yilin Lyu
  TITLE     Direct Submission
  JOURNAL   Exported Apr 15, 2024 from SnapGene 4.2.4
            http://www.snapgene.com
FEATURES             Location/Qualifiers
     source          1..9916
                     /organism="synthetic DNA construct"
                     /mol_type="other DNA"
     CDS             468..1328
                     /label=Lactamase-AmpC
                     /note="Lactamase-AmpC"
     misc_feature    2330..2704
                     /label=ScARSH4
                     /note="ScARSH4"
     misc_feature    2705..2821
                     /label=ScCEN6
                     /note="ScCEN6"
     terminator      2867..3101
                     /note="TEF terminator"
     CDS             complement(3102..4130)
                     /label=HphMX4
                     /note="HphMX4"
     misc_feature    4131..4509
                     /label=TEF Promoter
                     /note="TEF Promoter"
     promoter        4531..5116
                     /note="Promoter KmURA3"
     CDS             5117..5920
                     /codon_start=1
                     /label=KmURA3
```

```
                     /note="KmURA3"
                     /translation="MSTKSYSERAAAHRSPVAAKLLNLMEEKKSNLCASLDVRKTAELL
                     RLVEVLGPYICLLKTHVDILEDFSFENTIVPLKQLAEKHKFLIFEDRKFADIGNTVKLQ
                     YTSGVYRIAEWSDITNAHGVTGAGIVAGLKQGAEEVTKEPRGLLMLAELSSKGSLAHGE
                     YTRGTVEIAKSDKDFVIGFIAQNDMGGREEGYDWLIMTPGVGLDDKGDALGQQYRTVDE
                     VVAGGSDIIIVGRGLFAKGRDPVVEGERYRKAGWDAYLKRVGRSA"
     terminator      5921..6155
                     /note="TEF terminator"
     terminator      6172..6361
                     /note="ADH1 Terminator"
     CDS             complement(6378..7187)
                     /codon_start=1
                     /gene="aph(3')-Ia"
                     /product="aminoglycoside phosphotransferase"
                     /label=aph(3')-Ia
                     /note="KanR"
                     /note="confers resistance to kanamycin"
                     /translation="MGKEKTHVSRPRLNSNMDADLYGYKWARDNVGQSGATIYRLYGKP
                     DAPELFLKHGKGSVANDVTDEMVRLNWLTEFMPLPTIKHFIRTPDDAWLLTTAIPGKTA
                     FQVLEEYPDSGENIVDALAVFLRRLHSIPVCNCPFNSDRVFRLAQAQSRMNNGLVDASD
                     FDDERNGWPVEQVWKEMHKLLPFSPDSVVTHGDFSLDNLIFDEGKLIGCIDVGRVGIAD
                     RYQDLAILWNCLGEFSPSLQKRLFQKYGIDNPDMNKLQFHLMLDEFF"
     promoter        7194..8640
                     /note="ADH1 promoter"
     CDS             complement(8647..9892)
                     /label=kmARS1
                     /note="kmARS1"
     misc_feature    8842..9034
                     /label=KmCEN5
                     /note="KmCEN5"
     CDS             9505..9516
                     /label=ARS core sequence
                     /note="ARS core sequence"
ORIGIN
        1 tatagtgtca cctaaatcgt atgtgtatga tacataaggt tatgtattaa ttgtagccgc
       61 gttctaacga caatatgtcc atatggtgca ctctcagtac aatctgctct gatgccgcat
      121 agttaagcca gccccgacac ccgccaacac ccgctgacgc gccctgacgg gcttgtctgc
      181 tcccggcatc cgcttacaga caagctgtga ccgtctccgg gagctgcatg tgtcagaggt
      241 tttcaccgtc atcaccgaaa cgcgcgagac gaaagggcct cgtgatacgc ctatttttat
```

301 aggttaatgt catgataata atggtttctt agacgtcagg tggcactttt cggggaaatg
361 tgcgcggaac ccctatttgt ttatttttct aaatacattc aaatatgtat ccgctcatga
421 gacaataacc ctgataaatg cttcaataat attgaaaaag gaagagtatg agtattcaac
481 atttccgtgt cgcccttatt cccttttttg cggcattttg ccttcctgtt tttgctcacc
541 cagaaacgct ggtgaaagta aaagatgctg aagatcagtt gggtgcacga gtgggttaca
601 tcgaactgga tctcaacagc ggtaagatcc ttgagagttt tcgccccgaa gaacgttttc
661 caatgatgag cacttttaaa gttctgctat gtggcgcggt attatcccgt attgacgccg
721 ggcaagagca actcggtcgc cgcatacact attctcagaa tgacttggtt gagtactcac
781 cagtcacaga aaagcatctt acggatggca tgacagtaag agaattatgc agtgctgcca
841 taaccatgag tgataacact gcggccaact tacttctgac aacgatcgga ggaccgaagg
901 agctaaccgc ttttttgcac aacatggggg atcatgtaac tcgccttgat cgttgggaac
961 cggagctgaa tgaagccata ccaaacgacg agcgtgacac cacgatgcct gtagcaatgg
1021 caacaacgtt gcgcaaacta ttaactggcg aactacttac tctagcttcc cggcaacaat
1081 taatagactg gatggaggcg gataaagttg caggaccact tctgcgctcg gcccttccgg
1141 ctggctggtt tattgctgat aaatctggag ccggtgagcg tgggtctcgc ggtatcattg
1201 cagcactggg gccagatggt aagccctccc gtatcgtagt tatctacacg acggggagtc
1261 aggcaactat ggatgaacga aatagacaga tcgctgagat aggtgcctca ctgattaagc
1321 attggtaact gtcagaccaa gtttactcat atatacttta gattgattta aaacttcatt
1381 tttaatttaa aaggatctag gtgaagatcc tttttgataa tctcatgacc aaaatccctt
1441 aacgtgagtt ttcgttccac tgagcgtcag accccgtaga aaagatcaaa ggatcttctt
1501 gagatccttt ttttctgcgc gtaatctgct gcttgcaaac aaaaaaacca ccgctaccag
1561 cggtggtttg tttgccggat caagagctac caactctttt tccgaaggta actggcttca
1621 gcagagcgca gataccaaat actgtccttc tagtgtagcc gtagttaggc caccacttca
1681 agaactctgt agcaccgcct acatacctcg ctctgctaat cctgttacca gtggctgctg
1741 ccagtggcga taagtcgtgt cttaccgggt tggactcaag acgatagtta ccggataagg
1801 cgcagcggtc gggctgaacg gggggttcgt gcacacagcc cagcttggag cgaacgacct
1861 acaccgaact gagataccta cagcgtgagc attgagaaag cgccacgctt cccgaaggga
1921 gaaaggcgga caggtatccg gtaagcggca gggtcggaac aggagagcgc acgagggagc
1981 ttccaggggg aaacgcctgg tatctttata gtcctgtcgg gtttcgccac ctctgacttg
2041 agcgtcgatt tttgtgatgc tcgtcagggg ggcggagcct atggaaaaac gccagcaacg
2101 cggccttttt acggttcctg gccttttgct ggccttttgc tcacatgttc tttcctgcgt
2161 tatcccctga ttctgtggat aaccgtatta ccgcctttga gtgagctgat accgctcgcc
2221 gcagccgaac gaccgagcgc agcgagtcag tgagcgagga agcggaagag cgcccaatac
2281 gcaaaccgcc tctccccgcg cgttggccga ttcattaatg caggttaacg taacttacac
2341 gcgcctcgta tcttttaatg atggaataat ttgggaattt actctgtgtt tatttatttt
2401 tatgttttgt atttggattt tagaaagtaa ataaagaagg tagaagagtt acggaatgaa
2461 gaaaaaaaaa taaacaaagg tttaaaaaat ttcaacaaaa agcgtacttt acatatatat
2521 ttattagaca agaaaagcag attaaataga tatacattcg attaacgata agtaaaatgt
2581 aaaatcacag gattttcgtg tgtggtcttc tacacagaca agatgaaaca attcggcatt

2641 aatacctgag agcaggaaga gcaagataaa aggtagtatt tgttggcgat ccccctagag
2701 tcttttacat cttcggaaaa caaaaactat tttttcttta atttcttttt ttactttcta
2761 tttttaattt atatatttat attaaaaaat ttaaattata attattttta tagcacgtga
2821 tactagtgga tctgatatca tcgatgaatt cgagctcgtt ttcgacactg gatggcggcg
2881 ttagtatcga atcgacagca gtatagcgac cagcattcac atacgattga cgcatgatat
2941 tactttctgc gcacttaact tcgcatctgg gcagatgatg tcgaggcgaa aaaaaatata
3001 aatcacgcta acatttgatt aaaatagaac aactacaata taaaaaaact atacaaatga
3061 caagttcttg aaaacaagaa tctttttatt gtcagtactg attattcctt tgccctcgga
3121 cgagtgctgg ggcgtcggtt tccactatcg gcgagtactt ctacacagcc atcggtccag
3181 acggccgcgc ttctgcgggc gatttgtgta cgcccgacag tcccggctcc ggatcggacg
3241 attgcgtcgc atcgaccctg cgcccaagct gcatcatcga aattgccgtc aaccaagctc
3301 tgatagagtt ggtcaagacc aatgcggagc atatacgccc ggagccgcgg cgatcctgca
3361 agctccggat gcctccgctc gaagtagcgc gtctgctgct ccatacaagc caaccacggc
3421 ctccagaaga agatgttggc gacctcgtat tgggaatccc cgaacatcgc ctcgctccag
3481 tcaatgaccg ctgttatgcg gccattgtcc gtcaggacat tgttggagcc gaaatccgcg
3541 tgcacgaggt gccggacttc ggggcagtcc tcggcccaaa gcatcagctc atcgagagcc
3601 tgcgcgacgg acgcactgac ggtgtcgtcc atcacagttt gccagtgata cacatgggga
3661 tcagcaatcg cgcatatgaa atcacgccat gtagtgtatt gaccgattcc ttgcggtccg
3721 aatgggccga acccgctcgt ctggctaaga tcggccgcag cgatcgcatc catggcctcc
3781 gcgaccggct gcagaacagc gggcagttcg gtttcaggca ggtcttgcaa cgtgacaccc
3841 tgtgcacggc gggagatgca ataggtcagg ctctcgctga attccccaat gtcaagcact
3901 tccggaatcg ggagcgcggc cgatgcaaag tgccgataaa cataacgatc tttgtagaaa
3961 ccatcggcgc agctatttac ccgcaggaca tatccacgcc ctcctacatc gaagctgaaa
4021 gcacgagatt cttcgccctc cgagagctgc atcaggtcgg agacgctgtc gaacttttcg
4081 atcagaaact tctcgacaga cgtcgcggtg agttcaggct ttttacccat ggttgtttat
4141 gttcggatgt gatgtgagaa ctgtatccta gcaagatttt aaaaggaagt atatgaaaga
4201 agaacctcag tggcaaatcc taacctttta tatttctcta caggggcgcg gcgtggggac
4261 aattcaacgc gtctgtgagg ggagcgtttc cctgctcgca ggtctgcagc gaggagccgt
4321 aatttttgct tcgcgccgtg cggccatcaa aatgtatgga tgcaaatgat tatacatggg
4381 gatgtatggg ctaaatgtac gggcgacagt cacatcatgc ccctgagctg cgcacgtcaa
4441 gactgtcaag gagggtattc tgggcctcca tgtcgctggc cgggtgaccc ggcggggaca
4501 aggcaagcta aacagatcta tattaccctg cgaattctga ttggaaagac cattctgctt
4561 tacttttaga gcatcttggt cttctgagct cattatacct caatcaaaac tgaaattagg
4621 tgcctgtcac ggctcttttt ttactgtacc tttgacttcc tttcttattt ccaaggatgc
4681 tcatcacaat acgcttctag atctattatg cattataatt aatagttgta gctacaaaag
4741 gtaaaagaaa gtccggggca ggcaacaata gaaatcggca aaaaaaacta cagaaatact
4801 aagagcttct tccccattca gtcatcgcat ttcgaaacaa gaggggaatg gctctggcta
4861 gggaactaac caccatcgac tgactctatg cactaaccac gtgactacat atatgtgatc
4921 gtttttaaca tttttcaaag gctgtgtgtc tggctgtttc cattaatttt cactgattaa

4981 gcagtcatat tgaatctgag ctcatcacca acaagaaatt ctaccgtaaa agtgtaaaag
5041 ttcgtttaaa tcatttgtaa actggaacag caagaggaag tatcatcagc tagccccata
5101 aactaatcaa aggaggatgt cgactaagag ttactcggaa agagcagctg ctcatagaag
5161 tccagttgct gccaagcttt taaacttgat ggaagagaag aagtcaaact tatgtgcttc
5221 tcttgatgtt cgtaaaacag cagagttgtt aagattagtt gaggttttgg gtccatatat
5281 ctgtctattg aagacacatg tagatatctt ggaggatttc agctttgaga ataccattgt
5341 gccgttgaag caattagcag agaaacacaa gtttttgata tttgaagaca ggaagtttgc
5401 cgacattggg aacactgtta aattacaata cacgtctggt gtataccgta tcgccgaatg
5461 gtctgatatc accaatgcac acggtgtgac tggtgcgggc attgttgctg gtttgaagca
5521 aggtgccgag gaagttacaa aagaacctag agggttgtta atgcttgccg agttatcgtc
5581 caaggggtct ctagcgcacg gtgaatacac tcgtgggacc gtggaaattg ccaagagtga
5641 taaggacttt gttattggat ttattgctca aaacgatatg ggtggaagag aagagggcta
5701 cgattggttg atcatgacgc caggtgttgg tcttgatgac aaaggtgatg ctttgggaca
5761 acaatacaga actgtggatg aagttgttgc cggtggatca gacatcatta ttgttggtag
5821 aggtcttttc gcaaagggaa gagatcctgt agtggaaggt gagagataca gaaaggcggg
5881 atgggacgct tacttgaaga gagtaggcag atccgcttaa tcagtactga caataaaaag
5941 attcttgttt tcaagaactt gtcatttgta tagttttttt atattgtagt tgttctattt
6001 taatcaaatg ttagcgtgat ttatattttt tttcgcctcg acatcatctg cccagatgcg
6061 aagttaagtg cgcagaaagt aatatcatgc gtcaatcgta tgtgaatgct ggtcgctata
6121 ctgctgtcga ttcgatacta acgccgccat ccagtttatc cctagcggat ctgccggtag
6181 aggtgtggtc aataagagcg acctcatact atacctgaga aagcaacctg acctacagga
6241 aagagttact caagaataag aattttcgtt ttaaaaccta agagtcactt taaaatttgt
6301 atacacttat tttttttata acttatttaa taataaaaat cataaatcat aagaaattcg
6361 cctcgagtac cggtacctta gaaaaactca tcgagcatca aatgaaactg caatttattc
6421 atatcaggat tatcaatacc atatttttga aaaagccgtt tctgtaatga aggagaaaac
6481 tcaccgaggc agttccatag gatggcaaga tcctggtatc ggtctgcgat tccgactcgt
6541 ccaacatcaa tacaacctat taatttcccc tcgtcaaaaa taaggttatc aagtgagaaa
6601 tcaccatgag tgacgactga atccggtgag aatggcaaaa gcttatgcat ttctttccag
6661 acttgttcaa caggccagcc attacgctcg tcatcaaaat cactcgcatc aaccaaaccg
6721 ttattcattc gtgattgcgc ctgagcgaga cgaaatacgc gatcgctgtt aaaaggacaa
6781 ttacaaacag gaatcgaatg caaccggcgc aggaacactg ccagcgcatc aacaatattt
6841 tcacctgaat caggatattc ttctaatacc tggaatgctg ttttgccggg gatcgcagtg
6901 gtgagtaacc atgcatcatc aggagtacgg ataaaatgct tgatggtcgg aagaggcata
6961 aattccgtca gccagtttag tctgaccatc tcatctgtaa catcattggc aacgctacct
7021 ttgccatgtt tcagaaacaa ctctggcgca tcgggcttcc catacaatcg atagattgtc
7081 gcacctgatt gcccgacatt atcgcgagcc catttatacc catataaatc agcatccatg
7141 ttggaattta atcgcggcct cgaaacgtga gtcttttcct tacccatccc gggagttgat
7201 tgtatgcttg gtatagcttg aaatattgtg cagaaaaaga aacaaggaag aaagggaacg
7261 agaacaatga cgaggaaaca aaagattaat aattgcaggt ctatttatac ttgatagcaa

7321 gacagcaaac ttttttttat ttcaaattca agtaactgga aggaaggccg tataccgttg
7381 ctcattagag agtagtgtgc gtgaatgaag gaaggaaaaa gtttcgtgtg cttcgagata
7441 cccctcatca gctctggaac aacgacatct gttggtgctg tctttgtcgt taattttttc
7501 ctttagtgtc ttccatcatt ttttttgtca ttgcggatat ggtgagacaa caacggggga
7561 gagagaaaag aaaaaaaaag aaaagaagtt gcatgcgcct attattactt caatagatgg
7621 caaatggaaa aagggtagtg aaacttcgat atgatgatgg ctatcaagtc tagggctaca
7681 gtattagttc gttatgtacc accatcaatg aggcagtgta attggtgtag tcttgtttag
7741 cccattatgt cttgtctggt atctgttcta ttgtatatct cccctccgcc acctacatgt
7801 tagggagacc aacgaaggta ttataggaat cccgatgtat gggtttggtt gccagaaaag
7861 aggaagtcca tattgtacac ccggaaacaa caaaaggata tccgaaatat tccacggttt
7921 agaaaaaaat cggaaaagag cgcggagggg tgttaccccc cttctctact agcattggac
7981 tttaattaat atatgtgcat aggagaagtg taaagttccc ttccatattg taacataata
8041 aagtgcacac ccaaatgaat tgaaagcgta ctcaaacaga caaccatttc cagtgttgta
8101 tgtacctgtc tatttatact ggtagcaacc ctattgctgt ttcctcttca aagtactcta
8161 gcggttatgc gcgtctcacc ttcaaggtca tggtcgctct attgttcgca ccaccggcaa
8221 actcgcgtct cgcaagtctt ggctcattct tctagtatac tcatgttgca aatgcactca
8281 ggttctttcg gcaacttaaa taatgacacc agttgtcgtg gtcgtcatca tcgcaacccc
8341 aaccggcatt cttattgctt ctccaatctc gccccttagc gcagggtaaa ccttggaaaa
8401 tgcaggcgca aaaaactccg ccgggcacag cctcacgccc agcgttatcg ccgggccggc
8461 aagagcgcgg gtccgccaca gagtcagcat gattgtgcaa ttgcgtaaac tcgttttttc
8521 ggcgccgcaa agccaaatac atcatatcaa cacttttcac tttatttttc gttcgaccct
8581 tatatttgtc ttttgccttc atgctccttg atttcctatt tcatttacca tcatttcttc
8641 gtcgacgagc tcctttcatt tctgataaaa gtaaggcttc tctatttacc ttttaaccta
8701 catattcata gttggaagtt atccttctaa gtacgtatac aatattaatt caacgtaaaa
8761 acaaaactta ctgtaaatat gtgtaaaaaa aatctattaa attcatggca gtttcaagaa
8821 aagaaaacta ttatggtctg gtcacgtgta tacaaattat taattttaaa actatataat
8881 ttattatttt tttattttga agtttagagt aattttagta gtattttata ttttaaataa
8941 atatgcttta aatttttact taatatttta ttatttttaa atacaacgtt tttatttaaa
9001 acaaaattat aagttaaaaa gttgttccga aagtaaaata tattttatgg gttttacaaa
9061 aataaattat ttttaatgta tttttttaat tatatttttg tatgtaatta tatccacagg
9121 tattatgttg aatttagctg ttttagttta cctgtgtggt actatgattt ttttagaact
9181 ctcctcttag aaataggtgg tgttgcggtt gacttttaac gatatatcat tttcaattta
9241 tttattttaa agtgacatag agagattcct tttaattttt taatttttat tttcaataat
9301 tttaaaaatg ggggactttt aaattggaac aaaatgaaaa atatctgtta tacgtgcaac
9361 tgaattttac tgaccttaaa ggactatctc gaacttggtt cggaaatcct tgaaatgatt
9421 gatattttgg tggattttct ctgattttca aacaagtagt attttattta atatttatta
9481 tattttttac atttttttat atttttttat tgtttggaag gtaaagcaac aattactttc
9541 aaaatatata aatcaaactg aaatacttaa taagagacaa ataacattca agaatcaaat
9601 actgggttat taatcaaaag atctctctac atgcgcccaa attcactatt taaatttact

9661 ataccactga cagaatatat gaacccagat taagtagcca gaggctcttc cactatattg
9721 agtatatagc cttacatatt ttctgcgcat aatttactga tgtaaaataa acaaaaatag
9781 ttagtttgta gttatgaaaa aaggcttttg gaaaatgcga aatacgtgtt atttaaggtt
9841 aatcaacaaa acgcatatcc atagtggata gttggacaaa acttcaatcg ataagcttca
9901 gctggcggcc gcgttc

//

**LHZ1495 in genbank format.**

```
LOCUS       Exported               7057 bp ds-DNA     circular SYN 15-APR-2024
DEFINITION  synthetic circular DNA
ACCESSION   .
VERSION     .
KEYWORDS    LHZ1495
SOURCE      synthetic DNA construct
  ORGANISM  synthetic DNA construct
REFERENCE   1  (bases 1 to 7057)
  AUTHORS   Yao
  TITLE     Direct Submission
  JOURNAL   Exported Apr 15, 2024 from SnapGene 4.2.4
            http://www.snapgene.com
FEATURES             Location/Qualifiers
     source          1..7057
                     /organism="synthetic DNA construct"
                     /mol_type="other DNA"
     CDS             25..1270
                     /label=ARS1
     CDS             complement(401..412)
                     /label=ARS core sequence
     misc_feature    complement(883..1075)
                     /label=KmCEN5
     promoter        1277..2723
                     /label=ADH1 promoter
     terminator      2750..2939
                     /label=ADH1 Terminator
     misc_feature    2977..3355
                     /label=TEF Promoter
     CDS             3356..4384
                     /label=HphMX4
     terminator      4385..4619
                     /label=TEF terminator
     CDS             complement(5730..6590)
                     /label=Lactamase-AmpC
ORIGIN
        1 gaacgcggcc gccagctgaa gcttatcgat tgaagttttg tccaactatc cactatggat
       61 atgcgttttg ttgattaacc ttaaataaca cgtatttcgc attttccaaa agcctttttt
      121 cataactaca aactaactat ttttgtttat tttacatcag taaattatgc gcagaaaata
```

181 tgtaaggcta tatactcaat atagtggaag agcctctggc tacttaatct gggttcatat
241 attctgtcag tggtatagta aatttaaata gtgaatttgg gcgcatgtag agagatcttt
301 tgattaataa cccagtattt gattcttgaa tgttatttgt ctcttattaa gtatttcagt
361 ttgatttata tattttgaaa gtaattgttg ctttaccttc caaacaataa aaaaatataa
421 aaaaatgtaa aaaatataat aaatattaaa taaaatacta cttgtttgaa aatcagagaa
481 aatccaccaa aatatcaatc atttcaagga tttccgaacc aagttcgaga tagtccttta
541 aggtcagtaa aattcagttg cacgtataac agatattttt cattttgttc caatttaaaa
601 gtccccccatt tttaaaatta ttgaaaataa aaattaaaaa attaaaagga atctctctat
661 gtcactttaa aataaataaa ttgaaaatga tatatcgtta aaaggatccc gcaacaccac
721 ctatttctaa gaggagagtt ctaaaaaaat catagtacca cacaggtaaa ctaaaacagc
781 taaattcaac ataatacctg tggatataat tacatacaaa aatataatta aaaaaataca
841 ttaaaaataa tttattttg taaaacccat aaaatatatt ttactttcgg aacaactttt
901 taacttataa ttttgtttta aataaaaacg ttgtatttaa aaataataaa atattaagta
961 aaaatttaaa gcatatttat ttaaaatata aaatactact aaaattactc taaacttcaa
1021 aataaaaaaa taataaatta tatagtttta aaattaataa tttgtataca cgtgaccaga
1081 ccataatagt tttcttttct tgaaactgcc atgaatttaa tagatttttt ttacacatat
1141 ttacagtaag ttttgttttt acgttgaatt aatattgtat acgtacttag aaggataact
1201 tccaactatg aatatgtagg ttaaaaggta aatagagaag ccttactttt atcagaaatg
1261 aaaggagctc gtcgacgaag aaatgatggt aaatgaaata ggaaatcaag gagcatgaag
1321 gcaaaagaca aatataaggg tcgaacgaaa aataaagtga aaagtgttga tatgatgtat
1381 ttggctttgc ggcgccgaaa aaacgagttt acgcaattgc acaatcatgc tgactctgtg
1441 gcggacccgc gctcttgccg gcccggcgat aacgctgggc gtgaggctgt gcccggcgga
1501 gttttttgcg cctgcatttt ccaaggttta ccctgcgcta aggggcgaga ttggagaagc
1561 aataagaatg ccggttgggg ttgcgatgat gacgaccacg acaactggtg tcattattta
1621 agttgccgaa agaacctgag tgcatttgca acatgagtat actagaagaa tgagccaaga
1681 cttgcgagac gcgagtttgc cggtggtgcg aacaatagag cgaccatgac cttgaaggtg
1741 agacgcgcat aaccgctaga gtactttgaa gaggaaacag caatagggtt gctaccagta
1801 taaatagaca ggtacataca acactggaaa tggttgtctg tttgagtacg ctttcaattc
1861 atttgggtgt gcactttatt atgttacaat atggaaggga actttacact tctcctatgc
1921 acatatatta attaaagtcc aatgctagta gagaaggggg gtaacacccc tccgcgctct
1981 tttccgattt ttttctaaac cgtggaatat ttcggatatc cttttgttgt ttccgggtgt
2041 acaatatgga cttcctcttt tctggcaacc aaacccatac atcgggattc ctataatacc
2101 ttcgttggtc tccctaacat gtaggtggcg gaggggagat atacaataga acagatacca
2161 gacaagacat aatgggctaa acaagactac accaattaca ctgcctcatt gatggtggta
2221 cataacgaac taatactgta gccctagact tgatagccat catcatatcg aagtttcact
2281 accctttttc catttgccat ctattgaagt aataataggc gcatgcaact tcttttcttt
2341 ttttttcttt tctctctccc ccgttgttgt ctcaccatat ccgcaatgac aaaaaaaatg
2401 atggaagaca ctaaaggaaa aaattaacga caaagacagc accaacagat gtcgttgttc
2461 cagagctgat gaggggtatc tcgaagcaca cgaaactttt tccttccttc attcacgcac

2521 actactctct aatgagcaac ggtatacggc cttccttcca gttacttgaa tttgaaataa
2581 aaaaaagttt gctgtcttgc tatcaagtat aaatagacct gcaattatta atcttttgtt
2641 tcctcgtcat tgttctcgtt ccctttcttc cttgtttctt tttctgcaca atatttcaag
2701 ctataccaag catacaatca actcccgggt agcggtaccg gtactcgagg cgaatttctt
2761 atgatttatg atttttatta ttaaataagt tataaaaaaa ataagtgtat acaaatttta
2821 aagtgactct taggttttaa aacgaaaatt cttattcttg agtaactctt tcctgtaggt
2881 caggttgctt tctcaggtat agtatgaggt cgctcttatt gaccacacct ctaccggcag
2941 atccgctagg gataacaggg taatatagat ctgtttagct tgccttgtcc ccgccgggtc
3001 acccggccag cgacatggag gcccagaata ccctccttga cagtcttgac gtgcgcagct
3061 caggggcatg atgtgactgt cgcccgtaca tttagcccat acatccccat gtataatcat
3121 ttgcatccat acattttgat ggccgcacgg cgcgaagcaa aaattacggc tcctcgctgc
3181 agacctgcga gcagggaaac gctcccctca cagacgcgtt gaattgtccc cacgccgcgc
3241 ccctgtagag aaatataaaa ggttaggatt tgccactgag gttcttcttt catatacttc
3301 cttttaaaat cttgctagga tacagttctc acatcacatc cgaacataaa caaccatggg
3361 taaaaagcct gaactcaccg cgacgtctgt cgagaagttt ctgatcgaaa agttcgacag
3421 cgtctccgac ctgatgcagc tctcggaggg cgaagaatct cgtgctttca gcttcgatgt
3481 aggagggcgt ggatatgtcc tgcgggtaaa tagctgcgcc gatggtttct acaaagatcg
3541 ttatgtttat cggcactttg catcggccgc gctcccgatt ccggaagtgc ttgacattgg
3601 ggaattcagc gagagcctga cctattgcat ctcccgccgt gcacagggtg tcacgttgca
3661 agacctgcct gaaaccgaac tgcccgctgt tctgcagccg gtcgcggagg ccatggatgc
3721 gatcgctgcg gccgatctta gccagacgag cgggttcggc ccattcggac cgcaaggaat
3781 cggtcaatac actacatggc gtgatttcat atgcgcgatt gctgatcccc atgtgtatca
3841 ctggcaaact gtgatggacg acaccgtcag tgcgtccgtc gcgcaggctc tcgatgagct
3901 gatgctttgg gccgaggact gccccgaagt ccggcacctc gtgcacgcgg atttcggctc
3961 caacaatgtc ctgacggaca atggccgcat aacagcggtc attgactgga gcgaggcgat
4021 gttcggggat tcccaatacg aggtcgccaa catcttcttc tggaggccgt ggttggcttg
4081 tatggagcag cagacgcgct acttcgagcg gaggcatccg gagcttgcag gatcgccgcg
4141 gctccgggcg tatatgctcc gcattggtct tgaccaactc tatcagagct tggttgacgg
4201 caatttcgat gatgcagctt gggcgcaggg tcgatgcgac gcaatcgtcc gatccggagc
4261 cgggactgtc gggcgtacac aaatcgcccg cagaagcgcg gccgtctgga ccgatggctg
4321 tgtagaagta ctcgccgata gtggaaaccg acgccccagc actcgtccga gggcaaagga
4381 ataatcagta ctgacaataa aaagattctt gttttcaaga acttgtcatt tgtatagttt
4441 ttttatattg tagttgttct attttaatca aatgttagcg tgatttatat ttttttttcgc
4501 ctcgacatca tctgcccaga tgcgaagtta agtgcgcaga aagtaatatc atgcgtcaat
4561 cgtatgtgaa tgctggtcgc tatactgctg tcgattcgat actaacgccg ccatccagtg
4621 tcgaaaacga gctcgaattc atcgatgata tcagatccac tagtggccta tgcggccgcg
4681 gatctgccgg tctccctata gtgagtcgta ttaatttcga taagccaggt taacctgcat
4741 taatgaatcg gccaacgcgc ggggagaggc ggtttgcgta ttgggcgctc ttccgcttcc
4801 tcgctcactg actcgctgcg ctcggtcgtt cggctgcggc gagcggtatc agctcactca

4861 aaggcggtaa tacggttatc cacagaatca ggggataacg caggaaagaa catgtgagca
4921 aaaggccagc aaaaggccag gaaccgtaaa aaggccgcgt tgctggcgtt tttccatagg
4981 ctccgccccc ctgacgagca tcacaaaaat cgacgctcaa gtcagaggtg gcgaaacccg
5041 acaggactat aaagatacca ggcgtttccc cctggaagct ccctcgtgcg ctctcctgtt
5101 ccgaccctgc cgcttaccgg atacctgtcc gcctttctcc cttcgggaag cgtggcgctt
5161 tctcaatgct cacgctgtag gtatctcagt tcggtgtagg tcgttcgctc caagctgggc
5221 tgtgtgcacg aaccccccgt tcagcccgac cgctgcgcct tatccggtaa ctatcgtctt
5281 gagtccaacc cggtaagaca cgacttatcg ccactggcag cagccactgg taacaggatt
5341 agcagagcga ggtatgtagg cggtgctaca gagttcttga agtggtggcc taactacggc
5401 tacactagaa ggacagtatt tggtatctgc gctctgctga agccagttac cttcggaaaa
5461 agagttggta gctcttgatc cggcaaacaa accaccgctg gtagcggtgg tttttttgtt
5521 tgcaagcagc agattacgcg cagaaaaaaa ggatctcaag aagatccttt gatcttttct
5581 acggggtctg acgctcagtg gaacgaaaac tcacgttaag ggattttggt catgagatta
5641 tcaaaaagga tcttcaccta gatcctttta aattaaaaat gaagttttaa atcaatctaa
5701 agtatatatg agtaaacttg gtctgacagt taccaatgct taatcagtga ggcacctatc
5761 tcagcgatct gtctatttcg ttcatccata gttgcctgac tccccgtcgt gtagataact
5821 acgatacggg agggcttacc atctggcccc agtgctgcaa tgataccgcg agacccacgc
5881 tcaccggctc cagatttatc agcaataaac cagccagccg gaagggccga gcgcagaagt
5941 ggtcctgcaa ctttatccgc ctccatccag tctattaatt gttgccggga agctagagta
6001 agtagttcgc cagttaatag tttgcgcaac gttgttgcca ttgctacagg catcgtggtg
6061 tcacgctcgt cgtttggtat ggcttcattc agctccggtt cccaacgatc aaggcgagtt
6121 acatgatccc ccatgttgtg caaaaaagcg gttagctcct tcggtcctcc gatcgttgtc
6181 agaagtaagt tggccgcagt gttatcactc atggttatgg cagcactgca taattctctt
6241 actgtcatgc catccgtaag atgcttttct gtgactggtg agtactcaac caagtcattc
6301 tgagaatagt gtatgcggcg accgagttgc tcttgcccgg cgtcaatacg ggataatacc
6361 gcgccacata gcagaacttt aaaagtgctc atcattggaa aacgttcttc ggggcgaaaa
6421 ctctcaagga tcttaccgct gttgagatcc agttcgatgt aacccactcg tgcacccaac
6481 tgatcttcag catcttttac tttcaccagc gtttctgggt gagcaaaaac aggaaggcaa
6541 aatgccgcaa aaaagggaat aagggcgaca cggaaatgtt gaatactcat actcttcctt
6601 tttcaatatt attgaagcat ttatcagggt tattgtctca tgagcggata catatttgaa
6661 tgtatttaga aaaataaaca aataggggtt ccgcgcacat ttccccgaaa agtgccacct
6721 gacgtctaag aaaccattat tatcatgaca ttaacctata aaaataggcg tatcacgagg
6781 ccctttcgtc tcgcgcgttt cggtgatgac ggtgaaaacc tctgacacat gcagctcccg
6841 gagacggtca cagcttgtct gtaagcggat gccgggagca gacaagcccg tcagggcgcg
6901 tcagcgggtg ttggcgggtg tcggggctgg cttaactatg cggcatcaga gcagattgta
6961 ctgagagtgc accatatgga catattgtcg ttagaacgcg gctacaatta atacataacc
7021 ttatgtatca tacacatacg atttaggtga cactata
//

**pRS425-Cas9-2xSapI in genbank format.**

```
LOCUS       Exported              11990 bp ds-DNA     circular SYN 06-NOV-2021
DEFINITION  Complementary copy of pRS425-Cas9-2xSapI.
ACCESSION   .
VERSION     .
KEYWORDS    pRS425-Cas9-2xSapI
SOURCE      synthetic DNA construct
  ORGANISM  synthetic DNA construct
REFERENCE   1  (bases 1 to 11990)
  AUTHORS   .
  TITLE     Direct Submission
  JOURNAL   Exported Apr 15, 2024 from SnapGene 4.2.4
            http://www.snapgene.com
COMMENT     ORIGDB|GenBank
FEATURES             Location/Qualifiers
     source          1..11990
                     /organism="synthetic DNA construct"
                     /mol_type="other DNA"
     rep_origin      complement(69..1411)
                     /direction=LEFT
                     /label=2u origin
                     /note="2u ori yeast 2u plasmid origin of replication"
     protein_bind    401..448
                     /label=Protein_Bind_2
                     /bound_moiety="FLP recombinase from the Saccharomyces
                     cerevisiae 2u plasmid"
                     /note="FRT FLP-mediated recombination occurs in the 8-bp
                     core  sequence TCTAGAAA (Turan and Bode, 2011)."
     promoter        1438..1542
                     /gene="bla"
                     /label=AmpR promoter
                     /note="AmpR promoter"
     CDS             1543..2403
                     /codon_start=1
                     /gene="bla"
                     /product="beta-lactamase"
                     /label=Amp Resistance
                     /note="AmpR confers resistance to ampicillin,
                     carbenicillin, and related antibiotics"
```

/translation="MSIQHFRVALIPFFAAFCLPVFAHPETLVKVKDAEDQLGARVGYI
ELDLNSGKILESFRPEERFPMMSTFKVLLCGAVLSRIDAGQEQLGRRIHYSQNDLVEYS
PVTEKHLTDGMTVRELCSAAITMSDNTAANLLLTTIGGPKELTAFLHNMGDHVTRLDRW
EPELNEAIPNDERDTTMPVAMATTLRKLLTGELLTLASRQQLIDWMEADKVAGPLLRSA
LPAGWFIADKSGAGERGSRGIIAALGPDGKPSRIVVIYTTGSQATMDERNRQIAEIGAS
LIKHW"

primer_bind complement(1782..1800)
/label=Primer_Bind_8
/note="Amp-R"

rep_origin complement(2574..3162)
/direction=LEFT
/label=ColE1/pUC origin
/note="ori high-copy-number ColE1/pMB1/pBR322/pUC origin of
replication"

misc_feature complement(3340..3345)
/label=Misc_Feature_5
/note="Old SapI site used to be: ctcttc"

promoter 3486..3516
/label=Lac promoter
/note="lac promoter promoter for the E. coli lac operon"

protein_bind complement(3524..3540)
/label=Lac operator
/bound_moiety="lac repressor encoded by lacI"
/note="lac operator The lac repressor binds to the lac
operator to inhibit transcription in E. coli. This
inhibition can be relieved by adding lactose or
isopropyl-beta-D-thiogalactopyranoside (IPTG)."

primer_bind 3548..3564
/label=M13R primer
/note="M13 rev common sequencing primer, one of multiple
similar variants"

promoter 3585..3603
/label=T3 promoter
/note="T3 promoter promoter for bacteriophage T3 RNA
polymerase"

misc_feature 3622..4022
/label=TEF1 promoter
/note="TEF1 Promoter"

primer_bind 4023..4039

/label=SK primer
/note="SK primer common sequencing primer, one of multiple
similar  variants"
CDS  4051..8190
/codon_start=1
/label=Cas9 Streptococcus pyogenes
/note="Cas9 Streptococcus pyogenes Cas9 Gene (codon
optimized  for expression in human cells) ?"
/translation="MDKKYSIGLDIGTNSVGWAVITDEYKVPSKKFKVLGNTDRHSIKK
NLIGALLFDSGETAEATRLKRTARRRYTRRKNRICYLQEIFSNEMAKVDDSFFHRLEES
FLVEEDKKHERHPIFGNIVDEVAYHEKYPTIYHLRKKLVDSTDKADLRLIYLALAHMIK
FRGHFLIEGDLNPDNSDVDKLFIQLVQTYNQLFEENPINASGVDAKAILSARLSKSRRL
ENLIAQLPGEKKNGLFGNLIALSLGLTPNFKSNFDLAEDAKLQLSKDTYDDDLDNLLAQ
IGDQYADLFLAAKNLSDAILLSDILRVNTEITKAPLSASMIKRYDEHHQDLTLLKALVR
QQLPEKYKEIFFDQSKNGYAGYIDGGASQEEFYKFIKPILEKMDGTEELLVKLNREDLL
RKQRTFDNGSIPHQIHLGELHAILRRQEDFYPFLKDNREKIEKILTFRIPYYVGPLARG
NSRFAWMTRKSEETITPWNFEEVVDKGASAQSFIERMTNFDKNLPNEKVLPKHSLLYEY
FTVYNELTKVKYVTEGMRKPAFLSGEQKKAIVDLLFKTNRKVTVKQLKEDYFKKIECFD
SVEISGVEDRFNASLGTYHDLLKIIKDKDFLDNEENEDILEDIVLTLTLFEDREMIEER
LKTYAHLFDDKVMKQLKRRRYTGWGRLSRKLINGIRDKQSGKTILDFLKSDGFANRNFM
QLIHDDSLTFKEDIQKAQVSGQGDSLHEHIANLAGSPAIKKGILQTVKVVDELVKVMGR
HKPENIVIEMARENQTTQKGQKNSRERMKRIEEGIKELGSQILKEHPVENTQLQNEKLY
LYYLQNGRDMYVDQELDINRLSDYDVDHIVPQSFLKDDSIDNKVLTRSDKNRGKSDNVP
SEEVVKKMKNYWRQLLNAKLITQRKFDNLTKAERGGLSELDKAGFIKRQLVETRQITKH
VAQILDSRMNTKYDENDKLIREVKVITLKSKLVSDFRKDFQFYKVREINNYHHAHDAYL
NAVVGTALIKKYPKLESEFVYGDYKVYDVRKMIAKSEQEIGKATAKYFFYSNIMNFFKT
EITLANGEIRKRPLIETNGETGEIVWDKGRDFATVRKVLSMPQVNIVKKTEVQTGGFSK
ESILPKRNSDKLIARKKDWDPKKYGGFDSPTVAYSVLVVAKVEKGKSKKLKSVKELLGI
TIMERSSFEKNPIDFLEAKGYKEVKKDLIIKLPKYSLFELENGRKRMLASAGELQKGNE
LALPSKYVNFLYLASHYEKLKGSPEDNEQKQLFVEQHKHYLDEIIEQISEFSKRVILAD
ANLDKVLSAYNKHRDKPIREQAENIIHLFTLTNLGAPAAFKYFDTTIDRKRYTSTKEVL
DATLIHQSITGLYETRIDLSQLGGDSRADPKKKRKV"
CDS  6638..6649
/codon_start=1
/product="Factor Xa recognition and cleavage site"
/label=Factor Xa site (not in frame!)
/note="Factor Xa site (not in frame!)"
/translation="IEGR"
primer_bind  7465..7489

/label=YY161F
CDS 8167..8187
/codon_start=1
/product="nuclear localization signal of SV40 large T
antigen"
/label=SV40 NLS
/note="SV40 NLS"
/translation="PKKKRKV"
terminator complement(8203..8450)
/gene="S. cerevisiae CYC1"
/label=S. cerevisiae CYC1 terminator
/note="CYC1 terminator transcription terminator for CYC1"
misc_feature 8472..8740
/label=SNR52 Promoter
/note="SNR52 Promoter"
misc_feature complement(8762..8839)
/label=Structural guiding RNA
/note="Structural guiding RNA"
misc_feature complement(8840..8859)
/label=SUP4 Terminator
/note="SUP4 Terminator"
promoter complement(8882..8900)
/label=T7 promoter
/note="T7 promoter promoter for bacteriophage T7 RNA
polymerase"
primer_bind complement(8910..8926)
/label=M13F primer site
/note="M13 fwd common sequencing primer, one of multiple
similar variants"
primer_bind complement(8930..8953)
/label=M13F
misc_feature 9000..9019
/label=yy105f
rep_origin complement(9071..9526)
/direction=LEFT
/label=F1 origin
/note="f1 ori f1 bacteriophage origin of replication; arrow
indicates direction of (+) strand synthesis"
primer_bind 9527..9546

```
                /label=pRS-3B
                /note="prs-3b"
     promoter        9826..10233
                /gene="S. cerevisiae LEU2"
                /label=LEU2 promoter
                /note="LEU2 promoter"
     CDS             10234..11328
                /codon_start=1
                /gene="S. cerevisiae LEU2"
                /product="3-isopropylmalate dehydrogenase, required for
                leucine biosynthesis"
                /label=S. cerevisiae LEU2
                /note="LEU2 yeast auxotrophic marker"
                /translation="MSAPKKIVVLPGDHVGQEITAEAIKVLKAISDVRSNVKFDFENHL
                IGGAAIDATGVPLPDEALEASKKADAVLLGAVGGPKWGTGSVRPEQGLLKIRKELQLYA
                NLRPCNFASDSLLDLSPIKPQFAKGTDFVVVRELVGGIYFGKRKEDDGDGVAWDSEQYT
                VPEVQRITRMAAFMALQHEPPLPIWSLDKANLLASSRLWRKTVEETIKNEFPTLKVQHQ
                LIDSAAMILVKNPTHLNGIIITSNMFGDIISDEASVIPGSLGLLPSASLASLPDKNTAF
                GLYEPCHGSAPDLPKNKVDPIATILSAAMMLKLSLNLPEEGKAIEDAVKKVLDAGIRTG
                DLGGSNSTTEVGDAVAEEVKKILA"
     primer_bind     complement(11823..11845)
                /label=pRS-5
                /note="prs-5"
     primer_bind     11823..11845
                /label=PRS-5R
                /note="prs-5r"
     primer_bind     complement(11970..11990)
                /label=PRS-5B
                /note="prs-5b"
ORIGIN
        1 gacgaaaggg cctcgtgata cgcctatttt tataggttaa tgtcatgata ataatggttt
       61 cttagtatga tccaatatca aaggaaatga tagcattgaa ggatgagact aatccaattg
      121 aggagtggca gcatatagaa cagctaaagg gtagtgctga aggaagcata cgataccccg
      181 catggaatgg gataatatca caggaggtac tagactacct ttcatcctac ataaatagac
      241 gcatataagt acgcatttaa gcataaacac gcactatgcc gttcttctca tgtatatata
      301 tatacaggca acacgcagat ataggtgcga cgtgaacagt gagctgtatg tgcgcagctc
      361 gcgttgcatt ttcggaagcg ctcgttttcg gaaacgcttt gaagttccta ttccgaagtt
      421 cctattctct agaaagtata ggaacttcag agcgcttttg aaaaccaaaa gcgctctgaa
      481 gacgcacttt caaaaaacca aaaacgcacc ggactgtaac gagctactaa aatattgcga
```

541 ataccgcttc cacaaacatt gctcaaaagt atctctttgc tatatatctc tgtgctatat
601 ccctatataa cctacccatc cacctttcgc tccttgaact tgcatctaaa ctcgacctct
661 acatttttta tgtttatctc tagtattact ctttagacaa aaaaattgta gtaagaacta
721 ttcatagagt gaatcgaaaa caatacgaaa atgtaaacat ttcctatacg tagtatatag
781 agacaaaata gaagaaaccg ttcataattt tctgaccaat gaagaatcat caacgctatc
841 actttctgtt cacaaagtat gcgcaatcca catcggtata gaatataatc ggggatgcct
901 ttatcttgaa aaaatgcacc cgcagcttcg ctagtaatca gtaaacgcgg gaagtggagt
961 caggcttttt ttatggaaga gaaaatagac accaaagtag ccttcttcta accttaacgg
1021 acctacagtg caaaaagtta tcaagagact gcattataga gcgcacaaag gagaaaaaaa
1081 gtaatctaag atgctttgtt agaaaaatag cgctctcggg atgcattttt gtagaacaaa
1141 aaagaagtat agattctttg ttggtaaaat agcgctctcg cgttgcattt ctgttctgta
1201 aaaatgcagc tcagattctt tgtttgaaaa attagcgctc tcgcgttgca tttttgtttt
1261 acaaaaatga agcacagatt cttcgttggt aaaatagcgc tttcgcgttg catttctgtt
1321 ctgtaaaaat gcagctcaga ttctttgttt gaaaaattag cgctctcgcg ttgcattttt
1381 gttctacaaa atgaagcaca gatgcttcgt tcaggtggca cttttcgggg aaatgtgcgc
1441 ggaaccccta tttgtttatt tttctaaata cattcaaata tgtatccgct catgagacaa
1501 taaccctgat aaatgcttca ataatattga aaaaggaaga gtatgagtat tcaacatttc
1561 cgtgtcgccc ttattccctt ttttgcggca ttttgccttc ctgtttttgc tcacccagaa
1621 acgctggtga aagtaaaaga tgctgaagat cagttgggtg cacgagtggg ttacatcgaa
1681 ctggatctca acagcggtaa gatccttgag agttttcgcc ccgaagaacg ttttccaatg
1741 atgagcactt ttaaagttct gctatgtggc gcggtattat cccgtattga cgccgggcaa
1801 gagcaactcg gtcgccgcat acactattct cagaatgact tggttgagta ctcaccagtc
1861 acagaaaagc atcttacgga tggcatgaca gtaagagaat tatgcagtgc tgccataacc
1921 atgagtgata acactgcggc caacttactt ctgacaacga tcggaggacc gaaggagcta
1981 accgcttttt tgcacaacat gggggatcat gtaactcgcc ttgatcgttg ggaaccggag
2041 ctgaatgaag ccataccaaa cgacgagcgt gacaccacga tgcctgtagc aatggcaaca
2101 acgttgcgca aactattaac tggcgaacta cttactctag cttcccggca acaattaata
2161 gactggatgg aggcggataa agttgcagga ccacttctgc gctcggccct tccggctggc
2221 tggtttattg ctgataaatc tggagccggt gagcgtgggt ctcgcggtat cattgcagca
2281 ctggggccag atggtaagcc ctcccgtatc gtagttatct acacgacggg gagtcaggca
2341 actatggatg aacgaaatag acagatcgct gagataggtg cctcactgat taagcattgg
2401 taactgtcag accaagttta ctcatatata ctttagattg atttaaaact tcatttttaa
2461 tttaaaagga tctaggtgaa gatccttttt gataatctca tgaccaaaat cccttaacgt
2521 gagttttcgt tccactgagc gtcagacccc gtagaaaaga tcaaaggatc ttcttgagat
2581 cctttttttc tgcgcgtaat ctgctgcttg caaacaaaaa aaccaccgct accagcggtg
2641 gtttgtttgc cggatcaaga gctaccaact ctttttccga aggtaactgg cttcagcaga
2701 gcgcagatac caaatactgt ccttctagtg tagccgtagt taggccacca cttcaagaac
2761 tctgtagcac cgcctacata cctcgctctg ctaatcctgt taccagtggc tgctgccagt
2821 ggcgataagt cgtgtcttac cgggttggac tcaagacgat agttaccgga taaggcgcag

2881 cggtcgggct gaacgggggg ttcgtgcaca cagcccagct tggagcgaac gacctacacc
2941 gaactgagat acctacagcg tgagctatga gaaagcgcca cgcttcccga agggagaaag
3001 gcggacaggt atccggtaag cggcagggtc ggaacaggag agcgcacgag ggagcttcca
3061 gggggaaacg cctggtatct ttatagtcct gtcgggtttc gccacctctg acttgagcgt
3121 cgatttttgt gatgctcgtc aggggggcgg agcctatgga aaaacgccag caacgcggcc
3181 tttttacggt tcctggcctt ttgctggcct tttgctcaca tgttctttcc tgcgttatcc
3241 cctgattctg tggataaccg tattaccgcc tttgagtgag ctgataccgc tcgccgcagc
3301 cgaacgaccg agcgcagcga gtcagtgagc gaggaagcgg agagacgccc aatacgcaaa
3361 ccgcctctcc ccgcgcgttg gccgattcat taatgcagct ggcacgacag gtttcccgac
3421 tggaaagcgg gcagtgagcg caacgcaatt aatgtgagtt acctcactca ttaggcaccc
3481 caggctttac actttatgct tccggctcct atgttgtgtg gaattgtgag cggataacaa
3541 tttcacacag gaaacagcta tgaccatgat tacgccaagc gcgcaattaa ccctcactaa
3601 agggaacaaa agctggagct catagcttca aaatgtttct actccttttt tactcttcca
3661 gattttctcg gactccgcgc atcgccgtac cacttcaaaa cacccaagca cagcatacta
3721 aatttcccct ctttcttcct ctagggtgtc gttaattacc cgtactaaag gtttggaaaa
3781 gaaaaaagag accgcctcgt ttctttttct tcgtcgaaaa aggcaataaa aatttttatc
3841 acgtttcttt ttcttgaaaa tttttttttt gatttttttc tctttcgatg acctcccatt
3901 gatatttaag ttaataaacg gtcttcaatt tctcaagttt cagtttcatt tttcttgttc
3961 tattacaact ttttttactt cttgctcatt agaaagaaag catagcaatc taatctaagt
4021 tttctagaac tagtggatcc cccgggaaaa atggacaaga agtactccat tgggctcgat
4081 atcggcacaa acagcgtcgg ttgggccgtc attacggacg agtacaaggt gccgagcaaa
4141 aaattcaaag ttctgggcaa taccgatcgc cacagcataa agaagaacct cattggcgcc
4201 ctcctgttcg actccgggga gacggccgaa gccacgcggc tcaaaagaac agcacggcgc
4261 agatataccc gcagaaagaa tcggatctgc tacctgcagg agatctttag taatgagatg
4321 gctaaggtgg atgactcttt cttccatagg ctggaggagt cctttttggt ggaggaggat
4381 aaaaagcacg agcgccaccc aatctttggc aatatcgtgg acgaggtggc gtaccatgaa
4441 aagtacccaa ccatatatca tctgaggaag aagcttgtag acagtactga taaggctgac
4501 ttgcggttga tctatctcgc gctggcgcat atgatcaaat ttcggggaca cttcctcatc
4561 gagggggacc tgaacccaga caacagcgat gtcgacaaac tctttatcca actggttcag
4621 acttacaatc agcttttcga agagaacccg atcaacgcat ccggagttga cgccaaagca
4681 atcctgagcg ctaggctgtc caaatcccgg cggctcgaaa acctcatcgc acagctccct
4741 ggggagaaga agaacggcct gtttggtaat cttatcgccc tgtcactcgg gctgaccccc
4801 aactttaaat ctaacttcga cctggccgaa gatgccaagc ttcaactgag caaagacacc
4861 tacgatgatg atctcgacaa tctgctggcc cagatcggcg accagtacgc agaccttttt
4921 ttggcggcaa agaacctgtc agacgccatt ctgctgagtg atattctgcg agtgaacacg
4981 gagatcacca aagctccgct gagcgctagt atgatcaagc gctatgatga gcaccaccaa
5041 gacttgactt tgctgaaggc ccttgtcaga cagcaactgc ctgagaagta caaggaaatt
5101 ttcttcgatc agtctaaaaa tggctacgcc ggatacattg acggcggagc aagccaggag
5161 gaattttaca aatttattaa gcccatcttg gaaaaaatgg acggcaccga ggagctgctg

5221 gtaaagctta acagagaaga tctgttgcgc aaacagcgca ctttcgacaa tggaagcatc
5281 ccccaccaga ttcacctggg cgaactgcac gctatcctca ggcggcaaga ggatttctac
5341 ccctttttga aagataacag ggaaaagatt gagaaaatcc tcacatttcg gataccctac
5401 tatgtaggcc ccctcgcccg gggaaattcc agattcgcgt ggatgactcg caaatcagaa
5461 gagaccatca ctccctggaa cttcgaggaa gtcgtggata agggggcctc tgcccagtcc
5521 ttcatcgaaa ggatgactaa ctttgataaa aatctgccta acgaaaaggt gcttcctaaa
5581 cactctctgc tgtacgagta cttcacagtt tataacgagc tcaccaaggt caaatacgtc
5641 acagaaggga tgagaaagcc agcattcctg tctggagagc agaagaaagc tatcgtggac
5701 ctcctcttca agacgaaccg gaaagttacc gtgaaacagc tcaaagaaga ctatttcaaa
5761 aagattgaat gtttcgactc tgttgaaatc agcggagtgg aggatcgctt caacgcatcc
5821 ctgggaacgt atcacgatct cctgaaaatc attaaagaca aggacttcct ggacaatgag
5881 gagaacgagg acattcttga ggacattgtc ctcaccctta cgttgtttga agatagggag
5941 atgattgaag aacgcttgaa aacttacgct catctcttcg acgacaaagt catgaaacag
6001 ctcaagaggc gccgatatac aggatggggg cggctgtcaa gaaaactgat caatgggatc
6061 cgagacaagc agagtggaaa gacaatcctg gattttctta agtccgatgg atttgccaac
6121 cggaacttca tgcagttgat ccatgatgac tctctcacct ttaaggagga catccagaaa
6181 gcacaagttt ctggccaggg ggacagtctt cacgagcaca tcgctaatct tgcaggtagc
6241 ccagctatca aaaagggaat actgcagacc gttaaggtcg tggatgaact cgtcaaagta
6301 atgggaaggc ataagcccga gaatatcgtt atcgagatgg cccgagagaa ccaaactacc
6361 cagaagggac agaagaacag tagggaaagg atgaagagga ttgaagaggg tataaaagaa
6421 ctggggtccc aaatccttaa ggaacaccca gttgaaaaca cccagcttca gaatgagaag
6481 ctctacctgt actacctgca gaacggcagg gacatgtacg tggatcagga actggacatc
6541 aatcggctct ccgactacga cgtggatcat atcgtgcccc agtcttttct caaagatgat
6601 tctattgata ataaagtgtt gacaagatcc gataaaaata gagggaagag tgataacgtc
6661 ccctcagaag aagttgtcaa gaaaatgaaa aattattggc ggcagctgct gaacgccaaa
6721 ctgatcacac aacggaagtt cgataatctg actaaggctg aacgaggtgg cctgtctgag
6781 ttggataaag ccggcttcat caaaaggcag cttgttgaga cacgccagat caccaagcac
6841 gtggcccaaa ttctcgattc acgcatgaac accaagtacg atgaaaatga caaactgatt
6901 cgagaggtga aagttattac tctgaagtct aagctggtct cagatttcag aaaggacttt
6961 cagttttata aggtgagaga gatcaacaat taccaccatg cgcatgatgc ctacctgaat
7021 gcagtggtag gcactgcact tatcaaaaaa tatcccaagc ttgaatctga atttgtttac
7081 ggagactata aagtgtacga tgttaggaaa atgatcgcaa agtctgagca ggaaataggc
7141 aaggccaccg ctaagtactt cttttacagc aatattatga attttttcaa gaccgagatt
7201 acactggcca atggagagat tcggaagcga ccacttatcg aaacaaacgg agaaacagga
7261 gaaatcgtgt gggacaaggg tagggatttc gcgacagtcc ggaaggtcct gtccatgccg
7321 caggtgaaca tcgttaaaaa gaccgaagta cagaccggag gcttctccaa ggaaagtatc
7381 ctcccgaaaa ggaacagcga caagctgatc gcacgcaaaa aagattggga ccccaagaaa
7441 tacggcggat tcgattctcc tacagtcgct tacagtgtac tggttgtggc caaagtggag
7501 aaagggaagt ctaaaaaact caaaagcgtc aaggaactgc tgggcatcac aatcatggag

7561 cgatcaagct tcgaaaaaaa ccccatcgac tttctcgagg cgaaaggata taaagaggtc
7621 aaaaaagacc tcatcattaa gcttcccaag tactctctct ttgagcttga aaacggccgg
7681 aaacgaatgc tcgctagtgc gggcgagctg cagaaaggta acgagctggc actgccctct
7741 aaatacgtta atttcttgta tctggccagc cactatgaaa agctcaaagg gtctcccgaa
7801 gataatgagc agaagcagct gttcgtggaa caacacaaac actaccttga tgagatcatc
7861 gagcaaataa gcgaattctc caaaagagtg atcctcgccg acgctaacct cgataaggtg
7921 ctttctgctt acaataagca cagggataag cccatcaggg agcaggcaga aaacattatc
7981 cacttgttta ctctgaccaa cttgggcgcg cctgcagcct tcaagtactt cgacaccacc
8041 atagacagaa agcggtacac ctctacaaag gaggtcctgg acgccacact gattcatcag
8101 tcaattacgg ggctctatga aacaagaatc gacctctctc agctcggtgg agacagcagg
8161 gctgacccca agaagaagag gaaggtgtga tctcttctcg agtcatgtaa ttagttatgt
8221 cacgcttaca ttcacgccct ccccccacat ccgctctaac cgaaaaggaa ggagttagac
8281 aacctgaagt ctaggtccct atttattttt ttatagttat gttagtatta agaacgttat
8341 ttatatttca aatttttctt ttttttctgt acagacgcgt gtacgcatgt aacattatac
8401 tgaaaacctt gcttgagaag gttttgggac gctcgaaggc tttaatttgc ggccggtacc
8461 gcgggcccgc ttctttgaaa agataatgta tgattatgct ttcactcata tttatacaga
8521 aacttgatgt tttctttcga gtatatacaa ggtgattaca tgtacgtttg aagtacaact
8581 ctagattttg tagtgccctc ttgggctagc ggtaaaggtg cgcatttttt cacaccctac
8641 aatgttctgt tcaaaagatt ttggtcaaac gctgtagaag tgaaagttgg tgcgcatgtt
8701 tcggcgttcg aaacttctcc gcagtgaaag ataaatgatc agaagagcca tggctcttca
8761 gttttagagc tagaaatagc aagttaaaat aaggctagtc cgttatcaac ttgaaaaagt
8821 ggcaccgagt cggtggtgct ttttttgttt tttatgtctg cggccgcggt acccaattcg
8881 ccctatagtg agtcgtatta cgcgcgctca ctggccgtcg ttttacaacg tcgtgactgg
8941 gaaaaccctg gcgttaccca acttaatcgc cttgcagcac atcccccttt cgccagctgg
9001 cgtaatagcg aagaggcccg caccgatcgc ccttcccaac agttgcgcag cctgaatggc
9061 gaatggcgcg acgcgccctg tagcggcgca ttaagcgcgg cgggtgtggt ggttacgcgc
9121 agcgtgaccg ctacacttgc cagcgcccta gcgcccgctc ctttcgcttt cttcccttcc
9181 tttctcgcca cgttcgccgg ctttccccgt caagctctaa atcgggggct ccctttaggg
9241 ttccgattta gtgctttacg gcacctcgac cccaaaaaac ttgattaggg tgatggttca
9301 cgtagtgggc catcgccctg atagacggtt tttcgccctt tgacgttgga gtccacgttc
9361 tttaatagtg gactcttgtt ccaaactgga acaacactca accctatctc ggtctattct
9421 tttgatttat aagggatttt gccgatttcg gcctattggt taaaaaatga gctgatttaa
9481 caaaaattta acgcgaattt taacaaaata ttaacgttta caatttcctg atgcggtatt
9541 ttctccttac gcatctgtgc ggtatttcac accgcatatc gacggtcgag gagaacttct
9601 agtatatcca catacctaat attattgcct tattaaaaat ggaatcccaa caattacatc
9661 aaaatccaca ttctcttcaa aatcaattgt cctgtacttc cttgttcatg tgtgttcaaa
9721 aacgttatat ttataggata attatactct atttctcaac aagtaattgg ttgtttggcc
9781 gagcggtcta aggcgcctga ttcaagaaat atcttgaccg cagttaactg tgggaatact
9841 caggtatcgt aagatgcaag agttcgaatc tcttagcaac cattattttt ttcctcaaca

9901 taacgagaac acacaggggc gctatcgcac agaatcaaat tcgatgactg gaaatttttt
9961 gttaatttca gaggtcgcct gacgcatata ccttttcaa ctgaaaaatt gggagaaaaa
10021 ggaaaggtga gagcgccgga accggctttt catatagaat agagaagcgt tcatgactaa
10081 atgcttgcat cacaatactt gaagttgaca atattattta aggacctatt gtttttcca
10141 ataggtggtt agcaatcgtc ttactttcta acttttctta ccttttacat ttcagcaata
10201 tatatatata tatttcaagg atataccatt ctaatgtctg cccctaagaa gatcgtcgtt
10261 ttgccaggtg accacgttgg tcaagaaatc acagccgaag ccattaaggt tcttaaagct
10321 atttctgatg ttcgttccaa tgtcaagttc gatttcgaaa atcatttaat tggtggtgct
10381 gctatcgatg ctacaggtgt tccacttcca gatgaggcgc tggaagcctc caagaaggct
10441 gatgccgttt tgttaggtgc tgtgggtggt cctaaatggg gtaccggtag tgttagacct
10501 gaacaaggtt tactaaaaat ccgtaaagaa cttcaattgt acgccaactt aagaccatgt
10561 aactttgcat ccgactctct tttagactta tctccaatca agccacaatt tgctaaaggt
10621 actgacttcg ttgttgtcag agaattagtg ggaggtattt actttggtaa gagaaaggaa
10681 gacgatggtg atggtgtcgc ttgggatagt gaacaataca ccgttccaga agtgcaaaga
10741 atcacaagaa tggccgcttt catggcccta caacatgagc caccattgcc tatttggtcc
10801 ttggataaag ctaatctttt ggcctcttca agattatgga gaaaaactgt ggaggaaacc
10861 atcaagaacg aattccctac attgaaggtt caacatcaat tgattgattc tgccgccatg
10921 atcctagtta agaacccaac ccacctaaat ggtattataa tcaccagcaa catgtttggt
10981 gatatcatct ccgatgaagc ctccgttatc ccaggttcct tgggtttgtt gccatctgcg
11041 tccttggcct ctttgccaga caagaacacc gcatttggtt tgtacgaacc atgccacggt
11101 tctgctccag atttgccaaa gaataaggtt gaccctatcg ccactatctt gtctgctgca
11161 atgatgttga aattgtcatt gaacttgcct gaagaaggta aggccattga agatgcagtt
11221 aaaaaggttt tggatgcagg tatcagaact ggtgatttag gtggttccaa cagtaccacc
11281 gaagtcggtg atgctgtcgc cgaagaagtt aagaaaatcc ttgcttaaaa agattctctt
11341 tttttatgat atttgtacat aaactttata aatgaaattc ataatagaaa cgacacgaaa
11401 ttacaaaatg gaatatgttc atagggtaga cgaaactata tacgcaatct acatacattt
11461 atcaagaagg agaaaaagga ggatagtaaa ggaatacagg taagcaaatt gatactaatg
11521 gctcaacgtg ataaggaaaa agaattgcac tttaacatta atattgacaa ggaggagggc
11581 accacacaaa aagttaggtg taacagaaaa tcatgaaact acgattccta atttgatatt
11641 ggaggatttt ctctaaaaaa aaaaaaatac aacaaataaa aaacactcaa tgacctgacc
11701 atttgatgga gtttaagtca ataccttctt gaagcatttc ccataatggt gaaagttccc
11761 tcaagaattt tactctgtca gaaacggcct tacgacgtag tcgatatggt gcactctcag
11821 tacaatctgc tctgatgccg catagttaag ccagccccga cacccgccaa cacccgctga
11881 cgcgccctga cgggcttgtc tgctcccggc atccgcttac agacaagctg tgaccgtctc
11941 cgggagctgc atgtgtcaga ggttttcacc gtcatcaccg aaacgcgcga
//
