## Supplemental figures for "Intergeneric chromosomal transfer in yeast results in improved phenotypes and widespread transcriptional responses"

**Fig. S1**

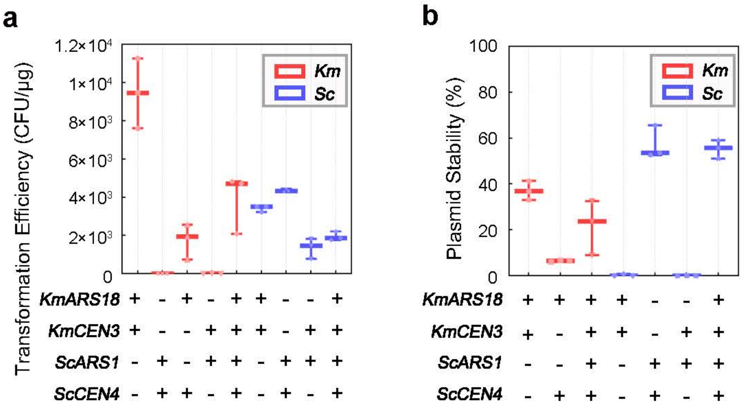


**Fig. S1. Transformation efficiency (a) and stability (b) of plasmids containing different combinations of *ARS* and *CEN* in *Km* (red) and *Sc* (blue).** Stability refers to the percentage of cells containing plasmid after being grown in the non-selective YPD medium for 24 h. Boxplots and whiskers indicate the median, maximum and minimum of three replicates.

**Fig. S2**

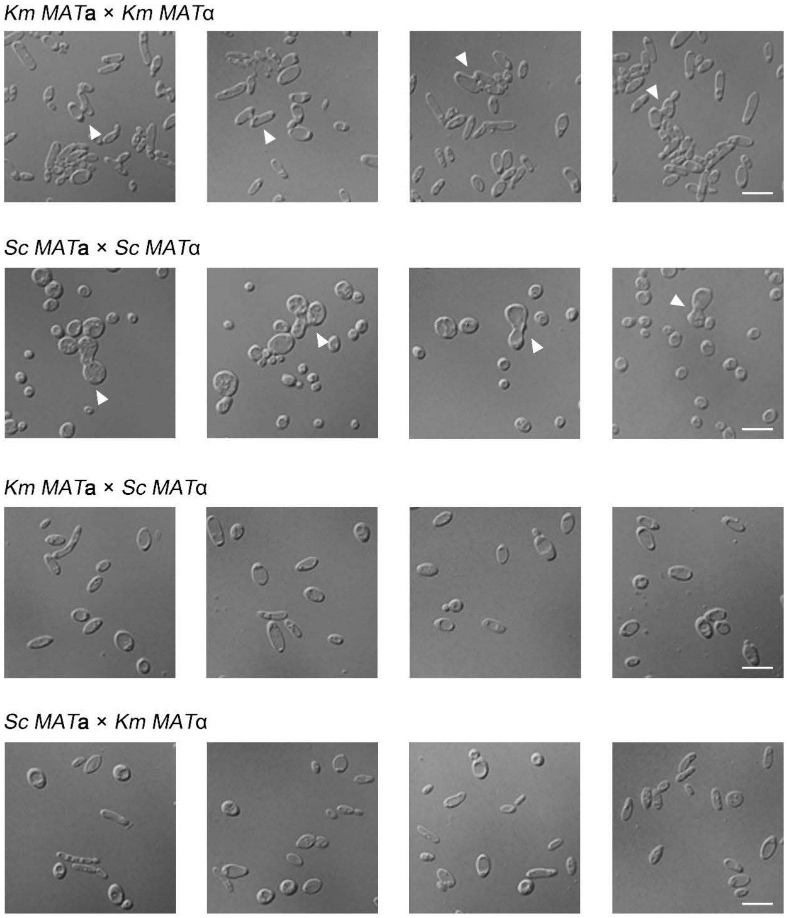


**Fig. S2. Zygotes were found in intraspecific but not interspecific crosses.** *MAT***a** and *MAT***α** cells from *Km* and *Sc* were mixed under the indicated combinations (see **Methods**). The mixture was incubated in ME medium for 24 h and subjected to microscopic observation. For each sample, more than 1500 cells were examined. Representative figures were shown. Zygotes were indicated by white arrows. Scale bar = 10 µm.

**Fig. S3**

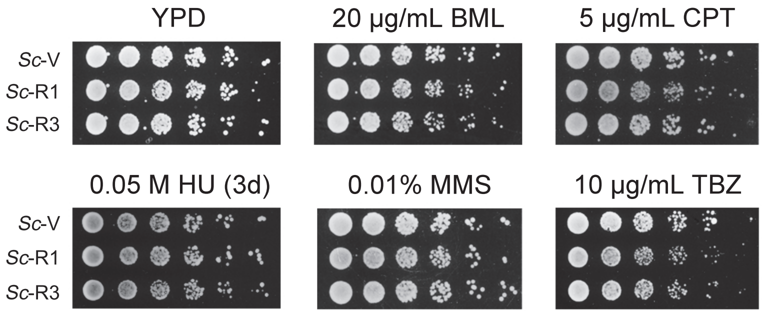


**Fig. S3. Growth of *Sc*-R1, *Sc*-R3 and *Sc*-Vector in the presence of microtubule-depolymerizing and DNA-damaging agents.** Spot assay in the presence of benomyl (BML), thiabendazole (TBZ), hydroxyurea (HU), methyl methane sulfonate (MMS) or camptothecin (CPT). The plates were incubated at 30 degrees for 1 day except for HU (3 days).

### Fig. S4

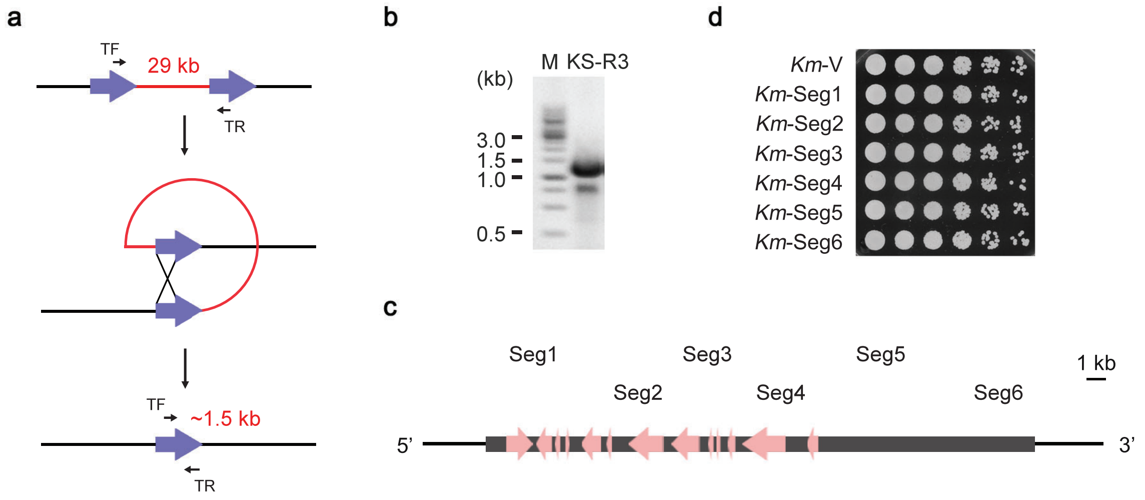


**Fig. S4. Identification of a 29 kb deletion in R3. (a)** Schematic representation of recombination between Ty elements. The positions of primers (TF & TR) used in **(b)** are indicated. **(b)** PCR-identification of a 29 kb deletion in R3, with the primer pair TF/TR. M: GeneRuler 1 kb DNA Ladder (Thermo). **(c)** The deleted region was divided into 6 segments: Seg1 (157,577-162,602 bp), Seg2 (162,580-168,195 bp), Seg3 (167,243-171,545 bp), Seg4 (170,602-175,731 bp), Seg5 (175,732-180,890 bp), and Seg6 (181,214-187,793 bp). ORFs in the segments are indicated with pink arrows. **(d)** Introducing *Sc* Seg1 to Seg6 did not affect *Km* growth. Seg1 to Seg6 were individually cloned into LHZ626 and transformed into *Km*. The transformants were subjected to spot assay on YPD alongside *Km*-Vector, at 30 degrees for 1 day.

**Fig. S5**

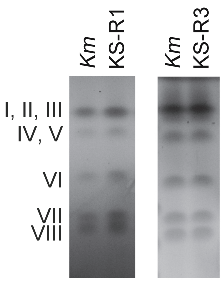


**Fig. S5. PFGE of *Km* chromosomes in KS-R1 and KS-R3.** *Km* chromosomes of Fim-1ΔU, KS-R1 and KS-R3 were separated in a 1% pulsed field certified agarose gel in 0.5×TBE at 14 °C. The running time was 24 h at 6.0 V/cm, with a 60~120 sec switch time ramp at included angle of 120 °.

**Fig. S6**

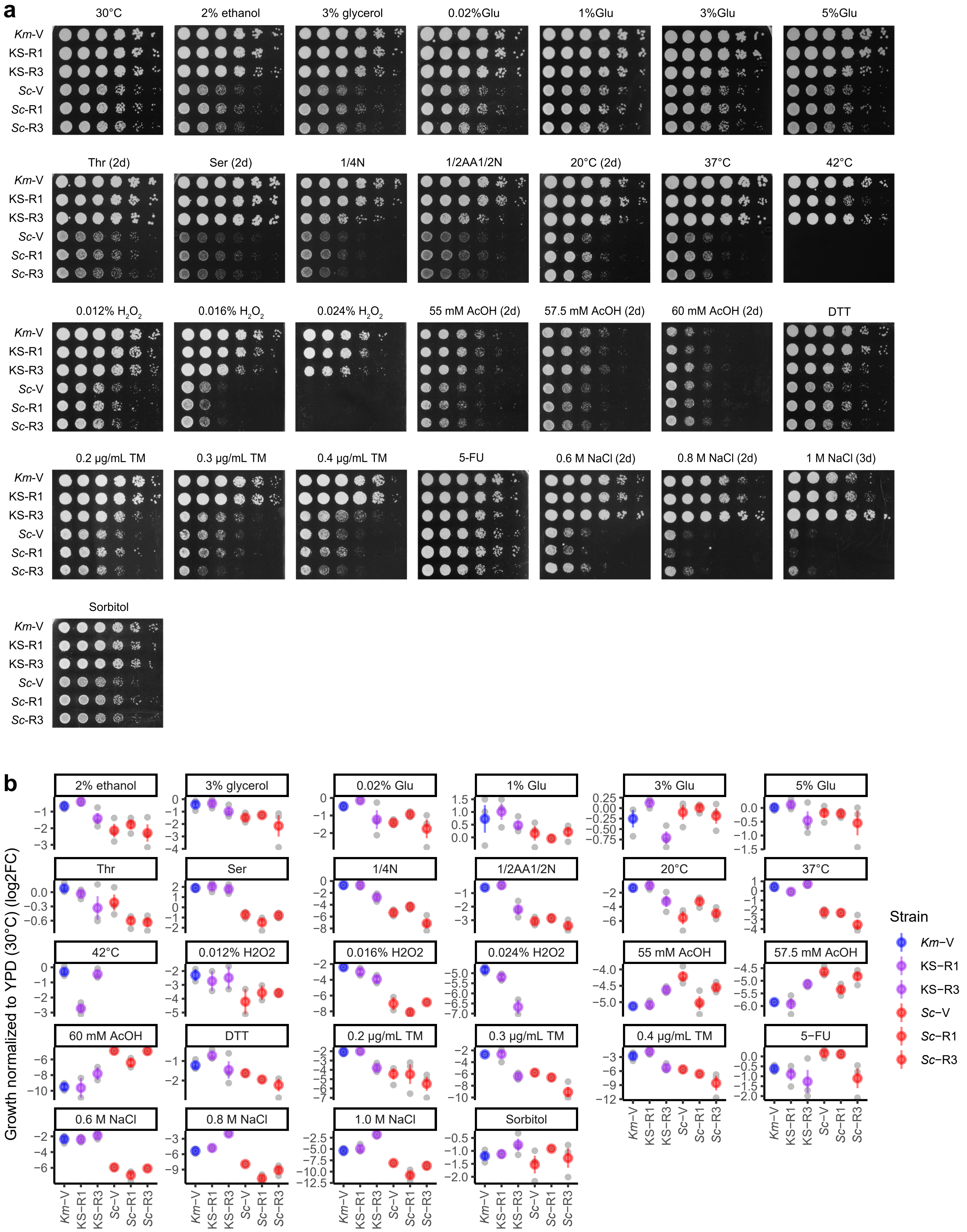


**Fig. S6. Growth of KS and their parental strains on YPD and under 28 environmental conditions.** **(a)** *Km* with an empty vector (*Km*-V), the synthetic strains with circularized *Sc* chrI (KS-R1) or chrIII (KS-R3), *Sc* with an empty vector (*Sc*-V), circularized chrI (*Sc*-R1) or chrIII (*Sc*-R3) were subjected to spot assays on YPD (30 °C) and under 28 other conditions. Unless otherwise indicated, the plates were incubated at 30°C for 1 day. Data for KS-R1 were from one representative transformant. See **Fig. S7** for all four independent transformants. **(b)** Quantification of growth. Growth under each condition was normalized to YPD growth at 30 °C for each strain, represented by log2 fold change. Grey points are data of biological replicates (n=2 or 3, see **Methods**). Colored points and error bars are mean ± standard error.

**Fig. S7**

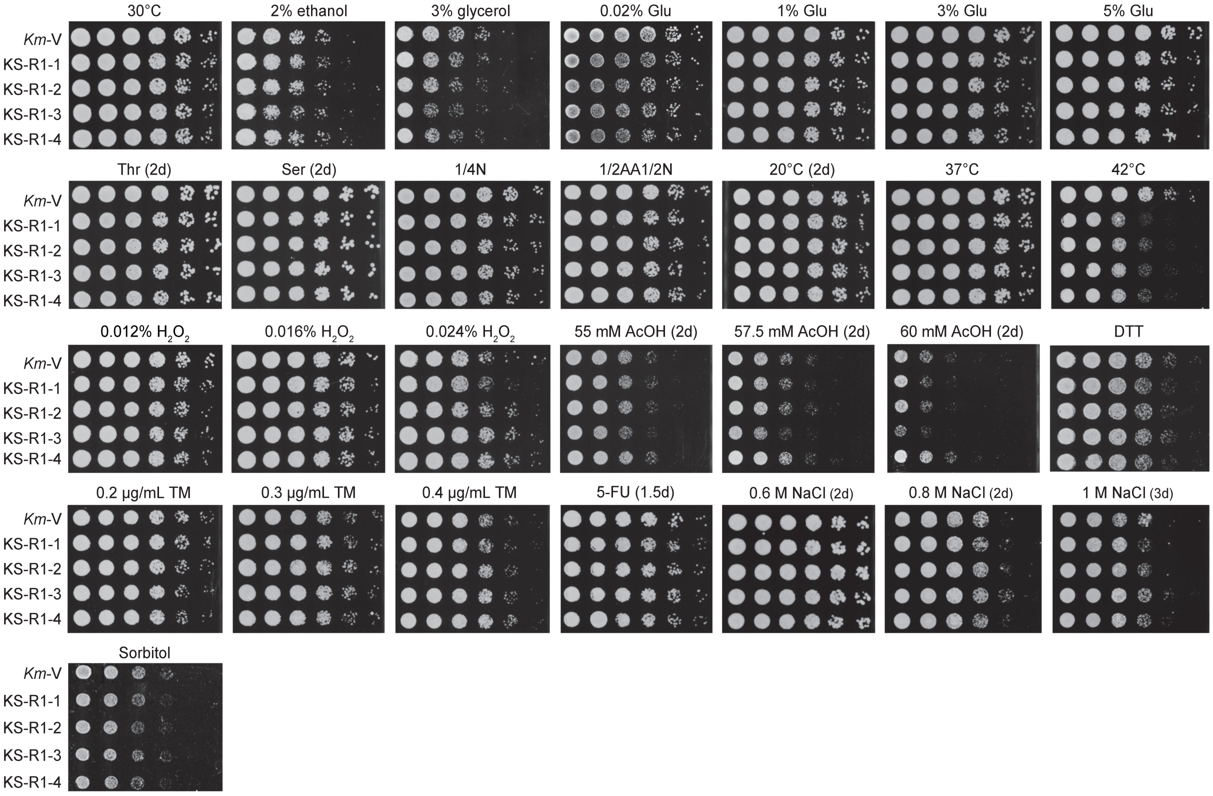


**Fig. S7. Four independent transformants of KS-R1 showed consistent phenotypes.** *Km*-V, *Km* with an empty vector. KS-R1-1 to 4, four independent transformant of circularized *Sc* chrI. Spot assay was performed as in **Fig. S6**. The four transformants showed consistent phenotypes except for KS-R1-3 under high concentration of acetic acid (AcOH), the cause of which remains to be investigated.

**Fig. S8**

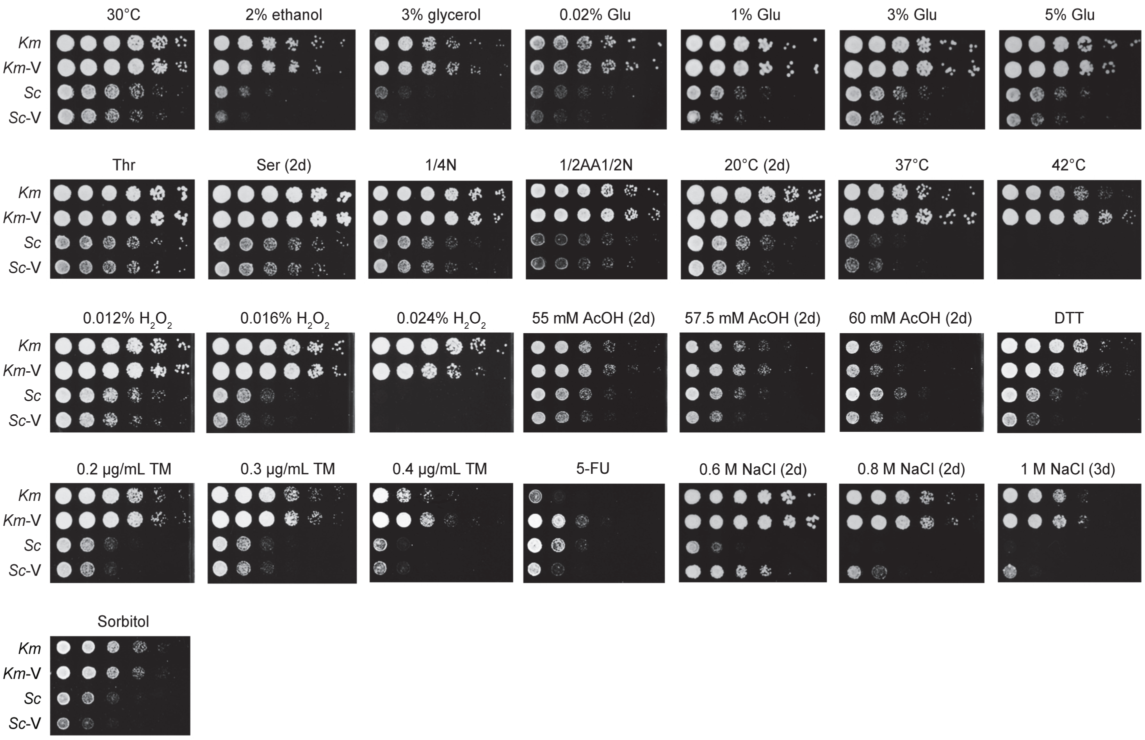


**Fig. S8. Effect of empty vector.** *Km* and *Sc* were compared to strains transformed with an empty vector, *Km*-V and *Sc*-V. The vector did not have any phenotype for most conditions, but caused differences under 42 °C, AcOH, TM (0.4 μg/mL), 5-FU, NaCl and sorbitol treatments.

**Fig. S9**

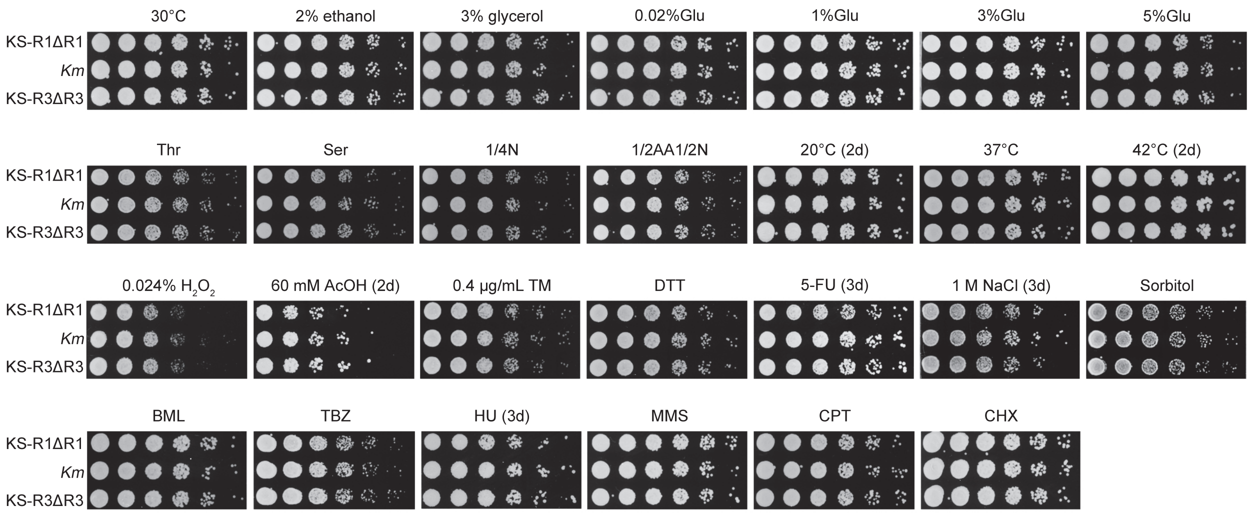


**Fig. S9. Growth of KS-R1 and KS-R3 after loss of *Sc* chromosomes.** KS-R1ΔR1 and KS-R3ΔR3 showed the same phenotype as *Km*, ruling out that the phenotypic differences in KS-R1 and KS-R3 were caused by mutations in the *Km* genome in these strains. Growth was after 1 day unless indicated otherwise.

**Fig. S10**

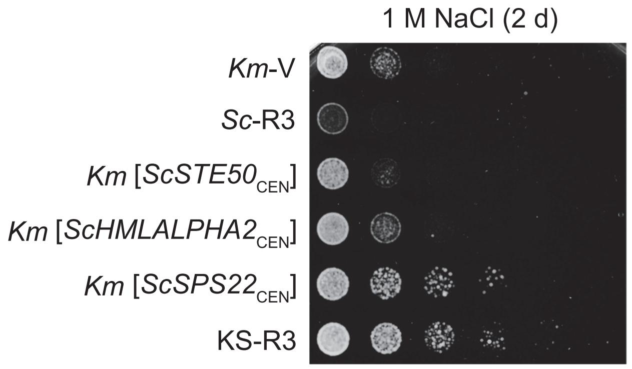


**Fig. S10. Screen three candidate genes.** *Sc STE50*, *HMLALPHA2* and *SPS22* were cloned into a centromeric vector, along with their native promoters, and transformed into *Km*. Only *ScSPS22* conferred the same level of salt resistance as R3.

**Fig. S11**

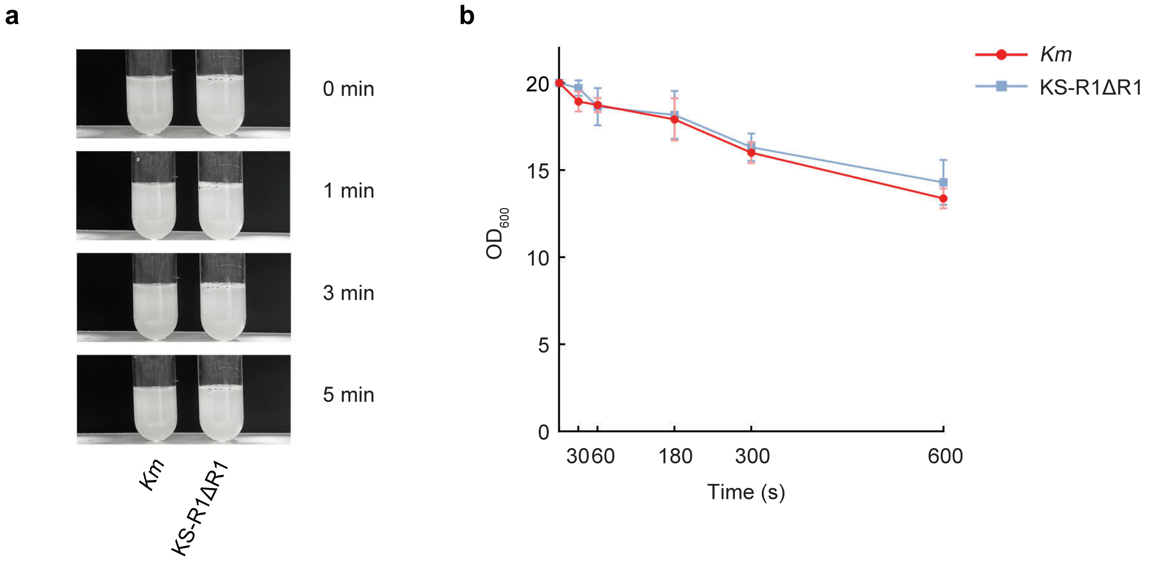


**Fig. S11. Flocculation of *Km* and KS-R1ΔR1.** A culture of 20 $OD_{600}$ of *Km* or KS-R1ΔR1was vortexed vigorously and then kept still. The cultures were imaged at designated time points after vortexing **(a)**, and the OD600 of the supernatant was monitored **(b)**. The values in **(b)** represent the mean ± SD (n=3).

### Fig. S12

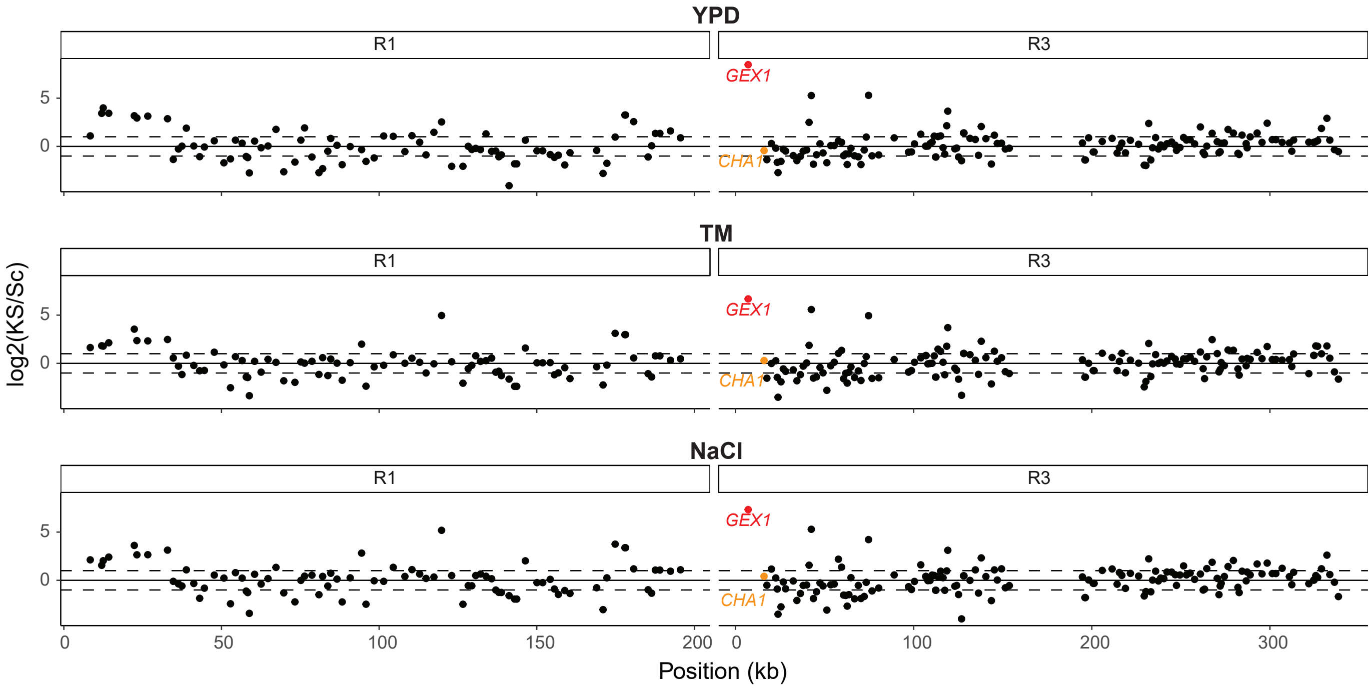


**Fig. S12. Expression differences [log2(KS/Sc)] along R1 and R3 chromosomes under YPD, TM and NaCl conditions.** *GEX1* is shown in red and *CHA1* is shown in orange. Dashed lines indicate two-fold differences.

### Fig. S13

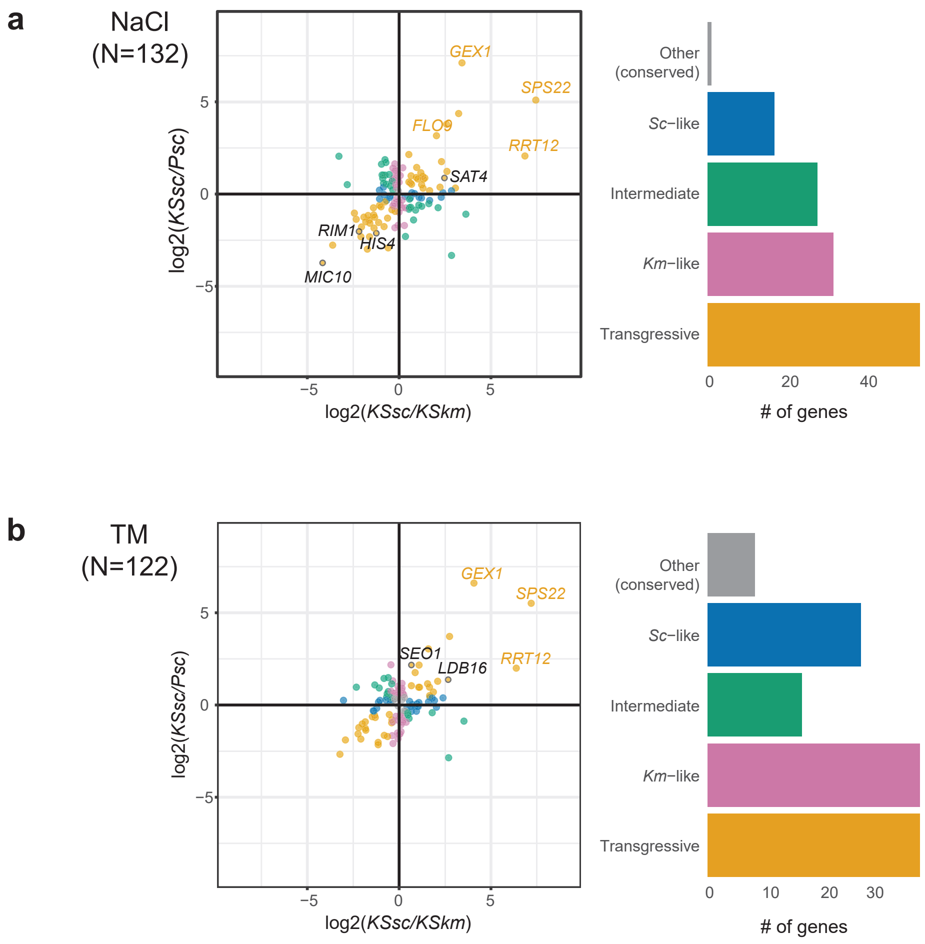


**Fig. S13. Divergence of gene expression under NaCl and TM.**
Genes showing divergent expression under NaCl (**a**, 132 genes) and TM (**b**, 122 genes) conditions were classified into five categories according to **Fig. 5c**. Log2 fold changes were extracted from DESeq models, adjusted with the ashr method. Genes showing the most prominent transgressive expression were labeled, with condition-specific transgressive expression labeled in black.

**Fig. S14**

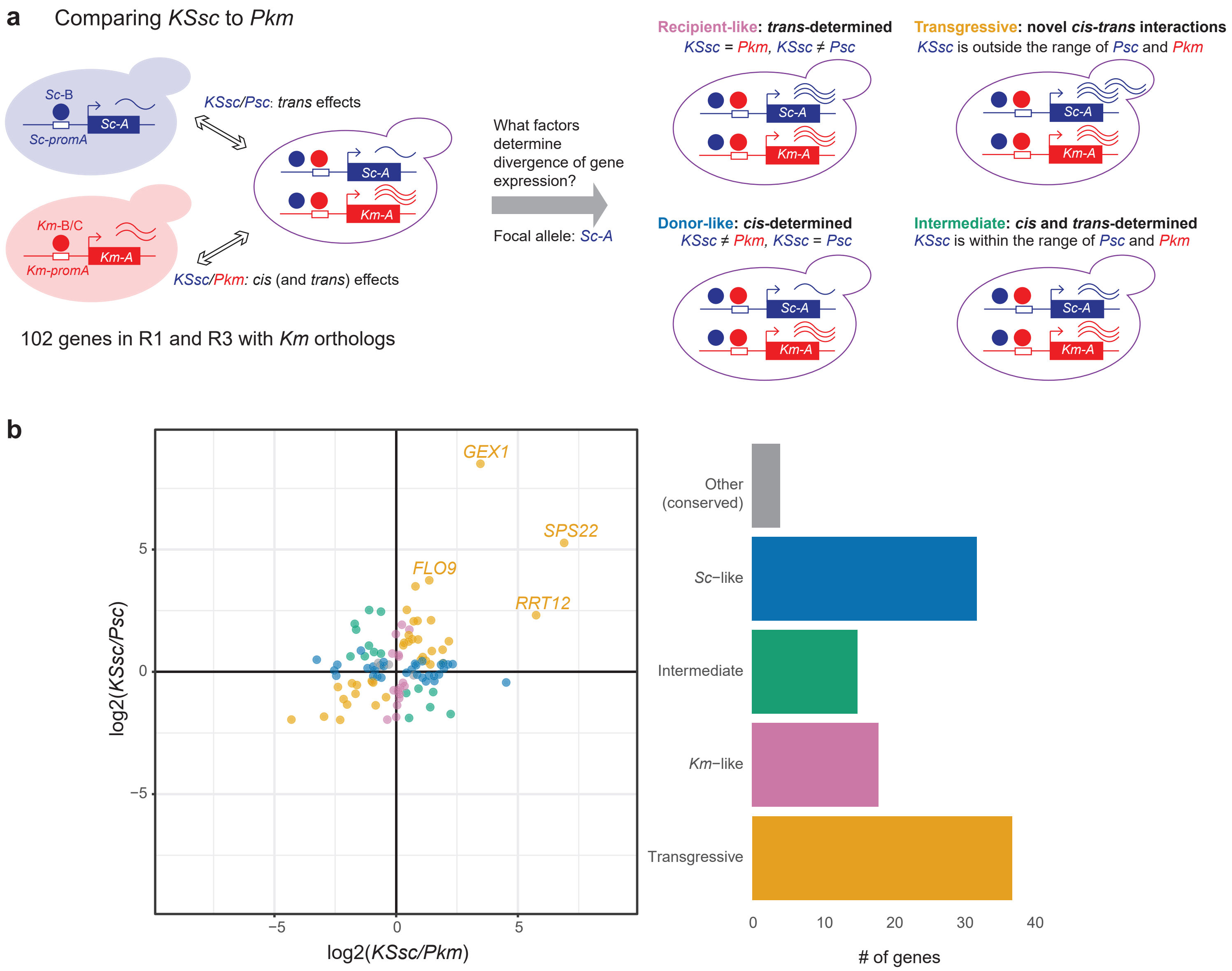


**Fig. S14. Comparing expression levels of transferred genes to donor and recipient.**
**(a)** Comparing allelic expression of the transferred genes (*KSsc*) to donor (*PSc*) and recipient (*Pkm*) to understand *cis* and *trans* evolution. Different from **Fig. 5**, here we take more into account scenarios where *Sc*-B is co-transferred into KS and may affect expression of *Km-A*. In the cases where *Km* alleles in KS show transgressive expression, transgressive expression of *Sc* is better defined by comparing *KSsc* levels to *Pkm* than to *KSkm*. **(b)** 102 genes showing divergent expression were classified into five categories according to **(a)**. Log2 fold changes were extracted from DESeq models, adjusted with the ashr method. Genes showing the most prominent transgressive expression were labeled. Data were from the YPD treatment.

Fig. S15

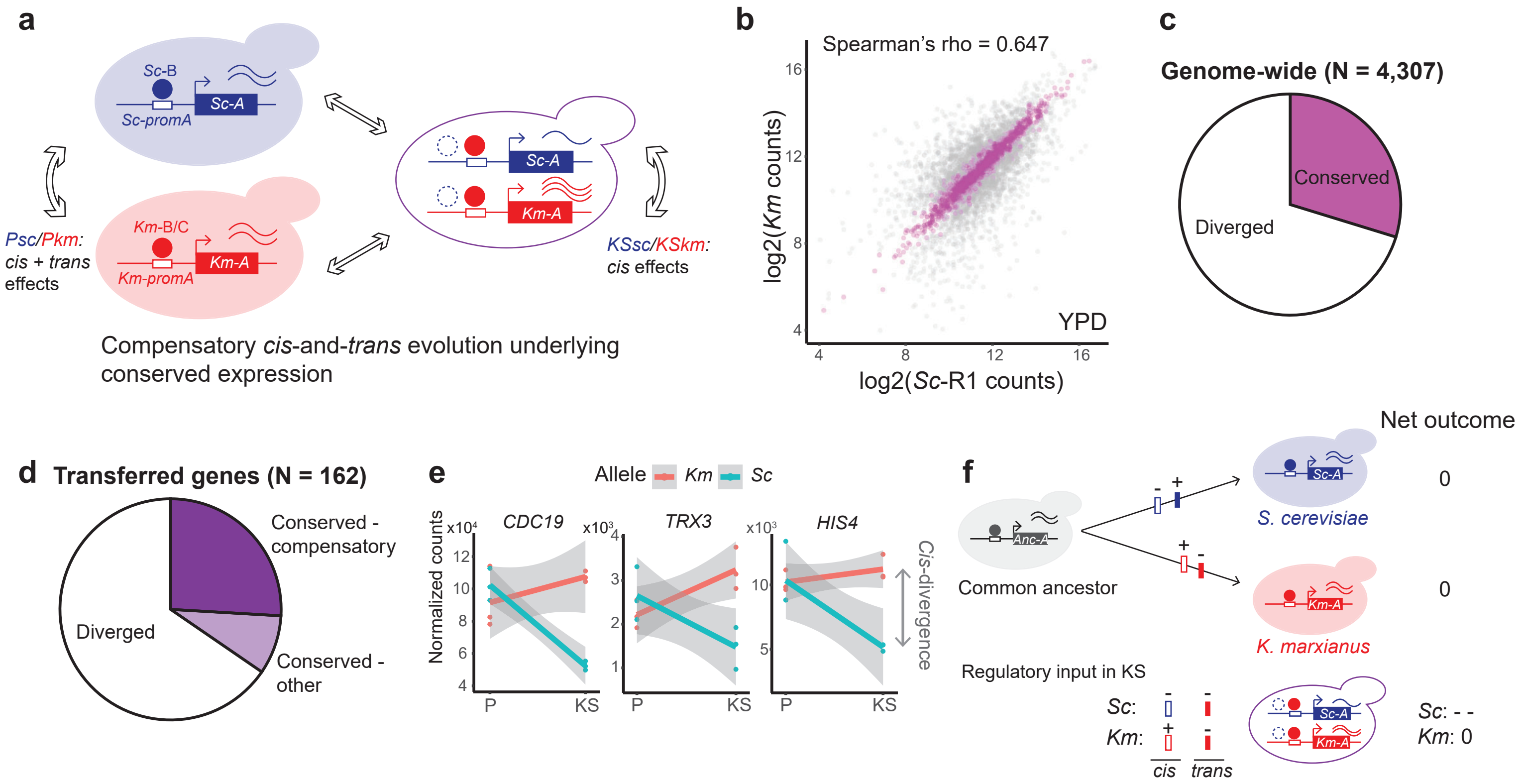


**Fig. S15. Compensatory *cis-trans* evolution underlying conserved expression levels.** **(a)** For genes showing conserved expression levels between species, a significant *cis*-effect in KS would indicate co-evolution of *cis* and *trans* changes with opposite effects (such that *cis* + *trans* effects = 0). **(b)** Correlation of gene expression, represented by log2 (normalized read counts) between *Sc*-R1 and *Km*. All data in this figure were from YPD treatment. N = 4,307 orthologs. Each point represents the average across three replicates. Pink shows genes with conserved expression levels (FDR-adjusted P > 0.05, Wald test). **(c)** Proportion of genes with conserved expression levels shown in **(b)**. **(d)** Proportion of genes with conserved expression levels among 162 transferred genes. Out of the conserved genes, 75% (dark purple) showed significant *cis*-effects in KS (FDR-adjusted P < 0.05, Wald test), consistent with compensatory evolution. **(e)** Examples of genes categorized as "conserved – compensatory". P, parents. KS, synthetic strains with *Sc* chr I or III transformed into *Km*. Differences in allelic expression in KS indicate *cis*-divergence (grey arrow). **(f)** Diagram showing that the *cis* and *trans* factors evolved in opposite directions, represented by "+" and "-", leading to conserved level of gene expression between species. The order and number of changes was arbitrary.

**Fig. S16**

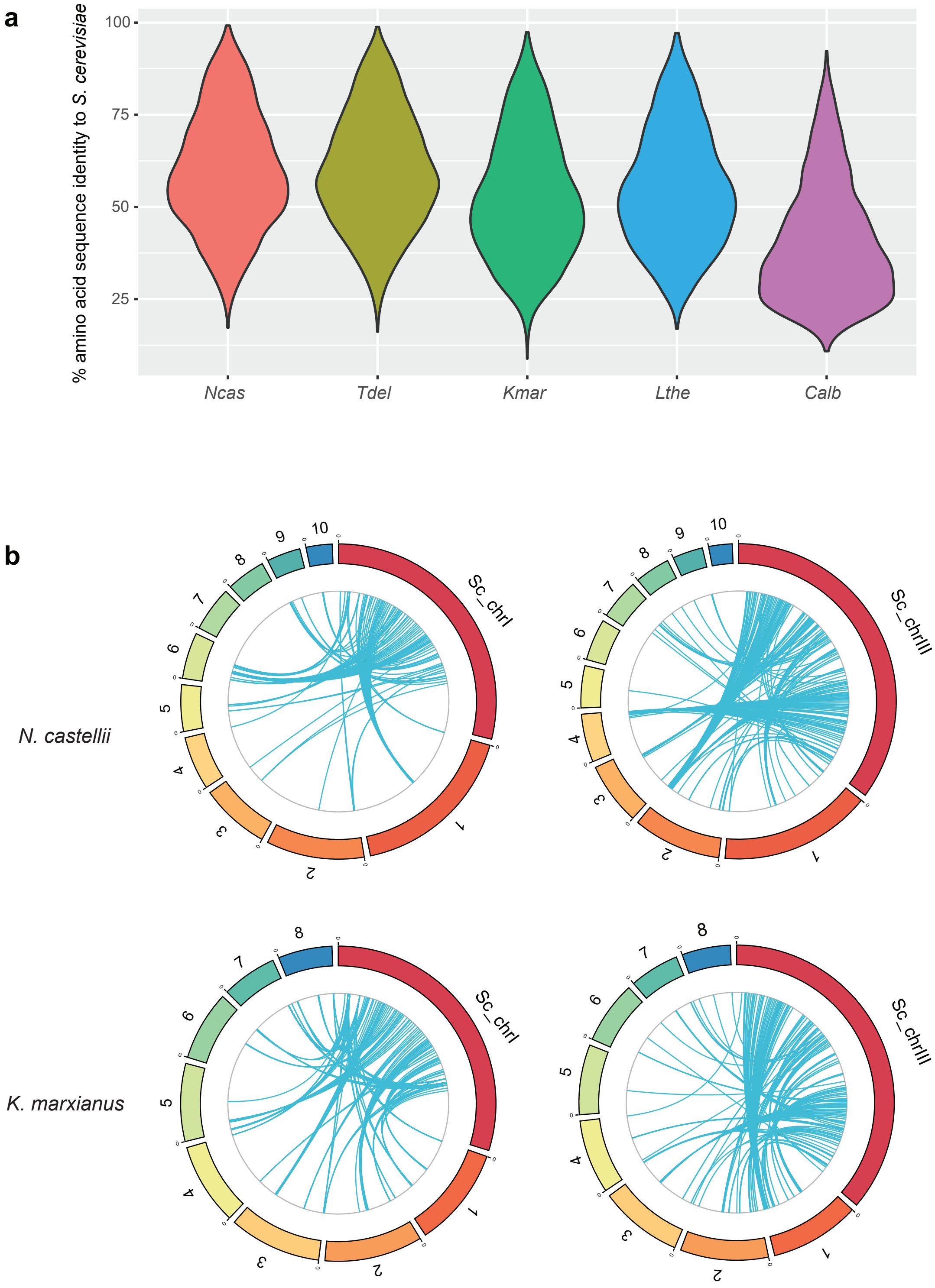


**Fig. S16. Divergence of orthologous genes.** (**a**) Amino-acid sequence similarity between *S. cerevisiae* and other species. Ncas, *N. castellii*; Tdel, *T. delbrueckii*; Kmar, *K. marxianus*; Lthe, *L. thermotolerans*; Calb, *C. albicans*. Violin plots show distribution of 2,847 single-copy orthologues. Similarity score was based on BLOSUM62 matrix. (**b**) Synteny between *Sc* chrI or chrIII and *N. castellii* and *K.marxianus* genomes. Note that the *Sc* chromosomes are not displayed to scale: they are enlarged 20x to better visualize the results. Blue lines connect orthologues. All possible gene pairs are shown in the cases of many-to-many, one-to-many or many-to-one orthologous relationships. Numbers denote chromosome IDs in *N. castellii* or *K. marxianus* genomes.
