## Supplemental data file1 for "Intergeneric chromosomal transfer in yeast results in improved phenotypes and widespread transcriptional responses"

**Supplementary Data File 1. Donor DNA sequences for engineering *Sc* chr1 and chr3.**

The promoter(s), and terminator(s) are labelled in pink and blue, respectively. Restriction sites were labelled in yellow. Donor was assembly from fragments by seamless ligation and sequence used for seamless ligation are underlined. Assembled donor was amplified by PCR and PCR product was transformed with CRISPR vector into Sc cells for chromosome engineering.

1. chr1- cir: donor used to cyclize Sc chr1

(1) Composition of the donor:

Left homologous arm (HA)-$T_{ADH1}$-*KmURA3*-$P_{KmUAR3}$-CRISPR site-directed repeat (DR)-AscI-right HA

DR was used to pop out *KmURA3* by recombination with another DR upstream of left HA. The design of DR followed a previous report: Shao, Y., Lu, N., Qin, Z., and Xue, X. (2018). CRISPR-Cas9 Facilitated Multiple-Chromosome Fusion in *Saccharomyces cerevisiae*. *ACS Synth Biol* 7, 2706–2708. https://doi.org/10.1021/acssynbio.8b00397.

(2) Sequence (5’-3’)：

CAGAGGTATTGTGGATCCTTCTACAGTACTTCTGAATACACCTAAAAGGTTGTTGGAT
GCTAAATTTAGCAAAAGTCTTTTTTAGCTCACTATTAGGCTTGTTAAAGTCTGAAATTG
TTGAAAGGCACTCAAAAGATAAATTGACAATTAGCATTAACGGCACAGTTGAAAGAG
TCACCCACTTGAAATTAGCTCGGTTATCAAATATAATTATCTCCGGTAAAGAGCTCTGC
AGCAGGGTTAATCTATTCCTATACTTACGCTGTAGGAACATTTTATTATTAGGATCCGAC
TACTGCCTACATATTTATTCGGAAGGCTTGATGTCGAAAATTTTTGAGCTTATAAAAAG
AACATATTTCACTCTTGCTCGTGCCGGTAGAGGTGTGGTCAATAAGAGCGACCTCATA
CTATACCTGAGAAAGCAACCTGACCTACAGGAAAGAGTTACTCAAGAATAAGAATTT
TCGTTTTAAAACCTAAGAGTCACTTTAAAATTTGTATACACTTATTTTTTTTATAACTTA
TTTAATAATAAAAATCATAAATCATAAGAAATTCGCTTAAGCGGATCTGCCTACTCTCT
TCAAGTAAGCGTCCCATCCCGCCTTTCTGTATCTCTCACCTTCCACTACAGGATCTCTT
CCCTTTGCGAAAAGACCTCTACCAACAATAATGATGTCTGATCCACCGGCAACAACTT
CATCCACAGTTCTGTATTGTTGTCCCAAAGCATCACCTTTGTCATCAAGACCAACACC
TGGCGTCATGATCAACCAATCGTAGCCCTCTTCTCTTCCACCCATATCGTTTTGAGCAA
TAAATCCAATAACAAAGTCCTTATCACTCTTGGCAATTTCCACGGTCCCACGAGTGTAT
TCACCGTGCGCTAGAGACCCCTTGGACGATAACTCGGCAAGCATTAACAACCCTCTA
GGTTCTTTTGTAACTTCCTCGGCACCTTGCTTCAAACCAGCAACAATGCCCGCACCAG
TCACACCGTGTGCATTGGTGATATCAGACCATTCGGCGATACGGTATACACCAGACGT
GTATTGTAATTTAACAGTGTTCCCAATGTCGGCAAACTTCCTGTCTTCAAATATCAAAA
ACTTGTGTTTCTCTGCTAATTGCTTCAACGGCACAATGGTATTCTCAAAGCTGAAATC
CTCCAAGATATCTACATGTGTCTTCAATAGACAGATATATGGACCCAAAACCTCAACTA
ATCTTAACAACTCTGCTGTTTTACGAACATCAAGAGAAGCACATAAGTTTGACTTCTT
CTCTTCCATCAAGTTTAAAAGCTTGGCAGCAACTGGACTTCTATGAGCAGCTGCTCTT
TCCGAGTAACTCTTAGTCGACATCCTCCTTTGATTAGTTTATGGGGCTAGCTGATGATA
CTTCCTCTTGCTGTTCCAGTTTACAAATGATTTAAACGAACTTTTACACTTTTACGGTA
GAATTTCTTGTTGGTGATGAGCTCAGATTCAATATGACTGCTTAATCAGTGAAAATTAA
TGGAAACAGCCAGACACACAGCCTTTGAAAAATGTTAAAAACGATCACATATATGTA
GTCACGTGGTTAGTGCATAGAGTCAGTCGATGGTGGTTAGTTCCCTAGCCAGAGCCAT
TCCCCTCTTGTTTCGAAATGCGATGACTGAATGGGGAAGAAGCTCTTAGTATTTCTGT
AGTTTTTTTTGCCGATTTCTATTGTTGCCTGCCCCGGACTTTCTTTTACCTTTTGTAGCT
ACAACTATTAATTATAATGCATAATAGATCTAGAAGCGTATTGTGATGAGCATCCTTGG 11
AAATAAGAAAGGAAGTCAAAGGTACAGTAAAAAAAGAGCCGTGACAGGCACCTAAT

TTCAGTTTTGATTGAGGTATAATGAGCTCAGAAGACCAAGATGCTCTAAAAGTAAAGC
AGAATGGTCTTTCCAATCAGAATTCGCTGATTACTAGCGAAGCTGCGGGAATATGGTG
GGGCCTTTGACTTCCAGAATGCTTTCACGGAATTATTTAGATGTACATTTAGCTCCAT
TTCCTGTGCCTCCGATAGGAAGGCATCATGGTACTACCGTGACGGAGAATACGTAGGC
TGACTTTTTCGTCAGTTTGTTGTCCGTTTACAAAATTGGTGAATGAATTCTAGCCTTTG
CTCATTAATTGCCCTCACAAGAATTTGGAAGTGCGTAGAACAAGTAAAAGGTTGCACT
AGGCGCGCCACGGATGCTATTTCAGAATATTTCGTACTTACACAGGCCATACATTAGAA
TAATATGTCACATCACTGTCGTAACACTCTTTATTCACCGAGCAATAATACGGTAGTGG
CTCAAACTCATGCGGGTGCTATGATACAATTATATCTTATTTCCATTCCCATATGCTAAC
CGCAATATCCTAAAAGCATAACTGATGCATCTTTAATCTTGTATGTGACACTACTCATAC
GAAGGGACTATATCTAGTCAAGACGATACTGTGATAGGTACGTTATTTAATAGGATCTA
TAACGAAATGTCAAATAATTTTACGGTAATATAACTTATCAGCGGCGTATACTAAAACG
GACGTTACGATATTGTCTCACTT

2. chr1-ARS1：donor used to insert *KmARS1* and *hyg*$^R$ into Sc chr1

(1) Composition of the donor:

Left HA-*KmARS1*-P$_{TEF}$- *hyg*$^R$ -T$_{TEF}$-Right HA

(2) Sequence (5'-3'):

TCCATACTCTTCAGGATATCATATAATAACAACTCTAATAATACCTCTAGTAGCGAGATC
TTCACACTTTTGGTAGAAAAAGTTTGGAATTTTGACGACTTGATAATGGCGATCAATT
CTAAAATTTCGAATACACATAATAACAACATTTCACCAATCACCAAGATCAAATATCAG
GACGAAGATGGGGATTTTGTTGTGTTAGGTAGCGATGAAGATTGGAATGTTGCTAAAG
AAATGTTGGCGGAAAACAATGAGAAATTCTTGAACATTCGTCTGTATTGATAAATAAA
ACTAGTATACAGCAAATACTAAATAATTCAAGAAAAAAACATTATCGATTGAAGTTTT
GTCCAACTATCCACTATGGATATGCGTTTTGTTGATTAACCTTAAATAACACGTATTTCG
CATTTTCCAAAAGCCTTTTTTCATAACTACAAACTAACTATTTTTGTTTATTTTACATCA
GTAAATTATGCGCAGAAAATATGTAAGGCTATATACTCAATATAGTGGAAGAGCCTCTG
GCTACTTAATCTGGGTTCATATATTCTGTCAGTGGTATAGTAAATTTAAATAGTGAATTT
GGGCGCATGTAGAGAGATCTTTTGATTAATAACCCAGTATTTGATTCTTGAATGTTATTT
GTCTCTTATTAAGTATTTCAGTTTGATTTATATATTTTGAAAGTAATTGTTGCTTTACCTT
CCAAACAATAAAAAAATATAAAAAAATGTAAAAAATATAATAAATATTAAATAAAATAC
TACTTGTTTGAAAATCAGAGAAAATCCACCAAAATATCAATCATTTCAAGGATTTCCG
AACCAAGTTCGAGATAGTCCTTTAAGGTCAGTAAAATTCAGTTGCACGTATAACAGAT
ATTTTTCATTTTGTTCCAATTTAAAAGTCCCCCATTTTTAAAATTATTGAAAATAAAAAT
TAAAAAATTAAAAGGAATCTCTCTATGTCACTTTAAAATAAATAAATTGAAAATGATAT
ATCGTTAAAAGAGCTTGCCTTGTCCCCGCCGGGTCACCCGGCCAGCGACATGGAGGC
CCAGAATACCCTCCTTGACAGTCTTGACGTGCGCAGCTCAGGGGCATGATGTGACTG
TCGCCCGTACATTTAGCCCATACATCCCCATGTATAATCATTTGCATCCATACATTTTGA
TGGCCGCACGGCGCGAAGCAAAAATTACGGCTCCTCGCTGCAGACCTGCGAGCAGG
GAAACGCTCCCCTCACAGACGCGTTGAATTGTCCCCACGCCGCGCCCCTGTAGAGAA
ATATAAAAGGTTAGGATTTGCCACTGAGGTTCTTCTTTCATATACTTCCTTTTAAAATCT
TGCTAGGATACAGTTCTCACATCACATCCGAACATAAACAACCATGGGTAAAAAGCCT
GAACTCACCGCGACGTCTGTCGAGAAGTTTCTGATCGAAAAGTTCGACAGCGTCTCC
GACCTGATGCAGCTCTCGGAGGGCGAAGAATCTCGTGCTTTCAGCTTCGATGTAGGA
GGGCGTGGATATGTCCTGCGGGTAAATAGCTGCGCCGATGGTTTCTACAAAGATCGTT
ATGTTTATCGGCACTTTGCATCGGCCGCGCTCCCGATTCCGGAAGTGCTTGACATTGG

GGAATTCAGCGAGAGCCTGACCTATTGCATCTCCCGCCGTGCACAGGGTGTCACGTT
GCAAGACCTGCCTGAAACCGAACTGCCCGCTGTTCTGCAGCCGGTCGCGGAGGCCAT
GGATGCGATCGCTGCGGCCGATCTTAGCCAGACGAGCGGGTTCGGCCCATTCGGACC
GCAAGGAATCGGTCAATACACTACATGGCGTGATTTCATATGCGCGATTGCTGATCCCC
ATGTGTATCACTGGCAAACTGTGATGGACGACACCGTCAGTGCGTCCGTCGCGCAGG
CTCTCGATGAGCTGATGCTTTGGGCCGAGGACTGCCCCGAAGTCCGGCACCTCGTGC
ACGCGGATTTCGGCTCCAACAATGTCCTGACGGACAATGGCCGCATAACAGCGGTCA
TTGACTGGAGCGAGGCGATGTTCGGGGATTCCCAATACGAGGTCGCCAACATCTTCTT
CTGGAGGCCGTGGTTGGCTTGTATGGAGCAGCAGACGCGCTACTTCGAGCGGAGGCA
TCCGGAGCTTGCAGGATCGCCGCGGCTCCGGGCGTATATGCTCCGCATTGGTCTTGAC
CAACTCTATCAGAGCTTGGTTGACGGCAATTTCGATGATGCAGCTTGGGCGCAGGGTC
GATGCGACGCAATCGTCCGATCCGGAGCCGGGACTGTCGGGCGTACACAAATCGCCC
GCAGAAGCGCGGCCGTCTGGACCGATGGCTGTGTAGAAGTACTCGCCGATAGTGGAA
ACCGACGCCCCAGCACTCGTCCGAGGGCAAAGGAATAATCAGTACTGACAATAAAAA
GATTCTTGTTTTCAAGAACTTGTCATTTGTATAGTTTTTTTATATTGTAGTTGTTCTATTT
TAATCAAATGTTAGCGTGATTTATATTTTTTTTCGCCTCGACATCATCTGCCCAGATGCG
AAGTTAAGTGCGCAGAAAGTAATATCATGCGTCAATCGTATGTGAATGCTGGTCGCTAT
ACTGCTGTCGATTCGATACTAACGCCGCCATCCAGTAGATAGAGAGGGGCAGATGTTC
AAGCTATACCCATTATATTGATCCACACTTAGTATTAAGATACGTCTGTGAAGGATGAA
AAAAAATGTATAATGTGACTAGAGGAAGTAAGGAGAAAAACGATAGTAATCGTATTT
TAGGTTGTGCGTTTTTATAATTTTTTTTTTTTTGTAATTCTATGCAAATGTAATATAAGTA
TATTTAAAGAAATAATGAGTCCTGTGAAAACAAAAAGAAAAAAAGATCATTAATGTAT
GTTAACGTATTTGCTTTGCAAATTTTAATTTATTTGTTGTTAAATGCATTTTTTTTTTGTC
GTTTCAGCGAGTTTTCTTGAGGTTGCTACTATCATTAAAATCACAATCCACAGAGGAA
GTTGATC

3. chr1-ARS1/CEN5: donor used to insert *KmARS1*/*KmCEN5* and $kan^R$ into Sc chr1

(1) Composition of the donor:

Left HA-*KmARS1*-*KmCEN5*-$P_{TEF}$-$kan^R$-$T_{TEF}$-Right HA

(2) Sequence (5'-3'):

GTTGATAGATATAAATTGGGACTTCTATAGTATGAGTATAGTGAGTAATGGTTTTGAATC
TGGGAACGTGTATGTGTGGTCTGTTGTTATTCCGCCAAAGTGGAGTGCTTTGGCGCCA
GATTTTGAAGAAGTAGAAGAGAATGTCGACTATTTGGAGAAGGAAGATGAATTTGAT
GAGGTCGATGAGGCAGAACAGCAGCAAGGACTAGAACAAGAGGAAGAAATAGCTAT
CGATCTTCGGACGAGAGAGCAATATGATGTTAGAGGTAATAACTTGCTTGTAGAACGG
TTCACAATCCCTACAGATTATACGAGGATAATTAAGATGCAGTCATCATAGGTTTCTCT
TCAAAAGGAGAAAGTTTAAATAGGCAACTATCGATTGAAGTTTTGTCCAACTATCCAC
TATGGATATGCGTTTTGTTGATTAACCTTAAATAACACGTATTTCGCATTTTCCAAAAGC
CTTTTTTCATAACTACAAACTAACTATTTTTGTTTATTTTACATCAGTAAATTATGCGCA
GAAAATATGTAAGGCTATATACTCAATATAGTGGAAGAGCCTCTGGCTACTTAATCTGG
GTTCATATATTCTGTCAGTGGTATAGTAAATTTAAATAGTGAATTTGGGCGCATGTAGA
GAGATCTTTTGATTAATAACCCAGTATTTGATTCTTGAATGTTATTTGTCTCTTATTAAGT
ATTTCAGTTTGATTTATATATTTTGAAAGTAATTGTTGCTTTACCTTCCAAACAATAAAA
AAATATAAAAAAATGTAAAAAATATAATAAATATTAAATAAAATACTACTTGTTTGAAA
ATCAGAGAAAATCCACCAAAATATCAATCATTTCAAGGATTTCCGAACCAAGTTCGAG
ATAGTCCTTTAAGGTCAGTAAAATTCAGTTGCACGTATAACAGATATTTTTCATTTTGTT

CCAATTTAAAAGTCCCCCATTTTTAAAATTATTGAAAATAAAAATTAAAAAATTAAAAG
GAATCTCTCTATGTCACTTTAAAATAAATAAATTGAAAATGATATATCGTTAAAAGGAT
CCCGCAACACCACCTATTTCTAAGAGGAGAGTTCTAAAAAAATCATAGTACCACACA
GGTAAACTAAAACAGCTAAATTCAACATAATACCTGTGGATATAATTACATACAAAAAT
ATAATTAAAAAAATACATTAAAAATAATTTATTTTTGTAAAACCCATAAAATATATTTTA
CTTTCGGAACAACTTTTTAACTTATAATTTTGTTTTAAATAAAAACGTTGTATTTAAAAA
TAATAAAATATTAAGTAAAAATTTAAAGCATATTTATTTAAAATATAAAATACTACTAAA
ATTACTCTAAACTTCAAAATAAAAAAATAATAAATTATATAGTTTTAAAATTAATAATTT
GTATACACGTGACCAGACCATAATAGTTTTCTTTTCTTGAAACTGCCATGAATTTAATA
GATTTTTTTTACACATATTTACAGTAAGTTTTGTTTTTACGTTGAATTAATATTGTATACG
TACTTAGAAGGATAACTTCCAACTATGAATATGTAGGTTAAAAGGTAAATAGAGAAGC
CTTACTTTTATCAGAAATGAAAGGAGCTCAGCTTGCCTCGTCCCCGCCGGGTCACCCG
GCCAGCGACATGGAGGCCCAGAATACCCTCCTTGACAGTCTTGACGTGCGCAGCTCA
GGGGCATGATGTGACTGTCGCCCGTACATTTAGCCCATACATCCCCATGTATAATCATT
TGCATCCATACATTTTGATGGCCGCACGGCGCGAAGCAAAAATTACGGCTCCTCGCTG
CAGACCTGCGAGCAGGGAAACGCTCCCCTCACAGACGCGTTGAATTGTCCCCACGCC
GCGCCCCTGTAGAGAAATATAAAAGGTTAGGATTTGCCACTGAGGTTCTTCTTTCATAT
ACTTCCTTTTAAAATCTTGCTAGGATACAGTTCTCACATCACATCCGAACATAAACAAC
CATGGGTAAGGAAAAGACTCACGTTTCGAGGCCGCGATTAAATTCCAACATGGATGC
TGATTTATATGGGTATAAATGGGCTCGCGATAATGTCGGGCAATCAGGTGCGACAATCT
ATCGATTGTATGGGAAGCCCGATGCGCCAGAGTTGTTTCTGAAACATGGCAAAGGTA
GCGTTGCCAATGATGTTACAGATGAGATGGTCAGACTAAACTGGCTGACGGAATTTAT
GCCTCTTCCGACCATCAAGCATTTTATCCGTACTCCTGATGATGCATGGTTACTCACCA
CTGCGATCCCCGGCAAAACAGCATTCCAGGTATTAGAAGAATATCCTGATTCAGGTGA
AAATATTGTTGATGCGCTGGCAGTGTTCCTGCGCCGGTTGCATTCGATTCCTGTTTGTA
ATTGTCCTTTTAACAGCGATCGCGTATTTCGTCTCGCTCAGGCGCAATCACGAATGAAT
AACGGTTTGGTTGATGCGAGTGATTTTGATGACGAGCGTAATGGCTGGCCTGTTGAAC
AAGTCTGGAAAGAAATGCATAAGCTTTTGCCATTCTCACCGGATTCAGTCGTCACTCA
TGGTGATTTCTCACTTGATAACCTTATTTTTGACGAGGGGAAATTAATAGGTTGTATTG
ATGTTGGACGAGTCGGAATCGCAGACCGATACCAGGATCTTGCCATCCTATGGAACTG
CCTCGGTGAGTTTTCTCCTTCATTACAGAAACGGCTTTTTCAAAAATATGGTATTGATA
ATCCTGATATGAATAAATTGCAGTTTCATTTGATGCTCGATGAGTTTTTCTAATCAGTAC
TGACAATAAAAAGATTCTTGTTTTCAAGAACTTGTCATTTGTATAGTTTTTTTATATTGT
AGTTGTTCTATTTTAATCAAATGTTAGCGTGATTTATATTTTTTTTCGCCTCGACATCATC
TGCCCAGATGCGAAGTTAAGTGCGCAGAAAGTAATATCATGCGTCAATCGTATGTGAA
TGCTGGTCGCTATACTGCTGTCGATTCGATACTAACGCCGCCATCCAGTGACACTGAA
GAGGTATAGTCATGCCTACCGCGATTTCTTTGACACAGAATTGAAAAATTTTGCATTTT
TTGGTAATTTCCTAATAATACGAAGTGCAATAATCTCACTTTGATAGGAGCACGTCATG
ATGGTAATTTCATACTACTGAACGTAAATGTTGAAGGTGAATTTGTAAAGCTTTGTTAT
TTGAAGCTGTACACCTAAAGGGCGGTAAATTGTTGGGTGTTAGTACCATGCCATTAATT
AGAATTTGTTAGCATTTTTATTCATTTGTGCATTATGGGTTCAAATTCATATGACTGAAC
GTGTAGTTTCATATCCAGTCATCAAGAGATGCTGAACCGCCCTTCAAAAACTTGACGA
AATAGA

4. chr3-cir: donor used to cyclize Sc chr3

(1) Composition of the donor:

Left HA-T$_{ADH1}$-*KmURA3*-P$_{KmUAR3}$-DR-AscI-Right HA

(2) Sequence (5'-3'):

GTTGGCGCCTCCGAACAAACAGATGCAGAAATCAAGATTAGTGAAGCATTGGCCAAG
ATTGCTGAGGAACATGGCACTGAGTCTGTTACTGCTATTGCTATTGCCTATGTTCGCTC
TAAGGCGAAAAATTTTTTCCGTCGGTTGAAGGAGGAAAAATTGAGGATCTCAAAGA
GAACATTAAGGCTCTCAGTATCGATCTAACGCCAGACAATATAAAATACTTAGAAAGT
ATAGTTCCTTTTGACATCGGATTTCCTAATAATTTTATCGTGTTAAATTCCTTGACTCAA
AAATATGGTACGAATAATGTTTAGATAATTTTTCAGTAATCAACTACGCAAGTAAAGCA
GTAAATACGTTACTGCTGGTATTAATGTCATGTATTGAGGCAATGATGCTATGCTCTTAA
CGACATGTAGCTTACAAAACGTTCCTATGGTTTTACTGCATGGTATTTACAAATTAGAT
AGCAGTTCCATCCGCCCTTGACATATTATTATTCAGACAAGGTGTATATGAGCATAAATA
TGTATATATGCACATGAGGTTTGTATTAACCTTGCAACTGTACCAAAATACAATCTTCTT
TGCTGCTATTTAACGGCGTATGTGCAGTTCATAAATGCGTGTTCTGTATAGTACATCCG
CGCACATTCTTCCTAGCGGAAGACATATTAACGTAGCCCGTACGCCCTGGTTAAGACT
TGGTATAGTTCCAATAATTGGAATATTACTGCATAGTGGTCCAATAGCAGGATTTAGCAT
AAACATGATAATATTTAGAGATGCTTTCATCCCTTGCCGGTAGAGGTGTGGTCAATAAG
AGCGACCTCATACTATACCTGAGAAAGCAACCTGACCTACAGGAAAGAGTTACTCAA
GAATAAGAATTTCGTTTTAAAACCTAAGAGTCACTTTAAAATTTGTATACACTTATTT
TTTTTATAACTTATTTAATAATAAAAATCATAAATCATAAGAAATTCGCTTAAGCGGATC
TGCCTACTCTCTTCAAGTAAGCGTCCCATCCCGCCTTTCTGTATCTCTCACCTTCCACT
ACAGGATCTCTTCCCTTTGCGAAAAGACCTCTACCAACAATAATGATGTCTGATCCAC
CGGCAACAACTTCATCCACAGTTCTGTATTGTTGTCCCAAAGCATCACCTTTGTCATC
AAGACCAACACCTGGCGTCATGATCAACCAATCGTAGCCCTCTTCTCTTCCACCCATA
TCGTTTTGAGCAATAAATCCAATAACAAAGTCCTTATCACTCTTGGCAATTTCCACGGT
CCCACGAGTGTATTCACCGTGCGCTAGAGACCCCTTGGACGATAACTCGGCAAGCATT
AACAACCCTCTAGGTTCTTTTGTAACTTCCTCGGCACCTTGCTTCAAACCAGCAACAA
TGCCCGCACCAGTCACACCGTGTGCATTGGTGATATCAGACCATTCGGCGATACGGTA
TACACCAGACGTGTATTGTAATTTAACAGTGTTCCCAATGTCGGCAAACTTCCTGTCTT
CAAATATCAAAAACTTGTGTTTCTCTGCTAATTGCTTCAACGGCACAATGGTATTCTCA
AAGCTGAAATCCTCCAAGATATCTACATGTGTCTTCAATAGACAGATATATGGACCCAA
AACCTCAACTAATCTTAACAACTCTGCTGTTTTACGAACATCAAGAGAAGCACATAAG
TTTGACTTCTTCTCTTCCATCAAGTTTAAAAGCTTGGCAGCAACTGGACTTCTATGAG
CAGCTGCTCTTTCCGAGTAACTCTTAGTCGACATCCTCCTTTGATTAGTTTATGGGGCT
AGCTGATGATACTTCCTCTTGCTGTTCCAGTTTACAAATGATTTAAACGAACTTTTACA
CTTTTACGGTAGAATTTCTTGTTGGTGATGAGCTCAGATTCAATATGACTGCTTAATCA
GTGAAAATTAATGGAAACAGCCAGACACACAGCCTTTGAAAAATGTTAAAAACGATC
ACATATATGTAGTCACGTGGTTAGTGCATAGAGTCAGTCGATGGTGGTTAGTTCCCTAG
CCAGAGCCATTCCCCTCTTGTTTCGAAATGCGATGACTGAATGGGGAAGAAGCTCTTA
GTATTTCTGTAGTTTTTTTGCCGATTTCTATTGTTGCCTGCCCCGGACTTTCTTTTACC
TTTTGTAGCTACAACTATTAATTATAATGCATAATAGATCTAGAAGCGTATTGTGATGAG
CATCCTTGGAAATAAGAAAGGAAGTCAAAGGTACAGTAAAAAAAGAGCCGTGACAG
GCACCTAATTTCAGTTTTGATTGAGGTATAATGAGCTCAGAAGACCAAGATGCTCTAA
AAGTAAAGCAGAATGGTCTTTCCAATCAGAATTCGGAAGCATATCCATCGTGCTTAAA
ATGATTGGGTCCGCGTCCGACTCACCTAGCAAGTTAGGACGCCTCCGATTTCTTTCTG
AAACTGCCGCTATTAAAGTATCCCCGTTAATCCTAGGAGAAGTCTCATACGATGGAGC
ACGTTCGGATTTTCTCAAATCAATGAACAAGAATCGAGCTTTTGAATTGCTTGATACTT

TTTACGAGGCAGGTGGAAATTTCATTGATGCCGCAAACAACTGCCAAAACGAGCAAT
CAGAAGAATGGATTGGTGAATGGATACAGTCCAGAAGGTTACGTGATCAAATTGTCAT
TGCAACCAAGTTTATAAAAAGCGATAAAAAGTATAAAGCAGGTGAAAGTAACACTGC
CAACTACTGTGGTAATCACAAGCGTAGTTTACATGTGAGTGTGAGGGATTCTCTCCGC
AAATTGCAAACTGATTGGATTGATATACTTTACGTTCACTGGTGGGATTATATGAGTTC
AATCGAAGAATTTATGGATAGTTTGCATATTCTGGTCCAGCAGGGCAAGGTCCTCTATT
TGGGTGTATCTGATACACCTGCTTGGGTTGTTTCTGCGGCAAATTACTACGCTACATCT
TATGGTAAAACTCCCTTTAGTATCTACCAAGGTAAATGGAACGTGTTGAACAGAGATT
TTGAGCGTGATATTATTCCAATGGCTAGGCATTTCGGTATGGCCCTCGCCCCATGGGAT
GTCATGGGAGGTGGAAGATTTCAGAGTAAAAAGCAATGGAGGAACGGAGGAAGAA
TGGAGAGGGTATTCGTTCTTTCGGCGCGCCTGTTGAATGTGGTAACCCAATAGCATGA
TATGAGTAATGCTTTAGTATTGTTTCAGAGTTGTTTCAGTAATGTTTTAGACAAGGAGA
ACATATAGTAGCAAACCTCTAATCCGGTAGTACTTAAGAAACTACAGTTTCTATGTACG
AAAGCAGTAACTATGTAATTATTACATTTACATGACATATAGGAAGGTCCAATAAACTT
ACTACATTATGACCTATAAGCTAGATCGTAATTCATTACGTCAACAGGTTATGAGCCCT
CAGAGCAATGCTTCTGAGAACATAATCAATCTATCTAGCCCCAACAATTATAAACAGT
GGCTGTACGGTATCGAGACCGCTGCTGAATATGCTAACGAATATATGAACGAATTCGTT
CATACCGGAGATATCCAATCAATGAAAAGGGATTACAATCTCAGCGCGAATGATGAAA
GCTTTGTCAAAACCGTATTTAACAGTTTCCTGGTAAAGCTCTACAAGAAAACTATCGT
GGGTGAAGCTGCATGTGAAATGAACTGGATATGTGATGATTCGCTTGGAAGGGTCTCT
GCTTATGATATTTTCTCGCACTTCGAAGAAAACTATAATGAAGTCACTATTGGATCCAG
GCTTACTCTTATAGAGGACCTACCAAATATATCCTCCAAGCCTGTAGATGAAATTGCTT
CCTTTTTGAAAACTCTATTCACGATGCTTGAAGACAATAGCGAAGAACAGGACAAAA
AAAAAAGACGCGACACCAATATCGCGTTGTTATTAATGACCTTCTTACCCGAGTTAAA
AGAATCATTCCA

5. chr3-ARS1/CEN5: donor used to insert *KmARS1*/*KmCEN5* and $hyg^R$ into Sc chr3

(1) Composition of the donor:

Left HA-*KmARS1*-*KmCEN5*-$P_{TEF}$-$hyg^R$-$T_{TEF}$-Right HA

(2) Sequence (5’-3’):

GTGCAGCTATGCATGGGAGTGATTTCACCTCTCTTCCGTGCCCTCGTGCAACTGCCTT
TATCAATTTTCGTATGGAATGGGTTTCAGCTTGTGGCATTGCCTATCAACATCCCATTG
AGGTTGTTCCTGGGGACGTCGCTGAGCCGTTTAGTTGCACAAACAAGCACATTAGAC
TTCTATGTGGTTTTGACGTTGTTTCAATATTTTGCTGTGCTTTGTGCTTTTGGCAGCATC
ATAGGACTCATCTTTGGATTTATATTGGGTGTGTTCCACTCAATCTGCGGGGTACCCAG
TGTATACATAAGTCTAGAATGGAAACGGTGGTTTGCTCCGATACGTACGGTCCTTGAA
CGTGCTTCCACTAGTATTGTCAACATTATGCGAGGACAAACTATTGCGCCAATACCCAT
GCCTAAGCCCAATCCCACGCATATATCAAAGCCTAACATGAAAAAATTCCATGATGAG
CCTGGAGCTGATGATATGACTATAACGCATGATGTGAACTGCTACATCACCCCTTGCCA
AACGCCTACTAACGAAAAAATTCAGCATTATAATAATGATTCATTCAACACGACCACAT
CGATTGAAGTTTTGTCCAACTATCCACTATGGATATGCGTTTTGTTGATTAACCTTAAAT
AACACGTATTTCGCATTTTCCAAAAGCCTTTTTTCATAACTACAAACTAACTATTTTTG
TTTATTTTACATCAGTAAATTATGCGCAGAAAATATGTAAGGCTATATACTCAATATAGT
GGAAGAGCCTCTGGCTACTTAATCTGGGTTCATATATTCTGTCAGTGGTATAGTAAATT
TAAATAGTGAATTTGGGCGCATGTAGAGAGATCTTTTGATTAATAACCCAGTATTTGAT
TCTTGAATGTTATTTGTCTCTTATTAAGTATTTCAGTTTGATTTATATATTTTGAAAGTAA

TTGTTGCTTTACCTTCCAAACAATAAAAAAATATAAAAAAATGTAAAAAATATAATAAA
TATTAAATAAAATACTACTTGTTTGAAAATCAGAGAAAATCCACCAAAATATCAATCAT
TTCAAGGATTTCCGAACCAAGTTCGAGATAGTCCTTTAAGGTCAGTAAAATTCAGTTG
CACGTATAACAGATATTTTTCATTTTGTTCCAATTTAAAAGTCCCCCATTTTTAAAATTA
TTGAAAATAAAAATTAAAAAATTAAAAGGAATCTCTCTATGTCACTTTAAAATAAATAA
ATTGAAAATGATATATCGTTAAAAGGATCCCGCAACACCACCTATTTCTAAGAGGAGA
GTTCTAAAAAAATCATAGTACCACACAGGTAAACTAAAACAGCTAAATTCAACATAAT
ACCTGTGGATATAATTACATACAAAAATATAATTAAAAAAATACATTAAAAATAATTTAT
TTTTGTAAAACCCATAAAATATATTTTACTTTCGGAACAACTTTTTAACTTATAATTTTG
TTTTAAATAAAAACGTTGTATTTAAAAATAATAAAATATTAAGTAAAAATTTAAAGCAT
ATTTATTTAAAATATAAAATACTACTAAAATTACTCTAAACTTCAAAATAAAAAAATAAT
AAATTATATAGTTTTAAAATTAATAATTTGTATACACGTGACCAGACCATAATAGTTTTC
TTTTCTTGAAACTGCCATGAATTTAATAGATTTTTTTTACACATATTTACAGTAAGTTTT
GTTTTTACGTTGAATTAATATTGTATACGTACTTAGAAGGATAACTTCCAACTATGAATA
TGTAGGTTAAAAGGTAAATAGAGAAGCCTTACTTTTATCAGAAATGAAAGGAGCTCA
GCTTGCCTTGTCCCCGCCGGGTCACCCGGCCAGCGACATGGAGGCCCAGAATACCCT
CCTTGACAGTCTTGACGTGCGCAGCTCAGGGGCATGATGTGACTGTCGCCCGTACATT
TAGCCCATACATCCCCATGTATAATCATTTGCATCCATACATTTTGATGGCCGCACGGCG
CGAAGCAAAAATTACGGCTCCTCGCTGCAGACCTGCGAGCAGGGAAACGCTCCCCTC
ACAGACGCGTTGAATTGTCCCCACGCCGCGCCCCTGTAGAGAAATATAAAAGGTTAG
GATTTGCCACTGAGGTTCTTCTTTCATATACTTCCTTTTAAAATCTTGCTAGGATACAGT
TCTCACATCACATCCGAACATAAACAACCATGGGTAAAAAGCCTGAACTCACCGCGA
CGTCTGTCGAGAAGTTTCTGATCGAAAAGTTCGACAGCGTCTCCGACCTGATGCAGC
TCTCGGAGGGCGAAGAATCTCGTGCTTTCAGCTTCGATGTAGGAGGGCGTGGATATGT
CCTGCGGGTAAATAGCTGCGCCGATGGTTTCTACAAAGATCGTTATGTTTATCGGCACT
TTGCATCGGCCGCGCTCCCGATTCCGGAAGTGCTTGACATTGGGGAATTCAGCGAGA
GCCTGACCTATTGCATCTCCCGCCGTGCACAGGGTGTCACGTTGCAAGACCTGCCTG
AAACCGAACTGCCCGCTGTTCTGCAGCCGGTCGCGGAGGCCATGGATGCGATCGCTG
CGGCCGATCTTAGCCAGACGAGCGGGTTCGGCCCATTCGGACCGCAAGGAATCGGTC
AATACACTACATGGCGTGATTTCATATGCGCGATTGCTGATCCCCATGTGTATCACTGG
CAAACTGTGATGGACGACACCGTCAGTGCGTCCGTCGCGCAGGCTCTCGATGAGCTG
ATGCTTTGGGCCGAGGACTGCCCCGAAGTCCGGCACCTCGTGCACGCGGATTTCGGC
TCCAACAATGTCCTGACGGACAATGGCCGCATAACAGCGGTCATTGACTGGAGCGAG
GCGATGTTCGGGGATTCCCAATACGAGGTCGCCAACATCTTCTTCTGGAGGCCGTGGT
TGGCTTGTATGGAGCAGCAGACGCGCTACTTCGAGCGGAGGCATCCGGAGCTTGCAG
GATCGCCGCGGCTCCGGGCGTATATGCTCCGCATTGGTCTTGACCAACTCTATCAGAG
CTTGGTTGACGGCAATTTCGATGATGCAGCTTGGGCGCAGGGTCGATGCGACGCAAT
CGTCCGATCCGGAGCCGGGACTGTCGGGCGTACACAAATCGCCCGCAGAAGCGCGG
CCGTCTGGACCGATGGCTGTGTAGAAGTACTCGCCGATAGTGGAAACCGACGCCCCA
GCACTCGTCCGAGGGCAAAGGAATAATCAGTACTGACAATAAAAAGATTCTTGTTTTC
AAGAACTTGTCATTTGTATAGTTTTTTTATATTGTAGTTGTTCTATTTTAATCAAATGTTA
GCGTGATTTATATTTTTTTTCGCCTCGACATCATCTGCCCAGATGCGAAGTTAAGTGCG
CAGAAAGTAATATCATGCGTCAATCGTATGTGAATGCTGGTCGCTATACTGCTGTCGAT
TCGATACTAACGCCGCCATCCAGTATAGGTCTGACACTTACCAAAACTCATTCGTCAC
CAATGAAACTTTGATGTCTCTTTCTAATAGAGCTAAGCTTCGAAGAAATGCCAGTGAT
GCGGACATCGTTAATATAAAGATTTTACGAAGGAATTCTAGGTGATATTGCAATTACTT

CTTCTCATGCACTAACAAGTGAATGATAGAAATATGTTGAGTTCCTAACTGCCTGATTT
TAAATAAGTTTCATATTATAATCTTTTAGCATATATATATATATATTGATCCTCTCTCTTCTT
TATTTTCTGCCAGTAACCCATGTGTGAAGAAGAAAACATAAATAAAAAAGCAGTAGC
ACATGGACACATTCACGCCCGAACACTCCTAAAAAGCAGCCCACACAAGAAAGTAG
ATATAATGTAGGACACCCAGCTTGTCCATAATTGCTAATAGCATACTCAGGATAACATAT
ATTAATGACGACTCGTTTGCTCCAACTCACTCGTCCTCATTACAGATTATTATCCCTACC
TCTCCAGAAACCCTTCAATATAAAAGGCAGATGTCCGCTGCGAACCCTTCTCCATTT
GGCAATTATTTGAACACGATCACTAAGTCCCTACAA

6. chr3-ARS1: donr used to insert *KmARS1* and $kan^R$ into Sc chr3
(1) Composition of the donor:
Left HA-KmARS1-$P_{TEF}$-*KanMX6*-$T_{TEF}$-Right HA
(2) Sequence (5'-3'):
TCTGTGGGTATTTCCGTGTATATGGTTAATAATAGTAGTATCTTGTCAGTTTTTTTATGT
TTTTCTTCGCGCGTCAACTTTCTACCAAGAGAAAAACAATATAAGGTCTCCTTACTCTA
TAGGAGAATAAAACAAACAAAAATAAAAAGCACATCGTAGCGCCAAGAAAATACTGC
AAATACCAAATACCACAATAATACTACAATTATCTATACACAAGTGTTTTGCCGCTTAA
AACTTCGATTTTCATAGTACGAAACTATACACCCTTGGTTTTTCTCTTTTCTAAATACAT
ATCTACCTTGTAAAGAATTTCCCGCACATCTTTGCGGGCATACAGTTCATGTATTGGCA
ACTAACGGAACTAAGGCAACATATCTTGCATATTGCAATGTTCACTATATAGATGAAAA
CTTATATCTAGGTTTCACGACACGAGAATAACTAAGAAGCACGATCCATGATATAGAA
AAATCAGTTACGACGAAGCACGGCAAATTAGCCGCCGAAGACCGATATTTTTGCCAC
CGGTCACAGTTTCTTTTCCACGGAGCTCTTCGCGGTTTTTTTGTTCCGGATTTTTTT
ACCGGCTCTTAGATCGATTGAAGTTTTGTCCAACTATCCACTATGGATATGCGTTTTGT
TGATTAACCTTAAATAACACGTATTTCGCATTTTCCAAAAGCCTTTTTTCATAACTACA
AACTAACTATTTTTGTTTATTTTACATCAGTAAATTATGCGCAGAAAATATGTAAGGCTA
TATACTCAATATAGTGGAAGAGCCTCTGGCTACTTAATCTGGGTTCATATATTCTGTCAG
TGGTATAGTAAATTTAAATAGTGAATTTGGGCGCATGTAGAGAGATCTTTTGATTAATA
ACCCAGTATTTGATTCTTGAATGTTATTTGTCTCTTATTAAGTATTTCAGTTTGATTTATA
TATTTTGAAAGTAATTGTTGCTTTACCTTCCAAACAATAAAAAAATATAAAAAAATGTA
AAAAATATAATAAATATTAAATAAAATACTACTTGTTTGAAAATCAGAGAAAATCCACC
AAAATATCAATCATTTCAAGGATTTCCGAACCAAGTTCGAGATAGTCCTTTAAGGTCA
GTAAAATTCAGTTGCACGTATAACAGATATTTTTCATTTTGTTCCAATTTAAAAGTCCC
CCATTTTTAAAATTATTGAAAATAAAAATTAAAAAATTAAAAGGAATCTCTCTATGTCA
CTTTAAAATAAATAAATTGAAAATGATATATCGTTAAAAGAGCTTGCCTCGTCCCCGCC
GGGTCACCCGGCCAGCGACATGGAGGCCCAGAATACCCTCCTTGACAGTCTTGACGT
GCGCAGCTCAGGGGCATGATGTGACTGTCGCCCGTACATTTAGCCCATACATCCCCAT
GTATAATCATTTGCATCCATACATTTTGATGGCCGCACGGCGCGAAGCAAAAATTACG
GCTCCTCGCTGCAGACCTGCGAGCAGGGAAACGCTCCCCTCACAGACGCGTTGAATT
GTCCCCACGCCGCGCCCCTGTAGAGAAATATAAAAGGTTAGGATTTGCCACTGAGGTT
CTTCTTTCATATACTTCCTTTTAAAATCTTGCTAGGATACAGTTCTCACATCACATCCGA
ACATAAACAACCATGGGTAAGGAAAAGACTCACGTTTCGAGGCCGCGATTAAATTCC
AACATGGATGCTGATTTATATGGGTATAAATGGGCTCGCGATAATGTCGGGCAATCAGG
TGCGACAATCTATCGATTGTATGGGAAGCCCGATGCGCCAGAGTTGTTTCTGAAACAT
GGCAAAGGTAGCGTTGCCAATGATGTTACAGATGAGATGGTCAGACTAAACTGGCTG
ACGGAATTTATGCCTCTTCCGACCATCAAGCATTTTATCCGTACTCCTGATGATGCATG

GTTACTCACCACTGCGATCCCCGGCAAAACAGCATTCCAGGTATTAGAAGAATATCCT
GATTCAGGTGAAAATATTGTTGATGCGCTGGCAGTGTTCCTGCGCCGGTTGCATTCGA
TTCCTGTTTGTAATTGTCCTTTTAACAGCGATCGCGTATTTCGTCTCGCTCAGGCGCAA
TCACGAATGAATAACGGTTTGGTTGATGCGAGTGATTTTGATGACGAGCGTAATGGCT
GGCCTGTTGAACAAGTCTGGAAAGAAATGCATAAGCTTTTGCCATTCTCACCGGATTC
AGTCGTCACTCATGGTGATTTCTCACTTGATAACCTTATTTTTGACGAGGGGAAATTAA
TAGGTTGTATTGATGTTGGACGAGTCGGAATCGCAGACCGATACCAGGATCTTGCCAT
CCTATGGAACTGCCTCGGTGAGTTTTCTCCTTCATTACAGAAACGGCTTTTTCAAAAA
TATGGTATTGATAATCCTGATATGAATAAATTGCAGTTTCATTTGATGCTCGATGAGTTT
TTCTAATCAGTACTGACAATAAAAAGATTCTTGTTTTCAAGAACTTGTCATTTGTATAG
TTTTTTTATATTGTAGTTGTTCTATTTTAATCAAATGTTAGCGTGATTTATATTTTTTTTCG
CCTCGACATCATCTGCCCAGATGCGAAGTTAAGTGCGCAGAAAGTAATATCATGCGTC
AATCGTATGTGAATGCTGGTCGCTATACTGCTGTCGATTCGATACTAACGCCGCCATCC
AGTTTATAAGGGGAGTGGCAGCGGCGGTAGACACTGCGCTCTATAAGAATACTTGCA
AGGGTCTTGTCTATTGTATAATTCGCTAGTATTTGTTTTGCATTGTACTCTTAATACCCC
AACCAAAAACAAAATAGTGAGAGTAATGGTGCGTTTTGTTTCAATTTTAAGTTTATTC
GGCTGCGCGGCGACGCTTGTCACGGCCCATGATGACATGGACATGGACATGGATATG
GACATGGATATGGACATGAATATCGATACGACAACGTCTCAATCCATAGATGTCTCATC
CACGGCTTCAATCGTCCCCGTGCCACATGAACCAAAACATTTGCATGGCCTTCCTATA
CTGCAATCGCCCTCGCTTACCCCTGCGGAGAGATTGTACTGGGAAAACTACAACACC
ACAACCTACTTTACTACACAGGCTGGGAATAGGTCTGCCCTTCGCTACCACATTATTAC
GCTGCTCTTGGTTGCATTTGTGCTCTACCCTGTGTCCCTGGCGCTAAGCGCCGCCCGT
TCTAGGTGGTACTTACCCCTGCTGTTTGTTAATCTATGCATTTGTATTTCGTCCGTAATG
GCATTGTCCGTGTTCAAAAA
